# Transcriptional profiling sheds light on the fibrotic aspects of idiopathic subglottic tracheal stenosis

**DOI:** 10.1101/2024.02.19.580975

**Authors:** Martin Direder, Maria Laggner, Dragan Copic, Katharina Klas, Daniel Bormann, Thomas Schweiger, Konrad Hoetzenecker, Clemens Aigner, Hendrik Jan Ankersmit, Michael Mildner

## Abstract

Idiopathic subglottic stenosis (ISGS) is a rare fibrotic disease of the upper trachea with an unknown pathomechanism. It typically affects adult Caucasian female patients, leading to severe airway constrictions caused by progressive scar formation and inflammation with clinical symptoms of dyspnoea, stridor and potential changes to the voice. Endoscopic treatment frequently leads to recurrence, whereas surgical resection and reconstruction provides excellent long-term functional outcome. This study aimed to identify so far unrecognized pathologic aspects of ISGS using single cell RNA sequencing. Our scRNAseq analysis uncovered the cellular composition of the subglottic scar tissue, including the presence of a pathologic, profibrotic fibroblast subtype and the presence of Schwann cells in a profibrotic state. In addition, a pathology-associated increase of plasma cells was identified. Using extended bioinformatics analyses, we decoded pathology-associated changes of factors of the extracellular matrix. Our data identified ongoing fibrotic processes in ISGS and provide novel insights on the contribution of fibroblasts, Schwann cells and plasma cells to the pathogenesis of ISGS. This knowledge could impact the development of novel approaches for diagnosis and therapy of ISGS.

## 2 Introduction

Idiopathic subglottic stenosis (ISGS) is a rare pathology of the subglottic larynx and upper trachea, occurring at a rate of 1 in 400,000 per year (1). It is characterized by fibrotic lesions with unknown etiology (1). These lesions typically exhibit a circumferential pattern, extending up to a height of 5 cm and are predominantly located within a region approximately 3 cm below the level of the vocal cords (2). Excessive deposition of extracellular matrix (ECM) in ISGS leads to symptoms including cough and wheezing, and a significant stridor and dyspnoea at rest in later stages of the disease (2). Due to significant recurrence rates of up to 50% following endoscopic treatment, repeated surgical interventions are often performed to prevent severe airway obstruction (1). Surgical resection with removal of all affected scar tissue provides the best long-term results with recurrence rates below 5% (3). Recent investigations into the pathomechanisms of ISGS revealed a high number of infiltrating T-cells accompanied by the activation of the IL-17A/IL23 pathway (4). Further research uncovered a direct influence of IL-17A on fibroblast proliferation in ISGS, along with an interplay with TGF-ß in regulating collagen production by fibroblasts (5). A potential association between a specific upper airway microbiota, especially members of the Moraxellaceae family, and ISGS has also been discussed (6). In contrast to patients with granulomatosis with polyangiitis a human leucocyte antigen (HLA) association with allele DPB1*04:01 or allele homozygosity was excluded in ISGS (7). Among ISGS patients, clonality of the TCR repertoire is driven by CD8+ T cells, and ISGS patients possess numerous TCRs targeting viral and intercellular pathogens. High frequency clonotypes do not map to known targets in public datasets (8).

There is a notably high incidence observed in Caucasian female perimenopausal individuals (1). Interestingly, premenopausal patients were shown to have a more aggressive disease variant than peri- and postmenopausal patients. However, it is unclear whether this is related to reduced estrogen in the peri- and postmenopausal state or the age-related physiology of wound healing and inflammation, regardless of estrogen (9).

At the pathophysiological level, numerous questions regarding the diseases are still unanswered. Recent studies found parallels of ISGS and other fibrotic diseases (10, 11). The application of newly developed methods such as single cell RNA sequencing (scRNAseq) provides important insights into the prevailing cellular and transcriptional situation of multiple pathologies (12). Using this method, research groups have decoded the pathologic airway epithelium of ISGS and identified S100A8/A9 as a key biomarker of ISGS macrophages (12-14). Fibrotic diseases are typically driven by multiple different cell types, with fibroblasts assuming a crucial role as the primary producers of the ECM in most cases (15). Transcriptional studies already identified pro-fibrotic subtypes of fibroblast in distinct fibrotic diseases (16-18). Beyond the well-established role of fibroblasts, recent discoveries have highlighted the involvement of Schwann cells in fibrotic processes (19, 20). These profibrotic Schwann cells exhibit a specific molecular pattern, contributing the matrix formation and influence other cell types, particularly macrophages (20, 21).

In this study, we performed single cell RNA sequencing utilizing resected specimens of patients suffering from ISGS, to unravel the cellular and transcriptional composition of ISGS compared to healthy, unaffected tracheal tissue. We hypothesize that an unbiased analysis of the transcriptional datasets may yield novel insights into the pathology of ISGS, providing new information on potential pathologic cellular states and disease-induced changes of the prevailing cellular environment. Our findings confirm the presence of pathologic, pro-fibrotic fibroblasts in the ISGS and unveil their transcriptional pattern. The results of this study further suggest a potential involvement of Schwann cell in this fibrotic disease and provide information on a pathological increase of plasma cells in ISGS.

## 3 Materials and Methods

### 3.1 Sample acquisition

Tissue samples of idiopathic subglottic stenosis and of unaffected, healthy tracheal areas were obtained from women who underwent open surgery with resection of the affected tracheal segment (donor information – Table S1). A total of six patients was included, whereby paired samples of diseased and healthy tissues were obtained from five. Tissue of healthy trachea has been obtained including cartilage, whereas the tissue of the stenosis was pure fibrotic tissue. For further processing, the cartilage was removed as much as possible using a scalpel. Patients with previous chemotherapeutic or radiation treatment were excluded. Only surplus tissue not required for pathologic examination was used in this study. Written informed consent was obtained from all donors.

## 3.2 Cell isolation and generation of cell suspension

Upon tracheal resection, tissues were cooled and immediately processed. Samples were washed with sterile Dulbecco’s phosphate-buffered saline (PBS, without Ca^2+^ and Mg^2+^, Gibco, Thermo Fisher Scientific, Waltham, MA, USA), mechanically minced, and enzymatically dissociated using MACS Miltenyi Multi Tissue Dissociation Kit 1 (Miltenyi Biotec, Bergisch Gladbach, Germany) in accordance with the manufacturer’s instructions. Cell aggregates were dissociated using gentleMACS OctoDissociator (Miltenyi) with the standard gentleMACs program’ 37C_tdk_1’. Afterwards, cell suspensions were passed through 70 and 40 µm cell strainers and washed twice with 0.04% bovine serum albumin (BSA, Sigma Aldrich, St. Louis, MO, USA) in PBS. Cell number and viability were determined using a LUNA-FL™ Dual Fluorescence Cell Counter (Logos Biosystems, Anyang-si, Gyeonggi-do, South Korea) and the Acridine Orange/Propidium Iodide (AO/PI) Cell Viability Kit (Logos Biosystems). Only cell suspensions displaying a viability >80% were further processed and cell counts were set to 0.7 - 1.0 x 10^6^ cells/ml. The isolation process was performed for all samples within 4 hours after resection and permanent cooling to minimize cell damage.

### 3.3 Single cell RNA sequencing

Single cell suspensions were further used for Gel Beads-in-emulsion (GEM) preparation, cDNA amplification and library preparation was performed using the Chromium Next GEM Single Cell 5’ Kit v2, the Dual Index Kit TT Set A (all 10x Genomics, Pleasanton, CA, USA), Chromium Next GEM Chips type K (10X Genomics) and the Chromium controller (10X Genomics), as described previously (20). RNA sequencing, demultiplexing and counting was performed by the Biomedical Sequencing Core Facility of the Center for Molecular Medicine (CeMM Research Center for Molecular Medicine, Vienna, Austria). Samples were sequenced (read length 50bp) on a NovaSeq 6000 (Illumina, San Diego, CA, USA). Raw reads were demultiplexed, aligned to the human reference genome (GrCh38-2020-A) and counted using the Cellranger pipeline (Cellranger v.6.1.2, 10x Genomics).

### 3.4 Bioinformatics analysis

For Bioinformatics analyses, R (R v.4.0.3, The R Foundation, Vienna, Austria), R-studio and Seurat (Seurat v.4.3.0.1, Satija wellLab) were used (22).First, all dataset features were proven for their annotation as NCBI symbol according to the Homo sapiens Ensemb ID and potential feature duplicates were removed. Datasets were converted into Seurat objects. To remove apoptotic cells and erythrocytes from analyses, only cells displaying <5% mitochondrial counts and <5% hemoglobin subunit beta (*HBB*) counts were included in the study. The purified data were pre-processed using sctransform-normalization with the glmGamPoi package and subsequently, all datasets were integrated according to the Seurat Vignette (23, 24). Next, PCA and UMAP were calculated. The “FindAllMarkers” command together with well-established marker genes were used for cluster annotation (marker genes information – Table S2). GO-Term analyses were performed applying Enrichr and Metascape [https://metascape.org; accessed on 2023-12-21] (25-28). For Metascape analysis, a p-value cutoff of 0.05 and a minimum enrichment score of 2 were defined as significant. All subset analyses were performed based on the raw data of the selected cell cluster. Subtype characterization was conducted using published subtype clustermarker (21, 29). The subtype annotation of plasma cells according to published subtype marker genes was not sufficient, therefore a consecutive numbering of the clusters was chosen. Potential cell-cell interactions were identified by CellChat (30). Monocle 3 (Monocle3, v0.2.3.0, Trapnell Lab) was used for pseudotime-trajectory analyses (31-35). The screening of the matrisome was performed according to the list published by Naba et al. (36).

### 3.5 Immunofluorescence and Hematoxylin & Eosin staining

In total 3 tissue samples of each condition were washed with PBS and fixed in 4.5% formaldehyde solution, neutral buffered (SAV Liquid Production GmbH, Flintsbach am Inn,Germany) for 24 h at 4 °C, directly following surgery. On the next day, the samples were washed with PBS overnight and then dehydrated by step-wise incubation with 10, 25, and 42% sucrose solution, each overnight at 4°C. The samples were snap-frozen using optimal cutting temperature compound (OCT compound, TissueTek, Sakura, Alphen aan den Rijn, The Netherlands) and preserved at -80 °C. Sections of 10 µm were cut with a cryotome (Leica, Wetzlar, Germany), dried for 30 min and immersed in PBS. Permeabilization and blocking of the sections was performed for 15 min using 1% BSA, 5% goat serum (DAKO, Glostrup, Denmark) and 0.3% Triton-X (Sigma Aldrich) in PBS. Details for antibodies, dilutions and incubation times can be found in Table S3. Antibodies not ready-to-use were diluted in antibody staining solution (1% BSA, 0,1% Triton-X in PBS). Incubation with secondary antibodies was performed for 1 h in combination with 50 µg/ml 4,6-diamidino-2-phenylindole (DAPI, Thermo Fisher Scientific). Next, sections were mounted with appropriate medium (Fluoromount-G, SouthernBiotech, Birmingham, AL, USA) and stored at 4°C. Imaging was performed with a confocal laser scanning microscope (TCS SP8X, Leica) equipped with a 20x (0.75 HC-Plan-Apochromat, Multimmersion), a 20x (0.75 HC-Plan-Apochromat) and a 63x (1.3 HC-Plan-Apochromat, Glycerol) objective using Leica application suite X version 1.8.1.13759 or LAS AF Lite software (both Leica) and an Olympus BX63 microscope (Olympus, Tokyo, Japan) with Olympus CellSens Dimension v2.3 (Olympus) software with standardized exposure time for all samples. A maximum projection of total z-stacks is depicted in the confocal images. Hematoxylin and eosin staining (H&E) of healthy trachea and ISGS tissue was performed according to standard protocol.

### 3.6 Fiber alignment examination

For fiber alignment analysis, H&E staining of three ISGS and three healthy trachea tissue slides were imaged and examined using Curvealign V4.0 Beta (MATLAB software, Cleve Moler, MathWorks, Natick, Massachusetts, USA). Fiber contrast, brightness and color were optimized by Adobe Photoshop CS6 (Adobe, Inc., San Jose, CA, USA). Three regions of interest (size 256 pixels, 256 pixels) were analyzed per image. For statistical evaluation, the coefficient of alignment was used.

### 3.7 Statistics

GraphPad Prism 8 software (GraphPad Software Inc., La Jolla, CA, USA) was used for statistical evaluations. Student’s t test was applied to compare two normally distributed groups. P-values were marked in figures by asterisks. * p<0.05, ** p<0.01, *** p<0.001 and **** p<0.0001.

### 3.8 Ethics approval statement

The study was conducted according to the principles of the Declaration of Helsinki. The use of resected tissue was approved by the ethics committee of the Medical University of Vienna (vote 1190/2020) in accordance with the guidelines of the Council for International Organizations of Medical Sciences (CIOMS).

## 4 Results

### 4.1 scRNAseq identifies the cellular composition of ISGS tissue

To elucidate the cellular composition of ISGS compared to the adjacent unaffected tissue and to identify previously uncharacterized cellular and transcriptional irregularities that may influence the progression of the fibrotic disease, scRNAseq was conducted. In line with previous publications, ISGS was characterized by areas with particularly high cellular infiltrations (Figure 1A) (37). Additionally, an augmented ECM featuring enhanced fiber alignment was detectable in ISGS (Figure 1A &B). Through scRNAseq, we obtained data from a total of 12723 cells from three ISGS and two healthy trachea samples for subsequent bioinformatics analyses. Following data pre-processing, 13 distinct cell clusters were identified (Figure 1C). These cell clusters were characterized, using well-established cell markers in combination with their identified cluster marker, as basal cells (Basal), secretory cells (Secretory), ciliated cells (Ciliated), T-cells (TC), B-cells (BC), plasma cells (PC), macrophages (Mac), mast cells (Mast), fibroblasts (FB), smooth muscle cells (SMC), endothelial cells (EC) (Figure 1D). One cluster expressed gene sets typical for chondrocytes and Schwann cells (CH_SC). (Figure 1D & S1).

**Figure 1:**
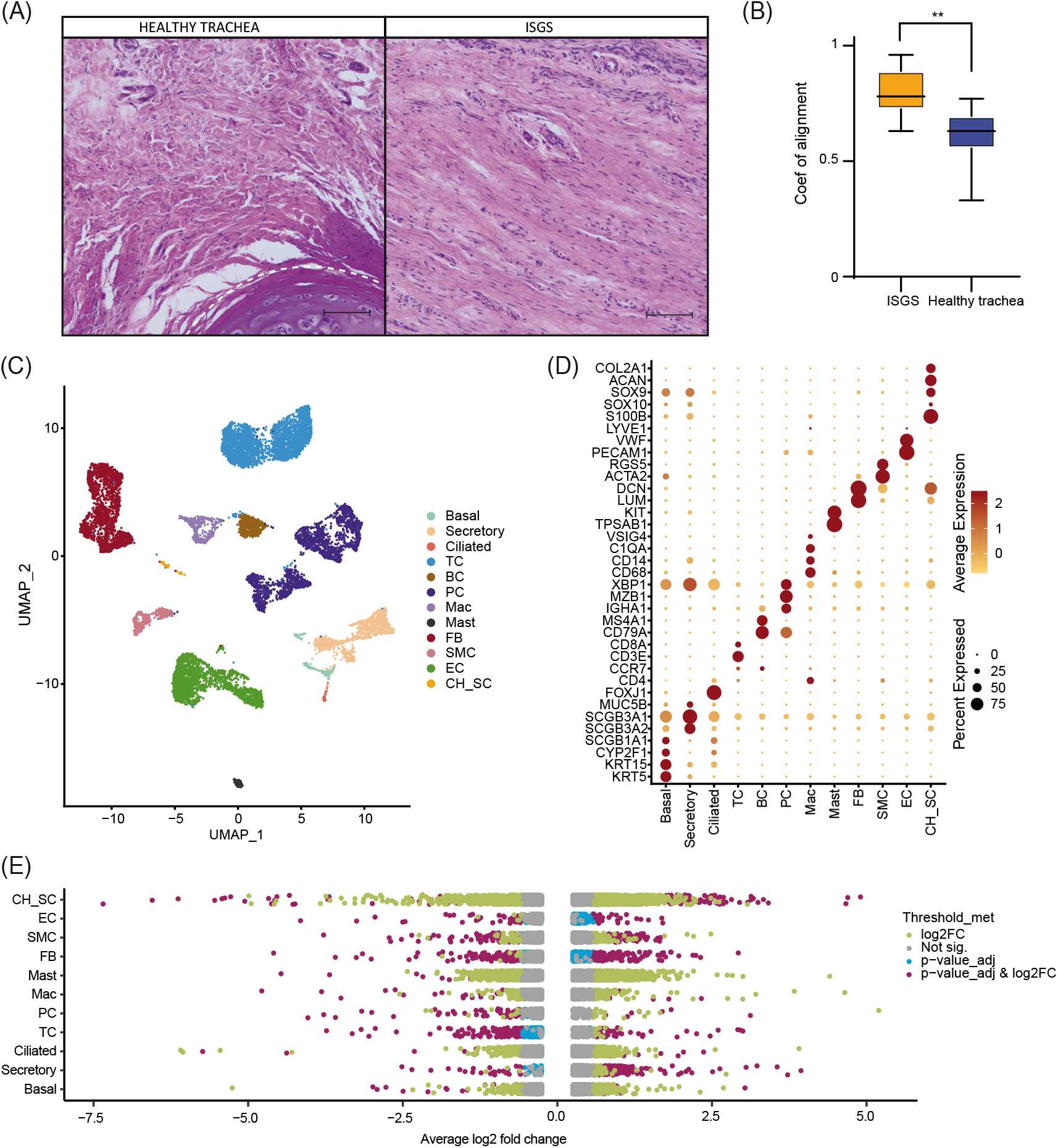
Single cell RNA sequencing of healthy trachea and ISGS. (A) Hematoxylin-eosin staining of healthy trachea and ISGS. Scale bars: 250 µm. Dashed line indicates soft tissue-cartilage border (B) Boxplot of the alignment coefficient evaluation of healthy trachea and ISGS. The middle line depicts the arithmetic mean. ** p<0.01 (C) UMAP-Plot after integration of all datasets. Clusters were characterized as basal cells (Basal), secretory cells (Secretory), ciliated cells (Ciliated), T-cells (TC), B-cells (BC), plasma cells (PC), macrophages (Mac), mast cells (Mast), fibroblasts (FB), smooth muscle cells (SMC), endothelial cells (EC), Chondrocyte-Schwann cells (CH_SC). (D) Dotplot depicting expression of well-known marker genes for cluster characterization. (E) Strip-Plots showing differentially expressed genes of all different clusters comparing ISGS with healthy trachea. Gene expression thresholds are colour coded: adjusted p-values ≤0.05 and log2(FC) ≥0.58 (purple), adjusted p-values ≤0.05 only (blue), log2(FC) ≥0.58 only (green), not significant (grey). Cell cluster with low cell number, leading to no meaningful results, were excluded.

Analysis of differentially expressed genes revealed substantial differences between ISGS and the normal trachea. The most pronounced genetic differences were observed in FB but also CH_SC (Figure 1E & S2). A comprehensive list of all identified differentially expressed genes (DEGs) is available in the supplements (Table S4).

### 4.2 Plasma cells with high subtype variety accumulate in ISGS

A comparison of the relative cellular distribution within the different samples revealed a remarkable cellular presence of PCs in ISGS (18,7%, 16.1%, 29.4%), whereas in the healthy trachea, the presence of PCs was below 9% in each sample (Figure 2A). IF staining of the PCs, using MZB1 as marker revealed spots of increased PC presence in the ISGS tissue, whereas in unaffected tissue, the abundance of PC was scarce (Figure 2B). GO-term analysis based on the top 30 differentially expressed genes of PCs from both conditions revealed an overrepresentation of biological pathways associated with intense protein processing and B-cell receptor signaling pathway in PCs from ISGS (Figure 2C) Subsetting of the PC cluster uncovered 6 transcriptionally distinguishable PC-subtypes (PC1-PC6) in ISGS (Figure 2D). A more precise annotation of the detected subclusters according to previously described PC subtypes was not feasible (Figures S3) (38, 39). PC2 appeared to constitute the predominant PC type in healthy tissue (Figure 2D). The cluster marker list indicated a distinct expression of genes crucial for immunoglobulin production, such as *IGHG1* in PC1, *IGHA1* in PC2, *IGKV3-20* in PC3 and *IGLC2* in PC4, among others (Figure 2E). Analysis of potential cell-cell interactions showed a more diverse interaction of PCs in ISGS with other cell types than in the healthy tissue (Figure 2F & G). A high probability of a crosstalk between PCs and BCs via MIF-(CD74+CXCR4) was exclusively detected in ISGS, a receptor pathway involved in wound-healing and recovery of multiple organs (Figure 2H) (40). In the healthy trachea, only PC2 revealed recognizable cell-cell interactions, mainly typical immunological interactions with TCs (Figure 2I). Our results indicate a highly heterogeneous population of PCs in ISGS. Their different expression patterns, especially of genes important for immunoglobulin production, suggest an involvement of specific immunoglobulins in the pathogenesis of ISGS.

**Figure 2:**
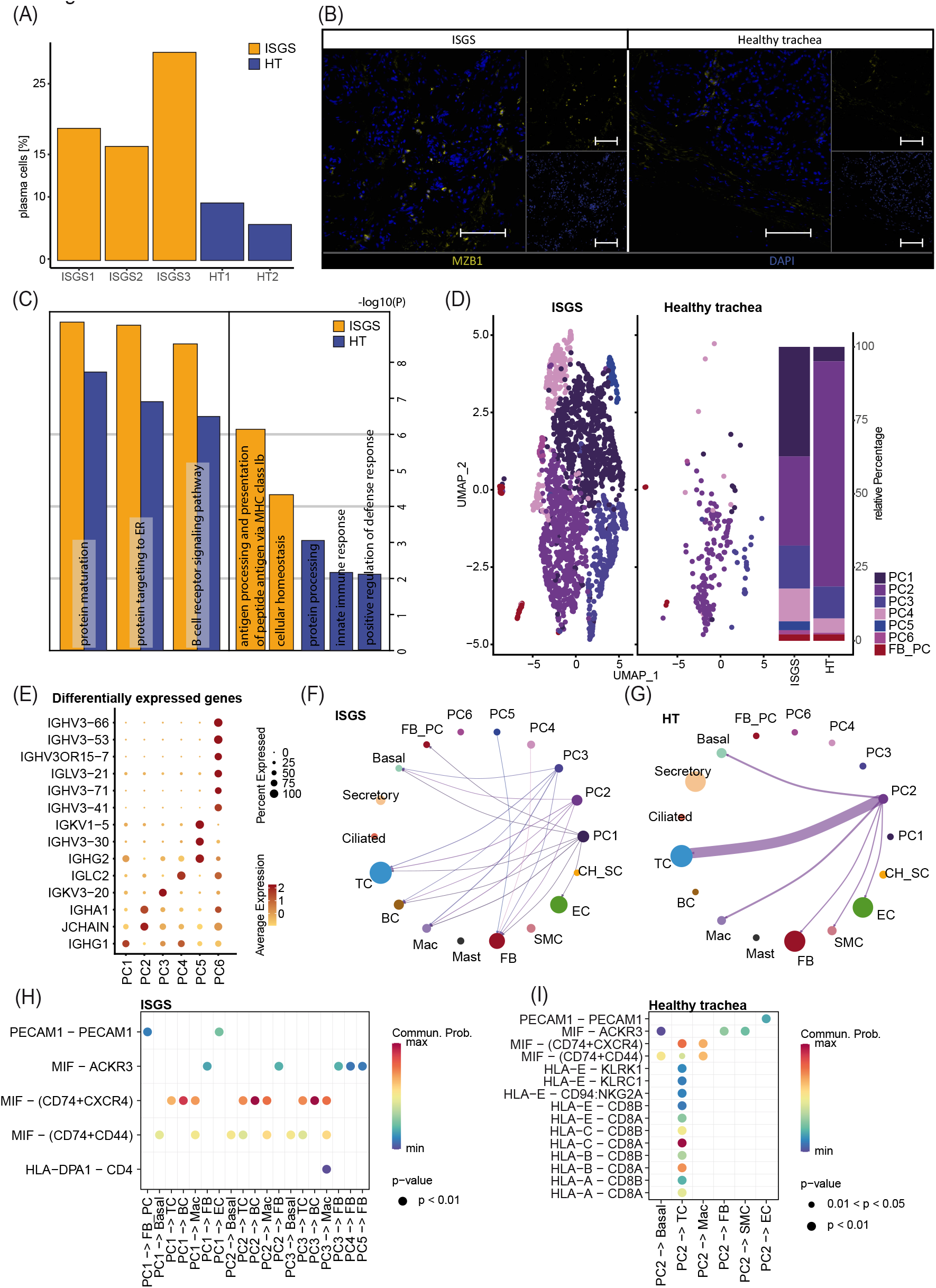
Transcriptional analysis reveals increased amount of plasma cells in ISGS tissue. (A) Barplot depicting relative amount of plasma cells in the distinct transcriptional datasets. (B) Representative immunofluorescence staining of Marginal zone B and B1 cell-specific protein (MZB1)-positive plasma cells in ISGS and healthy trachea, Tissues of n=3 donors per condition were stained. Scale bars: 100 µm (C) GO-Term enrichment of differentially expressed genes with average foldchange ≥1.5 comparing plasma cells of ISGS and healthy trachea with secretory cells as neutral reverence cluster respectively. Bar-length depicts statistical significance of the term. Identical relevant terms identified in both analysis (left) and relevant individual terms (right), ISGS results are colored in yellow, healthy trachea in blue. (D) UMAP-Plot of the subset of PC reveals distinct PC-subcluster: PC1-PC6, Barplot indicates relative amounts of subcluster within ISGS and healthy trachea. (E) Dotplot shows differentially expressed genes with foldchange ≥2 of the distinct PC subcluster. (F-G) Circle plots of identified interactions from distinct PC-subtypes with the remaining cell types in ISGS and healthy trachea. Bow thickness represents amount of interactions (H-I) Dotplots depict the detected receptor-ligand couples of PC-subtypes with remaining cell types in ISGS and healthy trachea tissue.

### 4.3 Fibrotic fibroblasts affect tissue environment in ISGS

To identify the cell types contributing most significantly to ECM production, we conducted module score analysis based on the recently published matrisome (36). This analysis revealed a substantial contribution of FB to ECM production, with potential matrix associated involvement of other cell clusters such as CH_SC (Figure 3A). A screening of ECM-associated genes unveiled multiple factors exhibiting upregulated expression in ISGS cells, including *CTHRC1, CTSG, MDK, MUC12, PLAT, SFRP1, TNC* (Figure 3B). Notably, the most pronounced differences in ECM gene expression were uncovered in the SC_CH cluster. Consequently, our data provide a transcriptional snapshot of the cellular landscape in ISGS, highlighting substantial genetic differences in matrix-associated gene expression.

**Figure 3:**
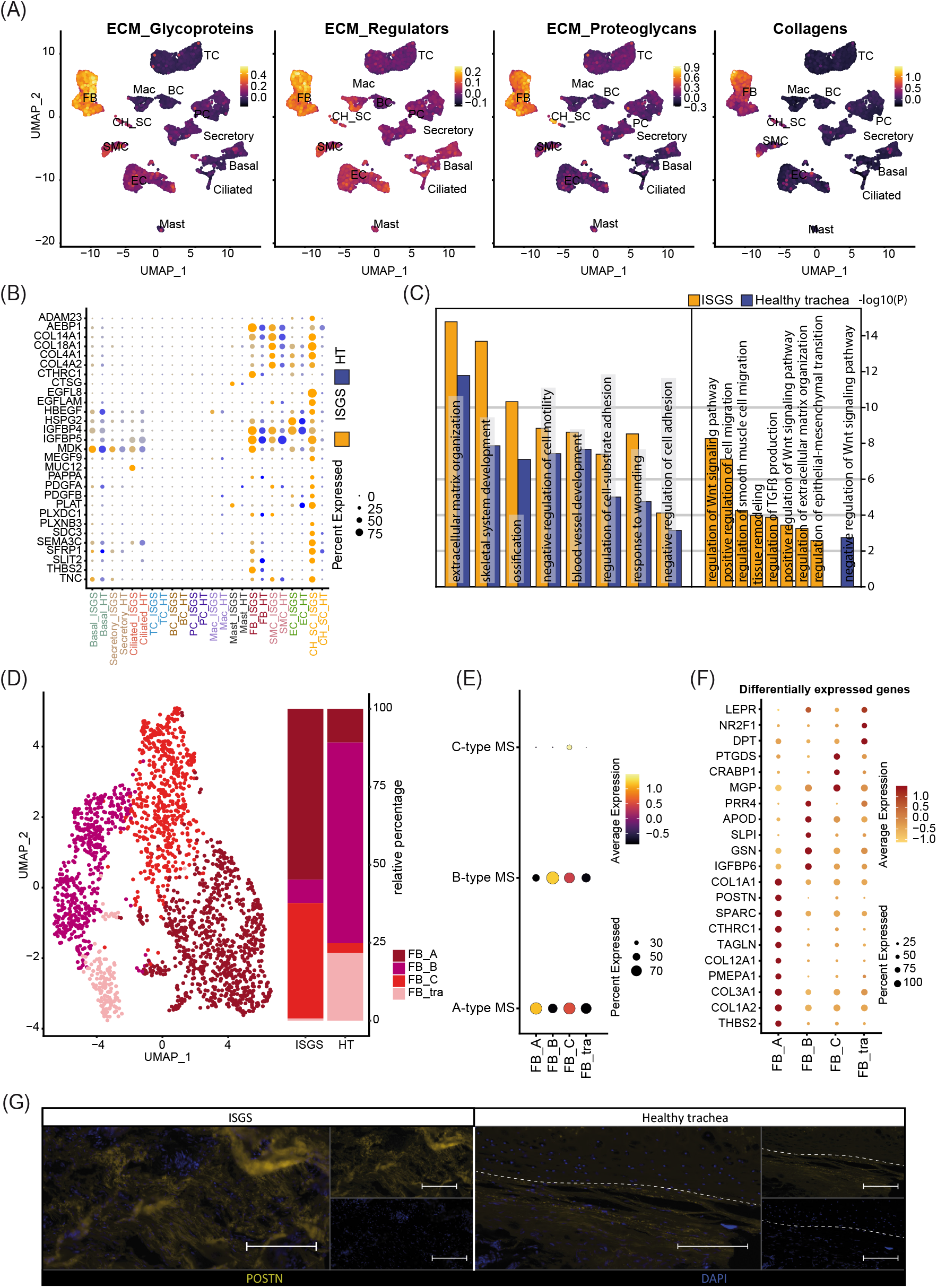
Impact of Fibroblasts on ISGS pathology. (A) Feature Plots showing module scores of extracellular matrix (ECM) -associated gene groups. (B) Dotplot depicting expression of matrix-associated genes by distinct cell types of healthy trachea and ISGS. (C) GO-Term enrichment of differentially expressed genes with average foldchange ≥2 comparing fibroblasts of ISGS with healthy controls. Identical relevant terms identified in both analysis (left) and relevant individual terms (right), ISGS results are colored in yellow, healthy trachea in blue. (D) UMAP-Plot of FB subclusters: FB-type A (FB_A), FB-type B (FB_B) and FB-type C (FB_C), tracheal specific FB subset (FB_tra). (E) Dot-Plot depicting module scores of published FB-subtype gene sets. (F) Dotplot shows differentially expressed genes with foldchange ≥2 of each FB subcluster. (G) Representative immunofluorescence image of periostin (POSTN) in ISGS and healthy trachea. Tissues of n=3 donors per condition were stained. Dashed line indicates soft tissue-cartilage border, Scale bars: 100 µm;

Fibroblasts are recognized as a pivotal factor in fibrotic diseases, being the primary producer of ECM and influencing the function of other cell-types (15). Given our data indicating significant alterations in gene expression associated with ECM formation in FBs in ISGS compared to the adjacent healthy tissue, we performed a detailed analysis of the FB cell cluster. To gain further insights into the functional role of the identified FBs, we conducted an enrichment analysis, revealing an enhanced involvement of ISGS-FBs in fibrotic features such as ECM organization, response to wounding and regulation of TGFß production (Figure 3C). Examination of FB cell-cell communication indicated only a slightly increased number of ligand-receptor interactions in ISGS compared to healthy trachea.

However, notable differences were observed in the specific interactions identified (Figure S4). Analyzing all resulting interactions indicated a high probability of communication between FBs, Mast and BCs. Notably, *COL4A2, MDK, NEGR1, POSTN, TNC, VEGFA* and *WNT5A* exclusively appeared in the FB interactions of the ISGS as communication partners. Subsetting of the identified FB cluster revealed several subclusters, which were classified according to the FB subtype scheme published by Ascension et al. (29) (Figure 3D & 3E). Cluster annotation revealed the presence of Type A (FB_A), Type B (FB_B) and Type C (FB_C) FBs in the samples. However, one FB cluster could not be explicitly assigned according to the Ascension nomenclature and was therefore designated as a tracheal specific FB subset (FB_tra) (Figure 3D & 3E). Comparing the cellular distribution in ISGS and healthy trachea, a pronounced prevalence of FB_A and FB_C was evident in the ISGS samples, whereas FB_B and FB_tra were predominant present in normal tissue (Figure 3D). Subsequent analysis revealed a strong expression of profibrotic genes, including *COL1A2, COL3A1, TAGLN, POSTN* and *COL1A1*, especially in the FB_A subset (Figure 3F). Immunofluorescence staining of periostin corroborated the increase of this pro-fibrotic factor in ISGS at the protein level (Figure 3G). These findings demonstrate the increased presence and the genetic profile of fibrotic FBs in ISGS.

### 4.4 In-depth analysis of Schwann cells suggests a pro-fibrotic impact on ISGS pathology

A recent study of our group demonstrated a significant involvement of Schwann cell to fibrotic processes in pathologic cutaneous scars (20). Therefore, we also conducted a detailed exploration of the SC_CH cluster with its distinctive expression pattern in our ISGS samples. Subset analysis of the SC_CH facilitated the segregation of the two cell types (Figure 4A). SCs were detected in both conditions, whereas CHs were only found in the unaffected tissue, as all fibrotic samples were obtained without any adherent cartilage (Figure 4A & 4B). Enrichr analysis of the two distinct clusters based on their cluster markers revealed cell type-specific features such as nervous system development and myelination for SCs and cartilage development and skeletal system development for CHs, supporting the annotation (Figure 4C).

**Figure 4:**
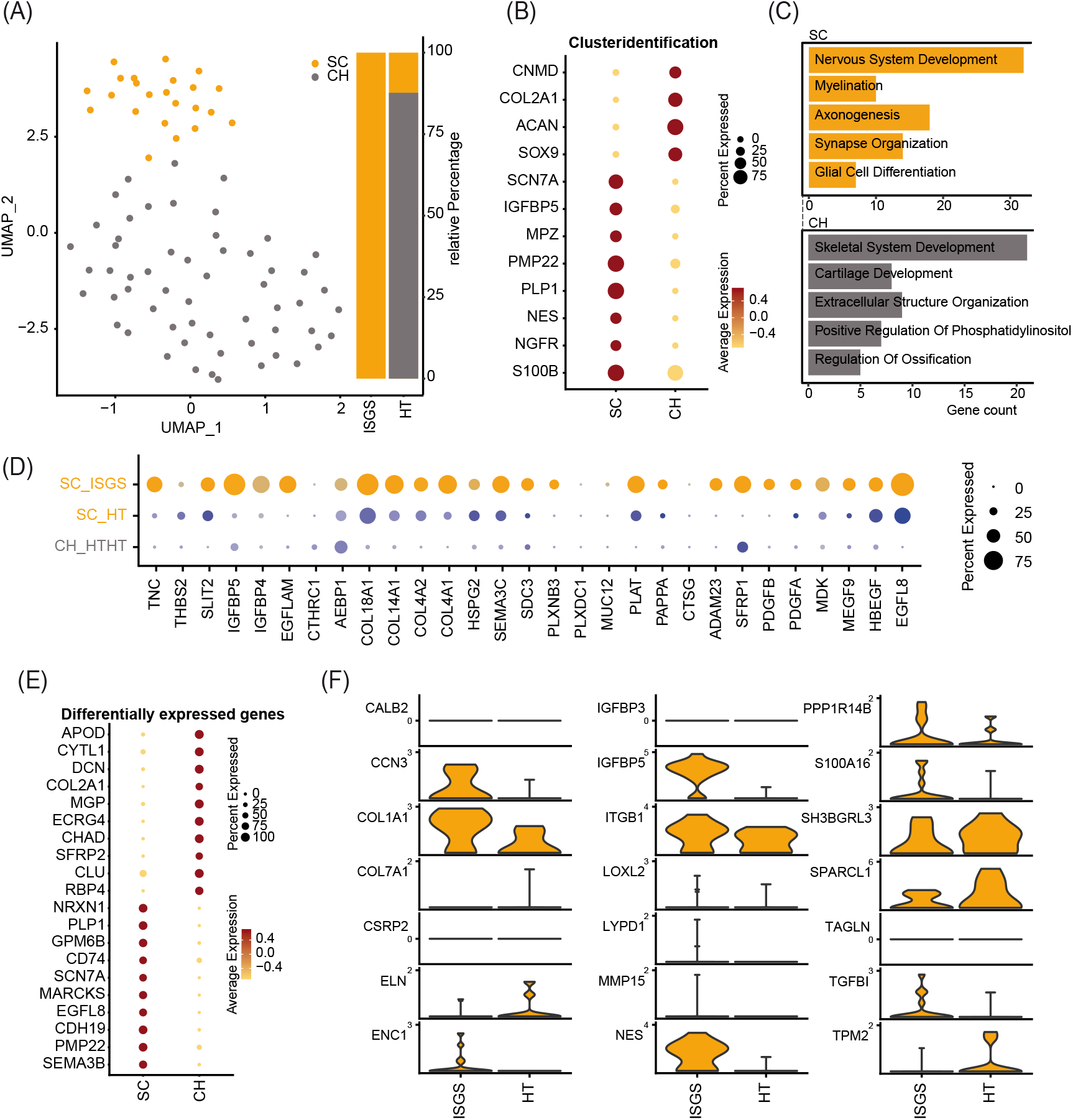
Transcriptional analysis reveals pro-fibrotic Schwann cells in ISGS. (A) UMAP-Plot of SC_CH-cluster. (B) Dotplot depicting expression of well-known marker genes for SC and CH. (C) Enrichment analysis of subset clustermarker with average foldchange ≥2. SC results depicted in yellow, CH in grey. (D) Dotplot shows differentially expressed genes comparing SCs of both conditions. (E) Dotplot of subset clustermarker genes with average foldchange ≥2. (F) Expression of pro-fibrotic SC-associated genes. Vertical lines depict maximum expression. Violin-width shows frequency of cells at the respective expression level.

Examination of the previously identified ECM-associated genes revealed a strong expression of these genes in SC of the ISGS, while SCs in healthy tissue and the detected CHs exhibit comparatively low expression levels (Figure 4D). The complete expression list of ECM-associated genes within the segregated SC_CH cluster is provided in the supplements (Figure S5 & S6). Genetic comparison of the two clusters showed a robust expression of characteristic SC genes (*PLP1, PMP22, SCN7A*) and CH genes (*DCN, COL2A1*). Interestingly, we also detected genes such as *EGFL8* and *CDH19*, which are known marker genes of SC precursors and repair SCs (Figure 4E) (41, 42). Recently, we identified the transcriptional pattern of profibrotic Schwann cells in pathologic scars (21).

Screening the detected SCs in ISGS and healthy tissue for these marker genes revealed the upregulation of 9 out of 21 genes in SCs of the ISGS (Figure 4F). Remarkably, among these 9 genes *IGFBP5, CCN3* and *NES* were included, three well-described main factors in profibrotic SCs. IF staining of SCs using S100B in combination with nestin uncovered double positive SCs in ISGS with a characteristic elongated form, comparable to those previously shown to be specific for pro-fibrotic SCs (Figure 5A & 5B) (20). In healthy tissue, S100 staining was only observed in chondrocytes and large nerve bundles (Figure 5A and 5D), while nestin staining was negative. In addition, IF-staining of S100 in combination with PGP9.5, an established axon marker, revealed that most SCs present in ISGS were not associated with axons (Figure 5C), while all SCs present in healthy tracheas were associated with axons in nerve bundles. (Figure 5D). These findings collectively suggest the presence of dedifferentiated, activated SCs with a profibrotic function in ISGS.

**Figure 5:**
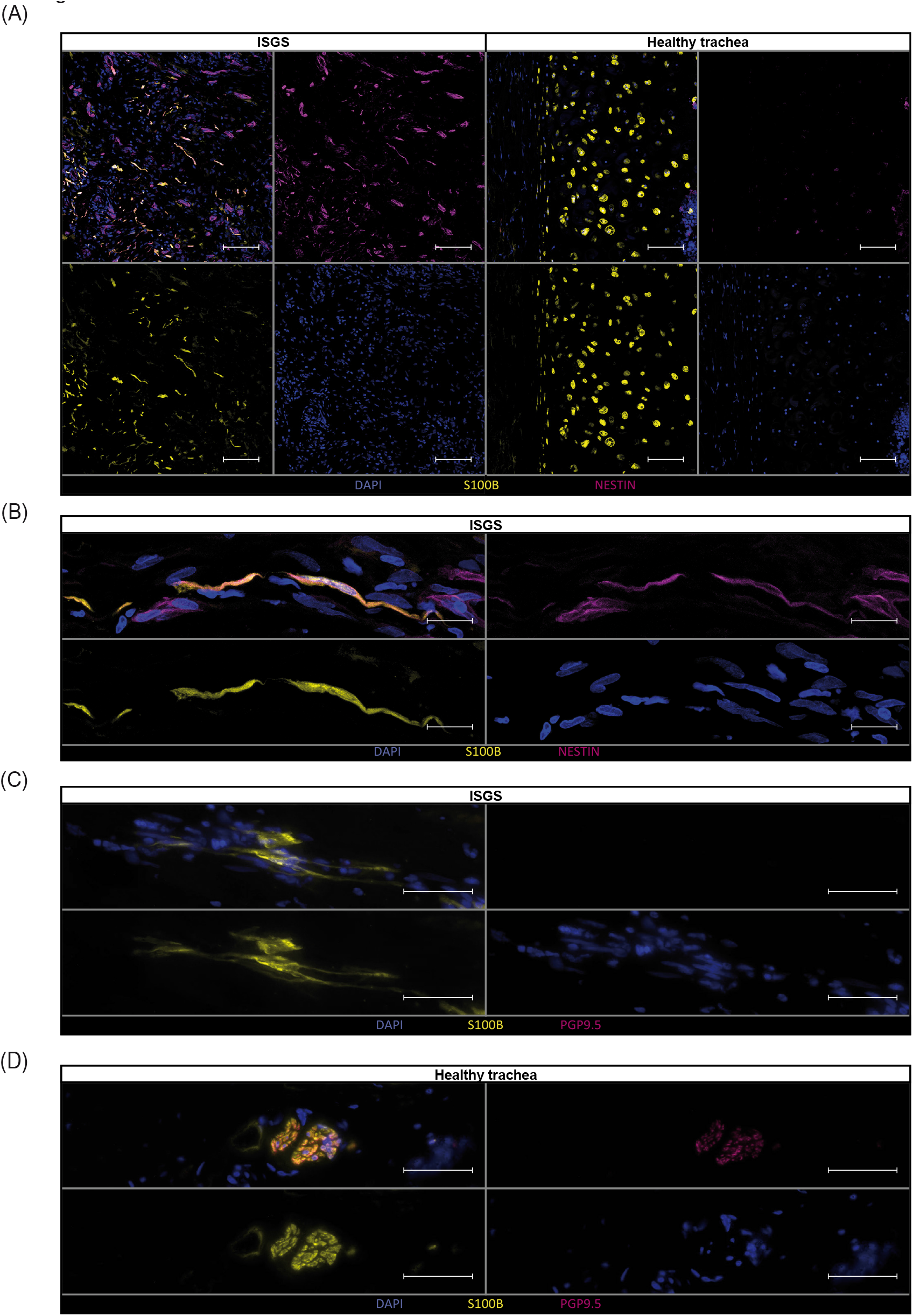
Verification of pro-fibrotic Schwann cells in ISGS tissue. Immunostainings of Schwann cells for (A-B) S100B and Nestin and (C-D) S100B and PGP9.5 shows double positive Schwann cells in ISGS compared to healthy trachea. In tissue of healthy trachea Schwann cells were only identified in nerve bundles (D). One representative micrograph of n=3 donors per condition is depicted. Scale bars in (A): 100µm, Scale bars in (B-D): 50µm.

## 5 Discussion

ISGS, characterized as a rare form of fibrotic disease, continues to elicit questions about its pathobiology. Specifically, the cellular drivers of this condition remain elusive. In this study, we employed single-cell RNA sequencing to elucidate the cellular composition of ISGS tissue, and compared it to normal, unaffected tracheal tissue. We aimed to identify cellular abnormalities and dysregulated gene expression that could potentially contribute to the disease pathology. On a histopathologic level, the disease is characterized by a dense fibrosis primarily constituted by concentrated cellular regions (37). The histologic examination of our samples was consistent with this characterization.

Our scRNAseq analysis revealed a substantial higher abundance of PCs in ISGS. These cells were observed in specific areas as dense cellular spots. The identified PCs expressed *IGHG1* and *IGHG2*, which encode crucial proteins of Immunoglobulin (Ig)G (43). Plasma cells have already been recognized for their pivotal crucial role in a specific fibro-inflammatory disorder, a IgG4-related disease (44). In this pathology, PCs often exhibit significant positivity for IgG4, and fibrotic manifestations can occur in nearly any organ (45). The presence of *IGHG1* and *IGHG2*-expressing PCs in ISGS suggests a disease-specific population of PCs, which differ from PCs commonly found in other fibrotic processes. The precise nature of these PCs and their contribution to ISGS requires further investigation in future studies.

In addition to IgG4-related fibrotic disease, an elevation of other IgG subclasses has also been reported in idiopathic pulmonary fibrosis, another fibrotic disorder in the thoracic region with an unsolved pathomechanism (46). In these cases, IgG accumulation is predominantly described as localized rather than systemic (47). Analysis of the broncho-alveolar lavage fluid from patients with idiopathic pulmonary fibrosis indicates a higher concentration of IgG1 and IgG3 compared to the healthy state (47). In addition, Prêle et al. recently reported the accumulation of PCs in fibrotic lungs, and depletion of PCs by the proteasome inhibitor bortezomib resulted in a reduced level of pulmonary fibrosis (48).

However, the mechanism leading to PC accumulation in fibrotic tissue and their precise function in pulmonary fibrosis still necessitate further research (49). These findings further support the previously postulated similarities between ISGS and pulmonary fibrosis (10) and suggest potential therapeutic targets for interventions that affect this pathologic cell type in the pulmonary fibrosis also in the context of ISGS. The transcriptional pattern of PCs in ISGS did not reveal specific direct involvement in fibrotic processes; however, GO-term analysis unveiled hyper-activated basal function of the PCs. Moreover, potential intercellular connections indicated a robust interaction between macrophage migration inhibitory factor (MIF)-positive cells and those expressing CD74 and CXCR4. This interaction with B-cells involves a receptor-ligand combination previously reported to be implicated in fibrotic processes such as wound healing. It is tempting to speculate that interventions affecting this interaction could represent an alternative treatment option for ISGS in the future(40).

Our scRNAseq analysis also demonstrated that while FB cell numbers did not exhibit a significantly increase, their transcriptional profile exhibited the highest deviation from the healthy condition. Our investigation contributes to a deeper understanding of the subsets of FBs present in ISGS and their role in the disease. A-type FBs present in ISGS have been recognized for their involvement in ECM homeostasis, and dysregulation of this cell type is thought to contribute to fibrotic pathologies (29). In our analyses, it is conceivable that the normal, tracheal FBs (FB_tra) are represented in the healthy control samples, while FB_A, characterized by a significant upregulation of ECM genes such as *COL1A1, COL3A1, POSTN*, and others, appears to be in the pro-fibrotic more pathologic-like state. Consistent with our findings, Tsukui et al. recently described this specific set of genes to be upregulated in a pathologic FB cell cluster associated with pulmonary fibrosis (50).

Collectively, our data suggest that specific FB subsets contribute to ISGS in a manner that parallels the common mechanisms observed in numerous other fibrotic organs. These data contribute to a deeper understanding of the subsets of FBs present in ISGS and their putative role to the disease.

Most notably, we identified a sub-population of SCs in ISGS exhibiting a pro-fibrotic phenotype. The pro-fibrotic role of SCs has recently been elucidated in keloids, a dermatological condition sharing hitherto unknown pathologic features (20). In keloids, SCs exist in an activated state, promoting synthesis of extracellular matrix (20). In our study, we detected the presence of SCs in ISGS with a specific gene expression pattern indicative of cell activation, featuring genes such as *IGFBP5* and *CCN3*, which are characteristic for pro-fibrotic SCs (21). Additionally, these SCs expressed *EGFL8*, a gene secreted by repair-related SCs, implicated in neurite growth and neuronal differentiation (50). Another upregulated factor was *CDH19*, a gene known to be elevated in SC precursors during SC development, subsequently downregulated in the inactive state of SCs, whether myelinating or non-myelinating (43). On a histologic level, we identified cells exhibiting a morphology similar to pro-fibrotic SCs found in keloids (20). Immunofluorescence staining indicated the presence of axon-independent SCs in ISGS. While the occurrence of SCs without contact to axons may not be as prominent as in keloids, our findings support the hypothesis of the involvement of axon free, pro-fibrotic SCs in diverse fibrotic diseases, including ISGS.

In summary, our study elucidates the cellular and the transcriptional landscape of ISGS. It delineates the FB subsets involved in this fibrotic disease and identifies the pathologic FB-subtype present in the tissue based on its transcriptional pattern. Apart from FBs, PCs and SCs also appear to play an important role in the pathological processes of ISGS, existing in an activated state and potentially contributing to fibrosis. These findings may aid in the development of novel therapeutic strategies to treat ISGS.

## 7 Acknowledgments

The authors acknowledge the core facilities of the Medical University of Vienna, a member of Vienna Life Science Instruments. We would especially like to thank the HP Haselsteiner and the CRISCAR Family Foundation for their support and trust in the Medical University/Aposcience AG public private partnership.

## 8 Data Availability Statement

scRNAseq datasets are publicly accessible on NCBI’s Gene Expression Omnibus (GEO) database (GEP series accession number GSE248105).

## 9 Author contributions

Conceptualization, M.D., K.H., M.M. and H.J.A.; methodology, M.D., M.L., D.C., K.K., D.B., T.S. and M.M.; software, M.D., D.C., and K.K.; validation, M.D. and M.M.; formal analysis, M.D.; investigation, M.D., K.H., M.M and H.J.A.; resources, K.H., C.A., H.J.A. and M.M.; data curation, M.D., D.B. and K.K.; writing original draft preparation, M.D., H.J.A. and M.M.; visualization, M.D.; supervision, H.J.A. and M.M.; project administration, M.D. and M.M.; funding acquisition, H.J.A. and M.M.; All authors reviewed the final version of the manuscript.

## 11 Data Availability Statement

The datasets generated for this study can be found in the on NCBI’s Gene Expression Omnibus (GEO) database (GEP series accession number GSE248105). [https://www.ncbi.nlm.nih.gov/geo/]

