## Supplementary Figures for "Transcriptional profiling sheds light on the fibrotic aspects of idiopathic subglottic tracheal stenosis"

### Figure S1

(A)

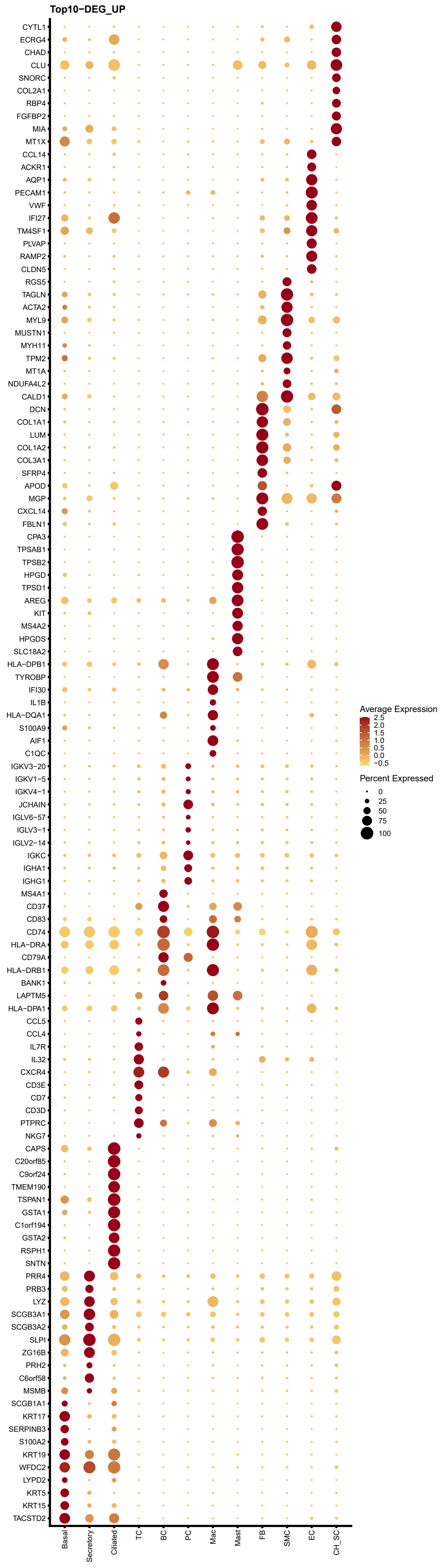
Top10-DEG_UP
CYTL1
ECRG4
CHAD
CLU
SNORC
COL2A1
RBP4
FGFBP2
MIA
MT1X
CCL14
ACKR1
AQP1
PECAM1
VWF
IFI27
TM4SF1
PLVAP
RAMP2
CLDN5
RGS5
TAGLN
ACTA2
MYL9
MUSTN1
MYH11
TPM2
MT1A
NDUFA4L2
CALD1
DCN
COL1A1
LUM
COL1A2
COL3A1
SFRP4
APOD
MGP
CXCL14
FBLN1
CPA3
TPSAB1
TPSB2
HPGD
TPSD1
AREG
KIT
MS4A2
HPGDS
SLC18A2
HLA-DPB1
TYROBP
IFI30
IL1B
HLA-DQA1
S100A9
AIF1
C1QC
IGKV3-20
IGKV1-5
IGKV4-1
JCHAIN
IGLV6-57
IGLV3-1
IGLV2-14
IGKC
IGHA1
IGHG1
MS4A1
CD37
CD83
CD74
HLA-DRA
CD79A
HLA-DRB1
BANK1
LAPTM5
HLA-DPA1
CCL5
CCL4
IL7R
IL32
CXCR4
CD3E
CD7
CD3D
PTPRC
NKG7
CAPS
C20orf85
C9orf24
TMEM190
TSPAN1
GSTA1
C1orf194
GSTA2
RSPH1
SNTN
PRR4
PRB3
LYZ
SCGB3A1
SCGB3A2
SLPI
ZG16B
PRH2
C6orf58
MSMB
SCGB1A1
KRT17
SERPINB3
S100A2
KRT19
WFDC2
LYPD2
KRT5
KRT15
TACSTD2
Basal
Secretory
Ciliated
TC
BC
PC
Mac
Mast
FB
SMC
EC
CH_SC
Average Expression
2.5
2.0
1.5
1.0
0.5
0.0
−0.5
Percent Expressed
0
25
50
75
100

### Figure S2

(A)

| | Basal | | Secretory | | Ciliated | | TC | | BC | | PC | | Mac | | Mast | | FB | | SMC | | EC | | CH_SC | | |
|---|---|---|---|---|---|---|---|---|---|---|---|---|---|---|---|---|---|---|---|---|---|---|---|---|---|
| **log2FC** | 279 | 186 | 67 | 64 | 350 | 189 | 15 | 15 | x | x | 26 | 17 | 160 | 94 | 1149 | 583 | 10 | 0 | 137 | 96 | 5 | 0 | 2049 | 1523 | UP |
| | | 93 | | 3 | | 161 | | 0 | | x | | 9 | | 66 | | 566 | | 10 | | 41 | | 5 | | 526 | DOWN |
| **Not sig.** | 1619 | 1064 | 1240 | 1225 | 2477 | 1242 | 238 | 226 | x | x | 261 | 76 | 1180 | 643 | 1134 | 480 | 356 | 6 | 1213 | 711 | 188 | 10 | 2414 | 1513 | UP |
| | | 555 | | 15 | | 1235 | | 12 | | x | | 185 | | 537 | | 654 | | 350 | | 502 | | 178 | | 901 | DOWN |
| **p-value_adj & log2FC** | 48 | 32 | 268 | 244 | 15 | 10 | 156 | 42 | x | x | 55 | 8 | 32 | 9 | 9 | 0 | 222 | 114 | 128 | 61 | 66 | 30 | 114 | 82 | UP |
| | | 16 | | 24 | | 5 | | 114 | | x | | 47 | | 23 | | 9 | | 108 | | 67 | | 36 | | 32 | DOWN |
| **p-value_adj** | 5 | 3 | 145 | 115 | 0 | 0 | 386 | 35 | x | x | 42 | 1 | 1 | 0 | 0 | 0 | 439 | 385 | 19 | 4 | 268 | 237 | 0 | 0 | UP |
| | | 2 | | 30 | | 0 | | 351 | | x | | 41 | | 1 | | 0 | | 54 | | 15 | | 31 | | 0 | DOWN |

Figure S3

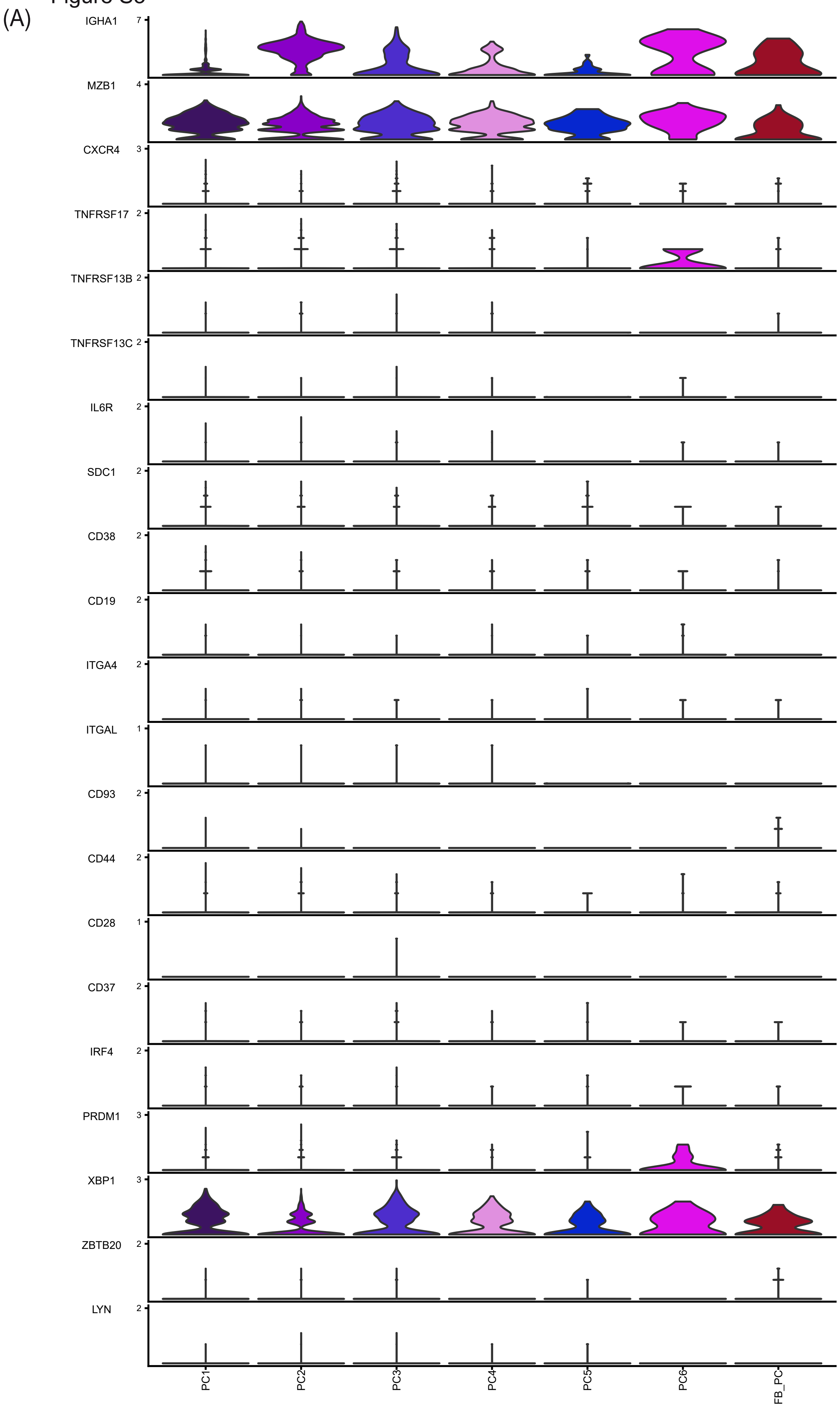
(A)
IGHA1
7
MZB1
4
CXCR4
3
TNFRSF17
2
TNFRSF13B
2
TNFRSF13C
2
IL6R
2
SDC1
2
CD38
2
CD19
2
ITGA4
2
ITGAL
1
CD93
2
CD44
2
CD28
1
CD37
2
IRF4
2
PRDM1
3
XBP1
3
ZBTB20
2
LYN
2
PC1
PC2
PC3
PC4
PC5
PC6
FB_PC

Figure S4

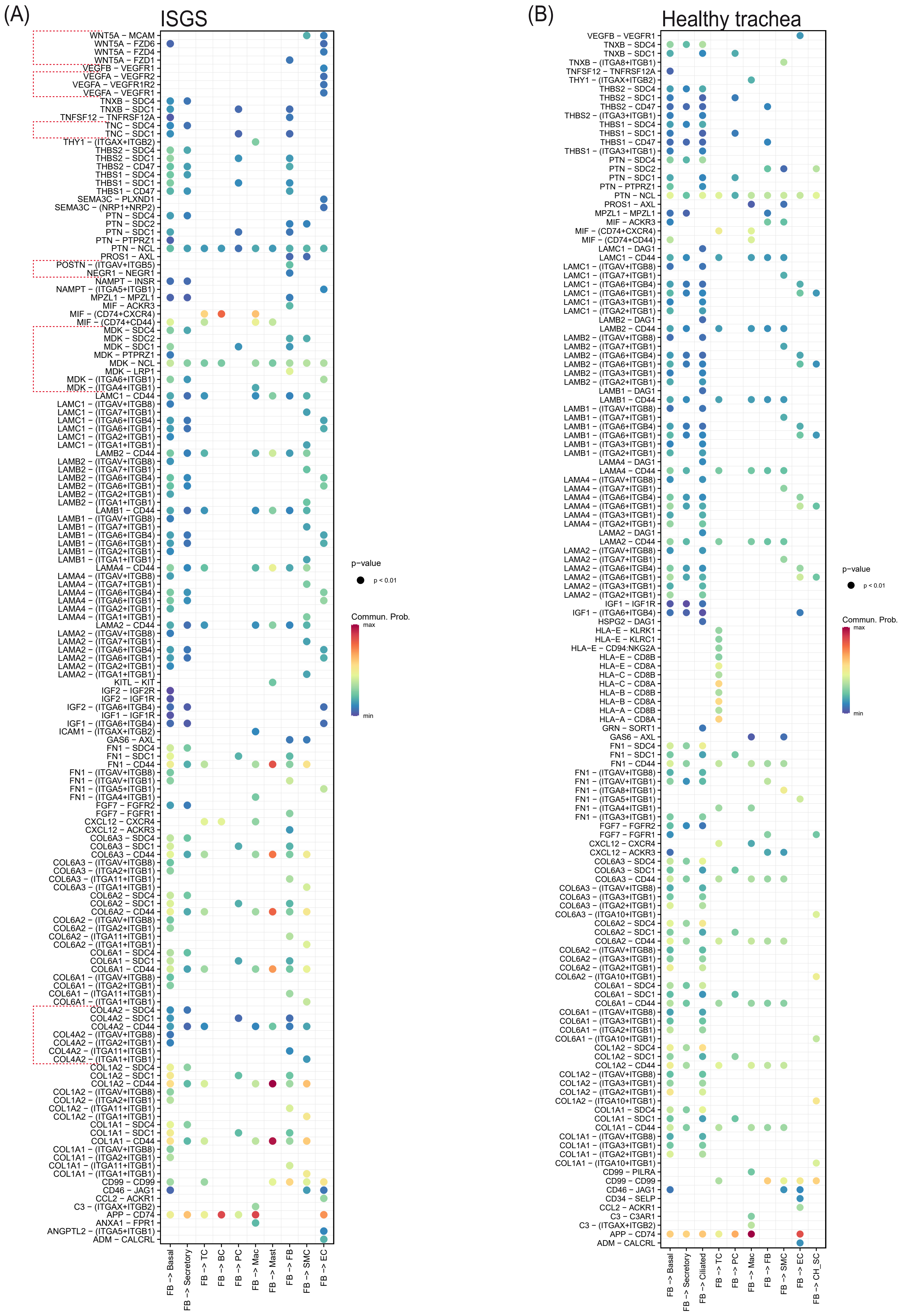
(A)
ISGS
WNT5A – MCAM
WNT5A – FZD6
WNT5A – FZD4
WNT5A – FZD1
VEGFB – VEGFR1
VEGFA – VEGFR2
VEGFA – VEGFR1R2
VEGFA – VEGFR1
TNXB – SDC4
TNXB – SDC1
TNFSF12 – TNFRSF12A
TNC – SDC4
TNC – SDC1
THY1 – (ITGAX+ITGB2)
THBS2 – SDC4
THBS2 – SDC1
THBS2 – CD47
THBS1 – SDC4
THBS1 – SDC1
THBS1 – CD47
SEMA3C – PLXND1
SEMA3C – (NRP1+NRP2)
PTN – SDC4
PTN – SDC2
PTN – SDC1
PTN – PTPRZ1
PTN – NCL
PROS1 – AXL
POSTN – (ITGAV+ITGB5)
NEGR1 – NEGR1
NAMPT – INSR
NAMPT – (ITGA5+ITGB1)
MPZL1 – MPZL1
MIF – ACKR3
MIF – (CD74+CXCR4)
MIF – (CD74+CD44)
MDK – SDC4
MDK – SDC2
MDK – SDC1
MDK – PTPRZ1
MDK – NCL
MDK – LRP1
MDK – (ITGA6+ITGB1)
MDK – (ITGA4+ITGB1)
LAMC1 – CD44
LAMC1 – (ITGAV+ITGB8)
LAMC1 – (ITGA7+ITGB1)
LAMC1 – (ITGA6+ITGB4)
LAMC1 – (ITGA6+ITGB1)
LAMC1 – (ITGA2+ITGB1)
LAMC1 – (ITGA1+ITGB1)
LAMB2 – CD44
LAMB2 – (ITGAV+ITGB8)
LAMB2 – (ITGA7+ITGB1)
LAMB2 – (ITGA6+ITGB4)
LAMB2 – (ITGA6+ITGB1)
LAMB2 – (ITGA2+ITGB1)
LAMB2 – (ITGA1+ITGB1)
LAMB1 – CD44
LAMB1 – (ITGAV+ITGB8)
LAMB1 – (ITGA7+ITGB1)
LAMB1 – (ITGA6+ITGB4)
LAMB1 – (ITGA6+ITGB1)
LAMB1 – (ITGA2+ITGB1)
LAMB1 – (ITGA1+ITGB1)
LAMA4 – CD44
LAMA4 – (ITGAV+ITGB8)
LAMA4 – (ITGA7+ITGB1)
LAMA4 – (ITGA6+ITGB4)
LAMA4 – (ITGA6+ITGB1)
LAMA4 – (ITGA2+ITGB1)
LAMA4 – (ITGA1+ITGB1)
LAMA2 – CD44
LAMA2 – (ITGAV+ITGB8)
LAMA2 – (ITGA7+ITGB1)
LAMA2 – (ITGA6+ITGB4)
LAMA2 – (ITGA6+ITGB1)
LAMA2 – (ITGA2+ITGB1)
LAMA2 – (ITGA1+ITGB1)
KITL – KIT
IGF2 – IGF2R
IGF2 – IGF1R
IGF2 – (ITGA6+ITGB4)
IGF1 – IGF1R
IGF1 – (ITGA6+ITGB4)
ICAM1 – (ITGAX+ITGB2)
GAS6 – AXL
FN1 – SDC4
FN1 – SDC1
FN1 – CD44
FN1 – (ITGAV+ITGB8)
FN1 – (ITGAV+ITGB1)
FN1 – (ITGA5+ITGB1)
FN1 – (ITGA4+ITGB1)
FGF7 – FGFR2
FGF7 – FGFR1
CXCL12 – CXCR4
CXCL12 – ACKR3
COL6A3 – SDC4
COL6A3 – SDC1
COL6A3 – CD44
COL6A3 – (ITGAV+ITGB8)
COL6A3 – (ITGA2+ITGB1)
COL6A3 – (ITGA11+ITGB1)
COL6A3 – (ITGA1+ITGB1)
COL6A2 – SDC4
COL6A2 – SDC1
COL6A2 – CD44
COL6A2 – (ITGAV+ITGB8)
COL6A2 – (ITGA2+ITGB1)
COL6A2 – (ITGA11+ITGB1)
COL6A2 – (ITGA1+ITGB1)
COL6A1 – SDC4
COL6A1 – SDC1
COL6A1 – CD44
COL6A1 – (ITGAV+ITGB8)
COL6A1 – (ITGA2+ITGB1)
COL6A1 – (ITGA11+ITGB1)
COL6A1 – (ITGA1+ITGB1)
COL4A2 – SDC4
COL4A2 – SDC1
COL4A2 – CD44
COL4A2 – (ITGAV+ITGB8)
COL4A2 – (ITGA2+ITGB1)
COL4A2 – (ITGA11+ITGB1)
COL4A2 – (ITGA1+ITGB1)
COL1A2 – SDC4
COL1A2 – SDC1
COL1A2 – CD44
COL1A2 – (ITGAV+ITGB8)
COL1A2 – (ITGA2+ITGB1)
COL1A2 – (ITGA11+ITGB1)
COL1A2 – (ITGA1+ITGB1)
COL1A1 – SDC4
COL1A1 – SDC1
COL1A1 – CD44
COL1A1 – (ITGAV+ITGB8)
COL1A1 – (ITGA2+ITGB1)
COL1A1 – (ITGA11+ITGB1)
COL1A1 – (ITGA1+ITGB1)
CD99 – CD99
CD46 – JAG1
CCL2 – ACKR1
C3 – (ITGAX+ITGB2)
APP – CD74
ANXA1 – FPR1
ANGPTL2 – (ITGA5+ITGB1)
ADM – CALCRL
FB -> Basal
FB -> Secretory
FB -> TC
FB -> BC
FB -> PC
FB -> Mac
FB -> Mast
FB -> FB
FB -> SMC
FB -> EC
p–value
p < 0.01
Commun. Prob.
max
min
(B)
Healthy trachea
VEGFB – VEGFR1
TNXB – SDC4
TNXB – SDC1
TNXB – (ITGA8+ITGB1)
TNFSF12 – TNFRSF12A
THY1 – (ITGAX+ITGB2)
THBS2 – SDC4
THBS2 – SDC1
THBS2 – CD47
THBS2 – (ITGA3+ITGB1)
THBS1 – SDC4
THBS1 – SDC1
THBS1 – CD47
THBS1 – (ITGA3+ITGB1)
PTN – SDC4
PTN – SDC2
PTN – SDC1
PTN – PTPRZ1
PTN – NCL
PROS1 – AXL
MPZL1 – MPZL1
MIF – ACKR3
MIF – (CD74+CXCR4)
MIF – (CD74+CD44)
LAMC1 – DAG1
LAMC1 – CD44
LAMC1 – (ITGAV+ITGB8)
LAMC1 – (ITGA7+ITGB1)
LAMC1 – (ITGA6+ITGB4)
LAMC1 – (ITGA6+ITGB1)
LAMC1 – (ITGA3+ITGB1)
LAMC1 – (ITGA2+ITGB1)
LAMB2 – DAG1
LAMB2 – CD44
LAMB2 – (ITGAV+ITGB8)
LAMB2 – (ITGA7+ITGB1)
LAMB2 – (ITGA6+ITGB4)
LAMB2 – (ITGA6+ITGB1)
LAMB2 – (ITGA3+ITGB1)
LAMB2 – (ITGA2+ITGB1)
LAMB1 – DAG1
LAMB1 – CD44
LAMB1 – (ITGAV+ITGB8)
LAMB1 – (ITGA7+ITGB1)
LAMB1 – (ITGA6+ITGB4)
LAMB1 – (ITGA6+ITGB1)
LAMB1 – (ITGA3+ITGB1)
LAMB1 – (ITGA2+ITGB1)
LAMA4 – DAG1
LAMA4 – CD44
LAMA4 – (ITGAV+ITGB8)
LAMA4 – (ITGA7+ITGB1)
LAMA4 – (ITGA6+ITGB4)
LAMA4 – (ITGA6+ITGB1)
LAMA4 – (ITGA3+ITGB1)
LAMA4 – (ITGA2+ITGB1)
LAMA2 – DAG1
LAMA2 – CD44
LAMA2 – (ITGAV+ITGB8)
LAMA2 – (ITGA7+ITGB1)
LAMA2 – (ITGA6+ITGB4)
LAMA2 – (ITGA6+ITGB1)
LAMA2 – (ITGA3+ITGB1)
LAMA2 – (ITGA2+ITGB1)
IGF1 – IGF1R
IGF1 – (ITGA6+ITGB4)
HSPG2 – DAG1
HLA–E – KLRK1
HLA–E – KLRC1
HLA–E – CD94:NKG2A
HLA–E – CD8B
HLA–E – CD8A
HLA–C – CD8B
HLA–C – CD8A
HLA–B – CD8B
HLA–B – CD8A
HLA–A – CD8B
HLA–A – CD8A
GRN – SORT1
GAS6 – AXL
FN1 – SDC4
FN1 – SDC1
FN1 – CD44
FN1 – (ITGAV+ITGB8)
FN1 – (ITGAV+ITGB1)
FN1 – (ITGA8+ITGB1)
FN1 – (ITGA5+ITGB1)
FN1 – (ITGA4+ITGB1)
FN1 – (ITGA3+ITGB1)
FGF7 – FGFR2
FGF7 – FGFR1
CXCL12 – CXCR4
CXCL12 – ACKR3
COL6A3 – SDC4
COL6A3 – SDC1
COL6A3 – CD44
COL6A3 – (ITGAV+ITGB8)
COL6A3 – (ITGA3+ITGB1)
COL6A3 – (ITGA2+ITGB1)
COL6A3 – (ITGA10+ITGB1)
COL6A2 – SDC4
COL6A2 – SDC1
COL6A2 – CD44
COL6A2 – (ITGAV+ITGB8)
COL6A2 – (ITGA3+ITGB1)
COL6A2 – (ITGA2+ITGB1)
COL6A2 – (ITGA10+ITGB1)
COL6A1 – SDC4
COL6A1 – SDC1
COL6A1 – CD44
COL6A1 – (ITGAV+ITGB8)
COL6A1 – (ITGA3+ITGB1)
COL6A1 – (ITGA2+ITGB1)
COL6A1 – (ITGA10+ITGB1)
COL1A2 – SDC4
COL1A2 – SDC1
COL1A2 – CD44
COL1A2 – (ITGAV+ITGB8)
COL1A2 – (ITGA3+ITGB1)
COL1A2 – (ITGA2+ITGB1)
COL1A2 – (ITGA10+ITGB1)
COL1A1 – SDC4
COL1A1 – SDC1
COL1A1 – CD44
COL1A1 – (ITGAV+ITGB8)
COL1A1 – (ITGA3+ITGB1)
COL1A1 – (ITGA2+ITGB1)
COL1A1 – (ITGA10+ITGB1)
CD99 – PILRA
CD99 – CD99
CD46 – JAG1
CD34 – SELP
CCL2 – ACKR1
C3 – C3AR1
C3 – (ITGAX+ITGB2)
APP – CD74
ADM – CALCRL
FB -> Basal
FB -> Secretory
FB -> Ciliated
FB -> TC
FB -> PC
FB -> Mac
FB -> FB
FB -> SMC
FB -> EC
FB -> CH_SC
p–value
p < 0.01
Commun. Prob.
max
min

### Figure S5

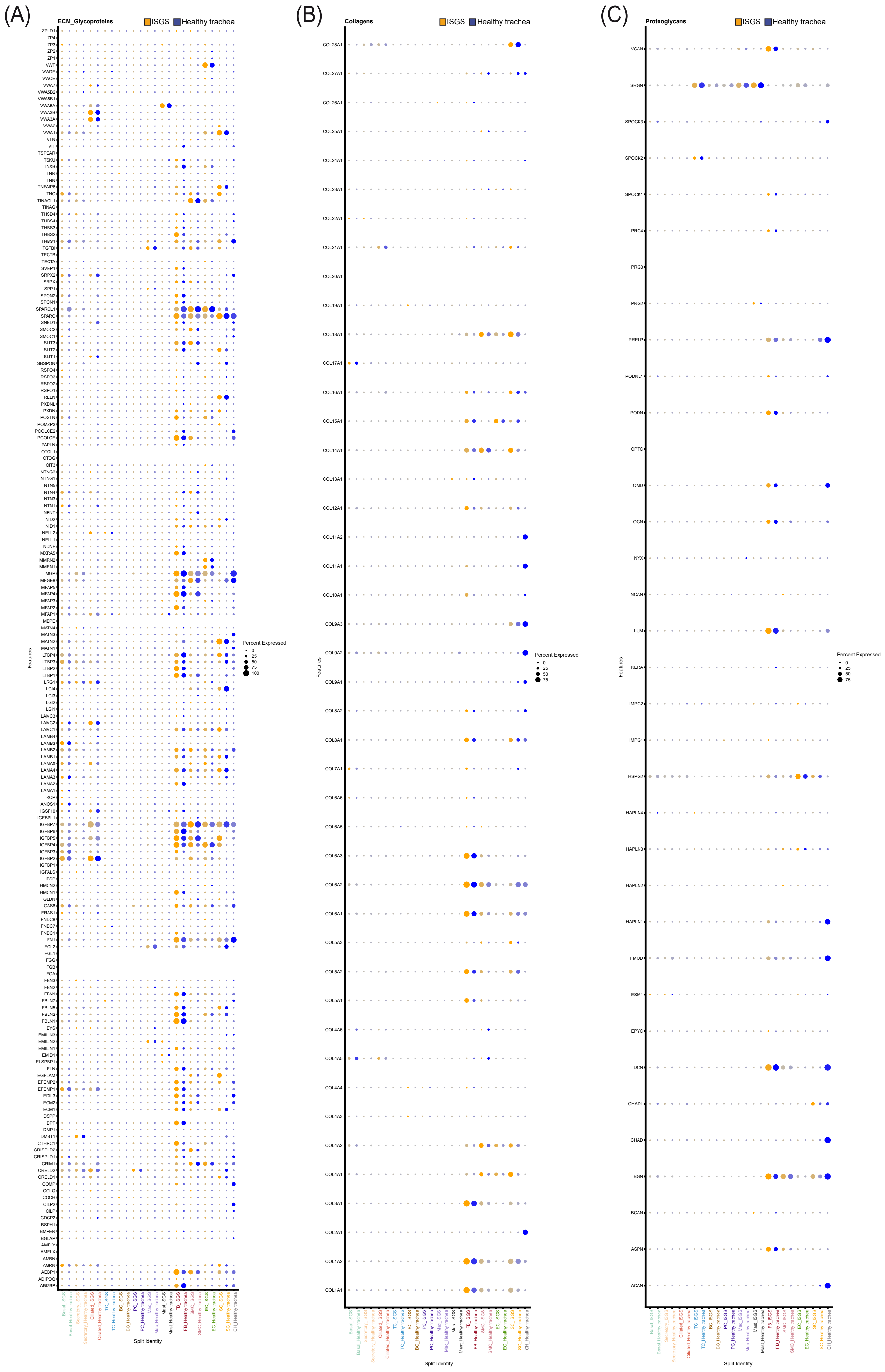
(A)
ECM_Glycoproteins
ISGS
Healthy trachea
Percent Expressed
Features
Split Identity
(B)
Collagens
ISGS
Healthy trachea
Percent Expressed
Features
Split Identity
(C)
Proteoglycans
ISGS
Healthy trachea
Percent Expressed
Features
Split Identity

### Figure S6

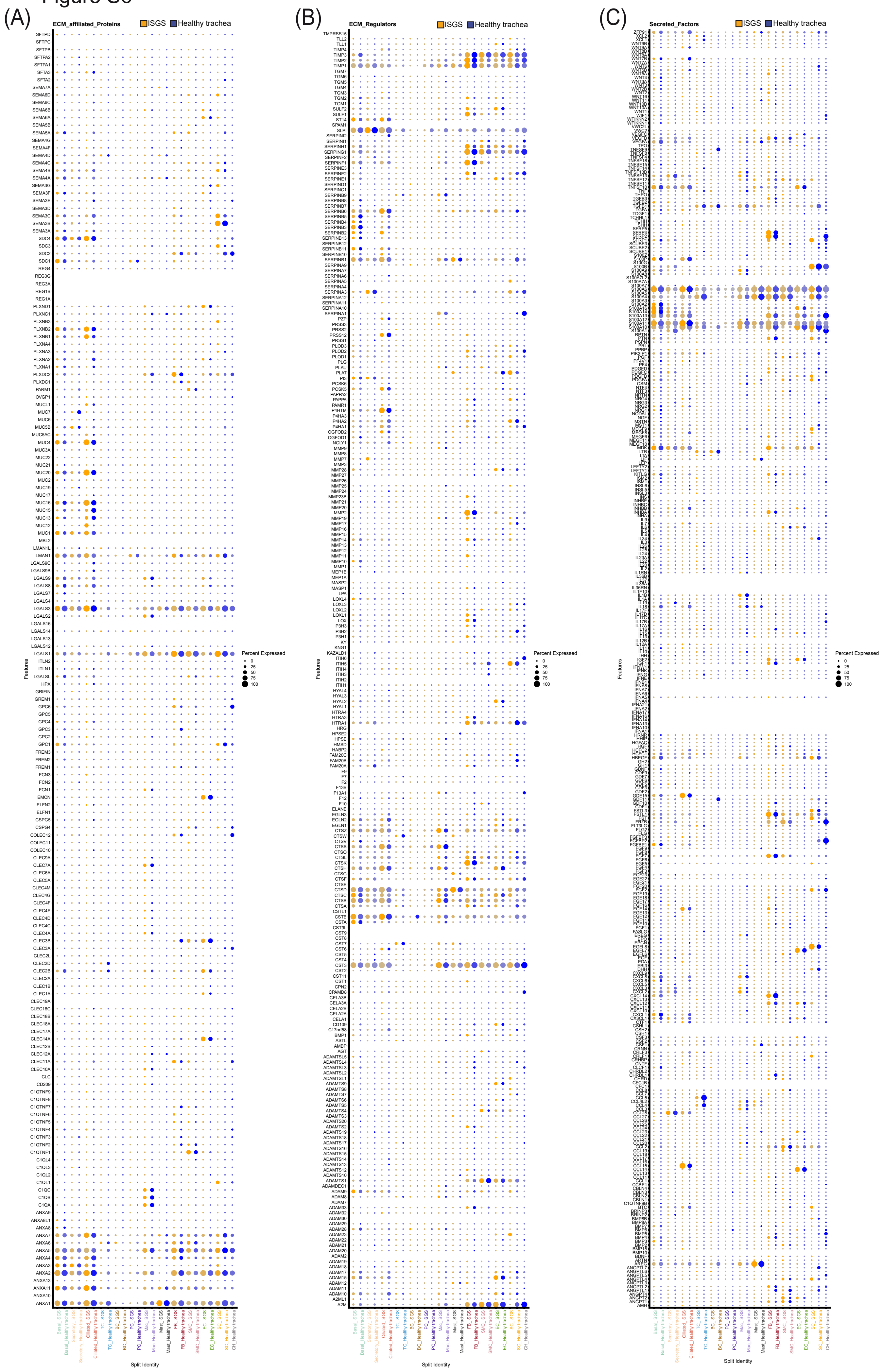

(A)
ECM_affiliated_Proteins
ISGS
Healthy trachea
(B)
ECM_Regulators
ISGS
Healthy trachea
(C)
Secreted_Factors
ISGS
Healthy trachea
Percent Expressed
0
25
50
75
100
Features
Split Identity
