## Supplementary Table 1 for "Transcriptional profiling sheds light on the fibrotic aspects of idiopathic subglottic tracheal stenosis"

### Supplementary Table 1 – donor information

| ID | sample | Age | Sex | Method |
|---|---|---|---|---|
| ISGS 1 | ISGS | 83 | Female | scRNAseq/Histo |
| ISGS 2 / HT 1 | ISGS + healthy trachea | 53 | Female | scRNAseq/Histo |
| ISGS 3 / HT 2 | ISGS + healthy trachea | 46 | Female | scRNAseq/Histo |
| ISGS 4 / HT 3 | ISGS + healthy trachea | 55 | Female | Histo |
| ISGS 5 / HT 4 | ISGS + healthy trachea | 31 | Female | Histo |
| ISGS 6 / HT 5 | ISGS + healthy trachea | 53 | Female | Histo |

**Supplementary Table 1. donor information.**
