## Supplementary Table 2 for "Transcriptional profiling sheds light on the fibrotic aspects of idiopathic subglottic tracheal stenosis"

Supplementary Table 2 - marker genes information

| celltype | abbreviation | marker gene | reference |
|---|---|---|---|
| basal cells | Basal | KRT5, KRT15, SCGB1A1, CYP2F1 | (1) |
| secretory cells | Secretory | MUC5B, SCGB3A1, SCGB3A2 | (2) |
| ciliated cells | Ciliated | FOXJ1 | (1) |
| T-cells | TC | CD3E, CD8A, CD4, CCR7 | (3) |
| B-cells | BC | CD79A, MS4A1 | (3, 4) |
| plasma cells | PC | IGHA1, MZB1, XBP1 | (3) |
| macrophages | Mac | CD68, CD14, C1QA, VSIG4 | (3) |
| Mast cells | Mast | TPSAB1, KIT | (3) |
| fibroblasts | FB | LUM, DCN | (5) |
| smooth muscle cells | SMC | ACTA2, RGS5 | (5) |
| endothelial cells | EC | PECAM1, VWF | (5) |
| Schwann cells | SC | S100B, SOX10 | (5) |
| chondrocytes | CH | SOX9, ACAN, COL2A1 | (6) |

**Supplementary Table 2. marker genes information.**
