## Supplementary Table 3 for "Transcriptional profiling sheds light on the fibrotic aspects of idiopathic subglottic tracheal stenosis"

Supplementary Table 3 – antibody information

| **1° Antibodies** | | | | | |
|---|---|---|---|---|---|
| Antigen | Species | catalog No | company | dilution | incubation |
| S100 | rabbit | #Z0311 | DAKO | ready to use | o.n., 4°C |
| Nestin | mouse | #MAB5326 | Millipore | 1:200 | o.n., 4°C |
| PGP9.5 | mouse | #7863-1004 | BioRad | 1:250 | o.n., 4°C |
| POSTN | rabbit | #EPR19934 | abcam | 1:2000 | o.n., 4°C |
| MZB1 | rabbit | #HPA052694 | merck | 1:100 | o.n., 4°C |
| **2° Antibodies** | | | | | |
| Antigen | Species | catalog No | company | dilution | incubation |
| α rb AF488 | goat | #A32731 | Sigma - Aldrich | 1:600 | 1 hr, RT |
| α m AF546 | goat | #A21123 | Thermo Fisher | 1:400 | 1 hr, RT |
| α g AF546 | donkey | #A11056 | Sigma – Aldrich | 1:400 | 1 hr, RT |
