## Supplementary Table 4 for "Transcriptional profiling sheds light on the fibrotic aspects of idiopathic subglottic tracheal stenosis"

| Supplementary Table 4 - clusermarker | | | | | | | | | |
|---|---|---|---|---|---|---|---|---|---|
| | **p_val** | **avg_log2FC** | **pct.1** | **pct.2** | **p_val_adj** | **celltype** | **Foldchange_UP** | **Foldchange_DOWN** | **Ratio_pct1_pct2** |
| PRR4 | 4,91324E-15 | -2,982882776 | 0,277 | 0,913 | 1,79761E-10 | Basal | 0,126491928 | 7,905642826 | 0,3033954 |
| GPX4 | 2,83403E-12 | 0,981880269 | 0,938 | 0,9 | 1,03689E-07 | Basal | 1,975037806 | 0,506319422 | 1,042222222 |
| MT-ATP8 | 4,78175E-11 | -1,535818791 | 0,738 | 0,867 | 1,7495E-06 | Basal | 0,344883547 | 2,89952945 | 0,851211073 |
| RPS27 | 2,80891E-10 | -0,677260192 | 1 | 1 | 1,0277E-05 | Basal | 0,625351747 | 1,599100035 | 1 |
| DDT | 2,97253E-10 | 1,095266961 | 0,615 | 0,26 | 1,08756E-05 | Basal | 2,13652613 | 0,468049506 | 2,365384615 |
| TYMP | 3,12007E-10 | 1,074687839 | 0,646 | 0,26 | 1,14154E-05 | Basal | 2,106266282 | 0,474773778 | 2,484615385 |
| RPS26 | 7,06349E-10 | -0,713836492 | 0,938 | 0,98 | 2,58432E-05 | Basal | 0,609696643 | 1,640159924 | 0,957142857 |
| FTH1 | 2,15667E-09 | 0,678369797 | 1 | 0,973 | 7,89059E-05 | Basal | 1,600330407 | 0,624870962 | 1,027749229 |
| PSMA7 | 2,5479E-09 | 0,784227924 | 0,846 | 0,68 | 9,32202E-05 | Basal | 1,722170433 | 0,580662623 | 1,244117647 |
| KRTCAP2 | 2,69831E-09 | 0,79612162 | 0,846 | 0,72 | 9,87232E-05 | Basal | 1,736426831 | 0,575895271 | 1,175 |
| PFN1 | 3,10699E-09 | 0,873920267 | 0,969 | 0,88 | 0,000113675 | Basal | 1,832636003 | 0,545662095 | 1,101136364 |
| RAC1 | 5,2609E-09 | 1,018590878 | 0,846 | 0,793 | 0,000192481 | Basal | 2,025939199 | 0,493598229 | 1,066834805 |
| ATP5F1D | 5,80979E-09 | 0,955044028 | 0,831 | 0,687 | 0,000212563 | Basal | 1,938638797 | 0,515825847 | 1,209606987 |
| SPINT2 | 5,82128E-09 | 0,755471901 | 0,892 | 0,88 | 0,000212983 | Basal | 1,688183701 | 0,592352597 | 1,013636364 |
| SCGB3A1 | 7,12374E-09 | -2,915549232 | 0,477 | 0,813 | 0,000260636 | Basal | 0,132535502 | 7,545148166 | 0,586715867 |
| FCGBP | 9,63949E-09 | 3,033894049 | 0,508 | 0,18 | 0,00035268 | Basal | 8,190173696 | 0,122097533 | 2,822222222 |
| SON | 9,74043E-09 | -0,933745697 | 0,677 | 0,893 | 0,000356373 | Basal | 0,52349741 | 1,910229128 | 0,758118701 |
| TSPO | 9,78357E-09 | 0,857885077 | 0,892 | 0,753 | 0,000357952 | Basal | 1,8123795 | 0,551760821 | 1,184594954 |
| SPARCL1 | 1,27093E-08 | -1,16983574 | 0,262 | 0,733 | 0,000464994 | Basal | 0,444471944 | 2,249860795 | 0,357435198 |
| HLA-DQB1 | 1,3859E-08 | -0,573600869 | 0,046 | 0,453 | 0,000507061 | Basal | 0,671937583 | 1,488233469 | 0,101545254 |
| RPS29 | 1,39194E-08 | -0,666428631 | 0,815 | 0,92 | 0,000509269 | Basal | 0,630064473 | 1,587139162 | 0,885869565 |
| MIF | 1,75025E-08 | 0,955654138 | 0,892 | 0,82 | 0,000640363 | Basal | 1,939458812 | 0,515607753 | 1,087804878 |
| IGKV3-11 | 1,79118E-08 | 0,586022141 | 0,2 | 0 | 0,000655338 | Basal | 1,501102135 | 0,666177189 | #DIV/0! |
| HINT1 | 2,63611E-08 | 0,751632607 | 0,985 | 0,853 | 0,000964473 | Basal | 1,683697086 | 0,593931063 | 1,154747948 |
| PPDPF | 4,40835E-08 | 1,039311019 | 0,938 | 0,913 | 0,001612885 | Basal | 2,055245904 | 0,486559782 | 1,027382256 |
| RPL18A | 4,41201E-08 | 0,563664045 | 1 | 0,98 | 0,001614221 | Basal | 1,47801821 | 0,67658165 | 1,020408163 |
| ETHE1 | 5,04266E-08 | 0,900036718 | 0,708 | 0,453 | 0,001844957 | Basal | 1,866113476 | 0,535873093 | 1,562913907 |
| ELOB | 7,65584E-08 | 0,79824727 | 0,908 | 0,773 | 0,002801043 | Basal | 1,738987147 | 0,575047378 | 1,174644243 |
| PGLS | 7,94411E-08 | 0,960396025 | 0,723 | 0,533 | 0,00290651 | Basal | 1,945843963 | 0,513915822 | 1,356472795 |
| MSMB | 1,11044E-07 | -2,092709261 | 0,185 | 0,573 | 0,004062784 | Basal | 0,234440015 | 4,265483437 | 0,322862129 |
| COX4I1 | 1,34415E-07 | 0,721589885 | 0,923 | 0,9 | 0,004917823 | Basal | 1,64899827 | 0,606428775 | 1,025555556 |

| Supplementary Table 4 - clusermarker | | | | | | | | | |
|---|---|---|---|---|---|---|---|---|---|
| | **p_val** | **avg_log2FC** | **pct.1** | **pct.2** | **p_val_adj** | **celltype** | **Foldchange_UP** | **Foldchange_DOWN** | **Ratio_pct1_pct2** |
| GSTA1 | 1,54932E-07 | -1,78568343 | 0,077 | 0,447 | 0,005668512 | Basal | 0,29003855 | 3,447817545 | 0,172259508 |
| LYZ | 1,71427E-07 | -2,495111805 | 0,4 | 0,793 | 0,006271995 | Basal | 0,177376673 | 5,637719911 | 0,504413619 |
| RPL13 | 2,5711E-07 | 0,519547914 | 1 | 1 | 0,009406882 | Basal | 1,433505971 | 0,697590398 | 1 |
| HSP90AB1 | 3,16409E-07 | -0,79034653 | 0,892 | 0,973 | 0,011576458 | Basal | 0,578205193 | 1,72948983 | 0,916752312 |
| SELENOW | 3,62534E-07 | 0,687145214 | 0,908 | 0,82 | 0,013264045 | Basal | 1,610094331 | 0,621081623 | 1,107317073 |
| PRB3 | 3,73212E-07 | -2,772435573 | 0,108 | 0,493 | 0,013654693 | Basal | 0,146357078 | 6,832604278 | 0,219066937 |
| MYL12B | 4,20173E-07 | 0,698478936 | 0,908 | 0,86 | 0,015372882 | Basal | 1,622792946 | 0,61622156 | 1,055813953 |
| ATP6V0E1 | 4,2064E-07 | 0,632071941 | 0,831 | 0,68 | 0,015389942 | Basal | 1,549789142 | 0,645249068 | 1,222058824 |
| HSPA8 | 5,09902E-07 | -1,028255999 | 0,815 | 0,9 | 0,018655771 | Basal | 0,490302492 | 2,039557243 | 0,905555556 |
| MRPL14 | 5,54901E-07 | 0,801787488 | 0,692 | 0,48 | 0,020302166 | Basal | 1,743259674 | 0,573638004 | 1,441666667 |
| TXNL4A | 5,608E-07 | 0,824948101 | 0,646 | 0,42 | 0,020517979 | Basal | 1,771471311 | 0,564502509 | 1,538095238 |
| ITM2B | 5,85539E-07 | 0,45920667 | 0,938 | 0,92 | 0,021423097 | Basal | 1,374785623 | 0,727386134 | 1,019565217 |
| CD9 | 6,74495E-07 | 1,000649134 | 0,923 | 0,893 | 0,024677753 | Basal | 2,000900093 | 0,499775078 | 1,033594625 |
| HSPA1A | 6,78501E-07 | -1,801033158 | 0,585 | 0,813 | 0,024824329 | Basal | 0,286969008 | 3,484696857 | 0,719557196 |
| GUK1 | 7,06448E-07 | 0,84310793 | 0,754 | 0,587 | 0,025846806 | Basal | 1,79391052 | 0,557441405 | 1,284497445 |
| COX8A | 8,61456E-07 | 0,691168147 | 0,846 | 0,733 | 0,031518081 | Basal | 1,61459032 | 0,619352159 | 1,154160982 |
| HSPA1B | 1,06977E-06 | -1,456919967 | 0,569 | 0,82 | 0,039139568 | Basal | 0,364269986 | 2,745216569 | 0,693902439 |
| CSTB | 1,14763E-06 | 0,819973776 | 0,892 | 0,833 | 0,041988213 | Basal | 1,765373903 | 0,566452239 | 1,070828331 |
| ATP6V0C | 1,15154E-06 | 0,785555047 | 0,785 | 0,72 | 0,042131526 | Basal | 1,723755371 | 0,580128722 | 1,090277778 |
| OAZ1 | 1,1711E-06 | 0,592595849 | 0,923 | 0,813 | 0,042846943 | Basal | 1,507957585 | 0,663148626 | 1,135301353 |
| CLIC3 | 1,18558E-06 | 0,850522212 | 0,446 | 0,16 | 0,043376894 | Basal | 1,803153494 | 0,554583957 | 2,7875 |
| RPL37 | 1,19506E-06 | -0,486456022 | 0,985 | 0,993 | 0,043723566 | Basal | 0,713776336 | 1,400999093 | 0,991943605 |
| METAP2 | 1,53751E-06 | -0,734473605 | 0,277 | 0,607 | 0,056252991 | Basal | 0,601037285 | 1,663790293 | 0,456342669 |
| COX6A1 | 1,68178E-06 | 0,532021289 | 0,908 | 0,847 | 0,06153142 | Basal | 1,445953631 | 0,691585109 | 1,07201889 |
| TMEM160 | 1,73748E-06 | 0,801519324 | 0,585 | 0,367 | 0,06356915 | Basal | 1,742935672 | 0,57374464 | 1,59400545 |
| GBP3 | 1,82703E-06 | -0,433417036 | 0,031 | 0,347 | 0,066845542 | Basal | 0,740505812 | 1,350428293 | 0,089337176 |
| PLAAT3 | 1,90202E-06 | 0,679308241 | 0,508 | 0,26 | 0,069589161 | Basal | 1,601371728 | 0,624464628 | 1,953846154 |
| RPL28 | 1,99619E-06 | 0,467687399 | 1 | 0,993 | 0,073034748 | Basal | 1,382890954 | 0,723122816 | 1,007049345 |
| CIAO2B | 2,24221E-06 | 0,664898508 | 0,646 | 0,44 | 0,082035804 | Basal | 1,585456734 | 0,630733074 | 1,468181818 |
| NME2 | 2,54643E-06 | 0,646322827 | 0,954 | 0,887 | 0,09316609 | Basal | 1,565173758 | 0,638906699 | 1,075535513 |
| DYNLL1 | 2,8973E-06 | 0,675873006 | 0,892 | 0,813 | 0,106003688 | Basal | 1,5975632 | 0,625953327 | 1,097170972 |

| Supplementary Table 4 - clusermarker | | | | | | | | | |
|---|---|---|---|---|---|---|---|---|---|
| | **p_val** | **avg_log2FC** | **pct.1** | **pct.2** | **p_val_adj** | **celltype** | **Foldchange_UP** | **Foldchange_DOWN** | **Ratio_pct1_pct2** |
| AURKAIP1 | 3,83358E-06 | 0,798045276 | 0,708 | 0,54 | 0,140259015 | Basal | 1,738743686 | 0,575127897 | 1,311111111 |
| GABARAP | 4,02604E-06 | 0,455325031 | 0,908 | 0,78 | 0,14730062 | Basal | 1,37109167 | 0,729345836 | 1,164102564 |
| TRNP1 | 4,2896E-06 | 0,963137492 | 0,492 | 0,247 | 0,156943681 | Basal | 1,949545049 | 0,512940186 | 1,991902834 |
| SNW1 | 4,38099E-06 | -0,561906226 | 0,169 | 0,5 | 0,16028726 | Basal | 0,677406518 | 1,47621845 | 0,338 |
| STATH | 4,56858E-06 | -2,33762514 | 0,031 | 0,32 | 0,167150763 | Basal | 0,197835723 | 5,054698846 | 0,096875 |
| CLTA | 5,11184E-06 | 0,808801514 | 0,815 | 0,68 | 0,187026999 | Basal | 1,751755608 | 0,570855886 | 1,198529412 |
| RPL22 | 5,74509E-06 | -0,529445272 | 0,938 | 0,967 | 0,210195525 | Basal | 0,692821078 | 1,443374099 | 0,970010341 |
| MGMT | 6,03136E-06 | 0,632178141 | 0,538 | 0,273 | 0,220669388 | Basal | 1,549903229 | 0,645201572 | 1,970695971 |
| S100A11 | 6,26553E-06 | 0,754944918 | 0,954 | 0,913 | 0,229236856 | Basal | 1,68756716 | 0,592569009 | 1,0449069 |
| BBLN | 6,27491E-06 | 0,934669722 | 0,738 | 0,607 | 0,229580006 | Basal | 1,911452994 | 0,523162224 | 1,215815486 |
| GSTP1 | 6,46355E-06 | 1,054674886 | 0,938 | 0,953 | 0,236481732 | Basal | 2,077250041 | 0,481405695 | 0,984260231 |
| FETUB | 7,36878E-06 | 0,333694359 | 0,154 | 0,007 | 0,269601607 | Basal | 1,260236377 | 0,793501932 | 22 |
| RHOD | 7,56235E-06 | 0,694271254 | 0,446 | 0,2 | 0,276683664 | Basal | 1,618066896 | 0,61802142 | 2,23 |
| SPRR3 | 7,64904E-06 | 1,609621406 | 0,4 | 0,147 | 0,279855583 | Basal | 3,051717477 | 0,327684331 | 2,721088435 |
| EFHD2 | 7,8797E-06 | 1,02393064 | 0,569 | 0,327 | 0,288294513 | Basal | 2,033451582 | 0,491774679 | 1,740061162 |
| RPL31 | 8,22086E-06 | -0,617228265 | 0,862 | 0,933 | 0,300776622 | Basal | 0,651922212 | 1,53392534 | 0,923901393 |
| SERINC2 | 1,01073E-05 | 0,839113345 | 0,646 | 0,487 | 0,369794195 | Basal | 1,788950346 | 0,558987007 | 1,326488706 |
| NDFIP1 | 1,04225E-05 | 0,93220534 | 0,708 | 0,547 | 0,381327965 | Basal | 1,908190676 | 0,524056643 | 1,294332724 |
| ASPG | 1,05039E-05 | 0,786844341 | 0,323 | 0,1 | 0,384305933 | Basal | 1,725296529 | 0,579610509 | 3,23 |
| AADAC | 1,13938E-05 | 0,421756461 | 0,185 | 0,02 | 0,416866332 | Basal | 1,339557455 | 0,746515199 | 9,25 |
| RPL8 | 1,21175E-05 | 0,594181763 | 0,985 | 0,993 | 0,443342549 | Basal | 1,509616151 | 0,662420046 | 0,991943605 |
| HEBP2 | 1,21549E-05 | 0,767780992 | 0,754 | 0,653 | 0,44470956 | Basal | 1,702648925 | 0,587320137 | 1,15467075 |
| SCGB1A1 | 1,23423E-05 | -5,259854643 | 0,2 | 0,467 | 0,451567896 | Basal | 0,026099127 | 38,31545793 | 0,428265525 |
| PDCD6 | 1,2716E-05 | 0,607993883 | 0,692 | 0,52 | 0,46524056 | Basal | 1,524138368 | 0,656108409 | 1,330769231 |
| SLC26A4 | 1,27668E-05 | 0,354899265 | 0,123 | 0 | 0,467098647 | Basal | 1,27889628 | 0,781924238 | #DIV/0! |
| IGLV3-21 | 1,27668E-05 | 0,630876758 | 0,123 | 0 | 0,467098647 | Basal | 1,548505769 | 0,645783839 | #DIV/0! |
| ALOX15 | 1,34591E-05 | 2,308643418 | 0,585 | 0,42 | 0,492429345 | Basal | 4,954170147 | 0,201850153 | 1,392857143 |
| ANTKMT | 1,37929E-05 | 0,616715874 | 0,462 | 0,24 | 0,50464108 | Basal | 1,533380645 | 0,652153791 | 1,925 |
| DUSP23 | 1,39071E-05 | 0,738292432 | 0,662 | 0,433 | 0,508820143 | Basal | 1,668200195 | 0,599448437 | 1,52886836 |
| RPL38 | 1,40818E-05 | -0,521938876 | 0,754 | 0,9 | 0,515209494 | Basal | 0,696435246 | 1,435883674 | 0,837777778 |
| CCDC60 | 1,51382E-05 | -0,312736425 | 0,015 | 0,273 | 0,553862097 | Basal | 0,805113215 | 1,242061341 | 0,054945055 |

| Supplementary Table 4 - clusermarker | | | | | | | | | |
|---|---|---|---|---|---|---|---|---|---|
| | **p_val** | **avg_log2FC** | **pct.1** | **pct.2** | **p_val_adj** | **celltype** | **Foldchange_UP** | **Foldchange_DOWN** | **Ratio_pct1_pct2** |
| MT-ND4L | 1,57999E-05 | -0,729607402 | 0,877 | 0,9 | 0,578071043 | Basal | 0,603068003 | 1,658187791 | 0,974444444 |
| ARL6IP4 | 1,82505E-05 | 0,679480662 | 0,769 | 0,68 | 0,667732641 | Basal | 1,601563125 | 0,62439 | 1,130882353 |
| TSPAN13 | 1,82775E-05 | 0,723061965 | 0,646 | 0,48 | 0,668717089 | Basal | 1,650681713 | 0,60581031 | 1,345833333 |
| GRN | 2,17497E-05 | 0,685864357 | 0,754 | 0,633 | 0,795756471 | Basal | 1,608665487 | 0,621633278 | 1,191153239 |
| CCDC190 | 2,34469E-05 | 0,377772332 | 0,138 | 0,007 | 0,857850647 | Basal | 1,299334001 | 0,769625053 | 19,71428571 |
| IGHG1 | 2,34469E-05 | 0,590185929 | 0,138 | 0,007 | 0,857850647 | Basal | 1,50544075 | 0,664257295 | 19,71428571 |
| S100A13 | 2,58307E-05 | 0,818775047 | 0,754 | 0,653 | 0,945066037 | Basal | 1,76390767 | 0,566923097 | 1,15467075 |
| RPS2 | 2,66869E-05 | 0,523586615 | 0,985 | 0,973 | 0,976393239 | Basal | 1,43752457 | 0,695640284 | 1,012332991 |
| TALDO1 | 2,67816E-05 | 0,735826344 | 0,738 | 0,573 | 0,979858278 | Basal | 1,665351072 | 0,600473988 | 1,287958115 |
| RPL34 | 2,68308E-05 | -0,541765419 | 1 | 0,993 | 0,981657517 | Basal | 0,686929802 | 1,455752826 | 1,007049345 |
| SRSF9 | 2,80556E-05 | 0,726825453 | 0,754 | 0,627 | 1 | Basal | 1,654993387 | 0,604232022 | 1,202551834 |
| IRX2 | 2,95988E-05 | -0,396733707 | 0,092 | 0,387 | 1 | Basal | 0,759576035 | 1,316523895 | 0,237726098 |
| CST3 | 3,01417E-05 | 0,326562076 | 0,892 | 0,86 | 1 | Basal | 1,254021494 | 0,797434498 | 1,037209302 |
| NRP2 | 3,37941E-05 | 0,599248272 | 0,538 | 0,287 | 1 | Basal | 1,514926995 | 0,660097815 | 1,87456446 |
| IGKC | 3,45318E-05 | 0,684189365 | 0,246 | 0,053 | 1 | Basal | 1,606798885 | 0,622355423 | 4,641509434 |
| VSTM2L | 3,93639E-05 | 0,54185857 | 0,262 | 0,067 | 1 | Basal | 1,455846823 | 0,68688545 | 3,910447761 |
| MESP1 | 4,30436E-05 | 0,362536624 | 0,215 | 0,04 | 1 | Basal | 1,285684471 | 0,777795814 | 5,375 |
| LSP1 | 4,45885E-05 | 0,602818386 | 0,369 | 0,147 | 1 | Basal | 1,518680498 | 0,658466347 | 2,510204082 |
| OCIAD2 | 4,62837E-05 | 0,760667922 | 0,615 | 0,44 | 1 | Basal | 1,694274839 | 0,590223013 | 1,397727273 |
| METTL7A | 4,76139E-05 | -0,610517758 | 0,215 | 0,533 | 1 | Basal | 0,654961605 | 1,526807055 | 0,403377111 |
| SLC25A6 | 4,78217E-05 | 0,525418897 | 0,923 | 0,807 | 1 | Basal | 1,439351446 | 0,694757353 | 1,143742255 |
| JTB | 4,79301E-05 | 0,655741847 | 0,692 | 0,533 | 1 | Basal | 1,575425843 | 0,634749014 | 1,298311445 |
| GNG5 | 4,89587E-05 | 0,757014964 | 0,8 | 0,74 | 1 | Basal | 1,689990297 | 0,591719373 | 1,081081081 |
| STAG2 | 4,91466E-05 | -0,318644966 | 0,185 | 0,487 | 1 | Basal | 0,801822627 | 1,24715862 | 0,379876797 |
| ANAPC11 | 5,02719E-05 | 0,889864597 | 0,723 | 0,627 | 1 | Basal | 1,853002203 | 0,539664766 | 1,153110048 |
| ARF5 | 5,18677E-05 | 0,674264066 | 0,723 | 0,627 | 1 | Basal | 1,595782539 | 0,6266518 | 1,153110048 |
| SSU72 | 5,33619E-05 | 0,557614659 | 0,692 | 0,54 | 1 | Basal | 1,471833685 | 0,679424591 | 1,281481481 |
| STMN1 | 5,36042E-05 | -0,959596845 | 0,246 | 0,54 | 1 | Basal | 0,514200584 | 1,944766363 | 0,455555556 |
| TAF10 | 5,52719E-05 | 0,847746553 | 0,6 | 0,453 | 1 | Basal | 1,79968767 | 0,55565197 | 1,324503311 |
| H4C3 | 5,62267E-05 | -0,896363938 | 0,508 | 0,733 | 1 | Basal | 0,537239044 | 1,86136881 | 0,693042292 |
| TENT5C | 5,62665E-05 | -0,507393682 | 0,231 | 0,52 | 1 | Basal | 0,703492193 | 1,421479883 | 0,444230769 |

| Supplementary Table 4 - clusermarker | | | | | | | | | |
|---|---|---|---|---|---|---|---|---|---|
| | **p_val** | **avg_log2FC** | **pct.1** | **pct.2** | **p_val_adj** | **celltype** | **Foldchange_UP** | **Foldchange_DOWN** | **Ratio_pct1_pct2** |
| VDAC2 | 5,70453E-05 | 0,778965788 | 0,785 | 0,653 | 1 | Basal | 1,715900369 | 0,582784419 | 1,202143951 |
| MRPS24 | 5,74477E-05 | 0,459152212 | 0,708 | 0,533 | 1 | Basal | 1,374733729 | 0,727413592 | 1,328330206 |
| OAT | 5,88328E-05 | -0,861329856 | 0,6 | 0,733 | 1 | Basal | 0,550444932 | 1,816712159 | 0,818553888 |
| RPL36A | 6,09114E-05 | -0,449212434 | 0,954 | 0,953 | 1 | Basal | 0,732442579 | 1,36529474 | 1,001049318 |
| RPL35A | 6,09623E-05 | -0,536004873 | 0,985 | 0,987 | 1 | Basal | 0,689678131 | 1,44995173 | 0,997973658 |
| IL18 | 6,73491E-05 | -0,566316728 | 0,169 | 0,447 | 1 | Basal | 0,675338763 | 1,480738342 | 0,378076063 |
| NENF | 6,81544E-05 | 0,622410677 | 0,723 | 0,573 | 1 | Basal | 1,539445376 | 0,649584594 | 1,261780105 |
| NDUFB9 | 7,05458E-05 | 0,43524264 | 0,846 | 0,733 | 1 | Basal | 1,352138223 | 0,73956936 | 1,154160982 |
| IL33 | 7,13919E-05 | -0,871463521 | 0,154 | 0,453 | 1 | Basal | 0,546592087 | 1,829517888 | 0,33995585 |
| CYBA | 7,3936E-05 | 1,145150522 | 0,723 | 0,633 | 1 | Basal | 2,211692049 | 0,452142513 | 1,142180095 |
| UBC | 7,45184E-05 | -0,596235576 | 0,923 | 0,987 | 1 | Basal | 0,661477699 | 1,511766763 | 0,935157042 |
| MRPL57 | 8,01579E-05 | 0,57463803 | 0,646 | 0,493 | 1 | Basal | 1,489303753 | 0,671454697 | 1,310344828 |
| NFIB | 8,03836E-05 | -0,747111464 | 0,462 | 0,733 | 1 | Basal | 0,595795254 | 1,678428946 | 0,630286494 |
| EDF1 | 8,12811E-05 | 0,39633692 | 0,846 | 0,8 | 1 | Basal | 1,316161859 | 0,759784971 | 1,0575 |
| PHPT1 | 8,17669E-05 | 0,56127934 | 0,692 | 0,593 | 1 | Basal | 1,475577137 | 0,677700932 | 1,166947723 |
| SH3BGRL3 | 8,28732E-05 | 0,736702963 | 0,738 | 0,613 | 1 | Basal | 1,66636329 | 0,600109235 | 1,203915171 |
| GNL3 | 9,12657E-05 | -0,479852652 | 0,185 | 0,46 | 1 | Basal | 0,717050856 | 1,394601223 | 0,402173913 |
| MYL12A | 9,21307E-05 | 0,547857984 | 0,877 | 0,78 | 1 | Basal | 1,461913534 | 0,684034984 | 1,124358974 |
| NRARP | 9,33004E-05 | -0,654988136 | 0,262 | 0,533 | 1 | Basal | 0,635080714 | 1,574603005 | 0,491557223 |
| RPS25 | 9,84083E-05 | -0,388044901 | 1 | 0,973 | 1 | Basal | 0,764164477 | 1,308618799 | 1,027749229 |
| TSPAN3 | 0,000104978 | 1,104991397 | 0,677 | 0,587 | 1 | Basal | 2,150975955 | 0,464905243 | 1,153321976 |
| SLC25A3 | 0,000107365 | 0,67393844 | 0,862 | 0,813 | 1 | Basal | 1,595422402 | 0,626793255 | 1,060270603 |
| PBX1 | 0,000108862 | -0,508553028 | 0,185 | 0,493 | 1 | Basal | 0,702927095 | 1,42262264 | 0,37525355 |
| LINC02185 | 0,000110772 | -0,454490857 | 0,062 | 0,293 | 1 | Basal | 0,72976767 | 1,370299126 | 0,211604096 |
| ZBTB20 | 0,000116971 | -0,341821626 | 0,154 | 0,473 | 1 | Basal | 0,789044392 | 1,267355818 | 0,325581395 |
| PRDX5 | 0,00011748 | 0,424673042 | 0,892 | 0,873 | 1 | Basal | 1,34226827 | 0,745007553 | 1,021764032 |
| ETFB | 0,000118246 | 0,602074147 | 0,662 | 0,493 | 1 | Basal | 1,517897263 | 0,658806116 | 1,342799189 |
| NAXE | 0,000119 | 0,290625314 | 0,6 | 0,42 | 1 | Basal | 1,223170327 | 0,817547628 | 1,428571429 |
| REEP5 | 0,000123432 | 0,55370751 | 0,8 | 0,66 | 1 | Basal | 1,467853016 | 0,681267122 | 1,212121212 |
| TACSTD2 | 0,000125551 | 0,676670165 | 0,862 | 0,813 | 1 | Basal | 1,598446176 | 0,625607553 | 1,060270603 |
| CHCHD2 | 0,00012729 | 0,365129953 | 0,892 | 0,847 | 1 | Basal | 1,287997643 | 0,776398937 | 1,053128689 |

| Supplementary Table 4 - clusermarker | | | | | | | | | |
|---|---|---|---|---|---|---|---|---|---|
| | **p_val** | **avg_log2FC** | **pct.1** | **pct.2** | **p_val_adj** | **celltype** | **Foldchange_UP** | **Foldchange_DOWN** | **Ratio_pct1_pct2** |
| CIB1 | 0,00012732 | 0,831353734 | 0,677 | 0,593 | 1 | Basal | 1,779354213 | 0,562001648 | 1,141652614 |
| NDUFA13 | 0,000127667 | 0,539879567 | 0,831 | 0,767 | 1 | Basal | 1,453851148 | 0,687828325 | 1,083441982 |
| CLDN3 | 0,000128547 | 1,23046461 | 0,508 | 0,327 | 1 | Basal | 2,346425427 | 0,426180175 | 1,55351682 |
| ST14 | 0,000133734 | 0,674513255 | 0,569 | 0,407 | 1 | Basal | 1,596058195 | 0,62654357 | 1,398034398 |
| HSPA2 | 0,000134977 | -0,71419059 | 0,262 | 0,567 | 1 | Basal | 0,609547016 | 1,640562537 | 0,462081129 |
| NDUFA11 | 0,000135251 | 0,51838891 | 0,862 | 0,74 | 1 | Basal | 1,432354812 | 0,698151039 | 1,164864865 |
| SMIM5 | 0,000135721 | 0,361637563 | 0,185 | 0,033 | 1 | Basal | 1,284883506 | 0,778280673 | 5,606060606 |
| SPCS1 | 0,000137514 | 0,470355924 | 0,769 | 0,673 | 1 | Basal | 1,385451227 | 0,721786506 | 1,142644874 |
| RBBP6 | 0,000140248 | -0,745627464 | 0,415 | 0,647 | 1 | Basal | 0,596408422 | 1,676703351 | 0,641421947 |
| CDKN2A | 0,00014144 | 0,441906574 | 0,185 | 0,033 | 1 | Basal | 1,358398315 | 0,736161102 | 5,606060606 |
| CHKA | 0,000156611 | -0,442605985 | 0,123 | 0,38 | 1 | Basal | 0,735804302 | 1,359057018 | 0,323684211 |
| GRAMD2B | 0,000158904 | -0,708478323 | 0,292 | 0,58 | 1 | Basal | 0,611965266 | 1,634079669 | 0,503448276 |
| HLA-B | 0,000160631 | 0,806254057 | 0,908 | 0,913 | 1 | Basal | 1,748665152 | 0,571864773 | 0,994523549 |
| PTPMT1 | 0,00016278 | 0,38392989 | 0,354 | 0,14 | 1 | Basal | 1,304891533 | 0,766347221 | 2,528571429 |
| GPX2 | 0,000162891 | 0,861097553 | 0,538 | 0,327 | 1 | Basal | 1,816419655 | 0,550533572 | 1,645259939 |
| NDUFC2 | 0,000164268 | 0,455852409 | 0,815 | 0,733 | 1 | Basal | 1,371592965 | 0,729079272 | 1,111869031 |
| FAM107A | 0,000169186 | -0,28012681 | 0,031 | 0,253 | 1 | Basal | 0,823518628 | 1,214301615 | 0,122529644 |
| RPS10 | 0,000176506 | 0,459337084 | 0,4 | 0,2 | 1 | Basal | 1,374909904 | 0,727320384 | 2 |
| GADD45B | 0,000177096 | -0,942229414 | 0,677 | 0,813 | 1 | Basal | 0,520428035 | 1,921495256 | 0,832718327 |
| CEACAM5 | 0,000181466 | 1,85082936 | 0,462 | 0,28 | 1 | Basal | 3,607074849 | 0,27723295 | 1,65 |
| P4HB | 0,000188489 | 0,588058584 | 0,754 | 0,673 | 1 | Basal | 1,503222519 | 0,665237506 | 1,120356612 |
| TUFM | 0,00019237 | 0,488865318 | 0,754 | 0,627 | 1 | Basal | 1,403340712 | 0,712585327 | 1,202551834 |
| ZNF429 | 0,000194614 | -0,350045156 | 0,031 | 0,253 | 1 | Basal | 0,784559541 | 1,274600522 | 0,122529644 |
| DNAJA1 | 0,000211865 | -0,702621418 | 0,692 | 0,86 | 1 | Basal | 0,61445471 | 1,627459248 | 0,804651163 |
| LYPD2 | 0,000217716 | 2,683852484 | 0,477 | 0,307 | 1 | Basal | 6,42569491 | 0,155625191 | 1,553745928 |
| FAM177A1 | 0,00021867 | 0,639180736 | 0,646 | 0,513 | 1 | Basal | 1,557444482 | 0,642077462 | 1,259259259 |
| EIF3F | 0,00021956 | 0,642265438 | 0,754 | 0,687 | 1 | Basal | 1,560778098 | 0,640706069 | 1,097525473 |
| COX5A | 0,000219686 | 0,674434598 | 0,738 | 0,68 | 1 | Basal | 1,595971178 | 0,626577731 | 1,085294118 |
| PSMD8 | 0,000223304 | 0,725359977 | 0,677 | 0,567 | 1 | Basal | 1,653313114 | 0,604846106 | 1,194003527 |
| SYNGR2 | 0,000225436 | 0,720535653 | 0,692 | 0,573 | 1 | Basal | 1,647793724 | 0,606872077 | 1,207678883 |
| CDC37 | 0,000227007 | 0,682546648 | 0,692 | 0,553 | 1 | Basal | 1,604970353 | 0,623064469 | 1,251356239 |

| Supplementary Table 4 - clusermarker | | | | | | | | | |
|---|---|---|---|---|---|---|---|---|---|
| | **p_val** | **avg_log2FC** | **pct.1** | **pct.2** | **p_val_adj** | **celltype** | **Foldchange_UP** | **Foldchange_DOWN** | **Ratio_pct1_pct2** |
| NAPRT | 0,000239669 | 0,571481082 | 0,523 | 0,327 | 1 | Basal | 1,486048377 | 0,672925603 | 1,599388379 |
| MAPK3 | 0,000240178 | 0,537399382 | 0,338 | 0,147 | 1 | Basal | 1,451353931 | 0,689011811 | 2,299319728 |
| DDX18 | 0,000242518 | -0,607989826 | 0,323 | 0,607 | 1 | Basal | 0,656110254 | 1,524134082 | 0,532125206 |
| MRPL41 | 0,000248584 | 0,539486079 | 0,646 | 0,547 | 1 | Basal | 1,453454672 | 0,688015952 | 1,180987203 |
| SHISA2 | 0,000251207 | -0,296586277 | 0,046 | 0,26 | 1 | Basal | 0,814176634 | 1,228234708 | 0,176923077 |
| ID2 | 0,000252893 | -0,77097907 | 0,415 | 0,667 | 1 | Basal | 0,586019644 | 1,706427439 | 0,622188906 |
| RPS16 | 0,000253725 | 0,387432628 | 0,985 | 0,927 | 1 | Basal | 1,308063545 | 0,764488853 | 1,062567422 |
| SDC1 | 0,000259764 | 0,601692338 | 0,646 | 0,493 | 1 | Basal | 1,517495605 | 0,658980492 | 1,310344828 |
| TMSB10 | 0,000264831 | 0,618478034 | 0,923 | 0,907 | 1 | Basal | 1,535254716 | 0,651357713 | 1,017640573 |
| UBE2E3 | 0,000275055 | 0,606961897 | 0,523 | 0,34 | 1 | Basal | 1,523048514 | 0,656577904 | 1,538235294 |
| TRAPPC5 | 0,000284448 | 0,472785629 | 0,692 | 0,513 | 1 | Basal | 1,387786491 | 0,720571937 | 1,348927875 |
| LPP | 0,000295642 | -0,728654157 | 0,446 | 0,72 | 1 | Basal | 0,603466606 | 1,657092523 | 0,619444444 |
| SLC52A2 | 0,00030587 | 0,479987319 | 0,354 | 0,167 | 1 | Basal | 1,394731407 | 0,716983926 | 2,119760479 |
| GSTM1 | 0,000314535 | -0,28122683 | 0,062 | 0,28 | 1 | Basal | 0,822890955 | 1,215227843 | 0,221428571 |
| CORO1B | 0,000320438 | 0,562587687 | 0,569 | 0,427 | 1 | Basal | 1,476915911 | 0,677086619 | 1,332552693 |
| FIS1 | 0,000323026 | 0,309834873 | 0,738 | 0,6 | 1 | Basal | 1,239565814 | 0,806734091 | 1,23 |
| IFITM3 | 0,000323271 | 0,913525588 | 0,908 | 0,887 | 1 | Basal | 1,883643034 | 0,530886151 | 1,02367531 |
| WSB2 | 0,00032356 | 0,287705276 | 0,308 | 0,12 | 1 | Basal | 1,220697115 | 0,819204033 | 2,566666667 |
| CSNK2A2 | 0,000326842 | -0,337237014 | 0,062 | 0,28 | 1 | Basal | 0,791555814 | 1,263334792 | 0,221428571 |
| TNFRSF6B | 0,000350152 | 0,269022146 | 0,169 | 0,033 | 1 | Basal | 1,204990812 | 0,829881847 | 5,121212121 |
| COX5B | 0,00036477 | 0,371502919 | 0,892 | 0,827 | 1 | Basal | 1,293699833 | 0,772976833 | 1,07859734 |
| PLEC | 0,000366489 | 0,728613972 | 0,662 | 0,5 | 1 | Basal | 1,657046367 | 0,603483415 | 1,324 |
| LAMTOR5 | 0,000368862 | 0,632749184 | 0,708 | 0,6 | 1 | Basal | 1,550516828 | 0,644946241 | 1,18 |
| RPL23 | 0,000368939 | -0,65039613 | 0,769 | 0,873 | 1 | Basal | 0,637105355 | 1,569599112 | 0,880870561 |
| MATR3.1 | 0,000387277 | -0,330700318 | 0,077 | 0,3 | 1 | Basal | 0,795150406 | 1,257623706 | 0,256666667 |
| CD24 | 0,000390994 | 1,074043903 | 0,662 | 0,567 | 1 | Basal | 2,105326376 | 0,474985737 | 1,167548501 |
| CHMP4B | 0,000411679 | 0,414814488 | 0,538 | 0,367 | 1 | Basal | 1,333127243 | 0,750115944 | 1,465940054 |
| PLEKHJ1 | 0,000418461 | 0,606827328 | 0,415 | 0,227 | 1 | Basal | 1,522906456 | 0,656639149 | 1,828193833 |
| UQCR10 | 0,000418924 | 0,521108791 | 0,8 | 0,72 | 1 | Basal | 1,435057745 | 0,69683607 | 1,111111111 |
| ATRX | 0,000437519 | -0,577053427 | 0,446 | 0,68 | 1 | Basal | 0,670331471 | 1,491799271 | 0,655882353 |
| TRPM4 | 0,000442815 | 0,523490642 | 0,338 | 0,16 | 1 | Basal | 1,437428945 | 0,695686562 | 2,1125 |

| Supplementary Table 4 - clusermarker | | | | | | | | | |
|---|---|---|---|---|---|---|---|---|---|
| | **p_val** | **avg_log2FC** | **pct.1** | **pct.2** | **p_val_adj** | **celltype** | **Foldchange_UP** | **Foldchange_DOWN** | **Ratio_pct1_pct2** |
| EPAS1 | 0,000444006 | -0,805633073 | 0,508 | 0,687 | 1 | Basal | 0,572110976 | 1,74791263 | 0,73944687 |
| TXNDC17 | 0,00044666 | 0,703163209 | 0,646 | 0,527 | 1 | Basal | 1,628070539 | 0,614224001 | 1,225806452 |
| PTPRZ1 | 0,000450277 | -0,589389882 | 0,292 | 0,513 | 1 | Basal | 0,664623918 | 1,50461031 | 0,56920078 |
| KIN | 0,000451642 | -0,291806 | 0,062 | 0,273 | 1 | Basal | 0,81687883 | 1,224171766 | 0,227106227 |
| NR1D2 | 0,000454367 | -0,380444884 | 0,308 | 0,567 | 1 | Basal | 0,768200664 | 1,301743212 | 0,543209877 |
| PSME2 | 0,000457285 | 0,396278888 | 0,769 | 0,627 | 1 | Basal | 1,316108918 | 0,759815534 | 1,226475279 |
| CHMP2A | 0,0004592 | 0,511964175 | 0,662 | 0,56 | 1 | Basal | 1,425990307 | 0,701267039 | 1,182142857 |
| CD177 | 0,000461169 | 0,629386358 | 0,154 | 0,027 | 1 | Basal | 1,546906886 | 0,646451321 | 5,703703704 |
| PNISR | 0,000465496 | -0,634490366 | 0,415 | 0,68 | 1 | Basal | 0,644168328 | 1,55238927 | 0,610294118 |
| AC073332.1 | 0,000478098 | 0,265821336 | 0,215 | 0,06 | 1 | Basal | 1,202320344 | 0,831725093 | 3,583333333 |
| THOC2 | 0,000481055 | -0,275544499 | 0,246 | 0,547 | 1 | Basal | 0,82613846 | 1,210450849 | 0,449725777 |
| DRAP1 | 0,000488062 | 0,522590242 | 0,631 | 0,547 | 1 | Basal | 1,436532111 | 0,696120882 | 1,153564899 |
| TMEM179B | 0,000490506 | 0,57899776 | 0,677 | 0,573 | 1 | Basal | 1,493811137 | 0,669428668 | 1,181500873 |
| HOXA5 | 0,000491019 | 0,274494739 | 0,154 | 0,027 | 1 | Basal | 1,209570399 | 0,826739808 | 5,703703704 |
| CRIP2 | 0,0004912 | 0,653878163 | 0,692 | 0,607 | 1 | Basal | 1,573392011 | 0,635569517 | 1,140032949 |
| ARHGAP5 | 0,000492723 | -0,395097259 | 0,277 | 0,547 | 1 | Basal | 0,76043811 | 1,315031409 | 0,506398537 |
| PAK1 | 0,000504881 | -0,399437333 | 0,215 | 0,473 | 1 | Basal | 0,758153914 | 1,318993389 | 0,454545455 |
| SAA2 | 0,000506942 | 0,506873793 | 0,138 | 0,02 | 1 | Basal | 1,420967732 | 0,703745749 | 6,9 |
| AKR1C1 | 0,000507936 | -0,45867567 | 0,354 | 0,62 | 1 | Basal | 0,727653906 | 1,374279711 | 0,570967742 |
| PPIG | 0,000518327 | -0,636006569 | 0,431 | 0,68 | 1 | Basal | 0,643491694 | 1,554021613 | 0,633823529 |
| NR4A2 | 0,000528565 | -0,630626506 | 0,431 | 0,653 | 1 | Basal | 0,645895867 | 1,548237187 | 0,660030628 |
| MRPS15 | 0,000537142 | 0,515795468 | 0,523 | 0,367 | 1 | Basal | 1,429782271 | 0,69940719 | 1,42506812 |
| MAFB | 0,000553246 | -0,761882599 | 0,308 | 0,533 | 1 | Basal | 0,589726284 | 1,695701934 | 0,577861163 |
| DDIT3 | 0,000563873 | -0,448307548 | 0,185 | 0,433 | 1 | Basal | 0,732902125 | 1,36443867 | 0,427251732 |
| MUC4 | 0,000570253 | 1,032181849 | 0,631 | 0,513 | 1 | Basal | 2,045114828 | 0,488970099 | 1,230019493 |
| CAPG | 0,000570294 | 0,64258006 | 0,646 | 0,48 | 1 | Basal | 1,561118508 | 0,64056636 | 1,345833333 |
| C6orf58 | 0,000571398 | -2,088536312 | 0,031 | 0,227 | 1 | Basal | 0,235119107 | 4,253163489 | 0,136563877 |
| KRT19 | 0,00057458 | 0,88866301 | 0,831 | 0,8 | 1 | Basal | 1,851459524 | 0,540114427 | 1,03875 |
| PIP | 0,000587695 | -1,999537781 | 0,046 | 0,24 | 1 | Basal | 0,250080109 | 3,998718662 | 0,191666667 |
| RPS19BP1 | 0,00059056 | 0,515984749 | 0,6 | 0,48 | 1 | Basal | 1,42996987 | 0,699315434 | 1,25 |
| AC008397.1 | 0,000590756 | 0,747895186 | 0,323 | 0,147 | 1 | Basal | 1,679340975 | 0,595471685 | 2,197278912 |

| Supplementary Table 4 - clusermarker | | | | | | | | | |
|---|---|---|---|---|---|---|---|---|---|
| | **p_val** | **avg_log2FC** | **pct.1** | **pct.2** | **p_val_adj** | **celltype** | **Foldchange_UP** | **Foldchange_DOWN** | **Ratio_pct1_pct2** |
| EPB41L2 | 0,000591536 | -0,592770974 | 0,062 | 0,267 | 1 | Basal | 0,663068133 | 1,508140643 | 0,232209738 |
| RBPJ | 0,000596307 | -0,676275053 | 0,246 | 0,5 | 1 | Basal | 0,625778912 | 1,598008467 | 0,492 |
| SCGB3A2 | 0,000597525 | -1,262115574 | 0,185 | 0,427 | 1 | Basal | 0,41693212 | 2,398471962 | 0,433255269 |
| ZBTB16 | 0,000661925 | -0,346953688 | 0,092 | 0,313 | 1 | Basal | 0,786242531 | 1,271872178 | 0,293929712 |
| OSBPL9 | 0,000663859 | -0,347627601 | 0,2 | 0,453 | 1 | Basal | 0,785875346 | 1,272466435 | 0,441501104 |
| AC005392.2 | 0,000670293 | 0,567002762 | 0,262 | 0,093 | 1 | Basal | 1,481442634 | 0,6750177 | 2,817204301 |
| NIBAN1 | 0,000671814 | -0,552656815 | 0,262 | 0,513 | 1 | Basal | 0,681763461 | 1,466784387 | 0,510721248 |
| BAZ2B | 0,000682908 | -0,338815895 | 0,292 | 0,547 | 1 | Basal | 0,790690011 | 1,264718139 | 0,533820841 |
| PPP1CA | 0,000684916 | 0,885968417 | 0,662 | 0,54 | 1 | Basal | 1,848004689 | 0,541124168 | 1,225925926 |
| KDM3A | 0,000718128 | -0,315608779 | 0,108 | 0,333 | 1 | Basal | 0,803511859 | 1,244536704 | 0,324324324 |
| CYCS | 0,000765499 | -0,616248901 | 0,431 | 0,647 | 1 | Basal | 0,652364915 | 1,532884399 | 0,666151468 |
| HLA-DRB5 | 0,000767407 | -0,614825201 | 0,123 | 0,347 | 1 | Basal | 0,653009008 | 1,531372443 | 0,354466859 |
| TRIM16 | 0,000768717 | 0,6118847 | 0,385 | 0,207 | 1 | Basal | 1,528254378 | 0,654341328 | 1,859903382 |
| PDLIM2 | 0,000771958 | 0,523670072 | 0,446 | 0,267 | 1 | Basal | 1,43760773 | 0,695600044 | 1,670411985 |
| ANKRD37 | 0,000793185 | -0,432889095 | 0,231 | 0,48 | 1 | Basal | 0,740776843 | 1,349934207 | 0,48125 |
| SERPINB5 | 0,000801532 | -0,951796442 | 0,354 | 0,587 | 1 | Basal | 0,516988308 | 1,934279721 | 0,60306644 |
| ZC3H13 | 0,000829113 | -0,555024708 | 0,215 | 0,453 | 1 | Basal | 0,680645401 | 1,469193795 | 0,474613687 |
| HNRNPA1 | 0,000837643 | -0,445636339 | 0,831 | 0,9 | 1 | Basal | 0,73426038 | 1,361914693 | 0,923333333 |
| MAL2 | 0,000854759 | 0,593794288 | 0,615 | 0,493 | 1 | Basal | 1,509210756 | 0,662597981 | 1,247464503 |
| TMEM219 | 0,000857786 | 0,577110642 | 0,708 | 0,647 | 1 | Basal | 1,491858435 | 0,670304887 | 1,094281298 |
| FAM3B | 0,00086281 | 0,987875072 | 0,554 | 0,44 | 1 | Basal | 1,983261717 | 0,504219888 | 1,259090909 |
| MARCKS | 0,000877086 | -0,479488758 | 0,446 | 0,707 | 1 | Basal | 0,717231741 | 1,394249505 | 0,630834512 |
| SLAMF7 | 0,00088844 | -0,262638335 | 0 | 0,153 | 1 | Basal | 0,833562144 | 1,199670602 | 0 |
| BTG2 | 0,00089119 | -0,769795648 | 0,569 | 0,7 | 1 | Basal | 0,586500544 | 1,705028256 | 0,812857143 |
| INSIG1 | 0,000908678 | -0,35904454 | 0,215 | 0,467 | 1 | Basal | 0,779680771 | 1,282576199 | 0,460385439 |
| IGFBP5 | 0,000921492 | -0,876469388 | 0,154 | 0,407 | 1 | Basal | 0,544698807 | 1,835876979 | 0,378378378 |
| RHBDL2 | 0,000932995 | 0,449101455 | 0,354 | 0,18 | 1 | Basal | 1,365189719 | 0,732498924 | 1,966666667 |
| AACS | 0,000938555 | 0,27213608 | 0,292 | 0,12 | 1 | Basal | 1,207594491 | 0,828092549 | 2,433333333 |
| BRD2 | 0,00093978 | -0,738665208 | 0,662 | 0,773 | 1 | Basal | 0,599293567 | 1,668631295 | 0,856403622 |
| MT-ND3 | 0,000943375 | -0,453202719 | 0,938 | 0,873 | 1 | Basal | 0,730419548 | 1,369076174 | 1,074455899 |
| CCN1 | 0,000975687 | -0,574954262 | 0,385 | 0,673 | 1 | Basal | 0,671307534 | 1,489630236 | 0,572065379 |

| Supplementary Table 4 - clusermarker | | | | | | | | | |
|---|---|---|---|---|---|---|---|---|---|
| | **p_val** | **avg_log2FC** | **pct.1** | **pct.2** | **p_val_adj** | **celltype** | **Foldchange_UP** | **Foldchange_DOWN** | **Ratio_pct1_pct2** |
| AP2S1 | 0,001000073 | 0,529806258 | 0,615 | 0,48 | 1 | Basal | 1,443735301 | 0,692647745 | 1,28125 |
| MMP28 | 0,001001666 | 0,4071888 | 0,262 | 0,107 | 1 | Basal | 1,326099291 | 0,754091347 | 2,448598131 |
| ARC | 0,001036789 | -0,536608968 | 0,062 | 0,26 | 1 | Basal | 0,689389405 | 1,450558991 | 0,238461538 |
| UQCRFS1 | 0,001039157 | 0,561056461 | 0,585 | 0,467 | 1 | Basal | 1,475349195 | 0,677805637 | 1,25267666 |
| TRADD | 0,001047192 | 0,455136041 | 0,477 | 0,28 | 1 | Basal | 1,370912071 | 0,729441385 | 1,703571429 |
| B4GALT4 | 0,001057481 | 0,790400215 | 0,538 | 0,42 | 1 | Basal | 1,729554188 | 0,578183677 | 1,280952381 |
| PGAP1 | 0,001070782 | -0,281946201 | 0,031 | 0,2 | 1 | Basal | 0,822480739 | 1,215833943 | 0,155 |
| DEPTOR | 0,001070962 | -0,403579471 | 0,092 | 0,3 | 1 | Basal | 0,755980292 | 1,322785807 | 0,306666667 |
| TMEM50A | 0,001078629 | 0,589604125 | 0,677 | 0,567 | 1 | Basal | 1,504833765 | 0,664525228 | 1,194003527 |
| CCND2 | 0,001079087 | -0,403262236 | 0,154 | 0,393 | 1 | Basal | 0,756146543 | 1,32249497 | 0,391857506 |
| GTF3C6 | 0,001080356 | 0,496826482 | 0,508 | 0,333 | 1 | Basal | 1,411106114 | 0,708663927 | 1,525525526 |
| RPL37A | 0,001106454 | -0,399814617 | 0,954 | 0,973 | 1 | Basal | 0,757955672 | 1,319338368 | 0,980472765 |
| NUDT8 | 0,001132715 | 0,405508316 | 0,415 | 0,247 | 1 | Basal | 1,324555519 | 0,754970241 | 1,680161943 |
| FXYD3 | 0,001157293 | 0,59402911 | 0,815 | 0,773 | 1 | Basal | 1,509456425 | 0,662490141 | 1,054333765 |
| ADIPOR2 | 0,001159204 | -0,33260656 | 0,123 | 0,333 | 1 | Basal | 0,794100462 | 1,259286511 | 0,369369369 |
| CD47 | 0,001168811 | 0,580945486 | 0,538 | 0,387 | 1 | Basal | 1,495829236 | 0,668525508 | 1,390180879 |
| CREB3 | 0,00117329 | 0,544391996 | 0,431 | 0,26 | 1 | Basal | 1,45840559 | 0,685680312 | 1,657692308 |
| RPS21 | 0,001173666 | -0,47604033 | 0,969 | 0,987 | 1 | Basal | 0,718948168 | 1,390920854 | 0,981762918 |
| ZNF428 | 0,001184914 | 0,503691376 | 0,631 | 0,52 | 1 | Basal | 1,417836696 | 0,705299843 | 1,213461538 |
| CLTB | 0,001196034 | 0,912357578 | 0,615 | 0,553 | 1 | Basal | 1,882118648 | 0,531316132 | 1,112115732 |
| TMED3 | 0,001200034 | 0,428119538 | 0,569 | 0,433 | 1 | Basal | 1,345478687 | 0,743229908 | 1,31408776 |
| GNG10 | 0,001214476 | 0,550851211 | 0,492 | 0,327 | 1 | Basal | 1,464949782 | 0,682617256 | 1,504587156 |
| HLF | 0,001217986 | -0,504031461 | 0,138 | 0,367 | 1 | Basal | 0,705133603 | 1,418170962 | 0,376021798 |
| CALR | 0,00122012 | 0,429842588 | 0,815 | 0,813 | 1 | Basal | 1,347086588 | 0,742342778 | 1,002460025 |
| MUC1 | 0,001235757 | 0,999984026 | 0,569 | 0,42 | 1 | Basal | 1,999977856 | 0,500005536 | 1,354761905 |
| OS9 | 0,001243405 | 0,800293746 | 0,615 | 0,527 | 1 | Basal | 1,741455667 | 0,574232247 | 1,166982922 |
| TMEM63B | 0,001246682 | 0,46175386 | 0,354 | 0,187 | 1 | Basal | 1,377215058 | 0,726103011 | 1,893048128 |
| PERP | 0,001251378 | 0,391459575 | 0,908 | 0,86 | 1 | Basal | 1,3117198 | 0,762357936 | 1,055813953 |
| PSMB8 | 0,00127113 | 0,635619351 | 0,677 | 0,527 | 1 | Basal | 1,553604572 | 0,643664429 | 1,284629981 |
| PRH2 | 0,001271138 | -0,593361319 | 0,015 | 0,18 | 1 | Basal | 0,662796864 | 1,508757893 | 0,083333333 |
| ZNF706 | 0,001274714 | 0,680635204 | 0,615 | 0,54 | 1 | Basal | 1,602845316 | 0,623890521 | 1,138888889 |

| Supplementary Table 4 - clusermarker | | | | | | | | | |
|---|---|---|---|---|---|---|---|---|---|
| | **p_val** | **avg_log2FC** | **pct.1** | **pct.2** | **p_val_adj** | **celltype** | **Foldchange_UP** | **Foldchange_DOWN** | **Ratio_pct1_pct2** |
| EIF4A1 | 0,001288484 | -0,619393913 | 0,785 | 0,893 | 1 | Basal | 0,650944337 | 1,536229664 | 0,879059351 |
| TMPRSS4 | 0,001298989 | 0,541231888 | 0,662 | 0,533 | 1 | Basal | 1,455214565 | 0,687183886 | 1,242026266 |
| PPP4C | 0,001311261 | 0,637175943 | 0,554 | 0,42 | 1 | Basal | 1,555281734 | 0,642970324 | 1,319047619 |
| ETS1 | 0,00136817 | -0,420457613 | 0,123 | 0,347 | 1 | Basal | 0,747187584 | 1,338352004 | 0,354466859 |
| NOS2 | 0,001370764 | 0,877659633 | 0,292 | 0,127 | 1 | Basal | 1,83739223 | 0,544249607 | 2,299212598 |
| RAB5IF | 0,001378152 | 0,518275641 | 0,554 | 0,413 | 1 | Basal | 1,432242359 | 0,698205854 | 1,341404358 |
| ITM2A | 0,001401361 | -0,430646659 | 0,092 | 0,3 | 1 | Basal | 0,741929156 | 1,347837583 | 0,306666667 |
| ADH1C | 0,001439773 | -0,793584392 | 0,4 | 0,593 | 1 | Basal | 0,576908973 | 1,733375709 | 0,674536256 |
| HMGCS1 | 0,001467281 | -0,370656588 | 0,154 | 0,373 | 1 | Basal | 0,773430419 | 1,29294113 | 0,412868633 |
| FXR1 | 0,001478609 | -0,558850625 | 0,308 | 0,553 | 1 | Basal | 0,678842773 | 1,473095156 | 0,556962025 |
| EGR2 | 0,001491451 | -0,568438488 | 0,215 | 0,433 | 1 | Basal | 0,674346277 | 1,482917655 | 0,496535797 |
| MDFI | 0,001505311 | 0,40956243 | 0,338 | 0,16 | 1 | Basal | 1,328282884 | 0,752851679 | 2,1125 |
| EMC6 | 0,001511168 | 0,45063453 | 0,492 | 0,353 | 1 | Basal | 1,366641204 | 0,73172095 | 1,393767705 |
| HSBP1 | 0,001514936 | 0,568245325 | 0,723 | 0,673 | 1 | Basal | 1,48271912 | 0,674436572 | 1,074294205 |
| SMAP1 | 0,001528784 | 0,618099549 | 0,569 | 0,433 | 1 | Basal | 1,534852 | 0,651528616 | 1,31408776 |
| ECHS1 | 0,001534904 | 0,490159928 | 0,692 | 0,507 | 1 | Basal | 1,404600572 | 0,711946172 | 1,364891519 |
| PLEKHG6 | 0,001539444 | 0,302386909 | 0,262 | 0,1 | 1 | Basal | 1,233183002 | 0,810909653 | 2,62 |
| RPS11 | 0,001589249 | -0,414349533 | 0,862 | 0,9 | 1 | Basal | 0,750357732 | 1,332697669 | 0,957777778 |
| CDH26 | 0,001593928 | 0,620657244 | 0,323 | 0,167 | 1 | Basal | 1,53757549 | 0,650374571 | 1,934131737 |
| FERMT2 | 0,001605032 | -0,507496919 | 0,215 | 0,44 | 1 | Basal | 0,703441854 | 1,421581605 | 0,488636364 |
| ALDOA | 0,001627959 | -0,541663776 | 0,692 | 0,827 | 1 | Basal | 0,6869782 | 1,455650267 | 0,836759371 |
| TRIR | 0,001631918 | 0,496073668 | 0,723 | 0,627 | 1 | Basal | 1,410369976 | 0,709033812 | 1,153110048 |
| PTRH1 | 0,001650209 | 0,280053492 | 0,292 | 0,127 | 1 | Basal | 1,214239905 | 0,823560481 | 2,299212598 |
| CES1 | 0,001653169 | -0,356152306 | 0,092 | 0,3 | 1 | Basal | 0,7812454 | 1,280007537 | 0,306666667 |
| S100A16 | 0,001654858 | 0,482723572 | 0,677 | 0,587 | 1 | Basal | 1,397379201 | 0,715625365 | 1,153321976 |
| TP53I3 | 0,001696239 | 0,605231651 | 0,523 | 0,353 | 1 | Basal | 1,521222994 | 0,65736582 | 1,481586402 |
| RPS6KB2 | 0,001705457 | 0,436902124 | 0,431 | 0,253 | 1 | Basal | 1,353694438 | 0,738719147 | 1,703557312 |
| ITGA2 | 0,001718334 | -0,452657949 | 0,308 | 0,6 | 1 | Basal | 0,73069541 | 1,368559301 | 0,513333333 |
| GSN | 0,001732651 | 0,916116343 | 0,846 | 0,833 | 1 | Basal | 1,887028671 | 0,529933655 | 1,015606242 |
| KDELR2 | 0,001737975 | 0,594065739 | 0,585 | 0,453 | 1 | Basal | 1,50949475 | 0,662473321 | 1,291390728 |
| CAPNS1 | 0,001741644 | 0,447137875 | 0,738 | 0,593 | 1 | Basal | 1,363332892 | 0,73349657 | 1,244519393 |

| Supplementary Table 4 - clusermarker | | | | | | | | | |
|---|---|---|---|---|---|---|---|---|---|
| | **p_val** | **avg_log2FC** | **pct.1** | **pct.2** | **p_val_adj** | **celltype** | **Foldchange_UP** | **Foldchange_DOWN** | **Ratio_pct1_pct2** |
| NEK6 | 0,001746492 | 0,533187418 | 0,323 | 0,16 | 1 | Basal | 1,447122867 | 0,691026327 | 2,01875 |
| HSPB8 | 0,001761351 | -0,327909108 | 0,246 | 0,48 | 1 | Basal | 0,796690287 | 1,25519291 | 0,5125 |
| PPIA | 0,001772289 | 0,252288558 | 0,954 | 0,96 | 1 | Basal | 1,19109506 | 0,839563553 | 0,99375 |
| UBL7 | 0,001808513 | 0,375506345 | 0,369 | 0,2 | 1 | Basal | 1,297294788 | 0,770834824 | 1,845 |
| UQCRHL | 0,001808713 | -0,306813656 | 0,138 | 0,347 | 1 | Basal | 0,808425282 | 1,236972696 | 0,397694524 |
| CD81 | 0,001827983 | -0,604819517 | 0,708 | 0,84 | 1 | Basal | 0,657553636 | 1,520788489 | 0,842857143 |
| ARPC1B | 0,001842408 | 0,611878036 | 0,646 | 0,553 | 1 | Basal | 1,528247319 | 0,654344351 | 1,168173599 |
| PAWR | 0,001846512 | 0,625509098 | 0,585 | 0,447 | 1 | Basal | 1,542755136 | 0,648191004 | 1,308724832 |
| TAGLN2 | 0,001849058 | 0,435918185 | 0,862 | 0,827 | 1 | Basal | 1,352771513 | 0,739223136 | 1,042321644 |
| NDUFA6 | 0,001858098 | 0,415700523 | 0,738 | 0,633 | 1 | Basal | 1,333946238 | 0,749655399 | 1,165876777 |
| MISP | 0,001866772 | 0,49928985 | 0,508 | 0,327 | 1 | Basal | 1,413517603 | 0,707454932 | 1,55351682 |
| CRB3 | 0,001884033 | 0,574487189 | 0,323 | 0,173 | 1 | Basal | 1,489148046 | 0,671524905 | 1,867052023 |
| ATP1B1 | 0,001904815 | 0,714758787 | 0,738 | 0,647 | 1 | Basal | 1,641208791 | 0,609306997 | 1,14064915 |
| RPL39 | 0,001906858 | -0,551287594 | 1 | 0,973 | 1 | Basal | 0,68241081 | 1,465392964 | 1,027749229 |
| ATP5ME | 0,001920518 | -0,450020955 | 0,554 | 0,727 | 1 | Basal | 0,732032215 | 1,366060098 | 0,762035763 |
| DPF2 | 0,001937282 | -0,343423875 | 0,092 | 0,287 | 1 | Basal | 0,78816857 | 1,268764117 | 0,320557491 |
| HMGN1 | 0,001971223 | 0,416424627 | 0,723 | 0,68 | 1 | Basal | 1,334615928 | 0,749279234 | 1,063235294 |
| PIAS4 | 0,001977956 | 0,331704987 | 0,169 | 0,047 | 1 | Basal | 1,2584998 | 0,794596869 | 3,595744681 |
| RBM22 | 0,002027868 | -0,443411464 | 0,185 | 0,42 | 1 | Basal | 0,735393605 | 1,359816013 | 0,44047619 |
| ANXA11 | 0,002058152 | 0,943026573 | 0,708 | 0,667 | 1 | Basal | 1,922557268 | 0,520140553 | 1,061469265 |
| FAM89B | 0,002061874 | 0,295383321 | 0,2 | 0,067 | 1 | Basal | 1,227211001 | 0,814855798 | 2,985074627 |
| ITGB4 | 0,002090688 | 0,442828065 | 0,646 | 0,56 | 1 | Basal | 1,35926624 | 0,735691044 | 1,153571429 |
| PACS1 | 0,002128035 | 0,347959122 | 0,446 | 0,267 | 1 | Basal | 1,272758872 | 0,785694779 | 1,670411985 |
| RNASEK | 0,002137265 | 0,325245562 | 0,815 | 0,8 | 1 | Basal | 1,252877673 | 0,798162519 | 1,01875 |
| PSMA6 | 0,002145433 | 0,472565875 | 0,677 | 0,56 | 1 | Basal | 1,387575117 | 0,720681704 | 1,208928571 |
| HBP1 | 0,002166392 | -0,39831219 | 0,308 | 0,54 | 1 | Basal | 0,758745421 | 1,31796512 | 0,57037037 |
| HEXIM1 | 0,002192843 | -0,507844814 | 0,385 | 0,58 | 1 | Basal | 0,703272245 | 1,421924451 | 0,663793103 |
| NDUFB1 | 0,002195565 | -0,616607439 | 0,415 | 0,613 | 1 | Basal | 0,65220281 | 1,533265398 | 0,676998369 |
| TNFRSF18 | 0,002230128 | 0,533912891 | 0,292 | 0,14 | 1 | Basal | 1,447850749 | 0,690678926 | 2,085714286 |
| NCOA7 | 0,002250511 | -0,538678585 | 0,431 | 0,647 | 1 | Basal | 0,688401151 | 1,452641383 | 0,666151468 |
| SEC62 | 0,002251206 | -0,577380207 | 0,615 | 0,773 | 1 | Basal | 0,670179654 | 1,492137212 | 0,795601552 |

| Supplementary Table 4 - clusermarker | | | | | | | | | |
|---|---|---|---|---|---|---|---|---|---|
| | **p_val** | **avg_log2FC** | **pct.1** | **pct.2** | **p_val_adj** | **celltype** | **Foldchange_UP** | **Foldchange_DOWN** | **Ratio_pct1_pct2** |
| NDUFV2 | 0,00226138 | 0,398436737 | 0,692 | 0,62 | 1 | Basal | 1,318078905 | 0,758679922 | 1,116129032 |
| EPS8L1 | 0,002263612 | 0,654148956 | 0,492 | 0,333 | 1 | Basal | 1,573687363 | 0,635450232 | 1,477477477 |
| ARL4A | 0,002272925 | -0,454135448 | 0,354 | 0,607 | 1 | Basal | 0,729947471 | 1,369961593 | 0,583196046 |
| TMBIM4 | 0,002298495 | 0,457442105 | 0,708 | 0,613 | 1 | Basal | 1,373105146 | 0,728276347 | 1,15497553 |
| NAT1 | 0,00232234 | 0,353943204 | 0,2 | 0,067 | 1 | Basal | 1,278049048 | 0,782442584 | 2,985074627 |
| MT1X | 0,00234712 | -0,489342499 | 0,6 | 0,84 | 1 | Basal | 0,712349674 | 1,403804953 | 0,714285714 |
| CDKN2AIP | 0,002349763 | -0,322771916 | 0,246 | 0,48 | 1 | Basal | 0,799532223 | 1,250731329 | 0,5125 |
| PRELID1 | 0,002351586 | 0,369512617 | 0,631 | 0,5 | 1 | Basal | 1,291916311 | 0,774043947 | 1,262 |
| ARG2 | 0,002362924 | -0,490008233 | 0,185 | 0,387 | 1 | Basal | 0,712021034 | 1,404452891 | 0,478036176 |
| HS3ST1 | 0,002371216 | 0,740410882 | 0,554 | 0,42 | 1 | Basal | 1,670651575 | 0,598568855 | 1,319047619 |
| CAPN5 | 0,002392308 | 0,392140535 | 0,292 | 0,14 | 1 | Basal | 1,312339085 | 0,761998184 | 2,085714286 |
| RBMXL1 | 0,00242351 | -0,34028369 | 0,108 | 0,293 | 1 | Basal | 0,789885974 | 1,266005515 | 0,368600683 |
| SLURP2 | 0,002427547 | 1,66868099 | 0,2 | 0,067 | 1 | Basal | 3,17923793 | 0,314540787 | 2,985074627 |
| CELF2 | 0,002441919 | -0,427289067 | 0,062 | 0,233 | 1 | Basal | 0,743657862 | 1,344704401 | 0,266094421 |
| AGR2 | 0,002487053 | -0,360530751 | 0,646 | 0,847 | 1 | Basal | 0,778877987 | 1,283898142 | 0,762691854 |
| GPBP1 | 0,002507553 | -0,454093773 | 0,308 | 0,54 | 1 | Basal | 0,729968557 | 1,36992202 | 0,57037037 |
| TLE4 | 0,002536238 | -0,48614833 | 0,323 | 0,553 | 1 | Basal | 0,713928584 | 1,400700325 | 0,584086799 |
| CXCL8 | 0,00254657 | -0,662316885 | 0,231 | 0,447 | 1 | Basal | 0,631862747 | 1,582622184 | 0,516778523 |
| RHOA | 0,00257551 | 0,25770159 | 0,877 | 0,833 | 1 | Basal | 1,195572476 | 0,836419389 | 1,052821128 |
| VPS28 | 0,002667064 | 0,353985646 | 0,692 | 0,6 | 1 | Basal | 1,278086647 | 0,782419566 | 1,153333333 |
| RPS19 | 0,002684595 | 0,420728507 | 0,985 | 0,987 | 1 | Basal | 1,338603329 | 0,747047298 | 0,997973658 |
| TMEM59 | 0,002691121 | 0,517291512 | 0,815 | 0,8 | 1 | Basal | 1,431265694 | 0,698682295 | 1,01875 |
| QKI | 0,002734803 | -0,296034974 | 0,185 | 0,413 | 1 | Basal | 0,814487818 | 1,227765447 | 0,447941889 |
| ATP6V0B | 0,002743235 | 0,687179334 | 0,646 | 0,62 | 1 | Basal | 1,61013241 | 0,621066934 | 1,041935484 |
| CLDN7 | 0,002764798 | 1,094189947 | 0,615 | 0,587 | 1 | Basal | 2,134931745 | 0,468399049 | 1,04770017 |
| TFRC | 0,002769938 | -0,311210631 | 0,154 | 0,353 | 1 | Basal | 0,805965154 | 1,240748431 | 0,436260623 |
| STXBP6 | 0,002779547 | -0,284689217 | 0,108 | 0,293 | 1 | Basal | 0,820918431 | 1,218147824 | 0,368600683 |
| PART1 | 0,002868219 | -0,256543636 | 0,108 | 0,293 | 1 | Basal | 0,837090996 | 1,194613256 | 0,368600683 |
| TECR | 0,002886876 | 0,531565963 | 0,723 | 0,627 | 1 | Basal | 1,445497349 | 0,691803413 | 1,153110048 |
| NSRP1 | 0,002887092 | -0,449387263 | 0,308 | 0,527 | 1 | Basal | 0,732353825 | 1,3654602 | 0,584440228 |
| H3-3A | 0,002917226 | 0,328395096 | 0,908 | 0,92 | 1 | Basal | 1,255615806 | 0,796421959 | 0,986956522 |

| Supplementary Table 4 - clusermarker | | | | | | | | | |
|---|---|---|---|---|---|---|---|---|---|
| | **p_val** | **avg_log2FC** | **pct.1** | **pct.2** | **p_val_adj** | **celltype** | **Foldchange_UP** | **Foldchange_DOWN** | **Ratio_pct1_pct2** |
| ZFAS1 | 0,002932613 | -0,536560755 | 0,8 | 0,853 | 1 | Basal | 0,689412444 | 1,450510516 | 0,937866354 |
| MROH6 | 0,002935837 | 0,386789034 | 0,246 | 0,107 | 1 | Basal | 1,307480141 | 0,764829972 | 2,299065421 |
| YTHDC1 | 0,002940613 | -0,421304367 | 0,354 | 0,607 | 1 | Basal | 0,746749169 | 1,339137747 | 0,583196046 |
| CDCP1 | 0,002988056 | 0,375013142 | 0,277 | 0,127 | 1 | Basal | 1,296851368 | 0,771098388 | 2,181102362 |
| SLC38A10 | 0,002989195 | 0,382011933 | 0,385 | 0,233 | 1 | Basal | 1,303157928 | 0,767366701 | 1,652360515 |
| PTOV1 | 0,002990746 | 0,567003627 | 0,569 | 0,473 | 1 | Basal | 1,481443522 | 0,675017296 | 1,202959831 |
| FN1 | 0,003012317 | -0,438093759 | 0,108 | 0,3 | 1 | Basal | 0,738109232 | 1,354813022 | 0,36 |
| LMTK3 | 0,003017158 | 0,429304452 | 0,323 | 0,173 | 1 | Basal | 1,346584209 | 0,742619729 | 1,867052023 |
| CHD9 | 0,00301986 | -0,385196741 | 0,338 | 0,593 | 1 | Basal | 0,765674576 | 1,30603788 | 0,569983137 |
| AUP1 | 0,003072762 | 0,471771282 | 0,569 | 0,48 | 1 | Basal | 1,386811094 | 0,721078743 | 1,185416667 |
| ST6GAL1 | 0,003077045 | 0,57839926 | 0,523 | 0,373 | 1 | Basal | 1,493191561 | 0,669706437 | 1,402144772 |
| MRPL35 | 0,003100036 | -0,374479167 | 0,123 | 0,313 | 1 | Basal | 0,771383843 | 1,296371462 | 0,392971246 |
| NUFIP2 | 0,00318875 | -0,349724221 | 0,369 | 0,587 | 1 | Basal | 0,78473409 | 1,274317011 | 0,628620102 |
| KRT13 | 0,003204465 | -1,04951033 | 0,369 | 0,187 | 1 | Basal | 0,483132118 | 2,069827201 | 1,973262032 |
| SNAI1 | 0,00320678 | -0,327207245 | 0,077 | 0,253 | 1 | Basal | 0,797077967 | 1,254582414 | 0,304347826 |
| BRI3 | 0,003269083 | 0,314398507 | 0,662 | 0,6 | 1 | Basal | 1,243493104 | 0,804186205 | 1,103333333 |
| SDF2L1 | 0,003282567 | 0,506467591 | 0,4 | 0,273 | 1 | Basal | 1,420567703 | 0,703943922 | 1,465201465 |
| IGFBP2 | 0,003291251 | 0,733478076 | 0,785 | 0,7 | 1 | Basal | 1,662642594 | 0,601452172 | 1,121428571 |
| KRT7 | 0,003320149 | 0,72778421 | 0,785 | 0,793 | 1 | Basal | 1,656093596 | 0,603830606 | 0,989911728 |
| YBX1 | 0,003359123 | 0,598062453 | 0,892 | 0,8 | 1 | Basal | 1,513682318 | 0,660640603 | 1,115 |
| PHYHD1 | 0,003366634 | -0,338002825 | 0,277 | 0,507 | 1 | Basal | 0,791135752 | 1,264005574 | 0,546351085 |
| PLPP2 | 0,003374286 | 0,529953379 | 0,662 | 0,6 | 1 | Basal | 1,443882536 | 0,692577114 | 1,103333333 |
| ANK3 | 0,003404576 | -0,438611629 | 0,354 | 0,56 | 1 | Basal | 0,737844328 | 1,355299434 | 0,632142857 |
| SULT1E1 | 0,003424518 | -0,365731559 | 0,031 | 0,18 | 1 | Basal | 0,776075245 | 1,288534851 | 0,172222222 |
| SPINT1 | 0,003430708 | 0,458652722 | 0,646 | 0,547 | 1 | Basal | 1,374257851 | 0,727665481 | 1,180987203 |
| SNAPC3 | 0,003439074 | 0,512856595 | 0,323 | 0,16 | 1 | Basal | 1,426872666 | 0,700833384 | 2,01875 |
| RPL41 | 0,003467409 | -0,272474591 | 1 | 1 | 1 | Basal | 0,82789827 | 1,207877871 | 1 |
| GLS | 0,003498061 | -0,254204497 | 0,231 | 0,453 | 1 | Basal | 0,838449329 | 1,192677918 | 0,509933775 |
| MYH14 | 0,003508736 | 0,806529112 | 0,554 | 0,467 | 1 | Basal | 1,748998572 | 0,571755755 | 1,186295503 |
| MEF2C | 0,003532968 | -0,430614693 | 0,062 | 0,227 | 1 | Basal | 0,741945595 | 1,347807718 | 0,273127753 |
| LTBP3 | 0,003543839 | 0,284602365 | 0,585 | 0,427 | 1 | Basal | 1,218074492 | 0,820967852 | 1,370023419 |

| Supplementary Table 4 - clusermarker | | | | | | | | | |
|---|---|---|---|---|---|---|---|---|---|
| | **p_val** | **avg_log2FC** | **pct.1** | **pct.2** | **p_val_adj** | **celltype** | **Foldchange_UP** | **Foldchange_DOWN** | **Ratio_pct1_pct2** |
| NDUFB2 | 0,003568184 | 0,568421147 | 0,815 | 0,713 | 1 | Basal | 1,48289983 | 0,674354383 | 1,143057504 |
| METTL26 | 0,003569932 | 0,459034633 | 0,523 | 0,387 | 1 | Basal | 1,374621694 | 0,727472878 | 1,351421189 |
| FGGY | 0,003588775 | -0,32339962 | 0,169 | 0,367 | 1 | Basal | 0,799184429 | 1,25127563 | 0,460490463 |
| ZDHHC9 | 0,003603994 | 0,422605637 | 0,277 | 0,127 | 1 | Basal | 1,340346155 | 0,746075927 | 2,181102362 |
| NDUFB11 | 0,003612043 | 0,355277465 | 0,769 | 0,707 | 1 | Basal | 1,279231584 | 0,781719285 | 1,087694484 |
| NOSIP | 0,003614629 | 0,462212252 | 0,508 | 0,373 | 1 | Basal | 1,377652715 | 0,72587234 | 1,361930295 |
| GPC1 | 0,003617226 | 0,455055833 | 0,446 | 0,293 | 1 | Basal | 1,370835856 | 0,72948194 | 1,5221843 |
| RBBP8 | 0,003632664 | -0,698476077 | 0,277 | 0,507 | 1 | Basal | 0,616222781 | 1,62278973 | 0,546351085 |
| SLC25A5 | 0,003635976 | 0,526597768 | 0,785 | 0,773 | 1 | Basal | 1,440528065 | 0,694189877 | 1,015523933 |
| SELENOH | 0,003641734 | 0,469865745 | 0,677 | 0,647 | 1 | Basal | 1,384980578 | 0,722031786 | 1,046367852 |
| DEDD | 0,003676009 | 0,355758872 | 0,231 | 0,093 | 1 | Basal | 1,279658518 | 0,78145848 | 2,483870968 |
| RAB5B | 0,003677547 | -0,331047448 | 0,215 | 0,44 | 1 | Basal | 0,794959105 | 1,257926343 | 0,488636364 |
| PDLIM4 | 0,00367898 | 0,305286196 | 0,554 | 0,38 | 1 | Basal | 1,235663739 | 0,809281659 | 1,457894737 |
| OST4 | 0,003716635 | 0,257542148 | 0,815 | 0,813 | 1 | Basal | 1,195440353 | 0,836511832 | 1,002460025 |
| CALM1 | 0,003724795 | 0,433747238 | 0,846 | 0,807 | 1 | Basal | 1,350737413 | 0,740336346 | 1,048327138 |
| FAM118B | 0,003735125 | -0,269679986 | 0,046 | 0,207 | 1 | Basal | 0,829503523 | 1,205540389 | 0,222222222 |
| CKS2 | 0,003736113 | -0,345054374 | 0,185 | 0,4 | 1 | Basal | 0,787278304 | 1,270198854 | 0,4625 |
| CD44 | 0,003783085 | 0,475724834 | 0,738 | 0,607 | 1 | Basal | 1,390616713 | 0,719105409 | 1,215815486 |
| LLGL1 | 0,003864065 | 0,381748692 | 0,123 | 0,027 | 1 | Basal | 1,30292017 | 0,767506731 | 4,555555556 |
| POLR2F | 0,00388012 | 0,649000374 | 0,554 | 0,44 | 1 | Basal | 1,568081314 | 0,637722031 | 1,259090909 |
| TIPARP | 0,003935983 | -0,535539156 | 0,262 | 0,46 | 1 | Basal | 0,689900803 | 1,449483746 | 0,569565217 |
| SNRPB2 | 0,003983868 | -0,346009515 | 0,277 | 0,493 | 1 | Basal | 0,786757256 | 1,271040072 | 0,561866126 |
| ZNF292 | 0,004000066 | -0,251955014 | 0,246 | 0,473 | 1 | Basal | 0,839757678 | 1,190819717 | 0,520084567 |
| MTRNR2L8 | 0,004027718 | -1,909128514 | 0,185 | 0,367 | 1 | Basal | 0,266253332 | 3,755821539 | 0,504087193 |
| NOL7 | 0,004065612 | 0,684163096 | 0,585 | 0,493 | 1 | Basal | 1,606769628 | 0,622366756 | 1,186612576 |
| YPEL3 | 0,004096032 | 0,620382975 | 0,569 | 0,52 | 1 | Basal | 1,537283212 | 0,650498225 | 1,094230769 |
| REXO2 | 0,004109401 | 0,40148919 | 0,646 | 0,533 | 1 | Basal | 1,320870647 | 0,757076404 | 1,212007505 |
| CAMTA1 | 0,004112683 | 0,479508992 | 0,585 | 0,487 | 1 | Basal | 1,394269059 | 0,717221682 | 1,201232033 |
| DST | 0,004177348 | -0,888446066 | 0,523 | 0,687 | 1 | Basal | 0,540195652 | 1,851181134 | 0,761280932 |
| PIP4P1 | 0,004190273 | 0,351530393 | 0,308 | 0,153 | 1 | Basal | 1,275913383 | 0,783752262 | 2,013071895 |
| CES2 | 0,004223591 | -0,273004958 | 0,154 | 0,367 | 1 | Basal | 0,827593972 | 1,208321996 | 0,419618529 |

| Supplementary Table 4 - clusermarker | | | | | | | | | |
|---|---|---|---|---|---|---|---|---|---|
| | **p_val** | **avg_log2FC** | **pct.1** | **pct.2** | **p_val_adj** | **celltype** | **Foldchange_UP** | **Foldchange_DOWN** | **Ratio_pct1_pct2** |
| ALPK1 | 0,004229104 | -0,259941927 | 0,077 | 0,24 | 1 | Basal | 0,835121535 | 1,197430503 | 0,320833333 |
| CYP1B1 | 0,004236273 | -0,370998108 | 0,046 | 0,2 | 1 | Basal | 0,773247351 | 1,293247236 | 0,23 |
| POLE4 | 0,004239505 | 0,582923877 | 0,446 | 0,32 | 1 | Basal | 1,497881897 | 0,667609377 | 1,39375 |
| UGT1A6 | 0,004247704 | -0,287874862 | 0,031 | 0,173 | 1 | Basal | 0,819107743 | 1,220840613 | 0,179190751 |
| RALA | 0,004289306 | 0,398111327 | 0,492 | 0,347 | 1 | Basal | 1,317781635 | 0,758851067 | 1,417867435 |
| DCXR | 0,004294611 | 0,513874773 | 0,662 | 0,533 | 1 | Basal | 1,427880033 | 0,700338948 | 1,242026266 |
| UNC93B1 | 0,004325223 | 0,521471959 | 0,508 | 0,38 | 1 | Basal | 1,435419037 | 0,696660678 | 1,336842105 |
| PDIA3 | 0,00438352 | 0,332557638 | 0,8 | 0,713 | 1 | Basal | 1,259243809 | 0,794127391 | 1,122019635 |
| DDAH2 | 0,004394605 | -0,37359753 | 0,215 | 0,453 | 1 | Basal | 0,771855383 | 1,295579486 | 0,474613687 |
| UBB | 0,004461816 | -0,391173278 | 0,877 | 0,9 | 1 | Basal | 0,762509239 | 1,31145952 | 0,974444444 |
| MLPH | 0,004470441 | 0,559682331 | 0,385 | 0,247 | 1 | Basal | 1,473944631 | 0,678451537 | 1,558704453 |
| MARK2 | 0,004568336 | 0,458920678 | 0,415 | 0,253 | 1 | Basal | 1,37451312 | 0,727530342 | 1,640316206 |
| SOBP | 0,004575067 | -0,371265307 | 0,062 | 0,213 | 1 | Basal | 0,773104153 | 1,293486778 | 0,291079812 |
| SERF2 | 0,004594495 | 0,49566232 | 0,877 | 0,867 | 1 | Basal | 1,409967902 | 0,709236004 | 1,011534025 |
| CAPS | 0,004617941 | -0,523689516 | 0,354 | 0,593 | 1 | Basal | 0,695590669 | 1,437627106 | 0,596964587 |
| RPS24 | 0,004622396 | -0,440623211 | 1 | 1 | 1 | Basal | 0,736816252 | 1,357190476 | 1 |
| CAST | 0,004646151 | 0,319270323 | 0,769 | 0,707 | 1 | Basal | 1,247699336 | 0,80147514 | 1,087694484 |
| BIK | 0,004677152 | 0,551059542 | 0,569 | 0,42 | 1 | Basal | 1,465161342 | 0,68251869 | 1,354761905 |
| PSMC6 | 0,004679381 | -0,369262581 | 0,292 | 0,52 | 1 | Basal | 0,774178109 | 1,291692426 | 0,561538462 |
| SIRT1 | 0,004703698 | -0,284269853 | 0,123 | 0,293 | 1 | Basal | 0,821157091 | 1,217793783 | 0,419795222 |
| RPL23A | 0,004707334 | -0,370835638 | 0,954 | 0,953 | 1 | Basal | 0,773334436 | 1,293101605 | 1,001049318 |
| HSP90AA1 | 0,004710266 | -0,587418688 | 0,877 | 0,92 | 1 | Basal | 0,665532633 | 1,502555924 | 0,95326087 |
| EGR1 | 0,004711591 | -0,430330508 | 0,723 | 0,82 | 1 | Basal | 0,742091759 | 1,347542251 | 0,881707317 |
| DUSP2 | 0,004740305 | -0,629398807 | 0,246 | 0,433 | 1 | Basal | 0,646445743 | 1,546920234 | 0,56812933 |
| PRRC2C | 0,004752337 | -0,441080296 | 0,631 | 0,767 | 1 | Basal | 0,736582846 | 1,357620539 | 0,822685789 |
| TAP1 | 0,004760988 | 0,543823632 | 0,4 | 0,26 | 1 | Basal | 1,45783115 | 0,685950496 | 1,538461538 |
| INO80C | 0,004774293 | 0,50376525 | 0,323 | 0,187 | 1 | Basal | 1,4179093 | 0,705263729 | 1,727272727 |
| SHOC2 | 0,004781017 | -0,320032936 | 0,215 | 0,447 | 1 | Basal | 0,80105159 | 1,248359048 | 0,48098434 |
| HNMT | 0,004960036 | -0,286482176 | 0,215 | 0,413 | 1 | Basal | 0,81989884 | 1,21966266 | 0,520581114 |
| LUC7L3 | 0,005005922 | -0,441294583 | 0,415 | 0,607 | 1 | Basal | 0,736473447 | 1,357822205 | 0,68369028 |
| WWTR1 | 0,005034786 | -0,375610929 | 0,292 | 0,52 | 1 | Basal | 0,770778946 | 1,297388836 | 0,561538462 |

| Supplementary Table 4 - clusermarker | | | | | | | | | |
|---|---|---|---|---|---|---|---|---|---|
| | **p_val** | **avg_log2FC** | **pct.1** | **pct.2** | **p_val_adj** | **celltype** | **Foldchange_UP** | **Foldchange_DOWN** | **Ratio_pct1_pct2** |
| GNB2 | 0,005046283 | 0,567185025 | 0,662 | 0,647 | 1 | Basal | 1,481629804 | 0,674932427 | 1,023183926 |
| ZSWIM6 | 0,005117499 | -0,373279443 | 0,2 | 0,38 | 1 | Basal | 0,772025581 | 1,295293866 | 0,526315789 |
| RIT1 | 0,005139315 | 0,322060706 | 0,323 | 0,18 | 1 | Basal | 1,250114904 | 0,799926468 | 1,794444444 |
| PCF11 | 0,005152419 | -0,543090632 | 0,292 | 0,48 | 1 | Basal | 0,6862991 | 1,457090648 | 0,608333333 |
| JAM3 | 0,00519405 | -0,499250251 | 0,062 | 0,22 | 1 | Basal | 0,70747435 | 1,413478806 | 0,281818182 |
| DAD1 | 0,005194407 | 0,369516302 | 0,723 | 0,687 | 1 | Basal | 1,291919611 | 0,774041969 | 1,052401747 |
| GSTK1 | 0,005220318 | 0,403327231 | 0,738 | 0,7 | 1 | Basal | 1,322554552 | 0,756112478 | 1,054285714 |
| FAM3C | 0,005235908 | 0,52170812 | 0,6 | 0,493 | 1 | Basal | 1,435654026 | 0,696546648 | 1,21703854 |
| SOX4 | 0,005307527 | -0,715628981 | 0,723 | 0,787 | 1 | Basal | 0,60893959 | 1,642199022 | 0,918678526 |
| ALDH7A1 | 0,005311037 | -0,495704975 | 0,246 | 0,48 | 1 | Basal | 0,709215035 | 1,41000959 | 0,5125 |
| SYNCRIP | 0,005315162 | -0,372734363 | 0,292 | 0,5 | 1 | Basal | 0,772317323 | 1,29480457 | 0,584 |
| TMEM141 | 0,00531986 | 0,425286655 | 0,6 | 0,487 | 1 | Basal | 1,342839291 | 0,744690751 | 1,232032854 |
| MVB12A | 0,005407224 | 0,374601954 | 0,446 | 0,307 | 1 | Basal | 1,2964818 | 0,771318194 | 1,45276873 |
| ALPL | 0,005407662 | 0,826838408 | 0,385 | 0,233 | 1 | Basal | 1,773793921 | 0,563763348 | 1,652360515 |
| PJA2 | 0,005427264 | -0,371120615 | 0,292 | 0,513 | 1 | Basal | 0,773181694 | 1,293357057 | 0,56920078 |
| CDC42EP2 | 0,005496335 | -0,317888004 | 0,077 | 0,233 | 1 | Basal | 0,802243442 | 1,246504425 | 0,330472103 |
| NFIA | 0,005523755 | -0,51186749 | 0,431 | 0,64 | 1 | Basal | 0,701314037 | 1,425894744 | 0,6734375 |
| SLC6A14 | 0,005559063 | 0,783844118 | 0,538 | 0,44 | 1 | Basal | 1,721712336 | 0,58081712 | 1,222727273 |
| LMOD1 | 0,005605411 | -0,683483466 | 0,077 | 0,24 | 1 | Basal | 0,622660011 | 1,606012884 | 0,320833333 |
| ARPC3 | 0,005624516 | 0,374478712 | 0,785 | 0,773 | 1 | Basal | 1,296371053 | 0,771384086 | 1,015523933 |
| DCAF6 | 0,005632622 | -0,260030381 | 0,169 | 0,36 | 1 | Basal | 0,835070334 | 1,197503922 | 0,469444444 |
| PRKAB1 | 0,005697109 | 0,275486607 | 0,262 | 0,12 | 1 | Basal | 1,210402277 | 0,826171612 | 2,183333333 |
| YAP1 | 0,005729527 | -0,425798473 | 0,277 | 0,48 | 1 | Basal | 0,744426608 | 1,343315768 | 0,577083333 |
| AL118516.1 | 0,005750417 | -0,266847636 | 0,138 | 0,32 | 1 | Basal | 0,831133634 | 1,203175951 | 0,43125 |
| ATXN10 | 0,00576384 | 0,27280687 | 0,477 | 0,34 | 1 | Basal | 1,2081561 | 0,827707612 | 1,402941176 |
| NBL1 | 0,005779418 | 0,535641096 | 0,569 | 0,467 | 1 | Basal | 1,44958617 | 0,689852056 | 1,218415418 |
| OSTM1 | 0,005783481 | 0,302077081 | 0,231 | 0,1 | 1 | Basal | 1,232918197 | 0,811083819 | 2,31 |
| KISS1 | 0,005826771 | -0,569574009 | 0,031 | 0,167 | 1 | Basal | 0,67381572 | 1,484085293 | 0,185628743 |
| ADI1 | 0,005850382 | 0,536547477 | 0,615 | 0,527 | 1 | Basal | 1,450497166 | 0,689418789 | 1,166982922 |
| JPT1 | 0,005863737 | 0,364746572 | 0,677 | 0,553 | 1 | Basal | 1,287655417 | 0,776605284 | 1,224231465 |
| SCAF1 | 0,005917383 | 0,384750625 | 0,323 | 0,18 | 1 | Basal | 1,305634084 | 0,765911378 | 1,794444444 |

| Supplementary Table 4 - clusermarker | | | | | | | | | |
|---|---|---|---|---|---|---|---|---|---|
| | **p_val** | **avg_log2FC** | **pct.1** | **pct.2** | **p_val_adj** | **celltype** | **Foldchange_UP** | **Foldchange_DOWN** | **Ratio_pct1_pct2** |
| CBR1 | 0,005924586 | 0,451942855 | 0,677 | 0,62 | 1 | Basal | 1,367881121 | 0,731057681 | 1,091935484 |
| LAGE3 | 0,005950994 | 0,435724514 | 0,492 | 0,32 | 1 | Basal | 1,352589926 | 0,739322378 | 1,5375 |
| CIR1 | 0,005999899 | -0,257864239 | 0,292 | 0,5 | 1 | Basal | 0,836325096 | 1,195707273 | 0,584 |
| CHIC2 | 0,006065068 | 0,467091225 | 0,431 | 0,3 | 1 | Basal | 1,382319611 | 0,723421698 | 1,436666667 |
| PRKAA2 | 0,006097894 | -0,310093968 | 0,108 | 0,287 | 1 | Basal | 0,806589221 | 1,239788449 | 0,37630662 |
| VAMP8 | 0,006115994 | 0,563963474 | 0,785 | 0,82 | 1 | Basal | 1,478325002 | 0,676441242 | 0,957317073 |
| HMGN5 | 0,00615906 | -0,397352145 | 0,092 | 0,26 | 1 | Basal | 0,759250498 | 1,317088368 | 0,353846154 |
| MICOS10 | 0,006161307 | 0,655105454 | 0,723 | 0,653 | 1 | Basal | 1,574731054 | 0,635029072 | 1,10719755 |
| TRAPPC1 | 0,006165542 | 0,475187386 | 0,6 | 0,48 | 1 | Basal | 1,390098762 | 0,719373348 | 1,25 |
| RAB14 | 0,006168222 | -0,529029727 | 0,462 | 0,66 | 1 | Basal | 0,693020663 | 1,442958419 | 0,7 |
| PSME1 | 0,006171841 | 0,40443069 | 0,785 | 0,713 | 1 | Basal | 1,323566507 | 0,75553438 | 1,100981767 |
| A4GALT | 0,006211208 | 0,444779302 | 0,538 | 0,427 | 1 | Basal | 1,361105884 | 0,734696699 | 1,259953162 |
| PLCG2 | 0,006344694 | -0,350500611 | 0,062 | 0,22 | 1 | Basal | 0,784311897 | 1,275002973 | 0,281818182 |
| CSK | 0,006344773 | 0,468522609 | 0,369 | 0,22 | 1 | Basal | 1,383691773 | 0,722704304 | 1,677272727 |
| IARS2 | 0,006352509 | -0,25638103 | 0,262 | 0,467 | 1 | Basal | 0,83718535 | 1,194478619 | 0,561027837 |
| SUGT1 | 0,00643059 | -0,399746995 | 0,246 | 0,473 | 1 | Basal | 0,7579912 | 1,319276529 | 0,520084567 |
| RAMP1 | 0,0064457 | 0,295999592 | 0,215 | 0,087 | 1 | Basal | 1,227735336 | 0,814507794 | 2,471264368 |
| HBEGF | 0,006525288 | -0,639032703 | 0,385 | 0,533 | 1 | Basal | 0,642143348 | 1,557284682 | 0,722326454 |
| NELFB | 0,006536521 | 0,402679105 | 0,323 | 0,18 | 1 | Basal | 1,321960532 | 0,756452236 | 1,794444444 |
| KDELR1 | 0,006543359 | 0,320603382 | 0,738 | 0,733 | 1 | Basal | 1,24885275 | 0,800734914 | 1,006821282 |
| CALM2 | 0,006589202 | -0,386695541 | 0,892 | 0,947 | 1 | Basal | 0,764879538 | 1,307395414 | 0,941921859 |
| ANAPC5 | 0,006745477 | 0,456660854 | 0,554 | 0,427 | 1 | Basal | 1,372361781 | 0,728670831 | 1,297423888 |
| TIMP3 | 0,006760288 | -0,623341834 | 0,354 | 0,587 | 1 | Basal | 0,649165469 | 1,540439299 | 0,60306644 |
| RIMKLB | 0,00678608 | -0,385831096 | 0,062 | 0,207 | 1 | Basal | 0,765337982 | 1,306612273 | 0,299516908 |
| SGMS1 | 0,006812644 | 0,526738107 | 0,415 | 0,267 | 1 | Basal | 1,4406682 | 0,694122353 | 1,554307116 |
| GTF2B | 0,006816981 | -0,388071301 | 0,292 | 0,5 | 1 | Basal | 0,764150494 | 1,308642746 | 0,584 |
| RPS5 | 0,006840083 | 0,361949168 | 1 | 0,98 | 1 | Basal | 1,285161055 | 0,778112592 | 1,020408163 |
| AKR1C2 | 0,006841987 | -0,474416263 | 0,323 | 0,54 | 1 | Basal | 0,719757957 | 1,389355951 | 0,598148148 |
| LEPROT | 0,006876755 | -0,551113987 | 0,4 | 0,647 | 1 | Basal | 0,682492933 | 1,465216636 | 0,618238022 |
| SLC38A2 | 0,006932042 | -0,4157912 | 0,6 | 0,76 | 1 | Basal | 0,749608283 | 1,334030083 | 0,789473684 |
| GNPTG | 0,006963255 | 0,396467414 | 0,4 | 0,273 | 1 | Basal | 1,316280913 | 0,75971625 | 1,465201465 |

| Supplementary Table 4 - clusermarker | | | | | | | | | |
|---|---|---|---|---|---|---|---|---|---|
| | p_val | avg_log2FC | pct.1 | pct.2 | p_val_adj | celltype | Foldchange_UP | Foldchange_DOWN | Ratio_pct1_pct2 |
| AGPAT2 | 0,007009974 | 0,606312236 | 0,523 | 0,42 | 1 | Basal | 1,522362824 | 0,656873634 | 1,245238095 |
| ARPC2 | 0,007024281 | 0,360753381 | 0,8 | 0,767 | 1 | Basal | 1,284096283 | 0,778757803 | 1,043024772 |
| PDZK1IP1 | 0,007056855 | 0,439321303 | 0,385 | 0,233 | 1 | Basal | 1,355966281 | 0,737481466 | 1,652360515 |
| LMAN2 | 0,007084313 | 0,336867698 | 0,6 | 0,553 | 1 | Basal | 1,263011432 | 0,791758471 | 1,084990958 |
| AC020656.1 | 0,00714364 | -1,17301077 | 0,015 | 0,133 | 1 | Basal | 0,443494842 | 2,254817656 | 0,112781955 |
| NPHP3 | 0,007151364 | -0,319793534 | 0,138 | 0,327 | 1 | Basal | 0,801184528 | 1,248151911 | 0,422018349 |
| ZFP36L1 | 0,00723838 | -0,558465788 | 0,877 | 0,907 | 1 | Basal | 0,679023877 | 1,472702262 | 0,966923925 |
| TPRKB | 0,007239007 | -0,342214771 | 0,262 | 0,487 | 1 | Basal | 0,788829401 | 1,267701228 | 0,53798768 |
| ERP29 | 0,007316883 | 0,391807442 | 0,692 | 0,627 | 1 | Basal | 1,312036124 | 0,762174136 | 1,103668262 |
| AQP5 | 0,007318746 | 0,975188512 | 0,677 | 0,673 | 1 | Basal | 1,965898057 | 0,508673375 | 1,005943536 |
| PKP4 | 0,007361865 | -0,341373001 | 0,277 | 0,5 | 1 | Basal | 0,789289794 | 1,266961778 | 0,554 |
| VAMP5 | 0,007390972 | 0,551762225 | 0,354 | 0,227 | 1 | Basal | 1,465875142 | 0,682186341 | 1,559471366 |
| SORBS1 | 0,007413215 | -0,415498029 | 0,062 | 0,213 | 1 | Basal | 0,749760627 | 1,333759021 | 0,291079812 |
| AMN | 0,007485779 | 0,448453978 | 0,277 | 0,147 | 1 | Basal | 1,364577163 | 0,732827741 | 1,884353741 |
| MRPL50 | 0,007515998 | -0,406096936 | 0,169 | 0,347 | 1 | Basal | 0,754662276 | 1,325096049 | 0,4870317 |
| ANKRD17 | 0,007553041 | -0,311757768 | 0,292 | 0,5 | 1 | Basal | 0,805659552 | 1,24121907 | 0,584 |
| PPP1R10 | 0,007630006 | -0,358908046 | 0,446 | 0,633 | 1 | Basal | 0,779754541 | 1,282454859 | 0,704581359 |
| FBXW5 | 0,007653089 | 0,382457223 | 0,492 | 0,387 | 1 | Basal | 1,303560211 | 0,767129889 | 1,271317829 |
| NEDD4L | 0,007738102 | -0,602613489 | 0,369 | 0,547 | 1 | Basal | 0,658559871 | 1,518464825 | 0,674588665 |
| SLC25A39 | 0,007752262 | 0,464259392 | 0,569 | 0,48 | 1 | Basal | 1,379608949 | 0,72484308 | 1,185416667 |
| NDUFB4 | 0,007793566 | 0,315083179 | 0,815 | 0,76 | 1 | Basal | 1,244083379 | 0,803804646 | 1,072368421 |
| UQCRC1 | 0,007793993 | 0,476426277 | 0,677 | 0,6 | 1 | Basal | 1,391293 | 0,718755862 | 1,128333333 |
| CCDC85B | 0,007813995 | 0,367955459 | 0,477 | 0,36 | 1 | Basal | 1,290522647 | 0,774879854 | 1,325 |
| MED22 | 0,007864712 | 0,364138337 | 0,154 | 0,047 | 1 | Basal | 1,28711266 | 0,776932767 | 3,276595745 |
| RABAC1 | 0,007868729 | 0,386364579 | 0,692 | 0,627 | 1 | Basal | 1,307095524 | 0,765055026 | 1,103668262 |
| HNRNPH3 | 0,007896483 | -0,375254383 | 0,369 | 0,58 | 1 | Basal | 0,770969459 | 1,29706824 | 0,636206897 |
| IFRD1 | 0,007916517 | -0,407664822 | 0,338 | 0,56 | 1 | Basal | 0,753842573 | 1,326536913 | 0,603571429 |
| NCOA4 | 0,007917868 | -0,409072219 | 0,415 | 0,6 | 1 | Basal | 0,753107533 | 1,327831625 | 0,691666667 |
| TSPAN15 | 0,007970342 | 0,51227285 | 0,292 | 0,16 | 1 | Basal | 1,426295441 | 0,701117014 | 1,825 |
| NIPBL | 0,007994086 | -0,344483778 | 0,385 | 0,607 | 1 | Basal | 0,78758974 | 1,269696582 | 0,634266886 |
| CIAO2A | 0,007995472 | 0,483836243 | 0,415 | 0,273 | 1 | Basal | 1,398457338 | 0,715073655 | 1,52014652 |

| Supplementary Table 4 - clusermarker | | | | | | | | | |
|---|---|---|---|---|---|---|---|---|---|
| | **p_val** | **avg_log2FC** | **pct.1** | **pct.2** | **p_val_adj** | **celltype** | **Foldchange_UP** | **Foldchange_DOWN** | **Ratio_pct1_pct2** |
| BLZF1 | 0,008080249 | -0,390750911 | 0,138 | 0,287 | 1 | Basal | 0,762732505 | 1,311075631 | 0,480836237 |
| KRT23 | 0,008106719 | 0,457247633 | 0,662 | 0,453 | 1 | Basal | 1,372920068 | 0,728374523 | 1,461368653 |
| ODC1 | 0,008122732 | -0,579112276 | 0,308 | 0,507 | 1 | Basal | 0,669375533 | 1,493929716 | 0,607495069 |
| GPC3 | 0,008270531 | -0,255120206 | 0,031 | 0,16 | 1 | Basal | 0,837917316 | 1,193435176 | 0,19375 |
| NASP | 0,00828444 | -0,517258653 | 0,354 | 0,553 | 1 | Basal | 0,698698209 | 1,431233095 | 0,640144665 |
| POLD4 | 0,008336315 | 0,506233427 | 0,677 | 0,573 | 1 | Basal | 1,420337149 | 0,704058188 | 1,181500873 |
| COX7B | 0,008428777 | -0,312405671 | 0,692 | 0,853 | 1 | Basal | 0,805297818 | 1,241776617 | 0,811254396 |
| SERP1 | 0,008448396 | 0,34597175 | 0,785 | 0,72 | 1 | Basal | 1,271006801 | 0,786777851 | 1,090277778 |
| TNPO1 | 0,008542506 | -0,27645094 | 0,215 | 0,4 | 1 | Basal | 0,825619563 | 1,21121161 | 0,5375 |
| VPS13C | 0,008612156 | -0,361323638 | 0,354 | 0,567 | 1 | Basal | 0,778450043 | 1,28460395 | 0,624338624 |
| TMEM87A | 0,008653338 | -0,370005263 | 0,385 | 0,587 | 1 | Basal | 0,773779674 | 1,292357545 | 0,655877342 |
| PELO | 0,008662622 | 0,389994439 | 0,231 | 0,1 | 1 | Basal | 1,310388352 | 0,763132546 | 2,31 |
| RAB11A | 0,008685472 | 0,568579482 | 0,585 | 0,527 | 1 | Basal | 1,483062586 | 0,674280377 | 1,110056926 |
| IER3IP1 | 0,008732964 | 0,311944522 | 0,554 | 0,467 | 1 | Basal | 1,241379754 | 0,805555268 | 1,186295503 |
| MAP1B | 0,00879693 | -0,854689061 | 0,138 | 0,307 | 1 | Basal | 0,552984497 | 1,808368962 | 0,449511401 |
| ROMO1 | 0,008861406 | 0,337828091 | 0,738 | 0,66 | 1 | Basal | 1,263852491 | 0,791231577 | 1,118181818 |
| SGCE | 0,008875828 | -0,254575194 | 0,077 | 0,24 | 1 | Basal | 0,838233919 | 1,192984414 | 0,320833333 |
| TMEM54 | 0,008879835 | 0,466918893 | 0,492 | 0,407 | 1 | Basal | 1,382154501 | 0,723508117 | 1,208845209 |
| LCN2 | 0,008967188 | 1,479014907 | 0,631 | 0,527 | 1 | Basal | 2,78758328 | 0,358733677 | 1,197343454 |
| AC036108.2 | 0,008974979 | -0,736866104 | 0,077 | 0,227 | 1 | Basal | 0,600041378 | 1,666551735 | 0,339207048 |
| TMSB4X | 0,009000012 | 0,577703176 | 0,862 | 0,9 | 1 | Basal | 1,492471287 | 0,670029641 | 0,957777778 |
| CHST6 | 0,009019478 | 0,26659513 | 0,2 | 0,087 | 1 | Basal | 1,202965385 | 0,831279115 | 2,298850575 |
| TMEM238 | 0,009083408 | 0,738988452 | 0,569 | 0,52 | 1 | Basal | 1,669005202 | 0,599159307 | 1,094230769 |
| PRPF40A | 0,009138858 | -0,406322651 | 0,369 | 0,573 | 1 | Basal | 0,754544216 | 1,325303381 | 0,643979058 |
| MDK | 0,009140551 | 0,463574871 | 0,723 | 0,713 | 1 | Basal | 1,378954516 | 0,725187081 | 1,014025245 |
| RPLP0 | 0,009142849 | 0,285646574 | 0,985 | 0,98 | 1 | Basal | 1,218956442 | 0,820373859 | 1,005102041 |
| EIF1AX | 0,009158003 | -0,540052524 | 0,477 | 0,66 | 1 | Basal | 0,68774587 | 1,454025453 | 0,722727273 |
| DBNDD2 | 0,009163042 | 0,563697052 | 0,477 | 0,36 | 1 | Basal | 1,478052026 | 0,676566171 | 1,325 |
| CRYAB | 0,009244631 | -1,002146678 | 0,138 | 0,3 | 1 | Basal | 0,499256571 | 2,002978143 | 0,46 |
| POLK | 0,009290698 | -0,354027799 | 0,108 | 0,26 | 1 | Basal | 0,782396706 | 1,278123991 | 0,415384615 |
| GADD45G | 0,009397217 | -0,640229247 | 0,231 | 0,407 | 1 | Basal | 0,641610987 | 1,558576801 | 0,567567568 |

| Supplementary Table 4 - clusermarker | | | | | | | | | |
|---|---|---|---|---|---|---|---|---|---|
| | **p_val** | **avg_log2FC** | **pct.1** | **pct.2** | **p_val_adj** | **celltype** | **Foldchange_UP** | **Foldchange_DOWN** | **Ratio_pct1_pct2** |
| TIMM10 | 0,009408006 | 0,375409466 | 0,4 | 0,253 | 1 | Basal | 1,297207676 | 0,770886589 | 1,581027668 |
| CNOT4 | 0,009443771 | -0,30968362 | 0,262 | 0,447 | 1 | Basal | 0,806818673 | 1,239435865 | 0,586129754 |
| UACA | 0,009449146 | -0,352696844 | 0,231 | 0,413 | 1 | Basal | 0,783118837 | 1,276945404 | 0,559322034 |
| AKR1A1 | 0,009480084 | 0,507060274 | 0,692 | 0,607 | 1 | Basal | 1,421151416 | 0,703654789 | 1,140032949 |
| RABL6 | 0,009539106 | 0,294739361 | 0,462 | 0,353 | 1 | Basal | 1,226663347 | 0,815219597 | 1,30878187 |
| FABP5 | 0,009574535 | -0,931613929 | 0,477 | 0,593 | 1 | Basal | 0,524271516 | 1,907408602 | 0,804384486 |
| ATP6V1F | 0,009587362 | 0,271944325 | 0,677 | 0,587 | 1 | Basal | 1,207433995 | 0,828202622 | 1,153321976 |
| FOXC1 | 0,009727978 | -0,279722327 | 0,215 | 0,427 | 1 | Basal | 0,823749548 | 1,213961213 | 0,503512881 |
| ISLR | 0,009835629 | -0,297483793 | 0,031 | 0,153 | 1 | Basal | 0,813670284 | 1,228999043 | 0,202614379 |
| ASH1L | 0,009845224 | -0,400055862 | 0,415 | 0,607 | 1 | Basal | 0,757828939 | 1,319559004 | 0,68369028 |
| PPP2R3A | 0,009850558 | -0,465698461 | 0,077 | 0,227 | 1 | Basal | 0,72412042 | 1,380985776 | 0,339207048 |
| MEG3 | 0,010006384 | -0,652594442 | 0,123 | 0,293 | 1 | Basal | 0,636135303 | 1,571992617 | 0,419795222 |
| CH25H | 0,01000968 | -0,283786956 | 0,108 | 0,273 | 1 | Basal | 0,821431993 | 1,217386233 | 0,395604396 |
| IFI35 | 0,010027157 | 0,356943332 | 0,369 | 0,233 | 1 | Basal | 1,280709555 | 0,780817162 | 1,583690987 |
| MYDGF | 0,010054654 | 0,586909467 | 0,569 | 0,453 | 1 | Basal | 1,502025668 | 0,665767584 | 1,25607064 |
| TCF25 | 0,010107384 | 0,337739842 | 0,662 | 0,627 | 1 | Basal | 1,263775184 | 0,791279978 | 1,055821372 |
| AL135905.2 | 0,010124979 | -0,255115797 | 0,138 | 0,3 | 1 | Basal | 0,837919877 | 1,193431529 | 0,46 |
| OSTF1 | 0,010179664 | 0,472448282 | 0,462 | 0,347 | 1 | Basal | 1,387462022 | 0,720740448 | 1,331412104 |
| EIF2D | 0,010216994 | -0,28460418 | 0,138 | 0,3 | 1 | Basal | 0,82096682 | 1,218076025 | 0,46 |
| CLN3 | 0,010335986 | 0,332028662 | 0,4 | 0,253 | 1 | Basal | 1,258782181 | 0,794418617 | 1,581027668 |
| TANK | 0,010372839 | -0,333308364 | 0,277 | 0,48 | 1 | Basal | 0,793714263 | 1,259899245 | 0,577083333 |
| UVRAG | 0,010415988 | 0,341201239 | 0,385 | 0,24 | 1 | Basal | 1,266810947 | 0,789383769 | 1,604166667 |
| CLEC2B | 0,010428606 | -0,494908026 | 0,277 | 0,447 | 1 | Basal | 0,709606915 | 1,409230912 | 0,619686801 |
| RUFY1 | 0,01043522 | 0,442823905 | 0,292 | 0,16 | 1 | Basal | 1,359262321 | 0,735693166 | 1,825 |
| PPIC | 0,010556543 | 0,524824813 | 0,477 | 0,367 | 1 | Basal | 1,43875886 | 0,695043504 | 1,29972752 |
| DUSP6 | 0,010566813 | -0,494235883 | 0,277 | 0,433 | 1 | Basal | 0,709937594 | 1,408574512 | 0,639722864 |
| SF3B5 | 0,010674973 | 0,397602182 | 0,662 | 0,6 | 1 | Basal | 1,317316657 | 0,759118922 | 1,103333333 |
| SNX9 | 0,010678014 | -0,26699883 | 0,215 | 0,407 | 1 | Basal | 0,831046536 | 1,20330205 | 0,528255528 |
| CCS | 0,010768541 | 0,530052612 | 0,415 | 0,287 | 1 | Basal | 1,443981853 | 0,692529479 | 1,445993031 |
| NPPC | 0,010778355 | -0,276615206 | 0,046 | 0,18 | 1 | Basal | 0,825525563 | 1,211349527 | 0,255555556 |
| TMEM147 | 0,010804605 | 0,417367256 | 0,477 | 0,373 | 1 | Basal | 1,335488225 | 0,748789829 | 1,278820375 |

| Supplementary Table 4 - clusermarker | | | | | | | | | |
|---|---|---|---|---|---|---|---|---|---|
| | **p_val** | **avg_log2FC** | **pct.1** | **pct.2** | **p_val_adj** | **celltype** | **Foldchange_UP** | **Foldchange_DOWN** | **Ratio_pct1_pct2** |
| ETFA | 0,010819765 | 0,338899452 | 0,508 | 0,393 | 1 | Basal | 1,26479139 | 0,790644218 | 1,292620865 |
| EIF6 | 0,010861653 | 0,387044342 | 0,6 | 0,547 | 1 | Basal | 1,307711541 | 0,764694635 | 1,096892139 |
| HNRNPU | 0,010957714 | 0,330404448 | 0,862 | 0,773 | 1 | Basal | 1,257365818 | 0,795313493 | 1,115135834 |
| RPL6 | 0,010977665 | -0,377246099 | 1 | 0,973 | 1 | Basal | 0,76990583 | 1,298860147 | 1,027749229 |
| AKAP9 | 0,011055332 | -0,379321145 | 0,538 | 0,747 | 1 | Basal | 0,768799261 | 1,300729658 | 0,72021419 |
| FAM174C | 0,011073205 | 0,407663284 | 0,354 | 0,227 | 1 | Basal | 1,326535499 | 0,753843377 | 1,559471366 |
| PLXNB2 | 0,011106915 | 0,560370232 | 0,554 | 0,46 | 1 | Basal | 1,4746476 | 0,678128117 | 1,204347826 |
| RAI1 | 0,011133904 | 0,421496064 | 0,369 | 0,253 | 1 | Basal | 1,339315696 | 0,746649952 | 1,458498024 |
| TMEM205 | 0,011207448 | 0,421345461 | 0,646 | 0,54 | 1 | Basal | 1,339175892 | 0,746727899 | 1,196296296 |
| CTSC | 0,011239919 | 0,467152119 | 0,692 | 0,56 | 1 | Basal | 1,382377957 | 0,723391164 | 1,235714286 |
| VPS37B | 0,01125418 | 0,459235429 | 0,492 | 0,38 | 1 | Basal | 1,374813029 | 0,727371634 | 1,294736842 |
| H2AZ1 | 0,011262148 | -0,500298531 | 0,769 | 0,813 | 1 | Basal | 0,706960478 | 1,41450623 | 0,945879459 |
| HSPE1 | 0,011286743 | -0,442555767 | 0,662 | 0,767 | 1 | Basal | 0,735829914 | 1,359009713 | 0,863102999 |
| BAG1 | 0,011313054 | 0,498078512 | 0,754 | 0,707 | 1 | Basal | 1,412331262 | 0,708049186 | 1,066478076 |
| CAVIN1 | 0,011386758 | -0,445922205 | 0,338 | 0,547 | 1 | Basal | 0,734114903 | 1,362184579 | 0,617915905 |
| ATAD1 | 0,011485494 | -0,390279713 | 0,246 | 0,427 | 1 | Basal | 0,762981662 | 1,310647491 | 0,576112412 |
| ERRFI1 | 0,011510643 | -0,9208949 | 0,508 | 0,653 | 1 | Basal | 0,528181289 | 1,893289331 | 0,777947933 |
| NDUFAB1 | 0,011551042 | 0,482864812 | 0,646 | 0,593 | 1 | Basal | 1,397516011 | 0,715555308 | 1,089376054 |
| VSIG2 | 0,011558062 | 0,941680507 | 0,477 | 0,387 | 1 | Basal | 1,920764317 | 0,520626082 | 1,23255814 |
| TBC1D22A | 0,011655308 | 0,424374511 | 0,277 | 0,14 | 1 | Basal | 1,341990549 | 0,745161731 | 1,978571429 |
| ARPC5L | 0,011682533 | 0,471140095 | 0,523 | 0,393 | 1 | Basal | 1,386204489 | 0,721394288 | 1,330788804 |
| NAA60 | 0,011733503 | 0,257084421 | 0,292 | 0,16 | 1 | Basal | 1,195061133 | 0,836777276 | 1,825 |
| NME3 | 0,011767644 | 0,330028363 | 0,631 | 0,553 | 1 | Basal | 1,257038088 | 0,795520844 | 1,141048825 |
| WSB1 | 0,011789999 | -0,466557518 | 0,554 | 0,68 | 1 | Basal | 0,723689368 | 1,381808334 | 0,814705882 |
| LIMA1 | 0,011844094 | -0,42456196 | 0,523 | 0,72 | 1 | Basal | 0,745064919 | 1,342164924 | 0,726388889 |
| RTN4 | 0,012005626 | -0,370390455 | 0,677 | 0,767 | 1 | Basal | 0,773573106 | 1,292702644 | 0,882659713 |
| RWDD4 | 0,012072728 | 0,355390761 | 0,369 | 0,24 | 1 | Basal | 1,279332048 | 0,781657899 | 1,5375 |
| TFDP2 | 0,01208759 | -0,269477833 | 0,231 | 0,4 | 1 | Basal | 0,829619763 | 1,205371479 | 0,5775 |
| FAM83D | 0,012114537 | 0,476913966 | 0,308 | 0,18 | 1 | Basal | 1,391763393 | 0,718512935 | 1,711111111 |
| TOMM5 | 0,012149424 | 0,612413023 | 0,569 | 0,487 | 1 | Basal | 1,528814136 | 0,654101749 | 1,168377823 |
| DNAJC15 | 0,012213198 | 0,327591702 | 0,6 | 0,54 | 1 | Basal | 1,254916786 | 0,796865586 | 1,111111111 |

| Supplementary Table 4 - clusermarker | | | | | | | | | |
|---|---|---|---|---|---|---|---|---|---|
| | **p_val** | **avg_log2FC** | **pct.1** | **pct.2** | **p_val_adj** | **celltype** | **Foldchange_UP** | **Foldchange_DOWN** | **Ratio_pct1_pct2** |
| PHLDA3 | 0,0122565 | 0,398310593 | 0,415 | 0,273 | 1 | Basal | 1,317963661 | 0,758746261 | 1,52014652 |
| SLC35A1 | 0,012292587 | -0,320248521 | 0,092 | 0,233 | 1 | Basal | 0,800931896 | 1,248545607 | 0,394849785 |
| IRX3 | 0,01231078 | -0,515795573 | 0,308 | 0,487 | 1 | Basal | 0,699407139 | 1,429782374 | 0,632443532 |
| PIK3R3 | 0,012321073 | -0,424003477 | 0,308 | 0,5 | 1 | Basal | 0,745353397 | 1,341645458 | 0,616 |
| PHGDH | 0,012373308 | -0,35336971 | 0,108 | 0,26 | 1 | Basal | 0,782753679 | 1,277541104 | 0,415384615 |
| CCDC124 | 0,012446214 | 0,368077243 | 0,492 | 0,387 | 1 | Basal | 1,29063159 | 0,774814445 | 1,271317829 |
| TGFB2 | 0,012460565 | 0,260347025 | 0,231 | 0,107 | 1 | Basal | 1,19776678 | 0,834887072 | 2,158878505 |
| CHST9 | 0,012474403 | -0,48459356 | 0,462 | 0,6 | 1 | Basal | 0,714698388 | 1,399191626 | 0,77 |
| PALM2AKAP2 | 0,012523417 | -0,25025762 | 0,077 | 0,22 | 1 | Basal | 0,840746271 | 1,189419489 | 0,35 |
| MBNL1 | 0,012577805 | -0,486370903 | 0,354 | 0,587 | 1 | Basal | 0,71381845 | 1,400916437 | 0,60306644 |
| TCF7L1 | 0,012618071 | -0,321323981 | 0,215 | 0,38 | 1 | Basal | 0,800335062 | 1,249476684 | 0,565789474 |
| MRPL4 | 0,012691307 | 0,316532266 | 0,477 | 0,36 | 1 | Basal | 1,245333602 | 0,802997685 | 1,325 |
| MBOAT2 | 0,012841755 | 0,525805004 | 0,354 | 0,227 | 1 | Basal | 1,439736709 | 0,694571441 | 1,559471366 |
| PTPRM | 0,012844572 | -0,272100987 | 0,077 | 0,213 | 1 | Basal | 0,828112692 | 1,207565117 | 0,361502347 |
| EIF4B | 0,012930955 | -0,46099487 | 0,6 | 0,78 | 1 | Basal | 0,726485108 | 1,376490707 | 0,769230769 |
| MYO9B | 0,012978751 | 0,315152906 | 0,246 | 0,127 | 1 | Basal | 1,244143509 | 0,803765798 | 1,937007874 |
| AREG | 0,012991002 | -0,812469972 | 0,446 | 0,56 | 1 | Basal | 0,569406169 | 1,756215607 | 0,796428571 |
| MMADHC | 0,013032534 | 0,495377136 | 0,662 | 0,58 | 1 | Basal | 1,409689215 | 0,709376215 | 1,14137931 |
| TICAM1 | 0,01330497 | 0,2724343 | 0,262 | 0,14 | 1 | Basal | 1,207844139 | 0,827921391 | 1,871428571 |
| HMGXB4 | 0,013353606 | -0,276451237 | 0,123 | 0,287 | 1 | Basal | 0,825619393 | 1,21121186 | 0,428571429 |
| AGRN | 0,013454242 | 0,68301799 | 0,431 | 0,32 | 1 | Basal | 1,605494797 | 0,622860941 | 1,346875 |
| RUSC1 | 0,013505206 | 0,378805624 | 0,369 | 0,247 | 1 | Basal | 1,300264948 | 0,769074027 | 1,493927126 |
| REX1BD | 0,013553628 | 0,463008309 | 0,569 | 0,527 | 1 | Basal | 1,378413092 | 0,725471926 | 1,079696395 |
| MXI1 | 0,013555667 | -0,276644953 | 0,169 | 0,36 | 1 | Basal | 0,825508541 | 1,211374505 | 0,469444444 |
| MIEN1 | 0,013589386 | 0,468393571 | 0,508 | 0,447 | 1 | Basal | 1,383568018 | 0,722768947 | 1,136465324 |
| XRN1 | 0,013623086 | -0,42960509 | 0,2 | 0,367 | 1 | Basal | 0,742464993 | 1,346864848 | 0,544959128 |
| IMPA2 | 0,013695205 | 0,282923036 | 0,323 | 0,193 | 1 | Basal | 1,216657451 | 0,821924034 | 1,67357513 |
| FOSB | 0,013731317 | -0,542635492 | 0,738 | 0,847 | 1 | Basal | 0,686515647 | 1,456631038 | 0,871310508 |
| CCNG2 | 0,013836362 | -0,461105094 | 0,185 | 0,353 | 1 | Basal | 0,726429606 | 1,376595877 | 0,52407932 |
| SAT1 | 0,013849041 | 0,723026656 | 0,862 | 0,853 | 1 | Basal | 1,650641314 | 0,605825137 | 1,010550996 |
| PSMB3 | 0,013959207 | 0,507763503 | 0,662 | 0,66 | 1 | Basal | 1,421844313 | 0,703311882 | 1,003030303 |

| Supplementary Table 4 - clusermarker | | | | | | | | | |
|---|---|---|---|---|---|---|---|---|---|
| | **p_val** | **avg_log2FC** | **pct.1** | **pct.2** | **p_val_adj** | **celltype** | **Foldchange_UP** | **Foldchange_DOWN** | **Ratio_pct1_pct2** |
| NIPSNAP2 | 0,013971633 | -0,308373993 | 0,215 | 0,4 | 1 | Basal | 0,807551407 | 1,23831126 | 0,5375 |
| CTSB | 0,014026448 | 0,465600529 | 0,738 | 0,74 | 1 | Basal | 1,380892037 | 0,724169576 | 0,997297297 |
| FAM118A | 0,014032335 | 0,307812387 | 0,215 | 0,1 | 1 | Basal | 1,237829309 | 0,807865828 | 2,15 |
| CDKN1A | 0,014040659 | -0,547612793 | 0,585 | 0,673 | 1 | Basal | 0,684151247 | 1,461665098 | 0,869242199 |
| NUP58 | 0,01406743 | -0,366193923 | 0,108 | 0,253 | 1 | Basal | 0,775826563 | 1,288947876 | 0,42687747 |
| ATP5F1B | 0,014074029 | 0,643732975 | 0,831 | 0,787 | 1 | Basal | 1,562366559 | 0,640054662 | 1,055908513 |
| PTTG1IP | 0,014167516 | 0,549667212 | 0,692 | 0,673 | 1 | Basal | 1,463748012 | 0,6831777 | 1,028231798 |
| RSRP1 | 0,014210956 | -0,515972503 | 0,569 | 0,713 | 1 | Basal | 0,69932137 | 1,429957732 | 0,798036466 |
| DHRS4 | 0,014315116 | 0,610731345 | 0,431 | 0,313 | 1 | Basal | 1,527033112 | 0,654864647 | 1,376996805 |
| CD200R1 | 0,014321298 | 0,269571536 | 0,169 | 0,067 | 1 | Basal | 1,20544977 | 0,829565881 | 2,52238806 |
| PPCS | 0,0143398 | 0,60470177 | 0,492 | 0,387 | 1 | Basal | 1,520664373 | 0,657607305 | 1,271317829 |
| STAU2 | 0,014373769 | -0,352367324 | 0,123 | 0,267 | 1 | Basal | 0,783297726 | 1,276653776 | 0,460674157 |
| ATP5IF1 | 0,014397047 | 0,272707899 | 0,769 | 0,7 | 1 | Basal | 1,208073221 | 0,827764396 | 1,098571429 |
| IER5 | 0,014431496 | -0,446884738 | 0,323 | 0,48 | 1 | Basal | 0,733625282 | 1,363093701 | 0,672916667 |
| RBIS | 0,014564265 | -0,30871154 | 0,523 | 0,707 | 1 | Basal | 0,807362487 | 1,238601021 | 0,739745403 |
| VPS25 | 0,014618282 | 0,327782042 | 0,4 | 0,267 | 1 | Basal | 1,255082363 | 0,796760459 | 1,498127341 |
| LRPAP1 | 0,014727438 | 0,773520195 | 0,538 | 0,413 | 1 | Basal | 1,709435744 | 0,584988353 | 1,302663438 |
| CBLC | 0,014749099 | 0,292858114 | 0,323 | 0,187 | 1 | Basal | 1,225064843 | 0,816283322 | 1,727272727 |
| ZNF24 | 0,014799003 | -0,456407722 | 0,292 | 0,487 | 1 | Basal | 0,728798693 | 1,372121011 | 0,599589322 |
| MCAM | 0,01490223 | -0,370555089 | 0,108 | 0,26 | 1 | Basal | 0,773484835 | 1,29285017 | 0,415384615 |
| KRT10 | 0,01499762 | 0,411715667 | 0,677 | 0,667 | 1 | Basal | 1,33026684 | 0,751728879 | 1,014992504 |
| THAP1 | 0,015002758 | 0,503079916 | 0,215 | 0,1 | 1 | Basal | 1,4172359 | 0,705598835 | 2,15 |
| IMP3 | 0,015074224 | 0,441202635 | 0,508 | 0,413 | 1 | Basal | 1,357735668 | 0,736520387 | 1,230024213 |
| RNF7 | 0,015082571 | 0,492961656 | 0,646 | 0,64 | 1 | Basal | 1,40733097 | 0,710564907 | 1,009375 |
| ASS1 | 0,01509111 | 0,70890854 | 0,662 | 0,62 | 1 | Basal | 1,63456703 | 0,611782803 | 1,067741935 |
| DDX3X | 0,015453208 | -0,483901064 | 0,585 | 0,753 | 1 | Basal | 0,715041527 | 1,398520173 | 0,77689243 |
| PRKCH | 0,01556133 | 0,270984729 | 0,215 | 0,107 | 1 | Basal | 1,206631147 | 0,828753677 | 2,009345794 |
| MDH1 | 0,015713105 | -0,375724074 | 0,492 | 0,68 | 1 | Basal | 0,7707185 | 1,297490589 | 0,723529412 |
| KDM4B | 0,015787354 | 0,299061949 | 0,246 | 0,127 | 1 | Basal | 1,230344175 | 0,8127807 | 1,937007874 |
| ADAM9 | 0,016015687 | 0,499053057 | 0,508 | 0,42 | 1 | Basal | 1,413285619 | 0,707571058 | 1,20952381 |
| HOXB2 | 0,016037306 | 0,315500896 | 0,431 | 0,293 | 1 | Basal | 1,244443642 | 0,803571946 | 1,470989761 |

| Supplementary Table 4 - clusermarker | | | | | | | | | |
|---|---|---|---|---|---|---|---|---|---|
| | **p_val** | **avg_log2FC** | **pct.1** | **pct.2** | **p_val_adj** | **celltype** | **Foldchange_UP** | **Foldchange_DOWN** | **Ratio_pct1_pct2** |
| SPECC1 | 0,016277865 | 0,32900972 | 0,292 | 0,167 | 1 | Basal | 1,256150844 | 0,796082735 | 1,748502994 |
| SNX8 | 0,016288117 | 0,400488687 | 0,246 | 0,133 | 1 | Basal | 1,319954946 | 0,757601616 | 1,84962406 |
| FIBP | 0,016320986 | 0,42676125 | 0,431 | 0,32 | 1 | Basal | 1,344212524 | 0,743929983 | 1,346875 |
| VEGFA | 0,01632611 | -0,532444603 | 0,415 | 0,58 | 1 | Basal | 0,691382215 | 1,446377964 | 0,715517241 |
| UBXN4 | 0,016569376 | 0,279564275 | 0,723 | 0,673 | 1 | Basal | 1,213828227 | 0,823839797 | 1,074294205 |
| MED16 | 0,016574106 | 0,335148502 | 0,338 | 0,213 | 1 | Basal | 1,261507253 | 0,792702537 | 1,58685446 |
| KIF1C | 0,016698736 | 0,458182507 | 0,431 | 0,32 | 1 | Basal | 1,373810015 | 0,727902686 | 1,346875 |
| S100A6 | 0,016710326 | 0,515352871 | 1 | 0,98 | 1 | Basal | 1,429343702 | 0,69962179 | 1,020408163 |
| CCT8 | 0,016754987 | -0,450141317 | 0,323 | 0,54 | 1 | Basal | 0,731971145 | 1,366174071 | 0,598148148 |
| SNRPD1 | 0,016764688 | 0,482054046 | 0,585 | 0,5 | 1 | Basal | 1,396730855 | 0,715957549 | 1,17 |
| YIF1A | 0,016860444 | 0,502756517 | 0,508 | 0,453 | 1 | Basal | 1,416918244 | 0,705757022 | 1,121412804 |
| CNPY2 | 0,016975868 | 0,411686304 | 0,6 | 0,487 | 1 | Basal | 1,330239765 | 0,751744179 | 1,232032854 |
| DUOXA1 | 0,017021337 | 0,446359106 | 0,354 | 0,247 | 1 | Basal | 1,362597161 | 0,73389262 | 1,433198381 |
| GADD45GIP1 | 0,017038428 | 0,419706713 | 0,554 | 0,493 | 1 | Basal | 1,337655593 | 0,747576585 | 1,123732252 |
| RPN1 | 0,017045077 | 0,39470874 | 0,6 | 0,56 | 1 | Basal | 1,314677318 | 0,760642924 | 1,071428571 |
| OGFRL1 | 0,017057545 | -0,316397114 | 0,246 | 0,433 | 1 | Basal | 0,803072914 | 1,245216944 | 0,56812933 |
| MFSD1 | 0,017203022 | 0,368198295 | 0,477 | 0,373 | 1 | Basal | 1,290739888 | 0,774749436 | 1,278820375 |
| DMKN | 0,0173019 | -0,309937754 | 0,323 | 0,52 | 1 | Basal | 0,806676563 | 1,239654213 | 0,621153846 |
| TMEM14C | 0,01753274 | 0,332924671 | 0,754 | 0,687 | 1 | Basal | 1,259564211 | 0,793925384 | 1,097525473 |
| MRPS11 | 0,017591006 | 0,398420344 | 0,369 | 0,247 | 1 | Basal | 1,318063927 | 0,758688543 | 1,493927126 |
| STOML2 | 0,017823403 | 0,254834176 | 0,508 | 0,42 | 1 | Basal | 1,193198588 | 0,838083459 | 1,20952381 |
| IDS | 0,017886328 | 0,627764054 | 0,431 | 0,347 | 1 | Basal | 1,545168374 | 0,647178662 | 1,242074928 |
| COBLL1 | 0,017992876 | -0,270033703 | 0,185 | 0,373 | 1 | Basal | 0,829300172 | 1,205835997 | 0,495978552 |
| BCAS1 | 0,01801272 | 0,525521521 | 0,369 | 0,253 | 1 | Basal | 1,439453836 | 0,694707934 | 1,458498024 |
| NME6 | 0,018067435 | 0,414786168 | 0,277 | 0,16 | 1 | Basal | 1,333101074 | 0,750130669 | 1,73125 |
| METTL9 | 0,01807746 | 0,522097402 | 0,492 | 0,427 | 1 | Basal | 1,43604146 | 0,696358725 | 1,152224824 |
| SORL1 | 0,018119514 | 0,706045823 | 0,492 | 0,407 | 1 | Basal | 1,6313268 | 0,61299796 | 1,208845209 |
| CRIP1 | 0,018146976 | 0,783254756 | 0,646 | 0,547 | 1 | Basal | 1,721009136 | 0,58105444 | 1,180987203 |
| CCT6A | 0,018171111 | 0,309669859 | 0,615 | 0,553 | 1 | Basal | 1,239424042 | 0,806826369 | 1,112115732 |
| MAGED2 | 0,018315505 | -0,421781323 | 0,523 | 0,68 | 1 | Basal | 0,746502335 | 1,339580539 | 0,769117647 |
| KANK2 | 0,018356481 | -0,273349751 | 0,077 | 0,207 | 1 | Basal | 0,827396207 | 1,20861081 | 0,371980676 |

| Supplementary Table 4 - clusermarker | | | | | | | | | |
|---|---|---|---|---|---|---|---|---|---|
| | p_val | avg_log2FC | pct.1 | pct.2 | p_val_adj | celltype | Foldchange_UP | Foldchange_DOWN | Ratio_pct1_pct2 |
| CSNK2B | 0,018511797 | 0,266058522 | 0,692 | 0,633 | 1 | Basal | 1,202518027 | 0,831588365 | 1,093206951 |
| RASD1 | 0,0185848 | -0,339834265 | 0,215 | 0,4 | 1 | Basal | 0,790132076 | 1,265611194 | 0,5375 |
| MT-ATP6 | 0,018818905 | 0,327867183 | 0,938 | 0,873 | 1 | Basal | 1,255156434 | 0,79671344 | 1,074455899 |
| CISD3 | 0,018868352 | 0,350383945 | 0,415 | 0,313 | 1 | Basal | 1,274899872 | 0,784375324 | 1,325878594 |
| ITGB1 | 0,018974467 | -0,399472365 | 0,569 | 0,747 | 1 | Basal | 0,758135505 | 1,319025417 | 0,761713521 |
| EXOSC8 | 0,019128041 | -0,257117349 | 0,138 | 0,293 | 1 | Basal | 0,836758178 | 1,19508841 | 0,470989761 |
| PPP1R14B | 0,019246319 | 0,60156295 | 0,569 | 0,527 | 1 | Basal | 1,517359514 | 0,659039595 | 1,079696395 |
| MGAT4B | 0,019290327 | 0,328391588 | 0,415 | 0,3 | 1 | Basal | 1,255612754 | 0,796423895 | 1,383333333 |
| CD63 | 0,019371243 | 0,313966801 | 0,877 | 0,867 | 1 | Basal | 1,243121062 | 0,804426882 | 1,011534025 |
| GGH | 0,019444997 | 0,357106342 | 0,338 | 0,227 | 1 | Basal | 1,280854271 | 0,780728942 | 1,488986784 |
| PDXK | 0,019505733 | 0,47330759 | 0,477 | 0,387 | 1 | Basal | 1,388288678 | 0,720311284 | 1,23255814 |
| ATP5PB | 0,019628595 | 0,399065752 | 0,692 | 0,667 | 1 | Basal | 1,318653711 | 0,75834921 | 1,037481259 |
| TCIRG1 | 0,019746942 | 0,362643318 | 0,385 | 0,26 | 1 | Basal | 1,285779557 | 0,777738295 | 1,480769231 |
| NXT1 | 0,019825421 | 0,314476494 | 0,308 | 0,193 | 1 | Basal | 1,243560324 | 0,804142735 | 1,595854922 |
| LONP2 | 0,019875583 | -0,457283425 | 0,308 | 0,473 | 1 | Basal | 0,728356453 | 1,372954129 | 0,651162791 |
| ZNF593 | 0,019879311 | 0,267616863 | 0,431 | 0,313 | 1 | Basal | 1,203817641 | 0,830690602 | 1,376996805 |
| ARMCX2 | 0,019913348 | -0,264592413 | 0,108 | 0,253 | 1 | Basal | 0,832433879 | 1,201296614 | 0,42687747 |
| MDC1 | 0,019990466 | -0,286312897 | 0,077 | 0,2 | 1 | Basal | 0,819995048 | 1,219519559 | 0,385 |
| PRMT7 | 0,020016171 | 0,272299894 | 0,277 | 0,153 | 1 | Basal | 1,207731617 | 0,827998527 | 1,810457516 |
| SUPT16H | 0,020266084 | -0,262895496 | 0,277 | 0,467 | 1 | Basal | 0,833413574 | 1,199884464 | 0,593147752 |
| FOXN3 | 0,020324708 | -0,36690694 | 0,385 | 0,54 | 1 | Basal | 0,775443224 | 1,289585065 | 0,712962963 |
| OIP5-AS1 | 0,020410995 | 0,290229281 | 0,492 | 0,373 | 1 | Basal | 1,222834602 | 0,817772083 | 1,319034853 |
| SEC14L1 | 0,020678761 | 0,448653148 | 0,354 | 0,247 | 1 | Basal | 1,364765563 | 0,732726578 | 1,433198381 |
| NFYB | 0,020712409 | -0,283879299 | 0,154 | 0,3 | 1 | Basal | 0,821379418 | 1,217464157 | 0,513333333 |
| ZNRD2 | 0,020764383 | 0,341195263 | 0,431 | 0,32 | 1 | Basal | 1,266805699 | 0,789387039 | 1,346875 |
| FADD | 0,020794115 | 0,333973236 | 0,215 | 0,1 | 1 | Basal | 1,260480007 | 0,793348561 | 2,15 |
| ANXA3 | 0,020834955 | 0,432058021 | 0,554 | 0,48 | 1 | Basal | 1,349156793 | 0,741203695 | 1,154166667 |
| CREB1 | 0,020837129 | -0,29112799 | 0,169 | 0,313 | 1 | Basal | 0,817262821 | 1,223596589 | 0,539936102 |
| IFI30 | 0,02086699 | 0,319926145 | 0,385 | 0,28 | 1 | Basal | 1,248266645 | 0,801110888 | 1,375 |
| ZFYVE21 | 0,020920124 | 0,453645883 | 0,538 | 0,447 | 1 | Basal | 1,369496789 | 0,730195213 | 1,203579418 |
| COA3 | 0,020920124 | 0,45832045 | 0,554 | 0,44 | 1 | Basal | 1,373941378 | 0,727833091 | 1,259090909 |

| Supplementary Table 4 - clusermarker | | | | | | | | | |
|---|---|---|---|---|---|---|---|---|---|
| | **p_val** | **avg_log2FC** | **pct.1** | **pct.2** | **p_val_adj** | **celltype** | **Foldchange_UP** | **Foldchange_DOWN** | **Ratio_pct1_pct2** |
| SPECC1L | 0,020987816 | -0,25289362 | 0,108 | 0,24 | 1 | Basal | 0,839211516 | 1,191594706 | 0,45 |
| TBCB | 0,021013876 | 0,584718388 | 0,554 | 0,487 | 1 | Basal | 1,499746213 | 0,66677948 | 1,137577002 |
| PLPP3 | 0,021203327 | -0,496353702 | 0,277 | 0,447 | 1 | Basal | 0,708896198 | 1,410643762 | 0,619686801 |
| KPNB1 | 0,021242197 | -0,324133128 | 0,415 | 0,62 | 1 | Basal | 0,798778204 | 1,251911976 | 0,669354839 |
| TOB1 | 0,021381365 | -0,38509771 | 0,538 | 0,693 | 1 | Basal | 0,765727136 | 1,305948233 | 0,776334776 |
| POMP | 0,021585152 | 0,3706231 | 0,708 | 0,64 | 1 | Basal | 1,292911118 | 0,773448372 | 1,10625 |
| STUB1 | 0,021667689 | 0,390988666 | 0,554 | 0,507 | 1 | Basal | 1,311291712 | 0,762606818 | 1,09270217 |
| MEA1 | 0,021684346 | 0,30866505 | 0,585 | 0,5 | 1 | Basal | 1,238561108 | 0,807388504 | 1,17 |
| TIMM17B | 0,021701926 | 0,257719733 | 0,477 | 0,367 | 1 | Basal | 1,195587512 | 0,83640887 | 1,29972752 |
| PPP6R3 | 0,021816449 | -0,371441504 | 0,231 | 0,4 | 1 | Basal | 0,773009739 | 1,293644762 | 0,5775 |
| UFC1 | 0,02181718 | 0,461468769 | 0,692 | 0,62 | 1 | Basal | 1,376942933 | 0,72624651 | 1,116129032 |
| CAPN15 | 0,021891343 | 0,256731981 | 0,215 | 0,107 | 1 | Basal | 1,194769224 | 0,83698172 | 2,009345794 |
| MAP2K2 | 0,021943161 | 0,420815932 | 0,554 | 0,44 | 1 | Basal | 1,338684449 | 0,747002029 | 1,259090909 |
| ANTXR1 | 0,021978288 | 0,284618656 | 0,446 | 0,307 | 1 | Basal | 1,218088247 | 0,820958582 | 1,45276873 |
| FBXO31 | 0,022107397 | 0,306201965 | 0,154 | 0,06 | 1 | Basal | 1,236448341 | 0,80876812 | 2,566666667 |
| SIVA1 | 0,02211307 | 0,304050562 | 0,631 | 0,587 | 1 | Basal | 1,234605875 | 0,809975086 | 1,074957411 |
| CIAO3 | 0,022120901 | 0,333547202 | 0,215 | 0,107 | 1 | Basal | 1,260107838 | 0,793582874 | 2,009345794 |
| UBE2E2 | 0,022198639 | -0,27382775 | 0,138 | 0,28 | 1 | Basal | 0,827122116 | 1,209011318 | 0,492857143 |
| DANCR | 0,022286893 | 0,360416078 | 0,6 | 0,513 | 1 | Basal | 1,283796095 | 0,778939899 | 1,169590643 |
| TXNDC15 | 0,022498697 | 0,301003552 | 0,385 | 0,287 | 1 | Basal | 1,232001106 | 0,811687583 | 1,341463415 |
| TMEM132A | 0,02252631 | 0,392731514 | 0,185 | 0,087 | 1 | Basal | 1,312876776 | 0,761686107 | 2,126436782 |
| ARF3 | 0,022554771 | 0,41712918 | 0,492 | 0,393 | 1 | Basal | 1,335267859 | 0,748913406 | 1,251908397 |
| IGKV3-20 | 0,022606638 | 0,811436767 | 0,123 | 0,04 | 1 | Basal | 1,75495832 | 0,569814102 | 3,075 |
| MED21 | 0,022703067 | -0,325504298 | 0,2 | 0,36 | 1 | Basal | 0,798019388 | 1,253102387 | 0,555555556 |
| HLA-C | 0,022818665 | 0,314378912 | 0,862 | 0,867 | 1 | Basal | 1,243476215 | 0,804197127 | 0,994232987 |
| RNF149 | 0,022849264 | 0,274801658 | 0,477 | 0,36 | 1 | Basal | 1,209827751 | 0,826563946 | 1,325 |
| MTDH | 0,022959048 | 0,403649583 | 0,692 | 0,68 | 1 | Basal | 1,322850094 | 0,755943553 | 1,017647059 |
| TTC9 | 0,023137839 | 0,391353692 | 0,308 | 0,193 | 1 | Basal | 1,311623533 | 0,76241389 | 1,595854922 |
| LOXL4 | 0,02320714 | 0,90928996 | 0,262 | 0,153 | 1 | Basal | 1,878120931 | 0,532447077 | 1,712418301 |
| TRAF4 | 0,023224059 | 0,397220347 | 0,523 | 0,413 | 1 | Basal | 1,316968051 | 0,759319863 | 1,266343826 |
| CTCF | 0,023271296 | 0,406132914 | 0,4 | 0,313 | 1 | Basal | 1,325129094 | 0,754643456 | 1,277955272 |

| Supplementary Table 4 - clusermarker | | | | | | | | | |
|---|---|---|---|---|---|---|---|---|---|
| | **p_val** | **avg_log2FC** | **pct.1** | **pct.2** | **p_val_adj** | **celltype** | **Foldchange_UP** | **Foldchange_DOWN** | **Ratio_pct1_pct2** |
| ARL1 | 0,023311082 | -0,329538041 | 0,354 | 0,573 | 1 | Basal | 0,79579126 | 1,256610937 | 0,617801047 |
| TSR3 | 0,023466979 | 0,357272092 | 0,338 | 0,227 | 1 | Basal | 1,281001435 | 0,78063925 | 1,488986784 |
| GABARAPL1 | 0,023494633 | -0,255972385 | 0,2 | 0,373 | 1 | Basal | 0,837422517 | 1,194140329 | 0,536193029 |
| ISG15 | 0,023562039 | 0,312638504 | 0,292 | 0,173 | 1 | Basal | 1,241977041 | 0,805167863 | 1,687861272 |
| TIMP1 | 0,02359609 | 0,357650992 | 0,723 | 0,66 | 1 | Basal | 1,281337913 | 0,780434255 | 1,095454545 |
| PABPC1 | 0,023838357 | -0,380520875 | 0,892 | 0,947 | 1 | Basal | 0,768160202 | 1,301811781 | 0,941921859 |
| C7orf50 | 0,023888483 | 0,389141292 | 0,415 | 0,293 | 1 | Basal | 1,309613675 | 0,763583963 | 1,416382253 |
| TRAPPC2L | 0,02395895 | 0,335368206 | 0,538 | 0,447 | 1 | Basal | 1,26169938 | 0,792581827 | 1,203579418 |
| SYTL2 | 0,024048265 | 0,42560246 | 0,462 | 0,36 | 1 | Basal | 1,34313327 | 0,744527757 | 1,283333333 |
| NUBP2 | 0,024066619 | 0,517560557 | 0,415 | 0,293 | 1 | Basal | 1,431532632 | 0,698552012 | 1,416382253 |
| DRAM2 | 0,024159627 | 0,290278122 | 0,308 | 0,187 | 1 | Basal | 1,222876001 | 0,817744399 | 1,647058824 |
| RPL26 | 0,024160437 | -0,254524739 | 1 | 1 | 1 | Basal | 0,838263235 | 1,192942692 | 1 |
| CPNE3 | 0,024497163 | 0,56542358 | 0,662 | 0,627 | 1 | Basal | 1,479821926 | 0,675756983 | 1,055821372 |
| ILK | 0,02451922 | 0,317973286 | 0,554 | 0,447 | 1 | Basal | 1,246578112 | 0,80219602 | 1,239373602 |
| COMTD1 | 0,024640132 | 0,369669269 | 0,446 | 0,347 | 1 | Basal | 1,292056599 | 0,773959903 | 1,285302594 |
| CEP170B | 0,024743066 | 0,263431066 | 0,262 | 0,153 | 1 | Basal | 1,200329978 | 0,833104245 | 1,712418301 |
| SH3BGR | 0,024794287 | -0,368362879 | 0,046 | 0,153 | 1 | Basal | 0,774661057 | 1,290887145 | 0,300653595 |
| ESD | 0,024863927 | -0,320695667 | 0,354 | 0,553 | 1 | Basal | 0,800683695 | 1,248932638 | 0,640144665 |
| VPS29 | 0,024906066 | 0,42266664 | 0,615 | 0,513 | 1 | Basal | 1,340402832 | 0,74604438 | 1,198830409 |
| KCNS3 | 0,025033246 | 0,320410059 | 0,246 | 0,133 | 1 | Basal | 1,248685414 | 0,800842221 | 1,84962406 |
| TPI1 | 0,025067669 | 0,423607322 | 0,892 | 0,833 | 1 | Basal | 1,341277102 | 0,745558094 | 1,070828331 |
| CYB5B | 0,025197 | -0,29694231 | 0,123 | 0,253 | 1 | Basal | 0,813975733 | 1,228537853 | 0,486166008 |
| TCN1 | 0,02543432 | -0,705736788 | 0,231 | 0,38 | 1 | Basal | 0,613129282 | 1,630977397 | 0,607894737 |
| UBXN1 | 0,025463259 | 0,360930953 | 0,646 | 0,627 | 1 | Basal | 1,284254344 | 0,778661957 | 1,03030303 |
| SERPINB1 | 0,0254913 | 0,273645704 | 0,662 | 0,587 | 1 | Basal | 1,208858769 | 0,827226493 | 1,127768313 |
| PLS3 | 0,025500799 | -0,274704355 | 0,431 | 0,613 | 1 | Basal | 0,826619696 | 1,209746156 | 0,703099511 |
| FAAP20 | 0,025513394 | 0,311411596 | 0,462 | 0,373 | 1 | Basal | 1,240921278 | 0,805852892 | 1,238605898 |
| SQOR | 0,025545646 | 0,280684866 | 0,369 | 0,247 | 1 | Basal | 1,214771415 | 0,823200141 | 1,493927126 |
| ZNF280D | 0,025751398 | -0,345051512 | 0,169 | 0,307 | 1 | Basal | 0,787279866 | 1,270196335 | 0,550488599 |
| COPE | 0,025831455 | 0,316877149 | 0,585 | 0,547 | 1 | Basal | 1,245631341 | 0,802805748 | 1,069469835 |
| RASSF9 | 0,026108282 | -0,323946278 | 0,215 | 0,353 | 1 | Basal | 0,798881664 | 1,251749846 | 0,609065156 |

| Supplementary Table 4 - clusermarker | | | | | | | | | |
|---|---|---|---|---|---|---|---|---|---|
| | **p_val** | **avg_log2FC** | **pct.1** | **pct.2** | **p_val_adj** | **celltype** | **Foldchange_UP** | **Foldchange_DOWN** | **Ratio_pct1_pct2** |
| OCIAD1 | 0,026178354 | -0,314518276 | 0,446 | 0,633 | 1 | Basal | 0,804119446 | 1,24359634 | 0,704581359 |
| FHL1 | 0,026204288 | -0,375943502 | 0,077 | 0,193 | 1 | Basal | 0,770601286 | 1,297687946 | 0,398963731 |
| GCHFR | 0,02620788 | 0,684875581 | 0,477 | 0,427 | 1 | Basal | 1,607563338 | 0,622059471 | 1,117096019 |
| TRMT112 | 0,026212338 | 0,394803098 | 0,692 | 0,627 | 1 | Basal | 1,314763306 | 0,760593177 | 1,103668262 |
| KCNMB1 | 0,026230785 | -0,607718156 | 0,108 | 0,24 | 1 | Basal | 0,656233816 | 1,523847103 | 0,45 |
| GABRP | 0,0263469 | 0,516724796 | 0,569 | 0,467 | 1 | Basal | 1,430703577 | 0,698956804 | 1,218415418 |
| PGD | 0,026622667 | 0,534320607 | 0,538 | 0,46 | 1 | Basal | 1,44825998 | 0,690483762 | 1,169565217 |
| RHOC | 0,02672245 | 0,379252969 | 0,8 | 0,7 | 1 | Basal | 1,300668192 | 0,768835592 | 1,142857143 |
| GCFC2 | 0,026810626 | 0,256077549 | 0,2 | 0,1 | 1 | Basal | 1,194227379 | 0,837361476 | 2 |
| KRT17 | 0,026831129 | -0,326144408 | 0,738 | 0,847 | 1 | Basal | 0,797665393 | 1,253658501 | 0,871310508 |
| ACY1 | 0,027290754 | 0,312740386 | 0,338 | 0,22 | 1 | Basal | 1,242064751 | 0,805111005 | 1,536363636 |
| LY75 | 0,027318779 | -0,402526464 | 0,138 | 0,28 | 1 | Basal | 0,756532275 | 1,321820673 | 0,492857143 |
| ARRDC3 | 0,027532644 | -0,276567776 | 0,154 | 0,307 | 1 | Basal | 0,825552703 | 1,211309704 | 0,501628664 |
| CCAR1 | 0,027582129 | -0,303530098 | 0,385 | 0,547 | 1 | Basal | 0,810267344 | 1,234160561 | 0,703839122 |
| ZNF703 | 0,027659283 | -0,326192455 | 0,277 | 0,44 | 1 | Basal | 0,797638828 | 1,253700252 | 0,629545455 |
| ADM | 0,027691908 | -0,606440934 | 0,354 | 0,487 | 1 | Basal | 0,656815039 | 1,522498635 | 0,726899384 |
| NUCB1 | 0,027947152 | 0,412788195 | 0,6 | 0,547 | 1 | Basal | 1,331256154 | 0,751170236 | 1,096892139 |
| UBR7 | 0,028155583 | -0,30372798 | 0,062 | 0,167 | 1 | Basal | 0,810156214 | 1,234329852 | 0,371257485 |
| FBP1 | 0,028192124 | 0,465054732 | 0,338 | 0,227 | 1 | Basal | 1,38036972 | 0,724443593 | 1,488986784 |
| SPSB3 | 0,02838377 | 0,518726416 | 0,477 | 0,4 | 1 | Basal | 1,432689938 | 0,697987732 | 1,1925 |
| PTRHD1 | 0,028429961 | 0,366737792 | 0,538 | 0,46 | 1 | Basal | 1,289433877 | 0,775534146 | 1,169565217 |
| PPP2R5E | 0,028790536 | 0,314159695 | 0,4 | 0,28 | 1 | Basal | 1,243287284 | 0,804319334 | 1,428571429 |
| CUL1 | 0,028923672 | -0,250814345 | 0,2 | 0,353 | 1 | Basal | 0,840421896 | 1,189878565 | 0,566572238 |
| ARMC10 | 0,028983863 | 0,387283086 | 0,415 | 0,327 | 1 | Basal | 1,307927965 | 0,764568101 | 1,26911315 |
| SFN | 0,028998083 | 0,477833553 | 0,677 | 0,593 | 1 | Basal | 1,392650798 | 0,718055094 | 1,141652614 |
| MYH11 | 0,029092575 | -1,127027129 | 0,169 | 0,3 | 1 | Basal | 0,457858233 | 2,184082161 | 0,563333333 |
| C1QBP | 0,029096102 | 0,395767726 | 0,631 | 0,6 | 1 | Basal | 1,31564269 | 0,760084792 | 1,051666667 |
| NEXN | 0,029122309 | -0,612519154 | 0,077 | 0,193 | 1 | Basal | 0,654053632 | 1,528926606 | 0,398963731 |
| TAX1BP3 | 0,029151815 | 0,438879651 | 0,523 | 0,447 | 1 | Basal | 1,355551243 | 0,737707265 | 1,170022371 |
| SLC16A3 | 0,029340024 | 0,564416153 | 0,385 | 0,293 | 1 | Basal | 1,478788934 | 0,676229026 | 1,313993174 |
| ITPR1 | 0,029634562 | -0,607271722 | 0,185 | 0,32 | 1 | Basal | 0,656436915 | 1,523375631 | 0,578125 |

| Supplementary Table 4 - clusermarker | | | | | | | | | |
|---|---|---|---|---|---|---|---|---|---|
| | **p_val** | **avg_log2FC** | **pct.1** | **pct.2** | **p_val_adj** | **celltype** | **Foldchange_UP** | **Foldchange_DOWN** | **Ratio_pct1_pct2** |
| GOLM2 | 0,029641656 | 0,334169782 | 0,523 | 0,44 | 1 | Basal | 1,260651741 | 0,793240486 | 1,188636364 |
| UQCRQ | 0,029654394 | 0,470118551 | 0,723 | 0,7 | 1 | Basal | 1,385223292 | 0,721905274 | 1,032857143 |
| SACM1L | 0,0296644 | -0,298940559 | 0,108 | 0,233 | 1 | Basal | 0,812849092 | 1,230240656 | 0,463519313 |
| TMUB1 | 0,030096227 | 0,295710369 | 0,385 | 0,3 | 1 | Basal | 1,227489232 | 0,814671097 | 1,283333333 |
| BPIFA1 | 0,030353155 | -0,473043786 | 0,062 | 0,173 | 1 | Basal | 0,720443009 | 1,388034845 | 0,358381503 |
| RPL7 | 0,030426482 | -0,282128494 | 0,908 | 0,92 | 1 | Basal | 0,82237682 | 1,215987581 | 0,986956522 |
| SLC25A1 | 0,030427037 | 0,409677701 | 0,385 | 0,293 | 1 | Basal | 1,328389018 | 0,752791529 | 1,313993174 |
| APOL3 | 0,030470557 | -0,302767969 | 0,015 | 0,1 | 1 | Basal | 0,810695495 | 1,233508767 | 0,15 |
| AL627171.2 | 0,030506177 | -0,410493926 | 0,169 | 0,32 | 1 | Basal | 0,752365747 | 1,329140785 | 0,528125 |
| COA5 | 0,030529067 | 0,651764846 | 0,323 | 0,213 | 1 | Basal | 1,571088931 | 0,636501206 | 1,516431925 |
| CREBBP | 0,030561738 | -0,431203035 | 0,323 | 0,467 | 1 | Basal | 0,741643086 | 1,348357477 | 0,691648822 |
| FGD5-AS1 | 0,030695426 | -0,37342772 | 0,277 | 0,447 | 1 | Basal | 0,771946238 | 1,295427001 | 0,619686801 |
| SRI | 0,03076185 | 0,385855201 | 0,708 | 0,653 | 1 | Basal | 1,306634105 | 0,765325194 | 1,084226646 |
| FUOM | 0,03081776 | 0,353346587 | 0,231 | 0,133 | 1 | Basal | 1,277520629 | 0,782766225 | 1,736842105 |
| NFE2L1 | 0,030818075 | 0,299245303 | 0,369 | 0,253 | 1 | Basal | 1,23050055 | 0,81267741 | 1,458498024 |
| CDK2AP1 | 0,030858704 | -0,3733757 | 0,385 | 0,56 | 1 | Basal | 0,771974073 | 1,295380292 | 0,6875 |
| SF3B2 | 0,030944098 | 0,313471723 | 0,646 | 0,573 | 1 | Basal | 1,242694543 | 0,804702978 | 1,127399651 |
| FKBP14 | 0,030982442 | -0,331401814 | 0,108 | 0,233 | 1 | Basal | 0,794763865 | 1,258235362 | 0,463519313 |
| CAPN1 | 0,031050919 | 0,382313975 | 0,615 | 0,593 | 1 | Basal | 1,303430785 | 0,767206062 | 1,037099494 |
| LSM4 | 0,031072096 | 0,399450833 | 0,492 | 0,42 | 1 | Basal | 1,319005731 | 0,75814682 | 1,171428571 |
| PHF20 | 0,031138299 | -0,271799036 | 0,2 | 0,353 | 1 | Basal | 0,828286031 | 1,207312404 | 0,566572238 |
| PYCARD | 0,031277682 | 0,481891463 | 0,492 | 0,42 | 1 | Basal | 1,396573461 | 0,716038238 | 1,171428571 |
| MAP4K3 | 0,031692061 | 0,364825668 | 0,354 | 0,24 | 1 | Basal | 1,287726014 | 0,776562707 | 1,475 |
| RHOG | 0,031692061 | 0,291699155 | 0,354 | 0,24 | 1 | Basal | 1,224081109 | 0,816939329 | 1,475 |
| MCRIP2 | 0,031731809 | 0,253565146 | 0,354 | 0,247 | 1 | Basal | 1,192149483 | 0,838820982 | 1,433198381 |
| PIN4 | 0,031852417 | -0,338706483 | 0,354 | 0,5 | 1 | Basal | 0,790749979 | 1,264622228 | 0,708 |
| SUMO1 | 0,031902824 | 0,258135983 | 0,631 | 0,573 | 1 | Basal | 1,195932516 | 0,836167582 | 1,10122164 |
| SERTAD2 | 0,031930099 | -0,251980457 | 0,231 | 0,393 | 1 | Basal | 0,839742868 | 1,190840718 | 0,58778626 |
| CITED2 | 0,03199949 | -0,78475615 | 0,492 | 0,593 | 1 | Basal | 0,580450059 | 1,7228011 | 0,829679595 |
| CSTA | 0,032355071 | 0,538930108 | 0,615 | 0,513 | 1 | Basal | 1,452894662 | 0,688281144 | 1,198830409 |
| IRF6 | 0,032397358 | -0,583154637 | 0,431 | 0,58 | 1 | Basal | 0,667502601 | 1,498121504 | 0,743103448 |

| Supplementary Table 4 - clusermarker | | | | | | | | | |
|---|---|---|---|---|---|---|---|---|---|
| | **p_val** | **avg_log2FC** | **pct.1** | **pct.2** | **p_val_adj** | **celltype** | **Foldchange_UP** | **Foldchange_DOWN** | **Ratio_pct1_pct2** |
| SLK | 0,032424934 | 0,35945222 | 0,646 | 0,547 | 1 | Basal | 1,282938684 | 0,779460478 | 1,180987203 |
| UQCC3 | 0,03254725 | 0,464662643 | 0,4 | 0,3 | 1 | Basal | 1,37999462 | 0,724640506 | 1,333333333 |
| HES4 | 0,032578553 | 0,350441788 | 0,492 | 0,433 | 1 | Basal | 1,274950988 | 0,784343876 | 1,136258661 |
| PSMB6 | 0,032674873 | 0,294958081 | 0,662 | 0,613 | 1 | Basal | 1,22684933 | 0,815096015 | 1,079934747 |
| CRNDE | 0,03268017 | 0,471837923 | 0,569 | 0,48 | 1 | Basal | 1,386875154 | 0,721045436 | 1,185416667 |
| PREPL | 0,032693084 | -0,286264984 | 0,169 | 0,3 | 1 | Basal | 0,820022281 | 1,219479059 | 0,563333333 |
| KLF3 | 0,032859371 | 0,419407362 | 0,631 | 0,56 | 1 | Basal | 1,337378067 | 0,747731718 | 1,126785714 |
| C1orf122 | 0,032862483 | 0,632906649 | 0,462 | 0,38 | 1 | Basal | 1,550686071 | 0,644875851 | 1,215789474 |
| SNF8 | 0,032899133 | 0,417669215 | 0,446 | 0,36 | 1 | Basal | 1,335767775 | 0,748633122 | 1,238888889 |
| CRABP2 | 0,033199622 | -0,313219215 | 0,169 | 0,327 | 1 | Basal | 0,804843834 | 1,242477059 | 0,516819572 |
| CDC42EP5 | 0,033409068 | 0,661547721 | 0,569 | 0,527 | 1 | Basal | 1,581778644 | 0,63219971 | 1,079696395 |
| ZNF331 | 0,033591346 | -0,291392704 | 0,123 | 0,24 | 1 | Basal | 0,817112879 | 1,223821122 | 0,5125 |
| ID4 | 0,033708585 | -0,482515189 | 0,508 | 0,673 | 1 | Basal | 0,715728737 | 1,397177378 | 0,754829123 |
| PDLIM7 | 0,033723784 | -0,476560114 | 0,138 | 0,273 | 1 | Basal | 0,718689187 | 1,391422075 | 0,505494505 |
| BCL7C | 0,033742619 | 0,569035592 | 0,462 | 0,367 | 1 | Basal | 1,483531533 | 0,674067236 | 1,258855586 |
| MZT2B | 0,033766599 | 0,362985613 | 0,692 | 0,673 | 1 | Basal | 1,286084658 | 0,77755379 | 1,028231798 |
| SLMAP | 0,034428227 | -0,29149855 | 0,277 | 0,447 | 1 | Basal | 0,817052932 | 1,223910914 | 0,619686801 |
| FOXD1 | 0,034488281 | 0,271488372 | 0,292 | 0,187 | 1 | Basal | 1,207052455 | 0,82846441 | 1,561497326 |
| GSTM2 | 0,034646824 | -0,26883434 | 0,108 | 0,233 | 1 | Basal | 0,829989885 | 1,20483396 | 0,463519313 |
| SLC25A12 | 0,034818684 | -0,335667917 | 0,138 | 0,273 | 1 | Basal | 0,79241719 | 1,261961518 | 0,505494505 |
| DDX5 | 0,034924703 | -0,338535855 | 0,892 | 0,887 | 1 | Basal | 0,790843506 | 1,26447267 | 1,005636979 |
| RPS3 | 0,035154761 | -0,315882981 | 1 | 0,993 | 1 | Basal | 0,803359155 | 1,244773266 | 1,007049345 |
| STN1 | 0,035259066 | 0,431098941 | 0,369 | 0,287 | 1 | Basal | 1,348260193 | 0,741696599 | 1,285714286 |
| HSPA13 | 0,0352797 | -0,250489193 | 0,092 | 0,207 | 1 | Basal | 0,84061133 | 1,189610423 | 0,444444444 |
| AP3S1 | 0,035324575 | 0,379467707 | 0,369 | 0,26 | 1 | Basal | 1,300861804 | 0,768721164 | 1,419230769 |
| PLAAT4 | 0,035473035 | 0,906655851 | 0,662 | 0,633 | 1 | Basal | 1,874694939 | 0,53342012 | 1,045813586 |
| SLC44A4 | 0,035514266 | 0,620939912 | 0,462 | 0,4 | 1 | Basal | 1,537876778 | 0,650247155 | 1,155 |
| DPP7 | 0,035753332 | 0,395500582 | 0,446 | 0,353 | 1 | Basal | 1,315399094 | 0,76022555 | 1,263456091 |
| HDHD5 | 0,035797478 | 0,375870057 | 0,215 | 0,12 | 1 | Basal | 1,297621885 | 0,770640517 | 1,791666667 |
| MGP | 0,036137162 | -1,164076033 | 0,077 | 0,193 | 1 | Basal | 0,446249967 | 2,240896525 | 0,398963731 |
| CMIP | 0,036282172 | 0,376936363 | 0,354 | 0,26 | 1 | Basal | 1,298581321 | 0,770071141 | 1,361538462 |

| Supplementary Table 4 - clusermarker | | | | | | | | | |
|---|---|---|---|---|---|---|---|---|---|
| | **p_val** | **avg_log2FC** | **pct.1** | **pct.2** | **p_val_adj** | **celltype** | **Foldchange_UP** | **Foldchange_DOWN** | **Ratio_pct1_pct2** |
| BZW2 | 0,036289677 | -0,348728814 | 0,308 | 0,453 | 1 | Basal | 0,785275714 | 1,273438083 | 0,6799117 |
| SMOC2 | 0,036562571 | -0,476193471 | 0,092 | 0,207 | 1 | Basal | 0,718871857 | 1,391068507 | 0,444444444 |
| UNC5B-AS1 | 0,036802334 | 0,299462433 | 0,4 | 0,28 | 1 | Basal | 1,230685759 | 0,812555108 | 1,428571429 |
| EMC10 | 0,03697946 | 0,317728614 | 0,462 | 0,393 | 1 | Basal | 1,246366718 | 0,802332079 | 1,175572519 |
| MTREX | 0,036992618 | -0,292998767 | 0,154 | 0,287 | 1 | Basal | 0,816203744 | 1,225184285 | 0,536585366 |
| FGFR1OP2 | 0,037174703 | 0,427939888 | 0,385 | 0,293 | 1 | Basal | 1,345311153 | 0,743322463 | 1,313993174 |
| CNN1 | 0,037187217 | -1,059613256 | 0,154 | 0,273 | 1 | Basal | 0,479760652 | 2,084372687 | 0,564102564 |
| LRRC8D | 0,037246732 | 0,284382467 | 0,323 | 0,207 | 1 | Basal | 1,217888845 | 0,821092995 | 1,560386473 |
| RDH14 | 0,037278514 | 0,36474045 | 0,308 | 0,193 | 1 | Basal | 1,287649952 | 0,776608579 | 1,595854922 |
| HIGD2A | 0,037596093 | 0,341287409 | 0,8 | 0,733 | 1 | Basal | 1,266886614 | 0,789336622 | 1,091405184 |
| SLC41A3 | 0,037669283 | -0,2588832 | 0,169 | 0,32 | 1 | Basal | 0,835734617 | 1,196552087 | 0,528125 |
| BAD | 0,037903999 | 0,354959962 | 0,492 | 0,427 | 1 | Basal | 1,278950087 | 0,781891342 | 1,152224824 |
| ZFP36L2 | 0,037937992 | -0,359524141 | 0,723 | 0,82 | 1 | Basal | 0,779421622 | 1,283002642 | 0,881707317 |
| CCNQ | 0,038014015 | 0,253885782 | 0,292 | 0,187 | 1 | Basal | 1,192414465 | 0,838634577 | 1,561497326 |
| GALNT15 | 0,038032787 | -0,374276547 | 0,062 | 0,167 | 1 | Basal | 0,771492188 | 1,296189405 | 0,371257485 |
| RFC2 | 0,038117203 | -0,260793714 | 0,108 | 0,227 | 1 | Basal | 0,834628614 | 1,198137691 | 0,475770925 |
| DCN | 0,038219718 | -0,56403409 | 0,031 | 0,12 | 1 | Basal | 0,676408132 | 1,478397364 | 0,258333333 |
| ARHGEF5 | 0,038298379 | 0,311694129 | 0,338 | 0,233 | 1 | Basal | 1,24116432 | 0,805695092 | 1,450643777 |
| NUP50 | 0,038310612 | -0,329193377 | 0,292 | 0,413 | 1 | Basal | 0,795981399 | 1,256310764 | 0,707021792 |
| PRPF8 | 0,038348561 | 0,395555212 | 0,554 | 0,487 | 1 | Basal | 1,315448905 | 0,760196763 | 1,137577002 |
| CENPC | 0,038361244 | -0,250230235 | 0,185 | 0,327 | 1 | Basal | 0,84076223 | 1,189396912 | 0,565749235 |
| KDSR | 0,03839558 | -0,381728516 | 0,292 | 0,453 | 1 | Basal | 0,767517464 | 1,302901949 | 0,644591611 |
| EGR3 | 0,038400312 | -0,267213442 | 0,231 | 0,373 | 1 | Basal | 0,83092292 | 1,203481064 | 0,619302949 |
| SH3BGRL | 0,038632046 | -0,306387362 | 0,462 | 0,633 | 1 | Basal | 0,808664194 | 1,236607244 | 0,72985782 |
| RLIM | 0,038660469 | -0,29953208 | 0,169 | 0,3 | 1 | Basal | 0,812515883 | 1,230745172 | 0,563333333 |
| RALY | 0,038665408 | 0,302841474 | 0,569 | 0,54 | 1 | Basal | 1,233571615 | 0,810654191 | 1,053703704 |
| GFER | 0,038671311 | 0,391369407 | 0,246 | 0,153 | 1 | Basal | 1,31163782 | 0,762405585 | 1,607843137 |
| CETN2 | 0,03868588 | -0,410503134 | 0,2 | 0,333 | 1 | Basal | 0,752360945 | 1,329149269 | 0,600600601 |
| COMMD1 | 0,039009969 | -0,576379865 | 0,338 | 0,5 | 1 | Basal | 0,670644507 | 1,491102946 | 0,676 |
| INPP5K | 0,039430697 | 0,294810239 | 0,262 | 0,16 | 1 | Basal | 1,226723613 | 0,815179548 | 1,6375 |
| PPM1G | 0,039534593 | 0,476960828 | 0,523 | 0,493 | 1 | Basal | 1,391808601 | 0,718489596 | 1,060851927 |

| Supplementary Table 4 - clusermarker | | | | | | | | | |
|---|---|---|---|---|---|---|---|---|---|
| | **p_val** | **avg_log2FC** | **pct.1** | **pct.2** | **p_val_adj** | **celltype** | **Foldchange_UP** | **Foldchange_DOWN** | **Ratio_pct1_pct2** |
| ANKRD9 | 0,039541862 | 0,34817849 | 0,323 | 0,233 | 1 | Basal | 1,272952415 | 0,785575319 | 1,386266094 |
| C17orf49 | 0,039852436 | 0,420990816 | 0,446 | 0,4 | 1 | Basal | 1,338846734 | 0,746911484 | 1,115 |
| HLA-E | 0,039925299 | 0,320150827 | 0,831 | 0,78 | 1 | Basal | 1,248461063 | 0,800986134 | 1,065384615 |
| CHCHD10 | 0,040056474 | 0,389975526 | 0,708 | 0,653 | 1 | Basal | 1,310371174 | 0,76314255 | 1,084226646 |
| HSD17B4 | 0,040082903 | -0,29206702 | 0,231 | 0,38 | 1 | Basal | 0,816731049 | 1,22439327 | 0,607894737 |
| DYM | 0,040184557 | 0,426713569 | 0,246 | 0,147 | 1 | Basal | 1,344168098 | 0,74395457 | 1,673469388 |
| ZNF503 | 0,040311622 | -0,321554279 | 0,277 | 0,427 | 1 | Basal | 0,800207314 | 1,249676155 | 0,648711944 |
| MFSD10 | 0,040385396 | 0,314193663 | 0,508 | 0,413 | 1 | Basal | 1,243316556 | 0,804300397 | 1,230024213 |
| PDCD10 | 0,040437654 | -0,275847433 | 0,354 | 0,533 | 1 | Basal | 0,825965007 | 1,210705043 | 0,664165103 |
| IGHA1 | 0,040510823 | 0,276823514 | 0,123 | 0,047 | 1 | Basal | 1,211524445 | 0,825406375 | 2,617021277 |
| H2AJ | 0,040655058 | 0,450263255 | 0,569 | 0,547 | 1 | Basal | 1,366289547 | 0,731909281 | 1,040219378 |
| TOMM7 | 0,040656289 | 0,297286938 | 0,908 | 0,847 | 1 | Basal | 1,228831359 | 0,813781316 | 1,07201889 |
| PIK3C2A | 0,040770989 | -0,329981036 | 0,2 | 0,347 | 1 | Basal | 0,795546941 | 1,256996851 | 0,576368876 |
| TMBIM1 | 0,041009899 | 0,351972818 | 0,6 | 0,547 | 1 | Basal | 1,276304722 | 0,783511949 | 1,096892139 |
| RPLP2 | 0,041016073 | 0,25041399 | 1 | 0,947 | 1 | Basal | 1,189548414 | 0,84065515 | 1,055966209 |
| LLGL2 | 0,041027103 | 0,45549982 | 0,462 | 0,393 | 1 | Basal | 1,371257793 | 0,729257478 | 1,175572519 |
| TMEM218 | 0,041243282 | 0,310166165 | 0,246 | 0,147 | 1 | Basal | 1,239850494 | 0,806548858 | 1,673469388 |
| LARS1 | 0,041612164 | -0,280378179 | 0,354 | 0,513 | 1 | Basal | 0,823375154 | 1,214513208 | 0,69005848 |
| ALDH9A1 | 0,04161758 | -0,371411728 | 0,246 | 0,407 | 1 | Basal | 0,773025694 | 1,293618063 | 0,604422604 |
| IMMP2L | 0,041757855 | -0,260369605 | 0,169 | 0,32 | 1 | Basal | 0,834874005 | 1,197785527 | 0,528125 |
| TCEAL4 | 0,041776576 | -0,452182015 | 0,338 | 0,513 | 1 | Basal | 0,730936501 | 1,368107898 | 0,658869396 |
| RBM3 | 0,041931426 | -0,424875192 | 0,692 | 0,827 | 1 | Basal | 0,744903171 | 1,342456361 | 0,836759371 |
| ATP5PF | 0,042022148 | 0,39187499 | 0,754 | 0,707 | 1 | Basal | 1,312097556 | 0,762138452 | 1,066478076 |
| ACBD6 | 0,042371538 | -0,320954043 | 0,169 | 0,32 | 1 | Basal | 0,800540311 | 1,249156333 | 0,528125 |
| YWHAH | 0,042463124 | 0,537851259 | 0,462 | 0,38 | 1 | Basal | 1,451808591 | 0,688796034 | 1,215789474 |
| NAA20 | 0,043503909 | 0,317013126 | 0,508 | 0,44 | 1 | Basal | 1,24574875 | 0,802730085 | 1,154545455 |
| NUDT14 | 0,043523222 | 0,380468903 | 0,446 | 0,393 | 1 | Basal | 1,301764885 | 0,768187874 | 1,134860051 |
| SERPINB4 | 0,043544509 | 0,521060086 | 0,354 | 0,253 | 1 | Basal | 1,435009299 | 0,696859596 | 1,399209486 |
| SEC13 | 0,043549346 | 0,277137204 | 0,523 | 0,447 | 1 | Basal | 1,211787899 | 0,825226924 | 1,170022371 |
| AL138963.4 | 0,043563764 | 0,641905093 | 0,492 | 0,427 | 1 | Basal | 1,560388307 | 0,64086612 | 1,152224824 |
| PFKP | 0,04357666 | 0,274347938 | 0,369 | 0,267 | 1 | Basal | 1,209447326 | 0,826823937 | 1,382022472 |

| Supplementary Table 4 - clusermarker | | | | | | | | | |
|---|---|---|---|---|---|---|---|---|---|
| | **p_val** | **avg_log2FC** | **pct.1** | **pct.2** | **p_val_adj** | **celltype** | **Foldchange_UP** | **Foldchange_DOWN** | **Ratio_pct1_pct2** |
| BBIP1 | 0,043603914 | -0,294429073 | 0,108 | 0,227 | 1 | Basal | 0,81539495 | 1,22639955 | 0,475770925 |
| ATP6AP1 | 0,043768488 | 0,301095862 | 0,477 | 0,36 | 1 | Basal | 1,232079938 | 0,811635649 | 1,325 |
| PITX1 | 0,043793308 | 0,387526205 | 0,523 | 0,473 | 1 | Basal | 1,308148392 | 0,764439269 | 1,105708245 |
| NOL3 | 0,043902171 | 0,314480826 | 0,385 | 0,273 | 1 | Basal | 1,243564059 | 0,80414032 | 1,41025641 |
| ZDHHC4 | 0,043945389 | 0,448890063 | 0,338 | 0,24 | 1 | Basal | 1,364989698 | 0,732606262 | 1,408333333 |
| AHCTF1 | 0,04409731 | -0,256408118 | 0,2 | 0,327 | 1 | Basal | 0,837169631 | 1,194501047 | 0,611620795 |
| RALGAPA2 | 0,044169682 | 0,411644901 | 0,446 | 0,38 | 1 | Basal | 1,33020159 | 0,751765753 | 1,173684211 |
| DTNA | 0,044717575 | -0,26627814 | 0,031 | 0,113 | 1 | Basal | 0,831461784 | 1,202701098 | 0,274336283 |
| LTF | 0,044828791 | -1,445419309 | 0,108 | 0,22 | 1 | Basal | 0,367185423 | 2,72341966 | 0,490909091 |
| OBI1 | 0,044845287 | -0,308121733 | 0,062 | 0,16 | 1 | Basal | 0,807692622 | 1,238094756 | 0,3875 |
| PRSS22 | 0,044940283 | 0,360687026 | 0,492 | 0,413 | 1 | Basal | 1,284037223 | 0,778793622 | 1,191283293 |
| SMDT1 | 0,045129331 | 0,274175722 | 0,677 | 0,567 | 1 | Basal | 1,209302961 | 0,826922642 | 1,194003527 |
| MPST | 0,045186593 | 0,290102193 | 0,508 | 0,433 | 1 | Basal | 1,222726886 | 0,817844125 | 1,173210162 |
| MTPN | 0,045231648 | 0,269084442 | 0,523 | 0,493 | 1 | Basal | 1,205042845 | 0,829846013 | 1,060851927 |
| CSRP1 | 0,045326787 | -0,789708078 | 0,415 | 0,573 | 1 | Basal | 0,578461129 | 1,728724628 | 0,72425829 |
| ACTG2 | 0,045454238 | -0,865876308 | 0,154 | 0,28 | 1 | Basal | 0,548713011 | 1,822446305 | 0,55 |
| PRR13 | 0,045782273 | 0,347359539 | 0,538 | 0,487 | 1 | Basal | 1,272230024 | 0,786021381 | 1,104722793 |
| BAX | 0,045880479 | 0,465759628 | 0,4 | 0,333 | 1 | Basal | 1,381044329 | 0,724089719 | 1,201201201 |
| CRYM | 0,045965779 | 0,268951677 | 0,262 | 0,153 | 1 | Basal | 1,204931955 | 0,829922384 | 1,712418301 |
| ARHGDIA | 0,046651504 | 0,566672736 | 0,492 | 0,427 | 1 | Basal | 1,481103783 | 0,675172133 | 1,152224824 |
| MT-ND4 | 0,04669044 | 0,308130479 | 0,985 | 0,88 | 1 | Basal | 1,238102261 | 0,807687726 | 1,119318182 |
| ERH | 0,046738696 | 0,297348751 | 0,662 | 0,64 | 1 | Basal | 1,22888401 | 0,81374645 | 1,034375 |
| TOMM6 | 0,04683718 | 0,414777727 | 0,677 | 0,607 | 1 | Basal | 1,333093275 | 0,750135057 | 1,115321252 |
| SLCO3A1 | 0,047157604 | 0,661798206 | 0,231 | 0,133 | 1 | Basal | 1,5820533 | 0,632089955 | 1,736842105 |
| PPP2R1B | 0,047208322 | 0,440535792 | 0,169 | 0,087 | 1 | Basal | 1,35710824 | 0,736860901 | 1,942528736 |
| PPIB | 0,047300964 | 0,292361516 | 0,862 | 0,8 | 1 | Basal | 1,224643229 | 0,816564348 | 1,0775 |
| TRPC1 | 0,047360875 | -0,333007798 | 0,031 | 0,113 | 1 | Basal | 0,79387964 | 1,259636789 | 0,274336283 |
| NFIL3 | 0,047492982 | -0,452050913 | 0,123 | 0,233 | 1 | Basal | 0,731002927 | 1,367983579 | 0,527896996 |
| PMVK | 0,047505991 | 0,622834085 | 0,415 | 0,347 | 1 | Basal | 1,539897245 | 0,649393979 | 1,195965418 |
| GABARAPL2 | 0,047589404 | 0,418807482 | 0,662 | 0,66 | 1 | Basal | 1,336822094 | 0,748042694 | 1,003030303 |
| REEP3 | 0,047793279 | 0,250664182 | 0,615 | 0,553 | 1 | Basal | 1,189754723 | 0,840509376 | 1,112115732 |

| Supplementary Table 4 - clusermarker | | | | | | | | | |
|---|---|---|---|---|---|---|---|---|---|
| | **p_val** | **avg_log2FC** | **pct.1** | **pct.2** | **p_val_adj** | **celltype** | **Foldchange_UP** | **Foldchange_DOWN** | **Ratio_pct1_pct2** |
| SYBU | 0,047871681 | 0,532963984 | 0,323 | 0,227 | 1 | Basal | 1,446898765 | 0,691133357 | 1,422907489 |
| MYO6 | 0,048403124 | 0,418076214 | 0,462 | 0,393 | 1 | Basal | 1,336144662 | 0,748421955 | 1,175572519 |
| KCMF1 | 0,04849247 | 0,602211106 | 0,446 | 0,373 | 1 | Basal | 1,518041368 | 0,658743576 | 1,195710456 |
| USP48 | 0,048574069 | 0,262919427 | 0,431 | 0,313 | 1 | Basal | 1,199904367 | 0,83339975 | 1,376996805 |
| PYURF | 0,048616918 | 0,433134846 | 0,554 | 0,493 | 1 | Basal | 1,350164176 | 0,740650669 | 1,123732252 |
| MRPS18B | 0,048700236 | -0,285339308 | 0,354 | 0,527 | 1 | Basal | 0,820548601 | 1,218696856 | 0,671726755 |
| RBX1 | 0,048854657 | 0,281396736 | 0,646 | 0,627 | 1 | Basal | 1,215370969 | 0,822794049 | 1,03030303 |
| EVPL | 0,04893351 | 0,422387564 | 0,277 | 0,173 | 1 | Basal | 1,340143569 | 0,746188709 | 1,601156069 |
| SGSM2 | 0,049013419 | 0,378455946 | 0,308 | 0,207 | 1 | Basal | 1,29994983 | 0,769260457 | 1,487922705 |
| USP7 | 0,049090813 | -0,431371473 | 0,246 | 0,387 | 1 | Basal | 0,741556502 | 1,34851491 | 0,635658915 |
| SRD5A3 | 0,049258 | 0,362999235 | 0,415 | 0,333 | 1 | Basal | 1,286096802 | 0,777546448 | 1,246246246 |
| ANKRD39 | 0,049692578 | 0,317358884 | 0,231 | 0,133 | 1 | Basal | 1,246047343 | 0,802537725 | 1,736842105 |
| RNF130 | 0,049700526 | 0,461997594 | 0,523 | 0,467 | 1 | Basal | 1,377447749 | 0,725980351 | 1,119914347 |
| SEC61B | 0,049997628 | 0,398226608 | 0,708 | 0,673 | 1 | Basal | 1,317886939 | 0,758790432 | 1,052005944 |
| MBTPS1 | 0,05003246 | 0,268628393 | 0,415 | 0,32 | 1 | Basal | 1,204661981 | 0,830108376 | 1,296875 |
| ACADSB | 0,050076465 | -0,290897007 | 0,108 | 0,213 | 1 | Basal | 0,81739368 | 1,2234007 | 0,507042254 |
| ULK3 | 0,050200142 | 0,259478984 | 0,231 | 0,14 | 1 | Basal | 1,197046324 | 0,835389558 | 1,65 |
| PRMT2 | 0,050430068 | 0,577885633 | 0,523 | 0,447 | 1 | Basal | 1,492660051 | 0,669944908 | 1,170022371 |
| ISOC2 | 0,05078204 | 0,357185816 | 0,462 | 0,4 | 1 | Basal | 1,280924831 | 0,780685935 | 1,155 |
| PRMT6 | 0,051064173 | 0,318270972 | 0,154 | 0,073 | 1 | Basal | 1,246835357 | 0,802030512 | 2,109589041 |
| ZNHIT1 | 0,051234801 | 0,254048103 | 0,585 | 0,493 | 1 | Basal | 1,192548634 | 0,838540225 | 1,186612576 |
| EPHB3 | 0,051263897 | -0,344699651 | 0,138 | 0,253 | 1 | Basal | 0,7874719 | 1,269886583 | 0,545454545 |
| EHMT1 | 0,05140669 | 0,309275171 | 0,385 | 0,3 | 1 | Basal | 1,239085011 | 0,807047128 | 1,283333333 |
| NDUFA2 | 0,051664634 | 0,282328931 | 0,662 | 0,627 | 1 | Basal | 1,216156533 | 0,822262573 | 1,055821372 |
| MICAL3 | 0,051667965 | -0,26815849 | 0,092 | 0,193 | 1 | Basal | 0,830378796 | 1,204269671 | 0,476683938 |
| LEF1 | 0,051814211 | -0,400585034 | 0,062 | 0,153 | 1 | Basal | 0,757551023 | 1,3200431 | 0,405228758 |
| NRDE2 | 0,051814211 | -0,360580235 | 0,062 | 0,153 | 1 | Basal | 0,778851272 | 1,28394218 | 0,405228758 |
| CALML5 | 0,051843537 | 0,529123227 | 0,108 | 0,04 | 1 | Basal | 1,443051939 | 0,69297575 | 2,7 |
| LGALS3 | 0,051968969 | 0,64710854 | 0,831 | 0,907 | 1 | Basal | 1,566026406 | 0,638558836 | 0,916207277 |
| DDR2 | 0,052120206 | -0,332461344 | 0,046 | 0,133 | 1 | Basal | 0,794180397 | 1,259159763 | 0,345864662 |
| INTS6 | 0,052147108 | -0,373772735 | 0,415 | 0,567 | 1 | Basal | 0,771761652 | 1,295736835 | 0,731922399 |

| Supplementary Table 4 - clusermarker | | | | | | | | | |
|---|---|---|---|---|---|---|---|---|---|
| | **p_val** | **avg_log2FC** | **pct.1** | **pct.2** | **p_val_adj** | **celltype** | **Foldchange_UP** | **Foldchange_DOWN** | **Ratio_pct1_pct2** |
| NOL8 | 0,05218835 | -0,331356523 | 0,169 | 0,287 | 1 | Basal | 0,794788816 | 1,258195862 | 0,588850174 |
| LDLR | 0,052224091 | -0,334953417 | 0,338 | 0,5 | 1 | Basal | 0,792809735 | 1,261336681 | 0,676 |
| MACC1 | 0,052559887 | 0,421483367 | 0,415 | 0,333 | 1 | Basal | 1,339303909 | 0,746656523 | 1,246246246 |
| CEP78 | 0,052609336 | -0,263457784 | 0,138 | 0,267 | 1 | Basal | 0,833088816 | 1,200352208 | 0,516853933 |
| MAP7D1 | 0,053288158 | 0,291461224 | 0,354 | 0,26 | 1 | Basal | 1,223879248 | 0,817074071 | 1,361538462 |
| COMT | 0,053339238 | 0,295805775 | 0,631 | 0,62 | 1 | Basal | 1,227570409 | 0,814617225 | 1,017741935 |
| ABRACL | 0,053744353 | 0,33426157 | 0,477 | 0,407 | 1 | Basal | 1,260731949 | 0,79319002 | 1,171990172 |
| LAMTOR1 | 0,053922314 | 0,371938365 | 0,554 | 0,547 | 1 | Basal | 1,294090367 | 0,772743562 | 1,012797075 |
| KDM5C | 0,054167223 | 0,362508028 | 0,323 | 0,233 | 1 | Basal | 1,285658988 | 0,777811231 | 1,386266094 |
| RPS20 | 0,054356878 | -0,326744112 | 0,769 | 0,82 | 1 | Basal | 0,797333885 | 1,254179734 | 0,937804878 |
| FER1L6 | 0,054662554 | 0,297383127 | 0,231 | 0,14 | 1 | Basal | 1,228913292 | 0,81372706 | 1,65 |
| CCNL1 | 0,054745964 | -0,254322443 | 0,523 | 0,68 | 1 | Basal | 0,838380785 | 1,192775428 | 0,769117647 |
| H1-2 | 0,055004615 | 0,607544223 | 0,277 | 0,187 | 1 | Basal | 1,523663398 | 0,656312937 | 1,481283422 |
| PRR15 | 0,055066008 | 0,333742863 | 0,338 | 0,253 | 1 | Basal | 1,260278748 | 0,793475254 | 1,335968379 |
| PSMB2 | 0,05548779 | 0,389190113 | 0,569 | 0,493 | 1 | Basal | 1,309657994 | 0,763558123 | 1,154158215 |
| COPS2 | 0,05555202 | -0,312058566 | 0,338 | 0,487 | 1 | Basal | 0,805491592 | 1,241477888 | 0,694045175 |
| LRP10 | 0,055736805 | 0,418107916 | 0,569 | 0,593 | 1 | Basal | 1,336174023 | 0,748405509 | 0,959527825 |
| GCNT3 | 0,05594912 | 0,736738343 | 0,323 | 0,253 | 1 | Basal | 1,666404156 | 0,600094519 | 1,276679842 |
| CCNB1IP1 | 0,056109306 | -0,262022904 | 0,2 | 0,333 | 1 | Basal | 0,833917804 | 1,199158951 | 0,600600601 |
| ENY2 | 0,056142695 | 0,321725445 | 0,631 | 0,513 | 1 | Basal | 1,249824429 | 0,800112381 | 1,230019493 |
| REC8 | 0,056711583 | -0,318449949 | 0,123 | 0,227 | 1 | Basal | 0,80193102 | 1,246990046 | 0,54185022 |
| NTRK2 | 0,056819075 | 0,328881941 | 0,308 | 0,18 | 1 | Basal | 1,256039592 | 0,796153247 | 1,711111111 |
| SMIM22 | 0,056826671 | 0,549374619 | 0,446 | 0,38 | 1 | Basal | 1,46345118 | 0,683316269 | 1,173684211 |
| TPM3 | 0,057067458 | 0,391046894 | 0,615 | 0,607 | 1 | Basal | 1,311344638 | 0,762576039 | 1,013179572 |
| CSNK1D | 0,05733134 | 0,42350195 | 0,462 | 0,387 | 1 | Basal | 1,341179141 | 0,745612551 | 1,19379845 |
| JUND | 0,057709243 | -0,32517474 | 0,6 | 0,733 | 1 | Basal | 0,798201702 | 1,252816171 | 0,818553888 |
| CS | 0,057834193 | -0,250770163 | 0,215 | 0,36 | 1 | Basal | 0,840447634 | 1,189842126 | 0,597222222 |
| MYO1B | 0,057913087 | 0,354822633 | 0,462 | 0,393 | 1 | Basal | 1,27882835 | 0,781965773 | 1,175572519 |
| GCLM | 0,058037037 | -0,393483455 | 0,292 | 0,46 | 1 | Basal | 0,761289215 | 1,313561234 | 0,634782609 |
| COG3 | 0,058169724 | 0,313637414 | 0,308 | 0,2 | 1 | Basal | 1,242837273 | 0,804610565 | 1,54 |
| BOD1L1 | 0,05832983 | -0,271763625 | 0,415 | 0,553 | 1 | Basal | 0,828306362 | 1,207282771 | 0,75045208 |

| Supplementary Table 4 - clusermarker | | | | | | | | | |
|---|---|---|---|---|---|---|---|---|---|
| | **p_val** | **avg_log2FC** | **pct.1** | **pct.2** | **p_val_adj** | **celltype** | **Foldchange_UP** | **Foldchange_DOWN** | **Ratio_pct1_pct2** |
| AOPEP | 0,058378907 | 0,532888881 | 0,538 | 0,473 | 1 | Basal | 1,446823445 | 0,691169336 | 1,137420719 |
| ATF4 | 0,058429793 | -0,354943151 | 0,538 | 0,653 | 1 | Basal | 0,781900453 | 1,278935185 | 0,82388974 |
| PUM1 | 0,058875839 | -0,495636295 | 0,446 | 0,607 | 1 | Basal | 0,709248798 | 1,409942467 | 0,73476112 |
| HCAR2 | 0,059326377 | -0,385914878 | 0,215 | 0,34 | 1 | Basal | 0,765293537 | 1,306688154 | 0,632352941 |
| ESYT2 | 0,059378957 | 0,250253024 | 0,385 | 0,287 | 1 | Basal | 1,1894157 | 0,840748949 | 1,341463415 |
| TM9SF2 | 0,059436184 | 0,272920519 | 0,708 | 0,667 | 1 | Basal | 1,208251276 | 0,827642412 | 1,061469265 |
| IL32 | 0,059564379 | 0,291502761 | 0,169 | 0,087 | 1 | Basal | 1,223914486 | 0,817050547 | 1,942528736 |
| ASAP2 | 0,059780228 | 0,353955075 | 0,385 | 0,313 | 1 | Basal | 1,278059564 | 0,782436146 | 1,230031949 |
| NRBP1 | 0,059792061 | 0,367286294 | 0,446 | 0,347 | 1 | Basal | 1,289924203 | 0,77523935 | 1,285302594 |
| CHADL | 0,060128698 | -0,286558426 | 0,046 | 0,133 | 1 | Basal | 0,819855507 | 1,219727125 | 0,345864662 |
| VCAN | 0,060238163 | -0,412751278 | 0,092 | 0,193 | 1 | Basal | 0,751189458 | 1,331222089 | 0,476683938 |
| S100A10 | 0,060342534 | 0,721018141 | 0,738 | 0,733 | 1 | Basal | 1,648344896 | 0,606669152 | 1,006821282 |
| SQSTM1 | 0,060449841 | 0,250977643 | 0,831 | 0,807 | 1 | Basal | 1,190013255 | 0,840326774 | 1,029739777 |
| UCHL3 | 0,060608813 | 0,269845727 | 0,415 | 0,327 | 1 | Basal | 1,205678893 | 0,829408233 | 1,26911315 |
| SMARCB1 | 0,060654223 | 0,360751034 | 0,431 | 0,367 | 1 | Basal | 1,284094194 | 0,77875907 | 1,174386921 |
| MRPS17 | 0,060714324 | 0,393797907 | 0,277 | 0,187 | 1 | Basal | 1,31384757 | 0,761123301 | 1,481283422 |
| ACTB | 0,06078887 | 0,393039693 | 1 | 0,987 | 1 | Basal | 1,313157254 | 0,761523418 | 1,013171226 |
| NECTIN2 | 0,060919592 | 0,273658513 | 0,508 | 0,467 | 1 | Basal | 1,208869501 | 0,827219149 | 1,087794433 |
| NUTF2 | 0,061085662 | 0,252853188 | 0,4 | 0,313 | 1 | Basal | 1,191561312 | 0,839235035 | 1,277955272 |
| MUC5B | 0,061461506 | -0,579330835 | 0,092 | 0,2 | 1 | Basal | 0,669274135 | 1,494156054 | 0,46 |
| TUBB6 | 0,061478387 | 0,26240257 | 0,292 | 0,187 | 1 | Basal | 1,199474569 | 0,833698376 | 1,561497326 |
| VDAC1 | 0,061684673 | 0,333365964 | 0,6 | 0,56 | 1 | Basal | 1,259949547 | 0,793682574 | 1,071428571 |
| MTCH1 | 0,061694915 | 0,33958169 | 0,585 | 0,573 | 1 | Basal | 1,26538964 | 0,790270418 | 1,020942408 |
| LYPLA2 | 0,061768238 | 0,363091652 | 0,4 | 0,333 | 1 | Basal | 1,286179189 | 0,777496641 | 1,201201201 |
| PRKCD | 0,062293148 | 0,289216976 | 0,462 | 0,373 | 1 | Basal | 1,221976869 | 0,818346096 | 1,238605898 |
| DNPH1 | 0,062353393 | 0,27070795 | 0,508 | 0,467 | 1 | Basal | 1,206399679 | 0,828912687 | 1,087794433 |
| MRPL36 | 0,062924815 | 0,287925478 | 0,354 | 0,267 | 1 | Basal | 1,220883447 | 0,819079006 | 1,325842697 |
| NUDT4 | 0,0629406 | -0,373750515 | 0,4 | 0,54 | 1 | Basal | 0,771773539 | 1,295716878 | 0,740740741 |
| MICALL2 | 0,063276691 | 0,302407242 | 0,262 | 0,167 | 1 | Basal | 1,233200383 | 0,810898224 | 1,568862275 |
| CHD2 | 0,063285215 | -0,294409821 | 0,385 | 0,553 | 1 | Basal | 0,815405831 | 1,226383185 | 0,696202532 |
| SF3A3 | 0,06344096 | -0,335609947 | 0,277 | 0,407 | 1 | Basal | 0,792449032 | 1,261910811 | 0,680589681 |

| Supplementary Table 4 - clusermarker | | | | | | | | | |
|---|---|---|---|---|---|---|---|---|---|
| | **p_val** | **avg_log2FC** | **pct.1** | **pct.2** | **p_val_adj** | **celltype** | **Foldchange_UP** | **Foldchange_DOWN** | **Ratio_pct1_pct2** |
| XPO6 | 0,063776563 | 0,261391912 | 0,262 | 0,173 | 1 | Basal | 1,198634589 | 0,834282615 | 1,514450867 |
| GSTZ1 | 0,063846947 | 0,435392644 | 0,308 | 0,207 | 1 | Basal | 1,352278819 | 0,739492467 | 1,487922705 |
| RAB2A | 0,064252295 | 0,390595038 | 0,692 | 0,653 | 1 | Basal | 1,310933986 | 0,762814917 | 1,059724349 |
| GTF3A | 0,064278464 | 0,388334681 | 0,446 | 0,387 | 1 | Basal | 1,308881674 | 0,764011002 | 1,15245478 |
| MTCH2 | 0,064718823 | 0,436712319 | 0,446 | 0,393 | 1 | Basal | 1,353516354 | 0,738816341 | 1,134860051 |
| RBM8A | 0,064910249 | 0,258881627 | 0,723 | 0,7 | 1 | Basal | 1,196550782 | 0,835735528 | 1,032857143 |
| GBP4 | 0,064923904 | 0,391691978 | 0,215 | 0,127 | 1 | Basal | 1,311931122 | 0,762235138 | 1,692913386 |
| FXN | 0,065619586 | 0,31448744 | 0,185 | 0,1 | 1 | Basal | 1,24356976 | 0,804136633 | 1,85 |
| SELENBP1 | 0,065865621 | -0,259136892 | 0,4 | 0,547 | 1 | Basal | 0,83558767 | 1,196762514 | 0,731261426 |
| CXCL1 | 0,065903263 | -0,732954823 | 0,4 | 0,527 | 1 | Basal | 0,601670353 | 1,662039677 | 0,759013283 |
| GFPT2 | 0,066389216 | -0,321240725 | 0,046 | 0,127 | 1 | Basal | 0,80038125 | 1,249404581 | 0,362204724 |
| UBL5 | 0,06639942 | 0,279010143 | 0,8 | 0,787 | 1 | Basal | 1,213362091 | 0,824156291 | 1,016518424 |
| STEAP4 | 0,066428525 | -0,39472163 | 0,2 | 0,307 | 1 | Basal | 0,760636128 | 1,314689065 | 0,651465798 |
| ERGIC1 | 0,066447257 | 0,258504239 | 0,4 | 0,32 | 1 | Basal | 1,196237823 | 0,835954173 | 1,25 |
| TMEM183A | 0,06663842 | 0,345552759 | 0,415 | 0,353 | 1 | Basal | 1,270637726 | 0,787006382 | 1,175637394 |
| NTAN1 | 0,067009499 | 0,323219568 | 0,169 | 0,087 | 1 | Basal | 1,251119477 | 0,799284176 | 1,942528736 |
| SNHG8 | 0,067059303 | -0,356396219 | 0,615 | 0,733 | 1 | Basal | 0,781113327 | 1,280223964 | 0,839017735 |
| MVP | 0,067208874 | 0,327731072 | 0,646 | 0,567 | 1 | Basal | 1,255038022 | 0,796788609 | 1,139329806 |
| NUCB2 | 0,067401374 | -0,321954482 | 0,477 | 0,6 | 1 | Basal | 0,799985368 | 1,250022863 | 0,795 |
| SOX2 | 0,067836183 | -0,284951652 | 0,6 | 0,693 | 1 | Basal | 0,820769114 | 1,218369432 | 0,865800866 |
| CXCL2 | 0,067917836 | 0,543724385 | 0,2 | 0,347 | 1 | Basal | 1,457730865 | 0,685997686 | 0,576368876 |
| PPL | 0,068018343 | 0,260145852 | 0,646 | 0,587 | 1 | Basal | 1,197599772 | 0,835003499 | 1,100511073 |
| FAM136A | 0,068476736 | 0,303268469 | 0,338 | 0,26 | 1 | Basal | 1,23393677 | 0,810414297 | 1,3 |
| RAB31 | 0,06894181 | 0,412709719 | 0,415 | 0,347 | 1 | Basal | 1,331183742 | 0,751211097 | 1,195965418 |
| HDAC7 | 0,069208017 | 0,267295309 | 0,4 | 0,307 | 1 | Basal | 1,203549358 | 0,83087577 | 1,302931596 |
| PCBP1 | 0,069249304 | 0,343469008 | 0,677 | 0,693 | 1 | Basal | 1,26880381 | 0,788143913 | 0,976911977 |
| ZNF226 | 0,069253871 | 0,315716542 | 0,2 | 0,113 | 1 | Basal | 1,244629669 | 0,803451842 | 1,769911504 |
| CTNNA1 | 0,069755389 | 0,291486468 | 0,692 | 0,667 | 1 | Basal | 1,223900664 | 0,817059774 | 1,037481259 |
| RXRB | 0,069849263 | -0,268660566 | 0,169 | 0,28 | 1 | Basal | 0,830089864 | 1,204688846 | 0,603571429 |
| DCTN6 | 0,069923432 | 0,25010355 | 0,262 | 0,167 | 1 | Basal | 1,189292474 | 0,840836062 | 1,568862275 |
| SLC44A1 | 0,070135578 | -0,327816116 | 0,262 | 0,427 | 1 | Basal | 0,796741641 | 1,255112006 | 0,613583138 |

| Supplementary Table 4 - clusermarker | | | | | | | | | |
|---|---|---|---|---|---|---|---|---|---|
| | **p_val** | **avg_log2FC** | **pct.1** | **pct.2** | **p_val_adj** | **celltype** | **Foldchange_UP** | **Foldchange_DOWN** | **Ratio_pct1_pct2** |
| GPAA1 | 0,070172312 | 0,263179794 | 0,477 | 0,427 | 1 | Basal | 1,200120937 | 0,833249358 | 1,117096019 |
| NEAT1 | 0,070298889 | 0,333811805 | 0,892 | 0,833 | 1 | Basal | 1,260338974 | 0,793437338 | 1,070828331 |
| ANKRD65 | 0,070719578 | -0,345183882 | 0,169 | 0,293 | 1 | Basal | 0,787207635 | 1,270312883 | 0,576791809 |
| GSK3B | 0,070874553 | -0,251316444 | 0,262 | 0,407 | 1 | Basal | 0,840129456 | 1,190292749 | 0,643734644 |
| ARID5A | 0,070922421 | -0,317032335 | 0,092 | 0,187 | 1 | Basal | 0,802719397 | 1,245765336 | 0,49197861 |
| TMEM125 | 0,071438078 | 0,302515408 | 0,277 | 0,2 | 1 | Basal | 1,233292845 | 0,810837429 | 1,385 |
| HNRNPA2B1 | 0,071793839 | -0,312511723 | 0,815 | 0,9 | 1 | Basal | 0,805238623 | 1,241867903 | 0,905555556 |
| NTM | 0,071942884 | -0,344309109 | 0,046 | 0,127 | 1 | Basal | 0,7876851 | 1,269542866 | 0,362204724 |
| CCDC50 | 0,072088318 | 0,396024879 | 0,369 | 0,293 | 1 | Basal | 1,315877217 | 0,759949323 | 1,259385666 |
| TXNIP | 0,072439877 | -0,444944076 | 0,723 | 0,78 | 1 | Basal | 0,734612792 | 1,361261348 | 0,926923077 |
| CTSZ | 0,072494742 | 0,458095542 | 0,538 | 0,487 | 1 | Basal | 1,373727205 | 0,727946565 | 1,104722793 |
| C8orf76 | 0,072531786 | 0,302138997 | 0,277 | 0,193 | 1 | Basal | 1,232971111 | 0,811049011 | 1,435233161 |
| ABI1 | 0,073176356 | -0,271309079 | 0,462 | 0,6 | 1 | Basal | 0,828567375 | 1,206902456 | 0,77 |
| CLIC1 | 0,073188055 | 0,314069452 | 0,846 | 0,813 | 1 | Basal | 1,243209516 | 0,804369647 | 1,040590406 |
| RMRP | 0,073302964 | -0,482145787 | 0,092 | 0,187 | 1 | Basal | 0,715912023 | 1,396819677 | 0,49197861 |
| ITPR3 | 0,073671899 | 0,37920576 | 0,4 | 0,307 | 1 | Basal | 1,300625631 | 0,768860751 | 1,302931596 |
| CLCA4 | 0,07371121 | -0,30777104 | 0,215 | 0,353 | 1 | Basal | 0,807888982 | 1,237793834 | 0,609065156 |
| EFNB2 | 0,073885208 | -0,288992984 | 0,292 | 0,407 | 1 | Basal | 0,818473162 | 1,22178716 | 0,717444717 |
| CLIC6 | 0,074125211 | 0,301049731 | 0,338 | 0,26 | 1 | Basal | 1,232040543 | 0,811661602 | 1,3 |
| GAS5 | 0,074130679 | -0,405493684 | 0,662 | 0,793 | 1 | Basal | 0,754977898 | 1,324542085 | 0,83480454 |
| EDN3 | 0,074517523 | -0,459472309 | 0,092 | 0,187 | 1 | Basal | 0,727252215 | 1,375038781 | 0,49197861 |
| MR1 | 0,074612908 | -0,271645868 | 0,123 | 0,22 | 1 | Basal | 0,828373974 | 1,207184233 | 0,559090909 |
| UGCG | 0,074651194 | 0,371145979 | 0,554 | 0,453 | 1 | Basal | 1,293379796 | 0,7731681 | 1,222958057 |
| GASK1B | 0,07524218 | -0,411999404 | 0,108 | 0,207 | 1 | Basal | 0,751581049 | 1,330528492 | 0,52173913 |
| PRRC2B | 0,075586228 | 0,333649083 | 0,554 | 0,5 | 1 | Basal | 1,260196828 | 0,793526835 | 1,108 |
| FAM184A | 0,075832694 | -0,253780733 | 0,108 | 0,207 | 1 | Basal | 0,838695643 | 1,192327643 | 0,52173913 |
| GIPC1 | 0,076282801 | 0,486324704 | 0,477 | 0,407 | 1 | Basal | 1,400871577 | 0,713841309 | 1,171990172 |
| VEGFB | 0,076653711 | -0,291243441 | 0,2 | 0,313 | 1 | Basal | 0,817197423 | 1,22369451 | 0,638977636 |
| PRPF31 | 0,076673785 | -0,280284179 | 0,231 | 0,367 | 1 | Basal | 0,823428804 | 1,214434078 | 0,629427793 |
| SYNPO2 | 0,07688379 | -0,402820771 | 0,138 | 0,247 | 1 | Basal | 0,756377959 | 1,322090349 | 0,558704453 |
| KMT2C | 0,076931515 | -0,297140177 | 0,415 | 0,553 | 1 | Basal | 0,813864104 | 1,22870636 | 0,75045208 |

| Supplementary Table 4 - clusermarker | | | | | | | | | |
|---|---|---|---|---|---|---|---|---|---|
| | **p_val** | **avg_log2FC** | **pct.1** | **pct.2** | **p_val_adj** | **celltype** | **Foldchange_UP** | **Foldchange_DOWN** | **Ratio_pct1_pct2** |
| MYLIP | 0,077557708 | -0,27963511 | 0,369 | 0,507 | 1 | Basal | 0,823799348 | 1,213887826 | 0,727810651 |
| SERTAD1 | 0,078065576 | -0,29877998 | 0,523 | 0,613 | 1 | Basal | 0,812939571 | 1,230103732 | 0,853181077 |
| EMP1 | 0,078193381 | -0,662443501 | 0,446 | 0,52 | 1 | Basal | 0,631807294 | 1,582761087 | 0,857692308 |
| RER1 | 0,07828961 | 0,364200921 | 0,523 | 0,5 | 1 | Basal | 1,287168496 | 0,776899064 | 1,046 |
| PPP1R14A | 0,078293835 | -0,538419046 | 0,123 | 0,227 | 1 | Basal | 0,688525005 | 1,452380078 | 0,54185022 |
| RAB11FIP1 | 0,079075279 | 0,656913841 | 0,523 | 0,507 | 1 | Basal | 1,576706183 | 0,634233576 | 1,031558185 |
| PLD5 | 0,079280252 | -0,252806532 | 0,031 | 0,1 | 1 | Basal | 0,839262176 | 1,191522778 | 0,31 |
| DYNLT1 | 0,079452992 | 0,573130756 | 0,631 | 0,6 | 1 | Basal | 1,487748595 | 0,672156575 | 1,051666667 |
| SDR16C5 | 0,080161462 | 0,35606903 | 0,415 | 0,347 | 1 | Basal | 1,279933655 | 0,781290496 | 1,195965418 |
| SEPHS2 | 0,080318019 | -0,253774623 | 0,323 | 0,473 | 1 | Basal | 0,838699195 | 1,192322594 | 0,682875264 |
| VMA21 | 0,080634925 | 0,374026336 | 0,277 | 0,2 | 1 | Basal | 1,295964623 | 0,771626002 | 1,385 |
| TPPP3 | 0,081299921 | 0,353645475 | 0,169 | 0,093 | 1 | Basal | 1,277785324 | 0,782604074 | 1,817204301 |
| CNBP | 0,081382994 | 0,251609262 | 0,769 | 0,78 | 1 | Basal | 1,190534363 | 0,839958956 | 0,985897436 |
| FBXO34 | 0,081470912 | 0,348155992 | 0,4 | 0,333 | 1 | Basal | 1,272932564 | 0,78558757 | 1,201201201 |
| PRKAR1A | 0,081546432 | -0,326601171 | 0,431 | 0,58 | 1 | Basal | 0,797412889 | 1,254055477 | 0,743103448 |
| YKT6 | 0,082581189 | 0,294320727 | 0,231 | 0,153 | 1 | Basal | 1,226307452 | 0,815456188 | 1,509803922 |
| SMARCA2 | 0,082771459 | -0,310504448 | 0,446 | 0,587 | 1 | Basal | 0,806359761 | 1,240141248 | 0,759795571 |
| MIIP | 0,082808785 | 0,252121444 | 0,169 | 0,093 | 1 | Basal | 1,190957098 | 0,839660809 | 1,817204301 |
| TMX4 | 0,082955816 | -0,269901183 | 0,231 | 0,373 | 1 | Basal | 0,829376352 | 1,205725239 | 0,619302949 |
| SERPINB2 | 0,082993729 | 0,327746695 | 0,354 | 0,233 | 1 | Basal | 1,255051613 | 0,796779981 | 1,519313305 |
| ICAM1 | 0,083076921 | 0,324531136 | 0,231 | 0,147 | 1 | Basal | 1,252257399 | 0,798557869 | 1,571428571 |
| SSH3 | 0,083174864 | 0,359080028 | 0,323 | 0,227 | 1 | Basal | 1,282607749 | 0,779661592 | 1,422907489 |
| UPF1 | 0,083222537 | 0,431521663 | 0,415 | 0,353 | 1 | Basal | 1,348655303 | 0,741479308 | 1,175637394 |
| EGFLAM | 0,083483491 | -0,290020326 | 0,046 | 0,12 | 1 | Basal | 0,817890535 | 1,222657504 | 0,383333333 |
| H2AC6 | 0,083789162 | 0,312626515 | 0,354 | 0,267 | 1 | Basal | 1,241966719 | 0,805174554 | 1,325842697 |
| FTSJ1 | 0,084037032 | 0,260126111 | 0,338 | 0,247 | 1 | Basal | 1,197583385 | 0,835014925 | 1,368421053 |
| HMGB2 | 0,084369356 | -0,26776797 | 0,277 | 0,4 | 1 | Basal | 0,8306036 | 1,203943734 | 0,6925 |
| ADIRF | 0,084421102 | 0,293953207 | 0,738 | 0,713 | 1 | Basal | 1,225995095 | 0,815663948 | 1,035063114 |
| UBE2V1 | 0,084436031 | 0,429488856 | 0,585 | 0,553 | 1 | Basal | 1,346756339 | 0,742524814 | 1,057866184 |
| IGFBP7 | 0,084524948 | 0,750935929 | 0,462 | 0,413 | 1 | Basal | 1,682884225 | 0,594217942 | 1,118644068 |
| TAOK1 | 0,084702226 | -0,252283525 | 0,277 | 0,413 | 1 | Basal | 0,839566481 | 1,191090905 | 0,670702179 |

| Supplementary Table 4 - clusermarker | | | | | | | | | |
|---|---|---|---|---|---|---|---|---|---|
| | **p_val** | **avg_log2FC** | **pct.1** | **pct.2** | **p_val_adj** | **celltype** | **Foldchange_UP** | **Foldchange_DOWN** | **Ratio_pct1_pct2** |
| SFPQ | 0,084812573 | -0,414788628 | 0,554 | 0,693 | 1 | Basal | 0,750129389 | 1,333103348 | 0,799422799 |
| SYNM | 0,085187724 | -0,592743303 | 0,185 | 0,293 | 1 | Basal | 0,663080851 | 1,508111716 | 0,631399317 |
| SETX | 0,086387421 | -0,273895863 | 0,246 | 0,373 | 1 | Basal | 0,827083067 | 1,2090684 | 0,659517426 |
| SPDEF | 0,086479584 | 0,446215089 | 0,277 | 0,2 | 1 | Basal | 1,362461147 | 0,733965884 | 1,385 |
| COIL | 0,086676757 | -0,257283072 | 0,123 | 0,22 | 1 | Basal | 0,836662065 | 1,195225698 | 0,559090909 |
| MRPS16 | 0,087405933 | 0,360510997 | 0,4 | 0,347 | 1 | Basal | 1,283880562 | 0,778888652 | 1,152737752 |
| PPP1R12B | 0,087425803 | -0,269753072 | 0,185 | 0,307 | 1 | Basal | 0,829461502 | 1,205601462 | 0,602605863 |
| C15orf48 | 0,087701647 | 0,668528569 | 0,323 | 0,227 | 1 | Basal | 1,58945103 | 0,62914804 | 1,422907489 |
| S100A1 | 0,087869943 | -0,388554823 | 0,062 | 0,14 | 1 | Basal | 0,76389443 | 1,309081413 | 0,442857143 |
| JDP2 | 0,088621729 | -0,436405737 | 0,323 | 0,453 | 1 | Basal | 0,738973361 | 1,353228753 | 0,713024283 |
| IREB2 | 0,088663208 | 0,341292937 | 0,277 | 0,187 | 1 | Basal | 1,266891468 | 0,789333597 | 1,481283422 |
| PRSS23 | 0,089186169 | 0,319089182 | 0,8 | 0,74 | 1 | Basal | 1,247542688 | 0,801575778 | 1,081081081 |
| C3 | 0,089243855 | -0,252146468 | 0,231 | 0,34 | 1 | Basal | 0,839646244 | 1,190977756 | 0,679411765 |
| CYTH2 | 0,089650417 | 0,264560544 | 0,431 | 0,353 | 1 | Basal | 1,201270078 | 0,832452267 | 1,220963173 |
| FUS | 0,089892351 | 0,252102004 | 0,8 | 0,78 | 1 | Basal | 1,19094105 | 0,839672123 | 1,025641026 |
| AC007255.1 | 0,090187012 | 0,296797408 | 0,2 | 0,127 | 1 | Basal | 1,228414467 | 0,814057492 | 1,57480315 |
| ARPC5 | 0,091594538 | 0,456621444 | 0,6 | 0,56 | 1 | Basal | 1,372324292 | 0,728690737 | 1,071428571 |
| CRAT | 0,091948722 | 0,254183723 | 0,292 | 0,213 | 1 | Basal | 1,192660744 | 0,838461402 | 1,370892019 |
| TRAPPC12 | 0,092582433 | 0,505837836 | 0,308 | 0,22 | 1 | Basal | 1,419947742 | 0,70425127 | 1,4 |
| AGPAT1 | 0,092661691 | -0,347750266 | 0,154 | 0,253 | 1 | Basal | 0,78580853 | 1,27257463 | 0,608695652 |
| NCBP2AS2 | 0,092676263 | 0,326939473 | 0,308 | 0,227 | 1 | Basal | 1,254349578 | 0,797225923 | 1,356828194 |
| NR4A1 | 0,093099155 | -0,354854024 | 0,723 | 0,773 | 1 | Basal | 0,781948759 | 1,278856176 | 0,935316947 |
| LSR | 0,093522909 | 0,30151394 | 0,538 | 0,587 | 1 | Basal | 1,232437034 | 0,811400479 | 0,916524702 |
| BLOC1S1 | 0,093576628 | 0,363564449 | 0,677 | 0,647 | 1 | Basal | 1,286600763 | 0,777241883 | 1,046367852 |
| SLC4A11 | 0,093681284 | 0,312358988 | 0,292 | 0,22 | 1 | Basal | 1,241736436 | 0,805323876 | 1,327272727 |
| CD99 | 0,093854433 | 0,449922577 | 0,554 | 0,52 | 1 | Basal | 1,36596695 | 0,732082134 | 1,065384615 |
| ASAP3 | 0,094432449 | 0,263104796 | 0,185 | 0,113 | 1 | Basal | 1,20005855 | 0,833292676 | 1,637168142 |
| SPPL2A | 0,094453691 | 0,364985345 | 0,4 | 0,333 | 1 | Basal | 1,287868547 | 0,776476762 | 1,201201201 |
| TMEM134 | 0,094616166 | 0,319420933 | 0,431 | 0,393 | 1 | Basal | 1,247829597 | 0,801391474 | 1,096692112 |
| NARS2 | 0,094642044 | -0,260147866 | 0,108 | 0,2 | 1 | Basal | 0,835002333 | 1,197601444 | 0,54 |
| CTTN | 0,095056497 | 0,35066319 | 0,508 | 0,493 | 1 | Basal | 1,275146662 | 0,784223517 | 1,030425963 |

| Supplementary Table 4 - clusermarker | | | | | | | | | |
|---|---|---|---|---|---|---|---|---|---|
| | **p_val** | **avg_log2FC** | **pct.1** | **pct.2** | **p_val_adj** | **celltype** | **Foldchange_UP** | **Foldchange_DOWN** | **Ratio_pct1_pct2** |
| SERINC1 | 0,095262012 | -0,341515587 | 0,477 | 0,64 | 1 | Basal | 0,78921179 | 1,267087001 | 0,7453125 |
| CYB5A | 0,095479705 | 0,351960222 | 0,831 | 0,8 | 1 | Basal | 1,276293578 | 0,78351879 | 1,03875 |
| MTA1 | 0,095525914 | 0,356729 | 0,338 | 0,267 | 1 | Basal | 1,280519302 | 0,780933172 | 1,265917603 |
| GTPBP6 | 0,095562694 | 0,34661672 | 0,292 | 0,22 | 1 | Basal | 1,271575143 | 0,786426194 | 1,327272727 |
| TMEM265 | 0,095641026 | 0,339170317 | 0,4 | 0,32 | 1 | Basal | 1,265028877 | 0,790495789 | 1,25 |
| TP53I11 | 0,095925752 | 0,312769514 | 0,385 | 0,32 | 1 | Basal | 1,242089829 | 0,80509475 | 1,203125 |
| DCUN1D5 | 0,095997745 | 0,367658694 | 0,292 | 0,227 | 1 | Basal | 1,290257211 | 0,775039264 | 1,286343612 |
| SLC25A26 | 0,096032959 | 0,327632674 | 0,262 | 0,18 | 1 | Basal | 1,254952426 | 0,796842955 | 1,455555556 |
| MUC5AC | 0,096262609 | 1,278430139 | 0,185 | 0,107 | 1 | Basal | 2,425748766 | 0,412243846 | 1,728971963 |
| PIBF1 | 0,09731987 | -0,279100532 | 0,215 | 0,32 | 1 | Basal | 0,824104657 | 1,213438114 | 0,671875 |
| TUBB | 0,097491557 | 0,424976049 | 0,677 | 0,653 | 1 | Basal | 1,342550214 | 0,744851097 | 1,036753446 |
| DENND5A | 0,097674331 | 0,391728714 | 0,215 | 0,14 | 1 | Basal | 1,311964528 | 0,76221573 | 1,535714286 |
| CHASERR | 0,098128518 | 0,306753963 | 0,662 | 0,62 | 1 | Basal | 1,236921517 | 0,808458731 | 1,067741935 |
| INTS11 | 0,0985739 | 0,299921288 | 0,385 | 0,293 | 1 | Basal | 1,231077245 | 0,812296713 | 1,313993174 |
| FAM120B | 0,098901019 | 0,318164785 | 0,323 | 0,233 | 1 | Basal | 1,24674359 | 0,802089546 | 1,386266094 |
| ABHD2 | 0,099067818 | 0,395277334 | 0,569 | 0,553 | 1 | Basal | 1,31519556 | 0,760343199 | 1,028933092 |
| RAB25 | 0,099672674 | 0,352500228 | 0,631 | 0,58 | 1 | Basal | 1,276771389 | 0,783225571 | 1,087931034 |
| MRPL27 | 0,099849995 | 0,272832286 | 0,477 | 0,407 | 1 | Basal | 1,208177384 | 0,82769303 | 1,171990172 |
| CALD1 | 0,099939235 | -1,010854146 | 0,308 | 0,407 | 1 | Basal | 0,496252355 | 2,015103787 | 0,756756757 |
| EIF4A2 | 0,100057344 | -0,311650345 | 0,769 | 0,807 | 1 | Basal | 0,805719543 | 1,241126653 | 0,95291202 |
| GOLGA3 | 0,100091918 | 0,283042133 | 0,308 | 0,247 | 1 | Basal | 1,216757893 | 0,821856185 | 1,246963563 |
| TRIB1 | 0,100261195 | -0,462941693 | 0,338 | 0,44 | 1 | Basal | 0,725505425 | 1,378349445 | 0,768181818 |
| ZC3H11A | 0,100711816 | -0,376813 | 0,046 | 0,113 | 1 | Basal | 0,770136992 | 1,298470286 | 0,407079646 |
| MRPS34 | 0,100893602 | 0,302113171 | 0,492 | 0,453 | 1 | Basal | 1,23294904 | 0,81106353 | 1,086092715 |
| CHMP5 | 0,100914651 | 0,343913505 | 0,569 | 0,507 | 1 | Basal | 1,269194791 | 0,787901122 | 1,122287968 |
| SNRPC | 0,101372714 | 0,403741932 | 0,585 | 0,553 | 1 | Basal | 1,322934774 | 0,755895166 | 1,057866184 |
| KDM1A | 0,101415717 | 0,270950936 | 0,354 | 0,287 | 1 | Basal | 1,206602884 | 0,828773089 | 1,233449477 |
| UQCR11 | 0,101609941 | 0,299135642 | 0,754 | 0,773 | 1 | Basal | 1,230407022 | 0,812739185 | 0,97542044 |
| STAT6 | 0,102060958 | -0,361866415 | 0,308 | 0,427 | 1 | Basal | 0,778157226 | 1,285087341 | 0,721311475 |
| DTYMK | 0,102526433 | 0,464616483 | 0,2 | 0,127 | 1 | Basal | 1,379950468 | 0,724663692 | 1,57480315 |
| ADRM1 | 0,102749094 | 0,370779238 | 0,477 | 0,44 | 1 | Basal | 1,293051053 | 0,773364669 | 1,084090909 |

| Supplementary Table 4 - clusermarker | | | | | | | | | |
|---|---|---|---|---|---|---|---|---|---|
| | **p_val** | **avg_log2FC** | **pct.1** | **pct.2** | **p_val_adj** | **celltype** | **Foldchange_UP** | **Foldchange_DOWN** | **Ratio_pct1_pct2** |
| PATZ1 | 0,103483891 | 0,284753245 | 0,231 | 0,153 | 1 | Basal | 1,218201887 | 0,820881999 | 1,509803922 |
| CALM3 | 0,10372712 | 0,331940076 | 0,477 | 0,407 | 1 | Basal | 1,25870489 | 0,794467399 | 1,171990172 |
| PIK3R1 | 0,10383394 | -0,29183947 | 0,431 | 0,567 | 1 | Basal | 0,816859879 | 1,224200167 | 0,760141093 |
| GPATCH2L | 0,104679634 | 0,275663934 | 0,277 | 0,2 | 1 | Basal | 1,210551061 | 0,82607007 | 1,385 |
| ADK | 0,105250597 | 0,352968166 | 0,446 | 0,4 | 1 | Basal | 1,277185577 | 0,782971573 | 1,115 |
| LPCAT4 | 0,105254299 | 0,41010409 | 0,492 | 0,46 | 1 | Basal | 1,328781682 | 0,752569074 | 1,069565217 |
| EIF4G2 | 0,105647718 | -0,300313314 | 0,754 | 0,833 | 1 | Basal | 0,812076017 | 1,231411813 | 0,905162065 |
| LRRFIP2 | 0,105838061 | 0,383068772 | 0,492 | 0,44 | 1 | Basal | 1,3041129 | 0,766804776 | 1,118181818 |
| LRRK1 | 0,105980135 | 0,374969666 | 0,169 | 0,1 | 1 | Basal | 1,296812288 | 0,771121626 | 1,69 |
| AZGP1 | 0,106114293 | -0,680268278 | 0,123 | 0,213 | 1 | Basal | 0,624049218 | 1,602437711 | 0,577464789 |
| CD55 | 0,106181342 | 0,666856038 | 0,692 | 0,687 | 1 | Basal | 1,587609431 | 0,62987784 | 1,00727802 |
| BRCC3 | 0,10624103 | -0,281155005 | 0,246 | 0,373 | 1 | Basal | 0,822931924 | 1,215167344 | 0,659517426 |
| ITGB8 | 0,106944201 | 0,351296249 | 0,508 | 0,48 | 1 | Basal | 1,275706324 | 0,783879472 | 1,058333333 |
| CHKB | 0,107393513 | 0,371019812 | 0,385 | 0,327 | 1 | Basal | 1,293266691 | 0,773235719 | 1,177370031 |
| SGCB | 0,107511524 | -0,265613327 | 0,154 | 0,253 | 1 | Basal | 0,831845021 | 1,202147005 | 0,608695652 |
| TPGS1 | 0,10753723 | 0,304350583 | 0,292 | 0,233 | 1 | Basal | 1,234862649 | 0,809806662 | 1,253218884 |
| TMEM14B | 0,107906658 | 0,47730165 | 0,585 | 0,593 | 1 | Basal | 1,392137441 | 0,718319881 | 0,986509275 |
| ARF6 | 0,108477117 | 0,470538978 | 0,538 | 0,52 | 1 | Basal | 1,38562703 | 0,721694928 | 1,034615385 |
| GOLGB1 | 0,109348345 | -0,378105033 | 0,6 | 0,687 | 1 | Basal | 0,76944759 | 1,299633677 | 0,873362445 |
| CPNE2 | 0,109440213 | 0,340813725 | 0,308 | 0,24 | 1 | Basal | 1,266470721 | 0,78959583 | 1,283333333 |
| MMUT | 0,109593402 | -0,301408812 | 0,169 | 0,273 | 1 | Basal | 0,811459607 | 1,232347231 | 0,619047619 |
| ARF4 | 0,109661905 | -0,27768403 | 0,6 | 0,707 | 1 | Basal | 0,824914196 | 1,212247292 | 0,848656294 |
| TMCO1 | 0,109707944 | -0,26487804 | 0,477 | 0,647 | 1 | Basal | 0,832269089 | 1,201534472 | 0,737248841 |
| EEF2 | 0,110262204 | 0,271915366 | 0,985 | 0,94 | 1 | Basal | 1,207409759 | 0,828219246 | 1,04787234 |
| VPS72 | 0,110593328 | 0,308711827 | 0,354 | 0,267 | 1 | Basal | 1,238601267 | 0,807362326 | 1,325842697 |
| AQP1 | 0,110755266 | -0,472306468 | 0,123 | 0,213 | 1 | Basal | 0,720811299 | 1,387325644 | 0,577464789 |
| A2M | 0,110817833 | -0,698353571 | 0,185 | 0,293 | 1 | Basal | 0,616275109 | 1,622651937 | 0,631399317 |
| HNRNPUL2 | 0,110954436 | -0,266733513 | 0,354 | 0,487 | 1 | Basal | 0,831199382 | 1,203080779 | 0,726899384 |
| PLEKHB2 | 0,112009994 | 0,262063091 | 0,446 | 0,38 | 1 | Basal | 1,199192355 | 0,833894576 | 1,173684211 |
| MMP24OS | 0,112384478 | 0,351120828 | 0,508 | 0,493 | 1 | Basal | 1,275551217 | 0,783974792 | 1,030425963 |
| CCND1 | 0,112561704 | 0,281026276 | 0,738 | 0,68 | 1 | Basal | 1,215058922 | 0,823005356 | 1,085294118 |

| Supplementary Table 4 - clusermarker | | | | | | | | | |
|---|---|---|---|---|---|---|---|---|---|
| | **p_val** | **avg_log2FC** | **pct.1** | **pct.2** | **p_val_adj** | **celltype** | **Foldchange_UP** | **Foldchange_DOWN** | **Ratio_pct1_pct2** |
| GRK6 | 0,112634283 | 0,394544059 | 0,2 | 0,127 | 1 | Basal | 1,314527259 | 0,760729755 | 1,57480315 |
| KIFAP3 | 0,112658611 | -0,360674047 | 0,215 | 0,32 | 1 | Basal | 0,778800628 | 1,284025672 | 0,671875 |
| ATP5F1C | 0,113341067 | 0,285519156 | 0,646 | 0,62 | 1 | Basal | 1,218848789 | 0,820446317 | 1,041935484 |
| RAD23A | 0,11385934 | 0,332386348 | 0,554 | 0,513 | 1 | Basal | 1,259094309 | 0,794221682 | 1,079922027 |
| GNA15 | 0,114463191 | 0,257541719 | 0,308 | 0,233 | 1 | Basal | 1,195439998 | 0,836512081 | 1,321888412 |
| CAB39 | 0,114535822 | -0,263932346 | 0,262 | 0,387 | 1 | Basal | 0,832814824 | 1,200747118 | 0,677002584 |
| MPZL1 | 0,114575347 | -0,410848232 | 0,323 | 0,42 | 1 | Basal | 0,752180999 | 1,329467244 | 0,769047619 |
| HSPA4 | 0,114951003 | 0,306975427 | 0,462 | 0,42 | 1 | Basal | 1,237111407 | 0,808334637 | 1,1 |
| SRSF3 | 0,115014564 | -0,285386343 | 0,708 | 0,767 | 1 | Basal | 0,82052185 | 1,218736588 | 0,923076923 |
| UBE2D2 | 0,115359272 | 0,469885336 | 0,538 | 0,527 | 1 | Basal | 1,384999385 | 0,722021981 | 1,020872865 |
| PER2 | 0,115443783 | 0,34243128 | 0,477 | 0,44 | 1 | Basal | 1,26789149 | 0,788711028 | 1,084090909 |
| TNIP1 | 0,116155761 | 0,260568679 | 0,4 | 0,327 | 1 | Basal | 1,197950818 | 0,834758811 | 1,22324159 |
| GLRX5 | 0,116333222 | 0,395899246 | 0,431 | 0,387 | 1 | Basal | 1,315762632 | 0,760015504 | 1,11369509 |
| NDUFS6 | 0,116681526 | 0,364369393 | 0,585 | 0,587 | 1 | Basal | 1,287318815 | 0,776808346 | 0,996592845 |
| NFKBIA | 0,117293986 | 0,526376004 | 0,8 | 0,753 | 1 | Basal | 1,440306651 | 0,694296593 | 1,062416999 |
| BCLAF1 | 0,117451973 | -0,33382732 | 0,492 | 0,66 | 1 | Basal | 0,793428805 | 1,260352527 | 0,745454545 |
| ORMDL2 | 0,117962638 | 0,298915016 | 0,4 | 0,333 | 1 | Basal | 1,230218875 | 0,812863483 | 1,201201201 |
| IFI27L2 | 0,118504869 | 0,293541881 | 0,369 | 0,3 | 1 | Basal | 1,225645602 | 0,815896535 | 1,23 |
| CHURC1 | 0,119883015 | -0,282887673 | 0,477 | 0,613 | 1 | Basal | 0,821944181 | 1,216627629 | 0,778140294 |
| NOLC1 | 0,1206551 | -0,308070123 | 0,369 | 0,493 | 1 | Basal | 0,807721517 | 1,238050465 | 0,748478702 |
| ATOX1 | 0,120955972 | 0,359942514 | 0,554 | 0,513 | 1 | Basal | 1,283374758 | 0,779195627 | 1,079922027 |
| ATP1A1 | 0,120993238 | 0,25013483 | 0,754 | 0,707 | 1 | Basal | 1,18931826 | 0,840817831 | 1,066478076 |
| CENPX | 0,121715197 | 0,378935965 | 0,462 | 0,413 | 1 | Basal | 1,300382427 | 0,769004547 | 1,118644068 |
| SURF1 | 0,121851669 | 0,33863394 | 0,431 | 0,38 | 1 | Basal | 1,264558641 | 0,790789741 | 1,134210526 |
| EHF | 0,122136908 | -0,392983435 | 0,554 | 0,633 | 1 | Basal | 0,761553114 | 1,313106048 | 0,875197472 |
| TSPAN1 | 0,12246015 | 0,625862107 | 0,615 | 0,593 | 1 | Basal | 1,543132675 | 0,648032419 | 1,037099494 |
| MYO1C | 0,122518724 | 0,36758438 | 0,369 | 0,32 | 1 | Basal | 1,290190751 | 0,775079188 | 1,153125 |
| RAB22A | 0,12304952 | 0,267663599 | 0,338 | 0,287 | 1 | Basal | 1,203856639 | 0,830663692 | 1,177700348 |
| MZT2A | 0,12307969 | 0,279355899 | 0,492 | 0,453 | 1 | Basal | 1,21365292 | 0,823958797 | 1,086092715 |
| PHLDA1 | 0,125666354 | -0,287697269 | 0,292 | 0,42 | 1 | Basal | 0,81920858 | 1,220690339 | 0,695238095 |
| APPL1 | 0,125765525 | -0,404536085 | 0,431 | 0,553 | 1 | Basal | 0,755479187 | 1,323663203 | 0,779385172 |

| Supplementary Table 4 - clusermarker | | | | | | | | | |
|---|---|---|---|---|---|---|---|---|---|
| | **p_val** | **avg_log2FC** | **pct.1** | **pct.2** | **p_val_adj** | **celltype** | **Foldchange_UP** | **Foldchange_DOWN** | **Ratio_pct1_pct2** |
| CMPK1 | 0,126225467 | 0,400084028 | 0,554 | 0,58 | 1 | Basal | 1,319584766 | 0,757814144 | 0,955172414 |
| RHOV | 0,126523094 | -0,413365662 | 0,262 | 0,347 | 1 | Basal | 0,750869626 | 1,331789123 | 0,755043228 |
| DSC2 | 0,127227638 | 0,621061961 | 0,215 | 0,147 | 1 | Basal | 1,538006885 | 0,650192148 | 1,462585034 |
| PHLDA2 | 0,127580677 | 0,286550929 | 0,646 | 0,593 | 1 | Basal | 1,219720786 | 0,819859767 | 1,089376054 |
| UROS | 0,127662387 | 0,454024379 | 0,308 | 0,24 | 1 | Basal | 1,369856128 | 0,73000367 | 1,283333333 |
| C22orf39 | 0,127985517 | 0,251219555 | 0,338 | 0,28 | 1 | Basal | 1,190212813 | 0,84018588 | 1,207142857 |
| SCNN1A | 0,128123616 | 0,452850254 | 0,538 | 0,533 | 1 | Basal | 1,368741736 | 0,730598019 | 1,009380863 |
| TMC5 | 0,12821805 | 0,512773196 | 0,369 | 0,313 | 1 | Basal | 1,426790184 | 0,700873899 | 1,178913738 |
| TM9SF3 | 0,128295613 | 0,257837071 | 0,615 | 0,56 | 1 | Basal | 1,195684756 | 0,836340846 | 1,098214286 |
| DLG5 | 0,128336909 | 0,279425481 | 0,338 | 0,267 | 1 | Basal | 1,213711456 | 0,823919058 | 1,265917603 |
| GLIPR1 | 0,128654749 | -0,298763476 | 0,154 | 0,24 | 1 | Basal | 0,812948871 | 1,23008966 | 0,641666667 |
| TNC | 0,128844429 | 0,704528857 | 0,385 | 0,34 | 1 | Basal | 1,629612393 | 0,613642854 | 1,132352941 |
| CCL2 | 0,12889385 | -0,50263445 | 0,154 | 0,247 | 1 | Basal | 0,705816739 | 1,416798363 | 0,623481781 |
| PPP1R12A | 0,128994469 | -0,319747977 | 0,462 | 0,593 | 1 | Basal | 0,801209828 | 1,248112498 | 0,779089376 |
| RAB21 | 0,129075952 | 0,370648857 | 0,415 | 0,367 | 1 | Basal | 1,292934202 | 0,773434564 | 1,130790191 |
| ZNF282 | 0,129101319 | 0,284993787 | 0,138 | 0,08 | 1 | Basal | 1,218405017 | 0,820745143 | 1,725 |
| MESD | 0,129353251 | 0,308432336 | 0,477 | 0,427 | 1 | Basal | 1,238361338 | 0,80751875 | 1,117096019 |
| KIAA1522 | 0,129376631 | 0,263981615 | 0,538 | 0,527 | 1 | Basal | 1,200788125 | 0,832786383 | 1,020872865 |
| PALLD | 0,129376631 | -0,425398524 | 0,446 | 0,567 | 1 | Basal | 0,744633009 | 1,342943421 | 0,78659612 |
| CLK1 | 0,130240388 | -0,343090039 | 0,369 | 0,5 | 1 | Basal | 0,788350971 | 1,268470562 | 0,738 |
| HLA-DRA | 0,130522509 | -0,349486796 | 0,462 | 0,6 | 1 | Basal | 0,784863244 | 1,274107314 | 0,77 |
| IL17RC | 0,130785284 | 0,311509661 | 0,308 | 0,24 | 1 | Basal | 1,241005631 | 0,805798117 | 1,283333333 |
| TWF2 | 0,130867297 | 0,367540085 | 0,354 | 0,293 | 1 | Basal | 1,290151139 | 0,775102986 | 1,208191126 |
| ATP2A2 | 0,131014768 | -0,283247831 | 0,6 | 0,667 | 1 | Basal | 0,821739014 | 1,216931389 | 0,899550225 |
| VILL | 0,13110706 | 0,331530159 | 0,369 | 0,313 | 1 | Basal | 1,258347302 | 0,794693165 | 1,178913738 |
| USP10 | 0,131155945 | 0,299219489 | 0,431 | 0,36 | 1 | Basal | 1,230478533 | 0,812691951 | 1,197222222 |
| CDK5RAP3 | 0,131457447 | -0,30162398 | 0,385 | 0,547 | 1 | Basal | 0,811338593 | 1,23253104 | 0,703839122 |
| LY6E | 0,131562863 | 0,267703092 | 0,785 | 0,753 | 1 | Basal | 1,203889594 | 0,830640953 | 1,04249668 |
| KRTCAP3 | 0,131899159 | 0,367413565 | 0,477 | 0,453 | 1 | Basal | 1,290038002 | 0,775170963 | 1,052980132 |
| MSRB3 | 0,132037789 | -0,583894672 | 0,154 | 0,24 | 1 | Basal | 0,667160291 | 1,498890167 | 0,641666667 |
| SLC9A3R1 | 0,132461124 | 0,259113082 | 0,6 | 0,58 | 1 | Basal | 1,196742763 | 0,83560146 | 1,034482759 |

| Supplementary Table 4 - clusermarker | | | | | | | | | |
|---|---|---|---|---|---|---|---|---|---|
| | **p_val** | **avg_log2FC** | **pct.1** | **pct.2** | **p_val_adj** | **celltype** | **Foldchange_UP** | **Foldchange_DOWN** | **Ratio_pct1_pct2** |
| MSRB2 | 0,132659344 | 0,333170611 | 0,369 | 0,307 | 1 | Basal | 1,25977895 | 0,793790053 | 1,201954397 |
| NNMT | 0,132893417 | 0,86513143 | 0,246 | 0,153 | 1 | Basal | 1,8215056 | 0,548996391 | 1,607843137 |
| CYP2F1 | 0,132941609 | -1,051641332 | 0,4 | 0,44 | 1 | Basal | 0,482419011 | 2,072886799 | 0,909090909 |
| MANF | 0,134570135 | 0,361871728 | 0,385 | 0,34 | 1 | Basal | 1,285092073 | 0,77815436 | 1,132352941 |
| ATP2C1 | 0,135373647 | 0,252089527 | 0,354 | 0,293 | 1 | Basal | 1,190930751 | 0,839679385 | 1,208191126 |
| LRRC41 | 0,135384284 | 0,377425403 | 0,323 | 0,267 | 1 | Basal | 1,299021585 | 0,769810149 | 1,209737828 |
| TACC1 | 0,13569617 | -0,262271587 | 0,338 | 0,467 | 1 | Basal | 0,833774071 | 1,199365673 | 0,723768737 |
| BGN | 0,136666966 | -0,319217754 | 0,154 | 0,253 | 1 | Basal | 0,801504345 | 1,247653873 | 0,608695652 |
| ILF3 | 0,137117439 | 0,309350786 | 0,554 | 0,547 | 1 | Basal | 1,239149956 | 0,80700483 | 1,012797075 |
| SARNP | 0,137141214 | -0,282180787 | 0,369 | 0,48 | 1 | Basal | 0,822347012 | 1,216031657 | 0,76875 |
| SDCBP2 | 0,137334414 | 0,360995968 | 0,231 | 0,167 | 1 | Basal | 1,28431222 | 0,778626867 | 1,383233533 |
| ECHDC2 | 0,137828869 | 0,34597503 | 0,431 | 0,367 | 1 | Basal | 1,271009691 | 0,786776063 | 1,174386921 |
| RSRC2 | 0,140096376 | -0,39476355 | 0,492 | 0,62 | 1 | Basal | 0,760614027 | 1,314727266 | 0,793548387 |
| ABCC1 | 0,140153457 | 0,3045644 | 0,277 | 0,2 | 1 | Basal | 1,235045678 | 0,809686652 | 1,385 |
| KLK11 | 0,141233177 | 0,414863077 | 0,538 | 0,52 | 1 | Basal | 1,333172142 | 0,750090681 | 1,034615385 |
| VKORC1 | 0,14255132 | 0,274806583 | 0,492 | 0,447 | 1 | Basal | 1,20983188 | 0,826561125 | 1,100671141 |
| PPP1R15A | 0,142582635 | -0,333983522 | 0,708 | 0,747 | 1 | Basal | 0,793342904 | 1,260488995 | 0,947791165 |
| SFRP1 | 0,142987685 | -0,518792637 | 0,2 | 0,3 | 1 | Basal | 0,697955694 | 1,432755701 | 0,666666667 |
| MXRA7 | 0,14303119 | -0,412871307 | 0,4 | 0,527 | 1 | Basal | 0,751126964 | 1,331332848 | 0,759013283 |
| COA4 | 0,143934155 | 0,250056071 | 0,508 | 0,473 | 1 | Basal | 1,189253335 | 0,840863734 | 1,073995772 |
| NIBAN2 | 0,144152743 | 0,340980532 | 0,431 | 0,38 | 1 | Basal | 1,266617162 | 0,78950454 | 1,134210526 |
| RNF113A | 0,144427657 | 0,434657011 | 0,2 | 0,133 | 1 | Basal | 1,351589466 | 0,739869631 | 1,503759398 |
| HMGN2 | 0,144758836 | -0,312685749 | 0,631 | 0,787 | 1 | Basal | 0,805141496 | 1,242017713 | 0,801778907 |
| PRXL2C | 0,144887514 | 0,265361425 | 0,262 | 0,18 | 1 | Basal | 1,201937122 | 0,831990278 | 1,455555556 |
| SIGIRR | 0,14489699 | 0,324899193 | 0,431 | 0,38 | 1 | Basal | 1,252576913 | 0,798354169 | 1,134210526 |
| METRN | 0,145989434 | 0,326909866 | 0,277 | 0,227 | 1 | Basal | 1,254323837 | 0,797242284 | 1,220264317 |
| PLS1 | 0,14614545 | 0,260313891 | 0,4 | 0,327 | 1 | Basal | 1,197739272 | 0,834906247 | 1,22324159 |
| KCNK6 | 0,147387689 | 0,345348586 | 0,246 | 0,18 | 1 | Basal | 1,270457916 | 0,787117769 | 1,366666667 |
| TIMM50 | 0,148057533 | 0,41942481 | 0,308 | 0,253 | 1 | Basal | 1,337394241 | 0,747722676 | 1,217391304 |
| S100A2 | 0,148088497 | 0,336754376 | 0,477 | 0,573 | 1 | Basal | 1,262912228 | 0,791820665 | 0,832460733 |
| RAB29 | 0,148915668 | -0,287447454 | 0,138 | 0,22 | 1 | Basal | 0,819350446 | 1,220478985 | 0,627272727 |

| Supplementary Table 4 - clusermarker | | | | | | | | | |
|---|---|---|---|---|---|---|---|---|---|
| | **p_val** | **avg_log2FC** | **pct.1** | **pct.2** | **p_val_adj** | **celltype** | **Foldchange_UP** | **Foldchange_DOWN** | **Ratio_pct1_pct2** |
| GALNT1 | 0,148994741 | 0,296379336 | 0,323 | 0,273 | 1 | Basal | 1,228058542 | 0,814293428 | 1,183150183 |
| DDX6 | 0,149118397 | -0,310566083 | 0,431 | 0,54 | 1 | Basal | 0,806325312 | 1,240194231 | 0,798148148 |
| NDUFB8 | 0,150116705 | 0,258790714 | 0,692 | 0,7 | 1 | Basal | 1,196475383 | 0,835788195 | 0,988571429 |
| EIF2AK1 | 0,150502682 | 0,278263534 | 0,354 | 0,313 | 1 | Basal | 1,212734326 | 0,82458291 | 1,130990415 |
| COA8 | 0,150525999 | 0,319092358 | 0,338 | 0,26 | 1 | Basal | 1,247545434 | 0,801574013 | 1,3 |
| DHPS | 0,150894194 | 0,350216545 | 0,338 | 0,293 | 1 | Basal | 1,27475195 | 0,784466342 | 1,153583618 |
| IGFBP4 | 0,151371006 | 0,624001279 | 0,554 | 0,527 | 1 | Basal | 1,541143583 | 0,648868808 | 1,051233397 |
| CALCOCO1 | 0,1515379 | -0,324232735 | 0,262 | 0,36 | 1 | Basal | 0,798723057 | 1,251998414 | 0,727777778 |
| B2M | 0,151722081 | 0,271234026 | 0,938 | 0,967 | 1 | Basal | 1,206839671 | 0,828610481 | 0,970010341 |
| BZW1 | 0,151732655 | -0,308034738 | 0,646 | 0,693 | 1 | Basal | 0,807741328 | 1,238020101 | 0,932178932 |
| NREP | 0,151974918 | -0,299448542 | 0,077 | 0,147 | 1 | Basal | 0,812562932 | 1,230673909 | 0,523809524 |
| S100A9 | 0,152510072 | 1,262182831 | 0,215 | 0,353 | 1 | Basal | 2,398583778 | 0,416912684 | 0,609065156 |
| ANXA1 | 0,152826329 | -0,469045597 | 0,846 | 0,827 | 1 | Basal | 0,722442366 | 1,384193463 | 1,022974607 |
| RIPOR1 | 0,152988802 | 0,265737005 | 0,2 | 0,133 | 1 | Basal | 1,202250066 | 0,831773712 | 1,503759398 |
| RBM43 | 0,153163849 | -0,281878915 | 0,092 | 0,16 | 1 | Basal | 0,8225191 | 1,215777239 | 0,575 |
| ERBB3 | 0,153409281 | 0,292337786 | 0,4 | 0,373 | 1 | Basal | 1,224623087 | 0,816577779 | 1,072386059 |
| AMD1 | 0,153968363 | -0,307966452 | 0,538 | 0,633 | 1 | Basal | 0,807779561 | 1,237961503 | 0,849921011 |
| CKS1B | 0,154605314 | -0,291947503 | 0,231 | 0,347 | 1 | Basal | 0,816798712 | 1,224291842 | 0,665706052 |
| TIMM23 | 0,154861786 | 0,389480053 | 0,323 | 0,253 | 1 | Basal | 1,309921224 | 0,763404686 | 1,276679842 |
| DAPL1 | 0,154994823 | -0,333399913 | 0,262 | 0,367 | 1 | Basal | 0,793663898 | 1,259979196 | 0,713896458 |
| SERPINB6 | 0,155031682 | 0,432996281 | 0,492 | 0,473 | 1 | Basal | 1,350034505 | 0,740721808 | 1,040169133 |
| HMOX2 | 0,155527049 | 0,263142218 | 0,323 | 0,26 | 1 | Basal | 1,200089679 | 0,833271061 | 1,242307692 |
| CD109 | 0,155799507 | -0,266371079 | 0,062 | 0,127 | 1 | Basal | 0,831408222 | 1,202778579 | 0,488188976 |
| IFITM1 | 0,155965902 | 0,653053073 | 0,754 | 0,72 | 1 | Basal | 1,572492431 | 0,635933109 | 1,047222222 |
| RRM1 | 0,157306494 | -0,301947415 | 0,246 | 0,353 | 1 | Basal | 0,811156721 | 1,23280739 | 0,696883853 |
| SLIRP | 0,15801892 | -0,309370719 | 0,492 | 0,64 | 1 | Basal | 0,80699368 | 1,239167077 | 0,76875 |
| PPP6R2 | 0,15841555 | 0,343220103 | 0,246 | 0,187 | 1 | Basal | 1,268584925 | 0,788279902 | 1,315508021 |
| IER5L | 0,159450357 | -0,303387205 | 0,292 | 0,387 | 1 | Basal | 0,810347601 | 1,234038329 | 0,754521964 |
| AP2M1 | 0,159490443 | 0,254734557 | 0,6 | 0,613 | 1 | Basal | 1,1931162 | 0,838141331 | 0,978792822 |
| SHISA5 | 0,160211854 | 0,358218339 | 0,477 | 0,44 | 1 | Basal | 1,281841905 | 0,780127406 | 1,084090909 |
| ST6GALNAC4 | 0,160319746 | 0,251390344 | 0,215 | 0,147 | 1 | Basal | 1,190353722 | 0,840086422 | 1,462585034 |

| Supplementary Table 4 - clusermarker | | | | | | | | | |
|---|---|---|---|---|---|---|---|---|---|
| | **p_val** | **avg_log2FC** | **pct.1** | **pct.2** | **p_val_adj** | **celltype** | **Foldchange_UP** | **Foldchange_DOWN** | **Ratio_pct1_pct2** |
| SLFN5 | 0,160886759 | 0,318651705 | 0,354 | 0,3 | 1 | Basal | 1,247164446 | 0,801818881 | 1,18 |
| AQP3 | 0,16106782 | -0,447751236 | 0,754 | 0,853 | 1 | Basal | 0,733184791 | 1,363912635 | 0,883939039 |
| CCNO | 0,161195471 | -0,373360835 | 0,169 | 0,26 | 1 | Basal | 0,771982027 | 1,295366945 | 0,65 |
| USP4 | 0,161699959 | -0,276219672 | 0,185 | 0,28 | 1 | Basal | 0,825751922 | 1,211017465 | 0,660714286 |
| MYLK | 0,162016637 | -1,147552188 | 0,246 | 0,307 | 1 | Basal | 0,451390453 | 2,215376937 | 0,801302932 |
| PFDN1 | 0,163617855 | -0,426167297 | 0,292 | 0,387 | 1 | Basal | 0,74423632 | 1,34365923 | 0,754521964 |
| CFB | 0,163995306 | 0,6790975 | 0,446 | 0,393 | 1 | Basal | 1,601137826 | 0,624555853 | 1,134860051 |
| SMARCE1 | 0,164095581 | -0,25855672 | 0,477 | 0,58 | 1 | Basal | 0,835923764 | 1,196281339 | 0,822413793 |
| CAV1 | 0,165336633 | -0,3140481 | 0,277 | 0,387 | 1 | Basal | 0,804381552 | 1,243191117 | 0,715762274 |
| ISG20 | 0,166189925 | 0,402050198 | 0,385 | 0,34 | 1 | Basal | 1,321384381 | 0,756782064 | 1,132352941 |
| ANXA4 | 0,166628323 | 0,281760429 | 0,585 | 0,587 | 1 | Basal | 1,215677394 | 0,822586654 | 0,996592845 |
| TTC1 | 0,166741858 | -0,305630531 | 0,262 | 0,36 | 1 | Basal | 0,809088527 | 1,235958696 | 0,727777778 |
| TGFBI | 0,167199053 | 0,550324285 | 0,2 | 0,14 | 1 | Basal | 1,464414826 | 0,682866618 | 1,428571429 |
| SUMO3 | 0,167628241 | 0,31327067 | 0,415 | 0,38 | 1 | Basal | 1,242521374 | 0,804815129 | 1,092105263 |
| MOB1A | 0,168939615 | 0,300745784 | 0,477 | 0,44 | 1 | Basal | 1,231781003 | 0,811832621 | 1,084090909 |
| VSIR | 0,1694834 | 0,282026313 | 0,369 | 0,3 | 1 | Basal | 1,215901459 | 0,822435069 | 1,23 |
| ACTA2 | 0,171636986 | -0,55990656 | 0,231 | 0,32 | 1 | Basal | 0,678346097 | 1,474173736 | 0,721875 |
| RUFY2 | 0,171868098 | -0,258993239 | 0,108 | 0,18 | 1 | Basal | 0,835670875 | 1,196643355 | 0,6 |
| H2AX | 0,172607709 | -0,290309115 | 0,431 | 0,567 | 1 | Basal | 0,817726832 | 1,222902271 | 0,760141093 |
| SRSF8 | 0,172983 | 0,300533857 | 0,4 | 0,347 | 1 | Basal | 1,231600072 | 0,811951885 | 1,152737752 |
| LGR4 | 0,173860329 | 0,499563131 | 0,323 | 0,26 | 1 | Basal | 1,413785383 | 0,707320936 | 1,242307692 |
| SLC22A18 | 0,173905533 | 0,334355891 | 0,231 | 0,173 | 1 | Basal | 1,260814377 | 0,793138164 | 1,335260116 |
| BSPRY | 0,174740726 | 0,250983676 | 0,277 | 0,22 | 1 | Basal | 1,190018231 | 0,84032326 | 1,259090909 |
| C11orf24 | 0,174862339 | 0,471477074 | 0,2 | 0,133 | 1 | Basal | 1,386528311 | 0,721225807 | 1,503759398 |
| AC127526.5 | 0,175202794 | 0,280920161 | 0,231 | 0,167 | 1 | Basal | 1,214969553 | 0,823065893 | 1,383233533 |
| ANP32A | 0,176196117 | 0,441355047 | 0,508 | 0,493 | 1 | Basal | 1,357879113 | 0,736442582 | 1,030425963 |
| MRPL34 | 0,177011066 | 0,307154129 | 0,415 | 0,38 | 1 | Basal | 1,237264653 | 0,808234517 | 1,092105263 |
| LRRC59 | 0,177184053 | 0,419669359 | 0,369 | 0,34 | 1 | Basal | 1,337620959 | 0,747595941 | 1,085294118 |
| PMEPA1 | 0,17767651 | 0,493313084 | 0,369 | 0,3 | 1 | Basal | 1,407673827 | 0,71039184 | 1,23 |
| ALDH16A1 | 0,177811172 | 0,256412467 | 0,277 | 0,22 | 1 | Basal | 1,194504648 | 0,837167107 | 1,259090909 |
| CLSTN1 | 0,179071318 | 0,251436196 | 0,523 | 0,487 | 1 | Basal | 1,190391554 | 0,840059723 | 1,073921971 |

| Supplementary Table 4 - clusermarker | | | | | | | | | |
|---|---|---|---|---|---|---|---|---|---|
| | p_val | avg_log2FC | pct.1 | pct.2 | p_val_adj | celltype | Foldchange_UP | Foldchange_DOWN | Ratio_pct1_pct2 |
| STAT2 | 0,179115851 | -0,332582289 | 0,246 | 0,34 | 1 | Basal | 0,794113822 | 1,259265325 | 0,723529412 |
| GAS1 | 0,179887983 | -0,361097072 | 0,062 | 0,12 | 1 | Basal | 0,778572303 | 1,284402227 | 0,516666667 |
| PEX16 | 0,180154953 | 0,329756945 | 0,292 | 0,24 | 1 | Basal | 1,25680162 | 0,795670521 | 1,216666667 |
| DUOX1 | 0,180356965 | 0,323982818 | 0,385 | 0,333 | 1 | Basal | 1,25178155 | 0,79886143 | 1,156156156 |
| BCL7B | 0,18040459 | 0,339839579 | 0,308 | 0,253 | 1 | Basal | 1,265615855 | 0,790129166 | 1,217391304 |
| ENDOG | 0,180707769 | 0,595941546 | 0,292 | 0,253 | 1 | Basal | 1,511458687 | 0,661612526 | 1,154150198 |
| FBXO32 | 0,181162567 | -0,524958948 | 0,462 | 0,58 | 1 | Basal | 0,694978885 | 1,438892635 | 0,796551724 |
| PEDS1 | 0,18136257 | 0,428623663 | 0,277 | 0,213 | 1 | Basal | 1,345948923 | 0,742970244 | 1,300469484 |
| TPD52L2 | 0,181629468 | 0,298924143 | 0,323 | 0,26 | 1 | Basal | 1,230226658 | 0,812858341 | 1,242307692 |
| GSTO2 | 0,182618444 | 0,481078793 | 0,2 | 0,313 | 1 | Basal | 1,395786993 | 0,716441695 | 0,638977636 |
| GOPC | 0,183089826 | 0,280090712 | 0,4 | 0,353 | 1 | Basal | 1,214271231 | 0,823539234 | 1,133144476 |
| NGDN | 0,184033626 | 0,352739134 | 0,323 | 0,253 | 1 | Basal | 1,276982836 | 0,783095882 | 1,276679842 |
| SEMA5A | 0,184054873 | -0,388412838 | 0,231 | 0,32 | 1 | Basal | 0,763969613 | 1,308952585 | 0,721875 |
| FGF7 | 0,186460207 | -0,39434061 | 0,108 | 0,18 | 1 | Basal | 0,760837041 | 1,314341897 | 0,6 |
| MUC13 | 0,186493219 | 0,35237426 | 0,292 | 0,233 | 1 | Basal | 1,276659913 | 0,783293961 | 1,253218884 |
| ZDHHC12 | 0,186895536 | 0,374777686 | 0,354 | 0,327 | 1 | Basal | 1,296639732 | 0,771224246 | 1,082568807 |
| NDUFS3 | 0,189234539 | 0,298312224 | 0,477 | 0,473 | 1 | Basal | 1,229704968 | 0,813203188 | 1,00845666 |
| HNRNPAB | 0,189810476 | -0,408898731 | 0,554 | 0,673 | 1 | Basal | 0,753198102 | 1,32767196 | 0,823179792 |
| ACOT2 | 0,190422203 | 0,37687876 | 0,138 | 0,087 | 1 | Basal | 1,298529474 | 0,770101889 | 1,586206897 |
| LYSMD2 | 0,190650166 | 0,360507278 | 0,215 | 0,16 | 1 | Basal | 1,283877253 | 0,77889066 | 1,34375 |
| TRIM7 | 0,191259121 | 0,340740044 | 0,292 | 0,247 | 1 | Basal | 1,266406042 | 0,789636157 | 1,182186235 |
| PCSK5 | 0,191581441 | 0,286386855 | 0,277 | 0,22 | 1 | Basal | 1,219582078 | 0,819953014 | 1,259090909 |
| CNOT2 | 0,191661846 | -0,306986861 | 0,246 | 0,34 | 1 | Basal | 0,80832823 | 1,237121212 | 0,723529412 |
| STK38L | 0,192624859 | -0,279121193 | 0,215 | 0,293 | 1 | Basal | 0,824092855 | 1,213455492 | 0,733788396 |
| FKBP1A | 0,192724701 | 0,368656194 | 0,462 | 0,46 | 1 | Basal | 1,291149622 | 0,774503576 | 1,004347826 |
| PNRC1 | 0,192881279 | 0,412706829 | 0,846 | 0,847 | 1 | Basal | 1,331181076 | 0,751212602 | 0,998819362 |
| AC016876.1 | 0,193802392 | 0,259174012 | 0,123 | 0,073 | 1 | Basal | 1,196793307 | 0,83556617 | 1,684931507 |
| MTX1 | 0,195649252 | 0,25983471 | 0,338 | 0,3 | 1 | Basal | 1,197341517 | 0,835183601 | 1,126666667 |
| MUC20 | 0,195958096 | 0,517130527 | 0,492 | 0,54 | 1 | Basal | 1,431105993 | 0,698760263 | 0,911111111 |
| PRDX4 | 0,198084223 | 0,411315445 | 0,492 | 0,467 | 1 | Basal | 1,329897858 | 0,751937447 | 1,053533191 |
| PURA | 0,198724067 | -0,305050561 | 0,385 | 0,5 | 1 | Basal | 0,809413849 | 1,235461935 | 0,77 |

| Supplementary Table 4 - clusermarker | | | | | | | | | |
|---|---|---|---|---|---|---|---|---|---|
| | **p_val** | **avg_log2FC** | **pct.1** | **pct.2** | **p_val_adj** | **celltype** | **Foldchange_UP** | **Foldchange_DOWN** | **Ratio_pct1_pct2** |
| TJAP1 | 0,199260644 | 0,283101584 | 0,169 | 0,113 | 1 | Basal | 1,216808034 | 0,821822319 | 1,495575221 |
| TOR1AIP1 | 0,201446319 | -0,286116206 | 0,292 | 0,4 | 1 | Basal | 0,820106851 | 1,219353306 | 0,73 |
| HIPK1 | 0,202203086 | 0,309866174 | 0,308 | 0,26 | 1 | Basal | 1,239592709 | 0,806716588 | 1,184615385 |
| KLHDC9 | 0,202249161 | 0,271895542 | 0,123 | 0,073 | 1 | Basal | 1,207393167 | 0,828230627 | 1,684931507 |
| HTRA1 | 0,203344466 | 0,333069113 | 0,277 | 0,213 | 1 | Basal | 1,259690324 | 0,793845901 | 1,300469484 |
| SDF4 | 0,203579915 | 0,25247412 | 0,523 | 0,52 | 1 | Basal | 1,191248271 | 0,839455573 | 1,005769231 |
| MYO1E | 0,20435973 | 0,337316647 | 0,431 | 0,42 | 1 | Basal | 1,263404527 | 0,791512123 | 1,026190476 |
| BACE2 | 0,204611024 | 0,285531838 | 0,431 | 0,4 | 1 | Basal | 1,218859504 | 0,820439105 | 1,0775 |
| ARPC1A | 0,204803784 | 0,352834416 | 0,554 | 0,553 | 1 | Basal | 1,277067176 | 0,783044164 | 1,001808318 |
| PIAS1 | 0,205172467 | -0,25907153 | 0,354 | 0,46 | 1 | Basal | 0,835625527 | 1,196708295 | 0,769565217 |
| POLR3C | 0,205665361 | 0,28754139 | 0,215 | 0,16 | 1 | Basal | 1,220558454 | 0,819297098 | 1,34375 |
| NDUFAF8 | 0,205760472 | 0,319042419 | 0,369 | 0,34 | 1 | Basal | 1,247502251 | 0,80160176 | 1,085294118 |
| SLC9A3R2 | 0,205927952 | 0,751667384 | 0,462 | 0,487 | 1 | Basal | 1,683737674 | 0,593916746 | 0,948665298 |
| AP001816.1 | 0,206249032 | 0,391591991 | 0,138 | 0,087 | 1 | Basal | 1,3118402 | 0,762287968 | 1,586206897 |
| C15orf40 | 0,206915158 | 0,276926365 | 0,323 | 0,267 | 1 | Basal | 1,211610818 | 0,825347533 | 1,209737828 |
| SLC35B2 | 0,208099193 | -0,327596961 | 0,277 | 0,387 | 1 | Basal | 0,796862681 | 1,25492136 | 0,715762274 |
| C19orf53 | 0,208335045 | 0,28133302 | 0,646 | 0,68 | 1 | Basal | 1,215317294 | 0,822830388 | 0,95 |
| GBF1 | 0,208376795 | 0,41074 | 0,323 | 0,273 | 1 | Basal | 1,329367511 | 0,75223743 | 1,183150183 |
| SERPINB3 | 0,208505133 | -1,04231972 | 0,631 | 0,607 | 1 | Basal | 0,485546132 | 2,059536537 | 1,039538715 |
| SOX15 | 0,209573411 | -0,383212101 | 0,369 | 0,467 | 1 | Basal | 0,766728599 | 1,304242467 | 0,790149893 |
| PLEKHA1 | 0,210764408 | 0,293906328 | 0,354 | 0,32 | 1 | Basal | 1,225955258 | 0,815690453 | 1,10625 |
| C19orf25 | 0,211157253 | 0,305947334 | 0,185 | 0,127 | 1 | Basal | 1,236230131 | 0,808910877 | 1,456692913 |
| MPRIP | 0,211929099 | -0,306056523 | 0,292 | 0,387 | 1 | Basal | 0,808849658 | 1,236323697 | 0,754521964 |
| R3HDM4 | 0,212207153 | 0,356715449 | 0,338 | 0,3 | 1 | Basal | 1,280507275 | 0,780940507 | 1,126666667 |
| GRINA | 0,212467289 | 0,302676248 | 0,492 | 0,473 | 1 | Basal | 1,233430347 | 0,810747038 | 1,040169133 |
| PYCR2 | 0,212542979 | -0,3244088 | 0,277 | 0,373 | 1 | Basal | 0,798625587 | 1,252151216 | 0,742627346 |
| PLCB1 | 0,212704915 | -0,371727941 | 0,123 | 0,187 | 1 | Basal | 0,772856279 | 1,293901631 | 0,657754011 |
| HES1 | 0,213367506 | -0,42771598 | 0,462 | 0,58 | 1 | Basal | 0,743437837 | 1,345102375 | 0,796551724 |
| NFIC | 0,213997724 | 0,278420896 | 0,646 | 0,633 | 1 | Basal | 1,212866613 | 0,824492973 | 1,020537125 |
| GALNT5 | 0,214310266 | 0,335077016 | 0,415 | 0,367 | 1 | Basal | 1,261444747 | 0,792741816 | 1,130790191 |
| PDE12 | 0,21481166 | 0,281583722 | 0,215 | 0,16 | 1 | Basal | 1,215528502 | 0,822687414 | 1,34375 |

| Supplementary Table 4 - clusermarker | | | | | | | | | |
|---|---|---|---|---|---|---|---|---|---|
| | **p_val** | **avg_log2FC** | **pct.1** | **pct.2** | **p_val_adj** | **celltype** | **Foldchange_UP** | **Foldchange_DOWN** | **Ratio_pct1_pct2** |
| DNAJC4 | 0,215434973 | 0,300349853 | 0,431 | 0,387 | 1 | Basal | 1,231443002 | 0,812055449 | 1,11369509 |
| CREB3L1 | 0,217452451 | 0,437315438 | 0,4 | 0,387 | 1 | Basal | 1,35408231 | 0,738507543 | 1,033591731 |
| PRH1 | 0,21788253 | 0,564047668 | 0,062 | 0,12 | 1 | Basal | 1,478411279 | 0,676401766 | 0,516666667 |
| SMARCD2 | 0,218140221 | 0,30421231 | 0,354 | 0,32 | 1 | Basal | 1,234744301 | 0,809884281 | 1,10625 |
| HSPA5 | 0,218573912 | 0,266471022 | 0,708 | 0,72 | 1 | Basal | 1,202861904 | 0,831350628 | 0,983333333 |
| NINJ1 | 0,219420279 | 0,371533994 | 0,446 | 0,413 | 1 | Basal | 1,293727699 | 0,772960184 | 1,079903148 |
| NR2F6 | 0,219447724 | 0,308163731 | 0,462 | 0,447 | 1 | Basal | 1,238130798 | 0,80766911 | 1,033557047 |
| CNN2 | 0,220278782 | 0,325853737 | 0,646 | 0,653 | 1 | Basal | 1,253405942 | 0,797826121 | 0,989280245 |
| RNF10 | 0,220283863 | 0,320033288 | 0,446 | 0,433 | 1 | Basal | 1,248359353 | 0,801051394 | 1,030023095 |
| NUDT1 | 0,220928345 | 0,361021871 | 0,185 | 0,133 | 1 | Basal | 1,284335279 | 0,778612888 | 1,390977444 |
| RNF181 | 0,221369908 | 0,325730441 | 0,569 | 0,58 | 1 | Basal | 1,253298827 | 0,797894308 | 0,981034483 |
| MRPL48 | 0,221521861 | 0,259836815 | 0,123 | 0,2 | 1 | Basal | 1,197343264 | 0,835182382 | 0,615 |
| ECHDC1 | 0,222711733 | -0,343494023 | 0,277 | 0,367 | 1 | Basal | 0,788130248 | 1,268825809 | 0,754768392 |
| DGCR6L | 0,223060039 | 0,301085591 | 0,385 | 0,327 | 1 | Basal | 1,232071167 | 0,811641427 | 1,177370031 |
| AC018521.5 | 0,223201912 | -0,272544324 | 0,185 | 0,273 | 1 | Basal | 0,827858254 | 1,207936256 | 0,677655678 |
| GPC4 | 0,223270914 | 0,287095453 | 0,154 | 0,1 | 1 | Basal | 1,220181238 | 0,819550382 | 1,54 |
| NCL | 0,224123538 | -0,265418606 | 0,831 | 0,867 | 1 | Basal | 0,831957302 | 1,201984762 | 0,958477509 |
| TMC4 | 0,224131875 | 0,313182975 | 0,369 | 0,333 | 1 | Basal | 1,242445849 | 0,804864051 | 1,108108108 |
| DDOST | 0,225365955 | 0,375100731 | 0,538 | 0,567 | 1 | Basal | 1,296930105 | 0,771051575 | 0,948853616 |
| GMPS | 0,22538551 | -0,252858315 | 0,185 | 0,267 | 1 | Basal | 0,839232053 | 1,191565547 | 0,692883895 |
| SLPI | 0,228075312 | -0,423387842 | 0,831 | 0,9 | 1 | Basal | 0,745671526 | 1,341073066 | 0,923333333 |
| PTPN18 | 0,228179131 | 0,294391 | 0,292 | 0,233 | 1 | Basal | 1,226367185 | 0,815416469 | 1,253218884 |
| BAIAP2 | 0,228472482 | 0,296798363 | 0,508 | 0,487 | 1 | Basal | 1,22841528 | 0,814056953 | 1,04312115 |
| UBE2F | 0,228888347 | 0,257014339 | 0,246 | 0,2 | 1 | Basal | 1,195003082 | 0,836817925 | 1,23 |
| CXCL3 | 0,228888347 | 0,39755942 | 0,154 | 0,24 | 1 | Basal | 1,317277611 | 0,759141423 | 0,641666667 |
| CSNK1E | 0,229334669 | 0,338029826 | 0,446 | 0,433 | 1 | Basal | 1,26402923 | 0,791120946 | 1,030023095 |
| TSC22D1 | 0,229873615 | -0,486907647 | 0,769 | 0,74 | 1 | Basal | 0,713552929 | 1,401437734 | 1,039189189 |
| PI3 | 0,231407937 | 0,637914678 | 0,338 | 0,293 | 1 | Basal | 1,556078323 | 0,642641174 | 1,153583618 |
| PLD3 | 0,231536861 | 0,283641781 | 0,585 | 0,56 | 1 | Basal | 1,217263736 | 0,821514656 | 1,044642857 |
| CARMN | 0,232419767 | -0,38136671 | 0,077 | 0,127 | 1 | Basal | 0,76770997 | 1,302575241 | 0,606299213 |
| BECN1 | 0,232568957 | 0,260277298 | 0,385 | 0,333 | 1 | Basal | 1,197708892 | 0,834927424 | 1,156156156 |

| Supplementary Table 4 - clusermarker | | | | | | | | | |
|---|---|---|---|---|---|---|---|---|---|
| | **p_val** | **avg_log2FC** | **pct.1** | **pct.2** | **p_val_adj** | **celltype** | **Foldchange_UP** | **Foldchange_DOWN** | **Ratio_pct1_pct2** |
| PIM3 | 0,23287625 | 0,324546155 | 0,508 | 0,527 | 1 | Basal | 1,252270436 | 0,798549555 | 0,963946869 |
| LAMB3 | 0,235065633 | 0,290996171 | 0,554 | 0,56 | 1 | Basal | 1,223484794 | 0,817337498 | 0,989285714 |
| XPNPEP1 | 0,235336564 | -0,273102098 | 0,169 | 0,24 | 1 | Basal | 0,82753825 | 1,208403358 | 0,704166667 |
| SPINK5 | 0,235630203 | 0,436073881 | 0,415 | 0,387 | 1 | Basal | 1,352917512 | 0,739143363 | 1,072351421 |
| EBLN3P | 0,235997613 | 0,2709051 | 0,323 | 0,3 | 1 | Basal | 1,20656455 | 0,828799421 | 1,076666667 |
| IKZF2 | 0,236640974 | 0,256795562 | 0,323 | 0,273 | 1 | Basal | 1,19482188 | 0,836944834 | 1,183150183 |
| USP22 | 0,238579691 | 0,309799498 | 0,477 | 0,48 | 1 | Basal | 1,23953542 | 0,806753872 | 0,99375 |
| ZFAND2B | 0,238952802 | 0,292246624 | 0,308 | 0,253 | 1 | Basal | 1,224545706 | 0,816629379 | 1,217391304 |
| PSMB9 | 0,239371839 | 0,393992048 | 0,462 | 0,433 | 1 | Basal | 1,314024384 | 0,761020885 | 1,066974596 |
| PAX9 | 0,239473991 | 0,511824277 | 0,446 | 0,427 | 1 | Basal | 1,425852035 | 0,701335044 | 1,044496487 |
| SMG7 | 0,2396884 | 0,270493188 | 0,308 | 0,267 | 1 | Basal | 1,206220106 | 0,82903609 | 1,153558052 |
| PLPP5 | 0,240443181 | 0,346502601 | 0,277 | 0,233 | 1 | Basal | 1,271474564 | 0,786488404 | 1,188841202 |
| EXOC7 | 0,241198945 | 0,253920199 | 0,338 | 0,287 | 1 | Basal | 1,192442912 | 0,83861457 | 1,177700348 |
| PTPRF | 0,242532749 | 0,292632432 | 0,462 | 0,433 | 1 | Basal | 1,22487322 | 0,816411024 | 1,066974596 |
| S100P | 0,242617927 | 0,723425493 | 0,477 | 0,5 | 1 | Basal | 1,651097702 | 0,605657678 | 0,954 |
| GINM1 | 0,243492037 | -0,301314257 | 0,385 | 0,487 | 1 | Basal | 0,811512793 | 1,232266464 | 0,790554415 |
| CYP2S1 | 0,244713399 | 0,293796798 | 0,246 | 0,2 | 1 | Basal | 1,225862187 | 0,815752383 | 1,23 |
| SREK1 | 0,244822799 | 0,336196658 | 0,338 | 0,46 | 1 | Basal | 1,262424105 | 0,792126827 | 0,734782609 |
| UNC13B | 0,245019554 | 0,261759284 | 0,323 | 0,273 | 1 | Basal | 1,198939852 | 0,834070198 | 1,183150183 |
| EBNA1BP2 | 0,246483213 | 0,336971375 | 0,415 | 0,387 | 1 | Basal | 1,2631022 | 0,791701574 | 1,072351421 |
| LGALS7 | 0,246650707 | -0,361611426 | 0,108 | 0,167 | 1 | Basal | 0,778294773 | 1,284860228 | 0,646706587 |
| TTC3 | 0,246702061 | -0,296926925 | 0,508 | 0,64 | 1 | Basal | 0,813984414 | 1,228524752 | 0,79375 |
| MBD2 | 0,247487125 | -0,30434219 | 0,385 | 0,473 | 1 | Basal | 0,809811373 | 1,234855465 | 0,813953488 |
| MAGEF1 | 0,247785006 | 0,260360917 | 0,431 | 0,427 | 1 | Basal | 1,197778313 | 0,834879033 | 1,009367681 |
| SPRY1 | 0,248878159 | -0,432560174 | 0,169 | 0,227 | 1 | Basal | 0,740945753 | 1,349626469 | 0,744493392 |
| HNRNPR | 0,249261711 | 0,356714035 | 0,554 | 0,533 | 1 | Basal | 1,28050602 | 0,780941272 | 1,039399625 |
| CD302 | 0,250055693 | 0,293295995 | 0,138 | 0,22 | 1 | Basal | 1,225436727 | 0,816035604 | 0,627272727 |
| DDB1 | 0,250058338 | 0,264022603 | 0,477 | 0,48 | 1 | Basal | 1,200822241 | 0,832762724 | 0,99375 |
| CNOT8 | 0,250963584 | 0,302011526 | 0,292 | 0,253 | 1 | Basal | 1,232862175 | 0,811120675 | 1,154150198 |
| SAMM50 | 0,251277016 | 0,359151359 | 0,292 | 0,24 | 1 | Basal | 1,282671166 | 0,779623045 | 1,216666667 |
| CAV2 | 0,251332541 | 0,261994993 | 0,538 | 0,54 | 1 | Basal | 1,199135752 | 0,833933938 | 0,996296296 |

| Supplementary Table 4 - clusermarker | | | | | | | | | |
|---|---|---|---|---|---|---|---|---|---|
| | **p_val** | **avg_log2FC** | **pct.1** | **pct.2** | **p_val_adj** | **celltype** | **Foldchange_UP** | **Foldchange_DOWN** | **Ratio_pct1_pct2** |
| HDGF | 0,252106664 | 0,285797136 | 0,554 | 0,527 | 1 | Basal | 1,219083661 | 0,820288247 | 1,051233397 |
| DNM2 | 0,252132632 | 0,289024058 | 0,369 | 0,333 | 1 | Basal | 1,221813476 | 0,818455533 | 1,108108108 |
| ADD3 | 0,252566637 | -0,299480319 | 0,462 | 0,553 | 1 | Basal | 0,812545035 | 1,230701016 | 0,835443038 |
| KRAS | 0,253102353 | 0,324803657 | 0,462 | 0,453 | 1 | Basal | 1,252493969 | 0,798407038 | 1,01986755 |
| DCTN3 | 0,255390999 | 0,316746614 | 0,477 | 0,46 | 1 | Basal | 1,245518641 | 0,802878389 | 1,036956522 |
| TRIM56 | 0,256318344 | 0,285023224 | 0,446 | 0,413 | 1 | Basal | 1,218429877 | 0,820728397 | 1,079903148 |
| MT-ND6 | 0,256481946 | 0,333136902 | 0,646 | 0,6 | 1 | Basal | 1,259749516 | 0,7938086 | 1,076666667 |
| EPCAM | 0,256976639 | 0,287977171 | 0,462 | 0,46 | 1 | Basal | 1,220927193 | 0,819049658 | 1,004347826 |
| TMEM259 | 0,261074264 | 0,280608484 | 0,431 | 0,4 | 1 | Basal | 1,214707102 | 0,823243725 | 1,0775 |
| MRPL21 | 0,261507389 | 0,333308021 | 0,431 | 0,387 | 1 | Basal | 1,259898944 | 0,793714452 | 1,11369509 |
| JMJD7 | 0,262689122 | 0,273749722 | 0,215 | 0,173 | 1 | Basal | 1,20894593 | 0,827166853 | 1,242774566 |
| AK2 | 0,263876418 | 0,38147688 | 0,492 | 0,493 | 1 | Basal | 1,302674715 | 0,767651347 | 0,997971602 |
| APEX2 | 0,264287081 | -0,379722278 | 0,092 | 0,147 | 1 | Basal | 0,768585531 | 1,301091368 | 0,62585034 |
| CLCA2 | 0,26437316 | -0,284722569 | 0,108 | 0,167 | 1 | Basal | 0,820899453 | 1,218175985 | 0,646706587 |
| DMPK | 0,265038507 | 0,371914974 | 0,138 | 0,093 | 1 | Basal | 1,294069386 | 0,772756091 | 1,483870968 |
| EPC1 | 0,265044725 | 0,305202242 | 0,538 | 0,513 | 1 | Basal | 1,235591834 | 0,809328754 | 1,048732943 |
| MAT2A | 0,269719833 | -0,256299305 | 0,538 | 0,593 | 1 | Basal | 0,837232775 | 1,194410957 | 0,907251265 |
| EMP2 | 0,270121211 | 0,348453121 | 0,585 | 0,62 | 1 | Basal | 1,273194757 | 0,785425792 | 0,943548387 |
| BOD1 | 0,271201596 | 0,446302175 | 0,308 | 0,267 | 1 | Basal | 1,362543391 | 0,733921581 | 1,153558052 |
| GOLGA5 | 0,272847796 | -0,287034079 | 0,215 | 0,28 | 1 | Basal | 0,819585247 | 1,220129332 | 0,767857143 |
| COX17 | 0,273821142 | 0,282748437 | 0,508 | 0,527 | 1 | Basal | 1,216510217 | 0,822023511 | 0,963946869 |
| LACTB | 0,276981974 | 0,349230148 | 0,215 | 0,167 | 1 | Basal | 1,273880677 | 0,78500288 | 1,28742515 |
| MPC2 | 0,278364491 | 0,56747666 | 0,492 | 0,493 | 1 | Basal | 1,48192934 | 0,674796006 | 0,997971602 |
| RAP1A | 0,280420625 | 0,364827052 | 0,415 | 0,427 | 1 | Basal | 1,287727249 | 0,776561963 | 0,971896956 |
| OSR1 | 0,2826038 | -0,263590084 | 0,169 | 0,24 | 1 | Basal | 0,833012423 | 1,200462289 | 0,704166667 |
| HSPH1 | 0,282751097 | -0,376701131 | 0,538 | 0,56 | 1 | Basal | 0,770196712 | 1,298369604 | 0,960714286 |
| TRMT11 | 0,283176782 | -0,279802588 | 0,123 | 0,18 | 1 | Basal | 0,823703722 | 1,214028751 | 0,683333333 |
| C9orf78 | 0,284387711 | 0,347286711 | 0,4 | 0,387 | 1 | Basal | 1,272165803 | 0,78606106 | 1,033591731 |
| TPM1 | 0,28473787 | -0,355061244 | 0,462 | 0,567 | 1 | Basal | 0,781836453 | 1,279039877 | 0,814814815 |
| DDR1 | 0,285247665 | 0,283386489 | 0,585 | 0,587 | 1 | Basal | 1,217048355 | 0,82166004 | 0,996592845 |
| RWDD1 | 0,285554959 | 0,26977929 | 0,6 | 0,56 | 1 | Basal | 1,205623372 | 0,829446428 | 1,071428571 |

| Supplementary Table 4 - clusermarker | | | | | | | | | |
|---|---|---|---|---|---|---|---|---|---|
| | **p_val** | **avg_log2FC** | **pct.1** | **pct.2** | **p_val_adj** | **celltype** | **Foldchange_UP** | **Foldchange_DOWN** | **Ratio_pct1_pct2** |
| TLE1 | 0,286317021 | 0,303296534 | 0,246 | 0,207 | 1 | Basal | 1,233960774 | 0,810398532 | 1,188405797 |
| LSM3 | 0,287258074 | 0,250034739 | 0,538 | 0,56 | 1 | Basal | 1,189235751 | 0,840876167 | 0,960714286 |
| CYSTM1 | 0,287575138 | -0,294579175 | 0,569 | 0,607 | 1 | Basal | 0,815310119 | 1,226527155 | 0,937397035 |
| MUC16 | 0,288723059 | -0,561483676 | 0,508 | 0,547 | 1 | Basal | 0,677604953 | 1,475786144 | 0,928702011 |
| BAZ2A | 0,288764264 | 0,282599137 | 0,385 | 0,367 | 1 | Basal | 1,216384331 | 0,822108584 | 1,049046322 |
| RBBP4 | 0,290033225 | -0,314607 | 0,415 | 0,52 | 1 | Basal | 0,804069995 | 1,243672822 | 0,798076923 |
| WDR1 | 0,294214605 | 0,424393609 | 0,554 | 0,593 | 1 | Basal | 1,342008314 | 0,745151867 | 0,934232715 |
| JUNB | 0,294483949 | -0,350087332 | 0,923 | 0,9 | 1 | Basal | 0,784536605 | 1,274637784 | 1,025555556 |
| HTATSF1 | 0,29451093 | 0,403986937 | 0,4 | 0,38 | 1 | Basal | 1,323159459 | 0,755766807 | 1,052631579 |
| WIPF2 | 0,295704163 | 0,25984969 | 0,338 | 0,287 | 1 | Basal | 1,197353949 | 0,835174929 | 1,177700348 |
| PPP1CB | 0,296017526 | 0,315338809 | 0,508 | 0,513 | 1 | Basal | 1,244303837 | 0,803662233 | 0,990253411 |
| PIGR | 0,298655318 | 0,414858508 | 0,538 | 0,52 | 1 | Basal | 1,333167921 | 0,750093056 | 1,034615385 |
| RAP1B | 0,299303826 | 0,293826978 | 0,477 | 0,5 | 1 | Basal | 1,225887831 | 0,815735318 | 0,954 |
| SBNO2 | 0,300093104 | 0,277062275 | 0,2 | 0,16 | 1 | Basal | 1,211724964 | 0,825269785 | 1,25 |
| MARCHF7 | 0,300558033 | 0,331123462 | 0,338 | 0,3 | 1 | Basal | 1,257992623 | 0,794917221 | 1,126666667 |
| CRLF3 | 0,300640431 | 0,325877092 | 0,123 | 0,08 | 1 | Basal | 1,253426232 | 0,797813205 | 1,5375 |
| ARHGEF12 | 0,301924957 | 0,328785023 | 0,523 | 0,507 | 1 | Basal | 1,255955216 | 0,796206733 | 1,031558185 |
| H2AZ2 | 0,302784072 | 0,37350907 | 0,492 | 0,54 | 1 | Basal | 1,295500049 | 0,771902711 | 0,911111111 |
| FAM3D | 0,302825541 | 0,352568649 | 0,554 | 0,553 | 1 | Basal | 1,276831943 | 0,783188426 | 1,001808318 |
| PLOD1 | 0,303198852 | 0,297955211 | 0,2 | 0,153 | 1 | Basal | 1,229400699 | 0,81340445 | 1,307189542 |
| CFD | 0,307147733 | -0,729933686 | 0,308 | 0,367 | 1 | Basal | 0,602931627 | 1,658562854 | 0,839237057 |
| GPS2 | 0,307937798 | 0,440586864 | 0,369 | 0,373 | 1 | Basal | 1,357156284 | 0,736834815 | 0,989276139 |
| CLPTM1 | 0,308113489 | 0,282270212 | 0,354 | 0,313 | 1 | Basal | 1,216107034 | 0,822296041 | 1,130990415 |
| OPTN | 0,312043087 | 0,369721981 | 0,415 | 0,413 | 1 | Basal | 1,292103808 | 0,773931625 | 1,004842615 |
| CCDC88C | 0,312570209 | 0,297779976 | 0,262 | 0,227 | 1 | Basal | 1,229251381 | 0,813503255 | 1,154185022 |
| GTF2A2 | 0,312597252 | 0,325730922 | 0,538 | 0,547 | 1 | Basal | 1,253299245 | 0,797894042 | 0,983546618 |
| TUBA1B | 0,312616815 | -0,264893874 | 0,723 | 0,793 | 1 | Basal | 0,832259954 | 1,201547659 | 0,911727617 |
| RPL27A | 0,312773194 | -0,285397076 | 0,846 | 0,893 | 1 | Basal | 0,820515746 | 1,218745655 | 0,947368421 |
| FJX1 | 0,313655322 | -0,353258778 | 0,092 | 0,14 | 1 | Basal | 0,78281387 | 1,277442875 | 0,657142857 |
| EIF3M | 0,318735265 | -0,276535065 | 0,462 | 0,62 | 1 | Basal | 0,825571421 | 1,21128224 | 0,74516129 |
| COL7A1 | 0,320565252 | 0,42409334 | 0,185 | 0,14 | 1 | Basal | 1,34172903 | 0,745306972 | 1,321428571 |

| Supplementary Table 4 - clusermarker | | | | | | | | | |
|---|---|---|---|---|---|---|---|---|---|
| | **p_val** | **avg_log2FC** | **pct.1** | **pct.2** | **p_val_adj** | **celltype** | **Foldchange_UP** | **Foldchange_DOWN** | **Ratio_pct1_pct2** |
| ZNF100 | 0,3208346 | -0,277262679 | 0,108 | 0,167 | 1 | Basal | 0,825155154 | 1,211893296 | 0,646706587 |
| B3GNT3 | 0,322202853 | 0,400859926 | 0,215 | 0,173 | 1 | Basal | 1,320294645 | 0,757406693 | 1,242774566 |
| RARRES1 | 0,322680251 | 0,588032213 | 0,231 | 0,307 | 1 | Basal | 1,503195041 | 0,665249667 | 0,752442997 |
| KHSRP | 0,322995359 | 0,402889533 | 0,262 | 0,22 | 1 | Basal | 1,322153364 | 0,756341909 | 1,190909091 |
| RNASET2 | 0,323435546 | 0,306746312 | 0,354 | 0,34 | 1 | Basal | 1,236914957 | 0,808463019 | 1,041176471 |
| WDR83OS | 0,326653286 | 0,308770495 | 0,708 | 0,78 | 1 | Basal | 1,238651636 | 0,807329495 | 0,907692308 |
| YME1L1 | 0,329502557 | 0,250356514 | 0,523 | 0,493 | 1 | Basal | 1,189501024 | 0,840688641 | 1,060851927 |
| CP | 0,330821739 | -0,398570179 | 0,446 | 0,493 | 1 | Basal | 0,758609751 | 1,318200825 | 0,904665314 |
| HTATIP2 | 0,332192022 | 0,259030013 | 0,277 | 0,253 | 1 | Basal | 1,196673858 | 0,835649574 | 1,09486166 |
| MRPL37 | 0,332551701 | 0,297872899 | 0,369 | 0,36 | 1 | Basal | 1,229330559 | 0,81345086 | 1,025 |
| COL9A2 | 0,332927718 | -0,256306736 | 0,2 | 0,253 | 1 | Basal | 0,837228463 | 1,194417109 | 0,790513834 |
| THAP7 | 0,333816832 | 0,2783928 | 0,308 | 0,273 | 1 | Basal | 1,212842993 | 0,82450903 | 1,128205128 |
| TSPAN31 | 0,33468653 | 0,263368688 | 0,338 | 0,32 | 1 | Basal | 1,20027808 | 0,833140267 | 1,05625 |
| CTSS | 0,335325778 | 0,293605323 | 0,369 | 0,36 | 1 | Basal | 1,225699501 | 0,815860657 | 1,025 |
| YIPF6 | 0,336052634 | 0,265716569 | 0,338 | 0,313 | 1 | Basal | 1,202233036 | 0,831785494 | 1,079872204 |
| WASL | 0,336189429 | 0,31234457 | 0,462 | 0,48 | 1 | Basal | 1,241724026 | 0,805331925 | 0,9625 |
| POLDIP3 | 0,336928405 | -0,285097446 | 0,169 | 0,233 | 1 | Basal | 0,820686174 | 1,218492563 | 0,725321888 |
| TBRG1 | 0,33704407 | 0,331092288 | 0,323 | 0,3 | 1 | Basal | 1,25796544 | 0,794934398 | 1,076666667 |
| CCDC80 | 0,337046206 | -0,311095076 | 0,415 | 0,46 | 1 | Basal | 0,806029711 | 1,240649056 | 0,902173913 |
| TSG101 | 0,33834582 | 0,331144691 | 0,431 | 0,38 | 1 | Basal | 1,258011134 | 0,794905524 | 1,134210526 |
| TMEM167A | 0,342255403 | 0,304492857 | 0,415 | 0,407 | 1 | Basal | 1,234984433 | 0,809726806 | 1,01965602 |
| BCAP31 | 0,342279652 | 0,333850226 | 0,692 | 0,68 | 1 | Basal | 1,260372539 | 0,793416208 | 1,017647059 |
| PLP2 | 0,34410254 | 0,518751041 | 0,523 | 0,547 | 1 | Basal | 1,432714392 | 0,697975818 | 0,956124314 |
| VMO1 | 0,34723608 | 0,638342663 | 0,554 | 0,507 | 1 | Basal | 1,556540012 | 0,642450558 | 1,09270217 |
| ZCRB1 | 0,348451614 | -0,293196621 | 0,446 | 0,533 | 1 | Basal | 0,816091815 | 1,225352321 | 0,836772983 |
| EDNRA | 0,349649823 | -0,328213747 | 0,138 | 0,193 | 1 | Basal | 0,796522076 | 1,255457984 | 0,715025907 |
| NUDT21 | 0,350378362 | -0,37213065 | 0,354 | 0,427 | 1 | Basal | 0,772640576 | 1,294262857 | 0,829039813 |
| GRTP1 | 0,350982675 | 0,323869776 | 0,262 | 0,233 | 1 | Basal | 1,251683471 | 0,798924028 | 1,124463519 |
| PLSCR1 | 0,351595164 | 0,291840933 | 0,385 | 0,367 | 1 | Basal | 1,224201408 | 0,81685905 | 1,049046322 |
| MAPRE3 | 0,351622669 | 0,357898922 | 0,123 | 0,087 | 1 | Basal | 1,281558132 | 0,780300148 | 1,413793103 |
| PAFAH1B3 | 0,352578913 | 0,619902844 | 0,369 | 0,353 | 1 | Basal | 1,536771686 | 0,650714748 | 1,045325779 |

| Supplementary Table 4 - clusermarker | | | | | | | | | |
|---|---|---|---|---|---|---|---|---|---|
| | **p_val** | **avg_log2FC** | **pct.1** | **pct.2** | **p_val_adj** | **celltype** | **Foldchange_UP** | **Foldchange_DOWN** | **Ratio_pct1_pct2** |
| FOSL1 | 0,354623957 | -0,382089805 | 0,262 | 0,313 | 1 | Basal | 0,767325282 | 1,30322827 | 0,837060703 |
| DESI1 | 0,354923076 | 0,365892675 | 0,246 | 0,213 | 1 | Basal | 1,28867876 | 0,775988579 | 1,154929577 |
| GRWD1 | 0,363621396 | 0,406362602 | 0,169 | 0,133 | 1 | Basal | 1,325340081 | 0,754523321 | 1,270676692 |
| KXD1 | 0,363706128 | 0,373312859 | 0,4 | 0,407 | 1 | Basal | 1,295323869 | 0,7720077 | 0,982800983 |
| ZBTB7A | 0,364757538 | 0,292462245 | 0,569 | 0,587 | 1 | Basal | 1,224728738 | 0,816507337 | 0,969335605 |
| ASPSCR1 | 0,365014027 | 0,296030579 | 0,231 | 0,193 | 1 | Basal | 1,227761707 | 0,814490299 | 1,196891192 |
| TFF3 | 0,366027379 | -0,349156185 | 0,292 | 0,38 | 1 | Basal | 0,785043126 | 1,27381537 | 0,768421053 |
| SYNGR1 | 0,366177794 | 0,277813855 | 0,277 | 0,26 | 1 | Basal | 1,212356383 | 0,824839968 | 1,065384615 |
| NIT2 | 0,371981054 | 0,325350729 | 0,338 | 0,307 | 1 | Basal | 1,252969007 | 0,798104338 | 1,100977199 |
| CDC20B | 0,372342609 | -0,585601358 | 0,077 | 0,12 | 1 | Basal | 0,666371517 | 1,50066438 | 0,641666667 |
| PCYOX1 | 0,374154215 | 0,27220666 | 0,446 | 0,433 | 1 | Basal | 1,20765357 | 0,828052038 | 1,030023095 |
| GFUS | 0,377587654 | 0,257396186 | 0,354 | 0,347 | 1 | Basal | 1,195319412 | 0,836596469 | 1,020172911 |
| GIHCG | 0,378603789 | 0,537494152 | 0,123 | 0,087 | 1 | Basal | 1,451449274 | 0,688966551 | 1,413793103 |
| UTP23 | 0,380533868 | 0,252751151 | 0,277 | 0,253 | 1 | Basal | 1,19147704 | 0,839294394 | 1,09486166 |
| DCP2 | 0,383214555 | 0,301805103 | 0,231 | 0,193 | 1 | Basal | 1,232685788 | 0,81123674 | 1,196891192 |
| NHP2 | 0,383398372 | 0,404588798 | 0,569 | 0,633 | 1 | Basal | 1,323711568 | 0,755451584 | 0,898894155 |
| NDUFA9 | 0,383571137 | 0,262868801 | 0,431 | 0,433 | 1 | Basal | 1,199862262 | 0,833428996 | 0,995381062 |
| PDHA1 | 0,38404844 | 0,253339828 | 0,338 | 0,327 | 1 | Basal | 1,191963309 | 0,838951998 | 1,033639144 |
| MYADM | 0,388628768 | -0,289933933 | 0,585 | 0,633 | 1 | Basal | 0,817939515 | 1,222584289 | 0,924170616 |
| DDX39A | 0,390190268 | -0,253896855 | 0,215 | 0,273 | 1 | Basal | 0,83862814 | 1,192423617 | 0,787545788 |
| CNDP2 | 0,39137861 | 0,258359565 | 0,446 | 0,433 | 1 | Basal | 1,19611787 | 0,836038007 | 1,030023095 |
| S100B | 0,391841534 | -0,301172282 | 0,138 | 0,193 | 1 | Basal | 0,811592657 | 1,232145204 | 0,715025907 |
| TMEM50B | 0,392175111 | 0,252111148 | 0,446 | 0,453 | 1 | Basal | 1,190948599 | 0,839666801 | 0,984547461 |
| AFDN | 0,392320685 | 0,334240795 | 0,477 | 0,493 | 1 | Basal | 1,260713796 | 0,793201442 | 0,967545639 |
| TMEM216 | 0,394790532 | 0,278464312 | 0,231 | 0,2 | 1 | Basal | 1,212903113 | 0,824468162 | 1,155 |
| CHST14 | 0,395395032 | 0,489334801 | 0,169 | 0,133 | 1 | Basal | 1,403797463 | 0,712353474 | 1,270676692 |
| THYN1 | 0,398543865 | 0,257886326 | 0,385 | 0,353 | 1 | Basal | 1,195725578 | 0,836312293 | 1,090651558 |
| SLC5A8 | 0,399831964 | 0,325362034 | 0,2 | 0,167 | 1 | Basal | 1,252978825 | 0,798098084 | 1,19760479 |
| NTS | 0,401072551 | 0,596122175 | 0,477 | 0,467 | 1 | Basal | 1,511647937 | 0,661529696 | 1,021413276 |
| DPH3 | 0,402784146 | 0,265007037 | 0,2 | 0,173 | 1 | Basal | 1,20164191 | 0,832194676 | 1,156069364 |
| RNF213 | 0,405052576 | 0,257533141 | 0,462 | 0,48 | 1 | Basal | 1,19543289 | 0,836517055 | 0,9625 |

| Supplementary Table 4 - clusermarker | | | | | | | | | |
|---|---|---|---|---|---|---|---|---|---|
| | **p_val** | **avg_log2FC** | **pct.1** | **pct.2** | **p_val_adj** | **celltype** | **Foldchange_UP** | **Foldchange_DOWN** | **Ratio_pct1_pct2** |
| NDUFB6 | 0,40697658 | 0,305357444 | 0,569 | 0,573 | 1 | Basal | 1,235724764 | 0,809241693 | 0,993019197 |
| HMGB3 | 0,407113376 | 0,317717085 | 0,415 | 0,393 | 1 | Basal | 1,246356757 | 0,802338491 | 1,055979644 |
| EIF4E | 0,408364486 | 0,446735177 | 0,4 | 0,487 | 1 | Basal | 1,362952399 | 0,733701339 | 0,821355236 |
| ATG4C | 0,409170813 | 0,278791165 | 0,077 | 0,12 | 1 | Basal | 1,213177936 | 0,824281394 | 0,641666667 |
| WBP1 | 0,411059701 | 0,268919057 | 0,369 | 0,34 | 1 | Basal | 1,204904712 | 0,829941148 | 1,085294118 |
| GNAS | 0,411318059 | -0,27449721 | 0,846 | 0,84 | 1 | Basal | 0,826738392 | 1,209572471 | 1,007142857 |
| RRAGA | 0,413203321 | 0,253374288 | 0,369 | 0,353 | 1 | Basal | 1,191991781 | 0,838931959 | 1,045325779 |
| TAGLN | 0,414214377 | -0,784833774 | 0,338 | 0,373 | 1 | Basal | 0,580418829 | 1,722893798 | 0,90616622 |
| LIN7C | 0,414280614 | -0,270646001 | 0,246 | 0,3 | 1 | Basal | 0,828948281 | 1,206347878 | 0,82 |
| EIF4H | 0,414883197 | 0,319512531 | 0,554 | 0,56 | 1 | Basal | 1,247908824 | 0,801340595 | 0,989285714 |
| PI4KB | 0,416032038 | 0,283406961 | 0,308 | 0,28 | 1 | Basal | 1,217065625 | 0,821648381 | 1,1 |
| DNAJB14 | 0,417391661 | 0,270621575 | 0,292 | 0,393 | 1 | Basal | 1,206327453 | 0,828962316 | 0,743002545 |
| PLAC8 | 0,419343161 | 0,390252285 | 0,492 | 0,493 | 1 | Basal | 1,310622573 | 0,762996167 | 0,997971602 |
| ADO | 0,419502719 | 0,336606061 | 0,108 | 0,08 | 1 | Basal | 1,262782403 | 0,791902071 | 1,35 |
| ZZEF1 | 0,420647184 | 0,286279783 | 0,277 | 0,26 | 1 | Basal | 1,219491568 | 0,82001387 | 1,065384615 |
| SRP68 | 0,421095286 | 0,332784381 | 0,246 | 0,227 | 1 | Basal | 1,259441735 | 0,79400259 | 1,083700441 |
| VEZF1 | 0,423229141 | 0,405340357 | 0,4 | 0,407 | 1 | Basal | 1,324401323 | 0,75505814 | 0,982800983 |
| MYL9 | 0,423943768 | -0,488181879 | 0,431 | 0,487 | 1 | Basal | 0,712922975 | 1,402676074 | 0,885010267 |
| TRAK1 | 0,42479956 | 0,356552358 | 0,446 | 0,5 | 1 | Basal | 1,280362526 | 0,781028794 | 0,892 |
| GAPVD1 | 0,426282547 | 0,301372058 | 0,338 | 0,32 | 1 | Basal | 1,232315836 | 0,81148028 | 1,05625 |
| HSPB1 | 0,427310114 | 0,278596752 | 0,938 | 0,933 | 1 | Basal | 1,213014463 | 0,824392479 | 1,005359057 |
| SMAGP | 0,427542518 | 0,28579562 | 0,338 | 0,327 | 1 | Basal | 1,21908238 | 0,82028911 | 1,033639144 |
| CHPT1 | 0,433288203 | 0,354948535 | 0,477 | 0,473 | 1 | Basal | 1,278939957 | 0,781897535 | 1,00845666 |
| COL6A2 | 0,435063176 | 0,291004038 | 0,231 | 0,193 | 1 | Basal | 1,223491465 | 0,817333041 | 1,196891192 |
| HSPD1 | 0,437423978 | -0,364481784 | 0,677 | 0,713 | 1 | Basal | 0,776747833 | 1,287419105 | 0,949509116 |
| LBH | 0,43963133 | 0,407887774 | 0,492 | 0,473 | 1 | Basal | 1,326741931 | 0,753726084 | 1,040169133 |
| BLVRA | 0,441167637 | 0,273212133 | 0,431 | 0,44 | 1 | Basal | 1,208495527 | 0,827475135 | 0,979545455 |
| DAP | 0,441492872 | 0,261769076 | 0,4 | 0,4 | 1 | Basal | 1,198947989 | 0,834064537 | 1 |
| SNHG5 | 0,442982641 | 0,413504828 | 0,615 | 0,633 | 1 | Basal | 1,331917597 | 0,750797198 | 0,971563981 |
| SMAD7 | 0,443201768 | 0,29605491 | 0,185 | 0,247 | 1 | Basal | 1,227782413 | 0,814476563 | 0,748987854 |
| PFKL | 0,44754746 | 0,291798003 | 0,369 | 0,38 | 1 | Basal | 1,224164981 | 0,816883358 | 0,971052632 |

| Supplementary Table 4 - clusermarker | | | | | | | | | |
|---|---|---|---|---|---|---|---|---|---|
| | p_val | avg_log2FC | pct.1 | pct.2 | p_val_adj | celltype | Foldchange_UP | Foldchange_DOWN | Ratio_pct1_pct2 |
| BAG3 | 0,450046569 | -0,284241402 | 0,569 | 0,62 | 1 | Basal | 0,821173285 | 1,217769767 | 0,917741935 |
| ALDH1A1 | 0,451492635 | -0,350094428 | 0,615 | 0,64 | 1 | Basal | 0,784532747 | 1,274644053 | 0,9609375 |
| SVIL | 0,456650142 | -0,268415197 | 0,323 | 0,393 | 1 | Basal | 0,830231055 | 1,204483973 | 0,821882952 |
| PRPF6 | 0,457131998 | 0,434314492 | 0,462 | 0,46 | 1 | Basal | 1,351268614 | 0,74004531 | 1,004347826 |
| ADH7 | 0,459972285 | -0,273963345 | 0,477 | 0,533 | 1 | Basal | 0,827044381 | 1,209124954 | 0,894934334 |
| SETD3 | 0,464379717 | 0,3106997 | 0,338 | 0,333 | 1 | Basal | 1,240309098 | 0,806250637 | 1,015015015 |
| VIM | 0,467704823 | 0,275142577 | 0,538 | 0,507 | 1 | Basal | 1,210113675 | 0,826368646 | 1,061143984 |
| CCNG1 | 0,46872268 | 0,258053457 | 0,185 | 0,247 | 1 | Basal | 1,195864107 | 0,836215414 | 0,748987854 |
| CHD1 | 0,469161085 | 0,254159815 | 0,338 | 0,427 | 1 | Basal | 1,19264098 | 0,838475297 | 0,791569087 |
| NRIP1 | 0,473484201 | 0,475523544 | 0,354 | 0,347 | 1 | Basal | 1,390422703 | 0,719205748 | 1,020172911 |
| ACYP2 | 0,481538054 | 0,306166895 | 0,292 | 0,28 | 1 | Basal | 1,236418285 | 0,80878778 | 1,042857143 |
| KRT14 | 0,481564698 | 1,29187005 | 0,385 | 0,427 | 1 | Basal | 2,448452232 | 0,408421282 | 0,901639344 |
| C1S | 0,48769625 | 0,503529183 | 0,231 | 0,207 | 1 | Basal | 1,417677308 | 0,70537914 | 1,115942029 |
| RGCC | 0,488120152 | 0,358524912 | 0,508 | 0,533 | 1 | Basal | 1,282114325 | 0,779961646 | 0,953095685 |
| ITGAV | 0,488225801 | 0,338332814 | 0,338 | 0,34 | 1 | Basal | 1,264294723 | 0,790954816 | 0,994117647 |
| PTPN3 | 0,490071208 | 0,336782624 | 0,323 | 0,313 | 1 | Basal | 1,262936957 | 0,791805161 | 1,031948882 |
| CISD1 | 0,490357185 | 0,283253725 | 0,415 | 0,433 | 1 | Basal | 1,216936361 | 0,821735657 | 0,958429561 |
| RC3H2 | 0,491304773 | 0,36304775 | 0,308 | 0,307 | 1 | Basal | 1,286140051 | 0,777520301 | 1,003257329 |
| ERN2 | 0,492408015 | 0,277832806 | 0,262 | 0,253 | 1 | Basal | 1,212372309 | 0,824829133 | 1,035573123 |
| SDHC | 0,493502166 | -0,301601829 | 0,523 | 0,613 | 1 | Basal | 0,81135105 | 1,232512116 | 0,853181077 |
| CCDC91 | 0,494772488 | -0,307370498 | 0,231 | 0,287 | 1 | Basal | 0,808113311 | 1,237450227 | 0,804878049 |
| IFNGR1 | 0,496251995 | 0,365171353 | 0,415 | 0,42 | 1 | Basal | 1,288034604 | 0,776376657 | 0,988095238 |
| SKIV2L | 0,496932725 | 0,257564115 | 0,169 | 0,147 | 1 | Basal | 1,195458555 | 0,836499095 | 1,149659864 |
| TM4SF1 | 0,498738024 | 0,319951213 | 0,6 | 0,607 | 1 | Basal | 1,248288336 | 0,801096967 | 0,988467875 |
| ITM2C | 0,501398009 | -0,268935566 | 0,292 | 0,34 | 1 | Basal | 0,829931651 | 1,2049185 | 0,858823529 |
| GOT2 | 0,501756116 | -0,263948165 | 0,262 | 0,313 | 1 | Basal | 0,832805693 | 1,200760284 | 0,837060703 |
| PAXX | 0,505054584 | 0,339378825 | 0,323 | 0,327 | 1 | Basal | 1,26521172 | 0,79038155 | 0,987767584 |
| ACAA1 | 0,505205678 | 0,267669645 | 0,354 | 0,347 | 1 | Basal | 1,203861683 | 0,830660211 | 1,020172911 |
| GORASP2 | 0,506517691 | 0,387011875 | 0,354 | 0,373 | 1 | Basal | 1,307682112 | 0,764711845 | 0,949061662 |
| TGIF2 | 0,50658776 | 0,416328543 | 0,2 | 0,253 | 1 | Basal | 1,334527046 | 0,749329138 | 0,790513834 |
| SEMA4C | 0,509655185 | 0,291039437 | 0,2 | 0,18 | 1 | Basal | 1,223521486 | 0,817312986 | 1,111111111 |

| Supplementary Table 4 - clusermarker | | | | | | | | | |
|---|---|---|---|---|---|---|---|---|---|
| | **p_val** | **avg_log2FC** | **pct.1** | **pct.2** | **p_val_adj** | **celltype** | **Foldchange_UP** | **Foldchange_DOWN** | **Ratio_pct1_pct2** |
| MCU | 0,509655185 | 0,347427118 | 0,2 | 0,18 | 1 | Basal | 1,27228962 | 0,785984562 | 1,111111111 |
| PCP4L1 | 0,511653966 | 0,57122481 | 0,169 | 0,227 | 1 | Basal | 1,485784427 | 0,673045148 | 0,744493392 |
| ECE1 | 0,512077982 | 0,255830182 | 0,477 | 0,48 | 1 | Basal | 1,194022632 | 0,837505063 | 0,99375 |
| IDO1 | 0,512169544 | 0,346754861 | 0,138 | 0,113 | 1 | Basal | 1,271696905 | 0,786350895 | 1,221238938 |
| PAIP2 | 0,515704813 | 0,330150864 | 0,431 | 0,473 | 1 | Basal | 1,257144828 | 0,795453298 | 0,911205074 |
| CEP63 | 0,515824195 | -0,252495624 | 0,246 | 0,287 | 1 | Basal | 0,839443061 | 1,191266027 | 0,857142857 |
| PRMT5 | 0,517077164 | -0,285534757 | 0,138 | 0,18 | 1 | Basal | 0,820437445 | 1,21886197 | 0,766666667 |
| ZG16B | 0,518396837 | -0,925023629 | 0,523 | 0,58 | 1 | Basal | 0,526671892 | 1,898715339 | 0,901724138 |
| ICE1 | 0,523863709 | 0,330345756 | 0,308 | 0,3 | 1 | Basal | 1,257314666 | 0,795345848 | 1,026666667 |
| NAA10 | 0,526306021 | 0,39034512 | 0,169 | 0,147 | 1 | Basal | 1,310706912 | 0,762947071 | 1,149659864 |
| PRRG4 | 0,527285876 | 0,361699511 | 0,323 | 0,333 | 1 | Basal | 1,284938679 | 0,778247255 | 0,96996997 |
| EMD | 0,527953565 | 0,321952733 | 0,462 | 0,487 | 1 | Basal | 1,250021347 | 0,799986338 | 0,948665298 |
| COASY | 0,529523563 | 0,368172597 | 0,215 | 0,193 | 1 | Basal | 1,290716896 | 0,774763237 | 1,113989637 |
| NDUFS2 | 0,531079014 | 0,277092968 | 0,415 | 0,453 | 1 | Basal | 1,211750744 | 0,825252227 | 0,91611479 |
| AKT1 | 0,532978198 | 0,324217338 | 0,338 | 0,433 | 1 | Basal | 1,251985053 | 0,798731581 | 0,780600462 |
| MIA | 0,53563219 | -0,316946739 | 0,246 | 0,287 | 1 | Basal | 0,802767024 | 1,245691427 | 0,857142857 |
| BAZ1A | 0,535924577 | 0,37676372 | 0,446 | 0,46 | 1 | Basal | 1,298425934 | 0,770163299 | 0,969565217 |
| HEXD | 0,536250548 | 0,262853518 | 0,185 | 0,16 | 1 | Basal | 1,199849551 | 0,833437825 | 1,15625 |
| UPK1B | 0,53647188 | 0,335435492 | 0,492 | 0,487 | 1 | Basal | 1,261758225 | 0,792544863 | 1,01026694 |
| LGALS7B | 0,549180963 | 0,412835918 | 0,323 | 0,36 | 1 | Basal | 1,331300191 | 0,751145389 | 0,897222222 |
| MBOAT7 | 0,551283652 | 0,288162467 | 0,231 | 0,213 | 1 | Basal | 1,221084016 | 0,818944468 | 1,084507042 |
| PKN2 | 0,551805158 | 0,25703298 | 0,4 | 0,52 | 1 | Basal | 1,195018522 | 0,836807113 | 0,769230769 |
| A1BG | 0,552171916 | 0,260470041 | 0,108 | 0,087 | 1 | Basal | 1,197868916 | 0,834815886 | 1,24137931 |
| MMP10 | 0,552570622 | 1,733598105 | 0,2 | 0,273 | 1 | Basal | 3,325561856 | 0,300701067 | 0,732600733 |
| HIF1AN | 0,554573702 | 0,294188228 | 0,215 | 0,2 | 1 | Basal | 1,226194831 | 0,815531084 | 1,075 |
| BPIFB1 | 0,555068437 | 0,958035779 | 0,462 | 0,46 | 1 | Basal | 1,942663168 | 0,514757276 | 1,004347826 |
| FUCA1 | 0,555160703 | 0,25586379 | 0,369 | 0,353 | 1 | Basal | 1,194050447 | 0,837485554 | 1,045325779 |
| S100A8 | 0,556993052 | 1,969209349 | 0,123 | 0,1 | 1 | Basal | 3,915534743 | 0,255392958 | 1,23 |
| UBA1 | 0,560369137 | 0,2851721 | 0,446 | 0,473 | 1 | Basal | 1,218555617 | 0,820643708 | 0,942917548 |
| MMP7 | 0,561714114 | 0,581824619 | 0,108 | 0,087 | 1 | Basal | 1,496741025 | 0,668118254 | 1,24137931 |
| PMPCA | 0,577507237 | 0,307289626 | 0,185 | 0,153 | 1 | Basal | 1,237380862 | 0,808158612 | 1,209150327 |

| Supplementary Table 4 - clusermarker | | | | | | | | | |
|---|---|---|---|---|---|---|---|---|---|
| | **p_val** | **avg_log2FC** | **pct.1** | **pct.2** | **p_val_adj** | **celltype** | **Foldchange_UP** | **Foldchange_DOWN** | **Ratio_pct1_pct2** |
| USF3 | 0,581890968 | 0,261136114 | 0,169 | 0,153 | 1 | Basal | 1,198422083 | 0,834430552 | 1,104575163 |
| PNKP | 0,582467599 | 0,382558279 | 0,246 | 0,227 | 1 | Basal | 1,303651525 | 0,767076156 | 1,083700441 |
| YES1 | 0,582738236 | 0,263344779 | 0,323 | 0,333 | 1 | Basal | 1,200258189 | 0,833154074 | 0,96996997 |
| POLR1H | 0,582995612 | 0,326287959 | 0,323 | 0,327 | 1 | Basal | 1,253783248 | 0,797586027 | 0,987767584 |
| UBE2I | 0,58624838 | 0,29111994 | 0,477 | 0,513 | 1 | Basal | 1,223589761 | 0,817267381 | 0,929824561 |
| NDUFS4 | 0,587511338 | 0,350812856 | 0,492 | 0,527 | 1 | Basal | 1,275278954 | 0,784142165 | 0,933586338 |
| LRP11 | 0,589364096 | 0,290333382 | 0,277 | 0,253 | 1 | Basal | 1,222922841 | 0,817713077 | 1,09486166 |
| LCMT1 | 0,589886618 | 0,258228552 | 0,262 | 0,32 | 1 | Basal | 1,196009254 | 0,836113932 | 0,81875 |
| DFFA | 0,591883494 | 0,263686457 | 0,246 | 0,22 | 1 | Basal | 1,200542484 | 0,832956779 | 1,118181818 |
| EPB41L1 | 0,592290307 | 0,41513526 | 0,385 | 0,427 | 1 | Basal | 1,333423686 | 0,74994918 | 0,901639344 |
| SERPINA3 | 0,595488004 | 0,969357875 | 0,169 | 0,147 | 1 | Basal | 1,957968935 | 0,510733333 | 1,149659864 |
| HRAS | 0,59693475 | 0,355529232 | 0,338 | 0,367 | 1 | Basal | 1,279454845 | 0,781582878 | 0,920980926 |
| MTERF1 | 0,597368152 | 0,323869958 | 0,108 | 0,087 | 1 | Basal | 1,251683629 | 0,798923927 | 1,24137931 |
| STRN3 | 0,604464334 | 0,262196566 | 0,292 | 0,293 | 1 | Basal | 1,199303307 | 0,833817429 | 0,996587031 |
| EYA2 | 0,605156332 | 0,25308161 | 0,323 | 0,32 | 1 | Basal | 1,191749987 | 0,83910217 | 1,009375 |
| ISCU | 0,606086028 | 0,356225661 | 0,538 | 0,567 | 1 | Basal | 1,280072623 | 0,781205677 | 0,948853616 |
| DSC3 | 0,607841908 | 0,298402973 | 0,277 | 0,273 | 1 | Basal | 1,229782321 | 0,813152037 | 1,014652015 |
| OGFOD1 | 0,608878649 | -0,282002603 | 0,169 | 0,193 | 1 | Basal | 0,822448585 | 1,215881477 | 0,875647668 |
| DNAJB1 | 0,61097365 | -0,404401815 | 0,754 | 0,753 | 1 | Basal | 0,755549501 | 1,323540017 | 1,001328021 |
| CGGBP1 | 0,612072018 | 0,321654969 | 0,385 | 0,4 | 1 | Basal | 1,249763377 | 0,800151468 | 0,9625 |
| SH3PXD2A | 0,616824121 | 0,321004815 | 0,369 | 0,407 | 1 | Basal | 1,249200295 | 0,800512139 | 0,906633907 |
| LDHA | 0,621045667 | 0,515620463 | 0,6 | 0,707 | 1 | Basal | 1,429608842 | 0,699492036 | 0,848656294 |
| MYG1 | 0,621055515 | 0,289343939 | 0,308 | 0,313 | 1 | Basal | 1,222084412 | 0,818274081 | 0,984025559 |
| IRF2 | 0,621386065 | 0,291970668 | 0,446 | 0,46 | 1 | Basal | 1,2243115 | 0,816785597 | 0,969565217 |
| CYP4B1 | 0,625135968 | -0,266364211 | 0,569 | 0,573 | 1 | Basal | 0,83141218 | 1,202772853 | 0,993019197 |
| SMIM26 | 0,632113957 | 0,253490719 | 0,554 | 0,567 | 1 | Basal | 1,192087983 | 0,838864257 | 0,97707231 |
| ACBD3 | 0,634328703 | 0,251437963 | 0,446 | 0,46 | 1 | Basal | 1,190393012 | 0,840058695 | 0,969565217 |
| KAT8 | 0,635726086 | 0,277995734 | 0,277 | 0,273 | 1 | Basal | 1,212509233 | 0,824735988 | 1,014652015 |
| PHYH | 0,636162304 | 0,305944132 | 0,292 | 0,287 | 1 | Basal | 1,236227388 | 0,808912672 | 1,017421603 |
| SPARC | 0,636940623 | -0,32860562 | 0,277 | 0,3 | 1 | Basal | 0,79630575 | 1,255799045 | 0,923333333 |
| NDRG2 | 0,637841345 | 0,268586363 | 0,523 | 0,573 | 1 | Basal | 1,204626885 | 0,83013256 | 0,912739965 |

| Supplementary Table 4 - clusermarker | | | | | | | | | |
|---|---|---|---|---|---|---|---|---|---|
| | **p_val** | **avg_log2FC** | **pct.1** | **pct.2** | **p_val_adj** | **celltype** | **Foldchange_UP** | **Foldchange_DOWN** | **Ratio_pct1_pct2** |
| C6orf132 | 0,640409458 | 0,263411484 | 0,354 | 0,347 | 1 | Basal | 1,200313686 | 0,833115553 | 1,020172911 |
| TDRP | 0,641933166 | 0,300484875 | 0,092 | 0,12 | 1 | Basal | 1,231558258 | 0,811979453 | 0,766666667 |
| NANS | 0,641947513 | 0,260765469 | 0,446 | 0,493 | 1 | Basal | 1,198114234 | 0,834644954 | 0,904665314 |
| PJA1 | 0,642739562 | 0,42228599 | 0,215 | 0,26 | 1 | Basal | 1,340049218 | 0,746241247 | 0,826923077 |
| EFHD1 | 0,650618651 | 0,370625234 | 0,169 | 0,153 | 1 | Basal | 1,292913031 | 0,773447228 | 1,104575163 |
| CAPN2 | 0,650880742 | 0,388602171 | 0,554 | 0,673 | 1 | Basal | 1,309124377 | 0,76386936 | 0,823179792 |
| FLYWCH2 | 0,654874804 | 0,260636053 | 0,338 | 0,34 | 1 | Basal | 1,198006763 | 0,834719828 | 0,994117647 |
| EFEMP1 | 0,656267879 | 0,659827885 | 0,523 | 0,593 | 1 | Basal | 1,57989413 | 0,632953804 | 0,881956155 |
| BCAM | 0,66625271 | 0,589243448 | 0,446 | 0,513 | 1 | Basal | 1,5044576 | 0,664691381 | 0,869395712 |
| APOD | 0,667542925 | -0,687562567 | 0,292 | 0,347 | 1 | Basal | 0,620901978 | 1,610560178 | 0,841498559 |
| AJUBA | 0,675874714 | 0,331300735 | 0,154 | 0,14 | 1 | Basal | 1,25814721 | 0,794819551 | 1,1 |
| SH3PXD2B | 0,677683474 | 0,273903568 | 0,262 | 0,253 | 1 | Basal | 1,209074857 | 0,82707865 | 1,035573123 |
| KLHDC3 | 0,681060365 | 0,287800558 | 0,2 | 0,24 | 1 | Basal | 1,220777737 | 0,819149932 | 0,833333333 |
| ATP2C2 | 0,681517613 | 0,251497499 | 0,277 | 0,293 | 1 | Basal | 1,190442137 | 0,840024028 | 0,945392491 |
| MRPS18C | 0,682030126 | -0,252399887 | 0,4 | 0,46 | 1 | Basal | 0,839498768 | 1,191186978 | 0,869565217 |
| DEF8 | 0,684278603 | 0,477768973 | 0,169 | 0,16 | 1 | Basal | 1,39258846 | 0,718087238 | 1,05625 |
| CTSF | 0,687544851 | 0,417502987 | 0,277 | 0,287 | 1 | Basal | 1,335613876 | 0,748719385 | 0,965156794 |
| MCRIP1 | 0,689112489 | 0,267890759 | 0,262 | 0,253 | 1 | Basal | 1,204046207 | 0,83053291 | 1,035573123 |
| MSLN | 0,69179533 | 0,346017148 | 0,492 | 0,507 | 1 | Basal | 1,271046797 | 0,786753094 | 0,970414201 |
| UBP1 | 0,694075815 | 0,266183462 | 0,246 | 0,253 | 1 | Basal | 1,202622172 | 0,831516351 | 0,972332016 |
| DMAC2 | 0,695369871 | 0,264656935 | 0,108 | 0,093 | 1 | Basal | 1,201350341 | 0,83239665 | 1,161290323 |
| ZXDB | 0,696555725 | -0,329962716 | 0,123 | 0,14 | 1 | Basal | 0,795557043 | 1,25698089 | 0,878571429 |
| KLF10 | 0,69696403 | 0,380396417 | 0,477 | 0,487 | 1 | Basal | 1,301699481 | 0,768226472 | 0,979466119 |
| STK39 | 0,703009879 | 0,260831448 | 0,354 | 0,407 | 1 | Basal | 1,198169029 | 0,834606784 | 0,86977887 |
| RGMA | 0,704719651 | 0,279898979 | 0,108 | 0,1 | 1 | Basal | 1,214109866 | 0,823648689 | 1,08 |
| SPAG9 | 0,709059272 | 0,407921088 | 0,369 | 0,4 | 1 | Basal | 1,326772568 | 0,75370868 | 0,9225 |
| SOD2 | 0,713806182 | 0,284405204 | 0,569 | 0,573 | 1 | Basal | 1,21790804 | 0,821080055 | 0,993019197 |
| NSF | 0,716270843 | 0,410763623 | 0,262 | 0,26 | 1 | Basal | 1,329389278 | 0,752225113 | 1,007692308 |
| MEX3D | 0,718684871 | 0,27939268 | 0,123 | 0,113 | 1 | Basal | 1,213683862 | 0,823937791 | 1,088495575 |
| CELSR2 | 0,719194075 | 0,260345588 | 0,215 | 0,24 | 1 | Basal | 1,197765587 | 0,834887904 | 0,895833333 |
| CYRIB | 0,728361073 | 0,340168646 | 0,277 | 0,3 | 1 | Basal | 1,265904565 | 0,789948964 | 0,923333333 |

| Supplementary Table 4 - clusermarker | | | | | | | | | |
|---|---|---|---|---|---|---|---|---|---|
| | **p_val** | **avg_log2FC** | **pct.1** | **pct.2** | **p_val_adj** | **celltype** | **Foldchange_UP** | **Foldchange_DOWN** | **Ratio_pct1_pct2** |
| AC244090.1 | 0,731550465 | 0,459769362 | 0,123 | 0,113 | 1 | Basal | 1,375321933 | 0,727102488 | 1,088495575 |
| SYT8 | 0,733173618 | 0,657711903 | 0,231 | 0,267 | 1 | Basal | 1,577578618 | 0,633882831 | 0,865168539 |
| KRT15 | 0,745386122 | 0,30991096 | 0,646 | 0,647 | 1 | Basal | 1,239631191 | 0,806691545 | 0,998454405 |
| POLR2K | 0,74699922 | 0,677630324 | 0,492 | 0,533 | 1 | Basal | 1,599510346 | 0,62519133 | 0,923076923 |
| INO80B | 0,747734458 | 0,253814629 | 0,123 | 0,14 | 1 | Basal | 1,192355658 | 0,838675938 | 0,878571429 |
| AC023157.3 | 0,751945393 | 0,333555949 | 0,2 | 0,2 | 1 | Basal | 1,260115478 | 0,793578063 | 1 |
| AK3 | 0,756478421 | 0,282272761 | 0,369 | 0,387 | 1 | Basal | 1,216109183 | 0,822294588 | 0,953488372 |
| SLC35A2 | 0,763222459 | 0,252490616 | 0,154 | 0,18 | 1 | Basal | 1,191261892 | 0,839445975 | 0,855555556 |
| STAT1 | 0,764525744 | 0,370849634 | 0,308 | 0,327 | 1 | Basal | 1,293114149 | 0,773326934 | 0,941896024 |
| SLAIN2 | 0,767503007 | 0,343194357 | 0,323 | 0,347 | 1 | Basal | 1,268562286 | 0,788293969 | 0,930835735 |
| MEN1 | 0,767842069 | 0,367579398 | 0,123 | 0,147 | 1 | Basal | 1,290186296 | 0,775081865 | 0,836734694 |
| FAAP100 | 0,775829437 | 0,260307151 | 0,092 | 0,113 | 1 | Basal | 1,197733676 | 0,834910148 | 0,814159292 |
| FHOD3 | 0,79297723 | -0,345280767 | 0,185 | 0,16 | 1 | Basal | 0,787154771 | 1,270398194 | 1,15625 |
| MBD1 | 0,793781918 | 0,28188834 | 0,2 | 0,2 | 1 | Basal | 1,215785182 | 0,822513726 | 1 |
| TMEM170A | 0,793805129 | 0,284820789 | 0,246 | 0,267 | 1 | Basal | 1,218258923 | 0,820843567 | 0,921348315 |
| ANXA8L1 | 0,796945282 | 0,308753126 | 0,123 | 0,14 | 1 | Basal | 1,238636724 | 0,807339215 | 0,878571429 |
| TRIOBP | 0,802801986 | 0,27341242 | 0,323 | 0,333 | 1 | Basal | 1,208663312 | 0,827360267 | 0,96996997 |
| HAUS4 | 0,810469178 | 0,264724824 | 0,338 | 0,373 | 1 | Basal | 1,201406874 | 0,832357481 | 0,90616622 |
| TFPI2 | 0,818549275 | 0,295803356 | 0,169 | 0,167 | 1 | Basal | 1,227568351 | 0,814618591 | 1,011976048 |
| ZBTB41 | 0,823966087 | -0,304042326 | 0,123 | 0,133 | 1 | Basal | 0,80997971 | 1,234598827 | 0,92481203 |
| VCL | 0,833418544 | 0,28003382 | 0,477 | 0,513 | 1 | Basal | 1,214223348 | 0,823571711 | 0,929824561 |
| KRT4 | 0,83380545 | 0,978960331 | 0,462 | 0,573 | 1 | Basal | 1,971044477 | 0,507345223 | 0,806282723 |
| ARHGAP18 | 0,836765798 | 0,480228719 | 0,292 | 0,313 | 1 | Basal | 1,394964801 | 0,716863966 | 0,932907348 |
| ESCO1 | 0,839024632 | 0,324848778 | 0,262 | 0,273 | 1 | Basal | 1,252533142 | 0,798382068 | 0,95970696 |
| IMMT | 0,839091565 | 0,289472871 | 0,292 | 0,333 | 1 | Basal | 1,222193633 | 0,818200957 | 0,876876877 |
| FLNA | 0,843021017 | 0,37360274 | 0,4 | 0,447 | 1 | Basal | 1,295584164 | 0,771852596 | 0,894854586 |
| ECPAS | 0,844309839 | 0,321814909 | 0,308 | 0,333 | 1 | Basal | 1,249901936 | 0,800062766 | 0,924924925 |
| ADD1 | 0,8451025 | 0,276914023 | 0,446 | 0,48 | 1 | Basal | 1,211600454 | 0,825354594 | 0,929166667 |
| SEC61A1 | 0,845115264 | 0,284551242 | 0,354 | 0,393 | 1 | Basal | 1,21803133 | 0,820996944 | 0,900763359 |
| NDUFA8 | 0,848086603 | 0,293389144 | 0,4 | 0,48 | 1 | Basal | 1,22551585 | 0,815982918 | 0,833333333 |
| ADSL | 0,861391114 | 0,269393547 | 0,277 | 0,293 | 1 | Basal | 1,20530106 | 0,829668233 | 0,945392491 |

| Supplementary Table 4 - clusermarker | | | | | | | | | |
|---|---|---|---|---|---|---|---|---|---|
| | **p_val** | **avg_log2FC** | **pct.1** | **pct.2** | **p_val_adj** | **celltype** | **Foldchange_UP** | **Foldchange_DOWN** | **Ratio_pct1_pct2** |
| MFGE8 | 0,86500734 | -0,303012173 | 0,231 | 0,213 | 1 | Basal | 0,810558281 | 1,23371758 | 1,084507042 |
| SLC34A2 | 0,867974246 | 0,442441907 | 0,277 | 0,3 | 1 | Basal | 1,358902462 | 0,735887989 | 0,923333333 |
| CLDN4 | 0,868212036 | -0,291156873 | 0,754 | 0,667 | 1 | Basal | 0,817246459 | 1,223621086 | 1,130434783 |
| NT5C | 0,878819089 | 0,367135418 | 0,431 | 0,467 | 1 | Basal | 1,289789311 | 0,775320428 | 0,922912206 |
| FXYD5 | 0,885632897 | 0,336172667 | 0,231 | 0,273 | 1 | Basal | 1,262403112 | 0,792139999 | 0,846153846 |
| TCEA1 | 0,885825775 | -0,310078009 | 0,492 | 0,587 | 1 | Basal | 0,806598144 | 1,239774735 | 0,838160136 |
| TMEM11 | 0,886604764 | 0,267706148 | 0,231 | 0,26 | 1 | Basal | 1,203892144 | 0,830639194 | 0,888461538 |
| VAT1 | 0,88807603 | 0,261882355 | 0,246 | 0,267 | 1 | Basal | 1,199042133 | 0,83399905 | 0,921348315 |
| GON4L | 0,888380114 | 0,266454703 | 0,277 | 0,333 | 1 | Basal | 1,202848298 | 0,831360032 | 0,831831832 |
| SAA1 | 0,889414269 | 1,089151747 | 0,308 | 0,333 | 1 | Basal | 2,12748911 | 0,470037659 | 0,924924925 |
| TEX30 | 0,896761267 | 0,345133433 | 0,092 | 0,1 | 1 | Basal | 1,270268462 | 0,787235163 | 0,92 |
| MRPL47 | 0,898126371 | 0,285153401 | 0,338 | 0,367 | 1 | Basal | 1,218539824 | 0,820654344 | 0,920980926 |
| MYO15B | 0,91632559 | 0,258665254 | 0,108 | 0,12 | 1 | Basal | 1,196371339 | 0,83586088 | 0,9 |
| MBTPS2 | 0,922476005 | 0,349108084 | 0,092 | 0,1 | 1 | Basal | 1,273772901 | 0,7850693 | 0,92 |
| TRAPPC2B | 0,926828474 | 0,28333932 | 0,123 | 0,133 | 1 | Basal | 1,217008564 | 0,821686905 | 0,92481203 |
| WDTC1 | 0,927815459 | 0,2930397 | 0,169 | 0,187 | 1 | Basal | 1,225219047 | 0,816180586 | 0,903743316 |
| CSTF1 | 0,92983918 | 0,299735841 | 0,092 | 0,1 | 1 | Basal | 1,23091901 | 0,812401134 | 0,92 |
| LIX1L | 0,932760945 | 0,301207088 | 0,169 | 0,18 | 1 | Basal | 1,232174931 | 0,811573077 | 0,938888889 |
| MED10 | 0,933586657 | 0,332638977 | 0,338 | 0,4 | 1 | Basal | 1,259314807 | 0,794082619 | 0,845 |
| VGLL4 | 0,938261343 | 0,332131228 | 0,462 | 0,533 | 1 | Basal | 1,258871676 | 0,794362141 | 0,866791745 |
| CEMIP | 0,947004747 | 0,259460509 | 0,108 | 0,12 | 1 | Basal | 1,197030995 | 0,835400256 | 0,9 |
| ARHGAP1 | 0,964226128 | -0,261552673 | 0,262 | 0,26 | 1 | Basal | 0,834189655 | 1,198768162 | 1,007692308 |
| NIPSNAP1 | 0,973537894 | 0,362138559 | 0,123 | 0,127 | 1 | Basal | 1,285329777 | 0,778010451 | 0,968503937 |
| CDH3 | 0,992315327 | 0,47664459 | 0,323 | 0,393 | 1 | Basal | 1,391503551 | 0,718647106 | 0,821882952 |
| LAP3 | 0,994550024 | 0,37858901 | 0,354 | 0,393 | 1 | Basal | 1,300069734 | 0,769189509 | 0,900763359 |
| SNRNP48 | 0,995323465 | 0,294596871 | 0,138 | 0,147 | 1 | Basal | 1,2265422 | 0,815300118 | 0,93877551 |
| BRI3BP | 0,99722169 | 0,311800951 | 0,185 | 0,193 | 1 | Basal | 1,241256223 | 0,805635437 | 0,958549223 |
| APOE | 0,998396113 | -0,465007975 | 0,138 | 0,133 | 1 | Basal | 0,724467073 | 1,380324984 | 1,037593985 |
| ITGBL1 | 1 | 0,356607017 | 0,154 | 0,16 | 1 | Basal | 1,280411037 | 0,780999204 | 0,9625 |
| PRB31 | 7,40938E-65 | -2,352828262 | 0,284 | 0,832 | 2,71087E-60 | Secretory | 0,195761876 | 5,108246915 | 0,341346154 |
| PRR41 | 6,57313E-56 | -1,358939115 | 0,67 | 0,99 | 2,40491E-51 | Secretory | 0,389868874 | 2,564964956 | 0,676767677 |

| Supplementary Table 4 - clusermarker | | | | | | | | | |
|---|---|---|---|---|---|---|---|---|---|
| | **p_val** | **avg_log2FC** | **pct.1** | **pct.2** | **p_val_adj** | **celltype** | **Foldchange_UP** | **Foldchange_DOWN** | **Ratio_pct1_pct2** |
| RPL131 | 9,58817E-56 | 1,103432069 | 0,95 | 0,94 | 3,50803E-51 | Secretory | 2,148652342 | 0,465408005 | 1,010638298 |
| C6orf581 | 4,99055E-51 | -1,493209414 | 0,45 | 0,882 | 1,82589E-46 | Secretory | 0,355221445 | 2,815145352 | 0,510204082 |
| RPL81 | 5,69541E-51 | 1,129618066 | 0,925 | 0,92 | 2,08378E-46 | Secretory | 2,18800808 | 0,457036704 | 1,005434783 |
| PIP1 | 6,86216E-50 | -1,763875248 | 0,526 | 0,889 | 2,51066E-45 | Secretory | 0,294456158 | 3,396091307 | 0,59167604 |
| MUC7 | 3,23943E-49 | -0,608629238 | 0,084 | 0,522 | 1,18521E-44 | Secretory | 0,655819526 | 1,524809738 | 0,16091954 |
| LYZ1 | 2,41227E-46 | -1,093955626 | 0,593 | 0,98 | 8,82578E-42 | Secretory | 0,468475132 | 2,13458502 | 0,605102041 |
| RPL18A1 | 5,26781E-43 | 1,133315586 | 0,897 | 0,902 | 1,92733E-38 | Secretory | 2,193622974 | 0,455866852 | 0,994456763 |
| RPS22 | 5,77628E-43 | 1,166227027 | 0,875 | 0,885 | 2,11337E-38 | Secretory | 2,244240095 | 0,445585124 | 0,988700565 |
| MMP71 | 9,12312E-43 | 3,052109164 | 0,412 | 0,073 | 3,33788E-38 | Secretory | 8,29423638 | 0,12056565 | 5,643835616 |
| RPLP01 | 2,26825E-42 | 1,123958262 | 0,903 | 0,915 | 8,29886E-38 | Secretory | 2,179441171 | 0,458833215 | 0,986885246 |
| RPS8 | 2,24353E-40 | 0,927841882 | 0,933 | 0,94 | 8,20841E-36 | Secretory | 1,902428037 | 0,525644062 | 0,992553191 |
| RPL10 | 5,98816E-39 | 0,968408305 | 0,948 | 0,972 | 2,19089E-34 | Secretory | 1,95668064 | 0,511069604 | 0,975308642 |
| ZG16B1 | 6,10889E-39 | -0,873536178 | 0,696 | 0,94 | 2,23506E-34 | Secretory | 0,545807386 | 1,832148165 | 0,740425532 |
| RPS51 | 3,09127E-38 | 1,033678435 | 0,873 | 0,877 | 1,131E-33 | Secretory | 2,047237437 | 0,488463127 | 0,995438997 |
| RPS12 | 3,51918E-37 | 0,911720997 | 0,944 | 0,947 | 1,28756E-32 | Secretory | 1,881288357 | 0,531550624 | 0,996832101 |
| RPL281 | 4,05842E-37 | 1,055992534 | 0,897 | 0,903 | 1,48485E-32 | Secretory | 2,079148111 | 0,480966216 | 0,993355482 |
| RPLP1 | 8,37599E-37 | 0,833245577 | 0,961 | 0,973 | 3,06452E-32 | Secretory | 1,781689056 | 0,561265164 | 0,987667009 |
| RPS191 | 1,52722E-36 | 1,29182484 | 0,849 | 0,85 | 5,58764E-32 | Secretory | 2,448375505 | 0,408434081 | 0,998823529 |
| RPS9 | 6,13507E-36 | 0,934525177 | 0,884 | 0,892 | 2,24464E-31 | Secretory | 1,911261492 | 0,523214643 | 0,99103139 |
| GNAS1 | 6,75983E-36 | -0,799230961 | 0,776 | 0,905 | 2,47322E-31 | Secretory | 0,57465542 | 1,740173268 | 0,857458564 |
| RPS23 | 9,99736E-36 | 0,855774381 | 0,935 | 0,92 | 3,65774E-31 | Secretory | 1,809729885 | 0,55256865 | 1,016304348 |
| RPL12 | 1,25746E-35 | 1,136419023 | 0,907 | 0,932 | 4,60068E-31 | Secretory | 2,19834684 | 0,454887273 | 0,973175966 |
| CD74 | 3,42295E-35 | 1,354694958 | 0,871 | 0,857 | 1,25235E-30 | Secretory | 2,557430365 | 0,391017489 | 1,016336056 |
| CLU | 5,89574E-34 | 2,509159524 | 0,692 | 0,449 | 2,15707E-29 | Secretory | 5,692883293 | 0,175657913 | 1,541202673 |
| FTH11 | 1,67039E-32 | 0,85656586 | 0,933 | 0,933 | 6,11146E-28 | Secretory | 1,810722996 | 0,552265588 | 1 |
| RACK1 | 2,62036E-32 | 0,973942361 | 0,851 | 0,864 | 9,58712E-28 | Secretory | 1,964200714 | 0,50911294 | 0,984953704 |
| KRT191 | 4,68482E-32 | 2,08009942 | 0,746 | 0,609 | 1,71403E-27 | Secretory | 4,22836354 | 0,236498113 | 1,224958949 |
| STATH1 | 4,82383E-32 | -2,045325589 | 0,144 | 0,501 | 1,76489E-27 | Secretory | 0,242267773 | 4,127664149 | 0,28742515 |
| RPL5 | 6,2803E-31 | 0,966707852 | 0,86 | 0,879 | 2,29778E-26 | Secretory | 1,954375728 | 0,511672339 | 0,978384528 |
| RPL18 | 6,36014E-31 | 0,751156458 | 0,899 | 0,917 | 2,32698E-26 | Secretory | 1,683141489 | 0,594127117 | 0,980370774 |
| RPL3 | 1,4099E-30 | 0,78709039 | 0,909 | 0,925 | 5,15841E-26 | Secretory | 1,7255908 | 0,579511666 | 0,982702703 |

| Supplementary Table 4 - clusermarker | | | | | | | | | |
|---|---|---|---|---|---|---|---|---|---|
| | **p_val** | **avg_log2FC** | **pct.1** | **pct.2** | **p_val_adj** | **celltype** | **Foldchange_UP** | **Foldchange_DOWN** | **Ratio_pct1_pct2** |
| SCGB3A21 | 1,56223E-30 | -1,013221939 | 0,425 | 0,81 | 5,71573E-26 | Secretory | 0,495438559 | 2,018413749 | 0,524691358 |
| RPS15 | 5,95261E-30 | 0,818692459 | 0,879 | 0,915 | 2,17788E-25 | Secretory | 1,763806698 | 0,566955552 | 0,960655738 |
| DUSP231 | 8,50606E-30 | 2,173957576 | 0,567 | 0,329 | 3,11211E-25 | Secretory | 4,512595849 | 0,221601941 | 1,723404255 |
| C31 | 9,86779E-30 | 1,101359314 | 0,373 | 0,093 | 3,61033E-25 | Secretory | 2,145567537 | 0,466077149 | 4,010752688 |
| NACA | 1,78602E-29 | 0,965028416 | 0,841 | 0,847 | 6,5345E-25 | Secretory | 1,952101971 | 0,512268321 | 0,992916175 |
| RPL10A | 2,18347E-29 | 0,936626904 | 0,853 | 0,872 | 7,98866E-25 | Secretory | 1,91404786 | 0,522452976 | 0,978211009 |
| PPDPF1 | 2,46488E-29 | 0,899010834 | 0,832 | 0,87 | 9,01825E-25 | Secretory | 1,864786976 | 0,536254281 | 0,956321839 |
| PTMA | 3,39838E-29 | 1,192561452 | 0,815 | 0,807 | 1,24337E-24 | Secretory | 2,285581797 | 0,437525361 | 1,009913259 |
| FMO2 | 4,05013E-29 | 1,371406727 | 0,358 | 0,092 | 1,48182E-24 | Secretory | 2,587227156 | 0,386514187 | 3,891304348 |
| TCN11 | 1,26233E-28 | -0,666106622 | 0,642 | 0,877 | 4,61849E-24 | Secretory | 0,630205119 | 1,586784953 | 0,732041049 |
| RPS18 | 2,73447E-28 | 0,842680725 | 0,922 | 0,935 | 1,00046E-23 | Secretory | 1,793379393 | 0,557606497 | 0,986096257 |
| AC020656.11 | 3,81148E-28 | -0,77931431 | 0,364 | 0,784 | 1,39451E-23 | Secretory | 0,582643649 | 1,716314942 | 0,464285714 |
| RPS161 | 9,09341E-28 | 0,721038196 | 0,843 | 0,835 | 3,32701E-23 | Secretory | 1,648367811 | 0,606660718 | 1,009580838 |
| RPS27A | 9,98209E-28 | 0,760112582 | 0,92 | 0,927 | 3,65215E-23 | Secretory | 1,693622784 | 0,590450252 | 0,992448759 |
| EEF21 | 1,33486E-27 | 0,94331929 | 0,83 | 0,84 | 4,88387E-23 | Secretory | 1,922947386 | 0,520035029 | 0,988095238 |
| RPL15 | 2,33731E-27 | 0,656273373 | 0,888 | 0,927 | 8,5515E-23 | Secretory | 1,576006377 | 0,634515199 | 0,957928803 |
| RARRES11 | 2,47757E-27 | 1,958979838 | 0,53 | 0,27 | 9,06468E-23 | Secretory | 3,887869617 | 0,257210272 | 1,962962963 |
| AP000851.1 | 2,60071E-27 | 0,845194317 | 0,272 | 0,042 | 9,51521E-23 | Secretory | 1,796506702 | 0,556635831 | 6,476190476 |
| KRT71 | 8,59793E-27 | 2,002285015 | 0,688 | 0,552 | 3,14572E-22 | Secretory | 4,006340426 | 0,249604351 | 1,246376812 |
| RPS4X | 1,44829E-26 | 0,829082174 | 0,905 | 0,925 | 5,29888E-22 | Secretory | 1,776554779 | 0,562887231 | 0,978378378 |
| PRH21 | 2,55439E-25 | -1,578121781 | 0,209 | 0,536 | 9,34574E-21 | Secretory | 0,334917628 | 2,985808794 | 0,389925373 |
| RPL14 | 5,606E-25 | 0,692687307 | 0,894 | 0,92 | 2,05107E-20 | Secretory | 1,616291381 | 0,618700323 | 0,97173913 |
| CA2 | 7,02678E-25 | -0,598550169 | 0,302 | 0,619 | 2,57089E-20 | Secretory | 0,660417306 | 1,514194118 | 0,487883683 |
| BTF3 | 7,65307E-25 | 1,098143781 | 0,741 | 0,747 | 2,80003E-20 | Secretory | 2,14079074 | 0,467117118 | 0,991967871 |
| RPS3A | 7,82194E-25 | 0,668707306 | 0,918 | 0,922 | 2,86181E-20 | Secretory | 1,589647962 | 0,629070099 | 0,995661605 |
| RPS7 | 1,81281E-24 | 0,781922931 | 0,858 | 0,887 | 6,63252E-20 | Secretory | 1,719421118 | 0,581591089 | 0,967305524 |
| TNFAIP2 | 1,9272E-24 | 0,97317459 | 0,284 | 0,06 | 7,05105E-20 | Secretory | 1,963155688 | 0,509383951 | 4,733333333 |
| PART11 | 2,22643E-24 | -0,484856026 | 0,308 | 0,647 | 8,14584E-20 | Secretory | 0,714568377 | 1,399446201 | 0,476043277 |
| GABRP1 | 2,25618E-24 | 0,767968976 | 0,319 | 0,08 | 8,2547E-20 | Secretory | 1,702870796 | 0,587243614 | 3,9875 |
| MTRNR2L12 | 3,38175E-24 | 0,978919605 | 0,832 | 0,787 | 1,23728E-19 | Secretory | 1,970988837 | 0,507359545 | 1,057179161 |
| HLA-B1 | 5,64622E-24 | 1,153448084 | 0,828 | 0,835 | 2,06578E-19 | Secretory | 2,224449095 | 0,44954951 | 0,991616766 |

| Supplementary Table 4 - clusermarker | | | | | | | | | |
|---|---|---|---|---|---|---|---|---|---|
| | **p_val** | **avg_log2FC** | **pct.1** | **pct.2** | **p_val_adj** | **celltype** | **Foldchange_UP** | **Foldchange_DOWN** | **Ratio_pct1_pct2** |
| RPL29 | 1,39859E-23 | 0,811816599 | 0,871 | 0,927 | 5,11701E-19 | Secretory | 1,755420425 | 0,569664102 | 0,939590076 |
| RPS13 | 2,15188E-23 | 0,693782728 | 0,894 | 0,925 | 7,8731E-19 | Secretory | 1,617519077 | 0,61823073 | 0,966486486 |
| IGKC1 | 2,46878E-23 | 3,93868938 | 0,263 | 0,053 | 9,03252E-19 | Secretory | 15,3342891 | 0,065213326 | 4,962264151 |
| RPL30 | 4,06141E-23 | 0,631810786 | 0,914 | 0,933 | 1,48595E-18 | Secretory | 1,549508625 | 0,645365882 | 0,979635584 |
| RPL32 | 4,10568E-23 | 0,707520582 | 0,905 | 0,913 | 1,50214E-18 | Secretory | 1,632995235 | 0,612371658 | 0,991237678 |
| EEF1D | 4,88866E-23 | 0,754044352 | 0,806 | 0,829 | 1,78861E-18 | Secretory | 1,686514067 | 0,592939021 | 0,97225573 |
| LCN21 | 5,32506E-23 | 1,7276513 | 0,634 | 0,401 | 1,94828E-18 | Secretory | 3,311882065 | 0,301943119 | 1,581047382 |
| PPIA1 | 6,71905E-23 | 1,044531004 | 0,744 | 0,734 | 2,4583E-18 | Secretory | 2,062695701 | 0,484802484 | 1,013623978 |
| RPL24 | 8,11843E-23 | 0,703360122 | 0,843 | 0,879 | 2,97029E-18 | Secretory | 1,62829277 | 0,614140171 | 0,959044369 |
| RPL19 | 8,66266E-23 | 0,578179801 | 0,92 | 0,953 | 3,16941E-18 | Secretory | 1,492964438 | 0,669808319 | 0,965372508 |
| ODAM | 1,07236E-22 | -0,941842944 | 0,138 | 0,421 | 3,92344E-18 | Secretory | 0,520567466 | 1,920980593 | 0,327790974 |
| RPS31 | 1,07461E-22 | 0,710768909 | 0,894 | 0,922 | 3,93169E-18 | Secretory | 1,636676179 | 0,610994412 | 0,969631236 |
| FAU | 1,50825E-22 | 0,844060623 | 0,836 | 0,874 | 5,51822E-18 | Secretory | 1,79509553 | 0,557073416 | 0,956521739 |
| PFN11 | 5,41424E-22 | 1,084173735 | 0,7 | 0,666 | 1,98091E-17 | Secretory | 2,12016087 | 0,471662323 | 1,051051051 |
| TTYH1 | 6,86455E-22 | -0,269929609 | 0,127 | 0,404 | 2,51153E-17 | Secretory | 0,829360011 | 1,205748996 | 0,314356436 |
| ATP2A3 | 8,70955E-22 | -0,456714839 | 0,239 | 0,559 | 3,18656E-17 | Secretory | 0,728643565 | 1,372413135 | 0,427549195 |
| MIF1 | 1,34699E-21 | 0,826278894 | 0,731 | 0,71 | 4,92824E-17 | Secretory | 1,773106132 | 0,563982032 | 1,029577465 |
| AZGP11 | 1,48229E-21 | -0,527104616 | 0,67 | 0,925 | 5,42325E-17 | Secretory | 0,693946037 | 1,44103424 | 0,724324324 |
| S100A61 | 1,531E-21 | 1,436306061 | 0,716 | 0,607 | 5,60145E-17 | Secretory | 2,706270524 | 0,369512209 | 1,179571664 |
| RPS14 | 2,1924E-21 | 0,62175641 | 0,933 | 0,935 | 8,02134E-17 | Secretory | 1,538747391 | 0,64987925 | 0,997860963 |
| ANPEP | 3,14394E-21 | 0,724487138 | 0,295 | 0,085 | 1,15027E-16 | Secretory | 1,652313153 | 0,605212153 | 3,470588235 |
| FTL | 3,62642E-21 | 0,648051289 | 0,862 | 0,879 | 1,3268E-16 | Secretory | 1,567050082 | 0,638141698 | 0,980659841 |
| IFITM31 | 3,72822E-21 | 1,858761764 | 0,634 | 0,521 | 1,36404E-16 | Secretory | 3,626962337 | 0,275712816 | 1,216890595 |
| RPL17 | 4,36198E-21 | 0,716380388 | 0,856 | 0,88 | 1,59592E-16 | Secretory | 1,643054561 | 0,608622516 | 0,972727273 |
| SLC25A61 | 4,43282E-21 | 0,877049113 | 0,748 | 0,742 | 1,62184E-16 | Secretory | 1,836614847 | 0,544479972 | 1,008086253 |
| RPL11 | 4,56527E-21 | 0,633069785 | 0,927 | 0,933 | 1,6703E-16 | Secretory | 1,550861429 | 0,644802934 | 0,993569132 |
| GAPDH | 1,18985E-20 | 0,704947717 | 0,845 | 0,862 | 4,35331E-16 | Secretory | 1,63008559 | 0,61346472 | 0,980278422 |
| EIF1 | 1,48567E-20 | 0,753294669 | 0,808 | 0,83 | 5,43562E-16 | Secretory | 1,685637913 | 0,593247216 | 0,973493976 |
| RPS6 | 1,77093E-20 | 0,560742116 | 0,918 | 0,95 | 6,47931E-16 | Secretory | 1,47502777 | 0,677953338 | 0,966315789 |
| S100A15 | 1,91145E-20 | -0,446719508 | 0,364 | 0,707 | 6,99343E-16 | Secretory | 0,733709307 | 1,362937596 | 0,514851485 |
| ACTB1 | 2,4777E-20 | 1,063478559 | 0,825 | 0,805 | 9,06514E-16 | Secretory | 2,089964677 | 0,478476986 | 1,02484472 |

| Supplementary Table 4 - clusermarker | | | | | | | | | |
|---|---|---|---|---|---|---|---|---|---|
| | **p_val** | **avg_log2FC** | **pct.1** | **pct.2** | **p_val_adj** | **celltype** | **Foldchange_UP** | **Foldchange_DOWN** | **Ratio_pct1_pct2** |
| LY6E1 | 2,92827E-20 | 1,492422988 | 0,537 | 0,361 | 1,07136E-15 | Secretory | 2,81361121 | 0,355415132 | 1,487534626 |
| S100A111 | 4,24845E-20 | 0,864844298 | 0,806 | 0,834 | 1,55438E-15 | Secretory | 1,821143111 | 0,549105665 | 0,966426859 |
| SLC12A2 | 5,59094E-20 | -0,614409549 | 0,506 | 0,794 | 2,04556E-15 | Secretory | 0,653197172 | 1,530931306 | 0,637279597 |
| RPSA | 1,36717E-19 | 0,590492817 | 0,851 | 0,86 | 5,00206E-15 | Secretory | 1,50576102 | 0,66411601 | 0,989534884 |
| INSIG11 | 1,41527E-19 | -0,342220036 | 0,14 | 0,389 | 5,17804E-15 | Secretory | 0,788826522 | 1,267705855 | 0,359897172 |
| COX4I11 | 1,611E-19 | 0,849933835 | 0,748 | 0,772 | 5,89418E-15 | Secretory | 1,802418261 | 0,55481018 | 0,968911917 |
| LPO | 1,98634E-19 | -0,342455388 | 0,287 | 0,612 | 7,26741E-15 | Secretory | 0,788697848 | 1,267912676 | 0,468954248 |
| TNFRSF19 | 2,27818E-19 | -0,326123188 | 0,196 | 0,473 | 8,33517E-15 | Secretory | 0,797677125 | 1,253640061 | 0,414376321 |
| BBOX1 | 2,4039E-19 | 0,506921097 | 0,179 | 0,022 | 8,79516E-15 | Secretory | 1,421014324 | 0,703722674 | 8,136363636 |
| GSTP11 | 2,67916E-19 | 0,979812692 | 0,748 | 0,719 | 9,80224E-15 | Secretory | 1,972209336 | 0,507045566 | 1,040333797 |
| MET | 3,71898E-19 | 0,932569523 | 0,312 | 0,111 | 1,36066E-14 | Secretory | 1,908672426 | 0,523924371 | 2,810810811 |
| NME21 | 5,24393E-19 | 0,854631159 | 0,737 | 0,767 | 1,9186E-14 | Secretory | 1,808296385 | 0,553006691 | 0,960886571 |
| IGHG11 | 6,19487E-19 | 1,663206846 | 0,134 | 0,003 | 2,26652E-14 | Secretory | 3,167197527 | 0,315736544 | 44,66666667 |
| EEF1A1 | 6,98124E-19 | 0,469061531 | 0,989 | 0,992 | 2,55423E-14 | Secretory | 1,384208751 | 0,722434386 | 0,996975806 |
| EEF1G | 7,7024E-19 | 0,722895402 | 0,793 | 0,822 | 2,81808E-14 | Secretory | 1,650491149 | 0,605880256 | 0,964720195 |
| RPL261 | 8,66595E-19 | 0,6825929 | 0,877 | 0,92 | 3,17061E-14 | Secretory | 1,605021808 | 0,623044494 | 0,95326087 |
| FXYD51 | 1,07146E-18 | 0,60667406 | 0,231 | 0,053 | 3,92014E-14 | Secretory | 1,522744676 | 0,656708913 | 4,358490566 |
| SAA21 | 1,62201E-18 | 1,741965598 | 0,284 | 0,093 | 5,93445E-14 | Secretory | 3,344905836 | 0,298962078 | 3,053763441 |
| SLPI1 | 3,14443E-18 | -0,588562053 | 0,974 | 1 | 1,15045E-13 | Secretory | 0,665005394 | 1,503747202 | 0,974 |
| LXN | 3,81912E-18 | 1,139322321 | 0,422 | 0,228 | 1,3973E-13 | Secretory | 2,202775276 | 0,453972773 | 1,850877193 |
| LINC01829 | 5,37152E-18 | -0,392598349 | 0,157 | 0,411 | 1,96528E-13 | Secretory | 0,761756416 | 1,312755599 | 0,381995134 |
| ETV1 | 5,90384E-18 | -0,320088195 | 0,08 | 0,291 | 2,16004E-13 | Secretory | 0,801020908 | 1,248406864 | 0,274914089 |
| SNHG29 | 1,05799E-17 | 0,993619938 | 0,731 | 0,759 | 3,87088E-13 | Secretory | 1,991174885 | 0,502216057 | 0,963109354 |
| CADM3 | 1,09695E-17 | 0,386637603 | 0,121 | 0,002 | 4,01341E-13 | Secretory | 1,30734291 | 0,764910256 | 60,5 |
| RPL7A | 1,36405E-17 | 0,533106944 | 0,907 | 0,947 | 4,99065E-13 | Secretory | 1,447042149 | 0,691064874 | 0,957761352 |
| SAA11 | 1,76304E-17 | 2,12304517 | 0,468 | 0,276 | 6,45045E-13 | Secretory | 4,356124446 | 0,229561853 | 1,695652174 |
| RPS28 | 1,78331E-17 | 0,453330802 | 0,916 | 0,932 | 6,52459E-13 | Secretory | 1,369197726 | 0,730354704 | 0,982832618 |
| DDIT4L | 2,6105E-17 | 0,337616782 | 0,136 | 0,008 | 9,55104E-13 | Secretory | 1,26366739 | 0,791347476 | 17 |
| LGALS31 | 3,52522E-17 | 1,07696868 | 0,709 | 0,691 | 1,28977E-12 | Secretory | 2,109598835 | 0,474023773 | 1,026049204 |
| F5 | 6,99792E-17 | -0,32305165 | 0,155 | 0,416 | 2,56033E-12 | Secretory | 0,799377211 | 1,250973866 | 0,372596154 |
| CP1 | 8,92406E-17 | 0,900653756 | 0,353 | 0,153 | 3,26505E-12 | Secretory | 1,866911781 | 0,53564395 | 2,307189542 |

| Supplementary Table 4 - clusermarker | | | | | | | | | |
|---|---|---|---|---|---|---|---|---|---|
| | **p_val** | **avg_log2FC** | **pct.1** | **pct.2** | **p_val_adj** | **celltype** | **Foldchange_UP** | **Foldchange_DOWN** | **Ratio_pct1_pct2** |
| NNMT1 | 1,04245E-16 | 1,017418099 | 0,278 | 0,098 | 3,81402E-12 | Secretory | 2,024292965 | 0,493999642 | 2,836734694 |
| ATP5F1D1 | 1,32405E-16 | 0,920949625 | 0,655 | 0,611 | 4,84429E-12 | Secretory | 1,89336115 | 0,528161254 | 1,072013093 |
| RPLP21 | 1,3979E-16 | 0,540525051 | 0,858 | 0,895 | 5,11451E-12 | Secretory | 1,454501769 | 0,687520649 | 0,958659218 |
| TAGLN21 | 3,19407E-16 | 1,213082431 | 0,631 | 0,576 | 1,16861E-11 | Secretory | 2,318324362 | 0,431346026 | 1,095486111 |
| ZFAS11 | 3,67202E-16 | 0,955842575 | 0,657 | 0,586 | 1,34348E-11 | Secretory | 1,93971215 | 0,515540411 | 1,12116041 |
| CHCHD21 | 4,54769E-16 | 0,832980999 | 0,716 | 0,74 | 1,66386E-11 | Secretory | 1,781362339 | 0,561368105 | 0,967567568 |
| H3-3B | 4,67329E-16 | 0,858739099 | 0,772 | 0,767 | 1,70982E-11 | Secretory | 1,813452679 | 0,551434295 | 1,006518905 |
| C19orf33 | 4,7821E-16 | 1,027607526 | 0,517 | 0,334 | 1,74963E-11 | Secretory | 2,038640694 | 0,490522927 | 1,547904192 |
| EDF11 | 4,96137E-16 | 0,840508025 | 0,694 | 0,687 | 1,81522E-11 | Secretory | 1,790680595 | 0,558446885 | 1,010189229 |
| ATP12A | 7,2054E-16 | -0,256490943 | 0,218 | 0,476 | 2,63624E-11 | Secretory | 0,83712157 | 1,194569625 | 0,457983193 |
| DEFB1 | 7,69024E-16 | 1,361179788 | 0,388 | 0,18 | 2,81363E-11 | Secretory | 2,568951739 | 0,389263833 | 2,155555556 |
| PTGER2 | 9,98603E-16 | 0,331267591 | 0,157 | 0,023 | 3,65359E-11 | Secretory | 1,258118306 | 0,794837811 | 6,826086957 |
| RPL411 | 1,39874E-15 | 0,418454259 | 0,978 | 0,978 | 5,11756E-11 | Secretory | 1,336494832 | 0,748225864 | 1 |
| TMEM2651 | 1,47204E-15 | 0,391268946 | 0,231 | 0,067 | 5,38576E-11 | Secretory | 1,311546489 | 0,762458676 | 3,447761194 |
| SOD3 | 1,90499E-15 | -0,293516581 | 0,196 | 0,434 | 6,96978E-11 | Secretory | 0,815910843 | 1,225624109 | 0,451612903 |
| HSPA1B1 | 2,28251E-15 | -0,830595161 | 0,459 | 0,669 | 8,35103E-11 | Secretory | 0,562297228 | 1,77841887 | 0,686098655 |
| H3-3A1 | 2,35258E-15 | 0,723647053 | 0,761 | 0,785 | 8,60737E-11 | Secretory | 1,651351287 | 0,605564672 | 0,969426752 |
| TACSTD21 | 2,61678E-15 | 1,064105357 | 0,651 | 0,587 | 9,574E-11 | Secretory | 2,090872886 | 0,478269151 | 1,109028961 |
| CIRBP | 2,66437E-15 | 0,893780968 | 0,685 | 0,686 | 9,74815E-11 | Secretory | 1,858039238 | 0,538201767 | 0,998542274 |
| S100A101 | 2,82024E-15 | 1,217994991 | 0,491 | 0,321 | 1,03184E-10 | Secretory | 2,326232006 | 0,429879736 | 1,529595016 |
| NCALD | 6,39865E-15 | -0,377051954 | 0,188 | 0,426 | 2,34108E-10 | Secretory | 0,770009444 | 1,29868537 | 0,441314554 |
| CLIC31 | 7,30291E-15 | 0,4084295 | 0,168 | 0,033 | 2,67192E-10 | Secretory | 1,32724021 | 0,753443116 | 5,090909091 |
| KRT141 | 7,60234E-15 | 3,228399869 | 0,306 | 0,14 | 2,78147E-10 | Secretory | 9,372278786 | 0,106697637 | 2,185714286 |
| ITGB2 | 8,18263E-15 | 0,28690147 | 0,162 | 0,028 | 2,99378E-10 | Secretory | 1,220017185 | 0,819660585 | 5,785714286 |
| TNC1 | 8,84983E-15 | 0,80963314 | 0,216 | 0,063 | 3,23789E-10 | Secretory | 1,752765679 | 0,570526917 | 3,428571429 |
| RPS15A | 9,32486E-15 | 0,547645339 | 0,899 | 0,938 | 3,41169E-10 | Secretory | 1,461698072 | 0,684135814 | 0,958422175 |
| COX6A11 | 1,04769E-14 | 0,894261117 | 0,69 | 0,7 | 3,8332E-10 | Secretory | 1,858657721 | 0,538022676 | 0,985714286 |
| RPL13A | 1,12161E-14 | 0,528729603 | 0,834 | 0,902 | 4,10363E-10 | Secretory | 1,442658271 | 0,693164847 | 0,924611973 |
| ASS11 | 1,14411E-14 | 1,206624833 | 0,612 | 0,559 | 4,18595E-10 | Secretory | 2,307970578 | 0,433281087 | 1,094812165 |
| RPL36 | 1,14457E-14 | 0,429406652 | 0,866 | 0,912 | 4,18762E-10 | Secretory | 1,346679604 | 0,742567124 | 0,949561404 |
| GPX41 | 1,25899E-14 | 1,019665817 | 0,664 | 0,671 | 4,60628E-10 | Secretory | 2,027449271 | 0,49323059 | 0,989567809 |

| Supplementary Table 4 - clusermarker | | | | | | | | | |
|---|---|---|---|---|---|---|---|---|---|
| | **p_val** | **avg_log2FC** | **pct.1** | **pct.2** | **p_val_adj** | **celltype** | **Foldchange_UP** | **Foldchange_DOWN** | **Ratio_pct1_pct2** |
| HINT11 | 1,36994E-14 | 0,797051887 | 0,722 | 0,769 | 5,0122E-10 | Secretory | 1,737546861 | 0,575524046 | 0,938881664 |
| CYBA1 | 1,56218E-14 | 1,147180876 | 0,534 | 0,419 | 5,71556E-10 | Secretory | 2,21480683 | 0,451506644 | 1,274463007 |
| IGKV4-1 | 1,82893E-14 | 0,836622341 | 0,103 | 0,003 | 6,69149E-10 | Secretory | 1,785864152 | 0,559953006 | 34,33333333 |
| PERP1 | 1,84318E-14 | 1,052722398 | 0,629 | 0,587 | 6,74365E-10 | Secretory | 2,074440672 | 0,482057652 | 1,071550256 |
| ADIRF1 | 1,94451E-14 | 1,430223426 | 0,601 | 0,542 | 7,11438E-10 | Secretory | 2,69488447 | 0,371073421 | 1,108856089 |
| RPL4 | 2,09001E-14 | 0,657334567 | 0,776 | 0,834 | 7,64672E-10 | Secretory | 1,577166057 | 0,634048644 | 0,930455635 |
| EEF1B2 | 2,58487E-14 | 0,632048828 | 0,774 | 0,785 | 9,45727E-10 | Secretory | 1,549764313 | 0,645259406 | 0,985987261 |
| ALOX5 | 3,09544E-14 | 0,335570933 | 0,151 | 0,025 | 1,13253E-09 | Secretory | 1,261876686 | 0,792470462 | 6,04 |
| SPINT21 | 3,20802E-14 | 0,592726927 | 0,744 | 0,777 | 1,17372E-09 | Secretory | 1,508094598 | 0,663088378 | 0,957528958 |
| UBA52 | 3,37141E-14 | 0,512450099 | 0,812 | 0,827 | 1,2335E-09 | Secretory | 1,426470685 | 0,70103088 | 0,981862152 |
| IGKV3-111 | 3,6863E-14 | 3,553823809 | 0,106 | 0,005 | 1,34871E-09 | Secretory | 11,74377078 | 0,085151526 | 21,2 |
| CSTB1 | 4,57896E-14 | 1,080786822 | 0,61 | 0,567 | 1,6753E-09 | Secretory | 2,115189356 | 0,472770912 | 1,075837743 |
| MYL12B1 | 4,74425E-14 | 1,109826015 | 0,64 | 0,629 | 1,73578E-09 | Secretory | 2,158196185 | 0,463349906 | 1,017488076 |
| PIGR1 | 7,61489E-14 | -0,255020098 | 0,873 | 0,952 | 2,78606E-09 | Secretory | 0,837975461 | 1,193352368 | 0,917016807 |
| RAC11 | 8,4043E-14 | 0,784097041 | 0,634 | 0,597 | 3,07488E-09 | Secretory | 1,722014201 | 0,580715304 | 1,061976549 |
| RPS211 | 1,0458E-13 | 0,507907075 | 0,841 | 0,87 | 3,82628E-09 | Secretory | 1,421985817 | 0,703241895 | 0,966666667 |
| SCARA3 | 1,08752E-13 | 0,559817764 | 0,291 | 0,126 | 3,9789E-09 | Secretory | 1,474083004 | 0,67838785 | 2,30952381 |
| IFITM2 | 1,36962E-13 | 1,076666923 | 0,45 | 0,301 | 5,01102E-09 | Secretory | 2,109157633 | 0,474122932 | 1,495016611 |
| HSPA1A1 | 1,66392E-13 | -1,080982919 | 0,53 | 0,692 | 6,08778E-09 | Secretory | 0,472706655 | 2,115476881 | 0,765895954 |
| PRB4 | 1,90704E-13 | -1,017722135 | 0,08 | 0,261 | 6,97729E-09 | Secretory | 0,493895546 | 2,024719614 | 0,30651341 |
| MUC11 | 2,15452E-13 | 0,847592454 | 0,429 | 0,27 | 7,88275E-09 | Secretory | 1,799495449 | 0,555711325 | 1,588888889 |
| ANXA12 | 2,16341E-13 | 0,633042965 | 0,575 | 0,404 | 7,91528E-09 | Secretory | 1,550832597 | 0,644814922 | 1,423267327 |
| BRK1 | 2,65506E-13 | 0,811426439 | 0,588 | 0,526 | 9,71408E-09 | Secretory | 1,754945756 | 0,569818182 | 1,117870722 |
| NPM1 | 3,45954E-13 | 0,819781028 | 0,683 | 0,674 | 1,26574E-08 | Secretory | 1,765138059 | 0,566527924 | 1,013353116 |
| SMR3A | 4,39707E-13 | -1,035840985 | 0,106 | 0,29 | 1,60876E-08 | Secretory | 0,487731486 | 2,050308476 | 0,365517241 |
| ACTG1 | 4,84229E-13 | 0,897229174 | 0,772 | 0,78 | 1,77165E-08 | Secretory | 1,862485475 | 0,536916939 | 0,98974359 |
| CES11 | 5,59624E-13 | -0,456637766 | 0,108 | 0,286 | 2,0475E-08 | Secretory | 0,728682493 | 1,372339819 | 0,377622378 |
| SNHG51 | 6,41546E-13 | 0,708551648 | 0,547 | 0,433 | 2,34722E-08 | Secretory | 1,634162723 | 0,611934164 | 1,263279446 |
| MT-ATP81 | 7,49113E-13 | -0,455804523 | 0,683 | 0,807 | 2,74078E-08 | Secretory | 0,729103472 | 1,371547439 | 0,846344486 |
| GABARAP1 | 7,62396E-13 | 0,688964932 | 0,703 | 0,745 | 2,78938E-08 | Secretory | 1,612126476 | 0,620298726 | 0,943624161 |
| AKR1C11 | 8,40339E-13 | 0,926820516 | 0,485 | 0,333 | 3,07455E-08 | Secretory | 1,901081676 | 0,526016327 | 1,456456456 |

| Supplementary Table 4 - clusermarker | | | | | | | | | |
|---|---|---|---|---|---|---|---|---|---|
| | **p_val** | **avg_log2FC** | **pct.1** | **pct.2** | **p_val_adj** | **celltype** | **Foldchange_UP** | **Foldchange_DOWN** | **Ratio_pct1_pct2** |
| EIF3F1 | 1,12343E-12 | 0,872057517 | 0,595 | 0,566 | 4,1103E-08 | Secretory | 1,830271303 | 0,546367087 | 1,051236749 |
| SKP1 | 1,47165E-12 | 0,797522712 | 0,666 | 0,672 | 5,38433E-08 | Secretory | 1,738114003 | 0,575336254 | 0,991071429 |
| ATP5MC2 | 1,52985E-12 | 0,65301117 | 0,769 | 0,812 | 5,59727E-08 | Secretory | 1,572446759 | 0,635951579 | 0,947044335 |
| MT-CO3 | 1,53297E-12 | 0,499539542 | 0,892 | 0,87 | 5,60869E-08 | Secretory | 1,413762267 | 0,707332501 | 1,025287356 |
| TPT1 | 1,8044E-12 | 0,349603859 | 0,97 | 0,977 | 6,60177E-08 | Secretory | 1,274210701 | 0,784799562 | 0,99283521 |
| CD91 | 1,95234E-12 | 0,902759351 | 0,688 | 0,697 | 7,14303E-08 | Secretory | 1,869638504 | 0,534862754 | 0,987087518 |
| HLA-DRB1 | 2,20008E-12 | 0,992765158 | 0,631 | 0,541 | 8,04942E-08 | Secretory | 1,989995485 | 0,502513703 | 1,166358595 |
| RNASE1 | 2,24833E-12 | -0,517250065 | 0,466 | 0,719 | 8,22596E-08 | Secretory | 0,698702368 | 1,431224575 | 0,648122392 |
| GALNT6 | 2,65225E-12 | -0,323502772 | 0,188 | 0,393 | 9,7038E-08 | Secretory | 0,79912729 | 1,251365098 | 0,478371501 |
| PNPLA2 | 2,87819E-12 | 0,763861121 | 0,502 | 0,389 | 1,05304E-07 | Secretory | 1,698029027 | 0,588918083 | 1,290488432 |
| ITM2B1 | 2,91669E-12 | 0,694482929 | 0,748 | 0,817 | 1,06713E-07 | Secretory | 1,618304319 | 0,617930749 | 0,915544676 |
| TMSB101 | 4,23169E-12 | 0,96258073 | 0,778 | 0,802 | 1,54825E-07 | Secretory | 1,94879283 | 0,513138177 | 0,970074813 |
| RPL35 | 4,33868E-12 | 0,440946735 | 0,808 | 0,86 | 1,58739E-07 | Secretory | 1,357494859 | 0,73665104 | 0,939534884 |
| ODF3B | 4,73184E-12 | 0,417917128 | 0,297 | 0,14 | 1,73124E-07 | Secretory | 1,335997333 | 0,748504488 | 2,121428571 |
| PRDX51 | 5,85239E-12 | 0,772333158 | 0,651 | 0,654 | 2,14121E-07 | Secretory | 1,708029814 | 0,585469874 | 0,995412844 |
| HES41 | 6,44822E-12 | 0,654163797 | 0,274 | 0,125 | 2,35921E-07 | Secretory | 1,573703552 | 0,635443695 | 2,192 |
| PRB1 | 6,87889E-12 | -1,370146091 | 0,071 | 0,228 | 2,51678E-07 | Secretory | 0,386852073 | 2,584967408 | 0,311403509 |
| TOMM71 | 8,38164E-12 | 0,520748676 | 0,722 | 0,722 | 3,06659E-07 | Secretory | 1,434699582 | 0,697010031 | 1 |
| CNBP1 | 9,03986E-12 | 0,846820787 | 0,601 | 0,589 | 3,30741E-07 | Secretory | 1,798533195 | 0,556008642 | 1,020373514 |
| PFDN5 | 1,44046E-11 | 0,496793197 | 0,789 | 0,839 | 5,2702E-07 | Secretory | 1,411073558 | 0,708680277 | 0,940405244 |
| FOXQ1 | 1,55365E-11 | 0,610001125 | 0,392 | 0,233 | 5,68434E-07 | Secretory | 1,526260399 | 0,655196191 | 1,682403433 |
| ARRDC2 | 1,60209E-11 | 0,286329962 | 0,175 | 0,052 | 5,86156E-07 | Secretory | 1,219533984 | 0,819985349 | 3,365384615 |
| RHOA1 | 1,99592E-11 | 0,850829288 | 0,629 | 0,629 | 7,30246E-07 | Secretory | 1,803537334 | 0,554465927 | 1 |
| APP | 2,01964E-11 | 0,996910362 | 0,631 | 0,611 | 7,38926E-07 | Secretory | 1,995721436 | 0,501071934 | 1,032733224 |
| SULF2 | 2,14452E-11 | 0,481537676 | 0,188 | 0,062 | 7,84617E-07 | Secretory | 1,396231026 | 0,716213851 | 3,032258065 |
| CLDN31 | 2,44646E-11 | 0,805063072 | 0,528 | 0,443 | 8,95086E-07 | Secretory | 1,747222176 | 0,572337058 | 1,191873589 |
| HLA-DPB1 | 3,65654E-11 | 0,803248428 | 0,429 | 0,275 | 1,33782E-06 | Secretory | 1,745025875 | 0,573057405 | 1,56 |
| HLA-DRA1 | 4,19039E-11 | 1,147269946 | 0,64 | 0,599 | 1,53314E-06 | Secretory | 2,214943573 | 0,45147877 | 1,068447412 |
| RPL61 | 4,42837E-11 | 0,415359638 | 0,894 | 0,923 | 1,62021E-06 | Secretory | 1,333631086 | 0,749832552 | 0,968580715 |
| RPS251 | 4,62431E-11 | 0,334804632 | 0,907 | 0,928 | 1,6919E-06 | Secretory | 1,261206607 | 0,792891501 | 0,97737069 |
| SLC34A21 | 4,75882E-11 | 0,976548543 | 0,399 | 0,265 | 1,74111E-06 | Secretory | 1,967752186 | 0,508194074 | 1,505660377 |

| Supplementary Table 4 - clusermarker | | | | | | | | | |
|---|---|---|---|---|---|---|---|---|---|
| | **p_val** | **avg_log2FC** | **pct.1** | **pct.2** | **p_val_adj** | **celltype** | **Foldchange_UP** | **Foldchange_DOWN** | **Ratio_pct1_pct2** |
| CFI | 4,7919E-11 | 0,288583055 | 0,175 | 0,055 | 1,75321E-06 | Secretory | 1,221440049 | 0,818705757 | 3,181818182 |
| MYL6 | 5,3228E-11 | 0,726530833 | 0,75 | 0,777 | 1,94745E-06 | Secretory | 1,654655448 | 0,604355427 | 0,965250965 |
| HMGB1 | 5,61318E-11 | 0,860028838 | 0,664 | 0,667 | 2,0537E-06 | Secretory | 1,815074592 | 0,550941545 | 0,995502249 |
| AHNAK | 5,6805E-11 | 1,052532315 | 0,545 | 0,463 | 2,07832E-06 | Secretory | 2,074167372 | 0,48212117 | 1,177105832 |
| CD151 | 6,32979E-11 | 0,910413066 | 0,612 | 0,592 | 2,31588E-06 | Secretory | 1,879583575 | 0,53203274 | 1,033783784 |
| APIP | 6,60303E-11 | -0,360913687 | 0,388 | 0,654 | 2,41585E-06 | Secretory | 0,778671276 | 1,284238973 | 0,593272171 |
| NDUFA131 | 7,2106E-11 | 0,770934998 | 0,655 | 0,671 | 2,63814E-06 | Secretory | 1,706375313 | 0,586037546 | 0,976154993 |
| NME31 | 7,32876E-11 | 0,653218153 | 0,534 | 0,464 | 2,68137E-06 | Secretory | 1,572672374 | 0,635860346 | 1,150862069 |
| SEC621 | 8,22256E-11 | -0,321424977 | 0,575 | 0,74 | 3,00839E-06 | Secretory | 0,800279036 | 1,249564158 | 0,777027027 |
| HEBP21 | 8,75779E-11 | 0,728852177 | 0,584 | 0,547 | 3,20421E-06 | Secretory | 1,657319986 | 0,603383781 | 1,067641682 |
| PGLS1 | 8,85722E-11 | 0,802694827 | 0,502 | 0,431 | 3,24059E-06 | Secretory | 1,744356389 | 0,573277345 | 1,164733179 |
| VIM1 | 9,2866E-11 | 1,228271675 | 0,562 | 0,441 | 3,39769E-06 | Secretory | 2,342861508 | 0,426828473 | 1,274376417 |
| FBLN2 | 1,04979E-10 | 0,389648531 | 0,183 | 0,063 | 3,84088E-06 | Secretory | 1,310074205 | 0,76331554 | 2,904761905 |
| NRP21 | 1,06112E-10 | 0,389923069 | 0,151 | 0,04 | 3,88233E-06 | Secretory | 1,31032353 | 0,763170299 | 3,775 |
| UBXN12 | 1,23496E-10 | 0,654187549 | 0,474 | 0,373 | 4,51835E-06 | Secretory | 1,57372946 | 0,635433234 | 1,27077748 |
| ARL6IP41 | 1,50057E-10 | 0,70811074 | 0,586 | 0,551 | 5,49014E-06 | Secretory | 1,633663376 | 0,612121209 | 1,063520871 |
| CTSD | 1,53783E-10 | 0,987661081 | 0,588 | 0,569 | 5,62645E-06 | Secretory | 1,982967566 | 0,504294683 | 1,033391916 |
| EIF3G | 1,72604E-10 | 0,846687425 | 0,584 | 0,564 | 6,31508E-06 | Secretory | 1,798366947 | 0,556060042 | 1,035460993 |
| HLA-A | 1,74871E-10 | 0,714398936 | 0,754 | 0,784 | 6,398E-06 | Secretory | 1,640799476 | 0,609458995 | 0,961734694 |
| KRTCAP31 | 1,9415E-10 | 0,683341364 | 0,494 | 0,404 | 7,10337E-06 | Secretory | 1,605854703 | 0,622721345 | 1,222772277 |
| PPIB1 | 1,96191E-10 | 0,468977001 | 0,746 | 0,787 | 7,17805E-06 | Secretory | 1,384127651 | 0,722476716 | 0,947903431 |
| PLP21 | 2,03532E-10 | 0,457131557 | 0,302 | 0,163 | 7,44661E-06 | Secretory | 1,37280961 | 0,728433129 | 1,852760736 |
| TSPO1 | 2,07391E-10 | 0,73425483 | 0,612 | 0,594 | 7,58783E-06 | Secretory | 1,663538009 | 0,601128435 | 1,03030303 |
| CLDN71 | 2,19919E-10 | 0,889971569 | 0,5 | 0,411 | 8,04618E-06 | Secretory | 1,853139604 | 0,539624753 | 1,216545012 |
| SNHG7 | 2,43906E-10 | 0,673955015 | 0,384 | 0,253 | 8,92379E-06 | Secretory | 1,595440732 | 0,626786054 | 1,517786561 |
| MGMT1 | 2,49439E-10 | 0,556163428 | 0,39 | 0,265 | 9,12623E-06 | Secretory | 1,470353888 | 0,680108379 | 1,471698113 |
| LSM2 | 3,16273E-10 | 0,493224694 | 0,332 | 0,196 | 1,15715E-05 | Secretory | 1,407587585 | 0,710435365 | 1,693877551 |
| PEBP1 | 3,2332E-10 | 0,536511932 | 0,709 | 0,735 | 1,18293E-05 | Secretory | 1,450461429 | 0,689435775 | 0,96462585 |
| TGM2 | 3,44723E-10 | 0,597278667 | 0,177 | 0,06 | 1,26124E-05 | Secretory | 1,512860189 | 0,660999613 | 2,95 |
| SLC66A3 | 3,52896E-10 | 0,423604106 | 0,289 | 0,153 | 1,29114E-05 | Secretory | 1,341274112 | 0,745559756 | 1,888888889 |
| ITGB41 | 3,90355E-10 | 0,749916064 | 0,489 | 0,401 | 1,42819E-05 | Secretory | 1,681694986 | 0,594638153 | 1,219451372 |

| Supplementary Table 4 - clusermarker | | | | | | | | | |
|---|---|---|---|---|---|---|---|---|---|
| | **p_val** | **avg_log2FC** | **pct.1** | **pct.2** | **p_val_adj** | **celltype** | **Foldchange_UP** | **Foldchange_DOWN** | **Ratio_pct1_pct2** |
| DDT1 | 3,9249E-10 | 0,598188105 | 0,427 | 0,314 | 1,436E-05 | Secretory | 1,513814158 | 0,660583067 | 1,359872611 |
| PROM1 | 4,01956E-10 | 0,707327273 | 0,498 | 0,396 | 1,47064E-05 | Secretory | 1,632776443 | 0,612453716 | 1,257575758 |
| GP2 | 4,20411E-10 | -0,359599443 | 0,071 | 0,208 | 1,53816E-05 | Secretory | 0,779380941 | 1,28306961 | 0,341346154 |
| SLC25A51 | 4,46917E-10 | 0,727628316 | 0,61 | 0,597 | 1,63514E-05 | Secretory | 1,655914651 | 0,603895859 | 1,021775544 |
| IGKV3-201 | 5,30205E-10 | 3,723881392 | 0,144 | 0,04 | 1,93986E-05 | Secretory | 13,21295632 | 0,07568329 | 3,6 |
| RERG | 5,30271E-10 | 0,343301152 | 0,179 | 0,062 | 1,9401E-05 | Secretory | 1,268656194 | 0,788235619 | 2,887096774 |
| MESP11 | 5,30314E-10 | 0,338791233 | 0,22 | 0,093 | 1,94026E-05 | Secretory | 1,26469652 | 0,790703528 | 2,365591398 |
| PHB | 5,92637E-10 | -0,259704206 | 0,558 | 0,75 | 2,16828E-05 | Secretory | 0,835259154 | 1,197233212 | 0,744 |
| OAT1 | 6,14862E-10 | -0,295561526 | 0,407 | 0,644 | 2,24959E-05 | Secretory | 0,814755152 | 1,227362598 | 0,631987578 |
| LIME1 | 6,24219E-10 | 0,377538733 | 0,22 | 0,093 | 2,28383E-05 | Secretory | 1,299123633 | 0,76974968 | 2,365591398 |
| PTTG1IP1 | 6,69516E-10 | 1,018355991 | 0,509 | 0,434 | 2,44956E-05 | Secretory | 2,02560938 | 0,493678598 | 1,17281106 |
| NUPR1 | 6,94409E-10 | -0,392809925 | 0,461 | 0,672 | 2,54064E-05 | Secretory | 0,76164471 | 1,312948133 | 0,686011905 |
| TFCP2L1 | 7,17999E-10 | 0,796112918 | 0,429 | 0,304 | 2,62694E-05 | Secretory | 1,736416357 | 0,575898745 | 1,411184211 |
| RBP1 | 7,22352E-10 | 0,913013328 | 0,351 | 0,231 | 2,64287E-05 | Secretory | 1,882974324 | 0,531074687 | 1,519480519 |
| RPL21 | 8,18199E-10 | 0,261179663 | 0,894 | 0,922 | 2,99354E-05 | Secretory | 1,19845826 | 0,834405364 | 0,969631236 |
| TSTD1 | 8,94683E-10 | 0,750031637 | 0,61 | 0,607 | 3,27338E-05 | Secretory | 1,681829711 | 0,594590519 | 1,004942339 |
| MUC41 | 1,1929E-09 | 0,869368898 | 0,147 | 0,042 | 4,36445E-05 | Secretory | 1,82686357 | 0,547386251 | 3,5 |
| NPC2 | 1,23883E-09 | 1,044750949 | 0,612 | 0,632 | 4,53251E-05 | Secretory | 2,063010192 | 0,48472858 | 0,96835443 |
| CYP27A1 | 1,27518E-09 | 0,354651255 | 0,235 | 0,105 | 4,6655E-05 | Secretory | 1,278676447 | 0,782058669 | 2,238095238 |
| CCNI | 1,30451E-09 | 0,671866476 | 0,659 | 0,686 | 4,77282E-05 | Secretory | 1,593132738 | 0,627694087 | 0,960641399 |
| GAS6 | 1,33416E-09 | 0,533814302 | 0,263 | 0,138 | 4,88128E-05 | Secretory | 1,447751812 | 0,690726126 | 1,905797101 |
| STEAP3 | 1,44789E-09 | 0,347198315 | 0,194 | 0,077 | 5,29738E-05 | Secretory | 1,272087858 | 0,786109225 | 2,519480519 |
| PLAAT31 | 1,58067E-09 | 0,66456077 | 0,364 | 0,245 | 5,7832E-05 | Secretory | 1,585085619 | 0,630880747 | 1,485714286 |
| CFB1 | 1,67137E-09 | 1,265028207 | 0,416 | 0,293 | 6,11506E-05 | Secretory | 2,403319087 | 0,416091232 | 1,419795222 |
| SLC25A31 | 1,70744E-09 | 0,716136128 | 0,621 | 0,631 | 6,247E-05 | Secretory | 1,642776401 | 0,608725569 | 0,984152139 |
| EIF3K | 2,10486E-09 | 0,776299516 | 0,647 | 0,652 | 7,70105E-05 | Secretory | 1,71273211 | 0,58386247 | 0,992331288 |
| CMTM7 | 2,14332E-09 | 0,430506377 | 0,312 | 0,178 | 7,84178E-05 | Secretory | 1,347706531 | 0,742001302 | 1,752808989 |
| PPA1 | 2,25277E-09 | 0,730564085 | 0,586 | 0,571 | 8,24221E-05 | Secretory | 1,659287736 | 0,602668228 | 1,026269702 |
| LTBP2 | 2,56367E-09 | 0,281302828 | 0,164 | 0,057 | 9,3797E-05 | Secretory | 1,215291861 | 0,822847608 | 2,877192982 |
| NDUFA111 | 2,86173E-09 | 0,663190166 | 0,638 | 0,667 | 0,000104702 | Secretory | 1,583580454 | 0,631480388 | 0,956521739 |
| CARHSP1 | 3,63992E-09 | 0,462030167 | 0,345 | 0,216 | 0,000133174 | Secretory | 1,377478849 | 0,72596396 | 1,597222222 |

| Supplementary Table 4 - clusermarker | | | | | | | | | |
|---|---|---|---|---|---|---|---|---|---|
| | **p_val** | **avg_log2FC** | **pct.1** | **pct.2** | **p_val_adj** | **celltype** | **Foldchange_UP** | **Foldchange_DOWN** | **Ratio_pct1_pct2** |
| RPL23A1 | 3,65475E-09 | 0,349319782 | 0,871 | 0,903 | 0,000133716 | Secretory | 1,273959824 | 0,78495411 | 0,964562569 |
| IFITM11 | 3,76117E-09 | 0,89022097 | 0,343 | 0,221 | 0,00013761 | Secretory | 1,853459986 | 0,539531475 | 1,552036199 |
| RPL36AL | 4,07398E-09 | 0,451579711 | 0,754 | 0,8 | 0,000149055 | Secretory | 1,367536852 | 0,73124172 | 0,9425 |
| CLDN1 | 4,13095E-09 | 0,288185249 | 0,149 | 0,047 | 0,000151139 | Secretory | 1,221103298 | 0,818931536 | 3,170212766 |
| H2AW | 4,14731E-09 | 0,280200809 | 0,159 | 0,055 | 0,000151738 | Secretory | 1,2143639 | 0,82347639 | 2,890909091 |
| CFH | 4,24869E-09 | 0,257311428 | 0,269 | 0,138 | 0,000155447 | Secretory | 1,19524919 | 0,83664562 | 1,949275362 |
| CXCL16 | 4,34431E-09 | 0,543442135 | 0,319 | 0,193 | 0,000158945 | Secretory | 1,457445701 | 0,686131908 | 1,652849741 |
| COL18A1 | 4,38005E-09 | 0,329182103 | 0,144 | 0,045 | 0,000160253 | Secretory | 1,256300947 | 0,79598762 | 3,2 |
| NDUFB91 | 4,38977E-09 | 0,638371891 | 0,588 | 0,576 | 0,000160609 | Secretory | 1,556571547 | 0,642437543 | 1,020833333 |
| ARF51 | 4,66357E-09 | 0,744802623 | 0,478 | 0,414 | 0,000170626 | Secretory | 1,675744993 | 0,596749508 | 1,154589372 |
| RNF150 | 4,81333E-09 | 0,259939457 | 0,159 | 0,053 | 0,000176105 | Secretory | 1,197428454 | 0,835122965 | 3 |
| XBP1 | 5,16766E-09 | -0,296575414 | 0,728 | 0,882 | 0,000189069 | Secretory | 0,814182764 | 1,22822546 | 0,825396825 |
| CLIC11 | 5,42206E-09 | 0,674549867 | 0,582 | 0,547 | 0,000198377 | Secretory | 1,596098698 | 0,626527671 | 1,063985375 |
| CTSB1 | 5,79623E-09 | 0,880333153 | 0,595 | 0,596 | 0,000212067 | Secretory | 1,840800337 | 0,543241969 | 0,998322148 |
| BBLN1 | 6,30303E-09 | 0,741924478 | 0,547 | 0,509 | 0,000230609 | Secretory | 1,672405251 | 0,597941198 | 1,074656189 |
| CX3CL1 | 6,36171E-09 | 0,504438214 | 0,274 | 0,148 | 0,000232756 | Secretory | 1,418570857 | 0,704934826 | 1,851351351 |
| FEZ2 | 7,85312E-09 | 0,311038411 | 0,25 | 0,128 | 0,000287322 | Secretory | 1,240600328 | 0,80606137 | 1,953125 |
| S100A14 | 8,01009E-09 | 0,492040026 | 0,373 | 0,241 | 0,000293065 | Secretory | 1,406432219 | 0,711018979 | 1,547717842 |
| SRP14 | 1,16723E-08 | 0,561531447 | 0,688 | 0,73 | 0,000427056 | Secretory | 1,475835012 | 0,677582516 | 0,942465753 |
| PDE4D | 1,28106E-08 | -0,299160396 | 0,136 | 0,285 | 0,000468703 | Secretory | 0,81272524 | 1,230428134 | 0,477192982 |
| PDXK1 | 1,32834E-08 | 0,603897918 | 0,412 | 0,306 | 0,000486001 | Secretory | 1,519817313 | 0,657973818 | 1,346405229 |
| PDLIM1 | 1,35098E-08 | 0,894676442 | 0,433 | 0,348 | 0,000494285 | Secretory | 1,859192871 | 0,537867811 | 1,244252874 |
| DYNC2I2 | 1,42309E-08 | 0,404345892 | 0,267 | 0,15 | 0,000520665 | Secretory | 1,323488713 | 0,75557879 | 1,78 |
| GSTA11 | 1,48003E-08 | 0,336889812 | 0,172 | 0,065 | 0,000541498 | Secretory | 1,263030792 | 0,791746334 | 2,646153846 |
| SPINT11 | 1,58542E-08 | 0,724721915 | 0,487 | 0,421 | 0,000580059 | Secretory | 1,652582064 | 0,605113671 | 1,156769596 |
| MUC161 | 1,76136E-08 | 1,027020126 | 0,297 | 0,16 | 0,00064443 | Secretory | 2,037810822 | 0,490722686 | 1,85625 |
| PDE7A | 2,01745E-08 | 0,373414097 | 0,231 | 0,116 | 0,000738126 | Secretory | 1,295414768 | 0,771953527 | 1,99137931 |
| RNASEK1 | 2,30891E-08 | 0,687126284 | 0,616 | 0,654 | 0,000844762 | Secretory | 1,610073204 | 0,621089772 | 0,941896024 |
| CTNNB1 | 2,32283E-08 | 0,849121535 | 0,517 | 0,459 | 0,000849855 | Secretory | 1,801403707 | 0,55512265 | 1,126361656 |
| OAZ11 | 2,40963E-08 | 0,466886359 | 0,679 | 0,722 | 0,000881611 | Secretory | 1,382123333 | 0,723524432 | 0,940443213 |
| TYMP1 | 2,56886E-08 | 0,541569799 | 0,265 | 0,148 | 0,00093987 | Secretory | 1,455555449 | 0,687022951 | 1,790540541 |

| Supplementary Table 4 - clusermarker | | | | | | | | | |
|---|---|---|---|---|---|---|---|---|---|
| | **p_val** | **avg_log2FC** | **pct.1** | **pct.2** | **p_val_adj** | **celltype** | **Foldchange_UP** | **Foldchange_DOWN** | **Ratio_pct1_pct2** |
| CAPG1 | 2,61172E-08 | 0,44775598 | 0,392 | 0,28 | 0,00095555 | Secretory | 1,36391712 | 0,73318238 | 1,4 |
| HLA-C1 | 2,69406E-08 | 0,513012289 | 0,782 | 0,827 | 0,000985677 | Secretory | 1,427026661 | 0,700757755 | 0,945586457 |
| ZFP36L11 | 3,01359E-08 | 0,677944033 | 0,666 | 0,664 | 0,001102582 | Secretory | 1,599858191 | 0,625055399 | 1,003012048 |
| GUK11 | 3,68827E-08 | 0,624577024 | 0,599 | 0,614 | 0,001349426 | Secretory | 1,541758739 | 0,648609912 | 0,975570033 |
| CAST1 | 3,69599E-08 | 0,840787016 | 0,504 | 0,441 | 0,001352253 | Secretory | 1,791026913 | 0,558338902 | 1,142857143 |
| ANXA2 | 3,87251E-08 | 0,694308536 | 0,698 | 0,712 | 0,001416834 | Secretory | 1,61810871 | 0,618005449 | 0,980337079 |
| MSMB1 | 3,92582E-08 | -1,234070141 | 0,235 | 0,399 | 0,001436339 | Secretory | 0,425116411 | 2,352296862 | 0,588972431 |
| RPL341 | 4,1622E-08 | 0,340730839 | 0,92 | 0,948 | 0,001522823 | Secretory | 1,266397962 | 0,789641195 | 0,970464135 |
| CRIP21 | 4,32154E-08 | 0,72309448 | 0,483 | 0,419 | 0,001581122 | Secretory | 1,650718917 | 0,605796656 | 1,15274463 |
| RPS241 | 4,40997E-08 | 0,343805754 | 0,897 | 0,912 | 0,001613476 | Secretory | 1,269100001 | 0,787959971 | 0,983552632 |
| TRIR1 | 4,62898E-08 | 0,660193949 | 0,541 | 0,514 | 0,001693604 | Secretory | 1,580295057 | 0,632793221 | 1,052529183 |
| MDK1 | 4,74234E-08 | 1,015320252 | 0,567 | 0,567 | 0,001735078 | Secretory | 2,021351546 | 0,494718498 | 1 |
| B2M1 | 4,7524E-08 | 0,326905301 | 0,942 | 0,982 | 0,001738762 | Secretory | 1,254319868 | 0,797244806 | 0,959266802 |
| RHOV1 | 5,12972E-08 | 0,326633921 | 0,17 | 0,07 | 0,00187681 | Secretory | 1,254083945 | 0,797394787 | 2,428571429 |
| YWHAB | 5,43329E-08 | 0,703071657 | 0,543 | 0,527 | 0,001987877 | Secretory | 1,627967227 | 0,61426298 | 1,030360531 |
| PSMA71 | 5,43345E-08 | 0,683995738 | 0,528 | 0,499 | 0,001987935 | Secretory | 1,606583247 | 0,622438957 | 1,058116232 |
| SLC4A111 | 6,26001E-08 | 0,405830947 | 0,267 | 0,146 | 0,00229035 | Secretory | 1,324851763 | 0,754801426 | 1,828767123 |
| YBX11 | 6,42264E-08 | 0,577329052 | 0,64 | 0,652 | 0,002349852 | Secretory | 1,492084304 | 0,670203417 | 0,981595092 |
| KRT5 | 6,91749E-08 | 0,980900541 | 0,272 | 0,165 | 0,002530902 | Secretory | 1,973697021 | 0,506663378 | 1,648484848 |
| CCND11 | 7,33983E-08 | 0,985266555 | 0,403 | 0,311 | 0,002685423 | Secretory | 1,979679048 | 0,505132386 | 1,295819936 |
| NUDT141 | 7,48703E-08 | 0,340306001 | 0,31 | 0,203 | 0,002739281 | Secretory | 1,266025094 | 0,789873759 | 1,527093596 |
| PSME11 | 7,51727E-08 | 0,761903008 | 0,573 | 0,584 | 0,002750343 | Secretory | 1,695725923 | 0,589717941 | 0,981164384 |
| AGRN1 | 7,81932E-08 | 0,385217304 | 0,284 | 0,168 | 0,002860856 | Secretory | 1,306056495 | 0,765663663 | 1,69047619 |
| CLTA1 | 7,9412E-08 | 0,510766868 | 0,558 | 0,551 | 0,002905445 | Secretory | 1,424807355 | 0,701849269 | 1,012704174 |
| SOD1 | 8,88522E-08 | 0,629889037 | 0,58 | 0,584 | 0,003250835 | Secretory | 1,54744597 | 0,646226117 | 0,993150685 |
| PRR15L | 8,99143E-08 | 0,369216912 | 0,304 | 0,188 | 0,003289694 | Secretory | 1,291651538 | 0,774202616 | 1,617021277 |
| NT5C1 | 9,08592E-08 | 0,434032538 | 0,317 | 0,203 | 0,003324265 | Secretory | 1,351004555 | 0,740189955 | 1,561576355 |
| SUMO2 | 9,33946E-08 | 0,732714725 | 0,61 | 0,651 | 0,00341703 | Secretory | 1,661763097 | 0,601770494 | 0,937019969 |
| CRABP21 | 9,35359E-08 | 0,538700596 | 0,235 | 0,123 | 0,003422198 | Secretory | 1,452663546 | 0,688390648 | 1,910569106 |
| VAMP2 | 1,06585E-07 | 0,737157966 | 0,509 | 0,469 | 0,00389964 | Secretory | 1,666888918 | 0,59992 | 1,085287846 |
| TMSB4X1 | 1,08945E-07 | 0,926616724 | 0,778 | 0,822 | 0,00398596 | Secretory | 1,900813153 | 0,526090636 | 0,946472019 |

| Supplementary Table 4 - clusermarker | | | | | | | | | |
|---|---|---|---|---|---|---|---|---|---|
| | **p_val** | **avg_log2FC** | **pct.1** | **pct.2** | **p_val_adj** | **celltype** | **Foldchange_UP** | **Foldchange_DOWN** | **Ratio_pct1_pct2** |
| TRMT1121 | 1,20545E-07 | 0,643808162 | 0,494 | 0,434 | 0,004410392 | Secretory | 1,562447984 | 0,640021306 | 1,138248848 |
| MRPS241 | 1,2386E-07 | 0,674923498 | 0,506 | 0,468 | 0,004531672 | Secretory | 1,596512112 | 0,626365433 | 1,081196581 |
| MYL12A1 | 1,30675E-07 | 0,733824876 | 0,573 | 0,564 | 0,004780992 | Secretory | 1,663042314 | 0,601307611 | 1,015957447 |
| SMR3B | 1,31859E-07 | -2,51415063 | 0,056 | 0,156 | 0,004824338 | Secretory | 0,175051262 | 5,712612336 | 0,358974359 |
| MGST1 | 1,32175E-07 | 0,877010289 | 0,502 | 0,456 | 0,004835875 | Secretory | 1,836565422 | 0,544494625 | 1,100877193 |
| PNRC11 | 1,33334E-07 | 0,627852459 | 0,606 | 0,604 | 0,004878298 | Secretory | 1,545263061 | 0,647139005 | 1,003311258 |
| ST13 | 1,3398E-07 | 0,650085253 | 0,552 | 0,547 | 0,004901911 | Secretory | 1,569260926 | 0,637242656 | 1,009140768 |
| RPL9 | 1,47352E-07 | 0,409674229 | 0,819 | 0,889 | 0,005391152 | Secretory | 1,328385821 | 0,752793341 | 0,921259843 |
| LAMB31 | 1,51568E-07 | 0,377760526 | 0,209 | 0,105 | 0,005545402 | Secretory | 1,299323369 | 0,769631351 | 1,99047619 |
| RPS291 | 1,62873E-07 | -0,253093338 | 0,718 | 0,847 | 0,005959033 | Secretory | 0,839095348 | 1,191759676 | 0,847697757 |
| MT-ATP61 | 1,6434E-07 | 0,446552675 | 0,821 | 0,834 | 0,006012714 | Secretory | 1,362779995 | 0,733794159 | 0,98441247 |
| MT-ND5 | 1,64444E-07 | 0,421161006 | 0,795 | 0,792 | 0,00601653 | Secretory | 1,339004683 | 0,746823378 | 1,003787879 |
| CRIP11 | 1,68634E-07 | 0,450090519 | 0,468 | 0,351 | 0,006169802 | Secretory | 1,366125969 | 0,731996919 | 1,333333333 |
| BLVRB | 1,82957E-07 | 0,566186432 | 0,431 | 0,343 | 0,006693843 | Secretory | 1,480604616 | 0,675399758 | 1,256559767 |
| KRTCAP21 | 1,83071E-07 | 0,434408423 | 0,657 | 0,694 | 0,006698033 | Secretory | 1,351356596 | 0,739997128 | 0,946685879 |
| SEC61B1 | 1,84905E-07 | 0,55422521 | 0,664 | 0,707 | 0,006765112 | Secretory | 1,468379838 | 0,681022699 | 0,939179632 |
| SERINC21 | 1,88479E-07 | 0,584501155 | 0,448 | 0,363 | 0,006895873 | Secretory | 1,499520406 | 0,666879888 | 1,23415978 |
| YPEL31 | 2,14676E-07 | 0,553753074 | 0,403 | 0,314 | 0,007854337 | Secretory | 1,467899375 | 0,681245606 | 1,28343949 |
| PDLIM41 | 2,3105E-07 | 0,505131179 | 0,269 | 0,166 | 0,008453433 | Secretory | 1,419252398 | 0,704596308 | 1,620481928 |
| ENO1 | 2,36807E-07 | 0,804253556 | 0,532 | 0,504 | 0,00866407 | Secretory | 1,74624206 | 0,572658295 | 1,055555556 |
| ATP5F1B1 | 2,70212E-07 | 0,614977278 | 0,575 | 0,581 | 0,009886257 | Secretory | 1,531533875 | 0,652940177 | 0,989672978 |
| WDR83OS1 | 2,71289E-07 | 0,625590056 | 0,608 | 0,644 | 0,009925664 | Secretory | 1,542841712 | 0,648154631 | 0,944099379 |
| PCBD1 | 2,82289E-07 | 0,60125585 | 0,47 | 0,413 | 0,01032809 | Secretory | 1,517036555 | 0,659179897 | 1,138014528 |
| ATP5MC3 | 2,87783E-07 | 0,562830936 | 0,593 | 0,617 | 0,010529117 | Secretory | 1,47716495 | 0,676972467 | 0,961102107 |
| EMP21 | 3,20092E-07 | 0,66285239 | 0,375 | 0,281 | 0,011711214 | Secretory | 1,583209737 | 0,631628253 | 1,334519573 |
| NDUFS7 | 3,27632E-07 | 0,553884627 | 0,485 | 0,439 | 0,011987062 | Secretory | 1,468033232 | 0,68118349 | 1,104783599 |
| TGFB21 | 3,29621E-07 | 0,276455654 | 0,114 | 0,035 | 0,012059835 | Secretory | 1,211215568 | 0,825616865 | 3,257142857 |
| HMGN11 | 3,34662E-07 | 0,593309441 | 0,541 | 0,542 | 0,012244294 | Secretory | 1,508703641 | 0,662820698 | 0,998154982 |
| PSMB91 | 3,81146E-07 | 0,523837484 | 0,263 | 0,156 | 0,01394499 | Secretory | 1,437774562 | 0,69551933 | 1,685897436 |
| PPL1 | 3,90047E-07 | 0,547750816 | 0,325 | 0,223 | 0,014270662 | Secretory | 1,461804942 | 0,684085798 | 1,457399103 |
| HLA-DPA1 | 4,13608E-07 | 0,801736849 | 0,409 | 0,301 | 0,015132658 | Secretory | 1,743198487 | 0,573658139 | 1,358803987 |

| Supplementary Table 4 - clusermarker | | | | | | | | | |
|---|---|---|---|---|---|---|---|---|---|
| | **p_val** | **avg_log2FC** | **pct.1** | **pct.2** | **p_val_adj** | **celltype** | **Foldchange_UP** | **Foldchange_DOWN** | **Ratio_pct1_pct2** |
| ATP6V0C1 | 4,15231E-07 | 0,615383475 | 0,562 | 0,566 | 0,015192073 | Secretory | 1,531965146 | 0,652756365 | 0,992932862 |
| RPL371 | 4,33935E-07 | 0,309249193 | 0,886 | 0,912 | 0,015876395 | Secretory | 1,239062699 | 0,807061661 | 0,971491228 |
| GNG51 | 4,89975E-07 | 0,600045988 | 0,591 | 0,606 | 0,017926702 | Secretory | 1,515764883 | 0,659732925 | 0,975247525 |
| MYLK1 | 5,19186E-07 | -0,299493405 | 0,052 | 0,146 | 0,018995472 | Secretory | 0,812537665 | 1,230712179 | 0,356164384 |
| CTSC1 | 5,19235E-07 | 0,763678582 | 0,377 | 0,288 | 0,018997245 | Secretory | 1,697814195 | 0,588992602 | 1,309027778 |
| PRXL2A | 5,26145E-07 | 0,286349472 | 0,213 | 0,11 | 0,019250064 | Secretory | 1,219550477 | 0,81997426 | 1,936363636 |
| GMDS | 5,38521E-07 | 0,717642185 | 0,569 | 0,589 | 0,019702867 | Secretory | 1,644492222 | 0,608090441 | 0,966044143 |
| ATP5PO | 5,7868E-07 | 0,628777245 | 0,578 | 0,606 | 0,02117217 | Secretory | 1,546253912 | 0,646724314 | 0,95379538 |
| CCL28 | 6,28066E-07 | 0,879012403 | 0,619 | 0,647 | 0,022979038 | Secretory | 1,839115903 | 0,543739521 | 0,956723338 |
| COX5B1 | 6,46145E-07 | 0,491694092 | 0,631 | 0,669 | 0,023640519 | Secretory | 1,406095021 | 0,71118949 | 0,943198804 |
| PLAAT41 | 7,37571E-07 | 0,734479032 | 0,446 | 0,369 | 0,026985517 | Secretory | 1,663796551 | 0,601035024 | 1,208672087 |
| TUFM1 | 7,87846E-07 | 0,613250736 | 0,485 | 0,458 | 0,028824939 | Secretory | 1,529702113 | 0,653722049 | 1,058951965 |
| HNRNPU1 | 7,93041E-07 | 0,8280033 | 0,562 | 0,537 | 0,029014988 | Secretory | 1,775226735 | 0,563308326 | 1,046554935 |
| EVA1C | 8,00256E-07 | 0,375177781 | 0,284 | 0,18 | 0,029278974 | Secretory | 1,296999372 | 0,771010397 | 1,577777778 |
| MVP1 | 8,07957E-07 | 0,476513322 | 0,31 | 0,208 | 0,029560726 | Secretory | 1,391376946 | 0,718712497 | 1,490384615 |
| TMEM541 | 8,11425E-07 | 0,392340839 | 0,347 | 0,25 | 0,029687607 | Secretory | 1,312521303 | 0,761892396 | 1,388 |
| CDC42EP1 | 8,15637E-07 | 0,404271517 | 0,289 | 0,183 | 0,029841706 | Secretory | 1,323420486 | 0,755617743 | 1,579234973 |
| PMEPA11 | 8,16718E-07 | 0,411099903 | 0,312 | 0,206 | 0,029881278 | Secretory | 1,329699183 | 0,752049796 | 1,514563107 |
| ST6GAL11 | 8,26025E-07 | 0,323646876 | 0,263 | 0,156 | 0,030221767 | Secretory | 1,251490098 | 0,799047473 | 1,685897436 |
| S100A91 | 8,41317E-07 | 0,704494711 | 0,155 | 0,065 | 0,030781262 | Secretory | 1,629573823 | 0,613657378 | 2,384615385 |
| REEP31 | 8,61064E-07 | 0,575304984 | 0,42 | 0,344 | 0,031503764 | Secretory | 1,489992412 | 0,671144357 | 1,220930233 |
| TUBB1 | 8,6264E-07 | 0,729942502 | 0,472 | 0,426 | 0,031561394 | Secretory | 1,658572988 | 0,602927943 | 1,107981221 |
| IGLV2-14 | 8,7983E-07 | 2,627017537 | 0,116 | 0,04 | 0,032190348 | Secretory | 6,177476155 | 0,161878407 | 2,9 |
| PLEC1 | 9,07821E-07 | 0,622282548 | 0,392 | 0,3 | 0,033214446 | Secretory | 1,53930866 | 0,649642288 | 1,306666667 |
| COMMD6 | 1,0082E-06 | 0,6880476 | 0,612 | 0,621 | 0,036886875 | Secretory | 1,611101738 | 0,620693266 | 0,985507246 |
| BANF1 | 1,02018E-06 | 0,611090969 | 0,487 | 0,443 | 0,037325219 | Secretory | 1,527413806 | 0,654701428 | 1,099322799 |
| TP53I31 | 1,06073E-06 | 0,270650388 | 0,224 | 0,123 | 0,038809017 | Secretory | 1,206351546 | 0,828945761 | 1,821138211 |
| MT-CO1 | 1,07128E-06 | 0,339129047 | 0,912 | 0,92 | 0,039195066 | Secretory | 1,264992689 | 0,790518403 | 0,991304348 |
| EPCAM1 | 1,07585E-06 | 0,624332434 | 0,584 | 0,601 | 0,039362242 | Secretory | 1,541497376 | 0,648719885 | 0,97171381 |
| SLIT3 | 1,07916E-06 | 0,310510673 | 0,181 | 0,088 | 0,039483177 | Secretory | 1,240146599 | 0,806356282 | 2,056818182 |
| POLE41 | 1,12754E-06 | 0,397147915 | 0,334 | 0,235 | 0,041253317 | Secretory | 1,316901933 | 0,759357986 | 1,421276596 |

| Supplementary Table 4 - clusermarker | | | | | | | | | |
|---|---|---|---|---|---|---|---|---|---|
| | **p_val** | **avg_log2FC** | **pct.1** | **pct.2** | **p_val_adj** | **celltype** | **Foldchange_UP** | **Foldchange_DOWN** | **Ratio_pct1_pct2** |
| DSTN | 1,15121E-06 | 0,544692424 | 0,675 | 0,724 | 0,042119352 | Secretory | 1,458709321 | 0,685537541 | 0,932320442 |
| TRAPPC51 | 1,18498E-06 | 0,591879112 | 0,502 | 0,481 | 0,043355044 | Secretory | 1,50720861 | 0,663478163 | 1,043659044 |
| ARPC1B1 | 1,19366E-06 | 0,499904957 | 0,306 | 0,21 | 0,04367226 | Secretory | 1,414120398 | 0,707153366 | 1,457142857 |
| RPL35A1 | 1,2111E-06 | 0,33705572 | 0,873 | 0,938 | 0,044310508 | Secretory | 1,263176047 | 0,79165529 | 0,930703625 |
| MVB12A1 | 1,29659E-06 | 0,37639742 | 0,282 | 0,181 | 0,047438216 | Secretory | 1,298096305 | 0,770358868 | 1,55801105 |
| ATP5MG | 1,30088E-06 | 0,415434158 | 0,683 | 0,724 | 0,047595478 | Secretory | 1,333699975 | 0,749793821 | 0,943370166 |
| MT-ND31 | 1,32744E-06 | -0,393634468 | 0,838 | 0,87 | 0,048566949 | Secretory | 0,761209532 | 1,313698736 | 0,963218391 |
| BAG11 | 1,35439E-06 | 0,664054854 | 0,489 | 0,438 | 0,049552906 | Secretory | 1,584529867 | 0,63110202 | 1,116438356 |
| SMDT11 | 1,37247E-06 | 0,677740157 | 0,578 | 0,609 | 0,050214408 | Secretory | 1,599632122 | 0,625143735 | 0,94909688 |
| NDUFB111 | 1,38061E-06 | 0,488369465 | 0,578 | 0,604 | 0,050512258 | Secretory | 1,402858467 | 0,712830284 | 0,956953642 |
| KRT18 | 1,47603E-06 | 0,388569862 | 0,726 | 0,78 | 0,054003625 | Secretory | 1,30909506 | 0,763886467 | 0,930769231 |
| RBPMS | 1,51605E-06 | 0,695137204 | 0,386 | 0,304 | 0,055467731 | Secretory | 1,6190384 | 0,617650576 | 1,269736842 |
| SPRY11 | 1,82795E-06 | 0,433705522 | 0,267 | 0,165 | 0,066879331 | Secretory | 1,350698357 | 0,740357753 | 1,618181818 |
| NDUFS8 | 1,8286E-06 | 0,610756355 | 0,526 | 0,506 | 0,066903082 | Secretory | 1,527059583 | 0,654853295 | 1,039525692 |
| CCT6A1 | 1,83033E-06 | 0,475670135 | 0,381 | 0,295 | 0,066966464 | Secretory | 1,39056399 | 0,719132674 | 1,291525424 |
| PPP1R13L | 1,93328E-06 | 0,294472805 | 0,22 | 0,125 | 0,070732737 | Secretory | 1,226436727 | 0,815370233 | 1,76 |
| TMEM179B1 | 2,02608E-06 | 0,606339987 | 0,455 | 0,416 | 0,074128098 | Secretory | 1,522392107 | 0,656860999 | 1,09375 |
| PHB2 | 2,0738E-06 | 0,610661448 | 0,487 | 0,468 | 0,075874086 | Secretory | 1,52695913 | 0,654896376 | 1,040598291 |
| NENF1 | 2,10045E-06 | 0,572663432 | 0,584 | 0,599 | 0,0768492 | Secretory | 1,487266755 | 0,672374338 | 0,974958264 |
| MCC | 2,17116E-06 | 0,363532755 | 0,228 | 0,131 | 0,079436318 | Secretory | 1,286572498 | 0,777258959 | 1,740458015 |
| S100A131 | 2,26162E-06 | 0,639382976 | 0,541 | 0,532 | 0,082745734 | Secretory | 1,557662822 | 0,641987461 | 1,016917293 |
| CUTA | 2,41762E-06 | 0,598024158 | 0,569 | 0,602 | 0,088453599 | Secretory | 1,513642139 | 0,66065814 | 0,945182724 |
| CD14 | 2,4357E-06 | 0,546711968 | 0,381 | 0,293 | 0,089114815 | Secretory | 1,460752712 | 0,684578568 | 1,300341297 |
| FIS11 | 2,61806E-06 | 0,560732562 | 0,541 | 0,537 | 0,095787017 | Secretory | 1,475018002 | 0,677957828 | 1,00744879 |
| TECR1 | 2,69518E-06 | 0,571153258 | 0,556 | 0,554 | 0,098608634 | Secretory | 1,48571074 | 0,673078529 | 1,003610108 |
| UNC93B11 | 2,70545E-06 | 0,399903483 | 0,254 | 0,156 | 0,098984227 | Secretory | 1,319419638 | 0,757908986 | 1,628205128 |
| HLA-F | 2,74986E-06 | 0,463737696 | 0,282 | 0,181 | 0,100609201 | Secretory | 1,379110156 | 0,72510524 | 1,55801105 |
| TMEM14C1 | 2,79545E-06 | 0,702829901 | 0,522 | 0,521 | 0,102277184 | Secretory | 1,627694448 | 0,614365922 | 1,001919386 |
| NDUFB81 | 2,87502E-06 | 0,545752735 | 0,53 | 0,512 | 0,105188523 | Secretory | 1,459781795 | 0,685033889 | 1,03515625 |
| CAPNS11 | 2,91426E-06 | 0,586580269 | 0,433 | 0,374 | 0,106624024 | Secretory | 1,501682971 | 0,665919518 | 1,157754011 |
| SELENOH1 | 3,00807E-06 | 0,624571688 | 0,539 | 0,531 | 0,110056201 | Secretory | 1,541753037 | 0,648612311 | 1,015065913 |

| Supplementary Table 4 - clusermarker | | | | | | | | | |
|---|---|---|---|---|---|---|---|---|---|
| | **p_val** | **avg_log2FC** | **pct.1** | **pct.2** | **p_val_adj** | **celltype** | **Foldchange_UP** | **Foldchange_DOWN** | **Ratio_pct1_pct2** |
| CDKN2A1 | 3,06733E-06 | 0,312325079 | 0,101 | 0,033 | 0,112224509 | Secretory | 1,241707251 | 0,805342805 | 3,060606061 |
| ALKBH7 | 3,09086E-06 | 0,626528476 | 0,547 | 0,556 | 0,113085289 | Secretory | 1,5438456 | 0,647733167 | 0,98381295 |
| SERF21 | 3,21207E-06 | 0,303745377 | 0,815 | 0,869 | 0,117519845 | Secretory | 1,234344737 | 0,810146445 | 0,937859609 |
| NDUFAF3 | 3,22661E-06 | 0,48246058 | 0,47 | 0,421 | 0,118051978 | Secretory | 1,397124493 | 0,715755829 | 1,116389549 |
| EMD1 | 3,52948E-06 | 0,476907442 | 0,422 | 0,353 | 0,12913303 | Secretory | 1,391757099 | 0,718516184 | 1,195467422 |
| TAX1BP31 | 3,80993E-06 | 0,37228789 | 0,304 | 0,208 | 0,139393764 | Secretory | 1,294403928 | 0,77255637 | 1,461538462 |
| BLVRA1 | 3,86926E-06 | 0,3587506 | 0,287 | 0,193 | 0,14156449 | Secretory | 1,282314909 | 0,779839643 | 1,487046632 |
| ATP1B3 | 3,89747E-06 | 0,41586571 | 0,287 | 0,19 | 0,142596699 | Secretory | 1,334098982 | 0,74956957 | 1,510526316 |
| FAM207A | 3,97264E-06 | 0,264098933 | 0,203 | 0,108 | 0,145346894 | Secretory | 1,200885775 | 0,832718665 | 1,87962963 |
| MRPL44 | 4,04333E-06 | 0,314758605 | 0,319 | 0,221 | 0,147933445 | Secretory | 1,24380352 | 0,803985504 | 1,443438914 |
| TFF31 | 4,12293E-06 | -1,152343049 | 0,222 | 0,354 | 0,150845644 | Secretory | 0,449893975 | 2,222745926 | 0,627118644 |
| ATP6V0B1 | 4,36204E-06 | 0,585560563 | 0,504 | 0,481 | 0,159593929 | Secretory | 1,500621947 | 0,66639036 | 1,047817048 |
| EPS8 | 4,46421E-06 | 0,484247837 | 0,28 | 0,188 | 0,163332034 | Secretory | 1,398856368 | 0,714869677 | 1,489361702 |
| TPI11 | 4,62161E-06 | 0,630199764 | 0,588 | 0,629 | 0,169090716 | Secretory | 1,547779293 | 0,646086948 | 0,93481717 |
| GCSH | 4,72007E-06 | 0,526518144 | 0,412 | 0,343 | 0,172693317 | Secretory | 1,440448563 | 0,694228191 | 1,201166181 |
| DAPK1 | 4,76286E-06 | 0,330461317 | 0,226 | 0,131 | 0,174258799 | Secretory | 1,257415382 | 0,795282143 | 1,72519084 |
| NEAT11 | 4,82767E-06 | 0,788757443 | 0,571 | 0,667 | 0,176630064 | Secretory | 1,727585896 | 0,57884242 | 0,856071964 |
| RHOC1 | 5,00902E-06 | 0,637184291 | 0,468 | 0,431 | 0,183264953 | Secretory | 1,555290734 | 0,642966603 | 1,085846868 |
| NCL1 | 5,02112E-06 | 0,664329182 | 0,565 | 0,571 | 0,183707571 | Secretory | 1,584831194 | 0,630982027 | 0,989492119 |
| PGGHG | 5,46019E-06 | 0,26007431 | 0,142 | 0,063 | 0,199772144 | Secretory | 1,197540386 | 0,835044907 | 2,253968254 |
| COX5A1 | 5,80683E-06 | 0,633953727 | 0,552 | 0,574 | 0,21245461 | Secretory | 1,551811935 | 0,644407984 | 0,961672474 |
| ANP32B | 5,89726E-06 | 0,503022658 | 0,425 | 0,371 | 0,215763105 | Secretory | 1,417179653 | 0,70562684 | 1,145552561 |
| SESN3 | 6,09854E-06 | 0,416016729 | 0,233 | 0,141 | 0,223127317 | Secretory | 1,334238641 | 0,74949111 | 1,65248227 |
| MUC201 | 6,16307E-06 | 0,492893676 | 0,284 | 0,185 | 0,225488347 | Secretory | 1,407264659 | 0,710598389 | 1,535135135 |
| IL321 | 6,65457E-06 | 0,340961188 | 0,17 | 0,083 | 0,243470875 | Secretory | 1,266600178 | 0,789515126 | 2,048192771 |
| ENC1 | 6,90292E-06 | 0,558414012 | 0,289 | 0,2 | 0,252557104 | Secretory | 1,47264941 | 0,679048247 | 1,445 |
| PABPC11 | 7,10244E-06 | 0,311545793 | 0,793 | 0,845 | 0,25985693 | Secretory | 1,241036712 | 0,805777936 | 0,938461538 |
| EID1 | 7,24645E-06 | 0,674876838 | 0,547 | 0,569 | 0,265125743 | Secretory | 1,596460478 | 0,626385691 | 0,961335677 |
| COX8A1 | 7,66814E-06 | 0,452015325 | 0,627 | 0,656 | 0,280554228 | Secretory | 1,367949835 | 0,731020959 | 0,955792683 |
| RPL391 | 7,82942E-06 | 0,340422869 | 0,877 | 0,897 | 0,286455158 | Secretory | 1,266127655 | 0,789809776 | 0,977703456 |
| AGPAT21 | 7,84656E-06 | 0,357912469 | 0,341 | 0,25 | 0,287082106 | Secretory | 1,281570166 | 0,780292821 | 1,364 |

| Supplementary Table 4 - clusermarker | | | | | | | | | |
|---|---|---|---|---|---|---|---|---|---|
| | **p_val** | **avg_log2FC** | **pct.1** | **pct.2** | **p_val_adj** | **celltype** | **Foldchange_UP** | **Foldchange_DOWN** | **Ratio_pct1_pct2** |
| HLA-E1 | 8,34965E-06 | 0,674207521 | 0,595 | 0,604 | 0,305488796 | Secretory | 1,595719995 | 0,626676361 | 0,985099338 |
| CD631 | 8,47757E-06 | 0,302400595 | 0,772 | 0,83 | 0,310168713 | Secretory | 1,233194701 | 0,81090196 | 0,930120482 |
| RANBP1 | 8,73495E-06 | 0,353101838 | 0,338 | 0,253 | 0,319585636 | Secretory | 1,277303919 | 0,78289903 | 1,335968379 |
| NOP53 | 8,84964E-06 | 0,612429103 | 0,591 | 0,626 | 0,323781653 | Secretory | 1,528831176 | 0,654094458 | 0,944089457 |
| DSG2 | 8,99059E-06 | 0,626713646 | 0,332 | 0,248 | 0,328938605 | Secretory | 1,544043765 | 0,647650036 | 1,338709677 |
| C12orf57 | 9,36112E-06 | 0,554424351 | 0,644 | 0,694 | 0,342495358 | Secretory | 1,468582538 | 0,680928701 | 0,92795389 |
| SLC25A15 | 9,36641E-06 | 0,39563682 | 0,336 | 0,253 | 0,342688854 | Secretory | 1,315523317 | 0,760153763 | 1,328063241 |
| KRT171 | 9,38332E-06 | 0,927770678 | 0,31 | 0,205 | 0,343307617 | Secretory | 1,902334145 | 0,525670005 | 1,512195122 |
| CCDC6 | 9,71916E-06 | 0,3746333 | 0,36 | 0,273 | 0,355594866 | Secretory | 1,29650997 | 0,771301435 | 1,318681319 |
| SMS | 9,73487E-06 | 0,52964237 | 0,356 | 0,278 | 0,35616977 | Secretory | 1,443571304 | 0,692726433 | 1,28057554 |
| COMTD11 | 1,00198E-05 | 0,396257047 | 0,317 | 0,23 | 0,366595024 | Secretory | 1,316088994 | 0,759827036 | 1,37826087 |
| SRI1 | 1,00876E-05 | 0,512740563 | 0,414 | 0,353 | 0,369075621 | Secretory | 1,426757911 | 0,700889753 | 1,172804533 |
| CYB5A1 | 1,07268E-05 | 0,548238132 | 0,539 | 0,521 | 0,392461264 | Secretory | 1,462298797 | 0,683854765 | 1,034548944 |
| TOMM20 | 1,0789E-05 | 0,554154753 | 0,554 | 0,572 | 0,394738942 | Secretory | 1,468308128 | 0,681055959 | 0,968531469 |
| SLC9A3R21 | 1,08755E-05 | 0,494255929 | 0,409 | 0,339 | 0,397900838 | Secretory | 1,408594084 | 0,709927729 | 1,206489676 |
| C1QBP1 | 1,10482E-05 | 0,418465914 | 0,425 | 0,368 | 0,40421903 | Secretory | 1,33650563 | 0,748219819 | 1,154891304 |
| HSPH11 | 1,2044E-05 | -0,406023174 | 0,235 | 0,358 | 0,440655507 | Secretory | 0,754700861 | 1,325028301 | 0,656424581 |
| SRSF91 | 1,25204E-05 | 0,532795097 | 0,515 | 0,521 | 0,458084172 | Secretory | 1,446729395 | 0,691214268 | 0,988483685 |
| BRI31 | 1,28774E-05 | 0,628242427 | 0,558 | 0,581 | 0,471145719 | Secretory | 1,54568081 | 0,646964104 | 0,960413081 |
| SMIM29 | 1,33347E-05 | 0,268848405 | 0,164 | 0,085 | 0,487875411 | Secretory | 1,204845706 | 0,829981794 | 1,929411765 |
| ANAPC111 | 1,41106E-05 | 0,549572799 | 0,578 | 0,614 | 0,516264583 | Secretory | 1,463652225 | 0,68322241 | 0,941368078 |
| ZFP36 | 1,44088E-05 | -0,379193781 | 0,595 | 0,696 | 0,527175552 | Secretory | 0,768867136 | 1,300614831 | 0,854885057 |
| ANXA111 | 1,44241E-05 | 0,549396459 | 0,453 | 0,388 | 0,52773298 | Secretory | 1,463473334 | 0,683305925 | 1,167525773 |
| FOS | 1,44702E-05 | 0,586180582 | 0,713 | 0,735 | 0,529422482 | Secretory | 1,501266999 | 0,666104031 | 0,970068027 |
| HNRNPA2B11 | 1,45853E-05 | 0,562100246 | 0,647 | 0,696 | 0,533633264 | Secretory | 1,476416991 | 0,677315424 | 0,929597701 |
| PSMB81 | 1,55057E-05 | 0,717186 | 0,435 | 0,399 | 0,567306692 | Secretory | 1,643972311 | 0,608282751 | 1,090225564 |
| ZBTB7C | 1,56949E-05 | 0,278445828 | 0,175 | 0,092 | 0,574230765 | Secretory | 1,212887573 | 0,824478725 | 1,902173913 |
| PRRC2B1 | 1,62154E-05 | 0,382421672 | 0,36 | 0,28 | 0,593271727 | Secretory | 1,30352809 | 0,767148792 | 1,285714286 |
| PRELID11 | 1,68887E-05 | 0,57043989 | 0,472 | 0,444 | 0,617908421 | Secretory | 1,484976284 | 0,673411428 | 1,063063063 |
| GTF2F1 | 1,71534E-05 | 0,425788504 | 0,336 | 0,255 | 0,627591393 | Secretory | 1,343306486 | 0,744431751 | 1,317647059 |
| ATP1A11 | 1,73136E-05 | 0,545629482 | 0,582 | 0,584 | 0,633451918 | Secretory | 1,459657088 | 0,685092415 | 0,996575342 |

| Supplementary Table 4 - clusermarker | | | | | | | | | |
|---|---|---|---|---|---|---|---|---|---|
| | **p_val** | **avg_log2FC** | **pct.1** | **pct.2** | **p_val_adj** | **celltype** | **Foldchange_UP** | **Foldchange_DOWN** | **Ratio_pct1_pct2** |
| S100A21 | 1,76208E-05 | 0,885836766 | 0,224 | 0,135 | 0,644692781 | Secretory | 1,847836059 | 0,54117355 | 1,659259259 |
| HSP90B1 | 1,76976E-05 | 0,387745052 | 0,709 | 0,767 | 0,647500731 | Secretory | 1,308346844 | 0,764323317 | 0,924380704 |
| DDAH21 | 1,8158E-05 | 0,539621506 | 0,386 | 0,318 | 0,664347231 | Secretory | 1,453591115 | 0,687951371 | 1,213836478 |
| HNRNPC | 1,83225E-05 | 0,71929067 | 0,502 | 0,496 | 0,670365892 | Secretory | 1,646372363 | 0,607396007 | 1,012096774 |
| COQ4 | 1,85855E-05 | 0,434330947 | 0,364 | 0,296 | 0,679988401 | Secretory | 1,351284027 | 0,740036869 | 1,22972973 |
| SIVA11 | 1,86147E-05 | 0,461748184 | 0,472 | 0,444 | 0,681055352 | Secretory | 1,37720964 | 0,726105867 | 1,063063063 |
| ARL6IP5 | 1,94812E-05 | 0,632049303 | 0,554 | 0,571 | 0,712759761 | Secretory | 1,549764823 | 0,645259194 | 0,970227671 |
| AURKAIP11 | 2,04679E-05 | 0,589214313 | 0,498 | 0,506 | 0,748858824 | Secretory | 1,504427218 | 0,664704804 | 0,984189723 |
| ITPA | 2,06123E-05 | 0,324509344 | 0,302 | 0,213 | 0,754142058 | Secretory | 1,252238484 | 0,798569931 | 1,417840376 |
| CD811 | 2,07293E-05 | 0,595103033 | 0,625 | 0,682 | 0,758422391 | Secretory | 1,510580463 | 0,661997175 | 0,916422287 |
| YWHAZ | 2,08573E-05 | 0,496897717 | 0,573 | 0,569 | 0,763105671 | Secretory | 1,411175791 | 0,708628936 | 1,007029877 |
| ANAPC16 | 2,09555E-05 | 0,554123885 | 0,582 | 0,612 | 0,766697338 | Secretory | 1,468276713 | 0,681070531 | 0,950980392 |
| SNRPD2 | 2,11785E-05 | 0,663435364 | 0,603 | 0,637 | 0,774857342 | Secretory | 1,58384962 | 0,631373072 | 0,946624804 |
| ELF3 | 2,13137E-05 | 0,609572958 | 0,558 | 0,531 | 0,779804016 | Secretory | 1,525807498 | 0,655390671 | 1,050847458 |
| RAN | 2,14884E-05 | 0,670964677 | 0,487 | 0,474 | 0,786194595 | Secretory | 1,592137215 | 0,628086569 | 1,02742616 |
| PRDX6 | 2,15949E-05 | 0,502360587 | 0,537 | 0,539 | 0,790092268 | Secretory | 1,416529441 | 0,705950735 | 0,996289425 |
| DMKN1 | 2,21371E-05 | 0,432997652 | 0,347 | 0,273 | 0,809929174 | Secretory | 1,350035788 | 0,740721105 | 1,271062271 |
| TRAP1 | 2,27136E-05 | 0,343960371 | 0,293 | 0,21 | 0,831023996 | Secretory | 1,269236022 | 0,787875528 | 1,395238095 |
| TSPAN4 | 2,34546E-05 | 0,280791336 | 0,241 | 0,155 | 0,858135105 | Secretory | 1,214861068 | 0,823139391 | 1,55483871 |
| VPS281 | 2,35005E-05 | 0,584983878 | 0,491 | 0,483 | 0,859811458 | Secretory | 1,500022226 | 0,666656788 | 1,016563147 |
| IFI27 | 2,51793E-05 | 0,526981755 | 0,183 | 0,101 | 0,921234856 | Secretory | 1,440911526 | 0,694005136 | 1,811881188 |
| LAMTOR4 | 2,53224E-05 | 0,483503575 | 0,571 | 0,601 | 0,926469997 | Secretory | 1,398134908 | 0,715238561 | 0,950083195 |
| REX1BD1 | 2,60665E-05 | 0,476294849 | 0,476 | 0,454 | 0,95369344 | Secretory | 1,39116626 | 0,718821343 | 1,04845815 |
| SERBP1 | 2,60738E-05 | 0,429271082 | 0,524 | 0,507 | 0,953962963 | Secretory | 1,346553063 | 0,742636906 | 1,033530572 |
| RAB5IF1 | 2,61511E-05 | 0,470554749 | 0,364 | 0,298 | 0,956790708 | Secretory | 1,385642177 | 0,721687039 | 1,22147651 |
| SPSB31 | 2,78842E-05 | 0,393425938 | 0,362 | 0,286 | 1 | Secretory | 1,313508866 | 0,761319566 | 1,265734266 |
| ENDOD1 | 2,85717E-05 | 0,454482969 | 0,263 | 0,18 | 1 | Secretory | 1,370291634 | 0,72977166 | 1,461111111 |
| TMEM50A1 | 2,86878E-05 | 0,590630423 | 0,504 | 0,496 | 1 | Secretory | 1,505904647 | 0,664052669 | 1,016129032 |
| THSD4 | 2,94414E-05 | 0,300041666 | 0,175 | 0,093 | 1 | Secretory | 1,231179971 | 0,812228938 | 1,88172043 |
| KHDRBS1 | 2,97334E-05 | 0,573670541 | 0,422 | 0,379 | 1 | Secretory | 1,488305342 | 0,671905134 | 1,113456464 |
| TRIP6 | 2,97698E-05 | 0,461688351 | 0,351 | 0,283 | 1 | Secretory | 1,377152524 | 0,726135982 | 1,240282686 |

| Supplementary Table 4 - clusermarker | | | | | | | | | |
|---|---|---|---|---|---|---|---|---|---|
| | **p_val** | **avg_log2FC** | **pct.1** | **pct.2** | **p_val_adj** | **celltype** | **Foldchange_UP** | **Foldchange_DOWN** | **Ratio_pct1_pct2** |
| ELOB1 | 3,07487E-05 | 0,337823925 | 0,692 | 0,754 | 1 | Secretory | 1,263848842 | 0,791233862 | 0,917771883 |
| PAFAH1B31 | 3,17521E-05 | 0,301425143 | 0,244 | 0,16 | 1 | Secretory | 1,23236118 | 0,811450422 | 1,525 |
| HNRNPA0 | 3,25541E-05 | 0,388792325 | 0,392 | 0,326 | 1 | Secretory | 1,309296937 | 0,763768685 | 1,202453988 |
| TMEM2381 | 3,29526E-05 | 0,566629632 | 0,494 | 0,484 | 1 | Secretory | 1,481059532 | 0,675192305 | 1,020661157 |
| LAPTM4A | 3,47413E-05 | 0,502031405 | 0,582 | 0,617 | 1 | Secretory | 1,416206266 | 0,706111831 | 0,943273906 |
| TTC19 | 3,58777E-05 | 0,566952264 | 0,351 | 0,28 | 1 | Secretory | 1,481390781 | 0,675041328 | 1,253571429 |
| CCDC24 | 3,69175E-05 | 0,267257176 | 0,211 | 0,125 | 1 | Secretory | 1,203517547 | 0,830897732 | 1,688 |
| CLTB1 | 3,92713E-05 | 0,560088405 | 0,392 | 0,334 | 1 | Secretory | 1,47435956 | 0,6782606 | 1,173652695 |
| TP53I13 | 4,04823E-05 | 0,257191264 | 0,233 | 0,151 | 1 | Secretory | 1,19514964 | 0,836715309 | 1,543046358 |
| NET1 | 4,17465E-05 | 0,648261099 | 0,494 | 0,476 | 1 | Secretory | 1,567277994 | 0,6380489 | 1,037815126 |
| MFSD101 | 4,60699E-05 | 0,277837527 | 0,323 | 0,238 | 1 | Secretory | 1,212376276 | 0,824826434 | 1,357142857 |
| HSBP1L1 | 4,67678E-05 | 0,380006696 | 0,353 | 0,283 | 1 | Secretory | 1,301347895 | 0,768434024 | 1,247349823 |
| RPL37A1 | 4,74684E-05 | 0,28989642 | 0,858 | 0,91 | 1 | Secretory | 1,2225525 | 0,817960783 | 0,942857143 |
| MYOF | 4,7751E-05 | 0,40335411 | 0,293 | 0,218 | 1 | Secretory | 1,322579193 | 0,756098391 | 1,344036697 |
| RPL71 | 4,8168E-05 | 0,35531924 | 0,731 | 0,804 | 1 | Secretory | 1,279268627 | 0,78169665 | 0,90920398 |
| MAP2K21 | 4,83184E-05 | 0,397921162 | 0,401 | 0,346 | 1 | Secretory | 1,317607947 | 0,758951099 | 1,158959538 |
| PRDX2 | 4,87883E-05 | 0,539728771 | 0,558 | 0,584 | 1 | Secretory | 1,453699194 | 0,687900223 | 0,955479452 |
| EIF3H | 5,00171E-05 | 0,497204771 | 0,582 | 0,627 | 1 | Secretory | 1,411476169 | 0,708478132 | 0,928229665 |
| SIGIRR1 | 5,02969E-05 | 0,36888353 | 0,353 | 0,283 | 1 | Secretory | 1,291353094 | 0,774381542 | 1,247349823 |
| NDUFC21 | 5,08123E-05 | 0,50791084 | 0,558 | 0,597 | 1 | Secretory | 1,421989527 | 0,70324006 | 0,934673367 |
| LTBP31 | 5,14683E-05 | 0,38728465 | 0,369 | 0,301 | 1 | Secretory | 1,307929383 | 0,764567271 | 1,225913621 |
| NUBP21 | 5,16261E-05 | 0,346836712 | 0,269 | 0,19 | 1 | Secretory | 1,271769057 | 0,786306283 | 1,415789474 |
| RAB11B | 5,23006E-05 | 0,466642597 | 0,431 | 0,383 | 1 | Secretory | 1,381889825 | 0,723646692 | 1,125326371 |
| CCT4 | 5,23198E-05 | 0,493908231 | 0,366 | 0,303 | 1 | Secretory | 1,408254646 | 0,710098847 | 1,207920792 |
| HSP90AB11 | 5,24254E-05 | 0,355749134 | 0,737 | 0,792 | 1 | Secretory | 1,27964988 | 0,781463755 | 0,930555556 |
| XRCC5 | 5,30095E-05 | 0,609097416 | 0,463 | 0,424 | 1 | Secretory | 1,525304644 | 0,655606737 | 1,091981132 |
| PGD1 | 5,32177E-05 | 0,387785054 | 0,358 | 0,291 | 1 | Secretory | 1,308383122 | 0,764302124 | 1,23024055 |
| CNIH4 | 5,42055E-05 | 0,337509455 | 0,291 | 0,213 | 1 | Secretory | 1,263573386 | 0,791406349 | 1,366197183 |
| VDAC21 | 5,46346E-05 | 0,625158415 | 0,489 | 0,493 | 1 | Secretory | 1,542380177 | 0,648348582 | 0,99188641 |
| CD241 | 5,53052E-05 | 0,475142078 | 0,726 | 0,76 | 1 | Secretory | 1,390055107 | 0,71939594 | 0,955263158 |
| FAAP201 | 5,74668E-05 | 0,352647849 | 0,351 | 0,285 | 1 | Secretory | 1,276902039 | 0,783145433 | 1,231578947 |

| Supplementary Table 4 - clusermarker | | | | | | | | | |
|---|---|---|---|---|---|---|---|---|---|
| | **p_val** | **avg_log2FC** | **pct.1** | **pct.2** | **p_val_adj** | **celltype** | **Foldchange_UP** | **Foldchange_DOWN** | **Ratio_pct1_pct2** |
| ADRM11 | 5,91073E-05 | 0,406892107 | 0,343 | 0,265 | 1 | Secretory | 1,325826604 | 0,754246443 | 1,294339623 |
| TUBA1A | 6,13442E-05 | 0,32444317 | 0,218 | 0,136 | 1 | Secretory | 1,252181047 | 0,798606561 | 1,602941176 |
| HSPA81 | 6,23534E-05 | 0,264450654 | 0,698 | 0,729 | 1 | Secretory | 1,20117858 | 0,832515678 | 0,957475995 |
| C15orf481 | 6,26024E-05 | 0,567536718 | 0,127 | 0,06 | 1 | Secretory | 1,481991032 | 0,674767916 | 2,116666667 |
| RPL27 | 6,2689E-05 | 0,293052299 | 0,746 | 0,817 | 1 | Secretory | 1,225229747 | 0,816173458 | 0,913096695 |
| TMEM8B | 6,35613E-05 | 0,253106056 | 0,172 | 0,097 | 1 | Secretory | 1,191770181 | 0,839087952 | 1,773195876 |
| RBMS2 | 6,38489E-05 | 0,280053832 | 0,181 | 0,101 | 1 | Secretory | 1,214240191 | 0,823560287 | 1,792079208 |
| CSNK2B1 | 6,43942E-05 | 0,452746954 | 0,504 | 0,506 | 1 | Secretory | 1,368643734 | 0,730650333 | 0,996047431 |
| ANKRD10 | 6,44608E-05 | 0,574874697 | 0,371 | 0,311 | 1 | Secretory | 1,489548085 | 0,671344558 | 1,192926045 |
| PPP1R14B1 | 6,93209E-05 | 0,491067235 | 0,422 | 0,384 | 1 | Secretory | 1,405484199 | 0,711498571 | 1,098958333 |
| ATP6V0E11 | 6,97984E-05 | 0,458676712 | 0,552 | 0,577 | 1 | Secretory | 1,374280704 | 0,727653381 | 0,956672444 |
| APLP2 | 7,17053E-05 | 0,519316462 | 0,515 | 0,511 | 1 | Secretory | 1,433276012 | 0,697702321 | 1,007827789 |
| ISCU1 | 7,22812E-05 | 0,493298666 | 0,483 | 0,481 | 1 | Secretory | 1,407659758 | 0,71039894 | 1,004158004 |
| PSMG2 | 7,95964E-05 | 0,443515042 | 0,353 | 0,301 | 1 | Secretory | 1,359913645 | 0,735340809 | 1,172757475 |
| EML4 | 8,51759E-05 | 0,42333343 | 0,44 | 0,389 | 1 | Secretory | 1,341022488 | 0,74569965 | 1,131105398 |
| STOM | 8,67785E-05 | 0,38726027 | 0,276 | 0,201 | 1 | Secretory | 1,307907281 | 0,764580192 | 1,373134328 |
| EIF3E | 8,88926E-05 | 0,613719109 | 0,593 | 0,641 | 1 | Secretory | 1,530198813 | 0,653509852 | 0,925117005 |
| EHF1 | 8,89419E-05 | 0,501865738 | 0,638 | 0,699 | 1 | Secretory | 1,416043651 | 0,70619292 | 0,912732475 |
| CFL1 | 8,9326E-05 | 0,553722028 | 0,629 | 0,689 | 1 | Secretory | 1,467867786 | 0,681260267 | 0,912917271 |
| GSTK11 | 9,0917E-05 | 0,579533715 | 0,541 | 0,572 | 1 | Secretory | 1,494366185 | 0,669180024 | 0,945804196 |
| H2AJ1 | 9,39651E-05 | 0,472925284 | 0,513 | 0,524 | 1 | Secretory | 1,387920838 | 0,720502187 | 0,979007634 |
| PLXNB21 | 9,58602E-05 | 0,346680286 | 0,31 | 0,236 | 1 | Secretory | 1,271631171 | 0,786391544 | 1,313559322 |
| RRAGA1 | 9,59839E-05 | 0,374069511 | 0,287 | 0,208 | 1 | Secretory | 1,296003408 | 0,771602909 | 1,379807692 |
| PKM | 9,66869E-05 | 0,631097039 | 0,56 | 0,589 | 1 | Secretory | 1,548742224 | 0,645685243 | 0,950764007 |
| TNFRSF14 | 0,000102904 | 0,432263158 | 0,323 | 0,258 | 1 | Secretory | 1,349348643 | 0,741098311 | 1,251937984 |
| HSD17B8 | 0,000103684 | 0,421910234 | 0,302 | 0,23 | 1 | Secretory | 1,339700242 | 0,746435634 | 1,313043478 |
| MSRB21 | 0,000104609 | 0,311140361 | 0,319 | 0,25 | 1 | Secretory | 1,240687999 | 0,806004411 | 1,276 |
| LAPTM4B | 0,000107345 | 0,405157511 | 0,336 | 0,28 | 1 | Secretory | 1,32423348 | 0,755153842 | 1,2 |
| FAM3C1 | 0,00010978 | 0,449489815 | 0,394 | 0,341 | 1 | Secretory | 1,365557265 | 0,732301768 | 1,15542522 |
| CAV21 | 0,000111047 | 0,401910219 | 0,254 | 0,176 | 1 | Secretory | 1,321256179 | 0,756855495 | 1,443181818 |
| RNH1 | 0,000117029 | 0,436747449 | 0,446 | 0,416 | 1 | Secretory | 1,353549313 | 0,738798351 | 1,072115385 |

| Supplementary Table 4 - clusermarker | | | | | | | | | |
|---|---|---|---|---|---|---|---|---|---|
| | **p_val** | **avg_log2FC** | **pct.1** | **pct.2** | **p_val_adj** | **celltype** | **Foldchange_UP** | **Foldchange_DOWN** | **Ratio_pct1_pct2** |
| HDAC2 | 0,000118419 | 0,366846763 | 0,321 | 0,251 | 1 | Secretory | 1,289531275 | 0,77547557 | 1,278884462 |
| MT-CYB | 0,000120027 | 0,251372243 | 0,871 | 0,887 | 1 | Secretory | 1,190338787 | 0,840096963 | 0,981961669 |
| GSDMD | 0,000122209 | 0,38564518 | 0,269 | 0,191 | 1 | Secretory | 1,306443904 | 0,765436615 | 1,408376963 |
| GPRC5C | 0,000124842 | 0,370116258 | 0,291 | 0,218 | 1 | Secretory | 1,292456978 | 0,773720145 | 1,334862385 |
| RNF71 | 0,000128509 | 0,437776441 | 0,433 | 0,409 | 1 | Secretory | 1,354515067 | 0,738271596 | 1,058679707 |
| MZT2B1 | 0,000131052 | 0,452491995 | 0,573 | 0,627 | 1 | Secretory | 1,368401884 | 0,730779467 | 0,913875598 |
| DCPS | 0,000133141 | 0,322462685 | 0,237 | 0,16 | 1 | Secretory | 1,250463273 | 0,799703615 | 1,48125 |
| MARCKSL1 | 0,000136293 | 0,508781241 | 0,571 | 0,601 | 1 | Secretory | 1,422847695 | 0,702815912 | 0,950083195 |
| CMTM6 | 0,000136885 | 0,363736504 | 0,401 | 0,341 | 1 | Secretory | 1,286754211 | 0,777149196 | 1,175953079 |
| TMEM1341 | 0,000138371 | 0,418723417 | 0,356 | 0,295 | 1 | Secretory | 1,3367442 | 0,748086283 | 1,206779661 |
| FCGRT | 0,000141588 | 0,579847028 | 0,446 | 0,428 | 1 | Secretory | 1,494690755 | 0,669034713 | 1,042056075 |
| CIAO2B1 | 0,00014695 | 0,45227154 | 0,379 | 0,329 | 1 | Secretory | 1,368192797 | 0,730891145 | 1,151975684 |
| RUNX1 | 0,000147115 | 0,675572975 | 0,384 | 0,329 | 1 | Secretory | 1,597230996 | 0,626083517 | 1,167173252 |
| PUF60 | 0,000148642 | 0,387882372 | 0,334 | 0,27 | 1 | Secretory | 1,308471383 | 0,76425057 | 1,237037037 |
| NOXO1 | 0,000153145 | 0,265499876 | 0,233 | 0,155 | 1 | Secretory | 1,202052473 | 0,831910438 | 1,503225806 |
| RTRAF | 0,00015358 | 0,436713941 | 0,478 | 0,469 | 1 | Secretory | 1,353517875 | 0,738815511 | 1,019189765 |
| PSAP | 0,000156 | 0,393392942 | 0,647 | 0,7 | 1 | Secretory | 1,313478825 | 0,761336979 | 0,924285714 |
| CXADR | 0,000159331 | 0,571476181 | 0,328 | 0,265 | 1 | Secretory | 1,486043329 | 0,672927889 | 1,237735849 |
| DLG51 | 0,000164399 | 0,331932722 | 0,19 | 0,115 | 1 | Secretory | 1,258698474 | 0,794471448 | 1,652173913 |
| SCGB2A1 | 0,000170411 | -0,283411886 | 0,06 | 0,128 | 1 | Secretory | 0,821645576 | 1,21706978 | 0,46875 |
| CISD31 | 0,000173162 | 0,397356085 | 0,317 | 0,251 | 1 | Secretory | 1,317091965 | 0,759248425 | 1,262948207 |
| BAX1 | 0,000174516 | 0,334070619 | 0,263 | 0,193 | 1 | Secretory | 1,260565094 | 0,793295011 | 1,362694301 |
| SERP11 | 0,000177034 | 0,324507823 | 0,72 | 0,79 | 1 | Secretory | 1,252237163 | 0,798570773 | 0,911392405 |
| HMGN21 | 0,000181378 | 0,475447664 | 0,556 | 0,601 | 1 | Secretory | 1,390349574 | 0,719243576 | 0,925124792 |
| RAB5C | 0,000183046 | 0,503446613 | 0,401 | 0,354 | 1 | Secretory | 1,417596171 | 0,705419512 | 1,132768362 |
| TMEM1411 | 0,000186978 | 0,425975972 | 0,414 | 0,376 | 1 | Secretory | 1,34348105 | 0,744335024 | 1,10106383 |
| RAB34 | 0,000187516 | 0,303958135 | 0,224 | 0,15 | 1 | Secretory | 1,234526782 | 0,810026979 | 1,493333333 |
| SYNGR21 | 0,000191618 | 0,433442902 | 0,547 | 0,579 | 1 | Secretory | 1,350452506 | 0,740492535 | 0,944732297 |
| FAAH2 | 0,00019301 | 0,273333446 | 0,162 | 0,092 | 1 | Secretory | 1,208597151 | 0,827405558 | 1,760869565 |
| IMPDH2 | 0,000195834 | 0,576418263 | 0,401 | 0,376 | 1 | Secretory | 1,491142633 | 0,670626657 | 1,066489362 |
| TAF101 | 0,000197704 | 0,402317421 | 0,489 | 0,491 | 1 | Secretory | 1,321629157 | 0,756641902 | 0,99592668 |

| Supplementary Table 4 - clusermarker | | | | | | | | | |
|---|---|---|---|---|---|---|---|---|---|
| | **p_val** | **avg_log2FC** | **pct.1** | **pct.2** | **p_val_adj** | **celltype** | **Foldchange_UP** | **Foldchange_DOWN** | **Ratio_pct1_pct2** |
| FAT1 | 0,000202629 | 0,333226287 | 0,188 | 0,115 | 1 | Secretory | 1,259827569 | 0,79375942 | 1,634782609 |
| NOL71 | 0,000202714 | 0,525977134 | 0,457 | 0,448 | 1 | Secretory | 1,439908497 | 0,694488575 | 1,020089286 |
| SYNE2 | 0,000203964 | 0,56981391 | 0,446 | 0,411 | 1 | Secretory | 1,484332098 | 0,673703682 | 1,085158151 |
| ERGIC3 | 0,000204457 | 0,454737074 | 0,606 | 0,656 | 1 | Secretory | 1,370533007 | 0,729643135 | 0,923780488 |
| EPS8L2 | 0,000204874 | 0,490686546 | 0,356 | 0,304 | 1 | Secretory | 1,405113378 | 0,711686342 | 1,171052632 |
| NEBL | 0,000206166 | 0,538181937 | 0,31 | 0,243 | 1 | Secretory | 1,452141397 | 0,688638174 | 1,275720165 |
| SELENOM | 0,000206287 | 0,432497963 | 0,653 | 0,722 | 1 | Secretory | 1,349568273 | 0,740977704 | 0,904432133 |
| CLMN | 0,000215869 | 0,337876011 | 0,228 | 0,153 | 1 | Secretory | 1,263894472 | 0,791205296 | 1,490196078 |
| PSMB1 | 0,000216871 | 0,448103638 | 0,513 | 0,539 | 1 | Secretory | 1,364245834 | 0,73300572 | 0,951762523 |
| DDX51 | 0,000223667 | 0,427224806 | 0,672 | 0,732 | 1 | Secretory | 1,344644505 | 0,743690987 | 0,918032787 |
| ZNHIT11 | 0,000230682 | 0,377177702 | 0,463 | 0,449 | 1 | Secretory | 1,298798571 | 0,769942332 | 1,031180401 |
| AFDN1 | 0,000234236 | 0,364444185 | 0,442 | 0,409 | 1 | Secretory | 1,287385554 | 0,776768076 | 1,080684597 |
| RBM42 | 0,00023976 | 0,290039255 | 0,306 | 0,231 | 1 | Secretory | 1,222673546 | 0,817879804 | 1,324675325 |
| TRIM22 | 0,000240473 | 0,381545033 | 0,252 | 0,178 | 1 | Secretory | 1,302736255 | 0,767615084 | 1,415730337 |
| GLRX51 | 0,000246863 | 0,425869966 | 0,328 | 0,265 | 1 | Secretory | 1,343382337 | 0,744389719 | 1,237735849 |
| FLNA1 | 0,000247544 | 0,420361195 | 0,213 | 0,143 | 1 | Secretory | 1,338262562 | 0,747237522 | 1,48951049 |
| SUPT4H1 | 0,000251579 | 0,321526265 | 0,302 | 0,235 | 1 | Secretory | 1,24965189 | 0,800222853 | 1,285106383 |
| PYCARD1 | 0,000261564 | 0,384752654 | 0,295 | 0,235 | 1 | Secretory | 1,30563592 | 0,7659103 | 1,255319149 |
| GADD45B1 | 0,000265913 | -0,30852386 | 0,578 | 0,664 | 1 | Secretory | 0,807467523 | 1,238439901 | 0,870481928 |
| MAD2L2 | 0,000266502 | 0,280898898 | 0,2 | 0,128 | 1 | Secretory | 1,214951647 | 0,823078023 | 1,5625 |
| TINAGL1 | 0,000269039 | 0,274171332 | 0,261 | 0,188 | 1 | Secretory | 1,209299282 | 0,826925158 | 1,388297872 |
| DAD11 | 0,000275729 | 0,416482572 | 0,588 | 0,646 | 1 | Secretory | 1,334669534 | 0,74924914 | 0,910216718 |
| LMNA | 0,000295572 | 0,589083811 | 0,522 | 0,517 | 1 | Secretory | 1,504291138 | 0,664764935 | 1,00967118 |
| APH1A | 0,000301599 | 0,362014803 | 0,386 | 0,343 | 1 | Secretory | 1,285219525 | 0,778077193 | 1,125364431 |
| MRPL12 | 0,000301681 | 0,332733861 | 0,3 | 0,231 | 1 | Secretory | 1,259397633 | 0,794030395 | 1,298701299 |
| SH3BP5 | 0,000306875 | 0,403044873 | 0,287 | 0,22 | 1 | Secretory | 1,322295732 | 0,756260476 | 1,304545455 |
| CNN3 | 0,000307491 | 0,527746633 | 0,455 | 0,416 | 1 | Secretory | 1,441675662 | 0,693637291 | 1,09375 |
| HIGD2A1 | 0,000320625 | 0,652446669 | 0,55 | 0,592 | 1 | Secretory | 1,571831609 | 0,636200465 | 0,929054054 |
| CAPN21 | 0,000324588 | 0,495716995 | 0,504 | 0,498 | 1 | Secretory | 1,410021338 | 0,709209126 | 1,012048193 |
| NAPRT1 | 0,000329715 | 0,328565456 | 0,302 | 0,231 | 1 | Secretory | 1,255764084 | 0,796327919 | 1,307359307 |
| DSC21 | 0,000334877 | 0,379507479 | 0,261 | 0,193 | 1 | Secretory | 1,300897667 | 0,768699972 | 1,352331606 |

| Supplementary Table 4 - clusermarker | | | | | | | | | |
|---|---|---|---|---|---|---|---|---|---|
| | **p_val** | **avg_log2FC** | **pct.1** | **pct.2** | **p_val_adj** | **celltype** | **Foldchange_UP** | **Foldchange_DOWN** | **Ratio_pct1_pct2** |
| ERH1 | 0,000335505 | 0,519750672 | 0,519 | 0,547 | 1 | Secretory | 1,433707452 | 0,697492364 | 0,9488117 |
| IL1R1 | 0,000337489 | 0,289399064 | 0,17 | 0,101 | 1 | Secretory | 1,222131108 | 0,818242816 | 1,683168317 |
| PPP4C1 | 0,000342848 | 0,308160806 | 0,312 | 0,245 | 1 | Secretory | 1,238128288 | 0,807670748 | 1,273469388 |
| EIF3I | 0,000355543 | 0,375754485 | 0,397 | 0,346 | 1 | Secretory | 1,297517939 | 0,770702254 | 1,147398844 |
| VAMP81 | 0,000359485 | 0,44643946 | 0,606 | 0,649 | 1 | Secretory | 1,362673056 | 0,733851745 | 0,933744222 |
| LTA4H | 0,000365812 | 0,351486285 | 0,302 | 0,233 | 1 | Secretory | 1,275874374 | 0,783776225 | 1,296137339 |
| ISCA2 | 0,000365963 | 0,294693875 | 0,228 | 0,153 | 1 | Secretory | 1,226624673 | 0,8152453 | 1,490196078 |
| TMEM591 | 0,000366707 | 0,311885153 | 0,731 | 0,774 | 1 | Secretory | 1,24132867 | 0,805588418 | 0,944444444 |
| NEK9 | 0,000378795 | 0,261453551 | 0,17 | 0,1 | 1 | Secretory | 1,198685802 | 0,834246971 | 1,7 |
| SCPEP1 | 0,000379983 | 0,482996192 | 0,366 | 0,314 | 1 | Secretory | 1,397643282 | 0,715490149 | 1,165605096 |
| TM4SF11 | 0,00038592 | 0,629325282 | 0,483 | 0,443 | 1 | Secretory | 1,546841399 | 0,646478689 | 1,090293454 |
| SELENOW1 | 0,000392953 | 0,370662003 | 0,636 | 0,702 | 1 | Secretory | 1,292945983 | 0,773427516 | 0,905982906 |
| TXNIP1 | 0,000394983 | 0,623185766 | 0,519 | 0,517 | 1 | Secretory | 1,540272666 | 0,649235698 | 1,003868472 |
| TMEM2191 | 0,000399939 | 0,471299854 | 0,554 | 0,606 | 1 | Secretory | 1,386358 | 0,721314408 | 0,914191419 |
| PLSCR11 | 0,000402958 | 0,385501214 | 0,323 | 0,261 | 1 | Secretory | 1,306313541 | 0,765513002 | 1,237547893 |
| EFEMP11 | 0,000403915 | 0,284446787 | 0,282 | 0,211 | 1 | Secretory | 1,217943144 | 0,821056389 | 1,336492891 |
| IGHA11 | 0,000404117 | 2,729576928 | 0,119 | 0,06 | 1 | Secretory | 6,63261106 | 0,150770186 | 1,983333333 |
| TCEA11 | 0,000410933 | 0,559586249 | 0,489 | 0,491 | 1 | Secretory | 1,473846472 | 0,678496722 | 0,99592668 |
| DSP | 0,000413297 | 0,691735799 | 0,448 | 0,429 | 1 | Secretory | 1,615225731 | 0,619108513 | 1,044289044 |
| SOX9 | 0,000423916 | 0,590176515 | 0,485 | 0,486 | 1 | Secretory | 1,505430927 | 0,664261629 | 0,997942387 |
| CD82 | 0,000427465 | 0,41870604 | 0,429 | 0,398 | 1 | Secretory | 1,336728099 | 0,748095294 | 1,077889447 |
| ALDH2 | 0,00043223 | 0,480033707 | 0,431 | 0,413 | 1 | Secretory | 1,394776253 | 0,716960873 | 1,043583535 |
| GNB21 | 0,000433137 | 0,390291411 | 0,519 | 0,544 | 1 | Secretory | 1,310658118 | 0,762975475 | 0,954044118 |
| BPIFB2 | 0,00044963 | -0,808608818 | 0,119 | 0,198 | 1 | Secretory | 0,570932139 | 1,751521647 | 0,601010101 |
| MGST3 | 0,000458211 | 0,433052049 | 0,606 | 0,652 | 1 | Secretory | 1,350086692 | 0,740693176 | 0,929447853 |
| DDOST1 | 0,000462043 | 0,494376 | 0,476 | 0,468 | 1 | Secretory | 1,408711322 | 0,709868647 | 1,017094017 |
| C11orf58 | 0,000465313 | 0,55800494 | 0,502 | 0,521 | 1 | Secretory | 1,472231903 | 0,679240817 | 0,96353167 |
| PSMB10 | 0,000472328 | 0,355515785 | 0,332 | 0,273 | 1 | Secretory | 1,27944292 | 0,781590163 | 1,216117216 |
| SRRM2 | 0,000474406 | 0,521880426 | 0,575 | 0,641 | 1 | Secretory | 1,435825501 | 0,696463463 | 0,897035881 |
| MIR205HG | 0,000486814 | 0,439314891 | 0,209 | 0,143 | 1 | Secretory | 1,355960254 | 0,737484743 | 1,461538462 |
| IFNGR11 | 0,000498763 | 0,327934394 | 0,278 | 0,208 | 1 | Secretory | 1,25521491 | 0,796676324 | 1,336538462 |

| Supplementary Table 4 - clusermarker | | | | | | | | | |
|---|---|---|---|---|---|---|---|---|---|
| | **p_val** | **avg_log2FC** | **pct.1** | **pct.2** | **p_val_adj** | **celltype** | **Foldchange_UP** | **Foldchange_DOWN** | **Ratio_pct1_pct2** |
| MRPS26 | 0,000507461 | 0,321308608 | 0,356 | 0,306 | 1 | Secretory | 1,249463371 | 0,80034359 | 1,163398693 |
| CBR3 | 0,000515104 | 0,546976114 | 0,422 | 0,396 | 1 | Secretory | 1,461020189 | 0,684453239 | 1,065656566 |
| TPM4 | 0,000522662 | 0,502606069 | 0,5 | 0,476 | 1 | Secretory | 1,416770492 | 0,705830624 | 1,050420168 |
| SLC39A4 | 0,000530257 | 0,264700059 | 0,276 | 0,205 | 1 | Secretory | 1,201386251 | 0,83237177 | 1,346341463 |
| TNFRSF12A | 0,000534993 | 0,440405354 | 0,2 | 0,135 | 1 | Secretory | 1,356985546 | 0,736927525 | 1,481481481 |
| RALY1 | 0,000535969 | 0,39610689 | 0,401 | 0,366 | 1 | Secretory | 1,315952021 | 0,759906124 | 1,095628415 |
| EIF5 | 0,000561827 | 0,570393087 | 0,506 | 0,521 | 1 | Secretory | 1,484928109 | 0,673433275 | 0,971209213 |
| CCS1 | 0,000572213 | 0,287593515 | 0,33 | 0,271 | 1 | Secretory | 1,220602555 | 0,819267497 | 1,217712177 |
| ATP5IF11 | 0,000576042 | 0,396335892 | 0,532 | 0,557 | 1 | Secretory | 1,316160922 | 0,759785512 | 0,955116697 |
| TUBA1B1 | 0,000578722 | 0,555099061 | 0,506 | 0,494 | 1 | Secretory | 1,469269515 | 0,680610323 | 1,024291498 |
| SNU13 | 0,00058055 | 0,491102512 | 0,506 | 0,526 | 1 | Secretory | 1,405518567 | 0,711481174 | 0,961977186 |
| ANXA41 | 0,000592487 | 0,349964014 | 0,321 | 0,26 | 1 | Secretory | 1,274528836 | 0,784603668 | 1,234615385 |
| TIMM17B1 | 0,000595202 | 0,30177935 | 0,293 | 0,231 | 1 | Secretory | 1,232663784 | 0,811251221 | 1,268398268 |
| FKBP4 | 0,000604537 | 0,270565187 | 0,338 | 0,28 | 1 | Secretory | 1,206280304 | 0,828994717 | 1,207142857 |
| TMEM1601 | 0,000612767 | 0,424870308 | 0,336 | 0,291 | 1 | Secretory | 1,342451817 | 0,744905692 | 1,154639175 |
| SQSTM11 | 0,000636524 | 0,35099594 | 0,623 | 0,647 | 1 | Secretory | 1,275440802 | 0,784042661 | 0,962905719 |
| LAMB2 | 0,00063742 | 0,318481172 | 0,248 | 0,183 | 1 | Secretory | 1,247017034 | 0,801913665 | 1,355191257 |
| SSU721 | 0,000640759 | 0,506738414 | 0,444 | 0,436 | 1 | Secretory | 1,420834397 | 0,70381179 | 1,018348624 |
| C19orf531 | 0,000660777 | 0,430496307 | 0,478 | 0,489 | 1 | Secretory | 1,347697124 | 0,742006481 | 0,977505112 |
| MTDH1 | 0,000674252 | 0,510304387 | 0,474 | 0,486 | 1 | Secretory | 1,424350681 | 0,702074295 | 0,975308642 |
| NDUFB21 | 0,000679326 | 0,453983162 | 0,606 | 0,659 | 1 | Secretory | 1,369816992 | 0,730024526 | 0,919575114 |
| ADAM15 | 0,00072632 | 0,319425142 | 0,254 | 0,188 | 1 | Secretory | 1,247833237 | 0,801389136 | 1,35106383 |
| EIF4B1 | 0,000728676 | 0,434484167 | 0,623 | 0,672 | 1 | Secretory | 1,351427546 | 0,739958278 | 0,927083333 |
| GAS51 | 0,000746808 | 0,556956993 | 0,61 | 0,684 | 1 | Secretory | 1,47116289 | 0,679734384 | 0,891812865 |
| RSL1D1 | 0,000753711 | 0,503715091 | 0,453 | 0,458 | 1 | Secretory | 1,417860003 | 0,70528825 | 0,989082969 |
| DCTN31 | 0,000756488 | 0,403389407 | 0,401 | 0,373 | 1 | Secretory | 1,322611551 | 0,756079893 | 1,075067024 |
| CDC42EP3 | 0,000778257 | 0,325381589 | 0,25 | 0,181 | 1 | Secretory | 1,252995809 | 0,798087267 | 1,38121547 |
| S100A161 | 0,000788219 | 0,355487088 | 0,347 | 0,288 | 1 | Secretory | 1,27941747 | 0,78160571 | 1,204861111 |
| KRT101 | 0,000796015 | 0,514570267 | 0,532 | 0,587 | 1 | Secretory | 1,428568551 | 0,70000141 | 0,906303237 |
| CD59 | 0,000804682 | 0,388035324 | 0,679 | 0,735 | 1 | Secretory | 1,308610112 | 0,76416955 | 0,923809524 |
| UQCR101 | 0,000816729 | 0,495271269 | 0,593 | 0,619 | 1 | Secretory | 1,409585774 | 0,709428272 | 0,957996769 |

| Supplementary Table 4 - clusermarker | | | | | | | | | |
|---|---|---|---|---|---|---|---|---|---|
| | **p_val** | **avg_log2FC** | **pct.1** | **pct.2** | **p_val_adj** | **celltype** | **Foldchange_UP** | **Foldchange_DOWN** | **Ratio_pct1_pct2** |
| UBE2I1 | 0,000829039 | 0,358716294 | 0,379 | 0,338 | 1 | Secretory | 1,282284417 | 0,779858187 | 1,121301775 |
| DCTN2 | 0,000845254 | 0,36939411 | 0,371 | 0,331 | 1 | Secretory | 1,291810194 | 0,774107531 | 1,120845921 |
| PLPP21 | 0,000871922 | 0,514521911 | 0,539 | 0,592 | 1 | Secretory | 1,42852067 | 0,700024873 | 0,910472973 |
| VAMP3 | 0,000875165 | 0,344239607 | 0,265 | 0,203 | 1 | Secretory | 1,269481707 | 0,787723048 | 1,305418719 |
| SERPINB61 | 0,000882273 | 0,525889398 | 0,371 | 0,331 | 1 | Secretory | 1,439820932 | 0,694530811 | 1,120845921 |
| JOSD2 | 0,000909422 | 0,306979911 | 0,22 | 0,16 | 1 | Secretory | 1,237115252 | 0,808332125 | 1,375 |
| YBX3 | 0,000909868 | 0,472433713 | 0,56 | 0,601 | 1 | Secretory | 1,387448011 | 0,720747727 | 0,931780366 |
| UQCRH | 0,00091746 | 0,422650881 | 0,578 | 0,619 | 1 | Secretory | 1,340388191 | 0,746052529 | 0,933764136 |
| CCT81 | 0,000941052 | 0,340988506 | 0,386 | 0,338 | 1 | Secretory | 1,266624163 | 0,789500176 | 1,142011834 |
| CSN3 | 0,000964486 | -0,313728951 | 0,06 | 0,118 | 1 | Secretory | 0,804559515 | 1,242916132 | 0,508474576 |
| YWHAH1 | 0,000977519 | 0,393610694 | 0,328 | 0,273 | 1 | Secretory | 1,313677088 | 0,761222076 | 1,201465201 |
| SLC25A11 | 0,000992351 | 0,289395527 | 0,248 | 0,186 | 1 | Secretory | 1,222128113 | 0,818244822 | 1,333333333 |
| ZNF4281 | 0,001006982 | 0,382378087 | 0,364 | 0,318 | 1 | Secretory | 1,303488709 | 0,767171969 | 1,144654088 |
| SAMM501 | 0,001025687 | 0,307910266 | 0,231 | 0,168 | 1 | Secretory | 1,237913292 | 0,807811021 | 1,375 |
| PPP2R1A | 0,00104559 | 0,494537209 | 0,388 | 0,351 | 1 | Secretory | 1,408868742 | 0,709789329 | 1,105413105 |
| CBX3 | 0,001050131 | 0,377626826 | 0,386 | 0,344 | 1 | Secretory | 1,299202961 | 0,769702679 | 1,122093023 |
| ZNF7061 | 0,001064619 | 0,482627028 | 0,405 | 0,376 | 1 | Secretory | 1,397285692 | 0,715673255 | 1,07712766 |
| PAXX1 | 0,00107092 | 0,390381759 | 0,278 | 0,22 | 1 | Secretory | 1,3107402 | 0,762927695 | 1,263636364 |
| ARPC31 | 0,001087525 | 0,525247142 | 0,588 | 0,657 | 1 | Secretory | 1,439180099 | 0,69484007 | 0,894977169 |
| PLEKHS1 | 0,001093094 | 0,531041406 | 0,429 | 0,401 | 1 | Secretory | 1,444971868 | 0,692054996 | 1,069825436 |
| HSBP11 | 0,001099677 | 0,479178932 | 0,547 | 0,596 | 1 | Secretory | 1,393950115 | 0,717385787 | 0,917785235 |
| CDC371 | 0,00113302 | 0,503692346 | 0,444 | 0,433 | 1 | Secretory | 1,41783765 | 0,705299369 | 1,025404157 |
| RCAN1 | 0,001169987 | -0,264715484 | 0,218 | 0,306 | 1 | Secretory | 0,83236287 | 1,201399097 | 0,712418301 |
| UQCR111 | 0,001181307 | 0,348469671 | 0,619 | 0,671 | 1 | Secretory | 1,273209363 | 0,785416781 | 0,922503726 |
| MRPL571 | 0,001194202 | 0,377904984 | 0,425 | 0,411 | 1 | Secretory | 1,299453477 | 0,769554291 | 1,03406326 |
| BIRC3 | 0,001204822 | 0,552273866 | 0,237 | 0,17 | 1 | Secretory | 1,466395096 | 0,681944452 | 1,394117647 |
| AHR | 0,001230767 | 0,459197596 | 0,259 | 0,203 | 1 | Secretory | 1,374776976 | 0,727390709 | 1,275862069 |
| SF3B51 | 0,001253342 | 0,483948716 | 0,494 | 0,519 | 1 | Secretory | 1,398566366 | 0,71501791 | 0,951830443 |
| GADD45GIP11 | 0,001268544 | 0,392405128 | 0,453 | 0,446 | 1 | Secretory | 1,312579792 | 0,761858445 | 1,015695067 |
| RAB251 | 0,001276387 | 0,528762014 | 0,42 | 0,411 | 1 | Secretory | 1,442690682 | 0,693149275 | 1,02189781 |
| DDIT4 | 0,001288876 | 0,624554823 | 0,459 | 0,463 | 1 | Secretory | 1,541735014 | 0,648619893 | 0,991360691 |

| Supplementary Table 4 - clusermarker | | | | | | | | | |
|---|---|---|---|---|---|---|---|---|---|
| | **p_val** | **avg_log2FC** | **pct.1** | **pct.2** | **p_val_adj** | **celltype** | **Foldchange_UP** | **Foldchange_DOWN** | **Ratio_pct1_pct2** |
| KRAS1 | 0,001297158 | 0,266859096 | 0,291 | 0,235 | 1 | Secretory | 1,203185509 | 0,831127031 | 1,238297872 |
| RNF2131 | 0,001299089 | 0,445149145 | 0,319 | 0,26 | 1 | Secretory | 1,361454855 | 0,73450838 | 1,226923077 |
| EIF4A21 | 0,001301346 | 0,479058404 | 0,541 | 0,581 | 1 | Secretory | 1,393833664 | 0,717445723 | 0,931153184 |
| AKR1C21 | 0,001327481 | 0,375108155 | 0,36 | 0,303 | 1 | Secretory | 1,296936779 | 0,771047607 | 1,188118812 |
| C5orf38 | 0,001330266 | 0,308437909 | 0,267 | 0,208 | 1 | Secretory | 1,238366122 | 0,807515631 | 1,283653846 |
| ATP5F1E | 0,00134042 | 0,286955265 | 0,746 | 0,795 | 1 | Secretory | 1,220062678 | 0,819630022 | 0,93836478 |
| PMVK1 | 0,001357265 | 0,324647657 | 0,276 | 0,22 | 1 | Secretory | 1,252358543 | 0,798493375 | 1,254545455 |
| TALDO11 | 0,001357684 | 0,399758734 | 0,489 | 0,514 | 1 | Secretory | 1,319287264 | 0,757985033 | 0,951361868 |
| HADH | 0,00135844 | 0,401768205 | 0,332 | 0,278 | 1 | Secretory | 1,321126125 | 0,756930001 | 1,194244604 |
| MRPL141 | 0,001372326 | 0,339758679 | 0,358 | 0,313 | 1 | Secretory | 1,265544887 | 0,790173474 | 1,143769968 |
| IGBP1 | 0,001375777 | 0,452622901 | 0,431 | 0,418 | 1 | Secretory | 1,368526054 | 0,730713162 | 1,031100478 |
| ACTN4 | 0,001385174 | 0,52585152 | 0,418 | 0,389 | 1 | Secretory | 1,43978313 | 0,694549046 | 1,074550129 |
| CLDN41 | 0,001385704 | 0,47732242 | 0,608 | 0,619 | 1 | Secretory | 1,392157482 | 0,718309539 | 0,982229402 |
| JAGN1 | 0,001390242 | 0,364025026 | 0,272 | 0,211 | 1 | Secretory | 1,287011572 | 0,77699379 | 1,289099526 |
| DUSP1 | 0,001391444 | 0,787693185 | 0,567 | 0,587 | 1 | Secretory | 1,726311947 | 0,579269582 | 0,96592845 |
| VAPA | 0,001407248 | 0,522973071 | 0,474 | 0,489 | 1 | Secretory | 1,436913355 | 0,695936186 | 0,969325153 |
| RPL36A1 | 0,001427954 | 0,283873919 | 0,828 | 0,874 | 1 | Secretory | 1,217459617 | 0,821382481 | 0,947368421 |
| CRTAP | 0,001430209 | 0,465875307 | 0,341 | 0,296 | 1 | Secretory | 1,381155069 | 0,724031662 | 1,152027027 |
| AL138963.41 | 0,00145349 | 0,667793457 | 0,498 | 0,498 | 1 | Secretory | 1,588641347 | 0,629468698 | 1 |
| MAP1LC3A | 0,001463624 | 0,374071977 | 0,308 | 0,26 | 1 | Secretory | 1,296005622 | 0,771601591 | 1,184615385 |
| PYGB | 0,001467327 | 0,379035865 | 0,379 | 0,341 | 1 | Secretory | 1,300472476 | 0,768951299 | 1,11143695 |
| MAGOH | 0,001478037 | 0,375993616 | 0,33 | 0,28 | 1 | Secretory | 1,297733025 | 0,770574518 | 1,178571429 |
| TPR | 0,001495877 | 0,407425283 | 0,379 | 0,344 | 1 | Secretory | 1,326316679 | 0,753967748 | 1,101744186 |
| CCL22 | 0,001499843 | -0,261157711 | 0,067 | 0,126 | 1 | Secretory | 0,83441806 | 1,198440024 | 0,531746032 |
| EGFR | 0,001507751 | 0,390512965 | 0,306 | 0,255 | 1 | Secretory | 1,310859411 | 0,762858314 | 1,2 |
| CHCHD5 | 0,001513389 | 0,451859044 | 0,341 | 0,303 | 1 | Secretory | 1,367801658 | 0,731100152 | 1,125412541 |
| MFGE81 | 0,001532151 | 0,373611321 | 0,252 | 0,191 | 1 | Secretory | 1,295591871 | 0,771848005 | 1,319371728 |
| IFT57 | 0,001565009 | 0,307084212 | 0,282 | 0,223 | 1 | Secretory | 1,237204694 | 0,808273687 | 1,264573991 |
| MACROH2A1 | 0,001575555 | 0,353965592 | 0,401 | 0,364 | 1 | Secretory | 1,278068881 | 0,782430443 | 1,101648352 |
| PTGR1 | 0,001589858 | 0,336664229 | 0,272 | 0,211 | 1 | Secretory | 1,262833318 | 0,791870143 | 1,289099526 |
| DUT | 0,001595418 | 0,379886808 | 0,47 | 0,488 | 1 | Secretory | 1,301239758 | 0,768497884 | 0,963114754 |

| Supplementary Table 4 - clusermarker | | | | | | | | | |
|---|---|---|---|---|---|---|---|---|---|
| | **p_val** | **avg_log2FC** | **pct.1** | **pct.2** | **p_val_adj** | **celltype** | **Foldchange_UP** | **Foldchange_DOWN** | **Ratio_pct1_pct2** |
| SCNM1 | 0,001614306 | 0,272864438 | 0,183 | 0,123 | 1 | Secretory | 1,20820431 | 0,827674584 | 1,487804878 |
| PHPT11 | 0,001665149 | 0,400274354 | 0,56 | 0,622 | 1 | Secretory | 1,319758863 | 0,757714177 | 0,900321543 |
| CHCHD101 | 0,00178264 | 0,451091943 | 0,545 | 0,596 | 1 | Secretory | 1,367074573 | 0,731488991 | 0,91442953 |
| UQCRC11 | 0,001787377 | 0,341879851 | 0,435 | 0,441 | 1 | Secretory | 1,267406967 | 0,789012548 | 0,986394558 |
| HUWE1 | 0,001799196 | 0,37905998 | 0,325 | 0,268 | 1 | Secretory | 1,300494213 | 0,768938446 | 1,212686567 |
| ETHE11 | 0,001828602 | 0,422835822 | 0,375 | 0,346 | 1 | Secretory | 1,340560028 | 0,745956898 | 1,083815029 |
| TRIM29 | 0,001829624 | 0,390591158 | 0,259 | 0,2 | 1 | Secretory | 1,31093046 | 0,762816969 | 1,295 |
| GCHFR1 | 0,001850033 | 0,360233673 | 0,336 | 0,291 | 1 | Secretory | 1,283633791 | 0,779038389 | 1,154639175 |
| C17orf491 | 0,001862794 | 0,362208738 | 0,295 | 0,243 | 1 | Secretory | 1,285392302 | 0,777972607 | 1,21399177 |
| TSC22D11 | 0,001884381 | 0,623153182 | 0,517 | 0,527 | 1 | Secretory | 1,540237879 | 0,649250362 | 0,981024668 |
| METTL261 | 0,001916462 | 0,46233018 | 0,418 | 0,418 | 1 | Secretory | 1,37776533 | 0,725813009 | 1 |
| CYB5R3 | 0,001924489 | 0,496485455 | 0,369 | 0,333 | 1 | Secretory | 1,410772594 | 0,708831462 | 1,108108108 |
| ARRDC1 | 0,001927835 | 0,415416365 | 0,33 | 0,285 | 1 | Secretory | 1,333683526 | 0,749803069 | 1,157894737 |
| TNFSF10 | 0,001930916 | 0,418539454 | 0,478 | 0,478 | 1 | Secretory | 1,336573758 | 0,74818168 | 1 |
| ZNF524 | 0,002084716 | 0,303999891 | 0,289 | 0,238 | 1 | Secretory | 1,234562513 | 0,810003535 | 1,214285714 |
| LAMTOR11 | 0,002103343 | 0,415237879 | 0,453 | 0,458 | 1 | Secretory | 1,333518536 | 0,749895838 | 0,989082969 |
| PSMD81 | 0,002113402 | 0,379867741 | 0,453 | 0,454 | 1 | Secretory | 1,301222561 | 0,76850804 | 0,997797357 |
| STUB11 | 0,002147824 | 0,388909007 | 0,429 | 0,411 | 1 | Secretory | 1,309402834 | 0,763706916 | 1,04379562 |
| SFRP11 | 0,002148856 | 0,378211022 | 0,287 | 0,215 | 1 | Secretory | 1,299729159 | 0,769391064 | 1,334883721 |
| MGP1 | 0,002189524 | 0,324864532 | 0,42 | 0,378 | 1 | Secretory | 1,25254682 | 0,79837335 | 1,111111111 |
| RNPS1 | 0,002231646 | 0,37899137 | 0,328 | 0,278 | 1 | Secretory | 1,300432367 | 0,768975016 | 1,179856115 |
| KLF31 | 0,002238667 | 0,415438148 | 0,334 | 0,293 | 1 | Secretory | 1,333703663 | 0,749791748 | 1,139931741 |
| SSNA1 | 0,002276881 | 0,379242993 | 0,356 | 0,313 | 1 | Secretory | 1,300659198 | 0,768840909 | 1,137380192 |
| APRT | 0,002281822 | 0,360594969 | 0,526 | 0,569 | 1 | Secretory | 1,283955293 | 0,778843318 | 0,924428822 |
| RBX11 | 0,002327206 | 0,400826754 | 0,44 | 0,438 | 1 | Secretory | 1,320264288 | 0,757424108 | 1,00456621 |
| MRPS21 | 0,002359775 | 0,471185032 | 0,509 | 0,519 | 1 | Secretory | 1,386247667 | 0,721371818 | 0,980732177 |
| HNRNPA3 | 0,002366615 | 0,456209976 | 0,459 | 0,451 | 1 | Secretory | 1,371932951 | 0,728898594 | 1,017738359 |
| STK38 | 0,002410839 | 0,250807802 | 0,237 | 0,173 | 1 | Secretory | 1,189873169 | 0,840425708 | 1,369942197 |
| MPV17 | 0,002433225 | 0,280099236 | 0,291 | 0,236 | 1 | Secretory | 1,214278406 | 0,823534369 | 1,233050847 |
| CTSZ1 | 0,002463838 | 0,524555947 | 0,489 | 0,512 | 1 | Secretory | 1,438490753 | 0,695173047 | 0,955078125 |
| TMCO3 | 0,002470132 | 0,290078342 | 0,235 | 0,178 | 1 | Secretory | 1,222706672 | 0,817857646 | 1,320224719 |

| Supplementary Table 4 - clusermarker | | | | | | | | | |
|---|---|---|---|---|---|---|---|---|---|
| | **p_val** | **avg_log2FC** | **pct.1** | **pct.2** | **p_val_adj** | **celltype** | **Foldchange_UP** | **Foldchange_DOWN** | **Ratio_pct1_pct2** |
| ARMC101 | 0,002470485 | 0,2755945 | 0,218 | 0,163 | 1 | Secretory | 1,210492801 | 0,826109828 | 1,337423313 |
| MEAF6 | 0,002471416 | 0,272658491 | 0,321 | 0,271 | 1 | Secretory | 1,208031849 | 0,827792745 | 1,184501845 |
| TCEA3 | 0,002553594 | 0,427324646 | 0,474 | 0,481 | 1 | Secretory | 1,344737563 | 0,743639523 | 0,985446985 |
| NUCKS1 | 0,00258733 | 0,527869476 | 0,509 | 0,544 | 1 | Secretory | 1,441798423 | 0,693578231 | 0,935661765 |
| HNRNPLL | 0,00260584 | 0,274424435 | 0,198 | 0,14 | 1 | Secretory | 1,209511457 | 0,826780098 | 1,414285714 |
| KDM1A1 | 0,002613079 | 0,269253542 | 0,218 | 0,158 | 1 | Secretory | 1,205184098 | 0,829748751 | 1,379746835 |
| UQCRQ1 | 0,002654432 | 0,416392213 | 0,647 | 0,704 | 1 | Secretory | 1,334585943 | 0,749296069 | 0,919034091 |
| GNAI2 | 0,002656497 | 0,37777713 | 0,33 | 0,283 | 1 | Secretory | 1,299338323 | 0,769622493 | 1,166077739 |
| CITED4 | 0,002660281 | 0,580667338 | 0,364 | 0,343 | 1 | Secretory | 1,495540871 | 0,668654411 | 1,06122449 |
| NDUFB10 | 0,002676557 | 0,41455536 | 0,524 | 0,586 | 1 | Secretory | 1,332887817 | 0,750250687 | 0,894197952 |
| RAD23A1 | 0,002680714 | 0,262344969 | 0,407 | 0,381 | 1 | Secretory | 1,19942668 | 0,833731663 | 1,06824147 |
| SRPK1 | 0,00270219 | 0,299391013 | 0,263 | 0,203 | 1 | Secretory | 1,230624835 | 0,812595335 | 1,295566502 |
| TXNL4A1 | 0,002742467 | 0,437680476 | 0,453 | 0,456 | 1 | Secretory | 1,35442497 | 0,738320706 | 0,993421053 |
| PCBP2 | 0,002766384 | 0,493968909 | 0,513 | 0,559 | 1 | Secretory | 1,408313877 | 0,710068982 | 0,917710197 |
| HADHA | 0,002778376 | 0,406667335 | 0,45 | 0,461 | 1 | Secretory | 1,325620055 | 0,754363964 | 0,976138829 |
| CCDC1241 | 0,002779106 | 0,302620246 | 0,315 | 0,271 | 1 | Secretory | 1,233382469 | 0,810778509 | 1,162361624 |
| PCBP11 | 0,002787054 | 0,493508501 | 0,496 | 0,512 | 1 | Secretory | 1,407864513 | 0,710295622 | 0,96875 |
| EIF3D | 0,002802589 | 0,513457482 | 0,425 | 0,428 | 1 | Secretory | 1,427467087 | 0,700541546 | 0,992990654 |
| MLST8 | 0,002887964 | 0,262042177 | 0,231 | 0,17 | 1 | Secretory | 1,199174971 | 0,833906664 | 1,358823529 |
| RAB13 | 0,003032438 | 0,479908385 | 0,448 | 0,453 | 1 | Secretory | 1,394655099 | 0,717023156 | 0,988962472 |
| DNAJC41 | 0,003039467 | 0,360816836 | 0,401 | 0,383 | 1 | Secretory | 1,284152763 | 0,778723551 | 1,046997389 |
| UQCRFS11 | 0,003062867 | 0,431259312 | 0,392 | 0,378 | 1 | Secretory | 1,348410075 | 0,741614156 | 1,037037037 |
| MID1 | 0,003073741 | 0,312523361 | 0,181 | 0,126 | 1 | Secretory | 1,241877921 | 0,805232127 | 1,436507937 |
| ERBB2 | 0,003151488 | 0,283234576 | 0,3 | 0,246 | 1 | Secretory | 1,216920209 | 0,821746564 | 1,219512195 |
| TRIM28 | 0,003230598 | 0,462390351 | 0,351 | 0,304 | 1 | Secretory | 1,377822794 | 0,725782738 | 1,154605263 |
| CD46 | 0,003263306 | 0,652561945 | 0,528 | 0,579 | 1 | Secretory | 1,571957209 | 0,636149632 | 0,911917098 |
| TSR2 | 0,003373344 | 0,321465325 | 0,259 | 0,206 | 1 | Secretory | 1,249599105 | 0,800256655 | 1,257281553 |
| SUGT11 | 0,003384581 | 0,279785095 | 0,282 | 0,228 | 1 | Secretory | 1,21401403 | 0,823713709 | 1,236842105 |
| TMEM14A | 0,003387235 | 0,350846946 | 0,308 | 0,266 | 1 | Secretory | 1,275309088 | 0,784123637 | 1,157894737 |
| EBPL | 0,003415746 | 0,380649189 | 0,343 | 0,306 | 1 | Secretory | 1,30192757 | 0,768091884 | 1,120915033 |
| SF3B6 | 0,003561741 | 0,400146339 | 0,44 | 0,446 | 1 | Secretory | 1,319641761 | 0,757781414 | 0,986547085 |

| Supplementary Table 4 - clusermarker | | | | | | | | | |
|---|---|---|---|---|---|---|---|---|---|
| | **p_val** | **avg_log2FC** | **pct.1** | **pct.2** | **p_val_adj** | **celltype** | **Foldchange_UP** | **Foldchange_DOWN** | **Ratio_pct1_pct2** |
| KDELR11 | 0,003568361 | 0,491972262 | 0,547 | 0,614 | 1 | Secretory | 1,40636616 | 0,711052376 | 0,890879479 |
| SP3 | 0,003587031 | 0,307693648 | 0,222 | 0,166 | 1 | Secretory | 1,237727435 | 0,807932321 | 1,337349398 |
| TJP1 | 0,003592411 | 0,385999241 | 0,332 | 0,283 | 1 | Secretory | 1,306764566 | 0,765248788 | 1,173144876 |
| TMEM18 | 0,003610293 | 0,307998681 | 0,353 | 0,316 | 1 | Secretory | 1,237989159 | 0,807761516 | 1,117088608 |
| PARD3 | 0,003628738 | 0,250279445 | 0,265 | 0,208 | 1 | Secretory | 1,189437482 | 0,840733552 | 1,274038462 |
| RNF169 | 0,003730427 | 0,291280603 | 0,155 | 0,101 | 1 | Secretory | 1,223726031 | 0,817176373 | 1,534653465 |
| PDIA6 | 0,003743777 | 0,41884851 | 0,582 | 0,639 | 1 | Secretory | 1,336860111 | 0,748021421 | 0,910798122 |
| MRPS151 | 0,003897573 | 0,353125216 | 0,334 | 0,293 | 1 | Secretory | 1,277324618 | 0,782886344 | 1,139931741 |
| EPB41L4A-AS1 | 0,003909837 | 0,334028737 | 0,356 | 0,328 | 1 | Secretory | 1,2605285 | 0,793318041 | 1,085365854 |
| ATP6AP2 | 0,003944241 | 0,438258231 | 0,345 | 0,311 | 1 | Secretory | 1,354967484 | 0,738025091 | 1,109324759 |
| IP6K2 | 0,00411542 | 0,304360011 | 0,246 | 0,188 | 1 | Secretory | 1,234870719 | 0,80980137 | 1,308510638 |
| TNIP11 | 0,004173552 | 0,304480665 | 0,228 | 0,175 | 1 | Secretory | 1,234973997 | 0,809733649 | 1,302857143 |
| TCF251 | 0,004219773 | 0,340311651 | 0,496 | 0,524 | 1 | Secretory | 1,266030053 | 0,789870665 | 0,946564885 |
| MRPL38 | 0,004221087 | 0,307608578 | 0,3 | 0,253 | 1 | Secretory | 1,237654453 | 0,807979963 | 1,185770751 |
| TUSC2 | 0,004240551 | 0,294103933 | 0,282 | 0,235 | 1 | Secretory | 1,226123188 | 0,815578736 | 1,2 |
| HNRNPL | 0,004259784 | 0,493415153 | 0,427 | 0,423 | 1 | Secretory | 1,407773421 | 0,710341583 | 1,009456265 |
| SARAF | 0,004279517 | 0,376908299 | 0,616 | 0,677 | 1 | Secretory | 1,29855606 | 0,770086121 | 0,909896603 |
| NFIB1 | 0,004303665 | 0,531993617 | 0,537 | 0,576 | 1 | Secretory | 1,445925897 | 0,691598374 | 0,932291667 |
| SPINT1-AS1 | 0,004453985 | 0,399395834 | 0,448 | 0,456 | 1 | Secretory | 1,318955448 | 0,758175723 | 0,98245614 |
| SLC25A37 | 0,004455362 | 0,466723514 | 0,42 | 0,404 | 1 | Secretory | 1,381967333 | 0,723606106 | 1,03960396 |
| SMIM19 | 0,004476322 | 0,387632291 | 0,353 | 0,318 | 1 | Secretory | 1,308244588 | 0,764383059 | 1,110062893 |
| CALD11 | 0,0044976 | 0,432209326 | 0,278 | 0,216 | 1 | Secretory | 1,349298295 | 0,741125964 | 1,287037037 |
| SEC11A | 0,004525148 | 0,367134684 | 0,446 | 0,458 | 1 | Secretory | 1,289788654 | 0,775320822 | 0,973799127 |
| CAPRIN1 | 0,004569678 | 0,407426404 | 0,319 | 0,28 | 1 | Secretory | 1,32631771 | 0,753967162 | 1,139285714 |
| CALM21 | 0,004579097 | 0,281460365 | 0,709 | 0,792 | 1 | Secretory | 1,215424573 | 0,822757761 | 0,89520202 |
| FAM174C1 | 0,004580237 | 0,336790165 | 0,312 | 0,268 | 1 | Secretory | 1,262943558 | 0,791801022 | 1,164179104 |
| SERPING1 | 0,00462585 | 0,443849375 | 0,33 | 0,291 | 1 | Secretory | 1,36022883 | 0,73517042 | 1,134020619 |
| STOML21 | 0,004628277 | 0,371709195 | 0,349 | 0,321 | 1 | Secretory | 1,293884819 | 0,772866321 | 1,087227414 |
| TLE5 | 0,004675238 | 0,517635305 | 0,571 | 0,629 | 1 | Secretory | 1,431606804 | 0,69851582 | 0,907790143 |
| CRB31 | 0,004734802 | 0,280022695 | 0,248 | 0,188 | 1 | Secretory | 1,214213985 | 0,823578061 | 1,319148936 |
| UBXN41 | 0,004737305 | 0,529474649 | 0,519 | 0,572 | 1 | Secretory | 1,44340349 | 0,692806971 | 0,907342657 |

| Supplementary Table 4 - clusermarker | | | | | | | | | |
|---|---|---|---|---|---|---|---|---|---|
| | **p_val** | **avg_log2FC** | **pct.1** | **pct.2** | **p_val_adj** | **celltype** | **Foldchange_UP** | **Foldchange_DOWN** | **Ratio_pct1_pct2** |
| NDUFAB11 | 0,004743452 | 0,439533403 | 0,433 | 0,449 | 1 | Secretory | 1,356165645 | 0,737373051 | 0,964365256 |
| APEX1 | 0,004880964 | 0,337136092 | 0,375 | 0,353 | 1 | Secretory | 1,26324642 | 0,791611188 | 1,062322946 |
| PRNP | 0,00492886 | 0,372140855 | 0,308 | 0,265 | 1 | Secretory | 1,294272013 | 0,772635111 | 1,162264151 |
| NEDD8 | 0,004932159 | 0,449362227 | 0,461 | 0,489 | 1 | Secretory | 1,365436504 | 0,732366534 | 0,942740286 |
| TMEM2051 | 0,004962536 | 0,31976827 | 0,427 | 0,423 | 1 | Secretory | 1,248130055 | 0,801198558 | 1,009456265 |
| C1orf43 | 0,00496461 | 0,462453517 | 0,401 | 0,394 | 1 | Secretory | 1,377883121 | 0,725750961 | 1,017766497 |
| SF1 | 0,004970933 | 0,294038276 | 0,453 | 0,449 | 1 | Secretory | 1,226067388 | 0,815615854 | 1,008908686 |
| CFAP36 | 0,004976835 | 0,267220271 | 0,263 | 0,218 | 1 | Secretory | 1,20348676 | 0,830918987 | 1,206422018 |
| POLR2E | 0,005184216 | 0,398582028 | 0,373 | 0,353 | 1 | Secretory | 1,318211652 | 0,75860352 | 1,056657224 |
| HAGH | 0,005190703 | 0,296375935 | 0,254 | 0,205 | 1 | Secretory | 1,228055647 | 0,814295348 | 1,23902439 |
| PGK1 | 0,005243439 | 0,309334108 | 0,502 | 0,521 | 1 | Secretory | 1,239135631 | 0,80701416 | 0,96353167 |
| LRIG3 | 0,00524519 | 0,31421439 | 0,282 | 0,23 | 1 | Secretory | 1,24333442 | 0,804288842 | 1,226086957 |
| EIF2B1 | 0,005292328 | 0,255760932 | 0,192 | 0,136 | 1 | Secretory | 1,193965319 | 0,837545265 | 1,411764706 |
| CEP57 | 0,005295416 | 0,253236079 | 0,267 | 0,22 | 1 | Secretory | 1,191877595 | 0,839012332 | 1,213636364 |
| MEA11 | 0,005352608 | 0,33299933 | 0,356 | 0,328 | 1 | Secretory | 1,259629395 | 0,7938843 | 1,085365854 |
| LSM8 | 0,00565476 | 0,404363084 | 0,356 | 0,328 | 1 | Secretory | 1,323504485 | 0,755569786 | 1,085365854 |
| ARID5B | 0,005667643 | 0,441599267 | 0,369 | 0,333 | 1 | Secretory | 1,358108995 | 0,736317927 | 1,108108108 |
| KIAA0232 | 0,00569632 | 0,269762578 | 0,151 | 0,1 | 1 | Secretory | 1,205609406 | 0,829456037 | 1,51 |
| ZKSCAN1 | 0,005702478 | 0,464970476 | 0,353 | 0,321 | 1 | Secretory | 1,380289106 | 0,724485904 | 1,099688474 |
| IMP31 | 0,00570984 | 0,3246378 | 0,347 | 0,313 | 1 | Secretory | 1,252349986 | 0,798498831 | 1,108626198 |
| SRRM1 | 0,005729034 | 0,514165446 | 0,509 | 0,536 | 1 | Secretory | 1,42816775 | 0,700197858 | 0,949626866 |
| MRTFB | 0,005810294 | 0,251361327 | 0,213 | 0,16 | 1 | Secretory | 1,19032978 | 0,84010332 | 1,33125 |
| LINC00511 | 0,005858982 | 0,261537822 | 0,101 | 0,058 | 1 | Secretory | 1,198755822 | 0,834198242 | 1,74137931 |
| ENY21 | 0,006021057 | 0,339431131 | 0,466 | 0,479 | 1 | Secretory | 1,265257592 | 0,790352895 | 0,972860125 |
| NPTN | 0,006032608 | 0,481544179 | 0,427 | 0,418 | 1 | Secretory | 1,39623732 | 0,716210622 | 1,0215311 |
| ANAPC51 | 0,006045488 | 0,349206303 | 0,381 | 0,368 | 1 | Secretory | 1,273859622 | 0,785015855 | 1,035326087 |
| PSMB61 | 0,00613051 | 0,450310187 | 0,468 | 0,488 | 1 | Secretory | 1,366333995 | 0,731885472 | 0,959016393 |
| SNHG81 | 0,006247112 | 0,51081831 | 0,506 | 0,556 | 1 | Secretory | 1,424858159 | 0,701824244 | 0,910071942 |
| BICDL2 | 0,006369437 | 0,376296692 | 0,289 | 0,246 | 1 | Secretory | 1,298005676 | 0,770412656 | 1,174796748 |
| SET | 0,006454991 | 0,488201878 | 0,476 | 0,519 | 1 | Secretory | 1,402695517 | 0,712913093 | 0,917148362 |
| ADD31 | 0,006569489 | 0,375716815 | 0,373 | 0,354 | 1 | Secretory | 1,29748406 | 0,770722378 | 1,053672316 |

| Supplementary Table 4 - clusermarker | | | | | | | | | |
|---|---|---|---|---|---|---|---|---|---|
| | **p_val** | **avg_log2FC** | **pct.1** | **pct.2** | **p_val_adj** | **celltype** | **Foldchange_UP** | **Foldchange_DOWN** | **Ratio_pct1_pct2** |
| AGR21 | 0,006653448 | -0,607058272 | 0,519 | 0,458 | 1 | Secretory | 0,656534044 | 1,52315026 | 1,133187773 |
| NDUFS5 | 0,006689568 | 0,488624928 | 0,569 | 0,651 | 1 | Secretory | 1,403106899 | 0,712704072 | 0,874039939 |
| TIMM13 | 0,006801367 | 0,296321607 | 0,334 | 0,304 | 1 | Secretory | 1,228009403 | 0,814326012 | 1,098684211 |
| TLN1 | 0,006884727 | 0,27381587 | 0,315 | 0,271 | 1 | Secretory | 1,209001362 | 0,827128928 | 1,162361624 |
| MDH2 | 0,00694589 | 0,311586938 | 0,478 | 0,506 | 1 | Secretory | 1,241072106 | 0,805754956 | 0,944664032 |
| EMC61 | 0,006989459 | 0,291531833 | 0,317 | 0,28 | 1 | Secretory | 1,223939149 | 0,817034083 | 1,132142857 |
| MRPL371 | 0,007105061 | 0,353496111 | 0,31 | 0,276 | 1 | Secretory | 1,27765304 | 0,782685102 | 1,123188406 |
| WASF2 | 0,007163776 | 0,320661315 | 0,418 | 0,409 | 1 | Secretory | 1,2489029 | 0,800702761 | 1,02200489 |
| ARPC21 | 0,007167638 | 0,392912603 | 0,429 | 0,428 | 1 | Secretory | 1,31304158 | 0,761590505 | 1,002336449 |
| ANXA5 | 0,007172243 | 0,375445685 | 0,58 | 0,629 | 1 | Secretory | 1,297240243 | 0,770867236 | 0,922098569 |
| HINT2 | 0,007182994 | 0,385495054 | 0,407 | 0,403 | 1 | Secretory | 1,306307964 | 0,76551627 | 1,009925558 |
| UBE3A | 0,007191122 | 0,364049134 | 0,321 | 0,28 | 1 | Secretory | 1,287033079 | 0,776980806 | 1,146428571 |
| RAB11A1 | 0,007258443 | 0,318670887 | 0,44 | 0,434 | 1 | Secretory | 1,247181028 | 0,80180822 | 1,013824885 |
| RASSF7 | 0,007277726 | 0,264390397 | 0,312 | 0,266 | 1 | Secretory | 1,201128412 | 0,83255045 | 1,172932331 |
| CHMP2A1 | 0,0072966 | 0,332916322 | 0,457 | 0,463 | 1 | Secretory | 1,259556922 | 0,793929979 | 0,987041037 |
| AAMDC | 0,007319712 | 0,34822464 | 0,31 | 0,275 | 1 | Secretory | 1,272993136 | 0,78555019 | 1,127272727 |
| OCIAD21 | 0,007339932 | 0,458467815 | 0,379 | 0,373 | 1 | Secretory | 1,374081727 | 0,72775875 | 1,016085791 |
| VPS721 | 0,007438592 | 0,269046593 | 0,209 | 0,158 | 1 | Secretory | 1,205011231 | 0,829867784 | 1,32278481 |
| REXO21 | 0,007458735 | 0,353938123 | 0,442 | 0,438 | 1 | Secretory | 1,278044547 | 0,78244534 | 1,00913242 |
| GLUD1 | 0,007467381 | 0,478528998 | 0,399 | 0,386 | 1 | Secretory | 1,393322282 | 0,717709042 | 1,033678756 |
| PSMB31 | 0,007478823 | 0,543723474 | 0,44 | 0,464 | 1 | Secretory | 1,457729944 | 0,685998119 | 0,948275862 |
| RCN2 | 0,007493556 | 0,348894782 | 0,293 | 0,256 | 1 | Secretory | 1,273584588 | 0,785185381 | 1,14453125 |
| ABHD14B | 0,007503075 | 0,377283958 | 0,349 | 0,328 | 1 | Secretory | 1,298894232 | 0,769885627 | 1,06402439 |
| IDH3G | 0,007521204 | 0,312597144 | 0,336 | 0,303 | 1 | Secretory | 1,241941435 | 0,805190947 | 1,108910891 |
| VWA1 | 0,007897163 | 0,287821853 | 0,28 | 0,241 | 1 | Secretory | 1,220795757 | 0,819137841 | 1,161825726 |
| UXT | 0,007901213 | 0,290713645 | 0,534 | 0,576 | 1 | Secretory | 1,22324522 | 0,817497574 | 0,927083333 |
| TMBIM41 | 0,00811591 | 0,444295022 | 0,513 | 0,567 | 1 | Secretory | 1,360649068 | 0,734943362 | 0,904761905 |
| SELENOF | 0,008119634 | 0,304548963 | 0,478 | 0,511 | 1 | Secretory | 1,235032462 | 0,809695316 | 0,935420744 |
| MRPL411 | 0,008251696 | 0,364236935 | 0,459 | 0,489 | 1 | Secretory | 1,287200628 | 0,776879671 | 0,938650307 |
| MT-ND61 | 0,008390968 | 0,3754402 | 0,558 | 0,559 | 1 | Secretory | 1,297235311 | 0,770870166 | 0,998211091 |
| KRT8 | 0,008423263 | 0,319320933 | 0,666 | 0,722 | 1 | Secretory | 1,247743106 | 0,801447025 | 0,922437673 |

| Supplementary Table 4 - clusermarker | | | | | | | | | |
|---|---|---|---|---|---|---|---|---|---|
| | **p_val** | **avg_log2FC** | **pct.1** | **pct.2** | **p_val_adj** | **celltype** | **Foldchange_UP** | **Foldchange_DOWN** | **Ratio_pct1_pct2** |
| AOPEP1 | 0,008529795 | 0,278160494 | 0,244 | 0,193 | 1 | Secretory | 1,212647714 | 0,824641805 | 1,264248705 |
| CHPT11 | 0,008607905 | 0,532075881 | 0,463 | 0,473 | 1 | Secretory | 1,446008348 | 0,69155894 | 0,978858351 |
| PML | 0,008622274 | 0,273350904 | 0,222 | 0,175 | 1 | Secretory | 1,208611776 | 0,827395546 | 1,268571429 |
| TPM31 | 0,008677988 | 0,52039615 | 0,446 | 0,453 | 1 | Secretory | 1,434349052 | 0,697180368 | 0,984547461 |
| COX6B1 | 0,008776391 | 0,343746528 | 0,662 | 0,722 | 1 | Secretory | 1,269047903 | 0,787992319 | 0,916897507 |
| OST41 | 0,008847362 | 0,299100716 | 0,657 | 0,734 | 1 | Secretory | 1,230377236 | 0,81275886 | 0,895095368 |
| ANXA31 | 0,008850063 | 0,514244194 | 0,502 | 0,537 | 1 | Secretory | 1,428245708 | 0,70015964 | 0,934823091 |
| HYI | 0,008940092 | 0,251352408 | 0,366 | 0,338 | 1 | Secretory | 1,190322421 | 0,840108514 | 1,082840237 |
| MIA1 | 0,008969447 | 0,390785306 | 0,565 | 0,572 | 1 | Secretory | 1,311106888 | 0,762714321 | 0,987762238 |
| CNN21 | 0,00898477 | 0,434887089 | 0,351 | 0,321 | 1 | Secretory | 1,351805031 | 0,739751648 | 1,093457944 |
| HBB | 0,009375991 | 0,324590489 | 0,144 | 0,095 | 1 | Secretory | 1,252308918 | 0,798525017 | 1,515789474 |
| PRPF81 | 0,009379737 | 0,407885961 | 0,371 | 0,343 | 1 | Secretory | 1,326740263 | 0,753727032 | 1,081632653 |
| FUS1 | 0,009532756 | 0,507759204 | 0,459 | 0,459 | 1 | Secretory | 1,421840075 | 0,703313978 | 1 |
| SH3BGRL31 | 0,009589787 | 0,294517146 | 0,55 | 0,599 | 1 | Secretory | 1,226474421 | 0,815345174 | 0,918196995 |
| MTPN1 | 0,009592856 | 0,344340783 | 0,353 | 0,329 | 1 | Secretory | 1,269570739 | 0,787667807 | 1,072948328 |
| SAP18 | 0,009966799 | 0,335643607 | 0,545 | 0,589 | 1 | Secretory | 1,261940253 | 0,792430543 | 0,925297114 |
| S100P1 | 0,009972549 | -0,408164511 | 0,062 | 0,108 | 1 | Secretory | 0,753581519 | 1,326996449 | 0,574074074 |
| CHASERR1 | 0,010171099 | 0,343852868 | 0,414 | 0,416 | 1 | Secretory | 1,269141447 | 0,787934239 | 0,995192308 |
| UBB1 | 0,010243897 | -0,336488639 | 0,778 | 0,877 | 1 | Secretory | 0,791966528 | 1,262679627 | 0,887115165 |
| CD471 | 0,010293804 | 0,518347468 | 0,429 | 0,438 | 1 | Secretory | 1,432313667 | 0,698171094 | 0,979452055 |
| ZFYVE211 | 0,010429595 | 0,309140252 | 0,317 | 0,285 | 1 | Secretory | 1,238969139 | 0,807122606 | 1,112280702 |
| C11orf1 | 0,010504781 | 0,361079094 | 0,315 | 0,28 | 1 | Secretory | 1,284386222 | 0,778582005 | 1,125 |
| MRPL43 | 0,010592013 | 0,389555401 | 0,39 | 0,391 | 1 | Secretory | 1,309989639 | 0,763364816 | 0,997442455 |
| MRPS12 | 0,010674695 | 0,318437172 | 0,284 | 0,246 | 1 | Secretory | 1,246979002 | 0,801938122 | 1,154471545 |
| RPS101 | 0,010852499 | 0,262790298 | 0,282 | 0,243 | 1 | Secretory | 1,199796974 | 0,833474348 | 1,160493827 |
| BCL2L1 | 0,010970827 | 0,289040859 | 0,246 | 0,201 | 1 | Secretory | 1,221827705 | 0,818446002 | 1,223880597 |
| SORD | 0,010984163 | 0,280829121 | 0,194 | 0,146 | 1 | Secretory | 1,214892886 | 0,823117833 | 1,328767123 |
| SNX3 | 0,010987429 | 0,372243029 | 0,491 | 0,532 | 1 | Secretory | 1,294363678 | 0,772580394 | 0,922932331 |
| GABARAPL21 | 0,011105183 | 0,341662037 | 0,461 | 0,489 | 1 | Secretory | 1,267215631 | 0,78913168 | 0,942740286 |
| PTPRK | 0,011173869 | 0,275654061 | 0,276 | 0,23 | 1 | Secretory | 1,210542777 | 0,826075723 | 1,2 |
| DYNLL11 | 0,011214156 | 0,462112891 | 0,526 | 0,576 | 1 | Secretory | 1,377557836 | 0,725922334 | 0,913194444 |

| Supplementary Table 4 - clusermarker | | | | | | | | | |
|---|---|---|---|---|---|---|---|---|---|
| | **p_val** | **avg_log2FC** | **pct.1** | **pct.2** | **p_val_adj** | **celltype** | **Foldchange_UP** | **Foldchange_DOWN** | **Ratio_pct1_pct2** |
| KTN1 | 0,011419154 | 0,438864113 | 0,504 | 0,537 | 1 | Secretory | 1,355536643 | 0,737715211 | 0,938547486 |
| NINJ11 | 0,011582204 | 0,397041812 | 0,323 | 0,301 | 1 | Secretory | 1,316805085 | 0,759413835 | 1,073089701 |
| CYC1 | 0,011598943 | 0,301872258 | 0,369 | 0,354 | 1 | Secretory | 1,232743168 | 0,811198979 | 1,042372881 |
| RPN11 | 0,011634446 | 0,319049751 | 0,463 | 0,494 | 1 | Secretory | 1,247508591 | 0,801597686 | 0,937246964 |
| COX6C | 0,011751214 | 0,362502523 | 0,649 | 0,72 | 1 | Secretory | 1,285654081 | 0,777814199 | 0,901388889 |
| NFKBIA1 | 0,011791562 | 0,417451376 | 0,554 | 0,564 | 1 | Secretory | 1,335566096 | 0,748746171 | 0,982269504 |
| DCXR1 | 0,011802315 | 0,401256395 | 0,539 | 0,604 | 1 | Secretory | 1,320657527 | 0,757198577 | 0,892384106 |
| ITGB81 | 0,011825176 | 0,308207306 | 0,289 | 0,246 | 1 | Secretory | 1,238168195 | 0,807644716 | 1,174796748 |
| PSMB5 | 0,011892367 | 0,349228643 | 0,412 | 0,404 | 1 | Secretory | 1,273879348 | 0,785003699 | 1,01980198 |
| TSEN34 | 0,01195269 | 0,266377648 | 0,317 | 0,28 | 1 | Secretory | 1,202784055 | 0,831404437 | 1,132142857 |
| NDUFA10 | 0,011953448 | 0,360766211 | 0,358 | 0,341 | 1 | Secretory | 1,284107703 | 0,778750877 | 1,049853372 |
| RPP21 | 0,011963865 | 0,313544949 | 0,345 | 0,313 | 1 | Secretory | 1,24275762 | 0,804662135 | 1,102236422 |
| RGL2 | 0,012023431 | 0,273458592 | 0,272 | 0,226 | 1 | Secretory | 1,208701994 | 0,827333788 | 1,203539823 |
| TPBG | 0,012030979 | 0,29581831 | 0,246 | 0,201 | 1 | Secretory | 1,227581075 | 0,814610147 | 1,223880597 |
| HLA-DMA | 0,012087216 | 0,420295371 | 0,487 | 0,499 | 1 | Secretory | 1,338201505 | 0,747271615 | 0,975951904 |
| NDUFB7 | 0,012327801 | 0,370412118 | 0,556 | 0,631 | 1 | Secretory | 1,292722055 | 0,773561491 | 0,881141046 |
| MYDGF1 | 0,012422269 | 0,370085595 | 0,552 | 0,594 | 1 | Secretory | 1,292429508 | 0,77373659 | 0,929292929 |
| REST | 0,012449173 | 0,362232262 | 0,274 | 0,231 | 1 | Secretory | 1,285413262 | 0,777959921 | 1,186147186 |
| EIF4EBP3 | 0,012530709 | 0,279895371 | 0,349 | 0,326 | 1 | Secretory | 1,21410683 | 0,823650749 | 1,070552147 |
| ATP2B1 | 0,012809342 | 0,320805958 | 0,218 | 0,171 | 1 | Secretory | 1,249028121 | 0,800622487 | 1,274853801 |
| ATL2 | 0,012851719 | 0,457249513 | 0,332 | 0,301 | 1 | Secretory | 1,372921857 | 0,728373574 | 1,102990033 |
| LAGE31 | 0,012926617 | 0,314264847 | 0,306 | 0,28 | 1 | Secretory | 1,243377905 | 0,804260713 | 1,092857143 |
| DRAP11 | 0,012975799 | 0,486145458 | 0,388 | 0,379 | 1 | Secretory | 1,400697537 | 0,713930005 | 1,023746702 |
| PMPCB | 0,012982266 | 0,272020895 | 0,3 | 0,258 | 1 | Secretory | 1,20749808 | 0,828158667 | 1,162790698 |
| HEBP1 | 0,01335534 | 0,303644304 | 0,312 | 0,285 | 1 | Secretory | 1,234258263 | 0,810203204 | 1,094736842 |
| RHEB | 0,01340258 | 0,311921747 | 0,362 | 0,354 | 1 | Secretory | 1,241360157 | 0,805567985 | 1,02259887 |
| PSMA1 | 0,013583166 | 0,343375432 | 0,377 | 0,373 | 1 | Secretory | 1,268721515 | 0,788195036 | 1,010723861 |
| NDUFC1 | 0,013630723 | 0,434753236 | 0,547 | 0,614 | 1 | Secretory | 1,351679617 | 0,739820285 | 0,890879479 |
| PRDX3 | 0,013630991 | 0,330890751 | 0,45 | 0,483 | 1 | Secretory | 1,257789721 | 0,795045454 | 0,931677019 |
| RNF1811 | 0,013747131 | 0,420107911 | 0,468 | 0,498 | 1 | Secretory | 1,338027634 | 0,74736872 | 0,939759036 |
| CFAP298 | 0,013787777 | 0,312743529 | 0,332 | 0,306 | 1 | Secretory | 1,242067457 | 0,805109251 | 1,08496732 |

| Supplementary Table 4 - clusermarker | | | | | | | | | |
|---|---|---|---|---|---|---|---|---|---|
| | **p_val** | **avg_log2FC** | **pct.1** | **pct.2** | **p_val_adj** | **celltype** | **Foldchange_UP** | **Foldchange_DOWN** | **Ratio_pct1_pct2** |
| CHKB1 | 0,013853084 | 0,328808595 | 0,248 | 0,203 | 1 | Secretory | 1,255975737 | 0,796193725 | 1,221674877 |
| PPP1CA1 | 0,013858046 | 0,39711585 | 0,433 | 0,444 | 1 | Secretory | 1,316872665 | 0,759374864 | 0,975225225 |
| NDUFV1 | 0,014000245 | 0,279863297 | 0,399 | 0,386 | 1 | Secretory | 1,214079838 | 0,823669061 | 1,033678756 |
| HNRNPM | 0,014096792 | 0,422713111 | 0,384 | 0,373 | 1 | Secretory | 1,340446009 | 0,746020349 | 1,029490617 |
| PCM1 | 0,014134272 | 0,344030787 | 0,39 | 0,369 | 1 | Secretory | 1,269297973 | 0,787837073 | 1,056910569 |
| LMAN21 | 0,014392251 | 0,384472723 | 0,463 | 0,493 | 1 | Secretory | 1,305382608 | 0,766058927 | 0,939148073 |
| STT3B | 0,014493544 | 0,294881358 | 0,321 | 0,278 | 1 | Secretory | 1,226784087 | 0,815139364 | 1,154676259 |
| TMEM98 | 0,014543067 | 0,378725721 | 0,353 | 0,334 | 1 | Secretory | 1,300192936 | 0,769116623 | 1,056886228 |
| NCOR2 | 0,014586682 | 0,314386941 | 0,278 | 0,241 | 1 | Secretory | 1,243483135 | 0,804192652 | 1,153526971 |
| TAF15 | 0,014628932 | 0,317042247 | 0,328 | 0,296 | 1 | Secretory | 1,245773896 | 0,802713882 | 1,108108108 |
| BSPRY1 | 0,014707151 | 0,259822244 | 0,228 | 0,186 | 1 | Secretory | 1,197331171 | 0,835190818 | 1,225806452 |
| AP2M11 | 0,014764969 | 0,396549803 | 0,414 | 0,423 | 1 | Secretory | 1,316356086 | 0,759672866 | 0,978723404 |
| SORBS3 | 0,014816669 | 0,267344727 | 0,308 | 0,27 | 1 | Secretory | 1,203590586 | 0,83084731 | 1,140740741 |
| CAVIN3 | 0,014948878 | 0,467504614 | 0,28 | 0,243 | 1 | Secretory | 1,382715757 | 0,723214439 | 1,152263374 |
| TBCB1 | 0,015123994 | 0,291180714 | 0,28 | 0,246 | 1 | Secretory | 1,223641306 | 0,817232955 | 1,138211382 |
| XRN2 | 0,015521066 | 0,259996041 | 0,315 | 0,286 | 1 | Secretory | 1,197475418 | 0,835090211 | 1,101398601 |
| ANKRD36B | 0,015636621 | 0,296928496 | 0,103 | 0,065 | 1 | Secretory | 1,22852609 | 0,813983527 | 1,584615385 |
| POLR2F1 | 0,015739286 | 0,297209085 | 0,388 | 0,376 | 1 | Secretory | 1,228765048 | 0,813825232 | 1,031914894 |
| SMCHD1 | 0,015757389 | 0,369782845 | 0,241 | 0,196 | 1 | Secretory | 1,29215832 | 0,773898976 | 1,229591837 |
| GLUL | 0,015829076 | 0,372446267 | 0,586 | 0,637 | 1 | Secretory | 1,294546033 | 0,772471565 | 0,919937206 |
| CFAP97 | 0,015992635 | 0,329471986 | 0,248 | 0,208 | 1 | Secretory | 1,256553403 | 0,795827696 | 1,192307692 |
| NDUFV21 | 0,015996717 | 0,393047414 | 0,444 | 0,458 | 1 | Secretory | 1,313164281 | 0,761519342 | 0,969432314 |
| HAUS41 | 0,015999021 | 0,294710323 | 0,218 | 0,178 | 1 | Secretory | 1,226638657 | 0,815236006 | 1,224719101 |
| SMIM261 | 0,016180783 | 0,404416957 | 0,489 | 0,536 | 1 | Secretory | 1,323553908 | 0,755541572 | 0,912313433 |
| LSM41 | 0,016225499 | 0,313580176 | 0,336 | 0,309 | 1 | Secretory | 1,242787965 | 0,804642488 | 1,087378641 |
| NDUFS31 | 0,016499935 | 0,365345805 | 0,362 | 0,354 | 1 | Secretory | 1,288190364 | 0,776282782 | 1,02259887 |
| HMG20B | 0,01690712 | 0,328847949 | 0,302 | 0,271 | 1 | Secretory | 1,256009998 | 0,796172006 | 1,114391144 |
| DDX39B | 0,01705955 | 0,39551552 | 0,42 | 0,424 | 1 | Secretory | 1,315412715 | 0,760217678 | 0,990566038 |
| GPAA11 | 0,017224479 | 0,265473766 | 0,345 | 0,328 | 1 | Secretory | 1,202030719 | 0,831925494 | 1,051829268 |
| RSBN1L | 0,017514161 | 0,287069802 | 0,328 | 0,303 | 1 | Secretory | 1,220159544 | 0,819564954 | 1,082508251 |
| OLA1 | 0,017554729 | 0,278976913 | 0,317 | 0,29 | 1 | Secretory | 1,213334143 | 0,824175274 | 1,093103448 |

| Supplementary Table 4 - clusermarker | | | | | | | | | |
|---|---|---|---|---|---|---|---|---|---|
| | **p_val** | **avg_log2FC** | **pct.1** | **pct.2** | **p_val_adj** | **celltype** | **Foldchange_UP** | **Foldchange_DOWN** | **Ratio_pct1_pct2** |
| IK | 0,017564094 | 0,32122793 | 0,36 | 0,338 | 1 | Secretory | 1,249393501 | 0,800388348 | 1,065088757 |
| CEBPB | 0,017761721 | 0,419559134 | 0,455 | 0,464 | 1 | Secretory | 1,337518767 | 0,747653061 | 0,980603448 |
| FLII | 0,018090625 | 0,266285524 | 0,289 | 0,253 | 1 | Secretory | 1,202707253 | 0,831457528 | 1,14229249 |
| WDR11 | 0,018099322 | 0,457869747 | 0,362 | 0,349 | 1 | Secretory | 1,373512221 | 0,728060504 | 1,037249284 |
| PTGES3 | 0,018131169 | 0,361572111 | 0,455 | 0,476 | 1 | Secretory | 1,284825214 | 0,778315983 | 0,955882353 |
| LUC7L31 | 0,018387253 | 0,523655849 | 0,366 | 0,338 | 1 | Secretory | 1,437593558 | 0,695606901 | 1,082840237 |
| CXCL81 | 0,018400809 | -0,49013237 | 0,088 | 0,133 | 1 | Secretory | 0,711959771 | 1,404573742 | 0,661654135 |
| MRPL54 | 0,018408212 | 0,290838207 | 0,377 | 0,376 | 1 | Secretory | 1,223350839 | 0,817426995 | 1,002659574 |
| ZYX | 0,018880062 | 0,27879114 | 0,237 | 0,195 | 1 | Secretory | 1,213177915 | 0,824281408 | 1,215384615 |
| IDS1 | 0,019154846 | 0,256517024 | 0,267 | 0,228 | 1 | Secretory | 1,19459122 | 0,837106437 | 1,171052632 |
| SNX6 | 0,019166667 | 0,427950544 | 0,384 | 0,378 | 1 | Secretory | 1,34532109 | 0,743316973 | 1,015873016 |
| ATP5F1C1 | 0,019189839 | 0,466508256 | 0,468 | 0,514 | 1 | Secretory | 1,381761152 | 0,723714079 | 0,910505837 |
| NDUFS21 | 0,019250935 | 0,324822734 | 0,319 | 0,296 | 1 | Secretory | 1,252510532 | 0,79839648 | 1,077702703 |
| SFPQ1 | 0,019417196 | 0,494131672 | 0,47 | 0,491 | 1 | Secretory | 1,408472769 | 0,709988877 | 0,957230143 |
| SPAG91 | 0,019497372 | 0,551243209 | 0,269 | 0,231 | 1 | Secretory | 1,465347882 | 0,682431805 | 1,164502165 |
| LDOC1 | 0,019566936 | 0,314063301 | 0,276 | 0,241 | 1 | Secretory | 1,243204216 | 0,804373077 | 1,145228216 |
| FCGBP1 | 0,019765758 | 1,51634739 | 0,435 | 0,443 | 1 | Secretory | 2,860658716 | 0,349569837 | 0,981941309 |
| UBA3 | 0,019865485 | 0,267067814 | 0,233 | 0,186 | 1 | Secretory | 1,203359589 | 0,831006799 | 1,252688172 |
| TUBB4B | 0,019867179 | 0,280601899 | 0,509 | 0,536 | 1 | Secretory | 1,214701558 | 0,823247483 | 0,949626866 |
| EIF4A11 | 0,019877128 | 0,384299182 | 0,582 | 0,631 | 1 | Secretory | 1,305225593 | 0,766151081 | 0,922345483 |
| SDHD | 0,020041524 | 0,457684201 | 0,401 | 0,413 | 1 | Secretory | 1,373335584 | 0,728154147 | 0,97094431 |
| SSB | 0,020452612 | 0,376814075 | 0,375 | 0,366 | 1 | Secretory | 1,298471254 | 0,770136418 | 1,024590164 |
| SNHG6 | 0,020516288 | 0,490748154 | 0,433 | 0,459 | 1 | Secretory | 1,405173382 | 0,711655951 | 0,94335512 |
| CDKN1C | 0,020644041 | 0,392460455 | 0,366 | 0,349 | 1 | Secretory | 1,312630131 | 0,761829229 | 1,048710602 |
| DNPH11 | 0,020645167 | 0,329618429 | 0,422 | 0,434 | 1 | Secretory | 1,256680957 | 0,795746919 | 0,97235023 |
| EPS8L11 | 0,020917043 | 0,258852053 | 0,196 | 0,153 | 1 | Secretory | 1,196526254 | 0,83575266 | 1,281045752 |
| DARS1 | 0,021116405 | 0,284834388 | 0,328 | 0,3 | 1 | Secretory | 1,218270406 | 0,82083583 | 1,093333333 |
| PABPC4 | 0,021376055 | 0,29688761 | 0,287 | 0,25 | 1 | Secretory | 1,228491274 | 0,814006596 | 1,148 |
| CCT3 | 0,021382587 | 0,435632804 | 0,364 | 0,359 | 1 | Secretory | 1,352503947 | 0,739369377 | 1,013927577 |
| SPCS11 | 0,021955464 | 0,305218178 | 0,593 | 0,684 | 1 | Secretory | 1,235605483 | 0,809319814 | 0,866959064 |
| DNAJA2 | 0,021990277 | 0,299476404 | 0,293 | 0,26 | 1 | Secretory | 1,230697676 | 0,81254724 | 1,126923077 |

| Supplementary Table 4 - clusermarker | | | | | | | | | |
|---|---|---|---|---|---|---|---|---|---|
| | **p_val** | **avg_log2FC** | **pct.1** | **pct.2** | **p_val_adj** | **celltype** | **Foldchange_UP** | **Foldchange_DOWN** | **Ratio_pct1_pct2** |
| SNRPF | 0,022184215 | 0,349534347 | 0,353 | 0,343 | 1 | Secretory | 1,274149309 | 0,784837376 | 1,029154519 |
| BLOC1S11 | 0,022474321 | 0,423734319 | 0,483 | 0,531 | 1 | Secretory | 1,341395176 | 0,745492468 | 0,90960452 |
| CALCOCO2 | 0,022507065 | 0,257973434 | 0,319 | 0,288 | 1 | Secretory | 1,195797777 | 0,836261799 | 1,107638889 |
| AHCY | 0,022625202 | 0,280655713 | 0,332 | 0,313 | 1 | Secretory | 1,214746868 | 0,823216776 | 1,060702875 |
| CRACR2B | 0,022933194 | 0,357880608 | 0,504 | 0,554 | 1 | Secretory | 1,281541864 | 0,780310053 | 0,909747292 |
| C1RL | 0,022958763 | 0,309824893 | 0,222 | 0,185 | 1 | Secretory | 1,239557239 | 0,806739671 | 1,2 |
| SRSF2 | 0,022982995 | 0,443854872 | 0,418 | 0,436 | 1 | Secretory | 1,360234013 | 0,735167619 | 0,958715596 |
| EIF61 | 0,023353341 | 0,321539875 | 0,356 | 0,344 | 1 | Secretory | 1,249663678 | 0,800215304 | 1,034883721 |
| SCAND1 | 0,023520466 | 0,365088406 | 0,435 | 0,464 | 1 | Secretory | 1,287960551 | 0,776421296 | 0,9375 |
| LDHA1 | 0,023567048 | 0,520694058 | 0,511 | 0,559 | 1 | Secretory | 1,434645268 | 0,697036419 | 0,914132379 |
| CIB11 | 0,023573955 | 0,294723774 | 0,476 | 0,509 | 1 | Secretory | 1,226650094 | 0,815228405 | 0,935166994 |
| LAMTOR51 | 0,023654915 | 0,364374562 | 0,397 | 0,406 | 1 | Secretory | 1,287323427 | 0,776805563 | 0,977832512 |
| DST1 | 0,023857312 | 0,4583962 | 0,39 | 0,368 | 1 | Secretory | 1,37401352 | 0,727794877 | 1,059782609 |
| TMEM123 | 0,024042522 | 0,431856785 | 0,53 | 0,607 | 1 | Secretory | 1,348968617 | 0,74130709 | 0,873146623 |
| PDCD61 | 0,024051816 | 0,334779561 | 0,399 | 0,401 | 1 | Secretory | 1,26118469 | 0,79290528 | 0,995012469 |
| PRPF61 | 0,024173713 | 0,33459102 | 0,315 | 0,29 | 1 | Secretory | 1,26101988 | 0,793008909 | 1,086206897 |
| NOA1 | 0,024237843 | 0,254755589 | 0,162 | 0,123 | 1 | Secretory | 1,193133594 | 0,838129112 | 1,317073171 |
| SF3A2 | 0,02441393 | 0,264415823 | 0,228 | 0,186 | 1 | Secretory | 1,201149581 | 0,832535777 | 1,225806452 |
| MRPS7 | 0,024633211 | 0,26365296 | 0,306 | 0,278 | 1 | Secretory | 1,200514609 | 0,832976119 | 1,100719424 |
| TAPBP | 0,025146508 | 0,413812738 | 0,397 | 0,401 | 1 | Secretory | 1,332201894 | 0,750636975 | 0,990024938 |
| PA2G4 | 0,025155965 | 0,386851826 | 0,394 | 0,399 | 1 | Secretory | 1,307537049 | 0,764796684 | 0,987468672 |
| YIF1A1 | 0,025217444 | 0,344430986 | 0,353 | 0,344 | 1 | Secretory | 1,26965012 | 0,787618561 | 1,026162791 |
| PNN | 0,02542614 | 0,303355462 | 0,358 | 0,341 | 1 | Secretory | 1,234011177 | 0,810365431 | 1,049853372 |
| PRDX1 | 0,025616959 | 0,277895204 | 0,623 | 0,699 | 1 | Secretory | 1,212424747 | 0,824793458 | 0,891273247 |
| TADA3 | 0,025677499 | 0,304694852 | 0,235 | 0,2 | 1 | Secretory | 1,235157359 | 0,809613442 | 1,175 |
| MLF2 | 0,026040059 | 0,351305479 | 0,431 | 0,456 | 1 | Secretory | 1,275714486 | 0,783874457 | 0,945175439 |
| TMA7 | 0,026187646 | 0,342832749 | 0,608 | 0,684 | 1 | Secretory | 1,268244363 | 0,788491578 | 0,888888889 |
| ADK1 | 0,026269201 | 0,262569088 | 0,228 | 0,193 | 1 | Secretory | 1,199613022 | 0,833602155 | 1,18134715 |
| NDUFA81 | 0,026388594 | 0,307519836 | 0,33 | 0,304 | 1 | Secretory | 1,237578326 | 0,808029665 | 1,085526316 |
| GGNBP2 | 0,026434354 | 0,336401042 | 0,3 | 0,27 | 1 | Secretory | 1,262602963 | 0,792014615 | 1,111111111 |
| ERP291 | 0,026597717 | 0,383655136 | 0,545 | 0,616 | 1 | Secretory | 1,304643046 | 0,766493182 | 0,88474026 |

| Supplementary Table 4 - clusermarker | | | | | | | | | |
|---|---|---|---|---|---|---|---|---|---|
| | **p_val** | **avg_log2FC** | **pct.1** | **pct.2** | **p_val_adj** | **celltype** | **Foldchange_UP** | **Foldchange_DOWN** | **Ratio_pct1_pct2** |
| SINHCAF | 0,027468228 | 0,34790048 | 0,237 | 0,201 | 1 | Secretory | 1,272707138 | 0,785726716 | 1,179104478 |
| POLR2L | 0,02793329 | 0,251442974 | 0,64 | 0,712 | 1 | Secretory | 1,190397147 | 0,840055777 | 0,898876404 |
| ACSL1 | 0,027945319 | 0,388522868 | 0,304 | 0,281 | 1 | Secretory | 1,309052418 | 0,76391135 | 1,081850534 |
| CFLAR | 0,028376668 | 0,34043371 | 0,349 | 0,333 | 1 | Secretory | 1,266137169 | 0,789803842 | 1,048048048 |
| SUPT16H1 | 0,028378834 | 0,294989494 | 0,246 | 0,208 | 1 | Secretory | 1,226876043 | 0,815078268 | 1,182692308 |
| NDFIP11 | 0,028590797 | 0,400251431 | 0,519 | 0,577 | 1 | Secretory | 1,319737893 | 0,757726216 | 0,899480069 |
| AKAP13 | 0,028995472 | 0,376718068 | 0,366 | 0,361 | 1 | Secretory | 1,298384848 | 0,77018767 | 1,013850416 |
| DEK | 0,029920367 | 0,556071476 | 0,416 | 0,446 | 1 | Secretory | 1,470260176 | 0,680151728 | 0,932735426 |
| JAK1 | 0,029928369 | 0,343599425 | 0,347 | 0,331 | 1 | Secretory | 1,268918513 | 0,78807267 | 1,048338369 |
| MYG11 | 0,030001665 | 0,258801168 | 0,304 | 0,276 | 1 | Secretory | 1,196484053 | 0,835782138 | 1,101449275 |
| SLTM | 0,030086635 | 0,372455259 | 0,362 | 0,354 | 1 | Secretory | 1,294554102 | 0,77246675 | 1,02259887 |
| PSMD9 | 0,030268756 | 0,293395129 | 0,332 | 0,316 | 1 | Secretory | 1,225520935 | 0,815979533 | 1,050632911 |
| NHP21 | 0,030517788 | 0,305259631 | 0,403 | 0,418 | 1 | Secretory | 1,235640986 | 0,80929656 | 0,964114833 |
| AK21 | 0,031192554 | 0,353001804 | 0,33 | 0,306 | 1 | Secretory | 1,277215356 | 0,782953317 | 1,078431373 |
| HOTAIRM1 | 0,031222154 | 0,354049361 | 0,302 | 0,283 | 1 | Secretory | 1,278143093 | 0,782385012 | 1,067137809 |
| MRPS341 | 0,031313095 | 0,396665539 | 0,412 | 0,434 | 1 | Secretory | 1,31646169 | 0,759611926 | 0,949308756 |
| ST6GALNAC1 | 0,031456941 | -0,288542367 | 0,095 | 0,136 | 1 | Secretory | 0,818728847 | 1,221405602 | 0,698529412 |
| PSMA5 | 0,031462106 | 0,366067419 | 0,323 | 0,304 | 1 | Secretory | 1,288834858 | 0,775894595 | 1,0625 |
| NDUFB5 | 0,031722977 | 0,316182149 | 0,366 | 0,359 | 1 | Secretory | 1,245031418 | 0,803192582 | 1,019498607 |
| RABAC11 | 0,032098415 | 0,379215521 | 0,55 | 0,611 | 1 | Secretory | 1,300634431 | 0,768855549 | 0,900163666 |
| UBE2D21 | 0,032238049 | 0,300176033 | 0,399 | 0,404 | 1 | Secretory | 1,231294643 | 0,812153294 | 0,987623762 |
| YWHAQ | 0,03270803 | 0,422545049 | 0,431 | 0,451 | 1 | Secretory | 1,340289867 | 0,74610726 | 0,955654102 |
| SPTAN1 | 0,032798919 | 0,448745786 | 0,364 | 0,356 | 1 | Secretory | 1,364853199 | 0,73267953 | 1,02247191 |
| IPO7 | 0,033091637 | 0,355304237 | 0,334 | 0,319 | 1 | Secretory | 1,279255324 | 0,781704779 | 1,047021944 |
| ADAR | 0,033250967 | 0,345352416 | 0,315 | 0,285 | 1 | Secretory | 1,270461288 | 0,78711568 | 1,105263158 |
| PSENEN | 0,033280247 | 0,379334423 | 0,325 | 0,311 | 1 | Secretory | 1,300741629 | 0,768792186 | 1,045016077 |
| CHMP1B | 0,033721369 | 0,298829188 | 0,263 | 0,225 | 1 | Secretory | 1,23014569 | 0,812911843 | 1,168888889 |
| SMARCA4 | 0,033808611 | 0,348151623 | 0,261 | 0,228 | 1 | Secretory | 1,272928709 | 0,785589949 | 1,144736842 |
| PRR131 | 0,034350615 | 0,44650119 | 0,418 | 0,434 | 1 | Secretory | 1,362731364 | 0,733820345 | 0,963133641 |
| RPS17 | 0,034441918 | 0,36663129 | 0,616 | 0,699 | 1 | Secretory | 1,289338692 | 0,775591399 | 0,881258941 |
| SLC66A2 | 0,035035656 | 0,290484666 | 0,235 | 0,206 | 1 | Secretory | 1,223051086 | 0,817627335 | 1,140776699 |

| Supplementary Table 4 - clusermarker | | | | | | | | | |
|---|---|---|---|---|---|---|---|---|---|
| | **p_val** | **avg_log2FC** | **pct.1** | **pct.2** | **p_val_adj** | **celltype** | **Foldchange_UP** | **Foldchange_DOWN** | **Ratio_pct1_pct2** |
| DYNLT11 | 0,035256046 | 0,409131102 | 0,457 | 0,484 | 1 | Secretory | 1,327885822 | 0,753076796 | 0,944214876 |
| BCAT2 | 0,035614066 | 0,262119203 | 0,239 | 0,205 | 1 | Secretory | 1,199238997 | 0,833862143 | 1,165853659 |
| RABGAP1L | 0,035878001 | 0,269522382 | 0,284 | 0,253 | 1 | Secretory | 1,2054087 | 0,829594145 | 1,122529644 |
| NUMA1 | 0,035973408 | 0,274498411 | 0,31 | 0,288 | 1 | Secretory | 1,209573478 | 0,826737704 | 1,076388889 |
| HNRNPA11 | 0,036234833 | 0,343844505 | 0,623 | 0,719 | 1 | Secretory | 1,26913409 | 0,787938806 | 0,866481224 |
| HNRNPF | 0,036929235 | 0,348957235 | 0,403 | 0,404 | 1 | Secretory | 1,273639721 | 0,785151392 | 0,997524752 |
| ARIH2 | 0,037170454 | 0,27333311 | 0,25 | 0,216 | 1 | Secretory | 1,208596869 | 0,827405751 | 1,157407407 |
| COMT1 | 0,037400537 | 0,423966228 | 0,466 | 0,511 | 1 | Secretory | 1,341610819 | 0,745372642 | 0,911937378 |
| RTL8A | 0,037676873 | 0,263189182 | 0,304 | 0,281 | 1 | Secretory | 1,200128746 | 0,833243936 | 1,081850534 |
| SELENBP11 | 0,037747812 | 0,432383759 | 0,405 | 0,413 | 1 | Secretory | 1,349461445 | 0,741036362 | 0,98062954 |
| AP2S11 | 0,037875162 | 0,319139366 | 0,407 | 0,418 | 1 | Secretory | 1,247586084 | 0,801547895 | 0,973684211 |
| SNRPD11 | 0,038564599 | 0,323885543 | 0,317 | 0,301 | 1 | Secretory | 1,251697151 | 0,798915296 | 1,053156146 |
| EIF3M1 | 0,039508144 | 0,302429695 | 0,384 | 0,389 | 1 | Secretory | 1,233219575 | 0,810885604 | 0,98714653 |
| BCAP311 | 0,039620449 | 0,296729555 | 0,478 | 0,529 | 1 | Secretory | 1,228356693 | 0,81409578 | 0,903591682 |
| PSME21 | 0,039747199 | 0,405304512 | 0,414 | 0,433 | 1 | Secretory | 1,324368418 | 0,755076901 | 0,956120092 |
| MALAT1 | 0,040110315 | 0,329087626 | 0,759 | 0,794 | 1 | Secretory | 1,256218679 | 0,796039747 | 0,955919395 |
| COPS6 | 0,040170055 | 0,274167518 | 0,36 | 0,354 | 1 | Secretory | 1,209296085 | 0,826927344 | 1,016949153 |
| KLHDC2 | 0,040197869 | 0,321690542 | 0,28 | 0,245 | 1 | Secretory | 1,249794193 | 0,800131738 | 1,142857143 |
| RALBP1 | 0,040199036 | 0,359339152 | 0,347 | 0,344 | 1 | Secretory | 1,28283814 | 0,779521569 | 1,00872093 |
| ILF31 | 0,040404939 | 0,415870352 | 0,332 | 0,316 | 1 | Secretory | 1,334103275 | 0,749567158 | 1,050632911 |
| FDPS | 0,040453017 | 0,359018334 | 0,407 | 0,419 | 1 | Secretory | 1,282552901 | 0,779694934 | 0,971360382 |
| KMT2E | 0,040513825 | 0,390996188 | 0,388 | 0,384 | 1 | Secretory | 1,31129855 | 0,762602841 | 1,010416667 |
| ZNF5931 | 0,040534717 | 0,291860511 | 0,289 | 0,26 | 1 | Secretory | 1,224218022 | 0,816847965 | 1,111538462 |
| PRSS221 | 0,04078469 | 0,401584283 | 0,254 | 0,225 | 1 | Secretory | 1,320957713 | 0,757026505 | 1,128888889 |
| MYH9 | 0,040873014 | 0,466795536 | 0,468 | 0,506 | 1 | Secretory | 1,382036325 | 0,723569983 | 0,924901186 |
| ACAA11 | 0,041087108 | 0,315148045 | 0,295 | 0,278 | 1 | Secretory | 1,244139317 | 0,803768506 | 1,061151079 |
| AKR7A2 | 0,041144687 | 0,344876993 | 0,321 | 0,314 | 1 | Secretory | 1,270042691 | 0,787375107 | 1,022292994 |
| SORBS2 | 0,041222238 | 0,438710901 | 0,42 | 0,441 | 1 | Secretory | 1,355392695 | 0,737793559 | 0,952380952 |
| MRPS161 | 0,04138439 | 0,260078461 | 0,304 | 0,278 | 1 | Secretory | 1,197543831 | 0,835042504 | 1,09352518 |
| B4GALT1 | 0,041421171 | 0,326160033 | 0,362 | 0,346 | 1 | Secretory | 1,253672078 | 0,797656754 | 1,046242775 |
| CCDC90B | 0,041527907 | 0,33387775 | 0,295 | 0,266 | 1 | Secretory | 1,260396585 | 0,793401071 | 1,109022556 |

| Supplementary Table 4 - clusermarker | | | | | | | | | |
|---|---|---|---|---|---|---|---|---|---|
| | **p_val** | **avg_log2FC** | **pct.1** | **pct.2** | **p_val_adj** | **celltype** | **Foldchange_UP** | **Foldchange_DOWN** | **Ratio_pct1_pct2** |
| ZMYM4 | 0,041923662 | 0,313414017 | 0,244 | 0,213 | 1 | Secretory | 1,242644839 | 0,804735166 | 1,145539906 |
| TMEM2591 | 0,041948812 | 0,358530555 | 0,3 | 0,278 | 1 | Secretory | 1,28211934 | 0,779958596 | 1,079136691 |
| NME4 | 0,041971343 | 0,389395607 | 0,427 | 0,448 | 1 | Secretory | 1,309844551 | 0,763449372 | 0,953125 |
| GLS1 | 0,042108239 | 0,271176847 | 0,216 | 0,181 | 1 | Secretory | 1,20679184 | 0,828643322 | 1,193370166 |
| ARPC5L1 | 0,042343963 | 0,261552464 | 0,306 | 0,285 | 1 | Secretory | 1,198767988 | 0,834189776 | 1,073684211 |
| DMBT1 | 0,042675058 | -0,514059111 | 0,371 | 0,418 | 1 | Secretory | 0,700249469 | 1,42806249 | 0,887559809 |
| PDGFC | 0,043195737 | 0,282620051 | 0,233 | 0,2 | 1 | Secretory | 1,216401964 | 0,822096667 | 1,165 |
| HNRNPDL | 0,043788468 | 0,321360918 | 0,457 | 0,488 | 1 | Secretory | 1,249508675 | 0,800314571 | 0,93647541 |
| ARGLU1 | 0,044228089 | 0,414745873 | 0,418 | 0,429 | 1 | Secretory | 1,33306384 | 0,75015162 | 0,974358974 |
| ANXA7 | 0,045194495 | 0,305273387 | 0,397 | 0,403 | 1 | Secretory | 1,235652768 | 0,809288844 | 0,985111663 |
| ARID1A | 0,045521762 | 0,258913172 | 0,278 | 0,255 | 1 | Secretory | 1,196576945 | 0,835717255 | 1,090196078 |
| MAML2 | 0,045856555 | 0,418727885 | 0,31 | 0,286 | 1 | Secretory | 1,33674834 | 0,748083966 | 1,083916084 |
| RER11 | 0,045958868 | 0,425699221 | 0,403 | 0,426 | 1 | Secretory | 1,343223355 | 0,744477823 | 0,94600939 |
| TOX4 | 0,045992364 | 0,281181986 | 0,312 | 0,291 | 1 | Secretory | 1,21519007 | 0,822916533 | 1,072164948 |
| TBC1D5 | 0,046270946 | 0,289782547 | 0,2 | 0,166 | 1 | Secretory | 1,222456006 | 0,818025348 | 1,204819277 |
| MTRNR2L81 | 0,046457231 | -0,309361617 | 0,412 | 0,329 | 1 | Secretory | 0,806998772 | 1,239159259 | 1,252279635 |
| SAT11 | 0,047599636 | 0,318837119 | 0,759 | 0,777 | 1 | Secretory | 1,24732474 | 0,801715838 | 0,976833977 |
| RBM39 | 0,048270825 | 0,383279324 | 0,491 | 0,532 | 1 | Secretory | 1,30430324 | 0,766692874 | 0,922932331 |
| ESYT21 | 0,048355581 | 0,306781885 | 0,248 | 0,218 | 1 | Secretory | 1,236945456 | 0,808443084 | 1,137614679 |
| HNRNPD | 0,048452034 | 0,470321051 | 0,466 | 0,507 | 1 | Secretory | 1,385417738 | 0,721803953 | 0,91913215 |
| CCT2 | 0,048551875 | 0,353725919 | 0,343 | 0,331 | 1 | Secretory | 1,277856575 | 0,782560437 | 1,036253776 |
| TMOD3 | 0,048782582 | 0,37285426 | 0,315 | 0,306 | 1 | Secretory | 1,294912181 | 0,772253141 | 1,029411765 |
| ZBTB7A1 | 0,049095262 | 0,344881559 | 0,334 | 0,319 | 1 | Secretory | 1,270046711 | 0,787372615 | 1,047021944 |
| FBXW51 | 0,049342193 | 0,335006907 | 0,328 | 0,314 | 1 | Secretory | 1,261383448 | 0,792780341 | 1,044585987 |
| UFC11 | 0,049679485 | 0,310279915 | 0,483 | 0,541 | 1 | Secretory | 1,239948255 | 0,806485268 | 0,892791128 |
| DBNL | 0,049695666 | 0,27651234 | 0,205 | 0,171 | 1 | Secretory | 1,21126316 | 0,825584425 | 1,198830409 |
| NRDC | 0,050003698 | 0,288201417 | 0,282 | 0,256 | 1 | Secretory | 1,221116983 | 0,818922359 | 1,1015625 |
| ILK1 | 0,05008665 | 0,356155905 | 0,323 | 0,313 | 1 | Secretory | 1,28001073 | 0,781243451 | 1,031948882 |
| RAB24 | 0,050134164 | 0,258278532 | 0,235 | 0,205 | 1 | Secretory | 1,196050688 | 0,836084967 | 1,146341463 |
| GPI | 0,050328998 | 0,355361827 | 0,366 | 0,373 | 1 | Secretory | 1,27930639 | 0,781673575 | 0,981233244 |
| NSD3 | 0,051623349 | 0,277861256 | 0,323 | 0,308 | 1 | Secretory | 1,212396217 | 0,824812867 | 1,048701299 |

| Supplementary Table 4 - clusermarker | | | | | | | | | |
|---|---|---|---|---|---|---|---|---|---|
| | **p_val** | **avg_log2FC** | **pct.1** | **pct.2** | **p_val_adj** | **celltype** | **Foldchange_UP** | **Foldchange_DOWN** | **Ratio_pct1_pct2** |
| PTRHD11 | 0,052306459 | 0,343722132 | 0,397 | 0,419 | 1 | Secretory | 1,269026444 | 0,788005644 | 0,947494033 |
| NDUFA61 | 0,052426309 | 0,281064001 | 0,442 | 0,471 | 1 | Secretory | 1,215090695 | 0,822983835 | 0,938428875 |
| ARL2 | 0,052611049 | 0,307222311 | 0,306 | 0,293 | 1 | Secretory | 1,237323128 | 0,808196321 | 1,044368601 |
| ADSL1 | 0,052915014 | 0,270272236 | 0,218 | 0,183 | 1 | Secretory | 1,206035384 | 0,829163069 | 1,191256831 |
| MRPL271 | 0,052948242 | 0,35157542 | 0,308 | 0,296 | 1 | Secretory | 1,275953205 | 0,783727801 | 1,040540541 |
| WASL1 | 0,054371534 | 0,353472301 | 0,315 | 0,309 | 1 | Secretory | 1,277631955 | 0,782698019 | 1,019417476 |
| RNF19A | 0,056256475 | 0,365748648 | 0,188 | 0,156 | 1 | Secretory | 1,288550115 | 0,776066052 | 1,205128205 |
| FRMD4B | 0,056261391 | 0,274158599 | 0,226 | 0,296 | 1 | Secretory | 1,209288609 | 0,826932457 | 0,763513514 |
| JMJD1C | 0,057341055 | 0,282918387 | 0,384 | 0,378 | 1 | Secretory | 1,216653531 | 0,821926682 | 1,015873016 |
| PSMC5 | 0,057862926 | 0,33220344 | 0,407 | 0,426 | 1 | Secretory | 1,258934688 | 0,794322382 | 0,955399061 |
| SEPTIN9 | 0,058139009 | 0,256619723 | 0,317 | 0,303 | 1 | Secretory | 1,194676261 | 0,837046849 | 1,04620462 |
| MSN | 0,058737785 | 0,334085099 | 0,246 | 0,22 | 1 | Secretory | 1,260577746 | 0,793287049 | 1,118181818 |
| CASC3 | 0,059106543 | 0,258589838 | 0,248 | 0,221 | 1 | Secretory | 1,196308801 | 0,835904575 | 1,122171946 |
| EIF3L | 0,059724878 | 0,338569758 | 0,528 | 0,596 | 1 | Secretory | 1,264502385 | 0,790824922 | 0,88590604 |
| AUP11 | 0,059825573 | 0,360523387 | 0,427 | 0,449 | 1 | Secretory | 1,283891589 | 0,778881962 | 0,951002227 |
| EIF1B | 0,059827924 | 0,306220177 | 0,349 | 0,346 | 1 | Secretory | 1,23646395 | 0,80875791 | 1,00867052 |
| IVNS1ABP | 0,060024556 | 0,361778875 | 0,438 | 0,446 | 1 | Secretory | 1,285009366 | 0,778204445 | 0,98206278 |
| RNF167 | 0,060207614 | 0,311394471 | 0,332 | 0,316 | 1 | Secretory | 1,240906548 | 0,805862457 | 1,050632911 |
| UBL51 | 0,060627321 | 0,292004307 | 0,61 | 0,719 | 1 | Secretory | 1,224340047 | 0,816766553 | 0,848400556 |
| PDZK1IP11 | 0,06090575 | 0,526004116 | 0,463 | 0,511 | 1 | Secretory | 1,439935426 | 0,694475587 | 0,906066536 |
| NDUFA12 | 0,062065679 | 0,364253926 | 0,381 | 0,399 | 1 | Secretory | 1,287215788 | 0,776870521 | 0,954887218 |
| SUMF1 | 0,062130531 | 0,271225037 | 0,252 | 0,225 | 1 | Secretory | 1,206832151 | 0,828615644 | 1,12 |
| HOOK2 | 0,062195507 | 0,393932735 | 0,416 | 0,433 | 1 | Secretory | 1,313970362 | 0,761052173 | 0,96073903 |
| CDC73 | 0,062226861 | 0,286090883 | 0,209 | 0,176 | 1 | Secretory | 1,219331904 | 0,820121246 | 1,1875 |
| PDK4 | 0,063227485 | 0,26301884 | 0,192 | 0,251 | 1 | Secretory | 1,199987053 | 0,833342324 | 0,764940239 |
| LSM14A | 0,063282792 | 0,307800508 | 0,3 | 0,278 | 1 | Secretory | 1,237819117 | 0,80787248 | 1,079136691 |
| ST141 | 0,063286904 | 0,258725006 | 0,306 | 0,283 | 1 | Secretory | 1,19642089 | 0,835826262 | 1,081272085 |
| SF3B21 | 0,063473521 | 0,319196248 | 0,442 | 0,484 | 1 | Secretory | 1,247635275 | 0,801516293 | 0,91322314 |
| GLG1 | 0,063573931 | 0,49470409 | 0,403 | 0,431 | 1 | Secretory | 1,40903172 | 0,709707231 | 0,935034803 |
| PTPN181 | 0,063971344 | 0,327094366 | 0,325 | 0,323 | 1 | Secretory | 1,254484257 | 0,797140335 | 1,00619195 |
| TAF1D | 0,065400552 | 0,410873141 | 0,356 | 0,359 | 1 | Secretory | 1,329490198 | 0,752168012 | 0,991643454 |

| Supplementary Table 4 - clusermarker | | | | | | | | | |
|---|---|---|---|---|---|---|---|---|---|
| | **p_val** | **avg_log2FC** | **pct.1** | **pct.2** | **p_val_adj** | **celltype** | **Foldchange_UP** | **Foldchange_DOWN** | **Ratio_pct1_pct2** |
| CSNK1D1 | 0,065465362 | 0,329272421 | 0,261 | 0,238 | 1 | Secretory | 1,256379598 | 0,795937789 | 1,096638655 |
| MRPL55 | 0,066349387 | 0,325845144 | 0,321 | 0,313 | 1 | Secretory | 1,253398476 | 0,797830873 | 1,025559105 |
| HNRNPR1 | 0,06648288 | 0,365984251 | 0,343 | 0,341 | 1 | Secretory | 1,288760562 | 0,775939325 | 1,005865103 |
| DAZAP2 | 0,066670336 | 0,390540257 | 0,466 | 0,519 | 1 | Secretory | 1,310884209 | 0,762843883 | 0,897880539 |
| LAP31 | 0,067014745 | 0,43553405 | 0,274 | 0,251 | 1 | Secretory | 1,352411369 | 0,739419989 | 1,091633466 |
| LSR1 | 0,067287034 | 0,257367833 | 0,377 | 0,376 | 1 | Secretory | 1,195295921 | 0,836612911 | 1,002659574 |
| NAXE1 | 0,067318551 | 0,40848375 | 0,369 | 0,386 | 1 | Secretory | 1,327290119 | 0,753414785 | 0,955958549 |
| GSN1 | 0,067602283 | 0,292537758 | 0,481 | 0,506 | 1 | Secretory | 1,224792843 | 0,816464601 | 0,950592885 |
| RALGAPA1 | 0,068155644 | 0,360540585 | 0,211 | 0,178 | 1 | Secretory | 1,283906894 | 0,778872677 | 1,185393258 |
| IRF9 | 0,068543396 | 0,277590126 | 0,25 | 0,228 | 1 | Secretory | 1,21216839 | 0,824967891 | 1,096491228 |
| HSPA51 | 0,068869456 | 0,319313734 | 0,586 | 0,672 | 1 | Secretory | 1,24773688 | 0,801451024 | 0,87202381 |
| TRIM8 | 0,068978115 | 0,313287671 | 0,332 | 0,323 | 1 | Secretory | 1,242536017 | 0,804805645 | 1,027863777 |
| RBM31 | 0,069229054 | 0,47160502 | 0,541 | 0,634 | 1 | Secretory | 1,386651281 | 0,721161848 | 0,853312303 |
| PYURF1 | 0,06924731 | 0,28698736 | 0,386 | 0,403 | 1 | Secretory | 1,220089821 | 0,819611788 | 0,957816377 |
| MPG | 0,069503485 | 0,332041805 | 0,36 | 0,364 | 1 | Secretory | 1,258793649 | 0,79441138 | 0,989010989 |
| PSMD6 | 0,070426701 | 0,317857629 | 0,3 | 0,278 | 1 | Secretory | 1,246478181 | 0,802260333 | 1,079136691 |
| ATP6V1F1 | 0,070445368 | 0,264240824 | 0,446 | 0,498 | 1 | Secretory | 1,20100389 | 0,83263677 | 0,895582329 |
| PTPRF1 | 0,070688049 | 0,313029221 | 0,409 | 0,431 | 1 | Secretory | 1,242313444 | 0,804949833 | 0,948955916 |
| RAB141 | 0,070797986 | 0,374674584 | 0,379 | 0,389 | 1 | Secretory | 1,296547071 | 0,771279364 | 0,974293059 |
| PRMT21 | 0,07098714 | 0,328445123 | 0,362 | 0,361 | 1 | Secretory | 1,255659348 | 0,796394342 | 1,002770083 |
| MZT2A1 | 0,071370368 | 0,354392351 | 0,405 | 0,434 | 1 | Secretory | 1,278446999 | 0,782199028 | 0,933179724 |
| DDX17 | 0,071927778 | 0,333300988 | 0,528 | 0,577 | 1 | Secretory | 1,259892803 | 0,793718321 | 0,91507799 |
| LAMTOR2 | 0,072135405 | 0,427338819 | 0,403 | 0,424 | 1 | Secretory | 1,344750774 | 0,743632217 | 0,950471698 |
| KIAA1217 | 0,072155359 | 0,39206239 | 0,282 | 0,266 | 1 | Secretory | 1,312268003 | 0,76203946 | 1,060150376 |
| MYL91 | 0,072357329 | 0,388095314 | 0,269 | 0,223 | 1 | Secretory | 1,308664527 | 0,764137775 | 1,206278027 |
| PSMG3 | 0,072567261 | 0,253366022 | 0,205 | 0,176 | 1 | Secretory | 1,191984951 | 0,838936766 | 1,164772727 |
| RAB6A | 0,072837597 | 0,331349136 | 0,317 | 0,301 | 1 | Secretory | 1,25818942 | 0,794792886 | 1,053156146 |
| UFM1 | 0,073200597 | 0,262272542 | 0,379 | 0,488 | 1 | Secretory | 1,199366467 | 0,833773519 | 0,776639344 |
| MPST1 | 0,074569271 | 0,274886714 | 0,356 | 0,359 | 1 | Secretory | 1,209899079 | 0,826515217 | 0,991643454 |
| ZNF5031 | 0,074697339 | 0,359375798 | 0,254 | 0,236 | 1 | Secretory | 1,282870726 | 0,779501769 | 1,076271186 |
| ARIH1 | 0,075030414 | 0,279558618 | 0,248 | 0,326 | 1 | Secretory | 1,213823467 | 0,823843028 | 0,760736196 |

| Supplementary Table 4 - clusermarker | | | | | | | | | |
|---|---|---|---|---|---|---|---|---|---|
| | **p_val** | **avg_log2FC** | **pct.1** | **pct.2** | **p_val_adj** | **celltype** | **Foldchange_UP** | **Foldchange_DOWN** | **Ratio_pct1_pct2** |
| HDDC2 | 0,075372221 | 0,337295833 | 0,353 | 0,358 | 1 | Secretory | 1,2633863 | 0,791523543 | 0,98603352 |
| ARPC1A1 | 0,075815868 | 0,395571836 | 0,375 | 0,379 | 1 | Secretory | 1,315464063 | 0,760188004 | 0,98944591 |
| KRT231 | 0,075896078 | 0,584986713 | 0,345 | 0,348 | 1 | Secretory | 1,500025174 | 0,666655478 | 0,99137931 |
| TMEM14B1 | 0,076258903 | 0,310406163 | 0,481 | 0,539 | 1 | Secretory | 1,240056765 | 0,806414697 | 0,892393321 |
| CSDE1 | 0,076472082 | 0,309812635 | 0,58 | 0,659 | 1 | Secretory | 1,239546708 | 0,806746526 | 0,880121396 |
| PPP1CB1 | 0,076833989 | 0,492238054 | 0,414 | 0,443 | 1 | Secretory | 1,406625283 | 0,710921389 | 0,934537246 |
| TMC41 | 0,077165231 | 0,275067161 | 0,328 | 0,309 | 1 | Secretory | 1,210050419 | 0,826411846 | 1,061488673 |
| CCSER2 | 0,077821826 | 0,363851707 | 0,254 | 0,231 | 1 | Secretory | 1,286856966 | 0,777087141 | 1,0995671 |
| HDAC71 | 0,078346417 | 0,280279619 | 0,213 | 0,185 | 1 | Secretory | 1,214430239 | 0,823431407 | 1,151351351 |
| EZR | 0,07867646 | 0,474064693 | 0,448 | 0,464 | 1 | Secretory | 1,38901742 | 0,719933376 | 0,965517241 |
| H1-0 | 0,079130309 | 0,403739644 | 0,386 | 0,406 | 1 | Secretory | 1,322932676 | 0,755896365 | 0,950738916 |
| ADI11 | 0,079521235 | 0,252475165 | 0,483 | 0,539 | 1 | Secretory | 1,191249133 | 0,839454965 | 0,896103896 |
| EMC4 | 0,079875669 | 0,386119178 | 0,412 | 0,433 | 1 | Secretory | 1,306873207 | 0,765185172 | 0,951501155 |
| AC020916.1 | 0,080292023 | 0,46844224 | 0,261 | 0,24 | 1 | Secretory | 1,383614694 | 0,722744565 | 1,0875 |
| TC2N | 0,080758895 | 0,484138997 | 0,338 | 0,341 | 1 | Secretory | 1,39875084 | 0,71492361 | 0,991202346 |
| UBA11 | 0,080918144 | 0,282385471 | 0,328 | 0,311 | 1 | Secretory | 1,216204195 | 0,822230349 | 1,054662379 |
| HMGA1 | 0,081428476 | 0,303889158 | 0,304 | 0,28 | 1 | Secretory | 1,23446776 | 0,810065708 | 1,085714286 |
| RNPEP | 0,081943658 | 0,270435285 | 0,259 | 0,238 | 1 | Secretory | 1,206171695 | 0,829069364 | 1,088235294 |
| DEPP1 | 0,082421349 | 0,275893287 | 0,172 | 0,143 | 1 | Secretory | 1,210743525 | 0,825938756 | 1,202797203 |
| PSMC3 | 0,082580676 | 0,251009435 | 0,369 | 0,374 | 1 | Secretory | 1,190039479 | 0,840308257 | 0,986631016 |
| PDAP1 | 0,082719782 | 0,279477877 | 0,336 | 0,331 | 1 | Secretory | 1,213755537 | 0,823889136 | 1,01510574 |
| ABLIM1 | 0,082788189 | 0,382502972 | 0,394 | 0,403 | 1 | Secretory | 1,30360155 | 0,767105563 | 0,977667494 |
| TXNL1 | 0,082891798 | 0,303567051 | 0,388 | 0,416 | 1 | Secretory | 1,234192173 | 0,81024659 | 0,932692308 |
| GPR108 | 0,083617584 | 0,375669987 | 0,284 | 0,273 | 1 | Secretory | 1,297441946 | 0,770747395 | 1,04029304 |
| MRPL361 | 0,083725917 | 0,324081391 | 0,25 | 0,228 | 1 | Secretory | 1,251867082 | 0,79880685 | 1,096491228 |
| ATP5F1A | 0,083813337 | 0,353892806 | 0,504 | 0,576 | 1 | Secretory | 1,278004402 | 0,782469918 | 0,875 |
| SLFN51 | 0,084379742 | 0,396807573 | 0,237 | 0,22 | 1 | Secretory | 1,316591303 | 0,759537145 | 1,077272727 |
| SCCPDH | 0,084537569 | 0,311008207 | 0,278 | 0,265 | 1 | Secretory | 1,240574355 | 0,806078246 | 1,049056604 |
| LSM5 | 0,084923301 | 0,297563892 | 0,347 | 0,349 | 1 | Secretory | 1,22906728 | 0,813625109 | 0,994269341 |
| PRMT1 | 0,085714129 | 0,321925678 | 0,31 | 0,303 | 1 | Secretory | 1,249997906 | 0,80000134 | 1,02310231 |
| RPL22L1 | 0,086454082 | 0,355937462 | 0,412 | 0,429 | 1 | Secretory | 1,279816935 | 0,78136175 | 0,96037296 |

| Supplementary Table 4 - clusermarker | | | | | | | | | |
|---|---|---|---|---|---|---|---|---|---|
| | **p_val** | **avg_log2FC** | **pct.1** | **pct.2** | **p_val_adj** | **celltype** | **Foldchange_UP** | **Foldchange_DOWN** | **Ratio_pct1_pct2** |
| SCARB2 | 0,0867772 | 0,370812428 | 0,369 | 0,378 | 1 | Secretory | 1,293080801 | 0,773346878 | 0,976190476 |
| ECHS11 | 0,086789511 | 0,372011598 | 0,466 | 0,524 | 1 | Secretory | 1,294156058 | 0,772704338 | 0,889312977 |
| JTB1 | 0,086873639 | 0,30813826 | 0,522 | 0,591 | 1 | Secretory | 1,238108938 | 0,80768337 | 0,883248731 |
| NBEAL1 | 0,087499234 | 0,256136556 | 0,261 | 0,238 | 1 | Secretory | 1,194276224 | 0,837327228 | 1,096638655 |
| KDM6B | 0,088497749 | 0,273234939 | 0,198 | 0,255 | 1 | Secretory | 1,208514631 | 0,827462055 | 0,776470588 |
| TAF7 | 0,089173554 | 0,342436156 | 0,302 | 0,293 | 1 | Secretory | 1,267895775 | 0,788708362 | 1,030716724 |
| SNRNP200 | 0,08931958 | 0,292570328 | 0,31 | 0,304 | 1 | Secretory | 1,224820495 | 0,816446169 | 1,019736842 |
| COG31 | 0,089808611 | 0,303756108 | 0,282 | 0,263 | 1 | Secretory | 1,234353918 | 0,810140419 | 1,072243346 |
| LEPROT1 | 0,09012068 | 0,255853198 | 0,377 | 0,389 | 1 | Secretory | 1,194041681 | 0,837491703 | 0,969151671 |
| ESD1 | 0,090143925 | 0,406806363 | 0,399 | 0,416 | 1 | Secretory | 1,325747808 | 0,754291272 | 0,959134615 |
| LGALS9 | 0,09016157 | 0,317679527 | 0,276 | 0,253 | 1 | Secretory | 1,246324311 | 0,802359379 | 1,090909091 |
| TRAF41 | 0,090171063 | 0,26462999 | 0,345 | 0,338 | 1 | Secretory | 1,201327904 | 0,832412197 | 1,020710059 |
| CORO1B1 | 0,090719082 | 0,317185478 | 0,369 | 0,371 | 1 | Secretory | 1,245897582 | 0,802634193 | 0,994609164 |
| CHD21 | 0,091930903 | 0,36330737 | 0,295 | 0,371 | 1 | Secretory | 1,286371519 | 0,777380395 | 0,795148248 |
| RWDD11 | 0,092122252 | 0,392956936 | 0,485 | 0,539 | 1 | Secretory | 1,31308193 | 0,761567102 | 0,899814471 |
| CISD11 | 0,094340943 | 0,272872951 | 0,28 | 0,265 | 1 | Secretory | 1,208211439 | 0,827669701 | 1,056603774 |
| PRH11 | 0,094605467 | 1,279682756 | 0,159 | 0,21 | 1 | Secretory | 2,427855833 | 0,411886071 | 0,757142857 |
| TWF1 | 0,094721042 | 0,339244905 | 0,323 | 0,314 | 1 | Secretory | 1,265094281 | 0,790454921 | 1,02866242 |
| NUCB11 | 0,094987039 | 0,324610762 | 0,44 | 0,478 | 1 | Secretory | 1,252326516 | 0,798513796 | 0,920502092 |
| MKRN1 | 0,095154705 | 0,276328606 | 0,278 | 0,265 | 1 | Secretory | 1,21110891 | 0,825689574 | 1,049056604 |
| PSMD12 | 0,09709455 | 0,333111027 | 0,289 | 0,275 | 1 | Secretory | 1,259726922 | 0,793822837 | 1,050909091 |
| FAM210B | 0,097941033 | 0,28249841 | 0,315 | 0,411 | 1 | Secretory | 1,216299408 | 0,822165984 | 0,766423358 |
| IER2 | 0,097945558 | 0,338779218 | 0,58 | 0,627 | 1 | Secretory | 1,264685987 | 0,790710113 | 0,925039872 |
| TMEM1471 | 0,0985093 | 0,300715761 | 0,373 | 0,386 | 1 | Secretory | 1,23175537 | 0,811849515 | 0,966321244 |
| PNKD | 0,098999538 | 0,29787655 | 0,45 | 0,498 | 1 | Secretory | 1,22933367 | 0,813448801 | 0,903614458 |
| TXN | 0,099008673 | 0,314739586 | 0,623 | 0,71 | 1 | Secretory | 1,243787123 | 0,803996103 | 0,877464789 |
| XPO1 | 0,0991459 | 0,299451813 | 0,239 | 0,223 | 1 | Secretory | 1,230676699 | 0,81256109 | 1,071748879 |
| PDIA31 | 0,099252249 | 0,33847728 | 0,642 | 0,752 | 1 | Secretory | 1,264421332 | 0,790875616 | 0,853723404 |
| CYP4X1 | 0,099307414 | 0,404649214 | 0,472 | 0,512 | 1 | Secretory | 1,323767002 | 0,755419948 | 0,921875 |
| PTOV11 | 0,09992467 | 0,290658994 | 0,373 | 0,386 | 1 | Secretory | 1,223198883 | 0,817528543 | 0,966321244 |
| MMP24OS1 | 0,100128119 | 0,271094757 | 0,42 | 0,451 | 1 | Secretory | 1,206723176 | 0,828690474 | 0,931263858 |

| Supplementary Table 4 - clusermarker | | | | | | | | | |
|---|---|---|---|---|---|---|---|---|---|
| | **p_val** | **avg_log2FC** | **pct.1** | **pct.2** | **p_val_adj** | **celltype** | **Foldchange_UP** | **Foldchange_DOWN** | **Ratio_pct1_pct2** |
| NUTF21 | 0,100420705 | 0,276797673 | 0,265 | 0,248 | 1 | Secretory | 1,211502745 | 0,825421159 | 1,068548387 |
| GPCPD1 | 0,100572279 | 0,563813094 | 0,36 | 0,368 | 1 | Secretory | 1,478170917 | 0,676511754 | 0,97826087 |
| PBRM1 | 0,101045193 | 0,338324645 | 0,222 | 0,198 | 1 | Secretory | 1,264287565 | 0,790959294 | 1,121212121 |
| LRP101 | 0,10362744 | 0,32216625 | 0,356 | 0,363 | 1 | Secretory | 1,250206363 | 0,79986795 | 0,980716253 |
| RHOB | 0,103735321 | 0,542558003 | 0,502 | 0,552 | 1 | Secretory | 1,456552803 | 0,686552522 | 0,90942029 |
| NUDC | 0,103747714 | 0,25300787 | 0,366 | 0,374 | 1 | Secretory | 1,191689075 | 0,839145059 | 0,978609626 |
| APPL11 | 0,103863166 | 0,300816259 | 0,358 | 0,366 | 1 | Secretory | 1,231841177 | 0,811792964 | 0,978142077 |
| DCUN1D51 | 0,104108809 | 0,277356817 | 0,17 | 0,145 | 1 | Secretory | 1,211972376 | 0,825101314 | 1,172413793 |
| DANCR1 | 0,104309197 | 0,321868953 | 0,401 | 0,433 | 1 | Secretory | 1,249948759 | 0,800032796 | 0,926096998 |
| TOMM22 | 0,104753908 | 0,313154043 | 0,336 | 0,346 | 1 | Secretory | 1,242420934 | 0,804880192 | 0,971098266 |
| ATP5PB1 | 0,104897176 | 0,326250475 | 0,506 | 0,582 | 1 | Secretory | 1,253750673 | 0,79760675 | 0,869415808 |
| ARPC51 | 0,105771417 | 0,375436543 | 0,306 | 0,3 | 1 | Secretory | 1,297232022 | 0,770872121 | 1,02 |
| PFDN2 | 0,108340458 | 0,268924075 | 0,289 | 0,278 | 1 | Secretory | 1,204908902 | 0,829938262 | 1,039568345 |
| SSBP1 | 0,108401369 | 0,448885382 | 0,412 | 0,453 | 1 | Secretory | 1,36498527 | 0,732608638 | 0,909492274 |
| LZTS3 | 0,108506571 | 0,258642534 | 0,207 | 0,265 | 1 | Secretory | 1,196352498 | 0,835874043 | 0,781132075 |
| ZC3H12A | 0,108870829 | 0,257509753 | 0,222 | 0,286 | 1 | Secretory | 1,19541351 | 0,836530616 | 0,776223776 |
| NDUFB41 | 0,112146059 | 0,352286494 | 0,526 | 0,626 | 1 | Secretory | 1,276582251 | 0,783341613 | 0,840255591 |
| CC2D1A | 0,112228939 | 0,258551355 | 0,166 | 0,141 | 1 | Secretory | 1,196276891 | 0,835926872 | 1,177304965 |
| NDUFA21 | 0,113174933 | 0,281607815 | 0,504 | 0,567 | 1 | Secretory | 1,215548801 | 0,822673675 | 0,888888889 |
| DPP71 | 0,113639755 | 0,265445254 | 0,433 | 0,469 | 1 | Secretory | 1,202006964 | 0,831941935 | 0,923240938 |
| REEP51 | 0,115235553 | 0,299207738 | 0,556 | 0,656 | 1 | Secretory | 1,230468511 | 0,812698571 | 0,847560976 |
| MMAB | 0,115359729 | 0,267936482 | 0,239 | 0,216 | 1 | Secretory | 1,204084367 | 0,830506588 | 1,106481481 |
| DDRGK1 | 0,116946589 | 0,259710304 | 0,405 | 0,438 | 1 | Secretory | 1,197238273 | 0,835255624 | 0,924657534 |
| UBTF | 0,118279188 | 0,266017177 | 0,224 | 0,201 | 1 | Secretory | 1,202483566 | 0,831612197 | 1,114427861 |
| TFAM | 0,118297003 | 0,314273459 | 0,252 | 0,236 | 1 | Secretory | 1,243385327 | 0,804255912 | 1,06779661 |
| PLEKHA5 | 0,118777195 | 0,395413253 | 0,228 | 0,208 | 1 | Secretory | 1,315319473 | 0,760271569 | 1,096153846 |
| EIF2S3 | 0,118818999 | 0,430497537 | 0,427 | 0,474 | 1 | Secretory | 1,347698273 | 0,742005848 | 0,900843882 |
| PPM1G1 | 0,119057642 | 0,332718255 | 0,319 | 0,316 | 1 | Secretory | 1,25938401 | 0,794038984 | 1,009493671 |
| GNPTG1 | 0,119353673 | 0,345146771 | 0,377 | 0,399 | 1 | Secretory | 1,270280207 | 0,787227885 | 0,944862155 |
| CHD4 | 0,11962665 | 0,318843279 | 0,33 | 0,326 | 1 | Secretory | 1,247330066 | 0,801712415 | 1,012269939 |
| COPE1 | 0,120125427 | 0,327267967 | 0,483 | 0,544 | 1 | Secretory | 1,254635219 | 0,79704442 | 0,887867647 |

| Supplementary Table 4 - clusermarker | | | | | | | | | |
|---|---|---|---|---|---|---|---|---|---|
| | **p_val** | **avg_log2FC** | **pct.1** | **pct.2** | **p_val_adj** | **celltype** | **Foldchange_UP** | **Foldchange_DOWN** | **Ratio_pct1_pct2** |
| GDI2 | 0,120199469 | 0,300250795 | 0,442 | 0,491 | 1 | Secretory | 1,231358451 | 0,812111209 | 0,900203666 |
| RPA3 | 0,120465501 | 0,279636467 | 0,22 | 0,198 | 1 | Secretory | 1,213888968 | 0,823798574 | 1,111111111 |
| VGLL41 | 0,120482891 | 0,345848407 | 0,349 | 0,361 | 1 | Secretory | 1,270898141 | 0,78684512 | 0,966759003 |
| TBCA | 0,12118536 | 0,297827154 | 0,442 | 0,499 | 1 | Secretory | 1,22929158 | 0,813476653 | 0,885771543 |
| RBCK1 | 0,121675482 | 0,341474994 | 0,284 | 0,28 | 1 | Secretory | 1,26705135 | 0,789233996 | 1,014285714 |
| STAT21 | 0,121752543 | 0,276993882 | 0,185 | 0,163 | 1 | Secretory | 1,211667522 | 0,825308908 | 1,134969325 |
| PPA2 | 0,122027002 | 0,410278587 | 0,317 | 0,313 | 1 | Secretory | 1,328942411 | 0,752478055 | 1,012779553 |
| CAPN11 | 0,122112882 | 0,368477771 | 0,358 | 0,368 | 1 | Secretory | 1,290989951 | 0,774599368 | 0,972826087 |
| NCKAP1 | 0,122182826 | 0,272833533 | 0,317 | 0,406 | 1 | Secretory | 1,208178428 | 0,827692315 | 0,780788177 |
| HMGN3 | 0,122628004 | 0,340128467 | 0,478 | 0,546 | 1 | Secretory | 1,26586931 | 0,789970965 | 0,875457875 |
| TOMM61 | 0,122969967 | 0,341022244 | 0,483 | 0,539 | 1 | Secretory | 1,266653783 | 0,789481714 | 0,896103896 |
| CACUL1 | 0,123651124 | 0,310410447 | 0,22 | 0,2 | 1 | Secretory | 1,240060447 | 0,806412302 | 1,1 |
| SURF11 | 0,123762155 | 0,277400442 | 0,341 | 0,346 | 1 | Secretory | 1,212009025 | 0,825076364 | 0,985549133 |
| RSF1 | 0,124720903 | 0,329162966 | 0,369 | 0,388 | 1 | Secretory | 1,256284282 | 0,795998178 | 0,951030928 |
| DHX16 | 0,125928984 | 0,300741641 | 0,125 | 0,1 | 1 | Secretory | 1,231777466 | 0,811834952 | 1,25 |
| PALLD1 | 0,126483733 | 0,401101039 | 0,375 | 0,383 | 1 | Secretory | 1,32051532 | 0,75728012 | 0,979112272 |
| GTF3A1 | 0,127703814 | 0,286251599 | 0,347 | 0,361 | 1 | Secretory | 1,219467745 | 0,82002989 | 0,961218837 |
| KRT151 | 0,129035149 | 0,662488166 | 0,259 | 0,231 | 1 | Secretory | 1,582810088 | 0,631787735 | 1,121212121 |
| TMEM101 | 0,129837935 | 0,273800451 | 0,224 | 0,201 | 1 | Secretory | 1,208988441 | 0,827137767 | 1,114427861 |
| LMAN1 | 0,130063257 | 0,349858884 | 0,483 | 0,537 | 1 | Secretory | 1,274435963 | 0,784660845 | 0,899441341 |
| TPM11 | 0,130294128 | 0,475305145 | 0,463 | 0,484 | 1 | Secretory | 1,390212233 | 0,719314631 | 0,95661157 |
| C18orf32 | 0,132511881 | 0,317269818 | 0,399 | 0,419 | 1 | Secretory | 1,245970419 | 0,802587272 | 0,952267303 |
| LRRFIP21 | 0,133812419 | 0,254210165 | 0,274 | 0,258 | 1 | Secretory | 1,192682604 | 0,838446035 | 1,062015504 |
| GET3 | 0,133982228 | 0,283211162 | 0,28 | 0,265 | 1 | Secretory | 1,216900459 | 0,8217599 | 1,056603774 |
| SEPTIN2 | 0,13438928 | 0,284485098 | 0,414 | 0,438 | 1 | Secretory | 1,217975487 | 0,821034586 | 0,945205479 |
| XRCC6 | 0,135244222 | 0,443603863 | 0,418 | 0,454 | 1 | Secretory | 1,359997371 | 0,735295539 | 0,920704846 |
| CPNE31 | 0,135585946 | 0,330060601 | 0,407 | 0,429 | 1 | Secretory | 1,257066177 | 0,795503068 | 0,948717949 |
| NDUFB61 | 0,137225472 | 0,298493876 | 0,427 | 0,461 | 1 | Secretory | 1,229859812 | 0,813100803 | 0,926247289 |
| CCDC85B1 | 0,137522678 | 0,295288982 | 0,284 | 0,275 | 1 | Secretory | 1,227130756 | 0,814909084 | 1,032727273 |
| MUC5B1 | 0,138504265 | -0,445342654 | 0,341 | 0,288 | 1 | Secretory | 0,734409866 | 1,36163748 | 1,184027778 |
| DNAJC151 | 0,139032897 | 0,391039343 | 0,399 | 0,449 | 1 | Secretory | 1,311337774 | 0,762580031 | 0,888641425 |

| Supplementary Table 4 - clusermarker | | | | | | | | | |
|---|---|---|---|---|---|---|---|---|---|
| | **p_val** | **avg_log2FC** | **pct.1** | **pct.2** | **p_val_adj** | **celltype** | **Foldchange_UP** | **Foldchange_DOWN** | **Ratio_pct1_pct2** |
| COX7A2 | 0,139255663 | 0,279181733 | 0,601 | 0,686 | 1 | Secretory | 1,213506413 | 0,824058274 | 0,876093294 |
| GRHPR | 0,140403742 | 0,366055094 | 0,338 | 0,348 | 1 | Secretory | 1,288823847 | 0,775901223 | 0,971264368 |
| FOSB1 | 0,141032607 | 0,422496445 | 0,578 | 0,611 | 1 | Secretory | 1,340244714 | 0,746132396 | 0,94599018 |
| ATP6V1G1 | 0,142276461 | 0,342259188 | 0,552 | 0,656 | 1 | Secretory | 1,267740258 | 0,788805115 | 0,841463415 |
| NONO | 0,14278639 | 0,39630636 | 0,388 | 0,416 | 1 | Secretory | 1,316133979 | 0,759801066 | 0,932692308 |
| CCDC57 | 0,143270135 | 0,25240663 | 0,293 | 0,278 | 1 | Secretory | 1,191192545 | 0,839494844 | 1,053956835 |
| MCM7 | 0,143700685 | 0,291830674 | 0,121 | 0,098 | 1 | Secretory | 1,224192703 | 0,816864859 | 1,234693878 |
| PGAM1 | 0,145262699 | 0,358917476 | 0,36 | 0,378 | 1 | Secretory | 1,282463242 | 0,779749444 | 0,952380952 |
| SNRPB | 0,149226718 | 0,286254327 | 0,343 | 0,353 | 1 | Secretory | 1,219470051 | 0,820028339 | 0,971671388 |
| NFE2L2 | 0,149849202 | 0,292983232 | 0,358 | 0,359 | 1 | Secretory | 1,225171092 | 0,816212533 | 0,997214485 |
| MIEN11 | 0,151118517 | 0,287656139 | 0,412 | 0,439 | 1 | Secretory | 1,22065554 | 0,819231935 | 0,938496583 |
| AGO2 | 0,151919364 | 0,269105241 | 0,149 | 0,125 | 1 | Secretory | 1,205060218 | 0,829834049 | 1,192 |
| TM9SF21 | 0,152770505 | 0,295596289 | 0,513 | 0,579 | 1 | Secretory | 1,227392174 | 0,814735519 | 0,886010363 |
| BTG3 | 0,153401927 | 0,298707203 | 0,263 | 0,248 | 1 | Secretory | 1,230041681 | 0,812980581 | 1,060483871 |
| RND3 | 0,155331879 | 0,31519054 | 0,237 | 0,215 | 1 | Secretory | 1,244175964 | 0,803744831 | 1,102325581 |
| STMP1 | 0,155704034 | 0,332931148 | 0,358 | 0,373 | 1 | Secretory | 1,259569866 | 0,79392182 | 0,959785523 |
| GNAQ | 0,15583931 | 0,409066951 | 0,409 | 0,441 | 1 | Secretory | 1,327826778 | 0,753110283 | 0,927437642 |
| ZRANB2 | 0,15620778 | 0,258294983 | 0,276 | 0,341 | 1 | Secretory | 1,196064327 | 0,836075433 | 0,809384164 |
| SPTBN1 | 0,157490073 | 0,374033714 | 0,399 | 0,424 | 1 | Secretory | 1,295971251 | 0,771622055 | 0,941037736 |
| SLC3A2 | 0,157556027 | 0,409465522 | 0,33 | 0,336 | 1 | Secretory | 1,328193665 | 0,752902251 | 0,982142857 |
| GRINA1 | 0,158873442 | 0,321880484 | 0,332 | 0,346 | 1 | Secretory | 1,249958749 | 0,800026401 | 0,959537572 |
| TRAPPC6A | 0,158884652 | 0,270260015 | 0,241 | 0,225 | 1 | Secretory | 1,206025168 | 0,829170092 | 1,071111111 |
| CREB11 | 0,159698423 | 0,316744475 | 0,177 | 0,151 | 1 | Secretory | 1,245516794 | 0,802879579 | 1,17218543 |
| CHMP4B1 | 0,160379406 | 0,278437898 | 0,272 | 0,256 | 1 | Secretory | 1,212880906 | 0,824483257 | 1,0625 |
| HDAC6 | 0,160601655 | 0,256399374 | 0,138 | 0,118 | 1 | Secretory | 1,194493807 | 0,837174704 | 1,169491525 |
| THYN11 | 0,16093143 | 0,259091988 | 0,216 | 0,196 | 1 | Secretory | 1,196725266 | 0,835613677 | 1,102040816 |
| HP1BP3 | 0,161230522 | 0,376378279 | 0,42 | 0,469 | 1 | Secretory | 1,298079083 | 0,770369089 | 0,895522388 |
| HNRNPK | 0,162659057 | 0,296316923 | 0,547 | 0,629 | 1 | Secretory | 1,228005415 | 0,814328657 | 0,86963434 |
| NEDD9 | 0,165682826 | 0,335872778 | 0,31 | 0,388 | 1 | Secretory | 1,262140727 | 0,792304676 | 0,798969072 |
| TUT4 | 0,165780692 | 0,274629217 | 0,228 | 0,208 | 1 | Secretory | 1,209683152 | 0,826662749 | 1,096153846 |
| EPC11 | 0,169449376 | 0,375648782 | 0,297 | 0,296 | 1 | Secretory | 1,297422876 | 0,770758724 | 1,003378378 |

| Supplementary Table 4 - clusermarker | | | | | | | | | |
|---|---|---|---|---|---|---|---|---|---|
| | **p_val** | **avg_log2FC** | **pct.1** | **pct.2** | **p_val_adj** | **celltype** | **Foldchange_UP** | **Foldchange_DOWN** | **Ratio_pct1_pct2** |
| TAX1BP1 | 0,171965668 | 0,382320687 | 0,487 | 0,567 | 1 | Secretory | 1,30343685 | 0,767202493 | 0,858906526 |
| MBTPS11 | 0,173789851 | 0,311156538 | 0,269 | 0,256 | 1 | Secretory | 1,240701911 | 0,805995373 | 1,05078125 |
| LUC7L2 | 0,173859634 | 0,344253972 | 0,351 | 0,361 | 1 | Secretory | 1,269494348 | 0,787715205 | 0,972299169 |
| NAALADL2 | 0,174807593 | 0,271067354 | 0,231 | 0,216 | 1 | Secretory | 1,206700255 | 0,828706214 | 1,069444444 |
| MARCKS1 | 0,17488642 | 0,269911764 | 0,211 | 0,196 | 1 | Secretory | 1,205734082 | 0,829370269 | 1,076530612 |
| TGFBR2 | 0,175054375 | 0,322259157 | 0,269 | 0,26 | 1 | Secretory | 1,250286877 | 0,799816441 | 1,034615385 |
| EMC2 | 0,176646657 | 0,345249267 | 0,259 | 0,246 | 1 | Secretory | 1,270370456 | 0,787171958 | 1,052845528 |
| TACC2 | 0,176885798 | 0,283456684 | 0,2 | 0,25 | 1 | Secretory | 1,217107572 | 0,821620063 | 0,8 |
| SUMO11 | 0,177110321 | 0,393226065 | 0,366 | 0,396 | 1 | Secretory | 1,313326903 | 0,761425048 | 0,924242424 |
| KIF5B | 0,177543182 | 0,311659983 | 0,409 | 0,453 | 1 | Secretory | 1,241134944 | 0,805714161 | 0,902869757 |
| KDM5B | 0,182250716 | 0,455633359 | 0,291 | 0,291 | 1 | Secretory | 1,371384726 | 0,729189979 | 1 |
| LNPEP | 0,183719249 | 0,270590872 | 0,233 | 0,213 | 1 | Secretory | 1,206301781 | 0,828979958 | 1,093896714 |
| SNRNP70 | 0,184892214 | 0,378546039 | 0,397 | 0,426 | 1 | Secretory | 1,300031012 | 0,769212419 | 0,931924883 |
| GTF2I | 0,185410219 | 0,395896486 | 0,422 | 0,461 | 1 | Secretory | 1,315760115 | 0,760016958 | 0,915401302 |
| RAD50 | 0,185641667 | 0,277421559 | 0,259 | 0,25 | 1 | Secretory | 1,212026766 | 0,825064287 | 1,036 |
| NIPBL1 | 0,187550185 | 0,299230587 | 0,312 | 0,309 | 1 | Secretory | 1,230487999 | 0,812685699 | 1,009708738 |
| CYBRD1 | 0,188119574 | 0,276781949 | 0,213 | 0,201 | 1 | Secretory | 1,211489541 | 0,825430156 | 1,059701493 |
| HCG18 | 0,188803578 | 0,267123121 | 0,157 | 0,135 | 1 | Secretory | 1,203405722 | 0,830974942 | 1,162962963 |
| BCAS2 | 0,189212361 | 0,287059306 | 0,25 | 0,238 | 1 | Secretory | 1,220150667 | 0,819570916 | 1,050420168 |
| SF3B1 | 0,189415371 | 0,502861334 | 0,459 | 0,517 | 1 | Secretory | 1,417021192 | 0,705705748 | 0,887814313 |
| AGAP1 | 0,190051052 | 0,253572654 | 0,177 | 0,158 | 1 | Secretory | 1,192155687 | 0,838816616 | 1,120253165 |
| ECSIT | 0,190155162 | 0,251322475 | 0,274 | 0,27 | 1 | Secretory | 1,190297725 | 0,840125944 | 1,014814815 |
| TRIM4 | 0,195371562 | 0,261608846 | 0,153 | 0,133 | 1 | Secretory | 1,198814838 | 0,834157176 | 1,15037594 |
| LPGAT1 | 0,196656761 | 0,284988452 | 0,235 | 0,225 | 1 | Secretory | 1,218400511 | 0,820748178 | 1,044444444 |
| ASH1L1 | 0,196898261 | 0,587326226 | 0,392 | 0,421 | 1 | Secretory | 1,502459629 | 0,665575288 | 0,93111639 |
| RBM6 | 0,200539953 | 0,271893042 | 0,282 | 0,278 | 1 | Secretory | 1,207391075 | 0,828232062 | 1,014388489 |
| GTF2A21 | 0,20061845 | 0,308497167 | 0,392 | 0,436 | 1 | Secretory | 1,238416988 | 0,807482463 | 0,899082569 |
| XIST | 0,200774783 | 0,581711493 | 0,433 | 0,552 | 1 | Secretory | 1,496623666 | 0,668170645 | 0,78442029 |
| ARPC4 | 0,203852978 | 0,348845963 | 0,282 | 0,273 | 1 | Secretory | 1,273541492 | 0,785211952 | 1,032967033 |
| ETV6 | 0,205636073 | 0,412707065 | 0,384 | 0,418 | 1 | Secretory | 1,331181293 | 0,75121248 | 0,918660287 |
| PSMD11 | 0,206524856 | 0,265659465 | 0,272 | 0,268 | 1 | Secretory | 1,202185451 | 0,831818418 | 1,014925373 |

| Supplementary Table 4 - clusermarker | | | | | | | | | |
|---|---|---|---|---|---|---|---|---|---|
| | **p_val** | **avg_log2FC** | **pct.1** | **pct.2** | **p_val_adj** | **celltype** | **Foldchange_UP** | **Foldchange_DOWN** | **Ratio_pct1_pct2** |
| MRPL32 | 0,206659264 | 0,273371641 | 0,276 | 0,273 | 1 | Secretory | 1,208629148 | 0,827383653 | 1,010989011 |
| ARHGDIB | 0,207240381 | 0,333947701 | 0,319 | 0,311 | 1 | Secretory | 1,260457698 | 0,793362603 | 1,025723473 |
| ABRACL1 | 0,210428749 | 0,283009413 | 0,295 | 0,295 | 1 | Secretory | 1,216730297 | 0,821874825 | 1 |
| MICOS13 | 0,211951932 | 0,369584734 | 0,433 | 0,493 | 1 | Secretory | 1,291980893 | 0,774005255 | 0,878296146 |
| PALMD | 0,212030577 | 0,315611893 | 0,295 | 0,293 | 1 | Secretory | 1,244539391 | 0,803510124 | 1,006825939 |
| MT1X1 | 0,213715482 | 0,513170767 | 0,364 | 0,358 | 1 | Secretory | 1,427183427 | 0,700680782 | 1,016759777 |
| NFIX | 0,214491715 | 0,398068501 | 0,438 | 0,483 | 1 | Secretory | 1,317742518 | 0,758873593 | 0,906832298 |
| CUL3 | 0,215842603 | 0,312390965 | 0,265 | 0,253 | 1 | Secretory | 1,241763959 | 0,805306026 | 1,04743083 |
| PAK2 | 0,216615188 | 0,447743406 | 0,319 | 0,324 | 1 | Secretory | 1,363905233 | 0,73318877 | 0,984567901 |
| LYRM2 | 0,221500234 | 0,330626239 | 0,269 | 0,258 | 1 | Secretory | 1,257559132 | 0,795191235 | 1,042635659 |
| CRNDE1 | 0,223241041 | 0,314368595 | 0,384 | 0,424 | 1 | Secretory | 1,243467323 | 0,804202878 | 0,905660377 |
| ACAP2 | 0,226252111 | 0,253062361 | 0,3 | 0,293 | 1 | Secretory | 1,191734087 | 0,839113365 | 1,023890785 |
| STK24 | 0,228491598 | 0,317927709 | 0,291 | 0,286 | 1 | Secretory | 1,246538731 | 0,802221363 | 1,017482517 |
| PCMT1 | 0,230233201 | 0,264783359 | 0,293 | 0,293 | 1 | Secretory | 1,20145562 | 0,832323711 | 1 |
| IGFBP71 | 0,230717391 | 0,645105082 | 0,203 | 0,185 | 1 | Secretory | 1,563853189 | 0,639446213 | 1,097297297 |
| SMARCC1 | 0,23130511 | 0,349656566 | 0,304 | 0,311 | 1 | Secretory | 1,274257254 | 0,78477089 | 0,977491961 |
| ILF2 | 0,231945792 | 0,326510884 | 0,366 | 0,391 | 1 | Secretory | 1,253976998 | 0,797462794 | 0,936061381 |
| ADD11 | 0,232236308 | 0,345383674 | 0,287 | 0,286 | 1 | Secretory | 1,270488814 | 0,787098626 | 1,003496503 |
| NORAD | 0,234504266 | 0,308005262 | 0,422 | 0,469 | 1 | Secretory | 1,237994806 | 0,807757832 | 0,89978678 |
| CDK16 | 0,236754612 | 0,265334177 | 0,233 | 0,218 | 1 | Secretory | 1,201914421 | 0,832005992 | 1,068807339 |
| REV3L | 0,240060001 | 0,277245327 | 0,19 | 0,176 | 1 | Secretory | 1,21187872 | 0,825165079 | 1,079545455 |
| IFI16 | 0,242707856 | 0,2623506 | 0,287 | 0,281 | 1 | Secretory | 1,199431361 | 0,833728409 | 1,021352313 |
| NFIC1 | 0,244983402 | 0,29796987 | 0,466 | 0,551 | 1 | Secretory | 1,229413191 | 0,813396185 | 0,845735027 |
| ATP5PD | 0,245650389 | 0,337785465 | 0,53 | 0,629 | 1 | Secretory | 1,26381515 | 0,791254955 | 0,842607313 |
| ECHDC11 | 0,245874211 | 0,300707191 | 0,33 | 0,354 | 1 | Secretory | 1,231748052 | 0,811854338 | 0,93220339 |
| ITPR2 | 0,248600081 | 0,299319888 | 0,362 | 0,383 | 1 | Secretory | 1,230564167 | 0,812635397 | 0,945169713 |
| POLR1D | 0,252112321 | 0,322270297 | 0,414 | 0,461 | 1 | Secretory | 1,25029653 | 0,799810266 | 0,898047722 |
| FLNB | 0,253112113 | 0,278750532 | 0,293 | 0,298 | 1 | Secretory | 1,213143768 | 0,82430461 | 0,983221477 |
| CDK2AP2 | 0,255553753 | 0,289900728 | 0,435 | 0,491 | 1 | Secretory | 1,22255615 | 0,817958341 | 0,885947047 |
| MAST4 | 0,256238475 | 0,421319577 | 0,295 | 0,295 | 1 | Secretory | 1,339151865 | 0,746741297 | 1 |
| DYNC1H1 | 0,258058102 | 0,382646304 | 0,407 | 0,439 | 1 | Secretory | 1,303731069 | 0,767029355 | 0,927107062 |

| Supplementary Table 4 - clusermarker | | | | | | | | | |
|---|---|---|---|---|---|---|---|---|---|
| | **p_val** | **avg_log2FC** | **pct.1** | **pct.2** | **p_val_adj** | **celltype** | **Foldchange_UP** | **Foldchange_DOWN** | **Ratio_pct1_pct2** |
| EGLN2 | 0,259447863 | 0,277427546 | 0,211 | 0,196 | 1 | Secretory | 1,212031796 | 0,825060863 | 1,076530612 |
| UBE2E31 | 0,260445978 | 0,332322648 | 0,328 | 0,343 | 1 | Secretory | 1,259038716 | 0,794256751 | 0,956268222 |
| FKBP8 | 0,26274469 | 0,290986286 | 0,42 | 0,479 | 1 | Secretory | 1,223476411 | 0,817343098 | 0,876826722 |
| PLEKHA3 | 0,264428513 | 0,272238131 | 0,22 | 0,215 | 1 | Secretory | 1,207679914 | 0,828033975 | 1,023255814 |
| SCAF11 | 0,266523996 | 0,357105248 | 0,388 | 0,421 | 1 | Secretory | 1,280853299 | 0,780729535 | 0,921615202 |
| TSHZ2 | 0,267634031 | 0,379025126 | 0,416 | 0,454 | 1 | Secretory | 1,300462795 | 0,768957024 | 0,916299559 |
| PHF14 | 0,268366961 | 0,260985224 | 0,259 | 0,256 | 1 | Secretory | 1,198296749 | 0,834517828 | 1,01171875 |
| SPG7 | 0,269651861 | 0,293062743 | 0,241 | 0,231 | 1 | Secretory | 1,225238616 | 0,81616755 | 1,043290043 |
| CHMP4A | 0,270622976 | 0,270236111 | 0,226 | 0,218 | 1 | Secretory | 1,206005186 | 0,829183831 | 1,036697248 |
| CSNK1A1 | 0,272235002 | 0,395095853 | 0,442 | 0,493 | 1 | Secretory | 1,315030129 | 0,760438851 | 0,896551724 |
| OTUB1 | 0,275326544 | 0,288247067 | 0,349 | 0,376 | 1 | Secretory | 1,221155622 | 0,818896447 | 0,928191489 |
| EMC3 | 0,275477426 | 0,367927641 | 0,319 | 0,333 | 1 | Secretory | 1,290497764 | 0,774894795 | 0,957957958 |
| LSM7 | 0,275880962 | 0,334330369 | 0,42 | 0,476 | 1 | Secretory | 1,260792073 | 0,793152195 | 0,882352941 |
| MYO61 | 0,276509896 | 0,381433705 | 0,274 | 0,275 | 1 | Secretory | 1,302635731 | 0,767674321 | 0,996363636 |
| MACF1 | 0,277568701 | 0,375532531 | 0,381 | 0,413 | 1 | Secretory | 1,297318335 | 0,770820833 | 0,92251816 |
| ZNF638 | 0,279069053 | 0,427417933 | 0,306 | 0,313 | 1 | Secretory | 1,344824519 | 0,74359144 | 0,977635783 |
| RASEF | 0,279946027 | 0,365489587 | 0,246 | 0,233 | 1 | Secretory | 1,288318754 | 0,77620542 | 1,055793991 |
| CENPC1 | 0,282678058 | 0,255012479 | 0,205 | 0,191 | 1 | Secretory | 1,193346065 | 0,837979887 | 1,073298429 |
| CAND1 | 0,285205689 | 0,315856678 | 0,209 | 0,2 | 1 | Secretory | 1,244750572 | 0,803373802 | 1,045 |
| SPTSSA | 0,285494511 | 0,258086185 | 0,356 | 0,443 | 1 | Secretory | 1,195891236 | 0,836196445 | 0,803611738 |
| EXOC71 | 0,287110791 | 0,270013155 | 0,213 | 0,205 | 1 | Secretory | 1,205818823 | 0,829311984 | 1,03902439 |
| CBR11 | 0,288250415 | 0,258124042 | 0,371 | 0,409 | 1 | Secretory | 1,195922617 | 0,836174503 | 0,907090465 |
| NCOR1 | 0,288322301 | 0,256303815 | 0,334 | 0,349 | 1 | Secretory | 1,19441469 | 0,837230158 | 0,957020057 |
| ENOSF1 | 0,288806437 | 0,288157641 | 0,278 | 0,276 | 1 | Secretory | 1,221079931 | 0,818947208 | 1,007246377 |
| TFDP21 | 0,290642381 | 0,280861665 | 0,256 | 0,25 | 1 | Secretory | 1,214920292 | 0,823099266 | 1,024 |
| LMBRD1 | 0,291480089 | 0,274286936 | 0,317 | 0,331 | 1 | Secretory | 1,209396187 | 0,826858899 | 0,957703927 |
| SMARCA21 | 0,292613616 | 0,32727887 | 0,414 | 0,466 | 1 | Secretory | 1,254644702 | 0,797038396 | 0,888412017 |
| WBP12 | 0,293446367 | 0,317047741 | 0,358 | 0,381 | 1 | Secretory | 1,24577864 | 0,802710825 | 0,939632546 |
| H1-10 | 0,294818338 | 0,28456992 | 0,369 | 0,393 | 1 | Secretory | 1,218047099 | 0,820986316 | 0,938931298 |
| STAT11 | 0,297121488 | 0,340762912 | 0,218 | 0,211 | 1 | Secretory | 1,266426116 | 0,78962364 | 1,033175355 |
| PSMA2 | 0,300498528 | 0,253279252 | 0,45 | 0,517 | 1 | Secretory | 1,191913262 | 0,838987225 | 0,87040619 |

| Supplementary Table 4 - clusermarker | | | | | | | | | |
|---|---|---|---|---|---|---|---|---|---|
| | **p_val** | **avg_log2FC** | **pct.1** | **pct.2** | **p_val_adj** | **celltype** | **Foldchange_UP** | **Foldchange_DOWN** | **Ratio_pct1_pct2** |
| PTEN | 0,301239343 | 0,252174258 | 0,252 | 0,246 | 1 | Secretory | 1,191000698 | 0,839630071 | 1,024390244 |
| AP3D1 | 0,303797563 | 0,263567905 | 0,297 | 0,3 | 1 | Secretory | 1,200443834 | 0,833025229 | 0,99 |
| CHD3 | 0,305228133 | 0,303537944 | 0,317 | 0,329 | 1 | Secretory | 1,234167273 | 0,810262937 | 0,963525836 |
| ISOC21 | 0,309319932 | 0,258054885 | 0,332 | 0,348 | 1 | Secretory | 1,19586529 | 0,836214587 | 0,954022989 |
| LRRFIP1 | 0,310469407 | 0,296805512 | 0,407 | 0,509 | 1 | Secretory | 1,228421367 | 0,814052919 | 0,799607073 |
| PTP4A2 | 0,312460864 | 0,364290477 | 0,414 | 0,463 | 1 | Secretory | 1,2872484 | 0,77685084 | 0,894168467 |
| FUT2 | 0,312851317 | 0,258165192 | 0,259 | 0,26 | 1 | Secretory | 1,195956729 | 0,836150653 | 0,996153846 |
| PPP6R1 | 0,314789723 | 0,282959144 | 0,112 | 0,097 | 1 | Secretory | 1,216687902 | 0,821903463 | 1,154639175 |
| FTX | 0,315492846 | 0,32141209 | 0,179 | 0,216 | 1 | Secretory | 1,249552996 | 0,800286185 | 0,828703704 |
| ACADVL | 0,316644146 | 0,308300157 | 0,47 | 0,544 | 1 | Secretory | 1,238247885 | 0,807592738 | 0,863970588 |
| POMP1 | 0,317199107 | 0,499365473 | 0,435 | 0,509 | 1 | Secretory | 1,413591699 | 0,707417849 | 0,854616896 |
| STK4 | 0,318112194 | 0,277795864 | 0,142 | 0,128 | 1 | Secretory | 1,212341265 | 0,824850254 | 1,109375 |
| IRF2BPL | 0,319207567 | 0,26557507 | 0,203 | 0,193 | 1 | Secretory | 1,202115127 | 0,83186708 | 1,051813472 |
| PNKP1 | 0,326198708 | 0,280600184 | 0,222 | 0,213 | 1 | Secretory | 1,214700114 | 0,823248462 | 1,042253521 |
| MRPL20 | 0,326741383 | 0,37227293 | 0,44 | 0,519 | 1 | Secretory | 1,294390505 | 0,772564382 | 0,8477842 |
| RBBP41 | 0,328108579 | 0,260325893 | 0,375 | 0,413 | 1 | Secretory | 1,197749236 | 0,834899301 | 0,907990315 |
| LONP21 | 0,328835776 | 0,349278614 | 0,321 | 0,338 | 1 | Secretory | 1,273923472 | 0,784976509 | 0,949704142 |
| EWSR1 | 0,332600522 | 0,413011145 | 0,36 | 0,389 | 1 | Secretory | 1,331461899 | 0,751054161 | 0,925449871 |
| SYPL1 | 0,33338622 | 0,326531809 | 0,468 | 0,557 | 1 | Secretory | 1,253995186 | 0,797451227 | 0,84021544 |
| FBL | 0,338167129 | 0,318669782 | 0,399 | 0,448 | 1 | Secretory | 1,247180072 | 0,801808834 | 0,890625 |
| PDS5A | 0,343840238 | 0,287382069 | 0,308 | 0,318 | 1 | Secretory | 1,220423672 | 0,819387581 | 0,968553459 |
| NDUFA5 | 0,34466887 | 0,262443818 | 0,36 | 0,441 | 1 | Secretory | 1,199508863 | 0,83367454 | 0,816326531 |
| CCNL11 | 0,349538804 | 0,463872529 | 0,405 | 0,494 | 1 | Secretory | 1,379239052 | 0,725037475 | 0,819838057 |
| N4BP2L2 | 0,35331178 | 0,387927472 | 0,397 | 0,434 | 1 | Secretory | 1,308512288 | 0,764226679 | 0,914746544 |
| BAG6 | 0,35628156 | 0,251999676 | 0,252 | 0,246 | 1 | Secretory | 1,190856582 | 0,839731681 | 1,024390244 |
| VPS291 | 0,357183839 | 0,261385172 | 0,371 | 0,403 | 1 | Secretory | 1,198628989 | 0,834286513 | 0,920595533 |
| HEXA | 0,360515277 | 0,305494186 | 0,345 | 0,361 | 1 | Secretory | 1,235841895 | 0,809164994 | 0,95567867 |
| DHCR24 | 0,363550711 | 0,295528129 | 0,345 | 0,374 | 1 | Secretory | 1,227334187 | 0,814774013 | 0,922459893 |
| EMP11 | 0,366312361 | 0,306948037 | 0,185 | 0,168 | 1 | Secretory | 1,23708792 | 0,808349984 | 1,101190476 |
| YPEL5 | 0,369205497 | 0,395727304 | 0,336 | 0,358 | 1 | Secretory | 1,315605828 | 0,760106089 | 0,938547486 |
| PI31 | 0,371313102 | 0,446823975 | 0,269 | 0,309 | 1 | Secretory | 1,363036291 | 0,733656181 | 0,870550162 |

| Supplementary Table 4 - clusermarker | | | | | | | | | |
|---|---|---|---|---|---|---|---|---|---|
| | **p_val** | **avg_log2FC** | **pct.1** | **pct.2** | **p_val_adj** | **celltype** | **Foldchange_UP** | **Foldchange_DOWN** | **Ratio_pct1_pct2** |
| MRPS36 | 0,374178437 | 0,28145477 | 0,31 | 0,324 | 1 | Secretory | 1,215419859 | 0,822760952 | 0,956790123 |
| CAPZA2 | 0,375088234 | 0,406927047 | 0,341 | 0,371 | 1 | Secretory | 1,325858713 | 0,754228177 | 0,919137466 |
| KANSL1 | 0,375485847 | 0,259762904 | 0,256 | 0,251 | 1 | Secretory | 1,197281924 | 0,835225171 | 1,019920319 |
| ARID4B | 0,377236544 | 0,292990414 | 0,328 | 0,343 | 1 | Secretory | 1,225177191 | 0,816208469 | 0,956268222 |
| CALR1 | 0,378413148 | 0,374583103 | 0,621 | 0,739 | 1 | Secretory | 1,296464859 | 0,771328272 | 0,840324763 |
| BSG | 0,378540744 | 0,31957643 | 0,491 | 0,569 | 1 | Secretory | 1,247964098 | 0,801305103 | 0,862917399 |
| SUMO31 | 0,384387267 | 0,291359873 | 0,343 | 0,376 | 1 | Secretory | 1,223793272 | 0,817131474 | 0,912234043 |
| TBL1XR1 | 0,386076445 | 0,276494851 | 0,25 | 0,248 | 1 | Secretory | 1,211248476 | 0,825594434 | 1,008064516 |
| EIF2A | 0,396502829 | 0,26846708 | 0,381 | 0,429 | 1 | Secretory | 1,20452729 | 0,830201199 | 0,888111888 |
| PSMD2 | 0,396835912 | 0,283424815 | 0,261 | 0,258 | 1 | Secretory | 1,217080686 | 0,821638213 | 1,011627907 |
| URI1 | 0,397886788 | 0,442700658 | 0,315 | 0,333 | 1 | Secretory | 1,359146206 | 0,735756018 | 0,945945946 |
| HNRNPUL1 | 0,398821576 | 0,261644762 | 0,336 | 0,354 | 1 | Secretory | 1,198844683 | 0,83413641 | 0,949152542 |
| TMX41 | 0,401479381 | 0,330423564 | 0,338 | 0,363 | 1 | Secretory | 1,257382478 | 0,795302955 | 0,931129477 |
| STX7 | 0,403443686 | 0,259964776 | 0,194 | 0,186 | 1 | Secretory | 1,197449468 | 0,835108309 | 1,043010753 |
| SPECC11 | 0,406404796 | 0,26925156 | 0,129 | 0,118 | 1 | Secretory | 1,205182442 | 0,829749891 | 1,093220339 |
| DCUN1D4 | 0,41069968 | 0,25021201 | 0,147 | 0,136 | 1 | Secretory | 1,189381887 | 0,840772851 | 1,080882353 |
| LGALS1 | 0,411873638 | 0,502457804 | 0,362 | 0,386 | 1 | Secretory | 1,416624898 | 0,705903166 | 0,937823834 |
| DCAF10 | 0,414715998 | 0,274779761 | 0,185 | 0,178 | 1 | Secretory | 1,209809388 | 0,826576492 | 1,039325843 |
| GOLM1 | 0,419750125 | -0,361030577 | 0,293 | 0,321 | 1 | Secretory | 0,778608189 | 1,28434303 | 0,912772586 |
| PRDM2 | 0,423281189 | 0,307903634 | 0,185 | 0,175 | 1 | Secretory | 1,237907601 | 0,807814734 | 1,057142857 |
| SBNO1 | 0,425827077 | 0,263435501 | 0,19 | 0,186 | 1 | Secretory | 1,200333668 | 0,833101684 | 1,021505376 |
| CADPS2 | 0,426200445 | 0,284407094 | 0,31 | 0,331 | 1 | Secretory | 1,217909635 | 0,82107898 | 0,936555891 |
| ARF61 | 0,426599539 | 0,323376558 | 0,317 | 0,339 | 1 | Secretory | 1,251255628 | 0,799197205 | 0,935103245 |
| OSTC | 0,427377946 | 0,280255151 | 0,433 | 0,519 | 1 | Secretory | 1,214409642 | 0,823445372 | 0,834296724 |
| HOMER2 | 0,434615672 | 0,251598856 | 0,304 | 0,318 | 1 | Secretory | 1,190525776 | 0,839965014 | 0,955974843 |
| BNIP3L | 0,440209175 | 0,264843412 | 0,334 | 0,351 | 1 | Secretory | 1,201505633 | 0,832289065 | 0,951566952 |
| CCT7 | 0,440647065 | 0,297332256 | 0,345 | 0,381 | 1 | Secretory | 1,228869959 | 0,813755754 | 0,905511811 |
| PIK3R11 | 0,441054242 | 0,34991012 | 0,282 | 0,298 | 1 | Secretory | 1,274481224 | 0,784632979 | 0,946308725 |
| PSMA61 | 0,441297395 | 0,309498061 | 0,405 | 0,473 | 1 | Secretory | 1,239276459 | 0,806922453 | 0,856236786 |
| BBX | 0,454077795 | 0,272234362 | 0,31 | 0,373 | 1 | Secretory | 1,207676759 | 0,828036138 | 0,831099196 |
| RAD21 | 0,455260016 | 0,325451403 | 0,308 | 0,333 | 1 | Secretory | 1,253056444 | 0,798048647 | 0,924924925 |

| Supplementary Table 4 - clusermarker | | | | | | | | | |
|---|---|---|---|---|---|---|---|---|---|
| | **p_val** | **avg_log2FC** | **pct.1** | **pct.2** | **p_val_adj** | **celltype** | **Foldchange_UP** | **Foldchange_DOWN** | **Ratio_pct1_pct2** |
| MPHOSPH8 | 0,459244199 | 0,287421553 | 0,442 | 0,509 | 1 | Secretory | 1,220457074 | 0,819365156 | 0,868369352 |
| KPNA4 | 0,46135474 | 0,260487301 | 0,216 | 0,211 | 1 | Secretory | 1,197883247 | 0,834805899 | 1,023696682 |
| CNPY21 | 0,461689191 | 0,273320839 | 0,494 | 0,587 | 1 | Secretory | 1,20858659 | 0,827412788 | 0,841567291 |
| PSMB4 | 0,467069441 | 0,25071724 | 0,353 | 0,388 | 1 | Secretory | 1,18979848 | 0,840478465 | 0,909793814 |
| CD320 | 0,473680104 | 0,278164592 | 0,235 | 0,24 | 1 | Secretory | 1,212651158 | 0,824639463 | 0,979166667 |
| CRYL1 | 0,479513101 | 0,334779808 | 0,261 | 0,313 | 1 | Secretory | 1,261184905 | 0,792905145 | 0,833865815 |
| MAP3K2 | 0,482245328 | 0,267305954 | 0,248 | 0,253 | 1 | Secretory | 1,203558239 | 0,830869639 | 0,980237154 |
| RRAS | 0,483718484 | 0,283432687 | 0,278 | 0,286 | 1 | Secretory | 1,217087327 | 0,82163373 | 0,972027972 |
| MPC21 | 0,484586497 | 0,309317217 | 0,403 | 0,459 | 1 | Secretory | 1,239121123 | 0,807023608 | 0,877995643 |
| CHURC11 | 0,487670601 | 0,300241133 | 0,349 | 0,388 | 1 | Secretory | 1,231350205 | 0,812116647 | 0,899484536 |
| VDAC11 | 0,489646243 | 0,254819471 | 0,427 | 0,489 | 1 | Secretory | 1,193186427 | 0,838092001 | 0,873210634 |
| PRKAR2B | 0,492255952 | 0,274050816 | 0,358 | 0,396 | 1 | Secretory | 1,209198267 | 0,826994239 | 0,904040404 |
| CD991 | 0,498551284 | 0,251092244 | 0,435 | 0,551 | 1 | Secretory | 1,190107788 | 0,840260026 | 0,789473684 |
| RERE | 0,49885751 | 0,271889635 | 0,369 | 0,394 | 1 | Secretory | 1,207388224 | 0,828234018 | 0,936548223 |
| MCL1 | 0,504339435 | 0,367697565 | 0,416 | 0,469 | 1 | Secretory | 1,290291975 | 0,775018383 | 0,886993603 |
| BRD4 | 0,507294436 | 0,341884688 | 0,293 | 0,313 | 1 | Secretory | 1,267411216 | 0,789009903 | 0,936102236 |
| PTBP3 | 0,513755146 | 0,328566019 | 0,231 | 0,24 | 1 | Secretory | 1,255764574 | 0,796327608 | 0,9625 |
| RBL2 | 0,515876667 | 0,264218338 | 0,272 | 0,285 | 1 | Secretory | 1,200985171 | 0,832649748 | 0,954385965 |
| SBDS | 0,517734712 | 0,262040246 | 0,394 | 0,502 | 1 | Secretory | 1,199173366 | 0,833907781 | 0,784860558 |
| ITGB11 | 0,517937035 | 0,315366948 | 0,422 | 0,484 | 1 | Secretory | 1,244328107 | 0,803646558 | 0,871900826 |
| NOP10 | 0,518906931 | 0,380040991 | 0,364 | 0,409 | 1 | Secretory | 1,301378831 | 0,768415758 | 0,88997555 |
| CALM31 | 0,519749386 | 0,273562867 | 0,274 | 0,28 | 1 | Secretory | 1,20878936 | 0,827273992 | 0,978571429 |
| PCNP | 0,526086925 | 0,366153716 | 0,315 | 0,333 | 1 | Secretory | 1,288911954 | 0,775848185 | 0,945945946 |
| SOD21 | 0,53148104 | 0,530951023 | 0,44 | 0,474 | 1 | Secretory | 1,444881346 | 0,692098353 | 0,928270042 |
| HSD17B10 | 0,532156662 | 0,304224354 | 0,295 | 0,319 | 1 | Secretory | 1,234754609 | 0,809877519 | 0,92476489 |
| UFD1 | 0,5369626 | 0,268412335 | 0,276 | 0,29 | 1 | Secretory | 1,204481584 | 0,830232702 | 0,951724138 |
| CCNG11 | 0,544043759 | 0,329625026 | 0,304 | 0,329 | 1 | Secretory | 1,256686704 | 0,79574328 | 0,924012158 |
| POLR2A | 0,544266734 | 0,254642892 | 0,291 | 0,306 | 1 | Secretory | 1,193040395 | 0,838194586 | 0,950980392 |
| CAPZB | 0,545138377 | 0,2514743 | 0,373 | 0,421 | 1 | Secretory | 1,190422995 | 0,840037536 | 0,885985748 |
| STAT3 | 0,546171879 | 0,263091682 | 0,422 | 0,481 | 1 | Secretory | 1,200047642 | 0,83330025 | 0,877338877 |
| PGRMC1 | 0,54635427 | 0,28474278 | 0,418 | 0,481 | 1 | Secretory | 1,218193051 | 0,820887953 | 0,869022869 |

| Supplementary Table 4 - clusermarker | | | | | | | | | |
|---|---|---|---|---|---|---|---|---|---|
| | **p_val** | **avg_log2FC** | **pct.1** | **pct.2** | **p_val_adj** | **celltype** | **Foldchange_UP** | **Foldchange_DOWN** | **Ratio_pct1_pct2** |
| SRSF5 | 0,546570055 | 0,275313224 | 0,47 | 0,539 | 1 | Secretory | 1,21025682 | 0,826270906 | 0,871985158 |
| NAP1L1 | 0,548968936 | 0,311041335 | 0,496 | 0,571 | 1 | Secretory | 1,240602842 | 0,806059736 | 0,868651489 |
| FOXC11 | 0,555300924 | 0,282217898 | 0,386 | 0,438 | 1 | Secretory | 1,216062938 | 0,822325859 | 0,881278539 |
| MPZL2 | 0,557653245 | 0,359159182 | 0,381 | 0,426 | 1 | Secretory | 1,282678122 | 0,779618817 | 0,894366197 |
| MAT2A1 | 0,562924875 | 0,32837751 | 0,31 | 0,328 | 1 | Secretory | 1,255600501 | 0,796431667 | 0,945121951 |
| WTAP | 0,563646198 | 0,39943205 | 0,315 | 0,334 | 1 | Secretory | 1,318988559 | 0,75815669 | 0,943113772 |
| ATP5MF | 0,563716956 | 0,250469286 | 0,534 | 0,656 | 1 | Secretory | 1,189594008 | 0,840622929 | 0,81402439 |
| HNRNPH1 | 0,569079228 | 0,31134353 | 0,401 | 0,494 | 1 | Secretory | 1,240862733 | 0,805890913 | 0,811740891 |
| MT1E | 0,570598742 | 0,299374944 | 0,317 | 0,384 | 1 | Secretory | 1,230611128 | 0,812604386 | 0,825520833 |
| ASAH1 | 0,574961712 | 0,28319293 | 0,435 | 0,507 | 1 | Secretory | 1,21688508 | 0,821770286 | 0,857988166 |
| PIK3C2A1 | 0,575647759 | 0,258655796 | 0,194 | 0,218 | 1 | Secretory | 1,196363496 | 0,835866359 | 0,889908257 |
| ENSA | 0,582186557 | 0,291476898 | 0,362 | 0,404 | 1 | Secretory | 1,223892545 | 0,817065195 | 0,896039604 |
| IL6ST | 0,583724892 | 0,259934991 | 0,384 | 0,471 | 1 | Secretory | 1,197424747 | 0,83512555 | 0,815286624 |
| SOX41 | 0,589602112 | 0,281016604 | 0,392 | 0,434 | 1 | Secretory | 1,215050776 | 0,823010873 | 0,903225806 |
| GOLM21 | 0,590938403 | 0,299784387 | 0,353 | 0,394 | 1 | Secretory | 1,230960431 | 0,812373798 | 0,895939086 |
| TRAPPC2L1 | 0,594675833 | 0,360655754 | 0,338 | 0,373 | 1 | Secretory | 1,284009391 | 0,778810504 | 0,90616622 |
| VPS13C1 | 0,602610423 | 0,321165747 | 0,332 | 0,364 | 1 | Secretory | 1,249339651 | 0,800422847 | 0,912087912 |
| CD551 | 0,606570407 | 0,651763037 | 0,474 | 0,567 | 1 | Secretory | 1,571086961 | 0,636502005 | 0,835978836 |
| CCDC59 | 0,609131786 | 0,261328988 | 0,256 | 0,275 | 1 | Secretory | 1,198582312 | 0,834319004 | 0,930909091 |
| RAB7A | 0,610052314 | 0,363243477 | 0,384 | 0,434 | 1 | Secretory | 1,286314551 | 0,777414824 | 0,884792627 |
| SPEN | 0,61047788 | 0,334312956 | 0,265 | 0,281 | 1 | Secretory | 1,260776855 | 0,793161768 | 0,943060498 |
| SNRPN | 0,611060609 | 0,26834277 | 0,246 | 0,261 | 1 | Secretory | 1,204423507 | 0,830272736 | 0,942528736 |
| MBNL11 | 0,615192839 | 0,335129476 | 0,289 | 0,309 | 1 | Secretory | 1,261490617 | 0,792712991 | 0,935275081 |
| CTSS1 | 0,616713514 | 0,26634138 | 0,261 | 0,275 | 1 | Secretory | 1,202753819 | 0,831425338 | 0,949090909 |
| ARFGAP3 | 0,619613732 | 0,282059298 | 0,392 | 0,484 | 1 | Secretory | 1,21592926 | 0,822416265 | 0,809917355 |
| SENP6 | 0,620895583 | 0,265738247 | 0,291 | 0,311 | 1 | Secretory | 1,2022511 | 0,831772996 | 0,935691318 |
| PPP4R3B | 0,623995159 | 0,253929203 | 0,25 | 0,263 | 1 | Secretory | 1,192450354 | 0,838609337 | 0,950570342 |
| MGST2 | 0,624308991 | 0,28363013 | 0,377 | 0,423 | 1 | Secretory | 1,217253906 | 0,82152129 | 0,891252955 |
| CENPX1 | 0,62646553 | 0,273025267 | 0,362 | 0,441 | 1 | Secretory | 1,208339006 | 0,827582322 | 0,820861678 |
| TLK1 | 0,626793243 | 0,257180275 | 0,231 | 0,245 | 1 | Secretory | 1,195140537 | 0,836721682 | 0,942857143 |
| MLEC | 0,632235931 | 0,252066912 | 0,418 | 0,511 | 1 | Secretory | 1,190912082 | 0,839692547 | 0,818003914 |

| Supplementary Table 4 - clusermarker | | | | | | | | | |
|---|---|---|---|---|---|---|---|---|---|
| | **p_val** | **avg_log2FC** | **pct.1** | **pct.2** | **p_val_adj** | **celltype** | **Foldchange_UP** | **Foldchange_DOWN** | **Ratio_pct1_pct2** |
| SDF41 | 0,632896219 | 0,301342656 | 0,386 | 0,446 | 1 | Secretory | 1,232290721 | 0,811496819 | 0,865470852 |
| SMARCE11 | 0,633130009 | 0,255295591 | 0,297 | 0,319 | 1 | Secretory | 1,193580268 | 0,837815459 | 0,931034483 |
| CGGBP11 | 0,637590687 | 0,294768065 | 0,213 | 0,248 | 1 | Secretory | 1,226687753 | 0,815203378 | 0,858870968 |
| ADIPOR1 | 0,644272599 | 0,251024558 | 0,379 | 0,431 | 1 | Secretory | 1,190051954 | 0,840299448 | 0,879350348 |
| BPIFB11 | 0,651428217 | 0,475796033 | 0,595 | 0,631 | 1 | Secretory | 1,390685344 | 0,719069921 | 0,942947702 |
| RBM8A1 | 0,659558642 | 0,347364209 | 0,381 | 0,438 | 1 | Secretory | 1,272234142 | 0,786018836 | 0,869863014 |
| ZBTB38 | 0,659954339 | 0,339659142 | 0,341 | 0,371 | 1 | Secretory | 1,265457576 | 0,790227993 | 0,919137466 |
| SRSF7 | 0,665833287 | 0,294289767 | 0,431 | 0,488 | 1 | Secretory | 1,226281136 | 0,815473688 | 0,883196721 |
| SKIL | 0,666953899 | 0,290760952 | 0,179 | 0,2 | 1 | Secretory | 1,223285332 | 0,817470768 | 0,895 |
| DYNC1LI2 | 0,672090266 | 0,286361875 | 0,347 | 0,414 | 1 | Secretory | 1,219560962 | 0,819967211 | 0,838164251 |
| CDC42 | 0,674322271 | 0,272231266 | 0,431 | 0,536 | 1 | Secretory | 1,207674168 | 0,828037915 | 0,804104478 |
| SERTAD4 | 0,677445058 | 0,270799642 | 0,211 | 0,211 | 1 | Secretory | 1,206476355 | 0,828860007 | 1 |
| CA8 | 0,679959058 | 0,307273141 | 0,269 | 0,306 | 1 | Secretory | 1,237366723 | 0,808167847 | 0,879084967 |
| ANK31 | 0,684268109 | 0,293840059 | 0,36 | 0,401 | 1 | Secretory | 1,225898947 | 0,815727922 | 0,897755611 |
| UBE2B | 0,688858381 | 0,263383207 | 0,319 | 0,358 | 1 | Secretory | 1,20029016 | 0,833131882 | 0,891061453 |
| NSA2 | 0,694426487 | 0,31532246 | 0,466 | 0,566 | 1 | Secretory | 1,244289736 | 0,80367134 | 0,823321555 |
| SDC4 | 0,695352463 | 0,287605021 | 0,448 | 0,506 | 1 | Secretory | 1,22061229 | 0,819260963 | 0,885375494 |
| EIF4G21 | 0,699080353 | 0,316363339 | 0,517 | 0,624 | 1 | Secretory | 1,245187794 | 0,803091714 | 0,828525641 |
| CCDC186 | 0,699164402 | 0,335441716 | 0,332 | 0,364 | 1 | Secretory | 1,26176367 | 0,792541444 | 0,912087912 |
| TXNDC151 | 0,701879927 | 0,303374211 | 0,287 | 0,318 | 1 | Secretory | 1,234027214 | 0,8103549 | 0,902515723 |
| SSR3 | 0,703377595 | 0,250785578 | 0,494 | 0,597 | 1 | Secretory | 1,18985484 | 0,840438654 | 0,827470687 |
| POLR2H | 0,705083834 | 0,272270001 | 0,284 | 0,338 | 1 | Secretory | 1,207706593 | 0,828015683 | 0,840236686 |
| HSPA9 | 0,705823188 | 0,297223827 | 0,338 | 0,371 | 1 | Secretory | 1,228777604 | 0,813816916 | 0,911051213 |
| BACE21 | 0,719284621 | 0,356913435 | 0,356 | 0,418 | 1 | Secretory | 1,280683015 | 0,780833343 | 0,851674641 |
| CLSTN11 | 0,723919369 | 0,253875402 | 0,325 | 0,384 | 1 | Secretory | 1,192405886 | 0,83864061 | 0,846354167 |
| SPPL2A1 | 0,725624393 | 0,267922467 | 0,246 | 0,255 | 1 | Secretory | 1,204072671 | 0,830514656 | 0,964705882 |
| HDGF1 | 0,725825976 | 0,329608582 | 0,276 | 0,319 | 1 | Secretory | 1,25667238 | 0,79575235 | 0,865203762 |
| F11R | 0,730502524 | 0,259123613 | 0,375 | 0,421 | 1 | Secretory | 1,196751499 | 0,83559536 | 0,890736342 |
| PPP4R3A | 0,731453426 | 0,270297263 | 0,216 | 0,236 | 1 | Secretory | 1,206056306 | 0,829148685 | 0,915254237 |
| OSBPL91 | 0,731846333 | 0,269449989 | 0,233 | 0,243 | 1 | Secretory | 1,205348215 | 0,829635775 | 0,958847737 |
| ALDH9A11 | 0,734688385 | 0,295835628 | 0,392 | 0,476 | 1 | Secretory | 1,227595811 | 0,814600369 | 0,823529412 |

| Supplementary Table 4 - clusermarker | | | | | | | | | |
|---|---|---|---|---|---|---|---|---|---|
| | **p_val** | **avg_log2FC** | **pct.1** | **pct.2** | **p_val_adj** | **celltype** | **Foldchange_UP** | **Foldchange_DOWN** | **Ratio_pct1_pct2** |
| PTK2 | 0,735483666 | 0,274204449 | 0,302 | 0,351 | 1 | Secretory | 1,209327041 | 0,826906177 | 0,86039886 |
| AKAP91 | 0,738891859 | 0,330850978 | 0,448 | 0,521 | 1 | Secretory | 1,257755046 | 0,795067373 | 0,859884837 |
| SRP9 | 0,739611633 | 0,289758216 | 0,466 | 0,547 | 1 | Secretory | 1,22243539 | 0,818039144 | 0,851919561 |
| GAMT | 0,739866407 | 0,387819303 | 0,263 | 0,308 | 1 | Secretory | 1,308414183 | 0,764283981 | 0,853896104 |
| USP34 | 0,743780172 | 0,306981906 | 0,28 | 0,304 | 1 | Secretory | 1,237116963 | 0,808331006 | 0,921052632 |
| YTHDC11 | 0,750203754 | 0,368612584 | 0,332 | 0,376 | 1 | Secretory | 1,291110593 | 0,774526989 | 0,882978723 |
| YWHAE | 0,756289526 | 0,25362118 | 0,494 | 0,581 | 1 | Secretory | 1,192195787 | 0,838788403 | 0,850258176 |
| TMC51 | 0,758084533 | 0,411887059 | 0,338 | 0,393 | 1 | Secretory | 1,330424885 | 0,751639579 | 0,860050891 |
| TRIP11 | 0,762053452 | 0,36544747 | 0,284 | 0,328 | 1 | Secretory | 1,288281144 | 0,776228081 | 0,865853659 |
| SPAG7 | 0,763553808 | 0,302879134 | 0,278 | 0,308 | 1 | Secretory | 1,233603817 | 0,81063303 | 0,902597403 |
| CTNNA11 | 0,76492053 | 0,312444378 | 0,414 | 0,476 | 1 | Secretory | 1,241809934 | 0,805276212 | 0,869747899 |
| ZFAND3 | 0,76693438 | 0,313053857 | 0,183 | 0,186 | 1 | Secretory | 1,242334658 | 0,804936088 | 0,983870968 |
| SON1 | 0,770560524 | 0,270126026 | 0,552 | 0,636 | 1 | Secretory | 1,205913165 | 0,829247104 | 0,867924528 |
| SMG1 | 0,780674796 | 0,3764397 | 0,306 | 0,328 | 1 | Secretory | 1,298134348 | 0,770336292 | 0,932926829 |
| DNAJC8 | 0,780750079 | 0,268066628 | 0,287 | 0,311 | 1 | Secretory | 1,204192993 | 0,830431672 | 0,922829582 |
| TFG | 0,782832864 | 0,253191833 | 0,302 | 0,344 | 1 | Secretory | 1,191841041 | 0,839038064 | 0,877906977 |
| THBS1 | 0,788791249 | 0,577894745 | 0,282 | 0,311 | 1 | Secretory | 1,492669479 | 0,669940676 | 0,906752412 |
| EIF5A | 0,789748471 | 0,264147758 | 0,388 | 0,441 | 1 | Secretory | 1,200926418 | 0,832690484 | 0,879818594 |
| DHX29 | 0,789817286 | 0,285442683 | 0,267 | 0,301 | 1 | Secretory | 1,218784184 | 0,820489807 | 0,887043189 |
| SERPINA31 | 0,790432535 | 0,660226379 | 0,547 | 0,624 | 1 | Secretory | 1,58033058 | 0,632778997 | 0,876602564 |
| CASP4 | 0,799608495 | 0,255270082 | 0,235 | 0,25 | 1 | Secretory | 1,193559164 | 0,837830273 | 0,94 |
| TSPYL1 | 0,80731819 | 0,317623244 | 0,3 | 0,324 | 1 | Secretory | 1,24627569 | 0,802390681 | 0,925925926 |
| PAFAH1B1 | 0,811525494 | 0,270025301 | 0,291 | 0,318 | 1 | Secretory | 1,205828975 | 0,829305002 | 0,91509434 |
| SAT2 | 0,81330982 | 0,300362235 | 0,364 | 0,446 | 1 | Secretory | 1,231453571 | 0,81204848 | 0,816143498 |
| UBE2D3 | 0,814170367 | 0,269793778 | 0,427 | 0,498 | 1 | Secretory | 1,205635479 | 0,829438099 | 0,857429719 |
| CHD91 | 0,814905414 | 0,275825211 | 0,334 | 0,398 | 1 | Secretory | 1,210686395 | 0,82597773 | 0,83919598 |
| MRPL40 | 0,815808619 | 0,324026811 | 0,366 | 0,436 | 1 | Secretory | 1,251819723 | 0,798837071 | 0,839449541 |
| FNBP1L | 0,816745138 | 0,254853419 | 0,388 | 0,451 | 1 | Secretory | 1,193214504 | 0,83807228 | 0,860310421 |
| ATOX11 | 0,820358111 | 0,29540647 | 0,429 | 0,493 | 1 | Secretory | 1,227230693 | 0,814842723 | 0,870182556 |
| EIF4A3 | 0,823775557 | 0,2991528 | 0,306 | 0,336 | 1 | Secretory | 1,230421655 | 0,812729519 | 0,910714286 |
| FAM177A11 | 0,830215315 | 0,286608668 | 0,341 | 0,391 | 1 | Secretory | 1,219769602 | 0,819826956 | 0,872122762 |

| Supplementary Table 4 - clusermarker | | | | | | | | | |
|---|---|---|---|---|---|---|---|---|---|
| | **p_val** | **avg_log2FC** | **pct.1** | **pct.2** | **p_val_adj** | **celltype** | **Foldchange_UP** | **Foldchange_DOWN** | **Ratio_pct1_pct2** |
| GTF2H5 | 0,835044689 | 0,294902875 | 0,328 | 0,384 | 1 | Secretory | 1,226802384 | 0,815127206 | 0,854166667 |
| RCN1 | 0,835431418 | 0,340085757 | 0,358 | 0,409 | 1 | Secretory | 1,265831835 | 0,789994352 | 0,875305623 |
| CAMTA11 | 0,83689322 | 0,331632889 | 0,33 | 0,379 | 1 | Secretory | 1,258436908 | 0,794636579 | 0,870712401 |
| EIF4E2 | 0,840076934 | 0,269512105 | 0,211 | 0,225 | 1 | Secretory | 1,205400113 | 0,829600055 | 0,937777778 |
| KLF4 | 0,84633337 | 0,338786858 | 0,351 | 0,403 | 1 | Secretory | 1,264692685 | 0,790705926 | 0,870967742 |
| TCF12 | 0,855143395 | 0,252624651 | 0,259 | 0,281 | 1 | Secretory | 1,191372572 | 0,839367989 | 0,921708185 |
| PDLIM5 | 0,863736017 | 0,276130906 | 0,397 | 0,458 | 1 | Secretory | 1,210942956 | 0,825802731 | 0,866812227 |
| MBD5 | 0,871719663 | 0,252954361 | 0,099 | 0,1 | 1 | Secretory | 1,191644877 | 0,839176183 | 0,99 |
| MAP4 | 0,871995159 | 0,25664649 | 0,261 | 0,286 | 1 | Secretory | 1,194698427 | 0,837031319 | 0,912587413 |
| LPIN1 | 0,873391121 | 0,283374749 | 0,19 | 0,205 | 1 | Secretory | 1,217038451 | 0,821666727 | 0,926829268 |
| SERPINB11 | 0,875349285 | 0,434869298 | 0,442 | 0,516 | 1 | Secretory | 1,351788362 | 0,739760771 | 0,856589147 |
| SH3YL1 | 0,890069871 | 0,303775034 | 0,422 | 0,498 | 1 | Secretory | 1,234370111 | 0,810129791 | 0,847389558 |
| ZCRB11 | 0,890781268 | 0,278264567 | 0,323 | 0,383 | 1 | Secretory | 1,212735195 | 0,824582319 | 0,843342037 |
| HBP11 | 0,891306622 | 0,257067701 | 0,278 | 0,316 | 1 | Secretory | 1,195047283 | 0,836786974 | 0,879746835 |
| WEE1 | 0,896532074 | 0,291989946 | 0,168 | 0,171 | 1 | Secretory | 1,22432786 | 0,816774683 | 0,98245614 |
| KMT2C1 | 0,899481445 | 0,254461311 | 0,362 | 0,406 | 1 | Secretory | 1,192890246 | 0,83830009 | 0,891625616 |
| MTCH11 | 0,907165559 | 0,260852027 | 0,461 | 0,554 | 1 | Secretory | 1,19818612 | 0,834594879 | 0,832129964 |
| PRPF4B | 0,91007278 | 0,333713626 | 0,278 | 0,308 | 1 | Secretory | 1,260253207 | 0,793491335 | 0,902597403 |
| UBE2N | 0,913583968 | 0,268565571 | 0,263 | 0,29 | 1 | Secretory | 1,204609524 | 0,830144524 | 0,906896552 |
| EMC7 | 0,917822987 | 0,298901203 | 0,291 | 0,329 | 1 | Secretory | 1,230207096 | 0,812871266 | 0,88449848 |
| SUB1 | 0,921116851 | 0,298081196 | 0,478 | 0,574 | 1 | Secretory | 1,229508063 | 0,813333422 | 0,832752613 |
| PKP41 | 0,921502541 | 0,30453606 | 0,328 | 0,376 | 1 | Secretory | 1,235021416 | 0,809702558 | 0,872340426 |
| TOMM51 | 0,922470189 | 0,335848573 | 0,36 | 0,423 | 1 | Secretory | 1,262119552 | 0,792317969 | 0,85106383 |
| DDX42 | 0,924206195 | 0,327232716 | 0,297 | 0,326 | 1 | Secretory | 1,254604564 | 0,797063894 | 0,911042945 |
| NANS1 | 0,924336418 | 0,285154055 | 0,466 | 0,559 | 1 | Secretory | 1,218540376 | 0,820653972 | 0,833631485 |
| MDM4 | 0,92725853 | 0,254728613 | 0,205 | 0,223 | 1 | Secretory | 1,193111285 | 0,838144784 | 0,919282511 |
| KPNB11 | 0,927538444 | 0,325178154 | 0,36 | 0,426 | 1 | Secretory | 1,252819135 | 0,798199813 | 0,845070423 |
| KRIT1 | 0,928449089 | 0,282590634 | 0,224 | 0,246 | 1 | Secretory | 1,216377162 | 0,822113429 | 0,910569106 |
| BTG1 | 0,949758471 | 0,271033261 | 0,489 | 0,547 | 1 | Secretory | 1,206671739 | 0,828725798 | 0,893967093 |
| TMEM106B | 0,950799033 | 0,258959735 | 0,308 | 0,341 | 1 | Secretory | 1,196615566 | 0,835690282 | 0,903225806 |
| YIPF3 | 0,95158218 | 0,34835325 | 0,425 | 0,507 | 1 | Secretory | 1,273106623 | 0,785480165 | 0,8382643 |

| Supplementary Table 4 - clusermarker | | | | | | | | | |
|---|---|---|---|---|---|---|---|---|---|
| | **p_val** | **avg_log2FC** | **pct.1** | **pct.2** | **p_val_adj** | **celltype** | **Foldchange_UP** | **Foldchange_DOWN** | **Ratio_pct1_pct2** |
| ZNF207 | 0,958478811 | 0,250191732 | 0,304 | 0,336 | 1 | Secretory | 1,189365169 | 0,840784669 | 0,904761905 |
| SMIM14 | 0,961863587 | 0,281401167 | 0,429 | 0,501 | 1 | Secretory | 1,215374701 | 0,822791522 | 0,856287425 |
| TTC32 | 0,969849476 | 0,274049248 | 0,478 | 0,567 | 1 | Secretory | 1,209196953 | 0,826995137 | 0,84303351 |
| MDH11 | 0,970939545 | 0,282050988 | 0,416 | 0,506 | 1 | Secretory | 1,215922255 | 0,822421002 | 0,822134387 |
| SH3GLB1 | 0,970978837 | 0,279823505 | 0,33 | 0,376 | 1 | Secretory | 1,214046353 | 0,823691779 | 0,877659574 |
| ZNHIT3 | 0,971289175 | 0,278740434 | 0,209 | 0,23 | 1 | Secretory | 1,213135276 | 0,82431038 | 0,908695652 |
| TFAP2C | 0,973319471 | 0,277540863 | 0,179 | 0,188 | 1 | Secretory | 1,212126999 | 0,824996061 | 0,95212766 |
| ARHGAP51 | 0,988817179 | 0,318092552 | 0,343 | 0,394 | 1 | Secretory | 1,246681169 | 0,802129707 | 0,870558376 |
| KCNQ1OT1 | 0,989007122 | 0,764041104 | 0,254 | 0,28 | 1 | Secretory | 1,698240877 | 0,588844618 | 0,907142857 |
| TIMM8B | 0,991250253 | 0,27785765 | 0,267 | 0,3 | 1 | Secretory | 1,212393187 | 0,824814929 | 0,89 |
| TXN2 | 0,997732458 | 0,310102447 | 0,36 | 0,426 | 1 | Secretory | 1,239795736 | 0,806584481 | 0,845070423 |
| DECR1 | 0,998366836 | 0,333526287 | 0,302 | 0,348 | 1 | Secretory | 1,260089569 | 0,79359438 | 0,867816092 |
| ZCCHC7 | 0,999299692 | 0,255988826 | 0,183 | 0,198 | 1 | Secretory | 1,194153938 | 0,837412973 | 0,924242424 |
| IGFBP72 | 5,26451E-10 | 1,952238753 | 1 | 0,756 | 1,92613E-05 | Ciliated | 3,869745673 | 0,258414915 | 1,322751323 |
| LYZ2 | 5,43877E-09 | -4,377761487 | 0,071 | 0,829 | 0,000198988 | Ciliated | 0,048101927 | 20,78918775 | 0,085645356 |
| ECRG4 | 5,54212E-09 | 2,038688231 | 0,964 | 0,707 | 0,000202769 | Ciliated | 4,108717761 | 0,243384934 | 1,363507779 |
| CYBA2 | 2,23912E-08 | 1,307347584 | 1 | 0,854 | 0,000819227 | Ciliated | 2,474861146 | 0,404063073 | 1,170960187 |
| RPS271 | 2,79233E-08 | -1,313496746 | 1 | 0,976 | 0,001021628 | Ciliated | 0,40234451 | 2,485432196 | 1,024590164 |
| TMEM190 | 4,49548E-08 | 1,672951311 | 1 | 0,854 | 0,001644761 | Ciliated | 3,188662292 | 0,313611135 | 1,170960187 |
| PRR42 | 8,20581E-08 | -5,735718868 | 0,214 | 0,878 | 0,003002259 | Ciliated | 0,01876621 | 53,28726447 | 0,243735763 |
| LCN22 | 1,96E-07 | 2,363243515 | 0,964 | 0,585 | 0,007171069 | Ciliated | 5,145258336 | 0,194353701 | 1,647863248 |
| BBLN2 | 2,46896E-07 | 1,111815678 | 0,929 | 0,707 | 0,009033186 | Ciliated | 2,161174669 | 0,462711327 | 1,314002829 |
| FTH12 | 2,95517E-07 | 0,854703788 | 1 | 1 | 0,010812097 | Ciliated | 1,808387422 | 0,552978852 | 1 |
| CCDC74B | 3,10914E-07 | 1,141458054 | 0,857 | 0,415 | 0,011375426 | Ciliated | 2,206038629 | 0,453301219 | 2,065060241 |
| GPX42 | 3,46541E-07 | 1,074398677 | 0,964 | 0,927 | 0,012678884 | Ciliated | 2,105844162 | 0,474868947 | 1,0399137 |
| GSTA12 | 5,60751E-07 | -1,499792797 | 0,893 | 1 | 0,020516206 | Ciliated | 0,353604172 | 2,828020929 | 0,893 |
| MGMT2 | 6,72761E-07 | 0,903682075 | 0,893 | 0,488 | 0,024614302 | Ciliated | 1,870834676 | 0,534520774 | 1,829918033 |
| GSTA2 | 1,18619E-06 | -2,312804654 | 0,643 | 0,951 | 0,043399006 | Ciliated | 0,201268785 | 4,96848033 | 0,676130389 |
| ST6GALNAC6 | 1,78155E-06 | 1,008143633 | 0,893 | 0,463 | 0,065181643 | Ciliated | 2,011321396 | 0,497185583 | 1,928725702 |
| CIB12 | 2,31821E-06 | 0,970106202 | 1 | 0,902 | 0,084816524 | Ciliated | 1,958984797 | 0,510468484 | 1,10864745 |
| ARL6IP42 | 2,3216E-06 | 0,87972469 | 1 | 0,707 | 0,08494034 | Ciliated | 1,840024135 | 0,543471132 | 1,414427157 |

| Supplementary Table 4 - clusermarker | | | | | | | | | |
|---|---|---|---|---|---|---|---|---|---|
| | **p_val** | **avg_log2FC** | **pct.1** | **pct.2** | **p_val_adj** | **celltype** | **Foldchange_UP** | **Foldchange_DOWN** | **Ratio_pct1_pct2** |
| GBP31 | 3,10052E-06 | -0,814713371 | 0,071 | 0,659 | 0,113438656 | Ciliated | 0,568521427 | 1,758948656 | 0,107738998 |
| SDF2L11 | 4,01457E-06 | 1,08563961 | 0,714 | 0,317 | 0,146881067 | Ciliated | 2,122316191 | 0,471183325 | 2,252365931 |
| MUC12 | 4,48391E-06 | 1,243756836 | 0,571 | 0,073 | 0,164052963 | Ciliated | 2,368144044 | 0,422271611 | 7,821917808 |
| CHMP2A2 | 5,08716E-06 | 0,961439685 | 0,964 | 0,756 | 0,186123855 | Ciliated | 1,947252115 | 0,513544185 | 1,275132275 |
| C9orf24 | 7,95452E-06 | 0,919346192 | 1 | 0,976 | 0,291031996 | Ciliated | 1,891258009 | 0,528748587 | 1,024590164 |
| BRI32 | 8,07589E-06 | 0,95084444 | 1 | 0,732 | 0,295472566 | Ciliated | 1,933003756 | 0,517329569 | 1,366120219 |
| CTXN1 | 8,98355E-06 | 1,107682957 | 0,964 | 0,659 | 0,328681169 | Ciliated | 2,154992662 | 0,464038703 | 1,462822458 |
| CSTB2 | 9,97633E-06 | 0,652254715 | 1 | 0,927 | 0,365003914 | Ciliated | 1,571622487 | 0,636285118 | 1,078748652 |
| RPS292 | 1,13489E-05 | -1,106025827 | 0,786 | 0,878 | 0,415222303 | Ciliated | 0,46457202 | 2,152518783 | 0,895216401 |
| JTB2 | 1,13736E-05 | 0,903565994 | 0,929 | 0,634 | 0,416125149 | Ciliated | 1,870684153 | 0,534563784 | 1,465299685 |
| SCGB3A11 | 1,14085E-05 | -6,0560617 | 0,393 | 0,805 | 0,417402708 | Ciliated | 0,015029474 | 66,53592937 | 0,488198758 |
| TMEM50A2 | 1,33604E-05 | 0,747323903 | 0,857 | 0,561 | 0,488816055 | Ciliated | 1,678676116 | 0,595707528 | 1,527629234 |
| GRIN3B | 1,35738E-05 | 0,929875115 | 0,679 | 0,22 | 0,496626442 | Ciliated | 1,905111075 | 0,524903778 | 3,086363636 |
| CLDN32 | 1,52846E-05 | 0,907134399 | 0,964 | 0,805 | 0,559216763 | Ciliated | 1,875316886 | 0,533243212 | 1,197515528 |
| HOTAIRM11 | 1,9254E-05 | 1,095224289 | 0,893 | 0,537 | 0,704444595 | Ciliated | 2,136462937 | 0,46806335 | 1,662942272 |
| JOSD21 | 1,98415E-05 | 0,704099618 | 0,929 | 0,585 | 0,725942058 | Ciliated | 1,629127613 | 0,613825456 | 1,588034188 |
| SERPINB21 | 2,06909E-05 | 1,25367757 | 0,5 | 0,073 | 0,757019019 | Ciliated | 2,384484774 | 0,419377809 | 6,849315068 |
| FAM27C | 2,80509E-05 | 1,005316765 | 0,714 | 0,39 | 1 | Ciliated | 2,007384199 | 0,498160741 | 1,830769231 |
| TMEM1472 | 2,9068E-05 | 0,619601496 | 0,786 | 0,463 | 1 | Ciliated | 1,536450721 | 0,650850682 | 1,69762419 |
| VSTM2L1 | 3,46684E-05 | 0,762727551 | 0,893 | 0,512 | 1 | Ciliated | 1,696695357 | 0,589380997 | 1,744140625 |
| CMBL | 3,7585E-05 | -0,376506987 | 0 | 0,463 | 1 | Ciliated | 0,770300364 | 1,298194894 | 0 |
| SAA12 | 4,29883E-05 | 2,550870126 | 0,5 | 0,098 | 1 | Ciliated | 5,859875957 | 0,170652076 | 5,102040816 |
| GABARAP2 | 4,33438E-05 | 0,742803019 | 1 | 0,854 | 1 | Ciliated | 1,673423986 | 0,597577188 | 1,170960187 |
| MAP6 | 4,42009E-05 | 0,816550109 | 0,857 | 0,585 | 1 | Ciliated | 1,761189452 | 0,567798086 | 1,464957265 |
| SAA22 | 4,47317E-05 | 1,586521378 | 0,357 | 0 | 1 | Ciliated | 3,003243347 | 0,332973351 | #DIV/0! |
| CDKN2A2 | 4,60188E-05 | 0,932725238 | 0,5 | 0,098 | 1 | Ciliated | 1,908878447 | 0,523867825 | 5,102040816 |
| DDT2 | 4,894E-05 | 1,028541129 | 0,607 | 0,195 | 1 | Ciliated | 2,039960375 | 0,4902056 | 3,112820513 |
| ATP5F1D2 | 5,24617E-05 | 0,976938241 | 0,857 | 0,634 | 1 | Ciliated | 1,968283783 | 0,50805682 | 1,351735016 |
| COX6A12 | 6,25938E-05 | 0,647920884 | 1 | 0,878 | 1 | Ciliated | 1,566908444 | 0,638199382 | 1,138952164 |
| RPGR | 6,36595E-05 | -1,005686 | 0,286 | 0,707 | 1 | Ciliated | 0,498033261 | 2,007898023 | 0,404526167 |
| CD92 | 6,43402E-05 | 0,649593009 | 0,964 | 0,927 | 1 | Ciliated | 1,568725588 | 0,637460119 | 1,0399137 |

| Supplementary Table 4 - clusermarker | | | | | | | | | |
|---|---|---|---|---|---|---|---|---|---|
| | **p_val** | **avg_log2FC** | **pct.1** | **pct.2** | **p_val_adj** | **celltype** | **Foldchange_UP** | **Foldchange_DOWN** | **Ratio_pct1_pct2** |
| PSMA72 | 6,79003E-05 | 0,899453478 | 0,893 | 0,585 | 1 | Ciliated | 1,865359213 | 0,536089774 | 1,526495726 |
| NDUFB101 | 7,19639E-05 | 0,716311639 | 0,929 | 0,707 | 1 | Ciliated | 1,642976265 | 0,608651519 | 1,314002829 |
| RPLP22 | 7,61509E-05 | 0,767648035 | 1 | 0,805 | 1 | Ciliated | 1,702492019 | 0,587374266 | 1,242236025 |
| RPS261 | 8,11612E-05 | -0,835455876 | 1 | 0,976 | 1 | Ciliated | 0,560405929 | 1,784420807 | 1,024590164 |
| KRT232 | 8,13446E-05 | 0,971202181 | 0,75 | 0,415 | 1 | Ciliated | 1,960473555 | 0,510080841 | 1,807228916 |
| NANS2 | 8,39262E-05 | 0,441195788 | 0,964 | 0,732 | 1 | Ciliated | 1,357729224 | 0,736523883 | 1,316939891 |
| PRDX52 | 8,53974E-05 | 0,651803829 | 1 | 1 | 1 | Ciliated | 1,571131384 | 0,636484008 | 1 |
| GSN2 | 8,60919E-05 | 1,15683175 | 0,929 | 0,756 | 1 | Ciliated | 2,229672394 | 0,448496381 | 1,228835979 |
| PSCA | 8,95757E-05 | -1,185787561 | 0,929 | 0,927 | 1 | Ciliated | 0,439584505 | 2,274875452 | 1,002157497 |
| ENDOG1 | 9,52757E-05 | 0,821736038 | 0,929 | 0,61 | 1 | Ciliated | 1,767531637 | 0,565760736 | 1,52295082 |
| USP40 | 9,7249E-05 | -0,611826935 | 0,071 | 0,561 | 1 | Ciliated | 0,654367529 | 1,528193189 | 0,126559715 |
| STMND1 | 0,000101336 | 0,599241142 | 0,786 | 0,512 | 1 | Ciliated | 1,514919509 | 0,660101077 | 1,53515625 |
| MRPS242 | 0,000110056 | 0,762226485 | 0,929 | 0,659 | 1 | Ciliated | 1,696106175 | 0,589585731 | 1,409711684 |
| TRMT1122 | 0,000117758 | 0,784644858 | 0,929 | 0,683 | 1 | Ciliated | 1,722668206 | 0,580494837 | 1,360175695 |
| C11orf97 | 0,000117775 | 0,810561325 | 0,964 | 0,659 | 1 | Ciliated | 1,753893717 | 0,570159976 | 1,462822458 |
| SH3BGRL1 | 0,000118895 | -0,986478079 | 0,214 | 0,707 | 1 | Ciliated | 0,504708371 | 1,981342211 | 0,302687412 |
| RAMP11 | 0,000128739 | 0,906962015 | 0,75 | 0,341 | 1 | Ciliated | 1,875092822 | 0,533306932 | 2,19941349 |
| COPRS | 0,00012901 | 0,652186697 | 0,929 | 0,61 | 1 | Ciliated | 1,571548392 | 0,636315118 | 1,52295082 |
| RSPH9 | 0,000130165 | 0,979444022 | 0,893 | 0,707 | 1 | Ciliated | 1,971705417 | 0,507175155 | 1,263083451 |
| CABCOCO1 | 0,000130779 | 1,073045941 | 0,857 | 0,439 | 1 | Ciliated | 2,103870553 | 0,475314415 | 1,952164009 |
| FOLR1 | 0,000150598 | -0,782140702 | 0,036 | 0,463 | 1 | Ciliated | 0,581503306 | 1,71968068 | 0,07775378 |
| GUK12 | 0,000153714 | 0,757384355 | 1 | 0,805 | 1 | Ciliated | 1,690423062 | 0,591567888 | 1,242236025 |
| LINC02166 | 0,000157252 | 0,834042944 | 0,893 | 0,512 | 1 | Ciliated | 1,782674055 | 0,560955042 | 1,744140625 |
| CLTB2 | 0,000159964 | 0,84840212 | 0,786 | 0,537 | 1 | Ciliated | 1,800505642 | 0,555399537 | 1,463687151 |
| CHCHD22 | 0,000168715 | 0,700679752 | 1 | 0,78 | 1 | Ciliated | 1,625270388 | 0,615282237 | 1,282051282 |
| ODF3B1 | 0,00017999 | 0,741019154 | 1 | 0,927 | 1 | Ciliated | 1,671356108 | 0,598316538 | 1,078748652 |
| U2AF2 | 0,000187185 | -0,397314867 | 0,036 | 0,488 | 1 | Ciliated | 0,759270117 | 1,317054337 | 0,073770492 |
| REX1BD2 | 0,000198784 | 0,484035351 | 0,821 | 0,537 | 1 | Ciliated | 1,398650354 | 0,714974974 | 1,52886406 |
| COX4I12 | 0,000208161 | 0,645328656 | 1 | 0,927 | 1 | Ciliated | 1,564095558 | 0,639347126 | 1,078748652 |
| RPS111 | 0,000218234 | -0,597094779 | 0,929 | 0,951 | 1 | Ciliated | 0,661083871 | 1,51266737 | 0,976866456 |
| TMEM8B1 | 0,000221443 | -0,38597006 | 0 | 0,39 | 1 | Ciliated | 0,765264266 | 1,306738136 | 0 |

| Supplementary Table 4 - clusermarker | | | | | | | | | |
|---|---|---|---|---|---|---|---|---|---|
| | **p_val** | **avg_log2FC** | **pct.1** | **pct.2** | **p_val_adj** | **celltype** | **Foldchange_UP** | **Foldchange_DOWN** | **Ratio_pct1_pct2** |
| SMARCD1 | 0,000221443 | -0,408176383 | 0 | 0,39 | 1 | Ciliated | 0,753575318 | 1,32700737 | 0 |
| HINT12 | 0,000264497 | 0,652067526 | 1 | 0,927 | 1 | Ciliated | 1,571418583 | 0,636367681 | 1,078748652 |
| SRSF92 | 0,00028384 | 0,711738843 | 0,929 | 0,683 | 1 | Ciliated | 1,637776898 | 0,610583774 | 1,360175695 |
| BCL7C1 | 0,000292221 | 0,73527433 | 0,607 | 0,268 | 1 | Ciliated | 1,664713987 | 0,600703789 | 2,264925373 |
| NGRN | 0,000293033 | 0,971687179 | 0,857 | 0,707 | 1 | Ciliated | 1,961132727 | 0,509909394 | 1,212164074 |
| ATP6V0E12 | 0,000301102 | 0,824505528 | 0,893 | 0,78 | 1 | Ciliated | 1,770927963 | 0,564675707 | 1,144871795 |
| RPL35A2 | 0,000305012 | -0,685628018 | 1 | 1 | 1 | Ciliated | 0,621735121 | 1,608401981 | 1 |
| ODAD2 | 0,000317656 | -1,031147276 | 0,321 | 0,756 | 1 | Ciliated | 0,489320871 | 2,043648779 | 0,424603175 |
| PTPMT11 | 0,000324585 | 0,908452348 | 0,643 | 0,317 | 1 | Ciliated | 1,877030832 | 0,532756299 | 2,028391167 |
| CCDC96 | 0,000327512 | 0,819343564 | 0,821 | 0,561 | 1 | Ciliated | 1,764602903 | 0,566699736 | 1,463458111 |
| S100A102 | 0,000341495 | 0,68426371 | 0,964 | 0,878 | 1 | Ciliated | 1,606881689 | 0,622323353 | 1,097949886 |
| IFT27 | 0,00034753 | 0,721414875 | 0,929 | 0,707 | 1 | Ciliated | 1,648798246 | 0,606502343 | 1,314002829 |
| GSTP12 | 0,000351105 | 0,464755589 | 1 | 0,976 | 1 | Ciliated | 1,38008353 | 0,724593822 | 1,024590164 |
| SRP14-DT | 0,000388615 | -0,430071292 | 0 | 0,366 | 1 | Ciliated | 0,742225107 | 1,347300153 | 0 |
| FABP51 | 0,000400699 | -1,132432684 | 0,143 | 0,585 | 1 | Ciliated | 0,456145919 | 2,192280929 | 0,244444444 |
| NDUFB112 | 0,00040085 | 0,557777365 | 0,786 | 0,561 | 1 | Ciliated | 1,471999688 | 0,67934797 | 1,401069519 |
| STUB12 | 0,000412465 | 0,582152631 | 0,857 | 0,61 | 1 | Ciliated | 1,497081363 | 0,667966367 | 1,404918033 |
| NDUFB12 | 0,000413436 | -0,826442884 | 0,643 | 0,878 | 1 | Ciliated | 0,563917929 | 1,773307691 | 0,732346241 |
| NAT14 | 0,000418109 | 0,595557446 | 0,75 | 0,366 | 1 | Ciliated | 1,511056333 | 0,661788696 | 2,049180328 |
| CIBAR2 | 0,0004408 | 0,765785638 | 1 | 0,854 | 1 | Ciliated | 1,700295664 | 0,588133006 | 1,170960187 |
| PFKP1 | 0,000458495 | 0,780163335 | 0,75 | 0,488 | 1 | Ciliated | 1,71732529 | 0,582300864 | 1,536885246 |
| OXTR | 0,000460066 | 0,538568665 | 0,75 | 0,39 | 1 | Ciliated | 1,452530709 | 0,688453603 | 1,923076923 |
| PRDX11 | 0,000463471 | 0,584793892 | 1 | 1 | 1 | Ciliated | 1,499824704 | 0,666744585 | 1 |
| RPL211 | 0,000463471 | -0,774148283 | 1 | 0,976 | 1 | Ciliated | 0,584733729 | 1,710180122 | 1,024590164 |
| CXXC4 | 0,000477467 | -0,643063972 | 0,036 | 0,439 | 1 | Ciliated | 0,640351536 | 1,56164223 | 0,082004556 |
| MSMB2 | 0,000479288 | -4,293208206 | 0,143 | 0,561 | 1 | Ciliated | 0,051005329 | 19,60579456 | 0,254901961 |
| BAX2 | 0,000494458 | 0,536119587 | 0,714 | 0,366 | 1 | Ciliated | 1,450067026 | 0,689623295 | 1,950819672 |
| KLHL6 | 0,00050681 | 0,730562257 | 0,786 | 0,561 | 1 | Ciliated | 1,659285634 | 0,602668992 | 1,401069519 |
| PRR29 | 0,000530986 | 0,763065403 | 0,893 | 0,634 | 1 | Ciliated | 1,697092737 | 0,589242991 | 1,40851735 |
| CRIP22 | 0,000543222 | 0,794128483 | 0,929 | 0,61 | 1 | Ciliated | 1,734029549 | 0,576691441 | 1,52295082 |
| FAM177A12 | 0,000543287 | 0,619091535 | 0,857 | 0,659 | 1 | Ciliated | 1,535907716 | 0,651080784 | 1,300455235 |

| Supplementary Table 4 - clusermarker | | | | | | | | | |
|---|---|---|---|---|---|---|---|---|---|
| | **p_val** | **avg_log2FC** | **pct.1** | **pct.2** | **p_val_adj** | **celltype** | **Foldchange_UP** | **Foldchange_DOWN** | **Ratio_pct1_pct2** |
| ARPC1B2 | 0,00055724 | 0,731799103 | 0,857 | 0,634 | 1 | Ciliated | 1,660708776 | 0,602152535 | 1,351735016 |
| COMT2 | 0,000579006 | 0,453990805 | 0,929 | 0,634 | 1 | Ciliated | 1,369824249 | 0,730020658 | 1,465299685 |
| AC244090.11 | 0,000595197 | 0,845311765 | 0,857 | 0,659 | 1 | Ciliated | 1,796652959 | 0,556590517 | 1,300455235 |
| RPL37A2 | 0,000604783 | -0,629855793 | 0,857 | 0,927 | 1 | Ciliated | 0,646241008 | 1,547410312 | 0,924487594 |
| MT1E1 | 0,000610689 | -1,298760163 | 0,536 | 0,829 | 1 | Ciliated | 0,406475369 | 2,46017367 | 0,646562123 |
| RPL372 | 0,000622477 | -0,765419415 | 0,964 | 0,976 | 1 | Ciliated | 0,588282321 | 1,699864104 | 0,987704918 |
| MRPL572 | 0,000629778 | 0,876720977 | 0,893 | 0,683 | 1 | Ciliated | 1,836197162 | 0,544603826 | 1,307467057 |
| TYMP2 | 0,000638832 | 0,631450065 | 0,893 | 0,61 | 1 | Ciliated | 1,549121246 | 0,645527264 | 1,463934426 |
| PSMG4 | 0,000655839 | 0,592634204 | 0,643 | 0,317 | 1 | Ciliated | 1,507997675 | 0,663130996 | 2,028391167 |
| PNKP2 | 0,000663064 | 0,547860127 | 0,607 | 0,293 | 1 | Ciliated | 1,461915705 | 0,684033968 | 2,071672355 |
| PRB32 | 0,000672929 | -1,801979246 | 0 | 0,341 | 1 | Ciliated | 0,286780882 | 3,486982795 | 0 |
| CSE1L | 0,000672929 | -0,288174947 | 0 | 0,341 | 1 | Ciliated | 0,818937384 | 1,221094578 | 0 |
| NDUFS81 | 0,000683178 | 0,576779782 | 0,893 | 0,707 | 1 | Ciliated | 1,491516339 | 0,670458629 | 1,263083451 |
| MORN1 | 0,000706312 | 0,662291497 | 0,679 | 0,439 | 1 | Ciliated | 1,582594334 | 0,631873866 | 1,546697039 |
| SSUH2 | 0,000716944 | 0,510310496 | 0,321 | 0,024 | 1 | Ciliated | 1,424356712 | 0,702071322 | 13,375 |
| INSR | 0,000721081 | -0,695011678 | 0,179 | 0,61 | 1 | Ciliated | 0,617704319 | 1,618897537 | 0,293442623 |
| DNMBP-AS1 | 0,000771435 | -0,575912892 | 0,036 | 0,415 | 1 | Ciliated | 0,670861617 | 1,490620383 | 0,086746988 |
| POLR2E1 | 0,000782873 | 0,389837713 | 0,857 | 0,61 | 1 | Ciliated | 1,310246007 | 0,763215453 | 1,404918033 |
| SNRPB21 | 0,000813403 | -0,845950772 | 0,25 | 0,659 | 1 | Ciliated | 0,556344044 | 1,797448919 | 0,379362671 |
| RSPH1 | 0,000831183 | 0,704932876 | 1 | 0,951 | 1 | Ciliated | 1,630068821 | 0,613471031 | 1,051524711 |
| RPL342 | 0,000831274 | -0,735672462 | 1 | 0,976 | 1 | Ciliated | 0,60053804 | 1,665173451 | 1,024590164 |
| MUC15 | 0,000843441 | -0,869692646 | 0,321 | 0,707 | 1 | Ciliated | 0,547263428 | 1,827273574 | 0,454031117 |
| REEP52 | 0,000875182 | 0,642163177 | 0,821 | 0,634 | 1 | Ciliated | 1,560667471 | 0,640751485 | 1,294952681 |
| PLA2G10 | 0,000893798 | 0,862159717 | 0,643 | 0,39 | 1 | Ciliated | 1,817757461 | 0,550128398 | 1,648717949 |
| WBP13 | 0,000896744 | 0,704903705 | 0,821 | 0,537 | 1 | Ciliated | 1,630035861 | 0,613483435 | 1,52886406 |
| OFD1 | 0,000897118 | -0,845397261 | 0,25 | 0,683 | 1 | Ciliated | 0,556557534 | 1,796759434 | 0,366032211 |
| PLAAT32 | 0,000910445 | 0,98807288 | 0,821 | 0,585 | 1 | Ciliated | 1,983533661 | 0,504150759 | 1,403418803 |
| CREBRF | 0,000927748 | -0,513128214 | 0,143 | 0,561 | 1 | Ciliated | 0,70070145 | 1,427141332 | 0,254901961 |
| ITM2B2 | 0,000947541 | 0,443077159 | 1 | 0,927 | 1 | Ciliated | 1,359500949 | 0,735564032 | 1,078748652 |
| TSPAN131 | 0,000969358 | 0,926287867 | 0,75 | 0,561 | 1 | Ciliated | 1,900379918 | 0,52621057 | 1,336898396 |
| PLXNA2 | 0,000980806 | -0,406099421 | 0,036 | 0,39 | 1 | Ciliated | 0,754660976 | 1,325098331 | 0,092307692 |

| Supplementary Table 4 - clusermarker | | | | | | | | | |
|---|---|---|---|---|---|---|---|---|---|
| | **p_val** | **avg_log2FC** | **pct.1** | **pct.2** | **p_val_adj** | **celltype** | **Foldchange_UP** | **Foldchange_DOWN** | **Ratio_pct1_pct2** |
| DNAJC1 | 0,000981506 | 0,781230068 | 0,643 | 0,39 | 1 | Ciliated | 1,718595555 | 0,581870468 | 1,648717949 |
| MIF2 | 0,00098812 | 0,462728497 | 0,964 | 0,902 | 1 | Ciliated | 1,378145773 | 0,725612645 | 1,068736142 |
| NDUFV22 | 0,001004225 | 0,52446418 | 0,929 | 0,659 | 1 | Ciliated | 1,438399257 | 0,695217267 | 1,409711684 |
| RPS162 | 0,001033716 | 0,547577549 | 1 | 0,927 | 1 | Ciliated | 1,46162939 | 0,684167961 | 1,078748652 |
| UBB2 | 0,001034156 | 0,30901651 | 1 | 0,976 | 1 | Ciliated | 1,238862875 | 0,807191837 | 1,024590164 |
| TRIR2 | 0,001062211 | 0,632236883 | 0,857 | 0,659 | 1 | Ciliated | 1,549966337 | 0,645175302 | 1,300455235 |
| MRPS10 | 0,001074657 | -0,546062947 | 0,107 | 0,512 | 1 | Ciliated | 0,684886607 | 1,460095715 | 0,208984375 |
| RHOQ | 0,001128874 | -0,644823398 | 0,071 | 0,439 | 1 | Ciliated | 0,639571076 | 1,563547879 | 0,161731207 |
| AL021026.1 | 0,001150513 | -0,619550176 | 0 | 0,317 | 1 | Ciliated | 0,650873835 | 1,536396068 | 0 |
| THAP12 | 0,001150513 | -0,314943075 | 0 | 0,317 | 1 | Ciliated | 0,803882709 | 1,243962569 | 0 |
| PPP3R1 | 0,001164216 | -0,65496367 | 0,036 | 0,39 | 1 | Ciliated | 0,635091484 | 1,574576302 | 0,092307692 |
| COX7C | 0,001175735 | -0,569175201 | 1 | 0,902 | 1 | Ciliated | 0,67400201 | 1,4836751 | 1,10864745 |
| PFN12 | 0,001192017 | 0,393911873 | 0,964 | 0,829 | 1 | Ciliated | 1,313951362 | 0,761063179 | 1,162846803 |
| TMED2 | 0,00122224 | 0,560148339 | 0,821 | 0,537 | 1 | Ciliated | 1,474420811 | 0,678232424 | 1,52886406 |
| TEKT4 | 0,001228592 | 0,883074431 | 0,75 | 0,463 | 1 | Ciliated | 1,844301384 | 0,54221073 | 1,61987041 |
| RSPH3 | 0,001274405 | 0,69397323 | 0,786 | 0,512 | 1 | Ciliated | 1,617732679 | 0,6181491 | 1,53515625 |
| MMP24OS2 | 0,001318154 | 0,835421701 | 0,857 | 0,732 | 1 | Ciliated | 1,784378538 | 0,560419204 | 1,170765027 |
| AC105446.1 | 0,001342287 | 0,629154791 | 0,75 | 0,39 | 1 | Ciliated | 1,546658612 | 0,646555091 | 1,923076923 |
| GRK61 | 0,001350529 | 0,287221567 | 0,286 | 0,024 | 1 | Ciliated | 1,220287906 | 0,819478744 | 11,91666667 |
| NELL2 | 0,001350529 | 0,602052331 | 0,286 | 0,024 | 1 | Ciliated | 1,51787431 | 0,658816078 | 11,91666667 |
| TPPP31 | 0,001395198 | 0,660226282 | 1 | 0,927 | 1 | Ciliated | 1,580330474 | 0,63277904 | 1,078748652 |
| TXNL4A2 | 0,001399079 | 0,875705321 | 0,786 | 0,488 | 1 | Ciliated | 1,834904937 | 0,544987361 | 1,610655738 |
| ALKBH71 | 0,001405954 | 0,417164483 | 0,893 | 0,634 | 1 | Ciliated | 1,335300533 | 0,74889508 | 1,40851735 |
| AC023300.2 | 0,001418111 | 0,540496713 | 0,679 | 0,366 | 1 | Ciliated | 1,454473199 | 0,687534154 | 1,855191257 |
| KRTCAP22 | 0,001435422 | 0,671148967 | 0,893 | 0,707 | 1 | Ciliated | 1,592340608 | 0,628006342 | 1,263083451 |
| EFHB | 0,001503212 | 0,797105667 | 0,786 | 0,537 | 1 | Ciliated | 1,737611633 | 0,575502593 | 1,463687151 |
| APRT1 | 0,001503683 | 0,690985654 | 0,786 | 0,61 | 1 | Ciliated | 1,614386095 | 0,619430509 | 1,28852459 |
| MAP1LC3A1 | 0,001507622 | 0,646043846 | 0,857 | 0,61 | 1 | Ciliated | 1,564871122 | 0,63903026 | 1,404918033 |
| SELENOH2 | 0,001513407 | 0,435926506 | 1 | 0,854 | 1 | Ciliated | 1,352779316 | 0,739218872 | 1,170960187 |
| ZNF5932 | 0,001552593 | 0,628056153 | 0,679 | 0,39 | 1 | Ciliated | 1,545481251 | 0,647047642 | 1,741025641 |
| UBC1 | 0,001583843 | -0,567888175 | 1 | 1 | 1 | Ciliated | 0,674603555 | 1,482352106 | 1 |

| Supplementary Table 4 - clusermarker | | | | | | | | | |
|---|---|---|---|---|---|---|---|---|---|
| | **p_val** | **avg_log2FC** | **pct.1** | **pct.2** | **p_val_adj** | **celltype** | **Foldchange_UP** | **Foldchange_DOWN** | **Ratio_pct1_pct2** |
| C20orf85 | 0,001583843 | 0,520301036 | 1 | 1 | 1 | Ciliated | 1,434254492 | 0,697226333 | 1 |
| AGO3 | 0,001635476 | -0,494268654 | 0,071 | 0,439 | 1 | Ciliated | 0,709921467 | 1,408606509 | 0,161731207 |
| PCNT | 0,001635476 | -0,470668467 | 0,071 | 0,439 | 1 | Ciliated | 0,721630156 | 1,385751402 | 0,161731207 |
| CERS4 | 0,001676374 | 0,650048893 | 0,643 | 0,341 | 1 | Ciliated | 1,569221375 | 0,637258717 | 1,885630499 |
| LSM42 | 0,001679235 | 0,560030127 | 0,857 | 0,707 | 1 | Ciliated | 1,474300004 | 0,678287999 | 1,212164074 |
| HOOK1 | 0,001683801 | -0,618478918 | 0,143 | 0,585 | 1 | Ciliated | 0,651357314 | 1,535255656 | 0,244444444 |
| TUFM2 | 0,001694325 | 0,463168873 | 0,929 | 0,805 | 1 | Ciliated | 1,37856651 | 0,725391189 | 1,154037267 |
| IDH2 | 0,001745133 | 0,524427401 | 0,821 | 0,585 | 1 | Ciliated | 1,438362587 | 0,695234991 | 1,403418803 |
| DYNLT4 | 0,001751132 | 0,794554124 | 0,929 | 0,659 | 1 | Ciliated | 1,734541218 | 0,576521324 | 1,409711684 |
| MGST21 | 0,001766074 | -0,672888639 | 0,286 | 0,683 | 1 | Ciliated | 0,627249518 | 1,594261888 | 0,418740849 |
| ABHD14A | 0,001778084 | 0,464742768 | 0,607 | 0,317 | 1 | Ciliated | 1,380071266 | 0,724600261 | 1,914826498 |
| GNAI21 | 0,001813284 | 0,914589733 | 0,714 | 0,537 | 1 | Ciliated | 1,885032939 | 0,530494709 | 1,329608939 |
| UBA31 | 0,001814655 | -0,426920157 | 0,071 | 0,439 | 1 | Ciliated | 0,743848047 | 1,344360591 | 0,161731207 |
| LRRCC1 | 0,001814655 | -0,637280175 | 0,071 | 0,439 | 1 | Ciliated | 0,642923872 | 1,555394104 | 0,161731207 |
| PLPP22 | 0,001819678 | 0,776236368 | 0,857 | 0,634 | 1 | Ciliated | 1,712657144 | 0,583888027 | 1,351735016 |
| IFT22 | 0,001819772 | 0,602414216 | 0,929 | 0,756 | 1 | Ciliated | 1,5182551 | 0,658650842 | 1,228835979 |
| ANAPC112 | 0,001882834 | 0,523609217 | 0,929 | 0,732 | 1 | Ciliated | 1,437547091 | 0,695629386 | 1,269125683 |
| CACHD1 | 0,001910854 | -0,440081399 | 0,071 | 0,439 | 1 | Ciliated | 0,73709302 | 1,356680871 | 0,161731207 |
| ZNF688 | 0,001928089 | 0,76754349 | 0,714 | 0,39 | 1 | Ciliated | 1,702368652 | 0,587416832 | 1,830769231 |
| PINLYP | 0,001934323 | 0,687655391 | 0,679 | 0,341 | 1 | Ciliated | 1,610663806 | 0,62086203 | 1,991202346 |
| STIL | 0,001943492 | -0,307596065 | 0 | 0,293 | 1 | Ciliated | 0,807986971 | 1,237643719 | 0 |
| H2AC19 | 0,001943492 | -0,509683891 | 0 | 0,293 | 1 | Ciliated | 0,702376319 | 1,423738206 | 0 |
| TAMM41 | 0,001943492 | -0,276627726 | 0 | 0,293 | 1 | Ciliated | 0,825518398 | 1,21136004 | 0 |
| CWF19L2 | 0,001943492 | -0,334619615 | 0 | 0,293 | 1 | Ciliated | 0,792993192 | 1,261044874 | 0 |
| RPL221 | 0,001949666 | -0,498080491 | 1 | 0,951 | 1 | Ciliated | 0,708048214 | 1,412333199 | 1,051524711 |
| TRAPPC52 | 0,001983446 | 0,71354479 | 0,786 | 0,634 | 1 | Ciliated | 1,63982833 | 0,609819932 | 1,239747634 |
| PRPSAP2 | 0,002011716 | -0,315299874 | 0,071 | 0,439 | 1 | Ciliated | 0,803683922 | 1,244270256 | 0,161731207 |
| TMEM542 | 0,00202793 | 0,681264104 | 0,571 | 0,366 | 1 | Ciliated | 1,603544181 | 0,623618614 | 1,56010929 |
| SRSF71 | 0,002045672 | -0,856611709 | 0,429 | 0,756 | 1 | Ciliated | 0,552248037 | 1,810780542 | 0,567460317 |
| OLA11 | 0,002069883 | 0,6485121 | 0,643 | 0,415 | 1 | Ciliated | 1,567550694 | 0,637937901 | 1,54939759 |
| HSPBP1 | 0,002110376 | 0,785447349 | 0,857 | 0,61 | 1 | Ciliated | 1,723626697 | 0,58017203 | 1,404918033 |

| Supplementary Table 4 - clusermarker | | | | | | | | | |
|---|---|---|---|---|---|---|---|---|---|
| | **p_val** | **avg_log2FC** | **pct.1** | **pct.2** | **p_val_adj** | **celltype** | **Foldchange_UP** | **Foldchange_DOWN** | **Ratio_pct1_pct2** |
| ADGRF1 | 0,002112837 | -0,776795521 | 0,5 | 0,732 | 1 | Ciliated | 0,58366177 | 1,713321057 | 0,683060109 |
| STOM1 | 0,002131058 | 0,783120212 | 0,75 | 0,585 | 1 | Ciliated | 1,720848644 | 0,581108631 | 1,282051282 |
| SLC25A4 | 0,002161774 | 0,733809128 | 0,714 | 0,488 | 1 | Ciliated | 1,663024161 | 0,601314174 | 1,463114754 |
| ZMAT5 | 0,002174666 | -0,537646853 | 0,036 | 0,366 | 1 | Ciliated | 0,688893632 | 1,451602909 | 0,098360656 |
| IGKV3-202 | 0,002192608 | 0,467453913 | 0,214 | 0 | 1 | Ciliated | 1,382667164 | 0,723239855 | #DIV/0! |
| RSPH10B2 | 0,002228244 | -0,3125035 | 0,071 | 0,439 | 1 | Ciliated | 0,805243212 | 1,241860825 | 0,161731207 |
| NDUFA121 | 0,002230123 | 0,26914561 | 0,893 | 0,634 | 1 | Ciliated | 1,205093938 | 0,829810829 | 1,40851735 |
| TLE51 | 0,002279073 | 0,460255732 | 1 | 0,805 | 1 | Ciliated | 1,375785668 | 0,726857404 | 1,242236025 |
| ALDH1A11 | 0,002296063 | -0,791861456 | 0,964 | 0,951 | 1 | Ciliated | 0,577598357 | 1,731306863 | 1,013669821 |
| TDRD3 | 0,002297917 | -0,365452272 | 0,036 | 0,366 | 1 | Ciliated | 0,776225497 | 1,288285432 | 0,098360656 |
| MGA | 0,002297917 | -0,250660427 | 0,036 | 0,366 | 1 | Ciliated | 0,840511564 | 1,189751626 | 0,098360656 |
| DBI | 0,002336497 | -0,651853027 | 0,893 | 0,927 | 1 | Ciliated | 0,636462303 | 1,571184963 | 0,963322546 |
| HACD4 | 0,002337611 | 0,724005394 | 0,786 | 0,488 | 1 | Ciliated | 1,651761506 | 0,605414278 | 1,610655738 |
| RAC12 | 0,002379796 | 0,46683479 | 0,964 | 0,854 | 1 | Ciliated | 1,38207393 | 0,723550295 | 1,128805621 |
| HSBP12 | 0,002391882 | 0,288371451 | 1 | 0,951 | 1 | Ciliated | 1,221260911 | 0,818825847 | 1,051524711 |
| GRAMD2B1 | 0,002437314 | -0,675868977 | 0,179 | 0,561 | 1 | Ciliated | 0,625955075 | 1,597558739 | 0,319073084 |
| ALDOA1 | 0,002449212 | -0,789760732 | 0,429 | 0,707 | 1 | Ciliated | 0,578440017 | 1,728787723 | 0,60678925 |
| MTCO2P12 | 0,002463682 | 0,636672447 | 0,964 | 0,805 | 1 | Ciliated | 1,55473904 | 0,643194758 | 1,197515528 |
| FAM204A | 0,002550617 | -0,501781289 | 0,179 | 0,561 | 1 | Ciliated | 0,706234258 | 1,415960764 | 0,319073084 |
| RECQL | 0,002575962 | -0,361573318 | 0,036 | 0,341 | 1 | Ciliated | 0,778315332 | 1,28482629 | 0,105571848 |
| POLR2I | 0,002588694 | 0,566150294 | 1 | 0,878 | 1 | Ciliated | 1,480567529 | 0,675416677 | 1,138952164 |
| NENF2 | 0,002626078 | 0,583110269 | 0,821 | 0,683 | 1 | Ciliated | 1,498075431 | 0,667523129 | 1,20204978 |
| NPTN1 | 0,002642579 | 0,605796571 | 0,857 | 0,707 | 1 | Ciliated | 1,52181878 | 0,657108463 | 1,212164074 |
| ALDH1L1 | 0,002647692 | 0,571426686 | 0,786 | 0,634 | 1 | Ciliated | 1,485992347 | 0,672950976 | 1,239747634 |
| CASP9 | 0,002664057 | -0,495130558 | 0,071 | 0,415 | 1 | Ciliated | 0,709497469 | 1,409448299 | 0,171084337 |
| ARID4A | 0,002664057 | -0,427238221 | 0,071 | 0,415 | 1 | Ciliated | 0,743684072 | 1,344657009 | 0,171084337 |
| DCAF5 | 0,002730764 | -0,568423992 | 0,179 | 0,512 | 1 | Ciliated | 0,674353053 | 1,482902754 | 0,349609375 |
| CBX4 | 0,002803213 | -0,425298125 | 0,071 | 0,415 | 1 | Ciliated | 0,74468483 | 1,342849967 | 0,171084337 |
| ST3GAL6 | 0,002839888 | -1,280177056 | 0,286 | 0,683 | 1 | Ciliated | 0,411744974 | 2,428687813 | 0,418740849 |
| PSMB92 | 0,002860529 | 0,881617742 | 0,75 | 0,61 | 1 | Ciliated | 1,842440133 | 0,542758477 | 1,229508197 |
| PSMA4 | 0,002927876 | -0,829546277 | 0,5 | 0,805 | 1 | Ciliated | 0,562706184 | 1,777126374 | 0,621118012 |

| Supplementary Table 4 - clusermarker | | | | | | | | | |
|---|---|---|---|---|---|---|---|---|---|
| | **p_val** | **avg_log2FC** | **pct.1** | **pct.2** | **p_val_adj** | **celltype** | **Foldchange_UP** | **Foldchange_DOWN** | **Ratio_pct1_pct2** |
| TANK1 | 0,002964592 | -0,519433902 | 0,179 | 0,537 | 1 | Ciliated | 0,697645528 | 1,43339269 | 0,333333333 |
| PPWD1 | 0,003041914 | -0,371226858 | 0,036 | 0,341 | 1 | Ciliated | 0,773124757 | 1,293452306 | 0,105571848 |
| ING5 | 0,003101628 | -0,414777347 | 0,071 | 0,415 | 1 | Ciliated | 0,750135255 | 1,333092923 | 0,171084337 |
| MYH141 | 0,003228891 | 0,481109987 | 0,857 | 0,707 | 1 | Ciliated | 1,395817174 | 0,716426205 | 1,212164074 |
| MRM2 | 0,003246098 | -0,284138584 | 0 | 0,268 | 1 | Ciliated | 0,82123181 | 1,217682982 | 0 |
| UQCRQ2 | 0,0032596 | 0,534660498 | 0,964 | 0,805 | 1 | Ciliated | 1,448601222 | 0,690321107 | 1,197515528 |
| DNAI7 | 0,003279592 | -0,882970364 | 0,429 | 0,805 | 1 | Ciliated | 0,542249843 | 1,844168352 | 0,532919255 |
| PPM1G2 | 0,003359862 | 0,321655197 | 0,929 | 0,659 | 1 | Ciliated | 1,249763574 | 0,800151341 | 1,409711684 |
| NDUFAB12 | 0,003408913 | 0,482226499 | 0,893 | 0,78 | 1 | Ciliated | 1,396897824 | 0,715871972 | 1,144871795 |
| SPINT22 | 0,003526618 | 0,510205587 | 0,964 | 0,805 | 1 | Ciliated | 1,42425314 | 0,702122377 | 1,197515528 |
| SCYL2 | 0,003583554 | -0,304449143 | 0,036 | 0,341 | 1 | Ciliated | 0,809751341 | 1,234947014 | 0,105571848 |
| C11orf53 | 0,003587575 | 0,440690465 | 0,25 | 0,024 | 1 | Ciliated | 1,357253745 | 0,736781905 | 10,41666667 |
| IGHA12 | 0,003587575 | 3,907122189 | 0,25 | 0,024 | 1 | Ciliated | 15,00240812 | 0,066655966 | 10,41666667 |
| AC004556.3 | 0,003587575 | 0,462616042 | 0,25 | 0,024 | 1 | Ciliated | 1,378038354 | 0,725669207 | 10,41666667 |
| ATF7IP2 | 0,003646431 | 0,820620884 | 0,821 | 0,634 | 1 | Ciliated | 1,766165923 | 0,566198219 | 1,294952681 |
| ANP32A1 | 0,003672839 | 0,480944646 | 0,75 | 0,512 | 1 | Ciliated | 1,395657214 | 0,716508316 | 1,46484375 |
| KIDINS220 | 0,003692324 | -0,435782894 | 0,107 | 0,463 | 1 | Ciliated | 0,739292461 | 1,352644661 | 0,231101512 |
| RPS61 | 0,003704971 | -0,581341608 | 1 | 0,976 | 1 | Ciliated | 0,668341976 | 1,496240003 | 1,024590164 |
| DEGS2 | 0,003760228 | 0,535666632 | 0,929 | 0,756 | 1 | Ciliated | 1,449611828 | 0,689839846 | 1,228835979 |
| SCGB1A11 | 0,003834966 | -6,090929142 | 0,143 | 0,463 | 1 | Ciliated | 0,014670592 | 68,16357706 | 0,308855292 |
| DNAI2 | 0,003942547 | 0,687336629 | 0,786 | 0,585 | 1 | Ciliated | 1,610307971 | 0,620999224 | 1,343589744 |
| BLVRB1 | 0,003944022 | 0,814129768 | 0,679 | 0,488 | 1 | Ciliated | 1,758237265 | 0,568751454 | 1,391393443 |
| ASRGL1 | 0,003950323 | 0,516406354 | 0,5 | 0,244 | 1 | Ciliated | 1,430387817 | 0,6991111 | 2,049180328 |
| STXBP5 | 0,003991917 | -0,374100203 | 0,036 | 0,341 | 1 | Ciliated | 0,771586495 | 1,296030978 | 0,105571848 |
| ARL14EP | 0,004018124 | 0,40744249 | 0,321 | 0,073 | 1 | Ciliated | 1,326332498 | 0,753958756 | 4,397260274 |
| TUSC3 | 0,004085786 | 0,647700054 | 0,893 | 0,732 | 1 | Ciliated | 1,566668619 | 0,638297077 | 1,219945355 |
| ISCA21 | 0,004183269 | 0,466139749 | 0,679 | 0,463 | 1 | Ciliated | 1,381408254 | 0,723898961 | 1,466522678 |
| MT-ATP82 | 0,004247592 | -1,339263194 | 0,893 | 0,732 | 1 | Ciliated | 0,395222451 | 2,530220635 | 1,219945355 |
| MRPL142 | 0,004265654 | 0,572330326 | 0,75 | 0,61 | 1 | Ciliated | 1,486923398 | 0,672529601 | 1,229508197 |
| METTL7A1 | 0,004265654 | -0,985340995 | 0,536 | 0,756 | 1 | Ciliated | 0,505106322 | 1,979781197 | 0,708994709 |
| MIEN12 | 0,004273822 | 0,44670267 | 0,821 | 0,61 | 1 | Ciliated | 1,362921689 | 0,733717871 | 1,345901639 |

| Supplementary Table 4 - clusermarker | | | | | | | | | |
|---|---|---|---|---|---|---|---|---|---|
| | **p_val** | **avg_log2FC** | **pct.1** | **pct.2** | **p_val_adj** | **celltype** | **Foldchange_UP** | **Foldchange_DOWN** | **Ratio_pct1_pct2** |
| KLHDC91 | 0,004275381 | 0,711691307 | 0,679 | 0,488 | 1 | Ciliated | 1,637722935 | 0,610603893 | 1,391393443 |
| DYDC2 | 0,004368484 | 0,468157461 | 0,857 | 0,683 | 1 | Ciliated | 1,383341603 | 0,722887245 | 1,254758419 |
| SERBP11 | 0,004417648 | 0,530868811 | 0,857 | 0,707 | 1 | Ciliated | 1,444799011 | 0,692137794 | 1,212164074 |
| WDR86-AS1 | 0,004418013 | 0,69107656 | 0,857 | 0,707 | 1 | Ciliated | 1,614487823 | 0,619391479 | 1,212164074 |
| PFN4 | 0,004453306 | 0,571937227 | 0,286 | 0,049 | 1 | Ciliated | 1,486518304 | 0,672712874 | 5,836734694 |
| OSCP1 | 0,004461613 | 0,710414462 | 0,893 | 0,707 | 1 | Ciliated | 1,636274124 | 0,611144542 | 1,263083451 |
| PIGQ | 0,00447065 | 0,501798038 | 0,643 | 0,366 | 1 | Ciliated | 1,415977203 | 0,706226059 | 1,756830601 |
| RAB11FIP11 | 0,004500655 | 0,619967392 | 0,929 | 0,707 | 1 | Ciliated | 1,536840445 | 0,650685635 | 1,314002829 |
| PPP1R16A | 0,004531234 | 0,660925589 | 0,75 | 0,537 | 1 | Ciliated | 1,581096681 | 0,632472392 | 1,396648045 |
| TSPO2 | 0,004636999 | 0,678486527 | 0,893 | 0,707 | 1 | Ciliated | 1,600459897 | 0,624820404 | 1,263083451 |
| BTF31 | 0,00465688 | 0,250370585 | 1 | 0,854 | 1 | Ciliated | 1,189512626 | 0,840680442 | 1,170960187 |
| IER51 | 0,004731158 | -0,349370378 | 0,071 | 0,39 | 1 | Ciliated | 0,784926582 | 1,274004504 | 0,182051282 |
| TBRG11 | 0,004794325 | -0,551027189 | 0,107 | 0,439 | 1 | Ciliated | 0,682533996 | 1,465128486 | 0,243735763 |
| CTTN1 | 0,004805599 | 0,582675799 | 0,714 | 0,561 | 1 | Ciliated | 1,497624352 | 0,667724185 | 1,272727273 |
| GOSR1 | 0,004822594 | -0,436171301 | 0,179 | 0,537 | 1 | Ciliated | 0,739093453 | 1,353008874 | 0,333333333 |
| PPDPF2 | 0,00485033 | 0,370525575 | 0,964 | 0,951 | 1 | Ciliated | 1,292823722 | 0,773500658 | 1,013669821 |
| UQCR112 | 0,004851885 | 0,427436006 | 1 | 0,976 | 1 | Ciliated | 1,344841366 | 0,743582125 | 1,024590164 |
| TNFRSF191 | 0,004900894 | 0,455880058 | 0,75 | 0,537 | 1 | Ciliated | 1,371619251 | 0,7290653 | 1,396648045 |
| CHCHD102 | 0,004903374 | 0,382215109 | 0,857 | 0,683 | 1 | Ciliated | 1,303341466 | 0,767258639 | 1,254758419 |
| SYNGR22 | 0,0049073 | 0,5328889 | 0,679 | 0,512 | 1 | Ciliated | 1,446823464 | 0,691169327 | 1,326171875 |
| MRPS27 | 0,004969448 | -0,40404074 | 0,036 | 0,317 | 1 | Ciliated | 0,755738623 | 1,323208805 | 0,113564669 |
| C1orf158 | 0,005160685 | -0,5952038 | 0,25 | 0,61 | 1 | Ciliated | 0,661950939 | 1,510685976 | 0,409836066 |
| TCEAL3 | 0,005160685 | 0,842997763 | 0,607 | 0,366 | 1 | Ciliated | 1,793773537 | 0,557483974 | 1,658469945 |
| TTC5 | 0,005176275 | -0,532416788 | 0,143 | 0,488 | 1 | Ciliated | 0,691395545 | 1,446350078 | 0,293032787 |
| CCDC74A | 0,005181055 | 0,719475555 | 0,786 | 0,585 | 1 | Ciliated | 1,646583364 | 0,607318173 | 1,343589744 |
| RBM10 | 0,005219911 | -0,299977491 | 0,071 | 0,39 | 1 | Ciliated | 0,812265069 | 1,231125205 | 0,182051282 |
| DYNLRB2 | 0,00523196 | 0,555423196 | 1 | 0,927 | 1 | Ciliated | 1,469599658 | 0,680457425 | 1,078748652 |
| GTF2IRD2B | 0,005244582 | -0,324100049 | 0,036 | 0,317 | 1 | Ciliated | 0,798796519 | 1,251883272 | 0,113564669 |
| LRRC61 | 0,005265358 | 0,690212503 | 0,679 | 0,415 | 1 | Ciliated | 1,613521166 | 0,619762555 | 1,636144578 |
| SMDT12 | 0,0052986 | 0,336589717 | 0,964 | 0,829 | 1 | Ciliated | 1,262768096 | 0,791911043 | 1,162846803 |
| IER5L1 | 0,005319225 | -0,275981394 | 0,071 | 0,366 | 1 | Ciliated | 0,825888316 | 1,210817468 | 0,193989071 |

| Supplementary Table 4 - clusermarker | | | | | | | | | |
|---|---|---|---|---|---|---|---|---|---|
| | p_val | avg_log2FC | pct.1 | pct.2 | p_val_adj | celltype | Foldchange_UP | Foldchange_DOWN | Ratio_pct1_pct2 |
| STATH2 | 0,005365129 | -0,648770494 | 0 | 0,244 | 1 | Ciliated | 0,637823654 | 1,567831476 | 0 |
| LINC00964 | 0,005365129 | -0,340482785 | 0 | 0,244 | 1 | Ciliated | 0,789776976 | 1,266180239 | 0 |
| SHTN1 | 0,005365129 | -0,328275774 | 0 | 0,244 | 1 | Ciliated | 0,796487832 | 1,255511962 | 0 |
| LRRC37A2 | 0,005365129 | -0,260438252 | 0 | 0,244 | 1 | Ciliated | 0,834834281 | 1,197842522 | 0 |
| RNF213-AS1 | 0,005365129 | -0,297267633 | 0 | 0,244 | 1 | Ciliated | 0,813792205 | 1,228814916 | 0 |
| NFS1 | 0,005365129 | -0,284027721 | 0 | 0,244 | 1 | Ciliated | 0,82129492 | 1,217589414 | 0 |
| FCGRT1 | 0,00538784 | 0,716442351 | 0,679 | 0,537 | 1 | Ciliated | 1,64312513 | 0,608596376 | 1,26443203 |
| FTL1 | 0,005430862 | 0,400183267 | 1 | 0,902 | 1 | Ciliated | 1,31967554 | 0,757762018 | 1,10864745 |
| TUBA1A1 | 0,005433412 | 0,542208838 | 1 | 0,927 | 1 | Ciliated | 1,456200326 | 0,686718703 | 1,078748652 |
| ZNF487 | 0,005520908 | 0,861268755 | 0,714 | 0,488 | 1 | Ciliated | 1,816635219 | 0,550468244 | 1,463114754 |
| MAPK9 | 0,005533406 | -0,429097014 | 0,036 | 0,317 | 1 | Ciliated | 0,742726514 | 1,346390604 | 0,113564669 |
| PEX3 | 0,005533406 | -0,258350687 | 0,036 | 0,317 | 1 | Ciliated | 0,836043152 | 1,196110509 | 0,113564669 |
| IGLV1-51 | 0,00555467 | 0,320773105 | 0,179 | 0 | 1 | Ciliated | 1,248999678 | 0,800640719 | #DIV/0! |
| METTL262 | 0,005594433 | 0,598313583 | 0,786 | 0,585 | 1 | Ciliated | 1,513945827 | 0,660525616 | 1,343589744 |
| UBR4 | 0,00559989 | -0,570048753 | 0,25 | 0,634 | 1 | Ciliated | 0,673594025 | 1,484573738 | 0,394321767 |
| CHST91 | 0,005634398 | -0,58195708 | 0,964 | 0,902 | 1 | Ciliated | 0,668056913 | 1,496878454 | 1,068736142 |
| RPS252 | 0,005641398 | -0,587914522 | 1 | 0,927 | 1 | Ciliated | 0,665303938 | 1,50307242 | 1,078748652 |
| C2CD5 | 0,005753407 | -0,427492177 | 0,143 | 0,512 | 1 | Ciliated | 0,743553174 | 1,344893728 | 0,279296875 |
| HOOK3 | 0,005753835 | -0,511202379 | 0,071 | 0,39 | 1 | Ciliated | 0,701637431 | 1,42523753 | 0,182051282 |
| RAB5B1 | 0,005770833 | -0,556781635 | 0,107 | 0,439 | 1 | Ciliated | 0,67981701 | 1,470984082 | 0,243735763 |
| NQO1 | 0,005804388 | 0,730929138 | 0,964 | 0,805 | 1 | Ciliated | 1,659707648 | 0,602515751 | 1,197515528 |
| ADGRA3 | 0,005836507 | -0,304067845 | 0,036 | 0,317 | 1 | Ciliated | 0,809965383 | 1,234620666 | 0,113564669 |
| ACTB2 | 0,005857892 | 0,468300638 | 1 | 0,951 | 1 | Ciliated | 1,383478897 | 0,722815507 | 1,051524711 |
| MICAL31 | 0,006038849 | -0,319085444 | 0,071 | 0,39 | 1 | Ciliated | 0,801577854 | 1,247539456 | 0,182051282 |
| MAOA | 0,006041447 | -0,261837985 | 0,107 | 0,439 | 1 | Ciliated | 0,8340247 | 1,199005258 | 0,243735763 |
| RASSF91 | 0,006071588 | -0,741282675 | 0,179 | 0,512 | 1 | Ciliated | 0,59820726 | 1,671661423 | 0,349609375 |
| CDC40 | 0,006100152 | -0,251453178 | 0,179 | 0,561 | 1 | Ciliated | 0,840049835 | 1,190405566 | 0,319073084 |
| PSMB41 | 0,006221794 | 0,704609933 | 0,679 | 0,463 | 1 | Ciliated | 1,629703976 | 0,61360837 | 1,466522678 |
| PLXNB1 | 0,006239934 | 0,55040628 | 0,679 | 0,39 | 1 | Ciliated | 1,464498057 | 0,682827809 | 1,741025641 |
| GTF2B1 | 0,006266588 | -0,442816424 | 0,214 | 0,585 | 1 | Ciliated | 0,735696981 | 1,359255272 | 0,365811966 |
| ZDHHC1 | 0,006276607 | 0,585133588 | 0,75 | 0,634 | 1 | Ciliated | 1,500177894 | 0,666587612 | 1,1829653 |

| Supplementary Table 4 - clusermarker | | | | | | | | | |
|---|---|---|---|---|---|---|---|---|---|
| | **p_val** | **avg_log2FC** | **pct.1** | **pct.2** | **p_val_adj** | **celltype** | **Foldchange_UP** | **Foldchange_DOWN** | **Ratio_pct1_pct2** |
| RAD9A | 0,00632583 | 0,533034026 | 0,607 | 0,366 | 1 | Ciliated | 1,446969013 | 0,691099803 | 1,658469945 |
| GAS52 | 0,006360328 | -0,498124568 | 0,179 | 0,561 | 1 | Ciliated | 0,708026582 | 1,41237635 | 0,319073084 |
| NME22 | 0,006371345 | 0,758216127 | 0,929 | 0,732 | 1 | Ciliated | 1,69139794 | 0,591226923 | 1,269125683 |
| ARF52 | 0,006433233 | 0,526139308 | 0,821 | 0,683 | 1 | Ciliated | 1,440070367 | 0,694410512 | 1,20204978 |
| SERPINB10 | 0,006442314 | 0,571737843 | 0,286 | 0,049 | 1 | Ciliated | 1,486312877 | 0,672805851 | 5,836734694 |
| TNKS | 0,006480749 | -0,311332936 | 0,071 | 0,366 | 1 | Ciliated | 0,805896831 | 1,240853621 | 0,193989071 |
| ASCC2 | 0,006480749 | -0,391447473 | 0,071 | 0,366 | 1 | Ciliated | 0,762364332 | 1,311708797 | 0,193989071 |
| UBE2D22 | 0,006591821 | 0,610857569 | 0,679 | 0,488 | 1 | Ciliated | 1,52716672 | 0,654807355 | 1,391393443 |
| PDLIM11 | 0,006684184 | 0,47810871 | 0,929 | 0,78 | 1 | Ciliated | 1,392916435 | 0,717918157 | 1,191025641 |
| CXCL17 | 0,006714415 | -0,758872973 | 0,571 | 0,829 | 1 | Ciliated | 0,590957804 | 1,692168194 | 0,688781665 |
| SPTLC2 | 0,006789729 | 0,493268867 | 0,679 | 0,537 | 1 | Ciliated | 1,407630683 | 0,710413613 | 1,26443203 |
| YBX12 | 0,006802787 | 0,560972613 | 0,857 | 0,659 | 1 | Ciliated | 1,475263451 | 0,677845031 | 1,300455235 |
| NUMBL | 0,006804693 | -0,420069821 | 0,071 | 0,366 | 1 | Ciliated | 0,747388453 | 1,337992307 | 0,193989071 |
| SLC25A62 | 0,007016902 | 0,601793053 | 0,786 | 0,634 | 1 | Ciliated | 1,517601545 | 0,65893449 | 1,239747634 |
| ELOB2 | 0,007043567 | 0,350901466 | 1 | 0,902 | 1 | Ciliated | 1,275357284 | 0,784094004 | 1,10864745 |
| HBB1 | 0,007216784 | -0,445603811 | 0,036 | 0,293 | 1 | Ciliated | 0,734276935 | 1,361883986 | 0,122866894 |
| IMMT1 | 0,007241879 | -0,303961051 | 0,107 | 0,439 | 1 | Ciliated | 0,810025342 | 1,234529277 | 0,243735763 |
| TUSC21 | 0,007403643 | 0,473771967 | 0,714 | 0,537 | 1 | Ciliated | 1,388735615 | 0,720079466 | 1,329608939 |
| POLR1D1 | 0,007408405 | 0,447972093 | 0,821 | 0,707 | 1 | Ciliated | 1,364121447 | 0,733072559 | 1,161244696 |
| CFAP221 | 0,007408955 | 0,418227977 | 0,929 | 0,634 | 1 | Ciliated | 1,336285224 | 0,74834323 | 1,465299685 |
| CEP350 | 0,007503781 | -0,310011634 | 0,214 | 0,537 | 1 | Ciliated | 0,806635254 | 1,239717697 | 0,398510242 |
| TSC22D12 | 0,007512597 | -0,436052735 | 0,429 | 0,78 | 1 | Ciliated | 0,739154197 | 1,352897682 | 0,55 |
| IDS2 | 0,007512597 | 0,700955444 | 0,75 | 0,561 | 1 | Ciliated | 1,625580999 | 0,615164671 | 1,336898396 |
| RPL311 | 0,007577561 | -0,567310221 | 0,929 | 0,951 | 1 | Ciliated | 0,67487386 | 1,481758385 | 0,976866456 |
| IRF5 | 0,00761317 | -0,267346661 | 0,036 | 0,293 | 1 | Ciliated | 0,830846196 | 1,203592199 | 0,122866894 |
| HLA-B2 | 0,007675902 | 0,641919734 | 0,964 | 0,829 | 1 | Ciliated | 1,560404143 | 0,640859616 | 1,162846803 |
| MTX11 | 0,007738504 | 0,461808702 | 0,643 | 0,415 | 1 | Ciliated | 1,377267412 | 0,726075409 | 1,54939759 |
| ADM1 | 0,007780589 | -0,44275665 | 0,107 | 0,415 | 1 | Ciliated | 0,735727463 | 1,359198956 | 0,257831325 |
| OAZ12 | 0,007864167 | 0,273817389 | 1 | 0,976 | 1 | Ciliated | 1,209002635 | 0,827128057 | 1,024590164 |
| SMURF1 | 0,007865629 | -0,378456025 | 0,071 | 0,366 | 1 | Ciliated | 0,769260414 | 1,299949902 | 0,193989071 |
| ATP6V0B2 | 0,007963297 | 0,398647158 | 0,786 | 0,659 | 1 | Ciliated | 1,318271163 | 0,758569274 | 1,192716237 |

| Supplementary Table 4 - clusermarker | | | | | | | | | |
|---|---|---|---|---|---|---|---|---|---|
| | **p_val** | **avg_log2FC** | **pct.1** | **pct.2** | **p_val_adj** | **celltype** | **Foldchange_UP** | **Foldchange_DOWN** | **Ratio_pct1_pct2** |
| CST31 | 0,007987806 | 0,385547068 | 0,964 | 0,878 | 1 | Ciliated | 1,306355062 | 0,765488671 | 1,097949886 |
| AP1B1 | 0,007999084 | -0,447926159 | 0,143 | 0,488 | 1 | Ciliated | 0,7330959 | 1,364078016 | 0,293032787 |
| AC005562.1 | 0,008028963 | -0,681132724 | 0,036 | 0,293 | 1 | Ciliated | 0,623675407 | 1,603398161 | 0,122866894 |
| FAM207A1 | 0,008121289 | 0,428847836 | 0,429 | 0,171 | 1 | Ciliated | 1,34615808 | 0,742854807 | 2,50877193 |
| UPF11 | 0,008133944 | 0,605446506 | 0,786 | 0,634 | 1 | Ciliated | 1,521449561 | 0,657267928 | 1,239747634 |
| FTO | 0,008140554 | -0,811215377 | 0,357 | 0,683 | 1 | Ciliated | 0,569901551 | 1,754689032 | 0,522693997 |
| SIX4 | 0,008141947 | -0,50546741 | 0,214 | 0,537 | 1 | Ciliated | 0,704432116 | 1,419583204 | 0,398510242 |
| PFN2 | 0,008182519 | 0,495433033 | 0,929 | 0,707 | 1 | Ciliated | 1,409743835 | 0,709348731 | 1,314002829 |
| MED211 | 0,00825088 | -0,311187937 | 0,071 | 0,366 | 1 | Ciliated | 0,805977832 | 1,240728914 | 0,193989071 |
| SLC25A14 | 0,00825088 | -0,58204098 | 0,071 | 0,366 | 1 | Ciliated | 0,668018064 | 1,496965508 | 0,193989071 |
| MIR22HG | 0,008270302 | -0,425529933 | 0,179 | 0,488 | 1 | Ciliated | 0,744565187 | 1,343065749 | 0,366803279 |
| PARP4 | 0,008336065 | -0,364654115 | 0,179 | 0,537 | 1 | Ciliated | 0,776655055 | 1,287572898 | 0,333333333 |
| XPOT | 0,008346204 | -0,401684675 | 0,143 | 0,488 | 1 | Ciliated | 0,756973828 | 1,321049637 | 0,293032787 |
| TMEM1602 | 0,008350522 | 0,969772429 | 0,536 | 0,39 | 1 | Ciliated | 1,958531631 | 0,510586597 | 1,374358974 |
| EGR11 | 0,008354102 | -1,051539763 | 0,5 | 0,659 | 1 | Ciliated | 0,482452976 | 2,072740868 | 0,758725341 |
| CCDC66 | 0,0083553 | -0,468276193 | 0,143 | 0,439 | 1 | Ciliated | 0,722827754 | 1,383455456 | 0,325740319 |
| PET100 | 0,008501579 | -0,746132963 | 0,679 | 0,829 | 1 | Ciliated | 0,596199486 | 1,677290946 | 0,819059107 |
| ARF41 | 0,008504532 | -0,645973672 | 0,5 | 0,707 | 1 | Ciliated | 0,639061344 | 1,564795007 | 0,707213579 |
| ATP5MC31 | 0,00853986 | 0,275051818 | 0,929 | 0,732 | 1 | Ciliated | 1,21003755 | 0,826420634 | 1,269125683 |
| TSTD11 | 0,008600556 | 0,254203756 | 0,964 | 0,927 | 1 | Ciliated | 1,192677305 | 0,83844976 | 1,0399137 |
| TMEM2192 | 0,008661179 | 0,508520086 | 0,929 | 0,805 | 1 | Ciliated | 1,422590157 | 0,702943146 | 1,154037267 |
| TFF32 | 0,008759965 | -0,868625976 | 0,964 | 0,951 | 1 | Ciliated | 0,547668202 | 1,825923061 | 1,013669821 |
| TC2N1 | 0,008761499 | 0,660683252 | 0,75 | 0,61 | 1 | Ciliated | 1,580831119 | 0,632578641 | 1,229508197 |
| RPL82 | 0,008761811 | 0,446557001 | 1 | 0,951 | 1 | Ciliated | 1,362784082 | 0,733791958 | 1,051524711 |
| RBMS3 | 0,008782893 | -0,251714391 | 0 | 0,22 | 1 | Ciliated | 0,83989775 | 1,19062112 | 0 |
| BLOC1S5 | 0,008782893 | -0,457549509 | 0 | 0,22 | 1 | Ciliated | 0,728222131 | 1,373207373 | 0 |
| LRRK11 | 0,008782893 | -0,285948003 | 0 | 0,22 | 1 | Ciliated | 0,820202472 | 1,219211151 | 0 |
| SLFN11 | 0,008782893 | -0,259832113 | 0 | 0,22 | 1 | Ciliated | 0,835185105 | 1,197339361 | 0 |
| AP000547.3 | 0,008782893 | -0,269260658 | 0 | 0,22 | 1 | Ciliated | 0,829744659 | 1,205190042 | 0 |
| NDUFV11 | 0,008824118 | 0,517586044 | 0,714 | 0,561 | 1 | Ciliated | 1,431557922 | 0,698539671 | 1,272727273 |
| SERPINI2 | 0,008922037 | -0,411848727 | 0,036 | 0,293 | 1 | Ciliated | 0,75165955 | 1,330389536 | 0,122866894 |

| Supplementary Table 4 - clusermarker | | | | | | | | | |
|---|---|---|---|---|---|---|---|---|---|
| | **p_val** | **avg_log2FC** | **pct.1** | **pct.2** | **p_val_adj** | **celltype** | **Foldchange_UP** | **Foldchange_DOWN** | **Ratio_pct1_pct2** |
| TMEM213 | 0,008922037 | -0,273948275 | 0,036 | 0,293 | 1 | Ciliated | 0,82705302 | 1,209112325 | 0,122866894 |
| ZBTB7A2 | 0,008951534 | 0,48568771 | 0,821 | 0,683 | 1 | Ciliated | 1,400253186 | 0,714156561 | 1,20204978 |
| SON2 | 0,00900429 | -0,785956269 | 0,679 | 0,756 | 1 | Ciliated | 0,579967407 | 1,724234825 | 0,898148148 |
| GK | 0,009023098 | -0,595953045 | 0,286 | 0,659 | 1 | Ciliated | 0,661607253 | 1,511470734 | 0,433990895 |
| NYNRIN | 0,009072401 | -0,354755963 | 0,071 | 0,366 | 1 | Ciliated | 0,78200191 | 1,278769255 | 0,193989071 |
| RPL392 | 0,009080975 | -0,836959078 | 1 | 0,976 | 1 | Ciliated | 0,559822323 | 1,786281037 | 1,024590164 |
| RPL231 | 0,009226191 | -0,486064498 | 0,786 | 0,854 | 1 | Ciliated | 0,71397007 | 1,400618936 | 0,920374707 |
| RPL62 | 0,009244417 | -0,421357897 | 0,964 | 0,976 | 1 | Ciliated | 0,746721463 | 1,339187435 | 0,987704918 |
| SLFN13 | 0,009254584 | -0,552874422 | 0,429 | 0,732 | 1 | Ciliated | 0,681660635 | 1,467005645 | 0,586065574 |
| IL15RA | 0,009312698 | 0,334130624 | 0,214 | 0,024 | 1 | Ciliated | 1,260617524 | 0,793262017 | 8,916666667 |
| H2AJ2 | 0,009333311 | 0,348013371 | 0,929 | 0,829 | 1 | Ciliated | 1,272806731 | 0,785665235 | 1,120627262 |
| CDK2AP11 | 0,009362971 | -0,477062033 | 0,214 | 0,585 | 1 | Ciliated | 0,718439196 | 1,391906239 | 0,365811966 |
| ASTN2 | 0,009401017 | -0,262184471 | 0,036 | 0,293 | 1 | Ciliated | 0,833824419 | 1,199293253 | 0,122866894 |
| NEK61 | 0,009401017 | 0,39500804 | 0,321 | 0,098 | 1 | Ciliated | 1,314950088 | 0,760485138 | 3,275510204 |
| STAT61 | 0,009428537 | -0,583646398 | 0,321 | 0,659 | 1 | Ciliated | 0,667275113 | 1,498632245 | 0,487101669 |
| CISD2 | 0,009469825 | 0,477944121 | 0,5 | 0,244 | 1 | Ciliated | 1,392757535 | 0,718000064 | 2,049180328 |
| VEZT | 0,009469825 | -0,408597916 | 0,143 | 0,488 | 1 | Ciliated | 0,753355166 | 1,327395158 | 0,293032787 |
| YWHAQ1 | 0,009470587 | 0,464623823 | 0,893 | 0,683 | 1 | Ciliated | 1,379957488 | 0,724660005 | 1,307467057 |
| RRAGA2 | 0,009510404 | 0,546483154 | 0,679 | 0,537 | 1 | Ciliated | 1,460521053 | 0,684687152 | 1,26443203 |
| KCNH3 | 0,009519363 | 0,460618472 | 0,464 | 0,244 | 1 | Ciliated | 1,376131628 | 0,726674672 | 1,901639344 |
| TXLNB | 0,009719707 | -0,513146276 | 0,25 | 0,634 | 1 | Ciliated | 0,700692677 | 1,4271592 | 0,394321767 |
| ROCK2 | 0,009732634 | -0,312058657 | 0,107 | 0,415 | 1 | Ciliated | 0,805491541 | 1,241477967 | 0,257831325 |
| ITGB42 | 0,009734197 | 0,416138355 | 0,643 | 0,415 | 1 | Ciliated | 1,334351128 | 0,749427927 | 1,54939759 |
| STAG21 | 0,009737622 | -0,64297136 | 0,25 | 0,561 | 1 | Ciliated | 0,640392643 | 1,561541986 | 0,445632799 |
| GSR | 0,009805717 | 0,56025349 | 0,536 | 0,293 | 1 | Ciliated | 1,474528278 | 0,678182992 | 1,829351536 |
| CMC1 | 0,009865967 | -0,311986656 | 0,107 | 0,439 | 1 | Ciliated | 0,805531742 | 1,241416009 | 0,243735763 |
| SAP30L | 0,009929813 | 0,319119867 | 0,464 | 0,22 | 1 | Ciliated | 1,247569223 | 0,801558729 | 2,109090909 |
| GNB22 | 0,009952035 | 0,494488103 | 0,857 | 0,829 | 1 | Ciliated | 1,408820789 | 0,709813489 | 1,033775633 |
| IQANK1 | 0,009954797 | 0,720032565 | 0,536 | 0,317 | 1 | Ciliated | 1,647219216 | 0,607083739 | 1,690851735 |
| WDR93 | 0,010002917 | -0,661909139 | 0,321 | 0,683 | 1 | Ciliated | 0,632041354 | 1,582174954 | 0,469985359 |
| MTMR12 | 0,01014151 | -0,256657382 | 0,143 | 0,463 | 1 | Ciliated | 0,837025 | 1,194707447 | 0,308855292 |

| Supplementary Table 4 - clusermarker | | | | | | | | | |
|---|---|---|---|---|---|---|---|---|---|
| | **p_val** | **avg_log2FC** | **pct.1** | **pct.2** | **p_val_adj** | **celltype** | **Foldchange_UP** | **Foldchange_DOWN** | **Ratio_pct1_pct2** |
| MSRB22 | 0,010207453 | 0,385404875 | 0,536 | 0,293 | 1 | Ciliated | 1,306226312 | 0,765564122 | 1,829351536 |
| EPPIN | 0,010352095 | -0,772299044 | 0,714 | 0,805 | 1 | Ciliated | 0,585483718 | 1,707989426 | 0,886956522 |
| ELAPOR1 | 0,010419164 | -0,339474955 | 0,107 | 0,39 | 1 | Ciliated | 0,790328886 | 1,265296027 | 0,274358974 |
| KRT172 | 0,010419164 | -0,948574563 | 0,107 | 0,39 | 1 | Ciliated | 0,518144155 | 1,929964837 | 0,274358974 |
| SH3BP5L | 0,010428316 | -0,434770764 | 0,036 | 0,293 | 1 | Ciliated | 0,739811297 | 1,351696039 | 0,122866894 |
| ASB8 | 0,010428316 | -0,468586289 | 0,036 | 0,293 | 1 | Ciliated | 0,722672404 | 1,383752851 | 0,122866894 |
| CD591 | 0,010450524 | 0,402566088 | 0,964 | 0,902 | 1 | Ciliated | 1,321856977 | 0,756511497 | 1,068736142 |
| MYL12B2 | 0,010461787 | 0,256806906 | 1 | 0,927 | 1 | Ciliated | 1,194831275 | 0,836938253 | 1,078748652 |
| HSD17B13 | 0,010495681 | 0,53779088 | 0,75 | 0,463 | 1 | Ciliated | 1,451747833 | 0,688824862 | 1,61987041 |
| TYK2 | 0,010595539 | 0,435523474 | 0,393 | 0,171 | 1 | Ciliated | 1,352401455 | 0,73942541 | 2,298245614 |
| UBXN13 | 0,010632549 | 0,382651919 | 0,929 | 0,756 | 1 | Ciliated | 1,303736143 | 0,767026369 | 1,228835979 |
| ALDH16A11 | 0,01067045 | 0,599795546 | 0,571 | 0,317 | 1 | Ciliated | 1,51550178 | 0,65984746 | 1,80126183 |
| TMEM14C2 | 0,010724665 | 0,373396188 | 0,857 | 0,707 | 1 | Ciliated | 1,295398688 | 0,77196311 | 1,212164074 |
| TSPAN32 | 0,010813094 | 0,469286879 | 0,929 | 0,902 | 1 | Ciliated | 1,38442498 | 0,722321552 | 1,029933481 |
| RPS242 | 0,010838385 | -0,608579437 | 1 | 1 | 1 | Ciliated | 0,655842165 | 1,524757104 | 1 |
| TMEM120B | 0,010943005 | 0,635273296 | 0,536 | 0,366 | 1 | Ciliated | 1,553231957 | 0,643818842 | 1,464480874 |
| NUDCD2 | 0,010976134 | 0,488513839 | 0,714 | 0,561 | 1 | Ciliated | 1,402998862 | 0,712758953 | 1,272727273 |
| KCNMB2 | 0,010978496 | -0,286221642 | 0,036 | 0,293 | 1 | Ciliated | 0,820046917 | 1,219442423 | 0,122866894 |
| DCP1A | 0,011027257 | -0,261082091 | 0,143 | 0,463 | 1 | Ciliated | 0,834461798 | 1,198377208 | 0,308855292 |
| RBM14 | 0,011095004 | -0,733551521 | 0,214 | 0,561 | 1 | Ciliated | 0,601421554 | 1,662727238 | 0,381461676 |
| CLDN72 | 0,011101834 | 0,502607204 | 0,929 | 0,805 | 1 | Ciliated | 1,416771606 | 0,705830069 | 1,154037267 |
| PFDN11 | 0,011103295 | -0,262169288 | 0,107 | 0,415 | 1 | Ciliated | 0,833833195 | 1,199280631 | 0,257831325 |
| RPL132 | 0,011223245 | 0,416656414 | 1 | 0,951 | 1 | Ciliated | 1,334830368 | 0,749158862 | 1,051524711 |
| BZW21 | 0,011265755 | -0,263214999 | 0,143 | 0,512 | 1 | Ciliated | 0,833229025 | 1,200150223 | 0,279296875 |
| PROS1 | 0,01127659 | -0,853657458 | 0,214 | 0,512 | 1 | Ciliated | 0,553380051 | 1,807076344 | 0,41796875 |
| EIF5B | 0,011361763 | -0,713577071 | 0,357 | 0,634 | 1 | Ciliated | 0,609806287 | 1,639865022 | 0,563091483 |
| CDC37L1 | 0,011454127 | -0,358398609 | 0,071 | 0,366 | 1 | Ciliated | 0,780029932 | 1,282002085 | 0,193989071 |
| LAMTOR21 | 0,011471528 | 0,267426827 | 0,786 | 0,61 | 1 | Ciliated | 1,203659081 | 0,83080003 | 1,28852459 |
| CCDC81 | 0,011507882 | -0,631438393 | 0,393 | 0,756 | 1 | Ciliated | 0,645532487 | 1,549108713 | 0,51984127 |
| HID1 | 0,011522736 | 0,570260852 | 0,25 | 0,049 | 1 | Ciliated | 1,48479201 | 0,673495003 | 5,102040816 |
| MTRNR2L82 | 0,011596881 | -5,452586103 | 0,143 | 0,39 | 1 | Ciliated | 0,022835369 | 43,79171674 | 0,366666667 |

| Supplementary Table 4 - clusermarker | | | | | | | | | |
|---|---|---|---|---|---|---|---|---|---|
| | **p_val** | **avg_log2FC** | **pct.1** | **pct.2** | **p_val_adj** | **celltype** | **Foldchange_UP** | **Foldchange_DOWN** | **Ratio_pct1_pct2** |
| ZMYND10 | 0,011614107 | 0,346742473 | 1 | 0,902 | 1 | Ciliated | 1,271685985 | 0,786357648 | 1,10864745 |
| MRPL551 | 0,01161706 | 0,278435845 | 0,714 | 0,512 | 1 | Ciliated | 1,21287918 | 0,824484431 | 1,39453125 |
| CDC372 | 0,011675286 | 0,335086758 | 0,75 | 0,634 | 1 | Ciliated | 1,261453265 | 0,792736463 | 1,1829653 |
| NAA40 | 0,011736359 | -0,287063029 | 0,071 | 0,341 | 1 | Ciliated | 0,819568801 | 1,220153816 | 0,208211144 |
| CLSTN12 | 0,011877272 | 0,650004604 | 0,786 | 0,561 | 1 | Ciliated | 1,569173203 | 0,63727828 | 1,401069519 |
| IRX31 | 0,011967667 | 0,468833493 | 0,893 | 0,61 | 1 | Ciliated | 1,383989975 | 0,722548586 | 1,463934426 |
| PCBP12 | 0,011968463 | 0,421005372 | 0,857 | 0,634 | 1 | Ciliated | 1,338860242 | 0,746903947 | 1,351735016 |
| PPP2CB | 0,012102157 | -0,635836325 | 0,5 | 0,78 | 1 | Ciliated | 0,643567633 | 1,553838243 | 0,641025641 |
| SEC22C | 0,01210984 | -0,304394972 | 0,107 | 0,415 | 1 | Ciliated | 0,809781746 | 1,234900644 | 0,257831325 |
| CNPY3 | 0,012132875 | 0,355203466 | 0,679 | 0,512 | 1 | Ciliated | 1,279165972 | 0,781759382 | 1,326171875 |
| RPL91 | 0,012233868 | -0,473162026 | 0,964 | 0,927 | 1 | Ciliated | 0,720383965 | 1,38814861 | 1,0399137 |
| UCHL31 | 0,012235072 | 0,621384511 | 0,5 | 0,293 | 1 | Ciliated | 1,538350782 | 0,650046798 | 1,706484642 |
| NEU1 | 0,012298929 | -0,359461919 | 0,071 | 0,341 | 1 | Ciliated | 0,779455238 | 1,282947308 | 0,208211144 |
| MCU1 | 0,012298929 | -0,407376963 | 0,071 | 0,341 | 1 | Ciliated | 0,753993001 | 1,326272258 | 0,208211144 |
| TCEAL1 | 0,012298929 | -0,298715171 | 0,071 | 0,341 | 1 | Ciliated | 0,812976091 | 1,230048474 | 0,208211144 |
| COTL1 | 0,012335982 | 0,331958341 | 0,286 | 0,073 | 1 | Ciliated | 1,258720826 | 0,79445734 | 3,917808219 |
| AUTS2 | 0,012422967 | -0,602764835 | 0,286 | 0,61 | 1 | Ciliated | 0,658490789 | 1,518624128 | 0,468852459 |
| SNX31 | 0,012439862 | 0,592726555 | 0,893 | 0,78 | 1 | Ciliated | 1,508094209 | 0,663088548 | 1,144871795 |
| DYNLL12 | 0,012454893 | 0,340561389 | 1 | 1 | 1 | Ciliated | 1,266249227 | 0,789733947 | 1 |
| ARHGEF28 | 0,0126324 | -0,268211097 | 0,286 | 0,659 | 1 | Ciliated | 0,830348517 | 1,204313585 | 0,433990895 |
| BHLHE41 | 0,012642808 | -0,285535313 | 0,107 | 0,415 | 1 | Ciliated | 0,820437129 | 1,218862439 | 0,257831325 |
| DNPH12 | 0,012844934 | 0,493322426 | 0,964 | 0,854 | 1 | Ciliated | 1,407682941 | 0,710387241 | 1,128805621 |
| HOATZ | 0,012873707 | -0,670374131 | 0,929 | 0,927 | 1 | Ciliated | 0,628343719 | 1,59148563 | 1,002157497 |
| CA5B | 0,01288526 | -0,392860927 | 0,071 | 0,341 | 1 | Ciliated | 0,761617785 | 1,312994549 | 0,208211144 |
| SOX151 | 0,013007337 | -0,307557743 | 0,143 | 0,463 | 1 | Ciliated | 0,808008434 | 1,237610844 | 0,308855292 |
| CFAP300 | 0,013034857 | 0,472268112 | 0,929 | 0,732 | 1 | Ciliated | 1,38728876 | 0,720830464 | 1,269125683 |
| LGMN | 0,013262679 | 0,516485422 | 0,393 | 0,171 | 1 | Ciliated | 1,430466213 | 0,699072785 | 2,298245614 |
| TTLL10 | 0,013401398 | 0,392457866 | 0,857 | 0,585 | 1 | Ciliated | 1,312627775 | 0,761830596 | 1,464957265 |
| LSP11 | 0,013496187 | 0,54375358 | 0,357 | 0,146 | 1 | Ciliated | 1,457760365 | 0,685983804 | 2,445205479 |
| NDUFB22 | 0,013497027 | 0,382362064 | 0,964 | 0,829 | 1 | Ciliated | 1,303474233 | 0,767180489 | 1,162846803 |
| SF11 | 0,01356521 | 0,322526908 | 0,821 | 0,707 | 1 | Ciliated | 1,25051894 | 0,799668016 | 1,161244696 |

| Supplementary Table 4 - clusermarker | | | | | | | | | |
|---|---|---|---|---|---|---|---|---|---|
| | **p_val** | **avg_log2FC** | **pct.1** | **pct.2** | **p_val_adj** | **celltype** | **Foldchange_UP** | **Foldchange_DOWN** | **Ratio_pct1_pct2** |
| AC007114.1 | 0,013569975 | -0,572229898 | 0,107 | 0,39 | 1 | Ciliated | 0,672576418 | 1,486819895 | 0,274358974 |
| PJA21 | 0,01360881 | -0,558663393 | 0,214 | 0,537 | 1 | Ciliated | 0,678930879 | 1,47290399 | 0,398510242 |
| ODAD1 | 0,013666183 | 0,555665207 | 0,893 | 0,634 | 1 | Ciliated | 1,469846203 | 0,680343289 | 1,40851735 |
| PGLS2 | 0,013667551 | 0,658915096 | 0,714 | 0,659 | 1 | Ciliated | 1,578894852 | 0,633354399 | 1,083459788 |
| DHPS1 | 0,01371418 | 0,541865553 | 0,571 | 0,39 | 1 | Ciliated | 1,455853869 | 0,686882126 | 1,464102564 |
| PRORP | 0,013738311 | 0,376060089 | 0,286 | 0,073 | 1 | Ciliated | 1,297792819 | 0,770539014 | 3,917808219 |
| ERCC4 | 0,013771391 | -0,254066764 | 0,107 | 0,415 | 1 | Ciliated | 0,838529379 | 1,19256406 | 0,257831325 |
| IDO11 | 0,013935985 | 0,266309452 | 0,143 | 0 | 1 | Ciliated | 1,202727201 | 0,831443738 | #DIV/0! |
| IGLV2-23 | 0,013935985 | 0,460285643 | 0,143 | 0 | 1 | Ciliated | 1,375814192 | 0,726842335 | #DIV/0! |
| S100A4 | 0,013991557 | 0,676340391 | 0,714 | 0,488 | 1 | Ciliated | 1,598080842 | 0,625750572 | 1,463114754 |
| COX5A2 | 0,014086684 | 0,518083113 | 0,857 | 0,829 | 1 | Ciliated | 1,432051238 | 0,698299037 | 1,033775633 |
| RIBC2 | 0,014129956 | 0,536070354 | 0,643 | 0,488 | 1 | Ciliated | 1,450017542 | 0,689646829 | 1,317622951 |
| SPATA7 | 0,014132568 | -0,510617769 | 0,071 | 0,341 | 1 | Ciliated | 0,701921807 | 1,424660112 | 0,208211144 |
| TSEN341 | 0,014140206 | 0,438314372 | 0,643 | 0,439 | 1 | Ciliated | 1,355020212 | 0,737996371 | 1,464692483 |
| COX19 | 0,014169996 | -0,280534708 | 0,107 | 0,39 | 1 | Ciliated | 0,823285825 | 1,214644986 | 0,274358974 |
| COPE2 | 0,014232172 | 0,309359334 | 0,821 | 0,659 | 1 | Ciliated | 1,239157298 | 0,807000049 | 1,245827011 |
| SULT2B1 | 0,014236596 | -0,330353964 | 0,036 | 0,268 | 1 | Ciliated | 0,795341324 | 1,25732182 | 0,134328358 |
| AGR22 | 0,014250327 | -0,921756794 | 0,929 | 0,902 | 1 | Ciliated | 0,527865838 | 1,894420757 | 1,029933481 |
| ZBTB40 | 0,014256176 | -0,391846927 | 0 | 0,195 | 1 | Ciliated | 0,762153277 | 1,312072034 | 0 |
| NCDN | 0,014256176 | -0,306626929 | 0 | 0,195 | 1 | Ciliated | 0,808529922 | 1,236812606 | 0 |
| AC021148.2 | 0,014256176 | -0,278302047 | 0 | 0,195 | 1 | Ciliated | 0,824560898 | 1,212766701 | 0 |
| AZGP12 | 0,014256176 | -0,686401955 | 0 | 0,195 | 1 | Ciliated | 0,621401679 | 1,609265043 | 0 |
| CPNE8 | 0,014256176 | -0,268174628 | 0 | 0,195 | 1 | Ciliated | 0,830369508 | 1,204283143 | 0 |
| INTS6-AS1 | 0,014256176 | -0,268141199 | 0 | 0,195 | 1 | Ciliated | 0,830388748 | 1,204255238 | 0 |
| MT1M | 0,014256176 | -0,449107117 | 0 | 0,195 | 1 | Ciliated | 0,732496049 | 1,365195077 | 0 |
| MT1G | 0,014256176 | -0,79667516 | 0 | 0,195 | 1 | Ciliated | 0,575674351 | 1,737093198 | 0 |
| EPB41L3 | 0,014256176 | -0,288801868 | 0 | 0,195 | 1 | Ciliated | 0,818581593 | 1,221625319 | 0 |
| MED26 | 0,014256176 | -0,387357476 | 0 | 0,195 | 1 | Ciliated | 0,764528678 | 1,307995408 | 0 |
| NCOA71 | 0,014349778 | -0,990695325 | 0,321 | 0,634 | 1 | Ciliated | 0,503235176 | 1,987142488 | 0,506309148 |
| CEBPD | 0,014395082 | 0,287666156 | 0,821 | 0,585 | 1 | Ciliated | 1,220664015 | 0,819226247 | 1,403418803 |
| MVP2 | 0,014404151 | 0,479707906 | 0,821 | 0,732 | 1 | Ciliated | 1,394461309 | 0,717122801 | 1,121584699 |

| Supplementary Table 4 - clusermarker | | | | | | | | | |
|---|---|---|---|---|---|---|---|---|---|
| | **p_val** | **avg_log2FC** | **pct.1** | **pct.2** | **p_val_adj** | **celltype** | **Foldchange_UP** | **Foldchange_DOWN** | **Ratio_pct1_pct2** |
| NCLN | 0,014485757 | 0,458978964 | 0,393 | 0,171 | 1 | Ciliated | 1,374568653 | 0,727500949 | 2,298245614 |
| CYP4B11 | 0,014494108 | -0,556895432 | 0,964 | 0,878 | 1 | Ciliated | 0,679763389 | 1,471100115 | 1,097949886 |
| TADA31 | 0,014641456 | 0,529551008 | 0,643 | 0,488 | 1 | Ciliated | 1,44347989 | 0,692770303 | 1,317622951 |
| TMEM87A1 | 0,014657464 | -0,508902426 | 0,321 | 0,659 | 1 | Ciliated | 0,702756878 | 1,422967218 | 0,487101669 |
| IFI27L21 | 0,014786765 | 0,560774661 | 0,536 | 0,341 | 1 | Ciliated | 1,475061045 | 0,677938044 | 1,571847507 |
| CEP95 | 0,014793307 | -0,310018472 | 0,107 | 0,39 | 1 | Ciliated | 0,806631431 | 1,239723573 | 0,274358974 |
| YWHAB1 | 0,014942393 | 0,451112951 | 0,929 | 0,732 | 1 | Ciliated | 1,36709448 | 0,731478339 | 1,269125683 |
| TKT | 0,015082513 | 0,68837744 | 0,714 | 0,561 | 1 | Ciliated | 1,611470122 | 0,620551375 | 1,272727273 |
| DPP72 | 0,015263633 | 0,571648959 | 0,679 | 0,561 | 1 | Ciliated | 1,486221308 | 0,672847304 | 1,210338681 |
| SDCBP | 0,015374775 | -0,473536787 | 0,393 | 0,683 | 1 | Ciliated | 0,720196859 | 1,388509249 | 0,575402635 |
| SGPL1 | 0,015394726 | 0,470069031 | 0,5 | 0,317 | 1 | Ciliated | 1,385175745 | 0,721930054 | 1,577287066 |
| MYH10 | 0,015430452 | -0,741264607 | 0,179 | 0,463 | 1 | Ciliated | 0,598214752 | 1,671640487 | 0,386609071 |
| ZNF143 | 0,015485235 | -0,506749146 | 0,071 | 0,341 | 1 | Ciliated | 0,703806554 | 1,420844967 | 0,208211144 |
| CAPN51 | 0,015485235 | 0,47861669 | 0,357 | 0,146 | 1 | Ciliated | 1,393406975 | 0,717665419 | 2,445205479 |
| IMPA21 | 0,015485235 | -0,500939766 | 0,071 | 0,341 | 1 | Ciliated | 0,706646324 | 1,415135076 | 0,208211144 |
| OMG | 0,015592724 | -1,061335334 | 0,821 | 0,854 | 1 | Ciliated | 0,479188326 | 2,086862193 | 0,961358314 |
| IGKC2 | 0,015605882 | 0,685213582 | 0,321 | 0,098 | 1 | Ciliated | 1,607940009 | 0,621913749 | 3,275510204 |
| HNMT1 | 0,01576817 | -0,250408102 | 0,036 | 0,268 | 1 | Ciliated | 0,840658581 | 1,189543559 | 0,134328358 |
| AC099518.4 | 0,01576817 | -0,436476422 | 0,036 | 0,268 | 1 | Ciliated | 0,738937156 | 1,353295057 | 0,134328358 |
| THAP6 | 0,015816978 | -0,490512365 | 0,071 | 0,317 | 1 | Ciliated | 0,711772271 | 1,404943745 | 0,223974763 |
| BRD8 | 0,015816978 | -0,356650359 | 0,071 | 0,317 | 1 | Ciliated | 0,780975741 | 1,280449504 | 0,223974763 |
| BBX1 | 0,015886286 | -0,291954476 | 0,25 | 0,61 | 1 | Ciliated | 0,816794765 | 1,224297759 | 0,409836066 |
| DHX40 | 0,015973252 | -0,456006064 | 0,25 | 0,561 | 1 | Ciliated | 0,729001625 | 1,371739054 | 0,445632799 |
| GLTP | 0,015994371 | 0,443504735 | 0,5 | 0,317 | 1 | Ciliated | 1,359903929 | 0,735346063 | 1,577287066 |
| PIR | 0,016059883 | 0,57005688 | 0,571 | 0,415 | 1 | Ciliated | 1,484582101 | 0,673590231 | 1,375903614 |
| ORC3 | 0,016112808 | -0,572029129 | 0,107 | 0,39 | 1 | Ciliated | 0,672670023 | 1,486613 | 0,274358974 |
| IFT43 | 0,016125314 | 0,378782589 | 0,893 | 0,805 | 1 | Ciliated | 1,300244187 | 0,769086307 | 1,10931677 |
| RIDA | 0,016203345 | -0,373649346 | 0,071 | 0,341 | 1 | Ciliated | 0,771827661 | 1,295626019 | 0,208211144 |
| NUDT142 | 0,016204708 | 0,542092217 | 0,821 | 0,683 | 1 | Ciliated | 1,456082619 | 0,686774217 | 1,20204978 |
| TTLL9 | 0,016238268 | -0,396495832 | 0,214 | 0,561 | 1 | Ciliated | 0,759701286 | 1,316306841 | 0,381461676 |
| ARFGAP31 | 0,016238268 | 0,432351346 | 0,571 | 0,317 | 1 | Ciliated | 1,349431127 | 0,741053011 | 1,80126183 |

| Supplementary Table 4 - clusermarker | | | | | | | | | |
|---|---|---|---|---|---|---|---|---|---|
| | **p_val** | **avg_log2FC** | **pct.1** | **pct.2** | **p_val_adj** | **celltype** | **Foldchange_UP** | **Foldchange_DOWN** | **Ratio_pct1_pct2** |
| SOX21 | 0,016394514 | -0,498660087 | 0,607 | 0,78 | 1 | Ciliated | 0,707763817 | 1,412900711 | 0,778205128 |
| ZNF217 | 0,016453564 | -0,347277459 | 0,214 | 0,537 | 1 | Ciliated | 0,786066101 | 1,272157644 | 0,398510242 |
| TOMM34 | 0,016474473 | 0,318523375 | 0,571 | 0,39 | 1 | Ciliated | 1,247053514 | 0,801890207 | 1,464102564 |
| TUBA4B | 0,016480039 | 0,550592896 | 0,857 | 0,659 | 1 | Ciliated | 1,464687507 | 0,682739489 | 1,300455235 |
| SCGB3A22 | 0,016567069 | -1,111447573 | 0,071 | 0,317 | 1 | Ciliated | 0,462829404 | 2,160623314 | 0,223974763 |
| CLDN11 | 0,016572644 | -0,495527307 | 0,25 | 0,561 | 1 | Ciliated | 0,70930238 | 1,409835958 | 0,445632799 |
| SLC39A10 | 0,016586949 | -0,392877689 | 0,036 | 0,268 | 1 | Ciliated | 0,761608936 | 1,313009805 | 0,134328358 |
| DDX27 | 0,016713239 | -0,307656195 | 0,143 | 0,415 | 1 | Ciliated | 0,807953296 | 1,237695303 | 0,344578313 |
| GTPBP10 | 0,016950559 | -0,370085676 | 0,071 | 0,341 | 1 | Ciliated | 0,773736546 | 1,292429581 | 0,208211144 |
| MIB1 | 0,01699053 | -0,304724612 | 0,107 | 0,415 | 1 | Ciliated | 0,809596741 | 1,235182837 | 0,257831325 |
| TMEM68 | 0,017024095 | -0,573009225 | 0,143 | 0,439 | 1 | Ciliated | 0,672213199 | 1,487623275 | 0,325740319 |
| SPOP | 0,017024095 | -0,42568547 | 0,143 | 0,439 | 1 | Ciliated | 0,744484919 | 1,343210553 | 0,325740319 |
| PTRHD12 | 0,017067871 | 0,57156158 | 0,679 | 0,61 | 1 | Ciliated | 1,486131296 | 0,672888057 | 1,113114754 |
| FUCA11 | 0,017257854 | 0,497007241 | 0,571 | 0,415 | 1 | Ciliated | 1,411282927 | 0,708575142 | 1,375903614 |
| PCNP1 | 0,017360915 | -0,298096694 | 0,179 | 0,512 | 1 | Ciliated | 0,813324684 | 1,229521271 | 0,349609375 |
| GYS1 | 0,017442833 | -0,349530882 | 0,036 | 0,268 | 1 | Ciliated | 0,784839261 | 1,274146249 | 0,134328358 |
| GOLGB11 | 0,017443158 | -0,576512848 | 0,679 | 0,756 | 1 | Ciliated | 0,670582692 | 1,491240398 | 0,898148148 |
| LSM51 | 0,01745141 | -0,619180321 | 0,357 | 0,683 | 1 | Ciliated | 0,651040717 | 1,536002241 | 0,522693997 |
| WDR54 | 0,017459084 | 0,532843054 | 0,893 | 0,78 | 1 | Ciliated | 1,446777487 | 0,691191292 | 1,144871795 |
| ANKRD45 | 0,017501812 | 0,641246222 | 0,607 | 0,463 | 1 | Ciliated | 1,559675849 | 0,641158867 | 1,311015119 |
| UBALD2 | 0,017501812 | -0,385875593 | 0,357 | 0,634 | 1 | Ciliated | 0,765314377 | 1,306652573 | 0,563091483 |
| TBL1X | 0,017534664 | -0,288665336 | 0,107 | 0,39 | 1 | Ciliated | 0,818659065 | 1,221509713 | 0,274358974 |
| MSRA | 0,017588132 | 0,365449693 | 0,321 | 0,122 | 1 | Ciliated | 1,288283129 | 0,776226885 | 2,631147541 |
| EMC41 | 0,017657893 | 0,478374268 | 0,75 | 0,634 | 1 | Ciliated | 1,393172855 | 0,717786021 | 1,1829653 |
| HELZ | 0,017736744 | -0,313900601 | 0,179 | 0,488 | 1 | Ciliated | 0,804463795 | 1,243064021 | 0,366803279 |
| RSRC21 | 0,017776246 | -0,475348596 | 0,321 | 0,634 | 1 | Ciliated | 0,719292967 | 1,390254104 | 0,506309148 |
| ATP12A1 | 0,017885714 | 0,817618723 | 0,571 | 0,415 | 1 | Ciliated | 1,76249446 | 0,56737767 | 1,375903614 |
| CLTA2 | 0,017925033 | 0,48121038 | 0,929 | 0,829 | 1 | Ciliated | 1,395914307 | 0,716376353 | 1,120627262 |
| PEX19 | 0,01798821 | -0,445048138 | 0,107 | 0,366 | 1 | Ciliated | 0,734559806 | 1,36135954 | 0,292349727 |
| FOS1 | 0,018030115 | -0,950518956 | 0,929 | 0,927 | 1 | Ciliated | 0,517446296 | 1,932567702 | 1,002157497 |
| SPCS2 | 0,018125318 | 0,548116718 | 0,679 | 0,561 | 1 | Ciliated | 1,462175737 | 0,683912319 | 1,210338681 |

| Supplementary Table 4 - clusermarker | | | | | | | | | |
|---|---|---|---|---|---|---|---|---|---|
| | **p_val** | **avg_log2FC** | **pct.1** | **pct.2** | **p_val_adj** | **celltype** | **Foldchange_UP** | **Foldchange_DOWN** | **Ratio_pct1_pct2** |
| TAPBPL | 0,018161488 | -0,637068837 | 0,071 | 0,317 | 1 | Ciliated | 0,64301806 | 1,555166274 | 0,223974763 |
| PCF111 | 0,018400548 | -0,327301309 | 0,214 | 0,537 | 1 | Ciliated | 0,797025999 | 1,254664215 | 0,398510242 |
| LAMTOR41 | 0,018418863 | 0,449045028 | 0,964 | 0,756 | 1 | Ciliated | 1,365136324 | 0,732527574 | 1,275132275 |
| MCUB | 0,018433994 | 0,375112137 | 0,321 | 0,122 | 1 | Ciliated | 1,296940358 | 0,771045479 | 2,631147541 |
| CES4A | 0,0184801 | 0,264113813 | 0,679 | 0,512 | 1 | Ciliated | 1,200898162 | 0,832710076 | 1,326171875 |
| NDUFAF31 | 0,018531649 | 0,46548699 | 0,857 | 0,707 | 1 | Ciliated | 1,380783367 | 0,724226569 | 1,212164074 |
| COQ10A | 0,018536191 | 0,386141012 | 0,357 | 0,146 | 1 | Ciliated | 1,306892986 | 0,765173592 | 2,445205479 |
| PRRC2B2 | 0,01856857 | 0,439750678 | 0,714 | 0,659 | 1 | Ciliated | 1,356369904 | 0,737262009 | 1,083459788 |
| TIMM17A | 0,018701511 | 0,499316387 | 0,643 | 0,488 | 1 | Ciliated | 1,413543604 | 0,707441919 | 1,317622951 |
| BUD23 | 0,019003905 | 0,470957957 | 0,821 | 0,707 | 1 | Ciliated | 1,386029493 | 0,721485369 | 1,161244696 |
| PPP1R13B | 0,019007947 | -0,294088507 | 0,071 | 0,317 | 1 | Ciliated | 0,815587457 | 1,226110078 | 0,223974763 |
| SMC5 | 0,019128932 | -0,578494465 | 0,25 | 0,585 | 1 | Ciliated | 0,669662244 | 1,493290102 | 0,427350427 |
| COX8A2 | 0,019215434 | 0,457450292 | 0,964 | 0,854 | 1 | Ciliated | 1,373112939 | 0,728272214 | 1,128805621 |
| GADD45B2 | 0,019379133 | -0,647093288 | 0,679 | 0,732 | 1 | Ciliated | 0,638565587 | 1,566009852 | 0,927595628 |
| JPT11 | 0,01955847 | 0,347570794 | 0,75 | 0,585 | 1 | Ciliated | 1,272416331 | 0,785906292 | 1,282051282 |
| AKR1A11 | 0,01964068 | 0,566525397 | 0,821 | 0,707 | 1 | Ciliated | 1,480952529 | 0,67524109 | 1,161244696 |
| DYNC1I2 | 0,019826234 | -0,361925161 | 0,25 | 0,585 | 1 | Ciliated | 0,778125541 | 1,285139669 | 0,427350427 |
| PBDC1 | 0,019873582 | -0,261432672 | 0,107 | 0,39 | 1 | Ciliated | 0,834259045 | 1,198668455 | 0,274358974 |
| CCNDBP1 | 0,019879277 | -0,474615807 | 0,25 | 0,537 | 1 | Ciliated | 0,719658411 | 1,389548131 | 0,465549348 |
| MED7 | 0,019888701 | -0,420198131 | 0,071 | 0,317 | 1 | Ciliated | 0,747321985 | 1,33811131 | 0,223974763 |
| HDHD2 | 0,019888701 | -0,266012605 | 0,071 | 0,317 | 1 | Ciliated | 0,831614833 | 1,202479755 | 0,223974763 |
| BAIAP3 | 0,02003274 | -0,462429279 | 0,214 | 0,512 | 1 | Ciliated | 0,725763154 | 1,377859972 | 0,41796875 |
| OAT2 | 0,020089907 | -0,679126937 | 0,357 | 0,61 | 1 | Ciliated | 0,624543109 | 1,601170496 | 0,585245902 |
| FIS12 | 0,020156337 | 0,496565049 | 0,893 | 0,756 | 1 | Ciliated | 1,410850429 | 0,708792356 | 1,181216931 |
| GON7 | 0,020156337 | 0,258110358 | 0,893 | 0,756 | 1 | Ciliated | 1,195911274 | 0,836182434 | 1,181216931 |
| NDUFS71 | 0,020177841 | 0,374765761 | 0,75 | 0,634 | 1 | Ciliated | 1,296629014 | 0,771230621 | 1,1829653 |
| AHSA1 | 0,020212051 | 0,527827529 | 0,786 | 0,585 | 1 | Ciliated | 1,441756503 | 0,693598398 | 1,343589744 |
| CBR31 | 0,020260234 | 0,467256006 | 0,464 | 0,268 | 1 | Ciliated | 1,382477505 | 0,723339075 | 1,731343284 |
| CIAO2B2 | 0,0204531 | 0,413892572 | 0,857 | 0,707 | 1 | Ciliated | 1,332275616 | 0,750595439 | 1,212164074 |
| COX7B1 | 0,02053377 | -0,711319726 | 0,929 | 0,902 | 1 | Ciliated | 0,610761181 | 1,637301177 | 1,029933481 |
| LARS11 | 0,020612877 | -0,459399226 | 0,25 | 0,537 | 1 | Ciliated | 0,727289057 | 1,374969128 | 0,465549348 |

| Supplementary Table 4 - clusermarker | | | | | | | | | |
|---|---|---|---|---|---|---|---|---|---|
| | **p_val** | **avg_log2FC** | **pct.1** | **pct.2** | **p_val_adj** | **celltype** | **Foldchange_UP** | **Foldchange_DOWN** | **Ratio_pct1_pct2** |
| EXPH5 | 0,020797466 | -0,464554639 | 0,357 | 0,61 | 1 | Ciliated | 0,724694757 | 1,379891314 | 0,585245902 |
| GPATCH4 | 0,020804875 | -0,41855903 | 0,071 | 0,317 | 1 | Ciliated | 0,748171528 | 1,336591894 | 0,223974763 |
| ARID2 | 0,020804875 | -0,535367765 | 0,071 | 0,317 | 1 | Ciliated | 0,689982767 | 1,449311559 | 0,223974763 |
| PLEKHA7 | 0,0210482 | -0,53727328 | 0,179 | 0,463 | 1 | Ciliated | 0,689072038 | 1,451227078 | 0,386609071 |
| SLC22A4 | 0,021200731 | 0,469645569 | 0,786 | 0,61 | 1 | Ciliated | 1,384769226 | 0,722141987 | 1,28852459 |
| CFAP44 | 0,021271218 | 0,304535721 | 0,821 | 0,659 | 1 | Ciliated | 1,235021126 | 0,809702748 | 1,245827011 |
| AL627171.21 | 0,021282276 | -0,553087863 | 0,143 | 0,415 | 1 | Ciliated | 0,681559794 | 1,467222699 | 0,344578313 |
| AKR1C12 | 0,021303607 | -0,283015517 | 0,107 | 0,366 | 1 | Ciliated | 0,821871347 | 1,216735445 | 0,292349727 |
| RCN21 | 0,021323763 | 0,543553084 | 0,679 | 0,537 | 1 | Ciliated | 1,457557789 | 0,686079144 | 1,26443203 |
| PUM2 | 0,021334309 | -0,43157408 | 0,286 | 0,61 | 1 | Ciliated | 0,741452368 | 1,348704304 | 0,468852459 |
| INPP4B | 0,021334309 | 0,633097773 | 0,571 | 0,415 | 1 | Ciliated | 1,550891514 | 0,644790426 | 1,375903614 |
| SMIM4 | 0,021369972 | -0,425138577 | 0,25 | 0,537 | 1 | Ciliated | 0,74476719 | 1,342701469 | 0,465549348 |
| ZSWIM61 | 0,021443778 | -0,425999874 | 0,179 | 0,488 | 1 | Ciliated | 0,744322693 | 1,343503308 | 0,366803279 |
| NDUFAF81 | 0,021446324 | 0,433225123 | 0,679 | 0,561 | 1 | Ciliated | 1,350248666 | 0,740604324 | 1,210338681 |
| SPATA33 | 0,021483249 | 0,511408341 | 0,786 | 0,634 | 1 | Ciliated | 1,425441014 | 0,701537272 | 1,239747634 |
| ZNF295-AS1 | 0,02156383 | 0,524978521 | 0,5 | 0,317 | 1 | Ciliated | 1,438912157 | 0,694969457 | 1,577287066 |
| SCAF8 | 0,021580215 | -0,367426226 | 0,107 | 0,39 | 1 | Ciliated | 0,77516416 | 1,290049323 | 0,274358974 |
| UFM11 | 0,02158481 | -0,671666051 | 0,5 | 0,732 | 1 | Ciliated | 0,627781295 | 1,592911429 | 0,683060109 |
| OCIAD22 | 0,021682759 | 0,543292346 | 0,607 | 0,537 | 1 | Ciliated | 1,457294388 | 0,68620315 | 1,130353818 |
| FIP1L1 | 0,02175752 | -0,46561025 | 0,214 | 0,561 | 1 | Ciliated | 0,724164696 | 1,380901342 | 0,381461676 |
| CYP4X11 | 0,021757623 | -0,297230283 | 0,071 | 0,317 | 1 | Ciliated | 0,813813274 | 1,228783103 | 0,223974763 |
| DPCD | 0,021830414 | 0,62481217 | 0,821 | 0,683 | 1 | Ciliated | 1,542010052 | 0,648504203 | 1,20204978 |
| LRRC23 | 0,021947916 | 0,4017238 | 1 | 0,951 | 1 | Ciliated | 1,321085463 | 0,756953299 | 1,051524711 |
| FGF14 | 0,022089625 | 0,566952203 | 0,571 | 0,415 | 1 | Ciliated | 1,481390719 | 0,675041356 | 1,375903614 |
| FBRS | 0,022089625 | 0,392846633 | 0,571 | 0,415 | 1 | Ciliated | 1,31298154 | 0,761625331 | 1,375903614 |
| C2CD3 | 0,022211299 | -0,273668856 | 0,107 | 0,366 | 1 | Ciliated | 0,827213218 | 1,208878168 | 0,292349727 |
| EXOC5 | 0,022211299 | -0,267466193 | 0,107 | 0,366 | 1 | Ciliated | 0,830777361 | 1,203691925 | 0,292349727 |
| GATC | 0,022328171 | 0,34393818 | 0,286 | 0,098 | 1 | Ciliated | 1,269216499 | 0,787887646 | 2,918367347 |
| BEX3 | 0,022480395 | 0,710974692 | 0,393 | 0,195 | 1 | Ciliated | 1,636909648 | 0,610907267 | 2,015384615 |
| AC107398.4 | 0,022497135 | -0,2562145 | 0,036 | 0,244 | 1 | Ciliated | 0,837281991 | 1,194340749 | 0,147540984 |
| MT1F | 0,022598809 | -0,464747282 | 0,143 | 0,39 | 1 | Ciliated | 0,724597994 | 1,380075583 | 0,366666667 |

| Supplementary Table 4 - clusermarker | | | | | | | | | |
|---|---|---|---|---|---|---|---|---|---|
| | **p_val** | **avg_log2FC** | **pct.1** | **pct.2** | **p_val_adj** | **celltype** | **Foldchange_UP** | **Foldchange_DOWN** | **Ratio_pct1_pct2** |
| RAB36 | 0,022791993 | 0,56706274 | 0,679 | 0,537 | 1 | Ciliated | 1,481504224 | 0,674989638 | 1,26443203 |
| LRRC40 | 0,02297475 | -0,538299435 | 0 | 0,171 | 1 | Ciliated | 0,688582091 | 1,452259669 | 0 |
| AC084198.2 | 0,02297475 | -0,411370098 | 0 | 0,171 | 1 | Ciliated | 0,751908962 | 1,32994824 | 0 |
| FBXL17 | 0,02297475 | -0,41018107 | 0 | 0,171 | 1 | Ciliated | 0,752528919 | 1,328852585 | 0 |
| FKTN | 0,02297475 | -0,296133698 | 0 | 0,171 | 1 | Ciliated | 0,814432084 | 1,227849466 | 0 |
| ZNF503-AS1 | 0,02297475 | -0,449154962 | 0 | 0,171 | 1 | Ciliated | 0,732471757 | 1,365240352 | 0 |
| EXOC6 | 0,02297475 | -0,262575638 | 0 | 0,171 | 1 | Ciliated | 0,83359837 | 1,199618469 | 0 |
| MTRF1 | 0,02297475 | -0,253256269 | 0 | 0,171 | 1 | Ciliated | 0,83900059 | 1,191894275 | 0 |
| SNAI11 | 0,02297475 | -0,266996855 | 0 | 0,171 | 1 | Ciliated | 0,831047673 | 1,203300403 | 0 |
| SH2D4A | 0,023105701 | -0,285475332 | 0,179 | 0,488 | 1 | Ciliated | 0,82047124 | 1,218811765 | 0,366803279 |
| IFIH1 | 0,023218673 | -0,469039718 | 0,071 | 0,293 | 1 | Ciliated | 0,72244531 | 1,384187823 | 0,242320819 |
| DYRK3 | 0,023330077 | -0,274361135 | 0,143 | 0,439 | 1 | Ciliated | 0,826816374 | 1,209458389 | 0,325740319 |
| CLDN12 | 0,023330077 | -0,356031077 | 0,143 | 0,439 | 1 | Ciliated | 0,78131105 | 1,279899984 | 0,325740319 |
| CEBPB1 | 0,023382727 | 0,631310023 | 0,714 | 0,659 | 1 | Ciliated | 1,548970881 | 0,645589928 | 1,083459788 |
| MAPRE31 | 0,023392951 | 0,68548787 | 0,643 | 0,537 | 1 | Ciliated | 1,608245743 | 0,621795521 | 1,197392924 |
| SMARCE12 | 0,023412163 | -0,494614577 | 0,321 | 0,683 | 1 | Ciliated | 0,709751266 | 1,408944299 | 0,469985359 |
| HSPA14.1 | 0,023651874 | 0,304499363 | 0,179 | 0,024 | 1 | Ciliated | 1,234990003 | 0,809723154 | 7,458333333 |
| LRRC26 | 0,023654643 | -0,266003781 | 0,036 | 0,244 | 1 | Ciliated | 0,831619919 | 1,202472401 | 0,147540984 |
| NTN1 | 0,023654643 | -0,270173209 | 0,036 | 0,244 | 1 | Ciliated | 0,829219984 | 1,205952605 | 0,147540984 |
| ZC3H131 | 0,023744708 | -0,543406875 | 0,25 | 0,512 | 1 | Ciliated | 0,686148678 | 1,45741008 | 0,48828125 |
| LINC02345 | 0,023756784 | -0,410069455 | 0,25 | 0,561 | 1 | Ciliated | 0,752587141 | 1,328749782 | 0,445632799 |
| CDC37L1-DT | 0,023777562 | -0,292322631 | 0,071 | 0,317 | 1 | Ciliated | 0,816586357 | 1,224610222 | 0,223974763 |
| FBXW9 | 0,023788892 | 0,49600326 | 0,571 | 0,439 | 1 | Ciliated | 1,410301147 | 0,709068416 | 1,300683371 |
| B3GNT5 | 0,023819213 | -0,573600533 | 0,5 | 0,732 | 1 | Ciliated | 0,67193774 | 1,488233122 | 0,683060109 |
| AGPAT22 | 0,023854852 | 0,551505014 | 0,607 | 0,463 | 1 | Ciliated | 1,465613822 | 0,682307976 | 1,311015119 |
| TMEM256 | 0,023951812 | 0,391726502 | 0,893 | 0,732 | 1 | Ciliated | 1,311962517 | 0,762216898 | 1,219945355 |
| FAM183A | 0,02415796 | 0,427974958 | 1 | 0,951 | 1 | Ciliated | 1,345343856 | 0,743304394 | 1,051524711 |
| BX284668.5 | 0,024295489 | -0,416968845 | 0,071 | 0,293 | 1 | Ciliated | 0,748996642 | 1,335119471 | 0,242320819 |
| CWC25 | 0,024295489 | -0,276177046 | 0,071 | 0,293 | 1 | Ciliated | 0,82577632 | 1,210981686 | 0,242320819 |
| TOB2 | 0,024347276 | 0,312155161 | 0,679 | 0,537 | 1 | Ciliated | 1,241561014 | 0,805437662 | 1,26443203 |
| ZNF160 | 0,024379087 | -0,84442506 | 0,107 | 0,39 | 1 | Ciliated | 0,556932713 | 1,795549045 | 0,274358974 |

| Supplementary Table 4 - clusermarker | | | | | | | | | |
|---|---|---|---|---|---|---|---|---|---|
| | **p_val** | **avg_log2FC** | **pct.1** | **pct.2** | **p_val_adj** | **celltype** | **Foldchange_UP** | **Foldchange_DOWN** | **Ratio_pct1_pct2** |
| CCHCR1 | 0,024473048 | -0,547502642 | 0,179 | 0,463 | 1 | Ciliated | 0,684203485 | 1,461553502 | 0,386609071 |
| IGFBP21 | 0,024532699 | 0,46338867 | 0,964 | 0,927 | 1 | Ciliated | 1,378776552 | 0,725280683 | 1,0399137 |
| ADAR1 | 0,024607333 | 0,462908955 | 0,714 | 0,585 | 1 | Ciliated | 1,378318168 | 0,725521888 | 1,220512821 |
| ANXA112 | 0,02464726 | 0,426655083 | 0,929 | 0,78 | 1 | Ciliated | 1,344113607 | 0,743984731 | 1,191025641 |
| UTRN | 0,024777109 | -0,441287377 | 0,393 | 0,683 | 1 | Ciliated | 0,736477126 | 1,357815423 | 0,575402635 |
| EFCAB7 | 0,024847179 | -0,375930294 | 0,071 | 0,317 | 1 | Ciliated | 0,77060834 | 1,297676067 | 0,223974763 |
| ZNF214 | 0,024847179 | -0,287080188 | 0,071 | 0,317 | 1 | Ciliated | 0,819559053 | 1,220168328 | 0,223974763 |
| STEAP41 | 0,024863477 | -0,26217338 | 0,036 | 0,244 | 1 | Ciliated | 0,83383083 | 1,199284032 | 0,147540984 |
| LGALS9C | 0,024863477 | -0,293625605 | 0,036 | 0,244 | 1 | Ciliated | 0,815849188 | 1,225716732 | 0,147540984 |
| CYP2S11 | 0,024863477 | -0,281693904 | 0,036 | 0,244 | 1 | Ciliated | 0,822624586 | 1,215621338 | 0,147540984 |
| OSBPL6 | 0,025084177 | -0,301998639 | 0,214 | 0,561 | 1 | Ciliated | 0,811127921 | 1,232851162 | 0,381461676 |
| BMPR1A | 0,025139038 | -0,385701448 | 0,107 | 0,366 | 1 | Ciliated | 0,765406762 | 1,306494859 | 0,292349727 |
| LAGE32 | 0,025180569 | 0,571554419 | 0,5 | 0,293 | 1 | Ciliated | 1,486123919 | 0,672891397 | 1,706484642 |
| PI4KA | 0,025192674 | -0,470592288 | 0,143 | 0,439 | 1 | Ciliated | 0,721668261 | 1,385678231 | 0,325740319 |
| RUVBL2 | 0,025247205 | 0,739924302 | 0,857 | 0,78 | 1 | Ciliated | 1,670088207 | 0,598770769 | 1,098717949 |
| PYGL | 0,025379491 | -0,428028353 | 0,107 | 0,39 | 1 | Ciliated | 0,743276884 | 1,345393649 | 0,274358974 |
| APAF1 | 0,025415327 | -0,576754886 | 0,071 | 0,293 | 1 | Ciliated | 0,670470199 | 1,491490601 | 0,242320819 |
| NDUFB51 | 0,025427027 | 0,383167751 | 0,75 | 0,585 | 1 | Ciliated | 1,304202374 | 0,766752169 | 1,282051282 |
| CS1 | 0,025771993 | -0,284346572 | 0,107 | 0,341 | 1 | Ciliated | 0,821113425 | 1,217858544 | 0,313782991 |
| PTPN182 | 0,025842827 | 0,450392787 | 0,5 | 0,317 | 1 | Ciliated | 1,366412224 | 0,73184357 | 1,577287066 |
| CCDC13 | 0,025844578 | -0,437044099 | 0,214 | 0,512 | 1 | Ciliated | 0,738646453 | 1,353827661 | 0,41796875 |
| ATP5MF1 | 0,025933723 | -0,77709299 | 0,786 | 0,805 | 1 | Ciliated | 0,583541437 | 1,713674362 | 0,976397516 |
| MYH91 | 0,026047343 | -0,485321888 | 0,286 | 0,585 | 1 | Ciliated | 0,714337672 | 1,39989817 | 0,488888889 |
| KIF26B-AS1 | 0,026053999 | 0,367270164 | 0,25 | 0,073 | 1 | Ciliated | 1,289909781 | 0,775248017 | 3,424657534 |
| STAG3 | 0,026053999 | 0,290653295 | 0,25 | 0,073 | 1 | Ciliated | 1,223194051 | 0,817531772 | 3,424657534 |
| MR11 | 0,026125449 | -0,347667178 | 0,036 | 0,244 | 1 | Ciliated | 0,785853788 | 1,272501343 | 0,147540984 |
| KBTBD2 | 0,026125449 | -0,262916633 | 0,036 | 0,244 | 1 | Ciliated | 0,833401364 | 1,199902043 | 0,147540984 |
| MALT1 | 0,026125449 | -0,30146806 | 0,036 | 0,244 | 1 | Ciliated | 0,811426284 | 1,232397841 | 0,147540984 |
| KDM7A | 0,026186528 | -0,261894127 | 0,107 | 0,366 | 1 | Ciliated | 0,833992244 | 1,199051918 | 0,292349727 |
| TMEM220 | 0,026186528 | 0,395729713 | 0,393 | 0,171 | 1 | Ciliated | 1,315608024 | 0,76010482 | 2,298245614 |
| TMEM111 | 0,026186528 | -0,446499143 | 0,107 | 0,366 | 1 | Ciliated | 0,733821387 | 1,36272943 | 0,292349727 |

| Supplementary Table 4 - clusermarker | | | | | | | | | |
|---|---|---|---|---|---|---|---|---|---|
| | **p_val** | **avg_log2FC** | **pct.1** | **pct.2** | **p_val_adj** | **celltype** | **Foldchange_UP** | **Foldchange_DOWN** | **Ratio_pct1_pct2** |
| U2SURP | 0,026196605 | -0,504202559 | 0,357 | 0,683 | 1 | Ciliated | 0,705049982 | 1,418339161 | 0,522693997 |
| BOLA3 | 0,026251562 | 0,613455247 | 0,714 | 0,585 | 1 | Ciliated | 1,529918972 | 0,653629387 | 1,220512821 |
| NR4A11 | 0,026336858 | -0,815635231 | 0,357 | 0,585 | 1 | Ciliated | 0,568158266 | 1,760072957 | 0,61025641 |
| VEZF11 | 0,026386347 | -0,254728833 | 0,286 | 0,659 | 1 | Ciliated | 0,838144657 | 1,193111466 | 0,433990895 |
| GNPTAB | 0,026413004 | 0,382437245 | 0,393 | 0,195 | 1 | Ciliated | 1,303542161 | 0,767140512 | 2,015384615 |
| ASH1L2 | 0,026418993 | -0,329442805 | 0,214 | 0,537 | 1 | Ciliated | 0,795843794 | 1,256527987 | 0,398510242 |
| MORF4L1 | 0,026482468 | -0,287609291 | 0,893 | 0,902 | 1 | Ciliated | 0,819258539 | 1,220615902 | 0,990022173 |
| ERG28 | 0,026767594 | 0,486070669 | 0,464 | 0,293 | 1 | Ciliated | 1,400624927 | 0,713967016 | 1,583617747 |
| EPHX1 | 0,026862076 | -0,568807819 | 0,893 | 0,878 | 1 | Ciliated | 0,674173666 | 1,483297331 | 1,017084282 |
| FBXW52 | 0,026951565 | 0,310115316 | 0,571 | 0,39 | 1 | Ciliated | 1,239806795 | 0,806577286 | 1,464102564 |
| RAB3B | 0,027177039 | 0,4379095 | 0,214 | 0,049 | 1 | Ciliated | 1,354639999 | 0,738203509 | 4,367346939 |
| RNFT2 | 0,027177039 | 0,329908717 | 0,214 | 0,049 | 1 | Ciliated | 1,256933842 | 0,795586821 | 4,367346939 |
| SLC10A3 | 0,027177039 | 0,283002418 | 0,214 | 0,049 | 1 | Ciliated | 1,216724398 | 0,82187881 | 4,367346939 |
| PRPF4B1 | 0,027267552 | -0,631063583 | 0,321 | 0,659 | 1 | Ciliated | 0,645700217 | 1,54870631 | 0,487101669 |
| PDE7A1 | 0,02727153 | -0,376724767 | 0,107 | 0,366 | 1 | Ciliated | 0,770184094 | 1,298390876 | 0,292349727 |
| C9orf72 | 0,027310919 | -0,366546203 | 0,25 | 0,561 | 1 | Ciliated | 0,775637143 | 1,289262652 | 0,445632799 |
| DYNLT2B | 0,027325496 | 0,532950018 | 0,964 | 0,854 | 1 | Ciliated | 1,446884758 | 0,691140047 | 1,128805621 |
| MAPK31 | 0,02736646 | 0,65179266 | 0,5 | 0,341 | 1 | Ciliated | 1,57111922 | 0,636488935 | 1,46627566 |
| CXCL82 | 0,027383314 | -1,368374244 | 0,107 | 0,317 | 1 | Ciliated | 0,387327477 | 2,581794627 | 0,337539432 |
| CALM11 | 0,027403094 | 0,302960227 | 1 | 0,951 | 1 | Ciliated | 1,233673159 | 0,810587466 | 1,051524711 |
| NEBL-AS1 | 0,027426372 | 0,419629423 | 0,357 | 0,171 | 1 | Ciliated | 1,337583933 | 0,747616636 | 2,087719298 |
| ZPR1 | 0,027426372 | 0,384698365 | 0,357 | 0,171 | 1 | Ciliated | 1,30558679 | 0,765939122 | 2,087719298 |
| NUDCD1 | 0,027442409 | -0,306234774 | 0,036 | 0,244 | 1 | Ciliated | 0,808749727 | 1,23647646 | 0,147540984 |
| SMYD2 | 0,027453406 | 0,39234661 | 0,786 | 0,61 | 1 | Ciliated | 1,312526553 | 0,761889348 | 1,28852459 |
| MIA3 | 0,027487279 | 0,92113862 | 0,393 | 0,195 | 1 | Ciliated | 1,8936092 | 0,528092069 | 2,015384615 |
| H2AZ21 | 0,027500335 | 0,480732826 | 0,821 | 0,659 | 1 | Ciliated | 1,395452315 | 0,716613523 | 1,245827011 |
| CUTA1 | 0,027618678 | 0,257830871 | 0,857 | 0,756 | 1 | Ciliated | 1,195679618 | 0,83634444 | 1,133597884 |
| SPRING1 | 0,027752361 | 0,280932278 | 0,5 | 0,317 | 1 | Ciliated | 1,214979758 | 0,82305898 | 1,577287066 |
| DDX52 | 0,027954881 | -0,270317995 | 0,143 | 0,415 | 1 | Ciliated | 0,82913677 | 1,206073638 | 0,344578313 |
| EFNB21 | 0,027997824 | -0,388157462 | 0,107 | 0,341 | 1 | Ciliated | 0,764104858 | 1,308720903 | 0,313782991 |
| TMEM254 | 0,028055064 | -0,478756383 | 0,214 | 0,488 | 1 | Ciliated | 0,717595932 | 1,393541902 | 0,43852459 |

| Supplementary Table 4 - clusermarker | | | | | | | | | |
|---|---|---|---|---|---|---|---|---|---|
| | **p_val** | **avg_log2FC** | **pct.1** | **pct.2** | **p_val_adj** | **celltype** | **Foldchange_UP** | **Foldchange_DOWN** | **Ratio_pct1_pct2** |
| BASP1 | 0,028191579 | 0,382071652 | 1 | 0,829 | 1 | Ciliated | 1,303211872 | 0,767334937 | 1,206272618 |
| ANXA13 | 0,028272293 | -0,465380997 | 1 | 1 | 1 | Ciliated | 0,724279779 | 1,380681926 | 1 |
| SMARCC2 | 0,028370685 | -0,269041757 | 0,179 | 0,463 | 1 | Ciliated | 0,829870565 | 1,205007192 | 0,386609071 |
| SLC41A1 | 0,028395106 | -0,451487143 | 0,107 | 0,366 | 1 | Ciliated | 0,73128864 | 1,367449109 | 0,292349727 |
| HLTF | 0,028395106 | -0,355943706 | 0,107 | 0,366 | 1 | Ciliated | 0,781358368 | 1,279822474 | 0,292349727 |
| HBS1L | 0,028395106 | 0,288091015 | 0,393 | 0,171 | 1 | Ciliated | 1,221023541 | 0,818985029 | 2,298245614 |
| PICK1 | 0,028395106 | -0,544586954 | 0,107 | 0,366 | 1 | Ciliated | 0,685587659 | 1,458602684 | 0,292349727 |
| REXO1 | 0,02857951 | 0,346419265 | 0,357 | 0,171 | 1 | Ciliated | 1,271401121 | 0,786533836 | 2,087719298 |
| ATP5ME1 | 0,028629133 | -0,652067676 | 0,786 | 0,829 | 1 | Ciliated | 0,636367615 | 1,571418746 | 0,948130277 |
| UBE2H | 0,028629133 | 0,340508709 | 0,893 | 0,756 | 1 | Ciliated | 1,266202991 | 0,789762784 | 1,181216931 |
| C6orf62 | 0,028841558 | -0,293678103 | 0,286 | 0,585 | 1 | Ciliated | 0,8158195 | 1,225761335 | 0,488888889 |
| CHD31 | 0,028841558 | -0,478461688 | 0,321 | 0,561 | 1 | Ciliated | 0,717742528 | 1,393257276 | 0,572192513 |
| NRARP1 | 0,028891964 | -0,315290322 | 0,179 | 0,439 | 1 | Ciliated | 0,803689244 | 1,244262018 | 0,407744875 |
| RABAC12 | 0,02889566 | 0,55249013 | 0,714 | 0,585 | 1 | Ciliated | 1,466614929 | 0,681842234 | 1,220512821 |
| TPRG1L | 0,029040975 | 0,374134711 | 0,643 | 0,463 | 1 | Ciliated | 1,296061979 | 0,771568039 | 1,388768898 |
| GEMIN8 | 0,029041989 | -0,268841148 | 0,143 | 0,415 | 1 | Ciliated | 0,829985968 | 1,204839646 | 0,344578313 |
| QKI1 | 0,029046778 | -0,289604165 | 0,071 | 0,293 | 1 | Ciliated | 0,818126499 | 1,222304866 | 0,242320819 |
| ZNF407-AS1 | 0,029046778 | -0,474895017 | 0,071 | 0,293 | 1 | Ciliated | 0,719519146 | 1,389817081 | 0,242320819 |
| MTDH2 | 0,029084504 | 0,40563881 | 0,821 | 0,683 | 1 | Ciliated | 1,324675333 | 0,754901956 | 1,20204978 |
| MRPL362 | 0,029126526 | 0,293864482 | 0,607 | 0,463 | 1 | Ciliated | 1,225919699 | 0,815714113 | 1,311015119 |
| OVOL2 | 0,029171575 | -0,359553926 | 0,107 | 0,341 | 1 | Ciliated | 0,77940553 | 1,283029131 | 0,313782991 |
| DNAAF11 | 0,029283934 | 0,487823257 | 0,821 | 0,659 | 1 | Ciliated | 1,402327443 | 0,713100214 | 1,245827011 |
| PIN1 | 0,029357799 | 0,580916664 | 0,714 | 0,61 | 1 | Ciliated | 1,495799353 | 0,668538864 | 1,170491803 |
| DENND10 | 0,029558337 | -0,250094376 | 0,107 | 0,366 | 1 | Ciliated | 0,840841409 | 1,189284911 | 0,292349727 |
| APTX | 0,029751211 | -0,390395983 | 0,143 | 0,39 | 1 | Ciliated | 0,762920174 | 1,310753123 | 0,366666667 |
| CCL15 | 0,029772404 | 0,423486426 | 0,929 | 0,634 | 1 | Ciliated | 1,341164709 | 0,745620574 | 1,465299685 |
| UROD | 0,029780368 | 0,377422199 | 0,5 | 0,317 | 1 | Ciliated | 1,2990187 | 0,769811859 | 1,577287066 |
| STOML22 | 0,029787216 | 0,551731286 | 0,679 | 0,585 | 1 | Ciliated | 1,465843706 | 0,682200971 | 1,160683761 |
| RPL36A2 | 0,029831255 | -0,620021759 | 0,857 | 0,854 | 1 | Ciliated | 0,650661114 | 1,536898361 | 1,003512881 |
| CYSTM11 | 0,030024967 | 0,38099383 | 1 | 0,854 | 1 | Ciliated | 1,30223862 | 0,767908419 | 1,170960187 |
| EIF11 | 0,030077591 | 0,273733305 | 1 | 0,951 | 1 | Ciliated | 1,208932173 | 0,827176265 | 1,051524711 |

| Supplementary Table 4 - clusermarker | | | | | | | | | |
|---|---|---|---|---|---|---|---|---|---|
| | **p_val** | **avg_log2FC** | **pct.1** | **pct.2** | **p_val_adj** | **celltype** | **Foldchange_UP** | **Foldchange_DOWN** | **Ratio_pct1_pct2** |
| MAPK6 | 0,030165288 | -0,574872036 | 0,143 | 0,415 | 1 | Ciliated | 0,671345796 | 1,489545338 | 0,344578313 |
| CIR11 | 0,030263211 | -0,748489419 | 0,5 | 0,707 | 1 | Ciliated | 0,595226466 | 1,680032822 | 0,707213579 |
| DPF21 | 0,03035264 | -0,445610926 | 0,071 | 0,293 | 1 | Ciliated | 0,734273314 | 1,361890703 | 0,242320819 |
| DCTN32 | 0,030360498 | 0,428090467 | 0,714 | 0,488 | 1 | Ciliated | 1,345451575 | 0,743244884 | 1,463114754 |
| SCCPDH1 | 0,030664353 | 0,448054095 | 0,714 | 0,537 | 1 | Ciliated | 1,364198986 | 0,733030893 | 1,329608939 |
| ERGIC31 | 0,03100377 | 0,432306362 | 1 | 0,902 | 1 | Ciliated | 1,349389052 | 0,741076118 | 1,10864745 |
| VILL1 | 0,031013255 | -0,368855518 | 0,286 | 0,61 | 1 | Ciliated | 0,774396578 | 1,291328021 | 0,468852459 |
| RPL121 | 0,031017589 | 0,353410282 | 1 | 0,951 | 1 | Ciliated | 1,277577032 | 0,782731667 | 1,051524711 |
| EMP22 | 0,031136845 | 0,483004513 | 0,714 | 0,659 | 1 | Ciliated | 1,397651344 | 0,715486022 | 1,083459788 |
| PRDX61 | 0,031241288 | 0,526409391 | 0,964 | 0,78 | 1 | Ciliated | 1,440339983 | 0,694280525 | 1,235897436 |
| RIMKLB1 | 0,031325722 | -0,31497511 | 0,143 | 0,415 | 1 | Ciliated | 0,803864859 | 1,243990192 | 0,344578313 |
| ISG151 | 0,031429726 | 0,253916371 | 0,536 | 0,341 | 1 | Ciliated | 1,192439748 | 0,838616795 | 1,571847507 |
| TSGA10 | 0,031505017 | -0,584040572 | 0,571 | 0,78 | 1 | Ciliated | 0,667092825 | 1,499041757 | 0,732051282 |
| CD164L2 | 0,031650385 | 0,675850167 | 0,679 | 0,561 | 1 | Ciliated | 1,59753791 | 0,625963236 | 1,210338681 |
| HNRNPA12 | 0,031758212 | -0,455405015 | 0,75 | 0,829 | 1 | Ciliated | 0,729305402 | 1,371167686 | 0,904704463 |
| MAGIX | 0,031890088 | 0,532789742 | 0,536 | 0,415 | 1 | Ciliated | 1,446724025 | 0,691216834 | 1,291566265 |
| TMEM94 | 0,031970414 | 0,459270215 | 0,5 | 0,366 | 1 | Ciliated | 1,374846178 | 0,727354097 | 1,366120219 |
| CDKL2 | 0,032008173 | -0,359579602 | 0,107 | 0,366 | 1 | Ciliated | 0,779391659 | 1,283051965 | 0,292349727 |
| LINC01571 | 0,032024471 | 0,41164011 | 0,714 | 0,512 | 1 | Ciliated | 1,330197173 | 0,751768249 | 1,39453125 |
| SMIM191 | 0,032082672 | 0,281124079 | 0,857 | 0,732 | 1 | Ciliated | 1,215141296 | 0,822949564 | 1,170765027 |
| SPRR31 | 0,03211099 | 0,383223695 | 0,214 | 0,049 | 1 | Ciliated | 1,304252949 | 0,766722438 | 4,367346939 |
| SMARCA5-AS1 | 0,03211099 | 0,265174069 | 0,214 | 0,049 | 1 | Ciliated | 1,201781043 | 0,832098331 | 4,367346939 |
| POLR2K1 | 0,032119102 | -0,445481613 | 0,357 | 0,634 | 1 | Ciliated | 0,734339132 | 1,361768638 | 0,563091483 |
| VAPB | 0,032253211 | -0,547161087 | 0,179 | 0,439 | 1 | Ciliated | 0,684365488 | 1,461207524 | 0,407744875 |
| ABRAXAS1 | 0,032324508 | -0,354111327 | 0,214 | 0,488 | 1 | Ciliated | 0,782351408 | 1,278197993 | 0,43852459 |
| C12orf75 | 0,032343455 | 0,496160794 | 0,964 | 0,829 | 1 | Ciliated | 1,410455152 | 0,708990994 | 1,162846803 |
| EXOC1 | 0,032791744 | -0,268397424 | 0,179 | 0,463 | 1 | Ciliated | 0,830241283 | 1,204469134 | 0,386609071 |
| SSH2 | 0,032791744 | 0,324103932 | 0,464 | 0,268 | 1 | Ciliated | 1,251886642 | 0,798794369 | 1,731343284 |
| CHPF | 0,032844816 | 0,356367799 | 0,429 | 0,268 | 1 | Ciliated | 1,280198744 | 0,781128715 | 1,600746269 |
| IRAK4 | 0,032867458 | 0,395237068 | 0,321 | 0,146 | 1 | Ciliated | 1,315158853 | 0,760364421 | 2,198630137 |
| SRI2 | 0,032919038 | 0,414841029 | 1 | 0,854 | 1 | Ciliated | 1,333151769 | 0,750102144 | 1,170960187 |

| Supplementary Table 4 - clusermarker | | | | | | | | | |
|---|---|---|---|---|---|---|---|---|---|
| | **p_val** | **avg_log2FC** | **pct.1** | **pct.2** | **p_val_adj** | **celltype** | **Foldchange_UP** | **Foldchange_DOWN** | **Ratio_pct1_pct2** |
| AL163051.1 | 0,032950173 | -0,330683661 | 0,107 | 0,341 | 1 | Ciliated | 0,795159586 | 1,257609186 | 0,313782991 |
| EDEM3 | 0,033060781 | -0,270479175 | 0,214 | 0,512 | 1 | Ciliated | 0,829044142 | 1,20620839 | 0,41796875 |
| FLYWCH21 | 0,033060781 | 0,452592803 | 0,5 | 0,317 | 1 | Ciliated | 1,368497503 | 0,730728407 | 1,577287066 |
| AC015813.8 | 0,033116225 | -0,592948713 | 0,071 | 0,293 | 1 | Ciliated | 0,662986449 | 1,508326455 | 0,242320819 |
| RABGAP1L1 | 0,033173508 | -0,534470945 | 0,357 | 0,634 | 1 | Ciliated | 0,690411813 | 1,448410906 | 0,563091483 |
| CFDP1 | 0,033251441 | 0,582503227 | 0,857 | 0,756 | 1 | Ciliated | 1,497445221 | 0,667804061 | 1,133597884 |
| GON4L1 | 0,033367659 | -0,50402852 | 0,143 | 0,39 | 1 | Ciliated | 0,705135041 | 1,41816807 | 0,366666667 |
| SS18L2 | 0,033367659 | -0,550367504 | 0,143 | 0,39 | 1 | Ciliated | 0,682846162 | 1,464458696 | 0,366666667 |
| ATP2A21 | 0,033394189 | 0,423462421 | 0,893 | 0,683 | 1 | Ciliated | 1,341142393 | 0,74563298 | 1,307467057 |
| PFDN21 | 0,033398897 | 0,38682292 | 0,714 | 0,488 | 1 | Ciliated | 1,307510852 | 0,764812008 | 1,463114754 |
| NUDT16 | 0,033475016 | -0,391238137 | 0,214 | 0,488 | 1 | Ciliated | 0,762474959 | 1,311518481 | 0,43852459 |
| SOX91 | 0,033535583 | -0,28664783 | 0,036 | 0,22 | 1 | Ciliated | 0,819804702 | 1,219802714 | 0,163636364 |
| RNASET21 | 0,033619041 | 0,387618922 | 0,357 | 0,171 | 1 | Ciliated | 1,308232465 | 0,764390142 | 2,087719298 |
| BANF11 | 0,033631683 | 0,262510223 | 0,929 | 0,78 | 1 | Ciliated | 1,199564076 | 0,833636168 | 1,191025641 |
| RPL23A2 | 0,033975612 | -0,463067636 | 1 | 0,902 | 1 | Ciliated | 0,725442093 | 1,378469777 | 1,10864745 |
| TFPT | 0,034055937 | 0,597898528 | 0,429 | 0,268 | 1 | Ciliated | 1,513510337 | 0,660715673 | 1,600746269 |
| COPS21 | 0,034220877 | -0,33563148 | 0,214 | 0,512 | 1 | Ciliated | 0,792437204 | 1,261929645 | 0,41796875 |
| PRRT3 | 0,034386996 | 0,425197882 | 0,464 | 0,293 | 1 | Ciliated | 1,342756665 | 0,744736575 | 1,583617747 |
| SSBP4 | 0,034389492 | 0,46009575 | 0,75 | 0,683 | 1 | Ciliated | 1,375633114 | 0,726938011 | 1,098096633 |
| FAM229B | 0,034452666 | 0,391086877 | 0,964 | 0,878 | 1 | Ciliated | 1,311380982 | 0,762554905 | 1,097949886 |
| WLS | 0,034466111 | 0,454716802 | 0,679 | 0,512 | 1 | Ciliated | 1,37051375 | 0,729653388 | 1,326171875 |
| CCDC78 | 0,034493881 | 0,359701186 | 0,964 | 0,927 | 1 | Ciliated | 1,283160099 | 0,779325979 | 1,0399137 |
| HMG20A | 0,034576968 | -0,422042699 | 0,071 | 0,293 | 1 | Ciliated | 0,746367101 | 1,339823256 | 0,242320819 |
| AXIN1 | 0,034576968 | -0,29607226 | 0,071 | 0,293 | 1 | Ciliated | 0,814466768 | 1,227797179 | 0,242320819 |
| YIPF2 | 0,034630031 | -0,357308362 | 0,107 | 0,366 | 1 | Ciliated | 0,780619625 | 1,28103364 | 0,292349727 |
| DYNLT2 | 0,034899728 | 0,261261174 | 0,107 | 0 | 1 | Ciliated | 1,198525974 | 0,834358222 | #DIV/0! |
| ANKRD24 | 0,034899728 | 0,253190328 | 0,107 | 0 | 1 | Ciliated | 1,191839798 | 0,839038939 | #DIV/0! |
| DCDC1 | 0,0349698 | -0,760768028 | 0,286 | 0,561 | 1 | Ciliated | 0,59018206 | 1,694392405 | 0,509803922 |
| RNF72 | 0,035192569 | 0,524330221 | 0,643 | 0,561 | 1 | Ciliated | 1,438265702 | 0,695281823 | 1,146167558 |
| LINC02832 | 0,035210797 | 0,493355947 | 0,571 | 0,39 | 1 | Ciliated | 1,407715649 | 0,710370735 | 1,464102564 |
| MALSU1 | 0,035210797 | 0,424983503 | 0,571 | 0,39 | 1 | Ciliated | 1,34255715 | 0,744847249 | 1,464102564 |

| Supplementary Table 4 - clusermarker | | | | | | | | | |
|---|---|---|---|---|---|---|---|---|---|
| | **p_val** | **avg_log2FC** | **pct.1** | **pct.2** | **p_val_adj** | **celltype** | **Foldchange_UP** | **Foldchange_DOWN** | **Ratio_pct1_pct2** |
| AC092683.1 | 0,035234491 | -0,292437561 | 0,036 | 0,22 | 1 | Ciliated | 0,816521308 | 1,224707783 | 0,163636364 |
| TRAPPC13 | 0,035234491 | -0,347454843 | 0,036 | 0,22 | 1 | Ciliated | 0,785969458 | 1,27231407 | 0,163636364 |
| SNF81 | 0,035329938 | 0,364029984 | 0,607 | 0,537 | 1 | Ciliated | 1,287015995 | 0,77699112 | 1,130353818 |
| IARS1 | 0,035366731 | -0,336444582 | 0,286 | 0,61 | 1 | Ciliated | 0,791990713 | 1,262641068 | 0,468852459 |
| PRPF311 | 0,035366731 | 0,409599226 | 0,571 | 0,415 | 1 | Ciliated | 1,328316763 | 0,752832478 | 1,375903614 |
| C12orf571 | 0,035401321 | 0,404455519 | 0,786 | 0,561 | 1 | Ciliated | 1,323589286 | 0,755521377 | 1,401069519 |
| NT5C2 | 0,035442348 | 0,454915131 | 0,571 | 0,439 | 1 | Ciliated | 1,370702168 | 0,729553088 | 1,300683371 |
| NTPCR | 0,035611823 | 0,422936341 | 0,464 | 0,293 | 1 | Ciliated | 1,340653434 | 0,745904926 | 1,583617747 |
| GAK | 0,035611823 | 0,419017126 | 0,464 | 0,293 | 1 | Ciliated | 1,337016367 | 0,747934 | 1,583617747 |
| HES11 | 0,035745476 | -0,602486947 | 0,464 | 0,683 | 1 | Ciliated | 0,658617638 | 1,518331643 | 0,679355783 |
| DRAM21 | 0,035812054 | 0,325071909 | 0,536 | 0,366 | 1 | Ciliated | 1,252726877 | 0,798258597 | 1,464480874 |
| MT-ND32 | 0,03590265 | -0,685686557 | 0,964 | 0,805 | 1 | Ciliated | 0,621709894 | 1,608467245 | 1,197515528 |
| ACTR3 | 0,036000826 | 0,295469258 | 0,786 | 0,659 | 1 | Ciliated | 1,227284105 | 0,814807261 | 1,192716237 |
| RPS210 | 0,036046929 | 0,376590749 | 1 | 0,854 | 1 | Ciliated | 1,298270269 | 0,770255642 | 1,170960187 |
| HDHD51 | 0,036092402 | -0,328577206 | 0,071 | 0,293 | 1 | Ciliated | 0,796321434 | 1,255774312 | 0,242320819 |
| PNRC12 | 0,036162312 | -0,457460231 | 0,75 | 0,878 | 1 | Ciliated | 0,728267197 | 1,373122399 | 0,854214123 |
| DPM3 | 0,036331461 | 0,431655122 | 0,821 | 0,659 | 1 | Ciliated | 1,348780068 | 0,741410719 | 1,245827011 |
| PRPF18 | 0,036358176 | -0,316885513 | 0,143 | 0,415 | 1 | Ciliated | 0,802801093 | 1,245638563 | 0,344578313 |
| TLCD2 | 0,036358176 | -0,271958003 | 0,143 | 0,415 | 1 | Ciliated | 0,82819477 | 1,207445442 | 0,344578313 |
| LARP6 | 0,03651619 | 0,370257716 | 0,643 | 0,439 | 1 | Ciliated | 1,292583711 | 0,773644284 | 1,464692483 |
| ZNHIT2 | 0,036556408 | 0,433754656 | 0,429 | 0,244 | 1 | Ciliated | 1,350744358 | 0,740332539 | 1,758196721 |
| MSI2 | 0,036738602 | -0,638354119 | 0,679 | 0,707 | 1 | Ciliated | 0,642445457 | 1,556552372 | 0,96039604 |
| CYP1B11 | 0,036823176 | -0,336858158 | 0 | 0,146 | 1 | Ciliated | 0,791763706 | 1,263003081 | 0 |
| LTF1 | 0,036823176 | -0,760609195 | 0 | 0,146 | 1 | Ciliated | 0,590247039 | 1,694205872 | 0 |
| GEM | 0,036823176 | -0,286493919 | 0 | 0,146 | 1 | Ciliated | 0,819892166 | 1,219672588 | 0 |
| PRH22 | 0,036823176 | -0,531522351 | 0 | 0,146 | 1 | Ciliated | 0,691824327 | 1,445453653 | 0 |
| PAQR4 | 0,036823176 | -0,323964004 | 0 | 0,146 | 1 | Ciliated | 0,798871849 | 1,251765226 | 0 |
| ZNF597 | 0,036823176 | -0,261662051 | 0 | 0,146 | 1 | Ciliated | 0,834126414 | 1,19885905 | 0 |
| ZNF816 | 0,036823176 | -0,362684695 | 0 | 0,146 | 1 | Ciliated | 0,777715989 | 1,285816434 | 0 |
| MAP4K4 | 0,036878356 | 0,385463874 | 0,464 | 0,293 | 1 | Ciliated | 1,306279731 | 0,765532815 | 1,583617747 |
| CRELD2 | 0,036947831 | 0,259977946 | 0,607 | 0,488 | 1 | Ciliated | 1,197460399 | 0,835100685 | 1,243852459 |

| Supplementary Table 4 - clusermarker | | | | | | | | | |
|---|---|---|---|---|---|---|---|---|---|
| | **p_val** | **avg_log2FC** | **pct.1** | **pct.2** | **p_val_adj** | **celltype** | **Foldchange_UP** | **Foldchange_DOWN** | **Ratio_pct1_pct2** |
| MT-ATP62 | 0,03699721 | 0,517825127 | 0,964 | 0,805 | 1 | Ciliated | 1,431795179 | 0,698423919 | 1,197515528 |
| PHF7 | 0,037006317 | -0,299874413 | 0,036 | 0,22 | 1 | Ciliated | 0,812323106 | 1,231037247 | 0,163636364 |
| SPARCL11 | 0,037006317 | -0,861688278 | 0,036 | 0,22 | 1 | Ciliated | 0,550308196 | 1,817163558 | 0,163636364 |
| OAS1 | 0,037006317 | -0,412483612 | 0,036 | 0,22 | 1 | Ciliated | 0,751328841 | 1,330975128 | 0,163636364 |
| RASD11 | 0,037006317 | -0,285004247 | 0,036 | 0,22 | 1 | Ciliated | 0,820739192 | 1,218413851 | 0,163636364 |
| DDX23 | 0,037018839 | 0,434125773 | 0,5 | 0,39 | 1 | Ciliated | 1,351091866 | 0,740142121 | 1,282051282 |
| SMS1 | 0,037110448 | 0,294114721 | 0,786 | 0,659 | 1 | Ciliated | 1,226132356 | 0,815572638 | 1,192716237 |
| SMIM221 | 0,037198069 | 0,427418823 | 1 | 0,927 | 1 | Ciliated | 1,344825348 | 0,743590981 | 1,078748652 |
| CUEDC2 | 0,037205589 | 0,481100485 | 0,536 | 0,341 | 1 | Ciliated | 1,39580798 | 0,716430923 | 1,571847507 |
| TMEM179B2 | 0,03744773 | 0,481774038 | 0,75 | 0,707 | 1 | Ciliated | 1,396459795 | 0,716096521 | 1,060820368 |
| HES42 | 0,037495402 | 0,448625563 | 0,393 | 0,195 | 1 | Ciliated | 1,364739468 | 0,732740588 | 2,015384615 |
| SEC61B2 | 0,037530145 | 0,309017473 | 0,857 | 0,756 | 1 | Ciliated | 1,238863702 | 0,807191298 | 1,133597884 |
| MRPS162 | 0,037584953 | 0,66816926 | 0,536 | 0,415 | 1 | Ciliated | 1,589055221 | 0,629304751 | 1,291566265 |
| PAN2 | 0,037664097 | -0,433714303 | 0,071 | 0,293 | 1 | Ciliated | 0,740353247 | 1,350706577 | 0,242320819 |
| SDCBP2-AS1 | 0,037664097 | -0,253609877 | 0,071 | 0,293 | 1 | Ciliated | 0,838794974 | 1,192186447 | 0,242320819 |
| CIAO2A1 | 0,037692388 | 0,537124518 | 0,607 | 0,463 | 1 | Ciliated | 1,451077443 | 0,689143095 | 1,311015119 |
| RPL72 | 0,037703665 | -0,409686296 | 0,964 | 0,878 | 1 | Ciliated | 0,752787044 | 1,328396932 | 1,097949886 |
| HHLA2 | 0,037718931 | -0,285812968 | 0,143 | 0,415 | 1 | Ciliated | 0,820279246 | 1,219097039 | 0,344578313 |
| ARFGEF3 | 0,038155452 | -0,495194699 | 0,464 | 0,707 | 1 | Ciliated | 0,709465925 | 1,409510963 | 0,656294201 |
| EEF1A11 | 0,038331947 | -0,427103796 | 1 | 1 | 1 | Ciliated | 0,743753369 | 1,344531724 | 1 |
| BAZ1A1 | 0,038385947 | -0,525886061 | 0,393 | 0,585 | 1 | Ciliated | 0,694532417 | 1,439817603 | 0,671794872 |
| CCNK | 0,038565847 | -0,436789293 | 0,214 | 0,537 | 1 | Ciliated | 0,738776923 | 1,353588571 | 0,398510242 |
| GPANK1 | 0,038853455 | -0,488075822 | 0,036 | 0,22 | 1 | Ciliated | 0,712975387 | 1,402572962 | 0,163636364 |
| PPIC1 | 0,038964828 | 0,512954248 | 0,571 | 0,415 | 1 | Ciliated | 1,426969252 | 0,700785948 | 1,375903614 |
| SYNE21 | 0,039181453 | -0,56940487 | 0,714 | 0,78 | 1 | Ciliated | 0,673894721 | 1,483911312 | 0,915384615 |
| HEBP22 | 0,039271754 | 0,303497199 | 0,893 | 0,854 | 1 | Ciliated | 1,234132418 | 0,810285821 | 1,045667447 |
| CFAP251 | 0,039768012 | 0,250230172 | 0,821 | 0,659 | 1 | Ciliated | 1,18939686 | 0,840762267 | 1,245827011 |
| USP3 | 0,03998924 | -0,501546616 | 0,179 | 0,439 | 1 | Ciliated | 0,706349146 | 1,415730458 | 0,407744875 |
| ATP5MJ | 0,040006003 | -0,473517009 | 0,821 | 0,805 | 1 | Ciliated | 0,720206732 | 1,388490214 | 1,019875776 |
| ZNF750 | 0,04017878 | -0,396728356 | 0,107 | 0,341 | 1 | Ciliated | 0,759578852 | 1,316519012 | 0,313782991 |
| NFAT5 | 0,040230716 | -0,347704004 | 0,143 | 0,39 | 1 | Ciliated | 0,785833729 | 1,272533824 | 0,366666667 |

| Supplementary Table 4 - clusermarker | | | | | | | | | |
|---|---|---|---|---|---|---|---|---|---|
| | **p_val** | **avg_log2FC** | **pct.1** | **pct.2** | **p_val_adj** | **celltype** | **Foldchange_UP** | **Foldchange_DOWN** | **Ratio_pct1_pct2** |
| GTF2IRD2 | 0,040778339 | -0,39960027 | 0,036 | 0,22 | 1 | Ciliated | 0,758068294 | 1,319142363 | 0,163636364 |
| FBXW10 | 0,040778339 | -0,456644623 | 0,036 | 0,22 | 1 | Ciliated | 0,728679029 | 1,372346341 | 0,163636364 |
| NLRP2 | 0,040778339 | -0,331977297 | 0,036 | 0,22 | 1 | Ciliated | 0,794446902 | 1,258737365 | 0,163636364 |
| ACYP1 | 0,040964984 | 0,395409574 | 0,643 | 0,488 | 1 | Ciliated | 1,315316119 | 0,760273508 | 1,317622951 |
| GOLGA7 | 0,040970559 | 0,497727909 | 0,357 | 0,171 | 1 | Ciliated | 1,411988079 | 0,708221276 | 2,087719298 |
| NEPRO | 0,040982679 | -0,317518121 | 0,071 | 0,293 | 1 | Ciliated | 0,80244915 | 1,246184883 | 0,242320819 |
| WIPI1 | 0,041087301 | -0,260190257 | 0,25 | 0,537 | 1 | Ciliated | 0,834977798 | 1,197636634 | 0,465549348 |
| RERE1 | 0,041248181 | 0,440442561 | 0,714 | 0,561 | 1 | Ciliated | 1,357020543 | 0,73690852 | 1,272727273 |
| HRAS1 | 0,041400867 | 0,348248835 | 0,571 | 0,39 | 1 | Ciliated | 1,273014485 | 0,785537016 | 1,464102564 |
| RFX3 | 0,041619503 | -0,470176235 | 0,607 | 0,732 | 1 | Ciliated | 0,72187641 | 1,385278679 | 0,829234973 |
| BMP2K | 0,041738273 | -0,263405135 | 0,143 | 0,39 | 1 | Ciliated | 0,833119219 | 1,200308403 | 0,366666667 |
| RB1 | 0,041738273 | -0,30086056 | 0,143 | 0,39 | 1 | Ciliated | 0,811768037 | 1,231879003 | 0,366666667 |
| UNC93B12 | 0,041842095 | 0,3129833 | 0,857 | 0,707 | 1 | Ciliated | 1,242273901 | 0,804975456 | 1,212164074 |
| PHB21 | 0,041843846 | 0,478836294 | 0,75 | 0,585 | 1 | Ciliated | 1,393619093 | 0,717556185 | 1,282051282 |
| SQLE | 0,041885539 | 0,646490565 | 0,714 | 0,585 | 1 | Ciliated | 1,565355747 | 0,63883242 | 1,220512821 |
| PDIA4 | 0,041910824 | 0,457758191 | 0,643 | 0,512 | 1 | Ciliated | 1,373406018 | 0,728116803 | 1,255859375 |
| SPATA18 | 0,042196031 | 0,503207652 | 0,964 | 0,78 | 1 | Ciliated | 1,417361387 | 0,705536364 | 1,235897436 |
| IL181 | 0,042550796 | -0,329088579 | 0,214 | 0,488 | 1 | Ciliated | 0,796039222 | 1,256219509 | 0,43852459 |
| FAM3B1 | 0,042641244 | 0,694821796 | 0,464 | 0,341 | 1 | Ciliated | 1,618684478 | 0,617785624 | 1,360703812 |
| SMIM7 | 0,042743656 | -0,496156206 | 0,286 | 0,585 | 1 | Ciliated | 0,708993248 | 1,410450667 | 0,488888889 |
| MUL1 | 0,042783442 | -0,346287363 | 0,036 | 0,22 | 1 | Ciliated | 0,78660575 | 1,271284884 | 0,163636364 |
| CNIH3 | 0,042783442 | -0,25534415 | 0,036 | 0,22 | 1 | Ciliated | 0,83778726 | 1,193620443 | 0,163636364 |
| AC022916.1 | 0,042783442 | -0,361661188 | 0,036 | 0,22 | 1 | Ciliated | 0,778267929 | 1,284904547 | 0,163636364 |
| CYB5A2 | 0,043132369 | -0,503813717 | 1 | 0,951 | 1 | Ciliated | 0,705240036 | 1,417956935 | 1,051524711 |
| HSF1 | 0,043181844 | 0,478762299 | 0,607 | 0,488 | 1 | Ciliated | 1,393547617 | 0,717592989 | 1,243852459 |
| CCDC88B | 0,043422058 | 0,348619889 | 0,464 | 0,268 | 1 | Ciliated | 1,27334194 | 0,785335006 | 1,731343284 |
| IRX5 | 0,043425652 | -0,30581493 | 0,107 | 0,341 | 1 | Ciliated | 0,808985119 | 1,236116681 | 0,313782991 |
| ARHGAP52 | 0,043564648 | -0,355369241 | 0,393 | 0,659 | 1 | Ciliated | 0,781669558 | 1,279312965 | 0,596358118 |
| ABHD12B | 0,043584752 | -0,34589033 | 0,25 | 0,561 | 1 | Ciliated | 0,786822255 | 1,270935073 | 0,445632799 |
| AC007255.11 | 0,043649191 | 0,360161236 | 0,429 | 0,268 | 1 | Ciliated | 1,283569342 | 0,779077505 | 1,600746269 |
| GFER1 | 0,043649191 | 0,397262179 | 0,429 | 0,268 | 1 | Ciliated | 1,317006238 | 0,759297846 | 1,600746269 |

| Supplementary Table 4 - clusermarker | | | | | | | | | |
|---|---|---|---|---|---|---|---|---|---|
| | **p_val** | **avg_log2FC** | **pct.1** | **pct.2** | **p_val_adj** | **celltype** | **Foldchange_UP** | **Foldchange_DOWN** | **Ratio_pct1_pct2** |
| MUC42 | 0,043705483 | 0,359375993 | 0,929 | 0,732 | 1 | Ciliated | 1,282870899 | 0,779501664 | 1,269125683 |
| HNRNPK1 | 0,04373285 | -0,385950373 | 0,75 | 0,78 | 1 | Ciliated | 0,765274709 | 1,306720303 | 0,961538462 |
| MSH3 | 0,043747163 | 0,359283551 | 0,75 | 0,683 | 1 | Ciliated | 1,282788701 | 0,779551612 | 1,098096633 |
| CTBP2 | 0,043851499 | -0,389842161 | 0,357 | 0,659 | 1 | Ciliated | 0,7632131 | 1,310250047 | 0,541729894 |
| ALG1L | 0,043861431 | 0,418381623 | 0,536 | 0,341 | 1 | Ciliated | 1,336427545 | 0,748263536 | 1,571847507 |
| ID21 | 0,043905951 | -0,320276438 | 0,286 | 0,561 | 1 | Ciliated | 0,800916397 | 1,248569768 | 0,509803922 |
| TMC42 | 0,043979773 | 0,461166995 | 0,786 | 0,634 | 1 | Ciliated | 1,376654943 | 0,726398438 | 1,239747634 |
| KLHL24 | 0,04400351 | -0,569869646 | 0,214 | 0,488 | 1 | Ciliated | 0,673677655 | 1,484389443 | 0,43852459 |
| RWDD41 | 0,04400351 | 0,46559555 | 0,464 | 0,317 | 1 | Ciliated | 1,380887272 | 0,724172074 | 1,463722397 |
| MTF1 | 0,044068009 | 0,4860146 | 0,571 | 0,488 | 1 | Ciliated | 1,400570494 | 0,713994764 | 1,170081967 |
| EFHD21 | 0,044171505 | 0,502983109 | 0,75 | 0,634 | 1 | Ciliated | 1,417140804 | 0,705646184 | 1,1829653 |
| MAP3K19 | 0,044397362 | -0,609027121 | 0,571 | 0,805 | 1 | Ciliated | 0,655638682 | 1,525230326 | 0,70931677 |
| PSMB82 | 0,044397362 | 0,321478367 | 0,786 | 0,659 | 1 | Ciliated | 1,249610401 | 0,800249421 | 1,192716237 |
| C22orf15 | 0,044460318 | 0,364241006 | 0,893 | 0,634 | 1 | Ciliated | 1,28720426 | 0,776877478 | 1,40851735 |
| MFSD102 | 0,044468711 | 0,378109993 | 0,679 | 0,585 | 1 | Ciliated | 1,299638144 | 0,769444945 | 1,160683761 |
| PCDH7 | 0,044535108 | -0,337083462 | 0,214 | 0,463 | 1 | Ciliated | 0,791640067 | 1,263200337 | 0,462203024 |
| SUMF2 | 0,044535108 | -0,483603319 | 0,214 | 0,463 | 1 | Ciliated | 0,715189113 | 1,398231575 | 0,462203024 |
| TSPAN151 | 0,044612137 | 0,554580381 | 0,536 | 0,39 | 1 | Ciliated | 1,468741377 | 0,680855061 | 1,374358974 |
| IPMK | 0,044897162 | -0,267620702 | 0,143 | 0,39 | 1 | Ciliated | 0,830688391 | 1,203820844 | 0,366666667 |
| ARL5A | 0,044936033 | -0,408922591 | 0,214 | 0,439 | 1 | Ciliated | 0,753185645 | 1,327693918 | 0,487471526 |
| C11orf71 | 0,045130514 | -0,286008466 | 0,107 | 0,341 | 1 | Ciliated | 0,820168098 | 1,219262249 | 0,313782991 |
| ZNF808 | 0,045130514 | -0,578083612 | 0,107 | 0,341 | 1 | Ciliated | 0,669852978 | 1,492864901 | 0,313782991 |
| BABAM1 | 0,045190418 | 0,527488943 | 0,429 | 0,268 | 1 | Ciliated | 1,441418177 | 0,693761197 | 1,600746269 |
| TM7SF2 | 0,045446822 | 0,270622471 | 0,607 | 0,463 | 1 | Ciliated | 1,206328203 | 0,828961801 | 1,311015119 |
| SPCS12 | 0,045480006 | 0,495498788 | 0,821 | 0,78 | 1 | Ciliated | 1,409808088 | 0,709316401 | 1,052564103 |
| EVA1C1 | 0,045496261 | 0,322940663 | 0,75 | 0,634 | 1 | Ciliated | 1,250877631 | 0,79943871 | 1,1829653 |
| PDE6D | 0,04581512 | 0,524705259 | 0,286 | 0,122 | 1 | Ciliated | 1,438639638 | 0,695101104 | 2,344262295 |
| HINT3 | 0,045978078 | -0,401071554 | 0,179 | 0,439 | 1 | Ciliated | 0,757295597 | 1,320488332 | 0,407744875 |
| ENAH | 0,046107896 | 0,360175478 | 0,821 | 0,659 | 1 | Ciliated | 1,283582013 | 0,779069814 | 1,245827011 |
| CCDC157 | 0,046316819 | 0,443414865 | 0,5 | 0,317 | 1 | Ciliated | 1,359819219 | 0,735391871 | 1,577287066 |
| DHRS3 | 0,046337055 | 0,404898533 | 0,643 | 0,561 | 1 | Ciliated | 1,323995788 | 0,755289412 | 1,146167558 |

| Supplementary Table 4 - clusermarker | | | | | | | | | |
|---|---|---|---|---|---|---|---|---|---|
| | **p_val** | **avg_log2FC** | **pct.1** | **pct.2** | **p_val_adj** | **celltype** | **Foldchange_UP** | **Foldchange_DOWN** | **Ratio_pct1_pct2** |
| TOR1AIP11 | 0,046550761 | -0,356850984 | 0,143 | 0,39 | 1 | Ciliated | 0,780867144 | 1,280627578 | 0,366666667 |
| PRR14L | 0,046550761 | -0,464752045 | 0,143 | 0,39 | 1 | Ciliated | 0,724595602 | 1,380080139 | 0,366666667 |
| UXT1 | 0,046767075 | 0,280602499 | 0,821 | 0,707 | 1 | Ciliated | 1,214702063 | 0,823247141 | 1,161244696 |
| TMEM107 | 0,046854344 | 0,618988463 | 0,821 | 0,585 | 1 | Ciliated | 1,535797988 | 0,651127302 | 1,403418803 |
| RAB4A | 0,046860958 | 0,353353093 | 0,643 | 0,537 | 1 | Ciliated | 1,27752639 | 0,782762695 | 1,197392924 |
| WASL2 | 0,046860958 | 0,555350134 | 0,607 | 0,561 | 1 | Ciliated | 1,469525235 | 0,680491887 | 1,081996435 |
| PART12 | 0,046891462 | -0,461981119 | 0,107 | 0,341 | 1 | Ciliated | 0,725988641 | 1,377432019 | 0,313782991 |
| PPP1R9A | 0,046891462 | -0,354361593 | 0,107 | 0,341 | 1 | Ciliated | 0,782215705 | 1,278419743 | 0,313782991 |
| AC105052.5 | 0,04689383 | -0,837133521 | 0,179 | 0,415 | 1 | Ciliated | 0,559754637 | 1,786497036 | 0,431325301 |
| BOD1L11 | 0,046994871 | -0,375838506 | 0,321 | 0,561 | 1 | Ciliated | 0,77065737 | 1,297593508 | 0,572192513 |
| RPP38 | 0,047026894 | 0,544911462 | 0,571 | 0,463 | 1 | Ciliated | 1,458930807 | 0,685433466 | 1,233261339 |
| PLAAT2 | 0,047037125 | 0,498215514 | 0,679 | 0,537 | 1 | Ciliated | 1,412465387 | 0,707981951 | 1,26443203 |
| NONO1 | 0,047108019 | -0,365545299 | 0,321 | 0,61 | 1 | Ciliated | 0,776175447 | 1,288368505 | 0,526229508 |
| COPB1 | 0,047724585 | -0,519603455 | 0,393 | 0,659 | 1 | Ciliated | 0,697563542 | 1,43356116 | 0,596358118 |
| TAF102 | 0,047729282 | 0,429774168 | 0,821 | 0,732 | 1 | Ciliated | 1,347022705 | 0,742377984 | 1,121584699 |
| DYNLT5 | 0,047903239 | 0,504907319 | 0,714 | 0,561 | 1 | Ciliated | 1,419032193 | 0,704705647 | 1,272727273 |
| CBX31 | 0,047977658 | -0,593149971 | 0,321 | 0,512 | 1 | Ciliated | 0,662893968 | 1,508536883 | 0,626953125 |
| PTOV12 | 0,048415888 | 0,355751963 | 0,75 | 0,61 | 1 | Ciliated | 1,279652389 | 0,781462223 | 1,229508197 |
| LCMT11 | 0,048554529 | 0,317756642 | 0,429 | 0,244 | 1 | Ciliated | 1,246390931 | 0,802316492 | 1,758196721 |
| CD461 | 0,048601648 | 0,546754148 | 0,714 | 0,585 | 1 | Ciliated | 1,46079542 | 0,684558554 | 1,220512821 |
| C6orf226 | 0,04864762 | 0,381838645 | 0,464 | 0,341 | 1 | Ciliated | 1,30300141 | 0,767458878 | 1,360703812 |
| NPM11 | 0,048894565 | -0,533339294 | 0,786 | 0,805 | 1 | Ciliated | 0,690953585 | 1,447275217 | 0,976397516 |
| IMPDH21 | 0,049238238 | 0,387306187 | 0,607 | 0,439 | 1 | Ciliated | 1,307948909 | 0,764555858 | 1,382687927 |
| TNIP12 | 0,049238238 | -0,428559442 | 0,321 | 0,634 | 1 | Ciliated | 0,743003318 | 1,34588901 | 0,506309148 |
| RBM4 | 0,049238238 | -0,491565052 | 0,321 | 0,634 | 1 | Ciliated | 0,711253104 | 1,40596926 | 0,506309148 |
| FCGBP2 | 0,049488652 | 1,279631107 | 0,321 | 0,146 | 1 | Ciliated | 2,427768916 | 0,411900817 | 2,198630137 |
| SMC6 | 0,049542581 | -0,293045901 | 0,143 | 0,366 | 1 | Ciliated | 0,816177078 | 1,225224314 | 0,390710383 |
| SLC30A5 | 0,049542581 | -0,317050845 | 0,143 | 0,366 | 1 | Ciliated | 0,802709099 | 1,24578132 | 0,390710383 |
| MTCH21 | 0,049800834 | 0,415002729 | 0,607 | 0,463 | 1 | Ciliated | 1,333301199 | 0,750018076 | 1,311015119 |
| SMIM41 | 0,049803178 | 0,274526925 | 0,679 | 0,415 | 1 | Ciliated | 1,209597384 | 0,826721365 | 1,636144578 |
| TUBB4B1 | 0,049843159 | 0,324319221 | 1 | 1 | 1 | Ciliated | 1,25207347 | 0,798675177 | 1 |

| Supplementary Table 4 - clusermarker | | | | | | | | | |
|---|---|---|---|---|---|---|---|---|---|
| | p_val | avg_log2FC | pct.1 | pct.2 | p_val_adj | celltype | Foldchange_UP | Foldchange_DOWN | Ratio_pct1_pct2 |
| ZNF552 | 0,049847494 | -0,313342019 | 0,071 | 0,268 | 1 | Ciliated | 0,804775327 | 1,242582826 | 0,264925373 |
| SNAP47 | 0,050012021 | -0,265344112 | 0,143 | 0,39 | 1 | Ciliated | 0,832000262 | 1,201922698 | 0,366666667 |
| MPG1 | 0,050022301 | 0,554903422 | 0,536 | 0,415 | 1 | Ciliated | 1,469070286 | 0,680702625 | 1,291566265 |
| MRPS261 | 0,050022301 | 0,415012281 | 0,536 | 0,415 | 1 | Ciliated | 1,333310027 | 0,75001311 | 1,291566265 |
| ENKD1 | 0,050098876 | 0,464829509 | 0,571 | 0,439 | 1 | Ciliated | 1,380154244 | 0,724556697 | 1,300683371 |
| PAX8-AS1 | 0,050213174 | -0,361710775 | 0,107 | 0,317 | 1 | Ciliated | 0,778241179 | 1,284948711 | 0,337539432 |
| PRKAA1 | 0,050239822 | -0,530273045 | 0,214 | 0,488 | 1 | Ciliated | 0,692423673 | 1,4442025 | 0,43852459 |
| ZNF444 | 0,050264339 | -0,318664501 | 0,179 | 0,415 | 1 | Ciliated | 0,801811769 | 1,247175508 | 0,431325301 |
| MAFB1 | 0,050264339 | -0,314397392 | 0,179 | 0,415 | 1 | Ciliated | 0,804186826 | 1,243492143 | 0,431325301 |
| DRAP12 | 0,050282413 | 0,494691761 | 0,607 | 0,488 | 1 | Ciliated | 1,409019679 | 0,709713296 | 1,243852459 |
| MAPK1IP1L | 0,050613911 | -0,308051522 | 0,25 | 0,585 | 1 | Ciliated | 0,807731931 | 1,238034503 | 0,427350427 |
| DLD | 0,05095063 | -0,460245768 | 0,179 | 0,439 | 1 | Ciliated | 0,726862425 | 1,375776166 | 0,407744875 |
| RALGPS1 | 0,05095063 | -0,277893607 | 0,179 | 0,439 | 1 | Ciliated | 0,824794372 | 1,212423404 | 0,407744875 |
| ST142 | 0,051043449 | 0,62776407 | 0,571 | 0,512 | 1 | Ciliated | 1,545168391 | 0,647178654 | 1,115234375 |
| MYADM1 | 0,051409643 | -0,420632204 | 0,286 | 0,561 | 1 | Ciliated | 0,747097167 | 1,338513977 | 0,509803922 |
| BCAP312 | 0,051552543 | 0,472081495 | 0,821 | 0,756 | 1 | Ciliated | 1,387109322 | 0,720923711 | 1,085978836 |
| PPL2 | 0,051596312 | 0,380467793 | 0,536 | 0,415 | 1 | Ciliated | 1,301763883 | 0,768188466 | 1,291566265 |
| TXN21 | 0,051653019 | -0,280218386 | 0,286 | 0,634 | 1 | Ciliated | 0,823466357 | 1,214378695 | 0,451104101 |
| SNHG82 | 0,051766014 | -0,590883963 | 0,25 | 0,512 | 1 | Ciliated | 0,663935978 | 1,506169319 | 0,48828125 |
| SF3B3 | 0,051822017 | -0,351093596 | 0,143 | 0,39 | 1 | Ciliated | 0,783989591 | 1,275527139 | 0,366666667 |
| RARRES2 | 0,051972773 | 0,518314421 | 0,286 | 0,122 | 1 | Ciliated | 1,432280858 | 0,698187087 | 2,344262295 |
| SNAP29 | 0,052024327 | -0,640479816 | 0,179 | 0,415 | 1 | Ciliated | 0,641499561 | 1,55884752 | 0,431325301 |
| HSPB12 | 0,052205941 | -0,419803534 | 0,5 | 0,756 | 1 | Ciliated | 0,747526415 | 1,337745368 | 0,661375661 |
| GTF3A2 | 0,052532653 | 0,347747929 | 0,679 | 0,488 | 1 | Ciliated | 1,272572569 | 0,785809803 | 1,391393443 |
| SLC27A1 | 0,052552315 | 0,442198979 | 0,357 | 0,195 | 1 | Ciliated | 1,358673662 | 0,736011912 | 1,830769231 |
| NSRP11 | 0,052736783 | -0,45145268 | 0,286 | 0,537 | 1 | Ciliated | 0,731306109 | 1,367416444 | 0,532588454 |
| GALNT3 | 0,052775348 | -0,510298094 | 0,429 | 0,683 | 1 | Ciliated | 0,702077358 | 1,424344467 | 0,628111274 |
| SYPL11 | 0,052825741 | 0,259329958 | 0,714 | 0,634 | 1 | Ciliated | 1,19692268 | 0,835475856 | 1,126182965 |
| TMEM1412 | 0,052897811 | 0,446489931 | 0,571 | 0,488 | 1 | Ciliated | 1,362720728 | 0,733826072 | 1,170081967 |
| COBL | 0,053032542 | -0,383109779 | 0,286 | 0,561 | 1 | Ciliated | 0,766782981 | 1,304149968 | 0,509803922 |
| EIF4EBP31 | 0,053149197 | -0,548443142 | 0,25 | 0,537 | 1 | Ciliated | 0,683757595 | 1,462506606 | 0,465549348 |

| Supplementary Table 4 - clusermarker | | | | | | | | | |
|---|---|---|---|---|---|---|---|---|---|
| | **p_val** | **avg_log2FC** | **pct.1** | **pct.2** | **p_val_adj** | **celltype** | **Foldchange_UP** | **Foldchange_DOWN** | **Ratio_pct1_pct2** |
| PPP1R14C | 0,053247304 | 0,47568609 | 0,571 | 0,439 | 1 | Ciliated | 1,390579369 | 0,719124721 | 1,300683371 |
| KLF2 | 0,053384647 | 0,397370016 | 0,607 | 0,488 | 1 | Ciliated | 1,317104684 | 0,759241093 | 1,243852459 |
| ZRANB21 | 0,053442456 | -0,316810356 | 0,25 | 0,512 | 1 | Ciliated | 0,802842916 | 1,245573673 | 0,48828125 |
| HMGCL | 0,053686499 | -0,380791903 | 0,143 | 0,39 | 1 | Ciliated | 0,768015907 | 1,302056365 | 0,366666667 |
| WDR38 | 0,054009467 | 0,407234156 | 0,893 | 0,854 | 1 | Ciliated | 1,326140981 | 0,75406764 | 1,045667447 |
| AP002387.2 | 0,054173328 | 0,435414678 | 0,286 | 0,122 | 1 | Ciliated | 1,352299473 | 0,739481173 | 2,344262295 |
| LMO4 | 0,054184708 | -0,548911625 | 0,429 | 0,61 | 1 | Ciliated | 0,683535596 | 1,4629816 | 0,703278689 |
| SARAF1 | 0,054196443 | 0,357707031 | 1 | 0,854 | 1 | Ciliated | 1,281387686 | 0,780403941 | 1,170960187 |
| COX6C1 | 0,05426466 | -0,417595246 | 0,964 | 0,951 | 1 | Ciliated | 0,748671507 | 1,33569929 | 1,013669821 |
| GCC2 | 0,054427619 | -0,737938485 | 0,679 | 0,78 | 1 | Ciliated | 0,599595522 | 1,667790973 | 0,870512821 |
| RAB311 | 0,054504415 | 0,499994124 | 0,571 | 0,488 | 1 | Ciliated | 1,414207803 | 0,707109661 | 1,170081967 |
| PSMD14 | 0,054509873 | -0,42647493 | 0,179 | 0,439 | 1 | Ciliated | 0,74407764 | 1,343945775 | 0,407744875 |
| GTF3C3 | 0,054509873 | -0,343207095 | 0,179 | 0,439 | 1 | Ciliated | 0,78828701 | 1,268573486 | 0,407744875 |
| BTG21 | 0,054515418 | -0,900251299 | 0,536 | 0,634 | 1 | Ciliated | 0,535793395 | 1,866391056 | 0,845425868 |
| ATF2 | 0,054523736 | -0,26218286 | 0,107 | 0,341 | 1 | Ciliated | 0,833825351 | 1,199291913 | 0,313782991 |
| DNAH7 | 0,054612581 | -0,582373367 | 0,5 | 0,683 | 1 | Ciliated | 0,667864175 | 1,497310438 | 0,732064422 |
| SEMA3E | 0,054741254 | -0,535480285 | 0,036 | 0,195 | 1 | Ciliated | 0,689928956 | 1,449424599 | 0,184615385 |
| MUC5B2 | 0,054741254 | -0,911902737 | 0,036 | 0,195 | 1 | Ciliated | 0,531483667 | 1,881525363 | 0,184615385 |
| DHRS7 | 0,054868834 | 0,500916755 | 0,536 | 0,415 | 1 | Ciliated | 1,415112505 | 0,706657596 | 1,291566265 |
| TMEM208 | 0,054868834 | 0,392118106 | 0,536 | 0,415 | 1 | Ciliated | 1,312318683 | 0,762010031 | 1,291566265 |
| ATP2A31 | 0,05498366 | 0,414227337 | 0,429 | 0,293 | 1 | Ciliated | 1,332584794 | 0,75042129 | 1,464163823 |
| AC097376.3 | 0,055169855 | 0,270673852 | 0,25 | 0,098 | 1 | Ciliated | 1,206371166 | 0,828932279 | 2,551020408 |
| HNRNPUL11 | 0,055398066 | -0,25128912 | 0,357 | 0,683 | 1 | Ciliated | 0,840145368 | 1,190270205 | 0,522693997 |
| EIF4A12 | 0,055552822 | -0,459634263 | 0,893 | 0,854 | 1 | Ciliated | 0,72717058 | 1,375193149 | 1,045667447 |
| ROPN1L | 0,055696024 | 0,303079118 | 0,964 | 0,854 | 1 | Ciliated | 1,233774828 | 0,81052067 | 1,128805621 |
| ATP6V0C2 | 0,055696024 | 0,286382923 | 1 | 0,829 | 1 | Ciliated | 1,219578754 | 0,819955248 | 1,206272618 |
| CRK | 0,055699155 | -0,409985884 | 0,179 | 0,415 | 1 | Ciliated | 0,752630738 | 1,328672814 | 0,431325301 |
| CTBP1 | 0,056180483 | -0,272228051 | 0,214 | 0,463 | 1 | Ciliated | 0,82803976 | 1,207671477 | 0,462203024 |
| ADAM10 | 0,056180483 | -0,49204541 | 0,214 | 0,463 | 1 | Ciliated | 0,711016325 | 1,406437468 | 0,462203024 |
| DDX171 | 0,056236661 | -0,578403706 | 0,893 | 0,829 | 1 | Ciliated | 0,669704373 | 1,493196163 | 1,077201448 |
| TSC22D4 | 0,05631755 | -0,486366264 | 0,107 | 0,317 | 1 | Ciliated | 0,713820746 | 1,400911932 | 0,337539432 |

| Supplementary Table 4 - clusermarker | | | | | | | | | |
|---|---|---|---|---|---|---|---|---|---|
| | **p_val** | **avg_log2FC** | **pct.1** | **pct.2** | **p_val_adj** | **celltype** | **Foldchange_UP** | **Foldchange_DOWN** | **Ratio_pct1_pct2** |
| WASHC4 | 0,05631755 | -0,290215978 | 0,107 | 0,317 | 1 | Ciliated | 0,817779624 | 1,222823326 | 0,337539432 |
| PSMG1 | 0,05631755 | 0,408634354 | 0,321 | 0,171 | 1 | Ciliated | 1,327428684 | 0,753336139 | 1,877192982 |
| ALCAM | 0,056371255 | -0,547944731 | 0,786 | 0,805 | 1 | Ciliated | 0,683993855 | 1,462001439 | 0,976397516 |
| CASP41 | 0,056401825 | -0,37872224 | 0,357 | 0,61 | 1 | Ciliated | 0,769118478 | 1,300189799 | 0,585245902 |
| PTAFR | 0,056451095 | 0,37826661 | 0,286 | 0,122 | 1 | Ciliated | 1,299779239 | 0,769361419 | 2,344262295 |
| CACFD1 | 0,056529496 | 0,342081825 | 0,357 | 0,195 | 1 | Ciliated | 1,267584413 | 0,788902096 | 1,830769231 |
| SMC3 | 0,05655933 | -0,536339052 | 0,321 | 0,61 | 1 | Ciliated | 0,689518396 | 1,450287629 | 0,526229508 |
| SIVA12 | 0,056577963 | 0,451938252 | 0,857 | 0,78 | 1 | Ciliated | 1,367876757 | 0,731060013 | 1,098717949 |
| NDUFC22 | 0,056755228 | 0,257167267 | 0,857 | 0,805 | 1 | Ciliated | 1,195129761 | 0,836729226 | 1,064596273 |
| PPP6R31 | 0,056828086 | -0,468182847 | 0,214 | 0,439 | 1 | Ciliated | 0,722874524 | 1,383365945 | 0,487471526 |
| CARS2 | 0,057198228 | -0,291334082 | 0,143 | 0,366 | 1 | Ciliated | 0,817146082 | 1,223771395 | 0,390710383 |
| RALGAPB | 0,057198228 | -0,324924369 | 0,143 | 0,366 | 1 | Ciliated | 0,798340237 | 1,252598771 | 0,390710383 |
| C2CD2L | 0,057200519 | -0,304236476 | 0,214 | 0,488 | 1 | Ciliated | 0,809870714 | 1,234764984 | 0,43852459 |
| PNLDC1 | 0,057648574 | 0,432228925 | 0,25 | 0,098 | 1 | Ciliated | 1,349316625 | 0,741115896 | 2,551020408 |
| CD992 | 0,057837829 | 0,62501147 | 0,714 | 0,585 | 1 | Ciliated | 1,542223086 | 0,648414622 | 1,220512821 |
| DDX181 | 0,057862271 | -0,333879606 | 0,429 | 0,732 | 1 | Ciliated | 0,79340005 | 1,260398206 | 0,586065574 |
| ADH71 | 0,057862271 | -0,781575108 | 0,536 | 0,659 | 1 | Ciliated | 0,581731323 | 1,719006628 | 0,813353566 |
| HIKESHI | 0,058106368 | 0,290975538 | 0,571 | 0,463 | 1 | Ciliated | 1,223467296 | 0,817349187 | 1,233261339 |
| LIMS1 | 0,058349718 | -0,451428309 | 0,357 | 0,561 | 1 | Ciliated | 0,731318464 | 1,367393345 | 0,636363636 |
| ACSL3 | 0,058349718 | 0,336896846 | 0,571 | 0,415 | 1 | Ciliated | 1,263036951 | 0,791742474 | 1,375903614 |
| SIK2 | 0,058484872 | -0,276479343 | 0,107 | 0,317 | 1 | Ciliated | 0,825603308 | 1,211235457 | 0,337539432 |
| NDUFA22 | 0,058518565 | 0,479032255 | 0,929 | 0,78 | 1 | Ciliated | 1,3938084 | 0,717458727 | 1,191025641 |
| ACADM | 0,058558792 | -0,577385646 | 0,5 | 0,756 | 1 | Ciliated | 0,670177127 | 1,492142837 | 0,661375661 |
| CP2 | 0,05874293 | -0,471005002 | 0,393 | 0,659 | 1 | Ciliated | 0,721461842 | 1,386074691 | 0,596358118 |
| KLF101 | 0,058745917 | -0,277153881 | 0,25 | 0,512 | 1 | Ciliated | 0,825217384 | 1,211801907 | 0,48828125 |
| ZBTB7B | 0,058808028 | 0,254374426 | 0,286 | 0,122 | 1 | Ciliated | 1,192818407 | 0,838350577 | 2,344262295 |
| RGS14 | 0,058808028 | 0,303525602 | 0,286 | 0,122 | 1 | Ciliated | 1,234156715 | 0,810269869 | 2,344262295 |
| TTC39A-AS1 | 0,058834531 | -0,433515557 | 0 | 0,122 | 1 | Ciliated | 0,740455245 | 1,350520517 | 0 |
| IGKV4-11 | 0,058834531 | 0,362271951 | 0,143 | 0,024 | 1 | Ciliated | 1,285448624 | 0,77793852 | 5,958333333 |
| MKRN2OS | 0,058834531 | -0,251104969 | 0 | 0,122 | 1 | Ciliated | 0,840252614 | 1,190118285 | 0 |
| AC011346.1 | 0,058834531 | -0,253534641 | 0 | 0,122 | 1 | Ciliated | 0,838838719 | 1,192124276 | 0 |

| Supplementary Table 4 - clusermarker | | | | | | | | | |
|---|---|---|---|---|---|---|---|---|---|
| | **p_val** | **avg_log2FC** | **pct.1** | **pct.2** | **p_val_adj** | **celltype** | **Foldchange_UP** | **Foldchange_DOWN** | **Ratio_pct1_pct2** |
| BX005040.1 | 0,058834531 | -0,324844649 | 0 | 0,122 | 1 | Ciliated | 0,798384352 | 1,252529558 | 0 |
| KLHDC1 | 0,058834531 | 0,269733506 | 0,143 | 0,024 | 1 | Ciliated | 1,205585112 | 0,829472752 | 5,958333333 |
| LINC021851 | 0,058834531 | -0,289258367 | 0 | 0,122 | 1 | Ciliated | 0,818322618 | 1,222011928 | 0 |
| THBD | 0,058834531 | -0,514661736 | 0 | 0,122 | 1 | Ciliated | 0,69995703 | 1,428659128 | 0 |
| SNHG17 | 0,058834531 | -0,407413492 | 0 | 0,122 | 1 | Ciliated | 0,75397391 | 1,32630584 | 0 |
| JADE3 | 0,058834531 | -0,394988152 | 0 | 0,122 | 1 | Ciliated | 0,760495622 | 1,314931961 | 0 |
| GSDMD1 | 0,058974562 | 0,62163679 | 0,536 | 0,488 | 1 | Ciliated | 1,538619811 | 0,649933137 | 1,098360656 |
| SLMAP1 | 0,058980861 | 0,382858001 | 0,464 | 0,293 | 1 | Ciliated | 1,303922389 | 0,766916811 | 1,583617747 |
| MIDEAS | 0,059258834 | -0,292758648 | 0,143 | 0,366 | 1 | Ciliated | 0,816339602 | 1,224980385 | 0,390710383 |
| DHX8 | 0,059258834 | -0,438639521 | 0,143 | 0,366 | 1 | Ciliated | 0,737830063 | 1,355325636 | 0,390710383 |
| OST42 | 0,059467613 | 0,368377869 | 0,964 | 0,78 | 1 | Ciliated | 1,290900557 | 0,774653008 | 1,235897436 |
| TMEM67 | 0,059853497 | -0,577951172 | 0,357 | 0,61 | 1 | Ciliated | 0,669914474 | 1,492727861 | 0,585245902 |
| UFD11 | 0,059853497 | 0,350098349 | 0,571 | 0,463 | 1 | Ciliated | 1,274647517 | 0,784530615 | 1,233261339 |
| EIF3I1 | 0,060041062 | -0,533056293 | 0,357 | 0,585 | 1 | Ciliated | 0,691089137 | 1,446991346 | 0,61025641 |
| SORD1 | 0,060041062 | -0,379097701 | 0,357 | 0,585 | 1 | Ciliated | 0,768918342 | 1,300528217 | 0,61025641 |
| ISCU2 | 0,0600721 | 0,438882 | 0,786 | 0,707 | 1 | Ciliated | 1,35555345 | 0,737706064 | 1,111739745 |
| KIAA0319L | 0,060125902 | -0,363343775 | 0,357 | 0,561 | 1 | Ciliated | 0,777360779 | 1,28640398 | 0,636363636 |
| TTC39C-AS1 | 0,060219181 | 0,269997165 | 0,25 | 0,098 | 1 | Ciliated | 1,205805458 | 0,829321176 | 2,551020408 |
| AC009237.14 | 0,060221959 | -0,339649404 | 0,036 | 0,195 | 1 | Ciliated | 0,790233327 | 1,265449034 | 0,184615385 |
| SPRY12 | 0,060221959 | -0,340407998 | 0,036 | 0,195 | 1 | Ciliated | 0,789817918 | 1,266114604 | 0,184615385 |
| C6orf582 | 0,060221959 | -1,442063448 | 0,036 | 0,195 | 1 | Ciliated | 0,368040529 | 2,717092061 | 0,184615385 |
| CCL14 | 0,060221959 | -0,702450034 | 0,036 | 0,195 | 1 | Ciliated | 0,614527708 | 1,627265926 | 0,184615385 |
| TET2 | 0,060235832 | -0,349965175 | 0,179 | 0,439 | 1 | Ciliated | 0,784603037 | 1,274529862 | 0,407744875 |
| NR4A21 | 0,060320269 | -0,561787159 | 0,5 | 0,659 | 1 | Ciliated | 0,677462427 | 1,476096621 | 0,758725341 |
| NUDT41 | 0,060416127 | 0,549437464 | 0,714 | 0,561 | 1 | Ciliated | 1,463514931 | 0,683286504 | 1,272727273 |
| CNBP2 | 0,060436632 | 0,263262184 | 0,821 | 0,707 | 1 | Ciliated | 1,200189476 | 0,833201774 | 1,161244696 |
| MRPL351 | 0,060721023 | -0,339705519 | 0,107 | 0,317 | 1 | Ciliated | 0,79020259 | 1,265498256 | 0,337539432 |
| NAA50 | 0,060721023 | -0,271152353 | 0,107 | 0,317 | 1 | Ciliated | 0,828657391 | 1,206771351 | 0,337539432 |
| UXS1 | 0,060754089 | 0,311877286 | 0,357 | 0,195 | 1 | Ciliated | 1,241321901 | 0,805592811 | 1,830769231 |
| AL035420.3 | 0,060754089 | 0,404137066 | 0,357 | 0,195 | 1 | Ciliated | 1,323297156 | 0,755688165 | 1,830769231 |
| NQO2 | 0,061380674 | -0,343589469 | 0,143 | 0,366 | 1 | Ciliated | 0,788078108 | 1,268909756 | 0,390710383 |

| Supplementary Table 4 - clusermarker | | | | | | | | | |
|---|---|---|---|---|---|---|---|---|---|
| | **p_val** | **avg_log2FC** | **pct.1** | **pct.2** | **p_val_adj** | **celltype** | **Foldchange_UP** | **Foldchange_DOWN** | **Ratio_pct1_pct2** |
| SLC11A2 | 0,061614935 | -0,287754733 | 0,179 | 0,415 | 1 | Ciliated | 0,819175951 | 1,220738962 | 0,431325301 |
| STRAP | 0,061643974 | 0,315864876 | 0,571 | 0,463 | 1 | Ciliated | 1,244757645 | 0,803369237 | 1,233261339 |
| ZNF506 | 0,061661049 | 0,269506805 | 0,214 | 0,073 | 1 | Ciliated | 1,205395684 | 0,829603103 | 2,931506849 |
| ANG | 0,061802123 | -0,264418738 | 0,286 | 0,561 | 1 | Ciliated | 0,832534095 | 1,201152007 | 0,509803922 |
| POC1B | 0,061896148 | -0,250988281 | 0,214 | 0,463 | 1 | Ciliated | 0,840320578 | 1,19002203 | 0,462203024 |
| LMAN11 | 0,061968493 | 0,311809164 | 0,714 | 0,61 | 1 | Ciliated | 1,24126329 | 0,805630851 | 1,170491803 |
| HIBADH | 0,062094765 | -0,341434696 | 0,25 | 0,537 | 1 | Ciliated | 0,789256042 | 1,267015959 | 0,465549348 |
| AC013470.2 | 0,06251887 | -0,287135726 | 0,25 | 0,512 | 1 | Ciliated | 0,819527505 | 1,2202153 | 0,48828125 |
| ZNF22 | 0,062884104 | 0,295620647 | 0,25 | 0,098 | 1 | Ciliated | 1,227412896 | 0,814721764 | 2,551020408 |
| ARRDC31 | 0,063027561 | -0,392175989 | 0,107 | 0,317 | 1 | Ciliated | 0,761979458 | 1,312371336 | 0,337539432 |
| TATDN1 | 0,063027561 | -0,281434476 | 0,107 | 0,317 | 1 | Ciliated | 0,822772525 | 1,215402762 | 0,337539432 |
| NOL31 | 0,063027561 | 0,483329853 | 0,321 | 0,171 | 1 | Ciliated | 1,397966562 | 0,715324692 | 1,877192982 |
| HSD17B11 | 0,063054127 | -0,842176952 | 0,286 | 0,512 | 1 | Ciliated | 0,557801241 | 1,792753273 | 0,55859375 |
| KLF16 | 0,063128489 | -0,503697709 | 0,036 | 0,195 | 1 | Ciliated | 0,705296747 | 1,417842921 | 0,184615385 |
| RGS22 | 0,06347692 | 0,46769512 | 0,643 | 0,512 | 1 | Ciliated | 1,382898355 | 0,723118945 | 1,255859375 |
| NCOA6 | 0,063565041 | -0,293834862 | 0,143 | 0,366 | 1 | Ciliated | 0,815730861 | 1,22589453 | 0,390710383 |
| EIF2AK4 | 0,063698899 | -0,290816811 | 0,179 | 0,415 | 1 | Ciliated | 0,817439118 | 1,223332696 | 0,431325301 |
| WDR90 | 0,063813526 | 0,483762506 | 0,536 | 0,415 | 1 | Ciliated | 1,398385864 | 0,715110204 | 1,291566265 |
| IFT20 | 0,063861378 | 0,328193296 | 0,643 | 0,561 | 1 | Ciliated | 1,255440187 | 0,796533368 | 1,146167558 |
| SEPTIN91 | 0,064171934 | 0,448064591 | 0,75 | 0,634 | 1 | Ciliated | 1,36420891 | 0,73302556 | 1,1829653 |
| YPEL2 | 0,064479226 | -0,266817778 | 0,25 | 0,512 | 1 | Ciliated | 0,831150835 | 1,20315105 | 0,48828125 |
| PDLIM71 | 0,064624676 | 0,298877894 | 0,214 | 0,073 | 1 | Ciliated | 1,23018722 | 0,8128844 | 2,931506849 |
| RTCB | 0,064790298 | -0,316380907 | 0,25 | 0,488 | 1 | Ciliated | 0,803081935 | 1,245202956 | 0,512295082 |
| ALDH3A1 | 0,064842261 | -0,965833408 | 0,536 | 0,683 | 1 | Ciliated | 0,511982567 | 1,953191505 | 0,78477306 |
| DYNC1H11 | 0,064844561 | -0,508268304 | 0,464 | 0,707 | 1 | Ciliated | 0,703065836 | 1,422341905 | 0,656294201 |
| MVB12A2 | 0,064942068 | 0,397828978 | 0,464 | 0,317 | 1 | Ciliated | 1,317523758 | 0,758999596 | 1,463722397 |
| ATG9B | 0,064986386 | 0,418187763 | 0,393 | 0,244 | 1 | Ciliated | 1,336247976 | 0,74836409 | 1,610655738 |
| TMEM592 | 0,06500074 | 0,419679637 | 0,929 | 0,732 | 1 | Ciliated | 1,337630489 | 0,747590615 | 1,269125683 |
| SDHD1 | 0,065009617 | -0,56023225 | 0,286 | 0,512 | 1 | Ciliated | 0,678192977 | 1,47450657 | 0,55859375 |
| TNFAIP21 | 0,065236995 | 0,454348484 | 0,357 | 0,195 | 1 | Ciliated | 1,370163904 | 0,729839691 | 1,830769231 |
| ASMTL | 0,065236995 | 0,355176685 | 0,357 | 0,195 | 1 | Ciliated | 1,279142226 | 0,781773895 | 1,830769231 |

| Supplementary Table 4 - clusermarker | | | | | | | | | |
|---|---|---|---|---|---|---|---|---|---|
| | **p_val** | **avg_log2FC** | **pct.1** | **pct.2** | **p_val_adj** | **celltype** | **Foldchange_UP** | **Foldchange_DOWN** | **Ratio_pct1_pct2** |
| TDRD7 | 0,065406051 | -0,26243269 | 0,107 | 0,317 | 1 | Ciliated | 0,833680971 | 1,199499611 | 0,337539432 |
| ADAM12 | 0,06541593 | 0,304495317 | 0,179 | 0,049 | 1 | Ciliated | 1,234986539 | 0,809725425 | 3,653061224 |
| DSCAML1 | 0,06541593 | 0,257419124 | 0,179 | 0,049 | 1 | Ciliated | 1,195338417 | 0,836583168 | 3,653061224 |
| C12orf56 | 0,06541593 | 0,270085103 | 0,179 | 0,049 | 1 | Ciliated | 1,205878959 | 0,829270627 | 3,653061224 |
| RHBDF1 | 0,06541593 | 0,28001091 | 0,179 | 0,049 | 1 | Ciliated | 1,214204066 | 0,823584789 | 3,653061224 |
| DEFB124 | 0,06541593 | 0,317839456 | 0,179 | 0,049 | 1 | Ciliated | 1,24646248 | 0,802270438 | 3,653061224 |
| SSU722 | 0,065540864 | 0,550009677 | 0,714 | 0,61 | 1 | Ciliated | 1,464095517 | 0,683015547 | 1,170491803 |
| ZMIZ1 | 0,065813235 | -0,358071804 | 0,143 | 0,366 | 1 | Ciliated | 0,780206648 | 1,281711714 | 0,390710383 |
| NCBP2 | 0,065840746 | -0,294329638 | 0,179 | 0,415 | 1 | Ciliated | 0,815451152 | 1,226315026 | 0,431325301 |
| CAPZA1 | 0,066005966 | -0,50385129 | 0,25 | 0,537 | 1 | Ciliated | 0,705221669 | 1,417993864 | 0,465549348 |
| MYCBP | 0,066012077 | 0,516675383 | 0,857 | 0,707 | 1 | Ciliated | 1,430654576 | 0,698980744 | 1,212164074 |
| DNMT3A | 0,066150006 | -0,329714934 | 0,036 | 0,195 | 1 | Ciliated | 0,795693691 | 1,256765023 | 0,184615385 |
| PLXNA1 | 0,066150006 | -0,326205197 | 0,036 | 0,195 | 1 | Ciliated | 0,797631783 | 1,253711325 | 0,184615385 |
| LINC02562 | 0,066150006 | -0,315422522 | 0,036 | 0,195 | 1 | Ciliated | 0,803615601 | 1,244376041 | 0,184615385 |
| LINS1 | 0,066150006 | -0,291003602 | 0,036 | 0,195 | 1 | Ciliated | 0,817333288 | 1,223491096 | 0,184615385 |
| RBM8A2 | 0,066193592 | 0,377671052 | 0,821 | 0,683 | 1 | Ciliated | 1,299242789 | 0,769679084 | 1,20204978 |
| GTF2H1 | 0,067011153 | -0,476921827 | 0,214 | 0,488 | 1 | Ciliated | 0,71850902 | 1,391770976 | 0,43852459 |
| FHAD1 | 0,067108656 | -0,494323337 | 0,643 | 0,805 | 1 | Ciliated | 0,70989456 | 1,4086599 | 0,798757764 |
| FASTK | 0,067578757 | 0,466496222 | 0,357 | 0,195 | 1 | Ciliated | 1,381749626 | 0,723720116 | 1,830769231 |
| MORN5 | 0,067628829 | 0,439596915 | 1 | 0,854 | 1 | Ciliated | 1,356225349 | 0,737340591 | 1,170960187 |
| NOS21 | 0,067704877 | 0,425482911 | 0,214 | 0,073 | 1 | Ciliated | 1,343021975 | 0,744589455 | 2,931506849 |
| AKR7A21 | 0,067832068 | 0,454215909 | 0,75 | 0,634 | 1 | Ciliated | 1,370038 | 0,729906761 | 1,1829653 |
| RAB20 | 0,067858068 | 0,270597024 | 0,321 | 0,171 | 1 | Ciliated | 1,206306925 | 0,828976423 | 1,877192982 |
| FMO5 | 0,068036424 | -0,276101263 | 0,25 | 0,537 | 1 | Ciliated | 0,825819698 | 1,210918076 | 0,465549348 |
| PRPF38B | 0,068041605 | -0,36953668 | 0,179 | 0,415 | 1 | Ciliated | 0,774031036 | 1,291937859 | 0,431325301 |
| SLC9A3R22 | 0,068386078 | 0,416741238 | 0,607 | 0,561 | 1 | Ciliated | 1,334908853 | 0,749114816 | 1,081996435 |
| CAPN13 | 0,068418464 | -0,474073175 | 0,071 | 0,244 | 1 | Ciliated | 0,719929143 | 1,389025587 | 0,290983607 |
| TBCEL | 0,068418464 | -0,350523839 | 0,071 | 0,244 | 1 | Ciliated | 0,784299269 | 1,275023501 | 0,290983607 |
| MAP2K6 | 0,068898704 | 0,273475641 | 0,179 | 0,049 | 1 | Ciliated | 1,208716278 | 0,827324012 | 3,653061224 |
| TRMT10C | 0,069005566 | -0,571768427 | 0,214 | 0,439 | 1 | Ciliated | 0,672791588 | 1,486344386 | 0,487471526 |
| SURF12 | 0,069129657 | 0,527326825 | 0,643 | 0,512 | 1 | Ciliated | 1,441256211 | 0,693839161 | 1,255859375 |

| Supplementary Table 4 - clusermarker | | | | | | | | | |
|---|---|---|---|---|---|---|---|---|---|
| | **p_val** | **avg_log2FC** | **pct.1** | **pct.2** | **p_val_adj** | **celltype** | **Foldchange_UP** | **Foldchange_DOWN** | **Ratio_pct1_pct2** |
| LMAN22 | 0,069374797 | 0,26367334 | 0,714 | 0,683 | 1 | Ciliated | 1,200531568 | 0,832964352 | 1,045387994 |
| TSR31 | 0,069434622 | 0,390717799 | 0,5 | 0,39 | 1 | Ciliated | 1,31104554 | 0,762750011 | 1,282051282 |
| SAT12 | 0,069572441 | 0,267083676 | 1 | 0,951 | 1 | Ciliated | 1,203372819 | 0,830997663 | 1,051524711 |
| TPM41 | 0,069736149 | -0,264259213 | 0,286 | 0,585 | 1 | Ciliated | 0,832626157 | 1,201019198 | 0,488888889 |
| OSBP | 0,069987765 | -0,256415494 | 0,214 | 0,512 | 1 | Ciliated | 0,837165351 | 1,194507154 | 0,41796875 |
| POLR1F | 0,069989243 | 0,380178321 | 0,357 | 0,195 | 1 | Ciliated | 1,301502715 | 0,768342616 | 1,830769231 |
| PSMA21 | 0,070026572 | -0,53306595 | 0,679 | 0,854 | 1 | Ciliated | 0,691084511 | 1,447001031 | 0,795081967 |
| GULP1 | 0,070257508 | -0,410844874 | 0,214 | 0,463 | 1 | Ciliated | 0,75218275 | 1,32946415 | 0,462203024 |
| SERINC11 | 0,070666692 | -0,286286257 | 0,25 | 0,512 | 1 | Ciliated | 0,82001019 | 1,219497041 | 0,48828125 |
| ARPC52 | 0,070689973 | 0,413733144 | 0,536 | 0,39 | 1 | Ciliated | 1,332128398 | 0,750678389 | 1,374358974 |
| MRPL23 | 0,071067271 | 0,35998327 | 0,464 | 0,341 | 1 | Ciliated | 1,283411015 | 0,779173615 | 1,360703812 |
| HLA-C2 | 0,071423229 | 0,328858199 | 1 | 0,878 | 1 | Ciliated | 1,256018922 | 0,79616635 | 1,138952164 |
| KLF3-AS1 | 0,071469295 | 0,291416753 | 0,25 | 0,098 | 1 | Ciliated | 1,223841523 | 0,817099258 | 2,551020408 |
| RPL412 | 0,071474991 | -0,292832022 | 1 | 1 | 1 | Ciliated | 0,816298085 | 1,225042688 | 1 |
| ETS2 | 0,071795585 | -0,477013884 | 0,429 | 0,634 | 1 | Ciliated | 0,718463174 | 1,391859786 | 0,676656151 |
| LKAAEAR1 | 0,071806467 | 0,597173219 | 0,393 | 0,244 | 1 | Ciliated | 1,512749617 | 0,661047928 | 1,610655738 |
| EXOSC4 | 0,071807907 | 0,372672968 | 0,536 | 0,415 | 1 | Ciliated | 1,29474947 | 0,772350191 | 1,291566265 |
| KRT102 | 0,071943987 | 0,292394448 | 0,821 | 0,756 | 1 | Ciliated | 1,224671184 | 0,816545709 | 1,085978836 |
| CHD8 | 0,072160391 | -0,288588104 | 0,214 | 0,512 | 1 | Ciliated | 0,818702892 | 1,221444324 | 0,41796875 |
| SSBP3 | 0,072200288 | -0,402890922 | 0,464 | 0,78 | 1 | Ciliated | 0,756341181 | 1,322154637 | 0,594871795 |
| DNAJA21 | 0,072327005 | 0,396446865 | 0,607 | 0,561 | 1 | Ciliated | 1,316262165 | 0,759727072 | 1,081996435 |
| GNPAT | 0,072486744 | -0,507051388 | 0,214 | 0,463 | 1 | Ciliated | 0,703659123 | 1,421142663 | 0,462203024 |
| HLA-DRB51 | 0,072486744 | -0,349864793 | 0,214 | 0,463 | 1 | Ciliated | 0,784657631 | 1,274441183 | 0,462203024 |
| PUF601 | 0,072507791 | 0,362500779 | 0,643 | 0,585 | 1 | Ciliated | 1,285652527 | 0,77781514 | 1,099145299 |
| PAMR1 | 0,072532402 | 0,260283273 | 0,179 | 0,049 | 1 | Ciliated | 1,197713853 | 0,834923966 | 3,653061224 |
| ANKRD66 | 0,072602705 | 0,338606615 | 0,893 | 0,683 | 1 | Ciliated | 1,26453469 | 0,790804719 | 1,307467057 |
| OAZ2 | 0,072624885 | -0,5085924 | 0,179 | 0,415 | 1 | Ciliated | 0,702907913 | 1,422661464 | 0,431325301 |
| RHPN1 | 0,072801149 | 0,382073345 | 0,536 | 0,39 | 1 | Ciliated | 1,303213401 | 0,767334037 | 1,374358974 |
| CTBS | 0,07292111 | 0,340986358 | 0,571 | 0,488 | 1 | Ciliated | 1,266622277 | 0,789501352 | 1,170081967 |
| CISD32 | 0,07292111 | 0,478241115 | 0,571 | 0,488 | 1 | Ciliated | 1,393044278 | 0,717852272 | 1,170081967 |
| C2orf49 | 0,07298902 | -0,252000377 | 0,107 | 0,317 | 1 | Ciliated | 0,839731274 | 1,19085716 | 0,337539432 |

| Supplementary Table 4 - clusermarker | | | | | | | | | |
|---|---|---|---|---|---|---|---|---|---|
| | **p_val** | **avg_log2FC** | **pct.1** | **pct.2** | **p_val_adj** | **celltype** | **Foldchange_UP** | **Foldchange_DOWN** | **Ratio_pct1_pct2** |
| EFNA1 | 0,073267241 | -0,390647107 | 0,25 | 0,488 | 1 | Ciliated | 0,762787387 | 1,3109813 | 0,512295082 |
| CHCHD3 | 0,07351274 | 0,359396414 | 0,429 | 0,293 | 1 | Ciliated | 1,282889058 | 0,77949063 | 1,464163823 |
| MBD6 | 0,073929819 | 0,321799751 | 0,536 | 0,415 | 1 | Ciliated | 1,249888804 | 0,800071172 | 1,291566265 |
| PLAC81 | 0,074179513 | -0,415861927 | 0,964 | 0,829 | 1 | Ciliated | 0,749571535 | 1,334095484 | 1,162846803 |
| IDH3B | 0,074206133 | 0,409168675 | 0,393 | 0,244 | 1 | Ciliated | 1,327920405 | 0,753057183 | 1,610655738 |
| CCDC911 | 0,074437878 | 0,789172607 | 0,464 | 0,39 | 1 | Ciliated | 1,728083113 | 0,578675871 | 1,18974359 |
| AHNAK2 | 0,074473733 | -0,61811841 | 0,429 | 0,61 | 1 | Ciliated | 0,651520099 | 1,534872067 | 0,703278689 |
| COPS9 | 0,074497912 | -0,487566385 | 0,464 | 0,732 | 1 | Ciliated | 0,713227193 | 1,402077781 | 0,633879781 |
| UBN1 | 0,074773555 | -0,263007927 | 0,214 | 0,463 | 1 | Ciliated | 0,833348628 | 1,199977976 | 0,462203024 |
| EVPL1 | 0,074773555 | 0,494886511 | 0,429 | 0,317 | 1 | Ciliated | 1,409209896 | 0,709617498 | 1,353312303 |
| TTC12 | 0,074924092 | -0,316081736 | 0,321 | 0,585 | 1 | Ciliated | 0,803248487 | 1,244944766 | 0,548717949 |
| MGAT4B1 | 0,075009584 | 0,510567248 | 0,393 | 0,268 | 1 | Ciliated | 1,424610222 | 0,701946388 | 1,46641791 |
| TPRKB1 | 0,075021975 | -0,393672456 | 0,143 | 0,341 | 1 | Ciliated | 0,761189488 | 1,313733329 | 0,419354839 |
| PARK7 | 0,075175876 | -0,373331825 | 0,929 | 0,854 | 1 | Ciliated | 0,77199755 | 1,295340898 | 1,087822014 |
| IFT571 | 0,075375908 | 0,374267214 | 1 | 0,902 | 1 | Ciliated | 1,29618102 | 0,771497179 | 1,10864745 |
| GAR1 | 0,07552261 | -0,419627464 | 0,25 | 0,488 | 1 | Ciliated | 0,747617651 | 1,337582117 | 0,512295082 |
| CD38 | 0,075855032 | -0,340820778 | 0,214 | 0,439 | 1 | Ciliated | 0,78959197 | 1,266476913 | 0,487471526 |
| PHKB | 0,07606441 | 0,459012034 | 0,5 | 0,415 | 1 | Ciliated | 1,374600161 | 0,727484274 | 1,204819277 |
| PGRMC11 | 0,076441797 | 0,394735184 | 0,679 | 0,61 | 1 | Ciliated | 1,314701416 | 0,760628982 | 1,113114754 |
| SP31 | 0,076678029 | -0,440419496 | 0,25 | 0,537 | 1 | Ciliated | 0,736920301 | 1,356998848 | 0,465549348 |
| NAP1L11 | 0,076836008 | -0,428692802 | 0,821 | 0,854 | 1 | Ciliated | 0,742934639 | 1,346013427 | 0,961358314 |
| EID11 | 0,076836008 | 0,317481495 | 0,929 | 0,78 | 1 | Ciliated | 1,246153246 | 0,802469523 | 1,191025641 |
| FAM50B | 0,077118957 | 0,494426 | 0,429 | 0,317 | 1 | Ciliated | 1,408760145 | 0,709844045 | 1,353312303 |
| CCDC241 | 0,077156617 | 0,566554107 | 0,571 | 0,488 | 1 | Ciliated | 1,480982001 | 0,675227652 | 1,170081967 |
| METTL27 | 0,077178323 | -0,590612796 | 0,321 | 0,537 | 1 | Ciliated | 0,664060782 | 1,505886249 | 0,597765363 |
| ABCA13 | 0,077222043 | -0,62225317 | 0,821 | 0,78 | 1 | Ciliated | 0,649655517 | 1,539277315 | 1,052564103 |
| ZNF862 | 0,077271608 | -0,261419907 | 0,071 | 0,244 | 1 | Ciliated | 0,834266426 | 1,198657849 | 0,290983607 |
| DICER1-AS1 | 0,077271608 | -0,447497258 | 0,071 | 0,244 | 1 | Ciliated | 0,733313875 | 1,363672547 | 0,290983607 |
| HMGXB41 | 0,077271608 | -0,339653685 | 0,071 | 0,244 | 1 | Ciliated | 0,790230982 | 1,265452789 | 0,290983607 |
| KRT81 | 0,077420632 | 0,371927868 | 0,964 | 0,951 | 1 | Ciliated | 1,294080951 | 0,772749185 | 1,013669821 |
| EPB41L11 | 0,077620708 | -0,309945194 | 0,357 | 0,61 | 1 | Ciliated | 0,806672403 | 1,239660606 | 0,585245902 |

| Supplementary Table 4 - clusermarker | | | | | | | | | |
|---|---|---|---|---|---|---|---|---|---|
| | **p_val** | **avg_log2FC** | **pct.1** | **pct.2** | **p_val_adj** | **celltype** | **Foldchange_UP** | **Foldchange_DOWN** | **Ratio_pct1_pct2** |
| CD812 | 0,077966819 | -0,498096086 | 0,75 | 0,805 | 1 | Ciliated | 0,708040561 | 1,412348466 | 0,931677019 |
| SETD2 | 0,07797613 | -0,385456452 | 0,357 | 0,585 | 1 | Ciliated | 0,765536753 | 1,306273012 | 0,61025641 |
| AMD11 | 0,07797613 | -0,416509857 | 0,357 | 0,585 | 1 | Ciliated | 0,74923497 | 1,334694775 | 0,61025641 |
| ORMDL21 | 0,078074505 | 0,547839082 | 0,607 | 0,512 | 1 | Ciliated | 1,46189438 | 0,684043946 | 1,185546875 |
| FOXA1 | 0,078074505 | -0,432031617 | 0,393 | 0,659 | 1 | Ciliated | 0,741217261 | 1,349132101 | 0,596358118 |
| FLII1 | 0,078074505 | 0,356341672 | 0,607 | 0,512 | 1 | Ciliated | 1,280175561 | 0,781142861 | 1,185546875 |
| ARMH4 | 0,078139214 | -0,263471281 | 0,286 | 0,537 | 1 | Ciliated | 0,833081023 | 1,200363438 | 0,532588454 |
| TTF2 | 0,078213344 | -0,362835122 | 0,107 | 0,293 | 1 | Ciliated | 0,777634903 | 1,28595051 | 0,365187713 |
| VTA1 | 0,078257863 | -0,434855414 | 0,214 | 0,439 | 1 | Ciliated | 0,73976789 | 1,351775352 | 0,487471526 |
| MTURN | 0,078305492 | 0,429313414 | 0,5 | 0,415 | 1 | Ciliated | 1,346592574 | 0,742615116 | 1,204819277 |
| PMPCA1 | 0,078433156 | 0,276666838 | 0,321 | 0,171 | 1 | Ciliated | 1,21139288 | 0,825496019 | 1,877192982 |
| C19orf251 | 0,078433156 | 0,381032489 | 0,321 | 0,171 | 1 | Ciliated | 1,302273516 | 0,767887842 | 1,877192982 |
| APLP21 | 0,078447076 | 0,462620031 | 0,75 | 0,707 | 1 | Ciliated | 1,378042164 | 0,725667201 | 1,060820368 |
| BCCIP | 0,078758625 | 0,384838688 | 0,643 | 0,463 | 1 | Ciliated | 1,305713783 | 0,765864627 | 1,388768898 |
| NEDD81 | 0,078964422 | 0,450179565 | 0,821 | 0,634 | 1 | Ciliated | 1,366210291 | 0,73195174 | 1,294952681 |
| FUCA2 | 0,079202091 | 0,353202495 | 0,393 | 0,244 | 1 | Ciliated | 1,27739304 | 0,782844409 | 1,610655738 |
| DOCK9 | 0,079202091 | -0,346566109 | 0,179 | 0,39 | 1 | Ciliated | 0,786453783 | 1,271530536 | 0,458974359 |
| HSD17B81 | 0,079216695 | 0,427747165 | 0,429 | 0,268 | 1 | Ciliated | 1,345131451 | 0,743421767 | 1,600746269 |
| TMEM1251 | 0,079348367 | 0,500701625 | 0,571 | 0,488 | 1 | Ciliated | 1,414901504 | 0,706762978 | 1,170081967 |
| MORN2 | 0,079477128 | 0,263416423 | 1 | 0,902 | 1 | Ciliated | 1,200317795 | 0,833112701 | 1,10864745 |
| DZANK1 | 0,080203344 | -0,476893049 | 0,25 | 0,488 | 1 | Ciliated | 0,718523352 | 1,391743215 | 0,512295082 |
| RRP7A | 0,080203344 | 0,417552522 | 0,464 | 0,341 | 1 | Ciliated | 1,335659735 | 0,748693678 | 1,360703812 |
| SOX42 | 0,080264255 | -0,952620672 | 0,464 | 0,61 | 1 | Ciliated | 0,51669303 | 1,935385116 | 0,760655738 |
| GPI1 | 0,080264255 | -0,304194735 | 0,393 | 0,659 | 1 | Ciliated | 0,809894147 | 1,23472926 | 0,596358118 |
| VKORC1L1 | 0,080460531 | 0,32652764 | 0,286 | 0,146 | 1 | Ciliated | 1,253991562 | 0,797453532 | 1,95890411 |
| CARD16 | 0,080460531 | 0,305391754 | 0,286 | 0,146 | 1 | Ciliated | 1,235754152 | 0,809222448 | 1,95890411 |
| DDX53 | 0,080461133 | -0,420563332 | 0,964 | 0,878 | 1 | Ciliated | 0,747132833 | 1,33845008 | 1,097949886 |
| CDKN1B | 0,080469917 | -0,484682611 | 0,179 | 0,439 | 1 | Ciliated | 0,714654274 | 1,399277995 | 0,407744875 |
| NBL11 | 0,080472931 | 0,287521227 | 0,571 | 0,415 | 1 | Ciliated | 1,220541397 | 0,819308548 | 1,375903614 |
| GMPR2 | 0,080604792 | -0,293769425 | 0,286 | 0,585 | 1 | Ciliated | 0,815767861 | 1,225838928 | 0,488888889 |
| PPP4C2 | 0,080831872 | 0,258429006 | 0,679 | 0,585 | 1 | Ciliated | 1,196175444 | 0,835997767 | 1,160683761 |

| Supplementary Table 4 - clusermarker | | | | | | | | | |
|---|---|---|---|---|---|---|---|---|---|
| | **p_val** | **avg_log2FC** | **pct.1** | **pct.2** | **p_val_adj** | **celltype** | **Foldchange_UP** | **Foldchange_DOWN** | **Ratio_pct1_pct2** |
| PIBF11 | 0,081767606 | -0,320698457 | 0,321 | 0,537 | 1 | Ciliated | 0,800682147 | 1,248935054 | 0,597765363 |
| ANAPC161 | 0,081864487 | 0,319097618 | 0,857 | 0,707 | 1 | Ciliated | 1,247549983 | 0,80157109 | 1,212164074 |
| SPG71 | 0,082055372 | -0,36627231 | 0,25 | 0,512 | 1 | Ciliated | 0,77578441 | 1,289017911 | 0,48828125 |
| SAR1B | 0,082102205 | 0,393415511 | 0,607 | 0,488 | 1 | Ciliated | 1,313499373 | 0,761325069 | 1,243852459 |
| B4GALT5 | 0,082519507 | -0,369932737 | 0,286 | 0,512 | 1 | Ciliated | 0,773818574 | 1,292292579 | 0,55859375 |
| ZMAT2 | 0,082997228 | -0,659906494 | 0,429 | 0,683 | 1 | Ciliated | 0,632919317 | 1,579980216 | 0,628111274 |
| PITX11 | 0,083041897 | 0,392559146 | 0,679 | 0,585 | 1 | Ciliated | 1,312719927 | 0,761777116 | 1,160683761 |
| DAZAP21 | 0,083164475 | -0,614632547 | 0,75 | 0,902 | 1 | Ciliated | 0,653096215 | 1,53116796 | 0,831485588 |
| RPS201 | 0,083447087 | -0,340336897 | 0,857 | 0,878 | 1 | Ciliated | 0,789856843 | 1,266052207 | 0,976082005 |
| RHBDD1 | 0,08367652 | -0,316213507 | 0,071 | 0,244 | 1 | Ciliated | 0,803175125 | 1,24505848 | 0,290983607 |
| C16orf89 | 0,08367652 | -0,25420062 | 0,071 | 0,244 | 1 | Ciliated | 0,838451582 | 1,192674714 | 0,290983607 |
| ALKBH6 | 0,08367652 | -0,333908843 | 0,071 | 0,244 | 1 | Ciliated | 0,793383972 | 1,260423749 | 0,290983607 |
| RPS15A1 | 0,083786221 | -0,420346291 | 1 | 0,976 | 1 | Ciliated | 0,747245241 | 1,338248737 | 1,024590164 |
| MUC162 | 0,084415706 | -0,594889639 | 0,964 | 0,78 | 1 | Ciliated | 0,662095101 | 1,510357044 | 1,235897436 |
| CCNB1IP11 | 0,084453451 | -0,479742035 | 0,036 | 0,171 | 1 | Ciliated | 0,717105836 | 1,394494298 | 0,210526316 |
| NR1H2 | 0,084516876 | -0,278960886 | 0,214 | 0,463 | 1 | Ciliated | 0,82418443 | 1,213320664 | 0,462203024 |
| SRGAP3 | 0,084845445 | -0,374581957 | 0,357 | 0,585 | 1 | Ciliated | 0,771328885 | 1,29646383 | 0,61025641 |
| MTIF3 | 0,084845445 | -0,425021764 | 0,357 | 0,585 | 1 | Ciliated | 0,744827495 | 1,342592757 | 0,61025641 |
| GDI1 | 0,084845445 | 0,486001466 | 0,571 | 0,439 | 1 | Ciliated | 1,400557744 | 0,714001264 | 1,300683371 |
| ACBD5 | 0,085195469 | -0,25088629 | 0,179 | 0,415 | 1 | Ciliated | 0,840379986 | 1,189937905 | 0,431325301 |
| SPAG92 | 0,085234476 | -0,254322248 | 0,286 | 0,537 | 1 | Ciliated | 0,838380898 | 1,192775267 | 0,532588454 |
| FBL1 | 0,085369953 | 0,343960277 | 0,607 | 0,585 | 1 | Ciliated | 1,269235938 | 0,787875579 | 1,037606838 |
| LSM31 | 0,085373012 | -0,487240094 | 0,464 | 0,683 | 1 | Ciliated | 0,713388521 | 1,401760711 | 0,679355783 |
| NDUFB71 | 0,085523132 | 0,256981316 | 0,929 | 0,805 | 1 | Ciliated | 1,194975729 | 0,83683708 | 1,154037267 |
| DDAH1 | 0,085556616 | -0,271123442 | 0,214 | 0,488 | 1 | Ciliated | 0,828673997 | 1,206747169 | 0,43852459 |
| PGP | 0,086254536 | 0,470072214 | 0,357 | 0,22 | 1 | Ciliated | 1,385178801 | 0,721928461 | 1,622727273 |
| THRA | 0,086254536 | 0,304914862 | 0,357 | 0,22 | 1 | Ciliated | 1,235345733 | 0,809489986 | 1,622727273 |
| PPP1R36 | 0,086351017 | -0,462431038 | 0,321 | 0,585 | 1 | Ciliated | 0,72576227 | 1,377861652 | 0,548717949 |
| ERRFI11 | 0,086575829 | -0,381511592 | 0,321 | 0,537 | 1 | Ciliated | 0,767632877 | 1,302706058 | 0,597765363 |
| PABPC12 | 0,086876032 | -0,398288633 | 0,893 | 0,878 | 1 | Ciliated | 0,75875781 | 1,3179436 | 1,017084282 |
| TRIM52 | 0,087036575 | -0,304027223 | 0,071 | 0,244 | 1 | Ciliated | 0,80998819 | 1,234585903 | 0,290983607 |

| Supplementary Table 4 - clusermarker | | | | | | | | | |
|---|---|---|---|---|---|---|---|---|---|
| | **p_val** | **avg_log2FC** | **pct.1** | **pct.2** | **p_val_adj** | **celltype** | **Foldchange_UP** | **Foldchange_DOWN** | **Ratio_pct1_pct2** |
| SESN31 | 0,087036575 | -0,324288088 | 0,071 | 0,244 | 1 | Ciliated | 0,798692412 | 1,252046451 | 0,290983607 |
| LRRC57 | 0,087036575 | -0,267117414 | 0,071 | 0,244 | 1 | Ciliated | 0,830978229 | 1,203400961 | 0,290983607 |
| HSPA82 | 0,08708958 | -0,358216648 | 0,964 | 0,927 | 1 | Ciliated | 0,78012832 | 1,281840402 | 1,0399137 |
| TIPARP1 | 0,087448088 | -0,280956429 | 0,286 | 0,512 | 1 | Ciliated | 0,823045202 | 1,215000097 | 0,55859375 |
| SEC14L11 | 0,08769275 | 0,488497445 | 0,643 | 0,585 | 1 | Ciliated | 1,402982919 | 0,712767053 | 1,099145299 |
| PRDX41 | 0,087696186 | 0,406653146 | 0,607 | 0,585 | 1 | Ciliated | 1,325607018 | 0,754371383 | 1,037606838 |
| CCDC69 | 0,087812894 | -0,32763638 | 0,286 | 0,561 | 1 | Ciliated | 0,796840908 | 1,25495565 | 0,509803922 |
| ST7 | 0,088487417 | -0,29735109 | 0,036 | 0,171 | 1 | Ciliated | 0,81374513 | 1,228886002 | 0,210526316 |
| LINC02453 | 0,088487417 | -0,391057856 | 0,036 | 0,171 | 1 | Ciliated | 0,762570245 | 1,311354602 | 0,210526316 |
| COG7 | 0,088495716 | -0,27025479 | 0,214 | 0,439 | 1 | Ciliated | 0,829173095 | 1,2060208 | 0,487471526 |
| SH2B1 | 0,088495716 | 0,33549043 | 0,429 | 0,293 | 1 | Ciliated | 1,261806275 | 0,792514683 | 1,464163823 |
| GFOD2 | 0,088716642 | 0,475825229 | 0,464 | 0,39 | 1 | Ciliated | 1,390713488 | 0,719055369 | 1,18974359 |
| FAM216B | 0,08891602 | -0,60452083 | 0,893 | 0,829 | 1 | Ciliated | 0,657689786 | 1,520473667 | 1,077201448 |
| YIPF31 | 0,089216612 | 0,46605984 | 0,571 | 0,463 | 1 | Ciliated | 1,381331741 | 0,723939058 | 1,233261339 |
| C1D | 0,089527095 | -0,491112554 | 0,393 | 0,659 | 1 | Ciliated | 0,711476222 | 1,40552835 | 0,596358118 |
| SRD5A31 | 0,089691116 | 0,416885822 | 0,536 | 0,463 | 1 | Ciliated | 1,335042641 | 0,749039746 | 1,157667387 |
| CD2AP | 0,089691116 | -0,300170074 | 0,357 | 0,585 | 1 | Ciliated | 0,812156649 | 1,231289557 | 0,61025641 |
| PDAP11 | 0,089691116 | 0,304058461 | 0,536 | 0,463 | 1 | Ciliated | 1,234612635 | 0,809970651 | 1,157667387 |
| UGCG1 | 0,08977425 | 0,748810492 | 0,714 | 0,585 | 1 | Ciliated | 1,680406756 | 0,595094013 | 1,220512821 |
| KCTD12 | 0,089989324 | -0,396173409 | 0,464 | 0,659 | 1 | Ciliated | 0,759871088 | 1,316012697 | 0,704097117 |
| INSIG12 | 0,090271455 | -0,257361163 | 0,25 | 0,488 | 1 | Ciliated | 0,836616779 | 1,195290395 | 0,512295082 |
| IFI301 | 0,090312259 | 0,38046796 | 0,321 | 0,171 | 1 | Ciliated | 1,301764034 | 0,768188377 | 1,877192982 |
| SELENOP | 0,090329786 | -0,291749155 | 0,286 | 0,561 | 1 | Ciliated | 0,816911017 | 1,224123532 | 0,509803922 |
| H2BC5 | 0,090377147 | -0,320893819 | 0,107 | 0,293 | 1 | Ciliated | 0,80057373 | 1,249104189 | 0,365187713 |
| NR4A3 | 0,090377147 | -0,297413819 | 0,107 | 0,293 | 1 | Ciliated | 0,813709749 | 1,228939436 | 0,365187713 |
| TBC1D1 | 0,09069273 | 0,370356867 | 0,393 | 0,268 | 1 | Ciliated | 1,292672549 | 0,773591116 | 1,46641791 |
| ATRAID | 0,090697497 | 0,395425411 | 0,643 | 0,659 | 1 | Ciliated | 1,315330558 | 0,760265162 | 0,975720789 |
| ANAPC10 | 0,091085276 | -0,296240285 | 0,179 | 0,439 | 1 | Ciliated | 0,814371916 | 1,227940184 | 0,407744875 |
| SUPT20H | 0,091217096 | -0,405314904 | 0,214 | 0,439 | 1 | Ciliated | 0,755071462 | 1,324377957 | 0,487471526 |
| SCP2 | 0,09123388 | -0,475079235 | 0,714 | 0,829 | 1 | Ciliated | 0,719427277 | 1,389994559 | 0,861278649 |
| TRAPPC11 | 0,091273119 | 0,328882716 | 0,643 | 0,561 | 1 | Ciliated | 1,256040267 | 0,796152819 | 1,146167558 |

| Supplementary Table 4 - clusermarker | | | | | | | | | |
|---|---|---|---|---|---|---|---|---|---|
| | **p_val** | **avg_log2FC** | **pct.1** | **pct.2** | **p_val_adj** | **celltype** | **Foldchange_UP** | **Foldchange_DOWN** | **Ratio_pct1_pct2** |
| RAB18 | 0,091298226 | -0,328817239 | 0,25 | 0,537 | 1 | Ciliated | 0,796188954 | 1,255983263 | 0,465549348 |
| XIAP | 0,091299123 | -0,255120816 | 0,321 | 0,585 | 1 | Ciliated | 0,837916962 | 1,193435681 | 0,548717949 |
| TMEM161B-DT | 0,091338402 | -0,300501948 | 0,143 | 0,317 | 1 | Ciliated | 0,811969843 | 1,231572833 | 0,451104101 |
| CDKN1A1 | 0,092168376 | -0,737350348 | 0,571 | 0,683 | 1 | Ciliated | 0,599840007 | 1,667111211 | 0,83601757 |
| TFG1 | 0,092196192 | -0,501404939 | 0,321 | 0,61 | 1 | Ciliated | 0,706418515 | 1,415591436 | 0,526229508 |
| EIF2AK11 | 0,092196192 | -0,525034878 | 0,393 | 0,561 | 1 | Ciliated | 0,694942309 | 1,438968367 | 0,700534759 |
| UBA5 | 0,092487956 | 0,397382743 | 0,429 | 0,268 | 1 | Ciliated | 1,317116303 | 0,759234395 | 1,600746269 |
| HSPB11 | 0,092550881 | 0,426215609 | 0,964 | 0,805 | 1 | Ciliated | 1,343704225 | 0,744211398 | 1,197515528 |
| SLC39A41 | 0,09260778 | 0,30821762 | 0,5 | 0,366 | 1 | Ciliated | 1,238177047 | 0,807638942 | 1,366120219 |
| DEFB11 | 0,092675275 | -0,29696549 | 0,036 | 0,171 | 1 | Ciliated | 0,813962655 | 1,228557592 | 0,210526316 |
| ALOX151 | 0,092849399 | 0,295960514 | 0,929 | 0,927 | 1 | Ciliated | 1,227702081 | 0,814529856 | 1,002157497 |
| C1orf116 | 0,092895013 | -0,344854971 | 0,179 | 0,39 | 1 | Ciliated | 0,787387126 | 1,270023305 | 0,458974359 |
| UQCC31 | 0,092941257 | 0,439187423 | 0,464 | 0,341 | 1 | Ciliated | 1,355840455 | 0,737549906 | 1,360703812 |
| ZKSCAN11 | 0,093034664 | -0,678822817 | 0,607 | 0,756 | 1 | Ciliated | 0,624674777 | 1,600833005 | 0,802910053 |
| TMX3 | 0,093377535 | -0,421749365 | 0,071 | 0,22 | 1 | Ciliated | 0,746518871 | 1,339550866 | 0,322727273 |
| TXNDC9 | 0,093501122 | -0,530158473 | 0,25 | 0,463 | 1 | Ciliated | 0,692478664 | 1,444087813 | 0,539956803 |
| MTA11 | 0,093546457 | 0,337625897 | 0,393 | 0,268 | 1 | Ciliated | 1,263675374 | 0,791342476 | 1,46641791 |
| BAG12 | 0,093576581 | 0,354746809 | 0,821 | 0,683 | 1 | Ciliated | 1,278761141 | 0,782006872 | 1,20204978 |
| HS1BP3 | 0,093644701 | -0,378634289 | 0,107 | 0,293 | 1 | Ciliated | 0,769165368 | 1,300110538 | 0,365187713 |
| CWC22 | 0,093644701 | -0,38062376 | 0,107 | 0,293 | 1 | Ciliated | 0,768105423 | 1,301904622 | 0,365187713 |
| PWWP2B | 0,094005356 | 0,476337781 | 0,429 | 0,293 | 1 | Ciliated | 1,391207659 | 0,718799953 | 1,464163823 |
| DNAJC27-AS1 | 0,094083146 | -0,288591414 | 0,071 | 0,244 | 1 | Ciliated | 0,818701013 | 1,221447126 | 0,290983607 |
| CCNB1 | 0,094083146 | -0,293901547 | 0,071 | 0,244 | 1 | Ciliated | 0,815693156 | 1,225951196 | 0,290983607 |
| SLC17A5 | 0,094083146 | 0,256906606 | 0,25 | 0,122 | 1 | Ciliated | 1,194913849 | 0,836880417 | 2,049180328 |
| P4HTM | 0,094208927 | 0,360007783 | 0,893 | 0,756 | 1 | Ciliated | 1,283432821 | 0,779160376 | 1,181216931 |
| CXCL11 | 0,094480403 | -0,868296698 | 0,143 | 0,317 | 1 | Ciliated | 0,547793215 | 1,825506364 | 0,451104101 |
| TMEM2382 | 0,094753598 | 0,287707319 | 0,714 | 0,659 | 1 | Ciliated | 1,220698843 | 0,819202874 | 1,083459788 |
| ABITRAM | 0,094889235 | -0,683732555 | 0,25 | 0,512 | 1 | Ciliated | 0,622552515 | 1,606290193 | 0,48828125 |
| PRKAG1 | 0,094904281 | -0,319304346 | 0,179 | 0,366 | 1 | Ciliated | 0,801456239 | 1,247728761 | 0,489071038 |
| PSIP1 | 0,094908687 | -0,41195206 | 0,429 | 0,683 | 1 | Ciliated | 0,751605714 | 1,330484829 | 0,628111274 |
| MT-CYB1 | 0,09494722 | 0,407170068 | 0,964 | 0,805 | 1 | Ciliated | 1,326082073 | 0,754101138 | 1,197515528 |

| Supplementary Table 4 - clusermarker | | | | | | | | | |
|---|---|---|---|---|---|---|---|---|---|
| | p_val | avg_log2FC | pct.1 | pct.2 | p_val_adj | celltype | Foldchange_UP | Foldchange_DOWN | Ratio_pct1_pct2 |
| PPIL3 | 0,095007661 | -0,411550124 | 0,143 | 0,341 | 1 | Ciliated | 0,751815141 | 1,330114207 | 0,419354839 |
| IK1 | 0,09505612 | -0,545441577 | 0,857 | 0,902 | 1 | Ciliated | 0,685181652 | 1,459466986 | 0,950110865 |
| SEMA3C | 0,095283475 | -0,255165292 | 0,286 | 0,512 | 1 | Ciliated | 0,83789113 | 1,193472473 | 0,55859375 |
| RPLP11 | 0,095310189 | 0,282303262 | 1 | 1 | 1 | Ciliated | 1,216134894 | 0,822277204 | 1 |
| UAP1 | 0,096231503 | -0,320855757 | 0,5 | 0,659 | 1 | Ciliated | 0,800594851 | 1,249071235 | 0,758725341 |
| SOX2-OT | 0,096471773 | 0,370815357 | 0,393 | 0,268 | 1 | Ciliated | 1,293083426 | 0,773345308 | 1,46641791 |
| FAHD1 | 0,096702599 | 0,441472651 | 0,286 | 0,146 | 1 | Ciliated | 1,357989807 | 0,736382552 | 1,95890411 |
| HSPA52 | 0,096715472 | 0,37925743 | 0,857 | 0,634 | 1 | Ciliated | 1,300672214 | 0,768833215 | 1,351735016 |
| BZW11 | 0,096810469 | -0,552199426 | 0,536 | 0,732 | 1 | Ciliated | 0,68197964 | 1,466319434 | 0,732240437 |
| ZG16B2 | 0,096861549 | -0,765531235 | 0,214 | 0,439 | 1 | Ciliated | 0,588236726 | 1,699995862 | 0,487471526 |
| VPS8 | 0,097006347 | -0,298004106 | 0,107 | 0,293 | 1 | Ciliated | 0,813376883 | 1,229442366 | 0,365187713 |
| SLC7A2 | 0,097020864 | -0,483009206 | 0,464 | 0,61 | 1 | Ciliated | 0,715483694 | 1,397655891 | 0,760655738 |
| SNX33 | 0,097182559 | -0,393086674 | 0,071 | 0,22 | 1 | Ciliated | 0,761498619 | 1,313200017 | 0,322727273 |
| PLIN5 | 0,097182559 | -0,306367041 | 0,071 | 0,22 | 1 | Ciliated | 0,808675584 | 1,236589826 | 0,322727273 |
| RABL2B | 0,097500081 | 0,255388829 | 0,679 | 0,537 | 1 | Ciliated | 1,193657409 | 0,837761314 | 1,26443203 |
| MAT2A2 | 0,097639588 | -0,343894998 | 0,25 | 0,512 | 1 | Ciliated | 0,787911229 | 1,26917851 | 0,48828125 |
| RBMS1 | 0,097639588 | -0,376362778 | 0,286 | 0,488 | 1 | Ciliated | 0,770377366 | 1,298065135 | 0,586065574 |
| WBP1L | 0,097960124 | 0,596361561 | 0,357 | 0,244 | 1 | Ciliated | 1,511898785 | 0,661419937 | 1,463114754 |
| FAM114A1 | 0,098090646 | -0,369418755 | 0,357 | 0,537 | 1 | Ciliated | 0,774094308 | 1,291832261 | 0,664804469 |
| BAZ2B1 | 0,098135035 | -0,37388921 | 0,393 | 0,585 | 1 | Ciliated | 0,771699347 | 1,295841449 | 0,671794872 |
| ZBED5 | 0,098878611 | -0,303178531 | 0,179 | 0,39 | 1 | Ciliated | 0,81046482 | 1,233859848 | 0,458974359 |
| ID3 | 0,099406855 | -0,492516009 | 0,321 | 0,488 | 1 | Ciliated | 0,710784434 | 1,406896314 | 0,657786885 |
| GNA11 | 0,099659644 | 0,368087352 | 0,679 | 0,61 | 1 | Ciliated | 1,290640633 | 0,774809017 | 1,113114754 |
| PLAAT42 | 0,099888912 | -0,414733792 | 0,821 | 0,902 | 1 | Ciliated | 0,750157902 | 1,333052678 | 0,910199557 |
| FLOT1 | 0,100035517 | 0,538439779 | 0,643 | 0,585 | 1 | Ciliated | 1,45240095 | 0,68851511 | 1,099145299 |
| EPS15L1 | 0,100246292 | 0,36753476 | 0,286 | 0,146 | 1 | Ciliated | 1,290146377 | 0,775105847 | 1,95890411 |
| UFL1 | 0,100453282 | -0,274599548 | 0,25 | 0,512 | 1 | Ciliated | 0,82667975 | 1,209658275 | 0,48828125 |
| HCAR21 | 0,100463803 | -0,263465514 | 0,107 | 0,293 | 1 | Ciliated | 0,833084353 | 1,200358639 | 0,365187713 |
| SSR31 | 0,100812531 | 0,319870215 | 0,5 | 0,366 | 1 | Ciliated | 1,248218254 | 0,801141945 | 1,366120219 |
| MDM2 | 0,101212628 | -0,317737184 | 0,429 | 0,659 | 1 | Ciliated | 0,802327313 | 1,246374122 | 0,650986343 |
| PHACTR4 | 0,101332525 | -0,300643933 | 0,25 | 0,488 | 1 | Ciliated | 0,811889936 | 1,231694046 | 0,512295082 |

| Supplementary Table 4 - clusermarker | | | | | | | | | |
|---|---|---|---|---|---|---|---|---|---|
| | **p_val** | **avg_log2FC** | **pct.1** | **pct.2** | **p_val_adj** | **celltype** | **Foldchange_UP** | **Foldchange_DOWN** | **Ratio_pct1_pct2** |
| IFI6 | 0,101396339 | 0,405393838 | 0,643 | 0,561 | 1 | Ciliated | 1,32445042 | 0,755030151 | 1,146167558 |
| AP002026.1 | 0,101441811 | -0,428868582 | 0,143 | 0,341 | 1 | Ciliated | 0,742844125 | 1,346177437 | 0,419354839 |
| AC025154.2 | 0,101441811 | -0,392446601 | 0,143 | 0,341 | 1 | Ciliated | 0,761836544 | 1,312617526 | 0,419354839 |
| AC008915.2 | 0,101441811 | -0,327469551 | 0,143 | 0,341 | 1 | Ciliated | 0,796933058 | 1,254810539 | 0,419354839 |
| CEP781 | 0,101983718 | -0,318288885 | 0,179 | 0,39 | 1 | Ciliated | 0,802020554 | 1,246850838 | 0,458974359 |
| NOLC11 | 0,102026867 | -0,360189936 | 0,25 | 0,463 | 1 | Ciliated | 0,779062006 | 1,283594877 | 0,539956803 |
| WAC | 0,102169017 | -0,305228775 | 0,357 | 0,61 | 1 | Ciliated | 0,809313869 | 1,235614559 | 0,585245902 |
| SELENBP12 | 0,102258719 | -0,526672272 | 0,571 | 0,683 | 1 | Ciliated | 0,694154028 | 1,440602459 | 0,83601757 |
| HSP90B11 | 0,102294091 | 0,296095899 | 0,929 | 0,805 | 1 | Ciliated | 1,227817297 | 0,814453423 | 1,154037267 |
| HNRNPA31 | 0,102506586 | -0,391779236 | 0,464 | 0,732 | 1 | Ciliated | 0,762189038 | 1,312010473 | 0,633879781 |
| DUSP18 | 0,102547765 | 0,423784205 | 0,607 | 0,537 | 1 | Ciliated | 1,34144156 | 0,74546669 | 1,130353818 |
| HADH1 | 0,102689203 | 0,446730552 | 0,607 | 0,561 | 1 | Ciliated | 1,362948029 | 0,733703691 | 1,081996435 |
| ARF31 | 0,102689203 | 0,347391408 | 0,607 | 0,561 | 1 | Ciliated | 1,272258128 | 0,786004018 | 1,081996435 |
| PSMB32 | 0,102815905 | 0,299104222 | 0,75 | 0,756 | 1 | Ciliated | 1,230380226 | 0,812756885 | 0,992063492 |
| DNAAF1 | 0,103632192 | 0,369382204 | 0,893 | 0,707 | 1 | Ciliated | 1,291799533 | 0,77411392 | 1,263083451 |
| SERTAD4-AS1 | 0,103671375 | 0,273078048 | 0,5 | 0,366 | 1 | Ciliated | 1,208383213 | 0,827552046 | 1,366120219 |
| DHRS9 | 0,103841975 | -0,52405597 | 0,893 | 0,878 | 1 | Ciliated | 0,695414006 | 1,43799232 | 1,017084282 |
| LINC00339 | 0,103892754 | -0,417222107 | 0,107 | 0,268 | 1 | Ciliated | 0,748865168 | 1,335353869 | 0,399253731 |
| CRELD1 | 0,103892754 | 0,287644292 | 0,286 | 0,146 | 1 | Ciliated | 1,220645516 | 0,819238663 | 1,95890411 |
| AC087477.2 | 0,103892754 | 0,30654526 | 0,286 | 0,146 | 1 | Ciliated | 1,236742594 | 0,808575693 | 1,95890411 |
| NUS1 | 0,10401878 | -0,28907131 | 0,107 | 0,293 | 1 | Ciliated | 0,818428727 | 1,221853494 | 0,365187713 |
| UGDH | 0,104163052 | -0,435594388 | 0,393 | 0,61 | 1 | Ciliated | 0,739389065 | 1,352467933 | 0,644262295 |
| TM2D1 | 0,104260051 | 0,321520582 | 0,464 | 0,341 | 1 | Ciliated | 1,249646967 | 0,800226005 | 1,360703812 |
| SYF2 | 0,104669814 | -0,312362427 | 0,321 | 0,585 | 1 | Ciliated | 0,805321957 | 1,241739396 | 0,548717949 |
| CHCHD6 | 0,104669814 | 0,516154093 | 0,536 | 0,439 | 1 | Ciliated | 1,43013773 | 0,699233353 | 1,22095672 |
| BCL7A | 0,104729322 | 0,418228445 | 0,357 | 0,22 | 1 | Ciliated | 1,336285657 | 0,748342987 | 1,622727273 |
| EEF1B21 | 0,104887179 | 0,281232016 | 0,929 | 0,805 | 1 | Ciliated | 1,215232211 | 0,822887997 | 1,154037267 |
| EIF3E1 | 0,104918165 | -0,664855281 | 0,5 | 0,732 | 1 | Ciliated | 0,630751973 | 1,585409231 | 0,683060109 |
| C1QBP2 | 0,104918165 | 0,422433315 | 0,679 | 0,61 | 1 | Ciliated | 1,340186068 | 0,746165047 | 1,113114754 |
| KIF3A | 0,105015141 | -0,267864017 | 0,393 | 0,659 | 1 | Ciliated | 0,830548305 | 1,204023889 | 0,596358118 |
| HINT21 | 0,105015141 | 0,394653624 | 0,643 | 0,488 | 1 | Ciliated | 1,314627094 | 0,760671984 | 1,317622951 |

| Supplementary Table 4 - clusermarker | | | | | | | | | |
|---|---|---|---|---|---|---|---|---|---|
| | **p_val** | **avg_log2FC** | **pct.1** | **pct.2** | **p_val_adj** | **celltype** | **Foldchange_UP** | **Foldchange_DOWN** | **Ratio_pct1_pct2** |
| DNAJA11 | 0,105033592 | -0,374660585 | 0,857 | 0,878 | 1 | Ciliated | 0,771286848 | 1,29653449 | 0,976082005 |
| SNRPN1 | 0,105165946 | 0,294415795 | 0,393 | 0,244 | 1 | Ciliated | 1,226388263 | 0,815402454 | 1,610655738 |
| ARPC5L2 | 0,105176194 | 0,31921412 | 0,679 | 0,585 | 1 | Ciliated | 1,247650731 | 0,801506363 | 1,160683761 |
| JAK11 | 0,105344968 | 0,319173074 | 0,679 | 0,561 | 1 | Ciliated | 1,247615234 | 0,801529167 | 1,210338681 |
| PRRG41 | 0,105392657 | -0,42574548 | 0,429 | 0,683 | 1 | Ciliated | 0,744453952 | 1,343266426 | 0,628111274 |
| RPS121 | 0,105415432 | 0,364333423 | 1 | 1 | 1 | Ciliated | 1,287286719 | 0,776827714 | 1 |
| GARS1 | 0,105502832 | 0,299049032 | 0,25 | 0,122 | 1 | Ciliated | 1,230333158 | 0,812787978 | 2,049180328 |
| SHARPIN | 0,105688662 | 0,47362191 | 0,393 | 0,268 | 1 | Ciliated | 1,388591177 | 0,720154367 | 1,46641791 |
| AC008014.1 | 0,105848211 | -0,584662626 | 0,214 | 0,439 | 1 | Ciliated | 0,666805252 | 1,499688247 | 0,487471526 |
| RAB8A | 0,105848211 | -0,381141513 | 0,214 | 0,439 | 1 | Ciliated | 0,767829815 | 1,302371932 | 0,487471526 |
| KIF1C1 | 0,105906661 | 0,594623396 | 0,393 | 0,293 | 1 | Ciliated | 1,51007834 | 0,662217299 | 1,341296928 |
| RBM7 | 0,106164778 | 0,348200414 | 0,321 | 0,195 | 1 | Ciliated | 1,27297176 | 0,785563381 | 1,646153846 |
| CD472 | 0,10645767 | 0,32279399 | 0,5 | 0,439 | 1 | Ciliated | 1,250750466 | 0,79951999 | 1,138952164 |
| C5orf49 | 0,106704251 | 0,334664585 | 0,964 | 0,927 | 1 | Ciliated | 1,261084183 | 0,792968474 | 1,0399137 |
| AP2M12 | 0,106951169 | 0,270114302 | 0,571 | 0,488 | 1 | Ciliated | 1,205903365 | 0,829253843 | 1,170081967 |
| DPY19L4 | 0,10767298 | -0,273427772 | 0,107 | 0,293 | 1 | Ciliated | 0,827351462 | 1,208676174 | 0,365187713 |
| FARP2 | 0,107923733 | -0,445743241 | 0,143 | 0,317 | 1 | Ciliated | 0,734205974 | 1,362015613 | 0,451104101 |
| PIK3C3 | 0,107923733 | -0,471907913 | 0,143 | 0,317 | 1 | Ciliated | 0,721010456 | 1,386942438 | 0,451104101 |
| LINC01765 | 0,108028964 | -0,556847545 | 0,464 | 0,634 | 1 | Ciliated | 0,679785953 | 1,471051286 | 0,731861199 |
| KLHDC31 | 0,108046618 | 0,333408037 | 0,464 | 0,317 | 1 | Ciliated | 1,259986291 | 0,793659429 | 1,463722397 |
| ABHD21 | 0,108113923 | 0,261100614 | 0,786 | 0,732 | 1 | Ciliated | 1,198392594 | 0,834451084 | 1,073770492 |
| PRELID12 | 0,108152281 | 0,496710168 | 0,607 | 0,561 | 1 | Ciliated | 1,410992352 | 0,708721063 | 1,081996435 |
| IQSEC1 | 0,108631269 | 0,441299018 | 0,464 | 0,293 | 1 | Ciliated | 1,357826378 | 0,736471184 | 1,583617747 |
| ICE2 | 0,108986603 | -0,290478574 | 0,214 | 0,439 | 1 | Ciliated | 0,817630787 | 1,223045922 | 0,487471526 |
| TMEM184A | 0,109088809 | 0,276720049 | 0,393 | 0,293 | 1 | Ciliated | 1,211437561 | 0,825465572 | 1,341296928 |
| AK1 | 0,10929013 | -0,350862345 | 0,786 | 0,805 | 1 | Ciliated | 0,784115267 | 1,2753227 | 0,976397516 |
| RNASEL | 0,109345323 | -0,252341412 | 0,071 | 0,22 | 1 | Ciliated | 0,839532795 | 1,191138697 | 0,322727273 |
| PALM2AKAP21 | 0,109345323 | -0,300645168 | 0,071 | 0,22 | 1 | Ciliated | 0,811889241 | 1,2316951 | 0,322727273 |
| AC011603.2 | 0,109345323 | -0,292072935 | 0,071 | 0,22 | 1 | Ciliated | 0,816727701 | 1,22439829 | 0,322727273 |
| NCK1 | 0,109544935 | 0,282680514 | 0,25 | 0,122 | 1 | Ciliated | 1,216452944 | 0,822062213 | 2,049180328 |
| EHF2 | 0,10957 | -0,438214725 | 0,643 | 0,683 | 1 | Ciliated | 0,738047346 | 1,354926625 | 0,941434846 |

| Supplementary Table 4 - clusermarker | | | | | | | | | |
|---|---|---|---|---|---|---|---|---|---|
| | **p_val** | **avg_log2FC** | **pct.1** | **pct.2** | **p_val_adj** | **celltype** | **Foldchange_UP** | **Foldchange_DOWN** | **Ratio_pct1_pct2** |
| PHPT12 | 0,110421188 | 0,427666236 | 0,714 | 0,659 | 1 | Ciliated | 1,345055997 | 0,743463471 | 1,083459788 |
| BCAS11 | 0,110855218 | -0,344757063 | 0,464 | 0,634 | 1 | Ciliated | 0,787440564 | 1,269937118 | 0,731861199 |
| PPP1R12A1 | 0,111160171 | -0,276999031 | 0,25 | 0,463 | 1 | Ciliated | 0,825305963 | 1,211671847 | 0,539956803 |
| ROGDI | 0,111341225 | 0,35600092 | 0,536 | 0,463 | 1 | Ciliated | 1,27987323 | 0,781327382 | 1,157667387 |
| ATF7 | 0,111542944 | 0,426238898 | 0,357 | 0,22 | 1 | Ciliated | 1,343725917 | 0,744199384 | 1,622727273 |
| LDLR1 | 0,111794764 | -0,373463709 | 0,25 | 0,439 | 1 | Ciliated | 0,771926982 | 1,295459317 | 0,569476082 |
| CCNL12 | 0,112095092 | -0,572068223 | 0,393 | 0,585 | 1 | Ciliated | 0,672651795 | 1,486653284 | 0,671794872 |
| PI4KB1 | 0,112198138 | -0,32862371 | 0,214 | 0,439 | 1 | Ciliated | 0,796295765 | 1,255814792 | 0,487471526 |
| NMRAL1 | 0,112198138 | -0,266871874 | 0,214 | 0,439 | 1 | Ciliated | 0,83111967 | 1,203196165 | 0,487471526 |
| MLLT10 | 0,112212016 | -0,453771366 | 0,179 | 0,415 | 1 | Ciliated | 0,730131705 | 1,36961591 | 0,431325301 |
| LONP1 | 0,112212016 | 0,362187769 | 0,393 | 0,268 | 1 | Ciliated | 1,28537362 | 0,777983914 | 1,46641791 |
| SERINC22 | 0,112343225 | 0,455453064 | 0,5 | 0,439 | 1 | Ciliated | 1,371213354 | 0,729281112 | 1,138952164 |
| MICU2 | 0,112343225 | -0,484094641 | 0,286 | 0,585 | 1 | Ciliated | 0,714945591 | 1,398707835 | 0,488888889 |
| TP53I131 | 0,1123461 | 0,337466796 | 0,393 | 0,293 | 1 | Ciliated | 1,263536023 | 0,791429751 | 1,341296928 |
| HDGF2 | 0,112540072 | 0,30284166 | 0,821 | 0,659 | 1 | Ciliated | 1,233571774 | 0,810654087 | 1,245827011 |
| PRXL2A1 | 0,112614219 | 0,53574675 | 0,5 | 0,415 | 1 | Ciliated | 1,449692332 | 0,689801538 | 1,204819277 |
| DHRS1 | 0,113339278 | 0,359007053 | 0,321 | 0,195 | 1 | Ciliated | 1,282542873 | 0,779701031 | 1,646153846 |
| PWWP2A | 0,113344024 | 0,341416558 | 0,286 | 0,171 | 1 | Ciliated | 1,267000029 | 0,789265964 | 1,67251462 |
| RNF40 | 0,113447066 | 0,304206009 | 0,464 | 0,341 | 1 | Ciliated | 1,234738908 | 0,809887818 | 1,360703812 |
| C1orf194 | 0,113538208 | -0,420318694 | 1 | 0,976 | 1 | Ciliated | 0,747259535 | 1,338223138 | 1,024590164 |
| GOLPH3L | 0,113573192 | -0,34064185 | 0,357 | 0,61 | 1 | Ciliated | 0,789689903 | 1,26631985 | 0,585245902 |
| CCDC113 | 0,113622395 | 0,440946982 | 0,821 | 0,707 | 1 | Ciliated | 1,357495092 | 0,736650914 | 1,161244696 |
| USP13 | 0,113657034 | -0,306453584 | 0,071 | 0,22 | 1 | Ciliated | 0,808627076 | 1,236664007 | 0,322727273 |
| TXNIP2 | 0,113739534 | -0,667960257 | 0,464 | 0,634 | 1 | Ciliated | 0,629395925 | 1,588825031 | 0,731861199 |
| PPM1A | 0,114003258 | -0,277074317 | 0,214 | 0,415 | 1 | Ciliated | 0,825262896 | 1,211735078 | 0,515662651 |
| SCAMP2 | 0,11431563 | -0,476434909 | 0,429 | 0,537 | 1 | Ciliated | 0,718751562 | 1,391301324 | 0,798882682 |
| RNF135 | 0,11443582 | 0,403139481 | 0,357 | 0,244 | 1 | Ciliated | 1,322382448 | 0,756210884 | 1,463114754 |
| HS3ST11 | 0,114921732 | 0,577190282 | 0,5 | 0,463 | 1 | Ciliated | 1,491940791 | 0,670267886 | 1,079913607 |
| PDHA11 | 0,115348121 | -0,407845376 | 0,143 | 0,341 | 1 | Ciliated | 0,753748235 | 1,326702941 | 0,419354839 |
| ZNRF1 | 0,115380707 | 0,492775048 | 0,536 | 0,415 | 1 | Ciliated | 1,407148949 | 0,710656822 | 1,291566265 |
| TCEANC2 | 0,115467145 | -0,290316799 | 0,107 | 0,268 | 1 | Ciliated | 0,817722477 | 1,222908785 | 0,399253731 |

| Supplementary Table 4 - clusermarker | | | | | | | | | |
|---|---|---|---|---|---|---|---|---|---|
| | **p_val** | **avg_log2FC** | **pct.1** | **pct.2** | **p_val_adj** | **celltype** | **Foldchange_UP** | **Foldchange_DOWN** | **Ratio_pct1_pct2** |
| RUFY3 | 0,115483842 | -0,554567418 | 0,214 | 0,439 | 1 | Ciliated | 0,680861179 | 1,46872818 | 0,487471526 |
| DBNL1 | 0,115590684 | 0,333418126 | 0,393 | 0,268 | 1 | Ciliated | 1,259995103 | 0,793653878 | 1,46641791 |
| TTLL3 | 0,115681067 | -0,364328706 | 0,286 | 0,561 | 1 | Ciliated | 0,776830254 | 1,287282511 | 0,509803922 |
| FAM210B1 | 0,115681067 | 0,32043146 | 0,5 | 0,415 | 1 | Ciliated | 1,248703937 | 0,800830341 | 1,204819277 |
| COMMD4 | 0,115810566 | -0,319986802 | 0,286 | 0,537 | 1 | Ciliated | 0,801077206 | 1,248319129 | 0,532588454 |
| CCDC89 | 0,116647309 | -0,322561609 | 0,25 | 0,488 | 1 | Ciliated | 0,799648782 | 1,250549018 | 0,512295082 |
| TMED1 | 0,116675753 | 0,319305472 | 0,429 | 0,317 | 1 | Ciliated | 1,247729734 | 0,801455614 | 1,353312303 |
| RPS6KA3 | 0,116675753 | -0,262406202 | 0,214 | 0,463 | 1 | Ciliated | 0,833696277 | 1,199477588 | 0,462203024 |
| PI32 | 0,117169485 | 0,265911424 | 0,214 | 0,098 | 1 | Ciliated | 1,202395424 | 0,831673158 | 2,183673469 |
| USO1 | 0,11723892 | -0,587893914 | 0,321 | 0,537 | 1 | Ciliated | 0,665313441 | 1,50305095 | 0,597765363 |
| FBXO9 | 0,117599417 | -0,458291801 | 0,25 | 0,463 | 1 | Ciliated | 0,727847544 | 1,373914094 | 0,539956803 |
| ICA1 | 0,117599417 | 0,302047519 | 0,464 | 0,317 | 1 | Ciliated | 1,232892933 | 0,811100439 | 1,463722397 |
| PKP2 | 0,117978905 | 0,410593151 | 0,357 | 0,244 | 1 | Ciliated | 1,329232204 | 0,752314003 | 1,463114754 |
| TMEM120A | 0,117995669 | 0,274799319 | 0,25 | 0,122 | 1 | Ciliated | 1,209825789 | 0,826565287 | 2,049180328 |
| MACROD1 | 0,117995669 | 0,453345206 | 0,25 | 0,122 | 1 | Ciliated | 1,369211397 | 0,730347412 | 2,049180328 |
| PPP2R1A1 | 0,118106287 | 0,338349521 | 0,75 | 0,61 | 1 | Ciliated | 1,264309365 | 0,790945656 | 1,229508197 |
| UBR5 | 0,118482399 | 0,342443083 | 0,536 | 0,415 | 1 | Ciliated | 1,267901862 | 0,788704575 | 1,291566265 |
| KPNA3 | 0,118482399 | -0,377601558 | 0,286 | 0,585 | 1 | Ciliated | 0,76971616 | 1,299180207 | 0,488888889 |
| ABCB7 | 0,118689612 | -0,255348568 | 0,179 | 0,39 | 1 | Ciliated | 0,837784694 | 1,193624099 | 0,458974359 |
| IFITM21 | 0,118716542 | -0,3794324 | 0,286 | 0,488 | 1 | Ciliated | 0,768739977 | 1,300829968 | 0,586065574 |
| KIAA1109 | 0,118813195 | -0,309457572 | 0,286 | 0,561 | 1 | Ciliated | 0,8069451 | 1,239241679 | 0,509803922 |
| TSPAN8 | 0,118966662 | -0,259328113 | 0,143 | 0,317 | 1 | Ciliated | 0,835476924 | 1,196921149 | 0,451104101 |
| PITPNA | 0,118971496 | 0,428920761 | 0,5 | 0,39 | 1 | Ciliated | 1,346226127 | 0,742817258 | 1,282051282 |
| TBCA1 | 0,119032927 | 0,287112185 | 0,964 | 0,854 | 1 | Ciliated | 1,220195389 | 0,819540877 | 1,128805621 |
| GAPDH1 | 0,119154391 | 0,47553888 | 0,964 | 0,902 | 1 | Ciliated | 1,390437484 | 0,719198103 | 1,068736142 |
| PIK3R31 | 0,119735676 | 0,302449683 | 0,679 | 0,537 | 1 | Ciliated | 1,233236661 | 0,810874369 | 1,26443203 |
| IFTAP | 0,119792133 | 0,251771144 | 0,786 | 0,634 | 1 | Ciliated | 1,190667958 | 0,839864711 | 1,239747634 |
| BTBD1 | 0,119918034 | -0,551579987 | 0,25 | 0,488 | 1 | Ciliated | 0,682272519 | 1,465689988 | 0,512295082 |
| ETNK1 | 0,120691801 | -0,475891412 | 0,429 | 0,61 | 1 | Ciliated | 0,719022383 | 1,390777288 | 0,703278689 |
| EGLN1 | 0,120890372 | 0,261539518 | 0,321 | 0,195 | 1 | Ciliated | 1,198757231 | 0,834197262 | 1,646153846 |
| AL022345.4 | 0,120890372 | 0,403595388 | 0,321 | 0,195 | 1 | Ciliated | 1,322800401 | 0,755971951 | 1,646153846 |

| Supplementary Table 4 - clusermarker | | | | | | | | | |
|---|---|---|---|---|---|---|---|---|---|
| | **p_val** | **avg_log2FC** | **pct.1** | **pct.2** | **p_val_adj** | **celltype** | **Foldchange_UP** | **Foldchange_DOWN** | **Ratio_pct1_pct2** |
| CFAP73 | 0,120899845 | 0,322840432 | 0,929 | 0,732 | 1 | Ciliated | 1,25079073 | 0,799494253 | 1,269125683 |
| SF3B22 | 0,120903263 | 0,320226717 | 0,857 | 0,78 | 1 | Ciliated | 1,248526737 | 0,800944001 | 1,098717949 |
| AC027644.3 | 0,121268388 | 0,27165061 | 0,286 | 0,171 | 1 | Ciliated | 1,207188201 | 0,828371251 | 1,67251462 |
| TUBG1 | 0,121607482 | 0,499696764 | 0,357 | 0,244 | 1 | Ciliated | 1,413916345 | 0,707255421 | 1,463114754 |
| FNDC11 | 0,121607482 | 0,564215551 | 0,357 | 0,244 | 1 | Ciliated | 1,478583328 | 0,67632306 | 1,463114754 |
| EI24 | 0,121645074 | 0,414338853 | 0,5 | 0,439 | 1 | Ciliated | 1,332687803 | 0,750363287 | 1,138952164 |
| SMARCA5 | 0,12178886 | -0,401864086 | 0,393 | 0,61 | 1 | Ciliated | 0,756879698 | 1,32121393 | 0,644262295 |
| AL121956.6 | 0,12178886 | -0,401691614 | 0,393 | 0,61 | 1 | Ciliated | 0,756970187 | 1,32105599 | 0,644262295 |
| ABAT | 0,121909834 | 0,314896704 | 0,214 | 0,098 | 1 | Ciliated | 1,243922586 | 0,803908548 | 2,183673469 |
| MPDZ | 0,122199654 | -0,278164555 | 0,286 | 0,537 | 1 | Ciliated | 0,824639484 | 1,212651127 | 0,532588454 |
| PIN41 | 0,122240578 | -0,413507808 | 0,393 | 0,634 | 1 | Ciliated | 0,750795648 | 1,331920347 | 0,619873817 |
| SLC25A23 | 0,12227503 | -0,383897739 | 0,179 | 0,39 | 1 | Ciliated | 0,766364299 | 1,304862453 | 0,458974359 |
| MAP3K2-DT | 0,122413613 | 0,387241014 | 0,357 | 0,22 | 1 | Ciliated | 1,307889824 | 0,764590397 | 1,622727273 |
| NME32 | 0,122622717 | 0,476578105 | 0,607 | 0,61 | 1 | Ciliated | 1,391439426 | 0,718680225 | 0,995081967 |
| ZNF740 | 0,122681244 | -0,384128076 | 0,071 | 0,22 | 1 | Ciliated | 0,766241953 | 1,305070801 | 0,322727273 |
| MT3 | 0,122838338 | -0,277755227 | 0,143 | 0,317 | 1 | Ciliated | 0,824873488 | 1,212307117 | 0,451104101 |
| IQCK | 0,123095771 | 0,365890426 | 0,714 | 0,61 | 1 | Ciliated | 1,28867675 | 0,775989789 | 1,170491803 |
| PGD2 | 0,123615051 | 0,334536397 | 0,536 | 0,463 | 1 | Ciliated | 1,260972136 | 0,793038935 | 1,157667387 |
| UMAD1 | 0,123730741 | -0,398151979 | 0,107 | 0,268 | 1 | Ciliated | 0,758829684 | 1,317818769 | 0,399253731 |
| NARS21 | 0,123730741 | -0,273171515 | 0,107 | 0,268 | 1 | Ciliated | 0,827498433 | 1,208461503 | 0,399253731 |
| PCM11 | 0,124135616 | -0,55444453 | 0,857 | 0,805 | 1 | Ciliated | 0,680919177 | 1,468603079 | 1,064596273 |
| TCERG1 | 0,124327813 | -0,281652182 | 0,25 | 0,463 | 1 | Ciliated | 0,822648376 | 1,215586184 | 0,539956803 |
| SLC25A391 | 0,125276547 | 0,411761127 | 0,5 | 0,415 | 1 | Ciliated | 1,330308758 | 0,751705192 | 1,204819277 |
| ZNF5241 | 0,125322751 | 0,535143418 | 0,357 | 0,244 | 1 | Ciliated | 1,449086201 | 0,690090071 | 1,463114754 |
| MRPS18C1 | 0,12538469 | -0,302465187 | 0,393 | 0,634 | 1 | Ciliated | 0,810865655 | 1,233249915 | 0,619873817 |
| PARVA | 0,125438558 | 0,499164997 | 0,679 | 0,585 | 1 | Ciliated | 1,413395281 | 0,707516159 | 1,160683761 |
| MUC14 | 0,125702668 | 0,429370024 | 0,679 | 0,561 | 1 | Ciliated | 1,346645414 | 0,742585976 | 1,210338681 |
| USP54 | 0,125944541 | -0,314167598 | 0,179 | 0,39 | 1 | Ciliated | 0,804314928 | 1,243294094 | 0,458974359 |
| CLASP1 | 0,126147911 | 0,385986469 | 0,393 | 0,293 | 1 | Ciliated | 1,306752998 | 0,765255562 | 1,341296928 |
| RFX2 | 0,126159452 | 0,415795825 | 0,714 | 0,61 | 1 | Ciliated | 1,334034359 | 0,74960588 | 1,170491803 |
| RTN41 | 0,126517844 | -0,358133603 | 0,607 | 0,732 | 1 | Ciliated | 0,780173228 | 1,281766619 | 0,829234973 |

| Supplementary Table 4 - clusermarker | | | | | | | | | |
|---|---|---|---|---|---|---|---|---|---|
| | **p_val** | **avg_log2FC** | **pct.1** | **pct.2** | **p_val_adj** | **celltype** | **Foldchange_UP** | **Foldchange_DOWN** | **Ratio_pct1_pct2** |
| CHIC21 | 0,126674558 | 0,368892556 | 0,464 | 0,341 | 1 | Ciliated | 1,291361174 | 0,774376697 | 1,360703812 |
| ABHD3 | 0,126729313 | -0,350841891 | 0,143 | 0,341 | 1 | Ciliated | 0,784126384 | 1,27530462 | 0,419354839 |
| TNPO2 | 0,126729313 | -0,36922422 | 0,143 | 0,341 | 1 | Ciliated | 0,774198694 | 1,291658081 | 0,419354839 |
| LDAF1 | 0,126807744 | -0,57644437 | 0,143 | 0,317 | 1 | Ciliated | 0,670614522 | 1,491169617 | 0,451104101 |
| STARD3 | 0,126807744 | -0,282370717 | 0,143 | 0,317 | 1 | Ciliated | 0,822238758 | 1,216191758 | 0,451104101 |
| KXD11 | 0,127214923 | 0,514437279 | 0,464 | 0,39 | 1 | Ciliated | 1,428436871 | 0,700065939 | 1,18974359 |
| ACTR3B | 0,127398528 | -0,26153496 | 0,071 | 0,22 | 1 | Ciliated | 0,834199898 | 1,198753444 | 0,322727273 |
| ANO6 | 0,128032103 | -0,282839845 | 0,107 | 0,268 | 1 | Ciliated | 0,82197143 | 1,216587297 | 0,399253731 |
| ELN-AS1 | 0,128156435 | -0,451026533 | 0,393 | 0,61 | 1 | Ciliated | 0,731522157 | 1,367012593 | 0,644262295 |
| ZBTB201 | 0,128180685 | -0,340403846 | 0,321 | 0,561 | 1 | Ciliated | 0,789820191 | 1,26611096 | 0,572192513 |
| ZNF7062 | 0,128185489 | 0,312271713 | 0,679 | 0,61 | 1 | Ciliated | 1,24166132 | 0,805372595 | 1,113114754 |
| P4HA1 | 0,128860967 | 0,354768427 | 0,464 | 0,415 | 1 | Ciliated | 1,278780302 | 0,781995154 | 1,118072289 |
| MRPL17 | 0,128889136 | 0,319720708 | 0,607 | 0,512 | 1 | Ciliated | 1,248088907 | 0,801224972 | 1,185546875 |
| USP1 | 0,128921577 | -0,272968132 | 0,286 | 0,512 | 1 | Ciliated | 0,827615097 | 1,208291153 | 0,55859375 |
| RTCA-AS1 | 0,129125898 | -0,474960159 | 0,179 | 0,366 | 1 | Ciliated | 0,719486659 | 1,389879837 | 0,489071038 |
| MAP4K3-DT | 0,129125898 | 0,367171835 | 0,357 | 0,244 | 1 | Ciliated | 1,289821868 | 0,775300857 | 1,463114754 |
| FBXL15 | 0,129125898 | 0,307631259 | 0,357 | 0,244 | 1 | Ciliated | 1,237673911 | 0,807967261 | 1,463114754 |
| CERS6 | 0,129211651 | 0,381462373 | 0,571 | 0,463 | 1 | Ciliated | 1,302661616 | 0,767659067 | 1,233261339 |
| CFL11 | 0,129279973 | -0,528924031 | 0,714 | 0,829 | 1 | Ciliated | 0,693071437 | 1,442852708 | 0,861278649 |
| IRF1-AS1 | 0,129619669 | 0,399414488 | 0,286 | 0,171 | 1 | Ciliated | 1,318972502 | 0,75816592 | 1,67251462 |
| CRB32 | 0,129910983 | 0,262200758 | 0,393 | 0,268 | 1 | Ciliated | 1,199306791 | 0,833815006 | 1,46641791 |
| NOL72 | 0,130289233 | 0,328316064 | 0,75 | 0,61 | 1 | Ciliated | 1,255547025 | 0,796465589 | 1,229508197 |
| TACSTD22 | 0,130411349 | 0,449305484 | 0,893 | 0,683 | 1 | Ciliated | 1,365382801 | 0,73239534 | 1,307467057 |
| WDR82 | 0,13071144 | -0,283184969 | 0,143 | 0,341 | 1 | Ciliated | 0,82177482 | 1,216878365 | 0,419354839 |
| ANKRD36 | 0,130876284 | -0,262117711 | 0,143 | 0,317 | 1 | Ciliated | 0,833863005 | 1,199237757 | 0,451104101 |
| SENP61 | 0,130918078 | -0,416417211 | 0,357 | 0,561 | 1 | Ciliated | 0,749283086 | 1,334609067 | 0,636363636 |
| METAP21 | 0,130918078 | -0,320763599 | 0,357 | 0,561 | 1 | Ciliated | 0,800645994 | 1,248991448 | 0,636363636 |
| SMAP11 | 0,131352808 | 0,430228588 | 0,429 | 0,341 | 1 | Ciliated | 1,347447056 | 0,742144187 | 1,258064516 |
| PPCS1 | 0,131547132 | 0,451401766 | 0,5 | 0,439 | 1 | Ciliated | 1,367368188 | 0,731331919 | 1,138952164 |
| DEUP1 | 0,131651641 | -0,378803828 | 0,214 | 0,415 | 1 | Ciliated | 0,769074984 | 1,30026333 | 0,515662651 |
| AC008969.1 | 0,131839052 | 0,257854592 | 0,214 | 0,098 | 1 | Ciliated | 1,195699277 | 0,836330689 | 2,183673469 |

| Supplementary Table 4 - clusermarker | | | | | | | | | |
|---|---|---|---|---|---|---|---|---|---|
| | **p_val** | **avg_log2FC** | **pct.1** | **pct.2** | **p_val_adj** | **celltype** | **Foldchange_UP** | **Foldchange_DOWN** | **Ratio_pct1_pct2** |
| TOB11 | 0,132054516 | -0,59686723 | 0,536 | 0,659 | 1 | Ciliated | 0,661188148 | 1,512428803 | 0,813353566 |
| PARP8 | 0,132272926 | -0,432290153 | 0,286 | 0,488 | 1 | Ciliated | 0,741084444 | 1,349373891 | 0,586065574 |
| PATJ | 0,132295757 | -0,254409753 | 0,286 | 0,537 | 1 | Ciliated | 0,838330049 | 1,192847616 | 0,532588454 |
| SLC38A21 | 0,132304354 | -0,430280921 | 0,464 | 0,634 | 1 | Ciliated | 0,742117266 | 1,347495935 | 0,731861199 |
| RABL61 | 0,132336013 | 0,275146778 | 0,643 | 0,537 | 1 | Ciliated | 1,210117198 | 0,826366241 | 1,197392924 |
| C4orf19 | 0,132448842 | -0,291688756 | 0,107 | 0,268 | 1 | Ciliated | 0,816945218 | 1,224072285 | 0,399253731 |
| RFX7 | 0,132448842 | -0,270668888 | 0,107 | 0,268 | 1 | Ciliated | 0,828935131 | 1,206367016 | 0,399253731 |
| FAM234A | 0,132448842 | -0,379878282 | 0,107 | 0,268 | 1 | Ciliated | 0,768502425 | 1,301232068 | 0,399253731 |
| UQCRH1 | 0,132519784 | 0,297630347 | 0,821 | 0,683 | 1 | Ciliated | 1,229123896 | 0,813587632 | 1,20204978 |
| RFXANK | 0,133060248 | 0,469887088 | 0,393 | 0,317 | 1 | Ciliated | 1,385001067 | 0,722021104 | 1,239747634 |
| C1orf141 | 0,133494712 | 0,334306135 | 0,536 | 0,463 | 1 | Ciliated | 1,260770894 | 0,793165519 | 1,157667387 |
| DANCR2 | 0,133516761 | 0,4258139 | 0,571 | 0,512 | 1 | Ciliated | 1,343330132 | 0,744418647 | 1,115234375 |
| CCDC187 | 0,133951665 | 0,377539298 | 0,643 | 0,512 | 1 | Ciliated | 1,299124141 | 0,769749378 | 1,255859375 |
| CFAP2981 | 0,134107793 | 0,325465579 | 1 | 0,805 | 1 | Ciliated | 1,253068757 | 0,798040805 | 1,242236025 |
| RB1CC1 | 0,134413515 | -0,306115671 | 0,571 | 0,756 | 1 | Ciliated | 0,808816497 | 1,236374386 | 0,755291005 |
| CLTC | 0,134587367 | -0,316041155 | 0,5 | 0,732 | 1 | Ciliated | 0,803271082 | 1,244909747 | 0,683060109 |
| RHOT1 | 0,135045346 | -0,31039429 | 0,143 | 0,317 | 1 | Ciliated | 0,806421333 | 1,24004656 | 0,451104101 |
| PRPS2 | 0,135045346 | -0,498402443 | 0,143 | 0,317 | 1 | Ciliated | 0,707890224 | 1,412648411 | 0,451104101 |
| SDF42 | 0,13547163 | 0,343784396 | 0,571 | 0,537 | 1 | Ciliated | 1,269081214 | 0,787971636 | 1,063314711 |
| SNRPA1 | 0,135481345 | 0,280621221 | 0,5 | 0,415 | 1 | Ciliated | 1,214717826 | 0,823236458 | 1,204819277 |
| TOMM221 | 0,135614797 | 0,253894568 | 0,607 | 0,537 | 1 | Ciliated | 1,192421727 | 0,838629469 | 1,130353818 |
| USP11 | 0,136039404 | 0,391462609 | 0,536 | 0,439 | 1 | Ciliated | 1,311722559 | 0,762356333 | 1,22095672 |
| DLG52 | 0,136123118 | -0,27488173 | 0,179 | 0,341 | 1 | Ciliated | 0,826518072 | 1,2098949 | 0,524926686 |
| DYNC2I21 | 0,136728507 | 0,454423808 | 0,786 | 0,683 | 1 | Ciliated | 1,370235443 | 0,729801586 | 1,150805271 |
| LGALS9B | 0,137255184 | -0,291330753 | 0,071 | 0,22 | 1 | Ciliated | 0,817147967 | 1,223768571 | 0,322727273 |
| CDK7 | 0,137569599 | -0,314405362 | 0,214 | 0,39 | 1 | Ciliated | 0,804182383 | 1,243499013 | 0,548717949 |
| RPSA1 | 0,137579916 | -0,366977039 | 0,929 | 0,951 | 1 | Ciliated | 0,775405547 | 1,289647726 | 0,976866456 |
| TBC1D9 | 0,137906161 | 0,29072027 | 0,179 | 0,073 | 1 | Ciliated | 1,223250837 | 0,81749382 | 2,452054795 |
| LINC01697 | 0,137906161 | 0,308834989 | 0,179 | 0,073 | 1 | Ciliated | 1,238707011 | 0,807293405 | 2,452054795 |
| FAAP202 | 0,138858807 | 0,621551737 | 0,607 | 0,512 | 1 | Ciliated | 1,538529106 | 0,649971454 | 1,185546875 |
| RPL381 | 0,138935312 | -0,308570302 | 0,964 | 0,829 | 1 | Ciliated | 0,80744153 | 1,238479769 | 1,162846803 |

| Supplementary Table 4 - clusermarker | | | | | | | | | |
|---|---|---|---|---|---|---|---|---|---|
| | **p_val** | **avg_log2FC** | **pct.1** | **pct.2** | **p_val_adj** | **celltype** | **Foldchange_UP** | **Foldchange_DOWN** | **Ratio_pct1_pct2** |
| AL138963.42 | 0,138966136 | 0,29072637 | 0,357 | 0,195 | 1 | Ciliated | 1,223256009 | 0,817490364 | 1,830769231 |
| NR2F61 | 0,139368164 | 0,405068857 | 0,571 | 0,463 | 1 | Ciliated | 1,324152108 | 0,755200248 | 1,233261339 |
| VPS36 | 0,139377623 | -0,402742495 | 0,321 | 0,512 | 1 | Ciliated | 0,756418999 | 1,322018618 | 0,626953125 |
| DNAJC81 | 0,139538643 | 0,448307801 | 0,536 | 0,439 | 1 | Ciliated | 1,364438908 | 0,732901996 | 1,22095672 |
| SYAP1 | 0,13956081 | -0,345001288 | 0,786 | 0,829 | 1 | Ciliated | 0,787307274 | 1,270152116 | 0,948130277 |
| XRN21 | 0,139992422 | 0,338700572 | 0,643 | 0,61 | 1 | Ciliated | 1,264617047 | 0,790753219 | 1,054098361 |
| CREB12 | 0,14107426 | -0,34517907 | 0,179 | 0,366 | 1 | Ciliated | 0,78721026 | 1,270308646 | 0,489071038 |
| RHOD1 | 0,14107426 | 0,456467112 | 0,357 | 0,244 | 1 | Ciliated | 1,372177496 | 0,728768693 | 1,463114754 |
| SERPINB12 | 0,141211216 | 0,370041951 | 0,821 | 0,707 | 1 | Ciliated | 1,292390411 | 0,773759997 | 1,161244696 |
| LY751 | 0,141370502 | -0,368675936 | 0,321 | 0,488 | 1 | Ciliated | 0,774492978 | 1,291167291 | 0,657786885 |
| INPP1 | 0,141635322 | -0,363983503 | 0,107 | 0,268 | 1 | Ciliated | 0,777016154 | 1,286974531 | 0,399253731 |
| FHOD31 | 0,141762367 | -0,261592714 | 0,036 | 0,146 | 1 | Ciliated | 0,834166504 | 1,198801433 | 0,246575342 |
| CKLF | 0,142058296 | 0,358135412 | 0,5 | 0,439 | 1 | Ciliated | 1,281768226 | 0,78017225 | 1,138952164 |
| TJP11 | 0,142059348 | -0,284742625 | 0,393 | 0,634 | 1 | Ciliated | 0,820888041 | 1,21819292 | 0,619873817 |
| SLC44A41 | 0,142233412 | 0,27777054 | 0,964 | 0,829 | 1 | Ciliated | 1,212319985 | 0,824864733 | 1,162846803 |
| GSTA3 | 0,142383772 | 0,339517119 | 0,214 | 0,098 | 1 | Ciliated | 1,265333006 | 0,790305789 | 2,183673469 |
| MICOS101 | 0,142412191 | 0,30385763 | 0,964 | 0,878 | 1 | Ciliated | 1,234440782 | 0,810083412 | 1,097949886 |
| WNK1 | 0,142427189 | -0,425172757 | 0,464 | 0,634 | 1 | Ciliated | 0,744749546 | 1,34273328 | 0,731861199 |
| INTS61 | 0,142427189 | -0,390778737 | 0,464 | 0,634 | 1 | Ciliated | 0,762717794 | 1,311100918 | 0,731861199 |
| DHX36 | 0,142633554 | -0,414581957 | 0,286 | 0,561 | 1 | Ciliated | 0,750236856 | 1,332912389 | 0,509803922 |
| MRPL45 | 0,143026281 | 0,510578262 | 0,5 | 0,39 | 1 | Ciliated | 1,424621099 | 0,701941029 | 1,282051282 |
| ANKMY2 | 0,14324126 | 0,313004807 | 0,321 | 0,22 | 1 | Ciliated | 1,242292421 | 0,804963456 | 1,459090909 |
| SLC16A1-AS1 | 0,143391525 | 0,314752257 | 0,107 | 0,024 | 1 | Ciliated | 1,243798047 | 0,803989042 | 4,458333333 |
| CLIC5 | 0,143391525 | 0,264586804 | 0,107 | 0,024 | 1 | Ciliated | 1,201291943 | 0,832437115 | 4,458333333 |
| SMIM30 | 0,143628621 | -0,465711729 | 0,143 | 0,293 | 1 | Ciliated | 0,72411376 | 1,380998477 | 0,488054608 |
| TAX1BP11 | 0,14417425 | -0,335848376 | 1 | 0,902 | 1 | Ciliated | 0,792318077 | 1,262119379 | 1,10864745 |
| SUMO21 | 0,14417425 | 0,271074533 | 1 | 0,902 | 1 | Ciliated | 1,20670626 | 0,82870209 | 1,10864745 |
| EIF4B2 | 0,144569672 | -0,271306432 | 0,714 | 0,78 | 1 | Ciliated | 0,828568896 | 1,206900241 | 0,915384615 |
| TGIF1 | 0,144671648 | -0,403300383 | 0,393 | 0,585 | 1 | Ciliated | 0,75612655 | 1,32252994 | 0,671794872 |
| TIMM101 | 0,145091851 | 0,324371434 | 0,464 | 0,39 | 1 | Ciliated | 1,252118785 | 0,798646272 | 1,18974359 |
| VPS292 | 0,145122176 | 0,376852487 | 0,679 | 0,585 | 1 | Ciliated | 1,298505826 | 0,770115913 | 1,160683761 |

| Supplementary Table 4 - clusermarker | | | | | | | | | |
|---|---|---|---|---|---|---|---|---|---|
| | **p_val** | **avg_log2FC** | **pct.1** | **pct.2** | **p_val_adj** | **celltype** | **Foldchange_UP** | **Foldchange_DOWN** | **Ratio_pct1_pct2** |
| KHDC4 | 0,145240498 | -0,286388147 | 0,179 | 0,366 | 1 | Ciliated | 0,819952279 | 1,21958317 | 0,489071038 |
| MT-CO11 | 0,145587557 | 0,314757071 | 0,964 | 0,829 | 1 | Ciliated | 1,243802197 | 0,803986359 | 1,162846803 |
| LAMP1 | 0,145701793 | 0,428242923 | 0,536 | 0,415 | 1 | Ciliated | 1,345593763 | 0,743166346 | 1,291566265 |
| C17orf97 | 0,145828383 | 0,475537511 | 0,607 | 0,512 | 1 | Ciliated | 1,390436164 | 0,719198785 | 1,185546875 |
| SNRNP2001 | 0,145930457 | 0,361483415 | 0,607 | 0,537 | 1 | Ciliated | 1,284746227 | 0,778363835 | 1,130353818 |
| SRGAP3-AS2 | 0,146049081 | -0,799939223 | 0,643 | 0,756 | 1 | Ciliated | 0,574373374 | 1,74102778 | 0,850529101 |
| DUSP11 | 0,146245903 | -0,45156458 | 0,857 | 0,78 | 1 | Ciliated | 0,731249389 | 1,367522509 | 1,098717949 |
| GIPC11 | 0,146316661 | 0,4036583 | 0,5 | 0,415 | 1 | Ciliated | 1,322858086 | 0,755938986 | 1,204819277 |
| SNHG10 | 0,146321607 | -0,55940192 | 0,25 | 0,463 | 1 | Ciliated | 0,678583417 | 1,473658175 | 0,539956803 |
| C16orf46 | 0,146321607 | 0,379920917 | 0,429 | 0,341 | 1 | Ciliated | 1,301270523 | 0,768479715 | 1,258064516 |
| SPECC1L1 | 0,146408429 | -0,264598369 | 0,107 | 0,268 | 1 | Ciliated | 0,832430442 | 1,201301573 | 0,399253731 |
| DNAAF8 | 0,14674779 | -0,353544512 | 0,321 | 0,512 | 1 | Ciliated | 0,782658844 | 1,277695905 | 0,626953125 |
| MAU2 | 0,146834801 | -0,42066922 | 0,107 | 0,244 | 1 | Ciliated | 0,747077999 | 1,33854832 | 0,43852459 |
| SCGB2A11 | 0,146863716 | 0,386823376 | 0,643 | 0,61 | 1 | Ciliated | 1,307511264 | 0,764811767 | 1,054098361 |
| RPL42 | 0,147153912 | -0,292186651 | 0,821 | 0,902 | 1 | Ciliated | 0,816663327 | 1,224494803 | 0,910199557 |
| MLEC1 | 0,147515398 | 0,330642165 | 0,643 | 0,561 | 1 | Ciliated | 1,257573014 | 0,795182458 | 1,146167558 |
| IDI1 | 0,147604653 | -0,290913528 | 0,357 | 0,585 | 1 | Ciliated | 0,817384319 | 1,22341471 | 0,61025641 |
| TCEAL41 | 0,147604653 | -0,384659095 | 0,357 | 0,585 | 1 | Ciliated | 0,765959971 | 1,305551252 | 0,61025641 |
| CPSF1 | 0,147654271 | 0,286524882 | 0,286 | 0,171 | 1 | Ciliated | 1,219698765 | 0,81987457 | 1,67251462 |
| ADRB1 | 0,147654271 | 0,315058466 | 0,286 | 0,171 | 1 | Ciliated | 1,244062068 | 0,803818415 | 1,67251462 |
| R3HDM41 | 0,147679714 | 0,322154422 | 0,357 | 0,268 | 1 | Ciliated | 1,250196113 | 0,799874508 | 1,332089552 |
| UACA1 | 0,147706643 | -0,323876839 | 0,429 | 0,659 | 1 | Ciliated | 0,798920116 | 1,251689599 | 0,650986343 |
| CFAP20 | 0,147706643 | -0,273243528 | 0,429 | 0,659 | 1 | Ciliated | 0,827457129 | 1,208521826 | 0,650986343 |
| BRD9 | 0,148169241 | -0,360443398 | 0,143 | 0,317 | 1 | Ciliated | 0,778925148 | 1,283820407 | 0,451104101 |
| APMAP | 0,148169241 | -0,253128052 | 0,143 | 0,317 | 1 | Ciliated | 0,839075158 | 1,191788352 | 0,451104101 |
| C3orf14 | 0,148194563 | -0,443203636 | 0,036 | 0,146 | 1 | Ciliated | 0,73549955 | 1,359620138 | 0,246575342 |
| CXCL6 | 0,148194563 | -0,285189527 | 0,036 | 0,146 | 1 | Ciliated | 0,820633795 | 1,218570337 | 0,246575342 |
| MOSPD3 | 0,148887775 | 0,395508453 | 0,464 | 0,39 | 1 | Ciliated | 1,315406271 | 0,760221402 | 1,18974359 |
| TP53RK | 0,148887775 | 0,545439476 | 0,464 | 0,39 | 1 | Ciliated | 1,459464861 | 0,685182649 | 1,18974359 |
| EAPP | 0,149187326 | -0,615324239 | 0,393 | 0,634 | 1 | Ciliated | 0,652783167 | 1,531902247 | 0,619873817 |
| PHIP | 0,149291417 | -0,401007168 | 0,536 | 0,634 | 1 | Ciliated | 0,757329395 | 1,320429402 | 0,845425868 |

| Supplementary Table 4 - clusermarker | | | | | | | | | |
|---|---|---|---|---|---|---|---|---|---|
| | **p_val** | **avg_log2FC** | **pct.1** | **pct.2** | **p_val_adj** | **celltype** | **Foldchange_UP** | **Foldchange_DOWN** | **Ratio_pct1_pct2** |
| SLC2A1 | 0,149416405 | 0,396470437 | 0,5 | 0,439 | 1 | Ciliated | 1,316283671 | 0,759714659 | 1,138952164 |
| WASF21 | 0,149578893 | 0,389295773 | 0,679 | 0,683 | 1 | Ciliated | 1,309753913 | 0,763502204 | 0,994143485 |
| RIN2 | 0,14978974 | -0,313290405 | 0,179 | 0,39 | 1 | Ciliated | 0,804804119 | 1,242538372 | 0,458974359 |
| CCDC181 | 0,150072103 | 0,504943274 | 0,5 | 0,415 | 1 | Ciliated | 1,419067558 | 0,704688085 | 1,204819277 |
| LAMA5 | 0,150260367 | 0,476777345 | 0,429 | 0,341 | 1 | Ciliated | 1,391631601 | 0,71858098 | 1,258064516 |
| SMKR1 | 0,150448238 | 0,285867129 | 0,321 | 0,195 | 1 | Ciliated | 1,219142807 | 0,820248452 | 1,646153846 |
| GEMIN5 | 0,150533128 | 0,262565186 | 0,143 | 0,049 | 1 | Ciliated | 1,199609777 | 0,83360441 | 2,918367347 |
| HSDL2 | 0,150542915 | -0,2969478 | 0,286 | 0,537 | 1 | Ciliated | 0,813972636 | 1,228542528 | 0,532588454 |
| CERKL | 0,150813947 | -0,301983959 | 0,286 | 0,512 | 1 | Ciliated | 0,811136174 | 1,232838618 | 0,55859375 |
| NOP58 | 0,150869091 | -0,4306753 | 0,286 | 0,488 | 1 | Ciliated | 0,741914427 | 1,347864341 | 0,586065574 |
| TIMM231 | 0,150869091 | 0,25048809 | 0,464 | 0,366 | 1 | Ciliated | 1,189609514 | 0,840611973 | 1,267759563 |
| RPS141 | 0,151045543 | -0,347853499 | 1 | 1 | 1 | Ciliated | 0,785752303 | 1,272665694 | 1 |
| MARCKS2 | 0,151270287 | -0,51483833 | 0,607 | 0,78 | 1 | Ciliated | 0,699871357 | 1,428834015 | 0,778205128 |
| RFC1 | 0,151304237 | 0,338398917 | 0,536 | 0,463 | 1 | Ciliated | 1,264352654 | 0,790918576 | 1,157667387 |
| OSBPL8 | 0,151304237 | 0,325118359 | 0,536 | 0,463 | 1 | Ciliated | 1,252767212 | 0,798232896 | 1,157667387 |
| ADGRG1 | 0,151304237 | -0,413636284 | 0,393 | 0,561 | 1 | Ciliated | 0,75072879 | 1,332038964 | 0,700534759 |
| ADRM12 | 0,151308517 | 0,403350026 | 0,571 | 0,561 | 1 | Ciliated | 1,322575449 | 0,756100532 | 1,017825312 |
| MLF21 | 0,152184271 | -0,588525098 | 0,536 | 0,683 | 1 | Ciliated | 0,665022428 | 1,503708684 | 0,78477306 |
| UQCR102 | 0,152477094 | -0,398110851 | 0,893 | 0,878 | 1 | Ciliated | 0,758851317 | 1,317781201 | 1,017084282 |
| HMGB21 | 0,152494752 | 0,365426204 | 0,571 | 0,488 | 1 | Ciliated | 1,288262154 | 0,776239523 | 1,170081967 |
| MISP3 | 0,152639574 | 0,376562619 | 0,393 | 0,317 | 1 | Ciliated | 1,298244955 | 0,770270661 | 1,239747634 |
| KRT181 | 0,152754675 | 0,25694026 | 0,964 | 0,927 | 1 | Ciliated | 1,194941723 | 0,836860895 | 1,0399137 |
| PSMA3 | 0,153062638 | -0,432914666 | 0,429 | 0,634 | 1 | Ciliated | 0,740763713 | 1,349958134 | 0,676656151 |
| TMEM231 | 0,153751309 | 0,456216067 | 0,821 | 0,732 | 1 | Ciliated | 1,371938743 | 0,728895517 | 1,121584699 |
| UNC119B | 0,154231505 | 0,302034987 | 0,536 | 0,439 | 1 | Ciliated | 1,232882224 | 0,811107485 | 1,22095672 |
| KCNE3 | 0,154412407 | 0,312113068 | 0,464 | 0,415 | 1 | Ciliated | 1,24152479 | 0,805461162 | 1,118072289 |
| MIDN | 0,154826679 | -0,326479737 | 0,679 | 0,732 | 1 | Ciliated | 0,797480011 | 1,253949925 | 0,927595628 |
| GPR137B | 0,154847352 | -0,315609597 | 0,036 | 0,146 | 1 | Ciliated | 0,803511403 | 1,244537409 | 0,246575342 |
| C11orf80 | 0,154847352 | -0,30278066 | 0,036 | 0,146 | 1 | Ciliated | 0,810688363 | 1,233519618 | 0,246575342 |
| CDK5RAP31 | 0,155074045 | -0,369925324 | 0,357 | 0,585 | 1 | Ciliated | 0,77382255 | 1,292285938 | 0,61025641 |
| ZMAT3 | 0,155542954 | -0,26313405 | 0,214 | 0,415 | 1 | Ciliated | 0,833275779 | 1,200082885 | 0,515662651 |

| Supplementary Table 4 - clusermarker | | | | | | | | | |
|---|---|---|---|---|---|---|---|---|---|
| | **p_val** | **avg_log2FC** | **pct.1** | **pct.2** | **p_val_adj** | **celltype** | **Foldchange_UP** | **Foldchange_DOWN** | **Ratio_pct1_pct2** |
| GCHFR2 | 0,155736471 | 0,350692027 | 0,536 | 0,488 | 1 | Ciliated | 1,275172151 | 0,784207841 | 1,098360656 |
| PLEKHG7 | 0,156241376 | -0,259155117 | 0,393 | 0,61 | 1 | Ciliated | 0,835577113 | 1,196777633 | 0,644262295 |
| IGF2BP3 | 0,156280037 | 0,587704828 | 0,179 | 0,073 | 1 | Ciliated | 1,502853966 | 0,665400646 | 2,452054795 |
| CYB561D2 | 0,156672646 | 0,321301512 | 0,321 | 0,22 | 1 | Ciliated | 1,249457226 | 0,800347527 | 1,459090909 |
| COPS61 | 0,156780627 | 0,368819274 | 0,607 | 0,512 | 1 | Ciliated | 1,29129558 | 0,774416032 | 1,185546875 |
| UBTD1 | 0,157061839 | 0,494952765 | 0,5 | 0,439 | 1 | Ciliated | 1,409274614 | 0,70958491 | 1,138952164 |
| PTPN23 | 0,157074232 | 0,283490652 | 0,25 | 0,146 | 1 | Ciliated | 1,217136229 | 0,821600718 | 1,712328767 |
| PROSER1 | 0,157074232 | 0,339756381 | 0,25 | 0,146 | 1 | Ciliated | 1,265542871 | 0,790174733 | 1,712328767 |
| FKBP15 | 0,157703441 | -0,35009391 | 0,143 | 0,293 | 1 | Ciliated | 0,784533028 | 1,274643596 | 0,488054608 |
| TRPC4AP | 0,157703441 | -0,419312048 | 0,143 | 0,293 | 1 | Ciliated | 0,74778112 | 1,337289713 | 0,488054608 |
| ERBB31 | 0,158082229 | -0,485815035 | 0,321 | 0,585 | 1 | Ciliated | 0,714093537 | 1,40037677 | 0,548717949 |
| ZFYVE16 | 0,158379725 | -0,291000149 | 0,25 | 0,463 | 1 | Ciliated | 0,817335244 | 1,223488167 | 0,539956803 |
| NDUFA3 | 0,158614075 | -0,461042854 | 0,75 | 0,756 | 1 | Ciliated | 0,726460946 | 1,37653649 | 0,992063492 |
| QRICH1 | 0,15873896 | 0,368241402 | 0,429 | 0,366 | 1 | Ciliated | 1,290778455 | 0,774726287 | 1,172131148 |
| IL13RA1 | 0,15873896 | -0,577858757 | 0,286 | 0,463 | 1 | Ciliated | 0,669957388 | 1,492632244 | 0,617710583 |
| MRPL121 | 0,158914752 | 0,270875566 | 0,536 | 0,463 | 1 | Ciliated | 1,20653985 | 0,828816388 | 1,157667387 |
| CFLAR1 | 0,159457167 | -0,413228262 | 0,643 | 0,732 | 1 | Ciliated | 0,750941141 | 1,331662291 | 0,878415301 |
| ACBD3-AS1 | 0,159781953 | 0,3528642 | 0,464 | 0,439 | 1 | Ciliated | 1,277093542 | 0,783027999 | 1,056947608 |
| EIF2S31 | 0,159781953 | -0,354545624 | 0,321 | 0,537 | 1 | Ciliated | 0,782115932 | 1,278582829 | 0,597765363 |
| CCDC25 | 0,16003469 | 0,334907872 | 0,5 | 0,463 | 1 | Ciliated | 1,261296862 | 0,792834764 | 1,079913607 |
| NFX1 | 0,160057417 | -0,412875471 | 0,357 | 0,634 | 1 | Ciliated | 0,751124795 | 1,331336692 | 0,563091483 |
| NFIA1 | 0,160511502 | -0,376994701 | 0,5 | 0,634 | 1 | Ciliated | 0,770040003 | 1,298633833 | 0,788643533 |
| FILIP1 | 0,160605622 | -0,448999622 | 0,464 | 0,634 | 1 | Ciliated | 0,732550629 | 1,36509336 | 0,731861199 |
| PIGU | 0,160686213 | 0,423912342 | 0,357 | 0,268 | 1 | Ciliated | 1,341560709 | 0,745400483 | 1,332089552 |
| PSME12 | 0,160971285 | -0,368078792 | 0,679 | 0,829 | 1 | Ciliated | 0,774813614 | 1,290632976 | 0,819059107 |
| NACC1 | 0,16135579 | 0,267570885 | 0,321 | 0,22 | 1 | Ciliated | 1,203779276 | 0,830717076 | 1,459090909 |
| SNX13 | 0,162342723 | -0,346044299 | 0,179 | 0,341 | 1 | Ciliated | 0,786738287 | 1,271070718 | 0,524926686 |
| BCL9 | 0,162394153 | 0,277728546 | 0,286 | 0,171 | 1 | Ciliated | 1,212284697 | 0,824888743 | 1,67251462 |
| RSL1D11 | 0,162561969 | 0,325060863 | 0,429 | 0,341 | 1 | Ciliated | 1,252717286 | 0,798264709 | 1,258064516 |
| MED13 | 0,162638683 | -0,297723507 | 0,214 | 0,39 | 1 | Ciliated | 0,813535097 | 1,229203268 | 0,548717949 |
| GOLM22 | 0,162688369 | 0,262086991 | 0,643 | 0,634 | 1 | Ciliated | 1,199212221 | 0,833880761 | 1,014195584 |

| Supplementary Table 4 - clusermarker | | | | | | | | | |
|---|---|---|---|---|---|---|---|---|---|
| | **p_val** | **avg_log2FC** | **pct.1** | **pct.2** | **p_val_adj** | **celltype** | **Foldchange_UP** | **Foldchange_DOWN** | **Ratio_pct1_pct2** |
| AGPAT5 | 0,162846127 | 0,329863085 | 0,357 | 0,244 | 1 | Ciliated | 1,256894087 | 0,795611986 | 1,463114754 |
| PURA1 | 0,163994228 | -0,410897033 | 0,357 | 0,561 | 1 | Ciliated | 0,752155556 | 1,329512216 | 0,636363636 |
| CYTH21 | 0,16412534 | 0,347365681 | 0,679 | 0,683 | 1 | Ciliated | 1,272235441 | 0,786018034 | 0,994143485 |
| VAMP31 | 0,164198259 | -0,297544962 | 0,214 | 0,415 | 1 | Ciliated | 0,813635785 | 1,229051153 | 0,515662651 |
| RHBDD2 | 0,165003511 | 0,357722714 | 0,643 | 0,683 | 1 | Ciliated | 1,281401615 | 0,780395458 | 0,941434846 |
| AL354766.2 | 0,165244969 | 0,383106842 | 0,143 | 0,049 | 1 | Ciliated | 1,304147314 | 0,766784542 | 2,918367347 |
| EML6 | 0,165400161 | 0,251651273 | 0,214 | 0,098 | 1 | Ciliated | 1,190569031 | 0,839934497 | 2,183673469 |
| SLC34A22 | 0,166016923 | 0,324017269 | 0,643 | 0,561 | 1 | Ciliated | 1,251811442 | 0,798842354 | 1,146167558 |
| RALBP11 | 0,166491445 | 0,539959264 | 0,643 | 0,634 | 1 | Ciliated | 1,453931464 | 0,687790329 | 1,014195584 |
| NDUFB82 | 0,166678121 | 0,295425232 | 0,964 | 0,78 | 1 | Ciliated | 1,227246653 | 0,814832127 | 1,235897436 |
| OXA1L | 0,166826993 | -0,392803672 | 0,25 | 0,463 | 1 | Ciliated | 0,761648011 | 1,312942443 | 0,539956803 |
| MAGED21 | 0,166938419 | -0,269372686 | 0,571 | 0,78 | 1 | Ciliated | 0,82968023 | 1,205283631 | 0,732051282 |
| MRFAP1L1 | 0,167130194 | -0,444941846 | 0,286 | 0,488 | 1 | Ciliated | 0,734613928 | 1,361259244 | 0,586065574 |
| AL357093.2 | 0,168991356 | -0,510549287 | 0,893 | 0,902 | 1 | Ciliated | 0,701955127 | 1,424592487 | 0,990022173 |
| NET11 | 0,169918171 | -0,344890074 | 0,5 | 0,659 | 1 | Ciliated | 0,787367968 | 1,270054207 | 0,758725341 |
| FLACC1 | 0,170868811 | -0,27228239 | 0,357 | 0,585 | 1 | Ciliated | 0,828008573 | 1,207716964 | 0,61025641 |
| ADI12 | 0,171394135 | 0,293164643 | 0,464 | 0,366 | 1 | Ciliated | 1,22532516 | 0,816109905 | 1,267759563 |
| SGSM3 | 0,171484391 | 0,435889127 | 0,536 | 0,488 | 1 | Ciliated | 1,352744266 | 0,739238025 | 1,098360656 |
| DRC3 | 0,17216677 | 0,285759089 | 0,893 | 0,805 | 1 | Ciliated | 1,219051511 | 0,820309881 | 1,10931677 |
| BPTF | 0,172324421 | -0,402694945 | 0,393 | 0,683 | 1 | Ciliated | 0,756443931 | 1,321975046 | 0,575402635 |
| BDH1 | 0,17233943 | 0,335380243 | 0,571 | 0,537 | 1 | Ciliated | 1,261709907 | 0,792575214 | 1,063314711 |
| SSBP11 | 0,172381204 | 0,321085122 | 0,75 | 0,707 | 1 | Ciliated | 1,249269833 | 0,80046758 | 1,060820368 |
| CCN11 | 0,173236923 | -0,596998086 | 0,357 | 0,537 | 1 | Ciliated | 0,661128179 | 1,512565991 | 0,664804469 |
| SIK1B | 0,173435645 | 0,287553855 | 0,25 | 0,146 | 1 | Ciliated | 1,220569001 | 0,819290019 | 1,712328767 |
| PRRC2C1 | 0,173553219 | -0,311967063 | 0,607 | 0,683 | 1 | Ciliated | 0,805542682 | 1,241399149 | 0,888726208 |
| LDLRAD1 | 0,17383367 | -0,288921155 | 0,821 | 0,829 | 1 | Ciliated | 0,818513913 | 1,221726331 | 0,990349819 |
| AUP12 | 0,174163984 | 0,331046495 | 0,5 | 0,415 | 1 | Ciliated | 1,257925511 | 0,794959631 | 1,204819277 |
| MTHFD2L | 0,174540227 | 0,47475483 | 0,357 | 0,268 | 1 | Ciliated | 1,389682039 | 0,719589066 | 1,332089552 |
| NTMT1 | 0,174540227 | 0,257737415 | 0,357 | 0,268 | 1 | Ciliated | 1,195602165 | 0,836398619 | 1,332089552 |
| VMP1 | 0,174936707 | -0,495074164 | 0,536 | 0,683 | 1 | Ciliated | 0,709525203 | 1,409393206 | 0,78477306 |
| CLPP | 0,174999596 | 0,300433466 | 0,5 | 0,488 | 1 | Ciliated | 1,231514374 | 0,812008387 | 1,024590164 |

| Supplementary Table 4 - clusermarker | | | | | | | | | |
|---|---|---|---|---|---|---|---|---|---|
| | **p_val** | **avg_log2FC** | **pct.1** | **pct.2** | **p_val_adj** | **celltype** | **Foldchange_UP** | **Foldchange_DOWN** | **Ratio_pct1_pct2** |
| HDHD3 | 0,175739 | -0,337628837 | 0,286 | 0,488 | 1 | Ciliated | 0,791340864 | 1,26367795 | 0,586065574 |
| VAMP51 | 0,176039409 | 0,32348521 | 0,321 | 0,22 | 1 | Ciliated | 1,251349866 | 0,799137018 | 1,459090909 |
| SNX19 | 0,176804013 | -0,278943705 | 0,179 | 0,341 | 1 | Ciliated | 0,824194245 | 1,213306216 | 0,524926686 |
| TRAF3IP1 | 0,176986719 | 0,251177109 | 0,786 | 0,707 | 1 | Ciliated | 1,190177797 | 0,840210599 | 1,111739745 |
| ZDHHC11B | 0,17708411 | -0,302459446 | 0,179 | 0,366 | 1 | Ciliated | 0,810868882 | 1,233245006 | 0,489071038 |
| NFE2L11 | 0,1776109 | -0,304975481 | 0,321 | 0,488 | 1 | Ciliated | 0,809455973 | 1,235397641 | 0,657786885 |
| ENC11 | 0,177854515 | -0,371788351 | 0,214 | 0,415 | 1 | Ciliated | 0,772823917 | 1,293955812 | 0,515662651 |
| NT5C3B | 0,178063965 | 0,315839434 | 0,286 | 0,195 | 1 | Ciliated | 1,244735693 | 0,803383405 | 1,466666667 |
| CFAP157 | 0,178228748 | 0,276513041 | 0,964 | 0,78 | 1 | Ciliated | 1,211263749 | 0,825584024 | 1,235897436 |
| TRIP13 | 0,17836118 | 0,261122334 | 0,571 | 0,512 | 1 | Ciliated | 1,198410637 | 0,834438521 | 1,115234375 |
| PPP1R15A1 | 0,178972177 | -0,410608237 | 0,75 | 0,732 | 1 | Ciliated | 0,752306136 | 1,329246103 | 1,024590164 |
| MOSMO | 0,179163247 | -0,320272196 | 0,107 | 0,244 | 1 | Ciliated | 0,800918753 | 1,248566096 | 0,43852459 |
| SGMS2 | 0,17952749 | -0,32053308 | 0,5 | 0,683 | 1 | Ciliated | 0,800773935 | 1,248791896 | 0,732064422 |
| MT-ND4L1 | 0,179726496 | -0,473832538 | 0,929 | 0,756 | 1 | Ciliated | 0,720049235 | 1,388793921 | 1,228835979 |
| CMTM8 | 0,180277683 | 0,268619477 | 0,429 | 0,366 | 1 | Ciliated | 1,204654535 | 0,830113506 | 1,172131148 |
| ANAPC13 | 0,180572439 | -0,452391847 | 0,357 | 0,561 | 1 | Ciliated | 0,730830198 | 1,368306896 | 0,636363636 |
| RACK11 | 0,180787966 | 0,279319052 | 1 | 0,878 | 1 | Ciliated | 1,213621923 | 0,823979842 | 1,138952164 |
| PEBP11 | 0,180787966 | 0,263166318 | 0,964 | 0,902 | 1 | Ciliated | 1,200109726 | 0,833257141 | 1,068736142 |
| HDAC21 | 0,181777159 | -0,264539252 | 0,321 | 0,561 | 1 | Ciliated | 0,832464553 | 1,201252349 | 0,572192513 |
| ATP2C21 | 0,182683279 | -0,480501742 | 0,607 | 0,756 | 1 | Ciliated | 0,716728316 | 1,395228816 | 0,802910053 |
| TEX26 | 0,182805034 | -0,661224382 | 0,357 | 0,512 | 1 | Ciliated | 0,632341416 | 1,581424172 | 0,697265625 |
| MNT | 0,183447231 | 0,316681525 | 0,286 | 0,195 | 1 | Ciliated | 1,245462449 | 0,802914613 | 1,466666667 |
| SYMPK | 0,183447231 | -0,297241841 | 0,143 | 0,293 | 1 | Ciliated | 0,813806754 | 1,228792948 | 0,488054608 |
| PPP1R32 | 0,183483986 | 0,272487614 | 0,571 | 0,439 | 1 | Ciliated | 1,207888775 | 0,827890797 | 1,300683371 |
| CEP126 | 0,183549965 | -0,492512278 | 0,643 | 0,756 | 1 | Ciliated | 0,710786271 | 1,406892677 | 0,850529101 |
| SCAP | 0,183634313 | 0,306380865 | 0,393 | 0,317 | 1 | Ciliated | 1,236601675 | 0,808667836 | 1,239747634 |
| TES | 0,183641373 | 0,261085114 | 0,5 | 0,39 | 1 | Ciliated | 1,198379719 | 0,83446005 | 1,282051282 |
| PPIA2 | 0,183999541 | 0,29209482 | 0,857 | 0,829 | 1 | Ciliated | 1,224416864 | 0,816715311 | 1,033775633 |
| SMPD2 | 0,184052697 | 0,457902253 | 0,536 | 0,488 | 1 | Ciliated | 1,373543169 | 0,7280441 | 1,098360656 |
| AHCYL2 | 0,184259687 | 0,35804942 | 0,357 | 0,268 | 1 | Ciliated | 1,281691828 | 0,780218753 | 1,332089552 |
| ALS2CL | 0,184338583 | 0,291684663 | 0,429 | 0,317 | 1 | Ciliated | 1,224068812 | 0,816947536 | 1,353312303 |

| Supplementary Table 4 - clusermarker | | | | | | | | | |
|---|---|---|---|---|---|---|---|---|---|
| | **p_val** | **avg_log2FC** | **pct.1** | **pct.2** | **p_val_adj** | **celltype** | **Foldchange_UP** | **Foldchange_DOWN** | **Ratio_pct1_pct2** |
| NSUN3 | 0,184422674 | -0,315473456 | 0,071 | 0,195 | 1 | Ciliated | 0,80358723 | 1,244419973 | 0,364102564 |
| TRIM21 | 0,184422674 | -0,29803706 | 0,071 | 0,195 | 1 | Ciliated | 0,813358304 | 1,22947045 | 0,364102564 |
| CYP4F11 | 0,184422674 | -0,291878932 | 0,071 | 0,195 | 1 | Ciliated | 0,816837535 | 1,224233653 | 0,364102564 |
| MRPS211 | 0,184658563 | -0,620576011 | 0,893 | 0,902 | 1 | Ciliated | 0,650411193 | 1,537488917 | 0,990022173 |
| TMEM50B1 | 0,184746222 | 0,274253496 | 0,571 | 0,537 | 1 | Ciliated | 1,209368156 | 0,826878065 | 1,063314711 |
| BPIFA11 | 0,184887616 | -0,793455789 | 0,071 | 0,171 | 1 | Ciliated | 0,576960402 | 1,733221201 | 0,415204678 |
| PPP2R3B | 0,185030325 | -0,252438913 | 0,107 | 0,244 | 1 | Ciliated | 0,839476059 | 1,191219201 | 0,43852459 |
| THUMPD3 | 0,186042962 | -0,430115585 | 0,214 | 0,39 | 1 | Ciliated | 0,74220232 | 1,347341518 | 0,548717949 |
| ILRUN | 0,186042962 | 0,409436832 | 0,357 | 0,293 | 1 | Ciliated | 1,328167251 | 0,752917224 | 1,218430034 |
| SMARCA22 | 0,186211798 | -0,303221635 | 0,5 | 0,659 | 1 | Ciliated | 0,810440606 | 1,233896714 | 0,758725341 |
| LINC00271 | 0,186530595 | 0,318184102 | 0,464 | 0,39 | 1 | Ciliated | 1,246760283 | 0,802078807 | 1,18974359 |
| AGAP3 | 0,186696371 | -0,265479539 | 0,25 | 0,439 | 1 | Ciliated | 0,831922165 | 1,202035529 | 0,569476082 |
| SPATA6 | 0,186791469 | 0,363698067 | 0,714 | 0,683 | 1 | Ciliated | 1,286719929 | 0,777169901 | 1,045387994 |
| SKA2 | 0,186964783 | 0,289340342 | 0,321 | 0,244 | 1 | Ciliated | 1,222081365 | 0,818276122 | 1,31557377 |
| PNN1 | 0,187242926 | -0,337164503 | 0,357 | 0,512 | 1 | Ciliated | 0,791595599 | 1,263271298 | 0,697265625 |
| MAPRE1 | 0,187417402 | -0,260970942 | 0,214 | 0,415 | 1 | Ciliated | 0,83452609 | 1,198284886 | 0,515662651 |
| KCNQ1OT11 | 0,188390157 | -0,444017529 | 0,393 | 0,585 | 1 | Ciliated | 0,735084737 | 1,360387382 | 0,671794872 |
| NIFK | 0,188418064 | 0,348377094 | 0,393 | 0,317 | 1 | Ciliated | 1,273127664 | 0,785467183 | 1,239747634 |
| WFS1 | 0,188950135 | 0,536646021 | 0,286 | 0,195 | 1 | Ciliated | 1,450596247 | 0,6893717 | 1,466666667 |
| BAMBI | 0,188950135 | -0,291623646 | 0,143 | 0,293 | 1 | Ciliated | 0,816982088 | 1,224017043 | 0,488054608 |
| APPL12 | 0,189267124 | 0,386284916 | 0,357 | 0,268 | 1 | Ciliated | 1,307023351 | 0,765097272 | 1,332089552 |
| RBCK11 | 0,18949148 | 0,458873978 | 0,464 | 0,439 | 1 | Ciliated | 1,374468628 | 0,727553892 | 1,056947608 |
| CHMP1A | 0,191008714 | 0,398234788 | 0,357 | 0,293 | 1 | Ciliated | 1,317894412 | 0,758786129 | 1,218430034 |
| TRIM14 | 0,19103835 | -0,327523426 | 0,107 | 0,244 | 1 | Ciliated | 0,796903298 | 1,254857399 | 0,43852459 |
| BET1L | 0,19103835 | 0,26953365 | 0,25 | 0,146 | 1 | Ciliated | 1,205418115 | 0,829587666 | 1,712328767 |
| CHST61 | 0,191085057 | 0,2865303 | 0,643 | 0,512 | 1 | Ciliated | 1,219703346 | 0,81987149 | 1,255859375 |
| MRPS23 | 0,191112648 | -0,263751451 | 0,321 | 0,488 | 1 | Ciliated | 0,832919255 | 1,20059657 | 0,657786885 |
| UBE2B1 | 0,191150841 | 0,40908873 | 0,571 | 0,512 | 1 | Ciliated | 1,327846822 | 0,753098914 | 1,115234375 |
| MKNK2 | 0,191150841 | 0,320557132 | 0,571 | 0,512 | 1 | Ciliated | 1,248812716 | 0,800760584 | 1,115234375 |
| IER21 | 0,191256505 | -0,595383326 | 0,857 | 0,756 | 1 | Ciliated | 0,661868573 | 1,510873973 | 1,133597884 |
| MARCHF71 | 0,191478649 | -0,288721657 | 0,25 | 0,439 | 1 | Ciliated | 0,818627106 | 1,221557401 | 0,569476082 |

| Supplementary Table 4 - clusermarker | | | | | | | | | |
|---|---|---|---|---|---|---|---|---|---|
| | **p_val** | **avg_log2FC** | **pct.1** | **pct.2** | **p_val_adj** | **celltype** | **Foldchange_UP** | **Foldchange_DOWN** | **Ratio_pct1_pct2** |
| HBEGF1 | 0,191697021 | -0,277209496 | 0,179 | 0,317 | 1 | Ciliated | 0,825185573 | 1,211848622 | 0,56466877 |
| TKFC | 0,192204059 | -0,537658633 | 0,179 | 0,341 | 1 | Ciliated | 0,688888007 | 1,451614762 | 0,524926686 |
| SH3BGRL2 | 0,192338857 | -0,313104853 | 0,214 | 0,415 | 1 | Ciliated | 0,804907636 | 1,242378573 | 0,515662651 |
| KMT2A | 0,192996412 | -0,327352891 | 0,429 | 0,683 | 1 | Ciliated | 0,796997503 | 1,254709076 | 0,628111274 |
| NUCB21 | 0,193095604 | -0,360241602 | 1 | 0,927 | 1 | Ciliated | 0,779034107 | 1,283640845 | 1,078748652 |
| COA6 | 0,193377047 | -0,399735481 | 0,393 | 0,659 | 1 | Ciliated | 0,75799725 | 1,319266001 | 0,596358118 |
| FARSB | 0,193389343 | -0,314285385 | 0,143 | 0,317 | 1 | Ciliated | 0,804249263 | 1,243395606 | 0,451104101 |
| PRELID3B | 0,193941409 | 0,393042176 | 0,429 | 0,39 | 1 | Ciliated | 1,313159514 | 0,761522107 | 1,1 |
| CDH261 | 0,193941409 | 0,33739106 | 0,464 | 0,366 | 1 | Ciliated | 1,263469695 | 0,791471299 | 1,267759563 |
| DUBR | 0,194573799 | -0,281557892 | 0,143 | 0,293 | 1 | Ciliated | 0,822702144 | 1,215506739 | 0,488054608 |
| GET4 | 0,194573799 | -0,365843333 | 0,143 | 0,293 | 1 | Ciliated | 0,77601512 | 1,288634686 | 0,488054608 |
| AP002008.4 | 0,194573799 | -0,275611833 | 0,143 | 0,293 | 1 | Ciliated | 0,826099903 | 1,210507345 | 0,488054608 |
| TNRC6B | 0,19520322 | -0,438628104 | 0,679 | 0,707 | 1 | Ciliated | 0,737835902 | 1,355314911 | 0,96039604 |
| AC092164.1 | 0,19554908 | 0,502919982 | 0,214 | 0,122 | 1 | Ciliated | 1,417078797 | 0,705677061 | 1,754098361 |
| RNF25 | 0,19554908 | 0,301116322 | 0,214 | 0,122 | 1 | Ciliated | 1,232097411 | 0,811624139 | 1,754098361 |
| NUDT81 | 0,19554908 | 0,337575721 | 0,214 | 0,122 | 1 | Ciliated | 1,263631425 | 0,791369999 | 1,754098361 |
| SYNGAP1 | 0,195777002 | 0,479150823 | 0,464 | 0,39 | 1 | Ciliated | 1,393922955 | 0,717399765 | 1,18974359 |
| BRD21 | 0,19583257 | -0,464853912 | 0,857 | 0,78 | 1 | Ciliated | 0,724544442 | 1,380177588 | 1,098717949 |
| LRBA | 0,196776282 | -0,389182784 | 0,357 | 0,585 | 1 | Ciliated | 0,763562003 | 1,30965134 | 0,61025641 |
| RHOBTB1 | 0,197188734 | -0,331971717 | 0,107 | 0,244 | 1 | Ciliated | 0,794449974 | 1,258732497 | 0,43852459 |
| MAPK10 | 0,19737964 | -0,472694406 | 0,464 | 0,659 | 1 | Ciliated | 0,720617501 | 1,387698743 | 0,704097117 |
| TMEM248 | 0,197550487 | -0,323361409 | 0,179 | 0,341 | 1 | Ciliated | 0,799205597 | 1,251242489 | 0,524926686 |
| SOWAHB | 0,197974952 | -0,280830279 | 0,071 | 0,195 | 1 | Ciliated | 0,823117172 | 1,214893861 | 0,364102564 |
| ARL6IP1 | 0,198228838 | -0,403543754 | 0,75 | 0,756 | 1 | Ciliated | 0,755999008 | 1,322753059 | 0,992063492 |
| WDR18 | 0,198260189 | 0,30188754 | 0,393 | 0,317 | 1 | Ciliated | 1,232756227 | 0,811190386 | 1,239747634 |
| TRAP11 | 0,198364089 | 0,413237469 | 0,5 | 0,463 | 1 | Ciliated | 1,33167079 | 0,750936348 | 1,079913607 |
| S100P2 | 0,198455494 | -0,760611602 | 0,607 | 0,683 | 1 | Ciliated | 0,590246054 | 1,694208699 | 0,888726208 |
| HSD11B1L | 0,198701012 | 0,434270022 | 0,429 | 0,39 | 1 | Ciliated | 1,351226963 | 0,740068121 | 1,1 |
| NAA38 | 0,19877951 | -0,508535814 | 0,821 | 0,878 | 1 | Ciliated | 0,702935483 | 1,422605665 | 0,935079727 |
| TBC1D2B | 0,198980713 | -0,257191275 | 0,143 | 0,317 | 1 | Ciliated | 0,836715302 | 1,19514965 | 0,451104101 |
| EFCAB11 | 0,199015805 | -0,340425797 | 0,286 | 0,463 | 1 | Ciliated | 0,789808174 | 1,266130224 | 0,617710583 |

| Supplementary Table 4 - clusermarker | | | | | | | | | |
|---|---|---|---|---|---|---|---|---|---|
| | **p_val** | **avg_log2FC** | **pct.1** | **pct.2** | **p_val_adj** | **celltype** | **Foldchange_UP** | **Foldchange_DOWN** | **Ratio_pct1_pct2** |
| NFKB1 | 0,199063331 | -0,29338925 | 0,25 | 0,463 | 1 | Ciliated | 0,815982858 | 1,225515941 | 0,539956803 |
| PIH1D2 | 0,199352623 | -0,492862924 | 0,536 | 0,707 | 1 | Ciliated | 0,710613537 | 1,407234662 | 0,758132956 |
| FMN1 | 0,199581598 | -0,343868765 | 0,214 | 0,366 | 1 | Ciliated | 0,787925556 | 1,269155432 | 0,584699454 |
| PRKDC | 0,199787674 | -0,421797937 | 0,357 | 0,537 | 1 | Ciliated | 0,746493738 | 1,339595966 | 0,664804469 |
| MPST2 | 0,200319304 | -0,364562778 | 0,143 | 0,293 | 1 | Ciliated | 0,776704227 | 1,287491384 | 0,488054608 |
| FXYD1 | 0,201360103 | 0,415576431 | 0,536 | 0,463 | 1 | Ciliated | 1,333831505 | 0,749719883 | 1,157667387 |
| RTRAF1 | 0,201391978 | 0,504402935 | 0,607 | 0,585 | 1 | Ciliated | 1,418536168 | 0,704952064 | 1,037606838 |
| TRA2B | 0,201854299 | -0,344085165 | 0,393 | 0,585 | 1 | Ciliated | 0,787807379 | 1,269345816 | 0,671794872 |
| MARCHF6 | 0,201854299 | -0,278569749 | 0,393 | 0,585 | 1 | Ciliated | 0,824407909 | 1,212991759 | 0,671794872 |
| CDKN1C1 | 0,201854299 | 0,334963715 | 0,536 | 0,488 | 1 | Ciliated | 1,261345684 | 0,792804076 | 1,098360656 |
| FSD1L | 0,202085819 | 0,472510723 | 0,464 | 0,415 | 1 | Ciliated | 1,387522074 | 0,720709255 | 1,118072289 |
| RALY2 | 0,202168905 | 0,29902739 | 0,536 | 0,512 | 1 | Ciliated | 1,230314702 | 0,812800171 | 1,046875 |
| PRKAB11 | 0,202252075 | 0,310639514 | 0,214 | 0,122 | 1 | Ciliated | 1,240257356 | 0,806284273 | 1,754098361 |
| SPPL3 | 0,202465543 | -0,257145317 | 0,214 | 0,415 | 1 | Ciliated | 0,836741957 | 1,195111578 | 0,515662651 |
| NCALD1 | 0,203005888 | 0,375195734 | 0,5 | 0,463 | 1 | Ciliated | 1,297015512 | 0,771000802 | 1,079913607 |
| MIS12 | 0,203482831 | -0,293757524 | 0,107 | 0,244 | 1 | Ciliated | 0,81577459 | 1,225828816 | 0,43852459 |
| HOOK21 | 0,20354545 | 0,264543996 | 0,464 | 0,366 | 1 | Ciliated | 1,201256299 | 0,832461816 | 1,267759563 |
| ZBTB8OS | 0,203827449 | -0,371457351 | 0,429 | 0,317 | 1 | Ciliated | 0,773001248 | 1,293658972 | 1,353312303 |
| DUSP232 | 0,203827449 | 0,358265404 | 0,429 | 0,317 | 1 | Ciliated | 1,281883723 | 0,780101956 | 1,353312303 |
| RANBP9 | 0,204017138 | -0,251612887 | 0,25 | 0,463 | 1 | Ciliated | 0,839956845 | 1,190537354 | 0,539956803 |
| MRPL42 | 0,204401371 | -0,539258319 | 0,429 | 0,659 | 1 | Ciliated | 0,688124579 | 1,453225231 | 0,650986343 |
| ORAI2 | 0,204487389 | 0,505617711 | 0,5 | 0,439 | 1 | Ciliated | 1,419731105 | 0,704358731 | 1,138952164 |
| MT-ND1 | 0,205041734 | -0,359954509 | 0,964 | 0,756 | 1 | Ciliated | 0,779189149 | 1,283385429 | 1,275132275 |
| EPS15 | 0,20535498 | -0,322697459 | 0,321 | 0,488 | 1 | Ciliated | 0,799573488 | 1,250666781 | 0,657786885 |
| DCXR2 | 0,20535498 | 0,354203922 | 0,464 | 0,39 | 1 | Ciliated | 1,278280033 | 0,782301197 | 1,18974359 |
| UBAC1 | 0,205585285 | 0,341739613 | 0,786 | 0,683 | 1 | Ciliated | 1,267283773 | 0,789089248 | 1,150805271 |
| BTG11 | 0,205585285 | -0,340383921 | 0,714 | 0,732 | 1 | Ciliated | 0,789831099 | 1,266093474 | 0,975409836 |
| SLC25A52 | 0,205722298 | 0,255197264 | 0,964 | 0,854 | 1 | Ciliated | 1,193498922 | 0,837872562 | 1,128805621 |
| MAK | 0,206365259 | -0,340787107 | 0,25 | 0,439 | 1 | Ciliated | 0,789610398 | 1,266447355 | 0,569476082 |
| MFSD14C | 0,20649021 | -0,384626629 | 0,214 | 0,39 | 1 | Ciliated | 0,765977209 | 1,305521873 | 0,548717949 |
| SSR2 | 0,207726298 | 0,262727635 | 0,357 | 0,561 | 1 | Ciliated | 1,199744862 | 0,83351055 | 0,636363636 |

| Supplementary Table 4 - clusermarker | | | | | | | | | |
|---|---|---|---|---|---|---|---|---|---|
| | **p_val** | **avg_log2FC** | **pct.1** | **pct.2** | **p_val_adj** | **celltype** | **Foldchange_UP** | **Foldchange_DOWN** | **Ratio_pct1_pct2** |
| UQCRFS12 | 0,208262954 | 0,253946 | 0,643 | 0,634 | 1 | Ciliated | 1,192464238 | 0,838599573 | 1,014195584 |
| CHMP51 | 0,208850048 | -0,439510353 | 0,786 | 0,829 | 1 | Ciliated | 0,737384833 | 1,356143977 | 0,948130277 |
| DENND2B | 0,209845332 | 0,263898382 | 0,607 | 0,61 | 1 | Ciliated | 1,20071885 | 0,832834431 | 0,995081967 |
| DDR11 | 0,210042312 | 0,292466591 | 0,75 | 0,707 | 1 | Ciliated | 1,224732427 | 0,816504877 | 1,060820368 |
| MRLN | 0,210116138 | -0,775010016 | 0,393 | 0,537 | 1 | Ciliated | 0,584384567 | 1,711201932 | 0,731843575 |
| PPP4R1 | 0,210765732 | 0,26861385 | 0,679 | 0,659 | 1 | Ciliated | 1,204649837 | 0,830116744 | 1,030349014 |
| PAK21 | 0,210951636 | -0,331735026 | 0,464 | 0,659 | 1 | Ciliated | 0,794580324 | 1,258526004 | 0,704097117 |
| SERTAD11 | 0,210951636 | -0,288305542 | 0,464 | 0,659 | 1 | Ciliated | 0,818863256 | 1,221205119 | 0,704097117 |
| SAT21 | 0,211482175 | 0,450815436 | 0,536 | 0,512 | 1 | Ciliated | 1,366812584 | 0,731629202 | 1,046875 |
| VWA3B | 0,211485535 | -0,481226373 | 0,75 | 0,683 | 1 | Ciliated | 0,716368411 | 1,395929782 | 1,098096633 |
| NEK7 | 0,211509557 | -0,409180031 | 0,25 | 0,439 | 1 | Ciliated | 0,753051256 | 1,327930858 | 0,569476082 |
| PITPNM1 | 0,211520151 | 0,511597802 | 0,571 | 0,537 | 1 | Ciliated | 1,425628222 | 0,701445148 | 1,063314711 |
| MRPL501 | 0,211847904 | -0,495364628 | 0,214 | 0,39 | 1 | Ciliated | 0,709382366 | 1,409676993 | 0,548717949 |
| PEX11B | 0,21217994 | 0,371077135 | 0,286 | 0,195 | 1 | Ciliated | 1,293318079 | 0,773204996 | 1,466666667 |
| ZNF385A | 0,21217994 | 0,327269779 | 0,286 | 0,195 | 1 | Ciliated | 1,254636795 | 0,797043418 | 1,466666667 |
| PIGH | 0,21217994 | 0,295597029 | 0,286 | 0,195 | 1 | Ciliated | 1,227392803 | 0,814735102 | 1,466666667 |
| AC010754.1 | 0,21217994 | -0,297268682 | 0,143 | 0,293 | 1 | Ciliated | 0,813791614 | 1,228815809 | 0,488054608 |
| CLUAP1 | 0,213025305 | -0,371214052 | 0,679 | 0,78 | 1 | Ciliated | 0,77313162 | 1,293440825 | 0,870512821 |
| FBXO311 | 0,213257662 | 0,286055164 | 0,286 | 0,171 | 1 | Ciliated | 1,219301715 | 0,820141551 | 1,67251462 |
| AKAP92 | 0,21332554 | -0,344316814 | 0,893 | 0,756 | 1 | Ciliated | 0,787680893 | 1,269549647 | 1,181216931 |
| YIPF61 | 0,213720303 | -0,380325937 | 0,25 | 0,415 | 1 | Ciliated | 0,768264003 | 1,301635891 | 0,602409639 |
| CSPP1 | 0,214138229 | 0,441399068 | 0,643 | 0,659 | 1 | Ciliated | 1,357920546 | 0,736420111 | 0,975720789 |
| NME5 | 0,21416204 | -0,277588609 | 0,786 | 0,902 | 1 | Ciliated | 0,824968759 | 1,212167115 | 0,871396896 |
| SLC16A5 | 0,214192655 | 0,519790315 | 0,393 | 0,366 | 1 | Ciliated | 1,433746849 | 0,697473198 | 1,073770492 |
| CFAP92 | 0,214241272 | -0,268809232 | 0,179 | 0,341 | 1 | Ciliated | 0,83000433 | 1,204812992 | 0,524926686 |
| KHNYN | 0,214241272 | 0,391920462 | 0,321 | 0,244 | 1 | Ciliated | 1,312138913 | 0,76211443 | 1,31557377 |
| TMC52 | 0,214619283 | -0,334501769 | 0,929 | 0,878 | 1 | Ciliated | 0,79305797 | 1,26094187 | 1,05808656 |
| COX171 | 0,215299272 | 0,309818674 | 0,857 | 0,659 | 1 | Ciliated | 1,239551896 | 0,806743149 | 1,300455235 |
| CD821 | 0,215577879 | 0,327207697 | 0,536 | 0,463 | 1 | Ciliated | 1,254582807 | 0,797077717 | 1,157667387 |
| DDX39B1 | 0,215622287 | -0,361297833 | 0,464 | 0,659 | 1 | Ciliated | 0,778463967 | 1,284580973 | 0,704097117 |
| TSC2 | 0,2156632 | 0,283400442 | 0,393 | 0,293 | 1 | Ciliated | 1,217060126 | 0,821652093 | 1,341296928 |

| Supplementary Table 4 - clusermarker | | | | | | | | | |
|---|---|---|---|---|---|---|---|---|---|
| | **p_val** | **avg_log2FC** | **pct.1** | **pct.2** | **p_val_adj** | **celltype** | **Foldchange_UP** | **Foldchange_DOWN** | **Ratio_pct1_pct2** |
| LINC00663 | 0,215853317 | -0,351376707 | 0,036 | 0,122 | 1 | Ciliated | 0,783835757 | 1,275777471 | 0,295081967 |
| FAM169A | 0,21615733 | 0,37107963 | 0,214 | 0,122 | 1 | Ciliated | 1,293320315 | 0,773203659 | 1,754098361 |
| HES6 | 0,216507266 | -0,267747322 | 0,107 | 0,244 | 1 | Ciliated | 0,830615488 | 1,203926503 | 0,43852459 |
| CCNG21 | 0,216507266 | 0,261272447 | 0,107 | 0,244 | 1 | Ciliated | 1,198535338 | 0,834351703 | 0,43852459 |
| RARA | 0,216507266 | -0,332601889 | 0,107 | 0,244 | 1 | Ciliated | 0,794103033 | 1,259282434 | 0,43852459 |
| C14orf119 | 0,216745925 | -0,255551284 | 0,25 | 0,439 | 1 | Ciliated | 0,837666983 | 1,193791829 | 0,569476082 |
| TALDO12 | 0,217039382 | 0,497571383 | 0,821 | 0,756 | 1 | Ciliated | 1,411834894 | 0,708298119 | 1,085978836 |
| TRNP11 | 0,217305345 | 0,420543341 | 0,357 | 0,293 | 1 | Ciliated | 1,338431534 | 0,747143186 | 1,218430034 |
| PRSS231 | 0,217455396 | 0,426943901 | 0,643 | 0,561 | 1 | Ciliated | 1,344382717 | 0,743835805 | 1,146167558 |
| UBN2 | 0,217800599 | -0,253700768 | 0,286 | 0,512 | 1 | Ciliated | 0,838742132 | 1,192261557 | 0,55859375 |
| MECOM | 0,219067933 | -0,313160107 | 0,357 | 0,537 | 1 | Ciliated | 0,80487681 | 1,242426155 | 0,664804469 |
| CCL281 | 0,219133603 | 0,348277788 | 0,429 | 0,366 | 1 | Ciliated | 1,273040033 | 0,785521251 | 1,172131148 |
| SCAF12 | 0,219139038 | 0,417062819 | 0,429 | 0,366 | 1 | Ciliated | 1,33520644 | 0,748947855 | 1,172131148 |
| SNTN | 0,219402048 | -0,345146639 | 1 | 0,951 | 1 | Ciliated | 0,787227957 | 1,27028009 | 1,051524711 |
| AHNAK1 | 0,219899563 | -0,34002897 | 0,714 | 0,756 | 1 | Ciliated | 0,790025447 | 1,265782012 | 0,944444444 |
| ULK2 | 0,220024719 | -0,349313173 | 0,179 | 0,341 | 1 | Ciliated | 0,784957705 | 1,273953989 | 0,524926686 |
| MAEA | 0,220029555 | -0,453618611 | 0,179 | 0,317 | 1 | Ciliated | 0,730209017 | 1,3694709 | 0,56466877 |
| CTNND1 | 0,220366967 | -0,3933267 | 0,464 | 0,659 | 1 | Ciliated | 0,761371936 | 1,313418517 | 0,704097117 |
| GPX21 | 0,220474156 | 0,602184669 | 0,536 | 0,463 | 1 | Ciliated | 1,51801355 | 0,658755648 | 1,157667387 |
| CLCN3 | 0,221101712 | -0,342280553 | 0,393 | 0,61 | 1 | Ciliated | 0,788793434 | 1,267759032 | 0,644262295 |
| C11orf581 | 0,2211365 | -0,409225048 | 0,714 | 0,805 | 1 | Ciliated | 0,753027758 | 1,327972294 | 0,886956522 |
| KIAA2012 | 0,221151844 | 0,278384703 | 0,607 | 0,488 | 1 | Ciliated | 1,212836186 | 0,824513658 | 1,243852459 |
| GTPBP61 | 0,22143053 | 0,250806526 | 0,357 | 0,268 | 1 | Ciliated | 1,189872117 | 0,840426451 | 1,332089552 |
| CLEC2B1 | 0,221912927 | -0,334443032 | 0,071 | 0,171 | 1 | Ciliated | 0,793090258 | 1,260890535 | 0,415204678 |
| CALCOCO21 | 0,222863177 | -0,350277421 | 0,214 | 0,39 | 1 | Ciliated | 0,784433242 | 1,274805741 | 0,548717949 |
| RAB40B | 0,222863177 | -0,325297636 | 0,214 | 0,39 | 1 | Ciliated | 0,79813371 | 1,252922897 | 0,548717949 |
| NUTF22 | 0,223592156 | 0,255625909 | 0,571 | 0,512 | 1 | Ciliated | 1,19385358 | 0,837623655 | 1,115234375 |
| H4C31 | 0,223911069 | -0,455698409 | 0,536 | 0,659 | 1 | Ciliated | 0,729157102 | 1,371446562 | 0,813353566 |
| RPAIN | 0,223915569 | 0,28960724 | 0,643 | 0,585 | 1 | Ciliated | 1,222307471 | 0,818124755 | 1,099145299 |
| SMARCA41 | 0,223915569 | 0,345759885 | 0,607 | 0,61 | 1 | Ciliated | 1,270820162 | 0,786893401 | 0,995081967 |
| PCYT1A | 0,224539785 | -0,273999398 | 0,143 | 0,293 | 1 | Ciliated | 0,827023714 | 1,209155171 | 0,488054608 |

| Supplementary Table 4 - clusermarker | | | | | | | | | |
|---|---|---|---|---|---|---|---|---|---|
| | **p_val** | **avg_log2FC** | **pct.1** | **pct.2** | **p_val_adj** | **celltype** | **Foldchange_UP** | **Foldchange_DOWN** | **Ratio_pct1_pct2** |
| FLCN | 0,224539785 | 0,259212377 | 0,286 | 0,195 | 1 | Ciliated | 1,196825133 | 0,835543951 | 1,466666667 |
| KIF3B | 0,224707419 | -0,395889311 | 0,571 | 0,732 | 1 | Ciliated | 0,760020738 | 1,315753571 | 0,780054645 |
| EPAS11 | 0,224729195 | -0,265605244 | 0,286 | 0,439 | 1 | Ciliated | 0,831849681 | 1,202140269 | 0,651480638 |
| CITED41 | 0,225181498 | 0,377399997 | 0,607 | 0,561 | 1 | Ciliated | 1,298998709 | 0,769823706 | 1,081996435 |
| HEY2 | 0,22527235 | -0,313725939 | 0,036 | 0,122 | 1 | Ciliated | 0,804561195 | 1,242913537 | 0,295081967 |
| ANKRD49 | 0,22527235 | -0,37818965 | 0,036 | 0,122 | 1 | Ciliated | 0,769402462 | 1,299709905 | 0,295081967 |
| SNX21 | 0,22527235 | -0,290442946 | 0,036 | 0,122 | 1 | Ciliated | 0,817650979 | 1,223015719 | 0,295081967 |
| KTN11 | 0,225663277 | 0,31434263 | 0,964 | 0,78 | 1 | Ciliated | 1,243444943 | 0,804217352 | 1,235897436 |
| SLIRP1 | 0,225874774 | -0,334505239 | 0,536 | 0,732 | 1 | Ciliated | 0,793056062 | 1,260944904 | 0,732240437 |
| DDX421 | 0,225913279 | -0,25343996 | 0,464 | 0,61 | 1 | Ciliated | 0,838893771 | 1,192046042 | 0,760655738 |
| ATP6AP11 | 0,225913279 | 0,343554049 | 0,607 | 0,512 | 1 | Ciliated | 1,268878603 | 0,788097457 | 1,185546875 |
| VEGFB1 | 0,225919339 | -0,363774639 | 0,179 | 0,341 | 1 | Ciliated | 0,777128653 | 1,286788225 | 0,524926686 |
| TM7SF3 | 0,226659077 | -0,338434825 | 0,25 | 0,39 | 1 | Ciliated | 0,79089889 | 1,264384124 | 0,641025641 |
| TPCN1 | 0,226728208 | 0,314846638 | 0,25 | 0,171 | 1 | Ciliated | 1,243879419 | 0,803936446 | 1,461988304 |
| MARK4 | 0,226728208 | 0,292081351 | 0,25 | 0,171 | 1 | Ciliated | 1,224405433 | 0,816722936 | 1,461988304 |
| MXD4 | 0,22699428 | 0,281069145 | 0,464 | 0,415 | 1 | Ciliated | 1,215095028 | 0,822980901 | 1,118072289 |
| WDR19 | 0,227403675 | -0,321275134 | 0,357 | 0,561 | 1 | Ciliated | 0,80036216 | 1,249434381 | 0,636363636 |
| GLRX52 | 0,227497134 | 0,275214976 | 0,393 | 0,341 | 1 | Ciliated | 1,210174404 | 0,826327178 | 1,152492669 |
| HIPK3 | 0,228073284 | -0,251637891 | 0,429 | 0,659 | 1 | Ciliated | 0,839942287 | 1,190557988 | 0,650986343 |
| MRPS30 | 0,228522011 | 0,508193498 | 0,357 | 0,293 | 1 | Ciliated | 1,422268157 | 0,703102292 | 1,218430034 |
| MROH1 | 0,228522011 | 0,326701054 | 0,357 | 0,293 | 1 | Ciliated | 1,254142302 | 0,797357683 | 1,218430034 |
| SSR4 | 0,228632703 | 0,358461374 | 0,964 | 0,902 | 1 | Ciliated | 1,282057861 | 0,779995998 | 1,068736142 |
| CCAR11 | 0,229194209 | -0,349202865 | 0,357 | 0,537 | 1 | Ciliated | 0,785017725 | 1,273856587 | 0,664804469 |
| GNPTG2 | 0,229731518 | 0,274560353 | 0,429 | 0,366 | 1 | Ciliated | 1,209625411 | 0,826702209 | 1,172131148 |
| GGA1 | 0,229731518 | 0,441561139 | 0,429 | 0,366 | 1 | Ciliated | 1,358073102 | 0,736337388 | 1,172131148 |
| NDUFS41 | 0,230860024 | -0,362518799 | 0,393 | 0,585 | 1 | Ciliated | 0,777805424 | 1,285668586 | 0,671794872 |
| LINC01184 | 0,230909114 | -0,295254793 | 0,143 | 0,293 | 1 | Ciliated | 0,814928396 | 1,227101675 | 0,488054608 |
| MRPS28 | 0,231030925 | -0,399555772 | 0,393 | 0,61 | 1 | Ciliated | 0,758091675 | 1,319101677 | 0,644262295 |
| MAP2K22 | 0,231030925 | 0,259960967 | 0,571 | 0,488 | 1 | Ciliated | 1,197446307 | 0,835110514 | 1,170081967 |
| KRT152 | 0,23120162 | -0,350065771 | 0,214 | 0,341 | 1 | Ciliated | 0,784548331 | 1,274618734 | 0,627565982 |
| THBS11 | 0,232773849 | -0,511095552 | 0,107 | 0,22 | 1 | Ciliated | 0,701689388 | 1,425131999 | 0,486363636 |

| Supplementary Table 4 - clusermarker | | | | | | | | | |
|---|---|---|---|---|---|---|---|---|---|
| | **p_val** | **avg_log2FC** | **pct.1** | **pct.2** | **p_val_adj** | **celltype** | **Foldchange_UP** | **Foldchange_DOWN** | **Ratio_pct1_pct2** |
| NACC2 | 0,232976168 | 0,386641173 | 0,357 | 0,268 | 1 | Ciliated | 1,307346145 | 0,764908363 | 1,332089552 |
| HDGFL2 | 0,233013039 | 0,280317146 | 0,393 | 0,341 | 1 | Ciliated | 1,214461829 | 0,823409988 | 1,152492669 |
| RPL27A1 | 0,234657265 | -0,266641058 | 0,821 | 0,829 | 1 | Ciliated | 0,831252651 | 1,203003682 | 0,990349819 |
| ZFAS12 | 0,23497723 | -0,393379494 | 0,679 | 0,732 | 1 | Ciliated | 0,761344076 | 1,313466581 | 0,927595628 |
| POLR3B | 0,234981315 | -0,28767427 | 0,036 | 0,122 | 1 | Ciliated | 0,81922164 | 1,22067088 | 0,295081967 |
| ASAH11 | 0,23525307 | -0,333572142 | 0,5 | 0,707 | 1 | Ciliated | 0,793569156 | 1,260129621 | 0,707213579 |
| OTUB11 | 0,235481191 | 0,256467165 | 0,714 | 0,659 | 1 | Ciliated | 1,194549937 | 0,837135367 | 1,083459788 |
| ELOC | 0,235589267 | 0,304513086 | 0,679 | 0,634 | 1 | Ciliated | 1,23500175 | 0,809715452 | 1,070977918 |
| LRRC591 | 0,235630973 | 0,351549176 | 0,429 | 0,341 | 1 | Ciliated | 1,275929995 | 0,783742058 | 1,258064516 |
| DAPL11 | 0,235660937 | 0,28032291 | 0,393 | 0,317 | 1 | Ciliated | 1,214466681 | 0,823406699 | 1,239747634 |
| SYTL21 | 0,236500579 | -0,480941835 | 0,643 | 0,732 | 1 | Ciliated | 0,716509712 | 1,395654494 | 0,878415301 |
| MFSD2A | 0,237198786 | 0,440402885 | 0,321 | 0,268 | 1 | Ciliated | 1,356983224 | 0,736928786 | 1,197761194 |
| DNAL4 | 0,237725976 | 0,250748838 | 0,5 | 0,463 | 1 | Ciliated | 1,189824539 | 0,840460057 | 1,079913607 |
| COPB2 | 0,237739646 | -0,364535794 | 0,536 | 0,683 | 1 | Ciliated | 0,776718754 | 1,287467304 | 0,78477306 |
| SPAG17 | 0,237811916 | -0,503617372 | 0,75 | 0,78 | 1 | Ciliated | 0,705336023 | 1,41776397 | 0,961538462 |
| MS4A8 | 0,23794679 | -0,303200211 | 0,857 | 0,902 | 1 | Ciliated | 0,810452641 | 1,23387839 | 0,950110865 |
| FOXP1 | 0,238063418 | -0,390714906 | 0,5 | 0,756 | 1 | Ciliated | 0,762751541 | 1,311042911 | 0,661375661 |
| HSPE11 | 0,238066974 | -0,428607639 | 0,929 | 0,878 | 1 | Ciliated | 0,742978496 | 1,345933974 | 1,05808656 |
| DOLK | 0,238120324 | -0,361950932 | 0,071 | 0,171 | 1 | Ciliated | 0,778111641 | 1,285162626 | 0,415204678 |
| DET1 | 0,238120324 | -0,33821043 | 0,071 | 0,171 | 1 | Ciliated | 0,791021915 | 1,264187478 | 0,415204678 |
| ZC2HC1A | 0,238257408 | -0,344865509 | 0,607 | 0,805 | 1 | Ciliated | 0,787381375 | 1,270032582 | 0,754037267 |
| MANF1 | 0,238623138 | 0,388537982 | 0,393 | 0,341 | 1 | Ciliated | 1,309066132 | 0,763903347 | 1,152492669 |
| MRPL11 | 0,238623138 | 0,35955186 | 0,393 | 0,341 | 1 | Ciliated | 1,283027293 | 0,779406647 | 1,152492669 |
| RPAP2 | 0,239227012 | -0,560827325 | 0,143 | 0,268 | 1 | Ciliated | 0,677913297 | 1,475114892 | 0,53358209 |
| KLHDC7A | 0,240377637 | 0,501868484 | 0,25 | 0,171 | 1 | Ciliated | 1,416046346 | 0,706191576 | 1,461988304 |
| TMEM65 | 0,240377637 | 0,286579886 | 0,25 | 0,171 | 1 | Ciliated | 1,219745267 | 0,819843312 | 1,461988304 |
| IQCA1 | 0,240615539 | 0,336415083 | 0,607 | 0,537 | 1 | Ciliated | 1,262615251 | 0,792006907 | 1,130353818 |
| INO80B1 | 0,240683226 | 0,382138078 | 0,429 | 0,366 | 1 | Ciliated | 1,303271877 | 0,767299608 | 1,172131148 |
| FNBP1L1 | 0,241386772 | 0,384671887 | 0,393 | 0,317 | 1 | Ciliated | 1,305562829 | 0,76595318 | 1,239747634 |
| PCYOX11 | 0,241386772 | 0,274006628 | 0,393 | 0,317 | 1 | Ciliated | 1,209161231 | 0,827019569 | 1,239747634 |
| GRHL2 | 0,241386772 | -0,281222832 | 0,25 | 0,415 | 1 | Ciliated | 0,822893235 | 1,215224476 | 0,602409639 |

| Supplementary Table 4 - clusermarker | | | | | | | | | |
|---|---|---|---|---|---|---|---|---|---|
| | **p_val** | **avg_log2FC** | **pct.1** | **pct.2** | **p_val_adj** | **celltype** | **Foldchange_UP** | **Foldchange_DOWN** | **Ratio_pct1_pct2** |
| FKBP3 | 0,241386772 | -0,286353486 | 0,25 | 0,415 | 1 | Ciliated | 0,819971979 | 1,21955387 | 0,602409639 |
| NECTIN21 | 0,241551287 | 0,355783796 | 0,464 | 0,39 | 1 | Ciliated | 1,279680625 | 0,78144498 | 1,18974359 |
| AQP31 | 0,241635732 | -0,358081846 | 0,821 | 0,805 | 1 | Ciliated | 0,780201217 | 1,281720636 | 1,019875776 |
| IFT74 | 0,241970638 | -0,453597553 | 0,5 | 0,659 | 1 | Ciliated | 0,730219675 | 1,369450912 | 0,758725341 |
| FOXD11 | 0,242299595 | 0,317073796 | 0,179 | 0,098 | 1 | Ciliated | 1,245801139 | 0,802696328 | 1,826530612 |
| AL354754.1 | 0,242299595 | 0,259934358 | 0,179 | 0,098 | 1 | Ciliated | 1,197424221 | 0,835125916 | 1,826530612 |
| MUC5AC1 | 0,24297232 | -1,565814367 | 0,214 | 0,098 | 1 | Ciliated | 0,337786982 | 2,960445647 | 2,183673469 |
| AP3D11 | 0,243009423 | 0,304370535 | 0,5 | 0,463 | 1 | Ciliated | 1,234879727 | 0,809795463 | 1,079913607 |
| RPL33 | 0,24320262 | -0,281996122 | 1 | 0,951 | 1 | Ciliated | 0,822452279 | 1,215876015 | 1,051524711 |
| DENND4C | 0,244276923 | 0,294028928 | 0,321 | 0,244 | 1 | Ciliated | 1,226059444 | 0,815621139 | 1,31557377 |
| SEC11C | 0,246172567 | 0,335103172 | 0,536 | 0,561 | 1 | Ciliated | 1,261467617 | 0,792727444 | 0,95543672 |
| GPAA12 | 0,246294886 | 0,406980817 | 0,429 | 0,366 | 1 | Ciliated | 1,32590813 | 0,754200067 | 1,172131148 |
| AC010894.2 | 0,246530194 | -0,256194232 | 0,071 | 0,171 | 1 | Ciliated | 0,837293754 | 1,194323969 | 0,415204678 |
| LNPK | 0,246530194 | -0,489009117 | 0,071 | 0,171 | 1 | Ciliated | 0,712514304 | 1,403480595 | 0,415204678 |
| BX255925.3 | 0,246530194 | -0,421989234 | 0,071 | 0,171 | 1 | Ciliated | 0,746394762 | 1,339773604 | 0,415204678 |
| MAZ | 0,246816791 | -0,460518649 | 0,357 | 0,512 | 1 | Ciliated | 0,726724954 | 1,376036415 | 0,697265625 |
| TRIM161 | 0,246852828 | 0,375253621 | 0,357 | 0,317 | 1 | Ciliated | 1,297067555 | 0,770969867 | 1,126182965 |
| SRPX2 | 0,246901765 | -0,254849373 | 0,286 | 0,439 | 1 | Ciliated | 0,838074631 | 1,193211157 | 0,651480638 |
| ALKBH3 | 0,247581313 | -0,36800889 | 0,107 | 0,22 | 1 | Ciliated | 0,774851156 | 1,290570443 | 0,486363636 |
| ZNF440 | 0,248370101 | -0,280234211 | 0,429 | 0,659 | 1 | Ciliated | 0,823457324 | 1,214392016 | 0,650986343 |
| STX2 | 0,248374873 | 0,33525905 | 0,5 | 0,463 | 1 | Ciliated | 1,261603922 | 0,792641797 | 1,079913607 |
| ATG3 | 0,248441641 | -0,269112672 | 0,321 | 0,512 | 1 | Ciliated | 0,829829775 | 1,205066425 | 0,626953125 |
| RAD211 | 0,2490588 | -0,473359329 | 0,429 | 0,61 | 1 | Ciliated | 0,720285452 | 1,388338466 | 0,703278689 |
| POLG2 | 0,250622708 | -0,44731074 | 0,179 | 0,341 | 1 | Ciliated | 0,733408687 | 1,363496257 | 0,524926686 |
| BID | 0,250622708 | 0,261620437 | 0,321 | 0,244 | 1 | Ciliated | 1,19882447 | 0,834150474 | 1,31557377 |
| SHC1 | 0,250784062 | -0,382730321 | 0,143 | 0,293 | 1 | Ciliated | 0,766984687 | 1,303806995 | 0,488054608 |
| NOP2 | 0,250784062 | 0,250086185 | 0,286 | 0,195 | 1 | Ciliated | 1,189278159 | 0,840846183 | 1,466666667 |
| TXNDC11 | 0,250784062 | -0,272351317 | 0,143 | 0,293 | 1 | Ciliated | 0,827969014 | 1,207774666 | 0,488054608 |
| PCED1A | 0,250784062 | -0,336507027 | 0,143 | 0,293 | 1 | Ciliated | 0,791956434 | 1,262695721 | 0,488054608 |
| ZFYVE27 | 0,250812284 | 0,33388006 | 0,179 | 0,098 | 1 | Ciliated | 1,260398603 | 0,7933998 | 1,826530612 |
| ARX | 0,250812284 | 0,299681603 | 0,179 | 0,098 | 1 | Ciliated | 1,230872735 | 0,812431677 | 1,826530612 |

| Supplementary Table 4 - clusermarker | | | | | | | | | |
|---|---|---|---|---|---|---|---|---|---|
| | **p_val** | **avg_log2FC** | **pct.1** | **pct.2** | **p_val_adj** | **celltype** | **Foldchange_UP** | **Foldchange_DOWN** | **Ratio_pct1_pct2** |
| LASP1 | 0,251712753 | 0,341476541 | 0,536 | 0,537 | 1 | Ciliated | 1,267052709 | 0,78923315 | 0,998137803 |
| SNRPA | 0,252178411 | -0,413605405 | 0,214 | 0,39 | 1 | Ciliated | 0,750744859 | 1,332010454 | 0,548717949 |
| TMEM131 | 0,252301775 | 0,307049196 | 0,643 | 0,561 | 1 | Ciliated | 1,237174666 | 0,808293305 | 1,146167558 |
| FAM104B | 0,253049009 | -0,410376433 | 0,536 | 0,683 | 1 | Ciliated | 0,752427022 | 1,329032545 | 0,78477306 |
| SERINC5 | 0,253130588 | 0,250394873 | 0,393 | 0,317 | 1 | Ciliated | 1,189532651 | 0,840666289 | 1,239747634 |
| HSPA1A2 | 0,253781982 | 0,255237363 | 0,821 | 0,78 | 1 | Ciliated | 1,193532096 | 0,837849274 | 1,052564103 |
| LIMA11 | 0,254477402 | -0,343030932 | 0,429 | 0,561 | 1 | Ciliated | 0,78838327 | 1,268418595 | 0,764705882 |
| IRF91 | 0,254539955 | 0,284328316 | 0,679 | 0,659 | 1 | Ciliated | 1,217843133 | 0,821123815 | 1,030349014 |
| PRSS222 | 0,255106766 | 0,384299143 | 0,464 | 0,439 | 1 | Ciliated | 1,305225558 | 0,766151102 | 1,056947608 |
| APBB2 | 0,255146021 | -0,309791902 | 0,071 | 0,171 | 1 | Ciliated | 0,806758119 | 1,239528895 | 0,415204678 |
| RNASE11 | 0,2552273 | -0,510972736 | 0,107 | 0,22 | 1 | Ciliated | 0,701749125 | 1,425010684 | 0,486363636 |
| ARMC3 | 0,255356286 | -0,27978535 | 0,714 | 0,805 | 1 | Ciliated | 0,823713563 | 1,214014245 | 0,886956522 |
| TENT2 | 0,255846966 | 0,401798309 | 0,321 | 0,268 | 1 | Ciliated | 1,321153694 | 0,756914207 | 1,197761194 |
| SELENOS | 0,255896602 | -0,374886006 | 0,536 | 0,732 | 1 | Ciliated | 0,771166344 | 1,296737089 | 0,732240437 |
| CREBBP1 | 0,25595483 | -0,288312066 | 0,357 | 0,537 | 1 | Ciliated | 0,818859553 | 1,221210642 | 0,664804469 |
| SF3B61 | 0,256343557 | -0,61586322 | 0,714 | 0,854 | 1 | Ciliated | 0,652539337 | 1,532474662 | 0,836065574 |
| SLC39A7 | 0,257333571 | -0,385182576 | 0,214 | 0,366 | 1 | Ciliated | 0,765682094 | 1,306025057 | 0,584699454 |
| ST8SIA4 | 0,257845291 | -0,295986048 | 0,179 | 0,317 | 1 | Ciliated | 0,81451544 | 1,227723811 | 0,56466877 |
| CCDC501 | 0,258066911 | -0,418450208 | 0,179 | 0,293 | 1 | Ciliated | 0,748227965 | 1,336491079 | 0,610921502 |
| ANKRD371 | 0,258085732 | -0,367667927 | 0,929 | 0,878 | 1 | Ciliated | 0,775034304 | 1,290265469 | 1,05808656 |
| ACVR1B | 0,258350224 | 0,27969658 | 0,357 | 0,293 | 1 | Ciliated | 1,213939548 | 0,823764249 | 1,218430034 |
| EEF1D1 | 0,258494522 | 0,453845917 | 0,75 | 0,707 | 1 | Ciliated | 1,369686686 | 0,730093977 | 1,060820368 |
| ARF1 | 0,259768271 | 0,271792187 | 0,714 | 0,634 | 1 | Ciliated | 1,207306673 | 0,828289964 | 1,126182965 |
| CRNKL1 | 0,260115271 | -0,264878127 | 0,143 | 0,268 | 1 | Ciliated | 0,832269038 | 1,201534545 | 0,53358209 |
| CPLANE1 | 0,261028401 | -0,325333183 | 0,536 | 0,61 | 1 | Ciliated | 0,798114045 | 1,252953768 | 0,878688525 |
| SPTAN11 | 0,261033106 | -0,27413024 | 0,536 | 0,61 | 1 | Ciliated | 0,826948712 | 1,209264838 | 0,878688525 |
| ACVR1 | 0,261482708 | 0,335588433 | 0,286 | 0,22 | 1 | Ciliated | 1,261891993 | 0,792460849 | 1,3 |
| RSU1 | 0,262014187 | -0,29920827 | 0,25 | 0,439 | 1 | Ciliated | 0,812698271 | 1,230468964 | 0,569476082 |
| ACOX1 | 0,262027656 | -0,332399738 | 0,25 | 0,39 | 1 | Ciliated | 0,794214311 | 1,259105995 | 0,641025641 |
| TMEM234 | 0,262283574 | 0,423926165 | 0,321 | 0,268 | 1 | Ciliated | 1,341573564 | 0,74539334 | 1,197761194 |
| MAPKAPK5 | 0,262283574 | 0,311242086 | 0,321 | 0,268 | 1 | Ciliated | 1,240775484 | 0,805947581 | 1,197761194 |

| Supplementary Table 4 - clusermarker | | | | | | | | | |
|---|---|---|---|---|---|---|---|---|---|
| | **p_val** | **avg_log2FC** | **pct.1** | **pct.2** | **p_val_adj** | **celltype** | **Foldchange_UP** | **Foldchange_DOWN** | **Ratio_pct1_pct2** |
| SLC22A181 | 0,262582568 | 0,323379019 | 0,429 | 0,39 | 1 | Ciliated | 1,251257762 | 0,799195841 | 1,1 |
| CDV3 | 0,262705953 | 0,288210752 | 0,536 | 0,512 | 1 | Ciliated | 1,221124884 | 0,81891706 | 1,046875 |
| FXR11 | 0,263624498 | -0,535396458 | 0,536 | 0,683 | 1 | Ciliated | 0,689969045 | 1,449340384 | 0,78477306 |
| ACTR3C | 0,263969001 | -0,291585841 | 0,071 | 0,171 | 1 | Ciliated | 0,817003497 | 1,223984969 | 0,415204678 |
| AL135925.1 | 0,263969001 | -0,280906875 | 0,071 | 0,171 | 1 | Ciliated | 0,823073472 | 1,214958365 | 0,415204678 |
| SARM1 | 0,263969001 | -0,292822054 | 0,071 | 0,171 | 1 | Ciliated | 0,816303725 | 1,225034224 | 0,415204678 |
| NECTIN4 | 0,264506973 | 0,408237898 | 0,393 | 0,366 | 1 | Ciliated | 1,327063953 | 0,753543186 | 1,073770492 |
| POLE42 | 0,264625972 | 0,31737546 | 0,357 | 0,293 | 1 | Ciliated | 1,24606166 | 0,802528504 | 1,218430034 |
| AP2B1 | 0,265278089 | 0,316162974 | 0,571 | 0,512 | 1 | Ciliated | 1,24501487 | 0,803203258 | 1,115234375 |
| CPNE32 | 0,266214756 | 0,264882581 | 0,643 | 0,707 | 1 | Ciliated | 1,201538254 | 0,832266469 | 0,909476662 |
| MT2A | 0,267512588 | -0,608200782 | 0,929 | 0,756 | 1 | Ciliated | 0,656014323 | 1,524356962 | 1,228835979 |
| CLNS1A | 0,268263605 | 0,343762225 | 0,5 | 0,537 | 1 | Ciliated | 1,269061711 | 0,787983745 | 0,931098696 |
| SUGP2 | 0,268301237 | -0,25572164 | 0,393 | 0,585 | 1 | Ciliated | 0,837568076 | 1,193932803 | 0,671794872 |
| GSS | 0,268407737 | 0,419947147 | 0,286 | 0,22 | 1 | Ciliated | 1,337878541 | 0,747452007 | 1,3 |
| TICAM11 | 0,269389107 | 0,443624756 | 0,25 | 0,171 | 1 | Ciliated | 1,360017067 | 0,735284891 | 1,461988304 |
| SLK1 | 0,27013324 | 0,332710715 | 0,607 | 0,61 | 1 | Ciliated | 1,259377428 | 0,794043134 | 0,995081967 |
| MCPH1 | 0,270153037 | -0,274659561 | 0,214 | 0,366 | 1 | Ciliated | 0,826645362 | 1,209708595 | 0,584699454 |
| KRR1 | 0,270153037 | -0,253040099 | 0,214 | 0,366 | 1 | Ciliated | 0,839126314 | 1,191715697 | 0,584699454 |
| PCYT2 | 0,270657732 | 0,328685056 | 0,5 | 0,463 | 1 | Ciliated | 1,255868192 | 0,796261906 | 1,079913607 |
| ITGA21 | 0,270662832 | -0,314469968 | 0,357 | 0,561 | 1 | Ciliated | 0,804146372 | 1,2435547 | 0,636363636 |
| DNAJC3-DT | 0,27100791 | -0,303434901 | 0,107 | 0,22 | 1 | Ciliated | 0,810320812 | 1,234079127 | 0,486363636 |
| ACSF3 | 0,271398227 | -0,313439 | 0,179 | 0,317 | 1 | Ciliated | 0,80472123 | 1,242666358 | 0,56466877 |
| ACAP1 | 0,271482583 | 0,28888327 | 0,393 | 0,317 | 1 | Ciliated | 1,221694249 | 0,818535407 | 1,239747634 |
| ARHGAP181 | 0,273560711 | -0,486921668 | 0,929 | 0,829 | 1 | Ciliated | 0,713545994 | 1,401451355 | 1,120627262 |
| PLEKHS11 | 0,273581899 | -0,356689815 | 0,643 | 0,756 | 1 | Ciliated | 0,780954383 | 1,280484523 | 0,850529101 |
| SSRP1 | 0,273901497 | -0,733059118 | 0,464 | 0,561 | 1 | Ciliated | 0,601626859 | 1,662159833 | 0,827094474 |
| PNPLA8 | 0,274622729 | -0,367412009 | 0,25 | 0,39 | 1 | Ciliated | 0,775171799 | 1,290036611 | 0,641025641 |
| FKBP11 | 0,275457705 | 0,365450992 | 0,286 | 0,22 | 1 | Ciliated | 1,288284289 | 0,776226186 | 1,3 |
| NUTM2B-AS1 | 0,275483304 | 0,309782165 | 0,321 | 0,268 | 1 | Ciliated | 1,239520528 | 0,806763565 | 1,197761194 |
| SYNJ2BP | 0,275972137 | -0,488046564 | 0,321 | 0,488 | 1 | Ciliated | 0,712989846 | 1,402544518 | 0,657786885 |
| ILVBL | 0,276713729 | 0,37063843 | 0,393 | 0,366 | 1 | Ciliated | 1,292924856 | 0,773440154 | 1,073770492 |

| Supplementary Table 4 - clusermarker | | | | | | | | | |
|---|---|---|---|---|---|---|---|---|---|
| | **p_val** | **avg_log2FC** | **pct.1** | **pct.2** | **p_val_adj** | **celltype** | **Foldchange_UP** | **Foldchange_DOWN** | **Ratio_pct1_pct2** |
| DHRS4L2 | 0,276724327 | 0,290364662 | 0,357 | 0,268 | 1 | Ciliated | 1,222949357 | 0,817695348 | 1,332089552 |
| FLAD1 | 0,277152248 | -0,279613364 | 0,179 | 0,341 | 1 | Ciliated | 0,823811766 | 1,213869529 | 0,524926686 |
| ITPA1 | 0,277191943 | -0,414243279 | 0,321 | 0,512 | 1 | Ciliated | 0,750412997 | 1,33259952 | 0,626953125 |
| ADIRF2 | 0,277490663 | -0,449077887 | 0,214 | 0,39 | 1 | Ciliated | 0,73251089 | 1,365167418 | 0,548717949 |
| RAI11 | 0,277835054 | 0,288406831 | 0,357 | 0,317 | 1 | Ciliated | 1,221290861 | 0,818805767 | 1,126182965 |
| TACC11 | 0,27835454 | -0,323410962 | 0,179 | 0,317 | 1 | Ciliated | 0,799178146 | 1,251285467 | 0,56466877 |
| TTN-AS1 | 0,278627404 | 0,284698501 | 0,214 | 0,122 | 1 | Ciliated | 1,218155663 | 0,820913148 | 1,754098361 |
| RAB1B | 0,278780538 | -0,550711473 | 0,5 | 0,732 | 1 | Ciliated | 0,682683376 | 1,464807896 | 0,683060109 |
| ELOVL5 | 0,278871567 | 0,262586675 | 0,5 | 0,439 | 1 | Ciliated | 1,199627646 | 0,833591993 | 1,138952164 |
| GPRC5A | 0,280556705 | -0,255275247 | 0,321 | 0,463 | 1 | Ciliated | 0,837827273 | 1,193563437 | 0,693304536 |
| CACYBP | 0,281128381 | 0,451464415 | 0,571 | 0,561 | 1 | Ciliated | 1,367427567 | 0,731300161 | 1,017825312 |
| URGCP | 0,282233109 | -0,313813994 | 0,143 | 0,268 | 1 | Ciliated | 0,804512089 | 1,242989401 | 0,53358209 |
| AC103957.2 | 0,282233109 | -0,279236843 | 0,143 | 0,268 | 1 | Ciliated | 0,824026796 | 1,21355277 | 0,53358209 |
| AGBL2 | 0,282283227 | 0,319878328 | 0,607 | 0,585 | 1 | Ciliated | 1,248225273 | 0,80113744 | 1,037606838 |
| ERP44 | 0,282958913 | -0,386067206 | 0,286 | 0,439 | 1 | Ciliated | 0,765212738 | 1,306826129 | 0,651480638 |
| F11R1 | 0,283416858 | -0,262755781 | 0,75 | 0,732 | 1 | Ciliated | 0,833494289 | 1,199768269 | 1,024590164 |
| C21orf58 | 0,283416858 | -0,379258193 | 0,679 | 0,78 | 1 | Ciliated | 0,768832808 | 1,300672902 | 0,870512821 |
| SNHG71 | 0,284073223 | 0,256501788 | 0,321 | 0,244 | 1 | Ciliated | 1,194578604 | 0,837115278 | 1,31557377 |
| DDX56 | 0,284080199 | -0,352784333 | 0,214 | 0,39 | 1 | Ciliated | 0,783071348 | 1,277022844 | 0,548717949 |
| SYTL1 | 0,284570407 | -0,388938801 | 0,643 | 0,756 | 1 | Ciliated | 0,763691144 | 1,309429876 | 0,850529101 |
| PITPNA-AS1 | 0,284761572 | 0,306840879 | 0,25 | 0,171 | 1 | Ciliated | 1,236996037 | 0,808410027 | 1,461988304 |
| ACAA12 | 0,286729569 | 0,269135338 | 0,429 | 0,39 | 1 | Ciliated | 1,205085358 | 0,829816737 | 1,1 |
| ANK32 | 0,286899914 | -0,50980268 | 0,571 | 0,634 | 1 | Ciliated | 0,702318489 | 1,423855439 | 0,900630915 |
| PIM1 | 0,287444414 | -0,279165546 | 0,107 | 0,22 | 1 | Ciliated | 0,82406752 | 1,213492798 | 0,486363636 |
| STAT5A | 0,287444414 | -0,283242422 | 0,107 | 0,22 | 1 | Ciliated | 0,821742095 | 1,216926827 | 0,486363636 |
| DYNC2LI1 | 0,287983834 | -0,391938341 | 0,464 | 0,683 | 1 | Ciliated | 0,762104986 | 1,312155174 | 0,679355783 |
| CAD | 0,288629919 | 0,253481995 | 0,143 | 0,073 | 1 | Ciliated | 1,192080774 | 0,83886933 | 1,95890411 |
| LINC00326 | 0,288629919 | 0,340902933 | 0,143 | 0,073 | 1 | Ciliated | 1,266549036 | 0,789547007 | 1,95890411 |
| EID3 | 0,288629919 | 0,257135296 | 0,143 | 0,073 | 1 | Ciliated | 1,195103277 | 0,836747769 | 1,95890411 |
| ARHGAP44 | 0,288629919 | 0,312266557 | 0,143 | 0,073 | 1 | Ciliated | 1,241656882 | 0,805375474 | 1,95890411 |
| AC244153.1 | 0,288629919 | 0,501947324 | 0,143 | 0,073 | 1 | Ciliated | 1,416123732 | 0,706152985 | 1,95890411 |

| Supplementary Table 4 - clusermarker | | | | | | | | | |
|---|---|---|---|---|---|---|---|---|---|
| | **p_val** | **avg_log2FC** | **pct.1** | **pct.2** | **p_val_adj** | **celltype** | **Foldchange_UP** | **Foldchange_DOWN** | **Ratio_pct1_pct2** |
| C19orf38 | 0,288629919 | 0,488295729 | 0,143 | 0,073 | 1 | Ciliated | 1,40278677 | 0,712866718 | 1,95890411 |
| LYRM1 | 0,289933924 | 0,257232559 | 0,286 | 0,22 | 1 | Ciliated | 1,19518385 | 0,836691359 | 1,3 |
| TAF11 | 0,290191497 | -0,305546108 | 0,214 | 0,366 | 1 | Ciliated | 0,809135874 | 1,235886373 | 0,584699454 |
| RPS181 | 0,290494097 | -0,316033155 | 1 | 0,951 | 1 | Ciliated | 0,803275536 | 1,244902845 | 1,051524711 |
| PRDM1 | 0,290725676 | -0,469481408 | 0,25 | 0,415 | 1 | Ciliated | 0,722224162 | 1,384611665 | 0,602409639 |
| MT1X2 | 0,290725676 | -0,605800636 | 0,286 | 0,39 | 1 | Ciliated | 0,657106612 | 1,521823068 | 0,733333333 |
| KIAA0930 | 0,290774857 | -0,466622186 | 0,214 | 0,39 | 1 | Ciliated | 0,72365693 | 1,381870274 | 0,548717949 |
| GMDS-DT | 0,290939532 | 0,327178415 | 0,357 | 0,317 | 1 | Ciliated | 1,254557343 | 0,797093896 | 1,126182965 |
| XIST1 | 0,29147574 | -0,329971054 | 0,393 | 0,585 | 1 | Ciliated | 0,795552445 | 1,256988154 | 0,671794872 |
| CAPN22 | 0,291800136 | -0,282775851 | 0,714 | 0,756 | 1 | Ciliated | 0,822007891 | 1,216533334 | 0,944444444 |
| EIF3A | 0,292485735 | -0,300381884 | 0,714 | 0,78 | 1 | Ciliated | 0,81203742 | 1,231470343 | 0,915384615 |
| TMEM251 | 0,292628531 | -0,386674946 | 0,179 | 0,317 | 1 | Ciliated | 0,764890457 | 1,30737675 | 0,56466877 |
| ADGRV1 | 0,292666169 | 0,301634943 | 0,25 | 0,171 | 1 | Ciliated | 1,232540406 | 0,811332428 | 1,461988304 |
| NDUFA51 | 0,293761576 | -0,29828648 | 0,464 | 0,683 | 1 | Ciliated | 0,813217699 | 1,229683024 | 0,679355783 |
| GLUD11 | 0,294284379 | 0,428614589 | 0,5 | 0,463 | 1 | Ciliated | 1,345940458 | 0,742974917 | 1,079913607 |
| SLC6A6 | 0,294493056 | 0,313232519 | 0,429 | 0,366 | 1 | Ciliated | 1,242488518 | 0,804836412 | 1,172131148 |
| TSN | 0,296138119 | -0,261471153 | 0,214 | 0,341 | 1 | Ciliated | 0,834236793 | 1,198700427 | 0,627565982 |
| STK40 | 0,296869553 | 0,434246351 | 0,536 | 0,537 | 1 | Ciliated | 1,351204794 | 0,740080264 | 0,998137803 |
| UBE2I2 | 0,296949239 | -0,282932642 | 0,393 | 0,512 | 1 | Ciliated | 0,821918561 | 1,216665552 | 0,767578125 |
| CEP250 | 0,29733915 | -0,266822496 | 0,25 | 0,415 | 1 | Ciliated | 0,831148117 | 1,203154985 | 0,602409639 |
| HIP1R | 0,297360757 | 0,292624456 | 0,286 | 0,22 | 1 | Ciliated | 1,224866449 | 0,816415537 | 1,3 |
| TMEM232 | 0,297501795 | -0,727292097 | 0,393 | 0,561 | 1 | Ciliated | 0,604036612 | 1,655528787 | 0,700534759 |
| CHID1 | 0,297501795 | 0,251152845 | 0,5 | 0,488 | 1 | Ciliated | 1,19015778 | 0,840224731 | 1,024590164 |
| DERL3 | 0,297645223 | 0,721788484 | 0,357 | 0,317 | 1 | Ciliated | 1,649225283 | 0,606345301 | 1,126182965 |
| AASDH | 0,29766761 | -0,256904155 | 0,143 | 0,268 | 1 | Ciliated | 0,836881839 | 1,194911819 | 0,53358209 |
| PIP2 | 0,29766761 | -1,304493712 | 0,143 | 0,268 | 1 | Ciliated | 0,404863163 | 2,469970331 | 0,53358209 |
| NCL2 | 0,297679358 | -0,279414421 | 0,821 | 0,829 | 1 | Ciliated | 0,823925374 | 1,213702152 | 0,990349819 |
| TRIP10 | 0,298263571 | 0,263757842 | 0,321 | 0,244 | 1 | Ciliated | 1,200601888 | 0,832915565 | 1,31557377 |
| SERPINB41 | 0,299358328 | 0,339735719 | 0,143 | 0,073 | 1 | Ciliated | 1,265524746 | 0,79018605 | 1,95890411 |
| STRN31 | 0,299946711 | -0,258228722 | 0,179 | 0,317 | 1 | Ciliated | 0,836113834 | 1,196009395 | 0,56466877 |
| GPR162 | 0,300102542 | 0,285608872 | 0,643 | 0,61 | 1 | Ciliated | 1,218924587 | 0,820395298 | 1,054098361 |

| Supplementary Table 4 - clusermarker | | | | | | | | | |
|---|---|---|---|---|---|---|---|---|---|
| | **p_val** | **avg_log2FC** | **pct.1** | **pct.2** | **p_val_adj** | **celltype** | **Foldchange_UP** | **Foldchange_DOWN** | **Ratio_pct1_pct2** |
| BIRC6 | 0,300696173 | -0,300705211 | 0,321 | 0,488 | 1 | Ciliated | 0,811855452 | 1,231746362 | 0,657786885 |
| TMX2 | 0,3010365 | -0,329521968 | 0,25 | 0,39 | 1 | Ciliated | 0,795800126 | 1,256596936 | 0,641025641 |
| VKORC11 | 0,302264056 | 0,318557284 | 0,393 | 0,366 | 1 | Ciliated | 1,247082824 | 0,80187136 | 1,073770492 |
| LRRC45 | 0,303245226 | 0,416618951 | 0,321 | 0,268 | 1 | Ciliated | 1,334795706 | 0,749178316 | 1,197761194 |
| AC008771.1 | 0,303300672 | -0,284570471 | 0,429 | 0,585 | 1 | Ciliated | 0,820986002 | 1,218047564 | 0,733333333 |
| DNAH5 | 0,303921584 | -0,465523579 | 0,75 | 0,756 | 1 | Ciliated | 0,724208202 | 1,380818385 | 0,992063492 |
| MBOAT71 | 0,304453418 | 0,362638658 | 0,357 | 0,317 | 1 | Ciliated | 1,285775404 | 0,777740807 | 1,126182965 |
| TAPT1 | 0,304540653 | -0,264945996 | 0,107 | 0,22 | 1 | Ciliated | 0,832229887 | 1,20159107 | 0,486363636 |
| ZNF335 | 0,304540653 | 0,309332334 | 0,214 | 0,146 | 1 | Ciliated | 1,239134107 | 0,807015152 | 1,465753425 |
| IGF2R | 0,304913692 | 0,267407688 | 0,286 | 0,22 | 1 | Ciliated | 1,203643113 | 0,830811051 | 1,3 |
| TM2D2 | 0,304913692 | 0,416701461 | 0,286 | 0,22 | 1 | Ciliated | 1,334872048 | 0,749135471 | 1,3 |
| STX17 | 0,30559255 | -0,260039681 | 0,143 | 0,268 | 1 | Ciliated | 0,835064951 | 1,197511641 | 0,53358209 |
| RSPH4A | 0,305940022 | -0,289363497 | 0,786 | 0,829 | 1 | Ciliated | 0,818262989 | 1,22210098 | 0,948130277 |
| TENT5A | 0,306598707 | -0,482738897 | 0,429 | 0,512 | 1 | Ciliated | 0,715617763 | 1,397394044 | 0,837890625 |
| PNMA1 | 0,306598707 | 0,395495026 | 0,5 | 0,463 | 1 | Ciliated | 1,315394029 | 0,760228478 | 1,079913607 |
| LRRC71 | 0,307543063 | -0,291205421 | 0,464 | 0,634 | 1 | Ciliated | 0,817218959 | 1,223662262 | 0,731861199 |
| SARNP1 | 0,308129238 | -0,276810537 | 0,321 | 0,512 | 1 | Ciliated | 0,825413799 | 1,211513547 | 0,626953125 |
| ELF2 | 0,308898871 | -0,303157001 | 0,321 | 0,537 | 1 | Ciliated | 0,810476915 | 1,233841435 | 0,597765363 |
| KMT5C | 0,308913592 | 0,261463234 | 0,25 | 0,171 | 1 | Ciliated | 1,198693847 | 0,834241372 | 1,461988304 |
| DGUOK | 0,309423325 | 0,364461715 | 0,536 | 0,512 | 1 | Ciliated | 1,287401196 | 0,776758638 | 1,046875 |
| CSNK1G2 | 0,30942716 | 0,655975107 | 0,464 | 0,463 | 1 | Ciliated | 1,575680584 | 0,634646394 | 1,002159827 |
| C2orf73 | 0,31032739 | 0,549008472 | 0,607 | 0,61 | 1 | Ciliated | 1,463079812 | 0,683489712 | 0,995081967 |
| STX4 | 0,310464806 | 0,353267794 | 0,321 | 0,268 | 1 | Ciliated | 1,277450859 | 0,782808977 | 1,197761194 |
| SMARCD21 | 0,310464806 | 0,286923838 | 0,321 | 0,268 | 1 | Ciliated | 1,220036101 | 0,819647877 | 1,197761194 |
| MLXIP | 0,310576286 | 0,45686242 | 0,571 | 0,561 | 1 | Ciliated | 1,372553534 | 0,728569032 | 1,017825312 |
| ERH2 | 0,310895206 | -0,408545604 | 0,75 | 0,805 | 1 | Ciliated | 0,753382484 | 1,327347027 | 0,931677019 |
| PLGRKT | 0,311364219 | 0,338530936 | 0,357 | 0,317 | 1 | Ciliated | 1,264468358 | 0,790846203 | 1,126182965 |
| MSRB1 | 0,311364219 | 0,279927151 | 0,357 | 0,317 | 1 | Ciliated | 1,214133575 | 0,823632606 | 1,126182965 |
| ATP13A1 | 0,311364219 | 0,526016221 | 0,357 | 0,317 | 1 | Ciliated | 1,439947508 | 0,69446976 | 1,126182965 |
| DCTPP1 | 0,312919664 | 0,27191997 | 0,321 | 0,244 | 1 | Ciliated | 1,207413611 | 0,828216604 | 1,31557377 |
| SPAG1 | 0,313333184 | -0,346470002 | 0,643 | 0,756 | 1 | Ciliated | 0,786506175 | 1,271445834 | 0,850529101 |

| Supplementary Table 4 - clusermarker | | | | | | | | | |
|---|---|---|---|---|---|---|---|---|---|
| | **p_val** | **avg_log2FC** | **pct.1** | **pct.2** | **p_val_adj** | **celltype** | **Foldchange_UP** | **Foldchange_DOWN** | **Ratio_pct1_pct2** |
| MUTYH | 0,313336916 | 0,378567633 | 0,214 | 0,146 | 1 | Ciliated | 1,300050471 | 0,769200906 | 1,465753425 |
| SPATA4 | 0,315121244 | 0,25872837 | 0,536 | 0,537 | 1 | Ciliated | 1,19642368 | 0,835824313 | 0,998137803 |
| GPRC5C1 | 0,315307942 | 0,298534188 | 0,464 | 0,439 | 1 | Ciliated | 1,229894177 | 0,813078083 | 1,056947608 |
| CTSC2 | 0,315794204 | 0,525182738 | 0,5 | 0,439 | 1 | Ciliated | 1,439115853 | 0,694871089 | 1,138952164 |
| ARID5B1 | 0,316055381 | -0,330571465 | 0,393 | 0,537 | 1 | Ciliated | 0,795221427 | 1,257511388 | 0,731843575 |
| AK22 | 0,316701772 | -0,427594328 | 0,429 | 0,659 | 1 | Ciliated | 0,743500528 | 1,344988958 | 0,650986343 |
| PYCARD2 | 0,316701772 | 0,283464282 | 0,571 | 0,561 | 1 | Ciliated | 1,217113982 | 0,821615736 | 1,017825312 |
| MYLIP1 | 0,316876368 | -0,273702622 | 0,357 | 0,463 | 1 | Ciliated | 0,827193857 | 1,208906463 | 0,771058315 |
| MRPL3 | 0,317446738 | 0,348714703 | 0,393 | 0,39 | 1 | Ciliated | 1,273425627 | 0,785283395 | 1,007692308 |
| TNFSF101 | 0,317888057 | -0,258217302 | 0,607 | 0,683 | 1 | Ciliated | 0,836120452 | 1,195999928 | 0,888726208 |
| YWHAH2 | 0,318354487 | 0,265374565 | 0,714 | 0,683 | 1 | Ciliated | 1,20194807 | 0,8319827 | 1,045387994 |
| ZCCHC17 | 0,318429617 | -0,387664442 | 0,214 | 0,366 | 1 | Ciliated | 0,764366024 | 1,308273744 | 0,584699454 |
| RPL111 | 0,319284767 | -0,330233118 | 1 | 0,976 | 1 | Ciliated | 0,795407947 | 1,257216506 | 1,024590164 |
| SELENOM1 | 0,320290524 | 0,266012881 | 0,393 | 0,341 | 1 | Ciliated | 1,202479985 | 0,831614673 | 1,152492669 |
| HEMK1 | 0,320398351 | 0,327146069 | 0,286 | 0,22 | 1 | Ciliated | 1,254529216 | 0,797111767 | 1,3 |
| EPS81 | 0,320718296 | -0,509576038 | 0,5 | 0,659 | 1 | Ciliated | 0,702428829 | 1,423631774 | 0,758725341 |
| LINC02541 | 0,320819105 | -0,313910076 | 0,286 | 0,463 | 1 | Ciliated | 0,804458512 | 1,243072185 | 0,617710583 |
| ATP5MK | 0,32099528 | -0,292094388 | 0,821 | 0,829 | 1 | Ciliated | 0,816715556 | 1,224416497 | 0,990349819 |
| PABPC1L | 0,32160392 | 0,268877045 | 0,143 | 0,073 | 1 | Ciliated | 1,204869624 | 0,829965317 | 1,95890411 |
| AC026704.1 | 0,322298785 | 0,301764781 | 0,214 | 0,146 | 1 | Ciliated | 1,232651336 | 0,811259414 | 1,465753425 |
| PARP9 | 0,322426083 | 0,519463126 | 0,393 | 0,366 | 1 | Ciliated | 1,433421726 | 0,697631396 | 1,073770492 |
| CYCS1 | 0,322473464 | -0,366314181 | 0,393 | 0,561 | 1 | Ciliated | 0,775761895 | 1,289055322 | 0,700534759 |
| CCDC801 | 0,324284521 | -0,269809255 | 0,893 | 0,805 | 1 | Ciliated | 0,829429201 | 1,205648413 | 1,10931677 |
| SART1 | 0,32479201 | 0,273392653 | 0,393 | 0,317 | 1 | Ciliated | 1,208646752 | 0,827371602 | 1,239747634 |
| TXNDC171 | 0,324991458 | 0,351008428 | 0,571 | 0,585 | 1 | Ciliated | 1,275451842 | 0,784035874 | 0,976068376 |
| BRMS1 | 0,325241628 | -0,304970945 | 0,214 | 0,341 | 1 | Ciliated | 0,809458518 | 1,235393757 | 0,627565982 |
| COQ9 | 0,325493826 | 0,262845352 | 0,357 | 0,317 | 1 | Ciliated | 1,19984276 | 0,833442542 | 1,126182965 |
| PYCR3 | 0,325746338 | 0,272599412 | 0,25 | 0,171 | 1 | Ciliated | 1,207982381 | 0,827826644 | 1,461988304 |
| ABHD12 | 0,326011803 | 0,266107098 | 0,25 | 0,195 | 1 | Ciliated | 1,202558517 | 0,831560366 | 1,282051282 |
| ZWILCH | 0,327730009 | 0,338617679 | 0,107 | 0,049 | 1 | Ciliated | 1,264544388 | 0,790798654 | 2,183673469 |
| ZNF562 | 0,328102305 | -0,259671378 | 0,107 | 0,195 | 1 | Ciliated | 0,835278161 | 1,19720597 | 0,548717949 |

| Supplementary Table 4 - clusermarker | | | | | | | | | |
|---|---|---|---|---|---|---|---|---|---|
| | **p_val** | **avg_log2FC** | **pct.1** | **pct.2** | **p_val_adj** | **celltype** | **Foldchange_UP** | **Foldchange_DOWN** | **Ratio_pct1_pct2** |
| USP101 | 0,328278011 | -0,338093084 | 0,464 | 0,561 | 1 | Ciliated | 0,791086258 | 1,264084656 | 0,827094474 |
| CEP128 | 0,328330153 | 0,272310161 | 0,286 | 0,22 | 1 | Ciliated | 1,207740212 | 0,827992634 | 1,3 |
| DYNLT12 | 0,331304939 | -0,318240499 | 1 | 0,976 | 1 | Ciliated | 0,802047453 | 1,246809022 | 1,024590164 |
| PRKAR1A1 | 0,331333511 | -0,252794589 | 0,821 | 0,732 | 1 | Ciliated | 0,839269124 | 1,191512915 | 1,121584699 |
| CFAP299 | 0,33186 | -0,436344555 | 0,536 | 0,707 | 1 | Ciliated | 0,7390047 | 1,353171367 | 0,758132956 |
| POLR3K | 0,332237555 | -0,293102382 | 0,357 | 0,561 | 1 | Ciliated | 0,816145126 | 1,225272281 | 0,636363636 |
| LINC00623 | 0,332343407 | -0,329191483 | 0,179 | 0,293 | 1 | Ciliated | 0,795982444 | 1,256309115 | 0,610921502 |
| SLC35A21 | 0,33249494 | 0,334906171 | 0,429 | 0,39 | 1 | Ciliated | 1,261295375 | 0,792835699 | 1,1 |
| CD552 | 0,332564121 | 0,290525842 | 0,75 | 0,634 | 1 | Ciliated | 1,223085994 | 0,817603999 | 1,1829653 |
| RESF1 | 0,332712618 | 0,2581188 | 0,357 | 0,317 | 1 | Ciliated | 1,195918272 | 0,836177541 | 1,126182965 |
| CEP631 | 0,333158045 | -0,485177918 | 0,25 | 0,366 | 1 | Ciliated | 0,714408961 | 1,399758478 | 0,683060109 |
| TUBB2 | 0,333617344 | -0,61275637 | 0,321 | 0,488 | 1 | Ciliated | 0,653946098 | 1,529178022 | 0,657786885 |
| VTI1B | 0,334311953 | 0,306643356 | 0,393 | 0,341 | 1 | Ciliated | 1,236826689 | 0,808520716 | 1,152492669 |
| CPEB4 | 0,334725127 | -0,259212175 | 0,393 | 0,61 | 1 | Ciliated | 0,835544068 | 1,196824965 | 0,644262295 |
| RBM38 | 0,336343973 | 0,631613472 | 0,393 | 0,366 | 1 | Ciliated | 1,549296718 | 0,645454153 | 1,073770492 |
| KRT192 | 0,3374057 | 0,339964623 | 0,964 | 0,951 | 1 | Ciliated | 1,265725556 | 0,790060685 | 1,013669821 |
| COPS5 | 0,33842269 | -0,289337152 | 0,5 | 0,415 | 1 | Ciliated | 0,818277931 | 1,222078663 | 1,204819277 |
| DMTN | 0,339112097 | 0,2782166 | 0,357 | 0,341 | 1 | Ciliated | 1,212694874 | 0,824609736 | 1,046920821 |
| DIAPH1 | 0,339398401 | -0,324991749 | 0,321 | 0,463 | 1 | Ciliated | 0,798302952 | 1,252657274 | 0,693304536 |
| BBOF1 | 0,339988757 | -0,435943283 | 0,75 | 0,756 | 1 | Ciliated | 0,739210276 | 1,352795047 | 0,992063492 |
| TMBIM42 | 0,340596923 | 0,253355712 | 0,786 | 0,756 | 1 | Ciliated | 1,191976433 | 0,838942761 | 1,03968254 |
| TMEM106C | 0,340719204 | 0,406014764 | 0,214 | 0,146 | 1 | Ciliated | 1,325020577 | 0,754705261 | 1,465753425 |
| TJP3 | 0,341340119 | -0,275985267 | 0,5 | 0,683 | 1 | Ciliated | 0,825886099 | 1,210820719 | 0,732064422 |
| FBLN21 | 0,341345345 | 0,259945332 | 0,107 | 0,049 | 1 | Ciliated | 1,19743333 | 0,835119564 | 2,183673469 |
| EMP12 | 0,342033992 | -0,427601983 | 0,179 | 0,268 | 1 | Ciliated | 0,743496583 | 1,344996094 | 0,667910448 |
| EPC12 | 0,342116435 | 0,257773529 | 0,5 | 0,439 | 1 | Ciliated | 1,195632095 | 0,836377682 | 1,138952164 |
| STN11 | 0,342116435 | 0,350422087 | 0,464 | 0,463 | 1 | Ciliated | 1,274933578 | 0,784354587 | 1,002159827 |
| PSMC31 | 0,342116435 | -0,288210859 | 0,393 | 0,512 | 1 | Ciliated | 0,818916999 | 1,221124975 | 0,767578125 |
| MARCHF10 | 0,342768248 | 0,273359015 | 0,607 | 0,585 | 1 | Ciliated | 1,208618571 | 0,827390894 | 1,037606838 |
| CNOT11 | 0,343445736 | 0,257436722 | 0,393 | 0,366 | 1 | Ciliated | 1,195352998 | 0,836572963 | 1,073770492 |
| ATP5PD1 | 0,346003868 | -0,351231506 | 0,893 | 0,829 | 1 | Ciliated | 0,783914651 | 1,275649076 | 1,077201448 |

| Supplementary Table 4 - clusermarker | | | | | | | | | |
|---|---|---|---|---|---|---|---|---|---|
| | **p_val** | **avg_log2FC** | **pct.1** | **pct.2** | **p_val_adj** | **celltype** | **Foldchange_UP** | **Foldchange_DOWN** | **Ratio_pct1_pct2** |
| CDK17 | 0,346399139 | 0,296657467 | 0,286 | 0,244 | 1 | Ciliated | 1,228295317 | 0,814136459 | 1,172131148 |
| GRN1 | 0,347457815 | 0,28063083 | 0,357 | 0,317 | 1 | Ciliated | 1,214725917 | 0,823230974 | 1,126182965 |
| IFT140 | 0,348160733 | 0,352590698 | 0,464 | 0,415 | 1 | Ciliated | 1,276851456 | 0,783176457 | 1,118072289 |
| CCDC12 | 0,348723076 | 0,250954597 | 0,393 | 0,341 | 1 | Ciliated | 1,189994246 | 0,840340198 | 1,152492669 |
| PTPN1 | 0,348910236 | -0,36561733 | 0,357 | 0,537 | 1 | Ciliated | 0,776136694 | 1,288432833 | 0,664804469 |
| UBLCP1 | 0,350177395 | 0,346350261 | 0,214 | 0,146 | 1 | Ciliated | 1,271340311 | 0,786571457 | 1,465753425 |
| TIMP31 | 0,350231862 | -0,479719129 | 0,071 | 0,146 | 1 | Ciliated | 0,717117223 | 1,394472157 | 0,48630137 |
| EIF1AX1 | 0,350586531 | -0,34313245 | 0,5 | 0,634 | 1 | Ciliated | 0,788327797 | 1,268507852 | 0,788643533 |
| RBM32 | 0,350586531 | -0,369447609 | 0,5 | 0,634 | 1 | Ciliated | 0,774078826 | 1,291858098 | 0,788643533 |
| NAA601 | 0,351208102 | -0,293575866 | 0,25 | 0,39 | 1 | Ciliated | 0,815877316 | 1,225674474 | 0,641025641 |
| STK32C | 0,351658902 | 0,348054413 | 0,25 | 0,195 | 1 | Ciliated | 1,272842941 | 0,785642885 | 1,282051282 |
| SAMM502 | 0,35221907 | 0,298814873 | 0,393 | 0,39 | 1 | Ciliated | 1,230133484 | 0,812919909 | 1,007692308 |
| ANKRD36C | 0,353478956 | -0,259693941 | 0,321 | 0,463 | 1 | Ciliated | 0,835265097 | 1,197224694 | 0,693304536 |
| COG1 | 0,354983939 | 0,423076025 | 0,357 | 0,317 | 1 | Ciliated | 1,340783244 | 0,74583271 | 1,126182965 |
| UBE2M | 0,355140664 | 0,3314731 | 0,464 | 0,415 | 1 | Ciliated | 1,258297535 | 0,794724596 | 1,118072289 |
| PIM31 | 0,355789116 | -0,357241597 | 0,357 | 0,537 | 1 | Ciliated | 0,780655752 | 1,280974358 | 0,664804469 |
| CEP120 | 0,356176139 | -0,258948395 | 0,214 | 0,366 | 1 | Ciliated | 0,835696851 | 1,19660616 | 0,584699454 |
| ZFP90 | 0,356176139 | -0,461051783 | 0,214 | 0,366 | 1 | Ciliated | 0,726456449 | 1,376545009 | 0,584699454 |
| DPY30 | 0,356396359 | 0,284792416 | 0,893 | 0,683 | 1 | Ciliated | 1,218234964 | 0,820859711 | 1,307467057 |
| CNOT1 | 0,357142884 | 0,380087794 | 0,571 | 0,512 | 1 | Ciliated | 1,30142105 | 0,76839083 | 1,115234375 |
| C2orf50 | 0,359684946 | 0,260204177 | 0,179 | 0,341 | 1 | Ciliated | 1,197648189 | 0,834969742 | 0,524926686 |
| UBE3D | 0,3604845 | 0,307066961 | 0,25 | 0,195 | 1 | Ciliated | 1,2371899 | 0,808283352 | 1,282051282 |
| CCDC39 | 0,362255362 | -0,274180245 | 0,714 | 0,683 | 1 | Ciliated | 0,826920049 | 1,209306753 | 1,045387994 |
| SHISA51 | 0,362421361 | 0,298687971 | 0,5 | 0,488 | 1 | Ciliated | 1,230025283 | 0,812991419 | 1,024590164 |
| BCAS4 | 0,36261215 | 0,284935972 | 0,357 | 0,317 | 1 | Ciliated | 1,218356191 | 0,820778035 | 1,126182965 |
| ASL | 0,362752914 | -0,29516993 | 0,357 | 0,537 | 1 | Ciliated | 0,814976334 | 1,227029496 | 0,664804469 |
| CASC31 | 0,362752914 | 0,380533173 | 0,464 | 0,463 | 1 | Ciliated | 1,301822877 | 0,768153654 | 1,002159827 |
| HMCES | 0,362850762 | 0,296101442 | 0,286 | 0,244 | 1 | Ciliated | 1,227822014 | 0,814450294 | 1,172131148 |
| IFITM12 | 0,363523391 | 0,355338648 | 0,357 | 0,293 | 1 | Ciliated | 1,279285837 | 0,781686134 | 1,218430034 |
| SLC44A11 | 0,364051003 | -0,265231947 | 0,214 | 0,366 | 1 | Ciliated | 0,83206495 | 1,201829256 | 0,584699454 |
| CRYBG1 | 0,365320683 | 0,32600739 | 0,393 | 0,366 | 1 | Ciliated | 1,253539441 | 0,797741154 | 1,073770492 |

| Supplementary Table 4 - clusermarker | | | | | | | | | |
|---|---|---|---|---|---|---|---|---|---|
| | **p_val** | **avg_log2FC** | **pct.1** | **pct.2** | **p_val_adj** | **celltype** | **Foldchange_UP** | **Foldchange_DOWN** | **Ratio_pct1_pct2** |
| LHPP | 0,365320683 | 0,285170211 | 0,393 | 0,366 | 1 | Ciliated | 1,218554022 | 0,820644782 | 1,073770492 |
| C19orf532 | 0,365375808 | 0,253645411 | 0,643 | 0,659 | 1 | Ciliated | 1,192215811 | 0,838774315 | 0,975720789 |
| MAML21 | 0,366177605 | -0,251990609 | 0,357 | 0,561 | 1 | Ciliated | 0,839736959 | 1,190849098 | 0,636363636 |
| SLC35F5 | 0,366778748 | -0,259357355 | 0,321 | 0,439 | 1 | Ciliated | 0,83545999 | 1,19694541 | 0,731207289 |
| RAN1 | 0,368083429 | -0,302123946 | 0,679 | 0,78 | 1 | Ciliated | 0,811057472 | 1,232958248 | 0,870512821 |
| TAPBP1 | 0,36817394 | 0,362993477 | 0,5 | 0,537 | 1 | Ciliated | 1,286091669 | 0,777549551 | 0,931098696 |
| MAP1LC3B | 0,36817394 | -0,279998012 | 0,464 | 0,561 | 1 | Ciliated | 0,823592152 | 1,214193211 | 0,827094474 |
| VIM-AS1 | 0,368966617 | 0,450231444 | 0,357 | 0,341 | 1 | Ciliated | 1,366259421 | 0,731925419 | 1,046920821 |
| MPND | 0,369447825 | 0,267600047 | 0,25 | 0,195 | 1 | Ciliated | 1,203803609 | 0,830700284 | 1,282051282 |
| MXRA71 | 0,369698823 | 0,303851439 | 0,464 | 0,439 | 1 | Ciliated | 1,234435485 | 0,810086888 | 1,056947608 |
| PCMT11 | 0,370342192 | 0,275708117 | 0,357 | 0,317 | 1 | Ciliated | 1,210588136 | 0,826044772 | 1,126182965 |
| ARHGEF18 | 0,370342192 | 0,443334265 | 0,357 | 0,317 | 1 | Ciliated | 1,359743251 | 0,735432957 | 1,126182965 |
| ATP6V1C2 | 0,370385423 | -0,288253075 | 0,143 | 0,244 | 1 | Ciliated | 0,818893037 | 1,221160707 | 0,586065574 |
| PDIA61 | 0,370598884 | 0,300515102 | 0,571 | 0,585 | 1 | Ciliated | 1,231584061 | 0,81196244 | 0,976068376 |
| PSMG31 | 0,370752 | 0,386262651 | 0,5 | 0,561 | 1 | Ciliated | 1,30700318 | 0,765109079 | 0,891265597 |
| TMEM63A | 0,371257008 | 0,273048849 | 0,286 | 0,244 | 1 | Ciliated | 1,208358757 | 0,827568794 | 1,172131148 |
| RO60 | 0,372268321 | -0,282544697 | 0,214 | 0,341 | 1 | Ciliated | 0,822139607 | 1,216338432 | 0,627565982 |
| ELF3-AS1 | 0,372416372 | 0,254327083 | 0,429 | 0,415 | 1 | Ciliated | 1,192779265 | 0,838378089 | 1,03373494 |
| PMAIP1 | 0,372416372 | -0,302986246 | 0,357 | 0,463 | 1 | Ciliated | 0,810572848 | 1,233695408 | 0,771058315 |
| GGNBP21 | 0,373146305 | 0,421569285 | 0,536 | 0,561 | 1 | Ciliated | 1,339383672 | 0,746612058 | 0,95543672 |
| TNFAIP1 | 0,373369168 | -0,284840214 | 0,071 | 0,146 | 1 | Ciliated | 0,820832515 | 1,218275326 | 0,48630137 |
| AFDN2 | 0,374680905 | -0,303771626 | 0,679 | 0,78 | 1 | Ciliated | 0,810131705 | 1,234367195 | 0,870512821 |
| GAS8 | 0,37510532 | 0,285850714 | 0,536 | 0,512 | 1 | Ciliated | 1,219128936 | 0,820257784 | 1,046875 |
| CFAP70 | 0,375891575 | -0,412401256 | 0,571 | 0,659 | 1 | Ciliated | 0,751371731 | 1,330899152 | 0,86646434 |
| FBXO321 | 0,376065635 | 0,307944635 | 0,321 | 0,293 | 1 | Ciliated | 1,237942782 | 0,807791777 | 1,09556314 |
| BUB3 | 0,376676693 | 0,376735093 | 0,357 | 0,341 | 1 | Ciliated | 1,298400169 | 0,770178581 | 1,046920821 |
| EIF5A1 | 0,37693444 | -0,406539554 | 0,393 | 0,512 | 1 | Ciliated | 0,754430782 | 1,32550265 | 0,767578125 |
| NRDC1 | 0,377424272 | 0,314338267 | 0,571 | 0,585 | 1 | Ciliated | 1,243441183 | 0,804219784 | 0,976068376 |
| GPBP11 | 0,377675692 | 0,415985085 | 0,571 | 0,512 | 1 | Ciliated | 1,334209376 | 0,74950755 | 1,115234375 |
| GPSM3 | 0,377740781 | -0,29513727 | 0,107 | 0,195 | 1 | Ciliated | 0,814994783 | 1,227001719 | 0,548717949 |
| ELMOD3 | 0,378548416 | 0,262148871 | 0,25 | 0,195 | 1 | Ciliated | 1,199263659 | 0,833844995 | 1,282051282 |

| Supplementary Table 4 - clusermarker | | | | | | | | | |
|---|---|---|---|---|---|---|---|---|---|
| | **p_val** | **avg_log2FC** | **pct.1** | **pct.2** | **p_val_adj** | **celltype** | **Foldchange_UP** | **Foldchange_DOWN** | **Ratio_pct1_pct2** |
| AC233723.1 | 0,378548416 | 0,269444241 | 0,25 | 0,195 | 1 | Ciliated | 1,205343412 | 0,82963908 | 1,282051282 |
| ENOPH1 | 0,37974925 | -0,348240645 | 0,143 | 0,244 | 1 | Ciliated | 0,785541475 | 1,273007259 | 0,586065574 |
| CGGBP12 | 0,380376555 | -0,305227086 | 0,286 | 0,439 | 1 | Ciliated | 0,809314817 | 1,235613112 | 0,651480638 |
| SAMD9 | 0,380497137 | -0,292207085 | 0,214 | 0,341 | 1 | Ciliated | 0,81665176 | 1,224512147 | 0,627565982 |
| CCDC65 | 0,382142012 | -0,280807265 | 0,607 | 0,683 | 1 | Ciliated | 0,823130303 | 1,214874482 | 0,888726208 |
| RHOB1 | 0,38263115 | 0,255171685 | 0,821 | 0,732 | 1 | Ciliated | 1,193477762 | 0,837887418 | 1,121584699 |
| JCHAIN | 0,38432644 | 1,594039414 | 0,107 | 0,049 | 1 | Ciliated | 3,018934413 | 0,331242705 | 2,183673469 |
| RCAN11 | 0,38432644 | -0,370410807 | 0,107 | 0,049 | 1 | Ciliated | 0,773562194 | 1,29272088 | 2,183673469 |
| PPP1R35 | 0,384485369 | 0,257778302 | 0,357 | 0,341 | 1 | Ciliated | 1,19563605 | 0,836374915 | 1,046920821 |
| THNSL1 | 0,385280639 | 0,380022611 | 0,179 | 0,122 | 1 | Ciliated | 1,301362251 | 0,768425547 | 1,467213115 |
| ANTKMT1 | 0,385280639 | 0,372652975 | 0,179 | 0,122 | 1 | Ciliated | 1,294731528 | 0,772360894 | 1,467213115 |
| SYCE3 | 0,385280639 | 0,361219685 | 0,179 | 0,122 | 1 | Ciliated | 1,284511392 | 0,778506136 | 1,467213115 |
| MAST4-AS1 | 0,385284646 | -0,26016214 | 0,071 | 0,146 | 1 | Ciliated | 0,834994072 | 1,197613293 | 0,48630137 |
| ATF3 | 0,385551014 | -0,583938572 | 0,714 | 0,61 | 1 | Ciliated | 0,66713999 | 1,498935778 | 1,170491803 |
| POP5 | 0,385759939 | 0,277976214 | 0,536 | 0,585 | 1 | Ciliated | 1,212492828 | 0,824747146 | 0,916239316 |
| DDX3X1 | 0,385991172 | -0,335597628 | 0,786 | 0,756 | 1 | Ciliated | 0,792455798 | 1,261900035 | 1,03968254 |
| MCFD2 | 0,386446892 | 0,285981064 | 0,393 | 0,341 | 1 | Ciliated | 1,219239091 | 0,820183676 | 1,152492669 |
| SDHA | 0,387057073 | 0,255248038 | 0,536 | 0,561 | 1 | Ciliated | 1,193540927 | 0,837843074 | 0,95543672 |
| HADHB | 0,38719196 | -0,420700852 | 0,357 | 0,463 | 1 | Ciliated | 0,747061618 | 1,338577669 | 0,771058315 |
| PGAP4 | 0,387785727 | 0,348034026 | 0,25 | 0,195 | 1 | Ciliated | 1,272824954 | 0,785653987 | 1,282051282 |
| FARSA | 0,388428405 | 0,261177825 | 0,286 | 0,244 | 1 | Ciliated | 1,198456733 | 0,834406427 | 1,172131148 |
| GRPEL1 | 0,388836755 | 0,262336044 | 0,321 | 0,268 | 1 | Ciliated | 1,19941926 | 0,833736821 | 1,197761194 |
| SFTA3 | 0,389257227 | -0,311382026 | 0,143 | 0,244 | 1 | Ciliated | 0,805869409 | 1,240895844 | 0,586065574 |
| CEACAM51 | 0,389257227 | 0,465941897 | 0,25 | 0,171 | 1 | Ciliated | 1,38121882 | 0,723998244 | 1,461988304 |
| TMEM165 | 0,389262882 | 0,301208564 | 0,571 | 0,561 | 1 | Ciliated | 1,23217619 | 0,811572247 | 1,017825312 |
| TMPRSS41 | 0,389694272 | 0,313274294 | 0,679 | 0,537 | 1 | Ciliated | 1,242524496 | 0,804813107 | 1,26443203 |
| OSTC1 | 0,391452853 | -0,319734247 | 0,357 | 0,537 | 1 | Ciliated | 0,801217453 | 1,24810062 | 0,664804469 |
| SLC25A24 | 0,391760135 | 0,401044027 | 0,5 | 0,561 | 1 | Ciliated | 1,320463137 | 0,757310047 | 0,891265597 |
| BPNT1 | 0,391827819 | -0,366625988 | 0,179 | 0,293 | 1 | Ciliated | 0,77559425 | 1,289333953 | 0,610921502 |
| ZNF518A | 0,392392273 | 0,265864287 | 0,357 | 0,341 | 1 | Ciliated | 1,202356139 | 0,831700332 | 1,046920821 |
| ZNF106 | 0,394912452 | 0,32408988 | 0,75 | 0,732 | 1 | Ciliated | 1,251874448 | 0,79880215 | 1,024590164 |

| Supplementary Table 4 - clusermarker | | | | | | | | | |
|---|---|---|---|---|---|---|---|---|---|
| | **p_val** | **avg_log2FC** | **pct.1** | **pct.2** | **p_val_adj** | **celltype** | **Foldchange_UP** | **Foldchange_DOWN** | **Ratio_pct1_pct2** |
| FKBP1A1 | 0,394937518 | -0,269188777 | 0,5 | 0,683 | 1 | Ciliated | 0,829786001 | 1,205129996 | 0,732064422 |
| C15orf482 | 0,396299367 | 0,364565191 | 0,179 | 0,122 | 1 | Ciliated | 1,287493538 | 0,776702928 | 1,467213115 |
| LYPD21 | 0,396310484 | 1,48157512 | 0,286 | 0,22 | 1 | Ciliated | 2,792534531 | 0,358097631 | 1,3 |
| CNIH2 | 0,397159122 | 0,349638071 | 0,25 | 0,195 | 1 | Ciliated | 1,274240919 | 0,784780951 | 1,282051282 |
| ODAD3 | 0,397264682 | 0,467407061 | 0,464 | 0,537 | 1 | Ciliated | 1,382622262 | 0,723263343 | 0,86405959 |
| EXOSC9 | 0,397429679 | -0,268463473 | 0,071 | 0,146 | 1 | Ciliated | 0,830203274 | 1,204524278 | 0,48630137 |
| HSD17B7 | 0,398828181 | -0,277578219 | 0,107 | 0,195 | 1 | Ciliated | 0,8249747 | 1,212158385 | 0,548717949 |
| H2AZ11 | 0,400608701 | -0,391930666 | 0,643 | 0,683 | 1 | Ciliated | 0,76210904 | 1,312148193 | 0,941434846 |
| SRSF31 | 0,400608701 | -0,351975436 | 0,643 | 0,683 | 1 | Ciliated | 0,783510527 | 1,276307038 | 0,941434846 |
| TRIM52-AS1 | 0,400818598 | 0,25763639 | 0,321 | 0,293 | 1 | Ciliated | 1,195518446 | 0,83645719 | 1,09556314 |
| UBE4A | 0,400818598 | 0,433467376 | 0,321 | 0,293 | 1 | Ciliated | 1,350475415 | 0,740479974 | 1,09556314 |
| SNRPE | 0,403793954 | -0,281019106 | 0,536 | 0,634 | 1 | Ciliated | 0,823009446 | 1,215052883 | 0,845425868 |
| CFAP52 | 0,404520786 | -0,31498417 | 0,643 | 0,78 | 1 | Ciliated | 0,803859811 | 1,243998003 | 0,824358974 |
| KDELR12 | 0,404566657 | 0,252327917 | 0,536 | 0,488 | 1 | Ciliated | 1,191127556 | 0,839540648 | 1,098360656 |
| ATN1 | 0,404789021 | 0,332307073 | 0,393 | 0,39 | 1 | Ciliated | 1,259025125 | 0,794265325 | 1,007692308 |
| KATNB1 | 0,404789021 | 0,347763744 | 0,393 | 0,39 | 1 | Ciliated | 1,272586519 | 0,785801189 | 1,007692308 |
| FIBP1 | 0,405333272 | 0,292681023 | 0,5 | 0,488 | 1 | Ciliated | 1,224914476 | 0,816383527 | 1,024590164 |
| RNASEH2C | 0,405846445 | -0,27691563 | 0,214 | 0,341 | 1 | Ciliated | 0,825353674 | 1,211601803 | 0,627565982 |
| SLC39A3 | 0,405846445 | 0,318080644 | 0,321 | 0,268 | 1 | Ciliated | 1,246670879 | 0,802136327 | 1,197761194 |
| EIF4E21 | 0,405917094 | 0,307701389 | 0,464 | 0,488 | 1 | Ciliated | 1,237734077 | 0,807927986 | 0,950819672 |
| C11orf16 | 0,406158706 | -0,274240644 | 0,429 | 0,61 | 1 | Ciliated | 0,826885431 | 1,209357382 | 0,703278689 |
| CHD6 | 0,406305665 | -0,2507527 | 0,393 | 0,512 | 1 | Ciliated | 0,840457807 | 1,189827724 | 0,767578125 |
| VPS37C | 0,40637832 | -0,27339798 | 0,25 | 0,39 | 1 | Ciliated | 0,827368548 | 1,208651215 | 0,641025641 |
| TPM12 | 0,406441052 | 0,269985952 | 0,429 | 0,439 | 1 | Ciliated | 1,205796086 | 0,829327621 | 0,977220957 |
| RAB40C | 0,406667881 | 0,329047533 | 0,25 | 0,195 | 1 | Ciliated | 1,256183769 | 0,79606187 | 1,282051282 |
| CEP290 | 0,407364832 | -0,38137025 | 0,571 | 0,61 | 1 | Ciliated | 0,767708087 | 1,302578438 | 0,936065574 |
| TSPYL5 | 0,407506224 | 0,46217203 | 0,179 | 0,122 | 1 | Ciliated | 1,377614306 | 0,725892578 | 1,467213115 |
| LINC01550 | 0,407506224 | 0,26586915 | 0,179 | 0,122 | 1 | Ciliated | 1,202360192 | 0,831697528 | 1,467213115 |
| ANKS1A | 0,408499043 | 0,327838284 | 0,393 | 0,317 | 1 | Ciliated | 1,255131292 | 0,796729399 | 1,239747634 |
| MCM9 | 0,408499043 | 0,291924433 | 0,357 | 0,341 | 1 | Ciliated | 1,224272265 | 0,816811774 | 1,046920821 |
| UBFD1 | 0,40870273 | -0,26861081 | 0,143 | 0,244 | 1 | Ciliated | 0,830118493 | 1,204647298 | 0,586065574 |

| Supplementary Table 4 - clusermarker | | | | | | | | | |
|---|---|---|---|---|---|---|---|---|---|
| | **p_val** | **avg_log2FC** | **pct.1** | **pct.2** | **p_val_adj** | **celltype** | **Foldchange_UP** | **Foldchange_DOWN** | **Ratio_pct1_pct2** |
| EPHX2 | 0,409282297 | 0,26455154 | 0,321 | 0,293 | 1 | Ciliated | 1,20126258 | 0,832457463 | 1,09556314 |
| SUOX | 0,409632493 | -0,276115727 | 0,107 | 0,195 | 1 | Ciliated | 0,825811419 | 1,210930216 | 0,548717949 |
| BPGM | 0,409943066 | -0,254609931 | 0,179 | 0,293 | 1 | Ciliated | 0,838213736 | 1,193013138 | 0,610921502 |
| SERPINB31 | 0,410612328 | -1,297808282 | 0,25 | 0,293 | 1 | Ciliated | 0,406743647 | 2,458550999 | 0,853242321 |
| IQCE | 0,410957397 | -0,40112788 | 0,429 | 0,585 | 1 | Ciliated | 0,757266031 | 1,320539888 | 0,733333333 |
| ZBED5-AS1 | 0,410957397 | 0,483954063 | 0,536 | 0,512 | 1 | Ciliated | 1,39857155 | 0,71501526 | 1,046875 |
| FLNB1 | 0,412278264 | -0,405441458 | 0,679 | 0,683 | 1 | Ciliated | 0,75500523 | 1,324494137 | 0,994143485 |
| MGST11 | 0,413856547 | -0,278995712 | 0,393 | 0,512 | 1 | Ciliated | 0,824164535 | 1,213349954 | 0,767578125 |
| MAGED1 | 0,41389461 | 0,285368724 | 0,393 | 0,439 | 1 | Ciliated | 1,218721705 | 0,820531871 | 0,895216401 |
| SIKE1 | 0,415096875 | 0,390888529 | 0,214 | 0,171 | 1 | Ciliated | 1,311200699 | 0,762659752 | 1,251461988 |
| EPB41L5 | 0,415096875 | 0,258731646 | 0,214 | 0,171 | 1 | Ciliated | 1,196426397 | 0,835822415 | 1,251461988 |
| RDH141 | 0,416697934 | 0,396884898 | 0,357 | 0,341 | 1 | Ciliated | 1,316661871 | 0,759496437 | 1,046920821 |
| DNAJB6 | 0,417060845 | 0,28655325 | 0,607 | 0,683 | 1 | Ciliated | 1,219722749 | 0,819858448 | 0,888726208 |
| EIF4G1 | 0,41718653 | 0,276986695 | 0,5 | 0,463 | 1 | Ciliated | 1,211661486 | 0,82531302 | 1,079913607 |
| MAF1 | 0,417851424 | 0,27202828 | 0,321 | 0,293 | 1 | Ciliated | 1,207504261 | 0,828154427 | 1,09556314 |
| SLC35D2 | 0,417851424 | 0,351573667 | 0,321 | 0,293 | 1 | Ciliated | 1,275951654 | 0,783728754 | 1,09556314 |
| SLC35A3 | 0,417865138 | -0,273923809 | 0,5 | 0,707 | 1 | Ciliated | 0,827067046 | 1,209091821 | 0,707213579 |
| USF1 | 0,41863856 | -0,278966639 | 0,143 | 0,244 | 1 | Ciliated | 0,824181143 | 1,213325503 | 0,586065574 |
| RAPGEF2 | 0,418840631 | 0,559434279 | 0,357 | 0,317 | 1 | Ciliated | 1,473691229 | 0,678568197 | 1,126182965 |
| FAM118A1 | 0,418899884 | 0,265164972 | 0,179 | 0,122 | 1 | Ciliated | 1,201773465 | 0,832103578 | 1,467213115 |
| RDH10 | 0,41940929 | -0,270085788 | 0,464 | 0,537 | 1 | Ciliated | 0,829270233 | 1,205879532 | 0,86405959 |
| ENPP4 | 0,41965123 | 0,274631591 | 0,393 | 0,366 | 1 | Ciliated | 1,209685143 | 0,826661389 | 1,073770492 |
| KIAA1191 | 0,420629954 | 0,287693579 | 0,393 | 0,39 | 1 | Ciliated | 1,220687217 | 0,819210675 | 1,007692308 |
| AQP51 | 0,421719582 | 0,25155057 | 0,321 | 0,488 | 1 | Ciliated | 1,19048593 | 0,839993128 | 0,657786885 |
| C11orf52 | 0,421791358 | -0,53682031 | 0,393 | 0,488 | 1 | Ciliated | 0,689288423 | 1,4507715 | 0,805327869 |
| ADGRE5 | 0,42244431 | -0,384658205 | 0,286 | 0,415 | 1 | Ciliated | 0,765960444 | 1,305550447 | 0,689156627 |
| ERICH3 | 0,422540072 | -0,295677106 | 0,857 | 0,829 | 1 | Ciliated | 0,814689881 | 1,227460931 | 1,033775633 |
| CDC42EP51 | 0,422880282 | 0,636828067 | 0,25 | 0,22 | 1 | Ciliated | 1,554906755 | 0,643125382 | 1,136363636 |
| UBE2F1 | 0,424191646 | 0,350208948 | 0,286 | 0,244 | 1 | Ciliated | 1,274745238 | 0,784470473 | 1,172131148 |
| COMTD12 | 0,424191646 | 0,331609822 | 0,286 | 0,244 | 1 | Ciliated | 1,258416788 | 0,794649284 | 1,172131148 |
| HDAC72 | 0,424993102 | 0,285054283 | 0,25 | 0,415 | 1 | Ciliated | 1,218456109 | 0,820710728 | 0,602409639 |

| Supplementary Table 4 - clusermarker | | | | | | | | | |
|---|---|---|---|---|---|---|---|---|---|
| | **p_val** | **avg_log2FC** | **pct.1** | **pct.2** | **p_val_adj** | **celltype** | **Foldchange_UP** | **Foldchange_DOWN** | **Ratio_pct1_pct2** |
| SFI1 | 0,424993102 | 0,290402662 | 0,357 | 0,341 | 1 | Ciliated | 1,222981569 | 0,817673811 | 1,046920821 |
| BMT2 | 0,42542435 | 0,257099515 | 0,214 | 0,171 | 1 | Ciliated | 1,195073636 | 0,836768522 | 1,251461988 |
| RREB1 | 0,427272347 | 0,287103625 | 0,357 | 0,317 | 1 | Ciliated | 1,22018815 | 0,81954574 | 1,126182965 |
| NEK5 | 0,427339888 | -0,256117305 | 0,536 | 0,61 | 1 | Ciliated | 0,837338401 | 1,194260288 | 0,878688525 |
| MORC3 | 0,427782931 | -0,263600174 | 0,321 | 0,415 | 1 | Ciliated | 0,833006597 | 1,200470685 | 0,773493976 |
| ZBTB381 | 0,428638343 | -0,253893854 | 0,393 | 0,537 | 1 | Ciliated | 0,838629884 | 1,192421137 | 0,731843575 |
| SIN3A | 0,428685035 | -0,303749644 | 0,321 | 0,439 | 1 | Ciliated | 0,810144049 | 1,234348388 | 0,731207289 |
| LBH1 | 0,429205121 | -0,356852227 | 0,393 | 0,512 | 1 | Ciliated | 0,780866471 | 1,280628682 | 0,767578125 |
| MUC202 | 0,429657626 | -0,304162664 | 0,964 | 0,756 | 1 | Ciliated | 0,809912151 | 1,234701812 | 1,275132275 |
| RPS3A1 | 0,430581947 | -0,390660981 | 1 | 0,951 | 1 | Ciliated | 0,762780051 | 1,310993908 | 1,051524711 |
| IQUB | 0,430649468 | -0,26275086 | 0,464 | 0,61 | 1 | Ciliated | 0,833497132 | 1,199764176 | 0,760655738 |
| PKN1 | 0,430702658 | -0,568351873 | 0,286 | 0,415 | 1 | Ciliated | 0,674386764 | 1,482828627 | 0,689156627 |
| FAM118B1 | 0,431756292 | -0,253648513 | 0,107 | 0,195 | 1 | Ciliated | 0,838772512 | 1,192218374 | 0,548717949 |
| SGCB1 | 0,432506426 | -0,267363231 | 0,179 | 0,268 | 1 | Ciliated | 0,830836654 | 1,203606023 | 0,667910448 |
| BCAR1 | 0,432506426 | 0,291386594 | 0,25 | 0,22 | 1 | Ciliated | 1,223815939 | 0,817116339 | 1,136363636 |
| CYB5B1 | 0,433037504 | 0,343075994 | 0,286 | 0,268 | 1 | Ciliated | 1,268458213 | 0,788358646 | 1,067164179 |
| CXXC1 | 0,433383944 | 0,320937643 | 0,357 | 0,341 | 1 | Ciliated | 1,249142133 | 0,800549412 | 1,046920821 |
| AHDC1 | 0,433424435 | 0,26047634 | 0,286 | 0,244 | 1 | Ciliated | 1,197874146 | 0,834812241 | 1,172131148 |
| RAB3IP | 0,433618277 | 0,36189825 | 0,429 | 0,415 | 1 | Ciliated | 1,285115698 | 0,778140055 | 1,03373494 |
| PHB1 | 0,43388563 | 0,287726089 | 0,643 | 0,634 | 1 | Ciliated | 1,220714725 | 0,819192216 | 1,014195584 |
| PAGR1 | 0,435303347 | 0,255788532 | 0,321 | 0,293 | 1 | Ciliated | 1,193988161 | 0,837529242 | 1,09556314 |
| SRSF11 | 0,437297536 | -0,278727092 | 0,679 | 0,78 | 1 | Ciliated | 0,824318003 | 1,213124057 | 0,870512821 |
| CRY2 | 0,437362234 | 0,266075602 | 0,429 | 0,39 | 1 | Ciliated | 1,202532264 | 0,83157852 | 1,1 |
| ZMYND8 | 0,438182236 | -0,361267703 | 0,5 | 0,585 | 1 | Ciliated | 0,778480225 | 1,284554145 | 0,854700855 |
| FUNDC2 | 0,438182236 | -0,277576146 | 0,5 | 0,585 | 1 | Ciliated | 0,824975885 | 1,212156644 | 0,854700855 |
| KIF27 | 0,438288075 | -0,32949172 | 0,571 | 0,634 | 1 | Ciliated | 0,795816811 | 1,256570591 | 0,900630915 |
| DNAJC27 | 0,438931388 | -0,376879468 | 0,143 | 0,244 | 1 | Ciliated | 0,770101511 | 1,298530111 | 0,586065574 |
| MYG12 | 0,441052436 | 0,431498455 | 0,5 | 0,537 | 1 | Ciliated | 1,348633607 | 0,741491236 | 0,931098696 |
| CYB561A3 | 0,441657436 | -0,304811673 | 0,357 | 0,463 | 1 | Ciliated | 0,809547887 | 1,235257378 | 0,771058315 |
| CYP2F11 | 0,442816845 | -0,789209374 | 0,214 | 0,268 | 1 | Ciliated | 0,578661123 | 1,728127155 | 0,798507463 |
| DSP1 | 0,443497169 | -0,37801612 | 0,929 | 0,732 | 1 | Ciliated | 0,769495012 | 1,299553583 | 1,269125683 |

| Supplementary Table 4 - clusermarker | | | | | | | | | |
|---|---|---|---|---|---|---|---|---|---|
| | **p_val** | **avg_log2FC** | **pct.1** | **pct.2** | **p_val_adj** | **celltype** | **Foldchange_UP** | **Foldchange_DOWN** | **Ratio_pct1_pct2** |
| MRPS18B1 | 0,443510182 | -0,302682642 | 0,429 | 0,61 | 1 | Ciliated | 0,810743444 | 1,233435814 | 0,703278689 |
| STIMATE | 0,444318739 | 0,360873164 | 0,393 | 0,366 | 1 | Ciliated | 1,284202902 | 0,778693148 | 1,073770492 |
| KIAA12171 | 0,445775456 | -0,453187711 | 0,536 | 0,659 | 1 | Ciliated | 0,730427146 | 1,369061932 | 0,813353566 |
| TIMM131 | 0,445790211 | 0,308754016 | 0,536 | 0,561 | 1 | Ciliated | 1,238637488 | 0,807338717 | 0,95543672 |
| LSM71 | 0,44586376 | -0,493966073 | 0,679 | 0,707 | 1 | Ciliated | 0,710070377 | 1,408311108 | 0,96039604 |
| PSMD61 | 0,446140679 | -0,320394092 | 0,429 | 0,561 | 1 | Ciliated | 0,800851084 | 1,248671594 | 0,764705882 |
| CDC421 | 0,446772014 | -0,264485043 | 0,821 | 0,78 | 1 | Ciliated | 0,832495833 | 1,201207213 | 1,052564103 |
| ANKUB1 | 0,4500402 | -0,357576598 | 0,571 | 0,707 | 1 | Ciliated | 0,7804745 | 1,281271842 | 0,807637907 |
| ZCCHC2 | 0,452234572 | -0,517134318 | 0,214 | 0,293 | 1 | Ciliated | 0,698758427 | 1,431109754 | 0,730375427 |
| RPL262 | 0,452251712 | -0,251862256 | 0,964 | 0,902 | 1 | Ciliated | 0,839811672 | 1,190743156 | 1,068736142 |
| C1orf1221 | 0,453168589 | 0,316317282 | 0,321 | 0,293 | 1 | Ciliated | 1,245148042 | 0,803117353 | 1,09556314 |
| KCTD1 | 0,456059158 | -0,286065307 | 0,429 | 0,512 | 1 | Ciliated | 0,820135785 | 1,219310288 | 0,837890625 |
| SHOC21 | 0,456561431 | -0,352946758 | 0,429 | 0,585 | 1 | Ciliated | 0,782983191 | 1,277166625 | 0,733333333 |
| TELO2 | 0,457292501 | 0,346967691 | 0,214 | 0,171 | 1 | Ciliated | 1,271884523 | 0,7862349 | 1,251461988 |
| PCSK1N | 0,457292501 | 0,495765545 | 0,214 | 0,171 | 1 | Ciliated | 1,410068789 | 0,70918526 | 1,251461988 |
| IQGAP2 | 0,457354478 | -0,302934137 | 0,179 | 0,293 | 1 | Ciliated | 0,810602125 | 1,233650849 | 0,610921502 |
| WRAP53 | 0,457989136 | 0,288414704 | 0,429 | 0,415 | 1 | Ciliated | 1,221297525 | 0,818801299 | 1,03373494 |
| JUNB1 | 0,458220969 | -0,526006744 | 0,857 | 0,78 | 1 | Ciliated | 0,694474321 | 1,43993805 | 1,098717949 |
| PTPRF2 | 0,458663327 | -0,259012014 | 0,607 | 0,634 | 1 | Ciliated | 0,83566 | 1,196658928 | 0,957413249 |
| STARD10 | 0,459303869 | 0,268814854 | 0,571 | 0,61 | 1 | Ciliated | 1,204817687 | 0,830001096 | 0,936065574 |
| CENPS | 0,4600554 | 0,318915794 | 0,464 | 0,488 | 1 | Ciliated | 1,247392763 | 0,801672119 | 0,950819672 |
| WDR83OS2 | 0,460957563 | 0,279109011 | 0,607 | 0,61 | 1 | Ciliated | 1,213445245 | 0,824099813 | 0,995081967 |
| ITPRID2 | 0,461789169 | 0,323921186 | 0,393 | 0,39 | 1 | Ciliated | 1,251728075 | 0,798895559 | 1,007692308 |
| B3GNT7 | 0,461789169 | -0,266836979 | 0,321 | 0,439 | 1 | Ciliated | 0,831139773 | 1,203167063 | 0,731207289 |
| RORC | 0,461809986 | 0,267992308 | 0,286 | 0,244 | 1 | Ciliated | 1,204130961 | 0,830474452 | 1,172131148 |
| SLC25A13 | 0,462134525 | 0,263319847 | 0,25 | 0,22 | 1 | Ciliated | 1,200237447 | 0,833168472 | 1,136363636 |
| TMEM175 | 0,462254155 | 0,283308875 | 0,321 | 0,293 | 1 | Ciliated | 1,216982882 | 0,821704245 | 1,09556314 |
| WAC-AS1 | 0,46455319 | -0,401659657 | 0,25 | 0,366 | 1 | Ciliated | 0,756986954 | 1,321026729 | 0,683060109 |
| TMEM138 | 0,466280379 | 0,250657063 | 0,429 | 0,415 | 1 | Ciliated | 1,189748852 | 0,840513524 | 1,03373494 |
| ANXA51 | 0,466952527 | -0,385827292 | 0,821 | 0,683 | 1 | Ciliated | 0,76534 | 1,306608828 | 1,20204978 |
| C2orf81 | 0,467890346 | 0,453176176 | 0,357 | 0,341 | 1 | Ciliated | 1,369050986 | 0,730432986 | 1,046920821 |

| Supplementary Table 4 - clusermarker | | | | | | | | | |
|---|---|---|---|---|---|---|---|---|---|
| | **p_val** | **avg_log2FC** | **pct.1** | **pct.2** | **p_val_adj** | **celltype** | **Foldchange_UP** | **Foldchange_DOWN** | **Ratio_pct1_pct2** |
| ARSA | 0,470887384 | 0,333367645 | 0,357 | 0,317 | 1 | Ciliated | 1,259951015 | 0,79368165 | 1,126182965 |
| AL121790.2 | 0,471440481 | -0,251099402 | 0,25 | 0,341 | 1 | Ciliated | 0,840255856 | 1,190113693 | 0,73313783 |
| AMFR | 0,471440481 | 0,276223157 | 0,321 | 0,293 | 1 | Ciliated | 1,211020391 | 0,825749928 | 1,09556314 |
| COX7A1 | 0,471440481 | -0,404091242 | 0,25 | 0,341 | 1 | Ciliated | 0,755712168 | 1,323255126 | 0,73313783 |
| TIMM17B2 | 0,472160926 | 0,298784871 | 0,5 | 0,463 | 1 | Ciliated | 1,230107902 | 0,812936815 | 1,079913607 |
| SCIN | 0,472256652 | 0,251956507 | 0,25 | 0,22 | 1 | Ciliated | 1,190820949 | 0,839756809 | 1,136363636 |
| ACTR1B | 0,47362413 | 0,397341645 | 0,321 | 0,317 | 1 | Ciliated | 1,317078783 | 0,759256024 | 1,012618297 |
| RAB4B | 0,47362413 | 0,293408409 | 0,321 | 0,317 | 1 | Ciliated | 1,225532216 | 0,815972022 | 1,012618297 |
| PHTF1 | 0,477845836 | 0,319011567 | 0,5 | 0,512 | 1 | Ciliated | 1,247475573 | 0,801618902 | 0,9765625 |
| PIH1D1 | 0,478743253 | -0,277441091 | 0,464 | 0,585 | 1 | Ciliated | 0,825053118 | 1,212043175 | 0,793162393 |
| SLC39A9 | 0,47886207 | 0,347858786 | 0,393 | 0,39 | 1 | Ciliated | 1,272670358 | 0,785749424 | 1,007692308 |
| PHLDA31 | 0,47926189 | -0,404570339 | 0,143 | 0,22 | 1 | Ciliated | 0,755461249 | 1,323694631 | 0,65 |
| ACBD61 | 0,480726593 | 0,452929469 | 0,321 | 0,293 | 1 | Ciliated | 1,368816892 | 0,730557904 | 1,09556314 |
| PTP4A21 | 0,480902547 | 0,281886697 | 0,536 | 0,61 | 1 | Ciliated | 1,215783797 | 0,822514663 | 0,878688525 |
| NDUFB3 | 0,481793703 | -0,333779089 | 0,857 | 0,878 | 1 | Ciliated | 0,793455331 | 1,260310393 | 0,976082005 |
| MZT2A2 | 0,482179316 | 0,268090182 | 0,357 | 0,366 | 1 | Ciliated | 1,204212654 | 0,830418113 | 0,975409836 |
| CAB391 | 0,483247265 | -0,367749531 | 0,429 | 0,537 | 1 | Ciliated | 0,774990467 | 1,290338453 | 0,798882682 |
| NARS1 | 0,484258962 | -0,518302401 | 0,536 | 0,659 | 1 | Ciliated | 0,698192904 | 1,432268925 | 0,813353566 |
| FOSB2 | 0,484448859 | -0,370543799 | 0,679 | 0,634 | 1 | Ciliated | 0,773490888 | 1,292840053 | 1,070977918 |
| CAMTA12 | 0,485460909 | 0,442342052 | 0,429 | 0,488 | 1 | Ciliated | 1,35880841 | 0,735938925 | 0,879098361 |
| HLA-DMA1 | 0,485460909 | -0,385856086 | 0,393 | 0,512 | 1 | Ciliated | 0,765324725 | 1,306634906 | 0,767578125 |
| PGAM11 | 0,485460909 | 0,328278503 | 0,464 | 0,463 | 1 | Ciliated | 1,255514337 | 0,796486325 | 1,002159827 |
| GNB1 | 0,48574704 | 0,331296414 | 0,571 | 0,659 | 1 | Ciliated | 1,258143442 | 0,794821931 | 0,86646434 |
| DNAI3 | 0,48574704 | 0,284470502 | 0,571 | 0,659 | 1 | Ciliated | 1,217963164 | 0,821042893 | 0,86646434 |
| VPS13C2 | 0,48574704 | -0,348148811 | 0,571 | 0,659 | 1 | Ciliated | 0,78559148 | 1,272926228 | 0,86646434 |
| MDK2 | 0,48660389 | 0,286769895 | 0,607 | 0,585 | 1 | Ciliated | 1,219905924 | 0,819735342 | 1,037606838 |
| GNG101 | 0,487526078 | 0,269079529 | 0,393 | 0,39 | 1 | Ciliated | 1,205038741 | 0,829848839 | 1,007692308 |
| DRC1 | 0,487685676 | -0,346069823 | 0,821 | 0,756 | 1 | Ciliated | 0,786724369 | 1,271093206 | 1,085978836 |
| MAP3K4 | 0,489656417 | 0,290501503 | 0,286 | 0,268 | 1 | Ciliated | 1,22306536 | 0,817617792 | 1,067164179 |
| SOCS4 | 0,490111463 | -0,359816424 | 0,321 | 0,293 | 1 | Ciliated | 0,779263731 | 1,283262599 | 1,09556314 |
| GALNS | 0,490499853 | 0,302561028 | 0,429 | 0,463 | 1 | Ciliated | 1,233331844 | 0,81081179 | 0,926565875 |

| Supplementary Table 4 - clusermarker | | | | | | | | | |
|---|---|---|---|---|---|---|---|---|---|
| | **p_val** | **avg_log2FC** | **pct.1** | **pct.2** | **p_val_adj** | **celltype** | **Foldchange_UP** | **Foldchange_DOWN** | **Ratio_pct1_pct2** |
| CNIH41 | 0,491072679 | 0,466917972 | 0,357 | 0,366 | 1 | Ciliated | 1,382153619 | 0,723508579 | 0,975409836 |
| FAM166B | 0,491480929 | -0,406596472 | 0,643 | 0,732 | 1 | Ciliated | 0,754401018 | 1,325554945 | 0,878415301 |
| TTC26 | 0,491647089 | -0,332016731 | 0,357 | 0,463 | 1 | Ciliated | 0,794425187 | 1,258771772 | 0,771058315 |
| C7orf26 | 0,492062052 | 0,260824492 | 0,143 | 0,244 | 1 | Ciliated | 1,198163252 | 0,834610808 | 0,586065574 |
| MIR100HG | 0,492862355 | 0,261792288 | 0,25 | 0,22 | 1 | Ciliated | 1,19896728 | 0,834051118 | 1,136363636 |
| ZFP361 | 0,493121302 | -0,415553857 | 0,893 | 0,829 | 1 | Ciliated | 0,749731614 | 1,333810635 | 1,077201448 |
| GTF3C2 | 0,493583736 | 0,285976547 | 0,321 | 0,341 | 1 | Ciliated | 1,219235273 | 0,820186245 | 0,941348974 |
| CALR2 | 0,493790281 | 0,351940984 | 0,786 | 0,707 | 1 | Ciliated | 1,276276559 | 0,783529238 | 1,111739745 |
| POLDIP2 | 0,494738187 | -0,289885412 | 0,286 | 0,39 | 1 | Ciliated | 0,817967024 | 1,222543172 | 0,733333333 |
| BCAS3 | 0,495945748 | -0,269038818 | 0,321 | 0,488 | 1 | Ciliated | 0,829872257 | 1,205004737 | 0,657786885 |
| MDN1 | 0,496055458 | -0,290049666 | 0,321 | 0,415 | 1 | Ciliated | 0,817873902 | 1,222682369 | 0,773493976 |
| PSME3 | 0,4962307 | -0,308267765 | 0,321 | 0,463 | 1 | Ciliated | 0,807610871 | 1,238220084 | 0,693304536 |
| NCOR21 | 0,496274007 | 0,255830287 | 0,393 | 0,39 | 1 | Ciliated | 1,194022719 | 0,837505002 | 1,007692308 |
| RNF2132 | 0,496851756 | 0,320205719 | 0,536 | 0,61 | 1 | Ciliated | 1,248508565 | 0,800955659 | 0,878688525 |
| SPAG6 | 0,497167168 | 0,265171936 | 0,893 | 0,854 | 1 | Ciliated | 1,201779265 | 0,832099562 | 1,045667447 |
| RTN3 | 0,499034909 | 0,304006187 | 0,536 | 0,585 | 1 | Ciliated | 1,234567901 | 0,81 | 0,916239316 |
| PDE6B | 0,499465773 | 0,289967265 | 0,286 | 0,268 | 1 | Ciliated | 1,222612536 | 0,817920617 | 1,067164179 |
| MAFG | 0,499465773 | 0,309833342 | 0,286 | 0,268 | 1 | Ciliated | 1,239564499 | 0,806734947 | 1,067164179 |
| TM9SF31 | 0,500148327 | -0,303982419 | 0,571 | 0,634 | 1 | Ciliated | 0,810013344 | 1,234547563 | 0,900630915 |
| TTC17 | 0,500264381 | 0,296801573 | 0,429 | 0,415 | 1 | Ciliated | 1,228418013 | 0,814055142 | 1,03373494 |
| LMNA1 | 0,500925504 | -0,258231322 | 0,679 | 0,756 | 1 | Ciliated | 0,836112327 | 1,19601155 | 0,898148148 |
| CSNK1D2 | 0,502237598 | 0,250567901 | 0,464 | 0,463 | 1 | Ciliated | 1,189675326 | 0,840565471 | 1,002159827 |
| COMMD7 | 0,502237598 | -0,339054265 | 0,393 | 0,512 | 1 | Ciliated | 0,79055938 | 1,26492712 | 0,767578125 |
| CD441 | 0,503343079 | 0,292213701 | 0,25 | 0,22 | 1 | Ciliated | 1,224517762 | 0,816648015 | 1,136363636 |
| SAFB | 0,504013646 | 0,369540222 | 0,429 | 0,439 | 1 | Ciliated | 1,291941031 | 0,774029136 | 0,977220957 |
| SCRN2 | 0,504972526 | 0,280273719 | 0,393 | 0,415 | 1 | Ciliated | 1,214425273 | 0,823434774 | 0,946987952 |
| AL022069.3 | 0,506264723 | -0,338667258 | 0,071 | 0,122 | 1 | Ciliated | 0,790771479 | 1,264587845 | 0,581967213 |
| MGP2 | 0,506264723 | -0,805608865 | 0,071 | 0,122 | 1 | Ciliated | 0,572120576 | 1,7478833 | 0,581967213 |
| CMIP1 | 0,507170403 | -0,328908395 | 0,5 | 0,561 | 1 | Ciliated | 0,796138649 | 1,256062624 | 0,891265597 |
| NUBPL | 0,508977587 | -0,355282161 | 0,179 | 0,244 | 1 | Ciliated | 0,781716741 | 1,279235749 | 0,733606557 |
| FSIP1 | 0,508977587 | -0,306513009 | 0,179 | 0,244 | 1 | Ciliated | 0,808593769 | 1,236714947 | 0,733606557 |

| Supplementary Table 4 - clusermarker | | | | | | | | | |
|---|---|---|---|---|---|---|---|---|---|
| | **p_val** | **avg_log2FC** | **pct.1** | **pct.2** | **p_val_adj** | **celltype** | **Foldchange_UP** | **Foldchange_DOWN** | **Ratio_pct1_pct2** |
| NDUFB92 | 0,51009026 | 0,25671209 | 0,75 | 0,756 | 1 | Ciliated | 1,194752752 | 0,83699326 | 0,992063492 |
| HADHA1 | 0,510282873 | -0,296515239 | 0,607 | 0,732 | 1 | Ciliated | 0,814216724 | 1,228174232 | 0,829234973 |
| C11orf49 | 0,510684584 | -0,317727741 | 0,536 | 0,634 | 1 | Ciliated | 0,802332565 | 1,246365963 | 0,845425868 |
| CDHR3 | 0,512407939 | -0,416724049 | 0,893 | 0,805 | 1 | Ciliated | 0,749123742 | 1,334892948 | 1,10931677 |
| APBB1 | 0,513084094 | 0,260549377 | 0,357 | 0,341 | 1 | Ciliated | 1,19793479 | 0,834769979 | 1,046920821 |
| SLC66A31 | 0,513268513 | 0,251687224 | 0,214 | 0,171 | 1 | Ciliated | 1,1905987 | 0,839913567 | 1,251461988 |
| NTN4 | 0,513940348 | 0,368979672 | 0,25 | 0,22 | 1 | Ciliated | 1,291439153 | 0,774329939 | 1,136363636 |
| ZNF296 | 0,513940348 | 0,311272322 | 0,25 | 0,22 | 1 | Ciliated | 1,240801488 | 0,80593069 | 1,136363636 |
| ITGAE | 0,514018026 | 0,25260656 | 0,393 | 0,39 | 1 | Ciliated | 1,191357633 | 0,839378515 | 1,007692308 |
| ZBBX | 0,51602306 | -0,251568718 | 0,643 | 0,756 | 1 | Ciliated | 0,839982562 | 1,190500905 | 0,850529101 |
| PYGO2 | 0,516557565 | -0,321793132 | 0,107 | 0,171 | 1 | Ciliated | 0,800074843 | 1,249883069 | 0,625730994 |
| ZFAND31 | 0,518273089 | 0,352879869 | 0,357 | 0,366 | 1 | Ciliated | 1,277107412 | 0,783019495 | 0,975409836 |
| EMB | 0,519496445 | -0,321252641 | 0,5 | 0,61 | 1 | Ciliated | 0,800374639 | 1,249414901 | 0,819672131 |
| AC013264.1 | 0,520073742 | -0,329389195 | 0,857 | 0,805 | 1 | Ciliated | 0,795873367 | 1,256481296 | 1,064596273 |
| RFK | 0,520406397 | 0,292712775 | 0,321 | 0,317 | 1 | Ciliated | 1,224941435 | 0,81636556 | 1,012618297 |
| SLPI2 | 0,521165931 | -0,399119322 | 1 | 0,976 | 1 | Ciliated | 0,758321051 | 1,318702677 | 1,024590164 |
| BOD11 | 0,521299524 | -0,263578045 | 0,071 | 0,122 | 1 | Ciliated | 0,833019374 | 1,200452271 | 0,581967213 |
| BOLA2B | 0,521299524 | -0,326355839 | 0,071 | 0,122 | 1 | Ciliated | 0,797548501 | 1,253842241 | 0,581967213 |
| LIPE-AS1 | 0,521299524 | -0,329570037 | 0,071 | 0,122 | 1 | Ciliated | 0,795773611 | 1,256638806 | 0,581967213 |
| TMEM212 | 0,522153298 | -0,285831991 | 0,393 | 0,439 | 1 | Ciliated | 0,82026843 | 1,219113114 | 0,895216401 |
| FAM174C2 | 0,522696676 | 0,349040224 | 0,393 | 0,415 | 1 | Ciliated | 1,273712987 | 0,785106229 | 0,946987952 |
| PSMB7 | 0,524484713 | -0,368386191 | 0,571 | 0,634 | 1 | Ciliated | 0,77464854 | 1,290908003 | 0,900630915 |
| MTRF1L | 0,524877038 | -0,3105361 | 0,143 | 0,22 | 1 | Ciliated | 0,80634207 | 1,240168456 | 0,65 |
| NUPR11 | 0,527012397 | -0,452157883 | 0,786 | 0,78 | 1 | Ciliated | 0,730948728 | 1,368085014 | 1,007692308 |
| LSR2 | 0,527966371 | -0,298273314 | 0,393 | 0,512 | 1 | Ciliated | 0,813225121 | 1,229671803 | 0,767578125 |
| GADD45A | 0,528620703 | 0,265946719 | 0,179 | 0,293 | 1 | Ciliated | 1,20242484 | 0,831652812 | 0,610921502 |
| NUDCD3 | 0,529508786 | 0,268989814 | 0,286 | 0,268 | 1 | Ciliated | 1,204963808 | 0,829900445 | 1,067164179 |
| DYM1 | 0,529508786 | 0,326831894 | 0,286 | 0,268 | 1 | Ciliated | 1,254256047 | 0,797285373 | 1,067164179 |
| HM13 | 0,530012355 | 0,318784335 | 0,464 | 0,561 | 1 | Ciliated | 1,247279106 | 0,801745171 | 0,827094474 |
| KLF6 | 0,53206709 | -0,345278828 | 0,893 | 0,78 | 1 | Ciliated | 0,787155829 | 1,270396487 | 1,144871795 |
| URI11 | 0,532086521 | 0,261239745 | 0,393 | 0,39 | 1 | Ciliated | 1,198508171 | 0,834370615 | 1,007692308 |

| Supplementary Table 4 - clusermarker | | | | | | | | | |
|---|---|---|---|---|---|---|---|---|---|
| | **p_val** | **avg_log2FC** | **pct.1** | **pct.2** | **p_val_adj** | **celltype** | **Foldchange_UP** | **Foldchange_DOWN** | **Ratio_pct1_pct2** |
| CYB5R1 | 0,533566531 | 0,42156718 | 0,429 | 0,463 | 1 | Ciliated | 1,339381717 | 0,746613148 | 0,926565875 |
| CAT | 0,535373856 | 0,268905167 | 0,607 | 0,585 | 1 | Ciliated | 1,204893111 | 0,829949139 | 1,037606838 |
| NDUFA82 | 0,536171968 | -0,332219786 | 0,5 | 0,61 | 1 | Ciliated | 0,794313382 | 1,258948953 | 0,819672131 |
| TIFA | 0,536560882 | -0,318508959 | 0,071 | 0,122 | 1 | Ciliated | 0,80189822 | 1,247041053 | 0,581967213 |
| PABIR1 | 0,536560882 | -0,257550081 | 0,071 | 0,122 | 1 | Ciliated | 0,836507233 | 1,195446926 | 0,581967213 |
| NPDC1 | 0,53661852 | 0,469895104 | 0,214 | 0,171 | 1 | Ciliated | 1,385008763 | 0,722017092 | 1,251461988 |
| GLT8D1 | 0,536685107 | 0,33851404 | 0,429 | 0,488 | 1 | Ciliated | 1,26445355 | 0,790855465 | 0,879098361 |
| ACBD31 | 0,536689505 | 0,271530617 | 0,464 | 0,463 | 1 | Ciliated | 1,2070878 | 0,828440152 | 1,002159827 |
| RIPOR2 | 0,536689505 | -0,304435148 | 0,393 | 0,512 | 1 | Ciliated | 0,809759196 | 1,234935034 | 0,767578125 |
| CD1511 | 0,536830047 | 0,54103089 | 0,357 | 0,366 | 1 | Ciliated | 1,455011837 | 0,687279632 | 0,975409836 |
| TOM1L1 | 0,539244281 | 0,311224406 | 0,25 | 0,244 | 1 | Ciliated | 1,240760279 | 0,805957458 | 1,024590164 |
| ACP1 | 0,539542488 | -0,467234365 | 0,571 | 0,61 | 1 | Ciliated | 0,723349926 | 1,382456767 | 0,936065574 |
| RMDN3 | 0,540677967 | 0,283156361 | 0,321 | 0,341 | 1 | Ciliated | 1,216854236 | 0,821791116 | 0,941348974 |
| COQ8B | 0,540735024 | 0,384268646 | 0,393 | 0,415 | 1 | Ciliated | 1,305197968 | 0,766167298 | 0,946987952 |
| SCARB21 | 0,541508742 | 0,347561866 | 0,357 | 0,39 | 1 | Ciliated | 1,272408457 | 0,785911155 | 0,915384615 |
| PRMT51 | 0,542346497 | -0,303395172 | 0,214 | 0,293 | 1 | Ciliated | 0,810343126 | 1,234045144 | 0,730375427 |
| USP16 | 0,545883742 | -0,354543415 | 0,607 | 0,707 | 1 | Ciliated | 0,782117129 | 1,278580872 | 0,858557284 |
| PRADC1 | 0,546232226 | -0,293569644 | 0,286 | 0,415 | 1 | Ciliated | 0,815880834 | 1,225669189 | 0,689156627 |
| ZNF394 | 0,546414724 | -0,30985686 | 0,179 | 0,268 | 1 | Ciliated | 0,806721796 | 1,239584705 | 0,667910448 |
| IFITM32 | 0,54658329 | 0,362690031 | 0,607 | 0,634 | 1 | Ciliated | 1,285821189 | 0,777713113 | 0,957413249 |
| ELP1 | 0,548430001 | -0,27778707 | 0,25 | 0,341 | 1 | Ciliated | 0,824855282 | 1,212333875 | 0,73313783 |
| HDAC10 | 0,548461829 | 0,357650643 | 0,214 | 0,195 | 1 | Ciliated | 1,281337604 | 0,780434444 | 1,097435897 |
| NLRC5 | 0,548490675 | -0,360983626 | 0,143 | 0,22 | 1 | Ciliated | 0,778633528 | 1,284301233 | 0,65 |
| FDX2 | 0,548490675 | -0,292324641 | 0,143 | 0,22 | 1 | Ciliated | 0,816585219 | 1,224611929 | 0,65 |
| UBE2A | 0,548853037 | 0,298068227 | 0,536 | 0,585 | 1 | Ciliated | 1,22949701 | 0,813340733 | 0,916239316 |
| AC127526.51 | 0,549577316 | -0,384352671 | 0,286 | 0,341 | 1 | Ciliated | 0,766122676 | 1,305273987 | 0,838709677 |
| C8orf34 | 0,550359365 | 0,511260681 | 0,321 | 0,341 | 1 | Ciliated | 1,425295128 | 0,701609078 | 0,941348974 |
| VSIG21 | 0,550359365 | 0,422147061 | 0,321 | 0,341 | 1 | Ciliated | 1,33992018 | 0,746313112 | 0,941348974 |
| PLAUR | 0,552220402 | 0,250788938 | 0,179 | 0,146 | 1 | Ciliated | 1,189857611 | 0,840436696 | 1,226027397 |
| TMEM2181 | 0,555715301 | -0,280513222 | 0,286 | 0,415 | 1 | Ciliated | 0,823298086 | 1,214626897 | 0,689156627 |
| CALML4 | 0,557189895 | -0,358888335 | 0,964 | 0,829 | 1 | Ciliated | 0,779765194 | 1,282437338 | 1,162846803 |

| Supplementary Table 4 - clusermarker | | | | | | | | | |
|---|---|---|---|---|---|---|---|---|---|
| | **p_val** | **avg_log2FC** | **pct.1** | **pct.2** | **p_val_adj** | **celltype** | **Foldchange_UP** | **Foldchange_DOWN** | **Ratio_pct1_pct2** |
| NTS1 | 0,558472724 | 0,281831419 | 0,321 | 0,293 | 1 | Ciliated | 1,215737214 | 0,822546179 | 1,09556314 |
| SMARCC11 | 0,55979295 | 0,299617094 | 0,464 | 0,512 | 1 | Ciliated | 1,230817699 | 0,812468005 | 0,90625 |
| RINT1 | 0,560113185 | 0,312975196 | 0,214 | 0,195 | 1 | Ciliated | 1,242266923 | 0,804979978 | 1,097435897 |
| RYK | 0,560255135 | 0,305397012 | 0,357 | 0,39 | 1 | Ciliated | 1,235758656 | 0,809219498 | 0,915384615 |
| SBDS1 | 0,561606456 | 0,278190241 | 0,464 | 0,488 | 1 | Ciliated | 1,212672718 | 0,824624802 | 0,950819672 |
| SEC23B | 0,563286385 | -0,393912041 | 0,393 | 0,512 | 1 | Ciliated | 0,76106309 | 1,313951515 | 0,767578125 |
| FAM162A | 0,564479797 | 0,2904972 | 0,5 | 0,537 | 1 | Ciliated | 1,223061712 | 0,817620231 | 0,931098696 |
| ARPC32 | 0,565151418 | -0,339123426 | 0,821 | 0,878 | 1 | Ciliated | 0,790521483 | 1,264987761 | 0,935079727 |
| TMEM9B | 0,56527804 | -0,413167528 | 0,286 | 0,415 | 1 | Ciliated | 0,750972754 | 1,331606233 | 0,689156627 |
| RAF1 | 0,566173063 | -0,267746318 | 0,357 | 0,488 | 1 | Ciliated | 0,830616066 | 1,203925665 | 0,731557377 |
| TSR21 | 0,56916331 | -0,304806827 | 0,321 | 0,439 | 1 | Ciliated | 0,809550606 | 1,235253229 | 0,731207289 |
| TMEM250 | 0,571743272 | 0,268690567 | 0,25 | 0,244 | 1 | Ciliated | 1,204713897 | 0,830072602 | 1,024590164 |
| THOP1 | 0,571743272 | 0,349971965 | 0,25 | 0,244 | 1 | Ciliated | 1,274535859 | 0,784599345 | 1,024590164 |
| CCDC88C1 | 0,571932723 | 0,297707804 | 0,607 | 0,634 | 1 | Ciliated | 1,229189889 | 0,813543952 | 0,957413249 |
| AC074117.1 | 0,573735181 | 0,345103748 | 0,107 | 0,073 | 1 | Ciliated | 1,270242326 | 0,787251361 | 1,465753425 |
| KRT131 | 0,573735181 | 0,253672762 | 0,107 | 0,073 | 1 | Ciliated | 1,192238413 | 0,838758414 | 1,465753425 |
| APOBEC4 | 0,576211343 | -0,434628409 | 0,571 | 0,659 | 1 | Ciliated | 0,7398843 | 1,351562669 | 0,86646434 |
| SOD22 | 0,576211343 | -0,394481015 | 0,571 | 0,659 | 1 | Ciliated | 0,760762999 | 1,314469817 | 0,86646434 |
| PAPSS1 | 0,57723847 | -0,294035705 | 0,464 | 0,61 | 1 | Ciliated | 0,815617307 | 1,226065204 | 0,760655738 |
| VAPA1 | 0,577438626 | 0,254874541 | 0,857 | 0,829 | 1 | Ciliated | 1,193231973 | 0,838060011 | 1,033775633 |
| AC008035.1 | 0,578152422 | 0,322117546 | 0,179 | 0,146 | 1 | Ciliated | 1,250164157 | 0,799894953 | 1,226027397 |
| RAB2B | 0,578152422 | 0,31881211 | 0,179 | 0,146 | 1 | Ciliated | 1,247303118 | 0,801729736 | 1,226027397 |
| CNOT10 | 0,579310531 | 0,400052201 | 0,357 | 0,39 | 1 | Ciliated | 1,319555655 | 0,757830862 | 0,915384615 |
| ABRACL2 | 0,579310531 | 0,295091017 | 0,357 | 0,39 | 1 | Ciliated | 1,226962382 | 0,815020912 | 0,915384615 |
| ARMH1 | 0,581363733 | -0,38040217 | 0,393 | 0,512 | 1 | Ciliated | 0,768223409 | 1,301704672 | 0,767578125 |
| PTK21 | 0,581363733 | -0,273508997 | 0,393 | 0,512 | 1 | Ciliated | 0,827304883 | 1,208744225 | 0,767578125 |
| RBBP42 | 0,586554572 | 0,271791651 | 0,5 | 0,488 | 1 | Ciliated | 1,207306224 | 0,828290272 | 1,024590164 |
| KARS1 | 0,587149547 | 0,297041799 | 0,393 | 0,415 | 1 | Ciliated | 1,228622577 | 0,813919603 | 0,946987952 |
| TMEM170A1 | 0,588950954 | 0,266285901 | 0,357 | 0,39 | 1 | Ciliated | 1,202707568 | 0,831457311 | 0,915384615 |
| BUD31 | 0,589743196 | -0,511992711 | 0,643 | 0,732 | 1 | Ciliated | 0,701253168 | 1,426018513 | 0,878415301 |
| C2 | 0,590883318 | 0,284961533 | 0,107 | 0,073 | 1 | Ciliated | 1,218377777 | 0,820763493 | 1,465753425 |

| Supplementary Table 4 - clusermarker | | | | | | | | | |
|---|---|---|---|---|---|---|---|---|---|
| | **p_val** | **avg_log2FC** | **pct.1** | **pct.2** | **p_val_adj** | **celltype** | **Foldchange_UP** | **Foldchange_DOWN** | **Ratio_pct1_pct2** |
| TOMM52 | 0,595605076 | 0,291805611 | 0,464 | 0,512 | 1 | Ciliated | 1,224171436 | 0,81687905 | 0,90625 |
| MX1 | 0,595605076 | 0,296270033 | 0,5 | 0,488 | 1 | Ciliated | 1,227965504 | 0,814355124 | 1,024590164 |
| SMAD2 | 0,599999213 | 0,278512455 | 0,429 | 0,585 | 1 | Ciliated | 1,212943588 | 0,82444065 | 0,733333333 |
| TCN12 | 0,601677088 | -0,564321701 | 0,107 | 0,146 | 1 | Ciliated | 0,676273299 | 1,478692122 | 0,732876712 |
| LY6G5C | 0,605145175 | 0,319448999 | 0,25 | 0,244 | 1 | Ciliated | 1,247853871 | 0,801375885 | 1,024590164 |
| INTS10 | 0,606909082 | -0,431920723 | 0,393 | 0,537 | 1 | Ciliated | 0,741274238 | 1,349028402 | 0,731843575 |
| SELENOF1 | 0,608438776 | -0,32722345 | 0,571 | 0,61 | 1 | Ciliated | 0,797069014 | 1,254596506 | 0,936065574 |
| USP25 | 0,608450602 | -0,2544101 | 0,321 | 0,415 | 1 | Ciliated | 0,838329847 | 1,192847903 | 0,773493976 |
| C1orf189 | 0,609384669 | -0,295725428 | 0,714 | 0,78 | 1 | Ciliated | 0,814662594 | 1,227502045 | 0,915384615 |
| COPG2 | 0,60972389 | 0,328211394 | 0,143 | 0,22 | 1 | Ciliated | 1,255455936 | 0,796523375 | 0,65 |
| CARMIL1 | 0,610929281 | -0,296856495 | 0,143 | 0,195 | 1 | Ciliated | 0,814024152 | 1,228464779 | 0,733333333 |
| PSME3IP1 | 0,613894443 | -0,262046437 | 0,393 | 0,561 | 1 | Ciliated | 0,833904202 | 1,199178512 | 0,700534759 |
| KRT72 | 0,61561595 | 0,32401556 | 0,357 | 0,512 | 1 | Ciliated | 1,25180996 | 0,7988433 | 0,697265625 |
| SCAMP4 | 0,615912903 | 0,285405113 | 0,5 | 0,634 | 1 | Ciliated | 1,218752445 | 0,820511175 | 0,788643533 |
| PPM1B | 0,616470364 | 0,267953655 | 0,25 | 0,244 | 1 | Ciliated | 1,2040987 | 0,830496703 | 1,024590164 |
| NUTM2A-AS1 | 0,619948933 | 0,269080661 | 0,286 | 0,39 | 1 | Ciliated | 1,205039687 | 0,829848187 | 0,733333333 |
| GLMP | 0,620100993 | 0,363275199 | 0,214 | 0,195 | 1 | Ciliated | 1,286342834 | 0,77739773 | 1,097435897 |
| MARK21 | 0,620533505 | 0,292276271 | 0,321 | 0,293 | 1 | Ciliated | 1,224570871 | 0,816612598 | 1,09556314 |
| ELP5 | 0,620533505 | -0,270185922 | 0,25 | 0,341 | 1 | Ciliated | 0,829212677 | 1,205963231 | 0,73313783 |
| FUZ | 0,62089816 | -0,277292099 | 0,607 | 0,659 | 1 | Ciliated | 0,825138328 | 1,21191801 | 0,921092564 |
| RNF101 | 0,623967258 | 0,401450597 | 0,393 | 0,439 | 1 | Ciliated | 1,320835313 | 0,757096657 | 0,895216401 |
| IRAK1BP1 | 0,624818707 | 0,261877928 | 0,286 | 0,268 | 1 | Ciliated | 1,199038454 | 0,834001609 | 1,067164179 |
| FNTA | 0,62555913 | -0,324927119 | 0,357 | 0,439 | 1 | Ciliated | 0,798338715 | 1,252601159 | 0,813211845 |
| METTL5 | 0,627281235 | -0,444076613 | 0,536 | 0,512 | 1 | Ciliated | 0,735054633 | 1,360443095 | 1,046875 |
| RNF170 | 0,62788757 | 0,389867157 | 0,25 | 0,244 | 1 | Ciliated | 1,310272749 | 0,763199876 | 1,024590164 |
| ZNF561 | 0,62788757 | 0,264206455 | 0,25 | 0,244 | 1 | Ciliated | 1,200975279 | 0,832656606 | 1,024590164 |
| SNHG32 | 0,628232664 | 0,336705227 | 0,357 | 0,39 | 1 | Ciliated | 1,262869205 | 0,79184764 | 0,915384615 |
| PIGS | 0,62933203 | -0,29598929 | 0,214 | 0,293 | 1 | Ciliated | 0,814513609 | 1,22772657 | 0,730375427 |
| LGALS3BP | 0,629765551 | -0,352135129 | 0,571 | 0,683 | 1 | Ciliated | 0,783423804 | 1,276448321 | 0,83601757 |
| MORN3 | 0,629903108 | 0,302683111 | 0,464 | 0,537 | 1 | Ciliated | 1,233436215 | 0,810743181 | 0,86405959 |
| AC006230.1 | 0,63073638 | 0,296180863 | 0,321 | 0,341 | 1 | Ciliated | 1,227889608 | 0,814405459 | 0,941348974 |

| Supplementary Table 4 - clusermarker | | | | | | | | | |
|---|---|---|---|---|---|---|---|---|---|
| | **p_val** | **avg_log2FC** | **pct.1** | **pct.2** | **p_val_adj** | **celltype** | **Foldchange_UP** | **Foldchange_DOWN** | **Ratio_pct1_pct2** |
| USP28 | 0,63073638 | 0,366010883 | 0,321 | 0,341 | 1 | Ciliated | 1,288784352 | 0,775925001 | 0,941348974 |
| WWP2 | 0,63073638 | 0,26855323 | 0,321 | 0,341 | 1 | Ciliated | 1,20459922 | 0,830151625 | 0,941348974 |
| NHLRC2 | 0,632426746 | 0,274894023 | 0,214 | 0,195 | 1 | Ciliated | 1,209905209 | 0,82651103 | 1,097435897 |
| DNAJC5 | 0,635851057 | 0,319871126 | 0,214 | 0,317 | 1 | Ciliated | 1,248219042 | 0,801141439 | 0,675078864 |
| CD200R1L | 0,637106953 | -0,255186436 | 0,143 | 0,195 | 1 | Ciliated | 0,837878851 | 1,193489965 | 0,733333333 |
| CFAP20DC | 0,641126557 | 0,282682392 | 0,25 | 0,39 | 1 | Ciliated | 1,216454528 | 0,822061143 | 0,641025641 |
| PTS | 0,641781929 | -0,283570803 | 0,429 | 0,537 | 1 | Ciliated | 0,821555075 | 1,21720385 | 0,798882682 |
| SMIM20 | 0,641801807 | -0,355503615 | 0,25 | 0,317 | 1 | Ciliated | 0,781596756 | 1,279432126 | 0,788643533 |
| MFF | 0,641826319 | 0,284831576 | 0,357 | 0,415 | 1 | Ciliated | 1,218268032 | 0,82083743 | 0,860240964 |
| FBXO7 | 0,644486199 | 0,348710658 | 0,357 | 0,366 | 1 | Ciliated | 1,273422057 | 0,785285597 | 0,975409836 |
| AK9 | 0,645511416 | -0,25996496 | 0,571 | 0,634 | 1 | Ciliated | 0,835108202 | 1,197449621 | 0,900630915 |
| ATP5PF1 | 0,646411885 | -0,491607821 | 0,893 | 0,854 | 1 | Ciliated | 0,711232019 | 1,406010941 | 1,045667447 |
| NUDT211 | 0,648285425 | -0,313074371 | 0,321 | 0,415 | 1 | Ciliated | 0,804924643 | 1,242352323 | 0,773493976 |
| RAB6A1 | 0,649018425 | 0,329462228 | 0,429 | 0,512 | 1 | Ciliated | 1,256544904 | 0,795833079 | 0,837890625 |
| RBPMS1 | 0,651587519 | 0,263946909 | 0,321 | 0,341 | 1 | Ciliated | 1,200759239 | 0,832806417 | 0,941348974 |
| TTC21B | 0,652648579 | -0,332086236 | 0,25 | 0,341 | 1 | Ciliated | 0,794386915 | 1,258832417 | 0,73313783 |
| FUS2 | 0,653146244 | -0,260936409 | 0,75 | 0,756 | 1 | Ciliated | 0,834546065 | 1,198256203 | 0,992063492 |
| WDR37 | 0,653495389 | 0,370547114 | 0,143 | 0,122 | 1 | Ciliated | 1,292843023 | 0,773489111 | 1,172131148 |
| PDCD101 | 0,656246738 | 0,284190119 | 0,429 | 0,488 | 1 | Ciliated | 1,217726481 | 0,821202475 | 0,879098361 |
| FOXP2 | 0,658066396 | 0,271299758 | 0,286 | 0,317 | 1 | Ciliated | 1,206894658 | 0,828572728 | 0,902208202 |
| LLPH | 0,658066396 | 0,418101242 | 0,286 | 0,317 | 1 | Ciliated | 1,336167842 | 0,748408971 | 0,902208202 |
| AGRN2 | 0,658157851 | 0,323502867 | 0,214 | 0,317 | 1 | Ciliated | 1,251365181 | 0,799127238 | 0,675078864 |
| THRB | 0,658157851 | 0,284487524 | 0,214 | 0,317 | 1 | Ciliated | 1,217977536 | 0,821033205 | 0,675078864 |
| BTG31 | 0,663796622 | -0,278980597 | 0,25 | 0,317 | 1 | Ciliated | 0,824173169 | 1,213337242 | 0,788643533 |
| TCEA12 | 0,664778205 | -0,280039339 | 0,571 | 0,659 | 1 | Ciliated | 0,82356856 | 1,214227993 | 0,86646434 |
| CDC123 | 0,670683235 | 0,33592639 | 0,393 | 0,463 | 1 | Ciliated | 1,26218763 | 0,792275234 | 0,848812095 |
| PTPA | 0,67271475 | 0,258243721 | 0,321 | 0,341 | 1 | Ciliated | 1,196021829 | 0,836105141 | 0,941348974 |
| NDUFAF2 | 0,674905163 | 0,322593439 | 0,286 | 0,293 | 1 | Ciliated | 1,25057661 | 0,79963114 | 0,976109215 |
| PPP1R14B2 | 0,682044156 | 0,381989192 | 0,321 | 0,366 | 1 | Ciliated | 1,303137387 | 0,767378797 | 0,87704918 |
| H1-101 | 0,684055713 | -0,295611798 | 0,607 | 0,659 | 1 | Ciliated | 0,814726761 | 1,227405367 | 0,921092564 |
| ERI3 | 0,687195119 | 0,25711088 | 0,393 | 0,39 | 1 | Ciliated | 1,195083051 | 0,83676193 | 1,007692308 |

| Supplementary Table 4 - clusermarker | | | | | | | | | |
|---|---|---|---|---|---|---|---|---|---|
| | **p_val** | **avg_log2FC** | **pct.1** | **pct.2** | **p_val_adj** | **celltype** | **Foldchange_UP** | **Foldchange_DOWN** | **Ratio_pct1_pct2** |
| KLHDC4 | 0,690815965 | 0,264768421 | 0,179 | 0,171 | 1 | Ciliated | 1,201443181 | 0,832332328 | 1,046783626 |
| B3GALT4 | 0,692620966 | 0,343943775 | 0,321 | 0,366 | 1 | Ciliated | 1,269221421 | 0,787884591 | 0,87704918 |
| DAZAP1 | 0,692907881 | 0,280731251 | 0,393 | 0,439 | 1 | Ciliated | 1,214810473 | 0,823173674 | 0,895216401 |
| NOP531 | 0,696195602 | -0,254558098 | 0,679 | 0,707 | 1 | Ciliated | 0,838243852 | 1,192970277 | 0,96039604 |
| DTHD1 | 0,698141077 | -0,258337186 | 0,75 | 0,756 | 1 | Ciliated | 0,836050976 | 1,196099316 | 0,992063492 |
| IRF1 | 0,701656743 | -0,331214136 | 0,643 | 0,61 | 1 | Ciliated | 0,794867262 | 1,258071691 | 1,054098361 |
| SAMD8 | 0,702036448 | 0,256014054 | 0,286 | 0,317 | 1 | Ciliated | 1,19417482 | 0,83739833 | 0,902208202 |
| NELFB1 | 0,70319018 | 0,261746598 | 0,214 | 0,22 | 1 | Ciliated | 1,19892931 | 0,834077532 | 0,972727273 |
| GBP6 | 0,704503654 | 0,336483812 | 0,179 | 0,171 | 1 | Ciliated | 1,262675403 | 0,791969177 | 1,046783626 |
| ABHD16A | 0,705417215 | 0,389670448 | 0,357 | 0,439 | 1 | Ciliated | 1,310094108 | 0,763303944 | 0,813211845 |
| ABCF3 | 0,707607295 | -0,255967745 | 0,25 | 0,341 | 1 | Ciliated | 0,83742521 | 1,194136489 | 0,73313783 |
| FGD5-AS11 | 0,710477021 | 0,295554918 | 0,393 | 0,463 | 1 | Ciliated | 1,227356977 | 0,814758883 | 0,848812095 |
| BORCS8 | 0,712909054 | 0,293113564 | 0,357 | 0,415 | 1 | Ciliated | 1,225281778 | 0,8161388 | 0,860240964 |
| LYRM21 | 0,715675252 | -0,298210404 | 0,536 | 0,683 | 1 | Ciliated | 0,813260583 | 1,229618183 | 0,78477306 |
| TSSC4 | 0,715738848 | 0,258987346 | 0,321 | 0,341 | 1 | Ciliated | 1,196638468 | 0,835674289 | 0,941348974 |
| TTC23L | 0,721420229 | -0,282104224 | 0,179 | 0,22 | 1 | Ciliated | 0,822390655 | 1,215967124 | 0,813636364 |
| ING2 | 0,721791143 | 0,342928244 | 0,25 | 0,268 | 1 | Ciliated | 1,268328315 | 0,788439388 | 0,932835821 |
| PTGR11 | 0,726652009 | -0,278218243 | 0,607 | 0,683 | 1 | Ciliated | 0,824608797 | 1,212696255 | 0,888726208 |
| TMEM106B1 | 0,727761956 | -0,417849673 | 0,393 | 0,488 | 1 | Ciliated | 0,748539486 | 1,335934869 | 0,805327869 |
| UVSSA | 0,728479098 | 0,290224191 | 0,214 | 0,22 | 1 | Ciliated | 1,222830288 | 0,817774968 | 0,972727273 |
| RSRP11 | 0,729046971 | -0,251027591 | 0,679 | 0,732 | 1 | Ciliated | 0,840297682 | 1,190054456 | 0,927595628 |
| SMIM141 | 0,734154437 | -0,256717413 | 0,714 | 0,732 | 1 | Ciliated | 0,836990171 | 1,19475716 | 0,975409836 |
| COA81 | 0,734390088 | 0,275407637 | 0,25 | 0,244 | 1 | Ciliated | 1,210336024 | 0,826216835 | 1,024590164 |
| COL28A1 | 0,734797643 | -0,252061723 | 0,25 | 0,22 | 1 | Ciliated | 0,839695568 | 1,190907799 | 1,136363636 |
| COX11 | 0,734849059 | 0,463523121 | 0,179 | 0,244 | 1 | Ciliated | 1,378905053 | 0,725213094 | 0,733606557 |
| CCDC28A | 0,741264855 | -0,253226276 | 0,464 | 0,561 | 1 | Ciliated | 0,839018033 | 1,191869496 | 0,827094474 |
| DYDC1 | 0,741529769 | 0,331140925 | 0,357 | 0,39 | 1 | Ciliated | 1,25800785 | 0,794907599 | 0,915384615 |
| PIGG | 0,747027365 | 0,300029437 | 0,286 | 0,317 | 1 | Ciliated | 1,231169534 | 0,812235823 | 0,902208202 |
| CDC14B | 0,747027365 | 0,400044144 | 0,286 | 0,317 | 1 | Ciliated | 1,319548286 | 0,757835095 | 0,902208202 |
| GOPC1 | 0,747962656 | 0,262680438 | 0,393 | 0,488 | 1 | Ciliated | 1,199705614 | 0,833537818 | 0,805327869 |
| MBP | 0,748165224 | -0,266932183 | 0,429 | 0,537 | 1 | Ciliated | 0,831084928 | 1,203246463 | 0,798882682 |

| Supplementary Table 4 - clusermarker | | | | | | | | | |
|---|---|---|---|---|---|---|---|---|---|
| | **p_val** | **avg_log2FC** | **pct.1** | **pct.2** | **p_val_adj** | **celltype** | **Foldchange_UP** | **Foldchange_DOWN** | **Ratio_pct1_pct2** |
| FAM110A | 0,751703879 | -0,311407151 | 0,143 | 0,171 | 1 | Ciliated | 0,805855374 | 1,240917455 | 0,83625731 |
| NUFIP21 | 0,752361754 | 0,25981005 | 0,5 | 0,561 | 1 | Ciliated | 1,197321051 | 0,835197877 | 0,891265597 |
| TTC21A | 0,753040616 | 0,310855106 | 0,429 | 0,463 | 1 | Ciliated | 1,24044271 | 0,806163793 | 0,926565875 |
| LINC00342 | 0,754124146 | -0,3075249 | 0,393 | 0,439 | 1 | Ciliated | 0,808026829 | 1,23758267 | 0,895216401 |
| PLTP | 0,757280989 | 0,329486936 | 0,286 | 0,39 | 1 | Ciliated | 1,256566423 | 0,79581945 | 0,733333333 |
| RUVBL1 | 0,757899711 | -0,372166877 | 0,75 | 0,732 | 1 | Ciliated | 0,772621175 | 1,294295357 | 1,024590164 |
| UBE2Q1 | 0,758418206 | 0,297928949 | 0,286 | 0,317 | 1 | Ciliated | 1,22937832 | 0,813419257 | 0,902208202 |
| DAPK3 | 0,766965072 | 0,287973728 | 0,214 | 0,22 | 1 | Ciliated | 1,220924279 | 0,819051613 | 0,972727273 |
| CALM32 | 0,769804545 | -0,307523222 | 0,607 | 0,707 | 1 | Ciliated | 0,808027768 | 1,237581231 | 0,858557284 |
| CERS5 | 0,770046124 | 0,25423817 | 0,107 | 0,146 | 1 | Ciliated | 1,192705756 | 0,83842976 | 0,732876712 |
| CENPB | 0,773438617 | 0,255666042 | 0,179 | 0,244 | 1 | Ciliated | 1,193886792 | 0,837600354 | 0,733606557 |
| YAE1 | 0,774304864 | 0,260074639 | 0,143 | 0,195 | 1 | Ciliated | 1,197540659 | 0,835044717 | 0,733333333 |
| PLAC4 | 0,774304864 | 0,267918775 | 0,143 | 0,195 | 1 | Ciliated | 1,204069589 | 0,830516782 | 0,733333333 |
| ARL13B | 0,775918366 | 0,322578745 | 0,357 | 0,415 | 1 | Ciliated | 1,250563873 | 0,799639284 | 0,860240964 |
| RAB11FIP4 | 0,776440807 | 0,296145523 | 0,321 | 0,39 | 1 | Ciliated | 1,22785953 | 0,814425409 | 0,823076923 |
| MTHFSD | 0,778039544 | -0,291069644 | 0,143 | 0,122 | 1 | Ciliated | 0,817295874 | 1,223547105 | 1,172131148 |
| ASXL2 | 0,778100518 | -0,267640921 | 0,357 | 0,439 | 1 | Ciliated | 0,830676749 | 1,203837715 | 0,813211845 |
| CEP571 | 0,77813814 | -0,311926876 | 0,464 | 0,512 | 1 | Ciliated | 0,805565121 | 1,241364571 | 0,90625 |
| SLC4A1AP | 0,779244168 | 0,307862317 | 0,321 | 0,366 | 1 | Ciliated | 1,23787215 | 0,807837869 | 0,87704918 |
| SYNE1 | 0,77985478 | -0,45155672 | 0,714 | 0,659 | 1 | Ciliated | 0,731253373 | 1,367515059 | 1,083459788 |
| C17orf75 | 0,77992796 | 0,254452895 | 0,214 | 0,22 | 1 | Ciliated | 1,192883287 | 0,83830498 | 0,972727273 |
| ABHD6 | 0,781355587 | 0,254805796 | 0,286 | 0,317 | 1 | Ciliated | 1,193175116 | 0,838099945 | 0,902208202 |
| NFU1 | 0,781991819 | -0,388086078 | 0,393 | 0,463 | 1 | Ciliated | 0,764142666 | 1,30865615 | 0,848812095 |
| TCTA | 0,784324985 | 0,280540177 | 0,357 | 0,39 | 1 | Ciliated | 1,214649591 | 0,823282705 | 0,915384615 |
| GBA2 | 0,786432674 | 0,303937035 | 0,179 | 0,244 | 1 | Ciliated | 1,234508726 | 0,810038826 | 0,733606557 |
| KIN1 | 0,786432674 | 0,305690808 | 0,179 | 0,244 | 1 | Ciliated | 1,236010336 | 0,809054723 | 0,733606557 |
| GABRP2 | 0,786577854 | 0,428342488 | 0,357 | 0,415 | 1 | Ciliated | 1,345686629 | 0,74311506 | 0,860240964 |
| CMAS | 0,78887449 | 0,395467732 | 0,393 | 0,488 | 1 | Ciliated | 1,315369143 | 0,760242861 | 0,805327869 |
| C20orf96 | 0,788910358 | -0,339835019 | 0,607 | 0,707 | 1 | Ciliated | 0,790131663 | 1,265611855 | 0,858557284 |
| POLR2J | 0,789485887 | -0,262298591 | 0,5 | 0,512 | 1 | Ciliated | 0,833758465 | 1,199388122 | 0,9765625 |
| CCDC47 | 0,790517452 | -0,25836796 | 0,536 | 0,634 | 1 | Ciliated | 0,836033142 | 1,19612483 | 0,845425868 |

| Supplementary Table 4 - clusermarker | | | | | | | | | |
|---|---|---|---|---|---|---|---|---|---|
| | **p_val** | **avg_log2FC** | **pct.1** | **pct.2** | **p_val_adj** | **celltype** | **Foldchange_UP** | **Foldchange_DOWN** | **Ratio_pct1_pct2** |
| IFT81 | 0,791962207 | 0,394295191 | 0,607 | 0,634 | 1 | Ciliated | 1,31430052 | 0,760860994 | 0,957413249 |
| USB1 | 0,792952328 | 0,306527854 | 0,214 | 0,22 | 1 | Ciliated | 1,236727673 | 0,808585449 | 0,972727273 |
| TARS2 | 0,795737569 | -0,657423143 | 0,143 | 0,171 | 1 | Ciliated | 0,634009717 | 1,577262892 | 0,83625731 |
| ARFIP2 | 0,796736588 | -0,317818308 | 0,536 | 0,537 | 1 | Ciliated | 0,802282199 | 1,246444208 | 0,998137803 |
| H2BC12 | 0,797535901 | 0,38524446 | 0,107 | 0,098 | 1 | Ciliated | 1,30608108 | 0,765649251 | 1,091836735 |
| HEBP11 | 0,800448116 | -0,310037397 | 0,107 | 0,122 | 1 | Ciliated | 0,80662085 | 1,239739836 | 0,87704918 |
| UNC5B-AS11 | 0,805835458 | 0,408348894 | 0,429 | 0,488 | 1 | Ciliated | 1,327166056 | 0,753485214 | 0,879098361 |
| UGDH-AS1 | 0,806034715 | 0,262688722 | 0,214 | 0,22 | 1 | Ciliated | 1,199712503 | 0,833533032 | 0,972727273 |
| KLHDC21 | 0,806365257 | -0,351635083 | 0,393 | 0,439 | 1 | Ciliated | 0,783695391 | 1,276005973 | 0,895216401 |
| KRT41 | 0,807189619 | 0,610996448 | 0,429 | 0,512 | 1 | Ciliated | 1,527313737 | 0,654744324 | 0,837890625 |
| COPZ1 | 0,807189619 | 0,266356868 | 0,429 | 0,512 | 1 | Ciliated | 1,202766731 | 0,831416412 | 0,837890625 |
| PELP1 | 0,809271266 | 0,442455749 | 0,321 | 0,39 | 1 | Ciliated | 1,358915499 | 0,735880929 | 0,823076923 |
| SIPA1L1 | 0,809558843 | -0,251081272 | 0,393 | 0,488 | 1 | Ciliated | 0,840266416 | 1,190098736 | 0,805327869 |
| ANAPC4 | 0,815860331 | 0,463586005 | 0,25 | 0,293 | 1 | Ciliated | 1,378965158 | 0,725181484 | 0,853242321 |
| GPR180 | 0,818507714 | 0,276110698 | 0,214 | 0,293 | 1 | Ciliated | 1,210925995 | 0,825814298 | 0,730375427 |
| PARP3 | 0,819171611 | 0,287335014 | 0,214 | 0,22 | 1 | Ciliated | 1,220383868 | 0,819414306 | 0,972727273 |
| PDS5A1 | 0,820292925 | 0,254726118 | 0,321 | 0,39 | 1 | Ciliated | 1,193109221 | 0,838146234 | 0,823076923 |
| MLX | 0,820570861 | -0,271448097 | 0,357 | 0,439 | 1 | Ciliated | 0,828487539 | 1,207018758 | 0,813211845 |
| TOGARAM1 | 0,828338426 | -0,414626736 | 0,536 | 0,585 | 1 | Ciliated | 0,75021357 | 1,332953761 | 0,916239316 |
| KLHDC10 | 0,8312765 | 0,293792278 | 0,357 | 0,439 | 1 | Ciliated | 1,225858346 | 0,815754939 | 0,813211845 |
| CFD1 | 0,834268256 | -0,299022172 | 0,464 | 0,415 | 1 | Ciliated | 0,81280311 | 1,230310253 | 1,118072289 |
| ZNF84 | 0,843190247 | 0,343474288 | 0,25 | 0,268 | 1 | Ciliated | 1,268808453 | 0,788141029 | 0,932835821 |
| CSTA1 | 0,845594668 | 0,286103031 | 0,214 | 0,22 | 1 | Ciliated | 1,219342171 | 0,82011434 | 0,972727273 |
| NAXE2 | 0,84722864 | -0,338567805 | 0,393 | 0,512 | 1 | Ciliated | 0,790825993 | 1,264500673 | 0,767578125 |
| CFAP126 | 0,849695688 | -0,303100302 | 0,964 | 0,854 | 1 | Ciliated | 0,810508768 | 1,233792945 | 1,128805621 |
| LMF2 | 0,852599621 | -0,44326272 | 0,214 | 0,268 | 1 | Ciliated | 0,735469429 | 1,359675822 | 0,798507463 |
| DDX61 | 0,853271494 | -0,35349601 | 0,5 | 0,61 | 1 | Ciliated | 0,782685157 | 1,277652951 | 0,819672131 |
| MALAT11 | 0,854516817 | -0,419720012 | 0,964 | 0,878 | 1 | Ciliated | 0,747569693 | 1,337667925 | 1,097949886 |
| CNTRL | 0,855000157 | -0,261519923 | 0,607 | 0,61 | 1 | Ciliated | 0,834208592 | 1,19874095 | 0,995081967 |
| PAAF1 | 0,855089148 | -0,29284099 | 0,179 | 0,22 | 1 | Ciliated | 0,816293011 | 1,225050302 | 0,813636364 |
| UPF3A | 0,856916277 | -0,252535985 | 0,536 | 0,537 | 1 | Ciliated | 0,839419577 | 1,191299354 | 0,998137803 |

| Supplementary Table 4 - clusermarker | | | | | | | | | |
|---|---|---|---|---|---|---|---|---|---|
| | **p_val** | **avg_log2FC** | **pct.1** | **pct.2** | **p_val_adj** | **celltype** | **Foldchange_UP** | **Foldchange_DOWN** | **Ratio_pct1_pct2** |
| LRRC10B | 0,857052425 | -0,29566117 | 0,643 | 0,707 | 1 | Ciliated | 0,81469888 | 1,227447373 | 0,909476662 |
| DNAH11 | 0,857359775 | -0,269192996 | 0,679 | 0,707 | 1 | Ciliated | 0,829783574 | 1,20513352 | 0,96039604 |
| CD164 | 0,858310378 | -0,315993324 | 0,75 | 0,756 | 1 | Ciliated | 0,803297714 | 1,244868475 | 0,992063492 |
| LINC02300 | 0,864402068 | 0,295773287 | 0,25 | 0,293 | 1 | Ciliated | 1,227542766 | 0,814635569 | 0,853242321 |
| GTF2H51 | 0,866815385 | -0,298792729 | 0,643 | 0,707 | 1 | Ciliated | 0,812932387 | 1,230114603 | 0,909476662 |
| QTRT1 | 0,868701897 | 0,312467246 | 0,286 | 0,39 | 1 | Ciliated | 1,241829618 | 0,805263448 | 0,733333333 |
| GPN2 | 0,869063486 | -0,362332987 | 0,107 | 0,122 | 1 | Ciliated | 0,777905608 | 1,285503009 | 0,87704918 |
| AC026304.1 | 0,871935643 | 0,383379411 | 0,179 | 0,195 | 1 | Ciliated | 1,304393729 | 0,766639687 | 0,917948718 |
| TSPYL11 | 0,872106767 | -0,266517795 | 0,5 | 0,561 | 1 | Ciliated | 0,831323676 | 1,202900902 | 0,891265597 |
| APOD1 | 0,873865546 | -0,80010758 | 0,607 | 0,561 | 1 | Ciliated | 0,574306351 | 1,741230963 | 1,081996435 |
| YWHAE1 | 0,878532814 | -0,404259948 | 0,857 | 0,927 | 1 | Ciliated | 0,755623802 | 1,323409873 | 0,924487594 |
| UBE2K | 0,882915091 | -0,254677592 | 0,393 | 0,488 | 1 | Ciliated | 0,838174426 | 1,193069091 | 0,805327869 |
| ZNHIT12 | 0,891307623 | -0,284244991 | 0,607 | 0,732 | 1 | Ciliated | 0,821171242 | 1,217772797 | 0,829234973 |
| ITGB1BP1 | 0,891379836 | 0,279898002 | 0,321 | 0,366 | 1 | Ciliated | 1,214109045 | 0,823649247 | 0,87704918 |
| TARDBP | 0,896101204 | 0,350295071 | 0,357 | 0,439 | 1 | Ciliated | 1,274821337 | 0,784423645 | 0,813211845 |
| COASY1 | 0,90005744 | 0,28944761 | 0,286 | 0,341 | 1 | Ciliated | 1,222172233 | 0,818215283 | 0,838709677 |
| ZFPM1 | 0,900224294 | 0,445843399 | 0,179 | 0,195 | 1 | Ciliated | 1,362110173 | 0,734155004 | 0,917948718 |
| KPNA1 | 0,905522017 | -0,27774894 | 0,25 | 0,268 | 1 | Ciliated | 0,824877083 | 1,212301834 | 0,932835821 |
| BIRC31 | 0,905522017 | 0,463313076 | 0,25 | 0,268 | 1 | Ciliated | 1,37870431 | 0,725318687 | 0,932835821 |
| WFDC2 | 0,90758133 | -0,537546065 | 1 | 0,976 | 1 | Ciliated | 0,68894176 | 1,451501503 | 1,024590164 |
| EMSY | 0,908929605 | 0,304005329 | 0,25 | 0,317 | 1 | Ciliated | 1,234567167 | 0,810000482 | 0,788643533 |
| LMTK2 | 0,913461158 | 0,252009392 | 0,25 | 0,293 | 1 | Ciliated | 1,190864602 | 0,839726026 | 0,853242321 |
| CAMSAP1 | 0,914146975 | -0,280652132 | 0,321 | 0,366 | 1 | Ciliated | 0,823218819 | 1,214743853 | 0,87704918 |
| ORAI1 | 0,914146975 | 0,258492069 | 0,286 | 0,39 | 1 | Ciliated | 1,196227732 | 0,835961225 | 0,733333333 |
| IFT122 | 0,917886922 | 0,363355751 | 0,357 | 0,439 | 1 | Ciliated | 1,286414658 | 0,777354326 | 0,813211845 |
| EPPK1 | 0,921029538 | 0,303285116 | 0,25 | 0,317 | 1 | Ciliated | 1,233951008 | 0,810404946 | 0,788643533 |
| FLVCR1-DT | 0,921210599 | 0,286521152 | 0,107 | 0,122 | 1 | Ciliated | 1,219695612 | 0,819876689 | 0,87704918 |
| PBRM11 | 0,923052143 | -0,291645048 | 0,5 | 0,561 | 1 | Ciliated | 0,816969969 | 1,224035201 | 0,891265597 |
| RPA31 | 0,926732894 | -0,316151893 | 0,857 | 0,78 | 1 | Ciliated | 0,803209427 | 1,245005308 | 1,098717949 |
| SLC6A141 | 0,927309695 | 0,513897778 | 0,321 | 0,415 | 1 | Ciliated | 1,427902803 | 0,70032778 | 0,773493976 |
| POMT1 | 0,92864027 | 0,261697532 | 0,179 | 0,195 | 1 | Ciliated | 1,198888535 | 0,8341059 | 0,917948718 |

| Supplementary Table 4 - clusermarker | | | | | | | | | |
|---|---|---|---|---|---|---|---|---|---|
| | **p_val** | **avg_log2FC** | **pct.1** | **pct.2** | **p_val_adj** | **celltype** | **Foldchange_UP** | **Foldchange_DOWN** | **Ratio_pct1_pct2** |
| ACADSB1 | 0,929191521 | 0,320740506 | 0,214 | 0,268 | 1 | Ciliated | 1,248971456 | 0,80065881 | 0,798507463 |
| TRAF7 | 0,934880572 | -0,294457385 | 0,393 | 0,439 | 1 | Ciliated | 0,815378948 | 1,226423618 | 0,895216401 |
| PRKD2 | 0,936145707 | 0,25196308 | 0,286 | 0,366 | 1 | Ciliated | 1,190826375 | 0,839752983 | 0,781420765 |
| RP1 | 0,936479587 | -0,390969802 | 0,857 | 0,78 | 1 | Ciliated | 0,762616789 | 1,311274567 | 1,098717949 |
| CCDC90B1 | 0,942040601 | 0,516172913 | 0,214 | 0,268 | 1 | Ciliated | 1,430156386 | 0,699224231 | 0,798507463 |
| RPS19BP11 | 0,943226613 | -0,268042806 | 0,464 | 0,561 | 1 | Ciliated | 0,830445384 | 1,20417311 | 0,827094474 |
| TUBB2A | 0,944312918 | -0,351632241 | 0,536 | 0,61 | 1 | Ciliated | 0,783696934 | 1,27600346 | 0,878688525 |
| GATAD2A | 0,945281536 | 0,29140378 | 0,25 | 0,317 | 1 | Ciliated | 1,223830517 | 0,817106606 | 0,788643533 |
| AZIN1 | 0,946011291 | -0,274479589 | 0,786 | 0,732 | 1 | Ciliated | 0,82674849 | 1,209557698 | 1,073770492 |
| PHGDH1 | 0,951457704 | 0,268874558 | 0,179 | 0,22 | 1 | Ciliated | 1,204867547 | 0,829966748 | 0,813636364 |
| NUP93 | 0,95387969 | 0,30329973 | 0,214 | 0,244 | 1 | Ciliated | 1,233963508 | 0,810396737 | 0,87704918 |
| MEIS2 | 0,957427999 | 0,26138631 | 0,25 | 0,317 | 1 | Ciliated | 1,198629935 | 0,834285855 | 0,788643533 |
| ARL16 | 0,957427999 | 0,253154491 | 0,25 | 0,317 | 1 | Ciliated | 1,191810193 | 0,839059782 | 0,788643533 |
| ZNF20 | 0,959874557 | 0,409567445 | 0,286 | 0,39 | 1 | Ciliated | 1,328287501 | 0,752849063 | 0,733333333 |
| STX16 | 0,963417327 | 0,420474137 | 0,429 | 0,537 | 1 | Ciliated | 1,338367333 | 0,747179026 | 0,798882682 |
| PRR151 | 0,965627609 | -0,345049464 | 0,679 | 0,805 | 1 | Ciliated | 0,787280983 | 1,270194532 | 0,843478261 |
| PRCC | 0,967413119 | -0,323509708 | 0,393 | 0,439 | 1 | Ciliated | 0,799123448 | 1,251371115 | 0,895216401 |
| TMEM70 | 0,971333048 | 0,331888749 | 0,321 | 0,366 | 1 | Ciliated | 1,25866011 | 0,794495664 | 0,87704918 |
| LINC02265 | 0,975229919 | 0,27360252 | 0,25 | 0,293 | 1 | Ciliated | 1,208822585 | 0,827251255 | 0,853242321 |
| AC009318.1 | 0,976116343 | -0,256936907 | 0,143 | 0,146 | 1 | Ciliated | 0,83686284 | 1,194938946 | 0,979452055 |
| CIZ1 | 0,980665071 | 0,312054826 | 0,214 | 0,268 | 1 | Ciliated | 1,24147467 | 0,80549368 | 0,798507463 |
| ZNF496 | 0,981747783 | 0,272307933 | 0,25 | 0,317 | 1 | Ciliated | 1,207738347 | 0,827993913 | 0,788643533 |
| ATP6V0A2 | 0,98204569 | -0,256381241 | 0,286 | 0,317 | 1 | Ciliated | 0,837185227 | 1,194478794 | 0,902208202 |
| MMAB1 | 0,983977215 | -0,266962388 | 0,429 | 0,463 | 1 | Ciliated | 0,831067528 | 1,203271655 | 0,926565875 |
| VDAC12 | 0,985015463 | -0,284758971 | 0,607 | 0,659 | 1 | Ciliated | 0,82087874 | 1,218206723 | 0,921092564 |
| RCAN3AS | 0,993407786 | 0,359959359 | 0,214 | 0,244 | 1 | Ciliated | 1,283389744 | 0,779186529 | 0,87704918 |
| TMEM223 | 0,993915456 | 0,431549842 | 0,25 | 0,317 | 1 | Ciliated | 1,348681645 | 0,741464825 | 0,788643533 |
| PHLDA21 | 0,994105621 | 0,29088221 | 0,286 | 0,341 | 1 | Ciliated | 1,223388152 | 0,817402063 | 0,838709677 |
| RARS1 | 0,994265425 | 0,250730844 | 0,321 | 0,366 | 1 | Ciliated | 1,189809699 | 0,84047054 | 0,87704918 |
| CCNO1 | 1 | -0,36094883 | 0,393 | 0,439 | 1 | Ciliated | 0,778652308 | 1,284270257 | 0,895216401 |
| CDYL | 1 | 0,272411382 | 0,321 | 0,39 | 1 | Ciliated | 1,207824952 | 0,827934544 | 0,823076923 |

| Supplementary Table 4 - clusermarker | | | | | | | | | |
|---|---|---|---|---|---|---|---|---|---|
| | **p_val** | **avg_log2FC** | **pct.1** | **pct.2** | **p_val_adj** | **celltype** | **Foldchange_UP** | **Foldchange_DOWN** | **Ratio_pct1_pct2** |
| SNTB1 | 1 | -0,268169285 | 0,5 | 0,512 | 1 | Ciliated | 0,830372583 | 1,204278682 | 0,9765625 |
| MAP4K5 | 1 | 0,268414767 | 0,321 | 0,39 | 1 | Ciliated | 1,204483614 | 0,830231303 | 0,823076923 |
| AC036176.1 | 1 | -0,356311002 | 0,321 | 0,341 | 1 | Ciliated | 0,781159467 | 1,280148346 | 0,941348974 |
| PRR43 | 1,3097E-268 | -3,733367931 | 0,166 | 0,852 | 4,7916E-264 | TC | 0,075187262 | 13,30012529 | 0,194835681 |
| LYZ3 | 4,1742E-220 | -2,944525874 | 0,109 | 0,688 | 1,5272E-215 | TC | 0,129900071 | 7,69822519 | 0,158430233 |
| PRB33 | 5,2041E-190 | -3,046733203 | 0,013 | 0,389 | 1,904E-185 | TC | 0,121015756 | 8,263386813 | 0,033419023 |
| RPS262 | 2,8473E-175 | -1,639727978 | 0,582 | 0,955 | 1,0417E-170 | TC | 0,320916978 | 3,116070723 | 0,609424084 |
| RPS272 | 4,7748E-150 | -0,920669805 | 0,981 | 0,998 | 1,747E-145 | TC | 0,528263705 | 1,892993956 | 0,982965932 |
| CCL5 | 4,2409E-145 | -1,764685421 | 0,385 | 0,86 | 1,5516E-140 | TC | 0,294290847 | 3,397998985 | 0,447674419 |
| MT-ATP83 | 5,8082E-144 | -1,541760191 | 0,39 | 0,873 | 2,125E-139 | TC | 0,343466146 | 2,9114951 | 0,446735395 |
| RPS293 | 2,1559E-116 | -1,212143167 | 0,642 | 0,923 | 7,8878E-112 | TC | 0,431626944 | 2,316815512 | 0,695557963 |
| MTRNR2L83 | 1,1765E-109 | -3,317249611 | 0,066 | 0,39 | 4,3045E-105 | TC | 0,100324814 | 9,967623747 | 0,169230769 |
| RPL133 | 1,9263E-107 | 0,911215696 | 0,992 | 0,973 | 7,0479E-103 | TC | 1,880629555 | 0,531736831 | 1,019527235 |
| SLPI3 | 4,5272E-102 | -2,059060133 | 0,118 | 0,495 | 1,65635E-97 | TC | 0,239972313 | 4,167147403 | 0,238383838 |
| LINC01871 | 7,92868E-85 | -1,441016812 | 0,046 | 0,304 | 2,90087E-80 | TC | 0,368307629 | 2,715121599 | 0,151315789 |
| KLRC2 | 3,38835E-84 | -1,61732142 | 0,024 | 0,244 | 1,2397E-79 | TC | 0,325940059 | 3,068048778 | 0,098360656 |
| SAMSN1 | 3,57267E-83 | -1,361035417 | 0,219 | 0,599 | 1,30713E-78 | TC | 0,389302788 | 2,568694676 | 0,365609349 |
| SYTL3 | 6,43777E-74 | -1,5419963 | 0,174 | 0,508 | 2,35539E-69 | TC | 0,343409939 | 2,91197163 | 0,342519685 |
| KLRD1 | 4,46856E-73 | -1,222917125 | 0,091 | 0,384 | 1,63491E-68 | TC | 0,428415587 | 2,334182114 | 0,236979167 |
| RPL122 | 1,38591E-70 | 0,805903017 | 0,968 | 0,934 | 5,07064E-66 | TC | 1,748239714 | 0,572003937 | 1,03640257 |
| KLRK1 | 1,53189E-69 | -1,215386258 | 0,116 | 0,422 | 5,60472E-65 | TC | 0,430657763 | 2,322029431 | 0,274881517 |
| RPLP12 | 2,05973E-68 | 0,609909008 | 0,997 | 0,992 | 7,53592E-64 | TC | 1,526162949 | 0,655238027 | 1,005040323 |
| RPL101 | 5,00359E-68 | 0,575379152 | 0,995 | 0,992 | 1,83066E-63 | TC | 1,490069014 | 0,671109855 | 1,003024194 |
| RPL83 | 2,36057E-66 | 0,820938672 | 0,942 | 0,88 | 8,63663E-62 | TC | 1,766555006 | 0,566073514 | 1,070454545 |
| KLRC1 | 5,29261E-66 | -1,492964941 | 0,059 | 0,297 | 1,93641E-61 | TC | 0,355281644 | 2,81466835 | 0,198653199 |
| METRNL | 5,44845E-65 | -1,279899109 | 0,056 | 0,285 | 1,99343E-60 | TC | 0,411824307 | 2,428219953 | 0,196491228 |
| RPL18A2 | 2,81209E-62 | 0,679414668 | 0,973 | 0,953 | 1,02886E-57 | TC | 1,601489865 | 0,624418563 | 1,020986359 |
| LDLRAD4 | 3,28205E-62 | -1,42767333 | 0,132 | 0,416 | 1,2008E-57 | TC | 0,371729908 | 2,690125222 | 0,317307692 |
| HOPX | 1,49303E-61 | -0,996767866 | 0,132 | 0,432 | 5,46255E-57 | TC | 0,501121428 | 1,995524326 | 0,305555556 |
| PTPN22 | 3,5195E-61 | -1,136747257 | 0,147 | 0,447 | 1,28768E-56 | TC | 0,454783792 | 2,198847054 | 0,32885906 |
| CAPG2 | 5,58256E-60 | -1,108538715 | 0,074 | 0,313 | 2,04249E-55 | TC | 0,463763533 | 2,156271311 | 0,236421725 |

| Supplementary Table 4 - clusermarker | | | | | | | | | |
|---|---|---|---|---|---|---|---|---|---|
| | **p_val** | **avg_log2FC** | **pct.1** | **pct.2** | **p_val_adj** | **celltype** | **Foldchange_UP** | **Foldchange_DOWN** | **Ratio_pct1_pct2** |
| STATH3 | 6,45939E-60 | -1,355870235 | 0,003 | 0,122 | 2,3633E-55 | TC | 0,39069908 | 2,559514599 | 0,024590164 |
| CD52 | 1,51822E-59 | -0,930374475 | 0,574 | 0,867 | 5,55473E-55 | TC | 0,524722124 | 1,905770606 | 0,662053057 |
| JUND1 | 3,01212E-58 | -0,984883956 | 0,399 | 0,739 | 1,10204E-53 | TC | 0,505266363 | 1,979154112 | 0,539918809 |
| GZMB | 1,23385E-57 | -1,139648223 | 0,123 | 0,401 | 4,51429E-53 | TC | 0,453870233 | 2,203272935 | 0,306733167 |
| KLRC4 | 4,19201E-57 | -0,972225377 | 0,02 | 0,181 | 1,53373E-52 | TC | 0,509719207 | 1,961864465 | 0,110497238 |
| IGKC3 | 5,88201E-56 | 2,469651833 | 0,348 | 0,041 | 2,15205E-51 | TC | 5,539100956 | 0,180534713 | 8,487804878 |
| LINC02446 | 2,09242E-53 | -1,361642409 | 0,034 | 0,208 | 7,65554E-49 | TC | 0,389139029 | 2,569775644 | 0,163461538 |
| RPS151 | 1,56754E-52 | 0,611096213 | 0,979 | 0,947 | 5,73516E-48 | TC | 1,527419357 | 0,654699049 | 1,033790919 |
| RPS122 | 3,02883E-52 | 0,564903766 | 0,991 | 0,986 | 1,10816E-47 | TC | 1,479288831 | 0,676000507 | 1,005070994 |
| ZNF683 | 2,14697E-50 | -1,032652988 | 0,056 | 0,255 | 7,85513E-46 | TC | 0,488810443 | 2,045782807 | 0,219607843 |
| NFKBIA2 | 2,33375E-50 | 1,313399422 | 0,694 | 0,453 | 8,53851E-46 | TC | 2,485264535 | 0,402371653 | 1,53200883 |
| ID22 | 3,14245E-50 | -1,119843433 | 0,259 | 0,563 | 1,14973E-45 | TC | 0,460143759 | 2,173233865 | 0,460035524 |
| SCGB3A23 | 5,60135E-50 | -1,910044609 | 0,078 | 0,288 | 2,04937E-45 | TC | 0,266084318 | 3,758207202 | 0,270833333 |
| S100A41 | 6,11033E-49 | -0,749367271 | 0,713 | 0,934 | 2,23559E-44 | TC | 0,594864393 | 1,681055401 | 0,763383298 |
| RPS81 | 7,13524E-49 | 0,580197786 | 0,981 | 0,977 | 2,61057E-44 | TC | 1,495054199 | 0,668872072 | 1,004094166 |
| AUTS21 | 7,68473E-49 | -1,039323673 | 0,078 | 0,295 | 2,81161E-44 | TC | 0,486555514 | 2,055263932 | 0,26440678 |
| HSPA1A3 | 6,58315E-48 | -3,172467612 | 0,127 | 0,361 | 2,40858E-43 | TC | 0,110915461 | 9,015875616 | 0,351800554 |
| ZG16B3 | 1,60007E-47 | -1,189175686 | 0,022 | 0,166 | 5,85418E-43 | TC | 0,438553366 | 2,280224206 | 0,13253012 |
| CD8A | 4,1391E-47 | -0,851546444 | 0,214 | 0,52 | 1,51437E-42 | TC | 0,554190373 | 1,804434086 | 0,411538462 |
| RPS212 | 1,39722E-46 | 0,664391876 | 0,941 | 0,901 | 5,11203E-42 | TC | 1,584900066 | 0,630954608 | 1,044395117 |
| CXCR6 | 4,68008E-46 | -1,203439201 | 0,091 | 0,307 | 1,7123E-41 | TC | 0,434238877 | 2,302879943 | 0,296416938 |
| THEMIS | 2,21276E-45 | -0,849992736 | 0,085 | 0,301 | 8,09584E-41 | TC | 0,554787529 | 1,80249185 | 0,282392027 |
| GABARAPL11 | 1,719E-44 | -0,952366322 | 0,112 | 0,345 | 6,28929E-40 | TC | 0,516784132 | 1,935043934 | 0,324637681 |
| C6orf583 | 2,10495E-44 | -1,247045982 | 0,011 | 0,125 | 7,70138E-40 | TC | 0,421309987 | 2,373549244 | 0,088 |
| ACTB3 | 5,25675E-44 | 0,732255746 | 0,983 | 0,967 | 1,92329E-39 | TC | 1,661234509 | 0,601961972 | 1,016546019 |
| PRH23 | 1,79939E-43 | -1,179596003 | 0,008 | 0,113 | 6,58343E-39 | TC | 0,441475107 | 2,265133377 | 0,07079646 |
| ANXA14 | 7,05663E-43 | -0,876939754 | 0,417 | 0,726 | 2,58181E-38 | TC | 0,544521246 | 1,836475634 | 0,574380165 |
| RPS52 | 3,02472E-41 | 0,656824632 | 0,922 | 0,866 | 1,10665E-36 | TC | 1,576608691 | 0,634272794 | 1,064665127 |
| HSPA1B2 | 4,479E-41 | -2,602274363 | 0,12 | 0,336 | 1,63873E-36 | TC | 0,164678673 | 6,072431721 | 0,357142857 |
| H4C32 | 8,39703E-41 | -0,799382162 | 0,317 | 0,611 | 3,07222E-36 | TC | 0,574595197 | 1,740355655 | 0,518821604 |
| MSMB3 | 1,16452E-40 | -0,867960076 | 0,032 | 0,176 | 4,26062E-36 | TC | 0,547921046 | 1,82508047 | 0,181818182 |

| Supplementary Table 4 - clusermarker | | | | | | | | | |
|---|---|---|---|---|---|---|---|---|---|
| | **p_val** | **avg_log2FC** | **pct.1** | **pct.2** | **p_val_adj** | **celltype** | **Foldchange_UP** | **Foldchange_DOWN** | **Ratio_pct1_pct2** |
| MT-ND33 | 2,30664E-40 | -0,580226445 | 0,788 | 0,911 | 8,4393E-36 | TC | 0,668858785 | 1,495083898 | 0,864983535 |
| SCGB3A12 | 3,44229E-40 | -1,605010479 | 0,293 | 0,602 | 1,25943E-35 | TC | 0,328733302 | 3,041979602 | 0,486710963 |
| ITGA1 | 8,58576E-40 | -0,857703305 | 0,059 | 0,232 | 3,14127E-35 | TC | 0,551830344 | 1,812151164 | 0,254310345 |
| NR4A31 | 1,91854E-39 | -1,087379027 | 0,182 | 0,422 | 7,01937E-35 | TC | 0,470615575 | 2,12487655 | 0,431279621 |
| HSP90AA11 | 2,44217E-39 | -1,174941769 | 0,632 | 0,816 | 8,93515E-35 | TC | 0,442901636 | 2,257837676 | 0,774509804 |
| PARP81 | 4,91819E-39 | -0,806269371 | 0,179 | 0,438 | 1,79942E-34 | TC | 0,571858703 | 1,748683714 | 0,408675799 |
| RPS163 | 4,24896E-38 | 0,641365613 | 0,887 | 0,821 | 1,55457E-33 | TC | 1,559804927 | 0,641105809 | 1,080389769 |
| CD2 | 4,64116E-38 | -0,811114511 | 0,383 | 0,655 | 1,69806E-33 | TC | 0,569941396 | 1,754566357 | 0,584732824 |
| RPL181 | 5,03856E-38 | 0,523589836 | 0,961 | 0,943 | 1,84346E-33 | TC | 1,43752778 | 0,695638731 | 1,019088017 |
| GPR65 | 5,65185E-38 | -0,91518052 | 0,139 | 0,361 | 2,06784E-33 | TC | 0,530277515 | 1,885805022 | 0,385041551 |
| ARHGDIB1 | 1,22336E-37 | -0,688917137 | 0,616 | 0,849 | 4,47591E-33 | TC | 0,620319277 | 1,612073069 | 0,725559482 |
| MT-ND4L2 | 3,5486E-37 | -0,579745008 | 0,691 | 0,884 | 1,29832E-32 | TC | 0,669082025 | 1,494585062 | 0,781674208 |
| SLA2 | 5,79406E-37 | -0,731165863 | 0,055 | 0,218 | 2,11987E-32 | TC | 0,602416895 | 1,659980004 | 0,252293578 |
| UBC2 | 7,59949E-37 | -0,533531025 | 0,892 | 0,956 | 2,78043E-32 | TC | 0,690861765 | 1,44746757 | 0,933054393 |
| XCL1 | 1,98568E-36 | -1,727626537 | 0,082 | 0,262 | 7,26499E-32 | TC | 0,301948301 | 3,31182522 | 0,312977099 |
| CD3G | 2,15142E-36 | -0,787269974 | 0,269 | 0,539 | 7,8714E-32 | TC | 0,579439534 | 1,725805612 | 0,499072356 |
| TRAV12-2 | 3,01435E-36 | -1,576154598 | 0,04 | 0,179 | 1,10286E-31 | TC | 0,335374616 | 2,981740278 | 0,223463687 |
| CCL4 | 2,07093E-35 | -1,216507004 | 0,265 | 0,53 | 7,57691E-31 | TC | 0,43032334 | 2,323833982 | 0,5 |
| AL627171.22 | 2,14855E-35 | -0,856004743 | 0,056 | 0,215 | 7,86091E-31 | TC | 0,552480426 | 1,810018876 | 0,260465116 |
| ITGAE1 | 2,97412E-35 | -0,789586852 | 0,106 | 0,306 | 1,08814E-30 | TC | 0,578509737 | 1,728579374 | 0,346405229 |
| ANKRD28 | 4,02623E-35 | -0,810372335 | 0,108 | 0,307 | 1,47308E-30 | TC | 0,570234671 | 1,753663975 | 0,351791531 |
| ALOX5AP | 1,48586E-34 | -0,72551813 | 0,309 | 0,586 | 5,4363E-30 | TC | 0,604779805 | 1,653494366 | 0,527303754 |
| CD813 | 6,04228E-34 | -0,727169042 | 0,144 | 0,366 | 2,21069E-29 | TC | 0,604088136 | 1,655387585 | 0,393442623 |
| ALDOA2 | 2,27717E-33 | -0,73545279 | 0,148 | 0,367 | 8,33146E-29 | TC | 0,600629488 | 1,664919923 | 0,403269755 |
| PFN13 | 7,32746E-33 | 0,745045155 | 0,875 | 0,815 | 2,6809E-28 | TC | 1,676026727 | 0,596649197 | 1,073619632 |
| RPLP02 | 7,50995E-33 | 0,606106171 | 0,925 | 0,867 | 2,74767E-28 | TC | 1,522145395 | 0,656967464 | 1,066897347 |
| RPS131 | 1,05521E-32 | 0,52966614 | 0,951 | 0,941 | 3,8607E-28 | TC | 1,443595089 | 0,69271502 | 1,010626993 |
| SYTL22 | 4,73664E-32 | -0,721417264 | 0,077 | 0,245 | 1,73299E-27 | TC | 0,606501339 | 1,648800976 | 0,314285714 |
| ZEB2 | 6,10494E-32 | -0,855842994 | 0,096 | 0,274 | 2,23361E-27 | TC | 0,552542371 | 1,809815956 | 0,350364964 |
| CD96 | 1,22236E-31 | -0,61434338 | 0,237 | 0,494 | 4,47223E-27 | TC | 0,653227132 | 1,530861091 | 0,479757085 |
| PTPRC | 1,47277E-31 | -0,558245005 | 0,684 | 0,877 | 5,38843E-27 | TC | 0,6791278 | 1,472476904 | 0,779931585 |

| Supplementary Table 4 - clusermarker | | | | | | | | | |
|---|---|---|---|---|---|---|---|---|---|
| | **p_val** | **avg_log2FC** | **pct.1** | **pct.2** | **p_val_adj** | **celltype** | **Foldchange_UP** | **Foldchange_DOWN** | **Ratio_pct1_pct2** |
| RUNX3 | 7,1097E-31 | -0,693200079 | 0,179 | 0,405 | 2,60123E-26 | TC | 0,61848046 | 1,616865956 | 0,441975309 |
| CLEC2B2 | 3,16013E-30 | -0,820692131 | 0,105 | 0,288 | 1,1562E-25 | TC | 0,566170258 | 1,766253147 | 0,364583333 |
| LGALS32 | 5,34042E-30 | -0,656168696 | 0,186 | 0,411 | 1,9539E-25 | TC | 0,634561239 | 1,575892032 | 0,452554745 |
| TUBA4A | 1,66292E-29 | -0,786744498 | 0,23 | 0,462 | 6,08414E-25 | TC | 0,579650623 | 1,725177132 | 0,497835498 |
| PTPRCAP | 1,72619E-29 | -0,59030308 | 0,067 | 0,218 | 6,31561E-25 | TC | 0,664203357 | 1,505563001 | 0,30733945 |
| HSPA83 | 5,07056E-29 | -0,672322519 | 0,788 | 0,913 | 1,85516E-24 | TC | 0,627495701 | 1,593636415 | 0,863088719 |
| ARL4C | 6,38989E-29 | -0,660488836 | 0,258 | 0,506 | 2,33787E-24 | TC | 0,632663892 | 1,580618103 | 0,509881423 |
| SPRY13 | 6,66124E-29 | -0,837327878 | 0,038 | 0,16 | 2,43715E-24 | TC | 0,559679233 | 1,786737727 | 0,2375 |
| IGKV4-12 | 7,34826E-29 | 2,998909816 | 0,169 | 0,003 | 2,68851E-24 | TC | 7,993957021 | 0,125094493 | 56,33333333 |
| CLEC2D | 8,47472E-29 | -0,658245932 | 0,134 | 0,331 | 3,10064E-24 | TC | 0,633648236 | 1,578162684 | 0,404833837 |
| OASL | 1,00571E-28 | -0,68427672 | 0,04 | 0,164 | 3,67961E-24 | TC | 0,622317741 | 1,606896179 | 0,243902439 |
| LCP1 | 1,10219E-28 | -0,63818513 | 0,404 | 0,67 | 4,03259E-24 | TC | 0,642520713 | 1,556370058 | 0,602985075 |
| PRF1 | 1,53662E-28 | -0,662817054 | 0,138 | 0,334 | 5,62203E-24 | TC | 0,631643724 | 1,583170959 | 0,413173653 |
| JAML | 1,58088E-28 | -0,707975842 | 0,084 | 0,245 | 5,78398E-24 | TC | 0,612178447 | 1,633510629 | 0,342857143 |
| AOAH | 2,29718E-28 | -0,549542221 | 0,083 | 0,247 | 8,40468E-24 | TC | 0,683236891 | 1,463621203 | 0,336032389 |
| GNLY | 6,35601E-28 | -0,783062937 | 0,074 | 0,227 | 2,32547E-23 | TC | 0,581131702 | 1,720780328 | 0,325991189 |
| GZMH | 1,32902E-27 | -0,693863062 | 0,091 | 0,256 | 4,86248E-23 | TC | 0,618196306 | 1,617609149 | 0,35546875 |
| IGKV1-5 | 4,54157E-27 | 2,916766456 | 0,17 | 0,009 | 1,66162E-22 | TC | 7,551516811 | 0,132423727 | 18,88888889 |
| TRBV19 | 4,97186E-27 | -1,655826473 | 0,034 | 0,143 | 1,81905E-22 | TC | 0,317355891 | 3,151036516 | 0,237762238 |
| HLA-C3 | 6,9403E-27 | -0,392890914 | 0,872 | 0,953 | 2,53925E-22 | TC | 0,761601954 | 1,313021841 | 0,915005247 |
| HIC1 | 1,49755E-26 | -0,474780481 | 0,021 | 0,114 | 5,47908E-22 | TC | 0,719576272 | 1,389706747 | 0,184210526 |
| IGKV3-203 | 1,88324E-26 | 2,960449623 | 0,208 | 0,036 | 6,89021E-22 | TC | 7,783665021 | 0,128474182 | 5,777777778 |
| KANSL1-AS1 | 2,24507E-26 | -0,359513217 | 0,038 | 0,154 | 8,21403E-22 | TC | 0,779427524 | 1,282992927 | 0,246753247 |
| IGHG12 | 3,00375E-26 | 1,935028666 | 0,156 | 0,003 | 1,09898E-21 | TC | 3,823857248 | 0,261516039 | 52 |
| GLIPR11 | 3,08509E-26 | -0,61016786 | 0,218 | 0,441 | 1,12874E-21 | TC | 0,655120473 | 1,526436803 | 0,494331066 |
| STAT4 | 4,68759E-26 | -0,664627043 | 0,148 | 0,339 | 1,71505E-21 | TC | 0,630851767 | 1,585158435 | 0,436578171 |
| SON3 | 4,70232E-26 | -0,526487584 | 0,426 | 0,685 | 1,72044E-21 | TC | 0,694242897 | 1,440418051 | 0,62189781 |
| PET1001 | 5,2152E-26 | -0,600404595 | 0,096 | 0,259 | 1,90809E-21 | TC | 0,659568958 | 1,516141699 | 0,370656371 |
| EGR12 | 5,82789E-26 | -0,481251059 | 0,11 | 0,286 | 2,13225E-21 | TC | 0,716356153 | 1,395953668 | 0,384615385 |
| RACK12 | 2,29187E-25 | 0,513477948 | 0,864 | 0,809 | 8,38526E-21 | TC | 1,427487338 | 0,700531608 | 1,067985167 |
| ZFP362 | 2,95197E-25 | -0,680514986 | 0,781 | 0,878 | 1,08004E-20 | TC | 0,623942512 | 1,602711758 | 0,88952164 |

| Supplementary Table 4 - clusermarker | | | | | | | | | |
|---|---|---|---|---|---|---|---|---|---|
| | **p_val** | **avg_log2FC** | **pct.1** | **pct.2** | **p_val_adj** | **celltype** | **Foldchange_UP** | **Foldchange_DOWN** | **Ratio_pct1_pct2** |
| RGS2 | 3,25616E-25 | -0,6645621 | 0,1 | 0,262 | 1,19133E-20 | TC | 0,630880166 | 1,58508708 | 0,381679389 |
| MXRA72 | 3,54468E-25 | -0,538622316 | 0,041 | 0,157 | 1,29689E-20 | TC | 0,688428001 | 1,452584727 | 0,261146497 |
| MYADM2 | 3,58679E-25 | -0,767499491 | 0,276 | 0,491 | 1,3123E-20 | TC | 0,587434747 | 1,702316734 | 0,562118126 |
| CALM22 | 3,65583E-25 | -0,6295978 | 0,465 | 0,702 | 1,33756E-20 | TC | 0,646356584 | 1,547133617 | 0,662393162 |
| SEC622 | 5,11949E-25 | -0,653480421 | 0,196 | 0,405 | 1,87307E-20 | TC | 0,635744763 | 1,572958296 | 0,483950617 |
| HSPH12 | 8,06572E-25 | -0,876370216 | 0,177 | 0,373 | 2,951E-20 | TC | 0,544736251 | 1,835750784 | 0,474530831 |
| CD8B | 1,21186E-24 | -0,535259474 | 0,132 | 0,315 | 4,43384E-20 | TC | 0,69003456 | 1,449202776 | 0,419047619 |
| RPL291 | 1,61907E-24 | 0,422585039 | 0,954 | 0,937 | 5,92369E-20 | TC | 1,340327019 | 0,746086579 | 1,01814301 |
| SUB11 | 1,6326E-24 | -0,545030242 | 0,353 | 0,604 | 5,9732E-20 | TC | 0,685377036 | 1,459050928 | 0,584437086 |
| RPL191 | 1,83278E-24 | 0,377176804 | 0,982 | 0,977 | 6,7056E-20 | TC | 1,298797762 | 0,769942811 | 1,005117707 |
| RPLP23 | 1,89089E-24 | 0,449874915 | 0,932 | 0,916 | 6,91821E-20 | TC | 1,365921823 | 0,73210632 | 1,017467249 |
| IGKV3-112 | 2,07771E-24 | 2,599202396 | 0,158 | 0,011 | 7,60173E-20 | TC | 6,059515294 | 0,165029701 | 14,36363636 |
| HSP90AB12 | 5,31268E-24 | -0,826265219 | 0,692 | 0,845 | 1,94375E-19 | TC | 0,563987378 | 1,773089326 | 0,818934911 |
| LMO41 | 9,84822E-24 | -0,511438888 | 0,04 | 0,151 | 3,60317E-19 | TC | 0,701522418 | 1,425471197 | 0,264900662 |
| RANBP2 | 1,97143E-23 | -0,685248485 | 0,118 | 0,28 | 7,21287E-19 | TC | 0,621898703 | 1,607978911 | 0,421428571 |
| NFE2L21 | 2,29141E-23 | -0,689023491 | 0,154 | 0,33 | 8,38359E-19 | TC | 0,620273549 | 1,612191914 | 0,466666667 |
| NDUFA1 | 2,44625E-23 | -0,562966325 | 0,212 | 0,422 | 8,9501E-19 | TC | 0,67690894 | 1,47730358 | 0,502369668 |
| CEBPZ | 5,22346E-23 | -0,656769883 | 0,068 | 0,197 | 1,91111E-18 | TC | 0,634296865 | 1,576548861 | 0,345177665 |
| CREM | 8,42219E-23 | -0,84452089 | 0,35 | 0,545 | 3,08143E-18 | TC | 0,55689572 | 1,795668317 | 0,642201835 |
| LTF2 | 1,18174E-22 | -0,691264316 | 0,021 | 0,104 | 4,32365E-18 | TC | 0,619310875 | 1,614697951 | 0,201923077 |
| HLA-B3 | 2,18954E-22 | 0,42694989 | 0,97 | 0,968 | 8,01086E-18 | TC | 1,344388298 | 0,743832717 | 1,002066116 |
| IFNG | 2,92058E-22 | -0,986132169 | 0,044 | 0,152 | 1,06855E-17 | TC | 0,504829398 | 1,980867208 | 0,289473684 |
| LFNG | 3,32501E-22 | -0,490136183 | 0,022 | 0,107 | 1,21652E-17 | TC | 0,71195789 | 1,404577454 | 0,205607477 |
| ANXA52 | 4,19007E-22 | -0,553166045 | 0,145 | 0,319 | 1,53302E-17 | TC | 0,68152286 | 1,467302211 | 0,454545455 |
| IDI11 | 6,19174E-22 | -0,427666688 | 0,12 | 0,288 | 2,26537E-17 | TC | 0,743463238 | 1,345056418 | 0,416666667 |
| RPL382 | 9,88461E-22 | -0,50715979 | 0,519 | 0,73 | 3,61648E-17 | TC | 0,703606253 | 1,421249449 | 0,710958904 |
| CLIC32 | 1,72359E-21 | -0,692889264 | 0,036 | 0,134 | 6,3061E-17 | TC | 0,61861372 | 1,616517655 | 0,268656716 |
| LTB | 1,81976E-21 | 1,375669547 | 0,369 | 0,202 | 6,65795E-17 | TC | 2,594883102 | 0,385373815 | 1,826732673 |
| EIF4A13 | 2,03028E-21 | -0,529681621 | 0,529 | 0,771 | 7,42818E-17 | TC | 0,692707587 | 1,443610579 | 0,68612192 |
| SRRM11 | 2,37965E-21 | -0,509224299 | 0,212 | 0,414 | 8,70643E-17 | TC | 0,702600107 | 1,423284725 | 0,512077295 |
| RPS4X1 | 3,20347E-21 | 0,327652712 | 0,981 | 0,983 | 1,17205E-16 | TC | 1,254969857 | 0,796831888 | 0,997965412 |

| Supplementary Table 4 - clusermarker | | | | | | | | | |
|---|---|---|---|---|---|---|---|---|---|
| | **p_val** | **avg_log2FC** | **pct.1** | **pct.2** | **p_val_adj** | **celltype** | **Foldchange_UP** | **Foldchange_DOWN** | **Ratio_pct1_pct2** |
| VPS37B1 | 4,22548E-21 | -0,575192363 | 0,129 | 0,289 | 1,54598E-16 | TC | 0,671196751 | 1,489876104 | 0,446366782 |
| WIPF1 | 5,87381E-21 | -0,594116378 | 0,189 | 0,372 | 2,14905E-16 | TC | 0,662450069 | 1,509547734 | 0,508064516 |
| JUN | 1,08295E-20 | -0,539306985 | 0,399 | 0,617 | 3,96219E-16 | TC | 0,688101367 | 1,453274252 | 0,646677472 |
| NBL12 | 1,45444E-20 | -0,544316183 | 0,038 | 0,134 | 5,32135E-16 | TC | 0,685716345 | 1,458328954 | 0,28358209 |
| IGLV3-211 | 1,92287E-20 | 2,252050854 | 0,121 | 0,002 | 7,03519E-16 | TC | 4,763595308 | 0,209925473 | 60,5 |
| EEF1B22 | 2,47017E-20 | 0,637979243 | 0,765 | 0,7 | 9,03761E-16 | TC | 1,556147963 | 0,642612414 | 1,092857143 |
| RPL37A3 | 2,71041E-20 | -0,426721015 | 0,735 | 0,872 | 9,91658E-16 | TC | 0,743950731 | 1,344175036 | 0,842889908 |
| PGK11 | 2,80179E-20 | -0,590726298 | 0,354 | 0,572 | 1,02509E-15 | TC | 0,66400854 | 1,506004727 | 0,618881119 |
| TNFSF14 | 3,2403E-20 | -0,495899257 | 0,022 | 0,101 | 1,18553E-15 | TC | 0,709119534 | 1,410199483 | 0,217821782 |
| DHX361 | 3,43654E-20 | -0,53956097 | 0,137 | 0,303 | 1,25733E-15 | TC | 0,687980238 | 1,453530123 | 0,452145215 |
| CD3E | 3,47607E-20 | -0,475381723 | 0,615 | 0,804 | 1,27179E-15 | TC | 0,719276451 | 1,390286027 | 0,764925373 |
| JMJD6 | 4,22248E-20 | -0,582428809 | 0,118 | 0,27 | 1,54488E-15 | TC | 0,66783851 | 1,49736798 | 0,437037037 |
| AC004687.1 | 5,14486E-20 | -0,411023913 | 0,057 | 0,172 | 1,88235E-15 | TC | 0,75208941 | 1,329629147 | 0,331395349 |
| DDX182 | 5,15574E-20 | -0,6371986 | 0,136 | 0,294 | 1,88633E-15 | TC | 0,642960226 | 1,555306159 | 0,462585034 |
| RPL373 | 7,90466E-20 | -0,302793647 | 0,911 | 0,962 | 2,89208E-15 | TC | 0,810681066 | 1,233530721 | 0,946985447 |
| ATP8A1 | 8,33209E-20 | -0,510891535 | 0,058 | 0,17 | 3,04846E-15 | TC | 0,701788623 | 1,424930481 | 0,341176471 |
| ARL4A1 | 1,1289E-19 | -0,491099794 | 0,069 | 0,19 | 4,13029E-15 | TC | 0,711482514 | 1,405515919 | 0,363157895 |
| MATR3.11 | 1,33586E-19 | -0,429278835 | 0,04 | 0,137 | 4,88753E-15 | TC | 0,742632915 | 1,346560299 | 0,291970803 |
| FAM102B | 1,98822E-19 | -0,409363339 | 0,034 | 0,125 | 7,27432E-15 | TC | 0,752955579 | 1,328099595 | 0,272 |
| RPL151 | 1,99262E-19 | 0,414431707 | 0,879 | 0,875 | 7,29038E-15 | TC | 1,33277358 | 0,750314994 | 1,004571429 |
| SELENOK | 2,27499E-19 | -0,560095511 | 0,331 | 0,541 | 8,3235E-15 | TC | 0,678257259 | 1,474366822 | 0,611829945 |
| IER5L2 | 2,75006E-19 | -0,487347123 | 0,098 | 0,236 | 1,00617E-14 | TC | 0,713335599 | 1,401864707 | 0,415254237 |
| ATP5ME2 | 3,04155E-19 | -0,371325665 | 0,092 | 0,229 | 1,11281E-14 | TC | 0,773071809 | 1,293540895 | 0,401746725 |
| SCML4 | 3,32501E-19 | -0,506692145 | 0,112 | 0,259 | 1,21652E-14 | TC | 0,703834362 | 1,42078883 | 0,432432432 |
| DDX24 | 3,64421E-19 | -0,531089614 | 0,32 | 0,538 | 1,33331E-14 | TC | 0,692031871 | 1,445020153 | 0,594795539 |
| RPL282 | 4,16924E-19 | 0,281260349 | 0,991 | 0,983 | 1,5254E-14 | TC | 1,215256077 | 0,822871836 | 1,008138352 |
| DBI1 | 4,91525E-19 | -0,651725492 | 0,187 | 0,364 | 1,79834E-14 | TC | 0,636518569 | 1,571046075 | 0,513736264 |
| NR4A22 | 5,78828E-19 | -0,66126233 | 0,447 | 0,62 | 2,11776E-14 | TC | 0,632324783 | 1,581465771 | 0,720967742 |
| SEPTIN7 | 6,01299E-19 | -0,48067363 | 0,304 | 0,517 | 2,19997E-14 | TC | 0,716642928 | 1,395395058 | 0,588007737 |
| RPL36A3 | 1,13592E-18 | -0,4071206 | 0,712 | 0,88 | 4,15598E-14 | TC | 0,754126995 | 1,326036604 | 0,809090909 |
| RPS71 | 1,19593E-18 | 0,349315405 | 0,959 | 0,944 | 4,37554E-14 | TC | 1,27395596 | 0,784956491 | 1,015889831 |

| Supplementary Table 4 - clusermarker | | | | | | | | | |
|---|---|---|---|---|---|---|---|---|---|
| | **p_val** | **avg_log2FC** | **pct.1** | **pct.2** | **p_val_adj** | **celltype** | **Foldchange_UP** | **Foldchange_DOWN** | **Ratio_pct1_pct2** |
| NUFIP22 | 1,34867E-18 | -0,483138956 | 0,109 | 0,252 | 4,93437E-14 | TC | 0,71541935 | 1,397781595 | 0,432539683 |
| COPB11 | 1,52552E-18 | -0,530374537 | 0,092 | 0,221 | 5,58142E-14 | TC | 0,692374964 | 1,444304101 | 0,416289593 |
| RPS281 | 1,642E-18 | 0,340153549 | 0,973 | 0,971 | 6,00759E-14 | TC | 1,265891318 | 0,78995723 | 1,002059732 |
| ANXA21 | 2,2747E-18 | -0,441907749 | 0,22 | 0,408 | 8,32243E-14 | TC | 0,736160502 | 1,358399421 | 0,539215686 |
| ATP5MK1 | 3,42391E-18 | -0,497253811 | 0,204 | 0,387 | 1,25271E-13 | TC | 0,70845405 | 1,411524149 | 0,527131783 |
| LPXN | 3,46553E-18 | -0,38985202 | 0,12 | 0,268 | 1,26793E-13 | TC | 0,763207884 | 1,310259001 | 0,447761194 |
| ARF42 | 3,50949E-18 | -0,403784652 | 0,18 | 0,358 | 1,28402E-13 | TC | 0,755872783 | 1,322973948 | 0,502793296 |
| TC2N2 | 5,16447E-18 | -0,403038087 | 0,134 | 0,289 | 1,88952E-13 | TC | 0,756264033 | 1,322289513 | 0,46366782 |
| CWF19L21 | 7,18639E-18 | -0,394341074 | 0,033 | 0,117 | 2,62928E-13 | TC | 0,760836796 | 1,31434232 | 0,282051282 |
| STK17B | 8,56007E-18 | -0,482016973 | 0,367 | 0,58 | 3,13187E-13 | TC | 0,715975947 | 1,396694964 | 0,632758621 |
| NDUFS51 | 9,5406E-18 | -0,496790433 | 0,303 | 0,514 | 3,49062E-13 | TC | 0,708681635 | 1,411070855 | 0,589494163 |
| SKIL1 | 1,5532E-17 | -0,505131073 | 0,117 | 0,256 | 5,6827E-13 | TC | 0,70459636 | 1,419252293 | 0,45703125 |
| PRNP1 | 2,51081E-17 | -0,514561146 | 0,139 | 0,289 | 9,18631E-13 | TC | 0,700005835 | 1,42855952 | 0,480968858 |
| HSPE12 | 3,49516E-17 | -0,901574788 | 0,254 | 0,434 | 1,27877E-12 | TC | 0,535302098 | 1,868104018 | 0,585253456 |
| IGHA13 | 3,8703E-17 | 0,971134717 | 0,142 | 0,024 | 1,41603E-12 | TC | 1,96038188 | 0,510104695 | 5,916666667 |
| SYAP11 | 4,37682E-17 | -0,703848292 | 0,136 | 0,276 | 1,60135E-12 | TC | 0,613932397 | 1,628843835 | 0,492753623 |
| ATP1B11 | 4,84807E-17 | -0,388975424 | 0,042 | 0,133 | 1,77376E-12 | TC | 0,763671758 | 1,309463117 | 0,315789474 |
| CBX41 | 5,53062E-17 | -0,438416664 | 0,047 | 0,142 | 2,02349E-12 | TC | 0,737944047 | 1,355116291 | 0,330985915 |
| LAG3 | 7,53585E-17 | -0,260994525 | 0,079 | 0,196 | 2,75714E-12 | TC | 0,834512448 | 1,198304474 | 0,403061224 |
| PMEPA12 | 8,98399E-17 | -0,381808781 | 0,05 | 0,146 | 3,28697E-12 | TC | 0,767474764 | 1,302974438 | 0,342465753 |
| KLF21 | 9,11849E-17 | 1,383372069 | 0,257 | 0,119 | 3,33618E-12 | TC | 2,608774183 | 0,383321794 | 2,159663866 |
| BZW12 | 9,61944E-17 | -0,442324482 | 0,227 | 0,408 | 3,51946E-12 | TC | 0,735947888 | 1,358791861 | 0,556372549 |
| APOBEC3G | 1,04786E-16 | -0,358436524 | 0,089 | 0,212 | 3,83382E-12 | TC | 0,780009433 | 1,282035778 | 0,419811321 |
| FOSL2 | 1,45192E-16 | -0,533143851 | 0,161 | 0,312 | 5,31213E-12 | TC | 0,691047196 | 1,447079166 | 0,516025641 |
| CDV31 | 1,59221E-16 | -0,410173939 | 0,156 | 0,312 | 5,82543E-12 | TC | 0,752532639 | 1,328846017 | 0,5 |
| RPS231 | 1,68992E-16 | 0,334518735 | 0,968 | 0,97 | 6,18291E-12 | TC | 1,2609567 | 0,793048643 | 0,997938144 |
| RPL35A3 | 1,777E-16 | -0,288269864 | 0,919 | 0,977 | 6,50153E-12 | TC | 0,818883507 | 1,221174919 | 0,940634596 |
| USP36 | 1,78717E-16 | -0,45606649 | 0,059 | 0,16 | 6,53871E-12 | TC | 0,728971092 | 1,37179651 | 0,36875 |
| LINC01138 | 1,79675E-16 | -0,420023164 | 0,038 | 0,122 | 6,57377E-12 | TC | 0,747412624 | 1,337949037 | 0,31147541 |
| ETV3 | 1,94519E-16 | -0,481785775 | 0,063 | 0,167 | 7,11688E-12 | TC | 0,716090695 | 1,396471156 | 0,377245509 |
| EEF22 | 2,2071E-16 | 0,577256413 | 0,707 | 0,669 | 8,0751E-12 | TC | 1,492009181 | 0,670237163 | 1,056801196 |

| Supplementary Table 4 - clusermarker | | | | | | | | | |
|---|---|---|---|---|---|---|---|---|---|
| | **p_val** | **avg_log2FC** | **pct.1** | **pct.2** | **p_val_adj** | **celltype** | **Foldchange_UP** | **Foldchange_DOWN** | **Ratio_pct1_pct2** |
| OAT3 | 2,42465E-16 | -0,421683143 | 0,106 | 0,235 | 8,87105E-12 | TC | 0,746553138 | 1,33948938 | 0,45106383 |
| ITGB7 | 2,72157E-16 | -0,383417372 | 0,078 | 0,193 | 9,9574E-12 | TC | 0,766619514 | 1,304428052 | 0,404145078 |
| COX4I13 | 2,84118E-16 | 0,591304281 | 0,682 | 0,646 | 1,0395E-11 | TC | 1,506608194 | 0,663742573 | 1,055727554 |
| WFDC21 | 3,4766E-16 | -0,698120866 | 0,031 | 0,108 | 1,27198E-11 | TC | 0,616374522 | 1,622390226 | 0,287037037 |
| RAB142 | 3,91502E-16 | -0,407476949 | 0,086 | 0,203 | 1,43239E-11 | TC | 0,753940748 | 1,326364178 | 0,42364532 |
| HNRNPH31 | 4,16703E-16 | -0,442658926 | 0,114 | 0,245 | 1,52459E-11 | TC | 0,735777301 | 1,359106891 | 0,465306122 |
| LY9 | 4,44741E-16 | -0,43148588 | 0,058 | 0,157 | 1,62717E-11 | TC | 0,741497699 | 1,348621853 | 0,369426752 |
| CELF21 | 6,00149E-16 | -0,461214772 | 0,235 | 0,404 | 2,19576E-11 | TC | 0,726374383 | 1,376700533 | 0,581683168 |
| RPL51 | 7,01304E-16 | 0,451100607 | 0,86 | 0,843 | 2,56586E-11 | TC | 1,367082783 | 0,731484598 | 1,020166074 |
| SRSF72 | 7,10798E-16 | -0,443455057 | 0,409 | 0,627 | 2,6006E-11 | TC | 0,735371384 | 1,359857103 | 0,6523126 |
| RPS142 | 7,37008E-16 | 0,273185473 | 0,987 | 0,983 | 2,69649E-11 | TC | 1,208473195 | 0,827490427 | 1,004069176 |
| PERP2 | 8,71744E-16 | -0,488691232 | 0,059 | 0,157 | 3,18945E-11 | TC | 0,712671318 | 1,403171385 | 0,375796178 |
| KIF21A | 9,96277E-16 | -0,309534684 | 0,038 | 0,122 | 3,64508E-11 | TC | 0,80690197 | 1,239307918 | 0,31147541 |
| CD632 | 9,98254E-16 | -0,342379684 | 0,214 | 0,392 | 3,65231E-11 | TC | 0,788739236 | 1,267846146 | 0,545918367 |
| TIPARP2 | 1,14434E-15 | -0,632624008 | 0,142 | 0,276 | 4,18679E-11 | TC | 0,645002202 | 1,550382303 | 0,514492754 |
| GPR174 | 1,28707E-15 | -0,390054409 | 0,052 | 0,146 | 4,70902E-11 | TC | 0,763100825 | 1,310442824 | 0,356164384 |
| ZNF2071 | 1,294E-15 | -0,418554763 | 0,138 | 0,28 | 4,73436E-11 | TC | 0,748173741 | 1,336587941 | 0,492857143 |
| BTG12 | 1,35925E-15 | -0,518102074 | 0,878 | 0,935 | 4,97309E-11 | TC | 0,698289859 | 1,43207006 | 0,939037433 |
| AMD12 | 1,52908E-15 | -0,413934733 | 0,16 | 0,31 | 5,59443E-11 | TC | 0,750573504 | 1,33231455 | 0,516129032 |
| PDE4B | 1,5546E-15 | -0,435219275 | 0,269 | 0,449 | 5,6878E-11 | TC | 0,739581337 | 1,352116326 | 0,599109131 |
| UQCRB | 1,56969E-15 | -0,381038573 | 0,391 | 0,608 | 5,74303E-11 | TC | 0,767884604 | 1,302279007 | 0,643092105 |
| GRAP2 | 1,71884E-15 | -0,365820534 | 0,054 | 0,148 | 6,28871E-11 | TC | 0,776027383 | 1,288614322 | 0,364864865 |
| EIF5B1 | 2,81733E-15 | -0,39099753 | 0,076 | 0,184 | 1,03078E-10 | TC | 0,762602132 | 1,311299769 | 0,413043478 |
| GZMK | 3,31297E-15 | 1,514502356 | 0,168 | 0,05 | 1,21212E-10 | TC | 2,857002614 | 0,350017181 | 3,36 |
| COX7B2 | 3,32885E-15 | -0,403536514 | 0,22 | 0,392 | 1,21793E-10 | TC | 0,756002801 | 1,322746421 | 0,56122449 |
| ZRANB22 | 3,50286E-15 | -0,364751495 | 0,068 | 0,172 | 1,28159E-10 | TC | 0,776602634 | 1,28765981 | 0,395348837 |
| SMDT13 | 3,76865E-15 | 1,00610696 | 0,274 | 0,142 | 1,37884E-10 | TC | 2,008483988 | 0,497887962 | 1,929577465 |
| JUNB2 | 4,95602E-15 | 0,497348864 | 0,884 | 0,866 | 1,81326E-10 | TC | 1,411617151 | 0,708407375 | 1,020785219 |
| RPL13A1 | 5,30421E-15 | 0,428905563 | 0,836 | 0,828 | 1,94065E-10 | TC | 1,346211945 | 0,742825083 | 1,009661836 |
| DDX271 | 5,85405E-15 | -0,359984978 | 0,05 | 0,14 | 2,14182E-10 | TC | 0,779172693 | 1,283412534 | 0,357142857 |
| ELF1 | 7,72695E-15 | -0,562563823 | 0,262 | 0,432 | 2,82706E-10 | TC | 0,677097819 | 1,47689148 | 0,606481481 |

| Supplementary Table 4 - clusermarker | | | | | | | | | |
|---|---|---|---|---|---|---|---|---|---|
| | **p_val** | **avg_log2FC** | **pct.1** | **pct.2** | **p_val_adj** | **celltype** | **Foldchange_UP** | **Foldchange_DOWN** | **Ratio_pct1_pct2** |
| ATP1B31 | 7,79181E-15 | -0,532198802 | 0,203 | 0,355 | 2,85079E-10 | TC | 0,69150002 | 1,446131557 | 0,571830986 |
| RPL312 | 8,37957E-15 | -0,37270437 | 0,591 | 0,774 | 3,06583E-10 | TC | 0,77233338 | 1,294777652 | 0,763565891 |
| DDX21 | 1,15915E-14 | -0,422276633 | 0,132 | 0,265 | 4,24099E-10 | TC | 0,746246087 | 1,340040527 | 0,498113208 |
| AL135905.21 | 1,22913E-14 | -0,381554544 | 0,052 | 0,142 | 4,497E-10 | TC | 0,767610024 | 1,302744843 | 0,366197183 |
| CD53 | 2,09817E-14 | -0,417921979 | 0,296 | 0,488 | 7,67658E-10 | TC | 0,748501971 | 1,336001826 | 0,606557377 |
| MT-ATP63 | 2,22506E-14 | 0,348513986 | 0,928 | 0,895 | 8,14082E-10 | TC | 1,273248472 | 0,785392657 | 1,036871508 |
| GZMA | 2,26238E-14 | -0,4048896 | 0,226 | 0,395 | 8,27738E-10 | TC | 0,755294088 | 1,323987591 | 0,572151899 |
| DOCK10 | 3,46821E-14 | -0,42984626 | 0,138 | 0,274 | 1,26891E-09 | TC | 0,742340888 | 1,347090017 | 0,503649635 |
| SPTY2D1 | 3,59798E-14 | -0,304622193 | 0,065 | 0,163 | 1,31639E-09 | TC | 0,809654217 | 1,235095153 | 0,398773006 |
| TMCO11 | 3,70138E-14 | -0,319616777 | 0,094 | 0,209 | 1,35422E-09 | TC | 0,801282694 | 1,247998999 | 0,449760766 |
| CXCR3 | 4,16313E-14 | -0,277361481 | 0,082 | 0,191 | 1,52316E-09 | TC | 0,825098646 | 1,211976294 | 0,429319372 |
| DBF4 | 4,24098E-14 | -0,34484763 | 0,056 | 0,146 | 1,55165E-09 | TC | 0,787391132 | 1,270016843 | 0,383561644 |
| FAM133B | 4,99768E-14 | -0,307415028 | 0,084 | 0,193 | 1,8285E-09 | TC | 0,808088368 | 1,237488422 | 0,435233161 |
| RNF125 | 5,04848E-14 | -0,438350588 | 0,129 | 0,256 | 1,84709E-09 | TC | 0,737977846 | 1,355054228 | 0,50390625 |
| IDS3 | 5,25402E-14 | -0,409782619 | 0,258 | 0,431 | 1,92229E-09 | TC | 0,752736785 | 1,328485626 | 0,598607889 |
| SRRT | 5,85405E-14 | -0,429300919 | 0,114 | 0,235 | 2,14182E-09 | TC | 0,742621548 | 1,346580911 | 0,485106383 |
| PDCL3 | 7,10957E-14 | -0,559295324 | 0,145 | 0,276 | 2,60118E-09 | TC | 0,678633557 | 1,473549295 | 0,525362319 |
| MPZL3 | 7,18007E-14 | -0,379663153 | 0,038 | 0,113 | 2,62697E-09 | TC | 0,76861703 | 1,301038048 | 0,336283186 |
| NIFK1 | 7,61333E-14 | -0,274206781 | 0,037 | 0,113 | 2,78549E-09 | TC | 0,82690484 | 1,209328996 | 0,327433628 |
| HCG181 | 7,68022E-14 | -0,370979594 | 0,085 | 0,193 | 2,80996E-09 | TC | 0,773257275 | 1,29323064 | 0,440414508 |
| SCAF111 | 7,98965E-14 | -0,348496638 | 0,161 | 0,304 | 2,92317E-09 | TC | 0,785402101 | 1,273233162 | 0,529605263 |
| TNFRSF181 | 9,42243E-14 | 1,039408742 | 0,129 | 0,029 | 3,44738E-09 | TC | 2,055385125 | 0,486526825 | 4,448275862 |
| CD226 | 9,48015E-14 | -0,410368383 | 0,059 | 0,149 | 3,4685E-09 | TC | 0,752431221 | 1,329025128 | 0,395973154 |
| KPNA2 | 9,50421E-14 | -0,418044702 | 0,126 | 0,25 | 3,4773E-09 | TC | 0,748438303 | 1,336115477 | 0,504 |
| CTSW | 1,00752E-13 | -0,310388232 | 0,224 | 0,396 | 3,68621E-09 | TC | 0,80642472 | 1,240041352 | 0,565656566 |
| YPEL51 | 1,04204E-13 | -0,388769298 | 0,293 | 0,468 | 3,81251E-09 | TC | 0,763780876 | 1,309276039 | 0,626068376 |
| MGAT4A | 1,15239E-13 | -0,322788565 | 0,118 | 0,244 | 4,21624E-09 | TC | 0,799522996 | 1,250745763 | 0,483606557 |
| HSPD11 | 1,16539E-13 | -0,819856986 | 0,242 | 0,399 | 4,26383E-09 | TC | 0,566498097 | 1,765230997 | 0,606516291 |
| NDUFA52 | 1,23544E-13 | -0,400556593 | 0,07 | 0,166 | 4,52012E-09 | TC | 0,757565957 | 1,320017076 | 0,421686747 |
| CYTOR | 1,28424E-13 | -0,357261941 | 0,167 | 0,31 | 4,69864E-09 | TC | 0,780644743 | 1,280992422 | 0,538709677 |
| CAP1 | 1,33211E-13 | -0,355576184 | 0,199 | 0,357 | 4,87379E-09 | TC | 0,781557442 | 1,279496485 | 0,557422969 |

| Supplementary Table 4 - clusermarker | | | | | | | | | |
|---|---|---|---|---|---|---|---|---|---|
| | **p_val** | **avg_log2FC** | **pct.1** | **pct.2** | **p_val_adj** | **celltype** | **Foldchange_UP** | **Foldchange_DOWN** | **Ratio_pct1_pct2** |
| YIPF5 | 1,33645E-13 | -0,375723804 | 0,054 | 0,14 | 4,88968E-09 | TC | 0,770718644 | 1,297490346 | 0,385714286 |
| TXN1 | 1,45193E-13 | -0,399277342 | 0,218 | 0,384 | 5,31219E-09 | TC | 0,758237996 | 1,318847123 | 0,567708333 |
| WBP11 | 1,51227E-13 | -0,424615394 | 0,07 | 0,166 | 5,53295E-09 | TC | 0,745037323 | 1,342214636 | 0,421686747 |
| RAB8A1 | 1,55342E-13 | -0,365506775 | 0,059 | 0,149 | 5,68351E-09 | TC | 0,776196173 | 1,288334102 | 0,395973154 |
| MRFAP1L11 | 1,83045E-13 | -0,375433544 | 0,062 | 0,154 | 6,69705E-09 | TC | 0,770873723 | 1,297229326 | 0,402597403 |
| MORF4L11 | 1,88465E-13 | -0,386565556 | 0,247 | 0,419 | 6,89537E-09 | TC | 0,764948456 | 1,307277624 | 0,589498807 |
| TLE41 | 1,94779E-13 | -0,346718766 | 0,078 | 0,179 | 7,12638E-09 | TC | 0,786370569 | 1,271665089 | 0,43575419 |
| PRKACB | 2,0316E-13 | -0,382927367 | 0,093 | 0,202 | 7,433E-09 | TC | 0,766879938 | 1,303985084 | 0,46039604 |
| FABP52 | 2,03745E-13 | -0,483908223 | 0,11 | 0,226 | 7,45441E-09 | TC | 0,715037979 | 1,398527113 | 0,486725664 |
| UBB3 | 2,16099E-13 | -0,440527207 | 0,589 | 0,748 | 7,90643E-09 | TC | 0,736865285 | 1,357100165 | 0,787433155 |
| EIF4G22 | 2,4838E-13 | -0,371650573 | 0,355 | 0,55 | 9,08747E-09 | TC | 0,772897726 | 1,293832245 | 0,645454545 |
| ARHGAP9 | 2,75963E-13 | -0,358583526 | 0,202 | 0,355 | 1,00967E-08 | TC | 0,779929959 | 1,282166416 | 0,569014085 |
| CAPZA11 | 3,10639E-13 | -0,371326335 | 0,157 | 0,292 | 1,13654E-08 | TC | 0,77307145 | 1,293541495 | 0,537671233 |
| DNAJB61 | 3,17198E-13 | -0,270986224 | 0,16 | 0,304 | 1,16053E-08 | TC | 0,828752818 | 1,206632398 | 0,526315789 |
| SHOC22 | 3,98439E-13 | -0,297283723 | 0,065 | 0,158 | 1,45777E-08 | TC | 0,813783129 | 1,22882862 | 0,411392405 |
| HNRNPA13 | 4,21955E-13 | -0,364870656 | 0,515 | 0,699 | 1,54381E-08 | TC | 0,776538492 | 1,287766171 | 0,73676681 |
| KMT2A1 | 4,2679E-13 | -0,40523593 | 0,186 | 0,333 | 1,5615E-08 | TC | 0,755112796 | 1,324305462 | 0,558558559 |
| AKAP131 | 4,63221E-13 | -0,505279881 | 0,311 | 0,483 | 1,69479E-08 | TC | 0,704523688 | 1,419398691 | 0,64389234 |
| VDAC3 | 4,72581E-13 | -0,392424129 | 0,072 | 0,167 | 1,72903E-08 | TC | 0,761848411 | 1,31259708 | 0,431137725 |
| PPP2R5C | 4,86663E-13 | -0,376869419 | 0,227 | 0,386 | 1,78055E-08 | TC | 0,770106875 | 1,298521066 | 0,588082902 |
| PHC3 | 4,92715E-13 | -0,332855503 | 0,071 | 0,167 | 1,8027E-08 | TC | 0,793963448 | 1,259503825 | 0,425149701 |
| TMBIM11 | 5,03035E-13 | -0,378196157 | 0,055 | 0,14 | 1,84045E-08 | TC | 0,769398991 | 1,299715767 | 0,392857143 |
| ETF1 | 5,26403E-13 | -0,320305328 | 0,081 | 0,182 | 1,92595E-08 | TC | 0,80090036 | 1,24859477 | 0,445054945 |
| CCDC591 | 5,30288E-13 | -0,343694742 | 0,079 | 0,179 | 1,94017E-08 | TC | 0,788020605 | 1,269002351 | 0,441340782 |
| SDCBP1 | 5,62146E-13 | -0,429396208 | 0,161 | 0,294 | 2,05672E-08 | TC | 0,742572499 | 1,346669855 | 0,547619048 |
| SELL | 5,91478E-13 | 0,941112006 | 0,167 | 0,059 | 2,16404E-08 | TC | 1,920007579 | 0,520831277 | 2,830508475 |
| TSC22D2 | 6,51922E-13 | -0,350240266 | 0,113 | 0,229 | 2,38519E-08 | TC | 0,784453444 | 1,27477291 | 0,493449782 |
| KMT2E1 | 6,74557E-13 | -0,416775537 | 0,253 | 0,416 | 2,468E-08 | TC | 0,749097007 | 1,334940589 | 0,608173077 |
| SREK1IP1 | 6,87687E-13 | -0,276196082 | 0,033 | 0,102 | 2,51604E-08 | TC | 0,825765425 | 1,210997664 | 0,323529412 |
| TRG-AS1 | 7,14177E-13 | -0,471189396 | 0,05 | 0,13 | 2,61296E-08 | TC | 0,721369636 | 1,38625186 | 0,384615385 |
| KIAA11911 | 7,74247E-13 | -0,356757869 | 0,042 | 0,117 | 2,83274E-08 | TC | 0,780917545 | 1,280544927 | 0,358974359 |

| Supplementary Table 4 - clusermarker | | | | | | | | | |
|---|---|---|---|---|---|---|---|---|---|
| | **p_val** | **avg_log2FC** | **pct.1** | **pct.2** | **p_val_adj** | **celltype** | **Foldchange_UP** | **Foldchange_DOWN** | **Ratio_pct1_pct2** |
| PRRC2C2 | 1,0926E-12 | -0,385564025 | 0,348 | 0,536 | 3,99749E-08 | TC | 0,765479674 | 1,306370416 | 0,649253731 |
| ARF11 | 1,126E-12 | -0,35015431 | 0,269 | 0,438 | 4,11971E-08 | TC | 0,784500184 | 1,274696961 | 0,614155251 |
| CXCR4 | 1,26869E-12 | -0,459558451 | 0,829 | 0,864 | 4,64174E-08 | TC | 0,727208793 | 1,375120886 | 0,959490741 |
| MAZ1 | 1,32902E-12 | -0,311517683 | 0,097 | 0,205 | 4,86248E-08 | TC | 0,805793636 | 1,241012531 | 0,473170732 |
| PRMT9 | 1,33855E-12 | -0,370837221 | 0,063 | 0,151 | 4,89736E-08 | TC | 0,773333588 | 1,293103023 | 0,417218543 |
| MBNL2 | 1,44808E-12 | -0,399891828 | 0,042 | 0,116 | 5,2981E-08 | TC | 0,757915109 | 1,319408979 | 0,362068966 |
| TMX42 | 1,54534E-12 | -0,319497141 | 0,172 | 0,313 | 5,65395E-08 | TC | 0,801349144 | 1,247895512 | 0,549520767 |
| IFITM22 | 1,77382E-12 | -0,32471605 | 0,41 | 0,602 | 6,48988E-08 | TC | 0,798455522 | 1,252417914 | 0,681063123 |
| LITAF | 2,22248E-12 | -0,505704913 | 0,169 | 0,301 | 8,13138E-08 | TC | 0,704316158 | 1,419816922 | 0,561461794 |
| PDE7A2 | 2,41891E-12 | -0,287646588 | 0,153 | 0,288 | 8,85005E-08 | TC | 0,819237359 | 1,220647459 | 0,53125 |
| CENPC2 | 2,47531E-12 | -0,301619042 | 0,067 | 0,157 | 9,05642E-08 | TC | 0,81134137 | 1,232526821 | 0,426751592 |
| PTGER4 | 2,7097E-12 | -0,296794102 | 0,233 | 0,392 | 9,91396E-08 | TC | 0,814059358 | 1,228411652 | 0,594387755 |
| HCST | 2,87387E-12 | -0,329929727 | 0,448 | 0,645 | 1,05146E-07 | TC | 0,795575235 | 1,256952148 | 0,694573643 |
| WDR47 | 3,23268E-12 | -0,348173941 | 0,049 | 0,127 | 1,18274E-07 | TC | 0,785577796 | 1,272948402 | 0,385826772 |
| ETS11 | 3,66039E-12 | 0,697310896 | 0,527 | 0,452 | 1,33923E-07 | TC | 1,621479626 | 0,616720669 | 1,165929204 |
| AP2B11 | 3,71943E-12 | -0,290710856 | 0,062 | 0,148 | 1,36083E-07 | TC | 0,817499155 | 1,223242855 | 0,418918919 |
| IK2 | 3,90368E-12 | -0,304789782 | 0,153 | 0,283 | 1,42824E-07 | TC | 0,809560171 | 1,235238635 | 0,540636042 |
| AKAP93 | 3,97982E-12 | -0,402799831 | 0,151 | 0,277 | 1,4561E-07 | TC | 0,756388938 | 1,32207116 | 0,545126354 |
| ASXL21 | 4,50784E-12 | -0,38842041 | 0,076 | 0,169 | 1,64928E-07 | TC | 0,763965604 | 1,308959454 | 0,449704142 |
| RSRC22 | 4,89287E-12 | -0,31129841 | 0,152 | 0,283 | 1,79015E-07 | TC | 0,805916117 | 1,240823926 | 0,537102473 |
| INTS62 | 4,95907E-12 | -0,328582014 | 0,164 | 0,297 | 1,81437E-07 | TC | 0,79631878 | 1,255778497 | 0,552188552 |
| USP111 | 5,09476E-12 | -0,29742508 | 0,054 | 0,134 | 1,86402E-07 | TC | 0,813703398 | 1,228949028 | 0,402985075 |
| MORF4L2 | 5,7386E-12 | -0,375315429 | 0,065 | 0,152 | 2,09958E-07 | TC | 0,770936837 | 1,297123125 | 0,427631579 |
| STOM2 | 6,50682E-12 | -0,269695494 | 0,103 | 0,212 | 2,38065E-07 | TC | 0,829494607 | 1,205553347 | 0,485849057 |
| MAPKAPK2 | 6,67204E-12 | -0,360770068 | 0,07 | 0,158 | 2,4411E-07 | TC | 0,778748796 | 1,284111135 | 0,443037975 |
| HMGB11 | 6,8142E-12 | -0,363545428 | 0,509 | 0,706 | 2,49311E-07 | TC | 0,777252131 | 1,286583799 | 0,720963173 |
| NDUFA31 | 7,23422E-12 | -0,278259688 | 0,12 | 0,236 | 2,64678E-07 | TC | 0,824585108 | 1,212731093 | 0,508474576 |
| MAGOH1 | 7,43791E-12 | -0,342246672 | 0,134 | 0,253 | 2,72131E-07 | TC | 0,788811958 | 1,26772926 | 0,529644269 |
| CDC42EP31 | 7,71576E-12 | -0,357427379 | 0,116 | 0,227 | 2,82297E-07 | TC | 0,78055523 | 1,281139325 | 0,511013216 |
| ARMCX3 | 8,7628E-12 | -0,356680765 | 0,047 | 0,122 | 3,20605E-07 | TC | 0,780959282 | 1,28047649 | 0,385245902 |
| CHCHD7 | 9,47691E-12 | -0,288047807 | 0,055 | 0,136 | 3,46732E-07 | TC | 0,819009558 | 1,220986972 | 0,404411765 |

| Supplementary Table 4 - clusermarker | | | | | | | | | |
|---|---|---|---|---|---|---|---|---|---|
| | **p_val** | **avg_log2FC** | **pct.1** | **pct.2** | **p_val_adj** | **celltype** | **Foldchange_UP** | **Foldchange_DOWN** | **Ratio_pct1_pct2** |
| PLEKHA2 | 9,58143E-12 | -0,385558126 | 0,085 | 0,181 | 3,50556E-07 | TC | 0,765482804 | 1,306365074 | 0,46961326 |
| UAP11 | 1,00787E-11 | -0,335369836 | 0,066 | 0,152 | 3,68748E-07 | TC | 0,792580932 | 1,261700806 | 0,434210526 |
| FXR12 | 1,03245E-11 | -0,38239809 | 0,071 | 0,16 | 3,77743E-07 | TC | 0,767161332 | 1,303506783 | 0,44375 |
| CCR7 | 1,05272E-11 | 0,860728875 | 0,155 | 0,056 | 3,8516E-07 | TC | 1,815955532 | 0,550674277 | 2,767857143 |
| RGS1 | 1,07293E-11 | -0,58564024 | 0,419 | 0,55 | 3,92554E-07 | TC | 0,666353558 | 1,500704825 | 0,761818182 |
| TPT11 | 1,07511E-11 | 0,288463318 | 0,979 | 0,974 | 3,93352E-07 | TC | 1,221338679 | 0,818773709 | 1,00513347 |
| GNAS2 | 1,15431E-11 | -0,388712746 | 0,342 | 0,517 | 4,22327E-07 | TC | 0,763810816 | 1,309224718 | 0,661508704 |
| KLF13 | 1,20803E-11 | -0,293378934 | 0,138 | 0,261 | 4,41984E-07 | TC | 0,815988693 | 1,225507178 | 0,528735632 |
| SYTL11 | 1,22224E-11 | -0,253520591 | 0,071 | 0,161 | 4,47181E-07 | TC | 0,838846888 | 1,192112666 | 0,440993789 |
| ARHGAP182 | 1,31338E-11 | -0,27114082 | 0,037 | 0,104 | 4,80528E-07 | TC | 0,828664016 | 1,206761704 | 0,355769231 |
| S100A62 | 1,34238E-11 | -0,339727231 | 0,57 | 0,744 | 4,91135E-07 | TC | 0,790190698 | 1,265517301 | 0,766129032 |
| ELL2 | 1,36335E-11 | -0,39905718 | 0,11 | 0,218 | 4,9881E-07 | TC | 0,758353715 | 1,318645877 | 0,504587156 |
| SNRPG | 1,47417E-11 | -0,338137325 | 0,174 | 0,306 | 5,39354E-07 | TC | 0,791061999 | 1,26412342 | 0,568627451 |
| ATP5MF2 | 1,62254E-11 | -0,307189136 | 0,178 | 0,313 | 5,9364E-07 | TC | 0,808214905 | 1,237294676 | 0,568690096 |
| CYTIP | 1,65444E-11 | -0,391712148 | 0,326 | 0,489 | 6,0531E-07 | TC | 0,762224482 | 1,311949464 | 0,666666667 |
| ENSA1 | 1,7456E-11 | -0,339880886 | 0,114 | 0,223 | 6,38661E-07 | TC | 0,790106543 | 1,265652093 | 0,511210762 |
| MIDN1 | 1,7609E-11 | -0,385356875 | 0,18 | 0,31 | 6,44261E-07 | TC | 0,765589594 | 1,306182853 | 0,580645161 |
| CCNY | 1,85033E-11 | -0,267649343 | 0,067 | 0,154 | 6,76979E-07 | TC | 0,8306719 | 1,203844743 | 0,435064935 |
| EVI2A | 1,86126E-11 | -0,401744697 | 0,066 | 0,151 | 6,80979E-07 | TC | 0,756942335 | 1,321104599 | 0,437086093 |
| TUBA1C | 2,03677E-11 | -0,358951086 | 0,105 | 0,208 | 7,45192E-07 | TC | 0,779731279 | 1,282493119 | 0,504807692 |
| TNFRSF4 | 2,0405E-11 | 1,181997737 | 0,112 | 0,027 | 7,46557E-07 | TC | 2,26890741 | 0,44074077 | 4,148148148 |
| GYG1 | 2,13675E-11 | -0,276503595 | 0,056 | 0,136 | 7,81772E-07 | TC | 0,82558943 | 1,211255818 | 0,411764706 |
| CKAP2 | 2,17207E-11 | -0,344415757 | 0,042 | 0,111 | 7,94695E-07 | TC | 0,787626874 | 1,269636718 | 0,378378378 |
| CCDC1861 | 2,50478E-11 | -0,282013161 | 0,051 | 0,127 | 9,16424E-07 | TC | 0,822442566 | 1,215890375 | 0,401574803 |
| FYN | 3,01017E-11 | -0,353045371 | 0,341 | 0,518 | 1,10133E-06 | TC | 0,782929674 | 1,277253927 | 0,658301158 |
| NASP1 | 3,04054E-11 | -0,330474872 | 0,181 | 0,315 | 1,11244E-06 | TC | 0,795274671 | 1,257427196 | 0,574603175 |
| G3BP2 | 3,31967E-11 | -0,362337783 | 0,224 | 0,369 | 1,21457E-06 | TC | 0,777903022 | 1,285507283 | 0,60704607 |
| ITGB12 | 3,41188E-11 | -0,345504374 | 0,153 | 0,276 | 1,2483E-06 | TC | 0,787032777 | 1,270595112 | 0,554347826 |
| RPS91 | 3,65548E-11 | 0,318234588 | 0,906 | 0,889 | 1,33743E-06 | TC | 1,246803913 | 0,802050739 | 1,01912261 |
| RTF1 | 4,16265E-11 | -0,309811745 | 0,065 | 0,148 | 1,52299E-06 | TC | 0,806747023 | 1,239545943 | 0,439189189 |
| SLC7A5 | 4,45094E-11 | -0,32216795 | 0,23 | 0,375 | 1,62846E-06 | TC | 0,799867007 | 1,250207836 | 0,613333333 |

| Supplementary Table 4 - clusermarker | | | | | | | | | |
|---|---|---|---|---|---|---|---|---|---|
| | **p_val** | **avg_log2FC** | **pct.1** | **pct.2** | **p_val_adj** | **celltype** | **Foldchange_UP** | **Foldchange_DOWN** | **Ratio_pct1_pct2** |
| CKLF1 | 4,76787E-11 | -0,335665725 | 0,288 | 0,449 | 1,74442E-06 | TC | 0,792418394 | 1,2619596 | 0,64142539 |
| TBCC | 4,83782E-11 | -0,284170749 | 0,062 | 0,143 | 1,77001E-06 | TC | 0,821213501 | 1,217710131 | 0,433566434 |
| CCNT1 | 4,98896E-11 | -0,272741154 | 0,065 | 0,148 | 1,82531E-06 | TC | 0,827745315 | 1,208101069 | 0,439189189 |
| TMBIM6 | 5,08445E-11 | -0,323720631 | 0,364 | 0,535 | 1,86025E-06 | TC | 0,799006624 | 1,251554079 | 0,680373832 |
| SMAD71 | 6,31193E-11 | -0,271076226 | 0,161 | 0,288 | 2,30934E-06 | TC | 0,828701118 | 1,206707676 | 0,559027778 |
| ATP5F1E1 | 6,46526E-11 | -0,313575898 | 0,588 | 0,753 | 2,36544E-06 | TC | 0,804644874 | 1,24278428 | 0,780876494 |
| SMURF2 | 6,51631E-11 | -0,280004057 | 0,059 | 0,139 | 2,38412E-06 | TC | 0,823588701 | 1,214198299 | 0,424460432 |
| SMIM3 | 7,9861E-11 | -0,511325085 | 0,053 | 0,125 | 2,92187E-06 | TC | 0,701577757 | 1,425358757 | 0,424 |
| GNG2 | 8,09149E-11 | -0,340262232 | 0,165 | 0,288 | 2,96043E-06 | TC | 0,789897723 | 1,265986685 | 0,572916667 |
| TNF | 8,1977E-11 | -0,549315533 | 0,118 | 0,223 | 2,99929E-06 | TC | 0,683344255 | 1,463391245 | 0,529147982 |
| MYLIP2 | 8,49646E-11 | -0,331054434 | 0,13 | 0,242 | 3,1086E-06 | TC | 0,794955256 | 1,257932434 | 0,537190083 |
| P2RY10 | 9,76792E-11 | -0,36604867 | 0,095 | 0,19 | 3,57379E-06 | TC | 0,775904678 | 1,288818109 | 0,5 |
| TAF71 | 9,87188E-11 | -0,28712053 | 0,098 | 0,196 | 3,61182E-06 | TC | 0,819536137 | 1,220202448 | 0,5 |
| CBLB | 1,0601E-10 | -0,311184566 | 0,192 | 0,327 | 3,8786E-06 | TC | 0,805979715 | 1,240726016 | 0,587155963 |
| SNRPB1 | 1,08192E-10 | -0,316128223 | 0,194 | 0,328 | 3,95842E-06 | TC | 0,803222605 | 1,244984881 | 0,591463415 |
| RBPJ1 | 1,11231E-10 | -0,284002666 | 0,134 | 0,247 | 4,06961E-06 | TC | 0,821309183 | 1,217568268 | 0,542510121 |
| MED1 | 1,11899E-10 | -0,285200406 | 0,041 | 0,107 | 4,09405E-06 | TC | 0,820627607 | 1,218579526 | 0,38317757 |
| LGALS11 | 1,14488E-10 | -0,298019835 | 0,278 | 0,426 | 4,18878E-06 | TC | 0,813368015 | 1,229455771 | 0,65258216 |
| EEF1D2 | 1,22027E-10 | 0,415236893 | 0,774 | 0,747 | 4,46461E-06 | TC | 1,333517625 | 0,74989635 | 1,036144578 |
| ACTG11 | 1,29927E-10 | 0,454143343 | 0,814 | 0,794 | 4,75364E-06 | TC | 1,36996909 | 0,729943476 | 1,025188917 |
| C11orf582 | 1,35375E-10 | -0,257095835 | 0,186 | 0,321 | 4,95295E-06 | TC | 0,836770656 | 1,195070588 | 0,579439252 |
| SLC38A22 | 1,3763E-10 | -0,367183952 | 0,349 | 0,52 | 5,03547E-06 | TC | 0,775294345 | 1,289832702 | 0,671153846 |
| MBD21 | 1,41358E-10 | -0,381885742 | 0,063 | 0,142 | 5,17188E-06 | TC | 0,767433825 | 1,303043947 | 0,443661972 |
| PIAS11 | 1,43752E-10 | -0,27947984 | 0,112 | 0,217 | 5,25946E-06 | TC | 0,823888014 | 1,213757188 | 0,516129032 |
| TCP1 | 1,51845E-10 | -0,326546193 | 0,094 | 0,19 | 5,55556E-06 | TC | 0,797443277 | 1,254007688 | 0,494736842 |
| RHOH | 1,54941E-10 | -0,369239559 | 0,23 | 0,366 | 5,66884E-06 | TC | 0,774190463 | 1,291671815 | 0,628415301 |
| TXNIP3 | 1,55321E-10 | 0,381240346 | 0,571 | 0,45 | 5,68275E-06 | TC | 1,302461155 | 0,767777216 | 1,268888889 |
| FBXO341 | 1,56343E-10 | -0,427913604 | 0,052 | 0,123 | 5,72012E-06 | TC | 0,743336006 | 1,345286644 | 0,422764228 |
| THOC21 | 1,58869E-10 | -0,290770022 | 0,085 | 0,175 | 5,81255E-06 | TC | 0,817465629 | 1,223293022 | 0,485714286 |
| PRPS1 | 1,65067E-10 | -0,284266203 | 0,061 | 0,139 | 6,0393E-06 | TC | 0,821159168 | 1,217790702 | 0,438848921 |
| HECA | 1,69066E-10 | -0,334256803 | 0,104 | 0,202 | 6,18563E-06 | TC | 0,79319264 | 1,260727784 | 0,514851485 |

| Supplementary Table 4 - clusermarker | | | | | | | | | |
|---|---|---|---|---|---|---|---|---|---|
| | **p_val** | **avg_log2FC** | **pct.1** | **pct.2** | **p_val_adj** | **celltype** | **Foldchange_UP** | **Foldchange_DOWN** | **Ratio_pct1_pct2** |
| SQSTM12 | 1,88836E-10 | -0,304787321 | 0,247 | 0,398 | 6,90893E-06 | TC | 0,809561552 | 1,235236528 | 0,620603015 |
| RBM221 | 1,91751E-10 | -0,324401518 | 0,067 | 0,148 | 7,01561E-06 | TC | 0,798629618 | 1,252144896 | 0,452702703 |
| MAP1LC3B1 | 1,98181E-10 | -0,29227097 | 0,19 | 0,324 | 7,25085E-06 | TC | 0,816615598 | 1,224566371 | 0,586419753 |
| SRSF51 | 2,18471E-10 | -0,25733962 | 0,214 | 0,357 | 7,99318E-06 | TC | 0,836629272 | 1,195272546 | 0,599439776 |
| MT-ND2 | 2,22264E-10 | -0,313268607 | 0,795 | 0,878 | 8,13196E-06 | TC | 0,804816279 | 1,242519598 | 0,90546697 |
| GLS2 | 2,26525E-10 | -0,261309566 | 0,19 | 0,322 | 8,28788E-06 | TC | 0,834330236 | 1,198566176 | 0,590062112 |
| MAT2A3 | 2,38984E-10 | -0,337735854 | 0,183 | 0,31 | 8,74371E-06 | TC | 0,791282165 | 1,263771691 | 0,590322581 |
| ZNF241 | 2,46465E-10 | -0,273230935 | 0,071 | 0,154 | 9,01743E-06 | TC | 0,827464351 | 1,208511277 | 0,461038961 |
| UQCRC2 | 2,87361E-10 | -0,256766142 | 0,078 | 0,164 | 1,05137E-05 | TC | 0,836961901 | 1,194797515 | 0,475609756 |
| COMMD11 | 3,00697E-10 | -0,506774979 | 0,063 | 0,137 | 1,10016E-05 | TC | 0,703793952 | 1,420870409 | 0,459854015 |
| DNPH13 | 3,02582E-10 | 0,947996624 | 0,132 | 0,048 | 1,10706E-05 | TC | 1,929191855 | 0,518351764 | 2,75 |
| SLC25A63 | 3,04007E-10 | 0,669487235 | 0,444 | 0,361 | 1,11227E-05 | TC | 1,590507567 | 0,628730112 | 1,229916898 |
| NR3C1 | 3,30635E-10 | -0,433168292 | 0,249 | 0,392 | 1,20969E-05 | TC | 0,740633498 | 1,350195477 | 0,635204082 |
| SAFB2 | 3,67537E-10 | -0,276650814 | 0,109 | 0,209 | 1,34471E-05 | TC | 0,825505188 | 1,211379426 | 0,5215311 |
| CDKN1A2 | 3,68443E-10 | -0,38224409 | 0,288 | 0,429 | 1,34802E-05 | TC | 0,767243227 | 1,303367648 | 0,671328671 |
| GPBP1L1 | 3,95784E-10 | -0,318252179 | 0,043 | 0,108 | 1,44806E-05 | TC | 0,80204096 | 1,246819115 | 0,398148148 |
| JOSD1 | 4,19164E-10 | -0,327610084 | 0,091 | 0,181 | 1,5336E-05 | TC | 0,796855432 | 1,254932776 | 0,502762431 |
| AC007384.1 | 4,28051E-10 | -0,293450982 | 0,066 | 0,145 | 1,56611E-05 | TC | 0,815947943 | 1,225568381 | 0,455172414 |
| LRRFIP11 | 4,49623E-10 | -0,347907957 | 0,273 | 0,42 | 1,64504E-05 | TC | 0,785722644 | 1,272713734 | 0,65 |
| PABPC13 | 4,65114E-10 | -0,341066642 | 0,58 | 0,718 | 1,70171E-05 | TC | 0,789457419 | 1,266692764 | 0,807799443 |
| RNF168 | 4,79255E-10 | -0,258866135 | 0,059 | 0,134 | 1,75345E-05 | TC | 0,835744503 | 1,196537933 | 0,440298507 |
| EZR1 | 4,94649E-10 | -0,350929409 | 0,506 | 0,667 | 1,80977E-05 | TC | 0,784078818 | 1,275381985 | 0,75862069 |
| CD3D | 5,04088E-10 | -0,331034364 | 0,533 | 0,712 | 1,84431E-05 | TC | 0,794966315 | 1,257914934 | 0,748595506 |
| FYB1 | 5,43053E-10 | -0,274381054 | 0,454 | 0,637 | 1,98687E-05 | TC | 0,826804958 | 1,209475088 | 0,712715856 |
| RBL21 | 5,81513E-10 | -0,250478219 | 0,152 | 0,27 | 2,12758E-05 | TC | 0,840617724 | 1,189601374 | 0,562962963 |
| POLR2L1 | 5,93184E-10 | -0,303328198 | 0,237 | 0,38 | 2,17028E-05 | TC | 0,810380746 | 1,233987857 | 0,623684211 |
| ALG13 | 6,52311E-10 | -0,28013069 | 0,083 | 0,17 | 2,38661E-05 | TC | 0,823516414 | 1,21430488 | 0,488235294 |
| DAZAP22 | 7,05876E-10 | -0,310132194 | 0,259 | 0,408 | 2,58259E-05 | TC | 0,80656785 | 1,2398213 | 0,634803922 |
| GPR171 | 7,06998E-10 | -0,374936895 | 0,047 | 0,113 | 2,58669E-05 | TC | 0,771139142 | 1,296782831 | 0,415929204 |
| PTBP31 | 7,52473E-10 | -0,316726898 | 0,118 | 0,22 | 2,75307E-05 | TC | 0,802889361 | 1,24550162 | 0,536363636 |
| IER52 | 7,53532E-10 | -0,341150433 | 0,116 | 0,215 | 2,75695E-05 | TC | 0,789411569 | 1,266766335 | 0,539534884 |

| Supplementary Table 4 - clusermarker | | | | | | | | | |
|---|---|---|---|---|---|---|---|---|---|
| | **p_val** | **avg_log2FC** | **pct.1** | **pct.2** | **p_val_adj** | **celltype** | **Foldchange_UP** | **Foldchange_DOWN** | **Ratio_pct1_pct2** |
| HNRNPAB1 | 8,30027E-10 | -0,313857583 | 0,199 | 0,33 | 3,03682E-05 | TC | 0,804487783 | 1,243026957 | 0,603030303 |
| OXNAD1 | 9,62165E-10 | -0,271387216 | 0,086 | 0,173 | 3,52027E-05 | TC | 0,828522501 | 1,206967824 | 0,497109827 |
| WAKMAR2 | 1,07835E-09 | -0,250476649 | 0,047 | 0,113 | 3,94536E-05 | TC | 0,840618639 | 1,189600079 | 0,415929204 |
| UPF2 | 1,08208E-09 | -0,341570016 | 0,058 | 0,13 | 3,95901E-05 | TC | 0,789182015 | 1,267134806 | 0,446153846 |
| LMNA2 | 1,10123E-09 | -0,433372442 | 0,269 | 0,408 | 4,02908E-05 | TC | 0,740528702 | 1,350386552 | 0,659313725 |
| DPYD | 1,16621E-09 | -0,271907949 | 0,05 | 0,117 | 4,2668E-05 | TC | 0,828223504 | 1,207403551 | 0,427350427 |
| CTSS2 | 1,20558E-09 | -0,325892611 | 0,076 | 0,157 | 4,41085E-05 | TC | 0,797804623 | 1,253439716 | 0,484076433 |
| SAMD3 | 1,22504E-09 | -0,291459653 | 0,043 | 0,107 | 4,48205E-05 | TC | 0,817074961 | 1,223877916 | 0,401869159 |
| OXCT1 | 1,27916E-09 | -0,268707513 | 0,046 | 0,11 | 4,68007E-05 | TC | 0,830062853 | 1,204728048 | 0,418181818 |
| CDC5L | 1,42517E-09 | -0,27564303 | 0,057 | 0,128 | 5,21429E-05 | TC | 0,826082039 | 1,210533521 | 0,4453125 |
| MBNL12 | 1,45906E-09 | -0,299975208 | 0,367 | 0,535 | 5,33826E-05 | TC | 0,812266355 | 1,231123257 | 0,685981308 |
| MRPL471 | 1,63557E-09 | -0,289903675 | 0,047 | 0,111 | 5,98405E-05 | TC | 0,817956669 | 1,222558648 | 0,423423423 |
| ABLIM11 | 1,87764E-09 | -0,458421388 | 0,096 | 0,182 | 6,86972E-05 | TC | 0,72778217 | 1,374037509 | 0,527472527 |
| TNFRSF1B | 2,02081E-09 | -0,371186553 | 0,182 | 0,3 | 7,39353E-05 | TC | 0,773146357 | 1,293416171 | 0,606666667 |
| SRGN | 2,15726E-09 | -0,318864603 | 0,831 | 0,899 | 7,89275E-05 | TC | 0,801700565 | 1,247348503 | 0,9243604 |
| TOB12 | 2,22557E-09 | -0,28788295 | 0,18 | 0,3 | 8,14269E-05 | TC | 0,819103151 | 1,220847458 | 0,6 |
| RPL27A2 | 2,2362E-09 | -0,308628504 | 0,477 | 0,631 | 8,18158E-05 | TC | 0,807408957 | 1,238529734 | 0,755942948 |
| ITGA4 | 2,33703E-09 | -0,252318698 | 0,228 | 0,367 | 8,55051E-05 | TC | 0,839546013 | 1,191119944 | 0,621253406 |
| RORA | 2,37757E-09 | -0,278635873 | 0,231 | 0,367 | 8,69882E-05 | TC | 0,824370124 | 1,213047357 | 0,629427793 |
| NR4A12 | 2,40501E-09 | -0,413489099 | 0,218 | 0,339 | 8,7992E-05 | TC | 0,750805384 | 1,331903075 | 0,643067847 |
| GPBP12 | 2,74572E-09 | -0,263935187 | 0,159 | 0,27 | 0,000100458 | TC | 0,832813184 | 1,200749483 | 0,588888889 |
| PRPF38B1 | 2,79944E-09 | -0,265020913 | 0,177 | 0,298 | 0,000102423 | TC | 0,832186672 | 1,201653468 | 0,593959732 |
| DNAJB9 | 2,89857E-09 | -0,292171555 | 0,094 | 0,181 | 0,00010605 | TC | 0,816671873 | 1,22448199 | 0,519337017 |
| CD27 | 2,93998E-09 | 0,816718614 | 0,201 | 0,11 | 0,000107565 | TC | 1,761395169 | 0,567731772 | 1,827272727 |
| HLA-DRB52 | 3,0371E-09 | -0,313630105 | 0,041 | 0,101 | 0,000111118 | TC | 0,804614641 | 1,242830977 | 0,405940594 |
| CDK161 | 3,4085E-09 | -0,321242362 | 0,051 | 0,116 | 0,000124707 | TC | 0,800380341 | 1,249405999 | 0,439655172 |
| PTTG1 | 3,64637E-09 | -0,293573852 | 0,09 | 0,175 | 0,00013341 | TC | 0,815878455 | 1,225672763 | 0,514285714 |
| METAP22 | 3,89779E-09 | -0,308694108 | 0,053 | 0,119 | 0,000142608 | TC | 0,807372242 | 1,238586055 | 0,445378151 |
| LEF11 | 4,06899E-09 | 0,723974652 | 0,1 | 0,03 | 0,000148872 | TC | 1,651726309 | 0,605427179 | 3,333333333 |
| BPTF1 | 4,60769E-09 | -0,269144487 | 0,154 | 0,264 | 0,000168581 | TC | 0,829811475 | 1,205093 | 0,583333333 |
| CNOT21 | 4,77064E-09 | -0,251485822 | 0,088 | 0,172 | 0,000174543 | TC | 0,840030828 | 1,190432502 | 0,511627907 |

| Supplementary Table 4 - clusermarker | | | | | | | | | |
|---|---|---|---|---|---|---|---|---|---|
| | **p_val** | **avg_log2FC** | **pct.1** | **pct.2** | **p_val_adj** | **celltype** | **Foldchange_UP** | **Foldchange_DOWN** | **Ratio_pct1_pct2** |
| BAG31 | 4,77246E-09 | -0,34544784 | 0,052 | 0,119 | 0,00017461 | TC | 0,787063619 | 1,270545323 | 0,43697479 |
| RBM27 | 5,01723E-09 | -0,256406131 | 0,076 | 0,155 | 0,000183565 | TC | 0,837170784 | 1,194499402 | 0,490322581 |
| PRPF4B2 | 5,14736E-09 | -0,26431317 | 0,096 | 0,182 | 0,000188327 | TC | 0,832595017 | 1,201064118 | 0,527472527 |
| C12orf751 | 5,63026E-09 | -0,333337846 | 0,088 | 0,17 | 0,000205994 | TC | 0,793698044 | 1,259924991 | 0,517647059 |
| DOK2 | 6,00384E-09 | -0,264673941 | 0,11 | 0,203 | 0,000219662 | TC | 0,832386839 | 1,201364502 | 0,541871921 |
| RSL24D1 | 6,59295E-09 | -0,290517852 | 0,156 | 0,264 | 0,000241216 | TC | 0,817608527 | 1,22307922 | 0,590909091 |
| MAPRE11 | 6,64826E-09 | -0,260223081 | 0,07 | 0,145 | 0,00024324 | TC | 0,834958801 | 1,197663883 | 0,482758621 |
| RMND5A | 6,74613E-09 | -0,266831013 | 0,049 | 0,113 | 0,000246821 | TC | 0,83114321 | 1,203162088 | 0,433628319 |
| CCL4L2 | 6,77012E-09 | -0,685681807 | 0,077 | 0,154 | 0,000247698 | TC | 0,621711941 | 1,608461949 | 0,5 |
| GOLGB12 | 7,75054E-09 | -0,359091264 | 0,125 | 0,22 | 0,000283569 | TC | 0,779655521 | 1,282617738 | 0,568181818 |
| GOLGA4 | 7,85111E-09 | -0,318637076 | 0,113 | 0,205 | 0,000287249 | TC | 0,801827011 | 1,2471518 | 0,551219512 |
| PDE4D1 | 8,26562E-09 | -0,338093836 | 0,156 | 0,259 | 0,000302414 | TC | 0,791085846 | 1,264085315 | 0,602316602 |
| ARHGEF7 | 8,53224E-09 | -0,317599337 | 0,063 | 0,133 | 0,000312169 | TC | 0,802403978 | 1,246255039 | 0,473684211 |
| SLC2A3 | 8,97667E-09 | -0,303670964 | 0,365 | 0,521 | 0,000328429 | TC | 0,810188232 | 1,234281072 | 0,700575816 |
| COX7C1 | 9,3301E-09 | -0,296989889 | 0,478 | 0,643 | 0,00034136 | TC | 0,81394889 | 1,22857837 | 0,743390358 |
| TGFB1 | 9,44255E-09 | -0,286109578 | 0,369 | 0,52 | 0,000345475 | TC | 0,820110618 | 1,219347705 | 0,709615385 |
| GSPT1 | 1,03396E-08 | -0,324092273 | 0,185 | 0,3 | 0,000378296 | TC | 0,798800825 | 1,251876524 | 0,616666667 |
| PDCD4 | 1,05637E-08 | -0,351247528 | 0,275 | 0,408 | 0,000386494 | TC | 0,783905945 | 1,275663243 | 0,674019608 |
| UBE2Z | 1,13874E-08 | -0,294870564 | 0,049 | 0,111 | 0,000416631 | TC | 0,815145463 | 1,226774908 | 0,441441441 |
| RPL36AL1 | 1,24523E-08 | -0,268304753 | 0,536 | 0,706 | 0,000455591 | TC | 0,830294615 | 1,204391769 | 0,759206799 |
| BHLHE40 | 1,28989E-08 | -0,250007785 | 0,143 | 0,247 | 0,000471931 | TC | 0,840891878 | 1,189213532 | 0,578947368 |
| CHORDC1 | 1,36697E-08 | -0,329401697 | 0,078 | 0,154 | 0,000500133 | TC | 0,795866471 | 1,256492184 | 0,506493506 |
| SCP21 | 1,39165E-08 | -0,309987588 | 0,118 | 0,211 | 0,000509164 | TC | 0,806648699 | 1,239697035 | 0,559241706 |
| BTN3A2 | 1,49951E-08 | -0,350033921 | 0,063 | 0,131 | 0,000548627 | TC | 0,784565651 | 1,274590596 | 0,480916031 |
| PSMA41 | 1,65941E-08 | -0,285042918 | 0,074 | 0,148 | 0,000607127 | TC | 0,820717193 | 1,21844651 | 0,5 |
| PRKAR1A2 | 1,86199E-08 | -0,297426394 | 0,077 | 0,152 | 0,000681247 | TC | 0,813702657 | 1,228950147 | 0,506578947 |
| CRTAM | 1,88591E-08 | -0,404658234 | 0,065 | 0,134 | 0,000689996 | TC | 0,755415225 | 1,323775278 | 0,485074627 |
| RBM391 | 1,98503E-08 | -0,264467739 | 0,368 | 0,527 | 0,000726262 | TC | 0,832505819 | 1,201192805 | 0,69829222 |
| ARNTL | 2,0379E-08 | -0,256988067 | 0,049 | 0,11 | 0,000745605 | TC | 0,836833164 | 1,194981321 | 0,445454545 |
| DHRS71 | 2,04486E-08 | -0,257602544 | 0,179 | 0,291 | 0,000748153 | TC | 0,836476814 | 1,195490399 | 0,615120275 |
| LINC00861 | 2,36996E-08 | -0,273492024 | 0,049 | 0,11 | 0,000867097 | TC | 0,827314616 | 1,208730005 | 0,445454545 |

| Supplementary Table 4 - clusermarker | | | | | | | | | |
|---|---|---|---|---|---|---|---|---|---|
| | **p_val** | **avg_log2FC** | **pct.1** | **pct.2** | **p_val_adj** | **celltype** | **Foldchange_UP** | **Foldchange_DOWN** | **Ratio_pct1_pct2** |
| DNAJA12 | 2,52345E-08 | -0,421353668 | 0,489 | 0,62 | 0,000923256 | TC | 0,746723651 | 1,339183509 | 0,788709677 |
| TENT5C1 | 2,80115E-08 | -0,313257717 | 0,222 | 0,337 | 0,001024856 | TC | 0,804822355 | 1,242510219 | 0,658753709 |
| CCNH | 2,83503E-08 | -0,353458148 | 0,249 | 0,373 | 0,001037253 | TC | 0,782705698 | 1,277619421 | 0,667560322 |
| DNAJC82 | 2,94224E-08 | -0,252280345 | 0,117 | 0,208 | 0,001076478 | TC | 0,839568332 | 1,19108828 | 0,5625 |
| ATP5F1D3 | 3,29599E-08 | 0,778807115 | 0,331 | 0,261 | 0,001205903 | TC | 1,715711658 | 0,58284852 | 1,268199234 |
| DUSP5 | 3,37406E-08 | -0,26682816 | 0,121 | 0,212 | 0,001234468 | TC | 0,831144854 | 1,203159709 | 0,570754717 |
| TPM42 | 3,47205E-08 | -0,265171835 | 0,208 | 0,324 | 0,00127032 | TC | 0,83209962 | 1,201779181 | 0,641975309 |
| APOBEC3C | 3,65142E-08 | -0,354234878 | 0,057 | 0,12 | 0,001335947 | TC | 0,782284411 | 1,278307461 | 0,475 |
| IGLV2-141 | 3,98204E-08 | 2,090819169 | 0,133 | 0,057 | 0,001456909 | TC | 4,259898834 | 0,234747359 | 2,333333333 |
| MSN1 | 4,04808E-08 | -0,265236203 | 0,349 | 0,492 | 0,001481072 | TC | 0,832062495 | 1,201832802 | 0,709349593 |
| PAG1 | 4,67645E-08 | -0,257143071 | 0,113 | 0,199 | 0,001710972 | TC | 0,836743259 | 1,195109717 | 0,567839196 |
| REL | 4,82906E-08 | -0,265179876 | 0,218 | 0,336 | 0,001766808 | TC | 0,832094982 | 1,201785879 | 0,648809524 |
| ICOS | 5,64638E-08 | -0,305590135 | 0,123 | 0,211 | 0,00206584 | TC | 0,809111182 | 1,235924089 | 0,582938389 |
| NCL3 | 6,32548E-08 | -0,277750996 | 0,47 | 0,636 | 0,002314303 | TC | 0,824875907 | 1,212303562 | 0,738993711 |
| JCHAIN1 | 7,60262E-08 | 0,526932119 | 0,106 | 0,039 | 0,00278157 | TC | 1,440861952 | 0,694029014 | 2,717948718 |
| NR1D21 | 8,08837E-08 | -0,271466607 | 0,083 | 0,157 | 0,002959292 | TC | 0,828476909 | 1,207034245 | 0,52866242 |
| BCAS21 | 8,12681E-08 | -0,304114414 | 0,086 | 0,161 | 0,002973358 | TC | 0,809939238 | 1,234660519 | 0,534161491 |
| DHX9 | 9,8498E-08 | -0,29115476 | 0,108 | 0,19 | 0,003603745 | TC | 0,817247657 | 1,223619293 | 0,568421053 |
| STK17A | 1,22412E-07 | -0,30005822 | 0,246 | 0,366 | 0,00447869 | TC | 0,812219619 | 1,231194097 | 0,672131148 |
| SBNO11 | 1,29303E-07 | -0,252200584 | 0,061 | 0,123 | 0,004730812 | TC | 0,839614749 | 1,191022431 | 0,495934959 |
| SAMD91 | 1,4174E-07 | -0,293529622 | 0,05 | 0,107 | 0,005185837 | TC | 0,815903468 | 1,225635188 | 0,46728972 |
| INSIG13 | 1,43634E-07 | -0,265000756 | 0,151 | 0,245 | 0,005255125 | TC | 0,832198299 | 1,201636679 | 0,616326531 |
| FXYD52 | 1,78826E-07 | 0,492659357 | 0,591 | 0,581 | 0,006542708 | TC | 1,407036113 | 0,710713812 | 1,017211704 |
| IL2RG | 1,79618E-07 | -0,269354412 | 0,449 | 0,607 | 0,006571691 | TC | 0,829690739 | 1,205268365 | 0,73970346 |
| SEPTIN11 | 1,95741E-07 | -0,281263742 | 0,051 | 0,108 | 0,007161592 | TC | 0,822869901 | 1,215258936 | 0,472222222 |
| RABGAP1L2 | 2,21283E-07 | -0,268286915 | 0,131 | 0,218 | 0,008096087 | TC | 0,830304881 | 1,204376877 | 0,600917431 |
| MYBL1 | 2,28286E-07 | -0,255784088 | 0,061 | 0,122 | 0,008352296 | TC | 0,837531822 | 1,193984483 | 0,5 |
| MT-ND41 | 2,40559E-07 | 0,26653476 | 0,837 | 0,816 | 0,008801343 | TC | 1,202915047 | 0,831313901 | 1,025735294 |
| HELZ1 | 2,59146E-07 | -0,28541544 | 0,079 | 0,148 | 0,00948136 | TC | 0,820505301 | 1,218761169 | 0,533783784 |
| RASGEF1B | 3,09506E-07 | -0,250638128 | 0,056 | 0,114 | 0,011323882 | TC | 0,840524555 | 1,189733237 | 0,49122807 |
| BTF32 | 3,237E-07 | 0,389622091 | 0,633 | 0,633 | 0,011843196 | TC | 1,310050196 | 0,76332953 | 1 |

| Supplementary Table 4 - clusermarker | | | | | | | | | |
|---|---|---|---|---|---|---|---|---|---|
| | **p_val** | **avg_log2FC** | **pct.1** | **pct.2** | **p_val_adj** | **celltype** | **Foldchange_UP** | **Foldchange_DOWN** | **Ratio_pct1_pct2** |
| ATXN7 | 3,40699E-07 | -0,296243609 | 0,088 | 0,158 | 0,012465167 | TC | 0,814370039 | 1,227943013 | 0,556962025 |
| EIF12 | 3,70556E-07 | 0,257746338 | 0,902 | 0,872 | 0,013557518 | TC | 1,195609561 | 0,836393446 | 1,03440367 |
| HINT13 | 4,21195E-07 | 0,562176571 | 0,511 | 0,48 | 0,015410266 | TC | 1,476495103 | 0,677279592 | 1,064583333 |
| CTLA4 | 4,27707E-07 | 0,916560514 | 0,175 | 0,099 | 0,015648511 | TC | 1,887609731 | 0,529770526 | 1,767676768 |
| TMEM50B2 | 4,62091E-07 | -0,257579685 | 0,068 | 0,131 | 0,016906525 | TC | 0,836490068 | 1,195471457 | 0,519083969 |
| POLR1E | 5,36815E-07 | -0,286194926 | 0,054 | 0,11 | 0,019640455 | TC | 0,820062103 | 1,219419842 | 0,490909091 |
| EIF1AX2 | 5,52629E-07 | -0,288550819 | 0,157 | 0,248 | 0,020219036 | TC | 0,81872405 | 1,221412757 | 0,633064516 |
| SH2D2A | 6,40974E-07 | -0,29611138 | 0,091 | 0,161 | 0,023451313 | TC | 0,814444683 | 1,227830472 | 0,565217391 |
| BATF | 6,71001E-07 | 0,795766452 | 0,193 | 0,117 | 0,024549926 | TC | 1,735999404 | 0,576037064 | 1,64957265 |
| SMU1 | 8,16573E-07 | -0,251035713 | 0,069 | 0,131 | 0,029875939 | TC | 0,840292951 | 1,190061155 | 0,526717557 |
| SLC20A1 | 8,51584E-07 | -0,25961977 | 0,086 | 0,154 | 0,031156914 | TC | 0,83530804 | 1,197163145 | 0,558441558 |
| RPL10A1 | 9,33834E-07 | 0,263637436 | 0,841 | 0,852 | 0,034166193 | TC | 1,200501691 | 0,832985082 | 0,987089202 |
| SFPQ2 | 9,84874E-07 | -0,25213866 | 0,411 | 0,548 | 0,036033584 | TC | 0,839650789 | 1,19097131 | 0,75 |
| CACYBP1 | 1,03877E-06 | -0,279865988 | 0,118 | 0,197 | 0,038005339 | TC | 0,823667524 | 1,214082103 | 0,598984772 |
| CD741 | 1,14948E-06 | 0,505145426 | 0,58 | 0,547 | 0,042055886 | TC | 1,419266414 | 0,70458935 | 1,060329068 |
| EEF1G1 | 1,17176E-06 | 0,375995035 | 0,613 | 0,599 | 0,042871269 | TC | 1,297734301 | 0,77057376 | 1,023372287 |
| RGCC1 | 1,25476E-06 | -0,393006854 | 0,504 | 0,596 | 0,045907924 | TC | 0,761540752 | 1,313127364 | 0,845637584 |
| SOCS1 | 1,3132E-06 | 0,790739175 | 0,311 | 0,248 | 0,048045952 | TC | 1,729960594 | 0,578047849 | 1,254032258 |
| C12orf572 | 2,27735E-06 | 0,546139129 | 0,505 | 0,485 | 0,083321361 | TC | 1,460172819 | 0,684850442 | 1,041237113 |
| COX8A3 | 2,32139E-06 | 0,679433882 | 0,353 | 0,303 | 0,084932634 | TC | 1,601511194 | 0,624410247 | 1,165016502 |
| PPIA3 | 2,97202E-06 | 0,343367562 | 0,731 | 0,748 | 0,108737477 | TC | 1,268714595 | 0,788199335 | 0,977272727 |
| PPIP5K2 | 3,07409E-06 | -0,270517341 | 0,053 | 0,104 | 0,112471796 | TC | 0,82902221 | 1,2062403 | 0,509615385 |
| SOCS3 | 3,12556E-06 | 0,739169982 | 0,152 | 0,089 | 0,114354945 | TC | 1,669215221 | 0,599083921 | 1,707865169 |
| PDCD102 | 3,25234E-06 | -0,259201829 | 0,084 | 0,148 | 0,118993277 | TC | 0,83555006 | 1,196816383 | 0,567567568 |
| MCL11 | 3,52494E-06 | -0,256958537 | 0,391 | 0,524 | 0,128966872 | TC | 0,836850293 | 1,194956862 | 0,746183206 |
| NUCB22 | 5,38548E-06 | -0,255797094 | 0,089 | 0,152 | 0,19703863 | TC | 0,837524272 | 1,193995247 | 0,585526316 |
| SC5D | 6,08927E-06 | -0,265177486 | 0,081 | 0,142 | 0,22278795 | TC | 0,832096361 | 1,201783888 | 0,570422535 |
| IFITM33 | 8,22285E-06 | 0,447914507 | 0,169 | 0,098 | 0,30084949 | TC | 1,364066999 | 0,73310182 | 1,724489796 |
| ST3GAL1 | 9,80901E-06 | -0,277304529 | 0,096 | 0,16 | 0,358882095 | TC | 0,825131219 | 1,211928451 | 0,6 |
| PGLS3 | 1,34458E-05 | 0,622637291 | 0,134 | 0,077 | 0,49194292 | TC | 1,539687206 | 0,649482568 | 1,74025974 |
| PNRC13 | 1,80523E-05 | 0,318732074 | 0,748 | 0,756 | 0,660479138 | TC | 1,247233924 | 0,801774215 | 0,989417989 |

| Supplementary Table 4 - clusermarker | | | | | | | | | |
|---|---|---|---|---|---|---|---|---|---|
| | **p_val** | **avg_log2FC** | **pct.1** | **pct.2** | **p_val_adj** | **celltype** | **Foldchange_UP** | **Foldchange_DOWN** | **Ratio_pct1_pct2** |
| COTL11 | 2,54001E-05 | 0,61407508 | 0,428 | 0,398 | 0,929313969 | TC | 1,530576421 | 0,653348625 | 1,075376884 |
| GABARAP3 | 2,72341E-05 | 0,580972231 | 0,376 | 0,339 | 0,99641492 | TC | 1,495856966 | 0,668513115 | 1,109144543 |
| ITGB21 | 3,54786E-05 | 0,590404733 | 0,298 | 0,244 | 1 | TC | 1,505669088 | 0,664156559 | 1,221311475 |
| SAMHD1 | 4,42641E-05 | 0,63072871 | 0,183 | 0,127 | 1 | TC | 1,548346871 | 0,645850112 | 1,440944882 |
| KLRB1 | 5,58743E-05 | -0,392952926 | 0,247 | 0,318 | 1 | TC | 0,761569219 | 1,31307828 | 0,77672956 |
| EIF3F2 | 7,36013E-05 | 0,56519607 | 0,388 | 0,361 | 1 | TC | 1,47958858 | 0,675863557 | 1,074792244 |
| CCR6 | 7,3966E-05 | 0,314605591 | 0,102 | 0,051 | 1 | TC | 1,243671608 | 0,80407078 | 2 |
| NME23 | 0,000105561 | 0,584336216 | 0,324 | 0,283 | 1 | TC | 1,49934898 | 0,666956135 | 1,144876325 |
| MCUB1 | 0,000125719 | 0,561790529 | 0,134 | 0,084 | 1 | TC | 1,476100068 | 0,677460845 | 1,595238095 |
| EIF3K1 | 0,000138885 | 0,432287769 | 0,487 | 0,48 | 1 | TC | 1,349371662 | 0,741085668 | 1,014583333 |
| DUSP4 | 0,0001406 | -0,260985699 | 0,213 | 0,288 | 1 | TC | 0,834517553 | 1,198297143 | 0,739583333 |
| MED4 | 0,000169685 | -0,261897121 | 0,067 | 0,111 | 1 | TC | 0,833990514 | 1,199054406 | 0,603603604 |
| DDX54 | 0,000250565 | 0,258839189 | 0,737 | 0,726 | 1 | TC | 1,196515585 | 0,835760112 | 1,015151515 |
| RPL43 | 0,000292072 | 0,327503205 | 0,661 | 0,676 | 1 | TC | 1,25483981 | 0,796914468 | 0,977810651 |
| CORO1B2 | 0,0002944 | 0,707948321 | 0,18 | 0,134 | 1 | TC | 1,633479468 | 0,612190125 | 1,343283582 |
| MIF3 | 0,000349823 | 0,407037766 | 0,549 | 0,548 | 1 | TC | 1,32596047 | 0,754170296 | 1,001824818 |
| TIGIT | 0,000356201 | 0,771523442 | 0,164 | 0,119 | 1 | TC | 1,707071447 | 0,585798563 | 1,378151261 |
| CYBA3 | 0,000356292 | 0,325385495 | 0,663 | 0,688 | 1 | TC | 1,252999201 | 0,798085106 | 0,963662791 |
| GLUL1 | 0,000368896 | -0,285569469 | 0,115 | 0,169 | 1 | TC | 0,820417705 | 1,218891297 | 0,680473373 |
| GYPC | 0,000383223 | 0,586856419 | 0,268 | 0,226 | 1 | TC | 1,501970439 | 0,665792065 | 1,185840708 |
| ATP5MC21 | 0,000454727 | 0,348640587 | 0,564 | 0,574 | 1 | TC | 1,273360208 | 0,785323739 | 0,982578397 |
| CHCHD23 | 0,000486056 | 0,409340927 | 0,478 | 0,485 | 1 | TC | 1,328078963 | 0,752967277 | 0,98556701 |
| LIMD2 | 0,000719638 | 0,602641521 | 0,258 | 0,22 | 1 | TC | 1,518494329 | 0,658547076 | 1,172727273 |
| ITM2A1 | 0,001115036 | 0,553316938 | 0,3 | 0,265 | 1 | TC | 1,467455686 | 0,681451583 | 1,132075472 |
| SSR41 | 0,001132288 | 0,479761229 | 0,448 | 0,455 | 1 | TC | 1,39451285 | 0,717096296 | 0,984615385 |
| FOSB3 | 0,001476306 | -0,297615327 | 0,469 | 0,53 | 1 | TC | 0,813596103 | 1,229111099 | 0,88490566 |
| SMAP2 | 0,001484497 | 0,379003088 | 0,355 | 0,319 | 1 | TC | 1,30044293 | 0,76896877 | 1,112852665 |
| CORO1A | 0,001569618 | 0,340090871 | 0,572 | 0,592 | 1 | TC | 1,265836322 | 0,789991551 | 0,966216216 |
| PRDM11 | 0,001570633 | 0,479704399 | 0,258 | 0,212 | 1 | TC | 1,39445792 | 0,717124544 | 1,216981132 |
| IL6ST1 | 0,001786889 | 0,665316096 | 0,169 | 0,131 | 1 | TC | 1,585915711 | 0,630550535 | 1,290076336 |
| ARPC1B3 | 0,002089243 | 0,532249567 | 0,389 | 0,386 | 1 | TC | 1,446182443 | 0,691475688 | 1,007772021 |

| Supplementary Table 4 - clusermarker | | | | | | | | | |
|---|---|---|---|---|---|---|---|---|---|
| | **p_val** | **avg_log2FC** | **pct.1** | **pct.2** | **p_val_adj** | **celltype** | **Foldchange_UP** | **Foldchange_DOWN** | **Ratio_pct1_pct2** |
| AREG1 | 0,00222946 | 1,023167632 | 0,255 | 0,218 | 1 | TC | 2,032376421 | 0,492034837 | 1,169724771 |
| NDUFV23 | 0,003027615 | 0,486462332 | 0,193 | 0,155 | 1 | TC | 1,401005221 | 0,713773215 | 1,24516129 |
| PPDPF3 | 0,003116662 | 0,33922469 | 0,519 | 0,548 | 1 | TC | 1,265076554 | 0,790465997 | 0,947080292 |
| STAT31 | 0,003724058 | 0,517675486 | 0,309 | 0,288 | 1 | TC | 1,431646677 | 0,698496365 | 1,072916667 |
| GPX43 | 0,003759986 | 0,471150386 | 0,346 | 0,331 | 1 | TC | 1,386214377 | 0,721389142 | 1,045317221 |
| IGFLR1 | 0,003966686 | 0,524145827 | 0,131 | 0,096 | 1 | TC | 1,438081886 | 0,695370695 | 1,364583333 |
| PIM2 | 0,004049409 | 0,593741861 | 0,171 | 0,137 | 1 | TC | 1,509155913 | 0,66262206 | 1,248175182 |
| BBLN3 | 0,00511344 | 0,497269657 | 0,267 | 0,238 | 1 | TC | 1,411539653 | 0,708446269 | 1,121848739 |
| SNHG291 | 0,00530878 | 0,386562431 | 0,401 | 0,398 | 1 | TC | 1,307274792 | 0,764950113 | 1,007537688 |
| UBXN14 | 0,00580167 | 0,567149651 | 0,258 | 0,235 | 1 | TC | 1,481593476 | 0,674948976 | 1,09787234 |
| TCF7 | 0,00590073 | 0,491461211 | 0,117 | 0,084 | 1 | TC | 1,405868066 | 0,711304299 | 1,392857143 |
| KLF9 | 0,006412994 | 0,39029777 | 0,116 | 0,081 | 1 | TC | 1,310663895 | 0,762972112 | 1,432098765 |
| CIRBP1 | 0,007372419 | 0,433496876 | 0,413 | 0,422 | 1 | TC | 1,350503029 | 0,740464833 | 0,978672986 |
| CTSB2 | 0,008348364 | 0,391832326 | 0,105 | 0,074 | 1 | TC | 1,312058755 | 0,76216099 | 1,418918919 |
| C17orf492 | 0,008469321 | 0,38606263 | 0,124 | 0,09 | 1 | TC | 1,306821984 | 0,765215165 | 1,377777778 |
| SH2D1A | 0,008863954 | 0,447361982 | 0,133 | 0,101 | 1 | TC | 1,363544687 | 0,733382638 | 1,316831683 |
| APRT2 | 0,012760394 | 0,515250616 | 0,358 | 0,369 | 1 | TC | 1,429242397 | 0,69967138 | 0,970189702 |
| FOS2 | 0,01283805 | 0,342557716 | 0,595 | 0,578 | 1 | TC | 1,268002611 | 0,788641909 | 1,029411765 |
| TLK11 | 0,014579253 | 0,499406395 | 0,119 | 0,092 | 1 | TC | 1,413631796 | 0,707397784 | 1,293478261 |
| CUTA2 | 0,014729256 | 0,498358968 | 0,275 | 0,261 | 1 | TC | 1,412605842 | 0,707911556 | 1,053639847 |
| TOMM72 | 0,017967552 | 0,25281273 | 0,546 | 0,557 | 1 | TC | 1,191527898 | 0,83925857 | 0,980251346 |
| PRDX21 | 0,018390731 | 0,415050048 | 0,192 | 0,163 | 1 | TC | 1,33334493 | 0,749993477 | 1,17791411 |
| PHTF2 | 0,018626544 | 0,376334036 | 0,129 | 0,099 | 1 | TC | 1,298039275 | 0,770392714 | 1,303030303 |
| TYMP3 | 0,020452349 | 0,55260264 | 0,112 | 0,086 | 1 | TC | 1,466729309 | 0,681789062 | 1,302325581 |
| JAK3 | 0,022900302 | 0,429749654 | 0,105 | 0,08 | 1 | TC | 1,346999817 | 0,742390599 | 1,3125 |
| ARID5B2 | 0,025327314 | 0,367278444 | 0,174 | 0,145 | 1 | TC | 1,289917184 | 0,775243568 | 1,2 |
| ANP32B1 | 0,027269077 | 0,50313167 | 0,265 | 0,256 | 1 | TC | 1,417286741 | 0,705573524 | 1,03515625 |
| NDUFA112 | 0,028253486 | 0,435311707 | 0,282 | 0,271 | 1 | TC | 1,352202957 | 0,739533955 | 1,040590406 |
| AL138963.43 | 0,028342171 | 0,489132331 | 0,234 | 0,215 | 1 | TC | 1,403600465 | 0,712453454 | 1,088372093 |
| DUSP21 | 0,034345764 | 0,283278222 | 0,554 | 0,554 | 1 | TC | 1,216957025 | 0,821721704 | 1 |
| COMMD61 | 0,034995958 | 0,297572596 | 0,473 | 0,506 | 1 | TC | 1,229074695 | 0,8136202 | 0,934782609 |

| Supplementary Table 4 - clusermarker | | | | | | | | | |
|---|---|---|---|---|---|---|---|---|---|
| | **p_val** | **avg_log2FC** | **pct.1** | **pct.2** | **p_val_adj** | **celltype** | **Foldchange_UP** | **Foldchange_DOWN** | **Ratio_pct1_pct2** |
| CD28 | 0,035282727 | 0,327145409 | 0,103 | 0,078 | 1 | TC | 1,254528642 | 0,797112132 | 1,320512821 |
| NDUFB102 | 0,036828087 | 0,512675136 | 0,212 | 0,196 | 1 | TC | 1,426693209 | 0,700921539 | 1,081632653 |
| FUS3 | 0,041882306 | 0,308407181 | 0,505 | 0,541 | 1 | TC | 1,238339746 | 0,80753283 | 0,933456562 |
| TNFRSF141 | 0,044805069 | 0,430259202 | 0,152 | 0,13 | 1 | TC | 1,347475649 | 0,742128439 | 1,169230769 |
| THEM4 | 0,045671544 | 0,392478069 | 0,107 | 0,084 | 1 | TC | 1,312646157 | 0,761819928 | 1,273809524 |
| WDR74 | 0,046346288 | 0,425357539 | 0,128 | 0,105 | 1 | TC | 1,34290527 | 0,744654163 | 1,219047619 |
| YWHAB2 | 0,050791386 | 0,311806759 | 0,462 | 0,489 | 1 | TC | 1,24126122 | 0,805632194 | 0,944785276 |
| SELENOM2 | 0,051686129 | 0,350970396 | 0,132 | 0,108 | 1 | TC | 1,27541822 | 0,784056543 | 1,222222222 |
| HIGD2A2 | 0,05895088 | 0,468846885 | 0,309 | 0,318 | 1 | TC | 1,384002822 | 0,722541879 | 0,971698113 |
| FKBP5 | 0,059824028 | 0,28749008 | 0,194 | 0,167 | 1 | TC | 1,220515046 | 0,819326237 | 1,161676647 |
| EDF12 | 0,060627293 | 0,440789649 | 0,325 | 0,342 | 1 | TC | 1,357347058 | 0,736731254 | 0,950292398 |
| CD473 | 0,06836218 | 0,434157522 | 0,209 | 0,196 | 1 | TC | 1,3511216 | 0,740125833 | 1,066326531 |
| PPP1CC | 0,071902766 | 0,395758455 | 0,126 | 0,107 | 1 | TC | 1,315634235 | 0,760089676 | 1,177570093 |
| LDHB | 0,090059475 | 0,36456348 | 0,36 | 0,373 | 1 | TC | 1,287492011 | 0,776703849 | 0,965147453 |
| ITK | 0,093082669 | 0,328254328 | 0,24 | 0,224 | 1 | TC | 1,255493299 | 0,796499672 | 1,071428571 |
| SPCS13 | 0,094319298 | 0,450874207 | 0,265 | 0,265 | 1 | TC | 1,366868265 | 0,731599398 | 1 |
| CSK1 | 0,09754611 | 0,524990357 | 0,156 | 0,143 | 1 | TC | 1,438923962 | 0,694963755 | 1,090909091 |
| TRIR3 | 0,102370893 | 0,393973683 | 0,384 | 0,417 | 1 | TC | 1,314007657 | 0,761030573 | 0,920863309 |
| LAMTOR42 | 0,109721138 | 0,42759292 | 0,224 | 0,215 | 1 | TC | 1,344987645 | 0,743501254 | 1,041860465 |
| CALM33 | 0,111390903 | 0,428508076 | 0,169 | 0,157 | 1 | TC | 1,345841092 | 0,743029772 | 1,076433121 |
| RNASEK2 | 0,117743176 | 0,372316333 | 0,405 | 0,438 | 1 | TC | 1,294429447 | 0,772541139 | 0,924657534 |
| CSNK2B2 | 0,121143304 | 0,375396626 | 0,21 | 0,199 | 1 | TC | 1,297196131 | 0,770893449 | 1,055276382 |
| NAP1L4 | 0,122356437 | 0,449819678 | 0,193 | 0,182 | 1 | TC | 1,365869527 | 0,732134351 | 1,06043956 |
| PDE3B | 0,125607373 | 0,299126078 | 0,142 | 0,125 | 1 | TC | 1,230398865 | 0,812744573 | 1,136 |
| AQP32 | 0,128963666 | 0,299892543 | 0,149 | 0,131 | 1 | TC | 1,231052717 | 0,812312898 | 1,13740458 |
| ALKBH72 | 0,13504026 | 0,410118521 | 0,134 | 0,12 | 1 | TC | 1,328794973 | 0,752561547 | 1,116666667 |
| DNAJB13 | 0,138415175 | -0,495590883 | 0,495 | 0,554 | 1 | TC | 0,709271123 | 1,409898087 | 0,893501805 |
| TSPO3 | 0,139117766 | 0,469888873 | 0,233 | 0,235 | 1 | TC | 1,385002781 | 0,722020211 | 0,991489362 |
| RNF19A1 | 0,141945063 | 0,349668975 | 0,25 | 0,241 | 1 | TC | 1,274268214 | 0,784764141 | 1,037344398 |
| PRELID13 | 0,14693739 | 0,268373628 | 0,149 | 0,188 | 1 | TC | 1,204449268 | 0,830254977 | 0,792553191 |
| TAF103 | 0,154541608 | 0,48459689 | 0,202 | 0,197 | 1 | TC | 1,399194856 | 0,714696738 | 1,025380711 |

| Supplementary Table 4 - clusermarker | | | | | | | | | |
|---|---|---|---|---|---|---|---|---|---|
| | **p_val** | **avg_log2FC** | **pct.1** | **pct.2** | **p_val_adj** | **celltype** | **Foldchange_UP** | **Foldchange_DOWN** | **Ratio_pct1_pct2** |
| SEPTIN6 | 0,159730169 | 0,379864614 | 0,307 | 0,318 | 1 | TC | 1,30121974 | 0,768509706 | 0,965408805 |
| SEPTIN92 | 0,167687326 | 0,323951886 | 0,298 | 0,306 | 1 | TC | 1,251754712 | 0,798878559 | 0,973856209 |
| VSIR1 | 0,16838344 | 0,419819407 | 0,146 | 0,136 | 1 | TC | 1,337760087 | 0,747518191 | 1,073529412 |
| JAK12 | 0,173130965 | 0,328629745 | 0,387 | 0,423 | 1 | TC | 1,255820044 | 0,796292434 | 0,914893617 |
| NEAT12 | 0,175279691 | 0,365864961 | 0,369 | 0,386 | 1 | TC | 1,288654004 | 0,776003486 | 0,955958549 |
| MRPS342 | 0,185891085 | 0,335492576 | 0,106 | 0,093 | 1 | TC | 1,261808152 | 0,792513504 | 1,139784946 |
| SRRM21 | 0,186215507 | 0,353718428 | 0,35 | 0,375 | 1 | TC | 1,27784994 | 0,782564501 | 0,933333333 |
| UCP2 | 0,187706934 | 0,285150592 | 0,16 | 0,145 | 1 | TC | 1,218537451 | 0,820655942 | 1,103448276 |
| ERP292 | 0,191651737 | 0,448044005 | 0,229 | 0,233 | 1 | TC | 1,364189444 | 0,733036019 | 0,982832618 |
| ARHGAP4 | 0,192363561 | 0,391746299 | 0,147 | 0,136 | 1 | TC | 1,31198052 | 0,762206439 | 1,080882353 |
| LAMP11 | 0,193833784 | 0,353534428 | 0,135 | 0,123 | 1 | TC | 1,277686975 | 0,782664314 | 1,097560976 |
| MALT11 | 0,196978068 | 0,345197228 | 0,122 | 0,11 | 1 | TC | 1,270324634 | 0,787200353 | 1,109090909 |
| ARL6IP43 | 0,202729834 | 0,399586373 | 0,229 | 0,23 | 1 | TC | 1,319129656 | 0,758075596 | 0,995652174 |
| FAM102A | 0,205230453 | 0,334815761 | 0,121 | 0,108 | 1 | TC | 1,261216336 | 0,792885385 | 1,12037037 |
| TUBB4 | 0,215217532 | 0,347438078 | 0,188 | 0,229 | 1 | TC | 1,272299285 | 0,785978592 | 0,820960699 |
| CIB13 | 0,222448031 | 0,379529151 | 0,251 | 0,255 | 1 | TC | 1,300917209 | 0,768688425 | 0,984313725 |
| SSH21 | 0,225152014 | 0,3372816 | 0,132 | 0,12 | 1 | TC | 1,263373836 | 0,791531352 | 1,1 |
| INPP4B1 | 0,22728137 | 0,300000823 | 0,121 | 0,108 | 1 | TC | 1,231145116 | 0,812251933 | 1,12037037 |
| CLPP1 | 0,22840202 | 0,319281238 | 0,132 | 0,12 | 1 | TC | 1,247708776 | 0,801469076 | 1,1 |
| ANAPC113 | 0,22850046 | 0,339061041 | 0,164 | 0,155 | 1 | TC | 1,264933061 | 0,790555667 | 1,058064516 |
| ANKRD101 | 0,232704884 | 0,412686128 | 0,136 | 0,128 | 1 | TC | 1,331161975 | 0,751223382 | 1,0625 |
| ZNF7063 | 0,253062629 | 0,343819011 | 0,135 | 0,125 | 1 | TC | 1,269111664 | 0,78795273 | 1,08 |
| LIME11 | 0,254110599 | 0,345922333 | 0,234 | 0,232 | 1 | TC | 1,270963265 | 0,786804802 | 1,00862069 |
| MLLT6 | 0,26139285 | 0,28571246 | 0,118 | 0,107 | 1 | TC | 1,219012111 | 0,820336394 | 1,102803738 |
| LMAN12 | 0,271438675 | 0,314145932 | 0,129 | 0,119 | 1 | TC | 1,243275423 | 0,804327007 | 1,084033613 |
| SYNGR23 | 0,285318958 | 0,420120498 | 0,126 | 0,119 | 1 | TC | 1,338039307 | 0,7473622 | 1,058823529 |
| SDF43 | 0,291965772 | 0,299788097 | 0,153 | 0,145 | 1 | TC | 1,230963596 | 0,812371709 | 1,055172414 |
| TLE52 | 0,298380686 | 0,342083508 | 0,339 | 0,363 | 1 | TC | 1,267585892 | 0,788901176 | 0,933884298 |
| PLEC2 | 0,300239293 | 0,258904766 | 0,153 | 0,184 | 1 | TC | 1,196569973 | 0,835722124 | 0,831521739 |
| HM131 | 0,301461167 | 0,329213256 | 0,114 | 0,105 | 1 | TC | 1,256328075 | 0,795970432 | 1,085714286 |
| CCT82 | 0,303022796 | 0,270597961 | 0,116 | 0,14 | 1 | TC | 1,206307709 | 0,828975885 | 0,828571429 |

| Supplementary Table 4 - clusermarker | | | | | | | | | |
|---|---|---|---|---|---|---|---|---|---|
| | **p_val** | **avg_log2FC** | **pct.1** | **pct.2** | **p_val_adj** | **celltype** | **Foldchange_UP** | **Foldchange_DOWN** | **Ratio_pct1_pct2** |
| TMEM243 | 0,306048212 | 0,314290813 | 0,14 | 0,133 | 1 | TC | 1,243400284 | 0,804246237 | 1,052631579 |
| SF12 | 0,311381219 | 0,321032643 | 0,378 | 0,414 | 1 | TC | 1,249224391 | 0,800496698 | 0,913043478 |
| MZT2B2 | 0,311770497 | 0,284874899 | 0,28 | 0,288 | 1 | TC | 1,218304615 | 0,820812781 | 0,972222222 |
| TSPYL2 | 0,321030732 | 0,305135143 | 0,297 | 0,301 | 1 | TC | 1,235534369 | 0,809366397 | 0,986710963 |
| EPB41 | 0,33065985 | 0,344280714 | 0,167 | 0,161 | 1 | TC | 1,26951788 | 0,787700603 | 1,037267081 |
| PHPT13 | 0,341875533 | 0,319059876 | 0,107 | 0,099 | 1 | TC | 1,247517347 | 0,80159206 | 1,080808081 |
| LAT | 0,359720734 | 0,261633055 | 0,22 | 0,268 | 1 | TC | 1,198834955 | 0,834143178 | 0,820895522 |
| NME33 | 0,360163695 | 0,381667605 | 0,133 | 0,128 | 1 | TC | 1,302846941 | 0,76754987 | 1,0390625 |
| AURKAIP12 | 0,36033464 | 0,293156788 | 0,144 | 0,137 | 1 | TC | 1,225318489 | 0,816114348 | 1,051094891 |
| PPP4C3 | 0,369109063 | 0,282992042 | 0,105 | 0,125 | 1 | TC | 1,216715647 | 0,821884721 | 0,84 |
| CMTM71 | 0,373767362 | 0,288035558 | 0,104 | 0,096 | 1 | TC | 1,220976606 | 0,819016511 | 1,083333333 |
| FIS13 | 0,375030661 | 0,376674815 | 0,16 | 0,158 | 1 | TC | 1,298345921 | 0,770210761 | 1,012658228 |
| SLC25A34 | 0,376496259 | 0,281166406 | 0,392 | 0,432 | 1 | TC | 1,215176947 | 0,82292542 | 0,907407407 |
| SEC11C1 | 0,377729526 | 0,363209372 | 0,16 | 0,158 | 1 | TC | 1,286284143 | 0,777433202 | 1,012658228 |
| HNRNPL1 | 0,379876085 | 0,374626879 | 0,253 | 0,265 | 1 | TC | 1,2965042 | 0,771304868 | 0,954716981 |
| BAX3 | 0,381419498 | 0,472039807 | 0,155 | 0,155 | 1 | TC | 1,387069241 | 0,720944543 | 1 |
| TNIP13 | 0,397221865 | 0,293595278 | 0,102 | 0,096 | 1 | TC | 1,225690966 | 0,815866338 | 1,0625 |
| ATP5IF12 | 0,39981689 | 0,256907451 | 0,235 | 0,236 | 1 | TC | 1,194914548 | 0,836879927 | 0,995762712 |
| TMED10 | 0,417831843 | 0,282340636 | 0,18 | 0,179 | 1 | TC | 1,216166399 | 0,822255902 | 1,005586592 |
| TBC1D10C | 0,418462958 | 0,419777488 | 0,185 | 0,19 | 1 | TC | 1,337721217 | 0,747539911 | 0,973684211 |
| FLNA2 | 0,419242056 | 0,324385748 | 0,3 | 0,322 | 1 | TC | 1,252131209 | 0,798638348 | 0,931677019 |
| CDKN1B1 | 0,42459721 | 0,308106907 | 0,368 | 0,402 | 1 | TC | 1,238082032 | 0,807700923 | 0,915422886 |
| ATRAID1 | 0,428629934 | 0,304194584 | 0,129 | 0,125 | 1 | TC | 1,23472913 | 0,809894232 | 1,032 |
| DENND2D | 0,429898337 | 0,258741356 | 0,094 | 0,11 | 1 | TC | 1,196434449 | 0,83581679 | 0,854545455 |
| ACTR31 | 0,430970407 | 0,2652753 | 0,301 | 0,322 | 1 | TC | 1,201865371 | 0,832039947 | 0,934782609 |
| PIK3IP1 | 0,437963052 | 0,338669127 | 0,152 | 0,149 | 1 | TC | 1,264589484 | 0,790770454 | 1,020134228 |
| MDH21 | 0,439640323 | 0,26168108 | 0,153 | 0,179 | 1 | TC | 1,198874863 | 0,834115411 | 0,854748603 |
| SELENOW2 | 0,442018032 | 0,399537669 | 0,21 | 0,215 | 1 | TC | 1,319085124 | 0,758101188 | 0,976744186 |
| SF3B52 | 0,4434219 | 0,316609131 | 0,176 | 0,178 | 1 | TC | 1,245399954 | 0,802954904 | 0,988764045 |
| FCMR | 0,449229528 | 0,405969226 | 0,179 | 0,184 | 1 | TC | 1,324978754 | 0,754729083 | 0,972826087 |
| C19orf533 | 0,465304908 | 0,346467022 | 0,202 | 0,209 | 1 | TC | 1,271443208 | 0,7865078 | 0,966507177 |

| Supplementary Table 4 - clusermarker | | | | | | | | | |
|---|---|---|---|---|---|---|---|---|---|
| | **p_val** | **avg_log2FC** | **pct.1** | **pct.2** | **p_val_adj** | **celltype** | **Foldchange_UP** | **Foldchange_DOWN** | **Ratio_pct1_pct2** |
| H2AZ22 | 0,470450347 | 0,307401967 | 0,259 | 0,274 | 1 | TC | 1,237477219 | 0,808095684 | 0,945255474 |
| PPP1R2 | 0,487303973 | 0,36323707 | 0,276 | 0,295 | 1 | TC | 1,286308838 | 0,777418277 | 0,93559322 |
| VPS282 | 0,489176504 | 0,382513525 | 0,196 | 0,203 | 1 | TC | 1,303611085 | 0,767099952 | 0,965517241 |
| GNB23 | 0,50983555 | 0,274992376 | 0,201 | 0,203 | 1 | TC | 1,209987695 | 0,826454686 | 0,990147783 |
| NDUFS82 | 0,524798437 | 0,279689674 | 0,127 | 0,123 | 1 | TC | 1,213933737 | 0,823768192 | 1,032520325 |
| CD5 | 0,525007705 | 0,311946993 | 0,131 | 0,13 | 1 | TC | 1,241381881 | 0,805553888 | 1,007692308 |
| REX1BD3 | 0,529582762 | 0,391641498 | 0,207 | 0,218 | 1 | TC | 1,311885217 | 0,76226181 | 0,949541284 |
| HLA-DRA2 | 0,5370966 | 0,282382134 | 0,126 | 0,12 | 1 | TC | 1,216201382 | 0,822232251 | 1,05 |
| COX5B2 | 0,54003579 | 0,282406819 | 0,21 | 0,244 | 1 | TC | 1,216222192 | 0,822218182 | 0,860655738 |
| CCDC85B2 | 0,543199501 | 0,327193903 | 0,156 | 0,16 | 1 | TC | 1,254570812 | 0,797085338 | 0,975 |
| WDR83OS3 | 0,561946491 | 0,350720376 | 0,236 | 0,252 | 1 | TC | 1,275197208 | 0,784192432 | 0,936507937 |
| PPM1G3 | 0,562530108 | 0,338288293 | 0,139 | 0,14 | 1 | TC | 1,264255709 | 0,790979224 | 0,992857143 |
| NOSIP1 | 0,565183452 | 0,336313008 | 0,13 | 0,13 | 1 | TC | 1,262525921 | 0,792062946 | 1 |
| ITM2C1 | 0,565853762 | 0,25092244 | 0,137 | 0,155 | 1 | TC | 1,189967721 | 0,840358929 | 0,883870968 |
| TGIF11 | 0,570077537 | 0,279584707 | 0,188 | 0,19 | 1 | TC | 1,213845418 | 0,823828129 | 0,989473684 |
| NDUFC23 | 0,57505547 | 0,28456013 | 0,175 | 0,202 | 1 | TC | 1,218038833 | 0,820991887 | 0,866336634 |
| GPSM31 | 0,577085385 | 0,280894443 | 0,239 | 0,282 | 1 | TC | 1,214947895 | 0,823080565 | 0,84751773 |
| ATP6AP21 | 0,579060586 | 0,272101113 | 0,128 | 0,146 | 1 | TC | 1,207565222 | 0,82811262 | 0,876712329 |
| DRAP13 | 0,57923467 | 0,339408077 | 0,241 | 0,258 | 1 | TC | 1,265237373 | 0,790365524 | 0,934108527 |
| H1-4 | 0,579314024 | 0,276476311 | 0,122 | 0,137 | 1 | TC | 1,211232911 | 0,825605043 | 0,890510949 |
| SYNRG | 0,580531516 | 0,270031653 | 0,142 | 0,163 | 1 | TC | 1,205834284 | 0,82930135 | 0,871165644 |
| CHCHD103 | 0,587696678 | 0,262448066 | 0,121 | 0,136 | 1 | TC | 1,199512396 | 0,833672085 | 0,889705882 |
| DPP73 | 0,588933778 | 0,372319375 | 0,176 | 0,184 | 1 | TC | 1,294432177 | 0,77253951 | 0,956521739 |
| ATP5PO1 | 0,591952907 | 0,345397386 | 0,245 | 0,265 | 1 | TC | 1,27050089 | 0,787091145 | 0,924528302 |
| RCSD1 | 0,597965108 | 0,42813943 | 0,145 | 0,149 | 1 | TC | 1,345497238 | 0,74321966 | 0,973154362 |
| YPEL32 | 0,604949395 | 0,319565237 | 0,221 | 0,259 | 1 | TC | 1,247954415 | 0,80131132 | 0,853281853 |
| LAMTOR12 | 0,61318068 | 0,25947036 | 0,089 | 0,101 | 1 | TC | 1,197039168 | 0,835394552 | 0,881188119 |
| TMEM2193 | 0,617154845 | 0,332964657 | 0,162 | 0,188 | 1 | TC | 1,259599122 | 0,79390338 | 0,861702128 |
| NDUFAB13 | 0,618273826 | 0,30776578 | 0,127 | 0,128 | 1 | TC | 1,237789321 | 0,807891927 | 0,9921875 |
| KIF2A | 0,630402778 | 0,27530381 | 0,122 | 0,12 | 1 | TC | 1,210248923 | 0,826276298 | 1,016666667 |
| ZFAND5 | 0,633210648 | 0,302471184 | 0,272 | 0,289 | 1 | TC | 1,233255041 | 0,810862285 | 0,941176471 |

| Supplementary Table 4 - clusermarker | | | | | | | | | |
|---|---|---|---|---|---|---|---|---|---|
| | **p_val** | **avg_log2FC** | **pct.1** | **pct.2** | **p_val_adj** | **celltype** | **Foldchange_UP** | **Foldchange_DOWN** | **Ratio_pct1_pct2** |
| NDUFA23 | 0,635248427 | 0,259636442 | 0,141 | 0,142 | 1 | TC | 1,197176979 | 0,835298387 | 0,992957746 |
| SRSF93 | 0,637946683 | 0,396645926 | 0,223 | 0,244 | 1 | TC | 1,316443794 | 0,759622253 | 0,913934426 |
| NDUFB93 | 0,647432776 | 0,2760192 | 0,169 | 0,193 | 1 | TC | 1,210849199 | 0,825866674 | 0,875647668 |
| PSMF1 | 0,649717506 | 0,298247297 | 0,097 | 0,11 | 1 | TC | 1,229649627 | 0,813239786 | 0,881818182 |
| PSME22 | 0,657895321 | 0,358059684 | 0,263 | 0,289 | 1 | TC | 1,281700947 | 0,780213202 | 0,910034602 |
| JTB3 | 0,660898903 | 0,365436048 | 0,206 | 0,221 | 1 | TC | 1,288270944 | 0,776234226 | 0,932126697 |
| PNPLA21 | 0,665186045 | 0,278660623 | 0,185 | 0,209 | 1 | TC | 1,213068166 | 0,824355982 | 0,885167464 |
| ERH3 | 0,669809207 | 0,255380943 | 0,165 | 0,185 | 1 | TC | 1,193650884 | 0,837765894 | 0,891891892 |
| ISCU3 | 0,672353636 | 0,267242518 | 0,201 | 0,23 | 1 | TC | 1,203505319 | 0,830906174 | 0,873913043 |
| RGS10 | 0,682886363 | 0,332890387 | 0,183 | 0,211 | 1 | TC | 1,259534279 | 0,793944251 | 0,867298578 |
| COX5A3 | 0,694393865 | 0,282962984 | 0,18 | 0,188 | 1 | TC | 1,216691141 | 0,821901275 | 0,957446809 |
| MAP4K1 | 0,696274986 | 0,332046807 | 0,11 | 0,111 | 1 | TC | 1,258798014 | 0,794408626 | 0,990990991 |
| RAP1A1 | 0,701592097 | 0,3094293 | 0,194 | 0,22 | 1 | TC | 1,239217395 | 0,806960913 | 0,881818182 |
| TRBC2 | 0,706897147 | 0,268120204 | 0,322 | 0,351 | 1 | TC | 1,204237713 | 0,830400833 | 0,917378917 |
| ERGIC32 | 0,707145894 | 0,319447387 | 0,147 | 0,152 | 1 | TC | 1,247852477 | 0,80137678 | 0,967105263 |
| PSAP1 | 0,709804274 | 0,264507322 | 0,213 | 0,242 | 1 | TC | 1,201225763 | 0,832482978 | 0,880165289 |
| CHASERR2 | 0,724080936 | 0,287234002 | 0,259 | 0,3 | 1 | TC | 1,220298424 | 0,81947168 | 0,863333333 |
| SELENOH3 | 0,730003986 | 0,286107261 | 0,165 | 0,187 | 1 | TC | 1,219345746 | 0,820111936 | 0,882352941 |
| OCIAD23 | 0,734205591 | 0,293053607 | 0,179 | 0,203 | 1 | TC | 1,225230858 | 0,816172718 | 0,881773399 |
| PLP22 | 0,739095645 | 0,304330237 | 0,245 | 0,282 | 1 | TC | 1,234845234 | 0,809818083 | 0,868794326 |
| LMAN23 | 0,747237361 | 0,305552165 | 0,164 | 0,184 | 1 | TC | 1,235891562 | 0,809132477 | 0,891304348 |
| DNAJC152 | 0,756570497 | 0,297722753 | 0,126 | 0,13 | 1 | TC | 1,229202625 | 0,813535523 | 0,969230769 |
| WDR14 | 0,756946661 | 0,290590575 | 0,183 | 0,206 | 1 | TC | 1,223140874 | 0,817567315 | 0,888349515 |
| CTSA | 0,764368618 | 0,260172637 | 0,12 | 0,133 | 1 | TC | 1,197622007 | 0,834987997 | 0,902255639 |
| DUT1 | 0,784703907 | 0,380648322 | 0,153 | 0,163 | 1 | TC | 1,301926787 | 0,768092346 | 0,938650307 |
| AC020916.11 | 0,787687588 | 0,308187078 | 0,23 | 0,245 | 1 | TC | 1,238150834 | 0,80765604 | 0,93877551 |
| CSRNP1 | 0,799466825 | 0,262971025 | 0,302 | 0,351 | 1 | TC | 1,199947282 | 0,833369944 | 0,86039886 |
| ARHGAP45 | 0,81609215 | 0,347574056 | 0,121 | 0,134 | 1 | TC | 1,272419209 | 0,785904514 | 0,902985075 |
| SNHG72 | 0,823322493 | 0,261740378 | 0,136 | 0,149 | 1 | TC | 1,19892414 | 0,834081129 | 0,912751678 |
| CUL31 | 0,826784319 | 0,276006975 | 0,129 | 0,134 | 1 | TC | 1,210838938 | 0,825873672 | 0,962686567 |
| TBCB2 | 0,833415303 | 0,34792426 | 0,175 | 0,196 | 1 | TC | 1,272728117 | 0,785713765 | 0,892857143 |

| Supplementary Table 4 - clusermarker | | | | | | | | | |
|---|---|---|---|---|---|---|---|---|---|
| | **p_val** | **avg_log2FC** | **pct.1** | **pct.2** | **p_val_adj** | **celltype** | **Foldchange_UP** | **Foldchange_DOWN** | **Ratio_pct1_pct2** |
| SEC11A1 | 0,833739936 | 0,286769406 | 0,162 | 0,179 | 1 | TC | 1,21990551 | 0,81973562 | 0,905027933 |
| PMAIP11 | 0,840614983 | 0,298366186 | 0,1 | 0,102 | 1 | TC | 1,229750964 | 0,813172772 | 0,980392157 |
| NUMA11 | 0,857705908 | 0,257956569 | 0,131 | 0,136 | 1 | TC | 1,195783798 | 0,836271575 | 0,963235294 |
| PKM1 | 0,859502938 | 0,262863883 | 0,296 | 0,325 | 1 | TC | 1,199858172 | 0,833431837 | 0,910769231 |
| ILF32 | 0,860857917 | 0,280763547 | 0,152 | 0,16 | 1 | TC | 1,214837668 | 0,823155247 | 0,95 |
| PCBP13 | 0,868843053 | 0,304605957 | 0,269 | 0,31 | 1 | TC | 1,235081253 | 0,80966333 | 0,867741935 |
| NFKBIZ | 0,874702341 | 0,265717576 | 0,145 | 0,151 | 1 | TC | 1,202233875 | 0,831784914 | 0,960264901 |
| ARHGEF1 | 0,89628565 | 0,309751454 | 0,291 | 0,336 | 1 | TC | 1,239494142 | 0,806780739 | 0,866071429 |
| UXT2 | 0,903428456 | 0,253555209 | 0,261 | 0,286 | 1 | TC | 1,192141272 | 0,83882676 | 0,912587413 |
| SORL11 | 0,91110507 | 0,289010322 | 0,118 | 0,123 | 1 | TC | 1,221801844 | 0,818463326 | 0,959349593 |
| C1QBP3 | 0,922746295 | 0,253354964 | 0,114 | 0,122 | 1 | TC | 1,191975815 | 0,838943196 | 0,93442623 |
| HMGN12 | 0,923429449 | 0,317511612 | 0,237 | 0,27 | 1 | TC | 1,24617926 | 0,80245277 | 0,877777778 |
| ACAP11 | 0,948375541 | 0,274213337 | 0,311 | 0,354 | 1 | TC | 1,209334491 | 0,826901082 | 0,878531073 |
| ZAP70 | 0,948496532 | 0,303732985 | 0,203 | 0,224 | 1 | TC | 1,234334135 | 0,810153403 | 0,90625 |
| UBE2D23 | 0,949540059 | 0,293495297 | 0,264 | 0,297 | 1 | TC | 1,225606027 | 0,815922881 | 0,888888889 |
| NDUFB72 | 0,953376853 | 0,2691487 | 0,151 | 0,161 | 1 | TC | 1,205096519 | 0,829809052 | 0,937888199 |
| NDUFS72 | 0,961259825 | 0,280104177 | 0,166 | 0,178 | 1 | TC | 1,214282565 | 0,823531548 | 0,93258427 |
| ID31 | 0,961696403 | 0,338938269 | 0,102 | 0,107 | 1 | TC | 1,264825421 | 0,790622946 | 0,953271028 |
| PEBP12 | 0,966573109 | 0,260296447 | 0,247 | 0,276 | 1 | TC | 1,19772479 | 0,834916342 | 0,894927536 |
| PDIA62 | 0,968745401 | 0,298690013 | 0,145 | 0,158 | 1 | TC | 1,230027025 | 0,812990268 | 0,917721519 |
| ICAM3 | 0,970564892 | 0,254251824 | 0,282 | 0,318 | 1 | TC | 1,192717044 | 0,838421824 | 0,886792453 |
| TUFM3 | 0,984929645 | 0,283676907 | 0,174 | 0,19 | 1 | TC | 1,217293374 | 0,821494655 | 0,915789474 |
| SEC61B3 | 0,990274315 | 0,260663148 | 0,296 | 0,33 | 1 | TC | 1,198029263 | 0,834704152 | 0,896969697 |
| PRB34 | 5,3248E-152 | -3,588107109 | 0,011 | 0,424 | 1,9482E-147 | PC | 0,083151891 | 12,02618464 | 0,025943396 |
| LYZ4 | 2,9506E-140 | -3,002034782 | 0,067 | 0,69 | 1,0795E-135 | PC | 0,124823824 | 8,011291188 | 0,097101449 |
| PRR44 | 3,4019E-135 | -4,034816272 | 0,127 | 0,853 | 1,2446E-130 | PC | 0,061009754 | 16,39082165 | 0,148886284 |
| PRH24 | 2,72009E-59 | -1,308321614 | 0,003 | 0,152 | 9,95199E-55 | PC | 0,403790363 | 2,476532604 | 0,019736842 |
| JCHAIN2 | 3,29262E-56 | -1,909821843 | 0,708 | 0,957 | 1,20467E-51 | PC | 0,266125407 | 3,757626942 | 0,739811912 |
| STATH4 | 4,45235E-55 | -3,778400501 | 0,003 | 0,147 | 1,62898E-50 | PC | 0,072876602 | 13,72182532 | 0,020408163 |
| C6orf584 | 4,59481E-55 | -1,197056875 | 0,007 | 0,179 | 1,6811E-50 | PC | 0,436164158 | 2,292714755 | 0,039106145 |
| PIP3 | 2,02236E-41 | -0,919243445 | 0,002 | 0,109 | 7,39922E-37 | PC | 0,528786245 | 1,891123321 | 0,018348624 |

| Supplementary Table 4 - clusermarker | | | | | | | | | |
|---|---|---|---|---|---|---|---|---|---|
| | **p_val** | **avg_log2FC** | **pct.1** | **pct.2** | **p_val_adj** | **celltype** | **Foldchange_UP** | **Foldchange_DOWN** | **Ratio_pct1_pct2** |
| IL2RG1 | 2,41253E-41 | -1,31497348 | 0,148 | 0,516 | 8,82673E-37 | PC | 0,401932884 | 2,487977572 | 0,286821705 |
| SLPI4 | 7,23406E-41 | -1,928165675 | 0,131 | 0,511 | 2,64673E-36 | PC | 0,262763051 | 3,805710117 | 0,256360078 |
| SCGB3A24 | 1,21997E-33 | -1,771970524 | 0,058 | 0,31 | 4,46351E-29 | PC | 0,292808528 | 3,41520108 | 0,187096774 |
| RRAD | 3,45784E-31 | -0,751076852 | 0,019 | 0,174 | 1,26512E-26 | PC | 0,594159901 | 1,683048618 | 0,109195402 |
| TXNDC5 | 6,46886E-31 | -1,237615805 | 0,526 | 0,832 | 2,36676E-26 | PC | 0,424072899 | 2,358085134 | 0,632211538 |
| ZG16B4 | 1,64276E-30 | -0,96725798 | 0,013 | 0,147 | 6,01038E-26 | PC | 0,511477265 | 1,955121113 | 0,088435374 |
| SCGB3A13 | 3,01451E-29 | -1,979579765 | 0,21 | 0,582 | 1,10292E-24 | PC | 0,253563719 | 3,943781886 | 0,360824742 |
| AMPD1 | 1,74309E-28 | -0,549768777 | 0,031 | 0,207 | 6,37744E-24 | PC | 0,683129606 | 1,463851063 | 0,149758454 |
| IGHG2 | 5,68245E-24 | 0,613852426 | 0,535 | 0,125 | 2,07904E-19 | PC | 1,530340223 | 0,653449465 | 4,28 |
| CD742 | 8,19432E-22 | -0,798760946 | 0,612 | 0,864 | 2,99806E-17 | PC | 0,574842668 | 1,739606429 | 0,708333333 |
| AL391056.1 | 1,17958E-21 | -0,255288837 | 0,009 | 0,103 | 4,31574E-17 | PC | 0,837819381 | 1,193574681 | 0,087378641 |
| IGHG13 | 3,02316E-21 | 1,84011796 | 0,58 | 0,185 | 1,10609E-16 | PC | 3,580393019 | 0,279298947 | 3,135135135 |
| RASGRP3 | 5,30228E-20 | -0,510126343 | 0,026 | 0,158 | 1,93994E-15 | PC | 0,702160944 | 1,424174912 | 0,164556962 |
| IGHD | 1,03393E-18 | -3,164095595 | 0,013 | 0,109 | 3,78285E-14 | PC | 0,111560979 | 8,963707642 | 0,119266055 |
| LTF3 | 1,91573E-18 | -0,807859214 | 0,021 | 0,136 | 7,00907E-14 | PC | 0,571228864 | 1,750611816 | 0,154411765 |
| SEPTIN10 | 1,25223E-17 | -0,294978666 | 0,013 | 0,103 | 4,58152E-13 | PC | 0,815084385 | 1,226866835 | 0,126213592 |
| QPCT | 3,76361E-17 | -0,40382011 | 0,026 | 0,147 | 1,37699E-12 | PC | 0,755854206 | 1,323006464 | 0,176870748 |
| BIRC32 | 4,21392E-17 | -0,918050243 | 0,126 | 0,342 | 1,54175E-12 | PC | 0,529223766 | 1,889559886 | 0,368421053 |
| HSPA1A4 | 4,89611E-16 | -1,49754003 | 0,148 | 0,364 | 1,79134E-11 | PC | 0,354156756 | 2,823608422 | 0,406593407 |
| FXYD53 | 5,34487E-16 | -0,756058807 | 0,233 | 0,484 | 1,95553E-11 | PC | 0,59211167 | 1,688870616 | 0,481404959 |
| CRIP12 | 4,51897E-15 | -1,659962827 | 0,304 | 0,522 | 1,65335E-10 | PC | 0,316447302 | 3,160083823 | 0,582375479 |
| RPS273 | 6,29667E-15 | -0,844321876 | 0,711 | 0,848 | 2,30376E-10 | PC | 0,556972547 | 1,795420629 | 0,838443396 |
| MSMB4 | 6,65963E-15 | -1,907514857 | 0,03 | 0,147 | 2,43656E-10 | PC | 0,266551304 | 3,751622994 | 0,204081633 |
| HSPA1B3 | 1,06131E-14 | -1,685422375 | 0,163 | 0,37 | 3,88303E-10 | PC | 0,310911875 | 3,216345468 | 0,440540541 |
| RPS294 | 1,12229E-14 | -0,960752836 | 0,377 | 0,625 | 4,10611E-10 | PC | 0,513788735 | 1,946325273 | 0,6032 |
| HLA-B4 | 4,73108E-14 | 1,133228419 | 0,766 | 0,685 | 1,73096E-09 | PC | 2,19349044 | 0,455894396 | 1,118248175 |
| HOMER3 | 9,20955E-14 | -0,306322947 | 0,017 | 0,103 | 3,3695E-09 | PC | 0,8087003 | 1,236552032 | 0,165048544 |
| MTRNR2L84 | 1,04805E-13 | -2,649850822 | 0,069 | 0,217 | 3,83451E-09 | PC | 0,159336553 | 6,276023796 | 0,31797235 |
| CTSH | 1,3751E-13 | -0,52129525 | 0,13 | 0,326 | 5,03108E-09 | PC | 0,696746015 | 1,435243229 | 0,398773006 |
| SLC22A17 | 1,97184E-13 | -0,270928658 | 0,017 | 0,103 | 7,21435E-09 | PC | 0,828785887 | 1,206584252 | 0,165048544 |
| IGHA14 | 4,64251E-12 | -1,179652645 | 0,548 | 0,766 | 1,69855E-07 | PC | 0,441457774 | 2,265222311 | 0,7154047 |

| Supplementary Table 4 - clusermarker | | | | | | | | | |
|---|---|---|---|---|---|---|---|---|---|
| | **p_val** | **avg_log2FC** | **pct.1** | **pct.2** | **p_val_adj** | **celltype** | **Foldchange_UP** | **Foldchange_DOWN** | **Ratio_pct1_pct2** |
| CDKN1C2 | 1,42893E-11 | -0,447547251 | 0,031 | 0,13 | 5,22803E-07 | PC | 0,733288464 | 1,363719803 | 0,238461538 |
| PERP3 | 1,61847E-11 | -0,408761371 | 0,026 | 0,12 | 5,92148E-07 | PC | 0,753269818 | 1,327545558 | 0,216666667 |
| CD48 | 2,01892E-11 | -0,427350985 | 0,062 | 0,196 | 7,38662E-07 | PC | 0,743625946 | 1,344762114 | 0,316326531 |
| IGHG4 | 2,88203E-11 | 3,120826476 | 0,345 | 0,109 | 1,05445E-06 | PC | 8,698860787 | 0,114957582 | 3,165137615 |
| RPL36A4 | 4,19741E-11 | -0,749078867 | 0,475 | 0,663 | 1,5357E-06 | PC | 0,594983321 | 1,680719381 | 0,716440422 |
| CD93 | 5,99284E-11 | -0,273198714 | 0,09 | 0,245 | 2,1926E-06 | PC | 0,827482833 | 1,208484286 | 0,367346939 |
| ANKRD372 | 6,01161E-11 | -0,791691839 | 0,114 | 0,272 | 2,19947E-06 | PC | 0,577666269 | 1,731103326 | 0,419117647 |
| TNFRSF13B | 1,0175E-10 | -0,357685189 | 0,068 | 0,201 | 3,72271E-06 | PC | 0,780415756 | 1,281368286 | 0,338308458 |
| HSPB13 | 1,07867E-10 | -0,598769729 | 0,115 | 0,277 | 3,94654E-06 | PC | 0,660316807 | 1,514424576 | 0,415162455 |
| RPL383 | 1,21148E-10 | -0,576734148 | 0,291 | 0,505 | 4,43246E-06 | PC | 0,670479837 | 1,491469161 | 0,576237624 |
| LINC02362 | 1,58516E-10 | -0,660570594 | 0,108 | 0,261 | 5,79964E-06 | PC | 0,632628039 | 1,580707679 | 0,413793103 |
| IGHG3 | 2,03279E-10 | 1,199522791 | 0,43 | 0,179 | 7,43736E-06 | PC | 2,296636912 | 0,435419284 | 2,402234637 |
| TXNIP4 | 2,30767E-10 | -1,138535974 | 0,235 | 0,429 | 8,44307E-06 | PC | 0,45422028 | 2,201574971 | 0,547785548 |
| ITGB71 | 2,45376E-10 | -0,335806734 | 0,095 | 0,245 | 8,97759E-06 | PC | 0,792340947 | 1,26208295 | 0,387755102 |
| SEC623 | 4,55669E-10 | -0,679431787 | 0,365 | 0,56 | 1,66716E-05 | PC | 0,624411153 | 1,601508869 | 0,651785714 |
| CCR10 | 4,98211E-10 | -0,358673371 | 0,032 | 0,125 | 1,8228E-05 | PC | 0,779881389 | 1,282246267 | 0,256 |
| RPL37A4 | 8,4963E-10 | -0,639511422 | 0,521 | 0,701 | 3,10854E-05 | PC | 0,641930306 | 1,55780151 | 0,743223966 |
| MT-ATP84 | 1,02927E-09 | -0,711933541 | 0,43 | 0,625 | 3,76579E-05 | PC | 0,610501379 | 1,637997938 | 0,688 |
| S100A42 | 3,44584E-09 | -0,545602726 | 0,192 | 0,37 | 0,000126073 | PC | 0,685105121 | 1,459630018 | 0,518918919 |
| TNFAIP3 | 3,56493E-09 | -0,789171307 | 0,121 | 0,266 | 0,00013043 | PC | 0,578676392 | 1,728081557 | 0,454887218 |
| MEF2C1 | 3,70745E-09 | -0,588150857 | 0,158 | 0,321 | 0,000135644 | PC | 0,66519496 | 1,503318666 | 0,492211838 |
| GADD45B3 | 4,50986E-09 | -1,175125218 | 0,237 | 0,402 | 0,000165002 | PC | 0,442845321 | 2,258124793 | 0,589552239 |
| IFNG-AS1 | 5,06694E-09 | -0,290098538 | 0,031 | 0,114 | 0,000185384 | PC | 0,817846197 | 1,222723788 | 0,271929825 |
| GNAS3 | 5,14463E-09 | -0,517590267 | 0,334 | 0,522 | 0,000188227 | PC | 0,698537627 | 1,431562112 | 0,639846743 |
| SLAMF1 | 5,60335E-09 | -0,415603071 | 0,044 | 0,141 | 0,00020501 | PC | 0,749706039 | 1,333856135 | 0,312056738 |
| IGLV3-212 | 6,73902E-09 | 1,695041426 | 0,241 | 0,054 | 0,000246561 | PC | 3,237861838 | 0,308845791 | 4,462962963 |
| RALY3 | 7,35869E-09 | -0,405066344 | 0,124 | 0,272 | 0,000269232 | PC | 0,755201563 | 1,324149802 | 0,455882353 |
| NFKB11 | 1,06023E-08 | -0,371801085 | 0,05 | 0,152 | 0,000387907 | PC | 0,772817096 | 1,293967234 | 0,328947368 |
| CIR12 | 1,81126E-08 | -0,267757395 | 0,055 | 0,158 | 0,000662687 | PC | 0,830609689 | 1,203934909 | 0,348101266 |
| IGKV1-51 | 3,65212E-08 | 1,324435906 | 0,299 | 0,109 | 0,001336201 | PC | 2,504349484 | 0,399305291 | 2,743119266 |
| ITGA41 | 5,18692E-08 | -0,396964525 | 0,059 | 0,163 | 0,001897739 | PC | 0,759454519 | 1,316734544 | 0,36196319 |

| Supplementary Table 4 - clusermarker | | | | | | | | | |
|---|---|---|---|---|---|---|---|---|---|
| | **p_val** | **avg_log2FC** | **pct.1** | **pct.2** | **p_val_adj** | **celltype** | **Foldchange_UP** | **Foldchange_DOWN** | **Ratio_pct1_pct2** |
| KLF61 | 5,45249E-08 | -0,807572996 | 0,257 | 0,429 | 0,001994901 | PC | 0,571342202 | 1,750264546 | 0,599067599 |
| EMB1 | 5,95773E-08 | -0,346337228 | 0,069 | 0,179 | 0,002179753 | PC | 0,786578562 | 1,271328826 | 0,38547486 |
| PABPC41 | 8,27532E-08 | -0,493420021 | 0,157 | 0,304 | 0,00302769 | PC | 0,710339186 | 1,407778171 | 0,516447368 |
| MIR155HG | 8,35903E-08 | -0,373467215 | 0,051 | 0,147 | 0,003058318 | PC | 0,771925106 | 1,295462465 | 0,346938776 |
| S100A63 | 8,43422E-08 | -0,290791904 | 0,311 | 0,5 | 0,003085826 | PC | 0,81745323 | 1,223311577 | 0,622 |
| SSR42 | 8,72064E-08 | 0,398350168 | 0,95 | 0,946 | 0,00319062 | PC | 1,317999815 | 0,758725448 | 1,00422833 |
| DNAJB15 | 1,10203E-07 | -0,765296683 | 0,182 | 0,326 | 0,004031994 | PC | 0,588332369 | 1,699719501 | 0,558282209 |
| RBM392 | 1,51776E-07 | -0,34932565 | 0,185 | 0,342 | 0,005553022 | PC | 0,784950917 | 1,273965006 | 0,540935673 |
| RPS263 | 1,78906E-07 | -0,61730243 | 0,558 | 0,707 | 0,006545637 | PC | 0,651888699 | 1,534004197 | 0,789250354 |
| NDUFA14 | 1,89511E-07 | -0,437755165 | 0,318 | 0,495 | 0,006933655 | PC | 0,738282484 | 1,354495091 | 0,642424242 |
| AREG2 | 1,97774E-07 | -0,596681955 | 0,085 | 0,201 | 0,007235974 | PC | 0,661273065 | 1,512234586 | 0,422885572 |
| MANEA | 2,31346E-07 | -0,360411133 | 0,123 | 0,25 | 0,008464263 | PC | 0,778942569 | 1,283791695 | 0,492 |
| DNAJA13 | 3,55456E-07 | -0,471225709 | 0,205 | 0,353 | 0,013005055 | PC | 0,72135148 | 1,386286752 | 0,580736544 |
| HSPH13 | 3,56E-07 | -0,324791252 | 0,071 | 0,174 | 0,013024987 | PC | 0,798413903 | 1,2524832 | 0,408045977 |
| RPS202 | 3,84156E-07 | -0,401474065 | 0,251 | 0,413 | 0,014055106 | PC | 0,757084342 | 1,320856799 | 0,607748184 |
| FOS3 | 4,53604E-07 | -0,667063321 | 0,323 | 0,484 | 0,016596001 | PC | 0,629787348 | 1,587837552 | 0,667355372 |
| UBE2J1 | 4,69812E-07 | -0,529845693 | 0,283 | 0,446 | 0,017188997 | PC | 0,692628812 | 1,443774765 | 0,634529148 |
| CD79A | 4,79715E-07 | 0,781861814 | 0,677 | 0,614 | 0,01755135 | PC | 1,71934828 | 0,581615727 | 1,102605863 |
| EVI2B | 5,29202E-07 | -0,381672367 | 0,203 | 0,353 | 0,01936193 | PC | 0,767547336 | 1,302851241 | 0,575070822 |
| HSP90AA12 | 5,45126E-07 | -1,222549352 | 0,292 | 0,435 | 0,01994452 | PC | 0,428524813 | 2,333587158 | 0,671264368 |
| LMNA3 | 5,71693E-07 | -0,86212044 | 0,199 | 0,337 | 0,020916521 | PC | 0,550143375 | 1,817707973 | 0,590504451 |
| TMEM258 | 5,86537E-07 | -0,48848567 | 0,436 | 0,587 | 0,021459646 | PC | 0,71277287 | 1,402971468 | 0,742759796 |
| MEI1 | 6,13215E-07 | -0,359258853 | 0,229 | 0,391 | 0,022435679 | PC | 0,779564958 | 1,28276674 | 0,585677749 |
| LITAF1 | 6,72996E-07 | -0,334462557 | 0,072 | 0,174 | 0,024622903 | PC | 0,793079525 | 1,260907599 | 0,413793103 |
| SRGN1 | 7,53364E-07 | -0,80200112 | 0,398 | 0,543 | 0,027563339 | PC | 0,573553067 | 1,743517833 | 0,732965009 |
| SON4 | 1,1032E-06 | -0,283942514 | 0,237 | 0,397 | 0,040362859 | PC | 0,821343428 | 1,217517504 | 0,59697733 |
| RPL343 | 1,11621E-06 | -0,545838838 | 0,752 | 0,821 | 0,040838606 | PC | 0,684993006 | 1,459868921 | 0,915956151 |
| GSTM21 | 1,5916E-06 | -0,309408131 | 0,048 | 0,13 | 0,058232037 | PC | 0,806972754 | 1,239199211 | 0,369230769 |
| RPL313 | 1,62063E-06 | -0,326529825 | 0,348 | 0,522 | 0,059293935 | PC | 0,797452324 | 1,253993461 | 0,666666667 |
| ITM2B3 | 1,93196E-06 | -0,328567487 | 0,342 | 0,538 | 0,070684607 | PC | 0,796326798 | 1,255765852 | 0,635687732 |
| NFKBIA3 | 2,25846E-06 | -0,647111367 | 0,327 | 0,484 | 0,0826304 | PC | 0,638557585 | 1,566029475 | 0,675619835 |

| Supplementary Table 4 - clusermarker | | | | | | | | | |
|---|---|---|---|---|---|---|---|---|---|
| | **p_val** | **avg_log2FC** | **pct.1** | **pct.2** | **p_val_adj** | **celltype** | **Foldchange_UP** | **Foldchange_DOWN** | **Ratio_pct1_pct2** |
| RGS12 | 2,51282E-06 | -0,571098977 | 0,19 | 0,326 | 0,091936397 | PC | 0,673103855 | 1,485654841 | 0,582822086 |
| SEC61G | 2,72123E-06 | -0,366556211 | 0,402 | 0,576 | 0,099561492 | PC | 0,775631763 | 1,289271595 | 0,697916667 |
| RPL232 | 3,05654E-06 | -0,38942535 | 0,239 | 0,386 | 0,111829567 | PC | 0,763433632 | 1,309871556 | 0,619170984 |
| RGS21 | 3,82578E-06 | -0,342669913 | 0,056 | 0,141 | 0,139973953 | PC | 0,78858058 | 1,268101226 | 0,397163121 |
| PTP4A3 | 4,75141E-06 | 0,507180038 | 0,103 | 0 | 0,173839807 | PC | 1,421269396 | 0,703596379 | #DIV/0! |
| LSR3 | 5,75342E-06 | -0,278570546 | 0,071 | 0,163 | 0,210500234 | PC | 0,824407454 | 1,21299243 | 0,435582822 |
| GTF2I1 | 6,44416E-06 | -0,441420741 | 0,061 | 0,147 | 0,235772321 | PC | 0,736409048 | 1,357940946 | 0,414965986 |
| PET1002 | 6,45286E-06 | -0,30305168 | 0,158 | 0,283 | 0,236090678 | PC | 0,810536085 | 1,233751364 | 0,558303887 |
| NT5DC2 | 7,09071E-06 | -0,314837973 | 0,044 | 0,12 | 0,259427724 | PC | 0,803941275 | 1,243871948 | 0,366666667 |
| VIM2 | 7,6021E-06 | -0,686289584 | 0,542 | 0,685 | 0,278137994 | PC | 0,621450082 | 1,609139703 | 0,791240876 |
| RPS15A2 | 7,7914E-06 | -0,434002977 | 0,752 | 0,815 | 0,285064067 | PC | 0,740205121 | 1,350976872 | 0,922699387 |
| RPL35A4 | 7,89435E-06 | -0,423850062 | 0,693 | 0,793 | 0,288830605 | PC | 0,745432662 | 1,341502796 | 0,873896595 |
| EEF1A12 | 7,9793E-06 | -0,539815715 | 0,879 | 0,913 | 0,291938753 | PC | 0,687858768 | 1,453786804 | 0,962760131 |
| TXN3 | 9,76114E-06 | -0,551104082 | 0,205 | 0,337 | 0,357130713 | PC | 0,682497619 | 1,465206577 | 0,608308605 |
| TMSB102 | 1,1255E-05 | -0,367464952 | 0,525 | 0,658 | 0,411788266 | PC | 0,775143353 | 1,290083952 | 0,79787234 |
| AQP33 | 1,17358E-05 | -0,524513569 | 0,092 | 0,19 | 0,429376694 | PC | 0,695193468 | 1,438448499 | 0,484210526 |
| HINT14 | 1,3128E-05 | -0,485993779 | 0,33 | 0,467 | 0,480313431 | PC | 0,714005069 | 1,400550282 | 0,706638116 |
| MAN1A2 | 1,345E-05 | -0,278936882 | 0,111 | 0,217 | 0,492095527 | PC | 0,824198143 | 1,213300477 | 0,511520737 |
| UBC3 | 1,49785E-05 | -0,450273886 | 0,505 | 0,668 | 0,548017594 | PC | 0,731903888 | 1,366299615 | 0,755988024 |
| HSP90B12 | 1,60419E-05 | -0,536035564 | 0,687 | 0,799 | 0,586924271 | PC | 0,68966346 | 1,449982576 | 0,859824781 |
| IGHV4-31 | 1,77189E-05 | -2,042688181 | 0,044 | 0,114 | 0,648282878 | PC | 0,242711071 | 4,120125208 | 0,385964912 |
| IGKV4-13 | 1,81584E-05 | 1,410391545 | 0,324 | 0,174 | 0,664361722 | PC | 2,658092933 | 0,37620957 | 1,862068966 |
| SEC11C2 | 2,11623E-05 | -0,364552967 | 0,578 | 0,75 | 0,774264831 | PC | 0,776709509 | 1,287482629 | 0,770666667 |
| RPS253 | 3,01932E-05 | -0,369592044 | 0,631 | 0,734 | 1 | PC | 0,774001333 | 1,291987439 | 0,859673025 |
| HSPA84 | 3,33358E-05 | -0,565216674 | 0,352 | 0,467 | 1 | PC | 0,675853904 | 1,479609711 | 0,753747323 |
| SLAMF71 | 3,73692E-05 | -0,275385769 | 0,272 | 0,413 | 1 | PC | 0,82622936 | 1,210317678 | 0,658595642 |
| RAPGEF21 | 3,74738E-05 | -0,256481495 | 0,048 | 0,12 | 1 | PC | 0,837127053 | 1,194561801 | 0,4 |
| TMCO12 | 3,83344E-05 | -0,250460822 | 0,12 | 0,223 | 1 | PC | 0,840627861 | 1,18958703 | 0,538116592 |
| GAS61 | 3,95174E-05 | 0,425504379 | 0,195 | 0,071 | 1 | PC | 1,34304196 | 0,744578375 | 2,746478873 |
| RTN42 | 3,9657E-05 | -0,348835868 | 0,067 | 0,147 | 1 | PC | 0,785217446 | 1,27353258 | 0,455782313 |
| CYBA4 | 4,07175E-05 | 0,403401087 | 0,809 | 0,793 | 1 | PC | 1,322622259 | 0,756073772 | 1,020176545 |

| Supplementary Table 4 - clusermarker | | | | | | | | | |
|---|---|---|---|---|---|---|---|---|---|
| | **p_val** | **avg_log2FC** | **pct.1** | **pct.2** | **p_val_adj** | **celltype** | **Foldchange_UP** | **Foldchange_DOWN** | **Ratio_pct1_pct2** |
| CSDE11 | 4,37039E-05 | -0,353483424 | 0,208 | 0,337 | 1 | PC | 0,782691985 | 1,277641805 | 0,617210682 |
| IGHV3-64 | 4,61043E-05 | 0,376076348 | 0,055 | 0,13 | 1 | PC | 1,297807445 | 0,770530331 | 0,423076923 |
| CTSS3 | 4,6146E-05 | -0,409703715 | 0,237 | 0,359 | 1 | PC | 0,752777955 | 1,328412971 | 0,660167131 |
| FTL2 | 4,81988E-05 | 0,486210283 | 0,857 | 0,826 | 1 | PC | 1,400760476 | 0,713897927 | 1,037530266 |
| TNFRSF17 | 4,89281E-05 | -0,396490347 | 0,323 | 0,462 | 1 | PC | 0,759704174 | 1,316301837 | 0,699134199 |
| RPL18A3 | 5,07376E-05 | 0,531062692 | 0,765 | 0,745 | 1 | PC | 1,444993188 | 0,692044785 | 1,026845638 |
| IGKV3-113 | 5,66487E-05 | 0,578976497 | 0,245 | 0,109 | 1 | PC | 1,493789121 | 0,669438534 | 2,247706422 |
| B2M2 | 7,0346E-05 | 0,319475013 | 0,971 | 0,967 | 1 | PC | 1,247876372 | 0,801361435 | 1,004136505 |
| ATP5ME3 | 7,40019E-05 | -0,347193029 | 0,133 | 0,239 | 1 | PC | 0,786112106 | 1,272083196 | 0,556485356 |
| CCNH1 | 7,40323E-05 | -0,385719569 | 0,108 | 0,201 | 1 | PC | 0,765397149 | 1,306511269 | 0,537313433 |
| EAF2 | 7,42876E-05 | -0,381151478 | 0,128 | 0,228 | 1 | PC | 0,767824512 | 1,302380928 | 0,561403509 |
| GLO1 | 7,79672E-05 | -0,284702799 | 0,061 | 0,136 | 1 | PC | 0,820910702 | 1,218159292 | 0,448529412 |
| RPL27A3 | 8,13543E-05 | -0,396299891 | 0,325 | 0,457 | 1 | PC | 0,759804472 | 1,316128078 | 0,711159737 |
| EIF4B3 | 9,21785E-05 | -0,364636872 | 0,213 | 0,332 | 1 | PC | 0,776664337 | 1,287557509 | 0,641566265 |
| BCAS22 | 9,32613E-05 | -0,350524445 | 0,065 | 0,141 | 1 | PC | 0,78429894 | 1,275024037 | 0,460992908 |
| DDX183 | 0,000110344 | -0,27306738 | 0,121 | 0,217 | 1 | PC | 0,827558165 | 1,208374278 | 0,557603687 |
| IGLV6-57 | 0,000111582 | 5,196059428 | 0,279 | 0,152 | 1 | PC | 36,65808273 | 0,027279114 | 1,835526316 |
| IGLV2-142 | 0,000113929 | 0,456579613 | 0,262 | 0,13 | 1 | PC | 1,372284503 | 0,728711866 | 2,015384615 |
| RPS143 | 0,000135261 | -0,439610857 | 0,803 | 0,821 | 1 | PC | 0,737333465 | 1,356238455 | 0,978075518 |
| RPL393 | 0,000136057 | -0,398993148 | 0,687 | 0,761 | 1 | PC | 0,758387374 | 1,318587352 | 0,902759527 |
| JUND2 | 0,000140239 | -0,362587397 | 0,305 | 0,429 | 1 | PC | 0,777768442 | 1,285729719 | 0,710955711 |
| RPL361 | 0,000142909 | -0,287184429 | 0,576 | 0,707 | 1 | PC | 0,819499839 | 1,220256493 | 0,814710042 |
| HDLBP | 0,000144814 | -0,305978842 | 0,285 | 0,413 | 1 | PC | 0,808893211 | 1,23625713 | 0,690072639 |
| HLA-DPA11 | 0,000148105 | 0,587450379 | 0,239 | 0,12 | 1 | PC | 1,502588931 | 0,665518013 | 1,991666667 |
| S100A103 | 0,000159335 | -0,414445604 | 0,152 | 0,261 | 1 | PC | 0,750307766 | 1,332786418 | 0,582375479 |
| CFLAR2 | 0,00016327 | -0,470127374 | 0,202 | 0,315 | 1 | PC | 0,721900859 | 1,385231764 | 0,641269841 |
| NSRP12 | 0,000165088 | -0,310025857 | 0,043 | 0,103 | 1 | PC | 0,806627302 | 1,239729919 | 0,417475728 |
| HLA-DRB11 | 0,000172322 | 0,617585729 | 0,11 | 0,022 | 1 | PC | 1,534305456 | 0,651760701 | 5 |
| EIF1AX3 | 0,000189146 | -0,271324156 | 0,118 | 0,207 | 1 | PC | 0,828558716 | 1,206915068 | 0,570048309 |
| RPL36AL2 | 0,000200713 | -0,338658617 | 0,5 | 0,636 | 1 | PC | 0,790776215 | 1,264580271 | 0,786163522 |
| RPS213 | 0,000217019 | -0,344745486 | 0,522 | 0,614 | 1 | PC | 0,787446882 | 1,269926928 | 0,850162866 |

| Supplementary Table 4 - clusermarker | | | | | | | | | |
|---|---|---|---|---|---|---|---|---|---|
| | **p_val** | **avg_log2FC** | **pct.1** | **pct.2** | **p_val_adj** | **celltype** | **Foldchange_UP** | **Foldchange_DOWN** | **Ratio_pct1_pct2** |
| RPL374 | 0,000218159 | -0,414544811 | 0,619 | 0,712 | 1 | PC | 0,750256173 | 1,332878071 | 0,869382022 |
| RPL271 | 0,000238381 | -0,341673239 | 0,412 | 0,543 | 1 | PC | 0,789125553 | 1,267225471 | 0,758747698 |
| SNHG61 | 0,000260159 | -0,426411367 | 0,128 | 0,217 | 1 | PC | 0,744110423 | 1,343886564 | 0,589861751 |
| IGKC4 | 0,000261275 | 0,700115126 | 0,751 | 0,647 | 1 | PC | 1,624634433 | 0,615523086 | 1,160741886 |
| RPL212 | 0,000266196 | -0,396818661 | 0,674 | 0,777 | 1 | PC | 0,759531308 | 1,316601422 | 0,867438867 |
| PLAAT43 | 0,000269599 | -0,260855121 | 0,065 | 0,136 | 1 | PC | 0,834593089 | 1,19818869 | 0,477941176 |
| HMGB12 | 0,000269741 | -0,262295099 | 0,291 | 0,418 | 1 | PC | 0,833760483 | 1,19938522 | 0,696172249 |
| RPS32 | 0,000275992 | -0,299624424 | 0,697 | 0,772 | 1 | PC | 0,812463877 | 1,230823952 | 0,902849741 |
| AC007384.11 | 0,000281424 | -0,339815374 | 0,082 | 0,158 | 1 | PC | 0,790142422 | 1,265594622 | 0,518987342 |
| HSP90AB13 | 0,000283278 | -0,566932351 | 0,354 | 0,457 | 1 | PC | 0,675050646 | 1,481370334 | 0,774617068 |
| BTG13 | 0,000304408 | -0,281472849 | 0,219 | 0,332 | 1 | PC | 0,822750641 | 1,21543509 | 0,659638554 |
| IGKV1-9 | 0,00031499 | 0,614678968 | 0,131 | 0,038 | 1 | PC | 1,531217229 | 0,653075201 | 3,447368421 |
| OS91 | 0,000329235 | -0,326009477 | 0,272 | 0,391 | 1 | PC | 0,79774 | 1,253541255 | 0,695652174 |
| EZR2 | 0,000334824 | -0,748118164 | 0,252 | 0,348 | 1 | PC | 0,595379658 | 1,679600548 | 0,724137931 |
| RPS243 | 0,000335148 | -0,307984878 | 0,585 | 0,717 | 1 | PC | 0,807769244 | 1,237977315 | 0,815899582 |
| RPS27A1 | 0,000354391 | -0,351841774 | 0,764 | 0,81 | 1 | PC | 0,783583121 | 1,276188797 | 0,943209877 |
| IGLV2-231 | 0,000354492 | 0,762618091 | 0,152 | 0,054 | 1 | PC | 1,69656663 | 0,589425715 | 2,814814815 |
| RPS62 | 0,000383729 | -0,328259604 | 0,667 | 0,745 | 1 | PC | 0,796496759 | 1,25549789 | 0,895302013 |
| ERLEC1 | 0,000398209 | -0,362393039 | 0,332 | 0,446 | 1 | PC | 0,777873229 | 1,285556519 | 0,744394619 |
| PRPF4B3 | 0,000402047 | -0,256594047 | 0,052 | 0,114 | 1 | PC | 0,837061746 | 1,194654999 | 0,456140351 |
| IGHV3-23 | 0,00041149 | 0,7032225 | 0,253 | 0,13 | 1 | PC | 1,62813745 | 0,614198758 | 1,946153846 |
| TAGLN22 | 0,00043033 | -0,258267293 | 0,238 | 0,353 | 1 | PC | 0,83609148 | 1,196041371 | 0,674220963 |
| ASH1L3 | 0,000438318 | -0,382969711 | 0,09 | 0,168 | 1 | PC | 0,766857429 | 1,304023358 | 0,535714286 |
| SEMA4A | 0,000439811 | -0,266093724 | 0,045 | 0,103 | 1 | PC | 0,831568074 | 1,202547369 | 0,436893204 |
| SELENOS1 | 0,000440904 | -0,372207066 | 0,328 | 0,44 | 1 | PC | 0,772599653 | 1,294331413 | 0,745454545 |
| SRSF52 | 0,000446531 | -0,250122982 | 0,132 | 0,223 | 1 | PC | 0,840824736 | 1,189308493 | 0,591928251 |
| CEP3501 | 0,000472656 | -0,446608455 | 0,092 | 0,168 | 1 | PC | 0,733765788 | 1,362832686 | 0,547619048 |
| ASAH12 | 0,000492783 | -0,299868719 | 0,12 | 0,207 | 1 | PC | 0,812326313 | 1,231032388 | 0,579710145 |
| UBXN42 | 0,000514895 | -0,345574252 | 0,233 | 0,342 | 1 | PC | 0,786994658 | 1,270656656 | 0,68128655 |
| RTF2 | 0,000526481 | -0,284404247 | 0,065 | 0,13 | 1 | PC | 0,821080599 | 1,217907232 | 0,5 |
| DUSP12 | 0,000546674 | -0,513197525 | 0,272 | 0,386 | 1 | PC | 0,700667787 | 1,427209897 | 0,704663212 |

| Supplementary Table 4 - clusermarker | | | | | | | | | |
|---|---|---|---|---|---|---|---|---|---|
| | **p_val** | **avg_log2FC** | **pct.1** | **pct.2** | **p_val_adj** | **celltype** | **Foldchange_UP** | **Foldchange_DOWN** | **Ratio_pct1_pct2** |
| SYAP12 | 0,000557793 | -0,269522946 | 0,064 | 0,13 | 1 | PC | 0,829593821 | 1,205409171 | 0,492307692 |
| H3-3B1 | 0,000572696 | -0,258572168 | 0,383 | 0,527 | 1 | PC | 0,835914813 | 1,196294149 | 0,726755218 |
| MBD22 | 0,000654706 | -0,389399485 | 0,091 | 0,163 | 1 | PC | 0,76344732 | 1,309848072 | 0,558282209 |
| MGP3 | 0,00072309 | 0,754752304 | 0,106 | 0,027 | 1 | PC | 1,687341868 | 0,592648128 | 3,925925926 |
| AL627171.23 | 0,000758971 | -0,316840168 | 0,081 | 0,152 | 1 | PC | 0,802826327 | 1,245599412 | 0,532894737 |
| VDAC22 | 0,000918915 | -0,302039287 | 0,142 | 0,228 | 1 | PC | 0,811105067 | 1,232885899 | 0,622807018 |
| MTCO2P121 | 0,000944992 | 0,358806157 | 0,831 | 0,679 | 1 | PC | 1,282364291 | 0,779809612 | 1,223858616 |
| RPL7A1 | 0,000967394 | -0,284165716 | 0,755 | 0,837 | 1 | PC | 0,821216366 | 1,217705883 | 0,902031063 |
| ENSA2 | 0,000997901 | -0,343293592 | 0,074 | 0,141 | 1 | PC | 0,788239749 | 1,268649546 | 0,524822695 |
| EGR13 | 0,001048692 | -0,449206645 | 0,118 | 0,201 | 1 | PC | 0,732445518 | 1,365289261 | 0,587064677 |
| ERN1 | 0,001091699 | -0,423032843 | 0,131 | 0,217 | 1 | PC | 0,745855034 | 1,340743113 | 0,603686636 |
| NR4A23 | 0,001159713 | -0,328080039 | 0,079 | 0,147 | 1 | PC | 0,796595901 | 1,255341634 | 0,537414966 |
| ALG131 | 0,001175825 | -0,274276543 | 0,053 | 0,109 | 1 | PC | 0,826864856 | 1,209387475 | 0,486238532 |
| IGHGP | 0,00123479 | 0,429874865 | 0,107 | 0,033 | 1 | PC | 1,347116727 | 0,74232617 | 3,242424242 |
| UBA521 | 0,00129848 | -0,38323203 | 0,46 | 0,571 | 1 | PC | 0,766718008 | 1,304260484 | 0,805604203 |
| U2SURP1 | 0,001385671 | -0,434381317 | 0,076 | 0,141 | 1 | PC | 0,740011032 | 1,351331206 | 0,539007092 |
| SQSTM13 | 0,001406187 | -0,511872272 | 0,278 | 0,375 | 1 | PC | 0,701311713 | 1,425899471 | 0,741333333 |
| PDIA63 | 0,001495659 | -0,256026746 | 0,405 | 0,527 | 1 | PC | 0,837390963 | 1,194185326 | 0,768500949 |
| UFM12 | 0,0015921 | -0,254254559 | 0,208 | 0,304 | 1 | PC | 0,838420235 | 1,192719306 | 0,684210526 |
| SF3B53 | 0,001680836 | -0,293927928 | 0,163 | 0,25 | 1 | PC | 0,815678241 | 1,225973613 | 0,652 |
| KMT2A2 | 0,001697949 | -0,347523338 | 0,095 | 0,163 | 1 | PC | 0,785932143 | 1,272374477 | 0,582822086 |
| CCNL13 | 0,001706639 | -0,386516452 | 0,175 | 0,261 | 1 | PC | 0,764974492 | 1,30723313 | 0,670498084 |
| HP1BP31 | 0,001759638 | -0,280152461 | 0,107 | 0,179 | 1 | PC | 0,823503987 | 1,214323205 | 0,597765363 |
| NCL4 | 0,001802491 | -0,344968014 | 0,262 | 0,364 | 1 | PC | 0,787325432 | 1,270122822 | 0,71978022 |
| MTRNR2L121 | 0,001832347 | 0,448112399 | 0,674 | 0,603 | 1 | PC | 1,364254118 | 0,733001269 | 1,11774461 |
| DDX241 | 0,002009024 | -0,359056647 | 0,155 | 0,239 | 1 | PC | 0,779674228 | 1,282586962 | 0,648535565 |
| NOP532 | 0,002203808 | -0,323180597 | 0,188 | 0,277 | 1 | PC | 0,799305767 | 1,251085682 | 0,678700361 |
| VPS37B2 | 0,002224433 | -0,424269472 | 0,07 | 0,13 | 1 | PC | 0,745215986 | 1,341892846 | 0,538461538 |
| DBI2 | 0,002332374 | -0,343583341 | 0,146 | 0,223 | 1 | PC | 0,788081456 | 1,268904365 | 0,65470852 |
| RPL13A2 | 0,002357612 | -0,336510692 | 0,499 | 0,62 | 1 | PC | 0,791954421 | 1,262698929 | 0,80483871 |
| IGHA2 | 0,002631952 | -0,87490608 | 0,246 | 0,337 | 1 | PC | 0,545289364 | 1,833888696 | 0,729970326 |

| Supplementary Table 4 - clusermarker | | | | | | | | | |
|---|---|---|---|---|---|---|---|---|---|
| | **p_val** | **avg_log2FC** | **pct.1** | **pct.2** | **p_val_adj** | **celltype** | **Foldchange_UP** | **Foldchange_DOWN** | **Ratio_pct1_pct2** |
| RPL73 | 0,002769369 | -0,270959993 | 0,383 | 0,484 | 1 | PC | 0,828767886 | 1,206610459 | 0,791322314 |
| TMEM593 | 0,002902783 | 0,582960917 | 0,584 | 0,538 | 1 | PC | 1,497920355 | 0,667592237 | 1,085501859 |
| DDX172 | 0,002952863 | -0,28183608 | 0,186 | 0,272 | 1 | PC | 0,822543521 | 1,215741142 | 0,683823529 |
| ZFAS13 | 0,003610253 | -0,260301304 | 0,126 | 0,201 | 1 | PC | 0,834913531 | 1,197728822 | 0,626865672 |
| FOSB4 | 0,003857792 | -0,415643921 | 0,304 | 0,402 | 1 | PC | 0,749684811 | 1,333893904 | 0,756218905 |
| ARF12 | 0,003879225 | -0,293412675 | 0,243 | 0,332 | 1 | PC | 0,815969609 | 1,225535839 | 0,731927711 |
| XBP11 | 0,003974815 | 0,627243807 | 0,626 | 0,641 | 1 | PC | 1,544611275 | 0,647412081 | 0,976599064 |
| MESD1 | 0,00465917 | -0,25914064 | 0,121 | 0,19 | 1 | PC | 0,835585498 | 1,196765623 | 0,636842105 |
| CLK11 | 0,004714781 | -0,279346355 | 0,147 | 0,223 | 1 | PC | 0,823964248 | 1,213644891 | 0,659192825 |
| EIF4E1 | 0,004875424 | -0,29591223 | 0,074 | 0,13 | 1 | PC | 0,814557117 | 1,227660994 | 0,569230769 |
| SLC1A5 | 0,004961714 | -0,268901065 | 0,106 | 0,174 | 1 | PC | 0,829951499 | 1,204889685 | 0,609195402 |
| IRF21 | 0,005068731 | -0,314825199 | 0,058 | 0,109 | 1 | PC | 0,803948393 | 1,243860935 | 0,532110092 |
| RPL112 | 0,005215194 | -0,295574422 | 0,725 | 0,793 | 1 | PC | 0,814747869 | 1,22737357 | 0,914249685 |
| BHLHE401 | 0,005371991 | -0,267371401 | 0,057 | 0,109 | 1 | PC | 0,830831948 | 1,203612839 | 0,52293578 |
| DDX55 | 0,005588819 | -0,481983708 | 0,322 | 0,413 | 1 | PC | 0,715992456 | 1,396662761 | 0,779661017 |
| UBL52 | 0,005589931 | -0,258358807 | 0,276 | 0,375 | 1 | PC | 0,836038446 | 1,196117242 | 0,736 |
| PABPN1 | 0,005590485 | -0,336559897 | 0,096 | 0,158 | 1 | PC | 0,791927412 | 1,262741996 | 0,607594937 |
| RPL263 | 0,005605336 | -0,257554559 | 0,684 | 0,75 | 1 | PC | 0,836504636 | 1,195450637 | 0,912 |
| CLPTM1L | 0,00569666 | -0,385230689 | 0,219 | 0,304 | 1 | PC | 0,765656559 | 1,306068613 | 0,720394737 |
| CD553 | 0,005745105 | -0,421471433 | 0,198 | 0,283 | 1 | PC | 0,7466627 | 1,33929283 | 0,699646643 |
| COX7C2 | 0,005874824 | -0,250454027 | 0,362 | 0,462 | 1 | PC | 0,84063182 | 1,189581427 | 0,783549784 |
| IGHV4-61 | 0,005954234 | -2,126872811 | 0,093 | 0,152 | 1 | PC | 0,228953605 | 4,367697108 | 0,611842105 |
| PHB3 | 0,006177312 | -0,309770376 | 0,081 | 0,136 | 1 | PC | 0,806770157 | 1,2395104 | 0,595588235 |
| RPL413 | 0,006290912 | -0,273313018 | 0,922 | 0,918 | 1 | PC | 0,827417274 | 1,208580038 | 1,004357298 |
| IGLV2-11 | 0,006528146 | 1,323930071 | 0,135 | 0,065 | 1 | PC | 2,503471569 | 0,399445319 | 2,076923077 |
| IGKV1-39 | 0,00673933 | -1,437989785 | 0,07 | 0,12 | 1 | PC | 0,369081215 | 2,70943077 | 0,583333333 |
| PABPC14 | 0,007228603 | -0,291918609 | 0,328 | 0,429 | 1 | PC | 0,816815071 | 1,224267322 | 0,764568765 |
| DNAJC83 | 0,007389278 | -0,258252949 | 0,076 | 0,13 | 1 | PC | 0,836099793 | 1,196029479 | 0,584615385 |
| IGKV3-15 | 0,007816312 | -0,268050827 | 0,181 | 0,098 | 1 | PC | 0,830440767 | 1,204179804 | 1,846938776 |
| TMSB4X2 | 0,007843204 | 0,440702862 | 0,59 | 0,527 | 1 | PC | 1,357265408 | 0,736775574 | 1,119544592 |
| JSRP1 | 0,007979891 | -0,40391121 | 0,138 | 0,207 | 1 | PC | 0,755806479 | 1,323090008 | 0,666666667 |

| Supplementary Table 4 - clusermarker | | | | | | | | | |
|---|---|---|---|---|---|---|---|---|---|
| | **p_val** | **avg_log2FC** | **pct.1** | **pct.2** | **p_val_adj** | **celltype** | **Foldchange_UP** | **Foldchange_DOWN** | **Ratio_pct1_pct2** |
| PELI1 | 0,008075896 | -0,395606308 | 0,163 | 0,234 | 1 | PC | 0,76016984 | 1,315495495 | 0,696581197 |
| IGHM | 0,008305131 | -0,480148618 | 0,079 | 0,136 | 1 | PC | 0,716903769 | 1,394887352 | 0,580882353 |
| EIF3D1 | 0,008383685 | -0,278574601 | 0,085 | 0,141 | 1 | PC | 0,824405137 | 1,212995838 | 0,602836879 |
| HNRNPA14 | 0,008518282 | -0,294328552 | 0,273 | 0,353 | 1 | PC | 0,815451765 | 1,226314103 | 0,773371105 |
| MARCKS3 | 0,008728957 | -0,318175366 | 0,149 | 0,217 | 1 | PC | 0,802083663 | 1,246752734 | 0,686635945 |
| USP481 | 0,008880876 | 0,257697078 | 0,121 | 0,06 | 1 | PC | 1,195568737 | 0,836422005 | 2,016666667 |
| ZFP363 | 0,008909336 | -0,550953497 | 0,326 | 0,408 | 1 | PC | 0,68256886 | 1,46505365 | 0,799019608 |
| UBB4 | 0,009026044 | -0,295715869 | 0,414 | 0,489 | 1 | PC | 0,814667992 | 1,227493912 | 0,846625767 |
| C11orf583 | 0,009518354 | -0,443210271 | 0,138 | 0,201 | 1 | PC | 0,735496168 | 1,359626391 | 0,686567164 |
| ARPC5L3 | 0,0095567 | 0,435902707 | 0,135 | 0,071 | 1 | PC | 1,352757 | 0,739231067 | 1,901408451 |
| GLA | 0,010037387 | -0,271688716 | 0,13 | 0,196 | 1 | PC | 0,828349372 | 1,207220086 | 0,663265306 |
| LTC4S | 0,010488648 | 0,328278621 | 0,1 | 0,043 | 1 | PC | 1,255514439 | 0,79648626 | 2,325581395 |
| MT-ND34 | 0,011224016 | -0,366809917 | 0,65 | 0,652 | 1 | PC | 0,775495376 | 1,289498341 | 0,996932515 |
| HSH2D | 0,011959008 | -0,275627867 | 0,112 | 0,168 | 1 | PC | 0,826090722 | 1,210520799 | 0,666666667 |
| PNISR1 | 0,012087081 | -0,425110051 | 0,103 | 0,163 | 1 | PC | 0,744781916 | 1,34267492 | 0,63190184 |
| UPF21 | 0,012096012 | -0,263706193 | 0,062 | 0,109 | 1 | PC | 0,832945384 | 1,200558907 | 0,568807339 |
| ATP6AP22 | 0,012099377 | 0,309700734 | 0,107 | 0,049 | 1 | PC | 1,239450567 | 0,806809103 | 2,183673469 |
| MT-ATP64 | 0,013635409 | 0,337045866 | 0,718 | 0,636 | 1 | PC | 1,26316742 | 0,791660697 | 1,128930818 |
| MT-CO31 | 0,014360409 | 0,267515989 | 0,812 | 0,658 | 1 | PC | 1,203733472 | 0,830748686 | 1,234042553 |
| TPD52 | 0,014422937 | -0,332582156 | 0,189 | 0,255 | 1 | PC | 0,794113895 | 1,25926521 | 0,741176471 |
| IGLV1-44 | 0,014555995 | 0,76676074 | 0,133 | 0,071 | 1 | PC | 1,701445263 | 0,587735628 | 1,873239437 |
| P4HB1 | 0,014590523 | -0,26458936 | 0,349 | 0,446 | 1 | PC | 0,83243564 | 1,201294072 | 0,782511211 |
| RPL310 | 0,014747236 | -0,261232677 | 0,758 | 0,799 | 1 | PC | 0,834374703 | 1,198502299 | 0,948685857 |
| HLA-DMA2 | 0,015955649 | 0,285951119 | 0,152 | 0,087 | 1 | PC | 1,219213784 | 0,8202007 | 1,747126437 |
| IGHV3-7 | 0,017510398 | 0,518454029 | 0,18 | 0,114 | 1 | PC | 1,432419465 | 0,698119527 | 1,578947368 |
| IFI302 | 0,017570357 | 0,30144739 | 0,154 | 0,092 | 1 | PC | 1,232380184 | 0,811437909 | 1,673913043 |
| VPS361 | 0,018439876 | -0,286100249 | 0,061 | 0,103 | 1 | PC | 0,820115921 | 1,21933982 | 0,59223301 |
| NIBAN11 | 0,018578334 | -0,273354752 | 0,064 | 0,109 | 1 | PC | 0,827393339 | 1,208615 | 0,587155963 |
| RPS182 | 0,0190044 | -0,274420009 | 0,813 | 0,87 | 1 | PC | 0,826782634 | 1,209507746 | 0,934482759 |
| NUCKS11 | 0,020222374 | -0,252862748 | 0,096 | 0,147 | 1 | PC | 0,839229474 | 1,191569208 | 0,653061224 |
| IGHV3-21 | 0,020363412 | 0,496592505 | 0,132 | 0,071 | 1 | PC | 1,410877279 | 0,708778868 | 1,85915493 |

| Supplementary Table 4 - clusermarker | | | | | | | | | |
|---|---|---|---|---|---|---|---|---|---|
| | **p_val** | **avg_log2FC** | **pct.1** | **pct.2** | **p_val_adj** | **celltype** | **Foldchange_UP** | **Foldchange_DOWN** | **Ratio_pct1_pct2** |
| WIPF11 | 0,021056584 | -0,287043777 | 0,065 | 0,109 | 1 | PC | 0,819579738 | 1,220137534 | 0,596330275 |
| UQCR103 | 0,021077982 | 0,326010855 | 0,174 | 0,109 | 1 | PC | 1,253542452 | 0,797739238 | 1,596330275 |
| RPS3A2 | 0,021174331 | -0,256275788 | 0,715 | 0,766 | 1 | PC | 0,837246423 | 1,194391487 | 0,933420366 |
| IGLC1 | 0,022341168 | 0,440255492 | 0,275 | 0,353 | 1 | PC | 1,356844594 | 0,737004079 | 0,779036827 |
| RGCC2 | 0,022930249 | -0,432244497 | 0,146 | 0,207 | 1 | PC | 0,741107897 | 1,34933119 | 0,70531401 |
| EMC42 | 0,025411807 | -0,279173404 | 0,15 | 0,212 | 1 | PC | 0,824063031 | 1,213499407 | 0,70754717 |
| TOP1 | 0,025502124 | -0,278037635 | 0,157 | 0,217 | 1 | PC | 0,824712034 | 1,21254445 | 0,723502304 |
| GNG52 | 0,027885645 | 0,319838419 | 0,268 | 0,201 | 1 | PC | 1,248190744 | 0,801159602 | 1,333333333 |
| TRAM1 | 0,028423634 | -0,308058542 | 0,337 | 0,413 | 1 | PC | 0,807728001 | 1,238040527 | 0,81598063 |
| PRDX42 | 0,031937417 | 0,475622758 | 0,558 | 0,543 | 1 | PC | 1,390518326 | 0,71915629 | 1,027624309 |
| KDELR21 | 0,034145591 | 0,44294086 | 0,264 | 0,201 | 1 | PC | 1,359372517 | 0,735633528 | 1,313432836 |
| SRSF94 | 0,036027705 | 0,367590293 | 0,265 | 0,201 | 1 | PC | 1,29019604 | 0,775076011 | 1,31840796 |
| MRPS361 | 0,036401402 | -0,386433246 | 0,074 | 0,114 | 1 | PC | 0,765018613 | 1,307157739 | 0,649122807 |
| BRD22 | 0,036702386 | -0,294586025 | 0,229 | 0,288 | 1 | PC | 0,815306248 | 1,226532978 | 0,795138889 |
| CDKN1A3 | 0,037652382 | -0,421741231 | 0,253 | 0,315 | 1 | PC | 0,74652308 | 1,339543314 | 0,803174603 |
| COPE3 | 0,037672589 | 0,306298957 | 0,307 | 0,245 | 1 | PC | 1,23653147 | 0,808713749 | 1,253061224 |
| TOMM53 | 0,038438663 | -0,264793314 | 0,115 | 0,163 | 1 | PC | 0,832317967 | 1,201463911 | 0,705521472 |
| CAST2 | 0,038830872 | -0,253371395 | 0,095 | 0,141 | 1 | PC | 0,838933641 | 1,191989391 | 0,673758865 |
| BOD1L12 | 0,038899809 | -0,340179367 | 0,078 | 0,12 | 1 | PC | 0,789943094 | 1,265913973 | 0,65 |
| MRPS243 | 0,042414319 | 0,286382575 | 0,231 | 0,168 | 1 | PC | 1,21957846 | 0,819955446 | 1,375 |
| RPS53 | 0,04751783 | 0,289933689 | 0,705 | 0,69 | 1 | PC | 1,222584082 | 0,817939653 | 1,02173913 |
| ATP2A32 | 0,047993325 | -0,325827041 | 0,095 | 0,136 | 1 | PC | 0,797840884 | 1,253382748 | 0,698529412 |
| PCBP21 | 0,050267806 | -0,290457872 | 0,123 | 0,168 | 1 | PC | 0,81764252 | 1,223028372 | 0,732142857 |
| IGLV2-8 | 0,051157306 | -1,062408553 | 0,146 | 0,087 | 1 | PC | 0,478831991 | 2,088415184 | 1,67816092 |
| NDUFB103 | 0,053706326 | 0,33092247 | 0,226 | 0,174 | 1 | PC | 1,257817376 | 0,795027974 | 1,298850575 |
| CSNK2B3 | 0,054823561 | 0,286775455 | 0,201 | 0,147 | 1 | PC | 1,219910625 | 0,819732183 | 1,367346939 |
| GLS3 | 0,059168602 | -0,259719152 | 0,117 | 0,163 | 1 | PC | 0,835250501 | 1,197245615 | 0,717791411 |
| NFE2L22 | 0,05970181 | -0,253079197 | 0,091 | 0,13 | 1 | PC | 0,839103573 | 1,191747994 | 0,7 |
| PCED1B-AS1 | 0,06217633 | 0,382231897 | 0,198 | 0,147 | 1 | PC | 1,303356632 | 0,767249711 | 1,346938776 |
| NPC21 | 0,062445608 | 0,628865562 | 0,381 | 0,353 | 1 | PC | 1,546348571 | 0,646684725 | 1,079320113 |
| GUSB | 0,069209517 | 0,259840172 | 0,145 | 0,098 | 1 | PC | 1,19734605 | 0,835180439 | 1,479591837 |

| Supplementary Table 4 - clusermarker | | | | | | | | | |
|---|---|---|---|---|---|---|---|---|---|
| | **p_val** | **avg_log2FC** | **pct.1** | **pct.2** | **p_val_adj** | **celltype** | **Foldchange_UP** | **Foldchange_DOWN** | **Ratio_pct1_pct2** |
| NDUFA7 | 0,074632984 | 0,255982164 | 0,151 | 0,103 | 1 | PC | 1,194148424 | 0,837416841 | 1,466019417 |
| PPA11 | 0,075416454 | 0,306879322 | 0,151 | 0,103 | 1 | PC | 1,237029 | 0,808388486 | 1,466019417 |
| LARP1B | 0,077889987 | -0,28880046 | 0,078 | 0,114 | 1 | PC | 0,818582392 | 1,221624127 | 0,684210526 |
| ATP6V0B3 | 0,07927557 | 0,507806256 | 0,232 | 0,19 | 1 | PC | 1,421886448 | 0,703291041 | 1,221052632 |
| PSME23 | 0,081071876 | 0,250181125 | 0,239 | 0,19 | 1 | PC | 1,189356425 | 0,84079085 | 1,257894737 |
| DANCR3 | 0,08738296 | 0,336739137 | 0,226 | 0,179 | 1 | PC | 1,262898888 | 0,791829029 | 1,262569832 |
| CLINT1 | 0,090255292 | -0,350000888 | 0,15 | 0,196 | 1 | PC | 0,784583615 | 1,274561412 | 0,765306122 |
| NAA381 | 0,092748419 | 0,290459077 | 0,139 | 0,098 | 1 | PC | 1,223029393 | 0,817641837 | 1,418367347 |
| SMDT14 | 0,098211772 | 0,350385963 | 0,3 | 0,245 | 1 | PC | 1,274901655 | 0,784374227 | 1,224489796 |
| MALAT12 | 0,102097397 | -0,385131527 | 0,529 | 0,587 | 1 | PC | 0,765709187 | 1,305978845 | 0,901192504 |
| SEC31A | 0,102949453 | -0,30213037 | 0,151 | 0,19 | 1 | PC | 0,811053861 | 1,232963738 | 0,794736842 |
| ORAI21 | 0,10388665 | -0,257250691 | 0,081 | 0,114 | 1 | PC | 0,836680844 | 1,195198871 | 0,710526316 |
| MT2A1 | 0,104350367 | 0,313036997 | 0,096 | 0,136 | 1 | PC | 1,24232014 | 0,804945495 | 0,705882353 |
| CTSZ2 | 0,10461801 | 0,286776921 | 0,191 | 0,141 | 1 | PC | 1,219911865 | 0,81973135 | 1,354609929 |
| SAT22 | 0,107872638 | -0,270109707 | 0,116 | 0,152 | 1 | PC | 0,829256484 | 1,205899525 | 0,763157895 |
| PIM32 | 0,109542829 | -0,250216179 | 0,121 | 0,158 | 1 | PC | 0,840770422 | 1,189385324 | 0,765822785 |
| RAB5IF2 | 0,111346212 | 0,275226249 | 0,118 | 0,082 | 1 | PC | 1,21018386 | 0,826320721 | 1,43902439 |
| IGHV1-2 | 0,114956507 | 0,420335871 | 0,103 | 0,065 | 1 | PC | 1,338239072 | 0,747250638 | 1,584615385 |
| PLTP1 | 0,118515666 | 0,28188265 | 0,125 | 0,087 | 1 | PC | 1,215780387 | 0,82251697 | 1,436781609 |
| IGHV1-46 | 0,119208589 | 1,661082947 | 0,107 | 0,071 | 1 | PC | 3,162538291 | 0,316201705 | 1,507042254 |
| MZB1 | 0,122011681 | 0,25320047 | 0,769 | 0,81 | 1 | PC | 1,191848177 | 0,839033041 | 0,949382716 |
| ANKHD1 | 0,126924864 | -0,28878514 | 0,107 | 0,141 | 1 | PC | 0,818591085 | 1,221611154 | 0,758865248 |
| LMO42 | 0,126958702 | -0,290537989 | 0,103 | 0,136 | 1 | PC | 0,817597115 | 1,223096292 | 0,757352941 |
| GPX44 | 0,141461302 | 0,542602495 | 0,287 | 0,255 | 1 | PC | 1,456597723 | 0,686531349 | 1,125490196 |
| NOPCHAP1 | 0,144509128 | -0,367392249 | 0,08 | 0,109 | 1 | PC | 0,775182416 | 1,290018942 | 0,733944954 |
| QTRT11 | 0,14458176 | -0,286861895 | 0,089 | 0,12 | 1 | PC | 0,81968307 | 1,219983719 | 0,741666667 |
| ATP5F1B2 | 0,144919631 | 0,325144562 | 0,241 | 0,201 | 1 | PC | 1,252789965 | 0,798218399 | 1,199004975 |
| KDELR13 | 0,151778437 | -0,293203884 | 0,358 | 0,402 | 1 | PC | 0,816087707 | 1,225358489 | 0,890547264 |
| FTH13 | 0,152688005 | 0,333875555 | 0,589 | 0,62 | 1 | PC | 1,260394667 | 0,793402278 | 0,95 |
| NEAT13 | 0,172694082 | -0,328919155 | 0,335 | 0,413 | 1 | PC | 0,796132711 | 1,256071992 | 0,811138015 |
| IGFLR11 | 0,176861644 | 0,429521285 | 0,241 | 0,207 | 1 | PC | 1,346786612 | 0,742508124 | 1,164251208 |

| Supplementary Table 4 - clusermarker | | | | | | | | | |
|---|---|---|---|---|---|---|---|---|---|
| | **p_val** | **avg_log2FC** | **pct.1** | **pct.2** | **p_val_adj** | **celltype** | **Foldchange_UP** | **Foldchange_DOWN** | **Ratio_pct1_pct2** |
| COX6A13 | 0,189889939 | 0,402740388 | 0,363 | 0,342 | 1 | PC | 1,322016687 | 0,756420104 | 1,061403509 |
| HINT22 | 0,19830704 | 0,252699303 | 0,127 | 0,098 | 1 | PC | 1,191434221 | 0,839324557 | 1,295918367 |
| HYOU1 | 0,199205121 | -0,369953273 | 0,152 | 0,185 | 1 | PC | 0,773807559 | 1,292310974 | 0,821621622 |
| ARL5A1 | 0,205272552 | -0,26443911 | 0,08 | 0,103 | 1 | PC | 0,832522339 | 1,201168969 | 0,776699029 |
| ARPC22 | 0,217192966 | 0,270274055 | 0,237 | 0,201 | 1 | PC | 1,206036905 | 0,829162023 | 1,179104478 |
| NDUFA113 | 0,231357388 | 0,293269417 | 0,317 | 0,283 | 1 | PC | 1,225414152 | 0,816050638 | 1,120141343 |
| TYMP4 | 0,234750012 | 0,267455284 | 0,143 | 0,114 | 1 | PC | 1,203682823 | 0,830783643 | 1,254385965 |
| IGHV3-30 | 0,23627978 | -0,287215314 | 0,168 | 0,13 | 1 | PC | 0,819482296 | 1,220282617 | 1,292307692 |
| YME1L11 | 0,247803283 | -0,296412957 | 0,091 | 0,114 | 1 | PC | 0,814274452 | 1,228087162 | 0,798245614 |
| JTB4 | 0,2509871 | 0,258485372 | 0,276 | 0,239 | 1 | PC | 1,19622218 | 0,835965105 | 1,154811715 |
| LIME12 | 0,267631297 | 0,313035711 | 0,302 | 0,272 | 1 | PC | 1,242319033 | 0,804946212 | 1,110294118 |
| C12orf573 | 0,276097765 | 0,250679176 | 0,283 | 0,255 | 1 | PC | 1,189767089 | 0,84050064 | 1,109803922 |
| CUTA3 | 0,278360612 | 0,345475608 | 0,395 | 0,38 | 1 | PC | 1,270569777 | 0,787048471 | 1,039473684 |
| FKBP111 | 0,279692705 | 0,272002724 | 0,635 | 0,663 | 1 | PC | 1,207482872 | 0,828169097 | 0,957767722 |
| TPI12 | 0,347305694 | 0,422059544 | 0,206 | 0,185 | 1 | PC | 1,3398389 | 0,746358387 | 1,113513514 |
| IGKV3D-20 | 0,422116543 | 1,468823639 | 0,114 | 0,098 | 1 | PC | 2,767961045 | 0,361276761 | 1,163265306 |
| SELENOM3 | 0,446717423 | 0,352641048 | 0,378 | 0,386 | 1 | PC | 1,276896019 | 0,783149125 | 0,979274611 |
| ACTB4 | 0,455695022 | 0,421428103 | 0,476 | 0,505 | 1 | PC | 1,339252606 | 0,746685126 | 0,942574257 |
| HLA-E2 | 0,479558555 | 0,336118907 | 0,464 | 0,489 | 1 | PC | 1,262356071 | 0,792169518 | 0,948875256 |
| PLD31 | 0,495293617 | -0,271989959 | 0,169 | 0,185 | 1 | PC | 0,828176425 | 1,207472188 | 0,913513514 |
| XRN11 | 0,500954774 | -0,264404342 | 0,108 | 0,12 | 1 | PC | 0,832542403 | 1,201140022 | 0,9 |
| COX7B3 | 0,530852009 | 0,289757669 | 0,196 | 0,179 | 1 | PC | 1,222434927 | 0,818039454 | 1,094972067 |
| CHCHD24 | 0,657132175 | 0,379918995 | 0,374 | 0,397 | 1 | PC | 1,301268789 | 0,768480738 | 0,942065491 |
| CHCHD104 | 0,660179472 | 0,286170875 | 0,242 | 0,239 | 1 | PC | 1,219399513 | 0,820075774 | 1,012552301 |
| ATP5F1D4 | 0,672868943 | 0,326982835 | 0,307 | 0,315 | 1 | PC | 1,25438728 | 0,797201962 | 0,974603175 |
| EEF23 | 0,689640967 | 0,262857212 | 0,52 | 0,56 | 1 | PC | 1,199852623 | 0,833435691 | 0,928571429 |
| IGLC3 | 0,698911083 | -0,471040341 | 0,26 | 0,245 | 1 | PC | 0,72144417 | 1,386108644 | 1,06122449 |
| PRDX12 | 0,761657692 | 0,264064097 | 0,403 | 0,418 | 1 | PC | 1,200856779 | 0,832738773 | 0,964114833 |
| IGHV3-11 | 0,774118443 | -0,645401898 | 0,124 | 0,114 | 1 | PC | 0,639314669 | 1,564174965 | 1,087719298 |
| MT-ND62 | 0,808603349 | 0,342362985 | 0,365 | 0,38 | 1 | PC | 1,267831471 | 0,788748365 | 0,960526316 |
| SH3BGRL32 | 0,858848429 | 0,289942098 | 0,248 | 0,261 | 1 | PC | 1,222591208 | 0,817934886 | 0,950191571 |

| Supplementary Table 4 - clusermarker | | | | | | | | | |
|---|---|---|---|---|---|---|---|---|---|
| | **p_val** | **avg_log2FC** | **pct.1** | **pct.2** | **p_val_adj** | **celltype** | **Foldchange_UP** | **Foldchange_DOWN** | **Ratio_pct1_pct2** |
| IGHV4-39 | 0,877315012 | 1,076463697 | 0,143 | 0,141 | 1 | PC | 2,108860547 | 0,474189724 | 1,014184397 |
| IGHV4-59 | 0,926214764 | -0,343926535 | 0,14 | 0,136 | 1 | PC | 0,787894006 | 1,269206254 | 1,029411765 |
| IGLV1-40 | 0,937166275 | -0,437153432 | 0,119 | 0,12 | 1 | PC | 0,738590478 | 1,353930263 | 0,991666667 |
| CRELD21 | 0,967269214 | 0,276257843 | 0,294 | 0,315 | 1 | PC | 1,211049507 | 0,825730075 | 0,933333333 |
| PRR45 | 1,04427E-44 | -4,780361567 | 0,182 | 0,883 | 3,82066E-40 | Mac | 0,036388804 | 27,48098036 | 0,206115515 |
| PRB35 | 9,96001E-28 | -3,813095335 | 0,024 | 0,398 | 3,64407E-23 | Mac | 0,071144925 | 14,05581629 | 0,060301508 |
| SLPI5 | 8,60344E-18 | -2,780479128 | 0,157 | 0,563 | 3,14774E-13 | Mac | 0,145543354 | 6,870804951 | 0,278863233 |
| SCGB3A25 | 2,86081E-16 | -3,500053653 | 0,07 | 0,379 | 1,04668E-11 | Mac | 0,088385061 | 11,31412925 | 0,18469657 |
| PRH25 | 7,75146E-16 | -1,483913245 | 0,005 | 0,194 | 2,83603E-11 | Mac | 0,357517745 | 2,797063963 | 0,025773196 |
| C6orf585 | 1,98373E-15 | -1,979937271 | 0,011 | 0,214 | 7,25788E-11 | Mac | 0,253500892 | 3,944759295 | 0,051401869 |
| SCGB3A14 | 7,19534E-15 | -1,237724669 | 0,287 | 0,699 | 2,63256E-10 | Mac | 0,424040901 | 2,358263079 | 0,410586552 |
| RPS295 | 1,06912E-14 | -1,248496291 | 0,721 | 0,922 | 3,9116E-10 | Mac | 0,420886666 | 2,37593652 | 0,781995662 |
| STATH5 | 2,62071E-14 | -1,521173213 | 0,003 | 0,165 | 9,58838E-10 | Mac | 0,348402477 | 2,870243655 | 0,018181818 |
| TYMP5 | 1,23563E-13 | 1,550492747 | 0,705 | 0,379 | 4,52081E-09 | Mac | 2,929171668 | 0,341393443 | 1,860158311 |
| PIP4 | 1,54736E-12 | -1,524289556 | 0,003 | 0,146 | 5,66133E-08 | Mac | 0,347650711 | 2,876450325 | 0,020547945 |
| ZG16B5 | 5,06218E-12 | -2,623178906 | 0,022 | 0,204 | 1,8521E-07 | Mac | 0,162309696 | 6,161061369 | 0,107843137 |
| HLA-DQB11 | 9,33014E-12 | -1,377785187 | 0,683 | 0,825 | 3,41362E-07 | Mac | 0,384809098 | 2,598691158 | 0,827878788 |
| RPS264 | 3,19384E-11 | -0,958510992 | 0,824 | 0,932 | 1,16853E-06 | Mac | 0,514587746 | 1,943303173 | 0,884120172 |
| MTRNR2L85 | 6,55434E-11 | -1,867862104 | 0,136 | 0,417 | 2,39804E-06 | Mac | 0,273979128 | 3,649913072 | 0,326139089 |
| PFN14 | 6,76529E-11 | 0,846116941 | 0,946 | 0,874 | 2,47522E-06 | Mac | 1,797655961 | 0,556279968 | 1,082379863 |
| SERPINE1 | 8,77307E-11 | -0,97574562 | 0,014 | 0,165 | 3,2098E-06 | Mac | 0,508476985 | 1,966657352 | 0,084848485 |
| RPS274 | 9,87946E-10 | -0,621795896 | 0,967 | 0,971 | 3,6146E-05 | Mac | 0,649861463 | 1,538789506 | 0,995880536 |
| MSMB5 | 1,22816E-09 | -1,237007389 | 0,024 | 0,184 | 4,49347E-05 | Mac | 0,424251778 | 2,357090888 | 0,130434783 |
| WFDC22 | 1,18709E-08 | -1,053737319 | 0,03 | 0,184 | 0,000434322 | Mac | 0,481718649 | 2,075900534 | 0,163043478 |
| C32 | 1,83164E-08 | -0,735602147 | 0,084 | 0,301 | 0,000670141 | Mac | 0,60056731 | 1,665092295 | 0,279069767 |
| MT-ATP85 | 2,5015E-08 | -0,884420964 | 0,512 | 0,728 | 0,000915224 | Mac | 0,541704896 | 1,846023558 | 0,703296703 |
| EEF1A13 | 3,02897E-08 | -0,435271744 | 0,989 | 0,981 | 0,001108209 | Mac | 0,73955444 | 1,352165501 | 1,008154944 |
| IGKC5 | 8,00285E-08 | 2,328486407 | 0,347 | 0,068 | 0,002928004 | Mac | 5,022781118 | 0,199092888 | 5,102941176 |
| FTH14 | 8,4164E-08 | 0,745217837 | 0,995 | 0,971 | 0,003079308 | Mac | 1,676227349 | 0,596577786 | 1,024716787 |
| MIF4 | 8,85702E-08 | 1,005513313 | 0,683 | 0,456 | 0,003240518 | Mac | 2,007657697 | 0,498092878 | 1,497807018 |
| HSPA1A5 | 2,88458E-07 | -2,489028302 | 0,293 | 0,505 | 0,010553822 | Mac | 0,178126207 | 5,613997035 | 0,58019802 |

| Supplementary Table 4 - clusermarker | | | | | | | | | |
|---|---|---|---|---|---|---|---|---|---|
| | **p_val** | **avg_log2FC** | **pct.1** | **pct.2** | **p_val_adj** | **celltype** | **Foldchange_UP** | **Foldchange_DOWN** | **Ratio_pct1_pct2** |
| S100A104 | 3,58344E-07 | 0,875371685 | 0,753 | 0,544 | 0,013110744 | Mac | 1,834480647 | 0,545113409 | 1,384191176 |
| AREG3 | 5,99248E-07 | 1,802586341 | 0,556 | 0,311 | 0,021924695 | Mac | 3,488450449 | 0,286660228 | 1,78778135 |
| RPL314 | 8,44454E-07 | -0,701567261 | 0,74 | 0,854 | 0,030896024 | Mac | 0,614903847 | 1,626270521 | 0,866510539 |
| ATP6V0C3 | 8,93507E-07 | 0,844744283 | 0,748 | 0,553 | 0,032690749 | Mac | 1,795946387 | 0,556809494 | 1,352622061 |
| PGLS4 | 1,30253E-06 | 0,798830327 | 0,482 | 0,243 | 0,04765552 | Mac | 1,739690091 | 0,574815023 | 1,983539095 |
| MT-ND35 | 1,36093E-06 | -0,594640655 | 0,881 | 0,845 | 0,049792264 | Mac | 0,662209377 | 1,510096406 | 1,04260355 |
| OAZ13 | 1,42285E-06 | 0,570250218 | 0,916 | 0,806 | 0,052057901 | Mac | 1,484781065 | 0,673499968 | 1,136476427 |
| RPL344 | 1,93161E-06 | -0,455813064 | 0,962 | 0,961 | 0,070671745 | Mac | 0,729099156 | 1,371555559 | 1,001040583 |
| TMSB103 | 1,98434E-06 | 0,374924819 | 0,997 | 0,951 | 0,072600951 | Mac | 1,296771976 | 0,771145597 | 1,048370137 |
| AP2S12 | 2,17433E-06 | 0,958960024 | 0,55 | 0,35 | 0,079552045 | Mac | 1,943908111 | 0,514427608 | 1,571428571 |
| RPL84 | 2,26616E-06 | 0,508250292 | 0,949 | 0,854 | 0,082912136 | Mac | 1,422324147 | 0,703074613 | 1,111241218 |
| SLC23A2 | 3,24109E-06 | -0,421070333 | 0,016 | 0,117 | 0,118581723 | Mac | 0,746870317 | 1,338920529 | 0,136752137 |
| RPS152 | 3,42811E-06 | 0,445011392 | 0,962 | 0,913 | 0,125424404 | Mac | 1,361324866 | 0,734578516 | 1,053669222 |
| PLP23 | 3,45354E-06 | 1,041272306 | 0,287 | 0,078 | 0,126354603 | Mac | 2,058041831 | 0,485898773 | 3,679487179 |
| MYL61 | 3,79572E-06 | 0,581660547 | 0,889 | 0,825 | 0,138874049 | Mac | 1,496570816 | 0,668194241 | 1,077575758 |
| HSPA1B4 | 4,30182E-06 | -2,550742434 | 0,165 | 0,35 | 0,157390555 | Mac | 0,170667181 | 5,859357327 | 0,471428571 |
| SLC16A10 | 4,52182E-06 | 1,070728701 | 0,241 | 0,039 | 0,165439875 | Mac | 2,100494052 | 0,476078473 | 6,179487179 |
| LTF4 | 4,5756E-06 | -1,182407998 | 0,03 | 0,146 | 0,167407373 | Mac | 0,440615454 | 2,269552715 | 0,205479452 |
| EIF4EBP1 | 4,80327E-06 | 0,80538387 | 0,274 | 0,058 | 0,175737236 | Mac | 1,747610731 | 0,572209807 | 4,724137931 |
| SERF22 | 8,31726E-06 | 0,548407835 | 0,902 | 0,786 | 0,304303462 | Mac | 1,462470816 | 0,683774328 | 1,147582697 |
| HLA-B5 | 1,30013E-05 | 0,527470591 | 0,924 | 0,883 | 0,475676939 | Mac | 1,44139984 | 0,693770023 | 1,046432616 |
| GNG53 | 1,30303E-05 | 0,617777854 | 0,675 | 0,485 | 0,476740119 | Mac | 1,534509794 | 0,651673912 | 1,391752577 |
| RPL375 | 1,32471E-05 | -0,545207856 | 0,962 | 0,981 | 0,484671707 | Mac | 0,685292662 | 1,459230567 | 0,980632008 |
| GABARAP4 | 1,36771E-05 | 0,480381044 | 0,892 | 0,786 | 0,500403352 | Mac | 1,395112095 | 0,716788281 | 1,134860051 |
| CD207 | 2,03924E-05 | -1,281820796 | 0,035 | 0,146 | 0,746095288 | Mac | 0,411276118 | 2,431456524 | 0,239726027 |
| DDT3 | 2,34352E-05 | 0,852004577 | 0,236 | 0,058 | 0,8574229 | Mac | 1,805007181 | 0,554014416 | 4,068965517 |
| RPS63 | 2,59598E-05 | -0,375310816 | 0,943 | 0,961 | 0,949791892 | Mac | 0,770939303 | 1,297118977 | 0,981269511 |
| HLA-DRB53 | 2,74827E-05 | -0,672400074 | 0,751 | 0,825 | 1 | Mac | 0,62746197 | 1,593722086 | 0,91030303 |
| RPL18A4 | 3,25732E-05 | 0,421534014 | 0,962 | 0,932 | 1 | Mac | 1,339350926 | 0,746630312 | 1,032188841 |
| IGKV4-14 | 3,28401E-05 | 4,647506853 | 0,149 | 0 | 1 | Mac | 25,06334128 | 0,03989891 | #DIV/0! |
| IGLV3-213 | 3,28401E-05 | 2,194167785 | 0,149 | 0 | 1 | Mac | 4,576256079 | 0,21851924 | #DIV/0! |

| Supplementary Table 4 - clusermarker | | | | | | | | | |
|---|---|---|---|---|---|---|---|---|---|
| | **p_val** | **avg_log2FC** | **pct.1** | **pct.2** | **p_val_adj** | **celltype** | **Foldchange_UP** | **Foldchange_DOWN** | **Ratio_pct1_pct2** |
| BLOC1S12 | 3,3541E-05 | 0,818755019 | 0,482 | 0,291 | 1 | Mac | 1,763883184 | 0,566930968 | 1,656357388 |
| IGHA15 | 3,63785E-05 | 1,934141399 | 0,179 | 0,019 | 1 | Mac | 3,821506273 | 0,261676922 | 9,421052632 |
| CRIP13 | 3,71455E-05 | 0,940750726 | 0,607 | 0,388 | 1 | Mac | 1,919526831 | 0,52096172 | 1,56443299 |
| CALM12 | 4,37748E-05 | 0,637528909 | 0,699 | 0,524 | 1 | Mac | 1,555662291 | 0,642813036 | 1,333969466 |
| LPXN1 | 4,60596E-05 | -0,662904281 | 0,206 | 0,388 | 1 | Mac | 0,631605535 | 1,583266682 | 0,530927835 |
| CTSZ3 | 4,67042E-05 | 0,786922586 | 0,748 | 0,544 | 1 | Mac | 1,725390103 | 0,579579075 | 1,375 |
| CWC221 | 4,67649E-05 | -0,625195582 | 0,038 | 0,146 | 1 | Mac | 0,648331879 | 1,542419912 | 0,260273973 |
| IGKV3-114 | 4,74927E-05 | 2,997251754 | 0,144 | 0 | 1 | Mac | 7,984774993 | 0,125238344 | #DIV/0! |
| ACTB5 | 5,41694E-05 | 0,596691988 | 0,978 | 0,99 | 1 | Mac | 1,512245103 | 0,661268467 | 0,987878788 |
| PRKX | 5,59394E-05 | -0,528681308 | 0,065 | 0,194 | 1 | Mac | 0,693188051 | 1,442609979 | 0,335051546 |
| IGHG14 | 6,214E-05 | 1,552929629 | 0,157 | 0,01 | 1 | Mac | 2,934123565 | 0,340817276 | 15,7 |
| RCBTB2 | 6,29128E-05 | -0,493630502 | 0,049 | 0,165 | 1 | Mac | 0,710235559 | 1,407983573 | 0,296969697 |
| S100A64 | 6,57774E-05 | 0,535490198 | 0,843 | 0,641 | 1 | Mac | 1,449434558 | 0,689924215 | 1,315132605 |
| PTPRC1 | 7,28776E-05 | -0,723859129 | 0,531 | 0,709 | 1 | Mac | 0,60547566 | 1,651594053 | 0,748942172 |
| IGKV1-52 | 8,22307E-05 | 1,088088208 | 0,136 | 0 | 1 | Mac | 2,125921327 | 0,470384293 | #DIV/0! |
| JAG1 | 8,5064E-05 | -0,354328157 | 0,03 | 0,126 | 1 | Mac | 0,782233834 | 1,278390114 | 0,238095238 |
| IGFLR12 | 8,64137E-05 | 0,77012944 | 0,393 | 0,194 | 1 | Mac | 1,705422789 | 0,586364863 | 2,025773196 |
| RPS164 | 9,55393E-05 | 0,494073017 | 0,916 | 0,806 | 1 | Mac | 1,408415507 | 0,710017743 | 1,136476427 |
| MT-ND4L3 | 0,000100026 | -0,542982894 | 0,743 | 0,786 | 1 | Mac | 0,686350353 | 1,456981839 | 0,945292621 |
| BHLHE411 | 0,000115597 | -0,665884485 | 0,057 | 0,175 | 1 | Mac | 0,630302161 | 1,586540649 | 0,325714286 |
| PPP4C4 | 0,00012979 | 0,621882161 | 0,344 | 0,165 | 1 | Mac | 1,538881519 | 0,649822607 | 2,084848485 |
| IGLV2-143 | 0,000131118 | 3,831310661 | 0,146 | 0,01 | 1 | Mac | 14,23440871 | 0,070252303 | 14,6 |
| EPB41L21 | 0,00013617 | -0,69818824 | 0,1 | 0,243 | 1 | Mac | 0,616345738 | 1,622465993 | 0,411522634 |
| VIM3 | 0,000140959 | 0,416013095 | 0,932 | 0,845 | 1 | Mac | 1,33423528 | 0,749492998 | 1,10295858 |
| PSMA73 | 0,000153218 | 0,681885245 | 0,602 | 0,398 | 1 | Mac | 1,604234724 | 0,623350178 | 1,512562814 |
| CYBA5 | 0,000163796 | 0,490353288 | 0,892 | 0,845 | 1 | Mac | 1,40478884 | 0,711850758 | 1,055621302 |
| CKB | 0,000171453 | -0,434528338 | 0,057 | 0,175 | 1 | Mac | 0,739935623 | 1,351468924 | 0,325714286 |
| ALDH1A12 | 0,000179564 | -0,42788089 | 0,054 | 0,165 | 1 | Mac | 0,743352862 | 1,345256138 | 0,327272727 |
| RPL134 | 0,000182868 | 0,369528769 | 0,973 | 0,942 | 1 | Mac | 1,291930775 | 0,77403528 | 1,032908705 |
| AURKAIP13 | 0,000196388 | 0,626574129 | 0,344 | 0,165 | 1 | Mac | 1,543894455 | 0,64771267 | 2,084848485 |
| H2AZ23 | 0,000206548 | 0,756393586 | 0,431 | 0,252 | 1 | Mac | 1,689262565 | 0,591974286 | 1,71031746 |

| Supplementary Table 4 - clusermarker | | | | | | | | | |
|---|---|---|---|---|---|---|---|---|---|
| | **p_val** | **avg_log2FC** | **pct.1** | **pct.2** | **p_val_adj** | **celltype** | **Foldchange_UP** | **Foldchange_DOWN** | **Ratio_pct1_pct2** |
| CTSS4 | 0,000213096 | -0,705823545 | 0,661 | 0,757 | 1 | Mac | 0,613092412 | 1,631075479 | 0,87318362 |
| GSTM11 | 0,000220315 | -0,707468115 | 0,03 | 0,117 | 1 | Mac | 0,612393929 | 1,632935849 | 0,256410256 |
| IFITM34 | 0,000243962 | 0,70535684 | 0,553 | 0,379 | 1 | Mac | 1,630547919 | 0,613290777 | 1,459102902 |
| HLA-A1 | 0,000246934 | -0,500988391 | 0,829 | 0,874 | 1 | Mac | 0,706622508 | 1,415182773 | 0,948512586 |
| CHCHD25 | 0,000258546 | 0,520242885 | 0,691 | 0,553 | 1 | Mac | 1,434196682 | 0,697254437 | 1,24954792 |
| C3AR1 | 0,000264224 | -0,562357789 | 0,165 | 0,32 | 1 | Mac | 0,677194523 | 1,476680578 | 0,515625 |
| JTB5 | 0,000272615 | 0,733373727 | 0,358 | 0,194 | 1 | Mac | 1,66252234 | 0,601495677 | 1,845360825 |
| GSTP13 | 0,000295803 | 0,659552949 | 0,783 | 0,68 | 1 | Mac | 1,579593076 | 0,633074439 | 1,151470588 |
| ELOB3 | 0,00031779 | 0,552455747 | 0,604 | 0,456 | 1 | Mac | 1,466579976 | 0,681858484 | 1,324561404 |
| LGALS12 | 0,00034739 | 0,628837201 | 0,802 | 0,709 | 1 | Mac | 1,546318172 | 0,646697438 | 1,131170663 |
| COX4I14 | 0,000371605 | 0,510901056 | 0,829 | 0,709 | 1 | Mac | 1,424939885 | 0,701783992 | 1,169252468 |
| MAPRE2 | 0,000381449 | -0,325271401 | 0,098 | 0,233 | 1 | Mac | 0,798148224 | 1,252900113 | 0,420600858 |
| IL7R | 0,000430003 | 1,162018889 | 0,203 | 0,058 | 1 | Mac | 2,2377035 | 0,44688673 | 3,5 |
| TMEM176A | 0,000432751 | 1,143076121 | 0,32 | 0,155 | 1 | Mac | 2,208514219 | 0,4527931 | 2,064516129 |
| PTMA1 | 0,000448845 | 0,392185863 | 0,943 | 0,903 | 1 | Mac | 1,312380318 | 0,761974244 | 1,044296788 |
| GRINA2 | 0,000449859 | 0,87860838 | 0,42 | 0,262 | 1 | Mac | 1,838600935 | 0,543891815 | 1,603053435 |
| SIVA13 | 0,000492484 | 0,582910421 | 0,268 | 0,107 | 1 | Mac | 1,497867927 | 0,667615603 | 2,504672897 |
| IGLV1-401 | 0,000494668 | 2,75285128 | 0,108 | 0 | 1 | Mac | 6,740479761 | 0,148357392 | #DIV/0! |
| TNF1 | 0,000511715 | -1,751720469 | 0,141 | 0,272 | 1 | Mac | 0,296947446 | 3,367599258 | 0,518382353 |
| S100A112 | 0,000535937 | 0,5520837 | 0,875 | 0,835 | 1 | Mac | 1,466201818 | 0,682034347 | 1,047904192 |
| PLXND1 | 0,000542762 | 0,575980238 | 0,16 | 0,029 | 1 | Mac | 1,490689968 | 0,670830301 | 5,517241379 |
| JDP21 | 0,000563783 | -0,488244739 | 0,127 | 0,262 | 1 | Mac | 0,712891913 | 1,402737191 | 0,484732824 |
| C4orf48 | 0,000574772 | 0,711561061 | 0,333 | 0,165 | 1 | Mac | 1,637575089 | 0,610659021 | 2,018181818 |
| RPL394 | 0,000575926 | -0,357622083 | 0,959 | 0,913 | 1 | Mac | 0,780449894 | 1,281312238 | 1,050383352 |
| IGLV3-1 | 0,00059028 | 1,234314973 | 0,106 | 0 | 1 | Mac | 2,352696092 | 0,425044273 | #DIV/0! |
| HECA1 | 0,000592563 | -0,429677592 | 0,114 | 0,252 | 1 | Mac | 0,742427682 | 1,346932535 | 0,452380952 |
| RPLP13 | 0,000602174 | 0,271565983 | 0,995 | 0,981 | 1 | Mac | 1,20711739 | 0,828419844 | 1,014271152 |
| LSM43 | 0,000610571 | 0,373802426 | 0,271 | 0,107 | 1 | Mac | 1,295763501 | 0,771745769 | 2,53271028 |
| KLHL15 | 0,000631293 | -0,297452352 | 0,033 | 0,117 | 1 | Mac | 0,813688016 | 1,22897226 | 0,282051282 |
| HLA-DQA2 | 0,00065337 | -0,725562815 | 0,344 | 0,485 | 1 | Mac | 0,604761073 | 1,653545582 | 0,709278351 |
| PPDPF4 | 0,00065454 | 0,63871028 | 0,542 | 0,408 | 1 | Mac | 1,556936689 | 0,642286875 | 1,328431373 |

| Supplementary Table 4 - clusermarker | | | | | | | | | |
|---|---|---|---|---|---|---|---|---|---|
| | **p_val** | **avg_log2FC** | **pct.1** | **pct.2** | **p_val_adj** | **celltype** | **Foldchange_UP** | **Foldchange_DOWN** | **Ratio_pct1_pct2** |
| PRELID14 | 0,000682186 | 0,703567994 | 0,496 | 0,359 | 1 | Mac | 1,628527401 | 0,614051688 | 1,381615599 |
| CEBPD1 | 0,000685195 | 0,56880732 | 0,645 | 0,495 | 1 | Mac | 1,483296818 | 0,6741739 | 1,303030303 |
| SRI3 | 0,000697384 | 0,538504804 | 0,268 | 0,117 | 1 | Mac | 1,452466414 | 0,688484078 | 2,290598291 |
| ATP5F1D5 | 0,00070019 | 0,596282697 | 0,528 | 0,369 | 1 | Mac | 1,51181614 | 0,661456095 | 1,430894309 |
| FUOM1 | 0,000706507 | 0,616371988 | 0,209 | 0,068 | 1 | Mac | 1,533015186 | 0,652309259 | 3,073529412 |
| S100B1 | 0,000718445 | 1,287043722 | 0,184 | 0,049 | 1 | Mac | 2,440274973 | 0,409789885 | 3,755102041 |
| MRC1 | 0,00073351 | 0,577544401 | 0,279 | 0,117 | 1 | Mac | 1,492307043 | 0,670103384 | 2,384615385 |
| IFI303 | 0,000743815 | 0,775122791 | 0,829 | 0,699 | 1 | Mac | 1,711335701 | 0,584338888 | 1,185979971 |
| TIMP11 | 0,000744597 | 0,838416246 | 0,561 | 0,388 | 1 | Mac | 1,788086149 | 0,55925717 | 1,445876289 |
| CST7 | 0,000775894 | 1,051834864 | 0,236 | 0,087 | 1 | Mac | 2,073164887 | 0,482354301 | 2,712643678 |
| TUBA1B2 | 0,000791256 | 0,627551208 | 0,71 | 0,583 | 1 | Mac | 1,544940426 | 0,647274149 | 1,217838765 |
| GLIPR2 | 0,00084282 | 0,676316842 | 0,233 | 0,087 | 1 | Mac | 1,598054756 | 0,625760786 | 2,67816092 |
| TUG1 | 0,000876223 | -0,259278376 | 0,079 | 0,194 | 1 | Mac | 0,835505728 | 1,196879885 | 0,407216495 |
| TLR7 | 0,000881247 | -0,263091727 | 0,054 | 0,155 | 1 | Mac | 0,833300224 | 1,200047679 | 0,348387097 |
| SPOP1 | 0,000891385 | -0,368702622 | 0,114 | 0,243 | 1 | Mac | 0,774478652 | 1,291191174 | 0,469135802 |
| FNIP2 | 0,000894545 | -0,369367146 | 0,171 | 0,32 | 1 | Mac | 0,774121999 | 1,291786051 | 0,534375 |
| JCHAIN3 | 0,000895623 | 1,835682416 | 0,184 | 0,049 | 1 | Mac | 3,569402057 | 0,280158969 | 3,755102041 |
| MTDH3 | 0,00093083 | 0,60773996 | 0,564 | 0,427 | 1 | Mac | 1,523870134 | 0,656223898 | 1,320843091 |
| EDF13 | 0,000979874 | 0,575374812 | 0,534 | 0,369 | 1 | Mac | 1,490064531 | 0,671111874 | 1,447154472 |
| TBXAS1 | 0,000980447 | -0,439466263 | 0,295 | 0,466 | 1 | Mac | 0,737407368 | 1,356102533 | 0,63304721 |
| CLEC12A | 0,000993732 | -0,588869402 | 0,07 | 0,175 | 1 | Mac | 0,664863737 | 1,504067592 | 0,4 |
| A2M1 | 0,001002699 | -0,78707515 | 0,26 | 0,408 | 1 | Mac | 0,579517788 | 1,725572572 | 0,637254902 |
| SDF2L12 | 0,001021503 | 0,572159432 | 0,271 | 0,117 | 1 | Mac | 1,486747276 | 0,67260927 | 2,316239316 |
| TENT5A1 | 0,001058899 | -0,394771629 | 0,089 | 0,204 | 1 | Mac | 0,760609768 | 1,314734628 | 0,43627451 |
| LARS12 | 0,001061692 | -0,306925761 | 0,111 | 0,243 | 1 | Mac | 0,808362465 | 1,23706882 | 0,456790123 |
| HSPA6 | 0,001081819 | -1,142202965 | 0,095 | 0,214 | 1 | Mac | 0,453067225 | 2,207177974 | 0,443925234 |
| STRN32 | 0,001083257 | -0,412728347 | 0,03 | 0,107 | 1 | Mac | 0,751201398 | 1,331200931 | 0,280373832 |
| BAG32 | 0,001113374 | -0,654133978 | 0,079 | 0,184 | 1 | Mac | 0,635456829 | 1,573671025 | 0,429347826 |
| FAM102B1 | 0,001125685 | -0,362999464 | 0,057 | 0,155 | 1 | Mac | 0,777546325 | 1,286097006 | 0,367741935 |
| SHOC23 | 0,001162198 | -0,605876875 | 0,16 | 0,291 | 1 | Mac | 0,657071888 | 1,52190349 | 0,549828179 |
| TSPO4 | 0,001204141 | 0,574954846 | 0,607 | 0,456 | 1 | Mac | 1,48963084 | 0,671307262 | 1,331140351 |

| Supplementary Table 4 - clusermarker | | | | | | | | | |
|---|---|---|---|---|---|---|---|---|---|
| | **p_val** | **avg_log2FC** | **pct.1** | **pct.2** | **p_val_adj** | **celltype** | **Foldchange_UP** | **Foldchange_DOWN** | **Ratio_pct1_pct2** |
| HSPH14 | 0,001259942 | -0,912198634 | 0,257 | 0,388 | 1 | Mac | 0,531374671 | 1,881911303 | 0,662371134 |
| HSP90B13 | 0,001263664 | 0,528664445 | 0,599 | 0,466 | 1 | Mac | 1,442593116 | 0,693196154 | 1,285407725 |
| FCER1A | 0,001266899 | -1,208856741 | 0,157 | 0,282 | 1 | Mac | 0,432611301 | 2,311543866 | 0,556737589 |
| BBLN4 | 0,001302437 | 0,745908695 | 0,339 | 0,184 | 1 | Mac | 1,677030231 | 0,596292173 | 1,842391304 |
| CFL12 | 0,00131548 | 0,322148393 | 0,897 | 0,835 | 1 | Mac | 1,250190889 | 0,79987785 | 1,074251497 |
| COX8A4 | 0,001336454 | 0,512398355 | 0,626 | 0,544 | 1 | Mac | 1,426419524 | 0,701056024 | 1,150735294 |
| ITGAL | 0,00135116 | -0,365098285 | 0,054 | 0,146 | 1 | Mac | 0,776415979 | 1,287969371 | 0,369863014 |
| ILF33 | 0,001378594 | 0,418991751 | 0,271 | 0,117 | 1 | Mac | 1,336992851 | 0,747947156 | 2,316239316 |
| ZNF2072 | 0,001386851 | -0,435167015 | 0,263 | 0,417 | 1 | Mac | 0,739608128 | 1,352067348 | 0,630695444 |
| CNBP3 | 0,001416152 | 0,371436293 | 0,509 | 0,34 | 1 | Mac | 1,293640089 | 0,773012531 | 1,497058824 |
| MARCKS4 | 0,001441282 | -0,614933824 | 0,439 | 0,583 | 1 | Mac | 0,652959844 | 1,531487747 | 0,753001715 |
| AL118516.11 | 0,001540534 | -0,672233968 | 0,1 | 0,214 | 1 | Mac | 0,627534217 | 1,593538603 | 0,46728972 |
| EIF3E2 | 0,001543486 | -0,588134901 | 0,428 | 0,553 | 1 | Mac | 0,665202317 | 1,503302039 | 0,773960217 |
| USPL1 | 0,001558734 | -0,387600686 | 0,054 | 0,146 | 1 | Mac | 0,764399804 | 1,308215929 | 0,369863014 |
| NDUFS83 | 0,001581092 | 0,598808774 | 0,341 | 0,194 | 1 | Mac | 1,514465564 | 0,660298936 | 1,757731959 |
| TXNL4A3 | 0,001606906 | 0,682407627 | 0,314 | 0,175 | 1 | Mac | 1,604815702 | 0,623124511 | 1,794285714 |
| TUBA1C1 | 0,00163834 | 0,819895094 | 0,379 | 0,252 | 1 | Mac | 1,765277626 | 0,566483133 | 1,503968254 |
| AC020656.12 | 0,001746421 | -0,563026671 | 0,087 | 0,194 | 1 | Mac | 0,676880626 | 1,477365375 | 0,448453608 |
| PABPC15 | 0,001747456 | -0,430020208 | 0,799 | 0,806 | 1 | Mac | 0,742251389 | 1,347252448 | 0,991315136 |
| RPL384 | 0,001774639 | -0,625196142 | 0,696 | 0,748 | 1 | Mac | 0,648331627 | 1,542420511 | 0,930481283 |
| AL136987.1 | 0,001900219 | -0,302509093 | 0,033 | 0,107 | 1 | Mac | 0,810840978 | 1,233287447 | 0,308411215 |
| GUK13 | 0,00192549 | 0,785043553 | 0,442 | 0,32 | 1 | Mac | 1,723144338 | 0,580334438 | 1,38125 |
| CD36 | 0,001964461 | -0,537185396 | 0,092 | 0,204 | 1 | Mac | 0,689114015 | 1,451138677 | 0,450980392 |
| PYCARD3 | 0,001979791 | 0,505442504 | 0,558 | 0,408 | 1 | Mac | 1,419558697 | 0,704444277 | 1,367647059 |
| CD993 | 0,00198607 | 0,54684566 | 0,363 | 0,214 | 1 | Mac | 1,460888083 | 0,684515132 | 1,696261682 |
| SH3BGRL33 | 0,002000351 | 0,3799697 | 0,848 | 0,738 | 1 | Mac | 1,301314525 | 0,76845373 | 1,149051491 |
| AMD13 | 0,00202912 | -0,545541566 | 0,179 | 0,311 | 1 | Mac | 0,685134165 | 1,459568141 | 0,575562701 |
| ZFAND2A | 0,002030623 | -0,470065685 | 0,095 | 0,204 | 1 | Mac | 0,721931728 | 1,385172533 | 0,465686275 |
| CRTC3 | 0,002033598 | 0,378301836 | 0,106 | 0,01 | 1 | Mac | 1,299810976 | 0,769342634 | 10,6 |
| MPZL11 | 0,002039717 | -0,331903931 | 0,054 | 0,146 | 1 | Mac | 0,794487303 | 1,258673356 | 0,369863014 |
| P4HB2 | 0,002060942 | 0,644299097 | 0,417 | 0,291 | 1 | Mac | 1,562979761 | 0,63980355 | 1,432989691 |

| Supplementary Table 4 - clusermarker | | | | | | | | | |
|---|---|---|---|---|---|---|---|---|---|
| | **p_val** | **avg_log2FC** | **pct.1** | **pct.2** | **p_val_adj** | **celltype** | **Foldchange_UP** | **Foldchange_DOWN** | **Ratio_pct1_pct2** |
| IFI27L22 | 0,00210319 | 0,705148315 | 0,225 | 0,097 | 1 | Mac | 1,630312259 | 0,613379427 | 2,319587629 |
| HDAC9 | 0,002120151 | -0,287573087 | 0,081 | 0,184 | 1 | Mac | 0,819279098 | 1,220585272 | 0,440217391 |
| RPL35A5 | 0,002144715 | -0,414958563 | 0,946 | 0,922 | 1 | Mac | 0,750041037 | 1,333260382 | 1,026030369 |
| MRPS121 | 0,002199703 | 0,43071453 | 0,119 | 0,019 | 1 | Mac | 1,347900993 | 0,741894253 | 6,263157895 |
| SSBP12 | 0,002229718 | 0,679907177 | 0,306 | 0,165 | 1 | Mac | 1,602036677 | 0,624205435 | 1,854545455 |
| GAS62 | 0,002246024 | 0,722304135 | 0,176 | 0,058 | 1 | Mac | 1,649814859 | 0,606128618 | 3,034482759 |
| CSNK2B4 | 0,002267673 | 0,535282698 | 0,491 | 0,359 | 1 | Mac | 1,449226104 | 0,690023452 | 1,367688022 |
| CALR3 | 0,002325361 | 0,59225854 | 0,615 | 0,495 | 1 | Mac | 1,507605057 | 0,663303692 | 1,242424242 |
| TNFRSF1A | 0,002411502 | -0,769249574 | 0,16 | 0,282 | 1 | Mac | 0,586722582 | 1,704383008 | 0,567375887 |
| FAM174C3 | 0,00241215 | 0,376411643 | 0,203 | 0,078 | 1 | Mac | 1,298109102 | 0,770351274 | 2,602564103 |
| CIAO2B3 | 0,00242739 | 0,524676652 | 0,36 | 0,223 | 1 | Mac | 1,438611111 | 0,695114887 | 1,614349776 |
| KLHL241 | 0,002431901 | -0,384964035 | 0,076 | 0,175 | 1 | Mac | 0,765798089 | 1,305827234 | 0,434285714 |
| RCC2 | 0,002465999 | -0,359419505 | 0,136 | 0,262 | 1 | Mac | 0,779478154 | 1,282909592 | 0,519083969 |
| GBP41 | 0,002546296 | -0,559556758 | 0,084 | 0,184 | 1 | Mac | 0,678510592 | 1,473816345 | 0,456521739 |
| CSF3R | 0,002585213 | -0,577736508 | 0,157 | 0,291 | 1 | Mac | 0,670014161 | 1,492505769 | 0,5395189 |
| CREM1 | 0,002604674 | 0,353686301 | 0,537 | 0,33 | 1 | Mac | 1,277821484 | 0,782581928 | 1,627272727 |
| SOCS31 | 0,00266976 | 0,666693961 | 0,407 | 0,262 | 1 | Mac | 1,587431084 | 0,629948607 | 1,553435115 |
| CD814 | 0,002702625 | -0,578165742 | 0,436 | 0,563 | 1 | Mac | 0,669814846 | 1,492949889 | 0,774422735 |
| SLC7A7 | 0,002705056 | -0,567256797 | 0,187 | 0,32 | 1 | Mac | 0,674898851 | 1,481703515 | 0,584375 |
| TNFSF102 | 0,002731007 | -0,424989071 | 0,087 | 0,194 | 1 | Mac | 0,744844374 | 1,342562332 | 0,448453608 |
| PSMB33 | 0,002756094 | 0,436493383 | 0,48 | 0,34 | 1 | Mac | 1,353310967 | 0,738928468 | 1,411764706 |
| FPR1 | 0,002761667 | 0,720777452 | 0,279 | 0,146 | 1 | Mac | 1,648069922 | 0,606770372 | 1,910958904 |
| RSRC23 | 0,002816045 | -0,578611692 | 0,176 | 0,301 | 1 | Mac | 0,669607832 | 1,493411444 | 0,584717608 |
| RPS54 | 0,002904534 | 0,40407722 | 0,894 | 0,864 | 1 | Mac | 1,323242264 | 0,755719513 | 1,034722222 |
| HLA-DRA3 | 0,002955654 | -0,391433639 | 0,976 | 0,971 | 1 | Mac | 0,762371642 | 1,311696219 | 1,005149331 |
| HPGDS | 0,003068878 | -0,377141209 | 0,051 | 0,136 | 1 | Mac | 0,769961808 | 1,298765717 | 0,375 |
| ST6GAL12 | 0,003264293 | -0,25885923 | 0,095 | 0,204 | 1 | Mac | 0,835748503 | 1,196532206 | 0,465686275 |
| CLTB3 | 0,003266243 | 0,308861462 | 0,257 | 0,126 | 1 | Mac | 1,23872974 | 0,807278592 | 2,03968254 |
| ARPC1B4 | 0,003283792 | 0,292056222 | 0,762 | 0,592 | 1 | Mac | 1,224384106 | 0,816737162 | 1,287162162 |
| UBE2M1 | 0,00331325 | -0,369337259 | 0,089 | 0,194 | 1 | Mac | 0,774138036 | 1,29175929 | 0,458762887 |
| KDELR22 | 0,003315675 | 0,727252826 | 0,257 | 0,136 | 1 | Mac | 1,655483723 | 0,604053055 | 1,889705882 |

| Supplementary Table 4 - clusermarker | | | | | | | | | |
|---|---|---|---|---|---|---|---|---|---|
| | **p_val** | **avg_log2FC** | **pct.1** | **pct.2** | **p_val_adj** | **celltype** | **Foldchange_UP** | **Foldchange_DOWN** | **Ratio_pct1_pct2** |
| LAMTOR13 | 0,003344576 | 0,416831663 | 0,442 | 0,301 | 1 | Mac | 1,334992524 | 0,749067865 | 1,468438538 |
| FAM118A2 | 0,003360349 | 0,408821569 | 0,114 | 0,019 | 1 | Mac | 1,327600952 | 0,753238387 | 6 |
| PSMA62 | 0,00337397 | 0,637056916 | 0,412 | 0,291 | 1 | Mac | 1,555153423 | 0,643023373 | 1,41580756 |
| COTL12 | 0,003425152 | 0,523517843 | 0,794 | 0,718 | 1 | Mac | 1,437456046 | 0,695673445 | 1,105849582 |
| NME24 | 0,003426501 | 0,292583529 | 0,764 | 0,641 | 1 | Mac | 1,224831702 | 0,816438698 | 1,191887676 |
| CXCL21 | 0,003436322 | 0,788478969 | 0,415 | 0,252 | 1 | Mac | 1,727252463 | 0,578954161 | 1,646825397 |
| F13A1 | 0,003436925 | 0,745180842 | 0,271 | 0,136 | 1 | Mac | 1,676184366 | 0,596593084 | 1,992647059 |
| CYP1B12 | 0,003464119 | -0,31870631 | 0,046 | 0,126 | 1 | Mac | 0,801788533 | 1,247211651 | 0,365079365 |
| CDC5L1 | 0,003469583 | 0,460348663 | 0,163 | 0,049 | 1 | Mac | 1,375874292 | 0,726810586 | 3,326530612 |
| MATR3.12 | 0,003573133 | -0,336698546 | 0,046 | 0,126 | 1 | Mac | 0,791851308 | 1,262863356 | 0,365079365 |
| TMEM1413 | 0,003677241 | 0,362312134 | 0,127 | 0,029 | 1 | Mac | 1,285484428 | 0,777916852 | 4,379310345 |
| GMPR21 | 0,004004792 | -0,515286286 | 0,081 | 0,175 | 1 | Mac | 0,699654081 | 1,429277735 | 0,462857143 |
| SCPEP11 | 0,004012036 | -0,370255561 | 0,171 | 0,301 | 1 | Mac | 0,77364544 | 1,29258178 | 0,568106312 |
| COX5A4 | 0,004103797 | 0,379388081 | 0,507 | 0,388 | 1 | Mac | 1,300790009 | 0,768763592 | 1,306701031 |
| NDUFA71 | 0,004185749 | 0,3996869 | 0,26 | 0,126 | 1 | Mac | 1,319221576 | 0,758022775 | 2,063492063 |
| NDUFA114 | 0,004258235 | 0,309935889 | 0,528 | 0,417 | 1 | Mac | 1,239652611 | 0,806677606 | 1,26618705 |
| PDGFB | 0,004274404 | -0,31235296 | 0,054 | 0,136 | 1 | Mac | 0,805327241 | 1,241731248 | 0,397058824 |
| TYROBP | 0,00427861 | 0,275351336 | 0,932 | 0,903 | 1 | Mac | 1,210288792 | 0,826249079 | 1,032115172 |
| ZFAND51 | 0,004314658 | 0,672622996 | 0,602 | 0,495 | 1 | Mac | 1,593968363 | 0,627365024 | 1,216161616 |
| MYL12A2 | 0,00433086 | 0,437589454 | 0,707 | 0,592 | 1 | Mac | 1,35433952 | 0,738367289 | 1,194256757 |
| CTSB3 | 0,004416998 | 0,933229655 | 0,759 | 0,718 | 1 | Mac | 1,909545975 | 0,523684694 | 1,057103064 |
| SYNGR24 | 0,004437063 | 0,621570519 | 0,518 | 0,388 | 1 | Mac | 1,538549136 | 0,649962992 | 1,335051546 |
| PUM11 | 0,004471388 | -0,280969506 | 0,136 | 0,252 | 1 | Mac | 0,823037741 | 1,21501111 | 0,53968254 |
| IGLC2 | 0,004539693 | 1,118204911 | 0,108 | 0,019 | 1 | Mac | 2,170767043 | 0,460666658 | 5,684210526 |
| PHPT14 | 0,004543886 | 0,521416959 | 0,295 | 0,165 | 1 | Mac | 1,435364315 | 0,696687238 | 1,787878788 |
| SEC11C3 | 0,004611451 | 0,5513275 | 0,247 | 0,126 | 1 | Mac | 1,465433499 | 0,682391935 | 1,96031746 |
| CCDC1242 | 0,004616032 | 0,525383699 | 0,165 | 0,058 | 1 | Mac | 1,439316329 | 0,694774303 | 2,844827586 |
| LDHA2 | 0,004680113 | 0,664004267 | 0,507 | 0,417 | 1 | Mac | 1,584474308 | 0,63112415 | 1,215827338 |
| RPS72 | 0,004707353 | 0,311325242 | 0,946 | 0,903 | 1 | Mac | 1,240847004 | 0,805901128 | 1,047619048 |
| TUT7 | 0,004717256 | -0,284312083 | 0,092 | 0,194 | 1 | Mac | 0,821133055 | 1,21782943 | 0,474226804 |
| MRPS343 | 0,004750941 | 0,424356481 | 0,192 | 0,078 | 1 | Mac | 1,341973777 | 0,745171044 | 2,461538462 |

| Supplementary Table 4 - clusermarker | | | | | | | | | |
|---|---|---|---|---|---|---|---|---|---|
| | **p_val** | **avg_log2FC** | **pct.1** | **pct.2** | **p_val_adj** | **celltype** | **Foldchange_UP** | **Foldchange_DOWN** | **Ratio_pct1_pct2** |
| TMEM50A3 | 0,004773247 | 0,319679985 | 0,344 | 0,204 | 1 | Mac | 1,248053678 | 0,801247589 | 1,68627451 |
| MBD61 | 0,004833454 | -0,397958683 | 0,049 | 0,126 | 1 | Mac | 0,758931361 | 1,317642215 | 0,388888889 |
| MBD23 | 0,004834831 | -0,292594176 | 0,152 | 0,272 | 1 | Mac | 0,816432673 | 1,224840741 | 0,558823529 |
| RPLP03 | 0,004901917 | 0,293469951 | 0,949 | 0,874 | 1 | Mac | 1,225584495 | 0,815937215 | 1,085812357 |
| RALA1 | 0,004923859 | 0,513550449 | 0,282 | 0,146 | 1 | Mac | 1,427559075 | 0,700496405 | 1,931506849 |
| GK1 | 0,005183784 | 0,733366703 | 0,339 | 0,214 | 1 | Mac | 1,662514246 | 0,601498605 | 1,58411215 |
| BPNT2 | 0,00524645 | 0,410249812 | 0,106 | 0,019 | 1 | Mac | 1,328915905 | 0,752493063 | 5,578947368 |
| PPIE | 0,005343165 | -0,348164901 | 0,049 | 0,126 | 1 | Mac | 0,785582719 | 1,272940425 | 0,388888889 |
| TMEM154 | 0,005406217 | -0,622718009 | 0,049 | 0,126 | 1 | Mac | 0,649446231 | 1,539773353 | 0,388888889 |
| AL627171.24 | 0,005481393 | -0,640431088 | 0,103 | 0,204 | 1 | Mac | 0,641521229 | 1,558794869 | 0,504901961 |
| ATP5MG1 | 0,005503531 | 0,320169015 | 0,726 | 0,641 | 1 | Mac | 1,248476802 | 0,800976036 | 1,132605304 |
| YBX13 | 0,005534338 | 0,260818983 | 0,783 | 0,67 | 1 | Mac | 1,198158677 | 0,834613995 | 1,168656716 |
| ATP10D | 0,005632225 | -0,250464913 | 0,049 | 0,126 | 1 | Mac | 0,840625478 | 1,189590402 | 0,388888889 |
| LTN1 | 0,005642645 | -0,490436882 | 0,057 | 0,136 | 1 | Mac | 0,711809513 | 1,404870239 | 0,419117647 |
| BAG61 | 0,005887002 | 0,452462178 | 0,163 | 0,058 | 1 | Mac | 1,368373603 | 0,730794571 | 2,810344828 |
| BTF33 | 0,005925967 | 0,475323608 | 0,745 | 0,689 | 1 | Mac | 1,390230024 | 0,719305426 | 1,081277213 |
| KANSL1-AS11 | 0,006011673 | -0,300693161 | 0,038 | 0,107 | 1 | Mac | 0,811862233 | 1,231736075 | 0,355140187 |
| B2M3 | 0,006169577 | -0,265455284 | 0,981 | 0,99 | 1 | Mac | 0,831936152 | 1,20201532 | 0,990909091 |
| DOK3 | 0,006227947 | -0,354130669 | 0,076 | 0,165 | 1 | Mac | 0,78234092 | 1,278215129 | 0,460606061 |
| SNX32 | 0,006238839 | 0,515852891 | 0,455 | 0,33 | 1 | Mac | 1,429839181 | 0,699379352 | 1,378787879 |
| RPL74 | 0,006378218 | -0,324044888 | 0,74 | 0,825 | 1 | Mac | 0,798827061 | 1,251835408 | 0,896969697 |
| DNPH14 | 0,006485301 | 0,599321865 | 0,22 | 0,107 | 1 | Mac | 1,515004275 | 0,660064144 | 2,056074766 |
| CTDP1 | 0,006491405 | 0,373133137 | 0,119 | 0,029 | 1 | Mac | 1,295162515 | 0,772103877 | 4,103448276 |
| MAILR | 0,006592188 | -0,589461279 | 0,046 | 0,117 | 1 | Mac | 0,664591028 | 1,504684774 | 0,393162393 |
| DNM21 | 0,006601374 | 0,498144687 | 0,293 | 0,165 | 1 | Mac | 1,412396045 | 0,708016709 | 1,775757576 |
| FGL2 | 0,006750729 | -0,461761252 | 0,466 | 0,583 | 1 | Mac | 0,72609929 | 1,377222115 | 0,799313894 |
| SRSF95 | 0,006820992 | 0,369223839 | 0,355 | 0,223 | 1 | Mac | 1,29165774 | 0,774198899 | 1,591928251 |
| RPL37A5 | 0,006879862 | -0,365364927 | 0,873 | 0,903 | 1 | Mac | 0,776272494 | 1,288207438 | 0,966777409 |
| AGO4 | 0,006979652 | -0,312013702 | 0,051 | 0,126 | 1 | Mac | 0,805516641 | 1,241439282 | 0,404761905 |
| CST32 | 0,007035943 | 0,368543809 | 0,962 | 0,883 | 1 | Mac | 1,291049047 | 0,774563912 | 1,089467724 |
| YIF1A2 | 0,007046178 | 0,557541404 | 0,171 | 0,068 | 1 | Mac | 1,471758953 | 0,679459091 | 2,514705882 |

| Supplementary Table 4 - clusermarker | | | | | | | | | |
|---|---|---|---|---|---|---|---|---|---|
| | **p_val** | **avg_log2FC** | **pct.1** | **pct.2** | **p_val_adj** | **celltype** | **Foldchange_UP** | **Foldchange_DOWN** | **Ratio_pct1_pct2** |
| TAF104 | 0,007046448 | 0,387019416 | 0,442 | 0,301 | 1 | Mac | 1,307688948 | 0,764707847 | 1,468438538 |
| NRIP11 | 0,007077299 | -0,342411412 | 0,133 | 0,243 | 1 | Mac | 0,78872189 | 1,267874029 | 0,547325103 |
| EIF13 | 0,007131943 | 0,261634367 | 0,924 | 0,854 | 1 | Mac | 1,198836046 | 0,83414242 | 1,081967213 |
| SLC40A1 | 0,007357031 | -0,566790346 | 0,168 | 0,291 | 1 | Mac | 0,675117094 | 1,481224529 | 0,577319588 |
| CHORDC11 | 0,007374392 | -0,49392989 | 0,106 | 0,204 | 1 | Mac | 0,710088186 | 1,408275788 | 0,519607843 |
| CTSH1 | 0,007455435 | 0,470915581 | 0,545 | 0,417 | 1 | Mac | 1,385988783 | 0,721506561 | 1,306954436 |
| BRI33 | 0,00775886 | 0,490493961 | 0,564 | 0,447 | 1 | Mac | 1,404925822 | 0,711781351 | 1,261744966 |
| PDIA64 | 0,007795986 | 0,494614308 | 0,304 | 0,175 | 1 | Mac | 1,408944035 | 0,709751399 | 1,737142857 |
| MARCKSL11 | 0,007980174 | 0,478480382 | 0,295 | 0,175 | 1 | Mac | 1,39327533 | 0,717733228 | 1,685714286 |
| ABI3 | 0,007992274 | -0,66863981 | 0,163 | 0,272 | 1 | Mac | 0,62909953 | 1,589573591 | 0,599264706 |
| TPM32 | 0,008061855 | 0,450039316 | 0,696 | 0,65 | 1 | Mac | 1,366077485 | 0,732022899 | 1,070769231 |
| SRSF73 | 0,008097603 | -0,296129453 | 0,301 | 0,447 | 1 | Mac | 0,81443448 | 1,227845854 | 0,673378076 |
| MANBA | 0,008099979 | -0,347821158 | 0,176 | 0,301 | 1 | Mac | 0,785769918 | 1,272637164 | 0,584717608 |
| NDUFB104 | 0,008104635 | 0,359578131 | 0,309 | 0,175 | 1 | Mac | 1,283050656 | 0,779392454 | 1,765714286 |
| CYFIP1 | 0,00815474 | -0,27904786 | 0,184 | 0,311 | 1 | Mac | 0,824134745 | 1,213393813 | 0,591639871 |
| SMDT15 | 0,008348138 | 0,358525327 | 0,425 | 0,291 | 1 | Mac | 1,282114695 | 0,779961422 | 1,4604811 |
| TRADD1 | 0,008471943 | 0,592293408 | 0,152 | 0,058 | 1 | Mac | 1,507641495 | 0,663287661 | 2,620689655 |
| EVI2B1 | 0,008526715 | -0,40905792 | 0,317 | 0,456 | 1 | Mac | 0,753114997 | 1,327818465 | 0,695175439 |
| AXL | 0,008652728 | -0,378086655 | 0,141 | 0,252 | 1 | Mac | 0,769457391 | 1,299617121 | 0,55952381 |
| TRAPPC53 | 0,008707368 | 0,745232265 | 0,385 | 0,272 | 1 | Mac | 1,676244113 | 0,596571819 | 1,415441176 |
| ARPC23 | 0,008785157 | 0,291006336 | 0,745 | 0,65 | 1 | Mac | 1,223493414 | 0,817331739 | 1,146153846 |
| IER53 | 0,008867692 | -0,600958661 | 0,279 | 0,388 | 1 | Mac | 0,659315699 | 1,516724085 | 0,719072165 |
| PIGR2 | 0,008930655 | -0,574161343 | 0,046 | 0,117 | 1 | Mac | 0,671676592 | 1,488811747 | 0,393162393 |
| RNF73 | 0,008956131 | 0,631070334 | 0,263 | 0,155 | 1 | Mac | 1,548713556 | 0,645697195 | 1,696774194 |
| MYDGF2 | 0,009070199 | 0,633641682 | 0,314 | 0,204 | 1 | Mac | 1,551476325 | 0,64454738 | 1,539215686 |
| YBX31 | 0,009129739 | 0,604963379 | 0,417 | 0,282 | 1 | Mac | 1,520940145 | 0,65748807 | 1,478723404 |
| PSMA51 | 0,009290943 | 0,439308373 | 0,255 | 0,136 | 1 | Mac | 1,355954129 | 0,737488075 | 1,875 |
| HSP90AA13 | 0,009383601 | -0,794776441 | 0,743 | 0,767 | 1 | Mac | 0,57643249 | 1,734808529 | 0,968709257 |
| UQCR113 | 0,009516528 | 0,355629014 | 0,61 | 0,515 | 1 | Mac | 1,27954334 | 0,781528823 | 1,184466019 |
| CCDC107 | 0,009554603 | 0,415523498 | 0,203 | 0,097 | 1 | Mac | 1,333782567 | 0,749747391 | 2,092783505 |
| PLA2G7 | 0,009563659 | 0,519162127 | 0,114 | 0,029 | 1 | Mac | 1,433122692 | 0,697776963 | 3,931034483 |

| Supplementary Table 4 - clusermarker | | | | | | | | | |
|---|---|---|---|---|---|---|---|---|---|
| | **p_val** | **avg_log2FC** | **pct.1** | **pct.2** | **p_val_adj** | **celltype** | **Foldchange_UP** | **Foldchange_DOWN** | **Ratio_pct1_pct2** |
| SH3PXD2B1 | 0,009567733 | 0,416531767 | 0,127 | 0,039 | 1 | Mac | 1,334715045 | 0,749223592 | 3,256410256 |
| EMC62 | 0,009690802 | 0,411837512 | 0,179 | 0,078 | 1 | Mac | 1,330379194 | 0,751665393 | 2,294871795 |
| VAMP82 | 0,009786557 | 0,353979924 | 0,688 | 0,583 | 1 | Mac | 1,278081578 | 0,782422669 | 1,180102916 |
| SEC624 | 0,010034915 | -0,386309071 | 0,293 | 0,417 | 1 | Mac | 0,765084462 | 1,307045235 | 0,70263789 |
| AFF4 | 0,010075211 | -0,35001921 | 0,268 | 0,388 | 1 | Mac | 0,784573651 | 1,274577599 | 0,690721649 |
| TCEAL8 | 0,010075636 | -0,322140884 | 0,06 | 0,136 | 1 | Mac | 0,799882013 | 1,250184381 | 0,441176471 |
| CX3CR1 | 0,010258181 | -0,371366538 | 0,146 | 0,252 | 1 | Mac | 0,773049908 | 1,293577542 | 0,579365079 |
| CHPT12 | 0,010302728 | 0,285764633 | 0,154 | 0,058 | 1 | Mac | 1,219056196 | 0,820306728 | 2,655172414 |
| MGMT3 | 0,010438145 | 0,380432775 | 0,141 | 0,049 | 1 | Mac | 1,301732286 | 0,768207112 | 2,87755102 |
| TRMT1123 | 0,010470484 | 0,575771734 | 0,398 | 0,301 | 1 | Mac | 1,490474542 | 0,67092726 | 1,322259136 |
| ACAT1 | 0,010767829 | -0,414975953 | 0,06 | 0,136 | 1 | Mac | 0,750031996 | 1,333276454 | 0,441176471 |
| HNRNPA15 | 0,011176693 | -0,463960678 | 0,604 | 0,68 | 1 | Mac | 0,724993177 | 1,379323326 | 0,888235294 |
| NR1D22 | 0,011250025 | -0,536033549 | 0,062 | 0,136 | 1 | Mac | 0,689664423 | 1,449980551 | 0,455882353 |
| ZNF3311 | 0,011333357 | 0,489260564 | 0,526 | 0,369 | 1 | Mac | 1,403725229 | 0,712390131 | 1,425474255 |
| ATP1B12 | 0,011370611 | -0,509084381 | 0,171 | 0,272 | 1 | Mac | 0,702668251 | 1,423146696 | 0,628676471 |
| NDUFA24 | 0,0113874 | 0,602628128 | 0,352 | 0,243 | 1 | Mac | 1,518480233 | 0,658553189 | 1,448559671 |
| DNAJA4 | 0,011462172 | -0,346235404 | 0,054 | 0,126 | 1 | Mac | 0,78663408 | 1,2712391 | 0,428571429 |
| RB11 | 0,011502531 | -0,347147926 | 0,282 | 0,398 | 1 | Mac | 0,786136682 | 1,272043428 | 0,708542714 |
| FCGR2A | 0,011580833 | -0,339996322 | 0,404 | 0,544 | 1 | Mac | 0,790043326 | 1,265753367 | 0,742647059 |
| PHF11 | 0,011612506 | -0,433218134 | 0,068 | 0,146 | 1 | Mac | 0,740607911 | 1,350242125 | 0,465753425 |
| HINT15 | 0,011613937 | 0,568170809 | 0,602 | 0,534 | 1 | Mac | 1,482642538 | 0,674471408 | 1,127340824 |
| IDH3G1 | 0,011648769 | 0,389260648 | 0,23 | 0,117 | 1 | Mac | 1,309722026 | 0,763520793 | 1,965811966 |
| PPP1R14B3 | 0,011730528 | 0,434731787 | 0,26 | 0,146 | 1 | Mac | 1,351659521 | 0,739831285 | 1,780821918 |
| GNL31 | 0,011750568 | -0,427537158 | 0,127 | 0,223 | 1 | Mac | 0,743529991 | 1,34493566 | 0,569506726 |
| WBP111 | 0,011765467 | -0,293601125 | 0,1 | 0,194 | 1 | Mac | 0,815863031 | 1,225695934 | 0,515463918 |
| FLNA3 | 0,011824308 | 0,318646951 | 0,474 | 0,33 | 1 | Mac | 1,247160336 | 0,801821523 | 1,436363636 |
| HLA-DOA | 0,011841808 | -0,647614011 | 0,182 | 0,291 | 1 | Mac | 0,638335146 | 1,566575185 | 0,625429553 |
| FLOT2 | 0,012056111 | 0,362745604 | 0,179 | 0,078 | 1 | Mac | 1,285870721 | 0,777683155 | 2,294871795 |
| FIS14 | 0,012338151 | 0,35521497 | 0,285 | 0,175 | 1 | Mac | 1,279176171 | 0,781753149 | 1,628571429 |
| DENR | 0,012490984 | -0,629241487 | 0,1 | 0,184 | 1 | Mac | 0,646516239 | 1,546751557 | 0,543478261 |
| SLC1A3 | 0,01252574 | -0,300200894 | 0,087 | 0,175 | 1 | Mac | 0,812139299 | 1,231315861 | 0,497142857 |

| Supplementary Table 4 - clusermarker | | | | | | | | | |
|---|---|---|---|---|---|---|---|---|---|
| | **p_val** | **avg_log2FC** | **pct.1** | **pct.2** | **p_val_adj** | **celltype** | **Foldchange_UP** | **Foldchange_DOWN** | **Ratio_pct1_pct2** |
| FGD2 | 0,012697894 | -0,369258109 | 0,108 | 0,204 | 1 | Mac | 0,774180509 | 1,291688423 | 0,529411765 |
| ZNF131 | 0,012786112 | 0,290690522 | 0,111 | 0,029 | 1 | Mac | 1,223225614 | 0,817510677 | 3,827586207 |
| CD1512 | 0,012982029 | 0,540900216 | 0,195 | 0,097 | 1 | Mac | 1,454880053 | 0,687341886 | 2,010309278 |
| PHTF21 | 0,013017363 | 0,388914623 | 0,122 | 0,039 | 1 | Mac | 1,309407931 | 0,763703943 | 3,128205128 |
| GSPT11 | 0,013042516 | 0,299750007 | 0,247 | 0,136 | 1 | Mac | 1,230931097 | 0,812393157 | 1,816176471 |
| IGKV3-204 | 0,013052253 | 3,208558953 | 0,152 | 0,058 | 1 | Mac | 9,244267149 | 0,108175152 | 2,620689655 |
| ERICH1 | 0,013172045 | -0,255257431 | 0,122 | 0,223 | 1 | Mac | 0,83783762 | 1,193548698 | 0,547085202 |
| BLVRA2 | 0,01321438 | 0,508626443 | 0,206 | 0,107 | 1 | Mac | 1,422695035 | 0,702891326 | 1,925233645 |
| SNRNP25 | 0,013227008 | 0,467075975 | 0,133 | 0,049 | 1 | Mac | 1,382305 | 0,723429345 | 2,714285714 |
| AL499604.1 | 0,013415671 | -0,396567106 | 0,043 | 0,107 | 1 | Mac | 0,759663755 | 1,316371873 | 0,401869159 |
| OFD11 | 0,013463428 | -0,306943222 | 0,068 | 0,146 | 1 | Mac | 0,808352681 | 1,237083792 | 0,465753425 |
| SIGIRR2 | 0,013551281 | 0,345045332 | 0,173 | 0,078 | 1 | Mac | 1,270190894 | 0,787283238 | 2,217948718 |
| SCLT1 | 0,013597432 | -0,364994346 | 0,111 | 0,204 | 1 | Mac | 0,776471918 | 1,287876582 | 0,544117647 |
| VAMP52 | 0,013657653 | 0,555219861 | 0,201 | 0,097 | 1 | Mac | 1,469392546 | 0,680553337 | 2,072164948 |
| CSRP11 | 0,013660268 | 0,325679529 | 0,16 | 0,068 | 1 | Mac | 1,253254599 | 0,797922466 | 2,352941176 |
| TBC1D91 | 0,01377312 | -0,273083073 | 0,117 | 0,214 | 1 | Mac | 0,827549163 | 1,208387422 | 0,546728972 |
| ODF3B2 | 0,013791956 | 0,384908466 | 0,211 | 0,107 | 1 | Mac | 1,305776938 | 0,765827586 | 1,971962617 |
| UBC4 | 0,013804629 | -0,478613475 | 0,875 | 0,883 | 1 | Mac | 0,717667018 | 1,39340387 | 0,990939977 |
| RGS18 | 0,014032282 | -0,608768931 | 0,084 | 0,165 | 1 | Mac | 0,655756028 | 1,524957389 | 0,509090909 |
| CLIC12 | 0,014145194 | 0,416999348 | 0,699 | 0,621 | 1 | Mac | 1,3351477 | 0,748980806 | 1,125603865 |
| SLC12A21 | 0,014196995 | -0,591789483 | 0,043 | 0,107 | 1 | Mac | 0,663519383 | 1,507114977 | 0,401869159 |
| GTF3C61 | 0,014245029 | 0,548853154 | 0,217 | 0,117 | 1 | Mac | 1,462922308 | 0,6835633 | 1,854700855 |
| HBEGF2 | 0,014263893 | 0,808297846 | 0,328 | 0,233 | 1 | Mac | 1,751144148 | 0,571055216 | 1,407725322 |
| PGM2 | 0,014415224 | -0,585347821 | 0,076 | 0,155 | 1 | Mac | 0,666488635 | 1,500400679 | 0,490322581 |
| ARF53 | 0,014450127 | 0,293421385 | 0,453 | 0,34 | 1 | Mac | 1,225543238 | 0,815964683 | 1,332352941 |
| ID32 | 0,014567682 | 0,703499375 | 0,187 | 0,087 | 1 | Mac | 1,628449945 | 0,614080895 | 2,149425287 |
| MS4A4E | 0,014734277 | -0,54714709 | 0,057 | 0,126 | 1 | Mac | 0,684372128 | 1,461193347 | 0,452380952 |
| NTAN11 | 0,01513315 | 0,366728712 | 0,119 | 0,039 | 1 | Mac | 1,289425761 | 0,775539027 | 3,051282051 |
| STOML23 | 0,015139264 | 0,425390364 | 0,157 | 0,068 | 1 | Mac | 1,342935825 | 0,744637221 | 2,308823529 |
| SYNRG1 | 0,015228411 | 0,313135064 | 0,106 | 0,029 | 1 | Mac | 1,242404589 | 0,804890781 | 3,655172414 |
| SQSTM14 | 0,015409263 | -0,585859285 | 0,545 | 0,612 | 1 | Mac | 0,666252393 | 1,500932695 | 0,890522876 |

| Supplementary Table 4 - clusermarker | | | | | | | | | |
|---|---|---|---|---|---|---|---|---|---|
| | **p_val** | **avg_log2FC** | **pct.1** | **pct.2** | **p_val_adj** | **celltype** | **Foldchange_UP** | **Foldchange_DOWN** | **Ratio_pct1_pct2** |
| PNRC2 | 0,015427584 | -0,367841556 | 0,22 | 0,33 | 1 | Mac | 0,774941034 | 1,290420762 | 0,666666667 |
| THUMPD3-AS1 | 0,015460283 | -0,359334824 | 0,136 | 0,233 | 1 | Mac | 0,779523908 | 1,282834291 | 0,583690987 |
| DHPS2 | 0,015477326 | 0,371404867 | 0,136 | 0,049 | 1 | Mac | 1,29361191 | 0,77302937 | 2,775510204 |
| BIN1 | 0,015491376 | -0,3133303 | 0,119 | 0,214 | 1 | Mac | 0,804781864 | 1,242572732 | 0,556074766 |
| PLXNC1 | 0,015639475 | -0,50386111 | 0,154 | 0,252 | 1 | Mac | 0,705216869 | 1,418003516 | 0,611111111 |
| HLA-DPA12 | 0,016092237 | -0,379471418 | 0,935 | 0,942 | 1 | Mac | 0,768719186 | 1,30086515 | 0,992569002 |
| TUSC22 | 0,016136572 | 0,385686047 | 0,119 | 0,039 | 1 | Mac | 1,306480913 | 0,765414933 | 3,051282051 |
| RAC13 | 0,016151864 | 0,325376633 | 0,718 | 0,621 | 1 | Mac | 1,252991505 | 0,798090008 | 1,156199678 |
| TMEM1342 | 0,016340015 | 0,469162551 | 0,13 | 0,049 | 1 | Mac | 1,38430568 | 0,722383802 | 2,653061224 |
| TFEC | 0,016439149 | -0,329650062 | 0,098 | 0,184 | 1 | Mac | 0,795729471 | 1,256708513 | 0,532608696 |
| HSPA21 | 0,016448815 | -0,298728509 | 0,079 | 0,155 | 1 | Mac | 0,812968575 | 1,230059846 | 0,509677419 |
| DGCR6L1 | 0,016524373 | 0,326404969 | 0,106 | 0,029 | 1 | Mac | 1,25388494 | 0,797521342 | 3,655172414 |
| GPX45 | 0,016635963 | 0,335763213 | 0,724 | 0,631 | 1 | Mac | 1,262044878 | 0,79236485 | 1,147385103 |
| TBCB3 | 0,016735374 | 0,431607591 | 0,298 | 0,194 | 1 | Mac | 1,348735632 | 0,741435146 | 1,536082474 |
| RNF1812 | 0,016845624 | 0,485836242 | 0,287 | 0,175 | 1 | Mac | 1,400397355 | 0,71408304 | 1,64 |
| PSME13 | 0,017144957 | 0,448533794 | 0,553 | 0,447 | 1 | Mac | 1,36465266 | 0,732787199 | 1,237136465 |
| PI4KB2 | 0,01730705 | -0,396894319 | 0,1 | 0,184 | 1 | Mac | 0,759491477 | 1,316670469 | 0,543478261 |
| AGGF1 | 0,017430114 | 0,383711742 | 0,117 | 0,039 | 1 | Mac | 1,304694237 | 0,766463108 | 3 |
| FKBP1A2 | 0,017644454 | 0,496828374 | 0,396 | 0,291 | 1 | Mac | 1,411107965 | 0,708662997 | 1,360824742 |
| THEMIS2 | 0,017885202 | 0,512827613 | 0,325 | 0,214 | 1 | Mac | 1,426844003 | 0,700847463 | 1,518691589 |
| APRT3 | 0,018422302 | 0,510137422 | 0,485 | 0,388 | 1 | Mac | 1,424185848 | 0,702155552 | 1,25 |
| MRPL341 | 0,01846997 | 0,439620073 | 0,141 | 0,058 | 1 | Mac | 1,356247119 | 0,737328755 | 2,431034483 |
| PRKRA | 0,018524625 | 0,424877834 | 0,103 | 0,029 | 1 | Mac | 1,34245882 | 0,744901806 | 3,551724138 |
| SSH1 | 0,018776244 | -0,252173454 | 0,07 | 0,146 | 1 | Mac | 0,839630539 | 1,191000034 | 0,479452055 |
| DRAP14 | 0,018847848 | 0,267609146 | 0,447 | 0,33 | 1 | Mac | 1,203811201 | 0,830695045 | 1,354545455 |
| DRAM22 | 0,018895643 | -0,420152432 | 0,144 | 0,243 | 1 | Mac | 0,747345657 | 1,338068925 | 0,592592593 |
| ITPK1 | 0,018920146 | 0,564648857 | 0,152 | 0,068 | 1 | Mac | 1,479027479 | 0,67611996 | 2,235294118 |
| NAP1L12 | 0,019062274 | -0,442638479 | 0,512 | 0,592 | 1 | Mac | 0,735787729 | 1,35908763 | 0,864864865 |
| PARL | 0,019405313 | 0,410657583 | 0,157 | 0,068 | 1 | Mac | 1,32929157 | 0,752280405 | 2,308823529 |
| C19orf534 | 0,019820318 | 0,490873216 | 0,32 | 0,214 | 1 | Mac | 1,405295198 | 0,711594263 | 1,495327103 |
| UQCRQ3 | 0,020018021 | 0,400622483 | 0,45 | 0,359 | 1 | Mac | 1,320077365 | 0,757531359 | 1,253481894 |

| Supplementary Table 4 - clusermarker | | | | | | | | | |
|---|---|---|---|---|---|---|---|---|---|
| | **p_val** | **avg_log2FC** | **pct.1** | **pct.2** | **p_val_adj** | **celltype** | **Foldchange_UP** | **Foldchange_DOWN** | **Ratio_pct1_pct2** |
| ATP8B4 | 0,020080306 | -0,574898278 | 0,046 | 0,107 | 1 | Mac | 0,671333584 | 1,489572432 | 0,429906542 |
| PSMB62 | 0,020255155 | 0,42566361 | 0,401 | 0,291 | 1 | Mac | 1,343190201 | 0,7444962 | 1,378006873 |
| TUBB5 | 0,020401584 | 0,779162733 | 0,507 | 0,408 | 1 | Mac | 1,716134626 | 0,582704867 | 1,242647059 |
| RPL27A4 | 0,020515094 | -0,392907013 | 0,653 | 0,709 | 1 | Mac | 0,761593456 | 1,313036493 | 0,921015515 |
| HTATIP21 | 0,020577846 | 0,354349706 | 0,114 | 0,039 | 1 | Mac | 1,278409209 | 0,78222215 | 2,923076923 |
| RPL233 | 0,020618704 | -0,343734789 | 0,637 | 0,728 | 1 | Mac | 0,787998731 | 1,269037577 | 0,875 |
| CCND3 | 0,020622892 | 0,445505019 | 0,179 | 0,087 | 1 | Mac | 1,361790732 | 0,734327218 | 2,057471264 |
| ARL5A2 | 0,020714068 | -0,300417858 | 0,187 | 0,282 | 1 | Mac | 0,812017172 | 1,23150105 | 0,663120567 |
| NR4A24 | 0,020718977 | 0,294093312 | 0,672 | 0,573 | 1 | Mac | 1,226114161 | 0,815584741 | 1,172774869 |
| NCOA2 | 0,021029993 | -0,498690318 | 0,06 | 0,126 | 1 | Mac | 0,707748986 | 1,412930318 | 0,476190476 |
| ANAPC114 | 0,021126915 | 0,461040621 | 0,404 | 0,301 | 1 | Mac | 1,376534359 | 0,72646207 | 1,342192691 |
| LSM72 | 0,021150627 | 0,381431947 | 0,347 | 0,243 | 1 | Mac | 1,302634144 | 0,767675256 | 1,427983539 |
| RNF1491 | 0,021343273 | 0,345155898 | 0,415 | 0,291 | 1 | Mac | 1,270288243 | 0,787222904 | 1,426116838 |
| RPL222 | 0,021353197 | -0,296282126 | 0,87 | 0,864 | 1 | Mac | 0,814348298 | 1,227975797 | 1,006944444 |
| HNRNPL2 | 0,021905136 | 0,450713908 | 0,352 | 0,252 | 1 | Mac | 1,3667164 | 0,731680691 | 1,396825397 |
| DEK1 | 0,022209291 | 0,61779646 | 0,388 | 0,282 | 1 | Mac | 1,534529584 | 0,651665507 | 1,375886525 |
| GNL3L | 0,022236818 | -0,423607457 | 0,054 | 0,117 | 1 | Mac | 0,745558025 | 1,341277227 | 0,461538462 |
| FAM120A | 0,022781992 | -0,335144813 | 0,168 | 0,262 | 1 | Mac | 0,792704563 | 1,261504028 | 0,641221374 |
| RGCC3 | 0,022849242 | 0,524313526 | 0,458 | 0,35 | 1 | Mac | 1,438249058 | 0,695289869 | 1,308571429 |
| PNPLA6 | 0,022992148 | 0,460827326 | 0,149 | 0,068 | 1 | Mac | 1,376330861 | 0,726569481 | 2,191176471 |
| PPIA4 | 0,023077966 | 0,352974878 | 0,851 | 0,854 | 1 | Mac | 1,277191519 | 0,78296793 | 0,996487119 |
| MYO9B1 | 0,023168933 | 0,309087993 | 0,344 | 0,223 | 1 | Mac | 1,23892426 | 0,807151843 | 1,542600897 |
| TNIP2 | 0,023320916 | 0,358201565 | 0,125 | 0,049 | 1 | Mac | 1,281827001 | 0,780136477 | 2,551020408 |
| NDUFA15 | 0,023578437 | -0,395932109 | 0,461 | 0,573 | 1 | Mac | 0,759998191 | 1,315792605 | 0,804537522 |
| DNAJB11 | 0,023617323 | 0,488552852 | 0,195 | 0,107 | 1 | Mac | 1,403036802 | 0,712739679 | 1,822429907 |
| NFKBIA4 | 0,023636067 | 0,265034924 | 0,767 | 0,621 | 1 | Mac | 1,201665138 | 0,83217859 | 1,23510467 |
| MARCHF72 | 0,023756309 | -0,309478143 | 0,087 | 0,165 | 1 | Mac | 0,806933593 | 1,23925935 | 0,527272727 |
| APOE1 | 0,023760811 | -0,380270463 | 0,182 | 0,282 | 1 | Mac | 0,768293545 | 1,301585842 | 0,645390071 |
| RPL264 | 0,023776891 | -0,25266008 | 0,967 | 0,961 | 1 | Mac | 0,839347376 | 1,19140183 | 1,006243496 |
| MLF22 | 0,023974458 | 0,533915814 | 0,279 | 0,184 | 1 | Mac | 1,447853683 | 0,690677526 | 1,516304348 |
| MRPL143 | 0,024027302 | 0,454169178 | 0,173 | 0,087 | 1 | Mac | 1,369993623 | 0,729930405 | 1,988505747 |

| Supplementary Table 4 - clusermarker | | | | | | | | | |
|---|---|---|---|---|---|---|---|---|---|
| | p_val | avg_log2FC | pct.1 | pct.2 | p_val_adj | celltype | Foldchange_UP | Foldchange_DOWN | Ratio_pct1_pct2 |
| SPTY2D11 | 0,024103779 | -0,453428371 | 0,114 | 0,194 | 1 | Mac | 0,730305312 | 1,369290328 | 0,587628866 |
| MICOS102 | 0,024162171 | 0,482456015 | 0,434 | 0,33 | 1 | Mac | 1,397120073 | 0,715758094 | 1,315151515 |
| SEPTIN111 | 0,024165306 | 0,404302323 | 0,125 | 0,049 | 1 | Mac | 1,323448745 | 0,755601608 | 2,551020408 |
| IRF4 | 0,024194924 | 0,81794325 | 0,157 | 0,078 | 1 | Mac | 1,762890969 | 0,567250055 | 2,012820513 |
| NAPSA | 0,024467066 | 0,372018595 | 0,16 | 0,078 | 1 | Mac | 1,294162335 | 0,77270059 | 2,051282051 |
| HNRNPAB2 | 0,024586424 | 0,381149475 | 0,379 | 0,262 | 1 | Mac | 1,30237912 | 0,767825578 | 1,446564885 |
| PDIA41 | 0,025095399 | 0,316574537 | 0,206 | 0,107 | 1 | Mac | 1,245370091 | 0,802974158 | 1,925233645 |
| ATP5PO2 | 0,025264162 | 0,356751953 | 0,482 | 0,369 | 1 | Mac | 1,280539675 | 0,780920747 | 1,306233062 |
| VDAC23 | 0,025267611 | 0,299254305 | 0,344 | 0,243 | 1 | Mac | 1,230508228 | 0,812672339 | 1,41563786 |
| MAD2L21 | 0,025298291 | 0,460380547 | 0,136 | 0,058 | 1 | Mac | 1,3759047 | 0,726794523 | 2,344827586 |
| ATXN71 | 0,025439051 | -0,315134286 | 0,081 | 0,155 | 1 | Mac | 0,803776172 | 1,244127451 | 0,522580645 |
| RP2 | 0,025651449 | -0,323506835 | 0,13 | 0,214 | 1 | Mac | 0,799125039 | 1,251368623 | 0,607476636 |
| DCTN33 | 0,02571005 | 0,465122533 | 0,217 | 0,126 | 1 | Mac | 1,380434594 | 0,724409548 | 1,722222222 |
| ARHGAP22 | 0,025753652 | -0,352621467 | 0,081 | 0,155 | 1 | Mac | 0,783159754 | 1,276878689 | 0,522580645 |
| CCND12 | 0,025795733 | -0,468318491 | 0,076 | 0,146 | 1 | Mac | 0,722806562 | 1,383496018 | 0,520547945 |
| DST2 | 0,025914969 | -0,346922421 | 0,13 | 0,214 | 1 | Mac | 0,786259571 | 1,271844613 | 0,607476636 |
| HIGD2A3 | 0,026345212 | 0,431609671 | 0,561 | 0,476 | 1 | Mac | 1,348737576 | 0,741434077 | 1,178571429 |
| SSR43 | 0,02650874 | 0,532268476 | 0,588 | 0,515 | 1 | Mac | 1,446201398 | 0,691466625 | 1,141747573 |
| CUEDC21 | 0,026687638 | 0,505653234 | 0,16 | 0,078 | 1 | Mac | 1,419766063 | 0,704341388 | 2,051282051 |
| CD141 | 0,026751149 | 0,774170467 | 0,472 | 0,388 | 1 | Mac | 1,710206419 | 0,584724738 | 1,216494845 |
| EIF4A14 | 0,026826065 | -0,294622984 | 0,791 | 0,816 | 1 | Mac | 0,815285361 | 1,226564401 | 0,969362745 |
| GLIPR12 | 0,026853581 | -0,611061275 | 0,425 | 0,495 | 1 | Mac | 0,654714903 | 1,527382369 | 0,858585859 |
| FBP11 | 0,0271667 | 0,74697744 | 0,255 | 0,165 | 1 | Mac | 1,678273031 | 0,595850605 | 1,545454545 |
| MIIP1 | 0,027358909 | 0,493251416 | 0,106 | 0,039 | 1 | Mac | 1,407613657 | 0,710422207 | 2,717948718 |
| HIF1A | 0,027386747 | 0,393580097 | 0,461 | 0,35 | 1 | Mac | 1,313649228 | 0,76123822 | 1,317142857 |
| ENO11 | 0,027559276 | 0,406272653 | 0,591 | 0,534 | 1 | Mac | 1,325257452 | 0,754570365 | 1,106741573 |
| TIMM17B3 | 0,027562605 | 0,370686888 | 0,173 | 0,087 | 1 | Mac | 1,292968285 | 0,773414175 | 1,988505747 |
| PNN2 | 0,027709755 | -0,422962137 | 0,152 | 0,243 | 1 | Mac | 0,745891589 | 1,340677405 | 0,625514403 |
| MRTO4 | 0,027855813 | -0,460243183 | 0,057 | 0,117 | 1 | Mac | 0,726863727 | 1,375773701 | 0,487179487 |
| HIKESHI1 | 0,028164879 | 0,288655329 | 0,146 | 0,068 | 1 | Mac | 1,221501241 | 0,818664743 | 2,147058824 |
| SNHG1 | 0,028585323 | -0,407878729 | 0,13 | 0,214 | 1 | Mac | 0,75373081 | 1,326733613 | 0,607476636 |

| Supplementary Table 4 - clusermarker | | | | | | | | | |
|---|---|---|---|---|---|---|---|---|---|
| | **p_val** | **avg_log2FC** | **pct.1** | **pct.2** | **p_val_adj** | **celltype** | **Foldchange_UP** | **Foldchange_DOWN** | **Ratio_pct1_pct2** |
| SERPINB62 | 0,028592208 | 0,386155629 | 0,279 | 0,184 | 1 | Mac | 1,306906227 | 0,765165839 | 1,516304348 |
| HLA-DQA1 | 0,028603955 | -0,320322792 | 0,78 | 0,806 | 1 | Mac | 0,800890664 | 1,248609885 | 0,967741935 |
| TPP1 | 0,028628826 | -0,38329543 | 0,331 | 0,437 | 1 | Mac | 0,766684315 | 1,304317801 | 0,757437071 |
| AL355881.1 | 0,02864382 | -0,452402534 | 0,076 | 0,146 | 1 | Mac | 0,730824784 | 1,368317032 | 0,520547945 |
| HOMER1 | 0,028734777 | -0,28668943 | 0,062 | 0,126 | 1 | Mac | 0,819781064 | 1,219837887 | 0,492063492 |
| GNPTAB1 | 0,028863604 | 0,422102898 | 0,144 | 0,068 | 1 | Mac | 1,339879163 | 0,746335959 | 2,117647059 |
| ADAP2 | 0,028982828 | -0,399254824 | 0,241 | 0,34 | 1 | Mac | 0,758249831 | 1,318826539 | 0,708823529 |
| LYZ5 | 0,02910383 | -0,931361792 | 0,827 | 0,874 | 1 | Mac | 0,52436315 | 1,907075277 | 0,946224256 |
| TPI13 | 0,029115323 | 0,329580086 | 0,634 | 0,563 | 1 | Mac | 1,256647559 | 0,795768068 | 1,126110124 |
| CRELD22 | 0,029143998 | 0,3472514 | 0,136 | 0,058 | 1 | Mac | 1,272134666 | 0,7860803 | 2,344827586 |
| ARL6IP44 | 0,029298449 | 0,425815134 | 0,447 | 0,359 | 1 | Mac | 1,343331282 | 0,74441801 | 1,245125348 |
| YIPF51 | 0,029521282 | -0,52782895 | 0,062 | 0,126 | 1 | Mac | 0,693597715 | 1,441757923 | 0,492063492 |
| HNRNPH32 | 0,029585384 | -0,409999615 | 0,19 | 0,291 | 1 | Mac | 0,752623575 | 1,32868546 | 0,652920962 |
| CD300E | 0,029630978 | -0,282398277 | 0,049 | 0,107 | 1 | Mac | 0,822223051 | 1,21621499 | 0,457943925 |
| ABRACL3 | 0,029808455 | 0,302357856 | 0,325 | 0,214 | 1 | Mac | 1,233158168 | 0,810925983 | 1,518691589 |
| TMEM50B3 | 0,029868185 | 0,363333415 | 0,171 | 0,087 | 1 | Mac | 1,286394742 | 0,777366361 | 1,965517241 |
| SPCS14 | 0,029912653 | 0,33546553 | 0,382 | 0,262 | 1 | Mac | 1,261784497 | 0,792528362 | 1,458015267 |
| OTULINL | 0,030221667 | -0,611797699 | 0,141 | 0,223 | 1 | Mac | 0,65438079 | 1,52816222 | 0,632286996 |
| SLAMF8 | 0,030264595 | 0,443986718 | 0,171 | 0,087 | 1 | Mac | 1,360358328 | 0,735100436 | 1,965517241 |
| SND1 | 0,030399917 | 0,606235521 | 0,285 | 0,194 | 1 | Mac | 1,522281874 | 0,656908564 | 1,469072165 |
| EIF62 | 0,030410252 | 0,29251906 | 0,268 | 0,165 | 1 | Mac | 1,22477697 | 0,816475182 | 1,624242424 |
| PLXDC2 | 0,030494307 | -0,383005028 | 0,333 | 0,437 | 1 | Mac | 0,766838657 | 1,30405528 | 0,76201373 |
| AC007384.12 | 0,030623593 | -0,272280159 | 0,106 | 0,184 | 1 | Mac | 0,828009853 | 1,207715097 | 0,576086957 |
| IRF51 | 0,030858521 | -0,257523527 | 0,076 | 0,146 | 1 | Mac | 0,836522629 | 1,195424923 | 0,520547945 |
| CHM | 0,03113378 | -0,258318031 | 0,049 | 0,107 | 1 | Mac | 0,836062076 | 1,196083435 | 0,457943925 |
| FAM126A | 0,031310268 | -0,465738886 | 0,057 | 0,117 | 1 | Mac | 0,72410013 | 1,381024473 | 0,487179487 |
| FBXW53 | 0,031485567 | 0,254730553 | 0,22 | 0,126 | 1 | Mac | 1,193112888 | 0,838143657 | 1,746031746 |
| SNX81 | 0,031599573 | 0,417446049 | 0,154 | 0,078 | 1 | Mac | 1,335561165 | 0,748748935 | 1,974358974 |
| TUBA1A2 | 0,031626482 | 0,485330044 | 0,39 | 0,291 | 1 | Mac | 1,399906084 | 0,714333634 | 1,340206186 |
| TGFBR1 | 0,031685897 | -0,271951019 | 0,138 | 0,223 | 1 | Mac | 0,828198779 | 1,207439597 | 0,618834081 |
| RAP1B1 | 0,031769113 | 0,327267275 | 0,398 | 0,291 | 1 | Mac | 1,254634618 | 0,797044802 | 1,367697595 |

| Supplementary Table 4 - clusermarker | | | | | | | | | |
|---|---|---|---|---|---|---|---|---|---|
| | **p_val** | **avg_log2FC** | **pct.1** | **pct.2** | **p_val_adj** | **celltype** | **Foldchange_UP** | **Foldchange_DOWN** | **Ratio_pct1_pct2** |
| EIF3L1 | 0,031930573 | -0,445246728 | 0,417 | 0,524 | 1 | Mac | 0,7344587 | 1,361546946 | 0,795801527 |
| RNASEH2B | 0,031994991 | -0,464619127 | 0,108 | 0,184 | 1 | Mac | 0,724662364 | 1,379952996 | 0,586956522 |
| UFC12 | 0,032000788 | 0,66316196 | 0,295 | 0,204 | 1 | Mac | 1,583549494 | 0,631492734 | 1,446078431 |
| PMP22 | 0,032012537 | 0,443921838 | 0,211 | 0,126 | 1 | Mac | 1,360297153 | 0,735133495 | 1,674603175 |
| NDUFB23 | 0,032386261 | 0,421559586 | 0,564 | 0,476 | 1 | Mac | 1,339374667 | 0,746617078 | 1,18487395 |
| RPN12 | 0,032624066 | 0,28613062 | 0,331 | 0,233 | 1 | Mac | 1,219365489 | 0,820098657 | 1,420600858 |
| CISD33 | 0,032676341 | 0,463424033 | 0,168 | 0,087 | 1 | Mac | 1,37881035 | 0,725262905 | 1,931034483 |
| FCGR3A | 0,032753195 | -0,830738775 | 0,225 | 0,301 | 1 | Mac | 0,562241256 | 1,778595912 | 0,747508306 |
| CTNND11 | 0,032825534 | -0,388238165 | 0,152 | 0,243 | 1 | Mac | 0,764062116 | 1,308794114 | 0,625514403 |
| CDC42EP32 | 0,033014659 | -0,405173182 | 0,127 | 0,204 | 1 | Mac | 0,755145639 | 1,324247865 | 0,62254902 |
| DUSP233 | 0,033248423 | 0,361892817 | 0,241 | 0,155 | 1 | Mac | 1,285110858 | 0,778142986 | 1,55483871 |
| CD633 | 0,033367302 | 0,45343019 | 0,648 | 0,602 | 1 | Mac | 1,369292054 | 0,730304391 | 1,07641196 |
| FNDC3B | 0,033561916 | 0,467000873 | 0,157 | 0,078 | 1 | Mac | 1,382233043 | 0,723467005 | 2,012820513 |
| NENF3 | 0,033569184 | 0,400160572 | 0,298 | 0,204 | 1 | Mac | 1,31965478 | 0,757773938 | 1,460784314 |
| MTIF31 | 0,033603156 | -0,526200252 | 0,1 | 0,175 | 1 | Mac | 0,694381178 | 1,4401312 | 0,571428571 |
| CYTH1 | 0,033633518 | 0,375356324 | 0,236 | 0,146 | 1 | Mac | 1,297159894 | 0,770914985 | 1,616438356 |
| TRAP12 | 0,033763615 | 0,333900215 | 0,106 | 0,039 | 1 | Mac | 1,260416211 | 0,793388716 | 2,717948718 |
| SF3B11 | 0,034127312 | -0,372182381 | 0,379 | 0,485 | 1 | Mac | 0,772612872 | 1,294309267 | 0,781443299 |
| TMEM1231 | 0,034150916 | -0,42546772 | 0,331 | 0,437 | 1 | Mac | 0,744597295 | 1,343007833 | 0,757437071 |
| PER1 | 0,03427214 | 0,473796265 | 0,336 | 0,252 | 1 | Mac | 1,388759004 | 0,720067339 | 1,333333333 |
| EMP3 | 0,034366042 | 0,424491802 | 0,58 | 0,495 | 1 | Mac | 1,342099657 | 0,745101152 | 1,171717172 |
| CXCL31 | 0,034541882 | 0,90701005 | 0,279 | 0,184 | 1 | Mac | 1,875155255 | 0,533289176 | 1,516304348 |
| NDUFAB14 | 0,034646898 | 0,553115072 | 0,271 | 0,184 | 1 | Mac | 1,46725037 | 0,68154694 | 1,472826087 |
| HERC4 | 0,034806765 | -0,305183626 | 0,057 | 0,117 | 1 | Mac | 0,809339197 | 1,235575891 | 0,487179487 |
| CIITA | 0,034942869 | 0,521735184 | 0,266 | 0,175 | 1 | Mac | 1,435680958 | 0,696533582 | 1,52 |
| GAS53 | 0,03498472 | -0,363455162 | 0,32 | 0,437 | 1 | Mac | 0,777300763 | 1,286503304 | 0,732265446 |
| ITGB13 | 0,03530476 | -0,369244805 | 0,266 | 0,379 | 1 | Mac | 0,774187648 | 1,291676511 | 0,701846966 |
| TPM43 | 0,035743914 | 0,35582572 | 0,512 | 0,427 | 1 | Mac | 1,279717813 | 0,781422271 | 1,199063232 |
| FABP53 | 0,035858338 | 0,389335456 | 0,333 | 0,233 | 1 | Mac | 1,309789941 | 0,763481203 | 1,429184549 |
| SLC43A3 | 0,036076672 | 0,558091751 | 0,173 | 0,097 | 1 | Mac | 1,472320494 | 0,679199946 | 1,783505155 |
| AKAP94 | 0,036237538 | -0,461878864 | 0,209 | 0,301 | 1 | Mac | 0,7260401 | 1,377334393 | 0,694352159 |

| Supplementary Table 4 - clusermarker | | | | | | | | | |
|---|---|---|---|---|---|---|---|---|---|
| | **p_val** | **avg_log2FC** | **pct.1** | **pct.2** | **p_val_adj** | **celltype** | **Foldchange_UP** | **Foldchange_DOWN** | **Ratio_pct1_pct2** |
| NAP1L41 | 0,036318313 | 0,416846514 | 0,165 | 0,087 | 1 | Mac | 1,335006266 | 0,749060154 | 1,896551724 |
| WDFY4 | 0,036390133 | 0,40679216 | 0,133 | 0,058 | 1 | Mac | 1,325734756 | 0,754298697 | 2,293103448 |
| DYNLL13 | 0,036401468 | 0,286579812 | 0,593 | 0,476 | 1 | Mac | 1,219745205 | 0,819843354 | 1,245798319 |
| ERP441 | 0,036647558 | -0,374223072 | 0,173 | 0,262 | 1 | Mac | 0,771520784 | 1,296141362 | 0,660305344 |
| C1QB | 0,036966094 | -0,314627359 | 0,436 | 0,553 | 1 | Mac | 0,804058648 | 1,243690373 | 0,788426763 |
| LACC1 | 0,036972494 | -0,353488226 | 0,065 | 0,126 | 1 | Mac | 0,78268938 | 1,277646058 | 0,515873016 |
| AP3S11 | 0,037173478 | 0,52814756 | 0,187 | 0,107 | 1 | Mac | 1,442076361 | 0,693444555 | 1,747663551 |
| CYBB | 0,037223136 | -0,429062025 | 0,512 | 0,602 | 1 | Mac | 0,742744527 | 1,346357952 | 0,850498339 |
| HSP90AB14 | 0,037284719 | -0,771705808 | 0,808 | 0,748 | 1 | Mac | 0,585724518 | 1,707287247 | 1,080213904 |
| ACOT13 | 0,038004306 | 0,361258444 | 0,117 | 0,049 | 1 | Mac | 1,284545901 | 0,778485221 | 2,387755102 |
| NSFL1C | 0,038296454 | -0,266599075 | 0,133 | 0,214 | 1 | Mac | 0,831276841 | 1,202968674 | 0,621495327 |
| CDKN1B2 | 0,038357739 | 0,378434595 | 0,222 | 0,136 | 1 | Mac | 1,299930592 | 0,769271841 | 1,632352941 |
| CD3021 | 0,038366999 | -0,347854114 | 0,328 | 0,437 | 1 | Mac | 0,785751968 | 1,272666237 | 0,750572082 |
| PIH1D11 | 0,038418947 | -0,306406542 | 0,114 | 0,194 | 1 | Mac | 0,808653443 | 1,236623684 | 0,587628866 |
| P2RY13 | 0,038478247 | -0,448647956 | 0,144 | 0,223 | 1 | Mac | 0,732729215 | 1,36476065 | 0,64573991 |
| AK23 | 0,038495659 | -0,611849551 | 0,154 | 0,233 | 1 | Mac | 0,654357271 | 1,528217145 | 0,660944206 |
| EIF4A22 | 0,038842363 | -0,335299491 | 0,32 | 0,417 | 1 | Mac | 0,792619579 | 1,261639287 | 0,767386091 |
| RABGGTB | 0,039023732 | -0,451276545 | 0,111 | 0,184 | 1 | Mac | 0,731395398 | 1,36724951 | 0,60326087 |
| SF13 | 0,039126877 | 0,396807814 | 0,363 | 0,272 | 1 | Mac | 1,316591523 | 0,759537018 | 1,334558824 |
| SNHG83 | 0,039247011 | -0,270405777 | 0,266 | 0,369 | 1 | Mac | 0,829086321 | 1,206147025 | 0,720867209 |
| SYS1 | 0,03950853 | -0,384593304 | 0,057 | 0,117 | 1 | Mac | 0,765994902 | 1,305491717 | 0,487179487 |
| PRKDC1 | 0,039520976 | -0,336883807 | 0,203 | 0,311 | 1 | Mac | 0,79174963 | 1,263025535 | 0,652733119 |
| NECTIN22 | 0,039557097 | 0,364475164 | 0,19 | 0,107 | 1 | Mac | 1,287413198 | 0,776751397 | 1,775700935 |
| PPM1M | 0,039683609 | 0,301152762 | 0,13 | 0,058 | 1 | Mac | 1,232128532 | 0,811603639 | 2,24137931 |
| IGFBP73 | 0,039896659 | 0,613567783 | 0,146 | 0,068 | 1 | Mac | 1,530038317 | 0,653578403 | 2,147058824 |
| C15orf483 | 0,040181321 | 0,939785479 | 0,306 | 0,223 | 1 | Mac | 1,918242985 | 0,521310391 | 1,372197309 |
| EMD2 | 0,040292889 | 0,356197504 | 0,285 | 0,184 | 1 | Mac | 1,28004764 | 0,781220924 | 1,548913043 |
| TAMALIN | 0,040307606 | 0,368429435 | 0,447 | 0,33 | 1 | Mac | 1,290946698 | 0,774625321 | 1,354545455 |
| SEC11A2 | 0,040833935 | -0,274940759 | 0,412 | 0,534 | 1 | Mac | 0,826484255 | 1,209944405 | 0,771535581 |
| KIF2A1 | 0,040835049 | 0,308904724 | 0,13 | 0,058 | 1 | Mac | 1,238766887 | 0,807254384 | 2,24137931 |
| RUFY11 | 0,040909199 | 0,387813186 | 0,138 | 0,068 | 1 | Mac | 1,308408635 | 0,764287221 | 2,029411765 |

| Supplementary Table 4 - clusermarker | | | | | | | | | |
|---|---|---|---|---|---|---|---|---|---|
| | **p_val** | **avg_log2FC** | **pct.1** | **pct.2** | **p_val_adj** | **celltype** | **Foldchange_UP** | **Foldchange_DOWN** | **Ratio_pct1_pct2** |
| UFM13 | 0,041158318 | -0,35259202 | 0,179 | 0,272 | 1 | Mac | 0,783175739 | 1,276852627 | 0,658088235 |
| CBX6 | 0,041298091 | 0,323475694 | 0,127 | 0,058 | 1 | Mac | 1,251341612 | 0,799142289 | 2,189655172 |
| COX6C2 | 0,04133897 | 0,255089854 | 0,585 | 0,485 | 1 | Mac | 1,193410069 | 0,837934944 | 1,206185567 |
| CXCR41 | 0,041343543 | 0,369698102 | 0,575 | 0,485 | 1 | Mac | 1,292082422 | 0,773944435 | 1,18556701 |
| SLC39A8 | 0,041853466 | 0,385156644 | 0,141 | 0,068 | 1 | Mac | 1,306001581 | 0,765695857 | 2,073529412 |
| BTG32 | 0,042276916 | 0,382172502 | 0,154 | 0,078 | 1 | Mac | 1,303302975 | 0,767281299 | 1,974358974 |
| APOC2 | 0,04257414 | -0,470938107 | 0,108 | 0,184 | 1 | Mac | 0,721495296 | 1,386010423 | 0,586956522 |
| TLE42 | 0,043158209 | -0,364328539 | 0,108 | 0,184 | 1 | Mac | 0,776830344 | 1,287282361 | 0,586956522 |
| CYRIA | 0,043175522 | 0,427813677 | 0,293 | 0,194 | 1 | Mac | 1,345193467 | 0,743387494 | 1,510309278 |
| MRPL122 | 0,043336739 | 0,36667994 | 0,1 | 0,039 | 1 | Mac | 1,289382171 | 0,775565246 | 2,564102564 |
| PPP1R21 | 0,043521901 | 0,43246436 | 0,263 | 0,175 | 1 | Mac | 1,349536839 | 0,740994963 | 1,502857143 |
| ZYX1 | 0,043655493 | 0,309179716 | 0,447 | 0,369 | 1 | Mac | 1,239003031 | 0,807100528 | 1,211382114 |
| CD300LF | 0,043685881 | -0,391841886 | 0,051 | 0,107 | 1 | Mac | 0,76215594 | 1,312067449 | 0,476635514 |
| LASP11 | 0,043793907 | -0,413324684 | 0,154 | 0,233 | 1 | Mac | 0,750890953 | 1,331751296 | 0,660944206 |
| FGD4 | 0,044052536 | -0,282129783 | 0,206 | 0,301 | 1 | Mac | 0,822376085 | 1,215988667 | 0,684385382 |
| FAM162A1 | 0,044080396 | 0,542168046 | 0,192 | 0,117 | 1 | Mac | 1,456159153 | 0,68673812 | 1,641025641 |
| RPL92 | 0,044108222 | -0,260955895 | 0,913 | 0,893 | 1 | Mac | 0,834534794 | 1,198272388 | 1,022396417 |
| AQR | 0,044387329 | -0,300538618 | 0,073 | 0,136 | 1 | Mac | 0,811949205 | 1,231604137 | 0,536764706 |
| CYB561D21 | 0,0445645 | 0,381008425 | 0,1 | 0,039 | 1 | Mac | 1,302251795 | 0,76790065 | 2,564102564 |
| RTN31 | 0,045114037 | -0,496648581 | 0,214 | 0,311 | 1 | Mac | 0,708751318 | 1,41093212 | 0,688102894 |
| UBQLN1 | 0,045141144 | 0,46984935 | 0,157 | 0,087 | 1 | Mac | 1,384964839 | 0,722039991 | 1,804597701 |
| CCNH2 | 0,045580809 | 0,428690549 | 0,309 | 0,214 | 1 | Mac | 1,346011325 | 0,742935799 | 1,443925234 |
| KRTCAP23 | 0,045723541 | 0,433242549 | 0,461 | 0,388 | 1 | Mac | 1,350264976 | 0,740595378 | 1,18814433 |
| ARPC5L4 | 0,045829923 | 0,313681261 | 0,152 | 0,078 | 1 | Mac | 1,242875047 | 0,804586111 | 1,948717949 |
| AFF1 | 0,04632675 | -0,356692992 | 0,103 | 0,175 | 1 | Mac | 0,780952663 | 1,280487342 | 0,588571429 |
| MRPL541 | 0,047149528 | 0,421556413 | 0,206 | 0,126 | 1 | Mac | 1,339371721 | 0,74661872 | 1,634920635 |
| WIPI2 | 0,047644021 | -0,478239656 | 0,114 | 0,184 | 1 | Mac | 0,717852998 | 1,393042869 | 0,619565217 |
| IRAG2 | 0,047770434 | -0,403463379 | 0,076 | 0,136 | 1 | Mac | 0,756041127 | 1,322679368 | 0,558823529 |
| ARHGAP91 | 0,048149986 | -0,327622259 | 0,127 | 0,204 | 1 | Mac | 0,796848708 | 1,254943366 | 0,62254902 |
| SSR32 | 0,048548567 | 0,508825434 | 0,385 | 0,301 | 1 | Mac | 1,422891281 | 0,702794383 | 1,279069767 |
| RFX5 | 0,04868857 | -0,492590929 | 0,06 | 0,117 | 1 | Mac | 0,710747523 | 1,406969378 | 0,512820513 |

| Supplementary Table 4 - clusermarker | | | | | | | | | |
|---|---|---|---|---|---|---|---|---|---|
| | **p_val** | **avg_log2FC** | **pct.1** | **pct.2** | **p_val_adj** | **celltype** | **Foldchange_UP** | **Foldchange_DOWN** | **Ratio_pct1_pct2** |
| APOL6 | 0,048708633 | -0,258101073 | 0,076 | 0,136 | 1 | Mac | 0,836187816 | 1,195903577 | 0,558823529 |
| SF3B62 | 0,049006941 | 0,303181348 | 0,271 | 0,175 | 1 | Mac | 1,233862258 | 0,810463237 | 1,548571429 |
| ARL4C1 | 0,04965725 | 0,317462838 | 0,355 | 0,243 | 1 | Mac | 1,246137131 | 0,8024799 | 1,46090535 |
| NCKAP1L | 0,04966528 | -0,341034988 | 0,266 | 0,359 | 1 | Mac | 0,78947474 | 1,266664972 | 0,740947075 |
| CCT6A2 | 0,049794838 | 0,523797896 | 0,282 | 0,204 | 1 | Mac | 1,437735109 | 0,695538416 | 1,382352941 |
| UBE2B2 | 0,049863851 | 0,450842483 | 0,301 | 0,214 | 1 | Mac | 1,366838209 | 0,731615485 | 1,406542056 |
| FMNL1 | 0,050005902 | 0,423453778 | 0,309 | 0,214 | 1 | Mac | 1,341134359 | 0,745637447 | 1,443925234 |
| MANF2 | 0,050032978 | 0,34522808 | 0,201 | 0,117 | 1 | Mac | 1,270351801 | 0,787183518 | 1,717948718 |
| ACSL31 | 0,050041005 | 0,420623332 | 0,168 | 0,097 | 1 | Mac | 1,338505746 | 0,747101761 | 1,731958763 |
| TIMM132 | 0,050095893 | 0,287268631 | 0,192 | 0,117 | 1 | Mac | 1,220327715 | 0,819452011 | 1,641025641 |
| HOTAIRM12 | 0,050166016 | -0,367284773 | 0,089 | 0,155 | 1 | Mac | 0,775240167 | 1,289922843 | 0,574193548 |
| IST1 | 0,050167693 | -0,365549213 | 0,13 | 0,204 | 1 | Mac | 0,776173341 | 1,288372 | 0,637254902 |
| HNRNPK2 | 0,050530183 | -0,398356177 | 0,499 | 0,583 | 1 | Mac | 0,758722288 | 1,318005305 | 0,855917667 |
| SERPING11 | 0,050654533 | -0,58244829 | 0,119 | 0,194 | 1 | Mac | 0,667829492 | 1,4973882 | 0,613402062 |
| NDUFAF32 | 0,050669239 | 0,376831755 | 0,203 | 0,126 | 1 | Mac | 1,298487166 | 0,77012698 | 1,611111111 |
| PARP1 | 0,050763551 | 0,264958282 | 0,209 | 0,126 | 1 | Mac | 1,201601303 | 0,832222799 | 1,658730159 |
| TMEM167A1 | 0,050767183 | 0,420024021 | 0,146 | 0,078 | 1 | Mac | 1,337949832 | 0,747412179 | 1,871794872 |
| PTGS2 | 0,050774758 | -0,776660022 | 0,157 | 0,233 | 1 | Mac | 0,58371659 | 1,713160148 | 0,673819742 |
| ADI13 | 0,051138425 | 0,257485355 | 0,154 | 0,078 | 1 | Mac | 1,195393294 | 0,836544763 | 1,974358974 |
| TRIO | 0,051138455 | 0,264621755 | 0,125 | 0,058 | 1 | Mac | 1,201321046 | 0,832416949 | 2,155172414 |
| ANAPC162 | 0,051223048 | 0,486786984 | 0,274 | 0,194 | 1 | Mac | 1,401320527 | 0,713612611 | 1,412371134 |
| BAZ1A2 | 0,052235176 | -0,268464486 | 0,276 | 0,369 | 1 | Mac | 0,830202692 | 1,204525124 | 0,74796748 |
| QSOX1 | 0,052242138 | 0,325775241 | 0,173 | 0,097 | 1 | Mac | 1,253337746 | 0,797869531 | 1,783505155 |
| MYO1E1 | 0,052529356 | 0,370969825 | 0,136 | 0,068 | 1 | Mac | 1,293221883 | 0,773262511 | 2 |
| UQCRC21 | 0,052555008 | -0,362249483 | 0,184 | 0,272 | 1 | Mac | 0,777950635 | 1,285428606 | 0,676470588 |
| UGCG2 | 0,052558759 | 0,570184436 | 0,19 | 0,117 | 1 | Mac | 1,484713366 | 0,673530678 | 1,623931624 |
| NDUFV24 | 0,052631574 | 0,359441011 | 0,339 | 0,262 | 1 | Mac | 1,282928716 | 0,779466535 | 1,29389313 |
| CYC11 | 0,05267217 | 0,296992349 | 0,252 | 0,165 | 1 | Mac | 1,228580465 | 0,813947502 | 1,527272727 |
| TXN4 | 0,052866083 | 0,559996683 | 0,493 | 0,388 | 1 | Mac | 1,474265827 | 0,678303723 | 1,270618557 |
| ACOT9 | 0,053175392 | 0,402913622 | 0,168 | 0,097 | 1 | Mac | 1,32217544 | 0,756329281 | 1,731958763 |
| TTC33 | 0,053206214 | -0,352057464 | 0,192 | 0,272 | 1 | Mac | 0,78346598 | 1,276379607 | 0,705882353 |

| Supplementary Table 4 - clusermarker | | | | | | | | | |
|---|---|---|---|---|---|---|---|---|---|
| | **p_val** | **avg_log2FC** | **pct.1** | **pct.2** | **p_val_adj** | **celltype** | **Foldchange_UP** | **Foldchange_DOWN** | **Ratio_pct1_pct2** |
| SRGAP2C | 0,053450777 | -0,311779676 | 0,054 | 0,107 | 1 | Mac | 0,805647318 | 1,241237919 | 0,504672897 |
| PET1003 | 0,053661202 | -0,313433589 | 0,257 | 0,35 | 1 | Mac | 0,804724249 | 1,242661696 | 0,734285714 |
| PARVB | 0,053861333 | 0,373214624 | 0,163 | 0,087 | 1 | Mac | 1,295235671 | 0,772060268 | 1,873563218 |
| MRPL412 | 0,053893807 | 0,318291542 | 0,247 | 0,165 | 1 | Mac | 1,246853135 | 0,802019077 | 1,496969697 |
| CRYBG11 | 0,054309744 | 0,503212041 | 0,247 | 0,165 | 1 | Mac | 1,417365699 | 0,705534218 | 1,496969697 |
| ILF3-DT | 0,0543703 | -0,258664551 | 0,068 | 0,126 | 1 | Mac | 0,835861287 | 1,196370756 | 0,53968254 |
| TSC22D3 | 0,055032714 | 0,371354257 | 0,588 | 0,553 | 1 | Mac | 1,293566531 | 0,773056489 | 1,063291139 |
| CYB5D1 | 0,055202373 | -0,358322403 | 0,092 | 0,155 | 1 | Mac | 0,780071136 | 1,28193437 | 0,593548387 |
| NRP1 | 0,055210814 | 0,369691671 | 0,149 | 0,078 | 1 | Mac | 1,292076662 | 0,773947885 | 1,91025641 |
| MRPL401 | 0,055397686 | 0,260544308 | 0,125 | 0,058 | 1 | Mac | 1,197930581 | 0,834772912 | 2,155172414 |
| GABPB1-AS1 | 0,055406815 | 0,339246784 | 0,136 | 0,068 | 1 | Mac | 1,265095928 | 0,790453892 | 2 |
| PSMB83 | 0,055816564 | 0,411265693 | 0,336 | 0,262 | 1 | Mac | 1,329851997 | 0,751963378 | 1,282442748 |
| LUCAT1 | 0,055983845 | -0,594965556 | 0,149 | 0,223 | 1 | Mac | 0,662060261 | 1,510436524 | 0,668161435 |
| EIF4B4 | 0,05624717 | -0,271678007 | 0,425 | 0,534 | 1 | Mac | 0,82835552 | 1,207211126 | 0,79588015 |
| HNRNPUL21 | 0,056526263 | -0,42674302 | 0,195 | 0,282 | 1 | Mac | 0,743939384 | 1,344195538 | 0,691489362 |
| LARP4 | 0,056750897 | -0,357504774 | 0,084 | 0,146 | 1 | Mac | 0,780513357 | 1,281208056 | 0,575342466 |
| LMNB1 | 0,056946797 | 0,278199717 | 0,125 | 0,058 | 1 | Mac | 1,212680683 | 0,824619386 | 2,155172414 |
| TMEM14C3 | 0,057032661 | 0,326534594 | 0,377 | 0,282 | 1 | Mac | 1,253997606 | 0,797449688 | 1,336879433 |
| DNAJC10 | 0,057184696 | -0,300751078 | 0,108 | 0,175 | 1 | Mac | 0,811829642 | 1,231785523 | 0,617142857 |
| LBR | 0,057255159 | 0,331374764 | 0,163 | 0,087 | 1 | Mac | 1,258211771 | 0,794778767 | 1,873563218 |
| PAXX2 | 0,057507854 | 0,323755878 | 0,23 | 0,146 | 1 | Mac | 1,251584657 | 0,798987103 | 1,575342466 |
| PPP1CC1 | 0,057534299 | 0,419636325 | 0,201 | 0,126 | 1 | Mac | 1,337590332 | 0,747613059 | 1,595238095 |
| PTP4A22 | 0,05761982 | 0,263101912 | 0,477 | 0,379 | 1 | Mac | 1,200056151 | 0,833294341 | 1,258575198 |
| SRSF10 | 0,057872593 | 0,33564876 | 0,19 | 0,107 | 1 | Mac | 1,26194476 | 0,792427713 | 1,775700935 |
| MAN2C1 | 0,058051346 | 0,291722615 | 0,108 | 0,049 | 1 | Mac | 1,224101014 | 0,816926045 | 2,204081633 |
| TRABD | 0,058255535 | 0,390943245 | 0,236 | 0,155 | 1 | Mac | 1,311250429 | 0,762630828 | 1,522580645 |
| TSC22D13 | 0,058632982 | -0,31845299 | 0,084 | 0,146 | 1 | Mac | 0,80192933 | 1,246992674 | 0,575342466 |
| NANS3 | 0,058759234 | 0,346164719 | 0,29 | 0,204 | 1 | Mac | 1,271176817 | 0,786672623 | 1,421568627 |
| PFDN12 | 0,058845266 | -0,251064357 | 0,084 | 0,146 | 1 | Mac | 0,840276267 | 1,190084784 | 0,575342466 |
| MICAL1 | 0,058960786 | 0,422791164 | 0,222 | 0,146 | 1 | Mac | 1,340518532 | 0,745979989 | 1,520547945 |
| HLA-DPB11 | 0,058981272 | -0,403705151 | 0,946 | 0,913 | 1 | Mac | 0,755914437 | 1,322901046 | 1,036144578 |

| Supplementary Table 4 - clusermarker | | | | | | | | | |
|---|---|---|---|---|---|---|---|---|---|
| | **p_val** | **avg_log2FC** | **pct.1** | **pct.2** | **p_val_adj** | **celltype** | **Foldchange_UP** | **Foldchange_DOWN** | **Ratio_pct1_pct2** |
| THBD1 | 0,05910593 | 0,347156078 | 0,271 | 0,175 | 1 | Mac | 1,272050616 | 0,78613224 | 1,548571429 |
| HSPE13 | 0,059235246 | -0,664569727 | 0,39 | 0,447 | 1 | Mac | 0,630876831 | 1,58509546 | 0,872483221 |
| KLF91 | 0,059825694 | 0,350350308 | 0,154 | 0,087 | 1 | Mac | 1,274870147 | 0,784393612 | 1,770114943 |
| AIDA | 0,05996845 | -0,30820581 | 0,106 | 0,175 | 1 | Mac | 0,807645553 | 1,238166911 | 0,605714286 |
| HCST1 | 0,060343867 | 0,302179682 | 0,488 | 0,417 | 1 | Mac | 1,233005882 | 0,811026139 | 1,170263789 |
| ATRAID2 | 0,060349891 | 0,326772693 | 0,241 | 0,155 | 1 | Mac | 1,25420458 | 0,79731809 | 1,55483871 |
| CYB561A31 | 0,060353945 | 0,365298226 | 0,133 | 0,068 | 1 | Mac | 1,288147881 | 0,776308384 | 1,955882353 |
| BTG22 | 0,061865076 | -0,265872905 | 0,355 | 0,447 | 1 | Mac | 0,831695364 | 1,202363322 | 0,794183445 |
| FTX1 | 0,061926898 | -0,311282551 | 0,062 | 0,117 | 1 | Mac | 0,805924976 | 1,240810286 | 0,52991453 |
| IL1A | 0,06214547 | -0,285150163 | 0,076 | 0,136 | 1 | Mac | 0,820656186 | 1,218537088 | 0,558823529 |
| KRCC1 | 0,06226287 | 0,355373008 | 0,106 | 0,049 | 1 | Mac | 1,279316305 | 0,781667517 | 2,163265306 |
| MIR155HG1 | 0,063181762 | -0,536518244 | 0,057 | 0,107 | 1 | Mac | 0,689432759 | 1,450467775 | 0,53271028 |
| SIPA1 | 0,063350971 | -0,440078895 | 0,1 | 0,165 | 1 | Mac | 0,737094299 | 1,356678516 | 0,606060606 |
| VMA211 | 0,063976843 | 0,525339069 | 0,209 | 0,136 | 1 | Mac | 1,439271804 | 0,694795797 | 1,536764706 |
| NCL5 | 0,065136979 | -0,279055888 | 0,528 | 0,592 | 1 | Mac | 0,824130159 | 1,213400564 | 0,891891892 |
| PRKACB1 | 0,065262035 | -0,467458721 | 0,125 | 0,194 | 1 | Mac | 0,723237445 | 1,382671773 | 0,644329897 |
| MRPL51 | 0,065401185 | 0,432337156 | 0,287 | 0,214 | 1 | Mac | 1,349417855 | 0,7410603 | 1,341121495 |
| UBE2J11 | 0,065541833 | 0,412605478 | 0,22 | 0,146 | 1 | Mac | 1,331087562 | 0,751265378 | 1,506849315 |
| HES12 | 0,065982634 | -0,25164645 | 0,084 | 0,146 | 1 | Mac | 0,839937305 | 1,190565051 | 0,575342466 |
| UPF22 | 0,066130823 | -0,448509891 | 0,13 | 0,204 | 1 | Mac | 0,73279934 | 1,364630051 | 0,637254902 |
| DDX1 | 0,066167446 | -0,289497374 | 0,057 | 0,107 | 1 | Mac | 0,81818706 | 1,222214392 | 0,53271028 |
| NDUFA132 | 0,066238114 | 0,318409052 | 0,528 | 0,437 | 1 | Mac | 1,246954697 | 0,801953754 | 1,208237986 |
| HCLS1 | 0,066698128 | -0,500360487 | 0,561 | 0,621 | 1 | Mac | 0,706930118 | 1,414566977 | 0,903381643 |
| BAX4 | 0,0667723 | 0,325313447 | 0,306 | 0,223 | 1 | Mac | 1,252936628 | 0,798124963 | 1,372197309 |
| GNA12 | 0,067614237 | 0,388888933 | 0,152 | 0,087 | 1 | Mac | 1,309384615 | 0,763717542 | 1,747126437 |
| FOXO3 | 0,067734252 | -0,445742539 | 0,165 | 0,243 | 1 | Mac | 0,734206332 | 1,36201495 | 0,679012346 |
| IL182 | 0,067755066 | -0,55669779 | 0,276 | 0,35 | 1 | Mac | 0,67985652 | 1,470898595 | 0,788571429 |
| ARPC53 | 0,067793696 | 0,28628302 | 0,577 | 0,515 | 1 | Mac | 1,219494305 | 0,82001203 | 1,12038835 |
| MEGF9 | 0,067809234 | -0,367576425 | 0,092 | 0,155 | 1 | Mac | 0,775083462 | 1,290183637 | 0,593548387 |
| TMEM2561 | 0,067825691 | 0,376554462 | 0,287 | 0,204 | 1 | Mac | 1,298237614 | 0,770275017 | 1,406862745 |
| TNFRSF12A1 | 0,068127261 | -0,341897967 | 0,057 | 0,107 | 1 | Mac | 0,78900264 | 1,267422882 | 0,53271028 |

| Supplementary Table 4 - clusermarker | | | | | | | | | |
|---|---|---|---|---|---|---|---|---|---|
| | **p_val** | **avg_log2FC** | **pct.1** | **pct.2** | **p_val_adj** | **celltype** | **Foldchange_UP** | **Foldchange_DOWN** | **Ratio_pct1_pct2** |
| UQCRH2 | 0,068309139 | 0,408864294 | 0,534 | 0,466 | 1 | Mac | 1,327640269 | 0,753216081 | 1,145922747 |
| UBE2I3 | 0,068714242 | 0,383205766 | 0,29 | 0,214 | 1 | Mac | 1,30423674 | 0,766731966 | 1,355140187 |
| CLPTM11 | 0,068797573 | 0,291928054 | 0,154 | 0,087 | 1 | Mac | 1,224275337 | 0,816809724 | 1,770114943 |
| EFHD22 | 0,068801454 | 0,323669395 | 0,491 | 0,388 | 1 | Mac | 1,251509633 | 0,799035 | 1,265463918 |
| COX7A21 | 0,068977405 | 0,259837281 | 0,485 | 0,427 | 1 | Mac | 1,197343651 | 0,835182113 | 1,135831382 |
| ANXA22 | 0,069265837 | 0,366956783 | 0,678 | 0,592 | 1 | Mac | 1,289629618 | 0,775416434 | 1,14527027 |
| RMND5A1 | 0,069306919 | -0,252929531 | 0,07 | 0,126 | 1 | Mac | 0,839190626 | 1,191624368 | 0,555555556 |
| PNRC14 | 0,069598977 | 0,263625955 | 0,851 | 0,757 | 1 | Mac | 1,200492138 | 0,832991711 | 1,124174373 |
| DENND1A | 0,069749707 | 0,273318191 | 0,108 | 0,049 | 1 | Mac | 1,208584371 | 0,827414307 | 2,204081633 |
| OAT4 | 0,069804737 | -0,301308421 | 0,133 | 0,204 | 1 | Mac | 0,811516076 | 1,232261479 | 0,651960784 |
| TIMM17A1 | 0,069851361 | 0,276733228 | 0,165 | 0,097 | 1 | Mac | 1,211448628 | 0,825458032 | 1,701030928 |
| ARHGAP451 | 0,069954475 | 0,307353741 | 0,247 | 0,165 | 1 | Mac | 1,237435854 | 0,808122697 | 1,496969697 |
| ID23 | 0,070068816 | -0,357609263 | 0,39 | 0,476 | 1 | Mac | 0,780456829 | 1,281300852 | 0,819327731 |
| SCIMP | 0,070090286 | -0,273439691 | 0,138 | 0,214 | 1 | Mac | 0,827344628 | 1,208686159 | 0,644859813 |
| SMIM201 | 0,070397414 | 0,386985262 | 0,106 | 0,049 | 1 | Mac | 1,30765799 | 0,764725951 | 2,163265306 |
| LGALS3BP1 | 0,070592404 | -0,48827005 | 0,065 | 0,117 | 1 | Mac | 0,712879406 | 1,402761802 | 0,555555556 |
| MEF2C2 | 0,071047681 | -0,402542173 | 0,271 | 0,359 | 1 | Mac | 0,756524037 | 1,321835065 | 0,754874652 |
| SFT2D1 | 0,071529814 | 0,484686046 | 0,257 | 0,184 | 1 | Mac | 1,399281326 | 0,714652573 | 1,39673913 |
| KCNAB2 | 0,071658619 | 0,389018861 | 0,198 | 0,126 | 1 | Mac | 1,309502543 | 0,763648765 | 1,571428571 |
| RPL315 | 0,071677049 | -0,334494282 | 0,924 | 0,903 | 1 | Mac | 0,793062086 | 1,260935327 | 1,023255814 |
| WRNIP1 | 0,071893494 | -0,330712509 | 0,057 | 0,107 | 1 | Mac | 0,795143686 | 1,257634334 | 0,53271028 |
| MSRB23 | 0,071919697 | 0,364723351 | 0,106 | 0,049 | 1 | Mac | 1,287634691 | 0,776617784 | 2,163265306 |
| TMEM2194 | 0,072274238 | 0,271767707 | 0,398 | 0,33 | 1 | Mac | 1,207286187 | 0,828304018 | 1,206060606 |
| REXO22 | 0,072416451 | 0,34816299 | 0,117 | 0,058 | 1 | Mac | 1,272938739 | 0,785583759 | 2,017241379 |
| SUMO12 | 0,074334209 | 0,3376759 | 0,233 | 0,155 | 1 | Mac | 1,263719173 | 0,79131505 | 1,503225806 |
| HAVCR2 | 0,074470353 | -0,283505159 | 0,23 | 0,32 | 1 | Mac | 0,821592457 | 1,217148468 | 0,71875 |
| MRPL372 | 0,074717137 | 0,304996992 | 0,144 | 0,078 | 1 | Mac | 1,235416062 | 0,809443904 | 1,846153846 |
| SEC61B4 | 0,074817755 | 0,25108803 | 0,583 | 0,495 | 1 | Mac | 1,190104312 | 0,84026248 | 1,177777778 |
| ADA | 0,074905242 | -0,273062728 | 0,057 | 0,107 | 1 | Mac | 0,827560833 | 1,208370382 | 0,53271028 |
| ARSA1 | 0,074945173 | -0,289689154 | 0,079 | 0,136 | 1 | Mac | 0,818078304 | 1,222376873 | 0,580882353 |
| TERF2IP | 0,076064944 | 0,3024179 | 0,255 | 0,175 | 1 | Mac | 1,233209493 | 0,810892234 | 1,457142857 |

| Supplementary Table 4 - clusermarker | | | | | | | | | |
|---|---|---|---|---|---|---|---|---|---|
| | **p_val** | **avg_log2FC** | **pct.1** | **pct.2** | **p_val_adj** | **celltype** | **Foldchange_UP** | **Foldchange_DOWN** | **Ratio_pct1_pct2** |
| TXNDC12 | 0,076155913 | -0,327481433 | 0,146 | 0,223 | 1 | Mac | 0,796926495 | 1,254820873 | 0,65470852 |
| LGALS91 | 0,076521092 | -0,367730665 | 0,341 | 0,427 | 1 | Mac | 0,775000601 | 1,29032158 | 0,798594848 |
| HLA-DMA3 | 0,076856415 | -0,389220458 | 0,718 | 0,699 | 1 | Mac | 0,763542064 | 1,30968554 | 1,027181688 |
| RAPGEF22 | 0,077160264 | -0,261758846 | 0,087 | 0,146 | 1 | Mac | 0,834070452 | 1,198939488 | 0,595890411 |
| SELENOW3 | 0,077366575 | 0,400180056 | 0,287 | 0,223 | 1 | Mac | 1,319672603 | 0,757763704 | 1,286995516 |
| DDX3X2 | 0,077519335 | -0,371212509 | 0,523 | 0,621 | 1 | Mac | 0,773132447 | 1,293439441 | 0,842190016 |
| RASSF5 | 0,077524237 | -0,269152602 | 0,244 | 0,33 | 1 | Mac | 0,829806808 | 1,205099779 | 0,739393939 |
| MCRIP11 | 0,077552536 | 0,431491278 | 0,127 | 0,068 | 1 | Mac | 1,348626898 | 0,741494924 | 1,867647059 |
| NFYC | 0,077600414 | -0,35618865 | 0,111 | 0,175 | 1 | Mac | 0,781225718 | 1,280039784 | 0,634285714 |
| HNRNPM1 | 0,077800155 | 0,312160212 | 0,331 | 0,252 | 1 | Mac | 1,24156536 | 0,805434842 | 1,313492063 |
| NXT11 | 0,077801278 | 0,261471063 | 0,127 | 0,068 | 1 | Mac | 1,198700352 | 0,834236845 | 1,867647059 |
| EMC43 | 0,078671687 | 0,398990239 | 0,171 | 0,107 | 1 | Mac | 1,318584693 | 0,758388904 | 1,598130841 |
| EIF3H1 | 0,079759226 | -0,399750546 | 0,439 | 0,524 | 1 | Mac | 0,757989335 | 1,319279776 | 0,83778626 |
| REX1BD4 | 0,079759527 | 0,488287469 | 0,358 | 0,282 | 1 | Mac | 1,402778738 | 0,712870799 | 1,269503546 |
| LAMTOR43 | 0,079803475 | 0,269868091 | 0,577 | 0,485 | 1 | Mac | 1,205697583 | 0,829395376 | 1,189690722 |
| DNAJC153 | 0,080031907 | 0,417445037 | 0,347 | 0,272 | 1 | Mac | 1,335560228 | 0,74874946 | 1,275735294 |
| CWC251 | 0,080261107 | -0,263416343 | 0,057 | 0,107 | 1 | Mac | 0,833112747 | 1,200317729 | 0,53271028 |
| EPS151 | 0,080279323 | 0,366317187 | 0,206 | 0,136 | 1 | Mac | 1,289058008 | 0,775760279 | 1,514705882 |
| FAF1 | 0,080290576 | -0,354409691 | 0,073 | 0,126 | 1 | Mac | 0,782189627 | 1,278462364 | 0,579365079 |
| CBX32 | 0,080421807 | 0,459085548 | 0,268 | 0,194 | 1 | Mac | 1,374670208 | 0,727447205 | 1,381443299 |
| ST8SIA41 | 0,08052916 | 0,254447188 | 0,217 | 0,146 | 1 | Mac | 1,192878568 | 0,838308296 | 1,48630137 |
| CUL32 | 0,080619721 | 0,322336507 | 0,152 | 0,087 | 1 | Mac | 1,250353913 | 0,79977356 | 1,747126437 |
| SOD11 | 0,080963123 | 0,511786908 | 0,325 | 0,262 | 1 | Mac | 1,425815103 | 0,70135321 | 1,240458015 |
| TBC1D9B | 0,081094436 | 0,299844716 | 0,127 | 0,068 | 1 | Mac | 1,231011907 | 0,812339827 | 1,867647059 |
| TOX41 | 0,081700913 | 0,407967013 | 0,149 | 0,087 | 1 | Mac | 1,326814803 | 0,753684687 | 1,712643678 |
| RIPOR11 | 0,08180943 | -0,455538163 | 0,119 | 0,184 | 1 | Mac | 0,729238096 | 1,371294239 | 0,64673913 |
| DUSP24 | 0,081899477 | 0,303605196 | 0,48 | 0,369 | 1 | Mac | 1,234224806 | 0,810225167 | 1,300813008 |
| LRP1 | 0,08233538 | -0,273002171 | 0,22 | 0,301 | 1 | Mac | 0,82759557 | 1,208319662 | 0,73089701 |
| PARP14 | 0,082791314 | -0,348386942 | 0,203 | 0,282 | 1 | Mac | 0,785461821 | 1,273136355 | 0,719858156 |
| FN11 | 0,083317103 | -1,383515181 | 0,084 | 0,136 | 1 | Mac | 0,383283771 | 2,609032979 | 0,617647059 |
| ARL8B | 0,083802333 | 0,390675596 | 0,317 | 0,243 | 1 | Mac | 1,311007189 | 0,762772324 | 1,304526749 |

| Supplementary Table 4 - clusermarker | | | | | | | | | |
|---|---|---|---|---|---|---|---|---|---|
| | **p_val** | **avg_log2FC** | **pct.1** | **pct.2** | **p_val_adj** | **celltype** | **Foldchange_UP** | **Foldchange_DOWN** | **Ratio_pct1_pct2** |
| NKAP | 0,084215455 | 0,253632381 | 0,106 | 0,049 | 1 | Mac | 1,192205043 | 0,83878189 | 2,163265306 |
| PHC2 | 0,084357631 | -0,495302252 | 0,103 | 0,165 | 1 | Mac | 0,709413037 | 1,409616046 | 0,624242424 |
| S100A12 | 0,08450049 | -0,438568487 | 0,079 | 0,136 | 1 | Mac | 0,737866393 | 1,355258906 | 0,580882353 |
| EHMT11 | 0,084741957 | 0,306132276 | 0,1 | 0,049 | 1 | Mac | 1,236388616 | 0,808807188 | 2,040816327 |
| ARHGEF2 | 0,084762884 | -0,354055254 | 0,119 | 0,184 | 1 | Mac | 0,782381817 | 1,278148314 | 0,64673913 |
| MLXIP1 | 0,084887492 | 0,280657799 | 0,141 | 0,078 | 1 | Mac | 1,214748625 | 0,823215585 | 1,807692308 |
| ARHGAP21 | 0,084995461 | -0,504857736 | 0,092 | 0,146 | 1 | Mac | 0,704729867 | 1,418983424 | 0,630136986 |
| GOLGA2 | 0,085170272 | -0,330050403 | 0,068 | 0,117 | 1 | Mac | 0,795508691 | 1,257057291 | 0,581196581 |
| FXR13 | 0,085650764 | -0,269937523 | 0,125 | 0,194 | 1 | Mac | 0,829355461 | 1,205755611 | 0,644329897 |
| MRPL211 | 0,085964294 | 0,375511058 | 0,125 | 0,068 | 1 | Mac | 1,297299027 | 0,770832306 | 1,838235294 |
| LILRB4 | 0,086150386 | -0,439793509 | 0,187 | 0,262 | 1 | Mac | 0,737240121 | 1,356410172 | 0,713740458 |
| TWF21 | 0,086171808 | 0,349579251 | 0,317 | 0,243 | 1 | Mac | 1,274188968 | 0,784812948 | 1,304526749 |
| JPT12 | 0,086245757 | 0,440791093 | 0,366 | 0,301 | 1 | Mac | 1,357348417 | 0,736730516 | 1,215946844 |
| GRPEL11 | 0,086489785 | 0,252335999 | 0,13 | 0,068 | 1 | Mac | 1,191134229 | 0,839535945 | 1,911764706 |
| UVRAG1 | 0,086655229 | 0,408030224 | 0,206 | 0,136 | 1 | Mac | 1,326872937 | 0,753651666 | 1,514705882 |
| SYNCRIP1 | 0,086669399 | 0,317000342 | 0,241 | 0,165 | 1 | Mac | 1,245737711 | 0,802737199 | 1,460606061 |
| PTPN2 | 0,08686237 | 0,270994708 | 0,236 | 0,165 | 1 | Mac | 1,206639494 | 0,828747944 | 1,43030303 |
| PXN | 0,087007914 | -0,339332545 | 0,06 | 0,107 | 1 | Mac | 0,790406905 | 1,265171134 | 0,560747664 |
| RBM393 | 0,087844153 | -0,29076212 | 0,444 | 0,524 | 1 | Mac | 0,817470107 | 1,223286322 | 0,847328244 |
| GTF2A22 | 0,087856703 | 0,467559002 | 0,23 | 0,165 | 1 | Mac | 1,382767885 | 0,723187175 | 1,393939394 |
| GNB11 | 0,088378103 | 0,266313048 | 0,401 | 0,32 | 1 | Mac | 1,202730199 | 0,831441666 | 1,253125 |
| BANF12 | 0,088509366 | 0,309289688 | 0,328 | 0,252 | 1 | Mac | 1,239097479 | 0,807039008 | 1,301587302 |
| DDX184 | 0,08864093 | -0,253124492 | 0,209 | 0,282 | 1 | Mac | 0,839077229 | 1,191785411 | 0,741134752 |
| DDIT31 | 0,088656692 | -0,251863051 | 0,089 | 0,146 | 1 | Mac | 0,839811209 | 1,190743812 | 0,609589041 |
| PPP1R18 | 0,088717033 | -0,513740151 | 0,252 | 0,32 | 1 | Mac | 0,700404301 | 1,4277468 | 0,7875 |
| LUC7L32 | 0,089381571 | -0,387585273 | 0,206 | 0,272 | 1 | Mac | 0,764407971 | 1,308201953 | 0,757352941 |
| ATOX12 | 0,089914786 | 0,300090743 | 0,328 | 0,252 | 1 | Mac | 1,231221853 | 0,812201309 | 1,301587302 |
| ST131 | 0,090725829 | 0,480741065 | 0,328 | 0,262 | 1 | Mac | 1,395460285 | 0,716609431 | 1,251908397 |
| NCF2 | 0,090872978 | -0,397473127 | 0,276 | 0,359 | 1 | Mac | 0,759186831 | 1,317198823 | 0,768802228 |
| ANXA113 | 0,091299289 | 0,319400779 | 0,344 | 0,262 | 1 | Mac | 1,247812165 | 0,80140267 | 1,312977099 |
| CCDC85B3 | 0,09144937 | 0,525072165 | 0,198 | 0,136 | 1 | Mac | 1,439005559 | 0,694924348 | 1,455882353 |

| Supplementary Table 4 - clusermarker | | | | | | | | | |
|---|---|---|---|---|---|---|---|---|---|
| | **p_val** | **avg_log2FC** | **pct.1** | **pct.2** | **p_val_adj** | **celltype** | **Foldchange_UP** | **Foldchange_DOWN** | **Ratio_pct1_pct2** |
| DAD12 | 0,0915049 | 0,379628896 | 0,347 | 0,272 | 1 | Mac | 1,301007154 | 0,768635281 | 1,275735294 |
| DNAJA14 | 0,091577829 | -0,349497204 | 0,534 | 0,583 | 1 | Mac | 0,784857582 | 1,274116505 | 0,915951973 |
| ESF1 | 0,092602568 | -0,280711468 | 0,06 | 0,107 | 1 | Mac | 0,823184962 | 1,214793815 | 0,560747664 |
| NEK71 | 0,092702929 | -0,452013368 | 0,1 | 0,155 | 1 | Mac | 0,731021951 | 1,367947979 | 0,64516129 |
| NME41 | 0,092990507 | 0,35425171 | 0,136 | 0,078 | 1 | Mac | 1,278322375 | 0,782275285 | 1,743589744 |
| UQCRFS13 | 0,09355159 | 0,280841534 | 0,257 | 0,184 | 1 | Mac | 1,214903339 | 0,823110751 | 1,39673913 |
| RFTN1 | 0,09361799 | 0,298375176 | 0,138 | 0,078 | 1 | Mac | 1,229758627 | 0,813167705 | 1,769230769 |
| PCGF5 | 0,09452543 | -0,273966964 | 0,146 | 0,214 | 1 | Mac | 0,827042307 | 1,209127988 | 0,682242991 |
| IVNS1ABP1 | 0,094576091 | -0,437546139 | 0,285 | 0,369 | 1 | Mac | 0,738389458 | 1,354298858 | 0,772357724 |
| SNRPC1 | 0,094850138 | 0,348700283 | 0,225 | 0,155 | 1 | Mac | 1,273412899 | 0,785291245 | 1,451612903 |
| MALAT13 | 0,094995615 | -0,323898182 | 0,846 | 0,748 | 1 | Mac | 0,798908297 | 1,251708117 | 1,131016043 |
| CD84 | 0,095216884 | -0,393658269 | 0,141 | 0,204 | 1 | Mac | 0,761196973 | 1,31372041 | 0,691176471 |
| MGP4 | 0,095626764 | -1,079132266 | 0,119 | 0,058 | 1 | Mac | 0,47331342 | 2,112764939 | 2,051724138 |
| JPX | 0,096073448 | -0,290311445 | 0,076 | 0,126 | 1 | Mac | 0,817725511 | 1,222904246 | 0,603174603 |
| TNFSF13 | 0,096122454 | 0,457996511 | 0,312 | 0,243 | 1 | Mac | 1,373632911 | 0,727996535 | 1,283950617 |
| UPF3B | 0,096243499 | -0,316524155 | 0,06 | 0,107 | 1 | Mac | 0,8030022 | 1,245326601 | 0,560747664 |
| RPS203 | 0,096690895 | -0,469420559 | 0,596 | 0,689 | 1 | Mac | 0,722254624 | 1,384553267 | 0,865021771 |
| HCFC1R1 | 0,097072619 | 0,2849807 | 0,163 | 0,097 | 1 | Mac | 1,218393964 | 0,820752589 | 1,680412371 |
| IL1B | 0,097539297 | -0,439484055 | 0,393 | 0,476 | 1 | Mac | 0,737398274 | 1,356119257 | 0,825630252 |
| FNBP1 | 0,097782902 | 0,299559885 | 0,287 | 0,214 | 1 | Mac | 1,230768892 | 0,812500223 | 1,341121495 |
| EIF1AX4 | 0,098189472 | -0,384301872 | 0,236 | 0,301 | 1 | Mac | 0,766149653 | 1,305228027 | 0,784053156 |
| TAGAP | 0,098234091 | -0,601100727 | 0,203 | 0,262 | 1 | Mac | 0,659250778 | 1,516873448 | 0,77480916 |
| LONP22 | 0,098636834 | -0,338338696 | 0,1 | 0,155 | 1 | Mac | 0,790951591 | 1,264299879 | 0,64516129 |
| NBEAL11 | 0,099191234 | 0,307475426 | 0,114 | 0,058 | 1 | Mac | 1,237540231 | 0,808054538 | 1,965517241 |
| ANKRD171 | 0,100119772 | 0,352523085 | 0,133 | 0,078 | 1 | Mac | 1,276791617 | 0,783213162 | 1,705128205 |
| MYD88 | 0,100207568 | 0,295757532 | 0,228 | 0,155 | 1 | Mac | 1,227529361 | 0,814644466 | 1,470967742 |
| MARCHF2 | 0,100671113 | 0,518053986 | 0,165 | 0,107 | 1 | Mac | 1,432022327 | 0,698313135 | 1,542056075 |
| SUCLG1 | 0,10112532 | 0,321918258 | 0,16 | 0,097 | 1 | Mac | 1,249991477 | 0,800005455 | 1,649484536 |
| UPP1 | 0,101845971 | 0,413779067 | 0,306 | 0,233 | 1 | Mac | 1,332170802 | 0,750654494 | 1,313304721 |
| TMEM167B | 0,102306331 | -0,296301566 | 0,06 | 0,107 | 1 | Mac | 0,814337325 | 1,227992344 | 0,560747664 |
| RNF13 | 0,103302454 | -0,311985307 | 0,244 | 0,32 | 1 | Mac | 0,805532495 | 1,241414848 | 0,7625 |

| Supplementary Table 4 - clusermarker | | | | | | | | | |
|---|---|---|---|---|---|---|---|---|---|
| | **p_val** | **avg_log2FC** | **pct.1** | **pct.2** | **p_val_adj** | **celltype** | **Foldchange_UP** | **Foldchange_DOWN** | **Ratio_pct1_pct2** |
| CD521 | 0,104592049 | -0,410857815 | 0,263 | 0,34 | 1 | Mac | 0,752176003 | 1,329476075 | 0,773529412 |
| INTS63 | 0,105010901 | -0,285240319 | 0,168 | 0,233 | 1 | Mac | 0,820604904 | 1,218613239 | 0,721030043 |
| GEM1 | 0,105115026 | -0,47639858 | 0,092 | 0,146 | 1 | Mac | 0,718769661 | 1,39126629 | 0,630136986 |
| ELL21 | 0,105388395 | -0,352826553 | 0,285 | 0,359 | 1 | Mac | 0,783048432 | 1,277060216 | 0,793871866 |
| TMEM1473 | 0,10690787 | 0,363217001 | 0,233 | 0,165 | 1 | Mac | 1,286290944 | 0,777429091 | 1,412121212 |
| FRMD4B1 | 0,107198104 | -0,299771516 | 0,187 | 0,262 | 1 | Mac | 0,812381045 | 1,230949449 | 0,713740458 |
| FLI1 | 0,107269874 | -0,476787792 | 0,103 | 0,155 | 1 | Mac | 0,718575777 | 1,391641679 | 0,664516129 |
| RAB7A1 | 0,107468498 | -0,325557605 | 0,379 | 0,476 | 1 | Mac | 0,797989902 | 1,25314869 | 0,796218487 |
| EGR31 | 0,108196221 | -0,500831185 | 0,079 | 0,126 | 1 | Mac | 0,706699511 | 1,415028573 | 0,626984127 |
| PIK3CD | 0,108809106 | -0,324610698 | 0,092 | 0,146 | 1 | Mac | 0,798513831 | 1,252326461 | 0,630136986 |
| RAC2 | 0,109324273 | 0,281111997 | 0,298 | 0,223 | 1 | Mac | 1,215131119 | 0,822956457 | 1,33632287 |
| NDUFB83 | 0,109981309 | 0,258620517 | 0,366 | 0,291 | 1 | Mac | 1,196334241 | 0,8358868 | 1,257731959 |
| DNAJC42 | 0,110320242 | 0,41480115 | 0,195 | 0,136 | 1 | Mac | 1,333114918 | 0,750122879 | 1,433823529 |
| CD531 | 0,111022395 | -0,449234605 | 0,412 | 0,466 | 1 | Mac | 0,732431323 | 1,365315722 | 0,884120172 |
| MAFB2 | 0,111761183 | -0,423321824 | 0,347 | 0,437 | 1 | Mac | 0,745705649 | 1,341011699 | 0,794050343 |
| XBP12 | 0,111976729 | 0,347550953 | 0,39 | 0,311 | 1 | Mac | 1,272398832 | 0,7859171 | 1,254019293 |
| PCBP14 | 0,112349952 | 0,257859854 | 0,542 | 0,476 | 1 | Mac | 1,195703638 | 0,836327638 | 1,138655462 |
| MAX | 0,112667669 | -0,440475389 | 0,171 | 0,233 | 1 | Mac | 0,736891752 | 1,357051422 | 0,733905579 |
| UQCR104 | 0,112869646 | 0,327141014 | 0,472 | 0,388 | 1 | Mac | 1,25452482 | 0,79711456 | 1,216494845 |
| ZBTB7A3 | 0,11313687 | 0,416406319 | 0,23 | 0,165 | 1 | Mac | 1,334598992 | 0,749288742 | 1,393939394 |
| TANK2 | 0,113149065 | -0,266938784 | 0,165 | 0,233 | 1 | Mac | 0,831081125 | 1,203251969 | 0,708154506 |
| RABAC13 | 0,113429169 | 0,293695581 | 0,306 | 0,233 | 1 | Mac | 1,225776186 | 0,815809617 | 1,313304721 |
| UBE2L6 | 0,113570661 | -0,397503425 | 0,211 | 0,282 | 1 | Mac | 0,759170888 | 1,317226485 | 0,74822695 |
| TNRC6B1 | 0,113760863 | -0,329704464 | 0,16 | 0,223 | 1 | Mac | 0,795699466 | 1,256755902 | 0,717488789 |
| ATP2B4 | 0,113982688 | 0,295456666 | 0,108 | 0,058 | 1 | Mac | 1,227273393 | 0,814814373 | 1,862068966 |
| EIF2S32 | 0,114316139 | -0,344339757 | 0,314 | 0,408 | 1 | Mac | 0,787668367 | 1,269569837 | 0,769607843 |
| RPP211 | 0,114453869 | 0,422162931 | 0,117 | 0,068 | 1 | Mac | 1,33993492 | 0,746304903 | 1,720588235 |
| SAP30 | 0,114858131 | 0,428952791 | 0,195 | 0,136 | 1 | Mac | 1,346256016 | 0,742800766 | 1,433823529 |
| PPIP5K21 | 0,115216836 | 0,258563578 | 0,152 | 0,097 | 1 | Mac | 1,196287026 | 0,83591979 | 1,567010309 |
| HNRNPA01 | 0,115548194 | 0,321678627 | 0,35 | 0,282 | 1 | Mac | 1,249783871 | 0,800138346 | 1,241134752 |
| MERTK | 0,115915483 | -0,403806616 | 0,103 | 0,155 | 1 | Mac | 0,755861276 | 1,322994089 | 0,664516129 |

| Supplementary Table 4 - clusermarker | | | | | | | | | |
|---|---|---|---|---|---|---|---|---|---|
| | **p_val** | **avg_log2FC** | **pct.1** | **pct.2** | **p_val_adj** | **celltype** | **Foldchange_UP** | **Foldchange_DOWN** | **Ratio_pct1_pct2** |
| RBM34 | 0,116692458 | 0,287751425 | 0,612 | 0,544 | 1 | Mac | 1,220736162 | 0,81917783 | 1,125 |
| LRRC592 | 0,11674769 | 0,362923154 | 0,173 | 0,117 | 1 | Mac | 1,286028981 | 0,777587453 | 1,478632479 |
| TMEM1603 | 0,117271865 | 0,374221633 | 0,182 | 0,126 | 1 | Mac | 1,296140068 | 0,771521554 | 1,444444444 |
| NME34 | 0,117536827 | 0,350777868 | 0,195 | 0,136 | 1 | Mac | 1,275248027 | 0,784161182 | 1,433823529 |
| GIMAP7 | 0,118425096 | -0,61605145 | 0,089 | 0,136 | 1 | Mac | 0,652454205 | 1,532674618 | 0,654411765 |
| HLA-DQB2 | 0,118456304 | -1,062608856 | 0,236 | 0,301 | 1 | Mac | 0,478765515 | 2,088705158 | 0,784053156 |
| C20orf27 | 0,118834912 | 0,296275422 | 0,263 | 0,194 | 1 | Mac | 1,227970091 | 0,814352082 | 1,355670103 |
| GSTO1 | 0,120498032 | 0,335930715 | 0,317 | 0,252 | 1 | Mac | 1,262191414 | 0,792272859 | 1,257936508 |
| GABARAPL22 | 0,121142954 | 0,302460206 | 0,355 | 0,291 | 1 | Mac | 1,233245656 | 0,810868455 | 1,219931271 |
| GLRX3 | 0,121299476 | -0,375564025 | 0,125 | 0,184 | 1 | Mac | 0,770804006 | 1,297346656 | 0,679347826 |
| ARHGAP53 | 0,122078564 | -0,258052583 | 0,108 | 0,165 | 1 | Mac | 0,836215921 | 1,195863382 | 0,654545455 |
| PLEKHA21 | 0,122866994 | -0,322755846 | 0,119 | 0,175 | 1 | Mac | 0,799541129 | 1,250717397 | 0,68 |
| SNX2 | 0,123195898 | 0,450155925 | 0,266 | 0,204 | 1 | Mac | 1,366187905 | 0,731963734 | 1,303921569 |
| ILK2 | 0,123560059 | 0,317602882 | 0,241 | 0,175 | 1 | Mac | 1,246258101 | 0,802402006 | 1,377142857 |
| ETHE12 | 0,123770212 | 0,347256367 | 0,176 | 0,117 | 1 | Mac | 1,272139046 | 0,786077594 | 1,504273504 |
| NFE2L23 | 0,124023108 | 0,378141284 | 0,496 | 0,427 | 1 | Mac | 1,299666334 | 0,769428256 | 1,161592506 |
| CCDC88B1 | 0,124243434 | -0,291438577 | 0,092 | 0,146 | 1 | Mac | 0,817086898 | 1,223860036 | 0,630136986 |
| RNMT | 0,124734826 | -0,251568826 | 0,127 | 0,184 | 1 | Mac | 0,839982499 | 1,190500994 | 0,690217391 |
| BIRC2 | 0,125640896 | -0,412210494 | 0,149 | 0,214 | 1 | Mac | 0,751471089 | 1,330723184 | 0,696261682 |
| SOX43 | 0,12643904 | -0,31673248 | 0,184 | 0,262 | 1 | Mac | 0,802886255 | 1,245506439 | 0,702290076 |
| SNAP23 | 0,127084292 | -0,329015027 | 0,195 | 0,262 | 1 | Mac | 0,796079807 | 1,256155465 | 0,744274809 |
| MYG13 | 0,127282354 | 0,328666148 | 0,106 | 0,058 | 1 | Mac | 1,255851732 | 0,796272342 | 1,827586207 |
| RNASE12 | 0,127513676 | 0,526009653 | 0,22 | 0,155 | 1 | Mac | 1,439940953 | 0,694472921 | 1,419354839 |
| SPEN1 | 0,127520208 | -0,325616667 | 0,146 | 0,204 | 1 | Mac | 0,797957234 | 1,253199993 | 0,715686275 |
| BST2 | 0,128025199 | -0,26883961 | 0,39 | 0,466 | 1 | Mac | 0,829986853 | 1,204838361 | 0,836909871 |
| MZT2A3 | 0,12809079 | 0,474294184 | 0,179 | 0,126 | 1 | Mac | 1,38923839 | 0,719818864 | 1,420634921 |
| TMED9 | 0,128652643 | 0,283126621 | 0,271 | 0,204 | 1 | Mac | 1,216829151 | 0,821808057 | 1,328431373 |
| AKAP11 | 0,128844515 | -0,389208111 | 0,079 | 0,126 | 1 | Mac | 0,763548598 | 1,309674332 | 0,626984127 |
| RGS19 | 0,128920735 | 0,305362607 | 0,271 | 0,204 | 1 | Mac | 1,235729187 | 0,809238797 | 1,328431373 |
| ZNF7064 | 0,128997168 | 0,342904257 | 0,358 | 0,301 | 1 | Mac | 1,268307227 | 0,788452497 | 1,189368771 |
| PMF1 | 0,129456359 | 0,320582358 | 0,127 | 0,078 | 1 | Mac | 1,248834551 | 0,800746583 | 1,628205128 |

| Supplementary Table 4 - clusermarker | | | | | | | | | |
|---|---|---|---|---|---|---|---|---|---|
| | **p_val** | **avg_log2FC** | **pct.1** | **pct.2** | **p_val_adj** | **celltype** | **Foldchange_UP** | **Foldchange_DOWN** | **Ratio_pct1_pct2** |
| LMO2 | 0,129461277 | -0,329648912 | 0,127 | 0,184 | 1 | Mac | 0,795730105 | 1,256707511 | 0,690217391 |
| MON2 | 0,129659165 | -0,350617592 | 0,087 | 0,136 | 1 | Mac | 0,784248303 | 1,275106361 | 0,639705882 |
| CPEB2 | 0,129686971 | -0,261343191 | 0,076 | 0,126 | 1 | Mac | 0,83431079 | 1,198594111 | 0,603174603 |
| CIRBP2 | 0,129775737 | -0,386076099 | 0,453 | 0,534 | 1 | Mac | 0,765208021 | 1,306834184 | 0,848314607 |
| TMEM179B3 | 0,130302944 | 0,334360914 | 0,236 | 0,175 | 1 | Mac | 1,260818767 | 0,793135402 | 1,348571429 |
| PLTP2 | 0,130597979 | 1,115573728 | 0,209 | 0,155 | 1 | Mac | 2,166811613 | 0,461507588 | 1,348387097 |
| RHBDD21 | 0,131027234 | 0,403758643 | 0,125 | 0,078 | 1 | Mac | 1,322950098 | 0,75588641 | 1,602564103 |
| WSB11 | 0,131199749 | -0,411043704 | 0,431 | 0,515 | 1 | Mac | 0,752079093 | 1,329647387 | 0,836893204 |
| NET12 | 0,13141304 | 0,315115204 | 0,119 | 0,068 | 1 | Mac | 1,244110996 | 0,803786803 | 1,75 |
| MBNL21 | 0,132087408 | -0,495163652 | 0,087 | 0,136 | 1 | Mac | 0,709481194 | 1,40948063 | 0,639705882 |
| MCUR1 | 0,132190238 | 0,315221395 | 0,119 | 0,068 | 1 | Mac | 1,244202573 | 0,803727642 | 1,75 |
| ATG31 | 0,133788375 | 0,296055559 | 0,339 | 0,282 | 1 | Mac | 1,227782966 | 0,814476196 | 1,20212766 |
| GNA151 | 0,133870794 | 0,420144062 | 0,298 | 0,233 | 1 | Mac | 1,338061162 | 0,747349993 | 1,278969957 |
| AKR7A22 | 0,135208963 | 0,303487419 | 0,152 | 0,097 | 1 | Mac | 1,234124052 | 0,810291314 | 1,567010309 |
| SKAP2 | 0,136223773 | -0,301283708 | 0,214 | 0,291 | 1 | Mac | 0,811529977 | 1,232240371 | 0,735395189 |
| WDR15 | 0,136578886 | 0,297382523 | 0,388 | 0,32 | 1 | Mac | 1,228912777 | 0,813727401 | 1,2125 |
| LDLRAD41 | 0,136631013 | 0,269392027 | 0,198 | 0,136 | 1 | Mac | 1,205299789 | 0,829669107 | 1,455882353 |
| IL1R2 | 0,136655128 | 0,255526493 | 0,133 | 0,078 | 1 | Mac | 1,193771315 | 0,837681378 | 1,705128205 |
| ANKRD373 | 0,136762989 | 0,455602218 | 0,138 | 0,087 | 1 | Mac | 1,371355124 | 0,72920572 | 1,586206897 |
| N4BP2L21 | 0,137525626 | -0,303345738 | 0,176 | 0,243 | 1 | Mac | 0,810370893 | 1,23400286 | 0,724279835 |
| HEXB | 0,137863668 | 0,265227747 | 0,339 | 0,272 | 1 | Mac | 1,201825757 | 0,832067372 | 1,246323529 |
| DPP8 | 0,137877046 | -0,330388565 | 0,065 | 0,107 | 1 | Mac | 0,795322249 | 1,257351975 | 0,607476636 |
| CREB13 | 0,138444112 | -0,399076595 | 0,119 | 0,175 | 1 | Mac | 0,75834351 | 1,318663622 | 0,68 |
| RNF1671 | 0,138606555 | -0,452738831 | 0,127 | 0,184 | 1 | Mac | 0,730654447 | 1,368636028 | 0,690217391 |
| BTK | 0,14018875 | -0,258317267 | 0,16 | 0,223 | 1 | Mac | 0,836062519 | 1,196082802 | 0,717488789 |
| HMGB22 | 0,140844192 | 0,693791007 | 0,214 | 0,165 | 1 | Mac | 1,61752836 | 0,618227182 | 1,296969697 |
| NINJ12 | 0,142782492 | 0,292920041 | 0,442 | 0,359 | 1 | Mac | 1,22511743 | 0,816248284 | 1,231197772 |
| NADK | 0,143012346 | 0,397648808 | 0,114 | 0,068 | 1 | Mac | 1,317359231 | 0,759094389 | 1,676470588 |
| YWHAQ2 | 0,14367926 | 0,432633266 | 0,263 | 0,214 | 1 | Mac | 1,349694848 | 0,740908215 | 1,228971963 |
| MAF11 | 0,144623537 | 0,383560056 | 0,198 | 0,146 | 1 | Mac | 1,304557068 | 0,766543699 | 1,356164384 |
| COPS7A | 0,144862001 | -0,587385328 | 0,073 | 0,117 | 1 | Mac | 0,665548022 | 1,502521181 | 0,623931624 |

| Supplementary Table 4 - clusermarker | | | | | | | | | |
|---|---|---|---|---|---|---|---|---|---|
| | **p_val** | **avg_log2FC** | **pct.1** | **pct.2** | **p_val_adj** | **celltype** | **Foldchange_UP** | **Foldchange_DOWN** | **Ratio_pct1_pct2** |
| PSME4 | 0,144940295 | -0,25581227 | 0,089 | 0,136 | 1 | Mac | 0,837515462 | 1,194007807 | 0,654411765 |
| LITAF2 | 0,145415604 | -0,450544976 | 0,575 | 0,612 | 1 | Mac | 0,731766372 | 1,366556374 | 0,939542484 |
| IER5L3 | 0,14553513 | -0,385609972 | 0,117 | 0,165 | 1 | Mac | 0,765455295 | 1,306412022 | 0,709090909 |
| ATP6V1G11 | 0,145672003 | 0,313603937 | 0,491 | 0,437 | 1 | Mac | 1,242808434 | 0,804629236 | 1,123569794 |
| STMP11 | 0,146338998 | 0,407423165 | 0,266 | 0,204 | 1 | Mac | 1,326314732 | 0,753968855 | 1,303921569 |
| APP1 | 0,146417039 | 0,653224545 | 0,222 | 0,175 | 1 | Mac | 1,572679342 | 0,635857529 | 1,268571429 |
| IDH21 | 0,146735152 | 0,372738575 | 0,225 | 0,165 | 1 | Mac | 1,29480835 | 0,772315069 | 1,363636364 |
| LINC-PINT | 0,14721085 | 0,339938616 | 0,125 | 0,078 | 1 | Mac | 1,26570274 | 0,790074927 | 1,602564103 |
| GNPTG3 | 0,14784665 | 0,401934901 | 0,23 | 0,175 | 1 | Mac | 1,321278784 | 0,756842547 | 1,314285714 |
| UQCRB1 | 0,149095861 | -0,293554293 | 0,607 | 0,641 | 1 | Mac | 0,815889516 | 1,225656147 | 0,946957878 |
| IGFBP41 | 0,149631371 | 0,350424911 | 0,103 | 0,058 | 1 | Mac | 1,274936074 | 0,784353051 | 1,775862069 |
| STK241 | 0,150099036 | -0,394081858 | 0,179 | 0,233 | 1 | Mac | 0,760973512 | 1,314106187 | 0,768240343 |
| SMIM71 | 0,150539714 | 0,332784858 | 0,125 | 0,078 | 1 | Mac | 1,259442151 | 0,794002328 | 1,602564103 |
| ATP2B11 | 0,150899313 | 0,351024562 | 0,371 | 0,301 | 1 | Mac | 1,275466106 | 0,784027106 | 1,23255814 |
| UTP6 | 0,150934276 | -0,281694442 | 0,095 | 0,146 | 1 | Mac | 0,822624279 | 1,215621791 | 0,650684932 |
| RASGRP2 | 0,152397028 | 0,284516671 | 0,103 | 0,058 | 1 | Mac | 1,218002143 | 0,821016618 | 1,775862069 |
| SEC31A1 | 0,152879764 | -0,349927693 | 0,198 | 0,262 | 1 | Mac | 0,784623422 | 1,274496749 | 0,755725191 |
| TET21 | 0,153087207 | -0,482019709 | 0,317 | 0,369 | 1 | Mac | 0,715974589 | 1,396697613 | 0,859078591 |
| LHFPL2 | 0,153562026 | 0,250266726 | 0,117 | 0,068 | 1 | Mac | 1,189426996 | 0,840740964 | 1,720588235 |
| RFC11 | 0,154015267 | 0,292729981 | 0,106 | 0,058 | 1 | Mac | 1,224956044 | 0,816355823 | 1,827586207 |
| DOK1 | 0,154038775 | -0,259312502 | 0,073 | 0,117 | 1 | Mac | 0,835485965 | 1,196908198 | 0,623931624 |
| USP12 | 0,154424687 | 0,290787132 | 0,141 | 0,087 | 1 | Mac | 1,22330753 | 0,817455934 | 1,620689655 |
| EMILIN2 | 0,155778108 | 0,280824511 | 0,306 | 0,243 | 1 | Mac | 1,214889004 | 0,823120463 | 1,259259259 |
| CDK2AP21 | 0,155884827 | 0,323588293 | 0,274 | 0,223 | 1 | Mac | 1,25143928 | 0,79907992 | 1,228699552 |
| SNRPG1 | 0,156979122 | 0,307282563 | 0,35 | 0,291 | 1 | Mac | 1,237374804 | 0,808162569 | 1,202749141 |
| LPCAT2 | 0,156986859 | -0,253933284 | 0,184 | 0,243 | 1 | Mac | 0,838606964 | 1,192453727 | 0,757201646 |
| FUCA12 | 0,157568203 | 0,253147731 | 0,103 | 0,058 | 1 | Mac | 1,191804609 | 0,839063713 | 1,775862069 |
| TXN22 | 0,157742473 | 0,442179893 | 0,165 | 0,117 | 1 | Mac | 1,358655688 | 0,736021649 | 1,41025641 |
| CENPW | 0,159798919 | -0,264376819 | 0,089 | 0,136 | 1 | Mac | 0,832558286 | 1,201117107 | 0,654411765 |
| SLC66A21 | 0,160177968 | 0,254640363 | 0,114 | 0,068 | 1 | Mac | 1,193038304 | 0,838196055 | 1,676470588 |
| CSDE12 | 0,160538852 | -0,439837367 | 0,379 | 0,447 | 1 | Mac | 0,737217709 | 1,356451408 | 0,84787472 |

| Supplementary Table 4 - clusermarker | | | | | | | | | |
|---|---|---|---|---|---|---|---|---|---|
| | p_val | avg_log2FC | pct.1 | pct.2 | p_val_adj | celltype | Foldchange_UP | Foldchange_DOWN | Ratio_pct1_pct2 |
| IGSF6 | 0,160612547 | -0,336782837 | 0,423 | 0,485 | 1 | Mac | 0,791805044 | 1,262937143 | 0,872164948 |
| SERPINA1 | 0,161456113 | 0,340200719 | 0,485 | 0,427 | 1 | Mac | 1,265932708 | 0,789931403 | 1,135831382 |
| KIAA0100 | 0,161687699 | 0,359171629 | 0,154 | 0,107 | 1 | Mac | 1,282689187 | 0,779612091 | 1,439252336 |
| TAOK3 | 0,162153581 | -0,287391742 | 0,268 | 0,33 | 1 | Mac | 0,819382087 | 1,220431855 | 0,812121212 |
| NUDT11 | 0,162536848 | 0,285474297 | 0,149 | 0,097 | 1 | Mac | 1,218810891 | 0,820471828 | 1,536082474 |
| POLDIP31 | 0,164149693 | 0,26336971 | 0,1 | 0,058 | 1 | Mac | 1,20027893 | 0,833139677 | 1,724137931 |
| IRAK2 | 0,164407822 | -0,261921825 | 0,1 | 0,146 | 1 | Mac | 0,833976233 | 1,199074938 | 0,684931507 |
| ANKRD13A | 0,165144273 | 0,309375302 | 0,1 | 0,058 | 1 | Mac | 1,239171013 | 0,806991117 | 1,724137931 |
| HSPD12 | 0,165222249 | -0,657724002 | 0,401 | 0,437 | 1 | Mac | 0,633877515 | 1,577591848 | 0,917620137 |
| SEC24D | 0,166563411 | -0,300981076 | 0,084 | 0,126 | 1 | Mac | 0,811700228 | 1,231981913 | 0,666666667 |
| SDCBP3 | 0,167282399 | -0,31934953 | 0,499 | 0,563 | 1 | Mac | 0,801431138 | 1,247767839 | 0,886323268 |
| IFITM23 | 0,167366947 | -0,563361346 | 0,453 | 0,398 | 1 | Mac | 0,676723622 | 1,477708133 | 1,138190955 |
| GTF2B2 | 0,167578335 | 0,436982786 | 0,317 | 0,272 | 1 | Mac | 1,353770126 | 0,738677845 | 1,165441176 |
| SDHC1 | 0,167728943 | -0,265536132 | 0,211 | 0,282 | 1 | Mac | 0,831889532 | 1,202082682 | 0,74822695 |
| PRDX43 | 0,167855081 | 0,262445812 | 0,146 | 0,097 | 1 | Mac | 1,199510521 | 0,833673388 | 1,505154639 |
| MRPS212 | 0,168254141 | -0,336606148 | 0,285 | 0,35 | 1 | Mac | 0,791902023 | 1,262782479 | 0,814285714 |
| ABCA1 | 0,168697463 | -0,328562639 | 0,136 | 0,194 | 1 | Mac | 0,796329474 | 1,255761632 | 0,701030928 |
| BUD231 | 0,170511463 | 0,348894376 | 0,152 | 0,107 | 1 | Mac | 1,273584229 | 0,785185602 | 1,420560748 |
| SPINT23 | 0,170710174 | 0,285497068 | 0,355 | 0,301 | 1 | Mac | 1,218830128 | 0,820458878 | 1,179401993 |
| TAOK11 | 0,171449552 | -0,25740785 | 0,165 | 0,223 | 1 | Mac | 0,836589705 | 1,195329076 | 0,739910314 |
| SPSB32 | 0,171749483 | 0,323962228 | 0,136 | 0,087 | 1 | Mac | 1,251763685 | 0,798872832 | 1,563218391 |
| TSTD12 | 0,171916609 | 0,319400938 | 0,111 | 0,068 | 1 | Mac | 1,247812302 | 0,801402581 | 1,632352941 |
| DUSP41 | 0,172560637 | 0,39875779 | 0,168 | 0,117 | 1 | Mac | 1,318372258 | 0,758511106 | 1,435897436 |
| AHCYL1 | 0,172601856 | -0,289331322 | 0,152 | 0,204 | 1 | Mac | 0,818281237 | 1,222073725 | 0,745098039 |
| RAPGEF1 | 0,173137173 | 0,273867197 | 0,244 | 0,184 | 1 | Mac | 1,209044376 | 0,827099501 | 1,326086957 |
| ERH4 | 0,173641157 | 0,29365168 | 0,298 | 0,252 | 1 | Mac | 1,225738885 | 0,815834442 | 1,182539683 |
| DDX39A1 | 0,173888748 | 0,326031335 | 0,222 | 0,165 | 1 | Mac | 1,253560247 | 0,797727913 | 1,345454545 |
| SET1 | 0,174027972 | 0,2531705 | 0,409 | 0,359 | 1 | Mac | 1,191823418 | 0,839050471 | 1,139275766 |
| SERINC12 | 0,17431501 | -0,362935375 | 0,241 | 0,301 | 1 | Mac | 0,777580867 | 1,286039874 | 0,800664452 |
| AC090559.1 | 0,174319095 | -0,269658565 | 0,098 | 0,146 | 1 | Mac | 0,82951584 | 1,205522489 | 0,671232877 |
| ZMAT21 | 0,174481529 | -0,359640307 | 0,133 | 0,184 | 1 | Mac | 0,779358866 | 1,283105953 | 0,722826087 |

| Supplementary Table 4 - clusermarker | | | | | | | | | |
|---|---|---|---|---|---|---|---|---|---|
| | **p_val** | **avg_log2FC** | **pct.1** | **pct.2** | **p_val_adj** | **celltype** | **Foldchange_UP** | **Foldchange_DOWN** | **Ratio_pct1_pct2** |
| MS4A6A | 0,174626723 | -0,286556917 | 0,607 | 0,65 | 1 | Mac | 0,819856365 | 1,219725848 | 0,933846154 |
| VASH1 | 0,174759984 | -0,323204508 | 0,154 | 0,214 | 1 | Mac | 0,799292519 | 1,251106417 | 0,719626168 |
| CHST11 | 0,174950043 | 0,343529682 | 0,214 | 0,165 | 1 | Mac | 1,268857171 | 0,788110768 | 1,296969697 |
| FTL3 | 0,175022935 | 0,475656247 | 0,995 | 0,99 | 1 | Mac | 1,390550604 | 0,719139596 | 1,005050505 |
| TPP2 | 0,175358982 | -0,542852724 | 0,108 | 0,155 | 1 | Mac | 0,686412284 | 1,456850385 | 0,696774194 |
| SNX91 | 0,175424893 | 0,325964857 | 0,263 | 0,204 | 1 | Mac | 1,253502486 | 0,797764672 | 1,289215686 |
| SOS2 | 0,175454337 | 0,274116179 | 0,122 | 0,078 | 1 | Mac | 1,209253052 | 0,826956772 | 1,564102564 |
| SMIM31 | 0,175559787 | 0,347526935 | 0,179 | 0,126 | 1 | Mac | 1,272377649 | 0,785930184 | 1,420634921 |
| ATP5PB2 | 0,176948451 | 0,277316774 | 0,358 | 0,32 | 1 | Mac | 1,211938738 | 0,825124215 | 1,11875 |
| POLR2I1 | 0,177606408 | 0,43460342 | 0,133 | 0,087 | 1 | Mac | 1,351539259 | 0,739897116 | 1,528735632 |
| SLC35F51 | 0,179567512 | -0,293012827 | 0,068 | 0,107 | 1 | Mac | 0,816195789 | 1,225196225 | 0,635514019 |
| SMAD4 | 0,179567512 | -0,318422269 | 0,068 | 0,107 | 1 | Mac | 0,801946407 | 1,246966121 | 0,635514019 |
| AHNAK3 | 0,180432826 | -0,357553008 | 0,526 | 0,553 | 1 | Mac | 0,780487262 | 1,281250891 | 0,951175407 |
| UBE2F2 | 0,180632943 | 0,267249864 | 0,209 | 0,155 | 1 | Mac | 1,203511447 | 0,830901943 | 1,348387097 |
| OGFR | 0,180965203 | -0,4196191 | 0,1 | 0,146 | 1 | Mac | 0,747621985 | 1,337574362 | 0,684931507 |
| HLA-DMB | 0,181185588 | -0,419742114 | 0,504 | 0,544 | 1 | Mac | 0,747558241 | 1,337688418 | 0,926470588 |
| CENPX2 | 0,18345705 | 0,307495671 | 0,19 | 0,136 | 1 | Mac | 1,237557597 | 0,808043199 | 1,397058824 |
| DANCR4 | 0,184336031 | 0,310315625 | 0,163 | 0,117 | 1 | Mac | 1,239978946 | 0,806465306 | 1,393162393 |
| TMEM183A1 | 0,184617452 | 0,275511066 | 0,133 | 0,087 | 1 | Mac | 1,210422798 | 0,826157605 | 1,528735632 |
| GBP1 | 0,185092037 | 0,436176671 | 0,133 | 0,087 | 1 | Mac | 1,35301391 | 0,739090702 | 1,528735632 |
| ITFG1 | 0,187080609 | 0,254771301 | 0,108 | 0,068 | 1 | Mac | 1,193146588 | 0,838119984 | 1,588235294 |
| LTB1 | 0,187120483 | 0,546578329 | 0,154 | 0,107 | 1 | Mac | 1,460617407 | 0,684641984 | 1,439252336 |
| BCAP313 | 0,187155929 | 0,258394208 | 0,352 | 0,291 | 1 | Mac | 1,196146592 | 0,836017932 | 1,209621993 |
| FBXL5 | 0,187420528 | -0,327736086 | 0,1 | 0,146 | 1 | Mac | 0,79678584 | 1,255042384 | 0,684931507 |
| PRXL2C1 | 0,188176178 | 0,297777231 | 0,119 | 0,078 | 1 | Mac | 1,229249042 | 0,813504803 | 1,525641026 |
| ITGAV1 | 0,189130251 | -0,276569749 | 0,141 | 0,194 | 1 | Mac | 0,825551574 | 1,21131136 | 0,726804124 |
| ARRB2 | 0,190575676 | -0,277903442 | 0,398 | 0,466 | 1 | Mac | 0,824788749 | 1,21243167 | 0,854077253 |
| FERMT3 | 0,190638745 | 0,326284895 | 0,347 | 0,291 | 1 | Mac | 1,253780585 | 0,797587721 | 1,192439863 |
| LPAR6 | 0,191393134 | -0,377111141 | 0,095 | 0,136 | 1 | Mac | 0,769977855 | 1,29873865 | 0,698529412 |
| HSPA85 | 0,191470898 | -0,385616615 | 0,764 | 0,738 | 1 | Mac | 0,765451771 | 1,306418037 | 1,035230352 |
| GPS21 | 0,191501664 | 0,259217653 | 0,206 | 0,155 | 1 | Mac | 1,19682951 | 0,835540895 | 1,329032258 |

| Supplementary Table 4 - clusermarker | | | | | | | | | |
|---|---|---|---|---|---|---|---|---|---|
| | **p_val** | **avg_log2FC** | **pct.1** | **pct.2** | **p_val_adj** | **celltype** | **Foldchange_UP** | **Foldchange_DOWN** | **Ratio_pct1_pct2** |
| SLC25A16 | 0,192390291 | 0,254844211 | 0,108 | 0,068 | 1 | Mac | 1,193206888 | 0,838077629 | 1,588235294 |
| CHMP4A1 | 0,194780692 | -0,500213132 | 0,176 | 0,233 | 1 | Mac | 0,707002327 | 1,414422502 | 0,755364807 |
| FCGR1A | 0,195422096 | 0,531362271 | 0,182 | 0,136 | 1 | Mac | 1,445293276 | 0,691901095 | 1,338235294 |
| FAM107B | 0,19576726 | -0,330193296 | 0,192 | 0,252 | 1 | Mac | 0,795429903 | 1,257181804 | 0,761904762 |
| CLDND1 | 0,195866677 | -0,252242037 | 0,149 | 0,204 | 1 | Mac | 0,839590625 | 1,191056653 | 0,730392157 |
| DYNLT13 | 0,196907768 | 0,263013027 | 0,211 | 0,155 | 1 | Mac | 1,199982218 | 0,833345682 | 1,361290323 |
| BCLAF11 | 0,197435065 | -0,468792754 | 0,238 | 0,291 | 1 | Mac | 0,72256899 | 1,383950895 | 0,817869416 |
| ZFYVE161 | 0,197727089 | -0,306870609 | 0,203 | 0,262 | 1 | Mac | 0,808393368 | 1,237021529 | 0,77480916 |
| CCDC691 | 0,197809927 | 0,31138081 | 0,108 | 0,068 | 1 | Mac | 1,240894798 | 0,805870088 | 1,588235294 |
| YIPF32 | 0,198081616 | 0,379296139 | 0,154 | 0,107 | 1 | Mac | 1,300707113 | 0,768812587 | 1,439252336 |
| PSMA11 | 0,198101034 | 0,312367189 | 0,26 | 0,204 | 1 | Mac | 1,241743495 | 0,805319298 | 1,274509804 |
| BIN2 | 0,198632389 | -0,310453435 | 0,144 | 0,194 | 1 | Mac | 0,806388274 | 1,240097398 | 0,742268041 |
| COX14 | 0,199033505 | 0,352088855 | 0,255 | 0,204 | 1 | Mac | 1,27640738 | 0,783448933 | 1,25 |
| NR4A32 | 0,200569047 | 0,250016609 | 0,512 | 0,417 | 1 | Mac | 1,189220805 | 0,840886735 | 1,227817746 |
| EMC31 | 0,200709719 | -0,254528434 | 0,108 | 0,155 | 1 | Mac | 0,838261088 | 1,192945747 | 0,696774194 |
| SIPA1L11 | 0,201825249 | 0,314381794 | 0,165 | 0,117 | 1 | Mac | 1,243478699 | 0,804195521 | 1,41025641 |
| GTF2H52 | 0,202104097 | 0,290295982 | 0,16 | 0,117 | 1 | Mac | 1,222891139 | 0,817734276 | 1,367521368 |
| COPS62 | 0,20223604 | 0,33898672 | 0,146 | 0,107 | 1 | Mac | 1,264867899 | 0,790596394 | 1,364485981 |
| SON5 | 0,202459368 | -0,270345298 | 0,507 | 0,553 | 1 | Mac | 0,829121078 | 1,206096463 | 0,91681736 |
| LGALS33 | 0,202920541 | 0,350245763 | 0,531 | 0,466 | 1 | Mac | 1,274777767 | 0,784450456 | 1,139484979 |
| PTAFR1 | 0,203752152 | 0,330213672 | 0,152 | 0,107 | 1 | Mac | 1,25719956 | 0,795418669 | 1,420560748 |
| RNF11 | 0,205104595 | -0,386533275 | 0,1 | 0,146 | 1 | Mac | 0,764965572 | 1,307248373 | 0,684931507 |
| SLC2A31 | 0,20567011 | 0,406683059 | 0,461 | 0,398 | 1 | Mac | 1,325634504 | 0,754355742 | 1,158291457 |
| S100A92 | 0,206233103 | -0,636467041 | 0,377 | 0,272 | 1 | Mac | 0,64328634 | 1,554517697 | 1,386029412 |
| JAGN11 | 0,206667822 | -0,511765181 | 0,079 | 0,117 | 1 | Mac | 0,701363773 | 1,425793631 | 0,675213675 |
| NSD1 | 0,207050904 | 0,313735153 | 0,13 | 0,087 | 1 | Mac | 1,242921475 | 0,804556056 | 1,494252874 |
| GADD45A1 | 0,207571023 | 0,318696036 | 0,13 | 0,087 | 1 | Mac | 1,247202769 | 0,801794243 | 1,494252874 |
| TRAPPC14 | 0,207609126 | 0,41145054 | 0,36 | 0,33 | 1 | Mac | 1,330022397 | 0,751867038 | 1,090909091 |
| IARS21 | 0,209553064 | 0,266966602 | 0,108 | 0,068 | 1 | Mac | 1,20327517 | 0,8310651 | 1,588235294 |
| TREM2 | 0,210035993 | -0,351114196 | 0,179 | 0,223 | 1 | Mac | 0,783978396 | 1,275545353 | 0,802690583 |
| TMEM2592 | 0,210137532 | 0,384704263 | 0,179 | 0,136 | 1 | Mac | 1,305592127 | 0,765935991 | 1,316176471 |

| Supplementary Table 4 - clusermarker | | | | | | | | | |
|---|---|---|---|---|---|---|---|---|---|
| | **p_val** | **avg_log2FC** | **pct.1** | **pct.2** | **p_val_adj** | **celltype** | **Foldchange_UP** | **Foldchange_DOWN** | **Ratio_pct1_pct2** |
| ATP6V1B2 | 0,210152742 | 0,266720196 | 0,352 | 0,301 | 1 | Mac | 1,203069673 | 0,831207055 | 1,169435216 |
| CREBRF1 | 0,210772424 | -0,450395002 | 0,125 | 0,175 | 1 | Mac | 0,731842446 | 1,366414322 | 0,714285714 |
| PPP6C | 0,210875094 | 0,263960237 | 0,127 | 0,087 | 1 | Mac | 1,200770332 | 0,832798724 | 1,459770115 |
| ISG152 | 0,212956787 | 0,301139544 | 0,176 | 0,126 | 1 | Mac | 1,232117244 | 0,811611074 | 1,396825397 |
| ARID5B3 | 0,213809258 | 0,258316789 | 0,106 | 0,068 | 1 | Mac | 1,196082405 | 0,836062796 | 1,558823529 |
| RBM8A3 | 0,215209093 | 0,296646944 | 0,388 | 0,34 | 1 | Mac | 1,228286358 | 0,814142397 | 1,141176471 |
| TDG | 0,217633959 | 0,295771328 | 0,13 | 0,087 | 1 | Mac | 1,227541099 | 0,814636676 | 1,494252874 |
| CCNDBP11 | 0,217701985 | -0,341654008 | 0,103 | 0,146 | 1 | Mac | 0,789136072 | 1,267208579 | 0,705479452 |
| ARRB1 | 0,218017023 | 0,343798156 | 0,119 | 0,078 | 1 | Mac | 1,269093318 | 0,787964121 | 1,525641026 |
| ZPR11 | 0,218146962 | -0,330141463 | 0,087 | 0,126 | 1 | Mac | 0,795458482 | 1,257136636 | 0,69047619 |
| ELOVL51 | 0,218351615 | -0,259911266 | 0,144 | 0,194 | 1 | Mac | 0,835139284 | 1,197405055 | 0,742268041 |
| CTSD1 | 0,218520355 | 0,722365038 | 0,528 | 0,495 | 1 | Mac | 1,649884507 | 0,606103031 | 1,066666667 |
| NOL73 | 0,219868892 | 0,377745133 | 0,225 | 0,175 | 1 | Mac | 1,299309506 | 0,769639563 | 1,285714286 |
| ALOX51 | 0,220298391 | -0,328103758 | 0,23 | 0,291 | 1 | Mac | 0,796582804 | 1,255362273 | 0,790378007 |
| RBCK12 | 0,220526304 | 0,337984235 | 0,176 | 0,136 | 1 | Mac | 1,263989286 | 0,791145946 | 1,294117647 |
| H2AX1 | 0,222963397 | 0,362799784 | 0,201 | 0,155 | 1 | Mac | 1,285919013 | 0,77765395 | 1,296774194 |
| DOCK4 | 0,223044157 | -0,52139519 | 0,144 | 0,184 | 1 | Mac | 0,69669775 | 1,435342657 | 0,782608696 |
| TPGS2 | 0,224641956 | -0,332230678 | 0,079 | 0,117 | 1 | Mac | 0,794307385 | 1,258958457 | 0,675213675 |
| SETDB2 | 0,226039064 | 0,3138136 | 0,103 | 0,068 | 1 | Mac | 1,242989061 | 0,804512309 | 1,514705882 |
| BEST1 | 0,226116107 | -0,419485074 | 0,149 | 0,204 | 1 | Mac | 0,747691442 | 1,337450108 | 0,730392157 |
| UBL71 | 0,226589609 | 0,262750242 | 0,133 | 0,087 | 1 | Mac | 1,199763663 | 0,833497489 | 1,528735632 |
| KCNK61 | 0,227232884 | 0,275321834 | 0,127 | 0,087 | 1 | Mac | 1,210264043 | 0,826265975 | 1,459770115 |
| HPS1 | 0,227708928 | -0,310435821 | 0,157 | 0,204 | 1 | Mac | 0,806398119 | 1,240082257 | 0,769607843 |
| TSPYL12 | 0,228112366 | -0,622260478 | 0,073 | 0,107 | 1 | Mac | 0,649652226 | 1,539285112 | 0,682242991 |
| KIF5B1 | 0,228378811 | 0,279937859 | 0,328 | 0,282 | 1 | Mac | 1,214142587 | 0,823626492 | 1,163120567 |
| TLN11 | 0,230585632 | 0,309109026 | 0,455 | 0,408 | 1 | Mac | 1,238942323 | 0,807140076 | 1,115196078 |
| CTSA1 | 0,230762194 | 0,398593934 | 0,222 | 0,184 | 1 | Mac | 1,318222531 | 0,75859726 | 1,206521739 |
| ZFAND32 | 0,23117652 | 0,261541745 | 0,127 | 0,087 | 1 | Mac | 1,198759082 | 0,834195974 | 1,459770115 |
| RPS19BP12 | 0,232193995 | 0,259534402 | 0,233 | 0,184 | 1 | Mac | 1,197092308 | 0,835357469 | 1,266304348 |
| LRRFIP12 | 0,232268172 | -0,251261115 | 0,512 | 0,553 | 1 | Mac | 0,840161677 | 1,190247101 | 0,925858951 |
| MOB1A1 | 0,233076584 | 0,313971509 | 0,404 | 0,359 | 1 | Mac | 1,243125119 | 0,804424257 | 1,125348189 |

| Supplementary Table 4 - clusermarker | | | | | | | | | |
|---|---|---|---|---|---|---|---|---|---|
| | **p_val** | **avg_log2FC** | **pct.1** | **pct.2** | **p_val_adj** | **celltype** | **Foldchange_UP** | **Foldchange_DOWN** | **Ratio_pct1_pct2** |
| ENPP2 | 0,233122866 | 0,328970972 | 0,106 | 0,068 | 1 | Mac | 1,256117107 | 0,796104117 | 1,558823529 |
| NAPA | 0,233399249 | -0,308125458 | 0,255 | 0,311 | 1 | Mac | 0,807690537 | 1,238097952 | 0,819935691 |
| AIP | 0,234048231 | 0,255216433 | 0,171 | 0,126 | 1 | Mac | 1,193514781 | 0,837861429 | 1,357142857 |
| GTF3A3 | 0,236155287 | 0,262280352 | 0,282 | 0,223 | 1 | Mac | 1,19937296 | 0,833769006 | 1,264573991 |
| ITM2B4 | 0,236222038 | -0,329996015 | 0,835 | 0,835 | 1 | Mac | 0,795538681 | 1,257009902 | 1 |
| TIMM8B1 | 0,237873052 | 0,298267674 | 0,203 | 0,165 | 1 | Mac | 1,229666996 | 0,8132283 | 1,23030303 |
| ATF6 | 0,238382417 | -0,313457032 | 0,16 | 0,214 | 1 | Mac | 0,804711172 | 1,24268189 | 0,747663551 |
| LY6E2 | 0,238557132 | 0,468728643 | 0,192 | 0,146 | 1 | Mac | 1,383889395 | 0,722601101 | 1,315068493 |
| TPRKB2 | 0,241226599 | -0,526182428 | 0,122 | 0,165 | 1 | Mac | 0,694389757 | 1,440113408 | 0,739393939 |
| TRMT1 | 0,241716089 | 0,278807414 | 0,149 | 0,107 | 1 | Mac | 1,2131916 | 0,82427211 | 1,392523364 |
| COMMD8 | 0,242268972 | -0,584765362 | 0,098 | 0,136 | 1 | Mac | 0,66675777 | 1,499795044 | 0,720588235 |
| SLC25A32 | 0,242268972 | 0,2828996 | 0,114 | 0,078 | 1 | Mac | 1,216637688 | 0,821937385 | 1,461538462 |
| ACTR10 | 0,24245572 | -0,28433303 | 0,073 | 0,107 | 1 | Mac | 0,821121132 | 1,217847112 | 0,682242991 |
| ACAA13 | 0,242492445 | 0,26670177 | 0,225 | 0,175 | 1 | Mac | 1,203054308 | 0,831217671 | 1,285714286 |
| GPR157 | 0,242589949 | 0,250105776 | 0,106 | 0,068 | 1 | Mac | 1,189294309 | 0,840834764 | 1,558823529 |
| SNHG292 | 0,242761538 | -0,337282972 | 0,572 | 0,66 | 1 | Mac | 0,791530599 | 1,263375038 | 0,866666667 |
| RCN22 | 0,243724482 | 0,25944402 | 0,103 | 0,068 | 1 | Mac | 1,197017314 | 0,835409804 | 1,514705882 |
| SORL12 | 0,243756835 | -0,293420104 | 0,179 | 0,233 | 1 | Mac | 0,815965407 | 1,225542151 | 0,768240343 |
| DICER1 | 0,244384403 | -0,336346134 | 0,103 | 0,146 | 1 | Mac | 0,792044759 | 1,26255491 | 0,705479452 |
| SIAH2 | 0,245551068 | 0,256779511 | 0,149 | 0,107 | 1 | Mac | 1,194808587 | 0,836954146 | 1,392523364 |
| SP1 | 0,24596233 | -0,287068453 | 0,073 | 0,107 | 1 | Mac | 0,81956572 | 1,220158403 | 0,682242991 |
| PTOV13 | 0,245984872 | 0,288510137 | 0,114 | 0,078 | 1 | Mac | 1,221378315 | 0,818747138 | 1,461538462 |
| ZC3HAV1 | 0,247291328 | 0,383102706 | 0,271 | 0,223 | 1 | Mac | 1,304143574 | 0,76678674 | 1,215246637 |
| PTBP1 | 0,248048273 | 0,292982937 | 0,209 | 0,165 | 1 | Mac | 1,225170842 | 0,816212699 | 1,266666667 |
| SF3B4 | 0,248144621 | -0,410492719 | 0,146 | 0,194 | 1 | Mac | 0,752366376 | 1,329139674 | 0,75257732 |
| GIMAP1 | 0,248398305 | -0,338936525 | 0,089 | 0,126 | 1 | Mac | 0,790623901 | 1,264823892 | 0,706349206 |
| SKP11 | 0,248850313 | 0,281939569 | 0,523 | 0,495 | 1 | Mac | 1,215828354 | 0,82248452 | 1,056565657 |
| C5AR1 | 0,248939719 | -0,643503094 | 0,407 | 0,437 | 1 | Mac | 0,640156657 | 1,562117629 | 0,931350114 |
| TCTN3 | 0,250217002 | -0,30261938 | 0,073 | 0,107 | 1 | Mac | 0,810778996 | 1,233381729 | 0,682242991 |
| BUD311 | 0,25184907 | 0,328820161 | 0,206 | 0,165 | 1 | Mac | 1,255985807 | 0,796187341 | 1,248484848 |
| GLUL2 | 0,253311533 | 0,29240275 | 0,599 | 0,544 | 1 | Mac | 1,224678232 | 0,81654101 | 1,101102941 |

| Supplementary Table 4 - clusermarker | | | | | | | | | |
|---|---|---|---|---|---|---|---|---|---|
| | **p_val** | **avg_log2FC** | **pct.1** | **pct.2** | **p_val_adj** | **celltype** | **Foldchange_UP** | **Foldchange_DOWN** | **Ratio_pct1_pct2** |
| BCL2L13 | 0,254541998 | 0,257950404 | 0,125 | 0,087 | 1 | Mac | 1,195778688 | 0,836275148 | 1,436781609 |
| CCNL14 | 0,254542699 | -0,285844656 | 0,341 | 0,388 | 1 | Mac | 0,820261229 | 1,219123816 | 0,878865979 |
| DPP74 | 0,257096246 | 0,392356546 | 0,268 | 0,223 | 1 | Mac | 1,312535593 | 0,761884101 | 1,201793722 |
| RPL362 | 0,258084356 | -0,297298296 | 0,892 | 0,922 | 1 | Mac | 0,813774909 | 1,228841033 | 0,967462039 |
| ATF41 | 0,258221305 | 0,271980828 | 0,35 | 0,301 | 1 | Mac | 1,207464545 | 0,828181667 | 1,162790698 |
| POMP2 | 0,259826035 | 0,281173908 | 0,434 | 0,417 | 1 | Mac | 1,215183266 | 0,822921141 | 1,040767386 |
| ALOX5AP1 | 0,260717988 | 0,626142161 | 0,358 | 0,32 | 1 | Mac | 1,543432255 | 0,647906636 | 1,11875 |
| FAM204A1 | 0,260728757 | -0,759980104 | 0,149 | 0,194 | 1 | Mac | 0,590504474 | 1,69346727 | 0,768041237 |
| CNPPD1 | 0,26202724 | 0,260958786 | 0,136 | 0,097 | 1 | Mac | 1,198274789 | 0,834533121 | 1,402061856 |
| PIM33 | 0,262626028 | 0,282641851 | 0,298 | 0,262 | 1 | Mac | 1,216420345 | 0,822084244 | 1,13740458 |
| PAFAH1B2 | 0,264378596 | -0,282950089 | 0,106 | 0,146 | 1 | Mac | 0,821908621 | 1,216680266 | 0,726027397 |
| MEF2A | 0,265616699 | -0,355107545 | 0,217 | 0,262 | 1 | Mac | 0,781811361 | 1,279080926 | 0,828244275 |
| AQP9 | 0,268979832 | -0,331984894 | 0,092 | 0,126 | 1 | Mac | 0,794442719 | 1,258743993 | 0,73015873 |
| LMAN13 | 0,26999086 | 0,275309568 | 0,19 | 0,146 | 1 | Mac | 1,210253753 | 0,826273001 | 1,301369863 |
| NAPRT2 | 0,274142992 | 0,272547675 | 0,103 | 0,068 | 1 | Mac | 1,207939062 | 0,827856331 | 1,514705882 |
| GPNMB | 0,275256348 | -0,48934497 | 0,136 | 0,175 | 1 | Mac | 0,712348454 | 1,403807357 | 0,777142857 |
| LEPROTL1 | 0,275906567 | 0,283647236 | 0,13 | 0,097 | 1 | Mac | 1,217268339 | 0,82151155 | 1,340206186 |
| LY86 | 0,276289337 | -0,396603334 | 0,249 | 0,291 | 1 | Mac | 0,759644679 | 1,31640493 | 0,855670103 |
| TMEM176B | 0,276966915 | 0,301433907 | 0,352 | 0,311 | 1 | Mac | 1,232368667 | 0,811445493 | 1,131832797 |
| RAB131 | 0,278440072 | 0,387303959 | 0,13 | 0,097 | 1 | Mac | 1,307946888 | 0,764557039 | 1,340206186 |
| NOP533 | 0,279683136 | 0,252273608 | 0,352 | 0,301 | 1 | Mac | 1,191082717 | 0,839572253 | 1,169435216 |
| TMEM2431 | 0,281138164 | 0,2586789 | 0,144 | 0,107 | 1 | Mac | 1,196382656 | 0,835852973 | 1,345794393 |
| CMTM3 | 0,284095878 | 0,42359172 | 0,238 | 0,204 | 1 | Mac | 1,341262596 | 0,745566157 | 1,166666667 |
| HEXA1 | 0,285030675 | -0,302721358 | 0,263 | 0,32 | 1 | Mac | 0,810721688 | 1,233468914 | 0,821875 |
| COMMD3 | 0,285662415 | 0,298611955 | 0,154 | 0,117 | 1 | Mac | 1,229960475 | 0,813034256 | 1,316239316 |
| JMY | 0,286671925 | -0,624122107 | 0,125 | 0,155 | 1 | Mac | 0,648814467 | 1,541272662 | 0,806451613 |
| HDLBP1 | 0,286960922 | -0,293479676 | 0,301 | 0,359 | 1 | Mac | 0,815931715 | 1,225592757 | 0,838440111 |
| SIRPA | 0,287415271 | -0,414912563 | 0,13 | 0,165 | 1 | Mac | 0,750064952 | 1,333217872 | 0,787878788 |
| GPR34 | 0,287612274 | -0,719522351 | 0,163 | 0,204 | 1 | Mac | 0,607298474 | 1,646636774 | 0,799019608 |
| ORAI22 | 0,288575521 | 0,273734922 | 0,119 | 0,087 | 1 | Mac | 1,208933528 | 0,827175338 | 1,367816092 |
| ELAVL1 | 0,288653 | -0,251421679 | 0,127 | 0,175 | 1 | Mac | 0,840068177 | 1,190379576 | 0,725714286 |

| Supplementary Table 4 - clusermarker | | | | | | | | | |
|---|---|---|---|---|---|---|---|---|---|
| | **p_val** | **avg_log2FC** | **pct.1** | **pct.2** | **p_val_adj** | **celltype** | **Foldchange_UP** | **Foldchange_DOWN** | **Ratio_pct1_pct2** |
| WASHC41 | 0,28953392 | -0,314921536 | 0,144 | 0,184 | 1 | Mac | 0,803894711 | 1,243943997 | 0,782608696 |
| XRN12 | 0,289543383 | -0,360804081 | 0,111 | 0,146 | 1 | Mac | 0,778730436 | 1,28414141 | 0,760273973 |
| PDE4A | 0,292007654 | 0,286193762 | 0,228 | 0,184 | 1 | Mac | 1,219418858 | 0,820062765 | 1,239130435 |
| SNX5 | 0,292016746 | -0,440334234 | 0,206 | 0,252 | 1 | Mac | 0,736963854 | 1,356918653 | 0,817460317 |
| BCKDHA | 0,294660155 | 0,251964343 | 0,111 | 0,078 | 1 | Mac | 1,190827417 | 0,839752248 | 1,423076923 |
| JUN1 | 0,297505662 | -0,434207223 | 0,407 | 0,437 | 1 | Mac | 0,740100336 | 1,351168147 | 0,931350114 |
| ME2 | 0,299261146 | -0,315794043 | 0,133 | 0,175 | 1 | Mac | 0,803408682 | 1,244696532 | 0,76 |
| STUB13 | 0,300434469 | 0,33264979 | 0,217 | 0,184 | 1 | Mac | 1,259324246 | 0,794076667 | 1,179347826 |
| LSM81 | 0,302221689 | -0,361971427 | 0,209 | 0,252 | 1 | Mac | 0,778100587 | 1,285180884 | 0,829365079 |
| HSPB14 | 0,303168614 | -0,680715047 | 0,379 | 0,408 | 1 | Mac | 0,623855994 | 1,602934025 | 0,928921569 |
| NCOA41 | 0,303728062 | -0,34310755 | 0,388 | 0,417 | 1 | Mac | 0,788341403 | 1,268485959 | 0,930455635 |
| HDAC22 | 0,308937306 | 0,261381378 | 0,152 | 0,117 | 1 | Mac | 1,198625837 | 0,834288707 | 1,299145299 |
| MFF1 | 0,309255493 | 0,289318827 | 0,157 | 0,126 | 1 | Mac | 1,22206314 | 0,818288325 | 1,246031746 |
| XRCC61 | 0,309406091 | -0,33057974 | 0,287 | 0,34 | 1 | Mac | 0,795216866 | 1,2575186 | 0,844117647 |
| NAF1 | 0,30976575 | -0,334896554 | 0,214 | 0,252 | 1 | Mac | 0,792840984 | 1,261286967 | 0,849206349 |
| ANKH | 0,310084122 | 0,393003648 | 0,119 | 0,087 | 1 | Mac | 1,313124446 | 0,761542444 | 1,367816092 |
| SUZ12 | 0,314054249 | 0,255042126 | 0,152 | 0,117 | 1 | Mac | 1,193370588 | 0,837962666 | 1,299145299 |
| ARHGDIB2 | 0,314236662 | -0,313612398 | 0,656 | 0,689 | 1 | Mac | 0,804624516 | 1,242815723 | 0,952104499 |
| MARK3 | 0,316547511 | -0,295917771 | 0,095 | 0,126 | 1 | Mac | 0,814553989 | 1,227665709 | 0,753968254 |
| DUSP51 | 0,321183386 | 0,395731885 | 0,201 | 0,165 | 1 | Mac | 1,315610005 | 0,760103675 | 1,218181818 |
| CPD | 0,321314107 | -0,276471513 | 0,114 | 0,146 | 1 | Mac | 0,825607789 | 1,211228883 | 0,780821918 |
| SCO2 | 0,322887517 | 0,335543827 | 0,157 | 0,126 | 1 | Mac | 1,261852978 | 0,792485351 | 1,246031746 |
| SPTLC1 | 0,322987272 | 0,252942935 | 0,108 | 0,078 | 1 | Mac | 1,191635439 | 0,83918283 | 1,384615385 |
| TFEB | 0,325993861 | -0,292031672 | 0,076 | 0,107 | 1 | Mac | 0,81675106 | 1,224363271 | 0,710280374 |
| LMO43 | 0,327208271 | -0,296687597 | 0,122 | 0,155 | 1 | Mac | 0,814119456 | 1,22832097 | 0,787096774 |
| HMGN22 | 0,327382533 | 0,29877825 | 0,439 | 0,408 | 1 | Mac | 1,230102257 | 0,812940546 | 1,075980392 |
| NECAP2 | 0,327749967 | 0,288342766 | 0,152 | 0,117 | 1 | Mac | 1,221236629 | 0,818842128 | 1,299145299 |
| LNPEP1 | 0,327771407 | -0,275695677 | 0,141 | 0,175 | 1 | Mac | 0,826051895 | 1,210577697 | 0,805714286 |
| NPEPPS | 0,329984067 | 0,291907383 | 0,136 | 0,107 | 1 | Mac | 1,224257796 | 0,816821427 | 1,271028037 |
| PTRHD13 | 0,331637792 | 0,262280156 | 0,23 | 0,194 | 1 | Mac | 1,199372797 | 0,833769119 | 1,18556701 |
| NIBAN12 | 0,331757627 | -0,282322274 | 0,217 | 0,262 | 1 | Mac | 0,822266368 | 1,216150921 | 0,828244275 |

| Supplementary Table 4 - clusermarker | | | | | | | | | |
|---|---|---|---|---|---|---|---|---|---|
| | **p_val** | **avg_log2FC** | **pct.1** | **pct.2** | **p_val_adj** | **celltype** | **Foldchange_UP** | **Foldchange_DOWN** | **Ratio_pct1_pct2** |
| EMB2 | 0,332435331 | -0,347951966 | 0,173 | 0,214 | 1 | Mac | 0,785698676 | 1,272752558 | 0,808411215 |
| TCEAL42 | 0,333079311 | -0,399485024 | 0,127 | 0,165 | 1 | Mac | 0,758128852 | 1,319036991 | 0,76969697 |
| PAK11 | 0,333539922 | -0,320093781 | 0,29 | 0,33 | 1 | Mac | 0,801017806 | 1,248411698 | 0,878787879 |
| DDRGK11 | 0,334156306 | 0,271475231 | 0,127 | 0,097 | 1 | Mac | 1,20704146 | 0,828471956 | 1,309278351 |
| SNHG73 | 0,334583507 | 0,302644146 | 0,211 | 0,175 | 1 | Mac | 1,233402902 | 0,810765078 | 1,205714286 |
| PTPRJ | 0,336290976 | -0,302672846 | 0,13 | 0,165 | 1 | Mac | 0,810748949 | 1,233427439 | 0,787878788 |
| VSIG4 | 0,337060363 | -0,396082724 | 0,184 | 0,223 | 1 | Mac | 0,759918853 | 1,315929978 | 0,825112108 |
| EIF3F3 | 0,337616927 | -0,275197195 | 0,569 | 0,621 | 1 | Mac | 0,826337362 | 1,210159489 | 0,91626409 |
| CYTH4 | 0,338441078 | -0,265496746 | 0,136 | 0,175 | 1 | Mac | 0,831912243 | 1,202049865 | 0,777142857 |
| UBE2S | 0,339664637 | 0,305757267 | 0,39 | 0,35 | 1 | Mac | 1,236067275 | 0,809017454 | 1,114285714 |
| DPEP2 | 0,340939835 | -0,306750257 | 0,079 | 0,107 | 1 | Mac | 0,808460808 | 1,236918339 | 0,738317757 |
| PHLDA22 | 0,342139303 | -0,385914814 | 0,16 | 0,194 | 1 | Mac | 0,765293571 | 1,306688097 | 0,824742268 |
| SKIL2 | 0,344005257 | -0,300892923 | 0,347 | 0,388 | 1 | Mac | 0,811749827 | 1,231906638 | 0,894329897 |
| IL10RA | 0,344467979 | 0,261047655 | 0,266 | 0,233 | 1 | Mac | 1,198348605 | 0,834481716 | 1,141630901 |
| PPIL4 | 0,346438937 | 0,267189523 | 0,114 | 0,087 | 1 | Mac | 1,203461111 | 0,830936696 | 1,310344828 |
| TXNIP5 | 0,346694977 | -0,271022044 | 0,369 | 0,408 | 1 | Mac | 0,828732242 | 1,206662357 | 0,904411765 |
| MNDA | 0,346882247 | -0,402060056 | 0,442 | 0,515 | 1 | Mac | 0,756776893 | 1,321393411 | 0,858252427 |
| MAFF | 0,349618973 | -0,269700755 | 0,233 | 0,272 | 1 | Mac | 0,829491582 | 1,205557744 | 0,856617647 |
| SLCO2B1 | 0,350364231 | -0,446584095 | 0,211 | 0,243 | 1 | Mac | 0,733778177 | 1,362809676 | 0,868312757 |
| IQGAP21 | 0,350980876 | 0,289748516 | 0,257 | 0,223 | 1 | Mac | 1,222427172 | 0,818044644 | 1,152466368 |
| DDOST2 | 0,351261129 | -0,301652722 | 0,279 | 0,33 | 1 | Mac | 0,811322429 | 1,232555596 | 0,845454545 |
| G3BP21 | 0,351473402 | 0,294499035 | 0,238 | 0,204 | 1 | Mac | 1,226459025 | 0,815355409 | 1,166666667 |
| SNHG52 | 0,352194207 | -0,26693708 | 0,369 | 0,398 | 1 | Mac | 0,831082107 | 1,203250548 | 0,927135678 |
| ZSWIM7 | 0,35354718 | 0,258328984 | 0,106 | 0,078 | 1 | Mac | 1,196092516 | 0,836055729 | 1,358974359 |
| EMC71 | 0,354376782 | 0,263752777 | 0,146 | 0,117 | 1 | Mac | 1,200597673 | 0,832918489 | 1,247863248 |
| HP1BP32 | 0,354585474 | 0,293426831 | 0,287 | 0,262 | 1 | Mac | 1,225547865 | 0,815961603 | 1,095419847 |
| GNAI3 | 0,355491959 | -0,372675374 | 0,222 | 0,262 | 1 | Mac | 0,772348902 | 1,29475163 | 0,847328244 |
| CBR12 | 0,357037898 | 0,250110216 | 0,127 | 0,097 | 1 | Mac | 1,189297969 | 0,840832177 | 1,309278351 |
| ABCE1 | 0,357117895 | -0,492413384 | 0,087 | 0,117 | 1 | Mac | 0,710834996 | 1,40679624 | 0,743589744 |
| FCGBP3 | 0,357483112 | -0,477019665 | 0,079 | 0,107 | 1 | Mac | 0,718460295 | 1,391865364 | 0,738317757 |
| STMN11 | 0,359128618 | 0,56890931 | 0,19 | 0,155 | 1 | Mac | 1,483401682 | 0,674126241 | 1,225806452 |

| Supplementary Table 4 - clusermarker | | | | | | | | | |
|---|---|---|---|---|---|---|---|---|---|
| | **p_val** | **avg_log2FC** | **pct.1** | **pct.2** | **p_val_adj** | **celltype** | **Foldchange_UP** | **Foldchange_DOWN** | **Ratio_pct1_pct2** |
| ZNF6381 | 0,360098227 | -0,30329176 | 0,141 | 0,175 | 1 | Mac | 0,810401214 | 1,233956691 | 0,805714286 |
| IFI61 | 0,360749275 | 0,3814503 | 0,146 | 0,117 | 1 | Mac | 1,302650715 | 0,76766549 | 1,247863248 |
| NDUFA53 | 0,362510095 | -0,265715907 | 0,098 | 0,126 | 1 | Mac | 0,831785876 | 1,202232484 | 0,777777778 |
| BRD23 | 0,363517459 | -0,279675972 | 0,431 | 0,466 | 1 | Mac | 0,823776016 | 1,213922208 | 0,924892704 |
| PLAAT44 | 0,364767824 | -0,272731232 | 0,157 | 0,194 | 1 | Mac | 0,827751008 | 1,20809276 | 0,809278351 |
| REL1 | 0,365120377 | -0,296358101 | 0,491 | 0,515 | 1 | Mac | 0,814305414 | 1,228040466 | 0,953398058 |
| ATP5PF2 | 0,365320066 | 0,260815982 | 0,415 | 0,388 | 1 | Mac | 1,198156185 | 0,834615731 | 1,069587629 |
| LRRK2 | 0,370766243 | -0,25077403 | 0,114 | 0,146 | 1 | Mac | 0,840445381 | 1,189845316 | 0,780821918 |
| IDS4 | 0,372234264 | 0,383459146 | 0,26 | 0,233 | 1 | Mac | 1,304465823 | 0,766597317 | 1,115879828 |
| TCIRG11 | 0,373363683 | 0,464707807 | 0,257 | 0,233 | 1 | Mac | 1,380037822 | 0,724617821 | 1,103004292 |
| SMIM291 | 0,375957045 | -0,302831924 | 0,106 | 0,136 | 1 | Mac | 0,810659557 | 1,23356345 | 0,779411765 |
| PHACTR41 | 0,376971527 | -0,250948815 | 0,114 | 0,146 | 1 | Mac | 0,840343566 | 1,189989476 | 0,780821918 |
| IER22 | 0,377391561 | -0,285610879 | 0,477 | 0,515 | 1 | Mac | 0,820394157 | 1,218926283 | 0,926213592 |
| B4GALT51 | 0,379533957 | -0,269325596 | 0,087 | 0,117 | 1 | Mac | 0,829707311 | 1,205244291 | 0,743589744 |
| FCGRT2 | 0,382424976 | 0,260382162 | 0,575 | 0,544 | 1 | Mac | 1,197795952 | 0,834866738 | 1,056985294 |
| NAA382 | 0,387059852 | 0,330112105 | 0,241 | 0,214 | 1 | Mac | 1,257111055 | 0,795474669 | 1,126168224 |
| LYL1 | 0,387422049 | 0,2791375 | 0,117 | 0,087 | 1 | Mac | 1,213469208 | 0,824083539 | 1,344827586 |
| CALM34 | 0,387599873 | 0,2709202 | 0,228 | 0,194 | 1 | Mac | 1,206577178 | 0,828790746 | 1,175257732 |
| CTBP11 | 0,388323685 | 0,286038429 | 0,111 | 0,087 | 1 | Mac | 1,219287572 | 0,820151065 | 1,275862069 |
| ATP5MK2 | 0,389927838 | -0,303151094 | 0,463 | 0,515 | 1 | Mac | 0,810480233 | 1,233836383 | 0,899029126 |
| RIPOR21 | 0,389957272 | -0,277900145 | 0,081 | 0,107 | 1 | Mac | 0,824790634 | 1,212428899 | 0,757009346 |
| SELENOS2 | 0,390874486 | 0,318318068 | 0,233 | 0,204 | 1 | Mac | 1,24687606 | 0,802004331 | 1,142156863 |
| NDUFA32 | 0,392206682 | -0,371551047 | 0,257 | 0,291 | 1 | Mac | 0,772951047 | 1,293742992 | 0,883161512 |
| HSD17B41 | 0,39553209 | 0,342623199 | 0,152 | 0,126 | 1 | Mac | 1,268060166 | 0,788606114 | 1,206349206 |
| HLA-C4 | 0,397033628 | -0,30581129 | 0,87 | 0,796 | 1 | Mac | 0,80898716 | 1,236113562 | 1,092964824 |
| PDHA12 | 0,397198078 | -0,267263725 | 0,089 | 0,117 | 1 | Mac | 0,83089396 | 1,20352301 | 0,760683761 |
| CREG1 | 0,397700121 | 0,483158278 | 0,282 | 0,262 | 1 | Mac | 1,397800316 | 0,715409768 | 1,076335878 |
| LILRA5 | 0,398281779 | 0,264293648 | 0,157 | 0,126 | 1 | Mac | 1,201047865 | 0,832606284 | 1,246031746 |
| ATP5MC1 | 0,399916787 | 0,270478147 | 0,252 | 0,223 | 1 | Mac | 1,20620753 | 0,829044733 | 1,130044843 |
| STT3B1 | 0,400271666 | 0,26433577 | 0,163 | 0,136 | 1 | Mac | 1,201082932 | 0,832581975 | 1,198529412 |
| PLIN3 | 0,406812225 | 0,270033299 | 0,163 | 0,136 | 1 | Mac | 1,205835659 | 0,829300405 | 1,198529412 |

| Supplementary Table 4 - clusermarker | | | | | | | | | |
|---|---|---|---|---|---|---|---|---|---|
| | **p_val** | **avg_log2FC** | **pct.1** | **pct.2** | **p_val_adj** | **celltype** | **Foldchange_UP** | **Foldchange_DOWN** | **Ratio_pct1_pct2** |
| APBB1IP | 0,40691285 | 0,3086418 | 0,309 | 0,282 | 1 | Mac | 1,238541148 | 0,807401515 | 1,095744681 |
| KLF62 | 0,410044375 | 0,296524011 | 0,621 | 0,563 | 1 | Mac | 1,228181699 | 0,814211774 | 1,103019538 |
| APOC1 | 0,410855214 | -0,423770463 | 0,182 | 0,223 | 1 | Mac | 0,745473791 | 1,341428783 | 0,816143498 |
| CTCF1 | 0,411439159 | 0,276063359 | 0,111 | 0,087 | 1 | Mac | 1,210886261 | 0,825841396 | 1,275862069 |
| AP2A2 | 0,414892673 | 0,35219182 | 0,122 | 0,097 | 1 | Mac | 1,27649848 | 0,78339302 | 1,257731959 |
| LSM12 | 0,415800844 | 0,323628607 | 0,152 | 0,126 | 1 | Mac | 1,25147425 | 0,799057591 | 1,206349206 |
| TMBIM43 | 0,41823873 | 0,374249685 | 0,333 | 0,301 | 1 | Mac | 1,296165271 | 0,771506552 | 1,106312292 |
| CFP | 0,418785288 | 0,351383499 | 0,23 | 0,204 | 1 | Mac | 1,275783477 | 0,783832067 | 1,12745098 |
| ERBIN | 0,418847467 | -0,270256442 | 0,127 | 0,097 | 1 | Mac | 0,829172145 | 1,206022182 | 1,309278351 |
| MMP19 | 0,418995025 | 0,557708572 | 0,192 | 0,165 | 1 | Mac | 1,471929499 | 0,679380365 | 1,163636364 |
| CD163 | 0,420303766 | 0,251467864 | 0,257 | 0,223 | 1 | Mac | 1,190417685 | 0,840041284 | 1,152466368 |
| PTEN1 | 0,424952433 | -0,322074329 | 0,22 | 0,252 | 1 | Mac | 0,799918915 | 1,250126708 | 0,873015873 |
| TNFAIP22 | 0,431283111 | -0,415902953 | 0,355 | 0,388 | 1 | Mac | 0,74955022 | 1,334133422 | 0,914948454 |
| RAB8A2 | 0,435344834 | -0,275834851 | 0,184 | 0,214 | 1 | Mac | 0,82597221 | 1,210694485 | 0,859813084 |
| PLIN2 | 0,435355473 | 0,557064195 | 0,339 | 0,32 | 1 | Mac | 1,471272211 | 0,679683877 | 1,059375 |
| FYB11 | 0,438096915 | -0,292771184 | 0,493 | 0,485 | 1 | Mac | 0,816332509 | 1,224991029 | 1,016494845 |
| PLCG21 | 0,439136147 | -0,383215785 | 0,117 | 0,146 | 1 | Mac | 0,766726641 | 1,304245798 | 0,801369863 |
| EIF2S2 | 0,439532391 | -0,257890346 | 0,198 | 0,233 | 1 | Mac | 0,836309962 | 1,19572891 | 0,849785408 |
| HSPA53 | 0,440967458 | 0,38694865 | 0,423 | 0,417 | 1 | Mac | 1,307624806 | 0,764745358 | 1,014388489 |
| CDK5RAP32 | 0,444550312 | 0,252189408 | 0,152 | 0,126 | 1 | Mac | 1,191013205 | 0,839621254 | 1,206349206 |
| PARVG | 0,445807057 | -0,367424077 | 0,26 | 0,291 | 1 | Mac | 0,775165315 | 1,290047401 | 0,89347079 |
| SGK1 | 0,446077177 | -0,644286534 | 0,561 | 0,524 | 1 | Mac | 0,639809122 | 1,562966151 | 1,070610687 |
| GM2A | 0,448725018 | 0,26029308 | 0,22 | 0,194 | 1 | Mac | 1,197721994 | 0,83491829 | 1,134020619 |
| DAB2 | 0,449941482 | -0,403721556 | 0,233 | 0,262 | 1 | Mac | 0,755905842 | 1,322916089 | 0,889312977 |
| RSL1D12 | 0,452514131 | -0,251534166 | 0,295 | 0,32 | 1 | Mac | 0,840002679 | 1,190472394 | 0,921875 |
| DNMT1 | 0,452567464 | 0,380327624 | 0,198 | 0,175 | 1 | Mac | 1,301637413 | 0,768263105 | 1,131428571 |
| SLC11A1 | 0,456310072 | 0,425107734 | 0,201 | 0,175 | 1 | Mac | 1,342672764 | 0,744783112 | 1,148571429 |
| PSMD13 | 0,458039051 | 0,288072279 | 0,141 | 0,117 | 1 | Mac | 1,221007684 | 0,818995665 | 1,205128205 |
| SIGLEC10 | 0,458444609 | -0,286028982 | 0,149 | 0,175 | 1 | Mac | 0,820156435 | 1,219279587 | 0,851428571 |
| PSMB11 | 0,464432855 | 0,253141832 | 0,417 | 0,417 | 1 | Mac | 1,191799735 | 0,839067144 | 1 |
| NCOR11 | 0,464798256 | 0,324544512 | 0,198 | 0,175 | 1 | Mac | 1,25226901 | 0,798550465 | 1,131428571 |

| Supplementary Table 4 - clusermarker | | | | | | | | | |
|---|---|---|---|---|---|---|---|---|---|
| | **p_val** | **avg_log2FC** | **pct.1** | **pct.2** | **p_val_adj** | **celltype** | **Foldchange_UP** | **Foldchange_DOWN** | **Ratio_pct1_pct2** |
| CHKB2 | 0,468824048 | 0,374695743 | 0,117 | 0,097 | 1 | Mac | 1,296566087 | 0,771268052 | 1,206185567 |
| NBDY | 0,471836249 | -0,478305789 | 0,209 | 0,243 | 1 | Mac | 0,717820092 | 1,393106728 | 0,860082305 |
| MMP24OS3 | 0,472300173 | -0,323796594 | 0,22 | 0,252 | 1 | Mac | 0,798964555 | 1,25161998 | 0,873015873 |
| FKBP81 | 0,481924969 | -0,271858928 | 0,282 | 0,252 | 1 | Mac | 0,828251647 | 1,207362525 | 1,119047619 |
| SAMD81 | 0,48199966 | -0,291443955 | 0,084 | 0,107 | 1 | Mac | 0,817083852 | 1,223864598 | 0,785046729 |
| RTF21 | 0,4825875 | 0,307922148 | 0,241 | 0,223 | 1 | Mac | 1,237923487 | 0,807804368 | 1,080717489 |
| CMC11 | 0,486382607 | 0,320987167 | 0,117 | 0,097 | 1 | Mac | 1,249185014 | 0,800521932 | 1,206185567 |
| SFT2D2 | 0,486428035 | -0,264468187 | 0,16 | 0,184 | 1 | Mac | 0,83250556 | 1,201193178 | 0,869565217 |
| COPS91 | 0,487773382 | 0,266012841 | 0,266 | 0,243 | 1 | Mac | 1,202479952 | 0,831614696 | 1,094650206 |
| PPP1R9B | 0,48787272 | 0,344929742 | 0,114 | 0,097 | 1 | Mac | 1,270089129 | 0,787346319 | 1,175257732 |
| COPB12 | 0,493539648 | 0,260514556 | 0,198 | 0,175 | 1 | Mac | 1,197905877 | 0,834790128 | 1,131428571 |
| PRRC2B3 | 0,499055761 | 0,363974459 | 0,152 | 0,136 | 1 | Mac | 1,286966463 | 0,777021025 | 1,117647059 |
| RPL52 | 0,501804416 | -0,269537194 | 0,837 | 0,864 | 1 | Mac | 0,829585628 | 1,205421075 | 0,96875 |
| HSPB111 | 0,502036694 | 0,318746028 | 0,154 | 0,136 | 1 | Mac | 1,247245987 | 0,80176646 | 1,132352941 |
| PSMC4 | 0,505218726 | -0,55110949 | 0,179 | 0,204 | 1 | Mac | 0,682495061 | 1,465212069 | 0,87745098 |
| NFKBIZ1 | 0,505535667 | 0,313034388 | 0,379 | 0,359 | 1 | Mac | 1,242317893 | 0,804946951 | 1,055710306 |
| CCL41 | 0,506638589 | -0,511788345 | 0,257 | 0,291 | 1 | Mac | 0,701352512 | 1,425816523 | 0,883161512 |
| GRSF1 | 0,515394057 | 0,250553339 | 0,176 | 0,155 | 1 | Mac | 1,189663317 | 0,840573955 | 1,135483871 |
| PAPSS2 | 0,520608942 | 0,291689213 | 0,125 | 0,107 | 1 | Mac | 1,224072673 | 0,816944959 | 1,168224299 |
| UBXN43 | 0,522760372 | -0,310719082 | 0,252 | 0,282 | 1 | Mac | 0,806239805 | 1,240325761 | 0,893617021 |
| RNPEPL1 | 0,52687822 | 0,319372904 | 0,144 | 0,126 | 1 | Mac | 1,247788055 | 0,801418154 | 1,142857143 |
| MTHFS | 0,535610593 | -0,293372854 | 0,111 | 0,087 | 1 | Mac | 0,815992131 | 1,225502013 | 1,275862069 |
| GASK1B1 | 0,537184505 | -0,481877363 | 0,108 | 0,126 | 1 | Mac | 0,716045236 | 1,396559812 | 0,857142857 |
| KHDRBS11 | 0,540656333 | 0,29440815 | 0,331 | 0,33 | 1 | Mac | 1,226381764 | 0,815406776 | 1,003030303 |
| TAPBP2 | 0,542279147 | 0,263870107 | 0,352 | 0,34 | 1 | Mac | 1,200695318 | 0,832850753 | 1,035294118 |
| WIPF12 | 0,543171972 | 0,326019164 | 0,274 | 0,252 | 1 | Mac | 1,253549671 | 0,797734643 | 1,087301587 |
| ARID4B1 | 0,546725728 | -0,251268646 | 0,228 | 0,252 | 1 | Mac | 0,840157291 | 1,190253314 | 0,904761905 |
| RGS13 | 0,548751367 | -0,371558073 | 0,496 | 0,417 | 1 | Mac | 0,772947283 | 1,293749292 | 1,189448441 |
| S100A81 | 0,551809302 | -1,216088712 | 0,225 | 0,233 | 1 | Mac | 0,430448125 | 2,323160312 | 0,965665236 |
| RAB10 | 0,554450469 | -0,335395266 | 0,295 | 0,252 | 1 | Mac | 0,792566961 | 1,261723045 | 1,170634921 |
| VPS26B | 0,555475094 | -0,320784303 | 0,098 | 0,117 | 1 | Mac | 0,800634504 | 1,249009373 | 0,837606838 |

| Supplementary Table 4 - clusermarker | | | | | | | | | |
|---|---|---|---|---|---|---|---|---|---|
| | **p_val** | **avg_log2FC** | **pct.1** | **pct.2** | **p_val_adj** | **celltype** | **Foldchange_UP** | **Foldchange_DOWN** | **Ratio_pct1_pct2** |
| VCAN1 | 0,555539674 | -0,262108258 | 0,236 | 0,194 | 1 | Mac | 0,833868469 | 1,199229899 | 1,216494845 |
| DNAJC2 | 0,556498118 | 0,254676357 | 0,136 | 0,117 | 1 | Mac | 1,193068069 | 0,838175143 | 1,162393162 |
| ISCU4 | 0,560007575 | 0,316405247 | 0,26 | 0,252 | 1 | Mac | 1,245223964 | 0,803068387 | 1,031746032 |
| ARHGDIA1 | 0,560057887 | -0,387208829 | 0,347 | 0,379 | 1 | Mac | 0,764607454 | 1,307860647 | 0,915567282 |
| VAMP32 | 0,560767377 | 0,252357383 | 0,16 | 0,136 | 1 | Mac | 1,191151884 | 0,839523501 | 1,176470588 |
| MRPL18 | 0,561724002 | 0,374323605 | 0,133 | 0,117 | 1 | Mac | 1,296231685 | 0,771467024 | 1,136752137 |
| CAT1 | 0,565901205 | 0,265668336 | 0,268 | 0,252 | 1 | Mac | 1,202192843 | 0,831813303 | 1,063492063 |
| HAUS42 | 0,57041154 | 0,253962034 | 0,157 | 0,136 | 1 | Mac | 1,192477491 | 0,838590252 | 1,154411765 |
| ZBTB382 | 0,573496366 | 0,254533586 | 0,138 | 0,126 | 1 | Mac | 1,192950008 | 0,838258094 | 1,095238095 |
| ELOC1 | 0,574436338 | 0,250885226 | 0,279 | 0,262 | 1 | Mac | 1,189937027 | 0,840380606 | 1,064885496 |
| BNIP3L1 | 0,576991526 | -0,498791785 | 0,32 | 0,359 | 1 | Mac | 0,707699211 | 1,413029695 | 0,891364903 |
| PSMG21 | 0,58272193 | 0,396594051 | 0,195 | 0,184 | 1 | Mac | 1,316396459 | 0,759649567 | 1,059782609 |
| SZRD1 | 0,583114941 | 0,317326046 | 0,111 | 0,097 | 1 | Mac | 1,246018981 | 0,802555992 | 1,144329897 |
| PPP1R101 | 0,584983645 | -0,278237465 | 0,412 | 0,447 | 1 | Mac | 0,82459781 | 1,212712413 | 0,921700224 |
| RBM421 | 0,593279253 | -0,332620557 | 0,154 | 0,175 | 1 | Mac | 0,794092758 | 1,259298729 | 0,88 |
| BCL2L11 | 0,595752359 | 0,252247479 | 0,149 | 0,136 | 1 | Mac | 1,191061146 | 0,839587458 | 1,095588235 |
| SMIM12 | 0,610298406 | 0,294653784 | 0,1 | 0,087 | 1 | Mac | 1,226590586 | 0,815267956 | 1,149425287 |
| PALD1 | 0,616519672 | -0,271338746 | 0,098 | 0,117 | 1 | Mac | 0,828550337 | 1,206927274 | 0,837606838 |
| PSMD1 | 0,618324173 | -0,252927813 | 0,144 | 0,165 | 1 | Mac | 0,839191626 | 1,191622949 | 0,872727273 |
| ABHD17A | 0,619642463 | 0,324834726 | 0,182 | 0,175 | 1 | Mac | 1,252520942 | 0,798389844 | 1,04 |
| MMP9 | 0,628265621 | 0,332410425 | 0,133 | 0,155 | 1 | Mac | 1,259115322 | 0,794208427 | 0,858064516 |
| GPAA13 | 0,641796891 | -0,252172992 | 0,144 | 0,165 | 1 | Mac | 0,839630808 | 1,190999652 | 0,872727273 |
| DEGS1 | 0,646293644 | -0,340093364 | 0,173 | 0,194 | 1 | Mac | 0,789990186 | 1,26583851 | 0,891752577 |
| VIM-AS11 | 0,649276142 | 0,29852393 | 0,136 | 0,126 | 1 | Mac | 1,229885432 | 0,813083865 | 1,079365079 |
| ARPIN | 0,659855205 | 0,328798771 | 0,154 | 0,146 | 1 | Mac | 1,255967185 | 0,796199146 | 1,054794521 |
| FLYWCH22 | 0,673367407 | -0,395897506 | 0,095 | 0,107 | 1 | Mac | 0,76001642 | 1,315761046 | 0,887850467 |
| TFRC1 | 0,680761818 | 0,286771083 | 0,238 | 0,272 | 1 | Mac | 1,219906928 | 0,819734667 | 0,875 |
| CXCL83 | 0,68357227 | -0,952890049 | 0,407 | 0,388 | 1 | Mac | 0,516596563 | 1,935746521 | 1,048969072 |
| DDX62 | 0,689821182 | -0,339324381 | 0,209 | 0,223 | 1 | Mac | 0,790411377 | 1,265163975 | 0,937219731 |
| ANKRD281 | 0,693821497 | -0,395865247 | 0,163 | 0,165 | 1 | Mac | 0,760033415 | 1,315731625 | 0,987878788 |
| DNAJB141 | 0,69390504 | 0,260427054 | 0,106 | 0,097 | 1 | Mac | 1,197833224 | 0,834840761 | 1,092783505 |

| Supplementary Table 4 - clusermarker | | | | | | | | | |
|---|---|---|---|---|---|---|---|---|---|
| | **p_val** | **avg_log2FC** | **pct.1** | **pct.2** | **p_val_adj** | **celltype** | **Foldchange_UP** | **Foldchange_DOWN** | **Ratio_pct1_pct2** |
| PECAM1 | 0,700800691 | 0,39184655 | 0,26 | 0,262 | 1 | Mac | 1,31207169 | 0,762153476 | 0,992366412 |
| CHMP1B1 | 0,702898514 | -0,346301628 | 0,444 | 0,408 | 1 | Mac | 0,786597972 | 1,271297455 | 1,088235294 |
| SLC38A1 | 0,703169935 | -0,285213105 | 0,095 | 0,107 | 1 | Mac | 0,820620384 | 1,218590252 | 0,887850467 |
| TRIM221 | 0,709459613 | -0,265515221 | 0,23 | 0,252 | 1 | Mac | 0,831901589 | 1,20206526 | 0,912698413 |
| CD1C | 0,709749369 | 0,380359415 | 0,163 | 0,155 | 1 | Mac | 1,301666096 | 0,768246176 | 1,051612903 |
| PITHD1 | 0,711076629 | 0,280127109 | 0,106 | 0,097 | 1 | Mac | 1,214301866 | 0,823518458 | 1,092783505 |
| CTSL | 0,71113755 | 0,430039449 | 0,29 | 0,32 | 1 | Mac | 1,347270416 | 0,742241489 | 0,90625 |
| BCL2A1 | 0,716647248 | -0,361096917 | 0,48 | 0,485 | 1 | Mac | 0,778572387 | 1,284402089 | 0,989690722 |
| CDC422 | 0,721878202 | -0,283428375 | 0,61 | 0,631 | 1 | Mac | 0,821636185 | 1,21708369 | 0,966719493 |
| ADGRE51 | 0,722519321 | 0,38018382 | 0,287 | 0,291 | 1 | Mac | 1,301507676 | 0,768339687 | 0,986254296 |
| SHISA52 | 0,727252853 | 0,27105089 | 0,114 | 0,107 | 1 | Mac | 1,206686484 | 0,828715671 | 1,065420561 |
| LTBR | 0,7286841 | -0,343498033 | 0,138 | 0,146 | 1 | Mac | 0,788128057 | 1,268829337 | 0,945205479 |
| SETX1 | 0,729141895 | -0,448578955 | 0,165 | 0,184 | 1 | Mac | 0,732764261 | 1,364695378 | 0,89673913 |
| AIG1 | 0,729555787 | 0,338895569 | 0,106 | 0,097 | 1 | Mac | 1,264787986 | 0,790646347 | 1,092783505 |
| DNAJB16 | 0,729829962 | -0,826313581 | 0,409 | 0,398 | 1 | Mac | 0,563968473 | 1,773148764 | 1,027638191 |
| RAB4A1 | 0,73360215 | 0,29256338 | 0,165 | 0,155 | 1 | Mac | 1,224814596 | 0,816450101 | 1,064516129 |
| TREM1 | 0,736669735 | 0,298407297 | 0,171 | 0,165 | 1 | Mac | 1,229786007 | 0,8131496 | 1,036363636 |
| FOS4 | 0,737330776 | -0,307129691 | 0,71 | 0,67 | 1 | Mac | 0,808248208 | 1,237243695 | 1,059701493 |
| LTC4S1 | 0,737916809 | -0,289913925 | 0,117 | 0,126 | 1 | Mac | 0,817950858 | 1,222567334 | 0,928571429 |
| MS4A4A | 0,739075164 | 0,262272838 | 0,217 | 0,214 | 1 | Mac | 1,199366713 | 0,833773348 | 1,014018692 |
| TMEM230 | 0,742394322 | -0,466811269 | 0,225 | 0,243 | 1 | Mac | 0,723562092 | 1,382051397 | 0,925925926 |
| ADD12 | 0,743789232 | 0,31995497 | 0,182 | 0,175 | 1 | Mac | 1,248291586 | 0,801094881 | 1,04 |
| MAT2B | 0,754439685 | -0,283740439 | 0,117 | 0,126 | 1 | Mac | 0,821458479 | 1,217346981 | 0,928571429 |
| MRPL431 | 0,756667579 | -0,376168446 | 0,16 | 0,175 | 1 | Mac | 0,770481143 | 1,297890297 | 0,914285714 |
| LCOR | 0,756868448 | -0,250866035 | 0,136 | 0,146 | 1 | Mac | 0,840391785 | 1,189921198 | 0,931506849 |
| PGD3 | 0,758781663 | 0,265889524 | 0,247 | 0,243 | 1 | Mac | 1,202377172 | 0,831685783 | 1,016460905 |
| PEPD | 0,759785449 | -0,253807302 | 0,203 | 0,184 | 1 | Mac | 0,838680198 | 1,192349602 | 1,10326087 |
| CHCHD31 | 0,762075697 | 0,286489109 | 0,13 | 0,126 | 1 | Mac | 1,219668522 | 0,819894899 | 1,031746032 |
| SCNM11 | 0,765077215 | -0,299348797 | 0,195 | 0,204 | 1 | Mac | 0,812619113 | 1,230588825 | 0,955882353 |
| VPS26A | 0,767060556 | -0,305240775 | 0,16 | 0,165 | 1 | Mac | 0,809307138 | 1,235624837 | 0,96969697 |
| SAMSN11 | 0,771961097 | -0,263755692 | 0,352 | 0,34 | 1 | Mac | 0,832916806 | 1,200600099 | 1,035294118 |

| Supplementary Table 4 - clusermarker | | | | | | | | | |
|---|---|---|---|---|---|---|---|---|---|
| | **p_val** | **avg_log2FC** | **pct.1** | **pct.2** | **p_val_adj** | **celltype** | **Foldchange_UP** | **Foldchange_DOWN** | **Ratio_pct1_pct2** |
| OSTM11 | 0,777679568 | 0,313416657 | 0,127 | 0,126 | 1 | Mac | 1,242647112 | 0,804733693 | 1,007936508 |
| LAP32 | 0,780166147 | -0,379748092 | 0,374 | 0,379 | 1 | Mac | 0,768571779 | 1,301114649 | 0,986807388 |
| POLD41 | 0,781517308 | -0,264932368 | 0,39 | 0,369 | 1 | Mac | 0,832237748 | 1,201579719 | 1,056910569 |
| POLR2K2 | 0,784141146 | -0,329384878 | 0,171 | 0,155 | 1 | Mac | 0,795875749 | 1,256477536 | 1,103225806 |
| METAP23 | 0,786204424 | 0,268446927 | 0,141 | 0,165 | 1 | Mac | 1,204510464 | 0,830212796 | 0,854545455 |
| SRGN2 | 0,789156519 | -0,358289305 | 0,843 | 0,786 | 1 | Mac | 0,780089032 | 1,28190496 | 1,072519084 |
| RERE2 | 0,79407882 | -0,384385107 | 0,114 | 0,126 | 1 | Mac | 0,766105452 | 1,305303334 | 0,904761905 |
| SPTAN12 | 0,795970878 | 0,25420647 | 0,187 | 0,184 | 1 | Mac | 1,19267955 | 0,838448182 | 1,016304348 |
| CCL24 | 0,796605464 | 0,345330045 | 0,106 | 0,117 | 1 | Mac | 1,270441588 | 0,787127885 | 0,905982906 |
| LAT2 | 0,797145768 | -0,338974092 | 0,146 | 0,155 | 1 | Mac | 0,790603314 | 1,264856827 | 0,941935484 |
| CAMLG | 0,807721007 | -0,306241209 | 0,184 | 0,175 | 1 | Mac | 0,80874612 | 1,236481976 | 1,051428571 |
| CD83 | 0,818703489 | -0,52447183 | 0,591 | 0,515 | 1 | Mac | 0,695213581 | 1,438406883 | 1,147572816 |
| ARFGEF1 | 0,820698265 | -0,510643728 | 0,119 | 0,126 | 1 | Mac | 0,701909177 | 1,424685746 | 0,944444444 |
| MLEC2 | 0,822627562 | -0,325025506 | 0,263 | 0,272 | 1 | Mac | 0,798284273 | 1,252686585 | 0,966911765 |
| POLR2A1 | 0,823424693 | 0,262113361 | 0,287 | 0,32 | 1 | Mac | 1,199234141 | 0,83386552 | 0,896875 |
| LDHB1 | 0,833516633 | -0,258817921 | 0,317 | 0,33 | 1 | Mac | 0,835772433 | 1,196497946 | 0,960606061 |
| VMO11 | 0,838466489 | 0,358279419 | 0,152 | 0,165 | 1 | Mac | 1,281896176 | 0,780094378 | 0,921212121 |
| ENSA3 | 0,842985008 | -0,329172952 | 0,23 | 0,233 | 1 | Mac | 0,795992669 | 1,256292978 | 0,987124464 |
| EIF2A1 | 0,851562525 | -0,295982874 | 0,214 | 0,194 | 1 | Mac | 0,814517232 | 1,227721109 | 1,103092784 |
| EREG | 0,852400178 | -0,428052958 | 0,187 | 0,175 | 1 | Mac | 0,743264208 | 1,345416594 | 1,068571429 |
| NDUFS22 | 0,853712084 | -0,343413853 | 0,168 | 0,175 | 1 | Mac | 0,788174045 | 1,268755304 | 0,96 |
| AZI2 | 0,858013256 | 0,262982978 | 0,106 | 0,117 | 1 | Mac | 1,199957225 | 0,83336304 | 0,905982906 |
| TNPO11 | 0,858204695 | -0,401210885 | 0,149 | 0,155 | 1 | Mac | 0,757222463 | 1,320615867 | 0,961290323 |
| EIF3M2 | 0,861214553 | -0,340488045 | 0,347 | 0,34 | 1 | Mac | 0,789774096 | 1,266184855 | 1,020588235 |
| BASP11 | 0,86315261 | -0,342977765 | 0,374 | 0,35 | 1 | Mac | 0,788412325 | 1,268371851 | 1,068571429 |
| DAPP1 | 0,864069741 | -0,275230923 | 0,133 | 0,126 | 1 | Mac | 0,826318044 | 1,21018778 | 1,055555556 |
| NDRG1 | 0,874279607 | 0,25669741 | 0,144 | 0,155 | 1 | Mac | 1,194740595 | 0,837001776 | 0,929032258 |
| GADD45B4 | 0,892417931 | -0,507553386 | 0,607 | 0,505 | 1 | Mac | 0,703414322 | 1,421637247 | 1,201980198 |
| CDC1231 | 0,893705983 | -0,31929146 | 0,13 | 0,136 | 1 | Mac | 0,801463398 | 1,247717616 | 0,955882353 |
| H1-41 | 0,90728244 | 0,44494263 | 0,125 | 0,136 | 1 | Mac | 1,361259984 | 0,734613529 | 0,919117647 |
| BMP2K1 | 0,907412307 | -0,309034683 | 0,225 | 0,223 | 1 | Mac | 0,807181669 | 1,23887848 | 1,00896861 |

| Supplementary Table 4 - clusermarker | | | | | | | | | |
|---|---|---|---|---|---|---|---|---|---|
| | **p_val** | **avg_log2FC** | **pct.1** | **pct.2** | **p_val_adj** | **celltype** | **Foldchange_UP** | **Foldchange_DOWN** | **Ratio_pct1_pct2** |
| DCTN21 | 0,918769029 | -0,305624731 | 0,173 | 0,165 | 1 | Mac | 0,809091779 | 1,235953727 | 1,048484848 |
| NUDT212 | 0,92276441 | 0,269480329 | 0,108 | 0,117 | 1 | Mac | 1,205373564 | 0,829618327 | 0,923076923 |
| SELENOK1 | 0,928472886 | -0,282450112 | 0,458 | 0,437 | 1 | Mac | 0,822193509 | 1,216258689 | 1,04805492 |
| PHACTR2 | 0,931544975 | -0,338253567 | 0,149 | 0,146 | 1 | Mac | 0,790998264 | 1,264225278 | 1,020547945 |
| NDUFA122 | 0,951129818 | -0,260453284 | 0,228 | 0,223 | 1 | Mac | 0,834825582 | 1,197855003 | 1,022421525 |
| AP3D12 | 0,973482786 | -0,386241894 | 0,146 | 0,146 | 1 | Mac | 0,765120088 | 1,306984375 | 1 |
| RSU11 | 0,976867883 | -0,299662784 | 0,184 | 0,184 | 1 | Mac | 0,812442275 | 1,230856679 | 1 |
| BUB31 | 0,977466631 | -0,324968629 | 0,16 | 0,155 | 1 | Mac | 0,798315746 | 1,2526372 | 1,032258065 |
| CCT71 | 0,980810684 | -0,252994375 | 0,187 | 0,184 | 1 | Mac | 0,839152909 | 1,191677929 | 1,016304348 |
| TCERG11 | 0,987698593 | -0,381601065 | 0,13 | 0,126 | 1 | Mac | 0,767585272 | 1,302786852 | 1,031746032 |
| ICAM31 | 0,989734033 | -0,268281524 | 0,179 | 0,175 | 1 | Mac | 0,830307984 | 1,204372377 | 1,022857143 |
| HTRA11 | 0,992864508 | 0,407062827 | 0,111 | 0,117 | 1 | Mac | 1,325983504 | 0,754157195 | 0,948717949 |
| ARPP19 | 1 | -0,281721058 | 0,127 | 0,126 | 1 | Mac | 0,822609103 | 1,215644218 | 1,007936508 |
| PRR46 | 7,56516E-13 | -4,46823359 | 0,056 | 0,733 | 2,76787E-08 | Mast | 0,045178069 | 22,13463364 | 0,076398363 |
| LYZ6 | 2,27312E-12 | -3,684205935 | 0,046 | 0,667 | 8,31668E-08 | Mast | 0,077793534 | 12,85453872 | 0,068965517 |
| C6orf586 | 2,10467E-11 | -1,74683688 | 0 | 0,4 | 7,70037E-07 | Mast | 0,29795433 | 3,356219052 | 0 |
| PRB36 | 2,10467E-11 | -2,277870524 | 0 | 0,4 | 7,70037E-07 | Mast | 0,206201892 | 4,849616024 | 0 |
| EAPP1 | 5,87206E-07 | -1,33009886 | 0,019 | 0,333 | 0,021484088 | Mast | 0,397740986 | 2,514199027 | 0,057057057 |
| MT-ATP86 | 8,38163E-07 | -2,175069477 | 0,259 | 0,8 | 0,030665859 | Mast | 0,221431216 | 4,516075097 | 0,32375 |
| RPL345 | 1,09817E-06 | -1,007030692 | 0,972 | 1 | 0,040178838 | Mast | 0,497569275 | 2,009770397 | 0,972 |
| RPS275 | 1,17162E-06 | -1,404549531 | 0,815 | 1 | 0,042866187 | Mast | 0,377736073 | 2,64735108 | 0,815 |
| RPS265 | 1,30491E-06 | -1,862714275 | 0,611 | 1 | 0,047742747 | Mast | 0,274958487 | 3,636912647 | 0,611 |
| RPS254 | 1,781E-06 | -1,121141132 | 0,806 | 1 | 0,065161463 | Mast | 0,459730048 | 2,175189559 | 0,806 |
| FARP21 | 2,99351E-06 | -1,004572052 | 0 | 0,2 | 0,109523653 | Mast | 0,498417956 | 2,006348264 | 0 |
| COX7A11 | 2,99351E-06 | -1,003234235 | 0 | 0,2 | 0,109523653 | Mast | 0,498880355 | 2,004488631 | 0 |
| MT-ND4L4 | 3,25282E-06 | -1,545509121 | 0,648 | 0,933 | 0,119011071 | Mast | 0,342574787 | 2,919070635 | 0,694533762 |
| RPL376 | 9,29321E-06 | -1,252257475 | 0,824 | 1 | 0,340010576 | Mast | 0,419790821 | 2,382138796 | 0,824 |
| MT-ND36 | 2,12852E-05 | -1,15060852 | 0,722 | 1 | 0,778762041 | Mast | 0,4504352 | 2,220075161 | 0,722 |
| CEBPZ1 | 2,5434E-05 | -1,521516948 | 0,056 | 0,4 | 0,930552955 | Mast | 0,348319477 | 2,870927599 | 0,14 |
| GTF2F11 | 2,85398E-05 | -1,550173562 | 0,083 | 0,467 | 1 | Mast | 0,341468982 | 2,928523685 | 0,177730193 |
| RPL37A6 | 4,43498E-05 | -1,256693724 | 0,722 | 1 | 1 | Mast | 0,418501958 | 2,389475083 | 0,722 |

| Supplementary Table 4 - clusermarker | | | | | | | | | |
|---|---|---|---|---|---|---|---|---|---|
| | **p_val** | **avg_log2FC** | **pct.1** | **pct.2** | **p_val_adj** | **celltype** | **Foldchange_UP** | **Foldchange_DOWN** | **Ratio_pct1_pct2** |
| TPSD1 | 6,79491E-05 | 1,82220691 | 0,917 | 0,667 | 1 | Mast | 3,536217249 | 0,282788056 | 1,374812594 |
| TMEM176B1 | 8,2742E-05 | 2,747523891 | 0,639 | 0,067 | 1 | Mast | 6,715635326 | 0,148906239 | 9,537313433 |
| RPS296 | 8,40926E-05 | -1,481681261 | 0,343 | 0,867 | 1 | Mast | 0,358071286 | 2,792739987 | 0,39561707 |
| MYSM1 | 9,18554E-05 | -1,093110558 | 0,009 | 0,2 | 1 | Mast | 0,468749625 | 2,133335039 | 0,045 |
| PELO1 | 9,18554E-05 | -1,025508052 | 0,009 | 0,2 | 1 | Mast | 0,491237277 | 2,035676131 | 0,045 |
| AL121944.1 | 0,000101908 | -1,318779555 | 0,009 | 0,2 | 1 | Mast | 0,400873914 | 2,494549946 | 0,045 |
| MAGI2 | 0,000101908 | -1,035995049 | 0,009 | 0,2 | 1 | Mast | 0,487679404 | 2,050527439 | 0,045 |
| QSER1 | 0,000101908 | -0,862872069 | 0,009 | 0,2 | 1 | Mast | 0,549856831 | 1,818655228 | 0,045 |
| LTA4H1 | 0,000101908 | -1,242121399 | 0,009 | 0,2 | 1 | Mast | 0,422750569 | 2,36546104 | 0,045 |
| PSMB42 | 0,000112732 | -1,471956215 | 0,074 | 0,4 | 1 | Mast | 0,360493158 | 2,773977748 | 0,185 |
| PRKG2 | 0,000125207 | -0,938748556 | 0,009 | 0,2 | 1 | Mast | 0,521685212 | 1,916864762 | 0,045 |
| WDR70 | 0,000125207 | -0,719281537 | 0,009 | 0,2 | 1 | Mast | 0,607399852 | 1,646361941 | 0,045 |
| ZFHX3 | 0,000125207 | -0,925358279 | 0,009 | 0,2 | 1 | Mast | 0,526549738 | 1,899155819 | 0,045 |
| RPL213 | 0,000145836 | -1,063114136 | 0,741 | 1 | 1 | Mast | 0,478597864 | 2,089436821 | 0,741 |
| RAMAC | 0,000148994 | -0,742489826 | 0 | 0,133 | 1 | Mast | 0,597706929 | 1,673060744 | 0 |
| AC103591.3 | 0,00014907 | -0,759738018 | 0 | 0,133 | 1 | Mast | 0,59060357 | 1,693183129 | 0 |
| DPH5 | 0,00014907 | -0,856934216 | 0 | 0,133 | 1 | Mast | 0,552124599 | 1,811185377 | 0 |
| ST3GAL5 | 0,00014907 | -0,967124508 | 0 | 0,133 | 1 | Mast | 0,511524587 | 1,954940241 | 0 |
| ATG7 | 0,00014907 | -0,678853306 | 0 | 0,133 | 1 | Mast | 0,624661575 | 1,600866836 | 0 |
| STATH6 | 0,00014907 | -2,427220782 | 0 | 0,133 | 1 | Mast | 0,185923265 | 5,378563019 | 0 |
| AC084724.1 | 0,00014907 | -0,730500937 | 0 | 0,133 | 1 | Mast | 0,602694608 | 1,659215109 | 0 |
| EXOC2 | 0,00014907 | -0,740662197 | 0 | 0,133 | 1 | Mast | 0,598464594 | 1,670942626 | 0 |
| KIAA1586 | 0,00014907 | -0,641933571 | 0 | 0,133 | 1 | Mast | 0,640853469 | 1,56041911 | 0 |
| ORC31 | 0,00014907 | -0,693090782 | 0 | 0,133 | 1 | Mast | 0,618527317 | 1,616743468 | 0 |
| PDGFA | 0,00014907 | -0,963581996 | 0 | 0,133 | 1 | Mast | 0,51278217 | 1,950145809 | 0 |
| AGFG2 | 0,00014907 | -0,959212545 | 0 | 0,133 | 1 | Mast | 0,514337574 | 1,944248391 | 0 |
| PIP5 | 0,00014907 | -1,379571631 | 0 | 0,133 | 1 | Mast | 0,384332896 | 2,601911029 | 0 |
| HPSE2 | 0,00014907 | -0,630282098 | 0 | 0,133 | 1 | Mast | 0,646050077 | 1,547867627 | 0 |
| MTRNR2L86 | 0,00014907 | -1,516859768 | 0 | 0,133 | 1 | Mast | 0,349445708 | 2,861674867 | 0 |
| ZDHHC13 | 0,00014907 | -0,751680168 | 0 | 0,133 | 1 | Mast | 0,593911483 | 1,683752594 | 0 |
| TCN13 | 0,00014907 | -0,804153512 | 0 | 0,133 | 1 | Mast | 0,572698007 | 1,74612097 | 0 |

| Supplementary Table 4 - clusermarker | | | | | | | | | |
|---|---|---|---|---|---|---|---|---|---|
| | p_val | avg_log2FC | pct.1 | pct.2 | p_val_adj | celltype | Foldchange_UP | Foldchange_DOWN | Ratio_pct1_pct2 |
| TTC121 | 0,00014907 | -0,849402723 | 0 | 0,133 | 1 | Mast | 0,555014465 | 1,801754843 | 0 |
| MORN31 | 0,00014907 | -0,892803718 | 0 | 0,133 | 1 | Mast | 0,538566456 | 1,856781069 | 0 |
| FRMD6-AS1 | 0,00014907 | -0,694215873 | 0 | 0,133 | 1 | Mast | 0,618045144 | 1,618004784 | 0 |
| AC007262.2 | 0,00014907 | -0,959212545 | 0 | 0,133 | 1 | Mast | 0,514337574 | 1,944248391 | 0 |
| MYO5C | 0,00014907 | -0,575585306 | 0 | 0,133 | 1 | Mast | 0,671013964 | 1,490281952 | 0 |
| SLC24A1 | 0,00014907 | -0,677864324 | 0 | 0,133 | 1 | Mast | 0,625089934 | 1,599769802 | 0 |
| S100B2 | 0,00014907 | -1,121719197 | 0 | 0,133 | 1 | Mast | 0,459545878 | 2,176061298 | 0 |
| RTL5 | 0,00014907 | -0,883955721 | 0 | 0,133 | 1 | Mast | 0,541879615 | 1,845428344 | 0 |
| ZDHHC15 | 0,00014907 | -0,750599026 | 0 | 0,133 | 1 | Mast | 0,594356721 | 1,682491278 | 0 |
| MTM1 | 0,00014907 | -0,678853306 | 0 | 0,133 | 1 | Mast | 0,624661575 | 1,600866836 | 0 |
| GNL32 | 0,000159828 | -1,00723623 | 0,046 | 0,333 | 1 | Mast | 0,497498393 | 2,010056745 | 0,138138138 |
| NFATC3 | 0,000167684 | -1,367982857 | 0,074 | 0,4 | 1 | Mast | 0,387432569 | 2,58109431 | 0,185 |
| PRDM8 | 0,000174207 | -1,11476234 | 0,028 | 0,267 | 1 | Mast | 0,461767218 | 2,165593316 | 0,104868914 |
| ELK3 | 0,000219021 | -1,030858058 | 0,028 | 0,267 | 1 | Mast | 0,489418975 | 2,043239128 | 0,104868914 |
| HMGCS11 | 0,000235569 | -1,083825946 | 0,046 | 0,333 | 1 | Mast | 0,471776039 | 2,119649826 | 0,138138138 |
| NDUFA101 | 0,000236222 | -1,009826667 | 0,028 | 0,267 | 1 | Mast | 0,496605909 | 2,013669153 | 0,104868914 |
| KLF41 | 0,000297379 | -1,941962628 | 0,176 | 0,533 | 1 | Mast | 0,260262141 | 3,842279923 | 0,330206379 |
| WBP2 | 0,000300705 | -1,237566611 | 0,056 | 0,333 | 1 | Mast | 0,42408736 | 2,358004728 | 0,168168168 |
| MT-ND21 | 0,000309465 | -0,946026324 | 0,889 | 1 | 1 | Mast | 0,519060164 | 1,92655894 | 0,889 |
| RPL93 | 0,000312635 | -0,914338224 | 0,62 | 0,933 | 1 | Mast | 0,530587199 | 1,884704345 | 0,664523044 |
| RPL27A5 | 0,000317169 | -1,313029398 | 0,241 | 0,667 | 1 | Mast | 0,402474867 | 2,484627193 | 0,36131934 |
| SAP301 | 0,000359955 | -1,374599307 | 0,056 | 0,333 | 1 | Mast | 0,385659804 | 2,592958844 | 0,168168168 |
| RPL385 | 0,000421789 | -1,216840494 | 0,343 | 0,8 | 1 | Mast | 0,430223879 | 2,324371215 | 0,42875 |
| SLC20A11 | 0,000428459 | -1,142398826 | 0,083 | 0,4 | 1 | Mast | 0,453005721 | 2,207477642 | 0,2075 |
| CRLF2 | 0,000467382 | -1,642322377 | 0,037 | 0,267 | 1 | Mast | 0,320340391 | 3,121679397 | 0,138576779 |
| MT-ND11 | 0,000505184 | -0,764989708 | 0,852 | 1 | 1 | Mast | 0,588457566 | 1,699357876 | 0,852 |
| PLPP1 | 0,000570684 | 3,001011109 | 0,5 | 0 | 1 | Mast | 8,005608746 | 0,124912425 | #DIV/0! |
| SLCO2B11 | 0,000611506 | -1,320717774 | 0,037 | 0,267 | 1 | Mast | 0,400335713 | 2,497903554 | 0,138576779 |
| DYNC1LI1 | 0,000653504 | -1,159621416 | 0,037 | 0,267 | 1 | Mast | 0,447629984 | 2,233987969 | 0,138576779 |
| RASGEF1B1 | 0,000662175 | -1,35981932 | 0,185 | 0,533 | 1 | Mast | 0,389631083 | 2,566530348 | 0,347091932 |
| EIF3E3 | 0,00070434 | -1,439986166 | 0,296 | 0,667 | 1 | Mast | 0,368570838 | 2,713182639 | 0,443778111 |

| Supplementary Table 4 - clusermarker | | | | | | | | | |
|---|---|---|---|---|---|---|---|---|---|
| | **p_val** | **avg_log2FC** | **pct.1** | **pct.2** | **p_val_adj** | **celltype** | **Foldchange_UP** | **Foldchange_DOWN** | **Ratio_pct1_pct2** |
| ALDH6A1 | 0,000732461 | -0,875536843 | 0,019 | 0,2 | 1 | Mast | 0,545051009 | 1,834690668 | 0,095 |
| FKBP51 | 0,000745583 | -0,926251119 | 0,037 | 0,267 | 1 | Mast | 0,526223974 | 1,900331513 | 0,138576779 |
| RPL234 | 0,000762995 | -1,230417702 | 0,398 | 0,8 | 1 | Mast | 0,426194033 | 2,346349136 | 0,4975 |
| ACTB6 | 0,000829423 | 1,031360302 | 0,991 | 1 | 1 | Mast | 2,043950563 | 0,489248624 | 0,991 |
| SLPI6 | 0,000830262 | -1,949919009 | 0,13 | 0,467 | 1 | Mast | 0,258830761 | 3,863528417 | 0,278372591 |
| ATP13A3 | 0,000837157 | -1,199767234 | 0,065 | 0,333 | 1 | Mast | 0,435345515 | 2,297026076 | 0,195195195 |
| RPL395 | 0,000852494 | -0,751901131 | 0,954 | 1 | 1 | Mast | 0,593820527 | 1,684010496 | 0,954 |
| PPP2R3C | 0,000935086 | -0,753850945 | 0,019 | 0,2 | 1 | Mast | 0,593018515 | 1,686287989 | 0,095 |
| PFN15 | 0,000980953 | 1,426262755 | 0,759 | 0,4 | 1 | Mast | 2,687496274 | 0,372093539 | 1,8975 |
| RRP15 | 0,001013424 | -0,912934 | 0,019 | 0,2 | 1 | Mast | 0,531103889 | 1,88287079 | 0,095 |
| VEZF12 | 0,001013424 | -0,991459384 | 0,019 | 0,2 | 1 | Mast | 0,50296873 | 1,988195169 | 0,095 |
| RPL35A6 | 0,00103027 | -0,729432803 | 0,898 | 1 | 1 | Mast | 0,603140993 | 1,657987125 | 0,898 |
| WDFY3 | 0,0010978 | -0,743982381 | 0,019 | 0,2 | 1 | Mast | 0,597088886 | 1,674792521 | 0,095 |
| RPL316 | 0,001115504 | -1,085034327 | 0,537 | 0,867 | 1 | Mast | 0,471381052 | 2,121425959 | 0,619377163 |
| MED12 | 0,001188632 | -0,61499526 | 0,019 | 0,2 | 1 | Mast | 0,652932039 | 1,531552965 | 0,095 |
| RPS64 | 0,001221541 | -0,765996841 | 0,815 | 1 | 1 | Mast | 0,588046913 | 1,700544597 | 0,815 |
| AL355881.11 | 0,001325909 | -1,386917685 | 0,139 | 0,467 | 1 | Mast | 0,382380886 | 2,615193479 | 0,29764454 |
| TMEM176A1 | 0,001340807 | 2,583246399 | 0,454 | 0 | 1 | Mast | 5,992867178 | 0,166865036 | #DIV/0! |
| SCGB3A26 | 0,001422304 | -1,535497975 | 0,065 | 0,333 | 1 | Mast | 0,344960248 | 2,898884744 | 0,195195195 |
| RPS244 | 0,001587852 | -0,705105303 | 0,907 | 1 | 1 | Mast | 0,613397715 | 1,630263654 | 0,907 |
| PPP4R3B1 | 0,001633643 | -1,231448939 | 0,046 | 0,267 | 1 | Mast | 0,425889498 | 2,348026903 | 0,172284644 |
| KLF7 | 0,001891175 | -0,942606474 | 0,074 | 0,333 | 1 | Mast | 0,520292035 | 1,921997519 | 0,222222222 |
| RPL75 | 0,001967877 | -1,169118953 | 0,648 | 1 | 1 | Mast | 0,444692829 | 2,248743254 | 0,648 |
| MT-ND42 | 0,002057473 | -0,546654251 | 0,88 | 1 | 1 | Mast | 0,684605956 | 1,460694274 | 0,88 |
| ABR | 0,002060759 | -0,888043304 | 0,046 | 0,267 | 1 | Mast | 0,540346482 | 1,850664405 | 0,172284644 |
| MAGI2-AS3 | 0,002310638 | -0,829531564 | 0,046 | 0,267 | 1 | Mast | 0,562711923 | 1,77710825 | 0,172284644 |
| TRA2B1 | 0,002355418 | -0,876666999 | 0,176 | 0,533 | 1 | Mast | 0,544624203 | 1,836128462 | 0,330206379 |
| CAPZA21 | 0,002391186 | -1,054104336 | 0,139 | 0,467 | 1 | Mast | 0,481596117 | 2,076428703 | 0,29764454 |
| PCF112 | 0,00275057 | -0,97890921 | 0,185 | 0,533 | 1 | Mast | 0,507363201 | 1,970974635 | 0,347091932 |
| NBEAL12 | 0,002767737 | -1,304608575 | 0,083 | 0,333 | 1 | Mast | 0,40483093 | 2,470166989 | 0,249249249 |
| METTL21A | 0,00290718 | -1,367024837 | 0,157 | 0,467 | 1 | Mast | 0,387689929 | 2,579380907 | 0,336188437 |

| Supplementary Table 4 - clusermarker | | | | | | | | | |
|---|---|---|---|---|---|---|---|---|---|
| | **p_val** | **avg_log2FC** | **pct.1** | **pct.2** | **p_val_adj** | **celltype** | **Foldchange_UP** | **Foldchange_DOWN** | **Ratio_pct1_pct2** |
| UBA7 | 0,00304804 | -1,019843129 | 0,102 | 0,4 | 1 | Mast | 0,493169974 | 2,027698467 | 0,255 |
| FER | 0,003170448 | -1,269172575 | 0,222 | 0,533 | 1 | Mast | 0,41489766 | 2,410232925 | 0,416510319 |
| EEF1A14 | 0,003191807 | -0,62582927 | 1 | 1 | 1 | Mast | 0,648047169 | 1,543097553 | 1 |
| RPL135 | 0,00326448 | 0,867354098 | 0,926 | 0,867 | 1 | Mast | 1,82431404 | 0,548151238 | 1,06805075 |
| IL4R | 0,003320913 | -1,081739021 | 0,083 | 0,333 | 1 | Mast | 0,472458979 | 2,116585871 | 0,249249249 |
| COL9A21 | 0,003389863 | -0,773859262 | 0,028 | 0,2 | 1 | Mast | 0,584850883 | 1,709837549 | 0,14 |
| USP24 | 0,003389863 | -0,876530475 | 0,028 | 0,2 | 1 | Mast | 0,544675744 | 1,835954715 | 0,14 |
| PARN | 0,003389863 | -0,931520283 | 0,028 | 0,2 | 1 | Mast | 0,524305548 | 1,907284796 | 0,14 |
| LTC4S2 | 0,003415019 | 1,206044463 | 0,806 | 0,667 | 1 | Mast | 2,307042311 | 0,433455423 | 1,208395802 |
| CPM | 0,00347096 | -1,155763688 | 0,333 | 0,667 | 1 | Mast | 0,448828537 | 2,228022324 | 0,499250375 |
| LINC01409 | 0,003485203 | -0,687943062 | 0,009 | 0,133 | 1 | Mast | 0,620738244 | 1,610985001 | 0,067669173 |
| GPR89B | 0,003485203 | -0,781429989 | 0,009 | 0,133 | 1 | Mast | 0,581789842 | 1,718833724 | 0,067669173 |
| C2orf491 | 0,003485203 | -0,747844473 | 0,009 | 0,133 | 1 | Mast | 0,595492617 | 1,679281944 | 0,067669173 |
| EXOC11 | 0,003485203 | -0,715660371 | 0,009 | 0,133 | 1 | Mast | 0,608926341 | 1,642234753 | 0,067669173 |
| CCSER1 | 0,003485203 | -0,700498819 | 0,009 | 0,133 | 1 | Mast | 0,615359406 | 1,625066571 | 0,067669173 |
| SLC22A41 | 0,003485203 | -0,749105536 | 0,009 | 0,133 | 1 | Mast | 0,594972323 | 1,68075045 | 0,067669173 |
| PGM3 | 0,003485203 | -0,640582849 | 0,009 | 0,133 | 1 | Mast | 0,641453749 | 1,558958852 | 0,067669173 |
| PEX2 | 0,003485203 | -0,793762831 | 0,009 | 0,133 | 1 | Mast | 0,576837623 | 1,733590114 | 0,067669173 |
| FABP54 | 0,003485203 | -1,421221435 | 0,009 | 0,133 | 1 | Mast | 0,373396048 | 2,67812154 | 0,067669173 |
| ZFAND1 | 0,003485203 | -0,701171214 | 0,009 | 0,133 | 1 | Mast | 0,615072673 | 1,62582414 | 0,067669173 |
| KLHL9 | 0,003485203 | -0,600684382 | 0,009 | 0,133 | 1 | Mast | 0,659441057 | 1,516435759 | 0,067669173 |
| STIM1 | 0,003485203 | -1,17292831 | 0,009 | 0,133 | 1 | Mast | 0,443520191 | 2,254688782 | 0,067669173 |
| LMBR1L | 0,003485203 | -1,015238671 | 0,009 | 0,133 | 1 | Mast | 0,494746473 | 2,021237247 | 0,067669173 |
| RAP2A | 0,003485203 | -0,652589506 | 0,009 | 0,133 | 1 | Mast | 0,636137479 | 1,571987239 | 0,067669173 |
| SHF | 0,003485203 | -0,65019117 | 0,009 | 0,133 | 1 | Mast | 0,637195874 | 1,569376139 | 0,067669173 |
| TLCD3A | 0,003485203 | -0,730236475 | 0,009 | 0,133 | 1 | Mast | 0,602805099 | 1,658910985 | 0,067669173 |
| MPDU1 | 0,003485203 | -0,732015385 | 0,009 | 0,133 | 1 | Mast | 0,60206227 | 1,66095776 | 0,067669173 |
| SPAG5 | 0,003485203 | -0,711055059 | 0,009 | 0,133 | 1 | Mast | 0,610873237 | 1,637000837 | 0,067669173 |
| ZNF519 | 0,003485203 | -0,702496725 | 0,009 | 0,133 | 1 | Mast | 0,61450782 | 1,627318591 | 0,067669173 |
| ZNF529-AS1 | 0,003485203 | -0,721042146 | 0,009 | 0,133 | 1 | Mast | 0,606659057 | 1,648372324 | 0,067669173 |
| GPR651 | 0,003493159 | -0,919026913 | 0,185 | 0,533 | 1 | Mast | 0,528865616 | 1,890839507 | 0,347091932 |

| Supplementary Table 4 - clusermarker | | | | | | | | | |
|---|---|---|---|---|---|---|---|---|---|
| | **p_val** | **avg_log2FC** | **pct.1** | **pct.2** | **p_val_adj** | **celltype** | **Foldchange_UP** | **Foldchange_DOWN** | **Ratio_pct1_pct2** |
| PPIB2 | 0,003585725 | 1,629512226 | 0,519 | 0,067 | 1 | Mast | 3,094083703 | 0,323197462 | 7,746268657 |
| MEF2A1 | 0,003603185 | -1,391446246 | 0,056 | 0,267 | 1 | Mast | 0,38118249 | 2,623415362 | 0,209737828 |
| BAIAP21 | 0,003622725 | -0,801282948 | 0,028 | 0,2 | 1 | Mast | 0,573838652 | 1,742650127 | 0,14 |
| NDUFB52 | 0,00363352 | -1,145701762 | 0,083 | 0,333 | 1 | Mast | 0,451969786 | 2,212537276 | 0,249249249 |
| PREPL1 | 0,003822819 | -0,925804747 | 0,009 | 0,133 | 1 | Mast | 0,526386813 | 1,899743639 | 0,067669173 |
| ZSWIM62 | 0,003822819 | -0,863334719 | 0,009 | 0,133 | 1 | Mast | 0,549680529 | 1,819238535 | 0,067669173 |
| SLC35B3 | 0,003822819 | -0,735804699 | 0,009 | 0,133 | 1 | Mast | 0,600482997 | 1,665326087 | 0,067669173 |
| B3GALT41 | 0,003822819 | -0,522715526 | 0,009 | 0,133 | 1 | Mast | 0,696060433 | 1,436656865 | 0,067669173 |
| LRWD1 | 0,003822819 | -0,713863281 | 0,009 | 0,133 | 1 | Mast | 0,609685322 | 1,64019038 | 0,067669173 |
| CCN3 | 0,003822819 | -1,350557448 | 0,009 | 0,133 | 1 | Mast | 0,392140499 | 2,550106408 | 0,067669173 |
| DDX58 | 0,003822819 | -0,519757171 | 0,009 | 0,133 | 1 | Mast | 0,697489222 | 1,43371391 | 0,067669173 |
| PPM1N | 0,003822819 | -0,76473331 | 0,009 | 0,133 | 1 | Mast | 0,588562157 | 1,69905589 | 0,067669173 |
| SLC25A17 | 0,003822819 | -0,773275415 | 0,009 | 0,133 | 1 | Mast | 0,585087615 | 1,709145732 | 0,067669173 |
| RRP12 | 0,003870045 | -1,196863809 | 0,028 | 0,2 | 1 | Mast | 0,436222531 | 2,292407957 | 0,14 |
| GLYR1 | 0,003870045 | -0,715492459 | 0,028 | 0,2 | 1 | Mast | 0,608997217 | 1,642043628 | 0,14 |
| MORF4L12 | 0,004088926 | -1,089092901 | 0,315 | 0,667 | 1 | Mast | 0,470056832 | 2,127402333 | 0,472263868 |
| CTSD2 | 0,004110339 | 1,125226934 | 0,898 | 0,8 | 1 | Mast | 2,181358564 | 0,458429905 | 1,1225 |
| RAP2B | 0,004132609 | -0,810369804 | 0,028 | 0,2 | 1 | Mast | 0,570235671 | 1,753660899 | 0,14 |
| PTCH2 | 0,004189894 | -0,629231464 | 0,009 | 0,133 | 1 | Mast | 0,646520731 | 1,546740812 | 0,067669173 |
| WNT2B | 0,004189894 | -0,596800004 | 0,009 | 0,133 | 1 | Mast | 0,661218959 | 1,51235833 | 0,067669173 |
| SLC19A2 | 0,004189894 | -0,540244583 | 0,009 | 0,133 | 1 | Mast | 0,68765432 | 1,454219033 | 0,067669173 |
| XPR1 | 0,004189894 | -0,501638585 | 0,009 | 0,133 | 1 | Mast | 0,706304119 | 1,415820711 | 0,067669173 |
| C3orf38 | 0,004189894 | -0,502040046 | 0,009 | 0,133 | 1 | Mast | 0,706107602 | 1,416214749 | 0,067669173 |
| LIN54 | 0,004189894 | -0,638370456 | 0,009 | 0,133 | 1 | Mast | 0,642438182 | 1,556569999 | 0,067669173 |
| RAB33B | 0,004189894 | -0,596519963 | 0,009 | 0,133 | 1 | Mast | 0,66134732 | 1,512064795 | 0,067669173 |
| TSPYL4 | 0,004189894 | -0,530591184 | 0,009 | 0,133 | 1 | Mast | 0,692270999 | 1,444521006 | 0,067669173 |
| GTF2IRD21 | 0,004189894 | -0,738333781 | 0,009 | 0,133 | 1 | Mast | 0,599431257 | 1,668248008 | 0,067669173 |
| IMPA1 | 0,004189894 | -0,610655428 | 0,009 | 0,133 | 1 | Mast | 0,654899108 | 1,526952758 | 0,067669173 |
| PTPDC1 | 0,004189894 | -0,780102703 | 0,009 | 0,133 | 1 | Mast | 0,582325337 | 1,717253117 | 0,067669173 |
| COQ10A1 | 0,004189894 | -0,767378694 | 0,009 | 0,133 | 1 | Mast | 0,587483935 | 1,702174205 | 0,067669173 |
| N4BP2L2-IT2 | 0,004189894 | -0,448438157 | 0,009 | 0,133 | 1 | Mast | 0,732835777 | 1,364562199 | 0,067669173 |

| Supplementary Table 4 - clusermarker | | | | | | | | | |
|---|---|---|---|---|---|---|---|---|---|
| | p_val | avg_log2FC | pct.1 | pct.2 | p_val_adj | celltype | Foldchange_UP | Foldchange_DOWN | Ratio_pct1_pct2 |
| ADAL | 0,004189894 | -0,601303035 | 0,009 | 0,133 | 1 | Mast | 0,659158338 | 1,517086172 | 0,067669173 |
| CCL23 | 0,004189894 | -0,667085683 | 0,009 | 0,133 | 1 | Mast | 0,629777586 | 1,587862163 | 0,067669173 |
| PECAM11 | 0,004189894 | -0,597534975 | 0,009 | 0,133 | 1 | Mast | 0,660882191 | 1,513128986 | 0,067669173 |
| IL27RA | 0,004189894 | -0,637110424 | 0,009 | 0,133 | 1 | Mast | 0,642999524 | 1,555211104 | 0,067669173 |
| ZNF5061 | 0,004189894 | -0,506572658 | 0,009 | 0,133 | 1 | Mast | 0,703892658 | 1,420671162 | 0,067669173 |
| SRGN3 | 0,004326511 | -0,751453253 | 0,898 | 0,933 | 1 | Mast | 0,594004904 | 1,683487784 | 0,962486602 |
| LYN | 0,004354614 | -0,941038723 | 0,194 | 0,533 | 1 | Mast | 0,520857734 | 1,919910053 | 0,363977486 |
| CERS61 | 0,004411236 | -0,726110229 | 0,028 | 0,2 | 1 | Mast | 0,604531647 | 1,654173119 | 0,14 |
| CREBBP2 | 0,004411236 | -0,672565392 | 0,028 | 0,2 | 1 | Mast | 0,627390073 | 1,593904721 | 0,14 |
| ESF11 | 0,004411236 | -0,795240373 | 0,028 | 0,2 | 1 | Mast | 0,576247154 | 1,735366487 | 0,14 |
| SGK11 | 0,004520919 | -1,330045555 | 0,481 | 0,8 | 1 | Mast | 0,397755682 | 2,514106134 | 0,60125 |
| MYB | 0,004663218 | -1,078172023 | 0,056 | 0,267 | 1 | Mast | 0,473628558 | 2,111359173 | 0,209737828 |
| AC025259.3 | 0,004706783 | -0,618368348 | 0,028 | 0,2 | 1 | Mast | 0,651407237 | 1,535137997 | 0,14 |
| RPL321 | 0,004961946 | -0,545758816 | 0,981 | 1 | 1 | Mast | 0,685031001 | 1,459787949 | 0,981 |
| CCNT11 | 0,00516105 | -0,821746053 | 0,056 | 0,267 | 1 | Mast | 0,565756809 | 1,767543907 | 0,209737828 |
| ATP6V0B4 | 0,005262112 | 1,977715202 | 0,435 | 0,067 | 1 | Mast | 3,938688171 | 0,25389164 | 6,492537313 |
| SH3BGRL34 | 0,005425547 | 0,905476684 | 0,907 | 0,733 | 1 | Mast | 1,873163308 | 0,533856282 | 1,237380628 |
| RPL23A3 | 0,005489875 | -0,736932438 | 0,648 | 0,933 | 1 | Mast | 0,600013789 | 1,666628363 | 0,694533762 |
| GNAS4 | 0,005626628 | -1,150793743 | 0,306 | 0,667 | 1 | Mast | 0,450377374 | 2,220360207 | 0,458770615 |
| CASP3 | 0,005706426 | -0,806348697 | 0,056 | 0,267 | 1 | Mast | 0,57182726 | 1,748779867 | 0,209737828 |
| JPH4 | 0,005706426 | -0,991700395 | 0,056 | 0,267 | 1 | Mast | 0,502884714 | 1,988527336 | 0,209737828 |
| WFDC23 | 0,005706426 | -0,971299132 | 0,056 | 0,267 | 1 | Mast | 0,510046564 | 1,960605305 | 0,209737828 |
| IFITM35 | 0,005862436 | 2,000965934 | 0,435 | 0,067 | 1 | Mast | 4,002679034 | 0,249832672 | 6,492537313 |
| EXOSC81 | 0,005894154 | -0,83873305 | 0,093 | 0,333 | 1 | Mast | 0,559134376 | 1,78847884 | 0,279279279 |
| ATP1B32 | 0,006055761 | -1,618622162 | 0,176 | 0,467 | 1 | Mast | 0,325646322 | 3,070816196 | 0,376873662 |
| MT-CO12 | 0,00614534 | 0,431796551 | 1 | 1 | 1 | Mast | 1,348912297 | 0,741338041 | 1 |
| CREM2 | 0,006523133 | -1,060724335 | 0,222 | 0,533 | 1 | Mast | 0,479391311 | 2,085978568 | 0,416510319 |
| LYST | 0,006574174 | -0,924383724 | 0,176 | 0,467 | 1 | Mast | 0,526905548 | 1,897873354 | 0,376873662 |
| LARP7 | 0,006622245 | -0,735165432 | 0,056 | 0,267 | 1 | Mast | 0,600749134 | 1,664588335 | 0,209737828 |
| MS4A2 | 0,006645435 | -0,668772857 | 0,759 | 1 | 1 | Mast | 0,629041517 | 1,589720191 | 0,759 |
| WDR33 | 0,006955683 | -0,750312937 | 0,056 | 0,267 | 1 | Mast | 0,594474595 | 1,68215767 | 0,209737828 |

| Supplementary Table 4 - clusermarker | | | | | | | | | |
|---|---|---|---|---|---|---|---|---|---|
| | **p_val** | **avg_log2FC** | **pct.1** | **pct.2** | **p_val_adj** | **celltype** | **Foldchange_UP** | **Foldchange_DOWN** | **Ratio_pct1_pct2** |
| HBP12 | 0,007239496 | -0,986502009 | 0,093 | 0,333 | 1 | Mast | 0,5047 | 1,981375075 | 0,279279279 |
| BAZ2A1 | 0,007304123 | -0,818059087 | 0,056 | 0,267 | 1 | Mast | 0,567204512 | 1,763032521 | 0,209737828 |
| RPL363 | 0,007569559 | -0,718271583 | 0,759 | 0,867 | 1 | Mast | 0,607825209 | 1,645209815 | 0,875432526 |
| SEPTIN21 | 0,007844323 | 1,766734139 | 0,435 | 0,067 | 1 | Mast | 3,402827784 | 0,293873232 | 6,492537313 |
| STAP1 | 0,007850665 | -0,946585531 | 0,093 | 0,333 | 1 | Mast | 0,518859009 | 1,927305843 | 0,279279279 |
| CALCOCO22 | 0,007850665 | -0,922471065 | 0,093 | 0,333 | 1 | Mast | 0,527604559 | 1,895358908 | 0,279279279 |
| TNFAIP31 | 0,008034301 | -0,974508437 | 0,639 | 0,933 | 1 | Mast | 0,508913216 | 1,964971567 | 0,68488746 |
| FMN11 | 0,008048346 | -0,780097665 | 0,056 | 0,267 | 1 | Mast | 0,582327371 | 1,71724712 | 0,209737828 |
| BIRC33 | 0,008165722 | -1,321305565 | 0,38 | 0,667 | 1 | Mast | 0,400172639 | 2,49892147 | 0,569715142 |
| SSBP13 | 0,008526521 | -1,089826107 | 0,185 | 0,467 | 1 | Mast | 0,469818 | 2,128483797 | 0,39614561 |
| MGA1 | 0,008598281 | -0,864729173 | 0,037 | 0,2 | 1 | Mast | 0,549149485 | 1,820997792 | 0,185 |
| TIMP12 | 0,008624582 | 1,42796242 | 0,676 | 0,467 | 1 | Mast | 2,690664328 | 0,371655427 | 1,447537473 |
| STMN12 | 0,008701939 | -0,869920639 | 0,343 | 0,733 | 1 | Mast | 0,547176949 | 1,827562366 | 0,467939973 |
| CD274 | 0,009097278 | -1,079227083 | 0,037 | 0,2 | 1 | Mast | 0,473282314 | 2,1129038 | 0,185 |
| FXYD11 | 0,009097278 | -1,269498632 | 0,037 | 0,2 | 1 | Mast | 0,414803901 | 2,410777712 | 0,185 |
| CPEB41 | 0,00924841 | -1,257114305 | 0,287 | 0,533 | 1 | Mast | 0,418379972 | 2,390171776 | 0,538461538 |
| DOK31 | 0,009621969 | -1,176845137 | 0,037 | 0,2 | 1 | Mast | 0,442317695 | 2,260818438 | 0,185 |
| DPYSL2 | 0,009621969 | -0,921751201 | 0,037 | 0,2 | 1 | Mast | 0,527867884 | 1,894413414 | 0,185 |
| ERGIC2 | 0,009621969 | -0,837164154 | 0,037 | 0,2 | 1 | Mast | 0,559742751 | 1,78653497 | 0,185 |
| ADRM13 | 0,009821816 | -0,86102014 | 0,139 | 0,4 | 1 | Mast | 0,550563113 | 1,816322192 | 0,3475 |
| BPTF2 | 0,00996273 | -0,90538171 | 0,185 | 0,467 | 1 | Mast | 0,533891428 | 1,873040001 | 0,39614561 |
| FOSB5 | 0,010073899 | -0,799521117 | 0,843 | 1 | 1 | Mast | 0,574539856 | 1,740523288 | 0,843 |
| HECA2 | 0,010173472 | -0,82977382 | 0,037 | 0,2 | 1 | Mast | 0,56261744 | 1,777406686 | 0,185 |
| IPO71 | 0,010173472 | -0,647254915 | 0,037 | 0,2 | 1 | Mast | 0,638494052 | 1,566185303 | 0,185 |
| CITED21 | 0,010255583 | -0,75516198 | 0,167 | 0,467 | 1 | Mast | 0,59247986 | 1,687821083 | 0,357601713 |
| ELOB4 | 0,010572583 | 1,754417908 | 0,407 | 0,067 | 1 | Mast | 3,373901624 | 0,296392756 | 6,074626866 |
| RPL414 | 0,01063623 | -0,444759717 | 1 | 1 | 1 | Mast | 0,734706673 | 1,361087407 | 1 |
| RUNX31 | 0,010752944 | -0,780143584 | 0,037 | 0,2 | 1 | Mast | 0,582308836 | 1,717301779 | 0,185 |
| PMPCB1 | 0,010752944 | -0,804690454 | 0,037 | 0,2 | 1 | Mast | 0,5724849 | 1,746770962 | 0,185 |
| RLF | 0,010924607 | -0,888435103 | 0,065 | 0,267 | 1 | Mast | 0,540199758 | 1,851167066 | 0,243445693 |
| CAVIN11 | 0,010950966 | -0,774458567 | 0,13 | 0,4 | 1 | Mast | 0,584607982 | 1,710547975 | 0,325 |

| Supplementary Table 4 - clusermarker | | | | | | | | | |
|---|---|---|---|---|---|---|---|---|---|
| | **p_val** | **avg_log2FC** | **pct.1** | **pct.2** | **p_val_adj** | **celltype** | **Foldchange_UP** | **Foldchange_DOWN** | **Ratio_pct1_pct2** |
| MYLIP3 | 0,011339953 | -0,838782934 | 0,102 | 0,333 | 1 | Mast | 0,559115043 | 1,788540682 | 0,306306306 |
| GLRX | 0,011361577 | -0,681760321 | 0,037 | 0,2 | 1 | Mast | 0,623404156 | 1,604095818 | 0,185 |
| PABPC16 | 0,011594789 | -0,911292768 | 0,593 | 0,933 | 1 | Mast | 0,531708425 | 1,880730025 | 0,635584137 |
| NR4A25 | 0,012183856 | -0,821588927 | 0,519 | 0,8 | 1 | Mast | 0,56581843 | 1,767351411 | 0,64875 |
| TTC11 | 0,012480502 | -0,663269439 | 0,065 | 0,267 | 1 | Mast | 0,631445691 | 1,583667471 | 0,243445693 |
| NUMA12 | 0,012480502 | -0,571882326 | 0,065 | 0,267 | 1 | Mast | 0,672738474 | 1,486461735 | 0,243445693 |
| SDF2L13 | 0,012569334 | 2,200652026 | 0,315 | 0 | 1 | Mast | 4,596870505 | 0,217539302 | #DIV/0! |
| TBC1D23 | 0,012671286 | -0,762327672 | 0,037 | 0,2 | 1 | Mast | 0,589544381 | 1,69622514 | 0,185 |
| LSM6 | 0,012671286 | -0,813262159 | 0,037 | 0,2 | 1 | Mast | 0,569093592 | 1,757180214 | 0,185 |
| UBA522 | 0,013313835 | -0,656988191 | 0,657 | 0,933 | 1 | Mast | 0,634200891 | 1,576787442 | 0,704180064 |
| BEST11 | 0,013374014 | -0,773259527 | 0,037 | 0,2 | 1 | Mast | 0,585094058 | 1,70912691 | 0,185 |
| RPL36A5 | 0,013494416 | -0,679634157 | 0,657 | 0,867 | 1 | Mast | 0,624323572 | 1,601733532 | 0,757785467 |
| RPS144 | 0,013532717 | -0,457797612 | 0,972 | 1 | 1 | Mast | 0,728096908 | 1,373443547 | 0,972 |
| PSMC1 | 0,01362387 | -0,690190083 | 0,065 | 0,267 | 1 | Mast | 0,619772186 | 1,613496092 | 0,243445693 |
| CHURC12 | 0,013627652 | -0,981732533 | 0,204 | 0,467 | 1 | Mast | 0,506371273 | 1,974835566 | 0,436830835 |
| CNBP4 | 0,014061537 | 1,411875105 | 0,519 | 0,2 | 1 | Mast | 2,660827723 | 0,375822903 | 2,595 |
| GSTM3 | 0,014112909 | -0,574614976 | 0,037 | 0,2 | 1 | Mast | 0,671465427 | 1,489279954 | 0,185 |
| RBM61 | 0,014229519 | -0,697438236 | 0,065 | 0,267 | 1 | Mast | 0,616666237 | 1,621622753 | 0,243445693 |
| HNRNPD1 | 0,014374211 | 1,667037248 | 0,306 | 0 | 1 | Mast | 3,175617711 | 0,314899365 | #DIV/0! |
| STK41 | 0,014381087 | -0,97516618 | 0,222 | 0,467 | 1 | Mast | 0,508681249 | 1,965867627 | 0,475374732 |
| KAT6B | 0,014708511 | -1,106012746 | 0,074 | 0,267 | 1 | Mast | 0,464576233 | 2,152499266 | 0,277153558 |
| RPL265 | 0,014739633 | -0,511163369 | 0,935 | 1 | 1 | Mast | 0,701656404 | 1,425198993 | 0,935 |
| ACLY | 0,01488658 | -0,633625776 | 0,037 | 0,2 | 1 | Mast | 0,644554486 | 1,55145922 | 0,185 |
| RPL63 | 0,015053147 | -0,455391121 | 0,926 | 1 | 1 | Mast | 0,729312425 | 1,371154481 | 0,926 |
| RPL223 | 0,01595373 | -0,626797456 | 0,778 | 0,933 | 1 | Mast | 0,647612413 | 1,544133465 | 0,833869239 |
| METRNL1 | 0,015976438 | -0,678467531 | 0,074 | 0,267 | 1 | Mast | 0,624828631 | 1,600438824 | 0,277153558 |
| SON6 | 0,01610136 | -0,695944518 | 0,333 | 0,667 | 1 | Mast | 0,617305043 | 1,619944648 | 0,499250375 |
| AKAP12 | 0,016411346 | 1,744922451 | 0,296 | 0 | 1 | Mast | 3,351768366 | 0,298349973 | #DIV/0! |
| TESC | 0,016411346 | 1,872006266 | 0,296 | 0 | 1 | Mast | 3,660412571 | 0,273193248 | #DIV/0! |
| ZFAS14 | 0,016423923 | -0,854659037 | 0,407 | 0,733 | 1 | Mast | 0,552996005 | 1,808331328 | 0,555252387 |
| B4GALT52 | 0,016645713 | -0,801835478 | 0,074 | 0,267 | 1 | Mast | 0,573618923 | 1,743317664 | 0,277153558 |

| Supplementary Table 4 - clusermarker | | | | | | | | | |
|---|---|---|---|---|---|---|---|---|---|
| | **p_val** | **avg_log2FC** | **pct.1** | **pct.2** | **p_val_adj** | **celltype** | **Foldchange_UP** | **Foldchange_DOWN** | **Ratio_pct1_pct2** |
| LXN1 | 0,016866555 | -1,008501883 | 0,111 | 0,333 | 1 | Mast | 0,497062137 | 2,011820909 | 0,333333333 |
| SYAP13 | 0,016929122 | -0,844283 | 0,25 | 0,533 | 1 | Mast | 0,556987555 | 1,795372249 | 0,469043152 |
| MYL62 | 0,017113601 | 0,799257508 | 0,843 | 0,933 | 1 | Mast | 1,740205289 | 0,574644846 | 0,903536977 |
| CCDC18 | 0,017459945 | -0,817418527 | 0,019 | 0,133 | 1 | Mast | 0,567456408 | 1,762249904 | 0,142857143 |
| ZCWPW2 | 0,017459945 | -0,742178109 | 0,019 | 0,133 | 1 | Mast | 0,597836087 | 1,672699292 | 0,142857143 |
| SLC39A81 | 0,017459945 | -0,622618119 | 0,019 | 0,133 | 1 | Mast | 0,649491199 | 1,539666746 | 0,142857143 |
| MIR3142HG | 0,017459945 | -1,122137473 | 0,019 | 0,133 | 1 | Mast | 0,459412663 | 2,176692288 | 0,142857143 |
| DMAC2L | 0,017459945 | -1,020514529 | 0,019 | 0,133 | 1 | Mast | 0,492940516 | 2,028642334 | 0,142857143 |
| TRIM69 | 0,017459945 | -1,07638781 | 0,019 | 0,133 | 1 | Mast | 0,474214667 | 2,108749623 | 0,142857143 |
| BBS4 | 0,017459945 | -0,653771775 | 0,019 | 0,133 | 1 | Mast | 0,635616387 | 1,573275989 | 0,142857143 |
| PAM16 | 0,017459945 | -1,213319046 | 0,019 | 0,133 | 1 | Mast | 0,431275287 | 2,318704619 | 0,142857143 |
| ZSCAN18 | 0,017459945 | -0,599868043 | 0,019 | 0,133 | 1 | Mast | 0,659814303 | 1,515577937 | 0,142857143 |
| WNK3 | 0,017459945 | -0,622621253 | 0,019 | 0,133 | 1 | Mast | 0,649489788 | 1,53967009 | 0,142857143 |
| LSM32 | 0,017467253 | -0,896501743 | 0,111 | 0,333 | 1 | Mast | 0,53718773 | 1,861546614 | 0,333333333 |
| HCLS11 | 0,01799128 | -0,98222865 | 0,148 | 0,4 | 1 | Mast | 0,506197171 | 1,975514794 | 0,37 |
| NSMCE1-DT | 0,018058481 | -1,070204404 | 0,074 | 0,267 | 1 | Mast | 0,476251518 | 2,09973084 | 0,277153558 |
| ALDH1A13 | 0,01840698 | -1,526075005 | 0,12 | 0,333 | 1 | Mast | 0,347220732 | 2,880012363 | 0,36036036 |
| ACTG12 | 0,018617399 | 1,004077804 | 0,759 | 0,533 | 1 | Mast | 2,005661034 | 0,498588736 | 1,424015009 |
| AMD14 | 0,018662885 | -1,290572871 | 0,157 | 0,4 | 1 | Mast | 0,408788674 | 2,44625173 | 0,3925 |
| ATAD3B | 0,018686629 | -0,72370898 | 0,019 | 0,133 | 1 | Mast | 0,605538679 | 1,651422172 | 0,142857143 |
| SKI | 0,018686629 | -0,93900365 | 0,019 | 0,133 | 1 | Mast | 0,521592977 | 1,917203728 | 0,142857143 |
| UBXN11 | 0,018686629 | -0,717120573 | 0,019 | 0,133 | 1 | Mast | 0,608310338 | 1,643897757 | 0,142857143 |
| BTBD19 | 0,018686629 | -0,799033622 | 0,019 | 0,133 | 1 | Mast | 0,57473403 | 1,739935254 | 0,142857143 |
| ARV1 | 0,018686629 | -0,822284695 | 0,019 | 0,133 | 1 | Mast | 0,565545618 | 1,768203957 | 0,142857143 |
| SYCP2L | 0,018686629 | -0,886853297 | 0,019 | 0,133 | 1 | Mast | 0,540792371 | 1,849138513 | 0,142857143 |
| SP4 | 0,018686629 | -0,726130605 | 0,019 | 0,133 | 1 | Mast | 0,604523109 | 1,654196482 | 0,142857143 |
| AP005019.1 | 0,018686629 | -0,768284115 | 0,019 | 0,133 | 1 | Mast | 0,587115352 | 1,703242808 | 0,142857143 |
| CSNK2A21 | 0,018686629 | -0,564507767 | 0,019 | 0,133 | 1 | Mast | 0,676186085 | 1,478882843 | 0,142857143 |
| CD79B | 0,018686629 | -0,806388524 | 0,019 | 0,133 | 1 | Mast | 0,571811474 | 1,748828145 | 0,142857143 |
| GRWD11 | 0,018686629 | -0,836811071 | 0,019 | 0,133 | 1 | Mast | 0,559879759 | 1,78609779 | 0,142857143 |
| PIGA | 0,018686629 | -0,802762502 | 0,019 | 0,133 | 1 | Mast | 0,573250454 | 1,744438217 | 0,142857143 |

| Supplementary Table 4 - clusermarker | | | | | | | | | |
|---|---|---|---|---|---|---|---|---|---|
| | **p_val** | **avg_log2FC** | **pct.1** | **pct.2** | **p_val_adj** | **celltype** | **Foldchange_UP** | **Foldchange_DOWN** | **Ratio_pct1_pct2** |
| CTSW1 | 0,018707018 | 1,721672394 | 0,287 | 0 | 1 | Mast | 3,298185161 | 0,303197047 | #DIV/0! |
| ELL22 | 0,018924617 | -0,852516253 | 0,481 | 0,733 | 1 | Mast | 0,553817961 | 1,805647471 | 0,656207367 |
| SEC625 | 0,018971736 | -0,638550513 | 0,324 | 0,667 | 1 | Mast | 0,642358007 | 1,55676428 | 0,485757121 |
| PPP1R9B1 | 0,01920311 | -0,937168433 | 0,046 | 0,2 | 1 | Mast | 0,522256905 | 1,91476645 | 0,23 |
| HMGN31 | 0,019739935 | -0,913660324 | 0,194 | 0,467 | 1 | Mast | 0,530836573 | 1,883818959 | 0,415417559 |
| CTNNBL1 | 0,019857264 | -0,773387931 | 0,241 | 0,533 | 1 | Mast | 0,585041986 | 1,709279034 | 0,452157598 |
| TPRG1L1 | 0,019988057 | -0,779922939 | 0,019 | 0,133 | 1 | Mast | 0,582397901 | 1,717039155 | 0,142857143 |
| LINC02801 | 0,019988057 | -0,643099185 | 0,019 | 0,133 | 1 | Mast | 0,640335906 | 1,561680347 | 0,142857143 |
| SORT1 | 0,019988057 | -0,596385952 | 0,019 | 0,133 | 1 | Mast | 0,661408755 | 1,511924346 | 0,142857143 |
| NRAS | 0,019988057 | -0,648047907 | 0,019 | 0,133 | 1 | Mast | 0,638143193 | 1,56704641 | 0,142857143 |
| ATP1A1-AS1 | 0,019988057 | -0,527482745 | 0,019 | 0,133 | 1 | Mast | 0,693764178 | 1,441411984 | 0,142857143 |
| STARD7-AS1 | 0,019988057 | -0,561899125 | 0,019 | 0,133 | 1 | Mast | 0,677409853 | 1,476211183 | 0,142857143 |
| CNNM4 | 0,019988057 | -0,539811259 | 0,019 | 0,133 | 1 | Mast | 0,687860893 | 1,453782314 | 0,142857143 |
| TOMM70 | 0,019988057 | -0,577126855 | 0,019 | 0,133 | 1 | Mast | 0,670297355 | 1,4918752 | 0,142857143 |
| PIGX | 0,019988057 | -0,54285496 | 0,019 | 0,133 | 1 | Mast | 0,68641122 | 1,456852643 | 0,142857143 |
| DCUN1D41 | 0,019988057 | -0,882916525 | 0,019 | 0,133 | 1 | Mast | 0,542270079 | 1,844099532 | 0,142857143 |
| MCUB2 | 0,019988057 | -0,64712655 | 0,019 | 0,133 | 1 | Mast | 0,638550865 | 1,566045957 | 0,142857143 |
| SSBP2 | 0,019988057 | -0,644951939 | 0,019 | 0,133 | 1 | Mast | 0,639514095 | 1,563687193 | 0,142857143 |
| TMEM64 | 0,019988057 | -0,566229875 | 0,019 | 0,133 | 1 | Mast | 0,67537942 | 1,480649202 | 0,142857143 |
| JAK2 | 0,019988057 | -0,524878156 | 0,019 | 0,133 | 1 | Mast | 0,695017806 | 1,438812059 | 0,142857143 |
| NCBP1 | 0,019988057 | -0,500846674 | 0,019 | 0,133 | 1 | Mast | 0,706691923 | 1,415043765 | 0,142857143 |
| MSMB6 | 0,019988057 | -1,012794362 | 0,019 | 0,133 | 1 | Mast | 0,495585416 | 2,017815633 | 0,142857143 |
| FBXW4 | 0,019988057 | -0,444223207 | 0,019 | 0,133 | 1 | Mast | 0,734979947 | 1,360581339 | 0,142857143 |
| AP5M1 | 0,019988057 | -0,520485736 | 0,019 | 0,133 | 1 | Mast | 0,697137077 | 1,434438122 | 0,142857143 |
| AQR1 | 0,019988057 | -0,647589636 | 0,019 | 0,133 | 1 | Mast | 0,638345931 | 1,566548717 | 0,142857143 |
| RFX1 | 0,019988057 | -0,683456566 | 0,019 | 0,133 | 1 | Mast | 0,622671621 | 1,605982939 | 0,142857143 |
| CD841 | 0,020019912 | -0,924070315 | 0,222 | 0,467 | 1 | Mast | 0,527020025 | 1,897461107 | 0,475374732 |
| ATF31 | 0,020061587 | -0,876112698 | 0,111 | 0,333 | 1 | Mast | 0,544833494 | 1,835423136 | 0,333333333 |
| PUM12 | 0,020153227 | -0,805635231 | 0,046 | 0,2 | 1 | Mast | 0,57211012 | 1,747915245 | 0,23 |
| APPBP2 | 0,020153227 | -0,775955083 | 0,046 | 0,2 | 1 | Mast | 0,58400188 | 1,712323256 | 0,23 |
| H2AZ12 | 0,020493158 | -0,826060101 | 0,5 | 0,8 | 1 | Mast | 0,56406757 | 1,772837251 | 0,625 |

| Supplementary Table 4 - clusermarker | | | | | | | | | |
|---|---|---|---|---|---|---|---|---|---|
| | **p_val** | **avg_log2FC** | **pct.1** | **pct.2** | **p_val_adj** | **celltype** | **Foldchange_UP** | **Foldchange_DOWN** | **Ratio_pct1_pct2** |
| NSA21 | 0,021001227 | -0,748435275 | 0,148 | 0,4 | 1 | Mast | 0,595248805 | 1,679969773 | 0,37 |
| TMED4 | 0,021144109 | -0,832899533 | 0,046 | 0,2 | 1 | Mast | 0,561399805 | 1,781261753 | 0,23 |
| DDX10 | 0,021144109 | -0,860103707 | 0,046 | 0,2 | 1 | Mast | 0,550912955 | 1,815168787 | 0,23 |
| UCKL1 | 0,021144109 | -0,890883989 | 0,046 | 0,2 | 1 | Mast | 0,539283579 | 1,854311976 | 0,23 |
| OFD12 | 0,021144109 | -0,74500622 | 0,046 | 0,2 | 1 | Mast | 0,596665299 | 1,675981495 | 0,23 |
| HCST2 | 0,02129157 | 1,724808294 | 0,278 | 0 | 1 | Mast | 3,305362025 | 0,302538721 | #DIV/0! |
| NUP133 | 0,02136791 | -0,449849517 | 0,019 | 0,133 | 1 | Mast | 0,732119209 | 1,365897777 | 0,142857143 |
| CALCOCO11 | 0,02136791 | -0,514838819 | 0,019 | 0,133 | 1 | Mast | 0,69987112 | 1,428834498 | 0,142857143 |
| DCN1 | 0,02136791 | -0,332063481 | 0,019 | 0,133 | 1 | Mast | 0,794399444 | 1,258812563 | 0,142857143 |
| ALDH21 | 0,02136791 | -0,713455159 | 0,019 | 0,133 | 1 | Mast | 0,609857819 | 1,639726455 | 0,142857143 |
| SNHG30 | 0,02136791 | -0,522204773 | 0,019 | 0,133 | 1 | Mast | 0,696306901 | 1,43614834 | 0,142857143 |
| STAT5B | 0,02136791 | -0,74288062 | 0,019 | 0,133 | 1 | Mast | 0,597545046 | 1,673514 | 0,142857143 |
| FCER1A1 | 0,022318312 | -1,001134299 | 0,398 | 0,667 | 1 | Mast | 0,499607036 | 2,001573091 | 0,596701649 |
| NDUFS52 | 0,02262356 | 1,550170208 | 0,361 | 0,067 | 1 | Mast | 2,928516876 | 0,341469775 | 5,388059701 |
| RPF1 | 0,02282999 | -0,565399844 | 0,019 | 0,133 | 1 | Mast | 0,675768101 | 1,47979758 | 0,142857143 |
| PER21 | 0,02282999 | -0,546626387 | 0,019 | 0,133 | 1 | Mast | 0,684619179 | 1,460666063 | 0,142857143 |
| NAALADL21 | 0,02282999 | -0,384194931 | 0,019 | 0,133 | 1 | Mast | 0,766206446 | 1,30513128 | 0,142857143 |
| RUFY12 | 0,02282999 | -0,456032918 | 0,019 | 0,133 | 1 | Mast | 0,728988056 | 1,371764588 | 0,142857143 |
| VCPIP1 | 0,02282999 | -0,497786368 | 0,019 | 0,133 | 1 | Mast | 0,70819258 | 1,412045295 | 0,142857143 |
| BMS1 | 0,02282999 | -0,585867657 | 0,019 | 0,133 | 1 | Mast | 0,666248527 | 1,500941405 | 0,142857143 |
| NRIP3 | 0,02282999 | -0,492114682 | 0,019 | 0,133 | 1 | Mast | 0,710982186 | 1,406505001 | 0,142857143 |
| KIAA0586 | 0,02282999 | -0,466775635 | 0,019 | 0,133 | 1 | Mast | 0,723579964 | 1,382017262 | 0,142857143 |
| GABPB1-IT1 | 0,02282999 | -0,549010906 | 0,019 | 0,133 | 1 | Mast | 0,683488559 | 1,463082281 | 0,142857143 |
| NOL11 | 0,02282999 | -0,498125023 | 0,019 | 0,133 | 1 | Mast | 0,708026359 | 1,412376795 | 0,142857143 |
| THOC1 | 0,02282999 | -0,53105395 | 0,019 | 0,133 | 1 | Mast | 0,692048978 | 1,444984433 | 0,142857143 |
| CCDC97 | 0,02282999 | -0,394222907 | 0,019 | 0,133 | 1 | Mast | 0,760899117 | 1,31423467 | 0,142857143 |
| PIGP | 0,02282999 | -0,541586718 | 0,019 | 0,133 | 1 | Mast | 0,687014894 | 1,455572518 | 0,142857143 |
| CHORDC12 | 0,022967035 | -1,105820191 | 0,083 | 0,267 | 1 | Mast | 0,464638243 | 2,152211993 | 0,310861423 |
| GCSAML | 0,023011239 | -0,782112603 | 0,148 | 0,4 | 1 | Mast | 0,581514632 | 1,719647186 | 0,37 |
| ARID5B4 | 0,023253848 | -0,639820089 | 0,046 | 0,2 | 1 | Mast | 0,641792979 | 1,55813484 | 0,23 |
| LCMT12 | 0,023253848 | -0,867463708 | 0,046 | 0,2 | 1 | Mast | 0,548109594 | 1,824452648 | 0,23 |

| Supplementary Table 4 - clusermarker | | | | | | | | | |
|---|---|---|---|---|---|---|---|---|---|
| | **p_val** | **avg_log2FC** | **pct.1** | **pct.2** | **p_val_adj** | **celltype** | **Foldchange_UP** | **Foldchange_DOWN** | **Ratio_pct1_pct2** |
| CYBA6 | 0,023533337 | 1,199710512 | 0,62 | 0,4 | 1 | Mast | 2,296935765 | 0,435362632 | 1,55 |
| DRAP15 | 0,023616207 | 1,396393521 | 0,435 | 0,133 | 1 | Mast | 2,632426994 | 0,379877582 | 3,270676692 |
| ACSL32 | 0,023862763 | -0,589756958 | 0,074 | 0,267 | 1 | Mast | 0,664454834 | 1,504993189 | 0,277153558 |
| SVIP | 0,023862763 | -0,431227015 | 0,074 | 0,267 | 1 | Mast | 0,741630758 | 1,348379889 | 0,277153558 |
| USP15 | 0,024023783 | -1,063040118 | 0,12 | 0,333 | 1 | Mast | 0,478622419 | 2,089329625 | 0,36036036 |
| PSMB63 | 0,024196411 | 1,429355506 | 0,269 | 0 | 1 | Mast | 2,693263726 | 0,371296725 | #DIV/0! |
| SMDT16 | 0,024196411 | 1,706959486 | 0,269 | 0 | 1 | Mast | 3,264720503 | 0,306304935 | #DIV/0! |
| S100A43 | 0,024246442 | -0,465579052 | 0,935 | 1 | 1 | Mast | 0,724180356 | 1,38087148 | 0,935 |
| SLC25A33 | 0,024375623 | -0,680178346 | 0,046 | 0,2 | 1 | Mast | 0,62408812 | 1,602337824 | 0,23 |
| STX5 | 0,024375623 | -0,680623726 | 0,046 | 0,2 | 1 | Mast | 0,623895485 | 1,602832564 | 0,23 |
| POLE3 | 0,024582423 | -0,739490303 | 0,111 | 0,333 | 1 | Mast | 0,598950921 | 1,669585878 | 0,333333333 |
| SAT13 | 0,024650735 | -0,683873007 | 0,583 | 0,933 | 1 | Mast | 0,62249191 | 1,60644658 | 0,624866024 |
| EIF3L2 | 0,025483632 | -0,771602302 | 0,269 | 0,533 | 1 | Mast | 0,585766543 | 1,707164761 | 0,504690432 |
| RBM5 | 0,025544001 | -0,845397214 | 0,046 | 0,2 | 1 | Mast | 0,556557552 | 1,796759375 | 0,23 |
| CARD8 | 0,025544001 | -0,676703235 | 0,046 | 0,2 | 1 | Mast | 0,625593213 | 1,598482816 | 0,23 |
| PIK3R6 | 0,025711396 | -0,710707538 | 0,083 | 0,267 | 1 | Mast | 0,611020404 | 1,636606558 | 0,310861423 |
| TANK3 | 0,025712234 | -1,131522128 | 0,083 | 0,267 | 1 | Mast | 0,456433906 | 2,19089771 | 0,310861423 |
| MT-ATP65 | 0,026189446 | -0,375204069 | 0,88 | 1 | 1 | Mast | 0,770996347 | 1,297023006 | 0,88 |
| COBL1 | 0,026760518 | -0,534546355 | 0,046 | 0,2 | 1 | Mast | 0,690375726 | 1,448486617 | 0,23 |
| DYNLL2 | 0,026760518 | -0,653397116 | 0,046 | 0,2 | 1 | Mast | 0,635781474 | 1,572867472 | 0,23 |
| SEC61B5 | 0,026955498 | 1,596494691 | 0,463 | 0,2 | 1 | Mast | 3,024076617 | 0,330679453 | 2,315 |
| NASP2 | 0,027283969 | -1,103251241 | 0,139 | 0,333 | 1 | Mast | 0,465466343 | 2,148383045 | 0,417417417 |
| PRKAR1A3 | 0,027457614 | 1,442999023 | 0,259 | 0 | 1 | Mast | 2,718854643 | 0,367801935 | #DIV/0! |
| ARIH11 | 0,027568218 | -1,164356844 | 0,185 | 0,4 | 1 | Mast | 0,446163116 | 2,241332743 | 0,4625 |
| PKM2 | 0,027693381 | 1,476893844 | 0,417 | 0,133 | 1 | Mast | 2,783487961 | 0,359261478 | 3,135338346 |
| RPL85 | 0,027884614 | 0,456400523 | 0,981 | 1 | 1 | Mast | 1,372114164 | 0,72880233 | 0,981 |
| ZNF433-AS1 | 0,028026745 | -0,553343221 | 0,046 | 0,2 | 1 | Mast | 0,681439168 | 1,467482421 | 0,23 |
| AKAP132 | 0,028197781 | -0,768394966 | 0,546 | 0,8 | 1 | Mast | 0,587070242 | 1,703373683 | 0,6825 |
| C12orf574 | 0,02833288 | -0,53354421 | 0,565 | 0,8 | 1 | Mast | 0,690855451 | 1,447480799 | 0,70625 |
| ALS2 | 0,028736774 | -0,9522204 | 0,083 | 0,267 | 1 | Mast | 0,516836405 | 1,934848223 | 0,310861423 |
| TCF252 | 0,029113337 | -0,531796748 | 0,241 | 0,533 | 1 | Mast | 0,691692756 | 1,445728601 | 0,452157598 |

| Supplementary Table 4 - clusermarker | | | | | | | | | |
|---|---|---|---|---|---|---|---|---|---|
| | **p_val** | **avg_log2FC** | **pct.1** | **pct.2** | **p_val_adj** | **celltype** | **Foldchange_UP** | **Foldchange_DOWN** | **Ratio_pct1_pct2** |
| TXLNG | 0,029344282 | -0,560351582 | 0,046 | 0,2 | 1 | Mast | 0,678136883 | 1,474628538 | 0,23 |
| PBDC11 | 0,029344282 | -0,531354916 | 0,046 | 0,2 | 1 | Mast | 0,691904622 | 1,445285907 | 0,23 |
| RPS192 | 0,029857015 | 0,927690733 | 0,861 | 0,733 | 1 | Mast | 1,902228733 | 0,525699135 | 1,174624829 |
| CD831 | 0,030295973 | -0,883063038 | 0,407 | 0,733 | 1 | Mast | 0,542215012 | 1,84428682 | 0,555252387 |
| ALOX5AP2 | 0,030315748 | -0,532540069 | 0,639 | 0,867 | 1 | Mast | 0,691336466 | 1,446473676 | 0,737024221 |
| ERN11 | 0,030919215 | -0,784074251 | 0,083 | 0,267 | 1 | Mast | 0,580724477 | 1,721986999 | 0,310861423 |
| PPHLN1 | 0,031070444 | -0,629149434 | 0,12 | 0,333 | 1 | Mast | 0,646557492 | 1,546652868 | 0,36036036 |
| NUDT143 | 0,03111411 | 1,666012556 | 0,25 | 0 | 1 | Mast | 3,173362991 | 0,315123105 | #DIV/0! |
| CLNK | 0,031114627 | 1,587032423 | 0,25 | 0 | 1 | Mast | 3,004307371 | 0,332855423 | #DIV/0! |
| KLF22 | 0,031114627 | 1,517414567 | 0,25 | 0 | 1 | Mast | 2,862775557 | 0,349311352 | #DIV/0! |
| ATRAID3 | 0,032065196 | -0,725272708 | 0,12 | 0,333 | 1 | Mast | 0,604882695 | 1,653213109 | 0,36036036 |
| ST8SIA6 | 0,032561895 | -1,228140761 | 0,093 | 0,267 | 1 | Mast | 0,426867207 | 2,342648919 | 0,348314607 |
| PSMD15 | 0,032989222 | -0,737136421 | 0,13 | 0,333 | 1 | Mast | 0,599928959 | 1,666864026 | 0,39039039 |
| PURA2 | 0,033068105 | -1,031238209 | 0,056 | 0,2 | 1 | Mast | 0,48929003 | 2,043777593 | 0,28 |
| PLAT | 0,033176654 | 1,383350709 | 0,352 | 0,067 | 1 | Mast | 2,608735559 | 0,383327469 | 5,253731343 |
| SEM1 | 0,033372086 | -1,091163983 | 0,185 | 0,4 | 1 | Mast | 0,469382519 | 2,13045855 | 0,4625 |
| SLIRP2 | 0,033718004 | -1,054903573 | 0,093 | 0,267 | 1 | Mast | 0,481329391 | 2,077579341 | 0,348314607 |
| EGR32 | 0,034034144 | 1,557865127 | 0,333 | 0,067 | 1 | Mast | 2,944178469 | 0,339653323 | 4,970149254 |
| MYL12B3 | 0,034096244 | 1,34750528 | 0,528 | 0,267 | 1 | Mast | 2,544717102 | 0,39297099 | 1,97752809 |
| RHOA2 | 0,034371849 | 1,152266312 | 0,574 | 0,333 | 1 | Mast | 2,222627702 | 0,449917905 | 1,723723724 |
| ABHD5 | 0,034908073 | -0,899483301 | 0,093 | 0,267 | 1 | Mast | 0,536078693 | 1,865397773 | 0,348314607 |
| DUSP10 | 0,035208193 | -0,887332784 | 0,167 | 0,4 | 1 | Mast | 0,540612665 | 1,849753186 | 0,4175 |
| ATP6V1F2 | 0,035694772 | -0,559148401 | 0,481 | 0,733 | 1 | Mast | 0,678702673 | 1,473399237 | 0,656207367 |
| PIM21 | 0,035991947 | -1,21051144 | 0,056 | 0,2 | 1 | Mast | 0,432115402 | 2,314196612 | 0,28 |
| H3-3A2 | 0,03682074 | 0,820447166 | 0,722 | 0,8 | 1 | Mast | 1,765953269 | 0,5662664 | 0,9025 |
| CD442 | 0,038022628 | -0,692719818 | 0,843 | 0,933 | 1 | Mast | 0,618686381 | 1,616327805 | 0,903536977 |
| PARP41 | 0,038339451 | -0,65197825 | 0,083 | 0,267 | 1 | Mast | 0,636407062 | 1,571321344 | 0,310861423 |
| KMT2C2 | 0,038699879 | -0,634777384 | 0,093 | 0,267 | 1 | Mast | 0,644040186 | 1,552698141 | 0,348314607 |
| RPS204 | 0,039016405 | -0,868125285 | 0,333 | 0,6 | 1 | Mast | 0,547858304 | 1,825289481 | 0,555 |
| ITM2B5 | 0,03911208 | 0,768140123 | 0,87 | 0,733 | 1 | Mast | 1,70307282 | 0,587173953 | 1,186903138 |
| KRCC11 | 0,03971287 | -0,503712294 | 0,083 | 0,267 | 1 | Mast | 0,705289617 | 1,417857255 | 0,310861423 |

| Supplementary Table 4 - clusermarker | | | | | | | | | |
|---|---|---|---|---|---|---|---|---|---|
| | **p_val** | **avg_log2FC** | **pct.1** | **pct.2** | **p_val_adj** | **celltype** | **Foldchange_UP** | **Foldchange_DOWN** | **Ratio_pct1_pct2** |
| C1orf162 | 0,039794285 | 1,561080705 | 0,231 | 0 | 1 | Mast | 2,950747976 | 0,338897123 | #DIV/0! |
| CDK2AP22 | 0,039794931 | 1,636030394 | 0,231 | 0 | 1 | Mast | 3,108094555 | 0,321740533 | #DIV/0! |
| RTL8C | 0,040038853 | -0,676340857 | 0,093 | 0,267 | 1 | Mast | 0,62575037 | 1,598081357 | 0,348314607 |
| CAP11 | 0,040087361 | 1,480706819 | 0,37 | 0,133 | 1 | Mast | 2,79085432 | 0,358313221 | 2,781954887 |
| MEPCE | 0,040789554 | -0,834221624 | 0,056 | 0,2 | 1 | Mast | 0,560885572 | 1,782894855 | 0,28 |
| RP21 | 0,040789554 | -0,849771182 | 0,056 | 0,2 | 1 | Mast | 0,554872734 | 1,802215063 | 0,28 |
| PRPS11 | 0,040789554 | -0,633178758 | 0,056 | 0,2 | 1 | Mast | 0,644754232 | 1,550978576 | 0,28 |
| KHDRBS12 | 0,040993732 | 1,487555372 | 0,315 | 0,067 | 1 | Mast | 2,804134154 | 0,356616319 | 4,701492537 |
| XIST2 | 0,041476058 | -0,654936004 | 0,25 | 0,533 | 1 | Mast | 0,635103663 | 1,574546107 | 0,469043152 |
| RPLP14 | 0,042472123 | 0,428189043 | 0,991 | 1 | 1 | Mast | 1,34554351 | 0,743194102 | 0,991 |
| SPTBN11 | 0,042585244 | -0,622372804 | 0,083 | 0,267 | 1 | Mast | 0,649601647 | 1,539404964 | 0,310861423 |
| TMEM2201 | 0,042585244 | -0,573935162 | 0,083 | 0,267 | 1 | Mast | 0,671781903 | 1,488578354 | 0,310861423 |
| TMEM87A2 | 0,042834947 | -0,644078741 | 0,093 | 0,267 | 1 | Mast | 0,639901281 | 1,562741051 | 0,348314607 |
| KMT2E2 | 0,043872166 | 1,517164599 | 0,38 | 0,133 | 1 | Mast | 2,862279582 | 0,34937188 | 2,857142857 |
| SNHG84 | 0,043960364 | -0,943312295 | 0,259 | 0,467 | 1 | Mast | 0,52003755 | 1,922938064 | 0,554603854 |
| NT5C3A | 0,0442805 | -0,769765157 | 0,056 | 0,2 | 1 | Mast | 0,58651294 | 1,704992221 | 0,28 |
| TM9SF32 | 0,0442805 | -0,59340546 | 0,056 | 0,2 | 1 | Mast | 0,662776585 | 1,508804056 | 0,28 |
| ARL11 | 0,0442805 | -0,865576259 | 0,056 | 0,2 | 1 | Mast | 0,548827144 | 1,822067315 | 0,28 |
| PFDN51 | 0,044402935 | -0,451677604 | 0,769 | 1 | 1 | Mast | 0,731192104 | 1,367629649 | 0,769 |
| TGFB11 | 0,044453689 | 1,371764895 | 0,444 | 0,2 | 1 | Mast | 2,58786955 | 0,386418241 | 2,22 |
| UBC5 | 0,044498196 | -0,746352469 | 0,833 | 0,933 | 1 | Mast | 0,596108781 | 1,677546165 | 0,892818864 |
| BRI34 | 0,044919642 | 1,300104199 | 0,222 | 0 | 1 | Mast | 2,462466673 | 0,406096867 | #DIV/0! |
| NDUFS84 | 0,044919642 | 1,387877515 | 0,222 | 0 | 1 | Mast | 2,616933956 | 0,382126571 | #DIV/0! |
| PRKCB | 0,044919642 | 1,198782831 | 0,222 | 0 | 1 | Mast | 2,295459265 | 0,435642669 | #DIV/0! |
| TRAPPC15 | 0,044919642 | 1,357922082 | 0,222 | 0 | 1 | Mast | 2,563157413 | 0,390143811 | #DIV/0! |
| PRDX13 | 0,045015352 | -0,850177394 | 0,667 | 0,867 | 1 | Mast | 0,554716524 | 1,802722575 | 0,769319493 |
| CD822 | 0,045161861 | -0,726038419 | 0,38 | 0,6 | 1 | Mast | 0,604561738 | 1,654090785 | 0,633333333 |
| CD815 | 0,045575124 | -0,777232506 | 0,306 | 0,533 | 1 | Mast | 0,583485008 | 1,713840091 | 0,574108818 |
| GRK2 | 0,045794367 | -0,580787753 | 0,093 | 0,267 | 1 | Mast | 0,668598604 | 1,495665702 | 0,348314607 |
| VPS37B3 | 0,046037042 | -1,245645993 | 0,222 | 0,4 | 1 | Mast | 0,421719023 | 2,371247074 | 0,555 |
| PRAG1 | 0,046118483 | -0,643934539 | 0,056 | 0,2 | 1 | Mast | 0,639965244 | 1,562584858 | 0,28 |

| Supplementary Table 4 - clusermarker | | | | | | | | | |
|---|---|---|---|---|---|---|---|---|---|
| | **p_val** | **avg_log2FC** | **pct.1** | **pct.2** | **p_val_adj** | **celltype** | **Foldchange_UP** | **Foldchange_DOWN** | **Ratio_pct1_pct2** |
| TBPL1 | 0,046291246 | -0,531115478 | 0,12 | 0,333 | 1 | Mast | 0,692019464 | 1,445046059 | 0,36036036 |
| NFE2L24 | 0,046803753 | -0,895376081 | 0,324 | 0,533 | 1 | Mast | 0,537607034 | 1,86009471 | 0,607879925 |
| LSM121 | 0,046822315 | -0,621523096 | 0,028 | 0,133 | 1 | Mast | 0,649984358 | 1,538498562 | 0,210526316 |
| KLF11 | 0,046825343 | -0,610473579 | 0,028 | 0,133 | 1 | Mast | 0,654981662 | 1,526760301 | 0,210526316 |
| HMGXB3 | 0,046825343 | -0,710704039 | 0,028 | 0,133 | 1 | Mast | 0,611021886 | 1,636602589 | 0,210526316 |
| RRM2B | 0,046825343 | -0,548827744 | 0,028 | 0,133 | 1 | Mast | 0,683575339 | 1,462896542 | 0,210526316 |
| CAAP1 | 0,046825343 | -0,729572686 | 0,028 | 0,133 | 1 | Mast | 0,603082515 | 1,658147889 | 0,210526316 |
| COMMD9 | 0,046825343 | -0,854714522 | 0,028 | 0,133 | 1 | Mast | 0,552974737 | 1,808400877 | 0,210526316 |
| SLC2A32 | 0,046895571 | -1,578390007 | 0,546 | 0,667 | 1 | Mast | 0,334855366 | 2,986363967 | 0,818590705 |
| CHMP3 | 0,047263798 | -1,148219428 | 0,204 | 0,4 | 1 | Mast | 0,451181735 | 2,216401776 | 0,51 |
| BATF1 | 0,047319691 | -0,556700567 | 0,333 | 0,6 | 1 | Mast | 0,679855211 | 1,470901427 | 0,555 |
| MRPL16 | 0,047336359 | -0,863625005 | 0,093 | 0,267 | 1 | Mast | 0,549569938 | 1,819604623 | 0,348314607 |
| HP1BP33 | 0,047360525 | -0,358915247 | 0,167 | 0,4 | 1 | Mast | 0,779750649 | 1,28246126 | 0,4175 |
| TUBA1B3 | 0,047638363 | -0,440105126 | 0,556 | 0,867 | 1 | Mast | 0,737080897 | 1,356703184 | 0,641291811 |
| CD743 | 0,048165604 | 1,945423375 | 0,324 | 0,067 | 1 | Mast | 3,851507875 | 0,259638571 | 4,835820896 |
| AL157895.1 | 0,048641026 | -0,616342434 | 0,352 | 0,6 | 1 | Mast | 0,652322622 | 1,532983781 | 0,586666667 |
| WWC2 | 0,048924195 | -0,686856775 | 0,093 | 0,267 | 1 | Mast | 0,621205809 | 1,609772455 | 0,348314607 |
| SSR44 | 0,049020718 | 1,14780931 | 0,63 | 0,4 | 1 | Mast | 2,215771804 | 0,451310012 | 1,575 |
| NSUN4 | 0,049381023 | -0,743819292 | 0,028 | 0,133 | 1 | Mast | 0,597156387 | 1,674603206 | 0,210526316 |
| TEC | 0,049381023 | -0,795778331 | 0,028 | 0,133 | 1 | Mast | 0,576032321 | 1,736013698 | 0,210526316 |
| MTREX1 | 0,049381023 | -0,650626702 | 0,028 | 0,133 | 1 | Mast | 0,637003541 | 1,569849986 | 0,210526316 |
| NOP16 | 0,049381023 | -0,896145404 | 0,028 | 0,133 | 1 | Mast | 0,537320429 | 1,861086878 | 0,210526316 |
| TRMT111 | 0,049381023 | -0,713367946 | 0,028 | 0,133 | 1 | Mast | 0,609894687 | 1,639627333 | 0,210526316 |
| ECHDC12 | 0,049381023 | -0,599128318 | 0,028 | 0,133 | 1 | Mast | 0,660152702 | 1,514801041 | 0,210526316 |
| TRAPPC9 | 0,049381023 | -0,722317154 | 0,028 | 0,133 | 1 | Mast | 0,606123148 | 1,649829747 | 0,210526316 |
| CHCHD1 | 0,049381023 | -0,944062171 | 0,028 | 0,133 | 1 | Mast | 0,519767319 | 1,923937817 | 0,210526316 |
| TFCP2 | 0,049381023 | -0,654435406 | 0,028 | 0,133 | 1 | Mast | 0,635324075 | 1,573999853 | 0,210526316 |
| MTMR6 | 0,049381023 | -0,689183244 | 0,028 | 0,133 | 1 | Mast | 0,620204869 | 1,612370445 | 0,210526316 |
| LSR4 | 0,049381023 | -0,719288907 | 0,028 | 0,133 | 1 | Mast | 0,607396749 | 1,646370352 | 0,210526316 |
| COX7C3 | 0,049801245 | -1,043628894 | 0,472 | 0,667 | 1 | Mast | 0,485105724 | 2,061406311 | 0,707646177 |
| CNPY31 | 0,049911991 | -0,606485183 | 0,167 | 0,4 | 1 | Mast | 0,656794894 | 1,522545331 | 0,4175 |

| Supplementary Table 4 - clusermarker | | | | | | | | | |
|---|---|---|---|---|---|---|---|---|---|
| | **p_val** | **avg_log2FC** | **pct.1** | **pct.2** | **p_val_adj** | **celltype** | **Foldchange_UP** | **Foldchange_DOWN** | **Ratio_pct1_pct2** |
| AP3S12 | 0,050644172 | 1,357136442 | 0,213 | 0 | 1 | Mast | 2,561761987 | 0,390356327 | #DIV/0! |
| AZIN11 | 0,050644172 | 1,301693809 | 0,213 | 0 | 1 | Mast | 2,465181396 | 0,405649662 | #DIV/0! |
| PSMD82 | 0,050644172 | 1,503626867 | 0,213 | 0 | 1 | Mast | 2,835546602 | 0,35266569 | #DIV/0! |
| APOC11 | 0,050644172 | 1,555301999 | 0,213 | 0 | 1 | Mast | 2,938952412 | 0,340257296 | #DIV/0! |
| RPS232 | 0,050985794 | -0,452362617 | 0,963 | 1 | 1 | Mast | 0,730845005 | 1,368279174 | 0,963 |
| ANKRD12 | 0,051230341 | -0,517056734 | 0,167 | 0,4 | 1 | Mast | 0,698796005 | 1,431032794 | 0,4175 |
| TAF151 | 0,051318253 | -0,679971283 | 0,065 | 0,2 | 1 | Mast | 0,624177699 | 1,602107865 | 0,325 |
| LUC7L21 | 0,051430146 | -0,648072334 | 0,13 | 0,333 | 1 | Mast | 0,638132389 | 1,567072942 | 0,39039039 |
| GPBP1L11 | 0,052052713 | -0,490307434 | 0,028 | 0,133 | 1 | Mast | 0,711873384 | 1,40474419 | 0,210526316 |
| MAN1A21 | 0,052052713 | -0,56242427 | 0,028 | 0,133 | 1 | Mast | 0,677163318 | 1,476748626 | 0,210526316 |
| NUP42 | 0,052052713 | -0,520789074 | 0,028 | 0,133 | 1 | Mast | 0,696990514 | 1,434739756 | 0,210526316 |
| TRAF1 | 0,052052713 | -0,648119442 | 0,028 | 0,133 | 1 | Mast | 0,638111552 | 1,567124112 | 0,210526316 |
| ARMH3 | 0,052052713 | -0,494639382 | 0,028 | 0,133 | 1 | Mast | 0,709739063 | 1,408968524 | 0,210526316 |
| CAT2 | 0,052052713 | -0,668590575 | 0,028 | 0,133 | 1 | Mast | 0,629121 | 1,589519345 | 0,210526316 |
| PDE8A | 0,052052713 | -0,492930763 | 0,028 | 0,133 | 1 | Mast | 0,710580123 | 1,407300835 | 0,210526316 |
| ALOX12-AS1 | 0,052052713 | -0,625167801 | 0,028 | 0,133 | 1 | Mast | 0,648344364 | 1,542390211 | 0,210526316 |
| NFIX1 | 0,052052713 | -0,681360221 | 0,028 | 0,133 | 1 | Mast | 0,623577068 | 1,603651019 | 0,210526316 |
| CNOT41 | 0,052230393 | -0,547141699 | 0,093 | 0,267 | 1 | Mast | 0,684374685 | 1,461187887 | 0,348314607 |
| HDC | 0,052238305 | 0,988104227 | 0,667 | 0,333 | 1 | Mast | 1,983576759 | 0,504139805 | 2,003003003 |
| CYCS2 | 0,052443205 | -0,941719133 | 0,148 | 0,333 | 1 | Mast | 0,520612143 | 1,920815743 | 0,444444444 |
| PABPN11 | 0,052653006 | -1,237440573 | 0,157 | 0,333 | 1 | Mast | 0,424124411 | 2,357798735 | 0,471471471 |
| DDX3X3 | 0,053208036 | -0,715804061 | 0,611 | 0,733 | 1 | Mast | 0,608865697 | 1,642398325 | 0,833560709 |
| PDE4DIP | 0,053299427 | -0,778302066 | 0,065 | 0,2 | 1 | Mast | 0,583052595 | 1,715111139 | 0,325 |
| CDK4 | 0,053513952 | -0,73287085 | 0,139 | 0,333 | 1 | Mast | 0,601705375 | 1,66194294 | 0,417417417 |
| ATP5PO3 | 0,053917659 | 1,289150228 | 0,296 | 0,067 | 1 | Mast | 2,443840668 | 0,409191979 | 4,417910448 |
| BMI1 | 0,054125202 | -0,519198338 | 0,056 | 0,2 | 1 | Mast | 0,697759449 | 1,433158664 | 0,28 |
| TNFAIP23 | 0,054125202 | -0,505961249 | 0,056 | 0,2 | 1 | Mast | 0,704191028 | 1,420069215 | 0,28 |
| ITM2C2 | 0,054154334 | 1,42476061 | 0,361 | 0,133 | 1 | Mast | 2,68469949 | 0,372481167 | 2,714285714 |
| TSC22D31 | 0,054583044 | -0,547270433 | 0,722 | 0,867 | 1 | Mast | 0,68431362 | 1,461318277 | 0,832756632 |
| TOMM73 | 0,054779677 | -1,117019719 | 0,407 | 0,667 | 1 | Mast | 0,461045257 | 2,168984463 | 0,610194903 |
| TMEM242 | 0,054844254 | -0,701698184 | 0,028 | 0,133 | 1 | Mast | 0,614848048 | 1,62641811 | 0,210526316 |

| Supplementary Table 4 - clusermarker | | | | | | | | | |
|---|---|---|---|---|---|---|---|---|---|
| | **p_val** | **avg_log2FC** | **pct.1** | **pct.2** | **p_val_adj** | **celltype** | **Foldchange_UP** | **Foldchange_DOWN** | **Ratio_pct1_pct2** |
| DUSP42 | 0,054844254 | -0,440684406 | 0,028 | 0,133 | 1 | Mast | 0,736784999 | 1,357248045 | 0,210526316 |
| DENND5A1 | 0,054844254 | -0,43284405 | 0,028 | 0,133 | 1 | Mast | 0,740799972 | 1,349892059 | 0,210526316 |
| FAR1 | 0,054844254 | -0,453543836 | 0,028 | 0,133 | 1 | Mast | 0,730246865 | 1,369399923 | 0,210526316 |
| CDK8 | 0,054844254 | -0,374625078 | 0,028 | 0,133 | 1 | Mast | 0,771305831 | 1,296502581 | 0,210526316 |
| VBP1 | 0,054844254 | -0,504288867 | 0,028 | 0,133 | 1 | Mast | 0,705007804 | 1,418424015 | 0,210526316 |
| NUP153 | 0,056298906 | -0,428935178 | 0,056 | 0,2 | 1 | Mast | 0,742809835 | 1,34623958 | 0,28 |
| UQCRB2 | 0,056991458 | -0,537853563 | 0,407 | 0,733 | 1 | Mast | 0,688794934 | 1,45181091 | 0,555252387 |
| RAB6A2 | 0,05703404 | 1,27934523 | 0,204 | 0 | 1 | Mast | 2,42728789 | 0,411982445 | #DIV/0! |
| TNFSF12 | 0,05703404 | 1,271138777 | 0,204 | 0 | 1 | Mast | 2,413519992 | 0,414332594 | #DIV/0! |
| APOE2 | 0,05703404 | 1,923945236 | 0,204 | 0 | 1 | Mast | 3,794593218 | 0,263532859 | #DIV/0! |
| SMARCA51 | 0,057450029 | -0,855786627 | 0,065 | 0,2 | 1 | Mast | 0,55256396 | 1,809745246 | 0,325 |
| FAM53C | 0,057540598 | -0,511990067 | 0,093 | 0,267 | 1 | Mast | 0,701254453 | 1,426015899 | 0,348314607 |
| IGKC6 | 0,057700145 | 2,828764375 | 0,315 | 0,067 | 1 | Mast | 7,104653919 | 0,14075281 | 4,701492537 |
| USP17 | 0,057759535 | -0,439663444 | 0,028 | 0,133 | 1 | Mast | 0,73730659 | 1,356287892 | 0,210526316 |
| MFSD14A | 0,057759535 | -0,416503091 | 0,028 | 0,133 | 1 | Mast | 0,749238484 | 1,334688516 | 0,210526316 |
| ANKRD36C1 | 0,057759535 | -0,658668618 | 0,028 | 0,133 | 1 | Mast | 0,633462614 | 1,578625127 | 0,210526316 |
| QRICH11 | 0,057759535 | -0,546443059 | 0,028 | 0,133 | 1 | Mast | 0,684706181 | 1,460480463 | 0,210526316 |
| IQCB1 | 0,057759535 | -0,381605687 | 0,028 | 0,133 | 1 | Mast | 0,767582813 | 1,302791026 | 0,210526316 |
| MRPL472 | 0,057759535 | -0,556122282 | 0,028 | 0,133 | 1 | Mast | 0,680127776 | 1,470311954 | 0,210526316 |
| HARS1 | 0,057759535 | -0,540544164 | 0,028 | 0,133 | 1 | Mast | 0,68751154 | 1,454521039 | 0,210526316 |
| GPNMB1 | 0,057759535 | -0,513108888 | 0,028 | 0,133 | 1 | Mast | 0,700710836 | 1,427122214 | 0,210526316 |
| LONRF1 | 0,057759535 | -0,582893032 | 0,028 | 0,133 | 1 | Mast | 0,667623651 | 1,497849872 | 0,210526316 |
| GOLGA71 | 0,057759535 | -0,487525138 | 0,028 | 0,133 | 1 | Mast | 0,713247585 | 1,402037695 | 0,210526316 |
| FBH1 | 0,057759535 | -0,427784388 | 0,028 | 0,133 | 1 | Mast | 0,743402586 | 1,345166157 | 0,210526316 |
| NSMCE3 | 0,057759535 | -0,513978801 | 0,028 | 0,133 | 1 | Mast | 0,700288451 | 1,427982996 | 0,210526316 |
| RBL22 | 0,057759535 | -0,46238147 | 0,028 | 0,133 | 1 | Mast | 0,725787206 | 1,377814313 | 0,210526316 |
| USP251 | 0,057759535 | -0,50745734 | 0,028 | 0,133 | 1 | Mast | 0,703461152 | 1,421542607 | 0,210526316 |
| ZNF844 | 0,058544805 | -0,524196155 | 0,056 | 0,2 | 1 | Mast | 0,695346437 | 1,438132054 | 0,28 |
| STXBP51 | 0,058740053 | -0,948626693 | 0,102 | 0,267 | 1 | Mast | 0,518125433 | 1,930034576 | 0,382022472 |
| TLE43 | 0,058740053 | -0,854189595 | 0,102 | 0,267 | 1 | Mast | 0,553175975 | 1,807743006 | 0,382022472 |
| TALDO13 | 0,058991856 | 1,171674211 | 0,37 | 0,133 | 1 | Mast | 2,252729688 | 0,4439059 | 2,781954887 |

| Supplementary Table 4 - clusermarker | | | | | | | | | |
|---|---|---|---|---|---|---|---|---|---|
| | **p_val** | **avg_log2FC** | **pct.1** | **pct.2** | **p_val_adj** | **celltype** | **Foldchange_UP** | **Foldchange_DOWN** | **Ratio_pct1_pct2** |
| GOSR11 | 0,059408841 | -0,483986006 | 0,093 | 0,267 | 1 | Mast | 0,714999429 | 1,398602516 | 0,348314607 |
| HDLBP2 | 0,060388356 | 1,1757426 | 0,296 | 0,067 | 1 | Mast | 2,259091335 | 0,442655852 | 4,417910448 |
| GTF2I2 | 0,06058174 | -0,752485011 | 0,102 | 0,267 | 1 | Mast | 0,593580247 | 1,684692179 | 0,382022472 |
| SNRNP481 | 0,060802488 | -0,295395315 | 0,028 | 0,133 | 1 | Mast | 0,814849024 | 1,227221204 | 0,210526316 |
| HNRNPA16 | 0,061278126 | -0,741905687 | 0,389 | 0,6 | 1 | Mast | 0,597948987 | 1,672383468 | 0,648333333 |
| P2RY14 | 0,061324371 | -0,543275245 | 0,093 | 0,267 | 1 | Mast | 0,686211284 | 1,457277115 | 0,348314607 |
| DDX46 | 0,061400238 | -0,721787483 | 0,139 | 0,333 | 1 | Mast | 0,606345721 | 1,64922414 | 0,417417417 |
| NCKAP1L1 | 0,063293587 | -0,4495072 | 0,093 | 0,267 | 1 | Mast | 0,732292944 | 1,365573721 | 0,348314607 |
| GNAQ1 | 0,063318741 | -0,757081639 | 0,148 | 0,333 | 1 | Mast | 0,591692028 | 1,690068403 | 0,444444444 |
| PRIMPOL | 0,063977084 | -0,504181847 | 0,028 | 0,133 | 1 | Mast | 0,705060104 | 1,418318799 | 0,210526316 |
| MRPS271 | 0,063977084 | -0,291114409 | 0,028 | 0,133 | 1 | Mast | 0,817270515 | 1,22358507 | 0,210526316 |
| TRIM521 | 0,063977084 | -0,504017349 | 0,028 | 0,133 | 1 | Mast | 0,705140501 | 1,418157089 | 0,210526316 |
| NDRG3 | 0,063977084 | -0,328053105 | 0,028 | 0,133 | 1 | Mast | 0,796610773 | 1,255318198 | 0,210526316 |
| YBEY | 0,063977084 | -0,479955225 | 0,028 | 0,133 | 1 | Mast | 0,716999876 | 1,39470038 | 0,210526316 |
| IGKV4-15 | 0,064160131 | 1,459395299 | 0,194 | 0 | 1 | Mast | 2,749930771 | 0,363645518 | #DIV/0! |
| ABHD14B1 | 0,064162129 | 1,094622572 | 0,194 | 0 | 1 | Mast | 2,13557205 | 0,46825861 | #DIV/0! |
| TMEM179B4 | 0,064162129 | 1,29454783 | 0,194 | 0 | 1 | Mast | 2,453001014 | 0,407663916 | #DIV/0! |
| TUFM4 | 0,064162129 | 1,103421592 | 0,194 | 0 | 1 | Mast | 2,148636738 | 0,465411385 | #DIV/0! |
| STXBP2 | 0,064162129 | 1,241672569 | 0,194 | 0 | 1 | Mast | 2,364725246 | 0,422882109 | #DIV/0! |
| APOC21 | 0,064162129 | 1,802104061 | 0,194 | 0 | 1 | Mast | 3,487284485 | 0,286756072 | #DIV/0! |
| BAX5 | 0,064162129 | 1,275284755 | 0,194 | 0 | 1 | Mast | 2,420465876 | 0,413143606 | #DIV/0! |
| SNRPA2 | 0,064177114 | -0,583019687 | 0,065 | 0,2 | 1 | Mast | 0,667565042 | 1,497981376 | 0,325 |
| UBXN15 | 0,064366734 | -0,520592383 | 0,222 | 0,467 | 1 | Mast | 0,697085545 | 1,434544163 | 0,475374732 |
| MTRNR2L122 | 0,065105534 | 0,891569946 | 0,676 | 0,467 | 1 | Mast | 1,855193854 | 0,539027228 | 1,447537473 |
| PDZD8 | 0,065111456 | -0,815007934 | 0,111 | 0,267 | 1 | Mast | 0,568405361 | 1,759307827 | 0,415730337 |
| TUBA1A3 | 0,066224289 | 1,431220022 | 0,444 | 0,2 | 1 | Mast | 2,696746707 | 0,370817177 | 2,22 |
| DDX185 | 0,066395887 | -0,505829091 | 0,102 | 0,267 | 1 | Mast | 0,704255538 | 1,419939135 | 0,382022472 |
| SAP30BP | 0,066557164 | -0,729947243 | 0,065 | 0,2 | 1 | Mast | 0,602925961 | 1,658578439 | 0,325 |
| RABGGTB1 | 0,066559116 | -0,782658161 | 0,065 | 0,2 | 1 | Mast | 0,581294772 | 1,720297597 | 0,325 |
| HMGA11 | 0,066559116 | -0,676029201 | 0,065 | 0,2 | 1 | Mast | 0,625885561 | 1,597736172 | 0,325 |
| TMEM120A1 | 0,066559116 | -0,787582039 | 0,065 | 0,2 | 1 | Mast | 0,579314211 | 1,726178956 | 0,325 |

| Supplementary Table 4 - clusermarker | | | | | | | | | |
|---|---|---|---|---|---|---|---|---|---|
| | **p_val** | **avg_log2FC** | **pct.1** | **pct.2** | **p_val_adj** | **celltype** | **Foldchange_UP** | **Foldchange_DOWN** | **Ratio_pct1_pct2** |
| SLC1A51 | 0,068032244 | -0,729816597 | 0,194 | 0,4 | 1 | Mast | 0,602980563 | 1,65842825 | 0,485 |
| GALNT11 | 0,069013461 | -0,594150152 | 0,065 | 0,2 | 1 | Mast | 0,662434561 | 1,509583074 | 0,325 |
| MTCO2P122 | 0,069916831 | -0,273308964 | 1 | 1 | 1 | Mast | 0,827419599 | 1,208576642 | 1 |
| OST43 | 0,07038394 | -0,646028225 | 0,5 | 0,733 | 1 | Mast | 0,639037179 | 1,564854178 | 0,68212824 |
| NUCB12 | 0,070772069 | 1,249743846 | 0,343 | 0,133 | 1 | Mast | 2,377991975 | 0,420522866 | 2,578947368 |
| ZFP364 | 0,071522724 | -0,545325026 | 0,796 | 1 | 1 | Mast | 0,685237007 | 1,459349085 | 0,796 |
| COPE4 | 0,072101738 | 1,223290503 | 0,287 | 0,067 | 1 | Mast | 2,334786292 | 0,428304725 | 4,28358209 |
| AGTRAP | 0,072109742 | 1,315686491 | 0,185 | 0 | 1 | Mast | 2,489207487 | 0,401734289 | #DIV/0! |
| PLK3 | 0,072109742 | 1,186562673 | 0,185 | 0 | 1 | Mast | 2,276097995 | 0,439348395 | #DIV/0! |
| SNX17 | 0,072109742 | 1,361935764 | 0,185 | 0 | 1 | Mast | 2,57029823 | 0,389059911 | #DIV/0! |
| NDUFB62 | 0,072109742 | 0,957717251 | 0,185 | 0 | 1 | Mast | 1,942234302 | 0,51487094 | #DIV/0! |
| AIP1 | 0,072109742 | 1,167259409 | 0,185 | 0 | 1 | Mast | 2,24584663 | 0,44526638 | #DIV/0! |
| HSP90B14 | 0,072164304 | 1,314928745 | 0,574 | 0,4 | 1 | Mast | 2,487900425 | 0,401945347 | 1,435 |
| VIM-AS12 | 0,073643594 | 1,287821975 | 0,278 | 0,067 | 1 | Mast | 2,441591721 | 0,409568886 | 4,149253731 |
| PLD32 | 0,073643594 | 1,331300568 | 0,278 | 0,067 | 1 | Mast | 2,516294129 | 0,397409821 | 4,149253731 |
| ARPC1B5 | 0,073682691 | 1,339023702 | 0,417 | 0,2 | 1 | Mast | 2,529800647 | 0,395288064 | 2,085 |
| PCNP2 | 0,074065934 | -0,524853174 | 0,148 | 0,333 | 1 | Mast | 0,695029841 | 1,438787145 | 0,444444444 |
| RPRD2 | 0,074145283 | -0,59565581 | 0,065 | 0,2 | 1 | Mast | 0,661743576 | 1,511159361 | 0,325 |
| SPART | 0,074145283 | -0,615704976 | 0,065 | 0,2 | 1 | Mast | 0,652610916 | 1,53230658 | 0,325 |
| GAS54 | 0,07442978 | -0,665500162 | 0,204 | 0,4 | 1 | Mast | 0,630470091 | 1,586118063 | 0,51 |
| EIF3G1 | 0,075291195 | 1,054032301 | 0,278 | 0,067 | 1 | Mast | 2,076325028 | 0,481620164 | 4,149253731 |
| EFCAB14 | 0,07669368 | 1,053471027 | 0,38 | 0,133 | 1 | Mast | 2,0755174 | 0,481807572 | 2,857142857 |
| PSMA31 | 0,076825826 | -0,627182467 | 0,065 | 0,2 | 1 | Mast | 0,647439608 | 1,544545603 | 0,325 |
| SLC9A1 | 0,077102517 | -0,680097868 | 0,102 | 0,267 | 1 | Mast | 0,624122935 | 1,602248443 | 0,382022472 |
| JPT13 | 0,077102517 | -0,546506281 | 0,102 | 0,267 | 1 | Mast | 0,684676176 | 1,460544466 | 0,382022472 |
| PPIA5 | 0,078186276 | 0,754641687 | 0,704 | 0,6 | 1 | Mast | 1,687212498 | 0,592693571 | 1,173333333 |
| GSTP14 | 0,078264452 | 1,188458318 | 0,426 | 0,2 | 1 | Mast | 2,279090666 | 0,438771487 | 2,13 |
| DDAH22 | 0,078363836 | 0,835758767 | 0,306 | 0,067 | 1 | Mast | 1,784795483 | 0,560288285 | 4,567164179 |
| PCMT12 | 0,078676678 | 1,250159377 | 0,278 | 0,067 | 1 | Mast | 2,378676992 | 0,420401762 | 4,149253731 |
| ETV31 | 0,079584843 | -0,48043782 | 0,065 | 0,2 | 1 | Mast | 0,716760073 | 1,395166998 | 0,325 |
| DYNC1I21 | 0,079584843 | -0,586669531 | 0,065 | 0,2 | 1 | Mast | 0,665878318 | 1,501775885 | 0,325 |

| Supplementary Table 4 - clusermarker | | | | | | | | | |
|---|---|---|---|---|---|---|---|---|---|
| | **p_val** | **avg_log2FC** | **pct.1** | **pct.2** | **p_val_adj** | **celltype** | **Foldchange_UP** | **Foldchange_DOWN** | **Ratio_pct1_pct2** |
| AREG4 | 0,079953985 | 0,814944865 | 0,926 | 0,933 | 1 | Mast | 1,759230919 | 0,56843021 | 0,99249732 |
| GSTK12 | 0,080397709 | 1,01566013 | 0,287 | 0,067 | 1 | Mast | 2,021827803 | 0,494601963 | 4,28358209 |
| ZMIZ11 | 0,080967865 | 0,882250934 | 0,176 | 0 | 1 | Mast | 1,843248948 | 0,542520315 | #DIV/0! |
| CD1513 | 0,080967865 | 1,163855508 | 0,176 | 0 | 1 | Mast | 2,240554017 | 0,446318184 | #DIV/0! |
| SIVA14 | 0,080967865 | 0,918268925 | 0,176 | 0 | 1 | Mast | 1,889846325 | 0,529143553 | #DIV/0! |
| IGHG15 | 0,080967865 | 1,406856436 | 0,176 | 0 | 1 | Mast | 2,651587645 | 0,377132546 | #DIV/0! |
| SKAP1 | 0,080967865 | 1,205893157 | 0,176 | 0 | 1 | Mast | 2,306800367 | 0,433500885 | #DIV/0! |
| NT5C3 | 0,080967865 | 1,155761188 | 0,176 | 0 | 1 | Mast | 2,228018463 | 0,448829315 | #DIV/0! |
| TMEM1604 | 0,080967865 | 0,955545136 | 0,176 | 0 | 1 | Mast | 1,939312283 | 0,515646711 | #DIV/0! |
| MT-CYB2 | 0,081343825 | -0,293733971 | 0,954 | 1 | 1 | Mast | 0,815787908 | 1,225808803 | 0,954 |
| DNAJA15 | 0,081859825 | -0,400405383 | 0,491 | 0,8 | 1 | Mast | 0,757645362 | 1,319878732 | 0,61375 |
| RTN32 | 0,082162078 | -0,575436439 | 0,139 | 0,333 | 1 | Mast | 0,671083207 | 1,490128183 | 0,417417417 |
| PET1004 | 0,082423874 | -0,342555404 | 0,065 | 0,2 | 1 | Mast | 0,788643173 | 1,268000579 | 0,325 |
| CALR4 | 0,082936397 | 0,95461361 | 0,38 | 0,133 | 1 | Mast | 1,938060503 | 0,515979763 | 2,857142857 |
| CTSG | 0,083130415 | 1,575833596 | 0,37 | 0,133 | 1 | Mast | 2,98107691 | 0,335449245 | 2,781954887 |
| ARPC54 | 0,083299597 | -0,545738624 | 0,296 | 0,533 | 1 | Mast | 0,685040589 | 1,459767518 | 0,555347092 |
| RPLP24 | 0,083520645 | -0,369555374 | 0,833 | 0,933 | 1 | Mast | 0,774021006 | 1,2919546 | 0,892818864 |
| HS3ST12 | 0,083728831 | 1,73681048 | 0,259 | 0,067 | 1 | Mast | 3,332974963 | 0,300032257 | 3,865671642 |
| NME25 | 0,08383574 | 0,949873614 | 0,611 | 0,533 | 1 | Mast | 1,931703426 | 0,517677811 | 1,146341463 |
| RPS112 | 0,08383574 | -0,521897095 | 0,574 | 0,8 | 1 | Mast | 0,696455416 | 1,435842091 | 0,7175 |
| ZFC3H1 | 0,08446261 | -0,810943529 | 0,074 | 0,2 | 1 | Mast | 0,570008947 | 1,754358426 | 0,37 |
| NDUFA33 | 0,08446261 | -1,391027931 | 0,074 | 0,2 | 1 | Mast | 0,381293031 | 2,622654804 | 0,37 |
| RARRES12 | 0,085344465 | -0,418420245 | 0,065 | 0,2 | 1 | Mast | 0,748243504 | 1,336463323 | 0,325 |
| MNAT1 | 0,085344465 | -0,5448965 | 0,065 | 0,2 | 1 | Mast | 0,685440575 | 1,458915677 | 0,325 |
| TMEM2052 | 0,085344465 | -0,511624815 | 0,065 | 0,2 | 1 | Mast | 0,701432015 | 1,425654916 | 0,325 |
| SUPT5H | 0,085344465 | -0,467734228 | 0,065 | 0,2 | 1 | Mast | 0,723099344 | 1,382935842 | 0,325 |
| SFPQ3 | 0,086032441 | -0,677676439 | 0,5 | 0,667 | 1 | Mast | 0,625171346 | 1,599561474 | 0,749625187 |
| CD532 | 0,086136789 | -0,369711519 | 0,185 | 0,4 | 1 | Mast | 0,773937238 | 1,292094438 | 0,4625 |
| RPL351 | 0,087544688 | -0,440129753 | 0,667 | 0,8 | 1 | Mast | 0,737068315 | 1,356726343 | 0,83375 |
| CD2AP1 | 0,088348159 | -0,379456766 | 0,065 | 0,2 | 1 | Mast | 0,768726993 | 1,300851939 | 0,325 |
| BTG14 | 0,088596229 | -0,468667268 | 0,648 | 0,867 | 1 | Mast | 0,722631842 | 1,383830523 | 0,747404844 |

| Supplementary Table 4 - clusermarker | | | | | | | | | |
|---|---|---|---|---|---|---|---|---|---|
| | **p_val** | **avg_log2FC** | **pct.1** | **pct.2** | **p_val_adj** | **celltype** | **Foldchange_UP** | **Foldchange_DOWN** | **Ratio_pct1_pct2** |
| TNPO12 | 0,089310087 | -0,623663501 | 0,111 | 0,267 | 1 | Mast | 0,649020746 | 1,540782797 | 0,415730337 |
| CHCHD26 | 0,089607848 | 0,771974451 | 0,63 | 0,467 | 1 | Mast | 1,707605188 | 0,585615461 | 1,349036403 |
| HSPA54 | 0,089860561 | 1,380952953 | 0,602 | 0,4 | 1 | Mast | 2,604403446 | 0,383965089 | 1,505 |
| EPAS12 | 0,090837342 | 0,924641013 | 0,167 | 0 | 1 | Mast | 1,89821185 | 0,526811589 | #DIV/0! |
| CCT41 | 0,090837342 | 0,9664756 | 0,167 | 0 | 1 | Mast | 1,95406113 | 0,511754717 | #DIV/0! |
| CLTB4 | 0,090837342 | 1,07968691 | 0,167 | 0 | 1 | Mast | 2,113577349 | 0,47313149 | #DIV/0! |
| NAP1L42 | 0,090837342 | 1,120688393 | 0,167 | 0 | 1 | Mast | 2,174507061 | 0,45987434 | #DIV/0! |
| PGLS5 | 0,090837342 | 1,222346379 | 0,167 | 0 | 1 | Mast | 2,333258869 | 0,428585106 | #DIV/0! |
| VAV3 | 0,09083872 | 0,904476748 | 0,167 | 0 | 1 | Mast | 1,871865463 | 0,534226428 | #DIV/0! |
| RHOC2 | 0,09083872 | 1,128549121 | 0,167 | 0 | 1 | Mast | 2,186387506 | 0,457375464 | #DIV/0! |
| IGKV3-205 | 0,09083872 | 2,106710901 | 0,167 | 0 | 1 | Mast | 4,307082329 | 0,232175734 | #DIV/0! |
| EIF4G11 | 0,09083872 | 0,836540711 | 0,167 | 0 | 1 | Mast | 1,785763107 | 0,55998469 | #DIV/0! |
| LSM52 | 0,09083872 | 0,945233285 | 0,167 | 0 | 1 | Mast | 1,925500216 | 0,519345566 | #DIV/0! |
| DDB11 | 0,09083872 | 0,861032064 | 0,167 | 0 | 1 | Mast | 1,816337204 | 0,550558563 | #DIV/0! |
| NME35 | 0,09083872 | 1,141471517 | 0,167 | 0 | 1 | Mast | 2,206059216 | 0,453296989 | #DIV/0! |
| YPEL33 | 0,09083872 | 0,982009975 | 0,167 | 0 | 1 | Mast | 1,97521538 | 0,506273903 | #DIV/0! |
| MIR22HG1 | 0,09083872 | 0,970607329 | 0,167 | 0 | 1 | Mast | 1,959665379 | 0,510291201 | #DIV/0! |
| LAT21 | 0,091360026 | -0,782239013 | 0,241 | 0,4 | 1 | Mast | 0,581463681 | 1,719797869 | 0,6025 |
| ACOT11 | 0,091487295 | -0,531737623 | 0,037 | 0,133 | 1 | Mast | 0,691721104 | 1,445669352 | 0,278195489 |
| CMC12 | 0,091491046 | -0,905357153 | 0,037 | 0,133 | 1 | Mast | 0,533900516 | 1,873008118 | 0,278195489 |
| ELMOD2 | 0,091491046 | -1,040855101 | 0,037 | 0,133 | 1 | Mast | 0,486039307 | 2,057446764 | 0,278195489 |
| BTN3A21 | 0,091491046 | -0,678878338 | 0,037 | 0,133 | 1 | Mast | 0,624650737 | 1,600894613 | 0,278195489 |
| TBC1D22B | 0,091491046 | -0,63303099 | 0,037 | 0,133 | 1 | Mast | 0,644820274 | 1,550819725 | 0,278195489 |
| PRXL2A2 | 0,091491046 | -0,612099611 | 0,037 | 0,133 | 1 | Mast | 0,654243862 | 1,528482051 | 0,278195489 |
| OPHN1 | 0,091491046 | -0,473639088 | 0,037 | 0,133 | 1 | Mast | 0,720145792 | 1,388607711 | 0,278195489 |
| HNRNPR2 | 0,091827763 | -0,513110024 | 0,111 | 0,267 | 1 | Mast | 0,700710284 | 1,427123338 | 0,415730337 |
| FTH15 | 0,092751997 | -0,582094761 | 1 | 1 | 1 | Mast | 0,667993162 | 1,497021313 | 1 |
| NPM12 | 0,09290935 | -0,629876735 | 0,472 | 0,667 | 1 | Mast | 0,646231628 | 1,547432774 | 0,707646177 |
| MAT2A4 | 0,092938909 | -0,691733798 | 0,148 | 0,333 | 1 | Mast | 0,619109371 | 1,615223492 | 0,444444444 |
| SRSF53 | 0,09337828 | -1,020930783 | 0,269 | 0,467 | 1 | Mast | 0,492798311 | 2,029227733 | 0,576017131 |
| H4C33 | 0,094001945 | -1,042135422 | 0,12 | 0,267 | 1 | Mast | 0,485608163 | 2,059273457 | 0,449438202 |

| Supplementary Table 4 - clusermarker | | | | | | | | | |
|---|---|---|---|---|---|---|---|---|---|
| | **p_val** | **avg_log2FC** | **pct.1** | **pct.2** | **p_val_adj** | **celltype** | **Foldchange_UP** | **Foldchange_DOWN** | **Ratio_pct1_pct2** |
| RPL7A2 | 0,094252282 | -0,336691983 | 0,954 | 1 | 1 | Mast | 0,79185491 | 1,262857611 | 0,954 |
| SMNDC1 | 0,094611025 | -0,38429324 | 0,065 | 0,2 | 1 | Mast | 0,766154237 | 1,305220217 | 0,325 |
| STX3 | 0,095271368 | -0,364906333 | 0,231 | 0,467 | 1 | Mast | 0,776519289 | 1,287798017 | 0,494646681 |
| UQCRHL1 | 0,095538708 | -0,695169733 | 0,037 | 0,133 | 1 | Mast | 0,617636649 | 1,619074906 | 0,278195489 |
| TDP2 | 0,095538708 | -0,61787778 | 0,037 | 0,133 | 1 | Mast | 0,651628776 | 1,534616083 | 0,278195489 |
| EXTL3 | 0,095538708 | -0,65235776 | 0,037 | 0,133 | 1 | Mast | 0,636239673 | 1,571734745 | 0,278195489 |
| RPS33 | 0,095792931 | -0,257549132 | 0,981 | 1 | 1 | Mast | 0,836507783 | 1,19544614 | 0,981 |
| ANKLE2 | 0,096331742 | -0,916420949 | 0,074 | 0,2 | 1 | Mast | 0,529821778 | 1,887427134 | 0,37 |
| HLA-B6 | 0,096582808 | 0,907717417 | 0,704 | 0,8 | 1 | Mast | 1,876074887 | 0,533027763 | 0,88 |
| CUTA4 | 0,097403708 | 0,978808227 | 0,352 | 0,133 | 1 | Mast | 1,97083668 | 0,507398716 | 2,646616541 |
| RPN13 | 0,097488956 | 1,091588854 | 0,333 | 0,133 | 1 | Mast | 2,131086058 | 0,469244307 | 2,503759398 |
| TFG2 | 0,099498568 | -0,585482594 | 0,074 | 0,2 | 1 | Mast | 0,666426376 | 1,500540849 | 0,37 |
| CCDC141 | 0,09972851 | -0,650026765 | 0,037 | 0,133 | 1 | Mast | 0,637268491 | 1,569197308 | 0,278195489 |
| MFSD6 | 0,09972851 | -0,534041847 | 0,037 | 0,133 | 1 | Mast | 0,690617192 | 1,447980172 | 0,278195489 |
| HDAC4 | 0,09972851 | -0,583882844 | 0,037 | 0,133 | 1 | Mast | 0,667165761 | 1,498877878 | 0,278195489 |
| SLMAP2 | 0,09972851 | -0,517925279 | 0,037 | 0,133 | 1 | Mast | 0,698375436 | 1,431894578 | 0,278195489 |
| EIF4E4 | 0,09972851 | -0,526088413 | 0,037 | 0,133 | 1 | Mast | 0,694435009 | 1,440019565 | 0,278195489 |
| C5orf51 | 0,09972851 | -0,823776465 | 0,037 | 0,133 | 1 | Mast | 0,564961137 | 1,770033254 | 0,278195489 |
| MICA | 0,09972851 | -0,73557903 | 0,037 | 0,133 | 1 | Mast | 0,600576933 | 1,665065615 | 0,278195489 |
| MYO62 | 0,09972851 | -0,44802968 | 0,037 | 0,133 | 1 | Mast | 0,733043298 | 1,364175899 | 0,278195489 |
| LETM2 | 0,09972851 | -0,50025773 | 0,037 | 0,133 | 1 | Mast | 0,706980471 | 1,414466227 | 0,278195489 |
| PRPF312 | 0,09972851 | -0,82563777 | 0,037 | 0,133 | 1 | Mast | 0,564232718 | 1,772318351 | 0,278195489 |
| RPL241 | 0,099972183 | -0,471250629 | 0,833 | 1 | 1 | Mast | 0,72133902 | 1,386310698 | 0,833 |
| LYL11 | 0,100590344 | -0,79285317 | 0,167 | 0,333 | 1 | Mast | 0,57720145 | 1,732497379 | 0,501501502 |
| DNAJC84 | 0,101635515 | -0,689717359 | 0,157 | 0,333 | 1 | Mast | 0,619975298 | 1,612967489 | 0,471471471 |
| CSF1 | 0,101688097 | -0,992309601 | 0,176 | 0,333 | 1 | Mast | 0,502672406 | 1,989367208 | 0,528528529 |
| MT2A2 | 0,101828891 | 1,471087762 | 0,269 | 0,067 | 1 | Mast | 2,77230841 | 0,360710228 | 4,014925373 |
| PSMB21 | 0,101837684 | 0,80473828 | 0,157 | 0 | 1 | Mast | 1,74682887 | 0,572465922 | #DIV/0! |
| RNF1813 | 0,101837684 | 1,110513666 | 0,157 | 0 | 1 | Mast | 2,15922512 | 0,463129106 | #DIV/0! |
| CAMLG1 | 0,101837684 | 1,097389798 | 0,157 | 0 | 1 | Mast | 2,13967221 | 0,467361307 | #DIV/0! |
| REPS1 | 0,101837684 | 0,88618731 | 0,157 | 0 | 1 | Mast | 1,848285099 | 0,541042072 | #DIV/0! |

| Supplementary Table 4 - clusermarker | | | | | | | | | |
|---|---|---|---|---|---|---|---|---|---|
| | **p_val** | **avg_log2FC** | **pct.1** | **pct.2** | **p_val_adj** | **celltype** | **Foldchange_UP** | **Foldchange_DOWN** | **Ratio_pct1_pct2** |
| RNASEH2B1 | 0,101837684 | 1,094650126 | 0,157 | 0 | 1 | Mast | 2,135612837 | 0,468249667 | #DIV/0! |
| PSMB101 | 0,101837684 | 1,130466129 | 0,157 | 0 | 1 | Mast | 2,18929464 | 0,456768121 | #DIV/0! |
| NCOR12 | 0,101837684 | 1,18631458 | 0,157 | 0 | 1 | Mast | 2,27570662 | 0,439423953 | #DIV/0! |
| AC011446.2 | 0,101837684 | 1,166852048 | 0,157 | 0 | 1 | Mast | 2,24521258 | 0,445392124 | #DIV/0! |
| RALY4 | 0,101837684 | 1,055148498 | 0,157 | 0 | 1 | Mast | 2,077932079 | 0,481247684 | #DIV/0! |
| PMEPA13 | 0,101837684 | 1,059021687 | 0,157 | 0 | 1 | Mast | 2,083518178 | 0,479957416 | #DIV/0! |
| PHYH1 | 0,102747902 | -0,705686654 | 0,074 | 0,2 | 1 | Mast | 0,613150589 | 1,630920721 | 0,37 |
| DYM2 | 0,102747902 | -0,855449102 | 0,074 | 0,2 | 1 | Mast | 0,55269325 | 1,809321898 | 0,37 |
| SNX34 | 0,103807968 | 1,031285212 | 0,278 | 0,067 | 1 | Mast | 2,043844181 | 0,489274089 | 4,149253731 |
| BCL10 | 0,104063584 | -0,43243532 | 0,037 | 0,133 | 1 | Mast | 0,741009878 | 1,349509675 | 0,278195489 |
| C1orf21 | 0,104063584 | -0,754856027 | 0,037 | 0,133 | 1 | Mast | 0,592605521 | 1,687463184 | 0,278195489 |
| ZC3H8 | 0,104063584 | -0,586356601 | 0,037 | 0,133 | 1 | Mast | 0,666022767 | 1,501450176 | 0,278195489 |
| TLK12 | 0,104063584 | -0,443435158 | 0,037 | 0,133 | 1 | Mast | 0,735381528 | 1,359838346 | 0,278195489 |
| XRN13 | 0,104063584 | -0,80246953 | 0,037 | 0,133 | 1 | Mast | 0,573366878 | 1,744084004 | 0,278195489 |
| KBTBD21 | 0,104063584 | -0,63778897 | 0,037 | 0,133 | 1 | Mast | 0,642697172 | 1,555942741 | 0,278195489 |
| HDHD31 | 0,104063584 | -0,651126722 | 0,037 | 0,133 | 1 | Mast | 0,636782802 | 1,57039417 | 0,278195489 |
| MRRF | 0,104063584 | -0,626309772 | 0,037 | 0,133 | 1 | Mast | 0,647831367 | 1,54361158 | 0,278195489 |
| PPP3CB-AS1 | 0,104063584 | -0,547955492 | 0,037 | 0,133 | 1 | Mast | 0,683988753 | 1,462012344 | 0,278195489 |
| TUBD1 | 0,104063584 | -0,493421632 | 0,037 | 0,133 | 1 | Mast | 0,710338393 | 1,407779743 | 0,278195489 |
| PPP6R11 | 0,104063584 | -0,422400868 | 0,037 | 0,133 | 1 | Mast | 0,746181828 | 1,340155927 | 0,278195489 |
| PITPNB | 0,104063584 | -0,509887805 | 0,037 | 0,133 | 1 | Mast | 0,70227705 | 1,423939455 | 0,278195489 |
| NUP501 | 0,104063584 | -0,46591064 | 0,037 | 0,133 | 1 | Mast | 0,72401393 | 1,381188895 | 0,278195489 |
| SEC22C1 | 0,104795129 | -0,431138849 | 0,194 | 0,4 | 1 | Mast | 0,741676082 | 1,34829749 | 0,485 |
| COX8A5 | 0,105267873 | 1,208435817 | 0,491 | 0,333 | 1 | Mast | 2,310869544 | 0,432737539 | 1,474474474 |
| PRKACB2 | 0,106081081 | -0,843658934 | 0,074 | 0,2 | 1 | Mast | 0,557228544 | 1,794595793 | 0,37 |
| NDFIP2 | 0,106194592 | 0,832238622 | 0,269 | 0,067 | 1 | Mast | 1,780445928 | 0,561657046 | 4,014925373 |
| ANXA53 | 0,107856463 | -0,658063373 | 0,167 | 0,333 | 1 | Mast | 0,633728423 | 1,577962995 | 0,501501502 |
| EIF2B3 | 0,108547046 | -0,410531568 | 0,037 | 0,133 | 1 | Mast | 0,752346117 | 1,329175465 | 0,278195489 |
| ARHGAP15 | 0,108547046 | -0,435837261 | 0,037 | 0,133 | 1 | Mast | 0,739264602 | 1,352695635 | 0,278195489 |
| CIR13 | 0,108547046 | -0,537624206 | 0,037 | 0,133 | 1 | Mast | 0,688904446 | 1,451580122 | 0,278195489 |
| NCBP21 | 0,108547046 | -0,483412197 | 0,037 | 0,133 | 1 | Mast | 0,715283865 | 1,398046355 | 0,278195489 |

| Supplementary Table 4 - clusermarker | | | | | | | | | |
|---|---|---|---|---|---|---|---|---|---|
| | **p_val** | **avg_log2FC** | **pct.1** | **pct.2** | **p_val_adj** | **celltype** | **Foldchange_UP** | **Foldchange_DOWN** | **Ratio_pct1_pct2** |
| SCOC | 0,108547046 | -0,427548735 | 0,037 | 0,133 | 1 | Mast | 0,743524025 | 1,344946452 | 0,278195489 |
| BTN2A1 | 0,108547046 | -0,452902591 | 0,037 | 0,133 | 1 | Mast | 0,730571515 | 1,368791391 | 0,278195489 |
| ZDHHC21 | 0,108547046 | -0,641887701 | 0,037 | 0,133 | 1 | Mast | 0,640873846 | 1,560369497 | 0,278195489 |
| PCID2 | 0,108547046 | -0,387354121 | 0,037 | 0,133 | 1 | Mast | 0,764530456 | 1,307992366 | 0,278195489 |
| MIS18BP1 | 0,108547046 | -0,473072038 | 0,037 | 0,133 | 1 | Mast | 0,7204289 | 1,388062027 | 0,278195489 |
| USP31 | 0,108547046 | -0,503958183 | 0,037 | 0,133 | 1 | Mast | 0,705169419 | 1,418098931 | 0,278195489 |
| TOB13 | 0,108547046 | -0,50448245 | 0,037 | 0,133 | 1 | Mast | 0,704913211 | 1,418614354 | 0,278195489 |
| MAVS | 0,108547046 | -0,367990877 | 0,037 | 0,133 | 1 | Mast | 0,774860831 | 1,29055433 | 0,278195489 |
| NRIP12 | 0,108938397 | -0,794467305 | 0,13 | 0,267 | 1 | Mast | 0,576556019 | 1,73443684 | 0,486891386 |
| POLR2A2 | 0,10939392 | -0,749764477 | 0,269 | 0,467 | 1 | Mast | 0,594700636 | 1,681518297 | 0,576017131 |
| ETV61 | 0,109497102 | -0,593982886 | 0,074 | 0,2 | 1 | Mast | 0,662511367 | 1,509408063 | 0,37 |
| NCOA72 | 0,109499437 | -0,554558508 | 0,074 | 0,2 | 1 | Mast | 0,680865384 | 1,468719109 | 0,37 |
| PSMA74 | 0,110753771 | 0,887349198 | 0,426 | 0,2 | 1 | Mast | 1,849774231 | 0,540606515 | 2,13 |
| GPBP13 | 0,111640307 | -0,583800771 | 0,167 | 0,333 | 1 | Mast | 0,667203716 | 1,498792612 | 0,501501502 |
| GLUL3 | 0,112056888 | -0,749511087 | 0,694 | 0,8 | 1 | Mast | 0,594805096 | 1,681222987 | 0,8675 |
| TPSAB1 | 0,112211437 | 0,439443191 | 0,963 | 1 | 1 | Mast | 1,356080846 | 0,737419161 | 0,963 |
| PHTF11 | 0,11300429 | -0,567675338 | 0,074 | 0,2 | 1 | Mast | 0,674703084 | 1,482133435 | 0,37 |
| DHX362 | 0,11300429 | -0,549189197 | 0,074 | 0,2 | 1 | Mast | 0,683404098 | 1,463263102 | 0,37 |
| LMNB11 | 0,11300429 | -0,379101785 | 0,074 | 0,2 | 1 | Mast | 0,768916165 | 1,300531899 | 0,37 |
| CHD7 | 0,11300429 | -0,584385601 | 0,074 | 0,2 | 1 | Mast | 0,666933304 | 1,499400305 | 0,37 |
| NGRN1 | 0,11300429 | -0,507840766 | 0,074 | 0,2 | 1 | Mast | 0,703274218 | 1,42192046 | 0,37 |
| CHKB3 | 0,11300429 | -0,582145594 | 0,074 | 0,2 | 1 | Mast | 0,667969626 | 1,497074061 | 0,37 |
| IPO11 | 0,113181985 | -0,324865692 | 0,037 | 0,133 | 1 | Mast | 0,798372708 | 1,252547827 | 0,278195489 |
| EPN2 | 0,113181985 | -0,593265525 | 0,037 | 0,133 | 1 | Mast | 0,662840875 | 1,508657716 | 0,278195489 |
| MRPS71 | 0,113181985 | -0,432204779 | 0,037 | 0,133 | 1 | Mast | 0,7411283 | 1,349294042 | 0,278195489 |
| ARPC24 | 0,113537875 | 0,797098745 | 0,528 | 0,4 | 1 | Mast | 1,737603296 | 0,575505354 | 1,32 |
| NUDC1 | 0,114093986 | 0,966344288 | 0,148 | 0 | 1 | Mast | 1,953883281 | 0,511801298 | #DIV/0! |
| KRT1 | 0,114093986 | 1,141834032 | 0,148 | 0 | 1 | Mast | 2,206613616 | 0,4531831 | #DIV/0! |
| FNDC3A | 0,114093986 | 0,979538651 | 0,148 | 0 | 1 | Mast | 1,97183475 | 0,507141889 | #DIV/0! |
| UBE2A1 | 0,114093986 | 1,304950453 | 0,148 | 0 | 1 | Mast | 2,470752419 | 0,404735008 | #DIV/0! |
| UBR41 | 0,114095693 | 0,918554398 | 0,148 | 0 | 1 | Mast | 1,890220316 | 0,529038859 | #DIV/0! |

| Supplementary Table 4 - clusermarker | | | | | | | | | |
|---|---|---|---|---|---|---|---|---|---|
| | **p_val** | **avg_log2FC** | **pct.1** | **pct.2** | **p_val_adj** | **celltype** | **Foldchange_UP** | **Foldchange_DOWN** | **Ratio_pct1_pct2** |
| PLEKHB21 | 0,114095693 | 0,916816378 | 0,148 | 0 | 1 | Mast | 1,887944531 | 0,529676579 | #DIV/0! |
| TADA32 | 0,114095693 | 0,925809627 | 0,148 | 0 | 1 | Mast | 1,899750065 | 0,526385033 | #DIV/0! |
| SELENOT | 0,114095693 | 0,842050481 | 0,148 | 0 | 1 | Mast | 1,792596123 | 0,557850141 | #DIV/0! |
| DNAJB111 | 0,114095693 | 1,073381576 | 0,148 | 0 | 1 | Mast | 2,104360063 | 0,475203848 | #DIV/0! |
| HINT16 | 0,114095693 | 1,128075916 | 0,148 | 0 | 1 | Mast | 2,185670487 | 0,457525508 | #DIV/0! |
| ZYX2 | 0,114095693 | 0,781927441 | 0,148 | 0 | 1 | Mast | 1,719426493 | 0,58158927 | #DIV/0! |
| CKAP5 | 0,114095693 | 0,879386818 | 0,148 | 0 | 1 | Mast | 1,839593261 | 0,543598425 | #DIV/0! |
| DNAJC43 | 0,114095693 | 0,962776295 | 0,148 | 0 | 1 | Mast | 1,949057017 | 0,513068623 | #DIV/0! |
| ARL21 | 0,114095693 | 1,00174065 | 0,148 | 0 | 1 | Mast | 2,002414509 | 0,499397101 | #DIV/0! |
| ITPR21 | 0,114095693 | 1,097725836 | 0,148 | 0 | 1 | Mast | 2,140170648 | 0,46725246 | #DIV/0! |
| ZBTB1 | 0,114095693 | 0,890714358 | 0,148 | 0 | 1 | Mast | 1,854093961 | 0,539346992 | #DIV/0! |
| ATP6V1E1 | 0,114095693 | 0,955163708 | 0,148 | 0 | 1 | Mast | 1,938799624 | 0,515783058 | #DIV/0! |
| AP1S2 | 0,114095693 | 1,036164065 | 0,148 | 0 | 1 | Mast | 2,050767678 | 0,487622275 | #DIV/0! |
| LAGE33 | 0,114095693 | 1,12173858 | 0,148 | 0 | 1 | Mast | 2,176090535 | 0,459539704 | #DIV/0! |
| ACTR32 | 0,11429126 | 0,970816231 | 0,398 | 0,2 | 1 | Mast | 1,959949159 | 0,510217316 | 1,99 |
| DOCK2 | 0,114602782 | -0,949845994 | 0,13 | 0,267 | 1 | Mast | 0,517687721 | 1,931666444 | 0,486891386 |
| HECW2-AS1 | 0,115778638 | 1,534905287 | 0,306 | 0,133 | 1 | Mast | 2,89769407 | 0,345101993 | 2,30075188 |
| ARL6IP11 | 0,116530204 | -0,508202928 | 0,398 | 0,6 | 1 | Mast | 0,703097696 | 1,422277453 | 0,663333333 |
| TLE3 | 0,116596953 | -0,497551101 | 0,074 | 0,2 | 1 | Mast | 0,708308077 | 1,411815046 | 0,37 |
| CMTM31 | 0,116596953 | -0,520119941 | 0,074 | 0,2 | 1 | Mast | 0,697313858 | 1,434074467 | 0,37 |
| RPS214 | 0,116767233 | -0,284453161 | 0,815 | 1 | 1 | Mast | 0,821052761 | 1,217948525 | 0,815 |
| SCML1 | 0,117967215 | -0,278916942 | 0,037 | 0,133 | 1 | Mast | 0,824209535 | 1,213283708 | 0,278195489 |
| ATP2A22 | 0,117971185 | 1,043152079 | 0,241 | 0,067 | 1 | Mast | 2,060725123 | 0,485266079 | 3,597014925 |
| ATPAF1 | 0,117971467 | -0,334448785 | 0,037 | 0,133 | 1 | Mast | 0,793087096 | 1,260895563 | 0,278195489 |
| MSX2 | 0,117971467 | -0,351891115 | 0,037 | 0,133 | 1 | Mast | 0,783556322 | 1,276232444 | 0,278195489 |
| LMTK21 | 0,117971467 | -0,362135846 | 0,037 | 0,133 | 1 | Mast | 0,778011915 | 1,28532736 | 0,278195489 |
| SEC23IP | 0,117971467 | -0,472069425 | 0,037 | 0,133 | 1 | Mast | 0,720929743 | 1,387097717 | 0,278195489 |
| SNAPC1 | 0,117971467 | -0,30020574 | 0,037 | 0,133 | 1 | Mast | 0,812136571 | 1,231319997 | 0,278195489 |
| ADPGK | 0,117971467 | -0,325003829 | 0,037 | 0,133 | 1 | Mast | 0,798296267 | 1,252667763 | 0,278195489 |
| PAF1 | 0,117971467 | -0,284081379 | 0,037 | 0,133 | 1 | Mast | 0,821264374 | 1,2176347 | 0,278195489 |
| ARF54 | 0,117992539 | 0,891307457 | 0,259 | 0,067 | 1 | Mast | 1,854856344 | 0,539125309 | 3,865671642 |

| Supplementary Table 4 - clusermarker | | | | | | | | | |
|---|---|---|---|---|---|---|---|---|---|
| | **p_val** | **avg_log2FC** | **pct.1** | **pct.2** | **p_val_adj** | **celltype** | **Foldchange_UP** | **Foldchange_DOWN** | **Ratio_pct1_pct2** |
| SET2 | 0,118979718 | -0,546223123 | 0,315 | 0,533 | 1 | Mast | 0,684810571 | 1,460257833 | 0,590994371 |
| CSNK1D3 | 0,119110266 | -0,728577752 | 0,176 | 0,333 | 1 | Mast | 0,603498566 | 1,657004766 | 0,528528529 |
| KTN12 | 0,119217514 | -0,545503731 | 0,213 | 0,4 | 1 | Mast | 0,685152133 | 1,459529864 | 0,5325 |
| RPS183 | 0,119342816 | -0,258527383 | 0,935 | 1 | 1 | Mast | 0,835940762 | 1,196257014 | 0,935 |
| UFM14 | 0,119709349 | -0,435432172 | 0,157 | 0,333 | 1 | Mast | 0,739472206 | 1,352315871 | 0,471471471 |
| CERT1 | 0,119710686 | -0,5242516 | 0,157 | 0,333 | 1 | Mast | 0,695319715 | 1,438187325 | 0,471471471 |
| C1D1 | 0,120241761 | -0,80683872 | 0,083 | 0,2 | 1 | Mast | 0,571633067 | 1,749373955 | 0,415 |
| ID24 | 0,122627634 | -1,198307338 | 0,287 | 0,467 | 1 | Mast | 0,435786274 | 2,294702839 | 0,614561028 |
| BST21 | 0,122955942 | -0,685112086 | 0,37 | 0,533 | 1 | Mast | 0,621957503 | 1,607826892 | 0,694183865 |
| BMP2K2 | 0,123169364 | -0,526375057 | 0,417 | 0,6 | 1 | Mast | 0,694297048 | 1,440305706 | 0,695 |
| GABARAP5 | 0,123503307 | 0,686604033 | 0,648 | 0,467 | 1 | Mast | 1,609490468 | 0,621314646 | 1,3875803 |
| ATP1A12 | 0,12353087 | -0,809175772 | 0,13 | 0,267 | 1 | Mast | 0,570707817 | 1,752210099 | 0,486891386 |
| AGAP11 | 0,124048435 | -0,47792418 | 0,074 | 0,2 | 1 | Mast | 0,718009989 | 1,392738284 | 0,37 |
| LY6E3 | 0,12405091 | -0,386532229 | 0,074 | 0,2 | 1 | Mast | 0,764966127 | 1,307247426 | 0,37 |
| GNG21 | 0,12405091 | -0,382613542 | 0,074 | 0,2 | 1 | Mast | 0,767046773 | 1,303701463 | 0,37 |
| ICAM32 | 0,12405091 | -0,358461683 | 0,074 | 0,2 | 1 | Mast | 0,77999583 | 1,282058135 | 0,37 |
| SYTL31 | 0,124854873 | -0,501137397 | 0,25 | 0,467 | 1 | Mast | 0,706549529 | 1,415328945 | 0,535331906 |
| MT-ND63 | 0,125118979 | -0,498064747 | 0,37 | 0,6 | 1 | Mast | 0,708055941 | 1,412317787 | 0,616666667 |
| ACSL4 | 0,125235865 | 0,667747814 | 0,537 | 0,333 | 1 | Mast | 1,588591087 | 0,629488613 | 1,612612613 |
| ROMO11 | 0,125680111 | -0,39816383 | 0,204 | 0,4 | 1 | Mast | 0,758823451 | 1,317829593 | 0,51 |
| ZNF3312 | 0,12728295 | -1,274264648 | 0,259 | 0,4 | 1 | Mast | 0,413435837 | 2,418755008 | 0,6475 |
| RAB32 | 0,127509159 | 1,035003398 | 0,315 | 0,133 | 1 | Mast | 2,049118473 | 0,488014731 | 2,368421053 |
| CD634 | 0,127576168 | 0,42560847 | 0,88 | 0,933 | 1 | Mast | 1,343138865 | 0,744524655 | 0,943193998 |
| TNF2 | 0,127760407 | 1,265539649 | 0,139 | 0 | 1 | Mast | 2,404171225 | 0,415943752 | #DIV/0! |
| UTS2 | 0,127762309 | 1,41720772 | 0,139 | 0 | 1 | Mast | 2,670681105 | 0,374436318 | #DIV/0! |
| NFYC1 | 0,127762309 | 0,755563245 | 0,139 | 0 | 1 | Mast | 1,688290593 | 0,592315093 | #DIV/0! |
| PSMA52 | 0,127762309 | 1,093024805 | 0,139 | 0 | 1 | Mast | 2,133208239 | 0,468777488 | #DIV/0! |
| ZBTB383 | 0,127762309 | 0,971534245 | 0,139 | 0 | 1 | Mast | 1,960924848 | 0,50996345 | #DIV/0! |
| GPAT3 | 0,127762309 | 1,068509562 | 0,139 | 0 | 1 | Mast | 2,097265577 | 0,476811335 | #DIV/0! |
| GMPR | 0,127762309 | 0,791499041 | 0,139 | 0 | 1 | Mast | 1,730872002 | 0,577743472 | #DIV/0! |
| PRRC2A | 0,127762309 | 0,894652101 | 0,139 | 0 | 1 | Mast | 1,859161503 | 0,537876886 | #DIV/0! |

| Supplementary Table 4 - clusermarker | | | | | | | | | |
|---|---|---|---|---|---|---|---|---|---|
| | **p_val** | **avg_log2FC** | **pct.1** | **pct.2** | **p_val_adj** | **celltype** | **Foldchange_UP** | **Foldchange_DOWN** | **Ratio_pct1_pct2** |
| FBXO91 | 0,127762309 | 0,898219938 | 0,139 | 0 | 1 | Mast | 1,863764967 | 0,536548341 | #DIV/0! |
| SMAP12 | 0,127762309 | 0,881686961 | 0,139 | 0 | 1 | Mast | 1,842528534 | 0,542732436 | #DIV/0! |
| FUCA21 | 0,127762309 | 0,796542558 | 0,139 | 0 | 1 | Mast | 1,736933545 | 0,575727265 | #DIV/0! |
| VWC2 | 0,127762309 | 0,924156391 | 0,139 | 0 | 1 | Mast | 1,89757432 | 0,526988582 | #DIV/0! |
| IKZF1 | 0,127762309 | 0,961679229 | 0,139 | 0 | 1 | Mast | 1,947575462 | 0,513458923 | #DIV/0! |
| YWHAG | 0,127762309 | 0,731740536 | 0,139 | 0 | 1 | Mast | 1,66064136 | 0,602176981 | #DIV/0! |
| PLEC3 | 0,127762309 | 0,908717676 | 0,139 | 0 | 1 | Mast | 1,877376071 | 0,532658329 | #DIV/0! |
| PSMD131 | 0,127762309 | 0,859942936 | 0,139 | 0 | 1 | Mast | 1,81496652 | 0,550974351 | #DIV/0! |
| TAF105 | 0,127762309 | 0,915140864 | 0,139 | 0 | 1 | Mast | 1,885753187 | 0,530292091 | #DIV/0! |
| POLR1D2 | 0,127762309 | 1,079889195 | 0,139 | 0 | 1 | Mast | 2,11387372 | 0,473065155 | #DIV/0! |
| HERC1 | 0,127762309 | 0,78187894 | 0,139 | 0 | 1 | Mast | 1,71936869 | 0,581608823 | #DIV/0! |
| BCL7C2 | 0,127762309 | 0,902389927 | 0,139 | 0 | 1 | Mast | 1,869159816 | 0,534999732 | #DIV/0! |
| SLC25A111 | 0,127762309 | 0,890485981 | 0,139 | 0 | 1 | Mast | 1,853800483 | 0,539432376 | #DIV/0! |
| TXNDC172 | 0,127762309 | 0,75266596 | 0,139 | 0 | 1 | Mast | 1,684903493 | 0,593505802 | #DIV/0! |
| NDUFA72 | 0,127762309 | 0,96628312 | 0,139 | 0 | 1 | Mast | 1,953800442 | 0,511822998 | #DIV/0! |
| LSM14A1 | 0,127762309 | 0,903474582 | 0,139 | 0 | 1 | Mast | 1,870565627 | 0,534597656 | #DIV/0! |
| FDPS1 | 0,127914771 | -0,335261449 | 0,074 | 0,2 | 1 | Mast | 0,792640479 | 1,26160602 | 0,37 |
| VTI1A | 0,128026146 | -0,399155665 | 0,037 | 0,133 | 1 | Mast | 0,758301948 | 1,318735897 | 0,278195489 |
| WNT11 | 0,128026146 | -0,379598383 | 0,037 | 0,133 | 1 | Mast | 0,768651538 | 1,300979639 | 0,278195489 |
| PIP5K1C | 0,128026146 | -0,341488633 | 0,037 | 0,133 | 1 | Mast | 0,789226535 | 1,267063329 | 0,278195489 |
| HPGD | 0,128227699 | 1,015291944 | 0,852 | 0,867 | 1 | Mast | 2,021311885 | 0,494728205 | 0,982698962 |
| BZW13 | 0,130167895 | -0,413699841 | 0,454 | 0,667 | 1 | Mast | 0,750695718 | 1,332097648 | 0,68065967 |
| SH3KBP1 | 0,131046056 | 0,936728081 | 0,241 | 0,067 | 1 | Mast | 1,914182098 | 0,522416337 | 3,597014925 |
| SNHG62 | 0,131051417 | -0,360443835 | 0,278 | 0,467 | 1 | Mast | 0,778924912 | 1,283820795 | 0,595289079 |
| PRNP2 | 0,131301877 | 0,711752197 | 0,352 | 0,133 | 1 | Mast | 1,637792058 | 0,610578123 | 2,646616541 |
| RANBP21 | 0,131762096 | -0,73489431 | 0,12 | 0,267 | 1 | Mast | 0,600862041 | 1,664275543 | 0,449438202 |
| SMIM262 | 0,131762096 | -0,394965007 | 0,12 | 0,267 | 1 | Mast | 0,760507823 | 1,314910866 | 0,449438202 |
| NDRG21 | 0,131871576 | -0,370579028 | 0,074 | 0,2 | 1 | Mast | 0,773472 | 1,292871623 | 0,37 |
| SERINC3 | 0,133439371 | 1,278424799 | 0,222 | 0,067 | 1 | Mast | 2,425739788 | 0,412245372 | 3,313432836 |
| PLAUR1 | 0,134490699 | -0,594026062 | 0,287 | 0,467 | 1 | Mast | 0,662491541 | 1,509453236 | 0,614561028 |
| GIHCG1 | 0,135251183 | -0,80167289 | 0,083 | 0,2 | 1 | Mast | 0,573683572 | 1,743121207 | 0,415 |

| Supplementary Table 4 - clusermarker | | | | | | | | | |
|---|---|---|---|---|---|---|---|---|---|
| | **p_val** | **avg_log2FC** | **pct.1** | **pct.2** | **p_val_adj** | **celltype** | **Foldchange_UP** | **Foldchange_DOWN** | **Ratio_pct1_pct2** |
| MED131 | 0,135251183 | -0,676678436 | 0,083 | 0,2 | 1 | Mast | 0,625603966 | 1,598455339 | 0,415 |
| NDUFB13 | 0,135833962 | -0,671029837 | 0,176 | 0,333 | 1 | Mast | 0,628058201 | 1,592209127 | 0,528528529 |
| ERCC5 | 0,135922574 | -0,459698399 | 0,074 | 0,2 | 1 | Mast | 0,727138254 | 1,375254286 | 0,37 |
| UBALD21 | 0,135922574 | -0,327714689 | 0,074 | 0,2 | 1 | Mast | 0,796797658 | 1,25502377 | 0,37 |
| DDX57 | 0,136253956 | -0,360232811 | 0,75 | 0,867 | 1 | Mast | 0,779038854 | 1,283633023 | 0,865051903 |
| DDIT41 | 0,13626479 | -0,607769559 | 0,426 | 0,667 | 1 | Mast | 0,656210435 | 1,523901398 | 0,63868066 |
| ANAPC163 | 0,1364172 | -0,403781555 | 0,204 | 0,4 | 1 | Mast | 0,755874406 | 1,322971108 | 0,51 |
| LDHB2 | 0,138091124 | -0,47693633 | 0,333 | 0,533 | 1 | Mast | 0,718501797 | 1,391784967 | 0,624765478 |
| ACOT7 | 0,138493208 | -0,472053805 | 0,12 | 0,267 | 1 | Mast | 0,720937548 | 1,387082699 | 0,449438202 |
| KDM6A | 0,139220836 | -0,637058147 | 0,083 | 0,2 | 1 | Mast | 0,643022824 | 1,55515475 | 0,415 |
| AGO31 | 0,139606254 | -0,74275522 | 0,13 | 0,267 | 1 | Mast | 0,597596987 | 1,673368543 | 0,486891386 |
| SRSF32 | 0,141040064 | -0,600255124 | 0,222 | 0,4 | 1 | Mast | 0,659637296 | 1,515984628 | 0,555 |
| RSBN1 | 0,141794393 | -0,531784333 | 0,176 | 0,333 | 1 | Mast | 0,691698708 | 1,44571616 | 0,528528529 |
| IFNGR2 | 0,142346541 | 1,013800038 | 0,231 | 0,067 | 1 | Mast | 2,019222705 | 0,495240073 | 3,447761194 |
| TMEM50A4 | 0,142564554 | 1,002313342 | 0,315 | 0,133 | 1 | Mast | 2,003209546 | 0,499198899 | 2,368421053 |
| UBR51 | 0,143007557 | 0,875943258 | 0,13 | 0 | 1 | Mast | 1,835207583 | 0,544897487 | #DIV/0! |
| OPTN1 | 0,143007557 | 0,748950765 | 0,13 | 0 | 1 | Mast | 1,68057015 | 0,595036155 | #DIV/0! |
| NDUFS32 | 0,143007557 | 0,811757329 | 0,13 | 0 | 1 | Mast | 1,755348309 | 0,569687506 | #DIV/0! |
| KPNB12 | 0,143007557 | 0,996356003 | 0,13 | 0 | 1 | Mast | 1,994954722 | 0,501264509 | #DIV/0! |
| CAMTA13 | 0,143009677 | 0,70359312 | 0,13 | 0 | 1 | Mast | 1,628555763 | 0,614040994 | #DIV/0! |
| DPYD1 | 0,143009677 | 0,85623594 | 0,13 | 0 | 1 | Mast | 1,810308961 | 0,552391896 | #DIV/0! |
| RNF115 | 0,143009677 | 0,998247057 | 0,13 | 0 | 1 | Mast | 1,99757138 | 0,500607893 | #DIV/0! |
| RGS16 | 0,143009677 | 0,962322649 | 0,13 | 0 | 1 | Mast | 1,948444245 | 0,513229979 | #DIV/0! |
| PYCR21 | 0,143009677 | 0,675958401 | 0,13 | 0 | 1 | Mast | 1,597657765 | 0,625916277 | #DIV/0! |
| SF3B63 | 0,143009677 | 0,706969446 | 0,13 | 0 | 1 | Mast | 1,632371521 | 0,61260564 | #DIV/0! |
| IGKV1-53 | 0,143009677 | 1,748139225 | 0,13 | 0 | 1 | Mast | 3,359250135 | 0,297685483 | #DIV/0! |
| UQCRC12 | 0,143009677 | 0,938343782 | 0,13 | 0 | 1 | Mast | 1,916327026 | 0,521831601 | #DIV/0! |
| TWF22 | 0,143009677 | 0,886793464 | 0,13 | 0 | 1 | Mast | 1,849061826 | 0,540814799 | #DIV/0! |
| JCHAIN4 | 0,143009677 | 1,187376916 | 0,13 | 0 | 1 | Mast | 2,277382966 | 0,4391005 | #DIV/0! |
| PSMB93 | 0,143009677 | 0,955097617 | 0,13 | 0 | 1 | Mast | 1,938710808 | 0,515806687 | #DIV/0! |
| TMEM14A1 | 0,143009677 | 1,176216414 | 0,13 | 0 | 1 | Mast | 2,259833394 | 0,442510498 | #DIV/0! |

| Supplementary Table 4 - clusermarker | | | | | | | | | |
|---|---|---|---|---|---|---|---|---|---|
| | **p_val** | **avg_log2FC** | **pct.1** | **pct.2** | **p_val_adj** | **celltype** | **Foldchange_UP** | **Foldchange_DOWN** | **Ratio_pct1_pct2** |
| STARD3NL | 0,143009677 | 0,781279596 | 0,13 | 0 | 1 | Mast | 1,718654554 | 0,581850493 | #DIV/0! |
| RSBN1L1 | 0,143009677 | 0,8352726 | 0,13 | 0 | 1 | Mast | 1,784194134 | 0,560477126 | #DIV/0! |
| GEM2 | 0,143009677 | 1,085636965 | 0,13 | 0 | 1 | Mast | 2,122312299 | 0,471184189 | #DIV/0! |
| UBE2R2 | 0,143009677 | 0,834417815 | 0,13 | 0 | 1 | Mast | 1,783137326 | 0,560809302 | #DIV/0! |
| GRHPR1 | 0,143009677 | 0,900922715 | 0,13 | 0 | 1 | Mast | 1,867259858 | 0,5355441 | #DIV/0! |
| PRRC2B4 | 0,143009677 | 0,914139749 | 0,13 | 0 | 1 | Mast | 1,884445078 | 0,530660199 | #DIV/0! |
| TMEM1414 | 0,143009677 | 0,712731888 | 0,13 | 0 | 1 | Mast | 1,638904611 | 0,610163638 | #DIV/0! |
| ILK3 | 0,143009677 | 0,838415812 | 0,13 | 0 | 1 | Mast | 1,78808561 | 0,559257339 | #DIV/0! |
| POLR2G | 0,143009677 | 0,884140184 | 0,13 | 0 | 1 | Mast | 1,845664316 | 0,541810334 | #DIV/0! |
| CHKA1 | 0,143009677 | 0,973516699 | 0,13 | 0 | 1 | Mast | 1,96362127 | 0,509263174 | #DIV/0! |
| CARD161 | 0,143009677 | 0,9530577 | 0,13 | 0 | 1 | Mast | 1,935971481 | 0,516536535 | #DIV/0! |
| MIA2 | 0,143009677 | 0,860732943 | 0,13 | 0 | 1 | Mast | 1,815960652 | 0,550672725 | #DIV/0! |
| PTGER21 | 0,143009677 | 0,860079122 | 0,13 | 0 | 1 | Mast | 1,815137856 | 0,550922343 | #DIV/0! |
| VPS13C3 | 0,143009677 | 0,814381074 | 0,13 | 0 | 1 | Mast | 1,758543563 | 0,56865239 | #DIV/0! |
| GNPTG4 | 0,143009677 | 0,940312069 | 0,13 | 0 | 1 | Mast | 1,91894328 | 0,521120145 | #DIV/0! |
| GSPT12 | 0,143009677 | 0,850830428 | 0,13 | 0 | 1 | Mast | 1,803538759 | 0,554465489 | #DIV/0! |
| CORO1A1 | 0,143009677 | 0,897414566 | 0,13 | 0 | 1 | Mast | 1,862724826 | 0,536847948 | #DIV/0! |
| ITGA2B | 0,143009677 | 0,875067753 | 0,13 | 0 | 1 | Mast | 1,834094219 | 0,54522826 | #DIV/0! |
| KXD12 | 0,143009677 | 0,896968735 | 0,13 | 0 | 1 | Mast | 1,862149285 | 0,537013873 | #DIV/0! |
| NKG7 | 0,143009677 | 1,139326333 | 0,13 | 0 | 1 | Mast | 2,202781402 | 0,45397151 | #DIV/0! |
| PFKL1 | 0,143009677 | 0,811719139 | 0,13 | 0 | 1 | Mast | 1,755301844 | 0,569702586 | #DIV/0! |
| IGLV3-214 | 0,143009677 | 1,400961671 | 0,13 | 0 | 1 | Mast | 2,640775522 | 0,378676639 | #DIV/0! |
| PDS5A2 | 0,143279594 | -0,548633374 | 0,083 | 0,2 | 1 | Mast | 0,683667442 | 1,462699464 | 0,415 |
| RSL24D11 | 0,144240527 | -0,48160117 | 0,167 | 0,333 | 1 | Mast | 0,71618233 | 1,396292477 | 0,501501502 |
| SNX131 | 0,144312055 | -0,277846509 | 0,074 | 0,2 | 1 | Mast | 0,824821298 | 1,212383825 | 0,37 |
| AKR1A12 | 0,145199157 | 1,084010559 | 0,222 | 0,067 | 1 | Mast | 2,119921083 | 0,471715673 | 3,313432836 |
| HNRNPDL1 | 0,145422297 | 0,949745757 | 0,38 | 0,2 | 1 | Mast | 1,931532239 | 0,517723691 | 1,9 |
| RBBP61 | 0,147179487 | -0,99566701 | 0,148 | 0,267 | 1 | Mast | 0,501503957 | 1,994002211 | 0,554307116 |
| WSB12 | 0,147388686 | -0,351564923 | 0,167 | 0,333 | 1 | Mast | 0,783733504 | 1,275943921 | 0,501501502 |
| IMP4 | 0,14742852 | -0,638355759 | 0,083 | 0,2 | 1 | Mast | 0,642444727 | 1,556554142 | 0,415 |
| CCNDBP12 | 0,14742852 | -0,764195174 | 0,083 | 0,2 | 1 | Mast | 0,588781736 | 1,698422248 | 0,415 |

| Supplementary Table 4 - clusermarker | | | | | | | | | |
|---|---|---|---|---|---|---|---|---|---|
| | **p_val** | **avg_log2FC** | **pct.1** | **pct.2** | **p_val_adj** | **celltype** | **Foldchange_UP** | **Foldchange_DOWN** | **Ratio_pct1_pct2** |
| ARID4B2 | 0,148329186 | -0,694183069 | 0,194 | 0,333 | 1 | Mast | 0,618059198 | 1,617967994 | 0,582582583 |
| USP482 | 0,14833047 | 0,978141223 | 0,231 | 0,067 | 1 | Mast | 1,969925709 | 0,507633357 | 3,447761194 |
| H1-102 | 0,14833047 | 1,071565586 | 0,231 | 0,067 | 1 | Mast | 2,101712871 | 0,475802387 | 3,447761194 |
| EXOC6B | 0,148809045 | -0,828420703 | 0,046 | 0,133 | 1 | Mast | 0,563145372 | 1,775740422 | 0,345864662 |
| CYTH3 | 0,148809045 | -0,664880716 | 0,046 | 0,133 | 1 | Mast | 0,630740853 | 1,585437182 | 0,345864662 |
| NDUFAF1 | 0,148809045 | -0,65960713 | 0,046 | 0,133 | 1 | Mast | 0,633050664 | 1,579652399 | 0,345864662 |
| CD69 | 0,14987886 | -0,918643564 | 0,713 | 0,867 | 1 | Mast | 0,529006163 | 1,890337145 | 0,822376009 |
| ANKRD11 | 0,149997121 | -0,659514719 | 0,13 | 0,267 | 1 | Mast | 0,633091215 | 1,579551219 | 0,486891386 |
| TMEM594 | 0,150285836 | 0,926547497 | 0,324 | 0,133 | 1 | Mast | 1,900721946 | 0,52611588 | 2,436090226 |
| EIF1B1 | 0,151369108 | -0,732624931 | 0,185 | 0,333 | 1 | Mast | 0,60180795 | 1,661659672 | 0,555555556 |
| BBLN5 | 0,153221684 | 0,868759918 | 0,389 | 0,2 | 1 | Mast | 1,826092591 | 0,547617358 | 1,945 |
| NFKB2 | 0,153991703 | -0,732246357 | 0,139 | 0,267 | 1 | Mast | 0,601965889 | 1,661223698 | 0,520599251 |
| MLLT3 | 0,154301515 | -0,747629095 | 0,046 | 0,133 | 1 | Mast | 0,595581524 | 1,679031266 | 0,345864662 |
| KIF22 | 0,154301515 | -0,708198981 | 0,046 | 0,133 | 1 | Mast | 0,61208377 | 1,6337633 | 0,345864662 |
| PRELID15 | 0,154528034 | 0,943963698 | 0,222 | 0,067 | 1 | Mast | 1,923806501 | 0,519802797 | 3,313432836 |
| SERF23 | 0,1558282 | 0,630857826 | 0,731 | 0,6 | 1 | Mast | 1,548485448 | 0,645792313 | 1,218333333 |
| IRF2BP2 | 0,15600105 | -0,604089053 | 0,083 | 0,2 | 1 | Mast | 0,657886652 | 1,520018679 | 0,415 |
| MT-ND51 | 0,156175838 | -0,273957235 | 0,75 | 1 | 1 | Mast | 0,827047884 | 1,209119834 | 0,75 |
| EIF4B5 | 0,156363326 | -0,668353383 | 0,269 | 0,4 | 1 | Mast | 0,629224442 | 1,589258035 | 0,6725 |
| SOD23 | 0,157245427 | -0,552417162 | 0,13 | 0,267 | 1 | Mast | 0,681876721 | 1,466540753 | 0,486891386 |
| IDI12 | 0,157245427 | -0,677722451 | 0,13 | 0,267 | 1 | Mast | 0,625151408 | 1,59961249 | 0,486891386 |
| MLEC3 | 0,157573213 | 1,0165402 | 0,213 | 0,067 | 1 | Mast | 2,023061531 | 0,494300339 | 3,179104478 |
| SOD12 | 0,157892664 | 0,877951015 | 0,426 | 0,267 | 1 | Mast | 1,837763366 | 0,544139696 | 1,595505618 |
| COA61 | 0,159946677 | -0,62648997 | 0,046 | 0,133 | 1 | Mast | 0,647750455 | 1,543804395 | 0,345864662 |
| UBP11 | 0,159946677 | -0,485228232 | 0,046 | 0,133 | 1 | Mast | 0,714384046 | 1,399807296 | 0,345864662 |
| PLEKHF2 | 0,159946677 | -0,520955238 | 0,046 | 0,133 | 1 | Mast | 0,696910242 | 1,434905014 | 0,345864662 |
| GANAB | 0,159946677 | -0,682836439 | 0,046 | 0,133 | 1 | Mast | 0,622939328 | 1,605292772 | 0,345864662 |
| COA82 | 0,159946677 | -0,808560234 | 0,046 | 0,133 | 1 | Mast | 0,570951366 | 1,751462664 | 0,345864662 |
| NSF1 | 0,159946677 | -0,444201578 | 0,046 | 0,133 | 1 | Mast | 0,734990966 | 1,360560941 | 0,345864662 |
| TMEM2481 | 0,160035375 | 0,802371249 | 0,12 | 0 | 1 | Mast | 1,743965196 | 0,573405939 | #DIV/0! |
| WASF22 | 0,16003774 | 0,784560832 | 0,12 | 0 | 1 | Mast | 1,722567876 | 0,580528648 | #DIV/0! |

| Supplementary Table 4 - clusermarker | | | | | | | | | |
|---|---|---|---|---|---|---|---|---|---|
| | **p_val** | **avg_log2FC** | **pct.1** | **pct.2** | **p_val_adj** | **celltype** | **Foldchange_UP** | **Foldchange_DOWN** | **Ratio_pct1_pct2** |
| C1orf1222 | 0,16003774 | 0,58660643 | 0,12 | 0 | 1 | Mast | 1,501710202 | 0,665907443 | #DIV/0! |
| CTSS5 | 0,16003774 | 0,736292849 | 0,12 | 0 | 1 | Mast | 1,665889662 | 0,600279852 | #DIV/0! |
| LGALS8 | 0,16003774 | 0,797365266 | 0,12 | 0 | 1 | Mast | 1,737924327 | 0,575399046 | #DIV/0! |
| AHCTF11 | 0,16003774 | 0,801505987 | 0,12 | 0 | 1 | Mast | 1,742919559 | 0,573749944 | #DIV/0! |
| RNF144A | 0,16003774 | 0,936503047 | 0,12 | 0 | 1 | Mast | 1,913883543 | 0,522497831 | #DIV/0! |
| ANKRD361 | 0,16003774 | 0,837021007 | 0,12 | 0 | 1 | Mast | 1,786357716 | 0,559798293 | #DIV/0! |
| DOCK101 | 0,16003774 | 0,734285449 | 0,12 | 0 | 1 | Mast | 1,663573316 | 0,601115677 | #DIV/0! |
| FBLN22 | 0,16003774 | 0,616365487 | 0,12 | 0 | 1 | Mast | 1,533008278 | 0,652312199 | #DIV/0! |
| REST1 | 0,16003774 | 0,773401139 | 0,12 | 0 | 1 | Mast | 1,709294682 | 0,58503663 | #DIV/0! |
| EIF2AK12 | 0,16003774 | 0,766392783 | 0,12 | 0 | 1 | Mast | 1,701011368 | 0,587885548 | #DIV/0! |
| SLC4A2 | 0,16003774 | 0,789179294 | 0,12 | 0 | 1 | Mast | 1,728091124 | 0,578673188 | #DIV/0! |
| FDFT1 | 0,16003774 | 0,861154097 | 0,12 | 0 | 1 | Mast | 1,816490848 | 0,550511995 | #DIV/0! |
| VDAC31 | 0,16003774 | 0,828357292 | 0,12 | 0 | 1 | Mast | 1,775662373 | 0,563170125 | #DIV/0! |
| PAXX3 | 0,16003774 | 0,814597647 | 0,12 | 0 | 1 | Mast | 1,75880757 | 0,568567032 | #DIV/0! |
| PCGF51 | 0,16003774 | 0,814553505 | 0,12 | 0 | 1 | Mast | 1,758753757 | 0,568584429 | #DIV/0! |
| HPS11 | 0,16003774 | 0,803715372 | 0,12 | 0 | 1 | Mast | 1,745590762 | 0,572871959 | #DIV/0! |
| MRPL432 | 0,16003774 | 0,861504639 | 0,12 | 0 | 1 | Mast | 1,816932267 | 0,550378249 | #DIV/0! |
| COPB13 | 0,16003774 | 0,835033598 | 0,12 | 0 | 1 | Mast | 1,783898583 | 0,560569984 | #DIV/0! |
| FNBP4 | 0,16003774 | 0,976801672 | 0,12 | 0 | 1 | Mast | 1,96809747 | 0,508104916 | #DIV/0! |
| FKBP31 | 0,16003774 | 0,665387233 | 0,12 | 0 | 1 | Mast | 1,585993912 | 0,630519444 | #DIV/0! |
| TMX1 | 0,16003774 | 0,723171488 | 0,12 | 0 | 1 | Mast | 1,650807031 | 0,605764321 | #DIV/0! |
| PPM1A1 | 0,16003774 | 0,796395033 | 0,12 | 0 | 1 | Mast | 1,736755942 | 0,57578614 | #DIV/0! |
| ZNF1061 | 0,16003774 | 0,736729618 | 0,12 | 0 | 1 | Mast | 1,666394078 | 0,600098148 | #DIV/0! |
| NME42 | 0,16003774 | 1,132605033 | 0,12 | 0 | 1 | Mast | 2,192542843 | 0,45609143 | #DIV/0! |
| SMG11 | 0,16003774 | 0,550637229 | 0,12 | 0 | 1 | Mast | 1,464732516 | 0,682718509 | #DIV/0! |
| DEF81 | 0,16003774 | 0,859122153 | 0,12 | 0 | 1 | Mast | 1,813934237 | 0,551287902 | #DIV/0! |
| COPS3 | 0,16003774 | 0,85467507 | 0,12 | 0 | 1 | Mast | 1,808351424 | 0,552989859 | #DIV/0! |
| NPEPPS1 | 0,16003774 | 0,792054894 | 0,12 | 0 | 1 | Mast | 1,731539015 | 0,577520917 | #DIV/0! |
| TMEM1474 | 0,16003774 | 0,772657072 | 0,12 | 0 | 1 | Mast | 1,708413344 | 0,585338439 | #DIV/0! |
| RANBP11 | 0,16003774 | 0,806867716 | 0,12 | 0 | 1 | Mast | 1,749409115 | 0,571621579 | #DIV/0! |
| POLR2F2 | 0,16003774 | 0,846858065 | 0,12 | 0 | 1 | Mast | 1,798579668 | 0,555994276 | #DIV/0! |

| Supplementary Table 4 - clusermarker | | | | | | | | | |
|---|---|---|---|---|---|---|---|---|---|
| | **p_val** | **avg_log2FC** | **pct.1** | **pct.2** | **p_val_adj** | **celltype** | **Foldchange_UP** | **Foldchange_DOWN** | **Ratio_pct1_pct2** |
| TDRD31 | 0,160426693 | -0,42009206 | 0,083 | 0,2 | 1 | Mast | 0,747376932 | 1,338012932 | 0,415 |
| RARA1 | 0,160872163 | -0,792366599 | 0,194 | 0,333 | 1 | Mast | 0,577396153 | 1,731913166 | 0,582582583 |
| CRBN | 0,160967693 | -0,951215049 | 0,13 | 0,267 | 1 | Mast | 0,517196691 | 1,933500382 | 0,486891386 |
| TUBB4B2 | 0,162412401 | -0,957428354 | 0,361 | 0,533 | 1 | Mast | 0,514974052 | 1,941845412 | 0,677298311 |
| RPS82 | 0,162990683 | 0,34566997 | 1 | 1 | 1 | Mast | 1,270740962 | 0,786942445 | 1 |
| LCP11 | 0,163045184 | 0,734903928 | 0,454 | 0,267 | 1 | Mast | 1,664286638 | 0,600858036 | 1,700374532 |
| CLINT11 | 0,164305372 | 0,742545897 | 0,222 | 0,067 | 1 | Mast | 1,673125769 | 0,5976837 | 3,313432836 |
| MRPL421 | 0,164944009 | -0,515971992 | 0,083 | 0,2 | 1 | Mast | 0,699321617 | 1,429957226 | 0,415 |
| SCIN1 | 0,164946578 | -0,357998892 | 0,083 | 0,2 | 1 | Mast | 0,78024608 | 1,28164694 | 0,415 |
| NARF | 0,164946578 | -0,46861043 | 0,083 | 0,2 | 1 | Mast | 0,722660312 | 1,383776006 | 0,415 |
| LRIF1 | 0,165746626 | -0,723493303 | 0,046 | 0,133 | 1 | Mast | 0,605629211 | 1,65117531 | 0,345864662 |
| CCDC93 | 0,165746626 | -0,514036823 | 0,046 | 0,133 | 1 | Mast | 0,700260287 | 1,428040428 | 0,345864662 |
| CDKN1C3 | 0,165746626 | -0,624445011 | 0,046 | 0,133 | 1 | Mast | 0,648669265 | 1,541617668 | 0,345864662 |
| FADS3 | 0,165746626 | -0,581831626 | 0,046 | 0,133 | 1 | Mast | 0,668115009 | 1,496748295 | 0,345864662 |
| CTDSP2 | 0,165746626 | -0,615612498 | 0,046 | 0,133 | 1 | Mast | 0,65265275 | 1,53220836 | 0,345864662 |
| GPN3 | 0,165746626 | -0,679378482 | 0,046 | 0,133 | 1 | Mast | 0,624434225 | 1,601449696 | 0,345864662 |
| AKT11 | 0,165746626 | -0,49734917 | 0,046 | 0,133 | 1 | Mast | 0,708407224 | 1,41161745 | 0,345864662 |
| TRADD2 | 0,165746626 | -0,411213964 | 0,046 | 0,133 | 1 | Mast | 0,751990341 | 1,329804315 | 0,345864662 |
| HSD17B101 | 0,165746626 | -0,578539705 | 0,046 | 0,133 | 1 | Mast | 0,669641245 | 1,493336928 | 0,345864662 |
| TPM44 | 0,166438609 | 0,680641604 | 0,407 | 0,2 | 1 | Mast | 1,602852427 | 0,623887754 | 2,035 |
| ARHGAP183 | 0,168490281 | -0,408491448 | 0,444 | 0,667 | 1 | Mast | 0,753410765 | 1,327297202 | 0,665667166 |
| CLDND11 | 0,168611341 | -0,443595288 | 0,13 | 0,267 | 1 | Mast | 0,735299909 | 1,359989288 | 0,486891386 |
| BAZ1A3 | 0,168693161 | -0,659805318 | 0,139 | 0,267 | 1 | Mast | 0,632963706 | 1,579869416 | 0,520599251 |
| SSR21 | 0,168979049 | -0,480089831 | 0,37 | 0,533 | 1 | Mast | 0,716932982 | 1,394830515 | 0,694183865 |
| RPS92 | 0,169065933 | 0,398113182 | 0,759 | 0,667 | 1 | Mast | 1,31778333 | 0,758850091 | 1,137931034 |
| DNAJB62 | 0,169364099 | -0,465101226 | 0,185 | 0,333 | 1 | Mast | 0,724420247 | 1,380414206 | 0,555555556 |
| MAX1 | 0,169561674 | -0,319243722 | 0,083 | 0,2 | 1 | Mast | 0,801489918 | 1,24767633 | 0,415 |
| MFAP1 | 0,169561674 | -0,327446025 | 0,083 | 0,2 | 1 | Mast | 0,796946054 | 1,254790077 | 0,415 |
| CIRBP3 | 0,169797849 | 0,73232205 | 0,574 | 0,4 | 1 | Mast | 1,661310858 | 0,601934307 | 1,435 |
| IL18R1 | 0,171241947 | -0,37344917 | 0,231 | 0,4 | 1 | Mast | 0,771934761 | 1,295446262 | 0,5775 |
| MED11 | 0,171699113 | -0,458119128 | 0,046 | 0,133 | 1 | Mast | 0,727934664 | 1,373749663 | 0,345864662 |

| Supplementary Table 4 - clusermarker | | | | | | | | | |
|---|---|---|---|---|---|---|---|---|---|
| | **p_val** | **avg_log2FC** | **pct.1** | **pct.2** | **p_val_adj** | **celltype** | **Foldchange_UP** | **Foldchange_DOWN** | **Ratio_pct1_pct2** |
| ATG4B | 0,171703399 | -0,435766532 | 0,046 | 0,133 | 1 | Mast | 0,739300845 | 1,35262932 | 0,345864662 |
| GAPT | 0,171703399 | -0,552809887 | 0,046 | 0,133 | 1 | Mast | 0,681691128 | 1,466940024 | 0,345864662 |
| FBXW11 | 0,171703399 | -0,436665664 | 0,046 | 0,133 | 1 | Mast | 0,738840234 | 1,353472583 | 0,345864662 |
| SMIM13 | 0,171703399 | -0,707096286 | 0,046 | 0,133 | 1 | Mast | 0,612551783 | 1,632515043 | 0,345864662 |
| PFDN6 | 0,171703399 | -0,454794969 | 0,046 | 0,133 | 1 | Mast | 0,729613855 | 1,370588008 | 0,345864662 |
| MRPL181 | 0,171703399 | -0,678746174 | 0,046 | 0,133 | 1 | Mast | 0,624707964 | 1,600747963 | 0,345864662 |
| VPS13B | 0,171703399 | -0,389014706 | 0,046 | 0,133 | 1 | Mast | 0,763650965 | 1,309498771 | 0,345864662 |
| TRPS1 | 0,171703399 | -0,271894102 | 0,046 | 0,133 | 1 | Mast | 0,828231454 | 1,207391962 | 0,345864662 |
| TASOR2 | 0,171703399 | -0,51895954 | 0,046 | 0,133 | 1 | Mast | 0,697874954 | 1,432921464 | 0,345864662 |
| MAPK8 | 0,171703399 | -0,671857309 | 0,046 | 0,133 | 1 | Mast | 0,627698076 | 1,593122616 | 0,345864662 |
| LDB1 | 0,171703399 | -0,433779407 | 0,046 | 0,133 | 1 | Mast | 0,740319838 | 1,350767532 | 0,345864662 |
| STARD101 | 0,171703399 | -0,345958567 | 0,046 | 0,133 | 1 | Mast | 0,786785041 | 1,270995187 | 0,345864662 |
| SUSD6 | 0,171703399 | -0,705780804 | 0,046 | 0,133 | 1 | Mast | 0,613110576 | 1,631027158 | 0,345864662 |
| WIPF21 | 0,171703399 | -0,346892729 | 0,046 | 0,133 | 1 | Mast | 0,786275753 | 1,271818438 | 0,345864662 |
| SPIRE1 | 0,171703399 | -0,400558301 | 0,046 | 0,133 | 1 | Mast | 0,75756506 | 1,320018639 | 0,345864662 |
| ZNF83 | 0,171703399 | -0,376758907 | 0,046 | 0,133 | 1 | Mast | 0,770165868 | 1,298421601 | 0,345864662 |
| POLDIP32 | 0,171703399 | -0,450709197 | 0,046 | 0,133 | 1 | Mast | 0,73168308 | 1,366711937 | 0,345864662 |
| CRTAP1 | 0,174154161 | 0,887716075 | 0,352 | 0,2 | 1 | Mast | 1,850244688 | 0,540469056 | 1,76 |
| ALG132 | 0,174272924 | -0,389674387 | 0,083 | 0,2 | 1 | Mast | 0,763301861 | 1,310097684 | 0,415 |
| NFKBID | 0,174540185 | -0,51600512 | 0,185 | 0,333 | 1 | Mast | 0,69930556 | 1,429990061 | 0,555555556 |
| RAP1A2 | 0,174647866 | 0,836388562 | 0,213 | 0,067 | 1 | Mast | 1,785574788 | 0,56004375 | 3,179104478 |
| GPX46 | 0,175492065 | 0,636273299 | 0,648 | 0,667 | 1 | Mast | 1,554308952 | 0,643372734 | 0,971514243 |
| ARRB11 | 0,175656389 | -0,436452328 | 0,093 | 0,2 | 1 | Mast | 0,738949497 | 1,353272456 | 0,465 |
| LAX1 | 0,177192079 | -0,774992131 | 0,269 | 0,4 | 1 | Mast | 0,584391812 | 1,711180718 | 0,6725 |
| HADHA2 | 0,177654616 | -0,701803926 | 0,194 | 0,333 | 1 | Mast | 0,614802985 | 1,626537321 | 0,582582583 |
| SFT2D21 | 0,177818969 | -0,380081917 | 0,046 | 0,133 | 1 | Mast | 0,768393959 | 1,301415749 | 0,345864662 |
| TBC1D51 | 0,177818969 | -0,477831373 | 0,046 | 0,133 | 1 | Mast | 0,718056179 | 1,392648694 | 0,345864662 |
| DIPK2A | 0,177818969 | -0,342076212 | 0,046 | 0,133 | 1 | Mast | 0,788905165 | 1,267579481 | 0,345864662 |
| HTT | 0,177818969 | -0,353921775 | 0,046 | 0,133 | 1 | Mast | 0,782454207 | 1,278030064 | 0,345864662 |
| LAMTOR3 | 0,177818969 | -0,470196194 | 0,046 | 0,133 | 1 | Mast | 0,721866424 | 1,385297843 | 0,345864662 |
| POLR1C | 0,177818969 | -0,335897003 | 0,046 | 0,133 | 1 | Mast | 0,792291372 | 1,262161921 | 0,345864662 |

| Supplementary Table 4 - clusermarker | | | | | | | | | |
|---|---|---|---|---|---|---|---|---|---|
| | **p_val** | **avg_log2FC** | **pct.1** | **pct.2** | **p_val_adj** | **celltype** | **Foldchange_UP** | **Foldchange_DOWN** | **Ratio_pct1_pct2** |
| SKAP21 | 0,177818969 | -0,379537542 | 0,046 | 0,133 | 1 | Mast | 0,768683954 | 1,300924775 | 0,345864662 |
| GALT | 0,177818969 | -0,460255999 | 0,046 | 0,133 | 1 | Mast | 0,72685727 | 1,375785922 | 0,345864662 |
| BORCS7 | 0,177818969 | -0,536410445 | 0,046 | 0,133 | 1 | Mast | 0,689484276 | 1,450359399 | 0,345864662 |
| SPHK1 | 0,177818969 | -0,418757957 | 0,046 | 0,133 | 1 | Mast | 0,748068373 | 1,336776204 | 0,345864662 |
| UBE2O | 0,177818969 | -0,548202344 | 0,046 | 0,133 | 1 | Mast | 0,683871729 | 1,462262523 | 0,345864662 |
| ESCO11 | 0,177818969 | -0,467537544 | 0,046 | 0,133 | 1 | Mast | 0,723197931 | 1,382747318 | 0,345864662 |
| SUGP21 | 0,177818969 | -0,541536266 | 0,046 | 0,133 | 1 | Mast | 0,68703892 | 1,455521617 | 0,345864662 |
| ZFAND6 | 0,179078602 | 0,817591373 | 0,111 | 0 | 1 | Mast | 1,762461048 | 0,567388426 | #DIV/0! |
| H6PD | 0,179081246 | 0,682811377 | 0,111 | 0 | 1 | Mast | 1,605264885 | 0,622950149 | #DIV/0! |
| MEAF61 | 0,179081246 | 0,918831506 | 0,111 | 0 | 1 | Mast | 1,890583417 | 0,528937253 | #DIV/0! |
| HS2ST1 | 0,179081246 | 0,650130528 | 0,111 | 0 | 1 | Mast | 1,569310173 | 0,637222658 | #DIV/0! |
| DGUOK1 | 0,179081246 | 0,819003313 | 0,111 | 0 | 1 | Mast | 1,764186782 | 0,566833405 | #DIV/0! |
| POLE43 | 0,179081246 | 0,847261114 | 0,111 | 0 | 1 | Mast | 1,799082212 | 0,555838968 | #DIV/0! |
| IGKV3-115 | 0,179081246 | 1,067037536 | 0,111 | 0 | 1 | Mast | 2,095126764 | 0,477298089 | #DIV/0! |
| BAZ2B2 | 0,179081246 | 0,82151188 | 0,111 | 0 | 1 | Mast | 1,767257029 | 0,565848648 | #DIV/0! |
| PRKRA1 | 0,179081246 | 0,778679965 | 0,111 | 0 | 1 | Mast | 1,715560454 | 0,58289989 | #DIV/0! |
| INPP5D | 0,179081246 | 0,549987462 | 0,111 | 0 | 1 | Mast | 1,464072972 | 0,683026064 | #DIV/0! |
| STT3B2 | 0,179081246 | 0,84636442 | 0,111 | 0 | 1 | Mast | 1,797964356 | 0,556184552 | #DIV/0! |
| COPB21 | 0,179081246 | 0,839899285 | 0,111 | 0 | 1 | Mast | 1,789925183 | 0,558682569 | #DIV/0! |
| ADD13 | 0,179081246 | 0,729495233 | 0,111 | 0 | 1 | Mast | 1,658058872 | 0,603114893 | #DIV/0! |
| HEXB1 | 0,179081246 | 0,796333526 | 0,111 | 0 | 1 | Mast | 1,736681899 | 0,575810688 | #DIV/0! |
| MACROH2A11 | 0,179081246 | 0,842822404 | 0,111 | 0 | 1 | Mast | 1,793555518 | 0,55755174 | #DIV/0! |
| TRIM27 | 0,179081246 | 0,76765173 | 0,111 | 0 | 1 | Mast | 1,70249638 | 0,587372761 | #DIV/0! |
| NELFE | 0,179081246 | 0,771437192 | 0,111 | 0 | 1 | Mast | 1,706969395 | 0,585833585 | #DIV/0! |
| PHF3 | 0,179081246 | 0,7611069 | 0,111 | 0 | 1 | Mast | 1,694790444 | 0,59004345 | #DIV/0! |
| POR | 0,179081246 | 0,635839292 | 0,111 | 0 | 1 | Mast | 1,553841439 | 0,643566309 | #DIV/0! |
| PSMB71 | 0,179081246 | 0,755193867 | 0,111 | 0 | 1 | Mast | 1,687858389 | 0,592466765 | #DIV/0! |
| BUB32 | 0,179081246 | 0,736044958 | 0,111 | 0 | 1 | Mast | 1,665603445 | 0,600383004 | #DIV/0! |
| PSMC32 | 0,179081246 | 0,643048279 | 0,111 | 0 | 1 | Mast | 1,561625244 | 0,640358501 | #DIV/0! |
| CUL5 | 0,179081246 | 0,790980146 | 0,111 | 0 | 1 | Mast | 1,73024957 | 0,577951307 | #DIV/0! |
| ZMYM2 | 0,179081246 | 0,775053435 | 0,111 | 0 | 1 | Mast | 1,711253432 | 0,58436698 | #DIV/0! |

| Supplementary Table 4 - clusermarker | | | | | | | | | |
|---|---|---|---|---|---|---|---|---|---|
| | **p_val** | **avg_log2FC** | **pct.1** | **pct.2** | **p_val_adj** | **celltype** | **Foldchange_UP** | **Foldchange_DOWN** | **Ratio_pct1_pct2** |
| ARHGAP54 | 0,179081246 | 0,88887479 | 0,111 | 0 | 1 | Mast | 1,851731328 | 0,540035147 | #DIV/0! |
| MVP3 | 0,179081246 | 0,82044731 | 0,111 | 0 | 1 | Mast | 1,765953444 | 0,566266344 | #DIV/0! |
| RANGRF | 0,179081246 | 0,875815045 | 0,111 | 0 | 1 | Mast | 1,835044494 | 0,544945914 | #DIV/0! |
| COA31 | 0,179081246 | 0,759600438 | 0,111 | 0 | 1 | Mast | 1,693021668 | 0,590659895 | #DIV/0! |
| TSEN54 | 0,179081246 | 0,944938847 | 0,111 | 0 | 1 | Mast | 1,925107283 | 0,51945157 | #DIV/0! |
| CYBC1 | 0,179081246 | 0,799048003 | 0,111 | 0 | 1 | Mast | 1,739952598 | 0,574728301 | #DIV/0! |
| SNRPD12 | 0,179081246 | 0,97561843 | 0,111 | 0 | 1 | Mast | 1,966483975 | 0,508521815 | #DIV/0! |
| KDSR1 | 0,179081246 | 0,547839138 | 0,111 | 0 | 1 | Mast | 1,461894436 | 0,68404392 | #DIV/0! |
| MAP2K23 | 0,179081246 | 0,741936494 | 0,111 | 0 | 1 | Mast | 1,67241918 | 0,597936218 | #DIV/0! |
| ARHGEF11 | 0,179081246 | 0,967868956 | 0,111 | 0 | 1 | Mast | 1,955949275 | 0,511260702 | #DIV/0! |
| TTC34 | 0,179081246 | 0,639675653 | 0,111 | 0 | 1 | Mast | 1,557978855 | 0,641857235 | #DIV/0! |
| ATXN101 | 0,179081246 | 0,716246532 | 0,111 | 0 | 1 | Mast | 1,642902121 | 0,608678988 | #DIV/0! |
| VMA212 | 0,179081246 | 0,685149206 | 0,111 | 0 | 1 | Mast | 1,607868261 | 0,621941501 | #DIV/0! |
| RBM394 | 0,179958133 | -0,324640295 | 0,472 | 0,667 | 1 | Mast | 0,79849745 | 1,252352152 | 0,707646177 |
| CDKN1A4 | 0,180095623 | -0,797811294 | 0,546 | 0,6 | 1 | Mast | 0,575221181 | 1,738461713 | 0,91 |
| LEPROT2 | 0,180321173 | -0,624516807 | 0,093 | 0,2 | 1 | Mast | 0,648636985 | 1,541694389 | 0,465 |
| PLXNC11 | 0,180321173 | -0,726356653 | 0,093 | 0,2 | 1 | Mast | 0,604428397 | 1,654455689 | 0,465 |
| EMP31 | 0,180472272 | -0,392242877 | 0,667 | 0,8 | 1 | Mast | 0,761944131 | 1,312432183 | 0,83375 |
| DDX39A2 | 0,181816026 | 0,820835266 | 0,204 | 0,067 | 1 | Mast | 1,766428392 | 0,566114089 | 3,044776119 |
| PNISR2 | 0,181851662 | 0,824706772 | 0,213 | 0,067 | 1 | Mast | 1,77117501 | 0,564596945 | 3,179104478 |
| REX1BD5 | 0,181851662 | 0,863747883 | 0,213 | 0,067 | 1 | Mast | 1,81975961 | 0,549523132 | 3,179104478 |
| RAN2 | 0,182294651 | 0,723243357 | 0,324 | 0,133 | 1 | Mast | 1,650889269 | 0,605734145 | 2,436090226 |
| RPS3A3 | 0,182448587 | -0,365072262 | 0,981 | 1 | 1 | Mast | 0,776429984 | 1,287946138 | 0,981 |
| JUN2 | 0,182456921 | 0,68493947 | 0,731 | 0,6 | 1 | Mast | 1,607634529 | 0,622031924 | 1,218333333 |
| CSTB3 | 0,183077503 | 0,945710897 | 0,352 | 0,2 | 1 | Mast | 1,926137768 | 0,519173663 | 1,76 |
| RPL18A5 | 0,18363339 | 0,401945681 | 0,935 | 0,867 | 1 | Mast | 1,321288657 | 0,756836892 | 1,078431373 |
| TAOK31 | 0,1839874 | -0,539034645 | 0,204 | 0,333 | 1 | Mast | 0,688231273 | 1,452999942 | 0,612612613 |
| PPM1H | 0,183987537 | -0,262808684 | 0,083 | 0,2 | 1 | Mast | 0,833463725 | 1,199812265 | 0,415 |
| CD300A | 0,184090857 | -0,407460887 | 0,046 | 0,133 | 1 | Mast | 0,753949141 | 1,326349412 | 0,345864662 |
| ACBD62 | 0,184095244 | -0,472604888 | 0,046 | 0,133 | 1 | Mast | 0,720662216 | 1,38761264 | 0,345864662 |
| TTN | 0,184095244 | -0,32656478 | 0,046 | 0,133 | 1 | Mast | 0,797433003 | 1,254023844 | 0,345864662 |

| Supplementary Table 4 - clusermarker | | | | | | | | | |
|---|---|---|---|---|---|---|---|---|---|
| | **p_val** | **avg_log2FC** | **pct.1** | **pct.2** | **p_val_adj** | **celltype** | **Foldchange_UP** | **Foldchange_DOWN** | **Ratio_pct1_pct2** |
| INO80D | 0,184095244 | -0,35561641 | 0,046 | 0,133 | 1 | Mast | 0,781535651 | 1,279532161 | 0,345864662 |
| WDR6 | 0,184095244 | -0,421450403 | 0,046 | 0,133 | 1 | Mast | 0,746673584 | 1,339273307 | 0,345864662 |
| ZNF654 | 0,184095244 | -0,345548053 | 0,046 | 0,133 | 1 | Mast | 0,78700895 | 1,270633581 | 0,345864662 |
| PARP141 | 0,184095244 | -0,455801343 | 0,046 | 0,133 | 1 | Mast | 0,729105079 | 1,371544416 | 0,345864662 |
| PPM1K | 0,184095244 | -0,387931454 | 0,046 | 0,133 | 1 | Mast | 0,76422457 | 1,308515899 | 0,345864662 |
| MED101 | 0,184095244 | -0,353846483 | 0,046 | 0,133 | 1 | Mast | 0,782495043 | 1,277963368 | 0,345864662 |
| TARS1 | 0,184095244 | -0,345000964 | 0,046 | 0,133 | 1 | Mast | 0,787307451 | 1,270151831 | 0,345864662 |
| SLU7 | 0,184095244 | -0,484893016 | 0,046 | 0,133 | 1 | Mast | 0,714550056 | 1,399482082 | 0,345864662 |
| BAG4 | 0,184095244 | -0,258951304 | 0,046 | 0,133 | 1 | Mast | 0,835695166 | 1,196608573 | 0,345864662 |
| CTSF1 | 0,184095244 | -0,323100558 | 0,046 | 0,133 | 1 | Mast | 0,799350113 | 1,251016274 | 0,345864662 |
| EXOC51 | 0,184095244 | -0,388795434 | 0,046 | 0,133 | 1 | Mast | 0,763767039 | 1,309299759 | 0,345864662 |
| PHB4 | 0,184095244 | -0,45812474 | 0,046 | 0,133 | 1 | Mast | 0,727931833 | 1,373755007 | 0,345864662 |
| PCNA | 0,184095244 | -0,372699801 | 0,046 | 0,133 | 1 | Mast | 0,772335825 | 1,294773552 | 0,345864662 |
| HLA-A2 | 0,184720322 | -0,504769125 | 0,546 | 0,8 | 1 | Mast | 0,704773154 | 1,418896272 | 0,6825 |
| APC | 0,185077361 | -0,576883165 | 0,093 | 0,2 | 1 | Mast | 0,670410586 | 1,491623224 | 0,465 |
| TMED7 | 0,185077361 | -0,464369592 | 0,093 | 0,2 | 1 | Mast | 0,724787716 | 1,379714334 | 0,465 |
| SUSD1 | 0,185077361 | -0,89041456 | 0,093 | 0,2 | 1 | Mast | 0,539459082 | 1,853708713 | 0,465 |
| AHR1 | 0,18657096 | 0,724508833 | 0,574 | 0,467 | 1 | Mast | 1,652338001 | 0,605203051 | 1,229122056 |
| HSPD13 | 0,187493032 | 0,943753944 | 0,287 | 0,133 | 1 | Mast | 1,923526818 | 0,519878377 | 2,157894737 |
| SF14 | 0,187493032 | -0,5873329 | 0,25 | 0,4 | 1 | Mast | 0,665572209 | 1,502466579 | 0,625 |
| NDUFA16 | 0,18765571 | 1,044149801 | 0,278 | 0,133 | 1 | Mast | 2,062150749 | 0,4849306 | 2,090225564 |
| BHLHE402 | 0,187946727 | -0,58209648 | 0,472 | 0,6 | 1 | Mast | 0,667992366 | 1,497023097 | 0,786666667 |
| AC020916.12 | 0,1888588 | -0,596871472 | 0,611 | 0,733 | 1 | Mast | 0,661186204 | 1,51243325 | 0,833560709 |
| PLK2 | 0,189925725 | -0,71771953 | 0,093 | 0,2 | 1 | Mast | 0,608057841 | 1,644580389 | 0,465 |
| SYNJ2 | 0,189925725 | -0,454522691 | 0,093 | 0,2 | 1 | Mast | 0,729751567 | 1,370329363 | 0,465 |
| DDIT32 | 0,189925725 | -0,561910475 | 0,093 | 0,2 | 1 | Mast | 0,677404523 | 1,476222797 | 0,465 |
| PBXIP1 | 0,19053406 | -0,306765111 | 0,046 | 0,133 | 1 | Mast | 0,808452485 | 1,236931074 | 0,345864662 |
| IRAK3 | 0,19053406 | -0,314105778 | 0,046 | 0,133 | 1 | Mast | 0,804349394 | 1,24324082 | 0,345864662 |
| MDM21 | 0,19053406 | -0,324727199 | 0,046 | 0,133 | 1 | Mast | 0,798449352 | 1,252427593 | 0,345864662 |
| UBN11 | 0,19053406 | -0,263195828 | 0,046 | 0,133 | 1 | Mast | 0,833240097 | 1,200134275 | 0,345864662 |
| CAVIN2 | 0,191536444 | -0,906124278 | 0,296 | 0,467 | 1 | Mast | 0,5336167 | 1,87400432 | 0,633832976 |

| Supplementary Table 4 - clusermarker | | | | | | | | | |
|---|---|---|---|---|---|---|---|---|---|
| | **p_val** | **avg_log2FC** | **pct.1** | **pct.2** | **p_val_adj** | **celltype** | **Foldchange_UP** | **Foldchange_DOWN** | **Ratio_pct1_pct2** |
| SELENOK2 | 0,1916819 | -0,28305894 | 0,574 | 0,8 | 1 | Mast | 0,821846611 | 1,216772068 | 0,7175 |
| SPCS15 | 0,192184958 | 0,887207499 | 0,472 | 0,333 | 1 | Mast | 1,849592559 | 0,540659614 | 1,417417417 |
| VEGFA1 | 0,192377367 | -1,110780589 | 0,222 | 0,333 | 1 | Mast | 0,463043428 | 2,15962465 | 0,666666667 |
| TMEM2562 | 0,193226106 | 0,860349988 | 0,204 | 0,067 | 1 | Mast | 1,81547868 | 0,550818917 | 3,044776119 |
| PSMD111 | 0,194097465 | -0,369209568 | 0,083 | 0,2 | 1 | Mast | 0,774206557 | 1,291644963 | 0,415 |
| TRAF5 | 0,194867017 | -0,485435632 | 0,093 | 0,2 | 1 | Mast | 0,714281355 | 1,400008544 | 0,465 |
| PLEKHA22 | 0,194867017 | -0,471440212 | 0,093 | 0,2 | 1 | Mast | 0,721244235 | 1,386492884 | 0,465 |
| RAB341 | 0,194867017 | -0,486474833 | 0,093 | 0,2 | 1 | Mast | 0,71376703 | 1,40101736 | 0,465 |
| TRIR4 | 0,195455366 | -0,368611782 | 0,417 | 0,6 | 1 | Mast | 0,774527419 | 1,291109876 | 0,695 |
| SOCS11 | 0,196754135 | 0,803037975 | 0,528 | 0,333 | 1 | Mast | 1,744771337 | 0,573141006 | 1,585585586 |
| TRIM44 | 0,197078959 | 0,888948845 | 0,194 | 0,067 | 1 | Mast | 1,851826381 | 0,540007427 | 2,895522388 |
| LUZP1 | 0,197137179 | -0,334564405 | 0,046 | 0,133 | 1 | Mast | 0,793023539 | 1,260996617 | 0,345864662 |
| ZBED6 | 0,197137179 | -0,283623758 | 0,046 | 0,133 | 1 | Mast | 0,821524919 | 1,217248529 | 0,345864662 |
| CEP170 | 0,197137179 | -0,340488367 | 0,046 | 0,133 | 1 | Mast | 0,78977392 | 1,266185138 | 0,345864662 |
| COPS4 | 0,197137179 | -0,286053853 | 0,046 | 0,133 | 1 | Mast | 0,820142296 | 1,219300607 | 0,345864662 |
| KRR11 | 0,197137179 | -0,335563252 | 0,046 | 0,133 | 1 | Mast | 0,792474681 | 1,261869968 | 0,345864662 |
| GCHFR3 | 0,197137179 | -0,325205431 | 0,046 | 0,133 | 1 | Mast | 0,798184722 | 1,252842823 | 0,345864662 |
| CCT72 | 0,197143989 | 0,850614996 | 0,204 | 0,067 | 1 | Mast | 1,803269464 | 0,554548291 | 3,044776119 |
| MYL12A3 | 0,197878157 | 0,571396575 | 0,657 | 0,6 | 1 | Mast | 1,485961333 | 0,672965021 | 1,095 |
| GTF2B3 | 0,197990876 | -0,342043272 | 0,222 | 0,4 | 1 | Mast | 0,788923178 | 1,26755054 | 0,555 |
| COX7A22 | 0,198955437 | -0,996480272 | 0,333 | 0,467 | 1 | Mast | 0,501221334 | 1,995126569 | 0,713062099 |
| PRPF38A | 0,199901959 | -0,668098504 | 0,093 | 0,2 | 1 | Mast | 0,629335615 | 1,588977289 | 0,465 |
| ETF11 | 0,199901959 | -0,432412423 | 0,093 | 0,2 | 1 | Mast | 0,741021639 | 1,349488257 | 0,465 |
| HNRNPH11 | 0,200265058 | -0,347662469 | 0,185 | 0,333 | 1 | Mast | 0,785856353 | 1,272497188 | 0,555555556 |
| NDUFA83 | 0,200416413 | 0,700825742 | 0,102 | 0 | 1 | Mast | 1,625434861 | 0,615219978 | #DIV/0! |
| CASP1 | 0,200416413 | 0,761949887 | 0,102 | 0 | 1 | Mast | 1,695781024 | 0,58969878 | #DIV/0! |
| MAP4K51 | 0,200416413 | 0,609383309 | 0,102 | 0 | 1 | Mast | 1,525606937 | 0,655476831 | #DIV/0! |
| SPG72 | 0,200416413 | 0,668177646 | 0,102 | 0 | 1 | Mast | 1,589064457 | 0,629301093 | #DIV/0! |
| MORC31 | 0,200416413 | 0,640709873 | 0,102 | 0 | 1 | Mast | 1,559096118 | 0,641397274 | #DIV/0! |
| FAAP203 | 0,200419375 | 0,59451692 | 0,102 | 0 | 1 | Mast | 1,509966895 | 0,662266175 | #DIV/0! |
| SDHB | 0,200419375 | 0,743298881 | 0,102 | 0 | 1 | Mast | 1,67399925 | 0,597371833 | #DIV/0! |

| Supplementary Table 4 - clusermarker | | | | | | | | | |
|---|---|---|---|---|---|---|---|---|---|
| | p_val | avg_log2FC | pct.1 | pct.2 | p_val_adj | celltype | Foldchange_UP | Foldchange_DOWN | Ratio_pct1_pct2 |
| MACO1 | 0,200419375 | 0,592484695 | 0,102 | 0 | 1 | Mast | 1,507841406 | 0,663199721 | #DIV/0! |
| KIAA15221 | 0,200419375 | 0,665680192 | 0,102 | 0 | 1 | Mast | 1,586316002 | 0,630391422 | #DIV/0! |
| LSM10 | 0,200419375 | 0,856102788 | 0,102 | 0 | 1 | Mast | 1,810141889 | 0,552442881 | #DIV/0! |
| ZMPSTE24 | 0,200419375 | 0,65066618 | 0,102 | 0 | 1 | Mast | 1,569892944 | 0,63698611 | #DIV/0! |
| RNF112 | 0,200419375 | 0,567825997 | 0,102 | 0 | 1 | Mast | 1,482288221 | 0,674632629 | #DIV/0! |
| NAXE3 | 0,200419375 | 0,833686739 | 0,102 | 0 | 1 | Mast | 1,782233962 | 0,561093561 | #DIV/0! |
| PRCC1 | 0,200419375 | 0,709360956 | 0,102 | 0 | 1 | Mast | 1,635079696 | 0,611590984 | #DIV/0! |
| SH2D2A1 | 0,200419375 | 0,778412173 | 0,102 | 0 | 1 | Mast | 1,715242042 | 0,583008098 | #DIV/0! |
| DUSP234 | 0,200419375 | 0,758524728 | 0,102 | 0 | 1 | Mast | 1,691759779 | 0,59110047 | #DIV/0! |
| ATP1A4 | 0,200419375 | 0,777713393 | 0,102 | 0 | 1 | Mast | 1,714411453 | 0,58329055 | #DIV/0! |
| MPC22 | 0,200419375 | 0,581074538 | 0,102 | 0 | 1 | Mast | 1,495963046 | 0,66846571 | #DIV/0! |
| UCHL5 | 0,200419375 | 0,517142886 | 0,102 | 0 | 1 | Mast | 1,431118252 | 0,698754277 | #DIV/0! |
| ARL8A | 0,200419375 | 0,824091438 | 0,102 | 0 | 1 | Mast | 1,770419735 | 0,564837807 | #DIV/0! |
| PCBP1-AS1 | 0,200419375 | 0,576277852 | 0,102 | 0 | 1 | Mast | 1,490997513 | 0,67069193 | #DIV/0! |
| SP110 | 0,200419375 | 0,746771077 | 0,102 | 0 | 1 | Mast | 1,678032988 | 0,595935841 | #DIV/0! |
| PPP1R7 | 0,200419375 | 0,500707974 | 0,102 | 0 | 1 | Mast | 1,41490773 | 0,706759868 | #DIV/0! |
| CCDC121 | 0,200419375 | 0,708069703 | 0,102 | 0 | 1 | Mast | 1,633616907 | 0,61213862 | #DIV/0! |
| ATG32 | 0,200419375 | 0,734784917 | 0,102 | 0 | 1 | Mast | 1,664149353 | 0,600907604 | #DIV/0! |
| NAA501 | 0,200419375 | 0,692531502 | 0,102 | 0 | 1 | Mast | 1,616116837 | 0,618767144 | #DIV/0! |
| LAP33 | 0,200419375 | 0,695705274 | 0,102 | 0 | 1 | Mast | 1,619676032 | 0,61740742 | #DIV/0! |
| TMEM1651 | 0,200419375 | 0,686080041 | 0,102 | 0 | 1 | Mast | 1,608906002 | 0,62154035 | #DIV/0! |
| GOLPH3 | 0,200419375 | 0,745347948 | 0,102 | 0 | 1 | Mast | 1,676378528 | 0,596523985 | #DIV/0! |
| PPWD11 | 0,200419375 | 0,747435739 | 0,102 | 0 | 1 | Mast | 1,67880625 | 0,595661352 | #DIV/0! |
| SPARC1 | 0,200419375 | 0,749958306 | 0,102 | 0 | 1 | Mast | 1,681744227 | 0,594620742 | #DIV/0! |
| CCNG12 | 0,200419375 | 0,744040523 | 0,102 | 0 | 1 | Mast | 1,674860019 | 0,597064823 | #DIV/0! |
| RAB241 | 0,200419375 | 0,587636882 | 0,102 | 0 | 1 | Mast | 1,502783189 | 0,665431985 | #DIV/0! |
| HLA-DRA4 | 0,200419375 | 0,84826679 | 0,102 | 0 | 1 | Mast | 1,800336756 | 0,555451638 | #DIV/0! |
| HDAC23 | 0,200419375 | 0,7731743 | 0,102 | 0 | 1 | Mast | 1,709025947 | 0,585128624 | #DIV/0! |
| GTF2H53 | 0,200419375 | 0,630268113 | 0,102 | 0 | 1 | Mast | 1,547852623 | 0,64605634 | #DIV/0! |
| VPS41 | 0,200419375 | 0,583544463 | 0,102 | 0 | 1 | Mast | 1,498526362 | 0,667322261 | #DIV/0! |
| BCL7B1 | 0,200419375 | 0,727092389 | 0,102 | 0 | 1 | Mast | 1,655299632 | 0,604120233 | #DIV/0! |

| Supplementary Table 4 - clusermarker | | | | | | | | | |
|---|---|---|---|---|---|---|---|---|---|
| | **p_val** | **avg_log2FC** | **pct.1** | **pct.2** | **p_val_adj** | **celltype** | **Foldchange_UP** | **Foldchange_DOWN** | **Ratio_pct1_pct2** |
| PTPRN2 | 0,200419375 | 0,650024207 | 0,102 | 0 | 1 | Mast | 1,569194525 | 0,637269621 | #DIV/0! |
| CYC12 | 0,200419375 | 0,686614845 | 0,102 | 0 | 1 | Mast | 1,609502531 | 0,621309989 | #DIV/0! |
| SPTAN13 | 0,200419375 | 0,853719397 | 0,102 | 0 | 1 | Mast | 1,807153929 | 0,553356294 | #DIV/0! |
| RSU12 | 0,200419375 | 0,671508519 | 0,102 | 0 | 1 | Mast | 1,592737504 | 0,627849848 | #DIV/0! |
| UNC5B | 0,200419375 | 0,530503473 | 0,102 | 0 | 1 | Mast | 1,444433187 | 0,692313088 | #DIV/0! |
| IFITM13 | 0,200419375 | 0,917101217 | 0,102 | 0 | 1 | Mast | 1,888317315 | 0,529572012 | #DIV/0! |
| PLAAT33 | 0,200419375 | 0,766002934 | 0,102 | 0 | 1 | Mast | 1,700551779 | 0,588044429 | #DIV/0! |
| CEP572 | 0,200419375 | 0,735714946 | 0,102 | 0 | 1 | Mast | 1,665222487 | 0,600520356 | #DIV/0! |
| STT3A | 0,200419375 | 0,75008996 | 0,102 | 0 | 1 | Mast | 1,681897703 | 0,594566482 | #DIV/0! |
| CBX5 | 0,200419375 | 0,590929206 | 0,102 | 0 | 1 | Mast | 1,506216554 | 0,663915157 | #DIV/0! |
| SHMT2 | 0,200419375 | 0,858036952 | 0,102 | 0 | 1 | Mast | 1,812570302 | 0,551702739 | #DIV/0! |
| CHPT13 | 0,200419375 | 0,698351598 | 0,102 | 0 | 1 | Mast | 1,622649718 | 0,616275952 | #DIV/0! |
| NEMF | 0,200419375 | 0,770487319 | 0,102 | 0 | 1 | Mast | 1,705845894 | 0,586219426 | #DIV/0! |
| IGHV3-231 | 0,200419375 | 4,404236884 | 0,102 | 0 | 1 | Mast | 21,17421946 | 0,047227243 | #DIV/0! |
| CSK2 | 0,200419375 | 0,806786155 | 0,102 | 0 | 1 | Mast | 1,749310217 | 0,571653895 | #DIV/0! |
| IMP32 | 0,200419375 | 0,832909925 | 0,102 | 0 | 1 | Mast | 1,781274583 | 0,561395761 | #DIV/0! |
| TMEM170A2 | 0,200419375 | 0,873772533 | 0,102 | 0 | 1 | Mast | 1,832448348 | 0,545717974 | #DIV/0! |
| CTDNEP1 | 0,200419375 | 0,766503998 | 0,102 | 0 | 1 | Mast | 1,701142502 | 0,58784023 | #DIV/0! |
| USP221 | 0,200419375 | 0,677133977 | 0,102 | 0 | 1 | Mast | 1,598960143 | 0,625406458 | #DIV/0! |
| TAOK12 | 0,200419375 | 0,766731316 | 0,102 | 0 | 1 | Mast | 1,701410563 | 0,587747615 | #DIV/0! |
| STAT5A1 | 0,200419375 | 0,685467175 | 0,102 | 0 | 1 | Mast | 1,608222673 | 0,621804441 | #DIV/0! |
| METTL23 | 0,200419375 | 0,725801886 | 0,102 | 0 | 1 | Mast | 1,653819615 | 0,604660866 | #DIV/0! |
| CCDC571 | 0,200419375 | 0,64201497 | 0,102 | 0 | 1 | Mast | 1,560507153 | 0,640817313 | #DIV/0! |
| TPGS21 | 0,200419375 | 0,586136563 | 0,102 | 0 | 1 | Mast | 1,501221193 | 0,666124356 | #DIV/0! |
| YIF1B | 0,200419375 | 0,858779116 | 0,102 | 0 | 1 | Mast | 1,81350298 | 0,551419 | #DIV/0! |
| ISOC22 | 0,200419375 | 0,730915036 | 0,102 | 0 | 1 | Mast | 1,659691425 | 0,602521641 | #DIV/0! |
| TRIM281 | 0,200419375 | 0,598280333 | 0,102 | 0 | 1 | Mast | 1,513910936 | 0,660540839 | #DIV/0! |
| PTPRA | 0,200419375 | 0,682016051 | 0,102 | 0 | 1 | Mast | 1,604380183 | 0,623293662 | #DIV/0! |
| CBR13 | 0,200419375 | 1,003123087 | 0,102 | 0 | 1 | Mast | 2,004334207 | 0,498918791 | #DIV/0! |
| DYRK1A | 0,200419375 | 0,733923093 | 0,102 | 0 | 1 | Mast | 1,663155535 | 0,601266676 | #DIV/0! |
| DOCK11 | 0,200419375 | 0,515347188 | 0,102 | 0 | 1 | Mast | 1,429338072 | 0,699624546 | #DIV/0! |

| Supplementary Table 4 - clusermarker | | | | | | | | | |
|---|---|---|---|---|---|---|---|---|---|
| | **p_val** | **avg_log2FC** | **pct.1** | **pct.2** | **p_val_adj** | **celltype** | **Foldchange_UP** | **Foldchange_DOWN** | **Ratio_pct1_pct2** |
| SELENOW4 | 0,20073767 | 1,055796011 | 0,324 | 0,2 | 1 | Mast | 2,07886491 | 0,481031738 | 1,62 |
| NTRK1 | 0,200821522 | 0,633129911 | 0,213 | 0,067 | 1 | Mast | 1,550926063 | 0,644776062 | 3,179104478 |
| ZNF6382 | 0,201119712 | 0,688928677 | 0,204 | 0,067 | 1 | Mast | 1,612085963 | 0,620314315 | 3,044776119 |
| EVI2B2 | 0,201119712 | 0,846670177 | 0,204 | 0,067 | 1 | Mast | 1,798345446 | 0,55606669 | 3,044776119 |
| HS6ST1 | 0,201214268 | -0,370610435 | 0,139 | 0,267 | 1 | Mast | 0,773455162 | 1,292899768 | 0,520599251 |
| SEC131 | 0,201214268 | -0,443311175 | 0,139 | 0,267 | 1 | Mast | 0,735444728 | 1,359721489 | 0,520599251 |
| COX6A14 | 0,202008973 | 0,742459241 | 0,417 | 0,267 | 1 | Mast | 1,673025275 | 0,597719601 | 1,561797753 |
| REL2 | 0,202258988 | -0,61133525 | 0,463 | 0,667 | 1 | Mast | 0,654590582 | 1,527672453 | 0,694152924 |
| PPP1R15B | 0,203296433 | -0,465089996 | 0,194 | 0,333 | 1 | Mast | 0,724425886 | 1,380403461 | 0,582582583 |
| SERP12 | 0,204420778 | 0,74773725 | 0,454 | 0,333 | 1 | Mast | 1,679157143 | 0,595536877 | 1,363363363 |
| GALNT2 | 0,205031252 | -0,436605419 | 0,093 | 0,2 | 1 | Mast | 0,738871087 | 1,353416065 | 0,465 |
| ABHD22 | 0,205031252 | -0,527512153 | 0,093 | 0,2 | 1 | Mast | 0,693750036 | 1,441441366 | 0,465 |
| C7orf501 | 0,206399557 | -0,255066483 | 0,13 | 0,267 | 1 | Mast | 0,837948519 | 1,193390736 | 0,486891386 |
| SRP141 | 0,206533967 | -0,366941438 | 0,565 | 0,733 | 1 | Mast | 0,775424682 | 1,289615901 | 0,770804911 |
| LGALS13 | 0,208349929 | 1,109297535 | 0,509 | 0,4 | 1 | Mast | 2,157405751 | 0,463519669 | 1,2725 |
| H2AZ24 | 0,208598071 | -0,343909449 | 0,241 | 0,4 | 1 | Mast | 0,787903337 | 1,269191223 | 0,6025 |
| NDUFB73 | 0,208800653 | 0,694495412 | 0,213 | 0,067 | 1 | Mast | 1,618318321 | 0,617925403 | 3,179104478 |
| ACAP21 | 0,209246004 | 0,931306446 | 0,204 | 0,067 | 1 | Mast | 1,907002118 | 0,524383267 | 3,044776119 |
| ASRGL11 | 0,209246004 | 0,717059144 | 0,204 | 0,067 | 1 | Mast | 1,643827763 | 0,60833624 | 3,044776119 |
| RALGAPA11 | 0,209246004 | 0,704191359 | 0,204 | 0,067 | 1 | Mast | 1,629231213 | 0,613786424 | 3,044776119 |
| SVOPL | 0,209405041 | 0,757442708 | 0,194 | 0,067 | 1 | Mast | 1,690491436 | 0,591543961 | 2,895522388 |
| THUMPD1 | 0,209684172 | -0,913375559 | 0,102 | 0,2 | 1 | Mast | 0,530941362 | 1,883447159 | 0,51 |
| KLF63 | 0,209799682 | 0,523624807 | 0,648 | 0,733 | 1 | Mast | 1,437562626 | 0,695621869 | 0,884038199 |
| RBBP43 | 0,210255567 | -0,286756175 | 0,093 | 0,2 | 1 | Mast | 0,819743138 | 1,219894323 | 0,465 |
| MLLT101 | 0,210255567 | -0,458412656 | 0,093 | 0,2 | 1 | Mast | 0,727786575 | 1,374029193 | 0,465 |
| CACUL11 | 0,210842982 | -0,292223589 | 0,046 | 0,133 | 1 | Mast | 0,816642418 | 1,224526155 | 0,345864662 |
| ELF11 | 0,211435408 | -0,590351629 | 0,5 | 0,667 | 1 | Mast | 0,664181006 | 1,505613667 | 0,749625187 |
| KRTCAP24 | 0,213074298 | 0,653268915 | 0,315 | 0,133 | 1 | Mast | 1,57272771 | 0,635837973 | 2,368421053 |
| GOLGB13 | 0,213108517 | -0,966549695 | 0,148 | 0,267 | 1 | Mast | 0,511728435 | 1,95416149 | 0,554307116 |
| RPLP04 | 0,213152249 | 0,474275473 | 0,898 | 0,933 | 1 | Mast | 1,389220372 | 0,7198282 | 0,962486602 |
| ATRX1 | 0,214520257 | -0,343955914 | 0,139 | 0,267 | 1 | Mast | 0,787877962 | 1,2692321 | 0,520599251 |

| Supplementary Table 4 - clusermarker | | | | | | | | | |
|---|---|---|---|---|---|---|---|---|---|
| | **p_val** | **avg_log2FC** | **pct.1** | **pct.2** | **p_val_adj** | **celltype** | **Foldchange_UP** | **Foldchange_DOWN** | **Ratio_pct1_pct2** |
| TMSB104 | 0,214666457 | 0,43179865 | 0,907 | 1 | 1 | Mast | 1,34891426 | 0,741336963 | 0,907 |
| POLD42 | 0,214784206 | 0,716340633 | 0,296 | 0,133 | 1 | Mast | 1,643009284 | 0,608639287 | 2,22556391 |
| TMBIM44 | 0,214784206 | 0,623146767 | 0,296 | 0,133 | 1 | Mast | 1,54023103 | 0,649253249 | 2,22556391 |
| NDUFA4 | 0,215198117 | 0,468910316 | 0,731 | 0,6 | 1 | Mast | 1,384063674 | 0,722510112 | 1,218333333 |
| DARS11 | 0,215498061 | -0,623537089 | 0,056 | 0,133 | 1 | Mast | 0,649077617 | 1,540647797 | 0,421052632 |
| ACO2 | 0,215498061 | -0,675354325 | 0,056 | 0,133 | 1 | Mast | 0,626178412 | 1,596988944 | 0,421052632 |
| APRT4 | 0,215941957 | 0,841808826 | 0,278 | 0,133 | 1 | Mast | 1,792295883 | 0,55794359 | 2,090225564 |
| OSBPL81 | 0,216240998 | 0,733992205 | 0,444 | 0,333 | 1 | Mast | 1,66323521 | 0,601237873 | 1,333333333 |
| CXCL161 | 0,216285493 | 0,738844182 | 0,269 | 0,133 | 1 | Mast | 1,66883831 | 0,599219226 | 2,022556391 |
| HPGDS1 | 0,217242749 | 0,330619069 | 0,889 | 0,8 | 1 | Mast | 1,257552882 | 0,795195188 | 1,11125 |
| TAGLN23 | 0,217522024 | 0,494877066 | 0,759 | 0,8 | 1 | Mast | 1,40920067 | 0,709622143 | 0,94875 |
| RPN2 | 0,217924538 | 0,822666195 | 0,194 | 0,067 | 1 | Mast | 1,768671595 | 0,565396087 | 2,895522388 |
| UBE2D24 | 0,219106907 | -0,38385887 | 0,194 | 0,333 | 1 | Mast | 0,766384947 | 1,304827298 | 0,582582583 |
| SLC25A44 | 0,22004732 | -0,82599932 | 0,102 | 0,2 | 1 | Mast | 0,564091335 | 1,772762563 | 0,51 |
| TXNL11 | 0,22026997 | -0,259629765 | 0,13 | 0,267 | 1 | Mast | 0,835302253 | 1,197171438 | 0,486891386 |
| UNC13D | 0,220991812 | -0,411799519 | 0,093 | 0,2 | 1 | Mast | 0,751685188 | 1,33034416 | 0,465 |
| C18orf321 | 0,221877235 | 0,683652614 | 0,204 | 0,067 | 1 | Mast | 1,60620119 | 0,622587012 | 3,044776119 |
| CD592 | 0,222084499 | -0,394293056 | 0,148 | 0,267 | 1 | Mast | 0,76086212 | 1,314298575 | 0,554307116 |
| TMEM2195 | 0,222186124 | 0,724997774 | 0,361 | 0,2 | 1 | Mast | 1,652898086 | 0,604997978 | 1,805 |
| WDR43 | 0,222272902 | -0,735246312 | 0,056 | 0,133 | 1 | Mast | 0,600715456 | 1,664681657 | 0,421052632 |
| YPEL52 | 0,222272902 | -0,804278631 | 0,056 | 0,133 | 1 | Mast | 0,572648341 | 1,746272412 | 0,421052632 |
| MTIF32 | 0,222272902 | -0,748996737 | 0,056 | 0,133 | 1 | Mast | 0,595017194 | 1,680623704 | 0,421052632 |
| GSS1 | 0,222272902 | -0,651843661 | 0,056 | 0,133 | 1 | Mast | 0,636466435 | 1,571174762 | 0,421052632 |
| DCXR3 | 0,222276518 | 0,812291002 | 0,194 | 0,067 | 1 | Mast | 1,755997757 | 0,569476809 | 2,895522388 |
| PSMG22 | 0,222276518 | 0,792655401 | 0,194 | 0,067 | 1 | Mast | 1,7322599 | 0,57728058 | 2,895522388 |
| PTPN221 | 0,225364551 | -0,343668464 | 0,102 | 0,2 | 1 | Mast | 0,788034958 | 1,268979237 | 0,51 |
| SEC31A2 | 0,225364551 | -0,54967373 | 0,102 | 0,2 | 1 | Mast | 0,683174613 | 1,463754626 | 0,51 |
| BOD1L13 | 0,226504943 | -0,338397599 | 0,093 | 0,2 | 1 | Mast | 0,790919298 | 1,264351499 | 0,465 |
| NAF11 | 0,226504943 | -0,39297133 | 0,093 | 0,2 | 1 | Mast | 0,761559504 | 1,313095031 | 0,465 |
| NDUFB105 | 0,226654558 | 0,881896709 | 0,287 | 0,133 | 1 | Mast | 1,842796432 | 0,542653536 | 2,157894737 |
| GM2A1 | 0,22685744 | 0,953727937 | 0,185 | 0,067 | 1 | Mast | 1,93687109 | 0,516296622 | 2,76119403 |

| Supplementary Table 4 - clusermarker | | | | | | | | | |
|---|---|---|---|---|---|---|---|---|---|
| | **p_val** | **avg_log2FC** | **pct.1** | **pct.2** | **p_val_adj** | **celltype** | **Foldchange_UP** | **Foldchange_DOWN** | **Ratio_pct1_pct2** |
| NFATC1 | 0,22844923 | -0,314205019 | 0,139 | 0,267 | 1 | Mast | 0,804294066 | 1,243326343 | 0,520599251 |
| LY752 | 0,229198913 | -0,624198599 | 0,056 | 0,133 | 1 | Mast | 0,648780067 | 1,541354383 | 0,421052632 |
| APMAP1 | 0,229198913 | -0,737817327 | 0,056 | 0,133 | 1 | Mast | 0,599645879 | 1,667650917 | 0,421052632 |
| SMAD72 | 0,230772909 | -0,61661063 | 0,102 | 0,2 | 1 | Mast | 0,652201367 | 1,533268789 | 0,51 |
| CD481 | 0,231166409 | 0,773713421 | 0,194 | 0,067 | 1 | Mast | 1,709664712 | 0,584910008 | 2,895522388 |
| LINC00963 | 0,231166409 | 0,784949291 | 0,194 | 0,067 | 1 | Mast | 1,723031756 | 0,580372356 | 2,895522388 |
| ERF | 0,231423909 | 0,719997463 | 0,185 | 0,067 | 1 | Mast | 1,647179138 | 0,60709851 | 2,76119403 |
| CSDE13 | 0,232617817 | -0,345727816 | 0,361 | 0,533 | 1 | Mast | 0,786910893 | 1,270791914 | 0,677298311 |
| CRIP14 | 0,233155257 | -0,262149715 | 0,491 | 0,733 | 1 | Mast | 0,833844508 | 1,19926436 | 0,669849932 |
| SAP181 | 0,234937686 | 0,819989948 | 0,389 | 0,267 | 1 | Mast | 1,765393693 | 0,566445889 | 1,456928839 |
| PPT1 | 0,235704868 | 0,708890293 | 0,194 | 0,067 | 1 | Mast | 1,634546357 | 0,611790541 | 2,895522388 |
| HM132 | 0,236050966 | -0,442713896 | 0,148 | 0,267 | 1 | Mast | 0,735749266 | 1,359158678 | 0,554307116 |
| TMED21 | 0,236053495 | 0,737185633 | 0,185 | 0,067 | 1 | Mast | 1,666920885 | 0,599908495 | 2,76119403 |
| ZBTB11 | 0,236277152 | -0,546431216 | 0,056 | 0,133 | 1 | Mast | 0,684711801 | 1,460468474 | 0,421052632 |
| SDAD1 | 0,236277152 | -0,829294989 | 0,056 | 0,133 | 1 | Mast | 0,562804204 | 1,776816861 | 0,421052632 |
| SERPINB9 | 0,236277152 | -0,39803527 | 0,056 | 0,133 | 1 | Mast | 0,758891073 | 1,317712166 | 0,421052632 |
| STK17A1 | 0,236277152 | -0,549454974 | 0,056 | 0,133 | 1 | Mast | 0,683278211 | 1,463532693 | 0,421052632 |
| C11orf11 | 0,236277152 | -0,622238811 | 0,056 | 0,133 | 1 | Mast | 0,649661983 | 1,539261995 | 0,421052632 |
| LRRFIP13 | 0,238462502 | 0,848432358 | 0,389 | 0,267 | 1 | Mast | 1,800543379 | 0,555387897 | 1,456928839 |
| ZEB21 | 0,239988628 | -0,643007264 | 0,324 | 0,467 | 1 | Mast | 0,640376706 | 1,561580848 | 0,69379015 |
| NR4A33 | 0,240433442 | -0,614714521 | 0,407 | 0,533 | 1 | Mast | 0,653059107 | 1,531254964 | 0,763602251 |
| PER11 | 0,240588359 | -0,728346066 | 0,213 | 0,333 | 1 | Mast | 0,603595491 | 1,656738685 | 0,63963964 |
| IDH3G2 | 0,240748211 | 0,84543982 | 0,185 | 0,067 | 1 | Mast | 1,796812438 | 0,556541116 | 2,76119403 |
| COTL13 | 0,241248509 | -0,393755042 | 0,259 | 0,4 | 1 | Mast | 0,761145916 | 1,313808535 | 0,6475 |
| SSR33 | 0,242586442 | -0,801840239 | 0,222 | 0,333 | 1 | Mast | 0,57361703 | 1,743323417 | 0,666666667 |
| YBX14 | 0,243088109 | 0,668394408 | 0,574 | 0,467 | 1 | Mast | 1,589303229 | 0,629206549 | 1,229122056 |
| EEF1B23 | 0,243482828 | -0,425943851 | 0,694 | 0,8 | 1 | Mast | 0,744351597 | 1,343451138 | 0,8675 |
| FOXN2 | 0,243508601 | -0,512683147 | 0,056 | 0,133 | 1 | Mast | 0,700917647 | 1,426701131 | 0,421052632 |
| PNO1 | 0,243508601 | -0,414677262 | 0,056 | 0,133 | 1 | Mast | 0,750187296 | 1,333000445 | 0,421052632 |
| TTC31 | 0,243508601 | -0,544508469 | 0,056 | 0,133 | 1 | Mast | 0,685624957 | 1,458523336 | 0,421052632 |
| KCMF11 | 0,243508601 | -0,454579491 | 0,056 | 0,133 | 1 | Mast | 0,729722837 | 1,370383315 | 0,421052632 |

| Supplementary Table 4 - clusermarker | | | | | | | | | |
|---|---|---|---|---|---|---|---|---|---|
| | **p_val** | **avg_log2FC** | **pct.1** | **pct.2** | **p_val_adj** | **celltype** | **Foldchange_UP** | **Foldchange_DOWN** | **Ratio_pct1_pct2** |
| ISY1 | 0,243508601 | -0,43633485 | 0,056 | 0,133 | 1 | Mast | 0,739009671 | 1,353162264 | 0,421052632 |
| SMIM192 | 0,243508601 | -0,403026701 | 0,056 | 0,133 | 1 | Mast | 0,756270002 | 1,322279077 | 0,421052632 |
| FAM120A1 | 0,243508601 | -0,448573641 | 0,056 | 0,133 | 1 | Mast | 0,73276696 | 1,364690352 | 0,421052632 |
| XPA | 0,243508601 | -0,452679168 | 0,056 | 0,133 | 1 | Mast | 0,730684664 | 1,368579429 | 0,421052632 |
| GOT1 | 0,243508601 | -0,449608712 | 0,056 | 0,133 | 1 | Mast | 0,73224142 | 1,365669809 | 0,421052632 |
| SPRY2 | 0,243508601 | -0,368941742 | 0,056 | 0,133 | 1 | Mast | 0,774350297 | 1,2914052 | 0,421052632 |
| MGRN1 | 0,243508601 | -0,375518562 | 0,056 | 0,133 | 1 | Mast | 0,770828297 | 1,297305774 | 0,421052632 |
| AFG3L2 | 0,243508601 | -0,62981292 | 0,056 | 0,133 | 1 | Mast | 0,646260213 | 1,547364328 | 0,421052632 |
| ZNF146 | 0,243508601 | -0,603780019 | 0,056 | 0,133 | 1 | Mast | 0,658027591 | 1,519693117 | 0,421052632 |
| RGS23 | 0,244199695 | -0,845407474 | 0,38 | 0,467 | 1 | Mast | 0,556553594 | 1,796772153 | 0,813704497 |
| ERGIC33 | 0,24497009 | 0,544394927 | 0,194 | 0,067 | 1 | Mast | 1,458408553 | 0,685678919 | 2,895522388 |
| SRP72 | 0,245508311 | 0,789315109 | 0,185 | 0,067 | 1 | Mast | 1,728253814 | 0,578618715 | 2,76119403 |
| ISCU5 | 0,245508311 | 0,822090172 | 0,185 | 0,067 | 1 | Mast | 1,76796556 | 0,565621878 | 2,76119403 |
| EIF4A15 | 0,246541492 | -0,401566732 | 0,472 | 0,667 | 1 | Mast | 0,757035714 | 1,320941643 | 0,707646177 |
| TXNIP6 | 0,2473933 | -0,471272386 | 0,407 | 0,533 | 1 | Mast | 0,721328141 | 1,386331605 | 0,763602251 |
| PPP3CA | 0,247549403 | -0,568515126 | 0,102 | 0,2 | 1 | Mast | 0,674310456 | 1,482996431 | 0,51 |
| VMP11 | 0,248863266 | -0,378088213 | 0,194 | 0,333 | 1 | Mast | 0,76945656 | 1,299618525 | 0,582582583 |
| HCFC1R11 | 0,250332257 | 0,782476356 | 0,185 | 0,067 | 1 | Mast | 1,720080824 | 0,58136803 | 2,76119403 |
| MRPS213 | 0,250334034 | 0,712040254 | 0,185 | 0,067 | 1 | Mast | 1,638119102 | 0,610456223 | 2,76119403 |
| TGFBR21 | 0,250334034 | 0,635840092 | 0,185 | 0,067 | 1 | Mast | 1,553842301 | 0,643565952 | 2,76119403 |
| NCF1 | 0,250334034 | 0,750793457 | 0,185 | 0,067 | 1 | Mast | 1,682718041 | 0,594276626 | 2,76119403 |
| C9orf781 | 0,250334034 | 0,837634372 | 0,185 | 0,067 | 1 | Mast | 1,787117351 | 0,559560344 | 2,76119403 |
| ERRFI12 | 0,25089416 | -0,458179772 | 0,056 | 0,133 | 1 | Mast | 0,727904066 | 1,373807411 | 0,421052632 |
| CREB14 | 0,25089416 | -0,57267399 | 0,056 | 0,133 | 1 | Mast | 0,672369417 | 1,48727764 | 0,421052632 |
| AC008875.3 | 0,25089416 | -0,329483129 | 0,056 | 0,133 | 1 | Mast | 0,79582155 | 1,256563108 | 0,421052632 |
| UHRF1BP1L | 0,25089416 | -0,585143898 | 0,056 | 0,133 | 1 | Mast | 0,666582848 | 1,500188615 | 0,421052632 |
| LINC005111 | 0,25089416 | -0,564872635 | 0,056 | 0,133 | 1 | Mast | 0,676015095 | 1,47925691 | 0,421052632 |
| RGS101 | 0,251979917 | 0,797332447 | 0,454 | 0,333 | 1 | Mast | 1,737884793 | 0,575412135 | 1,363363363 |
| GHITM | 0,252771576 | -0,61694777 | 0,157 | 0,267 | 1 | Mast | 0,652048973 | 1,533627137 | 0,588014981 |
| HEBP23 | 0,255225607 | 0,809178121 | 0,185 | 0,067 | 1 | Mast | 1,752212952 | 0,570706888 | 2,76119403 |
| SPAG93 | 0,255225607 | 0,866111094 | 0,185 | 0,067 | 1 | Mast | 1,822742916 | 0,548623721 | 2,76119403 |

| Supplementary Table 4 - clusermarker | | | | | | | | | |
|---|---|---|---|---|---|---|---|---|---|
| | **p_val** | **avg_log2FC** | **pct.1** | **pct.2** | **p_val_adj** | **celltype** | **Foldchange_UP** | **Foldchange_DOWN** | **Ratio_pct1_pct2** |
| ARMCX31 | 0,255225607 | 0,681450997 | 0,185 | 0,067 | 1 | Mast | 1,603751925 | 0,623537833 | 2,76119403 |
| FXR14 | 0,255623224 | 0,876895409 | 0,176 | 0,067 | 1 | Mast | 1,836419185 | 0,544537983 | 2,626865672 |
| RRAGC | 0,258434649 | -0,404313513 | 0,056 | 0,133 | 1 | Mast | 0,755595747 | 1,323459011 | 0,421052632 |
| DHRS91 | 0,258434649 | -0,29031645 | 0,056 | 0,133 | 1 | Mast | 0,817722674 | 1,222908489 | 0,421052632 |
| IL5RA | 0,258434649 | -0,278866115 | 0,056 | 0,133 | 1 | Mast | 0,824238572 | 1,213240964 | 0,421052632 |
| HACL1 | 0,258434649 | -0,333883019 | 0,056 | 0,133 | 1 | Mast | 0,793398173 | 1,260401187 | 0,421052632 |
| SHPRH | 0,258434649 | -0,342412279 | 0,056 | 0,133 | 1 | Mast | 0,788721416 | 1,267874791 | 0,421052632 |
| CRCP | 0,258434649 | -0,360103334 | 0,056 | 0,133 | 1 | Mast | 0,779108773 | 1,283517827 | 0,421052632 |
| TATDN11 | 0,258434649 | -0,383772823 | 0,056 | 0,133 | 1 | Mast | 0,766430658 | 1,304749476 | 0,421052632 |
| CAMK2G | 0,258434649 | -0,490215082 | 0,056 | 0,133 | 1 | Mast | 0,711918955 | 1,404654271 | 0,421052632 |
| RDX | 0,258434649 | -0,330584686 | 0,056 | 0,133 | 1 | Mast | 0,795214139 | 1,257522912 | 0,421052632 |
| RNF102 | 0,258434649 | -0,329409287 | 0,056 | 0,133 | 1 | Mast | 0,795862284 | 1,256498794 | 0,421052632 |
| SNW11 | 0,258434649 | -0,269940278 | 0,056 | 0,133 | 1 | Mast | 0,829353877 | 1,205757913 | 0,421052632 |
| PDXK2 | 0,258434649 | -0,254451967 | 0,056 | 0,133 | 1 | Mast | 0,83830552 | 1,192882519 | 0,421052632 |
| ARHGAP6 | 0,258434649 | -0,364082773 | 0,056 | 0,133 | 1 | Mast | 0,77696269 | 1,287063089 | 0,421052632 |
| ZMAT1 | 0,258434649 | -0,327424682 | 0,056 | 0,133 | 1 | Mast | 0,796957844 | 1,254771514 | 0,421052632 |
| MTDH4 | 0,258531624 | 0,531334289 | 0,5 | 0,333 | 1 | Mast | 1,445265243 | 0,691914515 | 1,501501502 |
| ABRACL4 | 0,258939489 | 0,752033226 | 0,315 | 0,2 | 1 | Mast | 1,684164693 | 0,593766158 | 1,575 |
| MARCKSL12 | 0,259197415 | -0,340115152 | 0,102 | 0,2 | 1 | Mast | 0,789978256 | 1,265857627 | 0,51 |
| SMIM32 | 0,260687677 | -0,324332204 | 0,148 | 0,267 | 1 | Mast | 0,798667989 | 1,252084738 | 0,554307116 |
| RSL1D13 | 0,260687677 | 0,811107188 | 0,176 | 0,067 | 1 | Mast | 1,754557451 | 0,56994429 | 2,626865672 |
| PSMC51 | 0,260687677 | 0,952644686 | 0,176 | 0,067 | 1 | Mast | 1,935417331 | 0,51668443 | 2,626865672 |
| RPL283 | 0,26072959 | 0,262120147 | 0,935 | 1 | 1 | Mast | 1,199239782 | 0,833861597 | 0,935 |
| HNRNPA32 | 0,261101865 | 0,422635272 | 0,389 | 0,2 | 1 | Mast | 1,340373688 | 0,746060601 | 1,945 |
| MAN2A1 | 0,262686885 | -0,512063701 | 0,157 | 0,267 | 1 | Mast | 0,701218663 | 1,426088683 | 0,588014981 |
| ARL5B | 0,265161618 | -0,337577421 | 0,102 | 0,2 | 1 | Mast | 0,791369067 | 1,263632914 | 0,51 |
| WAC1 | 0,265161618 | -0,359999379 | 0,102 | 0,2 | 1 | Mast | 0,779164915 | 1,283425345 | 0,51 |
| MRPS163 | 0,265161618 | -0,520304284 | 0,102 | 0,2 | 1 | Mast | 0,697224764 | 1,434257721 | 0,51 |
| DCAF7 | 0,265161618 | -0,418429429 | 0,102 | 0,2 | 1 | Mast | 0,748238741 | 1,33647183 | 0,51 |
| ATP5MJ1 | 0,265207118 | 0,546434455 | 0,185 | 0,067 | 1 | Mast | 1,460471753 | 0,684710264 | 2,76119403 |
| LMAN24 | 0,265819253 | -0,411610605 | 0,148 | 0,267 | 1 | Mast | 0,751783624 | 1,330169969 | 0,554307116 |

| Supplementary Table 4 - clusermarker | | | | | | | | | |
|---|---|---|---|---|---|---|---|---|---|
| | p_val | avg_log2FC | pct.1 | pct.2 | p_val_adj | celltype | Foldchange_UP | Foldchange_DOWN | Ratio_pct1_pct2 |
| KDELR23 | 0,26582112 | 0,92027475 | 0,176 | 0,067 | 1 | Mast | 1,892475667 | 0,528408379 | 2,626865672 |
| IFRD11 | 0,26582112 | -0,395529265 | 0,148 | 0,267 | 1 | Mast | 0,760210436 | 1,315425247 | 0,554307116 |
| SLC39A101 | 0,266130801 | -0,304377792 | 0,056 | 0,133 | 1 | Mast | 0,809791389 | 1,234885939 | 0,421052632 |
| IP6K21 | 0,266130801 | -0,264820528 | 0,056 | 0,133 | 1 | Mast | 0,832302267 | 1,201486575 | 0,421052632 |
| CCDC1071 | 0,266130801 | -0,308456522 | 0,056 | 0,133 | 1 | Mast | 0,807505213 | 1,238382099 | 0,421052632 |
| CMA1 | 0,266130801 | -0,265811346 | 0,056 | 0,133 | 1 | Mast | 0,831730852 | 1,202312019 | 0,421052632 |
| AC004687.11 | 0,266130801 | -0,299680728 | 0,056 | 0,133 | 1 | Mast | 0,81243217 | 1,230871988 | 0,421052632 |
| NFATC2 | 0,266130801 | -0,367673301 | 0,056 | 0,133 | 1 | Mast | 0,775031418 | 1,290270275 | 0,421052632 |
| GDI11 | 0,266741227 | -0,808654793 | 0,167 | 0,267 | 1 | Mast | 0,570913945 | 1,751577464 | 0,625468165 |
| C4orf3 | 0,270172868 | 0,70671697 | 0,259 | 0,133 | 1 | Mast | 1,632085876 | 0,612712857 | 1,947368421 |
| CLTA3 | 0,270172868 | 0,812102603 | 0,259 | 0,133 | 1 | Mast | 1,755768459 | 0,569551181 | 1,947368421 |
| RAB5A | 0,270297429 | 0,608759826 | 0,185 | 0,067 | 1 | Mast | 1,524947765 | 0,655760166 | 2,76119403 |
| ARHGEF121 | 0,270297429 | 0,776239889 | 0,185 | 0,067 | 1 | Mast | 1,712661325 | 0,583886601 | 2,76119403 |
| HNRNPAB3 | 0,270927249 | -0,483598603 | 0,278 | 0,4 | 1 | Mast | 0,715191451 | 1,398227004 | 0,695 |
| SNX29 | 0,271021868 | 0,784746726 | 0,176 | 0,067 | 1 | Mast | 1,722789846 | 0,58045385 | 2,626865672 |
| MRPL52 | 0,27102374 | 0,779272807 | 0,176 | 0,067 | 1 | Mast | 1,716265568 | 0,58266041 | 2,626865672 |
| RALBP12 | 0,27102374 | -0,346381328 | 0,148 | 0,267 | 1 | Mast | 0,786554518 | 1,271367689 | 0,554307116 |
| EIF4A31 | 0,27177768 | -0,27740468 | 0,194 | 0,333 | 1 | Mast | 0,82507394 | 1,212012586 | 0,582582583 |
| FOXP11 | 0,272282399 | 0,561704435 | 0,435 | 0,267 | 1 | Mast | 1,476011984 | 0,677501274 | 1,629213483 |
| PTAFR2 | 0,273983261 | -0,311599802 | 0,056 | 0,133 | 1 | Mast | 0,805747771 | 1,241083172 | 0,421052632 |
| EMC21 | 0,273983261 | -0,289570697 | 0,056 | 0,133 | 1 | Mast | 0,818145478 | 1,222276511 | 0,421052632 |
| TRAPPC4 | 0,273983261 | -0,295511751 | 0,056 | 0,133 | 1 | Mast | 0,814783262 | 1,227320254 | 0,421052632 |
| CLIP1 | 0,273983261 | -0,286105784 | 0,056 | 0,133 | 1 | Mast | 0,820112775 | 1,219344498 | 0,421052632 |
| RABEP1 | 0,273983261 | -0,277183993 | 0,056 | 0,133 | 1 | Mast | 0,82520016 | 1,2118272 | 0,421052632 |
| C4orf481 | 0,274515279 | 0,48746655 | 0,361 | 0,2 | 1 | Mast | 1,401980759 | 0,713276551 | 1,805 |
| UBE2I4 | 0,275454331 | 0,66482825 | 0,185 | 0,067 | 1 | Mast | 1,585379526 | 0,630763791 | 2,76119403 |
| RTRAF2 | 0,276295705 | 0,918183113 | 0,176 | 0,067 | 1 | Mast | 1,88973392 | 0,529175028 | 2,626865672 |
| NEDD91 | 0,276765678 | 0,723432943 | 0,167 | 0,067 | 1 | Mast | 1,651106228 | 0,60565455 | 2,492537313 |
| GPS22 | 0,276765678 | 0,862255296 | 0,167 | 0,067 | 1 | Mast | 1,817877892 | 0,550091953 | 2,492537313 |
| MON21 | 0,277371922 | -0,373685979 | 0,102 | 0,2 | 1 | Mast | 0,771808063 | 1,295658918 | 0,51 |
| NFAT51 | 0,277371922 | -0,348282151 | 0,102 | 0,2 | 1 | Mast | 0,785518876 | 1,273043882 | 0,51 |

| Supplementary Table 4 - clusermarker | | | | | | | | | |
|---|---|---|---|---|---|---|---|---|---|
| | **p_val** | **avg_log2FC** | **pct.1** | **pct.2** | **p_val_adj** | **celltype** | **Foldchange_UP** | **Foldchange_DOWN** | **Ratio_pct1_pct2** |
| PPP2CA | 0,278057799 | -0,282697696 | 0,157 | 0,267 | 1 | Mast | 0,822052423 | 1,216467432 | 0,588014981 |
| BEX4 | 0,280677968 | 0,579881243 | 0,185 | 0,067 | 1 | Mast | 1,494726204 | 0,669018846 | 2,76119403 |
| SRSF74 | 0,281202453 | -0,313577227 | 0,269 | 0,4 | 1 | Mast | 0,804644133 | 1,242785425 | 0,6725 |
| TMEM14C4 | 0,281396415 | -0,296550829 | 0,306 | 0,467 | 1 | Mast | 0,814196639 | 1,22820453 | 0,655246253 |
| CCNH3 | 0,281443974 | 0,445292099 | 0,278 | 0,133 | 1 | Mast | 1,361589766 | 0,734435602 | 2,090225564 |
| PNP | 0,281637167 | -0,386450585 | 0,148 | 0,267 | 1 | Mast | 0,765009419 | 1,307173449 | 0,554307116 |
| PTTG1IP2 | 0,281637167 | 0,702744124 | 0,176 | 0,067 | 1 | Mast | 1,627597674 | 0,614402451 | 2,626865672 |
| GMFG | 0,282346671 | -0,728508788 | 0,222 | 0,333 | 1 | Mast | 0,603527415 | 1,656925559 | 0,666666667 |
| PPP1R22 | 0,283313047 | -0,297205184 | 0,157 | 0,267 | 1 | Mast | 0,813827432 | 1,228761726 | 0,588014981 |
| MARCHF21 | 0,283618529 | -0,271784136 | 0,102 | 0,2 | 1 | Mast | 0,828294586 | 1,207299935 | 0,51 |
| FLOT11 | 0,283722824 | 0,660752372 | 0,333 | 0,2 | 1 | Mast | 1,580906859 | 0,632548334 | 1,665 |
| CDKN2AIP1 | 0,285397631 | 0,817335459 | 0,241 | 0,133 | 1 | Mast | 1,76214844 | 0,567489082 | 1,812030075 |
| NEK1 | 0,285768804 | -0,54053907 | 0,111 | 0,2 | 1 | Mast | 0,687513968 | 1,454515902 | 0,555 |
| ANXA6 | 0,285768804 | -0,314138914 | 0,111 | 0,2 | 1 | Mast | 0,80433092 | 1,243269375 | 0,555 |
| SF3B41 | 0,287766142 | 0,674576244 | 0,167 | 0,067 | 1 | Mast | 1,596127881 | 0,626516216 | 2,492537313 |
| RAB101 | 0,287766142 | 0,664779087 | 0,167 | 0,067 | 1 | Mast | 1,585325501 | 0,630785286 | 2,492537313 |
| KIF5B2 | 0,287766142 | 0,76392347 | 0,167 | 0,067 | 1 | Mast | 1,698102412 | 0,588892633 | 2,492537313 |
| FUBP1 | 0,288531741 | -0,523031581 | 0,065 | 0,133 | 1 | Mast | 0,695907962 | 1,436971632 | 0,488721805 |
| MFN1 | 0,288531741 | -0,614512564 | 0,065 | 0,133 | 1 | Mast | 0,653150533 | 1,531040625 | 0,488721805 |
| GLS4 | 0,28864951 | -0,250347478 | 0,194 | 0,333 | 1 | Mast | 0,840693907 | 1,189493574 | 0,582582583 |
| ZNF521 | 0,290159238 | -0,254684069 | 0,056 | 0,133 | 1 | Mast | 0,838170663 | 1,193074447 | 0,421052632 |
| OSTC2 | 0,291325946 | 0,468059719 | 0,185 | 0,067 | 1 | Mast | 1,383247886 | 0,722936221 | 2,76119403 |
| YWHAE2 | 0,291454409 | 0,56191685 | 0,352 | 0,2 | 1 | Mast | 1,47622932 | 0,67740153 | 1,76 |
| BCL3 | 0,291750847 | -0,64845785 | 0,222 | 0,333 | 1 | Mast | 0,63796189 | 1,56749175 | 0,666666667 |
| ANKRD102 | 0,292529094 | 0,845934185 | 0,176 | 0,067 | 1 | Mast | 1,797428253 | 0,55635044 | 2,626865672 |
| BRAF | 0,293181838 | -1,132950245 | 0,12 | 0,2 | 1 | Mast | 0,455982308 | 2,193067542 | 0,6 |
| FIBP2 | 0,293375519 | 0,767111791 | 0,167 | 0,067 | 1 | Mast | 1,701859327 | 0,587592631 | 2,492537313 |
| RNH11 | 0,294133004 | 0,813772412 | 0,25 | 0,133 | 1 | Mast | 1,757801804 | 0,568892351 | 1,879699248 |
| LMNA4 | 0,294332934 | -0,628605819 | 0,685 | 0,667 | 1 | Mast | 0,646801164 | 1,546070191 | 1,026986507 |
| PPDPF5 | 0,294475037 | 0,549576033 | 0,648 | 0,6 | 1 | Mast | 1,463655506 | 0,683220878 | 1,08 |
| STX171 | 0,296379191 | -0,838963735 | 0,065 | 0,133 | 1 | Mast | 0,559044978 | 1,788764838 | 0,488721805 |

| Supplementary Table 4 - clusermarker | | | | | | | | | |
|---|---|---|---|---|---|---|---|---|---|
| | **p_val** | **avg_log2FC** | **pct.1** | **pct.2** | **p_val_adj** | **celltype** | **Foldchange_UP** | **Foldchange_DOWN** | **Ratio_pct1_pct2** |
| DDX422 | 0,296379191 | -0,588903356 | 0,065 | 0,133 | 1 | Mast | 0,66484809 | 1,504102991 | 0,488721805 |
| MRTFA | 0,296379191 | -0,726082325 | 0,065 | 0,133 | 1 | Mast | 0,60454334 | 1,654141125 | 0,488721805 |
| PRPS21 | 0,29639566 | -0,284594258 | 0,102 | 0,2 | 1 | Mast | 0,820972466 | 1,218067648 | 0,51 |
| PA2G41 | 0,296750491 | 0,458484504 | 0,185 | 0,067 | 1 | Mast | 1,374097623 | 0,727750331 | 2,76119403 |
| SLC25A64 | 0,296759245 | 0,651902632 | 0,509 | 0,467 | 1 | Mast | 1,571238987 | 0,63644042 | 1,08993576 |
| DHRS72 | 0,296872122 | 0,806510452 | 0,315 | 0,2 | 1 | Mast | 1,748975952 | 0,57176315 | 1,575 |
| MARCHF3 | 0,298222116 | -0,512395626 | 0,111 | 0,2 | 1 | Mast | 0,70105735 | 1,426416826 | 0,555 |
| SARNP2 | 0,298222116 | -0,397908905 | 0,111 | 0,2 | 1 | Mast | 0,758957547 | 1,317596753 | 0,555 |
| MYDGF3 | 0,299057799 | 0,772033862 | 0,167 | 0,067 | 1 | Mast | 1,707675509 | 0,585591346 | 2,492537313 |
| TMEM2301 | 0,299057799 | 0,756276465 | 0,167 | 0,067 | 1 | Mast | 1,689125433 | 0,592022345 | 2,492537313 |
| TLN12 | 0,29997549 | 0,690823529 | 0,389 | 0,267 | 1 | Mast | 1,614204687 | 0,619500122 | 1,456928839 |
| OAZ14 | 0,3004419 | 0,337975095 | 0,75 | 0,8 | 1 | Mast | 1,263981278 | 0,791150959 | 0,9375 |
| RBX12 | 0,300483104 | -0,672457952 | 0,167 | 0,267 | 1 | Mast | 0,627436798 | 1,593786025 | 0,625468165 |
| EIF3K2 | 0,301129624 | 0,679148538 | 0,565 | 0,533 | 1 | Mast | 1,601194469 | 0,624533758 | 1,060037523 |
| TFDP22 | 0,302242181 | 0,301112586 | 0,185 | 0,067 | 1 | Mast | 1,232094221 | 0,81162624 | 2,76119403 |
| USF31 | 0,304368177 | -0,506224352 | 0,065 | 0,133 | 1 | Mast | 0,704062617 | 1,420328216 | 0,488721805 |
| RALB | 0,304582725 | -0,380507573 | 0,111 | 0,2 | 1 | Mast | 0,768167284 | 1,301799778 | 0,555 |
| PPIP5K22 | 0,304582725 | -0,445947836 | 0,111 | 0,2 | 1 | Mast | 0,734101861 | 1,36220878 | 0,555 |
| IK3 | 0,304582725 | -0,459458891 | 0,111 | 0,2 | 1 | Mast | 0,727258979 | 1,375025993 | 0,555 |
| MCTP2 | 0,304582725 | -0,510178746 | 0,111 | 0,2 | 1 | Mast | 0,70213544 | 1,424226643 | 0,555 |
| STRAP1 | 0,30481109 | 0,830058886 | 0,167 | 0,067 | 1 | Mast | 1,777757923 | 0,562506282 | 2,492537313 |
| EIF1AX5 | 0,30481147 | 0,802648808 | 0,241 | 0,133 | 1 | Mast | 1,744300749 | 0,573295632 | 1,812030075 |
| EML41 | 0,304813061 | 0,676270473 | 0,167 | 0,067 | 1 | Mast | 1,598003395 | 0,625780898 | 2,492537313 |
| YWHAH3 | 0,304813061 | 0,77257246 | 0,167 | 0,067 | 1 | Mast | 1,708313151 | 0,585372769 | 2,492537313 |
| CHMP52 | 0,309549727 | -0,278449298 | 0,102 | 0,2 | 1 | Mast | 0,824476742 | 1,21289049 | 0,51 |
| IFI161 | 0,311032753 | -0,37223955 | 0,111 | 0,2 | 1 | Mast | 0,772582257 | 1,294360556 | 0,555 |
| DIAPH11 | 0,311032753 | -0,441678284 | 0,111 | 0,2 | 1 | Mast | 0,7362776 | 1,358183381 | 0,555 |
| ZBTB202 | 0,311776969 | -0,824245587 | 0,12 | 0,2 | 1 | Mast | 0,564777458 | 1,770608911 | 0,6 |
| ALDOA3 | 0,312327005 | 0,545371454 | 0,259 | 0,133 | 1 | Mast | 1,45939605 | 0,685214956 | 1,947368421 |
| TDG1 | 0,312498868 | -0,690610453 | 0,065 | 0,133 | 1 | Mast | 0,619591624 | 1,613966298 | 0,488721805 |
| ETS21 | 0,312498868 | -0,549520363 | 0,065 | 0,133 | 1 | Mast | 0,683247242 | 1,463599029 | 0,488721805 |

| Supplementary Table 4 - clusermarker | | | | | | | | | |
|---|---|---|---|---|---|---|---|---|---|
| | **p_val** | **avg_log2FC** | **pct.1** | **pct.2** | **p_val_adj** | **celltype** | **Foldchange_UP** | **Foldchange_DOWN** | **Ratio_pct1_pct2** |
| SEC61G1 | 0,313672148 | 0,753112946 | 0,25 | 0,133 | 1 | Mast | 1,685425602 | 0,593321947 | 1,879699248 |
| JAK13 | 0,315151506 | 0,485434471 | 0,176 | 0,067 | 1 | Mast | 1,400007417 | 0,71428193 | 2,626865672 |
| TIPRL | 0,316540762 | 0,522093935 | 0,167 | 0,067 | 1 | Mast | 1,436038009 | 0,696360398 | 2,492537313 |
| PHLDA11 | 0,316542734 | 0,683803285 | 0,167 | 0,067 | 1 | Mast | 1,606368946 | 0,622521994 | 2,492537313 |
| TNS1 | 0,317549113 | 0,796775882 | 0,157 | 0,067 | 1 | Mast | 1,737214478 | 0,575634162 | 2,343283582 |
| PYCARD4 | 0,320599549 | 0,398388814 | 0,269 | 0,133 | 1 | Mast | 1,318035122 | 0,758705124 | 2,022556391 |
| SRM | 0,320771355 | -0,375262109 | 0,065 | 0,133 | 1 | Mast | 0,770965331 | 1,297075186 | 0,488721805 |
| IFI62 | 0,320771355 | -0,459795078 | 0,065 | 0,133 | 1 | Mast | 0,727089528 | 1,375346449 | 0,488721805 |
| PIK3R32 | 0,320771355 | -0,568634897 | 0,065 | 0,133 | 1 | Mast | 0,674254478 | 1,483119553 | 0,488721805 |
| CD58 | 0,320771355 | -0,400398398 | 0,065 | 0,133 | 1 | Mast | 0,757649031 | 1,319872342 | 0,488721805 |
| XPO11 | 0,320771355 | -0,303817675 | 0,065 | 0,133 | 1 | Mast | 0,810105847 | 1,234406595 | 0,488721805 |
| ANKRD374 | 0,320771355 | -0,514079212 | 0,065 | 0,133 | 1 | Mast | 0,700239712 | 1,428082387 | 0,488721805 |
| ZFR | 0,320771355 | -0,372480858 | 0,065 | 0,133 | 1 | Mast | 0,772453044 | 1,294577072 | 0,488721805 |
| STARD4 | 0,320771355 | -0,497169015 | 0,065 | 0,133 | 1 | Mast | 0,708495691 | 1,411441188 | 0,488721805 |
| PPA12 | 0,320771355 | -0,5286821 | 0,065 | 0,133 | 1 | Mast | 0,693187671 | 1,442610771 | 0,488721805 |
| RBBP81 | 0,320771355 | -0,670879401 | 0,065 | 0,133 | 1 | Mast | 0,628123695 | 1,592043109 | 0,488721805 |
| JOSD11 | 0,320771355 | -0,579576186 | 0,065 | 0,133 | 1 | Mast | 0,669160325 | 1,494410178 | 0,488721805 |
| OGT | 0,320771355 | -0,372309254 | 0,065 | 0,133 | 1 | Mast | 0,77254493 | 1,294423095 | 0,488721805 |
| NKAP1 | 0,320771355 | -0,506967331 | 0,065 | 0,133 | 1 | Mast | 0,703700122 | 1,421059864 | 0,488721805 |
| HIGD2A4 | 0,320982112 | 0,424323623 | 0,176 | 0,067 | 1 | Mast | 1,341943214 | 0,745188015 | 2,626865672 |
| DYNC1H12 | 0,321800019 | -0,371872816 | 0,287 | 0,4 | 1 | Mast | 0,772778673 | 1,294031571 | 0,7175 |
| PAK12 | 0,32195789 | -0,291328733 | 0,315 | 0,467 | 1 | Mast | 0,817149111 | 1,223766858 | 0,674518201 |
| SLC25A53 | 0,32294964 | 0,495984964 | 0,444 | 0,333 | 1 | Mast | 1,410283262 | 0,709077408 | 1,333333333 |
| VPS283 | 0,323286078 | 0,572126113 | 0,333 | 0,2 | 1 | Mast | 1,48671294 | 0,672624804 | 1,665 |
| AFF11 | 0,324201223 | -0,397134571 | 0,111 | 0,2 | 1 | Mast | 0,75936501 | 1,316889753 | 0,555 |
| CSNK2B5 | 0,324626155 | 0,600386631 | 0,38 | 0,267 | 1 | Mast | 1,516122821 | 0,65957717 | 1,423220974 |
| FUS4 | 0,324922316 | -0,466021697 | 0,481 | 0,533 | 1 | Mast | 0,723958198 | 1,381295221 | 0,902439024 |
| BTK1 | 0,326480904 | -0,255370699 | 0,315 | 0,467 | 1 | Mast | 0,837771842 | 1,193642409 | 0,674518201 |
| DDX242 | 0,326821363 | -0,306138695 | 0,269 | 0,4 | 1 | Mast | 0,808803589 | 1,236394117 | 0,6725 |
| JTB6 | 0,326882743 | 0,423146939 | 0,176 | 0,067 | 1 | Mast | 1,340849151 | 0,74579605 | 2,626865672 |
| ATP5MK3 | 0,32806239 | -0,373219142 | 0,241 | 0,333 | 1 | Mast | 0,77205785 | 1,295239728 | 0,723723724 |

| Supplementary Table 4 - clusermarker | | | | | | | | | |
|---|---|---|---|---|---|---|---|---|---|
| | **p_val** | **avg_log2FC** | **pct.1** | **pct.2** | **p_val_adj** | **celltype** | **Foldchange_UP** | **Foldchange_DOWN** | **Ratio_pct1_pct2** |
| ZC3H15 | 0,328294717 | -0,361609929 | 0,167 | 0,267 | 1 | Mast | 0,778295581 | 1,284858895 | 0,625468165 |
| EZR3 | 0,328294717 | -0,337494692 | 0,167 | 0,267 | 1 | Mast | 0,791414448 | 1,263560455 | 0,625468165 |
| PRKDC2 | 0,328564736 | 0,635702481 | 0,167 | 0,067 | 1 | Mast | 1,553694095 | 0,643627342 | 2,492537313 |
| CAPG3 | 0,328864933 | 0,350139454 | 0,778 | 0,667 | 1 | Mast | 1,274683835 | 0,784508262 | 1,166416792 |
| ZMYM6 | 0,329185643 | -0,384692731 | 0,065 | 0,133 | 1 | Mast | 0,765942114 | 1,305581691 | 0,488721805 |
| TMCO13 | 0,329185643 | -0,303142274 | 0,065 | 0,133 | 1 | Mast | 0,810485188 | 1,23382884 | 0,488721805 |
| GLB1 | 0,329185643 | -0,35173865 | 0,065 | 0,133 | 1 | Mast | 0,783639134 | 1,276097578 | 0,488721805 |
| MAML3 | 0,329185643 | -0,463641553 | 0,065 | 0,133 | 1 | Mast | 0,725153563 | 1,379018254 | 0,488721805 |
| NUDT3 | 0,329185643 | -0,413473238 | 0,065 | 0,133 | 1 | Mast | 0,750813638 | 1,331888433 | 0,488721805 |
| NANS4 | 0,329185643 | -0,482534333 | 0,065 | 0,133 | 1 | Mast | 0,715719239 | 1,397195918 | 0,488721805 |
| NXF1 | 0,329185643 | -0,403744858 | 0,065 | 0,133 | 1 | Mast | 0,755893633 | 1,322937456 | 0,488721805 |
| ATG101 | 0,329185643 | -0,497260988 | 0,065 | 0,133 | 1 | Mast | 0,708450526 | 1,411531171 | 0,488721805 |
| NDUFB24 | 0,329454644 | -0,275649911 | 0,417 | 0,6 | 1 | Mast | 0,826078099 | 1,210539295 | 0,695 |
| RNF145 | 0,329876709 | 0,69456039 | 0,157 | 0,067 | 1 | Mast | 1,618391211 | 0,617897572 | 2,343283582 |
| EPC13 | 0,330282071 | -0,251238462 | 0,213 | 0,333 | 1 | Mast | 0,840174869 | 1,190228412 | 0,63963964 |
| NAMPT | 0,330919663 | -0,295046634 | 0,111 | 0,2 | 1 | Mast | 0,815045986 | 1,226924636 | 0,555 |
| RAB11A2 | 0,330919663 | -0,41503274 | 0,111 | 0,2 | 1 | Mast | 0,750002474 | 1,333328935 | 0,555 |
| CAPNS12 | 0,330919663 | -0,394666198 | 0,111 | 0,2 | 1 | Mast | 0,760665355 | 1,314638552 | 0,555 |
| PRDX62 | 0,332459424 | -0,3983851 | 0,491 | 0,6 | 1 | Mast | 0,758707077 | 1,318031728 | 0,818333333 |
| CEP3502 | 0,334685326 | 0,414916306 | 0,167 | 0,067 | 1 | Mast | 1,333221331 | 0,750063006 | 2,492537313 |
| NONO2 | 0,334685326 | 0,421358597 | 0,167 | 0,067 | 1 | Mast | 1,339188085 | 0,7467211 | 2,492537313 |
| PRDX44 | 0,336153127 | 0,704296751 | 0,157 | 0,067 | 1 | Mast | 1,629350236 | 0,613741587 | 2,343283582 |
| S100A113 | 0,336623571 | 0,391139652 | 0,731 | 0,667 | 1 | Mast | 1,311428954 | 0,762527011 | 1,095952024 |
| KIAA1549 | 0,337727515 | -0,374032242 | 0,111 | 0,2 | 1 | Mast | 0,771622843 | 1,295969928 | 0,555 |
| PPP4C5 | 0,337727515 | -0,335358563 | 0,111 | 0,2 | 1 | Mast | 0,792587125 | 1,261690947 | 0,555 |
| PEA15 | 0,337737836 | -0,516271717 | 0,065 | 0,133 | 1 | Mast | 0,699176346 | 1,430254335 | 0,488721805 |
| CIAO1 | 0,337741657 | -0,443660452 | 0,065 | 0,133 | 1 | Mast | 0,735266698 | 1,360050718 | 0,488721805 |
| SMC51 | 0,337741657 | -0,371598179 | 0,065 | 0,133 | 1 | Mast | 0,772925796 | 1,293785258 | 0,488721805 |
| EEA1 | 0,337741657 | -0,56485761 | 0,065 | 0,133 | 1 | Mast | 0,676022135 | 1,479241505 | 0,488721805 |
| SUPT16H2 | 0,337741657 | -0,335865326 | 0,065 | 0,133 | 1 | Mast | 0,792308769 | 1,262134207 | 0,488721805 |
| SMARCD22 | 0,337741657 | -0,386586591 | 0,065 | 0,133 | 1 | Mast | 0,764937303 | 1,307296685 | 0,488721805 |

| Supplementary Table 4 - clusermarker | | | | | | | | | |
|---|---|---|---|---|---|---|---|---|---|
| | **p_val** | **avg_log2FC** | **pct.1** | **pct.2** | **p_val_adj** | **celltype** | **Foldchange_UP** | **Foldchange_DOWN** | **Ratio_pct1_pct2** |
| GATA1 | 0,337741657 | -0,301298597 | 0,065 | 0,133 | 1 | Mast | 0,811521602 | 1,232253088 | 0,488721805 |
| BEX31 | 0,337741657 | -0,47453757 | 0,065 | 0,133 | 1 | Mast | 0,719697439 | 1,389472778 | 0,488721805 |
| H2AX2 | 0,337756761 | -0,65655193 | 0,12 | 0,2 | 1 | Mast | 0,634392698 | 1,576310704 | 0,6 |
| NDUFAB15 | 0,337756761 | -0,549637572 | 0,12 | 0,2 | 1 | Mast | 0,683191736 | 1,46371794 | 0,6 |
| AL138963.44 | 0,340845374 | 0,602453893 | 0,241 | 0,133 | 1 | Mast | 1,518296857 | 0,658632728 | 1,812030075 |
| CHMP2A3 | 0,340878911 | 0,431406372 | 0,167 | 0,067 | 1 | Mast | 1,348547531 | 0,741538564 | 2,492537313 |
| WBP14 | 0,342510047 | 0,608423256 | 0,157 | 0,067 | 1 | Mast | 1,524592047 | 0,655913168 | 2,343283582 |
| KCTD20 | 0,342510047 | 0,69659663 | 0,157 | 0,067 | 1 | Mast | 1,620677043 | 0,617026078 | 2,343283582 |
| JMJD1C1 | 0,342510047 | 0,604389928 | 0,157 | 0,067 | 1 | Mast | 1,520335713 | 0,657749464 | 2,343283582 |
| SNAP231 | 0,342510047 | 0,608004492 | 0,157 | 0,067 | 1 | Mast | 1,524149576 | 0,656103585 | 2,343283582 |
| ST132 | 0,343182404 | 0,626929447 | 0,315 | 0,2 | 1 | Mast | 1,544274743 | 0,647553166 | 1,575 |
| ARGLU11 | 0,343437644 | 0,860073113 | 0,222 | 0,133 | 1 | Mast | 1,815130296 | 0,550924637 | 1,669172932 |
| ODC11 | 0,344001087 | 0,414767827 | 0,167 | 0,067 | 1 | Mast | 1,333084126 | 0,750140205 | 2,492537313 |
| CST33 | 0,345131869 | 0,525572947 | 0,778 | 0,8 | 1 | Mast | 1,439505147 | 0,694683171 | 0,9725 |
| SARAF2 | 0,346424962 | 0,515256722 | 0,657 | 0,533 | 1 | Mast | 1,429248447 | 0,699668418 | 1,232645403 |
| NRBP11 | 0,346439233 | -0,273913944 | 0,065 | 0,133 | 1 | Mast | 0,827072702 | 1,209083552 | 0,488721805 |
| TRNT1 | 0,346439233 | -0,479712184 | 0,065 | 0,133 | 1 | Mast | 0,717120674 | 1,394465445 | 0,488721805 |
| OARD1 | 0,346439233 | -0,366693318 | 0,065 | 0,133 | 1 | Mast | 0,775558054 | 1,289394127 | 0,488721805 |
| BAD1 | 0,346439233 | -0,333412116 | 0,065 | 0,133 | 1 | Mast | 0,793657185 | 1,259989854 | 0,488721805 |
| SART3 | 0,346439233 | -0,393791984 | 0,065 | 0,133 | 1 | Mast | 0,761126426 | 1,313842176 | 0,488721805 |
| CHMP1B2 | 0,346439233 | -0,580686024 | 0,065 | 0,133 | 1 | Mast | 0,66864575 | 1,495560242 | 0,488721805 |
| INTS64 | 0,347462761 | -0,448558022 | 0,296 | 0,4 | 1 | Mast | 0,732774893 | 1,364675578 | 0,74 |
| PNRC15 | 0,347499779 | 0,661645238 | 0,5 | 0,467 | 1 | Mast | 1,581885565 | 0,632156979 | 1,070663812 |
| GADD45GIP12 | 0,348941198 | 0,551672386 | 0,157 | 0,067 | 1 | Mast | 1,465783862 | 0,682228824 | 2,343283582 |
| ANP32B2 | 0,349872254 | 0,494592801 | 0,25 | 0,133 | 1 | Mast | 1,408923032 | 0,709761979 | 1,879699248 |
| GPR35 | 0,350333709 | 0,803832937 | 0,148 | 0,067 | 1 | Mast | 1,745733017 | 0,572825278 | 2,208955224 |
| HNRNPU2 | 0,350820835 | 0,609450352 | 0,472 | 0,4 | 1 | Mast | 1,525677834 | 0,655446371 | 1,18 |
| XRN22 | 0,351252461 | -0,387457806 | 0,12 | 0,2 | 1 | Mast | 0,764475512 | 1,308086374 | 0,6 |
| PROCR | 0,351611081 | -0,467040869 | 0,111 | 0,2 | 1 | Mast | 0,723446949 | 1,382271363 | 0,555 |
| TM9SF22 | 0,352968197 | 0,687775465 | 0,315 | 0,2 | 1 | Mast | 1,610797865 | 0,620810358 | 1,575 |
| TSPYL21 | 0,35348014 | -0,398598721 | 0,278 | 0,4 | 1 | Mast | 0,758594743 | 1,318226905 | 0,695 |

| Supplementary Table 4 - clusermarker | | | | | | | | | |
|---|---|---|---|---|---|---|---|---|---|
| | **p_val** | **avg_log2FC** | **pct.1** | **pct.2** | **p_val_adj** | **celltype** | **Foldchange_UP** | **Foldchange_DOWN** | **Ratio_pct1_pct2** |
| FARS2 | 0,355278124 | -0,269281129 | 0,065 | 0,133 | 1 | Mast | 0,829732885 | 1,205207144 | 0,488721805 |
| SUMF21 | 0,355278124 | -0,26856074 | 0,065 | 0,133 | 1 | Mast | 0,830147304 | 1,204605491 | 0,488721805 |
| CFDP11 | 0,355278124 | -0,323680595 | 0,065 | 0,133 | 1 | Mast | 0,799028798 | 1,251519348 | 0,488721805 |
| SUZ121 | 0,355278124 | -0,279430197 | 0,065 | 0,133 | 1 | Mast | 0,823916365 | 1,213715424 | 0,488721805 |
| TMEM1541 | 0,355448528 | 0,600155203 | 0,157 | 0,067 | 1 | Mast | 1,515879634 | 0,659682984 | 2,343283582 |
| MRPL413 | 0,355448528 | 0,60536466 | 0,157 | 0,067 | 1 | Mast | 1,521363249 | 0,657305217 | 2,343283582 |
| HPCAL1 | 0,357018978 | 0,694005896 | 0,148 | 0,067 | 1 | Mast | 1,617769309 | 0,618135104 | 2,208955224 |
| ACP5 | 0,357018978 | 0,635312257 | 0,148 | 0,067 | 1 | Mast | 1,553273904 | 0,643801455 | 2,208955224 |
| CIAO2B4 | 0,358757584 | 0,740842521 | 0,231 | 0,133 | 1 | Mast | 1,671151492 | 0,598389796 | 1,736842105 |
| CCL42 | 0,360683771 | 0,867485924 | 0,25 | 0,133 | 1 | Mast | 1,824480744 | 0,548101153 | 1,879699248 |
| PSMF11 | 0,362031899 | 0,465202083 | 0,157 | 0,067 | 1 | Mast | 1,380510712 | 0,724369605 | 2,343283582 |
| LSM73 | 0,362540075 | -0,531301233 | 0,185 | 0,267 | 1 | Mast | 0,691930369 | 1,445232129 | 0,692883895 |
| FERMT21 | 0,363784039 | 0,65286571 | 0,148 | 0,067 | 1 | Mast | 1,572288224 | 0,636015703 | 2,208955224 |
| SCAMP21 | 0,363784039 | 0,77890061 | 0,148 | 0,067 | 1 | Mast | 1,715822851 | 0,582810748 | 2,208955224 |
| SAMSN12 | 0,363991502 | -0,288956625 | 0,454 | 0,533 | 1 | Mast | 0,81849379 | 1,221756368 | 0,851782364 |
| HERPUD1 | 0,364023859 | 0,745257992 | 0,296 | 0,2 | 1 | Mast | 1,676274005 | 0,596561181 | 1,48 |
| TMEM1343 | 0,364257994 | -0,257388766 | 0,065 | 0,133 | 1 | Mast | 0,836600772 | 1,195313265 | 0,488721805 |
| SCAF112 | 0,364362165 | -0,400851499 | 0,241 | 0,333 | 1 | Mast | 0,757411117 | 1,320286933 | 0,723723724 |
| TOMM54 | 0,36508495 | -0,491353028 | 0,12 | 0,2 | 1 | Mast | 0,71135764 | 1,405762648 | 0,6 |
| SAR1B1 | 0,365418324 | -0,453224868 | 0,074 | 0,133 | 1 | Mast | 0,730408334 | 1,369097193 | 0,556390977 |
| MTPN2 | 0,365950164 | 0,695155627 | 0,222 | 0,133 | 1 | Mast | 1,619059075 | 0,617642688 | 1,669172932 |
| ITPRID21 | 0,366381342 | 0,371005626 | 0,167 | 0,067 | 1 | Mast | 1,293253975 | 0,773243322 | 2,492537313 |
| GNB24 | 0,368188301 | 0,589431527 | 0,241 | 0,133 | 1 | Mast | 1,504653743 | 0,664604733 | 1,812030075 |
| UBE2V11 | 0,368460708 | -0,702597351 | 0,185 | 0,267 | 1 | Mast | 0,61446496 | 1,627432099 | 0,692883895 |
| ZC3HAV11 | 0,369344696 | -0,36607267 | 0,176 | 0,267 | 1 | Mast | 0,775891771 | 1,288839549 | 0,65917603 |
| BLVRA3 | 0,36986154 | 1,030955818 | 0,269 | 0,2 | 1 | Mast | 2,043377587 | 0,489385812 | 1,345 |
| SF3B54 | 0,370044701 | 0,565501356 | 0,231 | 0,133 | 1 | Mast | 1,479901706 | 0,675720554 | 1,736842105 |
| EMD3 | 0,370044701 | 0,61777041 | 0,231 | 0,133 | 1 | Mast | 1,534501876 | 0,651677274 | 1,736842105 |
| RASA2 | 0,370626566 | 0,582631609 | 0,148 | 0,067 | 1 | Mast | 1,49757848 | 0,667744638 | 2,208955224 |
| UBE2H1 | 0,370628709 | 0,642383212 | 0,148 | 0,067 | 1 | Mast | 1,560905516 | 0,640653768 | 2,208955224 |
| ZNRF11 | 0,370628709 | 0,737316539 | 0,148 | 0,067 | 1 | Mast | 1,667072144 | 0,599854064 | 2,208955224 |

| Supplementary Table 4 - clusermarker | | | | | | | | | |
|---|---|---|---|---|---|---|---|---|---|
| | **p_val** | **avg_log2FC** | **pct.1** | **pct.2** | **p_val_adj** | **celltype** | **Foldchange_UP** | **Foldchange_DOWN** | **Ratio_pct1_pct2** |
| DCTN22 | 0,37212704 | -0,358594936 | 0,12 | 0,2 | 1 | Mast | 0,77992379 | 1,282176557 | 0,6 |
| SPI1 | 0,373101373 | -0,278596852 | 0,111 | 0,2 | 1 | Mast | 0,824392422 | 1,213014547 | 0,555 |
| IRAK41 | 0,373378418 | -0,28027416 | 0,065 | 0,133 | 1 | Mast | 0,823434522 | 1,214425644 | 0,488721805 |
| PANK2 | 0,373378418 | -0,268290647 | 0,065 | 0,133 | 1 | Mast | 0,830302734 | 1,204379992 | 0,488721805 |
| TSC22D21 | 0,37378956 | -0,280612224 | 0,278 | 0,4 | 1 | Mast | 0,823241591 | 1,214710251 | 0,695 |
| PHYKPL | 0,374120825 | -0,574857868 | 0,074 | 0,133 | 1 | Mast | 0,671352389 | 1,489530709 | 0,556390977 |
| IGFLR13 | 0,37728938 | 0,508077059 | 0,25 | 0,133 | 1 | Mast | 1,422153371 | 0,703159041 | 1,879699248 |
| NDUFB31 | 0,377544006 | 0,784561232 | 0,222 | 0,133 | 1 | Mast | 1,722568354 | 0,580528487 | 1,669172932 |
| RBM26 | 0,37755064 | 0,543376138 | 0,148 | 0,067 | 1 | Mast | 1,457379031 | 0,686163296 | 2,208955224 |
| ARMH11 | 0,377552776 | 0,572838598 | 0,148 | 0,067 | 1 | Mast | 1,487447344 | 0,672292706 | 2,208955224 |
| LSM82 | 0,377552776 | 0,640977725 | 0,148 | 0,067 | 1 | Mast | 1,559385609 | 0,641278202 | 2,208955224 |
| ETNK11 | 0,377552776 | 0,614151905 | 0,148 | 0,067 | 1 | Mast | 1,530657928 | 0,653313834 | 2,208955224 |
| HSBP13 | 0,377552776 | 0,640422796 | 0,148 | 0,067 | 1 | Mast | 1,55878591 | 0,641524916 | 2,208955224 |
| EMILIN21 | 0,377552776 | 0,652353526 | 0,148 | 0,067 | 1 | Mast | 1,571730132 | 0,63624154 | 2,208955224 |
| METTL263 | 0,379252722 | -0,363258893 | 0,12 | 0,2 | 1 | Mast | 0,777406517 | 1,286328295 | 0,6 |
| DNAJC7 | 0,379252722 | -0,304768486 | 0,12 | 0,2 | 1 | Mast | 0,809572121 | 1,235220401 | 0,6 |
| UXT3 | 0,380102109 | -0,486918797 | 0,222 | 0,333 | 1 | Mast | 0,713547414 | 1,401448565 | 0,666666667 |
| NCOA42 | 0,380467009 | 0,879910643 | 0,278 | 0,2 | 1 | Mast | 1,840261316 | 0,543401087 | 1,39 |
| MBOAT72 | 0,381546856 | 0,644851957 | 0,231 | 0,133 | 1 | Mast | 1,56357883 | 0,639558416 | 1,736842105 |
| ABCC11 | 0,382236486 | 0,392389613 | 0,157 | 0,067 | 1 | Mast | 1,312565677 | 0,761866639 | 2,343283582 |
| ZNF451 | 0,382950378 | -0,360210648 | 0,074 | 0,133 | 1 | Mast | 0,779050822 | 1,283613305 | 0,556390977 |
| CCDC82 | 0,382950378 | -0,447555019 | 0,074 | 0,133 | 1 | Mast | 0,733284516 | 1,363727145 | 0,556390977 |
| CDK171 | 0,382950378 | -0,396150757 | 0,074 | 0,133 | 1 | Mast | 0,759883019 | 1,315992034 | 0,556390977 |
| SCAND11 | 0,382950378 | -0,587650023 | 0,074 | 0,133 | 1 | Mast | 0,665425924 | 1,502796877 | 0,556390977 |
| HSPE14 | 0,384207877 | -0,266051746 | 0,278 | 0,4 | 1 | Mast | 0,831592271 | 1,202512379 | 0,695 |
| ARRB21 | 0,384553869 | 0,603045101 | 0,148 | 0,067 | 1 | Mast | 1,518919173 | 0,658362879 | 2,208955224 |
| GPSM32 | 0,384555998 | 0,651356929 | 0,148 | 0,067 | 1 | Mast | 1,570644773 | 0,636681201 | 2,208955224 |
| SND11 | 0,384555998 | 0,592824147 | 0,148 | 0,067 | 1 | Mast | 1,508196228 | 0,663043695 | 2,208955224 |
| FBRS1 | 0,384555998 | 0,546325092 | 0,148 | 0,067 | 1 | Mast | 1,460361046 | 0,684762171 | 2,208955224 |
| C1QBP4 | 0,384555998 | 0,70665766 | 0,148 | 0,067 | 1 | Mast | 1,632018782 | 0,612738046 | 2,208955224 |
| FKBP1A3 | 0,384555998 | 0,590415957 | 0,148 | 0,067 | 1 | Mast | 1,505680802 | 0,664151392 | 2,208955224 |

| Supplementary Table 4 - clusermarker | | | | | | | | | |
|---|---|---|---|---|---|---|---|---|---|
| | **p_val** | **avg_log2FC** | **pct.1** | **pct.2** | **p_val_adj** | **celltype** | **Foldchange_UP** | **Foldchange_DOWN** | **Ratio_pct1_pct2** |
| CTSZ4 | 0,384555998 | 0,574799089 | 0,148 | 0,067 | 1 | Mast | 1,489470023 | 0,671379742 | 2,208955224 |
| MBNL13 | 0,385985869 | -0,297729599 | 0,287 | 0,4 | 1 | Mast | 0,813531662 | 1,229208458 | 0,7175 |
| LARS13 | 0,386459489 | -0,277730668 | 0,12 | 0,2 | 1 | Mast | 0,82488753 | 1,21228648 | 0,6 |
| HMGB23 | 0,38646173 | -0,268667959 | 0,12 | 0,2 | 1 | Mast | 0,830085611 | 1,204695018 | 0,6 |
| RBM17 | 0,38646173 | -0,322590223 | 0,12 | 0,2 | 1 | Mast | 0,799632923 | 1,250573822 | 0,6 |
| UBE2J12 | 0,389120095 | 0,265003278 | 0,157 | 0,067 | 1 | Mast | 1,20163878 | 0,832196844 | 2,343283582 |
| RNF187 | 0,389122118 | 0,317842164 | 0,157 | 0,067 | 1 | Mast | 1,246464819 | 0,802268933 | 2,343283582 |
| SLC44A2 | 0,389122118 | 0,368110827 | 0,157 | 0,067 | 1 | Mast | 1,290661635 | 0,774796409 | 2,343283582 |
| G3BP1 | 0,389361514 | 0,711184651 | 0,222 | 0,133 | 1 | Mast | 1,637147889 | 0,610818367 | 1,669172932 |
| GSR1 | 0,391638105 | 0,594918678 | 0,148 | 0,067 | 1 | Mast | 1,510387446 | 0,662081774 | 2,208955224 |
| PDLIM72 | 0,391906454 | -0,517856698 | 0,074 | 0,133 | 1 | Mast | 0,698408635 | 1,431826512 | 0,556390977 |
| RGS17 | 0,391906454 | -0,676575156 | 0,074 | 0,133 | 1 | Mast | 0,625648754 | 1,598340913 | 0,556390977 |
| ENSA4 | 0,393267333 | 0,577605849 | 0,231 | 0,133 | 1 | Mast | 1,492370606 | 0,670074844 | 1,736842105 |
| ACTR2 | 0,393864484 | 0,480527531 | 0,315 | 0,2 | 1 | Mast | 1,395253756 | 0,716715505 | 1,575 |
| RSF11 | 0,396080693 | 0,377606028 | 0,157 | 0,067 | 1 | Mast | 1,299184232 | 0,769713775 | 2,343283582 |
| PRG2 | 0,396082709 | 0,395753247 | 0,157 | 0,067 | 1 | Mast | 1,315629485 | 0,760092421 | 2,343283582 |
| YWHAZ1 | 0,397220241 | -0,395647352 | 0,639 | 0,667 | 1 | Mast | 0,760148214 | 1,31553292 | 0,95802099 |
| ZFP36L21 | 0,397808924 | 0,676493229 | 0,63 | 0,6 | 1 | Mast | 1,598250149 | 0,625684284 | 1,05 |
| FLNB2 | 0,398798798 | 0,467658285 | 0,148 | 0,067 | 1 | Mast | 1,382863047 | 0,723137408 | 2,208955224 |
| PTBP32 | 0,398798798 | 0,571784153 | 0,148 | 0,067 | 1 | Mast | 1,486360588 | 0,672784255 | 2,208955224 |
| ADCYAP1 | 0,398798798 | 0,899901068 | 0,148 | 0,067 | 1 | Mast | 1,865938023 | 0,535923481 | 2,208955224 |
| CCDC85B4 | 0,399208477 | 0,433254844 | 0,231 | 0,133 | 1 | Mast | 1,350276483 | 0,740589067 | 1,736842105 |
| ATF42 | 0,399209926 | 0,411363459 | 0,231 | 0,133 | 1 | Mast | 1,329942119 | 0,751912422 | 1,736842105 |
| TRAPPC54 | 0,400414019 | -0,3500888 | 0,176 | 0,267 | 1 | Mast | 0,784535807 | 1,274639081 | 0,65917603 |
| ECH1 | 0,400414019 | -0,34900375 | 0,176 | 0,267 | 1 | Mast | 0,785126078 | 1,273680786 | 0,65917603 |
| TIAL1 | 0,400988451 | -0,727278546 | 0,074 | 0,133 | 1 | Mast | 0,604042286 | 1,655513236 | 0,556390977 |
| OIP5-AS11 | 0,400988451 | -0,550416961 | 0,074 | 0,133 | 1 | Mast | 0,682822753 | 1,464508901 | 0,556390977 |
| SMS2 | 0,400988451 | -0,627423747 | 0,074 | 0,133 | 1 | Mast | 0,647331338 | 1,544803938 | 0,556390977 |
| DEGS11 | 0,401128483 | 0,733626583 | 0,139 | 0,067 | 1 | Mast | 1,66281375 | 0,601390264 | 2,074626866 |
| CCDC88A | 0,401128483 | -0,420999226 | 0,12 | 0,2 | 1 | Mast | 0,746907129 | 1,338854539 | 0,6 |
| ANAPC115 | 0,402748168 | 0,391418353 | 0,241 | 0,133 | 1 | Mast | 1,311682321 | 0,762379719 | 1,812030075 |

| Supplementary Table 4 - clusermarker | | | | | | | | | |
|---|---|---|---|---|---|---|---|---|---|
| | **p_val** | **avg_log2FC** | **pct.1** | **pct.2** | **p_val_adj** | **celltype** | **Foldchange_UP** | **Foldchange_DOWN** | **Ratio_pct1_pct2** |
| MYADM3 | 0,403101025 | 0,310666784 | 0,574 | 0,467 | 1 | Mast | 1,2402808 | 0,806269032 | 1,229122056 |
| ATP5F1A1 | 0,406037746 | 0,404688246 | 0,148 | 0,067 | 1 | Mast | 1,323802817 | 0,755399511 | 2,208955224 |
| MYO9B2 | 0,406037746 | 0,399385022 | 0,148 | 0,067 | 1 | Mast | 1,318945564 | 0,758181405 | 2,208955224 |
| MEA12 | 0,407504172 | 0,59455547 | 0,222 | 0,133 | 1 | Mast | 1,510007244 | 0,662248479 | 1,669172932 |
| VWA5A | 0,408069786 | -0,281894389 | 0,731 | 0,733 | 1 | Mast | 0,822510278 | 1,215790279 | 0,997271487 |
| ARHGDIB3 | 0,408115122 | -0,279295932 | 0,593 | 0,8 | 1 | Mast | 0,823993047 | 1,213602474 | 0,74125 |
| ARID1B | 0,408583319 | -0,302625416 | 0,12 | 0,2 | 1 | Mast | 0,810775604 | 1,23338689 | 0,6 |
| TRIM24 | 0,408583319 | 0,559847402 | 0,139 | 0,067 | 1 | Mast | 1,474113288 | 0,678373913 | 2,074626866 |
| POLR3GL | 0,408585531 | -0,31554383 | 0,12 | 0,2 | 1 | Mast | 0,803548033 | 1,244480677 | 0,6 |
| PRDX31 | 0,408585531 | 0,6098102 | 0,139 | 0,067 | 1 | Mast | 1,526058428 | 0,655282905 | 2,074626866 |
| CLIC13 | 0,409567508 | 0,26188066 | 0,843 | 0,867 | 1 | Mast | 1,199040725 | 0,83400003 | 0,972318339 |
| LYPLA1 | 0,409628664 | 0,707394295 | 0,213 | 0,133 | 1 | Mast | 1,632852297 | 0,612425265 | 1,601503759 |
| AGO21 | 0,410195691 | -0,641522987 | 0,074 | 0,133 | 1 | Mast | 0,641035879 | 1,559975084 | 0,556390977 |
| EEF2K | 0,410195691 | -0,776525524 | 0,074 | 0,133 | 1 | Mast | 0,583771011 | 1,713000443 | 0,556390977 |
| UBA12 | 0,410195691 | -0,427731829 | 0,074 | 0,133 | 1 | Mast | 0,74342967 | 1,345117152 | 0,556390977 |
| MAGED22 | 0,411557885 | -0,391985554 | 0,185 | 0,267 | 1 | Mast | 0,762080046 | 1,312198115 | 0,692883895 |
| JUNB3 | 0,411769688 | -0,314486097 | 0,852 | 0,8 | 1 | Mast | 0,804137382 | 1,243568603 | 1,065 |
| RPL13A3 | 0,413845415 | 0,32402799 | 0,704 | 0,8 | 1 | Mast | 1,251820745 | 0,798836418 | 0,88 |
| AC020914.1 | 0,414809683 | -0,274719529 | 0,231 | 0,333 | 1 | Mast | 0,826611002 | 1,20975888 | 0,693693694 |
| CYRIB1 | 0,416124504 | 0,653995142 | 0,139 | 0,067 | 1 | Mast | 1,573519592 | 0,635517985 | 2,074626866 |
| EGFL7 | 0,416124504 | 0,525418238 | 0,139 | 0,067 | 1 | Mast | 1,439350788 | 0,69475767 | 2,074626866 |
| OTUB12 | 0,416124504 | 0,456455227 | 0,139 | 0,067 | 1 | Mast | 1,372166193 | 0,728774696 | 2,074626866 |
| RAB211 | 0,416124504 | -0,335755551 | 0,12 | 0,2 | 1 | Mast | 0,792369058 | 1,262038176 | 0,6 |
| RCOR1 | 0,416124504 | 0,64686122 | 0,139 | 0,067 | 1 | Mast | 1,565757968 | 0,638668313 | 2,074626866 |
| PDCD62 | 0,417354648 | 0,434353902 | 0,231 | 0,133 | 1 | Mast | 1,351305528 | 0,740025094 | 1,736842105 |
| SRSF96 | 0,418030415 | 0,306042604 | 0,333 | 0,2 | 1 | Mast | 1,236311769 | 0,808857462 | 1,665 |
| IER3 | 0,418687005 | 0,58309351 | 0,444 | 0,333 | 1 | Mast | 1,498058029 | 0,667530884 | 1,333333333 |
| BNIP3L2 | 0,419282853 | -0,313552047 | 0,259 | 0,333 | 1 | Mast | 0,804658176 | 1,242763734 | 0,777777778 |
| PLCG22 | 0,419524082 | -0,361261013 | 0,074 | 0,133 | 1 | Mast | 0,778483835 | 1,284548189 | 0,556390977 |
| ERO1B | 0,41952742 | -0,289988355 | 0,074 | 0,133 | 1 | Mast | 0,817908661 | 1,222630409 | 0,556390977 |
| COQ10B | 0,41952742 | -0,32571615 | 0,074 | 0,133 | 1 | Mast | 0,797902211 | 1,253286412 | 0,556390977 |

| Supplementary Table 4 - clusermarker | | | | | | | | | |
|---|---|---|---|---|---|---|---|---|---|
| | **p_val** | **avg_log2FC** | **pct.1** | **pct.2** | **p_val_adj** | **celltype** | **Foldchange_UP** | **Foldchange_DOWN** | **Ratio_pct1_pct2** |
| TTC37 | 0,41952742 | -0,27484218 | 0,074 | 0,133 | 1 | Mast | 0,826540731 | 1,209861732 | 0,556390977 |
| MAT2B1 | 0,41952742 | -0,397859672 | 0,074 | 0,133 | 1 | Mast | 0,758983447 | 1,31755179 | 0,556390977 |
| BLOC1S2 | 0,41952742 | -0,293965448 | 0,074 | 0,133 | 1 | Mast | 0,815657028 | 1,226005497 | 0,556390977 |
| BTBD10 | 0,41952742 | -0,25624103 | 0,074 | 0,133 | 1 | Mast | 0,837266594 | 1,194362712 | 0,556390977 |
| MADD | 0,41952742 | -0,384547181 | 0,074 | 0,133 | 1 | Mast | 0,766019391 | 1,305449981 | 0,556390977 |
| CWC15 | 0,41952742 | -0,531719342 | 0,074 | 0,133 | 1 | Mast | 0,691729869 | 1,445651034 | 0,556390977 |
| MED212 | 0,41952742 | -0,26237296 | 0,074 | 0,133 | 1 | Mast | 0,833715487 | 1,199449951 | 0,556390977 |
| ZCRB12 | 0,41952742 | -0,344294857 | 0,074 | 0,133 | 1 | Mast | 0,787692881 | 1,269530325 | 0,556390977 |
| PDXDC1 | 0,41952742 | -0,324535166 | 0,074 | 0,133 | 1 | Mast | 0,798555638 | 1,252260897 | 0,556390977 |
| APOBR | 0,41952742 | -0,328635892 | 0,074 | 0,133 | 1 | Mast | 0,796289041 | 1,255825395 | 0,556390977 |
| LTBP4 | 0,41952742 | -0,404713746 | 0,074 | 0,133 | 1 | Mast | 0,755386159 | 1,323826216 | 0,556390977 |
| LMO44 | 0,419766761 | 0,526370869 | 0,37 | 0,267 | 1 | Mast | 1,440301525 | 0,694299063 | 1,38576779 |
| TMOD31 | 0,419874501 | 0,493017747 | 0,222 | 0,133 | 1 | Mast | 1,407385688 | 0,710537281 | 1,669172932 |
| MIER1 | 0,420748943 | -0,463733179 | 0,13 | 0,2 | 1 | Mast | 0,72510751 | 1,379105838 | 0,65 |
| WTAP1 | 0,420748943 | 0,385541476 | 0,148 | 0,067 | 1 | Mast | 1,306349998 | 0,765491638 | 2,208955224 |
| P2RX1 | 0,420748943 | -0,4571014 | 0,13 | 0,2 | 1 | Mast | 0,728448356 | 1,372780914 | 0,65 |
| SUPT4H11 | 0,420748943 | 0,323524216 | 0,148 | 0,067 | 1 | Mast | 1,251383699 | 0,799115412 | 2,208955224 |
| C19orf535 | 0,420748943 | 0,326341182 | 0,148 | 0,067 | 1 | Mast | 1,253829503 | 0,797556604 | 2,208955224 |
| SUMO22 | 0,422762395 | 0,526580856 | 0,491 | 0,467 | 1 | Mast | 1,440511178 | 0,694198015 | 1,051391863 |
| PPP1R181 | 0,42374497 | 0,617711593 | 0,139 | 0,067 | 1 | Mast | 1,534439318 | 0,651703843 | 2,074626866 |
| FDX1 | 0,426338727 | 0,801311508 | 0,315 | 0,2 | 1 | Mast | 1,742684626 | 0,573827292 | 1,575 |
| ERH5 | 0,427634337 | 0,30426174 | 0,306 | 0,2 | 1 | Mast | 1,234786607 | 0,809856533 | 1,53 |
| TAPT11 | 0,428979491 | -0,34808804 | 0,074 | 0,133 | 1 | Mast | 0,785624573 | 1,27287261 | 0,556390977 |
| TPRKB3 | 0,428982804 | -0,310678431 | 0,074 | 0,133 | 1 | Mast | 0,806262523 | 1,240290812 | 0,556390977 |
| UBA51 | 0,428982804 | -0,299091385 | 0,074 | 0,133 | 1 | Mast | 0,812764117 | 1,230369277 | 0,556390977 |
| SCLT11 | 0,428982804 | -0,403614254 | 0,074 | 0,133 | 1 | Mast | 0,755962065 | 1,322817699 | 0,556390977 |
| SMIM15 | 0,428982804 | -0,274243201 | 0,074 | 0,133 | 1 | Mast | 0,826883966 | 1,209359525 | 0,556390977 |
| ATG12 | 0,428982804 | -0,513381048 | 0,074 | 0,133 | 1 | Mast | 0,700578662 | 1,427391462 | 0,556390977 |
| ATF71 | 0,428982804 | -0,426719193 | 0,074 | 0,133 | 1 | Mast | 0,74395167 | 1,344173338 | 0,556390977 |
| USP14 | 0,428982804 | -0,41541003 | 0,074 | 0,133 | 1 | Mast | 0,749806361 | 1,333677669 | 0,556390977 |
| EIF3D2 | 0,42971667 | 0,318655009 | 0,231 | 0,133 | 1 | Mast | 1,247167302 | 0,801817045 | 1,736842105 |

| Supplementary Table 4 - clusermarker | | | | | | | | | |
|---|---|---|---|---|---|---|---|---|---|
| | **p_val** | **avg_log2FC** | **pct.1** | **pct.2** | **p_val_adj** | **celltype** | **Foldchange_UP** | **Foldchange_DOWN** | **Ratio_pct1_pct2** |
| NDUFB94 | 0,430785555 | 0,602231531 | 0,278 | 0,2 | 1 | Mast | 1,51806286 | 0,65873425 | 1,39 |
| STMP12 | 0,431446464 | 0,595741499 | 0,139 | 0,067 | 1 | Mast | 1,511249119 | 0,661704273 | 2,074626866 |
| GTF3A4 | 0,431446464 | 0,656561577 | 0,139 | 0,067 | 1 | Mast | 1,576321245 | 0,634388456 | 2,074626866 |
| MESD2 | 0,431446464 | 0,735852004 | 0,139 | 0,067 | 1 | Mast | 1,665380694 | 0,600463308 | 2,074626866 |
| MPP1 | 0,432184035 | -0,285818437 | 0,241 | 0,333 | 1 | Mast | 0,820276136 | 1,219101661 | 0,723723724 |
| LAMTOR14 | 0,4324628 | 0,496602951 | 0,222 | 0,133 | 1 | Mast | 1,410887495 | 0,708773735 | 1,669172932 |
| TYROBP1 | 0,432824861 | 0,41328709 | 0,722 | 0,8 | 1 | Mast | 1,331716592 | 0,750910521 | 0,9025 |
| CMTM61 | 0,435053106 | 0,530942753 | 0,213 | 0,133 | 1 | Mast | 1,444873063 | 0,692102321 | 1,601503759 |
| ESD2 | 0,435054604 | 0,566905917 | 0,213 | 0,133 | 1 | Mast | 1,481343191 | 0,675063014 | 1,601503759 |
| CANX | 0,435390136 | 0,565920694 | 0,454 | 0,4 | 1 | Mast | 1,480331921 | 0,675524175 | 1,135 |
| HNRNPUL22 | 0,435768467 | 0,344953205 | 0,148 | 0,067 | 1 | Mast | 1,270109784 | 0,787333514 | 2,208955224 |
| MIDN2 | 0,435768467 | 0,262640405 | 0,148 | 0,067 | 1 | Mast | 1,199672324 | 0,833560948 | 2,208955224 |
| ATP5PF3 | 0,437114648 | -0,641307525 | 0,306 | 0,4 | 1 | Mast | 0,641131623 | 1,559742124 | 0,765 |
| ZMYM5 | 0,438557649 | -0,289220866 | 0,074 | 0,133 | 1 | Mast | 0,81834389 | 1,221980164 | 0,556390977 |
| SLC66A32 | 0,438560936 | -0,351463464 | 0,074 | 0,133 | 1 | Mast | 0,783788623 | 1,275854192 | 0,556390977 |
| HERC3 | 0,438560936 | -0,287314061 | 0,074 | 0,133 | 1 | Mast | 0,819426207 | 1,220366144 | 0,556390977 |
| CPT1A | 0,438560936 | -0,273235383 | 0,074 | 0,133 | 1 | Mast | 0,827461801 | 1,208515002 | 0,556390977 |
| KATNBL1 | 0,438560936 | -0,303281047 | 0,074 | 0,133 | 1 | Mast | 0,810407231 | 1,233947528 | 0,556390977 |
| M6PR | 0,43883802 | 0,498086823 | 0,222 | 0,133 | 1 | Mast | 1,412339398 | 0,708045107 | 1,669172932 |
| PCBP15 | 0,438973874 | 0,545837087 | 0,343 | 0,267 | 1 | Mast | 1,459867149 | 0,684993837 | 1,284644195 |
| ARPC41 | 0,439228488 | 0,462291038 | 0,139 | 0,067 | 1 | Mast | 1,377727951 | 0,725832701 | 2,074626866 |
| MRPS362 | 0,439228488 | 0,407027037 | 0,139 | 0,067 | 1 | Mast | 1,325950609 | 0,754175905 | 2,074626866 |
| LARP1 | 0,439228488 | 0,471423405 | 0,139 | 0,067 | 1 | Mast | 1,386476732 | 0,721252638 | 2,074626866 |
| VKORC12 | 0,439228488 | 0,460841094 | 0,139 | 0,067 | 1 | Mast | 1,376343996 | 0,726562548 | 2,074626866 |
| CDK5RAP33 | 0,439228488 | 0,527947412 | 0,139 | 0,067 | 1 | Mast | 1,441876312 | 0,693540765 | 2,074626866 |
| RAD23A2 | 0,439228488 | 0,398032654 | 0,139 | 0,067 | 1 | Mast | 1,317709776 | 0,75889245 | 2,074626866 |
| PARVB1 | 0,439228488 | 0,531145179 | 0,139 | 0,067 | 1 | Mast | 1,445075809 | 0,692005218 | 2,074626866 |
| TAX1BP12 | 0,442288796 | 0,344331438 | 0,231 | 0,133 | 1 | Mast | 1,269562516 | 0,787672909 | 1,736842105 |
| SENP62 | 0,442313821 | 0,646812044 | 0,13 | 0,067 | 1 | Mast | 1,565704597 | 0,638690084 | 1,940298507 |
| CHST12 | 0,442313821 | 0,647968172 | 0,13 | 0,067 | 1 | Mast | 1,566959804 | 0,638178464 | 1,940298507 |
| METTL91 | 0,442313821 | 0,6973018 | 0,13 | 0,067 | 1 | Mast | 1,621469402 | 0,616724558 | 1,940298507 |

| Supplementary Table 4 - clusermarker | | | | | | | | | |
|---|---|---|---|---|---|---|---|---|---|
| | **p_val** | **avg_log2FC** | **pct.1** | **pct.2** | **p_val_adj** | **celltype** | **Foldchange_UP** | **Foldchange_DOWN** | **Ratio_pct1_pct2** |
| COX172 | 0,44247174 | -0,608208812 | 0,259 | 0,333 | 1 | Mast | 0,656010671 | 1,524365447 | 0,777777778 |
| ELOC2 | 0,443392709 | -0,264533241 | 0,13 | 0,2 | 1 | Mast | 0,832468022 | 1,201247344 | 0,65 |
| ATP5MC32 | 0,444628325 | 0,430382478 | 0,306 | 0,2 | 1 | Mast | 1,347590794 | 0,742065028 | 1,53 |
| ELOVL1 | 0,447090507 | 0,367105406 | 0,139 | 0,067 | 1 | Mast | 1,28976248 | 0,775336557 | 2,074626866 |
| KIDINS2201 | 0,447090507 | 0,525519487 | 0,139 | 0,067 | 1 | Mast | 1,439451805 | 0,694708914 | 2,074626866 |
| COPS51 | 0,447090507 | 0,307425879 | 0,139 | 0,067 | 1 | Mast | 1,23749773 | 0,80808229 | 2,074626866 |
| SERPINB63 | 0,447254242 | 0,783594218 | 0,287 | 0,2 | 1 | Mast | 1,721414132 | 0,580917736 | 1,435 |
| RRBP1 | 0,448101748 | -0,353233996 | 0,194 | 0,267 | 1 | Mast | 0,782827316 | 1,277420932 | 0,72659176 |
| NBN | 0,448260828 | -0,257883749 | 0,074 | 0,133 | 1 | Mast | 0,836313786 | 1,195723443 | 0,556390977 |
| ANXA15 | 0,448723273 | 0,36670613 | 0,935 | 0,933 | 1 | Mast | 1,289405578 | 0,775551166 | 1,002143623 |
| PGK12 | 0,449949884 | -0,34379294 | 0,324 | 0,4 | 1 | Mast | 0,787966969 | 1,269088729 | 0,81 |
| MACF11 | 0,450774382 | -0,28497834 | 0,185 | 0,267 | 1 | Mast | 0,820753931 | 1,218391971 | 0,692883895 |
| NR3C11 | 0,451749138 | -0,379960662 | 0,204 | 0,267 | 1 | Mast | 0,768458544 | 1,301306373 | 0,764044944 |
| SECISBP2L | 0,453566388 | -0,606179683 | 0,083 | 0,133 | 1 | Mast | 0,656933989 | 1,522222957 | 0,62406015 |
| DPP75 | 0,45470778 | 0,507462994 | 0,213 | 0,133 | 1 | Mast | 1,421548178 | 0,703458395 | 1,601503759 |
| UBE2L3 | 0,455031957 | 0,4926315 | 0,139 | 0,067 | 1 | Mast | 1,407008944 | 0,710727536 | 2,074626866 |
| SYTL23 | 0,455068774 | 0,342758307 | 0,231 | 0,133 | 1 | Mast | 1,268178925 | 0,788532265 | 1,736842105 |
| PIK3R12 | 0,455399025 | 0,656025431 | 0,269 | 0,2 | 1 | Mast | 1,575735548 | 0,634624256 | 1,345 |
| SDCBP4 | 0,456892077 | -0,279112825 | 0,5 | 0,6 | 1 | Mast | 0,824097635 | 1,213448453 | 0,833333333 |
| CBX33 | 0,458282982 | 0,396678659 | 0,222 | 0,133 | 1 | Mast | 1,316473663 | 0,759605018 | 1,669172932 |
| MLPH1 | 0,458552316 | -0,26817552 | 0,296 | 0,4 | 1 | Mast | 0,830368994 | 1,204283887 | 0,74 |
| GSTO11 | 0,458635435 | 0,599894599 | 0,13 | 0,067 | 1 | Mast | 1,515605835 | 0,659802158 | 1,940298507 |
| DNAJA22 | 0,458865597 | -0,36302877 | 0,13 | 0,2 | 1 | Mast | 0,77753053 | 1,286123131 | 0,65 |
| PPP1CC2 | 0,458867609 | -0,250080409 | 0,148 | 0,067 | 1 | Mast | 0,840849549 | 1,189273397 | 2,208955224 |
| B2M4 | 0,460442903 | 0,298513208 | 1 | 1 | 1 | Mast | 1,229876292 | 0,813089907 | 1 |
| LSM44 | 0,46136833 | 0,46207336 | 0,213 | 0,133 | 1 | Mast | 1,37752009 | 0,725942225 | 1,601503759 |
| HNRNPUL12 | 0,46136833 | 0,426040337 | 0,213 | 0,133 | 1 | Mast | 1,34354099 | 0,744301817 | 1,601503759 |
| CST71 | 0,46136833 | 0,604232474 | 0,213 | 0,133 | 1 | Mast | 1,520169795 | 0,657821254 | 1,601503759 |
| EID12 | 0,461472106 | 0,575624505 | 0,269 | 0,2 | 1 | Mast | 1,490322446 | 0,670995732 | 1,345 |
| CSNK1A11 | 0,462001155 | 0,25486127 | 0,306 | 0,2 | 1 | Mast | 1,193220997 | 0,83806772 | 1,53 |
| RSRP12 | 0,46248506 | 0,709337406 | 0,417 | 0,4 | 1 | Mast | 1,635053005 | 0,611600967 | 1,0425 |

| Supplementary Table 4 - clusermarker | | | | | | | | | |
|---|---|---|---|---|---|---|---|---|---|
| | **p_val** | **avg_log2FC** | **pct.1** | **pct.2** | **p_val_adj** | **celltype** | **Foldchange_UP** | **Foldchange_DOWN** | **Ratio_pct1_pct2** |
| PHACTR21 | 0,463052237 | 0,323004826 | 0,139 | 0,067 | 1 | Mast | 1,250933265 | 0,799403156 | 2,074626866 |
| TRIM81 | 0,463052237 | 0,326983471 | 0,139 | 0,067 | 1 | Mast | 1,254387833 | 0,79720161 | 2,074626866 |
| NDUFV12 | 0,463052237 | 0,340870718 | 0,139 | 0,067 | 1 | Mast | 1,266520754 | 0,789564638 | 2,074626866 |
| DDX39B2 | 0,465035 | 0,424538476 | 0,287 | 0,2 | 1 | Mast | 1,342143077 | 0,745077047 | 1,435 |
| CALM23 | 0,46540646 | -0,272515553 | 0,5 | 0,533 | 1 | Mast | 0,827874764 | 1,207912167 | 0,938086304 |
| ADRB2 | 0,466376182 | 0,434095479 | 0,278 | 0,2 | 1 | Mast | 1,351063496 | 0,740157663 | 1,39 |
| EHD1 | 0,466717157 | -0,355175775 | 0,13 | 0,2 | 1 | Mast | 0,781774387 | 1,27914142 | 0,65 |
| DR1 | 0,466921195 | 0,47498939 | 0,13 | 0,067 | 1 | Mast | 1,389907998 | 0,719472081 | 1,940298507 |
| MANF3 | 0,466921195 | 0,661374619 | 0,13 | 0,067 | 1 | Mast | 1,581588865 | 0,632275569 | 1,940298507 |
| PICALM | 0,466921195 | 0,610401927 | 0,13 | 0,067 | 1 | Mast | 1,526684476 | 0,655014193 | 1,940298507 |
| SUGT12 | 0,466921195 | 0,583405059 | 0,13 | 0,067 | 1 | Mast | 1,498381569 | 0,667386746 | 1,940298507 |
| DAZAP23 | 0,468392604 | 0,686398066 | 0,38 | 0,333 | 1 | Mast | 1,609260705 | 0,621403354 | 1,141141141 |
| EPB41L22 | 0,469732769 | -0,494137718 | 0,139 | 0,2 | 1 | Mast | 0,709985902 | 1,408478672 | 0,695 |
| TAMALIN1 | 0,469732769 | -0,61211593 | 0,139 | 0,2 | 1 | Mast | 0,654236461 | 1,528499341 | 0,695 |
| ITSN2 | 0,471150714 | 0,416991025 | 0,139 | 0,067 | 1 | Mast | 1,335139997 | 0,748985127 | 2,074626866 |
| DENND4C1 | 0,471150714 | 0,341713925 | 0,139 | 0,067 | 1 | Mast | 1,267261209 | 0,789103298 | 2,074626866 |
| EIF2S21 | 0,471150714 | 0,405188805 | 0,139 | 0,067 | 1 | Mast | 1,324262205 | 0,755137462 | 2,074626866 |
| NDUFA62 | 0,471150714 | 0,295401801 | 0,139 | 0,067 | 1 | Mast | 1,227226721 | 0,81484536 | 2,074626866 |
| YWHAB3 | 0,472872922 | 0,62690688 | 0,352 | 0,267 | 1 | Mast | 1,544250588 | 0,647563296 | 1,31835206 |
| PITHD11 | 0,475289328 | 0,441345455 | 0,13 | 0,067 | 1 | Mast | 1,357870085 | 0,736447479 | 1,940298507 |
| PPCS2 | 0,475289328 | 0,492026039 | 0,13 | 0,067 | 1 | Mast | 1,406418584 | 0,711025872 | 1,940298507 |
| CAPZA12 | 0,475289328 | 0,454750235 | 0,13 | 0,067 | 1 | Mast | 1,37054551 | 0,729636479 | 1,940298507 |
| SF3B12 | 0,475522298 | -0,27446711 | 0,259 | 0,333 | 1 | Mast | 0,826755642 | 1,209547235 | 0,777777778 |
| ARPC33 | 0,476825412 | -0,275242552 | 0,556 | 0,733 | 1 | Mast | 0,826311383 | 1,210197536 | 0,758526603 |
| VAMP21 | 0,477492386 | -0,432580517 | 0,139 | 0,2 | 1 | Mast | 0,740935305 | 1,3496455 | 0,695 |
| COX5A5 | 0,478055577 | 0,733893998 | 0,204 | 0,133 | 1 | Mast | 1,663121994 | 0,601278802 | 1,533834586 |
| TNFSF103 | 0,481524793 | 0,440324525 | 0,296 | 0,2 | 1 | Mast | 1,356909522 | 0,736968813 | 1,48 |
| EIF5A2 | 0,481673391 | 0,478764589 | 0,213 | 0,133 | 1 | Mast | 1,393549829 | 0,71759185 | 1,601503759 |
| FNIP21 | 0,482283054 | -0,336860874 | 0,083 | 0,133 | 1 | Mast | 0,791762216 | 1,263005458 | 0,62406015 |
| OAZ21 | 0,482283054 | -0,5072214 | 0,083 | 0,133 | 1 | Mast | 0,703576207 | 1,421310144 | 0,62406015 |
| SLC43A31 | 0,482637446 | -0,283427462 | 0,13 | 0,2 | 1 | Mast | 0,821636705 | 1,21708292 | 0,65 |

| Supplementary Table 4 - clusermarker | | | | | | | | | |
|---|---|---|---|---|---|---|---|---|---|
| | **p_val** | **avg_log2FC** | **pct.1** | **pct.2** | **p_val_adj** | **celltype** | **Foldchange_UP** | **Foldchange_DOWN** | **Ratio_pct1_pct2** |
| CTDSP1 | 0,48373909 | 0,412871211 | 0,13 | 0,067 | 1 | Mast | 1,33133276 | 0,751127013 | 1,940298507 |
| RSRC24 | 0,48373909 | 0,362991551 | 0,13 | 0,067 | 1 | Mast | 1,286089952 | 0,777550589 | 1,940298507 |
| UBA2 | 0,48373909 | 0,366611723 | 0,13 | 0,067 | 1 | Mast | 1,289321205 | 0,775601918 | 1,940298507 |
| YME1L12 | 0,484553222 | -0,43320456 | 0,278 | 0,333 | 1 | Mast | 0,74061488 | 1,350229421 | 0,834834835 |
| EIF3F4 | 0,48477097 | 0,40757392 | 0,37 | 0,267 | 1 | Mast | 1,326453333 | 0,753890073 | 1,38576779 |
| MRPL33 | 0,484947388 | 0,302469693 | 0,222 | 0,133 | 1 | Mast | 1,233253766 | 0,810863123 | 1,669172932 |
| NDUFAF33 | 0,485013857 | 0,504651831 | 0,204 | 0,133 | 1 | Mast | 1,418780917 | 0,704830455 | 1,533834586 |
| HEXIM11 | 0,485013857 | 0,439518734 | 0,204 | 0,133 | 1 | Mast | 1,356151856 | 0,737380549 | 1,533834586 |
| CYB5R31 | 0,485013857 | 0,542443158 | 0,204 | 0,133 | 1 | Mast | 1,45643686 | 0,686607176 | 1,533834586 |
| ANKRD172 | 0,485321612 | -0,490290441 | 0,139 | 0,2 | 1 | Mast | 0,711881769 | 1,404727644 | 0,695 |
| SRSF21 | 0,485492136 | 0,631221263 | 0,324 | 0,267 | 1 | Mast | 1,548875585 | 0,645629649 | 1,213483146 |
| KLF102 | 0,486778797 | -0,28016241 | 0,287 | 0,4 | 1 | Mast | 0,823498307 | 1,214331579 | 0,7175 |
| DNAJC19 | 0,488548235 | 0,46092512 | 0,213 | 0,133 | 1 | Mast | 1,37642416 | 0,726520232 | 1,601503759 |
| RPS165 | 0,488840858 | 0,256603187 | 0,648 | 0,867 | 1 | Mast | 1,194662568 | 0,837056443 | 0,747404844 |
| SERPINB13 | 0,489740778 | 0,333355021 | 0,639 | 0,533 | 1 | Mast | 1,25993999 | 0,793688595 | 1,198874296 |
| COX5B3 | 0,489767926 | 0,266405979 | 0,315 | 0,2 | 1 | Mast | 1,202807675 | 0,83138811 | 1,575 |
| RHOH1 | 0,489956137 | -0,291718528 | 0,333 | 0,4 | 1 | Mast | 0,81692836 | 1,224097546 | 0,8325 |
| SRRM22 | 0,490512252 | 0,355237703 | 0,5 | 0,4 | 1 | Mast | 1,279196328 | 0,781740831 | 1,25 |
| NDST2 | 0,49098204 | 0,495484736 | 0,278 | 0,2 | 1 | Mast | 1,409794357 | 0,70932331 | 1,39 |
| TGOLN2 | 0,490983189 | -0,273772914 | 0,259 | 0,333 | 1 | Mast | 0,827153556 | 1,208965365 | 0,777777778 |
| SLC7A51 | 0,49202663 | 0,599555268 | 0,204 | 0,133 | 1 | Mast | 1,515249397 | 0,659957365 | 1,533834586 |
| PDCL31 | 0,492069377 | -0,381289946 | 0,083 | 0,133 | 1 | Mast | 0,767750821 | 1,302505934 | 0,62406015 |
| TMX21 | 0,492069377 | -0,438637213 | 0,083 | 0,133 | 1 | Mast | 0,737831244 | 1,355323467 | 0,62406015 |
| SPATA13 | 0,492069377 | -0,286976134 | 0,083 | 0,133 | 1 | Mast | 0,819618166 | 1,220080327 | 0,62406015 |
| EXOSC41 | 0,4922697 | 0,422465091 | 0,13 | 0,067 | 1 | Mast | 1,340215587 | 0,746148612 | 1,940298507 |
| NUTF23 | 0,4922697 | 0,579522882 | 0,13 | 0,067 | 1 | Mast | 1,494354964 | 0,669185049 | 1,940298507 |
| AP2S13 | 0,4922697 | 0,352867307 | 0,13 | 0,067 | 1 | Mast | 1,277096292 | 0,783026312 | 1,940298507 |
| TCEA13 | 0,494321701 | 0,660636081 | 0,324 | 0,267 | 1 | Mast | 1,580779432 | 0,632599324 | 1,213483146 |
| MRPS6 | 0,495367795 | 0,625404692 | 0,194 | 0,133 | 1 | Mast | 1,542643493 | 0,648237914 | 1,458646617 |
| ARF62 | 0,495474222 | -0,261787069 | 0,194 | 0,267 | 1 | Mast | 0,834054135 | 1,198962943 | 0,72659176 |
| NCF4 | 0,495475628 | -0,366819382 | 0,194 | 0,267 | 1 | Mast | 0,775490288 | 1,289506801 | 0,72659176 |

| Supplementary Table 4 - clusermarker | | | | | | | | | |
|---|---|---|---|---|---|---|---|---|---|
| | **p_val** | **avg_log2FC** | **pct.1** | **pct.2** | **p_val_adj** | **celltype** | **Foldchange_UP** | **Foldchange_DOWN** | **Ratio_pct1_pct2** |
| CD462 | 0,499093442 | 0,481536513 | 0,204 | 0,133 | 1 | Mast | 1,396229901 | 0,716214428 | 1,533834586 |
| STX11 | 0,499093442 | 0,565104409 | 0,204 | 0,133 | 1 | Mast | 1,479494577 | 0,675906499 | 1,533834586 |
| ZNF2921 | 0,500878203 | 0,359388803 | 0,13 | 0,067 | 1 | Mast | 1,28288229 | 0,779494742 | 1,940298507 |
| RNF1492 | 0,50088034 | 0,326071628 | 0,13 | 0,067 | 1 | Mast | 1,253595259 | 0,797705634 | 1,940298507 |
| BUD312 | 0,50088034 | 0,382489535 | 0,13 | 0,067 | 1 | Mast | 1,303589408 | 0,767112707 | 1,940298507 |
| GDI21 | 0,501526138 | 0,407295871 | 0,352 | 0,267 | 1 | Mast | 1,326197712 | 0,754035383 | 1,31835206 |
| EBPL1 | 0,501961242 | -0,385201169 | 0,083 | 0,133 | 1 | Mast | 0,765672226 | 1,306041889 | 0,62406015 |
| SLFN52 | 0,501961242 | -0,371186595 | 0,083 | 0,133 | 1 | Mast | 0,773146334 | 1,293416209 | 0,62406015 |
| RASGRP4 | 0,501961242 | -0,382338234 | 0,083 | 0,133 | 1 | Mast | 0,767193162 | 1,303452703 | 0,62406015 |
| EEF24 | 0,502301522 | 0,388248743 | 0,63 | 0,6 | 1 | Mast | 1,30880371 | 0,764056514 | 1,05 |
| GNAI31 | 0,505563716 | 0,456948285 | 0,12 | 0,067 | 1 | Mast | 1,372635227 | 0,728525671 | 1,791044776 |
| TMEM30A | 0,505563716 | 0,625286123 | 0,12 | 0,067 | 1 | Mast | 1,542516716 | 0,648291192 | 1,791044776 |
| COPS22 | 0,505563716 | 0,628036014 | 0,12 | 0,067 | 1 | Mast | 1,545459678 | 0,647056675 | 1,791044776 |
| SF3A1 | 0,505563716 | 0,690186115 | 0,12 | 0,067 | 1 | Mast | 1,613491653 | 0,619773891 | 1,791044776 |
| PDIA32 | 0,505723029 | 0,562835831 | 0,472 | 0,467 | 1 | Mast | 1,477169962 | 0,67697017 | 1,010706638 |
| CHD92 | 0,50621382 | 0,454223755 | 0,204 | 0,133 | 1 | Mast | 1,370045451 | 0,729902792 | 1,533834586 |
| ZFP36L12 | 0,507864753 | 0,373477179 | 0,509 | 0,4 | 1 | Mast | 1,295471412 | 0,771919774 | 1,2725 |
| CHD41 | 0,508214196 | -0,364439125 | 0,213 | 0,267 | 1 | Mast | 0,7767708 | 1,287381039 | 0,797752809 |
| MLF23 | 0,509486276 | 0,393374886 | 0,213 | 0,133 | 1 | Mast | 1,313462386 | 0,761346508 | 1,601503759 |
| MAP41 | 0,509570153 | 0,290446108 | 0,13 | 0,067 | 1 | Mast | 1,223018399 | 0,817649187 | 1,940298507 |
| OXA1L1 | 0,509570153 | 0,432699801 | 0,13 | 0,067 | 1 | Mast | 1,349757095 | 0,740874046 | 1,940298507 |
| SAT23 | 0,509570153 | 0,33346776 | 0,13 | 0,067 | 1 | Mast | 1,260038451 | 0,793626575 | 1,940298507 |
| NUCKS12 | 0,509851006 | 0,526678521 | 0,194 | 0,133 | 1 | Mast | 1,440608699 | 0,694151022 | 1,458646617 |
| MRPS111 | 0,511957352 | -0,371910761 | 0,083 | 0,133 | 1 | Mast | 0,772758348 | 1,294065607 | 0,62406015 |
| TRAPPC8 | 0,511957352 | -0,270492667 | 0,083 | 0,133 | 1 | Mast | 0,829036389 | 1,206219671 | 0,62406015 |
| QPCT1 | 0,514565543 | 0,560688657 | 0,12 | 0,067 | 1 | Mast | 1,474973114 | 0,67797846 | 1,791044776 |
| NFE2L3 | 0,514565543 | 0,460496455 | 0,12 | 0,067 | 1 | Mast | 1,376015246 | 0,726736134 | 1,791044776 |
| SYNRG2 | 0,514565543 | 0,484242016 | 0,12 | 0,067 | 1 | Mast | 1,398850724 | 0,714872561 | 1,791044776 |
| SAFB1 | 0,514565543 | 0,500463002 | 0,12 | 0,067 | 1 | Mast | 1,414667496 | 0,706879887 | 1,791044776 |
| IER54 | 0,517174839 | 0,436882331 | 0,194 | 0,133 | 1 | Mast | 1,353675866 | 0,738729281 | 1,458646617 |
| ARHGDIA2 | 0,517174839 | 0,571927313 | 0,194 | 0,133 | 1 | Mast | 1,486508089 | 0,672717497 | 1,458646617 |

| Supplementary Table 4 - clusermarker | | | | | | | | | |
|---|---|---|---|---|---|---|---|---|---|
| | **p_val** | **avg_log2FC** | **pct.1** | **pct.2** | **p_val_adj** | **celltype** | **Foldchange_UP** | **Foldchange_DOWN** | **Ratio_pct1_pct2** |
| STXBP61 | 0,518338245 | 0,34011983 | 0,13 | 0,067 | 1 | Mast | 1,265861732 | 0,789975694 | 1,940298507 |
| NMT1 | 0,522056341 | -0,403497059 | 0,083 | 0,133 | 1 | Mast | 0,756023477 | 1,322710247 | 0,62406015 |
| CFAP2982 | 0,522056341 | -0,352292363 | 0,083 | 0,133 | 1 | Mast | 0,783338427 | 1,276587444 | 0,62406015 |
| PBX11 | 0,523647774 | 0,340719401 | 0,12 | 0,067 | 1 | Mast | 1,266387922 | 0,789647456 | 1,791044776 |
| RASSF1 | 0,523649976 | 0,487699595 | 0,12 | 0,067 | 1 | Mast | 1,402207246 | 0,713161341 | 1,791044776 |
| KPNA41 | 0,523649976 | 0,393327366 | 0,12 | 0,067 | 1 | Mast | 1,313419124 | 0,761371585 | 1,791044776 |
| PSMD21 | 0,523649976 | 0,34093675 | 0,12 | 0,067 | 1 | Mast | 1,266578723 | 0,7895285 | 1,791044776 |
| BTN2A2 | 0,523649976 | 0,442706253 | 0,12 | 0,067 | 1 | Mast | 1,359151477 | 0,735753164 | 1,791044776 |
| SPIN1 | 0,523649976 | 0,54763767 | 0,12 | 0,067 | 1 | Mast | 1,461690302 | 0,684139451 | 1,791044776 |
| CCAR12 | 0,523649976 | 0,528764342 | 0,12 | 0,067 | 1 | Mast | 1,44269301 | 0,693148156 | 1,791044776 |
| FAM174C4 | 0,523649976 | 0,664081595 | 0,12 | 0,067 | 1 | Mast | 1,584559237 | 0,631090322 | 1,791044776 |
| USP112 | 0,523649976 | 0,484824442 | 0,12 | 0,067 | 1 | Mast | 1,399415564 | 0,714584021 | 1,791044776 |
| EIF3H2 | 0,524286639 | -0,813579115 | 0,222 | 0,267 | 1 | Mast | 0,568968578 | 1,757566303 | 0,831460674 |
| MSRA1 | 0,524552775 | 0,510353849 | 0,194 | 0,133 | 1 | Mast | 1,424399514 | 0,702050226 | 1,458646617 |
| KDM6B1 | 0,527320094 | -0,378187992 | 0,417 | 0,467 | 1 | Mast | 0,769403345 | 1,299708412 | 0,892933619 |
| LUC7L33 | 0,527678483 | -0,70012558 | 0,157 | 0,2 | 1 | Mast | 0,615518626 | 1,624646205 | 0,785 |
| RNASET22 | 0,527891372 | 0,399231575 | 0,204 | 0,133 | 1 | Mast | 1,318805286 | 0,75826205 | 1,533834586 |
| DBI3 | 0,529311957 | 0,519107556 | 0,509 | 0,4 | 1 | Mast | 1,433068484 | 0,697803358 | 1,2725 |
| RNF1301 | 0,531307543 | 0,425145523 | 0,444 | 0,4 | 1 | Mast | 1,342707934 | 0,744763604 | 1,11 |
| HTRA2 | 0,532256783 | -0,257351905 | 0,083 | 0,133 | 1 | Mast | 0,836622148 | 1,195282725 | 0,62406015 |
| HSPA22 | 0,532256783 | -0,446970483 | 0,083 | 0,133 | 1 | Mast | 0,733581681 | 1,363174717 | 0,62406015 |
| CNOT12 | 0,532256783 | -0,385387861 | 0,083 | 0,133 | 1 | Mast | 0,76557315 | 1,306210908 | 0,62406015 |
| STAU1 | 0,532256783 | -0,252026977 | 0,083 | 0,133 | 1 | Mast | 0,839715791 | 1,190879117 | 0,62406015 |
| N4BP2L22 | 0,532815971 | 0,526018558 | 0,12 | 0,067 | 1 | Mast | 1,439949841 | 0,694468634 | 1,791044776 |
| ZSWIM71 | 0,532815971 | 0,354983245 | 0,12 | 0,067 | 1 | Mast | 1,278970728 | 0,781878723 | 1,791044776 |
| CENPX3 | 0,532815971 | 0,510389443 | 0,12 | 0,067 | 1 | Mast | 1,424434657 | 0,702032905 | 1,791044776 |
| ZBTB7A4 | 0,532815971 | 0,363166727 | 0,12 | 0,067 | 1 | Mast | 1,286246122 | 0,777456183 | 1,791044776 |
| PNRC21 | 0,533281188 | -0,643751843 | 0,093 | 0,133 | 1 | Mast | 0,640046291 | 1,562386992 | 0,69924812 |
| ADH5 | 0,533281188 | -0,925323054 | 0,093 | 0,133 | 1 | Mast | 0,526562595 | 1,899109451 | 0,69924812 |
| AP2M13 | 0,535500259 | 0,47613285 | 0,398 | 0,333 | 1 | Mast | 1,391010057 | 0,718902063 | 1,195195195 |
| PRPF40A1 | 0,535958718 | -0,285396103 | 0,213 | 0,267 | 1 | Mast | 0,820516299 | 1,218744834 | 0,797752809 |

| Supplementary Table 4 - clusermarker | | | | | | | | | |
|---|---|---|---|---|---|---|---|---|---|
| | **p_val** | **avg_log2FC** | **pct.1** | **pct.2** | **p_val_adj** | **celltype** | **Foldchange_UP** | **Foldchange_DOWN** | **Ratio_pct1_pct2** |
| SLC38A23 | 0,536666104 | -0,298479805 | 0,361 | 0,4 | 1 | Mast | 0,813108733 | 1,229847816 | 0,9025 |
| SLC45A3 | 0,538117643 | 0,267721248 | 0,213 | 0,133 | 1 | Mast | 1,203904745 | 0,8306305 | 1,601503759 |
| PHF201 | 0,54070559 | -0,425379932 | 0,204 | 0,267 | 1 | Mast | 0,744642605 | 1,342926114 | 0,764044944 |
| RPL171 | 0,541890138 | 0,324549618 | 0,769 | 0,867 | 1 | Mast | 1,252273441 | 0,798547639 | 0,886966551 |
| H2AJ3 | 0,542060304 | 0,332044991 | 0,12 | 0,067 | 1 | Mast | 1,258796429 | 0,794409626 | 1,791044776 |
| CACYBP2 | 0,542062446 | 0,446769833 | 0,12 | 0,067 | 1 | Mast | 1,362985139 | 0,733683715 | 1,791044776 |
| CCDC61 | 0,542062446 | 0,410292682 | 0,12 | 0,067 | 1 | Mast | 1,328955395 | 0,752470703 | 1,791044776 |
| SPN | 0,542062446 | 0,40444034 | 0,12 | 0,067 | 1 | Mast | 1,32357536 | 0,755529326 | 1,791044776 |
| GNB12 | 0,542601483 | 0,405508421 | 0,204 | 0,133 | 1 | Mast | 1,324555615 | 0,754970187 | 1,533834586 |
| BRK11 | 0,543131615 | 0,370808931 | 0,343 | 0,267 | 1 | Mast | 1,293077666 | 0,773348752 | 1,284644195 |
| DAD13 | 0,543454888 | 0,484220948 | 0,435 | 0,4 | 1 | Mast | 1,398830297 | 0,714883001 | 1,0875 |
| PTP4A23 | 0,545056047 | 0,591493676 | 0,25 | 0,2 | 1 | Mast | 1,506805992 | 0,663655444 | 1,25 |
| SEC11A3 | 0,545851308 | 0,37616615 | 0,407 | 0,4 | 1 | Mast | 1,297888232 | 0,770482369 | 1,0175 |
| MRPL511 | 0,547004108 | 0,522721502 | 0,194 | 0,133 | 1 | Mast | 1,436662816 | 0,69605755 | 1,458646617 |
| CMPK11 | 0,547005505 | 0,495591288 | 0,194 | 0,133 | 1 | Mast | 1,409898483 | 0,709270924 | 1,458646617 |
| NPC22 | 0,547139332 | 0,299216754 | 0,398 | 0,333 | 1 | Mast | 1,2304762 | 0,812693492 | 1,195195195 |
| COPS63 | 0,547781398 | 0,581949205 | 0,111 | 0,067 | 1 | Mast | 1,496870283 | 0,66806056 | 1,656716418 |
| KLRG1 | 0,55003239 | 0,330381149 | 0,204 | 0,133 | 1 | Mast | 1,257345512 | 0,795326337 | 1,533834586 |
| RILPL2 | 0,551150584 | -0,46515689 | 0,148 | 0,2 | 1 | Mast | 0,724392297 | 1,380467468 | 0,74 |
| RPS6KA5 | 0,551150584 | -0,557477916 | 0,148 | 0,2 | 1 | Mast | 0,679488992 | 1,471694187 | 0,74 |
| PHLDA23 | 0,551317098 | 0,665332461 | 0,185 | 0,133 | 1 | Mast | 1,5859337 | 0,630543383 | 1,390977444 |
| JARID2 | 0,551388274 | 0,327085979 | 0,12 | 0,067 | 1 | Mast | 1,254476964 | 0,797144968 | 1,791044776 |
| PTMS | 0,551388274 | 0,327976946 | 0,12 | 0,067 | 1 | Mast | 1,255251932 | 0,796652827 | 1,791044776 |
| UBE2Z1 | 0,551388274 | 0,437864666 | 0,12 | 0,067 | 1 | Mast | 1,354597902 | 0,73822645 | 1,791044776 |
| HK2 | 0,553194047 | -0,462179895 | 0,093 | 0,133 | 1 | Mast | 0,725888621 | 1,377621816 | 0,69924812 |
| PGAM12 | 0,553194047 | -0,431512712 | 0,093 | 0,133 | 1 | Mast | 0,741483908 | 1,348646936 | 0,69924812 |
| NDUFA115 | 0,555145756 | 0,369386927 | 0,361 | 0,267 | 1 | Mast | 1,291803762 | 0,774111386 | 1,352059925 |
| CCND21 | 0,557471981 | 0,499348021 | 0,111 | 0,067 | 1 | Mast | 1,413574599 | 0,707426407 | 1,656716418 |
| DUSP25 | 0,557513085 | 0,673506733 | 0,204 | 0,133 | 1 | Mast | 1,594945064 | 0,626980842 | 1,533834586 |
| ATP5F1D6 | 0,55854632 | 0,287681139 | 0,454 | 0,4 | 1 | Mast | 1,220676692 | 0,81921774 | 1,135 |
| PPP1CA2 | 0,559073011 | 0,581882025 | 0,185 | 0,133 | 1 | Mast | 1,496800582 | 0,66809167 | 1,390977444 |

| Supplementary Table 4 - clusermarker | | | | | | | | | |
|---|---|---|---|---|---|---|---|---|---|
| | **p_val** | **avg_log2FC** | **pct.1** | **pct.2** | **p_val_adj** | **celltype** | **Foldchange_UP** | **Foldchange_DOWN** | **Ratio_pct1_pct2** |
| COX7A2L | 0,559134517 | -0,379905785 | 0,222 | 0,267 | 1 | Mast | 0,768487775 | 1,301256874 | 0,831460674 |
| CDV32 | 0,559390226 | -0,359908229 | 0,148 | 0,2 | 1 | Mast | 0,779214145 | 1,28334426 | 0,74 |
| MOB1A2 | 0,560054545 | 0,480535948 | 0,259 | 0,2 | 1 | Mast | 1,395261897 | 0,716711323 | 1,295 |
| NOP581 | 0,560792289 | 0,370697727 | 0,12 | 0,067 | 1 | Mast | 1,292977999 | 0,773408365 | 1,791044776 |
| GRSF11 | 0,560792289 | 0,274422065 | 0,12 | 0,067 | 1 | Mast | 1,20950947 | 0,826781455 | 1,791044776 |
| TMED91 | 0,56223376 | 0,456068104 | 0,194 | 0,133 | 1 | Mast | 1,371798045 | 0,728970277 | 1,458646617 |
| EEF1D3 | 0,562430996 | 0,296051542 | 0,769 | 0,8 | 1 | Mast | 1,227779547 | 0,814478464 | 0,96125 |
| EIF5B2 | 0,563285345 | -0,424311523 | 0,093 | 0,133 | 1 | Mast | 0,745194265 | 1,341931959 | 0,69924812 |
| CDC1232 | 0,563285345 | -0,415505845 | 0,093 | 0,133 | 1 | Mast | 0,749756565 | 1,333766247 | 0,69924812 |
| PIN42 | 0,563285345 | -0,419533831 | 0,093 | 0,133 | 1 | Mast | 0,747666174 | 1,337495308 | 0,69924812 |
| EIF2S33 | 0,564445889 | -0,30955273 | 0,213 | 0,267 | 1 | Mast | 0,806891876 | 1,23932342 | 0,797752809 |
| MTHFD2 | 0,56504296 | 0,279073614 | 0,204 | 0,133 | 1 | Mast | 1,213415473 | 0,824120033 | 1,533834586 |
| PPP1R14B4 | 0,56504296 | 0,330832355 | 0,204 | 0,133 | 1 | Mast | 1,257738811 | 0,795077636 | 1,533834586 |
| RASSF51 | 0,566185818 | -0,260350811 | 0,269 | 0,333 | 1 | Mast | 0,834884881 | 1,197769923 | 0,807807808 |
| IGBP11 | 0,56688012 | 0,509718422 | 0,185 | 0,133 | 1 | Mast | 1,423772284 | 0,702359508 | 1,390977444 |
| MACIR | 0,567242962 | 0,658593188 | 0,111 | 0,067 | 1 | Mast | 1,578542592 | 0,633495735 | 1,656716418 |
| PIP4K2A | 0,567242962 | 0,465126853 | 0,111 | 0,067 | 1 | Mast | 1,380438727 | 0,724407379 | 1,656716418 |
| NSMCE4A | 0,567242962 | 0,530467455 | 0,111 | 0,067 | 1 | Mast | 1,444397127 | 0,692330372 | 1,656716418 |
| IFI27L23 | 0,567242962 | 0,650189902 | 0,111 | 0,067 | 1 | Mast | 1,56937476 | 0,637196434 | 1,656716418 |
| EP300 | 0,567242962 | 0,407993557 | 0,111 | 0,067 | 1 | Mast | 1,326839215 | 0,75367082 | 1,656716418 |
| ACTN41 | 0,567929719 | 0,397753723 | 0,324 | 0,267 | 1 | Mast | 1,317455035 | 0,759039188 | 1,213483146 |
| SH2B3 | 0,570273283 | 0,258351118 | 0,12 | 0,067 | 1 | Mast | 1,196110867 | 0,836042902 | 1,791044776 |
| PTBP11 | 0,570273283 | 0,269831796 | 0,12 | 0,067 | 1 | Mast | 1,205667251 | 0,829416242 | 1,791044776 |
| POLR2E2 | 0,570273283 | 0,306687724 | 0,12 | 0,067 | 1 | Mast | 1,236864726 | 0,808495852 | 1,791044776 |
| PACSIN2 | 0,570273283 | 0,257987451 | 0,12 | 0,067 | 1 | Mast | 1,195809395 | 0,836253674 | 1,791044776 |
| RAPGEF23 | 0,572841659 | -0,312875347 | 0,269 | 0,333 | 1 | Mast | 0,805035692 | 1,242180949 | 0,807807808 |
| C6orf621 | 0,573464579 | -0,411096094 | 0,093 | 0,133 | 1 | Mast | 0,752051782 | 1,329695673 | 0,69924812 |
| HNRNPK3 | 0,574104275 | 0,443649043 | 0,343 | 0,267 | 1 | Mast | 1,360039963 | 0,735272512 | 1,284644195 |
| PHIP1 | 0,576046642 | -0,306382706 | 0,148 | 0,2 | 1 | Mast | 0,808666803 | 1,236603253 | 0,74 |
| G3BP22 | 0,576144466 | 0,546810904 | 0,25 | 0,2 | 1 | Mast | 1,46085289 | 0,684531623 | 1,25 |
| CLIC4 | 0,577095245 | 0,465047293 | 0,111 | 0,067 | 1 | Mast | 1,380362603 | 0,724447329 | 1,656716418 |

| Supplementary Table 4 - clusermarker | | | | | | | | | |
|---|---|---|---|---|---|---|---|---|---|
| | **p_val** | **avg_log2FC** | **pct.1** | **pct.2** | **p_val_adj** | **celltype** | **Foldchange_UP** | **Foldchange_DOWN** | **Ratio_pct1_pct2** |
| ST3GAL3 | 0,577095245 | 0,644188908 | 0,111 | 0,067 | 1 | Mast | 1,562860389 | 0,639852419 | 1,656716418 |
| PELI11 | 0,577095245 | 0,377717166 | 0,111 | 0,067 | 1 | Mast | 1,299284319 | 0,769654482 | 1,656716418 |
| UNC50 | 0,577095245 | 0,365560426 | 0,111 | 0,067 | 1 | Mast | 1,288382014 | 0,776167308 | 1,656716418 |
| SMU11 | 0,577095245 | 0,495872001 | 0,111 | 0,067 | 1 | Mast | 1,410172841 | 0,709132931 | 1,656716418 |
| ZNF267 | 0,577095245 | 0,459589556 | 0,111 | 0,067 | 1 | Mast | 1,375150535 | 0,727193114 | 1,656716418 |
| ATP5MC22 | 0,581443738 | 0,405597439 | 0,713 | 0,733 | 1 | Mast | 1,324637347 | 0,754923604 | 0,97271487 |
| IL1RL1 | 0,581878779 | 0,323384199 | 0,583 | 0,533 | 1 | Mast | 1,251262255 | 0,799192972 | 1,09380863 |
| NINJ13 | 0,583524747 | -0,566165292 | 0,167 | 0,2 | 1 | Mast | 0,675409655 | 1,480582921 | 0,835 |
| NRDC2 | 0,583730204 | -0,298475729 | 0,093 | 0,133 | 1 | Mast | 0,81311103 | 1,229844342 | 0,69924812 |
| YBX32 | 0,583730204 | -0,836396571 | 0,093 | 0,133 | 1 | Mast | 0,560040641 | 1,785584701 | 0,69924812 |
| PIAS12 | 0,583730204 | -0,35284276 | 0,093 | 0,133 | 1 | Mast | 0,783039635 | 1,277074563 | 0,69924812 |
| MIF5 | 0,584576863 | 0,422222619 | 0,556 | 0,533 | 1 | Mast | 1,339990357 | 0,746274027 | 1,04315197 |
| CISD34 | 0,587025306 | 0,47830228 | 0,111 | 0,067 | 1 | Mast | 1,393103339 | 0,717821838 | 1,656716418 |
| MARCHF73 | 0,587027423 | 0,352647325 | 0,111 | 0,067 | 1 | Mast | 1,276901575 | 0,783145717 | 1,656716418 |
| BTD | 0,587027423 | 0,343420213 | 0,111 | 0,067 | 1 | Mast | 1,268760897 | 0,788170571 | 1,656716418 |
| DSE | 0,587027423 | 0,573543472 | 0,111 | 0,067 | 1 | Mast | 1,488174261 | 0,671964316 | 1,656716418 |
| CTSC3 | 0,587027423 | 0,377981824 | 0,111 | 0,067 | 1 | Mast | 1,299522691 | 0,769513305 | 1,656716418 |
| VPS362 | 0,587027423 | 0,354181356 | 0,111 | 0,067 | 1 | Mast | 1,278260039 | 0,782313434 | 1,656716418 |
| SLC16A31 | 0,587027423 | 0,625004358 | 0,111 | 0,067 | 1 | Mast | 1,542215484 | 0,648417819 | 1,656716418 |
| SUMO32 | 0,587027423 | 0,439487871 | 0,111 | 0,067 | 1 | Mast | 1,356122845 | 0,737396324 | 1,656716418 |
| IDS5 | 0,588832476 | 0,274445434 | 0,398 | 0,333 | 1 | Mast | 1,209529062 | 0,826768063 | 1,195195195 |
| MAP3K8 | 0,58910869 | -1,102585422 | 0,231 | 0,267 | 1 | Mast | 0,46568121 | 2,147391773 | 0,865168539 |
| NFKB12 | 0,589646434 | -1,004221074 | 0,38 | 0,4 | 1 | Mast | 0,498539225 | 2,005860221 | 0,95 |
| PRMT22 | 0,593290524 | 0,448630109 | 0,194 | 0,133 | 1 | Mast | 1,364743768 | 0,732738279 | 1,458646617 |
| ATP5F1B3 | 0,59586356 | 0,608575469 | 0,343 | 0,333 | 1 | Mast | 1,52475291 | 0,655843969 | 1,03003003 |
| GABPB1-AS11 | 0,597035967 | 0,28137224 | 0,111 | 0,067 | 1 | Mast | 1,215350333 | 0,822808019 | 1,656716418 |
| CDC42SE1 | 0,597038044 | 0,370038168 | 0,111 | 0,067 | 1 | Mast | 1,292387021 | 0,773762026 | 1,656716418 |
| STARD7 | 0,597038044 | 0,447824485 | 0,111 | 0,067 | 1 | Mast | 1,363981886 | 0,733147566 | 1,656716418 |
| PIK3CA | 0,597038044 | 0,399434973 | 0,111 | 0,067 | 1 | Mast | 1,318991231 | 0,758155154 | 1,656716418 |
| FRYL | 0,597038044 | 0,324018681 | 0,111 | 0,067 | 1 | Mast | 1,251812667 | 0,798841573 | 1,656716418 |
| NUDT12 | 0,597038044 | 0,449791697 | 0,111 | 0,067 | 1 | Mast | 1,365843036 | 0,732148551 | 1,656716418 |

| Supplementary Table 4 - clusermarker | | | | | | | | | |
|---|---|---|---|---|---|---|---|---|---|
| | **p_val** | **avg_log2FC** | **pct.1** | **pct.2** | **p_val_adj** | **celltype** | **Foldchange_UP** | **Foldchange_DOWN** | **Ratio_pct1_pct2** |
| AC100810.1 | 0,597038044 | 0,407394355 | 0,111 | 0,067 | 1 | Mast | 1,326288246 | 0,753983912 | 1,656716418 |
| KRAS2 | 0,597038044 | 0,363899051 | 0,111 | 0,067 | 1 | Mast | 1,286899197 | 0,77706164 | 1,656716418 |
| TUBB61 | 0,597038044 | 0,271690352 | 0,111 | 0,067 | 1 | Mast | 1,207221456 | 0,828348432 | 1,656716418 |
| VASP | 0,597038044 | 0,285454284 | 0,111 | 0,067 | 1 | Mast | 1,218793984 | 0,82048321 | 1,656716418 |
| ATP6V0D1 | 0,59862734 | 0,395279022 | 0,185 | 0,133 | 1 | Mast | 1,315197099 | 0,76034231 | 1,390977444 |
| TNRC6B2 | 0,599818787 | -0,339404931 | 0,167 | 0,2 | 1 | Mast | 0,790367247 | 1,265234615 | 0,835 |
| TOMM62 | 0,600262325 | -0,392911515 | 0,287 | 0,333 | 1 | Mast | 0,761591079 | 1,31304059 | 0,861861862 |
| PARP11 | 0,600774243 | -0,26846648 | 0,157 | 0,2 | 1 | Mast | 0,830201544 | 1,204526789 | 0,785 |
| AHNAK4 | 0,602687557 | 0,327976437 | 0,722 | 0,6 | 1 | Mast | 1,25525149 | 0,796653108 | 1,203333333 |
| PRMT11 | 0,603923892 | 0,453889046 | 0,176 | 0,133 | 1 | Mast | 1,369727634 | 0,730072151 | 1,323308271 |
| CD94 | 0,60436121 | 0,344545364 | 0,75 | 0,733 | 1 | Mast | 1,269750783 | 0,78755612 | 1,02319236 |
| USP53 | 0,604514171 | -0,367545338 | 0,093 | 0,133 | 1 | Mast | 0,775100164 | 1,290155836 | 0,69924812 |
| LINC02207 | 0,604514171 | -0,291006012 | 0,093 | 0,133 | 1 | Mast | 0,817331923 | 1,223493139 | 0,69924812 |
| SSU723 | 0,606688033 | 0,346208797 | 0,185 | 0,133 | 1 | Mast | 1,271215656 | 0,786648588 | 1,390977444 |
| CSRP12 | 0,60712561 | 0,260768667 | 0,111 | 0,067 | 1 | Mast | 1,19811689 | 0,834643104 | 1,656716418 |
| MRPL542 | 0,60712561 | 0,326149476 | 0,111 | 0,067 | 1 | Mast | 1,253662905 | 0,79766259 | 1,656716418 |
| DDT4 | 0,60712561 | 0,306349456 | 0,111 | 0,067 | 1 | Mast | 1,236574753 | 0,808685441 | 1,656716418 |
| MAST41 | 0,607139846 | 0,598714482 | 0,241 | 0,2 | 1 | Mast | 1,514366583 | 0,660342093 | 1,205 |
| SIGLEC6 | 0,608551147 | 0,371501634 | 0,528 | 0,467 | 1 | Mast | 1,293698681 | 0,772977522 | 1,130620985 |
| MPG2 | 0,612171716 | 0,383233557 | 0,176 | 0,133 | 1 | Mast | 1,304261864 | 0,766717197 | 1,323308271 |
| PSMB84 | 0,614796875 | 0,315383448 | 0,185 | 0,133 | 1 | Mast | 1,244342338 | 0,803637367 | 1,390977444 |
| SETD5 | 0,615029161 | -0,271930781 | 0,093 | 0,133 | 1 | Mast | 0,828210397 | 1,207422659 | 0,69924812 |
| APEX11 | 0,615029161 | -0,261384799 | 0,093 | 0,133 | 1 | Mast | 0,834286728 | 1,19862868 | 0,69924812 |
| LIMS11 | 0,616314224 | -0,675929708 | 0,167 | 0,2 | 1 | Mast | 0,625928725 | 1,597625991 | 0,835 |
| SLC30A7 | 0,617288582 | 0,340826022 | 0,111 | 0,067 | 1 | Mast | 1,266481516 | 0,789589099 | 1,656716418 |
| TIMM17A2 | 0,617288582 | 0,360326942 | 0,111 | 0,067 | 1 | Mast | 1,283716779 | 0,778988026 | 1,656716418 |
| ELK4 | 0,617288582 | 0,325602947 | 0,111 | 0,067 | 1 | Mast | 1,253188075 | 0,797964822 | 1,656716418 |
| BIRC61 | 0,617288582 | 0,305627834 | 0,111 | 0,067 | 1 | Mast | 1,235956385 | 0,809090039 | 1,656716418 |
| PSMD62 | 0,617288582 | 0,293978569 | 0,111 | 0,067 | 1 | Mast | 1,226016648 | 0,81564961 | 1,656716418 |
| B3GNT51 | 0,617288582 | 0,313426099 | 0,111 | 0,067 | 1 | Mast | 1,242655245 | 0,804728426 | 1,656716418 |
| HES13 | 0,617288582 | 0,280494288 | 0,111 | 0,067 | 1 | Mast | 1,214610956 | 0,823308891 | 1,656716418 |

| Supplementary Table 4 - clusermarker | | | | | | | | | |
|---|---|---|---|---|---|---|---|---|---|
| | **p_val** | **avg_log2FC** | **pct.1** | **pct.2** | **p_val_adj** | **celltype** | **Foldchange_UP** | **Foldchange_DOWN** | **Ratio_pct1_pct2** |
| TPCN11 | 0,617288582 | 0,290979899 | 0,111 | 0,067 | 1 | Mast | 1,223470994 | 0,817346717 | 1,656716418 |
| SELENOM4 | 0,617288582 | 0,354414409 | 0,111 | 0,067 | 1 | Mast | 1,278466545 | 0,782187069 | 1,656716418 |
| THRAP3 | 0,620467676 | 0,515498271 | 0,176 | 0,133 | 1 | Mast | 1,429487764 | 0,699551284 | 1,323308271 |
| IFITM10 | 0,622397851 | -0,375255316 | 0,102 | 0,133 | 1 | Mast | 0,770968961 | 1,297069078 | 0,766917293 |
| SUPT6H | 0,622397851 | -0,650900155 | 0,102 | 0,133 | 1 | Mast | 0,636882813 | 1,570147568 | 0,766917293 |
| HNRNPH2 | 0,622397851 | -0,719534724 | 0,102 | 0,133 | 1 | Mast | 0,607293265 | 1,646650896 | 0,766917293 |
| ICAM11 | 0,622585204 | 0,628821728 | 0,398 | 0,333 | 1 | Mast | 1,546301589 | 0,646704373 | 1,195195195 |
| PHPT15 | 0,622953062 | 0,437236117 | 0,185 | 0,133 | 1 | Mast | 1,354007863 | 0,738548148 | 1,390977444 |
| TBCB4 | 0,622953062 | 0,290718823 | 0,185 | 0,133 | 1 | Mast | 1,22324961 | 0,81749464 | 1,390977444 |
| SIL1 | 0,625623834 | 0,25428778 | 0,102 | 0,067 | 1 | Mast | 1,19274677 | 0,838400929 | 1,52238806 |
| VAT11 | 0,626370279 | 0,323348542 | 0,324 | 0,267 | 1 | Mast | 1,25123133 | 0,799212725 | 1,213483146 |
| SRSF101 | 0,628814841 | 0,466191244 | 0,176 | 0,133 | 1 | Mast | 1,381457563 | 0,723873123 | 1,323308271 |
| FCER1G | 0,630180511 | -0,264961274 | 0,787 | 0,8 | 1 | Mast | 0,832221074 | 1,201603795 | 0,98375 |
| YY1 | 0,631155773 | 0,346352252 | 0,185 | 0,133 | 1 | Mast | 1,271342065 | 0,786570371 | 1,390977444 |
| HNRNPM2 | 0,631155773 | 0,358415354 | 0,185 | 0,133 | 1 | Mast | 1,282016966 | 0,780020879 | 1,390977444 |
| CSRNP11 | 0,631891593 | 0,287458558 | 0,37 | 0,333 | 1 | Mast | 1,220488379 | 0,819344139 | 1,111111111 |
| AK24 | 0,636296391 | 0,447006216 | 0,102 | 0,067 | 1 | Mast | 1,36320848 | 0,733563512 | 1,52238806 |
| PPM1G4 | 0,636296391 | 0,421424223 | 0,102 | 0,067 | 1 | Mast | 1,339249004 | 0,746687134 | 1,52238806 |
| GBE1 | 0,636296391 | 0,460635842 | 0,102 | 0,067 | 1 | Mast | 1,376148197 | 0,726665923 | 1,52238806 |
| MARCHF61 | 0,636296391 | 0,457519632 | 0,102 | 0,067 | 1 | Mast | 1,373178936 | 0,728237212 | 1,52238806 |
| UBE2D1 | 0,636296391 | 0,366815008 | 0,102 | 0,067 | 1 | Mast | 1,289502891 | 0,775492639 | 1,52238806 |
| CNPY22 | 0,636296391 | 0,46465863 | 0,102 | 0,067 | 1 | Mast | 1,379990782 | 0,724642522 | 1,52238806 |
| TAX1BP32 | 0,636296391 | 0,334674776 | 0,102 | 0,067 | 1 | Mast | 1,261093091 | 0,792962872 | 1,52238806 |
| PRDX22 | 0,636296391 | 0,439931109 | 0,102 | 0,067 | 1 | Mast | 1,356539549 | 0,737169809 | 1,52238806 |
| TRMT1124 | 0,640213414 | 0,325837955 | 0,315 | 0,267 | 1 | Mast | 1,25339223 | 0,797834849 | 1,179775281 |
| MAPK1 | 0,640640791 | 0,511578577 | 0,306 | 0,267 | 1 | Mast | 1,425609225 | 0,701454496 | 1,146067416 |
| LNPEP2 | 0,645648752 | 0,420799092 | 0,176 | 0,133 | 1 | Mast | 1,338668822 | 0,747010749 | 1,323308271 |
| CTR9 | 0,645648752 | 0,439105728 | 0,176 | 0,133 | 1 | Mast | 1,355763681 | 0,737591672 | 1,323308271 |
| ACADVL1 | 0,645648752 | 0,430149545 | 0,176 | 0,133 | 1 | Mast | 1,347373234 | 0,742184849 | 1,323308271 |
| FUNDC21 | 0,645648752 | 0,346200869 | 0,176 | 0,133 | 1 | Mast | 1,271208669 | 0,786652911 | 1,323308271 |
| ZNF5933 | 0,64704498 | 0,438999834 | 0,102 | 0,067 | 1 | Mast | 1,355664171 | 0,737645813 | 1,52238806 |

| Supplementary Table 4 - clusermarker | | | | | | | | | |
|---|---|---|---|---|---|---|---|---|---|
| | **p_val** | **avg_log2FC** | **pct.1** | **pct.2** | **p_val_adj** | **celltype** | **Foldchange_UP** | **Foldchange_DOWN** | **Ratio_pct1_pct2** |
| TAL1 | 0,64704498 | 0,341693 | 0,102 | 0,067 | 1 | Mast | 1,267242829 | 0,789114743 | 1,52238806 |
| THUMPD3-AS11 | 0,64704498 | 0,39567824 | 0,102 | 0,067 | 1 | Mast | 1,315561086 | 0,760131939 | 1,52238806 |
| ELF21 | 0,64704498 | 0,530995455 | 0,102 | 0,067 | 1 | Mast | 1,444925846 | 0,692077038 | 1,52238806 |
| PHF1 | 0,64704498 | 0,49460539 | 0,102 | 0,067 | 1 | Mast | 1,408935326 | 0,709755786 | 1,52238806 |
| OXR1 | 0,64704498 | 0,384579211 | 0,102 | 0,067 | 1 | Mast | 1,305478964 | 0,766002385 | 1,52238806 |
| AMZ2 | 0,64704498 | 0,442594874 | 0,102 | 0,067 | 1 | Mast | 1,359046552 | 0,735809968 | 1,52238806 |
| MALT12 | 0,64704498 | 0,312031271 | 0,102 | 0,067 | 1 | Mast | 1,2414544 | 0,805506831 | 1,52238806 |
| DHPS3 | 0,64704498 | 0,304124282 | 0,102 | 0,067 | 1 | Mast | 1,234668963 | 0,809933699 | 1,52238806 |
| ADSL2 | 0,64704498 | 0,406107125 | 0,102 | 0,067 | 1 | Mast | 1,325105407 | 0,754656947 | 1,52238806 |
| HTATSF11 | 0,64704498 | 0,338001713 | 0,102 | 0,067 | 1 | Mast | 1,2640046 | 0,791136362 | 1,52238806 |
| SPATS2L | 0,648065098 | 0,590130709 | 0,25 | 0,2 | 1 | Mast | 1,50538313 | 0,66428272 | 1,25 |
| DUSP61 | 0,648929994 | -0,601936747 | 0,352 | 0,4 | 1 | Mast | 0,658868862 | 1,517752708 | 0,88 |
| GTF2A23 | 0,651909103 | 0,634897345 | 0,167 | 0,133 | 1 | Mast | 1,552827254 | 0,643986636 | 1,255639098 |
| SRSF4 | 0,653429545 | -0,40376895 | 0,102 | 0,133 | 1 | Mast | 0,75588101 | 1,322959548 | 0,766917293 |
| NFIA2 | 0,653429545 | -0,427823414 | 0,102 | 0,133 | 1 | Mast | 0,743382477 | 1,345202546 | 0,766917293 |
| ESYT22 | 0,653429545 | -0,26576335 | 0,102 | 0,133 | 1 | Mast | 0,831758523 | 1,20227202 | 0,766917293 |
| OAT5 | 0,653429545 | -0,315014751 | 0,102 | 0,133 | 1 | Mast | 0,803842772 | 1,244024373 | 0,766917293 |
| CHD32 | 0,653429545 | -0,31327427 | 0,102 | 0,133 | 1 | Mast | 0,804813121 | 1,242524475 | 0,766917293 |
| ESYT1 | 0,654134988 | 0,370690885 | 0,176 | 0,133 | 1 | Mast | 1,292971867 | 0,773412033 | 1,323308271 |
| GRB2 | 0,654134988 | 0,54597459 | 0,176 | 0,133 | 1 | Mast | 1,460006295 | 0,684928554 | 1,323308271 |
| SYNGR25 | 0,655175171 | 0,469748345 | 0,287 | 0,267 | 1 | Mast | 1,384867879 | 0,722090544 | 1,074906367 |
| SMARCC12 | 0,657865756 | 0,294528732 | 0,102 | 0,067 | 1 | Mast | 1,226484271 | 0,815338626 | 1,52238806 |
| HES43 | 0,657867705 | 0,629503822 | 0,102 | 0,067 | 1 | Mast | 1,54703284 | 0,646398689 | 1,52238806 |
| HADHB1 | 0,657867705 | 0,39187864 | 0,102 | 0,067 | 1 | Mast | 1,312100876 | 0,762136524 | 1,52238806 |
| ZNF148 | 0,657867705 | 0,352644841 | 0,102 | 0,067 | 1 | Mast | 1,276899376 | 0,783147066 | 1,52238806 |
| RWDD42 | 0,657867705 | 0,337142335 | 0,102 | 0,067 | 1 | Mast | 1,263251887 | 0,791607763 | 1,52238806 |
| AKIRIN2 | 0,657867705 | 0,419688227 | 0,102 | 0,067 | 1 | Mast | 1,337638454 | 0,747586164 | 1,52238806 |
| PDAP12 | 0,657867705 | 0,389736854 | 0,102 | 0,067 | 1 | Mast | 1,310154412 | 0,763268811 | 1,52238806 |
| TSPAN41 | 0,657867705 | 0,284968865 | 0,102 | 0,067 | 1 | Mast | 1,218383969 | 0,820759322 | 1,52238806 |
| TCIRG12 | 0,657867705 | 0,319474011 | 0,102 | 0,067 | 1 | Mast | 1,247875506 | 0,801361991 | 1,52238806 |
| MYCBP2 | 0,657867705 | 0,34311457 | 0,102 | 0,067 | 1 | Mast | 1,268492131 | 0,788337567 | 1,52238806 |

| Supplementary Table 4 - clusermarker | | | | | | | | | |
|---|---|---|---|---|---|---|---|---|---|
| | **p_val** | **avg_log2FC** | **pct.1** | **pct.2** | **p_val_adj** | **celltype** | **Foldchange_UP** | **Foldchange_DOWN** | **Ratio_pct1_pct2** |
| PIM34 | 0,657867705 | 0,492732016 | 0,102 | 0,067 | 1 | Mast | 1,407106978 | 0,710678019 | 1,52238806 |
| SKIL3 | 0,659147991 | -0,489711115 | 0,324 | 0,333 | 1 | Mast | 0,712167688 | 1,404163678 | 0,972972973 |
| ASH1L4 | 0,660602138 | 0,470541186 | 0,167 | 0,133 | 1 | Mast | 1,38562915 | 0,721693824 | 1,255639098 |
| DNAJB91 | 0,66060338 | 0,595295317 | 0,167 | 0,133 | 1 | Mast | 1,510781808 | 0,66190895 | 1,255639098 |
| CHASERR3 | 0,662270875 | 0,581880341 | 0,231 | 0,2 | 1 | Mast | 1,496798836 | 0,668092449 | 1,155 |
| ACER3 | 0,662664982 | 0,266573972 | 0,176 | 0,133 | 1 | Mast | 1,202947743 | 0,831291306 | 1,323308271 |
| COX4I15 | 0,66325446 | 0,29317653 | 0,685 | 0,733 | 1 | Mast | 1,225335256 | 0,816103181 | 0,934515689 |
| RNASEK3 | 0,663440148 | 0,495091465 | 0,639 | 0,667 | 1 | Mast | 1,409410107 | 0,709516694 | 0,95802099 |
| TSNAX | 0,663911629 | -0,379681509 | 0,102 | 0,133 | 1 | Mast | 0,768607251 | 1,301054601 | 0,766917293 |
| TACC12 | 0,665604882 | 0,688101137 | 0,222 | 0,2 | 1 | Mast | 1,611161525 | 0,620670233 | 1,11 |
| PDE7B | 0,668760725 | 0,30023286 | 0,102 | 0,067 | 1 | Mast | 1,231343144 | 0,812121304 | 1,52238806 |
| TRAPPC3 | 0,66876262 | 0,256418711 | 0,102 | 0,067 | 1 | Mast | 1,194509817 | 0,837163484 | 1,52238806 |
| DESI2 | 0,66876262 | 0,359940133 | 0,102 | 0,067 | 1 | Mast | 1,28337264 | 0,779196913 | 1,52238806 |
| CGGBP13 | 0,66876262 | 0,254725971 | 0,102 | 0,067 | 1 | Mast | 1,1931091 | 0,838146319 | 1,52238806 |
| PPP1R11 | 0,66876262 | 0,322186001 | 0,102 | 0,067 | 1 | Mast | 1,250223479 | 0,799856999 | 1,52238806 |
| RAB143 | 0,66876262 | 0,295936768 | 0,102 | 0,067 | 1 | Mast | 1,227681875 | 0,814543263 | 1,52238806 |
| ROCK1 | 0,66876262 | 0,440759941 | 0,102 | 0,067 | 1 | Mast | 1,357319108 | 0,736746425 | 1,52238806 |
| NOSIP2 | 0,669177599 | -0,46618686 | 0,111 | 0,067 | 1 | Mast | 0,723875323 | 1,381453365 | 1,656716418 |
| LRPAP11 | 0,669342425 | 0,387421686 | 0,167 | 0,133 | 1 | Mast | 1,308053624 | 0,764494652 | 1,255639098 |
| DNAJC154 | 0,671241342 | 0,36892059 | 0,176 | 0,133 | 1 | Mast | 1,291386267 | 0,77436165 | 1,323308271 |
| BTG23 | 0,672264852 | 0,396617989 | 0,565 | 0,6 | 1 | Mast | 1,316418302 | 0,759636962 | 0,941666667 |
| CLK12 | 0,672837248 | -0,415726749 | 0,176 | 0,2 | 1 | Mast | 0,749641772 | 1,333970488 | 0,88 |
| SH3BGRL4 | 0,673531936 | 0,250599099 | 0,324 | 0,267 | 1 | Mast | 1,189701052 | 0,840547294 | 1,213483146 |
| RHEB1 | 0,67399118 | 0,350804346 | 0,241 | 0,2 | 1 | Mast | 1,275271432 | 0,78414679 | 1,205 |
| STOML24 | 0,678126479 | 0,500931355 | 0,167 | 0,133 | 1 | Mast | 1,415126826 | 0,706650444 | 1,255639098 |
| DAP3 | 0,679727736 | 0,278828767 | 0,102 | 0,067 | 1 | Mast | 1,213209556 | 0,82425991 | 1,52238806 |
| CDC731 | 0,679727736 | 0,309574814 | 0,102 | 0,067 | 1 | Mast | 1,239342391 | 0,806879525 | 1,52238806 |
| SREK11 | 0,679727736 | 0,307935806 | 0,102 | 0,067 | 1 | Mast | 1,237935207 | 0,80779672 | 1,52238806 |
| WDFY2 | 0,679727736 | 0,259383712 | 0,102 | 0,067 | 1 | Mast | 1,196967277 | 0,835444727 | 1,52238806 |
| C15orf61 | 0,679727736 | 0,291039948 | 0,102 | 0,067 | 1 | Mast | 1,22352192 | 0,817312697 | 1,52238806 |
| RAB5IF3 | 0,679727736 | 0,346687518 | 0,102 | 0,067 | 1 | Mast | 1,271637545 | 0,786387602 | 1,52238806 |

| Supplementary Table 4 - clusermarker | | | | | | | | | |
|---|---|---|---|---|---|---|---|---|---|
| | **p_val** | **avg_log2FC** | **pct.1** | **pct.2** | **p_val_adj** | **celltype** | **Foldchange_UP** | **Foldchange_DOWN** | **Ratio_pct1_pct2** |
| SMARCE13 | 0,679858399 | 0,279997361 | 0,176 | 0,133 | 1 | Mast | 1,214192663 | 0,823592524 | 1,323308271 |
| NFKBIA5 | 0,681852269 | 0,291717512 | 0,898 | 0,8 | 1 | Mast | 1,224096684 | 0,816928935 | 1,1225 |
| DDX173 | 0,684072707 | 0,463135639 | 0,343 | 0,333 | 1 | Mast | 1,378534753 | 0,7254079 | 1,03003003 |
| STX41 | 0,685072516 | -0,307563424 | 0,102 | 0,133 | 1 | Mast | 0,808005252 | 1,237615718 | 0,766917293 |
| FHL11 | 0,685072516 | -0,550084179 | 0,102 | 0,133 | 1 | Mast | 0,682980276 | 1,464171126 | 0,766917293 |
| S100A105 | 0,688930757 | 0,656471485 | 0,435 | 0,467 | 1 | Mast | 1,576222811 | 0,634428072 | 0,931477516 |
| SLC24A3 | 0,689804875 | -0,47520161 | 0,176 | 0,2 | 1 | Mast | 0,719366255 | 1,390112468 | 0,88 |
| TUT41 | 0,690761018 | 0,263818117 | 0,102 | 0,067 | 1 | Mast | 1,20065205 | 0,832880767 | 1,52238806 |
| TBC1D11 | 0,690761018 | 0,339890294 | 0,102 | 0,067 | 1 | Mast | 1,265660346 | 0,790101391 | 1,52238806 |
| CTNNA12 | 0,690761018 | 0,255424396 | 0,102 | 0,067 | 1 | Mast | 1,193686837 | 0,837740661 | 1,52238806 |
| NSD31 | 0,690761018 | 0,262720374 | 0,102 | 0,067 | 1 | Mast | 1,199738824 | 0,833514745 | 1,52238806 |
| SHOC24 | 0,690761018 | 0,267999507 | 0,102 | 0,067 | 1 | Mast | 1,20413697 | 0,830470308 | 1,52238806 |
| UBE2L61 | 0,690761018 | 0,300241577 | 0,102 | 0,067 | 1 | Mast | 1,231350584 | 0,812116397 | 1,52238806 |
| PAQR5 | 0,690761018 | 0,258360319 | 0,102 | 0,067 | 1 | Mast | 1,196118495 | 0,83603757 | 1,52238806 |
| TESPA1 | 0,69415737 | 0,259204315 | 0,324 | 0,267 | 1 | Mast | 1,196818445 | 0,83554862 | 1,213483146 |
| EGR14 | 0,694450123 | 0,572319754 | 0,546 | 0,6 | 1 | Mast | 1,486912503 | 0,672534529 | 0,91 |
| UBE2B3 | 0,697221005 | 0,321121263 | 0,176 | 0,133 | 1 | Mast | 1,249301129 | 0,800447528 | 1,323308271 |
| SERBP12 | 0,698493316 | 0,442820774 | 0,278 | 0,267 | 1 | Mast | 1,359259371 | 0,735694762 | 1,041198502 |
| CD22 | 0,699910916 | -0,360023976 | 0,111 | 0,133 | 1 | Mast | 0,779151631 | 1,283447227 | 0,834586466 |
| GPR183 | 0,700496431 | 0,296572702 | 0,25 | 0,2 | 1 | Mast | 1,228223151 | 0,814184295 | 1,25 |
| LDHA3 | 0,704096322 | 0,381251619 | 0,296 | 0,267 | 1 | Mast | 1,302471332 | 0,767771217 | 1,108614232 |
| STAG22 | 0,705962261 | 0,332058103 | 0,176 | 0,133 | 1 | Mast | 1,258807869 | 0,794402406 | 1,323308271 |
| COX7B4 | 0,709410907 | 0,59223339 | 0,278 | 0,267 | 1 | Mast | 1,507578776 | 0,663315255 | 1,041198502 |
| RTN43 | 0,710237511 | 0,430821317 | 0,306 | 0,267 | 1 | Mast | 1,348000766 | 0,741839341 | 1,146067416 |
| ITGB1BP11 | 0,710321233 | -0,643358197 | 0,111 | 0,133 | 1 | Mast | 0,640220955 | 1,561960745 | 0,834586466 |
| BRD24 | 0,711013686 | 0,309785034 | 0,444 | 0,4 | 1 | Mast | 1,239522993 | 0,80676196 | 1,11 |
| IER23 | 0,712476969 | 0,30754846 | 0,63 | 0,667 | 1 | Mast | 1,237602881 | 0,808013633 | 0,944527736 |
| BCLAF12 | 0,712753913 | 0,328618678 | 0,222 | 0,2 | 1 | Mast | 1,255810411 | 0,796298543 | 1,11 |
| BAG13 | 0,71369054 | 0,361054969 | 0,167 | 0,133 | 1 | Mast | 1,284364745 | 0,778595025 | 1,255639098 |
| DNAJC11 | 0,713691609 | 0,388294936 | 0,167 | 0,133 | 1 | Mast | 1,308845617 | 0,76403205 | 1,255639098 |
| NUTM2B-AS11 | 0,714742486 | 0,261999874 | 0,176 | 0,133 | 1 | Mast | 1,199139809 | 0,833931117 | 1,323308271 |

| Supplementary Table 4 - clusermarker | | | | | | | | | |
|---|---|---|---|---|---|---|---|---|---|
| | p_val | avg_log2FC | pct.1 | pct.2 | p_val_adj | celltype | Foldchange_UP | Foldchange_DOWN | Ratio_pct1_pct2 |
| RHBDD22 | 0,716525479 | 0,563930166 | 0,231 | 0,2 | 1 | Mast | 1,478290873 | 0,676456859 | 1,155 |
| YTHDC12 | 0,720785886 | -0,364885299 | 0,111 | 0,133 | 1 | Mast | 0,776530611 | 1,287779241 | 0,834586466 |
| DTNBP1 | 0,720785886 | -0,374890442 | 0,111 | 0,133 | 1 | Mast | 0,771163972 | 1,296741077 | 0,834586466 |
| STK242 | 0,720785886 | -0,48287338 | 0,111 | 0,133 | 1 | Mast | 0,715551058 | 1,397524311 | 0,834586466 |
| MAOB | 0,722802515 | 0,351108497 | 0,333 | 0,333 | 1 | Mast | 1,275540314 | 0,783981493 | 1 |
| RHOBTB3 | 0,723559633 | -0,404326249 | 0,176 | 0,133 | 1 | Mast | 0,755589077 | 1,323470694 | 1,323308271 |
| POLR2L2 | 0,723561458 | 0,275381126 | 0,37 | 0,333 | 1 | Mast | 1,210313783 | 0,826232018 | 1,111111111 |
| CLNS1A1 | 0,724204563 | -0,291023947 | 0,176 | 0,2 | 1 | Mast | 0,817321762 | 1,22350835 | 0,88 |
| SNU131 | 0,726810276 | 0,3516361 | 0,315 | 0,267 | 1 | Mast | 1,276006873 | 0,783694838 | 1,179775281 |
| PHACTR1 | 0,727983585 | -0,334930152 | 0,167 | 0,2 | 1 | Mast | 0,79282252 | 1,261316341 | 0,835 |
| PTGS1 | 0,73067645 | 0,396077046 | 0,546 | 0,6 | 1 | Mast | 1,315924799 | 0,759921844 | 0,91 |
| FYN1 | 0,731716205 | 0,256368237 | 0,167 | 0,133 | 1 | Mast | 1,194468027 | 0,837192773 | 1,255639098 |
| FAM204A2 | 0,731716205 | 0,382758921 | 0,167 | 0,133 | 1 | Mast | 1,303832842 | 0,766969482 | 1,255639098 |
| ATP6AP12 | 0,731716205 | 0,309751921 | 0,167 | 0,133 | 1 | Mast | 1,239494544 | 0,806780478 | 1,255639098 |
| TMEM2581 | 0,734014773 | 0,399114243 | 0,287 | 0,267 | 1 | Mast | 1,318698034 | 0,758323721 | 1,074906367 |
| PTGS21 | 0,739180529 | -0,456194562 | 0,5 | 0,6 | 1 | Mast | 0,728906382 | 1,371918293 | 0,833333333 |
| S100A65 | 0,748395428 | -0,382450519 | 0,972 | 1 | 1 | Mast | 0,767133453 | 1,303554155 | 0,972 |
| DEK2 | 0,749327503 | 0,324998335 | 0,157 | 0,133 | 1 | Mast | 1,252662993 | 0,798299307 | 1,180451128 |
| U2SURP2 | 0,750379476 | -0,305854298 | 0,176 | 0,2 | 1 | Mast | 0,808963044 | 1,236150412 | 0,88 |
| SLC11A21 | 0,752487884 | -0,318458329 | 0,111 | 0,133 | 1 | Mast | 0,801926362 | 1,24699729 | 0,834586466 |
| HSD17B12 | 0,756194295 | 0,456989938 | 0,231 | 0,2 | 1 | Mast | 1,372674857 | 0,728504638 | 1,155 |
| NFKBIZ2 | 0,766691148 | 0,346502825 | 0,648 | 0,6 | 1 | Mast | 1,271474761 | 0,786488282 | 1,08 |
| POMP3 | 0,767746094 | 0,273010343 | 0,306 | 0,267 | 1 | Mast | 1,208326506 | 0,827590883 | 1,146067416 |
| CCT5 | 0,768034145 | 0,363679225 | 0,157 | 0,133 | 1 | Mast | 1,286703124 | 0,777180051 | 1,180451128 |
| AC068587.4 | 0,769169887 | 0,376372566 | 0,222 | 0,2 | 1 | Mast | 1,298073942 | 0,770372139 | 1,11 |
| TMED101 | 0,769169887 | 0,520445523 | 0,222 | 0,2 | 1 | Mast | 1,43439814 | 0,697156509 | 1,11 |
| EMID1 | 0,773860772 | -0,253225397 | 0,111 | 0,133 | 1 | Mast | 0,839018544 | 1,19186877 | 0,834586466 |
| RHEX | 0,775541791 | 0,295193139 | 0,722 | 0,733 | 1 | Mast | 1,227049236 | 0,814963223 | 0,984993179 |
| SP100 | 0,777269316 | 0,417816981 | 0,148 | 0,133 | 1 | Mast | 1,335904596 | 0,748556449 | 1,112781955 |
| KMT2A3 | 0,777440035 | 0,330515335 | 0,157 | 0,133 | 1 | Mast | 1,257462464 | 0,795252367 | 1,180451128 |
| ARF43 | 0,780705215 | -0,380655976 | 0,185 | 0,2 | 1 | Mast | 0,768088271 | 1,301933695 | 0,925 |

| Supplementary Table 4 - clusermarker | | | | | | | | | |
|---|---|---|---|---|---|---|---|---|---|
| | **p_val** | **avg_log2FC** | **pct.1** | **pct.2** | **p_val_adj** | **celltype** | **Foldchange_UP** | **Foldchange_DOWN** | **Ratio_pct1_pct2** |
| GPI2 | 0,783166165 | 0,308807757 | 0,102 | 0,133 | 1 | Mast | 1,238683629 | 0,807308643 | 0,766917293 |
| GATA2 | 0,784108716 | -0,265960721 | 0,694 | 0,8 | 1 | Mast | 0,831644741 | 1,20243651 | 0,8675 |
| PPP1R15A2 | 0,784909913 | 0,283903988 | 0,731 | 0,8 | 1 | Mast | 1,217484991 | 0,821365361 | 0,91375 |
| CDK15 | 0,785651458 | -0,388416416 | 0,12 | 0,133 | 1 | Mast | 0,763967719 | 1,308955831 | 0,902255639 |
| GNAI1 | 0,79612529 | -0,389596492 | 0,12 | 0,133 | 1 | Mast | 0,763343074 | 1,310026951 | 0,902255639 |
| TSC22D14 | 0,803136548 | 0,270053997 | 0,231 | 0,267 | 1 | Mast | 1,20585296 | 0,829288507 | 0,865168539 |
| RER12 | 0,805846031 | 0,34666415 | 0,157 | 0,133 | 1 | Mast | 1,271616948 | 0,78640034 | 1,180451128 |
| DUT2 | 0,805846031 | 0,428033613 | 0,157 | 0,133 | 1 | Mast | 1,345398555 | 0,743274174 | 1,180451128 |
| CCL4L21 | 0,805846031 | 1,068304563 | 0,157 | 0,133 | 1 | Mast | 2,096967588 | 0,476879092 | 1,180451128 |
| IGHA16 | 0,806330101 | 0,492915457 | 0,148 | 0,133 | 1 | Mast | 1,407285905 | 0,710587661 | 1,112781955 |
| CHD22 | 0,806330911 | 0,487385455 | 0,148 | 0,133 | 1 | Mast | 1,401901955 | 0,713316645 | 1,112781955 |
| NOL74 | 0,807528388 | 0,382126934 | 0,213 | 0,2 | 1 | Mast | 1,30326181 | 0,767305535 | 1,065 |
| CDC373 | 0,810130062 | 0,315078326 | 0,278 | 0,267 | 1 | Mast | 1,244079195 | 0,80380735 | 1,041198502 |
| EIF3M3 | 0,812805391 | 0,284441325 | 0,222 | 0,267 | 1 | Mast | 1,217938533 | 0,821059497 | 0,831460674 |
| ACBD32 | 0,815373956 | 0,264176824 | 0,157 | 0,133 | 1 | Mast | 1,200950613 | 0,832673708 | 1,180451128 |
| WDR16 | 0,816080028 | 0,342815263 | 0,148 | 0,133 | 1 | Mast | 1,268228992 | 0,788501135 | 1,112781955 |
| SMARCA23 | 0,816080028 | 0,351624232 | 0,148 | 0,133 | 1 | Mast | 1,275996376 | 0,783701285 | 1,112781955 |
| USO11 | 0,817184711 | -0,593479573 | 0,12 | 0,133 | 1 | Mast | 0,662742538 | 1,508881568 | 0,902255639 |
| TNFRSF12A2 | 0,817184711 | -0,637182831 | 0,12 | 0,133 | 1 | Mast | 0,642967254 | 1,555289159 | 0,902255639 |
| RABAC14 | 0,818532342 | 0,267193579 | 0,222 | 0,2 | 1 | Mast | 1,203464495 | 0,83093436 | 1,11 |
| FIS15 | 0,818575524 | 0,450017541 | 0,315 | 0,333 | 1 | Mast | 1,366056866 | 0,732033947 | 0,945945946 |
| SUB12 | 0,819722657 | 0,366408675 | 0,417 | 0,467 | 1 | Mast | 1,289139756 | 0,775711086 | 0,892933619 |
| PTPRC2 | 0,821640413 | 0,317610853 | 0,259 | 0,267 | 1 | Mast | 1,246264987 | 0,802397573 | 0,970037453 |
| TSTD13 | 0,824343646 | 0,398971301 | 0,213 | 0,2 | 1 | Mast | 1,318567385 | 0,758398859 | 1,065 |
| SNRPB22 | 0,82492926 | 0,262861659 | 0,157 | 0,133 | 1 | Mast | 1,199856322 | 0,833433122 | 1,180451128 |
| UBE2N1 | 0,825857696 | 0,389593804 | 0,148 | 0,133 | 1 | Mast | 1,310024509 | 0,763344497 | 1,112781955 |
| RAB2A1 | 0,832782947 | 0,252457984 | 0,213 | 0,2 | 1 | Mast | 1,191234947 | 0,839464962 | 1,065 |
| TPI14 | 0,83482165 | 0,380224744 | 0,574 | 0,667 | 1 | Mast | 1,301544596 | 0,768317892 | 0,860569715 |
| AFF41 | 0,835662447 | 0,619423452 | 0,148 | 0,133 | 1 | Mast | 1,536261118 | 0,650931009 | 1,112781955 |
| LSM21 | 0,835662447 | 0,33545176 | 0,148 | 0,133 | 1 | Mast | 1,261772454 | 0,792535926 | 1,112781955 |
| RAD212 | 0,835662447 | 0,37356785 | 0,148 | 0,133 | 1 | Mast | 1,295552833 | 0,771871262 | 1,112781955 |

| Supplementary Table 4 - clusermarker | | | | | | | | | |
|---|---|---|---|---|---|---|---|---|---|
| | **p_val** | **avg_log2FC** | **pct.1** | **pct.2** | **p_val_adj** | **celltype** | **Foldchange_UP** | **Foldchange_DOWN** | **Ratio_pct1_pct2** |
| TASOR | 0,836703908 | 0,405576941 | 0,139 | 0,133 | 1 | Mast | 1,324618526 | 0,754934331 | 1,045112782 |
| IRS2 | 0,837616216 | -0,729047574 | 0,13 | 0,133 | 1 | Mast | 0,603302065 | 1,657544467 | 0,977443609 |
| ABCA11 | 0,838375938 | -0,395868286 | 0,12 | 0,133 | 1 | Mast | 0,760031814 | 1,315734397 | 0,902255639 |
| PSMB51 | 0,838375938 | -0,254425061 | 0,12 | 0,133 | 1 | Mast | 0,838321154 | 1,192860273 | 0,902255639 |
| TOP11 | 0,841142469 | -0,271482345 | 0,176 | 0,133 | 1 | Mast | 0,828467871 | 1,207047412 | 1,323308271 |
| BCAP314 | 0,841241014 | 0,299489688 | 0,213 | 0,2 | 1 | Mast | 1,230709009 | 0,812539758 | 1,065 |
| NR4A13 | 0,842738426 | 0,283037848 | 0,741 | 0,933 | 1 | Mast | 1,216754278 | 0,821858626 | 0,794212219 |
| BASP12 | 0,845492803 | 0,262782968 | 0,148 | 0,133 | 1 | Mast | 1,199790878 | 0,833478582 | 1,112781955 |
| TMEM14B2 | 0,845492803 | 0,397848569 | 0,148 | 0,133 | 1 | Mast | 1,31754165 | 0,758989289 | 1,112781955 |
| NDUFB113 | 0,848348573 | 0,318482923 | 0,278 | 0,267 | 1 | Mast | 1,247018547 | 0,801912692 | 1,041198502 |
| PLIN21 | 0,85256107 | 0,423935546 | 0,407 | 0,467 | 1 | Mast | 1,341582287 | 0,745388493 | 0,871520343 |
| SRSF111 | 0,862317199 | 0,277547177 | 0,259 | 0,333 | 1 | Mast | 1,212132304 | 0,824992451 | 0,777777778 |
| NBDY1 | 0,865224354 | 0,322900184 | 0,148 | 0,133 | 1 | Mast | 1,250842535 | 0,799461141 | 1,112781955 |
| P4HB3 | 0,866375305 | 0,282464466 | 0,287 | 0,333 | 1 | Mast | 1,21627079 | 0,822185329 | 0,861861862 |
| HIF1A1 | 0,869264702 | 0,44551045 | 0,37 | 0,4 | 1 | Mast | 1,361795857 | 0,734324454 | 0,925 |
| GNG54 | 0,871314497 | 0,546445052 | 0,481 | 0,533 | 1 | Mast | 1,460482481 | 0,684705235 | 0,902439024 |
| RNF2133 | 0,871948244 | 0,273858362 | 0,111 | 0,133 | 1 | Mast | 1,209036971 | 0,827104566 | 0,834586466 |
| PSMA22 | 0,874015383 | 0,311342312 | 0,269 | 0,267 | 1 | Mast | 1,240861685 | 0,805891593 | 1,007490637 |
| ACIN1 | 0,875122534 | 0,367192204 | 0,148 | 0,133 | 1 | Mast | 1,289840079 | 0,775289911 | 1,112781955 |
| BLOC1S6 | 0,87714965 | 0,414127212 | 0,139 | 0,133 | 1 | Mast | 1,332492314 | 0,750473372 | 1,045112782 |
| CERS41 | 0,87714965 | 0,505110775 | 0,139 | 0,133 | 1 | Mast | 1,419232326 | 0,704606273 | 1,045112782 |
| ENO12 | 0,877232287 | 0,284873924 | 0,315 | 0,333 | 1 | Mast | 1,218303793 | 0,820813336 | 0,945945946 |
| TMA71 | 0,880933274 | 0,260252649 | 0,352 | 0,333 | 1 | Mast | 1,197688429 | 0,834941689 | 1,057057057 |
| IKZF3 | 0,882012111 | 0,25644992 | 0,204 | 0,2 | 1 | Mast | 1,194535658 | 0,837145374 | 1,02 |
| TENT5C2 | 0,8829992 | 0,272374869 | 0,111 | 0,133 | 1 | Mast | 1,207794383 | 0,827955498 | 0,834586466 |
| IQGAP1 | 0,886370032 | -0,313465767 | 0,306 | 0,267 | 1 | Mast | 0,8047063 | 1,242689414 | 1,146067416 |
| SMARCC21 | 0,889582323 | -0,287890308 | 0,13 | 0,133 | 1 | Mast | 0,819098974 | 1,220853684 | 0,977443609 |
| NUCB23 | 0,894161398 | 0,275823811 | 0,176 | 0,2 | 1 | Mast | 1,21068522 | 0,825978532 | 0,88 |
| RAD23B | 0,894976091 | 0,430747202 | 0,148 | 0,133 | 1 | Mast | 1,347931518 | 0,741877452 | 1,112781955 |
| BACE22 | 0,89517844 | 0,260127166 | 0,25 | 0,267 | 1 | Mast | 1,19758426 | 0,835014314 | 0,936329588 |
| PTPN6 | 0,897769076 | 0,327326213 | 0,194 | 0,2 | 1 | Mast | 1,254685874 | 0,797012241 | 0,97 |

| Supplementary Table 4 - clusermarker | | | | | | | | | |
|---|---|---|---|---|---|---|---|---|---|
| | p_val | avg_log2FC | pct.1 | pct.2 | p_val_adj | celltype | Foldchange_UP | Foldchange_DOWN | Ratio_pct1_pct2 |
| XRCC51 | 0,909485621 | 0,320698904 | 0,213 | 0,2 | 1 | Mast | 1,248935441 | 0,800681899 | 1,065 |
| CUL33 | 0,917918771 | 0,314775933 | 0,139 | 0,133 | 1 | Mast | 1,243818459 | 0,803975848 | 1,045112782 |
| PSME14 | 0,919010151 | -0,32344774 | 0,389 | 0,333 | 1 | Mast | 0,799157774 | 1,251317366 | 1,168168168 |
| IL2RG2 | 0,921006278 | 0,394838996 | 0,13 | 0,133 | 1 | Mast | 1,314796021 | 0,760574252 | 0,977443609 |
| RAB5C1 | 0,924343546 | 0,317453582 | 0,194 | 0,2 | 1 | Mast | 1,246129136 | 0,802485049 | 0,97 |
| LGALS34 | 0,92737985 | -0,358140973 | 0,648 | 0,533 | 1 | Mast | 0,780169242 | 1,281773167 | 1,21575985 |
| PRKX1 | 0,927409818 | 0,347282327 | 0,111 | 0,133 | 1 | Mast | 1,272161937 | 0,786063449 | 0,834586466 |
| NUFIP23 | 0,92814916 | -0,503253357 | 0,139 | 0,133 | 1 | Mast | 0,705514013 | 1,417406291 | 1,045112782 |
| HSPH15 | 0,92893082 | 0,279617279 | 0,269 | 0,267 | 1 | Mast | 1,213872823 | 0,82380953 | 1,007490637 |
| MGST31 | 0,931445794 | -0,651215529 | 0,157 | 0,133 | 1 | Mast | 0,636743606 | 1,57049084 | 1,180451128 |
| FAM162A2 | 0,931510873 | 0,347046797 | 0,13 | 0,133 | 1 | Mast | 1,271954265 | 0,78619179 | 0,977443609 |
| MAP1LC3B2 | 0,933774704 | 0,429155581 | 0,231 | 0,267 | 1 | Mast | 1,346445263 | 0,742696363 | 0,865168539 |
| LAIR1 | 0,934868814 | -0,279352382 | 0,148 | 0,133 | 1 | Mast | 0,823960806 | 1,213649961 | 1,112781955 |
| OSTF11 | 0,935378374 | 0,279563065 | 0,25 | 0,267 | 1 | Mast | 1,213827209 | 0,823840488 | 0,936329588 |
| EIF4A23 | 0,940099183 | 0,312233689 | 0,259 | 0,267 | 1 | Mast | 1,241628595 | 0,805393822 | 0,970037453 |
| SMCHD11 | 0,942027139 | -0,321752215 | 0,13 | 0,133 | 1 | Mast | 0,800097534 | 1,249847621 | 0,977443609 |
| ENPP3 | 0,942109175 | 0,266620568 | 0,194 | 0,2 | 1 | Mast | 1,202986596 | 0,831264457 | 0,97 |
| ADAM121 | 0,945793254 | 0,252794956 | 0,12 | 0,133 | 1 | Mast | 1,191513218 | 0,83926891 | 0,902255639 |
| AHI1 | 0,948643921 | 0,29635058 | 0,139 | 0,133 | 1 | Mast | 1,228034064 | 0,814309659 | 1,045112782 |
| FBP12 | 0,948643921 | 0,268722882 | 0,139 | 0,133 | 1 | Mast | 1,204740882 | 0,83005401 | 1,045112782 |
| PAPOLA | 0,949266407 | 0,336440831 | 0,176 | 0,2 | 1 | Mast | 1,262637786 | 0,791992772 | 0,88 |
| EIF3A1 | 0,94926659 | 0,279594298 | 0,176 | 0,2 | 1 | Mast | 1,213853487 | 0,823822653 | 0,88 |
| PRPF38B2 | 0,952554068 | 0,436780976 | 0,13 | 0,133 | 1 | Mast | 1,353580768 | 0,738781182 | 0,977443609 |
| PJA22 | 0,952554068 | 0,275179317 | 0,13 | 0,133 | 1 | Mast | 1,210144492 | 0,826347602 | 0,977443609 |
| PEBP13 | 0,954399298 | -0,339926013 | 0,528 | 0,533 | 1 | Mast | 0,790081829 | 1,265691683 | 0,990619137 |
| NEK62 | 0,954883055 | -0,330617072 | 0,148 | 0,133 | 1 | Mast | 0,795196288 | 1,257551141 | 1,112781955 |
| SWAP70 | 0,955010893 | 0,284715834 | 0,241 | 0,267 | 1 | Mast | 1,218170299 | 0,820903285 | 0,902621723 |
| TKT1 | 0,956622579 | 0,274234682 | 0,12 | 0,133 | 1 | Mast | 1,209352385 | 0,826888848 | 0,902255639 |
| CAPRIN11 | 0,959218171 | 0,385000103 | 0,185 | 0,2 | 1 | Mast | 1,305859881 | 0,765778944 | 0,925 |
| RBMX | 0,960542484 | 0,409343867 | 0,204 | 0,2 | 1 | Mast | 1,32808167 | 0,752965742 | 1,02 |
| PLP24 | 0,964037866 | 0,345098187 | 0,259 | 0,267 | 1 | Mast | 1,270237429 | 0,787254396 | 0,970037453 |

| Supplementary Table 4 - clusermarker | | | | | | | | | |
|---|---|---|---|---|---|---|---|---|---|
| | **p_val** | **avg_log2FC** | **pct.1** | **pct.2** | **p_val_adj** | **celltype** | **Foldchange_UP** | **Foldchange_DOWN** | **Ratio_pct1_pct2** |
| PROS11 | 0,967459755 | 0,279264671 | 0,12 | 0,133 | 1 | Mast | 1,213576177 | 0,824010902 | 0,902255639 |
| RNF1672 | 0,973630398 | 0,332993694 | 0,13 | 0,133 | 1 | Mast | 1,259624475 | 0,793887401 | 0,977443609 |
| DCTN34 | 0,977336602 | 0,333170144 | 0,185 | 0,2 | 1 | Mast | 1,259778543 | 0,79379031 | 0,925 |
| UBE2S1 | 0,989806751 | 0,255404996 | 0,435 | 0,467 | 1 | Mast | 1,193670785 | 0,837751926 | 0,931477516 |
| TSPO5 | 0,996378409 | 0,521027912 | 0,343 | 0,4 | 1 | Mast | 1,434977298 | 0,696875136 | 0,8575 |
| ENC12 | 1 | 0,290594001 | 0,12 | 0,133 | 1 | Mast | 1,223143779 | 0,817565373 | 0,902255639 |
| SNX10 | 1 | 0,464881715 | 0,12 | 0,133 | 1 | Mast | 1,380204187 | 0,724530478 | 0,902255639 |
| FAM107B1 | 1 | 0,320843789 | 0,287 | 0,333 | 1 | Mast | 1,249060873 | 0,800601493 | 0,861861862 |
| NDUFB84 | 1 | 0,380745911 | 0,315 | 0,333 | 1 | Mast | 1,302014857 | 0,768040391 | 0,945945946 |
| RCN23 | 1 | 0,400208358 | 0,12 | 0,133 | 1 | Mast | 1,319698491 | 0,757748839 | 0,902255639 |
| NSMCE1 | 1 | 0,378829339 | 0,454 | 0,533 | 1 | Mast | 1,300286322 | 0,769061385 | 0,851782364 |
| PRR47 | 9,2862E-304 | -4,588491738 | 0,078 | 0,894 | 3,3976E-299 | FB | 0,041564863 | 24,05878262 | 0,087248322 |
| LYZ7 | 5,3789E-228 | -3,590700354 | 0,058 | 0,726 | 1,968E-223 | FB | 0,08300256 | 12,04782115 | 0,079889807 |
| PRB37 | 1,0594E-177 | -3,151643518 | 0,007 | 0,506 | 3,8761E-173 | FB | 0,112528043 | 8,886673712 | 0,013833992 |
| GSN3 | 9,2729E-129 | -1,853767627 | 0,866 | 0,968 | 3,3927E-124 | FB | 0,276668897 | 3,614428689 | 0,894628099 |
| SPARCL12 | 3,3253E-117 | -1,575818772 | 0,687 | 0,937 | 1,2166E-112 | FB | 0,335452692 | 2,981046278 | 0,733191035 |
| RARRES21 | 1,1118E-102 | 1,717547027 | 0,903 | 0,51 | 4,06757E-98 | FB | 3,288767515 | 0,304065275 | 1,770588235 |
| MYOC | 2,78038E-97 | -3,009808067 | 0,1 | 0,492 | 1,01726E-92 | FB | 0,124153078 | 8,054572764 | 0,203252033 |
| AEBP1 | 4,73038E-91 | 1,383552693 | 0,926 | 0,594 | 1,7307E-86 | FB | 2,60910082 | 0,383273805 | 1,558922559 |
| SLPI7 | 1,06458E-85 | -2,397920171 | 0,23 | 0,632 | 3,89499E-81 | FB | 0,189737905 | 5,270428168 | 0,363924051 |
| LEPR | 8,08724E-85 | -1,774232538 | 0,31 | 0,716 | 2,95888E-80 | FB | 0,29234979 | 3,420560004 | 0,432960894 |
| SCGB3A27 | 3,2946E-83 | -3,251644964 | 0,035 | 0,34 | 1,20539E-78 | FB | 0,104992271 | 9,524510597 | 0,102941176 |
| MDK3 | 2,13748E-82 | 1,459670229 | 0,714 | 0,244 | 7,82039E-78 | FB | 2,750454866 | 0,363576226 | 2,926229508 |
| CTHRC1 | 3,23573E-80 | 1,692547666 | 0,688 | 0,234 | 1,18386E-75 | FB | 3,232269889 | 0,309380106 | 2,94017094 |
| POSTN | 3,74884E-79 | 1,750334869 | 0,606 | 0,127 | 1,37159E-74 | FB | 3,364366485 | 0,297232779 | 4,771653543 |
| COL1A1 | 2,18563E-75 | 1,963433072 | 0,947 | 0,752 | 7,99657E-71 | FB | 3,899889026 | 0,256417553 | 1,259308511 |
| CLEC3B | 5,87618E-75 | -2,285796766 | 0,121 | 0,463 | 2,14992E-70 | FB | 0,205072115 | 4,876333392 | 0,261339093 |
| RPLP15 | 3,4137E-65 | 0,730062866 | 0,992 | 0,965 | 1,24897E-60 | FB | 1,658711369 | 0,602877643 | 1,027979275 |
| TCF21 | 3,27574E-62 | -1,246186761 | 0,008 | 0,213 | 1,19849E-57 | FB | 0,421560979 | 2,372136058 | 0,037558685 |
| SCGB3A15 | 5,56938E-62 | -1,692727324 | 0,163 | 0,494 | 2,03767E-57 | FB | 0,309341581 | 3,232672428 | 0,329959514 |
| SULF1 | 6,79089E-60 | 1,257241769 | 0,719 | 0,348 | 2,48458E-55 | FB | 2,39038296 | 0,418343009 | 2,066091954 |

| Supplementary Table 4 - clusermarker | | | | | | | | | |
|---|---|---|---|---|---|---|---|---|---|
| | **p_val** | **avg_log2FC** | **pct.1** | **pct.2** | **p_val_adj** | **celltype** | **Foldchange_UP** | **Foldchange_DOWN** | **Ratio_pct1_pct2** |
| SPARC2 | 1,3883E-57 | 1,452269802 | 0,927 | 0,766 | 5,07937E-53 | FB | 2,736382298 | 0,365446013 | 1,210182768 |
| ITGBL11 | 3,64929E-57 | 1,165894034 | 0,736 | 0,374 | 1,33517E-52 | FB | 2,243722154 | 0,445687982 | 1,967914439 |
| MFAP2 | 2,05696E-56 | 1,142912278 | 0,725 | 0,36 | 7,52578E-52 | FB | 2,208263419 | 0,452844525 | 2,013888889 |
| ABCA8 | 9,03063E-55 | -1,382253186 | 0,286 | 0,59 | 3,30404E-50 | FB | 0,383619195 | 2,606751731 | 0,484745763 |
| THBS2 | 2,90758E-54 | 1,23609663 | 0,664 | 0,295 | 1,0638E-49 | FB | 2,355603349 | 0,424519688 | 2,250847458 |
| FRZB | 1,67761E-53 | 1,171444994 | 0,489 | 0,123 | 6,13787E-49 | FB | 2,252371801 | 0,443976434 | 3,975609756 |
| PTGDS | 4,6183E-53 | 2,315833843 | 0,646 | 0,305 | 1,6897E-48 | FB | 4,978923481 | 0,20084663 | 2,118032787 |
| COL1A2 | 7,82516E-52 | 1,167525986 | 0,979 | 0,934 | 2,86299E-47 | FB | 2,24626165 | 0,445184113 | 1,048179872 |
| C6orf587 | 9,17974E-52 | -1,418794245 | 0,003 | 0,166 | 3,35859E-47 | FB | 0,374024779 | 2,673619656 | 0,018072289 |
| TPM13 | 1,32762E-51 | 1,294462857 | 0,703 | 0,345 | 4,85737E-47 | FB | 2,452856539 | 0,407687928 | 2,037681159 |
| STATH7 | 2,46255E-50 | -1,716919753 | 0,003 | 0,161 | 9,00973E-46 | FB | 0,30419751 | 3,287337892 | 0,01863354 |
| ADH1B | 6,72091E-50 | -2,257566357 | 0,133 | 0,403 | 2,45898E-45 | FB | 0,209124449 | 4,781841651 | 0,330024814 |
| ITM2A2 | 1,7003E-49 | -1,17779136 | 0,359 | 0,645 | 6,22088E-45 | FB | 0,442027686 | 2,262301732 | 0,556589147 |
| PRH26 | 6,02088E-49 | -1,315256376 | 0,003 | 0,155 | 2,20286E-44 | FB | 0,401854077 | 2,488465484 | 0,019354839 |
| CFH1 | 3,73333E-48 | -1,113005755 | 0,786 | 0,898 | 1,36591E-43 | FB | 0,462329795 | 2,162958155 | 0,875278396 |
| MYL92 | 3,7676E-48 | 1,197681517 | 0,765 | 0,439 | 1,37845E-43 | FB | 2,293707644 | 0,435975353 | 1,742596811 |
| RPL284 | 4,47708E-48 | 0,586808814 | 0,984 | 0,903 | 1,63803E-43 | FB | 1,501920879 | 0,665814034 | 1,089700997 |
| COL10A1 | 5,8335E-48 | 1,429790002 | 0,434 | 0,1 | 2,1343E-43 | FB | 2,694074977 | 0,371184918 | 4,34 |
| IGFBP6 | 6,86298E-46 | -1,470653509 | 0,796 | 0,873 | 2,51096E-41 | FB | 0,360818819 | 2,771474068 | 0,911798396 |
| PRSS232 | 9,16011E-46 | 1,217626666 | 0,607 | 0,263 | 3,35141E-41 | FB | 2,325638187 | 0,4299895 | 2,307984791 |
| GPC31 | 1,2279E-45 | -1,213342601 | 0,084 | 0,321 | 4,4925E-41 | FB | 0,431268245 | 2,318742477 | 0,261682243 |
| MMP11 | 1,49278E-45 | 2,191392851 | 0,379 | 0,069 | 5,46162E-41 | FB | 4,567462395 | 0,218939953 | 5,492753623 |
| DCN2 | 1,41173E-44 | -0,80442367 | 0,993 | 0,998 | 5,16509E-40 | FB | 0,572590774 | 1,746447979 | 0,99498998 |
| IGFBP22 | 4,31803E-44 | 1,264460872 | 0,332 | 0,042 | 1,57984E-39 | FB | 2,402374176 | 0,416254891 | 7,904761905 |
| TNXB | 6,91508E-44 | -1,389004996 | 0,279 | 0,537 | 2,53002E-39 | FB | 0,381828052 | 2,618979915 | 0,519553073 |
| TPM2 | 7,02561E-44 | 1,34011837 | 0,669 | 0,344 | 2,57046E-39 | FB | 2,531720902 | 0,394988247 | 1,944767442 |
| LUM | 8,69072E-44 | 0,987423027 | 0,964 | 0,921 | 3,17967E-39 | FB | 1,98264039 | 0,504377902 | 1,046688382 |
| TAGLN1 | 9,35536E-44 | 1,679729023 | 0,682 | 0,381 | 3,42285E-39 | FB | 3,203677717 | 0,31214126 | 1,790026247 |
| CFD2 | 1,83431E-43 | -1,913515478 | 0,169 | 0,434 | 6,71118E-39 | FB | 0,265444936 | 3,767259668 | 0,389400922 |
| COL3A1 | 2,2339E-43 | 0,991870522 | 0,943 | 0,816 | 8,17316E-39 | FB | 1,988761844 | 0,502825415 | 1,155637255 |
| CD34 | 2,93078E-43 | -1,197097809 | 0,139 | 0,387 | 1,07228E-38 | FB | 0,436151783 | 2,292779808 | 0,359173127 |

| Supplementary Table 4 - clusermarker | | | | | | | | | |
|---|---|---|---|---|---|---|---|---|---|
| | **p_val** | **avg_log2FC** | **pct.1** | **pct.2** | **p_val_adj** | **celltype** | **Foldchange_UP** | **Foldchange_DOWN** | **Ratio_pct1_pct2** |
| CHRDL1 | 2,20827E-42 | -1,368308224 | 0,146 | 0,402 | 8,07941E-38 | FB | 0,387345202 | 2,581676484 | 0,36318408 |
| MATN2 | 4,57136E-41 | -1,154337276 | 0,133 | 0,374 | 1,67252E-36 | FB | 0,449272519 | 2,225820538 | 0,355614973 |
| ZG16B6 | 1,02942E-40 | -1,222989759 | 0,008 | 0,147 | 3,76632E-36 | FB | 0,428394018 | 2,334299633 | 0,054421769 |
| ANGPTL1 | 2,2281E-40 | -0,987807021 | 0,083 | 0,303 | 8,15197E-36 | FB | 0,504243672 | 1,98316817 | 0,273927393 |
| LTBP41 | 3,26506E-40 | -1,02851555 | 0,647 | 0,766 | 1,19459E-35 | FB | 0,490214292 | 2,039924206 | 0,84464752 |
| ADIRF3 | 1,10021E-39 | -0,962877931 | 0,65 | 0,81 | 4,02535E-35 | FB | 0,513032479 | 1,949194331 | 0,802469136 |
| COL5A1 | 2,9516E-39 | 1,138298913 | 0,602 | 0,289 | 1,0799E-34 | FB | 2,201213242 | 0,454294923 | 2,083044983 |
| TIMP32 | 3,62807E-39 | -0,685252322 | 0,78 | 0,915 | 1,3274E-34 | FB | 0,621897049 | 1,607983188 | 0,852459016 |
| MIF6 | 4,56827E-39 | 0,759376281 | 0,831 | 0,519 | 1,67139E-34 | FB | 1,692758638 | 0,590751674 | 1,601156069 |
| LRRC15 | 9,05438E-39 | 0,854629212 | 0,309 | 0,042 | 3,31272E-34 | FB | 1,808293945 | 0,553007437 | 7,357142857 |
| SFRP12 | 1,00454E-38 | 1,376795079 | 0,432 | 0,129 | 3,67532E-34 | FB | 2,596908313 | 0,385073279 | 3,348837209 |
| MRPS61 | 3,11326E-38 | 1,13944642 | 0,43 | 0,132 | 1,13905E-33 | FB | 2,202964765 | 0,453933724 | 3,257575758 |
| MMP2 | 7,73841E-38 | 0,681776967 | 0,942 | 0,765 | 2,83125E-33 | FB | 1,604114326 | 0,623396963 | 1,231372549 |
| ABI3BP | 8,18817E-38 | -0,938227547 | 0,594 | 0,784 | 2,99581E-33 | FB | 0,521873645 | 1,916172639 | 0,757653061 |
| PLAC9 | 1,83591E-37 | -1,059615986 | 0,621 | 0,761 | 6,71704E-33 | FB | 0,479759744 | 2,084376633 | 0,816031537 |
| IGKC7 | 2,12573E-37 | 1,68889626 | 0,351 | 0,073 | 7,77742E-33 | FB | 3,224099481 | 0,310164127 | 4,808219178 |
| PPIB3 | 2,62918E-37 | 0,697460094 | 0,907 | 0,711 | 9,61938E-33 | FB | 1,621647321 | 0,616656894 | 1,275668073 |
| PIP6 | 6,88005E-37 | -0,871736509 | 0,003 | 0,119 | 2,5172E-32 | FB | 0,54648867 | 1,829864103 | 0,025210084 |
| CNN22 | 1,12275E-36 | 0,791406575 | 0,496 | 0,187 | 4,1078E-32 | FB | 1,730761069 | 0,577780502 | 2,652406417 |
| FLNA4 | 1,4196E-36 | 0,87899394 | 0,672 | 0,36 | 5,19389E-32 | FB | 1,839092367 | 0,543746479 | 1,866666667 |
| IGF2 | 2,23958E-36 | 0,95019515 | 0,467 | 0,158 | 8,19394E-32 | FB | 1,932133996 | 0,517562448 | 2,955696203 |
| APOD2 | 2,99515E-36 | -1,449286677 | 0,637 | 0,839 | 1,09583E-31 | FB | 0,366202443 | 2,730730004 | 0,759237187 |
| VCAM1 | 4,48241E-36 | 0,86379975 | 0,404 | 0,118 | 1,63998E-31 | FB | 1,819825034 | 0,549503376 | 3,423728814 |
| FHL2 | 5,0141E-36 | 0,838326961 | 0,457 | 0,152 | 1,83451E-31 | FB | 1,787975491 | 0,559291783 | 3,006578947 |
| IGFBP42 | 5,25126E-36 | 0,750735835 | 0,907 | 0,627 | 1,92128E-31 | FB | 1,682650834 | 0,594300362 | 1,446570973 |
| GREM1 | 1,04969E-35 | 1,058375927 | 0,246 | 0,016 | 3,84048E-31 | FB | 2,08258579 | 0,480172296 | 15,375 |
| MYLK2 | 1,93397E-35 | 0,768539076 | 0,455 | 0,152 | 7,0758E-31 | FB | 1,703543841 | 0,587011602 | 2,993421053 |
| CD816 | 5,27272E-35 | -0,627742499 | 0,864 | 0,898 | 1,92913E-30 | FB | 0,647188331 | 1,545145288 | 0,962138085 |
| ANXA16 | 2,61635E-34 | -0,678693248 | 0,864 | 0,908 | 9,57243E-30 | FB | 0,624730882 | 1,60068924 | 0,95154185 |
| CTSK | 3,02707E-34 | 0,662130167 | 0,935 | 0,768 | 1,10751E-29 | FB | 1,58241737 | 0,631944529 | 1,217447917 |
| TNFRSF12A3 | 6,05165E-34 | 0,846060873 | 0,342 | 0,081 | 2,21412E-29 | FB | 1,797586099 | 0,556301587 | 4,222222222 |

| Supplementary Table 4 - clusermarker | | | | | | | | | |
|---|---|---|---|---|---|---|---|---|---|
| | **p_val** | **avg_log2FC** | **pct.1** | **pct.2** | **p_val_adj** | **celltype** | **Foldchange_UP** | **Foldchange_DOWN** | **Ratio_pct1_pct2** |
| COL12A1 | 8,01543E-34 | 1,625673386 | 0,529 | 0,271 | 2,93261E-29 | FB | 3,085861658 | 0,324058597 | 1,95202952 |
| RPL136 | 9,43924E-34 | 0,638013416 | 0,984 | 0,915 | 3,45354E-29 | FB | 1,556184824 | 0,642597193 | 1,075409836 |
| VIT | 1,00456E-33 | -0,959596391 | 0,069 | 0,256 | 3,67538E-29 | FB | 0,514200746 | 1,944765751 | 0,26953125 |
| SELENOM5 | 3,16942E-33 | 0,732962535 | 0,907 | 0,668 | 1,1596E-28 | FB | 1,662048561 | 0,601667137 | 1,357784431 |
| LBH2 | 3,77447E-33 | 0,980954085 | 0,542 | 0,234 | 1,38096E-28 | FB | 1,973770274 | 0,506644574 | 2,316239316 |
| EDIL3 | 4,68151E-33 | 0,753665453 | 0,639 | 0,334 | 1,71282E-28 | FB | 1,686071191 | 0,593094767 | 1,913173653 |
| PALLD2 | 1,61227E-32 | 0,697069442 | 0,66 | 0,352 | 5,8988E-28 | FB | 1,621208273 | 0,616823894 | 1,875 |
| TFPI | 2,02039E-32 | -0,970567982 | 0,504 | 0,658 | 7,39202E-28 | FB | 0,510305119 | 1,959611934 | 0,765957447 |
| C1QTNF7 | 2,17044E-32 | -0,93305489 | 0,078 | 0,263 | 7,94098E-28 | FB | 0,523748136 | 1,90931467 | 0,296577947 |
| WNT5A | 2,94057E-32 | 1,044885451 | 0,297 | 0,061 | 1,07587E-27 | FB | 2,063202535 | 0,484683391 | 4,868852459 |
| HMCN1 | 8,4183E-32 | 0,631952862 | 0,644 | 0,321 | 3,08E-27 | FB | 1,549661228 | 0,645302329 | 2,00623053 |
| ACTA21 | 1,08151E-31 | 1,644745922 | 0,309 | 0,071 | 3,95691E-27 | FB | 3,126927831 | 0,319802712 | 4,352112676 |
| LGALS35 | 1,14912E-31 | -0,608214718 | 0,889 | 0,889 | 4,2043E-27 | FB | 0,656007985 | 1,524371688 | 1 |
| CCN2 | 1,17455E-31 | 0,807210412 | 0,852 | 0,635 | 4,29734E-27 | FB | 1,749824717 | 0,571485813 | 1,341732283 |
| THY1 | 1,53939E-31 | 0,847174416 | 0,661 | 0,377 | 5,63215E-27 | FB | 1,7989741 | 0,555872372 | 1,75331565 |
| SDC11 | 2,13686E-31 | 0,7567143 | 0,255 | 0,035 | 7,81811E-27 | FB | 1,689638133 | 0,591842703 | 7,285714286 |
| PMEPA14 | 2,93553E-31 | 0,896934717 | 0,56 | 0,265 | 1,07402E-26 | FB | 1,862105376 | 0,537026536 | 2,113207547 |
| SCN7A | 3,28818E-31 | -1,188132327 | 0,134 | 0,339 | 1,20305E-26 | FB | 0,438870643 | 2,278575741 | 0,395280236 |
| SERPING12 | 5,23941E-31 | -0,596368148 | 0,948 | 0,942 | 1,91694E-26 | FB | 0,661416918 | 1,511905688 | 1,006369427 |
| PI16 | 6,12612E-31 | -0,769000536 | 0,321 | 0,605 | 2,24136E-26 | FB | 0,586823871 | 1,704088823 | 0,530578512 |
| SLC5A3 | 6,47291E-31 | 0,998609801 | 0,297 | 0,065 | 2,36824E-26 | FB | 1,998073703 | 0,500482039 | 4,569230769 |
| VGLL3 | 7,04027E-31 | 0,689058664 | 0,351 | 0,098 | 2,57583E-26 | FB | 1,612231219 | 0,620258427 | 3,581632653 |
| CDH11 | 7,9754E-31 | 0,745708635 | 0,654 | 0,373 | 2,91796E-26 | FB | 1,67679769 | 0,596374867 | 1,753351206 |
| IFITM36 | 2,41409E-30 | 0,511241855 | 0,976 | 0,908 | 8,83244E-26 | FB | 1,425276529 | 0,701618233 | 1,074889868 |
| GPX47 | 3,93427E-30 | 0,640990611 | 0,838 | 0,563 | 1,43943E-25 | FB | 1,559399537 | 0,641272475 | 1,488454707 |
| RPS193 | 1,57911E-29 | 0,552667787 | 0,97 | 0,834 | 5,77749E-25 | FB | 1,466795543 | 0,681758276 | 1,163069544 |
| EFEMP2 | 2,25542E-29 | 0,592748472 | 0,605 | 0,306 | 8,25191E-25 | FB | 1,50811712 | 0,663078475 | 1,977124183 |
| RAB312 | 3,11151E-29 | 0,759993532 | 0,574 | 0,29 | 1,13841E-24 | FB | 1,693483032 | 0,590498978 | 1,979310345 |
| MEOX2 | 5,3283E-29 | -0,925682119 | 0,086 | 0,263 | 1,94947E-24 | FB | 0,526431558 | 1,899582168 | 0,326996198 |
| MSMB7 | 1,32828E-28 | -0,996523251 | 0,026 | 0,156 | 4,85978E-24 | FB | 0,501206402 | 1,995186005 | 0,166666667 |
| FTL4 | 2,19351E-28 | 0,442373906 | 0,991 | 0,958 | 8,0254E-24 | FB | 1,358838411 | 0,735922676 | 1,034446764 |

| Supplementary Table 4 - clusermarker | | | | | | | | | |
|---|---|---|---|---|---|---|---|---|---|
| | **p_val** | **avg_log2FC** | **pct.1** | **pct.2** | **p_val_adj** | **celltype** | **Foldchange_UP** | **Foldchange_DOWN** | **Ratio_pct1_pct2** |
| HLA-B7 | 2,48387E-28 | 0,744289002 | 0,936 | 0,773 | 9,08772E-24 | FB | 1,675148509 | 0,596961997 | 1,210866753 |
| SELENOP1 | 2,86628E-28 | -0,806762013 | 0,515 | 0,71 | 1,04869E-23 | FB | 0,571663461 | 1,749280945 | 0,725352113 |
| PFN16 | 2,94299E-28 | 0,676863339 | 0,859 | 0,639 | 1,07675E-23 | FB | 1,598660218 | 0,625523791 | 1,34428795 |
| TPPP32 | 3,5533E-28 | -0,704151637 | 0,037 | 0,177 | 1,30005E-23 | FB | 0,613803324 | 1,629186355 | 0,209039548 |
| SERPINH1 | 4,05913E-28 | 0,638632944 | 0,693 | 0,424 | 1,48511E-23 | FB | 1,556853231 | 0,642321306 | 1,634433962 |
| HIF1A2 | 8,18893E-28 | 0,523032785 | 0,423 | 0,16 | 2,99608E-23 | FB | 1,436972831 | 0,695907381 | 2,64375 |
| CYTOR1 | 8,78114E-28 | 0,618628406 | 0,319 | 0,089 | 3,21276E-23 | FB | 1,535414743 | 0,651289826 | 3,584269663 |
| ABLIM12 | 1,11123E-27 | -0,97678233 | 0,115 | 0,297 | 4,06565E-23 | FB | 0,508111728 | 1,968071084 | 0,387205387 |
| PLOD2 | 1,38466E-27 | 0,53069198 | 0,372 | 0,119 | 5,06605E-23 | FB | 1,444621934 | 0,692222634 | 3,12605042 |
| WNT2 | 1,5347E-27 | 0,725435942 | 0,23 | 0,032 | 5,61502E-23 | FB | 1,653400171 | 0,604814259 | 7,1875 |
| SPTBN12 | 1,60171E-27 | -0,876345578 | 0,63 | 0,716 | 5,86017E-23 | FB | 0,544745554 | 1,835719433 | 0,879888268 |
| SCARA5 | 1,83086E-27 | -1,067948273 | 0,161 | 0,358 | 6,69857E-23 | FB | 0,476996877 | 2,096449783 | 0,44972067 |
| NFIA3 | 2,05297E-27 | -0,844309343 | 0,349 | 0,545 | 7,51122E-23 | FB | 0,556977385 | 1,795405031 | 0,640366972 |
| TPM45 | 2,49445E-27 | 0,701843683 | 0,706 | 0,439 | 9,12643E-23 | FB | 1,626582145 | 0,614786043 | 1,608200456 |
| PCOLCE | 2,7386E-27 | 0,545367252 | 0,885 | 0,689 | 1,00197E-22 | FB | 1,4593918 | 0,685216952 | 1,284470247 |
| MT-ATP87 | 3,05563E-27 | -0,553823026 | 0,808 | 0,924 | 1,11796E-22 | FB | 0,681212576 | 1,46797055 | 0,874458874 |
| CPXM1 | 4,48314E-27 | 0,731698939 | 0,281 | 0,065 | 1,64025E-22 | FB | 1,66059348 | 0,602194343 | 4,323076923 |
| CTSZ5 | 5,89694E-27 | 0,573213122 | 0,588 | 0,297 | 2,15752E-22 | FB | 1,487833536 | 0,672118201 | 1,97979798 |
| C1R | 6,89673E-27 | 0,603896441 | 0,862 | 0,632 | 2,52331E-22 | FB | 1,519815758 | 0,657974491 | 1,363924051 |
| MAMDC2 | 1,23869E-26 | -0,858248712 | 0,094 | 0,263 | 4,53199E-22 | FB | 0,551621765 | 1,812836372 | 0,357414449 |
| FNDC1 | 1,64793E-26 | 0,728341165 | 0,308 | 0,084 | 6,02927E-22 | FB | 1,656733056 | 0,603597542 | 3,666666667 |
| THBS12 | 2,1649E-26 | 1,239754094 | 0,531 | 0,284 | 7,92071E-22 | FB | 2,361582759 | 0,423444826 | 1,86971831 |
| IGFBP3 | 2,43448E-26 | 1,493787995 | 0,4 | 0,163 | 8,90705E-22 | FB | 2,816274571 | 0,355079015 | 2,45398773 |
| SULF21 | 3,0178E-26 | 0,530809188 | 0,402 | 0,152 | 1,10412E-21 | FB | 1,444739303 | 0,692166398 | 2,644736842 |
| CEBPB2 | 4,52105E-26 | 0,548166518 | 0,5 | 0,223 | 1,65412E-21 | FB | 1,462226211 | 0,683888712 | 2,242152466 |
| VKORC13 | 5,99787E-26 | 0,499386141 | 0,675 | 0,376 | 2,19444E-21 | FB | 1,41361195 | 0,707407715 | 1,795212766 |
| GUCY1A1 | 7,69312E-26 | 0,463987442 | 0,4 | 0,147 | 2,81468E-21 | FB | 1,379348916 | 0,724979727 | 2,721088435 |
| NME26 | 8,58658E-26 | 0,577982963 | 0,861 | 0,616 | 3,14157E-21 | FB | 1,492760755 | 0,669899712 | 1,397727273 |
| IGLV3-215 | 8,63486E-26 | 0,372613676 | 0,191 | 0,016 | 3,15924E-21 | FB | 1,294696259 | 0,772381934 | 11,9375 |
| IRAG1 | 1,00638E-25 | 0,481509851 | 0,195 | 0,018 | 3,68204E-21 | FB | 1,396204098 | 0,716227665 | 10,83333333 |
| CST1 | 1,17017E-25 | 2,924941766 | 0,192 | 0,018 | 4,28128E-21 | FB | 7,594430413 | 0,131675444 | 10,66666667 |

| Supplementary Table 4 - clusermarker | | | | | | | | | |
|---|---|---|---|---|---|---|---|---|---|
| | **p_val** | **avg_log2FC** | **pct.1** | **pct.2** | **p_val_adj** | **celltype** | **Foldchange_UP** | **Foldchange_DOWN** | **Ratio_pct1_pct2** |
| LGALS14 | 1,44519E-25 | 0,539214466 | 0,98 | 0,926 | 5,28753E-21 | FB | 1,453181058 | 0,688145496 | 1,058315335 |
| KIF26B | 1,52472E-25 | 0,664871769 | 0,24 | 0,045 | 5,57849E-21 | FB | 1,58542735 | 0,630744764 | 5,333333333 |
| SON7 | 2,28417E-25 | -0,678534994 | 0,754 | 0,808 | 8,3571E-21 | FB | 0,624799414 | 1,600513664 | 0,933168317 |
| CHCHD105 | 3,73253E-25 | 0,655379411 | 0,325 | 0,105 | 1,36562E-20 | FB | 1,575030113 | 0,634908496 | 3,095238095 |
| B2M5 | 4,70386E-25 | 0,396485136 | 0,995 | 0,985 | 1,721E-20 | FB | 1,316297082 | 0,759706918 | 1,010152284 |
| HLA-A3 | 6,46173E-25 | 0,471959979 | 0,949 | 0,797 | 2,36415E-20 | FB | 1,386992493 | 0,720984436 | 1,190715182 |
| SEC61B6 | 1,55289E-24 | 0,572144339 | 0,682 | 0,411 | 5,68156E-20 | FB | 1,486731722 | 0,672616307 | 1,659367397 |
| MYH92 | 3,43874E-24 | 0,698265932 | 0,629 | 0,373 | 1,25813E-19 | FB | 1,622553368 | 0,616312548 | 1,686327078 |
| RPS123 | 4,10221E-24 | 0,416231947 | 0,988 | 0,939 | 1,50088E-19 | FB | 1,334437695 | 0,749379311 | 1,052183174 |
| C1orf1223 | 6,27028E-24 | 0,561468074 | 0,369 | 0,135 | 2,29411E-19 | FB | 1,475770185 | 0,67761228 | 2,733333333 |
| LOXL1 | 6,33578E-24 | 0,513220731 | 0,407 | 0,166 | 2,31807E-19 | FB | 1,427232854 | 0,700656517 | 2,451807229 |
| CRIP15 | 8,33662E-24 | -0,994708507 | 0,53 | 0,665 | 3,05012E-19 | FB | 0,501837259 | 1,99267787 | 0,796992481 |
| TPT12 | 8,75572E-24 | 0,434380908 | 0,981 | 0,934 | 3,20346E-19 | FB | 1,351330823 | 0,740011242 | 1,050321199 |
| NEAT14 | 8,80904E-24 | 0,555382113 | 0,826 | 0,634 | 3,22296E-19 | FB | 1,469557809 | 0,680476803 | 1,302839117 |
| NRP22 | 9,82547E-24 | 0,527198943 | 0,269 | 0,066 | 3,59485E-19 | FB | 1,441128462 | 0,693900666 | 4,075757576 |
| RPLP05 | 1,1966E-23 | 0,443794929 | 0,977 | 0,892 | 4,378E-19 | FB | 1,360177497 | 0,735198165 | 1,09529148 |
| ACTN1 | 1,20865E-23 | 0,583216204 | 0,519 | 0,265 | 4,42207E-19 | FB | 1,498185438 | 0,667474116 | 1,958490566 |
| TMSB105 | 1,4689E-23 | 0,473241216 | 0,977 | 0,908 | 5,37427E-19 | FB | 1,388224808 | 0,720344424 | 1,075991189 |
| CD95 | 1,55236E-23 | -0,752970849 | 0,686 | 0,755 | 5,67962E-19 | FB | 0,593380389 | 1,685259606 | 0,908609272 |
| BMP5 | 1,75779E-23 | -0,723303428 | 0,027 | 0,142 | 6,43124E-19 | FB | 0,605708924 | 1,65095801 | 0,190140845 |
| MEG31 | 1,7949E-23 | 0,561575483 | 0,642 | 0,371 | 6,567E-19 | FB | 1,475880059 | 0,677561834 | 1,730458221 |
| HTRA12 | 2,06602E-23 | 0,684914556 | 0,663 | 0,426 | 7,55896E-19 | FB | 1,607606768 | 0,622042666 | 1,556338028 |
| TCIM | 2,09263E-23 | 0,761826232 | 0,197 | 0,027 | 7,65631E-19 | FB | 1,695635683 | 0,589749325 | 7,296296296 |
| GJA1 | 2,60345E-23 | 0,560703834 | 0,318 | 0,105 | 9,52525E-19 | FB | 1,474988631 | 0,677971327 | 3,028571429 |
| PDPN | 4,39689E-23 | 0,496782369 | 0,293 | 0,085 | 1,60869E-18 | FB | 1,411062968 | 0,708685595 | 3,447058824 |
| NREP1 | 4,54172E-23 | 0,699004052 | 0,372 | 0,152 | 1,66168E-18 | FB | 1,623383721 | 0,615997307 | 2,447368421 |
| CALD12 | 4,84869E-23 | 0,626586252 | 0,931 | 0,79 | 1,77399E-18 | FB | 1,543907428 | 0,647707228 | 1,178481013 |
| COL11A1 | 6,91178E-23 | 1,009567226 | 0,18 | 0,019 | 2,52881E-18 | FB | 2,013307066 | 0,496695222 | 9,473684211 |
| MYL12A4 | 7,46739E-23 | 0,486878105 | 0,787 | 0,531 | 2,73209E-18 | FB | 1,401409038 | 0,71356754 | 1,482109228 |
| FTH16 | 7,46947E-23 | 0,566537403 | 0,99 | 0,95 | 2,73286E-18 | FB | 1,480964854 | 0,675235471 | 1,042105263 |
| ATP2A23 | 8,25874E-23 | 0,495730871 | 0,383 | 0,15 | 3,02162E-18 | FB | 1,4100349 | 0,709202304 | 2,553333333 |

| Supplementary Table 4 - clusermarker | | | | | | | | | |
|---|---|---|---|---|---|---|---|---|---|
| | **p_val** | **avg_log2FC** | **pct.1** | **pct.2** | **p_val_adj** | **celltype** | **Foldchange_UP** | **Foldchange_DOWN** | **Ratio_pct1_pct2** |
| PENK | 1,61522E-22 | -0,811980037 | 0,016 | 0,113 | 5,9096E-18 | FB | 0,56959957 | 1,755619302 | 0,14159292 |
| SEMA5A1 | 1,91937E-22 | 0,545558025 | 0,273 | 0,079 | 7,0224E-18 | FB | 1,459584792 | 0,685126349 | 3,455696203 |
| RRAD1 | 1,96525E-22 | 0,688719922 | 0,237 | 0,055 | 7,19027E-18 | FB | 1,611852715 | 0,62040408 | 4,309090909 |
| CHPF1 | 2,5277E-22 | 0,631777571 | 0,453 | 0,221 | 9,24811E-18 | FB | 1,549472952 | 0,64538074 | 2,049773756 |
| COL5A2 | 2,80932E-22 | 0,752837031 | 0,689 | 0,458 | 1,02785E-17 | FB | 1,685103297 | 0,59343543 | 1,504366812 |
| ABCA10 | 3,4735E-22 | -0,910000415 | 0,16 | 0,327 | 1,27085E-17 | FB | 0,532184938 | 1,879046039 | 0,489296636 |
| GAS63 | 4,74402E-22 | 0,925637767 | 0,487 | 0,255 | 1,73569E-17 | FB | 1,899523772 | 0,526447742 | 1,909803922 |
| KIAA12172 | 4,9131E-22 | 0,48794643 | 0,306 | 0,098 | 1,79755E-17 | FB | 1,402447175 | 0,713039334 | 3,12244898 |
| COPZ2 | 6,55465E-22 | 0,552671239 | 0,443 | 0,206 | 2,39815E-17 | FB | 1,466799052 | 0,681756644 | 2,150485437 |
| CCND13 | 7,1956E-22 | 0,512103147 | 0,368 | 0,145 | 2,63265E-17 | FB | 1,426127677 | 0,70119949 | 2,537931034 |
| NFKBIZ3 | 8,35386E-22 | 0,451493378 | 0,478 | 0,231 | 3,05643E-17 | FB | 1,367455019 | 0,73128548 | 2,069264069 |
| FHL12 | 9,14317E-22 | -1,136979024 | 0,478 | 0,595 | 3,34521E-17 | FB | 0,454710737 | 2,199200323 | 0,803361345 |
| RPL123 | 9,9491E-22 | 0,461061031 | 0,987 | 0,9 | 3,64008E-17 | FB | 1,376553833 | 0,726451793 | 1,096666667 |
| BASP13 | 1,16938E-21 | 0,51069175 | 0,393 | 0,165 | 4,27841E-17 | FB | 1,424733169 | 0,701885814 | 2,381818182 |
| SOD31 | 1,22465E-21 | -0,61344451 | 0,738 | 0,818 | 4,48062E-17 | FB | 0,653634251 | 1,529907586 | 0,902200489 |
| FOSB6 | 1,29335E-21 | 0,697244869 | 0,81 | 0,65 | 4,732E-17 | FB | 1,621405418 | 0,616748895 | 1,246153846 |
| PHPT16 | 1,60081E-21 | 0,571297671 | 0,638 | 0,394 | 5,85687E-17 | FB | 1,485859466 | 0,673011158 | 1,61928934 |
| TMEM176B2 | 1,89251E-21 | 0,694874553 | 0,442 | 0,218 | 6,92414E-17 | FB | 1,618743672 | 0,617763033 | 2,027522936 |
| PDK41 | 2,01954E-21 | -0,829369345 | 0,143 | 0,313 | 7,38887E-17 | FB | 0,562775198 | 1,776908441 | 0,45686901 |
| EDF14 | 2,07963E-21 | 0,432113613 | 0,755 | 0,473 | 7,60874E-17 | FB | 1,349208781 | 0,741175135 | 1,596194503 |
| CYBA7 | 2,2842E-21 | 0,497020694 | 0,669 | 0,395 | 8,35721E-17 | FB | 1,411296087 | 0,708568534 | 1,693670886 |
| CFI1 | 2,28754E-21 | 0,436000139 | 0,267 | 0,073 | 8,36942E-17 | FB | 1,352848362 | 0,739181144 | 3,657534247 |
| PTK7 | 2,53487E-21 | 0,487884782 | 0,315 | 0,11 | 9,27434E-17 | FB | 1,402387248 | 0,713069804 | 2,863636364 |
| MYH101 | 2,90459E-21 | 0,439722472 | 0,503 | 0,253 | 1,0627E-16 | FB | 1,356343385 | 0,737276423 | 1,988142292 |
| SOD24 | 3,75829E-21 | 0,870568346 | 0,561 | 0,329 | 1,37505E-16 | FB | 1,828383046 | 0,546931346 | 1,705167173 |
| RBP11 | 3,94646E-21 | 0,538391636 | 0,225 | 0,052 | 1,44389E-16 | FB | 1,452352484 | 0,688538086 | 4,326923077 |
| RUNX11 | 4,29171E-21 | 0,467301654 | 0,388 | 0,161 | 1,57021E-16 | FB | 1,382521249 | 0,723316189 | 2,409937888 |
| MARCKS5 | 4,68661E-21 | 0,547853144 | 0,773 | 0,54 | 1,71469E-16 | FB | 1,461908629 | 0,684037279 | 1,431481481 |
| NME43 | 4,72152E-21 | 0,472700333 | 0,354 | 0,14 | 1,72746E-16 | FB | 1,387704444 | 0,72061454 | 2,528571429 |
| SSR45 | 4,77203E-21 | 0,50182024 | 0,764 | 0,506 | 1,74594E-16 | FB | 1,415998994 | 0,706215191 | 1,509881423 |
| H19 | 5,75938E-21 | -0,728449918 | 0,043 | 0,165 | 2,10718E-16 | FB | 0,603552043 | 1,656857949 | 0,260606061 |

| Supplementary Table 4 - clusermarker | | | | | | | | | |
|---|---|---|---|---|---|---|---|---|---|
| | **p_val** | **avg_log2FC** | **pct.1** | **pct.2** | **p_val_adj** | **celltype** | **Foldchange_UP** | **Foldchange_DOWN** | **Ratio_pct1_pct2** |
| RPL192 | 9,31723E-21 | 0,334819836 | 0,977 | 0,916 | 3,4089E-16 | FB | 1,261219897 | 0,792883146 | 1,066593886 |
| TGFBI1 | 1,15008E-20 | -0,913706124 | 0,161 | 0,319 | 4,20779E-16 | FB | 0,530819721 | 1,883878764 | 0,504702194 |
| S100A106 | 1,32873E-20 | -0,708638397 | 0,866 | 0,861 | 4,86143E-16 | FB | 0,61189737 | 1,634260988 | 1,005807201 |
| PXDN | 1,58653E-20 | 0,488776151 | 0,306 | 0,106 | 5,80463E-16 | FB | 1,40325398 | 0,71262937 | 2,886792453 |
| TMEM176A2 | 2,23255E-20 | 0,659428289 | 0,433 | 0,211 | 8,16822E-16 | FB | 1,579456593 | 0,633129144 | 2,052132701 |
| IGHG16 | 2,7891E-20 | 1,051249971 | 0,155 | 0,015 | 1,02045E-15 | FB | 2,072324561 | 0,482549895 | 10,33333333 |
| IGKV1-54 | 3,77935E-20 | 2,238412749 | 0,169 | 0,023 | 1,38275E-15 | FB | 4,718776197 | 0,211919353 | 7,347826087 |
| LTBP21 | 3,90187E-20 | 0,408137992 | 0,778 | 0,505 | 1,42758E-15 | FB | 1,326972058 | 0,753595371 | 1,540594059 |
| ANTXR11 | 4,52809E-20 | 0,549139678 | 0,607 | 0,358 | 1,65669E-15 | FB | 1,463212878 | 0,683427555 | 1,695530726 |
| SERTAD12 | 6,67834E-20 | 0,551650723 | 0,426 | 0,205 | 2,4434E-15 | FB | 1,465761853 | 0,682239068 | 2,07804878 |
| CFL13 | 7,25613E-20 | 0,479986153 | 0,838 | 0,606 | 2,6548E-15 | FB | 1,39473028 | 0,716984506 | 1,382838284 |
| GPM6B | 7,72367E-20 | 0,468668575 | 0,189 | 0,034 | 2,82586E-15 | FB | 1,383831777 | 0,722631187 | 5,558823529 |
| CPXM2 | 8,45261E-20 | 0,474747738 | 0,442 | 0,205 | 3,09256E-15 | FB | 1,389675207 | 0,719592603 | 2,156097561 |
| FN12 | 8,85671E-20 | 0,712644509 | 0,897 | 0,753 | 3,24041E-15 | FB | 1,638805352 | 0,610200594 | 1,19123506 |
| IGKV4-16 | 8,99756E-20 | 0,980909932 | 0,167 | 0,023 | 3,29194E-15 | FB | 1,973709869 | 0,50666008 | 7,260869565 |
| CHN1 | 1,52395E-19 | 0,401455306 | 0,351 | 0,144 | 5,57569E-15 | FB | 1,320839625 | 0,757094186 | 2,4375 |
| GPNMB2 | 1,58845E-19 | -0,753556684 | 0,75 | 0,745 | 5,81165E-15 | FB | 0,593139484 | 1,685944078 | 1,006711409 |
| AC020916.13 | 1,67737E-19 | 0,526070561 | 0,375 | 0,163 | 6,13701E-15 | FB | 1,440001746 | 0,694443602 | 2,300613497 |
| MT-CO13 | 2,90164E-19 | 0,275701503 | 0,999 | 0,992 | 1,06162E-14 | FB | 1,210582586 | 0,826048559 | 1,007056452 |
| BGN1 | 3,18307E-19 | 0,453806953 | 0,941 | 0,763 | 1,16459E-14 | FB | 1,369649695 | 0,730113695 | 1,233289646 |
| MMP191 | 3,857E-19 | 0,495910464 | 0,36 | 0,152 | 1,41116E-14 | FB | 1,410210438 | 0,709114025 | 2,368421053 |
| C1QTNF6 | 6,04466E-19 | 0,394243272 | 0,186 | 0,035 | 2,21156E-14 | FB | 1,314253222 | 0,760888376 | 5,314285714 |
| RDH101 | 6,11012E-19 | 0,59911922 | 0,266 | 0,09 | 2,23551E-14 | FB | 1,514791488 | 0,660156865 | 2,955555556 |
| NNMT2 | 6,16718E-19 | 0,479270703 | 0,858 | 0,663 | 2,25639E-14 | FB | 1,394038787 | 0,717340155 | 1,294117647 |
| MIDN3 | 6,63824E-19 | 0,447667862 | 0,344 | 0,144 | 2,42873E-14 | FB | 1,363833816 | 0,733227163 | 2,388888889 |
| LSP12 | 6,71333E-19 | -0,883953746 | 0,241 | 0,398 | 2,45621E-14 | FB | 0,541880356 | 1,845425818 | 0,605527638 |
| VMP12 | 7,51294E-19 | 0,498496797 | 0,426 | 0,203 | 2,74876E-14 | FB | 1,412740803 | 0,707843929 | 2,098522167 |
| EMILIN1 | 8,38613E-19 | 0,467398186 | 0,384 | 0,173 | 3,06823E-14 | FB | 1,382613758 | 0,723267792 | 2,219653179 |
| GABARAP6 | 8,8301E-19 | 0,395775321 | 0,889 | 0,703 | 3,23067E-14 | FB | 1,315649615 | 0,760080791 | 1,26458037 |
| NEXN1 | 9,07368E-19 | 0,505696799 | 0,269 | 0,09 | 3,31979E-14 | FB | 1,419808936 | 0,70432012 | 2,988888889 |
| GAPDH2 | 9,23518E-19 | 0,383667878 | 0,95 | 0,842 | 3,37888E-14 | FB | 1,304654569 | 0,766486412 | 1,128266033 |

| Supplementary Table 4 - clusermarker | | | | | | | | | |
|---|---|---|---|---|---|---|---|---|---|
| | **p_val** | **avg_log2FC** | **pct.1** | **pct.2** | **p_val_adj** | **celltype** | **Foldchange_UP** | **Foldchange_DOWN** | **Ratio_pct1_pct2** |
| OSTC3 | 1,01083E-18 | 0,423587608 | 0,606 | 0,361 | 3,69832E-14 | FB | 1,341258773 | 0,745568282 | 1,67867036 |
| FGL21 | 1,0927E-18 | -0,957842927 | 0,308 | 0,469 | 3,99787E-14 | FB | 0,51482609 | 1,9424035 | 0,656716418 |
| C4orf482 | 1,19903E-18 | 0,479074889 | 0,278 | 0,095 | 4,38688E-14 | FB | 1,39384959 | 0,717437525 | 2,926315789 |
| UQCRQ4 | 1,71291E-18 | 0,461092291 | 0,623 | 0,366 | 6,26702E-14 | FB | 1,376583661 | 0,726436052 | 1,702185792 |
| RPS233 | 1,9661E-18 | 0,300822768 | 0,987 | 0,944 | 7,19337E-14 | FB | 1,231846734 | 0,811789302 | 1,045550847 |
| GPX3 | 2,19743E-18 | -0,975698518 | 0,237 | 0,389 | 8,03975E-14 | FB | 0,508493586 | 1,966593143 | 0,609254499 |
| ARID5B5 | 2,52002E-18 | 0,396785999 | 0,543 | 0,298 | 9,22001E-14 | FB | 1,316571615 | 0,759548503 | 1,822147651 |
| NEGR1 | 3,03748E-18 | 0,360256828 | 0,406 | 0,185 | 1,11132E-13 | FB | 1,283654393 | 0,779025885 | 2,194594595 |
| MYO1D | 3,89166E-18 | 0,365171274 | 0,276 | 0,094 | 1,42384E-13 | FB | 1,288034534 | 0,776376699 | 2,936170213 |
| MYL63 | 3,93257E-18 | 0,532074479 | 0,938 | 0,844 | 1,43881E-13 | FB | 1,446006942 | 0,691559612 | 1,111374408 |
| KDELR24 | 4,30154E-18 | 0,455603589 | 0,549 | 0,313 | 1,5738E-13 | FB | 1,371356428 | 0,729205026 | 1,75399361 |
| NME36 | 4,35471E-18 | 0,407000657 | 0,54 | 0,295 | 1,59326E-13 | FB | 1,325926364 | 0,754189695 | 1,830508475 |
| RARRES13 | 4,76098E-18 | -1,136446987 | 0,289 | 0,437 | 1,7419E-13 | FB | 0,454878456 | 2,198389452 | 0,661327231 |
| PDLIM51 | 4,91605E-18 | 0,400386333 | 0,421 | 0,208 | 1,79864E-13 | FB | 1,319861303 | 0,757655367 | 2,024038462 |
| CXCL14 | 6,72919E-18 | -0,527618252 | 0,648 | 0,798 | 2,46201E-13 | FB | 0,693699018 | 1,441547377 | 0,812030075 |
| MMP23B | 6,76916E-18 | 0,463947649 | 0,29 | 0,108 | 2,47663E-13 | FB | 1,37931087 | 0,724999724 | 2,685185185 |
| KCNQ1OT12 | 7,00954E-18 | 0,548343633 | 0,42 | 0,208 | 2,56458E-13 | FB | 1,462405735 | 0,683804758 | 2,019230769 |
| ITGB14 | 1,09392E-17 | 0,407569231 | 0,747 | 0,519 | 4,00233E-13 | FB | 1,326449022 | 0,753892523 | 1,439306358 |
| PCOLCE2 | 1,14822E-17 | -1,32425258 | 0,164 | 0,311 | 4,20101E-13 | FB | 0,399356035 | 2,504031273 | 0,52733119 |
| EEF1G2 | 1,2779E-17 | 0,386870012 | 0,852 | 0,615 | 4,67546E-13 | FB | 1,307553531 | 0,764787044 | 1,385365854 |
| VCAN2 | 1,42837E-17 | 0,429211351 | 0,874 | 0,71 | 5,22599E-13 | FB | 1,346497313 | 0,742667653 | 1,230985915 |
| BHLHE403 | 1,47742E-17 | 0,531316814 | 0,498 | 0,284 | 5,40542E-13 | FB | 1,445247737 | 0,691922896 | 1,753521127 |
| VASN | 1,78317E-17 | 0,372839192 | 0,386 | 0,174 | 6,52409E-13 | FB | 1,294898656 | 0,772261207 | 2,218390805 |
| RASGRF2 | 2,44423E-17 | 0,405270149 | 0,169 | 0,032 | 8,94271E-13 | FB | 1,324336873 | 0,755094886 | 5,28125 |
| SUGCT | 2,7516E-17 | 0,473155723 | 0,185 | 0,042 | 1,00673E-12 | FB | 1,388142546 | 0,720387112 | 4,404761905 |
| ABCA6 | 2,80246E-17 | -0,824499605 | 0,307 | 0,45 | 1,02534E-12 | FB | 0,564678026 | 1,770920692 | 0,682222222 |
| TUBB3 | 2,91197E-17 | 0,395855627 | 0,204 | 0,053 | 1,0654E-12 | FB | 1,315722852 | 0,760038483 | 3,849056604 |
| MSRB31 | 3,1113E-17 | 0,351469803 | 0,358 | 0,156 | 1,13833E-12 | FB | 1,275859799 | 0,783785179 | 2,294871795 |
| ADAM122 | 4,59027E-17 | 0,488113112 | 0,182 | 0,04 | 1,67944E-12 | FB | 1,402609216 | 0,712956958 | 4,55 |
| GOLM11 | 5,11743E-17 | 0,388910596 | 0,343 | 0,148 | 1,87231E-12 | FB | 1,309404277 | 0,763706074 | 2,317567568 |
| ITGA11 | 5,14419E-17 | 0,506155023 | 0,3 | 0,119 | 1,8821E-12 | FB | 1,420259963 | 0,704096451 | 2,521008403 |

| Supplementary Table 4 - clusermarker | | | | | | | | | |
|---|---|---|---|---|---|---|---|---|---|
| | **p_val** | **avg_log2FC** | **pct.1** | **pct.2** | **p_val_adj** | **celltype** | **Foldchange_UP** | **Foldchange_DOWN** | **Ratio_pct1_pct2** |
| RIN21 | 6,03445E-17 | 0,403304965 | 0,301 | 0,119 | 2,20782E-12 | FB | 1,322534141 | 0,756124148 | 2,529411765 |
| RPS215 | 6,27306E-17 | 0,442210206 | 0,948 | 0,787 | 2,29513E-12 | FB | 1,358684236 | 0,736006184 | 1,204574333 |
| PDLIM73 | 6,60325E-17 | 0,4908644 | 0,386 | 0,189 | 2,41593E-12 | FB | 1,40528661 | 0,711598611 | 2,042328042 |
| NDUFB114 | 6,65544E-17 | 0,420243043 | 0,508 | 0,282 | 2,43503E-12 | FB | 1,338152967 | 0,74729872 | 1,80141844 |
| NENF4 | 6,81486E-17 | 0,496567236 | 0,647 | 0,413 | 2,49335E-12 | FB | 1,410852567 | 0,708791282 | 1,566585956 |
| DUXAP8 | 7,06962E-17 | 0,3684837 | 0,195 | 0,048 | 2,58656E-12 | FB | 1,290995257 | 0,774596184 | 4,0625 |
| PTMA2 | 7,32036E-17 | 0,363369464 | 0,967 | 0,845 | 2,6783E-12 | FB | 1,286426886 | 0,777346937 | 1,144378698 |
| PLN | 7,53562E-17 | 0,379510785 | 0,143 | 0,019 | 2,75706E-12 | FB | 1,300900648 | 0,76869821 | 7,526315789 |
| PODNL1 | 1,12E-16 | 0,393318453 | 0,223 | 0,068 | 4,09775E-12 | FB | 1,313411009 | 0,761376289 | 3,279411765 |
| GLUL4 | 1,13847E-16 | -0,83834183 | 0,568 | 0,634 | 4,16533E-12 | FB | 0,559286018 | 1,787993919 | 0,895899054 |
| WDR17 | 1,21118E-16 | 0,367833589 | 0,316 | 0,132 | 4,43133E-12 | FB | 1,290413636 | 0,774945314 | 2,393939394 |
| LMNA5 | 1,49217E-16 | 0,390668803 | 0,901 | 0,734 | 5,45941E-12 | FB | 1,311001015 | 0,762775916 | 1,227520436 |
| TGM21 | 1,68939E-16 | 0,469907062 | 0,295 | 0,118 | 6,18097E-12 | FB | 1,385020243 | 0,722011108 | 2,5 |
| MYDGF4 | 2,00293E-16 | 0,386806433 | 0,53 | 0,303 | 7,32813E-12 | FB | 1,307495909 | 0,764820748 | 1,749174917 |
| TNFAIP32 | 2,11565E-16 | 0,709235102 | 0,285 | 0,115 | 7,74052E-12 | FB | 1,634937065 | 0,611644339 | 2,47826087 |
| AP2S14 | 2,33062E-16 | 0,354322783 | 0,363 | 0,168 | 8,52704E-12 | FB | 1,278385352 | 0,782236748 | 2,160714286 |
| MANF4 | 2,40983E-16 | 0,393726745 | 0,297 | 0,121 | 8,81685E-12 | FB | 1,313782765 | 0,761160845 | 2,454545455 |
| GEM3 | 2,4139E-16 | 0,597243227 | 0,642 | 0,442 | 8,83173E-12 | FB | 1,512823026 | 0,661015851 | 1,452488688 |
| NOX4 | 2,57941E-16 | 0,309710224 | 0,245 | 0,081 | 9,43729E-12 | FB | 1,239458721 | 0,806803795 | 3,024691358 |
| PCSK7 | 2,64659E-16 | 0,427048849 | 0,237 | 0,081 | 9,68307E-12 | FB | 1,344480517 | 0,743781697 | 2,925925926 |
| PRSS12 | 2,73457E-16 | 0,488205972 | 0,21 | 0,063 | 1,0005E-11 | FB | 1,402699499 | 0,71291107 | 3,333333333 |
| ATOX13 | 3,30159E-16 | 0,358685453 | 0,427 | 0,219 | 1,20795E-11 | FB | 1,282257005 | 0,779874858 | 1,949771689 |
| NDUFA41 | 3,7017E-16 | 0,393530146 | 0,798 | 0,589 | 1,35434E-11 | FB | 1,313603745 | 0,761264577 | 1,35483871 |
| FHOD32 | 4,46126E-16 | 0,316358364 | 0,152 | 0,026 | 1,63224E-11 | FB | 1,245183499 | 0,803094484 | 5,846153846 |
| LTF5 | 5,49076E-16 | -0,502962417 | 0,021 | 0,102 | 2,00891E-11 | FB | 0,705656304 | 1,417120479 | 0,205882353 |
| MMP14 | 5,5704E-16 | 0,48650633 | 0,432 | 0,235 | 2,03804E-11 | FB | 1,401047948 | 0,713751447 | 1,838297872 |
| IL1R11 | 6,57175E-16 | 0,470862809 | 0,411 | 0,213 | 2,40441E-11 | FB | 1,385938085 | 0,721532953 | 1,929577465 |
| C12orf752 | 6,98618E-16 | 0,409726519 | 0,216 | 0,066 | 2,55603E-11 | FB | 1,328433968 | 0,752766057 | 3,272727273 |
| RPL396 | 7,52215E-16 | 0,445474323 | 0,979 | 0,918 | 2,75213E-11 | FB | 1,361761757 | 0,734342843 | 1,066448802 |
| PSME24 | 7,75499E-16 | 0,36054119 | 0,451 | 0,24 | 2,83732E-11 | FB | 1,283907432 | 0,778872351 | 1,879166667 |
| SVIL1 | 8,42654E-16 | 0,435839999 | 0,513 | 0,292 | 3,08302E-11 | FB | 1,352698203 | 0,739263198 | 1,756849315 |

| Supplementary Table 4 - clusermarker | | | | | | | | | |
|---|---|---|---|---|---|---|---|---|---|
| | **p_val** | **avg_log2FC** | **pct.1** | **pct.2** | **p_val_adj** | **celltype** | **Foldchange_UP** | **Foldchange_DOWN** | **Ratio_pct1_pct2** |
| CEMIP1 | 8,62373E-16 | 0,489493891 | 0,134 | 0,018 | 3,15516E-11 | FB | 1,403952272 | 0,712274926 | 7,444444444 |
| RCN11 | 8,73422E-16 | 0,348556423 | 0,415 | 0,213 | 3,19559E-11 | FB | 1,273285925 | 0,785369555 | 1,948356808 |
| PDLIM12 | 8,90541E-16 | 0,293356244 | 0,386 | 0,185 | 3,25822E-11 | FB | 1,225487904 | 0,816001526 | 2,086486486 |
| ANXA23 | 9,16817E-16 | -0,515255772 | 0,917 | 0,889 | 3,35436E-11 | FB | 0,699668879 | 1,429247505 | 1,031496063 |
| GLIPR13 | 9,67528E-16 | 0,294846056 | 0,393 | 0,185 | 3,5399E-11 | FB | 1,226754068 | 0,81515931 | 2,124324324 |
| LMOD11 | 1,08219E-15 | 0,307326554 | 0,195 | 0,052 | 3,9594E-11 | FB | 1,237412535 | 0,808137926 | 3,75 |
| CDKN1A5 | 1,0971E-15 | 0,466976569 | 0,554 | 0,345 | 4,01397E-11 | FB | 1,382209758 | 0,723479193 | 1,605797101 |
| CRABP1 | 1,10576E-15 | 0,952089223 | 0,489 | 0,298 | 4,04565E-11 | FB | 1,934672305 | 0,516883401 | 1,640939597 |
| STMN13 | 1,1512E-15 | 0,395244359 | 0,263 | 0,098 | 4,21189E-11 | FB | 1,3151655 | 0,760360578 | 2,683673469 |
| ACTN42 | 1,16946E-15 | 0,391559286 | 0,41 | 0,213 | 4,27869E-11 | FB | 1,311810462 | 0,762305249 | 1,924882629 |
| NR4A34 | 1,28295E-15 | 0,508903092 | 0,188 | 0,052 | 4,69392E-11 | FB | 1,422967875 | 0,702756554 | 3,615384615 |
| MT-ND37 | 1,35812E-15 | -0,307105742 | 0,97 | 0,971 | 4,96895E-11 | FB | 0,808261625 | 1,237223157 | 0,998970134 |
| F13A11 | 1,39764E-15 | -0,861774942 | 0,039 | 0,132 | 5,11353E-11 | FB | 0,55027514 | 1,817272719 | 0,295454545 |
| CADM31 | 1,39914E-15 | -0,514029646 | 0,024 | 0,106 | 5,11904E-11 | FB | 0,70026377 | 1,428033324 | 0,226415094 |
| BRI35 | 1,60155E-15 | 0,407456286 | 0,593 | 0,373 | 5,8596E-11 | FB | 1,326345182 | 0,753951546 | 1,589812332 |
| RPS184 | 1,64689E-15 | 0,28802604 | 0,987 | 0,94 | 6,02547E-11 | FB | 1,22096855 | 0,819021915 | 1,05 |
| TYMP6 | 1,74657E-15 | 0,5062174 | 0,4 | 0,211 | 6,39016E-11 | FB | 1,420321371 | 0,70406601 | 1,895734597 |
| ENAH1 | 2,04331E-15 | 0,321432719 | 0,382 | 0,187 | 7,47585E-11 | FB | 1,249570863 | 0,800274742 | 2,042780749 |
| CALU | 2,08504E-15 | 0,411085779 | 0,668 | 0,452 | 7,62854E-11 | FB | 1,329686166 | 0,752057159 | 1,477876106 |
| SLC25A65 | 2,08909E-15 | 0,36593077 | 0,717 | 0,484 | 7,64335E-11 | FB | 1,288712788 | 0,775968089 | 1,481404959 |
| SLC44A12 | 2,12729E-15 | 0,338910282 | 0,447 | 0,237 | 7,78311E-11 | FB | 1,264800884 | 0,790638283 | 1,886075949 |
| RORB | 2,21171E-15 | 0,368279661 | 0,241 | 0,084 | 8,09198E-11 | FB | 1,290812685 | 0,774705743 | 2,869047619 |
| HSPA55 | 2,91685E-15 | 0,361621363 | 0,757 | 0,545 | 1,06719E-10 | FB | 1,284869078 | 0,778289413 | 1,388990826 |
| RGS3 | 3,20695E-15 | 0,391297401 | 0,199 | 0,06 | 1,17333E-10 | FB | 1,311572358 | 0,762443638 | 3,316666667 |
| NUAK1 | 3,9001E-15 | 0,400059957 | 0,178 | 0,047 | 1,42693E-10 | FB | 1,31956275 | 0,757826788 | 3,787234043 |
| CEMIP2 | 4,01726E-15 | 0,316532999 | 0,164 | 0,037 | 1,46979E-10 | FB | 1,245334235 | 0,802997277 | 4,432432432 |
| ADM2 | 4,07071E-15 | 0,424741619 | 0,448 | 0,252 | 1,48935E-10 | FB | 1,342332075 | 0,744972141 | 1,777777778 |
| DRAP16 | 4,18616E-15 | 0,374075551 | 0,439 | 0,237 | 1,53159E-10 | FB | 1,296008833 | 0,771599679 | 1,852320675 |
| FAP | 4,30489E-15 | 0,389837708 | 0,493 | 0,276 | 1,57503E-10 | FB | 1,310246003 | 0,763215455 | 1,786231884 |
| TNS3 | 4,5085E-15 | 0,380108585 | 0,253 | 0,095 | 1,64953E-10 | FB | 1,301439805 | 0,768379756 | 2,663157895 |
| CADM1 | 4,92654E-15 | 0,36735835 | 0,143 | 0,026 | 1,80247E-10 | FB | 1,28998863 | 0,775200631 | 5,5 |

| Supplementary Table 4 - clusermarker | | | | | | | | | |
|---|---|---|---|---|---|---|---|---|---|
| | **p_val** | **avg_log2FC** | **pct.1** | **pct.2** | **p_val_adj** | **celltype** | **Foldchange_UP** | **Foldchange_DOWN** | **Ratio_pct1_pct2** |
| DDT5 | 5,03264E-15 | 0,386563843 | 0,192 | 0,056 | 1,84129E-10 | FB | 1,307276072 | 0,764949364 | 3,428571429 |
| FAAP204 | 5,43925E-15 | 0,344446637 | 0,322 | 0,144 | 1,99006E-10 | FB | 1,269663894 | 0,787610016 | 2,236111111 |
| AL138963.45 | 5,53129E-15 | 0,455007037 | 0,302 | 0,131 | 2,02373E-10 | FB | 1,370789491 | 0,729506614 | 2,305343511 |
| C1orf198 | 5,63689E-15 | 0,364859605 | 0,248 | 0,092 | 2,06237E-10 | FB | 1,287756307 | 0,77654444 | 2,695652174 |
| MARCKSL13 | 5,7004E-15 | 0,355884531 | 0,173 | 0,044 | 2,08561E-10 | FB | 1,279769981 | 0,781390418 | 3,931818182 |
| TMEM2196 | 6,12929E-15 | 0,310038828 | 0,491 | 0,269 | 2,24252E-10 | FB | 1,239741065 | 0,80662005 | 1,82527881 |
| SCUBE2 | 6,28898E-15 | 0,296890794 | 0,11 | 0,008 | 2,30095E-10 | FB | 1,228493985 | 0,814004799 | 13,75 |
| EIF4A32 | 6,76786E-15 | 0,323090284 | 0,405 | 0,206 | 2,47616E-10 | FB | 1,251007366 | 0,799355805 | 1,966019417 |
| MAGED11 | 7,17774E-15 | 0,343178989 | 0,4 | 0,208 | 2,62612E-10 | FB | 1,268548773 | 0,788302367 | 1,923076923 |
| HEG1 | 7,30649E-15 | 0,418111443 | 0,411 | 0,218 | 2,67322E-10 | FB | 1,33617729 | 0,748403679 | 1,885321101 |
| LAMA2 | 8,95096E-15 | -0,709480143 | 0,472 | 0,556 | 3,27489E-10 | FB | 0,61154046 | 1,635214782 | 0,848920863 |
| SHOX2 | 9,12168E-15 | 0,314505188 | 0,151 | 0,031 | 3,33735E-10 | FB | 1,243585058 | 0,804126741 | 4,870967742 |
| RGCC4 | 9,82099E-15 | 0,614745731 | 0,29 | 0,129 | 3,59321E-10 | FB | 1,53128809 | 0,65304498 | 2,248062016 |
| ALDH1A14 | 1,0789E-14 | -0,88436759 | 0,305 | 0,429 | 3,94738E-10 | FB | 0,541724938 | 1,845955263 | 0,710955711 |
| EPAS13 | 1,09218E-14 | 0,372009213 | 0,29 | 0,124 | 3,99595E-10 | FB | 1,294153919 | 0,772705615 | 2,338709677 |
| SH3PXD2A1 | 1,22354E-14 | 0,374649407 | 0,332 | 0,156 | 4,47657E-10 | FB | 1,296524445 | 0,771292824 | 2,128205128 |
| NOVA1 | 1,50441E-14 | -0,6393145 | 0,089 | 0,203 | 5,50417E-10 | FB | 0,642017933 | 1,557588892 | 0,438423645 |
| CP3 | 1,72708E-14 | 0,592957915 | 0,179 | 0,052 | 6,31886E-10 | FB | 1,508336076 | 0,66298222 | 3,442307692 |
| POMP4 | 1,73679E-14 | 0,294723631 | 0,498 | 0,277 | 6,35439E-10 | FB | 1,226649972 | 0,815228486 | 1,797833935 |
| CERCAM | 2,17131E-14 | 0,392711652 | 0,487 | 0,284 | 7,94417E-10 | FB | 1,312858702 | 0,761696593 | 1,714788732 |
| ACKR3 | 2,7475E-14 | -1,20918045 | 0,328 | 0,442 | 1,00523E-09 | FB | 0,432514244 | 2,312062584 | 0,742081448 |
| GPR176 | 2,89809E-14 | 0,29160083 | 0,155 | 0,035 | 1,06032E-09 | FB | 1,223997685 | 0,816995009 | 4,428571429 |
| LMO7 | 2,89955E-14 | 0,253229879 | 0,308 | 0,137 | 1,06086E-09 | FB | 1,191872472 | 0,839015938 | 2,248175182 |
| PKM3 | 2,97743E-14 | 0,396133223 | 0,74 | 0,527 | 1,08935E-09 | FB | 1,315976041 | 0,759892254 | 1,404174573 |
| CXCL22 | 3,58796E-14 | 1,108432289 | 0,225 | 0,085 | 1,31273E-09 | FB | 2,156112251 | 0,463797745 | 2,647058824 |
| BBLN6 | 3,65077E-14 | 0,38321545 | 0,461 | 0,26 | 1,33571E-09 | FB | 1,304245495 | 0,766726819 | 1,773076923 |
| CRIM1 | 3,93457E-14 | 0,364645433 | 0,297 | 0,131 | 1,43954E-09 | FB | 1,287565149 | 0,776659729 | 2,267175573 |
| EGR21 | 4,02531E-14 | 0,470077555 | 0,28 | 0,123 | 1,47274E-09 | FB | 1,38518393 | 0,721925788 | 2,276422764 |
| ALKBH73 | 4,66296E-14 | 0,352056806 | 0,388 | 0,2 | 1,70604E-09 | FB | 1,276379025 | 0,783466338 | 1,94 |
| PLAUR2 | 4,81166E-14 | 0,343834621 | 0,277 | 0,119 | 1,76044E-09 | FB | 1,269125395 | 0,787944205 | 2,327731092 |
| HCFC1R12 | 5,04491E-14 | 0,376690288 | 0,647 | 0,431 | 1,84578E-09 | FB | 1,298359846 | 0,7702025 | 1,501160093 |

| Supplementary Table 4 - clusermarker | | | | | | | | | |
|---|---|---|---|---|---|---|---|---|---|
| | **p_val** | **avg_log2FC** | **pct.1** | **pct.2** | **p_val_adj** | **celltype** | **Foldchange_UP** | **Foldchange_DOWN** | **Ratio_pct1_pct2** |
| ATF43 | 5,2451E-14 | 0,318176925 | 0,474 | 0,266 | 1,91903E-09 | FB | 1,246754081 | 0,802082797 | 1,781954887 |
| ANAPC116 | 5,3511E-14 | 0,324051519 | 0,561 | 0,344 | 1,95781E-09 | FB | 1,251841161 | 0,79882339 | 1,630813953 |
| P3H1 | 5,80388E-14 | 0,333784556 | 0,292 | 0,129 | 2,12347E-09 | FB | 1,260315169 | 0,793452324 | 2,263565891 |
| TCEAL31 | 6,12431E-14 | 0,312851068 | 0,272 | 0,113 | 2,2407E-09 | FB | 1,242160044 | 0,80504924 | 2,407079646 |
| EGR33 | 6,84242E-14 | 0,305415734 | 0,274 | 0,115 | 2,50344E-09 | FB | 1,235774693 | 0,809208997 | 2,382608696 |
| ERGIC34 | 7,82586E-14 | 0,304879355 | 0,6 | 0,368 | 2,86325E-09 | FB | 1,23531533 | 0,809509909 | 1,630434783 |
| IGLV3-11 | 8,0509E-14 | 1,327325545 | 0,122 | 0,018 | 2,94558E-09 | FB | 2,509370587 | 0,398506305 | 6,777777778 |
| ETHE13 | 8,08124E-14 | 0,393402304 | 0,283 | 0,127 | 2,95668E-09 | FB | 1,313487348 | 0,761332039 | 2,228346457 |
| PDLIM21 | 8,40161E-14 | 0,390456594 | 0,607 | 0,389 | 3,0739E-09 | FB | 1,310808191 | 0,762888122 | 1,560411311 |
| RPS6KA2 | 9,98814E-14 | 0,291479781 | 0,161 | 0,04 | 3,65436E-09 | FB | 1,223894991 | 0,817063561 | 4,025 |
| FOXF1 | 1,06063E-13 | 0,252159179 | 0,232 | 0,085 | 3,88051E-09 | FB | 1,190988249 | 0,839638847 | 2,729411765 |
| NDUFS85 | 1,11901E-13 | 0,359970013 | 0,395 | 0,211 | 4,09412E-09 | FB | 1,283399222 | 0,779180775 | 1,872037915 |
| ADAMTS12 | 1,17919E-13 | 0,382077673 | 0,195 | 0,065 | 4,31429E-09 | FB | 1,303217311 | 0,767331735 | 3 |
| ITM2C3 | 1,25368E-13 | 0,289811598 | 0,277 | 0,116 | 4,58686E-09 | FB | 1,222480623 | 0,818008875 | 2,387931034 |
| SEPTIN112 | 1,28335E-13 | 0,35911708 | 0,593 | 0,377 | 4,69541E-09 | FB | 1,282640689 | 0,779641569 | 1,572944297 |
| DIO2 | 1,31724E-13 | 0,422713656 | 0,638 | 0,431 | 4,81939E-09 | FB | 1,340446516 | 0,746020067 | 1,480278422 |
| EEF1D4 | 1,31945E-13 | 0,325231788 | 0,878 | 0,679 | 4,82749E-09 | FB | 1,252865712 | 0,79817014 | 1,293078056 |
| TMEM1605 | 1,36808E-13 | 0,342973924 | 0,241 | 0,095 | 5,0054E-09 | FB | 1,268368474 | 0,788414424 | 2,536842105 |
| TMEM100 | 1,37493E-13 | -0,662007365 | 0,119 | 0,239 | 5,03045E-09 | FB | 0,631998323 | 1,58228268 | 0,49790795 |
| UBE2S2 | 1,53275E-13 | 0,324179626 | 0,406 | 0,213 | 5,60786E-09 | FB | 1,251952326 | 0,79875246 | 1,906103286 |
| MT-ND4L5 | 1,55127E-13 | -0,322949783 | 0,934 | 0,976 | 5,67562E-09 | FB | 0,799433657 | 1,250885538 | 0,956967213 |
| SSR34 | 1,57426E-13 | 0,269098345 | 0,558 | 0,339 | 5,75973E-09 | FB | 1,205054458 | 0,829838016 | 1,646017699 |
| FST | 1,69373E-13 | 0,30748388 | 0,246 | 0,097 | 6,19686E-09 | FB | 1,237547483 | 0,808049803 | 2,536082474 |
| HDLBP3 | 1,70242E-13 | 0,365172834 | 0,631 | 0,419 | 6,22863E-09 | FB | 1,288035926 | 0,77637586 | 1,505966587 |
| SOCS32 | 1,71087E-13 | 0,589377838 | 0,512 | 0,324 | 6,25956E-09 | FB | 1,50459775 | 0,664629467 | 1,580246914 |
| CDKN2A3 | 1,74217E-13 | 0,357581472 | 0,155 | 0,039 | 6,37408E-09 | FB | 1,28127617 | 0,780471863 | 3,974358974 |
| CNTN4 | 1,86841E-13 | -0,504933508 | 0,034 | 0,116 | 6,83594E-09 | FB | 0,704692855 | 1,419057952 | 0,293103448 |
| JCAD | 1,95692E-13 | 0,328771436 | 0,192 | 0,063 | 7,15979E-09 | FB | 1,255943388 | 0,796214232 | 3,047619048 |
| BCAT1 | 2,22585E-13 | 0,271027942 | 0,106 | 0,011 | 8,14371E-09 | FB | 1,20666729 | 0,828728854 | 9,636363636 |
| CAPN6 | 2,24345E-13 | -0,613303786 | 0,044 | 0,132 | 8,2081E-09 | FB | 0,653698011 | 1,529758363 | 0,333333333 |
| RPS297 | 2,26767E-13 | -0,492922389 | 0,77 | 0,734 | 8,29673E-09 | FB | 0,710584247 | 1,407292667 | 1,049046322 |

| Supplementary Table 4 - clusermarker | | | | | | | | | |
|---|---|---|---|---|---|---|---|---|---|
| | **p_val** | **avg_log2FC** | **pct.1** | **pct.2** | **p_val_adj** | **celltype** | **Foldchange_UP** | **Foldchange_DOWN** | **Ratio_pct1_pct2** |
| RPLP25 | 2,26902E-13 | 0,342071035 | 0,948 | 0,818 | 8,30165E-09 | FB | 1,267574933 | 0,788907996 | 1,158924205 |
| CYCS3 | 2,38467E-13 | 0,338100417 | 0,363 | 0,187 | 8,72478E-09 | FB | 1,264091081 | 0,791082237 | 1,941176471 |
| MIR4435-2HG | 2,40582E-13 | 0,338833687 | 0,271 | 0,116 | 8,80216E-09 | FB | 1,264733736 | 0,79068026 | 2,336206897 |
| SEMA4A1 | 2,52323E-13 | 0,291032759 | 0,16 | 0,042 | 9,23174E-09 | FB | 1,223515823 | 0,81731677 | 3,80952381 |
| ITM2B6 | 2,67155E-13 | -0,321882461 | 0,969 | 0,935 | 9,77438E-09 | FB | 0,800025305 | 1,249960462 | 1,036363636 |
| ZYX3 | 2,83234E-13 | 0,350016674 | 0,478 | 0,279 | 1,03627E-08 | FB | 1,274575358 | 0,78457503 | 1,713261649 |
| SERPINE11 | 2,98195E-13 | 0,65773593 | 0,274 | 0,123 | 1,09101E-08 | FB | 1,577604891 | 0,633872274 | 2,227642276 |
| PPIC2 | 3,05096E-13 | 0,31312303 | 0,598 | 0,377 | 1,11626E-08 | FB | 1,242394226 | 0,804897495 | 1,586206897 |
| AHNAK5 | 3,17695E-13 | -0,453811 | 0,914 | 0,881 | 1,16235E-08 | FB | 0,730111647 | 1,369653537 | 1,037457435 |
| COL18A11 | 3,37971E-13 | 0,455199118 | 0,404 | 0,229 | 1,23654E-08 | FB | 1,370972011 | 0,729409493 | 1,76419214 |
| FKBP10 | 3,4278E-13 | 0,280166302 | 0,488 | 0,277 | 1,25413E-08 | FB | 1,214334855 | 0,823496086 | 1,761732852 |
| CCL19 | 3,73249E-13 | 1,83096787 | 0,14 | 0,032 | 1,36561E-08 | FB | 3,557756739 | 0,281075991 | 4,375 |
| COL6A3 | 3,75901E-13 | 0,404014344 | 0,893 | 0,765 | 1,37531E-08 | FB | 1,323184595 | 0,75575245 | 1,167320261 |
| TIMP13 | 3,83404E-13 | 0,93313309 | 0,95 | 0,855 | 1,40276E-08 | FB | 1,909418165 | 0,523719748 | 1,111111111 |
| SPRY14 | 3,93046E-13 | -0,853925549 | 0,46 | 0,534 | 1,43804E-08 | FB | 0,553277228 | 1,807412179 | 0,861423221 |
| OLFML2B | 4,11842E-13 | 0,329379728 | 0,376 | 0,195 | 1,50681E-08 | FB | 1,256473051 | 0,79587859 | 1,928205128 |
| OST44 | 4,16414E-13 | 0,379177521 | 0,726 | 0,508 | 1,52353E-08 | FB | 1,300600173 | 0,768875801 | 1,429133858 |
| GPC6 | 4,22856E-13 | 0,272141747 | 0,373 | 0,19 | 1,5471E-08 | FB | 1,207599234 | 0,828089296 | 1,963157895 |
| COL8A1 | 4,44189E-13 | 0,397962225 | 0,633 | 0,448 | 1,62515E-08 | FB | 1,31764545 | 0,758929498 | 1,412946429 |
| IFITM24 | 4,47979E-13 | 0,355618036 | 0,861 | 0,663 | 1,63902E-08 | FB | 1,279533603 | 0,78153477 | 1,298642534 |
| RUNX2 | 4,64403E-13 | 0,326071782 | 0,185 | 0,06 | 1,69911E-08 | FB | 1,253595392 | 0,797705549 | 3,083333333 |
| ADAMTS2 | 4,73093E-13 | 0,328822144 | 0,308 | 0,15 | 1,7309E-08 | FB | 1,255987533 | 0,796186247 | 2,053333333 |
| BMP1 | 5,01102E-13 | 0,337182364 | 0,248 | 0,103 | 1,83338E-08 | FB | 1,263286938 | 0,791585799 | 2,40776699 |
| TNFRSF6B1 | 5,08555E-13 | 0,45667309 | 0,212 | 0,081 | 1,86065E-08 | FB | 1,372373421 | 0,728664651 | 2,617283951 |
| CLEC11A | 5,5065E-13 | 0,278778374 | 0,459 | 0,261 | 2,01466E-08 | FB | 1,21316718 | 0,824288702 | 1,75862069 |
| HSP90B15 | 5,87902E-13 | 0,333824014 | 0,85 | 0,674 | 2,15096E-08 | FB | 1,260349639 | 0,793430623 | 1,261127596 |
| RPL102 | 5,93022E-13 | 0,269350124 | 0,995 | 0,977 | 2,16969E-08 | FB | 1,205264783 | 0,829693205 | 1,018423746 |
| CISD12 | 5,97801E-13 | 0,268192565 | 0,323 | 0,155 | 2,18718E-08 | FB | 1,204298115 | 0,830359184 | 2,083870968 |
| SGIP1 | 6,12103E-13 | 0,353965777 | 0,227 | 0,09 | 2,2395E-08 | FB | 1,278069045 | 0,782430342 | 2,522222222 |
| CAB39L | 6,72657E-13 | -0,606430116 | 0,104 | 0,215 | 2,46105E-08 | FB | 0,656819964 | 1,522487218 | 0,48372093 |
| PTS1 | 6,73956E-13 | 0,302131082 | 0,186 | 0,061 | 2,4658E-08 | FB | 1,232964347 | 0,811053461 | 3,049180328 |

| Supplementary Table 4 - clusermarker | | | | | | | | | |
|---|---|---|---|---|---|---|---|---|---|
| | **p_val** | **avg_log2FC** | **pct.1** | **pct.2** | **p_val_adj** | **celltype** | **Foldchange_UP** | **Foldchange_DOWN** | **Ratio_pct1_pct2** |
| UNC5B1 | 7,06996E-13 | 0,294679324 | 0,197 | 0,068 | 2,58668E-08 | FB | 1,226612301 | 0,815253523 | 2,897058824 |
| MEGF6 | 7,26922E-13 | 0,333803485 | 0,205 | 0,074 | 2,65959E-08 | FB | 1,260331705 | 0,793441914 | 2,77027027 |
| ATP5F1D7 | 8,05891E-13 | 0,391064188 | 0,598 | 0,379 | 2,94851E-08 | FB | 1,311360358 | 0,762566898 | 1,577836412 |
| CCDC85B5 | 8,36712E-13 | 0,29844545 | 0,503 | 0,3 | 3,06128E-08 | FB | 1,229818531 | 0,813128096 | 1,676666667 |
| RGS161 | 9,13128E-13 | 0,501225751 | 0,377 | 0,202 | 3,34086E-08 | FB | 1,415415625 | 0,70650626 | 1,866336634 |
| HMGN13 | 9,84929E-13 | 0,267940235 | 0,443 | 0,244 | 3,60356E-08 | FB | 1,204087499 | 0,830504428 | 1,81557377 |
| FBLIM1 | 1,01266E-12 | 0,32659426 | 0,158 | 0,045 | 3,70501E-08 | FB | 1,254049469 | 0,797416709 | 3,511111111 |
| GLMP1 | 1,02354E-12 | 0,307391691 | 0,286 | 0,132 | 3,74482E-08 | FB | 1,237468405 | 0,80810144 | 2,166666667 |
| C11orf96 | 1,03263E-12 | 0,513614204 | 0,246 | 0,105 | 3,77809E-08 | FB | 1,427622163 | 0,700465449 | 2,342857143 |
| TLN13 | 1,12144E-12 | 0,346083635 | 0,546 | 0,344 | 4,10301E-08 | FB | 1,271105374 | 0,786716837 | 1,587209302 |
| COL7A11 | 1,12728E-12 | 0,542971458 | 0,145 | 0,035 | 4,12437E-08 | FB | 1,45697029 | 0,686355794 | 4,142857143 |
| WLS1 | 1,16013E-12 | 0,331054445 | 0,442 | 0,253 | 4,24455E-08 | FB | 1,257932444 | 0,79495525 | 1,747035573 |
| GASK1B2 | 1,214E-12 | 0,3388883 | 0,364 | 0,194 | 4,44165E-08 | FB | 1,264781613 | 0,79065033 | 1,87628866 |
| ARPC25 | 1,25605E-12 | 0,345693902 | 0,586 | 0,384 | 4,59549E-08 | FB | 1,270762042 | 0,786929391 | 1,526041667 |
| OSMR | 1,62474E-12 | 0,316783783 | 0,29 | 0,134 | 5,94443E-08 | FB | 1,24555073 | 0,802857704 | 2,164179104 |
| SEC626 | 1,7863E-12 | -0,517184546 | 0,724 | 0,735 | 6,53555E-08 | FB | 0,6987341 | 1,431159579 | 0,985034014 |
| COL6A1 | 1,80336E-12 | 0,410014035 | 0,886 | 0,716 | 6,59794E-08 | FB | 1,32869874 | 0,752616052 | 1,237430168 |
| STUB14 | 1,83408E-12 | 0,273979013 | 0,366 | 0,194 | 6,71035E-08 | FB | 1,209138086 | 0,8270354 | 1,886597938 |
| NEK63 | 1,99859E-12 | 0,258656165 | 0,129 | 0,027 | 7,31223E-08 | FB | 1,196363802 | 0,835866145 | 4,777777778 |
| PLEC4 | 2,013E-12 | 0,371470341 | 0,43 | 0,25 | 7,36497E-08 | FB | 1,29367062 | 0,772994288 | 1,72 |
| GALNT12 | 2,08049E-12 | 0,294097967 | 0,216 | 0,084 | 7,61188E-08 | FB | 1,226118117 | 0,815582109 | 2,571428571 |
| FIBP3 | 2,24081E-12 | 0,347060864 | 0,254 | 0,111 | 8,19847E-08 | FB | 1,271966667 | 0,786184124 | 2,288288288 |
| TUBA1C2 | 2,36201E-12 | 0,304125372 | 0,341 | 0,174 | 8,64188E-08 | FB | 1,234669896 | 0,809933087 | 1,959770115 |
| ABL2 | 2,48507E-12 | 0,319669141 | 0,206 | 0,076 | 9,09212E-08 | FB | 1,248044297 | 0,801253611 | 2,710526316 |
| EPHB6 | 2,54381E-12 | 0,34976008 | 0,141 | 0,035 | 9,30703E-08 | FB | 1,274348686 | 0,784714585 | 4,028571429 |
| C1S1 | 2,65735E-12 | 0,276848677 | 0,976 | 0,921 | 9,72246E-08 | FB | 1,211545575 | 0,825391979 | 1,059717698 |
| FILIP1L | 2,66975E-12 | 0,377783343 | 0,347 | 0,179 | 9,76782E-08 | FB | 1,299343918 | 0,769619179 | 1,938547486 |
| CCN4 | 2,73119E-12 | 0,290175501 | 0,253 | 0,106 | 9,99262E-08 | FB | 1,222789018 | 0,817802569 | 2,386792453 |
| CILP2 | 2,91166E-12 | 0,361091674 | 0,113 | 0,019 | 1,06529E-07 | FB | 1,284397421 | 0,778575216 | 5,947368421 |
| SPECC12 | 2,94953E-12 | 0,25875807 | 0,114 | 0,019 | 1,07914E-07 | FB | 1,19644831 | 0,835807106 | 6 |
| SERF24 | 3,01229E-12 | 0,370203602 | 0,927 | 0,805 | 1,10211E-07 | FB | 1,292535229 | 0,773673303 | 1,151552795 |

| Supplementary Table 4 - clusermarker | | | | | | | | | |
|---|---|---|---|---|---|---|---|---|---|
| | **p_val** | **avg_log2FC** | **pct.1** | **pct.2** | **p_val_adj** | **celltype** | **Foldchange_UP** | **Foldchange_DOWN** | **Ratio_pct1_pct2** |
| HLF1 | 3,49468E-12 | -0,568167916 | 0,154 | 0,271 | 1,2786E-07 | FB | 0,67447276 | 1,482639565 | 0,568265683 |
| FGF141 | 3,7519E-12 | 0,28551528 | 0,31 | 0,148 | 1,37271E-07 | FB | 1,218845514 | 0,820448521 | 2,094594595 |
| CDC42EP33 | 3,82465E-12 | 0,26881181 | 0,229 | 0,094 | 1,39933E-07 | FB | 1,204815145 | 0,830002847 | 2,436170213 |
| RPL18A6 | 4,07245E-12 | 0,332463213 | 0,974 | 0,856 | 1,48999E-07 | FB | 1,259161394 | 0,794179368 | 1,137850467 |
| NRG1 | 4,5921E-12 | 0,317211559 | 0,108 | 0,018 | 1,68011E-07 | FB | 1,245920106 | 0,802619682 | 6 |
| MICAL2 | 4,65391E-12 | 0,300877438 | 0,183 | 0,063 | 1,70273E-07 | FB | 1,231893415 | 0,81175854 | 2,904761905 |
| OSR11 | 5,13255E-12 | -0,489328441 | 0,052 | 0,139 | 1,87785E-07 | FB | 0,712356615 | 1,403791274 | 0,374100719 |
| C1QTNF3 | 5,32629E-12 | -0,789384573 | 0,045 | 0,127 | 1,94873E-07 | FB | 0,578590855 | 1,728337029 | 0,354330709 |
| SLIT2 | 5,44378E-12 | -0,719897934 | 0,321 | 0,432 | 1,99172E-07 | FB | 0,607140394 | 1,647065506 | 0,743055556 |
| NAMPT1 | 5,54006E-12 | 0,392647585 | 0,291 | 0,14 | 2,02694E-07 | FB | 1,312800401 | 0,761730419 | 2,078571429 |
| HES44 | 6,09186E-12 | 0,288272141 | 0,176 | 0,058 | 2,22883E-07 | FB | 1,221176846 | 0,818882215 | 3,034482759 |
| TSPAN9 | 6,10063E-12 | 0,278769689 | 0,199 | 0,076 | 2,23204E-07 | FB | 1,213159877 | 0,824293664 | 2,618421053 |
| SYTL24 | 6,82628E-12 | 0,299340637 | 0,2 | 0,076 | 2,49753E-07 | FB | 1,230581865 | 0,812623709 | 2,631578947 |
| GRN2 | 7,14614E-12 | 0,271242099 | 0,586 | 0,374 | 2,61456E-07 | FB | 1,206846424 | 0,828605844 | 1,56684492 |
| ZBTB161 | 7,7009E-12 | -0,646931506 | 0,17 | 0,284 | 2,81753E-07 | FB | 0,638637199 | 1,565834251 | 0,598591549 |
| MT-ND22 | 7,90298E-12 | -0,330638128 | 0,967 | 0,929 | 2,89146E-07 | FB | 0,795184682 | 1,257569496 | 1,040904198 |
| SLC12A8 | 7,97708E-12 | 0,270159622 | 0,12 | 0,024 | 2,91858E-07 | FB | 1,205941248 | 0,829227794 | 5 |
| PHLDA24 | 8,10319E-12 | 0,350984747 | 0,24 | 0,103 | 2,96471E-07 | FB | 1,275430907 | 0,784048743 | 2,330097087 |
| ARL4C2 | 8,50408E-12 | 0,268906496 | 0,142 | 0,037 | 3,11139E-07 | FB | 1,20489422 | 0,829948375 | 3,837837838 |
| RPS153 | 8,58669E-12 | 0,308419558 | 0,979 | 0,882 | 3,14161E-07 | FB | 1,23835037 | 0,807525902 | 1,109977324 |
| TAF106 | 9,88256E-12 | 0,312168634 | 0,358 | 0,192 | 3,61573E-07 | FB | 1,241572608 | 0,80543014 | 1,864583333 |
| ARL6IP51 | 1,03404E-11 | -0,486955364 | 0,782 | 0,761 | 3,78323E-07 | FB | 0,713529329 | 1,401484087 | 1,027595269 |
| LAMB21 | 1,04061E-11 | 0,25143013 | 0,53 | 0,318 | 3,80728E-07 | FB | 1,190386549 | 0,840063256 | 1,666666667 |
| NDUFA116 | 1,12423E-11 | 0,256102674 | 0,734 | 0,503 | 4,11324E-07 | FB | 1,194248176 | 0,837346893 | 1,459244533 |
| IRF11 | 1,19697E-11 | 0,357864425 | 0,45 | 0,266 | 4,37937E-07 | FB | 1,281527489 | 0,780318806 | 1,691729323 |
| RPS276 | 1,20996E-11 | -0,309986362 | 0,98 | 0,958 | 4,42686E-07 | FB | 0,806649385 | 1,239695981 | 1,022964509 |
| NFIB2 | 1,22818E-11 | -0,498578893 | 0,607 | 0,647 | 4,49353E-07 | FB | 0,70780365 | 1,412821197 | 0,938176198 |
| LRRN3 | 1,25266E-11 | 0,317012722 | 0,103 | 0,016 | 4,58312E-07 | FB | 1,2457484 | 0,80273031 | 6,4375 |
| GPC11 | 1,26901E-11 | 0,3186488 | 0,217 | 0,089 | 4,64294E-07 | FB | 1,247161934 | 0,801820495 | 2,438202247 |
| MCL12 | 1,28891E-11 | 0,313442825 | 0,642 | 0,437 | 4,71575E-07 | FB | 1,242669652 | 0,804719097 | 1,469107551 |
| FOSL21 | 1,29758E-11 | 0,251562818 | 0,248 | 0,11 | 4,74745E-07 | FB | 1,190496037 | 0,839985997 | 2,254545455 |

| Supplementary Table 4 - clusermarker | | | | | | | | | |
|---|---|---|---|---|---|---|---|---|---|
| | **p_val** | **avg_log2FC** | **pct.1** | **pct.2** | **p_val_adj** | **celltype** | **Foldchange_UP** | **Foldchange_DOWN** | **Ratio_pct1_pct2** |
| P4HB4 | 1,40923E-11 | 0,335030971 | 0,715 | 0,529 | 5,15593E-07 | FB | 1,261404487 | 0,792767118 | 1,351606805 |
| WFDC24 | 1,58E-11 | -0,61620672 | 0,049 | 0,131 | 5,78075E-07 | FB | 0,652383989 | 1,532839581 | 0,374045802 |
| LAP34 | 1,58651E-11 | 0,314727957 | 0,372 | 0,206 | 5,80456E-07 | FB | 1,243777097 | 0,804002584 | 1,805825243 |
| UGCG3 | 1,6497E-11 | 0,295213503 | 0,294 | 0,144 | 6,03575E-07 | FB | 1,227066556 | 0,814951719 | 2,041666667 |
| CNN31 | 1,66302E-11 | 0,315993786 | 0,696 | 0,484 | 6,0845E-07 | FB | 1,244868874 | 0,803297456 | 1,438016529 |
| COX4I16 | 1,67401E-11 | 0,337384884 | 0,828 | 0,606 | 6,1247E-07 | FB | 1,263464286 | 0,791474687 | 1,366336634 |
| STAT32 | 1,80392E-11 | 0,26766258 | 0,606 | 0,385 | 6,60001E-07 | FB | 1,203855788 | 0,830664279 | 1,574025974 |
| PLAAT34 | 1,87027E-11 | 0,319029848 | 0,183 | 0,066 | 6,84277E-07 | FB | 1,247491381 | 0,801608745 | 2,772727273 |
| NPM13 | 2,1554E-11 | 0,283653057 | 0,811 | 0,635 | 7,88596E-07 | FB | 1,217273251 | 0,821508235 | 1,277165354 |
| SSR22 | 2,2025E-11 | 0,253405891 | 0,755 | 0,539 | 8,0583E-07 | FB | 1,192017892 | 0,838913582 | 1,400742115 |
| ISG153 | 2,24648E-11 | 0,375976171 | 0,215 | 0,089 | 8,21919E-07 | FB | 1,297717332 | 0,770583836 | 2,415730337 |
| TRIB11 | 2,25723E-11 | 0,300043625 | 0,172 | 0,058 | 8,25854E-07 | FB | 1,231181642 | 0,812227835 | 2,965517241 |
| SPHK11 | 2,35767E-11 | 0,341172894 | 0,161 | 0,052 | 8,62601E-07 | FB | 1,266786058 | 0,789399278 | 3,096153846 |
| AIG11 | 2,37173E-11 | 0,278080547 | 0,288 | 0,142 | 8,67745E-07 | FB | 1,212580517 | 0,824687504 | 2,028169014 |
| ICAM12 | 2,43616E-11 | 0,316289938 | 0,272 | 0,134 | 8,91319E-07 | FB | 1,245124443 | 0,803132575 | 2,029850746 |
| PID1 | 2,51242E-11 | -0,60071458 | 0,164 | 0,273 | 9,19221E-07 | FB | 0,659427254 | 1,516467501 | 0,600732601 |
| A1BG1 | 2,51451E-11 | 0,310365649 | 0,243 | 0,11 | 9,19982E-07 | FB | 1,240021942 | 0,806437343 | 2,209090909 |
| LRRFIP14 | 2,64773E-11 | 0,305293606 | 0,307 | 0,156 | 9,68724E-07 | FB | 1,235670086 | 0,809277502 | 1,967948718 |
| IER31 | 2,71431E-11 | 0,581160259 | 0,625 | 0,445 | 9,93085E-07 | FB | 1,496051936 | 0,668425993 | 1,404494382 |
| COX6A15 | 2,76503E-11 | 0,290386842 | 0,694 | 0,481 | 1,01164E-06 | FB | 1,222968159 | 0,817682777 | 1,442827443 |
| ABCA9 | 2,91102E-11 | -0,57901692 | 0,125 | 0,232 | 1,06506E-06 | FB | 0,669419777 | 1,493830977 | 0,538793103 |
| CRABP22 | 2,91163E-11 | 0,381673356 | 0,656 | 0,463 | 1,06528E-06 | FB | 1,302852134 | 0,767546811 | 1,416846652 |
| UQCR114 | 3,04724E-11 | 0,284911081 | 0,63 | 0,415 | 1,1149E-06 | FB | 1,218335171 | 0,820792196 | 1,518072289 |
| JTB7 | 3,11824E-11 | 0,283399355 | 0,497 | 0,306 | 1,14087E-06 | FB | 1,217059208 | 0,821652713 | 1,624183007 |
| ITGA8 | 3,4535E-11 | -0,361179836 | 0,037 | 0,111 | 1,26353E-06 | FB | 0,778527639 | 1,284475913 | 0,333333333 |
| PTTG1IP3 | 3,54756E-11 | 0,327671919 | 0,533 | 0,339 | 1,29795E-06 | FB | 1,254986564 | 0,79682128 | 1,572271386 |
| SPRY21 | 3,622E-11 | -0,744503481 | 0,166 | 0,273 | 1,32518E-06 | FB | 0,596873257 | 1,675397564 | 0,608058608 |
| TTC35 | 3,71289E-11 | 0,275302809 | 0,621 | 0,413 | 1,35844E-06 | FB | 1,210248083 | 0,826276872 | 1,503631961 |
| TXNIP7 | 3,72631E-11 | -0,895769572 | 0,608 | 0,626 | 1,36334E-06 | FB | 0,537460423 | 1,860602116 | 0,971246006 |
| TPI15 | 3,93791E-11 | 0,283712749 | 0,716 | 0,485 | 1,44076E-06 | FB | 1,217323617 | 0,821474246 | 1,47628866 |
| PDGFC1 | 3,98684E-11 | 0,287842116 | 0,225 | 0,097 | 1,45867E-06 | FB | 1,220812904 | 0,819126335 | 2,319587629 |

| Supplementary Table 4 - clusermarker | | | | | | | | | |
|---|---|---|---|---|---|---|---|---|---|
| | **p_val** | **avg_log2FC** | **pct.1** | **pct.2** | **p_val_adj** | **celltype** | **Foldchange_UP** | **Foldchange_DOWN** | **Ratio_pct1_pct2** |
| MCTS1 | 4,04698E-11 | 0,26196867 | 0,186 | 0,071 | 1,48067E-06 | FB | 1,199113873 | 0,833949154 | 2,61971831 |
| RPL317 | 4,15932E-11 | 0,272356926 | 0,967 | 0,86 | 1,52177E-06 | FB | 1,207779362 | 0,827965796 | 1,124418605 |
| CXCL13 | 4,20745E-11 | 1,42438815 | 0,134 | 0,039 | 1,53938E-06 | FB | 2,684006472 | 0,372577343 | 3,435897436 |
| STAT12 | 4,52927E-11 | 0,327217095 | 0,342 | 0,184 | 1,65713E-06 | FB | 1,254590979 | 0,797072525 | 1,858695652 |
| IGKV3-116 | 4,60456E-11 | 0,920963849 | 0,105 | 0,019 | 1,68467E-06 | FB | 1,893379818 | 0,528156047 | 5,526315789 |
| STK17B1 | 4,60653E-11 | 0,296763896 | 0,186 | 0,069 | 1,68539E-06 | FB | 1,228385933 | 0,814076402 | 2,695652174 |
| COMP | 4,71602E-11 | 1,026278961 | 0,249 | 0,121 | 1,72545E-06 | FB | 2,036764193 | 0,490974853 | 2,05785124 |
| MEIS21 | 4,95041E-11 | 0,261204574 | 0,303 | 0,153 | 1,81121E-06 | FB | 1,198478953 | 0,834390956 | 1,980392157 |
| CYGB | 6,40799E-11 | -0,533129826 | 0,149 | 0,256 | 2,34449E-06 | FB | 0,691053914 | 1,447065099 | 0,58203125 |
| TMEM119 | 6,43854E-11 | 0,282166528 | 0,364 | 0,203 | 2,35567E-06 | FB | 1,216019638 | 0,82235514 | 1,793103448 |
| ARF44 | 6,57222E-11 | 0,276908836 | 0,735 | 0,544 | 2,40458E-06 | FB | 1,211596097 | 0,825357561 | 1,351102941 |
| SNHG293 | 6,6184E-11 | 0,297020491 | 0,816 | 0,585 | 2,42148E-06 | FB | 1,228604431 | 0,813931624 | 1,394871795 |
| PTN | 6,75761E-11 | -0,555552218 | 0,413 | 0,527 | 2,47241E-06 | FB | 0,680396574 | 1,469731092 | 0,783681214 |
| IGFBP51 | 7,25695E-11 | 0,408091588 | 0,895 | 0,734 | 2,6551E-06 | FB | 1,326929377 | 0,75361961 | 1,219346049 |
| KLF103 | 7,86885E-11 | 0,325645504 | 0,41 | 0,248 | 2,87898E-06 | FB | 1,253225043 | 0,797941284 | 1,653225806 |
| EPB41L23 | 8,10856E-11 | -0,749163031 | 0,429 | 0,49 | 2,96668E-06 | FB | 0,594948613 | 1,680817433 | 0,875510204 |
| CTSB4 | 8,24879E-11 | 0,329919566 | 0,687 | 0,487 | 3,01798E-06 | FB | 1,256943295 | 0,795580838 | 1,410677618 |
| ISLR1 | 9,85334E-11 | 0,31577831 | 0,766 | 0,573 | 3,60504E-06 | FB | 1,244682958 | 0,803417444 | 1,336823735 |
| SPINT24 | 9,88425E-11 | 0,30257141 | 0,142 | 0,042 | 3,61635E-06 | FB | 1,23334072 | 0,810805955 | 3,380952381 |
| PRRX1 | 1,02925E-10 | -0,589580056 | 0,698 | 0,711 | 3,76572E-06 | FB | 0,664536314 | 1,50480866 | 0,981715893 |
| COX8A6 | 1,21749E-10 | 0,254424395 | 0,562 | 0,363 | 4,45444E-06 | FB | 1,192859722 | 0,838321541 | 1,548209366 |
| RPS166 | 1,24634E-10 | 0,299461393 | 0,931 | 0,789 | 4,55998E-06 | FB | 1,230684871 | 0,812555694 | 1,179974651 |
| LDHA4 | 1,26142E-10 | 0,295462808 | 0,755 | 0,56 | 4,61517E-06 | FB | 1,227278618 | 0,814810904 | 1,348214286 |
| LMCD1 | 1,26205E-10 | 0,279931272 | 0,311 | 0,16 | 4,61746E-06 | FB | 1,214137043 | 0,823630253 | 1,94375 |
| NDUFB106 | 1,33809E-10 | 0,256796518 | 0,472 | 0,287 | 4,89566E-06 | FB | 1,194822672 | 0,836944279 | 1,644599303 |
| CFB2 | 1,36354E-10 | 0,320174605 | 0,356 | 0,197 | 4,98877E-06 | FB | 1,24848164 | 0,800972932 | 1,807106599 |
| LSM22 | 1,36454E-10 | 0,260980178 | 0,272 | 0,134 | 4,99243E-06 | FB | 1,198292557 | 0,834520747 | 2,029850746 |
| INHBA | 1,51834E-10 | 0,340519985 | 0,536 | 0,352 | 5,55515E-06 | FB | 1,266212888 | 0,789756612 | 1,522727273 |
| RPS55 | 1,75117E-10 | 0,262695398 | 0,968 | 0,84 | 6,40702E-06 | FB | 1,199718054 | 0,833529175 | 1,152380952 |
| MZT2B3 | 1,80034E-10 | 0,252936017 | 0,497 | 0,306 | 6,58689E-06 | FB | 1,191629725 | 0,839186854 | 1,624183007 |
| PDGFD | 1,82127E-10 | 0,252302895 | 0,187 | 0,073 | 6,66349E-06 | FB | 1,191106897 | 0,839555209 | 2,561643836 |

| Supplementary Table 4 - clusermarker | | | | | | | | | |
|---|---|---|---|---|---|---|---|---|---|
| | **p_val** | **avg_log2FC** | **pct.1** | **pct.2** | **p_val_adj** | **celltype** | **Foldchange_UP** | **Foldchange_DOWN** | **Ratio_pct1_pct2** |
| MIR22HG2 | 1,9896E-10 | 0,2543907 | 0,283 | 0,144 | 7,27936E-06 | FB | 1,192831863 | 0,83834112 | 1,965277778 |
| ELOB5 | 2,28711E-10 | 0,289584369 | 0,737 | 0,534 | 8,36784E-06 | FB | 1,222288093 | 0,818137725 | 1,380149813 |
| HSPA1A6 | 2,29674E-10 | -0,727130501 | 0,719 | 0,755 | 8,40309E-06 | FB | 0,604104274 | 1,655343361 | 0,952317881 |
| PDIA33 | 2,60672E-10 | 0,271944277 | 0,736 | 0,535 | 9,5372E-06 | FB | 1,207433954 | 0,82820265 | 1,375700935 |
| RAMP2 | 2,68814E-10 | -0,595704313 | 0,086 | 0,177 | 9,83511E-06 | FB | 0,661721329 | 1,511210167 | 0,485875706 |
| YBX15 | 2,7458E-10 | 0,270551754 | 0,733 | 0,516 | 1,0046E-05 | FB | 1,206269073 | 0,829002436 | 1,420542636 |
| NID2 | 2,7591E-10 | 0,273192237 | 0,207 | 0,09 | 1,00947E-05 | FB | 1,208478861 | 0,827486547 | 2,3 |
| VAMP53 | 2,78214E-10 | 0,32104571 | 0,453 | 0,282 | 1,0179E-05 | FB | 1,249235706 | 0,800489448 | 1,606382979 |
| NDUFAF82 | 3,17871E-10 | 0,274400948 | 0,209 | 0,092 | 1,16299E-05 | FB | 1,209491766 | 0,826793558 | 2,27173913 |
| HAPLN3 | 3,21807E-10 | 0,251273351 | 0,139 | 0,044 | 1,1774E-05 | FB | 1,190257196 | 0,840154551 | 3,159090909 |
| NCOR22 | 3,49883E-10 | 0,304293879 | 0,283 | 0,145 | 1,28012E-05 | FB | 1,234814115 | 0,809838492 | 1,951724138 |
| SMARCC13 | 3,51436E-10 | 0,25859494 | 0,192 | 0,077 | 1,2858E-05 | FB | 1,196313032 | 0,835901619 | 2,493506494 |
| C19orf536 | 3,67627E-10 | 0,284397438 | 0,488 | 0,313 | 1,34504E-05 | FB | 1,217901483 | 0,821084475 | 1,559105431 |
| CD635 | 3,85312E-10 | 0,265249155 | 0,958 | 0,848 | 1,40974E-05 | FB | 1,201843592 | 0,832055025 | 1,129716981 |
| PDGFRB | 4,05993E-10 | 0,283699765 | 0,607 | 0,405 | 1,48541E-05 | FB | 1,217312661 | 0,821481639 | 1,498765432 |
| NPC23 | 4,30118E-10 | 0,277106183 | 0,789 | 0,595 | 1,57367E-05 | FB | 1,211761843 | 0,825244668 | 1,32605042 |
| C12orf575 | 4,30169E-10 | 0,282898182 | 0,901 | 0,711 | 1,57386E-05 | FB | 1,216636491 | 0,821938194 | 1,267229255 |
| OSR2 | 4,42175E-10 | 0,284455491 | 0,443 | 0,276 | 1,61778E-05 | FB | 1,217950492 | 0,821051436 | 1,605072464 |
| IFI304 | 4,53994E-10 | 0,273464936 | 0,115 | 0,029 | 1,66103E-05 | FB | 1,208707309 | 0,82733015 | 3,965517241 |
| CYBRD11 | 4,6248E-10 | -0,575758518 | 0,699 | 0,69 | 1,69207E-05 | FB | 0,670933406 | 1,490460888 | 1,013043478 |
| C17orf493 | 4,81702E-10 | 0,258014924 | 0,248 | 0,121 | 1,7624E-05 | FB | 1,195832167 | 0,83623775 | 2,049586777 |
| PLXNB22 | 4,81961E-10 | 0,294037736 | 0,19 | 0,079 | 1,76335E-05 | FB | 1,226066929 | 0,815616159 | 2,405063291 |
| SAT14 | 5,27129E-10 | 0,307790099 | 0,654 | 0,466 | 1,92861E-05 | FB | 1,237810186 | 0,807878309 | 1,403433476 |
| NR3C12 | 5,43867E-10 | -0,611680035 | 0,382 | 0,455 | 1,98985E-05 | FB | 0,654434162 | 1,528037591 | 0,83956044 |
| COTL14 | 5,47592E-10 | 0,266482412 | 0,12 | 0,032 | 2,00348E-05 | FB | 1,2028714 | 0,831344065 | 3,75 |
| SLC38A24 | 5,88863E-10 | 0,271672613 | 0,734 | 0,553 | 2,15447E-05 | FB | 1,207206612 | 0,828358617 | 1,327305606 |
| PLPP31 | 6,16923E-10 | -0,492095143 | 0,67 | 0,689 | 2,25714E-05 | FB | 0,710991815 | 1,406485952 | 0,972423803 |
| HNRNPU3 | 6,41663E-10 | 0,294077213 | 0,663 | 0,471 | 2,34765E-05 | FB | 1,226100479 | 0,815593842 | 1,407643312 |
| ATP6V0B5 | 6,74388E-10 | 0,257672109 | 0,358 | 0,208 | 2,46738E-05 | FB | 1,195548046 | 0,836436481 | 1,721153846 |
| PARP142 | 6,77588E-10 | 0,27910928 | 0,281 | 0,144 | 2,47909E-05 | FB | 1,213445472 | 0,824099659 | 1,951388889 |
| PPP1R15A3 | 6,7873E-10 | 0,296538018 | 0,619 | 0,442 | 2,48327E-05 | FB | 1,228193623 | 0,814203869 | 1,400452489 |

| Supplementary Table 4 - clusermarker | | | | | | | | | |
|---|---|---|---|---|---|---|---|---|---|
| | **p_val** | **avg_log2FC** | **pct.1** | **pct.2** | **p_val_adj** | **celltype** | **Foldchange_UP** | **Foldchange_DOWN** | **Ratio_pct1_pct2** |
| GADD45G1 | 7,65783E-10 | -0,821239921 | 0,154 | 0,253 | 2,80177E-05 | FB | 0,565955325 | 1,766923919 | 0,608695652 |
| ATP5F1E2 | 7,82923E-10 | 0,297893764 | 0,818 | 0,587 | 2,86448E-05 | FB | 1,229348338 | 0,813439095 | 1,393526405 |
| DOCK91 | 7,90168E-10 | -0,578253274 | 0,079 | 0,165 | 2,89099E-05 | FB | 0,669774208 | 1,493040473 | 0,478787879 |
| WNT111 | 8,46881E-10 | 0,262164211 | 0,128 | 0,039 | 3,09848E-05 | FB | 1,19927641 | 0,833836129 | 3,282051282 |
| PPL3 | 8,80785E-10 | -0,454268897 | 0,086 | 0,176 | 3,22253E-05 | FB | 0,729879954 | 1,37008832 | 0,488636364 |
| CALR5 | 8,83831E-10 | 0,33251528 | 0,756 | 0,589 | 3,23367E-05 | FB | 1,259206837 | 0,794150707 | 1,283531409 |
| RABAC15 | 9,54835E-10 | 0,262099407 | 0,718 | 0,519 | 3,49345E-05 | FB | 1,199222542 | 0,833873585 | 1,383429672 |
| TPST1 | 1,0865E-09 | 0,266893015 | 0,302 | 0,163 | 3,97516E-05 | FB | 1,203213797 | 0,831107491 | 1,852760736 |
| CNN11 | 1,11144E-09 | 0,306937507 | 0,122 | 0,035 | 4,06644E-05 | FB | 1,237078891 | 0,808355883 | 3,485714286 |
| IFI63 | 1,14044E-09 | 0,290555601 | 0,411 | 0,245 | 4,17252E-05 | FB | 1,223111223 | 0,817587134 | 1,67755102 |
| SERPINA32 | 1,28318E-09 | -0,784026451 | 0,174 | 0,273 | 4,69478E-05 | FB | 0,580743718 | 1,721929947 | 0,637362637 |
| FOXD12 | 1,37378E-09 | -0,348510097 | 0,036 | 0,102 | 5,02626E-05 | FB | 0,785394774 | 1,27324504 | 0,352941176 |
| RPS266 | 1,4253E-09 | -0,318332219 | 0,911 | 0,871 | 5,21475E-05 | FB | 0,801996464 | 1,24688829 | 1,045924225 |
| NTM1 | 1,45555E-09 | 0,328804007 | 0,16 | 0,061 | 5,32542E-05 | FB | 1,255971743 | 0,796196256 | 2,62295082 |
| VIM-AS13 | 1,66769E-09 | 0,260214916 | 0,148 | 0,053 | 6,10156E-05 | FB | 1,197657105 | 0,834963527 | 2,79245283 |
| TAGLN24 | 1,67587E-09 | -0,443171408 | 0,8 | 0,774 | 6,13149E-05 | FB | 0,73551598 | 1,359589766 | 1,033591731 |
| SGK12 | 1,90244E-09 | 0,383527437 | 0,315 | 0,176 | 6,96044E-05 | FB | 1,304527573 | 0,76656103 | 1,789772727 |
| ARF55 | 2,08853E-09 | 0,263161338 | 0,461 | 0,287 | 7,64131E-05 | FB | 1,200105584 | 0,833260018 | 1,606271777 |
| NIBAN13 | 2,27436E-09 | -0,470764208 | 0,09 | 0,176 | 8,3212E-05 | FB | 0,721582268 | 1,385843367 | 0,511363636 |
| CILP | 2,34467E-09 | -0,629989922 | 0,04 | 0,106 | 8,57845E-05 | FB | 0,646180929 | 1,547554183 | 0,377358491 |
| CMTM32 | 2,79421E-09 | 0,295791051 | 0,386 | 0,234 | 0,000102232 | FB | 1,227557881 | 0,814625539 | 1,64957265 |
| ANXA42 | 4,21893E-09 | -0,630537863 | 0,432 | 0,481 | 0,000154358 | FB | 0,645935554 | 1,548142061 | 0,898128898 |
| NTN41 | 4,21914E-09 | -0,593197853 | 0,168 | 0,261 | 0,000154366 | FB | 0,662871967 | 1,508586952 | 0,643678161 |
| C7orf502 | 4,26396E-09 | 0,251913173 | 0,23 | 0,113 | 0,000156006 | FB | 1,190785181 | 0,839782033 | 2,03539823 |
| ADAMTSL3 | 4,51657E-09 | -0,488219932 | 0,092 | 0,179 | 0,000165248 | FB | 0,712904171 | 1,402713071 | 0,51396648 |
| IGLV2-144 | 4,55315E-09 | 1,839436494 | 0,165 | 0,066 | 0,000166586 | FB | 3,578702196 | 0,279430907 | 2,5 |
| PTHLH | 4,90885E-09 | -0,615888407 | 0,133 | 0,223 | 0,0001796 | FB | 0,652527945 | 1,532501417 | 0,596412556 |
| PPDPF6 | 4,93352E-09 | 0,280073561 | 0,883 | 0,719 | 0,000180503 | FB | 1,214256796 | 0,823549024 | 1,228094576 |
| MXRA5 | 5,0194E-09 | 0,322139477 | 0,682 | 0,506 | 0,000183645 | FB | 1,250183162 | 0,799882794 | 1,347826087 |
| HINT17 | 5,40596E-09 | 0,259857368 | 0,674 | 0,479 | 0,000197788 | FB | 1,197360322 | 0,835170484 | 1,407098121 |
| FKBP112 | 5,66735E-09 | 0,269249408 | 0,289 | 0,158 | 0,000207351 | FB | 1,205180645 | 0,829751129 | 1,829113924 |

| Supplementary Table 4 - clusermarker | | | | | | | | | |
|---|---|---|---|---|---|---|---|---|---|
| | **p_val** | **avg_log2FC** | **pct.1** | **pct.2** | **p_val_adj** | **celltype** | **Foldchange_UP** | **Foldchange_DOWN** | **Ratio_pct1_pct2** |
| DDIT42 | 7,46788E-09 | -0,675399958 | 0,398 | 0,477 | 0,000273227 | FB | 0,626158606 | 1,597039458 | 0,834381551 |
| OPHN11 | 7,70352E-09 | -0,5195397 | 0,075 | 0,152 | 0,000281849 | FB | 0,697594369 | 1,433497809 | 0,493421053 |
| CREB5 | 8,29798E-09 | -0,539221497 | 0,06 | 0,132 | 0,000303598 | FB | 0,688142142 | 1,453188141 | 0,454545455 |
| EMC101 | 9,52605E-09 | 0,272334364 | 0,269 | 0,15 | 0,000348529 | FB | 1,207760474 | 0,827978744 | 1,793333333 |
| ALDH22 | 1,28307E-08 | -0,501152131 | 0,366 | 0,452 | 0,000469438 | FB | 0,706542314 | 1,415343399 | 0,809734513 |
| CALCRL | 1,29792E-08 | -0,473548171 | 0,055 | 0,124 | 0,000474871 | FB | 0,720191176 | 1,388520206 | 0,443548387 |
| HSPG2 | 1,40999E-08 | 0,281933597 | 0,505 | 0,329 | 0,000515873 | FB | 1,215823321 | 0,822487924 | 1,534954407 |
| CADPS21 | 1,55608E-08 | 0,386124401 | 0,242 | 0,132 | 0,000569323 | FB | 1,306877939 | 0,765182402 | 1,833333333 |
| KCNK15 | 1,64941E-08 | 0,328902299 | 0,157 | 0,066 | 0,00060347 | FB | 1,256057316 | 0,796142013 | 2,378787879 |
| CNBP5 | 1,76779E-08 | -0,398041059 | 0,696 | 0,689 | 0,000646783 | FB | 0,758888028 | 1,317717454 | 1,010159652 |
| TCEAL7 | 1,7814E-08 | -0,332666591 | 0,045 | 0,111 | 0,000651761 | FB | 0,79406742 | 1,259338911 | 0,405405405 |
| PSMA75 | 2,22883E-08 | 0,264845599 | 0,614 | 0,431 | 0,000815463 | FB | 1,201507454 | 0,832287803 | 1,424593968 |
| CELF22 | 2,32739E-08 | -0,497058376 | 0,344 | 0,429 | 0,000851521 | FB | 0,708550027 | 1,411332949 | 0,801864802 |
| LAMP5 | 2,66573E-08 | 0,39406027 | 0,103 | 0,031 | 0,00097531 | FB | 1,314086523 | 0,760984899 | 3,322580645 |
| HLA-C5 | 3,12965E-08 | 0,254874656 | 0,931 | 0,829 | 0,001145045 | FB | 1,193232068 | 0,838059944 | 1,123039807 |
| CRIP23 | 4,65982E-08 | -0,503902078 | 0,358 | 0,44 | 0,00170489 | FB | 0,705196844 | 1,418043783 | 0,813636364 |
| CLEC14A | 4,80286E-08 | -0,517377318 | 0,073 | 0,145 | 0,001757221 | FB | 0,698640742 | 1,431350822 | 0,503448276 |
| EZR4 | 5,12032E-08 | -0,614150422 | 0,157 | 0,24 | 0,001873372 | FB | 0,653314506 | 1,530656355 | 0,654166667 |
| PLAU | 5,17309E-08 | 0,25736539 | 0,159 | 0,068 | 0,001892679 | FB | 1,195293897 | 0,836614328 | 2,338235294 |
| S100A44 | 5,85295E-08 | -0,354620714 | 0,956 | 0,937 | 0,00214142 | FB | 0,782075225 | 1,278649379 | 1,020277481 |
| ESD3 | 6,17252E-08 | -0,552556579 | 0,495 | 0,524 | 0,002258341 | FB | 0,68181083 | 1,466682482 | 0,944656489 |
| COL4A2 | 6,27729E-08 | 0,305116824 | 0,303 | 0,181 | 0,002296672 | FB | 1,235518681 | 0,809376674 | 1,674033149 |
| TSPO6 | 6,36991E-08 | 0,282996643 | 0,626 | 0,465 | 0,00233056 | FB | 1,216719527 | 0,8218821 | 1,346236559 |
| BAX6 | 7,96095E-08 | 0,267868105 | 0,265 | 0,152 | 0,002912671 | FB | 1,204027301 | 0,830545951 | 1,743421053 |
| GARS11 | 8,03787E-08 | 0,254817923 | 0,232 | 0,126 | 0,002940814 | FB | 1,193185146 | 0,8380929 | 1,841269841 |
| RPS93 | 8,22432E-08 | 0,254847386 | 0,961 | 0,852 | 0,003009031 | FB | 1,193209514 | 0,838075785 | 1,127934272 |
| IER5L4 | 8,6144E-08 | 0,265082173 | 0,383 | 0,247 | 0,00315175 | FB | 1,201704494 | 0,832151336 | 1,550607287 |
| LINC01140 | 8,74378E-08 | -0,439318587 | 0,08 | 0,153 | 0,003199086 | FB | 0,737482854 | 1,355963728 | 0,522875817 |
| EFEMP12 | 9,52177E-08 | 0,459144653 | 0,6 | 0,471 | 0,003483731 | FB | 1,374726527 | 0,727417403 | 1,27388535 |
| RPL86 | 9,93482E-08 | 0,285776729 | 0,964 | 0,865 | 0,003634854 | FB | 1,219066417 | 0,820299851 | 1,114450867 |
| APOE3 | 1,02981E-07 | 0,801240882 | 0,136 | 0,055 | 0,003767784 | FB | 1,742599316 | 0,573855384 | 2,472727273 |

| Supplementary Table 4 - clusermarker | | | | | | | | | |
|---|---|---|---|---|---|---|---|---|---|
| | **p_val** | **avg_log2FC** | **pct.1** | **pct.2** | **p_val_adj** | **celltype** | **Foldchange_UP** | **Foldchange_DOWN** | **Ratio_pct1_pct2** |
| LRRN4CL | 1,08582E-07 | -0,590393358 | 0,274 | 0,348 | 0,003972708 | FB | 0,664161795 | 1,505657216 | 0,787356322 |
| IGSF10 | 1,54871E-07 | -0,388512287 | 0,154 | 0,245 | 0,005666248 | FB | 0,763916953 | 1,309042817 | 0,628571429 |
| MGLL | 1,69349E-07 | -0,535098585 | 0,199 | 0,276 | 0,006195956 | FB | 0,690111517 | 1,44904117 | 0,721014493 |
| C1QTNF2 | 1,7036E-07 | -0,428065555 | 0,121 | 0,2 | 0,006232978 | FB | 0,743257718 | 1,345428343 | 0,605 |
| CKB1 | 1,79828E-07 | -0,523725648 | 0,174 | 0,258 | 0,006579355 | FB | 0,695573248 | 1,437663112 | 0,674418605 |
| NFKBIA6 | 1,82459E-07 | 0,269884641 | 0,801 | 0,687 | 0,006675612 | FB | 1,205711414 | 0,829385861 | 1,165938865 |
| IL331 | 1,94604E-07 | -0,458037963 | 0,248 | 0,344 | 0,007119974 | FB | 0,727975618 | 1,373672379 | 0,720930233 |
| ADRA2A | 2,34946E-07 | 0,250516894 | 0,112 | 0,04 | 0,00859598 | FB | 1,189633265 | 0,84059519 | 2,8 |
| METTL7A2 | 2,68642E-07 | -0,616702979 | 0,224 | 0,303 | 0,009828797 | FB | 0,65215962 | 1,533366939 | 0,739273927 |
| IL16 | 2,87932E-07 | -0,559799774 | 0,155 | 0,232 | 0,010534573 | FB | 0,678396309 | 1,474064624 | 0,668103448 |
| NDUFA4L2 | 3,20992E-07 | -0,547718059 | 0,102 | 0,177 | 0,011744123 | FB | 0,684101331 | 1,461771751 | 0,576271186 |
| LHFPL6 | 3,42094E-07 | -0,569505887 | 0,433 | 0,473 | 0,01251621 | FB | 0,673847537 | 1,484015219 | 0,915433404 |
| GDF10 | 4,98469E-07 | -0,600582169 | 0,165 | 0,242 | 0,0182375 | FB | 0,659487779 | 1,516328325 | 0,681818182 |
| SOX44 | 5,5494E-07 | 0,267907345 | 0,607 | 0,444 | 0,020303584 | FB | 1,20406005 | 0,830523361 | 1,367117117 |
| FXYD6 | 5,73401E-07 | -0,517952198 | 0,158 | 0,235 | 0,020979036 | FB | 0,698362405 | 1,431921295 | 0,672340426 |
| PLTP3 | 6,17475E-07 | 0,252998917 | 0,634 | 0,463 | 0,022591565 | FB | 1,19168168 | 0,839150267 | 1,369330454 |
| PIAS13 | 8,77149E-07 | -0,518140678 | 0,236 | 0,306 | 0,032092238 | FB | 0,698271174 | 1,43210838 | 0,77124183 |
| ITIH5 | 8,99012E-07 | -0,508416163 | 0,175 | 0,252 | 0,032892154 | FB | 0,702993784 | 1,422487685 | 0,694444444 |
| MFAP5 | 9,94751E-07 | -1,178491427 | 0,419 | 0,473 | 0,036394961 | FB | 0,441813245 | 2,263399779 | 0,885835095 |
| HSPA1B5 | 1,08249E-06 | -0,659623755 | 0,626 | 0,655 | 0,039604891 | FB | 0,633043369 | 1,579670603 | 0,955725191 |
| JUNB4 | 1,10523E-06 | 0,284097142 | 0,877 | 0,782 | 0,040437036 | FB | 1,217648005 | 0,821255401 | 1,121483376 |
| SRGN4 | 1,72884E-06 | 0,303977606 | 0,106 | 0,04 | 0,063252989 | FB | 1,234543444 | 0,810016047 | 2,65 |
| TMBIM61 | 1,78026E-06 | -0,453460859 | 0,712 | 0,679 | 0,065134363 | FB | 0,730288866 | 1,369321163 | 1,048600884 |
| OAF | 1,79188E-06 | -0,512223498 | 0,205 | 0,279 | 0,065559605 | FB | 0,701140998 | 1,42624665 | 0,734767025 |
| CCN31 | 1,91718E-06 | -0,926804364 | 0,106 | 0,174 | 0,070143877 | FB | 0,526022216 | 1,901060392 | 0,609195402 |
| CLU1 | 2,36507E-06 | -0,470806196 | 0,563 | 0,624 | 0,086530701 | FB | 0,721561268 | 1,385883701 | 0,90224359 |
| PLCB4 | 2,4562E-06 | -0,291251828 | 0,05 | 0,105 | 0,089865016 | FB | 0,817192672 | 1,223701624 | 0,476190476 |
| NR2F1 | 2,94361E-06 | -0,658935366 | 0,314 | 0,371 | 0,107697772 | FB | 0,633345501 | 1,578917035 | 0,846361186 |
| EGR15 | 3,03684E-06 | 0,308339806 | 0,808 | 0,732 | 0,111108768 | FB | 1,238281916 | 0,807570543 | 1,103825137 |
| PMP221 | 3,36145E-06 | -0,524651373 | 0,728 | 0,669 | 0,122985517 | FB | 0,695127067 | 1,438585904 | 1,08819133 |
| FAM102B2 | 3,84903E-06 | -0,419579666 | 0,113 | 0,179 | 0,14082456 | FB | 0,747642421 | 1,337537802 | 0,631284916 |

| Supplementary Table 4 - clusermarker | | | | | | | | | |
|---|---|---|---|---|---|---|---|---|---|
| | **p_val** | **avg_log2FC** | **pct.1** | **pct.2** | **p_val_adj** | **celltype** | **Foldchange_UP** | **Foldchange_DOWN** | **Ratio_pct1_pct2** |
| EBF1 | 4,53949E-06 | -0,630907058 | 0,167 | 0,234 | 0,16608622 | FB | 0,645770276 | 1,548538292 | 0,713675214 |
| C7 | 5,65053E-06 | -0,7678379 | 0,179 | 0,256 | 0,20673608 | FB | 0,58729697 | 1,702716089 | 0,69921875 |
| RBMS31 | 5,77103E-06 | -0,493871371 | 0,382 | 0,432 | 0,211144587 | FB | 0,71011699 | 1,408218666 | 0,884259259 |
| TSPAN81 | 5,90557E-06 | -0,588986615 | 0,143 | 0,211 | 0,21606709 | FB | 0,664809722 | 1,504189796 | 0,677725118 |
| AKR1C22 | 6,0859E-06 | -0,583491808 | 0,16 | 0,231 | 0,222664844 | FB | 0,667346617 | 1,49847167 | 0,692640693 |
| LAPTM4A1 | 6,36586E-06 | -0,290180565 | 0,894 | 0,866 | 0,232907739 | FB | 0,817799698 | 1,22279331 | 1,032332564 |
| ADAMTSL4 | 6,99863E-06 | -0,386031962 | 0,089 | 0,148 | 0,256059041 | FB | 0,765231431 | 1,306794205 | 0,601351351 |
| PHGDH2 | 8,84304E-06 | -0,402804843 | 0,115 | 0,182 | 0,323540365 | FB | 0,75638631 | 1,322075752 | 0,631868132 |
| SFRP4 | 9,67962E-06 | 0,456961258 | 0,758 | 0,64 | 0,354148285 | FB | 1,37264757 | 0,72851912 | 1,184375 |
| NRP11 | 1,11504E-05 | -0,573214796 | 0,287 | 0,344 | 0,407961383 | FB | 0,672117421 | 1,487835263 | 0,834302326 |
| TSC22D32 | 1,33627E-05 | -0,497461111 | 0,67 | 0,666 | 0,488899429 | FB | 0,70835226 | 1,411726984 | 1,006006006 |
| FGF71 | 1,37919E-05 | -0,54120441 | 0,642 | 0,634 | 0,504605656 | FB | 0,687196974 | 1,455186849 | 1,012618297 |
| RPL386 | 1,78815E-05 | -0,382167925 | 0,724 | 0,668 | 0,654230979 | FB | 0,767283734 | 1,30329884 | 1,083832335 |
| PCDH9 | 1,98009E-05 | -0,36216437 | 0,073 | 0,129 | 0,724455752 | FB | 0,777996533 | 1,285352772 | 0,565891473 |
| ANK2 | 2,00372E-05 | -0,48863508 | 0,287 | 0,347 | 0,733101224 | FB | 0,712699056 | 1,403116773 | 0,827089337 |
| RMRP1 | 2,26011E-05 | -0,437896868 | 0,068 | 0,121 | 0,826904721 | FB | 0,738209972 | 1,354628138 | 0,561983471 |
| ADAM33 | 2,29302E-05 | -0,604311474 | 0,263 | 0,319 | 0,838947414 | FB | 0,657785233 | 1,52025304 | 0,824451411 |
| GFRA1 | 2,99692E-05 | -0,459854007 | 0,289 | 0,348 | 1 | FB | 0,72705983 | 1,375402628 | 0,83045977 |
| EIF4A24 | 3,94433E-05 | -0,43337149 | 0,668 | 0,624 | 1 | FB | 0,74052919 | 1,350385661 | 1,070512821 |
| SDC2 | 5,06865E-05 | -0,477573961 | 0,388 | 0,423 | 1 | FB | 0,718184309 | 1,392400234 | 0,917257683 |
| EGFR1 | 5,17147E-05 | -0,533987337 | 0,171 | 0,227 | 1 | FB | 0,690643286 | 1,447925463 | 0,753303965 |
| ZBTB203 | 5,64154E-05 | -0,375973683 | 0,635 | 0,616 | 1 | FB | 0,770585165 | 1,297715094 | 1,030844156 |
| FGF18 | 5,66351E-05 | 0,525702312 | 0,152 | 0,087 | 1 | FB | 1,439634231 | 0,694620882 | 1,747126437 |
| CDO1 | 5,86904E-05 | -0,57861654 | 0,227 | 0,284 | 1 | FB | 0,669605582 | 1,493416462 | 0,799295775 |
| WBP21 | 8,09621E-05 | -0,469035265 | 0,216 | 0,271 | 1 | FB | 0,722447539 | 1,384183551 | 0,79704797 |
| RBP4 | 8,21548E-05 | -0,418575626 | 0,08 | 0,132 | 1 | FB | 0,748162922 | 1,336607269 | 0,606060606 |
| RHOJ | 8,59932E-05 | -0,360976908 | 0,12 | 0,179 | 1 | FB | 0,778637154 | 1,284295252 | 0,670391061 |
| FKBP52 | 0,000108925 | -0,348173452 | 0,118 | 0,173 | 1 | FB | 0,785578063 | 1,27294797 | 0,682080925 |
| PLP25 | 0,000114644 | -0,522621998 | 0,369 | 0,403 | 1 | FB | 0,696105559 | 1,436563732 | 0,915632754 |
| ZFP36L22 | 0,000129478 | -0,301366491 | 0,861 | 0,827 | 1 | FB | 0,811483412 | 1,23231108 | 1,041112455 |
| MYADM4 | 0,000139444 | -0,370460772 | 0,65 | 0,635 | 1 | FB | 0,773535403 | 1,292765652 | 1,023622047 |

| Supplementary Table 4 - clusermarker | | | | | | | | | |
|---|---|---|---|---|---|---|---|---|---|
| | **p_val** | **avg_log2FC** | **pct.1** | **pct.2** | **p_val_adj** | **celltype** | **Foldchange_UP** | **Foldchange_DOWN** | **Ratio_pct1_pct2** |
| SDCBP5 | 0,000144519 | -0,504614853 | 0,426 | 0,44 | 1 | FB | 0,704848521 | 1,418744553 | 0,968181818 |
| MBNL14 | 0,000180183 | -0,355354809 | 0,675 | 0,645 | 1 | FB | 0,781677378 | 1,279300167 | 1,046511628 |
| CAV11 | 0,000202649 | -0,544923588 | 0,507 | 0,513 | 1 | FB | 0,685427705 | 1,458943069 | 0,988304094 |
| SCARA31 | 0,000213844 | -0,409611914 | 0,13 | 0,184 | 1 | FB | 0,752825857 | 1,328328445 | 0,706521739 |
| EPHX11 | 0,0002235 | -0,370898846 | 0,571 | 0,581 | 1 | FB | 0,773300555 | 1,29315826 | 0,982788296 |
| GPSM2 | 0,000228869 | -0,307309548 | 0,064 | 0,11 | 1 | FB | 0,808147452 | 1,237397949 | 0,581818182 |
| C1orf211 | 0,000230708 | -0,406278307 | 0,188 | 0,242 | 1 | FB | 0,754567408 | 1,325262646 | 0,776859504 |
| AKR1C13 | 0,000252501 | -0,414423002 | 0,193 | 0,252 | 1 | FB | 0,750319521 | 1,332765538 | 0,765873016 |
| SEPTIN71 | 0,000280674 | -0,305534793 | 0,76 | 0,711 | 1 | FB | 0,80914222 | 1,23587668 | 1,068917018 |
| FXYD12 | 0,000334872 | -0,35865176 | 0,473 | 0,513 | 1 | FB | 0,779893072 | 1,282227059 | 0,92202729 |
| FMNL2 | 0,000364063 | -0,355630424 | 0,141 | 0,192 | 1 | FB | 0,781528059 | 1,27954459 | 0,734375 |
| MAFB3 | 0,000369203 | -0,407943441 | 0,441 | 0,471 | 1 | FB | 0,753697002 | 1,326793124 | 0,936305732 |
| DCLK1 | 0,000387184 | -0,441095746 | 0,149 | 0,203 | 1 | FB | 0,736574958 | 1,357635078 | 0,733990148 |
| TGFBR3 | 0,000420918 | -0,365021933 | 0,246 | 0,302 | 1 | FB | 0,77645707 | 1,287901209 | 0,814569536 |
| COL21A1 | 0,00046906 | -0,352168156 | 0,082 | 0,127 | 1 | FB | 0,78340587 | 1,276477542 | 0,645669291 |
| GNAI11 | 0,000478889 | -0,293489811 | 0,062 | 0,103 | 1 | FB | 0,815925983 | 1,225601366 | 0,601941748 |
| PRELP | 0,000503043 | -0,303849513 | 0,703 | 0,684 | 1 | FB | 0,810087969 | 1,234433837 | 1,027777778 |
| CCL25 | 0,000562687 | -0,323104729 | 0,417 | 0,305 | 1 | FB | 0,799347802 | 1,251019891 | 1,367213115 |
| PGF | 0,000641614 | -0,291733332 | 0,06 | 0,1 | 1 | FB | 0,816919977 | 1,224110107 | 0,6 |
| MXRA73 | 0,000667813 | -0,409313368 | 0,486 | 0,477 | 1 | FB | 0,75298166 | 1,328053594 | 1,018867925 |
| COX7C4 | 0,000668112 | -0,297392926 | 0,781 | 0,694 | 1 | FB | 0,813721533 | 1,228921638 | 1,125360231 |
| CCDC152 | 0,000673986 | -0,344815581 | 0,084 | 0,129 | 1 | FB | 0,787408624 | 1,26998863 | 0,651162791 |
| ARHGAP61 | 0,000718775 | -0,265216403 | 0,074 | 0,118 | 1 | FB | 0,832073915 | 1,201816307 | 0,627118644 |
| OLFML1 | 0,000740545 | -0,429956032 | 0,338 | 0,366 | 1 | FB | 0,742284407 | 1,347192519 | 0,923497268 |
| ADD32 | 0,000903899 | -0,360337258 | 0,596 | 0,563 | 1 | FB | 0,778982456 | 1,283725959 | 1,058614565 |
| RASL12 | 0,000951928 | -0,308343563 | 0,097 | 0,142 | 1 | FB | 0,807568441 | 1,23828514 | 0,683098592 |
| PLBD1 | 0,001036009 | -0,341345103 | 0,118 | 0,163 | 1 | FB | 0,789305057 | 1,266937278 | 0,72392638 |
| PLEKHA23 | 0,001080074 | -0,288510041 | 0,064 | 0,103 | 1 | FB | 0,818747192 | 1,221378234 | 0,621359223 |
| ATP2B41 | 0,001145789 | -0,477093054 | 0,307 | 0,335 | 1 | FB | 0,718423748 | 1,391936169 | 0,91641791 |
| PTGES | 0,001147669 | -0,279438748 | 0,11 | 0,156 | 1 | FB | 0,823911481 | 1,213722618 | 0,705128205 |
| C33 | 0,001297464 | 0,379860513 | 0,749 | 0,692 | 1 | FB | 1,301216042 | 0,76851189 | 1,082369942 |

| Supplementary Table 4 - clusermarker | | | | | | | | | |
|---|---|---|---|---|---|---|---|---|---|
| | **p_val** | **avg_log2FC** | **pct.1** | **pct.2** | **p_val_adj** | **celltype** | **Foldchange_UP** | **Foldchange_DOWN** | **Ratio_pct1_pct2** |
| TSHZ21 | 0,001337412 | -0,38208188 | 0,316 | 0,353 | 1 | FB | 0,767329497 | 1,303221111 | 0,895184136 |
| SLC43A32 | 0,00143198 | -0,33722355 | 0,099 | 0,142 | 1 | FB | 0,791563201 | 1,263323002 | 0,697183099 |
| TMEM109 | 0,00150635 | -0,46174162 | 0,45 | 0,447 | 1 | FB | 0,726109171 | 1,377203374 | 1,006711409 |
| SRP142 | 0,001791685 | -0,276388269 | 0,859 | 0,779 | 1 | FB | 0,825655428 | 1,211158996 | 1,102695764 |
| SYNE3 | 0,001792506 | -0,253287122 | 0,101 | 0,145 | 1 | FB | 0,838982648 | 1,191919764 | 0,696551724 |
| MALAT14 | 0,001854204 | -0,271197057 | 0,977 | 0,979 | 1 | FB | 0,828631715 | 1,206808745 | 0,997957099 |
| RNF131 | 0,001902922 | -0,455753143 | 0,289 | 0,321 | 1 | FB | 0,729129439 | 1,371498594 | 0,900311526 |
| IFIT1 | 0,001956593 | -0,367686445 | 0,125 | 0,168 | 1 | FB | 0,775024356 | 1,29028203 | 0,744047619 |
| CD554 | 0,002404713 | -0,318928606 | 0,551 | 0,402 | 1 | FB | 0,801665 | 1,24740384 | 1,370646766 |
| ACTB7 | 0,00243919 | 0,315787054 | 0,987 | 0,963 | 1 | FB | 1,244690501 | 0,803412574 | 1,024922118 |
| SLC9A9 | 0,002447276 | -0,391609391 | 0,108 | 0,148 | 1 | FB | 0,762278774 | 1,311856022 | 0,72972973 |
| NDRG22 | 0,002476453 | -0,314539324 | 0,131 | 0,176 | 1 | FB | 0,804107714 | 1,243614483 | 0,744318182 |
| GOLIM4 | 0,00263597 | -0,455847228 | 0,459 | 0,447 | 1 | FB | 0,72908189 | 1,371588038 | 1,026845638 |
| RABGAP1L3 | 0,00268997 | -0,414668111 | 0,184 | 0,224 | 1 | FB | 0,750192055 | 1,33299199 | 0,821428571 |
| PROCR1 | 0,002709426 | -0,331178756 | 0,069 | 0,105 | 1 | FB | 0,794886755 | 1,258040839 | 0,657142857 |
| U2SURP3 | 0,003081039 | -0,465665705 | 0,365 | 0,377 | 1 | FB | 0,72413686 | 1,380954423 | 0,968169761 |
| CEBPD2 | 0,003130004 | -0,314509811 | 0,719 | 0,695 | 1 | FB | 0,804124164 | 1,243589043 | 1,034532374 |
| SRSF54 | 0,003202749 | -0,415682094 | 0,579 | 0,545 | 1 | FB | 0,749664976 | 1,333929198 | 1,062385321 |
| SOBP1 | 0,003288419 | -0,274471065 | 0,1 | 0,14 | 1 | FB | 0,826753375 | 1,20955055 | 0,714285714 |
| SGCE1 | 0,003421468 | -0,443076181 | 0,359 | 0,377 | 1 | FB | 0,73556453 | 1,359500028 | 0,952254642 |
| CEP1261 | 0,003463261 | -0,350278581 | 0,138 | 0,177 | 1 | FB | 0,784432611 | 1,274806765 | 0,779661017 |
| GSTM5 | 0,004067751 | -0,371914745 | 0,17 | 0,211 | 1 | FB | 0,772756214 | 1,29406918 | 0,805687204 |
| CAPG4 | 0,004089101 | -0,438973272 | 0,332 | 0,352 | 1 | FB | 0,737659395 | 1,355639211 | 0,943181818 |
| FAXDC2 | 0,004391734 | -0,262784029 | 0,114 | 0,155 | 1 | FB | 0,833477969 | 1,199791761 | 0,735483871 |
| TACC13 | 0,004574218 | -0,366890805 | 0,542 | 0,51 | 1 | FB | 0,775451897 | 1,289570642 | 1,062745098 |
| SSPN | 0,004666394 | -0,378153685 | 0,557 | 0,523 | 1 | FB | 0,769421642 | 1,299677505 | 1,06500956 |
| SYNE11 | 0,004855946 | -0,443276388 | 0,477 | 0,453 | 1 | FB | 0,735462461 | 1,359688703 | 1,052980132 |
| ID25 | 0,004996894 | -0,406384161 | 0,562 | 0,55 | 1 | FB | 0,754512046 | 1,325359887 | 1,021818182 |
| RNF24 | 0,005123688 | -0,343950767 | 0,187 | 0,224 | 1 | FB | 0,787880773 | 1,269227572 | 0,834821429 |
| IGF1 | 0,005190912 | -0,442733472 | 0,418 | 0,447 | 1 | FB | 0,735739283 | 1,35917712 | 0,935123043 |
| HSD17B111 | 0,005775501 | -0,483046095 | 0,227 | 0,256 | 1 | FB | 0,7154654 | 1,397691628 | 0,88671875 |

| Supplementary Table 4 - clusermarker | | | | | | | | | |
|---|---|---|---|---|---|---|---|---|---|
| | **p_val** | **avg_log2FC** | **pct.1** | **pct.2** | **p_val_adj** | **celltype** | **Foldchange_UP** | **Foldchange_DOWN** | **Ratio_pct1_pct2** |
| ARHGAP211 | 0,006125327 | -0,455401147 | 0,357 | 0,368 | 1 | FB | 0,729307357 | 1,371164009 | 0,970108696 |
| PDGFRL | 0,006239649 | -0,539342223 | 0,498 | 0,477 | 1 | FB | 0,68808456 | 1,453309749 | 1,044025157 |
| PTGIS | 0,006382653 | -0,266008817 | 0,164 | 0,206 | 1 | FB | 0,831617016 | 1,202476598 | 0,796116505 |
| FMO21 | 0,006721628 | -0,613655392 | 0,323 | 0,35 | 1 | FB | 0,653538715 | 1,530131233 | 0,922857143 |
| GSTM12 | 0,007905301 | -0,279410513 | 0,131 | 0,168 | 1 | FB | 0,823927606 | 1,213698864 | 0,779761905 |
| DPT | 0,008556497 | -0,621650554 | 0,702 | 0,619 | 1 | FB | 0,649926936 | 1,538634491 | 1,134087237 |
| HAS2 | 0,008898023 | -0,262641766 | 0,09 | 0,124 | 1 | FB | 0,833560162 | 1,199673456 | 0,725806452 |
| CCDC1862 | 0,010105755 | -0,422862826 | 0,177 | 0,206 | 1 | FB | 0,745942936 | 1,34058512 | 0,859223301 |
| SOX152 | 0,010215185 | -0,335596184 | 0,098 | 0,132 | 1 | FB | 0,792456591 | 1,261898773 | 0,742424242 |
| AIMP1 | 0,010694312 | -0,425956347 | 0,276 | 0,297 | 1 | FB | 0,74434515 | 1,343462774 | 0,929292929 |
| IFITM14 | 0,011653548 | -0,253236483 | 0,741 | 0,697 | 1 | FB | 0,839012097 | 1,191877928 | 1,06312769 |
| SLC29A1 | 0,012080571 | -0,399233665 | 0,148 | 0,184 | 1 | FB | 0,758260951 | 1,318807197 | 0,804347826 |
| AXIN2 | 0,012681539 | -0,273178862 | 0,102 | 0,134 | 1 | FB | 0,827494219 | 1,208467657 | 0,76119403 |
| AQP11 | 0,013169497 | -0,536418692 | 0,172 | 0,203 | 1 | FB | 0,689480334 | 1,450367691 | 0,84729064 |
| H1-103 | 0,013396078 | -0,408526991 | 0,49 | 0,468 | 1 | FB | 0,753392203 | 1,327329903 | 1,047008547 |
| HSD17B121 | 0,013431305 | -0,427743886 | 0,227 | 0,252 | 1 | FB | 0,743423457 | 1,345128394 | 0,900793651 |
| RNH12 | 0,013594516 | -0,411998897 | 0,579 | 0,524 | 1 | FB | 0,751581314 | 1,330528024 | 1,104961832 |
| UAP12 | 0,013836496 | -0,418204391 | 0,22 | 0,245 | 1 | FB | 0,748355464 | 1,336263378 | 0,897959184 |
| BOC | 0,014056347 | -0,372129257 | 0,293 | 0,315 | 1 | FB | 0,772641322 | 1,294261608 | 0,93015873 |
| NDRG11 | 0,01414171 | -0,369395921 | 0,229 | 0,255 | 1 | FB | 0,77410656 | 1,291811815 | 0,898039216 |
| GNG22 | 0,014157418 | -0,302156471 | 0,095 | 0,126 | 1 | FB | 0,811039188 | 1,232986045 | 0,753968254 |
| CFL2 | 0,015863605 | -0,251680225 | 0,101 | 0,132 | 1 | FB | 0,839917641 | 1,190592924 | 0,765151515 |
| CYP27A11 | 0,018335645 | -0,371137849 | 0,195 | 0,221 | 1 | FB | 0,773172457 | 1,293372507 | 0,882352941 |
| RARG | 0,018507746 | -0,399312299 | 0,232 | 0,255 | 1 | FB | 0,758219624 | 1,31887908 | 0,909803922 |
| PNISR3 | 0,018818007 | -0,434215946 | 0,486 | 0,461 | 1 | FB | 0,740095861 | 1,351176317 | 1,054229935 |
| PHYHD11 | 0,019412758 | -0,278017389 | 0,107 | 0,137 | 1 | FB | 0,824723608 | 1,212527434 | 0,781021898 |
| AL627171.25 | 0,020015 | -0,539617387 | 0,159 | 0,19 | 1 | FB | 0,687953335 | 1,453586964 | 0,836842105 |
| SCPEP12 | 0,020332949 | -0,319255499 | 0,415 | 0,419 | 1 | FB | 0,801483375 | 1,247686516 | 0,990453461 |
| OTUD6B-AS1 | 0,020390462 | -0,405522991 | 0,22 | 0,244 | 1 | FB | 0,754962562 | 1,324568993 | 0,901639344 |
| ACVRL1 | 0,021079091 | -0,252897697 | 0,129 | 0,16 | 1 | FB | 0,839209144 | 1,191598074 | 0,80625 |
| CWF19L22 | 0,021279193 | -0,309406529 | 0,119 | 0,148 | 1 | FB | 0,80697365 | 1,239197835 | 0,804054054 |

| Supplementary Table 4 - clusermarker | | | | | | | | | |
|---|---|---|---|---|---|---|---|---|---|
| | **p_val** | **avg_log2FC** | **pct.1** | **pct.2** | **p_val_adj** | **celltype** | **Foldchange_UP** | **Foldchange_DOWN** | **Ratio_pct1_pct2** |
| JAG11 | 0,021560996 | -0,280206746 | 0,181 | 0,211 | 1 | FB | 0,823473001 | 1,214368898 | 0,857819905 |
| CAT3 | 0,022255774 | -0,37911741 | 0,199 | 0,224 | 1 | FB | 0,768907837 | 1,300545984 | 0,888392857 |
| SAMHD11 | 0,022400179 | -0,368301363 | 0,279 | 0,3 | 1 | FB | 0,774694089 | 1,290832103 | 0,93 |
| NTRK21 | 0,023998409 | -0,399013313 | 0,179 | 0,21 | 1 | FB | 0,758376775 | 1,318605782 | 0,852380952 |
| GNG11 | 0,024061492 | -0,321220825 | 0,401 | 0,398 | 1 | FB | 0,80039229 | 1,249387348 | 1,007537688 |
| GAS7 | 0,024551355 | -0,324772397 | 0,225 | 0,25 | 1 | FB | 0,798424338 | 1,252466831 | 0,9 |
| OLFML2A | 0,025152463 | -0,4484848 | 0,137 | 0,165 | 1 | FB | 0,732812085 | 1,364606317 | 0,83030303 |
| CD1091 | 0,025304058 | -0,345593038 | 0,129 | 0,156 | 1 | FB | 0,78698441 | 1,270673201 | 0,826923077 |
| EPHA3 | 0,025694602 | -0,325287839 | 0,216 | 0,248 | 1 | FB | 0,79813913 | 1,252914389 | 0,870967742 |
| ADH51 | 0,02581143 | -0,414276982 | 0,521 | 0,492 | 1 | FB | 0,750395467 | 1,332630652 | 1,058943089 |
| PIH1D12 | 0,026244723 | -0,425173938 | 0,225 | 0,247 | 1 | FB | 0,744748936 | 1,34273438 | 0,910931174 |
| VPS4B | 0,027156941 | -0,269716943 | 0,182 | 0,21 | 1 | FB | 0,829482274 | 1,205571271 | 0,866666667 |
| SELENBP13 | 0,027632267 | -0,341960922 | 0,169 | 0,198 | 1 | FB | 0,788968211 | 1,26747819 | 0,853535354 |
| MTRNR2L87 | 0,028051772 | -1,863946019 | 0,151 | 0,176 | 1 | FB | 0,274723834 | 3,640019095 | 0,857954545 |
| ZC2HC1A1 | 0,029720942 | -0,412102391 | 0,168 | 0,192 | 1 | FB | 0,7515274 | 1,330623475 | 0,875 |
| MGST32 | 0,029765698 | -0,308843137 | 0,773 | 0,677 | 1 | FB | 0,807288845 | 1,238714006 | 1,141802068 |
| PDLIM3 | 0,031431847 | -0,366719839 | 0,171 | 0,197 | 1 | FB | 0,775543797 | 1,289417831 | 0,868020305 |
| EIF1AX6 | 0,032457337 | -0,351075334 | 0,42 | 0,4 | 1 | FB | 0,783999514 | 1,275510994 | 1,05 |
| KLF23 | 0,033663705 | -0,312820822 | 0,508 | 0,394 | 1 | FB | 0,805066118 | 1,242134003 | 1,289340102 |
| DIAPH2 | 0,037353967 | -0,327519431 | 0,169 | 0,195 | 1 | FB | 0,796905505 | 1,254853924 | 0,866666667 |
| LRRC17 | 0,037780753 | -0,294992648 | 0,136 | 0,165 | 1 | FB | 0,815076486 | 1,226878725 | 0,824242424 |
| MFAP4 | 0,038414957 | -0,319644611 | 0,932 | 0,858 | 1 | FB | 0,801267235 | 1,248023077 | 1,086247086 |
| KIF221 | 0,039730554 | -0,305741699 | 0,143 | 0,169 | 1 | FB | 0,809026184 | 1,236053937 | 0,846153846 |
| SPON2 | 0,039828975 | -0,293150543 | 0,461 | 0,461 | 1 | FB | 0,816117881 | 1,225313185 | 1 |
| TCF7L2 | 0,041712377 | -0,422543848 | 0,291 | 0,302 | 1 | FB | 0,746107881 | 1,340288752 | 0,963576159 |
| BMERB1 | 0,042236887 | -0,364197857 | 0,218 | 0,237 | 1 | FB | 0,776900714 | 1,287165763 | 0,919831224 |
| H4C34 | 0,044316586 | -0,428240909 | 0,371 | 0,365 | 1 | FB | 0,743167384 | 1,345591884 | 1,016438356 |
| ZNF704 | 0,04438182 | -0,252883929 | 0,16 | 0,185 | 1 | FB | 0,839217153 | 1,191586702 | 0,864864865 |
| SWAP701 | 0,049011366 | -0,305651054 | 0,168 | 0,19 | 1 | FB | 0,809077017 | 1,235976278 | 0,884210526 |
| SMIM142 | 0,050084251 | -0,396880146 | 0,3 | 0,303 | 1 | FB | 0,759498939 | 1,316657534 | 0,99009901 |
| CRYAB1 | 0,050184192 | -0,328002362 | 0,349 | 0,35 | 1 | FB | 0,796638792 | 1,255274047 | 0,997142857 |

| Supplementary Table 4 - clusermarker | | | | | | | | | |
|---|---|---|---|---|---|---|---|---|---|
| | **p_val** | **avg_log2FC** | **pct.1** | **pct.2** | **p_val_adj** | **celltype** | **Foldchange_UP** | **Foldchange_DOWN** | **Ratio_pct1_pct2** |
| RAB3IL1 | 0,050301442 | -0,27113795 | 0,166 | 0,19 | 1 | FB | 0,828665664 | 1,206759304 | 0,873684211 |
| A4GALT1 | 0,050608603 | -0,349167814 | 0,22 | 0,237 | 1 | FB | 0,785036798 | 1,273825638 | 0,928270042 |
| IFI162 | 0,052651942 | -0,325305008 | 0,544 | 0,505 | 1 | FB | 0,798129631 | 1,2529293 | 1,077227723 |
| RFC12 | 0,053947996 | -0,366564136 | 0,227 | 0,245 | 1 | FB | 0,775627502 | 1,289278678 | 0,926530612 |
| RND31 | 0,054014974 | -0,50524306 | 0,476 | 0,442 | 1 | FB | 0,704541669 | 1,419362465 | 1,076923077 |
| EXOC12 | 0,05556775 | -0,305891214 | 0,136 | 0,158 | 1 | FB | 0,808942344 | 1,236182044 | 0,860759494 |
| TMX43 | 0,056098647 | -0,285702087 | 0,367 | 0,368 | 1 | FB | 0,820342292 | 1,219003347 | 0,997282609 |
| PROS12 | 0,056686926 | -0,330556164 | 0,264 | 0,281 | 1 | FB | 0,795229861 | 1,257498051 | 0,939501779 |
| TMBIM45 | 0,058171546 | -0,339174818 | 0,579 | 0,508 | 1 | FB | 0,790493323 | 1,265032823 | 1,13976378 |
| KLF92 | 0,058565716 | -0,300790032 | 0,462 | 0,437 | 1 | FB | 0,811807722 | 1,231818783 | 1,057208238 |
| MBNL22 | 0,060391845 | -0,36273944 | 0,347 | 0,344 | 1 | FB | 0,777686478 | 1,285865227 | 1,00872093 |
| COLEC12 | 0,062363778 | -0,378966944 | 0,316 | 0,319 | 1 | FB | 0,768988035 | 1,30041035 | 0,990595611 |
| TNFSF104 | 0,062896361 | -0,283811468 | 0,267 | 0,284 | 1 | FB | 0,821418037 | 1,217406917 | 0,940140845 |
| FAM13A | 0,064114826 | -0,362101493 | 0,216 | 0,232 | 1 | FB | 0,778030441 | 1,285296754 | 0,931034483 |
| ARL4A2 | 0,064800587 | -0,259968613 | 0,146 | 0,171 | 1 | FB | 0,835106087 | 1,197452653 | 0,85380117 |
| NFE2L25 | 0,066509311 | -0,304700407 | 0,556 | 0,513 | 1 | FB | 0,809610324 | 1,235162114 | 1,083820663 |
| TMBIM12 | 0,066858972 | -0,454749734 | 0,363 | 0,347 | 1 | FB | 0,729636732 | 1,370545034 | 1,04610951 |
| CPNE33 | 0,067591579 | -0,303376299 | 0,472 | 0,447 | 1 | FB | 0,810353727 | 1,234029001 | 1,055928412 |
| CPQ | 0,072604406 | -0,365457896 | 0,341 | 0,337 | 1 | FB | 0,776222471 | 1,288290454 | 1,011869436 |
| PTGS22 | 0,072648128 | -0,291714444 | 0,104 | 0,126 | 1 | FB | 0,816930672 | 1,224094081 | 0,825396825 |
| SPOP2 | 0,073168552 | -0,265936926 | 0,226 | 0,242 | 1 | FB | 0,831658458 | 1,202416678 | 0,933884298 |
| GSTM31 | 0,075420677 | -0,342157393 | 0,371 | 0,36 | 1 | FB | 0,788860774 | 1,267650811 | 1,030555556 |
| ZRANB23 | 0,075919505 | -0,508449761 | 0,358 | 0,348 | 1 | FB | 0,702977412 | 1,422520813 | 1,028735632 |
| SH3GLB2 | 0,076643883 | -0,252303256 | 0,145 | 0,168 | 1 | FB | 0,839554999 | 1,191107195 | 0,863095238 |
| EXPH51 | 0,07697687 | -0,315421689 | 0,125 | 0,145 | 1 | FB | 0,803616066 | 1,244375321 | 0,862068966 |
| FIBIN | 0,077321791 | -0,469325106 | 0,224 | 0,235 | 1 | FB | 0,722302413 | 1,384461664 | 0,953191489 |
| RAB144 | 0,077968728 | -0,35874079 | 0,402 | 0,387 | 1 | FB | 0,779844945 | 1,282306189 | 1,03875969 |
| NR1D23 | 0,078199398 | -0,282047127 | 0,302 | 0,305 | 1 | FB | 0,822423203 | 1,215919002 | 0,990163934 |
| PBX3 | 0,078717197 | -0,270979654 | 0,143 | 0,165 | 1 | FB | 0,828756592 | 1,206626903 | 0,866666667 |
| RASD12 | 0,079618628 | -0,447124751 | 0,341 | 0,345 | 1 | FB | 0,733503243 | 1,36332049 | 0,988405797 |
| ECI2 | 0,08018201 | -0,302717391 | 0,148 | 0,168 | 1 | FB | 0,810723917 | 1,233465523 | 0,880952381 |

| Supplementary Table 4 - clusermarker | | | | | | | | | |
|---|---|---|---|---|---|---|---|---|---|
| | p_val | avg_log2FC | pct.1 | pct.2 | p_val_adj | celltype | Foldchange_UP | Foldchange_DOWN | Ratio_pct1_pct2 |
| PCDH18 | 0,085019504 | -0,318151135 | 0,36 | 0,358 | 1 | FB | 0,802097135 | 1,246731794 | 1,005586592 |
| MTUS1 | 0,085115704 | -0,311256611 | 0,158 | 0,176 | 1 | FB | 0,805939467 | 1,240787976 | 0,897727273 |
| MRFAP1L12 | 0,090418175 | -0,28757615 | 0,196 | 0,211 | 1 | FB | 0,819277358 | 1,220587863 | 0,928909953 |
| MRPS31 | 0,090996431 | -0,28766411 | 0,105 | 0,124 | 1 | FB | 0,819227409 | 1,220662283 | 0,846774194 |
| ENSA5 | 0,092431431 | -0,371935801 | 0,262 | 0,265 | 1 | FB | 0,772744936 | 1,294088067 | 0,988679245 |
| DIAPH12 | 0,099233071 | -0,281053286 | 0,14 | 0,158 | 1 | FB | 0,822989948 | 1,21508167 | 0,886075949 |
| COIL1 | 0,108583405 | -0,264276931 | 0,099 | 0,118 | 1 | FB | 0,832615932 | 1,201033948 | 0,838983051 |
| MSX1 | 0,108717717 | -0,362696805 | 0,136 | 0,153 | 1 | FB | 0,777709461 | 1,285827227 | 0,888888889 |
| ZC3H6 | 0,11075706 | -0,335056968 | 0,139 | 0,156 | 1 | FB | 0,792752833 | 1,261427218 | 0,891025641 |
| HSPB6 | 0,112519282 | -0,261771725 | 0,263 | 0,274 | 1 | FB | 0,834063006 | 1,198950191 | 0,959854015 |
| ANKRD35 | 0,112599982 | -0,277340765 | 0,185 | 0,203 | 1 | FB | 0,825110494 | 1,211958892 | 0,911330049 |
| CRYBG3 | 0,1134141 | -0,372273515 | 0,237 | 0,244 | 1 | FB | 0,772564068 | 1,29439103 | 0,971311475 |
| FZD4 | 0,114036382 | -0,254079589 | 0,157 | 0,176 | 1 | FB | 0,838521924 | 1,192574661 | 0,892045455 |
| NR1D1 | 0,11529151 | -0,263543875 | 0,118 | 0,137 | 1 | FB | 0,833039105 | 1,200423839 | 0,861313869 |
| ZBED51 | 0,116010689 | -0,250075541 | 0,159 | 0,176 | 1 | FB | 0,840852386 | 1,189269385 | 0,903409091 |
| DBNDD21 | 0,11681257 | -0,310848259 | 0,173 | 0,189 | 1 | FB | 0,806167619 | 1,240436823 | 0,915343915 |
| RNPC3 | 0,119331628 | -0,269766591 | 0,147 | 0,166 | 1 | FB | 0,82945373 | 1,20561276 | 0,885542169 |
| ECM2 | 0,120217861 | -0,370284513 | 0,361 | 0,345 | 1 | FB | 0,773629915 | 1,292607719 | 1,046376812 |
| BHLHE412 | 0,12027813 | -0,331125157 | 0,19 | 0,205 | 1 | FB | 0,794916287 | 1,257994101 | 0,926829268 |
| A2M2 | 0,121510666 | -0,427270539 | 0,673 | 0,65 | 1 | FB | 0,743667413 | 1,344687131 | 1,035384615 |
| PALM | 0,123137953 | -0,30121837 | 0,134 | 0,153 | 1 | FB | 0,811566731 | 1,232184566 | 0,875816993 |
| NPEPPS2 | 0,12708056 | -0,303636809 | 0,256 | 0,266 | 1 | FB | 0,810207413 | 1,234251852 | 0,962406015 |
| MPHOSPH81 | 0,12749609 | -0,366615822 | 0,433 | 0,403 | 1 | FB | 0,775599715 | 1,289324868 | 1,074441687 |
| SLC27A3 | 0,12802036 | -0,260788698 | 0,112 | 0,129 | 1 | FB | 0,834631515 | 1,198133525 | 0,868217054 |
| GSTM22 | 0,128071275 | -0,298688879 | 0,206 | 0,221 | 1 | FB | 0,812990907 | 1,230026058 | 0,932126697 |
| ATP5ME4 | 0,132683221 | -0,289875171 | 0,335 | 0,327 | 1 | FB | 0,81797283 | 1,222534493 | 1,024464832 |
| LIFR | 0,134690992 | -0,277831922 | 0,115 | 0,132 | 1 | FB | 0,824829638 | 1,212371566 | 0,871212121 |
| KIFAP31 | 0,135138667 | -0,356964747 | 0,304 | 0,302 | 1 | FB | 0,780805572 | 1,280728566 | 1,006622517 |
| SNW12 | 0,144655654 | -0,414591673 | 0,236 | 0,239 | 1 | FB | 0,750231803 | 1,332921366 | 0,987447699 |
| MAPK14 | 0,14579293 | -0,290198198 | 0,124 | 0,14 | 1 | FB | 0,817789702 | 1,222808256 | 0,885714286 |
| SRSF75 | 0,148756121 | -0,378040492 | 0,549 | 0,492 | 1 | FB | 0,769482013 | 1,299575537 | 1,115853659 |

| Supplementary Table 4 - clusermarker | | | | | | | | | |
|---|---|---|---|---|---|---|---|---|---|
| | **p_val** | **avg_log2FC** | **pct.1** | **pct.2** | **p_val_adj** | **celltype** | **Foldchange_UP** | **Foldchange_DOWN** | **Ratio_pct1_pct2** |
| NFIX2 | 0,149610146 | -0,292653091 | 0,508 | 0,458 | 1 | FB | 0,816399333 | 1,224890761 | 1,109170306 |
| TMOD32 | 0,156494489 | -0,286139467 | 0,321 | 0,313 | 1 | FB | 0,820093628 | 1,219372966 | 1,025559105 |
| PPFIBP1 | 0,156667406 | -0,296339777 | 0,293 | 0,292 | 1 | FB | 0,814315757 | 1,228024869 | 1,003424658 |
| DDIT33 | 0,158256978 | -0,39914698 | 0,261 | 0,263 | 1 | FB | 0,758306514 | 1,318727958 | 0,992395437 |
| PTPRA1 | 0,158729003 | -0,341224825 | 0,227 | 0,232 | 1 | FB | 0,789370864 | 1,266831657 | 0,978448276 |
| NR2F2 | 0,167119928 | -0,289537452 | 0,469 | 0,445 | 1 | FB | 0,818164331 | 1,222248346 | 1,053932584 |
| EZH1 | 0,173543962 | -0,30340537 | 0,11 | 0,124 | 1 | FB | 0,810337398 | 1,234053867 | 0,887096774 |
| CA5B1 | 0,176368317 | -0,390407647 | 0,197 | 0,205 | 1 | FB | 0,762914005 | 1,31076372 | 0,96097561 |
| CDC401 | 0,181004182 | -0,309308676 | 0,198 | 0,206 | 1 | FB | 0,807028386 | 1,239113788 | 0,961165049 |
| UPF23 | 0,183406378 | -0,321470175 | 0,174 | 0,182 | 1 | FB | 0,800253965 | 1,249603306 | 0,956043956 |
| DDX186 | 0,183713112 | -0,323187595 | 0,382 | 0,361 | 1 | FB | 0,799301889 | 1,25109175 | 1,058171745 |
| AFDN3 | 0,18478438 | -0,251510047 | 0,128 | 0,142 | 1 | FB | 0,840016722 | 1,190452491 | 0,901408451 |
| C14orf132 | 0,186992117 | -0,270701617 | 0,163 | 0,174 | 1 | FB | 0,828916326 | 1,206394383 | 0,936781609 |
| RECK | 0,191499956 | -0,334350628 | 0,374 | 0,353 | 1 | FB | 0,793141057 | 1,260809778 | 1,059490085 |
| BRD7 | 0,195110788 | -0,283769946 | 0,19 | 0,2 | 1 | FB | 0,821441679 | 1,217371879 | 0,95 |
| CBR32 | 0,198106306 | -0,293966065 | 0,204 | 0,213 | 1 | FB | 0,815656679 | 1,226006022 | 0,957746479 |
| GALNT151 | 0,198224368 | -0,455429913 | 0,127 | 0,137 | 1 | FB | 0,729292816 | 1,371191349 | 0,927007299 |
| CEP1701 | 0,198396776 | -0,30774853 | 0,259 | 0,26 | 1 | FB | 0,807901587 | 1,237774521 | 0,996153846 |
| PLA2G2A | 0,199194478 | -0,397761218 | 0,245 | 0,211 | 1 | FB | 0,759035245 | 1,317461879 | 1,161137441 |
| METTL17 | 0,200819562 | -0,254154541 | 0,104 | 0,118 | 1 | FB | 0,838478362 | 1,192636621 | 0,881355932 |
| NOTCH2 | 0,20212361 | -0,339428844 | 0,307 | 0,3 | 1 | FB | 0,790354147 | 1,265255586 | 1,023333333 |
| SF3B13 | 0,204580137 | -0,319297477 | 0,579 | 0,502 | 1 | FB | 0,801460055 | 1,24772282 | 1,153386454 |
| ASH1L5 | 0,212369983 | -0,33293823 | 0,416 | 0,387 | 1 | FB | 0,793917922 | 1,259576049 | 1,074935401 |
| PPIG1 | 0,217049676 | -0,330722179 | 0,423 | 0,39 | 1 | FB | 0,795138356 | 1,257642764 | 1,084615385 |
| UBE4A1 | 0,220795279 | -0,281075842 | 0,181 | 0,187 | 1 | FB | 0,82297708 | 1,215100668 | 0,967914439 |
| MNAT11 | 0,224697546 | -0,274457571 | 0,188 | 0,152 | 1 | FB | 0,826761108 | 1,209539237 | 1,236842105 |
| TTC13 | 0,225176893 | -0,270928592 | 0,218 | 0,226 | 1 | FB | 0,828785925 | 1,206584197 | 0,96460177 |
| TAF72 | 0,226555965 | -0,27207325 | 0,276 | 0,279 | 1 | FB | 0,828128614 | 1,207541901 | 0,989247312 |
| LINC02802 | 0,226558466 | -0,271798479 | 0,173 | 0,184 | 1 | FB | 0,828286351 | 1,207311938 | 0,940217391 |
| DPYSL21 | 0,22685421 | -0,258853449 | 0,369 | 0,35 | 1 | FB | 0,835751851 | 1,196527412 | 1,054285714 |
| PHF31 | 0,228898019 | -0,285992233 | 0,354 | 0,339 | 1 | FB | 0,820177327 | 1,21924853 | 1,044247788 |

| Supplementary Table 4 - clusermarker | | | | | | | | | |
|---|---|---|---|---|---|---|---|---|---|
| | **p_val** | **avg_log2FC** | **pct.1** | **pct.2** | **p_val_adj** | **celltype** | **Foldchange_UP** | **Foldchange_DOWN** | **Ratio_pct1_pct2** |
| KLF5 | 0,229411874 | -0,258371292 | 0,125 | 0,139 | 1 | FB | 0,836031211 | 1,196127592 | 0,899280576 |
| MSN2 | 0,232690212 | -0,3399889 | 0,447 | 0,403 | 1 | FB | 0,79004739 | 1,265746855 | 1,109181141 |
| NSRP13 | 0,237819654 | -0,301736468 | 0,199 | 0,205 | 1 | FB | 0,811275335 | 1,232627145 | 0,970731707 |
| SH3BGRL35 | 0,242839313 | -0,446499638 | 0,613 | 0,539 | 1 | FB | 0,733821135 | 1,362729897 | 1,13729128 |
| SFT2D22 | 0,24294644 | -0,304752508 | 0,231 | 0,234 | 1 | FB | 0,809581087 | 1,235206721 | 0,987179487 |
| FMC1 | 0,248413926 | -0,274739164 | 0,166 | 0,174 | 1 | FB | 0,826599752 | 1,209775345 | 0,954022989 |
| PET1005 | 0,258482744 | -0,316013101 | 0,326 | 0,313 | 1 | FB | 0,803286702 | 1,24488554 | 1,041533546 |
| ARL12 | 0,26136808 | -0,315745434 | 0,362 | 0,345 | 1 | FB | 0,803435752 | 1,244654594 | 1,049275362 |
| GABPA | 0,261974832 | -0,276943758 | 0,124 | 0,134 | 1 | FB | 0,825337583 | 1,211625426 | 0,925373134 |
| TSPAN11 | 0,269288297 | -0,304113594 | 0,305 | 0,303 | 1 | FB | 0,809939699 | 1,234659817 | 1,00660066 |
| BOD1L14 | 0,271135187 | -0,313704146 | 0,334 | 0,316 | 1 | FB | 0,804573348 | 1,242894762 | 1,056962025 |
| SARS1 | 0,272561559 | -0,279174437 | 0,309 | 0,25 | 1 | FB | 0,824062441 | 1,213500277 | 1,236 |
| EMP32 | 0,272704293 | -0,258514152 | 0,755 | 0,666 | 1 | FB | 0,835948429 | 1,196246042 | 1,133633634 |
| USP531 | 0,274889655 | -0,297194794 | 0,209 | 0,213 | 1 | FB | 0,813833293 | 1,228752876 | 0,981220657 |
| GAP43 | 0,276637557 | -0,294200608 | 0,12 | 0,131 | 1 | FB | 0,815524086 | 1,226205353 | 0,916030534 |
| IL6ST2 | 0,289942164 | -0,255160469 | 0,586 | 0,524 | 1 | FB | 0,837893932 | 1,193468483 | 1,118320611 |
| GAS11 | 0,29036419 | -0,430169514 | 0,333 | 0,273 | 1 | FB | 0,742174576 | 1,347391883 | 1,21978022 |
| AR | 0,291340943 | -0,284660242 | 0,235 | 0,24 | 1 | FB | 0,820934918 | 1,218123359 | 0,979166667 |
| CLIC2 | 0,297919353 | -0,27893069 | 0,189 | 0,194 | 1 | FB | 0,82420168 | 1,21329527 | 0,974226804 |
| SRPK2 | 0,306379418 | -0,351354994 | 0,229 | 0,224 | 1 | FB | 0,783847554 | 1,27575827 | 1,022321429 |
| ZNF1062 | 0,309528726 | -0,279947511 | 0,372 | 0,345 | 1 | FB | 0,823620982 | 1,214150709 | 1,07826087 |
| PKN21 | 0,310154212 | -0,270126379 | 0,201 | 0,203 | 1 | FB | 0,829246901 | 1,205913461 | 0,990147783 |
| APCDD1 | 0,313458657 | -0,332751337 | 0,153 | 0,16 | 1 | FB | 0,794020777 | 1,259412888 | 0,95625 |
| SMC31 | 0,316466221 | -0,313757149 | 0,202 | 0,205 | 1 | FB | 0,80454379 | 1,242940425 | 0,985365854 |
| EPC14 | 0,321067408 | -0,277642326 | 0,351 | 0,329 | 1 | FB | 0,824938043 | 1,212212249 | 1,066869301 |
| PREPL2 | 0,321453578 | -0,253523595 | 0,139 | 0,147 | 1 | FB | 0,838845141 | 1,192115148 | 0,945578231 |
| HMG20A1 | 0,32400791 | -0,260381552 | 0,129 | 0,139 | 1 | FB | 0,834867091 | 1,197795446 | 0,928057554 |
| KMT2E3 | 0,32741516 | -0,333350382 | 0,442 | 0,394 | 1 | FB | 0,793691147 | 1,259935939 | 1,121827411 |
| THOC22 | 0,329333073 | -0,250315118 | 0,246 | 0,244 | 1 | FB | 0,840712764 | 1,189466894 | 1,008196721 |
| SH3D19 | 0,330980781 | -0,314618262 | 0,375 | 0,348 | 1 | FB | 0,804063719 | 1,24368253 | 1,077586207 |
| STAG23 | 0,335776496 | -0,26388449 | 0,245 | 0,247 | 1 | FB | 0,83284245 | 1,200707289 | 0,991902834 |

| Supplementary Table 4 - clusermarker | | | | | | | | | |
|---|---|---|---|---|---|---|---|---|---|
| | **p_val** | **avg_log2FC** | **pct.1** | **pct.2** | **p_val_adj** | **celltype** | **Foldchange_UP** | **Foldchange_DOWN** | **Ratio_pct1_pct2** |
| EIF4H1 | 0,339899863 | -0,285634992 | 0,344 | 0,319 | 1 | FB | 0,820380445 | 1,218946656 | 1,078369906 |
| KIAA09301 | 0,343910404 | -0,342838849 | 0,344 | 0,327 | 1 | FB | 0,788488244 | 1,268249726 | 1,051987768 |
| ABHD23 | 0,359070522 | -0,264775354 | 0,244 | 0,239 | 1 | FB | 0,832328329 | 1,201448954 | 1,020920502 |
| CLDND12 | 0,36872045 | -0,31808744 | 0,306 | 0,287 | 1 | FB | 0,802132548 | 1,246676752 | 1,066202091 |
| IFNGR12 | 0,380996884 | -0,26021975 | 0,304 | 0,295 | 1 | FB | 0,83496073 | 1,197661117 | 1,030508475 |
| PCMTD1 | 0,385752069 | -0,318094103 | 0,295 | 0,284 | 1 | FB | 0,802128844 | 1,24668251 | 1,038732394 |
| PFDN13 | 0,397445042 | -0,252483486 | 0,209 | 0,208 | 1 | FB | 0,839450123 | 1,191256005 | 1,004807692 |
| SSBP21 | 0,404900094 | -0,253920761 | 0,244 | 0,237 | 1 | FB | 0,838614243 | 1,192443377 | 1,029535865 |
| APPL13 | 0,409077401 | -0,280174245 | 0,353 | 0,326 | 1 | FB | 0,823491552 | 1,214341541 | 1,082822086 |
| LMBRD11 | 0,413135516 | -0,292635652 | 0,296 | 0,285 | 1 | FB | 0,816409202 | 1,224875954 | 1,038596491 |
| GPBP14 | 0,413688776 | -0,299767217 | 0,273 | 0,265 | 1 | FB | 0,812383466 | 1,23094578 | 1,030188679 |
| RNASE4 | 0,413853082 | -0,375220399 | 0,448 | 0,392 | 1 | FB | 0,770987621 | 1,297037687 | 1,142857143 |
| RERE3 | 0,416857064 | -0,251273526 | 0,204 | 0,205 | 1 | FB | 0,840154449 | 1,19025734 | 0,995121951 |
| SLC16A7 | 0,438113907 | -0,265340296 | 0,206 | 0,174 | 1 | FB | 0,832002463 | 1,201919519 | 1,183908046 |
| PTPN13 | 0,449508769 | -0,291010851 | 0,228 | 0,221 | 1 | FB | 0,817329181 | 1,223497243 | 1,031674208 |
| ZEB22 | 0,452820717 | -0,261404833 | 0,444 | 0,402 | 1 | FB | 0,834275143 | 1,198645325 | 1,104477612 |
| MRPS18B2 | 0,453771881 | -0,282207516 | 0,245 | 0,239 | 1 | FB | 0,822331777 | 1,216054186 | 1,025104603 |
| RHOB2 | 0,453944294 | -0,251555901 | 0,591 | 0,539 | 1 | FB | 0,839990024 | 1,190490329 | 1,096474954 |
| IFI271 | 0,456077203 | -0,279174518 | 0,327 | 0,321 | 1 | FB | 0,824062395 | 1,213500344 | 1,018691589 |
| DDAH23 | 0,463969572 | -0,278042142 | 0,574 | 0,5 | 1 | FB | 0,824709458 | 1,212548238 | 1,148 |
| PRPF4B4 | 0,467390962 | -0,299284421 | 0,225 | 0,216 | 1 | FB | 0,812655375 | 1,230533915 | 1,041666667 |
| SMIM72 | 0,468029714 | -0,258277857 | 0,288 | 0,247 | 1 | FB | 0,836085358 | 1,196050129 | 1,165991903 |
| CD463 | 0,469352533 | -0,287768705 | 0,343 | 0,324 | 1 | FB | 0,819168017 | 1,220750785 | 1,058641975 |
| ENG | 0,473817915 | -0,271155613 | 0,308 | 0,294 | 1 | FB | 0,828655519 | 1,206774078 | 1,047619048 |
| PLAGL1 | 0,490280949 | -0,320750512 | 0,266 | 0,255 | 1 | FB | 0,800653257 | 1,248980118 | 1,043137255 |
| MT1X3 | 0,491778406 | -0,448081983 | 0,302 | 0,261 | 1 | FB | 0,733016723 | 1,364225357 | 1,157088123 |
| SDAD11 | 0,498320623 | -0,332217237 | 0,173 | 0,173 | 1 | FB | 0,794314785 | 1,258946728 | 1 |
| RPL7L1 | 0,501262533 | -0,284885336 | 0,246 | 0,237 | 1 | FB | 0,820806843 | 1,218313429 | 1,037974684 |
| HMGN32 | 0,506253745 | -0,336180531 | 0,362 | 0,327 | 1 | FB | 0,792135681 | 1,262409993 | 1,107033639 |
| BMP4 | 0,508362811 | -0,323125335 | 0,253 | 0,245 | 1 | FB | 0,799336385 | 1,25103776 | 1,032653061 |
| MAGI2-AS31 | 0,510773547 | -0,262452226 | 0,303 | 0,261 | 1 | FB | 0,833669681 | 1,199515854 | 1,16091954 |

| Supplementary Table 4 - clusermarker | | | | | | | | | |
|---|---|---|---|---|---|---|---|---|---|
| | **p_val** | **avg_log2FC** | **pct.1** | **pct.2** | **p_val_adj** | **celltype** | **Foldchange_UP** | **Foldchange_DOWN** | **Ratio_pct1_pct2** |
| TWIST1 | 0,515014424 | -0,289158569 | 0,351 | 0,3 | 1 | FB | 0,818379227 | 1,221927399 | 1,17 |
| MAF | 0,527629992 | -0,284422688 | 0,13 | 0,132 | 1 | FB | 0,821070104 | 1,217922799 | 0,984848485 |
| N4BP2L23 | 0,530841571 | -0,360506459 | 0,455 | 0,406 | 1 | FB | 0,778891101 | 1,283876524 | 1,120689655 |
| GNPDA2 | 0,586856899 | -0,252840367 | 0,124 | 0,127 | 1 | FB | 0,839242494 | 1,191550723 | 0,976377953 |
| DST3 | 0,587516803 | -0,26222167 | 0,633 | 0,54 | 1 | FB | 0,83380292 | 1,199324176 | 1,172222222 |
| OMD | 0,606779229 | -0,342772141 | 0,581 | 0,505 | 1 | FB | 0,788524704 | 1,268191085 | 1,15049505 |
| ANKRD121 | 0,633545259 | -0,305526192 | 0,444 | 0,392 | 1 | FB | 0,809147044 | 1,235869312 | 1,132653061 |
| LUC7L34 | 0,636448527 | -0,340738104 | 0,382 | 0,345 | 1 | FB | 0,789637219 | 1,266404339 | 1,107246377 |
| DUSP62 | 0,648036055 | -0,390147626 | 0,182 | 0,177 | 1 | FB | 0,76305152 | 1,310527498 | 1,028248588 |
| MT1M1 | 0,661205817 | -0,360439077 | 0,255 | 0,244 | 1 | FB | 0,778927481 | 1,283816561 | 1,045081967 |
| JAM2 | 0,670283019 | -0,251569044 | 0,217 | 0,208 | 1 | FB | 0,839982372 | 1,190501175 | 1,043269231 |
| WASHC3 | 0,702261528 | -0,310059601 | 0,3 | 0,271 | 1 | FB | 0,806608435 | 1,239758917 | 1,10701107 |
| CIR14 | 0,711350755 | -0,335245217 | 0,281 | 0,256 | 1 | FB | 0,792649398 | 1,261591825 | 1,09765625 |
| ANTXR2 | 0,721900546 | -0,267583128 | 0,193 | 0,185 | 1 | FB | 0,830710026 | 1,203789492 | 1,043243243 |
| CHD93 | 0,722466041 | -0,353775007 | 0,414 | 0,365 | 1 | FB | 0,782533811 | 1,277900055 | 1,134246575 |
| DYNC1I22 | 0,726514155 | -0,344427937 | 0,302 | 0,269 | 1 | FB | 0,787620225 | 1,269647437 | 1,12267658 |
| LINC01436 | 0,736599916 | -0,336907533 | 0,241 | 0,213 | 1 | FB | 0,791736609 | 1,263046306 | 1,131455399 |
| PCNP3 | 0,737131042 | -0,27724143 | 0,35 | 0,313 | 1 | FB | 0,825167308 | 1,211875447 | 1,118210863 |
| C11orf584 | 0,741577247 | -0,262897403 | 0,603 | 0,51 | 1 | FB | 0,833412473 | 1,199886049 | 1,182352941 |
| LYPLAL1 | 0,752165056 | -0,266911713 | 0,104 | 0,103 | 1 | FB | 0,83109672 | 1,203229391 | 1,009708738 |
| OGT1 | 0,781859512 | -0,378890939 | 0,209 | 0,198 | 1 | FB | 0,769028548 | 1,300341843 | 1,055555556 |
| SMAD5 | 0,788647201 | -0,306278926 | 0,286 | 0,261 | 1 | FB | 0,808724977 | 1,236514301 | 1,095785441 |
| PHC31 | 0,795663004 | -0,255770355 | 0,185 | 0,176 | 1 | FB | 0,837539794 | 1,193973118 | 1,051136364 |
| SNRPB23 | 0,796845282 | -0,252181589 | 0,302 | 0,263 | 1 | FB | 0,839625805 | 1,191006749 | 1,148288973 |
| R3HDM2 | 0,797213293 | -0,264041837 | 0,163 | 0,156 | 1 | FB | 0,832751621 | 1,20083825 | 1,044871795 |
| RORA1 | 0,807211977 | -0,29085799 | 0,239 | 0,221 | 1 | FB | 0,817415786 | 1,223367614 | 1,081447964 |
| AHI11 | 0,80973833 | -0,250352928 | 0,262 | 0,235 | 1 | FB | 0,840690731 | 1,189498068 | 1,114893617 |
| PLEKHA51 | 0,810917279 | -0,266977666 | 0,395 | 0,348 | 1 | FB | 0,831058727 | 1,203284398 | 1,135057471 |
| TNRC6A | 0,823485848 | -0,286739616 | 0,184 | 0,174 | 1 | FB | 0,819752547 | 1,219880321 | 1,057471264 |
| PAPPA | 0,827419246 | -0,330382843 | 0,237 | 0,208 | 1 | FB | 0,795325403 | 1,257346988 | 1,139423077 |
| REV3L1 | 0,82783551 | -0,357384467 | 0,262 | 0,244 | 1 | FB | 0,780578447 | 1,28110122 | 1,073770492 |

| Supplementary Table 4 - clusermarker | | | | | | | | | |
|---|---|---|---|---|---|---|---|---|---|
| | **p_val** | **avg_log2FC** | **pct.1** | **pct.2** | **p_val_adj** | **celltype** | **Foldchange_UP** | **Foldchange_DOWN** | **Ratio_pct1_pct2** |
| JPX1 | 0,841423509 | -0,274167669 | 0,252 | 0,226 | 1 | FB | 0,826927258 | 1,209296211 | 1,115044248 |
| TLE53 | 0,849606417 | -0,271995219 | 0,44 | 0,384 | 1 | FB | 0,828173406 | 1,20747659 | 1,145833333 |
| SNHG12 | 0,864626073 | -0,272620621 | 0,173 | 0,163 | 1 | FB | 0,827814474 | 1,208000139 | 1,061349693 |
| NEMF1 | 0,874536813 | -0,251631679 | 0,213 | 0,194 | 1 | FB | 0,839945904 | 1,190552861 | 1,097938144 |
| PDCD103 | 0,883278329 | -0,279206862 | 0,244 | 0,223 | 1 | FB | 0,82404392 | 1,21352755 | 1,094170404 |
| MAN1A1 | 0,896680877 | -0,288669473 | 0,465 | 0,394 | 1 | FB | 0,818656717 | 1,221513217 | 1,180203046 |
| FAM204A3 | 0,897072751 | -0,260204055 | 0,276 | 0,245 | 1 | FB | 0,834969813 | 1,197648088 | 1,126530612 |
| PLK21 | 0,89793213 | -0,261838995 | 0,273 | 0,25 | 1 | FB | 0,834024116 | 1,199006097 | 1,092 |
| ITGB3BP | 0,917955079 | -0,255468838 | 0,108 | 0,105 | 1 | FB | 0,837714855 | 1,193723609 | 1,028571429 |
| ARGLU12 | 0,925784562 | -0,340623131 | 0,408 | 0,36 | 1 | FB | 0,789700149 | 1,26630342 | 1,133333333 |
| SESTD1 | 0,934260837 | -0,276658966 | 0,258 | 0,232 | 1 | FB | 0,825500523 | 1,211386271 | 1,112068966 |
| TIAL11 | 0,941924079 | -0,292435693 | 0,234 | 0,215 | 1 | FB | 0,816522365 | 1,224706197 | 1,088372093 |
| NKD2 | 0,94443041 | -0,252901963 | 0,156 | 0,147 | 1 | FB | 0,839206663 | 1,191601598 | 1,06122449 |
| SMU12 | 0,953055731 | -0,270243534 | 0,19 | 0,177 | 1 | FB | 0,829179564 | 1,206011391 | 1,073446328 |
| WIPI11 | 0,965314473 | -0,305537971 | 0,16 | 0,148 | 1 | FB | 0,809140437 | 1,235879402 | 1,081081081 |
| PRR48 | 6,13743E-69 | -3,780208098 | 0,074 | 0,914 | 2,2455E-64 | SMC | 0,07278535 | 13,73902859 | 0,080962801 |
| LYZ8 | 1,64921E-44 | -2,719784154 | 0,071 | 0,69 | 6,03397E-40 | SMC | 0,15179707 | 6,587742452 | 0,102898551 |
| PRB38 | 2,09342E-44 | -2,69695673 | 0 | 0,491 | 7,65919E-40 | SMC | 0,154218022 | 6,484326473 | 0 |
| SLPI8 | 2,17233E-21 | -2,521564319 | 0,131 | 0,534 | 7,94791E-17 | SMC | 0,17415402 | 5,742043736 | 0,245318352 |
| RERGL | 8,62777E-21 | -2,249499846 | 0,213 | 0,638 | 3,15664E-16 | SMC | 0,210296997 | 4,755179645 | 0,333855799 |
| PLN1 | 1,55087E-20 | -1,472720628 | 0,321 | 0,75 | 5,67416E-16 | SMC | 0,360302202 | 2,775447932 | 0,428 |
| RPL37A7 | 4,01575E-20 | -0,914646229 | 0,861 | 0,974 | 1,46924E-15 | SMC | 0,530473935 | 1,885106758 | 0,883983573 |
| B2M6 | 1,38967E-19 | 0,946610341 | 0,991 | 0,983 | 5,08437E-15 | SMC | 1,927338989 | 0,518850086 | 1,008138352 |
| IGFBP74 | 3,67832E-19 | 0,930983416 | 0,994 | 0,991 | 1,34579E-14 | SMC | 1,906575175 | 0,524500693 | 1,003027245 |
| PRH27 | 2,18937E-18 | -1,051516066 | 0 | 0,207 | 8,01024E-14 | SMC | 0,482460901 | 2,072706822 | 0 |
| TCF211 | 1,53848E-17 | -1,294281564 | 0,023 | 0,276 | 5,62883E-13 | SMC | 0,407739162 | 2,452548325 | 0,083333333 |
| STATH8 | 2,9726E-17 | -1,719702847 | 0,003 | 0,207 | 1,08759E-12 | SMC | 0,30361125 | 3,293685596 | 0,014492754 |
| HLA-B8 | 3,17548E-17 | 1,365114798 | 0,92 | 0,707 | 1,16181E-12 | SMC | 2,575968226 | 0,388203546 | 1,301272984 |
| SCGB3A28 | 4,03285E-17 | -1,305706786 | 0,048 | 0,345 | 1,4755E-12 | SMC | 0,404522881 | 2,472048051 | 0,139130435 |
| MYH111 | 1,19817E-16 | -1,704346239 | 0,551 | 0,81 | 4,38376E-12 | SMC | 0,306860267 | 3,258812255 | 0,680246914 |
| IGFBP43 | 3,8591E-16 | 1,410859665 | 0,778 | 0,448 | 1,41193E-11 | SMC | 2,658955559 | 0,376087519 | 1,736607143 |

| Supplementary Table 4 - clusermarker | | | | | | | | | |
|---|---|---|---|---|---|---|---|---|---|
| | **p_val** | **avg_log2FC** | **pct.1** | **pct.2** | **p_val_adj** | **celltype** | **Foldchange_UP** | **Foldchange_DOWN** | **Ratio_pct1_pct2** |
| C6orf588 | 4,91523E-16 | -1,041959867 | 0,011 | 0,224 | 1,79833E-11 | SMC | 0,485667258 | 2,059022889 | 0,049107143 |
| STEAP42 | 5,11129E-16 | 1,668635776 | 0,622 | 0,155 | 1,87007E-11 | SMC | 3,179138295 | 0,314550645 | 4,012903226 |
| CST34 | 1,0564E-15 | 1,112168661 | 0,909 | 0,716 | 3,86504E-11 | SMC | 2,161703507 | 0,46259813 | 1,269553073 |
| SORBS21 | 1,38759E-15 | -1,724772733 | 0,324 | 0,664 | 5,07679E-11 | SMC | 0,302546178 | 3,305280553 | 0,487951807 |
| MFAP41 | 2,46586E-15 | -1,076211556 | 0,366 | 0,776 | 9,02183E-11 | SMC | 0,474272605 | 2,108492012 | 0,471649485 |
| CNN12 | 2,51099E-15 | -1,538178274 | 0,295 | 0,647 | 9,18695E-11 | SMC | 0,344319961 | 2,90427542 | 0,455950541 |
| NDUFA42 | 3,82784E-15 | -1,178053645 | 0,497 | 0,802 | 1,40049E-10 | SMC | 0,441947332 | 2,26271306 | 0,619700748 |
| RPS277 | 4,31089E-15 | -0,658950146 | 0,974 | 1 | 1,57723E-10 | SMC | 0,633339012 | 1,578933211 | 0,974 |
| CLU2 | 6,62329E-15 | -1,922204328 | 0,224 | 0,586 | 2,42326E-10 | SMC | 0,263851058 | 3,790017022 | 0,38225256 |
| PIP7 | 9,90755E-15 | -1,083064676 | 0 | 0,164 | 3,62487E-10 | SMC | 0,472025048 | 2,118531641 | 0 |
| SCGB3A16 | 1,07454E-14 | -2,355342906 | 0,168 | 0,526 | 3,93141E-10 | SMC | 0,195420957 | 5,117158447 | 0,319391635 |
| EEF1A15 | 3,36367E-14 | -0,542249375 | 0,994 | 1 | 1,23067E-09 | SMC | 0,686699408 | 1,456241244 | 0,994 |
| IGFBP23 | 4,19422E-14 | 1,621288123 | 0,56 | 0,155 | 1,53454E-09 | SMC | 3,076496013 | 0,325045115 | 3,612903226 |
| RPS298 | 7,42726E-14 | -1,013647173 | 0,585 | 0,836 | 2,71741E-09 | SMC | 0,49529255 | 2,019008764 | 0,699760766 |
| HLA-A4 | 7,56793E-14 | 0,96712819 | 0,895 | 0,75 | 2,76888E-09 | SMC | 1,954945231 | 0,511523282 | 1,193333333 |
| SLC38A12 | 7,70388E-14 | -0,837734811 | 0,043 | 0,284 | 2,81862E-09 | SMC | 0,559521389 | 1,787241773 | 0,151408451 |
| ITGA81 | 8,62579E-14 | -1,241696681 | 0,159 | 0,466 | 3,15592E-09 | SMC | 0,422875041 | 2,36476477 | 0,341201717 |
| NET13 | 8,99462E-14 | -1,25031487 | 0,253 | 0,612 | 3,29086E-09 | SMC | 0,420356454 | 2,378933379 | 0,413398693 |
| COL1A21 | 2,14837E-13 | 1,653913151 | 0,699 | 0,397 | 7,86024E-09 | SMC | 3,146860336 | 0,317777052 | 1,76070529 |
| C12orf753 | 3,19604E-13 | -1,229687825 | 0,179 | 0,509 | 1,16933E-08 | SMC | 0,426409704 | 2,345162391 | 0,351669941 |
| TMSB106 | 3,45634E-13 | 0,993493305 | 0,92 | 0,784 | 1,26457E-08 | SMC | 1,991000115 | 0,502260142 | 1,173469388 |
| PCOLCE1 | 3,63411E-13 | 1,11852495 | 0,611 | 0,207 | 1,32961E-08 | SMC | 2,171248647 | 0,460564478 | 2,951690821 |
| KCNMA1 | 7,49295E-13 | -0,746030388 | 0,068 | 0,328 | 2,74145E-08 | SMC | 0,596241877 | 1,677171696 | 0,207317073 |
| IFITM15 | 1,93155E-12 | 0,998962067 | 0,824 | 0,474 | 7,06695E-08 | SMC | 1,998561637 | 0,50035985 | 1,738396624 |
| RPL387 | 2,6553E-12 | -0,881867241 | 0,631 | 0,871 | 9,71494E-08 | SMC | 0,542664621 | 1,842758791 | 0,72445465 |
| IFITM37 | 2,83726E-12 | 0,770117103 | 0,98 | 0,931 | 1,03807E-07 | SMC | 1,705408205 | 0,586369877 | 1,052631579 |
| C1S2 | 6,40759E-12 | 1,390661882 | 0,693 | 0,405 | 2,34435E-07 | SMC | 2,621989453 | 0,381389787 | 1,711111111 |
| ARHGDIB4 | 9,91334E-12 | 1,324977764 | 0,548 | 0,19 | 3,62699E-07 | SMC | 2,505290264 | 0,399155345 | 2,884210526 |
| NDUFA4L21 | 1,44748E-11 | 1,284928218 | 0,719 | 0,388 | 5,29591E-07 | SMC | 2,436699285 | 0,410391223 | 1,853092784 |
| RPS255 | 1,48612E-11 | -0,637029624 | 0,926 | 0,966 | 5,43728E-07 | SMC | 0,643035537 | 1,555124004 | 0,958592133 |
| DSTN1 | 1,82625E-11 | -0,940675696 | 0,759 | 0,862 | 6,6817E-07 | SMC | 0,520988815 | 1,919427005 | 0,880510441 |

| Supplementary Table 4 - clusermarker | | | | | | | | | |
|---|---|---|---|---|---|---|---|---|---|
| | **p_val** | **avg_log2FC** | **pct.1** | **pct.2** | **p_val_adj** | **celltype** | **Foldchange_UP** | **Foldchange_DOWN** | **Ratio_pct1_pct2** |
| PDGFRB1 | 2,55531E-11 | 1,109240737 | 0,744 | 0,414 | 9,34911E-07 | SMC | 2,157320818 | 0,463537918 | 1,797101449 |
| FTH17 | 2,60661E-11 | 0,548120783 | 0,989 | 0,966 | 9,53681E-07 | SMC | 1,462179857 | 0,683910392 | 1,023809524 |
| ITIH51 | 3,41165E-11 | -1,038379414 | 0,19 | 0,483 | 1,24822E-06 | SMC | 0,486874075 | 2,05391918 | 0,393374741 |
| SERPING13 | 3,52795E-11 | 0,831603703 | 0,892 | 0,776 | 1,29077E-06 | SMC | 1,779662539 | 0,561904281 | 1,149484536 |
| ACTG21 | 3,62295E-11 | -1,999516392 | 0,239 | 0,552 | 1,32553E-06 | SMC | 0,250083817 | 3,998659377 | 0,432971014 |
| FMO3 | 3,9855E-11 | 1,40827916 | 0,355 | 0,043 | 1,45817E-06 | SMC | 2,654203817 | 0,376760818 | 8,255813953 |
| KCNMB11 | 4,37896E-11 | -0,94841457 | 0,142 | 0,414 | 1,60213E-06 | SMC | 0,51820162 | 1,929750818 | 0,342995169 |
| HSPA1A7 | 4,40362E-11 | -1,738809094 | 0,608 | 0,802 | 1,61115E-06 | SMC | 0,2996169 | 3,337595444 | 0,758104738 |
| ZG16B7 | 4,71229E-11 | -0,919133525 | 0,011 | 0,164 | 1,72409E-06 | SMC | 0,528826535 | 1,89097924 | 0,067073171 |
| CFH2 | 5,09931E-11 | 0,936095902 | 0,605 | 0,224 | 1,86569E-06 | SMC | 1,9133435 | 0,522645307 | 2,700892857 |
| CASQ2 | 5,75328E-11 | -1,007999304 | 0,151 | 0,44 | 2,10495E-06 | SMC | 0,497235324 | 2,011120191 | 0,343181818 |
| COL4A1 | 5,81021E-11 | 1,457971144 | 0,56 | 0,233 | 2,12578E-06 | SMC | 2,747217519 | 0,364004668 | 2,403433476 |
| THY11 | 6,71901E-11 | 1,349895376 | 0,5 | 0,155 | 2,45829E-06 | SMC | 2,5489364 | 0,392320499 | 3,225806452 |
| KCNAB1 | 8,03062E-11 | -0,951777931 | 0,145 | 0,431 | 2,93816E-06 | SMC | 0,516994941 | 1,934254904 | 0,336426914 |
| C1R1 | 1,04506E-10 | 1,199599158 | 0,438 | 0,103 | 3,82356E-06 | SMC | 2,296758485 | 0,435396236 | 4,252427184 |
| GGT5 | 1,60425E-10 | 1,16597299 | 0,537 | 0,181 | 5,86946E-06 | SMC | 2,243844952 | 0,445663592 | 2,966850829 |
| CPED1 | 1,78941E-10 | -0,6962985 | 0,105 | 0,362 | 6,54691E-06 | SMC | 0,617153598 | 1,620342169 | 0,290055249 |
| AKAP6 | 2,06853E-10 | -0,711454787 | 0,04 | 0,233 | 7,56813E-06 | SMC | 0,610704006 | 1,637454463 | 0,17167382 |
| HOPX1 | 4,05861E-10 | 1,153629071 | 0,477 | 0,138 | 1,48492E-05 | SMC | 2,224728171 | 0,449493117 | 3,456521739 |
| PMP222 | 4,55266E-10 | 1,300382939 | 0,514 | 0,224 | 1,66568E-05 | SMC | 2,462942486 | 0,406018413 | 2,294642857 |
| RGS5 | 4,60605E-10 | 1,285358933 | 0,744 | 0,431 | 1,68521E-05 | SMC | 2,437426868 | 0,41026872 | 1,726218097 |
| COL4A21 | 4,67992E-10 | 1,329634101 | 0,67 | 0,397 | 1,71224E-05 | SMC | 2,513389218 | 0,397869137 | 1,687657431 |
| AC036108.21 | 5,14163E-10 | -0,70985781 | 0,026 | 0,19 | 1,88117E-05 | SMC | 0,611380393 | 1,635642902 | 0,136842105 |
| SBSPON | 5,63511E-10 | -0,887170456 | 0,128 | 0,379 | 2,06172E-05 | SMC | 0,540673497 | 1,849545069 | 0,337730871 |
| SYNM1 | 6,04398E-10 | -1,136027515 | 0,116 | 0,353 | 2,21131E-05 | SMC | 0,455010734 | 2,19775035 | 0,328611898 |
| RNF1501 | 6,07555E-10 | -0,606346989 | 0,051 | 0,25 | 2,22286E-05 | SMC | 0,656857811 | 1,522399496 | 0,204 |
| GADD45G2 | 6,27079E-10 | -1,279903997 | 0,125 | 0,371 | 2,2943E-05 | SMC | 0,411822912 | 2,428228179 | 0,336927224 |
| RPL346 | 7,82831E-10 | -0,539584744 | 0,972 | 0,991 | 2,86414E-05 | SMC | 0,687968901 | 1,453554076 | 0,980827447 |
| RPL318 | 1,42898E-09 | -0,702230447 | 0,761 | 0,914 | 5,22822E-05 | SMC | 0,61462125 | 1,627018265 | 0,832603939 |
| UQCRB3 | 1,69774E-09 | -0,814998711 | 0,634 | 0,81 | 6,21152E-05 | SMC | 0,568408994 | 1,75929658 | 0,782716049 |
| COL1A11 | 1,84388E-09 | 1,4365138 | 0,631 | 0,345 | 6,7462E-05 | SMC | 2,706660237 | 0,369459006 | 1,828985507 |

| Supplementary Table 4 - clusermarker | | | | | | | | | |
|---|---|---|---|---|---|---|---|---|---|
| | **p_val** | **avg_log2FC** | **pct.1** | **pct.2** | **p_val_adj** | **celltype** | **Foldchange_UP** | **Foldchange_DOWN** | **Ratio_pct1_pct2** |
| RPL76 | 2,34599E-09 | -0,66574249 | 0,81 | 0,931 | 8,58327E-05 | SMC | 0,630364201 | 1,586384503 | 0,870032223 |
| ACKR31 | 2,50745E-09 | -1,134946809 | 0,153 | 0,405 | 9,17402E-05 | SMC | 0,455351705 | 2,196104657 | 0,377777778 |
| RPL94 | 3,30047E-09 | -0,53392983 | 0,932 | 0,957 | 0,000120754 | SMC | 0,690670816 | 1,44786775 | 0,973876698 |
| ACTA22 | 3,8541E-09 | -0,91687561 | 0,864 | 0,914 | 0,00014101 | SMC | 0,529654833 | 1,888022045 | 0,945295405 |
| MT-CO14 | 5,21977E-09 | 0,359819466 | 0,983 | 1 | 0,000190976 | SMC | 1,283265304 | 0,779262088 | 0,983 |
| PPP1R1A | 5,57695E-09 | -0,503045165 | 0,017 | 0,155 | 0,000204044 | SMC | 0,705615831 | 1,417201763 | 0,109677419 |
| SPARC3 | 6,723E-09 | 1,180763064 | 0,835 | 0,75 | 0,000245975 | SMC | 2,266966488 | 0,441118122 | 1,113333333 |
| CLEC3B1 | 7,09577E-09 | 1,153163291 | 0,483 | 0,181 | 0,000259613 | SMC | 2,224010024 | 0,449638261 | 2,668508287 |
| RPS216 | 7,86625E-09 | -0,587826012 | 0,832 | 0,905 | 0,000287803 | SMC | 0,665344756 | 1,502980209 | 0,919337017 |
| RPL377 | 8,32214E-09 | -0,50333372 | 0,957 | 0,974 | 0,000304482 | SMC | 0,705474715 | 1,417485247 | 0,982546201 |
| NOTCH3 | 9,19968E-09 | 0,737478239 | 0,807 | 0,517 | 0,000336589 | SMC | 1,667259003 | 0,599786835 | 1,560928433 |
| LHFPL61 | 9,44388E-09 | 0,876248798 | 0,713 | 0,422 | 0,000345523 | SMC | 1,835596292 | 0,544782098 | 1,68957346 |
| COL3A11 | 1,1113E-08 | 1,591274123 | 0,571 | 0,31 | 0,00040659 | SMC | 3,013153401 | 0,331878224 | 1,841935484 |
| COL18A12 | 1,13047E-08 | 1,370397497 | 0,727 | 0,517 | 0,000413607 | SMC | 2,585417908 | 0,386784665 | 1,406189555 |
| DPYSL3 | 1,82902E-08 | -0,668964537 | 0,094 | 0,302 | 0,000669183 | SMC | 0,628957946 | 1,589931419 | 0,311258278 |
| PPIB4 | 1,85368E-08 | 0,794532961 | 0,724 | 0,474 | 0,000678207 | SMC | 1,734515774 | 0,576529781 | 1,52742616 |
| TMEM176A3 | 2,11766E-08 | 0,93391888 | 0,412 | 0,112 | 0,000774789 | SMC | 1,910458448 | 0,523434572 | 3,678571429 |
| ANGPTL11 | 2,14026E-08 | -0,799863576 | 0,153 | 0,388 | 0,000783058 | SMC | 0,574403492 | 1,740936492 | 0,394329897 |
| HSPA1B6 | 2,75904E-08 | -1,393874688 | 0,443 | 0,681 | 0,001009451 | SMC | 0,380541398 | 2,627834991 | 0,65051395 |
| EPS82 | 2,8032E-08 | 0,690775308 | 0,75 | 0,483 | 0,001025606 | SMC | 1,614150735 | 0,619520828 | 1,552795031 |
| PRSS233 | 2,85423E-08 | 1,105051765 | 0,31 | 0,052 | 0,001044277 | SMC | 2,151065961 | 0,464885791 | 5,961538462 |
| ADGRF5 | 3,04411E-08 | 0,907504451 | 0,409 | 0,112 | 0,001113748 | SMC | 1,875797967 | 0,533106453 | 3,651785714 |
| OLFML2B1 | 4,35168E-08 | 1,095576057 | 0,33 | 0,078 | 0,00159215 | SMC | 2,136983927 | 0,467949238 | 4,230769231 |
| ADIRF4 | 5,27035E-08 | -0,780459853 | 0,966 | 0,966 | 0,001928262 | SMC | 0,582181196 | 1,717678288 | 1 |
| PRRX11 | 5,35525E-08 | 0,946521691 | 0,599 | 0,302 | 0,001959325 | SMC | 1,927220562 | 0,518881969 | 1,983443709 |
| COL4A5 | 5,47598E-08 | -0,281129131 | 0,048 | 0,216 | 0,002003496 | SMC | 0,822946683 | 1,215145551 | 0,222222222 |
| FSTL1 | 6,88375E-08 | 1,000637115 | 0,443 | 0,181 | 0,002518557 | SMC | 2,000883424 | 0,499779241 | 2,447513812 |
| OGN | 7,3204E-08 | -0,737432247 | 0,043 | 0,198 | 0,002678313 | SMC | 0,599805956 | 1,667205852 | 0,217171717 |
| MSMB8 | 7,65033E-08 | -0,705760538 | 0,023 | 0,155 | 0,002799025 | SMC | 0,613119189 | 1,631004246 | 0,148387097 |
| WFDC25 | 8,03185E-08 | -0,721181554 | 0,043 | 0,198 | 0,002938613 | SMC | 0,606600438 | 1,648531614 | 0,217171717 |
| BCHE | 9,60574E-08 | -0,339671767 | 0,006 | 0,103 | 0,003514451 | SMC | 0,790221078 | 1,26546865 | 0,058252427 |

| Supplementary Table 4 - clusermarker | | | | | | | | | |
|---|---|---|---|---|---|---|---|---|---|
| | **p_val** | **avg_log2FC** | **pct.1** | **pct.2** | **p_val_adj** | **celltype** | **Foldchange_UP** | **Foldchange_DOWN** | **Ratio_pct1_pct2** |
| TIMP14 | 1,12072E-07 | 0,542845937 | 0,73 | 0,483 | 0,00410039 | SMC | 1,456843531 | 0,686415513 | 1,511387164 |
| IFITM25 | 1,14191E-07 | 0,604371041 | 0,898 | 0,793 | 0,00417792 | SMC | 1,52031581 | 0,657758075 | 1,132408575 |
| PHLDA12 | 1,17797E-07 | 0,851265928 | 0,52 | 0,224 | 0,004309848 | SMC | 1,804083268 | 0,55429814 | 2,321428571 |
| RPL137 | 1,33747E-07 | 0,499419836 | 0,986 | 0,966 | 0,004893398 | SMC | 1,413644966 | 0,707391194 | 1,020703934 |
| FHL21 | 1,52973E-07 | 1,255836029 | 0,469 | 0,224 | 0,005596835 | SMC | 2,388054942 | 0,418750835 | 2,09375 |
| IL322 | 1,61582E-07 | 1,222756493 | 0,287 | 0,06 | 0,005911783 | SMC | 2,333922237 | 0,42846329 | 4,783333333 |
| HLA-C6 | 1,65726E-07 | 0,650019937 | 0,866 | 0,75 | 0,006063422 | SMC | 1,56918988 | 0,637271507 | 1,154666667 |
| ANKRD103 | 1,85622E-07 | 0,844781893 | 0,455 | 0,198 | 0,006791357 | SMC | 1,795993206 | 0,556794979 | 2,297979798 |
| FRZB1 | 1,9066E-07 | 0,641119207 | 0,761 | 0,534 | 0,006975672 | SMC | 1,559538541 | 0,641215317 | 1,425093633 |
| ITM2B7 | 2,52225E-07 | 0,627255601 | 0,915 | 0,828 | 0,00922814 | SMC | 1,544623902 | 0,647406789 | 1,105072464 |
| CSRP13 | 2,53971E-07 | -0,930964136 | 0,548 | 0,716 | 0,009292039 | SMC | 0,524507702 | 1,906549696 | 0,765363128 |
| EPHX12 | 2,60004E-07 | 0,66491235 | 0,582 | 0,319 | 0,009512777 | SMC | 1,585471946 | 0,630727023 | 1,824451411 |
| DKK3 | 2,83838E-07 | 0,873389362 | 0,523 | 0,259 | 0,010384789 | SMC | 1,831961726 | 0,545862933 | 2,019305019 |
| MT-ATP88 | 3,3705E-07 | -0,558561003 | 0,79 | 0,957 | 0,012331633 | SMC | 0,678979065 | 1,47279946 | 0,825496343 |
| TSC22D15 | 3,71562E-07 | -0,634104032 | 0,662 | 0,828 | 0,013594335 | SMC | 0,644340851 | 1,551973616 | 0,799516908 |
| CTSC4 | 3,72809E-07 | 0,74465932 | 0,443 | 0,172 | 0,013639946 | SMC | 1,675578549 | 0,596808786 | 2,575581395 |
| CYBA8 | 4,72251E-07 | 1,037948282 | 0,364 | 0,121 | 0,017278262 | SMC | 2,053305482 | 0,487019593 | 3,008264463 |
| RPS145 | 4,74912E-07 | -0,405933587 | 0,977 | 0,966 | 0,017375603 | SMC | 0,754747727 | 1,324946023 | 1,011387164 |
| MT-ND38 | 4,94683E-07 | -0,420767587 | 0,935 | 0,957 | 0,018098983 | SMC | 0,747027062 | 1,338639589 | 0,977011494 |
| PLXDC1 | 5,03144E-07 | 1,082892573 | 0,256 | 0,043 | 0,018408546 | SMC | 2,11827893 | 0,472081361 | 5,953488372 |
| GADD45B5 | 5,04782E-07 | -1,150561026 | 0,821 | 0,862 | 0,018468469 | SMC | 0,450450029 | 2,220002077 | 0,952436195 |
| AZGP13 | 5,13417E-07 | -0,323762612 | 0,009 | 0,103 | 0,018784393 | SMC | 0,798983374 | 1,251590499 | 0,087378641 |
| COX7C5 | 5,29478E-07 | -0,71285728 | 0,699 | 0,802 | 0,019372004 | SMC | 0,610110608 | 1,639047063 | 0,871571072 |
| ISYNA1 | 6,64001E-07 | 0,811456834 | 0,636 | 0,397 | 0,024293804 | SMC | 1,75498273 | 0,569806177 | 1,602015113 |
| RASSF3 | 6,94832E-07 | -0,424281969 | 0,045 | 0,19 | 0,0254218 | SMC | 0,745209531 | 1,341904469 | 0,236842105 |
| MDK4 | 7,02671E-07 | 1,027666788 | 0,347 | 0,121 | 0,025708629 | SMC | 2,038724438 | 0,490502778 | 2,867768595 |
| SORBS11 | 7,08779E-07 | -0,834598317 | 0,179 | 0,388 | 0,025932097 | SMC | 0,560739141 | 1,783360437 | 0,461340206 |
| KRT182 | 7,22953E-07 | 0,862859878 | 0,202 | 0,009 | 0,026450683 | SMC | 1,81863986 | 0,549861477 | 22,44444444 |
| RHOB3 | 8,02602E-07 | -0,875528534 | 0,659 | 0,784 | 0,029364803 | SMC | 0,545054148 | 1,834680102 | 0,840561224 |
| ATP5ME5 | 8,03373E-07 | -0,563643867 | 0,347 | 0,586 | 0,029393023 | SMC | 0,676591113 | 1,477997538 | 0,592150171 |
| COL4A6 | 8,76416E-07 | -0,278261698 | 0,026 | 0,147 | 0,032065418 | SMC | 0,824583959 | 1,212732783 | 0,176870748 |

| Supplementary Table 4 - clusermarker | | | | | | | | | |
|---|---|---|---|---|---|---|---|---|---|
| | **p_val** | **avg_log2FC** | **pct.1** | **pct.2** | **p_val_adj** | **celltype** | **Foldchange_UP** | **Foldchange_DOWN** | **Ratio_pct1_pct2** |
| MYL93 | 9,54558E-07 | -0,694445137 | 0,986 | 0,983 | 0,034924397 | SMC | 0,617946936 | 1,618261927 | 1,003051882 |
| SUSD5 | 9,72049E-07 | -0,714664562 | 0,119 | 0,31 | 0,035564372 | SMC | 0,609346793 | 1,641101604 | 0,383870968 |
| ASAP1 | 1,0383E-06 | 0,846248898 | 0,327 | 0,103 | 0,037988449 | SMC | 1,797820391 | 0,55622909 | 3,174757282 |
| COL5A21 | 1,05233E-06 | 0,92115851 | 0,364 | 0,121 | 0,03850172 | SMC | 1,893635306 | 0,528084788 | 3,008264463 |
| LDOC11 | 1,11436E-06 | -0,680083618 | 0,139 | 0,336 | 0,040771058 | SMC | 0,624129099 | 1,602232617 | 0,413690476 |
| DES | 1,16201E-06 | -2,060038142 | 0,111 | 0,293 | 0,042514376 | SMC | 0,23980969 | 4,169973289 | 0,37883959 |
| TCEA31 | 1,21792E-06 | -0,478386733 | 0,065 | 0,224 | 0,044560133 | SMC | 0,71777982 | 1,393184891 | 0,290178571 |
| PGAM2 | 1,43689E-06 | -0,351081757 | 0,02 | 0,129 | 0,052571601 | SMC | 0,783996024 | 1,275516673 | 0,15503876 |
| MT-CO32 | 1,43817E-06 | 0,345595869 | 0,991 | 0,974 | 0,052618308 | SMC | 1,270675695 | 0,786982866 | 1,017453799 |
| RMRP2 | 1,50924E-06 | -0,75654897 | 0,048 | 0,19 | 0,055218674 | SMC | 0,591910531 | 1,689444515 | 0,252631579 |
| TAGLN3 | 1,52612E-06 | -0,55211911 | 0,977 | 0,983 | 0,055836158 | SMC | 0,682017607 | 1,466237806 | 0,993896236 |
| MYOCD | 1,54422E-06 | -0,828731886 | 0,153 | 0,353 | 0,05649837 | SMC | 0,563023917 | 1,776123481 | 0,433427762 |
| MSRB32 | 1,55582E-06 | -0,655903327 | 0,398 | 0,603 | 0,056922805 | SMC | 0,63467797 | 1,57560219 | 0,660033167 |
| LMOD12 | 1,59235E-06 | -0,85830335 | 0,625 | 0,776 | 0,058259408 | SMC | 0,551600875 | 1,812905029 | 0,805412371 |
| ID26 | 1,62875E-06 | -0,832640835 | 0,281 | 0,509 | 0,059591 | SMC | 0,561500482 | 1,780942373 | 0,552062868 |
| CX3CL11 | 1,63655E-06 | -0,381856827 | 0,043 | 0,181 | 0,059876311 | SMC | 0,767449206 | 1,303017832 | 0,237569061 |
| NR2F21 | 1,73278E-06 | 0,556855346 | 0,878 | 0,793 | 0,063397309 | SMC | 1,47105924 | 0,679782277 | 1,107187894 |
| KCNA5 | 1,81038E-06 | -0,539990647 | 0,045 | 0,181 | 0,0662363 | SMC | 0,687775368 | 1,453963091 | 0,248618785 |
| HOXA51 | 1,85171E-06 | -0,489744369 | 0,011 | 0,103 | 0,067748527 | SMC | 0,712151273 | 1,404196045 | 0,106796117 |
| FGL22 | 1,87802E-06 | -0,767891138 | 0,014 | 0,112 | 0,06871097 | SMC | 0,587275298 | 1,702778924 | 0,125 |
| PALLD3 | 2,30317E-06 | -0,968543268 | 0,506 | 0,647 | 0,084266199 | SMC | 0,511021796 | 1,956863694 | 0,782071097 |
| ADCY5 | 2,32988E-06 | -0,330658046 | 0,048 | 0,19 | 0,085243353 | SMC | 0,795173704 | 1,257586858 | 0,252631579 |
| TFPI1 | 2,64354E-06 | 0,704325211 | 0,534 | 0,293 | 0,096719204 | SMC | 1,629382378 | 0,61372948 | 1,822525597 |
| CD593 | 2,70756E-06 | 0,634964299 | 0,75 | 0,595 | 0,099061558 | SMC | 1,552899321 | 0,64395675 | 1,260504202 |
| FAU1 | 2,92798E-06 | -0,377523393 | 0,943 | 0,983 | 0,107125946 | SMC | 0,769757864 | 1,299109819 | 0,95930824 |
| SLIT31 | 3,07458E-06 | 0,709672297 | 0,645 | 0,422 | 0,1124895 | SMC | 1,635432593 | 0,611459014 | 1,528436019 |
| RPL285 | 3,10711E-06 | 0,456486706 | 0,96 | 0,914 | 0,113679844 | SMC | 1,372196133 | 0,728758795 | 1,050328228 |
| LAMB22 | 3,23009E-06 | 0,855381421 | 0,489 | 0,259 | 0,118179285 | SMC | 1,80923702 | 0,552719179 | 1,888030888 |
| ACTC1 | 3,48838E-06 | -0,408317854 | 0,023 | 0,129 | 0,127629312 | SMC | 0,753501426 | 1,327137502 | 0,178294574 |
| SASH1 | 3,53891E-06 | 0,803428412 | 0,295 | 0,086 | 0,129478107 | SMC | 1,745243589 | 0,572985918 | 3,430232558 |
| ACTB8 | 3,7334E-06 | -0,461838686 | 0,989 | 0,991 | 0,13659394 | SMC | 0,726060319 | 1,377296036 | 0,997981837 |

| Supplementary Table 4 - clusermarker | | | | | | | | | |
|---|---|---|---|---|---|---|---|---|---|
| | **p_val** | **avg_log2FC** | **pct.1** | **pct.2** | **p_val_adj** | **celltype** | **Foldchange_UP** | **Foldchange_DOWN** | **Ratio_pct1_pct2** |
| SERPINF1 | 3,87433E-06 | 0,707457681 | 0,469 | 0,216 | 0,141750275 | SMC | 1,632924039 | 0,612398358 | 2,171296296 |
| CDC42EP52 | 4,09983E-06 | 0,653004759 | 0,162 | 0 | 0,150000442 | SMC | 1,572439771 | 0,635954406 | #DIV/0! |
| MUSTN1 | 4,27089E-06 | -0,911347889 | 0,639 | 0,741 | 0,156259206 | SMC | 0,531688111 | 1,880801884 | 0,862348178 |
| SEMA5A2 | 4,69678E-06 | 0,828786212 | 0,23 | 0,043 | 0,171840993 | SMC | 1,776190364 | 0,563002716 | 5,348837209 |
| C20orf271 | 4,71535E-06 | 0,820589305 | 0,358 | 0,138 | 0,172520665 | SMC | 1,766127264 | 0,566210613 | 2,594202899 |
| RPL124 | 4,91473E-06 | 0,470256078 | 0,969 | 0,957 | 0,179815206 | SMC | 1,385355346 | 0,721836461 | 1,012539185 |
| PIGR3 | 5,05098E-06 | -0,486524245 | 0,02 | 0,121 | 0,184800086 | SMC | 0,713742584 | 1,401065346 | 0,165289256 |
| LGALS3BP2 | 5,16654E-06 | 0,925096261 | 0,432 | 0,233 | 0,189028103 | SMC | 1,898810932 | 0,526645378 | 1,854077253 |
| HSPB81 | 5,2414E-06 | -0,713915327 | 0,227 | 0,422 | 0,191767174 | SMC | 0,609663328 | 1,640249552 | 0,537914692 |
| ALDH1B1 | 5,49341E-06 | -0,313488205 | 0,02 | 0,121 | 0,200987405 | SMC | 0,804693785 | 1,242708741 | 0,165289256 |
| IGKV3-206 | 6,15498E-06 | 2,48159006 | 0,213 | 0,034 | 0,225192184 | SMC | 5,585126899 | 0,179046961 | 6,264705882 |
| DEPP11 | 6,5759E-06 | 1,211570014 | 0,477 | 0,267 | 0,240592324 | SMC | 2,315895272 | 0,431798455 | 1,786516854 |
| MYL64 | 6,63255E-06 | -0,525097753 | 0,957 | 0,957 | 0,242665286 | SMC | 0,694912023 | 1,439031082 | 1 |
| ITIH3 | 7,15717E-06 | -0,553991033 | 0,062 | 0,198 | 0,261859276 | SMC | 0,681133251 | 1,468141511 | 0,313131313 |
| PCP4L11 | 7,47057E-06 | -0,417225239 | 0,028 | 0,138 | 0,273325646 | SMC | 0,748863543 | 1,335356768 | 0,202898551 |
| IFI163 | 8,23076E-06 | 0,329227661 | 0,509 | 0,259 | 0,301138842 | SMC | 1,256340619 | 0,795962484 | 1,965250965 |
| SMTN | 8,34354E-06 | -0,864861084 | 0,193 | 0,371 | 0,30526525 | SMC | 0,549099277 | 1,821164301 | 0,520215633 |
| AHR2 | 8,74337E-06 | 0,83387965 | 0,412 | 0,198 | 0,319893777 | SMC | 1,782472291 | 0,561018539 | 2,080808081 |
| LINC01936 | 9,26006E-06 | -0,278171775 | 0,014 | 0,103 | 0,338797833 | SMC | 0,824635357 | 1,212657196 | 0,13592233 |
| PLPPR4 | 9,26148E-06 | 0,621212212 | 0,276 | 0,069 | 0,338849743 | SMC | 1,53816707 | 0,650124437 | 4 |
| H4C35 | 9,6058E-06 | -0,740271904 | 0,372 | 0,552 | 0,351447464 | SMC | 0,598626519 | 1,670490646 | 0,673913043 |
| PEAK1 | 1,02527E-05 | 0,81676355 | 0,364 | 0,155 | 0,375116585 | SMC | 1,761450033 | 0,567714089 | 2,348387097 |
| S100A114 | 1,05657E-05 | 0,561240671 | 0,795 | 0,621 | 0,386567741 | SMC | 1,475537586 | 0,677719097 | 1,280193237 |
| RPS267 | 1,14558E-05 | -0,369860875 | 0,844 | 0,957 | 0,419134875 | SMC | 0,77385712 | 1,29222821 | 0,881922675 |
| IGKV4-17 | 1,29736E-05 | 1,143518597 | 0,162 | 0,009 | 0,474665728 | SMC | 2,209191676 | 0,452654249 | 18 |
| COL14A1 | 1,30424E-05 | 0,557926843 | 0,79 | 0,638 | 0,477183356 | SMC | 1,472152209 | 0,679277587 | 1,238244514 |
| NEXN2 | 1,30427E-05 | -0,700586097 | 0,426 | 0,621 | 0,47719162 | SMC | 0,61532218 | 1,625164884 | 0,685990338 |
| SPEG | 1,33714E-05 | -0,49705162 | 0,105 | 0,267 | 0,489218483 | SMC | 0,708553346 | 1,41132634 | 0,393258427 |
| MYH93 | 1,34598E-05 | 0,760203434 | 0,759 | 0,664 | 0,492452745 | SMC | 1,69372944 | 0,590413071 | 1,143072289 |
| ITGA12 | 1,36622E-05 | 0,916224634 | 0,426 | 0,233 | 0,49985775 | SMC | 1,887170319 | 0,529893879 | 1,82832618 |
| RPL23A4 | 1,38051E-05 | -0,444755304 | 0,915 | 0,94 | 0,505085896 | SMC | 0,73470892 | 1,361083243 | 0,973404255 |

| Supplementary Table 4 - clusermarker | | | | | | | | | |
|---|---|---|---|---|---|---|---|---|---|
| | **p_val** | **avg_log2FC** | **pct.1** | **pct.2** | **p_val_adj** | **celltype** | **Foldchange_UP** | **Foldchange_DOWN** | **Ratio_pct1_pct2** |
| PRDM16 | 1,38267E-05 | -0,315295215 | 0,034 | 0,147 | 0,505878155 | SMC | 0,803686518 | 1,244266238 | 0,231292517 |
| RPL415 | 1,41506E-05 | -0,308977451 | 0,997 | 1 | 0,517727863 | SMC | 0,807213691 | 1,238829335 | 0,997 |
| PXDN1 | 1,50635E-05 | 0,740425131 | 0,188 | 0,026 | 0,551127668 | SMC | 1,670668076 | 0,598562943 | 7,230769231 |
| LIMCH1 | 1,52009E-05 | -0,320402855 | 0,034 | 0,147 | 0,556154803 | SMC | 0,80084622 | 1,248679178 | 0,231292517 |
| ZEB1 | 1,56117E-05 | -0,588198577 | 0,196 | 0,388 | 0,571184708 | SMC | 0,665172958 | 1,503368392 | 0,505154639 |
| FLNC | 1,78858E-05 | -0,557637162 | 0,045 | 0,164 | 0,65438911 | SMC | 0,679413994 | 1,471856643 | 0,274390244 |
| PRXL2A3 | 1,89062E-05 | 0,644231432 | 0,273 | 0,086 | 0,691720347 | SMC | 1,562906456 | 0,639833559 | 3,174418605 |
| IGLV3-216 | 1,96185E-05 | 1,732135446 | 0,156 | 0,009 | 0,71778199 | SMC | 3,322191982 | 0,301006084 | 17,33333333 |
| PRPH | 2,07347E-05 | -0,677968327 | 0,028 | 0,129 | 0,758621264 | SMC | 0,625044874 | 1,599885132 | 0,217054264 |
| SSTR2 | 2,27853E-05 | 0,543661701 | 0,261 | 0,069 | 0,833646205 | SMC | 1,457667529 | 0,686027493 | 3,782608696 |
| COL6A21 | 2,28337E-05 | 0,683768104 | 0,761 | 0,647 | 0,835414838 | SMC | 1,606329774 | 0,622537175 | 1,176197836 |
| CHSY3 | 2,30611E-05 | 0,557245497 | 0,139 | 0 | 0,843734724 | SMC | 1,471457116 | 0,679598467 | #DIV/0! |
| SLC38A25 | 2,35408E-05 | 0,623902361 | 0,753 | 0,629 | 0,861287321 | SMC | 1,541037918 | 0,648913299 | 1,197138315 |
| TMEM176B3 | 2,37745E-05 | 0,619243349 | 0,324 | 0,112 | 0,869837103 | SMC | 1,536069347 | 0,651012275 | 2,892857143 |
| FABP3 | 2,4398E-05 | -0,430397012 | 0,06 | 0,19 | 0,892647963 | SMC | 0,742057552 | 1,34760437 | 0,315789474 |
| SH3BGR1 | 2,45817E-05 | -0,431721672 | 0,045 | 0,164 | 0,899370662 | SMC | 0,741376519 | 1,348842288 | 0,274390244 |
| GNAS5 | 2,54803E-05 | -0,470270026 | 0,722 | 0,784 | 0,932249541 | SMC | 0,721829482 | 1,38536874 | 0,920918367 |
| ZBTB162 | 2,64225E-05 | -0,542486126 | 0,293 | 0,509 | 0,966720553 | SMC | 0,686586728 | 1,456480237 | 0,575638507 |
| RPL44 | 2,78591E-05 | -0,448767353 | 0,75 | 0,853 | 1 | SMC | 0,732668577 | 1,364873603 | 0,879249707 |
| SYTL25 | 2,78645E-05 | 0,702619491 | 0,5 | 0,284 | 1 | SMC | 1,627457073 | 0,614455531 | 1,76056338 |
| MYOM1 | 2,84432E-05 | -0,501483668 | 0,136 | 0,31 | 1 | SMC | 0,706379966 | 1,415668688 | 0,438709677 |
| ABCC9 | 2,99658E-05 | 0,59318049 | 0,426 | 0,207 | 1 | SMC | 1,508568796 | 0,662879945 | 2,057971014 |
| CHCHD106 | 3,03334E-05 | 0,757414153 | 0,537 | 0,362 | 1 | SMC | 1,690457977 | 0,591555669 | 1,483425414 |
| CPE | 3,22758E-05 | 0,568364076 | 0,835 | 0,793 | 1 | SMC | 1,48284117 | 0,67438106 | 1,05296343 |
| CCN32 | 3,25961E-05 | -0,523045321 | 0,08 | 0,224 | 1 | SMC | 0,695901335 | 1,436985317 | 0,357142857 |
| IGHG17 | 3,52619E-05 | 1,272021479 | 0,134 | 0 | 1 | SMC | 2,414997138 | 0,414079166 | #DIV/0! |
| NIBAN14 | 3,53271E-05 | -0,653281763 | 0,236 | 0,422 | 1 | SMC | 0,635832311 | 1,572741715 | 0,559241706 |
| SH3BGRL5 | 3,66587E-05 | -0,536614049 | 0,668 | 0,81 | 1 | SMC | 0,689386977 | 1,4505641 | 0,824691358 |
| TSPAN42 | 3,68359E-05 | 0,746121458 | 0,335 | 0,147 | 1 | SMC | 1,67727757 | 0,596204241 | 2,278911565 |
| EPAS14 | 3,69791E-05 | 0,60100011 | 0,739 | 0,586 | 1 | SMC | 1,516767661 | 0,659296757 | 1,26109215 |
| HSPB15 | 3,84333E-05 | -0,740310148 | 0,81 | 0,871 | 1 | SMC | 0,59861065 | 1,670534928 | 0,929965557 |

| Supplementary Table 4 - clusermarker | | | | | | | | | |
|---|---|---|---|---|---|---|---|---|---|
| | **p_val** | **avg_log2FC** | **pct.1** | **pct.2** | **p_val_adj** | **celltype** | **Foldchange_UP** | **Foldchange_DOWN** | **Ratio_pct1_pct2** |
| AC020916.14 | 3,86384E-05 | 0,667068146 | 0,23 | 0,06 | 1 | SMC | 1,587842863 | 0,629785241 | 3,833333333 |
| RPL36A6 | 3,88193E-05 | -0,440885072 | 0,793 | 0,897 | 1 | SMC | 0,736682526 | 1,35743684 | 0,884057971 |
| PPP1R3C | 3,8865E-05 | -0,426200456 | 0,062 | 0,19 | 1 | SMC | 0,744219214 | 1,343690113 | 0,326315789 |
| ROCK11 | 3,92605E-05 | -0,577769331 | 0,506 | 0,672 | 1 | SMC | 0,669998917 | 1,492539726 | 0,75297619 |
| BTC | 3,93231E-05 | -0,519012566 | 0,026 | 0,121 | 1 | SMC | 0,697849304 | 1,432974132 | 0,214876033 |
| ADAMTS4 | 3,97038E-05 | 0,651067517 | 0,406 | 0,19 | 1 | SMC | 1,570329726 | 0,636808935 | 2,136842105 |
| LUM1 | 4,05331E-05 | 1,246801517 | 0,239 | 0,06 | 1 | SMC | 2,37314708 | 0,421381384 | 3,983333333 |
| RBP5 | 4,18239E-05 | 0,688436982 | 0,165 | 0,017 | 1 | SMC | 1,611536631 | 0,620525765 | 9,705882353 |
| PMEPA15 | 4,23099E-05 | 0,840320722 | 0,472 | 0,284 | 1 | SMC | 1,790448127 | 0,558519392 | 1,661971831 |
| Z93241.1 | 4,30837E-05 | -0,286282107 | 0,026 | 0,121 | 1 | SMC | 0,820012549 | 1,219493533 | 0,214876033 |
| RPS113 | 4,3382E-05 | -0,488631365 | 0,79 | 0,888 | 1 | SMC | 0,712700892 | 1,403113159 | 0,88963964 |
| RPS245 | 4,41987E-05 | -0,349859451 | 0,955 | 0,983 | 1 | SMC | 0,784660537 | 1,274436464 | 0,971515768 |
| RYR2 | 4,54543E-05 | -0,297292165 | 0,034 | 0,138 | 1 | SMC | 0,813778367 | 1,228835811 | 0,246376812 |
| CRYAB2 | 4,56143E-05 | -1,001198547 | 0,554 | 0,69 | 1 | SMC | 0,499584788 | 2,001662229 | 0,802898551 |
| CD443 | 4,5656E-05 | 0,750266409 | 0,608 | 0,44 | 1 | SMC | 1,682103421 | 0,594493768 | 1,381818182 |
| NTRK22 | 4,56989E-05 | -0,497539305 | 0,267 | 0,474 | 1 | SMC | 0,708313869 | 1,411803502 | 0,563291139 |
| KLF24 | 4,6074E-05 | -0,829266196 | 0,438 | 0,586 | 1 | SMC | 0,562815437 | 1,776781401 | 0,747440273 |
| NCS1 | 4,6155E-05 | -0,402705456 | 0,068 | 0,198 | 1 | SMC | 0,756438419 | 1,321984678 | 0,343434343 |
| SVIL2 | 4,78625E-05 | -0,790879549 | 0,219 | 0,397 | 1 | SMC | 0,577991608 | 1,730128926 | 0,55163728 |
| LPP1 | 4,90644E-05 | -0,697345073 | 0,69 | 0,776 | 1 | SMC | 0,616706059 | 1,621518039 | 0,889175258 |
| FXYD13 | 5,3815E-05 | -0,541979906 | 0,523 | 0,698 | 1 | SMC | 0,686827683 | 1,45596927 | 0,749283668 |
| ASAH13 | 5,48459E-05 | 0,876369511 | 0,48 | 0,302 | 1 | SMC | 1,835749886 | 0,544736517 | 1,589403974 |
| LAPTM4A2 | 5,53941E-05 | 0,555601897 | 0,795 | 0,655 | 1 | SMC | 1,469781703 | 0,680373145 | 1,213740458 |
| CD636 | 5,55997E-05 | 0,433171543 | 0,878 | 0,836 | 1 | SMC | 1,350198521 | 0,740631829 | 1,050239234 |
| MTRNR2L123 | 5,76054E-05 | 0,436042549 | 0,938 | 0,905 | 1 | SMC | 1,352888131 | 0,739159415 | 1,036464088 |
| KCNJ8 | 5,81184E-05 | 0,67331038 | 0,29 | 0,103 | 1 | SMC | 1,594728004 | 0,627066181 | 2,815533981 |
| SRSF55 | 5,94469E-05 | -0,55428316 | 0,466 | 0,638 | 1 | SMC | 0,680995344 | 1,46843882 | 0,730407524 |
| PET1006 | 5,94733E-05 | -0,654387925 | 0,213 | 0,388 | 1 | SMC | 0,635344984 | 1,573948052 | 0,548969072 |
| NME27 | 6,05838E-05 | 0,566645066 | 0,761 | 0,629 | 1 | SMC | 1,481075377 | 0,675185082 | 1,209856916 |
| COX7A12 | 6,08321E-05 | -0,681139069 | 0,418 | 0,595 | 1 | SMC | 0,623672664 | 1,603405212 | 0,702521008 |
| RBPMS2 | 6,13615E-05 | -0,551187087 | 0,139 | 0,302 | 1 | SMC | 0,682458353 | 1,46529088 | 0,460264901 |

| Supplementary Table 4 - clusermarker | | | | | | | | | |
|---|---|---|---|---|---|---|---|---|---|
| | **p_val** | **avg_log2FC** | **pct.1** | **pct.2** | **p_val_adj** | **celltype** | **Foldchange_UP** | **Foldchange_DOWN** | **Ratio_pct1_pct2** |
| ARHGAP151 | 6,30534E-05 | 0,670442388 | 0,312 | 0,121 | 1 | SMC | 1,591560929 | 0,628313991 | 2,578512397 |
| NUPR12 | 6,47152E-05 | -0,67409581 | 0,457 | 0,612 | 1 | SMC | 0,626724888 | 1,59559644 | 0,746732026 |
| FBLN1 | 6,63111E-05 | 0,597381373 | 0,202 | 0,043 | 1 | SMC | 1,512967894 | 0,660952558 | 4,697674419 |
| NEAT15 | 6,64629E-05 | 0,653814613 | 0,727 | 0,56 | 1 | SMC | 1,573322705 | 0,635597514 | 1,298214286 |
| RPL27A6 | 6,82039E-05 | -0,506189061 | 0,639 | 0,784 | 1 | SMC | 0,70407984 | 1,420293471 | 0,81505102 |
| DYNLT14 | 6,89134E-05 | -0,544962045 | 0,241 | 0,431 | 1 | SMC | 0,685409434 | 1,458981961 | 0,559164733 |
| PRPSAP1 | 6,96404E-05 | -0,344019842 | 0,099 | 0,25 | 1 | SMC | 0,787843051 | 1,269288343 | 0,396 |
| KCNE4 | 6,98683E-05 | 0,742425976 | 0,386 | 0,198 | 1 | SMC | 1,6729867 | 0,597733383 | 1,949494949 |
| MIR4435-2HG1 | 7,23705E-05 | 0,55633688 | 0,256 | 0,078 | 1 | SMC | 1,470530676 | 0,680026617 | 3,282051282 |
| PPP1R12B1 | 7,29139E-05 | -0,638604826 | 0,264 | 0,44 | 1 | SMC | 0,642333824 | 1,556822889 | 0,6 |
| INMT | 7,33464E-05 | -0,464974819 | 0,091 | 0,233 | 1 | SMC | 0,724483722 | 1,380293262 | 0,39055794 |
| ARHGEF10L | 7,7752E-05 | -0,344053171 | 0,054 | 0,172 | 1 | SMC | 0,78782485 | 1,269317666 | 0,313953488 |
| CTSK1 | 7,95681E-05 | 0,805463105 | 0,287 | 0,112 | 1 | SMC | 1,747706716 | 0,572178381 | 2,5625 |
| RACK13 | 8,27413E-05 | 0,351092895 | 0,935 | 0,888 | 1 | SMC | 1,27552652 | 0,783989971 | 1,052927928 |
| DBI4 | 8,31761E-05 | -0,68493208 | 0,321 | 0,5 | 1 | SMC | 0,62203511 | 1,607626295 | 0,642 |
| PLEKHO1 | 8,53808E-05 | -0,517598054 | 0,116 | 0,267 | 1 | SMC | 0,698533856 | 1,431569839 | 0,434456929 |
| TAGLN25 | 8,73375E-05 | 0,558274851 | 0,739 | 0,569 | 1 | SMC | 1,472507366 | 0,67911375 | 1,298769772 |
| CCL191 | 8,97577E-05 | 1,155307185 | 0,307 | 0,121 | 1 | SMC | 2,227317436 | 0,44897058 | 2,537190083 |
| CSDE14 | 8,98233E-05 | -0,513370367 | 0,52 | 0,664 | 1 | SMC | 0,700583849 | 1,427380894 | 0,78313253 |
| COL6A11 | 9,07536E-05 | 0,634773506 | 0,534 | 0,353 | 1 | SMC | 1,552693968 | 0,644041917 | 1,512747875 |
| CPM1 | 9,1116E-05 | 0,647574367 | 0,259 | 0,086 | 1 | SMC | 1,566532137 | 0,638352688 | 3,011627907 |
| ECM1 | 9,53536E-05 | 0,335104093 | 0,213 | 0,052 | 1 | SMC | 1,261468423 | 0,792726938 | 4,096153846 |
| CTSD3 | 9,79957E-05 | 0,69796753 | 0,494 | 0,336 | 1 | SMC | 1,6222178 | 0,616440036 | 1,470238095 |
| EHD2 | 9,9127E-05 | 0,458670096 | 0,625 | 0,448 | 1 | SMC | 1,374274401 | 0,727656718 | 1,395089286 |
| CRISPLD2 | 0,000105629 | 0,65215561 | 0,548 | 0,345 | 1 | SMC | 1,571514529 | 0,636328829 | 1,588405797 |
| RNASEK4 | 0,0001077 | 0,555638853 | 0,716 | 0,578 | 1 | SMC | 1,469819353 | 0,680355717 | 1,238754325 |
| IGKV1-55 | 0,000109532 | 2,202288648 | 0,134 | 0,009 | 1 | SMC | 4,602088248 | 0,217292661 | 14,88888889 |
| FBLIM11 | 0,000112735 | 0,689779095 | 0,307 | 0,138 | 1 | SMC | 1,613036512 | 0,619948769 | 2,224637681 |
| RPL224 | 0,000113406 | -0,404381444 | 0,895 | 0,931 | 1 | SMC | 0,75556017 | 1,323521328 | 0,961331901 |
| RPL272 | 0,000123979 | -0,481467342 | 0,812 | 0,871 | 1 | SMC | 0,716248768 | 1,396162959 | 0,932261768 |
| SLC22A3 | 0,00013284 | -0,358130859 | 0,048 | 0,155 | 1 | SMC | 0,780174712 | 1,281764181 | 0,309677419 |

| Supplementary Table 4 - clusermarker | | | | | | | | | |
|---|---|---|---|---|---|---|---|---|---|
| | **p_val** | **avg_log2FC** | **pct.1** | **pct.2** | **p_val_adj** | **celltype** | **Foldchange_UP** | **Foldchange_DOWN** | **Ratio_pct1_pct2** |
| MAP3K20 | 0,000134454 | -0,510383523 | 0,446 | 0,629 | 1 | SMC | 0,702035786 | 1,424428812 | 0,709062003 |
| RPL214 | 0,000138587 | -0,370785872 | 0,943 | 0,948 | 1 | SMC | 0,773361113 | 1,293056999 | 0,994725738 |
| IGLV2-145 | 0,000139226 | 2,098903386 | 0,165 | 0,026 | 1 | SMC | 4,283836404 | 0,233435618 | 6,346153846 |
| SIGIRR3 | 0,000139287 | 0,659657259 | 0,23 | 0,078 | 1 | SMC | 1,579707288 | 0,633028668 | 2,948717949 |
| MAML31 | 0,000141606 | -0,275566625 | 0,026 | 0,112 | 1 | SMC | 0,82612579 | 1,210469414 | 0,232142857 |
| DUSP8 | 0,000143933 | -0,285340662 | 0,043 | 0,147 | 1 | SMC | 0,820547831 | 1,218698 | 0,292517007 |
| RPL35A7 | 0,000148379 | -0,288854923 | 0,969 | 0,983 | 1 | SMC | 0,818551491 | 1,221670245 | 0,985757884 |
| TRIM52-AS11 | 0,000149201 | -0,299531274 | 0,026 | 0,112 | 1 | SMC | 0,812516337 | 1,230744484 | 0,232142857 |
| UGCG4 | 0,000152162 | 0,821886945 | 0,321 | 0,155 | 1 | SMC | 1,767716532 | 0,56570156 | 2,070967742 |
| MIF7 | 0,000153135 | 0,443494661 | 0,787 | 0,698 | 1 | SMC | 1,359894433 | 0,735351198 | 1,127507163 |
| GEM4 | 0,000157831 | 0,539100113 | 0,577 | 0,397 | 1 | SMC | 1,453065879 | 0,688200043 | 1,453400504 |
| ID1 | 0,00015869 | -0,573667798 | 0,295 | 0,483 | 1 | SMC | 0,671906411 | 1,488302512 | 0,610766046 |
| YBX33 | 0,000159323 | -0,589741063 | 0,611 | 0,759 | 1 | SMC | 0,664462155 | 1,504976607 | 0,805006588 |
| TENT5B | 0,000159594 | -0,439678821 | 0,074 | 0,198 | 1 | SMC | 0,737298731 | 1,356302348 | 0,373737374 |
| SMIM301 | 0,000161512 | -0,387985016 | 0,074 | 0,198 | 1 | SMC | 0,764196197 | 1,308564481 | 0,373737374 |
| GJC1 | 0,000162292 | 0,761041521 | 0,264 | 0,103 | 1 | SMC | 1,694713643 | 0,590070189 | 2,563106796 |
| ZEB23 | 0,000163779 | 0,457680873 | 0,699 | 0,491 | 1 | SMC | 1,373332416 | 0,728155826 | 1,423625255 |
| TPM21 | 0,000165048 | -0,730524937 | 0,901 | 0,888 | 1 | SMC | 0,602684582 | 1,659242712 | 1,01463964 |
| NDUFB74 | 0,00016565 | -0,444177546 | 0,477 | 0,655 | 1 | SMC | 0,735003209 | 1,360538277 | 0,728244275 |
| MARCKS6 | 0,000171933 | 0,748086494 | 0,304 | 0,138 | 1 | SMC | 1,679563678 | 0,595392728 | 2,202898551 |
| CCL21 | 0,000184311 | 1,083561068 | 0,304 | 0,121 | 1 | SMC | 2,119260696 | 0,471862665 | 2,512396694 |
| ZBTB10 | 0,000194286 | -0,421846 | 0,077 | 0,198 | 1 | SMC | 0,746468869 | 1,339640595 | 0,388888889 |
| LRRFIP15 | 0,000196149 | -0,480620568 | 0,355 | 0,509 | 1 | SMC | 0,716669286 | 1,395343738 | 0,697445972 |
| HSP90AA14 | 0,000200269 | -0,516303274 | 0,815 | 0,862 | 1 | SMC | 0,699161053 | 1,43028562 | 0,945475638 |
| STOM3 | 0,000201249 | 0,430893181 | 0,588 | 0,405 | 1 | SMC | 1,348067915 | 0,741802389 | 1,451851852 |
| PROS13 | 0,000201484 | 0,711719958 | 0,307 | 0,138 | 1 | SMC | 1,63775546 | 0,610591767 | 2,224637681 |
| RPL235 | 0,00020829 | -0,439701518 | 0,602 | 0,759 | 1 | SMC | 0,737287131 | 1,356323686 | 0,79314888 |
| GAPDH3 | 0,000224267 | 0,342342191 | 0,952 | 0,914 | 1 | SMC | 1,267813197 | 0,788759734 | 1,041575492 |
| LINC014361 | 0,000230358 | 0,525332529 | 0,29 | 0,112 | 1 | SMC | 1,43926528 | 0,694798946 | 2,589285714 |
| GUCY1A2 | 0,000241499 | 0,48600665 | 0,239 | 0,078 | 1 | SMC | 1,400562777 | 0,713998699 | 3,064102564 |
| SUSD2 | 0,000243319 | 0,780051648 | 0,327 | 0,164 | 1 | SMC | 1,717192347 | 0,582345945 | 1,993902439 |

| Supplementary Table 4 - clusermarker | | | | | | | | | |
|---|---|---|---|---|---|---|---|---|---|
| | **p_val** | **avg_log2FC** | **pct.1** | **pct.2** | **p_val_adj** | **celltype** | **Foldchange_UP** | **Foldchange_DOWN** | **Ratio_pct1_pct2** |
| ANKRD29 | 0,000250893 | 0,559647074 | 0,239 | 0,086 | 1 | SMC | 1,473908611 | 0,678468117 | 2,779069767 |
| PLTP4 | 0,000252251 | 0,735682354 | 0,256 | 0,103 | 1 | SMC | 1,665184868 | 0,600533922 | 2,485436893 |
| SLC20A2 | 0,000255626 | -0,466079911 | 0,082 | 0,207 | 1 | SMC | 0,723928986 | 1,38135096 | 0,396135266 |
| CNN32 | 0,000256962 | 0,514660957 | 0,676 | 0,509 | 1 | SMC | 1,428658356 | 0,699957408 | 1,328094303 |
| ANO1 | 0,000263583 | 0,523725516 | 0,341 | 0,172 | 1 | SMC | 1,437662981 | 0,695573311 | 1,98255814 |
| F2R | 0,000266704 | 0,519749652 | 0,151 | 0,026 | 1 | SMC | 1,433706438 | 0,697492857 | 5,807692308 |
| CD4 | 0,000276713 | 0,466621329 | 0,196 | 0,052 | 1 | SMC | 1,381869453 | 0,72365736 | 3,769230769 |
| FILIP11 | 0,000284421 | -0,651099778 | 0,222 | 0,379 | 1 | SMC | 0,636794695 | 1,570364841 | 0,585751979 |
| CAMK2N1 | 0,000292063 | 0,65800811 | 0,335 | 0,172 | 1 | SMC | 1,577902552 | 0,633752698 | 1,947674419 |
| DAD14 | 0,000294191 | 0,577889746 | 0,571 | 0,397 | 1 | SMC | 1,492664306 | 0,669942998 | 1,438287154 |
| RAMP12 | 0,000302514 | -0,447560641 | 0,21 | 0,371 | 1 | SMC | 0,733281658 | 1,36373246 | 0,566037736 |
| NEURL1B | 0,000306712 | 0,521878015 | 0,403 | 0,224 | 1 | SMC | 1,435823102 | 0,696464626 | 1,799107143 |
| ARHGAP11 | 0,000310511 | 0,539626551 | 0,472 | 0,284 | 1 | SMC | 1,453596198 | 0,687948965 | 1,661971831 |
| AQP12 | 0,000316421 | -0,880505002 | 0,128 | 0,259 | 1 | SMC | 0,543177264 | 1,841019621 | 0,494208494 |
| SMOC21 | 0,000317016 | 0,393887749 | 0,474 | 0,267 | 1 | SMC | 1,313929391 | 0,761075905 | 1,775280899 |
| CD248 | 0,000317731 | 0,616749601 | 0,293 | 0,129 | 1 | SMC | 1,533416492 | 0,652138545 | 2,271317829 |
| RPS205 | 0,000318315 | -0,58260651 | 0,514 | 0,638 | 1 | SMC | 0,667756255 | 1,497552426 | 0,805642633 |
| CHORDC13 | 0,000320474 | -0,427581755 | 0,102 | 0,233 | 1 | SMC | 0,743507008 | 1,344977236 | 0,43776824 |
| RPL364 | 0,000325396 | -0,369783485 | 0,884 | 0,914 | 1 | SMC | 0,773898632 | 1,292158893 | 0,967177243 |
| COL6A31 | 0,000328446 | 0,556237538 | 0,361 | 0,181 | 1 | SMC | 1,470429421 | 0,680073443 | 1,994475138 |
| MT1F1 | 0,000328485 | 0,452160567 | 0,193 | 0,052 | 1 | SMC | 1,368087559 | 0,730947368 | 3,711538462 |
| COX173 | 0,000330654 | -0,486400809 | 0,213 | 0,371 | 1 | SMC | 0,713803654 | 1,400945477 | 0,574123989 |
| GABARAP7 | 0,00033368 | 0,424636036 | 0,807 | 0,664 | 1 | SMC | 1,342233841 | 0,745026663 | 1,215361446 |
| COL5A11 | 0,000337702 | 0,639961562 | 0,216 | 0,069 | 1 | SMC | 1,558287641 | 0,641730046 | 3,130434783 |
| COX6C3 | 0,000342829 | -0,431789134 | 0,71 | 0,828 | 1 | SMC | 0,741341852 | 1,348905362 | 0,857487923 |
| LAMB1 | 0,000344873 | 0,629277344 | 0,148 | 0,026 | 1 | SMC | 1,546790002 | 0,646500171 | 5,692307692 |
| MT-ATP66 | 0,000345637 | 0,292794858 | 0,977 | 0,957 | 1 | SMC | 1,225011131 | 0,816319113 | 1,020898642 |
| CPXM11 | 0,000348224 | 0,460545198 | 0,102 | 0 | 1 | SMC | 1,376061737 | 0,72671158 | #DIV/0! |
| AEBP11 | 0,000350812 | 0,79046865 | 0,688 | 0,569 | 1 | SMC | 1,729636232 | 0,578156251 | 1,20913884 |
| CD97 | 0,000351898 | -0,481541404 | 0,588 | 0,707 | 1 | SMC | 0,716212 | 1,396234635 | 0,831683168 |
| RPL242 | 0,000356677 | -0,345791028 | 0,929 | 0,948 | 1 | SMC | 0,786876415 | 1,270847596 | 0,979957806 |

| Supplementary Table 4 - clusermarker | | | | | | | | | |
|---|---|---|---|---|---|---|---|---|---|
| | **p_val** | **avg_log2FC** | **pct.1** | **pct.2** | **p_val_adj** | **celltype** | **Foldchange_UP** | **Foldchange_DOWN** | **Ratio_pct1_pct2** |
| GBP2 | 0,00035865 | -0,5398552 | 0,233 | 0,397 | 1 | SMC | 0,687839943 | 1,453826592 | 0,586901763 |
| EGFL6 | 0,000361705 | 0,540546175 | 0,207 | 0,06 | 1 | SMC | 1,454523066 | 0,687510582 | 3,45 |
| CEBPB3 | 0,000372065 | 0,548619797 | 0,537 | 0,371 | 1 | SMC | 1,462685698 | 0,683673875 | 1,447439353 |
| PLCG23 | 0,000375775 | -0,513611459 | 0,034 | 0,121 | 1 | SMC | 0,700466782 | 1,427619447 | 0,280991736 |
| ATP1A2 | 0,000394811 | -0,348991935 | 0,156 | 0,31 | 1 | SMC | 0,785132508 | 1,273670355 | 0,503225806 |
| TXNIP8 | 0,000404464 | -0,465681037 | 0,591 | 0,733 | 1 | SMC | 0,724129165 | 1,380969098 | 0,80627558 |
| BTF34 | 0,00040594 | 0,403291612 | 0,875 | 0,784 | 1 | SMC | 1,3225219 | 0,756131146 | 1,116071429 |
| SLC38A11 | 0,000412321 | 0,562706333 | 0,23 | 0,086 | 1 | SMC | 1,477037375 | 0,677030938 | 2,674418605 |
| MOB2 | 0,000421089 | -0,475438828 | 0,185 | 0,345 | 1 | SMC | 0,719247981 | 1,390341059 | 0,536231884 |
| RBM35 | 0,000422165 | -0,450317378 | 0,639 | 0,767 | 1 | SMC | 0,731881824 | 1,366340805 | 0,833116037 |
| IL34 | 0,000426518 | 0,500821934 | 0,145 | 0,026 | 1 | SMC | 1,4150195 | 0,706704042 | 5,576923077 |
| LOXL2 | 0,000429375 | 0,591449982 | 0,162 | 0,034 | 1 | SMC | 1,506760357 | 0,663675544 | 4,764705882 |
| LY6E4 | 0,000430982 | 0,663419421 | 0,312 | 0,164 | 1 | SMC | 1,583832117 | 0,631380049 | 1,902439024 |
| OAF1 | 0,000432515 | 0,515315505 | 0,21 | 0,069 | 1 | SMC | 1,429306682 | 0,699639911 | 3,043478261 |
| HNRNPA17 | 0,000442149 | -0,538043338 | 0,71 | 0,784 | 1 | SMC | 0,688704334 | 1,452001897 | 0,905612245 |
| ZNF7041 | 0,000449813 | -0,48139611 | 0,077 | 0,19 | 1 | SMC | 0,716284133 | 1,396094027 | 0,405263158 |
| BMP11 | 0,000457078 | 0,398670799 | 0,116 | 0,009 | 1 | SMC | 1,318292766 | 0,758556844 | 12,88888889 |
| CKB2 | 0,000462476 | -0,645949249 | 0,349 | 0,5 | 1 | SMC | 0,639072162 | 1,564768517 | 0,698 |
| SORBS31 | 0,000465263 | 0,521002355 | 0,568 | 0,414 | 1 | SMC | 1,434951877 | 0,696887482 | 1,371980676 |
| IFI272 | 0,000475464 | 0,52729948 | 0,409 | 0,233 | 1 | SMC | 1,441228893 | 0,693852312 | 1,755364807 |
| FN13 | 0,000504833 | 0,972159782 | 0,511 | 0,336 | 1 | SMC | 1,961775267 | 0,509742383 | 1,520833333 |
| PPP1R14A1 | 0,000512625 | -0,554773275 | 0,562 | 0,655 | 1 | SMC | 0,680764034 | 1,468937766 | 0,858015267 |
| PPP1R12A2 | 0,000539706 | -0,443831867 | 0,636 | 0,759 | 1 | SMC | 0,735179342 | 1,360212322 | 0,837944664 |
| HIGD1B | 0,00055101 | 0,748275723 | 0,276 | 0,121 | 1 | SMC | 1,679783989 | 0,595314639 | 2,280991736 |
| PDLIM13 | 0,000552097 | 0,499980488 | 0,477 | 0,31 | 1 | SMC | 1,414194436 | 0,707116345 | 1,538709677 |
| TUBB2A1 | 0,000554987 | -0,469974286 | 0,179 | 0,328 | 1 | SMC | 0,721977466 | 1,385084781 | 0,545731707 |
| PCSK51 | 0,000592931 | 0,526166242 | 0,111 | 0,009 | 1 | SMC | 1,440097252 | 0,694397548 | 12,33333333 |
| GPR4 | 0,000605222 | 0,450270805 | 0,128 | 0,017 | 1 | SMC | 1,366296697 | 0,731905451 | 7,529411765 |
| HNRNPA2B12 | 0,000617122 | -0,373806087 | 0,713 | 0,793 | 1 | SMC | 0,771743811 | 1,295766789 | 0,899117276 |
| FRMD3 | 0,000623308 | 0,468184735 | 0,17 | 0,043 | 1 | SMC | 1,383367756 | 0,722873578 | 3,953488372 |
| ADAMTS21 | 0,000627119 | 0,447666729 | 0,125 | 0,017 | 1 | SMC | 1,363832745 | 0,733227739 | 7,352941176 |

| Supplementary Table 4 - clusermarker | | | | | | | | | |
|---|---|---|---|---|---|---|---|---|---|
| | **p_val** | **avg_log2FC** | **pct.1** | **pct.2** | **p_val_adj** | **celltype** | **Foldchange_UP** | **Foldchange_DOWN** | **Ratio_pct1_pct2** |
| CYGB1 | 0,000642772 | 0,475336392 | 0,168 | 0,043 | 1 | SMC | 1,390242343 | 0,719299052 | 3,906976744 |
| MGST33 | 0,000646472 | -0,419884563 | 0,812 | 0,836 | 1 | SMC | 0,747484432 | 1,337820505 | 0,971291866 |
| VPS37B4 | 0,000663649 | -0,422296384 | 0,045 | 0,138 | 1 | SMC | 0,746235871 | 1,340058873 | 0,326086957 |
| TMEM1191 | 0,000666915 | 0,561624628 | 0,17 | 0,043 | 1 | SMC | 1,475930337 | 0,677538753 | 3,953488372 |
| SLCO3A11 | 0,000674595 | 0,308148393 | 0,318 | 0,147 | 1 | SMC | 1,238117635 | 0,807677697 | 2,163265306 |
| RAP2A1 | 0,000675643 | 0,531351078 | 0,224 | 0,086 | 1 | SMC | 1,445282063 | 0,691906463 | 2,604651163 |
| RFTN11 | 0,000679037 | 0,518962436 | 0,19 | 0,06 | 1 | SMC | 1,432924341 | 0,697873552 | 3,166666667 |
| KCNS31 | 0,000692878 | 0,323546302 | 0,111 | 0,009 | 1 | SMC | 1,251402856 | 0,799103179 | 12,33333333 |
| RPLP06 | 0,000716334 | 0,362926736 | 0,952 | 0,905 | 1 | SMC | 1,286032174 | 0,777585523 | 1,051933702 |
| SLC25A54 | 0,000733684 | 0,441095453 | 0,639 | 0,509 | 1 | SMC | 1,357634802 | 0,736575107 | 1,25540275 |
| USPL11 | 0,000750486 | -0,307628557 | 0,037 | 0,121 | 1 | SMC | 0,807968774 | 1,237671593 | 0,305785124 |
| HIF1A3 | 0,000751466 | 0,469699361 | 0,395 | 0,224 | 1 | SMC | 1,38482086 | 0,722115061 | 1,763392857 |
| LINC02202 | 0,000765432 | 0,586887188 | 0,196 | 0,069 | 1 | SMC | 1,502002473 | 0,665777865 | 2,84057971 |
| NID1 | 0,000774276 | 0,591692352 | 0,281 | 0,129 | 1 | SMC | 1,507013512 | 0,663564057 | 2,178294574 |
| HSP90AB15 | 0,000791662 | -0,524092676 | 0,892 | 0,922 | 1 | SMC | 0,695396314 | 1,438028906 | 0,967462039 |
| ASXL1 | 0,000795378 | 0,498645152 | 0,369 | 0,216 | 1 | SMC | 1,412886085 | 0,707771144 | 1,708333333 |
| EPHA4 | 0,00079897 | -0,382693231 | 0,028 | 0,103 | 1 | SMC | 0,767004405 | 1,303773477 | 0,27184466 |
| INAFM1 | 0,000837702 | 0,613483908 | 0,352 | 0,198 | 1 | SMC | 1,529949367 | 0,653616402 | 1,777777778 |
| PFDN14 | 0,000850902 | -0,417541206 | 0,108 | 0,233 | 1 | SMC | 0,748699551 | 1,335649258 | 0,463519313 |
| TPT13 | 0,000867622 | 0,30673774 | 0,974 | 0,983 | 1 | SMC | 1,236907607 | 0,808467823 | 0,990844354 |
| PSAP2 | 0,000867885 | 0,401990095 | 0,744 | 0,56 | 1 | SMC | 1,321329334 | 0,756813593 | 1,328571429 |
| KIRREL1 | 0,000868051 | 0,459957517 | 0,25 | 0,103 | 1 | SMC | 1,375501314 | 0,727007666 | 2,427184466 |
| TMEM2511 | 0,00087679 | -0,276854118 | 0,037 | 0,121 | 1 | SMC | 0,825388866 | 1,211550145 | 0,305785124 |
| HADH2 | 0,000878741 | -0,440515701 | 0,122 | 0,25 | 1 | SMC | 0,736871162 | 1,357089342 | 0,488 |
| EEF1G3 | 0,00091377 | -0,428039191 | 0,707 | 0,836 | 1 | SMC | 0,743271301 | 1,345403756 | 0,84569378 |
| DLGAP1-AS1 | 0,000914508 | -0,36114526 | 0,037 | 0,121 | 1 | SMC | 0,778546298 | 1,284445129 | 0,305785124 |
| LINC00702 | 0,000926129 | -0,428258151 | 0,111 | 0,233 | 1 | SMC | 0,743158502 | 1,345607965 | 0,47639485 |
| SEMA6D | 0,000955632 | 0,520800639 | 0,173 | 0,052 | 1 | SMC | 1,434751258 | 0,696984926 | 3,326923077 |
| SLMAP3 | 0,000971911 | -0,386582274 | 0,153 | 0,293 | 1 | SMC | 0,764939592 | 1,307292773 | 0,5221843 |
| XRCC62 | 0,000996295 | -0,382623551 | 0,398 | 0,552 | 1 | SMC | 0,767041451 | 1,303710508 | 0,721014493 |
| RUFY21 | 0,001011906 | -0,315754175 | 0,043 | 0,129 | 1 | SMC | 0,803430884 | 1,244662136 | 0,333333333 |

| Supplementary Table 4 - clusermarker | | | | | | | | | |
|---|---|---|---|---|---|---|---|---|---|
| | **p_val** | **avg_log2FC** | **pct.1** | **pct.2** | **p_val_adj** | **celltype** | **Foldchange_UP** | **Foldchange_DOWN** | **Ratio_pct1_pct2** |
| BEX41 | 0,001014055 | -0,562481306 | 0,077 | 0,181 | 1 | SMC | 0,677136547 | 1,47680701 | 0,425414365 |
| KANK1 | 0,001023386 | -0,367443993 | 0,06 | 0,155 | 1 | SMC | 0,775154614 | 1,290065211 | 0,387096774 |
| BTG24 | 0,00102584 | -0,689181437 | 0,452 | 0,569 | 1 | SMC | 0,620205645 | 1,612368425 | 0,794376098 |
| NNMT3 | 0,001029024 | 0,601890209 | 0,588 | 0,397 | 1 | SMC | 1,517703749 | 0,658890116 | 1,481108312 |
| CALM13 | 0,001045601 | -0,326026694 | 0,753 | 0,845 | 1 | SMC | 0,797730479 | 1,253556215 | 0,89112426 |
| LDB3 | 0,00112978 | -0,474155085 | 0,19 | 0,345 | 1 | SMC | 0,71988827 | 1,389104452 | 0,550724638 |
| CRABP23 | 0,001130169 | -0,316861201 | 0,06 | 0,155 | 1 | SMC | 0,802814622 | 1,245617572 | 0,387096774 |
| INPP5A | 0,001135726 | -0,368887222 | 0,094 | 0,207 | 1 | SMC | 0,77437956 | 1,291356399 | 0,45410628 |
| FHL13 | 0,001140573 | -0,533882714 | 0,514 | 0,655 | 1 | SMC | 0,690693373 | 1,447820465 | 0,784732824 |
| MED30 | 0,001157072 | -0,433014293 | 0,065 | 0,164 | 1 | SMC | 0,74071256 | 1,350051361 | 0,396341463 |
| GUCY1A11 | 0,001162837 | -0,618320786 | 0,389 | 0,526 | 1 | SMC | 0,651428712 | 1,535087388 | 0,739543726 |
| RPL64 | 0,001172985 | -0,279528429 | 0,946 | 0,957 | 1 | SMC | 0,823860267 | 1,213798068 | 0,988505747 |
| CYTH31 | 0,001188545 | 0,405928461 | 0,185 | 0,06 | 1 | SMC | 1,324941315 | 0,754750409 | 3,083333333 |
| ZNF7031 | 0,00119877 | 0,55306945 | 0,338 | 0,19 | 1 | SMC | 1,467203973 | 0,681568493 | 1,778947368 |
| VAMP54 | 0,001199216 | 0,605412011 | 0,392 | 0,25 | 1 | SMC | 1,521413183 | 0,657283643 | 1,568 |
| VPS284 | 0,001237711 | 0,421968819 | 0,503 | 0,345 | 1 | SMC | 1,339754646 | 0,746405324 | 1,457971014 |
| CYTOR2 | 0,001246121 | 0,625492438 | 0,327 | 0,181 | 1 | SMC | 1,542737321 | 0,648198489 | 1,806629834 |
| ISLR2 | 0,00125453 | 0,444658148 | 0,219 | 0,086 | 1 | SMC | 1,360991586 | 0,7347584 | 2,546511628 |
| YTHDC13 | 0,001266674 | -0,502718129 | 0,259 | 0,405 | 1 | SMC | 0,705775802 | 1,416880542 | 0,639506173 |
| GPC61 | 0,001336391 | -0,313641293 | 0,06 | 0,155 | 1 | SMC | 0,804608401 | 1,242840615 | 0,387096774 |
| EML42 | 0,001391563 | 0,585355741 | 0,233 | 0,103 | 1 | SMC | 1,500408916 | 0,666484976 | 2,262135922 |
| PSMB94 | 0,001415519 | 0,653270706 | 0,261 | 0,129 | 1 | SMC | 1,572729662 | 0,635837184 | 2,023255814 |
| RAMP3 | 0,001441612 | 0,556531319 | 0,114 | 0,017 | 1 | SMC | 1,47072888 | 0,679934972 | 6,705882353 |
| CERS62 | 0,001442621 | 0,396074632 | 0,134 | 0,026 | 1 | SMC | 1,315922597 | 0,759923116 | 5,153846154 |
| TIMP33 | 0,001446372 | 0,424594069 | 0,918 | 0,828 | 1 | SMC | 1,342194796 | 0,745048337 | 1,108695652 |
| CAMK2G1 | 0,001447159 | -0,494072172 | 0,159 | 0,293 | 1 | SMC | 0,710018159 | 1,408414682 | 0,542662116 |
| PHB5 | 0,001451664 | -0,482737242 | 0,241 | 0,371 | 1 | SMC | 0,715618584 | 1,397392441 | 0,649595687 |
| C21 | 0,001466383 | 0,444125165 | 0,131 | 0,026 | 1 | SMC | 1,36048888 | 0,735029896 | 5,038461538 |
| SLC35A4 | 0,001485992 | -0,289059774 | 0,065 | 0,164 | 1 | SMC | 0,818435271 | 1,221843725 | 0,396341463 |
| LTBP1 | 0,001509509 | -0,373442068 | 0,29 | 0,457 | 1 | SMC | 0,771938561 | 1,295439885 | 0,634573304 |
| TIMP2 | 0,001520864 | 0,576672341 | 0,614 | 0,483 | 1 | SMC | 1,491405267 | 0,670508562 | 1,271221532 |

| Supplementary Table 4 - clusermarker | | | | | | | | | |
|---|---|---|---|---|---|---|---|---|---|
| | **p_val** | **avg_log2FC** | **pct.1** | **pct.2** | **p_val_adj** | **celltype** | **Foldchange_UP** | **Foldchange_DOWN** | **Ratio_pct1_pct2** |
| TCF4 | 0,00152289 | 0,429430445 | 0,554 | 0,397 | 1 | SMC | 1,346701814 | 0,742554877 | 1,395465995 |
| TMED71 | 0,001563665 | -0,420646679 | 0,176 | 0,31 | 1 | SMC | 0,747089671 | 1,338527407 | 0,567741935 |
| ZNF189 | 0,001563936 | -0,313542346 | 0,031 | 0,103 | 1 | SMC | 0,804663587 | 1,242755377 | 0,300970874 |
| NSUN6 | 0,001576403 | -0,482547749 | 0,051 | 0,138 | 1 | SMC | 0,715712584 | 1,397208911 | 0,369565217 |
| TEAD1 | 0,001596834 | 0,512451634 | 0,23 | 0,103 | 1 | SMC | 1,426472203 | 0,701030134 | 2,233009709 |
| SNRPE1 | 0,00161064 | -0,438806197 | 0,168 | 0,302 | 1 | SMC | 0,737744826 | 1,355482228 | 0,556291391 |
| BST22 | 0,001638558 | 0,538438105 | 0,577 | 0,414 | 1 | SMC | 1,452399265 | 0,688515909 | 1,393719807 |
| TNFAIP24 | 0,001640088 | 0,538629101 | 0,139 | 0,034 | 1 | SMC | 1,452591558 | 0,688424764 | 4,088235294 |
| AP3S13 | 0,001653476 | 0,509966544 | 0,293 | 0,147 | 1 | SMC | 1,424017172 | 0,702238723 | 1,993197279 |
| RAC14 | 0,001684757 | 0,50505912 | 0,707 | 0,586 | 1 | SMC | 1,419181511 | 0,704631502 | 1,206484642 |
| STAT33 | 0,001685181 | 0,62106352 | 0,528 | 0,397 | 1 | SMC | 1,538008546 | 0,650191446 | 1,329974811 |
| PDLIM31 | 0,001691266 | -0,635182078 | 0,338 | 0,457 | 1 | SMC | 0,64385955 | 1,553133754 | 0,739606127 |
| PLSCR3 | 0,001710094 | 0,602895274 | 0,241 | 0,112 | 1 | SMC | 1,518761439 | 0,658431255 | 2,151785714 |
| RGS31 | 0,001716419 | -0,25275318 | 0,045 | 0,129 | 1 | SMC | 0,839293213 | 1,191478716 | 0,348837209 |
| LRRC47 | 0,001737737 | -0,252300586 | 0,077 | 0,181 | 1 | SMC | 0,839556553 | 1,191104991 | 0,425414365 |
| BTBD6 | 0,001793843 | -0,29330749 | 0,031 | 0,103 | 1 | SMC | 0,816029103 | 1,225446491 | 0,300970874 |
| VCAM11 | 0,001797322 | 0,541423631 | 0,182 | 0,06 | 1 | SMC | 1,455407985 | 0,687092561 | 3,033333333 |
| DAZAP24 | 0,001802112 | -0,430134428 | 0,42 | 0,543 | 1 | SMC | 0,742192626 | 1,347359115 | 0,773480663 |
| PTGER3 | 0,001804077 | 0,478563235 | 0,114 | 0,017 | 1 | SMC | 1,393355347 | 0,71769201 | 6,705882353 |
| AC092164.11 | 0,001815065 | -0,337137138 | 0,062 | 0,155 | 1 | SMC | 0,791610614 | 1,263247336 | 0,4 |
| CFI2 | 0,001847581 | 0,390381584 | 0,128 | 0,026 | 1 | SMC | 1,310740041 | 0,762927788 | 4,923076923 |
| UBA523 | 0,001855841 | -0,279362237 | 0,821 | 0,905 | 1 | SMC | 0,823955177 | 1,213658252 | 0,90718232 |
| AGAP12 | 0,001862818 | 0,298181948 | 0,139 | 0,034 | 1 | SMC | 1,22959393 | 0,813276624 | 4,088235294 |
| MYL12B4 | 0,001885384 | 0,462641916 | 0,688 | 0,603 | 1 | SMC | 1,378063069 | 0,725656193 | 1,140961857 |
| SEPTIN4 | 0,00200618 | 0,420802997 | 0,409 | 0,241 | 1 | SMC | 1,338672446 | 0,747008727 | 1,697095436 |
| TACC21 | 0,002017674 | -0,315679649 | 0,068 | 0,164 | 1 | SMC | 0,803472388 | 1,244597842 | 0,414634146 |
| COX7A23 | 0,002030658 | -0,463486118 | 0,631 | 0,681 | 1 | SMC | 0,725231695 | 1,378869687 | 0,926578561 |
| TPM46 | 0,002072544 | 0,404058836 | 0,67 | 0,56 | 1 | SMC | 1,323225403 | 0,755729143 | 1,196428571 |
| FRK | 0,002081883 | -0,267316547 | 0,057 | 0,147 | 1 | SMC | 0,830863539 | 1,203567076 | 0,387755102 |
| PRUNE2 | 0,002100533 | -0,412343715 | 0,077 | 0,172 | 1 | SMC | 0,7514017 | 1,330846071 | 0,447674419 |
| FEZ1 | 0,00221489 | 0,410280437 | 0,19 | 0,069 | 1 | SMC | 1,328944115 | 0,75247709 | 2,753623188 |

| Supplementary Table 4 - clusermarker | | | | | | | | | |
|---|---|---|---|---|---|---|---|---|---|
| | **p_val** | **avg_log2FC** | **pct.1** | **pct.2** | **p_val_adj** | **celltype** | **Foldchange_UP** | **Foldchange_DOWN** | **Ratio_pct1_pct2** |
| GSTP15 | 0,00224424 | -0,323975676 | 0,73 | 0,793 | 1 | SMC | 0,798865386 | 1,251775353 | 0,920554855 |
| CLDND13 | 0,002280389 | -0,511062827 | 0,27 | 0,405 | 1 | SMC | 0,701705304 | 1,425099673 | 0,666666667 |
| MAN2B1 | 0,002298065 | 0,335290036 | 0,108 | 0,017 | 1 | SMC | 1,261631019 | 0,792624773 | 6,352941176 |
| GPX48 | 0,002301398 | 0,471999369 | 0,619 | 0,5 | 1 | SMC | 1,387030362 | 0,720964751 | 1,238 |
| OAT6 | 0,00233703 | -0,428387274 | 0,364 | 0,509 | 1 | SMC | 0,743091992 | 1,345728404 | 0,715127701 |
| TUBB31 | 0,002368328 | 0,557510946 | 0,199 | 0,078 | 1 | SMC | 1,471727882 | 0,679473436 | 2,551282051 |
| HRH2 | 0,002376816 | -0,486093304 | 0,159 | 0,284 | 1 | SMC | 0,713955814 | 1,400646903 | 0,559859155 |
| SEC61G2 | 0,002380044 | -0,494450409 | 0,435 | 0,578 | 1 | SMC | 0,709832036 | 1,40878398 | 0,752595156 |
| NKD1 | 0,002399763 | -0,310361553 | 0,043 | 0,121 | 1 | SMC | 0,806439633 | 1,240018421 | 0,355371901 |
| TLE44 | 0,002401419 | -0,480444053 | 0,142 | 0,259 | 1 | SMC | 0,716756977 | 1,395173026 | 0,548262548 |
| PLPP11 | 0,002457088 | 0,481798072 | 0,378 | 0,233 | 1 | SMC | 1,396483059 | 0,716084591 | 1,622317597 |
| ATP5F1D8 | 0,002463698 | 0,440303978 | 0,574 | 0,431 | 1 | SMC | 1,356890196 | 0,73697931 | 1,331786543 |
| SOCS12 | 0,002485844 | 0,604622138 | 0,219 | 0,095 | 1 | SMC | 1,52058044 | 0,657643604 | 2,305263158 |
| PLPP51 | 0,002530262 | -0,271936844 | 0,048 | 0,129 | 1 | SMC | 0,828206916 | 1,207427733 | 0,372093023 |
| ARPC34 | 0,002530792 | 0,511999171 | 0,537 | 0,397 | 1 | SMC | 1,426024898 | 0,701250028 | 1,352644836 |
| ECE11 | 0,002540571 | 0,48782767 | 0,241 | 0,112 | 1 | SMC | 1,402331732 | 0,713098033 | 2,151785714 |
| GAS64 | 0,002585122 | 0,596113804 | 0,398 | 0,259 | 1 | SMC | 1,511639166 | 0,661533534 | 1,536679537 |
| POSTN1 | 0,002603776 | 0,763026389 | 0,134 | 0,034 | 1 | SMC | 1,697046845 | 0,589258925 | 3,941176471 |
| ATP2B42 | 0,002610589 | -0,366323513 | 0,327 | 0,466 | 1 | SMC | 0,775756878 | 1,28906366 | 0,701716738 |
| COX4I2 | 0,002664171 | 0,583641044 | 0,44 | 0,293 | 1 | SMC | 1,498626683 | 0,667277589 | 1,501706485 |
| CCNI1 | 0,002679044 | -0,340037832 | 0,699 | 0,759 | 1 | SMC | 0,790020595 | 1,265789787 | 0,920948617 |
| NTRK3 | 0,002695446 | -0,345743028 | 0,088 | 0,19 | 1 | SMC | 0,786902596 | 1,270805314 | 0,463157895 |
| ARC1 | 0,00269809 | -0,437104258 | 0,071 | 0,164 | 1 | SMC | 0,738615653 | 1,353884116 | 0,432926829 |
| TYMP7 | 0,002711536 | 0,410275734 | 0,239 | 0,112 | 1 | SMC | 1,328939782 | 0,752479543 | 2,133928571 |
| NDUFA34 | 0,002748487 | -0,511561214 | 0,176 | 0,293 | 1 | SMC | 0,701462938 | 1,425592067 | 0,600682594 |
| MYO1B1 | 0,002759476 | 0,570539362 | 0,46 | 0,31 | 1 | SMC | 1,485078674 | 0,673364999 | 1,483870968 |
| OSBPL82 | 0,002781813 | 0,3827367 | 0,435 | 0,276 | 1 | SMC | 1,30381276 | 0,766981295 | 1,576086957 |
| MAML22 | 0,002811854 | 0,643070908 | 0,247 | 0,129 | 1 | SMC | 1,561649738 | 0,640348457 | 1,914728682 |
| STK381 | 0,002812775 | -0,324710419 | 0,034 | 0,103 | 1 | SMC | 0,798458639 | 1,252413026 | 0,330097087 |
| TMEM30A1 | 0,002850165 | 0,467695822 | 0,372 | 0,224 | 1 | SMC | 1,382899028 | 0,723118594 | 1,660714286 |
| JAK14 | 0,002884872 | 0,405062124 | 0,537 | 0,405 | 1 | SMC | 1,324145928 | 0,755203772 | 1,325925926 |

| Supplementary Table 4 - clusermarker | | | | | | | | | |
|---|---|---|---|---|---|---|---|---|---|
| | **p_val** | **avg_log2FC** | **pct.1** | **pct.2** | **p_val_adj** | **celltype** | **Foldchange_UP** | **Foldchange_DOWN** | **Ratio_pct1_pct2** |
| CSDC2 | 0,002969065 | -0,339090466 | 0,034 | 0,103 | 1 | SMC | 0,790539543 | 1,26495886 | 0,330097087 |
| SSR23 | 0,002979444 | 0,446415209 | 0,577 | 0,448 | 1 | SMC | 1,36265015 | 0,733864081 | 1,287946429 |
| C4orf483 | 0,002991792 | 0,306382013 | 0,105 | 0,017 | 1 | SMC | 1,236602659 | 0,808667192 | 6,176470588 |
| PLXND11 | 0,002997574 | 0,448942099 | 0,205 | 0,086 | 1 | SMC | 1,365038933 | 0,732579838 | 2,38372093 |
| FAM162B | 0,003003184 | 0,46496667 | 0,395 | 0,241 | 1 | SMC | 1,380285465 | 0,724487815 | 1,639004149 |
| AL138963.46 | 0,003009163 | 0,622637668 | 0,267 | 0,138 | 1 | SMC | 1,539687609 | 0,649482398 | 1,934782609 |
| MCAM1 | 0,003017973 | -0,359575722 | 0,693 | 0,776 | 1 | SMC | 0,779393756 | 1,283048514 | 0,893041237 |
| TNFSF121 | 0,003042965 | 0,519720985 | 0,352 | 0,216 | 1 | SMC | 1,43367795 | 0,697506717 | 1,62962963 |
| ANGPT2 | 0,00306472 | 0,436894954 | 0,384 | 0,216 | 1 | SMC | 1,35368771 | 0,738722818 | 1,777777778 |
| DLC1 | 0,003128372 | 0,385541795 | 0,401 | 0,25 | 1 | SMC | 1,306350286 | 0,765491469 | 1,604 |
| UBC6 | 0,003153805 | -0,339955855 | 0,926 | 0,948 | 1 | SMC | 0,790065487 | 1,265717864 | 0,976793249 |
| NCKAP5 | 0,003170143 | 0,392850435 | 0,219 | 0,095 | 1 | SMC | 1,312985001 | 0,761623323 | 2,305263158 |
| ITPK11 | 0,003175763 | -0,356600863 | 0,091 | 0,19 | 1 | SMC | 0,781002535 | 1,280405574 | 0,478947368 |
| BST1 | 0,0031887 | 0,364587949 | 0,102 | 0,017 | 1 | SMC | 1,287513848 | 0,776690675 | 6 |
| MTIF33 | 0,003198258 | -0,357939445 | 0,151 | 0,276 | 1 | SMC | 0,78027823 | 1,28159413 | 0,547101449 |
| HTRA13 | 0,00319866 | 0,417607966 | 0,446 | 0,319 | 1 | SMC | 1,335711067 | 0,748664906 | 1,398119122 |
| CAPN23 | 0,003238207 | 0,473488642 | 0,364 | 0,241 | 1 | SMC | 1,388462913 | 0,720220894 | 1,510373444 |
| CAP2 | 0,003286793 | -0,29433512 | 0,131 | 0,25 | 1 | SMC | 0,815448053 | 1,226319686 | 0,524 |
| PDLIM52 | 0,003342985 | -0,409213456 | 0,358 | 0,491 | 1 | SMC | 0,753033809 | 1,327961624 | 0,729124236 |
| UBE2I5 | 0,003402255 | 0,448289345 | 0,273 | 0,147 | 1 | SMC | 1,364421454 | 0,732911372 | 1,857142857 |
| DNAJB12 | 0,003427812 | 0,401795782 | 0,128 | 0,034 | 1 | SMC | 1,321151379 | 0,756915533 | 3,764705882 |
| COMT3 | 0,00369755 | 0,406437768 | 0,429 | 0,284 | 1 | SMC | 1,325409135 | 0,754484011 | 1,51056338 |
| H2AJ4 | 0,003710311 | 0,505783351 | 0,486 | 0,345 | 1 | SMC | 1,419894117 | 0,704277867 | 1,408695652 |
| RPS65 | 0,0037557 | -0,302608401 | 0,938 | 0,94 | 1 | SMC | 0,810785166 | 1,233372343 | 0,99787234 |
| IFI27L24 | 0,003809766 | 0,427816511 | 0,256 | 0,129 | 1 | SMC | 1,345196109 | 0,743386034 | 1,984496124 |
| RASL11A | 0,003845289 | 0,495903871 | 0,54 | 0,405 | 1 | SMC | 1,410203993 | 0,709117266 | 1,333333333 |
| CSRNP3 | 0,003864003 | 0,376169023 | 0,261 | 0,129 | 1 | SMC | 1,297890816 | 0,770480835 | 2,023255814 |
| IGFBP52 | 0,003868742 | -0,267946202 | 0,668 | 0,871 | 1 | SMC | 0,830500993 | 1,20409248 | 0,766934558 |
| SELENOK3 | 0,00386902 | -0,506489139 | 0,429 | 0,552 | 1 | SMC | 0,703933408 | 1,420588921 | 0,777173913 |
| ASPN | 0,003924085 | 0,739142188 | 0,335 | 0,207 | 1 | SMC | 1,669183064 | 0,599095463 | 1,618357488 |
| MSC-AS1 | 0,003935133 | 0,436937584 | 0,128 | 0,034 | 1 | SMC | 1,353727711 | 0,73870099 | 3,764705882 |

| Supplementary Table 4 - clusermarker | | | | | | | | | |
|---|---|---|---|---|---|---|---|---|---|
| | **p_val** | **avg_log2FC** | **pct.1** | **pct.2** | **p_val_adj** | **celltype** | **Foldchange_UP** | **Foldchange_DOWN** | **Ratio_pct1_pct2** |
| CHMP31 | 0,003970763 | -0,397898815 | 0,301 | 0,431 | 1 | SMC | 0,758962855 | 1,317587538 | 0,69837587 |
| DDX561 | 0,004022686 | -0,363306324 | 0,085 | 0,181 | 1 | SMC | 0,777380959 | 1,286370586 | 0,46961326 |
| APOE4 | 0,004063545 | 0,36185076 | 0,222 | 0,103 | 1 | SMC | 1,285073396 | 0,77816567 | 2,155339806 |
| PRRC2B5 | 0,004112229 | 0,546547491 | 0,295 | 0,172 | 1 | SMC | 1,460586186 | 0,684656619 | 1,715116279 |
| ADAR2 | 0,004132433 | 0,413980324 | 0,298 | 0,181 | 1 | SMC | 1,332356654 | 0,750549785 | 1,64640884 |
| ARMCX6 | 0,004152001 | -0,316661439 | 0,08 | 0,172 | 1 | SMC | 0,802925791 | 1,245445109 | 0,465116279 |
| LIMA12 | 0,004203053 | 0,53570345 | 0,284 | 0,164 | 1 | SMC | 1,449648823 | 0,689822241 | 1,731707317 |
| FZD41 | 0,004207173 | 0,390729223 | 0,202 | 0,086 | 1 | SMC | 1,311055922 | 0,762743971 | 2,348837209 |
| CCDC102B | 0,004222967 | 0,337788796 | 0,25 | 0,121 | 1 | SMC | 1,263818067 | 0,791253129 | 2,066115702 |
| VEGFB2 | 0,004245063 | -0,472497055 | 0,222 | 0,345 | 1 | SMC | 0,720716083 | 1,387508928 | 0,643478261 |
| GPR20 | 0,004308751 | -0,277967547 | 0,062 | 0,147 | 1 | SMC | 0,824752101 | 1,212485544 | 0,421768707 |
| HSPB61 | 0,004357398 | -0,495678648 | 0,457 | 0,578 | 1 | SMC | 0,709227977 | 1,40998386 | 0,790657439 |
| SUPT16H3 | 0,004374036 | -0,34618758 | 0,185 | 0,31 | 1 | SMC | 0,786660157 | 1,271196961 | 0,596774194 |
| ITGB5 | 0,004377877 | 0,422705615 | 0,278 | 0,155 | 1 | SMC | 1,340439044 | 0,746024226 | 1,793548387 |
| MDH12 | 0,004440622 | -0,360938422 | 0,384 | 0,526 | 1 | SMC | 0,778657926 | 1,284260992 | 0,730038023 |
| EFEMP21 | 0,004559121 | 0,4186588 | 0,287 | 0,155 | 1 | SMC | 1,33668433 | 0,74811979 | 1,851612903 |
| PPP2R2B | 0,004581953 | -0,338667102 | 0,045 | 0,121 | 1 | SMC | 0,790771564 | 1,264587708 | 0,371900826 |
| GUCY1B1 | 0,004613883 | 0,385213679 | 0,526 | 0,388 | 1 | SMC | 1,306053213 | 0,765665587 | 1,355670103 |
| HDAC5 | 0,004638047 | -0,296489503 | 0,102 | 0,207 | 1 | SMC | 0,81423125 | 1,228152322 | 0,492753623 |
| ARHGAP55 | 0,004646553 | -0,313813437 | 0,168 | 0,284 | 1 | SMC | 0,8045124 | 1,24298892 | 0,591549296 |
| NFATC4 | 0,004653392 | 0,43729396 | 0,165 | 0,06 | 1 | SMC | 1,354062152 | 0,738518538 | 2,75 |
| OSTF12 | 0,00466124 | 0,403815691 | 0,159 | 0,06 | 1 | SMC | 1,323002411 | 0,755856521 | 2,65 |
| CBR14 | 0,00467244 | 0,530468656 | 0,315 | 0,198 | 1 | SMC | 1,444398329 | 0,692329796 | 1,590909091 |
| MGP5 | 0,004675698 | 0,356629535 | 0,855 | 0,828 | 1 | SMC | 1,280431021 | 0,780987014 | 1,032608696 |
| CALCOCO12 | 0,004709459 | -0,282941596 | 0,19 | 0,319 | 1 | SMC | 0,82191346 | 1,216673104 | 0,595611285 |
| VMP13 | 0,004720064 | 0,289812979 | 0,321 | 0,19 | 1 | SMC | 1,222481794 | 0,818008092 | 1,689473684 |
| VIRMA | 0,004733647 | -0,272483088 | 0,091 | 0,19 | 1 | SMC | 0,827893394 | 1,207884985 | 0,478947368 |
| ZFYVE212 | 0,004805323 | -0,294900562 | 0,213 | 0,353 | 1 | SMC | 0,815128514 | 1,226800417 | 0,603399433 |
| DYRK4 | 0,004813976 | -0,29679097 | 0,057 | 0,138 | 1 | SMC | 0,814061125 | 1,228408985 | 0,413043478 |
| HSPB7 | 0,00489516 | -0,36922179 | 0,185 | 0,302 | 1 | SMC | 0,774199998 | 1,291655906 | 0,612582781 |
| TNS31 | 0,004936271 | 0,370688679 | 0,162 | 0,06 | 1 | SMC | 1,29296989 | 0,773413215 | 2,7 |

| Supplementary Table 4 - clusermarker | | | | | | | | | |
|---|---|---|---|---|---|---|---|---|---|
| | **p_val** | **avg_log2FC** | **pct.1** | **pct.2** | **p_val_adj** | **celltype** | **Foldchange_UP** | **Foldchange_DOWN** | **Ratio_pct1_pct2** |
| EMP13 | 0,00493818 | 0,696890961 | 0,25 | 0,129 | 1 | SMC | 1,62100772 | 0,616900208 | 1,937984496 |
| TUSC31 | 0,004967419 | 0,390228884 | 0,136 | 0,043 | 1 | SMC | 1,310601315 | 0,763008543 | 3,162790698 |
| LGALS15 | 0,004983574 | 0,272881572 | 0,955 | 0,897 | 1 | SMC | 1,208218659 | 0,827664755 | 1,064659978 |
| SSR35 | 0,005039652 | 0,314600013 | 0,361 | 0,216 | 1 | SMC | 1,243666799 | 0,804073889 | 1,671296296 |
| TLE54 | 0,005052837 | 0,429800092 | 0,398 | 0,259 | 1 | SMC | 1,34704691 | 0,742364644 | 1,536679537 |
| AAMDC1 | 0,005055597 | -0,479753029 | 0,19 | 0,302 | 1 | SMC | 0,717100372 | 1,394504924 | 0,629139073 |
| C12orf29 | 0,005120137 | -0,293788919 | 0,051 | 0,129 | 1 | SMC | 0,815756838 | 1,225855492 | 0,395348837 |
| NME37 | 0,00514593 | 0,495607107 | 0,429 | 0,31 | 1 | SMC | 1,409913943 | 0,709263147 | 1,383870968 |
| SEMA4C1 | 0,005156584 | 0,486836033 | 0,148 | 0,052 | 1 | SMC | 1,40136817 | 0,71358835 | 2,846153846 |
| HSD17B112 | 0,005159854 | -0,322865436 | 0,156 | 0,276 | 1 | SMC | 0,799480397 | 1,250812408 | 0,565217391 |
| PGM5 | 0,005224261 | -0,517306147 | 0,185 | 0,302 | 1 | SMC | 0,698675208 | 1,431280213 | 0,612582781 |
| RPL18A7 | 0,005374721 | 0,253585498 | 0,972 | 0,983 | 1 | SMC | 1,1921663 | 0,838809149 | 0,988809766 |
| SGIP11 | 0,005383341 | 0,373569886 | 0,21 | 0,095 | 1 | SMC | 1,295554661 | 0,771870173 | 2,210526316 |
| ARHGAP42 | 0,005417132 | 0,301602651 | 0,207 | 0,095 | 1 | SMC | 1,232512819 | 0,811350588 | 2,178947368 |
| KLF42 | 0,005423804 | -0,525249505 | 0,185 | 0,302 | 1 | SMC | 0,694838932 | 1,439182456 | 0,612582781 |
| HSP90B16 | 0,005479516 | 0,328118581 | 0,676 | 0,534 | 1 | SMC | 1,255375172 | 0,79657462 | 1,265917603 |
| DDT6 | 0,005482564 | 0,476775451 | 0,156 | 0,06 | 1 | SMC | 1,391629774 | 0,718581924 | 2,6 |
| CKMT2 | 0,005490473 | -0,320204348 | 0,077 | 0,164 | 1 | SMC | 0,80095642 | 1,248507379 | 0,469512195 |
| CNTN41 | 0,005525868 | 0,339492994 | 0,125 | 0,034 | 1 | SMC | 1,265311848 | 0,790319004 | 3,676470588 |
| BCAM1 | 0,00552938 | -0,298864458 | 0,369 | 0,517 | 1 | SMC | 0,81289197 | 1,230175763 | 0,713733075 |
| TM4SF12 | 0,005561992 | 0,544765764 | 0,503 | 0,345 | 1 | SMC | 1,458783477 | 0,685502692 | 1,457971014 |
| METRNL2 | 0,005584943 | -0,297426932 | 0,043 | 0,112 | 1 | SMC | 0,813702353 | 1,228950606 | 0,383928571 |
| LINC00924 | 0,005604319 | 0,451829815 | 0,162 | 0,06 | 1 | SMC | 1,367773948 | 0,731114964 | 2,7 |
| CEBPD3 | 0,005867455 | 0,315875998 | 0,778 | 0,707 | 1 | SMC | 1,244767241 | 0,803363044 | 1,100424328 |
| TNFRSF1A1 | 0,00586866 | 0,480757618 | 0,375 | 0,25 | 1 | SMC | 1,395476296 | 0,716601208 | 1,5 |
| PLCB41 | 0,005940961 | -0,345231041 | 0,108 | 0,207 | 1 | SMC | 0,787181903 | 1,270354408 | 0,52173913 |
| SEC61B7 | 0,005946072 | 0,356990075 | 0,531 | 0,405 | 1 | SMC | 1,28075105 | 0,780791864 | 1,311111111 |
| EIF3L3 | 0,005948838 | -0,415934347 | 0,531 | 0,638 | 1 | SMC | 0,749533909 | 1,334162454 | 0,832288401 |
| POLR2K3 | 0,005949225 | -0,421296378 | 0,284 | 0,414 | 1 | SMC | 0,746753304 | 1,339130331 | 0,685990338 |
| ISG154 | 0,005971578 | 0,335465966 | 0,256 | 0,129 | 1 | SMC | 1,261784878 | 0,792528122 | 1,984496124 |
| ARPC1A2 | 0,005985827 | -0,346165079 | 0,369 | 0,5 | 1 | SMC | 0,786672426 | 1,271177134 | 0,738 |

| Supplementary Table 4 - clusermarker | | | | | | | | | |
|---|---|---|---|---|---|---|---|---|---|
| | **p_val** | **avg_log2FC** | **pct.1** | **pct.2** | **p_val_adj** | **celltype** | **Foldchange_UP** | **Foldchange_DOWN** | **Ratio_pct1_pct2** |
| COPRS1 | 0,006056395 | -0,277247197 | 0,128 | 0,241 | 1 | SMC | 0,825164009 | 1,211880291 | 0,531120332 |
| PSME15 | 0,006123152 | 0,422760288 | 0,577 | 0,448 | 1 | SMC | 1,340489843 | 0,745995955 | 1,287946429 |
| CDH6 | 0,006154226 | 0,490366619 | 0,321 | 0,198 | 1 | SMC | 1,40480182 | 0,71184418 | 1,621212121 |
| TTLL7 | 0,006173624 | -0,54949307 | 0,236 | 0,353 | 1 | SMC | 0,683260168 | 1,46357134 | 0,668555241 |
| CCDC3 | 0,006190568 | -0,317263347 | 0,301 | 0,431 | 1 | SMC | 0,802590872 | 1,245964831 | 0,69837587 |
| LIN7A | 0,006207469 | 0,305386357 | 0,165 | 0,06 | 1 | SMC | 1,235749529 | 0,809225475 | 2,75 |
| INPP4B2 | 0,006318333 | 0,535420814 | 0,483 | 0,345 | 1 | SMC | 1,449364852 | 0,689957396 | 1,4 |
| TNFAIP33 | 0,006318937 | 0,771472396 | 0,239 | 0,121 | 1 | SMC | 1,707011048 | 0,58581929 | 1,975206612 |
| LURAP1L | 0,006449639 | 0,450354376 | 0,276 | 0,155 | 1 | SMC | 1,366375845 | 0,731863055 | 1,780645161 |
| ACADM1 | 0,006451265 | -0,300317019 | 0,082 | 0,172 | 1 | SMC | 0,812073931 | 1,231414975 | 0,476744186 |
| CHN11 | 0,006456852 | 0,398068995 | 0,295 | 0,172 | 1 | SMC | 1,31774297 | 0,758873333 | 1,715116279 |
| CLEC11A1 | 0,006461172 | 0,51577379 | 0,179 | 0,078 | 1 | SMC | 1,429760786 | 0,699417699 | 2,294871795 |
| NPC24 | 0,00648702 | 0,475840405 | 0,389 | 0,259 | 1 | SMC | 1,390728117 | 0,719047805 | 1,501930502 |
| ECEL1 | 0,006511753 | 0,577017192 | 0,145 | 0,052 | 1 | SMC | 1,491761804 | 0,670348307 | 2,788461538 |
| ZNF512 | 0,006520801 | -0,25878745 | 0,048 | 0,121 | 1 | SMC | 0,835790086 | 1,196472676 | 0,396694215 |
| GABARAPL12 | 0,006525065 | -0,271607137 | 0,114 | 0,216 | 1 | SMC | 0,828396213 | 1,207151825 | 0,527777778 |
| TTC14 | 0,006557434 | -0,259889935 | 0,077 | 0,164 | 1 | SMC | 0,835151632 | 1,197387351 | 0,469512195 |
| DDIT34 | 0,006570718 | -0,492856908 | 0,261 | 0,388 | 1 | SMC | 0,7106165 | 1,407228794 | 0,672680412 |
| FAM13C | 0,006573407 | 0,359986183 | 0,429 | 0,302 | 1 | SMC | 1,283413606 | 0,779172042 | 1,420529801 |
| PPP3CA1 | 0,00661924 | 0,445852156 | 0,156 | 0,06 | 1 | SMC | 1,362118441 | 0,734150548 | 2,6 |
| GRK5 | 0,006620843 | 0,432416988 | 0,224 | 0,112 | 1 | SMC | 1,349492527 | 0,741019294 | 2 |
| TRPC6 | 0,006636828 | 0,393904105 | 0,196 | 0,086 | 1 | SMC | 1,313944288 | 0,761067276 | 2,279069767 |
| COX20 | 0,006693501 | -0,380774823 | 0,159 | 0,267 | 1 | SMC | 0,768025 | 1,30204095 | 0,595505618 |
| RPL172 | 0,006714964 | -0,261874178 | 0,938 | 0,966 | 1 | SMC | 0,834003777 | 1,199035338 | 0,971014493 |
| NIBAN21 | 0,006754762 | 0,358005073 | 0,156 | 0,06 | 1 | SMC | 1,281652431 | 0,780242736 | 2,6 |
| FILIP1L1 | 0,006839891 | 0,517824796 | 0,614 | 0,517 | 1 | SMC | 1,43179485 | 0,698424079 | 1,18762089 |
| MX2 | 0,006883441 | 0,354541113 | 0,119 | 0,034 | 1 | SMC | 1,278578831 | 0,782118377 | 3,5 |
| FLNA5 | 0,006883604 | -0,430607315 | 0,764 | 0,802 | 1 | SMC | 0,74194939 | 1,347800826 | 0,952618454 |
| MYL6B | 0,006918037 | 0,425979194 | 0,176 | 0,078 | 1 | SMC | 1,343484051 | 0,744333362 | 2,256410256 |
| PDIA34 | 0,006984933 | 0,405001428 | 0,489 | 0,362 | 1 | SMC | 1,324090221 | 0,755235545 | 1,350828729 |
| CXCL23 | 0,006992963 | 0,734145577 | 0,162 | 0,06 | 1 | SMC | 1,663412037 | 0,601173959 | 2,7 |

| Supplementary Table 4 - clusermarker | | | | | | | | | |
|---|---|---|---|---|---|---|---|---|---|
| | **p_val** | **avg_log2FC** | **pct.1** | **pct.2** | **p_val_adj** | **celltype** | **Foldchange_UP** | **Foldchange_DOWN** | **Ratio_pct1_pct2** |
| NFKBIA7 | 0,007018903 | 0,644898314 | 0,741 | 0,629 | 1 | SMC | 1,563629073 | 0,639537866 | 1,178060413 |
| WASHC42 | 0,007027103 | 0,285328869 | 0,19 | 0,086 | 1 | SMC | 1,218688038 | 0,820554538 | 2,209302326 |
| FAM3A | 0,007108794 | 0,333618079 | 0,119 | 0,034 | 1 | SMC | 1,260169746 | 0,793543888 | 3,5 |
| LRRC171 | 0,007109905 | 0,257113475 | 0,188 | 0,078 | 1 | SMC | 1,195085201 | 0,836760425 | 2,41025641 |
| RWDD12 | 0,007127759 | 0,399943823 | 0,457 | 0,328 | 1 | SMC | 1,319456531 | 0,757887794 | 1,393292683 |
| CUL34 | 0,007202374 | 0,363767278 | 0,193 | 0,086 | 1 | SMC | 1,286781659 | 0,777132618 | 2,244186047 |
| IL6ST3 | 0,007235034 | 0,499741007 | 0,509 | 0,397 | 1 | SMC | 1,413959705 | 0,707233733 | 1,282115869 |
| BRI36 | 0,007247105 | 0,498239697 | 0,432 | 0,319 | 1 | SMC | 1,412489064 | 0,707970083 | 1,354231975 |
| RELA | 0,007461818 | -0,318047897 | 0,128 | 0,233 | 1 | SMC | 0,802154535 | 1,246642581 | 0,549356223 |
| TAP11 | 0,007507884 | 0,303728714 | 0,159 | 0,06 | 1 | SMC | 1,23433048 | 0,810155802 | 2,65 |
| PDLIM74 | 0,007566187 | -0,442423389 | 0,335 | 0,448 | 1 | SMC | 0,735897435 | 1,358885019 | 0,747767857 |
| YY11 | 0,007617317 | 0,336716726 | 0,295 | 0,172 | 1 | SMC | 1,26287927 | 0,791841329 | 1,715116279 |
| PAG11 | 0,007633148 | 0,323853892 | 0,145 | 0,052 | 1 | SMC | 1,25166969 | 0,798932824 | 2,788461538 |
| CHSY1 | 0,007653384 | 0,362966757 | 0,182 | 0,078 | 1 | SMC | 1,28606785 | 0,777563952 | 2,333333333 |
| UACA2 | 0,007654577 | 0,617334159 | 0,349 | 0,233 | 1 | SMC | 1,534037934 | 0,651874362 | 1,497854077 |
| GPBP1L12 | 0,007663455 | -0,281966824 | 0,102 | 0,198 | 1 | SMC | 0,822468982 | 1,215851323 | 0,515151515 |
| CCDC88A1 | 0,007779371 | -0,32703971 | 0,151 | 0,259 | 1 | SMC | 0,797170534 | 1,254436732 | 0,583011583 |
| VPS13D | 0,007825104 | -0,455161112 | 0,125 | 0,224 | 1 | SMC | 0,729428709 | 1,370935895 | 0,558035714 |
| GULP11 | 0,007889662 | -0,310043664 | 0,111 | 0,207 | 1 | SMC | 0,806617346 | 1,239745221 | 0,536231884 |
| FXYD54 | 0,007945787 | 0,345427222 | 0,205 | 0,095 | 1 | SMC | 1,270527165 | 0,787074868 | 2,157894737 |
| NPNT | 0,00797312 | -0,430894453 | 0,168 | 0,276 | 1 | SMC | 0,741801735 | 1,348069104 | 0,608695652 |
| SYNE22 | 0,008016989 | -0,4094148 | 0,358 | 0,491 | 1 | SMC | 0,752928722 | 1,328146969 | 0,729124236 |
| ANPEP1 | 0,008098106 | 0,311972918 | 0,105 | 0,026 | 1 | SMC | 1,241404188 | 0,805539412 | 4,038461538 |
| LAMA51 | 0,008108004 | 0,5741054 | 0,224 | 0,121 | 1 | SMC | 1,488754016 | 0,671702638 | 1,851239669 |
| COL12A11 | 0,008127768 | 0,418414234 | 0,261 | 0,147 | 1 | SMC | 1,336457754 | 0,748246622 | 1,775510204 |
| MARCKSL14 | 0,008172846 | 0,450680647 | 0,188 | 0,086 | 1 | SMC | 1,366684891 | 0,73169756 | 2,186046512 |
| DDX39B3 | 0,00818333 | -0,353760677 | 0,42 | 0,56 | 1 | SMC | 0,782541584 | 1,277887362 | 0,75 |
| SLC7A21 | 0,008247004 | 0,378923596 | 0,514 | 0,388 | 1 | SMC | 1,300371278 | 0,769011141 | 1,324742268 |
| SLC25A41 | 0,008322936 | -0,261016506 | 0,17 | 0,293 | 1 | SMC | 0,834499734 | 1,198322731 | 0,580204778 |
| CLIC41 | 0,008448352 | -0,299564262 | 0,489 | 0,578 | 1 | SMC | 0,812497759 | 1,230772626 | 0,846020761 |
| NEDD82 | 0,008590019 | 0,508096731 | 0,477 | 0,379 | 1 | SMC | 1,422172763 | 0,703149453 | 1,258575198 |

| Supplementary Table 4 - clusermarker | | | | | | | | | |
|---|---|---|---|---|---|---|---|---|---|
| | **p_val** | **avg_log2FC** | **pct.1** | **pct.2** | **p_val_adj** | **celltype** | **Foldchange_UP** | **Foldchange_DOWN** | **Ratio_pct1_pct2** |
| CITED22 | 0,00861816 | 0,411614277 | 0,466 | 0,319 | 1 | SMC | 1,330173355 | 0,751781711 | 1,460815047 |
| ATE1 | 0,008663192 | 0,358227582 | 0,102 | 0,026 | 1 | SMC | 1,281850117 | 0,780122408 | 3,923076923 |
| ZMIZ12 | 0,008684594 | 0,452739503 | 0,21 | 0,112 | 1 | SMC | 1,368636666 | 0,730654106 | 1,875 |
| U2SURP4 | 0,00881773 | -0,370958735 | 0,278 | 0,397 | 1 | SMC | 0,773268455 | 1,293211942 | 0,700251889 |
| ADCY3 | 0,00887638 | 0,321681536 | 0,361 | 0,224 | 1 | SMC | 1,249786391 | 0,800136733 | 1,611607143 |
| HLA-F1 | 0,008929538 | 0,310468485 | 0,318 | 0,198 | 1 | SMC | 1,240110334 | 0,806379862 | 1,606060606 |
| CDC374 | 0,008933011 | 0,475829443 | 0,412 | 0,293 | 1 | SMC | 1,39071755 | 0,719053268 | 1,406143345 |
| ACOT91 | 0,008938841 | 0,478840674 | 0,148 | 0,06 | 1 | SMC | 1,393623324 | 0,717554007 | 2,466666667 |
| SH3GLB11 | 0,008960722 | 0,479434142 | 0,256 | 0,147 | 1 | SMC | 1,394196724 | 0,717258894 | 1,741496599 |
| VPS293 | 0,008962771 | -0,305943878 | 0,259 | 0,397 | 1 | SMC | 0,808912815 | 1,23622717 | 0,652392947 |
| CEP85L | 0,009166653 | 0,386122426 | 0,128 | 0,043 | 1 | SMC | 1,306876149 | 0,76518345 | 2,976744186 |
| CCL26 | 0,00921768 | 0,490449914 | 0,105 | 0,026 | 1 | SMC | 1,404882929 | 0,711803083 | 4,038461538 |
| PDCL32 | 0,009226087 | -0,253306026 | 0,085 | 0,172 | 1 | SMC | 0,838971655 | 1,191935382 | 0,494186047 |
| CALM35 | 0,009265638 | -0,474337406 | 0,281 | 0,388 | 1 | SMC | 0,719797299 | 1,389280012 | 0,724226804 |
| EIF4B6 | 0,009299619 | -0,393669778 | 0,457 | 0,569 | 1 | SMC | 0,761190902 | 1,313730889 | 0,803163445 |
| HIC11 | 0,009406457 | 0,39568769 | 0,128 | 0,043 | 1 | SMC | 1,315569704 | 0,76012696 | 2,976744186 |
| SCAF113 | 0,009492553 | -0,262586453 | 0,27 | 0,405 | 1 | SMC | 0,833592121 | 1,199627461 | 0,666666667 |
| MYO1C1 | 0,009552577 | 0,45377378 | 0,352 | 0,233 | 1 | SMC | 1,369618202 | 0,730130483 | 1,510729614 |
| RCSD11 | 0,009569621 | 0,414838261 | 0,259 | 0,147 | 1 | SMC | 1,333149211 | 0,750103583 | 1,761904762 |
| CDH111 | 0,009646975 | 0,393096059 | 0,114 | 0,034 | 1 | SMC | 1,31320856 | 0,761493666 | 3,352941176 |
| DNAJC191 | 0,009692984 | -0,283998785 | 0,151 | 0,259 | 1 | SMC | 0,821311392 | 1,217564994 | 0,583011583 |
| LACTB1 | 0,00977666 | 0,260581511 | 0,128 | 0,043 | 1 | SMC | 1,197961473 | 0,834751386 | 2,976744186 |
| COX7B5 | 0,00985365 | -0,304777117 | 0,531 | 0,629 | 1 | SMC | 0,809567277 | 1,235227792 | 0,844197138 |
| ABCE11 | 0,009923746 | -0,301121188 | 0,094 | 0,181 | 1 | SMC | 0,811621401 | 1,232101567 | 0,519337017 |
| DNAJB92 | 0,009924153 | -0,386447481 | 0,082 | 0,164 | 1 | SMC | 0,765011064 | 1,307170637 | 0,5 |
| PEG10 | 0,009954162 | -0,273929046 | 0,04 | 0,103 | 1 | SMC | 0,827064044 | 1,209096209 | 0,388349515 |
| RPL152 | 0,009973482 | -0,254404161 | 0,949 | 0,983 | 1 | SMC | 0,838333298 | 1,192842992 | 0,965412004 |
| GDF11 | 0,010002994 | 0,351719317 | 0,114 | 0,034 | 1 | SMC | 1,276080477 | 0,783649635 | 3,352941176 |
| ACAN | 0,010013172 | -0,42934672 | 0,045 | 0,112 | 1 | SMC | 0,742597972 | 1,346623661 | 0,401785714 |
| FTSJ3 | 0,010077238 | -0,283210543 | 0,057 | 0,129 | 1 | SMC | 0,821760253 | 1,216899937 | 0,441860465 |
| MFGE82 | 0,010105056 | 0,410495325 | 0,781 | 0,759 | 1 | SMC | 1,329142074 | 0,752365018 | 1,028985507 |

| Supplementary Table 4 - clusermarker | | | | | | | | | |
|---|---|---|---|---|---|---|---|---|---|
| | **p_val** | **avg_log2FC** | **pct.1** | **pct.2** | **p_val_adj** | **celltype** | **Foldchange_UP** | **Foldchange_DOWN** | **Ratio_pct1_pct2** |
| ZNF618 | 0,010159792 | 0,34614861 | 0,128 | 0,043 | 1 | SMC | 1,271162623 | 0,786681406 | 2,976744186 |
| CD40 | 0,010231622 | 0,354815967 | 0,17 | 0,078 | 1 | SMC | 1,278822442 | 0,781969386 | 2,179487179 |
| FAT11 | 0,010237416 | 0,347379683 | 0,304 | 0,181 | 1 | SMC | 1,272247788 | 0,786010406 | 1,679558011 |
| GNB25 | 0,01025458 | 0,397892439 | 0,435 | 0,31 | 1 | SMC | 1,317581715 | 0,75896621 | 1,403225806 |
| GOSR2 | 0,010319984 | 0,388063823 | 0,136 | 0,052 | 1 | SMC | 1,308635962 | 0,764154455 | 2,615384615 |
| SOCS33 | 0,010320844 | 0,467404741 | 0,594 | 0,466 | 1 | SMC | 1,382620039 | 0,723264506 | 1,274678112 |
| TPGS11 | 0,010422096 | 0,38152596 | 0,185 | 0,086 | 1 | SMC | 1,302719033 | 0,767625232 | 2,151162791 |
| EVA1B | 0,010423051 | 0,473207587 | 0,23 | 0,129 | 1 | SMC | 1,38819245 | 0,720361215 | 1,782945736 |
| ID41 | 0,010441179 | 0,408160194 | 0,685 | 0,586 | 1 | SMC | 1,326992479 | 0,753583774 | 1,16894198 |
| SH3KBP11 | 0,010496583 | 0,327317008 | 0,23 | 0,121 | 1 | SMC | 1,254677868 | 0,797017326 | 1,900826446 |
| NDUFB63 | 0,010566345 | -0,451776976 | 0,341 | 0,448 | 1 | SMC | 0,731141741 | 1,367723854 | 0,761160714 |
| NRIP2 | 0,010576173 | -0,311106596 | 0,099 | 0,19 | 1 | SMC | 0,806023275 | 1,240658962 | 0,521052632 |
| CSRP2 | 0,010604576 | -0,361773592 | 0,474 | 0,586 | 1 | SMC | 0,778207294 | 1,285004661 | 0,80887372 |
| RAB3GAP1 | 0,010677948 | -0,344733995 | 0,162 | 0,259 | 1 | SMC | 0,787453155 | 1,269916813 | 0,625482625 |
| SYNE12 | 0,010821094 | -0,252314475 | 0,281 | 0,414 | 1 | SMC | 0,83954847 | 1,191116458 | 0,678743961 |
| GLUL5 | 0,01084103 | -0,426305806 | 0,415 | 0,509 | 1 | SMC | 0,744164872 | 1,343788236 | 0,815324165 |
| MAN1A11 | 0,010972321 | 0,364090599 | 0,168 | 0,069 | 1 | SMC | 1,28707007 | 0,776958476 | 2,434782609 |
| MYOF1 | 0,011124608 | 0,43145659 | 0,324 | 0,207 | 1 | SMC | 1,348594473 | 0,741512753 | 1,565217391 |
| RANBP91 | 0,011192194 | 0,288277139 | 0,114 | 0,034 | 1 | SMC | 1,221181077 | 0,818879378 | 3,352941176 |
| GLG11 | 0,011348873 | 0,424418711 | 0,412 | 0,293 | 1 | SMC | 1,342031664 | 0,745138902 | 1,406143345 |
| AGT | 0,011604815 | 0,353493775 | 0,165 | 0,069 | 1 | SMC | 1,277650971 | 0,782686369 | 2,391304348 |
| LAMA4 | 0,011705339 | 0,401540676 | 0,415 | 0,284 | 1 | SMC | 1,320917786 | 0,757049387 | 1,461267606 |
| ARPC55 | 0,011750704 | 0,38818925 | 0,537 | 0,422 | 1 | SMC | 1,308749739 | 0,764088022 | 1,272511848 |
| RTL8C1 | 0,011771511 | -0,461193531 | 0,395 | 0,5 | 1 | SMC | 0,726385077 | 1,376680265 | 0,79 |
| ATP5IF13 | 0,011805901 | 0,421021731 | 0,477 | 0,371 | 1 | SMC | 1,338875424 | 0,746895478 | 1,285714286 |
| RAP1A3 | 0,011933625 | -0,291175138 | 0,273 | 0,388 | 1 | SMC | 0,817236113 | 1,223636578 | 0,703608247 |
| HTATIP22 | 0,012045639 | 0,344510662 | 0,122 | 0,043 | 1 | SMC | 1,269720242 | 0,787575064 | 2,837209302 |
| DACT3 | 0,012235583 | -0,320390008 | 0,102 | 0,19 | 1 | SMC | 0,800853351 | 1,248668059 | 0,536842105 |
| SOD32 | 0,012332467 | 0,322242147 | 0,906 | 0,862 | 1 | SMC | 1,250272135 | 0,799825871 | 1,051044084 |
| MVB12A3 | 0,012401421 | 0,386437058 | 0,168 | 0,078 | 1 | SMC | 1,307161193 | 0,765016591 | 2,153846154 |
| SLC9A3R23 | 0,012943781 | 0,47154958 | 0,239 | 0,138 | 1 | SMC | 1,386597996 | 0,721189561 | 1,731884058 |

| Supplementary Table 4 - clusermarker | | | | | | | | | |
|---|---|---|---|---|---|---|---|---|---|
| | **p_val** | **avg_log2FC** | **pct.1** | **pct.2** | **p_val_adj** | **celltype** | **Foldchange_UP** | **Foldchange_DOWN** | **Ratio_pct1_pct2** |
| PKP42 | 0,012981514 | -0,303058845 | 0,06 | 0,129 | 1 | SMC | 0,810532059 | 1,233757492 | 0,465116279 |
| AC018521.51 | 0,013071972 | -0,25214567 | 0,114 | 0,207 | 1 | SMC | 0,839646709 | 1,190977097 | 0,550724638 |
| TCEA2 | 0,013105199 | 0,465046197 | 0,193 | 0,103 | 1 | SMC | 1,380361554 | 0,724447879 | 1,873786408 |
| SELENOM6 | 0,013239599 | -0,416799124 | 0,849 | 0,81 | 1 | SMC | 0,74908476 | 1,334962414 | 1,048148148 |
| CDKN1A6 | 0,013293181 | -0,555953165 | 0,636 | 0,724 | 1 | SMC | 0,680207508 | 1,47013961 | 0,878453039 |
| SNAI2 | 0,013320744 | 0,306534524 | 0,233 | 0,121 | 1 | SMC | 1,23673339 | 0,80858171 | 1,925619835 |
| CYP7B1 | 0,013336798 | 0,314341892 | 0,227 | 0,121 | 1 | SMC | 1,243444308 | 0,804217763 | 1,876033058 |
| AKAP8L | 0,013416245 | 0,384609571 | 0,159 | 0,069 | 1 | SMC | 1,305506437 | 0,765986265 | 2,304347826 |
| TPGS22 | 0,013529705 | 0,403572339 | 0,168 | 0,078 | 1 | SMC | 1,322779268 | 0,755984029 | 2,153846154 |
| PPP4R3B2 | 0,013535815 | -0,35848818 | 0,131 | 0,224 | 1 | SMC | 0,779981505 | 1,282081683 | 0,584821429 |
| BCAR11 | 0,013581046 | 0,35690686 | 0,134 | 0,052 | 1 | SMC | 1,280677178 | 0,780836902 | 2,576923077 |
| C2CD2 | 0,013615752 | 0,368014107 | 0,145 | 0,06 | 1 | SMC | 1,29057511 | 0,774848355 | 2,416666667 |
| PTN1 | 0,013647207 | 0,504398734 | 0,179 | 0,086 | 1 | SMC | 1,418532037 | 0,704954117 | 2,081395349 |
| PSMB85 | 0,013799664 | 0,423695205 | 0,327 | 0,216 | 1 | SMC | 1,341358809 | 0,745512679 | 1,513888889 |
| SNX35 | 0,013871941 | 0,534786377 | 0,517 | 0,422 | 1 | SMC | 1,448727623 | 0,690260877 | 1,225118483 |
| FAM204A4 | 0,013919498 | -0,419084809 | 0,241 | 0,353 | 1 | SMC | 0,747898913 | 1,337079093 | 0,682719547 |
| GSDMD2 | 0,013922193 | 0,304730404 | 0,148 | 0,06 | 1 | SMC | 1,235187796 | 0,809593491 | 2,466666667 |
| TGFBI2 | 0,013934028 | 0,469457786 | 0,443 | 0,336 | 1 | SMC | 1,384588995 | 0,722235988 | 1,318452381 |
| LMCD11 | 0,013943551 | 0,274158955 | 0,199 | 0,095 | 1 | SMC | 1,209288907 | 0,826932253 | 2,094736842 |
| CCN5 | 0,013964206 | 0,743842634 | 0,122 | 0,043 | 1 | SMC | 1,674630301 | 0,597146725 | 2,837209302 |
| DNAAF2 | 0,014029722 | 0,387312522 | 0,108 | 0,034 | 1 | SMC | 1,307954652 | 0,764552501 | 3,176470588 |
| PSMB72 | 0,014137507 | -0,416870798 | 0,222 | 0,328 | 1 | SMC | 0,749047546 | 1,335028738 | 0,676829268 |
| RNF1673 | 0,014257105 | -0,397764237 | 0,196 | 0,293 | 1 | SMC | 0,759033656 | 1,317464636 | 0,66894198 |
| FRY | 0,014300914 | -0,327916703 | 0,247 | 0,362 | 1 | SMC | 0,796686094 | 1,255199517 | 0,682320442 |
| WDR110 | 0,014465128 | -0,495458089 | 0,352 | 0,457 | 1 | SMC | 0,709336411 | 1,409768319 | 0,7702407 |
| MSRB24 | 0,014703807 | -0,341110311 | 0,139 | 0,233 | 1 | SMC | 0,789433523 | 1,266731106 | 0,596566524 |
| SLC8A1 | 0,014724408 | -0,253864625 | 0,054 | 0,121 | 1 | SMC | 0,838646875 | 1,192396979 | 0,446280992 |
| GMFG1 | 0,014865852 | 0,317431948 | 0,122 | 0,043 | 1 | SMC | 1,24611045 | 0,802497082 | 2,837209302 |
| TMEM50A5 | 0,014877058 | 0,358347451 | 0,486 | 0,353 | 1 | SMC | 1,281956627 | 0,780057592 | 1,376770538 |
| FAM43A | 0,014926895 | 0,311668037 | 0,108 | 0,034 | 1 | SMC | 1,241141873 | 0,805709663 | 3,176470588 |
| MT2A3 | 0,014971091 | 0,252054834 | 0,889 | 0,853 | 1 | SMC | 1,190902113 | 0,839699577 | 1,042203986 |

| Supplementary Table 4 - clusermarker | | | | | | | | | |
|---|---|---|---|---|---|---|---|---|---|
| | **p_val** | **avg_log2FC** | **pct.1** | **pct.2** | **p_val_adj** | **celltype** | **Foldchange_UP** | **Foldchange_DOWN** | **Ratio_pct1_pct2** |
| TMEM35B | 0,015057883 | 0,47239474 | 0,301 | 0,207 | 1 | SMC | 1,38741053 | 0,720767198 | 1,45410628 |
| CEBPZ2 | 0,015071818 | -0,33753605 | 0,145 | 0,241 | 1 | SMC | 0,79139176 | 1,263596679 | 0,601659751 |
| DSE1 | 0,015115934 | 0,373298289 | 0,142 | 0,06 | 1 | SMC | 1,295310787 | 0,772015496 | 2,366666667 |
| GOLGA21 | 0,015204355 | -0,299941622 | 0,17 | 0,267 | 1 | SMC | 0,812285265 | 1,231094596 | 0,63670412 |
| SPOP3 | 0,015215115 | -0,304209341 | 0,168 | 0,267 | 1 | SMC | 0,809885947 | 1,23474176 | 0,629213483 |
| UGP2 | 0,015320288 | -0,462653817 | 0,23 | 0,336 | 1 | SMC | 0,725650207 | 1,378074436 | 0,68452381 |
| ECH11 | 0,015359862 | -0,339752329 | 0,29 | 0,397 | 1 | SMC | 0,790176952 | 1,265539317 | 0,730478589 |
| EXTL2 | 0,015467183 | -0,256919889 | 0,054 | 0,121 | 1 | SMC | 0,836872712 | 1,19492485 | 0,446280992 |
| GPI3 | 0,015483474 | 0,445624291 | 0,384 | 0,276 | 1 | SMC | 1,361903319 | 0,734266512 | 1,391304348 |
| CTDSP11 | 0,015512873 | -0,3329805 | 0,131 | 0,224 | 1 | SMC | 0,793894662 | 1,259612954 | 0,584821429 |
| EPHA2 | 0,01554725 | 0,398200389 | 0,131 | 0,052 | 1 | SMC | 1,317862989 | 0,758804222 | 2,519230769 |
| MTERF4 | 0,015629356 | -0,28572611 | 0,06 | 0,129 | 1 | SMC | 0,820328632 | 1,219023646 | 0,465116279 |
| SYNC | 0,015662999 | -0,293632751 | 0,116 | 0,207 | 1 | SMC | 0,815845146 | 1,225722804 | 0,560386473 |
| KDELR14 | 0,015714535 | 0,451384011 | 0,463 | 0,353 | 1 | SMC | 1,36735136 | 0,731340919 | 1,311614731 |
| TRIM222 | 0,015870438 | 0,534317037 | 0,196 | 0,103 | 1 | SMC | 1,448256397 | 0,690485471 | 1,902912621 |
| MPHOSPH82 | 0,015901099 | -0,289503064 | 0,315 | 0,431 | 1 | SMC | 0,818183833 | 1,222219212 | 0,730858469 |
| CRYZL1 | 0,016009675 | -0,451845241 | 0,108 | 0,19 | 1 | SMC | 0,731107146 | 1,367788572 | 0,568421053 |
| SLU71 | 0,016148948 | -0,419634673 | 0,165 | 0,259 | 1 | SMC | 0,747613915 | 1,337588801 | 0,637065637 |
| MRPS152 | 0,016278491 | 0,39795803 | 0,185 | 0,095 | 1 | SMC | 1,317641619 | 0,758931704 | 1,947368421 |
| CDK41 | 0,016411133 | 0,425369296 | 0,259 | 0,164 | 1 | SMC | 1,342916214 | 0,744648095 | 1,579268293 |
| MYC | 0,016442102 | 0,649058903 | 0,5 | 0,371 | 1 | SMC | 1,568144932 | 0,637696159 | 1,347708895 |
| HSD17B122 | 0,016558854 | -0,275536325 | 0,153 | 0,25 | 1 | SMC | 0,82614314 | 1,210443991 | 0,612 |
| REM1 | 0,016654627 | 0,385849541 | 0,219 | 0,121 | 1 | SMC | 1,306628979 | 0,765328197 | 1,809917355 |
| COTL15 | 0,016800843 | 0,256861494 | 0,108 | 0,034 | 1 | SMC | 1,194876485 | 0,836906586 | 3,176470588 |
| PIH1D13 | 0,016910038 | -0,309082688 | 0,131 | 0,224 | 1 | SMC | 0,807154811 | 1,238919704 | 0,584821429 |
| STOML25 | 0,017038302 | 0,463607944 | 0,227 | 0,138 | 1 | SMC | 1,378986128 | 0,725170457 | 1,644927536 |
| ALKBH74 | 0,017091033 | 0,436849469 | 0,381 | 0,267 | 1 | SMC | 1,353645032 | 0,738746109 | 1,426966292 |
| SEC627 | 0,01710125 | -0,433114499 | 0,534 | 0,603 | 1 | SMC | 0,740661114 | 1,350145135 | 0,885572139 |
| DDX39A3 | 0,017437691 | 0,385387787 | 0,128 | 0,052 | 1 | SMC | 1,306210841 | 0,76557319 | 2,461538462 |
| PIN43 | 0,017754713 | -0,453667611 | 0,165 | 0,259 | 1 | SMC | 0,730184217 | 1,369517414 | 0,637065637 |
| WBP22 | 0,017841783 | -0,324189215 | 0,199 | 0,302 | 1 | SMC | 0,798747151 | 1,251960648 | 0,658940397 |

| Supplementary Table 4 - clusermarker | | | | | | | | | |
|---|---|---|---|---|---|---|---|---|---|
| | **p_val** | **avg_log2FC** | **pct.1** | **pct.2** | **p_val_adj** | **celltype** | **Foldchange_UP** | **Foldchange_DOWN** | **Ratio_pct1_pct2** |
| ATP5MJ2 | 0,017990351 | -0,36471901 | 0,375 | 0,474 | 1 | SMC | 0,77662012 | 1,287630817 | 0,791139241 |
| TSPAN33 | 0,017994422 | 0,432336932 | 0,378 | 0,276 | 1 | SMC | 1,349417645 | 0,741060415 | 1,369565217 |
| LASP12 | 0,01808685 | 0,287917917 | 0,239 | 0,138 | 1 | SMC | 1,220877048 | 0,819083299 | 1,731884058 |
| ARHGAP24 | 0,018138 | 0,320918498 | 0,131 | 0,052 | 1 | SMC | 1,249125557 | 0,800560035 | 2,519230769 |
| ATP5MK4 | 0,0182328 | -0,470032086 | 0,523 | 0,586 | 1 | SMC | 0,721948541 | 1,385140273 | 0,892491468 |
| SNHG74 | 0,01849464 | 0,437352227 | 0,264 | 0,164 | 1 | SMC | 1,35411684 | 0,738488711 | 1,609756098 |
| LRP12 | 0,018579803 | 0,380166525 | 0,213 | 0,112 | 1 | SMC | 1,301492073 | 0,768348898 | 1,901785714 |
| ZFHX31 | 0,018757969 | -0,414411018 | 0,318 | 0,414 | 1 | SMC | 0,750325754 | 1,332754467 | 0,768115942 |
| PER3 | 0,018795759 | -0,273653408 | 0,17 | 0,267 | 1 | SMC | 0,827222076 | 1,208865224 | 0,63670412 |
| NABP1 | 0,018945649 | 0,367002265 | 0,153 | 0,069 | 1 | SMC | 1,289670275 | 0,775391989 | 2,217391304 |
| SPAG94 | 0,019097823 | 0,458305574 | 0,33 | 0,233 | 1 | SMC | 1,373927211 | 0,727840596 | 1,416309013 |
| PARM1 | 0,019159112 | 0,419854856 | 0,381 | 0,276 | 1 | SMC | 1,337792958 | 0,747499824 | 1,380434783 |
| PEAR1 | 0,019173676 | 0,406397277 | 0,179 | 0,095 | 1 | SMC | 1,325371936 | 0,754505187 | 1,884210526 |
| PDCD7 | 0,019325412 | -0,290572746 | 0,108 | 0,19 | 1 | SMC | 0,817577418 | 1,223125759 | 0,568421053 |
| BCL31 | 0,019364117 | 0,250325963 | 0,256 | 0,155 | 1 | SMC | 1,189475835 | 0,840706444 | 1,651612903 |
| AKAP1 | 0,019375115 | -0,372126838 | 0,193 | 0,293 | 1 | SMC | 0,772642618 | 1,294259438 | 0,658703072 |
| SAMD82 | 0,01944189 | 0,272698682 | 0,116 | 0,043 | 1 | SMC | 1,208065503 | 0,827769684 | 2,697674419 |
| SPTBN13 | 0,019531568 | 0,398489619 | 0,511 | 0,397 | 1 | SMC | 1,318127219 | 0,758652113 | 1,287153652 |
| P3H11 | 0,019611912 | 0,28347453 | 0,102 | 0,034 | 1 | SMC | 1,217122627 | 0,8216099 | 3 |
| SCYL21 | 0,019611912 | -0,297136109 | 0,068 | 0,138 | 1 | SMC | 0,813866398 | 1,228702895 | 0,492753623 |
| NFKBIZ4 | 0,019694055 | 0,480889779 | 0,386 | 0,284 | 1 | SMC | 1,395604137 | 0,716535566 | 1,35915493 |
| BRD25 | 0,019758371 | 0,330579989 | 0,636 | 0,56 | 1 | SMC | 1,257518818 | 0,795216728 | 1,135714286 |
| SPCS16 | 0,019777801 | 0,454697522 | 0,503 | 0,422 | 1 | SMC | 1,370495434 | 0,729663139 | 1,191943128 |
| IER5L5 | 0,019890076 | 0,401790502 | 0,426 | 0,328 | 1 | SMC | 1,321146544 | 0,756918303 | 1,298780488 |
| PTEN2 | 0,019967764 | 0,430479856 | 0,469 | 0,362 | 1 | SMC | 1,347681756 | 0,742014942 | 1,29558011 |
| ARPC42 | 0,020005302 | -0,378388789 | 0,236 | 0,336 | 1 | SMC | 0,769296266 | 1,299889319 | 0,702380952 |
| PFKFB3 | 0,020122694 | 0,276494623 | 0,188 | 0,095 | 1 | SMC | 1,211248285 | 0,825594564 | 1,978947368 |
| CFL21 | 0,02029141 | -0,268732703 | 0,162 | 0,259 | 1 | SMC | 0,83004836 | 1,204749083 | 0,625482625 |
| TMEM51 | 0,020313595 | -0,312600942 | 0,071 | 0,138 | 1 | SMC | 0,805188827 | 1,241944705 | 0,514492754 |
| ILK4 | 0,020440115 | -0,549185254 | 0,514 | 0,569 | 1 | SMC | 0,683405965 | 1,463259104 | 0,903339192 |
| FGF72 | 0,020621685 | 0,483344026 | 0,378 | 0,267 | 1 | SMC | 1,397980295 | 0,715317665 | 1,415730337 |

| Supplementary Table 4 - clusermarker | | | | | | | | | |
|---|---|---|---|---|---|---|---|---|---|
| | **p_val** | **avg_log2FC** | **pct.1** | **pct.2** | **p_val_adj** | **celltype** | **Foldchange_UP** | **Foldchange_DOWN** | **Ratio_pct1_pct2** |
| INTS111 | 0,020644746 | 0,39021162 | 0,128 | 0,052 | 1 | SMC | 1,310585632 | 0,763017674 | 2,461538462 |
| TASOR1 | 0,020668159 | 0,373654978 | 0,162 | 0,078 | 1 | SMC | 1,295631077 | 0,771824648 | 2,076923077 |
| ESD4 | 0,020691515 | -0,383023348 | 0,491 | 0,578 | 1 | SMC | 0,766828919 | 1,30407184 | 0,849480969 |
| ANXA17 | 0,020756068 | -0,353021736 | 0,591 | 0,638 | 1 | SMC | 0,7829425 | 1,277233002 | 0,926332288 |
| IFNGR13 | 0,021070027 | 0,388940189 | 0,222 | 0,129 | 1 | SMC | 1,309431136 | 0,763690409 | 1,720930233 |
| C2orf74 | 0,021106477 | -0,327196356 | 0,088 | 0,164 | 1 | SMC | 0,797083983 | 1,254572945 | 0,536585366 |
| DCAF71 | 0,02113349 | 0,368674638 | 0,222 | 0,129 | 1 | SMC | 1,291166129 | 0,774493675 | 1,720930233 |
| SH2D3C | 0,021179624 | 0,289762057 | 0,116 | 0,043 | 1 | SMC | 1,222438644 | 0,818036966 | 2,697674419 |
| CNGA1 | 0,021202333 | 0,298293997 | 0,105 | 0,034 | 1 | SMC | 1,229689431 | 0,813213462 | 3,088235294 |
| PIM11 | 0,021297249 | 0,366502033 | 0,102 | 0,034 | 1 | SMC | 1,28922318 | 0,775660891 | 3 |
| ANKLE21 | 0,021768003 | 0,341825241 | 0,114 | 0,043 | 1 | SMC | 1,267358993 | 0,789042415 | 2,651162791 |
| CTSZ6 | 0,021989247 | 0,340839282 | 0,409 | 0,293 | 1 | SMC | 1,266493157 | 0,789581842 | 1,395904437 |
| PDGFA1 | 0,022035285 | 0,334980495 | 0,301 | 0,198 | 1 | SMC | 1,261360355 | 0,792794855 | 1,52020202 |
| UBE2F3 | 0,022048521 | 0,312406208 | 0,136 | 0,06 | 1 | SMC | 1,24177708 | 0,805297518 | 2,266666667 |
| TGFB12 | 0,0220737 | 0,410543069 | 0,23 | 0,138 | 1 | SMC | 1,329186061 | 0,752340119 | 1,666666667 |
| DLX5 | 0,02230401 | 0,348142686 | 0,128 | 0,052 | 1 | SMC | 1,272920824 | 0,785594815 | 2,461538462 |
| FHL5 | 0,022326145 | -0,388910489 | 0,318 | 0,414 | 1 | SMC | 0,763706131 | 1,30940418 | 0,768115942 |
| CCNH4 | 0,022613353 | -0,315685614 | 0,168 | 0,267 | 1 | SMC | 0,803469066 | 1,244602987 | 0,629213483 |
| TMBIM13 | 0,022713239 | -0,338441218 | 0,216 | 0,319 | 1 | SMC | 0,790895385 | 1,264389726 | 0,677115987 |
| MTO1 | 0,022723171 | -0,322674561 | 0,062 | 0,129 | 1 | SMC | 0,799586178 | 1,250646931 | 0,480620155 |
| GNG55 | 0,02273784 | 0,39005985 | 0,506 | 0,405 | 1 | SMC | 1,310447767 | 0,763097947 | 1,249382716 |
| MAP2 | 0,022779533 | 0,326801419 | 0,159 | 0,078 | 1 | SMC | 1,254229553 | 0,797302214 | 2,038461538 |
| INTS12 | 0,022845596 | -0,295232311 | 0,051 | 0,112 | 1 | SMC | 0,814941095 | 1,227082553 | 0,455357143 |
| RUNX12 | 0,022904204 | 0,261950511 | 0,139 | 0,06 | 1 | SMC | 1,19909878 | 0,833959651 | 2,316666667 |
| WDR541 | 0,023324224 | 0,285711004 | 0,114 | 0,043 | 1 | SMC | 1,219010881 | 0,820337222 | 2,651162791 |
| STAT22 | 0,023375718 | 0,367734343 | 0,159 | 0,078 | 1 | SMC | 1,290324869 | 0,774998625 | 2,038461538 |
| PNISR4 | 0,023583 | -0,3755349 | 0,358 | 0,474 | 1 | SMC | 0,770819567 | 1,297320466 | 0,755274262 |
| PACSIN21 | 0,02404982 | 0,294124945 | 0,384 | 0,267 | 1 | SMC | 1,226141046 | 0,815566858 | 1,438202247 |
| BAX7 | 0,024171689 | 0,454710886 | 0,188 | 0,103 | 1 | SMC | 1,37050813 | 0,72965638 | 1,825242718 |
| SIGMAR1 | 0,024304815 | 0,441380015 | 0,196 | 0,112 | 1 | SMC | 1,357902613 | 0,736429837 | 1,75 |
| FAM107B2 | 0,024715 | -0,426337462 | 0,114 | 0,19 | 1 | SMC | 0,744148543 | 1,343817723 | 0,6 |

| Supplementary Table 4 - clusermarker | | | | | | | | | |
|---|---|---|---|---|---|---|---|---|---|
| | **p_val** | **avg_log2FC** | **pct.1** | **pct.2** | **p_val_adj** | **celltype** | **Foldchange_UP** | **Foldchange_DOWN** | **Ratio_pct1_pct2** |
| FAAP205 | 0,024786729 | 0,268667196 | 0,227 | 0,129 | 1 | SMC | 1,204694382 | 0,830086049 | 1,759689922 |
| POLR2H1 | 0,024817236 | 0,295740717 | 0,136 | 0,06 | 1 | SMC | 1,227515054 | 0,81465396 | 2,266666667 |
| TM9SF23 | 0,025030688 | 0,398158633 | 0,364 | 0,267 | 1 | SMC | 1,317824846 | 0,758826185 | 1,36329588 |
| CTNNAL1 | 0,025066713 | -0,344648857 | 0,077 | 0,147 | 1 | SMC | 0,787499626 | 1,269841873 | 0,523809524 |
| ECHDC21 | 0,0252306 | -0,31860219 | 0,128 | 0,216 | 1 | SMC | 0,801846401 | 1,247121642 | 0,592592593 |
| PCBP22 | 0,025238738 | 0,35022936 | 0,489 | 0,397 | 1 | SMC | 1,274763273 | 0,784459375 | 1,231738035 |
| IP6K22 | 0,025279072 | 0,25283847 | 0,139 | 0,06 | 1 | SMC | 1,191549156 | 0,839243597 | 2,316666667 |
| SGCA | 0,025308186 | -0,262728867 | 0,219 | 0,319 | 1 | SMC | 0,833509838 | 1,199745887 | 0,686520376 |
| MZT2B4 | 0,025373962 | -0,301088586 | 0,386 | 0,483 | 1 | SMC | 0,811639742 | 1,232073724 | 0,799171843 |
| PPP2CB1 | 0,025407063 | -0,3254357 | 0,176 | 0,267 | 1 | SMC | 0,798057333 | 1,253042806 | 0,65917603 |
| ARHGEF17 | 0,025641763 | 0,383767925 | 0,295 | 0,198 | 1 | SMC | 1,304745047 | 0,76643326 | 1,48989899 |
| AMOTL2 | 0,02576519 | 0,294130698 | 0,273 | 0,172 | 1 | SMC | 1,226145935 | 0,815563606 | 1,587209302 |
| LAPTM4B1 | 0,025809366 | 0,581196755 | 0,241 | 0,155 | 1 | SMC | 1,496089782 | 0,668409084 | 1,55483871 |
| SIRT2 | 0,026145313 | -0,348229694 | 0,102 | 0,181 | 1 | SMC | 0,785547438 | 1,272997595 | 0,563535912 |
| TOMM74 | 0,02618795 | -0,343465864 | 0,665 | 0,741 | 1 | SMC | 0,788145631 | 1,268801044 | 0,897435897 |
| AL627171.26 | 0,026271525 | -0,562239368 | 0,125 | 0,207 | 1 | SMC | 0,677250112 | 1,476559372 | 0,603864734 |
| CARHSP11 | 0,026282321 | -0,296544683 | 0,142 | 0,224 | 1 | SMC | 0,814200107 | 1,228199297 | 0,633928571 |
| CERS2 | 0,026290403 | -0,270792395 | 0,122 | 0,207 | 1 | SMC | 0,82886417 | 1,206470295 | 0,589371981 |
| SYPL12 | 0,026306427 | -0,270205166 | 0,347 | 0,457 | 1 | SMC | 0,829201617 | 1,205979318 | 0,759299781 |
| MIER11 | 0,026362608 | 0,36500492 | 0,307 | 0,216 | 1 | SMC | 1,287886022 | 0,776466227 | 1,421296296 |
| SLC2A33 | 0,026560284 | 0,592787522 | 0,432 | 0,336 | 1 | SMC | 1,508157941 | 0,663060528 | 1,285714286 |
| CAP12 | 0,0267122 | -0,297365848 | 0,48 | 0,578 | 1 | SMC | 0,813736806 | 1,228898573 | 0,830449827 |
| CDKN1B3 | 0,026893728 | 0,459219556 | 0,332 | 0,241 | 1 | SMC | 1,374797902 | 0,727379638 | 1,377593361 |
| RAP1B2 | 0,027037441 | 0,471700151 | 0,418 | 0,328 | 1 | SMC | 1,386742719 | 0,721114296 | 1,274390244 |
| CYB5R32 | 0,027137965 | 0,250192003 | 0,71 | 0,569 | 1 | SMC | 1,189365393 | 0,840784511 | 1,247803163 |
| TMTC3 | 0,027418917 | 0,281773718 | 0,122 | 0,052 | 1 | SMC | 1,215688591 | 0,822579078 | 2,346153846 |
| CDC42EP34 | 0,027570532 | -0,346102384 | 0,085 | 0,155 | 1 | SMC | 0,786706613 | 1,271121894 | 0,548387097 |
| ATF44 | 0,027954322 | 0,317450127 | 0,435 | 0,336 | 1 | SMC | 1,246126151 | 0,80248697 | 1,294642857 |
| SEPTIN93 | 0,027991614 | 0,436310706 | 0,241 | 0,155 | 1 | SMC | 1,353139618 | 0,739022039 | 1,55483871 |
| EIF2S22 | 0,028207223 | -0,492762615 | 0,224 | 0,31 | 1 | SMC | 0,710662946 | 1,407136822 | 0,722580645 |
| NR4A14 | 0,028477741 | -0,311730216 | 0,651 | 0,759 | 1 | SMC | 0,805674938 | 1,241195366 | 0,85770751 |

| Supplementary Table 4 - clusermarker | | | | | | | | | |
|---|---|---|---|---|---|---|---|---|---|
| | **p_val** | **avg_log2FC** | **pct.1** | **pct.2** | **p_val_adj** | **celltype** | **Foldchange_UP** | **Foldchange_DOWN** | **Ratio_pct1_pct2** |
| KAT81 | 0,028686914 | 0,435450474 | 0,159 | 0,086 | 1 | SMC | 1,352333026 | 0,739462825 | 1,848837209 |
| KIAA0040 | 0,028807274 | 0,364635014 | 0,321 | 0,216 | 1 | SMC | 1,28755585 | 0,776665338 | 1,486111111 |
| APPL14 | 0,028843113 | -0,272227929 | 0,241 | 0,353 | 1 | SMC | 0,82803983 | 1,207671374 | 0,682719547 |
| MOB1A3 | 0,028853323 | 0,34311921 | 0,236 | 0,147 | 1 | SMC | 1,268496211 | 0,788335031 | 1,605442177 |
| CLIC14 | 0,028900909 | 0,313017673 | 0,665 | 0,603 | 1 | SMC | 1,2423035 | 0,804956277 | 1,102819237 |
| RHOG1 | 0,029134165 | 0,269694481 | 0,122 | 0,052 | 1 | SMC | 1,205552501 | 0,829495189 | 2,346153846 |
| DEGS12 | 0,029363828 | 0,492119714 | 0,276 | 0,19 | 1 | SMC | 1,406509906 | 0,710979706 | 1,452631579 |
| CAMLG2 | 0,029632887 | 0,395585336 | 0,358 | 0,267 | 1 | SMC | 1,315476372 | 0,760180891 | 1,34082397 |
| S1PR3 | 0,029753228 | -0,303115092 | 0,105 | 0,181 | 1 | SMC | 0,810500459 | 1,233805594 | 0,580110497 |
| NOPCHAP11 | 0,029813852 | -0,292031419 | 0,068 | 0,129 | 1 | SMC | 0,816751204 | 1,224363056 | 0,527131783 |
| ZNF831 | 0,029870899 | 0,269029352 | 0,134 | 0,06 | 1 | SMC | 1,204996831 | 0,829877701 | 2,233333333 |
| HAX1 | 0,029876758 | -0,290447707 | 0,216 | 0,319 | 1 | SMC | 0,817648281 | 1,223019755 | 0,677115987 |
| ATF7IP | 0,029957835 | 0,34355168 | 0,23 | 0,138 | 1 | SMC | 1,26887652 | 0,788098751 | 1,666666667 |
| MTX2 | 0,029970016 | -0,261182112 | 0,06 | 0,121 | 1 | SMC | 0,834403948 | 1,198460294 | 0,495867769 |
| TWF23 | 0,030301609 | 0,293026442 | 0,168 | 0,086 | 1 | SMC | 1,225207787 | 0,816188087 | 1,953488372 |
| MAP2K1 | 0,030410167 | 0,317399698 | 0,162 | 0,086 | 1 | SMC | 1,246082594 | 0,802515022 | 1,88372093 |
| VDAC13 | 0,030458107 | -0,354701921 | 0,23 | 0,328 | 1 | SMC | 0,782031204 | 1,278721354 | 0,701219512 |
| PRDX45 | 0,030556208 | 0,375401842 | 0,261 | 0,164 | 1 | SMC | 1,297200821 | 0,770890662 | 1,591463415 |
| NR3C13 | 0,030659747 | 0,391879498 | 0,432 | 0,336 | 1 | SMC | 1,312101656 | 0,76213607 | 1,285714286 |
| MAP1A | 0,0306807 | 0,339072691 | 0,236 | 0,147 | 1 | SMC | 1,264943275 | 0,790549283 | 1,605442177 |
| OSBPL10 | 0,030709976 | 0,288461745 | 0,122 | 0,052 | 1 | SMC | 1,221337348 | 0,818774601 | 2,346153846 |
| COL5A3 | 0,030751986 | 0,378156843 | 0,131 | 0,06 | 1 | SMC | 1,29968035 | 0,769419958 | 2,183333333 |
| IRF92 | 0,030903253 | 0,535985057 | 0,241 | 0,164 | 1 | SMC | 1,449931815 | 0,689687604 | 1,469512195 |
| ALDH3A2 | 0,030933958 | -0,297403779 | 0,074 | 0,138 | 1 | SMC | 0,813715412 | 1,228930884 | 0,536231884 |
| PGLS6 | 0,030949338 | 0,399002324 | 0,261 | 0,172 | 1 | SMC | 1,318595739 | 0,758382551 | 1,51744186 |
| CD3022 | 0,031122121 | 0,397461809 | 0,304 | 0,207 | 1 | SMC | 1,317188489 | 0,759192787 | 1,468599034 |
| ANKRD50 | 0,031489814 | 0,291823079 | 0,108 | 0,043 | 1 | SMC | 1,224186259 | 0,816869159 | 2,511627907 |
| NSRP14 | 0,031625673 | -0,359097877 | 0,267 | 0,362 | 1 | SMC | 0,779651947 | 1,282623617 | 0,737569061 |
| IER24 | 0,031732344 | -0,302950387 | 0,645 | 0,724 | 1 | SMC | 0,810592995 | 1,233664745 | 0,890883978 |
| SPRY4 | 0,031738526 | 0,260279295 | 0,153 | 0,078 | 1 | SMC | 1,19771055 | 0,834926269 | 1,961538462 |
| PLAU1 | 0,031951486 | 0,266947536 | 0,182 | 0,095 | 1 | SMC | 1,203259269 | 0,831076083 | 1,915789474 |

| Supplementary Table 4 - clusermarker | | | | | | | | | |
|---|---|---|---|---|---|---|---|---|---|
| | **p_val** | **avg_log2FC** | **pct.1** | **pct.2** | **p_val_adj** | **celltype** | **Foldchange_UP** | **Foldchange_DOWN** | **Ratio_pct1_pct2** |
| MEOX21 | 0,031977981 | 0,397353952 | 0,173 | 0,095 | 1 | SMC | 1,317090018 | 0,759249547 | 1,821052632 |
| C1QTNF1 | 0,032299844 | 0,348265895 | 0,634 | 0,509 | 1 | SMC | 1,273029539 | 0,785527727 | 1,245579568 |
| MT1X4 | 0,032644159 | -0,568009971 | 0,366 | 0,457 | 1 | SMC | 0,674546605 | 1,482477255 | 0,800875274 |
| IGHA17 | 0,032660455 | -0,886571653 | 0,159 | 0,078 | 1 | SMC | 0,540897955 | 1,848777559 | 2,038461538 |
| PSMD83 | 0,032767932 | 0,481728901 | 0,347 | 0,267 | 1 | SMC | 1,396416105 | 0,716118925 | 1,299625468 |
| TWIST11 | 0,032896979 | 0,429739406 | 0,173 | 0,095 | 1 | SMC | 1,346990248 | 0,742395872 | 1,821052632 |
| MYCBP21 | 0,03325555 | 0,370964146 | 0,315 | 0,224 | 1 | SMC | 1,293216792 | 0,773265554 | 1,40625 |
| RABAC16 | 0,033401325 | 0,32223506 | 0,645 | 0,534 | 1 | SMC | 1,250265993 | 0,799829801 | 1,207865169 |
| RPN21 | 0,033429464 | 0,428843912 | 0,295 | 0,216 | 1 | SMC | 1,346154419 | 0,742856827 | 1,365740741 |
| RAB11A3 | 0,03344389 | 0,511344893 | 0,256 | 0,181 | 1 | SMC | 1,425378327 | 0,701568125 | 1,414364641 |
| EIF4G3 | 0,033496843 | 0,423299874 | 0,202 | 0,121 | 1 | SMC | 1,340991297 | 0,745716995 | 1,669421488 |
| FAM214A | 0,033571708 | -0,251323862 | 0,071 | 0,138 | 1 | SMC | 0,840125136 | 1,19029887 | 0,514492754 |
| UBL53 | 0,03360446 | -0,340525884 | 0,557 | 0,603 | 1 | SMC | 0,789753382 | 1,266218066 | 0,92371476 |
| TXNDC152 | 0,033649894 | 0,342330866 | 0,19 | 0,112 | 1 | SMC | 1,267803246 | 0,788765925 | 1,696428571 |
| RPS27L | 0,03377499 | 0,262169379 | 0,659 | 0,543 | 1 | SMC | 1,199280706 | 0,833833142 | 1,213627993 |
| ARL8B1 | 0,033810065 | 0,367182508 | 0,239 | 0,155 | 1 | SMC | 1,289831411 | 0,775295121 | 1,541935484 |
| ZCCHC24 | 0,033985265 | 0,259092988 | 0,196 | 0,112 | 1 | SMC | 1,196726095 | 0,835613098 | 1,75 |
| NAP1L13 | 0,03407381 | 0,295927826 | 0,568 | 0,491 | 1 | SMC | 1,227674266 | 0,814548311 | 1,156822811 |
| SLC25A231 | 0,034250931 | -0,308031342 | 0,097 | 0,164 | 1 | SMC | 0,807743229 | 1,238017186 | 0,591463415 |
| TINAGL11 | 0,034277813 | 0,329094739 | 0,798 | 0,75 | 1 | SMC | 1,256224873 | 0,796035823 | 1,064 |
| JUN3 | 0,034790234 | -0,371178329 | 0,812 | 0,836 | 1 | SMC | 0,773150764 | 1,293408798 | 0,971291866 |
| CSTB4 | 0,034801885 | 0,333095911 | 0,389 | 0,293 | 1 | SMC | 1,259713723 | 0,793831155 | 1,327645051 |
| FOXC12 | 0,034802998 | -0,284443469 | 0,134 | 0,216 | 1 | SMC | 0,821058278 | 1,217940343 | 0,62037037 |
| CLSTN2 | 0,034943475 | 0,286457795 | 0,134 | 0,06 | 1 | SMC | 1,219642049 | 0,819912696 | 2,233333333 |
| FIBP4 | 0,03520876 | 0,353564935 | 0,17 | 0,095 | 1 | SMC | 1,277713993 | 0,782647764 | 1,789473684 |
| EIF1AX7 | 0,035470949 | -0,369856592 | 0,378 | 0,474 | 1 | SMC | 0,773859417 | 1,292224373 | 0,797468354 |
| ARL4D | 0,035522408 | -0,269222181 | 0,094 | 0,164 | 1 | SMC | 0,829766788 | 1,2051579 | 0,573170732 |
| IGFBP61 | 0,035552849 | 0,385546873 | 0,386 | 0,276 | 1 | SMC | 1,306354885 | 0,765488775 | 1,398550725 |
| RPL22L11 | 0,035713393 | 0,413080396 | 0,293 | 0,207 | 1 | SMC | 1,331525812 | 0,751018111 | 1,415458937 |
| EIF2S34 | 0,03579733 | -0,279498177 | 0,386 | 0,474 | 1 | SMC | 0,823877543 | 1,213772616 | 0,814345992 |
| ATP5F1E3 | 0,035823634 | -0,291482651 | 0,693 | 0,716 | 1 | SMC | 0,817061936 | 1,223897426 | 0,967877095 |

| Supplementary Table 4 - clusermarker | | | | | | | | | |
|---|---|---|---|---|---|---|---|---|---|
| | **p_val** | **avg_log2FC** | **pct.1** | **pct.2** | **p_val_adj** | **celltype** | **Foldchange_UP** | **Foldchange_DOWN** | **Ratio_pct1_pct2** |
| SRPRA | 0,035896906 | 0,346845849 | 0,224 | 0,147 | 1 | SMC | 1,271777111 | 0,786301304 | 1,523809524 |
| LTBR1 | 0,036196943 | 0,379201657 | 0,21 | 0,129 | 1 | SMC | 1,300621933 | 0,768862938 | 1,627906977 |
| LEPROT3 | 0,036217174 | 0,309905182 | 0,531 | 0,422 | 1 | SMC | 1,239626226 | 0,806694776 | 1,258293839 |
| IRF3 | 0,036255191 | -0,310641603 | 0,074 | 0,138 | 1 | SMC | 0,806283105 | 1,240259152 | 0,536231884 |
| SON8 | 0,036413306 | -0,250767059 | 0,693 | 0,724 | 1 | SMC | 0,840449442 | 1,189839567 | 0,95718232 |
| TASP1 | 0,036768051 | -0,396670377 | 0,097 | 0,164 | 1 | SMC | 0,759609379 | 1,316466105 | 0,591463415 |
| GET1 | 0,036862511 | -0,28952245 | 0,114 | 0,19 | 1 | SMC | 0,818172839 | 1,222235635 | 0,6 |
| GOLGA31 | 0,036970388 | 0,327850794 | 0,116 | 0,052 | 1 | SMC | 1,255142176 | 0,79672249 | 2,230769231 |
| CTDNEP11 | 0,037024747 | 0,261555931 | 0,222 | 0,138 | 1 | SMC | 1,198770869 | 0,834187771 | 1,608695652 |
| SERPINH11 | 0,03724339 | 0,432716587 | 0,477 | 0,397 | 1 | SMC | 1,3497728 | 0,740865425 | 1,201511335 |
| CCND14 | 0,03724863 | 0,611325613 | 0,403 | 0,328 | 1 | SMC | 1,527662249 | 0,654594954 | 1,228658537 |
| SLC29A11 | 0,037571364 | -0,290800964 | 0,082 | 0,147 | 1 | SMC | 0,817448097 | 1,223319259 | 0,557823129 |
| AK11 | 0,038225043 | -0,327806252 | 0,165 | 0,25 | 1 | SMC | 0,796747089 | 1,255103425 | 0,66 |
| ADAMTS1 | 0,038511814 | -0,55323833 | 0,662 | 0,776 | 1 | SMC | 0,681488714 | 1,467375732 | 0,853092784 |
| PCDH1 | 0,03870814 | 0,298947042 | 0,139 | 0,069 | 1 | SMC | 1,230246184 | 0,812845439 | 2,014492754 |
| MYLK3 | 0,039375906 | -0,501762201 | 0,727 | 0,733 | 1 | SMC | 0,706243603 | 1,415942029 | 0,991814461 |
| MAP4K52 | 0,039572901 | 0,290059463 | 0,219 | 0,129 | 1 | SMC | 1,222690672 | 0,817868348 | 1,697674419 |
| DGKG | 0,039777345 | -0,273666503 | 0,08 | 0,147 | 1 | SMC | 0,827214567 | 1,208876197 | 0,544217687 |
| HSPD14 | 0,03980134 | -0,503843578 | 0,369 | 0,474 | 1 | SMC | 0,705225439 | 1,417986284 | 0,778481013 |
| ARL3 | 0,039814991 | -0,271529962 | 0,239 | 0,336 | 1 | SMC | 0,828440528 | 1,207087251 | 0,711309524 |
| ARHGAP10 | 0,03994462 | 0,393472095 | 0,429 | 0,345 | 1 | SMC | 1,31355089 | 0,761295209 | 1,243478261 |
| TSPAN14 | 0,040028833 | 0,418356335 | 0,199 | 0,121 | 1 | SMC | 1,33640412 | 0,748276652 | 1,644628099 |
| TRAPPC55 | 0,040151247 | 0,389537236 | 0,27 | 0,181 | 1 | SMC | 1,309973145 | 0,763374428 | 1,491712707 |
| ZNF580 | 0,040409751 | 0,273799494 | 0,202 | 0,121 | 1 | SMC | 1,208987639 | 0,827138316 | 1,669421488 |
| ZNFX1 | 0,040849918 | 0,372040752 | 0,261 | 0,172 | 1 | SMC | 1,29418221 | 0,772688723 | 1,51744186 |
| MED15 | 0,041227283 | 0,28379187 | 0,102 | 0,043 | 1 | SMC | 1,21739038 | 0,821429195 | 2,372093023 |
| CREM3 | 0,041601459 | -0,599397692 | 0,17 | 0,25 | 1 | SMC | 0,660029452 | 1,515083905 | 0,68 |
| PPM1G5 | 0,041670358 | 0,325724301 | 0,23 | 0,147 | 1 | SMC | 1,253293493 | 0,797897704 | 1,56462585 |
| ETFA1 | 0,04197978 | -0,257949173 | 0,151 | 0,233 | 1 | SMC | 0,836275862 | 1,195777668 | 0,64806867 |
| SAE1 | 0,042049601 | -0,266152967 | 0,108 | 0,181 | 1 | SMC | 0,831533927 | 1,202596752 | 0,596685083 |
| NUDT7 | 0,042472864 | -0,306808138 | 0,057 | 0,112 | 1 | SMC | 0,808428373 | 1,236967965 | 0,508928571 |

| Supplementary Table 4 - clusermarker | | | | | | | | | |
|---|---|---|---|---|---|---|---|---|---|
| | **p_val** | **avg_log2FC** | **pct.1** | **pct.2** | **p_val_adj** | **celltype** | **Foldchange_UP** | **Foldchange_DOWN** | **Ratio_pct1_pct2** |
| GPR1761 | 0,04252386 | 0,261288983 | 0,128 | 0,06 | 1 | SMC | 1,198549076 | 0,834342139 | 2,133333333 |
| PABPC42 | 0,042548362 | 0,325620867 | 0,233 | 0,155 | 1 | SMC | 1,253203642 | 0,797954911 | 1,503225806 |
| MRPL272 | 0,042619401 | 0,256664332 | 0,199 | 0,121 | 1 | SMC | 1,194713202 | 0,837020967 | 1,644628099 |
| TRIM82 | 0,042973699 | 0,453023662 | 0,241 | 0,164 | 1 | SMC | 1,368906264 | 0,730510208 | 1,469512195 |
| POLD43 | 0,042991226 | 0,30546872 | 0,241 | 0,155 | 1 | SMC | 1,23582008 | 0,809179278 | 1,55483871 |
| NDUFA73 | 0,043199619 | -0,322043018 | 0,216 | 0,31 | 1 | SMC | 0,799936276 | 1,250099577 | 0,696774194 |
| MR12 | 0,043205028 | -0,305091956 | 0,057 | 0,112 | 1 | SMC | 0,809390625 | 1,235497384 | 0,508928571 |
| GABARAPL23 | 0,043374708 | 0,310550222 | 0,423 | 0,345 | 1 | SMC | 1,240180595 | 0,806334177 | 1,226086957 |
| GFPT1 | 0,043391503 | 0,341618928 | 0,176 | 0,103 | 1 | SMC | 1,267177766 | 0,78915526 | 1,708737864 |
| HEBP24 | 0,043679484 | -0,302000791 | 0,233 | 0,319 | 1 | SMC | 0,811126711 | 1,232853001 | 0,730407524 |
| PSMC52 | 0,043680886 | -0,270211967 | 0,273 | 0,371 | 1 | SMC | 0,829197707 | 1,205985003 | 0,735849057 |
| TXNDC121 | 0,04403112 | 0,275323337 | 0,102 | 0,043 | 1 | SMC | 1,210265304 | 0,826265114 | 2,372093023 |
| RABL62 | 0,04403181 | 0,356456701 | 0,159 | 0,086 | 1 | SMC | 1,280277636 | 0,781080581 | 1,848837209 |
| EBPL2 | 0,044547082 | 0,281597463 | 0,136 | 0,069 | 1 | SMC | 1,215540079 | 0,822679579 | 1,971014493 |
| CDC42SE11 | 0,044583648 | 0,269787974 | 0,125 | 0,06 | 1 | SMC | 1,205630629 | 0,829441436 | 2,083333333 |
| ARFIP21 | 0,044928878 | -0,281206363 | 0,051 | 0,103 | 1 | SMC | 0,822902629 | 1,215210603 | 0,495145631 |
| RHOJ1 | 0,044960859 | 0,356267331 | 0,134 | 0,069 | 1 | SMC | 1,280109596 | 0,781183114 | 1,942028986 |
| ODF2L | 0,045441394 | 0,415644747 | 0,145 | 0,078 | 1 | SMC | 1,333894667 | 0,749684383 | 1,858974359 |
| HMG20B1 | 0,045502631 | 0,351441215 | 0,176 | 0,103 | 1 | SMC | 1,275834517 | 0,78380071 | 1,708737864 |
| ANK21 | 0,045533818 | 0,257063298 | 0,173 | 0,095 | 1 | SMC | 1,195043636 | 0,836789528 | 1,821052632 |
| MXRA8 | 0,045643298 | 0,359875277 | 0,278 | 0,19 | 1 | SMC | 1,283314949 | 0,779231942 | 1,463157895 |
| C1orf1981 | 0,045741371 | 0,360993803 | 0,205 | 0,129 | 1 | SMC | 1,284310292 | 0,778628036 | 1,589147287 |
| UAP13 | 0,045820882 | -0,270239925 | 0,142 | 0,224 | 1 | SMC | 0,829181638 | 1,206008375 | 0,633928571 |
| BTBD101 | 0,045932596 | -0,286516363 | 0,077 | 0,138 | 1 | SMC | 0,819879411 | 1,219691563 | 0,557971014 |
| AP2M14 | 0,046171851 | 0,402869332 | 0,46 | 0,362 | 1 | SMC | 1,322134852 | 0,7563525 | 1,270718232 |
| FOXS1 | 0,046273597 | 0,357393283 | 0,168 | 0,095 | 1 | SMC | 1,281109048 | 0,780573677 | 1,768421053 |
| CLIP11 | 0,046524217 | -0,32254077 | 0,156 | 0,233 | 1 | SMC | 0,799660333 | 1,250530955 | 0,669527897 |
| CAPRIN12 | 0,046649835 | 0,286643291 | 0,29 | 0,207 | 1 | SMC | 1,219798876 | 0,819807281 | 1,400966184 |
| SMIM263 | 0,046808917 | 0,312752301 | 0,341 | 0,25 | 1 | SMC | 1,242075009 | 0,805104356 | 1,364 |
| LINC02381 | 0,04698422 | -0,3714573 | 0,091 | 0,155 | 1 | SMC | 0,773001276 | 1,293658926 | 0,587096774 |
| ITGAV2 | 0,047105951 | 0,377255889 | 0,241 | 0,164 | 1 | SMC | 1,298868961 | 0,769900606 | 1,469512195 |

| Supplementary Table 4 - clusermarker | | | | | | | | | |
|---|---|---|---|---|---|---|---|---|---|
| | **p_val** | **avg_log2FC** | **pct.1** | **pct.2** | **p_val_adj** | **celltype** | **Foldchange_UP** | **Foldchange_DOWN** | **Ratio_pct1_pct2** |
| OAZ22 | 0,047341402 | 0,355668636 | 0,281 | 0,198 | 1 | SMC | 1,279578481 | 0,781507359 | 1,419191919 |
| MIDEAS1 | 0,047386461 | 0,286910472 | 0,145 | 0,078 | 1 | SMC | 1,220024798 | 0,819655471 | 1,858974359 |
| COQ10B1 | 0,047528974 | -0,496061707 | 0,114 | 0,181 | 1 | SMC | 0,70903969 | 1,410358283 | 0,629834254 |
| IQGAP11 | 0,04758209 | -0,330688139 | 0,429 | 0,491 | 1 | SMC | 0,795157118 | 1,25761309 | 0,873727088 |
| DDRGK12 | 0,047797464 | 0,295994399 | 0,182 | 0,103 | 1 | SMC | 1,227730917 | 0,814510725 | 1,766990291 |
| MAP42 | 0,048161074 | -0,36784926 | 0,44 | 0,517 | 1 | SMC | 0,774936896 | 1,290427653 | 0,85106383 |
| CLPP2 | 0,048311666 | 0,395839841 | 0,182 | 0,112 | 1 | SMC | 1,315708455 | 0,760046799 | 1,625 |
| LTBP42 | 0,048394331 | 0,407383538 | 0,312 | 0,233 | 1 | SMC | 1,326278302 | 0,753989565 | 1,339055794 |
| DNAJB17 | 0,048509107 | -0,532533702 | 0,571 | 0,612 | 1 | SMC | 0,691339517 | 1,446467293 | 0,933006536 |
| DYNC1I23 | 0,048793856 | -0,362791002 | 0,256 | 0,328 | 1 | SMC | 0,777658685 | 1,285911184 | 0,780487805 |
| CCDC821 | 0,049005357 | 0,257620019 | 0,19 | 0,112 | 1 | SMC | 1,19550488 | 0,836466682 | 1,696428571 |
| FGFR1 | 0,049633892 | -0,25637095 | 0,136 | 0,216 | 1 | SMC | 0,837191199 | 1,194470274 | 0,62962963 |
| TWSG1 | 0,04971337 | -0,30191296 | 0,278 | 0,362 | 1 | SMC | 0,811176093 | 1,232777948 | 0,767955801 |
| GPRC5C2 | 0,050009377 | -0,317803911 | 0,219 | 0,302 | 1 | SMC | 0,802290205 | 1,24643177 | 0,725165563 |
| PTGR12 | 0,050119479 | 0,365330251 | 0,193 | 0,121 | 1 | SMC | 1,288176476 | 0,776291152 | 1,595041322 |
| JAG12 | 0,050386326 | 0,255823717 | 0,534 | 0,44 | 1 | SMC | 1,194017281 | 0,837508817 | 1,213636364 |
| TMED31 | 0,050398792 | 0,255107311 | 0,21 | 0,129 | 1 | SMC | 1,193424509 | 0,837924806 | 1,627906977 |
| PPIG2 | 0,051075339 | -0,279241056 | 0,384 | 0,474 | 1 | SMC | 0,82402439 | 1,213556313 | 0,810126582 |
| GRINA3 | 0,051089157 | 0,360604026 | 0,293 | 0,216 | 1 | SMC | 1,283963354 | 0,778838428 | 1,356481481 |
| KIAA2026 | 0,051092815 | -0,331128372 | 0,108 | 0,172 | 1 | SMC | 0,794914516 | 1,257996904 | 0,627906977 |
| THBD2 | 0,051397504 | 0,313083686 | 0,199 | 0,121 | 1 | SMC | 1,242360345 | 0,804919446 | 1,644628099 |
| MRTFB1 | 0,051603371 | 0,255541628 | 0,239 | 0,155 | 1 | SMC | 1,193783839 | 0,83767259 | 1,541935484 |
| PRDX53 | 0,051699333 | 0,393014747 | 0,426 | 0,336 | 1 | SMC | 1,313134548 | 0,761536585 | 1,267857143 |
| HRC | 0,05218342 | -0,352811651 | 0,156 | 0,233 | 1 | SMC | 0,78305652 | 1,277047025 | 0,669527897 |
| CHRDL11 | 0,052197907 | 0,358476091 | 0,136 | 0,069 | 1 | SMC | 1,28207094 | 0,779988041 | 1,971014493 |
| SCAMP22 | 0,052382137 | 0,257052131 | 0,111 | 0,052 | 1 | SMC | 1,195034386 | 0,836796005 | 2,134615385 |
| GPX8 | 0,052567038 | 0,314229354 | 0,122 | 0,06 | 1 | SMC | 1,243347316 | 0,804280499 | 2,033333333 |
| SMG71 | 0,053005294 | 0,357910406 | 0,142 | 0,078 | 1 | SMC | 1,281568334 | 0,780293937 | 1,820512821 |
| SEPTIN113 | 0,053043788 | 0,311338623 | 0,267 | 0,181 | 1 | SMC | 1,240858512 | 0,805893654 | 1,475138122 |
| CNOT6L | 0,053247699 | -0,273779688 | 0,071 | 0,129 | 1 | SMC | 0,827149672 | 1,208971041 | 0,550387597 |
| AKR1A13 | 0,05341863 | 0,295573154 | 0,222 | 0,147 | 1 | SMC | 1,227372491 | 0,814748585 | 1,510204082 |

| Supplementary Table 4 - clusermarker | | | | | | | | | |
|---|---|---|---|---|---|---|---|---|---|
| | **p_val** | **avg_log2FC** | **pct.1** | **pct.2** | **p_val_adj** | **celltype** | **Foldchange_UP** | **Foldchange_DOWN** | **Ratio_pct1_pct2** |
| IMPDH22 | 0,053429191 | 0,474312444 | 0,358 | 0,276 | 1 | SMC | 1,389255974 | 0,719809753 | 1,297101449 |
| TRIR5 | 0,054272228 | 0,386010819 | 0,415 | 0,345 | 1 | SMC | 1,306775054 | 0,765242646 | 1,202898551 |
| CLK13 | 0,055598769 | 0,309468816 | 0,29 | 0,216 | 1 | SMC | 1,239251337 | 0,806938811 | 1,342592593 |
| PPP3CC | 0,055891243 | -0,329154176 | 0,119 | 0,19 | 1 | SMC | 0,796003028 | 1,256276628 | 0,626315789 |
| SMARCD3 | 0,055903272 | 0,356891372 | 0,261 | 0,181 | 1 | SMC | 1,28066343 | 0,780845284 | 1,44198895 |
| FLYWCH23 | 0,055994536 | 0,306594196 | 0,185 | 0,112 | 1 | SMC | 1,236784545 | 0,808548267 | 1,651785714 |
| PDCD63 | 0,056046061 | 0,33343169 | 0,267 | 0,19 | 1 | SMC | 1,260006949 | 0,793646417 | 1,405263158 |
| RGS162 | 0,056126813 | 0,436834229 | 0,497 | 0,397 | 1 | SMC | 1,353630733 | 0,738753913 | 1,251889169 |
| SMARCE14 | 0,056185818 | -0,276257345 | 0,267 | 0,353 | 1 | SMC | 0,82573036 | 1,211049089 | 0,756373938 |
| AIG12 | 0,056378523 | -0,32290603 | 0,219 | 0,31 | 1 | SMC | 0,799457901 | 1,250847603 | 0,706451613 |
| AOC3 | 0,0567102 | -0,358837068 | 0,364 | 0,422 | 1 | SMC | 0,779792904 | 1,282391767 | 0,862559242 |
| FAM91A1 | 0,056777387 | 0,265624355 | 0,108 | 0,052 | 1 | SMC | 1,202156194 | 0,831838662 | 2,076923077 |
| STX10 | 0,05691928 | 0,289973926 | 0,153 | 0,086 | 1 | SMC | 1,222618181 | 0,817916841 | 1,779069767 |
| MCL13 | 0,057051734 | 0,330795897 | 0,537 | 0,457 | 1 | SMC | 1,257707027 | 0,795097728 | 1,175054705 |
| NFIB3 | 0,057063783 | -0,251139625 | 0,44 | 0,543 | 1 | SMC | 0,84023243 | 1,190146874 | 0,810313076 |
| FABP4 | 0,057091426 | -0,873200439 | 0,23 | 0,31 | 1 | SMC | 0,545934419 | 1,831721843 | 0,741935484 |
| SUMO23 | 0,057195122 | 0,288514265 | 0,736 | 0,638 | 1 | SMC | 1,221381811 | 0,818744795 | 1,153605016 |
| TGIF12 | 0,057229784 | 0,252089985 | 0,122 | 0,06 | 1 | SMC | 1,190931129 | 0,839679118 | 2,033333333 |
| ENPEP | 0,057248398 | 0,282514359 | 0,199 | 0,121 | 1 | SMC | 1,216312853 | 0,822156896 | 1,644628099 |
| MAPRE12 | 0,057962981 | -0,252518891 | 0,119 | 0,19 | 1 | SMC | 0,839429523 | 1,191285239 | 0,626315789 |
| CPSF4 | 0,058214824 | 0,31706867 | 0,108 | 0,052 | 1 | SMC | 1,245796712 | 0,80269918 | 2,076923077 |
| GSN4 | 0,058555535 | -0,297514254 | 0,784 | 0,81 | 1 | SMC | 0,813653104 | 1,229024993 | 0,967901235 |
| TRIB2 | 0,058601018 | 0,30609686 | 0,119 | 0,06 | 1 | SMC | 1,236358265 | 0,808827043 | 1,983333333 |
| XIST3 | 0,058609361 | 0,578622352 | 0,381 | 0,302 | 1 | SMC | 1,49342248 | 0,669602884 | 1,261589404 |
| FAM117A | 0,059258906 | -0,2815271 | 0,074 | 0,129 | 1 | SMC | 0,822719703 | 1,215480796 | 0,573643411 |
| EWSR11 | 0,059736929 | -0,354364679 | 0,341 | 0,431 | 1 | SMC | 0,782214032 | 1,278422477 | 0,791183295 |
| LAMC1 | 0,059753054 | 0,418599911 | 0,386 | 0,319 | 1 | SMC | 1,336629769 | 0,748150328 | 1,210031348 |
| TAF9B | 0,059887511 | -0,251839526 | 0,091 | 0,155 | 1 | SMC | 0,839824903 | 1,190724395 | 0,587096774 |
| PGR | 0,060133906 | -0,25738045 | 0,074 | 0,129 | 1 | SMC | 0,836605594 | 1,195306375 | 0,573643411 |
| SNHG9 | 0,060718492 | -0,263919187 | 0,122 | 0,19 | 1 | SMC | 0,83282242 | 1,200736166 | 0,642105263 |
| NAA383 | 0,060878575 | -0,275965096 | 0,244 | 0,328 | 1 | SMC | 0,825897646 | 1,21080379 | 0,743902439 |

| Supplementary Table 4 - clusermarker | | | | | | | | | |
|---|---|---|---|---|---|---|---|---|---|
| | **p_val** | **avg_log2FC** | **pct.1** | **pct.2** | **p_val_adj** | **celltype** | **Foldchange_UP** | **Foldchange_DOWN** | **Ratio_pct1_pct2** |
| A2M3 | 0,060912484 | 0,27146028 | 0,898 | 0,871 | 1 | SMC | 1,207028951 | 0,828480542 | 1,030998852 |
| SMU13 | 0,060972008 | 0,280678396 | 0,224 | 0,147 | 1 | SMC | 1,214765967 | 0,823203833 | 1,523809524 |
| GRN3 | 0,061182079 | 0,259038603 | 0,375 | 0,293 | 1 | SMC | 1,196680983 | 0,835644599 | 1,279863481 |
| CLTA4 | 0,0613799 | 0,362446909 | 0,452 | 0,379 | 1 | SMC | 1,285604522 | 0,777844183 | 1,192612137 |
| CSNK1E1 | 0,061537644 | 0,311880681 | 0,185 | 0,112 | 1 | SMC | 1,241324823 | 0,805590915 | 1,651785714 |
| TNXB1 | 0,061646572 | 0,436894703 | 0,119 | 0,06 | 1 | SMC | 1,353687474 | 0,738722947 | 1,983333333 |
| RERE4 | 0,062039946 | -0,285122432 | 0,176 | 0,25 | 1 | SMC | 0,820671961 | 1,218513666 | 0,704 |
| WASF23 | 0,0626823 | 0,413484271 | 0,486 | 0,422 | 1 | SMC | 1,331898618 | 0,750807896 | 1,151658768 |
| GLT8D11 | 0,062742841 | 0,332898163 | 0,128 | 0,069 | 1 | SMC | 1,259541068 | 0,793939972 | 1,855072464 |
| RAI14 | 0,062883782 | 0,255314738 | 0,131 | 0,069 | 1 | SMC | 1,193596109 | 0,837804339 | 1,898550725 |
| METTL14 | 0,062930583 | 0,295992187 | 0,108 | 0,052 | 1 | SMC | 1,227729035 | 0,814511974 | 2,076923077 |
| TMEM245 | 0,063599425 | 0,27469823 | 0,185 | 0,112 | 1 | SMC | 1,20974102 | 0,826623206 | 1,651785714 |
| PCYOX12 | 0,063614122 | 0,395808301 | 0,392 | 0,319 | 1 | SMC | 1,315679691 | 0,760063415 | 1,228840125 |
| HEXB2 | 0,064003792 | 0,390942756 | 0,148 | 0,086 | 1 | SMC | 1,311249985 | 0,762631086 | 1,720930233 |
| TSEN342 | 0,064471739 | 0,392657917 | 0,216 | 0,147 | 1 | SMC | 1,312809803 | 0,761724964 | 1,469387755 |
| TERF1 | 0,064648154 | -0,29067266 | 0,165 | 0,241 | 1 | SMC | 0,817520799 | 1,22321047 | 0,684647303 |
| SRI4 | 0,064668566 | 0,388485795 | 0,318 | 0,25 | 1 | SMC | 1,30901878 | 0,76393098 | 1,272 |
| SNCG | 0,06475693 | -0,391421463 | 0,449 | 0,517 | 1 | SMC | 0,762378076 | 1,311685148 | 0,868471954 |
| LMO45 | 0,064891348 | -0,389913198 | 0,267 | 0,345 | 1 | SMC | 0,763175521 | 1,310314565 | 0,773913043 |
| BAZ2A2 | 0,065067147 | 0,320379779 | 0,233 | 0,155 | 1 | SMC | 1,248659206 | 0,800859029 | 1,503225806 |
| CACNA1H | 0,065087433 | -0,280927846 | 0,094 | 0,155 | 1 | SMC | 0,823061508 | 1,214976025 | 0,606451613 |
| GBF11 | 0,06514127 | 0,27963888 | 0,153 | 0,086 | 1 | SMC | 1,213890998 | 0,823797196 | 1,779069767 |
| TTC15 | 0,065199824 | -0,373894378 | 0,153 | 0,233 | 1 | SMC | 0,771696583 | 1,295846091 | 0,656652361 |
| RALGAPA12 | 0,065300737 | -0,251262703 | 0,094 | 0,155 | 1 | SMC | 0,840160752 | 1,190248411 | 0,606451613 |
| COPE5 | 0,065338081 | 0,298172265 | 0,352 | 0,276 | 1 | SMC | 1,229585677 | 0,813282082 | 1,275362319 |
| ADGRL3 | 0,065473006 | 0,271842884 | 0,119 | 0,06 | 1 | SMC | 1,207349099 | 0,828260858 | 1,983333333 |
| PSMD10 | 0,066021292 | -0,336891733 | 0,105 | 0,164 | 1 | SMC | 0,79174528 | 1,263032474 | 0,640243902 |
| EGFLAM1 | 0,066080307 | 0,44973855 | 0,281 | 0,207 | 1 | SMC | 1,365792721 | 0,732175523 | 1,357487923 |
| RASSF4 | 0,066353185 | 0,394875679 | 0,148 | 0,086 | 1 | SMC | 1,314829453 | 0,760554913 | 1,720930233 |
| CTSB5 | 0,066576918 | 0,319729102 | 0,398 | 0,31 | 1 | SMC | 1,248096169 | 0,80122031 | 1,283870968 |
| BMP2 | 0,066710936 | 0,293086965 | 0,182 | 0,112 | 1 | SMC | 1,225259188 | 0,816153847 | 1,625 |

| Supplementary Table 4 - clusermarker | | | | | | | | | |
|---|---|---|---|---|---|---|---|---|---|
| | **p_val** | **avg_log2FC** | **pct.1** | **pct.2** | **p_val_adj** | **celltype** | **Foldchange_UP** | **Foldchange_DOWN** | **Ratio_pct1_pct2** |
| EIF4A25 | 0,066751504 | -0,269970185 | 0,568 | 0,638 | 1 | SMC | 0,829336685 | 1,205782908 | 0,890282132 |
| PSME25 | 0,0667557 | 0,318052058 | 0,295 | 0,216 | 1 | SMC | 1,246646178 | 0,802152221 | 1,365740741 |
| KDELR25 | 0,066802734 | 0,309066029 | 0,216 | 0,147 | 1 | SMC | 1,238905399 | 0,807164131 | 1,469387755 |
| TBC1D12 | 0,066893733 | -0,298850274 | 0,256 | 0,345 | 1 | SMC | 0,812899962 | 1,230163669 | 0,742028986 |
| RPAIN1 | 0,066938426 | 0,308113226 | 0,219 | 0,147 | 1 | SMC | 1,238087455 | 0,807697385 | 1,489795918 |
| PPP6C1 | 0,066950689 | 0,284988659 | 0,168 | 0,103 | 1 | SMC | 1,218400686 | 0,820748061 | 1,631067961 |
| DICER11 | 0,066969725 | 0,263056746 | 0,19 | 0,121 | 1 | SMC | 1,200018582 | 0,833320429 | 1,570247934 |
| HNRNPH33 | 0,067015854 | -0,315708159 | 0,375 | 0,457 | 1 | SMC | 0,803456511 | 1,244622436 | 0,820568928 |
| SYDE1 | 0,068036058 | 0,268403343 | 0,151 | 0,086 | 1 | SMC | 1,204474077 | 0,830237877 | 1,755813953 |
| SOD13 | 0,068039071 | -0,32172778 | 0,651 | 0,69 | 1 | SMC | 0,800111086 | 1,249826452 | 0,943478261 |
| PRDX14 | 0,068332163 | -0,253274561 | 0,744 | 0,793 | 1 | SMC | 0,838989953 | 1,191909387 | 0,938209332 |
| EML2 | 0,068430182 | 0,287128238 | 0,128 | 0,069 | 1 | SMC | 1,220208968 | 0,819531758 | 1,855072464 |
| SYF21 | 0,06875241 | 0,299126434 | 0,332 | 0,25 | 1 | SMC | 1,230399169 | 0,812744372 | 1,328 |
| ADAP21 | 0,069109664 | 0,265716832 | 0,116 | 0,06 | 1 | SMC | 1,202233255 | 0,831785343 | 1,933333333 |
| GOLM23 | 0,069306622 | 0,357547909 | 0,224 | 0,155 | 1 | SMC | 1,281246362 | 0,780490021 | 1,44516129 |
| TERF2IP1 | 0,069436096 | -0,290162085 | 0,27 | 0,345 | 1 | SMC | 0,817810174 | 1,222777647 | 0,782608696 |
| ATF32 | 0,069994423 | -0,604067462 | 0,54 | 0,569 | 1 | SMC | 0,657896498 | 1,519995932 | 0,949033392 |
| NDUFB53 | 0,071024188 | 0,330475011 | 0,253 | 0,181 | 1 | SMC | 1,257427317 | 0,795274594 | 1,397790055 |
| SLIRP3 | 0,071072562 | -0,282020759 | 0,264 | 0,345 | 1 | SMC | 0,822438234 | 1,215896779 | 0,765217391 |
| LAMTOR44 | 0,071088124 | 0,399187593 | 0,528 | 0,448 | 1 | SMC | 1,318765082 | 0,758285167 | 1,178571429 |
| SECISBP2L1 | 0,071311973 | -0,268564741 | 0,213 | 0,293 | 1 | SMC | 0,830145001 | 1,204608832 | 0,726962457 |
| ARHGAP62 | 0,071624417 | 0,290820948 | 0,241 | 0,164 | 1 | SMC | 1,223336204 | 0,817436774 | 1,469512195 |
| SEPTIN8 | 0,072011585 | 0,259625239 | 0,202 | 0,129 | 1 | SMC | 1,197167683 | 0,835304874 | 1,565891473 |
| PLD33 | 0,072156258 | 0,304105154 | 0,315 | 0,241 | 1 | SMC | 1,234652594 | 0,809944437 | 1,307053942 |
| ZC3H132 | 0,072177251 | -0,38397874 | 0,256 | 0,328 | 1 | SMC | 0,766321273 | 1,304935718 | 0,780487805 |
| AC010642.2 | 0,072452193 | 0,325378069 | 0,128 | 0,069 | 1 | SMC | 1,252992752 | 0,798089214 | 1,855072464 |
| USP71 | 0,07257608 | -0,276939922 | 0,097 | 0,155 | 1 | SMC | 0,825339777 | 1,211622204 | 0,625806452 |
| PHIP2 | 0,072920978 | -0,262159357 | 0,247 | 0,319 | 1 | SMC | 0,833838935 | 1,199272375 | 0,774294671 |
| LARP71 | 0,072952829 | -0,330016013 | 0,227 | 0,302 | 1 | SMC | 0,795527654 | 1,257027326 | 0,751655629 |
| FERMT22 | 0,073336557 | 0,328728869 | 0,355 | 0,276 | 1 | SMC | 1,255906332 | 0,796237725 | 1,286231884 |
| INO80D1 | 0,073844821 | -0,334954192 | 0,085 | 0,138 | 1 | SMC | 0,792809309 | 1,261337358 | 0,615942029 |

| Supplementary Table 4 - clusermarker | | | | | | | | | |
|---|---|---|---|---|---|---|---|---|---|
| | **p_val** | **avg_log2FC** | **pct.1** | **pct.2** | **p_val_adj** | **celltype** | **Foldchange_UP** | **Foldchange_DOWN** | **Ratio_pct1_pct2** |
| GNA121 | 0,073975888 | -0,331714905 | 0,082 | 0,138 | 1 | SMC | 0,794591406 | 1,258508451 | 0,594202899 |
| EXOC72 | 0,07441555 | 0,282063412 | 0,139 | 0,078 | 1 | SMC | 1,215932727 | 0,82241392 | 1,782051282 |
| NCOA1 | 0,074443762 | -0,286224716 | 0,111 | 0,172 | 1 | SMC | 0,82004517 | 1,219445022 | 0,645348837 |
| COMMD2 | 0,074651712 | 0,34144522 | 0,128 | 0,069 | 1 | SMC | 1,267025201 | 0,789250284 | 1,855072464 |
| MBNL1-AS1 | 0,074888275 | -0,327378639 | 0,071 | 0,121 | 1 | SMC | 0,796983279 | 1,254731468 | 0,58677686 |
| SEPTIN22 | 0,075168447 | 0,302946434 | 0,483 | 0,414 | 1 | SMC | 1,233661364 | 0,810595216 | 1,166666667 |
| LDB2 | 0,075546053 | 0,27172874 | 0,139 | 0,078 | 1 | SMC | 1,207253579 | 0,828326391 | 1,782051282 |
| DNAJA16 | 0,076034224 | -0,46763737 | 0,537 | 0,569 | 1 | SMC | 0,723147892 | 1,382842999 | 0,943760984 |
| COX5A6 | 0,076097277 | 0,391368712 | 0,273 | 0,207 | 1 | SMC | 1,311637189 | 0,762405952 | 1,31884058 |
| SPRED1 | 0,076188124 | 0,285320383 | 0,136 | 0,078 | 1 | SMC | 1,218680869 | 0,820559365 | 1,743589744 |
| KATNBL11 | 0,076718545 | -0,256620904 | 0,102 | 0,164 | 1 | SMC | 0,837046164 | 1,194677239 | 0,62195122 |
| PRXL2C2 | 0,077027816 | 0,385721361 | 0,176 | 0,112 | 1 | SMC | 1,306512892 | 0,765396198 | 1,571428571 |
| ADAMTS5 | 0,077550233 | 0,298681409 | 0,116 | 0,06 | 1 | SMC | 1,230019689 | 0,812995116 | 1,933333333 |
| CD555 | 0,077757382 | -0,250171612 | 0,105 | 0,164 | 1 | SMC | 0,840796394 | 1,189348583 | 0,640243902 |
| SPRYD3 | 0,078549147 | -0,297642346 | 0,111 | 0,172 | 1 | SMC | 0,813580865 | 1,229134119 | 0,645348837 |
| MRPL144 | 0,078696308 | 0,308605614 | 0,145 | 0,086 | 1 | SMC | 1,238510083 | 0,807421767 | 1,686046512 |
| BAZ1A4 | 0,078706878 | 0,25030138 | 0,17 | 0,103 | 1 | SMC | 1,189455567 | 0,84072077 | 1,650485437 |
| CYP4X12 | 0,079759367 | 0,314759774 | 0,176 | 0,112 | 1 | SMC | 1,243804528 | 0,803984853 | 1,571428571 |
| GSTK13 | 0,079842657 | 0,311307605 | 0,432 | 0,371 | 1 | SMC | 1,240831834 | 0,805910981 | 1,164420485 |
| ATL3 | 0,080139769 | -0,312564504 | 0,318 | 0,405 | 1 | SMC | 0,805209163 | 1,241913338 | 0,785185185 |
| OSMR1 | 0,080685728 | 0,298541511 | 0,241 | 0,172 | 1 | SMC | 1,22990042 | 0,813073956 | 1,401162791 |
| SMG12 | 0,08254531 | 0,402941101 | 0,19 | 0,129 | 1 | SMC | 1,322200625 | 0,756314875 | 1,472868217 |
| SMC61 | 0,082600195 | 0,266406533 | 0,159 | 0,095 | 1 | SMC | 1,202808137 | 0,831387791 | 1,673684211 |
| SCN4B | 0,082652254 | 0,342998329 | 0,114 | 0,06 | 1 | SMC | 1,26838993 | 0,788401087 | 1,9 |
| USP113 | 0,083283167 | -0,288671673 | 0,176 | 0,25 | 1 | SMC | 0,818655469 | 1,221515079 | 0,704 |
| SRPX | 0,083294788 | 0,37787317 | 0,25 | 0,181 | 1 | SMC | 1,299424823 | 0,769571261 | 1,38121547 |
| HMGN51 | 0,083638928 | -0,33544389 | 0,071 | 0,121 | 1 | SMC | 0,792540249 | 1,261765571 | 0,58677686 |
| DNAJC3 | 0,083731094 | 0,390417391 | 0,273 | 0,207 | 1 | SMC | 1,310772573 | 0,762908853 | 1,31884058 |
| RMDN1 | 0,084004072 | 0,269231728 | 0,114 | 0,06 | 1 | SMC | 1,205165875 | 0,829761298 | 1,9 |
| FAM110B | 0,084079047 | 0,262129919 | 0,116 | 0,06 | 1 | SMC | 1,199247904 | 0,833855949 | 1,933333333 |
| CHURC13 | 0,084479257 | -0,266471537 | 0,597 | 0,664 | 1 | SMC | 0,831350331 | 1,202862334 | 0,899096386 |

| Supplementary Table 4 - clusermarker | | | | | | | | | |
|---|---|---|---|---|---|---|---|---|---|
| | **p_val** | **avg_log2FC** | **pct.1** | **pct.2** | **p_val_adj** | **celltype** | **Foldchange_UP** | **Foldchange_DOWN** | **Ratio_pct1_pct2** |
| MZT2A4 | 0,084784656 | 0,2736242 | 0,176 | 0,112 | 1 | SMC | 1,20884075 | 0,827238824 | 1,571428571 |
| TMEM1344 | 0,084823717 | 0,293121718 | 0,114 | 0,06 | 1 | SMC | 1,225288704 | 0,816134187 | 1,9 |
| IGF2R1 | 0,084934517 | 0,294750481 | 0,196 | 0,129 | 1 | SMC | 1,226672802 | 0,815213314 | 1,519379845 |
| ASPH | 0,084962963 | 0,277879273 | 0,412 | 0,345 | 1 | SMC | 1,212411358 | 0,824802567 | 1,194202899 |
| NHS | 0,085161665 | 0,256118867 | 0,125 | 0,069 | 1 | SMC | 1,194261581 | 0,837337495 | 1,811594203 |
| PITPNA1 | 0,085649819 | 0,27033188 | 0,114 | 0,06 | 1 | SMC | 1,206085246 | 0,82912879 | 1,9 |
| RBIS1 | 0,08578952 | -0,268072625 | 0,412 | 0,483 | 1 | SMC | 0,83042822 | 1,204197998 | 0,85300207 |
| CHRD | 0,085936087 | 0,27772768 | 0,102 | 0,052 | 1 | SMC | 1,21228397 | 0,824889238 | 1,961538462 |
| NF1 | 0,086634124 | 0,327117214 | 0,205 | 0,138 | 1 | SMC | 1,254504124 | 0,79712771 | 1,485507246 |
| BBIP11 | 0,086790238 | -0,311999343 | 0,094 | 0,147 | 1 | SMC | 0,805524658 | 1,241426927 | 0,639455782 |
| ITGA7 | 0,086929765 | 0,332954581 | 0,418 | 0,353 | 1 | SMC | 1,259590325 | 0,793908924 | 1,184135977 |
| PSMG32 | 0,087799467 | 0,257166022 | 0,111 | 0,06 | 1 | SMC | 1,195128729 | 0,836729948 | 1,85 |
| DTX3 | 0,088047549 | -0,285630364 | 0,188 | 0,267 | 1 | SMC | 0,820383076 | 1,218942746 | 0,70411985 |
| AUP13 | 0,088725397 | 0,355340295 | 0,207 | 0,147 | 1 | SMC | 1,279287297 | 0,781685242 | 1,408163265 |
| AC011603.21 | 0,089232658 | -0,293880793 | 0,085 | 0,138 | 1 | SMC | 0,81570489 | 1,22593356 | 0,615942029 |
| MAT2A5 | 0,089950129 | -0,321601688 | 0,46 | 0,543 | 1 | SMC | 0,800181019 | 1,249717222 | 0,847145488 |
| DMPK1 | 0,090161089 | -0,251455502 | 0,193 | 0,259 | 1 | SMC | 0,840048482 | 1,190407484 | 0,745173745 |
| CIAO11 | 0,090502011 | 0,261501729 | 0,188 | 0,121 | 1 | SMC | 1,198725832 | 0,834219113 | 1,553719008 |
| DDX187 | 0,091782975 | -0,330951296 | 0,304 | 0,371 | 1 | SMC | 0,795012089 | 1,257842508 | 0,819407008 |
| LAMP2 | 0,091839604 | 0,264495965 | 0,179 | 0,112 | 1 | SMC | 1,201216307 | 0,832489531 | 1,598214286 |
| WDFY31 | 0,092503156 | 0,295961902 | 0,142 | 0,086 | 1 | SMC | 1,227703263 | 0,814529073 | 1,651162791 |
| HSBP14 | 0,092566568 | 0,304595069 | 0,315 | 0,241 | 1 | SMC | 1,235071932 | 0,80966944 | 1,307053942 |
| ATXN2 | 0,09278641 | -0,277171951 | 0,142 | 0,207 | 1 | SMC | 0,825207048 | 1,211817085 | 0,685990338 |
| PDLIM22 | 0,092805499 | 0,28588266 | 0,321 | 0,241 | 1 | SMC | 1,219155931 | 0,820239622 | 1,331950207 |
| CCDC251 | 0,093311088 | 0,269353882 | 0,134 | 0,078 | 1 | SMC | 1,205267922 | 0,829691044 | 1,717948718 |
| KANK21 | 0,093377547 | -0,287575569 | 0,423 | 0,5 | 1 | SMC | 0,819277688 | 1,220587371 | 0,846 |
| SNTA1 | 0,093911808 | -0,326652435 | 0,176 | 0,241 | 1 | SMC | 0,797384554 | 1,254100039 | 0,730290456 |
| ZFAND52 | 0,094383905 | -0,341510502 | 0,52 | 0,586 | 1 | SMC | 0,789214571 | 1,267082536 | 0,887372014 |
| NCKAP11 | 0,0944176 | -0,380639611 | 0,321 | 0,379 | 1 | SMC | 0,768096983 | 1,301918926 | 0,846965699 |
| PPP1CB2 | 0,095063048 | -0,251996399 | 0,33 | 0,422 | 1 | SMC | 0,839733589 | 1,190853877 | 0,781990521 |
| CBLB1 | 0,095112422 | 0,313611817 | 0,119 | 0,069 | 1 | SMC | 1,242815222 | 0,804624841 | 1,724637681 |

| Supplementary Table 4 - clusermarker | | | | | | | | | |
|---|---|---|---|---|---|---|---|---|---|
| | **p_val** | **avg_log2FC** | **pct.1** | **pct.2** | **p_val_adj** | **celltype** | **Foldchange_UP** | **Foldchange_DOWN** | **Ratio_pct1_pct2** |
| TCAF1 | 0,095846115 | -0,416222656 | 0,094 | 0,147 | 1 | SMC | 0,749384137 | 1,334429101 | 0,639455782 |
| EIF1B2 | 0,096150968 | -0,259189782 | 0,381 | 0,457 | 1 | SMC | 0,835557037 | 1,196806389 | 0,833698031 |
| SF3B31 | 0,096572455 | 0,368240557 | 0,139 | 0,086 | 1 | SMC | 1,290777699 | 0,774726741 | 1,61627907 |
| ATXN1 | 0,096584332 | 0,322106627 | 0,119 | 0,069 | 1 | SMC | 1,250154696 | 0,799901007 | 1,724637681 |
| MTCH12 | 0,096735946 | 0,318375459 | 0,349 | 0,276 | 1 | SMC | 1,246925662 | 0,801972427 | 1,264492754 |
| TMX22 | 0,097571251 | 0,30248064 | 0,151 | 0,095 | 1 | SMC | 1,233263124 | 0,81085697 | 1,589473684 |
| DELE1 | 0,097647667 | -0,268764123 | 0,08 | 0,129 | 1 | SMC | 0,830030282 | 1,204775321 | 0,620155039 |
| ACADVL2 | 0,097674662 | 0,278102595 | 0,307 | 0,233 | 1 | SMC | 1,212599048 | 0,824674901 | 1,317596567 |
| MRPS18C2 | 0,097971514 | -0,322253671 | 0,213 | 0,284 | 1 | SMC | 0,799819482 | 1,250282122 | 0,75 |
| DNAJC44 | 0,098064392 | 0,275028751 | 0,134 | 0,078 | 1 | SMC | 1,210018203 | 0,826433848 | 1,717948718 |
| NDUFAF34 | 0,098700859 | 0,420580501 | 0,264 | 0,207 | 1 | SMC | 1,338466009 | 0,747123941 | 1,275362319 |
| RIMKLB2 | 0,099313964 | -0,264333495 | 0,065 | 0,112 | 1 | SMC | 0,832583288 | 1,201081039 | 0,580357143 |
| EPB41L24 | 0,099325714 | 0,254887608 | 0,332 | 0,259 | 1 | SMC | 1,19324278 | 0,83805242 | 1,281853282 |
| TMEM106C1 | 0,099511997 | -0,419682457 | 0,142 | 0,207 | 1 | SMC | 0,747589154 | 1,337633103 | 0,685990338 |
| PARP143 | 0,099568688 | 0,295995389 | 0,188 | 0,129 | 1 | SMC | 1,22773176 | 0,814510167 | 1,457364341 |
| RPL87 | 0,100291971 | 0,250566188 | 0,963 | 0,922 | 1 | SMC | 1,189673913 | 0,840566469 | 1,044468547 |
| DCTN35 | 0,100554979 | 0,376968828 | 0,295 | 0,241 | 1 | SMC | 1,298610543 | 0,770053813 | 1,22406639 |
| NR1D24 | 0,101069861 | 0,25615257 | 0,273 | 0,207 | 1 | SMC | 1,194289481 | 0,837317934 | 1,31884058 |
| UBR1 | 0,101585531 | -0,393830707 | 0,153 | 0,216 | 1 | SMC | 0,761105997 | 1,313877441 | 0,708333333 |
| SBDS2 | 0,101645598 | -0,275957149 | 0,56 | 0,595 | 1 | SMC | 0,825902196 | 1,21079712 | 0,941176471 |
| EIF3E4 | 0,102114354 | -0,297296354 | 0,582 | 0,647 | 1 | SMC | 0,813776004 | 1,228839379 | 0,899536321 |
| NEU11 | 0,102206868 | 0,260802007 | 0,139 | 0,086 | 1 | SMC | 1,198144578 | 0,834623816 | 1,61627907 |
| CCDC502 | 0,103575824 | 0,323499703 | 0,267 | 0,198 | 1 | SMC | 1,251362436 | 0,79912899 | 1,348484848 |
| MYL12A5 | 0,10358555 | 0,273573646 | 0,679 | 0,655 | 1 | SMC | 1,208798391 | 0,827267812 | 1,036641221 |
| HNRNPF1 | 0,103769473 | 0,316794182 | 0,347 | 0,284 | 1 | SMC | 1,245559709 | 0,802851917 | 1,221830986 |
| RBMS21 | 0,105120546 | 0,291565735 | 0,139 | 0,086 | 1 | SMC | 1,223967911 | 0,817014883 | 1,61627907 |
| SH3BGRL36 | 0,105158094 | 0,29494596 | 0,44 | 0,362 | 1 | SMC | 1,226839022 | 0,815102864 | 1,215469613 |
| NEK72 | 0,105165188 | -0,268125431 | 0,168 | 0,233 | 1 | SMC | 0,830397824 | 1,204242076 | 0,721030043 |
| SMIM10 | 0,105476442 | -0,266033646 | 0,193 | 0,259 | 1 | SMC | 0,831602704 | 1,202497293 | 0,745173745 |
| PLSCR4 | 0,10596495 | 0,34150286 | 0,179 | 0,121 | 1 | SMC | 1,267075824 | 0,789218752 | 1,479338843 |
| PSMD3 | 0,106168555 | 0,303468071 | 0,151 | 0,095 | 1 | SMC | 1,234107502 | 0,810302181 | 1,589473684 |

| Supplementary Table 4 - clusermarker | | | | | | | | | |
|---|---|---|---|---|---|---|---|---|---|
| | **p_val** | **avg_log2FC** | **pct.1** | **pct.2** | **p_val_adj** | **celltype** | **Foldchange_UP** | **Foldchange_DOWN** | **Ratio_pct1_pct2** |
| TBC1D52 | 0,106652072 | 0,282296186 | 0,108 | 0,06 | 1 | SMC | 1,216128929 | 0,822281237 | 1,8 |
| TM2D11 | 0,108511648 | 0,338176958 | 0,188 | 0,129 | 1 | SMC | 1,264158148 | 0,791040268 | 1,457364341 |
| BEX32 | 0,109096321 | 0,35270676 | 0,438 | 0,371 | 1 | SMC | 1,276954181 | 0,783113455 | 1,180592992 |
| NBEAL13 | 0,109153222 | -0,333755594 | 0,321 | 0,388 | 1 | SMC | 0,793468253 | 1,260289869 | 0,827319588 |
| PSMD4 | 0,10925869 | -0,264209859 | 0,327 | 0,397 | 1 | SMC | 0,832654642 | 1,200978113 | 0,823677582 |
| COL27A1 | 0,110985101 | -0,252097088 | 0,125 | 0,181 | 1 | SMC | 0,839674984 | 1,190936992 | 0,690607735 |
| RTRAF3 | 0,11115891 | 0,294587197 | 0,332 | 0,276 | 1 | SMC | 1,226533975 | 0,815305585 | 1,202898551 |
| RTN44 | 0,11130829 | 0,263238975 | 0,534 | 0,474 | 1 | SMC | 1,200170168 | 0,833215178 | 1,126582278 |
| COBLL11 | 0,111486118 | 0,352863671 | 0,315 | 0,259 | 1 | SMC | 1,277093073 | 0,783028286 | 1,216216216 |
| ETF12 | 0,111769665 | 0,381802987 | 0,196 | 0,138 | 1 | SMC | 1,302969205 | 0,767477847 | 1,420289855 |
| ADI14 | 0,111869385 | 0,37047762 | 0,372 | 0,302 | 1 | SMC | 1,292780749 | 0,77352637 | 1,231788079 |
| EBNA1BP21 | 0,113340024 | -0,28184536 | 0,134 | 0,19 | 1 | SMC | 0,82253823 | 1,215748962 | 0,705263158 |
| SNHG15 | 0,114218935 | 0,257058405 | 0,253 | 0,19 | 1 | SMC | 1,195039583 | 0,836792366 | 1,331578947 |
| TUBB7 | 0,114282313 | 0,271514356 | 0,616 | 0,517 | 1 | SMC | 1,207074194 | 0,828449489 | 1,191489362 |
| PKIG | 0,11465713 | 0,467087943 | 0,378 | 0,328 | 1 | SMC | 1,382316467 | 0,723423343 | 1,152439024 |
| MEF2C3 | 0,11466879 | -0,342582521 | 0,438 | 0,5 | 1 | SMC | 0,78862835 | 1,268024413 | 0,876 |
| TPRKB4 | 0,115480164 | -0,40656806 | 0,125 | 0,181 | 1 | SMC | 0,754415875 | 1,32552884 | 0,690607735 |
| COX5B4 | 0,115649016 | -0,273630416 | 0,534 | 0,578 | 1 | SMC | 0,827235259 | 1,208845959 | 0,923875433 |
| A4GALT2 | 0,115698357 | 0,258931754 | 0,162 | 0,103 | 1 | SMC | 1,196592358 | 0,835706491 | 1,572815534 |
| NHP22 | 0,11592956 | -0,278427601 | 0,247 | 0,319 | 1 | SMC | 0,824489141 | 1,21287225 | 0,774294671 |
| SGK13 | 0,116083778 | 0,255548134 | 0,182 | 0,121 | 1 | SMC | 1,193789222 | 0,837668812 | 1,504132231 |
| ARL13 | 0,116114005 | -0,251185818 | 0,156 | 0,224 | 1 | SMC | 0,840205528 | 1,190184981 | 0,696428571 |
| TRPC4AP1 | 0,117221925 | 0,327014656 | 0,105 | 0,06 | 1 | SMC | 1,254414948 | 0,797184378 | 1,75 |
| NKTR | 0,117261711 | 0,351195546 | 0,21 | 0,147 | 1 | SMC | 1,27561728 | 0,783934191 | 1,428571429 |
| CRBN1 | 0,117591313 | -0,271422774 | 0,148 | 0,207 | 1 | SMC | 0,828502081 | 1,206997572 | 0,714975845 |
| RNF1302 | 0,118018239 | 0,299057165 | 0,256 | 0,19 | 1 | SMC | 1,230340094 | 0,812783396 | 1,347368421 |
| JADE1 | 0,118559109 | 0,320419503 | 0,196 | 0,138 | 1 | SMC | 1,248693588 | 0,800836978 | 1,420289855 |
| TSC22D33 | 0,120336692 | 0,275239359 | 0,631 | 0,595 | 1 | SMC | 1,210194857 | 0,826313212 | 1,060504202 |
| GLIPR14 | 0,120642966 | -0,436434408 | 0,136 | 0,19 | 1 | SMC | 0,738958675 | 1,353255646 | 0,715789474 |
| VCP | 0,120748981 | 0,339467495 | 0,415 | 0,345 | 1 | SMC | 1,265289484 | 0,790332973 | 1,202898551 |
| FCHSD2 | 0,121903829 | -0,333554897 | 0,153 | 0,216 | 1 | SMC | 0,793578642 | 1,260114559 | 0,708333333 |

| Supplementary Table 4 - clusermarker | | | | | | | | | |
|---|---|---|---|---|---|---|---|---|---|
| | **p_val** | **avg_log2FC** | **pct.1** | **pct.2** | **p_val_adj** | **celltype** | **Foldchange_UP** | **Foldchange_DOWN** | **Ratio_pct1_pct2** |
| PGF1 | 0,121995361 | 0,300895996 | 0,247 | 0,181 | 1 | SMC | 1,231909262 | 0,811748098 | 1,364640884 |
| GTF2B4 | 0,122048817 | -0,263852999 | 0,188 | 0,25 | 1 | SMC | 0,83286063 | 1,200681079 | 0,752 |
| ATXN2L | 0,122122556 | 0,368039341 | 0,125 | 0,078 | 1 | SMC | 1,290597683 | 0,774834802 | 1,602564103 |
| MARCHF62 | 0,123040987 | -0,279057391 | 0,128 | 0,19 | 1 | SMC | 0,8241293 | 1,213401829 | 0,673684211 |
| ACAP22 | 0,123167961 | 0,325003316 | 0,293 | 0,233 | 1 | SMC | 1,252667318 | 0,798296551 | 1,25751073 |
| PSMB102 | 0,123427721 | 0,288222968 | 0,148 | 0,095 | 1 | SMC | 1,221135224 | 0,818910126 | 1,557894737 |
| AP1S1 | 0,123484587 | 0,344379435 | 0,114 | 0,069 | 1 | SMC | 1,269604753 | 0,787646705 | 1,652173913 |
| HEPH | 0,123822568 | 0,30399525 | 0,159 | 0,103 | 1 | SMC | 1,234558542 | 0,810006141 | 1,54368932 |
| SLC25A371 | 0,123991197 | 0,305578868 | 0,156 | 0,103 | 1 | SMC | 1,235914437 | 0,809117501 | 1,514563107 |
| HSPH16 | 0,124961767 | -0,372905181 | 0,193 | 0,259 | 1 | SMC | 0,772225885 | 1,294957887 | 0,745173745 |
| DST4 | 0,125032171 | -0,275105674 | 0,565 | 0,586 | 1 | SMC | 0,826389785 | 1,210082721 | 0,964163823 |
| RASD13 | 0,12516834 | 0,639405536 | 0,452 | 0,379 | 1 | SMC | 1,55768718 | 0,641977422 | 1,192612137 |
| HRAS2 | 0,125326736 | 0,303179504 | 0,114 | 0,069 | 1 | SMC | 1,233860681 | 0,810464273 | 1,652173913 |
| RARRES22 | 0,12634838 | 0,384110506 | 0,54 | 0,509 | 1 | SMC | 1,305054907 | 0,766251286 | 1,060903733 |
| NDUFS23 | 0,126817438 | 0,336015024 | 0,264 | 0,207 | 1 | SMC | 1,262265177 | 0,792226561 | 1,275362319 |
| ALDH9A12 | 0,127367317 | 0,280620543 | 0,153 | 0,103 | 1 | SMC | 1,214717255 | 0,823236844 | 1,485436893 |
| TTC36 | 0,127410393 | 0,274393528 | 0,321 | 0,259 | 1 | SMC | 1,209485546 | 0,82679781 | 1,239382239 |
| SIPA1L12 | 0,127565509 | -0,293895538 | 0,091 | 0,138 | 1 | SMC | 0,815696554 | 1,225946089 | 0,65942029 |
| GET31 | 0,12790882 | 0,310935598 | 0,202 | 0,147 | 1 | SMC | 1,24051192 | 0,806118816 | 1,37414966 |
| CDK2AP23 | 0,128262661 | 0,301950485 | 0,216 | 0,155 | 1 | SMC | 1,232810013 | 0,811154995 | 1,393548387 |
| TCEAL9 | 0,12853345 | -0,252447824 | 0,29 | 0,216 | 1 | SMC | 0,839470874 | 1,191226558 | 1,342592593 |
| KARS11 | 0,129201026 | 0,344257627 | 0,116 | 0,069 | 1 | SMC | 1,269497564 | 0,787713209 | 1,68115942 |
| B4GALT11 | 0,132529268 | 0,267115515 | 0,125 | 0,078 | 1 | SMC | 1,203399377 | 0,830979323 | 1,602564103 |
| S100A107 | 0,13299706 | 0,385804271 | 0,344 | 0,267 | 1 | SMC | 1,306587979 | 0,765352212 | 1,288389513 |
| MAP1LC3A2 | 0,13311519 | 0,313726238 | 0,33 | 0,267 | 1 | SMC | 1,242913795 | 0,804561028 | 1,235955056 |
| PWWP3A | 0,13329853 | 0,278983834 | 0,114 | 0,069 | 1 | SMC | 1,213339964 | 0,82417132 | 1,652173913 |
| GBP42 | 0,134430505 | 0,268765057 | 0,128 | 0,078 | 1 | SMC | 1,204776101 | 0,830029745 | 1,641025641 |
| PLEKHH2 | 0,13532604 | 0,26985387 | 0,116 | 0,069 | 1 | SMC | 1,205685698 | 0,829403552 | 1,68115942 |
| ANO61 | 0,135499112 | 0,313815837 | 0,327 | 0,267 | 1 | SMC | 1,242990988 | 0,804511062 | 1,224719101 |
| NR4A26 | 0,135938758 | -0,436652593 | 0,369 | 0,422 | 1 | SMC | 0,738846928 | 1,35346032 | 0,874407583 |
| NDUFS33 | 0,136819725 | 0,320571248 | 0,219 | 0,164 | 1 | SMC | 1,248824935 | 0,800752749 | 1,335365854 |

| Supplementary Table 4 - clusermarker | | | | | | | | | |
|---|---|---|---|---|---|---|---|---|---|
| | **p_val** | **avg_log2FC** | **pct.1** | **pct.2** | **p_val_adj** | **celltype** | **Foldchange_UP** | **Foldchange_DOWN** | **Ratio_pct1_pct2** |
| KANSL11 | 0,136843331 | 0,346604549 | 0,162 | 0,112 | 1 | SMC | 1,271564416 | 0,786432828 | 1,446428571 |
| MLX1 | 0,13686305 | 0,281254634 | 0,134 | 0,086 | 1 | SMC | 1,215251263 | 0,822875096 | 1,558139535 |
| MRPS62 | 0,137199136 | -0,257620754 | 0,224 | 0,293 | 1 | SMC | 0,836466256 | 1,195505489 | 0,764505119 |
| RBM51 | 0,137400542 | -0,275327953 | 0,105 | 0,155 | 1 | SMC | 0,826262471 | 1,210269176 | 0,677419355 |
| A1BG2 | 0,137587322 | 0,317279288 | 0,176 | 0,121 | 1 | SMC | 1,245978598 | 0,802582004 | 1,454545455 |
| AKR7A23 | 0,137661572 | 0,294159334 | 0,185 | 0,129 | 1 | SMC | 1,226170273 | 0,815547418 | 1,434108527 |
| SUMF22 | 0,13931873 | 0,425745493 | 0,165 | 0,121 | 1 | SMC | 1,343266438 | 0,744453946 | 1,363636364 |
| URM1 | 0,140039411 | -0,257484514 | 0,111 | 0,164 | 1 | SMC | 0,836545251 | 1,195392597 | 0,676829268 |
| PNN3 | 0,141189069 | 0,250023324 | 0,29 | 0,233 | 1 | SMC | 1,189226341 | 0,84088282 | 1,244635193 |
| PELI2 | 0,141667494 | 0,259650879 | 0,105 | 0,06 | 1 | SMC | 1,197188959 | 0,835290029 | 1,75 |
| PLSCR12 | 0,142093937 | 0,27882788 | 0,136 | 0,086 | 1 | SMC | 1,21320881 | 0,824260417 | 1,581395349 |
| ZNF37A | 0,143705544 | 0,302505803 | 0,114 | 0,069 | 1 | SMC | 1,233284634 | 0,810842828 | 1,652173913 |
| FOXO31 | 0,144080512 | 0,379427412 | 0,273 | 0,224 | 1 | SMC | 1,300825472 | 0,768742634 | 1,21875 |
| IDH3G3 | 0,145323468 | -0,278517254 | 0,227 | 0,293 | 1 | SMC | 0,824437907 | 1,212947623 | 0,774744027 |
| EFNA11 | 0,145925527 | -0,38891736 | 0,116 | 0,164 | 1 | SMC | 0,763702494 | 1,309410416 | 0,707317073 |
| RAB212 | 0,14626136 | -0,260124 | 0,196 | 0,259 | 1 | SMC | 0,835016147 | 1,197581633 | 0,756756757 |
| MAGED23 | 0,146524532 | -0,301522523 | 0,466 | 0,509 | 1 | SMC | 0,811395652 | 1,232444366 | 0,915520629 |
| STRN4 | 0,147321515 | -0,257158672 | 0,068 | 0,112 | 1 | SMC | 0,836734211 | 1,195122641 | 0,607142857 |
| STT3B3 | 0,14866292 | 0,252200528 | 0,162 | 0,112 | 1 | SMC | 1,191022384 | 0,839614782 | 1,446428571 |
| GATAD1 | 0,149983184 | 0,285306589 | 0,159 | 0,112 | 1 | SMC | 1,218669218 | 0,82056721 | 1,419642857 |
| THBS13 | 0,150874202 | -0,678499456 | 0,31 | 0,362 | 1 | SMC | 0,624814805 | 1,600474239 | 0,856353591 |
| PLXNB23 | 0,150900228 | 0,337627945 | 0,139 | 0,095 | 1 | SMC | 1,263677168 | 0,791341353 | 1,463157895 |
| CPQ1 | 0,153484425 | 0,319340117 | 0,182 | 0,129 | 1 | SMC | 1,247759698 | 0,801436368 | 1,410852713 |
| EHBP1L1 | 0,153536359 | 0,336955774 | 0,207 | 0,155 | 1 | SMC | 1,263088541 | 0,791710136 | 1,335483871 |
| C11orf961 | 0,154230259 | -0,395838141 | 0,795 | 0,741 | 1 | SMC | 0,760047695 | 1,315706905 | 1,072874494 |
| SPATS2L1 | 0,15528777 | 0,269891184 | 0,352 | 0,302 | 1 | SMC | 1,205716882 | 0,8293821 | 1,165562914 |
| RAB1A | 0,15534172 | -0,253319935 | 0,205 | 0,259 | 1 | SMC | 0,838963566 | 1,191946873 | 0,791505792 |
| IRAG11 | 0,155367035 | 0,302769865 | 0,52 | 0,474 | 1 | SMC | 1,233510387 | 0,81069443 | 1,097046414 |
| RCN12 | 0,155695713 | 0,339743586 | 0,227 | 0,181 | 1 | SMC | 1,265531648 | 0,790181741 | 1,254143646 |
| LSM33 | 0,156481733 | -0,255655137 | 0,23 | 0,293 | 1 | SMC | 0,837606686 | 1,193877768 | 0,784982935 |
| ATP1B2 | 0,156497058 | 0,322879854 | 0,312 | 0,259 | 1 | SMC | 1,250824908 | 0,799472407 | 1,204633205 |

| Supplementary Table 4 - clusermarker | | | | | | | | | |
|---|---|---|---|---|---|---|---|---|---|
| | **p_val** | **avg_log2FC** | **pct.1** | **pct.2** | **p_val_adj** | **celltype** | **Foldchange_UP** | **Foldchange_DOWN** | **Ratio_pct1_pct2** |
| ACYP21 | 0,157436534 | 0,271941998 | 0,111 | 0,069 | 1 | SMC | 1,207432047 | 0,828203958 | 1,608695652 |
| ZNF3313 | 0,157482518 | -0,437284671 | 0,236 | 0,302 | 1 | SMC | 0,738523293 | 1,354053433 | 0,781456954 |
| DUSP52 | 0,157672066 | 0,256034113 | 0,119 | 0,078 | 1 | SMC | 1,194191424 | 0,837386687 | 1,525641026 |
| BACE23 | 0,158592133 | 0,275438525 | 0,162 | 0,112 | 1 | SMC | 1,210361938 | 0,826199146 | 1,446428571 |
| NDUFV25 | 0,159715018 | 0,325673475 | 0,349 | 0,284 | 1 | SMC | 1,253249341 | 0,797925814 | 1,228873239 |
| PGRMC12 | 0,160814819 | -0,274514474 | 0,23 | 0,293 | 1 | SMC | 0,8267285 | 1,209586945 | 0,784982935 |
| CISD35 | 0,162224767 | -0,269259171 | 0,176 | 0,233 | 1 | SMC | 0,829745514 | 1,2051888 | 0,755364807 |
| ITPA2 | 0,163675443 | -0,329747023 | 0,111 | 0,155 | 1 | SMC | 0,795675994 | 1,256792976 | 0,716129032 |
| PNMA11 | 0,164229344 | 0,335596363 | 0,241 | 0,19 | 1 | SMC | 1,261898929 | 0,792456493 | 1,268421053 |
| MXRA74 | 0,164342697 | -0,318808328 | 0,344 | 0,397 | 1 | SMC | 0,801731838 | 1,247299848 | 0,866498741 |
| JCHAIN5 | 0,165394712 | -2,097739635 | 0,139 | 0,086 | 1 | SMC | 0,233623995 | 4,280382239 | 1,61627907 |
| SPSB33 | 0,165668582 | 0,311375118 | 0,216 | 0,164 | 1 | SMC | 1,240889902 | 0,805873268 | 1,317073171 |
| KDM2A | 0,1657663 | 0,293420403 | 0,139 | 0,095 | 1 | SMC | 1,225542404 | 0,815965238 | 1,463157895 |
| IER55 | 0,167812236 | -0,412290548 | 0,205 | 0,259 | 1 | SMC | 0,751429391 | 1,330797027 | 0,791505792 |
| MPRIP1 | 0,167952121 | 0,260122858 | 0,384 | 0,328 | 1 | SMC | 1,197580684 | 0,835016808 | 1,170731707 |
| APRT5 | 0,168576552 | 0,272413064 | 0,338 | 0,284 | 1 | SMC | 1,20782636 | 0,827933578 | 1,190140845 |
| PDK42 | 0,17006371 | -0,470290536 | 0,315 | 0,371 | 1 | SMC | 0,72181922 | 1,385388436 | 0,849056604 |
| EPHA31 | 0,170256257 | 0,295806334 | 0,176 | 0,129 | 1 | SMC | 1,227570885 | 0,814616909 | 1,364341085 |
| BRD41 | 0,170593782 | 0,261532879 | 0,298 | 0,241 | 1 | SMC | 1,198751715 | 0,834201101 | 1,236514523 |
| RPS19BP13 | 0,172511643 | 0,267799176 | 0,236 | 0,181 | 1 | SMC | 1,203969776 | 0,830585634 | 1,303867403 |
| PTBP12 | 0,172706415 | 0,277922006 | 0,239 | 0,181 | 1 | SMC | 1,21244727 | 0,824778136 | 1,320441989 |
| PLEC5 | 0,173155158 | 0,414130851 | 0,33 | 0,293 | 1 | SMC | 1,332495676 | 0,750471479 | 1,126279863 |
| FCGRT3 | 0,173956572 | 0,326620682 | 0,267 | 0,216 | 1 | SMC | 1,254072436 | 0,797402105 | 1,236111111 |
| RSRC1 | 0,175869608 | 0,299030654 | 0,156 | 0,112 | 1 | SMC | 1,230317486 | 0,812798332 | 1,392857143 |
| TMOD1 | 0,176023368 | -0,260177398 | 0,111 | 0,155 | 1 | SMC | 0,834985241 | 1,197625959 | 0,716129032 |
| ANAPC117 | 0,177211649 | 0,338006665 | 0,452 | 0,397 | 1 | SMC | 1,264008938 | 0,791133646 | 1,138539043 |
| CTCF2 | 0,17875465 | 0,266524344 | 0,136 | 0,095 | 1 | SMC | 1,202906363 | 0,831319902 | 1,431578947 |
| DHX30 | 0,178895658 | 0,258277994 | 0,119 | 0,078 | 1 | SMC | 1,196050242 | 0,836085278 | 1,525641026 |
| MAFB4 | 0,179670421 | -0,257771873 | 0,202 | 0,25 | 1 | SMC | 0,836378642 | 1,195630722 | 0,808 |
| PRKCSH | 0,179772143 | 0,316223177 | 0,23 | 0,181 | 1 | SMC | 1,245066825 | 0,803169741 | 1,270718232 |
| MRPL182 | 0,181999012 | -0,351954519 | 0,227 | 0,276 | 1 | SMC | 0,783521887 | 1,276288533 | 0,822463768 |

| Supplementary Table 4 - clusermarker | | | | | | | | | |
|---|---|---|---|---|---|---|---|---|---|
| | **p_val** | **avg_log2FC** | **pct.1** | **pct.2** | **p_val_adj** | **celltype** | **Foldchange_UP** | **Foldchange_DOWN** | **Ratio_pct1_pct2** |
| MAPRE21 | 0,182685361 | -0,25144412 | 0,304 | 0,362 | 1 | SMC | 0,840055109 | 1,190398093 | 0,839779006 |
| MRPL24 | 0,184189817 | -0,342759103 | 0,239 | 0,293 | 1 | SMC | 0,78853183 | 1,268179625 | 0,815699659 |
| DDX423 | 0,184598171 | -0,316453814 | 0,173 | 0,233 | 1 | SMC | 0,803041352 | 1,245265885 | 0,74248927 |
| ARF32 | 0,184668652 | 0,339190983 | 0,193 | 0,147 | 1 | SMC | 1,265046997 | 0,790484466 | 1,31292517 |
| KIAA11091 | 0,185058779 | -0,274464304 | 0,213 | 0,276 | 1 | SMC | 0,82675725 | 1,209544882 | 0,77173913 |
| HERC2 | 0,185576517 | 0,264591705 | 0,153 | 0,112 | 1 | SMC | 1,201296025 | 0,832434287 | 1,366071429 |
| BGN2 | 0,185751727 | 0,25266592 | 0,761 | 0,741 | 1 | SMC | 1,191406652 | 0,839343979 | 1,026990553 |
| JMJD8 | 0,185822064 | 0,31035749 | 0,145 | 0,103 | 1 | SMC | 1,240014929 | 0,806441904 | 1,40776699 |
| NREP2 | 0,187266153 | 0,266992642 | 0,156 | 0,112 | 1 | SMC | 1,203296889 | 0,8310501 | 1,392857143 |
| TCEA14 | 0,193611008 | -0,327869564 | 0,392 | 0,448 | 1 | SMC | 0,796712125 | 1,255158505 | 0,875 |
| NFASC | 0,19483665 | -0,279926489 | 0,162 | 0,216 | 1 | SMC | 0,823632983 | 1,214133018 | 0,75 |
| TNC2 | 0,195486594 | 0,441899781 | 0,369 | 0,319 | 1 | SMC | 1,358391918 | 0,736164568 | 1,156739812 |
| UROS1 | 0,195910297 | 0,25089648 | 0,116 | 0,078 | 1 | SMC | 1,189946309 | 0,840374051 | 1,487179487 |
| SLC35B21 | 0,196325488 | 0,303441891 | 0,125 | 0,086 | 1 | SMC | 1,234085107 | 0,810316886 | 1,453488372 |
| HUWE11 | 0,197666932 | 0,298359037 | 0,236 | 0,19 | 1 | SMC | 1,22974487 | 0,813176801 | 1,242105263 |
| C8orf33 | 0,197724988 | -0,260073564 | 0,077 | 0,112 | 1 | SMC | 0,835045339 | 1,197539766 | 0,6875 |
| NDUFA25 | 0,198467553 | 0,292171828 | 0,29 | 0,241 | 1 | SMC | 1,224482222 | 0,816671718 | 1,203319502 |
| COX8A7 | 0,198574734 | -0,459820108 | 0,534 | 0,569 | 1 | SMC | 0,727076913 | 1,37537031 | 0,938488576 |
| STUB15 | 0,199725687 | 0,252329277 | 0,27 | 0,233 | 1 | SMC | 1,191128679 | 0,839539857 | 1,158798283 |
| TCF121 | 0,20158134 | 0,280851731 | 0,193 | 0,147 | 1 | SMC | 1,214911926 | 0,823104934 | 1,31292517 |
| CSF11 | 0,20171235 | 0,270643439 | 0,188 | 0,138 | 1 | SMC | 1,206345735 | 0,828949754 | 1,362318841 |
| CIAO2B5 | 0,202705017 | 0,252826254 | 0,278 | 0,224 | 1 | SMC | 1,191539067 | 0,839250703 | 1,241071429 |
| SMAD41 | 0,202868748 | 0,262969405 | 0,173 | 0,129 | 1 | SMC | 1,199945935 | 0,83337088 | 1,341085271 |
| PSMA32 | 0,204573232 | 0,266726517 | 0,318 | 0,267 | 1 | SMC | 1,203074944 | 0,831203413 | 1,191011236 |
| KTN13 | 0,204598845 | -0,269690639 | 0,463 | 0,491 | 1 | SMC | 0,829497398 | 1,205549291 | 0,942973523 |
| ELP2 | 0,204857377 | 0,271942473 | 0,125 | 0,086 | 1 | SMC | 1,207432445 | 0,828203685 | 1,453488372 |
| TOX42 | 0,207652983 | 0,310548191 | 0,236 | 0,19 | 1 | SMC | 1,24017885 | 0,806335312 | 1,242105263 |
| PHACTR22 | 0,209097468 | 0,261253161 | 0,216 | 0,172 | 1 | SMC | 1,198519317 | 0,834362856 | 1,255813953 |
| PURA3 | 0,21042296 | -0,254331065 | 0,324 | 0,353 | 1 | SMC | 0,838375775 | 1,192782557 | 0,917847025 |
| BIRC62 | 0,210499351 | 0,295408185 | 0,213 | 0,172 | 1 | SMC | 1,227232151 | 0,814841755 | 1,238372093 |
| MCMBP | 0,210806443 | -0,285515493 | 0,091 | 0,129 | 1 | SMC | 0,8204484 | 1,218845694 | 0,705426357 |

| Supplementary Table 4 - clusermarker | | | | | | | | | |
|---|---|---|---|---|---|---|---|---|---|
| | **p_val** | **avg_log2FC** | **pct.1** | **pct.2** | **p_val_adj** | **celltype** | **Foldchange_UP** | **Foldchange_DOWN** | **Ratio_pct1_pct2** |
| TENT21 | 0,211965109 | 0,255647119 | 0,105 | 0,069 | 1 | SMC | 1,193871132 | 0,837611341 | 1,52173913 |
| UBE2J13 | 0,214474985 | 0,27652952 | 0,182 | 0,138 | 1 | SMC | 1,211277584 | 0,825574594 | 1,31884058 |
| RECQL1 | 0,216186106 | -0,312299349 | 0,185 | 0,233 | 1 | SMC | 0,805357168 | 1,241685105 | 0,793991416 |
| AK25 | 0,217041863 | 0,266412453 | 0,179 | 0,138 | 1 | SMC | 1,202813072 | 0,83138438 | 1,297101449 |
| FAM168B | 0,220565658 | 0,310391242 | 0,125 | 0,086 | 1 | SMC | 1,24004394 | 0,806423037 | 1,453488372 |
| LIMS2 | 0,220837421 | -0,28707995 | 0,196 | 0,25 | 1 | SMC | 0,819559189 | 1,220168126 | 0,784 |
| FBXO17 | 0,221900588 | 0,294566821 | 0,122 | 0,086 | 1 | SMC | 1,226516652 | 0,8153171 | 1,418604651 |
| NEK11 | 0,222214341 | 0,297838879 | 0,142 | 0,103 | 1 | SMC | 1,229301571 | 0,813470042 | 1,378640777 |
| TMEM2563 | 0,222341949 | 0,280764872 | 0,352 | 0,31 | 1 | SMC | 1,214838783 | 0,823154491 | 1,135483871 |
| SETD7 | 0,223068512 | 0,257081905 | 0,134 | 0,095 | 1 | SMC | 1,195059049 | 0,836778736 | 1,410526316 |
| SP1001 | 0,223706268 | 0,343951822 | 0,273 | 0,233 | 1 | SMC | 1,2692285 | 0,787880196 | 1,17167382 |
| ADAMTS10 | 0,227750276 | 0,334032028 | 0,102 | 0,069 | 1 | SMC | 1,260531376 | 0,793316231 | 1,47826087 |
| CNPY32 | 0,228353538 | 0,279748845 | 0,102 | 0,069 | 1 | SMC | 1,213983527 | 0,823734406 | 1,47826087 |
| UQCC32 | 0,233834142 | 0,25152857 | 0,102 | 0,069 | 1 | SMC | 1,190467776 | 0,840005937 | 1,47826087 |
| OXA1L2 | 0,235352824 | 0,293959095 | 0,295 | 0,259 | 1 | SMC | 1,226000099 | 0,81566062 | 1,138996139 |
| ILF21 | 0,235472882 | 0,264977092 | 0,227 | 0,19 | 1 | SMC | 1,20161697 | 0,832211949 | 1,194736842 |
| FDX11 | 0,236299657 | 0,370643481 | 0,102 | 0,069 | 1 | SMC | 1,292929384 | 0,773437446 | 1,47826087 |
| CSNK2B6 | 0,237086368 | 0,331398259 | 0,457 | 0,405 | 1 | SMC | 1,258232261 | 0,794765824 | 1,128395062 |
| ZFAND11 | 0,238695646 | -0,272728657 | 0,142 | 0,19 | 1 | SMC | 0,827752486 | 1,208090604 | 0,747368421 |
| NRP12 | 0,241243699 | 0,331275289 | 0,281 | 0,233 | 1 | SMC | 1,258125019 | 0,79483357 | 1,206008584 |
| PRR132 | 0,242256322 | 0,305008085 | 0,304 | 0,259 | 1 | SMC | 1,23542556 | 0,809437681 | 1,173745174 |
| LINC00989 | 0,243696423 | -0,455930846 | 0,102 | 0,138 | 1 | SMC | 0,729039634 | 1,371667538 | 0,739130435 |
| SOD25 | 0,244596985 | 0,538875823 | 0,364 | 0,328 | 1 | SMC | 1,452839994 | 0,688307043 | 1,109756098 |
| C15orf401 | 0,246051383 | 0,329819268 | 0,119 | 0,086 | 1 | SMC | 1,256855914 | 0,79563615 | 1,38372093 |
| GMPS1 | 0,247082074 | 0,274452174 | 0,122 | 0,086 | 1 | SMC | 1,209534712 | 0,826764201 | 1,418604651 |
| EMP23 | 0,24798742 | 0,316864488 | 0,389 | 0,353 | 1 | SMC | 1,245620409 | 0,802812793 | 1,101983003 |
| PTOV14 | 0,25198696 | 0,25363375 | 0,216 | 0,172 | 1 | SMC | 1,192206174 | 0,838781095 | 1,255813953 |
| SCAF4 | 0,253187439 | 0,264398751 | 0,111 | 0,078 | 1 | SMC | 1,201135367 | 0,832545629 | 1,423076923 |
| MARK31 | 0,253801166 | 0,325783566 | 0,145 | 0,112 | 1 | SMC | 1,253344979 | 0,797864927 | 1,294642857 |
| SNX61 | 0,255672907 | 0,289552747 | 0,355 | 0,31 | 1 | SMC | 1,222261303 | 0,818155657 | 1,14516129 |
| FUBP11 | 0,261770243 | 0,260088439 | 0,122 | 0,086 | 1 | SMC | 1,197552114 | 0,835036729 | 1,418604651 |

| Supplementary Table 4 - clusermarker | | | | | | | | | |
|---|---|---|---|---|---|---|---|---|---|
| | **p_val** | **avg_log2FC** | **pct.1** | **pct.2** | **p_val_adj** | **celltype** | **Foldchange_UP** | **Foldchange_DOWN** | **Ratio_pct1_pct2** |
| TAPBP3 | 0,263923265 | 0,327180429 | 0,224 | 0,19 | 1 | SMC | 1,254559095 | 0,797092783 | 1,178947368 |
| ZNF4282 | 0,26557564 | 0,31472387 | 0,276 | 0,233 | 1 | SMC | 1,243773574 | 0,804004862 | 1,184549356 |
| SAMHD12 | 0,269258353 | 0,302472125 | 0,273 | 0,233 | 1 | SMC | 1,233255845 | 0,810861756 | 1,17167382 |
| CRIP16 | 0,269323446 | -0,281210517 | 0,923 | 0,888 | 1 | SMC | 0,82290026 | 1,215214102 | 1,039414414 |
| PHLDB2 | 0,270763761 | -0,336701595 | 0,395 | 0,414 | 1 | SMC | 0,791849634 | 1,262866025 | 0,95410628 |
| UBE2R21 | 0,271224649 | 0,255054101 | 0,202 | 0,164 | 1 | SMC | 1,193380494 | 0,837955711 | 1,231707317 |
| TMX31 | 0,272120354 | 0,250721168 | 0,128 | 0,095 | 1 | SMC | 1,189801719 | 0,840476177 | 1,347368421 |
| SNHG121 | 0,273453404 | -0,259402028 | 0,142 | 0,181 | 1 | SMC | 0,835434121 | 1,196982473 | 0,784530387 |
| SNRNP27 | 0,274759458 | -0,303748192 | 0,074 | 0,103 | 1 | SMC | 0,810144864 | 1,234347145 | 0,718446602 |
| VPS4B1 | 0,275982356 | -0,254919183 | 0,173 | 0,216 | 1 | SMC | 0,838034078 | 1,193268897 | 0,800925926 |
| ZFYVE162 | 0,276642168 | 0,348138054 | 0,17 | 0,138 | 1 | SMC | 1,272916737 | 0,785597338 | 1,231884058 |
| USE1 | 0,277312174 | -0,284077032 | 0,108 | 0,147 | 1 | SMC | 0,821266848 | 1,217631032 | 0,734693878 |
| GYPC1 | 0,277349772 | 0,358240605 | 0,276 | 0,233 | 1 | SMC | 1,281861689 | 0,780115366 | 1,184549356 |
| SUMO33 | 0,283993392 | 0,270430122 | 0,188 | 0,155 | 1 | SMC | 1,206167378 | 0,829072331 | 1,212903226 |
| RER13 | 0,283997221 | 0,328913498 | 0,236 | 0,198 | 1 | SMC | 1,256067067 | 0,796135832 | 1,191919192 |
| ETHE14 | 0,28441439 | 0,255051872 | 0,136 | 0,103 | 1 | SMC | 1,19337865 | 0,837957005 | 1,32038835 |
| ADIPOR21 | 0,290479126 | -0,27465132 | 0,074 | 0,103 | 1 | SMC | 0,826650084 | 1,209701685 | 0,718446602 |
| PRKRA2 | 0,290903908 | 0,265858212 | 0,165 | 0,129 | 1 | SMC | 1,202351076 | 0,831703834 | 1,279069767 |
| DNM22 | 0,292868547 | 0,268516917 | 0,128 | 0,095 | 1 | SMC | 1,2045689 | 0,830172521 | 1,347368421 |
| MPZL12 | 0,293299379 | 0,307740611 | 0,17 | 0,138 | 1 | SMC | 1,237767727 | 0,807906022 | 1,231884058 |
| MLLT61 | 0,296588919 | 0,302718796 | 0,21 | 0,181 | 1 | SMC | 1,233466724 | 0,810723127 | 1,160220994 |
| HES45 | 0,304498837 | 0,272942518 | 0,48 | 0,431 | 1 | SMC | 1,208269701 | 0,827629791 | 1,113689095 |
| BPNT21 | 0,306278408 | 0,342462879 | 0,196 | 0,164 | 1 | SMC | 1,26791926 | 0,788693753 | 1,195121951 |
| PIN11 | 0,306704963 | 0,286610168 | 0,25 | 0,224 | 1 | SMC | 1,21977087 | 0,819826104 | 1,116071429 |
| RPP212 | 0,307557918 | 0,3053657 | 0,227 | 0,198 | 1 | SMC | 1,235731836 | 0,809237062 | 1,146464646 |
| HNRNPM3 | 0,308344385 | -0,33677805 | 0,278 | 0,336 | 1 | SMC | 0,791807671 | 1,262932952 | 0,827380952 |
| TXNDC173 | 0,3105416 | 0,308421129 | 0,216 | 0,181 | 1 | SMC | 1,238351718 | 0,807525023 | 1,193370166 |
| AL118516.12 | 0,316418126 | -0,305866152 | 0,102 | 0,138 | 1 | SMC | 0,808956397 | 1,236160569 | 0,739130435 |
| CD744 | 0,316834898 | 0,29818752 | 0,168 | 0,121 | 1 | SMC | 1,229598679 | 0,813273483 | 1,388429752 |
| NSA22 | 0,316944589 | -0,353045284 | 0,318 | 0,353 | 1 | SMC | 0,782929721 | 1,277253849 | 0,900849858 |
| MYSM11 | 0,317726376 | 0,250514479 | 0,105 | 0,078 | 1 | SMC | 1,189631274 | 0,840596597 | 1,346153846 |

| Supplementary Table 4 - clusermarker | | | | | | | | | |
|---|---|---|---|---|---|---|---|---|---|
| | **p_val** | **avg_log2FC** | **pct.1** | **pct.2** | **p_val_adj** | **celltype** | **Foldchange_UP** | **Foldchange_DOWN** | **Ratio_pct1_pct2** |
| APP2 | 0,322630713 | 0,280469094 | 0,634 | 0,569 | 1 | SMC | 1,214589746 | 0,823323269 | 1,114235501 |
| USP161 | 0,323446968 | 0,250080145 | 0,273 | 0,241 | 1 | SMC | 1,18927318 | 0,840849703 | 1,132780083 |
| MRPL414 | 0,32845023 | 0,271241441 | 0,213 | 0,181 | 1 | SMC | 1,206845873 | 0,828606222 | 1,17679558 |
| SPTSSA1 | 0,332368031 | 0,256491488 | 0,267 | 0,233 | 1 | SMC | 1,194570076 | 0,837121254 | 1,145922747 |
| MRPL46 | 0,343373923 | 0,376995182 | 0,159 | 0,138 | 1 | SMC | 1,298634266 | 0,770039746 | 1,152173913 |
| EIF3D3 | 0,347186724 | -0,331240672 | 0,338 | 0,371 | 1 | SMC | 0,794852641 | 1,258094831 | 0,911051213 |
| EBLN3P1 | 0,350341207 | 0,278423327 | 0,156 | 0,129 | 1 | SMC | 1,212868657 | 0,824491584 | 1,209302326 |
| ETFB1 | 0,351465708 | -0,253934908 | 0,244 | 0,284 | 1 | SMC | 0,83860602 | 1,19245507 | 0,85915493 |
| DIS3 | 0,357630217 | 0,309477156 | 0,111 | 0,086 | 1 | SMC | 1,239258502 | 0,806934145 | 1,290697674 |
| PTPN11 | 0,364440035 | -0,252429954 | 0,179 | 0,216 | 1 | SMC | 0,839481272 | 1,191211803 | 0,828703704 |
| HSPA86 | 0,364838892 | -0,288037073 | 0,764 | 0,776 | 1 | SMC | 0,819015651 | 1,220977888 | 0,984536082 |
| NFIL31 | 0,369082014 | -0,395090814 | 0,202 | 0,233 | 1 | SMC | 0,760441507 | 1,315025535 | 0,86695279 |
| SSU724 | 0,377909474 | 0,312054871 | 0,173 | 0,147 | 1 | SMC | 1,241474708 | 0,805493655 | 1,176870748 |
| CYCS4 | 0,379455933 | -0,431197509 | 0,406 | 0,414 | 1 | SMC | 0,741645927 | 1,348352312 | 0,980676329 |
| USP483 | 0,38123376 | 0,26742459 | 0,162 | 0,138 | 1 | SMC | 1,203657215 | 0,830801318 | 1,173913043 |
| POLR1H1 | 0,382445093 | -0,251083967 | 0,122 | 0,155 | 1 | SMC | 0,840264846 | 1,19010096 | 0,787096774 |
| SCRN1 | 0,383213656 | 0,304398465 | 0,116 | 0,095 | 1 | SMC | 1,234903634 | 0,809779786 | 1,221052632 |
| OLA12 | 0,386439538 | -0,481918072 | 0,173 | 0,207 | 1 | SMC | 0,716025031 | 1,39659922 | 0,835748792 |
| REEP32 | 0,387377439 | 0,329572987 | 0,236 | 0,216 | 1 | SMC | 1,256641375 | 0,795771984 | 1,092592593 |
| PLEKHH3 | 0,398943864 | 0,252228585 | 0,205 | 0,181 | 1 | SMC | 1,191045548 | 0,839598454 | 1,132596685 |
| CYHR1 | 0,404968477 | 0,260580608 | 0,179 | 0,155 | 1 | SMC | 1,197960723 | 0,834751908 | 1,15483871 |
| MTRNR2L88 | 0,406747112 | -2,236303423 | 0,148 | 0,172 | 1 | SMC | 0,212229421 | 4,711882043 | 0,860465116 |
| SIK3 | 0,409305132 | 0,258489263 | 0,119 | 0,095 | 1 | SMC | 1,196225405 | 0,835962851 | 1,252631579 |
| CYP1B13 | 0,413400946 | 0,318746215 | 0,119 | 0,095 | 1 | SMC | 1,247246149 | 0,801766356 | 1,252631579 |
| CTBP12 | 0,413741438 | 0,31106976 | 0,114 | 0,095 | 1 | SMC | 1,240627285 | 0,806043855 | 1,2 |
| MTHFR | 0,416431005 | 0,336127377 | 0,108 | 0,086 | 1 | SMC | 1,262363482 | 0,792164867 | 1,255813953 |
| ATP2B12 | 0,427262488 | 0,265842575 | 0,168 | 0,147 | 1 | SMC | 1,202338044 | 0,831712849 | 1,142857143 |
| PTP4A31 | 0,429053689 | 0,310216008 | 0,324 | 0,293 | 1 | SMC | 1,23989333 | 0,806520994 | 1,105802048 |
| EIF51 | 0,453470196 | -0,293789502 | 0,571 | 0,603 | 1 | SMC | 0,815756509 | 1,225855987 | 0,946932007 |
| COPS8 | 0,486806049 | 0,264364099 | 0,179 | 0,164 | 1 | SMC | 1,201106518 | 0,832565626 | 1,091463415 |
| SPG73 | 0,495189419 | 0,256622223 | 0,111 | 0,095 | 1 | SMC | 1,194678332 | 0,837045398 | 1,168421053 |

| Supplementary Table 4 - clusermarker | | | | | | | | | |
|---|---|---|---|---|---|---|---|---|---|
| | **p_val** | **avg_log2FC** | **pct.1** | **pct.2** | **p_val_adj** | **celltype** | **Foldchange_UP** | **Foldchange_DOWN** | **Ratio_pct1_pct2** |
| TXNL4A4 | 0,501913356 | 0,270788344 | 0,227 | 0,216 | 1 | SMC | 1,206466908 | 0,828866497 | 1,050925926 |
| TAF107 | 0,508008253 | 0,273946695 | 0,179 | 0,164 | 1 | SMC | 1,209111001 | 0,827053926 | 1,091463415 |
| SIRT11 | 0,51667018 | -0,268508621 | 0,088 | 0,103 | 1 | SMC | 0,830177294 | 1,204561974 | 0,854368932 |
| GTF3C62 | 0,532035266 | 0,258972259 | 0,179 | 0,164 | 1 | SMC | 1,196625953 | 0,835683028 | 1,091463415 |
| CCT51 | 0,560596601 | -0,359701671 | 0,213 | 0,233 | 1 | SMC | 0,779325717 | 1,283160531 | 0,91416309 |
| TUBB62 | 0,563189786 | 0,283354053 | 0,219 | 0,207 | 1 | SMC | 1,217020993 | 0,821678513 | 1,057971014 |
| DNAJB4 | 0,56375101 | -0,482127793 | 0,344 | 0,353 | 1 | SMC | 0,715920952 | 1,396802255 | 0,974504249 |
| EMP33 | 0,60859555 | -0,264050685 | 0,443 | 0,379 | 1 | SMC | 0,832746514 | 1,200845615 | 1,168865435 |
| TCEAL32 | 0,610997004 | 0,263445789 | 0,205 | 0,198 | 1 | SMC | 1,200342228 | 0,833095743 | 1,035353535 |
| GADD45A2 | 0,64279068 | 0,293486166 | 0,332 | 0,328 | 1 | SMC | 1,22559827 | 0,815928045 | 1,012195122 |
| MIF4GD | 0,643607913 | 0,265148316 | 0,131 | 0,121 | 1 | SMC | 1,20175959 | 0,832113185 | 1,082644628 |
| E2F4 | 0,683926475 | 0,26351098 | 0,134 | 0,129 | 1 | SMC | 1,200396469 | 0,833058099 | 1,03875969 |
| IL6 | 0,694823627 | 0,352642886 | 0,264 | 0,284 | 1 | SMC | 1,276897646 | 0,783148127 | 0,929577465 |
| CCL27 | 0,732902998 | -0,385575724 | 0,511 | 0,483 | 1 | SMC | 0,765473467 | 1,306381009 | 1,057971014 |
| LENG8 | 0,735257938 | 0,319452763 | 0,125 | 0,121 | 1 | SMC | 1,247857127 | 0,801373794 | 1,033057851 |
| ANKHD11 | 0,737692714 | -0,316830787 | 0,264 | 0,267 | 1 | SMC | 0,802831547 | 1,245591313 | 0,988764045 |
| MIR22HG3 | 0,749684488 | -0,33107336 | 0,25 | 0,241 | 1 | SMC | 0,794944828 | 1,257948936 | 1,037344398 |
| EIF4A33 | 0,842433812 | -0,29677633 | 0,386 | 0,345 | 1 | SMC | 0,814069386 | 1,22839652 | 1,11884058 |
| UBAC11 | 0,847802504 | -0,320993971 | 0,114 | 0,121 | 1 | SMC | 0,800518156 | 1,249190905 | 0,94214876 |
| SP1101 | 0,849110821 | -0,29186694 | 0,142 | 0,129 | 1 | SMC | 0,816844325 | 1,224223477 | 1,100775194 |
| PLRG1 | 0,920755447 | -0,347173437 | 0,128 | 0,129 | 1 | SMC | 0,786122781 | 1,272065922 | 0,992248062 |
| GLA1 | 0,940813183 | -0,385802095 | 0,227 | 0,216 | 1 | SMC | 0,765353367 | 1,306586008 | 1,050925926 |
| ATP1B33 | 0,959106773 | -0,275495216 | 0,281 | 0,259 | 1 | SMC | 0,826166681 | 1,2104095 | 1,084942085 |
| ALKAL2 | 0,959313498 | 0,302429625 | 0,139 | 0,147 | 1 | SMC | 1,233219515 | 0,810885643 | 0,945578231 |
| PRR49 | 7,552E-273 | -4,154002662 | 0,113 | 0,861 | 2,763E-268 | EC | 0,056172092 | 17,80243478 | 0,131242741 |
| LYZ9 | 1,5726E-191 | -3,247309965 | 0,095 | 0,684 | 5,7537E-187 | EC | 0,105308226 | 9,495934374 | 0,138888889 |
| PRB39 | 1,1378E-162 | -2,948247324 | 0,017 | 0,457 | 4,1628E-158 | EC | 0,129565424 | 7,718108491 | 0,037199125 |
| SLPI9 | 2,31852E-87 | -2,462951829 | 0,163 | 0,553 | 8,48278E-83 | EC | 0,181375082 | 5,513436516 | 0,294755877 |
| SCGB3A29 | 7,87852E-72 | -2,500901179 | 0,051 | 0,329 | 2,88252E-67 | EC | 0,176666306 | 5,660388905 | 0,155015198 |
| STATH9 | 6,43056E-69 | -1,310902986 | 0,005 | 0,197 | 2,35275E-64 | EC | 0,403068519 | 2,480967756 | 0,025380711 |
| RPS278 | 7,81745E-58 | -0,522198444 | 0,982 | 0,988 | 2,86017E-53 | EC | 0,696309956 | 1,43614204 | 0,993927126 |

| Supplementary Table 4 - clusermarker | | | | | | | | | |
|---|---|---|---|---|---|---|---|---|---|
| | **p_val** | **avg_log2FC** | **pct.1** | **pct.2** | **p_val_adj** | **celltype** | **Foldchange_UP** | **Foldchange_DOWN** | **Ratio_pct1_pct2** |
| TXNIP9 | 2,28995E-47 | -0,941973937 | 0,878 | 0,923 | 8,37825E-43 | EC | 0,520520202 | 1,921155022 | 0,951245937 |
| SCGB3A17 | 2,09979E-46 | -1,24012315 | 0,214 | 0,499 | 7,6825E-42 | EC | 0,423336518 | 2,362186953 | 0,428857715 |
| RPS299 | 5,47353E-45 | -0,674955901 | 0,792 | 0,867 | 2,0026E-40 | EC | 0,626351365 | 1,59654797 | 0,91349481 |
| RPLP07 | 1,76211E-43 | 0,568369361 | 0,984 | 0,934 | 6,44702E-39 | EC | 1,482846602 | 0,67437859 | 1,053533191 |
| PLAT1 | 2,25136E-43 | -1,273115205 | 0,237 | 0,496 | 8,23707E-39 | EC | 0,413765366 | 2,416828672 | 0,477822581 |
| C6orf589 | 1,68302E-41 | -1,370850265 | 0,01 | 0,145 | 6,15765E-37 | EC | 0,386663298 | 2,586229429 | 0,068965517 |
| GAPDH4 | 1,4884E-40 | 0,531157416 | 0,967 | 0,908 | 5,44562E-36 | EC | 1,445088067 | 0,691999348 | 1,064977974 |
| IGKC8 | 1,64893E-40 | 1,131348065 | 0,321 | 0,04 | 6,03296E-36 | EC | 2,190633391 | 0,456488979 | 8,025 |
| PRH28 | 2,54357E-40 | -1,045870287 | 0,007 | 0,129 | 9,30617E-36 | EC | 0,484352641 | 2,064611434 | 0,054263566 |
| SPARC4 | 2,89374E-40 | 0,954989333 | 0,772 | 0,469 | 1,05873E-35 | EC | 1,938565301 | 0,515845404 | 1,646055437 |
| RPL138 | 9,65067E-39 | 0,540594955 | 0,99 | 0,978 | 3,53089E-34 | EC | 1,454572247 | 0,687487337 | 1,012269939 |
| PIP8 | 1,07114E-38 | -0,871336411 | 0,002 | 0,108 | 3,91899E-34 | EC | 0,546640247 | 1,829356703 | 0,018518519 |
| MT-ATP89 | 2,43674E-38 | -0,702992351 | 0,765 | 0,916 | 8,9153E-34 | EC | 0,614296747 | 1,62787774 | 0,835152838 |
| RPS268 | 4,51588E-38 | -0,498706204 | 0,916 | 0,961 | 1,65223E-33 | EC | 0,707741193 | 1,412945877 | 0,953173777 |
| PFN17 | 1,30995E-37 | 0,684530647 | 0,874 | 0,677 | 4,7927E-33 | EC | 1,607179031 | 0,622208217 | 1,29098966 |
| ZG16B8 | 2,83073E-36 | -1,536354725 | 0,018 | 0,153 | 1,03568E-31 | EC | 0,344755453 | 2,900606769 | 0,117647059 |
| NME28 | 2,62402E-35 | 0,663541452 | 0,882 | 0,718 | 9,60051E-31 | EC | 1,583966093 | 0,631326646 | 1,228412256 |
| HSPA1A8 | 3,55073E-35 | -1,803388723 | 0,54 | 0,677 | 1,29911E-30 | EC | 0,286500841 | 3,490391155 | 0,797636632 |
| RPLP16 | 9,79102E-35 | 0,371195572 | 0,999 | 0,988 | 3,58224E-30 | EC | 1,293424256 | 0,773141523 | 1,011133603 |
| MIF8 | 6,24726E-34 | 0,728425764 | 0,808 | 0,565 | 2,28569E-29 | EC | 1,656830209 | 0,603562148 | 1,430088496 |
| RPL125 | 5,8778E-33 | 0,46276717 | 0,989 | 0,963 | 2,15051E-28 | EC | 1,378182717 | 0,725593195 | 1,026998962 |
| RPL286 | 6,00008E-33 | 0,399003641 | 0,993 | 0,966 | 2,19525E-28 | EC | 1,318596942 | 0,758381859 | 1,027950311 |
| ENG1 | 1,56423E-32 | 0,721669385 | 0,79 | 0,555 | 5,72304E-28 | EC | 1,649089141 | 0,606395358 | 1,423423423 |
| TPT14 | 4,6681E-32 | 0,418998448 | 0,994 | 0,97 | 1,70792E-27 | EC | 1,336999057 | 0,747943684 | 1,024742268 |
| SPARCL13 | 3,27735E-28 | -0,478031694 | 0,964 | 0,936 | 1,19908E-23 | EC | 0,717956482 | 1,39284208 | 1,02991453 |
| COX5A7 | 4,03976E-28 | 0,600008557 | 0,558 | 0,287 | 1,47803E-23 | EC | 1,515725557 | 0,659750042 | 1,944250871 |
| MSMB9 | 7,05639E-28 | -1,740598245 | 0,037 | 0,171 | 2,58172E-23 | EC | 0,299245562 | 3,341737115 | 0,216374269 |
| PPIA6 | 1,58935E-27 | 0,453567454 | 0,942 | 0,83 | 5,81494E-23 | EC | 1,369422341 | 0,73023491 | 1,134939759 |
| RPL103 | 2,38976E-27 | 0,348013077 | 0,998 | 0,987 | 8,74343E-23 | EC | 1,272806473 | 0,785665395 | 1,011144883 |
| RPL18A8 | 4,19361E-27 | 0,355456344 | 0,984 | 0,963 | 1,53431E-22 | EC | 1,279390205 | 0,781622367 | 1,021806854 |
| HSPA1B7 | 9,99069E-27 | -1,676586144 | 0,393 | 0,553 | 3,65529E-22 | EC | 0,312821994 | 3,196706172 | 0,710669078 |

| Supplementary Table 4 - clusermarker | | | | | | | | | |
|---|---|---|---|---|---|---|---|---|---|
| | **p_val** | **avg_log2FC** | **pct.1** | **pct.2** | **p_val_adj** | **celltype** | **Foldchange_UP** | **Foldchange_DOWN** | **Ratio_pct1_pct2** |
| RPS217 | 3,87357E-26 | 0,443543179 | 0,977 | 0,928 | 1,41722E-21 | EC | 1,359940167 | 0,735326468 | 1,052801724 |
| RPS56 | 4,30622E-26 | 0,382310332 | 0,981 | 0,941 | 1,57552E-21 | EC | 1,303427494 | 0,767207999 | 1,04250797 |
| LINC01235 | 5,7833E-26 | 0,576111833 | 0,2 | 0,018 | 2,11594E-21 | EC | 1,490825946 | 0,670769115 | 11,11111111 |
| HLA-E3 | 1,49743E-25 | -0,387325396 | 0,976 | 0,963 | 5,47865E-21 | EC | 0,764545678 | 1,307966323 | 1,013499481 |
| RPL88 | 2,04196E-25 | 0,427815948 | 0,986 | 0,963 | 7,4709E-21 | EC | 1,345195584 | 0,743386324 | 1,023883697 |
| PRSS234 | 3,80287E-25 | 0,667891478 | 0,6 | 0,358 | 1,39136E-20 | EC | 1,588749287 | 0,629425932 | 1,675977654 |
| GABARAP8 | 5,15528E-25 | 0,536229467 | 0,829 | 0,622 | 1,88616E-20 | EC | 1,450177472 | 0,689570773 | 1,332797428 |
| RPS124 | 7,01804E-25 | 0,315350363 | 0,995 | 0,982 | 2,56769E-20 | EC | 1,244313802 | 0,803655797 | 1,013238289 |
| ATP5F1D9 | 1,24601E-24 | 0,563145759 | 0,668 | 0,418 | 4,55879E-20 | EC | 1,47748733 | 0,676824755 | 1,598086124 |
| EEF25 | 1,31542E-24 | 0,453460948 | 0,925 | 0,813 | 4,81272E-20 | EC | 1,369321248 | 0,730288821 | 1,137761378 |
| NPC25 | 1,54058E-24 | 0,696923975 | 0,76 | 0,56 | 5,63651E-20 | EC | 1,621044814 | 0,616886092 | 1,357142857 |
| HSP90AA15 | 1,61265E-23 | -0,710394304 | 0,844 | 0,852 | 5,90019E-19 | EC | 0,611153081 | 1,636251261 | 0,990610329 |
| PTGDS1 | 1,71228E-23 | 1,472796757 | 0,259 | 0,067 | 6,26472E-19 | EC | 2,775594393 | 0,360283189 | 3,865671642 |
| RPL388 | 1,73531E-23 | -0,504091736 | 0,832 | 0,835 | 6,349E-19 | EC | 0,705104144 | 1,418230213 | 0,996407186 |
| AL138963.47 | 3,84295E-23 | 0,833685378 | 0,42 | 0,198 | 1,40602E-18 | EC | 1,782232282 | 0,56109409 | 2,121212121 |
| BRI37 | 7,4215E-23 | 0,569061297 | 0,663 | 0,424 | 2,7153E-18 | EC | 1,483557966 | 0,674055226 | 1,563679245 |
| EEF1B24 | 1,34998E-22 | 0,462831127 | 0,92 | 0,802 | 4,93919E-18 | EC | 1,378243815 | 0,725561028 | 1,14713217 |
| B2M7 | 3,01318E-22 | -0,300455072 | 0,999 | 0,997 | 1,10243E-17 | EC | 0,811996226 | 1,231532817 | 1,002006018 |
| MT-ATP67 | 4,20312E-22 | 0,345676442 | 0,973 | 0,929 | 1,53779E-17 | EC | 1,270746663 | 0,786938915 | 1,047362756 |
| HSPG21 | 7,26294E-22 | 0,653894422 | 0,808 | 0,612 | 2,65729E-17 | EC | 1,573409742 | 0,635562354 | 1,320261438 |
| GUK14 | 1,01091E-21 | 0,592904551 | 0,826 | 0,642 | 3,69861E-17 | EC | 1,508280285 | 0,663006743 | 1,286604361 |
| IGKV1-56 | 1,72914E-21 | 1,28238545 | 0,144 | 0,003 | 6,32639E-17 | EC | 2,432408356 | 0,41111518 | 48 |
| APRT6 | 2,44946E-21 | 0,447562589 | 0,553 | 0,314 | 8,96183E-17 | EC | 1,363734302 | 0,733280668 | 1,761146497 |
| SLC25A55 | 2,63134E-21 | 0,400284785 | 0,681 | 0,427 | 9,6273E-17 | EC | 1,319768404 | 0,757708699 | 1,594847775 |
| COL4A11 | 3,92758E-21 | 0,978630596 | 0,421 | 0,208 | 1,43698E-16 | EC | 1,970594037 | 0,507461193 | 2,024038462 |
| HLA-A5 | 4,81113E-21 | -0,363410907 | 0,966 | 0,963 | 1,76025E-16 | EC | 0,777324608 | 1,28646384 | 1,003115265 |
| DDT7 | 6,95941E-21 | 0,539615508 | 0,269 | 0,086 | 2,54624E-16 | EC | 1,453585071 | 0,687954231 | 3,127906977 |
| IGKV3-207 | 9,66372E-21 | 1,688759077 | 0,181 | 0,027 | 3,53567E-16 | EC | 3,223792923 | 0,310193621 | 6,703703704 |
| VAMP55 | 2,04258E-20 | 0,494100816 | 0,656 | 0,429 | 7,47318E-16 | EC | 1,408442646 | 0,710004062 | 1,529137529 |
| CAV12 | 2,87708E-20 | -0,634237297 | 0,873 | 0,894 | 1,05264E-15 | EC | 0,644281334 | 1,552116982 | 0,976510067 |
| AP2S15 | 2,99882E-20 | 0,453907485 | 0,436 | 0,213 | 1,09718E-15 | EC | 1,36974514 | 0,73006282 | 2,046948357 |

| Supplementary Table 4 - clusermarker | | | | | | | | | |
|---|---|---|---|---|---|---|---|---|---|
| | **p_val** | **avg_log2FC** | **pct.1** | **pct.2** | **p_val_adj** | **celltype** | **Foldchange_UP** | **Foldchange_DOWN** | **Ratio_pct1_pct2** |
| CCDC85B6 | 3,08526E-20 | 0,524288074 | 0,728 | 0,523 | 1,12881E-15 | EC | 1,438223685 | 0,695302136 | 1,391969407 |
| IGHG18 | 3,46928E-20 | 1,018757707 | 0,129 | 0 | 1,2693E-15 | EC | 2,026173487 | 0,493541153 | #DIV/0! |
| VWF | 4,3352E-20 | 0,59058646 | 0,854 | 0,713 | 1,58612E-15 | EC | 1,50585876 | 0,664072904 | 1,197755961 |
| RPS194 | 5,434E-20 | 0,322520112 | 0,978 | 0,951 | 1,98814E-15 | EC | 1,250513049 | 0,799671783 | 1,028391167 |
| IGKV4-18 | 7,45408E-20 | 1,116219365 | 0,141 | 0,007 | 2,72722E-15 | EC | 2,167781525 | 0,461301099 | 20,14285714 |
| BBLN7 | 9,77811E-20 | 0,457427225 | 0,576 | 0,353 | 3,57752E-15 | EC | 1,373090985 | 0,728283858 | 1,631728045 |
| RPL36A7 | 1,28531E-19 | -0,384940398 | 0,949 | 0,939 | 4,70255E-15 | EC | 0,765810636 | 1,305805839 | 1,010649627 |
| FTH18 | 1,34291E-19 | 0,376608799 | 0,975 | 0,913 | 4,91329E-15 | EC | 1,298286512 | 0,770246006 | 1,067907996 |
| AL583785.1 | 2,0663E-19 | 0,51456439 | 0,388 | 0,176 | 7,55998E-15 | EC | 1,428562731 | 0,700004262 | 2,204545455 |
| COL4A22 | 2,18747E-19 | 0,879085023 | 0,434 | 0,232 | 8,00329E-15 | EC | 1,83920848 | 0,543712152 | 1,870689655 |
| PDLIM14 | 4,13188E-19 | 0,496198478 | 0,761 | 0,561 | 1,51173E-14 | EC | 1,410491995 | 0,708972475 | 1,356506239 |
| IGLV3-217 | 5,08757E-19 | 1,350151999 | 0,129 | 0,003 | 1,86139E-14 | EC | 2,549389838 | 0,39225072 | 43 |
| IGFBP75 | 1,34411E-18 | 0,595369611 | 0,891 | 0,795 | 4,91771E-14 | EC | 1,510859611 | 0,661874864 | 1,120754717 |
| VWA11 | 1,50666E-18 | 0,630914615 | 0,426 | 0,222 | 5,51243E-14 | EC | 1,548546404 | 0,645766893 | 1,918918919 |
| BTF35 | 1,68085E-18 | 0,366980495 | 0,917 | 0,78 | 6,14972E-14 | EC | 1,289650815 | 0,77540369 | 1,175641026 |
| UBC7 | 2,12892E-18 | -0,478681361 | 0,963 | 0,956 | 7,78908E-14 | EC | 0,717633249 | 1,393469437 | 1,007322176 |
| RPS154 | 2,17407E-18 | 0,311526369 | 0,988 | 0,973 | 7,95429E-14 | EC | 1,241020003 | 0,805788785 | 1,015416238 |
| MARCKSL15 | 2,28895E-18 | 0,511987191 | 0,533 | 0,309 | 8,37459E-14 | EC | 1,426013056 | 0,701255852 | 1,724919094 |
| NUDT144 | 5,00677E-18 | 0,39017226 | 0,326 | 0,136 | 1,83183E-13 | EC | 1,310549876 | 0,763038491 | 2,397058824 |
| TNFRSF6B2 | 1,06265E-17 | 0,972099719 | 0,279 | 0,106 | 3,88792E-13 | EC | 1,961693595 | 0,509763606 | 2,632075472 |
| GNG56 | 1,17722E-17 | 0,399108206 | 0,792 | 0,6 | 4,30711E-13 | EC | 1,318692517 | 0,758326894 | 1,32 |
| RPS94 | 1,43175E-17 | 0,276779187 | 0,979 | 0,945 | 5,23836E-13 | EC | 1,211487221 | 0,825431736 | 1,035978836 |
| COL18A13 | 2,1445E-17 | 0,562739861 | 0,494 | 0,284 | 7,84607E-13 | EC | 1,477071702 | 0,677015204 | 1,73943662 |
| DUSP235 | 2,54802E-17 | 0,643195059 | 0,593 | 0,385 | 9,32243E-13 | EC | 1,561784132 | 0,640293354 | 1,54025974 |
| COL15A1 | 3,38169E-17 | 0,701003774 | 0,608 | 0,408 | 1,23726E-12 | EC | 1,625635457 | 0,615144063 | 1,490196078 |
| GPX49 | 3,43273E-17 | 0,420687299 | 0,708 | 0,476 | 1,25593E-12 | EC | 1,338565094 | 0,747068637 | 1,487394958 |
| SNHG75 | 5,11754E-17 | 0,424118695 | 0,712 | 0,494 | 1,87236E-12 | EC | 1,34175261 | 0,745293873 | 1,441295547 |
| POSTN2 | 5,16781E-17 | 0,635857051 | 0,461 | 0,25 | 1,89075E-12 | EC | 1,553860566 | 0,643558387 | 1,844 |
| C1orf1224 | 6,12521E-17 | 0,360576527 | 0,276 | 0,104 | 2,24103E-12 | EC | 1,283938881 | 0,778853273 | 2,653846154 |
| SNHG294 | 9,78906E-17 | 0,412678315 | 0,868 | 0,739 | 3,58152E-12 | EC | 1,331154766 | 0,75122745 | 1,174560217 |
| SH3BGRL37 | 1,04373E-16 | 0,40715583 | 0,849 | 0,681 | 3,81868E-12 | EC | 1,326068985 | 0,75410858 | 1,246696035 |

| Supplementary Table 4 - clusermarker | | | | | | | | | |
|---|---|---|---|---|---|---|---|---|---|
| | **p_val** | **avg_log2FC** | **pct.1** | **pct.2** | **p_val_adj** | **celltype** | **Foldchange_UP** | **Foldchange_DOWN** | **Ratio_pct1_pct2** |
| MT-CO33 | 1,11779E-16 | 0,272669309 | 0,982 | 0,958 | 4,08967E-12 | EC | 1,208040907 | 0,827786538 | 1,025052192 |
| EIF3F5 | 1,18447E-16 | 0,392795967 | 0,687 | 0,481 | 4,3336E-12 | EC | 1,312935431 | 0,761652079 | 1,428274428 |
| COX4I17 | 1,61307E-16 | 0,415337083 | 0,888 | 0,756 | 5,90176E-12 | EC | 1,333610236 | 0,749844274 | 1,174603175 |
| NPDC11 | 1,66649E-16 | 0,430713344 | 0,83 | 0,654 | 6,09718E-12 | EC | 1,347899885 | 0,741894863 | 1,26911315 |
| MT-ND39 | 2,70835E-16 | -0,30003814 | 0,966 | 0,951 | 9,90905E-12 | EC | 0,812230923 | 1,231176961 | 1,015772871 |
| PRDX54 | 5,62936E-16 | 0,318039859 | 0,579 | 0,37 | 2,05961E-11 | EC | 1,246635636 | 0,802159004 | 1,564864865 |
| MYL12B5 | 6,30293E-16 | 0,341605463 | 0,87 | 0,719 | 2,30605E-11 | EC | 1,26716594 | 0,789162626 | 1,210013908 |
| HINT18 | 6,72371E-16 | 0,388902121 | 0,822 | 0,649 | 2,46E-11 | EC | 1,309396584 | 0,763710561 | 1,266563945 |
| TIMP34 | 7,53266E-16 | -0,446191473 | 0,783 | 0,839 | 2,75598E-11 | EC | 0,733977899 | 1,362438844 | 0,933253874 |
| GADD45GIP13 | 9,68104E-16 | 0,404587192 | 0,372 | 0,182 | 3,542E-11 | EC | 1,323710094 | 0,755452424 | 2,043956044 |
| LTF6 | 1,12059E-15 | -0,766160751 | 0,037 | 0,126 | 4,09989E-11 | EC | 0,587980106 | 1,700737813 | 0,293650794 |
| MT-ND4L6 | 1,18667E-15 | -0,329770953 | 0,924 | 0,946 | 4,34167E-11 | EC | 0,795662796 | 1,256813822 | 0,976744186 |
| ADRM14 | 1,298E-15 | 0,306620098 | 0,378 | 0,19 | 4,749E-11 | EC | 1,23680675 | 0,80853375 | 1,989473684 |
| HLA-C7 | 1,50413E-15 | -0,393067783 | 0,954 | 0,948 | 5,50317E-11 | EC | 0,76150859 | 1,313182822 | 1,006329114 |
| EDF15 | 1,63774E-15 | 0,366776199 | 0,744 | 0,539 | 5,992E-11 | EC | 1,289468204 | 0,7755135 | 1,380333952 |
| NOVA11 | 1,64496E-15 | 0,30559413 | 0,135 | 0,018 | 6,01842E-11 | EC | 1,235927511 | 0,809108941 | 7,5 |
| RABAC17 | 1,71632E-15 | 0,299225948 | 0,664 | 0,454 | 6,27951E-11 | EC | 1,230484042 | 0,812688313 | 1,462555066 |
| NECTIN23 | 1,96372E-15 | 0,355866978 | 0,486 | 0,281 | 7,18467E-11 | EC | 1,27975441 | 0,781399925 | 1,729537367 |
| RACK14 | 2,84963E-15 | 0,285070102 | 0,962 | 0,926 | 1,04259E-10 | EC | 1,218469469 | 0,820701729 | 1,03887689 |
| MGMT4 | 2,85563E-15 | 0,407054861 | 0,333 | 0,16 | 1,04479E-10 | EC | 1,325976182 | 0,75416136 | 2,08125 |
| CLIC15 | 3,18455E-15 | 0,377919159 | 0,804 | 0,629 | 1,16513E-10 | EC | 1,299466245 | 0,76954673 | 1,278219396 |
| FAM174C5 | 3,40758E-15 | 0,361089905 | 0,254 | 0,097 | 1,24673E-10 | EC | 1,284395847 | 0,778576171 | 2,618556701 |
| TSPAN132 | 8,07278E-15 | 0,354890789 | 0,363 | 0,182 | 2,95359E-10 | EC | 1,278888767 | 0,781928832 | 1,994505495 |
| C71 | 1,24783E-14 | 0,451714971 | 0,281 | 0,121 | 4,56545E-10 | EC | 1,367665072 | 0,731173166 | 2,32231405 |
| ITM2A3 | 1,72008E-14 | -0,748300353 | 0,482 | 0,575 | 6,29325E-10 | EC | 0,595304476 | 1,679812668 | 0,83826087 |
| CAVIN21 | 2,02983E-14 | -0,764190548 | 0,566 | 0,639 | 7,42655E-10 | EC | 0,588783624 | 1,698416801 | 0,885758998 |
| CHCHD27 | 2,46626E-14 | 0,373499016 | 0,784 | 0,593 | 9,02331E-10 | EC | 1,295491021 | 0,77190809 | 1,322091062 |
| REXO23 | 2,90889E-14 | 0,315100052 | 0,419 | 0,235 | 1,06427E-09 | EC | 1,244097929 | 0,803795245 | 1,782978723 |
| AKR1C14 | 3,92859E-14 | -0,594379718 | 0,056 | 0,153 | 1,43735E-09 | EC | 0,66232916 | 1,509823302 | 0,366013072 |
| STC1 | 4,57159E-14 | -0,856244891 | 0,103 | 0,22 | 1,67261E-09 | EC | 0,552388469 | 1,810320194 | 0,468181818 |
| SLC29A12 | 5,22375E-14 | 0,360450284 | 0,409 | 0,227 | 1,91121E-09 | EC | 1,283826534 | 0,77892143 | 1,801762115 |

| Supplementary Table 4 - clusermarker | | | | | | | | | |
|---|---|---|---|---|---|---|---|---|---|
| | **p_val** | **avg_log2FC** | **pct.1** | **pct.2** | **p_val_adj** | **celltype** | **Foldchange_UP** | **Foldchange_DOWN** | **Ratio_pct1_pct2** |
| HTRA14 | 5,55422E-14 | 0,482444831 | 0,414 | 0,239 | 2,03212E-09 | EC | 1,397109241 | 0,715763643 | 1,732217573 |
| LDHB3 | 5,91172E-14 | 0,317121564 | 0,477 | 0,284 | 2,16292E-09 | EC | 1,245842388 | 0,802669751 | 1,679577465 |
| ACTG13 | 6,69014E-14 | 0,295829914 | 0,973 | 0,933 | 2,44772E-09 | EC | 1,227590949 | 0,814603595 | 1,042872454 |
| FKBP1A4 | 7,06607E-14 | 0,431373726 | 0,803 | 0,647 | 2,58526E-09 | EC | 1,348517016 | 0,741555344 | 1,241112828 |
| ZNF385D | 8,06011E-14 | 0,407607452 | 0,48 | 0,287 | 2,94895E-09 | EC | 1,326484163 | 0,753872551 | 1,672473868 |
| MMP21 | 9,25936E-14 | 0,573217732 | 0,301 | 0,141 | 3,38772E-09 | EC | 1,487838291 | 0,672116053 | 2,134751773 |
| COX6A16 | 9,99606E-14 | 0,351647822 | 0,741 | 0,538 | 3,65726E-09 | EC | 1,276017241 | 0,783688471 | 1,37732342 |
| TIMP15 | 1,01369E-13 | 0,388164398 | 0,753 | 0,555 | 3,7088E-09 | EC | 1,308727195 | 0,764101184 | 1,356756757 |
| SYNE23 | 1,18944E-13 | -0,603969285 | 0,64 | 0,684 | 4,35181E-09 | EC | 0,65794127 | 1,519892497 | 0,935672515 |
| HLA-DQB12 | 1,23295E-13 | -0,734894723 | 0,394 | 0,492 | 4,511E-09 | EC | 0,600861869 | 1,664276019 | 0,800813008 |
| BAX8 | 1,61331E-13 | 0,384323267 | 0,295 | 0,143 | 5,9026E-09 | EC | 1,305247384 | 0,766138291 | 2,062937063 |
| LAMTOR52 | 1,61697E-13 | 0,292085331 | 0,416 | 0,237 | 5,91601E-09 | EC | 1,22440881 | 0,816720683 | 1,755274262 |
| CLEC3B2 | 1,83076E-13 | -0,69227182 | 0,596 | 0,682 | 6,6982E-09 | EC | 0,618878531 | 1,615825967 | 0,873900293 |
| RPL37A8 | 2,19705E-13 | -0,288688278 | 0,951 | 0,928 | 8,03835E-09 | EC | 0,818646047 | 1,221529138 | 1,024784483 |
| CLTB5 | 2,36868E-13 | 0,331867974 | 0,363 | 0,195 | 8,66628E-09 | EC | 1,258641985 | 0,794507105 | 1,861538462 |
| ARPC5L5 | 2,37877E-13 | 0,361307665 | 0,345 | 0,185 | 8,7032E-09 | EC | 1,284589728 | 0,778458661 | 1,864864865 |
| CUTA5 | 2,49244E-13 | 0,358657064 | 0,462 | 0,276 | 9,11909E-09 | EC | 1,282231773 | 0,779890205 | 1,673913043 |
| HLA-DRB54 | 2,51138E-13 | -0,645989042 | 0,552 | 0,612 | 9,18839E-09 | EC | 0,639054535 | 1,564811678 | 0,901960784 |
| NDUFB25 | 2,58244E-13 | 0,313264069 | 0,571 | 0,366 | 9,44837E-09 | EC | 1,242515689 | 0,804818811 | 1,56010929 |
| ZNF7065 | 2,79896E-13 | 0,352677215 | 0,446 | 0,264 | 1,02405E-08 | EC | 1,27692803 | 0,783129492 | 1,689393939 |
| ARF56 | 3,65594E-13 | 0,342879125 | 0,491 | 0,301 | 1,3376E-08 | EC | 1,268285132 | 0,788466233 | 1,631229236 |
| S100A115 | 3,66656E-13 | 0,299623899 | 0,853 | 0,696 | 1,34149E-08 | EC | 1,230823504 | 0,812464173 | 1,225574713 |
| SMDT17 | 4,53946E-13 | 0,374354306 | 0,507 | 0,329 | 1,66085E-08 | EC | 1,29625927 | 0,771450606 | 1,541033435 |
| NDUFS86 | 5,13435E-13 | 0,293603028 | 0,36 | 0,195 | 1,8785E-08 | EC | 1,22569755 | 0,815861955 | 1,846153846 |
| BRK12 | 5,52933E-13 | 0,305054823 | 0,604 | 0,408 | 2,02301E-08 | EC | 1,235465584 | 0,809411458 | 1,480392157 |
| CTSB6 | 6,19162E-13 | 0,304546838 | 0,546 | 0,35 | 2,26533E-08 | EC | 1,235030643 | 0,809696509 | 1,56 |
| LDHA5 | 7,09502E-13 | 0,313691349 | 0,75 | 0,566 | 2,59586E-08 | EC | 1,242883737 | 0,804580485 | 1,325088339 |
| TUFM5 | 7,7955E-13 | 0,403221948 | 0,41 | 0,245 | 2,85214E-08 | EC | 1,32245804 | 0,756167659 | 1,673469388 |
| ABL21 | 8,31244E-13 | 0,456546301 | 0,408 | 0,232 | 3,04127E-08 | EC | 1,372252817 | 0,728728692 | 1,75862069 |
| ATP5MC23 | 9,92447E-13 | 0,288865607 | 0,873 | 0,739 | 3,63106E-08 | EC | 1,221679292 | 0,818545429 | 1,181326116 |
| TIMM133 | 1,00651E-12 | 0,334897708 | 0,306 | 0,155 | 3,6825E-08 | EC | 1,261287976 | 0,79284035 | 1,974193548 |

| Supplementary Table 4 - clusermarker | | | | | | | | | |
|---|---|---|---|---|---|---|---|---|---|
| | **p_val** | **avg_log2FC** | **pct.1** | **pct.2** | **p_val_adj** | **celltype** | **Foldchange_UP** | **Foldchange_DOWN** | **Ratio_pct1_pct2** |
| TRAPPC56 | 1,20917E-12 | 0,356937136 | 0,361 | 0,198 | 4,424E-08 | EC | 1,280704055 | 0,780820515 | 1,823232323 |
| PPP1R14B5 | 1,24322E-12 | 0,334072157 | 0,465 | 0,286 | 4,54857E-08 | EC | 1,260566438 | 0,793294165 | 1,625874126 |
| NDUFAB16 | 1,33875E-12 | 0,291681249 | 0,385 | 0,213 | 4,89809E-08 | EC | 1,224065916 | 0,816949469 | 1,807511737 |
| GIMAP71 | 1,53729E-12 | -0,690924695 | 0,694 | 0,709 | 5,62447E-08 | EC | 0,619456682 | 1,614317884 | 0,978843441 |
| SERPINB64 | 1,71267E-12 | 0,269756066 | 0,472 | 0,289 | 6,26613E-08 | EC | 1,205603965 | 0,829459781 | 1,633217993 |
| SEC61B8 | 1,86856E-12 | 0,261824288 | 0,637 | 0,44 | 6,83649E-08 | EC | 1,198993874 | 0,834032618 | 1,447727273 |
| SNCA | 1,95954E-12 | 0,305123729 | 0,197 | 0,071 | 7,16937E-08 | EC | 1,235524594 | 0,8093728 | 2,774647887 |
| PSMB52 | 2,06353E-12 | 0,319752692 | 0,438 | 0,262 | 7,54983E-08 | EC | 1,248116578 | 0,801207209 | 1,671755725 |
| PSMA76 | 2,15825E-12 | 0,309027697 | 0,666 | 0,467 | 7,89639E-08 | EC | 1,238872481 | 0,807185578 | 1,426124197 |
| TKT2 | 2,57406E-12 | 0,26084629 | 0,305 | 0,153 | 9,41771E-08 | EC | 1,198181356 | 0,834598197 | 1,993464052 |
| MZT2B5 | 2,59362E-12 | 0,327078248 | 0,537 | 0,35 | 9,48929E-08 | EC | 1,254470242 | 0,79714924 | 1,534285714 |
| PAXX4 | 2,61871E-12 | 0,341419996 | 0,331 | 0,176 | 9,58106E-08 | EC | 1,267003049 | 0,789264083 | 1,880681818 |
| EPN1 | 2,71891E-12 | 0,312171259 | 0,35 | 0,188 | 9,94767E-08 | EC | 1,241574867 | 0,805428675 | 1,861702128 |
| SERF25 | 2,81777E-12 | 0,273547005 | 0,924 | 0,825 | 1,03094E-07 | EC | 1,20877607 | 0,827283088 | 1,12 |
| CTSH2 | 2,94139E-12 | -0,648142041 | 0,563 | 0,618 | 1,07617E-07 | EC | 0,638101557 | 1,56714866 | 0,911003236 |
| HEBP12 | 3,66113E-12 | 0,309094214 | 0,327 | 0,171 | 1,3395E-07 | EC | 1,238929603 | 0,807148362 | 1,912280702 |
| NDUFA123 | 3,88013E-12 | 0,252190118 | 0,41 | 0,235 | 1,41962E-07 | EC | 1,19101379 | 0,839620841 | 1,744680851 |
| PARVB2 | 3,89699E-12 | 0,306931066 | 0,331 | 0,175 | 1,42579E-07 | EC | 1,237073368 | 0,808359493 | 1,891428571 |
| CSTB5 | 4,05839E-12 | 0,266469003 | 0,526 | 0,345 | 1,48484E-07 | EC | 1,202860221 | 0,831351792 | 1,524637681 |
| IGHA18 | 4,57084E-12 | 0,589909613 | 0,15 | 0,04 | 1,67233E-07 | EC | 1,505152445 | 0,66438453 | 3,75 |
| COL6A22 | 4,63153E-12 | 0,376951629 | 0,227 | 0,097 | 1,69454E-07 | EC | 1,298595062 | 0,770062993 | 2,340206186 |
| SON9 | 4,95653E-12 | -0,444670976 | 0,744 | 0,755 | 1,81345E-07 | EC | 0,734751867 | 1,361003687 | 0,985430464 |
| EIF3G2 | 5,02232E-12 | 0,293461299 | 0,659 | 0,455 | 1,83752E-07 | EC | 1,225577145 | 0,815942108 | 1,448351648 |
| TXNL4A5 | 5,36384E-12 | 0,281630172 | 0,32 | 0,166 | 1,96247E-07 | EC | 1,215567639 | 0,822660927 | 1,927710843 |
| ATP6V0C4 | 5,72985E-12 | 0,289314715 | 0,544 | 0,346 | 2,09638E-07 | EC | 1,222059658 | 0,818290657 | 1,572254335 |
| RTRAF4 | 5,77885E-12 | 0,284829231 | 0,521 | 0,338 | 2,11431E-07 | EC | 1,218266051 | 0,820838764 | 1,541420118 |
| VKORC14 | 6,3834E-12 | 0,333304024 | 0,489 | 0,318 | 2,33549E-07 | EC | 1,259895454 | 0,793716651 | 1,537735849 |
| IMPDH23 | 6,43638E-12 | 0,315896811 | 0,443 | 0,269 | 2,35488E-07 | EC | 1,244785199 | 0,803351455 | 1,646840149 |
| LIFR1 | 7,32314E-12 | -0,600706799 | 0,57 | 0,632 | 2,67932E-07 | EC | 0,659430811 | 1,516459322 | 0,901898734 |
| TFF33 | 7,38902E-12 | -1,410608576 | 0,036 | 0,111 | 2,70342E-07 | EC | 0,37615298 | 2,65849283 | 0,324324324 |
| NAMPT2 | 7,5616E-12 | 0,325095075 | 0,457 | 0,286 | 2,76656E-07 | EC | 1,252746994 | 0,798245779 | 1,597902098 |

| Supplementary Table 4 - clusermarker | | | | | | | | | |
|---|---|---|---|---|---|---|---|---|---|
| | **p_val** | **avg_log2FC** | **pct.1** | **pct.2** | **p_val_adj** | **celltype** | **Foldchange_UP** | **Foldchange_DOWN** | **Ratio_pct1_pct2** |
| NENF5 | 7,85976E-12 | 0,348032971 | 0,415 | 0,25 | 2,87565E-07 | EC | 1,272824024 | 0,785654561 | 1,66 |
| VIM-AS14 | 8,46953E-12 | 0,375345271 | 0,227 | 0,097 | 3,09875E-07 | EC | 1,297149956 | 0,770920891 | 2,340206186 |
| PSMG23 | 8,8521E-12 | 0,299721067 | 0,364 | 0,205 | 3,23872E-07 | EC | 1,230906405 | 0,812409454 | 1,775609756 |
| ATP5MC33 | 9,67393E-12 | 0,283492588 | 0,4 | 0,235 | 3,5394E-07 | EC | 1,217137862 | 0,821599616 | 1,70212766 |
| CD994 | 1,04179E-11 | 0,301061389 | 0,815 | 0,629 | 3,81158E-07 | EC | 1,232050498 | 0,811655043 | 1,295707472 |
| NXT12 | 1,04864E-11 | 0,262543297 | 0,214 | 0,086 | 3,83667E-07 | EC | 1,199591576 | 0,833617057 | 2,488372093 |
| RBPMS3 | 1,05608E-11 | 0,359489383 | 0,464 | 0,296 | 3,86388E-07 | EC | 1,282971731 | 0,779440401 | 1,567567568 |
| TRIR6 | 1,06809E-11 | 0,340672906 | 0,63 | 0,455 | 3,90781E-07 | EC | 1,26634711 | 0,789672904 | 1,384615385 |
| ATP5F1C2 | 1,24342E-11 | 0,283589252 | 0,476 | 0,299 | 4,54931E-07 | EC | 1,217219417 | 0,821544568 | 1,591973244 |
| PPP4C6 | 1,31613E-11 | 0,309014858 | 0,327 | 0,176 | 4,81533E-07 | EC | 1,238861457 | 0,807192761 | 1,857954545 |
| PHB22 | 1,34087E-11 | 0,278786353 | 0,514 | 0,333 | 4,90585E-07 | EC | 1,21317389 | 0,824284143 | 1,543543544 |
| EIF3K3 | 1,41389E-11 | 0,302692535 | 0,785 | 0,602 | 5,17301E-07 | EC | 1,233444272 | 0,810737885 | 1,303986711 |
| THBS14 | 1,41538E-11 | 0,475324743 | 0,288 | 0,146 | 5,17844E-07 | EC | 1,390231118 | 0,71930486 | 1,97260274 |
| LAMA52 | 1,66513E-11 | 0,348277271 | 0,37 | 0,212 | 6,09222E-07 | EC | 1,273039577 | 0,785521533 | 1,745283019 |
| HLA-DQA11 | 1,74365E-11 | -0,615766565 | 0,236 | 0,353 | 6,37948E-07 | EC | 0,652583056 | 1,532371996 | 0,668555241 |
| PCBP16 | 1,77002E-11 | 0,317731772 | 0,624 | 0,44 | 6,47598E-07 | EC | 1,246369446 | 0,802330323 | 1,418181818 |
| RNF74 | 2,31072E-11 | 0,309426524 | 0,451 | 0,286 | 8,45423E-07 | EC | 1,23921501 | 0,806962466 | 1,576923077 |
| ALKBH75 | 2,78769E-11 | 0,346236129 | 0,465 | 0,304 | 1,01993E-06 | EC | 1,271239739 | 0,786633685 | 1,529605263 |
| RAI141 | 2,99402E-11 | 0,335234887 | 0,405 | 0,247 | 1,09542E-06 | EC | 1,261582792 | 0,792655073 | 1,639676113 |
| MRPL373 | 3,08733E-11 | 0,272084105 | 0,245 | 0,113 | 1,12956E-06 | EC | 1,207550986 | 0,828122383 | 2,168141593 |
| SF3B55 | 3,24536E-11 | 0,302872655 | 0,464 | 0,296 | 1,18738E-06 | EC | 1,233598277 | 0,810636671 | 1,567567568 |
| BACE24 | 3,25533E-11 | 0,292110821 | 0,508 | 0,331 | 1,19103E-06 | EC | 1,224430444 | 0,816706253 | 1,534743202 |
| FAM241A | 3,30531E-11 | 0,27930504 | 0,527 | 0,341 | 1,20931E-06 | EC | 1,213610136 | 0,823987845 | 1,545454545 |
| TMEM2197 | 3,352E-11 | 0,292108876 | 0,631 | 0,442 | 1,22639E-06 | EC | 1,224428793 | 0,816707354 | 1,42760181 |
| SELENOH4 | 3,36467E-11 | 0,257595716 | 0,542 | 0,358 | 1,23103E-06 | EC | 1,195484741 | 0,836480773 | 1,51396648 |
| TPI16 | 3,56399E-11 | 0,325179318 | 0,706 | 0,523 | 1,30396E-06 | EC | 1,252820147 | 0,798199169 | 1,349904398 |
| RPS102 | 3,69208E-11 | 0,341465735 | 0,189 | 0,076 | 1,35082E-06 | EC | 1,267043219 | 0,789239061 | 2,486842105 |
| SPNS2 | 3,73108E-11 | 0,323569898 | 0,329 | 0,182 | 1,36509E-06 | EC | 1,251423323 | 0,799090109 | 1,807692308 |
| CD637 | 4,03709E-11 | 0,282025477 | 0,816 | 0,637 | 1,47705E-06 | EC | 1,215900755 | 0,822435545 | 1,28100471 |
| PYCARD5 | 4,92958E-11 | 0,277838174 | 0,269 | 0,131 | 1,80359E-06 | EC | 1,21237682 | 0,824826063 | 2,053435115 |
| COTL16 | 5,48633E-11 | 0,363439963 | 0,326 | 0,178 | 2,00728E-06 | EC | 1,28648975 | 0,777308952 | 1,831460674 |

| Supplementary Table 4 - clusermarker | | | | | | | | | |
|---|---|---|---|---|---|---|---|---|---|
| | **p_val** | **avg_log2FC** | **pct.1** | **pct.2** | **p_val_adj** | **celltype** | **Foldchange_UP** | **Foldchange_DOWN** | **Ratio_pct1_pct2** |
| MPZL21 | 5,48848E-11 | 0,2763815 | 0,288 | 0,145 | 2,00807E-06 | EC | 1,211153314 | 0,825659302 | 1,986206897 |
| PRELID16 | 5,57321E-11 | 0,311102642 | 0,385 | 0,227 | 2,03907E-06 | EC | 1,240655562 | 0,806025484 | 1,696035242 |
| CST35 | 6,07764E-11 | 0,433270008 | 0,921 | 0,82 | 2,22363E-06 | EC | 1,350290675 | 0,740581282 | 1,123170732 |
| ARHGDIB5 | 6,75684E-11 | 0,275665632 | 0,55 | 0,375 | 2,47212E-06 | EC | 1,210552487 | 0,826069097 | 1,466666667 |
| EBPL3 | 7,07937E-11 | 0,291855564 | 0,394 | 0,239 | 2,59013E-06 | EC | 1,224213824 | 0,816850766 | 1,648535565 |
| PRXL2A4 | 7,84052E-11 | 0,252385486 | 0,366 | 0,21 | 2,86861E-06 | EC | 1,191175087 | 0,839507148 | 1,742857143 |
| SELENOM7 | 7,86573E-11 | 0,331093339 | 0,658 | 0,474 | 2,87784E-06 | EC | 1,257966357 | 0,794933819 | 1,388185654 |
| FEZ21 | 7,88264E-11 | 0,307909423 | 0,384 | 0,23 | 2,88402E-06 | EC | 1,237912568 | 0,807811493 | 1,669565217 |
| PPIB5 | 8,26361E-11 | 0,296540399 | 0,713 | 0,565 | 3,02341E-06 | EC | 1,22819565 | 0,814202525 | 1,261946903 |
| ZYX4 | 8,52119E-11 | 0,255673154 | 0,36 | 0,203 | 3,11765E-06 | EC | 1,193892677 | 0,837596225 | 1,773399015 |
| AURKAIP14 | 8,54735E-11 | 0,265168651 | 0,376 | 0,224 | 3,12722E-06 | EC | 1,201776529 | 0,832101456 | 1,678571429 |
| SF3B64 | 9,38087E-11 | 0,294572614 | 0,5 | 0,334 | 3,43218E-06 | EC | 1,226521577 | 0,815313826 | 1,497005988 |
| EPB41L31 | 9,57619E-11 | 0,282380028 | 0,254 | 0,121 | 3,50364E-06 | EC | 1,216199607 | 0,822233451 | 2,099173554 |
| MEOX1 | 9,66098E-11 | 0,280116442 | 0,439 | 0,266 | 3,53466E-06 | EC | 1,214292888 | 0,823524547 | 1,65037594 |
| POLE44 | 1,09858E-10 | 0,301421092 | 0,44 | 0,276 | 4,01939E-06 | EC | 1,23235772 | 0,811452701 | 1,594202899 |
| MRPL415 | 1,15077E-10 | 0,295711851 | 0,303 | 0,163 | 4,21033E-06 | EC | 1,227490493 | 0,814670261 | 1,858895706 |
| FAM118A3 | 1,19528E-10 | 0,312996947 | 0,14 | 0,042 | 4,37318E-06 | EC | 1,242285653 | 0,804967841 | 3,333333333 |
| SERPING14 | 1,22328E-10 | -0,542825855 | 0,483 | 0,56 | 4,47562E-06 | EC | 0,686425068 | 1,456823253 | 0,8625 |
| COPE6 | 1,22828E-10 | 0,258772135 | 0,365 | 0,207 | 4,49392E-06 | EC | 1,196459975 | 0,835798958 | 1,763285024 |
| PON2 | 1,28024E-10 | 0,264017499 | 0,319 | 0,176 | 4,68402E-06 | EC | 1,200817993 | 0,83276567 | 1,8125 |
| ADIRF5 | 1,50116E-10 | -0,419529041 | 0,857 | 0,859 | 5,49231E-06 | EC | 0,747668657 | 1,337490867 | 0,997671711 |
| MDK5 | 1,52554E-10 | 0,289735672 | 0,504 | 0,333 | 5,58149E-06 | EC | 1,222416289 | 0,818051927 | 1,513513514 |
| EMD4 | 1,65017E-10 | 0,250950734 | 0,297 | 0,158 | 6,03747E-06 | EC | 1,189991059 | 0,840342448 | 1,879746835 |
| EVA1B1 | 1,67737E-10 | 0,278805393 | 0,26 | 0,131 | 6,13701E-06 | EC | 1,2131899 | 0,824273265 | 1,984732824 |
| NDUFB107 | 1,93039E-10 | 0,305609994 | 0,494 | 0,328 | 7,0627E-06 | EC | 1,235941102 | 0,809100044 | 1,506097561 |
| MT-ND23 | 2,28456E-10 | -0,285826348 | 0,954 | 0,943 | 8,35852E-06 | EC | 0,820271638 | 1,219108346 | 1,011664899 |
| GSN5 | 2,61428E-10 | -0,414688371 | 0,828 | 0,808 | 9,56485E-06 | EC | 0,75018152 | 1,33301071 | 1,024752475 |
| RPL22L12 | 2,84158E-10 | 0,256382996 | 0,45 | 0,287 | 1,03965E-05 | EC | 1,194480246 | 0,837184209 | 1,567944251 |
| UQCRH3 | 2,92856E-10 | 0,299188362 | 0,677 | 0,501 | 1,07147E-05 | EC | 1,230451985 | 0,812709486 | 1,351297405 |
| ACTB9 | 3,07805E-10 | 0,311425552 | 0,984 | 0,948 | 1,12617E-05 | EC | 1,240933282 | 0,805845096 | 1,037974684 |
| PHPT17 | 3,25298E-10 | 0,302119459 | 0,366 | 0,213 | 1,19017E-05 | EC | 1,232954413 | 0,811059995 | 1,718309859 |

| Supplementary Table 4 - clusermarker | | | | | | | | | |
|---|---|---|---|---|---|---|---|---|---|
| | **p_val** | **avg_log2FC** | **pct.1** | **pct.2** | **p_val_adj** | **celltype** | **Foldchange_UP** | **Foldchange_DOWN** | **Ratio_pct1_pct2** |
| LTBP22 | 3,29626E-10 | 0,261257852 | 0,102 | 0,02 | 1,206E-05 | EC | 1,198523214 | 0,834360143 | 5,1 |
| WFDC26 | 3,36246E-10 | -0,520109359 | 0,04 | 0,109 | 1,23022E-05 | EC | 0,697318973 | 1,434063949 | 0,366972477 |
| UPP11 | 3,60745E-10 | 0,329884162 | 0,387 | 0,239 | 1,31986E-05 | EC | 1,256912449 | 0,795600362 | 1,619246862 |
| PLEKHO11 | 4,37794E-10 | 0,259496227 | 0,173 | 0,067 | 1,60176E-05 | EC | 1,197060631 | 0,835379574 | 2,582089552 |
| TMUB11 | 4,61545E-10 | 0,258467764 | 0,188 | 0,079 | 1,68866E-05 | EC | 1,196207579 | 0,835975308 | 2,379746835 |
| GTF3A5 | 4,72227E-10 | 0,274215415 | 0,326 | 0,185 | 1,72774E-05 | EC | 1,209336233 | 0,826899891 | 1,762162162 |
| SRP19 | 4,88015E-10 | 0,259189511 | 0,256 | 0,128 | 1,7855E-05 | EC | 1,196806164 | 0,835557194 | 2 |
| LAMB11 | 5,43462E-10 | 0,325227669 | 0,366 | 0,217 | 1,98837E-05 | EC | 1,252862135 | 0,798172418 | 1,686635945 |
| CHMP2A4 | 5,59431E-10 | 0,252965563 | 0,426 | 0,266 | 2,04679E-05 | EC | 1,19165413 | 0,839169668 | 1,601503759 |
| PPM1G6 | 6,24038E-10 | 0,282356506 | 0,353 | 0,21 | 2,28317E-05 | EC | 1,216179778 | 0,822246857 | 1,680952381 |
| TSPAN152 | 6,34336E-10 | 0,262761934 | 0,156 | 0,057 | 2,32085E-05 | EC | 1,199773385 | 0,833490734 | 2,736842105 |
| SIGIRR4 | 6,44215E-10 | 0,304871562 | 0,395 | 0,249 | 2,35699E-05 | EC | 1,235308657 | 0,809514281 | 1,586345382 |
| EEF1G4 | 6,53668E-10 | 0,25924977 | 0,908 | 0,822 | 2,39158E-05 | EC | 1,196856153 | 0,835522295 | 1,104622871 |
| ENO13 | 6,57727E-10 | 0,286309354 | 0,627 | 0,467 | 2,40643E-05 | EC | 1,219516564 | 0,819997062 | 1,34261242 |
| TMEM2564 | 6,59427E-10 | 0,274353644 | 0,389 | 0,24 | 2,41265E-05 | EC | 1,209452109 | 0,826820667 | 1,620833333 |
| CTHRC11 | 6,87097E-10 | 0,262978223 | 0,301 | 0,166 | 2,51388E-05 | EC | 1,19995327 | 0,833365786 | 1,813253012 |
| HSP90AB16 | 7,99679E-10 | -0,395873366 | 0,95 | 0,924 | 2,92579E-05 | EC | 0,760029138 | 1,31573903 | 1,028138528 |
| TIMM102 | 8,57801E-10 | 0,302263529 | 0,289 | 0,156 | 3,13844E-05 | EC | 1,233077544 | 0,810979005 | 1,852564103 |
| NDUFA117 | 9,23962E-10 | 0,283571688 | 0,627 | 0,457 | 3,3805E-05 | EC | 1,217204597 | 0,821554571 | 1,371991247 |
| SRSF97 | 9,29128E-10 | 0,286650972 | 0,463 | 0,309 | 3,3994E-05 | EC | 1,219805369 | 0,819802917 | 1,498381877 |
| PCDH17 | 1,02353E-09 | 0,330432701 | 0,423 | 0,279 | 3,74479E-05 | EC | 1,257390442 | 0,795297918 | 1,516129032 |
| MALAT15 | 1,11965E-09 | -0,275231856 | 0,986 | 0,99 | 4,09648E-05 | EC | 0,82631751 | 1,210188563 | 0,995959596 |
| REX1BD6 | 1,15772E-09 | 0,277652488 | 0,421 | 0,271 | 4,23576E-05 | EC | 1,212220788 | 0,824932232 | 1,553505535 |
| LGALS16 | 1,18181E-09 | 0,513921223 | 0,509 | 0,351 | 4,32389E-05 | EC | 1,427926007 | 0,7003164 | 1,45014245 |
| GALNT18 | 1,21053E-09 | 0,253144663 | 0,326 | 0,187 | 4,42895E-05 | EC | 1,191802074 | 0,839065497 | 1,743315508 |
| RAMP31 | 1,36406E-09 | 0,449423952 | 0,635 | 0,482 | 4,99067E-05 | EC | 1,365494925 | 0,732335201 | 1,317427386 |
| AC020916.15 | 1,39546E-09 | 0,266279594 | 0,32 | 0,188 | 5,10558E-05 | EC | 1,20270231 | 0,831460946 | 1,70212766 |
| PRDX46 | 1,42376E-09 | 0,256097949 | 0,407 | 0,255 | 5,2091E-05 | EC | 1,194244265 | 0,837349636 | 1,596078431 |
| COX8A8 | 1,58103E-09 | 0,254760802 | 0,563 | 0,395 | 5,78451E-05 | EC | 1,193137905 | 0,838126084 | 1,425316456 |
| RGS163 | 1,64126E-09 | -0,923908169 | 0,175 | 0,274 | 6,00488E-05 | EC | 0,527079261 | 1,897247862 | 0,638686131 |
| RAC15 | 1,65624E-09 | 0,275011358 | 0,845 | 0,687 | 6,05968E-05 | EC | 1,210003615 | 0,826443812 | 1,229985444 |

| Supplementary Table 4 - clusermarker | | | | | | | | | |
|---|---|---|---|---|---|---|---|---|---|
| | **p_val** | **avg_log2FC** | **pct.1** | **pct.2** | **p_val_adj** | **celltype** | **Foldchange_UP** | **Foldchange_DOWN** | **Ratio_pct1_pct2** |
| SNCG1 | 1,77275E-09 | 0,306991355 | 0,595 | 0,44 | 6,48596E-05 | EC | 1,237125066 | 0,808325712 | 1,352272727 |
| IGLV2-146 | 1,88589E-09 | 1,704783612 | 0,143 | 0,049 | 6,89991E-05 | EC | 3,25980036 | 0,306767252 | 2,918367347 |
| RPL236 | 2,05969E-09 | -0,339830729 | 0,796 | 0,808 | 7,5358E-05 | EC | 0,790134013 | 1,265608092 | 0,985148515 |
| TMEM255B | 2,07018E-09 | 0,265951268 | 0,286 | 0,158 | 7,57419E-05 | EC | 1,202428632 | 0,83165019 | 1,810126582 |
| SNX36 | 2,35682E-09 | 0,2720311 | 0,615 | 0,439 | 8,62291E-05 | EC | 1,207506621 | 0,828152809 | 1,400911162 |
| NFKBIA8 | 2,52353E-09 | 0,295348687 | 0,869 | 0,723 | 9,23283E-05 | EC | 1,227181541 | 0,81487536 | 1,201936376 |
| CPXM21 | 2,89214E-09 | 0,259344745 | 0,434 | 0,281 | 0,000105815 | EC | 1,196934948 | 0,835467292 | 1,544483986 |
| PICALM1 | 2,90968E-09 | 0,275423874 | 0,441 | 0,287 | 0,000106457 | EC | 1,210349646 | 0,826207537 | 1,536585366 |
| PLVAP | 3,22038E-09 | 0,313383475 | 0,814 | 0,634 | 0,000117824 | EC | 1,242618532 | 0,804752202 | 1,283911672 |
| YWHAQ3 | 3,27077E-09 | 0,26825503 | 0,497 | 0,341 | 0,000119668 | EC | 1,204350259 | 0,830323232 | 1,457478006 |
| CCL231 | 3,31107E-09 | 0,578762245 | 0,265 | 0,146 | 0,000121142 | EC | 1,493567298 | 0,669537959 | 1,815068493 |
| CAVIN31 | 3,49864E-09 | 0,402302033 | 0,468 | 0,323 | 0,000128005 | EC | 1,321615061 | 0,756649973 | 1,448916409 |
| KIAA00401 | 3,93612E-09 | 0,262606012 | 0,224 | 0,111 | 0,000144011 | EC | 1,199643725 | 0,83358082 | 2,018018018 |
| ALDH1A2 | 4,52647E-09 | -0,321980154 | 0,062 | 0,138 | 0,00016561 | EC | 0,799971133 | 1,250045106 | 0,449275362 |
| STUB16 | 5,13966E-09 | 0,260367224 | 0,305 | 0,176 | 0,000188045 | EC | 1,19778355 | 0,834875383 | 1,732954545 |
| RHOJ2 | 5,81387E-09 | 0,383977205 | 0,438 | 0,297 | 0,000212712 | EC | 1,304934329 | 0,766322088 | 1,474747475 |
| VCAN3 | 5,92913E-09 | 0,411386739 | 0,237 | 0,123 | 0,000216929 | EC | 1,32996358 | 0,751900289 | 1,926829268 |
| PTOV15 | 6,37191E-09 | 0,305205329 | 0,27 | 0,153 | 0,000233129 | EC | 1,235594478 | 0,809327022 | 1,764705882 |
| RHOA3 | 6,51249E-09 | 0,262110284 | 0,879 | 0,746 | 0,000238272 | EC | 1,199231583 | 0,833867298 | 1,178284182 |
| PITPNC1 | 6,79511E-09 | 0,274903118 | 0,289 | 0,168 | 0,000248613 | EC | 1,209912837 | 0,826505819 | 1,720238095 |
| RELL1 | 6,85779E-09 | 0,303651644 | 0,305 | 0,178 | 0,000250906 | EC | 1,234264543 | 0,810199083 | 1,713483146 |
| YBX16 | 7,37601E-09 | 0,25031854 | 0,811 | 0,676 | 0,000269866 | EC | 1,189469715 | 0,84071077 | 1,199704142 |
| TUBB4B3 | 7,91406E-09 | 0,265862251 | 0,611 | 0,44 | 0,000289552 | EC | 1,202354442 | 0,831701506 | 1,388636364 |
| PHLDA32 | 1,27986E-08 | 0,293044534 | 0,141 | 0,054 | 0,000468263 | EC | 1,225223152 | 0,816177852 | 2,611111111 |
| ELOB6 | 1,34453E-08 | 0,251381154 | 0,681 | 0,492 | 0,000491922 | EC | 1,190346139 | 0,840091774 | 1,384146341 |
| SOX18 | 1,47349E-08 | 0,351982886 | 0,628 | 0,467 | 0,000539108 | EC | 1,276313628 | 0,783506481 | 1,344753747 |
| CNKSR3 | 1,60571E-08 | 0,37641123 | 0,339 | 0,21 | 0,000587479 | EC | 1,298108731 | 0,770351494 | 1,614285714 |
| SPTBN14 | 1,61887E-08 | -0,355629706 | 0,782 | 0,778 | 0,000592296 | EC | 0,781528448 | 1,279543954 | 1,005141388 |
| C1orf54 | 1,85729E-08 | 0,333439907 | 0,276 | 0,16 | 0,000679527 | EC | 1,260014125 | 0,793641897 | 1,725 |
| SLC6A61 | 2,33548E-08 | 0,381005054 | 0,17 | 0,076 | 0,000854483 | EC | 1,302248751 | 0,767902445 | 2,236842105 |
| CDC42EP11 | 2,91375E-08 | 0,271781707 | 0,185 | 0,087 | 0,001066056 | EC | 1,207297902 | 0,828295981 | 2,126436782 |

| Supplementary Table 4 - clusermarker | | | | | | | | | |
|---|---|---|---|---|---|---|---|---|---|
| | **p_val** | **avg_log2FC** | **pct.1** | **pct.2** | **p_val_adj** | **celltype** | **Foldchange_UP** | **Foldchange_DOWN** | **Ratio_pct1_pct2** |
| IL3RA | 3,83731E-08 | 0,285068486 | 0,439 | 0,291 | 0,001403958 | EC | 1,218468104 | 0,820702649 | 1,508591065 |
| TPM47 | 4,35031E-08 | 0,270820284 | 0,766 | 0,625 | 0,001591649 | EC | 1,206493618 | 0,828848147 | 1,2256 |
| CDH13 | 4,44104E-08 | -0,664533519 | 0,153 | 0,24 | 0,001624844 | EC | 0,630892664 | 1,585055679 | 0,6375 |
| ITGA6 | 5,48521E-08 | -0,433131047 | 0,635 | 0,657 | 0,002006873 | EC | 0,740652619 | 1,350160621 | 0,96651446 |
| CD79B1 | 6,85392E-08 | 0,25327779 | 0,229 | 0,124 | 0,002507643 | EC | 1,191912054 | 0,838988075 | 1,846774194 |
| SYNGR26 | 7,31723E-08 | 0,269440069 | 0,576 | 0,422 | 0,002677157 | EC | 1,205339927 | 0,829641479 | 1,36492891 |
| SERPINA33 | 7,96359E-08 | 0,367425422 | 0,142 | 0,057 | 0,00291364 | EC | 1,290048604 | 0,775164592 | 2,49122807 |
| SLC2A34 | 1,21615E-07 | 0,371290812 | 0,651 | 0,501 | 0,004449525 | EC | 1,293509646 | 0,773090486 | 1,299401198 |
| BGN3 | 1,23299E-07 | 0,324733186 | 0,219 | 0,116 | 0,004511124 | EC | 1,252432791 | 0,798446038 | 1,887931034 |
| JCHAIN6 | 1,40032E-07 | 0,41855822 | 0,129 | 0,05 | 0,005123334 | EC | 1,336591144 | 0,748171948 | 2,58 |
| SERPINE12 | 1,42279E-07 | 0,661674464 | 0,399 | 0,279 | 0,005205571 | EC | 1,581917612 | 0,632144173 | 1,430107527 |
| GNAS6 | 1,59255E-07 | -0,31903469 | 0,8 | 0,788 | 0,005826667 | EC | 0,801606054 | 1,247495568 | 1,015228426 |
| MYL94 | 1,91454E-07 | 0,366311758 | 0,478 | 0,341 | 0,007004728 | EC | 1,289053158 | 0,775763198 | 1,401759531 |
| GADD45B6 | 1,99759E-07 | -0,565522372 | 0,718 | 0,708 | 0,007308565 | EC | 0,675710711 | 1,479923264 | 1,014124294 |
| TACC14 | 2,80654E-07 | -0,407615893 | 0,676 | 0,671 | 0,010268274 | EC | 0,75386814 | 1,326491925 | 1,007451565 |
| EMCN | 7,70264E-07 | -0,313178005 | 0,673 | 0,699 | 0,028181654 | EC | 0,804866824 | 1,242441569 | 0,962804006 |
| GUCY1A12 | 8,55614E-07 | -0,400844571 | 0,092 | 0,161 | 0,031304334 | EC | 0,757414754 | 1,320280592 | 0,571428571 |
| H191 | 1,58387E-06 | 0,308823459 | 0,14 | 0,064 | 0,0579492 | EC | 1,23869711 | 0,807299857 | 2,1875 |
| HLA-DRA5 | 1,59216E-06 | -0,366267167 | 0,831 | 0,807 | 0,058252237 | EC | 0,775787176 | 1,289013316 | 1,029739777 |
| SEC628 | 1,87333E-06 | -0,390149346 | 0,682 | 0,661 | 0,068539353 | EC | 0,76305061 | 1,310529061 | 1,031770045 |
| TENT5A2 | 2,27611E-06 | -0,265250057 | 0,05 | 0,104 | 0,083276027 | EC | 0,832054505 | 1,201844343 | 0,480769231 |
| DEPP12 | 3,41988E-06 | -0,620984489 | 0,288 | 0,368 | 0,125123118 | EC | 0,650227064 | 1,537924296 | 0,782608696 |
| PLAC91 | 3,80401E-06 | -0,414841595 | 0,137 | 0,21 | 0,139177175 | EC | 0,75010185 | 1,333152292 | 0,652380952 |
| IGFBP44 | 4,48779E-06 | 0,251398771 | 0,873 | 0,768 | 0,164194644 | EC | 1,190360675 | 0,840081515 | 1,13671875 |
| AKR1C3 | 5,20592E-06 | -0,530848414 | 0,3 | 0,365 | 0,190469024 | EC | 0,692147579 | 1,444778585 | 0,821917808 |
| CRIP17 | 9,62547E-06 | -0,525475699 | 0,393 | 0,471 | 0,352167052 | EC | 0,694729999 | 1,439408117 | 0,834394904 |
| RMRP3 | 9,86845E-06 | -0,536728716 | 0,071 | 0,128 | 0,361056935 | EC | 0,689332186 | 1,450679397 | 0,5546875 |
| RBP7 | 1,03218E-05 | 0,400130403 | 0,413 | 0,296 | 0,377644591 | EC | 1,319627185 | 0,757789785 | 1,39527027 |
| TFPI3 | 1,28994E-05 | -0,64334105 | 0,481 | 0,521 | 0,471951045 | EC | 0,640228564 | 1,561942182 | 0,923224568 |
| BTG25 | 1,37694E-05 | -0,484697415 | 0,478 | 0,521 | 0,503780433 | EC | 0,714646942 | 1,399292353 | 0,917466411 |
| PGM51 | 1,56778E-05 | -0,478581249 | 0,25 | 0,318 | 0,573602788 | EC | 0,717683049 | 1,393372745 | 0,786163522 |

| Supplementary Table 4 - clusermarker | | | | | | | | | |
|---|---|---|---|---|---|---|---|---|---|
| | **p_val** | **avg_log2FC** | **pct.1** | **pct.2** | **p_val_adj** | **celltype** | **Foldchange_UP** | **Foldchange_DOWN** | **Ratio_pct1_pct2** |
| RPL27A7 | 1,60295E-05 | -0,253530658 | 0,787 | 0,761 | 0,586472398 | EC | 0,838841035 | 1,192120984 | 1,034165572 |
| CLIC21 | 1,62645E-05 | -0,468231263 | 0,385 | 0,437 | 0,595067526 | EC | 0,722850266 | 1,383412371 | 0,881006865 |
| CLU3 | 1,67084E-05 | -0,380267953 | 0,707 | 0,765 | 0,611311426 | EC | 0,768294881 | 1,301583577 | 0,924183007 |
| DDIT43 | 1,69809E-05 | 0,273864146 | 0,587 | 0,457 | 0,62128012 | EC | 1,209041819 | 0,82710125 | 1,284463895 |
| HLA-DQA21 | 1,90312E-05 | -0,32434987 | 0,069 | 0,123 | 0,696295162 | EC | 0,798658209 | 1,252100071 | 0,56097561 |
| FOS5 | 2,03639E-05 | -0,320548565 | 0,866 | 0,855 | 0,745052931 | EC | 0,80076534 | 1,248805299 | 1,012865497 |
| SLCO2A1 | 2,42352E-05 | 0,250754245 | 0,404 | 0,294 | 0,886693838 | EC | 1,189828998 | 0,840456907 | 1,37414966 |
| COL6A12 | 2,50243E-05 | 0,262231789 | 0,171 | 0,099 | 0,915565096 | EC | 1,199332588 | 0,833797072 | 1,727272727 |
| JUN4 | 2,93922E-05 | -0,416134565 | 0,828 | 0,787 | 1 | EC | 0,749429896 | 1,334347623 | 1,052096569 |
| PRRC2C3 | 3,33019E-05 | -0,337540172 | 0,619 | 0,617 | 1 | EC | 0,7913895 | 1,263600289 | 1,003241491 |
| CYTL1 | 3,56351E-05 | -0,551991507 | 0,28 | 0,188 | 1 | EC | 0,682077933 | 1,466108126 | 1,489361702 |
| CD98 | 3,816E-05 | -0,345736564 | 0,712 | 0,703 | 1 | EC | 0,786906121 | 1,27079962 | 1,012802276 |
| TSC22D16 | 3,84869E-05 | -0,512629556 | 0,578 | 0,6 | 1 | EC | 0,700943684 | 1,426648135 | 0,963333333 |
| AL627171.27 | 4,01967E-05 | -0,578998168 | 0,211 | 0,276 | 1 | EC | 0,669428479 | 1,49381156 | 0,764492754 |
| UBB5 | 4,80243E-05 | -0,253986008 | 0,824 | 0,778 | 1 | EC | 0,838576317 | 1,192497307 | 1,059125964 |
| AKAP121 | 6,47866E-05 | -0,426221529 | 0,154 | 0,218 | 1 | EC | 0,744208344 | 1,34370974 | 0,706422018 |
| TSPAN7 | 7,14803E-05 | -0,343829729 | 0,743 | 0,703 | 1 | EC | 0,787946876 | 1,269121092 | 1,056899004 |
| HES14 | 7,61127E-05 | -0,542603065 | 0,359 | 0,403 | 1 | EC | 0,686531078 | 1,456598299 | 0,890818859 |
| INMT1 | 7,91633E-05 | -0,260392583 | 0,06 | 0,108 | 1 | EC | 0,834860708 | 1,197804604 | 0,555555556 |
| HSPA87 | 8,59024E-05 | -0,317761511 | 0,88 | 0,852 | 1 | EC | 0,802313784 | 1,246395138 | 1,03286385 |
| ID11 | 8,88004E-05 | 0,284429372 | 0,715 | 0,605 | 1 | EC | 1,217928442 | 0,8210663 | 1,181818182 |
| TPM22 | 8,96313E-05 | 0,269149208 | 0,138 | 0,077 | 1 | EC | 1,205096943 | 0,82980876 | 1,792207792 |
| U2SURP5 | 0,000134215 | -0,51879668 | 0,371 | 0,413 | 1 | EC | 0,697953738 | 1,432759716 | 0,898305085 |
| THBD3 | 0,000150186 | -0,347513294 | 0,583 | 0,61 | 1 | EC | 0,785937615 | 1,272365619 | 0,955737705 |
| RPS206 | 0,000180594 | -0,282112511 | 0,729 | 0,704 | 1 | EC | 0,822385931 | 1,215974109 | 1,035511364 |
| MYCT1 | 0,000198733 | -0,420472012 | 0,441 | 0,464 | 1 | EC | 0,747180126 | 1,338365362 | 0,950431034 |
| ANXA32 | 0,000216366 | -0,339721324 | 0,066 | 0,111 | 1 | EC | 0,790193934 | 1,26551212 | 0,594594595 |
| MAOA1 | 0,000238837 | -0,341153175 | 0,121 | 0,175 | 1 | EC | 0,789410069 | 1,266768743 | 0,691428571 |
| DNAJB18 | 0,000248873 | -0,700234928 | 0,59 | 0,571 | 1 | EC | 0,615471975 | 1,624769348 | 1,033274956 |
| BTN3A22 | 0,000259824 | -0,385787557 | 0,166 | 0,218 | 1 | EC | 0,765361079 | 1,306572842 | 0,76146789 |
| CCDC802 | 0,000443339 | -0,343805512 | 0,13 | 0,183 | 1 | EC | 0,787960103 | 1,269099789 | 0,710382514 |

| Supplementary Table 4 - clusermarker | | | | | | | | | |
|---|---|---|---|---|---|---|---|---|---|
| | **p_val** | **avg_log2FC** | **pct.1** | **pct.2** | **p_val_adj** | **celltype** | **Foldchange_UP** | **Foldchange_DOWN** | **Ratio_pct1_pct2** |
| ATP2A33 | 0,000473333 | -0,335069297 | 0,076 | 0,121 | 1 | EC | 0,792746058 | 1,261437998 | 0,628099174 |
| MEF2C4 | 0,000795855 | -0,407530938 | 0,545 | 0,531 | 1 | EC | 0,753912534 | 1,326413815 | 1,026365348 |
| ADAMTS11 | 0,000844287 | -0,450633193 | 0,575 | 0,575 | 1 | EC | 0,731721628 | 1,366639938 | 1 |
| IFIT11 | 0,000910575 | -0,331350987 | 0,139 | 0,19 | 1 | EC | 0,794791866 | 1,258191034 | 0,731578947 |
| A2M4 | 0,000912987 | -0,299725745 | 0,873 | 0,861 | 1 | EC | 0,812406819 | 1,230910396 | 1,013937282 |
| PCAT19 | 0,001005033 | -0,358402016 | 0,453 | 0,476 | 1 | EC | 0,78002809 | 1,282005113 | 0,951680672 |
| PLK22 | 0,001246174 | -0,388243947 | 0,308 | 0,35 | 1 | EC | 0,764059054 | 1,30879936 | 0,88 |
| GAS65 | 0,001337822 | -0,476790343 | 0,282 | 0,328 | 1 | EC | 0,718574506 | 1,391644139 | 0,859756098 |
| SYT1 | 0,001540243 | -0,29351874 | 0,065 | 0,103 | 1 | EC | 0,815909622 | 1,225625943 | 0,631067961 |
| C19orf331 | 0,001781699 | -0,344350729 | 0,149 | 0,197 | 1 | EC | 0,787662377 | 1,269579492 | 0,756345178 |
| ZBTB204 | 0,001893686 | -0,386026417 | 0,477 | 0,479 | 1 | EC | 0,765234373 | 1,306789182 | 0,995824635 |
| TLE45 | 0,002112449 | -0,297658703 | 0,127 | 0,171 | 1 | EC | 0,813571641 | 1,229148054 | 0,742690058 |
| TLR4 | 0,002313711 | -0,281057671 | 0,109 | 0,153 | 1 | EC | 0,822987446 | 1,215085363 | 0,712418301 |
| H4C36 | 0,002566521 | -0,363579254 | 0,488 | 0,497 | 1 | EC | 0,777233907 | 1,286613966 | 0,981891348 |
| EIF4A26 | 0,002655065 | -0,273900501 | 0,619 | 0,61 | 1 | EC | 0,827080408 | 1,209072286 | 1,014754098 |
| CWF19L23 | 0,002996469 | -0,372744701 | 0,129 | 0,171 | 1 | EC | 0,772311789 | 1,294813849 | 0,754385965 |
| LINC021852 | 0,003742186 | -0,34402538 | 0,125 | 0,165 | 1 | EC | 0,787840026 | 1,269293215 | 0,757575758 |
| GAB1 | 0,003789627 | -0,30996989 | 0,09 | 0,126 | 1 | EC | 0,806658594 | 1,239681827 | 0,714285714 |
| ARL15 | 0,004032702 | -0,392003663 | 0,187 | 0,232 | 1 | EC | 0,76207048 | 1,312214586 | 0,806034483 |
| CALM24 | 0,005258413 | -0,296925055 | 0,719 | 0,689 | 1 | EC | 0,813985469 | 1,228523159 | 1,043541364 |
| FN14 | 0,005634918 | -0,288274033 | 0,373 | 0,271 | 1 | EC | 0,81888114 | 1,221178448 | 1,376383764 |
| DDX188 | 0,006564176 | -0,426091152 | 0,411 | 0,427 | 1 | EC | 0,744275601 | 1,343588313 | 0,962529274 |
| HLA-DPA13 | 0,006608973 | -0,30730319 | 0,742 | 0,704 | 1 | EC | 0,808151014 | 1,237392496 | 1,053977273 |
| FXYD61 | 0,007437852 | -0,368386936 | 0,074 | 0,106 | 1 | EC | 0,774648139 | 1,290908671 | 0,698113208 |
| GIMAP5 | 0,008616315 | -0,375644972 | 0,42 | 0,435 | 1 | EC | 0,770760759 | 1,29741945 | 0,965517241 |
| BOD1L15 | 0,009600961 | -0,350820774 | 0,346 | 0,366 | 1 | EC | 0,784137861 | 1,275285953 | 0,945355191 |
| MBNL15 | 0,010346284 | -0,251220097 | 0,622 | 0,585 | 1 | EC | 0,840185564 | 1,190213261 | 1,063247863 |
| UACA3 | 0,012327862 | -0,278641422 | 0,482 | 0,487 | 1 | EC | 0,824366954 | 1,213052022 | 0,98973306 |
| PLXDC21 | 0,012921552 | -0,379370866 | 0,244 | 0,274 | 1 | EC | 0,768772766 | 1,300774487 | 0,890510949 |
| COX174 | 0,013596992 | -0,35808408 | 0,308 | 0,334 | 1 | EC | 0,780200009 | 1,281722621 | 0,922155689 |
| KCNN3 | 0,014090359 | -0,382415752 | 0,215 | 0,249 | 1 | EC | 0,76715194 | 1,303522741 | 0,863453815 |

| Supplementary Table 4 - clusermarker | | | | | | | | | |
|---|---|---|---|---|---|---|---|---|---|
| | **p_val** | **avg_log2FC** | **pct.1** | **pct.2** | **p_val_adj** | **celltype** | **Foldchange_UP** | **Foldchange_DOWN** | **Ratio_pct1_pct2** |
| LIMCH11 | 0,014735634 | -0,382257799 | 0,248 | 0,279 | 1 | EC | 0,767235936 | 1,303380033 | 0,888888889 |
| SYNE13 | 0,014775724 | -0,294804823 | 0,234 | 0,271 | 1 | EC | 0,815182608 | 1,226719007 | 0,863468635 |
| PLCG24 | 0,017343685 | -0,389176885 | 0,268 | 0,289 | 1 | EC | 0,763565125 | 1,309645985 | 0,92733564 |
| HMOX1 | 0,017490501 | -0,311013103 | 0,111 | 0,143 | 1 | EC | 0,80607551 | 1,240578565 | 0,776223776 |
| TAF73 | 0,017906424 | -0,364790665 | 0,222 | 0,254 | 1 | EC | 0,776581549 | 1,287694771 | 0,874015748 |
| IFIT3 | 0,017965678 | -0,31279729 | 0,198 | 0,23 | 1 | EC | 0,80507925 | 1,242113743 | 0,860869565 |
| HBEGF3 | 0,018648066 | -0,282392327 | 0,295 | 0,324 | 1 | EC | 0,822226442 | 1,216209975 | 0,910493827 |
| BST23 | 0,018946519 | -0,490241991 | 0,503 | 0,484 | 1 | EC | 0,711905676 | 1,40468047 | 1,039256198 |
| METTL7A3 | 0,019570089 | -0,265184342 | 0,208 | 0,242 | 1 | EC | 0,832092406 | 1,2017896 | 0,859504132 |
| DDX272 | 0,020863262 | -0,269513219 | 0,112 | 0,143 | 1 | EC | 0,829599415 | 1,205401043 | 0,783216783 |
| GPRC5B | 0,020945022 | -0,355204488 | 0,19 | 0,222 | 1 | EC | 0,781758829 | 1,279166878 | 0,855855856 |
| ITGA9 | 0,022286361 | -0,294574601 | 0,138 | 0,173 | 1 | EC | 0,815312704 | 1,226523266 | 0,797687861 |
| FAM107A1 | 0,022707917 | -0,413879763 | 0,279 | 0,306 | 1 | EC | 0,750602103 | 1,332263787 | 0,911764706 |
| RBM52 | 0,026472902 | -0,322456296 | 0,21 | 0,239 | 1 | EC | 0,799707156 | 1,250457736 | 0,878661088 |
| ADGRA2 | 0,026855421 | -0,343995612 | 0,183 | 0,21 | 1 | EC | 0,787856282 | 1,269267026 | 0,871428571 |
| SRRM12 | 0,028140572 | -0,299938441 | 0,525 | 0,508 | 1 | EC | 0,812287055 | 1,231091883 | 1,033464567 |
| GBP11 | 0,028179583 | -0,272774093 | 0,244 | 0,272 | 1 | EC | 0,827726417 | 1,208128652 | 0,897058824 |
| NR4A15 | 0,028912469 | -0,435940338 | 0,537 | 0,513 | 1 | EC | 0,739211785 | 1,352792286 | 1,046783626 |
| ATF33 | 0,029420929 | -0,49432626 | 0,418 | 0,415 | 1 | EC | 0,709893121 | 1,408662755 | 1,007228916 |
| ITPR22 | 0,033823408 | -0,345652216 | 0,269 | 0,292 | 1 | EC | 0,786952129 | 1,270725324 | 0,921232877 |
| PET1007 | 0,035695168 | -0,32992222 | 0,289 | 0,311 | 1 | EC | 0,795579375 | 1,256945607 | 0,92926045 |
| UTRN1 | 0,037619054 | -0,329933594 | 0,553 | 0,539 | 1 | EC | 0,795573102 | 1,256955517 | 1,025974026 |
| ADGRF51 | 0,038299409 | -0,43096651 | 0,355 | 0,371 | 1 | EC | 0,741764686 | 1,348136436 | 0,956873315 |
| CALCRL1 | 0,042537263 | -0,259404312 | 0,613 | 0,598 | 1 | EC | 0,835432798 | 1,196984368 | 1,025083612 |
| GJA4 | 0,043483936 | 0,273073553 | 0,166 | 0,133 | 1 | EC | 1,208379449 | 0,827554624 | 1,248120301 |
| GIMAP8 | 0,047899117 | -0,407911549 | 0,415 | 0,408 | 1 | EC | 0,753713663 | 1,326763794 | 1,017156863 |
| HSD17B123 | 0,049276317 | -0,290000603 | 0,286 | 0,306 | 1 | EC | 0,817901717 | 1,222640789 | 0,934640523 |
| ATP5ME6 | 0,051126502 | -0,281320765 | 0,338 | 0,353 | 1 | EC | 0,822837377 | 1,21530697 | 0,957507082 |
| MT1E2 | 0,055987392 | -0,342398762 | 0,103 | 0,128 | 1 | EC | 0,788728806 | 1,267862912 | 0,8046875 |
| LY6E5 | 0,056258701 | -0,331176435 | 0,536 | 0,514 | 1 | EC | 0,794888034 | 1,258038815 | 1,042801556 |
| ITPRID22 | 0,061667446 | -0,309631283 | 0,497 | 0,476 | 1 | EC | 0,806847943 | 1,239390902 | 1,044117647 |

| Supplementary Table 4 - clusermarker | | | | | | | | | |
|---|---|---|---|---|---|---|---|---|---|
| | **p_val** | **avg_log2FC** | **pct.1** | **pct.2** | **p_val_adj** | **celltype** | **Foldchange_UP** | **Foldchange_DOWN** | **Ratio_pct1_pct2** |
| RBMS32 | 0,063023347 | -0,311568863 | 0,197 | 0,22 | 1 | EC | 0,805765051 | 1,241056557 | 0,895454545 |
| STING1 | 0,063404742 | -0,291904464 | 0,456 | 0,45 | 1 | EC | 0,81682308 | 1,224255319 | 1,013333333 |
| CD832 | 0,065139421 | -0,274097453 | 0,109 | 0,134 | 1 | EC | 0,826967506 | 1,209237356 | 0,813432836 |
| GIMAP6 | 0,066795062 | -0,398002366 | 0,465 | 0,455 | 1 | EC | 0,758908382 | 1,317682113 | 1,021978022 |
| VEGFC | 0,071827627 | -0,278033696 | 0,143 | 0,166 | 1 | EC | 0,824714286 | 1,212541139 | 0,861445783 |
| SEPTIN72 | 0,076869287 | -0,291985073 | 0,575 | 0,541 | 1 | EC | 0,816777442 | 1,224323724 | 1,06284658 |
| TAGLN4 | 0,079325689 | 0,370802563 | 0,195 | 0,161 | 1 | EC | 1,293071959 | 0,773352166 | 1,211180124 |
| MTUS11 | 0,086894704 | -0,284097385 | 0,602 | 0,573 | 1 | EC | 0,821255262 | 1,21764821 | 1,05061082 |
| DAAM1 | 0,089105718 | -0,375497383 | 0,342 | 0,348 | 1 | EC | 0,770839612 | 1,29728673 | 0,982758621 |
| CCND15 | 0,091838902 | -0,306631808 | 0,167 | 0,19 | 1 | EC | 0,808527188 | 1,236816789 | 0,878947368 |
| NDUFA17 | 0,095317585 | -0,266580077 | 0,575 | 0,541 | 1 | EC | 0,831287788 | 1,202952834 | 1,06284658 |
| EDNRB | 0,095541878 | -0,476884467 | 0,115 | 0,134 | 1 | EC | 0,718527627 | 1,391734935 | 0,858208955 |
| MT1X5 | 0,101556757 | -0,519727086 | 0,165 | 0,185 | 1 | EC | 0,697503767 | 1,433684013 | 0,891891892 |
| SF3B14 | 0,103853023 | -0,297527523 | 0,578 | 0,539 | 1 | EC | 0,81364562 | 1,229036297 | 1,072356215 |
| COBLL12 | 0,113541852 | -0,258086549 | 0,228 | 0,244 | 1 | EC | 0,836196234 | 1,195891538 | 0,93442623 |
| NSRP15 | 0,115047444 | -0,347930638 | 0,273 | 0,281 | 1 | EC | 0,785710291 | 1,272733743 | 0,971530249 |
| NDUFB14 | 0,116569573 | -0,291601418 | 0,326 | 0,329 | 1 | EC | 0,816994676 | 1,223998184 | 0,990881459 |
| PPFIBP11 | 0,123737477 | -0,335738377 | 0,387 | 0,382 | 1 | EC | 0,79237849 | 1,262023152 | 1,013089005 |
| CAV22 | 0,124411272 | -0,283725325 | 0,476 | 0,452 | 1 | EC | 0,821467085 | 1,217334228 | 1,053097345 |
| PNISR5 | 0,126495491 | -0,347024996 | 0,457 | 0,439 | 1 | EC | 0,78620367 | 1,271935044 | 1,041002278 |
| LINC01116 | 0,12676281 | -0,257281739 | 0,16 | 0,18 | 1 | EC | 0,836662838 | 1,195224593 | 0,888888889 |
| NDRG23 | 0,127942464 | -0,275204568 | 0,118 | 0,138 | 1 | EC | 0,826333139 | 1,210165674 | 0,855072464 |
| SMAD1 | 0,134770135 | -0,292180275 | 0,393 | 0,39 | 1 | EC | 0,816666937 | 1,224489391 | 1,007692308 |
| AKAP95 | 0,139608791 | -0,326089126 | 0,401 | 0,388 | 1 | EC | 0,797695959 | 1,253610463 | 1,033505155 |
| SELENOP2 | 0,139610631 | -0,298074448 | 0,39 | 0,382 | 1 | EC | 0,813337226 | 1,229502312 | 1,020942408 |
| SKAP22 | 0,146405496 | -0,282119208 | 0,214 | 0,23 | 1 | EC | 0,822382113 | 1,215979754 | 0,930434783 |
| HSPE15 | 0,148340037 | -0,335126054 | 0,584 | 0,534 | 1 | EC | 0,792714871 | 1,261487625 | 1,093632959 |
| ID27 | 0,152409594 | -0,403418901 | 0,407 | 0,393 | 1 | EC | 0,756064436 | 1,322638591 | 1,03562341 |
| DLL4 | 0,157750995 | -0,280016127 | 0,215 | 0,23 | 1 | EC | 0,823581811 | 1,214208458 | 0,934782609 |
| SEMA3G | 0,160124348 | -0,262473624 | 0,138 | 0,156 | 1 | EC | 0,833657317 | 1,199533645 | 0,884615385 |
| LYST1 | 0,162405168 | -0,370517984 | 0,347 | 0,343 | 1 | EC | 0,773504729 | 1,292816919 | 1,011661808 |

| Supplementary Table 4 - clusermarker | | | | | | | | | |
|---|---|---|---|---|---|---|---|---|---|
| | **p_val** | **avg_log2FC** | **pct.1** | **pct.2** | **p_val_adj** | **celltype** | **Foldchange_UP** | **Foldchange_DOWN** | **Ratio_pct1_pct2** |
| ZC3H133 | 0,163566759 | -0,263387111 | 0,288 | 0,297 | 1 | EC | 0,833129628 | 1,200293408 | 0,96969697 |
| JAG13 | 0,171697793 | -0,315253422 | 0,347 | 0,355 | 1 | EC | 0,803709799 | 1,244230194 | 0,977464789 |
| NFIA4 | 0,176256385 | -0,287793564 | 0,495 | 0,477 | 1 | EC | 0,819153903 | 1,220771819 | 1,037735849 |
| SPRY15 | 0,189701686 | -0,257367732 | 0,638 | 0,585 | 1 | EC | 0,836612969 | 1,195295837 | 1,090598291 |
| PTPN14 | 0,191043561 | -0,281419377 | 0,312 | 0,314 | 1 | EC | 0,822781137 | 1,215390042 | 0,993630573 |
| HSPH17 | 0,191321974 | -0,416568425 | 0,351 | 0,345 | 1 | EC | 0,749204554 | 1,33474896 | 1,017391304 |
| MGST22 | 0,201835078 | -0,251736849 | 0,656 | 0,605 | 1 | EC | 0,839884676 | 1,190639654 | 1,084297521 |
| PROX1 | 0,205207977 | -0,370750957 | 0,127 | 0,101 | 1 | EC | 0,773379829 | 1,293025706 | 1,257425743 |
| CIR15 | 0,207552565 | -0,327259364 | 0,254 | 0,261 | 1 | EC | 0,797049172 | 1,254627738 | 0,973180077 |
| DYNC1I24 | 0,211142807 | -0,267732631 | 0,289 | 0,294 | 1 | EC | 0,830623946 | 1,203914244 | 0,982993197 |
| SULF11 | 0,215953746 | -0,366721367 | 0,193 | 0,207 | 1 | EC | 0,775542975 | 1,289419196 | 0,93236715 |
| SCAPER | 0,230948336 | -0,26498477 | 0,147 | 0,16 | 1 | EC | 0,83220752 | 1,201623364 | 0,91875 |
| KMT2A4 | 0,234841294 | -0,304909215 | 0,45 | 0,422 | 1 | EC | 0,809493154 | 1,235340898 | 1,066350711 |
| BPTF3 | 0,241649923 | -0,254691943 | 0,444 | 0,422 | 1 | EC | 0,838166088 | 1,193080959 | 1,052132701 |
| SUN1 | 0,248060576 | -0,299880965 | 0,384 | 0,365 | 1 | EC | 0,812319417 | 1,231042837 | 1,052054795 |
| GLO11 | 0,255692626 | -0,253618876 | 0,176 | 0,187 | 1 | EC | 0,838789743 | 1,192193883 | 0,941176471 |
| CYP1B14 | 0,259121857 | -0,252048765 | 0,253 | 0,264 | 1 | EC | 0,839703109 | 1,190897103 | 0,958333333 |
| RBM431 | 0,27756988 | -0,269342751 | 0,112 | 0,124 | 1 | EC | 0,829697445 | 1,205258623 | 0,903225806 |
| PFDN15 | 0,278638857 | -0,395702114 | 0,245 | 0,247 | 1 | EC | 0,760119361 | 1,315582857 | 0,991902834 |
| DNAJA17 | 0,2833842 | -0,28658703 | 0,727 | 0,65 | 1 | EC | 0,819839252 | 1,219751308 | 1,118461538 |
| GBP43 | 0,288327079 | -0,278079572 | 0,318 | 0,309 | 1 | EC | 0,824688062 | 1,212579697 | 1,029126214 |
| AFDN4 | 0,295531201 | -0,348676464 | 0,451 | 0,425 | 1 | EC | 0,78530421 | 1,273391875 | 1,061176471 |
| ITPR11 | 0,316019031 | -0,251772879 | 0,26 | 0,266 | 1 | EC | 0,839863701 | 1,190669389 | 0,977443609 |
| HSPD15 | 0,321238408 | -0,37919966 | 0,523 | 0,486 | 1 | EC | 0,768864002 | 1,300620132 | 1,076131687 |
| MTRNR2L89 | 0,349101681 | -1,284648852 | 0,173 | 0,178 | 1 | EC | 0,4104707 | 2,436227484 | 0,971910112 |
| ZNF6383 | 0,351233679 | -0,369704428 | 0,34 | 0,323 | 1 | EC | 0,773941042 | 1,292088087 | 1,052631579 |
| ZRANB24 | 0,366563207 | -0,288078086 | 0,282 | 0,277 | 1 | EC | 0,818992369 | 1,221012598 | 1,018050542 |
| WNK11 | 0,383792914 | -0,298548502 | 0,546 | 0,492 | 1 | EC | 0,813070016 | 1,22990638 | 1,109756098 |
| MX11 | 0,409019959 | -0,322482626 | 0,366 | 0,351 | 1 | EC | 0,799692562 | 1,250480557 | 1,042735043 |
| YIPF52 | 0,432949588 | -0,261578157 | 0,251 | 0,245 | 1 | EC | 0,83417492 | 1,198789338 | 1,024489796 |
| DST5 | 0,441052581 | -0,297647149 | 0,376 | 0,355 | 1 | EC | 0,813578157 | 1,22913821 | 1,05915493 |

| Supplementary Table 4 - clusermarker | | | | | | | | | |
|---|---|---|---|---|---|---|---|---|---|
| | **p_val** | **avg_log2FC** | **pct.1** | **pct.2** | **p_val_adj** | **celltype** | **Foldchange_UP** | **Foldchange_DOWN** | **Ratio_pct1_pct2** |
| ABCB1 | 0,519153187 | -0,262449481 | 0,215 | 0,213 | 1 | EC | 0,833671267 | 1,199513572 | 1,009389671 |
| CFAP361 | 0,521963203 | -0,275736406 | 0,312 | 0,272 | 1 | EC | 0,826028574 | 1,210611874 | 1,147058824 |
| SOX45 | 0,53094666 | -0,30366522 | 0,454 | 0,439 | 1 | EC | 0,810191458 | 1,234276158 | 1,034168565 |
| LEPR1 | 0,604276284 | -0,33876537 | 0,285 | 0,271 | 1 | EC | 0,790717703 | 1,264673847 | 1,051660517 |
| LTBP43 | 0,613381275 | -0,35654745 | 0,297 | 0,262 | 1 | EC | 0,781031451 | 1,280358171 | 1,133587786 |
| MAN1A12 | 0,617857067 | -0,257373892 | 0,266 | 0,25 | 1 | EC | 0,836609397 | 1,195300941 | 1,064 |
| PARP144 | 0,635199904 | -0,345712729 | 0,34 | 0,318 | 1 | EC | 0,786919122 | 1,270778625 | 1,06918239 |
| IK4 | 0,644736514 | -0,355128874 | 0,391 | 0,356 | 1 | EC | 0,781799803 | 1,279099837 | 1,098314607 |
| TSHZ22 | 0,653042326 | -0,270559984 | 0,535 | 0,477 | 1 | EC | 0,828997707 | 1,206275955 | 1,121593291 |
| RCAN12 | 0,663502802 | -0,283205938 | 0,468 | 0,435 | 1 | EC | 0,821762876 | 1,216896052 | 1,075862069 |
| LMO21 | 0,694664829 | -0,31930388 | 0,446 | 0,407 | 1 | EC | 0,801456498 | 1,247728357 | 1,095823096 |
| SAT15 | 0,702870446 | -0,332879306 | 0,718 | 0,625 | 1 | EC | 0,793950349 | 1,259524605 | 1,1488 |
| ETS12 | 0,796574907 | -0,263888207 | 0,459 | 0,413 | 1 | EC | 0,832840304 | 1,200710382 | 1,111380145 |
| IER56 | 0,805300056 | -0,270638954 | 0,341 | 0,313 | 1 | EC | 0,828952331 | 1,206341985 | 1,089456869 |
| RIN22 | 0,809135311 | -0,276878821 | 0,215 | 0,205 | 1 | EC | 0,825374733 | 1,211570891 | 1,048780488 |
| ITGA13 | 0,811457329 | -0,277958385 | 0,175 | 0,17 | 1 | EC | 0,824757339 | 1,212477844 | 1,029411765 |
| IGFBP53 | 0,822364969 | -0,285756591 | 0,103 | 0,097 | 1 | EC | 0,820311301 | 1,2190494 | 1,06185567 |
| FAM204A5 | 0,842604501 | -0,266550064 | 0,294 | 0,274 | 1 | EC | 0,831305082 | 1,202927808 | 1,072992701 |
| IGFBP31 | 0,873546634 | -0,589060663 | 0,148 | 0,143 | 1 | EC | 0,664775601 | 1,504267002 | 1,034965035 |
| MT2A4 | 0,879020711 | -0,268164938 | 0,625 | 0,556 | 1 | EC | 0,830375085 | 1,204275053 | 1,124100719 |
| ENPP21 | 0,921205856 | -0,3152885 | 0,301 | 0,274 | 1 | EC | 0,803690259 | 1,244260446 | 1,098540146 |
| VPS35 | 0,948491614 | -0,25311482 | 0,288 | 0,266 | 1 | EC | 0,839082854 | 1,191777421 | 1,082706767 |
| XIST4 | 0,94904087 | -0,270573649 | 0,548 | 0,482 | 1 | EC | 0,828989855 | 1,20628738 | 1,136929461 |
| N4BP2L24 | 0,978162942 | -0,263541287 | 0,503 | 0,455 | 1 | EC | 0,833040599 | 1,200421686 | 1,105494505 |
| HLA-DRB12 | 1,15131E-17 | 4,901127409 | 1 | 0,015 | 4,21229E-13 | CH_SC | 29,88039696 | 0,033466758 | 66,66666667 |
| HLA-DPA14 | 2,64385E-13 | 3,132103973 | 0,8 | 0,015 | 9,67304E-09 | CH_SC | 8,767125941 | 0,114062465 | 53,33333333 |
| HLA-DRA6 | 6,27111E-13 | 4,665307237 | 0,8 | 0,031 | 2,29441E-08 | CH_SC | 25,37449555 | 0,03940965 | 25,80645161 |
| PMP2 | 3,14513E-12 | 2,68493701 | 0,8 | 0,031 | 1,15071E-07 | CH_SC | 6,430527154 | 0,155508246 | 25,80645161 |
| EGFL8 | 7,0802E-12 | 2,624475528 | 0,933 | 0,077 | 2,59043E-07 | CH_SC | 6,16660111 | 0,162163886 | 12,11688312 |
| PLP1 | 2,22917E-11 | 2,815691245 | 1 | 0,108 | 8,15587E-07 | CH_SC | 7,040565178 | 0,142034052 | 9,259259259 |
| COL4A12 | 2,42021E-11 | 2,533159774 | 0,733 | 0,031 | 8,85481E-07 | CH_SC | 5,788380552 | 0,172759892 | 23,64516129 |

| Supplementary Table 4 - clusermarker | | | | | | | | | |
|---|---|---|---|---|---|---|---|---|---|
| | **p_val** | **avg_log2FC** | **pct.1** | **pct.2** | **p_val_adj** | **celltype** | **Foldchange_UP** | **Foldchange_DOWN** | **Ratio_pct1_pct2** |
| NES | 3,35866E-11 | 2,686435233 | 0,667 | 0,015 | 1,22883E-06 | CH_SC | 6,437208657 | 0,155346836 | 44,46666667 |
| NRXN1 | 3,62332E-11 | 2,439145402 | 1 | 0,123 | 1,32566E-06 | CH_SC | 5,423203857 | 0,184392847 | 8,130081301 |
| SCN7A1 | 7,62665E-11 | 3,078118649 | 0,867 | 0,077 | 2,79036E-06 | CH_SC | 8,445124249 | 0,11841152 | 11,25974026 |
| CDH19 | 1,64792E-10 | 2,483155757 | 1 | 0,154 | 6,02925E-06 | CH_SC | 5,591191495 | 0,178852755 | 6,493506494 |
| ZEB24 | 2,08067E-10 | 2,72771044 | 0,867 | 0,123 | 7,61256E-06 | CH_SC | 6,624035662 | 0,150965371 | 7,048780488 |
| COL18A14 | 3,75547E-10 | 3,073240541 | 0,867 | 0,154 | 1,37401E-05 | CH_SC | 8,41661742 | 0,118812576 | 5,62987013 |
| EGFLAM2 | 3,84026E-10 | 1,707182728 | 0,667 | 0,031 | 1,40504E-05 | CH_SC | 3,265225724 | 0,306257541 | 21,51612903 |
| HLA-DMA4 | 4,9494E-10 | 2,629798179 | 0,667 | 0,031 | 1,81084E-05 | CH_SC | 6,189394069 | 0,161566704 | 21,51612903 |
| MARCKS7 | 5,33578E-10 | 2,678921655 | 0,867 | 0,108 | 1,9522E-05 | CH_SC | 6,40377072 | 0,156157996 | 8,027777778 |
| DST6 | 1,04025E-09 | 3,16852382 | 1 | 0,431 | 3,80597E-05 | CH_SC | 8,991263218 | 0,111219077 | 2,320185615 |
| HLA-DRB55 | 1,32741E-09 | 3,331487802 | 0,667 | 0,046 | 4,8566E-05 | CH_SC | 10,06648287 | 0,099339562 | 14,5 |
| GPR155 | 1,50201E-09 | 2,381571104 | 0,8 | 0,092 | 5,49541E-05 | CH_SC | 5,211039186 | 0,191900303 | 8,695652174 |
| ITGA14 | 2,13375E-09 | 1,973990792 | 0,667 | 0,046 | 7,80674E-05 | CH_SC | 3,928533306 | 0,254547925 | 14,5 |
| ITGB82 | 3,11917E-09 | 3,243953143 | 0,8 | 0,123 | 0,000114121 | CH_SC | 9,473865193 | 0,105553539 | 6,504065041 |
| AHR3 | 3,17195E-09 | 2,387837114 | 0,867 | 0,138 | 0,000116052 | CH_SC | 5,233721344 | 0,191068636 | 6,282608696 |
| NIBAN15 | 4,01986E-09 | 1,832221898 | 0,733 | 0,062 | 0,000147075 | CH_SC | 3,560850578 | 0,280831778 | 11,82258065 |
| PLAT2 | 4,30196E-09 | 2,099936057 | 0,667 | 0,046 | 0,000157396 | CH_SC | 4,286903842 | 0,233268587 | 14,5 |
| GFRA3 | 4,683E-09 | 2,542291958 | 0,733 | 0,077 | 0,000171337 | CH_SC | 5,825136909 | 0,171669785 | 9,519480519 |
| NTM2 | 4,79255E-09 | 2,548787162 | 0,667 | 0,062 | 0,000175345 | CH_SC | 5,851421572 | 0,170898642 | 10,75806452 |
| IGFBP45 | 5,35957E-09 | 2,370888964 | 0,667 | 0,062 | 0,000196091 | CH_SC | 5,172597608 | 0,193326463 | 10,75806452 |
| TCIM1 | 5,57385E-09 | 1,825829334 | 0,733 | 0,062 | 0,000203931 | CH_SC | 3,545107398 | 0,282078901 | 11,82258065 |
| CREB51 | 5,64603E-09 | 2,090661999 | 0,6 | 0,031 | 0,000206571 | CH_SC | 4,259434778 | 0,234772934 | 19,35483871 |
| CNP | 6,81136E-09 | 2,101375637 | 0,8 | 0,123 | 0,000249207 | CH_SC | 4,291183625 | 0,233035938 | 6,504065041 |
| SOD33 | 8,48483E-09 | -5,378687679 | 0 | 0,954 | 0,000310434 | CH_SC | 0,024035528 | 41,60507687 | 0 |
| ANTXR21 | 9,21609E-09 | 1,798684191 | 0,6 | 0,031 | 0,000337189 | CH_SC | 3,479027761 | 0,287436626 | 19,35483871 |
| SLC15A3 | 9,32705E-09 | 1,868216779 | 0,667 | 0,062 | 0,000341249 | CH_SC | 3,650810482 | 0,273911781 | 10,75806452 |
| ADAM23 | 1,1362E-08 | 1,420408016 | 0,467 | 0 | 0,000415702 | CH_SC | 2,676611989 | 0,373606636 | #DIV/0! |
| COL1A12 | 1,17049E-08 | 2,015188285 | 0,733 | 0,092 | 0,000428245 | CH_SC | 4,042333314 | 0,247381876 | 7,967391304 |
| CD745 | 1,20376E-08 | 4,694861463 | 0,933 | 0,4 | 0,00044042 | CH_SC | 25,89966375 | 0,03861054 | 2,3325 |
| COL14A11 | 1,20847E-08 | 3,411386811 | 0,733 | 0,108 | 0,000442141 | CH_SC | 10,63970917 | 0,093987531 | 6,787037037 |
| ENDOD11 | 1,21048E-08 | 1,901561575 | 0,8 | 0,123 | 0,000442879 | CH_SC | 3,736173818 | 0,2676535 | 6,504065041 |

| Supplementary Table 4 - clusermarker | | | | | | | | | |
|---|---|---|---|---|---|---|---|---|---|
| | p_val | avg_log2FC | pct.1 | pct.2 | p_val_adj | celltype | Foldchange_UP | Foldchange_DOWN | Ratio_pct1_pct2 |
| KCTD121 | 1,79327E-08 | 2,064814008 | 0,667 | 0,062 | 0,000656102 | CH_SC | 4,183800349 | 0,239017142 | 10,75806452 |
| HLA-DQB13 | 1,95998E-08 | 3,132625363 | 0,6 | 0,046 | 0,000717099 | CH_SC | 8,770294957 | 0,114021251 | 13,04347826 |
| SEMA3B | 2,13793E-08 | 2,057090929 | 0,867 | 0,154 | 0,000782206 | CH_SC | 4,161463342 | 0,240300086 | 5,62987013 |
| ERBB32 | 2,35939E-08 | 2,463275001 | 0,733 | 0,092 | 0,000863232 | CH_SC | 5,514671697 | 0,181334457 | 7,967391304 |
| RPS256 | 2,52283E-08 | -2,06809618 | 0,867 | 1 | 0,000923028 | CH_SC | 0,238473989 | 4,193329449 | 0,867 |
| L1CAM | 2,8751E-08 | 1,995710971 | 0,733 | 0,092 | 0,001051912 | CH_SC | 3,988125945 | 0,250744338 | 7,967391304 |
| IFITM38 | 2,91412E-08 | 2,08030823 | 1 | 0,8 | 0,00106619 | CH_SC | 4,228975581 | 0,236463886 | 1,25 |
| TNC3 | 3,09236E-08 | 1,935749443 | 0,6 | 0,046 | 0,0011314 | CH_SC | 3,825768143 | 0,261385417 | 13,04347826 |
| IGFBP54 | 3,50638E-08 | 2,946675776 | 0,867 | 0,231 | 0,001282878 | CH_SC | 7,709705625 | 0,129706638 | 3,753246753 |
| PRELP1 | 3,58882E-08 | -4,670829742 | 0,067 | 0,923 | 0,001313041 | CH_SC | 0,039259082 | 25,47181296 | 0,072589382 |
| MATN21 | 3,983E-08 | 2,144887809 | 0,933 | 0,262 | 0,001457259 | CH_SC | 4,422578679 | 0,226112427 | 3,561068702 |
| LGALS17 | 4,9851E-08 | 2,489386617 | 1 | 0,646 | 0,001823897 | CH_SC | 5,615391528 | 0,178081973 | 1,547987616 |
| TMSB4X3 | 5,05205E-08 | 1,722825046 | 0,933 | 0,262 | 0,001848392 | CH_SC | 3,300821322 | 0,302954902 | 3,561068702 |
| SYT11 | 6,05839E-08 | 1,796183694 | 0,6 | 0,046 | 0,002216582 | CH_SC | 3,473003088 | 0,287935246 | 13,04347826 |
| GPM6B1 | 6,16247E-08 | 2,090204442 | 0,933 | 0,185 | 0,002254663 | CH_SC | 4,258084092 | 0,234847405 | 5,043243243 |
| A2M5 | 6,21722E-08 | -4,623083282 | 0 | 0,892 | 0,002274694 | CH_SC | 0,040580114 | 24,64261207 | 0 |
| PRR410 | 6,21834E-08 | -3,690346978 | 0 | 0,892 | 0,002275105 | CH_SC | 0,077463099 | 12,90937257 | 0 |
| TUBA1A4 | 7,19556E-08 | 2,426039841 | 0,867 | 0,215 | 0,00263264 | CH_SC | 5,37416211 | 0,186075518 | 4,03255814 |
| TMEM176A4 | 7,21639E-08 | 2,60075574 | 0,733 | 0,123 | 0,002640261 | CH_SC | 6,066043063 | 0,16485211 | 5,959349593 |
| APOD3 | 7,72365E-08 | -6,55169533 | 0,133 | 0,908 | 0,002825853 | CH_SC | 0,010659656 | 93,81165909 | 0,146475771 |
| SGCB2 | 8,66719E-08 | 2,132456053 | 0,733 | 0,123 | 0,003171066 | CH_SC | 4,384632883 | 0,228069265 | 5,959349593 |
| VWA12 | 8,76795E-08 | 2,075599037 | 0,8 | 0,154 | 0,003207929 | CH_SC | 4,215194016 | 0,237237004 | 5,194805195 |
| NRXN3 | 9,41339E-08 | 1,830797008 | 0,6 | 0,046 | 0,003444078 | CH_SC | 3,557335411 | 0,281109281 | 13,04347826 |
| MAP1B1 | 9,49478E-08 | 2,314915311 | 0,733 | 0,123 | 0,003473856 | CH_SC | 4,975754518 | 0,200974545 | 5,959349593 |
| ECRG41 | 9,9819E-08 | -6,132924983 | 0 | 0,877 | 0,003652076 | CH_SC | 0,014249697 | 70,1769325 | 0 |
| CHAD | 9,9819E-08 | -5,424070948 | 0 | 0,877 | 0,003652076 | CH_SC | 0,023291204 | 42,93466256 | 0 |
| TMSB107 | 1,11719E-07 | 2,141931012 | 1 | 0,538 | 0,004087445 | CH_SC | 4,413523905 | 0,226576319 | 1,858736059 |
| SPTBN15 | 1,14015E-07 | 2,56340894 | 0,933 | 0,431 | 0,004171483 | CH_SC | 5,911027532 | 0,169175324 | 2,164733179 |
| MPZ | 1,1707E-07 | 2,585203449 | 0,6 | 0,046 | 0,004283231 | CH_SC | 6,001002161 | 0,166638834 | 13,04347826 |
| NLGN4X | 1,1808E-07 | 1,776993198 | 0,6 | 0,062 | 0,004320199 | CH_SC | 3,427111662 | 0,291790901 | 9,677419355 |
| ABLIM3 | 1,52274E-07 | 1,022719207 | 0,4 | 0 | 0,005571235 | CH_SC | 2,031744807 | 0,492187797 | #DIV/0! |

| Supplementary Table 4 - clusermarker | | | | | | | | | |
|---|---|---|---|---|---|---|---|---|---|
| | **p_val** | **avg_log2FC** | **pct.1** | **pct.2** | **p_val_adj** | **celltype** | **Foldchange_UP** | **Foldchange_DOWN** | **Ratio_pct1_pct2** |
| APOL61 | 1,52274E-07 | 1,292284287 | 0,4 | 0 | 0,005571235 | CH_SC | 2,449155349 | 0,40830403 | #DIV/0! |
| ASPA | 1,52281E-07 | 1,614794192 | 0,667 | 0,077 | 0,005571511 | CH_SC | 3,062679055 | 0,326511522 | 8,662337662 |
| SAMHD13 | 1,55772E-07 | 1,514092332 | 0,867 | 0,154 | 0,005699245 | CH_SC | 2,856190749 | 0,350116672 | 5,62987013 |
| THSD7A | 1,66984E-07 | 1,626971914 | 0,467 | 0,015 | 0,006109457 | CH_SC | 3,088640402 | 0,323767053 | 31,13333333 |
| KCNMB4 | 1,72786E-07 | 1,75805863 | 0,733 | 0,108 | 0,006321721 | CH_SC | 3,382426606 | 0,295645735 | 6,787037037 |
| MGP6 | 1,74579E-07 | -5,541592287 | 0,2 | 0,892 | 0,006387338 | CH_SC | 0,021469133 | 46,57850037 | 0,224215247 |
| GAS2L3 | 1,74961E-07 | 1,84878549 | 0,533 | 0,031 | 0,006401282 | CH_SC | 3,601968315 | 0,277625985 | 17,19354839 |
| HSPA12A | 1,95073E-07 | 1,637709767 | 0,733 | 0,123 | 0,007137138 | CH_SC | 3,111714647 | 0,321366228 | 5,959349593 |
| DCN3 | 2,28143E-07 | -5,283202716 | 0,133 | 0,877 | 0,008347083 | CH_SC | 0,025680147 | 38,94058671 | 0,151653364 |
| PLK23 | 3,28122E-07 | 2,115269261 | 0,733 | 0,138 | 0,012004987 | CH_SC | 4,332708752 | 0,230802497 | 5,311594203 |
| MYOT | 3,31595E-07 | 1,742172566 | 0,6 | 0,062 | 0,012132067 | CH_SC | 3,345385729 | 0,298919192 | 9,677419355 |
| GAS71 | 3,45161E-07 | 2,294393056 | 0,733 | 0,154 | 0,012628414 | CH_SC | 4,905475735 | 0,203853827 | 4,75974026 |
| GNG23 | 3,66943E-07 | 1,891891083 | 0,6 | 0,062 | 0,013425348 | CH_SC | 3,711213715 | 0,269453628 | 9,677419355 |
| ACAN1 | 3,89743E-07 | -4,242313715 | 0 | 0,831 | 0,01425951 | CH_SC | 0,052836777 | 18,92621108 | 0 |
| FMOD | 3,89806E-07 | -4,275305115 | 0 | 0,831 | 0,014261819 | CH_SC | 0,051642222 | 19,36400027 | 0 |
| FGFBP2 | 3,89806E-07 | -4,535367064 | 0 | 0,831 | 0,014261819 | CH_SC | 0,043123944 | 23,18897367 | 0 |
| XKR4 | 4,29884E-07 | 2,643084351 | 0,667 | 0,092 | 0,015728149 | CH_SC | 6,246657158 | 0,160085623 | 7,25 |
| RPL215 | 4,3253E-07 | -1,666398682 | 0,867 | 0,985 | 0,01582498 | CH_SC | 0,315038776 | 3,174212431 | 0,880203046 |
| CLU4 | 4,88098E-07 | -4,569685514 | 0,6 | 0,985 | 0,01785806 | CH_SC | 0,042110228 | 23,74720002 | 0,609137056 |
| RPL36A8 | 4,95667E-07 | -2,234568938 | 0,533 | 0,954 | 0,018134963 | CH_SC | 0,212484728 | 4,70622057 | 0,55870021 |
| SLIT21 | 5,06286E-07 | 1,815203641 | 0,533 | 0,046 | 0,018523469 | CH_SC | 3,519093 | 0,28416413 | 11,58695652 |
| ITIH52 | 5,33911E-07 | 2,253834194 | 0,733 | 0,169 | 0,019534192 | CH_SC | 4,76948731 | 0,209666141 | 4,337278107 |
| ADGRB3 | 6,26915E-07 | 2,296212626 | 0,533 | 0,046 | 0,022936955 | CH_SC | 4,911666568 | 0,203596882 | 11,58695652 |
| RPL65 | 6,30574E-07 | -1,894536411 | 0,733 | 0,954 | 0,023070794 | CH_SC | 0,26896001 | 3,718024846 | 0,768343816 |
| PMP223 | 6,46737E-07 | 2,43675903 | 0,933 | 0,431 | 0,023662171 | CH_SC | 5,414240714 | 0,184698105 | 2,164733179 |
| RPS125 | 6,7668E-07 | -1,760500126 | 0,867 | 0,969 | 0,024757707 | CH_SC | 0,295145832 | 3,388155586 | 0,894736842 |
| ARHGEF26 | 7,38244E-07 | 1,252986493 | 0,6 | 0,062 | 0,027010148 | CH_SC | 2,383342836 | 0,419578747 | 9,677419355 |
| ARHGAP152 | 8,14616E-07 | 1,803999822 | 0,6 | 0,062 | 0,029804365 | CH_SC | 3,491869934 | 0,28637951 | 9,677419355 |
| RPS4X2 | 8,17066E-07 | -1,936472085 | 0,8 | 0,969 | 0,029893989 | CH_SC | 0,261254522 | 3,827684941 | 0,825593395 |
| RPL347 | 8,7152E-07 | -2,180095708 | 0,867 | 0,969 | 0,031886313 | CH_SC | 0,22066111 | 4,531836173 | 0,894736842 |
| CCN21 | 8,90987E-07 | -3,711362871 | 0,2 | 0,892 | 0,032598556 | CH_SC | 0,076342864 | 13,09880114 | 0,224215247 |

| Supplementary Table 4 - clusermarker | | | | | | | | | |
|---|---|---|---|---|---|---|---|---|---|
| | **p_val** | **avg_log2FC** | **pct.1** | **pct.2** | **p_val_adj** | **celltype** | **Foldchange_UP** | **Foldchange_DOWN** | **Ratio_pct1_pct2** |
| RPL365 | 9,13234E-07 | -2,06800321 | 0,667 | 0,938 | 0,033412503 | CH_SC | 0,238489357 | 4,193059231 | 0,71108742 |
| RBP41 | 9,22425E-07 | -5,05482091 | 0 | 0,8 | 0,033748775 | CH_SC | 0,030084811 | 33,23936485 | 0 |
| SLPI10 | 9,22567E-07 | -2,992538511 | 0 | 0,8 | 0,033753944 | CH_SC | 0,125648163 | 7,958731528 | 0 |
| RPS218 | 9,38276E-07 | -2,363442354 | 0,4 | 0,938 | 0,034328694 | CH_SC | 0,194326916 | 5,145967529 | 0,426439232 |
| MT1X6 | 9,44393E-07 | -4,333089516 | 0,133 | 0,846 | 0,034552499 | CH_SC | 0,049614667 | 20,15533024 | 0,157210402 |
| CADM11 | 9,50046E-07 | 1,841404507 | 0,867 | 0,277 | 0,034759321 | CH_SC | 3,583587316 | 0,279049989 | 3,129963899 |
| EEF1A16 | 9,89058E-07 | -1,628824143 | 0,933 | 0,969 | 0,036186661 | CH_SC | 0,323351646 | 3,092608353 | 0,962848297 |
| CYTL11 | 1,08161E-06 | -7,346503572 | 0,467 | 0,877 | 0,039572707 | CH_SC | 0,006144437 | 162,7488537 | 0,532497149 |
| RPS234 | 1,12128E-06 | -1,67846174 | 0,867 | 1 | 0,04102439 | CH_SC | 0,31241557 | 3,200864799 | 0,867 |
| RPS3A4 | 1,26825E-06 | -1,923565907 | 0,733 | 1 | 0,046401492 | CH_SC | 0,263602159 | 3,793595633 | 0,733 |
| RPS279 | 1,27007E-06 | -1,659075813 | 1 | 1 | 0,046467888 | CH_SC | 0,316641924 | 3,158141501 | 1 |
| RPL416 | 1,35131E-06 | -1,772858913 | 0,933 | 1 | 0,049440335 | CH_SC | 0,292628276 | 3,417304755 | 0,933 |
| COL4A23 | 1,35366E-06 | 1,921031808 | 0,533 | 0,062 | 0,049526536 | CH_SC | 3,786938016 | 0,264065584 | 8,596774194 |
| DCPS1 | 1,36038E-06 | 1,633412838 | 0,467 | 0,031 | 0,049772157 | CH_SC | 3,10246049 | 0,322324814 | 15,06451613 |
| SNORC | 1,39991E-06 | -4,988006701 | 0 | 0,785 | 0,051218605 | CH_SC | 0,031510868 | 31,73508279 | 0 |
| RPL173 | 1,41329E-06 | -2,057092846 | 0,6 | 0,954 | 0,051708014 | CH_SC | 0,240299767 | 4,161468872 | 0,628930818 |
| RPL378 | 1,43672E-06 | -1,579770893 | 0,867 | 0,985 | 0,05256543 | CH_SC | 0,33453501 | 2,989223756 | 0,880203046 |
| SEMA3C1 | 1,52845E-06 | 2,479146819 | 0,667 | 0,138 | 0,055921518 | CH_SC | 5,575676347 | 0,179350439 | 4,833333333 |
| RPL266 | 1,62159E-06 | -1,706556459 | 0,8 | 0,969 | 0,059328962 | CH_SC | 0,306390515 | 3,263808607 | 0,825593395 |
| DAG1 | 1,65108E-06 | 1,664942012 | 0,533 | 0,062 | 0,060408108 | CH_SC | 3,171009088 | 0,315357027 | 8,596774194 |
| COL28A11 | 1,66322E-06 | 1,550363359 | 0,667 | 0,092 | 0,060852155 | CH_SC | 2,928908978 | 0,341424062 | 7,25 |
| CTSC5 | 1,70319E-06 | 1,68514891 | 0,467 | 0,031 | 0,062314617 | CH_SC | 3,215735862 | 0,310970814 | 15,06451613 |
| PDGFB1 | 1,7662E-06 | 1,258079147 | 0,4 | 0,015 | 0,064620137 | CH_SC | 2,391770803 | 0,418100262 | 26,66666667 |
| RPL193 | 1,83815E-06 | -1,326631578 | 0,867 | 1 | 0,067252444 | CH_SC | 0,398698041 | 2,508163815 | 0,867 |
| CADM32 | 1,88593E-06 | 1,327286944 | 0,6 | 0,077 | 0,06900039 | CH_SC | 2,509303446 | 0,398516968 | 7,792207792 |
| ANK33 | 1,91241E-06 | 2,20834006 | 0,733 | 0,154 | 0,0699692 | CH_SC | 4,621432336 | 0,216383131 | 4,75974026 |
| SEMA3G1 | 1,94179E-06 | 1,178775319 | 0,333 | 0 | 0,071044303 | CH_SC | 2,263845213 | 0,441726313 | #DIV/0! |
| PCDH11 | 1,94179E-06 | 1,29525482 | 0,333 | 0 | 0,071044303 | CH_SC | 2,454203397 | 0,40746419 | #DIV/0! |
| C22 | 1,94179E-06 | 1,751151384 | 0,333 | 0 | 0,071044303 | CH_SC | 3,366271139 | 0,297064603 | #DIV/0! |
| PLXNB3 | 1,94179E-06 | 1,522202822 | 0,333 | 0 | 0,071044303 | CH_SC | 2,872292795 | 0,348153921 | #DIV/0! |
| TSPAN111 | 1,95528E-06 | 2,61542875 | 0,667 | 0,154 | 0,07153765 | CH_SC | 6,128052894 | 0,16318397 | 4,331168831 |

| Supplementary Table 4 - clusermarker | | | | | | | | | |
|---|---|---|---|---|---|---|---|---|---|
| | **p_val** | **avg_log2FC** | **pct.1** | **pct.2** | **p_val_adj** | **celltype** | **Foldchange_UP** | **Foldchange_DOWN** | **Ratio_pct1_pct2** |
| SLC35F1 | 2,00088E-06 | 1,33836595 | 0,4 | 0,015 | 0,07320613 | CH_SC | 2,528647526 | 0,395468324 | 26,66666667 |
| CFH3 | 2,1061E-06 | -3,830859651 | 0 | 0,769 | 0,077055716 | CH_SC | 0,070274269 | 14,22995951 | 0 |
| SORCS1 | 2,12824E-06 | 1,1185561 | 0,467 | 0,031 | 0,077865885 | CH_SC | 2,171295529 | 0,460554534 | 15,06451613 |
| RXRG | 2,26686E-06 | 1,322010233 | 0,6 | 0,077 | 0,082937696 | CH_SC | 2,500142339 | 0,399977227 | 7,792207792 |
| EHBP1 | 2,34639E-06 | 1,826402664 | 0,733 | 0,185 | 0,085847433 | CH_SC | 3,546516511 | 0,281966825 | 3,962162162 |
| PRNP3 | 2,43002E-06 | 1,966239783 | 0,933 | 0,6 | 0,088907243 | CH_SC | 3,907483505 | 0,255919187 | 1,555 |
| TNFAIP6 | 2,48404E-06 | 1,29653527 | 0,6 | 0,077 | 0,090883392 | CH_SC | 2,456382569 | 0,40710271 | 7,792207792 |
| RPL37A9 | 2,60457E-06 | -1,543026238 | 0,667 | 0,938 | 0,095293543 | CH_SC | 0,343164867 | 2,914051225 | 0,71108742 |
| RPL35A8 | 2,64564E-06 | -1,38796657 | 0,867 | 0,985 | 0,096796148 | CH_SC | 0,382102984 | 2,617095499 | 0,880203046 |
| CNN33 | 2,77336E-06 | 1,655519548 | 0,933 | 0,462 | 0,101468793 | CH_SC | 3,150366222 | 0,317423413 | 2,019480519 |
| GNG111 | 3,26944E-06 | 1,578576478 | 0,533 | 0,062 | 0,119619048 | CH_SC | 2,986749986 | 0,334812088 | 8,596774194 |
| RASSF41 | 3,26944E-06 | 1,607515479 | 0,533 | 0,062 | 0,119619048 | CH_SC | 3,047266082 | 0,328163007 | 8,596774194 |
| RPL225 | 3,32831E-06 | -1,641456694 | 0,6 | 0,969 | 0,121772715 | CH_SC | 0,320532668 | 3,119806808 | 0,619195046 |
| RPS66 | 3,56297E-06 | -1,742625992 | 0,867 | 0,954 | 0,130358303 | CH_SC | 0,298825259 | 3,34643732 | 0,908805031 |
| RPS282 | 3,67549E-06 | -1,399899415 | 0,8 | 0,985 | 0,134475113 | CH_SC | 0,378955562 | 2,638831835 | 0,812182741 |
| RPS146 | 3,78942E-06 | -1,209678149 | 0,933 | 0,969 | 0,138643421 | CH_SC | 0,432365061 | 2,312860334 | 0,962848297 |
| ITGB43 | 4,26396E-06 | 1,616776639 | 0,6 | 0,092 | 0,156005571 | CH_SC | 3,06689046 | 0,326063162 | 6,52173913 |
| PRSS235 | 4,48287E-06 | 2,153127801 | 0,6 | 0,108 | 0,164014906 | CH_SC | 4,44791063 | 0,224824661 | 5,555555556 |
| PSME26 | 4,48287E-06 | 1,538010127 | 0,6 | 0,108 | 0,164014906 | CH_SC | 2,903936944 | 0,344360094 | 5,555555556 |
| FRZB2 | 4,64294E-06 | -4,547129598 | 0 | 0,738 | 0,16987115 | CH_SC | 0,042773777 | 23,37881009 | 0 |
| HAPLN1 | 4,64294E-06 | -3,440319183 | 0 | 0,738 | 0,16987115 | CH_SC | 0,092121443 | 10,85523598 | 0 |
| CRYAB3 | 4,65249E-06 | 2,091749319 | 0,933 | 0,646 | 0,170220569 | CH_SC | 4,262646208 | 0,234596059 | 1,444272446 |
| SCN9A | 4,79065E-06 | 1,593161592 | 0,533 | 0,062 | 0,175275666 | CH_SC | 3,01709807 | 0,331444314 | 8,596774194 |
| S100A66 | 4,80343E-06 | 1,302173547 | 1 | 1 | 0,175743028 | CH_SC | 2,466001278 | 0,405514794 | 1 |
| SOX22 | 5,26579E-06 | 1,284445007 | 0,533 | 0,062 | 0,192659536 | CH_SC | 2,435883282 | 0,410528701 | 8,596774194 |
| MFGE83 | 5,39946E-06 | -3,494865607 | 0,2 | 0,815 | 0,197549941 | CH_SC | 0,088703472 | 11,27351582 | 0,245398773 |
| RPL301 | 5,7244E-06 | -1,317794479 | 0,867 | 0,985 | 0,209438463 | CH_SC | 0,401147725 | 2,492847241 | 0,880203046 |
| RPS73 | 6,02889E-06 | -1,585846228 | 0,733 | 0,954 | 0,220578989 | CH_SC | 0,333129211 | 3,001838222 | 0,768343816 |
| GPX31 | 6,13024E-06 | -3,13305504 | 0,067 | 0,769 | 0,224287146 | CH_SC | 0,113987297 | 8,772907395 | 0,087126138 |
| LPAR1 | 6,33692E-06 | 1,778347831 | 0,667 | 0,154 | 0,231848996 | CH_SC | 3,430331093 | 0,29151705 | 4,331168831 |
| SLITRK6 | 6,35489E-06 | 1,176876569 | 0,533 | 0,062 | 0,232506376 | CH_SC | 2,260867696 | 0,442308058 | 8,596774194 |

| Supplementary Table 4 - clusermarker | | | | | | | | | |
|---|---|---|---|---|---|---|---|---|---|
| | **p_val** | **avg_log2FC** | **pct.1** | **pct.2** | **p_val_adj** | **celltype** | **Foldchange_UP** | **Foldchange_DOWN** | **Ratio_pct1_pct2** |
| RPL182 | 6,7491E-06 | -1,683990212 | 0,733 | 0,938 | 0,246929345 | CH_SC | 0,31122067 | 3,213154187 | 0,781449893 |
| C2orf88 | 6,97724E-06 | 1,330422295 | 0,533 | 0,062 | 0,255276263 | CH_SC | 2,514762744 | 0,397651827 | 8,596774194 |
| PMEPA16 | 7,22741E-06 | 3,074558357 | 0,8 | 0,431 | 0,264429293 | CH_SC | 8,424309013 | 0,118704098 | 1,856148492 |
| RND32 | 7,38075E-06 | 1,958915486 | 0,667 | 0,154 | 0,270039597 | CH_SC | 3,887696202 | 0,257221745 | 4,331168831 |
| CTNNA13 | 7,95776E-06 | 1,953612217 | 0,8 | 0,292 | 0,291150439 | CH_SC | 3,873431475 | 0,258169018 | 2,739726027 |
| CHL1 | 7,99441E-06 | 1,592680524 | 0,733 | 0,169 | 0,29249166 | CH_SC | 3,016092185 | 0,331554853 | 4,337278107 |
| RPS185 | 8,10522E-06 | -1,446135629 | 0,933 | 0,969 | 0,296545599 | CH_SC | 0,367003155 | 2,724772216 | 0,962848297 |
| COL9A22 | 8,26554E-06 | -3,274606356 | 0,067 | 0,738 | 0,302411249 | CH_SC | 0,103334481 | 9,677311867 | 0,090785908 |
| COTL17 | 8,95166E-06 | 1,289296659 | 0,467 | 0,046 | 0,327514348 | CH_SC | 2,444088726 | 0,409150449 | 10,15217391 |
| NGFR | 8,95166E-06 | 1,871932148 | 0,467 | 0,046 | 0,327514348 | CH_SC | 3,660224524 | 0,273207284 | 10,15217391 |
| FST1 | 9,29772E-06 | 1,733341418 | 0,533 | 0,077 | 0,34017574 | CH_SC | 3,324970218 | 0,300754574 | 6,922077922 |
| OLFML2A1 | 9,29772E-06 | 1,87214203 | 0,533 | 0,077 | 0,34017574 | CH_SC | 3,660757048 | 0,273167541 | 6,922077922 |
| AP1S21 | 9,47431E-06 | 1,918528934 | 0,6 | 0,108 | 0,346636461 | CH_SC | 3,780373905 | 0,264524099 | 5,555555556 |
| JUN5 | 9,6074E-06 | 1,63132674 | 1 | 0,723 | 0,351505877 | CH_SC | 3,097977659 | 0,322791224 | 1,383125864 |
| LYZ10 | 9,89616E-06 | -2,830082471 | 0 | 0,708 | 0,362070862 | CH_SC | 0,140624272 | 7,111147942 | 0 |
| RPLP17 | 9,90956E-06 | -1,402057996 | 0,933 | 0,985 | 0,362561091 | CH_SC | 0,378388987 | 2,642783049 | 0,947208122 |
| SSR36 | 1,05541E-05 | -2,018174126 | 0,333 | 0,862 | 0,386141275 | CH_SC | 0,246870418 | 4,050708099 | 0,386310905 |
| RPL113 | 1,07928E-05 | -1,18554442 | 1 | 0,969 | 0,394877672 | CH_SC | 0,439658596 | 2,274492094 | 1,031991744 |
| RPL53 | 1,09948E-05 | -1,436651847 | 0,667 | 0,954 | 0,402266892 | CH_SC | 0,369423655 | 2,706919243 | 0,699161426 |
| SORBS22 | 1,12862E-05 | 1,437164346 | 0,8 | 0,231 | 0,412929461 | CH_SC | 2,707881011 | 0,369292445 | 3,463203463 |
| RPS15A3 | 1,14229E-05 | -1,426731803 | 0,933 | 1 | 0,41793075 | CH_SC | 0,371972584 | 2,688370173 | 0,933 |
| DKK31 | 1,17633E-05 | 1,59170767 | 0,8 | 0,262 | 0,43038293 | CH_SC | 3,014059026 | 0,331778506 | 3,053435115 |
| LAP35 | 1,20491E-05 | 1,200375226 | 0,467 | 0,046 | 0,440840719 | CH_SC | 2,29799431 | 0,435162087 | 10,15217391 |
| CELF23 | 1,20491E-05 | 1,505921563 | 0,467 | 0,046 | 0,440840719 | CH_SC | 2,840060304 | 0,3521052 | 10,15217391 |
| IGKC9 | 1,21253E-05 | 1,736412792 | 0,533 | 0,077 | 0,443628625 | CH_SC | 3,332056333 | 0,300114974 | 6,922077922 |
| ALDH1A3 | 1,25081E-05 | 1,368341277 | 0,4 | 0,031 | 0,457634027 | CH_SC | 2,581735631 | 0,387336328 | 12,90322581 |
| SOX46 | 1,28633E-05 | 2,200275748 | 0,8 | 0,323 | 0,470629879 | CH_SC | 4,595671725 | 0,217596047 | 2,476780186 |
| LAMC11 | 1,30162E-05 | 1,797401135 | 0,6 | 0,123 | 0,476224383 | CH_SC | 3,475935074 | 0,28769237 | 4,87804878 |
| GFRA2 | 1,32919E-05 | 1,481577053 | 0,467 | 0,046 | 0,486312193 | CH_SC | 2,792538271 | 0,358097151 | 10,15217391 |
| PFN18 | 1,39266E-05 | 1,574101865 | 0,867 | 0,585 | 0,509533453 | CH_SC | 2,977500737 | 0,335852142 | 1,482051282 |
| SNAPC2 | 1,39364E-05 | 1,377744346 | 0,4 | 0,031 | 0,509892254 | CH_SC | 2,598617594 | 0,384819991 | 12,90322581 |

| Supplementary Table 4 - clusermarker | | | | | | | | | |
|---|---|---|---|---|---|---|---|---|---|
| | **p_val** | **avg_log2FC** | **pct.1** | **pct.2** | **p_val_adj** | **celltype** | **Foldchange_UP** | **Foldchange_DOWN** | **Ratio_pct1_pct2** |
| B2M8 | 1,43124E-05 | 1,555573391 | 1 | 0,969 | 0,523646488 | CH_SC | 2,939505324 | 0,340193294 | 1,031991744 |
| MAL | 1,44478E-05 | 1,771717333 | 0,533 | 0,077 | 0,528603175 | CH_SC | 3,41460177 | 0,29285992 | 6,922077922 |
| RDX1 | 1,47485E-05 | 1,654869034 | 0,667 | 0,169 | 0,539603616 | CH_SC | 3,148946035 | 0,317566573 | 3,946745562 |
| FRMD31 | 1,55195E-05 | 1,188912651 | 0,4 | 0,031 | 0,567812609 | CH_SC | 2,279808508 | 0,438633331 | 12,90322581 |
| FRMD4A | 1,72731E-05 | 1,456991249 | 0,4 | 0,031 | 0,6319726 | CH_SC | 2,745352209 | 0,364251988 | 12,90322581 |
| NPTX2 | 1,92146E-05 | 1,765915867 | 0,4 | 0,031 | 0,703004577 | CH_SC | 3,400898306 | 0,294039959 | 12,90322581 |
| RPS34 | 1,9943E-05 | -1,427187835 | 0,933 | 0,938 | 0,729654052 | CH_SC | 0,371855023 | 2,689220094 | 0,99466951 |
| IFI351 | 2,02522E-05 | 1,60598453 | 0,467 | 0,062 | 0,740966891 | CH_SC | 3,044034121 | 0,328511429 | 7,532258065 |
| RPS2910 | 2,05838E-05 | -1,789379559 | 0,4 | 0,862 | 0,753097699 | CH_SC | 0,289296433 | 3,456662045 | 0,464037123 |
| LMNA6 | 2,16339E-05 | 1,34593524 | 0,933 | 0,415 | 0,791519201 | CH_SC | 2,541949273 | 0,393398881 | 2,248192771 |
| TPT15 | 2,17075E-05 | -1,41794738 | 0,933 | 0,985 | 0,794211238 | CH_SC | 0,374244396 | 2,672050696 | 0,947208122 |
| TBX3 | 2,25903E-05 | 0,882421689 | 0,333 | 0,015 | 0,826510963 | CH_SC | 1,843467125 | 0,542456107 | 22,2 |
| RPGR1 | 2,25903E-05 | 1,307665995 | 0,333 | 0,015 | 0,826510963 | CH_SC | 2,475407422 | 0,403973904 | 22,2 |
| CTNNAL11 | 2,28786E-05 | 1,951307987 | 0,533 | 0,092 | 0,837061049 | CH_SC | 3,867249881 | 0,258581687 | 5,793478261 |
| PTPRF3 | 2,37202E-05 | 0,613363573 | 0,267 | 0 | 0,867851322 | CH_SC | 1,529821759 | 0,653670922 | #DIV/0! |
| CDK18 | 2,37202E-05 | 1,402541179 | 0,267 | 0 | 0,867851322 | CH_SC | 2,643668309 | 0,378262279 | #DIV/0! |
| NID11 | 2,37202E-05 | 0,876424209 | 0,267 | 0 | 0,867851322 | CH_SC | 1,835819488 | 0,544715865 | #DIV/0! |
| ASXL22 | 2,37202E-05 | 0,788803143 | 0,267 | 0 | 0,867851322 | CH_SC | 1,72764062 | 0,578824084 | #DIV/0! |
| CYP27C1 | 2,37202E-05 | 0,839918036 | 0,267 | 0 | 0,867851322 | CH_SC | 1,789948447 | 0,558675308 | #DIV/0! |
| GPR17 | 2,37202E-05 | 1,208038507 | 0,267 | 0 | 0,867851322 | CH_SC | 2,310233231 | 0,432856729 | #DIV/0! |
| ANXA33 | 2,37202E-05 | 1,053998445 | 0,267 | 0 | 0,867851322 | CH_SC | 2,076276303 | 0,481631466 | #DIV/0! |
| AC093849.2 | 2,37202E-05 | 0,667714709 | 0,267 | 0 | 0,867851322 | CH_SC | 1,588554634 | 0,629503058 | #DIV/0! |
| ENC13 | 2,37202E-05 | 1,080315964 | 0,267 | 0 | 0,867851322 | CH_SC | 2,114499126 | 0,472925237 | #DIV/0! |
| JAKMIP2 | 2,37202E-05 | 0,732587854 | 0,267 | 0 | 0,867851322 | CH_SC | 1,661616968 | 0,601823416 | #DIV/0! |
| PSMB8-AS1 | 2,37202E-05 | 0,940060819 | 0,267 | 0 | 0,867851322 | CH_SC | 1,918609118 | 0,521210908 | #DIV/0! |
| ADAMTSL1 | 2,37202E-05 | 0,730066347 | 0,267 | 0 | 0,867851322 | CH_SC | 1,658715372 | 0,602876188 | #DIV/0! |
| NEK64 | 2,37202E-05 | 0,451271298 | 0,267 | 0 | 0,867851322 | CH_SC | 1,367244537 | 0,731398059 | #DIV/0! |
| CXCL12 | 2,37202E-05 | 0,868748607 | 0,267 | 0 | 0,867851322 | CH_SC | 1,826078274 | 0,547621651 | #DIV/0! |
| B4GALNT4 | 2,37202E-05 | 1,443564654 | 0,267 | 0 | 0,867851322 | CH_SC | 2,719920822 | 0,367657761 | #DIV/0! |
| TMEM132B | 2,37202E-05 | 0,914627478 | 0,267 | 0 | 0,867851322 | CH_SC | 1,885082257 | 0,53048083 | #DIV/0! |
| SLFN53 | 2,37202E-05 | 0,60831236 | 0,267 | 0 | 0,867851322 | CH_SC | 1,524474861 | 0,655963588 | #DIV/0! |

| Supplementary Table 4 - clusermarker | | | | | | | | | |
|---|---|---|---|---|---|---|---|---|---|
| | **p_val** | **avg_log2FC** | **pct.1** | **pct.2** | **p_val_adj** | **celltype** | **Foldchange_UP** | **Foldchange_DOWN** | **Ratio_pct1_pct2** |
| LMNB2 | 2,37202E-05 | 1,303808346 | 0,267 | 0 | 0,867851322 | CH_SC | 2,468797227 | 0,405055543 | #DIV/0! |
| MFSD12 | 2,37202E-05 | 0,881935247 | 0,267 | 0 | 0,867851322 | CH_SC | 1,842845658 | 0,542639041 | #DIV/0! |
| ARNTL1 | 2,37384E-05 | 0,767778549 | 0,4 | 0,031 | 0,868517101 | CH_SC | 1,702646042 | 0,587321132 | 12,90322581 |
| LSM74 | 2,50289E-05 | 1,434590684 | 0,867 | 0,338 | 0,915733375 | CH_SC | 2,703054657 | 0,369951824 | 2,565088757 |
| FARP22 | 2,54662E-05 | 0,947036952 | 0,333 | 0,015 | 0,931732789 | CH_SC | 1,927908995 | 0,518696683 | 22,2 |
| ARRDC4 | 2,54662E-05 | 0,655487112 | 0,333 | 0,015 | 0,931732789 | CH_SC | 1,575147697 | 0,6348611 | 22,2 |
| DSTN2 | 2,60346E-05 | -2,44128321 | 0,067 | 0,708 | 0,952526734 | CH_SC | 0,184119813 | 5,431246001 | 0,094632768 |
| RPL139 | 2,62369E-05 | -1,344178337 | 0,8 | 0,969 | 0,959930551 | CH_SC | 0,393878251 | 2,53885559 | 0,825593395 |
| S100A45 | 2,66295E-05 | 2,181210179 | 0,867 | 0,446 | 0,974291947 | CH_SC | 4,535338333 | 0,220490717 | 1,943946188 |
| LGALS3BP3 | 2,66535E-05 | 1,891281493 | 0,467 | 0,062 | 0,975171002 | CH_SC | 3,709645927 | 0,269567506 | 7,532258065 |
| RPS246 | 2,75935E-05 | -1,436830896 | 0,733 | 0,954 | 1 | CH_SC | 0,369377809 | 2,707255212 | 0,768343816 |
| RPL322 | 2,77524E-05 | -1,339641796 | 0,867 | 0,985 | 1 | CH_SC | 0,395118747 | 2,530884721 | 0,880203046 |
| DPYD2 | 2,86883E-05 | 1,452993446 | 0,333 | 0,015 | 1 | CH_SC | 2,737755189 | 0,365262754 | 22,2 |
| GABRA2 | 2,86883E-05 | 1,088232095 | 0,333 | 0,015 | 1 | CH_SC | 2,126133366 | 0,470337382 | 22,2 |
| SFRP2 | 2,89874E-05 | -4,577705224 | 0 | 0,662 | 1 | CH_SC | 0,041876793 | 23,87957448 | 0 |
| COL11A2 | 2,89874E-05 | -2,753532088 | 0 | 0,662 | 1 | CH_SC | 0,148287399 | 6,743661345 | 0 |
| COL2A1 | 2,89874E-05 | -4,950439305 | 0 | 0,662 | 1 | CH_SC | 0,032342179 | 30,91937613 | 0 |
| SERPINA11 | 2,89874E-05 | -3,678721942 | 0 | 0,662 | 1 | CH_SC | 0,078089807 | 12,80576861 | 0 |
| BCL2 | 2,91852E-05 | 1,662501779 | 0,467 | 0,062 | 1 | CH_SC | 3,165650047 | 0,315890887 | 7,532258065 |
| RPL243 | 3,07577E-05 | -1,530785416 | 0,867 | 0,923 | 1 | CH_SC | 0,346088902 | 2,889430995 | 0,939328277 |
| PARP10 | 3,22955E-05 | 0,681375354 | 0,333 | 0,015 | 1 | CH_SC | 1,603667839 | 0,623570527 | 22,2 |
| GFRA11 | 3,22955E-05 | 1,28431863 | 0,333 | 0,015 | 1 | CH_SC | 2,435669914 | 0,410564664 | 22,2 |
| HTATIP23 | 3,22955E-05 | 0,801321409 | 0,333 | 0,015 | 1 | CH_SC | 1,742696585 | 0,573823354 | 22,2 |
| LINC005112 | 3,22955E-05 | 1,414015097 | 0,333 | 0,015 | 1 | CH_SC | 2,664777536 | 0,375265847 | 22,2 |
| SCAMP41 | 3,22955E-05 | 0,774361798 | 0,333 | 0,015 | 1 | CH_SC | 1,710433243 | 0,584647196 | 22,2 |
| RPS27A2 | 3,25893E-05 | -1,347480136 | 0,933 | 0,938 | 1 | CH_SC | 0,392977839 | 2,544672752 | 0,99466951 |
| FSTL3 | 3,49503E-05 | 1,855289876 | 0,467 | 0,062 | 1 | CH_SC | 3,618244438 | 0,276377126 | 7,532258065 |
| F3 | 3,6331E-05 | 0,853688188 | 0,333 | 0,015 | 1 | CH_SC | 1,807114837 | 0,553368264 | 22,2 |
| IL1RAP | 3,6331E-05 | 1,165657624 | 0,333 | 0,015 | 1 | CH_SC | 2,243354511 | 0,445761022 | 22,2 |
| HAND2-AS1 | 3,6331E-05 | 1,085537472 | 0,333 | 0,015 | 1 | CH_SC | 2,122165942 | 0,471216685 | 22,2 |
| NKAIN3 | 3,6331E-05 | 0,893791387 | 0,333 | 0,015 | 1 | CH_SC | 1,858052655 | 0,53819788 | 22,2 |

| Supplementary Table 4 - clusermarker | | | | | | | | | |
|---|---|---|---|---|---|---|---|---|---|
| | **p_val** | **avg_log2FC** | **pct.1** | **pct.2** | **p_val_adj** | **celltype** | **Foldchange_UP** | **Foldchange_DOWN** | **Ratio_pct1_pct2** |
| COL5A31 | 3,6331E-05 | 0,904108654 | 0,333 | 0,015 | 1 | CH_SC | 1,871387931 | 0,534362749 | 22,2 |
| ITM2A4 | 3,76186E-05 | -2,341034921 | 0,133 | 0,708 | 1 | CH_SC | 0,197368694 | 5,066659653 | 0,187853107 |
| VIM4 | 4,04697E-05 | 0,958624762 | 1 | 1 | 1 | CH_SC | 1,943456426 | 0,514547168 | 1 |
| SOX92 | 4,08377E-05 | -2,117757122 | 0 | 0,646 | 1 | CH_SC | 0,230404832 | 4,340186755 | 0 |
| EEF1G5 | 4,13542E-05 | -1,555468748 | 0,467 | 0,862 | 1 | CH_SC | 0,340217971 | 2,93929212 | 0,541763341 |
| RPS269 | 4,26654E-05 | -1,037394992 | 0,933 | 0,969 | 1 | CH_SC | 0,487206406 | 2,052518168 | 0,962848297 |
| RELN | 4,55967E-05 | 1,369582498 | 0,667 | 0,154 | 1 | CH_SC | 2,583957781 | 0,387003227 | 4,331168831 |
| RHOB4 | 4,8527E-05 | 1,638007314 | 0,867 | 0,723 | 1 | CH_SC | 3,112356486 | 0,321299955 | 1,199170124 |
| SOX10 | 4,98788E-05 | 1,561570253 | 0,467 | 0,062 | 1 | CH_SC | 2,95174942 | 0,338782145 | 7,532258065 |
| CLIC42 | 5,07758E-05 | 2,235704479 | 0,8 | 0,431 | 1 | CH_SC | 4,709926281 | 0,212317548 | 1,856148492 |
| PSME16 | 5,29521E-05 | 1,544637845 | 0,667 | 0,2 | 1 | CH_SC | 2,917308275 | 0,342781738 | 3,335 |
| IFI273 | 5,62349E-05 | 2,229224113 | 0,467 | 0,077 | 1 | CH_SC | 4,688817453 | 0,213273391 | 6,064935065 |
| RALA2 | 5,77408E-05 | 1,735473137 | 0,4 | 0,046 | 1 | CH_SC | 3,329886807 | 0,300310508 | 8,695652174 |
| MAPRE22 | 5,84961E-05 | 1,622192467 | 0,533 | 0,108 | 1 | CH_SC | 3,0784251 | 0,324841426 | 4,935185185 |
| AFAP1L2 | 5,94422E-05 | 0,744714128 | 0,467 | 0,062 | 1 | CH_SC | 1,675642205 | 0,596786114 | 7,532258065 |
| ATP1A21 | 6,03605E-05 | 1,371279208 | 0,533 | 0,092 | 1 | CH_SC | 2,586998483 | 0,386548352 | 5,793478261 |
| CAVIN32 | 6,168E-05 | 1,545674899 | 0,8 | 0,338 | 1 | CH_SC | 2,91940608 | 0,342535424 | 2,366863905 |
| SNHG295 | 6,54645E-05 | -1,842341544 | 0,333 | 0,8 | 1 | CH_SC | 0,278868803 | 3,585915627 | 0,41625 |
| CORO2B | 6,98133E-05 | 1,116009614 | 0,4 | 0,046 | 1 | CH_SC | 2,167466377 | 0,461368172 | 8,695652174 |
| SPINT25 | 7,02212E-05 | -1,738120221 | 0,067 | 0,738 | 1 | CH_SC | 0,299759998 | 3,336002156 | 0,090785908 |
| TSC22D17 | 7,03446E-05 | -2,028615253 | 0,133 | 0,769 | 1 | CH_SC | 0,245090207 | 4,080130378 | 0,172951886 |
| PEA151 | 7,35534E-05 | 1,558283021 | 0,533 | 0,108 | 1 | CH_SC | 2,945031409 | 0,339554952 | 4,935185185 |
| RPLP08 | 7,37544E-05 | -1,392754511 | 0,8 | 0,954 | 1 | CH_SC | 0,380836983 | 2,625795408 | 0,838574423 |
| RPL389 | 7,77725E-05 | -1,428366452 | 0,6 | 0,908 | 1 | CH_SC | 0,371551358 | 2,691417963 | 0,660792952 |
| ENPP1 | 7,9266E-05 | -2,194793543 | 0 | 0,615 | 1 | CH_SC | 0,218424479 | 4,578241424 | 0 |
| MEGF91 | 8,53483E-05 | 1,622359176 | 0,467 | 0,077 | 1 | CH_SC | 3,078780844 | 0,324803892 | 6,064935065 |
| CANX1 | 8,61624E-05 | 1,001963525 | 0,867 | 0,446 | 1 | CH_SC | 2,002723876 | 0,499319957 | 1,943946188 |
| RPL153 | 8,8289E-05 | -1,241719522 | 0,733 | 0,969 | 1 | CH_SC | 0,422868347 | 2,364802208 | 0,756449948 |
| RPL7A3 | 8,85926E-05 | -1,060734956 | 0,933 | 1 | 1 | CH_SC | 0,479387782 | 2,085993924 | 0,933 |
| AGAP13 | 9,24813E-05 | 1,430151458 | 0,4 | 0,046 | 1 | CH_SC | 2,69475004 | 0,371091932 | 8,695652174 |
| PPP2R2B1 | 9,24813E-05 | 1,928823498 | 0,4 | 0,046 | 1 | CH_SC | 3,807445796 | 0,262643266 | 8,695652174 |

| Supplementary Table 4 - clusermarker | | | | | | | | | |
|---|---|---|---|---|---|---|---|---|---|
| | **p_val** | **avg_log2FC** | **pct.1** | **pct.2** | **p_val_adj** | **celltype** | **Foldchange_UP** | **Foldchange_DOWN** | **Ratio_pct1_pct2** |
| FTH19 | 9,32261E-05 | 0,935807039 | 1 | 0,985 | 1 | CH_SC | 1,91296044 | 0,522749963 | 1,015228426 |
| ALCAM1 | 9,48722E-05 | 1,690633508 | 0,667 | 0,2 | 1 | CH_SC | 3,227984179 | 0,309790862 | 3,335 |
| SPARC5 | 0,000103024 | 0,937006924 | 1 | 0,923 | 1 | CH_SC | 1,914552105 | 0,522315375 | 1,083423619 |
| RPS132 | 0,000103217 | -1,017106194 | 0,933 | 1 | 1 | CH_SC | 0,494106454 | 2,023855369 | 0,933 |
| CD910 | 0,000108402 | 0,954458689 | 1 | 0,769 | 1 | CH_SC | 1,937852399 | 0,516035174 | 1,300390117 |
| SLC40A11 | 0,000109245 | -2,671718759 | 0 | 0,6 | 1 | CH_SC | 0,15693959 | 6,371878507 | 0 |
| ANGPTL4 | 0,000111281 | 1,451470723 | 0,4 | 0,046 | 1 | CH_SC | 2,734867092 | 0,365648482 | 8,695652174 |
| UQCRB4 | 0,000112269 | -1,440440664 | 0,533 | 0,892 | 1 | CH_SC | 0,368454745 | 2,714037517 | 0,597533632 |
| RPS57 | 0,000113612 | -1,234156299 | 0,667 | 0,969 | 1 | CH_SC | 0,425091024 | 2,352437345 | 0,688338493 |
| PLPP12 | 0,000113995 | 1,71175732 | 0,667 | 0,246 | 1 | CH_SC | 3,275595747 | 0,305287977 | 2,711382114 |
| CSRP14 | 0,000120458 | 1,267642195 | 0,8 | 0,4 | 1 | CH_SC | 2,407677556 | 0,415338008 | 2 |
| RPL23A5 | 0,000124905 | -1,284327799 | 0,733 | 0,923 | 1 | CH_SC | 0,410562055 | 2,435685395 | 0,794149512 |
| PRIMA1 | 0,000128296 | 1,057580505 | 0,467 | 0,062 | 1 | CH_SC | 2,081437884 | 0,480437109 | 7,532258065 |
| SIGIRR5 | 0,000128326 | 1,110969278 | 0,467 | 0,077 | 1 | CH_SC | 2,159907124 | 0,46298287 | 6,064935065 |
| HLA-B9 | 0,000129563 | 1,545312061 | 0,867 | 0,769 | 1 | CH_SC | 2,918671941 | 0,342621583 | 1,127438231 |
| COL11A11 | 0,000132256 | -2,496092337 | 0,067 | 0,631 | 1 | CH_SC | 0,177256159 | 5,641552906 | 0,106180666 |
| EEF1B25 | 0,000132641 | -1,658203942 | 0,533 | 0,877 | 1 | CH_SC | 0,316833339 | 3,156233503 | 0,607753706 |
| JAG14 | 0,000133646 | 0,983320813 | 0,4 | 0,046 | 1 | CH_SC | 1,977010884 | 0,50581411 | 8,695652174 |
| CMTM5 | 0,000139075 | 1,158231074 | 0,467 | 0,077 | 1 | CH_SC | 2,231836085 | 0,448061579 | 6,064935065 |
| ANKRD104 | 0,000140501 | 1,704352832 | 0,6 | 0,2 | 1 | CH_SC | 3,258827149 | 0,306858865 | 3 |
| FRMD6 | 0,000143785 | 1,275529287 | 0,533 | 0,108 | 1 | CH_SC | 2,420876171 | 0,413073585 | 4,935185185 |
| RPS95 | 0,00014595 | -1,178463419 | 0,667 | 0,969 | 1 | CH_SC | 0,441821822 | 2,263355838 | 0,688338493 |
| PRDM161 | 0,000146081 | 1,086990121 | 0,333 | 0,031 | 1 | CH_SC | 2,124303827 | 0,470742456 | 10,74193548 |
| SEMA6D1 | 0,000146081 | 1,431709804 | 0,333 | 0,031 | 1 | CH_SC | 2,697662383 | 0,370691309 | 10,74193548 |
| SNCA1 | 0,000146355 | 1,212440991 | 0,4 | 0,046 | 1 | CH_SC | 2,317293836 | 0,43153785 | 8,695652174 |
| RPL95 | 0,000146554 | -1,027706803 | 0,867 | 0,969 | 1 | CH_SC | 0,490489173 | 2,038780985 | 0,894736842 |
| AZGP14 | 0,000149511 | -3,059215423 | 0 | 0,585 | 1 | CH_SC | 0,119973242 | 8,33519195 | 0 |
| RPL13A4 | 0,000150256 | -1,386993472 | 0,6 | 0,908 | 1 | CH_SC | 0,3823608 | 2,615330863 | 0,660792952 |
| KHDRBS3 | 0,000150669 | 2,003865571 | 0,467 | 0,077 | 1 | CH_SC | 4,01073201 | 0,249331044 | 6,064935065 |
| PRRG42 | 0,000150669 | 1,120559983 | 0,467 | 0,077 | 1 | CH_SC | 2,174313523 | 0,459915274 | 6,064935065 |
| RPS195 | 0,000152507 | -1,281012329 | 0,6 | 0,954 | 1 | CH_SC | 0,411506656 | 2,430094351 | 0,628930818 |

| Supplementary Table 4 - clusermarker | | | | | | | | | |
|---|---|---|---|---|---|---|---|---|---|
| | **p_val** | **avg_log2FC** | **pct.1** | **pct.2** | **p_val_adj** | **celltype** | **Foldchange_UP** | **Foldchange_DOWN** | **Ratio_pct1_pct2** |
| HLA-DPB12 | 0,000153728 | 2,557932783 | 0,533 | 0,154 | 1 | CH_SC | 5,888633083 | 0,169818697 | 3,461038961 |
| IFNAR2 | 0,000160197 | 1,069713003 | 0,4 | 0,046 | 1 | CH_SC | 2,099015766 | 0,476413763 | 8,695652174 |
| ARHGAP12 | 0,000161627 | 1,112861784 | 0,333 | 0,031 | 1 | CH_SC | 2,162742317 | 0,462375934 | 10,74193548 |
| B4GALNT1 | 0,000161627 | 1,418578621 | 0,333 | 0,031 | 1 | CH_SC | 2,673220088 | 0,374080684 | 10,74193548 |
| CRLF1 | 0,000161627 | 1,235432897 | 0,333 | 0,031 | 1 | CH_SC | 2,354519869 | 0,42471504 | 10,74193548 |
| S100A108 | 0,000163444 | 1,46815334 | 0,867 | 0,815 | 1 | CH_SC | 2,766675304 | 0,361444655 | 1,063803681 |
| ABCA81 | 0,00016631 | 1,3854835 | 0,533 | 0,108 | 1 | CH_SC | 2,612595004 | 0,3827612 | 4,935185185 |
| RPS167 | 0,000175325 | -1,110676173 | 0,667 | 0,908 | 1 | CH_SC | 0,463076942 | 2,159468352 | 0,734581498 |
| NCAM1 | 0,000176638 | 1,420522303 | 0,467 | 0,077 | 1 | CH_SC | 2,676824033 | 0,37357704 | 6,064935065 |
| UCHL1 | 0,000178721 | 1,130082022 | 0,333 | 0,031 | 1 | CH_SC | 2,188711835 | 0,456889749 | 10,74193548 |
| RBM141 | 0,000178721 | 1,458177015 | 0,333 | 0,031 | 1 | CH_SC | 2,747609571 | 0,363952728 | 10,74193548 |
| PPME1 | 0,000178721 | 0,913829903 | 0,333 | 0,031 | 1 | CH_SC | 1,884040403 | 0,53077418 | 10,74193548 |
| GABPA1 | 0,000178721 | 1,336821735 | 0,333 | 0,031 | 1 | CH_SC | 2,52594239 | 0,395891848 | 10,74193548 |
| FN15 | 0,000187653 | -3,004282503 | 0,667 | 0,892 | 1 | CH_SC | 0,1246295 | 8,02378252 | 0,747757848 |
| FBLN5 | 0,000191149 | 1,149960696 | 0,467 | 0,077 | 1 | CH_SC | 2,219078489 | 0,450637508 | 6,064935065 |
| CCN33 | 0,000197386 | 1,437394979 | 0,4 | 0,062 | 1 | CH_SC | 2,708313935 | 0,369233414 | 6,451612903 |
| LINC01198 | 0,000197503 | 1,352668824 | 0,333 | 0,031 | 1 | CH_SC | 2,553841208 | 0,391567023 | 10,74193548 |
| MDK6 | 0,000203829 | 1,298710732 | 0,533 | 0,123 | 1 | CH_SC | 2,460089379 | 0,406489296 | 4,333333333 |
| RPL10A2 | 0,000215731 | -1,297964851 | 0,6 | 0,969 | 1 | CH_SC | 0,406699508 | 2,458817828 | 0,619195046 |
| MT-ND24 | 0,000216655 | 0,902905261 | 1 | 0,908 | 1 | CH_SC | 1,869827604 | 0,534808663 | 1,101321586 |
| DCBLD2 | 0,000218129 | 0,650742109 | 0,333 | 0,031 | 1 | CH_SC | 1,569975569 | 0,636952587 | 10,74193548 |
| ZNF622 | 0,000218129 | 0,775931515 | 0,333 | 0,031 | 1 | CH_SC | 1,712295284 | 0,58401142 | 10,74193548 |
| TNFRSF192 | 0,000218129 | 1,513267692 | 0,333 | 0,031 | 1 | CH_SC | 2,854558625 | 0,350316855 | 10,74193548 |
| SERPINA5 | 0,000218129 | 1,511079598 | 0,333 | 0,031 | 1 | CH_SC | 2,85023248 | 0,350848574 | 10,74193548 |
| CMTM33 | 0,000218129 | 0,762523929 | 0,333 | 0,031 | 1 | CH_SC | 1,696455903 | 0,589464187 | 10,74193548 |
| MT1E3 | 0,000220526 | -2,60874045 | 0,533 | 0,862 | 1 | CH_SC | 0,163942243 | 6,099709139 | 0,618329466 |
| ANXA24 | 0,000234467 | 1,300820032 | 0,867 | 0,769 | 1 | CH_SC | 2,463688797 | 0,40589542 | 1,127438231 |
| PRRX12 | 0,00023648 | -1,6365411 | 0,067 | 0,6 | 1 | CH_SC | 0,321626659 | 3,109194998 | 0,111666667 |
| EPB41L25 | 0,000240617 | 1,557324109 | 0,733 | 0,292 | 1 | CH_SC | 2,943074595 | 0,339780718 | 2,510273973 |
| CD817 | 0,000248696 | 0,787813183 | 1 | 0,862 | 1 | CH_SC | 1,726455541 | 0,579221402 | 1,160092807 |
| GAS55 | 0,000256642 | -1,720021685 | 0,333 | 0,769 | 1 | CH_SC | 0,303544158 | 3,294413587 | 0,433029909 |

| Supplementary Table 4 - clusermarker | | | | | | | | | |
|---|---|---|---|---|---|---|---|---|---|
| | **p_val** | **avg_log2FC** | **pct.1** | **pct.2** | **p_val_adj** | **celltype** | **Foldchange_UP** | **Foldchange_DOWN** | **Ratio_pct1_pct2** |
| BTC1 | 0,00026559 | 0,792852723 | 0,333 | 0,031 | 1 | CH_SC | 1,732496843 | 0,577201629 | 10,74193548 |
| SEMA4B | 0,00026559 | 0,898396769 | 0,333 | 0,031 | 1 | CH_SC | 1,863993422 | 0,53648258 | 10,74193548 |
| RHOC3 | 0,000268158 | 1,438798514 | 0,8 | 0,508 | 1 | CH_SC | 2,710950018 | 0,368874377 | 1,57480315 |
| AK5 | 0,00026949 | 0,640932677 | 0,267 | 0,015 | 1 | CH_SC | 1,559336917 | 0,641298227 | 17,8 |
| LBX2-AS1 | 0,00026949 | 0,774979343 | 0,267 | 0,015 | 1 | CH_SC | 1,71116555 | 0,584396992 | 17,8 |
| CCDC112 | 0,00026949 | 0,812601448 | 0,267 | 0,015 | 1 | CH_SC | 1,756375662 | 0,56935428 | 17,8 |
| HLA-DQA12 | 0,00026949 | 1,732303264 | 0,267 | 0,015 | 1 | CH_SC | 3,322578451 | 0,300971073 | 17,8 |
| AL139383.1 | 0,00026949 | 1,623353354 | 0,267 | 0,015 | 1 | CH_SC | 3,080903198 | 0,324580143 | 17,8 |
| PLEKHG3 | 0,00026949 | 1,43995648 | 0,267 | 0,015 | 1 | CH_SC | 2,71312681 | 0,368578423 | 17,8 |
| SHISA9 | 0,00026949 | 0,436805023 | 0,267 | 0,015 | 1 | CH_SC | 1,35360333 | 0,738768868 | 17,8 |
| CRKL | 0,00026949 | 0,574277144 | 0,267 | 0,015 | 1 | CH_SC | 1,488931254 | 0,671622681 | 17,8 |
| TMEM176B4 | 0,000270438 | 0,968868943 | 0,667 | 0,215 | 1 | CH_SC | 1,957305488 | 0,510906451 | 3,102325581 |
| NRN1 | 0,000274081 | 1,135000255 | 0,533 | 0,108 | 1 | CH_SC | 2,196186016 | 0,455334836 | 4,935185185 |
| OMD1 | 0,000274398 | -2,344180201 | 0 | 0,554 | 1 | CH_SC | 0,196938871 | 5,077717741 | 0 |
| ARL6IP6 | 0,000276346 | 1,063543 | 0,4 | 0,062 | 1 | CH_SC | 2,090058031 | 0,478455615 | 6,451612903 |
| AGL | 0,000280621 | 0,639892489 | 0,2 | 0 | 1 | CH_SC | 1,558213035 | 0,641760772 | #DIV/0! |
| CELSR21 | 0,000280621 | 0,454117569 | 0,2 | 0 | 1 | CH_SC | 1,369944616 | 0,729956517 | #DIV/0! |
| AC242426.2 | 0,000280621 | 0,555390318 | 0,2 | 0 | 1 | CH_SC | 1,469566167 | 0,680472933 | #DIV/0! |
| OLFML2B2 | 0,000280621 | 0,646116283 | 0,2 | 0 | 1 | CH_SC | 1,564949695 | 0,638998176 | #DIV/0! |
| CNTN2 | 0,000280621 | 0,76914774 | 0,2 | 0 | 1 | CH_SC | 1,704262707 | 0,586763998 | #DIV/0! |
| OSR12 | 0,000280621 | 0,488233764 | 0,2 | 0 | 1 | CH_SC | 1,40272652 | 0,712897336 | #DIV/0! |
| GPN1 | 0,000280621 | 0,599623129 | 0,2 | 0 | 1 | CH_SC | 1,515320672 | 0,659926324 | #DIV/0! |
| QPCT2 | 0,000280621 | 0,833726439 | 0,2 | 0 | 1 | CH_SC | 1,782283006 | 0,561078121 | #DIV/0! |
| KIF5C | 0,000280621 | 0,46334984 | 0,2 | 0 | 1 | CH_SC | 1,378739444 | 0,725300204 | #DIV/0! |
| FMNL21 | 0,000280621 | 1,297836779 | 0,2 | 0 | 1 | CH_SC | 2,458599562 | 0,406735613 | #DIV/0! |
| MFSD61 | 0,000280621 | 0,818353271 | 0,2 | 0 | 1 | CH_SC | 1,763392063 | 0,567088863 | #DIV/0! |
| IGSF11 | 0,000280621 | 0,744127688 | 0,2 | 0 | 1 | CH_SC | 1,674961213 | 0,59702875 | #DIV/0! |
| SPON21 | 0,000280621 | 0,496811189 | 0,2 | 0 | 1 | CH_SC | 1,411091156 | 0,708671439 | #DIV/0! |
| CASC15 | 0,000280621 | 0,492955563 | 0,2 | 0 | 1 | CH_SC | 1,407325027 | 0,710567908 | #DIV/0! |
| MANEA1 | 0,000280621 | 0,531647544 | 0,2 | 0 | 1 | CH_SC | 1,445579091 | 0,691764295 | #DIV/0! |
| KRBA1 | 0,000280621 | 0,747987165 | 0,2 | 0 | 1 | CH_SC | 1,679448044 | 0,595433722 | #DIV/0! |

| Supplementary Table 4 - clusermarker | | | | | | | | | |
|---|---|---|---|---|---|---|---|---|---|
| | **p_val** | **avg_log2FC** | **pct.1** | **pct.2** | **p_val_adj** | **celltype** | **Foldchange_UP** | **Foldchange_DOWN** | **Ratio_pct1_pct2** |
| NEFM | 0,000280621 | 0,647951021 | 0,2 | 0 | 1 | CH_SC | 1,566941176 | 0,63818605 | #DIV/0! |
| TRAF11 | 0,000280621 | 0,472454067 | 0,2 | 0 | 1 | CH_SC | 1,387467585 | 0,720737559 | #DIV/0! |
| NRP13 | 0,000280621 | 0,931089889 | 0,2 | 0 | 1 | CH_SC | 1,906715887 | 0,524461985 | #DIV/0! |
| USP541 | 0,000280621 | 0,907119195 | 0,2 | 0 | 1 | CH_SC | 1,875297123 | 0,533248832 | #DIV/0! |
| VWA2 | 0,000280621 | 0,821960216 | 0,2 | 0 | 1 | CH_SC | 1,767806312 | 0,56567283 | #DIV/0! |
| AC136475.1 | 0,000280621 | 0,531647544 | 0,2 | 0 | 1 | CH_SC | 1,445579091 | 0,691764295 | #DIV/0! |
| CYP2R1 | 0,000280621 | 0,531647544 | 0,2 | 0 | 1 | CH_SC | 1,445579091 | 0,691764295 | #DIV/0! |
| PRMT3 | 0,000280621 | 1,041690356 | 0,2 | 0 | 1 | CH_SC | 2,058638277 | 0,485757994 | #DIV/0! |
| ADAMTS8 | 0,000280621 | 0,706610711 | 0,2 | 0 | 1 | CH_SC | 1,631965673 | 0,612757987 | #DIV/0! |
| FREM2 | 0,000280621 | 0,548921456 | 0,2 | 0 | 1 | CH_SC | 1,46299157 | 0,683530938 | #DIV/0! |
| IGHA21 | 0,000280621 | 0,744057169 | 0,2 | 0 | 1 | CH_SC | 1,674879344 | 0,597057934 | #DIV/0! |
| NLGN2 | 0,000280621 | 0,585384087 | 0,2 | 0 | 1 | CH_SC | 1,500438396 | 0,666471881 | #DIV/0! |
| TEX2 | 0,000280621 | 0,72952797 | 0,2 | 0 | 1 | CH_SC | 1,658096497 | 0,603101208 | #DIV/0! |
| SIRT7 | 0,000280621 | 0,613372492 | 0,2 | 0 | 1 | CH_SC | 1,529831216 | 0,653666881 | #DIV/0! |
| GALR1 | 0,000280621 | 0,685407643 | 0,2 | 0 | 1 | CH_SC | 1,608156312 | 0,6218301 | #DIV/0! |
| ZNF253 | 0,000280621 | 0,559897692 | 0,2 | 0 | 1 | CH_SC | 1,474164674 | 0,678350267 | #DIV/0! |
| KCTD15 | 0,000280621 | 0,647982187 | 0,2 | 0 | 1 | CH_SC | 1,566975026 | 0,638172264 | #DIV/0! |
| HMGXB42 | 0,000280621 | 0,61413289 | 0,2 | 0 | 1 | CH_SC | 1,530637754 | 0,653322445 | #DIV/0! |
| SEPTIN3 | 0,000280621 | 0,488233764 | 0,2 | 0 | 1 | CH_SC | 1,40272652 | 0,712897336 | #DIV/0! |
| COL9A3 | 0,000280863 | -2,401156158 | 0,533 | 0,785 | 1 | CH_SC | 0,189312797 | 5,282263089 | 0,678980892 |
| SSRP11 | 0,000286831 | 1,340408137 | 0,467 | 0,092 | 1 | CH_SC | 2,532229451 | 0,394908921 | 5,076086957 |
| MT-CYB3 | 0,000289797 | 0,890502619 | 1 | 0,938 | 1 | CH_SC | 1,853821863 | 0,539426155 | 1,066098081 |
| SPATA61 | 0,000300271 | 1,405834865 | 0,4 | 0,062 | 1 | CH_SC | 2,649710723 | 0,377399688 | 6,451612903 |
| SRGAP2B | 0,000301962 | 0,599474747 | 0,267 | 0,015 | 1 | CH_SC | 1,515164829 | 0,659994201 | 17,8 |
| PPP1R12B2 | 0,000301962 | 0,670574257 | 0,267 | 0,015 | 1 | CH_SC | 1,591706411 | 0,628256564 | 17,8 |
| DTX3L | 0,000301962 | 1,40291909 | 0,267 | 0,015 | 1 | CH_SC | 2,644360904 | 0,378163207 | 17,8 |
| TGFBI3 | 0,000301962 | 1,306395296 | 0,267 | 0,015 | 1 | CH_SC | 2,473228089 | 0,404329873 | 17,8 |
| AC079781.5 | 0,000301962 | 1,039259602 | 0,267 | 0,015 | 1 | CH_SC | 2,055172658 | 0,486577124 | 17,8 |
| SLC9A3R11 | 0,000301962 | 0,952335374 | 0,267 | 0,015 | 1 | CH_SC | 1,935002425 | 0,516795218 | 17,8 |
| PCSK2 | 0,000304573 | 1,397124477 | 0,467 | 0,077 | 1 | CH_SC | 2,633761078 | 0,379685161 | 6,064935065 |
| CYBA9 | 0,000313837 | 1,370636087 | 0,6 | 0,2 | 1 | CH_SC | 2,585845515 | 0,386720705 | 3 |

| Supplementary Table 4 - clusermarker | | | | | | | | | |
|---|---|---|---|---|---|---|---|---|---|
| | **p_val** | **avg_log2FC** | **pct.1** | **pct.2** | **p_val_adj** | **celltype** | **Foldchange_UP** | **Foldchange_DOWN** | **Ratio_pct1_pct2** |
| WASHC1 | 0,000326126 | 1,040791795 | 0,4 | 0,062 | 1 | CH_SC | 2,057356483 | 0,486060636 | 6,451612903 |
| MAN2A2 | 0,000326126 | 1,14307383 | 0,4 | 0,062 | 1 | CH_SC | 2,208510713 | 0,452793819 | 6,451612903 |
| TMEM39B | 0,000338072 | 0,884277368 | 0,267 | 0,015 | 1 | CH_SC | 1,845839826 | 0,541758817 | 17,8 |
| TMEM651 | 0,000338072 | 0,732434755 | 0,267 | 0,015 | 1 | CH_SC | 1,661440646 | 0,601887285 | 17,8 |
| NELFB2 | 0,000338072 | 0,63044278 | 0,267 | 0,015 | 1 | CH_SC | 1,548040032 | 0,645978127 | 17,8 |
| MIF9 | 0,000346706 | 1,308793746 | 0,8 | 0,431 | 1 | CH_SC | 2,477343198 | 0,403658242 | 1,856148492 |
| STAT23 | 0,000357879 | 0,934101709 | 0,467 | 0,092 | 1 | CH_SC | 1,910700571 | 0,523368243 | 5,076086957 |
| RPS219 | 0,000361078 | -1,30712942 | 0,6 | 0,923 | 1 | CH_SC | 0,40412418 | 2,474486926 | 0,650054171 |
| KLF93 | 0,00036555 | 1,30436766 | 0,733 | 0,323 | 1 | CH_SC | 2,469754531 | 0,404898538 | 2,269349845 |
| FBXO2 | 0,00036809 | -1,755733731 | 0 | 0,538 | 1 | CH_SC | 0,296122551 | 3,376980224 | 0 |
| CST36 | 0,000376164 | -1,467745854 | 0,933 | 0,985 | 1 | CH_SC | 0,361546758 | 2,765893973 | 0,947208122 |
| RAP1GDS1 | 0,000378191 | 0,651958428 | 0,267 | 0,015 | 1 | CH_SC | 1,571299755 | 0,636415806 | 17,8 |
| RASA3 | 0,000378191 | 0,614096159 | 0,267 | 0,015 | 1 | CH_SC | 1,530598784 | 0,653339079 | 17,8 |
| RARA2 | 0,000378191 | 0,756839377 | 0,267 | 0,015 | 1 | CH_SC | 1,689784626 | 0,591791394 | 17,8 |
| GDI12 | 0,000387438 | 1,202659562 | 0,6 | 0,215 | 1 | CH_SC | 2,301635794 | 0,434473605 | 2,790697674 |
| UBE2D25 | 0,000388716 | 1,49457927 | 0,667 | 0,246 | 1 | CH_SC | 2,817819637 | 0,354884318 | 2,711382114 |
| COX7C6 | 0,000391691 | -1,261361179 | 0,467 | 0,846 | 1 | CH_SC | 0,417150194 | 2,397218111 | 0,552009456 |
| RPL397 | 0,000412744 | -1,166312201 | 0,933 | 0,954 | 1 | CH_SC | 0,445558818 | 2,244372594 | 0,977987421 |
| PDGFA2 | 0,000416741 | 0,911335711 | 0,4 | 0,062 | 1 | CH_SC | 1,880786008 | 0,531692599 | 6,451612903 |
| SIRT12 | 0,000422726 | 0,406679969 | 0,267 | 0,015 | 1 | CH_SC | 1,325631665 | 0,754357358 | 17,8 |
| FLNA6 | 0,00043141 | 1,09773285 | 0,6 | 0,185 | 1 | CH_SC | 2,140181053 | 0,467250188 | 3,243243243 |
| TJP12 | 0,00044513 | 1,075953063 | 0,467 | 0,092 | 1 | CH_SC | 2,10811426 | 0,47435759 | 5,076086957 |
| GRIK3 | 0,000451838 | 1,240683891 | 0,4 | 0,062 | 1 | CH_SC | 2,363105257 | 0,423172009 | 6,451612903 |
| OLFM1 | 0,000451838 | 0,933006932 | 0,4 | 0,062 | 1 | CH_SC | 1,909251202 | 0,523765547 | 6,451612903 |
| PFDN52 | 0,000456099 | -1,07492028 | 0,667 | 0,908 | 1 | CH_SC | 0,47469729 | 2,106605663 | 0,734581498 |
| HLA-A6 | 0,000472442 | 1,031211315 | 1 | 0,8 | 1 | CH_SC | 2,043739495 | 0,489299151 | 1,25 |
| RPS207 | 0,00047569 | -1,280282602 | 0,4 | 0,8 | 1 | CH_SC | 0,411714852 | 2,4288655 | 0,5 |
| COMT4 | 0,000479428 | 1,439875888 | 0,733 | 0,415 | 1 | CH_SC | 2,712975253 | 0,368599013 | 1,76626506 |
| PLEKHA4 | 0,000489679 | 0,821343831 | 0,4 | 0,062 | 1 | CH_SC | 1,767051186 | 0,565914563 | 6,451612903 |
| ISLR3 | 0,000490595 | -1,74980522 | 0 | 0,523 | 1 | CH_SC | 0,297341921 | 3,363131569 | 0 |
| BRD26 | 0,000503495 | 1,909765146 | 0,733 | 0,338 | 1 | CH_SC | 3,757479272 | 0,266135866 | 2,168639053 |

| Supplementary Table 4 - clusermarker | | | | | | | | | |
|---|---|---|---|---|---|---|---|---|---|
| | **p_val** | **avg_log2FC** | **pct.1** | **pct.2** | **p_val_adj** | **celltype** | **Foldchange_UP** | **Foldchange_DOWN** | **Ratio_pct1_pct2** |
| S100B3 | 0,000506561 | 1,019532871 | 0,933 | 0,908 | 1 | CH_SC | 2,027262447 | 0,493276044 | 1,02753304 |
| ADIRF6 | 0,000511311 | -1,787199854 | 0,4 | 0,846 | 1 | CH_SC | 0,289733849 | 3,451443468 | 0,472813239 |
| LAMA41 | 0,000516019 | 1,331776702 | 0,6 | 0,185 | 1 | CH_SC | 2,51712472 | 0,397278685 | 3,243243243 |
| RPL319 | 0,00053548 | -1,351398162 | 0,533 | 0,815 | 1 | CH_SC | 0,39191205 | 2,551592888 | 0,65398773 |
| HBEGF4 | 0,000554812 | 1,993873255 | 0,533 | 0,169 | 1 | CH_SC | 3,983049076 | 0,251063941 | 3,153846154 |
| LTBP44 | 0,000555852 | 1,581410211 | 0,667 | 0,308 | 1 | CH_SC | 2,992622308 | 0,334155098 | 2,165584416 |
| SGCE2 | 0,000561519 | 1,113960735 | 0,533 | 0,123 | 1 | CH_SC | 2,164390382 | 0,462023861 | 4,333333333 |
| SCCPDH2 | 0,000565219 | 1,510520572 | 0,533 | 0,154 | 1 | CH_SC | 2,849128264 | 0,350984549 | 3,461038961 |
| DMD | 0,0005739 | 1,587793366 | 0,467 | 0,108 | 1 | CH_SC | 3,005892397 | 0,332679906 | 4,324074074 |
| ITGA61 | 0,000588613 | 1,663273411 | 0,667 | 0,292 | 1 | CH_SC | 3,167343663 | 0,315721976 | 2,284246575 |
| PROS14 | 0,000589775 | 1,280742638 | 0,533 | 0,169 | 1 | CH_SC | 2,429640122 | 0,411583588 | 3,153846154 |
| MCAM2 | 0,000592528 | 1,362679315 | 0,467 | 0,092 | 1 | CH_SC | 2,571623279 | 0,388859445 | 5,076086957 |
| AGRN3 | 0,000596352 | 1,203578311 | 0,333 | 0,046 | 1 | CH_SC | 2,303102006 | 0,434197008 | 7,239130435 |
| PRR14 | 0,000596352 | 0,988769758 | 0,333 | 0,046 | 1 | CH_SC | 1,984492016 | 0,503907293 | 7,239130435 |
| ANKRD111 | 0,000614699 | 1,312754887 | 0,467 | 0,108 | 1 | CH_SC | 2,484154472 | 0,402551456 | 4,324074074 |
| RRBP11 | 0,000626393 | -1,744364036 | 0,067 | 0,554 | 1 | CH_SC | 0,298465475 | 3,350471269 | 0,120938628 |
| PSMB34 | 0,000639427 | 1,079754122 | 0,667 | 0,246 | 1 | CH_SC | 2,113675817 | 0,473109448 | 2,711382114 |
| DDIT44 | 0,000642692 | 1,107243521 | 0,867 | 0,415 | 1 | CH_SC | 2,154336365 | 0,464180068 | 2,089156627 |
| ZBTB384 | 0,000650483 | 1,595195597 | 0,6 | 0,215 | 1 | CH_SC | 3,02135477 | 0,330977352 | 2,790697674 |
| MICALL21 | 0,000650605 | 1,357921587 | 0,333 | 0,046 | 1 | CH_SC | 2,563156533 | 0,390143944 | 7,239130435 |
| P4HA2 | 0,000658185 | 1,190848221 | 0,467 | 0,108 | 1 | CH_SC | 2,282869232 | 0,43804524 | 4,324074074 |
| CASC32 | 0,000658185 | 1,270746743 | 0,467 | 0,108 | 1 | CH_SC | 2,412864237 | 0,414445199 | 4,324074074 |
| S100A132 | 0,000666378 | -1,597138884 | 0,4 | 0,862 | 1 | CH_SC | 0,33053183 | 3,025427229 | 0,464037123 |
| NR2F22 | 0,000674054 | 1,459730478 | 0,6 | 0,2 | 1 | CH_SC | 2,750569731 | 0,363561043 | 3 |
| B4GAT1 | 0,000709419 | 0,995549943 | 0,333 | 0,046 | 1 | CH_SC | 1,993840416 | 0,501544653 | 7,239130435 |
| MAN2B11 | 0,000709419 | 1,305410589 | 0,333 | 0,046 | 1 | CH_SC | 2,471540571 | 0,404605942 | 7,239130435 |
| VAPB1 | 0,000709419 | 1,328407846 | 0,333 | 0,046 | 1 | CH_SC | 2,511253806 | 0,39820746 | 7,239130435 |
| H3-3B2 | 0,000731459 | 1,038752448 | 1 | 1 | 1 | CH_SC | 2,054450326 | 0,486748201 | 1 |
| EIF4B7 | 0,000770438 | -1,790885581 | 0,2 | 0,615 | 1 | CH_SC | 0,288994596 | 3,460272321 | 0,325203252 |
| EVI5 | 0,000773142 | 1,140928331 | 0,333 | 0,046 | 1 | CH_SC | 2,205228775 | 0,453467691 | 7,239130435 |
| SOX13 | 0,000773142 | 0,998813787 | 0,333 | 0,046 | 1 | CH_SC | 1,998356235 | 0,500411279 | 7,239130435 |

| Supplementary Table 4 - clusermarker | | | | | | | | | |
|---|---|---|---|---|---|---|---|---|---|
| | p_val | avg_log2FC | pct.1 | pct.2 | p_val_adj | celltype | Foldchange_UP | Foldchange_DOWN | Ratio_pct1_pct2 |
| PXDN2 | 0,000773142 | 0,991174266 | 0,333 | 0,046 | 1 | CH_SC | 1,987802283 | 0,503068141 | 7,239130435 |
| ARHGAP241 | 0,000773142 | 0,949608894 | 0,333 | 0,046 | 1 | CH_SC | 1,93134901 | 0,517772808 | 7,239130435 |
| LRRCC11 | 0,000773142 | 0,950904856 | 0,333 | 0,046 | 1 | CH_SC | 1,933084705 | 0,517307905 | 7,239130435 |
| KIAA12173 | 0,000773142 | 1,475092591 | 0,333 | 0,046 | 1 | CH_SC | 2,780014854 | 0,359710308 | 7,239130435 |
| CEBPD4 | 0,000793467 | -1,934145752 | 0,4 | 0,754 | 1 | CH_SC | 0,261676133 | 3,821517805 | 0,530503979 |
| S100A116 | 0,000806188 | 1,164911913 | 0,867 | 0,738 | 1 | CH_SC | 2,242195249 | 0,44599149 | 1,174796748 |
| KIF5B3 | 0,00080692 | 1,269245938 | 0,667 | 0,277 | 1 | CH_SC | 2,410355491 | 0,414876562 | 2,407942238 |
| CD594 | 0,000809661 | 1,03047994 | 0,867 | 0,815 | 1 | CH_SC | 2,042703683 | 0,489547264 | 1,063803681 |
| PLEKHB1 | 0,000811045 | 1,300938544 | 0,467 | 0,123 | 1 | CH_SC | 2,463891187 | 0,405862079 | 3,796747967 |
| STMN14 | 0,000813952 | 1,67559484 | 0,667 | 0,338 | 1 | CH_SC | 3,194510407 | 0,313037014 | 1,973372781 |
| TNFRSF12A4 | 0,000815097 | 1,711661756 | 0,533 | 0,185 | 1 | CH_SC | 3,27537878 | 0,3053082 | 2,881081081 |
| ATP1B13 | 0,000842146 | 0,923861149 | 0,333 | 0,046 | 1 | CH_SC | 1,897186029 | 0,527096439 | 7,239130435 |
| PCBP4 | 0,000842146 | 1,204459656 | 0,333 | 0,046 | 1 | CH_SC | 2,304509406 | 0,433931837 | 7,239130435 |
| S1PR31 | 0,000842146 | 1,347760465 | 0,333 | 0,046 | 1 | CH_SC | 2,545167254 | 0,392901488 | 7,239130435 |
| PHF19 | 0,000842146 | 0,98161757 | 0,333 | 0,046 | 1 | CH_SC | 1,974678206 | 0,506411626 | 7,239130435 |
| AARS1 | 0,000842146 | 1,211029282 | 0,333 | 0,046 | 1 | CH_SC | 2,315027421 | 0,431960326 | 7,239130435 |
| MIA4 | 0,000845073 | -1,447554813 | 0,667 | 0,923 | 1 | CH_SC | 0,36664231 | 2,727453902 | 0,722643554 |
| TAGLN26 | 0,000855805 | 1,792577454 | 0,667 | 0,338 | 1 | CH_SC | 3,464332621 | 0,288655885 | 1,973372781 |
| CPE1 | 0,000855805 | -1,768452261 | 0 | 0,492 | 1 | CH_SC | 0,293523464 | 3,406882664 | 0 |
| TSPAN133 | 0,000855805 | -1,527475306 | 0 | 0,492 | 1 | CH_SC | 0,346883878 | 2,882809102 | 0 |
| CNMD | 0,000855805 | -2,373770322 | 0 | 0,492 | 1 | CH_SC | 0,192940736 | 5,182938668 | 0 |
| GPC62 | 0,000855805 | -1,743632326 | 0 | 0,492 | 1 | CH_SC | 0,29861689 | 3,3487724 | 0 |
| TBCB5 | 0,000866817 | 1,180156872 | 0,6 | 0,246 | 1 | CH_SC | 2,266014153 | 0,44130351 | 2,43902439 |
| RAB6A3 | 0,000869206 | 1,348446559 | 0,533 | 0,154 | 1 | CH_SC | 2,546377931 | 0,392714682 | 3,461038961 |
| C1orf1982 | 0,00087377 | 1,591636147 | 0,733 | 0,354 | 1 | CH_SC | 3,013909605 | 0,331794954 | 2,070621469 |
| ITGA10 | 0,000896684 | -1,582804466 | 0,067 | 0,538 | 1 | CH_SC | 0,333832318 | 2,995515847 | 0,124535316 |
| UTRN2 | 0,000898145 | 1,37077688 | 0,533 | 0,169 | 1 | CH_SC | 2,58609788 | 0,386682967 | 3,153846154 |
| RPS83 | 0,000912299 | -0,876420885 | 0,867 | 0,954 | 1 | CH_SC | 0,54471712 | 1,835815259 | 0,908805031 |
| PPT2 | 0,000916825 | 0,899030506 | 0,333 | 0,046 | 1 | CH_SC | 1,864812405 | 0,536246969 | 7,239130435 |
| PTPN12 | 0,000916825 | 0,758234235 | 0,333 | 0,046 | 1 | CH_SC | 1,69141917 | 0,591219502 | 7,239130435 |
| STK42 | 0,000916825 | 1,247369548 | 0,333 | 0,046 | 1 | CH_SC | 2,374081642 | 0,421215506 | 7,239130435 |

| Supplementary Table 4 - clusermarker | | | | | | | | | |
|---|---|---|---|---|---|---|---|---|---|
| | **p_val** | **avg_log2FC** | **pct.1** | **pct.2** | **p_val_adj** | **celltype** | **Foldchange_UP** | **Foldchange_DOWN** | **Ratio_pct1_pct2** |
| PLEC6 | 0,00091766 | 1,424275838 | 0,667 | 0,323 | 1 | CH_SC | 2,683797533 | 0,372606349 | 2,06501548 |
| ATP6AP23 | 0,000939892 | 1,091442395 | 0,667 | 0,292 | 1 | CH_SC | 2,130869726 | 0,469291946 | 2,284246575 |
| DLC11 | 0,000985115 | 0,888665302 | 0,467 | 0,108 | 1 | CH_SC | 1,851462465 | 0,540113569 | 4,324074074 |
| ZBTB44 | 0,000985115 | 0,863009719 | 0,467 | 0,108 | 1 | CH_SC | 1,818828756 | 0,549804371 | 4,324074074 |
| TSPAN153 | 0,000997602 | 0,854499115 | 0,333 | 0,046 | 1 | CH_SC | 1,808130886 | 0,553057308 | 7,239130435 |
| SEL1L | 0,000997602 | 1,033346535 | 0,333 | 0,046 | 1 | CH_SC | 2,046766513 | 0,488575513 | 7,239130435 |
| ITPR31 | 0,001084925 | 1,026947432 | 0,333 | 0,046 | 1 | CH_SC | 2,037708144 | 0,490747413 | 7,239130435 |
| S100A17 | 0,001085282 | -1,847132831 | 0,4 | 0,754 | 1 | CH_SC | 0,277944197 | 3,597844492 | 0,530503979 |
| VMP14 | 0,001120092 | 1,442117191 | 0,467 | 0,123 | 1 | CH_SC | 2,71719328 | 0,368026819 | 3,796747967 |
| GLUL6 | 0,001120147 | -1,310023437 | 0 | 0,477 | 1 | CH_SC | 0,403314328 | 2,47945568 | 0 |
| CHPF2 | 0,001120147 | -1,423523833 | 0 | 0,477 | 1 | CH_SC | 0,372800621 | 2,682398968 | 0 |
| PDGFRL1 | 0,001120245 | -1,345357443 | 0 | 0,477 | 1 | CH_SC | 0,393556468 | 2,540931431 | 0 |
| LUM2 | 0,001120245 | -2,86066146 | 0 | 0,477 | 1 | CH_SC | 0,137675003 | 7,263482706 | 0 |
| WWP21 | 0,001120245 | -1,917417918 | 0 | 0,477 | 1 | CH_SC | 0,264727887 | 3,777463769 | 0 |
| ITPK12 | 0,001141167 | 0,93926449 | 0,4 | 0,077 | 1 | CH_SC | 1,91755039 | 0,521498682 | 5,194805195 |
| NACA1 | 0,001155358 | -1,002854724 | 0,8 | 0,923 | 1 | CH_SC | 0,499011606 | 2,003961406 | 0,866738895 |
| BICD1 | 0,001179272 | 0,793211273 | 0,333 | 0,046 | 1 | CH_SC | 1,732927469 | 0,577058196 | 7,239130435 |
| TAMALIN2 | 0,001179272 | 0,692147638 | 0,333 | 0,046 | 1 | CH_SC | 1,615686888 | 0,618931804 | 7,239130435 |
| COL5A22 | 0,001199914 | 1,068745535 | 0,467 | 0,108 | 1 | CH_SC | 2,097608643 | 0,476733352 | 4,324074074 |
| SFSWAP | 0,001318091 | 0,825930514 | 0,4 | 0,077 | 1 | CH_SC | 1,772678017 | 0,564118238 | 5,194805195 |
| TMEM35B1 | 0,001319318 | 1,068267298 | 0,533 | 0,154 | 1 | CH_SC | 2,096913424 | 0,47689141 | 3,461038961 |
| PTPRZ11 | 0,001321142 | 2,046015109 | 0,467 | 0,138 | 1 | CH_SC | 4,129637391 | 0,242152011 | 3,384057971 |
| QKI2 | 0,001325006 | 1,656278114 | 0,6 | 0,246 | 1 | CH_SC | 3,152023113 | 0,317256557 | 2,43902439 |
| ITGB15 | 0,00134348 | 1,544360899 | 0,733 | 0,508 | 1 | CH_SC | 2,91674831 | 0,342847546 | 1,442913386 |
| SH3PXD2A2 | 0,001352488 | 1,187525832 | 0,4 | 0,092 | 1 | CH_SC | 2,277618051 | 0,439055179 | 4,347826087 |
| PTPDC11 | 0,001381649 | 1,220024743 | 0,267 | 0,031 | 1 | CH_SC | 2,329507124 | 0,429275356 | 8,612903226 |
| EDNRB1 | 0,001391101 | 0,598154828 | 0,333 | 0,046 | 1 | CH_SC | 1,513779241 | 0,660598304 | 7,239130435 |
| EMP24 | 0,001398915 | 1,199992317 | 0,667 | 0,338 | 1 | CH_SC | 2,297384475 | 0,4352776 | 1,973372781 |
| ZCCHC241 | 0,001404608 | 1,660602734 | 0,467 | 0,138 | 1 | CH_SC | 3,161485788 | 0,316306973 | 3,384057971 |
| RNF2134 | 0,001442181 | 0,918723768 | 0,467 | 0,123 | 1 | CH_SC | 1,890442237 | 0,528976755 | 3,796747967 |
| COMP1 | 0,001458022 | -3,139091385 | 0 | 0,462 | 1 | CH_SC | 0,113511362 | 8,809690799 | 0 |

| Supplementary Table 4 - clusermarker | | | | | | | | | |
|---|---|---|---|---|---|---|---|---|---|
| | **p_val** | **avg_log2FC** | **pct.1** | **pct.2** | **p_val_adj** | **celltype** | **Foldchange_UP** | **Foldchange_DOWN** | **Ratio_pct1_pct2** |
| ITIH6 | 0,001458022 | -1,470406073 | 0 | 0,462 | 1 | CH_SC | 0,360880708 | 2,770998774 | 0 |
| DDR21 | 0,001478956 | -1,672821409 | 0,133 | 0,554 | 1 | CH_SC | 0,313639374 | 3,188375193 | 0,240072202 |
| USP41 | 0,001516715 | 0,914034809 | 0,267 | 0,031 | 1 | CH_SC | 1,884308012 | 0,5306988 | 8,612903226 |
| LRRC4C | 0,001516715 | 1,537674847 | 0,267 | 0,031 | 1 | CH_SC | 2,903262151 | 0,344440132 | 8,612903226 |
| SSH11 | 0,001516715 | 0,94083624 | 0,267 | 0,031 | 1 | CH_SC | 1,919640612 | 0,520930842 | 8,612903226 |
| PGRMC13 | 0,001531046 | 1,389836507 | 0,667 | 0,308 | 1 | CH_SC | 2,620489825 | 0,381608045 | 2,165584416 |
| RPL45 | 0,001579294 | -1,112353228 | 0,6 | 0,923 | 1 | CH_SC | 0,462538953 | 2,161980076 | 0,650054171 |
| VCL1 | 0,001586307 | 1,722603014 | 0,467 | 0,138 | 1 | CH_SC | 3,300313363 | 0,30300153 | 3,384057971 |
| FADS31 | 0,001586307 | 1,332472647 | 0,467 | 0,138 | 1 | CH_SC | 2,518339254 | 0,397087088 | 3,384057971 |
| CCPG1 | 0,001612871 | -1,440355563 | 0,133 | 0,569 | 1 | CH_SC | 0,368476479 | 2,713877427 | 0,233743409 |
| ARPC26 | 0,001631336 | 1,144364753 | 0,733 | 0,354 | 1 | CH_SC | 2,210487771 | 0,452388841 | 2,070621469 |
| GSK3B1 | 0,001631353 | 0,658644244 | 0,4 | 0,077 | 1 | CH_SC | 1,578598457 | 0,633473317 | 5,194805195 |
| CCL29 | 0,001660442 | 1,281136921 | 0,4 | 0,092 | 1 | CH_SC | 2,430304223 | 0,41147112 | 4,347826087 |
| AGTRAP1 | 0,001663851 | 0,874642487 | 0,267 | 0,031 | 1 | CH_SC | 1,833553659 | 0,545389002 | 8,612903226 |
| CAMK2N11 | 0,001663851 | 0,754117926 | 0,267 | 0,031 | 1 | CH_SC | 1,686600077 | 0,592908783 | 8,612903226 |
| LBR1 | 0,001663851 | 0,759218179 | 0,267 | 0,031 | 1 | CH_SC | 1,692573142 | 0,590816418 | 8,612903226 |
| HAND2 | 0,001663851 | 1,417973652 | 0,267 | 0,031 | 1 | CH_SC | 2,672099356 | 0,374237581 | 8,612903226 |
| MCUR11 | 0,001663851 | 0,757981606 | 0,267 | 0,031 | 1 | CH_SC | 1,691123013 | 0,591323039 | 8,612903226 |
| PDPK1 | 0,001663851 | 0,737323595 | 0,267 | 0,031 | 1 | CH_SC | 1,667080297 | 0,59985113 | 8,612903226 |
| ERCC41 | 0,001663851 | 1,394220967 | 0,267 | 0,031 | 1 | CH_SC | 2,628465806 | 0,38045007 | 8,612903226 |
| ZDHHC7 | 0,001663851 | 0,788617539 | 0,267 | 0,031 | 1 | CH_SC | 1,727418373 | 0,578898555 | 8,612903226 |
| WDR181 | 0,001663851 | 0,600542394 | 0,267 | 0,031 | 1 | CH_SC | 1,516286521 | 0,659505961 | 8,612903226 |
| YBX34 | 0,001709493 | -1,819148814 | 0,067 | 0,508 | 1 | CH_SC | 0,28338812 | 3,528729431 | 0,131889764 |
| INSIG14 | 0,001737588 | 1,096078648 | 0,467 | 0,123 | 1 | CH_SC | 2,137728517 | 0,467786247 | 3,796747967 |
| VPS13C4 | 0,00176332 | 1,266182359 | 0,533 | 0,154 | 1 | CH_SC | 2,405242505 | 0,415758493 | 3,461038961 |
| ITSN21 | 0,001789454 | 1,319581908 | 0,333 | 0,062 | 1 | CH_SC | 2,495937671 | 0,40065103 | 5,370967742 |
| RPL126 | 0,001816379 | -0,91294629 | 0,867 | 0,985 | 1 | CH_SC | 0,531099365 | 1,88288683 | 0,880203046 |
| HEG11 | 0,001824019 | 0,515581115 | 0,267 | 0,031 | 1 | CH_SC | 1,429569852 | 0,699511114 | 8,612903226 |
| ARID5B6 | 0,001824019 | 0,91420555 | 0,267 | 0,031 | 1 | CH_SC | 1,88453103 | 0,530635996 | 8,612903226 |
| RGS11 | 0,001824019 | 0,858960202 | 0,267 | 0,031 | 1 | CH_SC | 1,813730624 | 0,551349791 | 8,612903226 |
| SORBS12 | 0,001824271 | 1,311165733 | 0,6 | 0,231 | 1 | CH_SC | 2,481419638 | 0,402995118 | 2,597402597 |

| Supplementary Table 4 - clusermarker | | | | | | | | | |
|---|---|---|---|---|---|---|---|---|---|
| | **p_val** | **avg_log2FC** | **pct.1** | **pct.2** | **p_val_adj** | **celltype** | **Foldchange_UP** | **Foldchange_DOWN** | **Ratio_pct1_pct2** |
| LRP13 | 0,001833226 | -1,248271713 | 0,2 | 0,631 | 1 | CH_SC | 0,420952188 | 2,375566697 | 0,316957211 |
| DHX15 | 0,001878604 | 1,482766932 | 0,4 | 0,108 | 1 | CH_SC | 2,7948424 | 0,357801928 | 3,703703704 |
| MPHOSPH6 | 0,001887109 | -1,351291719 | 0 | 0,446 | 1 | CH_SC | 0,391940967 | 2,551404636 | 0 |
| FGFR11 | 0,001887109 | -1,355761079 | 0 | 0,446 | 1 | CH_SC | 0,390728642 | 2,55932095 | 0 |
| PRB310 | 0,001887109 | -2,167221656 | 0 | 0,446 | 1 | CH_SC | 0,222639016 | 4,49157572 | 0 |
| COLEC121 | 0,001887109 | -1,392966913 | 0 | 0,446 | 1 | CH_SC | 0,380780918 | 2,626182021 | 0 |
| TCEAL2 | 0,001887109 | -1,317008947 | 0 | 0,446 | 1 | CH_SC | 0,401366205 | 2,491490282 | 0 |
| TENM3 | 0,001899714 | 1,783664596 | 0,467 | 0,138 | 1 | CH_SC | 3,442996217 | 0,290444699 | 3,384057971 |
| IAH1 | 0,001900286 | 1,171020598 | 0,4 | 0,092 | 1 | CH_SC | 2,251709321 | 0,444107057 | 4,347826087 |
| MARK32 | 0,001900286 | 1,032187256 | 0,4 | 0,092 | 1 | CH_SC | 2,045122493 | 0,488968266 | 4,347826087 |
| BACH1 | 0,001900286 | 1,021584985 | 0,4 | 0,092 | 1 | CH_SC | 2,030148112 | 0,492574898 | 4,347826087 |
| ZFP36L23 | 0,001924121 | 1,095278186 | 0,867 | 0,692 | 1 | CH_SC | 2,136542753 | 0,468045864 | 1,252890173 |
| BCKDHA1 | 0,001930961 | 0,999746501 | 0,333 | 0,062 | 1 | CH_SC | 1,999648607 | 0,500087864 | 5,370967742 |
| SERPING15 | 0,001931188 | -1,344347684 | 0,6 | 0,892 | 1 | CH_SC | 0,39383202 | 2,539153624 | 0,67264574 |
| MT1M2 | 0,001990191 | -3,093660524 | 0,2 | 0,6 | 1 | CH_SC | 0,117142742 | 8,536593747 | 0,333333333 |
| DLX2 | 0,001998247 | 1,084971715 | 0,267 | 0,031 | 1 | CH_SC | 2,121333891 | 0,47140151 | 8,612903226 |
| NAV2 | 0,001998247 | 0,641105294 | 0,267 | 0,031 | 1 | CH_SC | 1,559523502 | 0,6412215 | 8,612903226 |
| MGAT4C | 0,001998247 | 1,55095327 | 0,267 | 0,031 | 1 | CH_SC | 2,93010684 | 0,341284484 | 8,612903226 |
| PTPRS | 0,001998247 | 1,151179225 | 0,267 | 0,031 | 1 | CH_SC | 2,220953558 | 0,450257051 | 8,612903226 |
| TBC1D10A | 0,001998247 | 0,630121189 | 0,267 | 0,031 | 1 | CH_SC | 1,547694997 | 0,646122138 | 8,612903226 |
| TLN14 | 0,002010954 | 1,117900897 | 0,667 | 0,369 | 1 | CH_SC | 2,170309654 | 0,460763743 | 1,807588076 |
| EEA11 | 0,002021678 | 0,994580979 | 0,6 | 0,231 | 1 | CH_SC | 1,992501733 | 0,501881621 | 2,597402597 |
| SCGB3A18 | 0,002080166 | -1,649217196 | 0,067 | 0,492 | 1 | CH_SC | 0,318813097 | 3,136633998 | 0,136178862 |
| LYST2 | 0,002082677 | 0,941002386 | 0,333 | 0,062 | 1 | CH_SC | 1,919861697 | 0,520870853 | 5,370967742 |
| INO80E | 0,002082677 | 0,882142696 | 0,333 | 0,062 | 1 | CH_SC | 1,843110664 | 0,542561019 | 5,370967742 |
| ERBB21 | 0,002082677 | 1,050324632 | 0,333 | 0,062 | 1 | CH_SC | 2,070995806 | 0,4828595 | 5,370967742 |
| LSM45 | 0,002082677 | 0,914951116 | 0,333 | 0,062 | 1 | CH_SC | 1,885505183 | 0,530361841 | 5,370967742 |
| TEAD11 | 0,002120451 | 1,087332507 | 0,533 | 0,169 | 1 | CH_SC | 2,124808034 | 0,470630751 | 3,153846154 |
| METRN1 | 0,002126112 | 1,259368824 | 0,533 | 0,185 | 1 | CH_SC | 2,39390985 | 0,417726674 | 2,881081081 |
| UBB6 | 0,002147899 | -1,183607444 | 0,267 | 0,815 | 1 | CH_SC | 0,440249282 | 2,271440389 | 0,327607362 |
| METTL92 | 0,002171582 | 0,843935966 | 0,4 | 0,092 | 1 | CH_SC | 1,794940431 | 0,557121553 | 4,347826087 |

| Supplementary Table 4 - clusermarker | | | | | | | | | |
|---|---|---|---|---|---|---|---|---|---|
| | **p_val** | **avg_log2FC** | **pct.1** | **pct.2** | **p_val_adj** | **celltype** | **Foldchange_UP** | **Foldchange_DOWN** | **Ratio_pct1_pct2** |
| RPL22L13 | 0,00217862 | -1,403753069 | 0,067 | 0,492 | 1 | CH_SC | 0,377944665 | 2,645889972 | 0,136178862 |
| SDC21 | 0,002185976 | -1,598089096 | 0,133 | 0,585 | 1 | CH_SC | 0,330314202 | 3,027420543 | 0,227350427 |
| BGN4 | 0,002206128 | -1,183314424 | 0,667 | 0,908 | 1 | CH_SC | 0,440338708 | 2,270979093 | 0,734581498 |
| TPT1-AS1 | 0,002218146 | 1,109041882 | 0,467 | 0,123 | 1 | CH_SC | 2,157023482 | 0,463601814 | 3,796747967 |
| DOCK1 | 0,002245257 | 0,963993484 | 0,333 | 0,062 | 1 | CH_SC | 1,950702112 | 0,512635934 | 5,370967742 |
| PDE4DIP1 | 0,002273902 | -0,929050044 | 0,133 | 0,662 | 1 | CH_SC | 0,525204054 | 1,904021862 | 0,200906344 |
| SARAF3 | 0,002332011 | 1,24161188 | 0,667 | 0,4 | 1 | CH_SC | 2,364625773 | 0,422899899 | 1,6675 |
| AHNAK6 | 0,002347296 | 1,057090404 | 0,867 | 0,8 | 1 | CH_SC | 2,080730914 | 0,480600347 | 1,08375 |
| LAMB12 | 0,002355968 | 1,180127562 | 0,467 | 0,123 | 1 | CH_SC | 2,265968117 | 0,441312476 | 3,796747967 |
| RPL141 | 0,002372484 | -0,85001232 | 0,8 | 0,954 | 1 | CH_SC | 0,554779998 | 1,802516318 | 0,838574423 |
| MBD51 | 0,002393339 | 0,98466869 | 0,267 | 0,031 | 1 | CH_SC | 1,978858822 | 0,50534176 | 8,612903226 |
| SOX2-OT1 | 0,002393339 | 0,430739971 | 0,267 | 0,031 | 1 | CH_SC | 1,347924762 | 0,74188117 | 8,612903226 |
| TMEM1751 | 0,002393339 | 0,431997718 | 0,267 | 0,031 | 1 | CH_SC | 1,3491004 | 0,741234677 | 8,612903226 |
| COL21A11 | 0,002393339 | 1,213755733 | 0,267 | 0,031 | 1 | CH_SC | 2,319406571 | 0,431144765 | 8,612903226 |
| ZNRF2 | 0,002393339 | 0,709292469 | 0,267 | 0,031 | 1 | CH_SC | 1,635002078 | 0,611620018 | 8,612903226 |
| UPP12 | 0,002393339 | 0,511990931 | 0,267 | 0,031 | 1 | CH_SC | 1,426016754 | 0,701254033 | 8,612903226 |
| ADAM91 | 0,002393339 | 0,64027579 | 0,267 | 0,031 | 1 | CH_SC | 1,558627083 | 0,641590289 | 8,612903226 |
| HEY1 | 0,002393339 | 0,608223307 | 0,267 | 0,031 | 1 | CH_SC | 1,524380763 | 0,65600408 | 8,612903226 |
| FMNL3 | 0,002393339 | 0,605827697 | 0,267 | 0,031 | 1 | CH_SC | 1,521851613 | 0,657094287 | 8,612903226 |
| CERS42 | 0,002393339 | 0,563690776 | 0,267 | 0,031 | 1 | CH_SC | 1,478045597 | 0,676569114 | 8,612903226 |
| RBM432 | 0,002419391 | 1,363427733 | 0,333 | 0,062 | 1 | CH_SC | 2,572957688 | 0,388657771 | 5,370967742 |
| SH3BP2 | 0,002419391 | 1,045204389 | 0,333 | 0,062 | 1 | CH_SC | 2,063658699 | 0,484576253 | 5,370967742 |
| ACTB10 | 0,002425916 | 0,894939693 | 0,933 | 0,969 | 1 | CH_SC | 1,859532152 | 0,537769674 | 0,962848297 |
| NBL13 | 0,002429292 | -1,706270191 | 0 | 0,431 | 1 | CH_SC | 0,306451317 | 3,263161049 | 0 |
| PLA2G2A1 | 0,002429292 | -2,940168472 | 0 | 0,431 | 1 | CH_SC | 0,130293004 | 7,675009161 | 0 |
| TM4SF13 | 0,002429292 | -1,828183188 | 0 | 0,431 | 1 | CH_SC | 0,281619046 | 3,550896201 | 0 |
| FOXC13 | 0,002429292 | -1,40488243 | 0 | 0,431 | 1 | CH_SC | 0,377648921 | 2,647962022 | 0 |
| ASS12 | 0,002429292 | -1,4267429 | 0 | 0,431 | 1 | CH_SC | 0,371969723 | 2,688390852 | 0 |
| PPP1R3C1 | 0,002429292 | -1,652829481 | 0 | 0,431 | 1 | CH_SC | 0,318015837 | 3,144497484 | 0 |
| MTRNR2L124 | 0,002445582 | 1,208656123 | 0,933 | 0,785 | 1 | CH_SC | 2,31122245 | 0,432671464 | 1,188535032 |
| ADD33 | 0,002477971 | 1,107481758 | 0,4 | 0,092 | 1 | CH_SC | 2,154692147 | 0,464103423 | 4,347826087 |

| Supplementary Table 4 - clusermarker | | | | | | | | | |
|---|---|---|---|---|---|---|---|---|---|
| | **p_val** | **avg_log2FC** | **pct.1** | **pct.2** | **p_val_adj** | **celltype** | **Foldchange_UP** | **Foldchange_DOWN** | **Ratio_pct1_pct2** |
| SMIM51 | 0,00250159 | 1,185371391 | 0,467 | 0,123 | 1 | CH_SC | 2,27421932 | 0,439711329 | 3,796747967 |
| C1QL1 | 0,002550754 | 1,273268703 | 0,467 | 0,138 | 1 | CH_SC | 2,417085829 | 0,413721345 | 3,384057971 |
| SPTAN14 | 0,002560838 | 1,517144314 | 0,533 | 0,215 | 1 | CH_SC | 2,862239338 | 0,349376793 | 2,479069767 |
| OSGEP | 0,002605806 | 0,838270465 | 0,333 | 0,062 | 1 | CH_SC | 1,787905476 | 0,559313685 | 5,370967742 |
| RPL89 | 0,002619458 | -0,969789716 | 0,733 | 0,938 | 1 | CH_SC | 0,510580478 | 1,9585551 | 0,781449893 |
| ATP1B21 | 0,00264556 | 1,137273829 | 0,4 | 0,092 | 1 | CH_SC | 2,199649761 | 0,45461783 | 4,347826087 |
| RHOA4 | 0,002721621 | 1,464468037 | 0,733 | 0,585 | 1 | CH_SC | 2,75961697 | 0,36236913 | 1,252991453 |
| CD1514 | 0,002780993 | 1,353449876 | 0,667 | 0,385 | 1 | CH_SC | 2,55522419 | 0,391355093 | 1,732467532 |
| PAPPA1 | 0,002805267 | 0,810863554 | 0,333 | 0,062 | 1 | CH_SC | 1,754261177 | 0,570040546 | 5,370967742 |
| WARS1 | 0,002805267 | 0,805889859 | 0,333 | 0,062 | 1 | CH_SC | 1,74822377 | 0,572009154 | 5,370967742 |
| RHBDF11 | 0,002805267 | 1,096843405 | 0,333 | 0,062 | 1 | CH_SC | 2,138862004 | 0,467538344 | 5,370967742 |
| LPCAT21 | 0,002805267 | 1,163208285 | 0,333 | 0,062 | 1 | CH_SC | 2,239549083 | 0,446518457 | 5,370967742 |
| GRIPAP1 | 0,002805267 | 0,816311666 | 0,333 | 0,062 | 1 | CH_SC | 1,760898393 | 0,567891937 | 5,370967742 |
| SLC44A13 | 0,002817816 | 0,750997534 | 0,467 | 0,123 | 1 | CH_SC | 1,682956088 | 0,594192568 | 3,796747967 |
| PHC21 | 0,002977276 | 0,67684888 | 0,2 | 0,015 | 1 | CH_SC | 1,598644197 | 0,62553006 | 13,33333333 |
| FOXD3 | 0,002977276 | 0,336421801 | 0,2 | 0,015 | 1 | CH_SC | 1,262621131 | 0,792003219 | 13,33333333 |
| BCL91 | 0,002977276 | 1,059820487 | 0,2 | 0,015 | 1 | CH_SC | 2,084672112 | 0,479691743 | 13,33333333 |
| KCNS32 | 0,002977276 | 0,564230305 | 0,2 | 0,015 | 1 | CH_SC | 1,478598449 | 0,676316143 | 13,33333333 |
| ATP6V0E2 | 0,002977276 | 0,637813652 | 0,2 | 0,015 | 1 | CH_SC | 1,555969361 | 0,642686177 | 13,33333333 |
| HRAS3 | 0,002977276 | 0,535819028 | 0,2 | 0,015 | 1 | CH_SC | 1,449764963 | 0,68976698 | 13,33333333 |
| SLC2A35 | 0,002977276 | 0,777027898 | 0,2 | 0,015 | 1 | CH_SC | 1,713597045 | 0,583567766 | 13,33333333 |
| SLC39A91 | 0,002977276 | 0,549303159 | 0,2 | 0,015 | 1 | CH_SC | 1,463378694 | 0,683350116 | 13,33333333 |
| TMEM59L | 0,002977276 | 0,546705713 | 0,2 | 0,015 | 1 | CH_SC | 1,460746379 | 0,684581536 | 13,33333333 |
| RYR1 | 0,002977276 | 0,552086354 | 0,2 | 0,015 | 1 | CH_SC | 1,466204516 | 0,682033092 | 13,33333333 |
| IGLV1-511 | 0,002977276 | 1,396587687 | 0,2 | 0,015 | 1 | CH_SC | 2,632781305 | 0,379826459 | 13,33333333 |
| SHROOM4 | 0,002977276 | 1,067491726 | 0,2 | 0,015 | 1 | CH_SC | 2,095786458 | 0,477147849 | 13,33333333 |
| DUSP63 | 0,002989252 | 0,945299359 | 0,467 | 0,123 | 1 | CH_SC | 1,925588405 | 0,519321781 | 3,796747967 |
| NARF1 | 0,003012198 | 0,771001977 | 0,4 | 0,092 | 1 | CH_SC | 1,706454535 | 0,586010339 | 4,347826087 |
| RASGEF1C | 0,003018579 | 0,814950927 | 0,333 | 0,062 | 1 | CH_SC | 1,759238311 | 0,568427821 | 5,370967742 |
| PTPRJ1 | 0,003018579 | 0,675107411 | 0,333 | 0,062 | 1 | CH_SC | 1,596715646 | 0,62628559 | 5,370967742 |
| CDC42EP12 | 0,003018579 | 0,709935737 | 0,333 | 0,062 | 1 | CH_SC | 1,635731254 | 0,61134737 | 5,370967742 |

| Supplementary Table 4 - clusermarker | | | | | | | | | |
|---|---|---|---|---|---|---|---|---|---|
| | **p_val** | **avg_log2FC** | **pct.1** | **pct.2** | **p_val_adj** | **celltype** | **Foldchange_UP** | **Foldchange_DOWN** | **Ratio_pct1_pct2** |
| ZFP36L13 | 0,003019977 | 1,106244617 | 0,8 | 0,538 | 1 | CH_SC | 2,152845246 | 0,464501572 | 1,486988848 |
| SELENOP3 | 0,003067954 | -1,402269318 | 0,2 | 0,631 | 1 | CH_SC | 0,378333565 | 2,643170185 | 0,316957211 |
| DEGS13 | 0,003090926 | 1,011140423 | 0,533 | 0,185 | 1 | CH_SC | 2,015503687 | 0,496153893 | 2,881081081 |
| EDIL31 | 0,003110862 | -1,758341962 | 0 | 0,415 | 1 | CH_SC | 0,295587679 | 3,383090949 | 0 |
| ARF57 | 0,003110862 | 1,431597215 | 0,6 | 0,277 | 1 | CH_SC | 2,697451863 | 0,370720239 | 2,166064982 |
| APP3 | 0,003120407 | 1,008264111 | 0,8 | 0,6 | 1 | CH_SC | 2,011489366 | 0,497144065 | 1,333333333 |
| RAC16 | 0,003171956 | 1,374473221 | 0,733 | 0,538 | 1 | CH_SC | 2,592732238 | 0,385693511 | 1,362453532 |
| PTN2 | 0,003212099 | 0,906038603 | 0,467 | 0,138 | 1 | CH_SC | 1,873893034 | 0,53364839 | 3,384057971 |
| IQGAP22 | 0,003212391 | 1,521084736 | 0,4 | 0,092 | 1 | CH_SC | 2,870067636 | 0,348423845 | 4,347826087 |
| RPL292 | 0,00321817 | -0,770575979 | 1 | 0,969 | 1 | CH_SC | 0,586183401 | 1,705950728 | 1,031991744 |
| PSMB95 | 0,00324659 | 0,706253907 | 0,333 | 0,062 | 1 | CH_SC | 1,631562108 | 0,612909552 | 5,370967742 |
| CSNK1E2 | 0,00324659 | 0,594896515 | 0,333 | 0,062 | 1 | CH_SC | 1,510364242 | 0,662091946 | 5,370967742 |
| HLA-C8 | 0,003253027 | 1,106008868 | 0,8 | 0,892 | 1 | CH_SC | 2,152493481 | 0,464577481 | 0,896860987 |
| RPSA2 | 0,003276679 | -0,784345696 | 0,8 | 0,954 | 1 | CH_SC | 0,580615223 | 1,722311025 | 0,838574423 |
| EIF3E5 | 0,003280933 | -1,137802369 | 0,2 | 0,677 | 1 | CH_SC | 0,454451308 | 2,200455764 | 0,295420975 |
| FAM102B3 | 0,003285241 | 0,478454142 | 0,133 | 0 | 1 | CH_SC | 1,393249989 | 0,717746283 | #DIV/0! |
| KCNA2 | 0,003285241 | 0,735089105 | 0,133 | 0 | 1 | CH_SC | 1,66450027 | 0,600780918 | #DIV/0! |
| SYT14 | 0,003285241 | 0,587687061 | 0,133 | 0 | 1 | CH_SC | 1,502835459 | 0,665408841 | #DIV/0! |
| TP53BP2 | 0,003285241 | 0,53490553 | 0,133 | 0 | 1 | CH_SC | 1,448847279 | 0,690203871 | #DIV/0! |
| AL591848.4 | 0,003285241 | 0,495720851 | 0,133 | 0 | 1 | CH_SC | 1,410025106 | 0,70920723 | #DIV/0! |
| TRIM58 | 0,003285241 | 0,257450531 | 0,133 | 0 | 1 | CH_SC | 1,19536444 | 0,836564956 | #DIV/0! |
| EIPR1 | 0,003285241 | 1,035894699 | 0,133 | 0 | 1 | CH_SC | 2,050384815 | 0,487713327 | #DIV/0! |
| LRRTM4 | 0,003285241 | 0,472777577 | 0,133 | 0 | 1 | CH_SC | 1,387778746 | 0,720575958 | #DIV/0! |
| IGKV1-57 | 0,003285241 | 0,912617702 | 0,133 | 0 | 1 | CH_SC | 1,882458033 | 0,531220342 | #DIV/0! |
| SH3RF3 | 0,003285241 | 0,793634758 | 0,133 | 0 | 1 | CH_SC | 1,733436224 | 0,576888833 | #DIV/0! |
| LYPD1 | 0,003285241 | 0,499131584 | 0,133 | 0 | 1 | CH_SC | 1,413362547 | 0,707532545 | #DIV/0! |
| LINC011161 | 0,003285241 | 0,450405373 | 0,133 | 0 | 1 | CH_SC | 1,366424145 | 0,731837185 | #DIV/0! |
| CHROMR | 0,003285241 | 0,315853472 | 0,133 | 0 | 1 | CH_SC | 1,244747805 | 0,803375588 | #DIV/0! |
| ECEL11 | 0,003285241 | 0,484243334 | 0,133 | 0 | 1 | CH_SC | 1,398852002 | 0,714871908 | #DIV/0! |
| ATG71 | 0,003285241 | 0,495601528 | 0,133 | 0 | 1 | CH_SC | 1,409908491 | 0,70926589 | #DIV/0! |
| NCKIPSD | 0,003285241 | 0,250156809 | 0,133 | 0 | 1 | CH_SC | 1,189336379 | 0,840805022 | #DIV/0! |

| Supplementary Table 4 - clusermarker | | | | | | | | | |
|---|---|---|---|---|---|---|---|---|---|
| | **p_val** | **avg_log2FC** | **pct.1** | **pct.2** | **p_val_adj** | **celltype** | **Foldchange_UP** | **Foldchange_DOWN** | **Ratio_pct1_pct2** |
| RBM15B | 0,003285241 | 0,264412536 | 0,133 | 0 | 1 | CH_SC | 1,201146844 | 0,832537674 | #DIV/0! |
| C3orf141 | 0,003285241 | 0,426929563 | 0,133 | 0 | 1 | CH_SC | 1,344369356 | 0,743843197 | #DIV/0! |
| SORCS2 | 0,003285241 | 0,425658518 | 0,133 | 0 | 1 | CH_SC | 1,343185459 | 0,744498828 | #DIV/0! |
| NSUN2 | 0,003285241 | 0,274857983 | 0,133 | 0 | 1 | CH_SC | 1,209874985 | 0,826531677 | #DIV/0! |
| ARL151 | 0,003285241 | 1,019450091 | 0,133 | 0 | 1 | CH_SC | 2,02714613 | 0,493304348 | #DIV/0! |
| STARD4-AS1 | 0,003285241 | 1,066251643 | 0,133 | 0 | 1 | CH_SC | 2,093985776 | 0,477558163 | #DIV/0! |
| SEMA6A | 0,003285241 | 1,281385818 | 0,133 | 0 | 1 | CH_SC | 2,430723542 | 0,411400138 | #DIV/0! |
| AC106791.1 | 0,003285241 | 0,735089105 | 0,133 | 0 | 1 | CH_SC | 1,66450027 | 0,600780918 | #DIV/0! |
| AC005740.5 | 0,003285241 | 0,44841475 | 0,133 | 0 | 1 | CH_SC | 1,364540061 | 0,732847667 | #DIV/0! |
| ARSI | 0,003285241 | 0,486642661 | 0,133 | 0 | 1 | CH_SC | 1,40118035 | 0,713684002 | #DIV/0! |
| SMIM131 | 0,003285241 | 1,528036136 | 0,133 | 0 | 1 | CH_SC | 2,883929978 | 0,346749057 | #DIV/0! |
| TSBP1-AS1 | 0,003285241 | 0,250156809 | 0,133 | 0 | 1 | CH_SC | 1,189336379 | 0,840805022 | #DIV/0! |
| HLA-DQA22 | 0,003285241 | 0,62481019 | 0,133 | 0 | 1 | CH_SC | 1,542007936 | 0,648505093 | #DIV/0! |
| SAMD5 | 0,003285241 | 0,349381757 | 0,133 | 0 | 1 | CH_SC | 1,274014553 | 0,78492039 | #DIV/0! |
| PDE10A | 0,003285241 | 0,436265796 | 0,133 | 0 | 1 | CH_SC | 1,353097497 | 0,739045045 | #DIV/0! |
| NUP421 | 0,003285241 | 0,587687061 | 0,133 | 0 | 1 | CH_SC | 1,502835459 | 0,665408841 | #DIV/0! |
| PLEKHA8 | 0,003285241 | 0,52926142 | 0,133 | 0 | 1 | CH_SC | 1,443190173 | 0,692909374 | #DIV/0! |
| NOD1 | 0,003285241 | 0,44841475 | 0,133 | 0 | 1 | CH_SC | 1,364540061 | 0,732847667 | #DIV/0! |
| GRB10 | 0,003285241 | 0,609825057 | 0,133 | 0 | 1 | CH_SC | 1,526074144 | 0,655276157 | #DIV/0! |
| SAMD92 | 0,003285241 | 0,690813828 | 0,133 | 0 | 1 | CH_SC | 1,614193832 | 0,619504288 | #DIV/0! |
| TFPI21 | 0,003285241 | 0,288954746 | 0,133 | 0 | 1 | CH_SC | 1,221754778 | 0,818494855 | #DIV/0! |
| GIMAP2 | 0,003285241 | 0,49950539 | 0,133 | 0 | 1 | CH_SC | 1,4137288 | 0,707349245 | #DIV/0! |
| SOX7 | 0,003285241 | 0,463675399 | 0,133 | 0 | 1 | CH_SC | 1,379050605 | 0,725136551 | #DIV/0! |
| TRIM35 | 0,003285241 | 0,426929563 | 0,133 | 0 | 1 | CH_SC | 1,344369356 | 0,743843197 | #DIV/0! |
| UNC13B1 | 0,003285241 | 0,466172405 | 0,133 | 0 | 1 | CH_SC | 1,381439523 | 0,723882576 | #DIV/0! |
| SLC31A2 | 0,003285241 | 0,484243334 | 0,133 | 0 | 1 | CH_SC | 1,398852002 | 0,714871908 | #DIV/0! |
| MACORIS | 0,003285241 | 0,482020582 | 0,133 | 0 | 1 | CH_SC | 1,396698458 | 0,715974156 | #DIV/0! |
| LINC00844 | 0,003285241 | 0,82221413 | 0,133 | 0 | 1 | CH_SC | 1,768117473 | 0,565573281 | #DIV/0! |
| HKDC1 | 0,003285241 | 0,691141132 | 0,133 | 0 | 1 | CH_SC | 1,614560086 | 0,619363757 | #DIV/0! |
| ADAM123 | 0,003285241 | 0,426929563 | 0,133 | 0 | 1 | CH_SC | 1,344369356 | 0,743843197 | #DIV/0! |
| C10orf90 | 0,003285241 | 1,092715622 | 0,133 | 0 | 1 | CH_SC | 2,132751121 | 0,468877962 | #DIV/0! |

| Supplementary Table 4 - clusermarker | | | | | | | | | |
|---|---|---|---|---|---|---|---|---|---|
| | **p_val** | **avg_log2FC** | **pct.1** | **pct.2** | **p_val_adj** | **celltype** | **Foldchange_UP** | **Foldchange_DOWN** | **Ratio_pct1_pct2** |
| BBOX11 | 0,003285241 | 0,640183416 | 0,133 | 0 | 1 | CH_SC | 1,558527289 | 0,64163137 | #DIV/0! |
| FEN1 | 0,003285241 | 0,517406432 | 0,133 | 0 | 1 | CH_SC | 1,431379708 | 0,698626643 | #DIV/0! |
| GRK21 | 0,003285241 | 0,463675399 | 0,133 | 0 | 1 | CH_SC | 1,379050605 | 0,725136551 | #DIV/0! |
| TCIRG13 | 0,003285241 | 0,495601528 | 0,133 | 0 | 1 | CH_SC | 1,409908491 | 0,70926589 | #DIV/0! |
| NOX41 | 0,003285241 | 0,499131584 | 0,133 | 0 | 1 | CH_SC | 1,413362547 | 0,707532545 | #DIV/0! |
| GPR83 | 0,003285241 | 0,411273086 | 0,133 | 0 | 1 | CH_SC | 1,329858811 | 0,751959525 | #DIV/0! |
| RNF26 | 0,003285241 | 0,900875769 | 0,133 | 0 | 1 | CH_SC | 1,867199098 | 0,535561527 | #DIV/0! |
| PKP21 | 0,003285241 | 0,609825057 | 0,133 | 0 | 1 | CH_SC | 1,526074144 | 0,655276157 | #DIV/0! |
| AGAP2-AS1 | 0,003285241 | 0,425658518 | 0,133 | 0 | 1 | CH_SC | 1,343185459 | 0,744498828 | #DIV/0! |
| AC024909.1 | 0,003285241 | 0,587687061 | 0,133 | 0 | 1 | CH_SC | 1,502835459 | 0,665408841 | #DIV/0! |
| IGHG21 | 0,003285241 | 0,495601528 | 0,133 | 0 | 1 | CH_SC | 1,409908491 | 0,70926589 | #DIV/0! |
| IGHG19 | 0,003285241 | 1,587203302 | 0,133 | 0 | 1 | CH_SC | 3,004663236 | 0,332816 | #DIV/0! |
| INAFM2 | 0,003285241 | 0,912617702 | 0,133 | 0 | 1 | CH_SC | 1,882458033 | 0,531220342 | #DIV/0! |
| PEAK11 | 0,003285241 | 1,029740282 | 0,133 | 0 | 1 | CH_SC | 2,041656674 | 0,489798316 | #DIV/0! |
| AC015712.2 | 0,003285241 | 0,394623754 | 0,133 | 0 | 1 | CH_SC | 1,314599876 | 0,760687733 | #DIV/0! |
| E4F1 | 0,003285241 | 0,426929563 | 0,133 | 0 | 1 | CH_SC | 1,344369356 | 0,743843197 | #DIV/0! |
| TVP23A | 0,003285241 | 0,484621017 | 0,133 | 0 | 1 | CH_SC | 1,399218256 | 0,714684786 | #DIV/0! |
| GTF3C1 | 0,003285241 | 0,339543038 | 0,133 | 0 | 1 | CH_SC | 1,265355739 | 0,790291591 | #DIV/0! |
| EDC4 | 0,003285241 | 0,609825057 | 0,133 | 0 | 1 | CH_SC | 1,526074144 | 0,655276157 | #DIV/0! |
| PLA2G15 | 0,003285241 | 1,035894699 | 0,133 | 0 | 1 | CH_SC | 2,050384815 | 0,487713327 | #DIV/0! |
| MEOX11 | 0,003285241 | 0,484243334 | 0,133 | 0 | 1 | CH_SC | 1,398852002 | 0,714871908 | #DIV/0! |
| ABHD31 | 0,003285241 | 0,463675399 | 0,133 | 0 | 1 | CH_SC | 1,379050605 | 0,725136551 | #DIV/0! |
| DTNA1 | 0,003285241 | 0,388340962 | 0,133 | 0 | 1 | CH_SC | 1,308887373 | 0,764007676 | #DIV/0! |
| APC2 | 0,003285241 | 1,024810429 | 0,133 | 0 | 1 | CH_SC | 2,034692006 | 0,491474875 | #DIV/0! |
| SLC25A232 | 0,003285241 | 0,661280393 | 0,133 | 0 | 1 | CH_SC | 1,581485571 | 0,632316866 | #DIV/0! |
| AC011446.21 | 0,003285241 | 0,411273086 | 0,133 | 0 | 1 | CH_SC | 1,329858811 | 0,751959525 | #DIV/0! |
| ZNF850 | 0,003285241 | 0,679884626 | 0,133 | 0 | 1 | CH_SC | 1,602011635 | 0,624215192 | #DIV/0! |
| ZNF781 | 0,003285241 | 0,724089033 | 0,133 | 0 | 1 | CH_SC | 1,651857268 | 0,605379181 | #DIV/0! |
| ZNF2261 | 0,003285241 | 0,793634758 | 0,133 | 0 | 1 | CH_SC | 1,733436224 | 0,576888833 | #DIV/0! |
| APOC22 | 0,003285241 | 0,661895794 | 0,133 | 0 | 1 | CH_SC | 1,582160319 | 0,6320472 | #DIV/0! |
| AP5S1 | 0,003285241 | 0,394623754 | 0,133 | 0 | 1 | CH_SC | 1,314599876 | 0,760687733 | #DIV/0! |

| Supplementary Table 4 - clusermarker | | | | | | | | | |
|---|---|---|---|---|---|---|---|---|---|
| | **p_val** | **avg_log2FC** | **pct.1** | **pct.2** | **p_val_adj** | **celltype** | **Foldchange_UP** | **Foldchange_DOWN** | **Ratio_pct1_pct2** |
| KCNG1 | 0,003285241 | 0,436265796 | 0,133 | 0 | 1 | CH_SC | 1,353097497 | 0,739045045 | #DIV/0! |
| NKILA | 0,003285241 | 0,707813907 | 0,133 | 0 | 1 | CH_SC | 1,633327286 | 0,612247165 | #DIV/0! |
| NPEPL1 | 0,003285241 | 0,410460967 | 0,133 | 0 | 1 | CH_SC | 1,329110421 | 0,752382935 | #DIV/0! |
| DOP1B | 0,003285241 | 0,441160488 | 0,133 | 0 | 1 | CH_SC | 1,357696004 | 0,736541904 | #DIV/0! |
| PEX26 | 0,003285241 | 0,466172405 | 0,133 | 0 | 1 | CH_SC | 1,381439523 | 0,723882576 | #DIV/0! |
| PI4KA1 | 0,003285241 | 1,019450091 | 0,133 | 0 | 1 | CH_SC | 2,02714613 | 0,493304348 | #DIV/0! |
| MN1 | 0,003285241 | 0,49950539 | 0,133 | 0 | 1 | CH_SC | 1,4137288 | 0,707349245 | #DIV/0! |
| ODF3B3 | 0,003285241 | 0,426929563 | 0,133 | 0 | 1 | CH_SC | 1,344369356 | 0,743843197 | #DIV/0! |
| PRRG1 | 0,003285241 | 1,196876513 | 0,133 | 0 | 1 | CH_SC | 2,292428144 | 0,436218689 | #DIV/0! |
| PDZD4 | 0,003285241 | 0,539540453 | 0,133 | 0 | 1 | CH_SC | 1,453509452 | 0,687990022 | #DIV/0! |
| DNASE1L1 | 0,003285241 | 0,410460967 | 0,133 | 0 | 1 | CH_SC | 1,329110421 | 0,752382935 | #DIV/0! |
| TUBA1B4 | 0,003300533 | 1,386244555 | 0,8 | 0,738 | 1 | CH_SC | 2,613973572 | 0,382559338 | 1,08401084 |
| IFFO2 | 0,003309382 | 0,704145121 | 0,2 | 0,015 | 1 | CH_SC | 1,629178997 | 0,613806096 | 13,33333333 |
| SMAP21 | 0,003309382 | 0,91090694 | 0,2 | 0,015 | 1 | CH_SC | 1,880227119 | 0,531850642 | 13,33333333 |
| FCGR2B | 0,003309382 | 1,1008178 | 0,2 | 0,015 | 1 | CH_SC | 2,144762351 | 0,466252123 | 13,33333333 |
| CAMK1 | 0,003309382 | 0,481233767 | 0,2 | 0,015 | 1 | CH_SC | 1,395936937 | 0,716364739 | 13,33333333 |
| MKRN2 | 0,003309382 | 0,668857347 | 0,2 | 0,015 | 1 | CH_SC | 1,589813294 | 0,629004679 | 13,33333333 |
| HERC31 | 0,003309382 | 1,293478154 | 0,2 | 0,015 | 1 | CH_SC | 2,451182928 | 0,407966288 | 13,33333333 |
| DBN1 | 0,003309382 | 0,618076393 | 0,2 | 0,015 | 1 | CH_SC | 1,534827366 | 0,651539074 | 13,33333333 |
| FABP7 | 0,003309382 | 0,888143806 | 0,2 | 0,015 | 1 | CH_SC | 1,850793331 | 0,540308841 | 13,33333333 |
| TNFAIP34 | 0,003309382 | 1,018407044 | 0,2 | 0,015 | 1 | CH_SC | 2,025681062 | 0,493661129 | 13,33333333 |
| CRPPA | 0,003309382 | 0,72907219 | 0,2 | 0,015 | 1 | CH_SC | 1,657572749 | 0,603291771 | 13,33333333 |
| ZNF713 | 0,003309382 | 0,79217833 | 0,2 | 0,015 | 1 | CH_SC | 1,73168717 | 0,577471507 | 13,33333333 |
| DDHD2 | 0,003309382 | 1,237537605 | 0,2 | 0,015 | 1 | CH_SC | 2,35795732 | 0,424095886 | 13,33333333 |
| TMEM71 | 0,003309382 | 1,121189223 | 0,2 | 0,015 | 1 | CH_SC | 2,17526207 | 0,459714723 | 13,33333333 |
| TNKS1BP1 | 0,003309382 | 0,503871036 | 0,2 | 0,015 | 1 | CH_SC | 1,418013272 | 0,705212017 | 13,33333333 |
| KIRREL3 | 0,003309382 | 0,929980637 | 0,2 | 0,015 | 1 | CH_SC | 1,905250424 | 0,524865386 | 13,33333333 |
| R3HDM21 | 0,003309382 | 0,852724601 | 0,2 | 0,015 | 1 | CH_SC | 1,805908254 | 0,553737986 | 13,33333333 |
| AP5M11 | 0,003309382 | 0,661185244 | 0,2 | 0,015 | 1 | CH_SC | 1,581381271 | 0,63235857 | 13,33333333 |
| CRYZL11 | 0,003309382 | 0,422460688 | 0,2 | 0,015 | 1 | CH_SC | 1,340211497 | 0,746150889 | 13,33333333 |
| PCBP3 | 0,003309382 | 0,762173143 | 0,2 | 0,015 | 1 | CH_SC | 1,696043465 | 0,589607531 | 13,33333333 |

| Supplementary Table 4 - clusermarker | | | | | | | | | |
|---|---|---|---|---|---|---|---|---|---|
| | **p_val** | **avg_log2FC** | **pct.1** | **pct.2** | **p_val_adj** | **celltype** | **Foldchange_UP** | **Foldchange_DOWN** | **Ratio_pct1_pct2** |
| SKP12 | 0,00337812 | 1,075946484 | 0,8 | 0,815 | 1 | CH_SC | 2,108104646 | 0,474359753 | 0,981595092 |
| CAPS1 | 0,003400206 | 1,320389066 | 0,467 | 0,138 | 1 | CH_SC | 2,497334488 | 0,400426937 | 3,384057971 |
| HNRNPR3 | 0,00347734 | 1,243074056 | 0,533 | 0,215 | 1 | CH_SC | 2,367023544 | 0,422471505 | 2,479069767 |
| IER32 | 0,003550135 | 1,116192094 | 0,8 | 0,492 | 1 | CH_SC | 2,167740548 | 0,461309819 | 1,62601626 |
| MANBAL | 0,003558928 | 1,097668592 | 0,533 | 0,2 | 1 | CH_SC | 2,140085731 | 0,467271 | 2,665 |
| SDC3 | 0,00359829 | 1,072685108 | 0,467 | 0,138 | 1 | CH_SC | 2,103344419 | 0,47543331 | 3,384057971 |
| PNRC16 | 0,003647264 | 0,928100709 | 0,933 | 0,738 | 1 | CH_SC | 1,902769373 | 0,525549767 | 1,264227642 |
| YRDC | 0,003674901 | 0,453587961 | 0,2 | 0,015 | 1 | CH_SC | 1,369441806 | 0,73022453 | 13,33333333 |
| ROR1 | 0,003674901 | 0,760405689 | 0,2 | 0,015 | 1 | CH_SC | 1,693966905 | 0,590330305 | 13,33333333 |
| DENND1B | 0,003674901 | 0,502529762 | 0,2 | 0,015 | 1 | CH_SC | 1,416695558 | 0,705867958 | 13,33333333 |
| OGG1 | 0,003674901 | 0,520618923 | 0,2 | 0,015 | 1 | CH_SC | 1,434570554 | 0,697072722 | 13,33333333 |
| ARHGEF3 | 0,003674901 | 1,028341169 | 0,2 | 0,015 | 1 | CH_SC | 2,039677652 | 0,490273548 | 13,33333333 |
| PIK3CB | 0,003674901 | 0,417616116 | 0,2 | 0,015 | 1 | CH_SC | 1,335718612 | 0,748660677 | 13,33333333 |
| SPRY16 | 0,003674901 | 0,75947533 | 0,2 | 0,015 | 1 | CH_SC | 1,69287486 | 0,590711117 | 13,33333333 |
| CDH61 | 0,003674901 | 0,445733283 | 0,2 | 0,015 | 1 | CH_SC | 1,362006211 | 0,734211042 | 13,33333333 |
| WDR55 | 0,003674901 | 0,477556298 | 0,2 | 0,015 | 1 | CH_SC | 1,392383186 | 0,718193102 | 13,33333333 |
| PEX1 | 0,003674901 | 0,467139747 | 0,2 | 0,015 | 1 | CH_SC | 1,382366103 | 0,723397368 | 13,33333333 |
| PPP1R9A1 | 0,003674901 | 0,499514046 | 0,2 | 0,015 | 1 | CH_SC | 1,413737283 | 0,707345001 | 13,33333333 |
| ABCB8 | 0,003674901 | 0,457315131 | 0,2 | 0,015 | 1 | CH_SC | 1,372984303 | 0,728340446 | 13,33333333 |
| SNX16 | 0,003674901 | 0,455217483 | 0,2 | 0,015 | 1 | CH_SC | 1,370989463 | 0,729400208 | 13,33333333 |
| TMEM641 | 0,003674901 | 0,511498213 | 0,2 | 0,015 | 1 | CH_SC | 1,425529815 | 0,701493571 | 13,33333333 |
| INSC | 0,003674901 | 0,455332749 | 0,2 | 0,015 | 1 | CH_SC | 1,371099004 | 0,729341935 | 13,33333333 |
| TUBB32 | 0,003674901 | 0,403974693 | 0,2 | 0,015 | 1 | CH_SC | 1,32314823 | 0,755773221 | 13,33333333 |
| TEPSIN | 0,003674901 | 0,424658998 | 0,2 | 0,015 | 1 | CH_SC | 1,342255204 | 0,745014806 | 13,33333333 |
| B4GALT6 | 0,003674901 | 0,397866067 | 0,2 | 0,015 | 1 | CH_SC | 1,31755763 | 0,758980083 | 13,33333333 |
| CNN13 | 0,003674901 | 0,613237504 | 0,2 | 0,015 | 1 | CH_SC | 1,529688083 | 0,653728045 | 13,33333333 |
| LAMP51 | 0,003674901 | 0,482603229 | 0,2 | 0,015 | 1 | CH_SC | 1,397262643 | 0,715685061 | 13,33333333 |
| NDRG31 | 0,003674901 | 0,584475318 | 0,2 | 0,015 | 1 | CH_SC | 1,499493552 | 0,666891831 | 13,33333333 |
| URB1 | 0,003674901 | 0,673777019 | 0,2 | 0,015 | 1 | CH_SC | 1,595243901 | 0,62686339 | 13,33333333 |
| ASMTL1 | 0,003674901 | 0,553871166 | 0,2 | 0,015 | 1 | CH_SC | 1,468019534 | 0,681189846 | 13,33333333 |
| INTS6L | 0,003674901 | 0,666696578 | 0,2 | 0,015 | 1 | CH_SC | 1,587433964 | 0,629947464 | 13,33333333 |

| Supplementary Table 4 - clusermarker | | | | | | | | | |
|---|---|---|---|---|---|---|---|---|---|
| | **p_val** | **avg_log2FC** | **pct.1** | **pct.2** | **p_val_adj** | **celltype** | **Foldchange_UP** | **Foldchange_DOWN** | **Ratio_pct1_pct2** |
| CFL14 | 0,003687179 | 0,903085962 | 0,8 | 0,631 | 1 | CH_SC | 1,870061819 | 0,534741681 | 1,267828843 |
| HINT19 | 0,003703007 | -0,882770009 | 0,333 | 0,815 | 1 | CH_SC | 0,542325154 | 1,843912261 | 0,408588957 |
| ILF22 | 0,003744828 | 1,080373417 | 0,533 | 0,2 | 1 | CH_SC | 2,114583335 | 0,472906404 | 2,665 |
| PSMA12 | 0,003790782 | 0,934479915 | 0,467 | 0,154 | 1 | CH_SC | 1,911201531 | 0,523231058 | 3,032467532 |
| ARL8A1 | 0,003806815 | 0,799244436 | 0,467 | 0,138 | 1 | CH_SC | 1,740189521 | 0,574650053 | 3,384057971 |
| NUCKS13 | 0,003936379 | 0,962431159 | 0,8 | 0,569 | 1 | CH_SC | 1,9485908 | 0,513191379 | 1,405975395 |
| SQSTM15 | 0,003956571 | 0,858332308 | 0,933 | 0,785 | 1 | CH_SC | 1,812941419 | 0,551589803 | 1,188535032 |
| ALDH23 | 0,003963103 | -1,102844769 | 0 | 0,4 | 1 | CH_SC | 0,465597505 | 2,147777834 | 0 |
| PCOLCE21 | 0,003963409 | -1,633117077 | 0 | 0,4 | 1 | CH_SC | 0,322390899 | 3,101824533 | 0 |
| MELTF | 0,003963409 | -1,371717468 | 0 | 0,4 | 1 | CH_SC | 0,386430945 | 2,587784477 | 0 |
| FGFRL1 | 0,003963409 | -1,115253774 | 0 | 0,4 | 1 | CH_SC | 0,46160995 | 2,166331121 | 0 |
| PCDH20 | 0,004027936 | 0,727078423 | 0,333 | 0,062 | 1 | CH_SC | 1,655283609 | 0,604126081 | 5,370967742 |
| RPL18A9 | 0,004076485 | -0,801895301 | 0,867 | 0,985 | 1 | CH_SC | 0,573595137 | 1,743389953 | 0,880203046 |
| DGKG1 | 0,004076767 | 0,47970128 | 0,2 | 0,015 | 1 | CH_SC | 1,394454905 | 0,717126095 | 13,33333333 |
| F2R1 | 0,004076767 | 0,448536783 | 0,2 | 0,015 | 1 | CH_SC | 1,364655488 | 0,73278568 | 13,33333333 |
| TFEB1 | 0,004076767 | 0,269061801 | 0,2 | 0,015 | 1 | CH_SC | 1,205023934 | 0,829859036 | 13,33333333 |
| IL1RAPL1 | 0,004076767 | 0,389983833 | 0,2 | 0,015 | 1 | CH_SC | 1,31037872 | 0,763138156 | 13,33333333 |
| GSN6 | 0,004105789 | 0,879171898 | 0,867 | 0,723 | 1 | CH_SC | 1,839319235 | 0,543679412 | 1,199170124 |
| PRPSAP11 | 0,004155323 | 1,43368172 | 0,467 | 0,169 | 1 | CH_SC | 2,701352144 | 0,370184984 | 2,763313609 |
| MYL65 | 0,004190128 | 0,618289233 | 1 | 0,831 | 1 | CH_SC | 1,535053814 | 0,65144296 | 1,203369434 |
| COL3A12 | 0,004228383 | 1,293465312 | 0,467 | 0,154 | 1 | CH_SC | 2,451161108 | 0,407969919 | 3,032467532 |
| PON21 | 0,004228383 | 1,140928432 | 0,467 | 0,154 | 1 | CH_SC | 2,205228929 | 0,453467659 | 3,032467532 |
| SEPTIN94 | 0,004228383 | 1,351289941 | 0,467 | 0,154 | 1 | CH_SC | 2,551401491 | 0,39194145 | 3,032467532 |
| CIT | 0,004324937 | 1,204305308 | 0,333 | 0,077 | 1 | CH_SC | 2,304262869 | 0,433978264 | 4,324675325 |
| APPL15 | 0,004356761 | 1,203304244 | 0,6 | 0,292 | 1 | CH_SC | 2,302664531 | 0,434279499 | 2,054794521 |
| EIF52 | 0,004406837 | 0,978235667 | 0,867 | 0,6 | 1 | CH_SC | 1,970054672 | 0,507600126 | 1,445 |
| ELL23 | 0,004465905 | -1,423415156 | 0,133 | 0,538 | 1 | CH_SC | 0,372828704 | 2,682196912 | 0,247211896 |
| DLEU1 | 0,004483973 | 1,124571049 | 0,267 | 0,046 | 1 | CH_SC | 2,180367089 | 0,458638366 | 5,804347826 |
| RPL273 | 0,00449439 | -0,831934463 | 0,6 | 0,892 | 1 | CH_SC | 0,561775471 | 1,780070601 | 0,67264574 |
| PHLDA33 | 0,004499952 | 0,899001647 | 0,467 | 0,138 | 1 | CH_SC | 1,864775101 | 0,536257696 | 3,384057971 |
| TANC1 | 0,004529893 | 1,60636574 | 0,4 | 0,123 | 1 | CH_SC | 3,044838567 | 0,328424637 | 3,25203252 |

| Supplementary Table 4 - clusermarker | | | | | | | | | |
|---|---|---|---|---|---|---|---|---|---|
| | **p_val** | **avg_log2FC** | **pct.1** | **pct.2** | **p_val_adj** | **celltype** | **Foldchange_UP** | **Foldchange_DOWN** | **Ratio_pct1_pct2** |
| IFITM26 | 0,004574512 | 1,082191673 | 0,733 | 0,431 | 1 | CH_SC | 2,117250064 | 0,472310766 | 1,700696056 |
| CGGBP14 | 0,00461994 | 1,063157753 | 0,467 | 0,169 | 1 | CH_SC | 2,089499991 | 0,478583395 | 2,763313609 |
| DCTN1 | 0,004626065 | 1,065862945 | 0,333 | 0,077 | 1 | CH_SC | 2,093421682 | 0,477686846 | 4,324675325 |
| SFPQ4 | 0,004632308 | 1,102452394 | 0,733 | 0,462 | 1 | CH_SC | 2,147193775 | 0,465724152 | 1,586580087 |
| LPGAT11 | 0,004639856 | 0,45264284 | 0,333 | 0,062 | 1 | CH_SC | 1,368544968 | 0,730703063 | 5,370967742 |
| MTCO2P123 | 0,004755601 | 0,702272304 | 1 | 0,923 | 1 | CH_SC | 1,62706547 | 0,614603418 | 1,083423619 |
| MALAT16 | 0,004760266 | 0,624520323 | 1 | 0,969 | 1 | CH_SC | 1,541698146 | 0,648635404 | 1,031991744 |
| SLC20A21 | 0,004850601 | 1,265492728 | 0,267 | 0,046 | 1 | CH_SC | 2,404093035 | 0,41595728 | 5,804347826 |
| HDAC73 | 0,004850601 | 0,919674753 | 0,267 | 0,046 | 1 | CH_SC | 1,891688776 | 0,528628183 | 5,804347826 |
| GNB13 | 0,004917688 | 1,069439833 | 0,533 | 0,215 | 1 | CH_SC | 2,098618362 | 0,476503979 | 2,479069767 |
| PDLIM42 | 0,004917688 | 1,268685785 | 0,533 | 0,215 | 1 | CH_SC | 2,409419809 | 0,415037677 | 2,479069767 |
| BOC1 | 0,005024732 | -1,163965139 | 0 | 0,385 | 1 | CH_SC | 0,446284269 | 2,240724284 | 0 |
| COL9A1 | 0,005024732 | -2,122581645 | 0 | 0,385 | 1 | CH_SC | 0,229635621 | 4,354725088 | 0 |
| CILP21 | 0,005024732 | -1,444432536 | 0 | 0,385 | 1 | CH_SC | 0,367436656 | 2,721557535 | 0 |
| TMOD2 | 0,005082757 | 0,753638469 | 0,4 | 0,108 | 1 | CH_SC | 1,686039656 | 0,59310586 | 3,703703704 |
| CLIP4 | 0,005244142 | 0,995888335 | 0,267 | 0,046 | 1 | CH_SC | 1,994308137 | 0,501427027 | 5,804347826 |
| RNASEH2B2 | 0,005244142 | 0,847055856 | 0,267 | 0,046 | 1 | CH_SC | 1,798826267 | 0,555918055 | 5,804347826 |
| ARC2 | 0,005285934 | 1,154091967 | 0,333 | 0,077 | 1 | CH_SC | 2,2254421 | 0,449348918 | 4,324675325 |
| APIP1 | 0,005285934 | 1,437772013 | 0,333 | 0,077 | 1 | CH_SC | 2,709021819 | 0,369136931 | 4,324675325 |
| NUPR13 | 0,00545375 | -0,985908382 | 0,133 | 0,554 | 1 | CH_SC | 0,504907712 | 1,980559965 | 0,240072202 |
| PSMA77 | 0,005523076 | 1,307458634 | 0,6 | 0,369 | 1 | CH_SC | 2,475051653 | 0,404031972 | 1,62601626 |
| ARHGEF21 | 0,005666304 | 1,141598017 | 0,267 | 0,046 | 1 | CH_SC | 2,206252659 | 0,453257244 | 5,804347826 |
| NIT1 | 0,005666304 | 0,794424925 | 0,267 | 0,046 | 1 | CH_SC | 1,73438589 | 0,576572956 | 5,804347826 |
| FH | 0,005666304 | 0,583032235 | 0,267 | 0,046 | 1 | CH_SC | 1,497994405 | 0,667559236 | 5,804347826 |
| TSPAN5 | 0,005666304 | 1,637407674 | 0,267 | 0,046 | 1 | CH_SC | 3,111063138 | 0,321433528 | 5,804347826 |
| PPP3CC1 | 0,005666304 | 0,901295637 | 0,267 | 0,046 | 1 | CH_SC | 1,867742588 | 0,535405685 | 5,804347826 |
| ME3 | 0,005666304 | 1,365671145 | 0,267 | 0,046 | 1 | CH_SC | 2,576961789 | 0,388053872 | 5,804347826 |
| TNFAIP25 | 0,005666304 | 0,937233977 | 0,267 | 0,046 | 1 | CH_SC | 1,914853444 | 0,522233178 | 5,804347826 |
| XAF1 | 0,005666304 | 1,307632005 | 0,267 | 0,046 | 1 | CH_SC | 2,475349102 | 0,403983422 | 5,804347826 |
| NEDD4L1 | 0,005666304 | 0,66259296 | 0,267 | 0,046 | 1 | CH_SC | 1,582925064 | 0,631741845 | 5,804347826 |
| DNAAF9 | 0,005666304 | 0,848334389 | 0,267 | 0,046 | 1 | CH_SC | 1,800421114 | 0,555425612 | 5,804347826 |

| Supplementary Table 4 - clusermarker | | | | | | | | | |
|---|---|---|---|---|---|---|---|---|---|
| | **p_val** | **avg_log2FC** | **pct.1** | **pct.2** | **p_val_adj** | **celltype** | **Foldchange_UP** | **Foldchange_DOWN** | **Ratio_pct1_pct2** |
| RAP1A4 | 0,005692872 | 1,026767525 | 0,467 | 0,169 | 1 | CH_SC | 2,037454053 | 0,490808614 | 2,763313609 |
| POLK1 | 0,005720935 | 0,817027596 | 0,4 | 0,108 | 1 | CH_SC | 1,761772447 | 0,567610194 | 3,703703704 |
| MAN1A22 | 0,005912711 | 0,648217056 | 0,467 | 0,138 | 1 | CH_SC | 1,567230149 | 0,638068379 | 3,384057971 |
| GULP12 | 0,005912711 | 0,682031026 | 0,467 | 0,138 | 1 | CH_SC | 1,604396836 | 0,623287193 | 3,384057971 |
| ATP6V0A21 | 0,006029679 | 0,993290879 | 0,333 | 0,077 | 1 | CH_SC | 1,990720776 | 0,502330619 | 4,324675325 |
| ZNF462 | 0,006039674 | 1,42671291 | 0,4 | 0,123 | 1 | CH_SC | 2,688334969 | 0,371977455 | 3,25203252 |
| PALM2AKAP22 | 0,006039674 | 0,981204673 | 0,4 | 0,123 | 1 | CH_SC | 1,974113136 | 0,506556581 | 3,25203252 |
| VPS13D1 | 0,006118884 | 0,525911803 | 0,267 | 0,046 | 1 | CH_SC | 1,439843293 | 0,694520025 | 5,804347826 |
| SELENON | 0,006118884 | 0,761081687 | 0,267 | 0,046 | 1 | CH_SC | 1,694760826 | 0,590053761 | 5,804347826 |
| SMIM292 | 0,006118884 | 0,54198658 | 0,267 | 0,046 | 1 | CH_SC | 1,455976006 | 0,686824505 | 5,804347826 |
| LMBR1 | 0,006118884 | 0,692549838 | 0,267 | 0,046 | 1 | CH_SC | 1,616137378 | 0,61875928 | 5,804347826 |
| EXOC52 | 0,006118884 | 1,178465575 | 0,267 | 0,046 | 1 | CH_SC | 2,263359221 | 0,441821162 | 5,804347826 |
| RABEP11 | 0,006118884 | 0,730239796 | 0,267 | 0,046 | 1 | CH_SC | 1,658914803 | 0,602803711 | 5,804347826 |
| ZNF536 | 0,006118884 | 0,74461603 | 0,267 | 0,046 | 1 | CH_SC | 1,675528272 | 0,596826695 | 5,804347826 |
| LIMA13 | 0,006144411 | 1,057710766 | 0,467 | 0,154 | 1 | CH_SC | 2,081625826 | 0,480393732 | 3,032467532 |
| ARPC1B6 | 0,006231892 | 1,215843402 | 0,6 | 0,292 | 1 | CH_SC | 2,322765324 | 0,430521323 | 2,054794521 |
| EIF4G12 | 0,0062558 | 0,937054597 | 0,533 | 0,215 | 1 | CH_SC | 1,914615371 | 0,522298115 | 2,479069767 |
| TMEM50A6 | 0,006309517 | 0,774741779 | 0,467 | 0,169 | 1 | CH_SC | 1,710883801 | 0,584493231 | 2,763313609 |
| MATN3 | 0,006339897 | -1,841903668 | 0 | 0,369 | 1 | CH_SC | 0,278953456 | 3,584827424 | 0 |
| GALNT152 | 0,006339897 | -1,366180276 | 0 | 0,369 | 1 | CH_SC | 0,387916951 | 2,577871365 | 0 |
| SPOCK3 | 0,006339897 | -1,07066239 | 0 | 0,369 | 1 | CH_SC | 0,476100355 | 2,10039751 | 0 |
| BMP51 | 0,006339897 | -1,194036427 | 0 | 0,369 | 1 | CH_SC | 0,437078274 | 2,287919714 | 0 |
| NEBL1 | 0,006339897 | -1,301233357 | 0 | 0,369 | 1 | CH_SC | 0,40577915 | 2,464394732 | 0 |
| FXYD62 | 0,006339897 | -1,106852823 | 0 | 0,369 | 1 | CH_SC | 0,46430579 | 2,153753026 | 0 |
| CPAMD8 | 0,006339897 | -1,232585535 | 0 | 0,369 | 1 | CH_SC | 0,425554103 | 2,349877474 | 0 |
| DMKN2 | 0,006339897 | -1,08982788 | 0 | 0,369 | 1 | CH_SC | 0,469817422 | 2,128486412 | 0 |
| CALM14 | 0,006390474 | 1,12501589 | 0,8 | 0,738 | 1 | CH_SC | 2,181039488 | 0,458496971 | 1,08401084 |
| RPL352 | 0,006408213 | -0,833400956 | 0,667 | 0,938 | 1 | CH_SC | 0,561204718 | 1,781880955 | 0,71108742 |
| TRIO1 | 0,006435825 | 0,88465322 | 0,333 | 0,077 | 1 | CH_SC | 1,846320769 | 0,541617695 | 4,324675325 |
| WWP1 | 0,006435825 | 0,768767067 | 0,333 | 0,077 | 1 | CH_SC | 1,703813076 | 0,586918844 | 4,324675325 |
| NRBP2 | 0,006435825 | 0,873056088 | 0,333 | 0,077 | 1 | CH_SC | 1,831538576 | 0,545989046 | 4,324675325 |

| Supplementary Table 4 - clusermarker | | | | | | | | | |
|---|---|---|---|---|---|---|---|---|---|
| | **p_val** | **avg_log2FC** | **pct.1** | **pct.2** | **p_val_adj** | **celltype** | **Foldchange_UP** | **Foldchange_DOWN** | **Ratio_pct1_pct2** |
| CSDE15 | 0,006535121 | 0,837910605 | 0,733 | 0,538 | 1 | CH_SC | 1,787459564 | 0,559453215 | 1,362453532 |
| SCARB22 | 0,00656045 | 1,251677819 | 0,533 | 0,215 | 1 | CH_SC | 2,381181877 | 0,419959521 | 2,479069767 |
| SLC50A1 | 0,006603766 | 0,730541784 | 0,267 | 0,046 | 1 | CH_SC | 1,659262087 | 0,602677544 | 5,804347826 |
| ADAMTS41 | 0,006603766 | 0,662826427 | 0,267 | 0,046 | 1 | CH_SC | 1,583181245 | 0,63163962 | 5,804347826 |
| EXOSC91 | 0,006603766 | 0,628725377 | 0,267 | 0,046 | 1 | CH_SC | 1,546198322 | 0,646747565 | 5,804347826 |
| TBC1D9B1 | 0,006603766 | 0,731426452 | 0,267 | 0,046 | 1 | CH_SC | 1,660279867 | 0,602308093 | 5,804347826 |
| CSGALNACT2 | 0,006603766 | 0,668390816 | 0,267 | 0,046 | 1 | CH_SC | 1,589299272 | 0,629208116 | 5,804347826 |
| FAM111A | 0,006603766 | 0,642475542 | 0,267 | 0,046 | 1 | CH_SC | 1,561005415 | 0,640612768 | 5,804347826 |
| CPNE21 | 0,006603766 | 0,877521423 | 0,267 | 0,046 | 1 | CH_SC | 1,837216217 | 0,544301749 | 5,804347826 |
| CHCHD32 | 0,006761033 | 1,032590802 | 0,4 | 0,123 | 1 | CH_SC | 2,045694627 | 0,488831513 | 3,25203252 |
| TPM23 | 0,006769904 | -1,068058766 | 0,067 | 0,446 | 1 | CH_SC | 0,476960347 | 2,096610351 | 0,150224215 |
| STK391 | 0,006799945 | -1,429960386 | 0,067 | 0,431 | 1 | CH_SC | 0,371141083 | 2,69439317 | 0,155452436 |
| PCOLCE3 | 0,006799945 | -1,465015778 | 0,067 | 0,431 | 1 | CH_SC | 0,362231577 | 2,760664899 | 0,155452436 |
| ZNF5032 | 0,00681394 | 1,304007743 | 0,667 | 0,354 | 1 | CH_SC | 2,469138465 | 0,404999563 | 1,884180791 |
| NXPE3 | 0,007122926 | 1,012681162 | 0,267 | 0,046 | 1 | CH_SC | 2,017657313 | 0,495624303 | 5,804347826 |
| BRD3 | 0,007122926 | 0,501924467 | 0,267 | 0,046 | 1 | CH_SC | 1,416101296 | 0,706164173 | 5,804347826 |
| EHMT12 | 0,007122926 | 0,828167157 | 0,267 | 0,046 | 1 | CH_SC | 1,775428371 | 0,563244351 | 5,804347826 |
| ARHGAP19 | 0,007122926 | 0,799699947 | 0,267 | 0,046 | 1 | CH_SC | 1,740739049 | 0,574468644 | 5,804347826 |
| LYSMD21 | 0,007122926 | 0,624415554 | 0,267 | 0,046 | 1 | CH_SC | 1,541586191 | 0,64868251 | 5,804347826 |
| MT2A5 | 0,007198279 | -2,93093603 | 0,933 | 0,938 | 1 | CH_SC | 0,13112948 | 7,626050211 | 0,99466951 |
| SOD26 | 0,007311983 | -1,606260839 | 0,133 | 0,538 | 1 | CH_SC | 0,328448518 | 3,044617178 | 0,247211896 |
| LUZP11 | 0,007322725 | 0,704696676 | 0,333 | 0,077 | 1 | CH_SC | 1,629801966 | 0,613571477 | 4,324675325 |
| CCDC1521 | 0,007322725 | 0,786026808 | 0,333 | 0,077 | 1 | CH_SC | 1,724319131 | 0,579939051 | 4,324675325 |
| ZSWIM63 | 0,007322725 | 1,086057984 | 0,333 | 0,077 | 1 | CH_SC | 2,122931739 | 0,471046705 | 4,324675325 |
| VMA213 | 0,007322725 | 0,806025601 | 0,333 | 0,077 | 1 | CH_SC | 1,748388266 | 0,571955337 | 4,324675325 |
| ADAMTS13 | 0,007347531 | 1,487530461 | 0,467 | 0,169 | 1 | CH_SC | 2,804085735 | 0,356622477 | 2,763313609 |
| IFNAR1 | 0,007347531 | 1,218811149 | 0,467 | 0,169 | 1 | CH_SC | 2,32754837 | 0,429636614 | 2,763313609 |
| SF15 | 0,007366242 | 0,869360906 | 0,667 | 0,338 | 1 | CH_SC | 1,82685345 | 0,547389283 | 1,973372781 |
| BRI38 | 0,007617845 | -1,113010881 | 0,067 | 0,462 | 1 | CH_SC | 0,462328152 | 2,162965841 | 0,145021645 |
| CEP104 | 0,007678437 | 0,400114816 | 0,267 | 0,046 | 1 | CH_SC | 1,319612927 | 0,757797972 | 5,804347826 |
| PPFIBP2 | 0,007678437 | 0,527674174 | 0,267 | 0,046 | 1 | CH_SC | 1,441603256 | 0,693672129 | 5,804347826 |

| Supplementary Table 4 - clusermarker | | | | | | | | | |
|---|---|---|---|---|---|---|---|---|---|
| | **p_val** | **avg_log2FC** | **pct.1** | **pct.2** | **p_val_adj** | **celltype** | **Foldchange_UP** | **Foldchange_DOWN** | **Ratio_pct1_pct2** |
| PTPN21 | 0,007678437 | 0,521236219 | 0,267 | 0,046 | 1 | CH_SC | 1,435184504 | 0,696774524 | 5,804347826 |
| SLFN111 | 0,007678437 | 0,807277555 | 0,267 | 0,046 | 1 | CH_SC | 1,749906156 | 0,571459216 | 5,804347826 |
| CADM4 | 0,007678437 | 0,616044033 | 0,267 | 0,046 | 1 | CH_SC | 1,532666739 | 0,652457559 | 5,804347826 |
| PSMA63 | 0,007745643 | 0,93277932 | 0,6 | 0,292 | 1 | CH_SC | 1,908950006 | 0,523848187 | 2,054794521 |
| SDF2L14 | 0,007806054 | 0,616782638 | 0,333 | 0,077 | 1 | CH_SC | 1,533451607 | 0,652123612 | 4,324675325 |
| DYNC1I25 | 0,007818519 | 1,181800922 | 0,467 | 0,185 | 1 | CH_SC | 2,268597903 | 0,440800901 | 2,524324324 |
| VCP1 | 0,007962454 | 0,97434267 | 0,533 | 0,231 | 1 | CH_SC | 1,964745803 | 0,508971694 | 2,307359307 |
| NTRK23 | 0,007962454 | -1,417919334 | 0 | 0,354 | 1 | CH_SC | 0,374251672 | 2,671998753 | 0 |
| GPRC5C3 | 0,007962454 | -1,138381452 | 0 | 0,354 | 1 | CH_SC | 0,454268932 | 2,201339181 | 0 |
| TIPARP3 | 0,008068116 | -1,55257472 | 0,067 | 0,431 | 1 | CH_SC | 0,340901128 | 2,933401848 | 0,155452436 |
| RHOBTB31 | 0,008085267 | -1,414087179 | 0,067 | 0,415 | 1 | CH_SC | 0,375247098 | 2,664910681 | 0,161445783 |
| PTMA3 | 0,008187799 | 0,753509082 | 0,933 | 0,954 | 1 | CH_SC | 1,685888452 | 0,593159054 | 0,977987421 |
| CLPP3 | 0,008217684 | 1,01878409 | 0,4 | 0,138 | 1 | CH_SC | 2,02621054 | 0,493532128 | 2,898550725 |
| ILK5 | 0,008280387 | 1,022837023 | 0,533 | 0,262 | 1 | CH_SC | 2,031910734 | 0,492147604 | 2,034351145 |
| LRRC471 | 0,00831777 | 0,586653058 | 0,333 | 0,077 | 1 | CH_SC | 1,501758737 | 0,665885921 | 4,324675325 |
| PDE4D2 | 0,00831777 | 0,506873104 | 0,333 | 0,077 | 1 | CH_SC | 1,420967053 | 0,703746085 | 4,324675325 |
| DCAF72 | 0,00831777 | 0,740859145 | 0,333 | 0,077 | 1 | CH_SC | 1,671170748 | 0,598382901 | 4,324675325 |
| SUB13 | 0,008348848 | -0,457549084 | 0,2 | 0,631 | 1 | CH_SC | 0,728222346 | 1,373206969 | 0,316957211 |
| COL1A22 | 0,00835232 | 0,523296315 | 0,8 | 0,354 | 1 | CH_SC | 1,43723534 | 0,695780275 | 2,259887006 |
| RSL24D12 | 0,00835232 | -1,189436098 | 0,133 | 0,508 | 1 | CH_SC | 0,438474212 | 2,280635832 | 0,261811024 |
| PAM | 0,008438161 | -1,297520606 | 0,067 | 0,415 | 1 | CH_SC | 0,406824761 | 2,458060808 | 0,161445783 |
| ZNF832 | 0,008440203 | 0,870490147 | 0,4 | 0,123 | 1 | CH_SC | 1,828283944 | 0,546960992 | 3,25203252 |
| HMG20B2 | 0,008667699 | 1,00393621 | 0,4 | 0,138 | 1 | CH_SC | 2,005464196 | 0,498637673 | 2,898550725 |
| MRPS214 | 0,00873417 | -1,144269236 | 0,2 | 0,569 | 1 | CH_SC | 0,452418793 | 2,210341425 | 0,351493849 |
| SEPTIN73 | 0,008738444 | 0,931584465 | 0,8 | 0,708 | 1 | CH_SC | 1,907369648 | 0,524282224 | 1,129943503 |
| ECM11 | 0,008859293 | 0,684883689 | 0,333 | 0,077 | 1 | CH_SC | 1,607572373 | 0,622055975 | 4,324675325 |
| COL15A11 | 0,00890729 | 0,683702934 | 0,267 | 0,046 | 1 | CH_SC | 1,606257214 | 0,622565297 | 5,804347826 |
| TMEM1415 | 0,00890729 | 0,553902053 | 0,267 | 0,046 | 1 | CH_SC | 1,468050965 | 0,681175262 | 5,804347826 |
| USP222 | 0,008925181 | 0,859240043 | 0,333 | 0,092 | 1 | CH_SC | 1,814082469 | 0,551242855 | 3,619565217 |
| SMARCC22 | 0,008969089 | 0,961270378 | 0,467 | 0,169 | 1 | CH_SC | 1,947023609 | 0,513604455 | 2,763313609 |
| PHPT18 | 0,009043295 | 1,012567908 | 0,6 | 0,246 | 1 | CH_SC | 2,017498929 | 0,495663212 | 2,43902439 |

| Supplementary Table 4 - clusermarker | | | | | | | | | |
|---|---|---|---|---|---|---|---|---|---|
| | **p_val** | **avg_log2FC** | **pct.1** | **pct.2** | **p_val_adj** | **celltype** | **Foldchange_UP** | **Foldchange_DOWN** | **Ratio_pct1_pct2** |
| LMCD12 | 0,009185539 | -1,207463214 | 0,067 | 0,415 | 1 | CH_SC | 0,433029371 | 2,309312179 | 0,161445783 |
| ASAP21 | 0,009412281 | 1,302334689 | 0,4 | 0,123 | 1 | CH_SC | 2,466276733 | 0,405469503 | 3,25203252 |
| KIF2A2 | 0,009412281 | 1,132551152 | 0,4 | 0,123 | 1 | CH_SC | 2,192460957 | 0,456108464 | 3,25203252 |
| ZCCHC10 | 0,009477907 | 0,868474927 | 0,333 | 0,092 | 1 | CH_SC | 1,825731899 | 0,547725545 | 3,619565217 |
| ZFAS15 | 0,009482772 | -0,613111496 | 0,6 | 0,892 | 1 | CH_SC | 0,653785146 | 1,529554483 | 0,67264574 |
| CBR15 | 0,009513531 | 0,942076671 | 0,533 | 0,215 | 1 | CH_SC | 1,921291831 | 0,520483137 | 2,479069767 |
| DHH | 0,009585273 | 0,333908384 | 0,267 | 0,046 | 1 | CH_SC | 1,260423348 | 0,793384224 | 5,804347826 |
| ST6GALNAC2 | 0,009585273 | 0,610953369 | 0,267 | 0,046 | 1 | CH_SC | 1,527268133 | 0,654763874 | 5,804347826 |
| RPL104 | 0,00981008 | -0,56117006 | 0,933 | 0,985 | 1 | CH_SC | 0,677752268 | 1,475465369 | 0,947208122 |
| CALM36 | 0,009870502 | 1,413832577 | 0,467 | 0,215 | 1 | CH_SC | 2,664440428 | 0,375313326 | 2,172093023 |
| DOCK5 | 0,009934803 | 1,094690799 | 0,4 | 0,123 | 1 | CH_SC | 2,135673046 | 0,468236466 | 3,25203252 |
| VPS26A1 | 0,009934803 | 0,894905716 | 0,4 | 0,123 | 1 | CH_SC | 1,859488359 | 0,537782339 | 3,25203252 |
| SPPL2A2 | 0,009949103 | 0,966404957 | 0,533 | 0,246 | 1 | CH_SC | 1,953965449 | 0,511779776 | 2,166666667 |
| CDO11 | 0,009955846 | -1,178149037 | 0 | 0,338 | 1 | CH_SC | 0,441918111 | 2,262862677 | 0 |
| SCGB3A210 | 0,009955846 | -1,51757875 | 0 | 0,338 | 1 | CH_SC | 0,349271601 | 2,863101369 | 0 |
| PNP1 | 0,009955846 | -0,939254027 | 0 | 0,338 | 1 | CH_SC | 0,521502464 | 1,917536483 | 0 |
| ALDH1A15 | 0,0100377 | 0,678901579 | 0,333 | 0,077 | 1 | CH_SC | 1,600920403 | 0,624640674 | 4,324675325 |
| CEP1702 | 0,010060958 | 1,000476544 | 0,333 | 0,092 | 1 | CH_SC | 2,00066074 | 0,49983487 | 3,619565217 |
| CLCN31 | 0,010483011 | 1,067684033 | 0,4 | 0,123 | 1 | CH_SC | 2,096065838 | 0,477084251 | 3,25203252 |
| MORC32 | 0,010483011 | 0,788319768 | 0,4 | 0,123 | 1 | CH_SC | 1,727061872 | 0,579018052 | 3,25203252 |
| MYL95 | 0,010548639 | 1,489050083 | 0,667 | 0,446 | 1 | CH_SC | 2,807040896 | 0,356247036 | 1,495515695 |
| TGFBR22 | 0,010590936 | 1,240953102 | 0,667 | 0,385 | 1 | CH_SC | 2,36354626 | 0,423093052 | 1,732467532 |
| ARF13 | 0,010762412 | 0,835806483 | 0,667 | 0,492 | 1 | CH_SC | 1,784854514 | 0,560269754 | 1,355691057 |
| TSC22D34 | 0,010885829 | -0,919912654 | 0,333 | 0,754 | 1 | CH_SC | 0,528541019 | 1,892000741 | 0,441644562 |
| SRGN5 | 0,011017315 | -0,893345304 | 0,067 | 0,462 | 1 | CH_SC | 0,538364317 | 1,857478232 | 0,145021645 |
| CD638 | 0,011029804 | 0,588442191 | 0,867 | 0,877 | 1 | CH_SC | 1,503622273 | 0,665060646 | 0,988597491 |
| ATP5MJ3 | 0,011065024 | -0,988088187 | 0,267 | 0,615 | 1 | CH_SC | 0,50414541 | 1,983554706 | 0,434146341 |
| SFRP13 | 0,011066665 | 0,984111678 | 0,667 | 0,338 | 1 | CH_SC | 1,978094949 | 0,505536906 | 1,973372781 |
| VAMP22 | 0,01125228 | 0,993005727 | 0,667 | 0,415 | 1 | CH_SC | 1,990327344 | 0,502429916 | 1,607228916 |
| HSPG22 | 0,011264764 | 1,321665321 | 0,4 | 0,138 | 1 | CH_SC | 2,49954469 | 0,400072863 | 2,898550725 |
| CDC42BPB | 0,011264764 | 1,146755484 | 0,4 | 0,138 | 1 | CH_SC | 2,21415387 | 0,451639795 | 2,898550725 |

| Supplementary Table 4 - clusermarker | | | | | | | | | |
|---|---|---|---|---|---|---|---|---|---|
| | **p_val** | **avg_log2FC** | **pct.1** | **pct.2** | **p_val_adj** | **celltype** | **Foldchange_UP** | **Foldchange_DOWN** | **Ratio_pct1_pct2** |
| CXXC5 | 0,011323645 | 1,457846263 | 0,467 | 0,215 | 1 | CH_SC | 2,746979726 | 0,364036178 | 2,172093023 |
| SETD51 | 0,011323703 | 0,935434688 | 0,333 | 0,092 | 1 | CH_SC | 1,91246678 | 0,522884899 | 3,619565217 |
| PPP2R5C1 | 0,011323703 | 1,320248761 | 0,333 | 0,092 | 1 | CH_SC | 2,49709163 | 0,400465881 | 3,619565217 |
| IFITM16 | 0,011447827 | -1,168620053 | 0,133 | 0,508 | 1 | CH_SC | 0,444846636 | 2,247965748 | 0,261811024 |
| RASD14 | 0,011515106 | -2,228776032 | 0,133 | 0,462 | 1 | CH_SC | 0,213339641 | 4,6873614 | 0,287878788 |
| HSPB16 | 0,011594761 | 0,967877793 | 0,867 | 0,662 | 1 | CH_SC | 1,955961255 | 0,511257571 | 1,309667674 |
| FAM126A1 | 0,011660773 | 0,789718164 | 0,4 | 0,123 | 1 | CH_SC | 1,728736714 | 0,578457085 | 3,25203252 |
| SNRNP40 | 0,011679709 | 0,804853308 | 0,267 | 0,062 | 1 | CH_SC | 1,746968153 | 0,57242028 | 4,306451613 |
| ATP6V1A | 0,011679709 | 2,074251594 | 0,267 | 0,062 | 1 | CH_SC | 4,211258964 | 0,237458681 | 4,306451613 |
| WDFY32 | 0,011679709 | 1,116633408 | 0,267 | 0,062 | 1 | CH_SC | 2,168403752 | 0,461168728 | 4,306451613 |
| MAGI21 | 0,011679709 | 1,153060233 | 0,267 | 0,062 | 1 | CH_SC | 2,223851159 | 0,449670382 | 4,306451613 |
| USP72 | 0,011679709 | 0,755811456 | 0,267 | 0,062 | 1 | CH_SC | 1,688581082 | 0,592213196 | 4,306451613 |
| SLC25A56 | 0,011745316 | 0,960774785 | 0,6 | 0,354 | 1 | CH_SC | 1,946354884 | 0,513780918 | 1,694915254 |
| CCT21 | 0,012002432 | 0,815263862 | 0,467 | 0,169 | 1 | CH_SC | 1,759619949 | 0,568304537 | 2,763313609 |
| STX12 | 0,012006348 | 0,975322013 | 0,333 | 0,092 | 1 | CH_SC | 1,966079982 | 0,508626307 | 3,619565217 |
| OSMR2 | 0,012006348 | 1,292614228 | 0,333 | 0,092 | 1 | CH_SC | 2,449715531 | 0,408210663 | 3,619565217 |
| MGAT4B2 | 0,012006348 | 0,808088392 | 0,333 | 0,092 | 1 | CH_SC | 1,75088993 | 0,57113813 | 3,619565217 |
| RAPGEF5 | 0,012006348 | 1,317373995 | 0,333 | 0,092 | 1 | CH_SC | 2,492120789 | 0,40126466 | 3,619565217 |
| VIM-AS15 | 0,012006348 | 1,091678629 | 0,333 | 0,092 | 1 | CH_SC | 2,131218674 | 0,469215108 | 3,619565217 |
| ZMYM21 | 0,012006348 | 0,714910355 | 0,333 | 0,092 | 1 | CH_SC | 1,641381224 | 0,609242987 | 3,619565217 |
| GCC21 | 0,01203477 | 1,503542475 | 0,467 | 0,185 | 1 | CH_SC | 2,835380738 | 0,352686321 | 2,524324324 |
| ADK2 | 0,01203477 | 1,13436317 | 0,467 | 0,185 | 1 | CH_SC | 2,195216408 | 0,455535954 | 2,524324324 |
| NUCB24 | 0,012220716 | -1,237931811 | 0,2 | 0,538 | 1 | CH_SC | 0,423980021 | 2,358601702 | 0,371747212 |
| DCXR4 | 0,012365705 | -1,008469154 | 0,133 | 0,508 | 1 | CH_SC | 0,497073413 | 2,011775269 | 0,261811024 |
| CYBRD12 | 0,01238484 | -1,036547126 | 0,4 | 0,708 | 1 | CH_SC | 0,487492819 | 2,051312266 | 0,564971751 |
| FRMD4B2 | 0,012395047 | -1,079632838 | 0 | 0,323 | 1 | CH_SC | 0,473149223 | 2,113498132 | 0 |
| CLEC3A | 0,012395047 | -2,625058487 | 0 | 0,323 | 1 | CH_SC | 0,162098373 | 6,169093393 | 0 |
| GALNT13 | 0,012395047 | -0,838322442 | 0 | 0,323 | 1 | CH_SC | 0,559293535 | 1,78796989 | 0 |
| LDAH | 0,012517455 | 1,891983084 | 0,2 | 0,031 | 1 | CH_SC | 3,711450389 | 0,269436445 | 6,451612903 |
| GPD2 | 0,012517455 | 0,713500211 | 0,2 | 0,031 | 1 | CH_SC | 1,639777661 | 0,609838775 | 6,451612903 |
| PARD3B | 0,012517455 | 1,08153152 | 0,2 | 0,031 | 1 | CH_SC | 2,116281468 | 0,472526937 | 6,451612903 |

| Supplementary Table 4 - clusermarker | | | | | | | | | |
|---|---|---|---|---|---|---|---|---|---|
| | **p_val** | **avg_log2FC** | **pct.1** | **pct.2** | **p_val_adj** | **celltype** | **Foldchange_UP** | **Foldchange_DOWN** | **Ratio_pct1_pct2** |
| RBSN | 0,012517455 | 0,642170902 | 0,2 | 0,031 | 1 | CH_SC | 1,560675828 | 0,640748054 | 6,451612903 |
| SLC22A23 | 0,012517455 | 0,966167435 | 0,2 | 0,031 | 1 | CH_SC | 1,95364378 | 0,511864041 | 6,451612903 |
| TBL2 | 0,012517455 | 1,36509835 | 0,2 | 0,031 | 1 | CH_SC | 2,575938858 | 0,388207972 | 6,451612903 |
| ZNF25 | 0,012517455 | 0,744652627 | 0,2 | 0,031 | 1 | CH_SC | 1,675570776 | 0,596811555 | 6,451612903 |
| DLG53 | 0,012517455 | 0,496197534 | 0,2 | 0,031 | 1 | CH_SC | 1,410491072 | 0,708972939 | 6,451612903 |
| PGM2L1 | 0,012517455 | 1,052972425 | 0,2 | 0,031 | 1 | CH_SC | 2,074800216 | 0,481974116 | 6,451612903 |
| POLG | 0,012517455 | 0,949292944 | 0,2 | 0,031 | 1 | CH_SC | 1,930926091 | 0,517886213 | 6,451612903 |
| FBRS2 | 0,012517455 | 0,730416076 | 0,2 | 0,031 | 1 | CH_SC | 1,659117516 | 0,60273006 | 6,451612903 |
| MEX3C | 0,012517455 | 0,817553462 | 0,2 | 0,031 | 1 | CH_SC | 1,762414735 | 0,567403336 | 6,451612903 |
| GEM5 | 0,012660293 | -1,693016154 | 0,267 | 0,6 | 1 | CH_SC | 0,309279657 | 3,233319678 | 0,445 |
| RNF103 | 0,012725229 | 0,871232343 | 0,333 | 0,092 | 1 | CH_SC | 1,829224748 | 0,54667968 | 3,619565217 |
| DLG1 | 0,012725229 | 1,125078679 | 0,333 | 0,092 | 1 | CH_SC | 2,181134413 | 0,458477017 | 3,619565217 |
| LGI4 | 0,013083601 | 0,738101789 | 0,467 | 0,154 | 1 | CH_SC | 1,667979767 | 0,599527656 | 3,032467532 |
| ATF7IP1 | 0,013135719 | 0,892033408 | 0,4 | 0,138 | 1 | CH_SC | 1,855789926 | 0,538854095 | 2,898550725 |
| FTX2 | 0,013135719 | 1,265956806 | 0,4 | 0,138 | 1 | CH_SC | 2,404866495 | 0,415823499 | 2,898550725 |
| RPL237 | 0,013184699 | -1,06231792 | 0,467 | 0,738 | 1 | CH_SC | 0,478862073 | 2,08828399 | 0,632791328 |
| MAN1B1 | 0,01334897 | 0,792413128 | 0,267 | 0,062 | 1 | CH_SC | 1,731969025 | 0,577377531 | 4,306451613 |
| TSR32 | 0,01334897 | 0,833365534 | 0,267 | 0,062 | 1 | CH_SC | 1,781837206 | 0,561218498 | 4,306451613 |
| PAQR8 | 0,013593419 | 0,809323266 | 0,2 | 0,031 | 1 | CH_SC | 1,752389246 | 0,570649473 | 6,451612903 |
| ASTN21 | 0,013593419 | 0,783196119 | 0,2 | 0,031 | 1 | CH_SC | 1,720939188 | 0,581078057 | 6,451612903 |
| CBL | 0,013593419 | 1,156361222 | 0,2 | 0,031 | 1 | CH_SC | 2,228945316 | 0,44864268 | 6,451612903 |
| CSK3 | 0,013593419 | 0,741046165 | 0,2 | 0,031 | 1 | CH_SC | 1,6713874 | 0,598305336 | 6,451612903 |
| VPS35L | 0,013593419 | 0,676191701 | 0,2 | 0,031 | 1 | CH_SC | 1,597916145 | 0,625815067 | 6,451612903 |
| SNHG301 | 0,013593419 | 0,65326145 | 0,2 | 0,031 | 1 | CH_SC | 1,572719572 | 0,635841264 | 6,451612903 |
| GGA11 | 0,013593419 | 0,809510028 | 0,2 | 0,031 | 1 | CH_SC | 1,752616113 | 0,570575605 | 6,451612903 |
| ABLIM13 | 0,013647727 | 0,966256818 | 0,4 | 0,123 | 1 | CH_SC | 1,953764822 | 0,511832329 | 3,25203252 |
| CRTAP2 | 0,013720583 | -1,163001267 | 0,267 | 0,569 | 1 | CH_SC | 0,446582534 | 2,239227743 | 0,469244288 |
| LINC00632 | 0,013736351 | -1,291873245 | 0,333 | 0,677 | 1 | CH_SC | 0,408420378 | 2,448457655 | 0,491875923 |
| EIF3L4 | 0,013933675 | -1,189785385 | 0,333 | 0,662 | 1 | CH_SC | 0,438368067 | 2,281188057 | 0,503021148 |
| SNHG63 | 0,014113773 | -1,094492003 | 0,333 | 0,646 | 1 | CH_SC | 0,468300991 | 2,135378781 | 0,515479876 |
| BPTF4 | 0,014162784 | 1,456361717 | 0,533 | 0,308 | 1 | CH_SC | 2,744154514 | 0,364410967 | 1,730519481 |

| Supplementary Table 4 - clusermarker | | | | | | | | | |
|---|---|---|---|---|---|---|---|---|---|
| | **p_val** | **avg_log2FC** | **pct.1** | **pct.2** | **p_val_adj** | **celltype** | **Foldchange_UP** | **Foldchange_DOWN** | **Ratio_pct1_pct2** |
| LARP1B1 | 0,01426012 | 0,832319141 | 0,267 | 0,062 | 1 | CH_SC | 1,7805453 | 0,5616257 | 4,306451613 |
| GGCT | 0,01426012 | 0,723869628 | 0,267 | 0,062 | 1 | CH_SC | 1,651606072 | 0,605471254 | 4,306451613 |
| STX21 | 0,01426012 | 0,744589923 | 0,267 | 0,062 | 1 | CH_SC | 1,675497952 | 0,596837495 | 4,306451613 |
| MICU21 | 0,01426012 | 0,705370259 | 0,267 | 0,062 | 1 | CH_SC | 1,630563086 | 0,613285072 | 4,306451613 |
| ZNF532 | 0,01426012 | 0,836832779 | 0,267 | 0,062 | 1 | CH_SC | 1,786124665 | 0,559871335 | 4,306451613 |
| ID33 | 0,014260289 | 1,143702851 | 0,8 | 0,585 | 1 | CH_SC | 2,209473842 | 0,452596442 | 1,367521368 |
| TUBB2B | 0,014513786 | 1,62433178 | 0,533 | 0,292 | 1 | CH_SC | 3,082993355 | 0,324360089 | 1,825342466 |
| CNKSR31 | 0,014533485 | 1,397272202 | 0,4 | 0,154 | 1 | CH_SC | 2,634030776 | 0,379646286 | 2,597402597 |
| RPL320 | 0,014647994 | -0,666968953 | 0,8 | 0,969 | 1 | CH_SC | 0,629828544 | 1,587733693 | 0,825593395 |
| MTF11 | 0,014750284 | 0,52178128 | 0,2 | 0,031 | 1 | CH_SC | 1,435726831 | 0,696511327 | 6,451612903 |
| CDKN2C | 0,014750284 | 0,642970164 | 0,2 | 0,031 | 1 | CH_SC | 1,561540691 | 0,640393174 | 6,451612903 |
| ACTR1B1 | 0,014750284 | 0,779714508 | 0,2 | 0,031 | 1 | CH_SC | 1,716791107 | 0,582482048 | 6,451612903 |
| WDR12 | 0,014750284 | 0,491917709 | 0,2 | 0,031 | 1 | CH_SC | 1,406312982 | 0,711079264 | 6,451612903 |
| DYNC1LI11 | 0,014750284 | 0,872441387 | 0,2 | 0,031 | 1 | CH_SC | 1,830758365 | 0,546221729 | 6,451612903 |
| PHLDB21 | 0,014750284 | 1,186765633 | 0,2 | 0,031 | 1 | CH_SC | 2,276418222 | 0,439286591 | 6,451612903 |
| FRYL1 | 0,014750284 | 0,549587279 | 0,2 | 0,031 | 1 | CH_SC | 1,463666915 | 0,683215552 | 6,451612903 |
| TMEM184C | 0,014750284 | 0,484466047 | 0,2 | 0,031 | 1 | CH_SC | 1,399067963 | 0,71476156 | 6,451612903 |
| TAP2 | 0,014750284 | 0,745524733 | 0,2 | 0,031 | 1 | CH_SC | 1,676583961 | 0,596450893 | 6,451612903 |
| RNGTT | 0,014750284 | 0,771769719 | 0,2 | 0,031 | 1 | CH_SC | 1,70736288 | 0,585698572 | 6,451612903 |
| NFE2L31 | 0,014750284 | 0,557759693 | 0,2 | 0,031 | 1 | CH_SC | 1,471981657 | 0,679356292 | 6,451612903 |
| RUSC2 | 0,014750284 | 0,589179078 | 0,2 | 0,031 | 1 | CH_SC | 1,504390476 | 0,664721039 | 6,451612903 |
| DIP2C | 0,014750284 | 0,466630072 | 0,2 | 0,031 | 1 | CH_SC | 1,381877827 | 0,723652974 | 6,451612903 |
| ACTR1A | 0,014750284 | 0,460330287 | 0,2 | 0,031 | 1 | CH_SC | 1,375856768 | 0,726819843 | 6,451612903 |
| FADS2 | 0,014750284 | 0,727382083 | 0,2 | 0,031 | 1 | CH_SC | 1,655632052 | 0,603998937 | 6,451612903 |
| DPAGT1 | 0,014750284 | 0,64806053 | 0,2 | 0,031 | 1 | CH_SC | 1,56706012 | 0,63813761 | 6,451612903 |
| RNASE13 | 0,014750284 | 1,032412181 | 0,2 | 0,031 | 1 | CH_SC | 2,045441364 | 0,488892039 | 6,451612903 |
| SOS21 | 0,014750284 | 1,325652291 | 0,2 | 0,031 | 1 | CH_SC | 2,506461877 | 0,398968765 | 6,451612903 |
| LINS11 | 0,014750284 | 0,580781147 | 0,2 | 0,031 | 1 | CH_SC | 1,495658854 | 0,668601665 | 6,451612903 |
| RARA-AS1 | 0,014750284 | 0,876890823 | 0,2 | 0,031 | 1 | CH_SC | 1,836413348 | 0,544539715 | 6,451612903 |
| TANC2 | 0,014750284 | 0,538762254 | 0,2 | 0,031 | 1 | CH_SC | 1,452725631 | 0,688361229 | 6,451612903 |
| TTYH2 | 0,014750284 | 0,67511929 | 0,2 | 0,031 | 1 | CH_SC | 1,596728794 | 0,626280433 | 6,451612903 |

| Supplementary Table 4 - clusermarker | | | | | | | | | |
|---|---|---|---|---|---|---|---|---|---|
| | **p_val** | **avg_log2FC** | **pct.1** | **pct.2** | **p_val_adj** | **celltype** | **Foldchange_UP** | **Foldchange_DOWN** | **Ratio_pct1_pct2** |
| MIF4GD1 | 0,014750284 | 0,668281518 | 0,2 | 0,031 | 1 | CH_SC | 1,589178872 | 0,629255786 | 6,451612903 |
| CDH2 | 0,014750284 | 0,516447178 | 0,2 | 0,031 | 1 | CH_SC | 1,430428293 | 0,699091318 | 6,451612903 |
| GATAD2A1 | 0,014750284 | 0,73846574 | 0,2 | 0,031 | 1 | CH_SC | 1,668400604 | 0,599376431 | 6,451612903 |
| ZNF1001 | 0,014750284 | 1,126277842 | 0,2 | 0,031 | 1 | CH_SC | 2,182948118 | 0,458096091 | 6,451612903 |
| DMWD | 0,014750284 | 0,926533959 | 0,2 | 0,031 | 1 | CH_SC | 1,900704109 | 0,526120818 | 6,451612903 |
| VPS16 | 0,014750284 | 0,502044171 | 0,2 | 0,031 | 1 | CH_SC | 1,416218798 | 0,706105583 | 6,451612903 |
| FUS5 | 0,015077111 | 0,989373387 | 0,8 | 0,677 | 1 | CH_SC | 1,985322508 | 0,503696501 | 1,1816839 |
| RPLP26 | 0,015096169 | -0,669104607 | 0,667 | 0,938 | 1 | CH_SC | 0,628896884 | 1,590085792 | 0,71108742 |
| SGMS11 | 0,01511556 | 0,402839806 | 0,333 | 0,092 | 1 | CH_SC | 1,322107793 | 0,75636798 | 3,619565217 |
| ANKLE22 | 0,01511556 | 0,629550373 | 0,333 | 0,092 | 1 | CH_SC | 1,547082758 | 0,646377833 | 3,619565217 |
| EIF14 | 0,01515562 | 0,458464921 | 1 | 0,923 | 1 | CH_SC | 1,374078971 | 0,72776021 | 1,083423619 |
| SLC12A22 | 0,015225701 | 0,898797501 | 0,267 | 0,062 | 1 | CH_SC | 1,864511248 | 0,536333584 | 4,306451613 |
| CDK6 | 0,015225701 | 0,641872219 | 0,267 | 0,062 | 1 | CH_SC | 1,560352753 | 0,640880723 | 4,306451613 |
| DPY19L41 | 0,015225701 | 0,634808908 | 0,267 | 0,062 | 1 | CH_SC | 1,55273207 | 0,644026113 | 4,306451613 |
| MX12 | 0,015225701 | 0,777509313 | 0,267 | 0,062 | 1 | CH_SC | 1,714168954 | 0,583373067 | 4,306451613 |
| PEPD1 | 0,015259746 | 1,223567957 | 0,4 | 0,154 | 1 | CH_SC | 2,335235352 | 0,428222363 | 2,597402597 |
| YWHAZ2 | 0,015325562 | 0,763869788 | 0,733 | 0,538 | 1 | CH_SC | 1,698039228 | 0,588914545 | 1,362453532 |
| SNAI21 | 0,015368485 | -0,930263802 | 0 | 0,308 | 1 | CH_SC | 0,524762378 | 1,905624415 | 0 |
| ZBTB163 | 0,015368485 | -1,101486118 | 0 | 0,308 | 1 | CH_SC | 0,466036185 | 2,145756127 | 0 |
| GABARAPL13 | 0,015368485 | -1,347487941 | 0 | 0,308 | 1 | CH_SC | 0,392975713 | 2,54468652 | 0 |
| BEX42 | 0,015368485 | -0,951585017 | 0 | 0,308 | 1 | CH_SC | 0,517064077 | 1,933996276 | 0 |
| BIRC34 | 0,015485763 | 1,230043359 | 0,333 | 0,108 | 1 | CH_SC | 2,345740397 | 0,426304633 | 3,083333333 |
| GAS66 | 0,015635637 | -1,165373834 | 0,133 | 0,431 | 1 | CH_SC | 0,445848715 | 2,242913269 | 0,308584687 |
| RSL1D14 | 0,015635637 | -1,050977846 | 0,133 | 0,431 | 1 | CH_SC | 0,482640924 | 2,07193371 | 0,308584687 |
| YWHAB4 | 0,015734506 | 1,059359622 | 0,533 | 0,292 | 1 | CH_SC | 2,084006276 | 0,479845004 | 1,825342466 |
| C11orf962 | 0,015758505 | -2,483620386 | 0,067 | 0,385 | 1 | CH_SC | 0,178795163 | 5,592992461 | 0,174025974 |
| NPM14 | 0,015807186 | -0,950513474 | 0,6 | 0,846 | 1 | CH_SC | 0,517448262 | 1,932560359 | 0,709219858 |
| KDSR2 | 0,015865287 | 0,693842272 | 0,467 | 0,185 | 1 | CH_SC | 1,617585838 | 0,618205214 | 2,524324324 |
| PLPP32 | 0,015928008 | 0,625915147 | 0,4 | 0,123 | 1 | CH_SC | 1,543189409 | 0,648008595 | 3,25203252 |
| CD581 | 0,015993061 | 0,595709747 | 0,2 | 0,031 | 1 | CH_SC | 1,511215859 | 0,661718836 | 6,451612903 |
| SMC4 | 0,015993061 | 0,74439795 | 0,2 | 0,031 | 1 | CH_SC | 1,675275016 | 0,596916919 | 6,451612903 |

| Supplementary Table 4 - clusermarker | | | | | | | | | |
|---|---|---|---|---|---|---|---|---|---|
| | **p_val** | **avg_log2FC** | **pct.1** | **pct.2** | **p_val_adj** | **celltype** | **Foldchange_UP** | **Foldchange_DOWN** | **Ratio_pct1_pct2** |
| NAA15 | 0,015993061 | 0,524426066 | 0,2 | 0,031 | 1 | CH_SC | 1,438361256 | 0,695235634 | 6,451612903 |
| VCAN4 | 0,015993061 | 0,676690201 | 0,2 | 0,031 | 1 | CH_SC | 1,598468375 | 0,625598864 | 6,451612903 |
| AGPAT51 | 0,015993061 | 0,638007896 | 0,2 | 0,031 | 1 | CH_SC | 1,55617887 | 0,642599652 | 6,451612903 |
| DCAF101 | 0,015993061 | 0,267099458 | 0,2 | 0,031 | 1 | CH_SC | 1,203385983 | 0,830988572 | 6,451612903 |
| MMP17 | 0,015993061 | 0,566237488 | 0,2 | 0,031 | 1 | CH_SC | 1,480657015 | 0,675375857 | 6,451612903 |
| ATXN2L1 | 0,015993061 | 0,279491786 | 0,2 | 0,031 | 1 | CH_SC | 1,213767239 | 0,823881192 | 6,451612903 |
| SMIM101 | 0,015993061 | 0,469063445 | 0,2 | 0,031 | 1 | CH_SC | 1,384210588 | 0,722433428 | 6,451612903 |
| NME7 | 0,016248392 | 0,856208059 | 0,267 | 0,062 | 1 | CH_SC | 1,810273976 | 0,552402572 | 4,306451613 |
| NFKB21 | 0,016248392 | 0,808133196 | 0,267 | 0,062 | 1 | CH_SC | 1,750944307 | 0,571120393 | 4,306451613 |
| TBCD | 0,016248392 | 0,71944182 | 0,267 | 0,062 | 1 | CH_SC | 1,646544862 | 0,607332374 | 4,306451613 |
| SMARCB11 | 0,016248392 | 0,664633558 | 0,267 | 0,062 | 1 | CH_SC | 1,585165593 | 0,630848919 | 4,306451613 |
| BAMBI1 | 0,016410824 | -1,136888611 | 0,067 | 0,385 | 1 | CH_SC | 0,454739235 | 2,199062504 | 0,174025974 |
| XIST5 | 0,016471256 | 1,28067051 | 0,733 | 0,431 | 1 | CH_SC | 2,429518655 | 0,411604166 | 1,700696056 |
| APLP22 | 0,016721704 | 0,720193619 | 0,667 | 0,385 | 1 | CH_SC | 1,647403111 | 0,607015971 | 1,732467532 |
| SMAD21 | 0,016868801 | 0,887569757 | 0,4 | 0,138 | 1 | CH_SC | 1,850057047 | 0,540523873 | 2,898550725 |
| SCRG1 | 0,016965458 | -2,41041347 | 0,133 | 0,462 | 1 | CH_SC | 0,188101926 | 5,316266658 | 0,287878788 |
| TSPAN6 | 0,017074678 | -0,945585784 | 0,2 | 0,569 | 1 | CH_SC | 0,519218688 | 1,925970739 | 0,351493849 |
| C17orf494 | 0,01725965 | 1,193675039 | 0,333 | 0,108 | 1 | CH_SC | 2,287346673 | 0,437187774 | 3,083333333 |
| UBQLN4 | 0,017326977 | 0,469566168 | 0,2 | 0,031 | 1 | CH_SC | 1,384693015 | 0,722181732 | 6,451612903 |
| DNAJC271 | 0,017326977 | 0,370783669 | 0,2 | 0,031 | 1 | CH_SC | 1,293055025 | 0,773362294 | 6,451612903 |
| SEMA4C2 | 0,017326977 | 0,412726907 | 0,2 | 0,031 | 1 | CH_SC | 1,331199602 | 0,751202148 | 6,451612903 |
| SCD5 | 0,017326977 | 0,529150556 | 0,2 | 0,031 | 1 | CH_SC | 1,443079275 | 0,692962623 | 6,451612903 |
| FNIP1 | 0,017326977 | 0,606378854 | 0,2 | 0,031 | 1 | CH_SC | 1,522433121 | 0,656843303 | 6,451612903 |
| WASHC2A | 0,017326977 | 0,419538982 | 0,2 | 0,031 | 1 | CH_SC | 1,337500084 | 0,747663505 | 6,451612903 |
| RNF6 | 0,017326977 | 0,436161585 | 0,2 | 0,031 | 1 | CH_SC | 1,352999762 | 0,73909843 | 6,451612903 |
| GALK1 | 0,017326977 | 0,365912067 | 0,2 | 0,031 | 1 | CH_SC | 1,288696082 | 0,775978149 | 6,451612903 |
| ZNF574 | 0,017326977 | 0,674966993 | 0,2 | 0,031 | 1 | CH_SC | 1,596560245 | 0,626346549 | 6,451612903 |
| FLRT3 | 0,017326977 | 0,65448809 | 0,2 | 0,031 | 1 | CH_SC | 1,574057333 | 0,635300874 | 6,451612903 |
| SIPA1L2 | 0,017330963 | 0,665270294 | 0,267 | 0,062 | 1 | CH_SC | 1,585865363 | 0,630570554 | 4,306451613 |
| MMP141 | 0,017330963 | 0,6534417 | 0,267 | 0,062 | 1 | CH_SC | 1,572916079 | 0,635761827 | 4,306451613 |
| SH3BGRL38 | 0,017378999 | -0,775602453 | 0,4 | 0,754 | 1 | CH_SC | 0,584144641 | 1,711904774 | 0,530503979 |

| Supplementary Table 4 - clusermarker | | | | | | | | | |
|---|---|---|---|---|---|---|---|---|---|
| | **p_val** | **avg_log2FC** | **pct.1** | **pct.2** | **p_val_adj** | **celltype** | **Foldchange_UP** | **Foldchange_DOWN** | **Ratio_pct1_pct2** |
| IRF93 | 0,017718336 | 1,076320144 | 0,4 | 0,138 | 1 | CH_SC | 2,108650719 | 0,474236909 | 2,898550725 |
| RNF220 | 0,017892805 | 0,461680684 | 0,333 | 0,092 | 1 | CH_SC | 1,377145205 | 0,726139841 | 3,619565217 |
| NTHL1 | 0,017892805 | 0,465963248 | 0,333 | 0,092 | 1 | CH_SC | 1,381239261 | 0,723987529 | 3,619565217 |
| VPS4B2 | 0,017892805 | 0,417982902 | 0,333 | 0,092 | 1 | CH_SC | 1,336058245 | 0,748470364 | 3,619565217 |
| UBXN2A | 0,01821165 | 0,739256733 | 0,333 | 0,108 | 1 | CH_SC | 1,669315597 | 0,599047898 | 3,083333333 |
| FIGN | 0,01821165 | 0,905140759 | 0,333 | 0,108 | 1 | CH_SC | 1,872727202 | 0,533980603 | 3,083333333 |
| UBTF1 | 0,01821165 | 0,851674184 | 0,333 | 0,108 | 1 | CH_SC | 1,804593862 | 0,554141306 | 3,083333333 |
| POLR1A | 0,018476278 | 0,613295665 | 0,267 | 0,062 | 1 | CH_SC | 1,529749752 | 0,653701691 | 4,306451613 |
| ATP6V1B21 | 0,018476278 | 1,101304438 | 0,267 | 0,062 | 1 | CH_SC | 2,145485927 | 0,466094877 | 4,306451613 |
| ZNF516 | 0,018476278 | 0,447181521 | 0,267 | 0,062 | 1 | CH_SC | 1,363374138 | 0,73347438 | 4,306451613 |
| IGSF1 | 0,018476278 | 0,685959291 | 0,267 | 0,062 | 1 | CH_SC | 1,608771346 | 0,621592374 | 4,306451613 |
| SNRPE2 | 0,018504135 | 1,176950322 | 0,533 | 0,262 | 1 | CH_SC | 2,260983277 | 0,442285447 | 2,034351145 |
| VGLL31 | 0,018757481 | 0,398092436 | 0,2 | 0,031 | 1 | CH_SC | 1,31776438 | 0,758861003 | 6,451612903 |
| LMF1 | 0,018757481 | 0,318573812 | 0,2 | 0,031 | 1 | CH_SC | 1,247097111 | 0,801862174 | 6,451612903 |
| CPPED1 | 0,018757481 | 0,252788753 | 0,2 | 0,031 | 1 | CH_SC | 1,191508095 | 0,839272519 | 6,451612903 |
| MRPS301 | 0,018913111 | 0,576482003 | 0,333 | 0,092 | 1 | CH_SC | 1,491208515 | 0,670597029 | 3,619565217 |
| ARMC102 | 0,018913111 | 0,544547367 | 0,333 | 0,092 | 1 | CH_SC | 1,458562661 | 0,685606472 | 3,619565217 |
| KIAA00402 | 0,018980299 | -1,178241726 | 0 | 0,292 | 1 | CH_SC | 0,44188972 | 2,263008065 | 0 |
| CHI3L1 | 0,018980299 | -2,081074006 | 0 | 0,292 | 1 | CH_SC | 0,236338405 | 4,231220899 | 0 |
| ACAP23 | 0,018980299 | 0,865892966 | 0,467 | 0,185 | 1 | CH_SC | 1,822467347 | 0,548706676 | 2,524324324 |
| F13A12 | 0,018980299 | -1,671752205 | 0 | 0,292 | 1 | CH_SC | 0,313871903 | 3,186013113 | 0 |
| NT5E | 0,018980299 | -0,859733491 | 0 | 0,292 | 1 | CH_SC | 0,551054345 | 1,81470305 | 0 |
| AC005165.1 | 0,018980299 | -1,287223765 | 0 | 0,292 | 1 | CH_SC | 0,409738748 | 2,44057953 | 0 |
| SLC39A14 | 0,018980299 | -0,990992296 | 0 | 0,292 | 1 | CH_SC | 0,503131598 | 1,987551573 | 0 |
| TNFRSF11B | 0,018980299 | -1,278822107 | 0 | 0,292 | 1 | CH_SC | 0,412131858 | 2,426407911 | 0 |
| GLT8D2 | 0,018980299 | -0,864975471 | 0 | 0,292 | 1 | CH_SC | 0,549055742 | 1,8213087 | 0 |
| TPM14 | 0,018980299 | -0,88058163 | 0 | 0,292 | 1 | CH_SC | 0,543148414 | 1,841117407 | 0 |
| MT1A | 0,018980299 | -1,393219837 | 0 | 0,292 | 1 | CH_SC | 0,380714168 | 2,626642466 | 0 |
| SLC14A1 | 0,018980299 | -0,959376299 | 0 | 0,292 | 1 | CH_SC | 0,514279197 | 1,944469087 | 0 |
| HSPE16 | 0,019062739 | -0,959403738 | 0,333 | 0,662 | 1 | CH_SC | 0,514269416 | 1,94450607 | 0,503021148 |
| UQCRH4 | 0,019095521 | -0,939494362 | 0,267 | 0,631 | 1 | CH_SC | 0,521415595 | 1,917855948 | 0,423137876 |

| Supplementary Table 4 - clusermarker | | | | | | | | | |
|---|---|---|---|---|---|---|---|---|---|
| | **p_val** | **avg_log2FC** | **pct.1** | **pct.2** | **p_val_adj** | **celltype** | **Foldchange_UP** | **Foldchange_DOWN** | **Ratio_pct1_pct2** |
| ZNF644 | 0,019097349 | 1,314395779 | 0,4 | 0,169 | 1 | CH_SC | 2,486981505 | 0,402093863 | 2,366863905 |
| DDX3X4 | 0,019319565 | 1,107258763 | 0,733 | 0,554 | 1 | CH_SC | 2,154359126 | 0,464175164 | 1,323104693 |
| GTDC1 | 0,019687295 | 0,381576089 | 0,267 | 0,062 | 1 | CH_SC | 1,302764298 | 0,767598561 | 4,306451613 |
| SNHG151 | 0,019687295 | 0,58654853 | 0,267 | 0,062 | 1 | CH_SC | 1,501649934 | 0,665934168 | 4,306451613 |
| SECISBP2L2 | 0,019888648 | 1,213379915 | 0,6 | 0,385 | 1 | CH_SC | 2,31880245 | 0,431257091 | 1,558441558 |
| PTPRE | 0,019983937 | 0,504288416 | 0,333 | 0,092 | 1 | CH_SC | 1,418423571 | 0,705008025 | 3,619565217 |
| PRKDC3 | 0,020052293 | 0,839299973 | 0,533 | 0,262 | 1 | CH_SC | 1,789181781 | 0,558914701 | 2,034351145 |
| SDCBP6 | 0,020052293 | -1,201508336 | 0,067 | 0,369 | 1 | CH_SC | 0,43482044 | 2,299799892 | 0,181571816 |
| SCPEP13 | 0,020052293 | -0,853950563 | 0,067 | 0,369 | 1 | CH_SC | 0,553267635 | 1,807443517 | 0,181571816 |
| JAGN12 | 0,020254424 | 0,856776197 | 0,333 | 0,108 | 1 | CH_SC | 1,810987009 | 0,552185077 | 3,083333333 |
| IPO5 | 0,020254424 | 0,831714937 | 0,333 | 0,108 | 1 | CH_SC | 1,77979976 | 0,561860959 | 3,083333333 |
| DYNC1H13 | 0,020403723 | 1,25128891 | 0,467 | 0,231 | 1 | CH_SC | 2,380540065 | 0,420072745 | 2,021645022 |
| TAX1BP33 | 0,02043822 | 0,582967593 | 0,467 | 0,2 | 1 | CH_SC | 1,497927287 | 0,667589147 | 2,335 |
| ATP6V0E13 | 0,02083407 | 0,903969343 | 0,6 | 0,323 | 1 | CH_SC | 1,871207232 | 0,534414352 | 1,857585139 |
| RPS114 | 0,020887027 | -0,696305672 | 0,733 | 0,862 | 1 | CH_SC | 0,61715053 | 1,620350224 | 0,850348028 |
| FBLN23 | 0,020967065 | 0,504011543 | 0,267 | 0,062 | 1 | CH_SC | 1,418151382 | 0,705143339 | 4,306451613 |
| ITGA22 | 0,020967065 | 0,37168332 | 0,267 | 0,062 | 1 | CH_SC | 1,293861614 | 0,772880182 | 4,306451613 |
| ILF3-DT1 | 0,020967065 | 0,448754165 | 0,267 | 0,062 | 1 | CH_SC | 1,364861126 | 0,732675275 | 4,306451613 |
| B3GAT3 | 0,021107307 | 0,556231045 | 0,333 | 0,092 | 1 | CH_SC | 1,470422803 | 0,680076505 | 3,619565217 |
| TMEM2451 | 0,021348802 | 0,811021215 | 0,333 | 0,108 | 1 | CH_SC | 1,754452897 | 0,569978255 | 3,083333333 |
| ARHGEF122 | 0,021502584 | 0,526371236 | 0,4 | 0,138 | 1 | CH_SC | 1,440301891 | 0,694298887 | 2,898550725 |
| KHDRBS13 | 0,021678172 | -0,780958755 | 0,067 | 0,385 | 1 | CH_SC | 0,581979905 | 1,718272386 | 0,174025974 |
| PARP145 | 0,021918047 | 0,956003239 | 0,267 | 0,077 | 1 | CH_SC | 1,939928176 | 0,515483002 | 3,467532468 |
| FHL14 | 0,022020836 | -1,02054053 | 0,133 | 0,462 | 1 | CH_SC | 0,492931632 | 2,028678896 | 0,287878788 |
| ARID1A1 | 0,022318734 | 0,399575896 | 0,267 | 0,062 | 1 | CH_SC | 1,319120077 | 0,758081101 | 4,306451613 |
| POU3F1 | 0,022318734 | 0,785259321 | 0,267 | 0,062 | 1 | CH_SC | 1,723402069 | 0,58024765 | 4,306451613 |
| H1-42 | 0,022318734 | 0,555358788 | 0,267 | 0,062 | 1 | CH_SC | 1,46953405 | 0,680487805 | 4,306451613 |
| SP1002 | 0,022494322 | 1,163661043 | 0,333 | 0,108 | 1 | CH_SC | 2,240252026 | 0,446378349 | 3,083333333 |
| NCOR23 | 0,022494322 | 0,783033558 | 0,333 | 0,108 | 1 | CH_SC | 1,720745286 | 0,581143536 | 3,083333333 |
| DANCR5 | 0,022545135 | -0,903222914 | 0,067 | 0,354 | 1 | CH_SC | 0,534690921 | 1,870239349 | 0,189265537 |
| CYC13 | 0,022552069 | 0,65561684 | 0,4 | 0,138 | 1 | CH_SC | 1,575289342 | 0,634804016 | 2,898550725 |

| Supplementary Table 4 - clusermarker | | | | | | | | | |
|---|---|---|---|---|---|---|---|---|---|
| | **p_val** | **avg_log2FC** | **pct.1** | **pct.2** | **p_val_adj** | **celltype** | **Foldchange_UP** | **Foldchange_DOWN** | **Ratio_pct1_pct2** |
| NDRG24 | 0,022870881 | 0,741569481 | 0,733 | 0,508 | 1 | CH_SC | 1,67199378 | 0,598088349 | 1,442913386 |
| STARD13 | 0,023233646 | 0,906908097 | 0,467 | 0,2 | 1 | CH_SC | 1,875022746 | 0,533326863 | 2,335 |
| KAT6A | 0,023249079 | 0,641713131 | 0,267 | 0,077 | 1 | CH_SC | 1,560180699 | 0,640951398 | 3,467532468 |
| COL8A2 | 0,023353028 | -0,747660604 | 0 | 0,277 | 1 | CH_SC | 0,595568516 | 1,679067937 | 0 |
| SERPINI1 | 0,023353028 | -0,784360551 | 0 | 0,277 | 1 | CH_SC | 0,580609245 | 1,722328758 | 0 |
| SCIN2 | 0,023353028 | -0,985458797 | 0 | 0,277 | 1 | CH_SC | 0,50506508 | 1,979942862 | 0 |
| CSGALNACT1 | 0,023353028 | -1,021155258 | 0 | 0,277 | 1 | CH_SC | 0,49272164 | 2,029543494 | 0 |
| ECM21 | 0,023353028 | -0,98157315 | 0 | 0,277 | 1 | CH_SC | 0,506427218 | 1,974617407 | 0 |
| USE11 | 0,023353028 | -0,832184053 | 0 | 0,277 | 1 | CH_SC | 0,56167829 | 1,780378585 | 0 |
| ERCC1 | 0,023353028 | -0,820643279 | 0 | 0,277 | 1 | CH_SC | 0,56618943 | 1,766193339 | 0 |
| AR1 | 0,023353028 | -0,749891506 | 0 | 0,277 | 1 | CH_SC | 0,594648275 | 1,68166636 | 0 |
| BEX2 | 0,023353028 | -0,812948446 | 0 | 0,277 | 1 | CH_SC | 0,569217355 | 1,756798157 | 0 |
| SERPINB14 | 0,023459397 | -0,9709091 | 0,067 | 0,354 | 1 | CH_SC | 0,510184474 | 1,960075328 | 0,189265537 |
| TCEAL33 | 0,023460634 | -0,91209317 | 0,067 | 0,354 | 1 | CH_SC | 0,531413517 | 1,881773736 | 0,189265537 |
| PLOD21 | 0,023490691 | -1,289708807 | 0,067 | 0,369 | 1 | CH_SC | 0,40903358 | 2,44478705 | 0,181571816 |
| OSTF13 | 0,023548215 | 1,254972892 | 0,333 | 0,123 | 1 | CH_SC | 2,386626641 | 0,41900144 | 2,707317073 |
| PPP1R12A3 | 0,023711397 | 1,079418171 | 0,467 | 0,215 | 1 | CH_SC | 2,113183677 | 0,473219631 | 2,172093023 |
| ARHGAP56 | 0,023930364 | 0,685829162 | 0,533 | 0,246 | 1 | CH_SC | 1,608626244 | 0,621648443 | 2,166666667 |
| EGR34 | 0,023979187 | 1,284067551 | 0,4 | 0,169 | 1 | CH_SC | 2,435246059 | 0,410636123 | 2,366863905 |
| TSPAN43 | 0,024195211 | -0,729564385 | 0,067 | 0,4 | 1 | CH_SC | 0,603085985 | 1,658138349 | 0,1675 |
| ERGIC21 | 0,024236495 | 1,004756024 | 0,467 | 0,2 | 1 | CH_SC | 2,006604129 | 0,498354402 | 2,335 |
| CCT83 | 0,024236495 | 0,825529302 | 0,467 | 0,2 | 1 | CH_SC | 1,772185106 | 0,564275141 | 2,335 |
| SNRPC2 | 0,024713901 | 1,246450277 | 0,467 | 0,215 | 1 | CH_SC | 2,372569382 | 0,421483986 | 2,172093023 |
| FXYD14 | 0,024916365 | 1,046181396 | 0,733 | 0,631 | 1 | CH_SC | 2,065056702 | 0,484248204 | 1,161648177 |
| GRAMD2B2 | 0,024946502 | 0,544229125 | 0,333 | 0,108 | 1 | CH_SC | 1,458240955 | 0,685757725 | 3,083333333 |
| ATP6AP13 | 0,025388814 | -1,132039111 | 0,067 | 0,354 | 1 | CH_SC | 0,456270375 | 2,191682948 | 0,189265537 |
| YWHAE3 | 0,025844911 | 1,125915333 | 0,667 | 0,477 | 1 | CH_SC | 2,182399673 | 0,458211212 | 1,398322851 |
| DUSP13 | 0,025969946 | -0,989300887 | 0,533 | 0,785 | 1 | CH_SC | 0,503721814 | 1,985222742 | 0,678980892 |
| CCDC803 | 0,025993993 | -1,12813993 | 0,2 | 0,523 | 1 | CH_SC | 0,457505207 | 2,18576747 | 0,382409178 |
| PEX21 | 0,026122912 | 0,85503571 | 0,267 | 0,077 | 1 | CH_SC | 1,808803527 | 0,552851642 | 3,467532468 |
| INPP5F | 0,026122912 | 0,997179488 | 0,267 | 0,077 | 1 | CH_SC | 1,99609376 | 0,500978471 | 3,467532468 |

| Supplementary Table 4 - clusermarker | | | | | | | | | |
|---|---|---|---|---|---|---|---|---|---|
| | **p_val** | **avg_log2FC** | **pct.1** | **pct.2** | **p_val_adj** | **celltype** | **Foldchange_UP** | **Foldchange_DOWN** | **Ratio_pct1_pct2** |
| MTCH22 | 0,026122912 | 0,97160698 | 0,267 | 0,077 | 1 | CH_SC | 1,961023712 | 0,50993774 | 3,467532468 |
| ST133 | 0,026630329 | -0,945283954 | 0,333 | 0,662 | 1 | CH_SC | 0,519327327 | 1,925567843 | 0,503021148 |
| PTPN131 | 0,026800041 | 1,116484412 | 0,4 | 0,154 | 1 | CH_SC | 2,168179818 | 0,461216358 | 2,597402597 |
| RPL287 | 0,026847533 | -0,676048092 | 0,667 | 0,954 | 1 | CH_SC | 0,625877365 | 1,597757093 | 0,699161426 |
| DAD15 | 0,026977925 | 0,879674796 | 0,733 | 0,692 | 1 | CH_SC | 1,839960501 | 0,543489928 | 1,059248555 |
| CLDND14 | 0,026985082 | 0,649460801 | 0,6 | 0,323 | 1 | CH_SC | 1,568581837 | 0,637518538 | 1,857585139 |
| HSPA56 | 0,02726753 | 0,901736363 | 0,733 | 0,662 | 1 | CH_SC | 1,868313248 | 0,53524215 | 1,107250755 |
| MEF2A2 | 0,02736729 | -0,938538687 | 0,067 | 0,338 | 1 | CH_SC | 0,521761107 | 1,916585936 | 0,198224852 |
| MYO1D1 | 0,02736729 | -0,983645045 | 0,067 | 0,338 | 1 | CH_SC | 0,505700446 | 1,977455248 | 0,198224852 |
| C1orf1225 | 0,027429616 | 0,669210845 | 0,533 | 0,262 | 1 | CH_SC | 1,590202888 | 0,628850575 | 2,034351145 |
| TTYH11 | 0,027586966 | 0,324078588 | 0,333 | 0,092 | 1 | CH_SC | 1,25186465 | 0,798808402 | 3,619565217 |
| UQCC2 | 0,027626859 | 0,774840447 | 0,333 | 0,108 | 1 | CH_SC | 1,711000815 | 0,584453258 | 3,083333333 |
| PREB | 0,027671584 | 0,868424063 | 0,267 | 0,077 | 1 | CH_SC | 1,825667532 | 0,547744856 | 3,467532468 |
| PRICKLE2 | 0,027671584 | 0,921538839 | 0,267 | 0,077 | 1 | CH_SC | 1,894134579 | 0,527945591 | 3,467532468 |
| WAC-AS11 | 0,027671584 | 0,838884578 | 0,267 | 0,077 | 1 | CH_SC | 1,788666697 | 0,559075652 | 3,467532468 |
| ATP6V1D | 0,027671584 | 0,896036279 | 0,267 | 0,077 | 1 | CH_SC | 1,860946112 | 0,537361073 | 3,467532468 |
| PTPRM1 | 0,027671584 | 0,997037038 | 0,267 | 0,077 | 1 | CH_SC | 1,995896677 | 0,50102794 | 3,467532468 |
| FAM210B2 | 0,027794754 | 1,096908019 | 0,4 | 0,185 | 1 | CH_SC | 2,1389578 | 0,467517405 | 2,162162162 |
| PCMT13 | 0,027944714 | 1,462391635 | 0,467 | 0,215 | 1 | CH_SC | 2,755648041 | 0,362891046 | 2,172093023 |
| HEXIM12 | 0,028336147 | 1,165941772 | 0,533 | 0,308 | 1 | CH_SC | 2,243796398 | 0,445673235 | 1,730519481 |
| CTSB7 | 0,028608395 | -0,920559743 | 0,2 | 0,492 | 1 | CH_SC | 0,528304007 | 1,892849546 | 0,406504065 |
| SRSF22 | 0,028608395 | 1,003566894 | 0,6 | 0,4 | 1 | CH_SC | 2,004950882 | 0,498765336 | 1,5 |
| ORMDL3 | 0,02862902 | -0,809951752 | 0 | 0,262 | 1 | CH_SC | 0,570400934 | 1,753152811 | 0 |
| ZNF385D1 | 0,028630844 | -1,350734217 | 0 | 0,262 | 1 | CH_SC | 0,392092454 | 2,550418885 | 0 |
| PPIC3 | 0,028630844 | -0,63463518 | 0 | 0,262 | 1 | CH_SC | 0,644103671 | 1,552545102 | 0 |
| KCTD9 | 0,028630844 | 0,85203328 | 0,4 | 0,169 | 1 | CH_SC | 1,805043092 | 0,554003394 | 2,366863905 |
| CCDC1863 | 0,028630844 | -0,790336003 | 0 | 0,262 | 1 | CH_SC | 0,578209411 | 1,729477211 | 0 |
| SRSF81 | 0,028630844 | -0,624846071 | 0 | 0,262 | 1 | CH_SC | 0,648488964 | 1,542046288 | 0 |
| IGFBP62 | 0,028630844 | -1,283106036 | 0 | 0,262 | 1 | CH_SC | 0,410909891 | 2,433623578 | 0 |
| MT1F2 | 0,028630844 | -0,797525441 | 0 | 0,262 | 1 | CH_SC | 0,575335166 | 1,738117291 | 0 |
| CCNI2 | 0,028660978 | -0,743241491 | 0,533 | 0,831 | 1 | CH_SC | 0,597395597 | 1,673932659 | 0,641395909 |

| Supplementary Table 4 - clusermarker | | | | | | | | | |
|---|---|---|---|---|---|---|---|---|---|
| | **p_val** | **avg_log2FC** | **pct.1** | **pct.2** | **p_val_adj** | **celltype** | **Foldchange_UP** | **Foldchange_DOWN** | **Ratio_pct1_pct2** |
| MT-CO15 | 0,02866142 | 0,520353752 | 0,933 | 0,954 | 1 | CH_SC | 1,4343069 | 0,697200857 | 0,977987421 |
| HNRNPA33 | 0,028810694 | 1,19848237 | 0,6 | 0,431 | 1 | CH_SC | 2,294981255 | 0,435733407 | 1,392111369 |
| LUC7L | 0,02905777 | 0,807262792 | 0,333 | 0,108 | 1 | CH_SC | 1,749888249 | 0,571465064 | 3,083333333 |
| VKORC15 | 0,029194721 | -0,921582964 | 0,267 | 0,554 | 1 | CH_SC | 0,527929444 | 1,894192513 | 0,481949458 |
| RCAN13 | 0,029233688 | 0,910085238 | 0,667 | 0,4 | 1 | CH_SC | 1,87915652 | 0,532153649 | 1,6675 |
| DNAJB112 | 0,029298798 | 0,784824921 | 0,267 | 0,077 | 1 | CH_SC | 1,722883226 | 0,58042239 | 3,467532468 |
| MRPL161 | 0,029298798 | 0,87700232 | 0,267 | 0,077 | 1 | CH_SC | 1,836555278 | 0,544497632 | 3,467532468 |
| C1R2 | 0,029298798 | 0,939361791 | 0,267 | 0,077 | 1 | CH_SC | 1,917679721 | 0,521463511 | 3,467532468 |
| EGLN21 | 0,029298798 | 0,953310598 | 0,267 | 0,077 | 1 | CH_SC | 1,936310878 | 0,516445996 | 3,467532468 |
| PCDH91 | 0,029322129 | 1,278366403 | 0,4 | 0,154 | 1 | CH_SC | 2,425641604 | 0,412262058 | 2,597402597 |
| ANKH1 | 0,029446354 | -1,020063824 | 0,067 | 0,385 | 1 | CH_SC | 0,493094537 | 2,028008676 | 0,174025974 |
| IL1R12 | 0,02959999 | -1,272341078 | 0,067 | 0,338 | 1 | CH_SC | 0,413987445 | 2,415532188 | 0,198224852 |
| SPRY22 | 0,029601486 | -1,048455669 | 0,067 | 0,338 | 1 | CH_SC | 0,483485434 | 2,068314637 | 0,198224852 |
| FBXO44 | 0,029852105 | 0,785250255 | 0,133 | 0,015 | 1 | CH_SC | 1,723391239 | 0,580251296 | 8,866666667 |
| KIAA0319L1 | 0,029852105 | 0,477817312 | 0,133 | 0,015 | 1 | CH_SC | 1,39263512 | 0,718063178 | 8,866666667 |
| RETSAT | 0,029852105 | 0,608282729 | 0,133 | 0,015 | 1 | CH_SC | 1,52444355 | 0,655977061 | 8,866666667 |
| RGPD8 | 0,029852105 | 0,572624427 | 0,133 | 0,015 | 1 | CH_SC | 1,487226546 | 0,672392516 | 8,866666667 |
| EPB41L51 | 0,029852105 | 0,329810468 | 0,133 | 0,015 | 1 | CH_SC | 1,256848247 | 0,795641003 | 8,866666667 |
| GK5 | 0,029852105 | 0,68009342 | 0,133 | 0,015 | 1 | CH_SC | 1,602243503 | 0,624124859 | 8,866666667 |
| VPS81 | 0,029852105 | 0,736259939 | 0,133 | 0,015 | 1 | CH_SC | 1,665851662 | 0,600293545 | 8,866666667 |
| TNK2 | 0,029852105 | 0,455116636 | 0,133 | 0,015 | 1 | CH_SC | 1,370893632 | 0,729451197 | 8,866666667 |
| CFI3 | 0,029852105 | 0,493918707 | 0,133 | 0,015 | 1 | CH_SC | 1,408264872 | 0,710093691 | 8,866666667 |
| SYNPO21 | 0,029852105 | 0,458309695 | 0,133 | 0,015 | 1 | CH_SC | 1,373931135 | 0,727838517 | 8,866666667 |
| PRLR | 0,029852105 | 0,645635877 | 0,133 | 0,015 | 1 | CH_SC | 1,564428666 | 0,639210992 | 8,866666667 |
| PDLIM75 | 0,029852105 | 0,416351215 | 0,133 | 0,015 | 1 | CH_SC | 1,334548018 | 0,749317362 | 8,866666667 |
| BRPF3 | 0,029852105 | 0,4183708 | 0,133 | 0,015 | 1 | CH_SC | 1,336417519 | 0,748269149 | 8,866666667 |
| BTBD9 | 0,029852105 | 0,930298654 | 0,133 | 0,015 | 1 | CH_SC | 1,905670451 | 0,524749701 | 8,866666667 |
| HSF2 | 0,029852105 | 0,75165302 | 0,133 | 0,015 | 1 | CH_SC | 1,68372091 | 0,593922659 | 8,866666667 |
| PLEKHG1 | 0,029852105 | 0,746480898 | 0,133 | 0,015 | 1 | CH_SC | 1,677695507 | 0,596055718 | 8,866666667 |
| KDM4C | 0,029852105 | 0,627980976 | 0,133 | 0,015 | 1 | CH_SC | 1,54540072 | 0,64708136 | 8,866666667 |
| RASEF1 | 0,029852105 | 0,280771013 | 0,133 | 0,015 | 1 | CH_SC | 1,214843954 | 0,823150987 | 8,866666667 |

| Supplementary Table 4 - clusermarker | | | | | | | | | |
|---|---|---|---|---|---|---|---|---|---|
| | **p_val** | **avg_log2FC** | **pct.1** | **pct.2** | **p_val_adj** | **celltype** | **Foldchange_UP** | **Foldchange_DOWN** | **Ratio_pct1_pct2** |
| AUH | 0,029852105 | 0,573623752 | 0,133 | 0,015 | 1 | CH_SC | 1,488257074 | 0,671926925 | 8,866666667 |
| ZBTB43 | 0,029852105 | 0,45417726 | 0,133 | 0,015 | 1 | CH_SC | 1,370001298 | 0,729926316 | 8,866666667 |
| AIF1L | 0,029852105 | 0,81350923 | 0,133 | 0,015 | 1 | CH_SC | 1,757481168 | 0,56899614 | 8,866666667 |
| POLR3A | 0,029852105 | 0,453097051 | 0,133 | 0,015 | 1 | CH_SC | 1,368975902 | 0,730473048 | 8,866666667 |
| LGI1 | 0,029852105 | 0,486478537 | 0,133 | 0,015 | 1 | CH_SC | 1,401020958 | 0,713765197 | 8,866666667 |
| NANOS1 | 0,029852105 | 0,316373911 | 0,133 | 0,015 | 1 | CH_SC | 1,245196918 | 0,80308583 | 8,866666667 |
| ADCY6 | 0,029852105 | 0,873737074 | 0,133 | 0,015 | 1 | CH_SC | 1,83240331 | 0,545731387 | 8,866666667 |
| NOP9 | 0,029852105 | 0,554959013 | 0,133 | 0,015 | 1 | CH_SC | 1,469126895 | 0,680676396 | 8,866666667 |
| RPUSD2 | 0,029852105 | 0,46219923 | 0,133 | 0,015 | 1 | CH_SC | 1,377640279 | 0,725878892 | 8,866666667 |
| MCRIP21 | 0,029852105 | 0,374361177 | 0,133 | 0,015 | 1 | CH_SC | 1,296265443 | 0,771446933 | 8,866666667 |
| SETD6 | 0,029852105 | 0,402714323 | 0,133 | 0,015 | 1 | CH_SC | 1,321992803 | 0,75643377 | 8,866666667 |
| MYO18A | 0,029852105 | 0,310690915 | 0,133 | 0,015 | 1 | CH_SC | 1,240301545 | 0,806255546 | 8,866666667 |
| ACBD4 | 0,029852105 | 0,417099755 | 0,133 | 0,015 | 1 | CH_SC | 1,335240625 | 0,748928681 | 8,866666667 |
| CASKIN2 | 0,029852105 | 0,630894689 | 0,133 | 0,015 | 1 | CH_SC | 1,548525015 | 0,645775813 | 8,866666667 |
| DSEL | 0,029852105 | 0,835716297 | 0,133 | 0,015 | 1 | CH_SC | 1,784742943 | 0,560304779 | 8,866666667 |
| DOCK6 | 0,029852105 | 0,682582369 | 0,133 | 0,015 | 1 | CH_SC | 1,605010092 | 0,623049042 | 8,866666667 |
| MAP3K10 | 0,029852105 | 1,255070336 | 0,133 | 0,015 | 1 | CH_SC | 2,386787847 | 0,418973141 | 8,866666667 |
| BBC3 | 0,029852105 | 0,488553236 | 0,133 | 0,015 | 1 | CH_SC | 1,403037176 | 0,712739489 | 8,866666667 |
| ZNF544 | 0,029852105 | 0,62268947 | 0,133 | 0,015 | 1 | CH_SC | 1,539742894 | 0,649459078 | 8,866666667 |
| HIC2 | 0,029852105 | 0,276195434 | 0,133 | 0,015 | 1 | CH_SC | 1,21099712 | 0,825765795 | 8,866666667 |
| ZNF41 | 0,029852105 | 0,767440772 | 0,133 | 0,015 | 1 | CH_SC | 1,702247451 | 0,587458656 | 8,866666667 |
| PEBP14 | 0,029855398 | 0,50863746 | 0,8 | 0,8 | 1 | CH_SC | 1,4227059 | 0,702885958 | 1 |
| IFI64 | 0,029908513 | 1,547782919 | 0,4 | 0,169 | 1 | CH_SC | 2,923674942 | 0,342035288 | 2,366863905 |
| PLEKHB22 | 0,029908513 | 1,29224708 | 0,4 | 0,169 | 1 | CH_SC | 2,449092187 | 0,40831456 | 2,366863905 |
| ITPRID23 | 0,029908513 | 1,435067415 | 0,4 | 0,169 | 1 | CH_SC | 2,703948015 | 0,369829595 | 2,366863905 |
| NRIP13 | 0,030122851 | 0,886925118 | 0,333 | 0,123 | 1 | CH_SC | 1,849230571 | 0,540765449 | 2,707317073 |
| RAB5C2 | 0,030605734 | 0,807998956 | 0,467 | 0,231 | 1 | CH_SC | 1,750781393 | 0,571173537 | 2,021645022 |
| TIMP35 | 0,030711406 | 0,481769025 | 0,867 | 0,846 | 1 | CH_SC | 1,396454942 | 0,716099009 | 1,024822695 |
| AFAP1 | 0,031007674 | 0,684475212 | 0,267 | 0,077 | 1 | CH_SC | 1,607117278 | 0,622232126 | 3,467532468 |
| FNIP22 | 0,031007674 | 0,845696706 | 0,267 | 0,077 | 1 | CH_SC | 1,797132407 | 0,556442027 | 3,467532468 |
| UBLCP11 | 0,031007674 | 0,674791539 | 0,267 | 0,077 | 1 | CH_SC | 1,59636609 | 0,626422727 | 3,467532468 |

| Supplementary Table 4 - clusermarker | | | | | | | | | |
|---|---|---|---|---|---|---|---|---|---|
| | **p_val** | **avg_log2FC** | **pct.1** | **pct.2** | **p_val_adj** | **celltype** | **Foldchange_UP** | **Foldchange_DOWN** | **Ratio_pct1_pct2** |
| PAK1IP1 | 0,031007674 | 0,838950777 | 0,267 | 0,077 | 1 | CH_SC | 1,788748772 | 0,559049999 | 3,467532468 |
| BZW22 | 0,031007674 | 0,67831075 | 0,267 | 0,077 | 1 | CH_SC | 1,60026491 | 0,624896537 | 3,467532468 |
| NDEL1 | 0,031007674 | 0,668777591 | 0,267 | 0,077 | 1 | CH_SC | 1,589725408 | 0,629039452 | 3,467532468 |
| NAGA | 0,031007674 | 0,740674785 | 0,267 | 0,077 | 1 | CH_SC | 1,670957206 | 0,598459372 | 3,467532468 |
| PRPF82 | 0,031143072 | 0,606935688 | 0,533 | 0,277 | 1 | CH_SC | 1,523020845 | 0,656589832 | 1,924187726 |
| GOLPH31 | 0,031234893 | 0,751654028 | 0,4 | 0,169 | 1 | CH_SC | 1,683722086 | 0,593922244 | 2,366863905 |
| NDUFA43 | 0,031395447 | 0,810440249 | 0,8 | 0,723 | 1 | CH_SC | 1,753746529 | 0,570207828 | 1,106500692 |
| HGSNAT | 0,031612114 | 1,25915264 | 0,333 | 0,123 | 1 | CH_SC | 2,393551156 | 0,417789274 | 2,707317073 |
| MT-ATP810 | 0,03174464 | 0,485994407 | 1 | 0,831 | 1 | CH_SC | 1,400550891 | 0,714004758 | 1,203369434 |
| WLS2 | 0,031833618 | 0,914951356 | 0,467 | 0,231 | 1 | CH_SC | 1,885505496 | 0,530361753 | 2,021645022 |
| FAU2 | 0,031945477 | -0,587135639 | 0,867 | 0,954 | 1 | CH_SC | 0,665663219 | 1,50226116 | 0,908805031 |
| TNKS2 | 0,032045797 | 0,566748297 | 0,4 | 0,154 | 1 | CH_SC | 1,481181358 | 0,675136771 | 2,597402597 |
| RANBP12 | 0,032045797 | 0,702431522 | 0,4 | 0,154 | 1 | CH_SC | 1,627245046 | 0,614535593 | 2,597402597 |
| COL12A12 | 0,03211178 | 0,603392437 | 0,333 | 0,108 | 1 | CH_SC | 1,519284904 | 0,658204394 | 3,083333333 |
| FKBP1A5 | 0,03211178 | 0,525118527 | 0,333 | 0,108 | 1 | CH_SC | 1,439051803 | 0,694902017 | 3,083333333 |
| STAT34 | 0,03232026 | 1,040017521 | 0,533 | 0,277 | 1 | CH_SC | 2,056252626 | 0,486321567 | 1,924187726 |
| CD556 | 0,032371467 | -0,877954259 | 0,133 | 0,477 | 1 | CH_SC | 0,544138473 | 1,837767499 | 0,278825996 |
| PITPNA2 | 0,032611432 | 0,712151769 | 0,4 | 0,169 | 1 | CH_SC | 1,638245728 | 0,610409039 | 2,366863905 |
| PLEKHG5 | 0,032793854 | 0,412555741 | 0,133 | 0,015 | 1 | CH_SC | 1,331041673 | 0,751291278 | 8,866666667 |
| AL603840.1 | 0,032793854 | 0,704517743 | 0,133 | 0,015 | 1 | CH_SC | 1,62959984 | 0,613647581 | 8,866666667 |
| TTLL71 | 0,032793854 | 0,359513051 | 0,133 | 0,015 | 1 | CH_SC | 1,28299278 | 0,779427613 | 8,866666667 |
| AC105942.1 | 0,032793854 | 0,483126491 | 0,133 | 0,015 | 1 | CH_SC | 1,397769519 | 0,715425531 | 8,866666667 |
| KHK | 0,032793854 | 0,403416871 | 0,133 | 0,015 | 1 | CH_SC | 1,32263673 | 0,7560655 | 8,866666667 |
| WDPCP | 0,032793854 | 0,353365133 | 0,133 | 0,015 | 1 | CH_SC | 1,277537051 | 0,782756163 | 8,866666667 |
| TTC21B1 | 0,032793854 | 0,400152845 | 0,133 | 0,015 | 1 | CH_SC | 1,319647712 | 0,757777997 | 8,866666667 |
| HOXD-AS2 | 0,032793854 | 0,427449777 | 0,133 | 0,015 | 1 | CH_SC | 1,344854203 | 0,743575027 | 8,866666667 |
| KLF71 | 0,032793854 | 0,378344097 | 0,133 | 0,015 | 1 | CH_SC | 1,299849053 | 0,769320098 | 8,866666667 |
| LINC02610 | 0,032793854 | 0,387531243 | 0,133 | 0,015 | 1 | CH_SC | 1,308152961 | 0,764436599 | 8,866666667 |
| TRAF3IP11 | 0,032793854 | 0,384399896 | 0,133 | 0,015 | 1 | CH_SC | 1,305316714 | 0,766097599 | 8,866666667 |
| IRAK21 | 0,032793854 | 0,544168632 | 0,133 | 0,015 | 1 | CH_SC | 1,458179811 | 0,68578648 | 8,866666667 |
| UBP12 | 0,032793854 | 0,658353706 | 0,133 | 0,015 | 1 | CH_SC | 1,578280582 | 0,633600902 | 8,866666667 |

| Supplementary Table 4 - clusermarker | | | | | | | | | |
|---|---|---|---|---|---|---|---|---|---|
| | **p_val** | **avg_log2FC** | **pct.1** | **pct.2** | **p_val_adj** | **celltype** | **Foldchange_UP** | **Foldchange_DOWN** | **Ratio_pct1_pct2** |
| RRP9 | 0,032793854 | 0,343675839 | 0,133 | 0,015 | 1 | CH_SC | 1,268985724 | 0,78803093 | 8,866666667 |
| ST3GAL6-AS1 | 0,032793854 | 0,420522411 | 0,133 | 0,015 | 1 | CH_SC | 1,338412117 | 0,747154025 | 8,866666667 |
| RBP12 | 0,032793854 | 0,35831527 | 0,133 | 0,015 | 1 | CH_SC | 1,281928031 | 0,780074993 | 8,866666667 |
| AC093599.2 | 0,032793854 | 0,631557765 | 0,133 | 0,015 | 1 | CH_SC | 1,549236896 | 0,645479076 | 8,866666667 |
| NIM1K | 0,032793854 | 0,386322537 | 0,133 | 0,015 | 1 | CH_SC | 1,307057435 | 0,765077321 | 8,866666667 |
| TRAPPC131 | 0,032793854 | 0,440763578 | 0,133 | 0,015 | 1 | CH_SC | 1,35732253 | 0,736744567 | 8,866666667 |
| STARD41 | 0,032793854 | 0,891536454 | 0,133 | 0,015 | 1 | CH_SC | 1,855150786 | 0,539039741 | 8,866666667 |
| ZNF354C | 0,032793854 | 0,562796168 | 0,133 | 0,015 | 1 | CH_SC | 1,477129352 | 0,676988781 | 8,866666667 |
| HLA-DMB1 | 0,032793854 | 0,400361947 | 0,133 | 0,015 | 1 | CH_SC | 1,319838993 | 0,757668174 | 8,866666667 |
| TBCC1 | 0,032793854 | 0,3935768 | 0,133 | 0,015 | 1 | CH_SC | 1,313646226 | 0,76123996 | 8,866666667 |
| CNR1 | 0,032793854 | 0,980569463 | 0,133 | 0,015 | 1 | CH_SC | 1,973244137 | 0,506779664 | 8,866666667 |
| PREP | 0,032793854 | 0,402117899 | 0,133 | 0,015 | 1 | CH_SC | 1,321446392 | 0,756746552 | 8,866666667 |
| C6orf120 | 0,032793854 | 0,579722311 | 0,133 | 0,015 | 1 | CH_SC | 1,494561549 | 0,669092551 | 8,866666667 |
| SLC37A3 | 0,032793854 | 0,522455905 | 0,133 | 0,015 | 1 | CH_SC | 1,436398354 | 0,696185705 | 8,866666667 |
| GDAP1 | 0,032793854 | 0,391701796 | 0,133 | 0,015 | 1 | CH_SC | 1,31194005 | 0,762229951 | 8,866666667 |
| DGAT1 | 0,032793854 | 0,532272096 | 0,133 | 0,015 | 1 | CH_SC | 1,446205027 | 0,69146489 | 8,866666667 |
| AL162231.2 | 0,032793854 | 0,511912891 | 0,133 | 0,015 | 1 | CH_SC | 1,425939617 | 0,701291968 | 8,866666667 |
| MAMDC21 | 0,032793854 | 0,442327315 | 0,133 | 0,015 | 1 | CH_SC | 1,35879453 | 0,735946442 | 8,866666667 |
| VPS13A | 0,032793854 | 0,424447573 | 0,133 | 0,015 | 1 | CH_SC | 1,342058512 | 0,745123995 | 8,866666667 |
| SEMA4D | 0,032793854 | 0,449994585 | 0,133 | 0,015 | 1 | CH_SC | 1,36603513 | 0,732045596 | 8,866666667 |
| ZDHHC121 | 0,032793854 | 0,481821491 | 0,133 | 0,015 | 1 | CH_SC | 1,396505728 | 0,716072967 | 8,866666667 |
| HERC41 | 0,032793854 | 0,688230023 | 0,133 | 0,015 | 1 | CH_SC | 1,611305468 | 0,620614787 | 8,866666667 |
| CYB5R2 | 0,032793854 | 0,676501718 | 0,133 | 0,015 | 1 | CH_SC | 1,598259555 | 0,625680602 | 8,866666667 |
| UEVLD | 0,032793854 | 0,37433382 | 0,133 | 0,015 | 1 | CH_SC | 1,296240863 | 0,771461561 | 8,866666667 |
| CAPN52 | 0,032793854 | 0,528303175 | 0,133 | 0,015 | 1 | CH_SC | 1,442231918 | 0,693369761 | 8,866666667 |
| PANX1 | 0,032793854 | 0,560291482 | 0,133 | 0,015 | 1 | CH_SC | 1,474567109 | 0,678165133 | 8,866666667 |
| APOA1 | 0,032793854 | 0,376771234 | 0,133 | 0,015 | 1 | CH_SC | 1,298432696 | 0,770159287 | 8,866666667 |
| MMP192 | 0,032793854 | 0,93618369 | 0,133 | 0,015 | 1 | CH_SC | 1,91345993 | 0,522613505 | 8,866666667 |
| CDK81 | 0,032793854 | 0,337638947 | 0,133 | 0,015 | 1 | CH_SC | 1,263686805 | 0,791335318 | 8,866666667 |
| FGF14-AS2 | 0,032793854 | 0,436052648 | 0,133 | 0,015 | 1 | CH_SC | 1,352897601 | 0,739154241 | 8,866666667 |
| RABGGTA | 0,032793854 | 0,437375544 | 0,133 | 0,015 | 1 | CH_SC | 1,354138725 | 0,738476776 | 8,866666667 |

| Supplementary Table 4 - clusermarker | | | | | | | | | |
|---|---|---|---|---|---|---|---|---|---|
| | **p_val** | **avg_log2FC** | **pct.1** | **pct.2** | **p_val_adj** | **celltype** | **Foldchange_UP** | **Foldchange_DOWN** | **Ratio_pct1_pct2** |
| G2E3 | 0,032793854 | 0,456466826 | 0,133 | 0,015 | 1 | CH_SC | 1,372177225 | 0,728768837 | 8,866666667 |
| AL121603.2 | 0,032793854 | 0,401086388 | 0,133 | 0,015 | 1 | CH_SC | 1,32050191 | 0,75728781 | 8,866666667 |
| STARD9 | 0,032793854 | 0,999878802 | 0,133 | 0,015 | 1 | CH_SC | 1,999831991 | 0,500042006 | 8,866666667 |
| ADAMTS7 | 0,032793854 | 0,411342457 | 0,133 | 0,015 | 1 | CH_SC | 1,329922759 | 0,751923368 | 8,866666667 |
| CPEB1 | 0,032793854 | 0,525565986 | 0,133 | 0,015 | 1 | CH_SC | 1,439498201 | 0,694686523 | 8,866666667 |
| ALG1 | 0,032793854 | 0,412843106 | 0,133 | 0,015 | 1 | CH_SC | 1,331306825 | 0,751141646 | 8,866666667 |
| TBC1D10B | 0,032793854 | 0,968232715 | 0,133 | 0,015 | 1 | CH_SC | 1,956442507 | 0,51113181 | 8,866666667 |
| KARS12 | 0,032793854 | 0,274562659 | 0,133 | 0,015 | 1 | CH_SC | 1,209627345 | 0,826700888 | 8,866666667 |
| TSR1 | 0,032793854 | 0,375378508 | 0,133 | 0,015 | 1 | CH_SC | 1,29717984 | 0,770903131 | 8,866666667 |
| RAI12 | 0,032793854 | 0,600830296 | 0,133 | 0,015 | 1 | CH_SC | 1,516589139 | 0,659374365 | 8,866666667 |
| RNF1351 | 0,032793854 | 0,320979277 | 0,133 | 0,015 | 1 | CH_SC | 1,249178182 | 0,800526309 | 8,866666667 |
| RGS9 | 0,032793854 | 0,376212218 | 0,133 | 0,015 | 1 | CH_SC | 1,297929676 | 0,770457767 | 8,866666667 |
| FOXJ1 | 0,032793854 | 0,426933859 | 0,133 | 0,015 | 1 | CH_SC | 1,344373359 | 0,743840982 | 8,866666667 |
| PHLPP1 | 0,032793854 | 0,41368637 | 0,133 | 0,015 | 1 | CH_SC | 1,332085209 | 0,750702728 | 8,866666667 |
| PPAN | 0,032793854 | 0,679091154 | 0,133 | 0,015 | 1 | CH_SC | 1,601130782 | 0,6245586 | 8,866666667 |
| ZNF136 | 0,032793854 | 0,445082723 | 0,133 | 0,015 | 1 | CH_SC | 1,361392175 | 0,734542197 | 8,866666667 |
| C19orf44 | 0,032793854 | 0,684014933 | 0,133 | 0,015 | 1 | CH_SC | 1,606604623 | 0,622430675 | 8,866666667 |
| MPV17L2 | 0,032793854 | 0,370858911 | 0,133 | 0,015 | 1 | CH_SC | 1,293122464 | 0,773321961 | 8,866666667 |
| ZNF761 | 0,032793854 | 0,431953361 | 0,133 | 0,015 | 1 | CH_SC | 1,349058922 | 0,741257468 | 8,866666667 |
| EVA1C2 | 0,032793854 | 1,147265587 | 0,133 | 0,015 | 1 | CH_SC | 2,214936882 | 0,451480134 | 8,866666667 |
| ARVCF | 0,032793854 | 0,672609303 | 0,133 | 0,015 | 1 | CH_SC | 1,593953236 | 0,627370978 | 8,866666667 |
| STS | 0,032793854 | 0,510800267 | 0,133 | 0,015 | 1 | CH_SC | 1,424840339 | 0,701833021 | 8,866666667 |
| MPP11 | 0,032793854 | 0,408837775 | 0,133 | 0,015 | 1 | CH_SC | 1,327615865 | 0,753229926 | 8,866666667 |
| ECE12 | 0,032801405 | 0,829160639 | 0,267 | 0,077 | 1 | CH_SC | 1,776651405 | 0,562856617 | 3,467532468 |
| RBBP5 | 0,032801405 | 0,598684019 | 0,267 | 0,077 | 1 | CH_SC | 1,514334608 | 0,660356037 | 3,467532468 |
| PPP1R182 | 0,032801405 | 0,849681013 | 0,267 | 0,077 | 1 | CH_SC | 1,802102427 | 0,554907415 | 3,467532468 |
| CTXND1 | 0,032801405 | 0,732228104 | 0,267 | 0,077 | 1 | CH_SC | 1,66120268 | 0,601973505 | 3,467532468 |
| CLK3 | 0,032981782 | 0,487493933 | 0,2 | 0,046 | 1 | CH_SC | 1,402007369 | 0,713263013 | 4,347826087 |
| ROCK12 | 0,033143792 | 0,654352727 | 0,533 | 0,262 | 1 | CH_SC | 1,573909652 | 0,635360485 | 2,034351145 |
| MT-ATP68 | 0,033437783 | 0,5417199 | 0,933 | 0,923 | 1 | CH_SC | 1,455706895 | 0,686951476 | 1,010834236 |
| SH3GLB12 | 0,033487098 | 0,433265154 | 0,4 | 0,154 | 1 | CH_SC | 1,350286133 | 0,740583774 | 2,597402597 |

| Supplementary Table 4 - clusermarker | | | | | | | | | |
|---|---|---|---|---|---|---|---|---|---|
| | **p_val** | **avg_log2FC** | **pct.1** | **pct.2** | **p_val_adj** | **celltype** | **Foldchange_UP** | **Foldchange_DOWN** | **Ratio_pct1_pct2** |
| NCOR13 | 0,033487098 | 0,75267208 | 0,4 | 0,154 | 1 | CH_SC | 1,68491064 | 0,593503285 | 2,597402597 |
| SENP63 | 0,033697774 | 0,840950934 | 0,467 | 0,2 | 1 | CH_SC | 1,791230419 | 0,558275468 | 2,335 |
| DRAP17 | 0,033985994 | 0,878959064 | 0,533 | 0,308 | 1 | CH_SC | 1,839047909 | 0,543759624 | 1,730519481 |
| SNHG321 | 0,034041651 | -0,868589148 | 0,133 | 0,431 | 1 | CH_SC | 0,547682183 | 1,825876451 | 0,308584687 |
| ZNF2073 | 0,034139368 | 0,816151195 | 0,467 | 0,215 | 1 | CH_SC | 1,76070254 | 0,567955107 | 2,172093023 |
| H4C37 | 0,034416488 | -1,246029756 | 0,067 | 0,323 | 1 | CH_SC | 0,421606859 | 2,37187792 | 0,207430341 |
| FAM133B1 | 0,034416488 | 0,908059527 | 0,467 | 0,231 | 1 | CH_SC | 1,876519818 | 0,53290138 | 2,021645022 |
| LIX1L1 | 0,034683255 | 0,609783454 | 0,267 | 0,077 | 1 | CH_SC | 1,526030137 | 0,655295053 | 3,467532468 |
| UBE2J14 | 0,034683255 | 0,690582598 | 0,267 | 0,077 | 1 | CH_SC | 1,613935136 | 0,619603587 | 3,467532468 |
| LHPP1 | 0,034683255 | 0,683259779 | 0,267 | 0,077 | 1 | CH_SC | 1,605763893 | 0,622756561 | 3,467532468 |
| AGPAT11 | 0,034780804 | 0,971899332 | 0,333 | 0,123 | 1 | CH_SC | 1,96142114 | 0,509834415 | 2,707317073 |
| SNHG19 | 0,03498131 | -0,687176686 | 0 | 0,246 | 1 | CH_SC | 0,621068075 | 1,610129454 | 0 |
| MATN1 | 0,034983497 | -1,900332668 | 0 | 0,246 | 1 | CH_SC | 0,267881588 | 3,732992649 | 0 |
| GSTM51 | 0,034983497 | -0,723711901 | 0 | 0,246 | 1 | CH_SC | 0,605537453 | 1,651425516 | 0 |
| GLMP2 | 0,034983497 | -0,671814212 | 0 | 0,246 | 1 | CH_SC | 0,627716827 | 1,593075026 | 0 |
| SUSD51 | 0,034983497 | -0,794779452 | 0 | 0,246 | 1 | CH_SC | 0,576431287 | 1,73481215 | 0 |
| ADGRA31 | 0,034983497 | -0,90338708 | 0 | 0,246 | 1 | CH_SC | 0,534630082 | 1,870452176 | 0 |
| ZSCAN16-AS1 | 0,034983497 | -0,752148222 | 0 | 0,246 | 1 | CH_SC | 0,593718832 | 1,684298942 | 0 |
| SMOC22 | 0,034983497 | -0,673093736 | 0 | 0,246 | 1 | CH_SC | 0,627160353 | 1,594488548 | 0 |
| ZFAND12 | 0,034983497 | -0,8655545 | 0 | 0,246 | 1 | CH_SC | 0,548835421 | 1,822039834 | 0 |
| AKR1C15 | 0,034983497 | -1,323185684 | 0 | 0,246 | 1 | CH_SC | 0,399651474 | 2,502180185 | 0 |
| UCMA | 0,034983497 | -1,945079851 | 0 | 0,246 | 1 | CH_SC | 0,259700401 | 3,850590892 | 0 |
| ZBED52 | 0,034983497 | -0,917258873 | 0 | 0,246 | 1 | CH_SC | 0,529514145 | 1,888523678 | 0 |
| ATF1 | 0,034983497 | -0,672344005 | 0 | 0,246 | 1 | CH_SC | 0,627486356 | 1,593660149 | 0 |
| SPART1 | 0,034983497 | -0,817599932 | 0 | 0,246 | 1 | CH_SC | 0,56738506 | 1,762471504 | 0 |
| AL627171.28 | 0,034983497 | -1,574809068 | 0 | 0,246 | 1 | CH_SC | 0,335687549 | 2,978960652 | 0 |
| MAP2K61 | 0,034983497 | -0,95564928 | 0 | 0,246 | 1 | CH_SC | 0,515609489 | 1,939452281 | 0 |
| TCEAL6 | 0,034983497 | -0,652857137 | 0 | 0,246 | 1 | CH_SC | 0,636019482 | 1,572278881 | 0 |
| CTSK2 | 0,03509424 | -0,813564733 | 0,2 | 0,492 | 1 | CH_SC | 0,56897425 | 1,757548782 | 0,406504065 |
| KLF64 | 0,03509424 | 1,042925451 | 0,6 | 0,4 | 1 | CH_SC | 2,060401436 | 0,485342314 | 1,5 |
| OAF2 | 0,035428734 | 1,101227733 | 0,333 | 0,138 | 1 | CH_SC | 2,14537186 | 0,466119659 | 2,413043478 |

| Supplementary Table 4 - clusermarker | | | | | | | | | |
|---|---|---|---|---|---|---|---|---|---|
| | **p_val** | **avg_log2FC** | **pct.1** | **pct.2** | **p_val_adj** | **celltype** | **Foldchange_UP** | **Foldchange_DOWN** | **Ratio_pct1_pct2** |
| ABHD17A1 | 0,035436852 | 0,592017778 | 0,333 | 0,108 | 1 | CH_SC | 1,507353484 | 0,663414395 | 3,083333333 |
| KPNB13 | 0,035508248 | 0,761127022 | 0,467 | 0,215 | 1 | CH_SC | 1,694814084 | 0,59003522 | 2,172093023 |
| NME38 | 0,035510048 | 0,868399646 | 0,467 | 0,215 | 1 | CH_SC | 1,825636633 | 0,547754127 | 2,172093023 |
| C12orf576 | 0,035514299 | -0,751822419 | 0,533 | 0,877 | 1 | CH_SC | 0,593852926 | 1,683918621 | 0,607753706 |
| SRP143 | 0,035628317 | -0,577172549 | 0,6 | 0,877 | 1 | CH_SC | 0,670276125 | 1,491922453 | 0,684150513 |
| DNAJC85 | 0,035773719 | -0,801045084 | 0,067 | 0,323 | 1 | CH_SC | 0,573933271 | 1,742362832 | 0,207430341 |
| MEAF62 | 0,035815679 | 1,254430249 | 0,4 | 0,185 | 1 | CH_SC | 2,385729125 | 0,419159069 | 2,162162162 |
| FOS6 | 0,035943741 | 0,66694429 | 1 | 0,846 | 1 | CH_SC | 1,587706551 | 0,629839311 | 1,182033097 |
| BMP8B | 0,035980387 | 0,341838211 | 0,133 | 0,015 | 1 | CH_SC | 1,267370386 | 0,789035321 | 8,866666667 |
| LHFPL21 | 0,035980387 | 0,712820159 | 0,133 | 0,015 | 1 | CH_SC | 1,639004891 | 0,610126306 | 8,866666667 |
| DNAJC14 | 0,035980387 | 0,326623519 | 0,133 | 0,015 | 1 | CH_SC | 1,254074902 | 0,797400536 | 8,866666667 |
| PHF111 | 0,035980387 | 0,267581243 | 0,133 | 0,015 | 1 | CH_SC | 1,203787919 | 0,830711111 | 8,866666667 |
| TC2N3 | 0,035980387 | 0,338759009 | 0,133 | 0,015 | 1 | CH_SC | 1,264668272 | 0,790721189 | 8,866666667 |
| RILP | 0,035980387 | 0,329610227 | 0,133 | 0,015 | 1 | CH_SC | 1,256673813 | 0,795751443 | 8,866666667 |
| CDC732 | 0,036464358 | 0,70473465 | 0,333 | 0,123 | 1 | CH_SC | 1,629844865 | 0,613555328 | 2,707317073 |
| REEP33 | 0,036464358 | 1,010422667 | 0,333 | 0,123 | 1 | CH_SC | 2,014501202 | 0,496400796 | 2,707317073 |
| SRP681 | 0,036464358 | 0,716595457 | 0,333 | 0,123 | 1 | CH_SC | 1,643299516 | 0,608531792 | 2,707317073 |
| PDPN1 | 0,036551271 | -1,177775413 | 0,133 | 0,415 | 1 | CH_SC | 0,442032572 | 2,262276726 | 0,320481928 |
| CAP13 | 0,036552754 | 0,810401412 | 0,533 | 0,323 | 1 | CH_SC | 1,75369932 | 0,570223178 | 1,650154799 |
| STRN | 0,036656555 | 0,645157072 | 0,267 | 0,077 | 1 | CH_SC | 1,563909547 | 0,63942317 | 3,467532468 |
| NRP23 | 0,036656555 | 0,793006458 | 0,267 | 0,077 | 1 | CH_SC | 1,732681469 | 0,577140125 | 3,467532468 |
| CHP1 | 0,036656555 | 0,599702744 | 0,267 | 0,077 | 1 | CH_SC | 1,515404297 | 0,659889907 | 3,467532468 |
| SLC25A392 | 0,036656555 | 0,423881186 | 0,267 | 0,077 | 1 | CH_SC | 1,341531737 | 0,74541658 | 3,467532468 |
| URI12 | 0,036927406 | 0,650677172 | 0,467 | 0,215 | 1 | CH_SC | 1,569904904 | 0,636981258 | 2,172093023 |
| PPP2CB2 | 0,037056852 | 0,817211516 | 0,4 | 0,169 | 1 | CH_SC | 1,761997058 | 0,567537837 | 2,366863905 |
| STK32A | 0,037206748 | 0,48990433 | 0,333 | 0,108 | 1 | CH_SC | 1,404351745 | 0,712072316 | 3,083333333 |
| TUBB2A2 | 0,037636664 | 0,691448754 | 0,667 | 0,415 | 1 | CH_SC | 1,614904391 | 0,619231705 | 1,607228916 |
| ANO10 | 0,037679657 | 0,539875267 | 0,2 | 0,046 | 1 | CH_SC | 1,453846815 | 0,687830375 | 4,347826087 |
| DENND6A | 0,037679657 | 0,664711588 | 0,2 | 0,046 | 1 | CH_SC | 1,58525133 | 0,630814799 | 4,347826087 |
| ATP13A31 | 0,037679657 | 0,591511144 | 0,2 | 0,046 | 1 | CH_SC | 1,506824237 | 0,663647408 | 4,347826087 |
| ENOPH11 | 0,037679657 | 0,627063868 | 0,2 | 0,046 | 1 | CH_SC | 1,544418636 | 0,647492834 | 4,347826087 |

| Supplementary Table 4 - clusermarker | | | | | | | | | |
|---|---|---|---|---|---|---|---|---|---|
| | **p_val** | **avg_log2FC** | **pct.1** | **pct.2** | **p_val_adj** | **celltype** | **Foldchange_UP** | **Foldchange_DOWN** | **Ratio_pct1_pct2** |
| BICD2 | 0,037679657 | 0,553689971 | 0,2 | 0,046 | 1 | CH_SC | 1,46783517 | 0,681275405 | 4,347826087 |
| FBXO3 | 0,037679657 | 0,832906535 | 0,2 | 0,046 | 1 | CH_SC | 1,781270397 | 0,56139708 | 4,347826087 |
| FAM76B | 0,037679657 | 0,657180635 | 0,2 | 0,046 | 1 | CH_SC | 1,576997786 | 0,634116299 | 4,347826087 |
| TXNRD1 | 0,037679657 | 0,678493687 | 0,2 | 0,046 | 1 | CH_SC | 1,600467839 | 0,624817304 | 4,347826087 |
| MGAT2 | 0,037679657 | 0,749011725 | 0,2 | 0,046 | 1 | CH_SC | 1,680641164 | 0,595011012 | 4,347826087 |
| MAP2K11 | 0,037679657 | 0,788399273 | 0,2 | 0,046 | 1 | CH_SC | 1,727157051 | 0,578986144 | 4,347826087 |
| GINS3 | 0,037679657 | 0,707310466 | 0,2 | 0,046 | 1 | CH_SC | 1,632757421 | 0,612460851 | 4,347826087 |
| SMG9 | 0,037679657 | 0,579830406 | 0,2 | 0,046 | 1 | CH_SC | 1,494673534 | 0,669042421 | 4,347826087 |
| MCM3AP | 0,037679657 | 0,531806981 | 0,2 | 0,046 | 1 | CH_SC | 1,445738855 | 0,69168785 | 4,347826087 |
| MICAL32 | 0,037679657 | 0,635184102 | 0,2 | 0,046 | 1 | CH_SC | 1,553135932 | 0,643858647 | 4,347826087 |
| ZMAT31 | 0,038216863 | 0,862396663 | 0,333 | 0,123 | 1 | CH_SC | 1,818056031 | 0,550038053 | 2,707317073 |
| NSD11 | 0,038216863 | 1,060146993 | 0,333 | 0,123 | 1 | CH_SC | 2,085143961 | 0,479583194 | 2,707317073 |
| ITFG11 | 0,038216863 | 0,971749605 | 0,333 | 0,123 | 1 | CH_SC | 1,961217588 | 0,50988733 | 2,707317073 |
| COX111 | 0,038216863 | 0,607442593 | 0,333 | 0,123 | 1 | CH_SC | 1,523556068 | 0,656359173 | 2,707317073 |
| FTL5 | 0,038367674 | -0,39277235 | 0,933 | 0,985 | 1 | CH_SC | 0,761664547 | 1,312913938 | 0,947208122 |
| CCNY1 | 0,038419628 | 1,345938881 | 0,267 | 0,092 | 1 | CH_SC | 2,541955688 | 0,393397888 | 2,902173913 |
| NBDY2 | 0,038421676 | -0,655892649 | 0,067 | 0,369 | 1 | CH_SC | 0,634682668 | 1,575590528 | 0,181571816 |
| RARRES23 | 0,038620814 | 0,90540345 | 0,6 | 0,385 | 1 | CH_SC | 1,873068226 | 0,533883382 | 1,558441558 |
| NUDT31 | 0,038625504 | 0,821386807 | 0,467 | 0,231 | 1 | CH_SC | 1,767103825 | 0,565897706 | 2,021645022 |
| CUEDC22 | 0,038648992 | 0,753120503 | 0,4 | 0,169 | 1 | CH_SC | 1,685434431 | 0,593318839 | 2,366863905 |
| OTUB13 | 0,038648992 | 0,703300906 | 0,4 | 0,169 | 1 | CH_SC | 1,628225937 | 0,614165379 | 2,366863905 |
| PLEKHA52 | 0,038648992 | 0,581373554 | 0,4 | 0,169 | 1 | CH_SC | 1,496273135 | 0,668327177 | 2,366863905 |
| MAP43 | 0,038708538 | 1,291411037 | 0,533 | 0,369 | 1 | CH_SC | 2,447673348 | 0,408551248 | 1,444444444 |
| SNX7 | 0,038724705 | 0,52581326 | 0,267 | 0,077 | 1 | CH_SC | 1,439744949 | 0,694567465 | 3,467532468 |
| MAPKAP1 | 0,038724705 | 0,389856221 | 0,267 | 0,077 | 1 | CH_SC | 1,310262816 | 0,763205662 | 3,467532468 |
| SIAH1 | 0,038724705 | 0,827964172 | 0,267 | 0,077 | 1 | CH_SC | 1,775178589 | 0,563323604 | 3,467532468 |
| SSH22 | 0,038724705 | 0,52137403 | 0,267 | 0,077 | 1 | CH_SC | 1,435321605 | 0,696707969 | 3,467532468 |
| AATK | 0,038724705 | 1,026871451 | 0,267 | 0,077 | 1 | CH_SC | 2,037600827 | 0,49077326 | 3,467532468 |
| ST3GAL4 | 0,038810059 | 0,852545638 | 0,333 | 0,138 | 1 | CH_SC | 1,805684249 | 0,55380668 | 2,413043478 |
| MAGEH1 | 0,038810059 | 0,941574677 | 0,333 | 0,138 | 1 | CH_SC | 1,920623423 | 0,520664274 | 2,413043478 |
| EWSR12 | 0,039907044 | 1,265891229 | 0,467 | 0,215 | 1 | CH_SC | 2,404757185 | 0,4158424 | 2,172093023 |

| Supplementary Table 4 - clusermarker | | | | | | | | | |
|---|---|---|---|---|---|---|---|---|---|
| | **p_val** | **avg_log2FC** | **pct.1** | **pct.2** | **p_val_adj** | **celltype** | **Foldchange_UP** | **Foldchange_DOWN** | **Ratio_pct1_pct2** |
| SERTAD13 | 0,040083423 | 1,043163147 | 0,6 | 0,4 | 1 | CH_SC | 2,060740933 | 0,485262356 | 1,5 |
| MINDY1 | 0,040234686 | 0,660048326 | 0,2 | 0,046 | 1 | CH_SC | 1,580135553 | 0,632857098 | 4,347826087 |
| PDHB | 0,040234686 | 0,491973921 | 0,2 | 0,046 | 1 | CH_SC | 1,406367778 | 0,711051558 | 4,347826087 |
| SCHIP1 | 0,040234686 | 1,08391678 | 0,2 | 0,046 | 1 | CH_SC | 2,119783286 | 0,471746337 | 4,347826087 |
| SP41 | 0,040234686 | 0,851628943 | 0,2 | 0,046 | 1 | CH_SC | 1,804537273 | 0,554158684 | 4,347826087 |
| NAPEPLD | 0,040234686 | 0,619924195 | 0,2 | 0,046 | 1 | CH_SC | 1,536794429 | 0,650705118 | 4,347826087 |
| CPSF11 | 0,040234686 | 0,847576331 | 0,2 | 0,046 | 1 | CH_SC | 1,799475339 | 0,555717535 | 4,347826087 |
| INPPL1 | 0,040234686 | 0,478564345 | 0,2 | 0,046 | 1 | CH_SC | 1,393356419 | 0,717691458 | 4,347826087 |
| MX21 | 0,040234686 | 0,756160621 | 0,2 | 0,046 | 1 | CH_SC | 1,688989806 | 0,592069885 | 4,347826087 |
| TTC28 | 0,040234686 | 0,465603399 | 0,2 | 0,046 | 1 | CH_SC | 1,380894784 | 0,724168135 | 4,347826087 |
| WDR83OS4 | 0,040279413 | 0,736211639 | 0,8 | 0,646 | 1 | CH_SC | 1,665795891 | 0,600313643 | 1,238390093 |
| NUDT213 | 0,040298891 | 0,709086805 | 0,4 | 0,169 | 1 | CH_SC | 1,634769015 | 0,611707214 | 2,366863905 |
| FOXN21 | 0,040891172 | 0,942182377 | 0,267 | 0,077 | 1 | CH_SC | 1,921432609 | 0,520445003 | 3,467532468 |
| RETREG1 | 0,040891172 | 0,365739433 | 0,267 | 0,077 | 1 | CH_SC | 1,288541884 | 0,776071009 | 3,467532468 |
| FEM1C | 0,040891172 | 1,341972599 | 0,267 | 0,077 | 1 | CH_SC | 2,534976896 | 0,394480913 | 3,467532468 |
| GATM | 0,040891172 | 0,499504139 | 0,267 | 0,077 | 1 | CH_SC | 1,413727574 | 0,707349859 | 3,467532468 |
| BBS2 | 0,040891172 | 0,563714574 | 0,267 | 0,077 | 1 | CH_SC | 1,478069978 | 0,676557954 | 3,467532468 |
| PPDPF7 | 0,040983475 | 0,531726458 | 0,8 | 0,785 | 1 | CH_SC | 1,445658165 | 0,691726457 | 1,01910828 |
| ENHO | 0,041318179 | -0,658541082 | 0,067 | 0,369 | 1 | CH_SC | 0,633518616 | 1,578485581 | 0,181571816 |
| AC068888.1 | 0,041668208 | -0,756560299 | 0,067 | 0,323 | 1 | CH_SC | 0,591905883 | 1,689457781 | 0,207430341 |
| CCNB1IP12 | 0,041668208 | -0,732116405 | 0,067 | 0,323 | 1 | CH_SC | 0,602020114 | 1,661074068 | 0,207430341 |
| TENT5A3 | 0,041697361 | 0,394018508 | 0,533 | 0,231 | 1 | CH_SC | 1,314048485 | 0,761006927 | 2,307359307 |
| EI241 | 0,041934453 | 0,548710567 | 0,333 | 0,123 | 1 | CH_SC | 1,462777729 | 0,683630862 | 2,707317073 |
| MEF2C5 | 0,041937365 | 0,631149486 | 0,333 | 0,123 | 1 | CH_SC | 1,548798528 | 0,645661771 | 2,707317073 |
| MTURN1 | 0,041937365 | 0,587659414 | 0,333 | 0,123 | 1 | CH_SC | 1,50280666 | 0,665421592 | 2,707317073 |
| SPATA131 | 0,041937365 | 0,796531034 | 0,333 | 0,123 | 1 | CH_SC | 1,736919671 | 0,575731864 | 2,707317073 |
| ZMIZ13 | 0,042008135 | 0,636704473 | 0,4 | 0,169 | 1 | CH_SC | 1,554773553 | 0,64318048 | 2,366863905 |
| PNPLA81 | 0,042461714 | 0,879132166 | 0,333 | 0,138 | 1 | CH_SC | 1,839268581 | 0,543694385 | 2,413043478 |
| MYOC1 | 0,042611186 | -2,212592176 | 0 | 0,231 | 1 | CH_SC | 0,215746315 | 4,635073372 | 0 |
| COLGALT2 | 0,042611186 | -0,9750532 | 0 | 0,231 | 1 | CH_SC | 0,508721086 | 1,965713682 | 0 |
| RAMP13 | 0,042611186 | -0,596109044 | 0 | 0,231 | 1 | CH_SC | 0,661535717 | 1,511634179 | 0 |

| Supplementary Table 4 - clusermarker | | | | | | | | | |
|---|---|---|---|---|---|---|---|---|---|
| | **p_val** | **avg_log2FC** | **pct.1** | **pct.2** | **p_val_adj** | **celltype** | **Foldchange_UP** | **Foldchange_DOWN** | **Ratio_pct1_pct2** |
| GPX81 | 0,042611186 | -0,708997553 | 0 | 0,231 | 1 | CH_SC | 0,611745058 | 1,634667884 | 0 |
| IL17B | 0,042611186 | -1,362410781 | 0 | 0,231 | 1 | CH_SC | 0,388931831 | 2,571144657 | 0 |
| SOBP2 | 0,042611186 | -0,61882882 | 0 | 0,231 | 1 | CH_SC | 0,651199357 | 1,535628052 | 0 |
| PLAGL11 | 0,042611186 | -0,805533494 | 0 | 0,231 | 1 | CH_SC | 0,572150466 | 1,747791989 | 0 |
| CLIC33 | 0,042611186 | -0,929540572 | 0 | 0,231 | 1 | CH_SC | 0,52502551 | 1,904669355 | 0 |
| MRPL416 | 0,042611186 | -0,684046548 | 0 | 0,231 | 1 | CH_SC | 0,622417035 | 1,606639831 | 0 |
| IFITM101 | 0,042611186 | -0,672025761 | 0 | 0,231 | 1 | CH_SC | 0,627624789 | 1,593308642 | 0 |
| BCAT11 | 0,042611186 | -0,912103301 | 0 | 0,231 | 1 | CH_SC | 0,531409785 | 1,881786952 | 0 |
| IKBIP | 0,042611186 | -0,536316864 | 0 | 0,231 | 1 | CH_SC | 0,689529001 | 1,450265324 | 0 |
| MTHFS1 | 0,042611186 | -0,596609371 | 0 | 0,231 | 1 | CH_SC | 0,661306336 | 1,512158504 | 0 |
| FOXC2 | 0,042611186 | -0,668780151 | 0 | 0,231 | 1 | CH_SC | 0,629038336 | 1,589728228 | 0 |
| ABCA5 | 0,042611186 | -0,719573978 | 0 | 0,231 | 1 | CH_SC | 0,607276742 | 1,6466957 | 0 |
| CCDC68 | 0,042611186 | -0,688221412 | 0 | 0,231 | 1 | CH_SC | 0,620618491 | 1,611295851 | 0 |
| LENG81 | 0,042611186 | -0,788169634 | 0 | 0,231 | 1 | CH_SC | 0,57907831 | 1,726882154 | 0 |
| MZT2B6 | 0,042650738 | -0,750337538 | 0,2 | 0,477 | 1 | CH_SC | 0,594464458 | 1,682186354 | 0,419287212 |
| BTN2A11 | 0,042935162 | 0,536288577 | 0,2 | 0,046 | 1 | CH_SC | 1,45023689 | 0,68954252 | 4,347826087 |
| ZNF318 | 0,042935162 | 0,58693083 | 0,2 | 0,046 | 1 | CH_SC | 1,502047909 | 0,665757726 | 4,347826087 |
| ZNF664 | 0,042935162 | 0,447768807 | 0,2 | 0,046 | 1 | CH_SC | 1,363929246 | 0,733175861 | 4,347826087 |
| PHKB1 | 0,042935162 | 0,405865319 | 0,2 | 0,046 | 1 | CH_SC | 1,324883329 | 0,754783443 | 4,347826087 |
| NPC1 | 0,042935162 | 0,440692841 | 0,2 | 0,046 | 1 | CH_SC | 1,35725598 | 0,736780692 | 4,347826087 |
| SRPX21 | 0,043088221 | -1,117208889 | 0,067 | 0,308 | 1 | CH_SC | 0,460984808 | 2,169268885 | 0,217532468 |
| KCND3 | 0,043159482 | 0,548561334 | 0,267 | 0,077 | 1 | CH_SC | 1,462626427 | 0,683701581 | 3,467532468 |
| TCERG12 | 0,043159482 | 0,358993235 | 0,267 | 0,077 | 1 | CH_SC | 1,282530589 | 0,779708499 | 3,467532468 |
| WASHC5 | 0,043159482 | 0,426526204 | 0,267 | 0,077 | 1 | CH_SC | 1,34399354 | 0,744051195 | 3,467532468 |
| LHFPL62 | 0,043159482 | 0,438456007 | 0,267 | 0,077 | 1 | CH_SC | 1,355153246 | 0,737923923 | 3,467532468 |
| NSRP16 | 0,043264144 | -0,721652614 | 0,067 | 0,323 | 1 | CH_SC | 0,606402407 | 1,649069971 | 0,207430341 |
| IRF2BP21 | 0,043269436 | 0,886495943 | 0,467 | 0,246 | 1 | CH_SC | 1,848680541 | 0,54092634 | 1,898373984 |
| NNMT4 | 0,043379677 | -1,642365688 | 0,533 | 0,646 | 1 | CH_SC | 0,320330775 | 3,121773113 | 0,825077399 |
| CEP3503 | 0,043778334 | 0,636700058 | 0,4 | 0,169 | 1 | CH_SC | 1,554768796 | 0,643182448 | 2,366863905 |
| LPP2 | 0,043778334 | 0,742973957 | 0,4 | 0,169 | 1 | CH_SC | 1,673622273 | 0,597506388 | 2,366863905 |
| SIRT21 | 0,043909768 | 0,734444191 | 0,333 | 0,123 | 1 | CH_SC | 1,663756371 | 0,601049539 | 2,707317073 |

| Supplementary Table 4 - clusermarker | | | | | | | | | |
|---|---|---|---|---|---|---|---|---|---|
| | **p_val** | **avg_log2FC** | **pct.1** | **pct.2** | **p_val_adj** | **celltype** | **Foldchange_UP** | **Foldchange_DOWN** | **Ratio_pct1_pct2** |
| ABI3BP1 | 0,044737593 | -1,416176386 | 0,133 | 0,369 | 1 | CH_SC | 0,374704086 | 2,668772607 | 0,360433604 |
| CCDC88A2 | 0,04475756 | -0,803308732 | 0,067 | 0,308 | 1 | CH_SC | 0,573033452 | 1,745098817 | 0,217532468 |
| COBLL13 | 0,04475756 | -0,767407263 | 0,067 | 0,308 | 1 | CH_SC | 0,587472302 | 1,702207912 | 0,217532468 |
| SESN32 | 0,044839634 | -0,274694448 | 0,133 | 0,431 | 1 | CH_SC | 0,826625373 | 1,209737849 | 0,308584687 |
| RPL77 | 0,045124181 | -0,570019228 | 0,8 | 0,954 | 1 | CH_SC | 0,673607811 | 1,484543356 | 0,838574423 |
| UBE2F4 | 0,045533226 | 0,340260823 | 0,267 | 0,077 | 1 | CH_SC | 1,265985449 | 0,789898494 | 3,467532468 |
| CHCHD107 | 0,045533226 | 0,393748056 | 0,267 | 0,077 | 1 | CH_SC | 1,313802172 | 0,761149602 | 3,467532468 |
| TOPORS | 0,045611116 | 0,598038437 | 0,4 | 0,169 | 1 | CH_SC | 1,51365712 | 0,660651601 | 2,366863905 |
| PSMA23 | 0,045661717 | 0,902965005 | 0,6 | 0,4 | 1 | CH_SC | 1,869905037 | 0,534786516 | 1,5 |
| BAZ2A3 | 0,045732213 | 1,027088894 | 0,4 | 0,185 | 1 | CH_SC | 2,037907958 | 0,490699296 | 2,162162162 |
| ARHGAP31 | 0,045787286 | 0,470672777 | 0,2 | 0,046 | 1 | CH_SC | 1,385755541 | 0,721628 | 4,347826087 |
| TBRG4 | 0,045787286 | 0,580981771 | 0,2 | 0,046 | 1 | CH_SC | 1,495866858 | 0,668508694 | 4,347826087 |
| CHKA2 | 0,045787286 | 0,56674923 | 0,2 | 0,046 | 1 | CH_SC | 1,481182316 | 0,675136335 | 4,347826087 |
| GIT1 | 0,045787286 | 0,400983405 | 0,2 | 0,046 | 1 | CH_SC | 1,320407652 | 0,75734187 | 4,347826087 |
| ABCB71 | 0,045787286 | 0,30036527 | 0,2 | 0,046 | 1 | CH_SC | 1,231456161 | 0,812046772 | 4,347826087 |
| FADS1 | 0,045959928 | 0,743734428 | 0,333 | 0,123 | 1 | CH_SC | 1,674504703 | 0,597191515 | 2,707317073 |
| AMOTL1 | 0,045959928 | 0,806009868 | 0,333 | 0,123 | 1 | CH_SC | 1,7483692 | 0,571961574 | 2,707317073 |
| XYLT1 | 0,046481123 | -1,029547439 | 0,067 | 0,308 | 1 | CH_SC | 0,489863791 | 2,041383787 | 0,217532468 |
| NREP3 | 0,046611291 | -0,667793914 | 0,067 | 0,323 | 1 | CH_SC | 0,629468498 | 1,58864185 | 0,207430341 |
| TMEM981 | 0,04715612 | -0,476618751 | 0,2 | 0,523 | 1 | CH_SC | 0,718659978 | 1,391478629 | 0,382409178 |
| MRPS18B3 | 0,047427149 | 0,960765441 | 0,4 | 0,2 | 1 | CH_SC | 1,946342278 | 0,513784246 | 2 |
| DNAJC31 | 0,047427149 | 0,684739651 | 0,4 | 0,2 | 1 | CH_SC | 1,607411882 | 0,622118084 | 2 |
| ZNF1481 | 0,047508131 | 0,697165826 | 0,4 | 0,169 | 1 | CH_SC | 1,621316586 | 0,616782687 | 2,366863905 |
| C1QBP5 | 0,047591609 | 1,163051894 | 0,4 | 0,185 | 1 | CH_SC | 2,239306324 | 0,446566863 | 2,162162162 |
| CHMP1B3 | 0,048136161 | -1,209349797 | 0,133 | 0,415 | 1 | CH_SC | 0,432463477 | 2,312333995 | 0,320481928 |
| GTF2B5 | 0,048260219 | -0,96993553 | 0,067 | 0,308 | 1 | CH_SC | 0,510528876 | 1,958753062 | 0,217532468 |
| BLOC1S13 | 0,048289139 | 0,878138205 | 0,467 | 0,246 | 1 | CH_SC | 1,838001832 | 0,544069099 | 1,898373984 |
| UIMC1 | 0,048797381 | 0,315763616 | 0,2 | 0,046 | 1 | CH_SC | 1,244670281 | 0,803425626 | 4,347826087 |
| PLEKHF21 | 0,048797381 | 0,981860229 | 0,2 | 0,046 | 1 | CH_SC | 1,975010371 | 0,506326455 | 4,347826087 |
| APBB11 | 0,048797381 | 0,286753311 | 0,2 | 0,046 | 1 | CH_SC | 1,219891901 | 0,819744765 | 4,347826087 |
| GDF111 | 0,048797381 | 0,477864199 | 0,2 | 0,046 | 1 | CH_SC | 1,392680381 | 0,718039841 | 4,347826087 |

| Supplementary Table 4 - clusermarker | | | | | | | | | |
|---|---|---|---|---|---|---|---|---|---|
| | **p_val** | **avg_log2FC** | **pct.1** | **pct.2** | **p_val_adj** | **celltype** | **Foldchange_UP** | **Foldchange_DOWN** | **Ratio_pct1_pct2** |
| DDHD1 | 0,048797381 | 0,644391274 | 0,2 | 0,046 | 1 | CH_SC | 1,563079626 | 0,639762673 | 4,347826087 |
| GALK2 | 0,048797381 | 0,335864755 | 0,2 | 0,046 | 1 | CH_SC | 1,262133708 | 0,792309082 | 4,347826087 |
| PHF12 | 0,048797381 | 0,349909259 | 0,2 | 0,046 | 1 | CH_SC | 1,274480464 | 0,784633448 | 4,347826087 |
| ENDOV | 0,048797381 | 0,408199267 | 0,2 | 0,046 | 1 | CH_SC | 1,327028419 | 0,753563364 | 4,347826087 |
| ZNF4283 | 0,049304953 | 0,86027149 | 0,4 | 0,2 | 1 | CH_SC | 1,815379901 | 0,550848888 | 2 |
| IFI164 | 0,049514153 | 0,852172174 | 0,4 | 0,185 | 1 | CH_SC | 1,80521688 | 0,553950061 | 2,162162162 |
| RNF1493 | 0,04963717 | 0,720077336 | 0,267 | 0,092 | 1 | CH_SC | 1,647270335 | 0,607064899 | 2,902173913 |
| POP51 | 0,04963717 | 0,683079272 | 0,267 | 0,092 | 1 | CH_SC | 1,605562997 | 0,622834484 | 2,902173913 |
| CFAP201 | 0,04963717 | 0,804392253 | 0,267 | 0,092 | 1 | CH_SC | 1,746409947 | 0,572603243 | 2,902173913 |
| HELZ2 | 0,04963717 | 0,74905703 | 0,267 | 0,092 | 1 | CH_SC | 1,680693942 | 0,594992327 | 2,902173913 |
| MAPRE13 | 0,04963717 | 0,736077407 | 0,267 | 0,092 | 1 | CH_SC | 1,665640909 | 0,6003695 | 2,902173913 |
| FAM177A13 | 0,049780607 | -0,603915415 | 0,133 | 0,415 | 1 | CH_SC | 0,657965838 | 1,519835746 | 0,320481928 |
| TRIR7 | 0,049780607 | -0,67370438 | 0,133 | 0,415 | 1 | CH_SC | 0,626894953 | 1,595163584 | 0,320481928 |
| HNRNPL3 | 0,050067503 | 0,671660398 | 0,467 | 0,246 | 1 | CH_SC | 1,592905187 | 0,627783755 | 1,898373984 |
| XPO12 | 0,050302677 | 0,772455198 | 0,333 | 0,123 | 1 | CH_SC | 1,708174305 | 0,58542035 | 2,707317073 |
| IGFBP76 | 0,051267496 | 0,38920488 | 0,933 | 0,769 | 1 | CH_SC | 1,309671399 | 0,763550308 | 1,213263979 |
| CRIP18 | 0,051586732 | 0,830636404 | 0,533 | 0,308 | 1 | CH_SC | 1,778469711 | 0,562281153 | 1,730519481 |
| AKR7A24 | 0,051750705 | -0,790922026 | 0 | 0,215 | 1 | CH_SC | 0,577974591 | 1,730179866 | 0 |
| ZNF5934 | 0,051750705 | -0,396425083 | 0 | 0,215 | 1 | CH_SC | 0,759738542 | 1,316242292 | 0 |
| CRYZ | 0,051750705 | -0,63986277 | 0 | 0,215 | 1 | CH_SC | 0,641773992 | 1,558180937 | 0 |
| ARHGAP29 | 0,051750705 | -0,535115651 | 0 | 0,215 | 1 | CH_SC | 0,690103354 | 1,449058311 | 0 |
| ETV32 | 0,051750705 | -0,658651373 | 0 | 0,215 | 1 | CH_SC | 0,633470186 | 1,578606258 | 0 |
| CAPG5 | 0,051750705 | -0,880008457 | 0 | 0,215 | 1 | CH_SC | 0,543364246 | 1,840386089 | 0 |
| SUMF11 | 0,051750705 | -0,562214368 | 0 | 0,215 | 1 | CH_SC | 0,677261848 | 1,476533785 | 0 |
| FLNB3 | 0,051750705 | -0,682041164 | 0 | 0,215 | 1 | CH_SC | 0,623282813 | 1,60440811 | 0 |
| FGFR3 | 0,051750705 | -0,817433803 | 0 | 0,215 | 1 | CH_SC | 0,567450399 | 1,762268564 | 0 |
| TBC1D13 | 0,051750705 | -0,557864593 | 0 | 0,215 | 1 | CH_SC | 0,679306897 | 1,47208869 | 0 |
| CORIN | 0,051750705 | -0,689951558 | 0 | 0,215 | 1 | CH_SC | 0,619874663 | 1,613229349 | 0 |
| MAST42 | 0,051750705 | -0,811298278 | 0 | 0,215 | 1 | CH_SC | 0,569868804 | 1,754789863 | 0 |
| CCNG13 | 0,051750705 | -0,815428716 | 0 | 0,215 | 1 | CH_SC | 0,568239602 | 1,759821028 | 0 |
| TPBG1 | 0,051750705 | -0,607673577 | 0 | 0,215 | 1 | CH_SC | 0,656254094 | 1,523800018 | 0 |

| Supplementary Table 4 - clusermarker | | | | | | | | | |
|---|---|---|---|---|---|---|---|---|---|
| | **p_val** | **avg_log2FC** | **pct.1** | **pct.2** | **p_val_adj** | **celltype** | **Foldchange_UP** | **Foldchange_DOWN** | **Ratio_pct1_pct2** |
| TMEM60 | 0,051750705 | -0,610947107 | 0 | 0,215 | 1 | CH_SC | 0,654766717 | 1,527261503 | 0 |
| CRISPLD1 | 0,051750705 | -1,027712061 | 0 | 0,215 | 1 | CH_SC | 0,490487386 | 2,038788416 | 0 |
| TATDN12 | 0,051750705 | -0,496996643 | 0 | 0,215 | 1 | CH_SC | 0,708580347 | 1,41127256 | 0 |
| MSMB10 | 0,051750705 | -0,677970617 | 0 | 0,215 | 1 | CH_SC | 0,625043881 | 1,599887672 | 0 |
| PRH29 | 0,051750705 | -0,838438333 | 0 | 0,215 | 1 | CH_SC | 0,559248609 | 1,788113523 | 0 |
| PRR133 | 0,051750705 | -0,556607741 | 0 | 0,215 | 1 | CH_SC | 0,679898956 | 1,470806789 | 0 |
| CRYL11 | 0,051750705 | -0,554487245 | 0 | 0,215 | 1 | CH_SC | 0,680899017 | 1,468646562 | 0 |
| POLR1D3 | 0,051750705 | -0,62048547 | 0 | 0,215 | 1 | CH_SC | 0,650452012 | 1,53739243 | 0 |
| FLRT2 | 0,051750705 | -0,744052733 | 0 | 0,215 | 1 | CH_SC | 0,597059769 | 1,674874194 | 0 |
| ZG16B9 | 0,051750705 | -0,661326467 | 0 | 0,215 | 1 | CH_SC | 0,632296673 | 1,581536079 | 0 |
| SERPINF11 | 0,051750705 | -1,255139738 | 0 | 0,215 | 1 | CH_SC | 0,418952986 | 2,386902668 | 0 |
| MIR497HG | 0,051750705 | -0,669794715 | 0 | 0,215 | 1 | CH_SC | 0,628596126 | 1,590846586 | 0 |
| HS3ST3B1 | 0,051750705 | -0,849175614 | 0 | 0,215 | 1 | CH_SC | 0,555101842 | 1,801471233 | 0 |
| ACSF2 | 0,051750705 | -0,689471628 | 0 | 0,215 | 1 | CH_SC | 0,620080906 | 1,612692779 | 0 |
| NAA201 | 0,051750705 | -0,735337571 | 0 | 0,215 | 1 | CH_SC | 0,600677458 | 1,664786961 | 0 |
| PKIG1 | 0,051750705 | -0,619401519 | 0 | 0,215 | 1 | CH_SC | 0,650940905 | 1,536237764 | 0 |
| TRA2B2 | 0,051900446 | -0,544323172 | 0,067 | 0,338 | 1 | CH_SC | 0,685713024 | 1,458336017 | 0,198224852 |
| PER12 | 0,051900446 | 0,9782524 | 0,467 | 0,246 | 1 | CH_SC | 1,970077522 | 0,507594239 | 1,898373984 |
| CTCF3 | 0,051971883 | 0,251093246 | 0,2 | 0,046 | 1 | CH_SC | 1,190108614 | 0,840259442 | 4,347826087 |
| HEPH1 | 0,051971883 | 0,279297394 | 0,2 | 0,046 | 1 | CH_SC | 1,213603704 | 0,823992212 | 4,347826087 |
| TMEM126A | 0,052182744 | 0,681947036 | 0,267 | 0,092 | 1 | CH_SC | 1,604303434 | 0,62332348 | 2,902173913 |
| TBRG12 | 0,052182744 | 0,67655944 | 0,267 | 0,092 | 1 | CH_SC | 1,598323502 | 0,625655569 | 2,902173913 |
| HERC21 | 0,052182744 | 1,067535073 | 0,267 | 0,092 | 1 | CH_SC | 2,095849428 | 0,477133513 | 2,902173913 |
| PTPRA2 | 0,052182744 | 0,849097293 | 0,267 | 0,092 | 1 | CH_SC | 1,801373437 | 0,555131978 | 2,902173913 |
| NEGR11 | 0,052599911 | 1,067064302 | 0,333 | 0,123 | 1 | CH_SC | 2,095165636 | 0,477289233 | 2,707317073 |
| HNRNPA2B13 | 0,053076259 | 0,448135371 | 0,933 | 0,8 | 1 | CH_SC | 1,364275842 | 0,732989597 | 1,16625 |
| PLLP | 0,053323796 | 0,257316218 | 0,267 | 0,077 | 1 | CH_SC | 1,195253158 | 0,836642843 | 3,467532468 |
| PPP4R2 | 0,053555021 | 0,690749843 | 0,4 | 0,185 | 1 | CH_SC | 1,614122243 | 0,619531764 | 2,162162162 |
| WBP4 | 0,053555021 | 1,195199667 | 0,4 | 0,185 | 1 | CH_SC | 2,289765199 | 0,436726002 | 2,162162162 |
| ALDH3A21 | 0,053601347 | 0,719177296 | 0,4 | 0,169 | 1 | CH_SC | 1,646242988 | 0,607443741 | 2,366863905 |
| NPEPPS3 | 0,053601347 | 1,423079733 | 0,4 | 0,169 | 1 | CH_SC | 2,68157338 | 0,372915396 | 2,366863905 |

| Supplementary Table 4 - clusermarker | | | | | | | | | |
|---|---|---|---|---|---|---|---|---|---|
| | **p_val** | **avg_log2FC** | **pct.1** | **pct.2** | **p_val_adj** | **celltype** | **Foldchange_UP** | **Foldchange_DOWN** | **Ratio_pct1_pct2** |
| C12orf754 | 0,053714098 | -1,046054225 | 0,067 | 0,292 | 1 | CH_SC | 0,484290892 | 2,064874679 | 0,229452055 |
| SERPINE2 | 0,054368454 | -1,013743153 | 0,333 | 0,538 | 1 | CH_SC | 0,495259601 | 2,019143089 | 0,618959108 |
| MYL12A6 | 0,054693259 | 0,677499296 | 0,6 | 0,462 | 1 | CH_SC | 1,599365082 | 0,625248113 | 1,298701299 |
| DTD1 | 0,054836594 | 0,79814116 | 0,267 | 0,092 | 1 | CH_SC | 1,73885925 | 0,575089674 | 2,902173913 |
| DDT8 | 0,054836594 | 0,765549518 | 0,267 | 0,092 | 1 | CH_SC | 1,700017407 | 0,588229271 | 2,902173913 |
| SEPTIN101 | 0,054984195 | 0,452190567 | 0,333 | 0,123 | 1 | CH_SC | 1,368116008 | 0,730932168 | 2,707317073 |
| RAD501 | 0,055202349 | 0,645240507 | 0,333 | 0,138 | 1 | CH_SC | 1,563999994 | 0,639386192 | 2,413043478 |
| FGD5-AS12 | 0,055354021 | 0,760030068 | 0,467 | 0,262 | 1 | CH_SC | 1,69352592 | 0,590484024 | 1,782442748 |
| ATP1B34 | 0,057021154 | -0,880149624 | 0,133 | 0,4 | 1 | CH_SC | 0,543311081 | 1,840566179 | 0,3325 |
| CAPNS13 | 0,057021154 | 0,710857079 | 0,533 | 0,308 | 1 | CH_SC | 1,636776208 | 0,610957072 | 1,730519481 |
| NFAT52 | 0,057451112 | 1,318552261 | 0,4 | 0,2 | 1 | CH_SC | 2,494156965 | 0,400937076 | 2 |
| MICU1 | 0,057457916 | 0,924411429 | 0,333 | 0,123 | 1 | CH_SC | 1,8979098 | 0,52689543 | 2,707317073 |
| GTF3C63 | 0,05760205 | 0,553798854 | 0,267 | 0,092 | 1 | CH_SC | 1,467945955 | 0,68122399 | 2,902173913 |
| IFNGR14 | 0,05760205 | 0,614684669 | 0,267 | 0,092 | 1 | CH_SC | 1,531223279 | 0,653072621 | 2,902173913 |
| PTAR1 | 0,05760205 | 1,252815645 | 0,267 | 0,092 | 1 | CH_SC | 2,38306061 | 0,419628437 | 2,902173913 |
| PPP4C7 | 0,05760205 | 0,587369637 | 0,267 | 0,092 | 1 | CH_SC | 1,502504839 | 0,665555261 | 2,902173913 |
| GET32 | 0,05760205 | 0,752027719 | 0,267 | 0,092 | 1 | CH_SC | 1,684158265 | 0,593768425 | 2,902173913 |
| ARHGEF9 | 0,05760205 | 0,526294657 | 0,267 | 0,092 | 1 | CH_SC | 1,440225442 | 0,694335742 | 2,902173913 |
| ERLEC11 | 0,057739059 | -0,602681939 | 0,067 | 0,338 | 1 | CH_SC | 0,658528626 | 1,518536871 | 0,198224852 |
| ITPKC | 0,057865878 | -1,021611567 | 0,067 | 0,292 | 1 | CH_SC | 0,492565822 | 2,030185519 | 0,229452055 |
| AGO32 | 0,058015704 | 0,423598852 | 0,4 | 0,169 | 1 | CH_SC | 1,341269227 | 0,745562472 | 2,366863905 |
| SAP30BP1 | 0,058035463 | 0,425504199 | 0,467 | 0,215 | 1 | CH_SC | 1,343041792 | 0,744578468 | 2,172093023 |
| COL6A23 | 0,058306322 | -0,890888237 | 0,4 | 0,677 | 1 | CH_SC | 0,539281991 | 1,854317437 | 0,59084195 |
| HMGB13 | 0,058755732 | 0,548161141 | 0,8 | 0,846 | 1 | CH_SC | 1,462220761 | 0,683891261 | 0,945626478 |
| ABL22 | 0,059743796 | 0,866109051 | 0,333 | 0,154 | 1 | CH_SC | 1,822740334 | 0,548624498 | 2,162337662 |
| HSP90AA16 | 0,060006835 | 0,615846315 | 0,8 | 0,923 | 1 | CH_SC | 1,532456705 | 0,652546983 | 0,866738895 |
| TOX | 0,060039776 | -0,759162423 | 0,067 | 0,292 | 1 | CH_SC | 0,590839251 | 1,69250773 | 0,229452055 |
| COX8A9 | 0,060097222 | 0,709923809 | 0,6 | 0,554 | 1 | CH_SC | 1,63571773 | 0,611352425 | 1,083032491 |
| PPP1R111 | 0,06010107 | 0,856897722 | 0,333 | 0,138 | 1 | CH_SC | 1,811139563 | 0,552138565 | 2,413043478 |
| AKAP122 | 0,06010107 | 1,231967568 | 0,333 | 0,138 | 1 | CH_SC | 2,348871139 | 0,425736424 | 2,413043478 |
| OPTN2 | 0,06010107 | 1,144061909 | 0,333 | 0,138 | 1 | CH_SC | 2,210023804 | 0,452483814 | 2,413043478 |

| Supplementary Table 4 - clusermarker | | | | | | | | | |
|---|---|---|---|---|---|---|---|---|---|
| | **p_val** | **avg_log2FC** | **pct.1** | **pct.2** | **p_val_adj** | **celltype** | **Foldchange_UP** | **Foldchange_DOWN** | **Ratio_pct1_pct2** |
| MORF4L21 | 0,060311592 | -0,741706077 | 0,333 | 0,569 | 1 | CH_SC | 0,598031724 | 1,672152094 | 0,585237258 |
| KAT6B1 | 0,060333752 | 0,563654057 | 0,4 | 0,169 | 1 | CH_SC | 1,478007978 | 0,676586334 | 2,366863905 |
| ARL5A3 | 0,060482481 | 0,490252869 | 0,267 | 0,092 | 1 | CH_SC | 1,404691062 | 0,711900308 | 2,902173913 |
| RIC8A | 0,060482481 | 0,638578094 | 0,267 | 0,092 | 1 | CH_SC | 1,556794042 | 0,642345727 | 2,902173913 |
| ARHGEF40 | 0,060482481 | 0,511507352 | 0,267 | 0,092 | 1 | CH_SC | 1,425538845 | 0,701489127 | 2,902173913 |
| ERRFI13 | 0,060902395 | -0,813301869 | 0,133 | 0,4 | 1 | CH_SC | 0,569077928 | 1,75722858 | 0,3325 |
| LDHB4 | 0,061873328 | -0,431627726 | 0,333 | 0,615 | 1 | CH_SC | 0,741424798 | 1,348754456 | 0,541463415 |
| ANAPC164 | 0,0622387 | -0,697779867 | 0,267 | 0,554 | 1 | CH_SC | 0,616520227 | 1,622006799 | 0,481949458 |
| COPB14 | 0,062280965 | -0,746067856 | 0,067 | 0,292 | 1 | CH_SC | 0,596226393 | 1,677215253 | 0,229452055 |
| TPI17 | 0,062294716 | 0,721193141 | 0,6 | 0,431 | 1 | CH_SC | 1,648544854 | 0,606595567 | 1,392111369 |
| SF3B65 | 0,062455594 | -0,630352628 | 0,2 | 0,508 | 1 | CH_SC | 0,646018494 | 1,5479433 | 0,393700787 |
| FYB12 | 0,062679597 | -0,556581276 | 0 | 0,2 | 1 | CH_SC | 0,679911428 | 1,470779809 | 0 |
| SCAMP3 | 0,062682482 | 0,564163079 | 0,333 | 0,138 | 1 | CH_SC | 1,478529551 | 0,676347658 | 2,413043478 |
| SERTAD4-AS11 | 0,062683327 | -0,495689144 | 0 | 0,2 | 1 | CH_SC | 0,709222817 | 1,409994118 | 0 |
| GFPT11 | 0,062683327 | -0,61614832 | 0 | 0,2 | 1 | CH_SC | 0,652410398 | 1,532777534 | 0 |
| HNMT2 | 0,062683327 | -0,6148337 | 0 | 0,2 | 1 | CH_SC | 0,653005161 | 1,531381464 | 0 |
| KIF9 | 0,062683327 | -0,527943892 | 0 | 0,2 | 1 | CH_SC | 0,693542457 | 1,441872794 | 0 |
| TMEM45A | 0,062683327 | -0,579090759 | 0 | 0,2 | 1 | CH_SC | 0,669385517 | 1,493907434 | 0 |
| CD142 | 0,062683327 | -0,586670726 | 0 | 0,2 | 1 | CH_SC | 0,665877766 | 1,501777129 | 0 |
| PDGFRB2 | 0,062683327 | -0,603921334 | 0 | 0,2 | 1 | CH_SC | 0,657963138 | 1,519841981 | 0 |
| NQO21 | 0,062683327 | -0,549718001 | 0 | 0,2 | 1 | CH_SC | 0,683153649 | 1,463799544 | 0 |
| CCN6 | 0,062683327 | -0,498922989 | 0 | 0,2 | 1 | CH_SC | 0,707634853 | 1,413158208 | 0 |
| PERP4 | 0,062683327 | -0,720258771 | 0 | 0,2 | 1 | CH_SC | 0,606988559 | 1,64747751 | 0 |
| PCMTD11 | 0,062683327 | -0,49875094 | 0 | 0,2 | 1 | CH_SC | 0,707719247 | 1,412989691 | 0 |
| CEMIP21 | 0,062683327 | -0,731288887 | 0 | 0,2 | 1 | CH_SC | 0,602365527 | 1,660121563 | 0 |
| CISD13 | 0,062683327 | -0,660716743 | 0 | 0,2 | 1 | CH_SC | 0,632563956 | 1,580867817 | 0 |
| BICC1 | 0,062683327 | -0,694752153 | 0 | 0,2 | 1 | CH_SC | 0,617815447 | 1,618606341 | 0 |
| EIF4EBP2 | 0,062683327 | -0,505952958 | 0 | 0,2 | 1 | CH_SC | 0,704195075 | 1,420061054 | 0 |
| H192 | 0,062683327 | -0,700428662 | 0 | 0,2 | 1 | CH_SC | 0,615389331 | 1,624987547 | 0 |
| ANGPTL5 | 0,062683327 | -0,963839845 | 0 | 0,2 | 1 | CH_SC | 0,51269053 | 1,950494385 | 0 |
| HEBP13 | 0,062683327 | -0,575909862 | 0 | 0,2 | 1 | CH_SC | 0,670863026 | 1,490617252 | 0 |

| Supplementary Table 4 - clusermarker | | | | | | | | | |
|---|---|---|---|---|---|---|---|---|---|
| | p_val | avg_log2FC | pct.1 | pct.2 | p_val_adj | celltype | Foldchange_UP | Foldchange_DOWN | Ratio_pct1_pct2 |
| KRT82 | 0,062683327 | -0,538350912 | 0 | 0,2 | 1 | CH_SC | 0,688557523 | 1,452311488 | 0 |
| WIF1 | 0,062683327 | -2,402116385 | 0 | 0,2 | 1 | CH_SC | 0,189186836 | 5,285780021 | 0 |
| WSCD2 | 0,062683327 | -0,778858368 | 0 | 0,2 | 1 | CH_SC | 0,582827814 | 1,715772612 | 0 |
| SMAD9 | 0,062683327 | -0,466751975 | 0 | 0,2 | 1 | CH_SC | 0,723591831 | 1,381994597 | 0 |
| TOX43 | 0,062683327 | -0,506897254 | 0 | 0,2 | 1 | CH_SC | 0,703734304 | 1,42099084 | 0 |
| LTBP23 | 0,062683327 | -0,805458547 | 0 | 0,2 | 1 | CH_SC | 0,572180189 | 1,747701194 | 0 |
| TP53BP1 | 0,062683327 | -0,524761358 | 0 | 0,2 | 1 | CH_SC | 0,695074075 | 1,43869558 | 0 |
| RAB11A4 | 0,062683327 | -0,524513859 | 0 | 0,2 | 1 | CH_SC | 0,695193328 | 1,438448789 | 0 |
| MMP22 | 0,062683327 | -0,832778498 | 0 | 0,2 | 1 | CH_SC | 0,561446905 | 1,78111232 | 0 |
| DRG2 | 0,062683327 | -0,597414736 | 0 | 0,2 | 1 | CH_SC | 0,660937273 | 1,513002883 | 0 |
| AC091271.1 | 0,062683327 | -0,647682357 | 0 | 0,2 | 1 | CH_SC | 0,638304906 | 1,566649402 | 0 |
| ETS22 | 0,062683327 | -0,7016114 | 0 | 0,2 | 1 | CH_SC | 0,614885035 | 1,626320276 | 0 |
| ATP6V0B6 | 0,063427255 | 0,877355385 | 0,4 | 0,215 | 1 | CH_SC | 1,837004786 | 0,544364395 | 1,860465116 |
| SHC11 | 0,063481286 | 0,568162233 | 0,267 | 0,092 | 1 | CH_SC | 1,482633725 | 0,674475417 | 2,902173913 |
| SMAP13 | 0,063481286 | 0,653585059 | 0,267 | 0,092 | 1 | CH_SC | 1,573072386 | 0,635698655 | 2,902173913 |
| CREB31 | 0,063481286 | 0,398760535 | 0,267 | 0,092 | 1 | CH_SC | 1,318374766 | 0,758509663 | 2,902173913 |
| IVD | 0,063481286 | 0,778266752 | 0,267 | 0,092 | 1 | CH_SC | 1,715069158 | 0,583066867 | 2,902173913 |
| MPDU11 | 0,063481286 | 0,581444109 | 0,267 | 0,092 | 1 | CH_SC | 1,496346312 | 0,668294493 | 2,902173913 |
| SCAF13 | 0,063481286 | 0,474578832 | 0,267 | 0,092 | 1 | CH_SC | 1,389512518 | 0,719676856 | 2,902173913 |
| MFSD103 | 0,064823373 | 1,127323521 | 0,333 | 0,154 | 1 | CH_SC | 2,184530913 | 0,457764179 | 2,162337662 |
| RCN13 | 0,064823373 | 0,974269085 | 0,333 | 0,154 | 1 | CH_SC | 1,964645593 | 0,508997655 | 2,162337662 |
| C9orf782 | 0,064874693 | 0,476507056 | 0,4 | 0,185 | 1 | CH_SC | 1,391370903 | 0,718715619 | 2,162162162 |
| ATP5F1C3 | 0,064995074 | 0,718739892 | 0,467 | 0,277 | 1 | CH_SC | 1,645743947 | 0,607627937 | 1,685920578 |
| COX7A24 | 0,065839406 | 0,83912266 | 0,733 | 0,631 | 1 | CH_SC | 1,788961898 | 0,558983398 | 1,161648177 |
| UGCG5 | 0,066365875 | 0,985591904 | 0,2 | 0,062 | 1 | CH_SC | 1,980125545 | 0,505018484 | 3,225806452 |
| EXOC4 | 0,066601892 | 0,530614128 | 0,267 | 0,092 | 1 | CH_SC | 1,44454398 | 0,692259989 | 2,902173913 |
| DUSP3 | 0,066601892 | 0,512587396 | 0,267 | 0,092 | 1 | CH_SC | 1,426606444 | 0,700964168 | 2,902173913 |
| COPZ21 | 0,06668655 | -0,690710373 | 0,067 | 0,277 | 1 | CH_SC | 0,619548713 | 1,614078084 | 0,241877256 |
| RPAIN2 | 0,066782218 | -0,444463003 | 0,067 | 0,323 | 1 | CH_SC | 0,734857793 | 1,360807505 | 0,207430341 |
| ID12 | 0,067463956 | -0,573322256 | 0,333 | 0,615 | 1 | CH_SC | 0,67206736 | 1,487946089 | 0,541463415 |
| MXRA81 | 0,067468544 | 0,682927325 | 0,6 | 0,462 | 1 | CH_SC | 1,605393905 | 0,622900085 | 1,298701299 |

| Supplementary Table 4 - clusermarker | | | | | | | | | |
|---|---|---|---|---|---|---|---|---|---|
| | **p_val** | **avg_log2FC** | **pct.1** | **pct.2** | **p_val_adj** | **celltype** | **Foldchange_UP** | **Foldchange_DOWN** | **Ratio_pct1_pct2** |
| MT-ND4L7 | 0,067511829 | 0,5235486 | 0,933 | 0,815 | 1 | CH_SC | 1,437486692 | 0,695658614 | 1,144785276 |
| NOLC12 | 0,068024457 | 0,947036563 | 0,467 | 0,262 | 1 | CH_SC | 1,927908475 | 0,518696822 | 1,782442748 |
| RERE5 | 0,068120586 | 0,500963731 | 0,333 | 0,138 | 1 | CH_SC | 1,415158583 | 0,706634587 | 2,413043478 |
| NT5C4 | 0,068120586 | 0,776845115 | 0,333 | 0,138 | 1 | CH_SC | 1,713379955 | 0,583641706 | 2,413043478 |
| MAPK1IP1L1 | 0,068188041 | 0,774672886 | 0,4 | 0,215 | 1 | CH_SC | 1,710802103 | 0,584521143 | 1,860465116 |
| PTP4A24 | 0,068220244 | 1,00542108 | 0,467 | 0,323 | 1 | CH_SC | 2,00752935 | 0,498124722 | 1,445820433 |
| DDX243 | 0,06842972 | -0,748292003 | 0,267 | 0,569 | 1 | CH_SC | 0,595307922 | 1,679802945 | 0,469244288 |
| TSPYL22 | 0,068965754 | -0,786707844 | 0,2 | 0,4 | 1 | CH_SC | 0,57966535 | 1,725133302 | 0,5 |
| APBB21 | 0,069177285 | -1,022797299 | 0,067 | 0,277 | 1 | CH_SC | 0,492161156 | 2,031854787 | 0,241877256 |
| EBPL4 | 0,069177285 | -0,774890421 | 0,067 | 0,277 | 1 | CH_SC | 0,584433013 | 1,711060084 | 0,241877256 |
| ABHD24 | 0,069177285 | 0,64602837 | 0,4 | 0,2 | 1 | CH_SC | 1,564854335 | 0,639037115 | 2 |
| RFLNA | 0,069180181 | -0,923492808 | 0,133 | 0,338 | 1 | CH_SC | 0,527231032 | 1,89670171 | 0,393491124 |
| CSTB6 | 0,069479395 | -0,632774372 | 0,267 | 0,538 | 1 | CH_SC | 0,644934981 | 1,550543899 | 0,496282528 |
| AKR1A14 | 0,069479777 | -0,817882217 | 0,133 | 0,354 | 1 | CH_SC | 0,567274053 | 1,762816392 | 0,375706215 |
| CIAO12 | 0,069847754 | 0,569549371 | 0,267 | 0,092 | 1 | CH_SC | 1,484059949 | 0,673827227 | 2,902173913 |
| CDK172 | 0,069847754 | 0,398848651 | 0,267 | 0,092 | 1 | CH_SC | 1,318455292 | 0,758463337 | 2,902173913 |
| ROMO12 | 0,070015876 | -0,762745266 | 0,2 | 0,431 | 1 | CH_SC | 0,58937376 | 1,69671619 | 0,464037123 |
| PLOD3 | 0,070172666 | 1,147630186 | 0,2 | 0,062 | 1 | CH_SC | 2,215496713 | 0,45136605 | 3,225806452 |
| FAM13B | 0,070254099 | 0,651869706 | 0,333 | 0,154 | 1 | CH_SC | 1,571203127 | 0,636454945 | 2,162337662 |
| COMMD62 | 0,070456699 | -0,492021736 | 0,467 | 0,769 | 1 | CH_SC | 0,711027993 | 1,406414389 | 0,607282185 |
| NEDD92 | 0,071058347 | 0,728125958 | 0,467 | 0,246 | 1 | CH_SC | 1,65648594 | 0,603687587 | 1,898373984 |
| CCDC90B2 | 0,071882422 | -0,686385908 | 0,067 | 0,308 | 1 | CH_SC | 0,621408591 | 1,609247143 | 0,217532468 |
| DNAJA18 | 0,072495355 | 0,712475683 | 0,733 | 0,538 | 1 | CH_SC | 1,638613588 | 0,610272005 | 1,362453532 |
| HSPA88 | 0,072613647 | 0,807770569 | 0,8 | 0,738 | 1 | CH_SC | 1,750504256 | 0,571263964 | 1,08401084 |
| TMEM14C5 | 0,072770088 | -0,826277578 | 0,2 | 0,477 | 1 | CH_SC | 0,563982547 | 1,773104515 | 0,419287212 |
| DR11 | 0,073222345 | 0,384113918 | 0,267 | 0,092 | 1 | CH_SC | 1,305057993 | 0,766249473 | 2,902173913 |
| ARPC5L6 | 0,073222345 | 0,448579224 | 0,267 | 0,092 | 1 | CH_SC | 1,364695633 | 0,732764124 | 2,902173913 |
| STX8 | 0,073222345 | 0,435902479 | 0,267 | 0,092 | 1 | CH_SC | 1,352756787 | 0,739231183 | 2,902173913 |
| EIF4A16 | 0,07324833 | 0,991129959 | 0,8 | 0,723 | 1 | CH_SC | 1,987741236 | 0,503083592 | 1,106500692 |
| HEXA2 | 0,073965426 | -0,887663371 | 0,133 | 0,338 | 1 | CH_SC | 0,5404888 | 1,850177098 | 0,393491124 |
| MAGED12 | 0,073965426 | -0,882149323 | 0,133 | 0,338 | 1 | CH_SC | 0,542558527 | 1,843119131 | 0,393491124 |

| Supplementary Table 4 - clusermarker | | | | | | | | | |
|---|---|---|---|---|---|---|---|---|---|
| | **p_val** | **avg_log2FC** | **pct.1** | **pct.2** | **p_val_adj** | **celltype** | **Foldchange_UP** | **Foldchange_DOWN** | **Ratio_pct1_pct2** |
| HEBP25 | 0,074077666 | -0,388965585 | 0,067 | 0,323 | 1 | CH_SC | 0,763676966 | 1,309454186 | 0,207430341 |
| GPR89A | 0,074156823 | 0,794240701 | 0,2 | 0,062 | 1 | CH_SC | 1,734164432 | 0,576646586 | 3,225806452 |
| ATG4B1 | 0,074156823 | 0,575674166 | 0,2 | 0,062 | 1 | CH_SC | 1,490373746 | 0,670972635 | 3,225806452 |
| LSM1 | 0,074156823 | 0,693771058 | 0,2 | 0,062 | 1 | CH_SC | 1,617505993 | 0,618235731 | 3,225806452 |
| GIT2 | 0,074156823 | 0,77815745 | 0,2 | 0,062 | 1 | CH_SC | 1,714939225 | 0,583111043 | 3,225806452 |
| PIP4K2B | 0,074156823 | 0,722633873 | 0,2 | 0,062 | 1 | CH_SC | 1,650191978 | 0,605990099 | 3,225806452 |
| NAT141 | 0,074156823 | 0,739872972 | 0,2 | 0,062 | 1 | CH_SC | 1,670028788 | 0,598792073 | 3,225806452 |
| TFAP2C1 | 0,074156823 | 0,923086855 | 0,2 | 0,062 | 1 | CH_SC | 1,896168083 | 0,527379407 | 3,225806452 |
| RRAS1 | 0,074215505 | 0,714604931 | 0,467 | 0,277 | 1 | CH_SC | 1,641033774 | 0,60937198 | 1,685920578 |
| COMMD21 | 0,075740729 | -0,481343076 | 0 | 0,185 | 1 | CH_SC | 0,716310465 | 1,396042707 | 0 |
| ZNF33A | 0,075740729 | -0,451744475 | 0 | 0,185 | 1 | CH_SC | 0,731158213 | 1,367693042 | 0 |
| B4GALT41 | 0,075745174 | -0,417547852 | 0 | 0,185 | 1 | CH_SC | 0,748696102 | 1,335655412 | 0 |
| PFN21 | 0,075745174 | -0,582392574 | 0 | 0,185 | 1 | CH_SC | 0,667855283 | 1,497330372 | 0 |
| PLRG11 | 0,075745174 | -0,496107385 | 0 | 0,185 | 1 | CH_SC | 0,709017241 | 1,410402938 | 0 |
| LIFR2 | 0,075745174 | -0,580628784 | 0 | 0,185 | 1 | CH_SC | 0,66867228 | 1,495500905 | 0 |
| CREBRF2 | 0,075745174 | -0,588743274 | 0 | 0,185 | 1 | CH_SC | 0,664921865 | 1,503936104 | 0 |
| TES1 | 0,075745174 | -0,468192739 | 0 | 0,185 | 1 | CH_SC | 0,722869568 | 1,383375431 | 0 |
| OGN1 | 0,075745174 | -1,220834292 | 0 | 0,185 | 1 | CH_SC | 0,429034541 | 2,330814663 | 0 |
| GIHCG2 | 0,075745174 | -0,427366291 | 0 | 0,185 | 1 | CH_SC | 0,743618057 | 1,344776381 | 0 |
| CCDC592 | 0,075745174 | -0,545253263 | 0 | 0,185 | 1 | CH_SC | 0,685271094 | 1,459276496 | 0 |
| NUP581 | 0,075745174 | -0,513446295 | 0 | 0,185 | 1 | CH_SC | 0,700546978 | 1,427456018 | 0 |
| ANG1 | 0,075745174 | -0,44787532 | 0 | 0,185 | 1 | CH_SC | 0,733121733 | 1,364029948 | 0 |
| CDR1 | 0,075745174 | -0,762306664 | 0 | 0,185 | 1 | CH_SC | 0,589552965 | 1,696200441 | 0 |
| MPC1 | 0,075876676 | 1,077606551 | 0,4 | 0,215 | 1 | CH_SC | 2,110531776 | 0,473814235 | 1,860465116 |
| IRF12 | 0,076022153 | 0,848993305 | 0,467 | 0,246 | 1 | CH_SC | 1,801243601 | 0,555171993 | 1,898373984 |
| KIF1B | 0,076606565 | 0,707914068 | 0,267 | 0,108 | 1 | CH_SC | 1,633440686 | 0,61220466 | 2,472222222 |
| DDX59 | 0,076658163 | 0,511623282 | 0,8 | 0,738 | 1 | CH_SC | 1,425653401 | 0,70143276 | 1,08401084 |
| CLIP3 | 0,07699987 | 1,206903789 | 0,333 | 0,138 | 1 | CH_SC | 2,308416885 | 0,433197316 | 2,413043478 |
| SF3B15 | 0,077163408 | 0,935552392 | 0,6 | 0,4 | 1 | CH_SC | 1,912622817 | 0,522842241 | 1,5 |
| RSRC25 | 0,077541332 | 0,889399268 | 0,533 | 0,338 | 1 | CH_SC | 1,85240463 | 0,539838858 | 1,576923077 |
| RPL36AL3 | 0,07803557 | -0,432857895 | 0,6 | 0,8 | 1 | CH_SC | 0,740792863 | 1,349905013 | 0,75 |

| Supplementary Table 4 - clusermarker | | | | | | | | | |
|---|---|---|---|---|---|---|---|---|---|
| | **p_val** | **avg_log2FC** | **pct.1** | **pct.2** | **p_val_adj** | **celltype** | **Foldchange_UP** | **Foldchange_DOWN** | **Ratio_pct1_pct2** |
| PIGF | 0,078323972 | 0,45863078 | 0,2 | 0,062 | 1 | CH_SC | 1,374236951 | 0,727676548 | 3,225806452 |
| LRP1B | 0,078323972 | 0,636814397 | 0,2 | 0,062 | 1 | CH_SC | 1,554892021 | 0,643131476 | 3,225806452 |
| APEH | 0,078323972 | 0,648471327 | 0,2 | 0,062 | 1 | CH_SC | 1,567506393 | 0,637955931 | 3,225806452 |
| FBXO38 | 0,078323972 | 0,55220586 | 0,2 | 0,062 | 1 | CH_SC | 1,466325974 | 0,681976598 | 3,225806452 |
| TBC1D7 | 0,078323972 | 0,546070561 | 0,2 | 0,062 | 1 | CH_SC | 1,460103421 | 0,684882992 | 3,225806452 |
| RNF1701 | 0,078323972 | 0,827347294 | 0,2 | 0,062 | 1 | CH_SC | 1,774419708 | 0,563564525 | 3,225806452 |
| DDX581 | 0,078323972 | 0,71503201 | 0,2 | 0,062 | 1 | CH_SC | 1,641519639 | 0,609191615 | 3,225806452 |
| NTMT11 | 0,078323972 | 0,506700029 | 0,2 | 0,062 | 1 | CH_SC | 1,420796595 | 0,703830516 | 3,225806452 |
| FBXW54 | 0,078323972 | 0,317836944 | 0,2 | 0,062 | 1 | CH_SC | 1,246460309 | 0,802271835 | 3,225806452 |
| CAMKK2 | 0,078323972 | 0,601529876 | 0,2 | 0,062 | 1 | CH_SC | 1,517324729 | 0,659054704 | 3,225806452 |
| KATNAL1 | 0,078323972 | 0,601483613 | 0,2 | 0,062 | 1 | CH_SC | 1,517276074 | 0,659075838 | 3,225806452 |
| ARHGAP23 | 0,078323972 | 0,773361625 | 0,2 | 0,062 | 1 | CH_SC | 1,709247867 | 0,585052653 | 3,225806452 |
| TOMM40 | 0,078323972 | 0,438440253 | 0,2 | 0,062 | 1 | CH_SC | 1,355138449 | 0,737931981 | 3,225806452 |
| AL133415.1 | 0,078591564 | 0,986245168 | 0,4 | 0,215 | 1 | CH_SC | 1,981022365 | 0,504789859 | 1,860465116 |
| TYW3 | 0,079017898 | -0,753194079 | 0,133 | 0,338 | 1 | CH_SC | 0,593288581 | 1,685520389 | 0,393491124 |
| XRCC63 | 0,079151971 | 0,742159006 | 0,533 | 0,354 | 1 | CH_SC | 1,672677143 | 0,597844004 | 1,505649718 |
| TXLNA | 0,080095116 | 0,531740372 | 0,267 | 0,108 | 1 | CH_SC | 1,445672107 | 0,691719786 | 2,472222222 |
| UGGT2 | 0,080371688 | 0,487211201 | 0,267 | 0,092 | 1 | CH_SC | 1,401732638 | 0,713402808 | 2,902173913 |
| HSPA131 | 0,080371688 | 0,328609574 | 0,267 | 0,092 | 1 | CH_SC | 1,255802487 | 0,796303567 | 2,902173913 |
| MRPL512 | 0,080410917 | -0,627947643 | 0,267 | 0,492 | 1 | CH_SC | 0,64709631 | 1,545365016 | 0,542682927 |
| PNKD1 | 0,081581466 | -0,357142672 | 0,2 | 0,431 | 1 | CH_SC | 0,780709282 | 1,280886525 | 0,464037123 |
| KDELR26 | 0,081581466 | 0,83751101 | 0,533 | 0,354 | 1 | CH_SC | 1,786964545 | 0,559608193 | 1,505649718 |
| MT-ND310 | 0,082192956 | 0,491309539 | 1 | 0,954 | 1 | CH_SC | 1,405720274 | 0,711379083 | 1,048218029 |
| RALGAPA13 | 0,082235814 | 0,846089792 | 0,333 | 0,154 | 1 | CH_SC | 1,797622132 | 0,556290436 | 2,162337662 |
| NOP582 | 0,082469537 | -0,625219699 | 0,067 | 0,262 | 1 | CH_SC | 0,648321041 | 1,542445697 | 0,255725191 |
| ZMYM41 | 0,082679762 | 0,691363514 | 0,2 | 0,062 | 1 | CH_SC | 1,61480898 | 0,619268293 | 3,225806452 |
| TSEN15 | 0,082679762 | 0,785204674 | 0,2 | 0,062 | 1 | CH_SC | 1,72333679 | 0,580269629 | 3,225806452 |
| CCDC14 | 0,082679762 | 1,139873896 | 0,2 | 0,062 | 1 | CH_SC | 2,203617608 | 0,453799242 | 3,225806452 |
| ERAP1 | 0,082679762 | 0,562142037 | 0,2 | 0,062 | 1 | CH_SC | 1,47645976 | 0,677295804 | 3,225806452 |
| PHF21A | 0,082679762 | 0,657743211 | 0,2 | 0,062 | 1 | CH_SC | 1,577612853 | 0,633869075 | 3,225806452 |
| SYVN1 | 0,082679762 | 0,808469939 | 0,2 | 0,062 | 1 | CH_SC | 1,751353047 | 0,570987102 | 3,225806452 |

| Supplementary Table 4 - clusermarker | | | | | | | | | |
|---|---|---|---|---|---|---|---|---|---|
| | **p_val** | **avg_log2FC** | **pct.1** | **pct.2** | **p_val_adj** | **celltype** | **Foldchange_UP** | **Foldchange_DOWN** | **Ratio_pct1_pct2** |
| EMSY1 | 0,082679762 | 0,517383692 | 0,2 | 0,062 | 1 | CH_SC | 1,431357146 | 0,698637655 | 3,225806452 |
| CORO1C | 0,082679762 | 0,547320104 | 0,2 | 0,062 | 1 | CH_SC | 1,46136859 | 0,68429006 | 3,225806452 |
| AQR2 | 0,082679762 | 0,695375634 | 0,2 | 0,062 | 1 | CH_SC | 1,619305996 | 0,617548507 | 3,225806452 |
| FLOT21 | 0,082679762 | 0,660008621 | 0,2 | 0,062 | 1 | CH_SC | 1,580092066 | 0,632874515 | 3,225806452 |
| SHFL | 0,082679762 | 0,894874627 | 0,2 | 0,062 | 1 | CH_SC | 1,859448289 | 0,537793928 | 3,225806452 |
| MEGF8 | 0,082679762 | 0,450442073 | 0,2 | 0,062 | 1 | CH_SC | 1,366458905 | 0,731818569 | 3,225806452 |
| NECTIN24 | 0,082679762 | 0,533246604 | 0,2 | 0,062 | 1 | CH_SC | 1,447182236 | 0,690997979 | 3,225806452 |
| VGLL42 | 0,082801499 | 0,646258659 | 0,4 | 0,2 | 1 | CH_SC | 1,565104144 | 0,638935117 | 2 |
| TFDP23 | 0,083427429 | 0,542776675 | 0,333 | 0,138 | 1 | CH_SC | 1,456773592 | 0,686448468 | 2,413043478 |
| FDPS2 | 0,083711665 | 0,631917404 | 0,267 | 0,108 | 1 | CH_SC | 1,549623142 | 0,645318189 | 2,472222222 |
| TRIM282 | 0,083711665 | 0,770358254 | 0,267 | 0,108 | 1 | CH_SC | 1,705693293 | 0,586271872 | 2,472222222 |
| TCEAL71 | 0,083711665 | 1,195116426 | 0,267 | 0,108 | 1 | CH_SC | 2,289633088 | 0,436751201 | 2,472222222 |
| DDX461 | 0,08396962 | 0,430238983 | 0,4 | 0,185 | 1 | CH_SC | 1,347456765 | 0,74213884 | 2,162162162 |
| RXRB1 | 0,084153456 | 0,408580584 | 0,267 | 0,092 | 1 | CH_SC | 1,327379211 | 0,753364217 | 2,902173913 |
| CDC37L11 | 0,084153456 | 0,595345 | 0,267 | 0,092 | 1 | CH_SC | 1,510833837 | 0,661886155 | 2,902173913 |
| RSU13 | 0,084153456 | 0,253758596 | 0,267 | 0,092 | 1 | CH_SC | 1,192309348 | 0,838708513 | 2,902173913 |
| MRE11 | 0,084153456 | 0,431425108 | 0,267 | 0,092 | 1 | CH_SC | 1,348565045 | 0,741528934 | 2,902173913 |
| PLEKHJ11 | 0,084153456 | 0,402916965 | 0,267 | 0,092 | 1 | CH_SC | 1,322178504 | 0,756327528 | 2,902173913 |
| PTMS1 | 0,084330538 | 0,715198907 | 0,667 | 0,523 | 1 | CH_SC | 1,641709548 | 0,609121145 | 1,275334608 |
| OXA1L3 | 0,085392069 | -0,371357345 | 0,067 | 0,308 | 1 | CH_SC | 0,773054834 | 1,2935693 | 0,217532468 |
| DCTD | 0,085476905 | 1,075929803 | 0,333 | 0,154 | 1 | CH_SC | 2,108080271 | 0,474365238 | 2,162337662 |
| BDP1 | 0,085476905 | 1,731417861 | 0,333 | 0,154 | 1 | CH_SC | 3,320539963 | 0,301155839 | 2,162337662 |
| DIXDC1 | 0,085476905 | 0,655387306 | 0,333 | 0,154 | 1 | CH_SC | 1,575038731 | 0,634905022 | 2,162337662 |
| NUDT2 | 0,085488287 | -0,547917161 | 0,067 | 0,262 | 1 | CH_SC | 0,684006926 | 1,4619735 | 0,255725191 |
| ATF34 | 0,085721708 | 0,817483859 | 0,467 | 0,262 | 1 | CH_SC | 1,762329709 | 0,567430711 | 1,782442748 |
| PRDX47 | 0,085722984 | -0,770307634 | 0,2 | 0,4 | 1 | CH_SC | 0,586292443 | 1,705633447 | 0,5 |
| SRP91 | 0,086344373 | 0,636696077 | 0,533 | 0,308 | 1 | CH_SC | 1,554764506 | 0,643184223 | 1,730519481 |
| BEX33 | 0,086724503 | -0,527652693 | 0,4 | 0,631 | 1 | CH_SC | 0,693682458 | 1,441581791 | 0,633914422 |
| SPTSSA2 | 0,086798131 | 0,666199966 | 0,467 | 0,246 | 1 | CH_SC | 1,586887623 | 0,630164345 | 1,898373984 |
| KIDINS2202 | 0,086800498 | 1,37607903 | 0,333 | 0,138 | 1 | CH_SC | 2,595619717 | 0,385264449 | 2,413043478 |
| SMIM264 | 0,086844005 | -0,614918257 | 0,2 | 0,446 | 1 | CH_SC | 0,65296689 | 1,531471221 | 0,448430493 |

| Supplementary Table 4 - clusermarker | | | | | | | | | |
|---|---|---|---|---|---|---|---|---|---|
| | **p_val** | **avg_log2FC** | **pct.1** | **pct.2** | **p_val_adj** | **celltype** | **Foldchange_UP** | **Foldchange_DOWN** | **Ratio_pct1_pct2** |
| PAFAH1B11 | 0,087213443 | 0,715635451 | 0,4 | 0,215 | 1 | CH_SC | 1,642206386 | 0,60893686 | 1,860465116 |
| FOXD3-AS1 | 0,087229854 | 0,597833946 | 0,2 | 0,062 | 1 | CH_SC | 1,513442585 | 0,66074525 | 3,225806452 |
| NME61 | 0,087229854 | 0,893258453 | 0,2 | 0,062 | 1 | CH_SC | 1,857366415 | 0,538396728 | 3,225806452 |
| SLC33A1 | 0,087229854 | 0,461380931 | 0,2 | 0,062 | 1 | CH_SC | 1,376859101 | 0,726290729 | 3,225806452 |
| DVL3 | 0,087229854 | 0,377540909 | 0,2 | 0,062 | 1 | CH_SC | 1,299125592 | 0,769748519 | 3,225806452 |
| PPP1R351 | 0,087229854 | 0,475951291 | 0,2 | 0,062 | 1 | CH_SC | 1,390835012 | 0,718992541 | 3,225806452 |
| SHARPIN1 | 0,087229854 | 0,978461268 | 0,2 | 0,062 | 1 | CH_SC | 1,970362763 | 0,507520756 | 3,225806452 |
| SH3GLB21 | 0,087229854 | 0,462358433 | 0,2 | 0,062 | 1 | CH_SC | 1,377792312 | 0,725798795 | 3,225806452 |
| ARAP1 | 0,087229854 | 0,465983168 | 0,2 | 0,062 | 1 | CH_SC | 1,381258333 | 0,723977533 | 3,225806452 |
| NABP2 | 0,087229854 | 0,36479657 | 0,2 | 0,062 | 1 | CH_SC | 1,287700042 | 0,77657837 | 3,225806452 |
| SPATA20 | 0,087229854 | 0,519241507 | 0,2 | 0,062 | 1 | CH_SC | 1,433201549 | 0,697738571 | 3,225806452 |
| MAFG1 | 0,087229854 | 0,752267451 | 0,2 | 0,062 | 1 | CH_SC | 1,684438145 | 0,593669766 | 3,225806452 |
| TMX32 | 0,087229854 | 0,570697756 | 0,2 | 0,062 | 1 | CH_SC | 1,48524173 | 0,673291074 | 3,225806452 |
| SLC27A11 | 0,087229854 | 0,693041473 | 0,2 | 0,062 | 1 | CH_SC | 1,616688211 | 0,618548458 | 3,225806452 |
| TMEM255A | 0,087229854 | 0,624728773 | 0,2 | 0,062 | 1 | CH_SC | 1,541920916 | 0,648541692 | 3,225806452 |
| PHF20L1 | 0,087459319 | 0,607957857 | 0,267 | 0,108 | 1 | CH_SC | 1,524100309 | 0,656124793 | 2,472222222 |
| DLST | 0,087459319 | 0,54011574 | 0,267 | 0,108 | 1 | CH_SC | 1,454089166 | 0,687715735 | 2,472222222 |
| RTF22 | 0,088326145 | 0,759888214 | 0,4 | 0,231 | 1 | CH_SC | 1,693359411 | 0,590542087 | 1,731601732 |
| BCAS23 | 0,088602797 | -0,651932158 | 0,067 | 0,262 | 1 | CH_SC | 0,636427395 | 1,571271143 | 0,255725191 |
| AHI12 | 0,088602797 | -0,786862517 | 0,067 | 0,262 | 1 | CH_SC | 0,579603207 | 1,725318265 | 0,255725191 |
| AVPI1 | 0,088602797 | -0,65171213 | 0,067 | 0,262 | 1 | CH_SC | 0,636524465 | 1,571031524 | 0,255725191 |
| MFF2 | 0,088827311 | 0,472668721 | 0,4 | 0,2 | 1 | CH_SC | 1,387674038 | 0,72063033 | 2 |
| NDRG12 | 0,088827311 | 0,791370481 | 0,4 | 0,2 | 1 | CH_SC | 1,730717769 | 0,577794958 | 2 |
| TMEM1091 | 0,090024543 | 0,811253336 | 0,467 | 0,277 | 1 | CH_SC | 1,7547352 | 0,569886556 | 1,685920578 |
| SYPL13 | 0,090192209 | 0,309882323 | 0,4 | 0,185 | 1 | CH_SC | 1,239606584 | 0,806707558 | 2,162162162 |
| TLE55 | 0,09025153 | 0,955010644 | 0,4 | 0,215 | 1 | CH_SC | 1,938593936 | 0,515837784 | 1,860465116 |
| COBL2 | 0,09028282 | 0,331145038 | 0,333 | 0,138 | 1 | CH_SC | 1,258011437 | 0,794905333 | 2,413043478 |
| SELENOW5 | 0,090824539 | 0,648738319 | 0,6 | 0,492 | 1 | CH_SC | 1,56779651 | 0,637837879 | 1,219512195 |
| SAYSD1 | 0,091335889 | -0,444408693 | 0 | 0,169 | 1 | CH_SC | 0,734885457 | 1,360756279 | 0 |
| MAP3K5 | 0,091335889 | -0,407710355 | 0 | 0,169 | 1 | CH_SC | 0,753818781 | 1,326578781 | 0 |
| PKN22 | 0,091340958 | 1,115434358 | 0,4 | 0,231 | 1 | CH_SC | 2,166602301 | 0,461552173 | 1,731601732 |

| Supplementary Table 4 - clusermarker | | | | | | | | | |
|---|---|---|---|---|---|---|---|---|---|
| | **p_val** | **avg_log2FC** | **pct.1** | **pct.2** | **p_val_adj** | **celltype** | **Foldchange_UP** | **Foldchange_DOWN** | **Ratio_pct1_pct2** |
| FGGY1 | 0,091341185 | -0,581807191 | 0 | 0,169 | 1 | CH_SC | 0,668126325 | 1,496722944 | 0 |
| CYP1B15 | 0,091341185 | -0,814512413 | 0 | 0,169 | 1 | CH_SC | 0,568600624 | 1,758703663 | 0 |
| SLC1A4 | 0,091341185 | -0,585845547 | 0 | 0,169 | 1 | CH_SC | 0,666258737 | 1,500918403 | 0 |
| VAMP83 | 0,091341185 | -0,501268599 | 0 | 0,169 | 1 | CH_SC | 0,706485277 | 1,415457664 | 0 |
| ZNF385B | 0,091341185 | -0,623104079 | 0 | 0,169 | 1 | CH_SC | 0,64927246 | 1,540185456 | 0 |
| SNED1 | 0,091341185 | -0,713390158 | 0 | 0,169 | 1 | CH_SC | 0,609885298 | 1,639652577 | 0 |
| TADA33 | 0,091341185 | 0,721212954 | 0,267 | 0,108 | 1 | CH_SC | 1,648567494 | 0,606587236 | 2,472222222 |
| STATH10 | 0,091341185 | -1,082405395 | 0 | 0,169 | 1 | CH_SC | 0,472240803 | 2,117563738 | 0 |
| ADCY2 | 0,091341185 | -0,804664884 | 0 | 0,169 | 1 | CH_SC | 0,572495047 | 1,746740004 | 0 |
| DAB21 | 0,091341185 | -0,547191154 | 0 | 0,169 | 1 | CH_SC | 0,684351225 | 1,461237977 | 0 |
| ARHGEF281 | 0,091341185 | -0,550204694 | 0 | 0,169 | 1 | CH_SC | 0,682923226 | 1,46429344 | 0 |
| CXCL141 | 0,091341185 | -2,738957095 | 0 | 0,169 | 1 | CH_SC | 0,149793082 | 6,675875707 | 0 |
| SIL11 | 0,091341185 | 0,590772423 | 0,267 | 0,108 | 1 | CH_SC | 1,506052876 | 0,663987311 | 2,472222222 |
| GLO12 | 0,091341185 | -0,475246527 | 0 | 0,169 | 1 | CH_SC | 0,719343859 | 1,390155748 | 0 |
| SLC26A41 | 0,091341185 | -0,773545113 | 0 | 0,169 | 1 | CH_SC | 0,584978249 | 1,70946527 | 0 |
| UBE2V2 | 0,091341185 | -0,750829806 | 0 | 0,169 | 1 | CH_SC | 0,594261653 | 1,682760438 | 0 |
| SPTLC11 | 0,091341185 | -0,695484917 | 0 | 0,169 | 1 | CH_SC | 0,61750173 | 1,619428662 | 0 |
| NRBF2 | 0,091341185 | -0,648416165 | 0 | 0,169 | 1 | CH_SC | 0,637980324 | 1,56744646 | 0 |
| SCUBE21 | 0,091341185 | -0,641326456 | 0 | 0,169 | 1 | CH_SC | 0,64112321 | 1,559762591 | 0 |
| P3H3 | 0,091341185 | -0,465054975 | 0 | 0,169 | 1 | CH_SC | 0,724443471 | 1,380369952 | 0 |
| CHST111 | 0,091341185 | -0,623070731 | 0 | 0,169 | 1 | CH_SC | 0,649287468 | 1,540149856 | 0 |
| PRMT52 | 0,091341185 | 0,508339741 | 0,267 | 0,108 | 1 | CH_SC | 1,422412336 | 0,703031023 | 2,472222222 |
| SIPA1L13 | 0,091341185 | -0,599347731 | 0 | 0,169 | 1 | CH_SC | 0,66005231 | 1,515031438 | 0 |
| USP8 | 0,091341185 | 0,848494702 | 0,267 | 0,108 | 1 | CH_SC | 1,80062119 | 0,555363897 | 2,472222222 |
| ADAMTSL31 | 0,091341185 | -0,477877233 | 0 | 0,169 | 1 | CH_SC | 0,718033354 | 1,392692964 | 0 |
| WNK4 | 0,091341185 | -0,338659621 | 0 | 0,169 | 1 | CH_SC | 0,790775664 | 1,264581151 | 0 |
| SGTA | 0,091341185 | -0,588481627 | 0 | 0,169 | 1 | CH_SC | 0,665042467 | 1,503663375 | 0 |
| FBXO171 | 0,091341185 | -0,431209197 | 0 | 0,169 | 1 | CH_SC | 0,741639918 | 1,348363236 | 0 |
| DBP | 0,091341185 | -0,485375416 | 0 | 0,169 | 1 | CH_SC | 0,714311169 | 1,399950111 | 0 |
| GPCPD11 | 0,091341185 | -0,534341706 | 0 | 0,169 | 1 | CH_SC | 0,690473664 | 1,448281161 | 0 |
| DRG1 | 0,091341185 | -0,475389826 | 0 | 0,169 | 1 | CH_SC | 0,719272412 | 1,390293836 | 0 |

| Supplementary Table 4 - clusermarker | | | | | | | | | |
|---|---|---|---|---|---|---|---|---|---|
| | **p_val** | **avg_log2FC** | **pct.1** | **pct.2** | **p_val_adj** | **celltype** | **Foldchange_UP** | **Foldchange_DOWN** | **Ratio_pct1_pct2** |
| MID1IP1 | 0,091341185 | -0,512810641 | 0 | 0,169 | 1 | CH_SC | 0,700855708 | 1,426827217 | 0 |
| MAP4K53 | 0,091414796 | 0,602741337 | 0,467 | 0,262 | 1 | CH_SC | 1,518599394 | 0,658501514 | 1,782442748 |
| TIMP16 | 0,09146105 | 0,324310302 | 0,8 | 0,754 | 1 | CH_SC | 1,25206573 | 0,798680114 | 1,061007958 |
| TMBIM14 | 0,091804486 | -0,70698742 | 0,067 | 0,262 | 1 | CH_SC | 0,612598008 | 1,632391858 | 0,255725191 |
| PLPP52 | 0,091804486 | -0,559712067 | 0,067 | 0,262 | 1 | CH_SC | 0,678437553 | 1,473975012 | 0,255725191 |
| MAGT1 | 0,09197009 | 0,659025651 | 0,4 | 0,2 | 1 | CH_SC | 1,579015848 | 0,633305867 | 2 |
| CNST | 0,091979911 | 0,467960818 | 0,2 | 0,062 | 1 | CH_SC | 1,383153064 | 0,722985782 | 3,225806452 |
| IGFBP24 | 0,091979911 | 0,698788826 | 0,2 | 0,062 | 1 | CH_SC | 1,623141558 | 0,61608921 | 3,225806452 |
| COPS7B | 0,091979911 | 0,538354953 | 0,2 | 0,062 | 1 | CH_SC | 1,452315556 | 0,688555594 | 3,225806452 |
| KPNA11 | 0,091979911 | 0,388867353 | 0,2 | 0,062 | 1 | CH_SC | 1,309365029 | 0,763728966 | 3,225806452 |
| LEF12 | 0,091979911 | 0,555062505 | 0,2 | 0,062 | 1 | CH_SC | 1,469232287 | 0,680627569 | 3,225806452 |
| FBXO4 | 0,091979911 | 0,345942623 | 0,2 | 0,062 | 1 | CH_SC | 1,270981141 | 0,786793736 | 3,225806452 |
| CLK4 | 0,091979911 | 0,355688667 | 0,2 | 0,062 | 1 | CH_SC | 1,279596248 | 0,781496508 | 3,225806452 |
| KIF21A1 | 0,091979911 | 0,427535766 | 0,2 | 0,062 | 1 | CH_SC | 1,344934363 | 0,743530709 | 3,225806452 |
| MFSD5 | 0,091979911 | 0,419335357 | 0,2 | 0,062 | 1 | CH_SC | 1,337311319 | 0,747769039 | 3,225806452 |
| UNC45A | 0,091979911 | 0,328467998 | 0,2 | 0,062 | 1 | CH_SC | 1,255679257 | 0,796381715 | 3,225806452 |
| C16orf91 | 0,091979911 | 0,444250646 | 0,2 | 0,062 | 1 | CH_SC | 1,360607216 | 0,734965968 | 3,225806452 |
| BFAR | 0,091979911 | 0,470500892 | 0,2 | 0,062 | 1 | CH_SC | 1,385590451 | 0,721713981 | 3,225806452 |
| MED14 | 0,091979911 | 0,89696616 | 0,2 | 0,062 | 1 | CH_SC | 1,86214596 | 0,537014832 | 3,225806452 |
| STAT5A2 | 0,091979911 | 0,341773588 | 0,2 | 0,062 | 1 | CH_SC | 1,267313619 | 0,789070665 | 3,225806452 |
| DDX39A4 | 0,091979911 | 0,318264134 | 0,2 | 0,062 | 1 | CH_SC | 1,246829447 | 0,802034314 | 3,225806452 |
| UFD12 | 0,091979911 | 0,344084421 | 0,2 | 0,062 | 1 | CH_SC | 1,269345162 | 0,787807785 | 3,225806452 |
| FGL23 | 0,092148759 | -1,057992735 | 0,267 | 0,092 | 1 | CH_SC | 0,480299851 | 2,082032711 | 2,902173913 |
| VPS363 | 0,092268656 | 0,618098 | 0,333 | 0,154 | 1 | CH_SC | 1,534850352 | 0,651529316 | 2,162337662 |
| RBFOX2 | 0,092337701 | 0,445639382 | 0,533 | 0,292 | 1 | CH_SC | 1,361917565 | 0,734258832 | 1,825342466 |
| ACP6 | 0,092788888 | 0,632044488 | 0,133 | 0,031 | 1 | CH_SC | 1,549759651 | 0,645261347 | 4,290322581 |
| RPS6KC1 | 0,092788888 | 0,27944376 | 0,133 | 0,031 | 1 | CH_SC | 1,213726834 | 0,823908619 | 4,290322581 |
| STK16 | 0,092788888 | 0,400829267 | 0,133 | 0,031 | 1 | CH_SC | 1,320266588 | 0,757422788 | 4,290322581 |
| JCHAIN7 | 0,092788888 | 0,614242826 | 0,133 | 0,031 | 1 | CH_SC | 1,530754396 | 0,653272663 | 4,290322581 |
| GGH1 | 0,092788888 | 0,499716666 | 0,133 | 0,031 | 1 | CH_SC | 1,413935849 | 0,707245665 | 4,290322581 |
| VPS13B1 | 0,092788888 | 0,627526082 | 0,133 | 0,031 | 1 | CH_SC | 1,54491352 | 0,647285422 | 4,290322581 |

| Supplementary Table 4 - clusermarker | | | | | | | | | |
|---|---|---|---|---|---|---|---|---|---|
| | **p_val** | **avg_log2FC** | **pct.1** | **pct.2** | **p_val_adj** | **celltype** | **Foldchange_UP** | **Foldchange_DOWN** | **Ratio_pct1_pct2** |
| NOP21 | 0,092788888 | 0,686131158 | 0,133 | 0,031 | 1 | CH_SC | 1,608963009 | 0,621518329 | 4,290322581 |
| CAPS2 | 0,092788888 | 0,448684397 | 0,133 | 0,031 | 1 | CH_SC | 1,364795124 | 0,732710707 | 4,290322581 |
| NEDD1 | 0,092788888 | 0,554311628 | 0,133 | 0,031 | 1 | CH_SC | 1,468467797 | 0,680981907 | 4,290322581 |
| TCHP | 0,092788888 | 1,132816006 | 0,133 | 0,031 | 1 | CH_SC | 2,192863493 | 0,456024738 | 4,290322581 |
| NDUFAF11 | 0,092788888 | 0,630593627 | 0,133 | 0,031 | 1 | CH_SC | 1,548201902 | 0,645910587 | 4,290322581 |
| DMXL2 | 0,092788888 | 1,138708471 | 0,133 | 0,031 | 1 | CH_SC | 2,20183822 | 0,454165974 | 4,290322581 |
| AC093512.2 | 0,092788888 | 0,452505908 | 0,133 | 0,031 | 1 | CH_SC | 1,36841508 | 0,73077242 | 4,290322581 |
| ADAT1 | 0,092788888 | 0,392854272 | 0,133 | 0,031 | 1 | CH_SC | 1,312988493 | 0,761621298 | 4,290322581 |
| TP53 | 0,092788888 | 0,562431659 | 0,133 | 0,031 | 1 | CH_SC | 1,47675619 | 0,67715985 | 4,290322581 |
| TMEM1071 | 0,092788888 | 1,105617068 | 0,133 | 0,031 | 1 | CH_SC | 2,151908997 | 0,464703666 | 4,290322581 |
| HOXB4 | 0,092788888 | 0,684587691 | 0,133 | 0,031 | 1 | CH_SC | 1,60724258 | 0,622183616 | 4,290322581 |
| BRWD3 | 0,092788888 | 0,675527327 | 0,133 | 0,031 | 1 | CH_SC | 1,59718046 | 0,626103327 | 4,290322581 |
| FAM199X | 0,092788888 | 0,757629476 | 0,133 | 0,031 | 1 | CH_SC | 1,690710297 | 0,591467386 | 4,290322581 |
| UFC13 | 0,092905595 | -0,643095338 | 0,133 | 0,354 | 1 | CH_SC | 0,640337614 | 1,561676182 | 0,375706215 |
| ATRAID4 | 0,093376801 | 0,846125235 | 0,533 | 0,4 | 1 | CH_SC | 1,797666296 | 0,55627677 | 1,3325 |
| NDUFAB17 | 0,09433164 | -0,693279188 | 0,2 | 0,415 | 1 | CH_SC | 0,618446547 | 1,616954618 | 0,481927711 |
| NFE2L26 | 0,094371721 | 0,495640734 | 0,467 | 0,262 | 1 | CH_SC | 1,409946806 | 0,709246616 | 1,782442748 |
| CCND22 | 0,094371721 | 0,511531444 | 0,467 | 0,262 | 1 | CH_SC | 1,42556265 | 0,701477413 | 1,782442748 |
| RCN3 | 0,095098484 | -0,639472326 | 0,067 | 0,262 | 1 | CH_SC | 0,641947702 | 1,557759295 | 0,255725191 |
| TCP11 | 0,095201587 | 0,842153836 | 0,4 | 0,2 | 1 | CH_SC | 1,792724549 | 0,557810178 | 2 |
| HSPA23 | 0,095201587 | 0,604001713 | 0,4 | 0,2 | 1 | CH_SC | 1,519926661 | 0,657926481 | 2 |
| LRCH3 | 0,095360364 | 0,706314868 | 0,267 | 0,108 | 1 | CH_SC | 1,631631051 | 0,612883654 | 2,472222222 |
| SECISBP2 | 0,095360364 | 0,560101688 | 0,267 | 0,108 | 1 | CH_SC | 1,474373134 | 0,678254356 | 2,472222222 |
| FAR11 | 0,095360364 | 0,680877906 | 0,267 | 0,108 | 1 | CH_SC | 1,603114983 | 0,623785574 | 2,472222222 |
| ARHGAP13 | 0,095360364 | 0,650621867 | 0,267 | 0,108 | 1 | CH_SC | 1,569844724 | 0,637005676 | 2,472222222 |
| ATP5IF14 | 0,095456268 | -0,64059766 | 0,267 | 0,492 | 1 | CH_SC | 0,641447164 | 1,558974856 | 0,542682927 |
| KDELR15 | 0,095735516 | 0,726038015 | 0,533 | 0,338 | 1 | CH_SC | 1,654090322 | 0,604561907 | 1,576923077 |
| ATP5F1E4 | 0,095770786 | 0,759401046 | 0,867 | 0,877 | 1 | CH_SC | 1,692787696 | 0,590741534 | 0,988597491 |
| ADD14 | 0,095823534 | 0,67154299 | 0,333 | 0,154 | 1 | CH_SC | 1,59277556 | 0,627834847 | 2,162337662 |
| MED102 | 0,095823534 | 0,890351281 | 0,333 | 0,154 | 1 | CH_SC | 1,853627407 | 0,539482744 | 2,162337662 |
| H1-01 | 0,095860195 | 0,885574929 | 0,467 | 0,277 | 1 | CH_SC | 1,847500723 | 0,541271777 | 1,685920578 |

| Supplementary Table 4 - clusermarker | | | | | | | | | |
|---|---|---|---|---|---|---|---|---|---|
| | **p_val** | **avg_log2FC** | **pct.1** | **pct.2** | **p_val_adj** | **celltype** | **Foldchange_UP** | **Foldchange_DOWN** | **Ratio_pct1_pct2** |
| ARFGAP32 | 0,095860195 | 1,08493324 | 0,467 | 0,277 | 1 | CH_SC | 2,12127732 | 0,471414082 | 1,685920578 |
| KLF25 | 0,096680056 | -0,925718737 | 0,4 | 0,6 | 1 | CH_SC | 0,526418196 | 1,899630384 | 0,666666667 |
| PDE6D1 | 0,096935584 | 0,372186006 | 0,2 | 0,062 | 1 | CH_SC | 1,294312519 | 0,772610931 | 3,225806452 |
| CMTM62 | 0,096935584 | 0,251491757 | 0,2 | 0,062 | 1 | CH_SC | 1,1904374 | 0,840027372 | 3,225806452 |
| SLC35A5 | 0,096935584 | 0,342490953 | 0,2 | 0,062 | 1 | CH_SC | 1,267943933 | 0,788678406 | 3,225806452 |
| PTX3 | 0,096935584 | 0,315758275 | 0,2 | 0,062 | 1 | CH_SC | 1,244665673 | 0,803428601 | 3,225806452 |
| ECI21 | 0,096935584 | 0,446931651 | 0,2 | 0,062 | 1 | CH_SC | 1,363138026 | 0,733601426 | 3,225806452 |
| FAM102A1 | 0,096935584 | 0,578733859 | 0,2 | 0,062 | 1 | CH_SC | 1,493537912 | 0,669551132 | 3,225806452 |
| LRRC8A | 0,096935584 | 0,346766761 | 0,2 | 0,062 | 1 | CH_SC | 1,271707394 | 0,786344409 | 3,225806452 |
| XPOT1 | 0,096935584 | 0,413504329 | 0,2 | 0,062 | 1 | CH_SC | 1,331917135 | 0,750797458 | 3,225806452 |
| PIGB | 0,096935584 | 0,25823856 | 0,2 | 0,062 | 1 | CH_SC | 1,19601755 | 0,836108132 | 3,225806452 |
| POLR3H | 0,096935584 | 0,372395057 | 0,2 | 0,062 | 1 | CH_SC | 1,294500082 | 0,772498985 | 3,225806452 |
| CHD42 | 0,097112524 | 0,70447911 | 0,467 | 0,292 | 1 | CH_SC | 1,629556202 | 0,613664014 | 1,599315068 |
| TM9SF24 | 0,097453723 | -0,855151654 | 0,2 | 0,431 | 1 | CH_SC | 0,552807213 | 1,808948899 | 0,464037123 |
| NR4A16 | 0,098036853 | 0,630208198 | 0,867 | 0,708 | 1 | CH_SC | 1,547788341 | 0,646083171 | 1,224576271 |
| SNHG76 | 0,098143717 | -0,900759441 | 0,2 | 0,369 | 1 | CH_SC | 0,535604712 | 1,867048547 | 0,54200542 |
| TERF2IP2 | 0,098950337 | -0,631557219 | 0,133 | 0,338 | 1 | CH_SC | 0,64547932 | 1,549236309 | 0,393491124 |
| DVL1 | 0,099305869 | 0,678806739 | 0,133 | 0,031 | 1 | CH_SC | 1,600815165 | 0,624681738 | 4,290322581 |
| SNIP1 | 0,099305869 | 0,553315193 | 0,133 | 0,031 | 1 | CH_SC | 1,467453911 | 0,681452407 | 4,290322581 |
| CSF12 | 0,099305869 | 0,468015817 | 0,133 | 0,031 | 1 | CH_SC | 1,383205794 | 0,722958221 | 4,290322581 |
| BOLA1 | 0,099305869 | 0,427044476 | 0,133 | 0,031 | 1 | CH_SC | 1,344476442 | 0,743783951 | 4,290322581 |
| IGKV3-151 | 0,099305869 | 0,89414768 | 0,133 | 0,031 | 1 | CH_SC | 1,858511583 | 0,538064981 | 4,290322581 |
| AGPS | 0,099305869 | 0,406356763 | 0,133 | 0,031 | 1 | CH_SC | 1,325334717 | 0,754526375 | 4,290322581 |
| ITGA9-AS1 | 0,099305869 | 0,63904664 | 0,133 | 0,031 | 1 | CH_SC | 1,557299726 | 0,642137145 | 4,290322581 |
| ZNF662 | 0,099305869 | 0,587387575 | 0,133 | 0,031 | 1 | CH_SC | 1,50252352 | 0,665546986 | 4,290322581 |
| HTT1 | 0,099305869 | 0,439766059 | 0,133 | 0,031 | 1 | CH_SC | 1,356384364 | 0,737254149 | 4,290322581 |
| RAI142 | 0,099305869 | 0,402846068 | 0,133 | 0,031 | 1 | CH_SC | 1,322113532 | 0,756364696 | 4,290322581 |
| TMEM267 | 0,099305869 | 0,446738941 | 0,133 | 0,031 | 1 | CH_SC | 1,362955955 | 0,733699425 | 4,290322581 |
| PARP82 | 0,099305869 | 0,501139786 | 0,133 | 0,031 | 1 | CH_SC | 1,415331288 | 0,70654836 | 4,290322581 |
| RMND5B | 0,099305869 | 0,560938311 | 0,133 | 0,031 | 1 | CH_SC | 1,475228376 | 0,677861148 | 4,290322581 |
| VARS1 | 0,099305869 | 0,466886926 | 0,133 | 0,031 | 1 | CH_SC | 1,382123875 | 0,723524149 | 4,290322581 |

| Supplementary Table 4 - clusermarker | | | | | | | | | |
|---|---|---|---|---|---|---|---|---|---|
| | **p_val** | **avg_log2FC** | **pct.1** | **pct.2** | **p_val_adj** | **celltype** | **Foldchange_UP** | **Foldchange_DOWN** | **Ratio_pct1_pct2** |
| POU3F2 | 0,099305869 | 0,352062532 | 0,133 | 0,031 | 1 | CH_SC | 1,276384091 | 0,783463228 | 4,290322581 |
| MICAL11 | 0,099305869 | 0,529359246 | 0,133 | 0,031 | 1 | CH_SC | 1,443288036 | 0,692862391 | 4,290322581 |
| NCOA73 | 0,099305869 | 0,426581135 | 0,133 | 0,031 | 1 | CH_SC | 1,344044714 | 0,744022866 | 4,290322581 |
| NBN1 | 0,099305869 | 0,691302518 | 0,133 | 0,031 | 1 | CH_SC | 1,614740707 | 0,619294476 | 4,290322581 |
| MINDY3 | 0,099305869 | 0,376089099 | 0,133 | 0,031 | 1 | CH_SC | 1,297818916 | 0,77052352 | 4,290322581 |
| EID31 | 0,099305869 | 0,399493981 | 0,133 | 0,031 | 1 | CH_SC | 1,31904518 | 0,758124146 | 4,290322581 |
| PROSER11 | 0,099305869 | 0,354050433 | 0,133 | 0,031 | 1 | CH_SC | 1,278144043 | 0,782384431 | 4,290322581 |
| TMEM121 | 0,099305869 | 0,415980916 | 0,133 | 0,031 | 1 | CH_SC | 1,334205521 | 0,749509715 | 4,290322581 |
| SRP14-DT1 | 0,099305869 | 0,41174418 | 0,133 | 0,031 | 1 | CH_SC | 1,330293131 | 0,751714022 | 4,290322581 |
| TRAF71 | 0,099305869 | 0,58181111 | 0,133 | 0,031 | 1 | CH_SC | 1,49672701 | 0,66812451 | 4,290322581 |
| CORO7 | 0,099305869 | 0,498790785 | 0,133 | 0,031 | 1 | CH_SC | 1,413028716 | 0,707699701 | 4,290322581 |
| AC009133.1 | 0,099305869 | 0,586029136 | 0,133 | 0,031 | 1 | CH_SC | 1,501109413 | 0,666173959 | 4,290322581 |
| ZNF276 | 0,099305869 | 0,964066613 | 0,133 | 0,031 | 1 | CH_SC | 1,950800994 | 0,51260995 | 4,290322581 |
| MED111 | 0,099305869 | 0,374064754 | 0,133 | 0,031 | 1 | CH_SC | 1,295999134 | 0,771605454 | 4,290322581 |
| EPN21 | 0,099305869 | 0,470633616 | 0,133 | 0,031 | 1 | CH_SC | 1,385717927 | 0,721647588 | 4,290322581 |
| ATP9B | 0,099305869 | 0,509198601 | 0,133 | 0,031 | 1 | CH_SC | 1,423259373 | 0,702612622 | 4,290322581 |
| MED161 | 0,099305869 | 0,462800707 | 0,133 | 0,031 | 1 | CH_SC | 1,378214754 | 0,725576328 | 4,290322581 |
| ZNF414 | 0,099305869 | 0,413018919 | 0,133 | 0,031 | 1 | CH_SC | 1,331469073 | 0,751050114 | 4,290322581 |
| ZNF8441 | 0,099305869 | 0,722857276 | 0,133 | 0,031 | 1 | CH_SC | 1,650447532 | 0,605896268 | 4,290322581 |
| TSHZ3 | 0,099305869 | 0,418781408 | 0,133 | 0,031 | 1 | CH_SC | 1,336797933 | 0,748056213 | 4,290322581 |
| THAP8 | 0,099305869 | 0,381787994 | 0,133 | 0,031 | 1 | CH_SC | 1,302955664 | 0,767485823 | 4,290322581 |
| MIS18A | 0,099305869 | 0,658647682 | 0,133 | 0,031 | 1 | CH_SC | 1,578602219 | 0,633471807 | 4,290322581 |
| ZC3H7B | 0,099305869 | 0,489417722 | 0,133 | 0,031 | 1 | CH_SC | 1,40387815 | 0,712312532 | 4,290322581 |
| CHKB4 | 0,099305869 | 0,702499632 | 0,133 | 0,031 | 1 | CH_SC | 1,62732187 | 0,614506582 | 4,290322581 |
| NLGN3 | 0,099305869 | 0,585962761 | 0,133 | 0,031 | 1 | CH_SC | 1,501040352 | 0,666204608 | 4,290322581 |
| Z83843.1 | 0,099305869 | 0,563574379 | 0,133 | 0,031 | 1 | CH_SC | 1,477926352 | 0,676623702 | 4,290322581 |
| PLXNA3 | 0,099305869 | 0,421728001 | 0,133 | 0,031 | 1 | CH_SC | 1,33953103 | 0,746529926 | 4,290322581 |
| MED29 | 0,099487217 | 0,609197734 | 0,333 | 0,154 | 1 | CH_SC | 1,52541071 | 0,655561151 | 2,162337662 |
| TPP11 | 0,099519951 | 0,531831574 | 0,267 | 0,108 | 1 | CH_SC | 1,445763501 | 0,691676059 | 2,472222222 |
| SPIRE11 | 0,099519951 | 0,582376994 | 0,267 | 0,108 | 1 | CH_SC | 1,497314202 | 0,667862496 | 2,472222222 |
| NKAP2 | 0,099519951 | 0,628367636 | 0,267 | 0,108 | 1 | CH_SC | 1,545814963 | 0,646907957 | 2,472222222 |

| Supplementary Table 4 - clusermarker | | | | | | | | | |
|---|---|---|---|---|---|---|---|---|---|
| | **p_val** | **avg_log2FC** | **pct.1** | **pct.2** | **p_val_adj** | **celltype** | **Foldchange_UP** | **Foldchange_DOWN** | **Ratio_pct1_pct2** |
| ATP5MG2 | 0,101010338 | -0,263493177 | 0,467 | 0,738 | 1 | CH_SC | 0,833068379 | 1,200381655 | 0,632791328 |
| DUSP26 | 0,101611423 | -1,12344176 | 0,067 | 0,246 | 1 | CH_SC | 0,458997512 | 2,178661045 | 0,272357724 |
| TCTN1 | 0,101611423 | -0,694845281 | 0,067 | 0,246 | 1 | CH_SC | 0,617775567 | 1,618710829 | 0,272357724 |
| CTBS1 | 0,101970196 | -0,681847486 | 0,067 | 0,262 | 1 | CH_SC | 0,623366493 | 1,604192737 | 0,255725191 |
| ZNF362 | 0,102102505 | 0,434483913 | 0,2 | 0,062 | 1 | CH_SC | 1,351427308 | 0,739958408 | 3,225806452 |
| EFHC1 | 0,102102505 | 0,329360158 | 0,2 | 0,062 | 1 | CH_SC | 1,256456006 | 0,795889386 | 3,225806452 |
| FNBP11 | 0,102102505 | 0,288206906 | 0,2 | 0,062 | 1 | CH_SC | 1,221121629 | 0,818919243 | 3,225806452 |
| NEIL1 | 0,102102505 | 0,355498011 | 0,2 | 0,062 | 1 | CH_SC | 1,279427157 | 0,781599792 | 3,225806452 |
| ISYNA11 | 0,102102505 | 0,379698732 | 0,2 | 0,062 | 1 | CH_SC | 1,301070133 | 0,768598075 | 3,225806452 |
| DMAC21 | 0,102102505 | 0,285431922 | 0,2 | 0,062 | 1 | CH_SC | 1,218775092 | 0,820495928 | 3,225806452 |
| TSC22D41 | 0,103261799 | 0,669195466 | 0,333 | 0,154 | 1 | CH_SC | 1,590185936 | 0,628857279 | 2,162337662 |
| IFRD12 | 0,103264867 | 1,250994647 | 0,4 | 0,215 | 1 | CH_SC | 2,380054562 | 0,420158435 | 1,860465116 |
| MAP7D11 | 0,103823025 | 0,460498607 | 0,267 | 0,108 | 1 | CH_SC | 1,376017299 | 0,72673505 | 2,472222222 |
| ANTXR12 | 0,103823025 | 0,829840025 | 0,267 | 0,108 | 1 | CH_SC | 1,777488252 | 0,562591623 | 2,472222222 |
| PARL1 | 0,103823025 | 0,563005939 | 0,267 | 0,108 | 1 | CH_SC | 1,477344146 | 0,676890353 | 2,472222222 |
| CDC402 | 0,103823025 | 0,616265651 | 0,267 | 0,108 | 1 | CH_SC | 1,532902195 | 0,652357341 | 2,472222222 |
| ST6GALNAC61 | 0,103823025 | 0,734592736 | 0,267 | 0,108 | 1 | CH_SC | 1,663927686 | 0,600987656 | 2,472222222 |
| UPF3A1 | 0,103823025 | 0,36625274 | 0,267 | 0,108 | 1 | CH_SC | 1,289000426 | 0,775794934 | 2,472222222 |
| HUWE12 | 0,103823025 | 1,239770761 | 0,267 | 0,108 | 1 | CH_SC | 2,361610041 | 0,423439934 | 2,472222222 |
| SPEN2 | 0,104236251 | 0,656328785 | 0,4 | 0,231 | 1 | CH_SC | 1,576066912 | 0,634490828 | 1,731601732 |
| RWDD13 | 0,104846352 | -0,998150464 | 0,267 | 0,431 | 1 | CH_SC | 0,500641411 | 1,997437642 | 0,619489559 |
| ITGAV3 | 0,105239692 | 0,628064788 | 0,467 | 0,246 | 1 | CH_SC | 1,545490502 | 0,647043769 | 1,898373984 |
| ZFYVE163 | 0,10526265 | -0,699098836 | 0,067 | 0,246 | 1 | CH_SC | 0,615956837 | 1,623490381 | 0,272357724 |
| MYC1 | 0,10526265 | -0,839855804 | 0,067 | 0,246 | 1 | CH_SC | 0,558699408 | 1,789871237 | 0,272357724 |
| FRY1 | 0,10526265 | -0,590483914 | 0,067 | 0,246 | 1 | CH_SC | 0,664120108 | 1,505751727 | 0,272357724 |
| NBEAL14 | 0,105350426 | 0,323585686 | 0,333 | 0,138 | 1 | CH_SC | 1,251437019 | 0,799081364 | 2,413043478 |
| ADGRG6 | 0,105444334 | 0,518341753 | 0,4 | 0,2 | 1 | CH_SC | 1,432307994 | 0,69817386 | 2 |
| DNAJC6 | 0,106183001 | 0,464200744 | 0,133 | 0,031 | 1 | CH_SC | 1,379552867 | 0,724872547 | 4,290322581 |
| HS2ST11 | 0,106183001 | 0,455211532 | 0,133 | 0,031 | 1 | CH_SC | 1,370983807 | 0,729403217 | 4,290322581 |
| DENND4B | 0,106183001 | 0,274225682 | 0,133 | 0,031 | 1 | CH_SC | 1,20934484 | 0,826894006 | 4,290322581 |
| GPR137B1 | 0,106183001 | 0,412343043 | 0,133 | 0,031 | 1 | CH_SC | 1,330845451 | 0,75140205 | 4,290322581 |

| Supplementary Table 4 - clusermarker | | | | | | | | | |
|---|---|---|---|---|---|---|---|---|---|
| | **p_val** | **avg_log2FC** | **pct.1** | **pct.2** | **p_val_adj** | **celltype** | **Foldchange_UP** | **Foldchange_DOWN** | **Ratio_pct1_pct2** |
| HS1BP31 | 0,106183001 | 0,944203626 | 0,133 | 0,031 | 1 | CH_SC | 1,924126466 | 0,519716358 | 4,290322581 |
| SELENOI | 0,106183001 | 0,430623193 | 0,133 | 0,031 | 1 | CH_SC | 1,34781566 | 0,741941224 | 4,290322581 |
| TRMT61B | 0,106183001 | 0,289629144 | 0,133 | 0,031 | 1 | CH_SC | 1,222326029 | 0,818112334 | 4,290322581 |
| PRKCE | 0,106183001 | 0,900359439 | 0,133 | 0,031 | 1 | CH_SC | 1,86653096 | 0,535753235 | 4,290322581 |
| IGKV3-208 | 0,106183001 | 1,224212392 | 0,133 | 0,031 | 1 | CH_SC | 2,336278708 | 0,428031123 | 4,290322581 |
| LIPT1 | 0,106183001 | 0,337735779 | 0,133 | 0,031 | 1 | CH_SC | 1,263771625 | 0,791282206 | 4,290322581 |
| TWF24 | 0,106183001 | 0,267646415 | 0,133 | 0,031 | 1 | CH_SC | 1,2038423 | 0,830673586 | 4,290322581 |
| KCTD6 | 0,106183001 | 0,37652943 | 0,133 | 0,031 | 1 | CH_SC | 1,298215089 | 0,770288382 | 4,290322581 |
| CTBP1-DT | 0,106183001 | 0,300224645 | 0,133 | 0,031 | 1 | CH_SC | 1,231336132 | 0,812125929 | 4,290322581 |
| CKMT2-AS1 | 0,106183001 | 0,496899129 | 0,133 | 0,031 | 1 | CH_SC | 1,411177173 | 0,708628243 | 4,290322581 |
| IGIP | 0,106183001 | 0,351584596 | 0,133 | 0,031 | 1 | CH_SC | 1,275961321 | 0,783722816 | 4,290322581 |
| ARHGAP26 | 0,106183001 | 0,65815989 | 0,133 | 0,031 | 1 | CH_SC | 1,578068565 | 0,633686028 | 4,290322581 |
| SERPINB91 | 0,106183001 | 0,517179916 | 0,133 | 0,031 | 1 | CH_SC | 1,431154986 | 0,698736342 | 4,290322581 |
| ZSCAN26 | 0,106183001 | 0,642417734 | 0,133 | 0,031 | 1 | CH_SC | 1,560942867 | 0,640638438 | 4,290322581 |
| GRIK2 | 0,106183001 | 0,839969139 | 0,133 | 0,031 | 1 | CH_SC | 1,790011851 | 0,558655519 | 4,290322581 |
| AK91 | 0,106183001 | 0,339931359 | 0,133 | 0,031 | 1 | CH_SC | 1,265696373 | 0,790078901 | 4,290322581 |
| MAD1L1 | 0,106183001 | 0,343364216 | 0,133 | 0,031 | 1 | CH_SC | 1,268711652 | 0,788201164 | 4,290322581 |
| POMZP3 | 0,106183001 | 0,323577648 | 0,133 | 0,031 | 1 | CH_SC | 1,251430046 | 0,799085816 | 4,290322581 |
| BMP12 | 0,106183001 | 0,38088252 | 0,133 | 0,031 | 1 | CH_SC | 1,302138151 | 0,767967669 | 4,290322581 |
| AC090204.1 | 0,106183001 | 0,406135159 | 0,133 | 0,031 | 1 | CH_SC | 1,325131156 | 0,754642282 | 4,290322581 |
| TMEM671 | 0,106183001 | 0,409005993 | 0,133 | 0,031 | 1 | CH_SC | 1,327770674 | 0,753142105 | 4,290322581 |
| SIGMAR11 | 0,106183001 | 0,365138495 | 0,133 | 0,031 | 1 | CH_SC | 1,288005269 | 0,776394339 | 4,290322581 |
| AL158152.1 | 0,106183001 | 0,312369441 | 0,133 | 0,031 | 1 | CH_SC | 1,241745433 | 0,805318041 | 4,290322581 |
| NACC21 | 0,106183001 | 0,301319955 | 0,133 | 0,031 | 1 | CH_SC | 1,232271331 | 0,811509588 | 4,290322581 |
| CUBN | 0,106183001 | 0,410344028 | 0,133 | 0,031 | 1 | CH_SC | 1,329002693 | 0,752443923 | 4,290322581 |
| ELP4 | 0,106183001 | 0,948178036 | 0,133 | 0,031 | 1 | CH_SC | 1,929434457 | 0,518286587 | 4,290322581 |
| VPS37C1 | 0,106183001 | 0,701495751 | 0,133 | 0,031 | 1 | CH_SC | 1,626189913 | 0,614934327 | 4,290322581 |
| TSPAN91 | 0,106183001 | 0,410153223 | 0,133 | 0,031 | 1 | CH_SC | 1,328826936 | 0,752543445 | 4,290322581 |
| KPNA31 | 0,106183001 | 0,662801648 | 0,133 | 0,031 | 1 | CH_SC | 1,583154053 | 0,631650469 | 4,290322581 |
| TTLL5 | 0,106183001 | 0,503694058 | 0,133 | 0,031 | 1 | CH_SC | 1,417839332 | 0,705298532 | 4,290322581 |
| PSME3IP11 | 0,106183001 | 0,383220354 | 0,133 | 0,031 | 1 | CH_SC | 1,304249928 | 0,766724213 | 4,290322581 |

| Supplementary Table 4 - clusermarker | | | | | | | | | |
|---|---|---|---|---|---|---|---|---|---|
| | **p_val** | **avg_log2FC** | **pct.1** | **pct.2** | **p_val_adj** | **celltype** | **Foldchange_UP** | **Foldchange_DOWN** | **Ratio_pct1_pct2** |
| KIF1C2 | 0,106183001 | 0,337694439 | 0,133 | 0,031 | 1 | CH_SC | 1,263735412 | 0,791304881 | 4,290322581 |
| SLFN12 | 0,106183001 | 0,387602479 | 0,133 | 0,031 | 1 | CH_SC | 1,308217555 | 0,764398854 | 4,290322581 |
| STAT5B1 | 0,106183001 | 0,440791252 | 0,133 | 0,031 | 1 | CH_SC | 1,357348567 | 0,736730435 | 4,290322581 |
| AC005332.4 | 0,106183001 | 0,406805594 | 0,133 | 0,031 | 1 | CH_SC | 1,325747101 | 0,754291674 | 4,290322581 |
| RHBDF2 | 0,106183001 | 0,283304955 | 0,133 | 0,031 | 1 | CH_SC | 1,216979575 | 0,821706478 | 4,290322581 |
| FASN | 0,106183001 | 0,39358125 | 0,133 | 0,031 | 1 | CH_SC | 1,313650277 | 0,761237612 | 4,290322581 |
| FLT3LG | 0,106183001 | 0,338524273 | 0,133 | 0,031 | 1 | CH_SC | 1,264462519 | 0,790849855 | 4,290322581 |
| ZNF8081 | 0,106183001 | 0,437437405 | 0,133 | 0,031 | 1 | CH_SC | 1,35419679 | 0,738445112 | 4,290322581 |
| RBL1 | 0,106183001 | 0,381176812 | 0,133 | 0,031 | 1 | CH_SC | 1,302403798 | 0,767811029 | 4,290322581 |
| SRC | 0,106183001 | 0,350407376 | 0,133 | 0,031 | 1 | CH_SC | 1,274920578 | 0,784362585 | 4,290322581 |
| PES1 | 0,106183001 | 0,444338106 | 0,133 | 0,031 | 1 | CH_SC | 1,360689702 | 0,734921414 | 4,290322581 |
| TYMP8 | 0,106183001 | 0,278908792 | 0,133 | 0,031 | 1 | CH_SC | 1,213276854 | 0,82421419 | 4,290322581 |
| CSTF2 | 0,106183001 | 0,574920087 | 0,133 | 0,031 | 1 | CH_SC | 1,48959495 | 0,671323436 | 4,290322581 |
| RENBP | 0,106183001 | 0,257350713 | 0,133 | 0,031 | 1 | CH_SC | 1,195281737 | 0,836622839 | 4,290322581 |
| SNRNP2002 | 0,107149362 | 0,421117029 | 0,333 | 0,154 | 1 | CH_SC | 1,338963867 | 0,746846143 | 2,162337662 |
| FEZ11 | 0,107149362 | 0,400434594 | 0,333 | 0,154 | 1 | CH_SC | 1,319905456 | 0,757630022 | 2,162337662 |
| CREB3L2 | 0,107395484 | 0,701254129 | 0,467 | 0,308 | 1 | CH_SC | 1,625917583 | 0,615037325 | 1,516233766 |
| CASD1 | 0,107486274 | 0,259466885 | 0,2 | 0,062 | 1 | CH_SC | 1,197036286 | 0,835396564 | 3,225806452 |
| PRXL2C3 | 0,107486274 | 0,291914385 | 0,2 | 0,062 | 1 | CH_SC | 1,224263738 | 0,816817463 | 3,225806452 |
| SBDS3 | 0,107534461 | 0,590065649 | 0,6 | 0,431 | 1 | CH_SC | 1,505315245 | 0,664312677 | 1,392111369 |
| RBMS11 | 0,108284182 | 0,774974205 | 0,4 | 0,246 | 1 | CH_SC | 1,711159456 | 0,584399073 | 1,62601626 |
| PRRC2B6 | 0,10834565 | 0,777191779 | 0,333 | 0,169 | 1 | CH_SC | 1,71379171 | 0,58350148 | 1,970414201 |
| EPB41L4A-AS11 | 0,108437914 | -0,631754782 | 0,133 | 0,354 | 1 | CH_SC | 0,645390935 | 1,549448476 | 0,375706215 |
| SPTY2D12 | 0,109017231 | -0,651395665 | 0,067 | 0,246 | 1 | CH_SC | 0,636664106 | 1,570686945 | 0,272357724 |
| H2AX3 | 0,109017231 | -0,685099777 | 0,067 | 0,246 | 1 | CH_SC | 0,62196281 | 1,607813174 | 0,272357724 |
| MRFAP1 | 0,109366973 | 0,852012846 | 0,667 | 0,462 | 1 | CH_SC | 1,805017527 | 0,554011241 | 1,443722944 |
| PRKCZ | 0,109964501 | -0,460322848 | 0 | 0,154 | 1 | CH_SC | 0,726823591 | 1,375849673 | 0 |
| KAZN | 0,109964501 | -0,660814211 | 0 | 0,154 | 1 | CH_SC | 0,632521221 | 1,580974624 | 0 |
| MFAP21 | 0,109964501 | -0,488074355 | 0 | 0,154 | 1 | CH_SC | 0,712976112 | 1,402571536 | 0 |
| EXTL1 | 0,109964501 | -0,560161677 | 0 | 0,154 | 1 | CH_SC | 0,678226154 | 1,474434442 | 0 |
| SLC19A21 | 0,109964501 | -0,522612299 | 0 | 0,154 | 1 | CH_SC | 0,696110239 | 1,436554074 | 0 |

| Supplementary Table 4 - clusermarker | | | | | | | | | |
|---|---|---|---|---|---|---|---|---|---|
| | **p_val** | **avg_log2FC** | **pct.1** | **pct.2** | **p_val_adj** | **celltype** | **Foldchange_UP** | **Foldchange_DOWN** | **Ratio_pct1_pct2** |
| MDM41 | 0,109964501 | -0,480536446 | 0 | 0,154 | 1 | CH_SC | 0,716711076 | 1,395262379 | 0 |
| FAM98A | 0,109964501 | -0,393782167 | 0 | 0,154 | 1 | CH_SC | 0,761131605 | 1,313833236 | 0 |
| EFEMP13 | 0,109964501 | -0,554117048 | 0 | 0,154 | 1 | CH_SC | 0,681073758 | 1,468269755 | 0 |
| AFF3 | 0,109964501 | -0,455021762 | 0 | 0,154 | 1 | CH_SC | 0,729499168 | 1,370803482 | 0 |
| FBLN7 | 0,109964501 | -0,560127276 | 0 | 0,154 | 1 | CH_SC | 0,678242326 | 1,474399285 | 0 |
| GYPC2 | 0,109964501 | -0,649711411 | 0 | 0,154 | 1 | CH_SC | 0,637407805 | 1,568854339 | 0 |
| COL6A32 | 0,109964501 | -0,68278577 | 0 | 0,154 | 1 | CH_SC | 0,622961206 | 1,605236393 | 0 |
| NT5DC21 | 0,109964501 | -0,658294074 | 0 | 0,154 | 1 | CH_SC | 0,633627092 | 1,578215347 | 0 |
| SENP7 | 0,109964501 | -0,425631696 | 0 | 0,154 | 1 | CH_SC | 0,744512669 | 1,343160488 | 0 |
| AASDH1 | 0,109964501 | -0,638834673 | 0 | 0,154 | 1 | CH_SC | 0,642231497 | 1,557070938 | 0 |
| GRSF12 | 0,109964501 | -0,446209007 | 0 | 0,154 | 1 | CH_SC | 0,733968978 | 1,362455403 | 0 |
| MAPK101 | 0,109964501 | -0,512912437 | 0 | 0,154 | 1 | CH_SC | 0,700806258 | 1,426927897 | 0 |
| UNC5C | 0,109964501 | -0,603031619 | 0 | 0,154 | 1 | CH_SC | 0,658369032 | 1,518904979 | 0 |
| GAR11 | 0,109964501 | -0,264610704 | 0 | 0,154 | 1 | CH_SC | 0,832423325 | 1,201311844 | 0 |
| SERINC51 | 0,109964501 | -0,513420955 | 0 | 0,154 | 1 | CH_SC | 0,700559283 | 1,427430947 | 0 |
| AC008771.11 | 0,109964501 | -0,460371479 | 0 | 0,154 | 1 | CH_SC | 0,726799091 | 1,375896052 | 0 |
| SGCD | 0,109964501 | -0,613263655 | 0 | 0,154 | 1 | CH_SC | 0,653716196 | 1,52971581 | 0 |
| STC2 | 0,109964501 | -0,497202959 | 0 | 0,154 | 1 | CH_SC | 0,708479022 | 1,411474396 | 0 |
| PSMG41 | 0,109964501 | -0,327094828 | 0 | 0,154 | 1 | CH_SC | 0,797140079 | 1,254484659 | 0 |
| CCNC | 0,109964501 | -0,386499678 | 0 | 0,154 | 1 | CH_SC | 0,764983387 | 1,307217931 | 0 |
| AFDN5 | 0,109964501 | -0,45986089 | 0 | 0,154 | 1 | CH_SC | 0,727056361 | 1,37540919 | 0 |
| FZD9 | 0,109964501 | -0,482937717 | 0 | 0,154 | 1 | CH_SC | 0,715519149 | 1,397586635 | 0 |
| TMEM2432 | 0,109964501 | -0,427821036 | 0 | 0,154 | 1 | CH_SC | 0,743383702 | 1,345200328 | 0 |
| FDFT11 | 0,109964501 | -0,458088625 | 0 | 0,154 | 1 | CH_SC | 0,727950055 | 1,373720619 | 0 |
| R3HCC1 | 0,109964501 | -0,429264603 | 0 | 0,154 | 1 | CH_SC | 0,742640241 | 1,346547015 | 0 |
| SLC25A372 | 0,109964501 | -0,392464916 | 0 | 0,154 | 1 | CH_SC | 0,761826873 | 1,31263419 | 0 |
| C8orf82 | 0,109964501 | -0,50950687 | 0 | 0,154 | 1 | CH_SC | 0,702462507 | 1,423563522 | 0 |
| COQ41 | 0,109964501 | -0,444963727 | 0 | 0,154 | 1 | CH_SC | 0,734602786 | 1,36127989 | 0 |
| ZNF37A1 | 0,109964501 | -0,418782227 | 0 | 0,154 | 1 | CH_SC | 0,748055789 | 1,336798692 | 0 |
| DDX50 | 0,109964501 | -0,393683896 | 0 | 0,154 | 1 | CH_SC | 0,761183453 | 1,313743746 | 0 |
| BLOC1S21 | 0,109964501 | -0,46659897 | 0 | 0,154 | 1 | CH_SC | 0,723668575 | 1,381848037 | 0 |

| Supplementary Table 4 - clusermarker | | | | | | | | | |
|---|---|---|---|---|---|---|---|---|---|
| | **p_val** | **avg_log2FC** | **pct.1** | **pct.2** | **p_val_adj** | **celltype** | **Foldchange_UP** | **Foldchange_DOWN** | **Ratio_pct1_pct2** |
| NAALADL1 | 0,109964501 | -0,432883541 | 0 | 0,154 | 1 | CH_SC | 0,740779695 | 1,34992901 | 0 |
| BHLHE413 | 0,109964501 | -0,566475007 | 0 | 0,154 | 1 | CH_SC | 0,675264675 | 1,480900804 | 0 |
| STK38L1 | 0,109964501 | -0,766515512 | 0 | 0,154 | 1 | CH_SC | 0,587835539 | 1,701156079 | 0 |
| HSPB82 | 0,109964501 | -0,409679731 | 0 | 0,154 | 1 | CH_SC | 0,75279047 | 1,328390887 | 0 |
| SCARB1 | 0,109964501 | -0,369140752 | 0 | 0,154 | 1 | CH_SC | 0,774243488 | 1,291583353 | 0 |
| ARMH41 | 0,109964501 | -0,430800417 | 0 | 0,154 | 1 | CH_SC | 0,741850088 | 1,347981238 | 0 |
| FOXF11 | 0,109964501 | -0,455710443 | 0 | 0,154 | 1 | CH_SC | 0,72915102 | 1,371458001 | 0 |
| CYB5D2 | 0,109964501 | -0,419027893 | 0 | 0,154 | 1 | CH_SC | 0,747928419 | 1,337026345 | 0 |
| SOX9-AS1 | 0,109964501 | -0,678191708 | 0 | 0,154 | 1 | CH_SC | 0,624948101 | 1,600132871 | 0 |
| DLGAP1-AS11 | 0,109964501 | -0,420998406 | 0 | 0,154 | 1 | CH_SC | 0,746907554 | 1,338853778 | 0 |
| GNAL | 0,109964501 | -0,442047201 | 0 | 0,154 | 1 | CH_SC | 0,736089348 | 1,358530731 | 0 |
| XAB2 | 0,109964501 | -0,401811348 | 0 | 0,154 | 1 | CH_SC | 0,756907366 | 1,321165634 | 0 |
| TRAPPC6A1 | 0,109964501 | -0,428702158 | 0 | 0,154 | 1 | CH_SC | 0,742929821 | 1,346022157 | 0 |
| CD401 | 0,109964501 | -0,561306693 | 0 | 0,154 | 1 | CH_SC | 0,677688083 | 1,475605113 | 0 |
| ERG | 0,109964501 | -0,533540915 | 0 | 0,154 | 1 | CH_SC | 0,690857029 | 1,447477493 | 0 |
| JOSD12 | 0,109964501 | -0,362450011 | 0 | 0,154 | 1 | CH_SC | 0,777842511 | 1,285607286 | 0 |
| KLHL151 | 0,109964501 | -0,453561932 | 0 | 0,154 | 1 | CH_SC | 0,730237705 | 1,369417099 | 0 |
| STK26 | 0,109964501 | -0,583035937 | 0 | 0,154 | 1 | CH_SC | 0,667557523 | 1,497998249 | 0 |
| XPC | 0,11018973 | 0,992786429 | 0,267 | 0,123 | 1 | CH_SC | 1,990024827 | 0,502506294 | 2,170731707 |
| SNRPN2 | 0,11018973 | 0,740763618 | 0,267 | 0,123 | 1 | CH_SC | 1,671060096 | 0,598422524 | 2,170731707 |
| ALKBH76 | 0,111770485 | -0,995879637 | 0,133 | 0,308 | 1 | CH_SC | 0,50143005 | 1,994296113 | 0,431818182 |
| LRP102 | 0,112029769 | -0,569235256 | 0,133 | 0,338 | 1 | CH_SC | 0,673973954 | 1,483736863 | 0,393491124 |
| SLC2A8 | 0,112647068 | 0,873288422 | 0,2 | 0,077 | 1 | CH_SC | 1,831833554 | 0,545901126 | 2,597402597 |
| PDK2 | 0,112647068 | 1,321361128 | 0,2 | 0,077 | 1 | CH_SC | 2,499017714 | 0,400157228 | 2,597402597 |
| CRELD11 | 0,112871867 | 0,839324468 | 0,267 | 0,108 | 1 | CH_SC | 1,789212159 | 0,558905211 | 2,472222222 |
| MAP7D3 | 0,112871867 | 0,699464725 | 0,267 | 0,108 | 1 | CH_SC | 1,623902173 | 0,615800642 | 2,472222222 |
| FKBP53 | 0,112876946 | -0,587070627 | 0,067 | 0,246 | 1 | CH_SC | 0,665693217 | 1,502193465 | 0,272357724 |
| TMEM2341 | 0,113432461 | 0,342080464 | 0,133 | 0,031 | 1 | CH_SC | 1,267583217 | 0,78890284 | 4,290322581 |
| GAP431 | 0,113432461 | 0,473557078 | 0,133 | 0,031 | 1 | CH_SC | 1,388528778 | 0,72018673 | 4,290322581 |
| H4C8 | 0,113432461 | 0,299026429 | 0,133 | 0,031 | 1 | CH_SC | 1,230313883 | 0,812800712 | 4,290322581 |
| SUPT3H | 0,113432461 | 0,275114603 | 0,133 | 0,031 | 1 | CH_SC | 1,210090211 | 0,82638467 | 4,290322581 |

| Supplementary Table 4 - clusermarker | | | | | | | | | |
|---|---|---|---|---|---|---|---|---|---|
| | **p_val** | **avg_log2FC** | **pct.1** | **pct.2** | **p_val_adj** | **celltype** | **Foldchange_UP** | **Foldchange_DOWN** | **Ratio_pct1_pct2** |
| SUPV3L1 | 0,113432461 | 0,266274561 | 0,133 | 0,031 | 1 | CH_SC | 1,202698114 | 0,831463846 | 4,290322581 |
| FUNDC22 | 0,113918658 | 0,542845461 | 0,533 | 0,338 | 1 | CH_SC | 1,456843051 | 0,686415739 | 1,576923077 |
| SELENOM8 | 0,113957603 | 0,577720853 | 0,8 | 0,769 | 1 | CH_SC | 1,492489573 | 0,670021431 | 1,040312094 |
| ACTG14 | 0,114565873 | 0,267212717 | 0,933 | 0,938 | 1 | CH_SC | 1,203480459 | 0,830923338 | 0,99466951 |
| DENND5A2 | 0,114680937 | 0,818856379 | 0,267 | 0,123 | 1 | CH_SC | 1,764007113 | 0,566891138 | 2,170731707 |
| PHLDA13 | 0,11478329 | 0,684404468 | 0,6 | 0,4 | 1 | CH_SC | 1,607038473 | 0,622262638 | 1,5 |
| CRIM11 | 0,115198919 | -0,471693655 | 0,133 | 0,354 | 1 | CH_SC | 0,721117543 | 1,386736475 | 0,375706215 |
| ITCH | 0,115271695 | 0,434605764 | 0,333 | 0,154 | 1 | CH_SC | 1,351541456 | 0,739895913 | 2,162337662 |
| CAMTA14 | 0,115561273 | 1,063882972 | 0,4 | 0,262 | 1 | CH_SC | 2,090550613 | 0,478342879 | 1,526717557 |
| SUN11 | 0,116325791 | 0,597842418 | 0,333 | 0,169 | 1 | CH_SC | 1,513451474 | 0,66074137 | 1,970414201 |
| PUF602 | 0,116325791 | 0,522576968 | 0,333 | 0,169 | 1 | CH_SC | 1,436518894 | 0,696127287 | 1,970414201 |
| RAB102 | 0,116843562 | 0,923099964 | 0,333 | 0,185 | 1 | CH_SC | 1,896185313 | 0,527374615 | 1,8 |
| TUG11 | 0,116843562 | 1,180721465 | 0,333 | 0,185 | 1 | CH_SC | 2,266901123 | 0,441130842 | 1,8 |
| UBA524 | 0,11728813 | -0,402479458 | 0,867 | 0,908 | 1 | CH_SC | 0,756556924 | 1,321777606 | 0,954845815 |
| PXK | 0,11762369 | 0,553485199 | 0,267 | 0,108 | 1 | CH_SC | 1,467626845 | 0,68137211 | 2,472222222 |
| SMC1A | 0,117727738 | 0,727891455 | 0,4 | 0,215 | 1 | CH_SC | 1,656216708 | 0,603785721 | 1,860465116 |
| PLD34 | 0,118210692 | -0,562567775 | 0,467 | 0,692 | 1 | CH_SC | 0,677095964 | 1,476895526 | 0,674855491 |
| LAMTOR45 | 0,118423324 | -0,490250244 | 0,2 | 0,477 | 1 | CH_SC | 0,711901604 | 1,404688506 | 0,419287212 |
| MT-ND12 | 0,118560793 | 0,434324562 | 0,933 | 0,862 | 1 | CH_SC | 1,351278046 | 0,740040144 | 1,082366589 |
| SCP22 | 0,118701548 | 0,59231929 | 0,467 | 0,277 | 1 | CH_SC | 1,507668542 | 0,663275761 | 1,685920578 |
| RBL23 | 0,119062292 | -0,66703328 | 0,133 | 0,338 | 1 | CH_SC | 0,629800462 | 1,587804488 | 0,393491124 |
| SVBP | 0,119314947 | 0,860801663 | 0,267 | 0,123 | 1 | CH_SC | 1,816047154 | 0,550646495 | 2,170731707 |
| RBM222 | 0,119314947 | 0,645248383 | 0,267 | 0,123 | 1 | CH_SC | 1,564008532 | 0,639382701 | 2,170731707 |
| BZW14 | 0,119772605 | 0,584368523 | 0,6 | 0,431 | 1 | CH_SC | 1,499382556 | 0,666941199 | 1,392111369 |
| HCFC1R13 | 0,120918827 | 0,810102241 | 0,333 | 0,185 | 1 | CH_SC | 1,753335693 | 0,570341438 | 1,8 |
| CNBP6 | 0,121244847 | -0,625312705 | 0,4 | 0,662 | 1 | CH_SC | 0,648279247 | 1,542545137 | 0,604229607 |
| OAZ15 | 0,121871382 | 0,524785699 | 0,733 | 0,708 | 1 | CH_SC | 1,438719854 | 0,695062348 | 1,035310734 |
| GTF2A24 | 0,122705983 | -0,605239279 | 0,133 | 0,338 | 1 | CH_SC | 0,657362344 | 1,521231037 | 0,393491124 |
| MRPL402 | 0,122762392 | -0,58533348 | 0,133 | 0,308 | 1 | CH_SC | 0,66649526 | 1,500385765 | 0,431818182 |
| CAB39L1 | 0,122873419 | 0,627988965 | 0,467 | 0,246 | 1 | CH_SC | 1,545409279 | 0,647077776 | 1,898373984 |
| TPM48 | 0,123434855 | 0,551415012 | 0,467 | 0,292 | 1 | CH_SC | 1,465522392 | 0,682350543 | 1,599315068 |

| Supplementary Table 4 - clusermarker | | | | | | | | | |
|---|---|---|---|---|---|---|---|---|---|
| | **p_val** | **avg_log2FC** | **pct.1** | **pct.2** | **p_val_adj** | **celltype** | **Foldchange_UP** | **Foldchange_DOWN** | **Ratio_pct1_pct2** |
| UBR2 | 0,123791765 | 1,095019127 | 0,2 | 0,077 | 1 | CH_SC | 2,136159137 | 0,468129917 | 2,597402597 |
| RDH102 | 0,123870578 | 0,4043009 | 0,333 | 0,154 | 1 | CH_SC | 1,32344744 | 0,755602353 | 2,162337662 |
| PWP1 | 0,123870578 | 0,588307791 | 0,333 | 0,154 | 1 | CH_SC | 1,503482203 | 0,665122605 | 2,162337662 |
| ISG20L2 | 0,124094416 | 0,978559826 | 0,267 | 0,123 | 1 | CH_SC | 1,970497373 | 0,507486086 | 2,170731707 |
| TST | 0,124094416 | 0,510568127 | 0,267 | 0,123 | 1 | CH_SC | 1,42461109 | 0,70194596 | 2,170731707 |
| TXNL12 | 0,125104475 | 0,66730604 | 0,333 | 0,185 | 1 | CH_SC | 1,588104712 | 0,629681401 | 1,8 |
| SPCS21 | 0,125425311 | 0,857585814 | 0,6 | 0,554 | 1 | CH_SC | 1,81200359 | 0,551875286 | 1,083032491 |
| SNRPG2 | 0,125462381 | -0,683841961 | 0,2 | 0,4 | 1 | CH_SC | 0,622505306 | 1,606412011 | 0,5 |
| JOSD22 | 0,126243038 | 0,665370852 | 0,4 | 0,231 | 1 | CH_SC | 1,585975904 | 0,630526603 | 1,731601732 |
| PSMF12 | 0,126243038 | 0,624067137 | 0,4 | 0,231 | 1 | CH_SC | 1,541213937 | 0,648839188 | 1,731601732 |
| ARMCX32 | 0,126662564 | 0,77355928 | 0,4 | 0,262 | 1 | CH_SC | 1,709482057 | 0,584972504 | 1,526717557 |
| FKBP82 | 0,126715268 | 0,570731351 | 0,533 | 0,4 | 1 | CH_SC | 1,485276317 | 0,673275396 | 1,3325 |
| PBXIP11 | 0,127083993 | -0,734082316 | 0,2 | 0,354 | 1 | CH_SC | 0,601200321 | 1,663339099 | 0,564971751 |
| MMGT1 | 0,127597049 | 0,276601676 | 0,267 | 0,108 | 1 | CH_SC | 1,211338167 | 0,825533304 | 2,472222222 |
| PTGES31 | 0,128348473 | 0,813267672 | 0,6 | 0,492 | 1 | CH_SC | 1,757186928 | 0,569091418 | 1,219512195 |
| IER25 | 0,12895637 | 0,286269559 | 0,8 | 0,615 | 1 | CH_SC | 1,219482926 | 0,820019681 | 1,300813008 |
| CNOT6L1 | 0,129021972 | 0,606786706 | 0,267 | 0,123 | 1 | CH_SC | 1,522863577 | 0,656657638 | 2,170731707 |
| SCRN11 | 0,129021972 | 1,197850882 | 0,267 | 0,123 | 1 | CH_SC | 2,29397693 | 0,435924175 | 2,170731707 |
| PIK3C2A2 | 0,129021972 | 1,082389108 | 0,267 | 0,123 | 1 | CH_SC | 2,117539832 | 0,472246134 | 2,170731707 |
| KPNA21 | 0,129021972 | 0,980363301 | 0,267 | 0,123 | 1 | CH_SC | 1,97296218 | 0,506852088 | 2,170731707 |
| CEBPZ3 | 0,12915317 | -0,663504783 | 0,067 | 0,231 | 1 | CH_SC | 0,631342692 | 1,583925833 | 0,29004329 |
| TSPYL13 | 0,12915317 | -0,61198895 | 0,067 | 0,231 | 1 | CH_SC | 0,654294047 | 1,528364814 | 0,29004329 |
| MRPS33 | 0,12915317 | -0,65148511 | 0,067 | 0,231 | 1 | CH_SC | 0,636624635 | 1,570784329 | 0,29004329 |
| SLC43A33 | 0,12915317 | -0,529679581 | 0,067 | 0,231 | 1 | CH_SC | 0,692708566 | 1,443608538 | 0,29004329 |
| SAFB21 | 0,12915317 | -0,666204925 | 0,067 | 0,231 | 1 | CH_SC | 0,630162179 | 1,586893078 | 0,29004329 |
| ELF22 | 0,129175965 | 1,015959531 | 0,333 | 0,2 | 1 | CH_SC | 2,022247436 | 0,494499329 | 1,665 |
| CAMSAP2 | 0,12958284 | 0,55672642 | 0,4 | 0,215 | 1 | CH_SC | 1,470927785 | 0,679843028 | 1,860465116 |
| BCAR3 | 0,129677814 | 0,595360818 | 0,2 | 0,077 | 1 | CH_SC | 1,510850402 | 0,661878899 | 2,597402597 |
| VPS45 | 0,129677814 | 0,734366256 | 0,2 | 0,077 | 1 | CH_SC | 1,663666497 | 0,601082009 | 2,597402597 |
| R3HDM1 | 0,129677814 | 1,025889592 | 0,2 | 0,077 | 1 | CH_SC | 2,036214564 | 0,49110738 | 2,597402597 |
| TARS11 | 0,129677814 | 0,830199238 | 0,2 | 0,077 | 1 | CH_SC | 1,77793088 | 0,562451562 | 2,597402597 |

| Supplementary Table 4 - clusermarker | | | | | | | | | |
|---|---|---|---|---|---|---|---|---|---|
| | p_val | avg_log2FC | pct.1 | pct.2 | p_val_adj | celltype | Foldchange_UP | Foldchange_DOWN | Ratio_pct1_pct2 |
| RGMB | 0,129677814 | 0,588963064 | 0,2 | 0,077 | 1 | CH_SC | 1,504165241 | 0,664820575 | 2,597402597 |
| GSDMD3 | 0,129677814 | 0,589720265 | 0,2 | 0,077 | 1 | CH_SC | 1,504954913 | 0,664471734 | 2,597402597 |
| XRRA1 | 0,129677814 | 0,729154334 | 0,2 | 0,077 | 1 | CH_SC | 1,657667131 | 0,603257422 | 2,597402597 |
| SIK31 | 0,129677814 | 0,545384108 | 0,2 | 0,077 | 1 | CH_SC | 1,45940885 | 0,685208946 | 2,597402597 |
| BACE1 | 0,129677814 | 0,602093232 | 0,2 | 0,077 | 1 | CH_SC | 1,517917343 | 0,658797401 | 2,597402597 |
| OSBPL83 | 0,129711694 | 0,800890199 | 0,467 | 0,277 | 1 | CH_SC | 1,742175785 | 0,573994891 | 1,685920578 |
| CD1641 | 0,130521367 | -0,757693915 | 0,333 | 0,477 | 1 | CH_SC | 0,591440968 | 1,690785816 | 0,698113208 |
| SKIL4 | 0,130542058 | -0,803836085 | 0,133 | 0,323 | 1 | CH_SC | 0,572824028 | 1,745736825 | 0,411764706 |
| FNDC3A1 | 0,130547542 | 0,733092216 | 0,4 | 0,246 | 1 | CH_SC | 1,662197966 | 0,601613057 | 1,62601626 |
| MT-ND43 | 0,131014197 | 0,299802639 | 0,933 | 0,954 | 1 | CH_SC | 1,230976004 | 0,81236352 | 0,977987421 |
| MIIP2 | 0,132224288 | -0,438790836 | 0 | 0,138 | 1 | CH_SC | 0,737752681 | 1,355467795 | 0 |
| TCEA32 | 0,132224288 | -0,361281407 | 0 | 0,138 | 1 | CH_SC | 0,77847283 | 1,284566348 | 0 |
| CTH | 0,132224288 | -0,351187331 | 0 | 0,138 | 1 | CH_SC | 0,783938655 | 1,275610016 | 0 |
| TMEM167B1 | 0,132224288 | -0,549430347 | 0 | 0,138 | 1 | CH_SC | 0,683289874 | 1,463507711 | 0 |
| SLC25A441 | 0,132224288 | -0,344858151 | 0 | 0,138 | 1 | CH_SC | 0,78738539 | 1,270026105 | 0 |
| FMO22 | 0,132224288 | -0,83691068 | 0 | 0,138 | 1 | CH_SC | 0,559841104 | 1,786221113 | 0 |
| G0S2 | 0,132224288 | -0,649718201 | 0 | 0,138 | 1 | CH_SC | 0,637404805 | 1,568861723 | 0 |
| ATAD2B | 0,132224288 | -0,498522423 | 0 | 0,138 | 1 | CH_SC | 0,707831356 | 1,412765897 | 0 |
| GALNT51 | 0,132224288 | -0,302253092 | 0 | 0,138 | 1 | CH_SC | 0,810984872 | 1,233068624 | 0 |
| CTDSP12 | 0,132224288 | -0,388044791 | 0 | 0,138 | 1 | CH_SC | 0,764164535 | 1,308618699 | 0 |
| PTPRG | 0,132224288 | -0,40669659 | 0 | 0,138 | 1 | CH_SC | 0,754348667 | 1,325646937 | 0 |
| TRPC11 | 0,132224288 | -0,618476349 | 0 | 0,138 | 1 | CH_SC | 0,651358474 | 1,535252922 | 0 |
| SIAH21 | 0,132224288 | -0,499482859 | 0 | 0,138 | 1 | CH_SC | 0,707360293 | 1,413706721 | 0 |
| MAN2B2 | 0,132224288 | -0,439366076 | 0 | 0,138 | 1 | CH_SC | 0,737458579 | 1,356008363 | 0 |
| TNIP14 | 0,132224288 | -0,307369309 | 0 | 0,138 | 1 | CH_SC | 0,808113977 | 1,237449207 | 0 |
| ATXN11 | 0,132224288 | -0,512975894 | 0 | 0,138 | 1 | CH_SC | 0,700775434 | 1,426990662 | 0 |
| ZKSCAN8 | 0,132224288 | -0,341010747 | 0 | 0,138 | 1 | CH_SC | 0,789488006 | 1,266643689 | 0 |
| WDR46 | 0,132224288 | -0,501869966 | 0 | 0,138 | 1 | CH_SC | 0,70619085 | 1,4160478 | 0 |
| RARS2 | 0,132224288 | -0,283244082 | 0 | 0,138 | 1 | CH_SC | 0,82174115 | 1,216928227 | 0 |
| TPD52L1 | 0,132224288 | -0,301675799 | 0 | 0,138 | 1 | CH_SC | 0,811309451 | 1,232575312 | 0 |
| VTA11 | 0,132224288 | -0,517839913 | 0 | 0,138 | 1 | CH_SC | 0,698416761 | 1,431809853 | 0 |

| Supplementary Table 4 - clusermarker | | | | | | | | | |
|---|---|---|---|---|---|---|---|---|---|
| | **p_val** | **avg_log2FC** | **pct.1** | **pct.2** | **p_val_adj** | **celltype** | **Foldchange_UP** | **Foldchange_DOWN** | **Ratio_pct1_pct2** |
| HOTAIRM13 | 0,132224288 | -0,376519317 | 0 | 0,138 | 1 | CH_SC | 0,770293781 | 1,298205989 | 0 |
| HOXA7 | 0,132224288 | -0,329841296 | 0 | 0,138 | 1 | CH_SC | 0,795624002 | 1,256875104 | 0 |
| SLC7A22 | 0,132224288 | -0,490968354 | 0 | 0,138 | 1 | CH_SC | 0,711547338 | 1,405387873 | 0 |
| TNFRSF10D | 0,132224288 | -0,504731665 | 0 | 0,138 | 1 | CH_SC | 0,704791454 | 1,41885943 | 0 |
| KANK11 | 0,132224288 | -0,430803867 | 0 | 0,138 | 1 | CH_SC | 0,741848314 | 1,347984462 | 0 |
| TLE46 | 0,132224288 | -0,540897781 | 0 | 0,138 | 1 | CH_SC | 0,687343046 | 1,454877598 | 0 |
| AKR1C23 | 0,132224288 | -0,531745945 | 0 | 0,138 | 1 | CH_SC | 0,691717113 | 1,445677692 | 0 |
| ALDH18A1 | 0,132224288 | -0,355604058 | 0 | 0,138 | 1 | CH_SC | 0,781542342 | 1,279521206 | 0 |
| ATE11 | 0,132224288 | -0,510511368 | 0 | 0,138 | 1 | CH_SC | 0,701973577 | 1,424555044 | 0 |
| NUP98 | 0,132224288 | -0,35492667 | 0 | 0,138 | 1 | CH_SC | 0,781909385 | 1,278920574 | 0 |
| SLC36A4 | 0,132224288 | -0,393953612 | 0 | 0,138 | 1 | CH_SC | 0,76104116 | 1,313989377 | 0 |
| ARHGDIB6 | 0,132224288 | -0,292193655 | 0 | 0,138 | 1 | CH_SC | 0,816659362 | 1,224500748 | 0 |
| ETNK12 | 0,132224288 | -0,50569348 | 0 | 0,138 | 1 | CH_SC | 0,70432174 | 1,419805669 | 0 |
| TMTC1 | 0,132224288 | -0,517126 | 0 | 0,138 | 1 | CH_SC | 0,698762456 | 1,431101502 | 0 |
| STAT62 | 0,132224288 | -0,482059298 | 0 | 0,138 | 1 | CH_SC | 0,715954943 | 1,39673594 | 0 |
| SLC7A8 | 0,132224288 | -0,543299002 | 0 | 0,138 | 1 | CH_SC | 0,686199985 | 1,457301111 | 0 |
| CGRRF1 | 0,132224288 | -0,479634108 | 0 | 0,138 | 1 | CH_SC | 0,717159485 | 1,39438998 | 0 |
| SMOC1 | 0,132224288 | -0,4961155 | 0 | 0,138 | 1 | CH_SC | 0,709013253 | 1,410410871 | 0 |
| ZC3H14 | 0,132224288 | -0,474221675 | 0 | 0,138 | 1 | CH_SC | 0,719855043 | 1,38916857 | 0 |
| FGF73 | 0,132224288 | -0,813437384 | 0 | 0,138 | 1 | CH_SC | 0,569024476 | 1,757393648 | 0 |
| CILP1 | 0,132224288 | -2,032277149 | 0 | 0,138 | 1 | CH_SC | 0,2444689 | 4,090499854 | 0 |
| CEMIP3 | 0,132224288 | -0,556475532 | 0 | 0,138 | 1 | CH_SC | 0,679961265 | 1,47067201 | 0 |
| SNRNP251 | 0,132224288 | -0,413505603 | 0 | 0,138 | 1 | CH_SC | 0,750796795 | 1,331918312 | 0 |
| RUSF1 | 0,132224288 | -0,316293666 | 0 | 0,138 | 1 | CH_SC | 0,8031305 | 1,24512766 | 0 |
| CYB5D11 | 0,132224288 | -0,60213586 | 0 | 0,138 | 1 | CH_SC | 0,658777935 | 1,517962194 | 0 |
| SDK2 | 0,132224288 | -0,312826223 | 0 | 0,138 | 1 | CH_SC | 0,805063104 | 1,242138653 | 0 |
| PYCR1 | 0,132224288 | -0,349652576 | 0 | 0,138 | 1 | CH_SC | 0,784773061 | 1,27425373 | 0 |
| TMED11 | 0,132224288 | -0,446064778 | 0 | 0,138 | 1 | CH_SC | 0,734042358 | 1,362319202 | 0 |
| ELOF1 | 0,132224288 | -0,310619732 | 0 | 0,138 | 1 | CH_SC | 0,806295328 | 1,24024035 | 0 |
| CA11 | 0,132224288 | -0,375965716 | 0 | 0,138 | 1 | CH_SC | 0,77058942 | 1,297707928 | 0 |
| CHMP4B2 | 0,132224288 | -0,324141839 | 0 | 0,138 | 1 | CH_SC | 0,798773381 | 1,251919536 | 0 |

| Supplementary Table 4 - clusermarker | | | | | | | | | |
|---|---|---|---|---|---|---|---|---|---|
| | **p_val** | **avg_log2FC** | **pct.1** | **pct.2** | **p_val_adj** | **celltype** | **Foldchange_UP** | **Foldchange_DOWN** | **Ratio_pct1_pct2** |
| YTHDF1 | 0,132224288 | -0,392287416 | 0 | 0,138 | 1 | CH_SC | 0,761920609 | 1,312472702 | 0 |
| SLC5A31 | 0,132224288 | -0,358718429 | 0 | 0,138 | 1 | CH_SC | 0,779857032 | 1,282286315 | 0 |
| SIM2 | 0,132224288 | -0,37330748 | 0 | 0,138 | 1 | CH_SC | 0,772010578 | 1,295319039 | 0 |
| POFUT2 | 0,132224288 | -0,474333124 | 0 | 0,138 | 1 | CH_SC | 0,719799436 | 1,389275888 | 0 |
| PACSIN22 | 0,132224288 | -0,38805877 | 0 | 0,138 | 1 | CH_SC | 0,764157131 | 1,308631379 | 0 |
| GPRASP1 | 0,132224288 | -0,373300023 | 0 | 0,138 | 1 | CH_SC | 0,772014568 | 1,295312344 | 0 |
| ZDHHC91 | 0,132224288 | -0,430833566 | 0 | 0,138 | 1 | CH_SC | 0,741833042 | 1,348012212 | 0 |
| HNRNPM4 | 0,132281937 | 0,810233514 | 0,467 | 0,323 | 1 | CH_SC | 1,75349524 | 0,570289544 | 1,445820433 |
| UBR11 | 0,133484968 | -0,46569209 | 0,067 | 0,262 | 1 | CH_SC | 0,724123617 | 1,380979678 | 0,255725191 |
| CLEC11A2 | 0,133484968 | -0,435637374 | 0,067 | 0,262 | 1 | CH_SC | 0,739367034 | 1,352508231 | 0,255725191 |
| PLOD11 | 0,133664924 | 0,598631815 | 0,333 | 0,169 | 1 | CH_SC | 1,514279813 | 0,660379932 | 1,970414201 |
| EVA1B2 | 0,133664924 | -0,53067813 | 0,067 | 0,231 | 1 | CH_SC | 0,69222928 | 1,444608065 | 0,29004329 |
| VDAC32 | 0,133664924 | -0,533135953 | 0,067 | 0,231 | 1 | CH_SC | 0,691050979 | 1,447071245 | 0,29004329 |
| SSB1 | 0,133668666 | 0,797267983 | 0,533 | 0,4 | 1 | CH_SC | 1,737807141 | 0,575437847 | 1,3325 |
| SLC39A6 | 0,133734182 | 0,532553676 | 0,4 | 0,215 | 1 | CH_SC | 1,446487319 | 0,691329946 | 1,860465116 |
| EIF2AK2 | 0,133813753 | 0,705635924 | 0,333 | 0,185 | 1 | CH_SC | 1,630863373 | 0,613172149 | 1,8 |
| CDK42 | 0,133813753 | 0,703836936 | 0,333 | 0,185 | 1 | CH_SC | 1,628831013 | 0,61393723 | 1,8 |
| SUMO13 | 0,134157472 | -0,484070314 | 0,133 | 0,338 | 1 | CH_SC | 0,714957647 | 1,39868425 | 0,393491124 |
| SDF44 | 0,134317058 | -0,544212434 | 0,333 | 0,523 | 1 | CH_SC | 0,685765659 | 1,458224084 | 0,636711281 |
| PLSCR41 | 0,134333321 | 0,413080871 | 0,4 | 0,231 | 1 | CH_SC | 1,33152625 | 0,751017864 | 1,731601732 |
| FNTA1 | 0,134333321 | 0,658600125 | 0,4 | 0,231 | 1 | CH_SC | 1,578550183 | 0,633492689 | 1,731601732 |
| GOLIM41 | 0,134520808 | -0,643477061 | 0,267 | 0,431 | 1 | CH_SC | 0,640168209 | 1,562089441 | 0,619489559 |
| TUBB8 | 0,135063568 | 0,569922344 | 0,6 | 0,492 | 1 | CH_SC | 1,484443665 | 0,673653048 | 1,219512195 |
| GAPDH5 | 0,135547896 | 0,558047714 | 0,867 | 0,892 | 1 | CH_SC | 1,472275554 | 0,679220678 | 0,971973094 |
| MAD2L22 | 0,135778882 | 0,765339053 | 0,2 | 0,077 | 1 | CH_SC | 1,69976942 | 0,58831509 | 2,597402597 |
| BRIX1 | 0,135778882 | 0,439582932 | 0,2 | 0,077 | 1 | CH_SC | 1,356212205 | 0,737347737 | 2,597402597 |
| MOSPD31 | 0,135778882 | 0,403308748 | 0,2 | 0,077 | 1 | CH_SC | 1,322537608 | 0,756122165 | 2,597402597 |
| VPS37A | 0,135778882 | 0,421910465 | 0,2 | 0,077 | 1 | CH_SC | 1,339700456 | 0,746435515 | 2,597402597 |
| NUDT22 | 0,135778882 | 0,442255038 | 0,2 | 0,077 | 1 | CH_SC | 1,358726457 | 0,735983313 | 2,597402597 |
| GOSR21 | 0,135778882 | 0,546361447 | 0,2 | 0,077 | 1 | CH_SC | 1,460397847 | 0,684744915 | 2,597402597 |
| ZNF652 | 0,135778882 | 0,710873852 | 0,2 | 0,077 | 1 | CH_SC | 1,636795236 | 0,61094997 | 2,597402597 |

| Supplementary Table 4 - clusermarker | | | | | | | | | |
|---|---|---|---|---|---|---|---|---|---|
| | **p_val** | **avg_log2FC** | **pct.1** | **pct.2** | **p_val_adj** | **celltype** | **Foldchange_UP** | **Foldchange_DOWN** | **Ratio_pct1_pct2** |
| UBQLN2 | 0,138216123 | 0,343128485 | 0,267 | 0,108 | 1 | CH_SC | 1,268504366 | 0,788329963 | 2,472222222 |
| FOSB7 | 0,138250091 | 0,374884345 | 0,867 | 0,692 | 1 | CH_SC | 1,296735596 | 0,771167232 | 1,252890173 |
| PDLIM23 | 0,138297667 | 0,493755402 | 0,333 | 0,169 | 1 | CH_SC | 1,408105473 | 0,710174074 | 1,970414201 |
| FXYD55 | 0,138370584 | -0,498454239 | 0,267 | 0,477 | 1 | CH_SC | 0,70786481 | 1,412699129 | 0,559748428 |
| FLOT12 | 0,13871038 | 0,793169857 | 0,4 | 0,246 | 1 | CH_SC | 1,732877723 | 0,577074762 | 1,62601626 |
| PSMA33 | 0,13871038 | 1,059567124 | 0,4 | 0,246 | 1 | CH_SC | 2,084306038 | 0,479775993 | 1,62601626 |
| TAF152 | 0,13871038 | 0,989924122 | 0,4 | 0,246 | 1 | CH_SC | 1,986080531 | 0,503504256 | 1,62601626 |
| GUK15 | 0,138785606 | 0,676555682 | 0,533 | 0,538 | 1 | CH_SC | 1,598319338 | 0,625657199 | 0,99070632 |
| ATP5MF3 | 0,139024669 | 0,607028534 | 0,6 | 0,446 | 1 | CH_SC | 1,523118864 | 0,656547577 | 1,34529148 |
| LZIC | 0,139331652 | 0,809011205 | 0,267 | 0,123 | 1 | CH_SC | 1,752010237 | 0,570772921 | 2,170731707 |
| RAB9A | 0,139331652 | 0,73823391 | 0,267 | 0,123 | 1 | CH_SC | 1,668132527 | 0,599472754 | 2,170731707 |
| BANF13 | 0,140222416 | -0,251591341 | 0,2 | 0,4 | 1 | CH_SC | 0,83996939 | 1,190519574 | 0,5 |
| CDC423 | 0,140813408 | 1,102405815 | 0,533 | 0,462 | 1 | CH_SC | 2,147124452 | 0,465739189 | 1,153679654 |
| HSPA1B8 | 0,141363484 | -1,353588183 | 0,533 | 0,692 | 1 | CH_SC | 0,391317576 | 2,555469164 | 0,770231214 |
| WSB13 | 0,141363484 | 0,528073789 | 0,733 | 0,646 | 1 | CH_SC | 1,442002623 | 0,693480014 | 1,134674923 |
| AAK1 | 0,142099281 | 0,716740491 | 0,2 | 0,077 | 1 | CH_SC | 1,643464725 | 0,60847062 | 2,597402597 |
| SMARCC14 | 0,142099281 | 0,373678674 | 0,2 | 0,077 | 1 | CH_SC | 1,295652357 | 0,771811971 | 2,597402597 |
| CHST121 | 0,142099281 | 0,373190094 | 0,2 | 0,077 | 1 | CH_SC | 1,295213649 | 0,772073396 | 2,597402597 |
| MICU3 | 0,142099281 | 0,645040713 | 0,2 | 0,077 | 1 | CH_SC | 1,563783416 | 0,639474744 | 2,597402597 |
| KIF13B | 0,142099281 | 0,441748369 | 0,2 | 0,077 | 1 | CH_SC | 1,358249362 | 0,736241833 | 2,597402597 |
| TGFBR11 | 0,142099281 | 0,470804614 | 0,2 | 0,077 | 1 | CH_SC | 1,385882181 | 0,721562059 | 2,597402597 |
| ADIPOR22 | 0,142099281 | 0,503080383 | 0,2 | 0,077 | 1 | CH_SC | 1,417236359 | 0,705598606 | 2,597402597 |
| GNPTAB2 | 0,142099281 | 1,011557176 | 0,2 | 0,077 | 1 | CH_SC | 2,016085993 | 0,496010589 | 2,597402597 |
| PLBD2 | 0,142099281 | 0,413668189 | 0,2 | 0,077 | 1 | CH_SC | 1,332068422 | 0,750712188 | 2,597402597 |
| MSL1 | 0,142099281 | 0,623365274 | 0,2 | 0,077 | 1 | CH_SC | 1,540464327 | 0,649154922 | 2,597402597 |
| ME21 | 0,142099281 | 0,609343492 | 0,2 | 0,077 | 1 | CH_SC | 1,525564832 | 0,655494922 | 2,597402597 |
| GRAMD1A | 0,142099281 | 0,52246607 | 0,2 | 0,077 | 1 | CH_SC | 1,436408475 | 0,696180799 | 2,597402597 |
| MAP3K21 | 0,142344689 | 0,468080238 | 0,4 | 0,215 | 1 | CH_SC | 1,38326756 | 0,722925939 | 1,860465116 |
| C5orf24 | 0,142344689 | 0,287936299 | 0,4 | 0,215 | 1 | CH_SC | 1,220892604 | 0,819072862 | 1,860465116 |
| ALDH6A11 | 0,142435703 | -0,310174734 | 0,067 | 0,262 | 1 | CH_SC | 0,806544068 | 1,239857858 | 0,255725191 |
| COPA | 0,142818792 | 0,966158223 | 0,4 | 0,231 | 1 | CH_SC | 1,953631305 | 0,51186731 | 1,731601732 |

| Supplementary Table 4 - clusermarker | | | | | | | | | |
|---|---|---|---|---|---|---|---|---|---|
| | **p_val** | **avg_log2FC** | **pct.1** | **pct.2** | **p_val_adj** | **celltype** | **Foldchange_UP** | **Foldchange_DOWN** | **Ratio_pct1_pct2** |
| MPG3 | 0,142936546 | 0,623441279 | 0,4 | 0,246 | 1 | CH_SC | 1,540545485 | 0,649120723 | 1,62601626 |
| NDUFA74 | 0,143053162 | 0,42706921 | 0,333 | 0,169 | 1 | CH_SC | 1,344499492 | 0,7437712 | 1,970414201 |
| FIBIN1 | 0,143290631 | -1,061487585 | 0,2 | 0,369 | 1 | CH_SC | 0,479137759 | 2,087082435 | 0,54200542 |
| MSN3 | 0,143373661 | 0,517440186 | 0,533 | 0,385 | 1 | CH_SC | 1,431413197 | 0,698610298 | 1,384415584 |
| NENF6 | 0,143554139 | -0,581621077 | 0,333 | 0,538 | 1 | CH_SC | 0,668212522 | 1,496529872 | 0,618959108 |
| CDKN1A7 | 0,1441088 | -0,712226641 | 0,467 | 0,646 | 1 | CH_SC | 0,610377361 | 1,63833075 | 0,722910217 |
| GPR107 | 0,144718826 | 0,363181391 | 0,267 | 0,123 | 1 | CH_SC | 1,286259196 | 0,777448281 | 2,170731707 |
| IFT431 | 0,144718826 | 0,715135852 | 0,267 | 0,123 | 1 | CH_SC | 1,641637796 | 0,609147768 | 2,170731707 |
| PRKCA | 0,144718826 | 0,392399899 | 0,267 | 0,123 | 1 | CH_SC | 1,312575036 | 0,761861206 | 2,170731707 |
| SET3 | 0,145366541 | 0,756350672 | 0,533 | 0,492 | 1 | CH_SC | 1,689212318 | 0,591991894 | 1,083333333 |
| TXNIP10 | 0,145381953 | 0,393857625 | 0,667 | 0,554 | 1 | CH_SC | 1,313901956 | 0,761091796 | 1,203971119 |
| JTB8 | 0,14575355 | -0,27747217 | 0,2 | 0,492 | 1 | CH_SC | 0,825035344 | 1,212069286 | 0,406504065 |
| RNF5 | 0,146806567 | 0,522331331 | 0,4 | 0,215 | 1 | CH_SC | 1,436274329 | 0,696245821 | 1,860465116 |
| PAPSS21 | 0,146987475 | -0,634669912 | 0,133 | 0,323 | 1 | CH_SC | 0,644088165 | 1,552582479 | 0,411764706 |
| ADRM15 | 0,14708049 | 0,702879232 | 0,333 | 0,2 | 1 | CH_SC | 1,627750106 | 0,614344915 | 1,665 |
| MYL12B6 | 0,14716331 | 1,134423502 | 0,6 | 0,569 | 1 | CH_SC | 2,195308211 | 0,455516904 | 1,054481547 |
| LSM14A2 | 0,1475569 | 0,418140028 | 0,333 | 0,154 | 1 | CH_SC | 1,336203764 | 0,748388851 | 2,162337662 |
| MPZL13 | 0,147933142 | -0,565406293 | 0,067 | 0,231 | 1 | CH_SC | 0,67576508 | 1,479804194 | 0,29004329 |
| SHC4 | 0,147933142 | 0,390271968 | 0,333 | 0,169 | 1 | CH_SC | 1,310640455 | 0,762985757 | 1,970414201 |
| RNF1303 | 0,147998517 | 1,044409553 | 0,467 | 0,369 | 1 | CH_SC | 2,062522064 | 0,484843298 | 1,265582656 |
| YARS1 | 0,148643224 | 0,282466717 | 0,2 | 0,077 | 1 | CH_SC | 1,216272688 | 0,822184046 | 2,597402597 |
| ERO1B1 | 0,148643224 | 0,517450338 | 0,2 | 0,077 | 1 | CH_SC | 1,43142327 | 0,698605382 | 2,597402597 |
| PDZRN3 | 0,148643224 | 0,386385139 | 0,2 | 0,077 | 1 | CH_SC | 1,307114152 | 0,765044123 | 2,597402597 |
| RNF4 | 0,148643224 | 0,476276747 | 0,2 | 0,077 | 1 | CH_SC | 1,391148805 | 0,718830363 | 2,597402597 |
| PPID | 0,148643224 | 1,429107735 | 0,2 | 0,077 | 1 | CH_SC | 2,692801219 | 0,371360497 | 2,597402597 |
| ZDHHC41 | 0,148643224 | 0,354318058 | 0,2 | 0,077 | 1 | CH_SC | 1,278381165 | 0,782239309 | 2,597402597 |
| CLDN121 | 0,148643224 | 0,334309513 | 0,2 | 0,077 | 1 | CH_SC | 1,260773847 | 0,793163661 | 2,597402597 |
| FBH11 | 0,148643224 | 0,356485087 | 0,2 | 0,077 | 1 | CH_SC | 1,280302827 | 0,781065213 | 2,597402597 |
| ATAD11 | 0,148643224 | 0,356921012 | 0,2 | 0,077 | 1 | CH_SC | 1,280689742 | 0,780829242 | 2,597402597 |
| IFIT31 | 0,148643224 | 0,58427257 | 0,2 | 0,077 | 1 | CH_SC | 1,499282836 | 0,666985559 | 2,597402597 |
| TULP3 | 0,148643224 | 0,387464891 | 0,2 | 0,077 | 1 | CH_SC | 1,308092797 | 0,764471758 | 2,597402597 |

| Supplementary Table 4 - clusermarker | | | | | | | | | |
|---|---|---|---|---|---|---|---|---|---|
| | **p_val** | **avg_log2FC** | **pct.1** | **pct.2** | **p_val_adj** | **celltype** | **Foldchange_UP** | **Foldchange_DOWN** | **Ratio_pct1_pct2** |
| C12orf43 | 0,148643224 | 0,529918936 | 0,2 | 0,077 | 1 | CH_SC | 1,443848064 | 0,69259365 | 2,597402597 |
| RMDN31 | 0,148643224 | 0,498516383 | 0,2 | 0,077 | 1 | CH_SC | 1,412759982 | 0,707834319 | 2,597402597 |
| Z93241.11 | 0,148643224 | 0,487613223 | 0,2 | 0,077 | 1 | CH_SC | 1,4021233 | 0,713204038 | 2,597402597 |
| KDM5C1 | 0,148643224 | 0,503747774 | 0,2 | 0,077 | 1 | CH_SC | 1,417892124 | 0,705272272 | 2,597402597 |
| TAF1 | 0,148643224 | 0,326561294 | 0,2 | 0,077 | 1 | CH_SC | 1,254020814 | 0,79743493 | 2,597402597 |
| SELENOK4 | 0,149405484 | -0,581774318 | 0,4 | 0,585 | 1 | CH_SC | 0,668141549 | 1,49668884 | 0,683760684 |
| EIF4A34 | 0,149633877 | 0,58466527 | 0,533 | 0,369 | 1 | CH_SC | 1,499690995 | 0,666804031 | 1,444444444 |
| MRPS14 | 0,150264175 | 0,397962638 | 0,267 | 0,123 | 1 | CH_SC | 1,317645828 | 0,75892928 | 2,170731707 |
| TMEM33 | 0,150264175 | 0,444272994 | 0,267 | 0,123 | 1 | CH_SC | 1,360628293 | 0,734954583 | 2,170731707 |
| DHRS41 | 0,150264175 | 0,509713784 | 0,267 | 0,123 | 1 | CH_SC | 1,423767707 | 0,702361765 | 2,170731707 |
| POLDIP33 | 0,150264175 | 0,517041204 | 0,267 | 0,123 | 1 | CH_SC | 1,43101739 | 0,698803527 | 2,170731707 |
| SEC61G3 | 0,151236062 | -0,383763255 | 0,333 | 0,6 | 1 | CH_SC | 0,766435741 | 1,304740824 | 0,555 |
| COX175 | 0,151465758 | 0,633281755 | 0,467 | 0,308 | 1 | CH_SC | 1,551089307 | 0,644708203 | 1,516233766 |
| USP9X | 0,151709729 | -0,66572848 | 0,133 | 0,292 | 1 | CH_SC | 0,630370322 | 1,586369098 | 0,455479452 |
| FAM120A2 | 0,151840214 | 0,711780847 | 0,333 | 0,2 | 1 | CH_SC | 1,637824583 | 0,610565997 | 1,665 |
| PUM13 | 0,152630734 | 0,733109599 | 0,333 | 0,185 | 1 | CH_SC | 1,662217995 | 0,601605808 | 1,8 |
| FUBP12 | 0,152684756 | -0,496937122 | 0,067 | 0,215 | 1 | CH_SC | 0,708609582 | 1,411214336 | 0,311627907 |
| DNAJB142 | 0,152684756 | -0,576724586 | 0,067 | 0,215 | 1 | CH_SC | 0,670484281 | 1,491459276 | 0,311627907 |
| CLUAP11 | 0,152684756 | -0,482061792 | 0,067 | 0,215 | 1 | CH_SC | 0,715953705 | 1,396738355 | 0,311627907 |
| CHID11 | 0,154092019 | -0,398678816 | 0,133 | 0,354 | 1 | CH_SC | 0,758552629 | 1,318300091 | 0,375706215 |
| RPS27L1 | 0,154956472 | -0,611639991 | 0,333 | 0,6 | 1 | CH_SC | 0,654452327 | 1,527995179 | 0,555 |
| NFIA5 | 0,155022215 | -0,653162694 | 0,2 | 0,354 | 1 | CH_SC | 0,63588479 | 1,572611919 | 0,564971751 |
| DSTYK | 0,155414821 | 0,354254307 | 0,2 | 0,077 | 1 | CH_SC | 1,278324676 | 0,782273876 | 2,597402597 |
| SPOPL | 0,155414821 | 0,2570513 | 0,2 | 0,077 | 1 | CH_SC | 1,195033698 | 0,836796487 | 2,597402597 |
| MAPK141 | 0,155414821 | 0,890329119 | 0,2 | 0,077 | 1 | CH_SC | 1,853598934 | 0,539491031 | 2,597402597 |
| RASSF8 | 0,155414821 | 0,918554214 | 0,2 | 0,077 | 1 | CH_SC | 1,890220074 | 0,529038927 | 2,597402597 |
| EXOSC82 | 0,155414821 | 0,398189722 | 0,2 | 0,077 | 1 | CH_SC | 1,317853245 | 0,758809833 | 2,597402597 |
| DHRS12 | 0,155414821 | 0,257332504 | 0,2 | 0,077 | 1 | CH_SC | 1,195266652 | 0,836633398 | 2,597402597 |
| NID21 | 0,155414821 | 0,354285106 | 0,2 | 0,077 | 1 | CH_SC | 1,278351967 | 0,782257176 | 2,597402597 |
| CHMP1A1 | 0,155414821 | 0,346917965 | 0,2 | 0,077 | 1 | CH_SC | 1,271840685 | 0,786262 | 2,597402597 |
| CPD1 | 0,155414821 | 0,351308909 | 0,2 | 0,077 | 1 | CH_SC | 1,275717518 | 0,783872594 | 2,597402597 |

| Supplementary Table 4 - clusermarker | | | | | | | | | |
|---|---|---|---|---|---|---|---|---|---|
| | **p_val** | **avg_log2FC** | **pct.1** | **pct.2** | **p_val_adj** | **celltype** | **Foldchange_UP** | **Foldchange_DOWN** | **Ratio_pct1_pct2** |
| NF11 | 0,155414821 | 0,418543443 | 0,2 | 0,077 | 1 | CH_SC | 1,336577454 | 0,748179612 | 2,597402597 |
| SMARCA42 | 0,155414821 | 0,303948281 | 0,2 | 0,077 | 1 | CH_SC | 1,23451835 | 0,810032512 | 2,597402597 |
| RAB22A1 | 0,155414821 | 0,34233175 | 0,2 | 0,077 | 1 | CH_SC | 1,267804022 | 0,788765442 | 2,597402597 |
| RBM36 | 0,155431674 | 0,461167167 | 0,667 | 0,585 | 1 | CH_SC | 1,376655108 | 0,726398351 | 1,14017094 |
| LRRC58 | 0,155970095 | 0,29910137 | 0,267 | 0,123 | 1 | CH_SC | 1,230377794 | 0,812758492 | 2,170731707 |
| ZMAT22 | 0,155970095 | 1,022113259 | 0,267 | 0,123 | 1 | CH_SC | 2,03089163 | 0,492394565 | 2,170731707 |
| ZNF1063 | 0,156208513 | 0,635823645 | 0,4 | 0,246 | 1 | CH_SC | 1,553824587 | 0,643573289 | 1,62601626 |
| AHCTF12 | 0,156716335 | 0,786054414 | 0,333 | 0,2 | 1 | CH_SC | 1,724352126 | 0,579927954 | 1,665 |
| SCARA32 | 0,157388404 | -0,550825688 | 0,267 | 0,477 | 1 | CH_SC | 0,682629332 | 1,464923866 | 0,559748428 |
| ANXA54 | 0,157944868 | 0,347947362 | 0,8 | 0,785 | 1 | CH_SC | 1,272748497 | 0,785701183 | 1,01910828 |
| CWC222 | 0,157947095 | -0,568149227 | 0,067 | 0,215 | 1 | CH_SC | 0,674481498 | 1,482620359 | 0,311627907 |
| TFRC2 | 0,157947095 | -0,726836785 | 0,067 | 0,215 | 1 | CH_SC | 0,604227275 | 1,655006388 | 0,311627907 |
| ZNF7042 | 0,157947095 | -0,5491762 | 0,067 | 0,215 | 1 | CH_SC | 0,683410254 | 1,46324992 | 0,311627907 |
| FKBP32 | 0,157947095 | 0,351232661 | 0,333 | 0,154 | 1 | CH_SC | 1,275650097 | 0,783914023 | 2,162337662 |
| MSI21 | 0,157947095 | -0,590507609 | 0,067 | 0,215 | 1 | CH_SC | 0,664109201 | 1,505776458 | 0,311627907 |
| BCL6 | 0,158073321 | -0,620861876 | 0,067 | 0,231 | 1 | CH_SC | 0,650282329 | 1,537793595 | 0,29004329 |
| NDUFB32 | 0,158307749 | 0,502866694 | 0,533 | 0,431 | 1 | CH_SC | 1,417026456 | 0,705703126 | 1,236658933 |
| OCIAD11 | 0,158547447 | -0,545415195 | 0,333 | 0,538 | 1 | CH_SC | 0,685194181 | 1,459440298 | 0,618959108 |
| ANXA18 | 0,158617834 | 0,931089593 | 0,733 | 0,708 | 1 | CH_SC | 1,906715496 | 0,524462093 | 1,035310734 |
| TRIM41 | 0,158878233 | -0,294609667 | 0 | 0,123 | 1 | CH_SC | 0,815292887 | 1,226553079 | 0 |
| FUCA13 | 0,158887242 | -0,447086071 | 0 | 0,123 | 1 | CH_SC | 0,733522909 | 1,363283938 | 0 |
| CIART | 0,158887242 | -0,481984225 | 0 | 0,123 | 1 | CH_SC | 0,715992199 | 1,396663261 | 0 |
| FAM20B | 0,158887242 | -0,333951841 | 0 | 0,123 | 1 | CH_SC | 0,793360326 | 1,260461315 | 0 |
| KCNT2 | 0,158887242 | -0,429853699 | 0 | 0,123 | 1 | CH_SC | 0,742337061 | 1,347096963 | 0 |
| CYS1 | 0,158887242 | -0,334843528 | 0 | 0,123 | 1 | CH_SC | 0,792870125 | 1,26124061 | 0 |
| VIT1 | 0,158887242 | -0,427800057 | 0 | 0,123 | 1 | CH_SC | 0,743394512 | 1,345180767 | 0 |
| HNRNPLL1 | 0,158887242 | -0,412465736 | 0 | 0,123 | 1 | CH_SC | 0,75133815 | 1,330958637 | 0 |
| ATOH8 | 0,158887242 | -0,376960479 | 0 | 0,123 | 1 | CH_SC | 0,770058269 | 1,298603029 | 0 |
| TXNDC91 | 0,158887242 | -0,37324969 | 0 | 0,123 | 1 | CH_SC | 0,772041503 | 1,295267154 | 0 |
| POU3F3 | 0,158887242 | -0,330231824 | 0 | 0,123 | 1 | CH_SC | 0,79540866 | 1,257215378 | 0 |
| SLC38A3 | 0,158887242 | -0,47054086 | 0 | 0,123 | 1 | CH_SC | 0,721693987 | 1,385628837 | 0 |

| Supplementary Table 4 - clusermarker | | | | | | | | | |
|---|---|---|---|---|---|---|---|---|---|
| | p_val | avg_log2FC | pct.1 | pct.2 | p_val_adj | celltype | Foldchange_UP | Foldchange_DOWN | Ratio_pct1_pct2 |
| MGLL1 | 0,158887242 | -0,393223993 | 0 | 0,123 | 1 | CH_SC | 0,761426142 | 1,313325016 | 0 |
| CP4 | 0,158887242 | -0,639202323 | 0 | 0,123 | 1 | CH_SC | 0,642067855 | 1,557467786 | 0 |
| DCUN1D1 | 0,158887242 | -0,314882995 | 0 | 0,123 | 1 | CH_SC | 0,803916187 | 1,243910766 | 0 |
| ETV5 | 0,158887242 | -0,478432274 | 0 | 0,123 | 1 | CH_SC | 0,717757162 | 1,39322887 | 0 |
| PGM21 | 0,158887242 | -0,286061024 | 0 | 0,123 | 1 | CH_SC | 0,82013822 | 1,219306668 | 0 |
| PDS5A3 | 0,158887242 | -0,535569476 | 0 | 0,123 | 1 | CH_SC | 0,689886304 | 1,449514209 | 0 |
| PDGFRA | 0,158887242 | -0,411750197 | 0 | 0,123 | 1 | CH_SC | 0,751710886 | 1,33029868 | 0 |
| BMP3 | 0,158887242 | -0,396985326 | 0 | 0,123 | 1 | CH_SC | 0,75944357 | 1,316753529 | 0 |
| SEC24D1 | 0,158887242 | -0,379946385 | 0 | 0,123 | 1 | CH_SC | 0,768466149 | 1,301293494 | 0 |
| RAD1 | 0,158887242 | -0,285255319 | 0 | 0,123 | 1 | CH_SC | 0,820596372 | 1,218625909 | 0 |
| RICTOR | 0,158887242 | -0,322172677 | 0 | 0,123 | 1 | CH_SC | 0,799864386 | 1,250211933 | 0 |
| SNX18 | 0,158887242 | -0,398330563 | 0 | 0,123 | 1 | CH_SC | 0,758735758 | 1,317981905 | 0 |
| PCDHB16 | 0,158887242 | -0,56415428 | 0 | 0,123 | 1 | CH_SC | 0,676351783 | 1,478520534 | 0 |
| DAXX | 0,158887242 | -0,335783892 | 0 | 0,123 | 1 | CH_SC | 0,792353492 | 1,262062968 | 0 |
| OARD11 | 0,158887242 | -0,299121533 | 0 | 0,123 | 1 | CH_SC | 0,812747133 | 1,230394989 | 0 |
| MMUT1 | 0,158887242 | -0,407926698 | 0 | 0,123 | 1 | CH_SC | 0,753705749 | 1,326777726 | 0 |
| UST | 0,158887242 | -0,395437059 | 0 | 0,123 | 1 | CH_SC | 0,760259024 | 1,315341177 | 0 |
| OSBPL3 | 0,158887242 | -0,352703904 | 0 | 0,123 | 1 | CH_SC | 0,783115005 | 1,276951653 | 0 |
| PHTF22 | 0,158887242 | -0,607235517 | 0 | 0,123 | 1 | CH_SC | 0,656453389 | 1,523337401 | 0 |
| SEMA3E1 | 0,158887242 | -0,365214629 | 0 | 0,123 | 1 | CH_SC | 0,776353368 | 1,288073242 | 0 |
| ZYX5 | 0,158887242 | -0,352190169 | 0 | 0,123 | 1 | CH_SC | 0,783393917 | 1,276497019 | 0 |
| TRIQK | 0,158887242 | -0,359457198 | 0 | 0,123 | 1 | CH_SC | 0,779457789 | 1,28294311 | 0 |
| CTHRC12 | 0,158887242 | -0,412680753 | 0 | 0,123 | 1 | CH_SC | 0,75122618 | 1,331157015 | 0 |
| BNC2 | 0,158887242 | -0,42129238 | 0 | 0,123 | 1 | CH_SC | 0,746755374 | 1,33912662 | 0 |
| ZNF483 | 0,158887242 | -0,410403373 | 0 | 0,123 | 1 | CH_SC | 0,752412972 | 1,329057363 | 0 |
| RGS32 | 0,158887242 | -0,407795627 | 0 | 0,123 | 1 | CH_SC | 0,753774227 | 1,326657192 | 0 |
| PTPA1 | 0,158887242 | -0,342969965 | 0 | 0,123 | 1 | CH_SC | 0,788416588 | 1,268364993 | 0 |
| AKR1C31 | 0,158887242 | -0,440819603 | 0 | 0,123 | 1 | CH_SC | 0,736715957 | 1,357375241 | 0 |
| PROSER2 | 0,158887242 | -0,319882835 | 0 | 0,123 | 1 | CH_SC | 0,801134937 | 1,248229173 | 0 |
| RUFY22 | 0,158887242 | -0,347420217 | 0 | 0,123 | 1 | CH_SC | 0,785988322 | 1,272283534 | 0 |
| PALD11 | 0,158887242 | -0,253660277 | 0 | 0,123 | 1 | CH_SC | 0,838765672 | 1,192228095 | 0 |

| Supplementary Table 4 - clusermarker | | | | | | | | | |
|---|---|---|---|---|---|---|---|---|---|
| | **p_val** | **avg_log2FC** | **pct.1** | **pct.2** | **p_val_adj** | **celltype** | **Foldchange_UP** | **Foldchange_DOWN** | **Ratio_pct1_pct2** |
| DENND101 | 0,158887242 | -0,296696614 | 0 | 0,123 | 1 | CH_SC | 0,814114368 | 1,228328646 | 0 |
| FGFR2 | 0,158887242 | -0,397850885 | 0 | 0,123 | 1 | CH_SC | 0,75898807 | 1,317543765 | 0 |
| CPXM22 | 0,158887242 | -0,541076641 | 0 | 0,123 | 1 | CH_SC | 0,687257837 | 1,455057979 | 0 |
| SVIP1 | 0,158887242 | -0,478337116 | 0 | 0,123 | 1 | CH_SC | 0,717804506 | 1,393136978 | 0 |
| IFTAP1 | 0,158887242 | -0,338805273 | 0 | 0,123 | 1 | CH_SC | 0,790695833 | 1,264708827 | 0 |
| RAB3IL11 | 0,158887242 | -0,316030079 | 0 | 0,123 | 1 | CH_SC | 0,803277249 | 1,24490019 | 0 |
| FCHSD21 | 0,158887242 | -0,36245473 | 0 | 0,123 | 1 | CH_SC | 0,777839967 | 1,285611492 | 0 |
| ANKRD42 | 0,158887242 | -0,419499404 | 0 | 0,123 | 1 | CH_SC | 0,747684016 | 1,337463392 | 0 |
| LAYN | 0,158887242 | -0,409619029 | 0 | 0,123 | 1 | CH_SC | 0,752822145 | 1,328334995 | 0 |
| THYN12 | 0,158887242 | -0,366746603 | 0 | 0,123 | 1 | CH_SC | 0,77552941 | 1,289441751 | 0 |
| SCNN1A1 | 0,158887242 | -0,302152495 | 0 | 0,123 | 1 | CH_SC | 0,811041423 | 1,232982647 | 0 |
| ING4 | 0,158887242 | -0,496004179 | 0 | 0,123 | 1 | CH_SC | 0,709067964 | 1,410302045 | 0 |
| PPHLN11 | 0,158887242 | -0,415798279 | 0 | 0,123 | 1 | CH_SC | 0,749604605 | 1,334036629 | 0 |
| SLC38A13 | 0,158887242 | -0,385671837 | 0 | 0,123 | 1 | CH_SC | 0,765422472 | 1,306468044 | 0 |
| AAAS | 0,158887242 | -0,482122176 | 0 | 0,123 | 1 | CH_SC | 0,715923739 | 1,396796816 | 0 |
| TBC1D15 | 0,158887242 | -0,302900449 | 0 | 0,123 | 1 | CH_SC | 0,810621054 | 1,233622043 | 0 |
| N4BP2L1 | 0,158887242 | -0,376231448 | 0 | 0,123 | 1 | CH_SC | 0,770447498 | 1,297946976 | 0 |
| COG32 | 0,158887242 | -0,290024661 | 0 | 0,123 | 1 | CH_SC | 0,817888078 | 1,222661177 | 0 |
| TPP21 | 0,158887242 | -0,350219341 | 0 | 0,123 | 1 | CH_SC | 0,784464822 | 1,274754421 | 0 |
| KLHDC11 | 0,158887242 | -0,3863594 | 0 | 0,123 | 1 | CH_SC | 0,765057772 | 1,307090832 | 0 |
| BMP41 | 0,158887242 | -0,445025876 | 0 | 0,123 | 1 | CH_SC | 0,734571141 | 1,361338533 | 0 |
| SIX1 | 0,158887242 | -0,453071265 | 0 | 0,123 | 1 | CH_SC | 0,730486104 | 1,368951434 | 0 |
| SGPP1 | 0,158887242 | -0,393612646 | 0 | 0,123 | 1 | CH_SC | 0,761221046 | 1,313678866 | 0 |
| GREM11 | 0,158887242 | -0,558223575 | 0 | 0,123 | 1 | CH_SC | 0,679137888 | 1,472455031 | 0 |
| ZNF770 | 0,158887242 | -0,276874649 | 0 | 0,123 | 1 | CH_SC | 0,82537712 | 1,211567387 | 0 |
| MFAP11 | 0,158887242 | -0,297570041 | 0 | 0,123 | 1 | CH_SC | 0,813621642 | 1,229072518 | 0 |
| NEDD4 | 0,158887242 | -0,286053141 | 0 | 0,123 | 1 | CH_SC | 0,820142701 | 1,219300006 | 0 |
| THSD41 | 0,158887242 | -0,471759883 | 0 | 0,123 | 1 | CH_SC | 0,72108444 | 1,386800136 | 0 |
| HAPLN31 | 0,158887242 | -0,31466885 | 0 | 0,123 | 1 | CH_SC | 0,804035524 | 1,243726141 | 0 |
| NUDT16L1 | 0,158887242 | -0,447776432 | 0 | 0,123 | 1 | CH_SC | 0,733171986 | 1,363936455 | 0 |
| COQ7 | 0,158887242 | -0,27613905 | 0 | 0,123 | 1 | CH_SC | 0,825798069 | 1,210949792 | 0 |

| Supplementary Table 4 - clusermarker | | | | | | | | | |
|---|---|---|---|---|---|---|---|---|---|
| | **p_val** | **avg_log2FC** | **pct.1** | **pct.2** | **p_val_adj** | **celltype** | **Foldchange_UP** | **Foldchange_DOWN** | **Ratio_pct1_pct2** |
| PAGR11 | 0,158887242 | -0,30321959 | 0 | 0,123 | 1 | CH_SC | 0,810441755 | 1,233894964 | 0 |
| NKD11 | 0,158887242 | -0,463463149 | 0 | 0,123 | 1 | CH_SC | 0,725243241 | 1,378847734 | 0 |
| B3GNT9 | 0,158887242 | -0,373415131 | 0 | 0,123 | 1 | CH_SC | 0,771952974 | 1,295415697 | 0 |
| SYCE1L | 0,158887242 | -0,367478469 | 0 | 0,123 | 1 | CH_SC | 0,77513609 | 1,29009604 | 0 |
| DDX521 | 0,158887242 | -0,316926711 | 0 | 0,123 | 1 | CH_SC | 0,802778169 | 1,245674134 | 0 |
| FN3KRP | 0,158887242 | -0,479192953 | 0 | 0,123 | 1 | CH_SC | 0,717378815 | 1,393963661 | 0 |
| RNF152 | 0,158887242 | -0,355528076 | 0 | 0,123 | 1 | CH_SC | 0,781583504 | 1,27945382 | 0 |
| TPGS12 | 0,158887242 | -0,329940613 | 0 | 0,123 | 1 | CH_SC | 0,795569232 | 1,256961632 | 0 |
| FDX21 | 0,158887242 | -0,320126486 | 0 | 0,123 | 1 | CH_SC | 0,800999648 | 1,248439999 | 0 |
| PODNL11 | 0,158887242 | -0,361560997 | 0 | 0,123 | 1 | CH_SC | 0,778321979 | 1,284815317 | 0 |
| RFXANK1 | 0,158887242 | -0,299008631 | 0 | 0,123 | 1 | CH_SC | 0,81281074 | 1,230298705 | 0 |
| FOXA3 | 0,158887242 | -0,477482649 | 0 | 0,123 | 1 | CH_SC | 0,718229767 | 1,392312107 | 0 |
| CRNKL11 | 0,158887242 | -0,426737261 | 0 | 0,123 | 1 | CH_SC | 0,743942354 | 1,344190172 | 0 |
| LNCNEF | 0,158887242 | -0,299782008 | 0 | 0,123 | 1 | CH_SC | 0,812375137 | 1,230958401 | 0 |
| NXT13 | 0,158887242 | -0,304801527 | 0 | 0,123 | 1 | CH_SC | 0,80955358 | 1,235248691 | 0 |
| GZF1 | 0,158887242 | -0,387372954 | 0 | 0,123 | 1 | CH_SC | 0,764520476 | 1,308009441 | 0 |
| JAM21 | 0,158887242 | -0,357421861 | 0 | 0,123 | 1 | CH_SC | 0,780558215 | 1,281134425 | 0 |
| BEX5 | 0,158887242 | -0,311387497 | 0 | 0,123 | 1 | CH_SC | 0,805866353 | 1,240900549 | 0 |
| TCEAL11 | 0,158887242 | -0,320132833 | 0 | 0,123 | 1 | CH_SC | 0,800996124 | 1,248445491 | 0 |
| INTS65 | 0,159091851 | 0,599029612 | 0,533 | 0,385 | 1 | CH_SC | 1,514697405 | 0,66019787 | 1,384415584 |
| BCAP315 | 0,159094915 | 0,339515884 | 0,533 | 0,385 | 1 | CH_SC | 1,265331923 | 0,790306466 | 1,384415584 |
| SNRPD21 | 0,159785625 | -0,57995753 | 0,467 | 0,662 | 1 | CH_SC | 0,668983471 | 1,494805244 | 0,705438066 |
| EIF4G23 | 0,160160863 | 0,919890672 | 0,533 | 0,477 | 1 | CH_SC | 1,891971914 | 0,528549072 | 1,117400419 |
| DNAJC192 | 0,160323919 | -0,361135564 | 0,133 | 0,323 | 1 | CH_SC | 0,77855153 | 1,284436497 | 0,411764706 |
| SSR1 | 0,160327758 | -0,350466752 | 0,133 | 0,323 | 1 | CH_SC | 0,784330304 | 1,27497305 | 0,411764706 |
| PAPOLA1 | 0,160327758 | 1,340177736 | 0,4 | 0,262 | 1 | CH_SC | 2,531825083 | 0,394971993 | 1,526717557 |
| ARL6IP12 | 0,160790922 | 0,39607655 | 0,667 | 0,508 | 1 | CH_SC | 1,315924347 | 0,759922105 | 1,312992126 |
| TIMMDC1 | 0,160834504 | -0,61112438 | 0,133 | 0,277 | 1 | CH_SC | 0,654686266 | 1,527449179 | 0,480144404 |
| ARPC1A3 | 0,161016057 | 0,375142253 | 0,4 | 0,231 | 1 | CH_SC | 1,296967432 | 0,771029384 | 1,731601732 |
| PURB | 0,161252712 | 0,655932972 | 0,267 | 0,138 | 1 | CH_SC | 1,575634566 | 0,634664929 | 1,934782609 |
| TGFB1I1 | 0,161252712 | 0,730916536 | 0,267 | 0,138 | 1 | CH_SC | 1,659693151 | 0,602521014 | 1,934782609 |

| Supplementary Table 4 - clusermarker | | | | | | | | | |
|---|---|---|---|---|---|---|---|---|---|
| | **p_val** | **avg_log2FC** | **pct.1** | **pct.2** | **p_val_adj** | **celltype** | **Foldchange_UP** | **Foldchange_DOWN** | **Ratio_pct1_pct2** |
| CCDC115 | 0,161838926 | 0,446536214 | 0,267 | 0,123 | 1 | CH_SC | 1,362764447 | 0,733802531 | 2,170731707 |
| UBA21 | 0,161838926 | 0,410999417 | 0,267 | 0,123 | 1 | CH_SC | 1,329606571 | 0,75210218 | 2,170731707 |
| ZBTB401 | 0,162418065 | 0,539290565 | 0,2 | 0,077 | 1 | CH_SC | 1,453257712 | 0,688109199 | 2,597402597 |
| HPF1 | 0,162418065 | 0,386749854 | 0,2 | 0,077 | 1 | CH_SC | 1,307444633 | 0,764850744 | 2,597402597 |
| LOXL21 | 0,162418065 | 0,492788482 | 0,2 | 0,077 | 1 | CH_SC | 1,407162052 | 0,710650204 | 2,597402597 |
| ARL5B1 | 0,162418065 | 0,25995368 | 0,2 | 0,077 | 1 | CH_SC | 1,197440258 | 0,835114732 | 2,597402597 |
| GTF2H3 | 0,162418065 | 0,47981637 | 0,2 | 0,077 | 1 | CH_SC | 1,394566151 | 0,717068889 | 2,597402597 |
| N4BP1 | 0,162418065 | 0,345945169 | 0,2 | 0,077 | 1 | CH_SC | 1,270983384 | 0,786792347 | 2,597402597 |
| MLX2 | 0,162418065 | 0,452843388 | 0,2 | 0,077 | 1 | CH_SC | 1,368735222 | 0,730601496 | 2,597402597 |
| AP1M1 | 0,162418065 | 0,360686669 | 0,2 | 0,077 | 1 | CH_SC | 1,284036906 | 0,778793815 | 2,597402597 |
| KMT2B | 0,162418065 | 0,282484887 | 0,2 | 0,077 | 1 | CH_SC | 1,216288006 | 0,822173692 | 2,597402597 |
| CAPZB1 | 0,162764 | 0,456347629 | 0,333 | 0,185 | 1 | CH_SC | 1,372063858 | 0,728829051 | 1,8 |
| SLC25A36 | 0,163221902 | 0,858769294 | 0,533 | 0,385 | 1 | CH_SC | 1,813490634 | 0,551422754 | 1,384415584 |
| ZCCHC71 | 0,16333681 | 0,573838666 | 0,333 | 0,169 | 1 | CH_SC | 1,488478792 | 0,671826838 | 1,970414201 |
| G3BP23 | 0,163345677 | -0,479995373 | 0,067 | 0,215 | 1 | CH_SC | 0,716979923 | 1,394739193 | 0,311627907 |
| TUBB63 | 0,163345677 | -0,569494513 | 0,067 | 0,215 | 1 | CH_SC | 0,67385285 | 1,484003519 | 0,311627907 |
| LMAN14 | 0,164388045 | 0,550934152 | 0,467 | 0,308 | 1 | CH_SC | 1,465034006 | 0,682578013 | 1,516233766 |
| COL8A11 | 0,16446305 | 0,486198251 | 0,533 | 0,4 | 1 | CH_SC | 1,400748795 | 0,71390388 | 1,3325 |
| SEC629 | 0,164558044 | -0,517331847 | 0,6 | 0,8 | 1 | CH_SC | 0,698662762 | 1,431305709 | 0,75 |
| NDUFB95 | 0,164701693 | 0,824272819 | 0,6 | 0,523 | 1 | CH_SC | 1,770642332 | 0,564766798 | 1,147227533 |
| NET14 | 0,164819648 | -0,734232956 | 0,2 | 0,4 | 1 | CH_SC | 0,601137549 | 1,663512787 | 0,5 |
| GSTK14 | 0,164968234 | 0,982789728 | 0,4 | 0,262 | 1 | CH_SC | 1,97628324 | 0,506000344 | 1,526717557 |
| HSPA91 | 0,164972113 | 0,844762322 | 0,4 | 0,262 | 1 | CH_SC | 1,795968843 | 0,556802533 | 1,526717557 |
| SPARCL14 | 0,16522051 | -0,667745566 | 0,4 | 0,2 | 1 | CH_SC | 0,629489594 | 1,588588611 | 2 |
| WDR111 | 0,165562689 | 0,413270868 | 0,4 | 0,246 | 1 | CH_SC | 1,331701619 | 0,750918964 | 1,62601626 |
| TMEM1652 | 0,165827987 | 0,455116474 | 0,4 | 0,231 | 1 | CH_SC | 1,370893478 | 0,729451279 | 1,731601732 |
| LSM23 | 0,165827987 | 0,431074388 | 0,4 | 0,231 | 1 | CH_SC | 1,348237247 | 0,741709222 | 1,731601732 |
| MAP2K24 | 0,165827987 | 0,34329106 | 0,4 | 0,231 | 1 | CH_SC | 1,26864732 | 0,788241132 | 1,731601732 |
| COX7B6 | 0,166731679 | -0,485678294 | 0,333 | 0,569 | 1 | CH_SC | 0,714161222 | 1,400244046 | 0,585237258 |
| KIAA11092 | 0,16682344 | 0,853911922 | 0,333 | 0,2 | 1 | CH_SC | 1,807395107 | 0,553282454 | 1,665 |
| HBP13 | 0,16682344 | 0,712640142 | 0,333 | 0,2 | 1 | CH_SC | 1,638800391 | 0,610202442 | 1,665 |

| Supplementary Table 4 - clusermarker | | | | | | | | | |
|---|---|---|---|---|---|---|---|---|---|
| | **p_val** | **avg_log2FC** | **pct.1** | **pct.2** | **p_val_adj** | **celltype** | **Foldchange_UP** | **Foldchange_DOWN** | **Ratio_pct1_pct2** |
| NELFE1 | 0,167069735 | 0,641672115 | 0,267 | 0,138 | 1 | CH_SC | 1,560136344 | 0,64096962 | 1,934782609 |
| BTBD7 | 0,167872942 | 0,695135606 | 0,267 | 0,123 | 1 | CH_SC | 1,619036608 | 0,61765126 | 2,170731707 |
| EPB41L32 | 0,167872942 | 0,882017905 | 0,267 | 0,123 | 1 | CH_SC | 1,842951245 | 0,542607952 | 2,170731707 |
| MPV171 | 0,168017165 | 0,889835096 | 0,333 | 0,185 | 1 | CH_SC | 1,852964313 | 0,539675801 | 1,8 |
| MZT2A5 | 0,168017165 | -0,30450413 | 0,067 | 0,246 | 1 | CH_SC | 0,809720478 | 1,234994083 | 0,272357724 |
| ARF45 | 0,168244478 | 0,429313234 | 0,667 | 0,538 | 1 | CH_SC | 1,346592406 | 0,742615208 | 1,239776952 |
| ARL161 | 0,168731361 | 0,320268891 | 0,333 | 0,169 | 1 | CH_SC | 1,248563235 | 0,800920587 | 1,970414201 |
| TWSG11 | 0,168731361 | -0,401517486 | 0,067 | 0,231 | 1 | CH_SC | 0,757061556 | 1,320896554 | 0,29004329 |
| GNG12 | 0,168748239 | 0,827035541 | 0,4 | 0,277 | 1 | CH_SC | 1,774036313 | 0,563686319 | 1,444043321 |
| PITPNA-AS11 | 0,168882213 | -0,424423059 | 0,067 | 0,215 | 1 | CH_SC | 0,745136656 | 1,342035709 | 0,311627907 |
| TPM33 | 0,169162992 | 0,52118383 | 0,467 | 0,323 | 1 | CH_SC | 1,435132389 | 0,696799827 | 1,445820433 |
| NR4A27 | 0,16925921 | 0,546194142 | 0,733 | 0,692 | 1 | CH_SC | 1,460228498 | 0,684824328 | 1,059248555 |
| ILKAP | 0,169656821 | 0,296262913 | 0,2 | 0,077 | 1 | CH_SC | 1,227959444 | 0,814359143 | 2,597402597 |
| CYP2U1 | 0,169656821 | 0,283154461 | 0,2 | 0,077 | 1 | CH_SC | 1,216852633 | 0,821792198 | 2,597402597 |
| DYNC2I1 | 0,169656821 | 0,325932475 | 0,2 | 0,077 | 1 | CH_SC | 1,253474351 | 0,797782579 | 2,597402597 |
| ACIN11 | 0,169717034 | 0,966963546 | 0,4 | 0,262 | 1 | CH_SC | 1,95472214 | 0,511581661 | 1,526717557 |
| SERINC13 | 0,170523834 | 0,690402091 | 0,6 | 0,538 | 1 | CH_SC | 1,613733217 | 0,619681116 | 1,115241636 |
| SLC38A26 | 0,170549052 | 0,506112645 | 0,6 | 0,477 | 1 | CH_SC | 1,420218244 | 0,704117134 | 1,257861635 |
| ACBD33 | 0,170734354 | -0,74297769 | 0,133 | 0,277 | 1 | CH_SC | 0,597504842 | 1,673626604 | 0,480144404 |
| H3-3A3 | 0,171448014 | 0,458774813 | 0,8 | 0,8 | 1 | CH_SC | 1,374374156 | 0,727603903 | 1 |
| RAD23B1 | 0,172057175 | 0,652952579 | 0,333 | 0,2 | 1 | CH_SC | 1,572382899 | 0,635977408 | 1,665 |
| SSNA11 | 0,173043052 | 0,45640889 | 0,267 | 0,138 | 1 | CH_SC | 1,372122122 | 0,728798103 | 1,934782609 |
| TNFSF122 | 0,173043052 | 0,555826648 | 0,267 | 0,138 | 1 | CH_SC | 1,470010691 | 0,680267161 | 1,934782609 |
| LAMTOR22 | 0,173396619 | 0,467418583 | 0,333 | 0,185 | 1 | CH_SC | 1,382633305 | 0,723257567 | 1,8 |
| FGD41 | 0,173396619 | 0,359247124 | 0,067 | 0,246 | 1 | CH_SC | 1,282756312 | 0,779571296 | 0,272357724 |
| SLC22A171 | 0,173453336 | 0,65529513 | 0,467 | 0,308 | 1 | CH_SC | 1,574938103 | 0,634945588 | 1,516233766 |
| ADI15 | 0,17369527 | 0,582156312 | 0,467 | 0,323 | 1 | CH_SC | 1,497085183 | 0,667964663 | 1,445820433 |
| TDP21 | 0,17407435 | 0,255806803 | 0,267 | 0,123 | 1 | CH_SC | 1,194003283 | 0,837518636 | 2,170731707 |
| PNMA12 | 0,17407435 | 0,325658246 | 0,267 | 0,123 | 1 | CH_SC | 1,253236111 | 0,797934237 | 2,170731707 |
| VBP11 | 0,17407435 | 0,43856669 | 0,267 | 0,123 | 1 | CH_SC | 1,355257218 | 0,737867312 | 2,170731707 |
| RECK1 | 0,174258514 | -0,342033391 | 0,067 | 0,231 | 1 | CH_SC | 0,788928581 | 1,267541859 | 0,29004329 |

| Supplementary Table 4 - clusermarker | | | | | | | | | |
|---|---|---|---|---|---|---|---|---|---|
| | **p_val** | **avg_log2FC** | **pct.1** | **pct.2** | **p_val_adj** | **celltype** | **Foldchange_UP** | **Foldchange_DOWN** | **Ratio_pct1_pct2** |
| MRPS164 | 0,174558368 | -0,404855545 | 0,067 | 0,215 | 1 | CH_SC | 0,755311917 | 1,323956338 | 0,311627907 |
| BUB33 | 0,174558368 | -0,578326188 | 0,067 | 0,215 | 1 | CH_SC | 0,669740358 | 1,493115933 | 0,311627907 |
| HSD17B102 | 0,174558368 | -0,494978958 | 0,067 | 0,215 | 1 | CH_SC | 0,709572027 | 1,4093002 | 0,311627907 |
| PKM4 | 0,175414924 | -0,530632718 | 0,467 | 0,615 | 1 | CH_SC | 0,692251069 | 1,444562593 | 0,759349593 |
| XBP13 | 0,176525849 | -0,541191713 | 0,333 | 0,523 | 1 | CH_SC | 0,687203023 | 1,455174042 | 0,636711281 |
| ANKRD44 | 0,177134815 | 0,259824929 | 0,2 | 0,077 | 1 | CH_SC | 1,197333399 | 0,835189264 | 2,597402597 |
| GLB11 | 0,177134815 | 0,273784166 | 0,2 | 0,077 | 1 | CH_SC | 1,208974794 | 0,827147104 | 2,597402597 |
| LSM61 | 0,177134815 | 0,291677088 | 0,2 | 0,077 | 1 | CH_SC | 1,224062386 | 0,816951825 | 2,597402597 |
| KIF3A1 | 0,177134815 | 0,267035766 | 0,2 | 0,077 | 1 | CH_SC | 1,203332858 | 0,831025259 | 2,597402597 |
| LAMA21 | 0,177134815 | 0,282517995 | 0,2 | 0,077 | 1 | CH_SC | 1,21631592 | 0,822154823 | 2,597402597 |
| WSB21 | 0,177134815 | 0,318313444 | 0,2 | 0,077 | 1 | CH_SC | 1,246872064 | 0,802006901 | 2,597402597 |
| CSNK2A22 | 0,177134815 | 0,279253707 | 0,2 | 0,077 | 1 | CH_SC | 1,213566955 | 0,824017163 | 2,597402597 |
| TMPO | 0,177623365 | 1,134737914 | 0,2 | 0,092 | 1 | CH_SC | 2,195786696 | 0,455417642 | 2,173913043 |
| USP341 | 0,178903764 | 0,302129996 | 0,333 | 0,185 | 1 | CH_SC | 1,232963418 | 0,811054071 | 1,8 |
| GBP21 | 0,179174604 | 0,498879756 | 0,267 | 0,138 | 1 | CH_SC | 1,41311586 | 0,707656059 | 1,934782609 |
| MRPL241 | 0,179174604 | 0,608119428 | 0,267 | 0,138 | 1 | CH_SC | 1,524271006 | 0,656051316 | 1,934782609 |
| YPEL53 | 0,179174604 | 0,6771353 | 0,267 | 0,138 | 1 | CH_SC | 1,598961608 | 0,625405885 | 1,934782609 |
| CCNK1 | 0,179174604 | 0,931996237 | 0,267 | 0,138 | 1 | CH_SC | 1,907914125 | 0,524132605 | 1,934782609 |
| ESD5 | 0,179179967 | 0,651763299 | 0,533 | 0,431 | 1 | CH_SC | 1,571087246 | 0,636501889 | 1,236658933 |
| ASH1L6 | 0,179512643 | 0,706757815 | 0,4 | 0,262 | 1 | CH_SC | 1,632132083 | 0,612695511 | 1,526717557 |
| UBR42 | 0,179919768 | 0,678017433 | 0,333 | 0,169 | 1 | CH_SC | 1,599939589 | 0,625023599 | 1,970414201 |
| GSTM32 | 0,179919768 | -0,303359537 | 0,067 | 0,231 | 1 | CH_SC | 0,810363143 | 1,234014663 | 0,29004329 |
| COL16A1 | 0,180375414 | 0,503307131 | 0,4 | 0,246 | 1 | CH_SC | 1,417459123 | 0,705487717 | 1,62601626 |
| PTRHD14 | 0,180375756 | -0,449125666 | 0,067 | 0,215 | 1 | CH_SC | 0,732486631 | 1,36521263 | 0,311627907 |
| CUL11 | 0,180445286 | 0,77310517 | 0,267 | 0,123 | 1 | CH_SC | 1,708944056 | 0,585156662 | 2,170731707 |
| EIF2AK41 | 0,180445286 | 0,259308846 | 0,267 | 0,123 | 1 | CH_SC | 1,196905164 | 0,835488082 | 2,170731707 |
| CAVIN12 | 0,181179224 | 0,719427865 | 0,533 | 0,523 | 1 | CH_SC | 1,646528935 | 0,607338249 | 1,019120459 |
| LAMB23 | 0,181412615 | -0,340710831 | 0,2 | 0,446 | 1 | CH_SC | 0,789652146 | 1,2663804 | 0,448430493 |
| DYNC1LI21 | 0,182891613 | 0,755082549 | 0,333 | 0,2 | 1 | CH_SC | 1,687728159 | 0,592512482 | 1,665 |
| TMEM2582 | 0,183182181 | 0,429817362 | 0,8 | 0,723 | 1 | CH_SC | 1,347063034 | 0,742355758 | 1,106500692 |
| NME29 | 0,183182181 | -0,305866227 | 0,533 | 0,785 | 1 | CH_SC | 0,808956355 | 1,236160634 | 0,678980892 |

| Supplementary Table 4 - clusermarker | | | | | | | | | |
|---|---|---|---|---|---|---|---|---|---|
| | **p_val** | **avg_log2FC** | **pct.1** | **pct.2** | **p_val_adj** | **celltype** | **Foldchange_UP** | **Foldchange_DOWN** | **Ratio_pct1_pct2** |
| UFM15 | 0,183315401 | 0,69691248 | 0,4 | 0,277 | 1 | CH_SC | 1,621031898 | 0,616891007 | 1,444043321 |
| CTSD4 | 0,183320111 | 0,542817061 | 0,6 | 0,508 | 1 | CH_SC | 1,456814373 | 0,686429252 | 1,181102362 |
| ATP5F1B4 | 0,183589497 | 0,844714184 | 0,533 | 0,431 | 1 | CH_SC | 1,795908918 | 0,556821112 | 1,236658933 |
| CDV33 | 0,184539955 | 0,412041575 | 0,333 | 0,185 | 1 | CH_SC | 1,330567384 | 0,751559081 | 1,8 |
| ZNF3314 | 0,184565297 | -0,660118077 | 0,133 | 0,323 | 1 | CH_SC | 0,632826501 | 1,58021195 | 0,411764706 |
| MAP4K41 | 0,184958838 | 0,490125023 | 0,2 | 0,092 | 1 | CH_SC | 1,404566589 | 0,711963397 | 2,173913043 |
| RASA21 | 0,185466264 | 0,47697832 | 0,267 | 0,138 | 1 | CH_SC | 1,391825477 | 0,718480885 | 1,934782609 |
| TSSC41 | 0,185466264 | 0,866111629 | 0,267 | 0,138 | 1 | CH_SC | 1,822743591 | 0,548623517 | 1,934782609 |
| NDUFV13 | 0,185466264 | 0,524499167 | 0,267 | 0,138 | 1 | CH_SC | 1,43843414 | 0,695200407 | 1,934782609 |
| KANK22 | 0,185466264 | 0,489798026 | 0,267 | 0,138 | 1 | CH_SC | 1,40424827 | 0,712124787 | 1,934782609 |
| GALNT21 | 0,185525566 | 0,5047036 | 0,4 | 0,246 | 1 | CH_SC | 1,418831829 | 0,704805164 | 1,62601626 |
| SLC16A4 | 0,186150594 | 0,680053196 | 0,133 | 0,046 | 1 | CH_SC | 1,602198832 | 0,62414226 | 2,891304348 |
| RIT11 | 0,186150594 | 0,53994841 | 0,133 | 0,046 | 1 | CH_SC | 1,453920525 | 0,687795504 | 2,891304348 |
| KCTD3 | 0,186150594 | 0,526206941 | 0,133 | 0,046 | 1 | CH_SC | 1,440137878 | 0,694377959 | 2,891304348 |
| TMEM87B | 0,186150594 | 0,823641067 | 0,133 | 0,046 | 1 | CH_SC | 1,769867143 | 0,565014161 | 2,891304348 |
| RELL11 | 0,186150594 | 0,6114244 | 0,133 | 0,046 | 1 | CH_SC | 1,527766858 | 0,654550133 | 2,891304348 |
| ANKRD501 | 0,186150594 | 0,444561832 | 0,133 | 0,046 | 1 | CH_SC | 1,360900728 | 0,734807455 | 2,891304348 |
| FHDC1 | 0,186150594 | 0,457029672 | 0,133 | 0,046 | 1 | CH_SC | 1,372712664 | 0,728484574 | 2,891304348 |
| PCSK52 | 0,186150594 | 0,524260789 | 0,133 | 0,046 | 1 | CH_SC | 1,438196485 | 0,695315286 | 2,891304348 |
| NCBP11 | 0,186150594 | 1,149945789 | 0,133 | 0,046 | 1 | CH_SC | 2,219055559 | 0,450642164 | 2,891304348 |
| PLPP7 | 0,186150594 | 0,707113388 | 0,133 | 0,046 | 1 | CH_SC | 1,632534395 | 0,612544522 | 2,891304348 |
| LINC00863 | 0,186150594 | 0,621040414 | 0,133 | 0,046 | 1 | CH_SC | 1,537983914 | 0,650201859 | 2,891304348 |
| ABCD4 | 0,186150594 | 0,536243254 | 0,133 | 0,046 | 1 | CH_SC | 1,45019133 | 0,689564183 | 2,891304348 |
| ATG2B | 0,186150594 | 0,563176692 | 0,133 | 0,046 | 1 | CH_SC | 1,47751901 | 0,676810243 | 2,891304348 |
| RNF111 | 0,186150594 | 0,598412786 | 0,133 | 0,046 | 1 | CH_SC | 1,514049933 | 0,660480198 | 2,891304348 |
| TMEM2311 | 0,186150594 | 1,064688586 | 0,133 | 0,046 | 1 | CH_SC | 2,091718322 | 0,478075843 | 2,891304348 |
| RND2 | 0,186150594 | 0,540267953 | 0,133 | 0,046 | 1 | CH_SC | 1,45424259 | 0,687643181 | 2,891304348 |
| FAM118A4 | 0,186150594 | 0,569004638 | 0,133 | 0,046 | 1 | CH_SC | 1,483499703 | 0,674081699 | 2,891304348 |
| SUCLG11 | 0,186294618 | -0,513448014 | 0,067 | 0,2 | 1 | CH_SC | 0,700546144 | 1,427457719 | 0,335 |
| NUDT5 | 0,186294618 | -0,477112599 | 0,067 | 0,2 | 1 | CH_SC | 0,718414016 | 1,391955026 | 0,335 |
| GANAB1 | 0,186294618 | -0,576990476 | 0,067 | 0,2 | 1 | CH_SC | 0,670360721 | 1,491734179 | 0,335 |

| Supplementary Table 4 - clusermarker | | | | | | | | | |
|---|---|---|---|---|---|---|---|---|---|
| | **p_val** | **avg_log2FC** | **pct.1** | **pct.2** | **p_val_adj** | **celltype** | **Foldchange_UP** | **Foldchange_DOWN** | **Ratio_pct1_pct2** |
| NDUFV3 | 0,186294618 | -0,507290234 | 0,067 | 0,2 | 1 | CH_SC | 0,703542638 | 1,42137796 | 0,335 |
| PCNX4 | 0,186330374 | -0,435664872 | 0,067 | 0,215 | 1 | CH_SC | 0,739352942 | 1,35253401 | 0,311627907 |
| DNTTIP2 | 0,186335939 | -0,49068204 | 0,067 | 0,215 | 1 | CH_SC | 0,711688565 | 1,40510899 | 0,311627907 |
| POLR2G1 | 0,186335939 | -0,523604995 | 0,067 | 0,215 | 1 | CH_SC | 0,695631421 | 1,437542884 | 0,311627907 |
| GTF2F12 | 0,186335939 | -0,470561061 | 0,067 | 0,215 | 1 | CH_SC | 0,721683882 | 1,385648239 | 0,311627907 |
| PLS31 | 0,186765278 | 1,116752633 | 0,4 | 0,292 | 1 | CH_SC | 2,168582957 | 0,461130618 | 1,369863014 |
| TTC371 | 0,186987807 | 0,395149296 | 0,267 | 0,123 | 1 | CH_SC | 1,315078843 | 0,760410682 | 2,170731707 |
| CYB5B2 | 0,186987807 | 0,335665876 | 0,267 | 0,123 | 1 | CH_SC | 1,261959732 | 0,792418311 | 2,170731707 |
| TPGS23 | 0,186987807 | 0,44872474 | 0,267 | 0,123 | 1 | CH_SC | 1,364833289 | 0,732690218 | 2,170731707 |
| RPS171 | 0,187791932 | -0,651369376 | 0,467 | 0,662 | 1 | CH_SC | 0,636675708 | 1,570658324 | 0,705438066 |
| IER5L6 | 0,188494838 | 1,016850565 | 0,333 | 0,2 | 1 | CH_SC | 2,023496797 | 0,494194012 | 1,665 |
| SYNE24 | 0,188494838 | 0,73273303 | 0,333 | 0,2 | 1 | CH_SC | 1,661784182 | 0,601762859 | 1,665 |
| PSMC11 | 0,190306505 | 0,425554813 | 0,333 | 0,185 | 1 | CH_SC | 1,343088911 | 0,744552346 | 1,8 |
| STAG24 | 0,19078366 | 0,778083628 | 0,4 | 0,246 | 1 | CH_SC | 1,714851474 | 0,583140881 | 1,62601626 |
| SMIM121 | 0,190942499 | -0,297212763 | 0 | 0,108 | 1 | CH_SC | 0,813823157 | 1,228768181 | 0 |
| RRAGC1 | 0,190942499 | -0,399209839 | 0 | 0,108 | 1 | CH_SC | 0,758273474 | 1,318785417 | 0 |
| ZMPSTE241 | 0,190942499 | -0,560371122 | 0 | 0,108 | 1 | CH_SC | 0,678127698 | 1,47464851 | 0 |
| TACSTD23 | 0,190942499 | -0,363752716 | 0 | 0,108 | 1 | CH_SC | 0,777140462 | 1,286768671 | 0 |
| PALMD1 | 0,190942499 | -0,41816063 | 0 | 0,108 | 1 | CH_SC | 0,748378164 | 1,336222845 | 0 |
| GPSM21 | 0,190942499 | -0,561866513 | 0 | 0,108 | 1 | CH_SC | 0,677425166 | 1,476177814 | 0 |
| PIAS3 | 0,190942499 | -0,251906768 | 0 | 0,108 | 1 | CH_SC | 0,839785761 | 1,190779895 | 0 |
| LYPLAL11 | 0,190942499 | -0,485351426 | 0 | 0,108 | 1 | CH_SC | 0,714323047 | 1,399926832 | 0 |
| PLEKHH21 | 0,190942499 | -0,278645627 | 0 | 0,108 | 1 | CH_SC | 0,824364551 | 1,213055558 | 0 |
| MCEE | 0,190942499 | -0,666964217 | 0 | 0,108 | 1 | CH_SC | 0,629830611 | 1,587728482 | 0 |
| POLE45 | 0,190942499 | -0,352473957 | 0 | 0,108 | 1 | CH_SC | 0,783239833 | 1,27674814 | 0 |
| PANTR1 | 0,190942499 | -0,347951469 | 0 | 0,108 | 1 | CH_SC | 0,785698946 | 1,27275212 | 0 |
| MRPS9 | 0,190942499 | -0,315167425 | 0 | 0,108 | 1 | CH_SC | 0,803757709 | 1,244156029 | 0 |
| FHL22 | 0,190942499 | -0,433902682 | 0 | 0,108 | 1 | CH_SC | 0,740256582 | 1,350882956 | 0 |
| RALB1 | 0,190942499 | -0,327881946 | 0 | 0,108 | 1 | CH_SC | 0,796705287 | 1,255169278 | 0 |
| PLA2R1 | 0,190942499 | -0,262038868 | 0 | 0,108 | 1 | CH_SC | 0,833908577 | 1,19917222 | 0 |
| PMS1 | 0,190942499 | -0,521971013 | 0 | 0,108 | 1 | CH_SC | 0,696419733 | 1,43591566 | 0 |

| Supplementary Table 4 - clusermarker | | | | | | | | | |
|---|---|---|---|---|---|---|---|---|---|
| | **p_val** | **avg_log2FC** | **pct.1** | **pct.2** | **p_val_adj** | **celltype** | **Foldchange_UP** | **Foldchange_DOWN** | **Ratio_pct1_pct2** |
| ICA1L | 0,190942499 | -0,365547678 | 0 | 0,108 | 1 | CH_SC | 0,776174167 | 1,28837063 | 0 |
| CCNYL1 | 0,190942499 | -0,279441512 | 0 | 0,108 | 1 | CH_SC | 0,823909903 | 1,213724943 | 0 |
| PIKFYVE | 0,190942499 | -0,29414217 | 0 | 0,108 | 1 | CH_SC | 0,81555712 | 1,226155686 | 0 |
| MAP21 | 0,190942499 | -0,429546633 | 0 | 0,108 | 1 | CH_SC | 0,742495078 | 1,346810275 | 0 |
| THRB1 | 0,190942499 | -0,413120559 | 0 | 0,108 | 1 | CH_SC | 0,750997204 | 1,331562881 | 0 |
| SLC6A20 | 0,190942499 | -0,342459924 | 0 | 0,108 | 1 | CH_SC | 0,788695368 | 1,267916664 | 0 |
| LTF7 | 0,190942499 | -0,359329239 | 0 | 0,108 | 1 | CH_SC | 0,779526926 | 1,282829325 | 0 |
| IL17RB | 0,190942499 | -0,300092285 | 0 | 0,108 | 1 | CH_SC | 0,812200441 | 1,231223168 | 0 |
| TBC1D231 | 0,190942499 | -0,531390211 | 0 | 0,108 | 1 | CH_SC | 0,691887695 | 1,445321267 | 0 |
| TRMT10C1 | 0,190942499 | -0,290834541 | 0 | 0,108 | 1 | CH_SC | 0,817429072 | 1,223347731 | 0 |
| ALDH1L11 | 0,190942499 | -0,321336565 | 0 | 0,108 | 1 | CH_SC | 0,800328081 | 1,249487583 | 0 |
| YEATS2 | 0,190942499 | -0,370669679 | 0 | 0,108 | 1 | CH_SC | 0,773423401 | 1,292952862 | 0 |
| DYNLT2B1 | 0,190942499 | -0,331651294 | 0 | 0,108 | 1 | CH_SC | 0,794626442 | 1,258452963 | 0 |
| UTP3 | 0,190942499 | -0,590102337 | 0 | 0,108 | 1 | CH_SC | 0,664295784 | 1,505353525 | 0 |
| RWDD43 | 0,190942499 | -0,317931357 | 0 | 0,108 | 1 | CH_SC | 0,802219335 | 1,246541883 | 0 |
| DROSHA | 0,190942499 | -0,317656521 | 0 | 0,108 | 1 | CH_SC | 0,802372173 | 1,246304437 | 0 |
| DUSP22 | 0,190942499 | -0,320047917 | 0 | 0,108 | 1 | CH_SC | 0,801043272 | 1,248372011 | 0 |
| BMP6 | 0,190942499 | -0,388372601 | 0 | 0,108 | 1 | CH_SC | 0,763990921 | 1,308916078 | 0 |
| HMGN4 | 0,190942499 | -0,274344827 | 0 | 0,108 | 1 | CH_SC | 0,82682572 | 1,209444718 | 0 |
| MOCS1 | 0,190942499 | -0,362171356 | 0 | 0,108 | 1 | CH_SC | 0,777992765 | 1,285358997 | 0 |
| MRPS101 | 0,190942499 | -0,355810774 | 0 | 0,108 | 1 | CH_SC | 0,781430367 | 1,279704555 | 0 |
| PGM31 | 0,190942499 | -0,409682408 | 0 | 0,108 | 1 | CH_SC | 0,752789073 | 1,328393352 | 0 |
| NT5DC1 | 0,190942499 | -0,307416337 | 0 | 0,108 | 1 | CH_SC | 0,808087635 | 1,237489546 | 0 |
| COL10A11 | 0,190942499 | -3,741934495 | 0 | 0,108 | 1 | CH_SC | 0,074742131 | 13,3793349 | 0 |
| C6orf5810 | 0,190942499 | -0,340666435 | 0 | 0,108 | 1 | CH_SC | 0,789676447 | 1,266341429 | 0 |
| FKBP141 | 0,190942499 | -0,282020931 | 0 | 0,108 | 1 | CH_SC | 0,822438136 | 1,215896924 | 0 |
| PMPCB2 | 0,190942499 | -0,430988958 | 0 | 0,108 | 1 | CH_SC | 0,741753144 | 1,348157413 | 0 |
| LINC-PINT1 | 0,190942499 | -0,376507214 | 0 | 0,108 | 1 | CH_SC | 0,770300243 | 1,298195098 | 0 |
| PIP9 | 0,190942499 | -0,351710589 | 0 | 0,108 | 1 | CH_SC | 0,783654376 | 1,276072758 | 0 |
| SCARA51 | 0,190942499 | -0,405210412 | 0 | 0,108 | 1 | CH_SC | 0,755126152 | 1,324282038 | 0 |
| TGS1 | 0,190942499 | -0,251269904 | 0 | 0,108 | 1 | CH_SC | 0,840156559 | 1,190254352 | 0 |

| Supplementary Table 4 - clusermarker | | | | | | | | | |
|---|---|---|---|---|---|---|---|---|---|
| | **p_val** | **avg_log2FC** | **pct.1** | **pct.2** | **p_val_adj** | **celltype** | **Foldchange_UP** | **Foldchange_DOWN** | **Ratio_pct1_pct2** |
| MSMP | 0,190942499 | -2,839538601 | 0 | 0,108 | 1 | CH_SC | 0,139705565 | 7,157910978 | 0 |
| HRCT1 | 0,190942499 | -0,417669201 | 0 | 0,108 | 1 | CH_SC | 0,74863313 | 1,335767762 | 0 |
| GKAP1 | 0,190942499 | -0,320605369 | 0 | 0,108 | 1 | CH_SC | 0,800733811 | 1,248854471 | 0 |
| SURF2 | 0,190942499 | -0,368438409 | 0 | 0,108 | 1 | CH_SC | 0,774620502 | 1,290954729 | 0 |
| SVIL3 | 0,190942499 | -0,359386495 | 0 | 0,108 | 1 | CH_SC | 0,779495989 | 1,282880237 | 0 |
| GDF101 | 0,190942499 | -0,47513721 | 0 | 0,108 | 1 | CH_SC | 0,719398367 | 1,390050417 | 0 |
| EPS8L21 | 0,190942499 | -0,32024575 | 0 | 0,108 | 1 | CH_SC | 0,800933434 | 1,248543209 | 0 |
| CHST1 | 0,190942499 | -0,263879126 | 0 | 0,108 | 1 | CH_SC | 0,832845547 | 1,200702824 | 0 |
| CORO1B3 | 0,190942499 | -0,373537887 | 0 | 0,108 | 1 | CH_SC | 0,771887293 | 1,295525926 | 0 |
| C11orf711 | 0,190942499 | -0,356971574 | 0 | 0,108 | 1 | CH_SC | 0,780801877 | 1,280734626 | 0 |
| TAGLN5 | 0,190942499 | -1,099083399 | 0 | 0,108 | 1 | CH_SC | 0,466812986 | 2,142185478 | 0 |
| CENATAC | 0,190942499 | -0,316362523 | 0 | 0,108 | 1 | CH_SC | 0,803092169 | 1,245187089 | 0 |
| RHNO1 | 0,190942499 | -0,324584053 | 0 | 0,108 | 1 | CH_SC | 0,798528579 | 1,252303332 | 0 |
| C12orf4 | 0,190942499 | -0,260219012 | 0 | 0,108 | 1 | CH_SC | 0,834961156 | 1,197660505 | 0 |
| ZNF384 | 0,190942499 | -0,279762015 | 0 | 0,108 | 1 | CH_SC | 0,823726887 | 1,213994609 | 0 |
| GPRC5A1 | 0,190942499 | -0,325233112 | 0 | 0,108 | 1 | CH_SC | 0,798169407 | 1,252866862 | 0 |
| DIP2B | 0,190942499 | -0,367325613 | 0 | 0,108 | 1 | CH_SC | 0,775218221 | 1,289959359 | 0 |
| NDUFA4L22 | 0,190942499 | -0,38781505 | 0 | 0,108 | 1 | CH_SC | 0,764286234 | 1,308410326 | 0 |
| SLC16A71 | 0,190942499 | -0,345846702 | 0 | 0,108 | 1 | CH_SC | 0,78684605 | 1,270896639 | 0 |
| TAFA2 | 0,190942499 | -0,487149007 | 0 | 0,108 | 1 | CH_SC | 0,713433563 | 1,401672212 | 0 |
| HECTD4 | 0,190942499 | -0,300822269 | 0 | 0,108 | 1 | CH_SC | 0,811789582 | 1,231846308 | 0 |
| CCDC92 | 0,190942499 | -0,351324175 | 0 | 0,108 | 1 | CH_SC | 0,783864299 | 1,275731017 | 0 |
| MRPS311 | 0,190942499 | -0,417638298 | 0 | 0,108 | 1 | CH_SC | 0,748649166 | 1,33573915 | 0 |
| DZIP1 | 0,190942499 | -0,318511272 | 0 | 0,108 | 1 | CH_SC | 0,801896934 | 1,247043052 | 0 |
| ITGBL12 | 0,190942499 | -0,407239666 | 0 | 0,108 | 1 | CH_SC | 0,75406476 | 1,326146046 | 0 |
| GCH1 | 0,190942499 | -0,373780686 | 0 | 0,108 | 1 | CH_SC | 0,771757399 | 1,295743975 | 0 |
| TMEM30B | 0,190942499 | -0,442004452 | 0 | 0,108 | 1 | CH_SC | 0,736111159 | 1,358490477 | 0 |
| MTMR10 | 0,190942499 | -0,272969997 | 0 | 0,108 | 1 | CH_SC | 0,827614028 | 1,208292715 | 0 |
| POLR2M | 0,190942499 | -0,34706558 | 0 | 0,108 | 1 | CH_SC | 0,786181554 | 1,271970825 | 0 |
| SCAPER1 | 0,190942499 | -0,293506553 | 0 | 0,108 | 1 | CH_SC | 0,815916515 | 1,225615589 | 0 |
| IL161 | 0,190942499 | -0,324518457 | 0 | 0,108 | 1 | CH_SC | 0,798564887 | 1,252246393 | 0 |

| Supplementary Table 4 - clusermarker | | | | | | | | | |
|---|---|---|---|---|---|---|---|---|---|
| | **p_val** | **avg_log2FC** | **pct.1** | **pct.2** | **p_val_adj** | **celltype** | **Foldchange_UP** | **Foldchange_DOWN** | **Ratio_pct1_pct2** |
| DEXI | 0,190942499 | -0,473628058 | 0 | 0,108 | 1 | CH_SC | 0,720151298 | 1,388597095 | 0 |
| LCMT13 | 0,190942499 | -0,298129719 | 0 | 0,108 | 1 | CH_SC | 0,813306067 | 1,229549417 | 0 |
| MAPK32 | 0,190942499 | -0,362634377 | 0 | 0,108 | 1 | CH_SC | 0,777743115 | 1,285771589 | 0 |
| PYCARD6 | 0,190942499 | -0,348208653 | 0 | 0,108 | 1 | CH_SC | 0,785558895 | 1,27297903 | 0 |
| ATP6V0D11 | 0,190942499 | -0,327652469 | 0 | 0,108 | 1 | CH_SC | 0,796832022 | 1,254969645 | 0 |
| ATMIN | 0,190942499 | -0,347299513 | 0 | 0,108 | 1 | CH_SC | 0,786054085 | 1,272177091 | 0 |
| TLCD21 | 0,190942499 | -0,303327602 | 0 | 0,108 | 1 | CH_SC | 0,810381081 | 1,233987347 | 0 |
| PELP11 | 0,190942499 | -0,479893593 | 0 | 0,108 | 1 | CH_SC | 0,717030507 | 1,3946408 | 0 |
| MYH102 | 0,190942499 | -0,329654758 | 0 | 0,108 | 1 | CH_SC | 0,795726881 | 1,256712603 | 0 |
| RAB11FIP41 | 0,190942499 | -0,316252205 | 0 | 0,108 | 1 | CH_SC | 0,803153581 | 1,245091878 | 0 |
| CWC252 | 0,190942499 | -0,355965833 | 0 | 0,108 | 1 | CH_SC | 0,781346384 | 1,279842103 | 0 |
| PGAP3 | 0,190942499 | -0,283113803 | 0 | 0,108 | 1 | CH_SC | 0,821815358 | 1,216818341 | 0 |
| JUP | 0,190942499 | -0,437494776 | 0 | 0,108 | 1 | CH_SC | 0,738415747 | 1,354250643 | 0 |
| ZNF385C | 0,190942499 | -0,342992081 | 0 | 0,108 | 1 | CH_SC | 0,788404502 | 1,268384437 | 0 |
| ATP6V0A1 | 0,190942499 | -0,406888371 | 0 | 0,108 | 1 | CH_SC | 0,754248396 | 1,32582317 | 0 |
| HLF2 | 0,190942499 | -0,314574705 | 0 | 0,108 | 1 | CH_SC | 0,804087995 | 1,243644983 | 0 |
| MRPS231 | 0,190942499 | -0,308952213 | 0 | 0,108 | 1 | CH_SC | 0,807227812 | 1,238807664 | 0 |
| TRIM47 | 0,190942499 | -0,351816161 | 0 | 0,108 | 1 | CH_SC | 0,783597032 | 1,27616614 | 0 |
| LPIN2 | 0,190942499 | -0,372791076 | 0 | 0,108 | 1 | CH_SC | 0,772286964 | 1,294855471 | 0 |
| PIEZO2 | 0,190942499 | -0,392032313 | 0 | 0,108 | 1 | CH_SC | 0,762055347 | 1,312240645 | 0 |
| LDLRAD42 | 0,190942499 | -0,456890254 | 0 | 0,108 | 1 | CH_SC | 0,728554976 | 1,372580015 | 0 |
| TAF4B | 0,190942499 | -0,36756931 | 0 | 0,108 | 1 | CH_SC | 0,775087284 | 1,290177274 | 0 |
| CCDC159 | 0,190942499 | -0,263285956 | 0 | 0,108 | 1 | CH_SC | 0,833188045 | 1,200209252 | 0 |
| RBM422 | 0,190942499 | -0,265200134 | 0 | 0,108 | 1 | CH_SC | 0,832083298 | 1,201802755 | 0 |
| LIN37 | 0,190942499 | -0,3039645 | 0 | 0,108 | 1 | CH_SC | 0,810023406 | 1,234532229 | 0 |
| XRCC1 | 0,190942499 | -0,266048675 | 0 | 0,108 | 1 | CH_SC | 0,831594041 | 1,20250982 | 0 |
| SIRPA1 | 0,190942499 | -0,332681563 | 0 | 0,108 | 1 | CH_SC | 0,794059179 | 1,25935198 | 0 |
| AAR2 | 0,190942499 | -0,273308028 | 0 | 0,108 | 1 | CH_SC | 0,827420136 | 1,208575857 | 0 |
| BCL2L131 | 0,190942499 | -0,36870508 | 0 | 0,108 | 1 | CH_SC | 0,774477333 | 1,291193374 | 0 |
| ZMAT51 | 0,190942499 | -0,268493212 | 0 | 0,108 | 1 | CH_SC | 0,830186161 | 1,204549108 | 0 |
| PNPLA4 | 0,190942499 | -0,336508856 | 0 | 0,108 | 1 | CH_SC | 0,791955429 | 1,262697323 | 0 |

| Supplementary Table 4 - clusermarker | | | | | | | | | |
|---|---|---|---|---|---|---|---|---|---|
| | **p_val** | **avg_log2FC** | **pct.1** | **pct.2** | **p_val_adj** | **celltype** | **Foldchange_UP** | **Foldchange_DOWN** | **Ratio_pct1_pct2** |
| MAOB1 | 0,190942499 | -0,268215022 | 0 | 0,108 | 1 | CH_SC | 0,830346258 | 1,204316862 | 0 |
| KDM6A1 | 0,190942499 | -0,335037095 | 0 | 0,108 | 1 | CH_SC | 0,792763753 | 1,261409842 | 0 |
| LPAR4 | 0,190942499 | -0,280505248 | 0 | 0,108 | 1 | CH_SC | 0,823302637 | 1,214620184 | 0 |
| BEX1 | 0,190942499 | -0,269903112 | 0 | 0,108 | 1 | CH_SC | 0,829375243 | 1,205726851 | 0 |
| ZC3H12A1 | 0,191520392 | -0,741455398 | 0,133 | 0,292 | 1 | CH_SC | 0,598135646 | 1,671861571 | 0,455479452 |
| ABHD14A1 | 0,191650318 | -0,290582598 | 0,067 | 0,231 | 1 | CH_SC | 0,817571835 | 1,223134112 | 0,29004329 |
| SEC31A3 | 0,191842642 | 0,792430746 | 0,533 | 0,415 | 1 | CH_SC | 1,731990175 | 0,577370481 | 1,284337349 |
| DPY301 | 0,191891784 | -0,689401143 | 0,133 | 0,277 | 1 | CH_SC | 0,620111202 | 1,61261399 | 0,480144404 |
| PHYHD12 | 0,191891784 | -0,578193398 | 0,133 | 0,277 | 1 | CH_SC | 0,669802006 | 1,492978508 | 0,480144404 |
| DPYSL22 | 0,191919836 | 0,467414381 | 0,267 | 0,138 | 1 | CH_SC | 1,382629278 | 0,723259673 | 1,934782609 |
| SERPINH12 | 0,192399769 | 0,584754087 | 0,467 | 0,292 | 1 | CH_SC | 1,499783324 | 0,666762981 | 1,599315068 |
| ELOVL52 | 0,192516075 | 0,748732752 | 0,2 | 0,092 | 1 | CH_SC | 1,68031621 | 0,59512608 | 2,173913043 |
| CRTC31 | 0,192516075 | 0,744535239 | 0,2 | 0,092 | 1 | CH_SC | 1,675434445 | 0,596860118 | 2,173913043 |
| EDEM31 | 0,192572735 | -0,589929712 | 0,067 | 0,2 | 1 | CH_SC | 0,664375275 | 1,505173413 | 0,335 |
| ARF33 | 0,192572735 | -0,439895396 | 0,067 | 0,2 | 1 | CH_SC | 0,737188057 | 1,356505969 | 0,335 |
| SDF2 | 0,192572735 | -0,395134962 | 0,067 | 0,2 | 1 | CH_SC | 0,760418237 | 1,315065777 | 0,335 |
| SUPT5H1 | 0,192572735 | -0,436142644 | 0,067 | 0,2 | 1 | CH_SC | 0,739108134 | 1,352981998 | 0,335 |
| AKAP96 | 0,192695934 | -0,490596734 | 0,2 | 0,369 | 1 | CH_SC | 0,711730648 | 1,405025908 | 0,54200542 |
| CCDC471 | 0,192695934 | 0,488249485 | 0,467 | 0,308 | 1 | CH_SC | 1,402741805 | 0,712889568 | 1,516233766 |
| HNRNPD2 | 0,193531779 | -0,356706343 | 0,133 | 0,338 | 1 | CH_SC | 0,780945436 | 1,280499192 | 0,393491124 |
| MT-CO34 | 0,195532849 | 0,267967439 | 1 | 0,938 | 1 | CH_SC | 1,204110205 | 0,830488767 | 1,066098081 |
| ISG155 | 0,196123911 | 0,89064968 | 0,133 | 0,046 | 1 | CH_SC | 1,854010841 | 0,539371172 | 2,891304348 |
| IFT172 | 0,196123911 | 0,541294304 | 0,133 | 0,046 | 1 | CH_SC | 1,455277524 | 0,687154157 | 2,891304348 |
| ALAS1 | 0,196123911 | 0,426894803 | 0,133 | 0,046 | 1 | CH_SC | 1,344336965 | 0,74386112 | 2,891304348 |
| TAP12 | 0,196123911 | 0,319101903 | 0,133 | 0,046 | 1 | CH_SC | 1,247553688 | 0,80156871 | 2,891304348 |
| TRAM2-AS1 | 0,196123911 | 0,399915131 | 0,133 | 0,046 | 1 | CH_SC | 1,319430291 | 0,757902867 | 2,891304348 |
| UNC5B2 | 0,196123911 | 0,445774725 | 0,133 | 0,046 | 1 | CH_SC | 1,362045336 | 0,734189952 | 2,891304348 |
| DUSP53 | 0,196123911 | 0,36468327 | 0,133 | 0,046 | 1 | CH_SC | 1,287598918 | 0,77663936 | 2,891304348 |
| COMMD91 | 0,196123911 | 0,420650456 | 0,133 | 0,046 | 1 | CH_SC | 1,338530911 | 0,747087715 | 2,891304348 |
| ARID21 | 0,196123911 | 0,682076884 | 0,133 | 0,046 | 1 | CH_SC | 1,604447835 | 0,623267381 | 2,891304348 |
| CUL4A | 0,196123911 | 0,777056871 | 0,133 | 0,046 | 1 | CH_SC | 1,71363146 | 0,583556047 | 2,891304348 |

| Supplementary Table 4 - clusermarker | | | | | | | | | |
|---|---|---|---|---|---|---|---|---|---|
| | **p_val** | **avg_log2FC** | **pct.1** | **pct.2** | **p_val_adj** | **celltype** | **Foldchange_UP** | **Foldchange_DOWN** | **Ratio_pct1_pct2** |
| DMAC2L1 | 0,196123911 | 0,49681216 | 0,133 | 0,046 | 1 | CH_SC | 1,411092106 | 0,708670962 | 2,891304348 |
| GOT21 | 0,196123911 | 0,369487763 | 0,133 | 0,046 | 1 | CH_SC | 1,291894055 | 0,774057281 | 2,891304348 |
| MAP1S | 0,196123911 | 0,922466739 | 0,133 | 0,046 | 1 | CH_SC | 1,895353225 | 0,527606141 | 2,891304348 |
| CDC375 | 0,196150644 | 0,51491731 | 0,4 | 0,246 | 1 | CH_SC | 1,428912237 | 0,699833044 | 1,62601626 |
| MAGEF11 | 0,196204676 | 0,406606878 | 0,333 | 0,185 | 1 | CH_SC | 1,325564506 | 0,754395577 | 1,8 |
| SAP182 | 0,196359054 | 0,711922551 | 0,733 | 0,662 | 1 | CH_SC | 1,637985461 | 0,610506029 | 1,107250755 |
| FOXO1 | 0,196988434 | 0,767841044 | 0,333 | 0,246 | 1 | CH_SC | 1,7027198 | 0,58729569 | 1,353658537 |
| MTUS12 | 0,196992817 | -0,450167563 | 0,133 | 0,292 | 1 | CH_SC | 0,731957829 | 1,366198926 | 0,455479452 |
| HNRNPH21 | 0,196992817 | -0,582209381 | 0,133 | 0,292 | 1 | CH_SC | 0,667940093 | 1,497140254 | 0,455479452 |
| SEPTIN23 | 0,197299781 | 0,612824548 | 0,533 | 0,431 | 1 | CH_SC | 1,529250288 | 0,653915195 | 1,236658933 |
| PDLIM53 | 0,197502368 | -0,537344891 | 0,2 | 0,354 | 1 | CH_SC | 0,689037835 | 1,451299115 | 0,564971751 |
| EAPP2 | 0,197536255 | 0,823529945 | 0,333 | 0,215 | 1 | CH_SC | 1,769730826 | 0,565057683 | 1,548837209 |
| S100A162 | 0,198537051 | 0,410273656 | 0,267 | 0,138 | 1 | CH_SC | 1,328937868 | 0,752480627 | 1,934782609 |
| SLU72 | 0,198537051 | 0,862987434 | 0,267 | 0,138 | 1 | CH_SC | 1,818800662 | 0,549812864 | 1,934782609 |
| CD823 | 0,198537051 | 0,504448477 | 0,267 | 0,138 | 1 | CH_SC | 1,418580948 | 0,704929811 | 1,934782609 |
| HIP1 | 0,198690645 | -0,455110208 | 0,067 | 0,215 | 1 | CH_SC | 0,729454447 | 1,370887524 | 0,311627907 |
| BMI11 | 0,198690645 | -0,350029466 | 0,067 | 0,215 | 1 | CH_SC | 0,784568074 | 1,274586659 | 0,311627907 |
| IMP33 | 0,198794079 | 0,729655429 | 0,4 | 0,277 | 1 | CH_SC | 1,658242992 | 0,603047928 | 1,444043321 |
| ECHDC22 | 0,199004983 | -0,49469657 | 0,067 | 0,2 | 1 | CH_SC | 0,70971093 | 1,409024376 | 0,335 |
| PRRC1 | 0,199004983 | -0,412563242 | 0,067 | 0,2 | 1 | CH_SC | 0,751287372 | 1,331048594 | 0,335 |
| CYHR11 | 0,199004983 | -0,413363669 | 0,067 | 0,2 | 1 | CH_SC | 0,750870663 | 1,331787283 | 0,335 |
| TMEM2161 | 0,199004983 | -0,459108342 | 0,067 | 0,2 | 1 | CH_SC | 0,727435712 | 1,374691926 | 0,335 |
| CARHSP12 | 0,199004983 | -0,863743722 | 0,067 | 0,2 | 1 | CH_SC | 0,549524717 | 1,819754362 | 0,335 |
| IL13RA11 | 0,199004983 | -0,499920037 | 0,067 | 0,2 | 1 | CH_SC | 0,707145974 | 1,41413518 | 0,335 |
| RNF1871 | 0,20007928 | 0,68343158 | 0,333 | 0,2 | 1 | CH_SC | 1,605955124 | 0,622682406 | 1,665 |
| GOLGA72 | 0,20007928 | 1,01531786 | 0,333 | 0,2 | 1 | CH_SC | 2,021348195 | 0,494719318 | 1,665 |
| MIR4435-2HG2 | 0,200297852 | 0,665912804 | 0,2 | 0,092 | 1 | CH_SC | 1,586571792 | 0,630289789 | 2,173913043 |
| USF32 | 0,200297852 | 0,502934028 | 0,2 | 0,092 | 1 | CH_SC | 1,417092593 | 0,70567019 | 2,173913043 |
| DPM2 | 0,200595424 | 0,278566183 | 0,267 | 0,123 | 1 | CH_SC | 1,212988761 | 0,824409947 | 2,170731707 |
| ERH6 | 0,202032423 | 0,306192058 | 0,533 | 0,431 | 1 | CH_SC | 1,23643985 | 0,808773674 | 1,236658933 |
| PDCD41 | 0,202703305 | -0,573170943 | 0,2 | 0,354 | 1 | CH_SC | 0,672137852 | 1,487790038 | 0,564971751 |

| Supplementary Table 4 - clusermarker | | | | | | | | | |
|---|---|---|---|---|---|---|---|---|---|
| | **p_val** | **avg_log2FC** | **pct.1** | **pct.2** | **p_val_adj** | **celltype** | **Foldchange_UP** | **Foldchange_DOWN** | **Ratio_pct1_pct2** |
| GOLGA41 | 0,202808773 | -0,413102701 | 0,2 | 0,385 | 1 | CH_SC | 0,7510065 | 1,331546399 | 0,519480519 |
| ARRDC32 | 0,203933977 | 0,350046146 | 0,067 | 0,231 | 1 | CH_SC | 1,274601396 | 0,784559003 | 0,29004329 |
| AHCYL11 | 0,205319559 | 0,535742432 | 0,267 | 0,138 | 1 | CH_SC | 1,449687993 | 0,689803602 | 1,934782609 |
| RPRD21 | 0,205319559 | 0,48418837 | 0,267 | 0,138 | 1 | CH_SC | 1,398798709 | 0,714899144 | 1,934782609 |
| SYNM2 | 0,205319559 | 0,640914708 | 0,267 | 0,138 | 1 | CH_SC | 1,559317496 | 0,641306214 | 1,934782609 |
| CDC27 | 0,205319559 | 0,478584427 | 0,267 | 0,138 | 1 | CH_SC | 1,393375814 | 0,717681468 | 1,934782609 |
| PDCD71 | 0,205592862 | -0,564014014 | 0,067 | 0,2 | 1 | CH_SC | 0,676417545 | 1,478376792 | 0,335 |
| PSENEN1 | 0,205592862 | -0,377217184 | 0,067 | 0,2 | 1 | CH_SC | 0,769921261 | 1,298834115 | 0,335 |
| YIPF1 | 0,206483244 | 0,474254626 | 0,133 | 0,046 | 1 | CH_SC | 1,389200299 | 0,719838601 | 2,891304348 |
| ZBTB411 | 0,206483244 | 0,344328816 | 0,133 | 0,046 | 1 | CH_SC | 1,269560208 | 0,787674341 | 2,891304348 |
| ELF31 | 0,206483244 | 0,376973499 | 0,133 | 0,046 | 1 | CH_SC | 1,298614748 | 0,770051319 | 2,891304348 |
| NOL10 | 0,206483244 | 0,560135251 | 0,133 | 0,046 | 1 | CH_SC | 1,474407435 | 0,678238577 | 2,891304348 |
| IWS1 | 0,206483244 | 0,372702895 | 0,133 | 0,046 | 1 | CH_SC | 1,294776329 | 0,772334169 | 2,891304348 |
| LIMS21 | 0,206483244 | 0,627514651 | 0,133 | 0,046 | 1 | CH_SC | 1,544901279 | 0,647290551 | 2,891304348 |
| LYAR | 0,206483244 | 0,394878305 | 0,133 | 0,046 | 1 | CH_SC | 1,314831846 | 0,760553529 | 2,891304348 |
| MSX11 | 0,206483244 | 0,730197647 | 0,133 | 0,046 | 1 | CH_SC | 1,658866338 | 0,602821323 | 2,891304348 |
| GALNT7 | 0,206483244 | 0,393272347 | 0,133 | 0,046 | 1 | CH_SC | 1,313369036 | 0,761400622 | 2,891304348 |
| CDK71 | 0,206483244 | 0,537075653 | 0,133 | 0,046 | 1 | CH_SC | 1,451028296 | 0,689166436 | 2,891304348 |
| ELOVL4 | 0,206483244 | 0,38299506 | 0,133 | 0,046 | 1 | CH_SC | 1,30404627 | 0,766843956 | 2,891304348 |
| STEAP1B | 0,206483244 | 0,457628411 | 0,133 | 0,046 | 1 | CH_SC | 1,373282477 | 0,728182305 | 2,891304348 |
| MCM71 | 0,206483244 | 0,338682167 | 0,133 | 0,046 | 1 | CH_SC | 1,264600913 | 0,790763307 | 2,891304348 |
| DEPTOR1 | 0,206483244 | 0,307189585 | 0,133 | 0,046 | 1 | CH_SC | 1,237295061 | 0,808214654 | 2,891304348 |
| NAPRT3 | 0,206483244 | 0,382783814 | 0,133 | 0,046 | 1 | CH_SC | 1,303855339 | 0,766956249 | 2,891304348 |
| AP3M1 | 0,206483244 | 0,299547225 | 0,133 | 0,046 | 1 | CH_SC | 1,230758092 | 0,812507354 | 2,891304348 |
| C10orf82 | 0,206483244 | 0,491768686 | 0,133 | 0,046 | 1 | CH_SC | 1,406167724 | 0,711152719 | 2,891304348 |
| C11orf491 | 0,206483244 | 0,693469453 | 0,133 | 0,046 | 1 | CH_SC | 1,617167878 | 0,618364991 | 2,891304348 |
| PPP2R5B | 0,206483244 | 0,283331223 | 0,133 | 0,046 | 1 | CH_SC | 1,217001733 | 0,821691517 | 2,891304348 |
| SNX15 | 0,206483244 | 0,352865892 | 0,133 | 0,046 | 1 | CH_SC | 1,277095039 | 0,78302708 | 2,891304348 |
| ETV62 | 0,206483244 | 0,362543365 | 0,133 | 0,046 | 1 | CH_SC | 1,285690479 | 0,77779218 | 2,891304348 |
| WASHC43 | 0,206483244 | 0,328497527 | 0,133 | 0,046 | 1 | CH_SC | 1,255704958 | 0,796365415 | 2,891304348 |
| SYNE31 | 0,206483244 | 0,350622055 | 0,133 | 0,046 | 1 | CH_SC | 1,275110306 | 0,784245877 | 2,891304348 |

| Supplementary Table 4 - clusermarker | | | | | | | | | |
|---|---|---|---|---|---|---|---|---|---|
| | **p_val** | **avg_log2FC** | **pct.1** | **pct.2** | **p_val_adj** | **celltype** | **Foldchange_UP** | **Foldchange_DOWN** | **Ratio_pct1_pct2** |
| EFTUD2 | 0,206483244 | 0,329288687 | 0,133 | 0,046 | 1 | CH_SC | 1,256393763 | 0,795928816 | 2,891304348 |
| MRPL49 | 0,206483244 | 0,410493773 | 0,133 | 0,046 | 1 | CH_SC | 1,329140644 | 0,752365827 | 2,891304348 |
| APOBEC3C1 | 0,206483244 | 0,335244149 | 0,133 | 0,046 | 1 | CH_SC | 1,261590891 | 0,792649984 | 2,891304348 |
| RPN22 | 0,206847283 | 0,295696789 | 0,533 | 0,431 | 1 | CH_SC | 1,227477678 | 0,814678766 | 1,236658933 |
| SERPINE13 | 0,20766421 | 0,269797173 | 0,267 | 0,123 | 1 | CH_SC | 1,205638316 | 0,829436147 | 2,170731707 |
| NDUFAF35 | 0,207987951 | 0,731416991 | 0,467 | 0,323 | 1 | CH_SC | 1,66026898 | 0,602312042 | 1,445820433 |
| ALDOA4 | 0,208125787 | -0,514747679 | 0,2 | 0,369 | 1 | CH_SC | 0,699915334 | 1,428744236 | 0,54200542 |
| NOC2L | 0,208306797 | 0,643961853 | 0,2 | 0,092 | 1 | CH_SC | 1,562614442 | 0,639953128 | 2,173913043 |
| MEF2D | 0,208306797 | 0,5362094 | 0,2 | 0,092 | 1 | CH_SC | 1,450157301 | 0,689580364 | 2,173913043 |
| WDR41 | 0,208306797 | 0,358878306 | 0,2 | 0,092 | 1 | CH_SC | 1,282428423 | 0,779770615 | 2,173913043 |
| CCNDBP13 | 0,208306797 | 0,528420613 | 0,2 | 0,092 | 1 | CH_SC | 1,442349322 | 0,693313322 | 2,173913043 |
| C15orf402 | 0,208306797 | 0,526987583 | 0,2 | 0,092 | 1 | CH_SC | 1,440917347 | 0,694002332 | 2,173913043 |
| IRS21 | 0,208400684 | 0,576155704 | 0,333 | 0,185 | 1 | CH_SC | 1,490871281 | 0,670748718 | 1,8 |
| DNAJC155 | 0,20927497 | 0,73156352 | 0,333 | 0,215 | 1 | CH_SC | 1,660437615 | 0,602250871 | 1,548837209 |
| TMED102 | 0,210299073 | -0,473703643 | 0,4 | 0,631 | 1 | CH_SC | 0,720113569 | 1,388669847 | 0,633914422 |
| PNPLA22 | 0,211413667 | 0,621114678 | 0,4 | 0,262 | 1 | CH_SC | 1,538063085 | 0,65016839 | 1,526717557 |
| NDUFB15 | 0,211413667 | 0,853419025 | 0,4 | 0,262 | 1 | CH_SC | 1,806777715 | 0,553471515 | 1,526717557 |
| SLC26A2 | 0,211633791 | -0,588206312 | 0,067 | 0,215 | 1 | CH_SC | 0,665169391 | 1,503376453 | 0,311627907 |
| LRP6 | 0,21226893 | 0,552138491 | 0,267 | 0,138 | 1 | CH_SC | 1,466257504 | 0,682008445 | 1,934782609 |
| MRPL2 | 0,212337801 | -0,561861394 | 0,067 | 0,2 | 1 | CH_SC | 0,677427569 | 1,476172576 | 0,335 |
| UBL3 | 0,212337801 | -0,487482907 | 0,067 | 0,2 | 1 | CH_SC | 0,713268464 | 1,401996655 | 0,335 |
| RHEB2 | 0,213463017 | -0,615798138 | 0,2 | 0,369 | 1 | CH_SC | 0,652568775 | 1,532405531 | 0,54200542 |
| FMC11 | 0,21409541 | -0,364068137 | 0,133 | 0,292 | 1 | CH_SC | 0,776970572 | 1,287050032 | 0,455479452 |
| WDR61 | 0,21409541 | -0,460866483 | 0,133 | 0,292 | 1 | CH_SC | 0,726549762 | 1,376368217 | 0,455479452 |
| PRDX15 | 0,214611041 | -0,443356238 | 0,6 | 0,8 | 1 | CH_SC | 0,735421756 | 1,359763961 | 0,75 |
| AKIRIN21 | 0,214700787 | 0,361699044 | 0,333 | 0,185 | 1 | CH_SC | 1,284938262 | 0,778247508 | 1,8 |
| PSAP3 | 0,215314165 | 0,283651639 | 0,933 | 0,923 | 1 | CH_SC | 1,217272054 | 0,821509043 | 1,010834236 |
| MTRES1 | 0,21654538 | 0,969533355 | 0,2 | 0,092 | 1 | CH_SC | 1,958207104 | 0,510671215 | 2,173913043 |
| GLRX31 | 0,21654538 | 0,327760235 | 0,2 | 0,092 | 1 | CH_SC | 1,255063392 | 0,796772503 | 2,173913043 |
| SCO1 | 0,21654538 | 0,68585814 | 0,2 | 0,092 | 1 | CH_SC | 1,608658555 | 0,621635957 | 2,173913043 |
| NOL111 | 0,21654538 | 0,419066944 | 0,2 | 0,092 | 1 | CH_SC | 1,337062536 | 0,747908174 | 2,173913043 |

| Supplementary Table 4 - clusermarker | | | | | | | | | |
|---|---|---|---|---|---|---|---|---|---|
| | **p_val** | **avg_log2FC** | **pct.1** | **pct.2** | **p_val_adj** | **celltype** | **Foldchange_UP** | **Foldchange_DOWN** | **Ratio_pct1_pct2** |
| SETDB1 | 0,217234252 | 0,255550291 | 0,133 | 0,046 | 1 | CH_SC | 1,193791007 | 0,83766756 | 2,891304348 |
| ASB1 | 0,217234252 | 0,368469053 | 0,133 | 0,046 | 1 | CH_SC | 1,29098215 | 0,774604049 | 2,891304348 |
| NEK4 | 0,217234252 | 0,315310955 | 0,133 | 0,046 | 1 | CH_SC | 1,244279814 | 0,803677749 | 2,891304348 |
| HACD2 | 0,217234252 | 0,404786413 | 0,133 | 0,046 | 1 | CH_SC | 1,323892897 | 0,755348112 | 2,891304348 |
| CEP70 | 0,217234252 | 0,25366491 | 0,133 | 0,046 | 1 | CH_SC | 1,192231924 | 0,838762979 | 2,891304348 |
| CDKAL1 | 0,217234252 | 0,305329421 | 0,133 | 0,046 | 1 | CH_SC | 1,235700761 | 0,809257412 | 2,891304348 |
| AL391807.1 | 0,217234252 | 0,428404462 | 0,133 | 0,046 | 1 | CH_SC | 1,345744438 | 0,743083138 | 2,891304348 |
| RBM28 | 0,217234252 | 0,35064161 | 0,133 | 0,046 | 1 | CH_SC | 1,275127589 | 0,784235248 | 2,891304348 |
| CLN8 | 0,217234252 | 0,345022869 | 0,133 | 0,046 | 1 | CH_SC | 1,270171117 | 0,787295496 | 2,891304348 |
| QSOX2 | 0,217234252 | 0,460313653 | 0,133 | 0,046 | 1 | CH_SC | 1,375840904 | 0,726828223 | 2,891304348 |
| NMT2 | 0,217234252 | 0,813246423 | 0,133 | 0,046 | 1 | CH_SC | 1,757161047 | 0,5690998 | 2,891304348 |
| SLC35E3 | 0,217234252 | 0,649407109 | 0,133 | 0,046 | 1 | CH_SC | 1,568523462 | 0,637542264 | 2,891304348 |
| RNF34 | 0,217234252 | 0,315311621 | 0,133 | 0,046 | 1 | CH_SC | 1,244280388 | 0,803677378 | 2,891304348 |
| NBEA | 0,217234252 | 0,327113913 | 0,133 | 0,046 | 1 | CH_SC | 1,254501254 | 0,797129534 | 2,891304348 |
| ERCC51 | 0,217234252 | 0,373812867 | 0,133 | 0,046 | 1 | CH_SC | 1,295772879 | 0,771740184 | 2,891304348 |
| METTL171 | 0,217234252 | 0,359328848 | 0,133 | 0,046 | 1 | CH_SC | 1,282828978 | 0,779527137 | 2,891304348 |
| MIA21 | 0,217234252 | 0,378513644 | 0,133 | 0,046 | 1 | CH_SC | 1,300001821 | 0,769229692 | 2,891304348 |
| FNTB | 0,217234252 | 0,264459585 | 0,133 | 0,046 | 1 | CH_SC | 1,201186016 | 0,832510524 | 2,891304348 |
| AC127502.2 | 0,217234252 | 0,311373808 | 0,133 | 0,046 | 1 | CH_SC | 1,240888775 | 0,805873999 | 2,891304348 |
| LETR1 | 0,217234252 | 0,808541074 | 0,133 | 0,046 | 1 | CH_SC | 1,751439403 | 0,570958949 | 2,891304348 |
| BECN11 | 0,217234252 | 0,289892476 | 0,133 | 0,046 | 1 | CH_SC | 1,222549158 | 0,817963019 | 2,891304348 |
| BAHCC1 | 0,217234252 | 0,408318426 | 0,133 | 0,046 | 1 | CH_SC | 1,327138029 | 0,753501127 | 2,891304348 |
| SPPL2B | 0,217234252 | 0,327547771 | 0,133 | 0,046 | 1 | CH_SC | 1,254878574 | 0,796889851 | 2,891304348 |
| ZNF260 | 0,217234252 | 0,434769058 | 0,133 | 0,046 | 1 | CH_SC | 1,35169444 | 0,739812172 | 2,891304348 |
| TASP11 | 0,217234252 | 0,733442693 | 0,133 | 0,046 | 1 | CH_SC | 1,662601817 | 0,601466924 | 2,891304348 |
| APMAP2 | 0,217234252 | 0,42378477 | 0,133 | 0,046 | 1 | CH_SC | 1,341442085 | 0,745466398 | 2,891304348 |
| FUNDC1 | 0,217234252 | 0,334431966 | 0,133 | 0,046 | 1 | CH_SC | 1,260880863 | 0,793096342 | 2,891304348 |
| RBM101 | 0,217234252 | 0,592569311 | 0,133 | 0,046 | 1 | CH_SC | 1,507929846 | 0,663160824 | 2,891304348 |
| LITAF3 | 0,21785633 | 0,774986378 | 0,6 | 0,492 | 1 | CH_SC | 1,711173894 | 0,584394142 | 1,219512195 |
| COX201 | 0,218034929 | -0,52386596 | 0,2 | 0,323 | 1 | CH_SC | 0,695505602 | 1,437802941 | 0,619195046 |
| KDM2A1 | 0,218417474 | 0,447740135 | 0,333 | 0,2 | 1 | CH_SC | 1,36390214 | 0,733190433 | 1,665 |

| Supplementary Table 4 - clusermarker | | | | | | | | | |
|---|---|---|---|---|---|---|---|---|---|
| | **p_val** | **avg_log2FC** | **pct.1** | **pct.2** | **p_val_adj** | **celltype** | **Foldchange_UP** | **Foldchange_DOWN** | **Ratio_pct1_pct2** |
| ASXL11 | 0,218417474 | 0,744222913 | 0,333 | 0,2 | 1 | CH_SC | 1,675071773 | 0,596989345 | 1,665 |
| CCT73 | 0,218903314 | 0,362558093 | 0,467 | 0,292 | 1 | CH_SC | 1,285703604 | 0,77778424 | 1,599315068 |
| CDR2 | 0,219241152 | -0,388796325 | 0,067 | 0,2 | 1 | CH_SC | 0,763766568 | 1,309300567 | 0,335 |
| ORMDL1 | 0,219386648 | 0,492954139 | 0,267 | 0,138 | 1 | CH_SC | 1,407323638 | 0,710568609 | 1,934782609 |
| RAB1B1 | 0,219386648 | 0,883026366 | 0,267 | 0,138 | 1 | CH_SC | 1,84423994 | 0,542228795 | 1,934782609 |
| WDR13 | 0,219386648 | 0,375527412 | 0,267 | 0,138 | 1 | CH_SC | 1,297313732 | 0,770823568 | 1,934782609 |
| WTAP2 | 0,22002762 | -0,343328153 | 0,133 | 0,292 | 1 | CH_SC | 0,788220866 | 1,268679938 | 0,455479452 |
| TMEM106B2 | 0,22002762 | -0,312549799 | 0,133 | 0,292 | 1 | CH_SC | 0,805217371 | 1,241900679 | 0,455479452 |
| TXNL4A6 | 0,22002762 | 0,314622327 | 0,4 | 0,231 | 1 | CH_SC | 1,243686035 | 0,804061453 | 1,731601732 |
| BTF3L4 | 0,22097221 | -0,70012005 | 0,133 | 0,277 | 1 | CH_SC | 0,615520986 | 1,624639977 | 0,480144404 |
| NPHP31 | 0,22097221 | -0,616364071 | 0,133 | 0,277 | 1 | CH_SC | 0,652312839 | 1,533006773 | 0,480144404 |
| ITPR12 | 0,22113704 | 0,499440123 | 0,333 | 0,185 | 1 | CH_SC | 1,413664845 | 0,707381246 | 1,8 |
| RBM261 | 0,22113704 | 0,388063992 | 0,333 | 0,185 | 1 | CH_SC | 1,308636116 | 0,764154365 | 1,8 |
| IGBP12 | 0,22113704 | 0,495835549 | 0,333 | 0,185 | 1 | CH_SC | 1,410137211 | 0,709150849 | 1,8 |
| BIRC63 | 0,222340259 | 0,723487382 | 0,267 | 0,123 | 1 | CH_SC | 1,651168533 | 0,605631696 | 2,170731707 |
| PRRC2C4 | 0,222642158 | 0,567999853 | 0,533 | 0,462 | 1 | CH_SC | 1,482466858 | 0,674551336 | 1,153679654 |
| ATOX14 | 0,222734521 | 0,670367146 | 0,467 | 0,354 | 1 | CH_SC | 1,591477926 | 0,628346761 | 1,31920904 |
| SDAD12 | 0,222752633 | 1,013363842 | 0,267 | 0,154 | 1 | CH_SC | 2,01861229 | 0,49538983 | 1,733766234 |
| ATP5F1D10 | 0,222908277 | -0,297959437 | 0,133 | 0,323 | 1 | CH_SC | 0,813402068 | 1,229404301 | 0,411764706 |
| IK5 | 0,223523439 | -0,545320248 | 0,267 | 0,385 | 1 | CH_SC | 0,685239277 | 1,459344252 | 0,693506494 |
| HTRA15 | 0,223701784 | -0,532442538 | 0,4 | 0,646 | 1 | CH_SC | 0,691383204 | 1,446375893 | 0,619195046 |
| LARS14 | 0,224790675 | 0,35122856 | 0,333 | 0,2 | 1 | CH_SC | 1,275646471 | 0,783916252 | 1,665 |
| WAC2 | 0,224790675 | 0,434452701 | 0,333 | 0,2 | 1 | CH_SC | 1,351398071 | 0,739974417 | 1,665 |
| BSCL2 | 0,224790675 | 0,527549564 | 0,333 | 0,2 | 1 | CH_SC | 1,441478745 | 0,693732046 | 1,665 |
| TNFRSF142 | 0,22501591 | 0,440113288 | 0,2 | 0,092 | 1 | CH_SC | 1,356710859 | 0,737076727 | 2,173913043 |
| GNL2 | 0,22501591 | 0,336903909 | 0,2 | 0,092 | 1 | CH_SC | 1,263043133 | 0,791738598 | 2,173913043 |
| ANAPC131 | 0,22501591 | 0,490285253 | 0,2 | 0,092 | 1 | CH_SC | 1,404722593 | 0,711884328 | 2,173913043 |
| CTBP13 | 0,22501591 | 0,508127219 | 0,2 | 0,092 | 1 | CH_SC | 1,422202817 | 0,703134594 | 2,173913043 |
| CTBP21 | 0,22501591 | 0,511256943 | 0,2 | 0,092 | 1 | CH_SC | 1,425291435 | 0,701610896 | 2,173913043 |
| SNX191 | 0,22501591 | 0,3774913 | 0,2 | 0,092 | 1 | CH_SC | 1,29908092 | 0,769774988 | 2,173913043 |
| IREB21 | 0,22501591 | 0,391429766 | 0,2 | 0,092 | 1 | CH_SC | 1,311692697 | 0,762373689 | 2,173913043 |

| Supplementary Table 4 - clusermarker | | | | | | | | | |
|---|---|---|---|---|---|---|---|---|---|
| | **p_val** | **avg_log2FC** | **pct.1** | **pct.2** | **p_val_adj** | **celltype** | **Foldchange_UP** | **Foldchange_DOWN** | **Ratio_pct1_pct2** |
| FLYWCH24 | 0,22501591 | 0,343813804 | 0,2 | 0,092 | 1 | CH_SC | 1,269107083 | 0,787955574 | 2,173913043 |
| DDRGK13 | 0,22501591 | 0,304083229 | 0,2 | 0,092 | 1 | CH_SC | 1,234633831 | 0,809956746 | 2,173913043 |
| RTN45 | 0,225451337 | -0,597965243 | 0,267 | 0,462 | 1 | CH_SC | 0,66068512 | 1,513580328 | 0,577922078 |
| BBIP12 | 0,226076853 | -0,331226365 | 0,133 | 0,292 | 1 | CH_SC | 0,794860524 | 1,258082355 | 0,455479452 |
| FOXJ3 | 0,226674106 | 0,80457348 | 0,267 | 0,138 | 1 | CH_SC | 1,746629341 | 0,572531319 | 1,934782609 |
| SMIM42 | 0,226674106 | -0,471143158 | 0,067 | 0,185 | 1 | CH_SC | 0,721392757 | 1,386207431 | 0,362162162 |
| SRPRB | 0,226674106 | -0,426354725 | 0,067 | 0,185 | 1 | CH_SC | 0,744139638 | 1,343833803 | 0,362162162 |
| PPP3CA2 | 0,226674106 | 0,425660117 | 0,267 | 0,138 | 1 | CH_SC | 1,343186949 | 0,744498002 | 1,934782609 |
| DENR1 | 0,226674106 | 0,360101576 | 0,267 | 0,138 | 1 | CH_SC | 1,283516263 | 0,779109723 | 1,934782609 |
| SF3A21 | 0,226674106 | -0,486182168 | 0,067 | 0,185 | 1 | CH_SC | 0,713911839 | 1,400733179 | 0,362162162 |
| ARHGDIA3 | 0,226678342 | 0,475074086 | 0,4 | 0,277 | 1 | CH_SC | 1,389989598 | 0,719429844 | 1,444043321 |
| RIOK3 | 0,227148898 | -0,598143964 | 0,133 | 0,277 | 1 | CH_SC | 0,660603279 | 1,513767842 | 0,480144404 |
| NONO3 | 0,227817515 | 0,667556516 | 0,333 | 0,215 | 1 | CH_SC | 1,588380458 | 0,629572087 | 1,548837209 |
| EGR16 | 0,227860459 | 0,256607802 | 0,8 | 0,677 | 1 | CH_SC | 1,19466639 | 0,837053766 | 1,1816839 |
| PCM12 | 0,228160367 | 0,677185132 | 0,467 | 0,354 | 1 | CH_SC | 1,59901684 | 0,625384283 | 1,31920904 |
| TUBB4B4 | 0,228160367 | 1,207343246 | 0,467 | 0,354 | 1 | CH_SC | 2,309120155 | 0,433065381 | 1,31920904 |
| KDM1A2 | 0,228382117 | 0,258593374 | 0,133 | 0,046 | 1 | CH_SC | 1,196311734 | 0,835902526 | 2,891304348 |
| OSBPL101 | 0,228382117 | 0,280048385 | 0,133 | 0,046 | 1 | CH_SC | 1,214235607 | 0,823563396 | 2,891304348 |
| QTRT2 | 0,228382117 | 0,457912572 | 0,133 | 0,046 | 1 | CH_SC | 1,373552993 | 0,728038893 | 2,891304348 |
| NUDT9 | 0,228382117 | 0,261113474 | 0,133 | 0,046 | 1 | CH_SC | 1,198403277 | 0,834443646 | 2,891304348 |
| DENND5B | 0,228382117 | 0,31668287 | 0,133 | 0,046 | 1 | CH_SC | 1,24546361 | 0,802913864 | 2,891304348 |
| GTF2A1 | 0,228382117 | 0,280375451 | 0,133 | 0,046 | 1 | CH_SC | 1,214510911 | 0,823376711 | 2,891304348 |
| TUBGCP4 | 0,228382117 | 0,253245443 | 0,133 | 0,046 | 1 | CH_SC | 1,191885331 | 0,839006886 | 2,891304348 |
| A1BG3 | 0,228382117 | 0,256414774 | 0,133 | 0,046 | 1 | CH_SC | 1,194506558 | 0,837165768 | 2,891304348 |
| GOLM24 | 0,229021726 | -0,69148791 | 0,267 | 0,385 | 1 | CH_SC | 0,619214899 | 1,614948222 | 0,693506494 |
| GADD45B7 | 0,229338386 | -0,626493414 | 0,667 | 0,738 | 1 | CH_SC | 0,647748909 | 1,543808081 | 0,903794038 |
| TMBIM62 | 0,229435664 | 0,434046085 | 0,733 | 0,677 | 1 | CH_SC | 1,35101724 | 0,740183005 | 1,082717873 |
| RAD213 | 0,229663201 | 0,647538495 | 0,467 | 0,323 | 1 | CH_SC | 1,566493187 | 0,63836856 | 1,445820433 |
| UBE2B4 | 0,229733618 | 0,583663005 | 0,467 | 0,4 | 1 | CH_SC | 1,498649495 | 0,667267432 | 1,1675 |
| HIGD2A5 | 0,229895962 | 0,433905499 | 0,267 | 0,154 | 1 | CH_SC | 1,350885594 | 0,740255137 | 1,733766234 |
| RB1CC11 | 0,229895962 | 0,689838111 | 0,267 | 0,154 | 1 | CH_SC | 1,613102498 | 0,619923409 | 1,733766234 |

| Supplementary Table 4 - clusermarker | | | | | | | | | |
|---|---|---|---|---|---|---|---|---|---|
| | **p_val** | **avg_log2FC** | **pct.1** | **pct.2** | **p_val_adj** | **celltype** | **Foldchange_UP** | **Foldchange_DOWN** | **Ratio_pct1_pct2** |
| PPP1R14B6 | 0,229895962 | 0,539610084 | 0,267 | 0,154 | 1 | CH_SC | 1,453579606 | 0,687956818 | 1,733766234 |
| FKBP101 | 0,23006559 | -0,672261404 | 0,2 | 0,338 | 1 | CH_SC | 0,627522284 | 1,593568907 | 0,591715976 |
| HNRNPUL13 | 0,230282991 | 0,601049409 | 0,333 | 0,231 | 1 | CH_SC | 1,516819493 | 0,659274228 | 1,441558442 |
| RSF12 | 0,231295648 | 0,33276734 | 0,333 | 0,2 | 1 | CH_SC | 1,259426859 | 0,794011969 | 1,665 |
| UXT4 | 0,232172834 | 0,510827008 | 0,533 | 0,431 | 1 | CH_SC | 1,42486675 | 0,701820012 | 1,236658933 |
| STUB17 | 0,232243852 | -0,445438914 | 0,133 | 0,292 | 1 | CH_SC | 0,734360866 | 1,361728335 | 0,455479452 |
| HNRNPK4 | 0,232848364 | 0,444762104 | 0,6 | 0,569 | 1 | CH_SC | 1,361089658 | 0,734705457 | 1,054481547 |
| EMC1 | 0,233720521 | 0,4734518 | 0,2 | 0,092 | 1 | CH_SC | 1,388427456 | 0,720239286 | 2,173913043 |
| AHDC11 | 0,233720521 | 0,334905675 | 0,2 | 0,092 | 1 | CH_SC | 1,261294941 | 0,792835972 | 2,173913043 |
| SUPT7L | 0,233720521 | 0,393354529 | 0,2 | 0,092 | 1 | CH_SC | 1,313443853 | 0,76135725 | 2,173913043 |
| CLASP2 | 0,233720521 | 0,382566006 | 0,2 | 0,092 | 1 | CH_SC | 1,303658507 | 0,767072047 | 2,173913043 |
| WDR48 | 0,233720521 | 0,300444092 | 0,2 | 0,092 | 1 | CH_SC | 1,231523444 | 0,812002406 | 2,173913043 |
| RASA1 | 0,233720521 | 0,442637212 | 0,2 | 0,092 | 1 | CH_SC | 1,359086436 | 0,735788375 | 2,173913043 |
| NT5C3A1 | 0,233720521 | 0,275045371 | 0,2 | 0,092 | 1 | CH_SC | 1,210032143 | 0,826424327 | 2,173913043 |
| DLD1 | 0,233720521 | 0,339220052 | 0,2 | 0,092 | 1 | CH_SC | 1,265072487 | 0,790468539 | 2,173913043 |
| ZDHHC17 | 0,233720521 | 0,274812372 | 0,2 | 0,092 | 1 | CH_SC | 1,209836735 | 0,826557808 | 2,173913043 |
| MLXIP2 | 0,233720521 | 0,275877226 | 0,2 | 0,092 | 1 | CH_SC | 1,210730046 | 0,82594795 | 2,173913043 |
| MLLT62 | 0,233720521 | 0,362666993 | 0,2 | 0,092 | 1 | CH_SC | 1,285800657 | 0,777725532 | 2,173913043 |
| KDM6B2 | 0,234132605 | -0,7483805 | 0,067 | 0,185 | 1 | CH_SC | 0,595271406 | 1,67990599 | 0,362162162 |
| ZNF667-AS1 | 0,234132605 | -0,398023668 | 0,067 | 0,185 | 1 | CH_SC | 0,758897177 | 1,317701569 | 0,362162162 |
| ERVK3-1 | 0,234132605 | -0,590452312 | 0,067 | 0,185 | 1 | CH_SC | 0,664134656 | 1,505718745 | 0,362162162 |
| DDX211 | 0,235563422 | -0,57524925 | 0,2 | 0,323 | 1 | CH_SC | 0,671170285 | 1,489934852 | 0,619195046 |
| KCNQ1OT13 | 0,235943303 | -0,727160575 | 0,2 | 0,338 | 1 | CH_SC | 0,604091681 | 1,655377869 | 0,591715976 |
| MBOAT21 | 0,237194462 | 0,383477196 | 0,267 | 0,154 | 1 | CH_SC | 1,304482144 | 0,766587726 | 1,733766234 |
| NEK12 | 0,237194462 | 0,754311424 | 0,267 | 0,154 | 1 | CH_SC | 1,686826304 | 0,592829266 | 1,733766234 |
| HECTD1 | 0,23720073 | 0,655297306 | 0,267 | 0,154 | 1 | CH_SC | 1,574940479 | 0,63494463 | 1,733766234 |
| SNW13 | 0,23720073 | 0,5384668 | 0,267 | 0,154 | 1 | CH_SC | 1,452428153 | 0,688502215 | 1,733766234 |
| SYNGR11 | 0,23720073 | 0,482689469 | 0,267 | 0,154 | 1 | CH_SC | 1,39734617 | 0,715642281 | 1,733766234 |
| HNRNPA18 | 0,237861592 | -0,330707657 | 0,667 | 0,862 | 1 | CH_SC | 0,795146361 | 1,257630104 | 0,773781903 |
| CREBBP3 | 0,237933231 | 0,603565166 | 0,333 | 0,2 | 1 | CH_SC | 1,519466814 | 0,658125594 | 1,665 |
| USP114 | 0,237933231 | 0,511189329 | 0,333 | 0,2 | 1 | CH_SC | 1,425224638 | 0,701643778 | 1,665 |

| Supplementary Table 4 - clusermarker | | | | | | | | | |
|---|---|---|---|---|---|---|---|---|---|
| | p_val | avg_log2FC | pct.1 | pct.2 | p_val_adj | celltype | Foldchange_UP | Foldchange_DOWN | Ratio_pct1_pct2 |
| AKAP133 | 0,238003421 | 0,860904805 | 0,2 | 0,108 | 1 | CH_SC | 1,816176993 | 0,550607129 | 1,851851852 |
| JADE11 | 0,238529315 | -0,371363267 | 0,133 | 0,292 | 1 | CH_SC | 0,773051661 | 1,293574609 | 0,455479452 |
| LNPEP3 | 0,238529315 | 0,77015266 | 0,4 | 0,231 | 1 | CH_SC | 1,705450237 | 0,586355426 | 1,731601732 |
| SELENOF2 | 0,239293043 | 0,497134028 | 0,467 | 0,354 | 1 | CH_SC | 1,411406959 | 0,708512873 | 1,31920904 |
| HMGN23 | 0,240880141 | 0,444982848 | 0,6 | 0,477 | 1 | CH_SC | 1,361297932 | 0,73459305 | 1,257861635 |
| EPHX13 | 0,240913988 | -0,557157942 | 0,067 | 0,2 | 1 | CH_SC | 0,679639712 | 1,471367818 | 0,335 |
| ETF13 | 0,240913988 | -0,370828943 | 0,067 | 0,2 | 1 | CH_SC | 0,773338025 | 1,293095604 | 0,335 |
| POLD44 | 0,240913988 | -0,328123867 | 0,067 | 0,2 | 1 | CH_SC | 0,796571701 | 1,255379771 | 0,335 |
| COX4I18 | 0,24091604 | 0,541088778 | 0,733 | 0,708 | 1 | CH_SC | 1,45507022 | 0,687252056 | 1,035310734 |
| IER3IP11 | 0,241622066 | -0,538340414 | 0,2 | 0,323 | 1 | CH_SC | 0,688562533 | 1,45230092 | 0,619195046 |
| CYP27A12 | 0,24176335 | -0,3836238 | 0,067 | 0,185 | 1 | CH_SC | 0,766509831 | 1,30461471 | 0,362162162 |
| LRMDA | 0,24176335 | -0,568313014 | 0,067 | 0,185 | 1 | CH_SC | 0,674404929 | 1,482788688 | 0,362162162 |
| SLF2 | 0,24176335 | 1,198032084 | 0,267 | 0,138 | 1 | CH_SC | 2,294265071 | 0,435869426 | 1,934782609 |
| GRB21 | 0,24176335 | 0,76735954 | 0,267 | 0,138 | 1 | CH_SC | 1,702151607 | 0,587491735 | 1,934782609 |
| UCKL11 | 0,24176335 | -0,53445048 | 0,067 | 0,185 | 1 | CH_SC | 0,690421607 | 1,44839036 | 0,362162162 |
| YY1AP1 | 0,242661167 | 0,257416768 | 0,2 | 0,092 | 1 | CH_SC | 1,195336465 | 0,836584534 | 2,173913043 |
| ELK41 | 0,242661167 | 0,54796996 | 0,2 | 0,092 | 1 | CH_SC | 1,462027005 | 0,683981894 | 2,173913043 |
| USP39 | 0,242661167 | 0,40007698 | 0,2 | 0,092 | 1 | CH_SC | 1,319578319 | 0,757817846 | 2,173913043 |
| DHX301 | 0,242661167 | 0,348085143 | 0,2 | 0,092 | 1 | CH_SC | 1,272870053 | 0,78562615 | 2,173913043 |
| ENG2 | 0,242661167 | 0,408007073 | 0,2 | 0,092 | 1 | CH_SC | 1,326851645 | 0,75366376 | 2,173913043 |
| SURF13 | 0,242661167 | 0,390708724 | 0,2 | 0,092 | 1 | CH_SC | 1,311037293 | 0,762754809 | 2,173913043 |
| RESF11 | 0,242661167 | 0,512700354 | 0,2 | 0,092 | 1 | CH_SC | 1,426718147 | 0,700909287 | 2,173913043 |
| ADAM101 | 0,242661167 | 0,310862888 | 0,2 | 0,092 | 1 | CH_SC | 1,240449401 | 0,806159444 | 2,173913043 |
| RAB8B | 0,242661167 | 0,337011534 | 0,2 | 0,092 | 1 | CH_SC | 1,263137361 | 0,791679536 | 2,173913043 |
| LRPAP12 | 0,243107879 | -0,324751487 | 0,133 | 0,308 | 1 | CH_SC | 0,79843591 | 1,252448678 | 0,431818182 |
| DDX63 | 0,244000633 | 0,944385214 | 0,4 | 0,338 | 1 | CH_SC | 1,924368666 | 0,519650947 | 1,183431953 |
| YIPF21 | 0,244667997 | 0,456788325 | 0,267 | 0,154 | 1 | CH_SC | 1,372483043 | 0,728606452 | 1,733766234 |
| ZNF6384 | 0,244687965 | 0,823463268 | 0,4 | 0,277 | 1 | CH_SC | 1,769649036 | 0,565083799 | 1,444043321 |
| SRSF1 | 0,244692173 | 0,702288021 | 0,4 | 0,277 | 1 | CH_SC | 1,627083197 | 0,614596723 | 1,444043321 |
| ARPC43 | 0,244704202 | 0,3916762 | 0,333 | 0,2 | 1 | CH_SC | 1,311916773 | 0,762243475 | 1,665 |
| KTN14 | 0,246199492 | 1,263450441 | 0,467 | 0,4 | 1 | CH_SC | 2,400692197 | 0,416546528 | 1,1675 |

| Supplementary Table 4 - clusermarker | | | | | | | | | |
|---|---|---|---|---|---|---|---|---|---|
| | **p_val** | **avg_log2FC** | **pct.1** | **pct.2** | **p_val_adj** | **celltype** | **Foldchange_UP** | **Foldchange_DOWN** | **Ratio_pct1_pct2** |
| CHMP2B | 0,246414548 | -0,397116588 | 0,133 | 0,277 | 1 | CH_SC | 0,759374475 | 1,316873338 | 0,480144404 |
| SRRM13 | 0,246525773 | -0,553724921 | 0,533 | 0,646 | 1 | CH_SC | 0,681258901 | 1,46787073 | 0,825077399 |
| NINJ14 | 0,247224071 | 0,577586151 | 0,4 | 0,262 | 1 | CH_SC | 1,492350229 | 0,670083993 | 1,526717557 |
| NOTCH21 | 0,247501926 | -0,407615493 | 0,133 | 0,262 | 1 | CH_SC | 0,753868349 | 1,326491557 | 0,507633588 |
| CTDNEP12 | 0,247501926 | -0,581044296 | 0,133 | 0,262 | 1 | CH_SC | 0,668479723 | 1,495931688 | 0,507633588 |
| PHB23 | 0,247890236 | -0,475123988 | 0,267 | 0,4 | 1 | CH_SC | 0,71940496 | 1,390037678 | 0,6675 |
| DYNLT15 | 0,247932271 | -0,396870866 | 0,2 | 0,354 | 1 | CH_SC | 0,759503824 | 1,316649065 | 0,564971751 |
| NDUFA124 | 0,248262427 | 0,273332564 | 0,333 | 0,185 | 1 | CH_SC | 1,208596412 | 0,827406064 | 1,8 |
| SLC25A35 | 0,24826736 | 0,43149856 | 0,6 | 0,569 | 1 | CH_SC | 1,348633706 | 0,741491182 | 1,054481547 |
| PDXK3 | 0,248462874 | -0,338159249 | 0,067 | 0,2 | 1 | CH_SC | 0,791049978 | 1,26414263 | 0,335 |
| NDUFS42 | 0,249447711 | 0,529138008 | 0,467 | 0,369 | 1 | CH_SC | 1,443066724 | 0,69296865 | 1,265582656 |
| ANKRD351 | 0,249567443 | -0,353465815 | 0,067 | 0,185 | 1 | CH_SC | 0,782701538 | 1,27762621 | 0,362162162 |
| MAT2B2 | 0,249567443 | -0,355378818 | 0,067 | 0,185 | 1 | CH_SC | 0,781664369 | 1,279321457 | 0,362162162 |
| SNX92 | 0,249567443 | 0,304175532 | 0,267 | 0,138 | 1 | CH_SC | 1,234712825 | 0,809904927 | 1,934782609 |
| RAN3 | 0,249928009 | 0,712310867 | 0,467 | 0,415 | 1 | CH_SC | 1,6384264 | 0,610341728 | 1,125301205 |
| SSR46 | 0,250905088 | -0,35734647 | 0,4 | 0,615 | 1 | CH_SC | 0,780599006 | 1,281067478 | 0,650406504 |
| NCKAP12 | 0,251458186 | -0,409177379 | 0,133 | 0,292 | 1 | CH_SC | 0,75305264 | 1,327928416 | 0,455479452 |
| GET11 | 0,251458186 | -0,295919955 | 0,133 | 0,292 | 1 | CH_SC | 0,814552756 | 1,227667567 | 0,455479452 |
| SMAD51 | 0,251609279 | 0,362535345 | 0,333 | 0,2 | 1 | CH_SC | 1,285683331 | 0,777796504 | 1,665 |
| CIB14 | 0,251609279 | 0,37756524 | 0,333 | 0,2 | 1 | CH_SC | 1,299147502 | 0,769735537 | 1,665 |
| JUND3 | 0,251727095 | 0,84853792 | 0,667 | 0,615 | 1 | CH_SC | 1,800675131 | 0,55534726 | 1,084552846 |
| RNF1151 | 0,251839615 | 0,456067468 | 0,2 | 0,092 | 1 | CH_SC | 1,371797439 | 0,728970598 | 2,173913043 |
| PRKD3 | 0,251839615 | 0,270022911 | 0,2 | 0,092 | 1 | CH_SC | 1,205826977 | 0,829306376 | 2,173913043 |
| SNX37 | 0,251959643 | 0,557927837 | 0,467 | 0,338 | 1 | CH_SC | 1,472153224 | 0,679277119 | 1,381656805 |
| CBX51 | 0,253651975 | 0,672076171 | 0,467 | 0,385 | 1 | CH_SC | 1,593364316 | 0,627602859 | 1,212987013 |
| MYCBP22 | 0,254066401 | -0,669516726 | 0,2 | 0,323 | 1 | CH_SC | 0,62871726 | 1,59054008 | 0,619195046 |
| PXYLP1 | 0,254320571 | -0,556833803 | 0,133 | 0,262 | 1 | CH_SC | 0,679792428 | 1,471037274 | 0,507633588 |
| CWC151 | 0,255112315 | -0,535054229 | 0,133 | 0,246 | 1 | CH_SC | 0,690132735 | 1,448996619 | 0,540650407 |
| PABPC43 | 0,25617568 | -0,278163226 | 0,067 | 0,2 | 1 | CH_SC | 0,824640244 | 1,21265001 | 0,335 |
| AP2B12 | 0,25617568 | -0,285258337 | 0,067 | 0,2 | 1 | CH_SC | 0,820594656 | 1,218628458 | 0,335 |
| APRT7 | 0,256465276 | 0,538555195 | 0,333 | 0,231 | 1 | CH_SC | 1,452517148 | 0,688460031 | 1,441558442 |

| Supplementary Table 4 - clusermarker | | | | | | | | | |
|---|---|---|---|---|---|---|---|---|---|
| | **p_val** | **avg_log2FC** | **pct.1** | **pct.2** | **p_val_adj** | **celltype** | **Foldchange_UP** | **Foldchange_DOWN** | **Ratio_pct1_pct2** |
| NDUFB75 | 0,257109326 | 0,371317863 | 0,6 | 0,477 | 1 | CH_SC | 1,293533899 | 0,77307599 | 1,257861635 |
| TMEM237 | 0,257545885 | -0,391395084 | 0,067 | 0,185 | 1 | CH_SC | 0,762392016 | 1,311661165 | 0,362162162 |
| APC1 | 0,257545885 | 0,25329179 | 0,267 | 0,138 | 1 | CH_SC | 1,191923621 | 0,838979933 | 1,934782609 |
| ZBTB101 | 0,257545885 | -0,330424493 | 0,067 | 0,185 | 1 | CH_SC | 0,795302442 | 1,257383288 | 0,362162162 |
| BCLAF13 | 0,257936201 | 0,813925647 | 0,4 | 0,354 | 1 | CH_SC | 1,757988517 | 0,56883193 | 1,129943503 |
| ATP5PF4 | 0,258297713 | 0,303270753 | 0,667 | 0,6 | 1 | CH_SC | 1,233938724 | 0,810413014 | 1,111666667 |
| CTDSPL | 0,258649117 | 0,432083641 | 0,333 | 0,2 | 1 | CH_SC | 1,349180752 | 0,741190533 | 1,665 |
| PET1008 | 0,259624666 | -0,264017955 | 0,267 | 0,492 | 1 | CH_SC | 0,832765407 | 1,200818372 | 0,542682927 |
| ATP5MC24 | 0,259687544 | -0,374150889 | 0,733 | 0,8 | 1 | CH_SC | 0,771559387 | 1,296076512 | 0,91625 |
| CSNK2B7 | 0,259927627 | 0,633503136 | 0,533 | 0,446 | 1 | CH_SC | 1,551327339 | 0,64460928 | 1,195067265 |
| FEZ22 | 0,260093816 | 0,47442463 | 0,267 | 0,154 | 1 | CH_SC | 1,389364009 | 0,719753782 | 1,733766234 |
| SHISA53 | 0,260093816 | 0,430786044 | 0,267 | 0,154 | 1 | CH_SC | 1,347967809 | 0,741857479 | 1,733766234 |
| SASH11 | 0,260093816 | 0,860115672 | 0,267 | 0,154 | 1 | CH_SC | 1,815183843 | 0,550908385 | 1,733766234 |
| UBR52 | 0,260093816 | 0,82809465 | 0,267 | 0,154 | 1 | CH_SC | 1,775339144 | 0,563272659 | 1,733766234 |
| MRPS344 | 0,260093816 | 0,525249354 | 0,267 | 0,154 | 1 | CH_SC | 1,439182305 | 0,694839004 | 1,733766234 |
| SRPRA1 | 0,260513593 | 0,746175314 | 0,4 | 0,292 | 1 | CH_SC | 1,677340184 | 0,596181985 | 1,369863014 |
| NRDC3 | 0,261257433 | 0,262951096 | 0,2 | 0,092 | 1 | CH_SC | 1,199930707 | 0,833381456 | 2,173913043 |
| MRPL552 | 0,261257433 | 0,284579575 | 0,2 | 0,092 | 1 | CH_SC | 1,21805525 | 0,820980821 | 2,173913043 |
| POLR2F3 | 0,261268839 | 0,624012141 | 0,333 | 0,215 | 1 | CH_SC | 1,541155186 | 0,648863923 | 1,548837209 |
| UBE2D31 | 0,261381937 | 0,755588453 | 0,467 | 0,415 | 1 | CH_SC | 1,688320091 | 0,592304744 | 1,125301205 |
| DOCK7 | 0,261706552 | 0,522951801 | 0,267 | 0,169 | 1 | CH_SC | 1,436892171 | 0,695946446 | 1,579881657 |
| MMP24OS4 | 0,262219758 | -0,524446146 | 0,133 | 0,246 | 1 | CH_SC | 0,695225958 | 1,438381276 | 0,540650407 |
| NDUFS73 | 0,263687698 | 0,480343016 | 0,4 | 0,277 | 1 | CH_SC | 1,395075321 | 0,716807175 | 1,444043321 |
| N4BP2L25 | 0,264297789 | -0,431097644 | 0,267 | 0,462 | 1 | CH_SC | 0,741697266 | 1,348258981 | 0,577922078 |
| IGSF8 | 0,264581083 | 0,534023941 | 0,2 | 0,108 | 1 | CH_SC | 1,4479622 | 0,690625763 | 1,851851852 |
| ITGB1BP12 | 0,264581083 | 0,447848306 | 0,2 | 0,108 | 1 | CH_SC | 1,364004407 | 0,733135461 | 1,851851852 |
| DCTN4 | 0,264581083 | 0,437010565 | 0,2 | 0,108 | 1 | CH_SC | 1,353796193 | 0,738663623 | 1,851851852 |
| STX51 | 0,264581083 | 0,690802514 | 0,2 | 0,108 | 1 | CH_SC | 1,614181174 | 0,619509146 | 1,851851852 |
| TRMT1125 | 0,265164609 | 0,962007411 | 0,533 | 0,508 | 1 | CH_SC | 1,948018544 | 0,513342136 | 1,049212598 |
| KDM3B | 0,265699567 | -0,391211113 | 0,067 | 0,185 | 1 | CH_SC | 0,762489242 | 1,311493914 | 0,362162162 |
| PAF11 | 0,265699567 | 0,501195643 | 0,267 | 0,138 | 1 | CH_SC | 1,415386087 | 0,706521004 | 1,934782609 |

| Supplementary Table 4 - clusermarker | | | | | | | | | |
|---|---|---|---|---|---|---|---|---|---|
| | **p_val** | **avg_log2FC** | **pct.1** | **pct.2** | **p_val_adj** | **celltype** | **Foldchange_UP** | **Foldchange_DOWN** | **Ratio_pct1_pct2** |
| GPI4 | 0,265824305 | 0,435264979 | 0,333 | 0,2 | 1 | CH_SC | 1,352159161 | 0,739557908 | 1,665 |
| PPIP5K23 | 0,266640035 | 0,644574125 | 0,4 | 0,262 | 1 | CH_SC | 1,563277748 | 0,639681593 | 1,526717557 |
| IST11 | 0,266797196 | -0,270385205 | 0,133 | 0,277 | 1 | CH_SC | 0,829098144 | 1,206129826 | 0,480144404 |
| TCEAL91 | 0,266923333 | -0,41099268 | 0,2 | 0,338 | 1 | CH_SC | 0,752105692 | 1,329600362 | 0,591715976 |
| MAN2A11 | 0,268054027 | 0,51279476 | 0,267 | 0,154 | 1 | CH_SC | 1,42681151 | 0,700863424 | 1,733766234 |
| RNF1451 | 0,268054027 | 0,540319569 | 0,267 | 0,154 | 1 | CH_SC | 1,454294621 | 0,687618579 | 1,733766234 |
| PPP1R16A1 | 0,268054027 | 0,43552236 | 0,267 | 0,154 | 1 | CH_SC | 1,352400411 | 0,73942598 | 1,733766234 |
| BRD71 | 0,268054027 | 0,44923251 | 0,267 | 0,154 | 1 | CH_SC | 1,365313739 | 0,732432386 | 1,733766234 |
| TFG3 | 0,268347261 | -0,558157484 | 0,133 | 0,262 | 1 | CH_SC | 0,679169 | 1,472387579 | 0,507633588 |
| TNS32 | 0,268347261 | 0,364694732 | 0,333 | 0,215 | 1 | CH_SC | 1,287609148 | 0,77663319 | 1,548837209 |
| PYURF2 | 0,26878419 | 0,561146964 | 0,467 | 0,354 | 1 | CH_SC | 1,475441749 | 0,677763118 | 1,31920904 |
| CDC42BPA | 0,269462822 | -0,475603941 | 0,133 | 0,246 | 1 | CH_SC | 0,71916567 | 1,390500189 | 0,540650407 |
| TLE11 | 0,269478625 | 0,568216045 | 0,267 | 0,169 | 1 | CH_SC | 1,482689027 | 0,67445026 | 1,579881657 |
| MICOS103 | 0,270239846 | 0,425198239 | 0,4 | 0,277 | 1 | CH_SC | 1,342756996 | 0,744736391 | 1,444043321 |
| PSMB12 | 0,270827857 | 0,263752052 | 0,533 | 0,508 | 1 | CH_SC | 1,20059707 | 0,832918908 | 1,049212598 |
| P4HB5 | 0,270827857 | 0,588659897 | 0,533 | 0,508 | 1 | CH_SC | 1,50384919 | 0,664960294 | 1,049212598 |
| CALU1 | 0,271436345 | 0,530475651 | 0,467 | 0,385 | 1 | CH_SC | 1,444405332 | 0,692326439 | 1,212987013 |
| ATP5MC34 | 0,271445728 | 0,513285254 | 0,533 | 0,446 | 1 | CH_SC | 1,427296687 | 0,700625181 | 1,195067265 |
| AL118516.13 | 0,272096272 | -0,296006858 | 0,067 | 0,2 | 1 | CH_SC | 0,814503691 | 1,22774152 | 0,335 |
| CHST3 | 0,273019386 | 0,423588292 | 0,467 | 0,292 | 1 | CH_SC | 1,34125941 | 0,745567929 | 1,599315068 |
| MBNL16 | 0,273292485 | 0,716135683 | 0,467 | 0,369 | 1 | CH_SC | 1,642775894 | 0,608725757 | 1,265582656 |
| PSMC53 | 0,273292485 | -0,366622764 | 0,267 | 0,415 | 1 | CH_SC | 0,775595983 | 1,289331072 | 0,643373494 |
| DCTN23 | 0,273339143 | 0,397084301 | 0,4 | 0,262 | 1 | CH_SC | 1,316843867 | 0,75939147 | 1,526717557 |
| HS3ST3A1 | 0,273842156 | -0,788157209 | 0,133 | 0,277 | 1 | CH_SC | 0,579083298 | 1,726867282 | 0,480144404 |
| AK31 | 0,273878521 | 0,703949756 | 0,2 | 0,108 | 1 | CH_SC | 1,628958394 | 0,613889221 | 1,851851852 |
| ASB81 | 0,273878521 | 0,500048705 | 0,2 | 0,108 | 1 | CH_SC | 1,414261306 | 0,70708291 | 1,851851852 |
| DMTF1 | 0,27402927 | -0,270860008 | 0,067 | 0,185 | 1 | CH_SC | 0,828825326 | 1,206526839 | 0,362162162 |
| MYL6B1 | 0,27402927 | -0,297852043 | 0,067 | 0,185 | 1 | CH_SC | 0,813462619 | 1,229312787 | 0,362162162 |
| MRPL381 | 0,27402927 | -0,382005017 | 0,067 | 0,185 | 1 | CH_SC | 0,76737038 | 1,303151681 | 0,362162162 |
| RNASET23 | 0,275118486 | -0,344068656 | 0,067 | 0,169 | 1 | CH_SC | 0,787816394 | 1,269331291 | 0,396449704 |
| CHMP4A2 | 0,275118486 | -0,409404486 | 0,067 | 0,169 | 1 | CH_SC | 0,752934105 | 1,328137474 | 0,396449704 |

| Supplementary Table 4 - clusermarker | | | | | | | | | |
|---|---|---|---|---|---|---|---|---|---|
| | **p_val** | **avg_log2FC** | **pct.1** | **pct.2** | **p_val_adj** | **celltype** | **Foldchange_UP** | **Foldchange_DOWN** | **Ratio_pct1_pct2** |
| DNAJC71 | 0,275118486 | -0,497053312 | 0,067 | 0,169 | 1 | CH_SC | 0,708552514 | 1,411327996 | 0,396449704 |
| PSMA42 | 0,275556309 | 0,726684069 | 0,333 | 0,215 | 1 | CH_SC | 1,654831207 | 0,604291239 | 1,548837209 |
| CNIH42 | 0,276180048 | 0,567067019 | 0,267 | 0,154 | 1 | CH_SC | 1,481508619 | 0,674987636 | 1,733766234 |
| SH3YL11 | 0,276180048 | 0,763212773 | 0,267 | 0,154 | 1 | CH_SC | 1,697266103 | 0,589182803 | 1,733766234 |
| SETD71 | 0,276180048 | 0,454122783 | 0,267 | 0,154 | 1 | CH_SC | 1,369949567 | 0,729953879 | 1,733766234 |
| PGM1 | 0,276841998 | -0,476236154 | 0,133 | 0,246 | 1 | CH_SC | 0,718850589 | 1,391109663 | 0,540650407 |
| MT-ND64 | 0,277442938 | 0,690454524 | 0,6 | 0,538 | 1 | CH_SC | 1,613791867 | 0,619658594 | 1,115241636 |
| UBXN44 | 0,279234457 | 0,823303417 | 0,467 | 0,415 | 1 | CH_SC | 1,769452969 | 0,565146414 | 1,125301205 |
| DHX363 | 0,279553428 | -0,337613935 | 0,2 | 0,354 | 1 | CH_SC | 0,791349038 | 1,263664897 | 0,564971751 |
| MGST34 | 0,279901518 | -0,388914162 | 0,333 | 0,538 | 1 | CH_SC | 0,763704187 | 1,309407513 | 0,618959108 |
| PAPSS11 | 0,280066072 | -0,341534846 | 0,2 | 0,338 | 1 | CH_SC | 0,789201254 | 1,267103916 | 0,591715976 |
| SARS11 | 0,280152427 | 0,522429757 | 0,4 | 0,262 | 1 | CH_SC | 1,43637232 | 0,696198323 | 1,526717557 |
| SNX51 | 0,280152427 | -0,530305226 | 0,2 | 0,308 | 1 | CH_SC | 0,692408228 | 1,444234715 | 0,649350649 |
| NAA384 | 0,280275872 | -0,368810177 | 0,2 | 0,323 | 1 | CH_SC | 0,774420916 | 1,291287437 | 0,619195046 |
| TMEM192 | 0,280305422 | 0,37908534 | 0,067 | 0,2 | 1 | CH_SC | 1,300517074 | 0,76892493 | 0,335 |
| SHOC25 | 0,280582762 | 0,261217817 | 0,333 | 0,2 | 1 | CH_SC | 1,198489955 | 0,834383297 | 1,665 |
| PMVK2 | 0,282535661 | 0,759571136 | 0,267 | 0,138 | 1 | CH_SC | 1,692987283 | 0,590671891 | 1,934782609 |
| CIR16 | 0,282535661 | -0,286190655 | 0,067 | 0,185 | 1 | CH_SC | 0,820064531 | 1,219416232 | 0,362162162 |
| GTF2I3 | 0,282859996 | 0,492854001 | 0,4 | 0,308 | 1 | CH_SC | 1,407225959 | 0,710617932 | 1,298701299 |
| SNHG16 | 0,282859996 | 0,596284059 | 0,4 | 0,308 | 1 | CH_SC | 1,511817568 | 0,66145547 | 1,298701299 |
| PRMT23 | 0,28289639 | 0,435276932 | 0,333 | 0,215 | 1 | CH_SC | 1,352170363 | 0,739551781 | 1,548837209 |
| CEBPZOS | 0,283396789 | 0,425619318 | 0,2 | 0,108 | 1 | CH_SC | 1,343148964 | 0,744519057 | 1,851851852 |
| WIPI21 | 0,283396789 | 0,500822855 | 0,2 | 0,108 | 1 | CH_SC | 1,415020402 | 0,706703591 | 1,851851852 |
| C8orf331 | 0,283396789 | 0,377521699 | 0,2 | 0,108 | 1 | CH_SC | 1,299108294 | 0,769758768 | 1,851851852 |
| UBE4A2 | 0,283396789 | 0,554666353 | 0,2 | 0,108 | 1 | CH_SC | 1,468828903 | 0,68081449 | 1,851851852 |
| PALLD4 | 0,283756515 | 0,433026661 | 0,467 | 0,385 | 1 | CH_SC | 1,350062934 | 0,740706211 | 1,212987013 |
| EFNA12 | 0,283940982 | -0,316508911 | 0,067 | 0,169 | 1 | CH_SC | 0,803010685 | 1,245313442 | 0,396449704 |
| NDUFS1 | 0,283940982 | -0,414548193 | 0,067 | 0,169 | 1 | CH_SC | 0,750254414 | 1,332881195 | 0,396449704 |
| GMDS1 | 0,283940982 | -0,543835484 | 0,067 | 0,169 | 1 | CH_SC | 0,685944861 | 1,457843126 | 0,396449704 |
| RING1 | 0,283940982 | -0,483058855 | 0,067 | 0,169 | 1 | CH_SC | 0,715459072 | 1,39770399 | 0,396449704 |
| REV3L2 | 0,283940982 | -0,554165715 | 0,067 | 0,169 | 1 | CH_SC | 0,681050784 | 1,468319285 | 0,396449704 |

| Supplementary Table 4 - clusermarker | | | | | | | | | |
|---|---|---|---|---|---|---|---|---|---|
| | **p_val** | **avg_log2FC** | **pct.1** | **pct.2** | **p_val_adj** | **celltype** | **Foldchange_UP** | **Foldchange_DOWN** | **Ratio_pct1_pct2** |
| LDLRAD3 | 0,283940982 | -0,29500129 | 0,067 | 0,169 | 1 | CH_SC | 0,815071604 | 1,226886074 | 0,396449704 |
| ERG281 | 0,283940982 | -0,377401544 | 0,067 | 0,169 | 1 | CH_SC | 0,76982288 | 1,299000102 | 0,396449704 |
| RNF114 | 0,283940982 | -0,3866832 | 0,067 | 0,169 | 1 | CH_SC | 0,764886081 | 1,307384229 | 0,396449704 |
| RALGPS2 | 0,284357706 | -0,482606789 | 0,133 | 0,246 | 1 | CH_SC | 0,715683295 | 1,397266091 | 0,540650407 |
| NGRN2 | 0,284429256 | 0,758629893 | 0,4 | 0,369 | 1 | CH_SC | 1,691883105 | 0,591057383 | 1,08401084 |
| CDC5L2 | 0,284472458 | 0,327932003 | 0,267 | 0,154 | 1 | CH_SC | 1,255212829 | 0,796677644 | 1,733766234 |
| CIAO2A2 | 0,284472458 | 0,375198891 | 0,267 | 0,154 | 1 | CH_SC | 1,29701835 | 0,770999115 | 1,733766234 |
| ANP32A2 | 0,284472458 | 0,381793007 | 0,267 | 0,154 | 1 | CH_SC | 1,302960192 | 0,767483156 | 1,733766234 |
| SNF82 | 0,284472458 | 0,259299571 | 0,267 | 0,154 | 1 | CH_SC | 1,196897469 | 0,835493454 | 1,733766234 |
| COQ10B2 | 0,285494059 | 0,656881119 | 0,267 | 0,169 | 1 | CH_SC | 1,576670422 | 0,634247961 | 1,579881657 |
| EPRS1 | 0,287080184 | -0,566224703 | 0,2 | 0,308 | 1 | CH_SC | 0,675381842 | 1,480643894 | 0,649350649 |
| CHMP32 | 0,287080184 | 0,349698679 | 0,4 | 0,262 | 1 | CH_SC | 1,27429445 | 0,784747983 | 1,526717557 |
| PRPF40A2 | 0,287080184 | 0,367914716 | 0,4 | 0,262 | 1 | CH_SC | 1,290486202 | 0,774901737 | 1,526717557 |
| HSBP15 | 0,287630769 | 0,351685231 | 0,467 | 0,354 | 1 | CH_SC | 1,276050329 | 0,78366815 | 1,31920904 |
| TRAM11 | 0,28894797 | 0,391804436 | 0,533 | 0,415 | 1 | CH_SC | 1,312033391 | 0,762175724 | 1,284337349 |
| BTG15 | 0,289176086 | 0,418723833 | 0,667 | 0,677 | 1 | CH_SC | 1,336744585 | 0,748086067 | 0,985228951 |
| XRN14 | 0,290367844 | -0,382325907 | 0,133 | 0,262 | 1 | CH_SC | 0,767199717 | 1,303441566 | 0,507633588 |
| RNF132 | 0,290367844 | 0,882687864 | 0,333 | 0,215 | 1 | CH_SC | 1,843807274 | 0,542356034 | 1,548837209 |
| NEDD83 | 0,292885655 | 0,591520727 | 0,533 | 0,538 | 1 | CH_SC | 1,506834246 | 0,663643 | 0,99070632 |
| NFKB13 | 0,292931734 | 0,260332593 | 0,267 | 0,154 | 1 | CH_SC | 1,197754798 | 0,834895424 | 1,733766234 |
| RABGAP1 | 0,292931734 | 0,266943416 | 0,267 | 0,154 | 1 | CH_SC | 1,203255832 | 0,831078457 | 1,733766234 |
| TMOD33 | 0,292931734 | 0,660803587 | 0,267 | 0,154 | 1 | CH_SC | 1,580962981 | 0,632525879 | 1,733766234 |
| GLA2 | 0,292931734 | 0,345796734 | 0,267 | 0,154 | 1 | CH_SC | 1,270852622 | 0,786873303 | 1,733766234 |
| PRPF38B3 | 0,292953062 | -0,283626771 | 0,067 | 0,169 | 1 | CH_SC | 0,821523204 | 1,217251072 | 0,396449704 |
| ETV11 | 0,292953062 | -0,472498405 | 0,067 | 0,169 | 1 | CH_SC | 0,720715409 | 1,387510227 | 0,396449704 |
| FASTK1 | 0,292953062 | -0,373794277 | 0,067 | 0,169 | 1 | CH_SC | 0,771750128 | 1,295756182 | 0,396449704 |
| ESF12 | 0,292953062 | -0,344173523 | 0,067 | 0,169 | 1 | CH_SC | 0,787759131 | 1,26942356 | 0,396449704 |
| GID8 | 0,292953062 | -0,365354397 | 0,067 | 0,169 | 1 | CH_SC | 0,776278159 | 1,288198036 | 0,396449704 |
| HS3ST13 | 0,293136646 | 0,715612134 | 0,2 | 0,108 | 1 | CH_SC | 1,642179845 | 0,608946701 | 1,851851852 |
| SAP30L1 | 0,293136646 | 0,419058816 | 0,2 | 0,108 | 1 | CH_SC | 1,337055003 | 0,747912388 | 1,851851852 |
| PSMB86 | 0,293136646 | 0,618468127 | 0,2 | 0,108 | 1 | CH_SC | 1,535244172 | 0,651362186 | 1,851851852 |

| Supplementary Table 4 - clusermarker | | | | | | | | | |
|---|---|---|---|---|---|---|---|---|---|
| | **p_val** | **avg_log2FC** | **pct.1** | **pct.2** | **p_val_adj** | **celltype** | **Foldchange_UP** | **Foldchange_DOWN** | **Ratio_pct1_pct2** |
| MED19 | 0,293136646 | 0,63742105 | 0,2 | 0,108 | 1 | CH_SC | 1,555545991 | 0,642861096 | 1,851851852 |
| MAF2 | 0,294046624 | -0,364261735 | 0,2 | 0,323 | 1 | CH_SC | 0,776866316 | 1,287222755 | 0,619195046 |
| PRRC2A1 | 0,295888015 | 0,579137642 | 0,333 | 0,2 | 1 | CH_SC | 1,493955983 | 0,669363764 | 1,665 |
| MAP1LC3B3 | 0,297376981 | -0,569541466 | 0,267 | 0,4 | 1 | CH_SC | 0,673830919 | 1,484051817 | 0,6675 |
| DAZAP25 | 0,297559032 | 0,322978343 | 0,4 | 0,277 | 1 | CH_SC | 1,250910302 | 0,799417831 | 1,444043321 |
| SNRNP701 | 0,297559032 | 0,80009177 | 0,4 | 0,277 | 1 | CH_SC | 1,741211882 | 0,574312644 | 1,444043321 |
| SNHG14 | 0,29797095 | -0,353239206 | 0,133 | 0,262 | 1 | CH_SC | 0,78282449 | 1,277425545 | 0,507633588 |
| C1orf174 | 0,298729728 | 0,523202916 | 0,133 | 0,062 | 1 | CH_SC | 1,437142298 | 0,695825321 | 2,14516129 |
| BEND5 | 0,298729728 | 0,591871072 | 0,133 | 0,062 | 1 | CH_SC | 1,507200211 | 0,663481861 | 2,14516129 |
| ABCD3 | 0,298729728 | 0,488319122 | 0,133 | 0,062 | 1 | CH_SC | 1,402809516 | 0,712855159 | 2,14516129 |
| POLR3C1 | 0,298729728 | 1,118965465 | 0,133 | 0,062 | 1 | CH_SC | 2,171911721 | 0,46042387 | 2,14516129 |
| PLEKHO12 | 0,298729728 | 0,556690699 | 0,133 | 0,062 | 1 | CH_SC | 1,470891366 | 0,679859861 | 2,14516129 |
| MOGS | 0,298729728 | 0,865329763 | 0,133 | 0,062 | 1 | CH_SC | 1,821756027 | 0,548920923 | 2,14516129 |
| ANKRD362 | 0,298729728 | 1,023913285 | 0,133 | 0,062 | 1 | CH_SC | 2,033427121 | 0,491780595 | 2,14516129 |
| ZDHHC3 | 0,298729728 | 0,502218815 | 0,133 | 0,062 | 1 | CH_SC | 1,416390247 | 0,706020111 | 2,14516129 |
| MAML32 | 0,298729728 | 0,561941525 | 0,133 | 0,062 | 1 | CH_SC | 1,47625457 | 0,677389944 | 2,14516129 |
| ZNF1311 | 0,298729728 | 1,092710595 | 0,133 | 0,062 | 1 | CH_SC | 2,13274369 | 0,468879596 | 2,14516129 |
| DCTN61 | 0,298729728 | 0,550291282 | 0,133 | 0,062 | 1 | CH_SC | 1,464381326 | 0,68288224 | 2,14516129 |
| STK3 | 0,298729728 | 0,805660223 | 0,133 | 0,062 | 1 | CH_SC | 1,747945524 | 0,572100209 | 2,14516129 |
| JCAD1 | 0,298729728 | 0,458986121 | 0,133 | 0,062 | 1 | CH_SC | 1,374575472 | 0,72749734 | 2,14516129 |
| TMEM1381 | 0,298729728 | 0,549847343 | 0,133 | 0,062 | 1 | CH_SC | 1,463930784 | 0,683092405 | 2,14516129 |
| BEST12 | 0,298729728 | 0,525266511 | 0,133 | 0,062 | 1 | CH_SC | 1,439199421 | 0,694830741 | 2,14516129 |
| TBK1 | 0,298729728 | 0,486687742 | 0,133 | 0,062 | 1 | CH_SC | 1,401224134 | 0,713661702 | 2,14516129 |
| ATXN3 | 0,298729728 | 0,850753964 | 0,133 | 0,062 | 1 | CH_SC | 1,803443173 | 0,554494877 | 2,14516129 |
| CIAPIN1 | 0,298729728 | 0,490567874 | 0,133 | 0,062 | 1 | CH_SC | 1,404997803 | 0,711744885 | 2,14516129 |
| PLEKHG4 | 0,298729728 | 0,519596177 | 0,133 | 0,062 | 1 | CH_SC | 1,433553928 | 0,697567061 | 2,14516129 |
| USP102 | 0,298729728 | 0,700485332 | 0,133 | 0,062 | 1 | CH_SC | 1,625051378 | 0,615365159 | 2,14516129 |
| TSIX | 0,298729728 | 0,6453315 | 0,133 | 0,062 | 1 | CH_SC | 1,56409864 | 0,639345866 | 2,14516129 |
| CMYA5 | 0,299494505 | 0,855781762 | 0,333 | 0,231 | 1 | CH_SC | 1,809739144 | 0,552565823 | 1,441558442 |
| DIS31 | 0,299800052 | -0,419374531 | 0,133 | 0,246 | 1 | CH_SC | 0,747748734 | 1,337347633 | 0,540650407 |
| MAFB5 | 0,299800052 | -0,925666446 | 0,133 | 0,246 | 1 | CH_SC | 0,526437277 | 1,899561532 | 0,540650407 |

| Supplementary Table 4 - clusermarker | | | | | | | | | |
|---|---|---|---|---|---|---|---|---|---|
| | **p_val** | **avg_log2FC** | **pct.1** | **pct.2** | **p_val_adj** | **celltype** | **Foldchange_UP** | **Foldchange_DOWN** | **Ratio_pct1_pct2** |
| ACTR101 | 0,300520013 | 0,646349494 | 0,333 | 0,246 | 1 | CH_SC | 1,565202689 | 0,63889489 | 1,353658537 |
| EIF3A2 | 0,300592419 | 0,623817755 | 0,4 | 0,292 | 1 | CH_SC | 1,540947547 | 0,648951356 | 1,369863014 |
| SLC3A21 | 0,300592419 | 0,622365966 | 0,4 | 0,292 | 1 | CH_SC | 1,539397667 | 0,649604726 | 1,369863014 |
| CKAP4 | 0,300766134 | -0,448511827 | 0,267 | 0,369 | 1 | CH_SC | 0,732798356 | 1,364631882 | 0,723577236 |
| SRSF33 | 0,301359389 | 0,520685927 | 0,533 | 0,477 | 1 | CH_SC | 1,434637182 | 0,697040347 | 1,117400419 |
| TPRKB5 | 0,301558243 | 0,386188413 | 0,267 | 0,154 | 1 | CH_SC | 1,306935926 | 0,765148452 | 1,733766234 |
| DNAJB2 | 0,302140862 | 0,451477573 | 0,267 | 0,169 | 1 | CH_SC | 1,367440039 | 0,731293491 | 1,579881657 |
| CTNND12 | 0,302140862 | 0,613231693 | 0,267 | 0,169 | 1 | CH_SC | 1,529681921 | 0,653730679 | 1,579881657 |
| COA51 | 0,302155175 | -0,504242297 | 0,067 | 0,169 | 1 | CH_SC | 0,705030562 | 1,418378229 | 0,396449704 |
| ARHGAP212 | 0,302155175 | -0,406229398 | 0,067 | 0,169 | 1 | CH_SC | 0,754592989 | 1,325217719 | 0,396449704 |
| RAPGEF24 | 0,303098667 | 0,877395949 | 0,2 | 0,108 | 1 | CH_SC | 1,837056437 | 0,54434909 | 1,851851852 |
| ATP6V1C1 | 0,303098667 | 0,378223878 | 0,2 | 0,108 | 1 | CH_SC | 1,299740741 | 0,769384207 | 1,851851852 |
| ISCA1 | 0,303098667 | 0,399116159 | 0,2 | 0,108 | 1 | CH_SC | 1,318699786 | 0,758322714 | 1,851851852 |
| NOC3L | 0,303098667 | 0,874213738 | 0,2 | 0,108 | 1 | CH_SC | 1,833008833 | 0,545551108 | 1,851851852 |
| MTA12 | 0,303098667 | 1,044099402 | 0,2 | 0,108 | 1 | CH_SC | 2,062078711 | 0,484947541 | 1,851851852 |
| HMOX21 | 0,303098667 | 0,412443942 | 0,2 | 0,108 | 1 | CH_SC | 1,330938531 | 0,7513495 | 1,851851852 |
| TALDO14 | 0,30415895 | 0,477726342 | 0,467 | 0,323 | 1 | CH_SC | 1,39254731 | 0,718108457 | 1,445820433 |
| PCF113 | 0,305705914 | -0,597483944 | 0,133 | 0,262 | 1 | CH_SC | 0,660905568 | 1,513075465 | 0,507633588 |
| HMGN14 | 0,305705914 | 0,465064314 | 0,333 | 0,215 | 1 | CH_SC | 1,380378888 | 0,724438782 | 1,548837209 |
| MDH13 | 0,306980302 | -0,442535854 | 0,267 | 0,446 | 1 | CH_SC | 0,73584007 | 1,358990955 | 0,598654709 |
| ZNF221 | 0,307109109 | 0,534680955 | 0,333 | 0,231 | 1 | CH_SC | 1,448621763 | 0,690311319 | 1,441558442 |
| IL6ST4 | 0,307139496 | 0,632474032 | 0,533 | 0,492 | 1 | CH_SC | 1,55022114 | 0,645069258 | 1,083333333 |
| REXO24 | 0,307896167 | -0,250804317 | 0,333 | 0,462 | 1 | CH_SC | 0,840427738 | 1,189870295 | 0,720779221 |
| CALM25 | 0,308708787 | 0,372748856 | 0,8 | 0,892 | 1 | CH_SC | 1,294817578 | 0,772309565 | 0,896860987 |
| KLF32 | 0,309119854 | -0,288548906 | 0,067 | 0,185 | 1 | CH_SC | 0,818725136 | 1,221411137 | 0,362162162 |
| LINC02256 | 0,309286756 | -0,525000673 | 0,133 | 0,231 | 1 | CH_SC | 0,694958786 | 1,438934252 | 0,575757576 |
| ATG33 | 0,310352245 | 0,281309444 | 0,267 | 0,154 | 1 | CH_SC | 1,215297434 | 0,822843834 | 1,733766234 |
| AKT12 | 0,310352245 | 0,367304746 | 0,267 | 0,154 | 1 | CH_SC | 1,289940701 | 0,775229435 | 1,733766234 |
| ACTR33 | 0,310701783 | 0,461199055 | 0,267 | 0,169 | 1 | CH_SC | 1,376685536 | 0,726382296 | 1,579881657 |
| YTHDF2 | 0,311456733 | 0,437100009 | 0,133 | 0,062 | 1 | CH_SC | 1,353880128 | 0,738617828 | 2,14516129 |
| UCHL51 | 0,311456733 | 0,385556162 | 0,133 | 0,062 | 1 | CH_SC | 1,306363296 | 0,765483846 | 2,14516129 |

| Supplementary Table 4 - clusermarker | | | | | | | | | |
|---|---|---|---|---|---|---|---|---|---|
| | **p_val** | **avg_log2FC** | **pct.1** | **pct.2** | **p_val_adj** | **celltype** | **Foldchange_UP** | **Foldchange_DOWN** | **Ratio_pct1_pct2** |
| ABHD61 | 0,311456733 | 0,371030825 | 0,133 | 0,062 | 1 | CH_SC | 1,293276565 | 0,773229816 | 2,14516129 |
| LSG1 | 0,311456733 | 0,570290689 | 0,133 | 0,062 | 1 | CH_SC | 1,484822718 | 0,673481075 | 2,14516129 |
| MRPL502 | 0,311456733 | 0,523270133 | 0,133 | 0,062 | 1 | CH_SC | 1,437209258 | 0,695792902 | 2,14516129 |
| PITRM1 | 0,311456733 | 0,707691859 | 0,133 | 0,062 | 1 | CH_SC | 1,633189117 | 0,612298961 | 2,14516129 |
| PRKD21 | 0,311456733 | 0,493553789 | 0,133 | 0,062 | 1 | CH_SC | 1,407908708 | 0,710273325 | 2,14516129 |
| VAMP7 | 0,311456733 | 0,401996244 | 0,133 | 0,062 | 1 | CH_SC | 1,321334965 | 0,756810367 | 2,14516129 |
| SMPD1 | 0,311547629 | -0,425134953 | 0,067 | 0,169 | 1 | CH_SC | 0,744769061 | 1,342698096 | 0,396449704 |
| ALG133 | 0,311547629 | -0,323261023 | 0,067 | 0,169 | 1 | CH_SC | 0,799261209 | 1,251155428 | 0,396449704 |
| SAP302 | 0,311742288 | 0,349296952 | 0,333 | 0,2 | 1 | CH_SC | 1,273939665 | 0,784966531 | 1,665 |
| LMBRD12 | 0,311888107 | 0,287411673 | 0,4 | 0,277 | 1 | CH_SC | 1,220448716 | 0,819370767 | 1,444043321 |
| SNX62 | 0,311888107 | 0,376930311 | 0,4 | 0,277 | 1 | CH_SC | 1,298575874 | 0,770074371 | 1,444043321 |
| SEC11A4 | 0,312018166 | -0,316393865 | 0,4 | 0,554 | 1 | CH_SC | 0,803074722 | 1,245214141 | 0,722021661 |
| CYTH22 | 0,313022098 | 0,638131746 | 0,2 | 0,123 | 1 | CH_SC | 1,556312468 | 0,642544489 | 1,62601626 |
| SNHG171 | 0,313022098 | 0,640028462 | 0,2 | 0,123 | 1 | CH_SC | 1,558359903 | 0,641700289 | 1,62601626 |
| PDIA35 | 0,313215571 | 0,735437817 | 0,6 | 0,538 | 1 | CH_SC | 1,664902644 | 0,600635721 | 1,115241636 |
| RYBP | 0,313283229 | 0,373651001 | 0,2 | 0,108 | 1 | CH_SC | 1,295627505 | 0,771826776 | 1,851851852 |
| CTDSP21 | 0,313283229 | 0,346903364 | 0,2 | 0,108 | 1 | CH_SC | 1,271827813 | 0,786269957 | 1,851851852 |
| ISG201 | 0,313283229 | 0,484228127 | 0,2 | 0,108 | 1 | CH_SC | 1,398837257 | 0,714879443 | 1,851851852 |
| LINC00667 | 0,313283229 | 0,414100596 | 0,2 | 0,108 | 1 | CH_SC | 1,332467732 | 0,750487217 | 1,851851852 |
| NDUFS61 | 0,313572876 | 0,417046991 | 0,333 | 0,215 | 1 | CH_SC | 1,335191792 | 0,748956072 | 1,548837209 |
| METTL7A4 | 0,313572876 | -0,251672419 | 0,133 | 0,262 | 1 | CH_SC | 0,839922186 | 1,190586482 | 0,507633588 |
| NDUFAF83 | 0,313572876 | -0,257646866 | 0,133 | 0,262 | 1 | CH_SC | 0,836451116 | 1,195527127 | 0,507633588 |
| NDUFV26 | 0,313572876 | 0,446393112 | 0,333 | 0,215 | 1 | CH_SC | 1,362629279 | 0,733875321 | 1,548837209 |
| ARL14 | 0,314820267 | -0,369781002 | 0,2 | 0,338 | 1 | CH_SC | 0,773899964 | 1,292156669 | 0,591715976 |
| ROCK21 | 0,314850831 | 0,574703894 | 0,333 | 0,231 | 1 | CH_SC | 1,489371745 | 0,671424044 | 1,441558442 |
| COL27A11 | 0,315791789 | -0,43500732 | 0,133 | 0,246 | 1 | CH_SC | 0,739690001 | 1,351917693 | 0,540650407 |
| ARL22 | 0,317381398 | 0,561419536 | 0,4 | 0,308 | 1 | CH_SC | 1,475720534 | 0,677635078 | 1,298701299 |
| RNF216 | 0,318337275 | -0,250509444 | 0,067 | 0,185 | 1 | CH_SC | 0,84059953 | 1,189627122 | 0,362162162 |
| SNX22 | 0,319313885 | 0,457751345 | 0,267 | 0,154 | 1 | CH_SC | 1,373399501 | 0,728120259 | 1,733766234 |
| FBXL3 | 0,319313885 | 0,415292613 | 0,267 | 0,154 | 1 | CH_SC | 1,333569129 | 0,749867388 | 1,733766234 |
| SRP54 | 0,319313885 | 0,388746813 | 0,267 | 0,154 | 1 | CH_SC | 1,309255634 | 0,76379278 | 1,733766234 |

| Supplementary Table 4 - clusermarker | | | | | | | | | |
|---|---|---|---|---|---|---|---|---|---|
| | **p_val** | **avg_log2FC** | **pct.1** | **pct.2** | **p_val_adj** | **celltype** | **Foldchange_UP** | **Foldchange_DOWN** | **Ratio_pct1_pct2** |
| DDX424 | 0,319313885 | 0,262952271 | 0,267 | 0,154 | 1 | CH_SC | 1,199931684 | 0,833380778 | 1,733766234 |
| UBE2A2 | 0,319313885 | 0,253515545 | 0,267 | 0,154 | 1 | CH_SC | 1,192108497 | 0,838849822 | 1,733766234 |
| KDM5B1 | 0,319421258 | 0,592042492 | 0,267 | 0,169 | 1 | CH_SC | 1,507379306 | 0,663403031 | 1,579881657 |
| PHF141 | 0,319421258 | 0,731150947 | 0,267 | 0,169 | 1 | CH_SC | 1,659962841 | 0,602423124 | 1,579881657 |
| UBA13 | 0,319421258 | 0,448345944 | 0,267 | 0,169 | 1 | CH_SC | 1,364474983 | 0,732882619 | 1,579881657 |
| ZEB11 | 0,321130588 | -0,39158498 | 0,067 | 0,169 | 1 | CH_SC | 0,762291672 | 1,311833825 | 0,396449704 |
| ZSCAN181 | 0,321130588 | -0,3168128 | 0,067 | 0,169 | 1 | CH_SC | 0,802841556 | 1,245575783 | 0,396449704 |
| MICOS131 | 0,321502991 | 0,943838568 | 0,4 | 0,338 | 1 | CH_SC | 1,92363965 | 0,519847883 | 1,183431953 |
| PRPF4B5 | 0,321571906 | 0,631077814 | 0,333 | 0,215 | 1 | CH_SC | 1,548721587 | 0,645693848 | 1,548837209 |
| NDUFA18 | 0,321827963 | -0,414712529 | 0,533 | 0,631 | 1 | CH_SC | 0,750168958 | 1,333033031 | 0,844690967 |
| IVNS1ABP2 | 0,322097541 | -0,519988494 | 0,2 | 0,338 | 1 | CH_SC | 0,697377395 | 1,433943812 | 0,591715976 |
| APPBP21 | 0,323000252 | 1,067050417 | 0,2 | 0,123 | 1 | CH_SC | 2,095145471 | 0,477293827 | 1,62601626 |
| CDC42EP53 | 0,323112394 | 0,524406284 | 0,4 | 0,354 | 1 | CH_SC | 1,438341534 | 0,695245167 | 1,129943503 |
| UBE2V12 | 0,323380465 | 0,64261038 | 0,333 | 0,246 | 1 | CH_SC | 1,561151318 | 0,640552898 | 1,353658537 |
| SZRD11 | 0,323690512 | 0,333448902 | 0,2 | 0,108 | 1 | CH_SC | 1,260021981 | 0,793636948 | 1,851851852 |
| USP33 | 0,323690512 | 0,262608928 | 0,2 | 0,108 | 1 | CH_SC | 1,199646149 | 0,833579135 | 1,851851852 |
| IFI44 | 0,323690512 | 0,425007216 | 0,2 | 0,108 | 1 | CH_SC | 1,342579218 | 0,744835006 | 1,851851852 |
| GALT1 | 0,323690512 | 0,327484123 | 0,2 | 0,108 | 1 | CH_SC | 1,254823213 | 0,796925009 | 1,851851852 |
| RELA1 | 0,323690512 | 0,363715171 | 0,2 | 0,108 | 1 | CH_SC | 1,286735184 | 0,777160687 | 1,851851852 |
| YIF1B1 | 0,323690512 | 0,548754336 | 0,2 | 0,108 | 1 | CH_SC | 1,462822108 | 0,683610122 | 1,851851852 |
| ZFX | 0,323690512 | 0,341844671 | 0,2 | 0,108 | 1 | CH_SC | 1,267376061 | 0,789031788 | 1,851851852 |
| TMA72 | 0,324337743 | -0,337753986 | 0,667 | 0,8 | 1 | CH_SC | 0,79127222 | 1,263787574 | 0,83375 |
| KLF104 | 0,324449407 | -0,589807629 | 0,267 | 0,4 | 1 | CH_SC | 0,664431497 | 1,505046049 | 0,6675 |
| NAV1 | 0,324536208 | 0,306436917 | 0,133 | 0,062 | 1 | CH_SC | 1,236649721 | 0,808636417 | 2,14516129 |
| RABIF | 0,324536208 | 0,30799918 | 0,133 | 0,062 | 1 | CH_SC | 1,237989587 | 0,807761237 | 2,14516129 |
| FAIM | 0,324536208 | 0,290700931 | 0,133 | 0,062 | 1 | CH_SC | 1,22323444 | 0,817504779 | 2,14516129 |
| AC093323.1 | 0,324536208 | 0,307066424 | 0,133 | 0,062 | 1 | CH_SC | 1,237189439 | 0,808283654 | 2,14516129 |
| IRF1-AS11 | 0,324536208 | 0,303895202 | 0,133 | 0,062 | 1 | CH_SC | 1,234472931 | 0,810062315 | 2,14516129 |
| FAM20C | 0,324536208 | 0,366376408 | 0,133 | 0,062 | 1 | CH_SC | 1,289110923 | 0,775728436 | 2,14516129 |
| IMPDH1 | 0,324536208 | 0,605239245 | 0,133 | 0,062 | 1 | CH_SC | 1,521231001 | 0,657362359 | 2,14516129 |
| SYBU1 | 0,324536208 | 0,323829168 | 0,133 | 0,062 | 1 | CH_SC | 1,25164824 | 0,798946515 | 2,14516129 |

| Supplementary Table 4 - clusermarker | | | | | | | | | |
|---|---|---|---|---|---|---|---|---|---|
| | **p_val** | **avg_log2FC** | **pct.1** | **pct.2** | **p_val_adj** | **celltype** | **Foldchange_UP** | **Foldchange_DOWN** | **Ratio_pct1_pct2** |
| MDP1 | 0,324536208 | 0,338097638 | 0,133 | 0,062 | 1 | CH_SC | 1,264088646 | 0,791083761 | 2,14516129 |
| SPTLC21 | 0,324536208 | 0,904316708 | 0,133 | 0,062 | 1 | CH_SC | 1,871657827 | 0,534285693 | 2,14516129 |
| ZNF397 | 0,324536208 | 0,46689743 | 0,133 | 0,062 | 1 | CH_SC | 1,382133939 | 0,72351888 | 2,14516129 |
| DNMT11 | 0,324536208 | 0,366253517 | 0,133 | 0,062 | 1 | CH_SC | 1,28900112 | 0,775794516 | 2,14516129 |
| ZNF43 | 0,324536208 | 0,375825446 | 0,133 | 0,062 | 1 | CH_SC | 1,297581761 | 0,770664347 | 2,14516129 |
| SLC35A22 | 0,324536208 | 0,477766357 | 0,133 | 0,062 | 1 | CH_SC | 1,392585935 | 0,718088539 | 2,14516129 |
| ARHGEF6 | 0,324536208 | 0,434237454 | 0,133 | 0,062 | 1 | CH_SC | 1,35119646 | 0,740084828 | 2,14516129 |
| ANXA114 | 0,325974387 | -0,595707795 | 0,133 | 0,231 | 1 | CH_SC | 0,661719732 | 1,511213814 | 0,575757576 |
| TRPS11 | 0,326701861 | -0,332090989 | 0,133 | 0,277 | 1 | CH_SC | 0,794384298 | 1,258836564 | 0,480144404 |
| PGLS7 | 0,327439459 | 0,509104504 | 0,267 | 0,185 | 1 | CH_SC | 1,423166547 | 0,70265845 | 1,443243243 |
| KDM5A | 0,328299312 | 0,292154291 | 0,267 | 0,169 | 1 | CH_SC | 1,224467338 | 0,816681645 | 1,579881657 |
| ZC3H134 | 0,328299312 | 0,461474739 | 0,267 | 0,169 | 1 | CH_SC | 1,376948631 | 0,726243505 | 1,579881657 |
| EFS | 0,328299312 | 0,496247153 | 0,267 | 0,169 | 1 | CH_SC | 1,410539584 | 0,708948555 | 1,579881657 |
| C6orf622 | 0,328443196 | 0,27354579 | 0,267 | 0,154 | 1 | CH_SC | 1,208775052 | 0,827283785 | 1,733766234 |
| RBM8A4 | 0,329483723 | 1,085470976 | 0,4 | 0,292 | 1 | CH_SC | 2,12206813 | 0,471238405 | 1,369863014 |
| KAT2B | 0,329703002 | 0,286513116 | 0,333 | 0,215 | 1 | CH_SC | 1,219688818 | 0,819881256 | 1,548837209 |
| LBH3 | 0,332333648 | -0,359886881 | 0,133 | 0,246 | 1 | CH_SC | 0,779225675 | 1,28332527 | 0,540650407 |
| CNOT42 | 0,332333648 | -0,384929436 | 0,133 | 0,246 | 1 | CH_SC | 0,765816455 | 1,305795917 | 0,540650407 |
| CAST3 | 0,332488337 | 0,330828768 | 0,467 | 0,369 | 1 | CH_SC | 1,257735683 | 0,795079613 | 1,265582656 |
| CALR6 | 0,332616402 | 0,469827523 | 0,667 | 0,615 | 1 | CH_SC | 1,384943886 | 0,722050915 | 1,084552846 |
| OXCT11 | 0,333183526 | -0,3606329 | 0,067 | 0,154 | 1 | CH_SC | 0,778822841 | 1,283989051 | 0,435064935 |
| TRIB12 | 0,333183526 | -0,526151619 | 0,067 | 0,154 | 1 | CH_SC | 0,694404586 | 1,440082655 | 0,435064935 |
| LARP61 | 0,333183526 | -0,304197824 | 0,067 | 0,154 | 1 | CH_SC | 0,809892412 | 1,234731904 | 0,435064935 |
| HOXB21 | 0,333183526 | -0,570424069 | 0,067 | 0,154 | 1 | CH_SC | 0,673418813 | 1,484959999 | 0,435064935 |
| DKC1 | 0,333183526 | -0,384951462 | 0,067 | 0,154 | 1 | CH_SC | 0,765804763 | 1,305815854 | 0,435064935 |
| CD200 | 0,334320494 | 0,312871839 | 0,2 | 0,108 | 1 | CH_SC | 1,242177929 | 0,80503765 | 1,851851852 |
| C5orf511 | 0,334320494 | 0,516285186 | 0,2 | 0,108 | 1 | CH_SC | 1,430267688 | 0,699169819 | 1,851851852 |
| MED13L | 0,334320494 | 0,284790013 | 0,2 | 0,108 | 1 | CH_SC | 1,218232935 | 0,820861078 | 1,851851852 |
| QTRT12 | 0,334320494 | 0,346566893 | 0,2 | 0,108 | 1 | CH_SC | 1,271531227 | 0,786453355 | 1,851851852 |
| U2AF21 | 0,334320494 | 0,443337182 | 0,2 | 0,108 | 1 | CH_SC | 1,359746001 | 0,73543147 | 1,851851852 |
| MXI11 | 0,334533713 | -0,521228254 | 0,133 | 0,231 | 1 | CH_SC | 0,696778371 | 1,435176581 | 0,575757576 |

| Supplementary Table 4 - clusermarker | | | | | | | | | |
|---|---|---|---|---|---|---|---|---|---|
| | **p_val** | **avg_log2FC** | **pct.1** | **pct.2** | **p_val_adj** | **celltype** | **Foldchange_UP** | **Foldchange_DOWN** | **Ratio_pct1_pct2** |
| NFIX3 | 0,336978753 | -0,598763452 | 0,2 | 0,338 | 1 | CH_SC | 0,660319679 | 1,514417988 | 0,591715976 |
| SAT16 | 0,337265473 | -0,6590483 | 0,667 | 0,677 | 1 | CH_SC | 0,633295925 | 1,579040637 | 0,985228951 |
| NCOA43 | 0,337335868 | 0,424998958 | 0,267 | 0,169 | 1 | CH_SC | 1,342571533 | 0,74483927 | 1,579881657 |
| KRT103 | 0,337798393 | -0,432997568 | 0,333 | 0,477 | 1 | CH_SC | 0,740721148 | 1,350035709 | 0,698113208 |
| ATXN21 | 0,337966087 | 0,285457655 | 0,333 | 0,215 | 1 | CH_SC | 1,218796832 | 0,820481293 | 1,548837209 |
| UBE2J2 | 0,337968125 | 0,307454602 | 0,133 | 0,062 | 1 | CH_SC | 1,237522368 | 0,808066202 | 2,14516129 |
| CDK11B | 0,337968125 | 0,274819002 | 0,133 | 0,062 | 1 | CH_SC | 1,209842295 | 0,82655401 | 2,14516129 |
| TRIM33 | 0,337968125 | 0,306063596 | 0,133 | 0,062 | 1 | CH_SC | 1,236329759 | 0,808845692 | 2,14516129 |
| SLC30A1 | 0,337968125 | 0,343892015 | 0,133 | 0,062 | 1 | CH_SC | 1,269175885 | 0,787912859 | 2,14516129 |
| SMYD3 | 0,337968125 | 0,266630734 | 0,133 | 0,062 | 1 | CH_SC | 1,202995073 | 0,8312586 | 2,14516129 |
| GTF3C21 | 0,337968125 | 0,276238489 | 0,133 | 0,062 | 1 | CH_SC | 1,21103326 | 0,825741152 | 2,14516129 |
| MAP4K31 | 0,337968125 | 0,815450432 | 0,133 | 0,062 | 1 | CH_SC | 1,759847518 | 0,568231048 | 2,14516129 |
| MTMR14 | 0,337968125 | 0,448669592 | 0,133 | 0,062 | 1 | CH_SC | 1,364781118 | 0,732718226 | 2,14516129 |
| PCYT1A1 | 0,337968125 | 0,458957655 | 0,133 | 0,062 | 1 | CH_SC | 1,37454835 | 0,727511695 | 2,14516129 |
| CXCL24 | 0,337968125 | 0,254516889 | 0,133 | 0,062 | 1 | CH_SC | 1,192936201 | 0,838267796 | 2,14516129 |
| TRIM522 | 0,337968125 | 0,567033509 | 0,133 | 0,062 | 1 | CH_SC | 1,481474207 | 0,675003314 | 2,14516129 |
| CDK14 | 0,337968125 | 0,259726181 | 0,133 | 0,062 | 1 | CH_SC | 1,197251448 | 0,835246432 | 2,14516129 |
| AC058791.1 | 0,337968125 | 0,272931844 | 0,133 | 0,062 | 1 | CH_SC | 1,208260762 | 0,827635914 | 2,14516129 |
| FHIP2B | 0,337968125 | 0,285375384 | 0,133 | 0,062 | 1 | CH_SC | 1,218727331 | 0,820528083 | 2,14516129 |
| RIPK2 | 0,337968125 | 0,280373527 | 0,133 | 0,062 | 1 | CH_SC | 1,214509292 | 0,823377809 | 2,14516129 |
| FBXO322 | 0,337968125 | 0,353111764 | 0,133 | 0,062 | 1 | CH_SC | 1,277312707 | 0,782893644 | 2,14516129 |
| OTUD1 | 0,337968125 | 0,261469486 | 0,133 | 0,062 | 1 | CH_SC | 1,198699042 | 0,834237757 | 2,14516129 |
| ZSWIM8 | 0,337968125 | 0,323643825 | 0,133 | 0,062 | 1 | CH_SC | 1,251487451 | 0,799049163 | 2,14516129 |
| STIM11 | 0,337968125 | 0,401356151 | 0,133 | 0,062 | 1 | CH_SC | 1,320748848 | 0,757146222 | 2,14516129 |
| FOXJ2 | 0,337968125 | 0,282692751 | 0,133 | 0,062 | 1 | CH_SC | 1,216463262 | 0,822055241 | 2,14516129 |
| ANO62 | 0,337968125 | 0,585602517 | 0,133 | 0,062 | 1 | CH_SC | 1,500665586 | 0,666370982 | 2,14516129 |
| NOC4L | 0,337968125 | 0,537686248 | 0,133 | 0,062 | 1 | CH_SC | 1,451642548 | 0,688874821 | 2,14516129 |
| DLEU2 | 0,337968125 | 0,346139952 | 0,133 | 0,062 | 1 | CH_SC | 1,271154995 | 0,786686127 | 2,14516129 |
| SOX8 | 0,337968125 | 0,319799191 | 0,133 | 0,062 | 1 | CH_SC | 1,248156806 | 0,801181386 | 2,14516129 |
| GLYR11 | 0,337968125 | 0,314876612 | 0,133 | 0,062 | 1 | CH_SC | 1,243905263 | 0,803919744 | 2,14516129 |
| XPO61 | 0,337968125 | 0,285637827 | 0,133 | 0,062 | 1 | CH_SC | 1,218949052 | 0,820378832 | 2,14516129 |

| Supplementary Table 4 - clusermarker | | | | | | | | | |
|---|---|---|---|---|---|---|---|---|---|
| | **p_val** | **avg_log2FC** | **pct.1** | **pct.2** | **p_val_adj** | **celltype** | **Foldchange_UP** | **Foldchange_DOWN** | **Ratio_pct1_pct2** |
| FTO1 | 0,337968125 | 0,570714582 | 0,133 | 0,062 | 1 | CH_SC | 1,485259053 | 0,673283221 | 2,14516129 |
| SREBF1 | 0,337968125 | 0,297966979 | 0,133 | 0,062 | 1 | CH_SC | 1,229410727 | 0,813397815 | 2,14516129 |
| MAP2K3 | 0,337968125 | 0,306666438 | 0,133 | 0,062 | 1 | CH_SC | 1,236846478 | 0,80850778 | 2,14516129 |
| CCDC43 | 0,337968125 | 0,543297326 | 0,133 | 0,062 | 1 | CH_SC | 1,457299418 | 0,686200782 | 2,14516129 |
| NSF2 | 0,337968125 | 0,292242739 | 0,133 | 0,062 | 1 | CH_SC | 1,224542409 | 0,816631578 | 2,14516129 |
| EXOC73 | 0,337968125 | 0,313570423 | 0,133 | 0,062 | 1 | CH_SC | 1,242779563 | 0,804647927 | 2,14516129 |
| ZNF4441 | 0,337968125 | 0,29444631 | 0,133 | 0,062 | 1 | CH_SC | 1,226414203 | 0,815385208 | 2,14516129 |
| NRSN2 | 0,337968125 | 0,34755108 | 0,133 | 0,062 | 1 | CH_SC | 1,272398945 | 0,78591703 | 2,14516129 |
| KIZ | 0,337968125 | 0,453728387 | 0,133 | 0,062 | 1 | CH_SC | 1,369575109 | 0,730153457 | 2,14516129 |
| MIEF1 | 0,337968125 | 0,316783753 | 0,133 | 0,062 | 1 | CH_SC | 1,245550705 | 0,802857721 | 2,14516129 |
| CASK | 0,337968125 | 0,336413647 | 0,133 | 0,062 | 1 | CH_SC | 1,262613995 | 0,792007695 | 2,14516129 |
| RNF113A1 | 0,337968125 | 0,275781432 | 0,133 | 0,062 | 1 | CH_SC | 1,210649657 | 0,826002794 | 2,14516129 |
| KLF43 | 0,33867524 | -0,772826799 | 0,467 | 0,569 | 1 | CH_SC | 0,58526958 | 1,708614344 | 0,820738137 |
| GPC12 | 0,33923445 | 0,485865786 | 0,333 | 0,246 | 1 | CH_SC | 1,400426034 | 0,714068416 | 1,353658537 |
| TNRC6A1 | 0,33923445 | 0,855036138 | 0,333 | 0,246 | 1 | CH_SC | 1,808804063 | 0,552851478 | 1,353658537 |
| PPP1R102 | 0,340816503 | 1,046170397 | 0,4 | 0,415 | 1 | CH_SC | 2,065040959 | 0,484251896 | 0,963855422 |
| CAV13 | 0,34086793 | -0,584147283 | 0,067 | 0,169 | 1 | CH_SC | 0,667043484 | 1,499152641 | 0,396449704 |
| NFATC11 | 0,34086793 | -0,747506222 | 0,067 | 0,169 | 1 | CH_SC | 0,595632251 | 1,67888827 | 0,396449704 |
| CDK2AP12 | 0,340988821 | 0,685607807 | 0,533 | 0,446 | 1 | CH_SC | 1,608379448 | 0,621743831 | 1,195067265 |
| ASAH14 | 0,341416555 | 0,548679401 | 0,4 | 0,323 | 1 | CH_SC | 1,46274613 | 0,68364563 | 1,238390093 |
| PGAM13 | 0,341416555 | -0,533285925 | 0,267 | 0,354 | 1 | CH_SC | 0,690979146 | 1,447221679 | 0,754237288 |
| ATP2B43 | 0,341888734 | 0,431377903 | 0,4 | 0,277 | 1 | CH_SC | 1,348520921 | 0,741553197 | 1,444043321 |
| PPA13 | 0,343151772 | -0,258339964 | 0,267 | 0,369 | 1 | CH_SC | 0,836049366 | 1,196101619 | 0,723577236 |
| RAB3GAP11 | 0,343571731 | -0,373298801 | 0,067 | 0,154 | 1 | CH_SC | 0,772015222 | 1,295311247 | 0,435064935 |
| ANKRD282 | 0,343571731 | -0,463172396 | 0,067 | 0,154 | 1 | CH_SC | 0,725389418 | 1,378569877 | 0,435064935 |
| C3orf70 | 0,343571731 | -0,470355111 | 0,067 | 0,154 | 1 | CH_SC | 0,721786912 | 1,385450447 | 0,435064935 |
| RNF217 | 0,343571731 | -0,453332142 | 0,067 | 0,154 | 1 | CH_SC | 0,730354026 | 1,369198998 | 0,435064935 |
| NSUN5 | 0,343571731 | -0,358769802 | 0,067 | 0,154 | 1 | CH_SC | 0,779829263 | 1,282331976 | 0,435064935 |
| SAMD83 | 0,343571731 | 0,59252392 | 0,2 | 0,123 | 1 | CH_SC | 1,507882403 | 0,66318169 | 1,62601626 |
| RIN23 | 0,343571731 | -0,581034756 | 0,067 | 0,154 | 1 | CH_SC | 0,668484143 | 1,495921796 | 0,435064935 |
| SEPTIN5 | 0,343571731 | -0,388856963 | 0,067 | 0,154 | 1 | CH_SC | 0,763734466 | 1,309355599 | 0,435064935 |

| Supplementary Table 4 - clusermarker | | | | | | | | | |
|---|---|---|---|---|---|---|---|---|---|
| | **p_val** | **avg_log2FC** | **pct.1** | **pct.2** | **p_val_adj** | **celltype** | **Foldchange_UP** | **Foldchange_DOWN** | **Ratio_pct1_pct2** |
| PDHA13 | 0,343571731 | -0,46565751 | 0,067 | 0,154 | 1 | CH_SC | 0,724140974 | 1,380946578 | 0,435064935 |
| CFLAR3 | 0,345172943 | 0,793696912 | 0,2 | 0,108 | 1 | CH_SC | 1,733510905 | 0,57686398 | 1,851851852 |
| ENPP22 | 0,345172943 | 0,327200929 | 0,2 | 0,108 | 1 | CH_SC | 1,254576921 | 0,797081457 | 1,851851852 |
| GSK3A | 0,345172943 | 0,403666476 | 0,2 | 0,108 | 1 | CH_SC | 1,322865583 | 0,755934702 | 1,851851852 |
| MKLN1 | 0,346361014 | 0,378245245 | 0,333 | 0,215 | 1 | CH_SC | 1,299759991 | 0,769372812 | 1,548837209 |
| COA41 | 0,346530742 | 0,50715442 | 0,267 | 0,169 | 1 | CH_SC | 1,421244159 | 0,703608872 | 1,579881657 |
| TOMM222 | 0,346530742 | 0,636049254 | 0,267 | 0,169 | 1 | CH_SC | 1,554067593 | 0,643472655 | 1,579881657 |
| YTHDC14 | 0,346908372 | 0,582393448 | 0,4 | 0,308 | 1 | CH_SC | 1,497331279 | 0,667854879 | 1,298701299 |
| PLAC92 | 0,349024578 | 0,702280363 | 0,6 | 0,569 | 1 | CH_SC | 1,627074559 | 0,614599985 | 1,054481547 |
| ZNF2922 | 0,34942511 | -0,326471114 | 0,133 | 0,246 | 1 | CH_SC | 0,797484778 | 1,25394243 | 0,540650407 |
| PICALM2 | 0,34942511 | -0,406934554 | 0,133 | 0,246 | 1 | CH_SC | 0,754224252 | 1,325865613 | 0,540650407 |
| CARD19 | 0,351021887 | -0,308164351 | 0,067 | 0,169 | 1 | CH_SC | 0,807668763 | 1,23813133 | 0,396449704 |
| ANGPTL2 | 0,351021887 | -0,251368749 | 0,067 | 0,169 | 1 | CH_SC | 0,840098998 | 1,190335904 | 0,396449704 |
| LDAF11 | 0,351021887 | -0,253159625 | 0,067 | 0,169 | 1 | CH_SC | 0,839056796 | 1,191814434 | 0,396449704 |
| AC245297.3 | 0,351751936 | 0,264854326 | 0,133 | 0,062 | 1 | CH_SC | 1,201514722 | 0,832282769 | 2,14516129 |
| KLHL18 | 0,351751936 | 0,339516306 | 0,133 | 0,062 | 1 | CH_SC | 1,265332293 | 0,790306234 | 2,14516129 |
| NUP205 | 0,351751936 | 0,267928135 | 0,133 | 0,062 | 1 | CH_SC | 1,204077401 | 0,830511393 | 2,14516129 |
| PAG12 | 0,351751936 | 0,26602698 | 0,133 | 0,062 | 1 | CH_SC | 1,202491737 | 0,831606546 | 2,14516129 |
| PLIN22 | 0,351751936 | 0,277382361 | 0,133 | 0,062 | 1 | CH_SC | 1,211993835 | 0,825086705 | 2,14516129 |
| COMTD13 | 0,351751936 | 0,288164008 | 0,133 | 0,062 | 1 | CH_SC | 1,22108532 | 0,818943594 | 2,14516129 |
| IFIT2 | 0,351751936 | 0,914654939 | 0,133 | 0,062 | 1 | CH_SC | 1,88511814 | 0,530470732 | 2,14516129 |
| GATD1 | 0,351751936 | 0,293314634 | 0,133 | 0,062 | 1 | CH_SC | 1,225452559 | 0,816025062 | 2,14516129 |
| RAB30 | 0,351751936 | 0,82107397 | 0,133 | 0,062 | 1 | CH_SC | 1,766720684 | 0,566020429 | 2,14516129 |
| ARHGAP421 | 0,351751936 | 0,276806672 | 0,133 | 0,062 | 1 | CH_SC | 1,211510301 | 0,825416011 | 2,14516129 |
| B4GALNT3 | 0,351751936 | 0,281056423 | 0,133 | 0,062 | 1 | CH_SC | 1,215084313 | 0,822988158 | 2,14516129 |
| TAPBPL1 | 0,351751936 | 0,555821808 | 0,133 | 0,062 | 1 | CH_SC | 1,47000576 | 0,680269443 | 2,14516129 |
| PUS7L | 0,351751936 | 0,281700994 | 0,133 | 0,062 | 1 | CH_SC | 1,215627312 | 0,822620543 | 2,14516129 |
| AHNAK21 | 0,351751936 | 0,256684791 | 0,133 | 0,062 | 1 | CH_SC | 1,194730145 | 0,837009097 | 2,14516129 |
| USP32 | 0,351751936 | 0,474265828 | 0,133 | 0,062 | 1 | CH_SC | 1,389211085 | 0,719833012 | 2,14516129 |
| CHSY11 | 0,351751936 | 0,403737681 | 0,133 | 0,062 | 1 | CH_SC | 1,322930875 | 0,755897393 | 2,14516129 |
| TENT4B | 0,351751936 | 0,284730801 | 0,133 | 0,062 | 1 | CH_SC | 1,218182937 | 0,820894769 | 2,14516129 |

| Supplementary Table 4 - clusermarker | | | | | | | | | |
|---|---|---|---|---|---|---|---|---|---|
| | **p_val** | **avg_log2FC** | **pct.1** | **pct.2** | **p_val_adj** | **celltype** | **Foldchange_UP** | **Foldchange_DOWN** | **Ratio_pct1_pct2** |
| SERTAD3 | 0,351751936 | 0,273740094 | 0,133 | 0,062 | 1 | CH_SC | 1,208937862 | 0,827172373 | 2,14516129 |
| ABHD121 | 0,351751936 | 0,26989606 | 0,133 | 0,062 | 1 | CH_SC | 1,205720957 | 0,829379297 | 2,14516129 |
| SAMM503 | 0,351751936 | 0,382257978 | 0,133 | 0,062 | 1 | CH_SC | 1,303380195 | 0,767235841 | 2,14516129 |
| ERGIC35 | 0,353073336 | 0,752621563 | 0,4 | 0,369 | 1 | CH_SC | 1,684851644 | 0,593524067 | 1,08401084 |
| EPM2AIP1 | 0,354164501 | -0,376590519 | 0,067 | 0,154 | 1 | CH_SC | 0,770255765 | 1,298270062 | 0,435064935 |
| WDR62 | 0,354164501 | -0,318811695 | 0,067 | 0,154 | 1 | CH_SC | 0,801729967 | 1,247302759 | 0,435064935 |
| ALDH7A11 | 0,354164501 | -0,392340172 | 0,067 | 0,154 | 1 | CH_SC | 0,761892748 | 1,312520697 | 0,435064935 |
| FOXP4 | 0,354164501 | -0,313076382 | 0,067 | 0,154 | 1 | CH_SC | 0,804923521 | 1,242354055 | 0,435064935 |
| TMEM8B2 | 0,354164501 | -0,322361348 | 0,067 | 0,154 | 1 | CH_SC | 0,799759789 | 1,250375442 | 0,435064935 |
| PTGR13 | 0,354164501 | -0,381023806 | 0,067 | 0,154 | 1 | CH_SC | 0,767892463 | 1,302265679 | 0,435064935 |
| NUP214 | 0,354164501 | -0,283067505 | 0,067 | 0,154 | 1 | CH_SC | 0,821841732 | 1,216779291 | 0,435064935 |
| TRAF42 | 0,354164501 | -0,327827168 | 0,067 | 0,154 | 1 | CH_SC | 0,796735538 | 1,255121621 | 0,435064935 |
| PHF202 | 0,354164501 | -0,443390657 | 0,067 | 0,154 | 1 | CH_SC | 0,735404211 | 1,359796402 | 0,435064935 |
| MAX2 | 0,354264046 | 0,521945591 | 0,267 | 0,185 | 1 | CH_SC | 1,435890358 | 0,696432005 | 1,443243243 |
| MRPS63 | 0,354264046 | -0,730354272 | 0,133 | 0,215 | 1 | CH_SC | 0,602755881 | 1,659046442 | 0,618604651 |
| LAMP21 | 0,354554969 | 0,655834615 | 0,4 | 0,308 | 1 | CH_SC | 1,57552715 | 0,634708199 | 1,298701299 |
| PDK43 | 0,355236747 | -0,658310977 | 0,333 | 0,462 | 1 | CH_SC | 0,633619668 | 1,578233838 | 0,720779221 |
| PSMB73 | 0,355465761 | 0,549093943 | 0,333 | 0,231 | 1 | CH_SC | 1,463166493 | 0,683449221 | 1,441558442 |
| ACTN43 | 0,355578476 | 0,8176896 | 0,333 | 0,246 | 1 | CH_SC | 1,762581051 | 0,567349796 | 1,353658537 |
| ITSN1 | 0,355883642 | 0,276953418 | 0,267 | 0,169 | 1 | CH_SC | 1,211633538 | 0,825332057 | 1,579881657 |
| COA62 | 0,356247417 | 0,362004077 | 0,2 | 0,108 | 1 | CH_SC | 1,285209969 | 0,778082978 | 1,851851852 |
| KPNA5 | 0,356247417 | 0,253429178 | 0,2 | 0,108 | 1 | CH_SC | 1,192037133 | 0,838900041 | 1,851851852 |
| SMIM193 | 0,356247417 | 0,262991814 | 0,2 | 0,108 | 1 | CH_SC | 1,199964574 | 0,833357936 | 1,851851852 |
| KLHL21 | 0,35817643 | -0,306457527 | 0,133 | 0,246 | 1 | CH_SC | 0,808624865 | 1,236667388 | 0,540650407 |
| SF3B56 | 0,358203126 | -0,383426289 | 0,333 | 0,415 | 1 | CH_SC | 0,766614776 | 1,304436114 | 0,802409639 |
| ATP5F1A2 | 0,359414125 | 0,79293126 | 0,4 | 0,354 | 1 | CH_SC | 1,732591158 | 0,577170209 | 1,129943503 |
| TMEM2053 | 0,361071878 | -0,366129336 | 0,133 | 0,231 | 1 | CH_SC | 0,775861296 | 1,288890173 | 0,575757576 |
| OSTC4 | 0,362356411 | -0,266794463 | 0,333 | 0,462 | 1 | CH_SC | 0,831164267 | 1,203131607 | 0,720779221 |
| NDUFB42 | 0,363360925 | 0,573218506 | 0,533 | 0,538 | 1 | CH_SC | 1,487839089 | 0,672115693 | 0,99070632 |
| HINT23 | 0,363505665 | -0,369819925 | 0,133 | 0,215 | 1 | CH_SC | 0,773879085 | 1,292191531 | 0,618604651 |
| GTF3A6 | 0,363505665 | 0,448674496 | 0,267 | 0,185 | 1 | CH_SC | 1,364785757 | 0,732715735 | 1,443243243 |

| Supplementary Table 4 - clusermarker | | | | | | | | | |
|---|---|---|---|---|---|---|---|---|---|
| | **p_val** | **avg_log2FC** | **pct.1** | **pct.2** | **p_val_adj** | **celltype** | **Foldchange_UP** | **Foldchange_DOWN** | **Ratio_pct1_pct2** |
| SMARCE15 | 0,363545419 | 0,410113715 | 0,333 | 0,215 | 1 | CH_SC | 1,328790547 | 0,752564053 | 1,548837209 |
| PIP4P2 | 0,364953444 | -0,353923029 | 0,067 | 0,154 | 1 | CH_SC | 0,782453526 | 1,278031176 | 0,435064935 |
| FAM136A1 | 0,364961285 | -0,39179568 | 0,067 | 0,154 | 1 | CH_SC | 0,76218035 | 1,312025428 | 0,435064935 |
| FBXL51 | 0,364961285 | -0,335943088 | 0,067 | 0,154 | 1 | CH_SC | 0,792266064 | 1,262202239 | 0,435064935 |
| GTF2H11 | 0,364961285 | -0,430092252 | 0,067 | 0,154 | 1 | CH_SC | 0,742214324 | 1,347319727 | 0,435064935 |
| DTX31 | 0,364961285 | 0,530372246 | 0,2 | 0,123 | 1 | CH_SC | 1,444301808 | 0,692376063 | 1,62601626 |
| GOLGA32 | 0,364961285 | -0,381585414 | 0,067 | 0,154 | 1 | CH_SC | 0,767593599 | 1,302772719 | 0,435064935 |
| THUMPD11 | 0,364961285 | 0,684911116 | 0,2 | 0,123 | 1 | CH_SC | 1,607602934 | 0,622044149 | 1,62601626 |
| AHCY1 | 0,364961285 | -0,470293072 | 0,067 | 0,154 | 1 | CH_SC | 0,721817951 | 1,385390871 | 0,435064935 |
| KPNA42 | 0,365388106 | 0,750139423 | 0,267 | 0,169 | 1 | CH_SC | 1,681955368 | 0,594546097 | 1,579881657 |
| RBBP82 | 0,36588656 | -0,277058171 | 0,133 | 0,062 | 1 | CH_SC | 0,825272132 | 1,211721517 | 2,14516129 |
| NDUFS24 | 0,366633659 | 0,407707355 | 0,267 | 0,154 | 1 | CH_SC | 1,326576022 | 0,753820349 | 1,733766234 |
| ARMCX1 | 0,366633659 | 0,269359353 | 0,267 | 0,154 | 1 | CH_SC | 1,205272493 | 0,829687897 | 1,733766234 |
| CHASERR4 | 0,366970815 | 0,706626395 | 0,4 | 0,354 | 1 | CH_SC | 1,631983413 | 0,612751326 | 1,129943503 |
| RBBP44 | 0,367064372 | -0,292330354 | 0,133 | 0,246 | 1 | CH_SC | 0,816581986 | 1,224616778 | 0,540650407 |
| AP2A1 | 0,367543258 | 0,415442152 | 0,2 | 0,108 | 1 | CH_SC | 1,333707364 | 0,749789667 | 1,851851852 |
| CHURC14 | 0,367577833 | 0,646784247 | 0,533 | 0,477 | 1 | CH_SC | 1,565674431 | 0,638702389 | 1,117400419 |
| TMEM595 | 0,367730992 | -0,423628445 | 0,667 | 0,769 | 1 | CH_SC | 0,745547179 | 1,34129674 | 0,867360208 |
| NORAD1 | 0,369186953 | -0,301077483 | 0,467 | 0,6 | 1 | CH_SC | 0,811645989 | 1,232064243 | 0,778333333 |
| UBE2S3 | 0,371750033 | 0,498991988 | 0,467 | 0,354 | 1 | CH_SC | 1,413225796 | 0,70760101 | 1,31920904 |
| TUSC32 | 0,372334221 | -0,304649416 | 0,133 | 0,262 | 1 | CH_SC | 0,80963894 | 1,235118459 | 0,507633588 |
| HM133 | 0,372598232 | 0,263845531 | 0,333 | 0,231 | 1 | CH_SC | 1,200674865 | 0,832864941 | 1,441558442 |
| C1S3 | 0,372616432 | 0,460044927 | 0,667 | 0,554 | 1 | CH_SC | 1,375584655 | 0,72696362 | 1,203971119 |
| VAMP33 | 0,372896654 | 0,624714285 | 0,267 | 0,185 | 1 | CH_SC | 1,541905433 | 0,648548205 | 1,443243243 |
| PSMC61 | 0,372896654 | 0,379351258 | 0,267 | 0,185 | 1 | CH_SC | 1,300756808 | 0,768783214 | 1,443243243 |
| CD3201 | 0,372896654 | -0,410893333 | 0,133 | 0,215 | 1 | CH_SC | 0,752157485 | 1,329508806 | 0,618604651 |
| SUMO34 | 0,372896654 | 0,415819132 | 0,267 | 0,185 | 1 | CH_SC | 1,334055911 | 0,74959377 | 1,443243243 |
| TXN23 | 0,372896654 | -0,695183774 | 0,133 | 0,215 | 1 | CH_SC | 0,617630638 | 1,619090664 | 0,618604651 |
| KMT2A5 | 0,373674353 | 0,78627986 | 0,4 | 0,277 | 1 | CH_SC | 1,724621607 | 0,579837337 | 1,444043321 |
| PFDN16 | 0,375061804 | 0,267050442 | 0,267 | 0,169 | 1 | CH_SC | 1,203345099 | 0,831016806 | 1,579881657 |
| SND12 | 0,375061804 | 0,510961658 | 0,267 | 0,169 | 1 | CH_SC | 1,424999742 | 0,701754513 | 1,579881657 |

| Supplementary Table 4 - clusermarker | | | | | | | | | |
|---|---|---|---|---|---|---|---|---|---|
| | **p_val** | **avg_log2FC** | **pct.1** | **pct.2** | **p_val_adj** | **celltype** | **Foldchange_UP** | **Foldchange_DOWN** | **Ratio_pct1_pct2** |
| ISOC23 | 0,375061804 | 0,296585323 | 0,267 | 0,169 | 1 | CH_SC | 1,228233896 | 0,814177172 | 1,579881657 |
| SLC20A12 | 0,375961348 | 0,436138526 | 0,2 | 0,123 | 1 | CH_SC | 1,352978137 | 0,739110243 | 1,62601626 |
| TNS11 | 0,375961348 | 0,470287337 | 0,2 | 0,123 | 1 | CH_SC | 1,385385364 | 0,721820821 | 1,62601626 |
| AMOTL21 | 0,375961348 | 0,395426428 | 0,2 | 0,123 | 1 | CH_SC | 1,315331485 | 0,760264627 | 1,62601626 |
| SFXN3 | 0,375961348 | 0,432347639 | 0,2 | 0,123 | 1 | CH_SC | 1,34942766 | 0,741054915 | 1,62601626 |
| VPS26B1 | 0,375961348 | 0,441848891 | 0,2 | 0,123 | 1 | CH_SC | 1,358344003 | 0,736190536 | 1,62601626 |
| DAP1 | 0,377772315 | -0,441511363 | 0,2 | 0,323 | 1 | CH_SC | 0,736362793 | 1,358026246 | 0,619195046 |
| PIK3R33 | 0,379059589 | 0,743216248 | 0,2 | 0,108 | 1 | CH_SC | 1,67390337 | 0,59740605 | 1,851851852 |
| RAB181 | 0,379477479 | -0,330202479 | 0,133 | 0,231 | 1 | CH_SC | 0,79542484 | 1,257189806 | 0,575757576 |
| HNRNPA02 | 0,379841841 | 0,474566786 | 0,4 | 0,323 | 1 | CH_SC | 1,389500916 | 0,719682865 | 1,238390093 |
| RNF113 | 0,380270038 | -0,597801102 | 0,2 | 0,292 | 1 | CH_SC | 0,660760293 | 1,513408132 | 0,684931507 |
| TMCO14 | 0,380270038 | 0,915656024 | 0,333 | 0,262 | 1 | CH_SC | 1,886426676 | 0,530102767 | 1,270992366 |
| CYSTM12 | 0,380270038 | -0,587049664 | 0,2 | 0,292 | 1 | CH_SC | 0,66570289 | 1,502171637 | 0,684931507 |
| NAPA1 | 0,380270038 | 0,517256291 | 0,333 | 0,262 | 1 | CH_SC | 1,431230752 | 0,698699353 | 1,270992366 |
| SNHG85 | 0,380814826 | -0,388098229 | 0,533 | 0,631 | 1 | CH_SC | 0,764136231 | 1,308667171 | 0,844690967 |
| ODC12 | 0,381253511 | 0,592946128 | 0,333 | 0,215 | 1 | CH_SC | 1,508323753 | 0,662987636 | 1,548837209 |
| NDUFA54 | 0,382378652 | -0,299044119 | 0,267 | 0,385 | 1 | CH_SC | 0,812790746 | 1,230328969 | 0,693506494 |
| PDAP13 | 0,382436456 | 0,362994696 | 0,267 | 0,185 | 1 | CH_SC | 1,286092755 | 0,777548895 | 1,443243243 |
| POLR2C | 0,382436456 | -0,447609011 | 0,133 | 0,215 | 1 | CH_SC | 0,733257074 | 1,363778184 | 0,618604651 |
| PLAUR3 | 0,382436456 | -0,515934595 | 0,133 | 0,215 | 1 | CH_SC | 0,699339745 | 1,429920159 | 0,618604651 |
| MAP3K13 | 0,382619059 | 0,525657195 | 0,067 | 0,169 | 1 | CH_SC | 1,439589211 | 0,694642605 | 0,396449704 |
| POMP5 | 0,383878565 | 0,282620298 | 0,533 | 0,431 | 1 | CH_SC | 1,216402172 | 0,822096526 | 1,236658933 |
| RACK15 | 0,384593655 | -0,27081703 | 0,8 | 0,923 | 1 | CH_SC | 0,828850017 | 1,206490897 | 0,866738895 |
| SPCS17 | 0,386931415 | -0,296556895 | 0,4 | 0,538 | 1 | CH_SC | 0,814193216 | 1,228209694 | 0,743494424 |
| TASOR3 | 0,387163768 | 0,631941301 | 0,2 | 0,123 | 1 | CH_SC | 1,54964881 | 0,6453075 | 1,62601626 |
| ARFIP1 | 0,387163768 | 0,429814485 | 0,2 | 0,123 | 1 | CH_SC | 1,347060349 | 0,742357238 | 1,62601626 |
| SREK1IP11 | 0,387163768 | -0,311553478 | 0,067 | 0,154 | 1 | CH_SC | 0,805773644 | 1,241043323 | 0,435064935 |
| FIBP5 | 0,387163768 | -0,314572379 | 0,067 | 0,154 | 1 | CH_SC | 0,804089291 | 1,243642977 | 0,435064935 |
| C11orf241 | 0,387163768 | 0,554203723 | 0,2 | 0,123 | 1 | CH_SC | 1,468357969 | 0,681032842 | 1,62601626 |
| MAPKAPK5-AS1 | 0,387163768 | 0,389616249 | 0,2 | 0,123 | 1 | CH_SC | 1,310044891 | 0,763332621 | 1,62601626 |
| TARS3 | 0,387163768 | 0,621481615 | 0,2 | 0,123 | 1 | CH_SC | 1,538454328 | 0,650003046 | 1,62601626 |

| Supplementary Table 4 - clusermarker | | | | | | | | | |
|---|---|---|---|---|---|---|---|---|---|
| | **p_val** | **avg_log2FC** | **pct.1** | **pct.2** | **p_val_adj** | **celltype** | **Foldchange_UP** | **Foldchange_DOWN** | **Ratio_pct1_pct2** |
| GLOD4 | 0,387163768 | -0,261591444 | 0,067 | 0,154 | 1 | CH_SC | 0,834167238 | 1,198800378 | 0,435064935 |
| TSEN343 | 0,387163768 | 0,612992648 | 0,2 | 0,123 | 1 | CH_SC | 1,529428484 | 0,653839006 | 1,62601626 |
| THBS15 | 0,38775571 | -1,055219417 | 0,6 | 0,615 | 1 | CH_SC | 0,481224027 | 2,078034227 | 0,975609756 |
| EIF2S35 | 0,387834436 | -0,417438981 | 0,267 | 0,354 | 1 | CH_SC | 0,748752603 | 1,335554622 | 0,754237288 |
| RIF1 | 0,388835555 | 0,579311762 | 0,333 | 0,262 | 1 | CH_SC | 1,494136301 | 0,669282983 | 1,270992366 |
| TTC171 | 0,388892848 | 0,869857716 | 0,267 | 0,2 | 1 | CH_SC | 1,827482658 | 0,547200815 | 1,335 |
| PRMT12 | 0,388892848 | 0,420071549 | 0,267 | 0,2 | 1 | CH_SC | 1,337993909 | 0,747387558 | 1,335 |
| PPIB6 | 0,389712898 | 0,304539936 | 0,8 | 0,8 | 1 | CH_SC | 1,235024734 | 0,809700383 | 1 |
| PKD2 | 0,389727818 | -0,402287164 | 0,2 | 0,277 | 1 | CH_SC | 0,756657771 | 1,32160144 | 0,722021661 |
| SLTM1 | 0,389727818 | 0,720260285 | 0,333 | 0,246 | 1 | CH_SC | 1,647479239 | 0,606987922 | 1,353658537 |
| PSMC2 | 0,39023316 | 0,479167083 | 0,333 | 0,231 | 1 | CH_SC | 1,393938666 | 0,717391679 | 1,441558442 |
| RAB342 | 0,39023316 | 0,383865187 | 0,333 | 0,231 | 1 | CH_SC | 1,304833012 | 0,766381591 | 1,441558442 |
| DDX11 | 0,392124413 | -0,3634056 | 0,133 | 0,215 | 1 | CH_SC | 0,777327467 | 1,286459109 | 0,618604651 |
| BIRC21 | 0,39247624 | -0,293646516 | 0,2 | 0,338 | 1 | CH_SC | 0,815837362 | 1,225734498 | 0,591715976 |
| SUMO24 | 0,392670362 | 0,341999538 | 0,6 | 0,677 | 1 | CH_SC | 1,267512116 | 0,788947094 | 0,886262925 |
| GLTP1 | 0,394865807 | 0,275794583 | 0,267 | 0,169 | 1 | CH_SC | 1,210660693 | 0,825995265 | 1,579881657 |
| SSU725 | 0,397517502 | -0,380844268 | 0,2 | 0,292 | 1 | CH_SC | 0,767988031 | 1,302103626 | 0,684931507 |
| NDUFA26 | 0,398176793 | -0,396112492 | 0,267 | 0,385 | 1 | CH_SC | 0,759903173 | 1,315957131 | 0,693506494 |
| TUT42 | 0,398449057 | -0,33801253 | 0,133 | 0,231 | 1 | CH_SC | 0,79113043 | 1,264014077 | 0,575757576 |
| RAPH1 | 0,398449057 | -0,3130729 | 0,133 | 0,231 | 1 | CH_SC | 0,804925463 | 1,242351057 | 0,575757576 |
| BLVRB2 | 0,398449057 | 0,600260358 | 0,267 | 0,2 | 1 | CH_SC | 1,515990127 | 0,659634903 | 1,335 |
| RRAGA3 | 0,398567102 | 0,290524578 | 0,333 | 0,246 | 1 | CH_SC | 1,223084923 | 0,817604715 | 1,353658537 |
| RABGAP1L4 | 0,398567432 | 0,572175739 | 0,2 | 0,123 | 1 | CH_SC | 1,48676408 | 0,672601668 | 1,62601626 |
| NUB1 | 0,398567432 | 0,923461793 | 0,2 | 0,123 | 1 | CH_SC | 1,896660936 | 0,527242366 | 1,62601626 |
| URM11 | 0,398567432 | 0,554731899 | 0,2 | 0,123 | 1 | CH_SC | 1,468895638 | 0,680783559 | 1,62601626 |
| RASSF8-AS1 | 0,398567432 | 0,369141848 | 0,2 | 0,123 | 1 | CH_SC | 1,291584335 | 0,774242899 | 1,62601626 |
| ORMDL22 | 0,398567432 | 0,680440083 | 0,2 | 0,123 | 1 | CH_SC | 1,60262855 | 0,623974907 | 1,62601626 |
| ATL1 | 0,398567432 | 0,390631854 | 0,2 | 0,123 | 1 | CH_SC | 1,31096744 | 0,762795451 | 1,62601626 |
| SMOX | 0,398567432 | 0,392311576 | 0,2 | 0,123 | 1 | CH_SC | 1,312494681 | 0,76190785 | 1,62601626 |
| CDK5RAP34 | 0,401954111 | -0,546893675 | 0,133 | 0,215 | 1 | CH_SC | 0,684492351 | 1,460936705 | 0,618604651 |
| SAR1B2 | 0,401959763 | 0,407920496 | 0,267 | 0,185 | 1 | CH_SC | 1,326772023 | 0,753708989 | 1,443243243 |

| Supplementary Table 4 - clusermarker | | | | | | | | | |
|---|---|---|---|---|---|---|---|---|---|
| | **p_val** | **avg_log2FC** | **pct.1** | **pct.2** | **p_val_adj** | **celltype** | **Foldchange_UP** | **Foldchange_DOWN** | **Ratio_pct1_pct2** |
| MCTS11 | 0,401959763 | 0,406401547 | 0,267 | 0,185 | 1 | CH_SC | 1,325375859 | 0,754502953 | 1,443243243 |
| RBIS2 | 0,40272428 | 0,609770946 | 0,533 | 0,492 | 1 | CH_SC | 1,526016907 | 0,655300734 | 1,083333333 |
| ZDHHC2 | 0,402740747 | -0,358525983 | 0,067 | 0,138 | 1 | CH_SC | 0,779961067 | 1,282115277 | 0,485507246 |
| UBE4B | 0,402749099 | -0,399019789 | 0,067 | 0,138 | 1 | CH_SC | 0,75837337 | 1,318611701 | 0,485507246 |
| MIGA1 | 0,402749099 | -0,443622802 | 0,067 | 0,138 | 1 | CH_SC | 0,735285886 | 1,360015225 | 0,485507246 |
| NSL1 | 0,402749099 | -0,317376086 | 0,067 | 0,138 | 1 | CH_SC | 0,802528156 | 1,246062201 | 0,485507246 |
| HDAC24 | 0,402749099 | -0,346278414 | 0,067 | 0,138 | 1 | CH_SC | 0,786610629 | 1,271276999 | 0,485507246 |
| VPS411 | 0,402749099 | -0,323987953 | 0,067 | 0,138 | 1 | CH_SC | 0,798858587 | 1,251786006 | 0,485507246 |
| ABCF2 | 0,402749099 | -0,51688901 | 0,067 | 0,138 | 1 | CH_SC | 0,69887725 | 1,430866435 | 0,485507246 |
| PPP6R32 | 0,402749099 | -0,445648784 | 0,067 | 0,138 | 1 | CH_SC | 0,734254046 | 1,361926441 | 0,485507246 |
| LETMD1 | 0,402749099 | -0,497365533 | 0,067 | 0,138 | 1 | CH_SC | 0,708399189 | 1,411633462 | 0,485507246 |
| CLIP12 | 0,402749099 | -0,408268006 | 0,067 | 0,138 | 1 | CH_SC | 0,753527461 | 1,327091649 | 0,485507246 |
| OAZ23 | 0,402749099 | -0,402750541 | 0,067 | 0,138 | 1 | CH_SC | 0,756414781 | 1,322025991 | 0,485507246 |
| TRPC4AP2 | 0,402749099 | -0,308643816 | 0,067 | 0,138 | 1 | CH_SC | 0,807400387 | 1,238542879 | 0,485507246 |
| DDX273 | 0,402749099 | -0,322061394 | 0,067 | 0,138 | 1 | CH_SC | 0,799926087 | 1,2501155 | 0,485507246 |
| SUN2 | 0,402937118 | 0,637415512 | 0,333 | 0,292 | 1 | CH_SC | 1,55554002 | 0,642863563 | 1,140410959 |
| BOD1L16 | 0,404774176 | -0,349660572 | 0,2 | 0,308 | 1 | CH_SC | 0,784768711 | 1,274260792 | 0,649350649 |
| TMEM50B4 | 0,404774176 | 0,842419667 | 0,333 | 0,277 | 1 | CH_SC | 1,793054907 | 0,557707405 | 1,202166065 |
| EIF2S23 | 0,405306234 | -0,430062118 | 0,267 | 0,369 | 1 | CH_SC | 0,742229826 | 1,347291586 | 0,723577236 |
| MEG32 | 0,40692763 | -0,474216246 | 0,333 | 0,462 | 1 | CH_SC | 0,719857752 | 1,389163342 | 0,720779221 |
| JUNB5 | 0,407324709 | 0,50739013 | 0,8 | 0,815 | 1 | CH_SC | 1,421476383 | 0,703493925 | 0,981595092 |
| CCT31 | 0,409297466 | 0,925757641 | 0,2 | 0,338 | 1 | CH_SC | 1,89968161 | 0,526404001 | 0,591715976 |
| WDR471 | 0,410171035 | 0,571078226 | 0,2 | 0,123 | 1 | CH_SC | 1,485633472 | 0,673113536 | 1,62601626 |
| LRPPRC | 0,410171035 | 0,312781679 | 0,2 | 0,123 | 1 | CH_SC | 1,242100303 | 0,805087961 | 1,62601626 |
| POLR1H2 | 0,410171035 | -0,267711872 | 0,067 | 0,154 | 1 | CH_SC | 0,830635898 | 1,203896921 | 0,435064935 |
| PIM12 | 0,410171035 | -0,424468248 | 0,067 | 0,154 | 1 | CH_SC | 0,745113316 | 1,342077746 | 0,435064935 |
| NXF11 | 0,410171035 | 0,510186758 | 0,2 | 0,123 | 1 | CH_SC | 1,424234552 | 0,70213154 | 1,62601626 |
| HSPB2 | 0,410171035 | 0,279505829 | 0,2 | 0,123 | 1 | CH_SC | 1,213779054 | 0,823873173 | 1,62601626 |
| JAM31 | 0,410171035 | 0,280042597 | 0,2 | 0,123 | 1 | CH_SC | 1,214230735 | 0,8235667 | 1,62601626 |
| APPL2 | 0,410171035 | 0,456310717 | 0,2 | 0,123 | 1 | CH_SC | 1,372028754 | 0,728847699 | 1,62601626 |
| ITGAE2 | 0,410171035 | 0,296374431 | 0,2 | 0,123 | 1 | CH_SC | 1,228054367 | 0,814296197 | 1,62601626 |

| Supplementary Table 4 - clusermarker | | | | | | | | | |
|---|---|---|---|---|---|---|---|---|---|
| | **p_val** | **avg_log2FC** | **pct.1** | **pct.2** | **p_val_adj** | **celltype** | **Foldchange_UP** | **Foldchange_DOWN** | **Ratio_pct1_pct2** |
| ATRN | 0,410171035 | 0,329418005 | 0,2 | 0,123 | 1 | CH_SC | 1,256506387 | 0,795857475 | 1,62601626 |
| SPHK12 | 0,410376259 | -0,28395263 | 0,133 | 0,062 | 1 | CH_SC | 0,821337669 | 1,217526041 | 2,14516129 |
| HIF1A4 | 0,411543662 | -0,489368406 | 0,2 | 0,323 | 1 | CH_SC | 0,712336882 | 1,403830162 | 0,619195046 |
| YIF1A3 | 0,411941641 | 0,414340953 | 0,267 | 0,185 | 1 | CH_SC | 1,332689743 | 0,750362194 | 1,443243243 |
| CPNE34 | 0,412420283 | 0,480952229 | 0,4 | 0,323 | 1 | CH_SC | 1,39566455 | 0,71650455 | 1,238390093 |
| CD995 | 0,412757576 | -0,429591342 | 0,6 | 0,8 | 1 | CH_SC | 0,742472068 | 1,346852014 | 0,75 |
| BRK13 | 0,414661922 | 0,721473322 | 0,467 | 0,431 | 1 | CH_SC | 1,648865044 | 0,606477773 | 1,083526682 |
| GSTM23 | 0,414919403 | -0,323730033 | 0,067 | 0,138 | 1 | CH_SC | 0,799001417 | 1,251562236 | 0,485507246 |
| SUCO | 0,414919403 | -0,385291427 | 0,067 | 0,138 | 1 | CH_SC | 0,765624325 | 1,3061236 | 0,485507246 |
| LTBP11 | 0,414919403 | -0,404671313 | 0,067 | 0,138 | 1 | CH_SC | 0,755408377 | 1,32378728 | 0,485507246 |
| RBMS33 | 0,414919403 | -0,377333033 | 0,067 | 0,138 | 1 | CH_SC | 0,769859438 | 1,298938417 | 0,485507246 |
| ABHD14B2 | 0,414919403 | -0,420227647 | 0,067 | 0,138 | 1 | CH_SC | 0,747306695 | 1,338138687 | 0,485507246 |
| ALG3 | 0,414919403 | -0,281192508 | 0,067 | 0,138 | 1 | CH_SC | 0,822910532 | 1,215198933 | 0,485507246 |
| MARCHF5 | 0,414919403 | -0,407981413 | 0,067 | 0,138 | 1 | CH_SC | 0,753677165 | 1,326828046 | 0,485507246 |
| PRPS12 | 0,414919403 | -0,314363031 | 0,067 | 0,138 | 1 | CH_SC | 0,80420598 | 1,243462527 | 0,485507246 |
| C19orf537 | 0,414945076 | 0,620686876 | 0,4 | 0,369 | 1 | CH_SC | 1,537607072 | 0,650361213 | 1,08401084 |
| BROX | 0,415289286 | 0,334161795 | 0,267 | 0,169 | 1 | CH_SC | 1,260644762 | 0,793244878 | 1,579881657 |
| SNAPC11 | 0,415901343 | 0,738889279 | 0,2 | 0,138 | 1 | CH_SC | 1,668890477 | 0,599200495 | 1,449275362 |
| MRPS18C3 | 0,417617851 | -0,5538928 | 0,2 | 0,262 | 1 | CH_SC | 0,681179631 | 1,468041549 | 0,763358779 |
| TXNDC153 | 0,417617851 | 0,296096289 | 0,333 | 0,231 | 1 | CH_SC | 1,227817629 | 0,814453203 | 1,441558442 |
| POLR2L3 | 0,418492748 | -0,314553041 | 0,533 | 0,615 | 1 | CH_SC | 0,804100069 | 1,243626308 | 0,866666667 |
| SEC63 | 0,419499918 | 0,364071797 | 0,4 | 0,308 | 1 | CH_SC | 1,287053297 | 0,776968601 | 1,298701299 |
| ATL21 | 0,42093651 | 0,527365124 | 0,133 | 0,077 | 1 | CH_SC | 1,441294472 | 0,693820742 | 1,727272727 |
| TIA1 | 0,42093651 | 0,348235507 | 0,133 | 0,077 | 1 | CH_SC | 1,273002724 | 0,785544273 | 1,727272727 |
| INSIG2 | 0,42093651 | 0,968500177 | 0,133 | 0,077 | 1 | CH_SC | 1,956805247 | 0,51103706 | 1,727272727 |
| TENT22 | 0,42093651 | 0,617404592 | 0,133 | 0,077 | 1 | CH_SC | 1,534112829 | 0,651842538 | 1,727272727 |
| IFIT12 | 0,42093651 | 0,616670101 | 0,133 | 0,077 | 1 | CH_SC | 1,533331995 | 0,652174482 | 1,727272727 |
| PLCE1 | 0,42093651 | 0,582433137 | 0,133 | 0,077 | 1 | CH_SC | 1,497372472 | 0,667836506 | 1,727272727 |
| TMEM2182 | 0,42093651 | 0,547508797 | 0,133 | 0,077 | 1 | CH_SC | 1,461559737 | 0,684200566 | 1,727272727 |
| NECAP1 | 0,42093651 | 1,333058991 | 0,133 | 0,077 | 1 | CH_SC | 2,519362973 | 0,396925735 | 1,727272727 |
| GOLT1B | 0,42093651 | 0,594578187 | 0,133 | 0,077 | 1 | CH_SC | 1,51003102 | 0,662238051 | 1,727272727 |

| Supplementary Table 4 - clusermarker | | | | | | | | | |
|---|---|---|---|---|---|---|---|---|---|
| | **p_val** | **avg_log2FC** | **pct.1** | **pct.2** | **p_val_adj** | **celltype** | **Foldchange_UP** | **Foldchange_DOWN** | **Ratio_pct1_pct2** |
| PRKAG11 | 0,42093651 | 0,437646207 | 0,133 | 0,077 | 1 | CH_SC | 1,354392798 | 0,738338244 | 1,727272727 |
| C16orf72 | 0,42093651 | 0,61415027 | 0,133 | 0,077 | 1 | CH_SC | 1,530656193 | 0,653314575 | 1,727272727 |
| OGFOD3 | 0,42093651 | 0,445559081 | 0,133 | 0,077 | 1 | CH_SC | 1,361841763 | 0,734299701 | 1,727272727 |
| EZR5 | 0,421841791 | 0,427127625 | 0,4 | 0,338 | 1 | CH_SC | 1,344553932 | 0,743741085 | 1,183431953 |
| TMED5 | 0,421973078 | 0,693903582 | 0,2 | 0,123 | 1 | CH_SC | 1,617654583 | 0,618178943 | 1,62601626 |
| SNX14 | 0,421973078 | 0,264321273 | 0,2 | 0,123 | 1 | CH_SC | 1,201070864 | 0,832590341 | 1,62601626 |
| AC100810.11 | 0,421973078 | 0,575858102 | 0,2 | 0,123 | 1 | CH_SC | 1,490563774 | 0,670887095 | 1,62601626 |
| RRAD2 | 0,421973078 | 0,493777014 | 0,2 | 0,123 | 1 | CH_SC | 1,408126567 | 0,710163435 | 1,62601626 |
| TRIP12 | 0,422069075 | 0,480275837 | 0,267 | 0,185 | 1 | CH_SC | 1,395010361 | 0,716840554 | 1,443243243 |
| LASP13 | 0,422069075 | 0,31394476 | 0,267 | 0,185 | 1 | CH_SC | 1,243102071 | 0,804439172 | 1,443243243 |
| NUDCD21 | 0,423230891 | 0,506612764 | 0,267 | 0,215 | 1 | CH_SC | 1,420710657 | 0,70387309 | 1,241860465 |
| SELENOS3 | 0,423382764 | 0,254495475 | 0,467 | 0,369 | 1 | CH_SC | 1,192918494 | 0,838280238 | 1,265582656 |
| DDOST3 | 0,423445009 | -0,250801384 | 0,267 | 0,431 | 1 | CH_SC | 0,840429446 | 1,189867876 | 0,619489559 |
| SPCS3 | 0,424255815 | 0,375870046 | 0,333 | 0,262 | 1 | CH_SC | 1,297621875 | 0,770640523 | 1,270992366 |
| EBNA1BP22 | 0,425724988 | -0,526163765 | 0,133 | 0,2 | 1 | CH_SC | 0,69439874 | 1,440094779 | 0,665 |
| MRPL543 | 0,425730837 | -0,461205453 | 0,133 | 0,2 | 1 | CH_SC | 0,726379075 | 1,376691641 | 0,665 |
| ERP293 | 0,426991862 | -0,482456003 | 0,2 | 0,262 | 1 | CH_SC | 0,7157581 | 1,397120061 | 0,763358779 |
| BSDC1 | 0,427304443 | -0,270413806 | 0,067 | 0,138 | 1 | CH_SC | 0,829081708 | 1,206153737 | 0,485507246 |
| RRP151 | 0,427304443 | -0,339862994 | 0,067 | 0,138 | 1 | CH_SC | 0,790116342 | 1,265636397 | 0,485507246 |
| MORN21 | 0,427304443 | -0,254222546 | 0,067 | 0,138 | 1 | CH_SC | 0,838438839 | 1,19269284 | 0,485507246 |
| MFSD11 | 0,427304443 | -0,271003097 | 0,067 | 0,138 | 1 | CH_SC | 0,828743126 | 1,20664651 | 0,485507246 |
| RAB28 | 0,427304443 | -0,296173621 | 0,067 | 0,138 | 1 | CH_SC | 0,814409547 | 1,227883445 | 0,485507246 |
| CFB3 | 0,427304443 | -0,360233055 | 0,067 | 0,138 | 1 | CH_SC | 0,779038722 | 1,283633241 | 0,485507246 |
| SRPK11 | 0,427304443 | -0,379340783 | 0,067 | 0,138 | 1 | CH_SC | 0,768788797 | 1,300747363 | 0,485507246 |
| KLHDC32 | 0,427304443 | -0,329600214 | 0,067 | 0,138 | 1 | CH_SC | 0,795756966 | 1,256665091 | 0,485507246 |
| RNF146 | 0,427304443 | -0,295546944 | 0,067 | 0,138 | 1 | CH_SC | 0,814763386 | 1,227350193 | 0,485507246 |
| IGF2R2 | 0,427304443 | -0,392148675 | 0,067 | 0,138 | 1 | CH_SC | 0,761993885 | 1,31234649 | 0,485507246 |
| TRA2A | 0,427304443 | -0,470786887 | 0,067 | 0,138 | 1 | CH_SC | 0,721570925 | 1,385865152 | 0,485507246 |
| MRPL171 | 0,427304443 | -0,434885923 | 0,067 | 0,138 | 1 | CH_SC | 0,739752246 | 1,351803938 | 0,485507246 |
| CFL22 | 0,427304443 | -0,34285945 | 0,067 | 0,138 | 1 | CH_SC | 0,788476985 | 1,268267836 | 0,485507246 |
| NCBP3 | 0,427304443 | -0,50008131 | 0,067 | 0,138 | 1 | CH_SC | 0,70706693 | 1,41429327 | 0,485507246 |

| Supplementary Table 4 - clusermarker | | | | | | | | | |
|---|---|---|---|---|---|---|---|---|---|
| | **p_val** | **avg_log2FC** | **pct.1** | **pct.2** | **p_val_adj** | **celltype** | **Foldchange_UP** | **Foldchange_DOWN** | **Ratio_pct1_pct2** |
| FAM104B1 | 0,427304443 | -0,373435381 | 0,067 | 0,138 | 1 | CH_SC | 0,771942139 | 1,29543388 | 0,485507246 |
| SERP13 | 0,427760744 | -0,459585325 | 0,4 | 0,492 | 1 | CH_SC | 0,727195247 | 1,375146502 | 0,81300813 |
| SELENBP14 | 0,427954511 | -0,279538363 | 0,133 | 0,231 | 1 | CH_SC | 0,823854594 | 1,213806426 | 0,575757576 |
| ATP2A24 | 0,427954511 | -0,433912118 | 0,133 | 0,231 | 1 | CH_SC | 0,74025174 | 1,350891793 | 0,575757576 |
| PRDX23 | 0,42808289 | -0,448164042 | 0,267 | 0,338 | 1 | CH_SC | 0,732975031 | 1,364302955 | 0,789940828 |
| TIAL12 | 0,432340991 | -0,326587148 | 0,133 | 0,215 | 1 | CH_SC | 0,797420639 | 1,254043287 | 0,618604651 |
| C12orf76 | 0,432340991 | 0,447792859 | 0,267 | 0,185 | 1 | CH_SC | 1,363951985 | 0,733163638 | 1,443243243 |
| SLC9A3R24 | 0,432340991 | 0,481271822 | 0,267 | 0,185 | 1 | CH_SC | 1,395973759 | 0,716345844 | 1,443243243 |
| BTG26 | 0,433296982 | -1,035248632 | 0,533 | 0,615 | 1 | CH_SC | 0,487931784 | 2,049466818 | 0,866666667 |
| GRHPR2 | 0,433971867 | 0,482906484 | 0,2 | 0,123 | 1 | CH_SC | 1,397556379 | 0,71553464 | 1,62601626 |
| CISD36 | 0,433971867 | 0,305159979 | 0,2 | 0,123 | 1 | CH_SC | 1,235555639 | 0,809352464 | 1,62601626 |
| WASF24 | 0,434971378 | 0,286786595 | 0,467 | 0,415 | 1 | CH_SC | 1,219920045 | 0,819725853 | 1,125301205 |
| SRP721 | 0,435113597 | 0,274421922 | 0,333 | 0,246 | 1 | CH_SC | 1,20950935 | 0,826781538 | 1,353658537 |
| EPS83 | 0,435113597 | -0,409772015 | 0,2 | 0,277 | 1 | CH_SC | 0,752742318 | 1,328475862 | 0,722021661 |
| RNPS11 | 0,435113597 | -0,44617027 | 0,2 | 0,277 | 1 | CH_SC | 0,733988686 | 1,362418821 | 0,722021661 |
| KIN2 | 0,435598308 | 0,637145762 | 0,133 | 0,077 | 1 | CH_SC | 1,555249197 | 0,642983775 | 1,727272727 |
| TUBG2 | 0,435598308 | 0,405342532 | 0,133 | 0,077 | 1 | CH_SC | 1,32440332 | 0,755057002 | 1,727272727 |
| ZNF426 | 0,435598308 | 0,859743697 | 0,133 | 0,077 | 1 | CH_SC | 1,814715887 | 0,551050446 | 1,727272727 |
| RHOQ1 | 0,436324032 | -0,398495811 | 0,133 | 0,2 | 1 | CH_SC | 0,758648857 | 1,318132877 | 0,665 |
| ARID4A1 | 0,436324032 | -0,520807382 | 0,133 | 0,2 | 1 | CH_SC | 0,696981669 | 1,434757963 | 0,665 |
| BCL7C3 | 0,436324032 | -0,356487059 | 0,133 | 0,2 | 1 | CH_SC | 0,781064146 | 1,280304576 | 0,665 |
| CDC26 | 0,436487362 | 0,400829854 | 0,333 | 0,231 | 1 | CH_SC | 1,320267124 | 0,75742248 | 1,441558442 |
| ANXA61 | 0,438065391 | 0,518179174 | 0,267 | 0,2 | 1 | CH_SC | 1,432146595 | 0,698252542 | 1,335 |
| AASS | 0,438065391 | 0,786940028 | 0,267 | 0,2 | 1 | CH_SC | 1,725410964 | 0,579572068 | 1,335 |
| CPSF6 | 0,439009423 | 1,029581625 | 0,2 | 0,138 | 1 | CH_SC | 2,041432159 | 0,489852183 | 1,449275362 |
| AC007952.4 | 0,439009423 | 0,450338111 | 0,2 | 0,138 | 1 | CH_SC | 1,366360441 | 0,731871306 | 1,449275362 |
| FAM114A11 | 0,439706481 | 0,278702273 | 0,4 | 0,369 | 1 | CH_SC | 1,213103188 | 0,824332184 | 1,08401084 |
| BTBD61 | 0,439902209 | -0,285027607 | 0,067 | 0,138 | 1 | CH_SC | 0,820725904 | 1,218433579 | 0,485507246 |
| NFIC2 | 0,44158171 | 0,582656465 | 0,467 | 0,446 | 1 | CH_SC | 1,497604282 | 0,667733133 | 1,047085202 |
| RTN33 | 0,442651179 | 0,896085167 | 0,333 | 0,262 | 1 | CH_SC | 1,861009174 | 0,537342864 | 1,270992366 |
| DPM31 | 0,442756206 | -0,295853497 | 0,133 | 0,215 | 1 | CH_SC | 0,814590279 | 1,227611016 | 0,618604651 |

| Supplementary Table 4 - clusermarker | | | | | | | | | |
|---|---|---|---|---|---|---|---|---|---|
| | **p_val** | **avg_log2FC** | **pct.1** | **pct.2** | **p_val_adj** | **celltype** | **Foldchange_UP** | **Foldchange_DOWN** | **Ratio_pct1_pct2** |
| BTF36 | 0,442892779 | 0,257532571 | 0,733 | 0,815 | 1 | CH_SC | 1,195432417 | 0,836517385 | 0,899386503 |
| PCYOX13 | 0,444543617 | -0,336947103 | 0,2 | 0,277 | 1 | CH_SC | 0,791714894 | 1,26308095 | 0,722021661 |
| EIF2A2 | 0,444543617 | -0,492322713 | 0,2 | 0,277 | 1 | CH_SC | 0,710879673 | 1,406707828 | 0,722021661 |
| PTOV16 | 0,444543617 | 0,300189376 | 0,333 | 0,246 | 1 | CH_SC | 1,231306031 | 0,812145782 | 1,353658537 |
| ZBTB205 | 0,445071578 | 0,65154436 | 0,667 | 0,692 | 1 | CH_SC | 1,570848841 | 0,63659849 | 0,963872832 |
| SNX171 | 0,446103401 | 0,570822149 | 0,333 | 0,231 | 1 | CH_SC | 1,485369797 | 0,673233024 | 1,441558442 |
| SCNM12 | 0,446165513 | 0,273488591 | 0,067 | 0,154 | 1 | CH_SC | 1,208727128 | 0,827316585 | 0,435064935 |
| DNAJC21 | 0,446165513 | 0,58884652 | 0,067 | 0,154 | 1 | CH_SC | 1,504043737 | 0,664874282 | 0,435064935 |
| MRPL22 | 0,446165513 | 0,644117594 | 0,2 | 0,123 | 1 | CH_SC | 1,562783138 | 0,639884048 | 1,62601626 |
| FAM107B3 | 0,446165513 | 0,270625457 | 0,2 | 0,123 | 1 | CH_SC | 1,2063307 | 0,828960086 | 1,62601626 |
| SDHB1 | 0,447067553 | -0,29562248 | 0,133 | 0,2 | 1 | CH_SC | 0,814720729 | 1,227414456 | 0,665 |
| ARL6IP45 | 0,448034575 | 0,559275137 | 0,4 | 0,4 | 1 | CH_SC | 1,473528677 | 0,678643053 | 1 |
| SEC132 | 0,449658853 | -0,293895749 | 0,2 | 0,308 | 1 | CH_SC | 0,815696435 | 1,225946268 | 0,649350649 |
| CC2D1B | 0,450550418 | 0,560841943 | 0,133 | 0,077 | 1 | CH_SC | 1,475129837 | 0,677906429 | 1,727272727 |
| RUSC11 | 0,450550418 | 0,307820994 | 0,133 | 0,077 | 1 | CH_SC | 1,237836693 | 0,807861009 | 1,727272727 |
| EIF2AK3 | 0,450550418 | 0,325273525 | 0,133 | 0,077 | 1 | CH_SC | 1,252901958 | 0,798147049 | 1,727272727 |
| MRPL363 | 0,450550418 | 0,368010177 | 0,133 | 0,077 | 1 | CH_SC | 1,290571594 | 0,774850465 | 1,727272727 |
| CASP8AP2 | 0,450550418 | 0,290019235 | 0,133 | 0,077 | 1 | CH_SC | 1,222656579 | 0,817891154 | 1,727272727 |
| KATNA1 | 0,450550418 | 0,360760357 | 0,133 | 0,077 | 1 | CH_SC | 1,284102492 | 0,778754038 | 1,727272727 |
| EBAG9 | 0,450550418 | 0,365025742 | 0,133 | 0,077 | 1 | CH_SC | 1,287904609 | 0,776455021 | 1,727272727 |
| SART11 | 0,450550418 | 0,622395028 | 0,133 | 0,077 | 1 | CH_SC | 1,539428677 | 0,649591641 | 1,727272727 |
| NAA16 | 0,450550418 | 0,361212267 | 0,133 | 0,077 | 1 | CH_SC | 1,284504788 | 0,778510138 | 1,727272727 |
| YJU2B | 0,450550418 | 0,427617841 | 0,133 | 0,077 | 1 | CH_SC | 1,345010878 | 0,74348841 | 1,727272727 |
| ATF21 | 0,450834634 | 0,986157108 | 0,2 | 0,138 | 1 | CH_SC | 1,980901451 | 0,504820671 | 1,449275362 |
| LARP72 | 0,450834634 | 0,785605378 | 0,2 | 0,138 | 1 | CH_SC | 1,723815509 | 0,580108483 | 1,449275362 |
| PPRC1 | 0,450834634 | 0,594799775 | 0,2 | 0,138 | 1 | CH_SC | 1,510262969 | 0,662136344 | 1,449275362 |
| COA32 | 0,451601077 | -0,518885619 | 0,267 | 0,308 | 1 | CH_SC | 0,697910712 | 1,432848045 | 0,866883117 |
| CCT52 | 0,452017289 | 0,426099049 | 0,333 | 0,262 | 1 | CH_SC | 1,343595668 | 0,744271528 | 1,270992366 |
| ENY22 | 0,452210289 | -0,298518058 | 0,333 | 0,415 | 1 | CH_SC | 0,813087174 | 1,229880426 | 0,802409639 |
| LUC7L22 | 0,452398551 | 0,299534443 | 0,467 | 0,354 | 1 | CH_SC | 1,230747188 | 0,812514552 | 1,31920904 |
| AARD | 0,452710456 | -0,372434838 | 0,067 | 0,138 | 1 | CH_SC | 0,772477685 | 1,294535778 | 0,485507246 |

| Supplementary Table 4 - clusermarker | | | | | | | | | |
|---|---|---|---|---|---|---|---|---|---|
| | **p_val** | **avg_log2FC** | **pct.1** | **pct.2** | **p_val_adj** | **celltype** | **Foldchange_UP** | **Foldchange_DOWN** | **Ratio_pct1_pct2** |
| TECR2 | 0,454436897 | 0,379692561 | 0,4 | 0,308 | 1 | CH_SC | 1,301064568 | 0,768601363 | 1,298701299 |
| PAIP21 | 0,456488379 | 0,45984477 | 0,4 | 0,354 | 1 | CH_SC | 1,375393821 | 0,727064485 | 1,129943503 |
| SRM1 | 0,457959963 | -0,467473544 | 0,133 | 0,2 | 1 | CH_SC | 0,723230014 | 1,382685979 | 0,665 |
| ZSWIM72 | 0,457959963 | -0,42378615 | 0,133 | 0,2 | 1 | CH_SC | 0,745465685 | 1,341443369 | 0,665 |
| ARID5A1 | 0,458551926 | 0,333485443 | 0,2 | 0,123 | 1 | CH_SC | 1,260053896 | 0,793616847 | 1,62601626 |
| RAB242 | 0,458551926 | 0,282604478 | 0,2 | 0,123 | 1 | CH_SC | 1,216388834 | 0,822105541 | 1,62601626 |
| EIF4G31 | 0,458693951 | 0,837750296 | 0,267 | 0,2 | 1 | CH_SC | 1,787260956 | 0,559515384 | 1,335 |
| PBX12 | 0,458693951 | 0,78056723 | 0,267 | 0,2 | 1 | CH_SC | 1,717806136 | 0,582137867 | 1,335 |
| PARVA1 | 0,461494411 | 0,529890771 | 0,333 | 0,262 | 1 | CH_SC | 1,443819877 | 0,692607171 | 1,270992366 |
| MRPL521 | 0,461494411 | 0,267730787 | 0,333 | 0,262 | 1 | CH_SC | 1,203912705 | 0,830625008 | 1,270992366 |
| FBXO92 | 0,462838097 | 0,473807926 | 0,2 | 0,138 | 1 | CH_SC | 1,388770228 | 0,720061519 | 1,449275362 |
| SLC39A71 | 0,46342065 | 0,476729389 | 0,4 | 0,308 | 1 | CH_SC | 1,391585343 | 0,718604867 | 1,298701299 |
| SMC52 | 0,464011274 | -0,536344959 | 0,133 | 0,215 | 1 | CH_SC | 0,689515573 | 1,450293567 | 0,618604651 |
| SNHG53 | 0,465181656 | 0,344088551 | 0,467 | 0,369 | 1 | CH_SC | 1,269348795 | 0,78780553 | 1,265582656 |
| JPX2 | 0,4656929 | -0,501598761 | 0,2 | 0,262 | 1 | CH_SC | 0,706323616 | 1,41578163 | 0,763358779 |
| MYSM12 | 0,465788947 | 0,425965761 | 0,133 | 0,077 | 1 | CH_SC | 1,343471541 | 0,744340293 | 1,727272727 |
| AC118549.1 | 0,465788947 | 0,342739375 | 0,133 | 0,077 | 1 | CH_SC | 1,268162283 | 0,788542613 | 1,727272727 |
| FAM114A2 | 0,465788947 | 0,378280495 | 0,133 | 0,077 | 1 | CH_SC | 1,299791749 | 0,769354015 | 1,727272727 |
| BCKDHB | 0,465788947 | 0,266330604 | 0,133 | 0,077 | 1 | CH_SC | 1,202744835 | 0,831431548 | 1,727272727 |
| NUS11 | 0,465788947 | 0,818367012 | 0,133 | 0,077 | 1 | CH_SC | 1,763408858 | 0,567083462 | 1,727272727 |
| THAP11 | 0,465788947 | 0,339405775 | 0,133 | 0,077 | 1 | CH_SC | 1,265235355 | 0,790366785 | 1,727272727 |
| NANS5 | 0,465788947 | 0,318026404 | 0,133 | 0,077 | 1 | CH_SC | 1,24662401 | 0,802166485 | 1,727272727 |
| PPP4R3A1 | 0,465788947 | 0,265759181 | 0,133 | 0,077 | 1 | CH_SC | 1,202268546 | 0,831760927 | 1,727272727 |
| CPSF2 | 0,465788947 | 0,396160915 | 0,133 | 0,077 | 1 | CH_SC | 1,316001301 | 0,759877668 | 1,727272727 |
| CTDSPL2 | 0,465788947 | 0,320729613 | 0,133 | 0,077 | 1 | CH_SC | 1,248962026 | 0,800664856 | 1,727272727 |
| DENND4A | 0,465788947 | 0,311396683 | 0,133 | 0,077 | 1 | CH_SC | 1,24090845 | 0,805861222 | 1,727272727 |
| BMERB11 | 0,465788947 | 0,257761784 | 0,133 | 0,077 | 1 | CH_SC | 1,195622361 | 0,836384491 | 1,727272727 |
| ZNF830 | 0,465788947 | 0,259231338 | 0,133 | 0,077 | 1 | CH_SC | 1,196840863 | 0,835532969 | 1,727272727 |
| CXXC11 | 0,465788947 | 0,314547989 | 0,133 | 0,077 | 1 | CH_SC | 1,243621952 | 0,804102885 | 1,727272727 |
| RALGAPB1 | 0,465788947 | 0,339822876 | 0,133 | 0,077 | 1 | CH_SC | 1,265601202 | 0,790138314 | 1,727272727 |
| RRP7A1 | 0,465788947 | 0,330469229 | 0,133 | 0,077 | 1 | CH_SC | 1,257422278 | 0,795277782 | 1,727272727 |

| Supplementary Table 4 - clusermarker | | | | | | | | | |
|---|---|---|---|---|---|---|---|---|---|
| | **p_val** | **avg_log2FC** | **pct.1** | **pct.2** | **p_val_adj** | **celltype** | **Foldchange_UP** | **Foldchange_DOWN** | **Ratio_pct1_pct2** |
| CNOT22 | 0,469209026 | 0,310022988 | 0,267 | 0,2 | 1 | CH_SC | 1,239727454 | 0,806628906 | 1,335 |
| FSTL11 | 0,469638886 | 0,268217739 | 0,467 | 0,354 | 1 | CH_SC | 1,20431913 | 0,830344695 | 1,31920904 |
| RTF11 | 0,471081507 | 0,289621867 | 0,333 | 0,262 | 1 | CH_SC | 1,222319864 | 0,81811646 | 1,270992366 |
| PPP3R11 | 0,471128822 | 0,381731573 | 0,2 | 0,123 | 1 | CH_SC | 1,302904709 | 0,767515838 | 1,62601626 |
| AFF42 | 0,473255274 | 0,662536669 | 0,4 | 0,323 | 1 | CH_SC | 1,582863303 | 0,631766494 | 1,238390093 |
| CLK14 | 0,473747882 | 0,600081543 | 0,4 | 0,338 | 1 | CH_SC | 1,515802239 | 0,659716666 | 1,183431953 |
| RAB132 | 0,473808492 | 0,764405103 | 0,4 | 0,385 | 1 | CH_SC | 1,698669406 | 0,588696068 | 1,038961039 |
| GOLGA22 | 0,473990366 | 0,524035551 | 0,4 | 0,354 | 1 | CH_SC | 1,437971967 | 0,695423849 | 1,129943503 |
| REEP53 | 0,474288889 | -0,265755419 | 0,333 | 0,477 | 1 | CH_SC | 0,831763096 | 1,20226541 | 0,698113208 |
| ATP6V1G12 | 0,474288889 | 1,024321543 | 0,467 | 0,446 | 1 | CH_SC | 2,034002628 | 0,491641449 | 1,047085202 |
| MXD41 | 0,47484823 | 0,380889777 | 0,267 | 0,185 | 1 | CH_SC | 1,302144701 | 0,767963806 | 1,443243243 |
| GPBP15 | 0,47484823 | 0,261177152 | 0,267 | 0,185 | 1 | CH_SC | 1,198456174 | 0,834406816 | 1,443243243 |
| PIAS14 | 0,47484823 | 0,648392574 | 0,267 | 0,185 | 1 | CH_SC | 1,567420829 | 0,637990756 | 1,443243243 |
| PSMD112 | 0,47484823 | -0,253088338 | 0,133 | 0,215 | 1 | CH_SC | 0,839098257 | 1,191755545 | 0,618604651 |
| LRRC411 | 0,475017702 | 0,424429054 | 0,2 | 0,138 | 1 | CH_SC | 1,342041285 | 0,74513356 | 1,449275362 |
| CYTOR3 | 0,475017702 | 0,454957859 | 0,2 | 0,138 | 1 | CH_SC | 1,370742765 | 0,729531481 | 1,449275362 |
| BRD42 | 0,477466338 | 0,694459044 | 0,267 | 0,231 | 1 | CH_SC | 1,618277526 | 0,61794098 | 1,155844156 |
| MED213 | 0,477685281 | 0,782435686 | 0,2 | 0,154 | 1 | CH_SC | 1,720032334 | 0,581384419 | 1,298701299 |
| SMARCA24 | 0,477890711 | 0,505910115 | 0,333 | 0,277 | 1 | CH_SC | 1,420018884 | 0,704215987 | 1,202166065 |
| EIF3G3 | 0,47839339 | -0,565537848 | 0,333 | 0,385 | 1 | CH_SC | 0,675703462 | 1,479939139 | 0,864935065 |
| EP400 | 0,478948261 | -0,464614654 | 0,067 | 0,138 | 1 | CH_SC | 0,724664611 | 1,379948718 | 0,485507246 |
| LINC014362 | 0,478948261 | -0,48642147 | 0,067 | 0,138 | 1 | CH_SC | 0,713793431 | 1,40096554 | 0,485507246 |
| CENPC3 | 0,479856066 | 0,459087146 | 0,267 | 0,2 | 1 | CH_SC | 1,37467173 | 0,727446399 | 1,335 |
| RTL8A1 | 0,480185111 | -0,290819816 | 0,133 | 0,2 | 1 | CH_SC | 0,817437415 | 1,223335245 | 0,665 |
| AL138963.48 | 0,48077332 | -0,315177149 | 0,2 | 0,292 | 1 | CH_SC | 0,803752292 | 1,244164415 | 0,684931507 |
| DNPEP | 0,48130956 | 0,511564421 | 0,133 | 0,077 | 1 | CH_SC | 1,425595236 | 0,701461379 | 1,727272727 |
| BLOC1S4 | 0,48130956 | 0,280294507 | 0,133 | 0,077 | 1 | CH_SC | 1,214442771 | 0,823422909 | 1,727272727 |
| SMAD11 | 0,48130956 | 0,301304022 | 0,133 | 0,077 | 1 | CH_SC | 1,232257722 | 0,81151855 | 1,727272727 |
| JMY1 | 0,48130956 | 0,397922494 | 0,133 | 0,077 | 1 | CH_SC | 1,317609164 | 0,758950398 | 1,727272727 |
| LINC011841 | 0,48130956 | 0,80018891 | 0,133 | 0,077 | 1 | CH_SC | 1,741329125 | 0,574273976 | 1,727272727 |
| WRNIP11 | 0,48130956 | 0,377644307 | 0,133 | 0,077 | 1 | CH_SC | 1,299218704 | 0,769693352 | 1,727272727 |

| Supplementary Table 4 - clusermarker | | | | | | | | | |
|---|---|---|---|---|---|---|---|---|---|
| | **p_val** | **avg_log2FC** | **pct.1** | **pct.2** | **p_val_adj** | **celltype** | **Foldchange_UP** | **Foldchange_DOWN** | **Ratio_pct1_pct2** |
| SEC23IP1 | 0,48130956 | 0,533459215 | 0,133 | 0,077 | 1 | CH_SC | 1,447395523 | 0,690896154 | 1,727272727 |
| GXYLT1 | 0,48130956 | 0,454619505 | 0,133 | 0,077 | 1 | CH_SC | 1,370421324 | 0,729702598 | 1,727272727 |
| PARN1 | 0,48130956 | 0,790732956 | 0,133 | 0,077 | 1 | CH_SC | 1,729953135 | 0,578050341 | 1,727272727 |
| ZNF699 | 0,48130956 | 0,364577964 | 0,133 | 0,077 | 1 | CH_SC | 1,287504936 | 0,776696051 | 1,727272727 |
| KIF3B1 | 0,48130956 | 0,538600048 | 0,133 | 0,077 | 1 | CH_SC | 1,452562307 | 0,688438627 | 1,727272727 |
| PAXBP1 | 0,48130956 | 0,266414192 | 0,133 | 0,077 | 1 | CH_SC | 1,202814523 | 0,831383377 | 1,727272727 |
| DYRK1A1 | 0,48130956 | 0,6389418 | 0,133 | 0,077 | 1 | CH_SC | 1,557186562 | 0,64218381 | 1,727272727 |
| NDFIP12 | 0,48168152 | 0,311759581 | 0,4 | 0,308 | 1 | CH_SC | 1,24122063 | 0,80565854 | 1,298701299 |
| VAMP56 | 0,482336109 | 0,285211202 | 0,4 | 0,323 | 1 | CH_SC | 1,218588645 | 0,820621466 | 1,238390093 |
| TCEAL43 | 0,482668018 | 0,566648583 | 0,467 | 0,446 | 1 | CH_SC | 1,481078987 | 0,675183436 | 1,047085202 |
| LDHA6 | 0,483266776 | -0,84659736 | 0,667 | 0,692 | 1 | CH_SC | 0,556094757 | 1,798254681 | 0,963872832 |
| CRIPT | 0,483893718 | 0,300288926 | 0,2 | 0,123 | 1 | CH_SC | 1,231390997 | 0,812089744 | 1,62601626 |
| KRAS3 | 0,485817456 | 0,300461517 | 0,267 | 0,185 | 1 | CH_SC | 1,231538318 | 0,811992599 | 1,443243243 |
| C2orf741 | 0,486101407 | -0,484090384 | 0,067 | 0,123 | 1 | CH_SC | 0,714947701 | 1,398703708 | 0,544715447 |
| CEP85L1 | 0,486101407 | -0,541790649 | 0,067 | 0,123 | 1 | CH_SC | 0,686917789 | 1,455778284 | 0,544715447 |
| TMEM25 | 0,486101407 | -0,274867451 | 0,067 | 0,123 | 1 | CH_SC | 0,826526252 | 1,209882925 | 0,544715447 |
| CES12 | 0,486101407 | -0,324518794 | 0,067 | 0,123 | 1 | CH_SC | 0,7985647 | 1,252246687 | 0,544715447 |
| PCNA1 | 0,486101407 | -0,37783513 | 0,067 | 0,123 | 1 | CH_SC | 0,769591553 | 1,29939056 | 0,544715447 |
| GART | 0,486101407 | -0,39549607 | 0,067 | 0,123 | 1 | CH_SC | 0,760227928 | 1,31539498 | 0,544715447 |
| CBX7 | 0,486101407 | -0,344691557 | 0,067 | 0,123 | 1 | CH_SC | 0,787476318 | 1,269879458 | 0,544715447 |
| IARS22 | 0,487371171 | 0,560675764 | 0,2 | 0,138 | 1 | CH_SC | 1,474959932 | 0,677984519 | 1,449275362 |
| IBTK | 0,487371171 | 0,484179936 | 0,2 | 0,138 | 1 | CH_SC | 1,398790532 | 0,714903323 | 1,449275362 |
| JDP22 | 0,487371171 | 0,566719013 | 0,2 | 0,138 | 1 | CH_SC | 1,481151293 | 0,675150476 | 1,449275362 |
| ADAR3 | 0,49058114 | 0,464818778 | 0,333 | 0,262 | 1 | CH_SC | 1,380143978 | 0,724562086 | 1,270992366 |
| GADD45GIP14 | 0,49058114 | 0,586552633 | 0,333 | 0,262 | 1 | CH_SC | 1,501654205 | 0,665932274 | 1,270992366 |
| YWHAH4 | 0,49058114 | 0,308573486 | 0,333 | 0,262 | 1 | CH_SC | 1,238482502 | 0,807439748 | 1,270992366 |
| ATRX2 | 0,491126936 | 0,344157949 | 0,467 | 0,446 | 1 | CH_SC | 1,269409855 | 0,787767635 | 1,047085202 |
| CCDC122 | 0,491514385 | -0,434325546 | 0,133 | 0,2 | 1 | CH_SC | 0,740039639 | 1,351278968 | 0,665 |
| RPA32 | 0,491514385 | -0,285472717 | 0,133 | 0,2 | 1 | CH_SC | 0,820472727 | 1,218809556 | 0,665 |
| CCDC252 | 0,491514385 | -0,362247962 | 0,133 | 0,2 | 1 | CH_SC | 0,777951455 | 1,28542725 | 0,665 |
| PMM1 | 0,491514385 | -0,523416366 | 0,133 | 0,2 | 1 | CH_SC | 0,695722379 | 1,437354942 | 0,665 |

| Supplementary Table 4 - clusermarker | | | | | | | | | |
|---|---|---|---|---|---|---|---|---|---|
| | **p_val** | **avg_log2FC** | **pct.1** | **pct.2** | **p_val_adj** | **celltype** | **Foldchange_UP** | **Foldchange_DOWN** | **Ratio_pct1_pct2** |
| RTL8C2 | 0,49179975 | 0,371558384 | 0,4 | 0,338 | 1 | CH_SC | 1,293749571 | 0,772947116 | 1,183431953 |
| COX6C4 | 0,492615757 | -0,264014074 | 0,533 | 0,754 | 1 | CH_SC | 0,832767647 | 1,200815141 | 0,706896552 |
| SWAP702 | 0,493775147 | 0,431224771 | 0,2 | 0,323 | 1 | CH_SC | 1,348377792 | 0,741631912 | 0,619195046 |
| PPIA7 | 0,496870672 | 0,28892591 | 0,667 | 0,877 | 1 | CH_SC | 1,221730358 | 0,818511215 | 0,76054732 |
| RBM62 | 0,49693295 | 0,674029623 | 0,267 | 0,185 | 1 | CH_SC | 1,595523241 | 0,626753641 | 1,443243243 |
| KRIT11 | 0,49693295 | -0,278860781 | 0,133 | 0,215 | 1 | CH_SC | 0,82424162 | 1,213236479 | 0,618604651 |
| NFYC2 | 0,497107486 | 0,306913363 | 0,133 | 0,077 | 1 | CH_SC | 1,237058188 | 0,808369412 | 1,727272727 |
| NAALADL22 | 0,497107486 | 0,516650036 | 0,133 | 0,077 | 1 | CH_SC | 1,430629441 | 0,698993025 | 1,727272727 |
| CCDC126 | 0,497107486 | 0,304920828 | 0,133 | 0,077 | 1 | CH_SC | 1,235350842 | 0,809486638 | 1,727272727 |
| DPY19L1 | 0,497107486 | 0,39626006 | 0,133 | 0,077 | 1 | CH_SC | 1,316091742 | 0,75982545 | 1,727272727 |
| KLHDC101 | 0,497107486 | 0,393281097 | 0,133 | 0,077 | 1 | CH_SC | 1,313377001 | 0,761396004 | 1,727272727 |
| SBSPON1 | 0,497107486 | 0,460839617 | 0,133 | 0,077 | 1 | CH_SC | 1,376342587 | 0,726563291 | 1,727272727 |
| VIRMA1 | 0,497107486 | 0,291752297 | 0,133 | 0,077 | 1 | CH_SC | 1,224126199 | 0,816909238 | 1,727272727 |
| RNF19A2 | 0,497107486 | 0,407223278 | 0,133 | 0,077 | 1 | CH_SC | 1,326130982 | 0,754073326 | 1,727272727 |
| UBE2L62 | 0,497107486 | 0,291987774 | 0,133 | 0,077 | 1 | CH_SC | 1,224326017 | 0,816775913 | 1,727272727 |
| MCRS1 | 0,497107486 | 0,329854257 | 0,133 | 0,077 | 1 | CH_SC | 1,256886396 | 0,795616854 | 1,727272727 |
| TTBK2 | 0,497107486 | 0,257286593 | 0,133 | 0,077 | 1 | CH_SC | 1,195228615 | 0,836660023 | 1,727272727 |
| PRDX32 | 0,497237549 | 0,415725845 | 0,333 | 0,277 | 1 | CH_SC | 1,333969652 | 0,749642242 | 1,202166065 |
| RBM271 | 0,498188716 | -0,477842188 | 0,2 | 0,246 | 1 | CH_SC | 0,718050797 | 1,392659133 | 0,81300813 |
| RAF11 | 0,499896051 | 0,28883155 | 0,2 | 0,138 | 1 | CH_SC | 1,221650452 | 0,818564752 | 1,449275362 |
| SH3BP51 | 0,499896051 | 0,301743056 | 0,2 | 0,138 | 1 | CH_SC | 1,232632774 | 0,81127163 | 1,449275362 |
| SPRED11 | 0,499896051 | 0,48346106 | 0,2 | 0,138 | 1 | CH_SC | 1,398093707 | 0,715259639 | 1,449275362 |
| TPRG1L2 | 0,500276703 | -0,270462776 | 0,067 | 0,123 | 1 | CH_SC | 0,829053566 | 1,206194679 | 0,544715447 |
| ITGB3BP1 | 0,500276703 | -0,263976946 | 0,067 | 0,123 | 1 | CH_SC | 0,832789078 | 1,200784239 | 0,544715447 |
| NAF12 | 0,500276703 | -0,439832152 | 0,067 | 0,123 | 1 | CH_SC | 0,737220375 | 1,356446504 | 0,544715447 |
| C1QTNF4 | 0,500276703 | -0,408465643 | 0,067 | 0,123 | 1 | CH_SC | 0,753424241 | 1,327273461 | 0,544715447 |
| VEZT1 | 0,500276703 | -0,305133116 | 0,067 | 0,123 | 1 | CH_SC | 0,809367534 | 1,235532633 | 0,544715447 |
| POLR3K1 | 0,500276703 | -0,332934619 | 0,067 | 0,123 | 1 | CH_SC | 0,79391991 | 1,259572896 | 0,544715447 |
| BANP | 0,500276703 | -0,28185767 | 0,067 | 0,123 | 1 | CH_SC | 0,822531212 | 1,215759336 | 0,544715447 |
| PHF23 | 0,500276703 | -0,265153966 | 0,067 | 0,123 | 1 | CH_SC | 0,832109926 | 1,201764296 | 0,544715447 |
| KANSL12 | 0,500276703 | -0,384815364 | 0,067 | 0,123 | 1 | CH_SC | 0,765877009 | 1,305692674 | 0,544715447 |

| Supplementary Table 4 - clusermarker | | | | | | | | | |
|---|---|---|---|---|---|---|---|---|---|
| | **p_val** | **avg_log2FC** | **pct.1** | **pct.2** | **p_val_adj** | **celltype** | **Foldchange_UP** | **Foldchange_DOWN** | **Ratio_pct1_pct2** |
| UBE2Z2 | 0,500276703 | -0,267901431 | 0,067 | 0,123 | 1 | CH_SC | 0,830526766 | 1,204055114 | 0,544715447 |
| GPR1081 | 0,500276703 | -0,252501204 | 0,067 | 0,123 | 1 | CH_SC | 0,839439814 | 1,191270634 | 0,544715447 |
| KEAP1 | 0,500276703 | -0,419885111 | 0,067 | 0,123 | 1 | CH_SC | 0,747484148 | 1,337821013 | 0,544715447 |
| SLC27A5 | 0,500276703 | -0,261115826 | 0,067 | 0,123 | 1 | CH_SC | 0,834442286 | 1,198405231 | 0,544715447 |
| RPS6KA31 | 0,500276703 | -0,257127208 | 0,067 | 0,123 | 1 | CH_SC | 0,83675246 | 1,195096576 | 0,544715447 |
| GLS5 | 0,500920892 | -0,258086128 | 0,267 | 0,385 | 1 | CH_SC | 0,836196478 | 1,195891189 | 0,693506494 |
| FNDC3B1 | 0,500920892 | 0,472400131 | 0,4 | 0,354 | 1 | CH_SC | 1,387415715 | 0,720764504 | 1,129943503 |
| ATP6V1F3 | 0,500962234 | 0,310249253 | 0,4 | 0,338 | 1 | CH_SC | 1,239921901 | 0,806502409 | 1,183431953 |
| SRSF98 | 0,500962234 | 0,352546624 | 0,4 | 0,338 | 1 | CH_SC | 1,27681245 | 0,783200383 | 1,183431953 |
| PFKFB31 | 0,50298562 | -0,287067421 | 0,267 | 0,169 | 1 | CH_SC | 0,819566306 | 1,22015753 | 1,579881657 |
| C14orf1191 | 0,50298562 | -0,358932406 | 0,133 | 0,2 | 1 | CH_SC | 0,779741374 | 1,282476514 | 0,665 |
| OFD13 | 0,50298562 | -0,340907227 | 0,133 | 0,2 | 1 | CH_SC | 0,789544657 | 1,266552805 | 0,665 |
| CALD13 | 0,50310171 | 0,638279946 | 0,533 | 0,569 | 1 | CH_SC | 1,556472348 | 0,642478488 | 0,936731107 |
| PRELID17 | 0,50342547 | 0,465421465 | 0,333 | 0,292 | 1 | CH_SC | 1,380720654 | 0,724259463 | 1,140410959 |
| CLTC1 | 0,503511248 | -0,445709786 | 0,2 | 0,277 | 1 | CH_SC | 0,734223 | 1,361984029 | 0,722021661 |
| GPAA14 | 0,506267998 | -0,528118887 | 0,2 | 0,262 | 1 | CH_SC | 0,693458337 | 1,4420477 | 0,763358779 |
| CSNK1A12 | 0,507065163 | 0,320279503 | 0,333 | 0,277 | 1 | CH_SC | 1,24857242 | 0,800914696 | 1,202166065 |
| CDKN2AIP2 | 0,508177176 | 0,313891782 | 0,267 | 0,185 | 1 | CH_SC | 1,243056423 | 0,804468712 | 1,443243243 |
| CCDC85B7 | 0,508726548 | 0,625672703 | 0,267 | 0,231 | 1 | CH_SC | 1,542930098 | 0,648117501 | 1,155844156 |
| FBN1 | 0,508734071 | -0,538394979 | 0,133 | 0,185 | 1 | CH_SC | 0,688536491 | 1,452355849 | 0,718918919 |
| SPATS2L2 | 0,5097913 | -0,286314586 | 0,267 | 0,338 | 1 | CH_SC | 0,819994089 | 1,219520987 | 0,789940828 |
| GLUD12 | 0,5097913 | 0,378194126 | 0,4 | 0,308 | 1 | CH_SC | 1,299713937 | 0,769400074 | 1,298701299 |
| C1orf431 | 0,510506557 | 0,353454885 | 0,333 | 0,262 | 1 | CH_SC | 1,277616531 | 0,782707468 | 1,270992366 |
| BUD313 | 0,51257305 | 0,832315112 | 0,267 | 0,2 | 1 | CH_SC | 1,780540328 | 0,561627268 | 1,335 |
| COPG1 | 0,512589724 | 0,478955842 | 0,2 | 0,138 | 1 | CH_SC | 1,393734578 | 0,717496728 | 1,449275362 |
| GNL1 | 0,512589724 | 0,282889191 | 0,2 | 0,138 | 1 | CH_SC | 1,216628909 | 0,821943316 | 1,449275362 |
| RLIM1 | 0,512589724 | 0,28730908 | 0,2 | 0,138 | 1 | CH_SC | 1,22036193 | 0,819429036 | 1,449275362 |
| ARMCX21 | 0,512589724 | 0,565234151 | 0,2 | 0,138 | 1 | CH_SC | 1,479627635 | 0,675845717 | 1,449275362 |
| LIMCH12 | 0,513170729 | 0,3806615 | 0,333 | 0,292 | 1 | CH_SC | 1,301938679 | 0,76808533 | 1,140410959 |
| SMG13 | 0,514104228 | 0,446416318 | 0,2 | 0,154 | 1 | CH_SC | 1,362651197 | 0,733863517 | 1,298701299 |
| MPRIP2 | 0,514104228 | 0,569871606 | 0,2 | 0,154 | 1 | CH_SC | 1,48439146 | 0,67367674 | 1,298701299 |

| Supplementary Table 4 - clusermarker | | | | | | | | | |
|---|---|---|---|---|---|---|---|---|---|
| | **p_val** | **avg_log2FC** | **pct.1** | **pct.2** | **p_val_adj** | **celltype** | **Foldchange_UP** | **Foldchange_DOWN** | **Ratio_pct1_pct2** |
| TBC1D53 | 0,514668308 | -0,348404872 | 0,067 | 0,123 | 1 | CH_SC | 0,78545206 | 1,273152177 | 0,544715447 |
| ARHGAP101 | 0,514668308 | -0,354243169 | 0,067 | 0,123 | 1 | CH_SC | 0,782279916 | 1,278314807 | 0,544715447 |
| CD1092 | 0,514668308 | -0,361186222 | 0,067 | 0,123 | 1 | CH_SC | 0,778524193 | 1,284481598 | 0,544715447 |
| GPNMB3 | 0,514668308 | -0,459994365 | 0,067 | 0,123 | 1 | CH_SC | 0,726989098 | 1,375536445 | 0,544715447 |
| TINF2 | 0,514668308 | -0,44716749 | 0,067 | 0,123 | 1 | CH_SC | 0,733481513 | 1,363360878 | 0,544715447 |
| TMUB2 | 0,514668308 | -0,271838492 | 0,067 | 0,123 | 1 | CH_SC | 0,828263379 | 1,207345423 | 0,544715447 |
| ICAM13 | 0,514668308 | -0,253124072 | 0,067 | 0,123 | 1 | CH_SC | 0,839077473 | 1,191785064 | 0,544715447 |
| ZNF585A | 0,514668308 | -0,262819723 | 0,067 | 0,123 | 1 | CH_SC | 0,833457348 | 1,199821445 | 0,544715447 |
| CDK162 | 0,514668308 | -0,289764419 | 0,067 | 0,123 | 1 | CH_SC | 0,818035626 | 1,222440647 | 0,544715447 |
| IDH3G4 | 0,514668308 | -0,256213254 | 0,067 | 0,123 | 1 | CH_SC | 0,837282714 | 1,194339717 | 0,544715447 |
| CCND31 | 0,5162785 | 0,632942825 | 0,267 | 0,215 | 1 | CH_SC | 1,550724955 | 0,644859681 | 1,241860465 |
| EIF3K4 | 0,517130525 | 0,396060883 | 0,6 | 0,615 | 1 | CH_SC | 1,315910056 | 0,759930358 | 0,975609756 |
| RAB1A1 | 0,518857407 | -0,33494719 | 0,267 | 0,323 | 1 | CH_SC | 0,792813157 | 1,261331237 | 0,826625387 |
| CHADL1 | 0,519569165 | 0,714354984 | 0,4 | 0,323 | 1 | CH_SC | 1,640749489 | 0,609477563 | 1,238390093 |
| VAPA2 | 0,519572615 | 0,350281504 | 0,4 | 0,323 | 1 | CH_SC | 1,274809349 | 0,784431022 | 1,238390093 |
| CAPZA13 | 0,519814116 | 0,481036741 | 0,067 | 0,138 | 1 | CH_SC | 1,395746309 | 0,716462579 | 0,485507246 |
| MRPS22 | 0,519814116 | 0,304157136 | 0,067 | 0,138 | 1 | CH_SC | 1,234697081 | 0,809915254 | 0,485507246 |
| TRIM23 | 0,519814116 | 0,469331988 | 0,067 | 0,138 | 1 | CH_SC | 1,384468268 | 0,722298967 | 0,485507246 |
| AGO22 | 0,519814116 | 1,227190182 | 0,067 | 0,138 | 1 | CH_SC | 2,341105877 | 0,427148558 | 0,485507246 |
| GTPBP4 | 0,519814116 | 0,313425153 | 0,067 | 0,138 | 1 | CH_SC | 1,24265443 | 0,804728954 | 0,485507246 |
| BTAF1 | 0,519814116 | 0,603337282 | 0,067 | 0,138 | 1 | CH_SC | 1,519226823 | 0,658229558 | 0,485507246 |
| GSTP16 | 0,520411979 | -0,295321574 | 0,733 | 0,846 | 1 | CH_SC | 0,814890675 | 1,227158478 | 0,86643026 |
| PDCD104 | 0,520746071 | -0,507428624 | 0,133 | 0,185 | 1 | CH_SC | 0,703475155 | 1,421514312 | 0,718918919 |
| CRIP24 | 0,524767348 | 0,338001674 | 0,533 | 0,477 | 1 | CH_SC | 1,264004565 | 0,791136383 | 1,117400419 |
| GTF2F2 | 0,525449399 | 0,390187491 | 0,2 | 0,138 | 1 | CH_SC | 1,310563712 | 0,763030435 | 1,449275362 |
| SBNO12 | 0,526557104 | 0,448157457 | 0,2 | 0,154 | 1 | CH_SC | 1,364296727 | 0,732978376 | 1,298701299 |
| TMEM2081 | 0,526557104 | 0,648172569 | 0,2 | 0,154 | 1 | CH_SC | 1,567181822 | 0,638088054 | 1,298701299 |
| TUFM6 | 0,527022354 | 0,311086249 | 0,333 | 0,277 | 1 | CH_SC | 1,240641465 | 0,806034643 | 1,202166065 |
| NDUFA102 | 0,527234578 | 0,364992298 | 0,267 | 0,215 | 1 | CH_SC | 1,287874754 | 0,77647302 | 1,241860465 |
| POLR2K4 | 0,527369868 | 0,715903249 | 0,4 | 0,385 | 1 | CH_SC | 1,642511246 | 0,608823837 | 1,038961039 |
| SAT24 | 0,528997289 | 0,258499153 | 0,4 | 0,308 | 1 | CH_SC | 1,196233606 | 0,83595712 | 1,298701299 |

| Supplementary Table 4 - clusermarker | | | | | | | | | |
|---|---|---|---|---|---|---|---|---|---|
| | **p_val** | **avg_log2FC** | **pct.1** | **pct.2** | **p_val_adj** | **celltype** | **Foldchange_UP** | **Foldchange_DOWN** | **Ratio_pct1_pct2** |
| ANKHD12 | 0,529108238 | 0,811130529 | 0,4 | 0,323 | 1 | CH_SC | 1,754585838 | 0,569935069 | 1,238390093 |
| STRIP1 | 0,529272149 | -0,293385067 | 0,067 | 0,123 | 1 | CH_SC | 0,815985224 | 1,225512387 | 0,544715447 |
| AC245407.2 | 0,529272149 | -0,275797674 | 0,067 | 0,123 | 1 | CH_SC | 0,825993495 | 1,210663287 | 0,544715447 |
| MR13 | 0,529272149 | -0,253256266 | 0,067 | 0,123 | 1 | CH_SC | 0,839000592 | 1,191894272 | 0,544715447 |
| HMCN11 | 0,529272149 | -0,267635552 | 0,067 | 0,123 | 1 | CH_SC | 0,830679841 | 1,203833235 | 0,544715447 |
| GNL33 | 0,530147553 | -0,369255078 | 0,2 | 0,246 | 1 | CH_SC | 0,774182135 | 1,291685708 | 0,81300813 |
| RARG1 | 0,530147553 | -0,389157346 | 0,2 | 0,246 | 1 | CH_SC | 0,763575466 | 1,309628248 | 0,81300813 |
| CLSTN13 | 0,530847077 | 0,380991358 | 0,333 | 0,262 | 1 | CH_SC | 1,302236389 | 0,767909735 | 1,270992366 |
| CLPTM1L1 | 0,532902521 | -0,43188216 | 0,133 | 0,185 | 1 | CH_SC | 0,741294052 | 1,348992343 | 0,718918919 |
| NFIL32 | 0,532902521 | -0,527417382 | 0,133 | 0,185 | 1 | CH_SC | 0,69379561 | 1,441346681 | 0,718918919 |
| ZNRD21 | 0,532902521 | -0,424643465 | 0,133 | 0,185 | 1 | CH_SC | 0,745022827 | 1,342240753 | 0,718918919 |
| CD474 | 0,534473743 | 0,640499528 | 0,333 | 0,246 | 1 | CH_SC | 1,558868819 | 0,641490796 | 1,353658537 |
| CFDP12 | 0,534473743 | -0,35322665 | 0,2 | 0,277 | 1 | CH_SC | 0,782831302 | 1,277414427 | 0,722021661 |
| NDUFAF21 | 0,535013305 | 0,265359083 | 0,267 | 0,2 | 1 | CH_SC | 1,201935171 | 0,831991629 | 1,335 |
| MAGI2-AS32 | 0,535013305 | 0,281712172 | 0,267 | 0,2 | 1 | CH_SC | 1,215636731 | 0,82261417 | 1,335 |
| CHMP2A5 | 0,538308061 | 0,279191637 | 0,267 | 0,215 | 1 | CH_SC | 1,213514744 | 0,824052617 | 1,241860465 |
| PLSCR13 | 0,538472119 | 0,357594285 | 0,2 | 0,138 | 1 | CH_SC | 1,281287549 | 0,780464932 | 1,449275362 |
| CWF19L24 | 0,538472119 | 0,347843153 | 0,2 | 0,138 | 1 | CH_SC | 1,272656567 | 0,785757938 | 1,449275362 |
| CD99L2 | 0,538472119 | 0,25354179 | 0,2 | 0,138 | 1 | CH_SC | 1,192130184 | 0,838834562 | 1,449275362 |
| EIF4H2 | 0,538490443 | -0,314136216 | 0,267 | 0,323 | 1 | CH_SC | 0,804332424 | 1,243267049 | 0,826625387 |
| NCL6 | 0,53856639 | 0,61703941 | 0,667 | 0,769 | 1 | CH_SC | 1,533724556 | 0,652007556 | 0,867360208 |
| RAD23A3 | 0,541169374 | -0,347065961 | 0,2 | 0,292 | 1 | CH_SC | 0,786181347 | 1,271971161 | 0,684931507 |
| PNISR6 | 0,5416559 | 0,594340807 | 0,533 | 0,538 | 1 | CH_SC | 1,509782582 | 0,662347024 | 0,99070632 |
| HDGF3 | 0,545200896 | -0,307323836 | 0,133 | 0,185 | 1 | CH_SC | 0,808139448 | 1,237410204 | 0,718918919 |
| AUP14 | 0,545200896 | -0,375145348 | 0,133 | 0,185 | 1 | CH_SC | 0,77102773 | 1,296970215 | 0,718918919 |
| TCAF11 | 0,545200896 | -0,377157156 | 0,133 | 0,185 | 1 | CH_SC | 0,769953297 | 1,298780074 | 0,718918919 |
| TSPAN311 | 0,545200896 | -0,531157145 | 0,133 | 0,185 | 1 | CH_SC | 0,691999478 | 1,445087795 | 0,718918919 |
| TENT5C3 | 0,545988803 | 0,432242373 | 0,133 | 0,092 | 1 | CH_SC | 1,349329203 | 0,741108988 | 1,445652174 |
| PIP5K1A | 0,545988803 | 0,944193904 | 0,133 | 0,092 | 1 | CH_SC | 1,9241135 | 0,519719861 | 1,445652174 |
| COPS81 | 0,545988803 | 0,51868271 | 0,133 | 0,092 | 1 | CH_SC | 1,432646536 | 0,698008877 | 1,445652174 |
| NR2F1-AS1 | 0,545988803 | 0,559898029 | 0,133 | 0,092 | 1 | CH_SC | 1,474165018 | 0,678350108 | 1,445652174 |

| Supplementary Table 4 - clusermarker | | | | | | | | | |
|---|---|---|---|---|---|---|---|---|---|
| | **p_val** | **avg_log2FC** | **pct.1** | **pct.2** | **p_val_adj** | **celltype** | **Foldchange_UP** | **Foldchange_DOWN** | **Ratio_pct1_pct2** |
| GEMIN51 | 0,545988803 | 0,998244049 | 0,133 | 0,092 | 1 | CH_SC | 1,997567216 | 0,500608937 | 1,445652174 |
| TSPYL41 | 0,545988803 | 0,412538293 | 0,133 | 0,092 | 1 | CH_SC | 1,331025576 | 0,751300364 | 1,445652174 |
| PSD3 | 0,545988803 | 0,450888823 | 0,133 | 0,092 | 1 | CH_SC | 1,366882113 | 0,731591986 | 1,445652174 |
| CD2BP2 | 0,545988803 | 0,355783509 | 0,133 | 0,092 | 1 | CH_SC | 1,279680371 | 0,781445135 | 1,445652174 |
| TK2 | 0,545988803 | 0,660043892 | 0,133 | 0,092 | 1 | CH_SC | 1,580130697 | 0,632859043 | 1,445652174 |
| DHX81 | 0,545988803 | 1,354185588 | 0,133 | 0,092 | 1 | CH_SC | 2,556527576 | 0,39115557 | 1,445652174 |
| RAB8A3 | 0,545988803 | 0,469906846 | 0,133 | 0,092 | 1 | CH_SC | 1,385020035 | 0,722011216 | 1,445652174 |
| AP1B11 | 0,545988803 | 0,60822468 | 0,133 | 0,092 | 1 | CH_SC | 1,524382213 | 0,656003456 | 1,445652174 |
| MARCKSL16 | 0,546416324 | 0,323382852 | 0,267 | 0,2 | 1 | CH_SC | 1,251261087 | 0,799193718 | 1,335 |
| HCG182 | 0,546416324 | 0,412929697 | 0,267 | 0,2 | 1 | CH_SC | 1,331386732 | 0,751096564 | 1,335 |
| CDC42EP35 | 0,547372874 | 0,410807121 | 0,333 | 0,277 | 1 | CH_SC | 1,32942936 | 0,752202433 | 1,202166065 |
| SF3B23 | 0,547546073 | 0,580817068 | 0,4 | 0,354 | 1 | CH_SC | 1,495696094 | 0,668585018 | 1,129943503 |
| GNB26 | 0,548442952 | 0,416065418 | 0,333 | 0,308 | 1 | CH_SC | 1,334283671 | 0,749465816 | 1,081168831 |
| COPRS2 | 0,548629468 | -0,260774347 | 0,2 | 0,262 | 1 | CH_SC | 0,834639818 | 1,198121607 | 0,763358779 |
| COL6A13 | 0,548659483 | 0,44862353 | 0,467 | 0,477 | 1 | CH_SC | 1,364737544 | 0,732741621 | 0,979035639 |
| RGCC5 | 0,548794007 | -0,387238055 | 0,4 | 0,523 | 1 | CH_SC | 0,764591966 | 1,307887141 | 0,764818356 |
| DYNLL14 | 0,549894455 | 0,440895922 | 0,6 | 0,754 | 1 | CH_SC | 1,357447048 | 0,736676986 | 0,795755968 |
| PPP1R15A4 | 0,55099951 | 0,334700865 | 0,467 | 0,492 | 1 | CH_SC | 1,261115897 | 0,792948533 | 0,949186992 |
| CDKN1C4 | 0,552007198 | -0,728899455 | 0,467 | 0,569 | 1 | CH_SC | 0,603364008 | 1,657374298 | 0,820738137 |
| RBM25 | 0,552501999 | 0,363535716 | 0,467 | 0,446 | 1 | CH_SC | 1,286575138 | 0,777257363 | 1,047085202 |
| DCAF8 | 0,557638542 | -0,284468053 | 0,133 | 0,185 | 1 | CH_SC | 0,821044286 | 1,217961097 | 0,718918919 |
| HADH3 | 0,557638542 | -0,258146074 | 0,133 | 0,185 | 1 | CH_SC | 0,836161733 | 1,195940881 | 0,718918919 |
| USP47 | 0,557938476 | 0,534123342 | 0,267 | 0,2 | 1 | CH_SC | 1,448061968 | 0,690578181 | 1,335 |
| NKTR1 | 0,559486848 | -0,321388702 | 0,2 | 0,262 | 1 | CH_SC | 0,800299159 | 1,249532739 | 0,763358779 |
| DHX91 | 0,561844511 | 0,416618611 | 0,133 | 0,092 | 1 | CH_SC | 1,334795392 | 0,749178493 | 1,445652174 |
| LPIN11 | 0,561844511 | 0,490089859 | 0,133 | 0,092 | 1 | CH_SC | 1,404532355 | 0,71198075 | 1,445652174 |
| CADM2 | 0,561844511 | 0,519040255 | 0,133 | 0,092 | 1 | CH_SC | 1,433001635 | 0,69783591 | 1,445652174 |
| SUDS3 | 0,561844511 | 0,974420909 | 0,133 | 0,092 | 1 | CH_SC | 1,964852357 | 0,508944093 | 1,445652174 |
| DUSP14 | 0,561844511 | 0,613276752 | 0,133 | 0,092 | 1 | CH_SC | 1,529729698 | 0,653710261 | 1,445652174 |
| HDAC51 | 0,561844511 | 0,32259312 | 0,133 | 0,092 | 1 | CH_SC | 1,250576334 | 0,799631317 | 1,445652174 |
| ZNF5801 | 0,561844511 | 0,327327467 | 0,133 | 0,092 | 1 | CH_SC | 1,254686965 | 0,797011548 | 1,445652174 |

| Supplementary Table 4 - clusermarker | | | | | | | | | |
|---|---|---|---|---|---|---|---|---|---|
| | **p_val** | **avg_log2FC** | **pct.1** | **pct.2** | **p_val_adj** | **celltype** | **Foldchange_UP** | **Foldchange_DOWN** | **Ratio_pct1_pct2** |
| MFAP42 | 0,562396181 | -0,34300216 | 0,133 | 0,2 | 1 | CH_SC | 0,788398994 | 1,268393298 | 0,665 |
| COPS92 | 0,564719319 | 0,488886055 | 0,467 | 0,369 | 1 | CH_SC | 1,403360883 | 0,712575085 | 1,265582656 |
| STARD71 | 0,564817753 | 0,268773725 | 0,2 | 0,154 | 1 | CH_SC | 1,204783339 | 0,830024758 | 1,298701299 |
| TIMM134 | 0,564817753 | 0,285680669 | 0,2 | 0,154 | 1 | CH_SC | 1,21898525 | 0,820354471 | 1,298701299 |
| SLC30A9 | 0,564994033 | 0,864801492 | 0,2 | 0,138 | 1 | CH_SC | 1,821089077 | 0,549121958 | 1,449275362 |
| SAMD4B | 0,564994033 | 0,69135575 | 0,2 | 0,138 | 1 | CH_SC | 1,614800289 | 0,619271625 | 1,449275362 |
| SYNCRIP2 | 0,566368893 | 0,298250586 | 0,333 | 0,246 | 1 | CH_SC | 1,229652431 | 0,813237932 | 1,353658537 |
| AAMDC2 | 0,566368893 | 0,60300172 | 0,267 | 0,262 | 1 | CH_SC | 1,518873501 | 0,658382676 | 1,019083969 |
| ATP5MK5 | 0,56650007 | 0,397266094 | 0,533 | 0,492 | 1 | CH_SC | 1,317009812 | 0,759295786 | 1,083333333 |
| TMEM2302 | 0,56810473 | -0,399583798 | 0,267 | 0,292 | 1 | CH_SC | 0,758076949 | 1,319127302 | 0,914383562 |
| RTRAF5 | 0,568483175 | 0,462272973 | 0,4 | 0,308 | 1 | CH_SC | 1,377710699 | 0,72584179 | 1,298701299 |
| PRPF38A1 | 0,56957755 | 0,284856977 | 0,267 | 0,2 | 1 | CH_SC | 1,218289481 | 0,820822978 | 1,335 |
| BPNT22 | 0,572213281 | 0,25762464 | 0,267 | 0,215 | 1 | CH_SC | 1,195508709 | 0,836464003 | 1,241860465 |
| CAPZA22 | 0,572725469 | 0,305876811 | 0,2 | 0,292 | 1 | CH_SC | 1,236169702 | 0,80895042 | 0,684931507 |
| ID28 | 0,573243484 | 0,678462155 | 0,467 | 0,462 | 1 | CH_SC | 1,600432859 | 0,62483096 | 1,010822511 |
| SLC44A21 | 0,574321388 | 0,591396364 | 0,267 | 0,231 | 1 | CH_SC | 1,50670436 | 0,66370021 | 1,155844156 |
| UBE2M2 | 0,574321388 | 0,563218134 | 0,267 | 0,231 | 1 | CH_SC | 1,477561452 | 0,676790802 | 1,155844156 |
| SNRPB24 | 0,575464244 | 0,467046693 | 0,267 | 0,4 | 1 | CH_SC | 1,382276943 | 0,723444028 | 0,6675 |
| RBPJ2 | 0,577856641 | 0,789117076 | 0,2 | 0,154 | 1 | CH_SC | 1,728016599 | 0,578698145 | 1,298701299 |
| MRPL374 | 0,577862043 | 0,521391861 | 0,2 | 0,154 | 1 | CH_SC | 1,435339344 | 0,696699358 | 1,298701299 |
| TXNDC174 | 0,577862043 | 0,518705656 | 0,2 | 0,154 | 1 | CH_SC | 1,432669322 | 0,697997775 | 1,298701299 |
| MTFR1L | 0,577919497 | 0,328222497 | 0,133 | 0,092 | 1 | CH_SC | 1,255465599 | 0,796517245 | 1,445652174 |
| GRPEL12 | 0,577919497 | 0,505689399 | 0,133 | 0,092 | 1 | CH_SC | 1,419801654 | 0,704323732 | 1,445652174 |
| PYGL1 | 0,577919497 | 0,788624784 | 0,133 | 0,092 | 1 | CH_SC | 1,727427047 | 0,578895648 | 1,445652174 |
| DYM3 | 0,577919497 | 0,412461932 | 0,133 | 0,092 | 1 | CH_SC | 1,330955127 | 0,751340131 | 1,445652174 |
| RNMT1 | 0,578486486 | 0,309370131 | 0,2 | 0,138 | 1 | CH_SC | 1,239166572 | 0,806994009 | 1,449275362 |
| AOPEP2 | 0,580180362 | 0,414830819 | 0,533 | 0,462 | 1 | CH_SC | 1,333142334 | 0,750107452 | 1,153679654 |
| UQCRC22 | 0,581509299 | -0,33626534 | 0,2 | 0,262 | 1 | CH_SC | 0,792089117 | 1,262484207 | 0,763358779 |
| NPC26 | 0,583792747 | -0,353630394 | 0,467 | 0,631 | 1 | CH_SC | 0,782612254 | 1,277771967 | 0,740095087 |
| NCK2 | 0,586037326 | -0,259890172 | 0,067 | 0,108 | 1 | CH_SC | 0,835151494 | 1,197387548 | 0,62037037 |
| SENP2 | 0,586037326 | -0,252668701 | 0,067 | 0,108 | 1 | CH_SC | 0,839342361 | 1,191408949 | 0,62037037 |

| Supplementary Table 4 - clusermarker | | | | | | | | | |
|---|---|---|---|---|---|---|---|---|---|
| | **p_val** | **avg_log2FC** | **pct.1** | **pct.2** | **p_val_adj** | **celltype** | **Foldchange_UP** | **Foldchange_DOWN** | **Ratio_pct1_pct2** |
| WDR191 | 0,586037326 | -0,283770316 | 0,067 | 0,108 | 1 | CH_SC | 0,821441468 | 1,217372192 | 0,62037037 |
| BBS7 | 0,586037326 | -0,379419772 | 0,067 | 0,108 | 1 | CH_SC | 0,768746706 | 1,300818583 | 0,62037037 |
| PAIP1 | 0,586037326 | -0,287667216 | 0,067 | 0,108 | 1 | CH_SC | 0,819225645 | 1,220664912 | 0,62037037 |
| RNF14 | 0,586037326 | -0,288912639 | 0,067 | 0,108 | 1 | CH_SC | 0,818518745 | 1,22171912 | 0,62037037 |
| METTL15 | 0,586037326 | -0,363236669 | 0,067 | 0,108 | 1 | CH_SC | 0,777418493 | 1,28630848 | 0,62037037 |
| CREBZF | 0,586037326 | -0,324653715 | 0,067 | 0,108 | 1 | CH_SC | 0,798490022 | 1,252363802 | 0,62037037 |
| C11orf12 | 0,586037326 | -0,27998781 | 0,067 | 0,108 | 1 | CH_SC | 0,823597976 | 1,214184625 | 0,62037037 |
| RSPRY1 | 0,586037326 | -0,267652562 | 0,067 | 0,108 | 1 | CH_SC | 0,830670047 | 1,203847429 | 0,62037037 |
| PCBP17 | 0,586949498 | 0,660719254 | 0,467 | 0,492 | 1 | CH_SC | 1,580870569 | 0,632562855 | 0,949186992 |
| LAGE34 | 0,587074263 | 0,253485072 | 0,133 | 0,2 | 1 | CH_SC | 1,192083317 | 0,838867541 | 0,665 |
| MARVELD1 | 0,58732054 | 0,682921398 | 0,4 | 0,338 | 1 | CH_SC | 1,605387309 | 0,622902644 | 1,183431953 |
| ZKSCAN12 | 0,588740895 | -0,304657049 | 0,267 | 0,338 | 1 | CH_SC | 0,809634657 | 1,235124994 | 0,789940828 |
| YME1L13 | 0,589138095 | 0,459145005 | 0,333 | 0,308 | 1 | CH_SC | 1,374726862 | 0,727417226 | 1,081168831 |
| AIMP11 | 0,589187552 | 0,458605049 | 0,467 | 0,446 | 1 | CH_SC | 1,374212441 | 0,727689526 | 1,047085202 |
| SDHC2 | 0,589204967 | 0,388874476 | 0,333 | 0,277 | 1 | CH_SC | 1,309371494 | 0,763725195 | 1,202166065 |
| ZNF242 | 0,589300542 | -0,263296469 | 0,267 | 0,308 | 1 | CH_SC | 0,833181974 | 1,200217998 | 0,866883117 |
| USP484 | 0,589323418 | 0,624662891 | 0,2 | 0,169 | 1 | CH_SC | 1,541850505 | 0,648571309 | 1,183431953 |
| HMGCS12 | 0,589323418 | 0,553593777 | 0,2 | 0,169 | 1 | CH_SC | 1,467737304 | 0,681320831 | 1,183431953 |
| CUX1 | 0,591046381 | 0,562145221 | 0,2 | 0,154 | 1 | CH_SC | 1,476463019 | 0,677294309 | 1,298701299 |
| DNM1L | 0,592128506 | 0,271140622 | 0,2 | 0,138 | 1 | CH_SC | 1,206761539 | 0,828664129 | 1,449275362 |
| NGDN1 | 0,592128506 | 0,377404541 | 0,2 | 0,138 | 1 | CH_SC | 1,2990028 | 0,769821281 | 1,449275362 |
| PHF13 | 0,594207661 | 0,56087833 | 0,133 | 0,092 | 1 | CH_SC | 1,475167043 | 0,677889331 | 1,445652174 |
| INTS3 | 0,594207661 | 0,407963943 | 0,133 | 0,092 | 1 | CH_SC | 1,326811979 | 0,753686291 | 1,445652174 |
| ADAM151 | 0,594207661 | 0,268848138 | 0,133 | 0,092 | 1 | CH_SC | 1,204845483 | 0,829981947 | 1,445652174 |
| LGALSL | 0,594207661 | 0,347010198 | 0,133 | 0,092 | 1 | CH_SC | 1,271921997 | 0,786211735 | 1,445652174 |
| GMPPA | 0,594207661 | 0,288573991 | 0,133 | 0,092 | 1 | CH_SC | 1,221432375 | 0,818710901 | 1,445652174 |
| DAAM2 | 0,594207661 | 0,269664239 | 0,133 | 0,092 | 1 | CH_SC | 1,205527231 | 0,829512577 | 1,445652174 |
| FBXO25 | 0,594207661 | 0,374116123 | 0,133 | 0,092 | 1 | CH_SC | 1,296045281 | 0,77157798 | 1,445652174 |
| ACBD51 | 0,594207661 | 0,334614494 | 0,133 | 0,092 | 1 | CH_SC | 1,261040398 | 0,792996007 | 1,445652174 |
| NFYB1 | 0,594207661 | 0,385613286 | 0,133 | 0,092 | 1 | CH_SC | 1,306415023 | 0,765453537 | 1,445652174 |
| SPPL31 | 0,594207661 | 0,47033856 | 0,133 | 0,092 | 1 | CH_SC | 1,385434552 | 0,721795193 | 1,445652174 |

| Supplementary Table 4 - clusermarker | | | | | | | | | |
|---|---|---|---|---|---|---|---|---|---|
| | **p_val** | **avg_log2FC** | **pct.1** | **pct.2** | **p_val_adj** | **celltype** | **Foldchange_UP** | **Foldchange_DOWN** | **Ratio_pct1_pct2** |
| SPG21 | 0,594207661 | 0,294161233 | 0,133 | 0,092 | 1 | CH_SC | 1,226171887 | 0,815546345 | 1,445652174 |
| PLSCR31 | 0,594207661 | 0,303633134 | 0,133 | 0,092 | 1 | CH_SC | 1,234248707 | 0,810209478 | 1,445652174 |
| TMEM126B | 0,595758683 | -0,305622688 | 0,133 | 0,185 | 1 | CH_SC | 0,809092925 | 1,235951977 | 0,718918919 |
| TMEM106C2 | 0,597043061 | 0,40382417 | 0,267 | 0,231 | 1 | CH_SC | 1,323010187 | 0,755852079 | 1,155844156 |
| SNRPB3 | 0,597043061 | 0,255996434 | 0,267 | 0,231 | 1 | CH_SC | 1,194160236 | 0,837408557 | 1,155844156 |
| VPS294 | 0,598287607 | 0,526139432 | 0,267 | 0,246 | 1 | CH_SC | 1,44007049 | 0,694410452 | 1,085365854 |
| SERTAD21 | 0,599831477 | -0,312792767 | 0,267 | 0,308 | 1 | CH_SC | 0,805081774 | 1,242109849 | 0,866883117 |
| MCFD21 | 0,601220609 | -0,512195853 | 0,2 | 0,231 | 1 | CH_SC | 0,701154434 | 1,426219321 | 0,865800866 |
| DNAJC12 | 0,601220609 | -0,32701332 | 0,2 | 0,231 | 1 | CH_SC | 0,797185116 | 1,254413786 | 0,865800866 |
| ANKIB1 | 0,6022258 | 0,525464501 | 0,2 | 0,169 | 1 | CH_SC | 1,439396945 | 0,694735392 | 1,183431953 |
| AL135905.22 | 0,60242969 | -0,276640499 | 0,067 | 0,108 | 1 | CH_SC | 0,82551109 | 1,211370765 | 0,62037037 |
| SURF6 | 0,60242969 | -0,344756985 | 0,067 | 0,108 | 1 | CH_SC | 0,787440606 | 1,26993705 | 0,62037037 |
| PRKG1 | 0,60242969 | -0,323511055 | 0,067 | 0,108 | 1 | CH_SC | 0,799122702 | 1,251372283 | 0,62037037 |
| ITPRIP | 0,60242969 | -0,273598322 | 0,067 | 0,108 | 1 | CH_SC | 0,827253662 | 1,208819067 | 0,62037037 |
| MGMT5 | 0,60242969 | -0,264967715 | 0,067 | 0,108 | 1 | CH_SC | 0,832217358 | 1,201609159 | 0,62037037 |
| EHBP1L11 | 0,60242969 | -0,270638168 | 0,067 | 0,108 | 1 | CH_SC | 0,828952782 | 1,206341328 | 0,62037037 |
| SIK21 | 0,60242969 | -0,482549903 | 0,067 | 0,108 | 1 | CH_SC | 0,715711515 | 1,397210997 | 0,62037037 |
| CD27-AS1 | 0,60242969 | -0,287649327 | 0,067 | 0,108 | 1 | CH_SC | 0,819235804 | 1,220649776 | 0,62037037 |
| DCAF51 | 0,60242969 | -0,30141438 | 0,067 | 0,108 | 1 | CH_SC | 0,811456476 | 1,232351986 | 0,62037037 |
| IGF1R | 0,60242969 | -0,263967762 | 0,067 | 0,108 | 1 | CH_SC | 0,83279438 | 1,200776595 | 0,62037037 |
| POLI | 0,60242969 | -0,443021171 | 0,067 | 0,108 | 1 | CH_SC | 0,735592579 | 1,359448191 | 0,62037037 |
| LONP11 | 0,60242969 | -0,292606294 | 0,067 | 0,108 | 1 | CH_SC | 0,816425815 | 1,224851029 | 0,62037037 |
| NR1H21 | 0,60242969 | -0,280091601 | 0,067 | 0,108 | 1 | CH_SC | 0,823538727 | 1,21427198 | 0,62037037 |
| NFIB4 | 0,603296495 | 0,666957562 | 0,4 | 0,385 | 1 | CH_SC | 1,587721157 | 0,629833517 | 1,038961039 |
| EMILIN11 | 0,604367364 | 0,314921651 | 0,2 | 0,154 | 1 | CH_SC | 1,243944096 | 0,803894647 | 1,298701299 |
| BCAR12 | 0,604367364 | 0,50637757 | 0,2 | 0,154 | 1 | CH_SC | 1,420479066 | 0,703987847 | 1,298701299 |
| PPP1R1B | 0,604367364 | 0,409419736 | 0,2 | 0,154 | 1 | CH_SC | 1,328151513 | 0,752926146 | 1,298701299 |
| PFKL2 | 0,604367364 | 0,251591851 | 0,2 | 0,154 | 1 | CH_SC | 1,190519994 | 0,839969093 | 1,298701299 |
| SCARF2 | 0,604367364 | -0,304355119 | 0,133 | 0,169 | 1 | CH_SC | 0,809804116 | 1,234866532 | 0,786982249 |
| FABP55 | 0,605172803 | 0,255961335 | 0,267 | 0,2 | 1 | CH_SC | 1,194131183 | 0,837428931 | 1,335 |
| RBBP7 | 0,605172803 | 0,312553545 | 0,267 | 0,2 | 1 | CH_SC | 1,241903904 | 0,80521528 | 1,335 |

| Supplementary Table 4 - clusermarker | | | | | | | | | |
|---|---|---|---|---|---|---|---|---|---|
| | **p_val** | **avg_log2FC** | **pct.1** | **pct.2** | **p_val_adj** | **celltype** | **Foldchange_UP** | **Foldchange_DOWN** | **Ratio_pct1_pct2** |
| TET22 | 0,605313184 | 0,694946789 | 0,067 | 0,123 | 1 | CH_SC | 1,618824725 | 0,617732102 | 0,544715447 |
| PELO2 | 0,605313184 | 0,699706481 | 0,067 | 0,123 | 1 | CH_SC | 1,624174318 | 0,615697459 | 0,544715447 |
| GTF3C5 | 0,605313184 | 0,288986493 | 0,067 | 0,123 | 1 | CH_SC | 1,221781663 | 0,818476844 | 0,544715447 |
| PTGR2 | 0,605313184 | 0,283432187 | 0,067 | 0,123 | 1 | CH_SC | 1,217086905 | 0,821634014 | 0,544715447 |
| WDR73 | 0,605313184 | 0,260175136 | 0,067 | 0,123 | 1 | CH_SC | 1,197624081 | 0,83498655 | 0,544715447 |
| PREX1 | 0,605313184 | 0,31033249 | 0,067 | 0,123 | 1 | CH_SC | 1,239993441 | 0,806455879 | 0,544715447 |
| SNAP291 | 0,605313184 | 0,427842517 | 0,067 | 0,123 | 1 | CH_SC | 1,345220358 | 0,743372633 | 0,544715447 |
| MYADM5 | 0,60635596 | 0,259057002 | 0,533 | 0,523 | 1 | CH_SC | 1,196696245 | 0,835633942 | 1,019120459 |
| PSMC33 | 0,607100811 | 0,383965293 | 0,267 | 0,215 | 1 | CH_SC | 1,304923555 | 0,766328416 | 1,241860465 |
| DDAH24 | 0,60855516 | -0,384292022 | 0,2 | 0,246 | 1 | CH_SC | 0,766154884 | 1,305219115 | 0,81300813 |
| TMEM263 | 0,610659697 | 0,457521918 | 0,333 | 0,277 | 1 | CH_SC | 1,373181111 | 0,728236058 | 1,202166065 |
| PIGK | 0,61070258 | 0,259505157 | 0,133 | 0,092 | 1 | CH_SC | 1,197068041 | 0,835374403 | 1,445652174 |
| KLHL12 | 0,61070258 | 0,322063501 | 0,133 | 0,092 | 1 | CH_SC | 1,250117326 | 0,799924918 | 1,445652174 |
| GGCX | 0,61070258 | 0,379034947 | 0,133 | 0,092 | 1 | CH_SC | 1,300471648 | 0,768951789 | 1,445652174 |
| WDR821 | 0,61070258 | 0,271935537 | 0,133 | 0,092 | 1 | CH_SC | 1,20742664 | 0,828207667 | 1,445652174 |
| LRIG1 | 0,61070258 | 0,803425553 | 0,133 | 0,092 | 1 | CH_SC | 1,74524013 | 0,572987054 | 1,445652174 |
| KIF13A | 0,61070258 | 0,539908619 | 0,133 | 0,092 | 1 | CH_SC | 1,453880425 | 0,687814474 | 1,445652174 |
| RASL11A1 | 0,61070258 | 0,296652752 | 0,133 | 0,092 | 1 | CH_SC | 1,228291303 | 0,81413912 | 1,445652174 |
| RBMX2 | 0,61070258 | 0,516649673 | 0,133 | 0,092 | 1 | CH_SC | 1,430629081 | 0,6989932 | 1,445652174 |
| KRTCAP25 | 0,612524988 | 0,287764736 | 0,467 | 0,477 | 1 | CH_SC | 1,220747426 | 0,819170271 | 0,979035639 |
| MARCHF74 | 0,617821453 | 0,450257154 | 0,2 | 0,154 | 1 | CH_SC | 1,366283769 | 0,731912376 | 1,298701299 |
| AZI21 | 0,617821453 | 0,720790529 | 0,2 | 0,154 | 1 | CH_SC | 1,64808486 | 0,606764872 | 1,298701299 |
| UBA32 | 0,617821453 | 0,363909745 | 0,2 | 0,154 | 1 | CH_SC | 1,286908736 | 0,77705588 | 1,298701299 |
| MTRNR2L810 | 0,617821453 | -3,276431219 | 0,133 | 0,169 | 1 | CH_SC | 0,103203856 | 9,689560432 | 0,786982249 |
| PDS5B | 0,617821453 | 0,318012032 | 0,2 | 0,154 | 1 | CH_SC | 1,246611591 | 0,802174476 | 1,298701299 |
| PKD1 | 0,617821453 | 0,272778441 | 0,2 | 0,154 | 1 | CH_SC | 1,208132293 | 0,827723922 | 1,298701299 |
| PIN12 | 0,617821453 | -0,464962531 | 0,133 | 0,169 | 1 | CH_SC | 0,724489893 | 1,380281505 | 0,786982249 |
| NSFL1C1 | 0,617821453 | 0,272383158 | 0,2 | 0,154 | 1 | CH_SC | 1,207801323 | 0,827950741 | 1,298701299 |
| NIPAL3 | 0,61902548 | -0,267533234 | 0,067 | 0,108 | 1 | CH_SC | 0,830738756 | 1,203747861 | 0,62037037 |
| HERPUD2 | 0,61902548 | -0,284866947 | 0,067 | 0,108 | 1 | CH_SC | 0,820817305 | 1,218297901 | 0,62037037 |
| MED27 | 0,61902548 | -0,301565438 | 0,067 | 0,108 | 1 | CH_SC | 0,811371517 | 1,232481027 | 0,62037037 |

| Supplementary Table 4 - clusermarker | | | | | | | | | |
|---|---|---|---|---|---|---|---|---|---|
| | **p_val** | **avg_log2FC** | **pct.1** | **pct.2** | **p_val_adj** | **celltype** | **Foldchange_UP** | **Foldchange_DOWN** | **Ratio_pct1_pct2** |
| PACSIN3 | 0,61902548 | -0,404232095 | 0,067 | 0,108 | 1 | CH_SC | 0,75563839 | 1,323384324 | 0,62037037 |
| GNS | 0,61902548 | -0,261769201 | 0,067 | 0,108 | 1 | CH_SC | 0,834064465 | 1,198948093 | 0,62037037 |
| HERC11 | 0,61902548 | -0,321073877 | 0,067 | 0,108 | 1 | CH_SC | 0,800473819 | 1,249260096 | 0,62037037 |
| ZHX3 | 0,61902548 | -0,289780556 | 0,067 | 0,108 | 1 | CH_SC | 0,818026477 | 1,22245432 | 0,62037037 |
| SSBP31 | 0,619846002 | 0,402271866 | 0,2 | 0,138 | 1 | CH_SC | 1,321587426 | 0,756665795 | 1,449275362 |
| SNRPA3 | 0,620165072 | -0,311447671 | 0,2 | 0,246 | 1 | CH_SC | 0,805832741 | 1,240952308 | 0,81300813 |
| ARL2BP | 0,621144607 | -0,323075398 | 0,267 | 0,308 | 1 | CH_SC | 0,799364053 | 1,250994457 | 0,866883117 |
| CNPY23 | 0,624537092 | -0,360570587 | 0,4 | 0,446 | 1 | CH_SC | 0,77885648 | 1,283933594 | 0,896860987 |
| MLEC4 | 0,624971064 | 0,684053754 | 0,333 | 0,323 | 1 | CH_SC | 1,606647855 | 0,622413926 | 1,030959752 |
| SLC25A66 | 0,625860825 | 0,343229553 | 0,467 | 0,508 | 1 | CH_SC | 1,268593234 | 0,788274738 | 0,919291339 |
| MEPCE1 | 0,627397517 | 0,298496476 | 0,133 | 0,092 | 1 | CH_SC | 1,229862028 | 0,813099337 | 1,445652174 |
| ERLIN2 | 0,627397517 | 0,274524822 | 0,133 | 0,092 | 1 | CH_SC | 1,209595621 | 0,826722569 | 1,445652174 |
| ADGRA21 | 0,627397517 | 0,265659291 | 0,133 | 0,092 | 1 | CH_SC | 1,202185305 | 0,831818519 | 1,445652174 |
| CHRAC1 | 0,627397517 | 0,733318668 | 0,133 | 0,092 | 1 | CH_SC | 1,662458893 | 0,601518633 | 1,445652174 |
| DPM1 | 0,627397517 | 0,46284911 | 0,133 | 0,092 | 1 | CH_SC | 1,378260995 | 0,725551985 | 1,445652174 |
| GINM11 | 0,627795138 | 0,386398095 | 0,4 | 0,338 | 1 | CH_SC | 1,30712589 | 0,765037253 | 1,183431953 |
| STT3A1 | 0,630873931 | 0,530513542 | 0,267 | 0,215 | 1 | CH_SC | 1,444443268 | 0,692308256 | 1,241860465 |
| CHD94 | 0,631031652 | 0,292289482 | 0,4 | 0,4 | 1 | CH_SC | 1,224582085 | 0,81660512 | 1 |
| EFEMP22 | 0,631173376 | 0,287016717 | 0,333 | 0,308 | 1 | CH_SC | 1,220114648 | 0,819595111 | 1,081168831 |
| UBAC2 | 0,631404975 | 0,288541905 | 0,2 | 0,154 | 1 | CH_SC | 1,22140521 | 0,818729109 | 1,298701299 |
| TIMM9 | 0,631404975 | -0,286120469 | 0,133 | 0,169 | 1 | CH_SC | 0,820104427 | 1,21935691 | 0,786982249 |
| TXNDC122 | 0,631870432 | 0,706653045 | 0,267 | 0,231 | 1 | CH_SC | 1,63201356 | 0,612740007 | 1,155844156 |
| LTBP32 | 0,631923153 | -0,35456925 | 0,267 | 0,308 | 1 | CH_SC | 0,782103123 | 1,278603767 | 0,866883117 |
| ASPH1 | 0,634191926 | 0,262256872 | 0,467 | 0,431 | 1 | CH_SC | 1,19935344 | 0,833782575 | 1,083526682 |
| SPRED2 | 0,635817802 | -0,299375889 | 0,067 | 0,108 | 1 | CH_SC | 0,812603853 | 1,230611935 | 0,62037037 |
| IDE | 0,635817802 | -0,320011069 | 0,067 | 0,108 | 1 | CH_SC | 0,801063731 | 1,248340127 | 0,62037037 |
| NHP23 | 0,636526376 | -0,254799864 | 0,333 | 0,369 | 1 | CH_SC | 0,838103391 | 1,193170211 | 0,902439024 |
| RNH13 | 0,639522308 | 0,957870117 | 0,333 | 0,338 | 1 | CH_SC | 1,942440109 | 0,514816388 | 0,985207101 |
| EMC32 | 0,640080491 | 0,315417239 | 0,133 | 0,215 | 1 | CH_SC | 1,244371483 | 0,803618544 | 0,618604651 |
| LDOC12 | 0,640080491 | 0,259603941 | 0,133 | 0,215 | 1 | CH_SC | 1,197150009 | 0,835317205 | 0,618604651 |
| NUCB13 | 0,640866356 | 0,58246706 | 0,467 | 0,477 | 1 | CH_SC | 1,497407681 | 0,667820803 | 0,979035639 |

| Supplementary Table 4 - clusermarker | | | | | | | | | |
|---|---|---|---|---|---|---|---|---|---|
| | **p_val** | **avg_log2FC** | **pct.1** | **pct.2** | **p_val_adj** | **celltype** | **Foldchange_UP** | **Foldchange_DOWN** | **Ratio_pct1_pct2** |
| KLC1 | 0,641674202 | 0,448098987 | 0,2 | 0,169 | 1 | CH_SC | 1,364241436 | 0,733008083 | 1,183431953 |
| IPO72 | 0,643473838 | 0,270879246 | 0,333 | 0,277 | 1 | CH_SC | 1,206542928 | 0,828814273 | 1,202166065 |
| BSG1 | 0,643909419 | -0,382041487 | 0,4 | 0,446 | 1 | CH_SC | 0,767350981 | 1,303184624 | 0,896860987 |
| EEFSEC | 0,644285426 | 0,327084993 | 0,133 | 0,092 | 1 | CH_SC | 1,254476107 | 0,797145513 | 1,445652174 |
| CISD21 | 0,644285426 | 0,34646227 | 0,133 | 0,092 | 1 | CH_SC | 1,27143902 | 0,78651039 | 1,445652174 |
| KBTBD22 | 0,644285426 | 0,367181435 | 0,133 | 0,092 | 1 | CH_SC | 1,289830451 | 0,775295698 | 1,445652174 |
| TP53TG1 | 0,644285426 | 0,72387364 | 0,133 | 0,092 | 1 | CH_SC | 1,651610665 | 0,60546957 | 1,445652174 |
| GMFB | 0,644285426 | 0,268451879 | 0,133 | 0,092 | 1 | CH_SC | 1,204514598 | 0,830209946 | 1,445652174 |
| GPT2 | 0,644285426 | 0,44567544 | 0,133 | 0,092 | 1 | CH_SC | 1,361951605 | 0,73424048 | 1,445652174 |
| RTCB1 | 0,644285426 | 0,298579233 | 0,133 | 0,092 | 1 | CH_SC | 1,229932578 | 0,813052697 | 1,445652174 |
| PCID21 | 0,645109432 | -0,386676806 | 0,133 | 0,169 | 1 | CH_SC | 0,764889471 | 1,307378435 | 0,786982249 |
| GORASP21 | 0,645114126 | 0,28819569 | 0,2 | 0,154 | 1 | CH_SC | 1,221112135 | 0,81892561 | 1,298701299 |
| DYNLL21 | 0,645114126 | -0,357306919 | 0,133 | 0,169 | 1 | CH_SC | 0,780620405 | 1,28103236 | 0,786982249 |
| PARK71 | 0,645378781 | 0,392885023 | 0,533 | 0,6 | 1 | CH_SC | 1,31301648 | 0,761605064 | 0,888333333 |
| TCF41 | 0,64748928 | 0,800840359 | 0,6 | 0,615 | 1 | CH_SC | 1,7421156 | 0,574014721 | 0,975609756 |
| DHRS73 | 0,648959732 | -0,544364309 | 0,2 | 0,231 | 1 | CH_SC | 0,685693471 | 1,458377602 | 0,865800866 |
| SMU14 | 0,650860406 | 1,051615242 | 0,2 | 0,185 | 1 | CH_SC | 2,072849312 | 0,482427736 | 1,081081081 |
| CKS21 | 0,650860406 | 0,606414937 | 0,2 | 0,185 | 1 | CH_SC | 1,522471199 | 0,656826875 | 1,081081081 |
| SPRYD7 | 0,650860406 | 0,940650036 | 0,2 | 0,185 | 1 | CH_SC | 1,919392865 | 0,520998081 | 1,081081081 |
| PAK22 | 0,652659088 | 0,339718988 | 0,333 | 0,308 | 1 | CH_SC | 1,265510071 | 0,790195213 | 1,081168831 |
| HTRA21 | 0,652799462 | -0,294515535 | 0,067 | 0,108 | 1 | CH_SC | 0,815346084 | 1,226473051 | 0,62037037 |
| CRK1 | 0,655059264 | 0,67374281 | 0,2 | 0,169 | 1 | CH_SC | 1,595206076 | 0,626878254 | 1,183431953 |
| BMPR2 | 0,655557683 | -0,28614347 | 0,2 | 0,246 | 1 | CH_SC | 0,820091352 | 1,21937635 | 0,81300813 |
| TRAPPC57 | 0,655557683 | 0,273070401 | 0,267 | 0,231 | 1 | CH_SC | 1,208376809 | 0,827556432 | 1,155844156 |
| CPQ2 | 0,655735566 | 0,579558492 | 0,267 | 0,246 | 1 | CH_SC | 1,49439185 | 0,669168532 | 1,085365854 |
| ACSL33 | 0,658944974 | -0,445187467 | 0,133 | 0,169 | 1 | CH_SC | 0,734488869 | 1,36149102 | 0,786982249 |
| CCDC503 | 0,658944974 | -0,281417909 | 0,133 | 0,169 | 1 | CH_SC | 0,822781973 | 1,215388806 | 0,786982249 |
| CAMK2D | 0,658944974 | 0,441193797 | 0,2 | 0,154 | 1 | CH_SC | 1,357727351 | 0,736524899 | 1,298701299 |
| CEP112 | 0,661358964 | 0,280814822 | 0,133 | 0,092 | 1 | CH_SC | 1,214880845 | 0,823125991 | 1,445652174 |
| TMED92 | 0,66543454 | 0,266743035 | 0,4 | 0,369 | 1 | CH_SC | 1,203088719 | 0,831193896 | 1,08401084 |
| TIMM17A3 | 0,667254927 | 0,327257438 | 0,267 | 0,215 | 1 | CH_SC | 1,254626063 | 0,797050236 | 1,241860465 |

| Supplementary Table 4 - clusermarker | | | | | | | | | |
|---|---|---|---|---|---|---|---|---|---|
| | **p_val** | **avg_log2FC** | **pct.1** | **pct.2** | **p_val_adj** | **celltype** | **Foldchange_UP** | **Foldchange_DOWN** | **Ratio_pct1_pct2** |
| PSMD22 | 0,667534494 | 0,421852029 | 0,267 | 0,231 | 1 | CH_SC | 1,339646194 | 0,746465749 | 1,155844156 |
| MOXD1 | 0,668556096 | 0,34355863 | 0,2 | 0,169 | 1 | CH_SC | 1,268882632 | 0,788094954 | 1,183431953 |
| OGA | 0,668556096 | 0,860929264 | 0,2 | 0,169 | 1 | CH_SC | 1,816207784 | 0,550597794 | 1,183431953 |
| SLC25A331 | 0,669451331 | 0,497936455 | 0,133 | 0,108 | 1 | CH_SC | 1,412192201 | 0,708118908 | 1,231481481 |
| GON4L2 | 0,669451331 | 0,480579583 | 0,133 | 0,108 | 1 | CH_SC | 1,395304098 | 0,716689646 | 1,231481481 |
| ALDH9A13 | 0,669451331 | 0,561199425 | 0,133 | 0,108 | 1 | CH_SC | 1,475495402 | 0,677738472 | 1,231481481 |
| STAG1 | 0,669451331 | 0,53865371 | 0,133 | 0,108 | 1 | CH_SC | 1,452616336 | 0,688413021 | 1,231481481 |
| TLR3 | 0,669451331 | 0,463733486 | 0,133 | 0,108 | 1 | CH_SC | 1,379106131 | 0,725107356 | 1,231481481 |
| MINCR | 0,669451331 | 0,438866326 | 0,133 | 0,108 | 1 | CH_SC | 1,355538723 | 0,737714079 | 1,231481481 |
| GNAQ2 | 0,669451331 | 0,742094461 | 0,133 | 0,108 | 1 | CH_SC | 1,672602311 | 0,597870751 | 1,231481481 |
| MARCHF9 | 0,669451331 | 0,555832372 | 0,133 | 0,108 | 1 | CH_SC | 1,470016524 | 0,680264462 | 1,231481481 |
| PEG3 | 0,669451331 | 0,623449158 | 0,133 | 0,108 | 1 | CH_SC | 1,540553898 | 0,649117179 | 1,231481481 |
| ZRSR2 | 0,669451331 | 0,541556622 | 0,133 | 0,108 | 1 | CH_SC | 1,455542154 | 0,687029226 | 1,231481481 |
| CUTA6 | 0,671559044 | 0,360632859 | 0,467 | 0,492 | 1 | CH_SC | 1,283989014 | 0,778822863 | 0,949186992 |
| TRAPPC31 | 0,672893462 | -0,376263859 | 0,133 | 0,169 | 1 | CH_SC | 0,770430189 | 1,297976137 | 0,786982249 |
| HAT1 | 0,672893462 | -0,397508866 | 0,133 | 0,169 | 1 | CH_SC | 0,759168025 | 1,317231453 | 0,786982249 |
| PHF15 | 0,672893462 | -0,25536315 | 0,133 | 0,169 | 1 | CH_SC | 0,837776226 | 1,193636163 | 0,786982249 |
| ACADVL3 | 0,674436895 | 0,310108528 | 0,333 | 0,308 | 1 | CH_SC | 1,239800961 | 0,806581081 | 1,081168831 |
| SCAND12 | 0,674436895 | 0,25984883 | 0,333 | 0,308 | 1 | CH_SC | 1,197353236 | 0,835175427 | 1,081168831 |
| CTSO | 0,677145244 | 0,93172496 | 0,2 | 0,185 | 1 | CH_SC | 1,907555404 | 0,524231169 | 1,081081081 |
| TMEM39A | 0,682160916 | 0,277985028 | 0,2 | 0,169 | 1 | CH_SC | 1,212500236 | 0,824742108 | 1,183431953 |
| RBM71 | 0,682160916 | 0,609996528 | 0,2 | 0,169 | 1 | CH_SC | 1,526255536 | 0,655198279 | 1,183431953 |
| UBE2Q2 | 0,682160916 | 0,440055516 | 0,2 | 0,169 | 1 | CH_SC | 1,356656532 | 0,737106244 | 1,183431953 |
| NCSTN | 0,685894584 | 0,414452453 | 0,133 | 0,108 | 1 | CH_SC | 1,332792745 | 0,750304204 | 1,231481481 |
| CREB15 | 0,685894584 | 0,858758509 | 0,133 | 0,108 | 1 | CH_SC | 1,813477077 | 0,551426877 | 1,231481481 |
| NSUN61 | 0,685894584 | 0,781959529 | 0,133 | 0,108 | 1 | CH_SC | 1,719464736 | 0,581576335 | 1,231481481 |
| MLLT1 | 0,685894584 | 0,330365661 | 0,133 | 0,108 | 1 | CH_SC | 1,257332014 | 0,795334875 | 1,231481481 |
| HSPH18 | 0,686958116 | -0,297674958 | 0,267 | 0,308 | 1 | CH_SC | 0,813562475 | 1,229161903 | 0,866883117 |
| UBQLN11 | 0,690439165 | 0,836099547 | 0,2 | 0,185 | 1 | CH_SC | 1,78521712 | 0,560155955 | 1,081081081 |
| DAAM11 | 0,690439165 | 0,408681543 | 0,2 | 0,185 | 1 | CH_SC | 1,327472103 | 0,753311499 | 1,081081081 |
| SMCHD12 | 0,690483082 | 0,394155292 | 0,2 | 0,277 | 1 | CH_SC | 1,314173078 | 0,760934779 | 0,722021661 |

| Supplementary Table 4 - clusermarker | | | | | | | | | |
|---|---|---|---|---|---|---|---|---|---|
| | **p_val** | **avg_log2FC** | **pct.1** | **pct.2** | **p_val_adj** | **celltype** | **Foldchange_UP** | **Foldchange_DOWN** | **Ratio_pct1_pct2** |
| DNPH15 | 0,691741275 | 0,598434513 | 0,267 | 0,231 | 1 | CH_SC | 1,514072735 | 0,660470251 | 1,155844156 |
| ATG1011 | 0,691848036 | -0,384843445 | 0,2 | 0,215 | 1 | CH_SC | 0,765862102 | 1,305718088 | 0,930232558 |
| C1orf212 | 0,695869838 | 0,702146208 | 0,2 | 0,169 | 1 | CH_SC | 1,626923267 | 0,614657139 | 1,183431953 |
| NOSIP3 | 0,695869838 | 0,309041355 | 0,2 | 0,169 | 1 | CH_SC | 1,23888421 | 0,807177936 | 1,183431953 |
| RAB7A2 | 0,695974234 | -0,44891159 | 0,333 | 0,354 | 1 | CH_SC | 0,73259533 | 1,365010066 | 0,940677966 |
| UGDH1 | 0,698186524 | -0,269457775 | 0,2 | 0,231 | 1 | CH_SC | 0,829631297 | 1,20535472 | 0,865800866 |
| NDUFB108 | 0,699094323 | 0,507429284 | 0,4 | 0,415 | 1 | CH_SC | 1,421514962 | 0,703474833 | 0,963855422 |
| ARPC35 | 0,700786929 | 0,35151989 | 0,467 | 0,492 | 1 | CH_SC | 1,275904094 | 0,783757968 | 0,949186992 |
| SLC39A1 | 0,701126525 | 0,548950118 | 0,2 | 0,154 | 1 | CH_SC | 1,463020635 | 0,683517359 | 1,298701299 |
| HOOK31 | 0,701126525 | 0,301081662 | 0,2 | 0,154 | 1 | CH_SC | 1,232067811 | 0,811643638 | 1,298701299 |
| NDUFS34 | 0,701126525 | -0,323362884 | 0,133 | 0,169 | 1 | CH_SC | 0,799204779 | 1,251243768 | 0,786982249 |
| FBXO21 | 0,701126525 | -0,257204698 | 0,133 | 0,169 | 1 | CH_SC | 0,836707517 | 1,195160769 | 0,786982249 |
| RFC13 | 0,702259927 | 0,324012195 | 0,267 | 0,262 | 1 | CH_SC | 1,25180704 | 0,798845164 | 1,019083969 |
| RBM33 | 0,702487932 | 0,604822251 | 0,133 | 0,108 | 1 | CH_SC | 1,52079137 | 0,65755239 | 1,231481481 |
| RANGRF1 | 0,702487932 | 0,38852033 | 0,133 | 0,108 | 1 | CH_SC | 1,309050115 | 0,763912694 | 1,231481481 |
| POLR2E3 | 0,703965768 | 0,585332095 | 0,267 | 0,231 | 1 | CH_SC | 1,500384324 | 0,6664959 | 1,155844156 |
| RALY5 | 0,703965768 | 0,304047522 | 0,267 | 0,231 | 1 | CH_SC | 1,234603274 | 0,809976793 | 1,155844156 |
| HNRNPF2 | 0,704527754 | 0,306096359 | 0,4 | 0,385 | 1 | CH_SC | 1,236357835 | 0,808827324 | 1,038961039 |
| DMAP1 | 0,704804153 | 0,387358189 | 0,067 | 0,108 | 1 | CH_SC | 1,307996054 | 0,7645283 | 0,62037037 |
| GFM1 | 0,704804153 | 0,793550004 | 0,067 | 0,108 | 1 | CH_SC | 1,733334392 | 0,576922724 | 0,62037037 |
| MCCC1 | 0,704804153 | 0,286895896 | 0,067 | 0,108 | 1 | CH_SC | 1,220012472 | 0,819663752 | 0,62037037 |
| KLHL91 | 0,704804153 | 0,642592482 | 0,067 | 0,108 | 1 | CH_SC | 1,56113195 | 0,640560844 | 0,62037037 |
| UBAC12 | 0,704804153 | 0,419249754 | 0,067 | 0,108 | 1 | CH_SC | 1,337231972 | 0,747813409 | 0,62037037 |
| SPRYD31 | 0,704804153 | 0,29992845 | 0,067 | 0,108 | 1 | CH_SC | 1,231083357 | 0,812292681 | 0,62037037 |
| ALKBH2 | 0,704804153 | 0,396004449 | 0,067 | 0,108 | 1 | CH_SC | 1,315858583 | 0,759960084 | 0,62037037 |
| TRPV4 | 0,704804153 | 0,393995881 | 0,067 | 0,108 | 1 | CH_SC | 1,314027875 | 0,761018863 | 0,62037037 |
| MIPOL1 | 0,704804153 | 0,38746268 | 0,067 | 0,108 | 1 | CH_SC | 1,308090793 | 0,764472929 | 0,62037037 |
| SMARCD23 | 0,704804153 | 0,281716556 | 0,067 | 0,108 | 1 | CH_SC | 1,215640424 | 0,82261167 | 0,62037037 |
| CDK5RAP2 | 0,709678871 | 0,263461689 | 0,2 | 0,169 | 1 | CH_SC | 1,200355457 | 0,833086561 | 1,183431953 |
| BRD81 | 0,710704508 | -0,302024459 | 0,2 | 0,231 | 1 | CH_SC | 0,811113404 | 1,232873227 | 0,865800866 |
| DDX189 | 0,711146964 | 0,604436588 | 0,333 | 0,323 | 1 | CH_SC | 1,520384885 | 0,657728191 | 1,030959752 |

| Supplementary Table 4 - clusermarker | | | | | | | | | |
|---|---|---|---|---|---|---|---|---|---|
| | **p_val** | **avg_log2FC** | **pct.1** | **pct.2** | **p_val_adj** | **celltype** | **Foldchange_UP** | **Foldchange_DOWN** | **Ratio_pct1_pct2** |
| NUMA13 | 0,714108695 | 0,283966629 | 0,267 | 0,262 | 1 | CH_SC | 1,217537856 | 0,821329699 | 1,019083969 |
| CEP573 | 0,714108695 | 0,69258432 | 0,267 | 0,262 | 1 | CH_SC | 1,616176006 | 0,618744491 | 1,019083969 |
| BBX2 | 0,71529622 | 0,288796526 | 0,267 | 0,246 | 1 | CH_SC | 1,221620795 | 0,818584625 | 1,085365854 |
| SNHG11 | 0,71529622 | 0,703429487 | 0,267 | 0,246 | 1 | CH_SC | 1,62837106 | 0,614110644 | 1,085365854 |
| ZNF302 | 0,715402394 | 0,29524781 | 0,2 | 0,154 | 1 | CH_SC | 1,227095736 | 0,814932341 | 1,298701299 |
| UQCR105 | 0,715477746 | 0,274303827 | 0,467 | 0,446 | 1 | CH_SC | 1,209410347 | 0,826849218 | 1,047085202 |
| HLTF1 | 0,716267237 | 0,766842503 | 0,267 | 0,231 | 1 | CH_SC | 1,701541694 | 0,587702319 | 1,155844156 |
| ATIC | 0,719224293 | 0,899586838 | 0,133 | 0,108 | 1 | CH_SC | 1,865531652 | 0,536040222 | 1,231481481 |
| COIL2 | 0,719224293 | 0,409947942 | 0,133 | 0,108 | 1 | CH_SC | 1,32863787 | 0,752650532 | 1,231481481 |
| SH3D191 | 0,722702281 | 0,576411915 | 0,267 | 0,292 | 1 | CH_SC | 1,491136072 | 0,670629608 | 0,914383562 |
| TMEM43 | 0,723583929 | 0,331936194 | 0,2 | 0,169 | 1 | CH_SC | 1,258701503 | 0,794469537 | 1,183431953 |
| FAM200B | 0,724443581 | 0,364008934 | 0,267 | 0,277 | 1 | CH_SC | 1,286997217 | 0,777002457 | 0,963898917 |
| SUGT13 | 0,726026804 | 0,472104029 | 0,267 | 0,262 | 1 | CH_SC | 1,387130988 | 0,720912451 | 1,019083969 |
| EIF4E3 | 0,72840499 | -0,523877807 | 0,133 | 0,154 | 1 | CH_SC | 0,69549989 | 1,437814748 | 0,863636364 |
| ACD | 0,72840499 | -0,274630081 | 0,133 | 0,154 | 1 | CH_SC | 0,826662254 | 1,209683876 | 0,863636364 |
| GPS1 | 0,72840499 | -0,383219847 | 0,133 | 0,154 | 1 | CH_SC | 0,766724482 | 1,30424947 | 0,863636364 |
| SEC23A | 0,729778495 | -0,277033349 | 0,133 | 0,169 | 1 | CH_SC | 0,825286331 | 1,21170067 | 0,786982249 |
| PDCD6IP | 0,730356932 | -0,271996014 | 0,2 | 0,215 | 1 | CH_SC | 0,828172949 | 1,207477256 | 0,930232558 |
| ANGPTL12 | 0,731352594 | -0,31586554 | 0,133 | 0,092 | 1 | CH_SC | 0,803368867 | 1,244758218 | 1,445652174 |
| OSBP1 | 0,733528515 | 0,495003632 | 0,2 | 0,262 | 1 | CH_SC | 1,409324303 | 0,709559892 | 0,763358779 |
| DEK3 | 0,734479426 | 0,261578056 | 0,467 | 0,538 | 1 | CH_SC | 1,198789254 | 0,834174979 | 0,86802974 |
| TAF12 | 0,73609638 | 0,280292084 | 0,133 | 0,108 | 1 | CH_SC | 1,214440731 | 0,823424292 | 1,231481481 |
| UHMK1 | 0,73609638 | 0,73857911 | 0,133 | 0,108 | 1 | CH_SC | 1,668531716 | 0,599329333 | 1,231481481 |
| SLC39A102 | 0,73609638 | 0,407183535 | 0,133 | 0,108 | 1 | CH_SC | 1,326094451 | 0,754094099 | 1,231481481 |
| EEF1E1 | 0,73609638 | 0,291975335 | 0,133 | 0,108 | 1 | CH_SC | 1,224315461 | 0,816782955 | 1,231481481 |
| HAX11 | 0,736389327 | 0,252799555 | 0,333 | 0,338 | 1 | CH_SC | 1,191517017 | 0,839266235 | 0,985207101 |
| OBSL1 | 0,73758083 | 0,320281679 | 0,2 | 0,169 | 1 | CH_SC | 1,248574303 | 0,800913488 | 1,183431953 |
| TMEM14B3 | 0,741360499 | 0,619244813 | 0,333 | 0,308 | 1 | CH_SC | 1,536070906 | 0,651011614 | 1,081168831 |
| NAMPT3 | 0,741360499 | 0,396273978 | 0,333 | 0,308 | 1 | CH_SC | 1,316104439 | 0,75981812 | 1,081168831 |
| KRCC12 | 0,743361369 | -0,349397538 | 0,133 | 0,154 | 1 | CH_SC | 0,784911805 | 1,274028489 | 0,863636364 |
| CRISPLD21 | 0,743361369 | -0,334228285 | 0,133 | 0,154 | 1 | CH_SC | 0,79320832 | 1,260702864 | 0,863636364 |

| Supplementary Table 4 - clusermarker | | | | | | | | | |
|---|---|---|---|---|---|---|---|---|---|
| | **p_val** | **avg_log2FC** | **pct.1** | **pct.2** | **p_val_adj** | **celltype** | **Foldchange_UP** | **Foldchange_DOWN** | **Ratio_pct1_pct2** |
| TOB21 | 0,744539058 | 0,855229626 | 0,2 | 0,185 | 1 | CH_SC | 1,809046669 | 0,552777337 | 1,081081081 |
| LRIG31 | 0,744612388 | 0,590611093 | 0,133 | 0,185 | 1 | CH_SC | 1,505884471 | 0,664061566 | 0,718918919 |
| NDUFA35 | 0,75006045 | 0,639541275 | 0,267 | 0,262 | 1 | CH_SC | 1,557833746 | 0,641917023 | 1,019083969 |
| REST2 | 0,751665297 | 0,413886034 | 0,2 | 0,169 | 1 | CH_SC | 1,332269578 | 0,75059884 | 1,183431953 |
| HNRNPH12 | 0,752730752 | 0,803083596 | 0,333 | 0,369 | 1 | CH_SC | 1,744826512 | 0,573122883 | 0,902439024 |
| ZMYM61 | 0,753096711 | 0,651289082 | 0,133 | 0,108 | 1 | CH_SC | 1,570570911 | 0,636711143 | 1,231481481 |
| NFKBIL1 | 0,753096711 | 0,321750818 | 0,133 | 0,108 | 1 | CH_SC | 1,249846411 | 0,800098309 | 1,231481481 |
| SCYL1 | 0,753096711 | 0,656089499 | 0,133 | 0,108 | 1 | CH_SC | 1,575805526 | 0,634596074 | 1,231481481 |
| CTTN2 | 0,753096711 | 0,522658961 | 0,133 | 0,108 | 1 | CH_SC | 1,436600538 | 0,696087725 | 1,231481481 |
| FAM168A | 0,753096711 | 0,439240327 | 0,133 | 0,108 | 1 | CH_SC | 1,355890175 | 0,73752286 | 1,231481481 |
| AC018521.52 | 0,753096711 | 0,317756494 | 0,133 | 0,108 | 1 | CH_SC | 1,246390804 | 0,802316574 | 1,231481481 |
| SCAF114 | 0,753604286 | 0,51564515 | 0,267 | 0,231 | 1 | CH_SC | 1,429633306 | 0,699480066 | 1,155844156 |
| VPS285 | 0,754111231 | 0,292540858 | 0,4 | 0,4 | 1 | CH_SC | 1,224795475 | 0,816462846 | 1 |
| FRA10AC1 | 0,758275834 | 0,585433223 | 0,2 | 0,185 | 1 | CH_SC | 1,500489499 | 0,666449182 | 1,081081081 |
| SEC22B | 0,758415227 | -0,344831988 | 0,133 | 0,154 | 1 | CH_SC | 0,787399669 | 1,270003074 | 0,863636364 |
| UQCC33 | 0,758415227 | -0,411314244 | 0,133 | 0,154 | 1 | CH_SC | 0,751938073 | 1,329896752 | 0,863636364 |
| ARMCX61 | 0,758415227 | -0,338062996 | 0,133 | 0,154 | 1 | CH_SC | 0,791102757 | 1,264058293 | 0,863636364 |
| TMBIM46 | 0,761346849 | 0,338930226 | 0,333 | 0,431 | 1 | CH_SC | 1,26481837 | 0,790627353 | 0,77262181 |
| TCEAL81 | 0,762170573 | 0,526649474 | 0,267 | 0,262 | 1 | CH_SC | 1,440579694 | 0,694164998 | 1,019083969 |
| CIAO2B6 | 0,764247675 | 0,535803066 | 0,267 | 0,246 | 1 | CH_SC | 1,449748923 | 0,689774612 | 1,085365854 |
| MOB4 | 0,765832971 | 0,259895281 | 0,2 | 0,169 | 1 | CH_SC | 1,197391788 | 0,835148537 | 1,183431953 |
| LGALS36 | 0,766747392 | 0,287476062 | 0,733 | 0,846 | 1 | CH_SC | 1,220503187 | 0,819334198 | 0,86643026 |
| MAGOH2 | 0,767961713 | 0,448495114 | 0,267 | 0,215 | 1 | CH_SC | 1,364616073 | 0,732806846 | 1,241860465 |
| TNPO13 | 0,767961713 | 0,522498502 | 0,267 | 0,215 | 1 | CH_SC | 1,436440765 | 0,696165149 | 1,241860465 |
| EIF3I2 | 0,769729112 | 0,256999768 | 0,267 | 0,354 | 1 | CH_SC | 1,194991013 | 0,836826377 | 0,754237288 |
| BOLA31 | 0,770217615 | 0,256271044 | 0,133 | 0,108 | 1 | CH_SC | 1,19438756 | 0,837249176 | 1,231481481 |
| MRPL13 | 0,770217615 | 0,402742327 | 0,133 | 0,108 | 1 | CH_SC | 1,322018465 | 0,756419087 | 1,231481481 |
| IKZF5 | 0,770217615 | 0,412059514 | 0,133 | 0,108 | 1 | CH_SC | 1,33058393 | 0,751549735 | 1,231481481 |
| TIMM21 | 0,770217615 | 0,412519378 | 0,133 | 0,108 | 1 | CH_SC | 1,331008125 | 0,751310215 | 1,231481481 |
| ZMYND81 | 0,770217615 | 0,341811763 | 0,133 | 0,108 | 1 | CH_SC | 1,267347153 | 0,789049786 | 1,231481481 |
| UBE2G2 | 0,770217615 | 0,457861335 | 0,133 | 0,108 | 1 | CH_SC | 1,373504213 | 0,728064749 | 1,231481481 |

| Supplementary Table 4 - clusermarker | | | | | | | | | |
|---|---|---|---|---|---|---|---|---|---|
| | **p_val** | **avg_log2FC** | **pct.1** | **pct.2** | **p_val_adj** | **celltype** | **Foldchange_UP** | **Foldchange_DOWN** | **Ratio_pct1_pct2** |
| TUBA1C3 | 0,770438061 | 0,337938495 | 0,2 | 0,262 | 1 | CH_SC | 1,263949212 | 0,79117103 | 0,763358779 |
| TMEM167A2 | 0,77208935 | 0,421793129 | 0,2 | 0,185 | 1 | CH_SC | 1,339591502 | 0,746496225 | 1,081081081 |
| GADD45G3 | 0,773561235 | -0,423899529 | 0,133 | 0,154 | 1 | CH_SC | 0,745407103 | 1,341548794 | 0,863636364 |
| UACA4 | 0,781236236 | 0,648763673 | 0,267 | 0,292 | 1 | CH_SC | 1,567824063 | 0,637826669 | 0,914383562 |
| NEU12 | 0,785975483 | 0,817076016 | 0,2 | 0,185 | 1 | CH_SC | 1,761831577 | 0,567591144 | 1,081081081 |
| ARIH12 | 0,785975483 | 0,302830487 | 0,2 | 0,185 | 1 | CH_SC | 1,233562221 | 0,810660365 | 1,081081081 |
| UAP14 | 0,786563544 | 0,369375633 | 0,267 | 0,262 | 1 | CH_SC | 1,29179365 | 0,774117445 | 1,019083969 |
| CETN21 | 0,787451246 | -0,255438404 | 0,133 | 0,108 | 1 | CH_SC | 0,837732527 | 1,193698427 | 1,231481481 |
| SUCLA2 | 0,788191218 | 0,418654166 | 0,133 | 0,169 | 1 | CH_SC | 1,336680036 | 0,748122193 | 0,786982249 |
| CHCHD11 | 0,788560779 | 0,467009916 | 0,133 | 0,123 | 1 | CH_SC | 1,382241707 | 0,72346247 | 1,081300813 |
| MIR100HG1 | 0,788560779 | 0,886425337 | 0,133 | 0,123 | 1 | CH_SC | 1,848590068 | 0,540952814 | 1,081300813 |
| MAPK61 | 0,788560779 | 0,443743724 | 0,133 | 0,123 | 1 | CH_SC | 1,360129222 | 0,73522426 | 1,081300813 |
| CAPRIN13 | 0,789755898 | 0,668928534 | 0,333 | 0,323 | 1 | CH_SC | 1,589891742 | 0,628973642 | 1,030959752 |
| MIR22HG4 | 0,791526606 | 0,278645994 | 0,267 | 0,231 | 1 | CH_SC | 1,213055867 | 0,824364341 | 1,155844156 |
| NUFIP24 | 0,792198883 | 0,532855766 | 0,333 | 0,338 | 1 | CH_SC | 1,446790235 | 0,691185201 | 0,985207101 |
| OS92 | 0,795088682 | 0,444498022 | 0,533 | 0,538 | 1 | CH_SC | 1,360840537 | 0,734839956 | 0,99070632 |
| NAP1L43 | 0,795342443 | 0,498463713 | 0,2 | 0,262 | 1 | CH_SC | 1,412708406 | 0,707860161 | 0,763358779 |
| MYLIP4 | 0,798840712 | 0,340255888 | 0,2 | 0,277 | 1 | CH_SC | 1,265981119 | 0,789901196 | 0,722021661 |
| ANP32B3 | 0,798840712 | 0,277142049 | 0,267 | 0,262 | 1 | CH_SC | 1,211791968 | 0,825224152 | 1,019083969 |
| GIGYF2 | 0,799930041 | 0,403223078 | 0,2 | 0,185 | 1 | CH_SC | 1,322459076 | 0,756167067 | 1,081081081 |
| TMX23 | 0,799930041 | 0,591286582 | 0,2 | 0,185 | 1 | CH_SC | 1,506589711 | 0,663750716 | 1,081081081 |
| C1D2 | 0,800350717 | 1,050546893 | 0,2 | 0,231 | 1 | CH_SC | 2,071314887 | 0,482785117 | 0,865800866 |
| CERS21 | 0,801184092 | 0,852737107 | 0,333 | 0,323 | 1 | CH_SC | 1,805923908 | 0,553733186 | 1,030959752 |
| MPC23 | 0,801602358 | 0,383893383 | 0,267 | 0,246 | 1 | CH_SC | 1,304858514 | 0,766366613 | 1,085365854 |
| IRF2BPL1 | 0,802997134 | -0,390828937 | 0,2 | 0,154 | 1 | CH_SC | 0,762691255 | 1,31114654 | 1,298701299 |
| LSM122 | 0,802997134 | 0,250090974 | 0,133 | 0,169 | 1 | CH_SC | 1,189282107 | 0,840843391 | 0,786982249 |
| NR4A35 | 0,803500127 | -0,853386506 | 0,333 | 0,338 | 1 | CH_SC | 0,553483991 | 1,80673699 | 0,985207101 |
| CHD11 | 0,804282806 | 0,610514237 | 0,267 | 0,231 | 1 | CH_SC | 1,526803329 | 0,654963204 | 1,155844156 |
| TGIF13 | 0,805022856 | -0,288718221 | 0,267 | 0,292 | 1 | CH_SC | 0,818629056 | 1,221554491 | 0,914383562 |
| ZFC3H11 | 0,805120603 | 0,854505874 | 0,133 | 0,123 | 1 | CH_SC | 1,808139357 | 0,553054717 | 1,081300813 |
| PFDN61 | 0,808790378 | 0,391321609 | 0,2 | 0,169 | 1 | CH_SC | 1,311594366 | 0,762430845 | 1,183431953 |

| Supplementary Table 4 - clusermarker | | | | | | | | | |
|---|---|---|---|---|---|---|---|---|---|
| | **p_val** | **avg_log2FC** | **pct.1** | **pct.2** | **p_val_adj** | **celltype** | **Foldchange_UP** | **Foldchange_DOWN** | **Ratio_pct1_pct2** |
| YAE11 | 0,808790378 | 0,277551005 | 0,2 | 0,169 | 1 | CH_SC | 1,21213552 | 0,824990262 | 1,183431953 |
| MESD3 | 0,809357067 | -0,306253857 | 0,2 | 0,215 | 1 | CH_SC | 0,80873903 | 1,236492816 | 0,930232558 |
| IDH1 | 0,816010187 | 0,624119748 | 0,133 | 0,185 | 1 | CH_SC | 1,541270142 | 0,648815527 | 0,718918919 |
| SLC35A11 | 0,816010187 | 0,488375896 | 0,133 | 0,185 | 1 | CH_SC | 1,402864721 | 0,712827107 | 0,718918919 |
| POGLUT3 | 0,816010187 | 0,337225037 | 0,133 | 0,185 | 1 | CH_SC | 1,263324304 | 0,791562386 | 0,718918919 |
| ABI11 | 0,817874098 | -0,325885507 | 0,2 | 0,154 | 1 | CH_SC | 0,797808552 | 1,253433543 | 1,298701299 |
| UVRAG2 | 0,817874098 | 0,319811686 | 0,133 | 0,169 | 1 | CH_SC | 1,248167616 | 0,801174448 | 0,786982249 |
| ING1 | 0,818572072 | -0,364111037 | 0,2 | 0,2 | 1 | CH_SC | 0,776947469 | 1,287088304 | 1 |
| GSPT13 | 0,818572072 | 0,622620454 | 0,2 | 0,2 | 1 | CH_SC | 1,539669237 | 0,649490148 | 1 |
| ZNF7066 | 0,818775253 | 0,634259342 | 0,267 | 0,385 | 1 | CH_SC | 1,5521407 | 0,644271489 | 0,693506494 |
| AK6 | 0,81949732 | -0,318371916 | 0,133 | 0,154 | 1 | CH_SC | 0,801974397 | 1,2469226 | 0,863636364 |
| DNAJB19 | 0,820587368 | 0,336944316 | 0,6 | 0,662 | 1 | CH_SC | 1,26307851 | 0,791716423 | 0,906344411 |
| PHF5A | 0,820821901 | -0,264275254 | 0,133 | 0,138 | 1 | CH_SC | 0,832616899 | 1,201032553 | 0,963768116 |
| SYNPO | 0,821768312 | 0,328977436 | 0,133 | 0,123 | 1 | CH_SC | 1,256122735 | 0,79610055 | 1,081300813 |
| DNAJB121 | 0,821768312 | 0,830558986 | 0,133 | 0,123 | 1 | CH_SC | 1,778374277 | 0,562311327 | 1,081300813 |
| EPN11 | 0,821768312 | 0,419301458 | 0,133 | 0,123 | 1 | CH_SC | 1,337279898 | 0,747786609 | 1,081300813 |
| CCNL15 | 0,821779772 | 0,375641072 | 0,467 | 0,492 | 1 | CH_SC | 1,297415943 | 0,770762843 | 0,949186992 |
| NUTM2A-AS11 | 0,82222448 | -0,26751268 | 0,133 | 0,108 | 1 | CH_SC | 0,830750592 | 1,203730711 | 1,231481481 |
| ARFRP1 | 0,82222448 | -0,266572364 | 0,133 | 0,108 | 1 | CH_SC | 0,831292232 | 1,202946402 | 1,231481481 |
| PLEKHA11 | 0,82802734 | 0,663378961 | 0,2 | 0,185 | 1 | CH_SC | 1,5837877 | 0,631397756 | 1,081081081 |
| NDUFA63 | 0,829936235 | 0,254057738 | 0,267 | 0,338 | 1 | CH_SC | 1,192556599 | 0,838534625 | 0,789940828 |
| TMX44 | 0,831873683 | 0,458555298 | 0,4 | 0,431 | 1 | CH_SC | 1,374165053 | 0,727714621 | 0,928074246 |
| TMEM1232 | 0,8323101 | 0,892850229 | 0,2 | 0,2 | 1 | CH_SC | 1,85684093 | 0,538549094 | 1 |
| SMARCA52 | 0,832312084 | 0,329635852 | 0,2 | 0,2 | 1 | CH_SC | 1,256696134 | 0,795737309 | 1 |
| GHITM1 | 0,833140616 | 0,5875589 | 0,4 | 0,385 | 1 | CH_SC | 1,502701961 | 0,665467954 | 1,038961039 |
| CITED23 | 0,835949895 | 0,414821618 | 0,4 | 0,415 | 1 | CH_SC | 1,333133832 | 0,750112236 | 0,963855422 |
| ZMYND11 | 0,836173505 | 0,622389169 | 0,2 | 0,215 | 1 | CH_SC | 1,539422426 | 0,649594278 | 0,930232558 |
| ARL8B2 | 0,836870454 | -0,362968349 | 0,133 | 0,138 | 1 | CH_SC | 0,777563094 | 1,286069269 | 0,963768116 |
| APCDD11 | 0,836870454 | -0,460221441 | 0,133 | 0,138 | 1 | CH_SC | 0,726874681 | 1,375752968 | 0,963768116 |
| HIPK2 | 0,838496672 | 0,769786186 | 0,133 | 0,123 | 1 | CH_SC | 1,705017073 | 0,586504391 | 1,081300813 |
| NRDE21 | 0,838496672 | 0,261460967 | 0,133 | 0,123 | 1 | CH_SC | 1,198691964 | 0,834242683 | 1,081300813 |

| Supplementary Table 4 - clusermarker | | | | | | | | | |
|---|---|---|---|---|---|---|---|---|---|
| | **p_val** | **avg_log2FC** | **pct.1** | **pct.2** | **p_val_adj** | **celltype** | **Foldchange_UP** | **Foldchange_DOWN** | **Ratio_pct1_pct2** |
| MAPK11 | 0,838496672 | 0,80412024 | 0,133 | 0,123 | 1 | CH_SC | 1,746080701 | 0,572711215 | 1,081300813 |
| AMD15 | 0,84110485 | 0,674623913 | 0,333 | 0,369 | 1 | CH_SC | 1,59618062 | 0,626495515 | 0,902439024 |
| MED41 | 0,842161385 | 0,307398595 | 0,2 | 0,185 | 1 | CH_SC | 1,237474327 | 0,808097573 | 1,081081081 |
| CLINT12 | 0,842852157 | 0,51818952 | 0,2 | 0,246 | 1 | CH_SC | 1,432156865 | 0,698247535 | 0,81300813 |
| HES15 | 0,844729172 | 0,37245421 | 0,267 | 0,277 | 1 | CH_SC | 1,29455316 | 0,772467312 | 0,963898917 |
| PSMD31 | 0,846103524 | 0,360117653 | 0,2 | 0,2 | 1 | CH_SC | 1,283530566 | 0,779101041 | 1 |
| PRPF313 | 0,846103524 | 0,278162302 | 0,2 | 0,2 | 1 | CH_SC | 1,212649233 | 0,824640772 | 1 |
| RC3H1 | 0,847820452 | -0,438159695 | 0,2 | 0,154 | 1 | CH_SC | 0,738075499 | 1,354874943 | 1,298701299 |
| PSIP11 | 0,848419205 | -0,308889068 | 0,267 | 0,262 | 1 | CH_SC | 0,807263144 | 1,238753444 | 1,019083969 |
| TMEM179B5 | 0,848419205 | 0,265886539 | 0,2 | 0,277 | 1 | CH_SC | 1,202374685 | 0,831687504 | 0,722021661 |
| UFL11 | 0,852108338 | 0,532775145 | 0,267 | 0,246 | 1 | CH_SC | 1,446709388 | 0,691223827 | 1,085365854 |
| AP2M15 | 0,852318966 | 0,341177099 | 0,4 | 0,354 | 1 | CH_SC | 1,26678975 | 0,789396978 | 1,129943503 |
| ABCE12 | 0,852987186 | -0,410645197 | 0,133 | 0,138 | 1 | CH_SC | 0,752286863 | 1,329280157 | 0,963768116 |
| IRF22 | 0,852987186 | -0,302289443 | 0,133 | 0,138 | 1 | CH_SC | 0,810964438 | 1,233099693 | 0,963768116 |
| STAU11 | 0,853120081 | 0,267333048 | 0,267 | 0,292 | 1 | CH_SC | 1,203580842 | 0,830854036 | 0,914383562 |
| PITHD12 | 0,855298339 | 0,384868413 | 0,133 | 0,123 | 1 | CH_SC | 1,305740686 | 0,765848848 | 1,081300813 |
| OTUD4 | 0,857350526 | -0,25784483 | 0,133 | 0,108 | 1 | CH_SC | 0,836336348 | 1,195691187 | 1,231481481 |
| RANBP22 | 0,858917951 | 0,437157996 | 0,333 | 0,323 | 1 | CH_SC | 1,353934546 | 0,738588141 | 1,030959752 |
| COPB22 | 0,858917951 | 0,466265342 | 0,333 | 0,323 | 1 | CH_SC | 1,381528517 | 0,723835945 | 1,030959752 |
| PPP2CA1 | 0,860916347 | 0,329857011 | 0,267 | 0,262 | 1 | CH_SC | 1,256888795 | 0,795615335 | 1,019083969 |
| MTR | 0,862879305 | 0,405470962 | 0,133 | 0,169 | 1 | CH_SC | 1,324521225 | 0,754989789 | 0,786982249 |
| POLR3GL1 | 0,863181728 | 0,478992115 | 0,2 | 0,215 | 1 | CH_SC | 1,393769621 | 0,717478689 | 0,930232558 |
| TMF1 | 0,863181728 | 0,3570987 | 0,2 | 0,215 | 1 | CH_SC | 1,280847486 | 0,780733078 | 0,930232558 |
| WASHC31 | 0,866061465 | 0,411154316 | 0,133 | 0,154 | 1 | CH_SC | 1,329749336 | 0,752021432 | 0,863636364 |
| SON10 | 0,867483699 | 0,347192255 | 0,667 | 0,862 | 1 | CH_SC | 1,272082515 | 0,786112527 | 0,773781903 |
| TMEM14A2 | 0,872092545 | 0,265863082 | 0,333 | 0,338 | 1 | CH_SC | 1,202355135 | 0,831701027 | 0,985207101 |
| DSE2 | 0,872165868 | 0,339593898 | 0,133 | 0,123 | 1 | CH_SC | 1,265400348 | 0,790263731 | 1,081300813 |
| FKBP41 | 0,872165868 | 0,301671626 | 0,133 | 0,123 | 1 | CH_SC | 1,232571746 | 0,811311799 | 1,081300813 |
| HAUS43 | 0,872165868 | 0,259969218 | 0,133 | 0,123 | 1 | CH_SC | 1,197453155 | 0,835105737 | 1,081300813 |
| CNOT13 | 0,872165868 | 0,303256946 | 0,133 | 0,123 | 1 | CH_SC | 1,233926914 | 0,81042077 | 1,081300813 |
| RAB5B2 | 0,873824179 | 0,316043312 | 0,2 | 0,2 | 1 | CH_SC | 1,244911609 | 0,803269881 | 1 |

| Supplementary Table 4 - clusermarker | | | | | | | | | |
|---|---|---|---|---|---|---|---|---|---|
| | **p_val** | **avg_log2FC** | **pct.1** | **pct.2** | **p_val_adj** | **celltype** | **Foldchange_UP** | **Foldchange_DOWN** | **Ratio_pct1_pct2** |
| SPSB34 | 0,873824179 | 0,395782324 | 0,2 | 0,2 | 1 | CH_SC | 1,315656002 | 0,760077101 | 1 |
| GBF12 | 0,875024679 | -0,370167331 | 0,133 | 0,108 | 1 | CH_SC | 0,773692755 | 1,292502732 | 1,231481481 |
| TIMP4 | 0,87798815 | 0,502665482 | 0,133 | 0,169 | 1 | CH_SC | 1,416828838 | 0,705801557 | 0,786982249 |
| TXN5 | 0,878085083 | -0,427021823 | 0,6 | 0,585 | 1 | CH_SC | 0,74379563 | 1,344455331 | 1,025641026 |
| SRPK21 | 0,879395208 | 0,259650904 | 0,2 | 0,231 | 1 | CH_SC | 1,19718898 | 0,835290014 | 0,865800866 |
| RUNX13 | 0,879395208 | 0,262088351 | 0,2 | 0,231 | 1 | CH_SC | 1,199213351 | 0,833879975 | 0,865800866 |
| SIVA15 | 0,880745886 | 0,439280347 | 0,333 | 0,308 | 1 | CH_SC | 1,355927787 | 0,737502402 | 1,081168831 |
| PIBF12 | 0,881695208 | 0,532523524 | 0,133 | 0,154 | 1 | CH_SC | 1,446457089 | 0,691344394 | 0,863636364 |
| COX5B5 | 0,884749134 | 0,358391267 | 0,467 | 0,508 | 1 | CH_SC | 1,281995562 | 0,780033902 | 0,919291339 |
| TKT3 | 0,884851798 | 0,621749793 | 0,2 | 0,185 | 1 | CH_SC | 1,538740333 | 0,649882231 | 1,081081081 |
| SP1102 | 0,885398826 | -0,337140766 | 0,133 | 0,138 | 1 | CH_SC | 0,791608623 | 1,263250514 | 0,963768116 |
| CELF1 | 0,885398826 | -0,282092569 | 0,133 | 0,138 | 1 | CH_SC | 0,822397299 | 1,215957302 | 0,963768116 |
| UBE2H2 | 0,887745033 | 0,370250359 | 0,2 | 0,2 | 1 | CH_SC | 1,292577119 | 0,77364823 | 1 |
| ZNHIT31 | 0,889091725 | 0,400343113 | 0,133 | 0,123 | 1 | CH_SC | 1,319821764 | 0,757678065 | 1,081300813 |
| SCRN21 | 0,889091725 | 0,353633395 | 0,133 | 0,123 | 1 | CH_SC | 1,277774625 | 0,782610626 | 1,081300813 |
| ARGLU13 | 0,889565376 | 0,370066977 | 0,267 | 0,292 | 1 | CH_SC | 1,292412829 | 0,773746575 | 0,914383562 |
| ATP2B13 | 0,89035082 | 0,41294441 | 0,2 | 0,215 | 1 | CH_SC | 1,331400311 | 0,751088904 | 0,930232558 |
| CKB3 | 0,892594982 | 0,294546013 | 0,267 | 0,323 | 1 | CH_SC | 1,226498962 | 0,815328859 | 0,826625387 |
| TAOK13 | 0,892709605 | 0,404220855 | 0,2 | 0,231 | 1 | CH_SC | 1,323374013 | 0,755644277 | 0,865800866 |
| TMEM1345 | 0,893141571 | 0,290975074 | 0,133 | 0,169 | 1 | CH_SC | 1,223466903 | 0,81734945 | 0,786982249 |
| PIGH1 | 0,893141571 | 0,334906998 | 0,133 | 0,169 | 1 | CH_SC | 1,261296098 | 0,792835244 | 0,786982249 |
| UBE2R22 | 0,894053708 | -0,267285996 | 0,267 | 0,277 | 1 | CH_SC | 0,830881134 | 1,203541589 | 0,963898917 |
| COPS23 | 0,894053708 | -0,333722614 | 0,267 | 0,277 | 1 | CH_SC | 0,793486392 | 1,260261059 | 0,963898917 |
| TIMM8B2 | 0,897375184 | 0,281955876 | 0,133 | 0,154 | 1 | CH_SC | 1,215842096 | 0,822475224 | 0,863636364 |
| PIGT | 0,89860493 | -0,300500634 | 0,267 | 0,262 | 1 | CH_SC | 0,811970583 | 1,231571711 | 1,019083969 |
| TTC191 | 0,899162948 | 0,566107693 | 0,2 | 0,185 | 1 | CH_SC | 1,480523811 | 0,675436621 | 1,081081081 |
| DNAJC101 | 0,90168037 | -0,290609803 | 0,133 | 0,138 | 1 | CH_SC | 0,817556418 | 1,223157176 | 0,963768116 |
| TOP2B | 0,90168037 | 0,468117179 | 0,133 | 0,138 | 1 | CH_SC | 1,383302979 | 0,722907429 | 0,963768116 |
| CAND11 | 0,90168037 | 0,434231482 | 0,133 | 0,138 | 1 | CH_SC | 1,351190867 | 0,740087892 | 0,963768116 |
| ELP51 | 0,90168037 | 0,432580202 | 0,133 | 0,138 | 1 | CH_SC | 1,349645206 | 0,740935466 | 0,963768116 |
| AC005670.3 | 0,90168037 | -0,339307963 | 0,133 | 0,138 | 1 | CH_SC | 0,790420372 | 1,265149577 | 0,963768116 |

| Supplementary Table 4 - clusermarker | | | | | | | | | |
|---|---|---|---|---|---|---|---|---|---|
| | **p_val** | **avg_log2FC** | **pct.1** | **pct.2** | **p_val_adj** | **celltype** | **Foldchange_UP** | **Foldchange_DOWN** | **Ratio_pct1_pct2** |
| MRPL273 | 0,90168037 | 0,38985702 | 0,133 | 0,138 | 1 | CH_SC | 1,310263543 | 0,763205239 | 0,963768116 |
| OSBPL92 | 0,901700602 | 0,300311707 | 0,2 | 0,2 | 1 | CH_SC | 1,231410442 | 0,812076921 | 1 |
| TOP12 | 0,901768918 | 0,326592297 | 0,267 | 0,292 | 1 | CH_SC | 1,254047763 | 0,797417793 | 0,914383562 |
| H2AZ13 | 0,903001127 | 0,316379097 | 0,533 | 0,585 | 1 | CH_SC | 1,245201394 | 0,803082943 | 0,911111111 |
| SEC22C2 | 0,906068297 | 0,405450977 | 0,133 | 0,123 | 1 | CH_SC | 1,324502877 | 0,755000248 | 1,081300813 |
| NHLRC21 | 0,906068297 | 0,711549033 | 0,133 | 0,123 | 1 | CH_SC | 1,637561436 | 0,610664112 | 1,081300813 |
| ZFAND61 | 0,906068297 | 0,426687129 | 0,133 | 0,123 | 1 | CH_SC | 1,344143464 | 0,743968205 | 1,081300813 |
| ACOX11 | 0,906068297 | 0,668771563 | 0,133 | 0,123 | 1 | CH_SC | 1,589718765 | 0,629042081 | 1,081300813 |
| CZIB | 0,908334103 | 0,602588999 | 0,133 | 0,169 | 1 | CH_SC | 1,518439049 | 0,658571051 | 0,786982249 |
| ENSA6 | 0,910551919 | -0,273957627 | 0,133 | 0,108 | 1 | CH_SC | 0,827047659 | 1,209120163 | 1,231481481 |
| SGK3 | 0,910551919 | -0,274624976 | 0,133 | 0,108 | 1 | CH_SC | 0,826665179 | 1,209679596 | 1,231481481 |
| ECHDC3 | 0,910551919 | -0,278828907 | 0,133 | 0,108 | 1 | CH_SC | 0,82425983 | 1,213209674 | 1,231481481 |
| WBP15 | 0,915686617 | 0,282314955 | 0,2 | 0,2 | 1 | CH_SC | 1,216144751 | 0,822270539 | 1 |
| RAP1B3 | 0,915686617 | 0,288343674 | 0,2 | 0,2 | 1 | CH_SC | 1,221237397 | 0,818841613 | 1 |
| SOX153 | 0,915686617 | 0,31823095 | 0,2 | 0,2 | 1 | CH_SC | 1,246800769 | 0,802052762 | 1 |
| PCNX1 | 0,91800338 | -0,258388418 | 0,133 | 0,138 | 1 | CH_SC | 0,836021287 | 1,196141792 | 0,963768116 |
| PCBP23 | 0,91818093 | 0,72167513 | 0,4 | 0,431 | 1 | CH_SC | 1,649095707 | 0,606392943 | 0,928074246 |
| EIF1AX8 | 0,91818093 | 0,322120704 | 0,4 | 0,431 | 1 | CH_SC | 1,250166894 | 0,799893202 | 0,928074246 |
| CAPN24 | 0,918882931 | 0,277570966 | 0,267 | 0,277 | 1 | CH_SC | 1,212152291 | 0,824978848 | 0,963898917 |
| SSBP22 | 0,921040386 | -0,306973773 | 0,267 | 0,231 | 1 | CH_SC | 0,808335564 | 1,237109989 | 1,155844156 |
| UPF24 | 0,921260736 | 0,591080958 | 0,267 | 0,308 | 1 | CH_SC | 1,506374996 | 0,663845326 | 0,866883117 |
| TACC15 | 0,923015601 | 0,255830787 | 0,333 | 0,431 | 1 | CH_SC | 1,194023133 | 0,837504712 | 0,77262181 |
| PPP1CC3 | 0,925846479 | -0,334196649 | 0,2 | 0,169 | 1 | CH_SC | 0,793225714 | 1,260675218 | 1,183431953 |
| C15orf611 | 0,928849573 | 0,325582513 | 0,133 | 0,154 | 1 | CH_SC | 1,253170325 | 0,797976125 | 0,863636364 |
| RGS164 | 0,933343682 | 0,507030616 | 0,267 | 0,308 | 1 | CH_SC | 1,421122201 | 0,703669255 | 0,866883117 |
| WDR331 | 0,934361025 | 0,457240228 | 0,133 | 0,138 | 1 | CH_SC | 1,372913021 | 0,728378262 | 0,963768116 |
| MTREX2 | 0,934361025 | 0,253404197 | 0,133 | 0,138 | 1 | CH_SC | 1,192016493 | 0,838914567 | 0,963768116 |
| RAB213 | 0,936523649 | -0,284567521 | 0,267 | 0,262 | 1 | CH_SC | 0,820987681 | 1,218045073 | 1,019083969 |
| TAF1D1 | 0,938511207 | 0,353272618 | 0,267 | 0,292 | 1 | CH_SC | 1,27745513 | 0,78280636 | 0,914383562 |
| ZCRB13 | 0,940205268 | 0,634589185 | 0,267 | 0,323 | 1 | CH_SC | 1,552495606 | 0,644124206 | 0,826625387 |
| CCN12 | 0,940806265 | -0,253103141 | 0,733 | 0,815 | 1 | CH_SC | 0,839089647 | 1,191767773 | 0,899386503 |

| Supplementary Table 4 - clusermarker | | | | | | | | | |
|---|---|---|---|---|---|---|---|---|---|
| | **p_val** | **avg_log2FC** | **pct.1** | **pct.2** | **p_val_adj** | **celltype** | **Foldchange_UP** | **Foldchange_DOWN** | **Ratio_pct1_pct2** |
| VPS351 | 0,94379151 | 0,268842973 | 0,267 | 0,277 | 1 | CH_SC | 1,20484117 | 0,829984919 | 0,963898917 |
| TIMP21 | 0,950405471 | -0,445998084 | 0,733 | 0,738 | 1 | CH_SC | 0,734076293 | 1,362256226 | 0,993224932 |
| RYK1 | 0,950746431 | -0,274510319 | 0,133 | 0,138 | 1 | CH_SC | 0,82673088 | 1,209583461 | 0,963768116 |
| CNIH1 | 0,956267982 | 0,268069882 | 0,267 | 0,277 | 1 | CH_SC | 1,204195709 | 0,830429798 | 0,963898917 |
| ZC3H61 | 0,957225176 | -0,474981027 | 0,133 | 0,123 | 1 | CH_SC | 0,719476252 | 1,389899941 | 1,081300813 |
| FAF2 | 0,957225176 | -0,311793827 | 0,133 | 0,123 | 1 | CH_SC | 0,805639415 | 1,241250094 | 1,081300813 |
| HSPA1A9 | 0,960134327 | -0,592642273 | 0,733 | 0,692 | 1 | CH_SC | 0,663127287 | 1,508006109 | 1,059248555 |
| RECQL2 | 0,967152689 | 0,378345094 | 0,133 | 0,138 | 1 | CH_SC | 1,29984995 | 0,769319566 | 0,963768116 |
| SH3KBP12 | 0,967152689 | 0,669933508 | 0,133 | 0,138 | 1 | CH_SC | 1,590999638 | 0,628535655 | 0,963768116 |
| LTBR2 | 0,971848873 | 0,277154127 | 0,2 | 0,2 | 1 | CH_SC | 1,211802113 | 0,825217244 | 1 |
| PPIL41 | 0,97310115 | 0,343182028 | 0,2 | 0,231 | 1 | CH_SC | 1,268551445 | 0,788300706 | 0,865800866 |
| USP162 | 0,973641817 | -0,397785378 | 0,267 | 0,231 | 1 | CH_SC | 0,759022534 | 1,317483942 | 1,155844156 |
| SPSB1 | 0,974327231 | -0,298740868 | 0,133 | 0,123 | 1 | CH_SC | 0,81296161 | 1,230070384 | 1,081300813 |
| CBX1 | 0,974327231 | -0,452450736 | 0,133 | 0,123 | 1 | CH_SC | 0,730800367 | 1,36836275 | 1,081300813 |
| FOSL22 | 0,975383993 | 0,297426388 | 0,267 | 0,292 | 1 | CH_SC | 1,228950143 | 0,81370266 | 0,914383562 |
| ATP1A13 | 0,977590333 | -0,251510089 | 0,4 | 0,385 | 1 | CH_SC | 0,840016698 | 1,190452526 | 1,038961039 |
| POLR2B | 0,983572595 | 0,661315178 | 0,133 | 0,138 | 1 | CH_SC | 1,581523702 | 0,632301621 | 0,963768116 |
| MGAT5 | 0,983572864 | 0,286138779 | 0,133 | 0,138 | 1 | CH_SC | 1,219372385 | 0,820094019 | 0,963768116 |
| HSDL21 | 0,983572864 | 0,653349479 | 0,133 | 0,138 | 1 | CH_SC | 1,572815538 | 0,635802468 | 0,963768116 |
| CPNE1 | 0,983572864 | 0,289491402 | 0,133 | 0,138 | 1 | CH_SC | 1,222209332 | 0,818190447 | 0,963768116 |
| RPL7L11 | 0,987690532 | 0,25185015 | 0,267 | 0,292 | 1 | CH_SC | 1,190733164 | 0,839818719 | 0,914383562 |
| SRGAP2 | 0,991441097 | -0,292992928 | 0,133 | 0,123 | 1 | CH_SC | 0,816207047 | 1,225179327 | 1,081300813 |
| HSD17B124 | 0,991441097 | -0,395187733 | 0,133 | 0,123 | 1 | CH_SC | 0,760390423 | 1,315113881 | 1,081300813 |
| SP11 | 0,991441097 | -0,371979752 | 0,133 | 0,123 | 1 | CH_SC | 0,772721395 | 1,294127491 | 1,081300813 |
| SMAD42 | 0,991441097 | -0,292910285 | 0,133 | 0,123 | 1 | CH_SC | 0,816253804 | 1,225109146 | 1,081300813 |
| PPM1B1 | 0,99208402 | 0,564701177 | 0,133 | 0,154 | 1 | CH_SC | 1,479081118 | 0,67609544 | 0,863636364 |
| SP32 | 0,99208402 | 0,298602904 | 0,133 | 0,154 | 1 | CH_SC | 1,229952758 | 0,813039357 | 0,863636364 |
| LNPK1 | 0,99208402 | 0,372882729 | 0,133 | 0,154 | 1 | CH_SC | 1,294937734 | 0,772237903 | 0,863636364 |
| PTRH2 | 0,99208402 | 0,805134997 | 0,133 | 0,154 | 1 | CH_SC | 1,747309285 | 0,572308525 | 0,863636364 |
| IQSEC11 | 0,993274093 | -0,28405002 | 0,267 | 0,215 | 1 | CH_SC | 0,821282225 | 1,217608234 | 1,241860465 |
| ARPC56 | 0,993933327 | 0,904820937 | 0,267 | 0,308 | 1 | CH_SC | 1,872312095 | 0,534098991 | 0,866883117 |

| Supplementary Table 4 - clusermarker | | | | | | | | | |
|---|---|---|---|---|---|---|---|---|---|
| | **p_val** | **avg_log2FC** | **pct.1** | **pct.2** | **p_val_adj** | **celltype** | **Foldchange_UP** | **Foldchange_DOWN** | **Ratio_pct1_pct2** |
| GYG11 | 1 | -0,371664236 | 0,133 | 0,123 | 1 | CH_SC | 0,772890406 | 1,293844498 | 1,081300813 |
| MEA13 | 1 | 0,514687305 | 0,267 | 0,292 | 1 | CH_SC | 1,428684448 | 0,699944625 | 0,914383562 |
| GARS12 | 1 | -0,308410946 | 0,2 | 0,185 | 1 | CH_SC | 0,807530723 | 1,238342978 | 1,081081081 |
| STMP13 | 1 | -0,370528788 | 0,2 | 0,185 | 1 | CH_SC | 0,773498936 | 1,292826601 | 1,081081081 |
| PDIA42 | 1 | 0,396794148 | 0,2 | 0,246 | 1 | CH_SC | 1,316579051 | 0,759544213 | 0,81300813 |
| IMPA11 | 1 | -0,388924226 | 0,133 | 0,123 | 1 | CH_SC | 0,763698859 | 1,309416647 | 1,081300813 |
| EBLN3P2 | 1 | 0,414849755 | 0,133 | 0,154 | 1 | CH_SC | 1,333159832 | 0,750097607 | 0,863636364 |
| GOLM12 | 1 | 0,449014772 | 0,133 | 0,154 | 1 | CH_SC | 1,365107695 | 0,732542937 | 0,863636364 |
| TNS2 | 1 | 0,26061973 | 0,2 | 0,231 | 1 | CH_SC | 1,197993209 | 0,834729273 | 0,865800866 |
| CLPX | 1 | 0,318945043 | 0,133 | 0,138 | 1 | CH_SC | 1,247418053 | 0,801655867 | 0,963768116 |
| COX5A8 | 1 | 0,268850698 | 0,267 | 0,292 | 1 | CH_SC | 1,204847621 | 0,829980474 | 0,914383562 |
| PQBP1 | 1 | -0,296320794 | 0,2 | 0,185 | 1 | CH_SC | 0,814326471 | 1,228008711 | 1,081081081 |
| PBDC12 | 1 | 0,277020727 | 0,133 | 0,138 | 1 | CH_SC | 1,211690068 | 0,825293552 | 0,963768116 |
